# SETDB1 Fuels the Lung Cancer Phenotype by Modulating Epigenome, 3D Genome Organization and Chromatin Mechanical Properties

**DOI:** 10.1101/2021.09.06.459062

**Authors:** Vlada V. Zakharova, Mikhail D. Magnitov, Laurence Del-Maestro, Sergey V. Ulianov, Alexandros Glentis, Burhan Ulyanik, Alice Williart, Anna Karpukhina, Oleg Demidov, Veronique Joliot, Yegor S. Vassetzky, René-Marc Mège, Matthieu Piel, Sergey V. Razin, Slimane Ait-Si-Ali

## Abstract

Imbalance in the finely orchestrated system of chromatin-modifying enzymes is a hallmark of many pathologies such as cancers, since causing the affection of the epigenome and transcriptional reprogramming. Here, we demonstrate that a loss-of-function mutation (LOF) of the major histone lysine methyltransferase SETDB1 possessing oncogenic activity in lung cancer cells leads to broad changes in the overall architecture and mechanical properties of the nucleus through genome-wide redistribution of heterochromatin, which perturbs chromatin spatial compartmentalization. Together with the enforced activation of the epithelial expression program, cytoskeleton remodeling, reduced proliferation rate and restricted cellular migration, this leads to the reversed oncogenic potential of lung adenocarcinoma cells. These results emphasize an essential role of chromatin architecture in the determination of oncogenic programs and illustrate a relationship between gene expression, epigenome, 3D genome and nuclear mechanics.

## Introduction

The eukaryotic nucleus is separated from the cytoplasm by the nuclear envelope (NE) supported by a network of intermediate filament proteins referred to as lamins^1^. Inside the nucleus, the genome is packaged and non-randomly organized to ensure efficient gene expression regulation. High-resolution C-techniques revealed that chromatin fibers fold into a hierarchical structure, including loops, topologically associated domains (TADs), and spatially segregated transcriptionally active (A) and repressive (B) compartments^2^. The organized chromatin occupies specific positions relative to the nuclear periphery, with euchromatin shifting away from the nuclear periphery where heterochromatin is enriched^3^. Perturbations in the higher-order chromatin organization and disruption of the peripheral heterochromatin pattern promote instability of the overall nuclear organization and are common for many pathologies, including cancer^4–6^.

The methylation of histone 3 at lysine positions 9 (H3K9) and 27 (H3K27) is essential for the assembly of constitutive and facultative heterochromatin, respectively. H3K27 methylation is mediated by EZH (enhancer-of-zeste) lysine methyltransferases (KMTs)^7^, which belong to the polycomb repressive complex 2 (PRC2), whereas H3K9 is mainly methylated by the KMTs of the SUV39 family composed of SUV39H1/2, G9A, GLP^8, 9^ and SETDB1^10^. Although in some cases these KMTs can substitute for each other, their functions are not redundant. In contrast to the SUV39H KMTs, which are key in the heterochromatin protein 1 (HP1)-dependent assembly of constitutive pericentromeric heterochromatin and heterochromatin tethering to the nuclear lamina^11^, SETDB1 catalyzes mostly H3K9 di- and tri-methylation within the euchromatic regions^10^ to repress gene expression and silence retrotransposons and other repetitive elements^12^. SETDB1 is essential for the early embryo identity, pluripotency, self-renewal, and terminal differentiation of many progenitor cell types^13, 14^.

Imbalance in the functions of epigenetic modifiers can affect the global epigenetic status of the cells, causing widespread cascade consequences. The overexpression of *SETDB1* is a common feature of different epithelial cancers^15–17^. Pro-oncogenic properties of SETDB1 involve the promotion of cell cycle and cell proliferation, migration, and invasive potential of cancer cells^18, 19^.

In addition to H3K9 methylation, SETDB1 might also impact the cancer phenotype by methylation of non-histone proteins, including the tumor suppressor p53 and the kinase AKT^17, 20^.

A decrease in SETDB1 levels remodels heterochromatin patterns^21, 22^ and affects the major characteristics of cancer progression, such as proliferation and migration, in various epithelial cancers^17, 23^. However, only certain issues have been addressed previously, which makes it difficult to draw a comprehensive model of the outcome of SETDB1 inactivation and gain insights into the underlying molecular mechanisms. Here, we analyzed the consequences of SETDB1 inactivation (LOF mutation) in the A549 lung adenocarcinoma cell line. SETDB1 LOF triggers a widespread decrease of H3K9me3 at euchromatic regions with a substantial modulation of the gene expression program, resulting in reduced proliferation and migration of cancer cells. This was accompanied by an unanticipated increase of H3K9me3 in lamina-associated domains (LADs) and overall remodeling of the heterochromatin patterns at the nuclear periphery that led to the enforcement of chromatin compartment spatial segregation and acquisition of non-malignant structural and mechanical properties of the cell nucleus. These results demonstrate an important role of SETDB1 as an organizer of 2D and 3D chromatin architecture, the disbalance of which serves as a trigger for oncogenesis.

## Results

### SETDB1 LOF reduces proliferation and migration velocity of A549 lung adenocarcinoma cells

To impair the H3K9 methylation function of SETDB1 in A549 lung adenocarcinoma cells, we performed CRISPR/Cas9-induced deletion in the catalytic SET domain of SETDB1 (Fig. 1b and Fig. S1a). The A549 cell line with a stable LOF mutation of *SETDB1* (SETDB1^LOF^) did not show an increase in a fraction of apoptotic cells (Fig. S1b). However, SETDB1^LOF^ cells displayed a reduced mitotic index (Fig. S1c) without significant changes in the duration of mitotic phases (Fig. S1c); a decreased proliferation rate (Fig. 1c), and an increased G1/S ratio (55.7/31 % in SETDB1^LOF^ *vs.* 49.8/34.6 % in the control cells; Fig. S1d).

**Figure 1.**
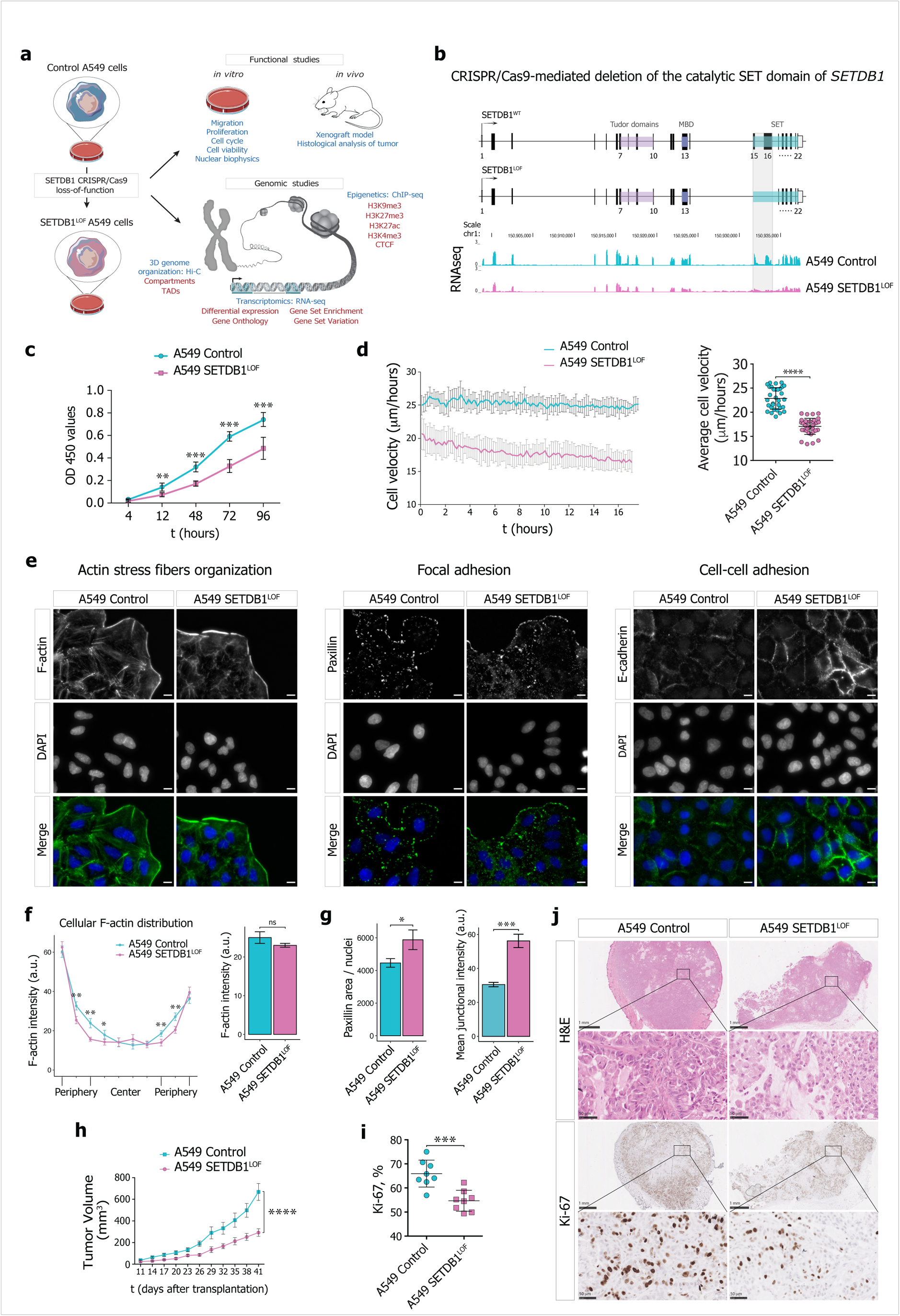
SETDB1 loss-of-function reduces A549 lung adenocarcinoma cell proliferation and migration velocity. a. Overall experimental strategy design.
b. *Upper*: diagram of the strategy used to generate *SETDB1* loss-of-function mutation (SETDB1^LOF^) in A549 cells by CRISPR/Cas9-mediated deletion in the catalytic SET domain (15^th^ and 16^th^ exons). *Lower*: RNA-seq data (CPM-normalized coverage) for *SETDB1* mRNA levels in control (cyan) and SETDB1^LOF^ (pink) A549 cells. Deleted region where the designed guide RNAs (gRNAs) target the 15^th^ (forward gRNA) and the 16^th^ (reverse gRNA) exons of the *SETDB1* gene (see Methods) is highlighted.
c. Cell proliferation curve for control (cyan) and SETDB1^LOF^ (pink) A549 cells at the indicated time points (t, hours) (N ≥ 3 independent experiments; mean ± SEM of 8 wells; **, *P-*value < 0.01; ***, *P*-value < 0.001, Mann-Whitney U-test).
d. Wound-healing assay. *Left*: representative cell velocity measured by wound-healing assay for control (cyan) and SETDB1^LOF^ (pink) A549 cells. *Right*: average cell velocity for control (cyan) and SETDB1^LOF^ (pink) A549 cells. Each point represents a migration rate in one location (N = 3 independent experiments, on average of 32 measurements for the control and 35 measurements for SETDB1^LOF^ cells; mean ± SEM; ****, *P-*value < 0.0001, t-test).
e. Representative immunofluorescence images of control and SETDB1^LOF^ A549 cells stained with phalloidin-TRITC (F-actin), anti-paxillin, or anti-E-cadherin antibodies. DNA was labeled with DAPI. Scale bar, 10 μm.
f. Quantification of F-actin cellular signal. *Left*: representation of F-actin cellular distribution stained with phalloidin-TRITC in control and SETDB1^LOF^ A549 cells; each point represents the mean of the F-actin signal intensity in one of the 10 bins along the cellular axis (N = 3 independent experiments; mean ± SEM, 92-104 of cells analyzed; *, *P-*value < 0.05; **, *P-*value < 0.01, Mann-Whitney U-test). *Right*: quantification of overall F-actin intensity (mean ± SEM; ns, not significant, Mann-Whitney U-test). A corresponding representative image is presented in Fig. 1e.
g. Quantification of paxillin and E-cadherin signals. *Left*: quantification of paxillin area normalized to the number of nuclei. *Right*: quantification of overall E-cadherin intensity at cellular junctions. Corresponding representative images are presented in Fig. 1e (N ≥ 3 independent experiments; mean ± SEM; *, *P-*value < 0.05; ***, *P-*value < 0.001, Mann-Whitney U-test).
h. Volume of tumors in NSG mice injected with control (N = 8 mice, cyan) or SETDB1^LOF^ A549 (N = 7 mice, pink) A549 cells at the indicated post-injection time in days (****, *P-*value < 0.0001, two-way ANOVA test).
i. Percentage of Ki-67-positive nuclei in control (cyan) or SETDB1^LOF^ (pink) A549 tumor sections (mean ± SEM; ***, *P-*value < 0.001, Mann-Whitney U-test). Corresponding representative images are presented in Fig. 1j.
j. Representative images of control and SETDB1^LOF^ A549 tumor sections stained with hematoxylin and eosin (H&E) or with an anti-Ki-67 antibody. Scale bar, 1 mm and 50 μm.

We next asked whether SETDB1 LOF affects cell migration using a live imaging wound-healing assay. To avoid bias by cell proliferation rates, cells were pre-treated with a mitosis-blocking agent Mitomycin C (as described in Methods). SETDB1^LOF^ cells showed a reduced overall displacement (Fig. 1d) and cell detachment (Movie S1), displaying collective migration, whereas some control cells in the leading front exhibited elongated shapes (Fig. S1e and Movie S1), often detached from the cohort (Movie S1), and demonstrated lamellipodia-like migration (Fig. S1e, Movie S1 and Movie S2). Furthermore, F-actin stained with phalloidin-TRITC formed thinner and poorly oriented stress fibers, as compared to control cells (Fig. 1e, 1f and Fig. S1f). The focal adhesion complex, which ensures the correct communication between the cell and extracellular matrix during migration^24^, was also affected in SETDB1^LOF^ cells, as highlighted by a markedly increased number of focal adhesion points revealed by paxillin immunostaining (Fig. 1e, 1g and Fig. S1f). Moreover, SETDB1 LOF promoted the formation of strong E-cadherin-based cell-cell junctions (Fig. 1e, 1g and Fig. S1f). To discriminate the effects of cytoskeleton remodeling and altered cell-cell adhesion in the suppression of cell migration, we impaired cell-cell junctions by chelation of divalent ions with EGTA^25^ and measured the cellular migration velocity. In the presence of 2.5 mM EGTA in the growth media, the SETDB1^LOF^ cells did not show significant changes in the average cellular velocity (Fig. S1g). A progressive decrease in cell velocity was observed at 3 mM EGTA due to the disruption of cell-cell contacts and detachment of cells from the surface. Thus, we conclude that SETDB1 LOF-mediated migration suppression is primarily determined by cytoskeleton remodeling, in particular by F-actin stress fiber reorganization.

To check whether SETDB1 LOF affected cancer cell proliferation *in vivo*, we performed a xenograft assay by subcutaneous injection of control and SETDB1^LOF^ cells into immunocompromised Nod-SCID-Gamma (NSG) mice. Tumor growth of SETDB1^LOF^ cells was significantly delayed compared to the control cells (Fig. 1h). Hematoxylin and Eosin (H&E) staining revealed a decreased cell density in SETDB1^LOF^ xenograft tumors compared to control (Fig. 1j). Apoptosis or necrosis was not microscopically observed upon SETDB1 LOF (Fig. 1j). The cell proliferation antigen Ki-67 immunostaining of xenografts (Fig. 1i, 1j) showed a significantly lower number of Ki-67-positive cells in SETDB1^LOF^ xenografts as compared to the control, indicating that SETDB1 LOF reduced tumor aggressiveness.

Collectively, these data suggest that SETDB1 LOF slows the proliferation and migration of lung adenocarcinoma cells. In agreement with these and previous results^26^, we found that lower *SETDB1* mRNA expression was associated with higher survival of NSCLC patients, as revealed by Kaplan-Meier analysis of publicly available data^27^ (Fig. S1h).

### Global transcriptional changes induced by SETDB1 LOF correlate with malignant phenotype attenuation

To obtain insights into the molecular mechanisms behind the aforementioned SETDB1 LOF-induced cellular phenotype, we performed differential gene expression analysis of RNA-seq data and identified 2848 differentially expressed genes (DEGs), with 1887 up- and 961 down-regulated ones (Fig. 2a and Fig. S2a). Analysis of transcriptomic data revealed the activation of 179 genes encoding KRüppel-associated box (KRAB)-containing zinc finger transcription factors, or KRAB-ZNFs (Fig. S2b), which function as a DNA-binding recognition scaffold for the silencing of endogenous retroviruses through heterochromatin establishment by SETDB1 in association with and its cofactor KRAB-associated protein 1 (KAP1)^12^. Another group of genes that was massively de-repressed upon SETDB1 LOF was the clustered *Protocadherin (PCDH)* genes (Fig. S2b) involved in the regulation of cell-cell adhesion^28^, cell death, and proliferation^29^, and are known to be regulated by SETDB1^30^.

**Figure 2.**
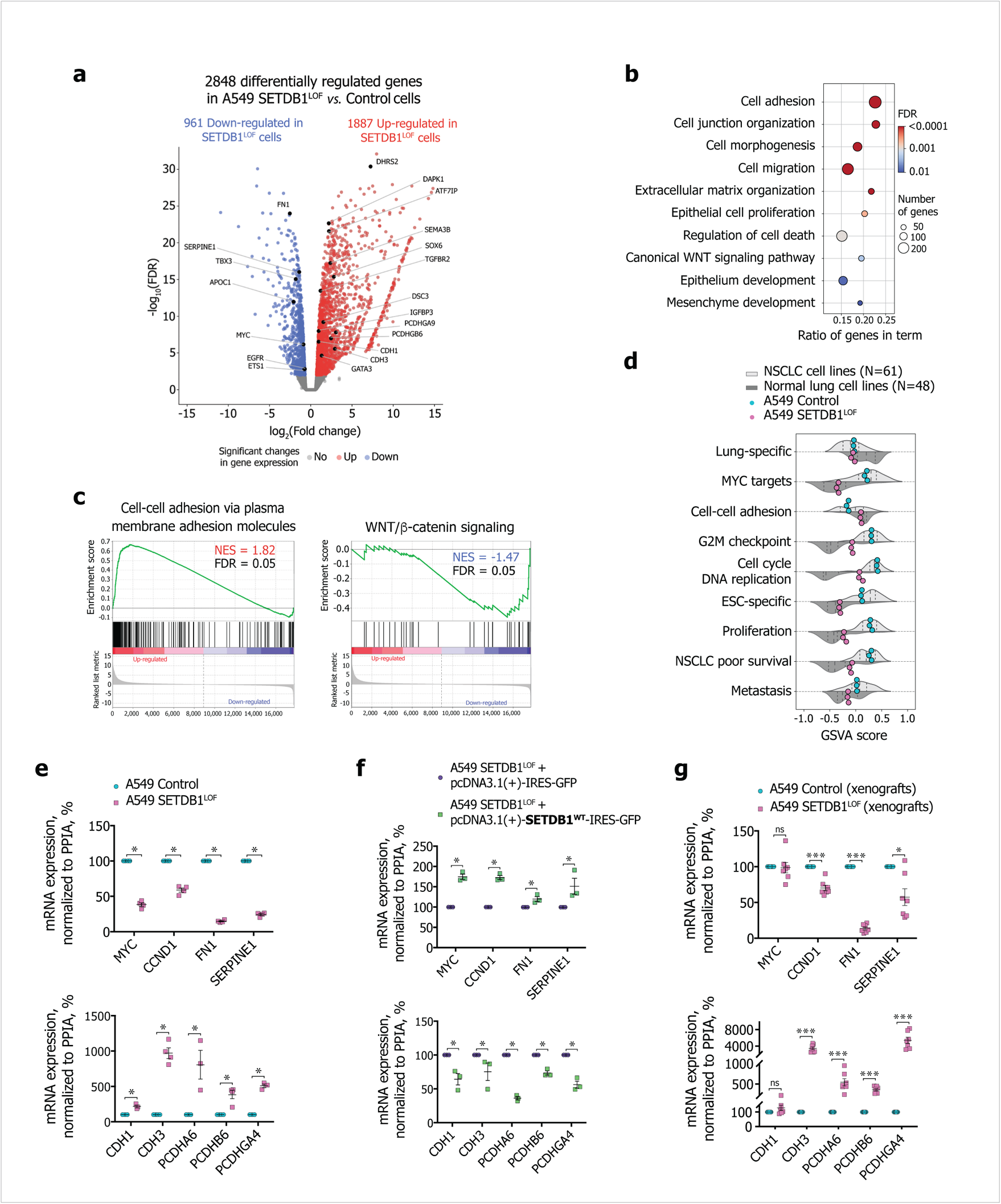
Global transcriptional changes induced by SETDB1 LOF correlate with malignant phenotype attenuation. a. Volcano plot showing significant SETDB1 LOF-induced changes in gene expression in A549 cells (N = 3 independent experiments; FDR < 0.01, t-test relative to a threshold). Up-regulated genes (red): 1887 (log_2_(fold change) > 0). Down-regulated genes (blue): 961 (log_2_(fold change) < 0).
b. Gene ontology terms over-represented in the differentially expressed genes in SETDB1^LOF^ *vs.* control A549 cells.
c. Gene set enrichment analysis in SETDB1^LOF^ *vs.* control A549 cells. Normalized enrichment score (NES) and FDR are indicated.
d. Gene set variation analysis scores for each gene set in the normal lung (N = 48), NSCLC (N = 61), control (cyan), and SETDB1^LOF^ (pink) A549 cells. Dashed vertical lines correspond to the median, 25^th^, and 75^th^ percentiles.
e. mRNA expression of selected mesenchymal and epithelial markers, and genes related to the WNT/β-catenin signaling pathway in control (cyan) and SETDB1^LOF^ (pink) A549 cells. RT-qPCR data were normalized to the *PPIA* mRNA expression values in control A549 cells that were set as 100% (N ≥ 3 independent experiments; mean ± SEM; *, *P-*value < 0.05, Mann-Whitney U-test).
f. Effect of rescued expression of *SETDB1* on mRNA expression of selected mesenchymal and epithelial markers, and genes related to the WNT/β-catenin signaling pathway in A549 SETDB1^LOF^ transfected with pcDNA3.1(+)-IRES-GFP or pcDNA3.1(+)-SETDB1^WT^-IRES-GFP. RT-qPCR data were normalized to the *PPIA* mRNA expression values in A549 SETDB1^LOF^ transfected with pcDNA3.1(+)-IRES-GFP that were set as 100% (N = 3 independent experiments; mean ± SEM; *, *P-*value < 0.05, Mann-Whitney U-test).
g. mRNA expression of selected mesenchymal and epithelial markers, and genes related to the WNT/β-catenin signaling pathway in control (N = 8 mice, cyan) and SETDB1^LOF^ A549 xenografts (N = 7 mice, pink) (see Figure 1h). RT-qPCR data were normalized to the *PPIA* expression values in control A549 xenografts that were set as 100% (mean ± SEM; ns, not significant, ***, *P-*value < 0.001, Mann-Whitney U-test).

Gene ontology (GO) enrichment analysis revealed that DEGs upon SETDB1 LOF were significantly enriched in terms of cell adhesion, migration, proliferation, and epithelium and mesenchyme development (Fig. 2b). Analysis of previously published RNA-seq data from *SETDB1* knockout cells (cervical adenocarcinoma cell line HeLa^21^; mouse postnatal forebrain neurons^30^; retinal pigment epithelial cells RPE1^23^) showed similar GO terms, such as cell adhesion, proliferation, migration, canonical WNT/β-catenin signaling pathway, and cell death regulation (Fig. S2c), thus demostrating that SETDB1 LOF-mediated effects on the transcription program are similar in non-related cancer and normal cell types.

To determine the gene signature transcriptional changes, we applied gene set enrichment analysis (GSEA). We found that in SETDB1^LOF^ cells, genes controlling cell-cell adhesion were significantly up-regulated (Fig. 2c); whereas NSCLC-associated signatures were down-regulated (Fig. 2c and Fig. S2d), such as WNT/β-catenin^31^, mTORC1^32^, and Hedgehog^33^ signaling pathways were down-regulated (Fig. 2c and Fig. S2d); MYC targets and proteins providing the G2M checkpoint breakthrough, including effector cyclin-dependent kinases^34^ were also down-regulated (Fig. 2c and Fig. S2d). Together, these data suggest that SETDB1 LOF attenuates the NSCLC-like phenotype of A549 cells.

To assess the overall trend of gene expression program changes in SETDB1^LOF^ cells, we re-analyzed public transcriptomic data for 48 normal lung and 61 NSCLC cell lines datasets and compared the expression levels of several relevant gene sets using gene set variation analysis (GSVA). The apparent feature of SETDB1^LOF^ cells was a correlation with normal epithelial cell lines across several gene sets, including MYC-targets, cell-cell adhesion, G2M-checkpoint, DNA replication, cell proliferation, embryonic stem cell (ESC)-specific genes, and genes associated with poor survival of NSCLC patients (Fig. 2d).

We confirmed RNA-seq data by RT-qPCR and found a reduction in mRNA levels of mesenchymal genes and genes related to WNT/β-catenin signaling (*MYC*, *CCND1*, *FN1*, *SERPINE1*) and an increase in mRNA levels of epithelial genes (*CDH1*, *CDH3*, *PCDHA6*, *PCDHB6*, *PCDHGA4*) (Fig. 2e). We checked the reproducibility of the observed effects in several SETDB1^LOF^ clones (Fig. S2e). All SETDB1 LOF clones exhibited the same trend, as compared to the controls (Fig. S2f), and this trend was reversed after the rescue of SETDB1 expression in SETDB1^LOF^ cells (Fig. 2f). Finally, we confirmed similar trends for mRNA expression of selected genes on the xenograft tumors from the NSG mice (Fig. 2g). Therefore, we concluded that SETDB1^LOF^ cells (re)acquired features of a normal epithelial phenotype underpinned by the gene expression program changes.

### Global transcriptional changes induced by SETDB1 LOF are linked to genome-wide modulation of enhancer and super-enhancer activities

To elucidate the mechanism of SETDB1 LOF-induced transcriptional changes, we examined the epigenetic status of potential regulatory elements by ChIP-seq assay for H3K27ac and H3K4me3 (Fig. S3a), which mark active enhancers and promoters, respectively. Increased and decreased H3K27ac peaks were predominantly located in intronic and intergenic regions, suggesting a possible association with enhancers (Fig. 3a, 3b). As expected, an increase of H3K27ac levels was accompanied by the up-regulation of associated genes, and *vice versa* (Fig. 3c). Moreover, genes associated with increased H3K27ac regions contributed to the GO terms associated with the epithelial phenotype, such as cell adhesion and cell-cell junction organization (Fig. 3d), whereas genes associated with down-regulated H3K27ac regions were involved in mesenchymal development and epithelial-to-mesenchymal (EMT) transition, cell migration, stress fiber assembly, and other terms, typical for cancer progression (Fig. 3d).

**Figure 3.**
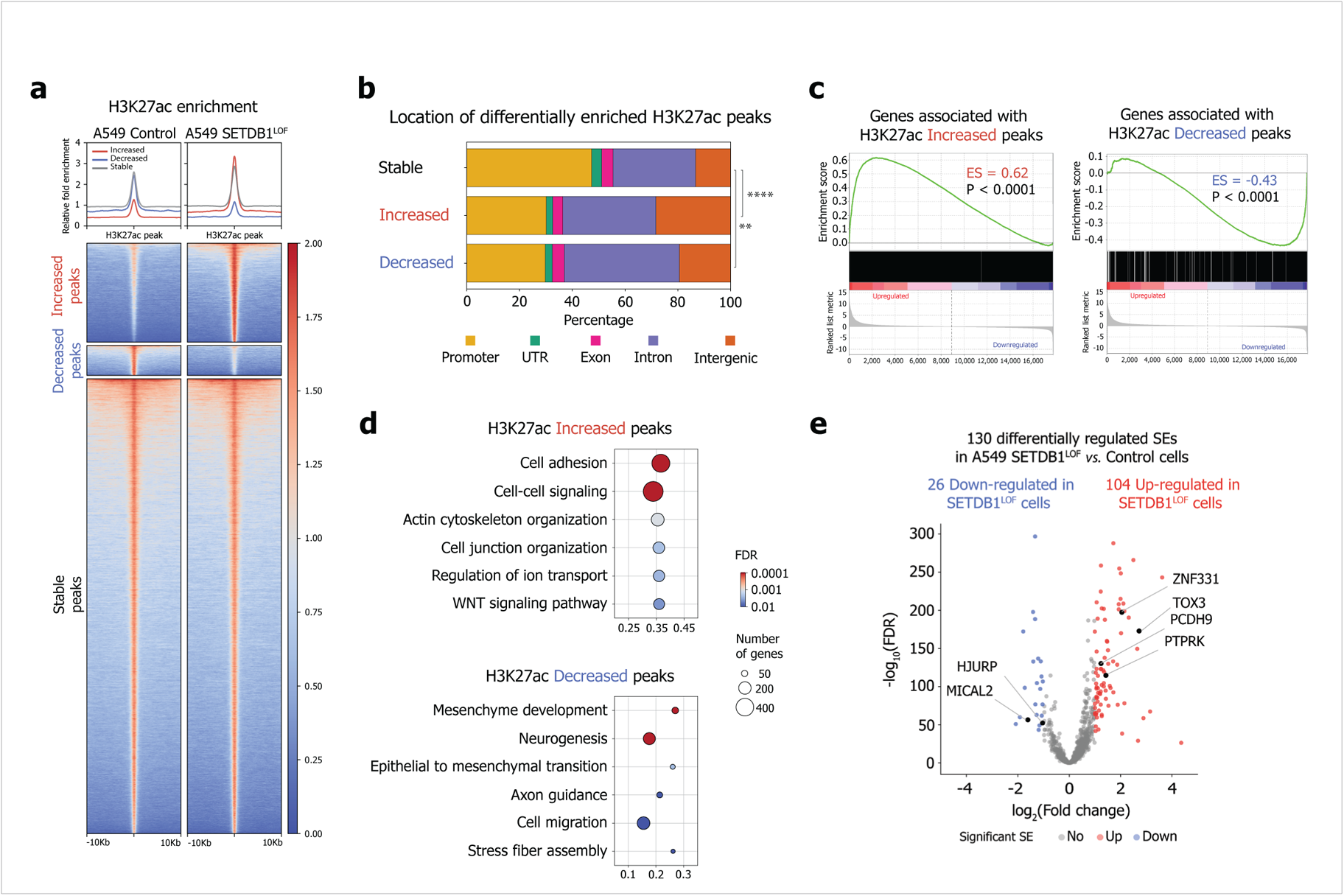
Global transcriptional changes induced by SETDB1 LOF are linked to genome-wide modulation of enhancer activity. a. Pile-ups of the H3K27ac enrichment at the differentially enriched H3K27ac peaks in control and SETDB1^LOF^ A549 cells.
b. Genomic locations of the differentially enriched H3K27ac peaks (**, *P-*value < 0.01; ****, *P-*value < 0.0001, chi-squared test).
c. Gene expression changes associated with increased and decreased H3K27ac enrichments revealed by gene set enrichment analysis in SETDB1^LOF^ *vs.* control A549 cells. Enrichment score (ES) and *P-*values are indicated.
d. Gene ontology terms overrepresented in genes associated with differentially enriched H3K27ac peaks.
e. Volcano plot showing differential (FDR < 0.01, absolute log_2_(fold change) > 1, Wald test) super-enhancers (SEs) in control and SETDB1^LOF^ A549 cells (N = 2 independent experiments). Increased SEs (red): 104; decreased SEs (blue): 26. Genes associated with SEs linked to NSCLC progression are indicated.

Since super-enhancers (SEs) are responsible for cell identity and play an important role in oncogenesis^35^, we asked whether SETDB1 LOF affected SEs genome-wide (i.e. those with an altered level of H3K27ac). First, to distinguish SEs from typical enhancers, we used the rank ordering of SE (ROSE) approach^36^ and annotated 832 and 883 SEs in the control and SETDB1^LOF^ cells (Fig. S3b and S3c). Next, we assessed the H3K27ac level at SEs and identified 130 differentially regulated SEs (Fig. 3e). Among 104 SEs up-regulated in SETDB1^LOF^ cells, we found SEs associated with *ZNF331, TOX3, PCDH9*, *PTPRK*, and *LIMA1* genes involved in the suppression of EMT, cell proliferation, cell migration, invasion, and metastasis^37–41^ (Fig. 3e and Fig. S3d). Among the 26 SEs down-regulated in SETDB1^LOF^, we found those associated with *MICAL2*, *RSPO3*, and *HJURP* genes involved in cancer progression^42–44^ (Fig. 3e and Fig. S3d). Finally, we applied HOMER^45^ to identify transcription factor (TF) binding motifs enriched in ATAC-seq peaks within differentially regulated SEs (ENCODE ATAC-seq for A549 cell line, as described in Methods) (Fig. S3e). We found that motifs of TFs involved in the regulation of proliferation, migration, and invasion of lung cancer cells, including FOSL1, ATF3, BATF, AP-1, SIX4, and NFE2L2 were enriched in differential SEs (Fig. S3e), denoting a potential role of these factors in the observed phenotype. Together, these data suggest that SETDB1 LOF-induced changes in the transcriptional pattern of A549 cells are mediated by the modulation of enhancer and SE activities.

### SETDB1 LOF induces H3K9me3 and H3K27me3 redistribution within the nuclear space and along the genome

To investigate how SETDB1 LOF affects heterochromatin profiles, we first assayed the total abundance of H3K9me3 and H3K27me3 marks by western blot analysis. A moderate, but significant decrease of both marks was observed in SETDB1^LOF^ cells (on average, by 25% for H3K9me3 and by 10% for H3K27me3; Fig. S4a). Immunofluorescence imaging showed an accumulation of H3K9me3 at the nuclear periphery in SETDB1^LOF^ nuclei (Fig. 4a). To quantitatively estimate the H3K9me3 shifting, we calculated the distribution of the immunofluorescence signal intensity along the nuclear radius. In control cells, H3K9me3 signal monotonically radially increased (Fig. 4b), whereas, in SETDB1^LOF^ cells, the distribution of H3K9me3 had a pronounced peak at the nuclear edge, co-localizing with the nuclear lamina as revealed by anti-Lamin-A/C staining (Fig. 4a, 4b and Fig. S4b, S4c). Notably, in normal human bronchial epithelial (NHBE) cells, H3K9me3 distribution demonstrated a pattern similar to that in SETDB1^LOF^ cells, with a well-defined peak at the nuclear lamina (Fig. S4b, S4c). Concerning the H3K27me3 pattern, we observed an opposite and less pronounced trend, with a slight shifting of H3K27me3 signal intensity from the nuclear lamina towards the nuclear interior in SETDB1^LOF^ cells, without detectable changes in the curve shape (Fig. 4a, 4b).

**Figure 4.**
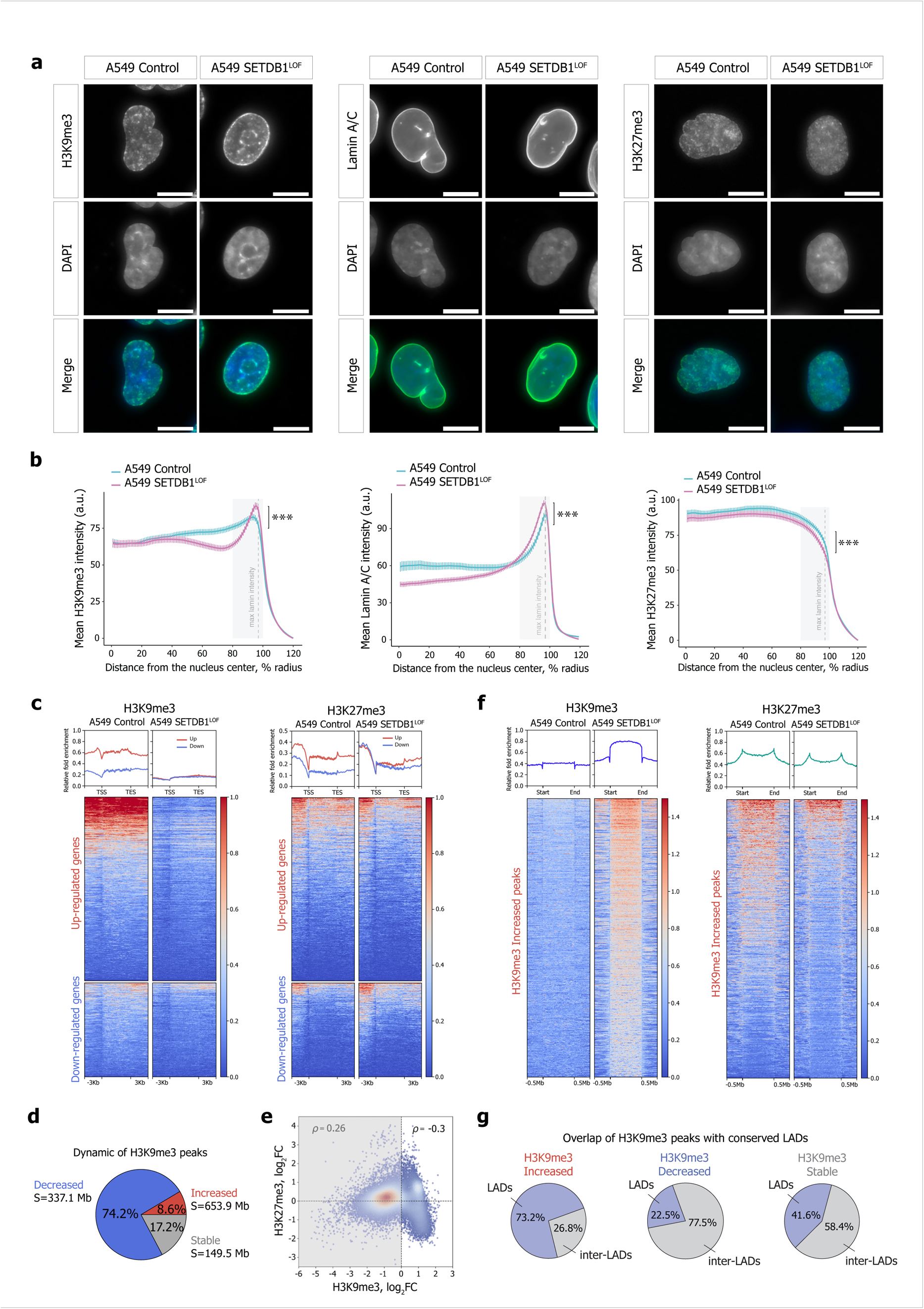
SETDB1 LOF induces H3K9me3 and H3K27me3 redistribution within the nuclear space and along the genome. a. Representative immunofluorescence images of control and SETDB1^LOF^ A549 cells stained with anti-H3K9me3, anti-Lamin A/C, and anti-H3K27me3 antibodies. DNA was labeled with DAPI. Scale bar, 10 μm.
b. Nuclear distribution of H3K9me3, Lamin A/C, and H3K27me3 in control (cyan) and SETDB1^LOF^ (cyan) A549 cells. Intensities are plotted as a function of distance from the nuclear center (N = 3 independent experiments; mean intensity ± SEM; 92-104 of cells analyzed; ***, *P-*value < 0.001, Kolmogorov-Smirnov test). The region with maximum lamin intensity is indicated with a grey dashed line.
c. Pile-ups of the H3K9me3 and H3K27me3 enrichments at the differentially expressed genes in control and SETDB1^LOF^ A549 cells. Up- and down-regulated genes are shown in red and blue, respectively.
d. Pie chart of the genome-wide dynamics of H3K9me3-enriched regions in SETDB1^LOF^ *vs.* control A549 cells. The perscentages indicate the fraction of changed H3K9me3 peaks from the total peaks number. S – the total length of changed H3K9me3 regions.
e. Correlation between H3K27me3 and H3K9me3 enrichment in 100-kb genomic bins in control and SETDB1^LOF^ A549 cells. Spearman’s correlation coefficients are indicated for bins with increased and decreased H3K9me3 levels separately.
f. Pile-ups of the H3K9me3 and H3K27me3 enrichment at increased H3K9me3 regions in control and SETDB1^LOF^ A549 cells.
g. Pie charts of the overlap between increased, decreased, and stable H3K9me3 enriched regions with constitutive LADs from Lenain et al. (2017).

Next, to characterize genome-wide the SETDB1 LOF-driven changes in the distributions of constitutive and facultative heterochromatin, we profiled H3K9me3 and H3K27me3 marks using ChIP-seq. In SETDB1^LOF^ cells we observed dramatic changes in H3K9me3 and H3K27me3 peak positions (Fig. S4d) concomitant to an overall alteration in levels of these marks across large segments of the genome, as compared to the control cells (Fig. S4f). In particular, H3K9me3 levels were substantially reduced along all genes, including the down-regulated ones (Fig. 4c and Fig. S4e). In contrast, H3K27me3 levels decreased and increased at up-regulated and down-regulated genes, respectively (Fig. 4c and Fig. S4e). Hence, we suggest that the up-regulation of gene expression upon SETDB1 LOF is associated with the loss of both constitutive and facultative heterochromatin, whereas transcription repression is not linked to the H3K9me3 (Fig. S4g), but appears to be mediated by other mechanisms, including Polycomb repressive^46^.

Surprisingly, the overall decrease in H3K9me3 levels evidenced by western blot in SETDB1^LOF^ cells (Fig. S4a) was accompanied by an increase in H3K9me3 levels at specific genomic regions (Fig. 4d, 4f and Fig. S4f), with a simultaneous decrease in H3K27me3 levels (Fig. 4e, 4f and Fig. S4f), as revealed by differential enrichment analysis. Visual inspection of the ChIP-seq profiles in the genome browser revealed that these regions are typically gene-poor, heterochromatin rich (Fig. S4f), and relatively long (up to 7.4 Mb in length, 202 kb on average, Fig. S4h). The majority of regions with increased H3K9me3 in SETDB1^LOF^ overlapped with constitutive LADs (cLADs) from Lenain et al. (2017) that are conserved in several non-related cell types, whereas regions with decreased H3K9me3 coincided with inter-LADs (Fig. 4g and Fig. S4i, S4k). These ChIP-seq results were validated by ChIP-qPCR for several genes, confirming a significant increase in H3K9me3 occupancy in cLAD regions in SETDB1^LOF^ cells (Fig. S4j). The predominant location of regions with increased H3K9me3 (Fig. S4k) and decreased H3K27me3 (Fig. S4l) levels within LADs was also confirmed by the comparison with LAD positions in fibroblast cell lines: IMR90, TIG-3, and NHDF. Moreover, analysis of published H3K9me3 ChIP-seq data in HeLa and RPE1 cells after *SETDB1* knockout^21, 23, 47^ showed an increase in H3K9me3 occupancy at constitutive LADs, as compared to the corresponding control cells (Fig. S4m), confirming that the observed increase in H3K9me3 upon SETDB1 LOF is not an artifact and not cell-type specific.

The enrichment in H3K9me3 at LADs upon SETDB1 LOF could be established by other H3K9-specific methyltransferases, SUV39H1, which co-exists with SETDB1 in the common complex(es)^48^, and/or its homolog SUV39H2, associated with H3K9me3 establishment in lamina-associated chromatin^49^. Indeed, we found that the knockdown of both *SUV39H1* and *SUV39H2* in SETDB1^LOF^ cells abolished the H3K9me3 enrichment at cLADs at several tested loci (Fig. S4n).

Collectively, these results demonstrate that SETDB1 LOF triggers a widespread decrease in H3K9me3 at euchromatic regions, accompanied by a SUV39H1/H2-mediated increase in H3K9me3 within LADs that is coupled with a slight decrease in H3K27me3.

### SETDB1 LOF leads to H3K9me3 redistribution between A/B compartments and affects both large-scale and local chromatin topology

To test whether the SETDB1 LOF-induced global heterochromatin re-distribution along the genome and in the nuclear space affected the 3D genome organization, we performed *in situ* Hi-C and generated contact maps at a resolution of up to 20 kb. The dependence of relative contact probability on the genomic distance between pairs of loci was essentially the same in SETDB1^LOF^ and control cells (Fig. S5a). Thus, SETDB1 LOF did not affect the general properties of the 3D genome. At the same time, visual inspection of heatmaps revealed that SETDB1 LOF substantially disturbed contact profiles at a subset of genome loci, as manifested by changes in the “plaid pattern”^50^, reflecting genome compartmentalization changes (Fig. 5a). To analyze the chromatin contact profile systematically, we annotated A (active) and B (inactive) chromatin compartments using principal component analysis (PCA)^50^ (Fig. 5a) and used publicly available profiles of histone marks to validate compartment segmentation (Fig. S5b). The first principal component (PC1) was highly correlated between SETDB1^LOF^ and control cells (Pearson’s *r* = 0.89; Fig. 5b and Fig. S5c), and the proportion of A and B compartments was similar in the two cell lines (Fig. S5d), suggesting that the overall compartmentalization was preserved upon SETDB1 LOF. Nonetheless, we found that 5.5% (N = 1454) and 3.8% (N = 1009) of 100-kb genomic bins switched from B to A and from A to B compartment, respectively (Fig. 5b, 5c).

**Figure 5.**
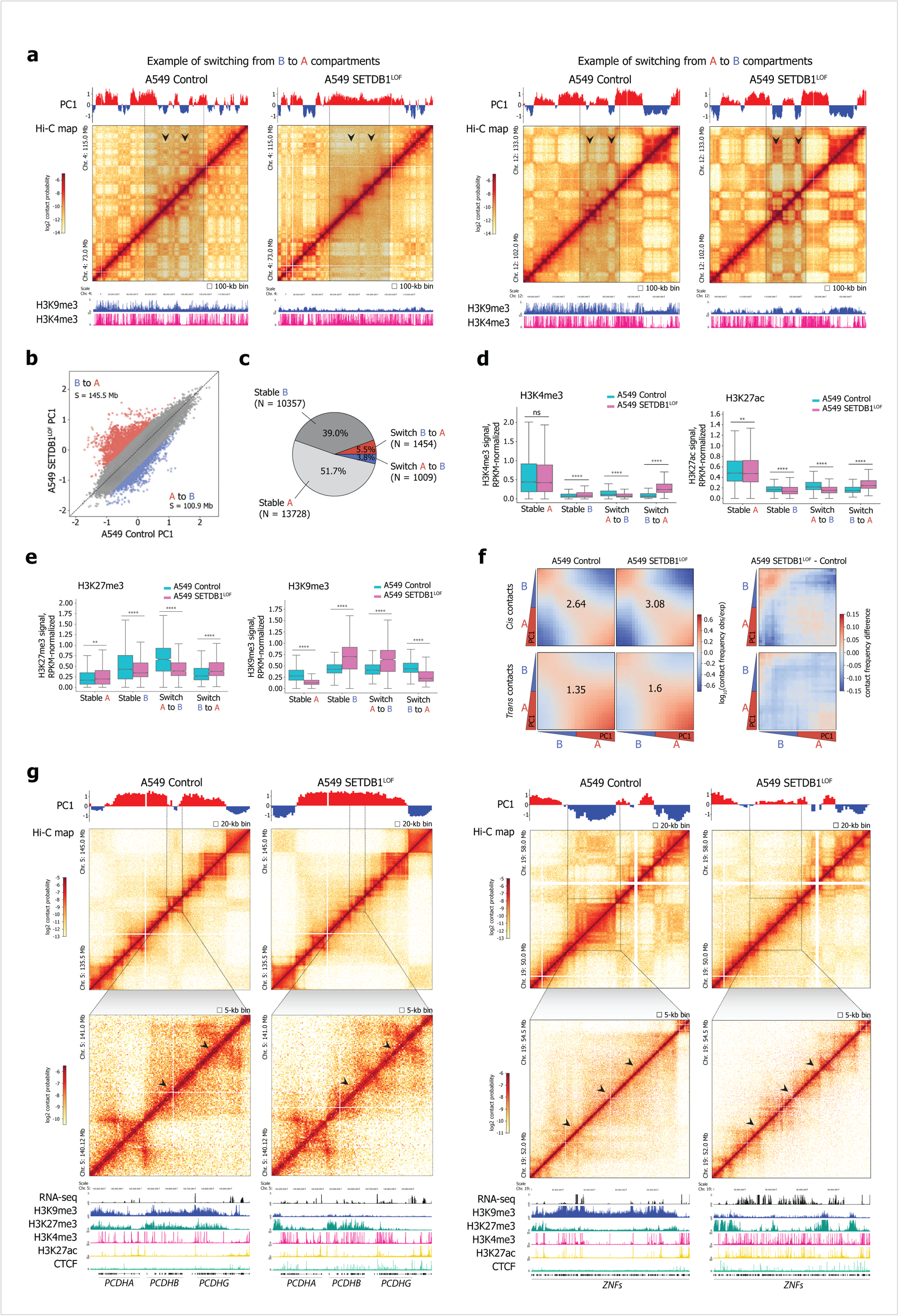
SETDB1 LOF leads to H3K9me3 redistribution between A/B compartments and affects both large-scale and local chromatin topology. a. Hi-C interaction matrices at 100-kb resolution showing examples of chromatin compartment switches in control and SETDB1^LOF^ A549 cells for Chr. 4: 73.0 – 115.0 Mb (left panel) and Chr. 12: 102.0 – 133.0 Mb (right panel). Switching regions are highlighted with grey boxes and black arrows. Eigenvector (PC1 track) for compartments A (red) and B (blue) is shown. *Lower panels:* ChIP-seq profiles (RPKM- and input-normalized ratio) for H3K9me3 and H3K4me3 in control and SETDB1^LOF^ A549 cells are shown according to the UCSC genome browser.
b. Scatter plot of PC1 values at 100-kb resolution for the control and SETDB1^LOF^ A549 cells. S – total length of the regions that switched compartment state (absolute z-score of ΔPC1 > 1.645) from B to A (red) or A to B (blue).
c. Pie chart showing the dynamics of compartment bins in SETDB1^LOF^ *vs.* control A549 cells. N – number of 100-kb bins.
d. Distribution of RPKM-normalized H3K4me3 and H3K27ac ChIP-seq signals within stable A and B compartment bins and in switched from B to A, or A to B compartment bins in control (cyan) and SETDB1^LOF^ (pink) A549 cell lines (ns, not significant; **, *P-*value < 0.01; ****, *P-*value < 0.0001, Mann-Whitney U-test).
e. Distribution of RPKM-normalized H3K27me3 and H3K9me3 ChIP-seq signals within stable A and B compartment bins and in switched from B to A, or A to B compartment bins in control (cyan) and SETDB1^LOF^ (pink) A549 cell lines (**, *P-*value < 0.01, ****, *P-*value < 0.0001, Mann-Whitney U-test).
f. *Left:* saddle plots of contact enrichments in *cis* and in *trans* between 100-kb genomic bins belonging to A and B compartments in control and SETDB1^LOF^ A549 cells. Numbers represent the compartment strength values calculated as described in Nora et al. (2017). *Right*: subtraction of A549 SETDB1^LOF^ from control saddle plots for *cis* and *trans* contacts.
g. Hi-C interaction matrices at 20- (top) and 5-kb (bottom) resolutions showing the *PCDH* gene cluster on Chr. 5: 135.5 – 145.0 Mb (left panels) and *ZNF* genes on Chr. 19: 50.0 – 58.0 Mb (right panels). Strengthened TAD boundaries in the SETDB1^LOF^ as compared to control A549 cells are highlighted with black arrows. Eigenvector (PC1 track) for compartments A and B is shown in red and blue, respectively. Genes, RNA-seq profile (CPM-normalized coverage), ChIP-seq profiles (RPKM- and input-normalized ratio) for H3K9me3, H3K27me3, H3K4me3, H3K27ac, and CTCF in control and SETDB1^LOF^ A549 cells are shown according to the UCSC genome browser.

To further analyze these compartment changes, we divided all 100-kb genomic bins into four groups according to their compartment state (Fig. 5c): bins belonging to A and to B compartments in both cell lines (“stable A” and “stable B”, respectively), and bins that switched their compartment state in SETDB1^LOF^ (“A-to-B” and “B-to-A”). As expected, the H3K4me3 and H3K27ac histone marks and transcription were substantially enriched within “stable A” bins, as compared to “stable B” bins; the decrease and increase of these marks correlated with the “A-to-B” and “B-to-A” switched bins, respectively (Fig. 5d and Fig. S5e). The changes in the H3K27me3 mark in the switched bins mirrored those of the active chromatin (Fig. 5e), probably due to colocalization of H3K4me3 and H3K27me3 within the so-called bivalent domains^51^. However, we observed a remarkably distinct trend for the H3K9me3 mark. In control cells, in contrast to the active marks displaying high differences between compartments, H3K9me3 abundance was only 1.4-fold enriched in the B as compared to the A compartment (Fig. 5e). In SETDB1^LOF^ cells, H3K9me3 increased in “stable B” and in “A-to-B” bins and decreased in the two other bin groups, resulting in a 4.2-fold overall enrichment of H3K9me3 in the B *vs.* A compartment. This finding agrees with the above-described increase of this mark in LADs (Fig. 4g), which mostly reside in the B compartment. As a result, we observed a gain of spatial contacts within the B compartment in *cis* and a decrease of inter-compartment interactions in SETDB1^LOF^ cells, suggesting an increased density of the B compartment folding and further segregation of compartments within the nuclear space (Fig. 5f). Consequently, the compartment strength was augmented upon SETDB1 LOF (Fig. S5f), in agreement with previously published Hi-C data from mouse postnatal forebrain neurons after *SETDB1* knockout^30^ (Fig. S5g).

Notably, the aforementioned changes in compartmentalization upon SETDB1 LOF displayed a remarkably stronger correlation with differential enrichment in H3K4me3 (Spearman’s *ρ* = 0.5), H3K27ac (Spearman’s *ρ* = 0.41), and H3K9me3 (Spearman’s *ρ* = −0.33) marks, as compared to changes in transcription level (Spearman’s *ρ* = 0.24; Fig. S5h, S5i). Thus, we suggest that SETDB1 LOF-induced shifts in compartment state and chromatin compaction are mostly driven by changes in the epigenetic profiles.

Next, we asked whether SETDB1 LOF impacted the organization of TADs, and found that TAD boundary positions were largely unaffected in SETDB1^LOF^ cells (Fig. S5j). A minor increase of TAD boundary insulation (Fig. S5k, S5l) and no changes in intra-TAD interactions (Fig. S5m) were detected. In parallel, we observed substantial alterations in the TAD profiles within several loci. One such example is the *PCDH* locus, containing 53 genes within three clusters: *PCDH-alpha (PCDHA), PCDH-beta (PCDHB)*, and *PCDH-gamma (PCDHG)* (Fig. 5g). In A549 control cells and many other cancer cells, the major part of the *PCDH* locus is transcriptionally repressed and marked with H3K9me3^29^. SETDB1 LOF resulted in a global reactivation of the *PCDH* locus (Fig. S2b), with a concomitant decrease in H3K9me3 and an increase in active histone marks (Fig. 5g). This change was coupled with the partition of the entire *PCDH* locus into several contact domains, the boundaries of which demarcated multi-gene clusters from each other (Fig. 5g). Interestingly, the up-regulation of the *PCDH* locus was correlated with the emergence of numerous CTCF ChIP-seq peaks within the locus, and the majority of them did not coincide with the positions of new TAD boundaries in SETDB1^LOF^ cells (Fig. 5g). An increase in CTCF binding strength (Fig. S5n) and in the number of CTCF-occupied sites (including promoters (Fig. S5q) and TAD boundaries (Fig. S5r)) without an increase in the total CTCF protein level (Fig. S5n, S5o, S5p) turned out to be a general trend in SETDB1 LOF condition. This is in agreement with published data, demonstrating that SETDB1 is required for shielding the genome from excess CTCF binding^30^.

Another example of TAD profile changes upon SETDB1 LOF is a *ZNF* locus on chromosome 19, which contains numerous *KRAB-ZNF* genes, including the tumor suppressor *ZNF331*^52, 53^ (Fig. 5g). Here, the entire region, located in a large virtually unstructured TAD in the control cells, was partitioned in SETDB1^LOF^ cells into several TADs with well-defined boundaries. Similar to the *PCDH* locus, this region lost H3K9me3 methylation (Fig. 5g) and was transcriptionally up-regulated (Fig. S2b) upon SETDB1 LOF. Thus, local changes in TAD profiles upon SETDB1 LOF occur within specific multi-gene loci, where alterations in H3K9me3 and active epigenetic marks coincide with large-scale perturbations of the transcription profile.

### SETDB1 LOF changes A549 cell nuclei shape and mechanical properties

To understand how changes in the epigenome and 3D genome organization upon SETDB1 LOF contribute to suppression of the malignant phenotype of A549 cells, we studied characteristics of nuclei and chromatin, which potentially reflect mechanical properties and were previously shown to influence cell migration^54, 55^. Examination under an optical microscope revealed the differences in the nuclear shape between the control and SETDB1^LOF^ cells (Fig. 4a, 6a and Fig. S4b), manifested by more invaginations and protrusions in the control cells. We assessed the overall nuclear shape by quantification of the nuclear irregularity index and found that SETDB1^LOF^ cells have an average nuclear irregularity index of 0.029, which perfectly matches that of normal primary NHBE cells (Fig. 6a, 6b). In contrast, A549 control cells have an abnormal nuclear morphology, with a significantly higher irregularity index (0.038; Fig. 6a, 6b). Correspondingly, the circularity index and the average nucleus area (Fig. 6b) of SETDB1^LOF^ cells were more similar to NHBE, rather than to the control A549.

**Figure 6.**
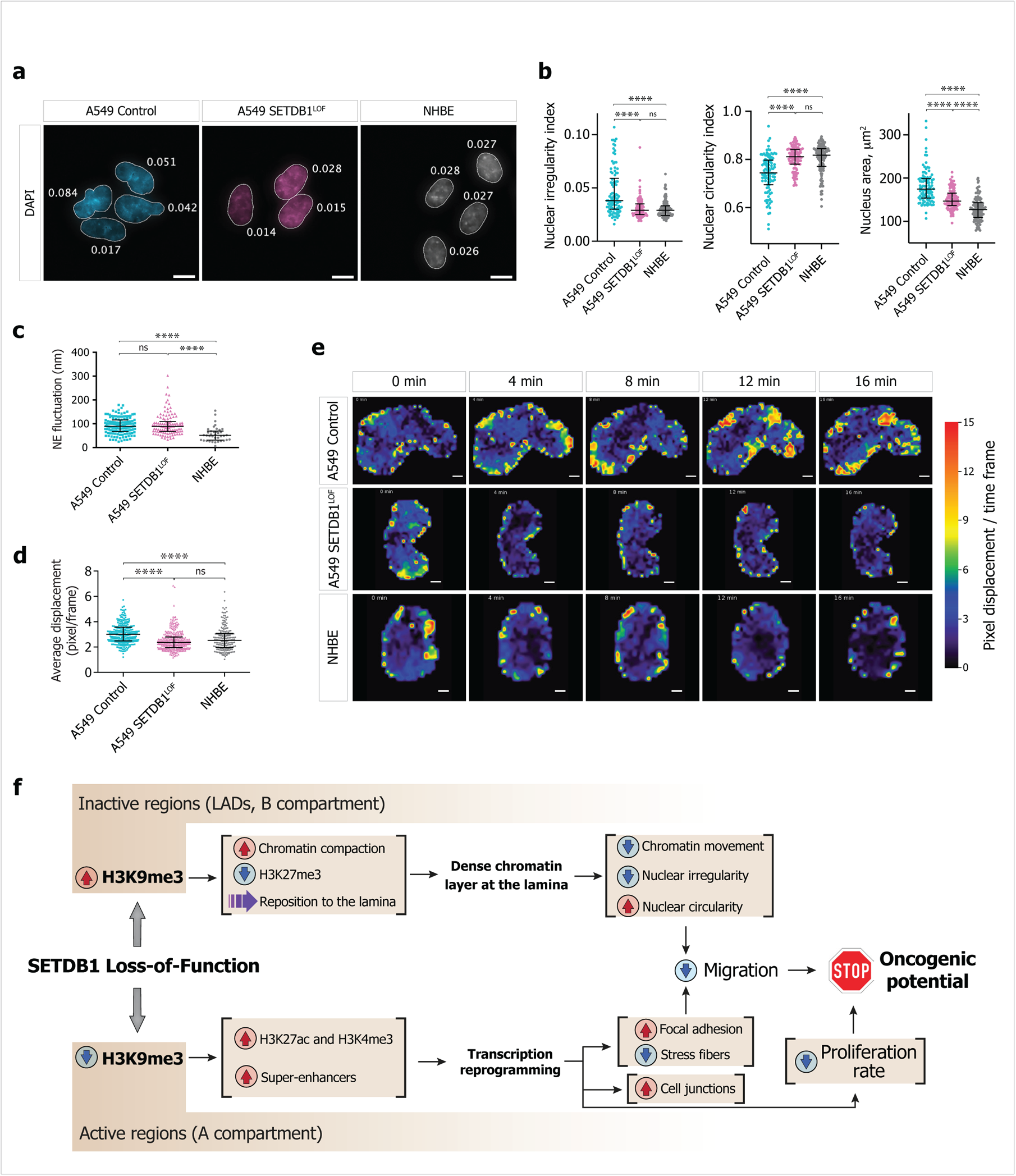
SETDB1 LOF changes the shape and mechanical properties of the A549 cell nuclei. a. Representative DAPI-stained nuclei of control (cyan) and SETDB1^LOF^ A549 (pink), and NHBE (grey) cells showing different nuclear morphology. Numbers denote the nuclear irregularity index of the corresponding nucleus. Bar, 10 μm.
b. Quantification of irregularity index (left panel), circularity index (middle panel), and nucleus area (right panel) for control (cyan), SETDB1^LOF^ A549 (pink), and NHBE cells (grey). Each point represents the measurement for individual nuclei, horizontal lines correspond to median, error bars represent the interquartile range (ns, not significant; ****, *P-*value < 0.0001, Kruskall-Wallis test with Dunn’s correction for multiple comparisons).
c. Quantification of NE fluctuations for control (cyan) and SETDB1^LOF^ (pink) A549, and NHBE cells (grey). Each point represents the average for each individual cell on 2 to 4 kymographs of the square root of the NE mean square displacement relative to its mean position; horizontal lines correspond to the median, error bars represent the interquartile range (ns, not significant; ****, *P-*value < 0.0001, one-way ANOVA-Kruskal-Wallis test).
d. Quantification of DNA displacement for control (cyan) and SETDB1^LOF^ (pink) A549, and NHBE nuclei (grey). Each point represents the average for each individual cell on 10 timeframes of the displacement between two consecutive frames; horizontal lines correspond to the median, error bars represent the interquartile range (ns, not significant; ****, *P-*value < 0.0001, one-way ANOVA-Kruskal-Wallis test).
e. Representative particle image velocimetry (PIV) of DNA within nuclei in control and SETDB1^LOF^ A549, and NHBE cells showing the overall flow of DNA displacement. Scale bar, 5 μm.
f. Graphical representation of the SETDB1 LOF-induced effects in lung adenocarcinoma cells.

The nuclear shape is primarily determined by the nuclear envelope’s (NE) mechanical properties, consisting of the nuclear membrane and nuclear lamina^56^. Therefore, we assayed the NE dynamics *in vivo* by measuring the NE fluctuations constrained by the cytoskeleton from the cytoplasm on one side, and chromatin and lamins from the nucleoplasm, on the other side. Surprisingly, we found that the NE fluctuation magnitude was the same between control and SETDB1^LOF^ cells, and, on average, two-fold higher than in the normal NHBE cells (Fig. 6c). Another factor influencing the overall mechanical properties of the nucleus is the state of the bulk chromatin, particularly the constitutive heterochromatin positioned near the nuclear lamina^55^. Thus, to follow the spatiotemporal dynamics of chromatin, we measured the chromatin movement by *in vivo* imaging of Hoechst 33342-stained nuclei. Particle image velocimetry (PIV) analysis showed that the average chromatin displacement significantly decreased from 3.044 pixels displacement/time frame in the control cells to 2.659 in SETDB1^LOF^ cells, which was similar to that in NHBE cells (Fig. 6d). Moreover, the observed decrease of chromatin mobility was more prominent at the nuclear periphery (Fig. 6e). Together with the above-described H3K9me3-marked heterochromatin redistribution towards the nuclear periphery and strengthening of the B compartments, these results show an increase in the density of tightly compacted chromatin at the periphery upon SETDB1 LOF. This increase could make nuclei more robust, preventing changes in their shape and restricting cell mobility.

## Discussion

Here, we sought to disambiguate the multiple roles of SETDB1 in cancer and their consequences on various scales, which fuel the oncogenic process. To this end, we have studied a plethora of effects triggered by SETDB1 loss of function in NSCLC cells, ranging from gene expression, and epigenome analyses, the 3D genome organization, and the mechanical properties of chromatin and nuclei, to xenograft tumor assays.

SETDB1 target loci are predominantly located within euchromatin gene-rich chromosome segments, where SETDB1 is involved in gene repression and silencing of repetitive elements^57^. We suppose that the decrease of H3K9me3 level at these loci is the first outcome of SETDB1 LOF, preceding and then directly or indirectly promoting all the other observed effects (proposed model is described in Fig. 6f). Writing and erasing epigenetic enzyme complexes for both activating and repressive histone modifications typically compete within these genomic regions in determining their epigenetic state^58, 59^. Our observations suggest that SETDB1 LOF shifts the equilibrium to the benefit of histone lysine acetyltransferases in hundreds of loci genome-wide. This shift results in a drastic increase of H3K27 acetylation responsible for an apparent activation of numerous super-enhancers, and an overall transcriptional reprogramming leading to massive up-regulation of gene repertoire underpinning the reacquired normal epithelial phenotype of SETDB1^LOF^ cells. These changes, together with increased focal adhesion and stress fiber reorganization, restrict cell migration.

Surprisingly, we have also found that SETDB1 LOF is followed by a SUV39H1/H2-dependent increase of H3K9me3 levels within LADs. Since H3K9me3 is an anchor for chromatin attachment to the lamina through the recruitment of HP1-beta interacting with the Lamin B receptor^60–62^, the increase in H3K9me3 levels could be responsible for the observed deposition of H3K9me3-marked chromatin at the nuclear periphery. H3K9me3 occupancy positively correlates with chromatin compaction driven by the oligomerization of H3K9me3 binding HP1^63–65^ and the formation of liquid condensates^66, 67^. Moreover, attachment to the lamina *per se* is sufficient for chromatin compaction^68^. These facts could explain the increased density of chromatin packaging in the B compartment, which is enriched in LADs, upon SETDB1 LOF, and tighter H3K9me3-driven association of LADs with the nuclear lamina, as revealed by immunofluorescence analysis. Thus, in SETDB1^LOF^ cells, tightly packed H3K9me3-marked chromatin forms a clearly visible layer adjacent to the nuclear lamina, similarly to non-cancer cells. The highly compact state and low mobility of these chromatin regions imply that they could serve as a tough carcass for the NE. The presence of this layer is associated with a more spherical shape of the NE, although it does not affect its fluctuations. Since the ability of nuclei to change their shape is an essential property of migrating cancer cells, suppression of this ability upon SETDB1 LOF, ensured by the tightly packed heterochromatin located at the lamina, may contribute to the reduction of cell migration, and, along with the decreased proliferation rate, restrict their oncogenic potential.

Taken together, our findings demonstrate that the loss of H3K9 methylation activity of SETDB1 reshapes the epigenetic profiles within both active and inactive chromatin. Ultimately, this causes perturbations in the spatial genome organization, affecting both the gene expression patterns and mechanical properties of the cell nuclei. Collectively, these changes result in the suppression of the lung adenocarcinoma cell’s malignant phenotype.

## Methods

### Cell lines

A549 human male lung carcinoma cells (CCL-185) were obtained from ATCC and cultured in Dulbecco’s modified Eagle Medium DMEM (Gibco; Cat#: 31966-021) supplemented with 10% Fœtal Bovine Serum (FBS; Gibco; Cat#: 10270-106) and 1% penicillin/streptomycin (Gibco; Cat#: 15140-122). Normal human bronchial epithelial (NHBE)-Bronchial Epi Cells for B-ALI from Lonza (CC-2540S) were cultured in PneumaCult-Ex Plus Medium (StemCell; Cat#: 05040) supplemented with 500 μl of Hydrocortisone (StemCell; Cat#: 07925). Cells were maintained at 37 °C in the presence of 5 % CO2 and were periodically screened for *Mycoplasma* contamination.

### Mice

All animals were bred and maintained in specific pathogen-free facilities in accordance with FELASA and Animal Experimental Ethics Committee guidelines (University of Burgundy, France). This study was complied with all relevant ethical regulations for animal testing and research and received ethical approval from the Animal Experimental Ethics Committee (University of Burgundy, France). Animals had water ad libitum and were fed regular chow. Experiments were performed in 9 to 13-weeks-old extremely immunodeficient Nod/SCID-Gamma (NSG) mice (JAX stock #005557; Jackson Laboratory; USA) ^69^. Littermate animals from different cages were randomly assigned into experimental groups and were either co-housed or systematically exposed to other groups’ bedding to ensure equal exposure to common microbiota.

### Cell lines establishment

To generate *SETDB1* loss-of-function (SETDB1^LOF^) mutation in A549 cells, optimal sgRNA target sequences, closest to the genomic target sites, were designed using CRISPOR tool^70^. The sgRNA were synthesized (Sigma-Aldrich) as oligonucleotide DNA sequences and cloned into pSpCas9(BB)-2A-GFP (PX458) (Addgene; Cat#: 48138) to construct two active CRISPR vectors; one construct targeted *SETDB1* gene at exon 15 (SETDB1_sgRNA_ex15: 5’-ACAAACCGGTTGGTGCAACA-3’) and the other one targeted *SETDB1* gene at exon 16 (SETDB1_sgRNA_ex16: 5’-GGCGGAGCTATGCTA CCCGG-3’). Cloning was performed according to a protocol described by^71^. For transfection, A549 cells were seeded into 10 cm dishes at 50-60% confluency and transfected with 10 μg of the appropriate sgRNA-containing PX458 plasmids, using jetOPTIMUS (Polyplus; Cat#: 117-01). The transfection was performed according to the manufacturer’s recommended protocol, using a 1:1 ratio of jetOPTIMUS transfection reagent/DNA. At 48 hours after transfection, single GFP-positive cells were isolated using FACSDiva version 6.1.3 into a 96-well plate. Genotyping PCRs were performed with Phusion High-Fidelity DNA Polymerase (NEB; Cat#: M0530S), using primers surrounding the genomic target site.

For the rescue experiments, full-length human *SETDB1* cDNA was cloned into the NotI sites of the pcDNA3.1(+)-IRES-GFP (Addgene; Cat#: 51406). For transfection, A549 SETDB1^LOF^ cells were seeded into 10 cm dishes at 50-60 % confluency and transfected with 10 μg of the pcDNA3.1 (+)-SETDB1^WT^-IRES-GFP or pcDNA3.1(+)-IRES-GFP (control) plasmids, using jetOPTIMUS. The transfection was performed according to the manufacturer’s recommended protocol, using a 1:1 ratio of jetOPTIMUS transfection reagent/DNA. At 48 hours after transfection, selected reagent Neomycin (G418) (700 μg/ml; Gibco; Cat#: 10131-035) was added for 2 weeks, and stable cell lines were isolated.

### Gene silencing by siRNA

siRNAs targeting human SUV39H1/H2 and negative control siRNAs^72^ were from Sigma-Aldrich. Transfections were performed using Lipofectamine RNAiMax (Thermo Fisher; Cat#: 13778150) according to the manufacturer’s instructions. Knockdown efficiency after 72 hours of RNAi treatment was examined using RT-qPCR.

### Immunofluorescence

Cells were seeded on glass coverslips at 60 - 70 % confluence, fixed in 4 % paraformaldehyde, incubated with 50 mM NH4Cl to quench formaldehyde, and permeabilized with 0.5 % Triton X-100. Samples were subsequently incubated overnight with primary antibody against SETDB1 (Thermo Fisher; Cat#: MA5-15722), H3K9me3 (Abcam; Cat#: ab8898), H3K27me3 (Diagenode; Cat#: C15410069), Lamin A/C (Sigma-Aldrich; Cat#: SAB4200236), E-cadherin (Abcam; Cat#: ab40772), phalloidin-TRITC (Sigma-Aldrich; Cat#: P1951), paxillin (Millipore; Cat#: 05-417) overnight at 4 °C, followed by incubation with appropriate fluorophore-conjugated secondary antibodies (Donkey F(ab’)2; Jackson ImmunoResearch Laboratories) for an additional 1 hour. Cell nuclei were stained with DAPI (Life Technologies; Cat#: 62248). Coverslips were mounted with Vectashield mounting media (Clinisciences). Microscopy was performed using an inverted microscope Leica DMI-6000, using 40x, 63x or 100x immersion objectives. Images were taken with the HQ2 Coolsnap motorized by MetaMorph 7.10.2.240 software. All images were processed with ImageJ (Fiji) software.

### Western blot

Cell lysis was carried out with RIPA lysis buffer (50 mM Tris pH 7.5, 150 mM NaCl, 1 % NP-40, 0.5 % Na-deoxycholate, 0.1 % SDS) supplemented with FPIC (Fast Protease Inhibitors cocktail; Sigma-Aldrich; Cat#: S8830-20TAB). The lysates were reduced in NuPAGE LDS Sample Buffer (Thermo Fisher; Cat#: NP0007) supplemented with NuPAGE Sample Reducing Agent (Thermo Fisher; Cat#: NP0009), separated by polyacrylamide gel in NuPAGE MOPS SDS Running Buffer (Thermo Fisher; Cat#: NP0001) and transferred into nitrocellulose membrane (Amersham; Cat#: 10600007). Membranes were blocked in 5 % milk powder in PBST Buffer (1 X PBS, 0.2 % Tween 20) and incubated overnight at 4 °C with the primary antibodies against SETDB1 (Abcam; Cat#: ab107225), H3 (Santa Cruz Biotechnology; Cat#: sc-8654), H3K9me3 (Abcam; Cat#: ab8898), H3K27me3 (Diagenode; Cat#: C15410069), Lamin A/C (Proteintech; Cat#: 10298-1-AP), Lamin B1 (Abcam; Cat#: ab16048), E-cadherin (BD Transduction Laboratory; Cat#: 610181), Paxillin (Millipore; Cat#: 05-417), β-actin (Sigma-Aldrich; Cat#: A5441). Membranes were incubated with the appropriate LI-COR IRDye secondary antibody (LI-COR Biosciences GmbH) and revealed by Odyssey Fc imaging system (LI-COR Biosciences GmbH). The images obtained from the slot blot assay were analyzed with the Image Studio Lite 5.2.5 software.

### Cell proliferation

Cell proliferation analysis was performed with Cell Counting Kit-8 (WST-8 / CCK8) (Abcam, Cat#: ab228554) according to the manufacturer’s instructions. Cells were seeded in 96-well plates and cultured for 24, 48, 72 and 96 hours, respectively. At indicated time points, 10 μl of CCK-8 reagent was added to the wells and incubated for another 1 hour. The absorbance at OD490 was measured with the Gen5 microplate reader (BioTek).

### Cell cycle analysis

Cells were seeded on 6-well plates at 60 - 70 % confluence. After 48 hours, cells were harvested and fixed with 70 % cold ethanol overnight at 4 °C. Samples were stained with 10 μg/ml propidium iodide (PI) staining solution from FITC Annexin V Apoptosis Detection Kit (BD Pharmingen; Cat#: 556547) and 0.2 mg/mL RNase A (Sigma-Aldrich; Cat#: R4642). Samples of at least 10000 cells were acquired using a BD FACSCalibur flow cytometer. Subsequent analysis was done with FlowJo 10.7.1 software.

### Mitotic index

Mitotic index was calculated as the number of positive cells stained with phospho-H3 (Ser10) (Millipore; Cat#: 05-598) in prometaphase, metaphase, anaphase, or telophase divided by the total number of nuclei stained with DAPI (Life Technologies; Cat#: 62248) × 100. Microscopy analysis was performed using an inverted microscope Leica DMI-6000, using 40x immersion objectives. Images were taken with the HQ2 Coolsnap motorized by MetaMorph 7.10.2.240 software. All images were processed with ImageJ (Fiji) software.

### Apoptosis assay

Apoptosis detection was performed by using Annexin V-FITC Apoptosis Detection Kit (BD Pharmingen; Cat#: 556547) according to the manufacturer’s instructions. Cells were seeded on 6-well plates at 80 - 90 % confluence. After 48 hours, cells were harvested, washed twice in cold PBS and then resuspended in 100 μl of a 1X Binding Buffer (10 mM HEPES (pH 7.4), 150 mM NaCl, 2.5 mM CaCl_2_), stained with 5 µl of FITC Annexin V and 5 µl PI and incubated in the dark for 15 min at room temperature. Then, 400 μl binding buffer was added to the samples and at least 10000 cells were acquired using a flow cytometer using a BD FACSCalibur flow cytometer. Subsequent analysis was done with FlowJo 10.7.1 software.

### Wound-healing and cell migration

Cells were seeded on a glass-bottom petri-dish (Fluorodish; Cat#: FD35-100) with the polydimethylsiloxane (PDMS, Sylgard 184; Dow Corning; Cat#: VAR0000625) coating for live microscopy. PDMS was prepared by mixing base and curing agent in a ratio of 1∶10. PDMS was homogeneously distributed on the surface of the dish to form a thin layer and then incubated at 80 °C for 2 hours. Some PDMS was poured over a square-shape mold, degassed, and cured at 80 °C for 2 hours to form square blocks. After curing, PDMS blocks were cut to form hollow 0.5 mm^2^ square-shape stencils. To allow cell adhesion and migration on the substrate, the Fluorodish surface was coated with 20 µg/ml of Fibronectin (Sigma-Aldrich; Cat#: F0556) diluted in PBS overnight at 4 °C. Excess of Fibronectin washed away with PBS and the liquid solution was removed to allow the surface to dry. Once the surface was dried, a hollow square PDMS stencil was placed in the center of the petri dish. Cells were seeded inside the PDMS stencils. A549 control and SETDB1^LOF^ cells were prepared at a 2/3 ratio due to their different proliferation rates, seeded in 2 separate dishes within concentrated drops of media, and incubated at 37 °C for 45 min to allow cells to attach on the surface. The medium was then removed, to wash off unattached cells, and cells incubated with fresh medium overnight. After cells had reached confluence inside of the PDMS stencils (usually 24 hours after seeding), samples were treated with mitosis-ceasing agent Mitomycin C (20 µg/ml, 1 hour at 37 °C; Sigma-Aldrich; Cat#: M4287) in order to block proliferation and record only active cell migration. PDMS stencils were then removed to allow cells to migrate away from their pool, and samples were transferred to a Biostation microscope (Nikon). To track cell movements, we used phase-contrast images and recorded multi-positional time-lapse movies. 10 min time step was applied between images, and movies were recorded over 16 to 25 hours.

For cell-cell adhesion disruption experiments, we used plastic-bottom petri dishes (Corning; Cat#: 353001), suitable for live microscopy. Petri-dish were functionalized with 20 µg/ml of Fibronectin (Sigma-Aldrich; Cat#: F0556) diluted in PBS overnight at 4 °C, and A549 SETDB1^LOF^ cells were plated and treated as described above. EGTA was added to the sample media to disrupt Ca^2+^-dependent cell-cell adhesions at concentrations as mentioned in Figure S1G.

Image processing was performed in the ImageJ (Fiji) program. Data analysis from processed files was performed using particle image velocimetry (PIV) from the MatPIV package (PIVlab v1.41) and implemented in the MATLAB program.

### Xenograft studies in NSG mice

A549 control or SETDB1^LOF^ cells (2 x 10^6^) were suspended in 200 µL of mixture Dulbecco-modified phosphate buffered saline (DPBS, Sigma-Aldrich, Cat#: 59331C) and BD Matrigel Matrix (BD Biosciences; Cat#: 356234) in a ratio of 1∶1, and injected subcutaneously into the right flanks of NSG mice. Tumor growth was monitored every third day using digital calipers. Tumor size was calculated as *Tumor_volume = (length x width^2^)/2*. Mice were euthanized at the end of the experiment, tumors were dissected and fixed in 10 % formalin for subsequent immunohistochemical analysis.

### Immunohistochemistry

Formalin-fixed paraffin-embedded (FFPE) tumor sections (5 µm) from A549 tumors were used for hematoxylin and eosin (H&E) staining and proliferation analysis with anti-Ki67 antibody (Leica Biosystems; Cat#: NCL-Ki-67p). Briefly, sections were deparaffinized, incubated overnight at 4 °C with primary antibodies, washed, and incubated with secondary antibodies HRP Labelled Polymer Anti-Rabbit (Dako EnVision+ System; Cat#: K4002) following the manufacturer’s protocol. Slides were counterstained with hematoxylin counterstain and coverslips were mounted using non-aqueous mounting media. Images were scanned at 20 x magnification using the Aperio Scanscope system.

### RNA and quantitative reverse transcription-PCR (qRT-PCR)

Total RNA was extracted from the cell lines or frozen tissue samples using RNeasy Mini Kit (Qiagen; Cat#: 74104). DNase treatment was performed using RNase-Free DNase Set (Qiagen; Cat#: 79254) to remove residual DNA. cDNA was prepared with a High-Capacity cDNA Reverse Transcription Kit (Thermo Fisher; Cat#: 4368814). Real-time quantitative PCR (RT-qPCR) was performed with GoTaq qPCR Master Mix (Promega; Cat#: A6002). Gene expression analysis was calculated following normalization to *PPAI* using the comparative Ct (cycle threshold) method. The primers used are provided in Table S1.

### RNA-seq

Total RNA was isolated using RNeasy Mini Kit (Qiagen; Cat#: 74104) followed by Turbo DNA-free Kit (Thermo Fisher; Cat#: AM1907). To construct libraries, 1 µg of high-quality total RNA (RIN > 9.9) was processed with Truseq Stranded Total RNA Library Prep Kit (Illumina; Cat#: 20020596) according to the manufacturer’s instructions. Briefly, after removal of ribosomal RNA using Ribo-zero rRNA Kit (Illumina; Cat#: 20037135), confirmed by quality control on the Agilent 2100 Bioanalyzer, total RNA molecules were fragmented and reverse-transcribed using random primers. Replacement of dTTP by dUTP during the second strand synthesis permitted to achieve strand specificity. The addition of a single A base to the cDNA was followed by ligation of adapters. Libraries were quantified by qPCR using the KAPA Library Quantification Kit for Illumina Libraries (KapaBiosystems; Cat#: KR0405) and library profiles were assessed using the DNA High Sensitivity HS Kit (Agilent; Cat#: 5067-4626) on an Agilent Bioanalyzer 2100. Libraries were sequenced on an Illumina Nextseq 500 in a paired-end mode in three independent biological replicates at Platform GENOM’IC - Institute Cochin. Demultiplexing and quality of sequences were performed with the Aozan software v2.2.1.

### Chromatin immunoprecipitation (ChIP)

#### Native ChIP (NChIP)

Around 1 x 10^7^ cells were collected, washed with PBS and resuspended in 250 μl of ice-cold Douncing buffer (10 mM Tris-HCl (pH 7.5), 4 mM MgCl_2_, 1 mM CaCl_2_, 0.2 % Triton X100), followed by homogenization with syringe with 26 gauge X1/2’’ needles (BD Microlance; Cat#: BD 303800). To obtain mononucleosomes, cells were digested with 1.25 μl of micrococcal nuclease (Mnase) (0.5 U/μl; Sigma-Aldrich; Cat#: N3755-500UN) for 5 min at 37 °C with 600 rpm. The reaction was stopped with 10 mM EDTA, pH 8. Nuclei were swollen to release chromatin after addition of Hypotonization buffer (0.2 mM EDTA (pH 8), FPIC (Fast Protease Inhibitors cocktail; Sigma-Aldrich; Cat#: S8830-20TAB), DTT). Chromatin corresponding to 8 μg of DNA was incubated with anti-H3K9me3 (Diagenode; Cat#: C15410193), anti-H3K27me3 (Diagenode; Cat#: C15410069), anti-H3K27ac (Diagenode; Cat#: C15410196) or anti-H3K4me3 (Diagenode; Cat#: C15410003) antibodies overnight at 4 °C. The DNA-protein-antibody complexes were captured by DiaMag protein A-coated magnetic beads (Diagenode; Cat#: C03010020) by incubating at 4 °C for 2 hours. Magnetic beads were then washed with Low-Salt buffer (20 mM Tris-HCl (pH 8), 0.1 % SDS, 1 % Triton X100, 2 mM EDTA (pH 8), 150 mM NaCI and FPIC) and High-Salt buffer (20 mM Tris-HCl (pH 8), 0.1 % SDS, 1 % Triton X100, 2 mM EDTA (pH 8), 500 mM NaCI and FPIC). DNA was eluted with TE buffer (100 mM Tris-HCl (pH 8), 1 % SDS, 1 mM EDTA (pH 8)) from the beads overnight, inputs corresponding to 1 % of IP were treated as well, followed by incubation with 1 μl of RNase A (10 μg/ml; Sigma-Aldrich; Cat#: R4642) for 1 hour. DNA was purified using MinElute PCR Purification Kit (Qiagen; Cat#: 28004).

#### Cross-linked ChIP (XChIP)

Around 1 x 10^7^ cells were cross-linked directly in the culture plate with culture medium supplemented with 1 % of formaldehyde (Sigma-Aldrich; Cat#: F8775) and 15 mM NaCl, 0.15 mM EDTA (pH 8), 0.075 mM EGTA and 0.015 mM HEPES (pH 8) during 10 min at RT. Formaldehyde was quenched with 0.125 M glycine and cells were washed in PBS. Fixed cells were prepared with buffers from Chromatin Shearing Optimization Kit, low SDS (Diagenode; Cat#: C01020013) for sonication for 5 min (30 sec ON, 30 sec OFF) (Bioruptor Diagenode), yielding genomic DNA fragments with a bulk size of 150-600 bp. 1 % of chromatin extracts were taken aside for inputs. Chromatin corresponding to 8 μg of DNA was incubated with anti-CTCF (Active Motif; Cat#: 61311) antibody overnight at 4 °C. The DNA-protein-antibody complexes were captured by DiaMag protein A-coated magnetic beads (Diagenode; Cat#: C03010020) by incubating at 4 °C for 2 hours. Magnetic beads were then washed with Low-Salt buffer (20 mM Tris-HCl (pH 8), 0.1 % SDS, 1 % Triton X100, 2 mM EDTA (pH 8), 150 mM NaCI and FPIC) and High-Salt buffer (20 mM Tris-HCl (pH 8), 0.1 % SDS, 1 % Triton X100, 2 mM EDTA (pH 8), 500 mM NaCI and FPIC). DNA was eluted with TE buffer (100 mM Tris-HCl (pH 8), 1 % SDS, 1 mM EDTA (pH 8)) from the beads overnight, and then reverse cross-linked with 1 μl of RNase A (10 μg/ml; Sigma-Aldrich; Cat#: R4642), followed by Proteinase K digestion (5 μl of 20 mg/mL; Sigma-Aldrich; Cat#: P2308) during 2 hours; inputs corresponding to 1 % of IP were reverse cross-linked in the same conditions. DNA was purified using MinElute PCR Purification Kit (Qiagen; Cat#: 28004).

### ChIP-seq

Libraries were prepared using the MicroPlex V2 library preparation Kit (Diagenode; Cat#: C05010012), following the manufacturer’s instructions, 10 µl of DNA was used as starting material. After end repair of the double-stranded DNA templates, cleavable stem-loop adaptors were ligated, and then adaptor enriched DNA was amplified with 10 PCR cycles to add Illumina compatible indexes. The libraries were then purified with the Agencourt AMPure XP bead-based purification system. Final libraries were quantified with Qubit dsDNA HS assay (Thermo Fisher; Cat#: Q32851), and size distribution was monitored by capillary electrophoresis on Agilent 2100 Bioanalyzer using DNA High Sensitivity HS Kit. Libraries were then normalized to 10 pM and pooled before sequencing. Libraries were sequenced on an Illumina Nextseq 500 in a paired-end mode in two independent biological replicates at Platform GENOM’IC - Institute Cochin. Demultiplexing and quality of sequences were performed with the Aozan software v2.2.1.

### Hi-C

Hi-C libraries were performed as described in^73^. 10 million cells were cross-linked with fresh 1 % formaldehyde (Sigma-Aldrich; Cat#: F8775) for 10 min at room temperature. Excess of formaldehyde was quenched with 125 mM glycine for 5 min. Cells were centrifuged (1,000 × g, 10 min, 4 °C), resuspended in 50 μl of 1 × PBS, snap-frozen in liquid nitrogen, and stored at −80 °C. Defrosted cells were lysed in 1.5 ml of Isotonic buffer (50 mM Tris-HCl pH 8.0, 150 mM NaCl, 0.5 % (v/v) NP-40 substitute, 1 % (v/v) Triton-X100, 1 × Protease Inhibitor Cocktail (Bimake; Cat#: B14001)) on ice for 15 min. Cells were centrifuged at 2,500 × g for 5 min, resuspended in 100 μl of 1.1 × DpnII buffer (NEB; Cat#: B0543S), and pelleted again. The pellet was resuspended in 200 μl of 0.3 % SDS in 1.1 × DpnII buffer and incubated at 37 °C for 1 hour. Then, 330 μl of 1.1 × DpnII buffer and 53 μl of 20 % Triton X-100 were added, and the suspension was incubated at 37 °C for 1 hour. Next, 600 U of DpnII enzyme (NEB; Cat#: R0543M) were added, and the chromatin was digested overnight (14 - 16 hours) at 37 °C with shaking (1,400 rpm). On the following day, 200 U of DpnII enzyme were added, and the cells were incubated for an additional 2 hours. DpnII was then inactivated by incubation at 65 °C for 20 min. After DpnII inactivation, the nuclei were harvested for 10 min at 5,000 × g, washed with 100 μl of 1 × NEBuffer 2.1 (NEB; Cat#: B7202S), and resuspended in 50 μl of 1 × NEBuffer 2.1. Cohesive DNA ends were biotinylated by the addition of 7.6 μl of the biotin fill-in mixture prepared in 1 × NEBuffer 2.1 (0.025 mM dCTP (Thermo Fisher; Cat#: R0151), 0.025 mM dGTP (Thermo Fisher; Cat#: R0161), 0.025 mM dTTP (Thermo Fisher; Cat#: R0171), 0.025 mM biotin-14-dATP (Invitrogen; Cat#: 19524-016), and 0.8 U/μl Klenow enzyme (NEB; Cat#: M0210L). The samples were incubated at 37 °C for 75 min with shaking (1,400 rpm). Nuclei were centrifuged at 3,000 × g for 5 min, resuspended in 100 μl of 1× T4 DNA ligase buffer (Thermo Fisher; Cat#: EL0011), and pelleted again. The pellet was resuspended in 300 μl of 1 × T4 DNA ligase buffer, and 75 U of T4 DNA ligase (Thermo Fisher; Cat#: EL0011) were added. Chromatin fragments were ligated at 20 °C for 6 hours. The cross-links were reversed by overnight incubation at 65 °C in the presence of Proteinase K (100 μg/ml; Sigma-Aldrich; Cat#: P2308). After cross-link reversal, the DNA was purified by single phenol-chloroform extraction followed by ethanol precipitation with 20 μg/ml glycogen (Thermo Scientific; Cat#: R0561) as the co-precipitator. After precipitation, the pellets were dissolved in 100 μl of 10 mM Tris-HCl (pH 8.0). To remove residual RNA, samples were treated with 50 μg of RNase A (Thermo Fisher; Cat#: R1253) for 45 min at 37 °C. To remove residual salts and DTT, the DNA was additionally purified using Agencourt AMPure XP beads (Beckman Coulter; Cat#: A63881). Biotinylated nucleotides from the non-ligated DNA ends were removed by incubating the Hi-C libraries (2 μg) in the presence of 6 U of T4 DNA polymerase (NEB; Cat#: M0203L) in NEBuffer 2.1 supplied with 0.025 mM dATP (Thermo Fisher; Cat#: R0141) and 0.025 mM dGTP at 20 °C for 4 hours. Next, the DNA was purified using Agencourt AMPure XP beads. The DNA was then dissolved in 500 μl of sonication buffer (50 mM Tris-HCl (pH 8.0), 10 mM EDTA, 0.1 % SDS) and sheared to a size of approximately 100 - 500 bp using a VirSonic 100 (VerTis). The samples were concentrated (and simultaneously purified) using AMICON Ultra Centrifugal Filter Units (Millipore; Cat#: UFC503096) to a total volume of approximately 50 μl. The DNA ends were repaired by adding 62.5 μl MQ water, 14 μl of 10 × T4 DNA ligase reaction buffer, 3.5 μl of 10 mM dNTP mix (Thermo Fisher; Cat#: R0191), 5 μl of 3 U/μl T4 DNA polymerase, 5 μl of 10 U/μl T4 polynucleotide kinase (NEB; Cat#: M0201L), 1 μl of 5 U/μl Klenow DNA polymerase, and then incubating at 25 °C for 30 min. The DNA was purified with Agencourt AMPure XP beads and eluted with 50 μl of 10 mM Tris-HCl (pH 8.0). To perform an A-tailing reaction, the DNA samples were supplemented with 6 μl 10 × NEBuffer 2.1, 1.2 μl of 10 mM dATP, 1 μl of MQ water, and 3.6 μl of 5 U/μl Klenow (exo-) (NEB; Cat#: M0212S). The reactions were carried out for 45 min at 37 °C in a PCR machine, and the enzyme was then heat-inactivated by incubation at 65 °C for 20 min. The DNA was purified using Agencourt AMPure XP beads and eluted with 100 μl of 10 mM Tris-HCl (pH 8.0). Biotin pulldown of the ligation junctions: 15 μl of MyOne Dynabeads Streptavidin C1 (Invitrogen; Cat#: 65001) beads washed with the TWB buffer (5 mM Tris-HCl (pH 8.0), 0.5 mM EDTA (pH 8.0), 1 M NaCl, 0.05 % Tween-20), resuspended in 200 μl of 2 × Binding buffer (10 mM Tris-HCl (pH 8.0), 1 mM EDTA, 2 M NaCl) and added to 200 μl of DNA. Biotin pulldown was performed for 30 min at 25 °C with shaking. Next, beads with captured ligation junctions were washed once with TWB buffer, once with 1 × T4 DNA ligase buffer, and then resuspended in 50 μl of adapter ligation mixture comprising 41.5 μl MQ water, 5 μl 10 × T4 DNA ligase reaction buffer, 2.5 μl of Illumina TruSeq adapters, and 1 μl of 5 U/μl T4 DNA ligase. Adapter ligation was performed at 22 °C for 2.5 hours, and the beads were sequentially washed twice with 100 μl of TWB, once with 100 μl of CWB (10 mM Tris-HCl (pH 8.0) and 50 mM NaCl), and then resuspended in 25 μl of MQ water. Test PCR reactions containing 4 μl of the streptavidin-bound Hi-C library were performed to determine the optimal number of PCR cycles required to generate sufficient PCR products for sequencing. The PCR reactions were performed using KAPA High Fidelity DNA Polymerase (Kapa Biosystems; Cat#: 08201595001) and Illumina PE1.0 and PE2.0 PCR primers (10 pmol each). The temperature profile was 5 min at 98 °C, followed by 6, 9, 12, 15, and 18 cycles of 20 s at 98 °C, 15 s at 65 °C, and 20 s at 72 °C. The PCR reactions were separated on a 2 % agarose gel containing ethidium bromide, and the number of PCR cycles necessary to obtain a sufficient amount of DNA was determined based on the visual inspection of gels (typically 8 - 12 cycles). Four preparative PCR reactions were performed for each sample. The PCR mixtures were combined, and the products were purified with Agencourt AMPure XP beads. Libraries were sequenced on an Illumina Novaseq 6000 in a paired-end mode in two independent biological replicates at Genetico.

### Chromatin mobility

Cells were plated in Fluorodishes (Worldprecision; Cat#: FD35100) at the rate of 2 x 10^5^ cells per dish the day before performing the live imaging. 30 min before live imaging, the medium was replaced with 1 mL of fresh medium with 2 drops of NucBlue LiveProbe reagent (Thermo Fisher; Cat#: R37605). Live imaging was performed on a Leica DMI8 microscope, equipped with a CSU-X1 Yokogawa spinning disk module. The acquisition was realized with a 40x dry objective (N.A. 1.40) and collected by the Hamamatsu Orca flash 4.0 camera. The microscope was controlled by the Metamorph software v7.10.2.240 from Molecular Devices. Images were acquired in brightfield (one plane, exposure time 50 ms) with a 405 nm laser at 15 % power (Z stack, 7 planes centered on the middle plane of the nuclei, Z step of 0.5 µm, exposure time 150 ms) every 2 min for 1.5 to 2 hours. Analysis was performed on the best Z plane chosen for each stage position, using a set of custom-written Fiji scripts. Briefly, the nuclei were segmented and registered for XY translation and rotation using the MultiStackReg plugin. The chromatin flow was then measured using the PIV plugin on the first 10 time points, restricted to the nuclei. The PIV magnitude value was obtained by averaging the values measured for the entire nucleus over the 10 time points.

### Nuclear envelope fluctuations

Cells were transfected with the pEGFP-LAP2b (Euroscarf plasmid bank; Cat#: P30463) plasmid 24 hours prior to imaging using jetOPTIMUS. The transfection was performed according to the manufacturer’s recommended protocol, using a 1:1 ratio of jetOPTIMUS transfection reagent/DNA. 30 min before live imaging, the medium was replaced with 1 mL of fresh medium with 2 drops of NucBlue LiveProbe reagent. Live imaging was performed on a Leica DMI8 microscope, equipped with CSU-X1 Yokogawa spinning disk module. The acquisition was realized with a 63x oil objective (N.A. 1.40) and collected by the Hamamatsu Orca flash 4.0 camera. The microscope was controlled by the Metamorph software v7.10.2.240 from Molecular Devices. Snapshots were first acquired in brightfield (one plane, exposure time 50 ms) with a 488 nm laser at 2% (one plane, exposure time 200 ms) and with a 405 nm laser at 15 % power (one plane, exposure time 150 ms). Movies at high temporal resolution were then acquired with a 488 nm laser at 2% (exposure time 200 ms) at the rate of 1 frame every 250 ms for 700 timeframes (1.75 min). Analysis was performed with Fiji and Python with custom-written scripts. Briefly, the nuclei in the 488 nm channel were registered for XY translation and rotation using the StackReg plugin. For each nucleus, 4 lines were hand-drawn on the nuclear envelope (NE) and resliced over time to obtain 4 kymographs, one for each individual point in the NE. A custom-written Python script was used to determine the position of the NE at each timepoint of each kymograph. For each timepoint, a parabola was fitted on the pixel with the highest intensity and its 10 neighboring pixels. The maximum of the parabola was assigned as the NE position. For each kymograph, the average NE position was determined by averaging the successive NE positions. The square root of the Mean Square Displacement was calculated using the displacements calculated for each time point as the distance to the average NE position. Timepoints with displacements larger than twice the average displacement were interpreted as NE positioning errors and excluded from the analysis. The values per cell were obtained by averaging the values of the corresponding kymographs.

### Immunofluorescence analysis

#### E-cadherin

Images were processed using the “rolling ball” background subtraction method implemented in Fiji (rolling ball radius set to 400). Mean junctional E-cadherin intensity was calculated by measuring the average intensity inside fixed-width regions of interest (ROIs) placed around all visible cellular junctions in the field (~20-50 junctions per image), excluding nodes. At least 4 fields were analyzed for each biological replicate. Images from three independent experiments were analyzed.

#### Paxillin

Images were processed using the “rolling ball” background subtraction method implemented in Fiji (rolling ball radius set to 50). Normalized paxillin area was calculated by dividing the total paxillin-positive area by the number of nuclei in the field. To measure the total paxillin-positive area, the paxillin-stained images were thresholded above the intensity value, similar for all of the images in the batch, so that no cytosolic background was present. The nuclei were counted using the corresponding images with the DAPI-staining. The measurements were performed for at least 4 fields per biological replicate and the mean values for all of the fields for each replicate were retained. Image processing was looped using a custom-written macro.

#### E-actin

The Phalloidin (F-actin) distribution quantification was performed using the method adopted from^74^. Images were processed using the “rolling ball” background subtraction method implemented in Fiji (rolling ball radius set to 50), and the ROIs around all fully visible cells in the field were manually outlined. The mean Phalloidin intensity was measured inside of each ROI. Each of the cells was then treated as a separate image with the rest of the pixels outside of the ROI set to zero. Using the *Fit Ellipse* function, implemented in Fiji, the center of mass of each cell was detected and the line, passing through the center of mass and the frontier of the cell with the maximum Phalloidin intensity, was drawn. Using the macro, adapted from Zonderland et al. (2019), the Phalloidin intensity profile along the line was obtained with the Fiji *getProfile* function, the line was trimmed on both sides by the first non-zero intensity value and then divided into 10 equal-size bins. The average intensity for all values inside each bin was calculated and the process was repeated for all of the cells in at least 4 fields for each biological replicate.

#### LaminA/C and H3K9me3 co-immunofluorescence

The images were subjected to background subtraction (rolling ball radius 50) and the ROIs around all fully visible nuclei were outlined. All ROIs were then analysed using the Clock Scan protocol^75^ implemented as a Fiji Plugin. Briefly, the plugin sequentially collected multiple radial pixel-intensity profiles, scanned from the ROI center to the predetermined distance outside of the ROI border. The profiles were scaled according to the measured ROI radius, so that the distance from the center of the ROI to its border was represented in % of radius. The individual profiles were averaged into one integral radial pixel-intensity profile for each nucleus. The intensity determined outside of the ROI border (at a distance 100-120% of radius) was used for background correction.

The intensities were measured separately for the H3K9me3, H3K27me3, and Lamin A/C channels. The measurements were performed for 30-35 nuclei per biological replicate (92-104 nuclei per condition in total). The H3K9me3, H3K27me3 and Lamin A/C distributions were plotted using the means for all the measured intensity values at a given distance from the nucleus center.

#### Nuclear morphology assessment

Nuclear morphology was assessed using the images in the DAPI channel. Fully visible nuclei were outlined, and their area, perimeter, circularity and solidity were measured with Fiji. Circularity was defined as *4π*Area/Perimeter^2^*, with a value of 1 indicating a perfect circle, and the values approaching 0 indicating a largely elongated shape. Solidity was defined as a ratio *Area/Convex_Area*, with a value of 1 indicating a convex object, and values less than 1 indicating an invaginated object with an irregular boundary. To quantify the deviation of the nuclear shape from the ellipse, the irregularity index (defined as *1-solidity*) was used^76^, with a value of 0 indicating a regular-shaped elliptical nuclei, and higher for nuclei with invaginated borders. The measurements were performed for 30-35 nuclei per biological replicate (92-104 nuclei per condition in total).

### Immunohistochemistry analysis

The percentage of Ki-67 positive nuclei was determined using digital images of the tumor section by examining 8 fixed-area (500000 mkm^2^) hot-spot fields. Hot spots were defined as areas in which Ki-67 staining was particularly higher relative to the adjacent areas. For each field the ratio of Ki-67-positive nuclei to the total number of nuclei was calculated. The fields contained 1500-2500 cells in total.

### RNA-seq analysis

#### RNA-seq reads mapping and quantification

RNA-seq reads were mapped to the hg19 reference human genome assembly using STAR v2.6.1c^77^ with GENCODE v33 (GRCh37) gene annotation^77^. Read counts per gene were obtained using the ‘--quantMode’ parameter in STAR. Uniquely mapped reads with MAPQ>30 were selected using SAMtools v1.5^78^. Reads overlapping with the hg19 blacklist regions^79^ were discarded using bedtools v2.25.0^80^. The bigwig files with the CPM-normalized transcription signal in 50 bp bins were generated for merged replicates using *bamCoverage* function from deepTools v3.3.0^81^. Visualization of the signal profiles was created in the UCSC Genome Browser^82^. Statistics of the RNA-seq data processing can be found in Table S2.

#### Differential gene expression

Differentially expressed genes were identified with edgeR v3.30.0 R package^83^. Genes with low expression counts were filtered (*filterByExp* function), and the remaining genes were normalized using the trimmed mean of M values (TMM) method^84^. To check the reproducibility between the replicates, a multidimensional scaling (MDS) plot of distances between the gene expression profiles was generated^85^. Quasi-likelihood negative binomial generalized log-linear model (*glmQLFit* function)^86^ and t-test relative to a fold change threshold of 1.5 (*glmTreat* function)^87^ were used to calculate *P*-values per gene. The obtained *P*-values were adjusted with Benjamini– Hochberg correction for multiple testing (FDR). FDR threshold of 0.01 was used to define differentially expressed genes (Table S3a).

#### Gene ontology analysis

Gene ontology (GO) analysis of differentially expressed genes was performed using GO term over-representation test from g:Profiler^88^. The annotation of genes with GO terms from Ensembl (release 100) was utilized, and electronic GO annotations were removed prior to the analysis. FDR threshold of 0.05 was applied, and significant GO terms of size between 25 and 1500 were extracted (Table S3b).

#### Gene set enrichment analysis

Gene set enrichment analysis (GSEA) was performed with GSEA v4.0.3^89^ in pre-ranked mode with 10000 permutations (Table S3C). Genes were pre-ranked by fold change derived from the differential expression analysis. Gene sets of size between 25 and 1500 from MSigDB v7.1^90^ were used.

#### Gene set variation analysis

Gene set variation analysis (GSVA) was performed using the GSVA v1.36.2 R package^91^. Publicly available RNA-seq datasets of normal lung (N=48) and NSCLC (N=61) cell lines were used as a background. The datasets were downloaded from the GEO database (GEO accession numbers: GSE63900, GSE93526, GSE85447, GSE101993, GSE97036, GSE42006, GSE152446, GSE113185, GSE80386, GSE148729, GSE109821, GSE123769, GSE141666, GSE123631, GSE113493, GSE150809, GSE74866, GSE99015, GSE79051, GSE107637, GSE86337, GSE78531, GSE78628, GSE61955 and processed as described above (section “*RNA-seq Reads Mapping and Quantification”*) to obtain read counts per gene. Lung-specific gene set was extracted from the Human Protein Atlas v20.0 database^92^, other gene sets were retrieved from the MSigDB v7.1 database. Ensembl gene identifiers were converted to Entrez using Biomart^93^ with Ensembl genes (release 100). GSVA scores were estimated for log-transformed CPM gene counts for each sample (Table S3d).

### ChIP-seq analysis

#### ChIP-seq reads mapping and normalization

ChIP-seq reads were mapped to the hg19 reference human genome assembly using Bowtie v2.2.3 with ‘--no-discordant’ and ‘--no-mixed’ options. Uniquely mapped reads with MAPQ>30 were selected using SAMtools v1.5 for further analysis. PCR duplicates were filtered out using the Picard v2.22 *MarkDuplicates* function. Reads overlapping with the hg19 blacklist regions were discarded using bedtools v2.25.0. The bigWig files with the ratio of RPKM-normalized ChIP-seq signal to the input in 50 bp bins were generated using deepTools v3.3.0 with pseudocount set to ‘0 1’ and smoothing window of 3 bins. Visualization of the signal profiles were created in the UCSC Genome Browser. Statistics of the ChIP-seq data processing can be found in Table S2.

#### Peak calling

Peak calling was performed with MACS2 v2.2.7.1^94^. For histone modifications (H3K4me3, H3K27ac, H3K9me3, H3K27me3) ChIP-seq, peaks were called both for replicates and pooled data. For H3K4me3 and H3K27ac, MACS2 was executed in a narrowPeak mode with tag size of 40 or 41 bp, mappable genome size set to ‘hs’, and q-value cutoff of 0.05. For H3K9me3 and H3K27me3, MACS2 was executed in a broadPeak mode with tag size of 40 or 41 bp, mappable genome size set to ‘hs’, and broad-cutoff of 0.1. The consensus peak list was obtained by overlapping the peaks, annotated for pooled replicates, with peaks from both replicates. Only the pooled peaks from canonical chromosomes that had an overlap of at least 50 % with peaks from both replicates were retained. For CTCF ChIP-seq, peaks were called using the pooled data in a narrowPeak mode with tag size of 40 bp, mappable genome size set to ‘hs’, and p-value cutoff of 0.001. No additional peak calling filters were applied to the obtained CTCF peaks.

#### Super-enhancers annotation

To identify the super-enhancers, we used rank ordering of super-enhancers (ROSE) approach^36^. For ROSE, H3K27ac peaks within 12.5 kb from one another were stitched into continuous enhancer clusters, within which the density of input-normalized H3K27ac ChIP-seq signal was calculated. These stitched regions were then ranked based on the total signal density, and plotted to geometrically define the cutoff. The cutoff was set at a point where a line with a slope of 1 was tangent to the curve, and regions above the curve were defined as super-enhancers (Table S4a).

#### Differential enrichment analysis

DiffBind v2.16.0^95^ was utilized to analyze the differential enrichment of the H3K9me3 and H3K27ac ChIP-seq peaks, and differentially regulated super-enhancers. To analyze heterochromatin domains and continuous enhancer clusters, peaks within 12.5 kb from one another for H3K9me3 and H3K27ac data were stitched together^96^. Stitched ChIP-seq peaks and annotated super-enhancers were pooled together for A549 control and A549 SETDB1^LOF^ conditions and used for differential enrichment analysis. DiffBind pipeline was executed with default parameters. FDR threshold of 0.01 and absolute log_2_FC of 1 were used to define differentially regulated regions (Table S4b, S4c and S4d).

Differential enrichment of CTCF was determined based on the fold change of the ChIP-seq signal density at the pooled peaks level. Absolute fold change threshold of 1.5 was used to define differentially regulated CTCF peaks (Table S4e).

Genomic location of differential H3K27ac and CTCF peaks was assessed using ChIPseeker v1.24.0^97^ with GENCODE v33 (GRCh37) gene annotation.

#### Lamina-associated domains

For the pile-ups generation, genomic regions of the constitutive^98^, HeLa^99, 100^, IMR90 ^101–103^, TIG-3^104^ and NHDF^100, 105^ lamina-associated domains (LADs) were used. Overlaps of differentially regulated H3K9me3 domains with conserved LADs and inter-LADs were calculated with *intersect* function from bedtools v2.25.0. Jaccard similarity coefficients between differentially regulated H3K9me3 domains and LADs were calculated with *jaccard* function from bedtools v2.25.0.

#### Pile-ups generation

Pile-ups of the ChIP-seq signals were calculated using the *computeMatrix* function from deepTools v3.3.0. Pile-ups at the genes were generated in *scale-regions* mode with parameters ‘--beforeRegionStartLength’ 3000, ‘--regionBodyLength’ 5000, ‘--afterRegionStartLength’ 3000, ‘--missingDataAsZero’, and ‘--skipZeros’. Pile-ups at the H3K9me3 domains and LADs were generated in *scale-regions* mode with parameters ‘--beforeRegionStartLength’ 500000, ‘--regionBodyLength’ 1000000, ‘--afterRegionStartLength’ 500000, ‘-bs’ 10000, ‘--unscaled5prime’ 10000, ‘--unscaled3prime’ 10000, ‘--missingDataAsZero’, and ‘--skipZeros’. Pile-ups of the H3K27ac and CTCF peaks were generated in *reference-point* mode with parameters ‘--referencePoint’ center, ‘-a’ 10000 (‘-a’ 2500 for CTCF), ‘-b’ 10000 (‘-b’ 2500 for CTCF), ‘--missingDataAsZero’, and ‘--skipZeros’. Pile-ups were visualized using *plotHeatmap* function from deepTools v3.3.0.

#### Association between enhancers and gene expression

To assess the degree of association between differentially regulated enhancers and gene expression changes, Genomic Regions Enrichment of Annotations Tool (GREAT) v4.0.4^106^ was utilized. GREAT was run in basal extension mode excluding regions 5 kb upstream and 1 kb downstream of TSS, and searching for H3K27ac peaks up to distal 500 kb. Genes associated with differential H3K27ac peaks were extracted and analyzed with g:Profiler to calculate GO terms enrichment, as described above for the differentially expressed genes (section “*Gene Ontology Analysis”*, Table S4f, S4g). To associate enhancer-linked genes with gene expression, fgsea v1.14.0 R package^107^ was used. It was run in pre-ranked mode with 10000 permutations, and parameter scoreType set to ‘pos’ and ‘neg’ for genes linked with increased and decreased H3K27ac peaks respectively.

#### Motifs search

HOMER v4.11^45^ was used to identify the motifs present in the ATAC-seq peaks within the annotated super-enhancers. ATAC-seq peaks for the A549 cell line from the ENCODE Project Consortium were used (ENCODE accession numbers ENCSR032RGS and ENCSR220ASC). Peak coordinates were converted from hg38 to hg19 reference human genome assembly with UCSC liftOver tool. HOMER motif discovery algorithm (findMotifsGenome.pl) was executed with default parameters, and known motifs with *P*-values < 0.01 were used for the analysis. The obtained motifs were filtered for transcription factor (TFs) motifs only using the Human Transcription Factor Database v1.01 and converted to the orthologous human gene names if necessary. In case more than one motif for TF was present, the motif with higher fold enrichment was retained.

### Hi-C analysis

#### Hi-C reads mapping and filtering

Hi-C reads were mapped to the reference human genome hg19 assembly using Bowtie v2.2.3 with the iterative mapping procedure implemented in hiclib Python package^109^. The mapping was performed in the ‘--very-sensitive’ mode. Minimal read size was set to 25 bp, and the read was extended by 5 bp during the iterative mapping procedure, until a maximal read length was reached. Non-uniquely mapped reads, ‘same fragment’ and ‘dangling end’ reads, PCR duplicates, reads from restriction fragments shorter than 100 bp and longer than 100 kb, and reads from the top 0.5 % of restriction fragments with the greatest number of reads were discarded. Read pairs were then aggregated into genomic bins to produce contact matrices. Low coverage bins removal and iterative correction of the matrices were performed using *balance* function from cooler v0.8.5^110^. Statistics of the Hi-C data processing can be found in Table S2.

#### Annotation of chromatin compartments

A/B-compartments were annotated using cooltools v0.3.2 *call-compartments* function for 100 kb resolution contact matrices. The orientation of the eigenvectors (PC1) was selected such that it correlates positively with GC content. Consequently, B compartment bins were assigned with negative eigenvector values, and A compartment bins were assigned with positive. To validate the compartment annotation, histone modification ChIP-seq signals from the ENCODE Project Consortium were used (ENCODE accession numbers ENCSR000AUL, ENCSR000AUN, ENCSR000AUI, ENCSR000ATP, ENCSR000AUK, ENCSR000AVI, ENCSR000ASH, ENCSR000ASV, and ENCSR000AUM). ChIP-seq signal fold change was calculated for each compartment, and was defined as the mean signal in the compartment divided by the mean signal across the whole genome.

#### Identification of compartment switches

Compartment switches were identified using the genome-wide eigenvector difference (ΔPC1) between A549 SETDB1^LOF^ and A549 control conditions. Only genomic bins where both conditions had annotated A/B compartment state were considered. Genomic bins with ΔPC1 difference between conditions greater than 1.645 standard deviations from the mean ΔPC1 were annotated as compartment switches. The annotated compartment switches were then classified into Stable A, Stable B, A to B, and B to A compartments based on the sign of the eigenvector difference (Table S5).

#### Saddle plots

Saddle plots were generated using cooltools v0.3.2 *compute-saddle* function using 100 kb observed-over-expected *cis* and *trans* matrices, respectively. The expected contact matrices were obtained using cooltools v0.3.2 *compute-expected* function with default parameters. To exclude the extreme eigenvector values from the analysis, top and bottom 2.5 percentiles of the genome-wide eigenvector were clipped. The obtained saddle plots were zoomed into 25 equally sized bins by quantiles.

#### Compartment strength

Compartment strength was calculated from the compartmentalization saddle plots using two metrics as described in^111^ and ^112^ (see Fig. S5f). AA stands for the average contact enrichment within top 20% of the genome-wide eigenvector values, BB stands for the average contact enrichment within bottom 20% of the genome-wide eigenvector values, and AB/BA stands for the average contact depletion between AA and BB.

#### Domains detection

Topologically associating domains (TADs) were annotated using the Armatus^113^ algorithm implementation from the lavaburst Python package ^112^ for 20 kb resolution contact matrices. In this algorithm, the average size and the number of TADs were controlled by the scaling parameter γ. To find the optimal γ for TAD partition, the domains at γ values from 0 to 2.5 were called. Median TAD size, number of TADs and genome coverage with TADs were then calculated for each annotation and plotted as curves. Based on these curves, γ_Control_=0.46 and γ_SETDB1-KO_=0.52 were selected. TADs smaller than 60 kb were dropped out due to their poor resolution. The remaining domains were subjected to manual refinement in order to increase the annotation quality.

#### Domain analysis

Insulation score was calculated using the insulation score algorithm^114^ implemented in cooltools v0.3.2 *diamond-insulation* function for 20 kb resolution contact matrices with window size of 360 kb. The average TAD and average TAD boundary were calculated using coolpup.py v0.9.5^115^ from 20 and 5 kb observed-over-expected contact matrices respectively. The expected contact matrices were obtained using cooltools v0.3.2 *compute-expected* function with ‘--ignore-diags’ set to 0. For the average TAD pile-up, coolpup.py was run with options ‘--local’, ‘--rescale’, and ‘--rescale_size’ set to 99 pixels. For the average TAD boundary, coolpup.py was run with options ‘--local’ and ‘--pad’ set to 200 kb.

#### P(s) curves

P(s) curves were computed using hiclib and the range between 20 kb and 100 Mb was extracted.

## Statistical tests

Statistical analyses were performed using Graphpad Prism v8.4.1 software, MATLAB, R and Python. Statistical significance was determined by the specific tests indicated in the corresponding figure legends.

## Data and code availability

Raw and processed data for Hi-C, RNA-seq, and ChIP-seq are available in the GEO repository under the accession number GSE168233. All custom analysis scripts, macro and data that support the conclusions are available from the authors on request.

## Acknowledgments

We thank Ekaterina Boyarchuk (UMR7216, Université de Paris, Paris, France) for technical help and Sonja Boland (Institut Jacques Monod, Paris, France) for providing the NHBE cell line, advice, and help with their culture. We thank Natalia Klimenko (Institute of Gene Biology, Russian Academy of Science, Moscow, Russia) for advice on the ChIP-seq data analysis. We also thank Mathieu Maurin (U932, Institute Curie, INSERM, Paris, France) for help with image analysis. Work in Ait-Si-Ali’s laboratory was supported by the Fondation pour la Recherche Medicale (FRM, « Equipe FRM » grant # DEQ20160334922); Association Française contre les Myopathies Telethon (AFM-Telethon, grant # 22480); Agence Nationale de la Recherche (ANR, « MuSIC » grant # ANR-17-CE12-0010-01). Work in Razin’s laboratory was supported by the Russian Science Foundation (RSF) grant 21-64-00001 and by the Interdisciplinary Scientific and Educational School of Moscow University “Molecular Technologies of the Living Systems and Synthetic Biology”. Hi-C experiments were performed using the equipment of IGB RAS facilities, supported by the Ministry of Science and Higher Education of the Russian Federation and facilities of the Center for Precision Genome Editing and Genetic Technologies for Biomedicine, IGB RAS.

## Author contributions

V.V.Z. and S.A. conceived the project; V.V.Z. engineered and cultured cell lines, performed functional and validation studies, and immunofluorescence imaging; V.V.Z. and L.D.-M. prepared ChIP-seq and RNA-seq libraries; S.V.U. prepared Hi-C libraries; M.D.M. performed computational analysis of NGS and publicly available data; V. J. engineered constructs for the rescue experiments; A.G. and R.-M.M performed *in vitro* migration assays; B.U. and O.D. performed *in vivo* studies; A.W. and M.P. performed mechanical studies; A.K. and Y.S.V. performed the analysis of immunofluorescence images; V.V.Z., S.V.U., S.V.R., Y.S.V., M.D.M., and S.A wrote the manuscript with input from all authors.

## Author approvals

All authors have seen and approved the manuscript. The manuscript hasn’t been accepted or published elsewhere.

---

**Supplementary Figure 1.**
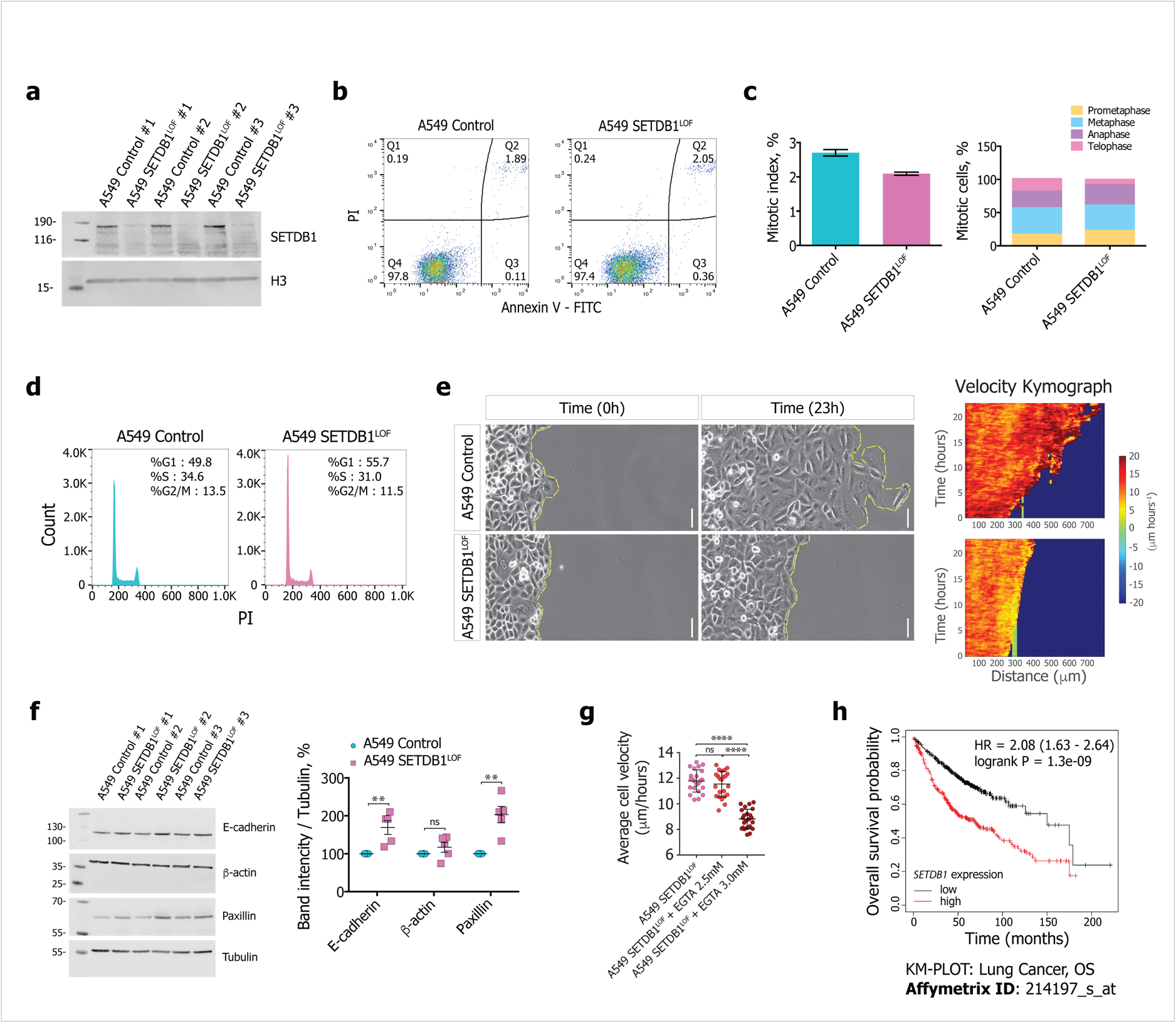
Analysis of SETDB1 LOF-induced phenotype. a. Representative SETDB1 western blot (histone H3 is used as a loading control) in control and SETDB1^LOF^ A549 cells (N = 3 independent experiments). Molecular weight marker is indicated on the left (in kDa).
b. Representative panels of cell apoptosis analysis of control and SETDB1^LOF^ A549 cells assayed by flow cytometry after Annexin V/PI double staining; Q1 (Annexin V-, PI+)-dead cells, Q2 (Annexin V+, PI+) - late-stage apoptosis cells, Q3 (Annexin V+, PI-) - early-stage apoptosis cells and Q4 (Annexin V-, PI-) - live cells.
c. *Left*: mitotic index of control (cyan, the total number of analyzed cells = 1622) and SETDB1^LOF^ (pink, the total number of analyzed cells = 1294) A549 cells. *Right*: percentage of control and SETDB1^LOF^ A549 cells in prometaphase, metaphase, anaphase, and telophase.
d. Representative FACs panels of cell cycle analysis for control (cyan) and SETDB1^LOF^ (pink) A549 cells. Indicated percentages of cells in G1, S, and G2/M phases were determined by flow cytometry after propidium iodide (PI) staining.
e. *Left*: representative images of cellular migration assessment of control and SETDB1^LOF^ A549 cells by wound healing assay at the indicated times (zero and 23 hours). Scale bar, 50 μm. *Right*: space-time diagram (“kymograph”) of cell migration for control (top) and SETDB1^LOF^ (bottom) A549 cells at the indicated time. Corresponding live migration assessment is presented in Supplementary Movies 1 and 2.
f. *Left*: representative western blots for E-cadherin, paxillin, and β-actin in control and SETDB1^LOF^ A549 cells; tubulin served as a loading control. A molecular weight marker is indicated on the left (in kDa). *Right*: quantification of western blot signals normalized to values of control cells (N = 5 independent experiments; mean ± SEM; ns, not significant; **, *P-*value < 0.01, Mann-Whitney U-test).
g. Average cell velocity for SETDB1^LOF^ A549 cells under EGTA treatment. Each point represents a migration rate in one location (N = 2 independent experiments, on average of 23 measurements; mean ± SEM; ns, not significant; ****, *P-*value < 0.0001, t-test).
h. Kaplan-Meier analysis of overall survival (OS) of lung cancer patients (histology type: adenocarcinoma) with high (red) and low (black) *SETDB1* expression (total patient number is 719, number of samples with *SETDB1* high expression = 357, number of samples with *SETDB1* low expression = 362; *P-*value = 1.3e-9, log-rank test; Affymetrix ID: 214197-s-at).

**Supplementary Figure 2.**
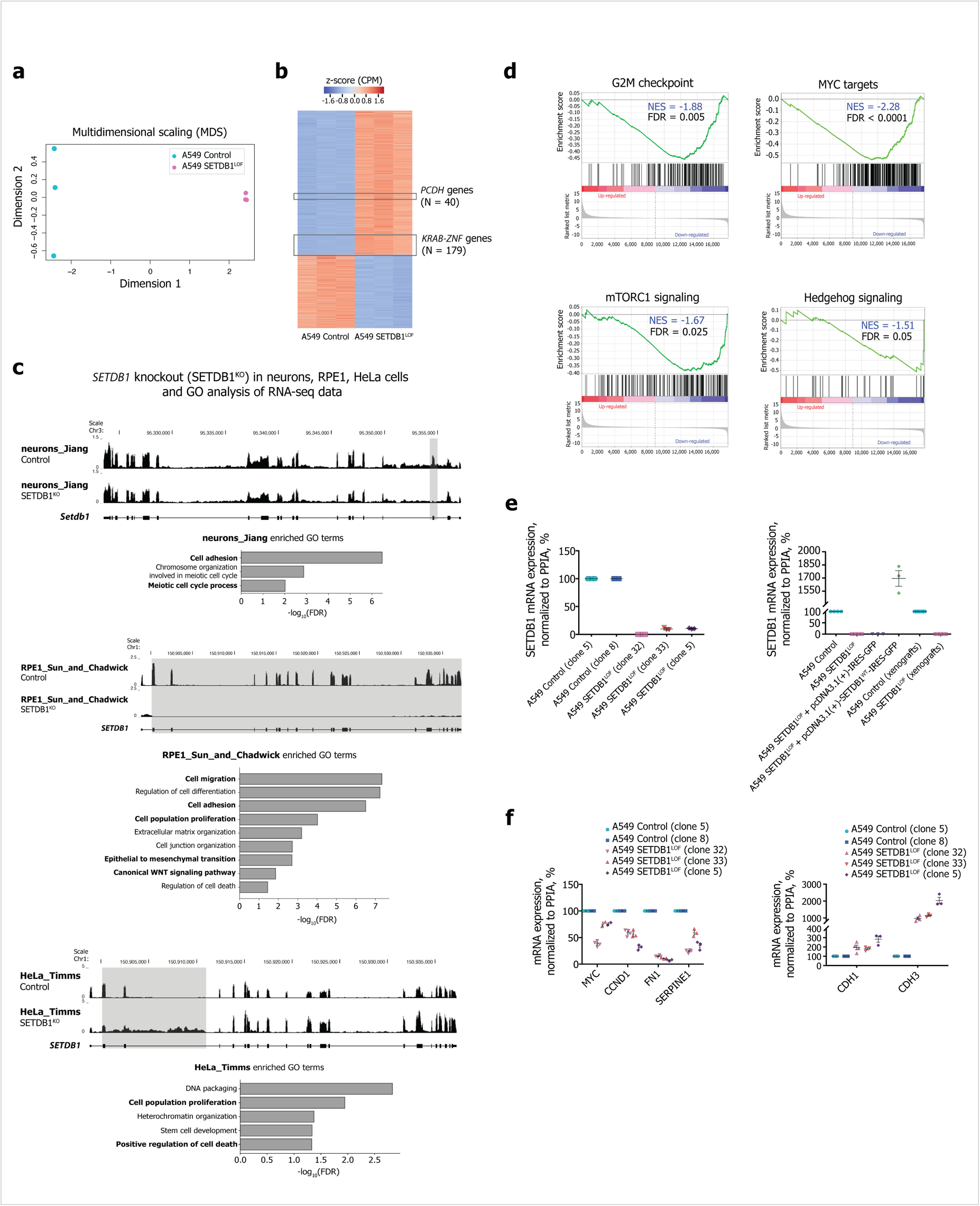
Analysis of transcription changes in lung adenocarcinoma cells with SETDB1 LOF. a. Multidimensional scaling (MDS) plot of the RNA-seq replicates in control (cyan) and SETDB1^LOF^ (pink) A549 cells (N = 3 independent experiments).
b. Heatmap of z-transformed CPM expression counts of differentially expressed genes in SETDB1^LOF^ and control A549 cells. Genes located within the *PCDH* (N=40) and *KRAB-ZNF* (N=179) clusters are highlighted in boxes.
c. UCSC genome browser screenshots of RNA-seq data (CPM-normalized coverage) for *SETDB1* mRNA in cells with *SETDB1* knockout (SETDB1^KO^) and corresponding controls: mouse postnatal forebrain neurons^30^, HeLa^21^, and RPE1 cells^23^. Gene ontology (GO) terms overrepresented in differentially expressed genes (DEGs) are shown under each UCSC genome browser screenshot.
d. Gene set enrichment analysis in SETDB1^LOF^ *vs.* control A549 cells. Normalized enrichment score (NES) and FDR are indicated.
e. *Left*: *SETDB1* mRNA levels in different clones of control and SETDB1^LOF^ A549 cells using primers for validation of the deletion, as measured by RT-qPCR. Data were normalized to the *PPIA* mRNA expression values in control clones that were set as 100 % (N = 3 independent experiments; mean ± SEM). *Right*: *SETDB1* mRNA levels in control and SETDB1^LOF^ A549 cells, control and SETDB1^LOF^ A549 xenografts, and in A549 SETDB1^LOF^ cells transfected with pcDNA3.1(+)-IRES-GFP or pcDNA3.1(+) SETDB1^WT^-IRES-GFP plasmids. RT-qPCR data were normalized to the *PPIA* mRNA expression values in control A549 cells, control xenografts or SETDB1^LOF^ A549 cells transfected with pcDNA3.1(+)-IRES-GFP (control for pcDNA3.1(+)-SETDB1^WT^-IRES-GFP plasmid) that were set as 100 % (N = 3 independent experiments; mean ± SEM).
f. mRNA levels of selected mesenchymal and epithelial markers, and genes related to the WNT/β-catenin signaling pathway in different clones of control and SETDB1^LOF^ A549 clones. RT-qPCR data were normalized to the *PPIA* mRNA expression values in control A549 cells that were set as 100 % (N ≥ 2 independent experiments; mean ± SEM).

**Supplementary Figure 3.**
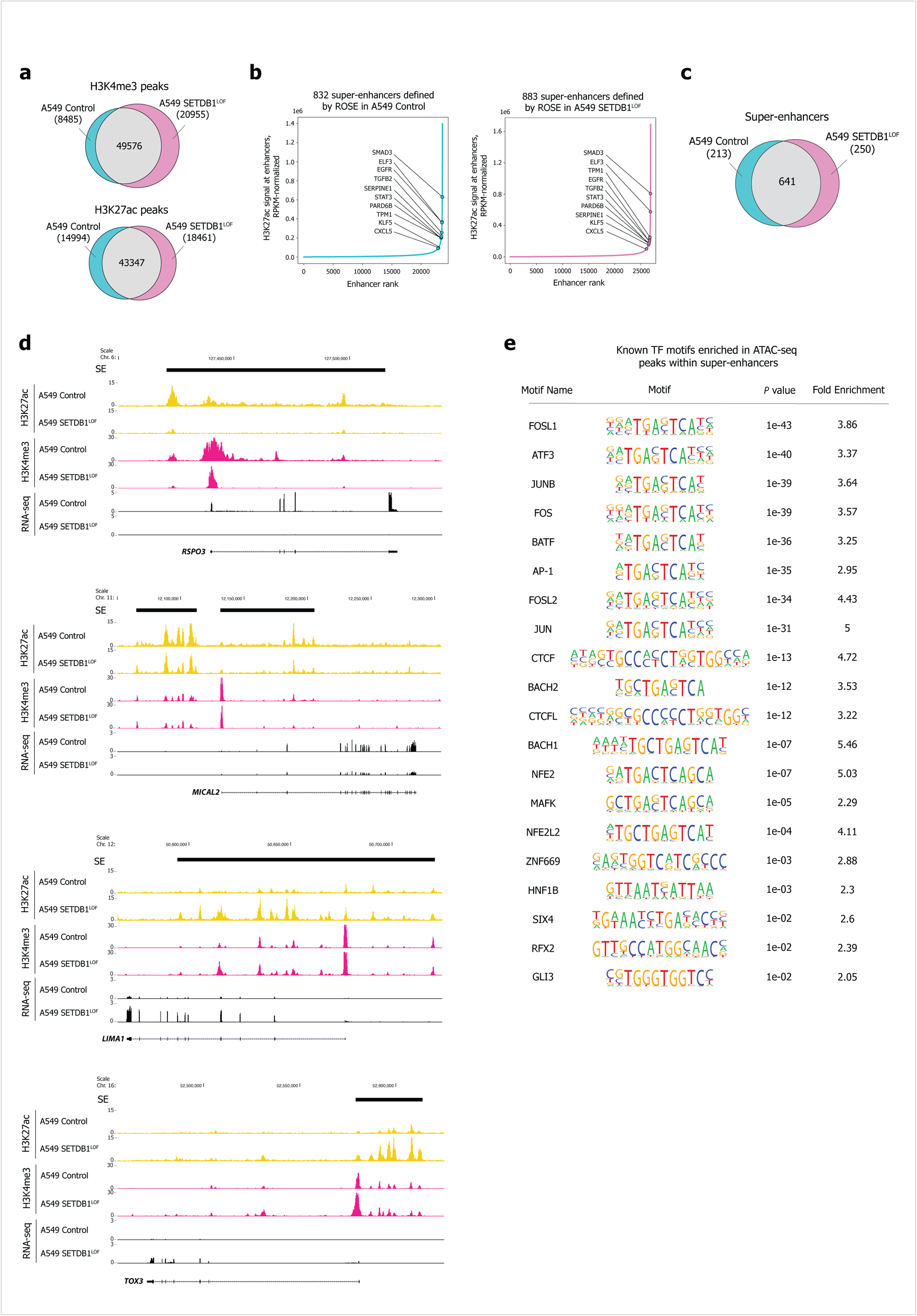
SETDB1 LOF drives changes in the super-enhancers’ landscape. a. *Upper:* venn diagram showing the H3K4me3 ChIP-seq peaks overlap between control and SETDB1^LOF^ A549 cell lines. The number of peaks and overlaps are indicated. *Bottom:* venn diagram showing overlap of H3K27ac ChIP-seq peaks between control (cyan) and SETDB1^LOF^ (pink) A549 cell lines. The numbers of peaks and overlaps are indicated.
b. Super-enhancers (SEs) defined by Rank Ordering of SE approach based on H3K27ac ChIP-seq signals in control (N = 832) and SETDB1^LOF^ (N = 883) A549 cells. Known SEs for the A549 cell line are highlighted.
c. Venn diagram showing the overlap of SEs between control and SETDB1^LOF^ A549 cells. The number of peaks and overlaps are indicated.
d. UCSC genome browser screenshots of H3K27ac (yellow), H3K4me3 (magenta) ChIP-seq, and RNA-seq (black) tracks for the down-regulated *RSPO3* and *MICAL2* and the up-regulated *LIMA1* and *TOX3* SEs in control and SETDB1^LOF^ A549 cells. Annotated coordinates of SEs are indicated in black boxes.
e. Transcription factor motifs identified in differentially regulated SEs (fold enrichment > 2) based on ATAC-seq peaks (data for A549 cells from ENCODE Project Consortium).

**Supplementary Figure 4.**
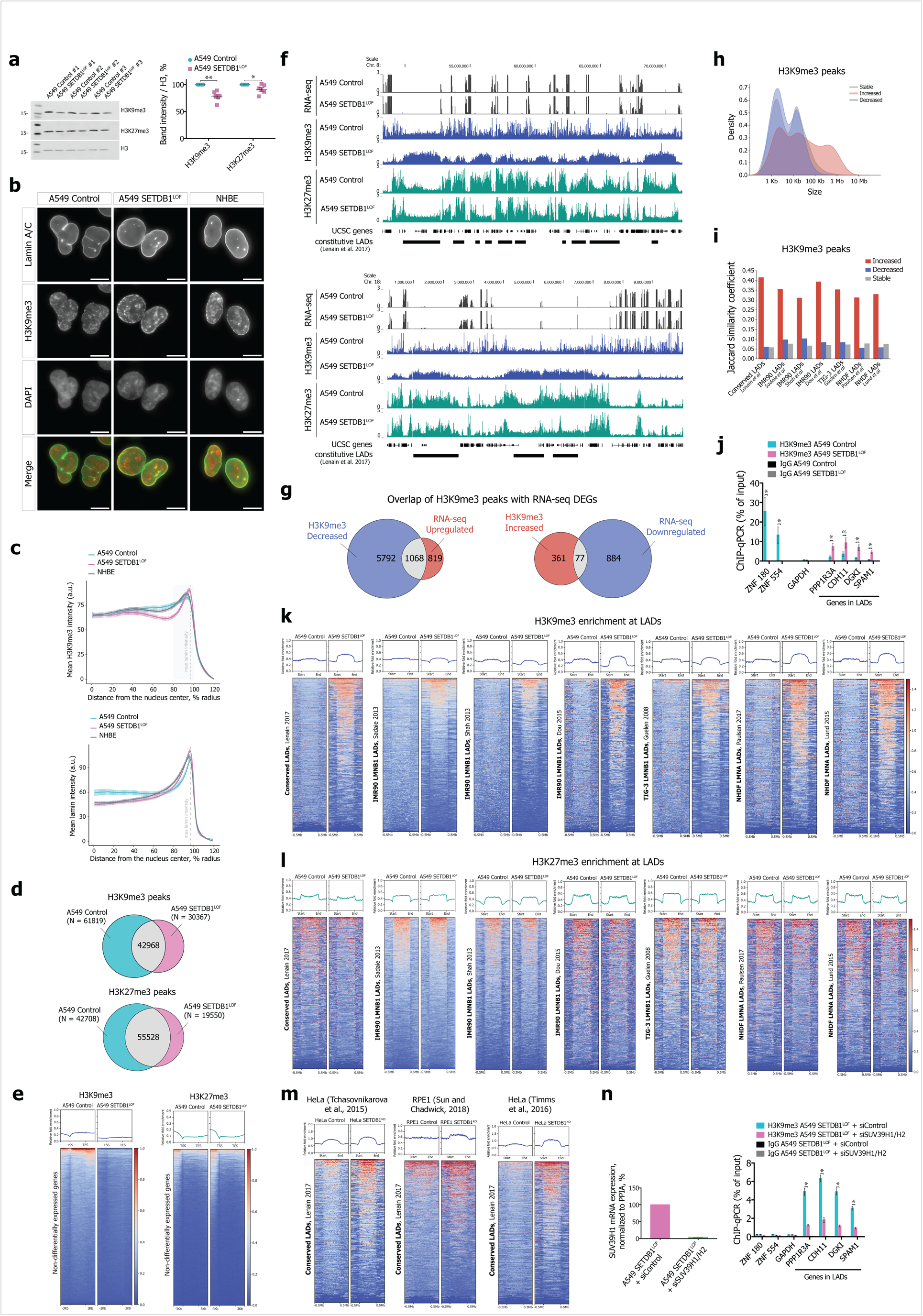
SETDB1 LOF triggered SUV39H1/H2-mediated increase in H3K9me3 within LADs. a. *Left*: representative western blots of H3K9me3 and H3K27me3 total levels in control and SETDB1^LOF^ A549 cells; histone H3 used as a loading control. A molecular weight marker is indicated on the left (in kDa). *Right*: quantification of western blot signals. Data were normalized to H3 values in control A549 cells that were set as 100 % (N = 6 independent experiments; mean ± SEM; *, *P-*value < 0.05; **, *P-*value < 0.01, Mann-Whitney U-test).
b. Representative images of Lamin A/C and H3K9me3 co-immunofluorescence in control and SETDB1^LOF^ A549, and NHBE cells. DNA was labeled with DAPI. Scale bar, 10 μm.
c. Nuclear distribution of H3K9me3 (upper panel) and Lamin A/C (bottom panel) in control and SETDB1^LOF^ A549 cells, and NHBE cells. Intensities are plotted as a function of distance from the nuclear center (N = 3 independent experiments; mean intensity ± SEM for 92-104 cells analyzed). The region with maximum lamin intensity is indicated with a grey dashed line.
d. Venn diagrams showing overlap of H3K9me3 and H3K27me3 ChIP-seq peaks between control and SETDB1^LOF^ A549 cells. The number of peaks and overlaps are indicated.
e. Pile-ups of the H3K9me3 and H3K27me3 enrichment at non-differentially expressed genes in control and SETDB1^LOF^ A549 cells.
f. UCSC genome browser screenshots of RNA-seq, H3K9me3, and H3K27me3 ChIP-seq tracks on Chr. 8: 48.5 – 71.4 Mb and Chr. 18: 0.0 – 10.0 Mb in control and SETDB1^LOF^ A549 cells. Constitutive LADs from Lenain et al. (2017) are indicated.
g. Venn diagram showing the number of up- and down-regulated DEGs in A549 SETDB1^LOF^ *vs.* control cells overlapping with decreased and increased H3K9me3 regions, respectively.
h. Size distribution of increased (red), decreased (blue), and stable (grey) H3K9me3 regions in SETDB1^LOF^ *vs.* control A549 cells.
i. Jaccard’s similarity coefficient between increased H3K9me3 regions and constitutive LADs from Lenain et al. (2017), LADs from IMR90^101–103^, TIG-3^104^, and NHDF^99, 100^ cells.
j. H3K9me3 ChIP-qPCR showing decreased enrichment at selected *ZNF* genes and increased enrichment occupancy at selected genes located at constitutive LADs from Lenain et al. (2017) in control and SETDB1^LOF^ A549 cells (N = 3 independent experiments; mean ± SEM; ns, not significant; *, *P-*value < 0.05, Mann-Whitney U-test).
k. Pile-ups of the H3K9me3 enrichment at constitutive LADs from Lenain et al. (2017), LADs from IMR90^101–103^, TIG-3^104^, and NHDF^99, 100^ cells in control and SETDB1^LOF^ A549 cells.
l. Pile-ups of the H3K27me3 enrichment at constitutive LADs from Lenain et al., (2017), LADs from IMR90^101–103^, TIG-3^104^, and NHDF^99, 100^ cells in control and SETDB1^LOF^ A549 cells.
m. Pile-ups of the H3K9me3 enrichments at constitutive LADs from Lenain et al. (2017) in control and *SETDB1* knockout (SETDB1^KO^) HeLa^21, 47^, and RPE1^23^ cells.
n. *Left*: *SUV39H1* mRNA levels in SETDB1^LOF^ A549 cells upon RNAi-mediated depletion of SUV39H1/H2 (N = 3 independent experiments; mean ± SEM), as measured by RT-qPCR. RT-qPCR data were normalized to the *PPIA* mRNA expression values in SETDB1^LOF^ A549 cells that were set as 100%. *Right*: ChIP-qPCR showing decreased H3K9me3 enrichment at selected genes located at constitutive LADs from Lenain et al. (2017) in SETDB1^LOF^ A549 cells upon RNAi-mediated depletion of SUV39H1/H2 (N = 3 independent experiments; mean ± SEM; *, *P-*value < 0.05, Mann-Whitney U-test).

**Supplementary Figure 5.**
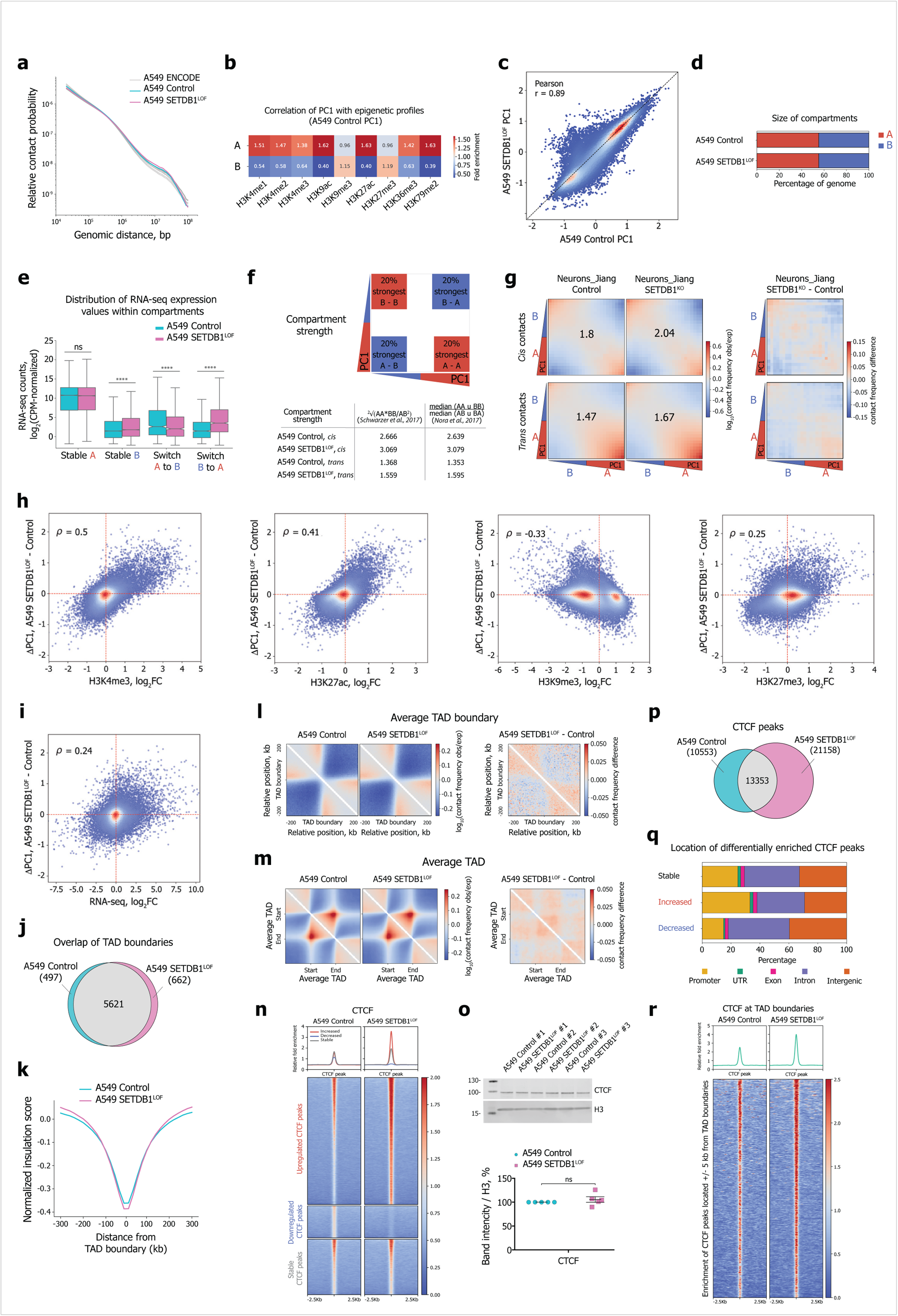
SETDB1 LOF drives A/B compartment switching and minor changes at TAD boundaries. a. Genome-wide relative contact probability scaling for A549 control (cyan) and SETDB1^LOF^ (pink), and A549 cells from ENCODE (grey).
b. Fold enrichment of epigenetic marks in the A and B compartments annotated in control A549 cells.
c. Correlation of PC1 profiles at 100-kb resolution for the control and SETDB1^LOF^ A549 cells. Pearson’s correlation coefficient is indicated.
d. Total size of A (red) and B (blue) compartments in control and SETDB1^LOF^ A549 cells.
e. Distribution of CPM-normalized RNA-seq expression values within stable A and B compartment bins and in switched from B to A, or from A to B compartment bins in control (cyan) and SETDB1^LOF^ (pink) A549 cells (ns, not significant, ****, *P-*value < 0.0001, Mann-Whitney U-test).
f. Compartment strength values for control and SETDB1^LOF^ A549 cells calculated as described in^111, 112^.
g. *Left*: saddle plots of contact enrichments in *cis* and in *trans* between 100-kb genomic bins belonging to A and B compartments in mouse postnatal forebrain control and SETDB1^KO^ neurons^30^. *Right*: subtraction of A549 SETDB1^KO^ and control saddle plots for *cis* and *trans* contacts. Numbers represent compartment strength values calculated as described in^111^.
h. Correlation between H3K4me3, H3K27ac, H3K9me3, H3K27me3 log_2_(fold change), and PC1 values difference in SETDB1^LOF^ *vs.* control A549 cells. Spearman’s correlation coefficients are indicated.
i. Correlation between RNA-seq and PC1 values in SETDB1^LOF^ *vs.* control A549 cells. Spearman’s correlation coefficient is indicated.
j. Venn diagram showing overlap of TAD boundaries between control and SETDB1^LOF^ A549 cells.
k. Average insulation score around TAD boundaries in control and SETDB1^LOF^ A549 cells.
l. *Left:* average TAD boundary in control and SETDB1^LOF^ A549 cells. *Right:* subtraction of SETDB1^LOF^ and control A549 average TAD boundary.
m. *Left:* average TAD in control and SETDB1^LOF^ A549 cells. *Right:* subtraction of SETDB1^LOF^ and control A549 average TADs.
n. Pile-ups of CTCF enrichment at differential CTCF peaks in control and SETDB1^LOF^ A549 cells.
o. *Upper*: representative western blot for CTCF in control and SETDB1^LOF^ A549 cell lines; H3 served as a loading control (N = 3 independent experiments). A molecular weight marker is indicated on the left (in kDa). *Bottom:* quantification of western blot signals. Data normalized to H3 values in control cells that were set as 100% (N = 5 independent experiments; ns, not significant, mean ± SEM).
p. Venn diagram showing overlap of CTCF ChIP-seq peaks between control and SETDB1^LOF^ A549 cells. Numbers of peaks and overlaps are indicated.
q. Genomic locations of differential and stable CTCF ChIP-seq peaks.
r. Pile-ups of CTCF enrichment at TAD boundaries (CTCF peaks from ± 5 kb around TAD boundaries; 1377 peaks for control and 2287 peaks for SETDB1^LOF^) in control and SETDB1^LOF^ A549 cells.

