## Supplemental Table 1 for "SETDB1 Fuels the Lung Cancer Phenotype by Modulating Epigenome, 3D Genome Organization and Chromatin Mechanical Properties"

**Supplementary Table S1. Primer sequences used in the study**

| **qPCR primers** | | |
|---|---|---|
| **Gene** | **Forward** | **Reverse** |
| *MYC* | TGCTGCCAAGAGGGTCAAGT | GTGTGTTCGCCTCTTGACATTC |
| *CCND1* | ACGAAGGTCTGCGCGTGTT | CCGCTGGCCATGAACTACCT |
| *FN1* | CGGTGGCTGTCAGTCAAAG | AAACCTCGGCTTCCTCCATAA |
| *SERPINE1* | CACAAATCAGACGGCAGCACT | CATCGGGCGTGGTGAACTC |
| *PPIA* | GTCAACCCCACCGTGTTCTT | CTGCTGTCTTTGGGACCTTGT- |
| *CDH1* | CCCGCCTTATGATTCTCTGCTCGTG- | TCCGTACATGTCAGCCAGCTTCTTG- |
| *CDH3* | TGACCACAAGCCCAAGTTTAC | TAAGCAACCACCCCATTGTAG |
| *PCDHA6* | CTGTTGAATGATGGCGGACG | GGGAGGAGCAGACATTGCTT |
| *PCDHB6* | AGCCATTCATACTTATGCATATTT | GCTCACAAACTAATGTGTTGACA |
| *PCDHGA4* | CCAACCCAGCTATGCAGACA | TGAGAGACATTTACACAACTCTTTG |
| *SETDB1* | CGCTGCAAATGTGACCCAAA | CTGCTGTCAGAGGAACAGGG |
| *SUV39H1* | GTCATGGAGTACGTGGGAGAG | CCTGACGGTCGTAGATCTGG |

| **ChIP primers** | | |
|---|---|---|
| **Target** | **Forward** | **Reverse** |
| *ZNF180* | TGATGCACAATAAGTCGAGCA | TGCAGTCAATGTGGGAAGTC |
| *ZNF554* | TGATGCACAATAAGTCGAGCA | TGCAGTCAATGTGGGAAGTC |
| *GAPDH* | GTAGGAGGGACTTAGAGAAGG | CTCAAAGGGCAGGAGTAAAG |
| *PPP1R3A* | AGCCTGCTAAGGAGCAACAG | GGGACGCGAATAAGTTTGCT |
| *CDH11* | ATCTGTGGTGGCCGACTCTA | CAACAGCGTGGATGTCGATG |
| *DGKI* | CCATCAGCTTCTTCCAGGGG | ACCCTGGTATTCGGGCAAAG |
| *SPAM1* | AGGAAGCCCCCGAATAAACG | GGGGATTCCTCCATTCACAGT |
