## Supplemental Table 2 for "SETDB1 Fuels the Lung Cancer Phenotype by Modulating Epigenome, 3D Genome Organization and Chromatin Mechanical Properties"

**Supplementary Table 2A. RNA-seq processing statistics**

| | Initial reads | Read pair uniquely mapped |
|---|---|---|
| **RNAseq_Control_rep1** | 34 448 512 | 30 740 862 |
| **RNAseq_Control_rep2** | 25 804 495 | 23 329 514 |
| **RNAseq_Control_rep3** | 41 313 462 | 37 371 810 |
| **RNAseq_SETDB1_LOF_rep1** | 31 209 511 | 28 448 598 |
| **RNAseq_SETDB1_LOF_rep2** | 66 034 540 | 60 697 286 |
| **RNAseq_SETDB1_LOF_rep3** | 30 698 765 | 27 979 051 |

### Supplementary Table 2B, ChIP-seq processing statistics

| | Initial reads | Read pair uniquely mapped with MAPQ>=30 | Read pair is not a duplicate | Read pair is not from blacklist regions |
|---|---|---|---|---|
| **Input_histones_Control_rep1** | 77 783 271 | 60 351 426 | 59 335 436 | 59 050 716 |
| **Input_histones_Control_rep2** | 54 945 495 | 43 625 535 | 42 767 259 | 42 563 137 |
| **Input_histones_Control_rep3** | 56 404 518 | 44 756 173 | 43 826 029 | 43 617 537 |
| **Input_histones_SETDB1_LOF_rep1** | 77 772 272 | 59 857 907 | 58 808 612 | 58 503 672 |
| **Input_histones_SETDB1_LOF_rep2** | 46 435 285 | 36 530 936 | 35 862 227 | 35 675 691 |
| **Input_histones_SETDB1_LOF_rep3** | 53 665 403 | 42 411 998 | 41 579 780 | 41 364 749 |
| **ChIPseq_H3K9me3_Control_rep1** | 91 068 904 | 63 350 153 | 60 909 113 | 60 471 583 |
| **ChIPseq_H3K9me3_Control_rep2** | 83 608 620 | 53 167 462 | 48 533 048 | 48 177 335 |
| **ChIPseq_H3K9me3_SETDB1_LOF_rep1** | 60 207 830 | 39 199 395 | 37 816 325 | 37 542 928 |
| **ChIPseq_H3K9me3_SETDB1_LOF_rep2** | 70 110 882 | 42 793 022 | 39 445 191 | 39 167 486 |
| **ChIPseq_H3K27me3_Control_rep1** | 33 369 314 | 26 289 515 | 25 222 160 | 25 106 656 |
| **ChIPseq_H3K27me3_Control_rep2** | 34 731 938 | 27 810 367 | 26 827 368 | 26 704 608 |
| **ChIPseq_H3K27me3_SETDB1_LOF_rep1** | 31 286 858 | 25 054 585 | 24 049 073 | 23 926 413 |
| **ChIPseq_H3K27me3_SETDB1_LOF_rep2** | 30 103 407 | 24 130 283 | 23 187 718 | 23 069 901 |
| **ChIPseq_H3K4me3_Control_rep1** | 64 801 780 | 55 551 664 | 50 034 307 | 49 748 082 |
| **ChIPseq_H3K4me3_Control_rep2** | 36 955 964 | 31 000 979 | 28 989 432 | 28 819 024 |
| **ChIPseq_H3K4me3_SETDB1_LOF_rep1** | 35 781 741 | 30 299 991 | 28 725 822 | 28 563 519 |
| **ChIPseq_H3K4me3_SETDB1_LOF_rep2** | 42 894 985 | 36 316 951 | 34 021 847 | 33 822 621 |
| **ChIPseq_H3K27ac_Control_rep1** | 33 369 314 | 26 289 515 | 25 222 160 | 25 106 656 |
| **ChIPseq_H3K27ac_Control_rep2** | 34 731 938 | 27 810 367 | 26 827 368 | 26 704 608 |
| **ChIPseq_H3K27ac_SETDB1_LOF_rep1** | 31 286 858 | 25 054 585 | 24 049 073 | 23 926 413 |
| **ChIPseq_H3K27ac_SETDB1_LOF_rep2** | 30 103 407 | 24 130 283 | 23 187 718 | 23 069 901 |

### Supplementary Table 2C, Hi-C processing statistics

| Replicate | Initial reads | Total DS+SS reads | Total DS reads | Same fragment: Total | Same fragment: Self-circles | Same fragment: Dangling ends | Extra Dangling Ends | Valid Pairs | Duplicates | Too Large or Small fragments | Overrepresented fragments | Reads after all filters | cis Reads | trans Reads |
|---|---|---|---|---|---|---|---|---|---|---|---|---|---|---|
| HiC_Control_rep1 | 309 889 029 | 278 203 380 | 192 648 068 | 20 488 586 | 294 778 | 19 694 239 | 17 344 432 | 154 815 050 | 10 167 253 | 6 448 411 | 6 580 128 | 131 619 258 | 103 445 115 | 28 174 143 |
| HiC_Control_rep2 | 285 383 970 | 259 447 734 | 175 767 996 | 2 583 424 | 292 710 | 1 786 573 | 8 124 060 | 165 060 512 | 7 550 508 | 6 974 466 | 7 684 356 | 142 851 182 | 114 300 482 | 28 550 700 |
| HiC_SETDB1_LOF_rep1 | 316 639 388 | 287 249 556 | 194 323 487 | 2 480 192 | 328 728 | 1 535 555 | 9 171 012 | 182 672 283 | 10 777 377 | 7 949 331 | 9 177 289 | 154 768 286 | 123 104 729 | 31 663 557 |
| HiC_SETDB1_LOF_rep2 | 291 295 678 | 263 467 295 | 177 675 577 | 2 109 460 | 307 537 | 1 298 336 | 9 429 711 | 166 136 406 | 8 364 544 | 7 804 620 | 8 323 816 | 141 643 426 | 111 375 377 | 30 268 049 |
