## Supplemental Table 3 for "SETDB1 Fuels the Lung Cancer Phenotype by Modulating Epigenome, 3D Genome Organization and Chromatin Mechanical Properties"

### Supplementary Table 3A, Differentially expression analysis results, Related to Figure 2A

| geneID | gene_name | PValue | FDR | logFC | logCPM | chrom | start | end | strand | gene_type |
|---|---|---|---|---|---|---|---|---|---|---|
| ENSG00000256229.8_3 | ZNF486 | 1.93E-31 | 4.03E-28 | 14.820 | 6.073 | chr19 | 20278023 | 20311297 | + | protein_coding |
| ENSG00000213793.5_5 | ZNF888 | 1.01E-30 | 1.38E-27 | 14.619 | 5.875 | chr19 | 53407670 | 53426734 | - | protein_coding |
| ENSG00000214652.6_4 | ZNF727 | 9.56E-30 | 9.43E-27 | 14.261 | 5.528 | chr7 | 53407670 | 63545717 | + | protein_coding |
| ENSG00000160321.15_6 | ZNF208 | 1.41E-26 | 5.83E-24 | 13.223 | 4.519 | chr19 | 22115760 | 22193751 | - | protein_coding |
| ENSG00000197172.6 | MAGEA6 | 3.02E-25 | 8.39E-23 | 12.595 | 3.914 | chrX | 151867214 | 151870825 | + | protein_coding |
| ENSG00000196109.9_6 | ZNF676 | 1.60E-23 | 2.74E-21 | 12.283 | 3.607 | chr19 | 22361891 | 22398603 | - | protein_coding |
| ENSG00000182986.14_5 | ZNF320 | 6.96E-32 | 1.88E-28 | 12.247 | 5.554 | chr19 | 53364104 | 53400946 | - | protein_coding |
| ENSG00000256771.4_5 | ZNF253 | 1.13E-23 | 2.01E-21 | 12.149 | 3.483 | chr19 | 19976691 | 20005483 | + | protein_coding |
| ENSG00000269825.1_7 | AC022150.4 | 2.41E-23 | 3.99E-21 | 12.021 | 3.361 | chr19 | 53153690 | 53156537 | - | lncRNA |
| ENSG00000197134.13_6 | ZNF257 | 4.27E-23 | 6.55E-21 | 11.990 | 3.329 | chr19 | 22235232 | 22274282 | + | protein_coding |
| ENSG00000198429.10_3 | ZNF69 | 2.52E-23 | 4.11E-21 | 11.952 | 3.295 | chr19 | 11998597 | 12025144 | + | protein_coding |
| ENSG00000188171.16_6 | ZNF626 | 5.04E-23 | 7.47E-21 | 11.892 | 3.235 | chr19 | 20802745 | 20844402 | - | protein_coding |
| ENSG00000233836.7_5 | AC139769.1 | 9.44E-23 | 1.34E-20 | 11.820 | 3.169 | chr19 | 23945746 | 24016116 | + | transcribed_unprocessed_pseudogene |
| ENSG00000184635.16_7 | ZNF93 | 5.55E-22 | 6.37E-20 | 11.659 | 3.009 | chr19 | 20011722 | 20074273 | + | protein_coding |
| ENSG00000213988.11_4 | ZNF90 | 4.42E-22 | 5.13E-20 | 11.644 | 2.999 | chr19 | 20188803 | 20237885 | + | protein_coding |
| ENSG00000105750.15_8 | ZNF85 | 1.18E-21 | 1.28E-19 | 11.597 | 2.948 | chr19 | 21106028 | 21133503 | + | protein_coding |
| ENSG00000269834.6_7 | ZNF528-AS1 | 8.82E-22 | 9.80E-20 | 11.520 | 2.878 | chr19 | 52892095 | 52901036 | - | lncRNA |
| ENSG00000176293.19_2 | ZNF135 | 3.88E-21 | 3.68E-19 | 11.404 | 2.769 | chr19 | 58570607 | 58597677 | + | protein_coding |
| ENSG00000182636.8_7 | NDN | 1.84E-20 | 1.54E-18 | 11.248 | 2.622 | chr15 | 23930547 | 23932452 | - | protein_coding |
| ENSG00000196172.9_4 | ZNF681 | 3.00E-28 | 2.05E-25 | 11.244 | 4.582 | chr19 | 23921997 | 23941693 | - | protein_coding |
| ENSG00000183850.14_4 | ZNF730 | 9.43E-21 | 8.33E-19 | 11.218 | 2.587 | chr19 | 23258012 | 23330023 | + | protein_coding |
| ENSG00000170954.11_6 | ZNF415 | 2.04E-20 | 1.69E-18 | 11.149 | 2.518 | chr19 | 53611132 | 53636330 | - | protein_coding |
| ENSG00000241149.3_5 | AC115220.1 | 3.12E-20 | 2.45E-18 | 11.109 | 2.490 | chr7 | 63459252 | 63476508 | + | protein_coding |
| ENSG00000189190.10_5 | ZNF600 | 2.61E-28 | 1.86E-25 | 11.080 | 4.422 | chr19 | 53253195 | 53290060 | - | protein_coding |
| ENSG00000215146.5_6 | BX322639.1 | 5.26E-20 | 4.06E-18 | 11.073 | 2.449 | chr10 | 42827314 | 42863422 | - | transcribed_unprocessed_pseudogene |
| ENSG00000213801.4_3 | ZNF321P | 3.15E-20 | 2.46E-18 | 10.994 | 2.374 | chr19 | 53430388 | 53445854 | - | transcribed_processed_pseudogene |
| ENSG00000198482.13_6 | ZNF808 | 1.44E-27 | 8.01E-25 | 10.845 | 4.195 | chr19 | 53030905 | 53067717 | + | protein_coding |
| ENSG00000198028.4_4 | ZNF560 | 3.20E-19 | 2.16E-17 | 10.763 | 2.146 | chr19 | 9577031 | 9609292 | - | protein_coding |
| ENSG00000197013.10_7 | ZNF429 | 2.57E-19 | 1.75E-17 | 10.734 | 2.125 | chr19 | 21679484 | 21739072 | + | protein_coding |
| ENSG00000197044.11_3 | ZNF441 | 6.88E-19 | 4.37E-17 | 10.685 | 2.079 | chr19 | 11877815 | 11894893 | + | protein_coding |
| ENSG00000269001.2_6 | AC092070.2 | 1.98E-17 | 1.04E-15 | 10.649 | 2.035 | chr19 | 53700364 | 53717775 | + | transcribed_unprocessed_pseudogene |
| ENSG00000169851.15_4 | PCDH7 | 1.54E-18 | 9.32E-17 | 10.635 | 2.036 | chr4 | 30722037 | 31148427 | + | protein_coding |
| ENSG00000167555.14_5 | ZNF528 | 1.75E-22 | 2.21E-20 | 10.491 | 5.136 | chr19 | 52901102 | 52921665 | + | protein_coding |
| ENSG00000167766.18_4 | ZNF83 | 1.36E-29 | 1.27E-26 | 10.367 | 5.019 | chr19 | 53097313 | 53193749 | - | protein_coding |
| ENSG00000123870.10_2 | ZNF137P | 1.64E-17 | 8.64E-16 | 10.204 | 1.614 | chr19 | 53091758 | 53100598 | + | transcribed_unprocessed_pseudogene |
| ENSG00000286949.1_2 | AC008969.2 | 9.16E-17 | 4.33E-15 | 10.136 | 1.554 | chr19 | 58528063 | 58536957 | - | lncRNA |
| ENSG00000197937.12_5 | ZNF347 | 1.17E-22 | 1.59E-20 | 10.104 | 3.474 | chr19 | 53627325 | 53662328 | - | protein_coding |
| ENSG00000182632.15_4 | CCNYL2 | 1.51E-16 | 6.82E-15 | 10.101 | 1.520 | chr10 | 42903616 | 42970797 | - | transcribed_unprocessed_pseudogene |
| ENSG00000170955.10_7 | CAVIN3 | 1.53E-22 | 2.00E-20 | 10.099 | 3.470 | chr11 | 6340176 | 6341762 | - | protein_coding |
| ENSG00000268696.3_8 | ZNF723 | 6.26E-17 | 3.06E-15 | 10.089 | 1.507 | chr19 | 23015093 | 23041469 | + | protein_coding |
| ENSG00000127324.9_4 | TSPAN8 | 4.17E-17 | 2.10E-15 | 10.087 | 1.498 | chr12 | 71518865 | 71835678 | - | protein_coding |
| ENSG00000235974.1_3 | VN2R19P | 5.82E-17 | 2.87E-15 | 10.023 | 1.442 | chr19 | 58523957 | 58537294 | - | unitary_pseudogene |
| ENSG00000229676.3_3 | ZNF492 | 8.17E-17 | 3.89E-15 | 9.997 | 1.419 | chr19 | 22817126 | 22850473 | + | protein_coding |
| ENSG00000269526.1_2 | ERVV-1 | 2.86E-11 | 5.56E-10 | 9.963 | 1.367 | chr19 | 53517174 | 53519375 | + | protein_coding |
| ENSG00000197497.11_6 | ZNF665 | 4.18E-16 | 1.74E-14 | 9.956 | 1.371 | chr19 | 53662466 | 53696639 | - | protein_coding |
| ENSG00000188868.14_6 | ZNF563 | 2.67E-16 | 1.15E-14 | 9.948 | 1.360 | chr19 | 12428291 | 12444534 | - | protein_coding |
| ENSG00000185037.9_5 | ZNF733P | 2.45E-16 | 1.06E-14 | 9.900 | 1.325 | chr7 | 62751896 | 62764434 | - | transcribed_unprocessed_pseudogene |
| ENSG00000167232.14_5 | ZNF91 | 2.73E-32 | 1.21E-28 | 9.895 | 6.969 | chr19 | 23487793 | 23578273 | - | protein_coding |
| ENSG00000267886.1_7 | AC074135.1 | 6.02E-16 | 2.43E-14 | 9.892 | 1.310 | chr19 | 23258003 | 23283163 | + | lncRNA |
| ENSG00000185177.8 | ZNF479 | 3.05E-16 | 1.30E-14 | 9.831 | 1.260 | chr7 | 57187321 | 57207571 | - | protein_coding |
| ENSG00000268964.3_4 | ERVV-2 | 3.70E-16 | 1.55E-14 | 9.793 | 1.217 | chr19 | 53547993 | 53554933 | + | protein_coding |
| ENSG00000269416.6_6 | LINC01224 | 8.24E-24 | 1.54E-21 | 9.792 | 3.180 | chr19 | 23582035 | 23598877 | - | lncRNA |
| ENSG00000011677.13_4 | GABRA3 | 9.05E-16 | 3.57E-14 | 9.693 | 1.127 | chrX | 151334706 | 151619787 | - | protein_coding |

| | | | | | | | | | | |
|---|---|---|---|---|---|---|---|---|---|---|
| ENSG00000171295.13_5 | ZNF440 | 3.22E-23 | 5.06E-21 | 9.636 | 3.032 | chr19 | 11925088 | 11946031 | + | protein_coding |
| ENSG00000197124.12_7 | ZNF682 | 3.14E-15 | 1.15E-13 | 9.584 | 1.029 | chr19 | 20107867 | 20150314 | - | protein_coding |
| ENSG00000234444.10_4 | ZNF736 | 1.30E-27 | 7.47E-25 | 9.577 | 4.252 | chr7 | 63767837 | 63817012 | + | protein_coding |
| ENSG00000180257.13_4 | ZNF816 | 2.63E-27 | 1.26E-24 | 9.569 | 4.244 | chr19 | 53452632 | 53466164 | - | protein_coding |
| ENSG00000269067.2_6 | ZNF728 | 1.10E-13 | 3.12E-12 | 9.528 | 0.952 | chr19 | 23157685 | 23185978 | - | protein_coding |
| ENSG00000085276.19_8 | MECOM | 4.61E-23 | 7.00E-21 | 9.513 | 2.910 | chr3 | 168801287 | 169381563 | - | protein_coding |
| ENSG00000270619.1_5 | AL391358.1 | 4.52E-15 | 1.60E-13 | 9.509 | 0.943 | chrX | 117412122 | 117417330 | + | processed_pseudogene |
| ENSG00000197647.11_6 | ZNF433 | 9.37E-15 | 3.21E-13 | 9.473 | 0.919 | chr19 | 12125547 | 12146556 | - | protein_coding |
| ENSG00000268100.1_5 | ZNF725P | 4.85E-15 | 1.72E-13 | 9.454 | 0.895 | chr19 | 23675230 | 23695458 | - | unprocessed_pseudogene |
| ENSG00000060718.22_6 | COL11A1 | 7.10E-13 | 1.77E-11 | 9.267 | 0.734 | chr1 | 103342023 | 103574428 | - | protein_coding |
| ENSG00000156755.10_3 | IGKV1OR-2 | 1.83E-14 | 5.99E-13 | 9.264 | 0.716 | chr9 | 69777472 | 69777953 | - | IG_V_pseudogene |
| ENSG00000085563.15_6 | ABCB1 | 8.73E-12 | 1.83E-10 | 9.237 | 0.675 | chr7 | 87132333 | 87342639 | - | protein_coding |
| ENSG00000197990.6_4 | ZNF734P | 4.87E-14 | 1.48E-12 | 9.237 | 0.682 | chr7 | 62910197 | 62917065 | + | unprocessed_pseudogene |
| ENSG00000287326.1_1 | AL589787.1 | 5.27E-14 | 1.59E-12 | 9.188 | 0.638 | chr10 | 128114499 | 128149526 | + | lncRNA |
| ENSG00000128573.26_6 | FOXP2 | 1.90E-13 | 5.12E-12 | 9.172 | 0.637 | chr7 | 113726382 | 114333827 | + | protein_coding |
| ENSG00000182568.17_6 | SATB1 | 7.12E-14 | 2.10E-12 | 9.138 | 0.604 | chr3 | 18386869 | 18487080 | - | protein_coding |
| ENSG00000170927.15_4 | PKHD1 | 5.45E-14 | 1.64E-12 | 9.136 | 0.600 | chr6 | 51480097 | 51952411 | - | protein_coding |
| ENSG00000267383.7_8 | AC011447.3 | 1.03E-13 | 2.97E-12 | 9.133 | 0.599 | chr19 | 20233378 | 20432114 | - | lncRNA |
| ENSG00000213973.9_4 | ZNF99 | 7.44E-14 | 2.18E-12 | 9.129 | 0.590 | chr19 | 22934985 | 22966953 | - | protein_coding |
| ENSG00000267254.7_7 | ZNF790-AS1 | 1.39E-24 | 3.13E-22 | 9.103 | 3.300 | chr19 | 37288404 | 37322498 | + | lncRNA |
| ENSG00000242779.7_5 | ZNF702P | 1.65E-21 | 1.68E-19 | 9.048 | 2.466 | chr19 | 53470149 | 53541151 | - | transcribed_processed_pseudogene |
| ENSG00000267260.2_7 | AC020928.1 | 1.24E-13 | 3.48E-12 | 9.020 | 0.485 | chr19 | 37264014 | 37267980 | + | lncRNA |
| ENSG00000182111.9_3 | ZNF716 | 4.41E-21 | 4.12E-19 | 8.967 | 2.387 | chr7 | 57509883 | 57533265 | + | protein_coding |
| ENSG00000197360.9_7 | ZNF98 | 3.16E-13 | 8.26E-12 | 8.934 | 0.399 | chr19 | 22573821 | 22715287 | - | protein_coding |
| ENSG00000269646.1_7 | AC010487.1 | 3.94E-12 | 8.61E-11 | 8.929 | 0.398 | chr19 | 53426635 | 53427328 | + | lncRNA |
| ENSG00000251247.11_8 | ZNF345 | 1.44E-20 | 1.24E-18 | 8.877 | 2.299 | chr19 | 37341263 | 37403931 | + | protein_coding |
| ENSG00000105388.16_4 | CEACAM5 | 1.22E-12 | 2.93E-11 | 8.841 | 0.307 | chr19 | 42212504 | 42234353 | + | protein_coding |
| ENSG00000188227.13_7 | ZNF793 | 6.67E-28 | 4.23E-25 | 8.837 | 4.439 | chr19 | 37997841 | 38039664 | + | protein_coding |
| ENSG00000198155.5_2 | ZNF876P | 8.18E-13 | 2.02E-11 | 8.820 | 0.292 | chr4 | 206399 | 249774 | + | transcribed_unprocessed_pseudogene |
| ENSG00000197123.9_3 | ZNF679 | 9.79E-13 | 2.36E-11 | 8.811 | 0.299 | chr7 | 63688852 | 63727309 | + | protein_coding |
| ENSG00000248874.5_4 | C5orf17 | 1.23E-12 | 2.95E-11 | 8.750 | 0.232 | chr5 | 23951457 | 24178372 | + | lncRNA |
| ENSG00000177599.13_3 | ZNF491 | 1.45E-12 | 3.46E-11 | 8.681 | 0.171 | chr19 | 11908482 | 11920437 | + | protein_coding |
| ENSG00000286132.1_2 | AC022415.3 | 4.39E-12 | 9.50E-11 | 8.611 | 0.096 | chr19 | 12186142 | 12224140 | + | protein_coding |
| ENSG00000226686.8_5 | LINC01535 | 1.20E-21 | 1.29E-19 | 8.606 | 2.823 | chr19 | 37742787 | 37756437 | + | lncRNA |
| ENSG00000069535.14_4 | MAOB | 9.01E-12 | 1.88E-10 | 8.596 | 0.093 | chrX | 43625857 | 43741696 | - | protein_coding |
| ENSG00000219665.8_7 | ZNF433-AS1 | 1.56E-11 | 3.15E-10 | 8.573 | 0.069 | chr19 | 12098432 | 12157090 | + | lncRNA |
| ENSG00000287394.1_2 | AC244102.4 | 9.41E-12 | 1.96E-10 | 8.550 | 0.052 | chrX | 151743149 | 151867178 | - | lncRNA |
| ENSG00000287158.1_1 | AC117464.1 | 4.90E-12 | 1.05E-10 | 8.539 | 0.043 | chr3 | 84002790 | 84100570 | + | lncRNA |
| ENSG00000161551.14_3 | ZNF577 | 1.50E-21 | 1.56E-19 | 8.478 | 3.193 | chr19 | 52308069 | 52394203 | - | protein_coding |
| ENSG00000274813.1_5 | AC126915.2 | 7.10E-12 | 1.50E-10 | 8.470 | -0.024 | chr5 | 178252383 | 178252572 | + | unprocessed_pseudogene |
| ENSG00000154080.14_5 | CHST9 | 1.75E-11 | 3.50E-10 | 8.460 | -0.040 | chr18 | 24486445 | 24765272 | - | protein_coding |
| ENSG00000164946.19_3 | FREM1 | 2.07E-11 | 4.09E-10 | 8.393 | -0.089 | chr9 | 14734664 | 14910993 | - | protein_coding |
| ENSG00000197863.9_6 | ZNF790 | 1.68E-25 | 5.06E-23 | 8.368 | 3.745 | chr19 | 37308330 | 37341689 | - | protein_coding |
| ENSG00000178222.13_6 | RNF212 | 1.61E-25 | 4.92E-23 | 8.329 | 3.709 | chr4 | 1050038 | 1107352 | - | protein_coding |
| ENSG00000267605.7_8 | AC016590.1 | 1.80E-11 | 3.60E-10 | 8.325 | -0.160 | chr19 | 37726409 | 37795297 | - | lncRNA |
| ENSG00000237440.9_8 | ZNF737 | 2.21E-26 | 8.55E-24 | 8.309 | 3.929 | chr19 | 20718631 | 20748615 | - | protein_coding |
| ENSG00000160282.14_4 | FTCD | 9.85E-11 | 1.78E-09 | 8.237 | -0.232 | chr21 | 47556176 | 47575493 | - | protein_coding |
| ENSG00000286447.1_2 | AC109782.1 | 4.40E-11 | 8.27E-10 | 8.234 | -0.241 | chr3 | 96529682 | 96533251 | - | lncRNA |
| ENSG00000267053.8_7 | AC092296.2 | 6.54E-08 | 7.97E-07 | 8.208 | -0.254 | chr19 | 36795466 | 36812176 | + | lncRNA |
| ENSG00000237268.2_6 | AC092447.7 | 4.48E-11 | 8.42E-10 | 8.200 | -0.271 | chr7 | 56489550 | 56516068 | - | transcribed_unprocessed_pseudogene |
| ENSG00000283236.1_5 | AC074141.1 | 9.23E-18 | 4.99E-16 | 8.144 | 1.598 | chr19 | 52363495 | 52366489 | - | transcribed_unprocessed_pseudogene |
| ENSG00000104447.12_5 | TRPS1 | 1.60E-27 | 8.61E-25 | 8.044 | 4.360 | chr8 | 116420724 | 116821899 | - | protein_coding |
| ENSG00000267565.1_7 | AC011477.2 | 3.02E-10 | 5.13E-09 | 8.027 | -0.426 | chr19 | 20003242 | 20006656 | - | lncRNA |
| ENSG00000185904.12_5 | LINC00839 | 2.99E-23 | 4.83E-21 | 8.011 | 3.116 | chr10 | 42970928 | 42990785 | + | lncRNA |
| ENSG00000007216.14_5 | SLC13A2 | 3.34E-09 | 4.95E-08 | 7.991 | -0.478 | chr17 | 26800311 | 26824799 | + | protein_coding |
| ENSG00000086548.9_5 | CEACAM6 | 4.62E-37 | 8.20E-33 | 7.982 | 8.569 | chr19 | 42254885 | 42276116 | + | protein_coding |
| ENSG00000145794.17_4 | MEGF10 | 2.87E-10 | 4.89E-09 | 7.951 | -0.496 | chr5 | 126626488 | 126801429 | + | protein_coding |

| | | | | | | | | | | |
|---|---|---|---|---|---|---|---|---|---|---|
| ENSG00000267641.1_5 | BNIP3P16 | 3.31E-10 | 5.58E-09 | 7.896 | -0.545 | chr19 | 20285822 | 20287872 | - | processed_pseudogene |
| ENSG00000267934.1_6 | AC010300.1 | 1.08E-09 | 1.71E-08 | 7.883 | -0.548 | chr19 | 23506770 | 23511903 | + | lncRNA |
| ENSG00000198153.8_2 | ZNF849P | 7.63E-10 | 1.23E-08 | 7.854 | -0.584 | chr19 | 22867969 | 22869534 | + | unprocessed_pseudogene |
| ENSG00000268992.1_4 | CDC42EP3P1 | 4.71E-10 | 7.76E-09 | 7.845 | -0.595 | chr19 | 23567332 | 23568095 | + | processed_pseudogene |
| ENSG00000221923.9_3 | ZNF880 | 8.34E-28 | 5.11E-25 | 7.822 | 4.388 | chr19 | 52873170 | 52889048 | + | protein_coding |
| ENSG00000223547.10_4 | ZNF844 | 6.49E-17 | 3.14E-15 | 7.794 | 1.265 | chr19 | 12175546 | 12192380 | + | protein_coding |
| ENSG00000204604.12_6 | ZNF468 | 7.65E-31 | 1.24E-27 | 7.741 | 5.321 | chr19 | 53341263 | 53360902 | - | protein_coding |
| ENSG00000226919.4_7 | AL365184.1 | 1.47E-09 | 2.28E-08 | 7.675 | -0.746 | chr1 | 240926634 | 240940124 | + | lncRNA |
| ENSG00000268970.1_8 | AC022150.2 | 1.59E-09 | 2.46E-08 | 7.666 | -0.748 | chr19 | 53100952 | 53102140 | - | lncRNA |
| ENSG00000138115.14_4 | CYP2C8 | 1.83E-09 | 2.79E-08 | 7.656 | -0.761 | chr10 | 96796529 | 96829254 | - | protein_coding |
| ENSG00000144290.17_5 | SLC4A10 | 2.06E-09 | 3.12E-08 | 7.640 | -0.776 | chr2 | 162280843 | 162841792 | + | protein_coding |
| ENSG00000223558.1_2 | TRIM60P17 | 3.19E-09 | 4.73E-08 | 7.613 | -0.804 | chr7 | 63545938 | 63546954 | + | processed_pseudogene |
| ENSG00000171051.8_4 | FPR1 | 3.63E-09 | 5.36E-08 | 7.574 | -0.836 | chr19 | 52248425 | 52307363 | - | protein_coding |
| ENSG00000164794.9_4 | KCNV1 | 4.31E-09 | 6.24E-08 | 7.542 | -0.854 | chr8 | 110975865 | 110988000 | - | protein_coding |
| ENSG00000167562.12_3 | ZNF701 | 2.00E-26 | 7.92E-24 | 7.507 | 4.089 | chr19 | 53058710 | 53090427 | + | protein_coding |
| ENSG00000154655.15_8 | L3MBTL4 | 3.06E-08 | 3.86E-07 | 7.492 | -0.901 | chr18 | 5954705 | 6415236 | - | protein_coding |
| ENSG00000196757.8_5 | ZNF700 | 1.01E-22 | 1.41E-20 | 7.474 | 2.887 | chr19 | 12035883 | 12061588 | + | protein_coding |
| ENSG00000270325.1_4 | BNIP3P9 | 1.08E-08 | 1.45E-07 | 7.469 | -0.930 | chr19 | 19967784 | 19968336 | + | processed_pseudogene |
| ENSG00000185988.15_8 | PLK5 | 7.28E-09 | 1.01E-07 | 7.468 | -0.918 | chr19 | 1508022 | 1536045 | + | protein_coding |
| ENSG00000160229.13_5 | ZNF66 | 5.86E-22 | 6.63E-20 | 7.468 | 2.593 | chr19 | 20959110 | 20992801 | + | protein_coding |
| ENSG00000196917.6_5 | HCAR1 | 8.93E-09 | 1.22E-07 | 7.436 | -0.950 | chr12 | 123210623 | 123215391 | - | protein_coding |
| ENSG00000251273.4_8 | LINC02228 | 8.77E-09 | 1.20E-07 | 7.434 | -0.950 | chr5 | 25087559 | 25191065 | - | lncRNA |
| ENSG00000206075.14_4 | SERPINB5 | 1.09E-08 | 1.47E-07 | 7.428 | -0.964 | chr18 | 61144191 | 61172318 | + | protein_coding |
| ENSG00000257446.4_5 | ZNF878 | 3.66E-08 | 4.58E-07 | 7.426 | -0.965 | chr19 | 12154620 | 12163776 | - | protein_coding |
| ENSG00000268658.5_7 | LINC00664 | 8.09E-09 | 1.12E-07 | 7.415 | -0.967 | chr19 | 21666176 | 21686040 | + | lncRNA |
| ENSG00000134343.14_6 | ANO3 | 9.73E-09 | 1.32E-07 | 7.411 | -0.968 | chr11 | 26210389 | 26684836 | + | protein_coding |
| ENSG00000286625.1_1 | AC113370.1 | 7.85E-09 | 1.09E-07 | 7.400 | -0.983 | chr5 | 25963948 | 25990796 | + | lncRNA |
| ENSG00000197083.11_5 | ZNF300P1 | 8.99E-09 | 1.23E-07 | 7.379 | -1.000 | chr5 | 150310207 | 150325851 | - | transcribed_unprocessed_pseudogene |
| ENSG00000130303.13_5 | BST2 | 4.88E-08 | 6.01E-07 | 7.377 | -1.014 | chr19 | 17513748 | 17516439 | - | protein_coding |
| ENSG00000287985.1_1 | AC073270.2 | 1.17E-07 | 1.37E-06 | 7.377 | -1.014 | chr7 | 63829619 | 63835962 | + | lncRNA |
| ENSG00000287550.1_2 | AL450345.2 | 9.91E-09 | 1.35E-07 | 7.360 | -1.018 | chr6 | 165204783 | 165241172 | - | lncRNA |
| ENSG00000271225.1_2 | BNIP3P4 | 1.02E-08 | 1.38E-07 | 7.358 | -1.018 | chr9 | 69823283 | 69823847 | - | processed_pseudogene |
| ENSG00000197020.11_7 | ZNF100 | 1.73E-14 | 5.67E-13 | 7.336 | 0.826 | chr19 | 21905573 | 21950381 | - | protein_coding |
| ENSG00000283285.1_6 | AC011460.1 | 1.46E-08 | 1.93E-07 | 7.296 | -1.071 | chr19 | 52417928 | 52420956 | - | unprocessed_pseudogene |
| ENSG00000239893.2_6 | ZNF736P9Y | 5.03E-14 | 1.52E-12 | 7.295 | 0.791 | chrY | 7936486 | 7994334 | - | transcribed_processed_pseudogene |
| ENSG00000172000.7_3 | ZNF556 | 6.66E-15 | 2.33E-13 | 7.293 | 0.787 | chr19 | 2867333 | 2883443 | + | protein_coding |
| ENSG00000253363.7_9 | AC090809.1 | 3.23E-08 | 4.06E-07 | 7.284 | -1.074 | chr8 | 36235587 | 36636727 | - | lncRNA |
| ENSG00000269345.1_2 | VN1R85P | 9.25E-08 | 1.10E-06 | 7.278 | -1.075 | chr19 | 22357568 | 22357993 | - | unprocessed_pseudogene |
| ENSG00000100867.14_3 | DHRS2 | 4.74E-35 | 4.21E-31 | 7.250 | 7.561 | chr14 | 24099324 | 24114848 | + | protein_coding |
| ENSG00000150471.16_4 | ADGRL3 | 6.18E-08 | 7.54E-07 | 7.240 | -1.125 | chr4 | 62066976 | 62944053 | + | protein_coding |
| ENSG00000169248.13_6 | CXCL11 | 1.56E-14 | 5.17E-13 | 7.223 | 0.728 | chr4 | 76954835 | 76962568 | - | protein_coding |
| ENSG00000198342.5 | ZNF442 | 2.59E-14 | 8.32E-13 | 7.151 | 0.649 | chr19 | 12460185 | 12476719 | - | protein_coding |
| ENSG00000145147.20_8 | SLIT2 | 2.92E-13 | 7.67E-12 | 7.116 | 0.608 | chr4 | 20253528 | 20622184 | + | protein_coding |
| ENSG00000167554.15_4 | ZNF610 | 1.35E-16 | 6.15E-15 | 7.107 | 1.391 | chr19 | 52839496 | 52871031 | + | protein_coding |
| ENSG00000196350.9_3 | ZNF729 | 3.22E-07 | 3.53E-06 | 7.105 | -1.242 | chr19 | 22469243 | 22499978 | + | protein_coding |
| ENSG00000249267.7_6 | LINC00939 | 1.43E-07 | 1.66E-06 | 7.098 | -1.243 | chr12 | 126443234 | 126468211 | - | lncRNA |
| ENSG00000224127.1_8 | AC099786.1 | 7.20E-08 | 8.71E-07 | 7.092 | -1.245 | chr1 | 75593515 | 75598743 | - | lncRNA |
| ENSG00000287407.1_3 | AL596442.3 | 7.55E-08 | 9.10E-07 | 7.090 | -1.245 | chr6 | 170473952 | 170475695 | - | lncRNA |
| ENSG00000271095.1_4 | BNIP3P28 | 7.19E-08 | 8.70E-07 | 7.070 | -1.265 | chr19 | 22180450 | 22181642 | + | processed_pseudogene |
| ENSG00000151883.18_4 | PARP8 | 3.71E-22 | 4.37E-20 | 7.064 | 2.732 | chr5 | 49961733 | 50142353 | + | protein_coding |
| ENSG00000276476.3_4 | AL136962.1 | 6.77E-08 | 8.22E-07 | 7.064 | -1.266 | chr13 | 22615111 | 22850665 | + | lncRNA |
| ENSG00000266916.7_9 | ZNF793-AS1 | 6.11E-16 | 2.45E-14 | 7.064 | 1.348 | chr19 | 37988061 | 37998024 | - | lncRNA |
| ENSG00000163638.13_5 | ADAMTS9 | 2.95E-13 | 7.74E-12 | 7.061 | 0.565 | chr3 | 64501330 | 64673676 | - | protein_coding |
| ENSG00000259129.6_6 | LINC00648 | 5.75E-14 | 1.72E-12 | 7.046 | 0.552 | chr14 | 48234157 | 48264505 | - | lncRNA |
| ENSG00000124491.16_3 | F13A1 | 7.61E-08 | 9.17E-07 | 7.043 | -1.286 | chr6 | 6144317 | 6321246 | - | protein_coding |

| | | | | | | | | | | |
|---|---|---|---|---|---|---|---|---|---|---|
| ENSG00000232677.8_8 | LINC00665 | 2.04E-31 | 4.03E-28 | 7.039 | 5.458 | chr19 | 36803969 | 36822672 | - | lncRNA |
| ENSG00000286393.1_2 | Z97192.4 | 9.00E-14 | 2.61E-12 | 7.031 | 0.543 | chr22 | 50055960 | 50058616 | - | lncRNA |
| ENSG00000282556.3_7 | AC068733.3 | 2.03E-07 | 2.31E-06 | 6.993 | -1.328 | chr11 | 6340370 | 6381684 | + | lncRNA |
| ENSG00000231205.7 | ZNF826P | 9.72E-17 | 4.56E-15 | 6.992 | 1.283 | chr19 | 20451078 | 20607775 | - | pseudogene |
| ENSG00000268833.1_7 | AC243967.2 | 2.95E-07 | 3.26E-06 | 6.941 | -1.372 | chr19 | 42261806 | 42290801 | - | lncRNA |
| ENSG00000268223.5_4 | ARL14EPL | 4.23E-16 | 1.76E-14 | 6.926 | 1.218 | chr5 | 115368021 | 115395815 | + | protein_coding |
| ENSG00000162624.14_3 | LHX8 | 3.69E-18 | 2.10E-16 | 6.921 | 1.704 | chr1 | 75594119 | 75627218 | + | protein_coding |
| ENSG00000244694.7_4 | PTCHD4 | 2.66E-07 | 2.97E-06 | 6.914 | -1.395 | chr6 | 47845764 | 48036425 | - | protein_coding |
| ENSG00000249790.3_6 | AC092490.1 | 1.86E-07 | 2.12E-06 | 6.908 | -1.396 | chr12 | 8940849 | 8948385 | + | lncRNA |
| ENSG00000105278.11_4 | ZFR2 | 2.84E-14 | 9.02E-13 | 6.863 | 1.152 | chr19 | 3804022 | 3869036 | - | protein_coding |
| ENSG00000168743.12_7 | NPNT | 3.21E-16 | 1.36E-14 | 6.860 | 1.156 | chr4 | 106815932 | 106925184 | + | protein_coding |
| ENSG00000135063.19_6 | FAM189A2 | 1.79E-13 | 4.85E-12 | 6.858 | 0.382 | chr9 | 71939483 | 72007474 | + | protein_coding |
| ENSG00000240225.10_4 | ZNF542P | 2.24E-24 | 4.91E-22 | 6.847 | 4.704 | chr19 | 56879468 | 56891197 | + | transcribed_unprocessed_pseudogene |
| ENSG00000080224.17_5 | EPHA6 | 1.32E-17 | 7.03E-16 | 6.844 | 1.636 | chr3 | 96533425 | 97471304 | + | protein_coding |
| ENSG00000258342.2_8 | AL133304.3 | 3.64E-07 | 3.97E-06 | 6.793 | -1.490 | chr14 | 36367142 | 36537467 | + | lncRNA |
| ENSG00000198093.11_6 | ZNF649 | 5.08E-24 | 1.01E-21 | 6.776 | 3.388 | chr19 | 52392488 | 52408293 | - | protein_coding |
| ENSG00000058091.17_6 | CDK14 | 6.89E-18 | 3.79E-16 | 6.771 | 2.201 | chr7 | 90095738 | 90839905 | + | protein_coding |
| ENSG00000176571.12_6 | CNBD1 | 1.23E-12 | 2.95E-11 | 6.769 | 0.286 | chr8 | 87878643 | 88627447 | + | protein_coding |
| ENSG00000166148.3_3 | AVPR1A | 3.18E-17 | 1.62E-15 | 6.711 | 1.507 | chr12 | 63536539 | 63544722 | - | protein_coding |
| ENSG00000268455.1_5 | AC073544.1 | 5.92E-07 | 6.29E-06 | 6.662 | -1.592 | chr19 | 23778924 | 23780961 | + | unprocessed_pseudogene |
| ENSG00000121413.12_3 | ZSCAN18 | 1.03E-26 | 4.46E-24 | 6.643 | 4.212 | chr19 | 58595205 | 58629794 | - | protein_coding |
| ENSG00000275713.2_3 | H2BC9 | 1.70E-24 | 3.77E-22 | 6.624 | 3.543 | chr6 | 26251879 | 26253938 | + | protein_coding |
| ENSG00000255980.1_7 | AP000439.2 | 2.23E-12 | 5.15E-11 | 6.581 | 0.119 | chr11 | 69240458 | 69244389 | + | lncRNA |
| ENSG00000196263.8_4 | ZNF471 | 2.11E-22 | 2.58E-20 | 6.572 | 2.947 | chr19 | 57019219 | 57041590 | + | protein_coding |
| ENSG00000164176.13_6 | EDIL3 | 1.70E-14 | 5.60E-13 | 6.555 | 0.866 | chr5 | 83236373 | 83680698 | - | protein_coding |
| ENSG00000242242.6_8 | NECTIN3-AS1 | 2.17E-30 | 2.41E-27 | 6.460 | 5.302 | chr3 | 110607231 | 110790400 | - | lncRNA |
| ENSG00000230392.1_6 | AC004835.1 | 1.10E-11 | 2.26E-10 | 6.443 | -0.012 | chrX | 118425492 | 118469573 | + | lncRNA |
| ENSG00000287498.1_2 | AC007749.2 | 1.07E-13 | 3.06E-12 | 6.419 | 0.726 | chr2 | 217925790 | 217929541 | - | lncRNA |
| ENSG00000101441.4_3 | CST4 | 6.64E-14 | 1.97E-12 | 6.389 | 0.716 | chr20 | 23666277 | 23669677 | - | protein_coding |
| ENSG00000244675.2_5 | AC108676.1 | 1.55E-11 | 3.13E-10 | 6.381 | -0.066 | chr3 | 194217046 | 194222232 | - | lncRNA |
| ENSG00000196268.11_3 | ZNF493 | 5.54E-24 | 1.08E-21 | 6.357 | 3.376 | chr19 | 21579921 | 21610375 | + | protein_coding |
| ENSG00000146039.11_4 | SLC17A4 | 4.04E-11 | 7.69E-10 | 6.286 | -0.157 | chr6 | 25754927 | 25781427 | + | protein_coding |
| ENSG00000125999.11_4 | BPIFB1 | 1.15E-16 | 5.30E-15 | 6.259 | 1.715 | chr20 | 31861286 | 31897677 | + | protein_coding |
| ENSG00000198939.8_4 | ZFP2 | 2.67E-13 | 7.06E-12 | 6.247 | 0.566 | chr5 | 178322899 | 178360213 | + | protein_coding |
| ENSG00000120324.9_3 | PCDHB10 | 8.17E-11 | 1.49E-09 | 6.231 | -0.199 | chr5 | 140571926 | 140575220 | + | protein_coding |
| ENSG00000184305.15_4 | CCSER1 | 3.33E-11 | 6.41E-10 | 6.225 | -0.208 | chr4 | 91048686 | 92523064 | + | protein_coding |
| ENSG00000205809.9_5 | KLRC2 | 1.69E-14 | 5.58E-13 | 6.187 | 1.006 | chr12 | 10579453 | 10594899 | - | protein_coding |
| ENSG00000161298.18_5 | ZNF382 | 6.77E-15 | 2.37E-13 | 6.175 | 1.005 | chr19 | 37095719 | 37125016 | + | protein_coding |
| ENSG00000118596.12_5 | SLC16A7 | 4.36E-28 | 2.87E-25 | 6.147 | 4.506 | chr12 | 59989810 | 60183636 | + | protein_coding |
| ENSG00000117148.8_3 | ACTL8 | 6.71E-24 | 1.29E-21 | 6.142 | 3.260 | chr1 | 18081828 | 18153558 | + | protein_coding |
| ENSG00000134827.8_4 | TCN1 | 5.65E-22 | 6.44E-20 | 6.120 | 3.060 | chr11 | 59620273 | 59633962 | - | protein_coding |
| ENSG00000177025.3_4 | C19orf18 | 1.67E-14 | 5.51E-13 | 6.106 | 0.935 | chr19 | 58469805 | 58485902 | - | protein_coding |
| ENSG00000285996.1_3 | AC006116.12 | 1.24E-13 | 3.48E-12 | 6.017 | 0.855 | chr19 | 56891599 | 56895293 | + | lncRNA |
| ENSG00000228933.8_8 | AC107419.1 | 1.98E-10 | 3.45E-09 | 6.003 | -0.402 | chrX | 27193037 | 27417123 | - | lncRNA |
| ENSG00000267041.6_4 | ZNF850 | 2.52E-31 | 4.47E-28 | 5.994 | 5.857 | chr19 | 37205285 | 37263727 | - | protein_coding |
| ENSG00000231056.1_7 | LINC02522 | 5.42E-12 | 1.16E-10 | 5.979 | 0.339 | chr6 | 10474733 | 10478735 | - | lncRNA |
| ENSG00000187094.12_4 | CCK | 6.31E-10 | 1.02E-08 | 5.977 | 0.343 | chr3 | 42299317 | 42307677 | - | protein_coding |
| ENSG00000231290.6_6 | APCDD1L-DT | 5.51E-15 | 1.94E-13 | 5.976 | 0.807 | chr20 | 57090435 | 57194944 | + | lncRNA |
| ENSG00000265972.6_4 | TXNIP | 3.49E-25 | 9.40E-23 | 5.936 | 3.643 | chr1 | 145438490 | 145442635 | + | protein_coding |
| ENSG00000160505.16_8 | NLRP4 | 3.31E-10 | 5.58E-09 | 5.929 | -0.470 | chr19 | 56347906 | 56393221 | + | protein_coding |
| ENSG00000117228.10_4 | GBP1 | 7.86E-15 | 2.73E-13 | 5.922 | 0.761 | chr1 | 89518002 | 89530913 | - | protein_coding |
| ENSG00000134258.12 | VTCN1 | 2.56E-16 | 1.10E-14 | 5.910 | 1.646 | chr1 | 117686209 | 117753556 | - | protein_coding |
| ENSG00000125207.7_4 | PIWIL1 | 3.36E-10 | 5.65E-09 | 5.910 | -0.484 | chr12 | 130822432 | 130857182 | + | protein_coding |
| ENSG00000255085.8_5 | AF186192.2 | 4.39E-10 | 7.28E-09 | 5.906 | -0.484 | chr8 | 145925738 | 145933902 | + | transcribed_unprocessed_pseudogene |
| ENSG00000169562.12_6 | GJB1 | 6.17E-12 | 1.31E-10 | 5.897 | 0.248 | chrX | 70435044 | 70445366 | + | protein_coding |
| ENSG00000170373.8_3 | CST1 | 1.22E-24 | 2.77E-22 | 5.894 | 3.602 | chr20 | 23728190 | 23731905 | - | protein_coding |
| ENSG00000250584.3_7 | LINC01511 | 7.66E-14 | 2.24E-12 | 5.877 | 0.713 | chr5 | 1354011 | 1380222 | - | lncRNA |

| | | | | | | | | | | |
|---|---|---|---|---|---|---|---|---|---|---|
| ENSG00000203326.12_5 | ZNF525 | 2.22E-25 | 6.36E-23 | 5.814 | 4.047 | chr19 | 53868946 | 53895470 | + | protein_coding |
| ENSG00000018869.16_6 | ZNF582 | 9.48E-23 | 1.34E-20 | 5.789 | 2.998 | chr19 | 56887215 | 56904914 | - | protein_coding |
| ENSG00000151882.11_4 | CCL28 | 5.66E-09 | 8.02E-08 | 5.789 | -0.585 | chr5 | 43376747 | 43412493 | - | protein_coding |
| ENSG00000196542.8_3 | SPTSSB | 1.01E-11 | 2.09E-10 | 5.783 | 0.154 | chr3 | 161062580 | 161090668 | - | protein_coding |
| ENSG00000174607.11_5 | UGT8 | 2.72E-09 | 4.06E-08 | 5.774 | -0.598 | chr4 | 115519926 | 115599381 | + | protein_coding |
| ENSG00000136237.19_7 | RAPGEF5 | 6.40E-12 | 1.35E-10 | 5.762 | 0.114 | chr7 | 22157854 | 22396773 | - | protein_coding |
| ENSG00000133687.16_7 | TMTC1 | 1.66E-13 | 4.55E-12 | 5.740 | 0.591 | chr12 | 29653773 | 29937692 | - | protein_coding |
| ENSG00000173467.9_5 | AGR3 | 1.25E-11 | 2.56E-10 | 5.693 | 0.061 | chr7 | 16899036 | 16921611 | - | protein_coding |
| ENSG00000233539.5_7 | AC011294.1 | 1.18E-08 | 1.58E-07 | 5.651 | -0.706 | chr7 | 46713383 | 46799449 | - | lncRNA |
| ENSG00000087116.16_6 | ADAMTS2 | 1.79E-08 | 2.33E-07 | 5.628 | -0.733 | chr5 | 178537854 | 178772462 | - | protein_coding |
| ENSG00000169306.10_4 | IL1RAPL1 | 7.78E-09 | 1.08E-07 | 5.622 | -0.745 | chrX | 28605563 | 29974835 | + | protein_coding |
| ENSG00000081803.16_7 | CADPS2 | 2.17E-19 | 1.52E-17 | 5.619 | 2.396 | chr7 | 121958465 | 122526813 | - | protein_coding |
| ENSG00000267454.6_6 | ZNF582-AS1 | 9.40E-18 | 5.06E-16 | 5.618 | 1.737 | chr19 | 56905025 | 56910541 | + | lncRNA |
| ENSG00000214189.9_5 | ZNF788P | 4.79E-09 | 6.88E-08 | 5.606 | -0.748 | chr19 | 12203078 | 12248050 | + | transcribed_unprocessed_pseudogene |
| ENSG00000154764.6_4 | WNT7A | 3.45E-11 | 6.63E-10 | 5.569 | -0.050 | chr3 | 13857755 | 13921568 | - | protein_coding |
| ENSG00000197550.3_5 | AL359955.1 | 4.12E-11 | 7.82E-10 | 5.568 | -0.057 | chr9 | 69830288 | 69848759 | + | unprocessed_pseudogene |
| ENSG00000249378.6_6 | LINC01060 | 1.82E-16 | 8.03E-15 | 5.566 | 1.844 | chr4 | 189321890 | 189602205 | + | lncRNA |
| ENSG00000145335.16_4 | SNCA | 9.81E-09 | 1.33E-07 | 5.516 | -0.824 | chr4 | 90621496 | 90759466 | - | protein_coding |
| ENSG00000164761.9_4 | TNFRSF11B | 4.62E-16 | 1.90E-14 | 5.507 | 1.260 | chr8 | 119935796 | 119964124 | - | protein_coding |
| ENSG00000120327.6_3 | PCDHB14 | 2.53E-15 | 9.42E-14 | 5.462 | 0.976 | chr5 | 140602504 | 140607331 | + | protein_coding |
| ENSG00000196684.12_6 | HSH2D | 1.10E-10 | 1.98E-09 | 5.444 | -0.162 | chr19 | 16244838 | 16269386 | + | protein_coding |
| ENSG00000012504.15_6 | NR1H4 | 5.33E-21 | 4.91E-19 | 5.442 | 2.745 | chr12 | 100867486 | 100958192 | + | protein_coding |
| ENSG00000261780.3_6 | LINC02582 | 1.13E-28 | 9.11E-26 | 5.421 | 5.062 | chr18 | 70992176 | 71017114 | + | lncRNA |
| ENSG00000213401.10_5 | MAGEA12 | 2.40E-10 | 4.12E-09 | 5.415 | -0.183 | chrX | 151899293 | 151903206 | - | protein_coding |
| ENSG00000215367.10_4 | TMED11P | 2.38E-08 | 3.05E-07 | 5.375 | -0.950 | chr4 | 1108985 | 1147514 | - | transcribed_unitary_pseudogene |
| ENSG00000250722.6_8 | SELENOP | 3.49E-08 | 4.38E-07 | 5.371 | -0.951 | chr5 | 42799982 | 42887494 | - | protein_coding |
| ENSG00000113248.6_4 | PCDHB15 | 2.34E-16 | 1.02E-14 | 5.363 | 1.497 | chr5 | 140624963 | 140628933 | + | protein_coding |
| ENSG00000078596.11_4 | ITM2A | 3.15E-15 | 1.15E-13 | 5.344 | 1.629 | chrX | 78615881 | 78623164 | - | protein_coding |
| ENSG00000246082.2_3 | NUDT16P1 | 1.16E-14 | 3.92E-13 | 5.339 | 0.862 | chr3 | 131080742 | 131083966 | + | transcribed_unitary_pseudogene |
| ENSG00000162645.13_5 | GBP2 | 1.60E-21 | 1.64E-19 | 5.305 | 3.042 | chr1 | 89571815 | 89616139 | - | protein_coding |
| ENSG00000179869.15_5 | ABCA13 | 2.64E-08 | 3.35E-07 | 5.295 | -1.018 | chr7 | 48211055 | 48687093 | + | protein_coding |
| ENSG00000132010.16_5 | ZNF20 | 3.22E-08 | 4.05E-07 | 5.281 | -1.034 | chr19 | 12203658 | 12251170 | - | protein_coding |
| ENSG00000127249.15_5 | ATP13A4 | 1.24E-09 | 1.96E-08 | 5.278 | -0.321 | chr3 | 193116756 | 193310900 | - | protein_coding |
| ENSG00000224265.1_7 | LINC02633 | 4.31E-08 | 5.35E-07 | 5.256 | -1.053 | chr10 | 43819113 | 43823380 | + | lncRNA |
| ENSG00000106571.14_6 | GLI3 | 8.86E-08 | 1.06E-06 | 5.241 | -1.069 | chr7 | 42000547 | 42277469 | - | protein_coding |
| ENSG00000253159.3_4 | PCDHGA12 | 3.09E-10 | 5.23E-09 | 5.211 | -0.368 | chr5 | 140810074 | 140892542 | + | protein_coding |
| ENSG00000224271.7_7 | AL117329.1 | 4.66E-10 | 7.70E-09 | 5.196 | -0.388 | chr22 | 48026576 | 48418772 | + | lncRNA |
| ENSG00000181143.15_5 | MUC16 | 1.29E-14 | 4.31E-13 | 5.192 | 0.965 | chr19 | 8959520 | 9092018 | - | protein_coding |
| ENSG00000260160.1_8 | AC011468.1 | 2.77E-13 | 7.31E-12 | 5.189 | 0.441 | chr19 | 52561743 | 52566956 | - | lncRNA |
| ENSG00000272674.3_6 | PCDHB16 | 2.45E-17 | 1.26E-15 | 5.170 | 1.618 | chr5 | 140560980 | 140565974 | + | protein_coding |
| ENSG00000146001.5_5 | PCDHB18P | 1.59E-07 | 1.84E-06 | 5.162 | -1.128 | chr5 | 140613905 | 140617101 | + | transcribed_unprocessed_pseudogene |
| ENSG00000173175.14_8 | ADCY5 | 9.38E-27 | 4.27E-24 | 5.155 | 4.332 | chr3 | 123001143 | 123168605 | - | protein_coding |
| ENSG00000171817.17_5 | ZNF540 | 1.44E-19 | 1.05E-17 | 5.153 | 2.239 | chr19 | 38042308 | 38105080 | + | protein_coding |
| ENSG00000119125.16_2 | GDA | 3.75E-26 | 1.36E-23 | 5.139 | 4.152 | chr9 | 74729511 | 74872109 | + | protein_coding |
| ENSG00000256615.1_7 | AC010197.1 | 5.94E-18 | 3.30E-16 | 5.135 | 1.942 | chr12 | 21815247 | 21912966 | + | lncRNA |
| ENSG00000268095.1_6 | ZNF649-AS1 | 2.80E-07 | 3.11E-06 | 5.128 | -1.149 | chr19 | 52391278 | 52403716 | + | lncRNA |
| ENSG00000274373.1_6 | AC148476.1 | 4.84E-11 | 9.04E-10 | 5.106 | 0.017 | chr12 | 133001090 | 133001794 | - | lncRNA |
| ENSG00000177839.7_7 | PCDHB9 | 1.97E-09 | 2.99E-08 | 5.047 | -0.510 | chr5 | 140566701 | 140571114 | + | protein_coding |
| ENSG00000154493.19_7 | C10orf90 | 4.75E-11 | 8.88E-10 | 5.016 | -0.082 | chr10 | 128113566 | 128487277 | - | protein_coding |
| ENSG00000257743.8_2 | MGAM2 | 9.61E-17 | 4.52E-15 | 5.006 | 1.701 | chr7 | 141811549 | 141922124 | + | protein_coding |
| ENSG00000069431.11_4 | ABCC9 | 7.75E-17 | 3.71E-15 | 4.948 | 1.535 | chr12 | 21950335 | 22095463 | - | protein_coding |
| ENSG00000170298.15_4 | LGALS9B | 6.11E-08 | 7.46E-07 | 4.945 | -0.598 | chr17 | 20352708 | 20370852 | - | protein_coding |
| ENSG00000203635.2_7 | AC144450.1 | 1.13E-08 | 1.52E-07 | 4.936 | -0.610 | chr2 | 1624282 | 1629191 | - | lncRNA |
| ENSG00000150540.14_3 | HNMT | 2.81E-22 | 3.37E-20 | 4.919 | 3.374 | chr2 | 138721590 | 138773934 | + | protein_coding |
| ENSG00000148053.16_3 | NTRK2 | 1.74E-10 | 3.06E-09 | 4.915 | -0.169 | chr9 | 87283466 | 87641985 | + | protein_coding |
| ENSG00000106560.11_3 | GIMAP2 | 7.58E-09 | 1.05E-07 | 4.905 | -0.647 | chr7 | 150382785 | 150390729 | + | protein_coding |
| ENSG00000153993.13_3 | SEMA3D | 2.18E-10 | 3.75E-09 | 4.873 | -0.208 | chr7 | 84624869 | 84816171 | - | protein_coding |
| ENSG00000198734.12_4 | F5 | 3.23E-15 | 1.17E-13 | 4.872 | 1.333 | chr1 | 169481189 | 169555826 | - | protein_coding |

| | | | | | | | | | | |
|---|---|---|---|---|---|---|---|---|---|---|
| ENSG00000114200.10_5 | BCHE | 1.04E-17 | 5.58E-16 | 4.869 | 1.880 | chr3 | 165490692 | 165555250 | - | protein_coding |
| ENSG00000229401.1_5 | MIR5689HG | 4.27E-09 | 6.20E-08 | 4.863 | -0.676 | chr6 | 10434549 | 10457014 | + | lncRNA |
| ENSG00000180787.6_3 | ZFP3 | 1.02E-24 | 2.44E-22 | 4.860 | 4.235 | chr17 | 4981762 | 4999669 | + | protein_coding |
| ENSG00000130508.11_5 | PXDN | 1.84E-29 | 1.64E-26 | 4.849 | 5.488 | chr2 | 1635659 | 1748624 | - | protein_coding |
| ENSG00000183668.18_5 | PSG9 | 5.34E-11 | 9.94E-10 | 4.849 | 0.138 | chr19 | 43715943 | 43773682 | - | protein_coding |
| ENSG00000148346.12_4 | LCN2 | 1.53E-22 | 2.00E-20 | 4.820 | 3.516 | chr9 | 130911350 | 130915732 | + | protein_coding |
| ENSG00000255644.1_7 | AC010185.1 | 4.62E-08 | 5.71E-07 | 4.780 | -0.758 | chr12 | 21912914 | 21919639 | + | lncRNA |
| ENSG00000126467.11_6 | TSKS | 8.35E-07 | 8.71E-06 | 4.780 | -1.442 | chr19 | 50243010 | 50266563 | - | protein_coding |
| ENSG00000231535.7_6 | LINC00278 | 1.75E-09 | 2.68E-08 | 4.778 | -0.291 | chrY | 2870928 | 3068550 | + | lncRNA |
| ENSG00000276170.4_7 | AC244153.1 | 4.25E-10 | 7.07E-09 | 4.777 | -0.290 | chr17 | 36606638 | 36608688 | - | lncRNA |
| ENSG00000285407.1_3 | AL033530.1 | 1.40E-08 | 1.85E-07 | 4.762 | -0.762 | chr1 | 69144885 | 69409135 | + | lncRNA |
| ENSG00000286339.1_1 | Z99496.2 | 9.30E-10 | 1.48E-08 | 4.754 | -0.303 | chr6 | 118886823 | 118898003 | + | lncRNA |
| ENSG00000188488.14_5 | SERPINA5 | 2.64E-09 | 3.95E-08 | 4.746 | 1.769 | chr14 | 95027779 | 95059455 | + | protein_coding |
| ENSG00000066468.22_6 | FGFR2 | 5.22E-13 | 1.33E-11 | 4.738 | 0.544 | chr10 | 123237848 | 123357972 | - | protein_coding |
| ENSG00000133636.11_3 | NTS | 1.85E-20 | 1.54E-18 | 4.725 | 2.644 | chr12 | 86268073 | 86276770 | + | protein_coding |
| ENSG00000187372.11_3 | PCDHB13 | 9.34E-19 | 5.85E-17 | 4.690 | 2.109 | chr5 | 140593491 | 140598551 | + | protein_coding |
| ENSG00000012779.11_3 | ALOX5 | 6.32E-17 | 3.09E-15 | 4.676 | 1.786 | chr10 | 45869624 | 45941567 | + | protein_coding |
| ENSG00000196267.12_5 | ZNF836 | 2.49E-19 | 1.71E-17 | 4.670 | 2.599 | chr19 | 52657117 | 52674896 | - | protein_coding |
| ENSG00000159871.15_6 | LYPD5 | 8.88E-10 | 1.42E-08 | 4.667 | -0.027 | chr19 | 44290026 | 44331358 | - | protein_coding |
| ENSG00000250376.1_5 | AC093720.1 | 2.17E-08 | 2.80E-07 | 4.626 | -0.883 | chr4 | 69650336 | 69653192 | + | unprocessed_pseudogene |
| ENSG00000250420.8_5 | AACSP1 | 5.17E-21 | 4.79E-19 | 4.621 | 2.895 | chr5 | 178191862 | 178245436 | - | transcribed_unprocessed_pseudogene |
| ENSG00000198346.11_5 | ZNF813 | 7.57E-26 | 2.56E-23 | 4.607 | 4.482 | chr19 | 53970987 | 53999509 | + | protein_coding |
| ENSG00000146090.16_4 | RASGEF1C | 2.97E-15 | 1.09E-13 | 4.597 | 1.216 | chr5 | 179527795 | 179636211 | - | protein_coding |
| ENSG00000137965.11_3 | IFI44 | 1.32E-07 | 1.54E-06 | 4.587 | -0.915 | chr1 | 79115481 | 79129763 | + | protein_coding |
| ENSG00000101276.17_7 | SLC52A3 | 3.96E-21 | 3.74E-19 | 4.582 | 2.995 | chr20 | 740724 | 756658 | - | protein_coding |
| ENSG00000140835.9_2 | CHST4 | 4.66E-09 | 6.72E-08 | 4.580 | -0.468 | chr16 | 71559136 | 71572649 | + | protein_coding |
| ENSG00000104267.10_3 | CA2 | 3.02E-16 | 1.29E-14 | 4.568 | 1.507 | chr8 | 86376197 | 86393722 | + | protein_coding |
| ENSG00000140450.9_3 | ARRDC4 | 1.97E-18 | 1.17E-16 | 4.549 | 2.042 | chr15 | 98503933 | 98517063 | + | protein_coding |
| ENSG00000240280.7_3 | TCAM1P | 1.97E-12 | 4.58E-11 | 4.540 | 0.558 | chr17 | 61926652 | 61941739 | + | transcribed_unitary_pseudogene |
| ENSG00000165591.6_2 | FAAH2 | 5.42E-19 | 3.53E-17 | 4.535 | 2.379 | chrX | 57313139 | 57515629 | + | protein_coding |
| ENSG00000267781.1_4 | SLC25A36P1 | 2.65E-07 | 2.96E-06 | 4.518 | -0.980 | chr19 | 56897595 | 56898059 | + | processed_pseudogene |
| ENSG00000113361.13_4 | CDH6 | 1.47E-21 | 1.54E-19 | 4.517 | 3.149 | chr5 | 31193793 | 31329253 | + | protein_coding |
| ENSG00000250208.6_7 | FZD10-AS1 | 2.01E-10 | 3.49E-09 | 4.496 | 0.087 | chr12 | 130628860 | 130646801 | - | lncRNA |
| ENSG00000236078.2_5 | LINC01447 | 1.85E-22 | 2.31E-20 | 4.495 | 3.295 | chr7 | 47648063 | 47669491 | + | lncRNA |
| ENSG00000120322.4_4 | PCDHB8 | 2.21E-18 | 1.31E-16 | 4.479 | 2.217 | chr5 | 140557371 | 140560120 | + | protein_coding |
| ENSG00000114812.13_3 | VIPR1 | 1.88E-13 | 5.09E-12 | 4.469 | 0.825 | chr3 | 42530791 | 42579065 | + | protein_coding |
| ENSG00000213976.4_5 | AC010615.1 | 1.80E-10 | 3.16E-09 | 4.468 | 0.297 | chr19 | 21565667 | 21569979 | + | unprocessed_pseudogene |
| ENSG00000204740.11_4 | MALRD1 | 2.08E-08 | 2.68E-07 | 4.462 | -0.569 | chr10 | 19337730 | 20079330 | + | protein_coding |
| ENSG00000087495.17_5 | PHACTR3 | 2.04E-07 | 2.32E-06 | 4.461 | -1.018 | chr20 | 58152564 | 58422766 | + | protein_coding |
| ENSG00000263567.1_6 | AC007923.1 | 9.93E-08 | 1.18E-06 | 4.460 | -1.018 | chr17 | 30089459 | 30096067 | + | lncRNA |
| ENSG00000283108.1_7 | AC011451.3 | 1.69E-07 | 1.94E-06 | 4.453 | -1.033 | chr19 | 9494257 | 9517731 | - | lncRNA |
| ENSG00000242516.2_3 | LINC00960 | 1.86E-10 | 3.26E-09 | 4.436 | 0.024 | chr3 | 75721451 | 75791199 | + | lncRNA |
| ENSG00000163565.18_5 | IFI16 | 1.08E-13 | 3.08E-12 | 4.421 | 1.354 | chr1 | 158969758 | 159024945 | + | protein_coding |
| ENSG00000269837.1_4 | IPO5P1 | 6.00E-23 | 8.74E-21 | 4.387 | 3.647 | chr19 | 23437855 | 23440741 | - | transcribed_processed_pseudogene |
| ENSG00000197608.11_4 | ZNF841 | 1.91E-28 | 1.48E-25 | 4.365 | 5.214 | chr19 | 52567719 | 52599018 | - | protein_coding |
| ENSG00000164749.13_6 | HNF4G | 3.17E-22 | 3.76E-20 | 4.363 | 3.248 | chr8 | 76320149 | 76479069 | + | protein_coding |
| ENSG00000172296.13_4 | SPTLC3 | 4.05E-21 | 3.80E-19 | 4.362 | 2.884 | chr20 | 12989620 | 13149750 | + | protein_coding |
| ENSG00000185432.12_3 | METTL7A | 1.10E-14 | 3.73E-13 | 4.361 | 1.211 | chr12 | 51317255 | 51326293 | + | protein_coding |
| ENSG00000204611.7_5 | ZNF616 | 9.46E-26 | 3.06E-23 | 4.348 | 4.405 | chr19 | 52616344 | 52643191 | - | protein_coding |
| ENSG00000114279.13_5 | FGF12 | 1.31E-20 | 1.13E-18 | 4.343 | 2.962 | chr3 | 191857184 | 192485553 | - | protein_coding |
| ENSG00000165511.6_6 | C10orf25 | 1.01E-06 | 1.05E-05 | 4.339 | -1.124 | chr10 | 45493146 | 45496336 | - | lncRNA |
| ENSG00000245648.2_6 | AC022075.1 | 1.84E-10 | 3.23E-09 | 4.322 | -0.060 | chr12 | 10516237 | 10551105 | + | lncRNA |
| ENSG00000284428.1_7 | AC092329.4 | 1.89E-21 | 1.92E-19 | 4.319 | 3.229 | chr19 | 23442708 | 23457053 | - | lncRNA |
| ENSG00000198788.7 | MUC2 | 4.19E-12 | 9.13E-11 | 4.309 | 0.522 | chr11 | 1074875 | 1104419 | + | protein_coding |
| ENSG00000276122.1_6 | AC026336.3 | 2.71E-07 | 3.02E-06 | 4.308 | -1.145 | chr12 | 130623238 | 130625313 | + | lncRNA |
| ENSG00000235180.1_6 | LINC00601 | 1.43E-06 | 1.45E-05 | 4.301 | -1.161 | chr10 | 128102438 | 128110448 | - | lncRNA |
| ENSG00000285722.1_3 | AC207130.1 | 4.18E-22 | 4.88E-20 | 4.293 | 3.469 | chr22 | 49768827 | 49773011 | + | lncRNA |
| ENSG00000257114.3_7 | LINC02450 | 1.66E-10 | 2.92E-09 | 4.287 | -0.095 | chr12 | 43080997 | 43110922 | + | lncRNA |

| | | | | | | | | | | |
|---|---|---|---|---|---|---|---|---|---|---|
| ENSG00000196074.13_7 | SYCP2 | 3.26E-25 | 8.90E-23 | 4.261 | 5.388 | chr20 | 58438619 | 58508710 | - | protein_coding |
| ENSG00000176593.8_7 | AC008969.1 | 4.45E-22 | 5.13E-20 | 4.257 | 3.659 | chr19 | 58513429 | 58522600 | + | lncRNA |
| ENSG00000137968.16_3 | SLC44A5 | 9.39E-20 | 7.04E-18 | 4.257 | 3.313 | chr1 | 75667816 | 76076801 | - | protein_coding |
| ENSG00000254245.2_3 | PCDHGA3 | 5.91E-09 | 8.35E-08 | 4.247 | -0.403 | chr5 | 140723396 | 140892546 | + | protein_coding |
| ENSG00000231453.1_5 | LINC01305 | 1.70E-13 | 4.65E-12 | 4.244 | 0.758 | chr2 | 175190755 | 175195371 | + | lncRNA |
| ENSG00000038427.16_5 | VCAN | 9.55E-29 | 8.08E-26 | 4.242 | 6.778 | chr5 | 82767437 | 82878122 | + | protein_coding |
| ENSG00000224149.1_8 | AC099786.2 | 1.44E-10 | 2.56E-09 | 4.240 | 0.093 | chr1 | 75595659 | 75598261 | - | lncRNA |
| ENSG00000148513.18_4 | ANKRD30A | 2.94E-13 | 7.72E-12 | 4.216 | 0.863 | chr10 | 37414653 | 37673039 | + | protein_coding |
| ENSG00000081665.14_7 | ZNF506 | 3.91E-24 | 8.07E-22 | 4.215 | 3.913 | chr19 | 19896648 | 19932560 | - | protein_coding |
| ENSG00000170848.15_6 | PSG6 | 1.93E-11 | 3.84E-10 | 4.209 | 0.578 | chr19 | 43406231 | 43423715 | - | protein_coding |
| ENSG00000137251.16_3 | TINAG | 2.50E-13 | 6.65E-12 | 4.208 | 0.852 | chr6 | 54172657 | 54254940 | + | protein_coding |
| ENSG00000006210.7_4 | CX3CL1 | 3.68E-12 | 8.11E-11 | 4.191 | 0.717 | chr16 | 57406389 | 57418956 | + | protein_coding |
| ENSG00000259033.2_6 | AL356804.1 | 6.57E-10 | 1.07E-08 | 4.187 | -0.196 | chr14 | 70722346 | 70810105 | + | lncRNA |
| ENSG00000244468.1_6 | AC093001.1 | 1.93E-08 | 2.50E-07 | 4.185 | -0.183 | chr3 | 149002569 | 149051440 | + | lncRNA |
| ENSG00000197249.14_6 | SERPINA1 | 4.74E-12 | 1.02E-10 | 4.177 | 0.404 | chr14 | 94843084 | 94857030 | - | protein_coding |
| ENSG00000110723.12_8 | EXPH5 | 1.44E-21 | 1.52E-19 | 4.172 | 3.464 | chr11 | 108376162 | 108464495 | - | protein_coding |
| ENSG00000187808.5_3 | SOWAHD | 1.16E-07 | 1.36E-06 | 4.167 | -0.808 | chrX | 118892551 | 118894165 | + | protein_coding |
| ENSG00000213013.4_4 | RPS15AP36 | 1.35E-06 | 1.37E-05 | 4.161 | -1.265 | chr19 | 58727546 | 58727927 | + | processed_pseudogene |
| ENSG00000156096.14_5 | UGT2B4 | 1.18E-06 | 1.21E-05 | 4.151 | -1.267 | chr4 | 70345883 | 70391732 | - | protein_coding |
| ENSG00000258175.1_7 | LINC02300 | 9.33E-08 | 1.11E-06 | 4.141 | -0.836 | chr14 | 29061429 | 29082829 | - | lncRNA |
| ENSG00000157613.11_6 | CREB3L1 | 3.55E-15 | 1.28E-13 | 4.136 | 1.440 | chr11 | 46299213 | 46342959 | + | protein_coding |
| ENSG00000162009.8_3 | SSTR5 | 8.84E-13 | 2.16E-11 | 4.125 | 0.777 | chr16 | 1128781 | 1130142 | + | protein_coding |
| ENSG00000121361.5_5 | KCNJ8 | 5.46E-12 | 1.17E-10 | 4.124 | 0.506 | chr12 | 21917889 | 21928534 | - | protein_coding |
| ENSG00000180479.14_5 | ZNF571 | 3.08E-23 | 4.92E-21 | 4.119 | 3.696 | chr19 | 38045684 | 38085693 | - | protein_coding |
| ENSG00000286802.1_1 | AC104459.1 | 3.77E-08 | 4.71E-07 | 4.110 | -0.867 | chr1 | 89594115 | 89610378 | + | lncRNA |
| ENSG00000162595.7_6 | DIRAS3 | 3.85E-21 | 3.68E-19 | 4.109 | 3.170 | chr1 | 68511569 | 68517314 | - | protein_coding |
| ENSG00000178662.16_4 | CSRNP3 | 9.52E-13 | 2.30E-11 | 4.103 | 0.757 | chr2 | 166326157 | 166545917 | + | protein_coding |
| ENSG00000109511.12_4 | ANXA10 | 2.52E-15 | 9.40E-14 | 4.095 | 1.479 | chr4 | 169013688 | 169108887 | + | protein_coding |
| ENSG00000042980.13_3 | ADAM28 | 3.31E-12 | 7.34E-11 | 4.083 | 0.469 | chr8 | 24151582 | 24216527 | + | protein_coding |
| ENSG00000196872.12_6 | KIAA1211L | 4.50E-14 | 1.38E-12 | 4.083 | 1.626 | chr2 | 99410309 | 99552722 | - | protein_coding |
| ENSG00000004799.8_4 | PDK4 | 2.74E-25 | 7.73E-23 | 4.076 | 4.468 | chr7 | 95212811 | 95225828 | - | protein_coding |
| ENSG00000151623.15_5 | NR3C2 | 1.78E-14 | 5.83E-13 | 4.073 | 1.320 | chr4 | 148999913 | 149365850 | - | protein_coding |
| ENSG00000262576.3_4 | PCDHGA4 | 2.49E-19 | 1.71E-17 | 4.070 | 2.649 | chr5 | 140734588 | 140892542 | + | protein_coding |
| ENSG00000268751.1_6 | SCGB1B2P | 2.78E-07 | 3.09E-06 | 4.066 | -0.900 | chr19 | 35067638 | 35068606 | - | lncRNA |
| ENSG00000185869.14_4 | ZNF829 | 1.32E-25 | 4.19E-23 | 4.060 | 4.494 | chr19 | 37379026 | 37407193 | - | protein_coding |
| ENSG00000128422.17_8 | KRT17 | 1.70E-09 | 2.61E-08 | 4.059 | -0.309 | chr17 | 39775694 | 39781094 | - | protein_coding |
| ENSG00000269289.6_7 | AC011503.1 | 3.43E-07 | 3.75E-06 | 4.041 | -0.917 | chr19 | 24101883 | 24140837 | - | lncRNA |
| ENSG00000276547.1_3 | PCDHGB5 | 3.29E-21 | 3.21E-19 | 4.039 | 3.424 | chr5 | 140777554 | 140892546 | + | protein_coding |
| ENSG00000187550.8_3 | SBK2 | 2.07E-20 | 1.70E-18 | 4.037 | 3.198 | chr19 | 56041100 | 56048456 | - | protein_coding |
| ENSG00000117707.16_6 | PROX1 | 2.45E-08 | 3.13E-07 | 4.011 | -0.360 | chr1 | 214156524 | 214214853 | + | protein_coding |
| ENSG00000198930.13_4 | CSAG1 | 2.36E-07 | 2.66E-06 | 4.000 | -0.950 | chrX | 151903228 | 151909518 | + | protein_coding |
| ENSG00000198768.11_6 | APCDD1L | 5.00E-09 | 7.15E-08 | 3.990 | -0.357 | chr20 | 57034157 | 57090455 | - | protein_coding |
| ENSG00000266066.1_4 | POLRMTP1 | 1.63E-19 | 1.16E-17 | 3.988 | 2.719 | chr17 | 60214333 | 60218000 | - | processed_pseudogene |
| ENSG00000102349.18_4 | KLF8 | 2.13E-07 | 2.42E-06 | 3.987 | -0.966 | chrX | 56258789 | 56317964 | + | protein_coding |
| ENSG00000167895.15_5 | TMC8 | 2.20E-15 | 8.28E-14 | 3.983 | 1.381 | chr17 | 76126851 | 76139049 | + | protein_coding |
| ENSG00000166432.15_4 | ZMAT1 | 8.38E-13 | 2.06E-11 | 3.980 | 1.141 | chrX | 101137260 | 101187052 | - | protein_coding |
| ENSG00000251191.8_6 | LINC00589 | 2.93E-11 | 5.69E-10 | 3.958 | 0.204 | chr8 | 29531438 | 29605625 | - | lncRNA |
| ENSG00000111348.9_5 | ARHGDIB | 3.23E-18 | 1.87E-16 | 3.951 | 2.273 | chr12 | 15094965 | 15114662 | - | protein_coding |
| ENSG00000179776.19_5 | CDH5 | 1.12E-14 | 3.77E-13 | 3.946 | 1.494 | chr16 | 66400525 | 66438687 | + | protein_coding |
| ENSG00000228283.3_4 | KATNBL1P6 | 1.25E-11 | 2.56E-10 | 3.891 | 0.665 | chr6 | 147123495 | 147124960 | - | transcribed_processed_pseudogene |
| ENSG00000163082.10_3 | SGPP2 | 5.94E-21 | 5.41E-19 | 3.890 | 3.334 | chr2 | 223289262 | 223427340 | + | protein_coding |
| ENSG00000003096.14_5 | KLHL13 | 1.70E-23 | 2.88E-21 | 3.882 | 3.825 | chrX | 117031776 | 117251303 | - | protein_coding |
| ENSG00000277075.2_3 | H2AC8 | 4.14E-07 | 4.48E-06 | 3.878 | -1.054 | chr6 | 26217203 | 26217711 | + | protein_coding |
| ENSG00000146233.8_4 | CYP39A1 | 1.83E-06 | 1.82E-05 | 3.872 | -1.068 | chr6 | 46517317 | 46620567 | - | protein_coding |
| ENSG00000176024.18_6 | ZNF613 | 1.36E-25 | 4.25E-23 | 3.862 | 4.700 | chr19 | 52430400 | 52452012 | + | protein_coding |
| ENSG00000230006.7_5 | ANKRD36BP2 | 1.11E-06 | 1.14E-05 | 3.860 | -1.070 | chr2 | 89065324 | 89106126 | + | transcribed_unprocessed_pseudogene |
| ENSG00000117226.12_5 | GBP3 | 2.26E-21 | 2.28E-19 | 3.850 | 3.357 | chr1 | 89472362 | 89488577 | - | protein_coding |
| ENSG00000156218.13_4 | ADAMTSL3 | 5.37E-09 | 7.63E-08 | 3.843 | -0.066 | chr15 | 84322840 | 84708594 | + | protein_coding |

| | | | | | | | | | | |
|---|---|---|---|---|---|---|---|---|---|---|
| ENSG00000104059.4_4 | FAM189A1 | 5.77E-17 | 2.85E-15 | 3.841 | 2.214 | chr15 | 29412457 | 29862927 | - | protein_coding |
| ENSG00000261713.6_6 | SSTR5-AS1 | 3.47E-16 | 1.46E-14 | 3.836 | 1.735 | chr16 | 1114093 | 1128731 | - | lncRNA |
| ENSG00000241119.2_4 | UGT1A9 | 1.17E-16 | 5.38E-15 | 3.823 | 2.070 | chr2 | 234580544 | 234681946 | + | protein_coding |
| ENSG00000170099.6_5 | SERPINA6 | 2.91E-20 | 2.33E-18 | 3.817 | 3.005 | chr14 | 94770585 | 94789726 | - | protein_coding |
| ENSG00000137460.8_2 | FHDC1 | 1.07E-14 | 3.61E-13 | 3.799 | 1.603 | chr4 | 153857504 | 153900848 | + | protein_coding |
| ENSG00000116106.12_7 | EPHA4 | 3.04E-20 | 2.41E-18 | 3.787 | 3.233 | chr2 | 222282747 | 222438922 | - | protein_coding |
| ENSG00000255622.3_4 | PCDHB17P | 6.52E-13 | 1.64E-11 | 3.779 | 1.135 | chr5 | 140535577 | 140538642 | + | transcribed_unprocessed_pseudogene |
| ENSG00000117983.17_5 | MUC5B | 1.95E-22 | 2.42E-20 | 3.772 | 6.229 | chr11 | 1244296 | 1283402 | + | protein_coding |
| ENSG00000114115.10_5 | RBP1 | 3.08E-12 | 6.89E-11 | 3.768 | 0.776 | chr3 | 139236276 | 139258671 | - | protein_coding |
| ENSG00000133321.11_4 | PLAAT4 | 8.51E-17 | 4.03E-15 | 3.768 | 2.067 | chr11 | 63304280 | 63313934 | + | protein_coding |
| ENSG00000197415.12_8 | VEPH1 | 1.21E-12 | 2.90E-11 | 3.760 | 0.869 | chr3 | 156977531 | 157251408 | - | protein_coding |
| ENSG00000150394.14_5 | CDH8 | 9.11E-07 | 9.48E-06 | 3.759 | -1.147 | chr16 | 61681146 | 62070939 | - | protein_coding |
| ENSG00000267470.6_8 | ZNF571-AS1 | 4.88E-14 | 1.48E-12 | 3.758 | 1.178 | chr19 | 38039816 | 38078249 | + | lncRNA |
| ENSG00000180861.9_6 | LINC01559 | 3.65E-23 | 5.70E-21 | 3.752 | 3.876 | chr12 | 13524023 | 13540101 | - | lncRNA |
| ENSG00000249884.8_4 | RNF103-CHMP3 | 5.35E-12 | 1.15E-10 | 3.729 | 0.643 | chr2 | 86732791 | 86948245 | - | protein_coding |
| ENSG00000123838.11_4 | C4BPA | 9.49E-19 | 5.92E-17 | 3.723 | 2.677 | chr1 | 207277578 | 207318317 | + | protein_coding |
| ENSG00000073910.22_7 | FRY | 3.40E-14 | 1.06E-12 | 3.721 | 1.347 | chr13 | 32420850 | 32873262 | + | protein_coding |
| ENSG00000197372.10_4 | ZNF675 | 8.67E-31 | 1.28E-27 | 3.721 | 6.706 | chr19 | 23708433 | 23870022 | - | protein_coding |
| ENSG00000159618.16_3 | ADGRG5 | 1.58E-16 | 7.07E-15 | 3.717 | 1.932 | chr16 | 57576555 | 57625593 | + | protein_coding |
| ENSG00000061918.13_4 | GUCY1B1 | 1.56E-12 | 3.68E-11 | 3.700 | 0.802 | chr4 | 156680144 | 156728926 | + | protein_coding |
| ENSG00000230234.1_6 | AL162582.1 | 2.14E-06 | 2.11E-05 | 3.691 | -1.205 | chr6 | 160693649 | 160697162 | + | lncRNA |
| ENSG00000169507.9_6 | SLC38A11 | 2.58E-08 | 3.28E-07 | 3.687 | -0.621 | chr2 | 165752696 | 165812035 | - | protein_coding |
| ENSG00000215018.10_6 | COL28A1 | 2.86E-12 | 6.45E-11 | 3.673 | 0.789 | chr7 | 7395834 | 7575504 | - | protein_coding |
| ENSG00000248771.5_8 | SMIM31 | 5.71E-13 | 1.45E-11 | 3.672 | 0.867 | chr4 | 165675216 | 165724947 | + | protein_coding |
| ENSG00000196620.10_5 | UGT2B15 | 2.63E-06 | 2.55E-05 | 3.650 | -1.243 | chr4 | 69512315 | 69536370 | - | protein_coding |
| ENSG00000171431.4_4 | KRT20 | 5.85E-16 | 2.36E-14 | 3.631 | 1.929 | chr17 | 39032141 | 39041494 | - | protein_coding |
| ENSG00000257588.1_8 | AC025154.2 | 2.28E-08 | 2.93E-07 | 3.616 | -0.458 | chr12 | 50345295 | 50356707 | - | lncRNA |
| ENSG00000178947.9_3 | SMIM10L2A | 9.42E-16 | 3.68E-14 | 3.607 | 1.761 | chrX | 134555869 | 134562000 | + | protein_coding |
| ENSG00000254122.2_3 | PCDHGB7 | 2.79E-08 | 3.54E-07 | 3.599 | -0.263 | chr5 | 140797212 | 140892546 | + | protein_coding |
| ENSG00000143816.8_4 | WNT9A | 1.41E-22 | 1.88E-20 | 3.594 | 3.743 | chr1 | 228106357 | 228135633 | - | protein_coding |
| ENSG00000256751.6_6 | PLBD1-AS1 | 1.36E-11 | 2.76E-10 | 3.591 | 0.794 | chr12 | 14720327 | 14777125 | + | lncRNA |
| ENSG00000253873.6_6 | PCDHGA11 | 2.52E-07 | 2.82E-06 | 3.588 | -0.704 | chr5 | 140800614 | 140892542 | + | protein_coding |
| ENSG00000136869.16_5 | TLR4 | 7.04E-11 | 1.29E-09 | 3.573 | 0.503 | chr9 | 120466453 | 120487013 | + | protein_coding |
| ENSG00000173702.7_7 | MUC13 | 1.39E-26 | 5.83E-24 | 3.565 | 5.192 | chr3 | 124624289 | 124672663 | - | protein_coding |
| ENSG00000198453.13_5 | ZNF568 | 9.78E-24 | 1.77E-21 | 3.552 | 4.342 | chr19 | 37407231 | 37489602 | + | protein_coding |
| ENSG00000125845.7_4 | BMP2 | 1.54E-09 | 2.38E-08 | 3.543 | 0.001 | chr20 | 6748333 | 6760893 | + | protein_coding |
| ENSG00000196104.11_8 | SPOCK3 | 1.11E-11 | 2.29E-10 | 3.541 | 0.753 | chr4 | 167654535 | 168155947 | - | protein_coding |
| ENSG00000160255.18_8 | ITGB2 | 2.73E-18 | 1.60E-16 | 3.535 | 2.439 | chr21 | 46305868 | 46351904 | - | protein_coding |
| ENSG00000081842.13 | PCDHA6 | 1.29E-09 | 2.02E-08 | 3.532 | -0.014 | chr5 | 140207563 | 140391929 | + | protein_coding |
| ENSG00000070526.15_5 | ST6GALNAC1 | 3.73E-15 | 1.34E-13 | 3.513 | 1.858 | chr17 | 74620843 | 74639868 | - | protein_coding |
| ENSG00000251095.7_7 | AC093866.1 | 3.26E-06 | 3.14E-05 | 3.503 | -1.351 | chr4 | 90332111 | 90647903 | + | lncRNA |
| ENSG00000152926.15_7 | ZNF117 | 8.10E-21 | 7.23E-19 | 3.491 | 3.201 | chr7 | 64432150 | 64467062 | - | protein_coding |
| ENSG00000162078.11_4 | ZG16B | 1.09E-13 | 3.09E-12 | 3.480 | 1.240 | chr16 | 2880170 | 2889586 | + | protein_coding |
| ENSG00000134709.12_4 | HOOK1 | 1.00E-21 | 1.11E-19 | 3.465 | 3.822 | chr1 | 60280458 | 60341994 | + | protein_coding |
| ENSG00000261502.4_9 | AC040174.1 | 8.91E-07 | 9.29E-06 | 3.465 | -0.591 | chr16 | 63090690 | 63163558 | - | lncRNA |
| ENSG00000103460.17_4 | TOX3 | 1.75E-19 | 1.23E-17 | 3.462 | 3.234 | chr16 | 52470329 | 52581714 | - | protein_coding |
| ENSG00000277806.1_7 | AC006213.4 | 6.42E-09 | 9.04E-08 | 3.461 | -0.082 | chr19 | 44480967 | 44481600 | + | lncRNA |
| ENSG00000196482.18_7 | ESRRG | 5.37E-06 | 5.01E-05 | 3.460 | -1.069 | chr1 | 216676588 | 217311097 | - | protein_coding |
| ENSG00000264063.1 | MIR3687 | 0.00459746 | 0.02640904 | 3.459 | 3.100 | chr21 | 9826203 | 9826263 | + | miRNA |
| ENSG00000245680.10_7 | ZNF585B | 9.63E-27 | 4.28E-24 | 3.457 | 5.110 | chr19 | 37672481 | 37709055 | - | protein_coding |
| ENSG00000264462.1 | MIR3648 | 0.00782699 | 0.0428411 | 3.453 | 3.784 | chr21 | 9825832 | 9826011 | + | miRNA |
| ENSG00000117472.10_3 | TSPAN1 | 9.36E-17 | 4.41E-15 | 3.452 | 2.581 | chr1 | 46640745 | 46651634 | + | protein_coding |
| ENSG00000135549.10 | PKIB | 1.41E-11 | 2.86E-10 | 3.443 | 0.580 | chr6 | 122793076 | 123047518 | + | protein_coding |
| ENSG00000236345.1_7 | SCAT8 | 9.09E-10 | 1.45E-08 | 3.440 | 0.045 | chr6 | 64516729 | 64532535 | + | lncRNA |
| ENSG00000181894.15_4 | ZNF329 | 1.93E-26 | 7.77E-24 | 3.437 | 5.279 | chr19 | 58637615 | 58666477 | - | protein_coding |
| ENSG00000233820.2_5 | AL589843.2 | 1.01E-06 | 1.04E-05 | 3.435 | -0.834 | chr9 | 99488172 | 99489751 | + | transcribed_processed_pseudogene |
| ENSG00000168421.13_4 | RHOH | 2.67E-07 | 2.98E-06 | 3.435 | -0.608 | chr4 | 40192673 | 40248587 | + | protein_coding |
| ENSG00000233008.6_6 | LINC01725 | 2.54E-06 | 2.47E-05 | 3.434 | -1.089 | chr1 | 84041459 | 84326706 | - | lncRNA |

| | | | | | | | | | | |
|---|---|---|---|---|---|---|---|---|---|---|
| ENSG00000177694.16_5 | NAALADL2 | 2.18E-07 | 2.47E-06 | 3.432 | -0.619 | chr3 | 174156363 | 175528336 | + | protein_coding |
| ENSG00000168143.9_3 | FAM83B | 1.75E-12 | 4.09E-11 | 3.424 | 1.010 | chr6 | 54711569 | 54809897 | + | protein_coding |
| ENSG00000148848.14_6 | ADAM12 | 4.21E-19 | 2.75E-17 | 3.418 | 3.152 | chr10 | 127700950 | 128077024 | - | protein_coding |
| ENSG00000214691.9_5 | LINC01913 | 1.48E-14 | 4.92E-13 | 3.409 | 1.491 | chr2 | 42087295 | 42121190 | + | lncRNA |
| ENSG00000198680.4_2 | TUSC1 | 6.57E-21 | 5.90E-19 | 3.407 | 3.554 | chr9 | 25676387 | 25678438 | - | protein_coding |
| ENSG00000248144.6_6 | ADH1C | 4.31E-09 | 6.24E-08 | 3.400 | -0.131 | chr4 | 100257654 | 100273903 | - | protein_coding |
| ENSG00000134207.16_8 | SYT6 | 2.23E-09 | 3.36E-08 | 3.399 | -0.003 | chr1 | 114631913 | 114696541 | - | protein_coding |
| ENSG00000118160.14_5 | SLC8A2 | 1.62E-07 | 1.87E-06 | 3.398 | -0.636 | chr19 | 47931274 | 47975150 | - | protein_coding |
| ENSG00000184574.10_4 | LPAR5 | 5.38E-13 | 1.36E-11 | 3.389 | 1.507 | chr12 | 6728001 | 6745126 | - | protein_coding |
| ENSG00000175920.18_4 | DOK7 | 4.56E-14 | 1.40E-12 | 3.369 | 2.207 | chr4 | 3465033 | 3503200 | + | protein_coding |
| ENSG00000267421.6_7 | AC005498.2 | 4.50E-06 | 4.24E-05 | 3.368 | -0.895 | chr19 | 57047525 | 57049944 | - | lncRNA |
| ENSG00000148702.15_4 | HABP2 | 4.40E-18 | 2.47E-16 | 3.367 | 2.535 | chr10 | 115310596 | 115349361 | + | protein_coding |
| ENSG00000134152.11_5 | KATNBL1 | 4.26E-30 | 4.45E-27 | 3.367 | 8.435 | chr15 | 34432875 | 34502297 | - | protein_coding |
| ENSG00000015413.9_5 | DPEP1 | 3.17E-14 | 9.94E-13 | 3.351 | 1.390 | chr16 | 89679716 | 89704864 | + | protein_coding |
| ENSG00000162782.16_6 | TDRD5 | 4.24E-12 | 9.22E-11 | 3.334 | 0.991 | chr1 | 179560748 | 179660407 | + | protein_coding |
| ENSG00000171827.10_2 | ZNF570 | 1.76E-25 | 5.20E-23 | 3.326 | 5.188 | chr19 | 37958487 | 37979554 | + | protein_coding |
| ENSG00000286261.1_1 | AC022137.3 | 7.75E-07 | 8.12E-06 | 3.306 | -0.932 | chr19 | 53935237 | 53965116 | + | protein_coding |
| ENSG00000169255.15_7 | B3GALNT1 | 5.68E-08 | 6.95E-07 | 3.305 | -0.530 | chr3 | 160801671 | 160823199 | - | protein_coding |
| ENSG00000171234.14_4 | UGT2B7 | 1.23E-08 | 1.64E-07 | 3.301 | -0.217 | chr4 | 69917081 | 69978705 | + | protein_coding |
| ENSG00000124749.17_7 | COL21A1 | 2.69E-21 | 2.67E-19 | 3.300 | 3.769 | chr6 | 55921388 | 56258892 | - | protein_coding |
| ENSG00000249196.6_6 | TMEM132D-AS1 | 5.02E-09 | 7.18E-08 | 3.297 | -0.077 | chr12 | 129594174 | 129597839 | + | lncRNA |
| ENSG00000160182.3_4 | TFF1 | 7.58E-13 | 1.88E-11 | 3.288 | 1.176 | chr21 | 43782391 | 43786644 | - | protein_coding |
| ENSG00000278921.2_5 | EPB41L4A-DT | 9.73E-09 | 1.32E-07 | 3.286 | 0.777 | chr5 | 111755280 | 111756675 | + | lncRNA |
| ENSG00000229489.2 | ATP5A1P5 | 9.62E-06 | 8.65E-05 | 3.280 | -1.205 | chr9 | 43399987 | 43401315 | + | pseudogene |
| ENSG00000164684.13_5 | ZNF704 | 1.34E-23 | 2.32E-21 | 3.267 | 4.432 | chr8 | 81540686 | 81787016 | - | protein_coding |
| ENSG00000171502.15_5 | COL24A1 | 5.54E-10 | 9.06E-09 | 3.250 | 0.668 | chr1 | 86194916 | 86622626 | - | protein_coding |
| ENSG00000198758.10_4 | EPS8L3 | 9.75E-17 | 4.56E-15 | 3.250 | 2.236 | chr1 | 110292702 | 110306649 | - | protein_coding |
| ENSG00000198354.7_4 | DCAF12L2 | 4.03E-06 | 3.82E-05 | 3.240 | -0.984 | chrX | 125297482 | 125300272 | - | protein_coding |
| ENSG00000235750.10_4 | KIAA0040 | 2.88E-16 | 1.23E-14 | 3.229 | 2.774 | chr1 | 175126122 | 175162135 | - | protein_coding |
| ENSG00000152433.14_3 | ZNF547 | 5.27E-15 | 1.86E-13 | 3.228 | 2.032 | chr19 | 57874845 | 57890933 | + | protein_coding |
| ENSG00000137393.9_2 | RNF144B | 1.21E-20 | 1.05E-18 | 3.211 | 3.700 | chr6 | 18387581 | 18469105 | + | protein_coding |
| ENSG00000204969.7_5 | PCDHA2 | 1.08E-10 | 1.95E-09 | 3.202 | 0.538 | chr5 | 140174437 | 140391932 | + | protein_coding |
| ENSG00000132821.12_3 | VSTM2L | 1.65E-15 | 6.29E-14 | 3.197 | 2.007 | chr20 | 36531513 | 36573752 | + | protein_coding |
| ENSG00000135960.10_5 | EDAR | 6.52E-10 | 1.06E-08 | 3.194 | 0.365 | chr2 | 109510927 | 109605828 | - | protein_coding |
| ENSG00000158486.13_3 | DNAH3 | 1.45E-15 | 5.54E-14 | 3.187 | 1.848 | chr16 | 20944433 | 21170762 | - | protein_coding |
| ENSG00000250731.1_4 | TPM3P6 | 1.78E-07 | 2.04E-06 | 3.181 | -0.468 | chr19 | 53982604 | 53983345 | + | processed_pseudogene |
| ENSG00000099260.11_4 | PALMD | 2.58E-12 | 5.85E-11 | 3.177 | 0.973 | chr1 | 100111669 | 100160097 | + | protein_coding |
| ENSG00000078725.13_4 | BRINP1 | 9.80E-06 | 8.79E-05 | 3.157 | -1.307 | chr9 | 121915736 | 122131713 | - | protein_coding |
| ENSG00000019549.13_7 | SNAI2 | 5.52E-11 | 1.03E-09 | 3.156 | 0.832 | chr8 | 49830157 | 49834299 | - | protein_coding |
| ENSG00000137726.17_6 | FXYD6 | 1.65E-06 | 1.65E-05 | 3.154 | -1.053 | chr11 | 117707691 | 117748201 | - | protein_coding |
| ENSG00000081853.15_4 | PCDHGA2 | 8.41E-15 | 2.91E-13 | 3.154 | 3.058 | chr5 | 140718327 | 140892542 | + | protein_coding |
| ENSG00000228623.6_3 | ZNF883 | 9.80E-12 | 2.03E-10 | 3.139 | 1.104 | chr9 | 115720002 | 115774507 | - | transcribed_unprocessed_pseudogene |
| ENSG00000171346.16_6 | KRT15 | 5.10E-21 | 4.75E-19 | 3.136 | 3.678 | chr17 | 39669997 | 39678781 | - | protein_coding |
| ENSG00000251985.1 | RNU6-1161P | 2.41E-08 | 3.09E-07 | 3.129 | -0.220 | chr22 | 46020411 | 46020512 | - | snRNA |
| ENSG00000170775.3_4 | GPR37 | 1.24E-18 | 7.65E-17 | 3.127 | 2.986 | chr7 | 124383939 | 124405846 | - | protein_coding |
| ENSG00000182175.14_5 | RGMA | 3.40E-08 | 4.27E-07 | 3.125 | -0.103 | chr15 | 93578503 | 93632433 | - | protein_coding |
| ENSG00000254497.1_6 | AC103681.1 | 5.43E-06 | 5.07E-05 | 3.124 | -1.072 | chr11 | 45275435 | 45276200 | + | lncRNA |
| ENSG00000108679.13_5 | LGALS3BP | 6.74E-24 | 1.29E-21 | 3.122 | 7.810 | chr17 | 76967320 | 76976029 | - | protein_coding |
| ENSG00000219891.2_2 | ZSCAN12P1 | 3.01E-07 | 3.33E-06 | 3.121 | -0.676 | chr6 | 28058932 | 28061442 | + | transcribed_unprocessed_pseudogene |
| ENSG00000248367.1_6 | AC008610.1 | 1.59E-13 | 4.39E-12 | 3.120 | 1.559 | chr5 | 179720245 | 179722253 | + | lncRNA |
| ENSG00000133962.8_9 | CATSPERB | 3.74E-09 | 5.51E-08 | 3.098 | 0.535 | chr14 | 92047040 | 92247051 | - | protein_coding |
| ENSG00000235269.2_7 | LINC02331 | 2.67E-18 | 1.57E-16 | 3.089 | 3.014 | chr14 | 54151857 | 54317600 | - | lncRNA |
| ENSG00000139174.12_7 | PRICKLE1 | 5.74E-15 | 2.02E-13 | 3.088 | 1.881 | chr12 | 42850559 | 42984157 | - | protein_coding |
| ENSG00000213020.10_7 | ZNF611 | 1.09E-24 | 2.54E-22 | 3.084 | 4.746 | chr19 | 53206066 | 53238326 | - | protein_coding |
| ENSG00000047365.12_5 | ARAP2 | 3.31E-19 | 2.22E-17 | 3.076 | 3.490 | chr4 | 35949843 | 36246136 | - | protein_coding |
| ENSG00000287294.1_1 | AL035694.1 | 1.92E-07 | 2.19E-06 | 3.074 | -0.555 | chr6 | 85478728 | 85479937 | + | lncRNA |
| ENSG00000087245.13_6 | MMP2 | 1.11E-16 | 5.14E-15 | 3.071 | 2.492 | chr16 | 55423612 | 55540603 | + | protein_coding |

| | | | | | | | | | | |
|---|---|---|---|---|---|---|---|---|---|---|
| ENSG00000173535.14_3 | TNFRSF10C | 1.07E-08 | 1.44E-07 | 3.070 | 0.167 | chr8 | 22960103 | 22974950 | + | protein_coding |
| ENSG00000153253.18_7 | SCN3A | 9.40E-08 | 1.12E-06 | 3.063 | -0.420 | chr2 | 165944032 | 166060577 | - | protein_coding |
| ENSG00000233251.8_6 | AC007743.1 | 3.83E-08 | 4.77E-07 | 3.054 | -0.281 | chr2 | 56400669 | 56413005 | - | lncRNA |
| ENSG00000114378.17_4 | HYAL1 | 1.29E-13 | 3.58E-12 | 3.049 | 2.361 | chr3 | 50337321 | 50349812 | - | protein_coding |
| ENSG00000261934.2_3 | PCDHGA9 | 9.73E-10 | 1.55E-08 | 3.048 | 0.323 | chr5 | 140782499 | 140892546 | + | protein_coding |
| ENSG00000198650.11_4 | TAT | 6.35E-07 | 6.71E-06 | 3.046 | -0.155 | chr16 | 71599563 | 71610995 | - | protein_coding |
| ENSG00000253910.2_4 | PCDHGB2 | 1.39E-11 | 2.83E-10 | 3.042 | 1.011 | chr5 | 140739609 | 140892546 | + | protein_coding |
| ENSG00000241635.8_4 | UGT1A1 | 5.48E-06 | 5.10E-05 | 3.038 | -1.145 | chr2 | 234668916 | 234681946 | + | protein_coding |
| ENSG00000178826.11_3 | TMEM139 | 3.77E-09 | 5.55E-08 | 3.025 | 0.298 | chr7 | 142977050 | 142985141 | + | protein_coding |
| ENSG00000125735.10_4 | TNFSF14 | 1.76E-05 | 0.00015307 | 3.023 | -1.162 | chr19 | 6661264 | 6670599 | - | protein_coding |
| ENSG00000146950.13_3 | SHROOM2 | 2.95E-14 | 9.31E-13 | 3.017 | 1.791 | chrX | 9754469 | 9917483 | + | protein_coding |
| ENSG00000116774.12_3 | OLFML3 | 4.57E-13 | 1.18E-11 | 3.011 | 1.497 | chr1 | 114522013 | 114578194 | + | protein_coding |
| ENSG00000105289.15_4 | TJP3 | 1.05E-22 | 1.45E-20 | 3.004 | 4.530 | chr19 | 3708360 | 3750811 | + | protein_coding |
| ENSG00000213958.2_4 | KRT18P29 | 1.73E-06 | 1.72E-05 | 3.003 | -0.774 | chr2 | 182825939 | 182827205 | - | processed_pseudogene |
| ENSG00000196967.10_4 | ZNF585A | 3.27E-21 | 3.21E-19 | 2.991 | 3.796 | chr19 | 37597636 | 37663643 | - | protein_coding |
| ENSG00000166704.11_4 | ZNF606 | 2.27E-24 | 4.93E-22 | 2.990 | 4.997 | chr19 | 58488421 | 58514717 | - | protein_coding |
| ENSG00000232306.2_5 | AC012485.2 | 4.64E-06 | 4.37E-05 | 2.987 | -0.790 | chr2 | 239204033 | 239209261 | - | lncRNA |
| ENSG00000130600.19_6 | H19 | 6.04E-17 | 2.96E-15 | 2.985 | 2.693 | chr11 | 2016406 | 2022700 | - | lncRNA |
| ENSG00000267481.1_8 | AC011477.1 | 2.42E-08 | 3.09E-07 | 2.971 | -0.009 | chr19 | 19899564 | 19901340 | - | lncRNA |
| ENSG00000213468.7_9 | FIRRE | 3.16E-10 | 5.33E-09 | 2.966 | 0.609 | chrX | 130822807 | 130964890 | - | lncRNA |
| ENSG00000157064.11_4 | NMNAT2 | 3.86E-13 | 1.00E-11 | 2.964 | 1.526 | chr1 | 183217372 | 183387515 | - | protein_coding |
| ENSG00000176532.4_3 | PRR15 | 6.44E-20 | 4.95E-18 | 2.963 | 4.300 | chr7 | 29603451 | 29606909 | + | protein_coding |
| ENSG00000054179.12_5 | ENTPD2 | 1.74E-07 | 2.00E-06 | 2.960 | -0.495 | chr9 | 139942559 | 139948513 | - | protein_coding |
| ENSG00000233030.2_8 | AC243772.2 | 5.46E-06 | 5.09E-05 | 2.957 | -0.998 | chr1 | 149757215 | 149764576 | - | lncRNA |
| ENSG00000072041.17_5 | SLC6A15 | 1.66E-11 | 3.33E-10 | 2.957 | 1.084 | chr12 | 85253270 | 85307394 | - | protein_coding |
| ENSG00000168350.8_4 | DEGS2 | 5.60E-14 | 1.68E-12 | 2.955 | 1.797 | chr14 | 100610294 | 100625982 | - | protein_coding |
| ENSG00000124785.9_4 | NRN1 | 8.39E-10 | 1.35E-08 | 2.950 | 0.532 | chr6 | 5998232 | 6007838 | - | protein_coding |
| ENSG00000253846.2_3 | PCDHGA10 | 7.76E-08 | 9.33E-07 | 2.949 | -0.242 | chr5 | 140792554 | 140892546 | + | protein_coding |
| ENSG00000163624.6_3 | CDS1 | 1.12E-15 | 4.35E-14 | 2.940 | 2.391 | chr4 | 85504280 | 85572487 | + | protein_coding |
| ENSG00000137440.5_4 | FGFBP1 | 7.65E-19 | 4.82E-17 | 2.926 | 3.037 | chr4 | 15937200 | 15940363 | - | protein_coding |
| ENSG00000062038.14_4 | CDH3 | 2.39E-07 | 2.68E-06 | 2.920 | -0.399 | chr16 | 68670092 | 68756519 | + | protein_coding |
| ENSG00000198515.13_6 | CNGA1 | 2.30E-05 | 0.00019716 | 2.905 | -1.034 | chr4 | 47937994 | 48018689 | - | protein_coding |
| ENSG00000236699.9_4 | ARHGEF38 | 1.46E-10 | 2.59E-09 | 2.902 | 0.614 | chr4 | 106473777 | 106629250 | + | protein_coding |
| ENSG00000172367.16_4 | PDZD3 | 5.28E-10 | 8.66E-09 | 2.900 | 0.619 | chr11 | 119056166 | 119060932 | + | protein_coding |
| ENSG00000149212.12_5 | SESN3 | 3.40E-18 | 1.96E-16 | 2.894 | 4.095 | chr11 | 94898677 | 94965705 | - | protein_coding |
| ENSG00000155066.16_4 | PROM2 | 8.24E-06 | 7.49E-05 | 2.890 | -1.051 | chr2 | 95940197 | 95957056 | + | protein_coding |
| ENSG00000176658.17_5 | MYO1D | 4.92E-23 | 7.34E-21 | 2.884 | 4.500 | chr17 | 30819540 | 31204195 | - | protein_coding |
| ENSG00000280623.1_3 | PCAT14 | 1.49E-05 | 0.00013058 | 2.883 | -1.264 | chr22 | 23879068 | 23889984 | + | lncRNA |
| ENSG00000114638.8_5 | UPK1B | 1.63E-19 | 1.16E-17 | 2.873 | 3.925 | chr3 | 118892364 | 118923990 | + | protein_coding |
| ENSG00000165449.11_3 | SLC16A9 | 6.54E-07 | 6.90E-06 | 2.872 | -0.579 | chr10 | 61410519 | 61495760 | - | protein_coding |
| ENSG00000197128.11_4 | ZNF772 | 5.25E-23 | 7.72E-21 | 2.858 | 4.584 | chr19 | 57978031 | 57988938 | - | protein_coding |
| ENSG00000136883.14_5 | KIF12 | 7.36E-25 | 1.84E-22 | 2.853 | 5.257 | chr9 | 116848406 | 116861571 | - | protein_coding |
| ENSG00000228653.2_4 | HNRNPCP7 | 2.68E-06 | 2.60E-05 | 2.851 | -0.732 | chr7 | 63961203 | 63962107 | + | processed_pseudogene |
| ENSG00000147689.16_4 | FAM83A | 2.05E-09 | 3.11E-08 | 2.850 | 0.234 | chr8 | 124191200 | 124222319 | + | protein_coding |
| ENSG00000120675.6_4 | DNAJC15 | 2.56E-06 | 2.49E-05 | 2.848 | -0.743 | chr13 | 43597339 | 43688349 | + | protein_coding |
| ENSG00000181007.9_4 | ZFP82 | 1.61E-22 | 2.08E-20 | 2.841 | 4.399 | chr19 | 36874022 | 36909546 | - | protein_coding |
| ENSG00000127831.11_4 | VIL1 | 1.42E-14 | 4.74E-13 | 2.836 | 1.870 | chr2 | 219283844 | 219318018 | + | protein_coding |
| ENSG00000280734.3_4 | LINC01232 | 2.85E-12 | 6.43E-11 | 2.836 | 1.269 | chr13 | 100139216 | 100151560 | - | lncRNA |
| ENSG00000155465.18_4 | SLC7A7 | 1.47E-21 | 1.54E-19 | 2.832 | 4.306 | chr14 | 23242431 | 23299029 | - | protein_coding |
| ENSG00000257335.8_3 | MGAM | 6.46E-21 | 5.83E-19 | 2.831 | 4.719 | chr7 | 141607613 | 141806547 | + | protein_coding |
| ENSG00000196646.12_4 | ZNF136 | 1.27E-15 | 4.89E-14 | 2.831 | 2.548 | chr19 | 12273879 | 12300686 | + | protein_coding |
| ENSG00000234284.7_7 | ZNF879 | 1.09E-19 | 8.15E-18 | 2.816 | 3.592 | chr5 | 178450805 | 178462065 | + | protein_coding |
| ENSG00000177409.12_5 | SAMD9L | 2.16E-16 | 9.43E-15 | 2.814 | 2.629 | chr7 | 92759369 | 92777698 | - | protein_coding |
| ENSG00000159713.11_4 | TPPP3 | 2.86E-21 | 2.82E-19 | 2.814 | 4.144 | chr16 | 67423712 | 67427421 | - | protein_coding |
| ENSG00000050405.13_6 | LIMA1 | 2.42E-23 | 3.99E-21 | 2.814 | 5.010 | chr12 | 50569571 | 50677329 | - | protein_coding |
| ENSG00000178201.4_3 | VN1R1 | 6.88E-07 | 7.23E-06 | 2.810 | -0.267 | chr19 | 57966158 | 57968510 | - | protein_coding |
| ENSG00000057704.13_6 | TMCC3 | 6.61E-09 | 9.27E-08 | 2.808 | 0.114 | chr12 | 94960898 | 95044333 | - | protein_coding |
| ENSG00000110693.18_7 | SOX6 | 7.76E-18 | 4.23E-16 | 2.803 | 2.978 | chr11 | 15987995 | 16761138 | - | protein_coding |
| ENSG00000196724.12_5 | ZNF418 | 5.69E-21 | 5.21E-19 | 2.802 | 3.907 | chr19 | 58433252 | 58446761 | - | protein_coding |
| ENSG00000181690.8_4 | PLAG1 | 8.29E-13 | 2.04E-11 | 2.797 | 1.567 | chr8 | 57073468 | 57123883 | - | protein_coding |

| | | | | | | | | | | |
|---|---|---|---|---|---|---|---|---|---|---|
| ENSG00000271366.1_7 | AC002128.2 | 0.00011008 | 0.0008452 | 2.797 | -1.123 | chr19 | 35742343 | 35744542 | - | lncRNA |
| ENSG00000162591.16_4 | MEGF6 | 1.06E-21 | 1.17E-19 | 2.788 | 5.152 | chr1 | 3404515 | 3528072 | - | protein_coding |
| ENSG00000197635.10_6 | DPP4 | 5.53E-12 | 1.18E-10 | 2.787 | 1.272 | chr2 | 162848755 | 162930725 | - | protein_coding |
| ENSG00000142619.4_3 | PADI3 | 5.08E-24 | 1.01E-21 | 2.786 | 5.207 | chr1 | 17575593 | 17610728 | + | protein_coding |
| ENSG00000260963.2_6 | AC026462.3 | 1.30E-08 | 1.72E-07 | 2.773 | 0.177 | chr16 | 52640918 | 52648552 | + | lncRNA |
| ENSG00000236039.3_8 | AC019117.2 | 8.86E-13 | 2.16E-11 | 2.773 | 1.539 | chr7 | 17445103 | 17598533 | - | lncRNA |
| ENSG00000115461.5_6 | IGFBP5 | 2.17E-10 | 3.74E-09 | 2.771 | 0.681 | chr2 | 217536828 | 217560272 | - | protein_coding |
| ENSG00000111335.12_4 | OAS2 | 6.92E-05 | 0.00054897 | 2.762 | -1.352 | chr12 | 113416200 | 113449528 | + | protein_coding |
| ENSG00000267058.1_9 | AC006213.2 | 2.10E-05 | 0.00018096 | 2.757 | -1.147 | chr19 | 44395956 | 44405957 | - | lncRNA |
| ENSG00000166446.15_5 | CDYL2 | 2.01E-09 | 3.05E-08 | 2.754 | 0.424 | chr16 | 80631804 | 80838495 | - | protein_coding |
| ENSG00000197479.7_4 | PCDHB11 | 4.92E-10 | 8.10E-09 | 2.752 | 0.618 | chr5 | 140579183 | 140583352 | + | protein_coding |
| ENSG00000184731.6_4 | FAM110C | 3.12E-19 | 2.11E-17 | 2.743 | 3.390 | chr2 | 38814 | 46870 | - | protein_coding |
| ENSG00000117480.16_3 | FAAH | 2.28E-11 | 4.49E-10 | 2.739 | 1.182 | chr1 | 46859989 | 46879520 | + | protein_coding |
| ENSG00000280217.1_5 | AC126564.1 | 1.84E-13 | 4.98E-12 | 2.738 | 1.726 | chr12 | 131720056 | 131722068 | + | TEC |
| ENSG00000112299.8_4 | VNN1 | 4.23E-05 | 0.00034882 | 2.736 | -1.165 | chr6 | 133001988 | 133035194 | - | protein_coding |
| ENSG00000204603.7_6 | LINC01257 | 1.20E-08 | 1.60E-07 | 2.735 | 0.357 | chr12 | 131649556 | 131717485 | + | lncRNA |
| ENSG00000151150.22_6 | ANK3 | 2.48E-21 | 2.49E-19 | 2.730 | 4.128 | chr10 | 61786056 | 62493248 | - | protein_coding |
| ENSG00000141668.10_4 | CBLN2 | 7.21E-13 | 1.79E-11 | 2.730 | 1.669 | chr18 | 70203915 | 70305756 | - | protein_coding |
| ENSG00000169302.16_4 | STK32A | 5.18E-09 | 7.37E-08 | 2.727 | 0.469 | chr5 | 146614526 | 146767418 | + | protein_coding |
| ENSG00000197050.11_6 | ZNF420 | 3.10E-23 | 4.92E-21 | 2.724 | 4.778 | chr19 | 37498759 | 37621270 | + | protein_coding |
| ENSG00000107954.10_3 | NEURL1 | 4.80E-09 | 6.89E-08 | 2.720 | 0.405 | chr10 | 105253736 | 105352309 | + | protein_coding |
| ENSG00000147041.12_5 | SYTL5 | 1.82E-16 | 8.03E-15 | 2.717 | 3.061 | chrX | 37865806 | 37988072 | + | protein_coding |
| ENSG00000183729.3_3 | NPBWR1 | 4.13E-19 | 2.71E-17 | 2.714 | 3.669 | chr8 | 53850991 | 53853677 | + | protein_coding |
| ENSG00000197857.14_3 | ZNF44 | 1.58E-17 | 8.34E-16 | 2.714 | 3.263 | chr19 | 12335501 | 12405702 | - | protein_coding |
| ENSG00000269397.1_7 | AC011503.2 | 6.83E-09 | 9.55E-08 | 2.712 | 0.878 | chr19 | 24110590 | 24112089 | + | lncRNA |
| ENSG00000169752.17_5 | NRG4 | 1.57E-14 | 5.22E-13 | 2.702 | 1.993 | chr15 | 76228310 | 76352136 | - | protein_coding |
| ENSG00000185101.13_4 | ANO9 | 4.36E-13 | 1.12E-11 | 2.697 | 1.958 | chr11 | 417933 | 442011 | - | protein_coding |
| ENSG00000104760.17_5 | FGL1 | 1.75E-22 | 2.21E-20 | 2.693 | 5.179 | chr8 | 17721889 | 17767874 | - | protein_coding |
| ENSG00000229689.3_5 | AC009237.3 | 4.64E-07 | 4.99E-06 | 2.693 | -0.355 | chr2 | 96191093 | 96198153 | + | transcribed_unprocessed_pseudogene |
| ENSG00000275234.1_7 | AC010503.5 | 6.54E-09 | 9.19E-08 | 2.691 | 0.309 | chr19 | 6469476 | 6470163 | + | lncRNA |
| ENSG00000047457.14_5 | CP | 6.36E-25 | 1.62E-22 | 2.686 | 5.712 | chr3 | 148880197 | 148939616 | - | protein_coding |
| ENSG00000213967.11_5 | ZNF726 | 3.89E-16 | 1.62E-14 | 2.684 | 2.639 | chr19 | 24097678 | 24127961 | + | protein_coding |
| ENSG00000100968.14_4 | NFATC4 | 4.34E-08 | 5.39E-07 | 2.682 | 0.174 | chr14 | 24834879 | 24848810 | + | protein_coding |
| ENSG00000009709.12_3 | PAX7 | 2.04E-15 | 7.71E-14 | 2.680 | 2.321 | chr1 | 18957340 | 19075360 | + | protein_coding |
| ENSG00000105855.10_3 | ITGB8 | 9.03E-24 | 1.65E-21 | 2.679 | 5.114 | chr7 | 20370325 | 20455377 | + | protein_coding |
| ENSG00000267345.1_7 | AC010632.1 | 8.73E-05 | 0.00068202 | 2.679 | -1.418 | chr19 | 37566015 | 37569507 | - | lncRNA |
| ENSG00000084453.16_6 | SLCO1A2 | 4.96E-06 | 4.64E-05 | 2.669 | -0.605 | chr12 | 21417534 | 21572528 | - | protein_coding |
| ENSG00000165164.14_9 | CFAP47 | 4.00E-06 | 3.79E-05 | 2.657 | -0.745 | chrX | 35937851 | 36403432 | + | protein_coding |
| ENSG00000182218.9_2 | HHIPL1 | 2.52E-07 | 2.82E-06 | 2.656 | -0.086 | chr14 | 100111447 | 100146906 | + | protein_coding |
| ENSG00000198440.9_6 | ZNF583 | 5.68E-19 | 3.68E-17 | 2.652 | 3.462 | chr19 | 56909335 | 56947404 | + | protein_coding |
| ENSG00000197119.13_5 | SLC25A29 | 3.97E-20 | 3.08E-18 | 2.651 | 4.717 | chr14 | 100757453 | 100772884 | - | protein_coding |
| ENSG00000113140.11_6 | SPARC | 8.14E-13 | 2.01E-11 | 2.646 | 1.628 | chr5 | 151040657 | 151066536 | - | protein_coding |
| ENSG00000019505.8_5 | SYT13 | 2.36E-28 | 1.75E-25 | 2.646 | 7.863 | chr11 | 45261853 | 45307892 | - | protein_coding |
| ENSG00000120262.10_3 | CCDC170 | 0.00018623 | 0.00138325 | 2.638 | -0.758 | chr6 | 151815152 | 151942328 | + | protein_coding |
| ENSG00000125730.17_6 | C3 | 7.63E-26 | 2.56E-23 | 2.637 | 6.094 | chr19 | 6677715 | 6730573 | - | protein_coding |
| ENSG00000253485.2_4 | PCDHGA5 | 2.76E-08 | 3.51E-07 | 2.634 | 0.121 | chr5 | 140743799 | 140892546 | + | protein_coding |
| ENSG00000166589.13_7 | CDH16 | 1.71E-06 | 1.71E-05 | 2.626 | -0.209 | chr16 | 66942025 | 66952820 | - | protein_coding |
| ENSG00000169122.11_2 | FAM110B | 8.88E-08 | 1.06E-06 | 2.625 | -0.114 | chr8 | 58907068 | 59116838 | + | protein_coding |
| ENSG00000163520.14_5 | FBLN2 | 4.08E-12 | 8.89E-11 | 2.622 | 3.336 | chr3 | 13590625 | 13679922 | + | protein_coding |
| ENSG00000000971.16_3 | CFH | 3.06E-26 | 1.13E-23 | 2.620 | 7.433 | chr1 | 196621173 | 196716634 | + | protein_coding |
| ENSG00000164197.12_4 | RNF180 | 2.91E-09 | 4.34E-08 | 2.619 | 0.439 | chr5 | 63461670 | 63668696 | + | protein_coding |
| ENSG00000258314.3_6 | AL589182.1 | 2.33E-07 | 2.63E-06 | 2.617 | -0.130 | chr14 | 19611768 | 19643232 | + | lncRNA |
| ENSG00000159905.14_5 | ZNF221 | 1.29E-15 | 4.97E-14 | 2.607 | 3.187 | chr19 | 44455375 | 44471861 | + | protein_coding |
| ENSG00000006747.15_4 | SCIN | 2.12E-09 | 3.20E-08 | 2.604 | 0.431 | chr7 | 12610203 | 12699807 | + | protein_coding |
| ENSG00000258512.1_3 | LINC00239 | 1.25E-06 | 1.28E-05 | 2.601 | -0.424 | chr14 | 102196774 | 102198859 | + | lncRNA |
| ENSG00000114315.4_3 | HES1 | 7.34E-20 | 5.62E-18 | 2.601 | 3.822 | chr3 | 193853937 | 193856521 | + | protein_coding |
| ENSG00000143341.12_3 | HMCN1 | 1.38E-07 | 1.61E-06 | 2.601 | -0.228 | chr1 | 185703523 | 186160081 | + | protein_coding |
| ENSG00000122728.6_4 | TAF1L | 3.77E-05 | 0.00031344 | 2.600 | -0.943 | chr9 | 32629452 | 32635667 | - | protein_coding |
| ENSG00000175311.7_3 | ANKS4B | 6.11E-18 | 3.38E-16 | 2.597 | 3.222 | chr16 | 21245020 | 21265171 | + | protein_coding |
| ENSG00000164199.18_9 | ADGRV1 | 5.57E-09 | 7.91E-08 | 2.592 | 0.420 | chr5 | 89825161 | 90460254 | + | protein_coding |

| | | | | | | | | | | |
|---|---|---|---|---|---|---|---|---|---|---|
| ENSG00000162878.13_3 | PKDCC | 1.03E-24 | 2.44E-22 | 2.592 | 5.780 | chr2 | 42275161 | 42285657 | + | protein_coding |
| ENSG00000151655.19_5 | ITIH2 | 3.11E-08 | 3.92E-07 | 2.588 | 0.227 | chr10 | 7745279 | 7791483 | + | protein_coding |
| ENSG00000231419.6_3 | LINC00689 | 9.64E-09 | 1.31E-07 | 2.588 | 0.227 | chr7 | 158799213 | 158822886 | + | lncRNA |
| ENSG00000255652.4_8 | AC140847.2 | 0.00014154 | 0.00107422 | 2.587 | -1.284 | chr12 | 34343406 | 34372181 | - | lncRNA |
| ENSG00000205413.8_5 | SAMD9 | 2.65E-22 | 3.20E-20 | 2.586 | 5.074 | chr7 | 92728826 | 92747336 | - | protein_coding |
| ENSG00000176222.9_6 | ZNF404 | 2.79E-08 | 3.53E-07 | 2.584 | 1.425 | chr19 | 44376517 | 44405537 | - | protein_coding |
| ENSG00000083814.13_5 | ZNF671 | 6.82E-17 | 3.28E-15 | 2.580 | 2.897 | chr19 | 58231119 | 58238992 | - | protein_coding |
| ENSG00000198774.5_5 | RASSF9 | 7.48E-22 | 8.36E-20 | 2.577 | 4.652 | chr12 | 86194481 | 86230187 | - | protein_coding |
| ENSG00000121858.11_4 | TNFSF10 | 4.39E-10 | 7.28E-09 | 2.576 | 0.711 | chr3 | 172223298 | 172241265 | - | protein_coding |
| ENSG00000145242.13_5 | EPHA5 | 0.0001874 | 0.00138964 | 2.576 | -1.489 | chr4 | 66185281 | 66536213 | - | protein_coding |
| ENSG00000151715.8_3 | TMEM45B | 6.14E-10 | 1.00E-08 | 2.574 | 0.813 | chr11 | 129685743 | 129729898 | + | protein_coding |
| ENSG00000256087.7_5 | ZNF432 | 3.51E-21 | 3.38E-19 | 2.573 | 4.545 | chr19 | 52534631 | 52598991 | - | protein_coding |
| ENSG00000156959.9_4 | LHFPL4 | 7.18E-09 | 1.00E-07 | 2.565 | 0.277 | chr3 | 9540045 | 9595506 | - | protein_coding |
| ENSG00000178685.14_7 | PARP10 | 3.26E-14 | 1.02E-12 | 2.563 | 2.270 | chr8 | 145051321 | 145086940 | - | protein_coding |
| ENSG00000198521.7 | ZNF43 | 7.70E-24 | 1.45E-21 | 2.563 | 5.265 | chr19 | 21987752 | 22034927 | - | protein_coding |
| ENSG00000268686.2_7 | AC010643.1 | 2.40E-13 | 6.41E-12 | 2.562 | 1.816 | chr19 | 49871815 | 49891348 | - | lncRNA |
| ENSG00000267390.1_6 | AC036176.1 | 6.94E-07 | 7.29E-06 | 2.560 | -0.355 | chr18 | 61034561 | 61048862 | + | lncRNA |
| ENSG00000187510.9_7 | PLEKHG7 | 1.14E-05 | 0.00010126 | 2.555 | -0.965 | chr12 | 93096619 | 93166231 | + | protein_coding |
| ENSG00000085117.12_5 | CD82 | 1.87E-15 | 7.10E-14 | 2.554 | 2.645 | chr11 | 44585977 | 44641908 | + | protein_coding |
| ENSG00000159674.12_7 | SPON2 | 3.07E-07 | 3.39E-06 | 2.551 | 0.209 | chr4 | 1160720 | 1202750 | - | protein_coding |
| ENSG00000100033.16_5 | PRODH | 1.50E-20 | 1.27E-18 | 2.550 | 4.246 | chr22 | 18900290 | 18924066 | - | protein_coding |
| ENSG00000172794.20_5 | RAB37 | 1.21E-22 | 1.64E-20 | 2.548 | 4.880 | chr17 | 72666717 | 72743474 | + | protein_coding |
| ENSG00000137731.14_6 | FXYD2 | 2.23E-16 | 9.69E-15 | 2.536 | 2.795 | chr11 | 117671559 | 117699413 | - | protein_coding |
| ENSG00000146151.13_4 | HMGCLL1 | 3.92E-11 | 7.48E-10 | 2.535 | 1.216 | chr6 | 55299167 | 55444012 | - | protein_coding |
| ENSG00000188396.3_4 | TCTEX1D4 | 1.95E-05 | 0.00016903 | 2.530 | -0.983 | chr1 | 45271585 | 45272347 | - | protein_coding |
| ENSG00000162039.15_5 | MEIOB | 1.32E-19 | 9.69E-18 | 2.522 | 3.940 | chr16 | 1883988 | 1934295 | - | protein_coding |
| ENSG00000217236.2_4 | SP9 | 3.99E-12 | 8.71E-11 | 2.522 | 1.496 | chr2 | 175199682 | 175203228 | + | protein_coding |
| ENSG00000157227.13_3 | MMP14 | 3.38E-14 | 1.06E-12 | 2.513 | 2.503 | chr14 | 23305769 | 23318236 | + | protein_coding |
| ENSG00000129204.17_3 | USP6 | 3.32E-10 | 5.59E-09 | 2.510 | 0.792 | chr17 | 5019327 | 5078329 | + | protein_coding |
| ENSG00000179299.17_5 | NSUN7 | 2.39E-13 | 6.37E-12 | 2.505 | 1.804 | chr4 | 40751942 | 40813201 | + | protein_coding |
| ENSG00000205336.12_7 | ADGRG1 | 4.18E-26 | 1.48E-23 | 2.502 | 6.893 | chr16 | 57644564 | 57699492 | + | protein_coding |
| ENSG00000065618.21_7 | COL17A1 | 6.65E-14 | 1.97E-12 | 2.502 | 2.204 | chr10 | 105791044 | 105845638 | - | protein_coding |
| ENSG00000134755.17_8 | DSC2 | 3.76E-19 | 2.48E-17 | 2.497 | 4.128 | chr18 | 28638806 | 28682485 | - | protein_coding |
| ENSG00000162777.17_5 | DENND2D | 6.06E-16 | 2.44E-14 | 2.492 | 2.775 | chr1 | 111728591 | 111747157 | - | protein_coding |
| ENSG00000171345.13_5 | KRT19 | 4.57E-25 | 1.19E-22 | 2.492 | 7.947 | chr17 | 39679869 | 39684560 | - | protein_coding |
| ENSG00000107282.8_6 | APBA1 | 3.33E-12 | 7.37E-11 | 2.492 | 1.592 | chr9 | 72042448 | 72287287 | - | protein_coding |
| ENSG00000109255.11_5 | NMU | 3.02E-06 | 2.92E-05 | 2.478 | -0.424 | chr4 | 56461396 | 56502865 | - | protein_coding |
| ENSG00000126217.21_5 | MCF2L | 1.19E-16 | 5.43E-15 | 2.471 | 3.288 | chr13 | 113548692 | 113754056 | + | protein_coding |
| ENSG00000230666.5_3 | CEACAM22P | 1.60E-06 | 1.60E-05 | 2.471 | -0.341 | chr19 | 45041045 | 45124119 | - | transcribed_unprocessed_pseudogene |
| ENSG00000125266.8_6 | EFNB2 | 3.64E-11 | 6.97E-10 | 2.468 | 1.339 | chr13 | 107142093 | 107188010 | - | protein_coding |
| ENSG00000077935.17_4 | SMC1B | 1.17E-08 | 1.57E-07 | 2.467 | 0.529 | chr22 | 45739944 | 45809500 | - | protein_coding |
| ENSG00000267309.1_6 | AC092295.2 | 8.88E-08 | 1.06E-06 | 2.464 | 0.316 | chr19 | 36980551 | 36981942 | + | lncRNA |
| ENSG00000275055.1_6 | AC011468.5 | 4.02E-05 | 0.00033281 | 2.463 | -1.034 | chr19 | 52552260 | 52553007 | + | lncRNA |
| ENSG00000256043.3_4 | CTSO | 2.40E-12 | 5.50E-11 | 2.462 | 1.917 | chr4 | 156845270 | 156875018 | - | protein_coding |
| ENSG00000225255.2 | LA16c-83F12.6 | 0.00011591 | 0.00088728 | 2.460 | -1.372 | chr22 | 16199666 | 16231333 | - | lincRNA |
| ENSG00000271133.5_8 | AC004130.2 | 2.32E-05 | 0.00019926 | 2.459 | -1.035 | chr7 | 20367922 | 20371370 | - | lncRNA |
| ENSG00000253305.2_3 | PCDHGB6 | 7.18E-09 | 1.00E-07 | 2.459 | 0.424 | chr5 | 140787588 | 140892546 | + | protein_coding |
| ENSG00000104324.16_6 | CPQ | 3.27E-11 | 6.31E-10 | 2.454 | 1.533 | chr8 | 97657470 | 98161882 | + | protein_coding |
| ENSG00000167165.19_6 | UGT1A6 | 8.52E-07 | 8.89E-06 | 2.452 | -0.258 | chr2 | 234600253 | 234681946 | + | protein_coding |
| ENSG00000169418.10_3 | NPR1 | 1.68E-13 | 4.60E-12 | 2.452 | 2.042 | chr1 | 153651164 | 153666468 | + | protein_coding |
| ENSG00000134538.3_4 | SLCO1B1 | 2.02E-13 | 5.43E-12 | 2.451 | 2.100 | chr12 | 21284128 | 21392730 | + | protein_coding |
| ENSG00000090339.9_3 | ICAM1 | 2.03E-08 | 2.62E-07 | 2.447 | 0.518 | chr19 | 10381769 | 10397291 | + | protein_coding |
| ENSG00000162383.12_6 | SLC1A7 | 3.74E-05 | 0.00031093 | 2.447 | -1.051 | chr1 | 53552851 | 53608304 | - | protein_coding |
| ENSG00000076641.4_4 | PAG1 | 2.46E-06 | 2.40E-05 | 2.442 | -0.456 | chr8 | 81880045 | 82024303 | - | protein_coding |
| ENSG00000167136.7_3 | ENDOG | 1.29E-07 | 1.51E-06 | 2.441 | 0.179 | chr9 | 131580779 | 131584955 | + | protein_coding |
| ENSG00000280414.1_5 | AC018470.1 | 7.62E-07 | 8.00E-06 | 2.438 | -0.276 | chr2 | 175199166 | 175202294 | - | TEC |
| ENSG00000081818.4_3 | PCDHB4 | 0.00017698 | 0.00132061 | 2.437 | -1.205 | chr5 | 140501400 | 140505205 | + | protein_coding |
| ENSG00000079112.10_6 | CDH17 | 1.42E-21 | 1.51E-19 | 2.437 | 4.961 | chr8 | 95139390 | 95229531 | - | protein_coding |
| ENSG00000196110.8_4 | ZNF699 | 2.31E-17 | 1.19E-15 | 2.435 | 3.498 | chr19 | 9401816 | 9420514 | - | protein_coding |

| | | | | | | | | | | |
|---|---|---|---|---|---|---|---|---|---|---|
| ENSG00000278318.5_6 | ZNF229 | 2.38E-20 | 1.93E-18 | 2.431 | 4.287 | chr19 | 44921685 | 44952766 | - | protein_coding |
| ENSG00000184357.4_3 | H1-5 | 2.50E-24 | 5.29E-22 | 2.431 | 6.095 | chr6 | 27834627 | 27835307 | - | protein_coding |
| ENSG00000184454.7_4 | NCMAP | 1.03E-13 | 2.97E-12 | 2.429 | 2.319 | chr1 | 24882578 | 24935819 | + | protein_coding |
| ENSG00000188785.12_4 | ZNF548 | 1.38E-18 | 8.37E-17 | 2.425 | 3.785 | chr19 | 57901218 | 57914360 | + | protein_coding |
| ENSG00000196793.13_3 | ZNF239 | 5.66E-17 | 2.80E-15 | 2.421 | 3.173 | chr10 | 44051792 | 44070066 | - | protein_coding |
| ENSG00000278917.1_7 | AC006213.6 | 9.72E-06 | 8.73E-05 | 2.410 | -0.805 | chr19 | 44395385 | 44399563 | + | TEC |
| ENSG00000132016.11_4 | C19orf57 | 5.74E-09 | 8.13E-08 | 2.407 | 0.711 | chr19 | 13993161 | 14017265 | - | protein_coding |
| ENSG00000267475.1_9 | AC008736.1 | 7.65E-13 | 1.90E-11 | 2.404 | 1.723 | chr19 | 33177995 | 33182656 | - | lncRNA |
| ENSG00000269235.2_6 | ZNF350-AS1 | 4.13E-11 | 7.82E-10 | 2.400 | 1.473 | chr19 | 52452387 | 52484620 | + | lncRNA |
| ENSG00000256683.7_7 | ZNF350 | 9.33E-20 | 7.02E-18 | 2.400 | 4.333 | chr19 | 52467593 | 52490109 | - | protein_coding |
| ENSG00000267319.1_5 | SELENOKP1 | 8.28E-05 | 0.00064825 | 2.399 | -1.243 | chr19 | 38184181 | 38184465 | - | processed_pseudogene |
| ENSG00000233527.9_7 | ZNF529-AS1 | 9.54E-15 | 3.26E-13 | 2.394 | 2.511 | chr19 | 37063972 | 37085610 | + | lncRNA |
| ENSG00000196081.9_8 | ZNF724 | 9.85E-17 | 4.59E-15 | 2.390 | 3.378 | chr19 | 23404401 | 23433192 | - | protein_coding |
| ENSG00000137672.13_5 | TRPC6 | 7.00E-18 | 3.84E-16 | 2.381 | 3.524 | chr11 | 101322295 | 101743293 | - | protein_coding |
| ENSG00000139865.16_2 | TTC6 | 1.56E-13 | 4.31E-12 | 2.380 | 2.138 | chr14 | 38065052 | 38510647 | + | protein_coding |
| ENSG00000136155.17_3 | SCEL | 4.70E-16 | 1.92E-14 | 2.375 | 3.121 | chr13 | 78109809 | 78219398 | + | protein_coding |
| ENSG00000204941.14_4 | PSG5 | 4.23E-11 | 7.99E-10 | 2.372 | 1.203 | chr19 | 43670408 | 43690688 | - | protein_coding |
| ENSG00000223542.1_7 | AL024497.1 | 0.0001158 | 0.00088684 | 2.368 | -1.264 | chr6 | 133756215 | 133760602 | - | lncRNA |
| ENSG00000012171.20_6 | SEMA3B | 7.56E-20 | 5.77E-18 | 2.367 | 5.125 | chr3 | 50304990 | 50314977 | + | protein_coding |
| ENSG00000170477.13_4 | KRT4 | 1.13E-20 | 9.92E-19 | 2.365 | 5.454 | chr12 | 53200333 | 53207900 | - | protein_coding |
| ENSG00000131845.14_5 | AC005261.1 | 1.55E-19 | 1.12E-17 | 2.365 | 4.101 | chr19 | 57862675 | 57871266 | + | protein_coding |
| ENSG00000079385.22_6 | CEACAM1 | 8.40E-13 | 2.06E-11 | 2.361 | 1.802 | chr19 | 43011456 | 43065386 | - | protein_coding |
| ENSG00000005471.17_8 | ABCB4 | 6.55E-07 | 6.91E-06 | 2.359 | -0.100 | chr7 | 87031013 | 87109751 | - | protein_coding |
| ENSG00000204956.5_3 | PCDHGA1 | 6.68E-13 | 1.68E-11 | 2.359 | 1.975 | chr5 | 140710138 | 140892548 | + | protein_coding |
| ENSG00000231160.10_7 | KLF3-AS1 | 1.78E-10 | 3.13E-09 | 2.358 | 1.139 | chr4 | 38604059 | 38666535 | - | lncRNA |
| ENSG00000272070.1_6 | AC005618.1 | 5.22E-07 | 5.59E-06 | 2.357 | -0.029 | chr5 | 140705777 | 140708924 | + | lncRNA |
| ENSG00000139192.12_4 | TAPBPL | 9.10E-16 | 3.58E-14 | 2.355 | 2.898 | chr12 | 6560856 | 6575683 | + | protein_coding |
| ENSG00000121905.10_3 | HPCA | 5.39E-05 | 0.00043631 | 2.354 | -0.980 | chr1 | 33351595 | 33364042 | + | protein_coding |
| ENSG00000233493.4_4 | TMEM238 | 7.49E-08 | 9.04E-07 | 2.352 | 0.626 | chr19 | 55890612 | 55895660 | - | protein_coding |
| ENSG00000204514.10_6 | ZNF814 | 2.47E-20 | 1.99E-18 | 2.351 | 4.341 | chr19 | 58360099 | 58400442 | - | protein_coding |
| ENSG00000205488.9_9 | CALML3-AS1 | 2.84E-05 | 0.00024074 | 2.348 | -0.851 | chr10 | 5551999 | 5568209 | - | lncRNA |
| ENSG00000267370.1_6 | AC008752.4 | 4.88E-05 | 0.00039845 | 2.345 | -1.125 | chr19 | 9866828 | 9867539 | - | unprocessed_pseudogene |
| ENSG00000149243.15_3 | KLHL35 | 9.66E-12 | 2.01E-10 | 2.339 | 1.743 | chr11 | 75133438 | 75141674 | - | protein_coding |
| ENSG00000238120.1_7 | LINC01589 | 1.31E-13 | 3.64E-12 | 2.332 | 2.551 | chr22 | 46000312 | 46001501 | - | lncRNA |
| ENSG00000006047.13_4 | YBX2 | 2.45E-13 | 6.51E-12 | 2.331 | 2.211 | chr17 | 7191582 | 7197958 | - | protein_coding |
| ENSG00000138792.10_4 | ENPEP | 7.14E-09 | 9.95E-08 | 2.328 | 0.644 | chr4 | 111286889 | 111486441 | + | protein_coding |
| ENSG00000129295.9_6 | LRRC6 | 2.00E-12 | 4.63E-11 | 2.325 | 2.037 | chr8 | 133582664 | 133687838 | - | protein_coding |
| ENSG00000139725.8_5 | RHOF | 3.30E-07 | 3.62E-06 | 2.323 | 3.118 | chr12 | 122215660 | 122241309 | - | protein_coding |
| ENSG00000132274.16_4 | TRIM22 | 1.44E-11 | 2.92E-10 | 2.312 | 1.641 | chr11 | 5710927 | 5758319 | + | protein_coding |
| ENSG00000174473.16_3 | GALNTL6 | 1.18E-09 | 1.86E-08 | 2.312 | 0.927 | chr4 | 172733405 | 173962710 | + | protein_coding |
| ENSG00000087266.17_9 | SH3BP2 | 3.54E-21 | 3.40E-19 | 2.307 | 4.870 | chr4 | 2794798 | 2842825 | + | protein_coding |
| ENSG00000260711.2_7 | AL121839.2 | 4.03E-08 | 5.02E-07 | 2.302 | 0.342 | chr14 | 92219200 | 92226142 | - | lncRNA |
| ENSG00000226652.3_4 | PSMD10P2 | 3.34E-07 | 3.67E-06 | 2.300 | 0.049 | chr3 | 186478482 | 186479839 | + | transcribed_processed_pseudogene |
| ENSG00000129595.13_5 | EPB41L4A | 6.39E-19 | 4.10E-17 | 2.299 | 3.859 | chr5 | 111478138 | 111755010 | - | protein_coding |
| ENSG00000155324.10_7 | GRAMD2B | 1.05E-11 | 2.16E-10 | 2.285 | 1.663 | chr5 | 125695824 | 125832186 | + | protein_coding |
| ENSG00000140022.13_8 | STON2 | 1.08E-16 | 4.99E-15 | 2.285 | 3.238 | chr14 | 81727000 | 81902809 | - | protein_coding |
| ENSG00000152784.16_3 | PRDM8 | 6.86E-06 | 6.31E-05 | 2.280 | -0.482 | chr4 | 81105033 | 81125483 | + | protein_coding |
| ENSG00000254872.3_6 | LINC02688 | 2.22E-10 | 3.81E-09 | 2.280 | 1.265 | chr11 | 1049880 | 1055749 | + | lncRNA |
| ENSG00000267414.1_7 | AC120049.1 | 1.21E-05 | 0.00010711 | 2.278 | -0.676 | chr18 | 42256892 | 42259682 | - | lncRNA |
| ENSG00000227039.6_6 | ITGB2-AS1 | 9.02E-10 | 1.44E-08 | 2.276 | 1.121 | chr21 | 46340966 | 46349593 | + | lncRNA |
| ENSG00000285712.1_4 | AC068707.1 | 0.00050038 | 0.00344714 | 2.274 | -1.033 | chr10 | 43821281 | 43846240 | - | lncRNA |
| ENSG00000132470.14_5 | ITGB4 | 2.54E-22 | 3.09E-20 | 2.273 | 8.393 | chr17 | 73717408 | 73753899 | + | protein_coding |
| ENSG00000132688.11_4 | NES | 6.62E-17 | 3.20E-15 | 2.271 | 3.614 | chr1 | 156638555 | 156647199 | - | protein_coding |
| ENSG00000003147.19_7 | ICA1 | 8.43E-15 | 2.91E-13 | 2.271 | 2.863 | chr7 | 8152814 | 8302317 | - | protein_coding |
| ENSG00000177932.7_3 | ZNF354C | 5.74E-19 | 3.71E-17 | 2.270 | 4.207 | chr5 | 178487374 | 178510978 | + | protein_coding |
| ENSG00000275832.5_3 | ARHGAP23 | 4.91E-23 | 7.34E-21 | 2.270 | 6.193 | chr17 | 36575523 | 36668621 | + | protein_coding |
| ENSG00000261150.3_3 | EPPK1 | 2.63E-06 | 2.55E-05 | 2.269 | -0.322 | chr8 | 144939497 | 144952635 | - | protein_coding |
| ENSG00000226314.8_4 | ZNF192P1 | 3.80E-06 | 3.62E-05 | 2.268 | -0.493 | chr6 | 28129547 | 28137372 | + | transcribed_unprocessed_pseudogene |
| ENSG00000101986.12_3 | ABCD1 | 4.05E-15 | 1.44E-13 | 2.263 | 3.935 | chrX | 152990311 | 153010209 | + | protein_coding |

| | | | | | | | | | | |
|---|---|---|---|---|---|---|---|---|---|---|
| ENSG00000187546.14_4 | AGMO | 7.34E-15 | 2.55E-13 | 2.261 | 2.686 | chr7 | 15239942 | 15601640 | - | protein_coding |
| ENSG00000145358.6_6 | DDIT4L | 1.81E-12 | 4.23E-11 | 2.257 | 2.151 | chr4 | 101107027 | 101111939 | - | protein_coding |
| ENSG00000111319.13_7 | SCNN1A | 1.78E-13 | 4.84E-12 | 2.256 | 4.180 | chr12 | 6456009 | 6486896 | - | protein_coding |
| ENSG00000274414.1_7 | AL121772.1 | 1.70E-08 | 2.22E-07 | 2.249 | 1.539 | chr20 | 25219643 | 25225865 | - | lncRNA |
| ENSG00000130545.16_4 | CRB3 | 7.75E-12 | 1.63E-10 | 2.247 | 1.911 | chr19 | 6463788 | 6467232 | + | protein_coding |
| ENSG00000115380.20_5 | EFEMP1 | 1.46E-22 | 1.94E-20 | 2.244 | 5.928 | chr2 | 56093102 | 56151274 | - | protein_coding |
| ENSG00000152137.6_3 | HSPB8 | 2.20E-18 | 1.31E-16 | 2.243 | 3.800 | chr12 | 119616447 | 119658936 | + | protein_coding |
| ENSG00000165168.8_4 | CYBB | 5.73E-05 | 0.00046213 | 2.241 | -0.932 | chrX | 37639312 | 37672714 | + | protein_coding |
| ENSG00000100346.18_3 | CACNA1I | 3.45E-05 | 0.00028816 | 2.239 | -0.607 | chr22 | 39966758 | 40085740 | + | protein_coding |
| ENSG00000161609.10_6 | CCDC155 | 8.97E-06 | 8.11E-05 | 2.236 | -0.618 | chr19 | 49891476 | 49921247 | + | protein_coding |
| ENSG00000234745.11_4 | HLA-B | 1.73E-05 | 0.0001507 | 2.235 | -0.619 | chr6 | 31237268 | 31324965 | - | protein_coding |
| ENSG00000259925.1_8 | AC130456.2 | 0.00046594 | 0.00323123 | 2.230 | -1.204 | chr16 | 19354969 | 19413015 | - | lncRNA |
| ENSG00000223813.3_7 | AC007255.1 | 1.17E-08 | 1.56E-07 | 2.224 | 0.676 | chr7 | 29553840 | 29603286 | - | lncRNA |
| ENSG00000170369.4_4 | CST2 | 7.37E-05 | 0.00058218 | 2.223 | -0.621 | chr20 | 23804406 | 23807366 | - | protein_coding |
| ENSG00000121316.11_4 | PLBD1 | 1.67E-16 | 7.43E-15 | 2.222 | 3.329 | chr12 | 14656595 | 14720817 | - | protein_coding |
| ENSG00000253767.2_4 | PCDHGA8 | 1.09E-07 | 1.29E-06 | 2.219 | 0.327 | chr5 | 140769724 | 140892546 | + | protein_coding |
| ENSG00000188039.15_6 | NWD1 | 9.40E-05 | 0.00073043 | 2.218 | -0.949 | chr19 | 16830786 | 16928774 | + | protein_coding |
| ENSG00000164125.15_6 | GASK1B | 4.74E-14 | 1.45E-12 | 2.215 | 2.744 | chr4 | 159045626 | 159094470 | - | protein_coding |
| ENSG00000113209.9_4 | PCDHB5 | 1.63E-15 | 6.21E-14 | 2.214 | 3.265 | chr5 | 140514788 | 140518197 | + | protein_coding |
| ENSG00000251209.9_6 | LINC00923 | 1.57E-09 | 2.43E-08 | 2.214 | 0.951 | chr15 | 98115415 | 98418880 | - | lncRNA |
| ENSG00000153896.19_4 | ZNF599 | 6.52E-13 | 1.64E-11 | 2.212 | 2.263 | chr19 | 35248978 | 35264134 | - | protein_coding |
| ENSG00000110852.4_4 | CLEC2B | 4.52E-11 | 8.47E-10 | 2.212 | 1.576 | chr12 | 10005583 | 10022735 | - | protein_coding |
| ENSG00000231924.9_6 | PSG1 | 2.26E-05 | 0.00019442 | 2.211 | -0.732 | chr19 | 43370616 | 43383974 | - | protein_coding |
| ENSG00000241269.1_7 | AC093620.1 | 8.74E-05 | 0.00068218 | 2.208 | -1.087 | chr7 | 5459458 | 5460398 | + | lncRNA |
| ENSG00000114698.15_5 | PLSCR4 | 8.10E-20 | 6.15E-18 | 2.206 | 4.703 | chr3 | 145910122 | 145968966 | - | protein_coding |
| ENSG00000171681.12_8 | ATF7IP | 1.12E-24 | 2.60E-22 | 2.206 | 7.529 | chr12 | 14518566 | 14655869 | + | protein_coding |
| ENSG00000203706.8_6 | SERTAD4-AS1 | 5.22E-05 | 0.00042379 | 2.200 | -0.850 | chr1 | 210404801 | 210407392 | - | lncRNA |
| ENSG00000140993.11_4 | TIGD7 | 2.82E-11 | 5.49E-10 | 2.196 | 1.711 | chr16 | 3348808 | 3355729 | - | protein_coding |
| ENSG00000128242.13_5 | GAL3ST1 | 1.93E-16 | 8.47E-15 | 2.195 | 4.466 | chr22 | 30950622 | 30970652 | - | protein_coding |
| ENSG00000183778.17_4 | B3GALT5 | 2.95E-14 | 9.31E-13 | 2.193 | 2.501 | chr21 | 40928369 | 41045064 | + | protein_coding |
| ENSG00000227954.7_7 | TARID | 1.04E-16 | 4.85E-15 | 2.192 | 3.518 | chr6 | 133823390 | 134213940 | - | lncRNA |
| ENSG00000273983.1_3 | H3C8 | 7.93E-06 | 7.22E-05 | 2.190 | -0.387 | chr6 | 26269633 | 26272043 | - | protein_coding |
| ENSG00000196440.11_2 | ARMCX4 | 1.30E-05 | 0.00011494 | 2.184 | -0.395 | chrX | 100673275 | 100788446 | + | protein_coding |
| ENSG00000196730.13_4 | DAPK1 | 6.29E-26 | 2.19E-23 | 2.183 | 8.710 | chr9 | 90112143 | 90323549 | + | protein_coding |
| ENSG00000169247.13_6 | SH3TC2 | 7.20E-10 | 1.16E-08 | 2.182 | 1.216 | chr5 | 148303202 | 148442726 | - | protein_coding |
| ENSG00000271447.6_6 | MMP28 | 5.23E-08 | 6.43E-07 | 2.182 | 0.486 | chr17 | 34083268 | 34122711 | - | protein_coding |
| ENSG00000129946.10_5 | SHC2 | 3.38E-07 | 3.70E-06 | 2.180 | 0.437 | chr19 | 416583 | 460996 | - | protein_coding |
| ENSG00000018236.15_6 | CNTN1 | 8.44E-26 | 2.78E-23 | 2.179 | 9.648 | chr12 | 41086241 | 41466217 | + | protein_coding |
| ENSG00000156299.13_5 | TIAM1 | 5.96E-14 | 1.78E-12 | 2.178 | 2.568 | chr21 | 32490734 | 32932290 | - | protein_coding |
| ENSG00000159915.12_4 | ZNF233 | 3.13E-14 | 9.83E-13 | 2.176 | 2.801 | chr19 | 44764033 | 44779470 | + | protein_coding |
| ENSG00000177103.15_6 | DSCAML1 | 2.73E-14 | 8.70E-13 | 2.174 | 2.884 | chr11 | 117298488 | 117688240 | - | protein_coding |
| ENSG00000184471.8_4 | C1QTNF8 | 5.93E-06 | 5.50E-05 | 2.165 | -0.241 | chr16 | 1138226 | 1146244 | - | protein_coding |
| ENSG00000129514.7_5 | FOXA1 | 2.90E-22 | 3.46E-20 | 2.163 | 5.535 | chr14 | 38058757 | 38065264 | - | protein_coding |
| ENSG00000102271.14_5 | KLHL4 | 4.53E-17 | 2.27E-15 | 2.158 | 3.575 | chrX | 86772412 | 86925050 | + | protein_coding |
| ENSG00000180938.6_3 | ZNF572 | 1.01E-14 | 3.43E-13 | 2.157 | 2.760 | chr8 | 125985537 | 125991631 | + | protein_coding |
| ENSG00000204616.11_4 | TRIM31 | 0.00026852 | 0.00194728 | 2.152 | -0.883 | chr6 | 30070674 | 30080867 | - | protein_coding |
| ENSG00000172346.15_3 | CSDC2 | 4.21E-06 | 3.97E-05 | 2.142 | -0.129 | chr22 | 41957014 | 41973745 | + | protein_coding |
| ENSG00000204396.11_4 | VWA7 | 7.66E-08 | 9.22E-07 | 2.137 | 0.530 | chr6 | 31733367 | 31745105 | - | protein_coding |
| ENSG00000112852.7_4 | PCDHB2 | 1.23E-11 | 2.53E-10 | 2.136 | 1.767 | chr5 | 140474191 | 140478287 | + | protein_coding |
| ENSG00000076356.7_4 | PLXNA2 | 1.60E-18 | 9.58E-17 | 2.135 | 4.237 | chr1 | 208195587 | 208417729 | - | protein_coding |
| ENSG00000172264.17_6 | MACROD2 | 1.80E-08 | 2.35E-07 | 2.134 | 0.854 | chr20 | 13976015 | 16033842 | + | protein_coding |
| ENSG00000050438.17_3 | SLC4A8 | 9.63E-15 | 3.28E-13 | 2.132 | 2.954 | chr12 | 51785101 | 51909547 | + | protein_coding |
| ENSG00000100156.10_6 | SLC16A8 | 1.16E-06 | 1.19E-05 | 2.130 | 0.097 | chr22 | 38474141 | 38480100 | - | protein_coding |
| ENSG00000241859.7_5 | ANOS2P | 6.79E-08 | 8.24E-07 | 2.125 | 0.517 | chrY | 15815782 | 16027704 | + | transcribed_unprocessed_pseudogene |
| ENSG00000286389.1_1 | AL731702.1 | 1.85E-06 | 1.83E-05 | 2.121 | -0.090 | chr1 | 227974454 | 227979880 | + | lncRNA |
| ENSG00000082497.12_3 | SERTAD4 | 2.76E-08 | 3.51E-07 | 2.110 | 0.732 | chr1 | 210406141 | 210419976 | + | protein_coding |
| ENSG00000060140.9_7 | STYK1 | 6.48E-05 | 0.00051824 | 2.108 | -0.915 | chr12 | 10771522 | 10826917 | - | protein_coding |
| ENSG00000188959.9_3 | C9orf152 | 5.35E-10 | 8.77E-09 | 2.104 | 1.612 | chr9 | 112952328 | 112970469 | - | protein_coding |
| ENSG00000065717.15_6 | TLE2 | 2.94E-12 | 6.60E-11 | 2.102 | 2.665 | chr19 | 2997637 | 3047633 | - | protein_coding |

| | | | | | | | | | | |
|---|---|---|---|---|---|---|---|---|---|---|
| ENSG00000168070.12_7 | MAJIN | 0.00015121 | 0.00114226 | 2.101 | -0.916 | chr11 | 64704989 | 64739580 | - | protein_coding |
| ENSG00000171606.18_4 | ZNF274 | 1.74E-22 | 2.21E-20 | 2.097 | 5.732 | chr19 | 58694396 | 58724928 | + | protein_coding |
| ENSG00000068078.18_5 | FGFR3 | 3.53E-14 | 1.10E-12 | 2.094 | 3.069 | chr4 | 1795020 | 1810599 | + | protein_coding |
| ENSG00000112837.17_7 | TBX18 | 8.43E-24 | 1.56E-21 | 2.088 | 6.684 | chr6 | 85397069 | 85474316 | - | protein_coding |
| ENSG00000104537.17_7 | ANXA13 | 2.28E-19 | 1.59E-17 | 2.080 | 4.392 | chr8 | 124693034 | 124749642 | - | protein_coding |
| ENSG00000240875.6_7 | LINC00886 | 4.71E-13 | 1.21E-11 | 2.079 | 2.568 | chr3 | 156465132 | 156534851 | - | lncRNA |
| ENSG00000258429.2_5 | PDF | 0.00062549 | 0.00423036 | 2.078 | 0.650 | chr16 | 69360816 | 69364491 | - | protein_coding |
| ENSG00000177875.5_3 | CCDC184 | 8.48E-10 | 1.36E-08 | 2.077 | 1.686 | chr12 | 48577427 | 48579709 | + | protein_coding |
| ENSG00000166311.10_6 | SMPD1 | 3.08E-15 | 1.13E-13 | 2.076 | 3.458 | chr11 | 6411670 | 6416228 | + | protein_coding |
| ENSG00000124613.8_5 | ZNF391 | 8.10E-09 | 1.12E-07 | 2.071 | 1.093 | chr6 | 27342394 | 27371683 | + | protein_coding |
| ENSG00000265962.1_7 | GACAT2 | 1.16E-19 | 8.55E-18 | 2.071 | 4.533 | chr18 | 8695854 | 8707619 | - | lncRNA |
| ENSG00000287263.1_2 | AC008875.3 | 7.14E-16 | 2.84E-14 | 2.070 | 3.240 | chr5 | 42983738 | 42993378 | - | lncRNA |
| ENSG00000004848.8_6 | ARX | 2.14E-10 | 3.71E-09 | 2.070 | 1.636 | chrX | 25021811 | 25034537 | - | protein_coding |
| ENSG00000005108.16_6 | THSD7A | 1.22E-23 | 2.14E-21 | 2.070 | 6.847 | chr7 | 11409992 | 11871824 | - | protein_coding |
| ENSG00000197619.14_6 | ZNF615 | 3.66E-19 | 2.43E-17 | 2.058 | 4.663 | chr19 | 52494585 | 52511483 | - | protein_coding |
| ENSG00000186020.13_7 | ZNF529 | 6.78E-19 | 4.32E-17 | 2.058 | 4.489 | chr19 | 37025676 | 37096178 | - | protein_coding |
| ENSG00000256223.6_7 | ZNF10 | 1.00E-13 | 2.90E-12 | 2.058 | 2.661 | chr12 | 133707161 | 133736051 | + | protein_coding |
| ENSG00000008441.16_5 | NFIX | 1.55E-12 | 3.65E-11 | 2.057 | 2.296 | chr19 | 13106422 | 13209610 | + | protein_coding |
| ENSG00000184005.11_4 | ST6GALNAC3 | 3.52E-10 | 5.90E-09 | 2.057 | 1.393 | chr1 | 76540431 | 77100288 | + | protein_coding |
| ENSG00000239857.7_6 | GET4 | 0.00042683 | 0.00298092 | 2.056 | -0.839 | chr7 | 916191 | 936073 | + | protein_coding |
| ENSG00000138735.16_5 | PDE5A | 2.20E-17 | 1.14E-15 | 2.045 | 3.867 | chr4 | 120415552 | 120549959 | - | protein_coding |
| ENSG00000260704.1_3 | LINC00543 | 9.73E-08 | 1.16E-06 | 2.045 | 0.537 | chr13 | 28527663 | 28529507 | + | lncRNA |
| ENSG00000267106.7_6 | ZNF561-AS1 | 1.96E-09 | 2.98E-08 | 2.043 | 1.251 | chr19 | 9731925 | 9757879 | + | lncRNA |
| ENSG00000132744.8_4 | ACY3 | 1.48E-08 | 1.95E-07 | 2.040 | 0.962 | chr11 | 67410026 | 67418201 | - | protein_coding |
| ENSG00000182809.10_2 | CRIP2 | 4.15E-07 | 4.49E-06 | 2.037 | 5.554 | chr14 | 105939299 | 105946507 | + | protein_coding |
| ENSG00000263935.1_4 | TOMM20P3 | 6.68E-05 | 0.00053274 | 2.037 | -0.761 | chr18 | 8688020 | 8688408 | - | processed_pseudogene |
| ENSG00000287064.1_1 | AL606500.1 | 3.11E-05 | 0.00026227 | 2.036 | -0.346 | chr1 | 154644069 | 154650821 | + | lncRNA |
| ENSG00000188037.12_5 | CLCN1 | 0.0007087 | 0.00474432 | 2.035 | -1.349 | chr7 | 143013204 | 143049176 | + | protein_coding |
| ENSG00000249746.1_7 | AC099509.1 | 0.00010112 | 0.00078037 | 2.034 | -0.761 | chr5 | 95549050 | 95550779 | - | lncRNA |
| ENSG00000245750.9_5 | DRAIC | 5.87E-09 | 8.30E-08 | 2.028 | 1.282 | chr15 | 69755260 | 70135459 | + | lncRNA |
| ENSG00000188732.11_3 | FAM221A | 4.79E-09 | 6.88E-08 | 2.027 | 1.007 | chr7 | 23719749 | 23742868 | + | protein_coding |
| ENSG00000196437.12_6 | ZNF569 | 1.44E-19 | 1.05E-17 | 2.019 | 4.862 | chr19 | 37902057 | 37960177 | - | protein_coding |
| ENSG00000114541.15_5 | FRMD4B | 3.84E-16 | 1.61E-14 | 2.016 | 3.515 | chr3 | 69217933 | 69591734 | - | protein_coding |
| ENSG00000204282.4_7 | TNRC6C-AS1 | 2.12E-05 | 0.00018265 | 2.016 | -0.365 | chr17 | 76103479 | 76107880 | - | lncRNA |
| ENSG00000196605.7_4 | ZNF846 | 3.53E-11 | 6.78E-10 | 2.014 | 1.814 | chr19 | 9862669 | 9903856 | - | protein_coding |
| ENSG00000287067.1_2 | AC106791.3 | 0.00035477 | 0.00251625 | 2.010 | -0.982 | chr5 | 137091429 | 137096429 | - | transcribed_processed_pseudogene |
| ENSG00000006606.8_3 | CCL26 | 6.02E-09 | 8.49E-08 | 2.009 | 1.068 | chr7 | 75398851 | 75419214 | - | protein_coding |
| ENSG00000176855.11_4 | KRT18P28 | 1.14E-06 | 1.17E-05 | 2.007 | 0.304 | chr1 | 182928209 | 182929540 | - | processed_pseudogene |
| ENSG00000111052.7_4 | LIN7A | 0.00016978 | 0.00127013 | 2.004 | -0.788 | chr12 | 81186299 | 81331704 | - | protein_coding |
| ENSG00000198478.8_3 | SH3BGRL2 | 7.28E-14 | 2.14E-12 | 2.003 | 2.783 | chr6 | 80341046 | 80413372 | + | protein_coding |
| ENSG00000253953.2_4 | PCDHGB4 | 4.15E-13 | 1.07E-11 | 1.999 | 2.497 | chr5 | 140767265 | 140892546 | + | protein_coding |
| ENSG00000102313.9_4 | ITIH6 | 0.00035036 | 0.00248895 | 1.998 | -1.108 | chrX | 54775351 | 54824688 | - | protein_coding |
| ENSG00000189431.8_4 | RASSF10 | 9.16E-16 | 3.59E-14 | 1.997 | 5.407 | chr11 | 13030863 | 13033666 | + | protein_coding |
| ENSG00000249464.6_6 | LINC01091 | 6.27E-10 | 1.02E-08 | 1.994 | 1.487 | chr4 | 124460181 | 124883853 | + | lncRNA |
| ENSG00000282849.1_7 | AL359834.1 | 0.00039446 | 0.00277448 | 1.991 | -1.243 | chr1 | 200447148 | 200452732 | - | lncRNA |
| ENSG00000275395.6_6 | FCGBP | 9.85E-19 | 6.12E-17 | 1.986 | 5.739 | chr19 | 40353963 | 40440533 | - | protein_coding |
| ENSG00000231752.6_3 | EMBP1 | 3.04E-09 | 4.52E-08 | 1.983 | 1.246 | chr1 | 121260910 | 121317500 | + | transcribed_unprocessed_pseudogene |
| ENSG00000204866.8_4 | IGFL2 | 1.88E-06 | 1.87E-05 | 1.979 | 0.401 | chr19 | 46646363 | 46664556 | + | protein_coding |
| ENSG00000243566.6_4 | UPK3B | 4.81E-16 | 1.97E-14 | 1.978 | 3.902 | chr7 | 76139745 | 76145838 | + | protein_coding |
| ENSG00000137486.17_8 | ARRB1 | 2.67E-21 | 2.66E-19 | 1.969 | 6.197 | chr11 | 74971166 | 75062749 | - | protein_coding |
| ENSG00000267549.6_8 | AC006116.9 | 0.00018366 | 0.0013659 | 1.969 | -0.555 | chr19 | 56826072 | 56876685 | + | lncRNA |
| ENSG00000254221.2_3 | PCDHGB1 | 1.44E-11 | 2.91E-10 | 1.967 | 2.358 | chr5 | 140729669 | 140892546 | + | protein_coding |
| ENSG00000269994.3_7 | AL513318.2 | 5.47E-07 | 5.84E-06 | 1.964 | 0.648 | chr9 | 89623355 | 89657334 | - | lncRNA |
| ENSG00000163293.12_5 | NIPAL1 | 1.94E-11 | 3.86E-10 | 1.963 | 2.262 | chr4 | 47916159 | 48042190 | + | protein_coding |
| ENSG00000110002.16_4 | VWA5A | 2.57E-05 | 0.00021908 | 1.961 | -0.269 | chr11 | 123986111 | 124018428 | + | protein_coding |
| ENSG00000109654.15_5 | TRIM2 | 3.68E-09 | 5.42E-08 | 1.961 | 3.016 | chr4 | 154073494 | 154260469 | + | protein_coding |
| ENSG00000125462.18_8 | C1orf61 | 1.71E-14 | 5.60E-13 | 1.956 | 3.178 | chr1 | 156374042 | 156426555 | - | protein_coding |

| | | | | | | | | | | |
|---|---|---|---|---|---|---|---|---|---|---|
| ENSG00000153291.16_3 | SLC25A27 | 2.88E-08 | 3.64E-07 | 1.955 | 1.036 | chr6 | 46620652 | 46645927 | + | protein_coding |
| ENSG00000139304.8 | PTPRQ | 0.00053833 | 0.00368715 | 1.951 | -1.145 | chr12 | 80799774 | 81072802 | + | protein_coding |
| ENSG00000286010.1_3 | AC108925.1 | 2.51E-06 | 2.44E-05 | 1.951 | 0.256 | chr8 | 128675525 | 128683235 | - | lncRNA |
| ENSG00000239887.6_6 | C1orf226 | 1.48E-08 | 1.95E-07 | 1.951 | 1.306 | chr1 | 162348631 | 162356602 | + | protein_coding |
| ENSG00000221994.11_8 | ZNF630 | 3.65E-09 | 5.39E-08 | 1.950 | 1.305 | chrX | 47842756 | 47931025 | - | protein_coding |
| ENSG00000152454.4_5 | AC010326.1 | 2.34E-13 | 6.27E-12 | 1.947 | 2.893 | chr19 | 58452201 | 58459074 | - | protein_coding |
| ENSG00000285867.1_3 | BX470102.2 | 2.44E-07 | 2.75E-06 | 1.942 | 0.864 | chr1 | 153559289 | 153578759 | - | lncRNA |
| ENSG00000227755.1_5 | AP000344.1 | 0.0002586 | 0.00187763 | 1.942 | -0.834 | chr22 | 23788785 | 23803020 | + | transcribed_processed_pseudogene |
| ENSG00000164520.11_4 | RAET1E | 7.32E-06 | 6.71E-05 | 1.940 | 0.192 | chr6 | 150204511 | 150219238 | - | protein_coding |
| ENSG00000138798.13_5 | EGF | 5.46E-07 | 5.83E-06 | 1.937 | 0.584 | chr4 | 110834039 | 110934922 | + | protein_coding |
| ENSG00000286538.1_2 | AC008798.2 | 0.00093428 | 0.00611168 | 1.936 | -0.945 | chr19 | 30329439 | 30332725 | - | lncRNA |
| ENSG00000198182.13_7 | ZNF607 | 2.91E-19 | 1.98E-17 | 1.936 | 5.019 | chr19 | 38187272 | 38210662 | - | protein_coding |
| ENSG00000236617.2_6 | AC127070.1 | 2.78E-05 | 0.0002355 | 1.936 | -0.288 | chr12 | 133464428 | 133465169 | + | lncRNA |
| ENSG00000254166.3_7 | CASC19 | 4.83E-14 | 1.47E-12 | 1.931 | 3.007 | chr8 | 128084939 | 128239786 | - | lncRNA |
| ENSG00000162415.7_5 | ZSWIM5 | 1.02E-14 | 3.47E-13 | 1.931 | 3.465 | chr1 | 45482071 | 45771881 | - | protein_coding |
| ENSG00000188001.10_3 | TPRG1 | 1.90E-07 | 2.17E-06 | 1.929 | 0.928 | chr3 | 188665003 | 189043093 | + | protein_coding |
| ENSG00000283635.1_6 | AC012485.3 | 8.30E-07 | 8.67E-06 | 1.928 | 0.562 | chr2 | 239197768 | 239208826 | + | lncRNA |
| ENSG00000163485.17_4 | ADORA1 | 1.42E-07 | 1.65E-06 | 1.927 | 0.750 | chr1 | 203059782 | 203136533 | + | protein_coding |
| ENSG00000168952.15_3 | STXBP6 | 1.39E-09 | 2.17E-08 | 1.927 | 1.484 | chr14 | 25278862 | 25519503 | - | protein_coding |
| ENSG00000272168.8_7 | CASC15 | 2.02E-13 | 5.43E-12 | 1.924 | 2.858 | chr6 | 21664416 | 22654684 | + | lncRNA |
| ENSG00000102109.9_4 | PCSK1N | 0.00016312 | 0.00122648 | 1.921 | 3.241 | chrX | 48689506 | 48694012 | - | protein_coding |
| ENSG00000095539.15_4 | SEMA4G | 4.01E-18 | 2.26E-16 | 1.918 | 4.663 | chr10 | 102729275 | 102745628 | + | protein_coding |
| ENSG00000162086.15_4 | ZNF75A | 8.30E-20 | 6.27E-18 | 1.916 | 5.491 | chr16 | 3355406 | 3368852 | + | protein_coding |
| ENSG00000196878.15_5 | LAMB3 | 1.38E-19 | 1.01E-17 | 1.916 | 5.247 | chr1 | 209788215 | 209825770 | - | protein_coding |
| ENSG00000030419.16_5 | IKZF2 | 4.71E-09 | 6.77E-08 | 1.915 | 1.477 | chr2 | 213864415 | 214017151 | - | protein_coding |
| ENSG00000172164.15_6 | SNTB1 | 1.51E-12 | 3.57E-11 | 1.912 | 2.468 | chr8 | 121547985 | 121825513 | - | protein_coding |
| ENSG00000286472.1_3 | AL035252.6 | 0.00018512 | 0.00137615 | 1.907 | -0.677 | chr20 | 25165294 | 25168485 | - | lncRNA |
| ENSG00000214212.9_4 | C19orf38 | 0.00072983 | 0.00486561 | 1.901 | -1.182 | chr19 | 10947251 | 10980466 | + | protein_coding |
| ENSG00000197016.12_4 | ZNF470 | 7.24E-17 | 3.47E-15 | 1.901 | 4.119 | chr19 | 57078837 | 57100279 | + | protein_coding |
| ENSG00000250708.2_6 | LINC02269 | 2.70E-05 | 0.00022939 | 1.900 | -0.189 | chr4 | 174620233 | 174922639 | + | lncRNA |
| ENSG00000174599.5_4 | TRAM1L1 | 3.64E-08 | 4.57E-07 | 1.892 | 1.159 | chr4 | 118004710 | 118006732 | - | protein_coding |
| ENSG00000099139.14_6 | PCSK5 | 4.22E-14 | 1.30E-12 | 1.887 | 3.019 | chr9 | 78505560 | 78977891 | + | protein_coding |
| ENSG00000188761.13_5 | BCL2L15 | 3.11E-10 | 5.25E-09 | 1.886 | 1.897 | chr1 | 114419438 | 114430203 | - | protein_coding |
| ENSG00000182795.13_4 | C1orf116 | 2.35E-19 | 1.63E-17 | 1.886 | 4.955 | chr1 | 207191867 | 207206101 | - | protein_coding |
| ENSG00000100079.7_4 | LGALS2 | 3.18E-06 | 3.06E-05 | 1.886 | 0.326 | chr22 | 37966255 | 37978623 | - | protein_coding |
| ENSG00000267419.2_6 | ZNF56 | 0.00015142 | 0.00114337 | 1.883 | -0.533 | chr19 | 19887411 | 19946882 | + | transcribed_unprocessed_pseudogene |
| ENSG00000110492.15_5 | MDK | 1.99E-13 | 5.36E-12 | 1.878 | 2.879 | chr11 | 46402306 | 46405387 | + | protein_coding |
| ENSG00000181449.4_6 | SOX2 | 7.91E-17 | 3.78E-15 | 1.876 | 4.088 | chr3 | 181429713 | 181432224 | + | protein_coding |
| ENSG00000188112.9_4 | C6orf132 | 1.64E-07 | 1.89E-06 | 1.872 | 0.889 | chr6 | 42068857 | 42110358 | - | protein_coding |
| ENSG00000168477.19_9 | TNXB | 1.94E-05 | 0.00016825 | 1.872 | -0.048 | chr6 | 32008930 | 32083111 | - | protein_coding |
| ENSG00000106541.12_4 | AGR2 | 1.91E-19 | 1.34E-17 | 1.870 | 9.287 | chr7 | 16831435 | 16873057 | - | protein_coding |
| ENSG00000171219.9_5 | CDC42BPG | 1.76E-12 | 4.11E-11 | 1.868 | 2.659 | chr11 | 64590524 | 64612125 | - | protein_coding |
| ENSG00000166816.14_6 | LDHD | 7.41E-11 | 1.35E-09 | 1.865 | 2.357 | chr16 | 75145758 | 75150669 | - | protein_coding |
| ENSG00000170786.12_3 | SDR16C5 | 1.16E-07 | 1.36E-06 | 1.864 | 0.931 | chr8 | 57212569 | 57233335 | - | protein_coding |
| ENSG00000141933.9_3 | TPGS1 | 6.69E-08 | 8.14E-07 | 1.863 | 1.342 | chr19 | 507497 | 519654 | + | protein_coding |
| ENSG00000142556.19_5 | ZNF614 | 1.48E-19 | 1.07E-17 | 1.858 | 5.260 | chr19 | 52516018 | 52533493 | - | protein_coding |
| ENSG00000224848.2_6 | AL589843.1 | 3.43E-06 | 3.29E-05 | 1.855 | 0.415 | chr9 | 99449332 | 99536600 | + | lncRNA |
| ENSG00000184545.11_4 | DUSP8 | 7.62E-08 | 9.18E-07 | 1.853 | 1.008 | chr11 | 1575281 | 1593501 | - | protein_coding |
| ENSG00000154620.6_3 | TMSB4Y | 8.40E-06 | 7.62E-05 | 1.847 | 0.639 | chrY | 15815779 | 15817904 | + | protein_coding |
| ENSG00000162687.18_5 | KCNT2 | 2.42E-09 | 3.64E-08 | 1.846 | 1.530 | chr1 | 196194909 | 196578355 | - | protein_coding |
| ENSG00000166035.11_4 | LIPC | 0.00014458 | 0.00109634 | 1.843 | -0.647 | chr15 | 58702768 | 58862043 | + | protein_coding |
| ENSG00000100342.20_2 | APOL1 | 2.03E-10 | 3.52E-09 | 1.837 | 1.982 | chr22 | 36649056 | 36663576 | + | protein_coding |
| ENSG00000245552.7_6 | AP000787.1 | 0.00125011 | 0.00799809 | 1.837 | -1.349 | chr11 | 94883703 | 94967555 | + | lncRNA |
| ENSG00000286681.1_2 | AF235103.3 | 0.00018716 | 0.00138844 | 1.834 | -0.732 | chr8 | 146080260 | 146094874 | - | lncRNA |
| ENSG00000113211.5_2 | PCDHB6 | 2.49E-10 | 4.27E-09 | 1.834 | 2.059 | chr5 | 140529603 | 140532868 | + | protein_coding |
| ENSG00000178386.13_8 | ZNF223 | 2.02E-06 | 1.99E-05 | 1.833 | 0.549 | chr19 | 44555520 | 44572152 | + | protein_coding |
| ENSG00000185404.16_2 | SP140L | 1.02E-09 | 1.62E-08 | 1.833 | 2.257 | chr2 | 231191899 | 231268447 | + | protein_coding |
| ENSG00000158955.11_5 | WNT9B | 0.00106641 | 0.0069099 | 1.831 | -1.122 | chr17 | 44910567 | 44964096 | + | protein_coding |
| ENSG00000065054.14_5 | SLC9A3R2 | 1.68E-11 | 3.37E-10 | 1.830 | 6.477 | chr16 | 2075357 | 2089027 | + | protein_coding |
| ENSG00000164736.6_3 | SOX17 | 5.34E-07 | 5.72E-06 | 1.829 | 0.788 | chr8 | 55370495 | 55373452 | + | protein_coding |

| | | | | | | | | | | |
|---|---|---|---|---|---|---|---|---|---|---|
| ENSG00000247774.7_7 | PCED1B-AS1 | 5.52E-05 | 0.00044662 | 1.826 | -0.311 | chr12 | 47599681 | 47610239 | - | lncRNA |
| ENSG00000269388.1_5 | AC018755.3 | 0.00044393 | 0.00308821 | 1.825 | -0.916 | chr19 | 52097814 | 52109403 | + | transcribed_processed_pseudogene |
| ENSG00000159208.16_4 | CIART | 0.00020691 | 0.00152349 | 1.822 | -0.508 | chr1 | 150254953 | 150259505 | + | protein_coding |
| ENSG00000169933.15_4 | FRMPD4 | 3.06E-07 | 3.38E-06 | 1.822 | 0.999 | chrX | 11840542 | 12742642 | + | protein_coding |
| ENSG00000152475.7_4 | ZNF837 | 5.01E-09 | 7.17E-08 | 1.821 | 1.807 | chr19 | 58878985 | 58892397 | - | protein_coding |
| ENSG00000246982.7_7 | Z84485.1 | 0.00015522 | 0.00117008 | 1.821 | -0.583 | chr6 | 36114475 | 36164982 | - | lncRNA |
| ENSG00000269696.2_7 | AC005498.3 | 3.34E-06 | 3.21E-05 | 1.819 | 0.693 | chr19 | 57056935 | 57078780 | - | lncRNA |
| ENSG00000236548.2_7 | RNF217-AS1 | 2.90E-06 | 2.81E-05 | 1.817 | 0.360 | chr6 | 124965580 | 125284311 | - | lncRNA |
| ENSG00000012124.17_4 | CD22 | 3.48E-06 | 3.34E-05 | 1.816 | 0.284 | chr19 | 35810164 | 35838264 | + | protein_coding |
| ENSG00000288541.1_1 | AL353732.2 | 0.00023067 | 0.00168791 | 1.815 | -0.745 | chr9 | 21410968 | 21422959 | - | lncRNA |
| ENSG00000168237.18_3 | GLYCTK | 1.99E-14 | 6.46E-13 | 1.811 | 3.404 | chr3 | 52321105 | 52329273 | + | protein_coding |
| ENSG00000214207.2_4 | KRT18P10 | 1.52E-08 | 2.01E-07 | 1.810 | 1.545 | chrY | 5441186 | 5442472 | + | processed_pseudogene |
| ENSG00000227502.3_6 | MROCKI | 5.53E-07 | 5.90E-06 | 1.807 | 0.779 | chr6 | 114189178 | 114194516 | - | lncRNA |
| ENSG00000215217.7_4 | C5orf49 | 0.00208102 | 0.01276706 | 1.807 | -1.243 | chr5 | 7830491 | 7851264 | - | protein_coding |
| ENSG00000109794.13_6 | FAM149A | 2.42E-12 | 5.54E-11 | 1.805 | 2.996 | chr4 | 187025573 | 187093821 | + | protein_coding |
| ENSG00000092421.16_6 | SEMA6A | 1.36E-05 | 0.00011979 | 1.801 | 0.055 | chr5 | 115779312 | 115910630 | - | protein_coding |
| ENSG00000213638.6_6 | ADAT3 | 2.64E-06 | 2.56E-05 | 1.800 | 1.294 | chr19 | 1905398 | 1913446 | + | protein_coding |
| ENSG00000164342.12_5 | TLR3 | 8.58E-10 | 1.37E-08 | 1.798 | 1.827 | chr4 | 186990306 | 187009223 | + | protein_coding |
| ENSG00000172006.12_3 | ZNF554 | 1.50E-09 | 2.32E-08 | 1.797 | 1.686 | chr19 | 2819866 | 2836733 | + | protein_coding |
| ENSG00000255087.4_7 | AP001993.1 | 7.33E-06 | 6.72E-05 | 1.796 | 0.649 | chr11 | 126891631 | 127042258 | + | lncRNA |
| ENSG00000258181.1_8 | AC008083.3 | 6.67E-06 | 6.14E-05 | 1.796 | 0.581 | chr12 | 47641907 | 47651322 | + | lncRNA |
| ENSG00000139832.5_4 | RAB20 | 6.61E-09 | 9.27E-08 | 1.794 | 1.680 | chr13 | 111175413 | 111214069 | - | protein_coding |
| ENSG00000164626.9_4 | KCNK5 | 3.11E-17 | 1.59E-15 | 1.791 | 4.578 | chr6 | 39156747 | 39197251 | - | protein_coding |
| ENSG00000165929.13_5 | TC2N | 1.11E-19 | 8.24E-18 | 1.791 | 5.527 | chr14 | 92246090 | 92333880 | - | protein_coding |
| ENSG00000170801.10_3 | HTRA3 | 3.84E-14 | 1.19E-12 | 1.790 | 3.441 | chr4 | 8271481 | 8308825 | + | protein_coding |
| ENSG00000149582.16_5 | TMEM25 | 1.14E-05 | 0.00010126 | 1.789 | 0.040 | chr11 | 118401756 | 118417995 | + | protein_coding |
| ENSG00000171533.11_5 | MAP6 | 5.95E-11 | 1.10E-09 | 1.786 | 2.321 | chr11 | 75297963 | 75380165 | - | protein_coding |
| ENSG00000133083.14_4 | DCLK1 | 0.00057023 | 0.00388917 | 1.786 | -0.946 | chr13 | 36342789 | 36705443 | - | protein_coding |
| ENSG00000185339.9_4 | TCN2 | 4.15E-14 | 1.28E-12 | 1.783 | 3.329 | chr22 | 31002990 | 31023258 | + | protein_coding |
| ENSG00000139737.22_7 | SLAIN1 | 8.48E-13 | 2.08E-11 | 1.783 | 2.803 | chr13 | 78271822 | 78338377 | + | protein_coding |
| ENSG00000186832.9_7 | KRT16 | 0.00196613 | 0.01211661 | 1.782 | -1.264 | chr17 | 39766030 | 39772151 | - | protein_coding |
| ENSG00000131781.13_5 | FMO5 | 2.23E-11 | 4.40E-10 | 1.780 | 2.480 | chr1 | 146646930 | 146714700 | - | protein_coding |
| ENSG00000134326.11_5 | CMPK2 | 0.00019771 | 0.00146119 | 1.777 | -0.343 | chr2 | 6980701 | 7006766 | - | protein_coding |
| ENSG00000064787.13_5 | BCAS1 | 4.46E-17 | 2.24E-15 | 1.776 | 4.590 | chr20 | 52553316 | 52687133 | - | protein_coding |
| ENSG00000137860.12_3 | SLC28A2 | 8.36E-05 | 0.00065451 | 1.775 | -0.235 | chr15 | 45544432 | 45570044 | + | protein_coding |
| ENSG00000240990.10_6 | HOXA11-AS | 0.00016532 | 0.00124092 | 1.774 | -0.420 | chr7 | 27224126 | 27228917 | + | lncRNA |
| ENSG00000159784.17_4 | FAM131B | 5.80E-08 | 7.08E-07 | 1.774 | 1.147 | chr7 | 143050493 | 143059863 | - | protein_coding |
| ENSG00000196511.14_5 | TPK1 | 1.83E-08 | 2.38E-07 | 1.773 | 1.480 | chr7 | 144149034 | 144533488 | - | protein_coding |
| ENSG00000160336.15_5 | ZNF761 | 3.48E-19 | 2.31E-17 | 1.772 | 5.528 | chr19 | 53948254 | 53961515 | + | protein_coding |
| ENSG00000205209.7_4 | SCGB2B2 | 0.00261419 | 0.01572849 | 1.771 | -1.161 | chr19 | 35084234 | 35166604 | - | protein_coding |
| ENSG00000267123.7_7 | SCAT1 | 0.00071958 | 0.0048099 | 1.770 | -0.963 | chr17 | 76601315 | 76628237 | - | lncRNA |
| ENSG00000196653.12_5 | ZNF502 | 3.76E-13 | 9.79E-12 | 1.768 | 3.292 | chr3 | 44754135 | 44765323 | + | protein_coding |
| ENSG00000244122.2_5 | UGT1A7 | 0.00077758 | 0.00516221 | 1.768 | -0.962 | chr2 | 234590584 | 234681945 | + | protein_coding |
| ENSG00000267152.1_7 | AC093227.1 | 0.00080136 | 0.00529877 | 1.767 | -0.868 | chr19 | 38219487 | 38221544 | - | lncRNA |
| ENSG00000254154.8_8 | CRYZL2P-SEC16B | 0.00074368 | 0.00495054 | 1.767 | -0.963 | chr1 | 177897923 | 178007142 | - | lncRNA |
| ENSG00000133328.4_5 | PLAAT2 | 0.00052553 | 0.00360365 | 1.765 | -0.787 | chr11 | 63320242 | 63330851 | - | protein_coding |
| ENSG00000242841.1_4 | KRT8P35 | 0.00103193 | 0.00670114 | 1.763 | -0.640 | chr3 | 63082050 | 63083387 | - | processed_pseudogene |
| ENSG00000131849.12_4 | ZNF132 | 1.51E-10 | 2.67E-09 | 1.761 | 2.150 | chr19 | 58944181 | 58951520 | - | protein_coding |
| ENSG00000139132.14_8 | FGD4 | 6.04E-19 | 3.89E-17 | 1.760 | 5.228 | chr12 | 32552463 | 32798984 | + | protein_coding |
| ENSG00000147003.7_6 | CLTRN | 3.85E-09 | 5.65E-08 | 1.760 | 1.941 | chrX | 15645441 | 15693135 | - | protein_coding |
| ENSG00000267422.1_5 | AC016582.2 | 0.00026972 | 0.00195439 | 1.759 | -0.363 | chr19 | 38270326 | 38283505 | + | transcribed_processed_pseudogene |
| ENSG00000109472.14_3 | CPE | 3.90E-09 | 5.71E-08 | 1.758 | 1.664 | chr4 | 166282346 | 166419699 | + | protein_coding |
| ENSG00000206159.11_4 | GYG2P1 | 8.05E-06 | 7.33E-05 | 1.758 | 0.319 | chrY | 14208716 | 14533386 | - | transcribed_unprocessed_pseudogene |
| ENSG00000106789.13_4 | CORO2A | 1.13E-13 | 3.17E-12 | 1.754 | 3.454 | chr9 | 100883257 | 100954919 | - | protein_coding |
| ENSG00000147642.17_8 | SYBU | 3.18E-15 | 1.16E-13 | 1.754 | 3.871 | chr8 | 110586207 | 110704020 | - | protein_coding |
| ENSG00000255408.4_4 | PCDHA3 | 1.42E-11 | 2.89E-10 | 1.753 | 2.530 | chr5 | 140180613 | 140391932 | + | protein_coding |
| ENSG00000108771.13_6 | DHX58 | 4.18E-05 | 0.00034523 | 1.752 | -0.138 | chr17 | 40253422 | 40264732 | - | protein_coding |

| | | | | | | | | | | |
|---|---|---|---|---|---|---|---|---|---|---|
| ENSG00000116833.14_4 | NR5A2 | 1.55E-21 | 1.60E-19 | 1.749 | 6.661 | chr1 | 199996742 | 200146548 | + | protein_coding |
| ENSG00000102174.9_4 | PHEX | 3.25E-10 | 5.48E-09 | 1.742 | 2.694 | chrX | 22050443 | 22269427 | + | protein_coding |
| ENSG00000248712.7_3 | CCDC153 | 0.00026998 | 0.00195546 | 1.742 | -0.577 | chr11 | 119060347 | 119067479 | - | protein_coding |
| ENSG00000168282.6_3 | MGAT2 | 0.00045478 | 0.00315996 | 1.738 | -0.804 | chr14 | 50087517 | 50090199 | + | protein_coding |
| ENSG00000255433.5_7 | LINC02735 | 6.96E-11 | 1.28E-09 | 1.734 | 2.365 | chr11 | 56615954 | 56645554 | + | lncRNA |
| ENSG00000253438.4_4 | PCAT1 | 0.00127363 | 0.00812516 | 1.733 | -1.184 | chr8 | 127568568 | 128431295 | + | lncRNA |
| ENSG00000163993.7_3 | S100P | 2.83E-15 | 1.05E-13 | 1.732 | 3.880 | chr4 | 6695605 | 6698897 | + | protein_coding |
| ENSG00000100557.9_5 | CCDC198 | 3.16E-16 | 1.34E-14 | 1.732 | 4.500 | chr14 | 57936019 | 57960585 | - | protein_coding |
| ENSG00000111057.11_6 | KRT18 | 1.37E-22 | 1.85E-20 | 1.730 | 10.716 | chr12 | 53342655 | 53346690 | + | protein_coding |
| ENSG00000262209.3_5 | PCDHGB3 | 1.85E-08 | 2.41E-07 | 1.729 | 1.403 | chr5 | 140749809 | 140892542 | + | protein_coding |
| ENSG00000127329.15_6 | PTPRB | 4.60E-08 | 5.69E-07 | 1.725 | 1.663 | chr12 | 70909646 | 71031220 | - | protein_coding |
| ENSG00000135373.12_4 | EHF | 8.61E-18 | 4.66E-16 | 1.725 | 4.968 | chr11 | 34642640 | 34682604 | + | protein_coding |
| ENSG00000224417.2_8 | AL606970.1 | 1.40E-06 | 1.42E-05 | 1.723 | 0.785 | chr6 | 168698749 | 168702794 | - | lncRNA |
| ENSG00000268201.1_9 | AC020915.2 | 2.24E-05 | 0.00019281 | 1.723 | 0.082 | chr19 | 58778001 | 58779051 | + | lncRNA |
| ENSG00000146555.19_4 | SDK1 | 4.25E-12 | 9.22E-11 | 1.721 | 2.919 | chr7 | 3340884 | 4308632 | + | protein_coding |
| ENSG00000214012.4_4 | KRT18P38 | 7.24E-05 | 0.00057293 | 1.720 | -0.276 | chr6 | 19612986 | 19614267 | + | processed_pseudogene |
| ENSG00000186493.12_4 | C5orf38 | 4.49E-05 | 0.00036897 | 1.718 | -0.058 | chr5 | 2752245 | 2755511 | + | protein_coding |
| ENSG00000130818.12_5 | ZNF426 | 2.25E-14 | 7.27E-13 | 1.717 | 3.802 | chr19 | 9633899 | 9649321 | - | protein_coding |
| ENSG00000257660.5_7 | ADCY6-DT | 0.00040196 | 0.00282164 | 1.715 | -0.602 | chr12 | 49182930 | 49184318 | + | lncRNA |
| ENSG00000041982.16_5 | TNC | 1.84E-11 | 3.67E-10 | 1.712 | 2.631 | chr9 | 117781854 | 117880536 | - | protein_coding |
| ENSG00000082458.12_4 | DLG3 | 1.54E-16 | 6.94E-15 | 1.711 | 4.356 | chrX | 69664685 | 69725340 | + | protein_coding |
| ENSG00000090376.11_4 | IRAK3 | 7.44E-06 | 6.80E-05 | 1.710 | 0.387 | chr12 | 66582975 | 66648402 | + | protein_coding |
| ENSG00000130748.7_4 | TMEM160 | 0.01399129 | 0.07232566 | 1.708 | 3.312 | chr19 | 47549167 | 47551882 | - | protein_coding |
| ENSG00000253944.1_7 | AC027117.2 | 8.29E-09 | 1.14E-07 | 1.706 | 1.808 | chr8 | 17658854 | 17718578 | + | lncRNA |
| ENSG00000182621.18_5 | PLCB1 | 3.95E-14 | 1.22E-12 | 1.705 | 3.839 | chr20 | 8057898 | 8949007 | + | protein_coding |
| ENSG00000165215.6_4 | CLDN3 | 9.77E-08 | 1.16E-06 | 1.705 | 1.728 | chr7 | 73183327 | 73184600 | - | protein_coding |
| ENSG00000131848.9_7 | ZSCAN5A | 1.93E-12 | 4.49E-11 | 1.705 | 3.226 | chr19 | 56731039 | 56879752 | - | protein_coding |
| ENSG00000172687.13_3 | ZNF738 | 4.60E-10 | 7.61E-09 | 1.704 | 2.462 | chr19 | 21541732 | 21562104 | + | protein_coding |
| ENSG00000213462.5_5 | ERV3-1 | 9.71E-12 | 2.01E-10 | 1.704 | 2.949 | chr7 | 64450734 | 64467121 | - | protein_coding |
| ENSG00000235904.3_9 | RBMS3-AS3 | 0.00063877 | 0.00431683 | 1.702 | -0.914 | chr3 | 29096061 | 29332217 | - | lncRNA |
| ENSG00000006611.16_5 | USH1C | 2.05E-07 | 2.33E-06 | 1.701 | 1.086 | chr11 | 17515442 | 17565963 | - | protein_coding |
| ENSG00000130762.15_5 | ARHGEF16 | 1.87E-13 | 5.06E-12 | 1.700 | 3.682 | chr1 | 3371229 | 3397677 | + | protein_coding |
| ENSG00000124107.5_4 | SLPI | 4.18E-07 | 4.52E-06 | 1.699 | 1.114 | chr20 | 43880880 | 43883205 | - | protein_coding |
| ENSG00000171747.9_4 | LGALS4 | 8.65E-05 | 0.00067591 | 1.697 | -0.022 | chr19 | 39292311 | 39303585 | - | protein_coding |
| ENSG00000205744.10_5 | DENND1C | 3.91E-09 | 5.72E-08 | 1.695 | 1.836 | chr19 | 6467218 | 6482568 | - | protein_coding |
| ENSG00000150347.16_4 | ARID5B | 3.86E-11 | 7.37E-10 | 1.694 | 2.797 | chr10 | 63661443 | 63856703 | + | protein_coding |
| ENSG00000123080.12_5 | CDKN2C | 3.19E-17 | 1.62E-15 | 1.693 | 4.866 | chr1 | 51426417 | 51440306 | + | protein_coding |
| ENSG00000134917.10_4 | ADAMTS8 | 0.00037844 | 0.00266813 | 1.692 | -0.685 | chr11 | 130274818 | 130298504 | - | protein_coding |
| ENSG00000143772.9_4 | ITPKB | 7.15E-14 | 2.11E-12 | 1.692 | 3.526 | chr1 | 226819391 | 226927024 | - | protein_coding |
| ENSG00000166592.12_7 | RRAD | 9.60E-05 | 0.00074357 | 1.690 | 0.068 | chr16 | 66955582 | 66959439 | - | protein_coding |
| ENSG00000235947.1_4 | EGOT | 0.00207792 | 0.0127525 | 1.690 | -1.106 | chr3 | 4790876 | 4793274 | - | lncRNA |
| ENSG00000188290.10_4 | HES4 | 1.74E-06 | 1.73E-05 | 1.689 | 1.537 | chr1 | 934342 | 935552 | - | protein_coding |
| ENSG00000105699.16_5 | LSR | 1.97E-14 | 6.43E-13 | 1.687 | 4.428 | chr19 | 35739233 | 35758867 | + | protein_coding |
| ENSG00000139508.14_3 | SLC46A3 | 1.43E-15 | 5.48E-14 | 1.686 | 4.059 | chr13 | 29274201 | 29293107 | - | protein_coding |
| ENSG00000256955.2_6 | AC131009.2 | 0.00218037 | 0.01332139 | 1.684 | -0.928 | chr12 | 132341965 | 132349083 | + | lncRNA |
| ENSG00000166165.13_5 | CKB | 2.16E-14 | 6.99E-13 | 1.679 | 5.373 | chr14 | 103986004 | 103989170 | - | protein_coding |
| ENSG00000136999.5_3 | CCN3 | 1.43E-05 | 0.00012587 | 1.677 | 0.283 | chr8 | 120428686 | 120436674 | + | protein_coding |
| ENSG00000205754.11_4 | SLCO1B7 | 9.47E-05 | 0.00073557 | 1.676 | -0.106 | chr12 | 21115702 | 21245679 | + | protein_coding |
| ENSG00000234840.2_6 | LINC01239 | 0.00020587 | 0.00151644 | 1.672 | -0.317 | chr9 | 22646199 | 22824212 | + | lncRNA |
| ENSG00000130768.15_4 | SMPDL3B | 4.35E-08 | 5.39E-07 | 1.672 | 1.409 | chr1 | 28261511 | 28285668 | + | protein_coding |
| ENSG00000111490.14_5 | TBC1D30 | 4.36E-12 | 9.46E-11 | 1.669 | 3.018 | chr12 | 65174296 | 65274813 | + | protein_coding |
| ENSG00000262294.1_7 | AC129507.2 | 0.00171873 | 0.01068091 | 1.667 | -0.773 | chr17 | 202429 | 207372 | + | lncRNA |
| ENSG00000288095.1_2 | AC005618.4 | 0.00058395 | 0.00397052 | 1.667 | -0.502 | chr5 | 140700547 | 140705552 | + | lncRNA |
| ENSG00000159708.18_4 | LRRC36 | 0.00024098 | 0.00175685 | 1.666 | -0.432 | chr16 | 67360701 | 67419107 | + | protein_coding |
| ENSG00000250221.1_4 | KRT8P32 | 0.00102745 | 0.00667445 | 1.666 | 0.017 | chr5 | 97727774 | 97729234 | - | processed_pseudogene |
| ENSG00000139112.11_4 | GABARAPL1 | 8.21E-17 | 3.90E-15 | 1.662 | 4.621 | chr12 | 10365057 | 10375727 | + | protein_coding |
| ENSG00000087303.18_5 | NID2 | 2.99E-11 | 5.79E-10 | 1.662 | 2.631 | chr14 | 52471521 | 52535946 | - | protein_coding |
| ENSG00000138172.11_4 | CALHM2 | 1.48E-09 | 2.30E-08 | 1.657 | 2.442 | chr10 | 105206543 | 105212159 | - | protein_coding |
| ENSG00000140678.16_2 | ITGAX | 9.66E-07 | 1.00E-05 | 1.656 | 1.037 | chr16 | 31366455 | 31394318 | + | protein_coding |
| ENSG00000286081.1_1 | AC016682.1 | 3.51E-05 | 0.0002929 | 1.655 | 0.189 | chr2 | 151007147 | 151022780 | - | lncRNA |

| | | | | | | | | | | |
|---|---|---|---|---|---|---|---|---|---|---|
| ENSG00000248727.6_6 | LINC01948 | 9.20E-07 | 9.58E-06 | 1.652 | 0.982 | chr5 | 55747454 | 55777773 | - | lncRNA |
| ENSG00000250303.4_8 | LINC02762 | 4.13E-06 | 3.91E-05 | 1.652 | 0.665 | chr11 | 112141472 | 112233257 | + | lncRNA |
| ENSG00000087253.13_4 | LPCAT2 | 4.27E-16 | 1.77E-14 | 1.651 | 4.873 | chr16 | 55542984 | 55620578 | + | protein_coding |
| ENSG00000256060.3_7 | TRAPPC2B | 8.91E-10 | 1.42E-08 | 1.649 | 2.107 | chr19 | 57874919 | 57876773 | + | protein_coding |
| ENSG00000106852.15_5 | LHX6 | 0.00040685 | 0.00285374 | 1.649 | -0.514 | chr9 | 124964856 | 124991905 | - | protein_coding |
| ENSG00000113924.12_5 | HGD | 2.28E-15 | 8.57E-14 | 1.647 | 4.661 | chr3 | 120347019 | 120401116 | - | protein_coding |
| ENSG00000221947.8_5 | XKR9 | 6.90E-05 | 0.00054797 | 1.645 | -0.079 | chr8 | 71581574 | 71702606 | + | protein_coding |
| ENSG00000283511.1_5 | AC137936.2 | 1.18E-05 | 0.00010455 | 1.642 | 0.467 | chr3 | 58165736 | 58173222 | + | transcribed_unitary_pseudogene |
| ENSG00000251381.8_6 | LINC00958 | 0.00279927 | 0.01672876 | 1.642 | -1.144 | chr11 | 12983088 | 13011144 | - | lncRNA |
| ENSG00000274150.1_3 | FADS2B | 0.00036154 | 0.00256017 | 1.641 | -0.386 | chr11 | 56658089 | 56695360 | + | unprocessed_pseudogene |
| ENSG00000163131.11_5 | CTSS | 2.69E-16 | 1.16E-14 | 1.641 | 4.667 | chr1 | 150702664 | 150738268 | - | protein_coding |
| ENSG00000196387.9_2 | ZNF140 | 6.26E-13 | 1.58E-11 | 1.633 | 3.727 | chr12 | 133656424 | 133684130 | + | protein_coding |
| ENSG00000185467.7_3 | KPNA7 | 7.75E-05 | 0.00060974 | 1.633 | 0.002 | chr7 | 98771197 | 98805129 | - | protein_coding |
| ENSG00000223586.6_5 | LINC01312 | 8.10E-05 | 0.00063544 | 1.632 | 0.144 | chr6 | 134142285 | 134194060 | + | lncRNA |
| ENSG00000184860.10_4 | SDR42E1 | 2.38E-11 | 4.67E-10 | 1.626 | 2.781 | chr16 | 82022460 | 82045086 | - | protein_coding |
| ENSG00000177570.14_5 | SAMD12 | 4.93E-08 | 6.07E-07 | 1.621 | 1.535 | chr8 | 119201694 | 119634351 | - | protein_coding |
| ENSG00000111261.14_6 | MANSC1 | 8.90E-14 | 2.58E-12 | 1.620 | 4.163 | chr12 | 12478990 | 12503176 | - | protein_coding |
| ENSG00000174028.6_2 | FAM3C2 | 2.36E-06 | 2.31E-05 | 1.619 | 0.874 | chrX | 23093875 | 23094884 | + | processed_pseudogene |
| ENSG00000181634.8_4 | TNFSF15 | 2.51E-17 | 1.29E-15 | 1.618 | 5.765 | chr9 | 117546932 | 117568319 | - | protein_coding |
| ENSG00000174652.19_5 | ZNF266 | 2.38E-15 | 8.92E-14 | 1.618 | 4.263 | chr19 | 9523137 | 9546247 | - | protein_coding |
| ENSG00000169594.13_4 | BNC1 | 0.00056855 | 0.00388071 | 1.618 | -0.660 | chr15 | 83924655 | 83953466 | - | protein_coding |
| ENSG00000243449.6_3 | C4orf48 | 5.88E-05 | 0.00047354 | 1.617 | 2.038 | chr4 | 2043720 | 2045697 | + | protein_coding |
| ENSG00000132970.14_6 | WASF3 | 0.00216941 | 0.01325902 | 1.615 | -1.163 | chr13 | 27131820 | 27263085 | + | protein_coding |
| ENSG00000004468.13_5 | CD38 | 8.70E-16 | 3.44E-14 | 1.613 | 4.842 | chr4 | 15779898 | 15854855 | + | protein_coding |
| ENSG00000163453.11_4 | IGFBP7 | 2.91E-18 | 1.69E-16 | 1.613 | 5.546 | chr4 | 57896939 | 57976551 | - | protein_coding |
| ENSG00000101084.18_5 | RAB5IF | 0.00766286 | 0.04200741 | 1.612 | 3.081 | chr20 | 35234182 | 35240960 | + | protein_coding |
| ENSG00000019991.18_7 | HGF | 0.0007778 | 0.00516221 | 1.609 | -0.672 | chr7 | 81328326 | 81399754 | - | protein_coding |
| ENSG00000161249.21_7 | DMKN | 6.02E-13 | 1.52E-11 | 1.608 | 3.504 | chr19 | 35988122 | 36004560 | - | protein_coding |
| ENSG00000185046.18_6 | ANKS1B | 3.67E-10 | 6.15E-09 | 1.606 | 2.356 | chr12 | 99120235 | 100378432 | - | protein_coding |
| ENSG00000253764.3_7 | AC019257.1 | 0.00071278 | 0.00476803 | 1.605 | -0.532 | chr8 | 1919319 | 1922803 | - | lncRNA |
| ENSG00000260588.1_8 | AC027702.2 | 2.71E-07 | 3.02E-06 | 1.603 | 1.253 | chr8 | 41685683 | 41687452 | - | lncRNA |
| ENSG00000198131.14_5 | ZNF544 | 3.67E-18 | 2.09E-16 | 1.597 | 5.505 | chr19 | 58739960 | 58788861 | + | protein_coding |
| ENSG00000267191.2_7 | AC006213.3 | 6.09E-06 | 5.64E-05 | 1.590 | 0.848 | chr19 | 44406061 | 44431919 | + | lncRNA |
| ENSG00000255236.3_6 | AP002992.1 | 0.00058043 | 0.00395114 | 1.589 | -0.431 | chr11 | 67889091 | 67897985 | + | lncRNA |
| ENSG00000198959.12_4 | TGM2 | 1.46E-20 | 1.25E-18 | 1.588 | 8.559 | chr20 | 36755787 | 36794980 | - | protein_coding |
| ENSG00000146678.10_3 | IGFBP1 | 1.29E-09 | 2.02E-08 | 1.587 | 2.307 | chr7 | 45927959 | 45933259 | + | protein_coding |
| ENSG00000164093.17_6 | PITX2 | 8.53E-08 | 1.02E-06 | 1.586 | 1.787 | chr4 | 111538579 | 111563279 | - | protein_coding |
| ENSG00000254343.2_6 | AC091563.1 | 0.00019792 | 0.00146211 | 1.584 | -0.215 | chr8 | 121064419 | 121068440 | + | lncRNA |
| ENSG00000197558.13_5 | SSPO | 3.63E-06 | 3.47E-05 | 1.584 | 1.931 | chr7 | 149473131 | 149531068 | + | transcribed_unitary_pseudogene |
| ENSG00000134215.16_5 | VAV3 | 1.01E-15 | 3.93E-14 | 1.582 | 4.752 | chr1 | 108113783 | 108507802 | - | protein_coding |
| ENSG00000157388.19_8 | CACNA1D | 2.60E-12 | 5.88E-11 | 1.580 | 3.483 | chr3 | 53362990 | 53847760 | + | protein_coding |
| ENSG00000286613.1_2 | AC006213.7 | 0.00306705 | 0.01818835 | 1.579 | -0.915 | chr19 | 44401363 | 44402685 | - | lncRNA |
| ENSG00000196557.13_2 | CACNA1H | 3.88E-09 | 5.68E-08 | 1.579 | 5.535 | chr16 | 1203106 | 1271771 | + | protein_coding |
| ENSG00000204920.11_4 | ZNF155 | 1.46E-09 | 2.27E-08 | 1.575 | 2.755 | chr19 | 44472014 | 44502478 | + | protein_coding |
| ENSG00000131015.5_3 | ULBP2 | 7.38E-06 | 6.75E-05 | 1.575 | 0.629 | chr6 | 150263150 | 150270371 | + | protein_coding |
| ENSG00000189164.15_3 | ZNF527 | 2.62E-15 | 9.74E-14 | 1.574 | 4.315 | chr19 | 37861963 | 37883968 | + | protein_coding |
| ENSG00000250312.8_6 | ZNF718 | 4.72E-08 | 5.83E-07 | 1.571 | 1.832 | chr4 | 124383 | 196092 | + | protein_coding |
| ENSG00000213214.4_3 | ARHGEF35 | 0.00220842 | 0.01346962 | 1.570 | -0.772 | chr7 | 143883176 | 143892748 | - | protein_coding |
| ENSG00000260253.1_6 | AC084262.1 | 0.00290185 | 0.01728367 | 1.570 | -0.916 | chr8 | 29209937 | 29210687 | - | lncRNA |
| ENSG00000196275.9 | GTF2IRD2 | 6.98E-06 | 6.41E-05 | 1.569 | 0.857 | chr7 | 74210483 | 74267847 | - | protein_coding |
| ENSG00000164488.12_7 | DACT2 | 3.75E-07 | 4.08E-06 | 1.566 | 1.540 | chr6 | 168693510 | 168720457 | - | protein_coding |
| ENSG00000167642.13_6 | SPINT2 | 8.77E-06 | 7.94E-05 | 1.565 | 6.972 | chr19 | 38734675 | 38783255 | + | protein_coding |
| ENSG00000174502.19_5 | SLC26A9 | 8.27E-07 | 8.64E-06 | 1.564 | 1.107 | chr1 | 205882176 | 205912588 | - | protein_coding |
| ENSG00000121406.9_3 | ZNF549 | 1.44E-11 | 2.92E-10 | 1.561 | 3.125 | chr19 | 58038693 | 58068910 | + | protein_coding |
| ENSG00000170381.14_5 | SEMA3E | 9.25E-15 | 3.19E-13 | 1.561 | 4.355 | chr7 | 82992554 | 83278455 | - | protein_coding |
| ENSG00000073849.15_4 | ST6GAL1 | 2.32E-12 | 5.33E-11 | 1.560 | 3.464 | chr3 | 186648291 | 186796341 | + | protein_coding |
| ENSG00000266680.1_6 | AL135905.1 | 7.10E-05 | 0.00056254 | 1.556 | 0.156 | chr6 | 64280910 | 64282313 | - | lncRNA |
| ENSG00000131981.16_3 | LGALS3 | 2.78E-18 | 1.62E-16 | 1.555 | 6.103 | chr14 | 55590828 | 55612141 | + | protein_coding |
| ENSG00000231483.1_8 | LINC02678 | 0.00010852 | 0.00083398 | 1.553 | 0.081 | chr10 | 5650438 | 5652756 | - | lncRNA |

| | | | | | | | | | | |
|---|---|---|---|---|---|---|---|---|---|---|
| ENSG00000145945.7_6 | FAM50B | 2.03E-08 | 2.62E-07 | 1.552 | 2.175 | chr6 | 3849607 | 3851554 | + | protein_coding |
| ENSG00000261115.6_6 | TMEM178B | 1.11E-07 | 1.32E-06 | 1.549 | 1.867 | chr7 | 140773864 | 141180180 | + | protein_coding |
| ENSG00000214029.5_5 | ZNF891 | 4.51E-14 | 1.38E-12 | 1.549 | 4.096 | chr12 | 133683403 | 133707059 | - | protein_coding |
| ENSG00000241388.5_5 | HNF1A-AS1 | 1.54E-13 | 4.24E-12 | 1.548 | 3.958 | chr12 | 121379531 | 121418768 | - | lncRNA |
| ENSG00000244219.7_5 | TMEM225B | 0.0008104 | 0.00535255 | 1.542 | -0.043 | chr7 | 99195890 | 99208668 | + | protein_coding |
| ENSG00000163297.17_4 | ANTXR2 | 8.92E-09 | 1.22E-07 | 1.542 | 2.402 | chr4 | 80822300 | 81046608 | - | protein_coding |
| ENSG00000106258.15_5 | CYP3A5 | 5.71E-16 | 2.31E-14 | 1.538 | 4.760 | chr7 | 99245817 | 99277621 | - | protein_coding |
| ENSG00000170421.12_7 | KRT8 | 3.42E-19 | 2.29E-17 | 1.538 | 12.114 | chr12 | 53290971 | 53343738 | - | protein_coding |
| ENSG00000050327.15_5 | ARHGEF5 | 2.21E-09 | 3.34E-08 | 1.536 | 2.402 | chr7 | 144052381 | 144077725 | + | protein_coding |
| ENSG00000186448.15_5 | ZNF197 | 4.60E-13 | 1.18E-11 | 1.533 | 3.772 | chr3 | 44626380 | 44689963 | + | protein_coding |
| ENSG00000167996.16_7 | FTH1 | 0.00631752 | 0.03521915 | 1.532 | 6.709 | chr11 | 61727190 | 61735106 | - | protein_coding |
| ENSG00000235257.9_9 | ITGA9-AS1 | 0.00032026 | 0.00228423 | 1.531 | -0.018 | chr3 | 37735146 | 37903293 | - | lncRNA |
| ENSG00000215807.4_4 | KRT18P65 | 0.00613625 | 0.03427312 | 1.529 | -1.222 | chr6 | 112682639 | 112683924 | - | processed_pseudogene |
| ENSG00000180626.10_5 | ZNF594 | 1.70E-15 | 6.44E-14 | 1.528 | 4.620 | chr17 | 5082830 | 5095163 | - | protein_coding |
| ENSG00000133256.12_2 | PDE6B | 0.00492101 | 0.02804993 | 1.528 | -1.221 | chr4 | 619373 | 664571 | + | protein_coding |
| ENSG00000174586.11_6 | ZNF497 | 9.20E-08 | 1.10E-06 | 1.527 | 1.908 | chr19 | 58865723 | 58874214 | - | protein_coding |
| ENSG00000188910.8_3 | GJB3 | 2.63E-07 | 2.94E-06 | 1.526 | 1.412 | chr1 | 35246815 | 35251970 | + | protein_coding |
| ENSG00000134762.17_7 | DSC3 | 3.19E-11 | 6.15E-10 | 1.525 | 3.169 | chr18 | 28569331 | 28622781 | - | protein_coding |
| ENSG00000047597.7_5 | XK | 1.54E-08 | 2.03E-07 | 1.519 | 2.067 | chrX | 37545044 | 37591383 | + | protein_coding |
| ENSG00000246695.8_7 | RASSF8-AS1 | 3.38E-10 | 5.67E-09 | 1.517 | 2.711 | chr12 | 26089616 | 26112698 | - | lncRNA |
| ENSG00000234156.3_8 | AL359636.2 | 1.34E-11 | 2.73E-10 | 1.516 | 3.437 | chr9 | 125165686 | 125401964 | + | lncRNA |
| ENSG00000179104.9_3 | TMTC2 | 5.94E-14 | 1.77E-12 | 1.516 | 4.578 | chr12 | 83080659 | 83528649 | + | protein_coding |
| ENSG00000263327.6_7 | TAPT1-AS1 | 0.00061872 | 0.00418932 | 1.515 | -0.071 | chr4 | 16228308 | 16321763 | + | lncRNA |
| ENSG00000255366.1_6 | AC120036.4 | 0.00325721 | 0.01920705 | 1.515 | -1.050 | chr8 | 48102395 | 48104885 | + | lncRNA |
| ENSG00000276900.1_6 | AC023157.3 | 1.69E-08 | 2.22E-07 | 1.512 | 2.076 | chr12 | 31882051 | 31884138 | + | lncRNA |
| ENSG00000154822.18_5 | PLCL2 | 1.34E-15 | 5.17E-14 | 1.511 | 4.882 | chr3 | 16844159 | 17132096 | + | protein_coding |
| ENSG00000213762.12_6 | ZNF134 | 1.80E-16 | 7.97E-15 | 1.510 | 5.103 | chr19 | 58125601 | 58136092 | + | protein_coding |
| ENSG00000215105.4_4 | TTC3P1 | 1.40E-10 | 2.50E-09 | 1.508 | 3.007 | chrX | 74960666 | 74966746 | - | processed_pseudogene |
| ENSG00000178038.17_4 | ALS2CL | 3.97E-14 | 1.23E-12 | 1.507 | 4.231 | chr3 | 46710485 | 46735194 | - | protein_coding |
| ENSG00000106025.9_4 | TSPAN12 | 2.31E-06 | 2.27E-05 | 1.507 | 1.080 | chr7 | 120427374 | 120498456 | - | protein_coding |
| ENSG00000118898.16_6 | PPL | 8.65E-14 | 2.52E-12 | 1.504 | 5.113 | chr16 | 4932508 | 5010742 | - | protein_coding |
| ENSG00000270344.3_6 | POC1B-AS1 | 2.32E-09 | 3.50E-08 | 1.504 | 2.351 | chr12 | 89918371 | 89941782 | + | lncRNA |
| ENSG00000188596.11_6 | CFAP54 | 6.31E-05 | 0.00050542 | 1.504 | 1.419 | chr12 | 96883349 | 97269333 | + | protein_coding |
| ENSG00000180318.4_3 | ALX1 | 2.40E-10 | 4.12E-09 | 1.502 | 2.961 | chr12 | 85673998 | 85695562 | + | protein_coding |
| ENSG00000150593.18_5 | PDCD4 | 2.06E-16 | 9.02E-15 | 1.501 | 5.367 | chr10 | 112631553 | 112659764 | + | protein_coding |
| ENSG00000152217.19_7 | SETBP1 | 1.07E-13 | 3.06E-12 | 1.501 | 4.146 | chr18 | 42260138 | 42648475 | + | protein_coding |
| ENSG00000119922.10_5 | IFIT2 | 1.61E-08 | 2.12E-07 | 1.501 | 2.255 | chr10 | 91043451 | 91069028 | + | protein_coding |
| ENSG00000130270.16_3 | ATP8B3 | 1.17E-14 | 3.94E-13 | 1.500 | 4.515 | chr19 | 1782074 | 1812275 | - | protein_coding |
| ENSG00000104361.10_4 | NIPAL2 | 2.14E-12 | 4.96E-11 | 1.498 | 3.739 | chr8 | 99202054 | 99306621 | - | protein_coding |
| ENSG00000196850.6_4 | PPTC7 | 6.80E-15 | 2.37E-13 | 1.496 | 4.647 | chr12 | 110971050 | 111021123 | - | protein_coding |
| ENSG00000090612.21_7 | ZNF268 | 4.31E-16 | 1.78E-14 | 1.493 | 5.566 | chr12 | 133757995 | 133791417 | + | protein_coding |
| ENSG00000183647.10_2 | ZNF530 | 2.28E-12 | 5.26E-11 | 1.492 | 3.741 | chr19 | 58111253 | 58124090 | + | protein_coding |
| ENSG00000179674.4_4 | ARL14 | 1.91E-07 | 2.19E-06 | 1.491 | 1.587 | chr3 | 160394948 | 160396236 | + | protein_coding |
| ENSG00000250120.8_6 | PCDHA10 | 4.23E-07 | 4.57E-06 | 1.490 | 1.513 | chr5 | 140235468 | 140391932 | + | protein_coding |
| ENSG00000197099.8_9 | AC068631.1 | 0.00311875 | 0.01845798 | 1.487 | -0.981 | chr3 | 186297265 | 186490775 | - | lncRNA |
| ENSG00000179954.16_4 | SSC5D | 7.73E-05 | 0.00060818 | 1.486 | 0.252 | chr19 | 55999771 | 56030466 | + | protein_coding |
| ENSG00000177879.16_3 | AP3S1 | 3.74E-18 | 2.12E-16 | 1.486 | 6.238 | chr5 | 115177289 | 115249778 | + | protein_coding |
| ENSG00000145687.16_5 | SSBP2 | 1.15E-10 | 2.07E-09 | 1.483 | 3.102 | chr5 | 80708840 | 81047616 | - | protein_coding |
| ENSG00000204580.13_3 | DDR1 | 2.81E-14 | 8.93E-13 | 1.483 | 5.809 | chr6 | 30844198 | 30867933 | + | protein_coding |
| ENSG00000181444.13_4 | ZNF467 | 2.49E-07 | 2.79E-06 | 1.481 | 1.813 | chr7 | 149461271 | 149470677 | - | protein_coding |
| ENSG00000167840.13_3 | ZNF232 | 3.87E-12 | 8.49E-11 | 1.479 | 3.974 | chr17 | 5008836 | 5026411 | - | protein_coding |
| ENSG00000111424.12_7 | VDR | 6.84E-09 | 9.56E-08 | 1.475 | 2.377 | chr12 | 48235320 | 48336831 | - | polymorphic_pseudogene |
| ENSG00000151062.15_7 | CACNA2D4 | 0.00473534 | 0.02711737 | 1.475 | -1.163 | chr12 | 1901129 | 2027832 | - | protein_coding |
| ENSG00000105519.17_5 | CAPS | 3.30E-06 | 3.17E-05 | 1.474 | 1.119 | chr19 | 5912350 | 5916222 | + | protein_coding |
| ENSG00000204052.4_3 | LRRC73 | 0.00015973 | 0.00120299 | 1.474 | 0.166 | chr6 | 43474707 | 43478424 | - | protein_coding |
| ENSG00000167105.8_3 | TMEM92 | 2.50E-11 | 4.91E-10 | 1.473 | 3.587 | chr17 | 48348767 | 48358846 | + | protein_coding |
| ENSG00000173041.12_4 | ZNF680 | 3.51E-15 | 1.27E-13 | 1.472 | 4.737 | chr7 | 63980256 | 64023453 | - | protein_coding |
| ENSG00000185522.9_5 | LMNTD2 | 1.33E-09 | 2.08E-08 | 1.470 | 2.706 | chr11 | 554850 | 560738 | - | protein_coding |
| ENSG00000141524.15_4 | TMC6 | 2.86E-08 | 3.62E-07 | 1.470 | 4.375 | chr17 | 76106539 | 76128488 | - | protein_coding |

| | | | | | | | | | | |
|---|---|---|---|---|---|---|---|---|---|---|
| ENSG00000213943.3_4 | KRT18P17 | 0.00047214 | 0.00327036 | 1.468 | 0.206 | chr3 | 12828892 | 12830170 | - | processed_pseudogene |
| ENSG00000059377.17_4 | TBXAS1 | 2.36E-14 | 7.60E-13 | 1.468 | 4.465 | chr7 | 139476850 | 139720125 | + | protein_coding |
| ENSG00000233143.2_5 | DIRC3-AS1 | 0.00263687 | 0.01584354 | 1.462 | -0.701 | chr2 | 218126422 | 218200843 | + | lncRNA |
| ENSG00000221869.5_3 | CEBPD | 2.16E-06 | 2.13E-05 | 1.460 | 4.420 | chr8 | 48649475 | 48650726 | - | protein_coding |
| ENSG00000122779.18_4 | TRIM24 | 1.43E-17 | 7.57E-16 | 1.460 | 6.656 | chr7 | 138145004 | 138274741 | + | protein_coding |
| ENSG00000251138.7_9 | AC090015.1 | 5.38E-07 | 5.76E-06 | 1.459 | 1.642 | chr12 | 74214095 | 74796380 | - | lncRNA |
| ENSG00000225269.3_6 | LINC00705 | 0.0001921 | 0.00142269 | 1.456 | 0.256 | chr10 | 4697619 | 4724914 | + | lncRNA |
| ENSG00000185561.10_4 | TLCD2 | 3.81E-12 | 8.37E-11 | 1.456 | 3.605 | chr17 | 1606110 | 1613671 | - | protein_coding |
| ENSG00000143036.17_4 | SLC44A3 | 4.08E-08 | 5.07E-07 | 1.455 | 2.643 | chr1 | 95285898 | 95360802 | + | protein_coding |
| ENSG00000159712.10_5 | ANKRD18CP | 0.00023429 | 0.00171162 | 1.453 | 0.488 | chr9 | 99918852 | 99983517 | - | unprocessed_pseudogene |
| ENSG00000135324.6_3 | MRAP2 | 2.82E-09 | 4.21E-08 | 1.451 | 2.779 | chr6 | 84743491 | 84800600 | + | protein_coding |
| ENSG00000179044.16_5 | EXOC3L1 | 0.00223913 | 0.01364288 | 1.450 | -0.325 | chr16 | 67218282 | 67224088 | - | protein_coding |
| ENSG00000251664.5_5 | PCDHA12 | 5.06E-09 | 7.22E-08 | 1.449 | 2.602 | chr5 | 140254887 | 140391932 | + | protein_coding |
| ENSG00000151491.14_9 | EPS8 | 2.22E-18 | 1.31E-16 | 1.448 | 6.961 | chr12 | 15773068 | 16035263 | - | protein_coding |
| ENSG00000145431.11_5 | PDGFC | 1.62E-12 | 3.82E-11 | 1.447 | 3.829 | chr4 | 157681606 | 157892951 | - | protein_coding |
| ENSG00000106003.13_4 | LFNG | 7.18E-13 | 1.79E-11 | 1.442 | 4.223 | chr7 | 2552163 | 2568811 | + | protein_coding |
| ENSG00000134138.20_6 | MEIS2 | 2.30E-14 | 7.41E-13 | 1.442 | 4.783 | chr15 | 37181405 | 37393500 | - | protein_coding |
| ENSG00000116176.6_3 | TPSG1 | 0.00113488 | 0.00731616 | 1.439 | -0.224 | chr16 | 1271651 | 1275257 | - | protein_coding |
| ENSG00000142530.11_6 | FAM71E1 | 0.00020818 | 0.00153093 | 1.437 | 0.316 | chr19 | 50970042 | 50980105 | - | protein_coding |
| ENSG00000237487.1_4 | VN1R48P | 0.00319993 | 0.01890063 | 1.437 | -0.788 | chr9 | 38504444 | 38505479 | - | processed_pseudogene |
| ENSG00000114631.11_4 | PODXL2 | 4.94E-13 | 1.26E-11 | 1.436 | 4.409 | chr3 | 127348028 | 127391645 | + | protein_coding |
| ENSG00000272144.1_6 | AC025171.4 | 0.00144048 | 0.00909164 | 1.436 | -0.318 | chr5 | 43061497 | 43062543 | + | lncRNA |
| ENSG00000232006.9_7 | AC005537.2 | 0.0009616 | 0.00627657 | 1.435 | -0.233 | chr7 | 42993734 | 43153530 | - | lncRNA |
| ENSG00000204044.6_7 | SLC12A5-AS1 | 0.00775084 | 0.04245042 | 1.434 | -1.284 | chr20 | 44642139 | 44650712 | - | lncRNA |
| ENSG00000139044.12_5 | B4GALNT3 | 0.00026405 | 0.0019164 | 1.434 | 0.147 | chr12 | 569105 | 672675 | + | protein_coding |
| ENSG00000171236.10_5 | LRG1 | 0.00017142 | 0.00128127 | 1.432 | 0.214 | chr19 | 4536414 | 4540048 | - | protein_coding |
| ENSG00000269729.1_7 | AC006262.2 | 6.76E-05 | 0.00053833 | 1.429 | 0.600 | chr19 | 46667150 | 46683904 | - | lncRNA |
| ENSG00000263155.6_7 | MYZAP | 0.00385961 | 0.02252748 | 1.427 | -0.866 | chr15 | 57884102 | 57977562 | + | protein_coding |
| ENSG00000248858.7_8 | AC104211.1 | 0.00597446 | 0.03345359 | 1.426 | -1.105 | chr8 | 93711970 | 93891730 | - | lncRNA |
| ENSG00000257732.1_7 | AC089983.1 | 6.73E-05 | 0.00053684 | 1.423 | 0.571 | chr12 | 104656092 | 104674500 | - | lncRNA |
| ENSG00000188321.13_4 | ZNF559 | 9.13E-11 | 1.66E-09 | 1.423 | 3.296 | chr19 | 9434448 | 9461838 | + | protein_coding |
| ENSG00000197405.8_4 | C5AR1 | 3.32E-05 | 0.00027813 | 1.422 | 0.747 | chr19 | 47793280 | 47825323 | + | protein_coding |
| ENSG00000152439.13_5 | ZNF773 | 3.53E-11 | 6.77E-10 | 1.421 | 3.389 | chr19 | 58011283 | 58029772 | + | protein_coding |
| ENSG00000287725.1_1 | AP003071.5 | 0.00232225 | 0.01408654 | 1.420 | -0.483 | chr11 | 68840383 | 68929908 | + | protein_coding |
| ENSG00000168542.15_4 | COL3A1 | 0.00262756 | 0.01579293 | 1.419 | -0.492 | chr2 | 189839099 | 189877472 | + | protein_coding |
| ENSG00000136274.9_4 | NACAD | 0.00557201 | 0.03140798 | 1.418 | -0.946 | chr7 | 45120036 | 45128568 | - | protein_coding |
| ENSG00000180481.11_3 | GLIPR1L2 | 0.00089972 | 0.00590301 | 1.418 | -0.331 | chr12 | 75784869 | 75826468 | + | protein_coding |
| ENSG00000254290.1_7 | AC124067.4 | 9.57E-06 | 8.61E-05 | 1.418 | 1.249 | chr8 | 37454998 | 37457376 | - | lncRNA |
| ENSG00000178665.16_5 | ZNF713 | 2.19E-07 | 2.48E-06 | 1.418 | 1.825 | chr7 | 55955149 | 56010223 | + | protein_coding |
| ENSG00000274180.1_4 | NATD1 | 1.61E-07 | 1.86E-06 | 1.417 | 2.249 | chr17 | 21142183 | 21156722 | - | protein_coding |
| ENSG00000119681.12_7 | LTBP2 | 4.63E-16 | 1.90E-14 | 1.417 | 5.588 | chr14 | 74964886 | 75079081 | - | protein_coding |
| ENSG00000006468.14_6 | ETV1 | 2.88E-14 | 9.11E-13 | 1.414 | 4.659 | chr7 | 13930854 | 14031050 | - | protein_coding |
| ENSG00000120784.16_7 | ZFP30 | 5.53E-17 | 2.74E-15 | 1.414 | 5.964 | chr19 | 38104650 | 38183238 | - | protein_coding |
| ENSG00000119888.11_3 | EPCAM | 8.64E-10 | 1.38E-08 | 1.413 | 2.955 | chr2 | 47572297 | 47614740 | + | protein_coding |
| ENSG00000198203.10_5 | SULT1C2 | 0.00413778 | 0.02400913 | 1.412 | -0.879 | chr2 | 108905095 | 108926371 | + | protein_coding |
| ENSG00000266921.2_8 | AC006213.1 | 0.00215855 | 0.01320627 | 1.412 | -0.494 | chr19 | 44501048 | 44507450 | - | lncRNA |
| ENSG00000198108.4_4 | CHSY3 | 0.0031243 | 0.0184775 | 1.410 | -0.673 | chr5 | 129240158 | 129522327 | + | protein_coding |
| ENSG00000173156.7_3 | RHOD | 5.48E-12 | 1.17E-10 | 1.409 | 4.124 | chr11 | 66824318 | 66839488 | + | protein_coding |
| ENSG00000261997.1_7 | AC007336.1 | 0.00512012 | 0.0290822 | 1.409 | -1.032 | chr16 | 55572112 | 55575939 | + | lncRNA |
| ENSG00000196867.8_4 | ZFP28 | 7.88E-13 | 1.95E-11 | 1.409 | 4.095 | chr19 | 57050317 | 57068177 | + | protein_coding |
| ENSG00000186026.7_4 | ZNF284 | 2.06E-06 | 2.04E-05 | 1.409 | 1.706 | chr19 | 44576312 | 44593766 | + | protein_coding |
| ENSG00000215182.6 | MUC5AC | 7.61E-06 | 6.94E-05 | 1.409 | 6.256 | chr11 | 1151580 | 1222364 | + | protein_coding |
| ENSG00000139266.6_4 | MARCHF9 | 8.76E-12 | 1.83E-10 | 1.408 | 3.687 | chr12 | 58148886 | 58154194 | + | protein_coding |
| ENSG00000163235.16_7 | TGFA | 3.11E-15 | 1.14E-13 | 1.408 | 5.110 | chr2 | 70674416 | 70781325 | - | protein_coding |
| ENSG00000008311.15_4 | AASS | 0.00049541 | 0.00341677 | 1.408 | 0.085 | chr7 | 121713603 | 121784309 | - | protein_coding |
| ENSG00000172361.6_5 | CFAP53 | 0.00084449 | 0.00556529 | 1.408 | -0.197 | chr18 | 47753563 | 47792865 | - | protein_coding |
| ENSG00000068079.7_2 | IFI35 | 3.71E-11 | 7.10E-10 | 1.407 | 3.699 | chr17 | 41158742 | 41166473 | + | protein_coding |
| ENSG00000276071.1_6 | AC074138.1 | 5.55E-05 | 0.00044833 | 1.406 | 0.708 | chr19 | 37159004 | 37160306 | + | lncRNA |

| | | | | | | | | | | |
|---|---|---|---|---|---|---|---|---|---|---|
| ENSG00000279198.1_5 | AC008894.3 | 0.00036306 | 0.00256786 | 1.405 | 0.122 | chr19 | 16185103 | 16188205 | - | TEC |
| ENSG00000203849.6 | RP11-417J8.6 | 0.00067942 | 0.00457417 | 1.404 | -0.110 | chr1 | 142620645 | 142729967 | - | lincRNA |
| ENSG00000142920.17_3 | AZIN2 | 1.64E-05 | 0.00014311 | 1.404 | 1.197 | chr1 | 33546705 | 33589093 | + | protein_coding |
| ENSG00000268912.1_6 | AC012313.5 | 3.03E-05 | 0.0002563 | 1.404 | 1.255 | chr19 | 58939999 | 58942515 | - | lncRNA |
| ENSG00000176728.9_8 | TTTY14 | 0.00329741 | 0.01942036 | 1.402 | -0.745 | chrY | 20934592 | 21239449 | - | lncRNA |
| ENSG00000152503.9_5 | TRIM36 | 0.00030615 | 0.00219773 | 1.401 | 0.575 | chr5 | 114460459 | 114516243 | - | protein_coding |
| ENSG00000267733.5_5 | AP005264.5 | 0.00032124 | 0.00228941 | 1.401 | 0.123 | chr18 | 12201981 | 12223538 | + | transcribed_processed_pseudogene |
| ENSG00000187650.4_4 | VMAC | 6.92E-07 | 7.27E-06 | 1.400 | 1.907 | chr19 | 5904883 | 5910864 | + | protein_coding |
| ENSG00000090776.6_3 | EFNB1 | 6.11E-09 | 8.61E-08 | 1.400 | 3.692 | chrX | 68048864 | 68062003 | + | protein_coding |
| ENSG00000159885.14_4 | ZNF222 | 1.12E-08 | 1.50E-07 | 1.396 | 2.852 | chr19 | 44529494 | 44537263 | + | protein_coding |
| ENSG00000229404.6_7 | LINC00858 | 1.15E-07 | 1.35E-06 | 1.395 | 2.078 | chr10 | 86027503 | 86054415 | + | lncRNA |
| ENSG00000164778.4_3 | EN2 | 0.00061391 | 0.0041599 | 1.395 | 0.078 | chr7 | 155250824 | 155257526 | + | protein_coding |
| ENSG00000224383.7_3 | PRR29 | 0.00206691 | 0.01269803 | 1.394 | -0.219 | chr17 | 62075711 | 62081664 | + | protein_coding |
| ENSG00000156265.16_5 | MAP3K7CL | 2.84E-07 | 3.16E-06 | 1.389 | 2.169 | chr21 | 30449792 | 30548210 | + | protein_coding |
| ENSG00000173080.5_2 | RXFP4 | 0.00235028 | 0.01424195 | 1.388 | -0.514 | chr1 | 155911501 | 155912740 | + | protein_coding |
| ENSG00000273987.1_7 | AC121761.2 | 0.00636199 | 0.03544478 | 1.387 | -1.050 | chr12 | 75727578 | 75728266 | - | lncRNA |
| ENSG00000139910.20_6 | NOVA1 | 4.58E-10 | 7.58E-09 | 1.386 | 3.405 | chr14 | 26912296 | 27067239 | - | protein_coding |
| ENSG00000260793.2_6 | AC003102.1 | 0.00016923 | 0.00126707 | 1.385 | 0.877 | chr17 | 42298769 | 42301078 | + | lncRNA |
| ENSG00000174567.8_4 | GOLT1A | 8.13E-07 | 8.51E-06 | 1.383 | 1.920 | chr1 | 204167291 | 204183116 | - | protein_coding |
| ENSG00000171368.12_4 | TPPP | 1.55E-11 | 3.13E-10 | 1.381 | 3.657 | chr5 | 659977 | 693467 | - | protein_coding |
| ENSG00000203497.2_7 | PDCD4-AS1 | 0.00010625 | 0.00081793 | 1.381 | 0.511 | chr10 | 112629626 | 112631991 | - | lncRNA |
| ENSG00000237172.4_4 | B3GNT9 | 6.82E-12 | 1.44E-10 | 1.380 | 3.947 | chr16 | 67182007 | 67184901 | - | protein_coding |
| ENSG00000230524.8_3 | COL6A4P1 | 0.00312517 | 0.0184775 | 1.380 | -0.579 | chr3 | 15193340 | 15247466 | - | transcribed_unprocessed_pseudogene |
| ENSG00000172748.14_6 | ZNF596 | 2.08E-06 | 2.06E-05 | 1.379 | 1.772 | chr8 | 182137 | 214703 | + | protein_coding |
| ENSG00000254285.3_4 | KRT8P3 | 4.16E-09 | 6.05E-08 | 1.379 | 4.238 | chr8 | 62490779 | 62492227 | + | processed_pseudogene |
| ENSG00000165512.5_3 | ZNF22 | 1.21E-10 | 2.17E-09 | 1.377 | 3.627 | chr10 | 45496371 | 45500774 | + | protein_coding |
| ENSG00000168056.16_7 | LTBP3 | 9.84E-09 | 1.34E-07 | 1.374 | 6.664 | chr11 | 65306276 | 65326401 | - | protein_coding |
| ENSG00000085465.13_4 | OVGP1 | 0.00011818 | 0.00090351 | 1.372 | 0.550 | chr1 | 111956941 | 111970357 | - | protein_coding |
| ENSG00000104490.18_7 | NCALD | 8.95E-13 | 2.18E-11 | 1.372 | 4.496 | chr8 | 102698770 | 103137135 | - | protein_coding |
| ENSG00000231203.1_4 | KRT8P10 | 5.00E-05 | 0.00040698 | 1.371 | 1.342 | chr2 | 183935768 | 183937214 | - | processed_pseudogene |
| ENSG00000119938.9_4 | PPP1R3C | 2.08E-15 | 7.84E-14 | 1.371 | 5.469 | chr10 | 93388199 | 93392828 | - | protein_coding |
| ENSG00000104870.13_7 | FCGRT | 4.86E-14 | 1.48E-12 | 1.370 | 4.830 | chr19 | 50010073 | 50029685 | + | protein_coding |
| ENSG00000120149.9_3 | MSX2 | 8.38E-08 | 1.01E-06 | 1.370 | 2.373 | chr5 | 174151585 | 174157899 | + | protein_coding |
| ENSG00000235034.7_4 | C19orf81 | 0.00020275 | 0.00149596 | 1.370 | 0.707 | chr19 | 51152702 | 51162567 | + | protein_coding |
| ENSG00000162913.10_9 | OBSCN-AS1 | 2.03E-06 | 2.00E-05 | 1.370 | 1.483 | chr1 | 228391204 | 228401365 | - | lncRNA |
| ENSG00000143416.21_7 | SELENBP1 | 5.01E-13 | 1.28E-11 | 1.366 | 4.555 | chr1 | 151336780 | 151345183 | - | protein_coding |
| ENSG00000188833.9_4 | ENTPD8 | 6.42E-05 | 0.00051368 | 1.366 | 0.925 | chr9 | 140328816 | 140336268 | - | protein_coding |
| ENSG00000246523.8_8 | FZD4-DT | 0.01007664 | 0.05389275 | 1.365 | -0.983 | chr11 | 86666658 | 86732951 | + | lncRNA |
| ENSG00000112297.16_8 | CRYBG1 | 3.16E-07 | 3.49E-06 | 1.364 | 2.226 | chr6 | 106808592 | 107018335 | + | protein_coding |
| ENSG00000197808.12_5 | ZNF461 | 5.60E-11 | 1.04E-09 | 1.363 | 3.865 | chr19 | 37127520 | 37157755 | - | protein_coding |
| ENSG00000188641.14_5 | DPYD | 1.31E-11 | 2.68E-10 | 1.362 | 4.009 | chr1 | 97543299 | 98460556 | - | protein_coding |
| ENSG00000273706.5_4 | LHX1 | 7.70E-06 | 7.02E-05 | 1.362 | 1.480 | chr17 | 35294084 | 35301912 | + | protein_coding |
| ENSG00000135148.12_5 | TRAFD1 | 6.55E-14 | 1.94E-12 | 1.360 | 4.947 | chr12 | 112563342 | 112591408 | + | protein_coding |
| ENSG00000172594.13_3 | SMPDL3A | 6.94E-08 | 8.41E-07 | 1.360 | 2.455 | chr6 | 123110194 | 123130865 | + | protein_coding |
| ENSG00000231156.1_4 | AC093702.1 | 0.00183809 | 0.01137889 | 1.358 | -0.194 | chr2 | 45240353 | 45240807 | - | lncRNA |
| ENSG00000171766.16_5 | GATM | 3.55E-12 | 7.84E-11 | 1.356 | 4.098 | chr15 | 45653322 | 45694525 | - | protein_coding |
| ENSG00000170571.12_5 | EMB | 7.49E-16 | 2.97E-14 | 1.355 | 5.941 | chr5 | 49692026 | 49739082 | - | protein_coding |
| ENSG00000139946.10_3 | PELI2 | 2.48E-12 | 5.66E-11 | 1.355 | 4.306 | chr14 | 56584532 | 56768242 | + | protein_coding |
| ENSG00000133985.3_3 | TTC9 | 3.94E-10 | 6.57E-09 | 1.354 | 3.538 | chr14 | 71108633 | 71142083 | + | protein_coding |
| ENSG00000154760.14_5 | SLFN13 | 6.13E-13 | 1.55E-11 | 1.354 | 4.411 | chr17 | 33762115 | 33775856 | - | protein_coding |
| ENSG00000102359.7_3 | SRPX2 | 1.69E-08 | 2.21E-07 | 1.354 | 2.729 | chrX | 99899192 | 99930785 | + | protein_coding |
| ENSG00000142173.16_5 | COL6A2 | 9.44E-13 | 2.29E-11 | 1.353 | 4.872 | chr21 | 47518026 | 47552762 | + | protein_coding |
| ENSG00000157240.4_3 | FZD1 | 3.01E-09 | 4.49E-08 | 1.350 | 3.292 | chr7 | 90893748 | 90900641 | + | protein_coding |
| ENSG00000233369.7_5 | GTF2IP4 | 5.20E-09 | 7.39E-08 | 1.349 | 3.026 | chr7 | 72569026 | 72621323 | + | transcribed_processed_pseudogene |
| ENSG00000160190.14_7 | SLC37A1 | 4.44E-09 | 6.42E-08 | 1.348 | 2.834 | chr21 | 43916118 | 44001550 | + | protein_coding |
| ENSG00000089127.13_6 | OAS1 | 2.66E-14 | 8.51E-13 | 1.348 | 5.105 | chr12 | 113344588 | 113371027 | + | protein_coding |
| ENSG00000063438.17_5 | AHRR | 0.00513292 | 0.02914562 | 1.347 | -0.649 | chr5 | 321874 | 435225 | + | protein_coding |
| ENSG00000272325.2_5 | NUDT3 | 0.00295089 | 0.01754633 | 1.347 | -0.377 | chr6 | 34247456 | 34360446 | - | protein_coding |

| | | | | | | | | | | |
|---|---|---|---|---|---|---|---|---|---|---|
| ENSG00000099953.10_6 | MMP11 | 0.00549089 | 0.03099004 | 1.345 | -0.717 | chr22 | 24110413 | 24126503 | + | protein_coding |
| ENSG00000160191.18_4 | PDE9A | 1.75E-08 | 2.29E-07 | 1.344 | 2.788 | chr21 | 44073731 | 44195619 | + | protein_coding |
| ENSG00000183111.12_3 | ARHGEF37 | 1.33E-08 | 1.77E-07 | 1.343 | 2.740 | chr5 | 148931510 | 149014531 | + | protein_coding |
| ENSG00000169435.14_5 | RASSF6 | 6.15E-10 | 1.00E-08 | 1.342 | 3.438 | chr4 | 74437267 | 74486348 | - | protein_coding |
| ENSG00000196954.14_7 | CASP4 | 4.23E-09 | 6.15E-08 | 1.342 | 2.954 | chr11 | 104813593 | 104840093 | - | protein_coding |
| ENSG00000181649.8_6 | PHLDA2 | 1.59E-08 | 2.09E-07 | 1.342 | 2.939 | chr11 | 2949503 | 2950650 | - | protein_coding |
| ENSG00000071242.12_6 | RPS6KA2 | 0.0104776 | 0.05576857 | 1.342 | -1.162 | chr6 | 166822852 | 167319939 | - | protein_coding |
| ENSG00000112773.16_9 | TENT5A | 6.91E-13 | 1.73E-11 | 1.340 | 4.496 | chr6 | 82201156 | 82462491 | - | protein_coding |
| ENSG00000215580.11_4 | BCORP1 | 0.00093094 | 0.00609207 | 1.336 | -0.044 | chrY | 21617317 | 21728978 | - | transcribed_unprocessed_pseudogene |
| ENSG00000160588.10_4 | MPZL3 | 4.89E-05 | 0.00039915 | 1.335 | 0.972 | chr11 | 118097405 | 118123080 | - | protein_coding |
| ENSG00000198722.14_5 | UNC13B | 1.17E-16 | 5.38E-15 | 1.334 | 6.341 | chr9 | 35161989 | 35405335 | + | protein_coding |
| ENSG00000167566.17_4 | NCKAP5L | 8.46E-14 | 2.47E-12 | 1.334 | 5.181 | chr12 | 50184929 | 50222533 | - | protein_coding |
| ENSG00000092929.11_6 | UNC13D | 3.62E-11 | 6.93E-10 | 1.331 | 4.315 | chr17 | 73823306 | 73840798 | - | protein_coding |
| ENSG00000186204.15_5 | CYP4F12 | 7.60E-10 | 1.23E-08 | 1.329 | 3.515 | chr19 | 15783828 | 15807984 | + | protein_coding |
| ENSG00000168772.11_4 | CXXC4 | 3.77E-08 | 4.71E-07 | 1.328 | 2.892 | chr4 | 105389465 | 105416058 | - | protein_coding |
| ENSG00000279329.1_6 | AC020910.6 | 4.24E-07 | 4.58E-06 | 1.327 | 2.100 | chr19 | 35166622 | 35168486 | - | TEC |
| ENSG00000168447.11_4 | SCNN1B | 0.00477643 | 0.02732223 | 1.327 | -0.730 | chr16 | 23289552 | 23392615 | + | protein_coding |
| ENSG00000140832.9_3 | MARVELD3 | 0.00078212 | 0.00518701 | 1.327 | 0.477 | chr16 | 71660064 | 71676017 | + | protein_coding |
| ENSG00000223855.2_7 | AC147651.1 | 5.27E-06 | 4.92E-05 | 1.326 | 1.721 | chr7 | 560028 | 564875 | + | lncRNA |
| ENSG00000139438.6_6 | FAM222A | 1.32E-08 | 1.75E-07 | 1.326 | 2.684 | chr12 | 110151630 | 110208300 | + | protein_coding |
| ENSG00000135643.5_3 | KCNMB4 | 3.16E-05 | 0.00026595 | 1.326 | 1.006 | chr12 | 70760070 | 70828072 | + | protein_coding |
| ENSG00000239912.1_2 | RPL39P36 | 0.00329776 | 0.01942036 | 1.325 | -0.399 | chr19 | 53916443 | 53916598 | + | processed_pseudogene |
| ENSG00000166046.11_5 | TCP11L2 | 6.20E-12 | 1.31E-10 | 1.325 | 4.236 | chr12 | 106695707 | 106740781 | + | protein_coding |
| ENSG00000083817.9_4 | ZNF416 | 3.24E-09 | 4.79E-08 | 1.324 | 3.288 | chr19 | 58082934 | 58090279 | - | protein_coding |
| ENSG00000167740.9_3 | CYB5D2 | 1.47E-13 | 4.09E-12 | 1.323 | 4.782 | chr17 | 4046462 | 4090605 | + | protein_coding |
| ENSG00000008277.14_2 | ADAM22 | 6.07E-07 | 6.44E-06 | 1.323 | 1.953 | chr7 | 87563458 | 87832204 | + | protein_coding |
| ENSG00000121075.11_5 | TBX4 | 0.01460059 | 0.07516912 | 1.322 | -1.013 | chr17 | 59529765 | 59562471 | + | protein_coding |
| ENSG00000278922.1_6 | AC002310.5 | 7.35E-05 | 0.0005814 | 1.321 | 1.060 | chr16 | 30538239 | 30539615 | + | TEC |
| ENSG00000107485.18_5 | GATA3 | 2.32E-06 | 2.27E-05 | 1.321 | 2.048 | chr10 | 8087341 | 8117161 | + | protein_coding |
| ENSG00000234928.1_5 | LINC01659 | 0.00129601 | 0.00824718 | 1.320 | -0.175 | chr22 | 23775751 | 23777258 | + | lncRNA |
| ENSG00000006756.16_4 | ARSD | 9.28E-15 | 3.19E-13 | 1.317 | 5.427 | chrX | 2822013 | 2847390 | - | protein_coding |
| ENSG00000049192.15_5 | ADAMTS6 | 0.00060266 | 0.00408678 | 1.316 | 0.224 | chr5 | 64444565 | 64777747 | - | protein_coding |
| ENSG00000130940.15_4 | CASZ1 | 1.32E-05 | 0.00011704 | 1.316 | 1.335 | chr1 | 10696661 | 10856707 | - | protein_coding |
| ENSG00000269386.5_7 | RAB11B-AS1 | 0.00128071 | 0.00816741 | 1.315 | 0.420 | chr19 | 8439257 | 8455569 | - | lncRNA |
| ENSG00000105717.14_5 | PBX4 | 0.00128941 | 0.00821108 | 1.310 | -0.018 | chr19 | 19672516 | 19729502 | - | protein_coding |
| ENSG00000234327.8_7 | AC012146.1 | 9.63E-07 | 9.99E-06 | 1.309 | 1.900 | chr17 | 5014763 | 5018299 | + | lncRNA |
| ENSG00000168490.14_6 | PHYHIP | 0.00152555 | 0.00957075 | 1.309 | 0.150 | chr8 | 22077216 | 22089614 | - | protein_coding |
| ENSG00000226937.9_7 | CEP164P1 | 0.01069225 | 0.05674108 | 1.309 | -1.104 | chr10 | 45497670 | 45571514 | - | transcribed_processed_pseudogene |
| ENSG00000072071.16_6 | ADGRL1 | 3.18E-09 | 4.73E-08 | 1.308 | 3.902 | chr19 | 14258555 | 14316999 | - | protein_coding |
| ENSG00000279611.1_6 | AC012313.10 | 0.00497414 | 0.02830736 | 1.307 | -0.576 | chr19 | 58862161 | 58863086 | + | TEC |
| ENSG00000130203.10_4 | APOE | 3.64E-05 | 0.00030328 | 1.306 | 1.187 | chr19 | 45409048 | 45412650 | + | protein_coding |
| ENSG00000181333.12_4 | HEPHL1 | 0.01185406 | 0.06221893 | 1.303 | -1.105 | chr11 | 93754520 | 93847374 | + | protein_coding |
| ENSG00000249867.6_8 | LINC02742 | 9.90E-05 | 0.0007657 | 1.302 | 0.907 | chr11 | 28724154 | 29085869 | + | lncRNA |
| ENSG00000189144.13_5 | ZNF573 | 9.22E-11 | 1.67E-09 | 1.302 | 4.005 | chr19 | 38226734 | 38307940 | - | protein_coding |
| ENSG00000142089.16_7 | IFITM3 | 6.96E-08 | 8.43E-07 | 1.301 | 2.641 | chr11 | 319676 | 327537 | - | protein_coding |
| ENSG00000141441.16_5 | GAREM1 | 2.33E-09 | 3.51E-08 | 1.300 | 3.368 | chr18 | 29704840 | 30050845 | - | protein_coding |
| ENSG00000148344.11_4 | PTGES | 1.50E-14 | 4.99E-13 | 1.298 | 6.063 | chr9 | 132500610 | 132515321 | - | protein_coding |
| ENSG00000198040.10_5 | ZNF84 | 1.23E-07 | 1.45E-06 | 1.298 | 2.498 | chr12 | 133613878 | 133639890 | + | protein_coding |
| ENSG00000107159.13_3 | CA9 | 0.00238945 | 0.01444973 | 1.298 | -0.276 | chr9 | 35673915 | 35681156 | + | protein_coding |
| ENSG00000012817.15_8 | KDM5D | 3.88E-15 | 1.39E-13 | 1.297 | 5.716 | chrY | 21865751 | 21906825 | - | protein_coding |
| ENSG00000162076.13_6 | FLYWCH2 | 1.28E-12 | 3.06E-11 | 1.295 | 4.891 | chr16 | 2933214 | 2949383 | + | protein_coding |
| ENSG00000021300.14_6 | PLEKHB1 | 0.00042359 | 0.00295944 | 1.295 | 0.645 | chr11 | 73357223 | 73373864 | + | protein_coding |
| ENSG00000189143.9_3 | CLDN4 | 2.95E-08 | 3.72E-07 | 1.294 | 2.703 | chr7 | 73213872 | 73247023 | + | protein_coding |
| ENSG00000105136.20_3 | ZNF419 | 3.04E-11 | 5.87E-10 | 1.294 | 4.129 | chr19 | 57999079 | 58007465 | + | protein_coding |
| ENSG00000259674.1_6 | AC092868.2 | 6.65E-09 | 9.32E-08 | 1.293 | 3.222 | chr15 | 59699162 | 59700353 | - | processed_pseudogene |
| ENSG00000285601.1_2 | AC084262.2 | 0.00203602 | 0.01251692 | 1.293 | 0.282 | chr8 | 29208809 | 29217739 | + | lncRNA |
| ENSG00000166126.10_5 | AMN | 0.00042866 | 0.00299134 | 1.292 | 0.624 | chr14 | 103388993 | 103399933 | + | protein_coding |
| ENSG00000103089.9_4 | FA2H | 3.40E-12 | 7.52E-11 | 1.290 | 4.885 | chr16 | 74746853 | 74808729 | - | protein_coding |

| | | | | | | | | | | |
|---|---|---|---|---|---|---|---|---|---|---|
| ENSG00000283162.1_6 | AL390726.5 | 6.72E-06 | 6.18E-05 | 1.289 | 1.545 | chr9 | 38433693 | 38543877 | + | transcribed_unprocessed_pseudogene |
| ENSG00000228630.5_6 | HOTAIR | 0.00357579 | 0.02097424 | 1.285 | -0.422 | chr12 | 54356092 | 54368740 | - | lncRNA |
| ENSG00000133519.12_5 | ZDHHC8P1 | 2.23E-08 | 2.87E-07 | 1.284 | 2.880 | chr22 | 23732793 | 23744913 | - | transcribed_unprocessed_pseudogene |
| ENSG00000260230.3_4 | FRRS1L | 3.55E-06 | 3.40E-05 | 1.284 | 1.858 | chr9 | 111892573 | 111929575 | - | protein_coding |
| ENSG00000234618.1_4 | RPSAP9 | 0.00636 | 0.03544478 | 1.284 | -0.644 | chr9 | 79013615 | 79014502 | + | processed_pseudogene |
| ENSG00000221949.5_5 | LINC01465 | 1.99E-05 | 0.00017168 | 1.282 | 3.599 | chr12 | 62995531 | 62997470 | - | lncRNA |
| ENSG00000135766.9_4 | EGLN1 | 0.00180694 | 0.01119771 | 1.281 | 4.824 | chr1 | 231499497 | 231558033 | - | protein_coding |
| ENSG00000261468.1_6 | AC096921.2 | 3.30E-05 | 0.00027657 | 1.280 | 1.429 | chr3 | 30739153 | 30741068 | + | lncRNA |
| ENSG00000174827.14_3 | PDZK1 | 7.47E-05 | 0.00058886 | 1.280 | 0.934 | chr1 | 145726918 | 145764207 | + | protein_coding |
| ENSG00000170965.10_4 | PLAC1 | 0.0032055 | 0.018921 | 1.279 | -0.245 | chrX | 133699868 | 133898352 | - | protein_coding |
| ENSG00000074966.11_4 | TXK | 0.00308648 | 0.01828525 | 1.276 | -0.080 | chr4 | 48068410 | 48136267 | - | protein_coding |
| ENSG00000181982.18_3 | CCDC149 | 1.35E-06 | 1.37E-05 | 1.275 | 2.592 | chr4 | 24807739 | 24981826 | - | protein_coding |
| ENSG00000165246.14_7 | NLGN4Y | 2.37E-08 | 3.03E-07 | 1.274 | 3.093 | chrY | 16634453 | 16957530 | + | protein_coding |
| ENSG00000253616.5_7 | AC107959.3 | 3.88E-05 | 0.00032187 | 1.274 | 1.180 | chr8 | 22928890 | 22932001 | - | lncRNA |
| ENSG00000187140.6_3 | FOXD3 | 0.00211372 | 0.01294979 | 1.273 | -0.003 | chr1 | 63788238 | 63790799 | + | protein_coding |
| ENSG00000172458.4_3 | IL17D | 0.0065082 | 0.03614602 | 1.272 | -0.646 | chr13 | 21276266 | 21297237 | + | protein_coding |
| ENSG00000086159.13_3 | AQP6 | 0.00550916 | 0.03106354 | 1.272 | -0.539 | chr12 | 50360977 | 50370922 | + | protein_coding |
| ENSG00000261534.1_6 | AL596244.1 | 0.01325838 | 0.06883741 | 1.271 | -1.125 | chr9 | 124577953 | 124581731 | - | lncRNA |
| ENSG00000108187.16_5 | PBLD | 1.94E-10 | 3.39E-09 | 1.271 | 3.718 | chr10 | 70042417 | 70092806 | - | protein_coding |
| ENSG00000281357.3_3 | ARRDC3-AS1 | 0.00148518 | 0.00934724 | 1.271 | 0.336 | chr5 | 90676166 | 90908383 | + | lncRNA |
| ENSG00000115267.8_6 | IFIH1 | 5.93E-07 | 6.29E-06 | 1.271 | 2.269 | chr2 | 163123584 | 163175194 | - | protein_coding |
| ENSG00000155093.19_6 | PTPRN2 | 2.18E-06 | 2.14E-05 | 1.270 | 2.152 | chr7 | 157331750 | 158380515 | - | protein_coding |
| ENSG00000123388.4_3 | HOXC11 | 0.00103257 | 0.00670285 | 1.270 | 0.283 | chr12 | 54366910 | 54371427 | + | protein_coding |
| ENSG00000166578.10_4 | IQCD | 8.31E-06 | 7.55E-05 | 1.268 | 1.601 | chr12 | 113633246 | 113658899 | - | protein_coding |
| ENSG00000185499.16_6 | MUC1 | 0.002204 | 0.01344726 | 1.267 | 0.435 | chr1 | 155158300 | 155162707 | - | protein_coding |
| ENSG00000196458.11_5 | ZNF605 | 3.77E-09 | 5.54E-08 | 1.267 | 4.627 | chr12 | 133494892 | 133532892 | - | protein_coding |
| ENSG00000170476.16_4 | MZB1 | 9.31E-05 | 0.00072334 | 1.265 | 1.188 | chr5 | 138723156 | 138725770 | - | protein_coding |
| ENSG00000205593.12_5 | DENND6B | 4.57E-06 | 4.31E-05 | 1.265 | 1.852 | chr22 | 50747459 | 50765441 | - | protein_coding |
| ENSG00000100065.15_6 | CARD10 | 8.93E-14 | 2.59E-12 | 1.263 | 6.050 | chr22 | 37886400 | 37915549 | - | protein_coding |
| ENSG00000080493.17_6 | SLC4A4 | 4.11E-11 | 7.81E-10 | 1.263 | 4.095 | chr4 | 71928384 | 72437804 | + | protein_coding |
| ENSG00000124664.11_5 | SPDEF | 2.89E-12 | 6.51E-11 | 1.262 | 5.290 | chr6 | 34505579 | 34524110 | - | protein_coding |
| ENSG00000109072.14_6 | VTN | 2.01E-08 | 2.61E-07 | 1.262 | 3.177 | chr17 | 26694305 | 26700110 | - | protein_coding |
| ENSG00000155792.10_3 | DEPTOR | 1.07E-08 | 1.44E-07 | 1.260 | 3.150 | chr8 | 120885957 | 121063157 | + | protein_coding |
| ENSG00000102934.10_4 | PLLP | 4.77E-09 | 6.86E-08 | 1.260 | 3.210 | chr16 | 57282459 | 57318584 | - | protein_coding |
| ENSG00000283064.1_7 | AL353759.1 | 0.00362244 | 0.02122686 | 1.259 | -0.299 | chr6 | 26160993 | 26173030 | - | lncRNA |
| ENSG00000169740.14_4 | ZNF32 | 4.13E-11 | 7.82E-10 | 1.259 | 4.083 | chr10 | 44139308 | 44144329 | - | protein_coding |
| ENSG00000253731.2_4 | PCDHGA6 | 4.21E-06 | 3.97E-05 | 1.258 | 2.002 | chr5 | 140753481 | 140892546 | + | protein_coding |
| ENSG00000047346.12_6 | FAM214A | 1.70E-10 | 2.99E-09 | 1.257 | 4.617 | chr15 | 52873514 | 53002014 | - | protein_coding |
| ENSG00000064989.13_4 | CALCRL | 0.00879374 | 0.04754665 | 1.257 | -0.716 | chr2 | 188206691 | 188313187 | - | protein_coding |
| ENSG00000138315.13_3 | OIT3 | 0.0134494 | 0.06970697 | 1.257 | -0.980 | chr10 | 74653342 | 74692794 | + | protein_coding |
| ENSG00000171557.17_6 | FGG | 1.50E-10 | 2.65E-09 | 1.256 | 4.402 | chr4 | 155525286 | 155534119 | - | protein_coding |
| ENSG00000256151.1_7 | ADGRD1-AS1 | 2.09E-05 | 0.00018024 | 1.255 | 1.401 | chr12 | 131474683 | 131478521 | - | lncRNA |
| ENSG00000184903.10_5 | IMMP2L | 1.01E-11 | 2.09E-10 | 1.255 | 4.330 | chr7 | 110302700 | 111202573 | - | protein_coding |
| ENSG00000005238.19_3 | FAM214B | 3.04E-10 | 5.15E-09 | 1.254 | 3.857 | chr9 | 35104109 | 35116338 | - | protein_coding |
| ENSG00000157343.8_3 | ARMC12 | 0.00505717 | 0.02875223 | 1.253 | -0.397 | chr6 | 35704809 | 35716856 | + | protein_coding |
| ENSG00000126709.15_4 | IFI6 | 1.02E-06 | 1.05E-05 | 1.253 | 2.275 | chr1 | 27992575 | 27998709 | - | protein_coding |
| ENSG00000272369.1_7 | AC008035.1 | 0.00549793 | 0.03101007 | 1.252 | -0.075 | chr12 | 46931285 | 47046333 | + | lncRNA |
| ENSG00000083844.10_5 | ZNF264 | 9.56E-13 | 2.31E-11 | 1.251 | 4.983 | chr19 | 57702868 | 57734214 | + | protein_coding |
| ENSG00000171843.16_4 | MLLT3 | 5.34E-14 | 1.61E-12 | 1.251 | 5.974 | chr9 | 20341667 | 20622517 | - | protein_coding |
| ENSG00000204965.9_5 | PCDHA5 | 1.92E-05 | 0.00016637 | 1.250 | 1.525 | chr5 | 140201189 | 140391932 | + | protein_coding |
| ENSG00000087494.16_5 | PTHLH | 1.17E-08 | 1.57E-07 | 1.247 | 3.271 | chr12 | 28111017 | 28125666 | - | protein_coding |
| ENSG00000176244.7_4 | ACBD7 | 6.66E-07 | 7.01E-06 | 1.246 | 2.210 | chr10 | 15117474 | 15130775 | - | protein_coding |
| ENSG00000109738.11_3 | GLRB | 1.07E-07 | 1.27E-06 | 1.246 | 2.666 | chr4 | 157997277 | 158093242 | + | protein_coding |
| ENSG00000104964.14_4 | TLE5 | 4.98E-09 | 7.13E-08 | 1.244 | 6.757 | chr19 | 3052908 | 3063105 | - | protein_coding |
| ENSG00000203780.11_7 | FANK1 | 7.71E-05 | 0.00060719 | 1.244 | 1.200 | chr10 | 127585133 | 127698161 | + | protein_coding |
| ENSG00000182459.5_3 | TEX19 | 0.00294684 | 0.01752813 | 1.243 | -0.185 | chr17 | 80317123 | 80321651 | + | protein_coding |
| ENSG00000215089.3_4 | KRT18P11 | 0.0006462 | 0.00436207 | 1.243 | 0.993 | chrX | 91714669 | 91715951 | + | processed_pseudogene |

| | | | | | | | | | | |
|---|---|---|---|---|---|---|---|---|---|---|
| ENSG00000152518.8_5 | ZFP36L2 | 1.68E-09 | 2.58E-08 | 1.241 | 4.726 | chr2 | 43449541 | 43453745 | - | protein_coding |
| ENSG00000124406.16_6 | ATP8A1 | 2.54E-05 | 0.00021683 | 1.241 | 1.514 | chr4 | 42410390 | 42659122 | - | protein_coding |
| ENSG00000171564.11_2 | FGB | 1.30E-10 | 2.33E-09 | 1.239 | 4.295 | chr4 | 155484108 | 155492238 | + | protein_coding |
| ENSG00000272473.1_6 | AC006273.1 | 1.16E-05 | 0.00010289 | 1.237 | 1.711 | chr19 | 781002 | 781862 | - | lncRNA |
| ENSG00000138496.16_4 | PARP9 | 1.42E-11 | 2.88E-10 | 1.237 | 4.494 | chr3 | 122246771 | 122283424 | - | protein_coding |
| ENSG00000154556.18_8 | SORBS2 | 0.00578141 | 0.03249552 | 1.236 | -0.318 | chr4 | 186506598 | 186877806 | - | protein_coding |
| ENSG00000139410.15_8 | SDSL | 2.95E-08 | 3.72E-07 | 1.236 | 3.085 | chr12 | 113860185 | 113876081 | + | protein_coding |
| ENSG00000104814.13_5 | MAP4K1 | 0.00015271 | 0.00115259 | 1.235 | 1.075 | chr19 | 39078281 | 39109522 | - | protein_coding |
| ENSG00000161860.8_6 | SYCE2 | 0.00528749 | 0.02997537 | 1.234 | -0.409 | chr19 | 13009600 | 13030107 | - | protein_coding |
| ENSG00000197748.13_6 | CFAP43 | 0.00114629 | 0.0073817 | 1.233 | 0.394 | chr10 | 105889646 | 105992122 | - | protein_coding |
| ENSG00000118849.10_4 | RARRES1 | 6.21E-10 | 1.01E-08 | 1.231 | 3.766 | chr3 | 158414681 | 158450278 | - | protein_coding |
| ENSG00000021826.17_6 | CPS1 | 6.16E-06 | 5.70E-05 | 1.231 | 1.903 | chr2 | 211342406 | 211543831 | + | protein_coding |
| ENSG00000052126.14_5 | PLEKHA5 | 2.71E-14 | 8.66E-13 | 1.230 | 5.938 | chr12 | 19282686 | 19529334 | + | protein_coding |
| ENSG00000130363.12_3 | RSPH3 | 6.62E-10 | 1.07E-08 | 1.228 | 3.732 | chr6 | 159393903 | 159421234 | - | protein_coding |
| ENSG00000177788.6_9 | AL162595.1 | 4.05E-10 | 6.76E-09 | 1.228 | 4.145 | chr1 | 229392639 | 229406989 | - | lncRNA |
| ENSG00000246228.6_7 | CASC8 | 1.86E-05 | 0.00016094 | 1.227 | 1.570 | chr8 | 128302062 | 128494384 | - | lncRNA |
| ENSG00000166510.14_6 | CCDC68 | 9.12E-11 | 1.66E-09 | 1.226 | 4.417 | chr18 | 52568740 | 52626692 | - | protein_coding |
| ENSG00000144802.11_4 | NFKBIZ | 2.35E-12 | 5.39E-11 | 1.226 | 4.931 | chr3 | 101546835 | 101579866 | + | protein_coding |
| ENSG00000141738.14_5 | GRB7 | 1.26E-11 | 2.58E-10 | 1.225 | 4.709 | chr17 | 37894180 | 37903544 | + | protein_coding |
| ENSG00000188051.7_5 | TMEM221 | 0.00110986 | 0.00716792 | 1.223 | 0.288 | chr19 | 17546318 | 17559477 | - | protein_coding |
| ENSG00000174021.11_5 | GNG5 | 0.00050206 | 0.00345387 | 1.223 | 0.819 | chr1 | 84964008 | 84972264 | - | protein_coding |
| ENSG00000188033.10_5 | ZNF490 | 1.47E-05 | 0.00012871 | 1.223 | 1.866 | chr19 | 12686914 | 12750912 | - | protein_coding |
| ENSG00000131459.13_5 | GFPT2 | 1.11E-13 | 3.12E-12 | 1.222 | 5.579 | chr5 | 179727698 | 179780336 | - | protein_coding |
| ENSG00000286193.2_3 | AC025449.2 | 0.00152236 | 0.00956083 | 1.222 | 0.194 | chr5 | 37249128 | 37252719 | + | lncRNA |
| ENSG00000016391.11_6 | CHDH | 1.39E-09 | 2.17E-08 | 1.221 | 4.371 | chr3 | 53846362 | 53880446 | - | protein_coding |
| ENSG00000130653.16_5 | PNPLA7 | 6.87E-05 | 0.00054579 | 1.221 | 1.516 | chr9 | 140354404 | 140444854 | - | protein_coding |
| ENSG00000214290.8_6 | COLCA2 | 0.00039554 | 0.00278098 | 1.220 | 1.060 | chr11 | 111169280 | 111179460 | + | protein_coding |
| ENSG00000102854.16_6 | MSLN | 0.00014762 | 0.00111698 | 1.219 | 1.171 | chr16 | 810734 | 818865 | + | protein_coding |
| ENSG00000126705.15_6 | AHDC1 | 1.12E-07 | 1.32E-06 | 1.218 | 3.787 | chr1 | 27860546 | 27930942 | - | protein_coding |
| ENSG00000168016.14_7 | TRANK1 | 4.26E-08 | 5.30E-07 | 1.218 | 3.016 | chr3 | 36868311 | 36986589 | - | protein_coding |
| ENSG00000250799.10_4 | PRODH2 | 0.0195238 | 0.09765746 | 1.218 | -0.945 | chr19 | 36290890 | 36304201 | - | protein_coding |
| ENSG00000171823.7_4 | FBXL14 | 3.58E-09 | 5.29E-08 | 1.218 | 3.970 | chr12 | 1675159 | 1703747 | - | protein_coding |
| ENSG00000136002.19_5 | ARHGEF4 | 2.45E-07 | 2.75E-06 | 1.217 | 3.134 | chr2 | 131594489 | 131804836 | + | protein_coding |
| ENSG00000101846.7_3 | STS | 2.58E-09 | 3.86E-08 | 1.217 | 3.676 | chrX | 7065278 | 7772399 | + | protein_coding |
| ENSG00000181085.15_3 | MAPK15 | 0.0004685 | 0.0032464 | 1.216 | 0.687 | chr8 | 144798510 | 144804628 | + | protein_coding |
| ENSG00000198948.12_6 | MFAP3L | 5.02E-10 | 8.26E-09 | 1.215 | 3.965 | chr4 | 170907748 | 170954182 | - | protein_coding |
| ENSG00000123094.15_7 | RASSF8 | 3.80E-12 | 8.37E-11 | 1.214 | 5.195 | chr12 | 26111962 | 26232825 | + | protein_coding |
| ENSG00000171604.12_4 | CXXC5 | 9.89E-12 | 2.05E-10 | 1.213 | 4.805 | chr5 | 139026884 | 139063467 | + | protein_coding |
| ENSG00000104783.14_7 | KCNN4 | 8.21E-13 | 2.02E-11 | 1.213 | 5.295 | chr19 | 44270685 | 44285409 | - | protein_coding |
| ENSG00000130518.17_6 | IQCN | 0.00030745 | 0.00220438 | 1.211 | 1.757 | chr19 | 18367907 | 18385319 | - | protein_coding |
| ENSG00000174428.18_5 | GTF2IRD2B | 7.44E-05 | 0.00058708 | 1.211 | 1.520 | chr7 | 74508364 | 74565623 | + | protein_coding |
| ENSG00000167711.13_2 | SERPINF2 | 1.07E-05 | 9.61E-05 | 1.211 | 2.467 | chr17 | 1646130 | 1658562 | + | protein_coding |
| ENSG00000241015.2_4 | TPM3P9 | 7.09E-11 | 1.30E-09 | 1.209 | 4.967 | chr19 | 53935237 | 53947923 | + | transcribed_processed_pseudogene |
| ENSG00000168676.11_5 | KCTD19 | 4.59E-05 | 0.00037689 | 1.209 | 1.723 | chr16 | 67323331 | 67360663 | - | protein_coding |
| ENSG00000139722.7_7 | VPS37B | 4.94E-10 | 8.13E-09 | 1.208 | 3.926 | chr12 | 123349877 | 123380674 | - | protein_coding |
| ENSG00000073067.14_4 | CYP2W1 | 0.00464054 | 0.02661344 | 1.207 | 0.018 | chr7 | 1022817 | 1029276 | + | protein_coding |
| ENSG00000119917.14_3 | IFIT3 | 2.92E-07 | 3.23E-06 | 1.207 | 2.847 | chr10 | 91087754 | 91100728 | + | protein_coding |
| ENSG00000183508.5_4 | TENT5C | 0.01426426 | 0.07360823 | 1.206 | -0.881 | chr1 | 118148670 | 118171011 | + | protein_coding |
| ENSG00000215165.3_4 | TCEA1P3 | 0.0175015 | 0.08864001 | 1.205 | -1.087 | chr9 | 38478468 | 38479347 | + | processed_pseudogene |
| ENSG00000156042.18_6 | CFAP70 | 0.00038189 | 0.00268929 | 1.204 | 0.991 | chr10 | 75013517 | 75118617 | - | protein_coding |
| ENSG00000248099.4_5 | INSL3 | 0.01919441 | 0.09622659 | 1.204 | -0.816 | chr19 | 17927321 | 17932383 | - | protein_coding |
| ENSG00000124459.12_5 | ZNF45 | 9.49E-13 | 2.30E-11 | 1.202 | 5.441 | chr19 | 44416776 | 44439434 | - | protein_coding |
| ENSG00000167380.16_6 | ZNF226 | 2.95E-12 | 6.61E-11 | 1.202 | 4.958 | chr19 | 44669226 | 44682534 | + | protein_coding |
| ENSG00000163739.5_3 | CXCL1 | 1.27E-05 | 0.00011268 | 1.201 | 2.044 | chr4 | 74735110 | 74737025 | + | protein_coding |
| ENSG00000261879.7_7 | AC087500.1 | 0.00141024 | 0.00891345 | 1.201 | 0.434 | chr17 | 5095322 | 5151477 | + | lncRNA |
| ENSG00000168916.15_4 | ZNF608 | 2.81E-09 | 4.20E-08 | 1.200 | 3.735 | chr5 | 123972606 | 124084500 | - | protein_coding |
| ENSG00000128298.17_5 | BAIAP2L2 | 2.51E-08 | 3.20E-07 | 1.200 | 3.457 | chr22 | 38480907 | 38506691 | - | protein_coding |
| ENSG00000139318.8_5 | DUSP6 | 7.89E-10 | 1.27E-08 | 1.200 | 3.934 | chr12 | 89741012 | 89746278 | - | protein_coding |
| ENSG00000229656.7_6 | ITGB1-DT | 2.25E-10 | 3.86E-09 | 1.199 | 4.291 | chr10 | 33247365 | 33371030 | + | lncRNA |
| ENSG00000196417.13_4 | ZNF765 | 1.67E-12 | 3.91E-11 | 1.199 | 5.251 | chr19 | 53893046 | 53933666 | + | protein_coding |

| | | | | | | | | | | |
|---|---|---|---|---|---|---|---|---|---|---|
| ENSG00000152049.6_2 | KCNE4 | 0.00071807 | 0.00480165 | 1.198 | 0.693 | chr2 | 223916532 | 224063117 | + | protein_coding |
| ENSG00000268756.1_6 | AC104534.1 | 0.00069465 | 0.00466607 | 1.196 | 0.582 | chr19 | 39310806 | 39313863 | + | lncRNA |
| ENSG00000256546.2_4 | AC156455.1 | 0.00010045 | 0.00077594 | 1.196 | 1.393 | chr12 | 122501212 | 122506522 | + | lncRNA |
| ENSG00000148400.12_6 | NOTCH1 | 0.00016184 | 0.00121733 | 1.196 | 6.130 | chr9 | 139388885 | 139440500 | - | protein_coding |
| ENSG00000225342.2_7 | LRRK2-DT | 0.00200871 | 0.01235762 | 1.194 | 0.345 | chr12 | 40579811 | 40618717 | - | lncRNA |
| ENSG00000229873.1_7 | OGFR-AS1 | 0.01062014 | 0.05644273 | 1.194 | -0.759 | chr20 | 61431979 | 61436939 | - | lncRNA |
| ENSG00000255874.2_4 | LINC00346 | 0.01290509 | 0.0672784 | 1.193 | -0.759 | chr13 | 111516334 | 111522598 | - | lncRNA |
| ENSG00000188283.11_6 | ZNF383 | 7.82E-11 | 1.43E-09 | 1.193 | 4.337 | chr19 | 37708828 | 37739640 | + | protein_coding |
| ENSG00000261308.2_4 | FIGNL2 | 0.00143704 | 0.00907315 | 1.192 | 0.606 | chr12 | 52211624 | 52242550 | - | protein_coding |
| ENSG00000134780.10_3 | DAGLA | 7.31E-09 | 1.02E-07 | 1.191 | 3.727 | chr11 | 61447863 | 61514473 | + | protein_coding |
| ENSG00000022267.19_8 | FHL1 | 2.84E-12 | 6.42E-11 | 1.190 | 5.502 | chrX | 135228861 | 135293518 | + | protein_coding |
| ENSG00000242193.11_6 | CRYZL2P | 4.92E-11 | 9.18E-10 | 1.190 | 4.564 | chr1 | 177975271 | 178007085 | - | transcribed_unitary_pseudogene |
| ENSG00000180537.13_7 | RNF182 | 6.97E-06 | 6.40E-05 | 1.189 | 2.359 | chr6 | 13924677 | 13980541 | + | protein_coding |
| ENSG00000105499.14_5 | PLA2G4C | 0.00025753 | 0.00187061 | 1.189 | 1.239 | chr19 | 48551100 | 48614074 | - | protein_coding |
| ENSG00000183628.14_4 | DGCR6 | 1.61E-07 | 1.86E-06 | 1.188 | 2.900 | chr22 | 18893541 | 18901751 | + | protein_coding |
| ENSG00000242797.3_6 | GLYCTK-AS1 | 0.01786316 | 0.09031724 | 1.188 | -0.960 | chr3 | 52322596 | 52333083 | - | lncRNA |
| ENSG00000146021.15_6 | KLHL3 | 7.71E-05 | 0.00060719 | 1.187 | 1.573 | chr5 | 136953189 | 137071778 | - | protein_coding |
| ENSG00000119669.5_3 | IRF2BPL | 9.88E-07 | 1.02E-05 | 1.185 | 5.521 | chr14 | 77490886 | 77495051 | - | protein_coding |
| ENSG00000254413.8_4 | CHKB-CPT1B | 0.0284052 | 0.13614114 | 1.184 | -1.181 | chr22 | 51007298 | 51021394 | - | protein_coding |
| ENSG00000204257.15_5 | HLA-DMA | 0.00064234 | 0.0043377 | 1.184 | 0.701 | chr6 | 32916390 | 32936871 | - | protein_coding |
| ENSG00000118985.16_4 | ELL2 | 1.11E-13 | 3.12E-12 | 1.183 | 5.982 | chr5 | 95220802 | 95297555 | - | protein_coding |
| ENSG00000160323.18_3 | ADAMTS13 | 0.00036778 | 0.00259814 | 1.183 | 1.484 | chr9 | 136279478 | 136324524 | + | protein_coding |
| ENSG00000206077.11_4 | ZDHHC11B | 0.00092011 | 0.0060279 | 1.183 | 0.792 | chr5 | 710470 | 784844 | - | protein_coding |
| ENSG00000167637.17_4 | ZNF283 | 1.60E-10 | 2.82E-09 | 1.182 | 4.730 | chr19 | 44331473 | 44356169 | + | protein_coding |
| ENSG00000253537.3_5 | PCDHGA7 | 2.62E-05 | 0.00022336 | 1.181 | 1.731 | chr5 | 140762306 | 140892542 | + | protein_coding |
| ENSG00000168040.4_2 | FADD | 0.00055045 | 0.00376622 | 1.181 | 3.092 | chr11 | 70049269 | 70053496 | + | protein_coding |
| ENSG00000168405.17_5 | CMAHP | 0.01716178 | 0.08701864 | 1.180 | -1.032 | chr6 | 25061854 | 25452491 | - | transcribed_unitary_pseudogene |
| ENSG00000180264.11_7 | ADGRD2 | 0.0075541 | 0.04144955 | 1.180 | 0.037 | chr9 | 127213704 | 127240868 | + | protein_coding |
| ENSG00000174718.12_6 | RESF1 | 2.18E-11 | 4.32E-10 | 1.179 | 6.910 | chr12 | 32112304 | 32146041 | + | protein_coding |
| ENSG00000138764.15_5 | CCNG2 | 5.45E-08 | 6.67E-07 | 1.178 | 3.492 | chr4 | 78078360 | 78354542 | + | protein_coding |
| ENSG00000213433.5_4 | RPLP1P6 | 0.01436532 | 0.07408667 | 1.178 | -0.713 | chr5 | 151145420 | 151145939 | - | transcribed_processed_pseudogene |
| ENSG00000163513.19_5 | TGFBR2 | 8.58E-16 | 3.40E-14 | 1.177 | 8.334 | chr3 | 30648093 | 30735634 | + | protein_coding |
| ENSG00000261324.2_6 | AC010168.2 | 6.45E-06 | 5.94E-05 | 1.176 | 2.244 | chr12 | 14915438 | 14920865 | - | lncRNA |
| ENSG00000205220.12_5 | PSMB10 | 3.08E-05 | 0.00026037 | 1.172 | 1.878 | chr16 | 67968409 | 67970767 | - | protein_coding |
| ENSG00000204525.16_3 | HLA-C | 0.00155791 | 0.00974759 | 1.168 | 0.463 | chr6 | 31236526 | 31239907 | - | protein_coding |
| ENSG00000138642.14_3 | HERC6 | 5.58E-09 | 7.92E-08 | 1.167 | 3.915 | chr4 | 89299891 | 89364263 | + | protein_coding |
| ENSG00000177675.8_4 | CD163L1 | 0.00082704 | 0.00545841 | 1.167 | 0.668 | chr12 | 7499281 | 7632493 | - | protein_coding |
| ENSG00000214562.14_4 | NUTM2D | 0.00496746 | 0.02828744 | 1.166 | -0.118 | chr10 | 89117425 | 89130452 | + | protein_coding |
| ENSG00000128039.11_3 | SRD5A3 | 9.54E-08 | 1.13E-06 | 1.165 | 3.095 | chr4 | 56212409 | 56239267 | + | protein_coding |
| ENSG00000165309.14_5 | ARMC3 | 0.02342518 | 0.11478082 | 1.165 | -0.978 | chr10 | 23216953 | 23327452 | + | protein_coding |
| ENSG00000151023.17_4 | ENKUR | 0.00548392 | 0.03097035 | 1.162 | -0.022 | chr10 | 25270908 | 25351208 | - | protein_coding |
| ENSG00000250579.2_7 | AC022424.1 | 0.01014864 | 0.05424516 | 1.161 | -0.321 | chr5 | 5132214 | 5140232 | - | lncRNA |
| ENSG00000286169.1_1 | AHRR | 0.00014553 | 0.00110217 | 1.161 | 1.430 | chr5 | 271785 | 438406 | + | protein_coding |
| ENSG00000175938.7_5 | ORAI3 | 1.08E-06 | 1.11E-05 | 1.160 | 2.844 | chr16 | 30960389 | 30967782 | + | protein_coding |
| ENSG00000224520.2_4 | KRT8P45 | 0.00013206 | 0.00100483 | 1.160 | 2.080 | chr1 | 157043049 | 157044495 | + | processed_pseudogene |
| ENSG00000180525.12_4 | PRR26 | 9.02E-05 | 0.00070279 | 1.159 | 1.757 | chr10 | 695888 | 715521 | + | protein_coding |
| ENSG00000132801.7_3 | ZSWIM3 | 6.64E-05 | 0.00052983 | 1.159 | 1.657 | chr20 | 44486253 | 44507761 | + | protein_coding |
| ENSG00000136052.9_6 | SLC41A2 | 1.84E-11 | 3.67E-10 | 1.159 | 6.081 | chr12 | 105196331 | 105352522 | - | protein_coding |
| ENSG00000187244.12_4 | BCAM | 6.24E-06 | 5.76E-05 | 1.159 | 6.138 | chr19 | 45312328 | 45324678 | + | protein_coding |
| ENSG00000012822.16_6 | CALCOCO1 | 1.42E-12 | 3.38E-11 | 1.159 | 5.586 | chr12 | 54102301 | 54121529 | - | protein_coding |
| ENSG00000174233.11_4 | ADCY6 | 2.37E-12 | 5.43E-11 | 1.158 | 5.746 | chr12 | 49159977 | 49182820 | - | protein_coding |
| ENSG00000187554.14_7 | TLR5 | 0.02280008 | 0.11218196 | 1.157 | -1.049 | chr1 | 223282746 | 223316590 | - | protein_coding |
| ENSG00000114374.13_4 | USP9Y | 3.24E-12 | 7.20E-11 | 1.156 | 5.879 | chrY | 14649585 | 14972764 | + | protein_coding |
| ENSG00000240583.12_5 | AQP1 | 0.02271964 | 0.11194116 | 1.156 | -1.123 | chr7 | 30951309 | 30965132 | + | protein_coding |
| ENSG00000164976.9_5 | MYORG | 6.30E-09 | 8.87E-08 | 1.156 | 3.799 | chr9 | 34366664 | 34376896 | - | protein_coding |
| ENSG00000288009.1_2 | AL353135.2 | 0.02603752 | 0.12594842 | 1.154 | -1.201 | chr6 | 89840606 | 89855704 | - | lncRNA |
| ENSG00000069011.16_4 | PITX1 | 4.47E-08 | 5.54E-07 | 1.154 | 4.177 | chr5 | 134363424 | 134370503 | - | protein_coding |
| ENSG00000091583.11_5 | APOH | 2.34E-07 | 2.64E-06 | 1.153 | 3.889 | chr17 | 64208151 | 64252643 | - | protein_coding |

| | | | | | | | | | | |
|---|---|---|---|---|---|---|---|---|---|---|
| ENSG00000175170.16_8 | FAM182B | 0.00077108 | 0.00512336 | 1.152 | 0.990 | chr20 | 25743020 | 25848861 | - | lncRNA |
| ENSG00000235890.2_6 | TSPEAR-AS1 | 6.99E-05 | 0.00055462 | 1.152 | 1.551 | chr21 | 45926690 | 45936458 | + | lncRNA |
| ENSG00000182749.5_4 | PAQR7 | 3.62E-06 | 3.46E-05 | 1.151 | 2.688 | chr1 | 26187701 | 26197744 | - | protein_coding |
| ENSG00000251533.2_6 | LINC00605 | 0.00210375 | 0.01289318 | 1.151 | 0.367 | chr14 | 103653558 | 103655365 | - | lncRNA |
| ENSG00000088808.18_8 | PPP1R13B | 1.91E-08 | 2.49E-07 | 1.150 | 3.933 | chr14 | 104199532 | 104313912 | - | protein_coding |
| ENSG00000205133.12_7 | TRIQK | 4.38E-11 | 8.25E-10 | 1.149 | 5.009 | chr8 | 93895760 | 94029901 | - | protein_coding |
| ENSG00000080823.23_5 | MOK | 1.46E-09 | 2.27E-08 | 1.149 | 4.283 | chr14 | 102690837 | 102771527 | - | protein_coding |
| ENSG00000186814.14_7 | ZSCAN30 | 6.13E-07 | 6.49E-06 | 1.148 | 3.006 | chr18 | 32831022 | 32870209 | - | protein_coding |
| ENSG00000142623.11_4 | PADI1 | 1.89E-06 | 1.88E-05 | 1.148 | 2.555 | chr1 | 17531623 | 17572502 | + | protein_coding |
| ENSG00000226137.5_5 | BAIAP2-DT | 5.97E-09 | 8.43E-08 | 1.148 | 3.895 | chr17 | 79002930 | 79008681 | - | lncRNA |
| ENSG00000257354.2_7 | AC048341.1 | 5.13E-08 | 6.30E-07 | 1.147 | 4.075 | chr12 | 62996532 | 63015993 | + | lncRNA |
| ENSG00000131188.12_3 | PRR7 | 0.00079348 | 0.00525644 | 1.146 | 2.852 | chr5 | 176873446 | 176883287 | + | protein_coding |
| ENSG00000197766.8_4 | CFD | 0.00031324 | 0.00223954 | 1.146 | 2.713 | chr19 | 859664 | 863641 | + | protein_coding |
| ENSG00000162576.16_3 | MXRA8 | 1.14E-05 | 0.00010109 | 1.146 | 2.839 | chr1 | 1288069 | 1297157 | - | protein_coding |
| ENSG00000137673.9_4 | MMP7 | 5.18E-09 | 7.37E-08 | 1.145 | 4.236 | chr11 | 102391239 | 102401481 | - | protein_coding |
| ENSG00000089335.21_5 | ZNF302 | 8.53E-14 | 2.49E-12 | 1.145 | 6.385 | chr19 | 35168544 | 35177302 | + | protein_coding |
| ENSG00000285588.1_2 | AL160254.1 | 0.00013861 | 0.00105246 | 1.145 | 1.538 | chr13 | 67695213 | 67730756 | + | lncRNA |
| ENSG00000100060.18_7 | MFNG | 9.60E-05 | 0.00074383 | 1.144 | 1.669 | chr22 | 37865101 | 37882431 | - | protein_coding |
| ENSG00000268357.1_2 | VN1R81P | 0.00685988 | 0.03790963 | 1.144 | -0.105 | chr19 | 21306620 | 21306991 | + | unprocessed_pseudogene |
| ENSG00000100290.3_3 | BIK | 0.00254068 | 0.01531219 | 1.143 | 0.344 | chr22 | 43506756 | 43525718 | + | protein_coding |
| ENSG00000049540.17_8 | ELN | 0.00655797 | 0.03639969 | 1.143 | 0.324 | chr7 | 73442119 | 73484237 | + | protein_coding |
| ENSG00000010438.16_4 | PRSS3 | 0.00015522 | 0.00117008 | 1.142 | 1.511 | chr9 | 33750464 | 33799229 | + | protein_coding |
| ENSG00000250091.3_6 | DNAH10OS | 6.45E-05 | 0.00051597 | 1.142 | 1.833 | chr12 | 124410008 | 124419531 | - | lncRNA |
| ENSG00000001561.7_3 | ENPP4 | 7.36E-05 | 0.00058151 | 1.140 | 1.973 | chr6 | 46097726 | 46114425 | + | protein_coding |
| ENSG00000146166.17_5 | LGSN | 9.53E-11 | 1.72E-09 | 1.139 | 5.147 | chr6 | 63985856 | 64029888 | - | protein_coding |
| ENSG00000142459.8_6 | EVI5L | 1.08E-06 | 1.11E-05 | 1.139 | 2.982 | chr19 | 7895119 | 7929862 | + | protein_coding |
| ENSG00000156050.10_6 | FAM161B | 1.76E-05 | 0.00015307 | 1.139 | 2.101 | chr14 | 74398204 | 74417117 | - | protein_coding |
| ENSG00000135821.18_5 | GLUL | 2.94E-12 | 6.60E-11 | 1.138 | 5.713 | chr1 | 182350839 | 182361341 | - | protein_coding |
| ENSG00000099840.13_5 | IZUMO4 | 0.00446095 | 0.02570797 | 1.137 | 0.132 | chr19 | 2096428 | 2099592 | + | protein_coding |
| ENSG00000106537.8_3 | TSPAN13 | 5.95E-12 | 1.27E-10 | 1.136 | 5.497 | chr7 | 16793380 | 16824161 | + | protein_coding |
| ENSG00000257261.6_8 | AC008014.1 | 1.94E-05 | 0.00016825 | 1.135 | 1.991 | chr12 | 46777435 | 47270567 | + | lncRNA |
| ENSG00000168672.4_5 | LRATD2 | 7.12E-12 | 1.50E-10 | 1.134 | 5.359 | chr8 | 127564688 | 127570723 | - | protein_coding |
| ENSG00000225096.3_6 | AL445250.1 | 0.02785911 | 0.13366805 | 1.134 | -1.066 | chr6 | 58287716 | 58764642 | + | lncRNA |
| ENSG00000235162.8_4 | C12orf75 | 1.04E-13 | 2.98E-12 | 1.133 | 7.337 | chr12 | 105629068 | 105789875 | + | protein_coding |
| ENSG00000174837.15_5 | ADGRE1 | 1.41E-07 | 1.64E-06 | 1.132 | 3.287 | chr19 | 6887577 | 6940470 | + | protein_coding |
| ENSG00000079931.15_4 | MOXD1 | 8.12E-05 | 0.00063698 | 1.131 | 1.896 | chr6 | 132617194 | 132722614 | - | protein_coding |
| ENSG00000067048.17_5 | DDX3Y | 1.21E-13 | 3.38E-12 | 1.131 | 6.733 | chrY | 15016019 | 15032390 | + | protein_coding |
| ENSG00000078804.13_4 | TP53INP2 | 1.43E-05 | 0.00012566 | 1.131 | 2.680 | chr20 | 33292143 | 33301243 | + | protein_coding |
| ENSG00000254419.1_7 | AL139349.1 | 0.00952804 | 0.05117433 | 1.130 | -0.395 | chr20 | 57285851 | 57286689 | + | lncRNA |
| ENSG00000180155.20_7 | LYNX1 | 1.29E-07 | 1.50E-06 | 1.128 | 3.451 | chr8 | 143852615 | 143859226 | - | protein_coding |
| ENSG00000164442.10_5 | CITED2 | 1.10E-08 | 1.48E-07 | 1.128 | 3.783 | chr6 | 139692944 | 139695785 | - | protein_coding |
| ENSG00000249395.3_6 | CASC9 | 8.98E-06 | 8.11E-05 | 1.127 | 2.864 | chr8 | 76135362 | 76236976 | - | lncRNA |
| ENSG00000076864.19_7 | RAP1GAP | 7.00E-13 | 1.75E-11 | 1.127 | 6.779 | chr1 | 21922708 | 21995856 | - | protein_coding |
| ENSG00000117643.14_3 | MAN1C1 | 0.0064564 | 0.0359145 | 1.126 | 0.131 | chr1 | 25943959 | 26112698 | + | protein_coding |
| ENSG00000204852.16_5 | TCTN1 | 1.05E-11 | 2.16E-10 | 1.126 | 5.562 | chr12 | 111051832 | 111087235 | + | protein_coding |
| ENSG00000183779.7_3 | ZNF703 | 3.81E-05 | 0.00031616 | 1.125 | 2.130 | chr8 | 37553300 | 37557537 | + | protein_coding |
| ENSG00000267924.1_7 | AC139769.2 | 0.00088234 | 0.0057911 | 1.125 | 1.097 | chr19 | 24000401 | 24057503 | - | lncRNA |
| ENSG00000100106.22_8 | TRIOBP | 3.14E-12 | 7.01E-11 | 1.125 | 6.614 | chr22 | 38093055 | 38172563 | + | protein_coding |
| ENSG00000171813.14_3 | PWWP2B | 2.64E-06 | 2.56E-05 | 1.124 | 4.049 | chr10 | 134210672 | 134231363 | + | protein_coding |
| ENSG00000171889.5_7 | MIR31HG | 8.75E-09 | 1.20E-07 | 1.123 | 3.904 | chr9 | 21453801 | 21559899 | - | lncRNA |
| ENSG00000248121.8_2 | SMURF2P1 | 0.0209212 | 0.10397397 | 1.123 | -0.866 | chr17 | 28927814 | 28942998 | + | transcribed_unprocessed_pseudogene |
| ENSG00000138152.10_4 | BTBD16 | 0.00216337 | 0.01322664 | 1.120 | 0.643 | chr10 | 124030811 | 124097674 | + | protein_coding |
| ENSG00000130522.6_5 | JUND | 0.01005433 | 0.05378963 | 1.118 | 5.373 | chr19 | 18390504 | 18392432 | - | protein_coding |
| ENSG00000197279.4_3 | ZNF165 | 0.00023675 | 0.00172744 | 1.118 | 1.368 | chr6 | 28048346 | 28057341 | + | protein_coding |
| ENSG00000178764.8_4 | ZHX2 | 8.79E-06 | 7.96E-05 | 1.117 | 2.597 | chr8 | 123793894 | 123986750 | + | protein_coding |
| ENSG00000107742.13_4 | SPOCK2 | 0.00155812 | 0.00974759 | 1.116 | 0.653 | chr10 | 73818792 | 73848790 | - | protein_coding |
| ENSG00000116785.13_2 | CFHR3 | 0.00040395 | 0.00283449 | 1.114 | 1.270 | chr1 | 196743925 | 196764536 | + | protein_coding |
| ENSG00000081760.17_4 | AACS | 2.05E-10 | 3.56E-09 | 1.114 | 5.341 | chr12 | 125549980 | 125627879 | + | protein_coding |
| ENSG00000171222.10_3 | SCAND1 | 0.03051091 | 0.14544872 | 1.114 | 4.719 | chr20 | 34541539 | 34547394 | - | protein_coding |

| | | | | | | | | | | |
|---|---|---|---|---|---|---|---|---|---|---|
| ENSG00000254815.6_7 | LMNTD2-AS1 | 0.002644 | 0.01587561 | 1.112 | 0.490 | chr11 | 557595 | 560107 | + | lncRNA |
| ENSG00000279799.1_6 | AC006077.2 | 0.015133 | 0.07757302 | 1.110 | -0.444 | chr5 | 134240925 | 134241381 | + | TEC |
| ENSG00000287824.1_2 | AL627443.3 | 0.0058123 | 0.03264137 | 1.110 | 0.052 | chr6 | 166511599 | 166513077 | + | lncRNA |
| ENSG00000198910.14_6 | L1CAM | 2.80E-07 | 3.11E-06 | 1.110 | 3.261 | chrX | 153126969 | 153151627 | - | protein_coding |
| ENSG00000101222.12_4 | SPEF1 | 0.00792317 | 0.04331415 | 1.110 | 0.201 | chr20 | 3758151 | 3762095 | - | protein_coding |
| ENSG00000218891.5_4 | ZNF579 | 0.00522706 | 0.0296612 | 1.110 | 4.035 | chr19 | 56088140 | 56092214 | - | protein_coding |
| ENSG00000145819.18_7 | ARHGAP26 | 2.60E-13 | 6.88E-12 | 1.109 | 6.575 | chr5 | 142149942 | 142608576 | + | protein_coding |
| ENSG00000197580.13_6 | BCO2 | 0.00218992 | 0.01337515 | 1.109 | 0.633 | chr11 | 112046233 | 112095422 | + | protein_coding |
| ENSG00000165810.17_3 | BTNL9 | 0.00034673 | 0.00246514 | 1.107 | 1.427 | chr5 | 180467225 | 180488521 | + | protein_coding |
| ENSG00000003436.16_6 | TFPI | 2.68E-12 | 6.05E-11 | 1.107 | 7.332 | chr2 | 188328957 | 188430487 | - | protein_coding |
| ENSG00000179455.9_4 | MKRN3 | 0.00011429 | 0.00087599 | 1.107 | 1.872 | chr15 | 23810821 | 23875222 | + | protein_coding |
| ENSG00000113205.5_4 | PCDHB3 | 0.02276743 | 0.11208338 | 1.106 | -0.758 | chr5 | 140480057 | 140483411 | + | protein_coding |
| ENSG00000133640.20_5 | LRRIQ1 | 0.00022451 | 0.00164628 | 1.105 | 1.640 | chr12 | 85430092 | 85658235 | + | protein_coding |
| ENSG00000215039.7_5 | CD27-AS1 | 0.00481901 | 0.0275392 | 1.102 | 0.317 | chr12 | 6548167 | 6560733 | - | lncRNA |
| ENSG00000122971.9_3 | ACADS | 1.27E-06 | 1.29E-05 | 1.102 | 4.130 | chr12 | 121163577 | 121177811 | + | protein_coding |
| ENSG00000179222.17_4 | MAGED1 | 3.32E-13 | 8.64E-12 | 1.100 | 6.948 | chrX | 51546103 | 51645453 | + | protein_coding |
| ENSG00000204967.6 | PCDHA4 | 3.22E-06 | 3.10E-05 | 1.099 | 3.018 | chr5 | 140186659 | 140391929 | + | protein_coding |
| ENSG00000173077.16_5 | DEC1 | 0.0142289 | 0.07344708 | 1.098 | -0.409 | chr9 | 117613248 | 118164923 | + | lncRNA |
| ENSG00000118640.11_3 | VAMP8 | 8.14E-06 | 7.40E-05 | 1.097 | 2.696 | chr2 | 85788685 | 85809154 | + | protein_coding |
| ENSG00000230555.2_6 | AL450326.1 | 0.00585575 | 0.03287175 | 1.097 | 0.287 | chr10 | 43916186 | 43917548 | + | lncRNA |
| ENSG00000076826.9_2 | CAMSAP3 | 5.76E-06 | 5.35E-05 | 1.097 | 2.799 | chr19 | 7660788 | 7683190 | + | protein_coding |
| ENSG00000277053.4_4 | GTF2IP1 | 0.01033106 | 0.05510126 | 1.097 | 0.605 | chr7 | 74601106 | 74653445 | - | transcribed_unprocessed_pseudogene |
| ENSG00000145246.14_6 | ATP10D | 3.81E-07 | 4.14E-06 | 1.096 | 3.372 | chr4 | 47487292 | 47595503 | + | protein_coding |
| ENSG00000184588.18_6 | PDE4B | 1.51E-12 | 3.58E-11 | 1.095 | 6.470 | chr1 | 66258197 | 66840262 | + | protein_coding |
| ENSG00000236204.6_6 | LINC01376 | 0.00013625 | 0.00103543 | 1.092 | 2.012 | chr2 | 19167729 | 19547828 | - | lncRNA |
| ENSG00000283413.1_1 | AC008703.1 | 0.00058078 | 0.00395146 | 1.091 | 1.442 | chr5 | 158911138 | 158938695 | - | lncRNA |
| ENSG00000109758.8_3 | HGFAC | 0.03964482 | 0.18347733 | 1.091 | -1.015 | chr4 | 3443614 | 3451222 | + | protein_coding |
| ENSG00000168487.19_5 | BMP1 | 4.12E-11 | 7.82E-10 | 1.090 | 6.179 | chr8 | 22022653 | 22069839 | + | protein_coding |
| ENSG00000286315.1_2 | AC087175.1 | 0.00280433 | 0.01674771 | 1.089 | 0.501 | chr7 | 47291737 | 47294156 | - | lncRNA |
| ENSG00000167700.9_3 | MFSD3 | 0.00091562 | 0.00600068 | 1.087 | 5.092 | chr8 | 145734453 | 145736596 | + | protein_coding |
| ENSG00000182472.9_7 | CAPN12 | 5.37E-05 | 0.00043548 | 1.087 | 2.030 | chr19 | 39220827 | 39260544 | - | protein_coding |
| ENSG00000268895.6_6 | A1BG-AS1 | 0.00050987 | 0.00350034 | 1.085 | 1.443 | chr19 | 58859084 | 58866821 | + | lncRNA |
| ENSG00000170190.16_6 | SLC16A5 | 4.23E-10 | 7.04E-09 | 1.084 | 6.034 | chr17 | 73083822 | 73102257 | + | protein_coding |
| ENSG00000262558.1_7 | AC129507.3 | 0.00363539 | 0.02129572 | 1.084 | 0.456 | chr17 | 204528 | 206673 | + | lncRNA |
| ENSG00000132141.14_6 | CCT6B | 0.00162157 | 0.01010895 | 1.084 | 1.155 | chr17 | 33254878 | 33308097 | - | protein_coding |
| ENSG00000162337.12_4 | LRP5 | 1.84E-07 | 2.10E-06 | 1.084 | 7.164 | chr11 | 68080059 | 68216743 | + | protein_coding |
| ENSG00000182196.13_6 | ARL6IP4 | 0.0270155 | 0.13025356 | 1.084 | -0.833 | chr12 | 123464607 | 123467460 | + | protein_coding |
| ENSG00000050748.17_8 | MAPK9 | 6.61E-11 | 1.22E-09 | 1.084 | 6.525 | chr5 | 179660143 | 179719099 | - | protein_coding |
| ENSG00000237356.7_8 | AL365295.1 | 0.00033073 | 0.00235418 | 1.084 | 1.651 | chr14 | 53635728 | 54350458 | + | lncRNA |
| ENSG00000178226.11_5 | PRSS36 | 0.02201461 | 0.10870863 | 1.083 | -0.773 | chr16 | 31150247 | 31161404 | - | protein_coding |
| ENSG00000064687.13_3 | ABCA7 | 3.40E-09 | 5.02E-08 | 1.083 | 4.905 | chr19 | 1039996 | 1065571 | + | protein_coding |
| ENSG00000100219.16_7 | XBP1 | 3.10E-12 | 6.92E-11 | 1.082 | 6.537 | chr22 | 29190543 | 29196585 | - | protein_coding |
| ENSG00000158246.8_5 | TENT5B | 0.00097984 | 0.00638154 | 1.082 | 1.449 | chr1 | 27331511 | 27339341 | - | protein_coding |
| ENSG00000149633.12_9 | KIAA1755 | 0.02867505 | 0.13732336 | 1.082 | -0.832 | chr20 | 36838905 | 36889174 | - | protein_coding |
| ENSG00000130700.7_4 | GATA5 | 0.00128356 | 0.00817675 | 1.082 | 1.124 | chr20 | 61038553 | 61051051 | - | protein_coding |
| ENSG00000224081.9_6 | SLC44A3-AS1 | 0.00145756 | 0.00918312 | 1.082 | 1.009 | chr1 | 95079370 | 95320982 | - | transcribed_processed_pseudogene |
| ENSG00000185507.21_6 | IRF7 | 1.15E-05 | 0.00010228 | 1.081 | 2.741 | chr11 | 612553 | 615983 | - | protein_coding |
| ENSG00000111727.12_5 | HCFC2 | 6.86E-09 | 9.57E-08 | 1.081 | 4.280 | chr12 | 104458309 | 104500302 | + | protein_coding |
| ENSG00000239305.7_5 | RNF103 | 2.89E-07 | 3.20E-06 | 1.080 | 5.162 | chr2 | 86830521 | 86850989 | - | protein_coding |
| ENSG00000221963.6_3 | APOL6 | 1.33E-08 | 1.77E-07 | 1.079 | 4.441 | chr22 | 36044493 | 36064451 | + | protein_coding |
| ENSG00000187609.16_5 | EXD3 | 3.81E-05 | 0.00031671 | 1.079 | 2.460 | chr9 | 140201348 | 140317663 | - | protein_coding |
| ENSG00000204961.6_3 | PCDHA9 | 0.02325023 | 0.11408085 | 1.079 | -0.774 | chr5 | 140227048 | 140391929 | + | protein_coding |
| ENSG00000287808.1_2 | AL355999.1 | 0.02242157 | 0.11059522 | 1.078 | -0.524 | chr1 | 182806320 | 182808696 | - | lncRNA |
| ENSG00000103227.18_7 | LMF1 | 4.11E-07 | 4.45E-06 | 1.076 | 3.646 | chr16 | 903634 | 1031318 | - | protein_coding |
| ENSG00000196466.10_5 | ZNF799 | 2.47E-06 | 2.41E-05 | 1.076 | 3.079 | chr19 | 12500830 | 12512085 | - | protein_coding |
| ENSG00000013619.14_7 | MAMLD1 | 2.22E-05 | 0.00019108 | 1.076 | 2.442 | chrX | 149529689 | 149682448 | + | protein_coding |
| ENSG00000100344.11_4 | PNPLA3 | 0.0001603 | 0.00120678 | 1.076 | 1.780 | chr22 | 44319672 | 44360368 | + | protein_coding |
| ENSG00000265660.1 | MIR4664 | 0.00509392 | 0.02894268 | 1.075 | 0.696 | chr8 | 144815253 | 144815323 | - | miRNA |

| | | | | | | | | | | |
|---|---|---|---|---|---|---|---|---|---|---|
| ENSG00000120658.13_4 | ENOX1 | 0.00670726 | 0.03713551 | 1.074 | 0.454 | chr13 | 43787654 | 44361044 | - | protein_coding |
| ENSG00000204060.7_4 | FOXO6 | 0.02021527 | 0.10062909 | 1.074 | -0.422 | chr1 | 41827594 | 41849262 | + | protein_coding |
| ENSG00000105641.4_3 | SLC5A5 | 0.01958449 | 0.09790593 | 1.074 | -0.619 | chr19 | 17982754 | 18005983 | + | protein_coding |
| ENSG00000226823.1_5 | SUGT1P1 | 0.0088497 | 0.04780491 | 1.073 | 0.111 | chr9 | 33503653 | 33510709 | - | unprocessed_pseudogene |
| ENSG00000197077.13_4 | KIAA1671 | 6.90E-10 | 1.12E-08 | 1.073 | 5.010 | chr22 | 25348697 | 25593415 | + | protein_coding |
| ENSG00000109452.12_7 | INPP4B | 2.38E-11 | 4.67E-10 | 1.073 | 5.781 | chr4 | 142944313 | 143768585 | - | protein_coding |
| ENSG00000178163.8_5 | ZNF518B | 6.64E-11 | 1.22E-09 | 1.073 | 5.558 | chr4 | 10441504 | 10459050 | - | protein_coding |
| ENSG00000234737.1_4 | KRT18P15 | 0.0193258 | 0.09677607 | 1.073 | -0.527 | chr3 | 32300432 | 32301722 | + | processed_pseudogene |
| ENSG00000105204.14_6 | DYRK1B | 4.65E-10 | 7.69E-09 | 1.072 | 5.020 | chr19 | 40315990 | 40324841 | - | protein_coding |
| ENSG00000198929.13_6 | NOS1AP | 0.02193702 | 0.10835558 | 1.071 | -0.431 | chr1 | 162039481 | 162340265 | + | protein_coding |
| ENSG00000184271.17_9 | POU6F1 | 0.00228524 | 0.01387623 | 1.070 | 1.035 | chr12 | 51580719 | 51611492 | - | protein_coding |
| ENSG00000256139.2_7 | AC007637.1 | 0.00860597 | 0.04661653 | 1.070 | 0.179 | chr12 | 109549023 | 109563399 | + | lncRNA |
| ENSG00000119242.8_6 | CCDC92 | 1.02E-07 | 1.21E-06 | 1.069 | 4.346 | chr12 | 124403207 | 124457378 | - | protein_coding |
| ENSG00000130584.11_5 | ZBTB46 | 0.00080215 | 0.00530203 | 1.068 | 1.485 | chr20 | 62375019 | 62463391 | - | protein_coding |
| ENSG00000225828.1_6 | FAM229A | 0.00414268 | 0.02402972 | 1.068 | 0.751 | chr1 | 32826871 | 32829879 | - | protein_coding |
| ENSG00000285999.1_2 | AC025442.2 | 0.02357789 | 0.11540177 | 1.067 | -0.673 | chr5 | 65381233 | 65389765 | + | lncRNA |
| ENSG00000159640.16_6 | ACE | 0.02360538 | 0.11550451 | 1.066 | -0.620 | chr17 | 61554422 | 61575741 | + | protein_coding |
| ENSG00000280002.1_5 | AC078925.4 | 0.00027515 | 0.00198966 | 1.066 | 1.634 | chr12 | 131472857 | 131474573 | - | TEC |
| ENSG00000263711.6_8 | LINC02864 | 1.40E-10 | 2.49E-09 | 1.066 | 5.499 | chr18 | 70818476 | 70931715 | - | lncRNA |
| ENSG00000185442.13_7 | FAM174B | 4.58E-07 | 4.93E-06 | 1.065 | 3.469 | chr15 | 93160678 | 93353114 | - | protein_coding |
| ENSG00000111907.21_8 | TPD52L1 | 1.02E-09 | 1.62E-08 | 1.065 | 5.029 | chr6 | 125440195 | 125585553 | + | protein_coding |
| ENSG00000152192.8_5 | POU4F1 | 7.34E-05 | 0.00058091 | 1.064 | 2.188 | chr13 | 79172497 | 79177687 | - | protein_coding |
| ENSG00000143554.14_7 | SLC27A3 | 0.00018113 | 0.00134935 | 1.064 | 1.954 | chr1 | 153746830 | 153752633 | + | protein_coding |
| ENSG00000127415.13_5 | IDUA | 1.60E-06 | 1.61E-05 | 1.064 | 3.339 | chr4 | 980785 | 998352 | + | protein_coding |
| ENSG00000173480.11_7 | ZNF417 | 1.04E-08 | 1.40E-07 | 1.063 | 4.462 | chr19 | 58411664 | 58427978 | - | protein_coding |
| ENSG00000273472.1_7 | AC096733.3 | 0.01213275 | 0.06353154 | 1.063 | -0.051 | chr4 | 141677682 | 141679075 | + | lncRNA |
| ENSG00000230850.3 | RP11-782C8.1 | 0.00155522 | 0.00973975 | 1.062 | 1.094 | chr1 | 143119061 | 143233417 | + | lincRNA |
| ENSG00000140297.13_6 | GCNT3 | 2.55E-12 | 5.78E-11 | 1.062 | 7.560 | chr15 | 59887074 | 59932438 | + | protein_coding |
| ENSG00000135404.11_5 | CD63 | 3.96E-13 | 1.03E-11 | 1.062 | 8.092 | chr12 | 56119107 | 56123491 | - | protein_coding |
| ENSG00000275202.1_7 | AL161421.1 | 0.02310695 | 0.11344052 | 1.060 | -0.396 | chr13 | 49549103 | 49551003 | - | lncRNA |
| ENSG00000284052.1_1 | AC006460.2 | 0.00603444 | 0.03375755 | 1.060 | 0.333 | chr2 | 191472386 | 191514566 | - | lncRNA |
| ENSG00000105997.22_5 | HOXA3 | 9.14E-05 | 0.00071163 | 1.059 | 2.039 | chr7 | 27145803 | 27192200 | - | protein_coding |
| ENSG00000258096.1_6 | AC025031.2 | 0.00840103 | 0.0455759 | 1.058 | 0.349 | chr12 | 46765246 | 46767561 | + | lncRNA |
| ENSG00000135318.12_4 | NT5E | 3.19E-12 | 7.11E-11 | 1.058 | 6.570 | chr6 | 86159302 | 86205509 | + | protein_coding |
| ENSG00000140986.8_5 | RPL3L | 0.01924267 | 0.09638695 | 1.058 | -0.529 | chr16 | 1993975 | 2007607 | - | protein_coding |
| ENSG00000248932.6_8 | AC046134.2 | 0.00020308 | 0.00149774 | 1.056 | 1.785 | chr3 | 139108603 | 139501541 | + | lncRNA |
| ENSG00000040608.14_6 | RTN4R | 0.00027022 | 0.00195638 | 1.056 | 2.196 | chr22 | 20228938 | 20270769 | - | protein_coding |
| ENSG00000170873.18_5 | MTSS1 | 0.00029758 | 0.00213965 | 1.055 | 2.034 | chr8 | 125563031 | 125740670 | - | protein_coding |
| ENSG00000144452.15_5 | ABCA12 | 5.96E-06 | 5.53E-05 | 1.054 | 3.146 | chr2 | 215796266 | 216003349 | - | protein_coding |
| ENSG00000273062.1_6 | AL449106.1 | 0.02355461 | 0.11531959 | 1.053 | -0.539 | chr1 | 178693441 | 178695420 | - | lncRNA |
| ENSG00000247317.3_7 | LY6E-DT | 0.01152731 | 0.06064724 | 1.053 | 0.097 | chr8 | 144063155 | 144099854 | - | lncRNA |
| ENSG00000239389.8_5 | PCDHA13 | 0.0433782 | 0.19827645 | 1.052 | -1.263 | chr5 | 140261709 | 140391932 | + | protein_coding |
| ENSG00000214274.9_3 | ANG | 0.00016629 | 0.00124663 | 1.051 | 2.172 | chr14 | 21152336 | 21167130 | + | protein_coding |
| ENSG00000106952.7_3 | TNFSF8 | 0.02497033 | 0.12144755 | 1.051 | -0.530 | chr9 | 117655623 | 117692875 | - | protein_coding |
| ENSG00000245025.3_8 | AC107959.1 | 0.0004793 | 0.00331353 | 1.050 | 1.635 | chr8 | 22842109 | 22876848 | - | lncRNA |
| ENSG00000108691.9_3 | CCL2 | 1.80E-07 | 2.06E-06 | 1.050 | 4.222 | chr17 | 32582237 | 32584222 | + | protein_coding |
| ENSG00000125398.8_5 | SOX9 | 2.15E-10 | 3.72E-09 | 1.049 | 5.475 | chr17 | 70117161 | 70122557 | + | protein_coding |
| ENSG00000171469.11_7 | ZNF561 | 2.50E-08 | 3.19E-07 | 1.049 | 4.370 | chr19 | 9715356 | 9731912 | - | protein_coding |
| ENSG00000184381.20_7 | PLA2G6 | 6.18E-06 | 5.71E-05 | 1.047 | 3.082 | chr22 | 38507502 | 38610785 | - | protein_coding |
| ENSG00000268230.5_8 | AC012313.3 | 0.00101341 | 0.00658567 | 1.047 | 1.234 | chr19 | 58858220 | 58874117 | - | lncRNA |
| ENSG00000232909.1_7 | AL157823.2 | 0.02881852 | 0.1378989 | 1.047 | -0.743 | chr6 | 35701644 | 35704724 | - | lncRNA |
| ENSG00000165731.20_6 | RET | 0.01507435 | 0.07731696 | 1.046 | -0.092 | chr10 | 43572512 | 43625799 | + | protein_coding |
| ENSG00000141401.12_3 | IMPA2 | 1.73E-10 | 3.05E-09 | 1.045 | 5.512 | chr18 | 11981024 | 12030876 | + | protein_coding |
| ENSG00000185015.8_3 | CA13 | 0.0109858 | 0.05805614 | 1.045 | -0.025 | chr8 | 86132816 | 86196302 | + | protein_coding |
| ENSG00000212916.4_2 | MAP10 | 9.71E-06 | 8.73E-05 | 1.044 | 2.806 | chr1 | 232940638 | 232944153 | + | protein_coding |
| ENSG00000072858.10_3 | SIDT1 | 0.0144164 | 0.074307 | 1.044 | -0.063 | chr3 | 113251143 | 113348425 | + | protein_coding |
| ENSG00000185565.12_5 | LSAMP | 0.00828773 | 0.04503878 | 1.043 | 0.516 | chr3 | 115521210 | 116858236 | - | protein_coding |
| ENSG00000285796.1_1 | AL162458.1 | 3.79E-05 | 0.00031521 | 1.043 | 2.645 | chr20 | 44555157 | 44563694 | - | lncRNA |
| ENSG00000150967.18_8 | ABCB9 | 6.10E-06 | 5.64E-05 | 1.041 | 3.332 | chr12 | 123405498 | 123466196 | - | protein_coding |

| | | | | | | | | | | |
|---|---|---|---|---|---|---|---|---|---|---|
| ENSG00000196937.11_5 | FAM3C | 3.71E-08 | 4.65E-07 | 1.041 | 4.380 | chr7 | 120988932 | 121036418 | - | protein_coding |
| ENSG00000176809.10_5 | LRRC37A3 | 7.13E-05 | 0.00056523 | 1.041 | 2.207 | chr17 | 62850430 | 62915598 | - | protein_coding |
| ENSG00000113916.18_5 | BCL6 | 2.55E-08 | 3.24E-07 | 1.040 | 4.681 | chr3 | 187439165 | 187463513 | - | protein_coding |
| ENSG00000099814.16_4 | CEP170B | 1.83E-07 | 2.10E-06 | 1.039 | 5.827 | chr14 | 105331605 | 105363107 | + | protein_coding |
| ENSG00000287390.1_2 | AC006525.1 | 0.01937558 | 0.09699799 | 1.037 | -0.129 | chr5 | 129991076 | 130279871 | - | lncRNA |
| ENSG00000261512.2_7 | AC092368.3 | 0.0409256 | 0.18881463 | 1.037 | -1.123 | chr16 | 46656773 | 46658363 | - | lncRNA |
| ENSG00000196167.10_7 | COLCA1 | 0.01003651 | 0.05371044 | 1.035 | 0.209 | chr11 | 111161512 | 111176223 | - | lncRNA |
| ENSG00000099849.15_5 | RASSF7 | 2.52E-09 | 3.79E-08 | 1.035 | 5.704 | chr11 | 560404 | 564025 | + | protein_coding |
| ENSG00000218048.2_5 | AL121949.1 | 4.88E-05 | 0.00039889 | 1.035 | 2.458 | chr6 | 64150671 | 64153485 | + | unprocessed_pseudogene |
| ENSG00000186472.20_4 | PCLO | 3.87E-07 | 4.20E-06 | 1.034 | 3.937 | chr7 | 82383328 | 82792246 | - | protein_coding |
| ENSG00000259605.3_7 | AC074212.1 | 0.03906218 | 0.18120568 | 1.034 | -0.746 | chr19 | 46268043 | 46273064 | + | lncRNA |
| ENSG00000237686.7_7 | AL109615.3 | 0.00076354 | 0.00507512 | 1.033 | 1.422 | chr6 | 43963460 | 44042389 | - | lncRNA |
| ENSG00000180855.16_5 | ZNF443 | 4.78E-06 | 4.50E-05 | 1.033 | 3.135 | chr19 | 12540520 | 12551835 | - | protein_coding |
| ENSG00000130733.10_5 | YIPF2 | 1.13E-05 | 0.00010108 | 1.032 | 3.364 | chr19 | 11032861 | 11039357 | - | protein_coding |
| ENSG00000177406.5_7 | NINJ2-AS1 | 3.99E-05 | 0.00033048 | 1.032 | 2.684 | chr12 | 740024 | 773362 | + | lncRNA |
| ENSG00000259032.2_4 | ENSAP2 | 0.03288694 | 0.15544089 | 1.030 | -0.700 | chr14 | 83159048 | 83159399 | - | processed_pseudogene |
| ENSG00000172137.19_6 | CALB2 | 1.30E-06 | 1.32E-05 | 1.030 | 3.732 | chr16 | 71392616 | 71424339 | + | protein_coding |
| ENSG00000151065.14_4 | DCP1B | 2.36E-07 | 2.66E-06 | 1.029 | 3.836 | chr12 | 2055219 | 2113701 | - | protein_coding |
| ENSG00000181666.18_6 | ZNF875 | 1.35E-09 | 2.12E-08 | 1.029 | 5.427 | chr19 | 37803739 | 37860267 | + | protein_coding |
| ENSG00000165102.15_5 | HGSNAT | 2.17E-10 | 3.74E-09 | 1.027 | 5.725 | chr8 | 42995607 | 43057998 | + | protein_coding |
| ENSG00000108639.7_5 | SYNGR2 | 5.97E-11 | 1.10E-09 | 1.026 | 7.058 | chr17 | 76164639 | 76169608 | + | protein_coding |
| ENSG00000145908.12_3 | ZNF300 | 2.47E-08 | 3.15E-07 | 1.026 | 4.883 | chr5 | 150273954 | 150284545 | - | protein_coding |
| ENSG00000120549.18_6 | KIAA1217 | 1.71E-07 | 1.97E-06 | 1.025 | 4.065 | chr10 | 23983675 | 24836777 | + | protein_coding |
| ENSG00000249158.6_4 | PCDHA11 | 0.00010311 | 0.00079476 | 1.025 | 2.200 | chr5 | 140247768 | 140391929 | + | protein_coding |
| ENSG00000197363.9_4 | ZNF517 | 2.87E-06 | 2.77E-05 | 1.025 | 3.476 | chr8 | 146024261 | 146036554 | + | protein_coding |
| ENSG00000227124.11_7 | ZNF717 | 0.00036479 | 0.00257807 | 1.025 | 1.854 | chr3 | 75727811 | 75834734 | - | protein_coding |
| ENSG00000176153.12_4 | GPX2 | 4.31E-12 | 9.36E-11 | 1.024 | 9.096 | chr14 | 65405870 | 65409623 | - | protein_coding |
| ENSG00000177990.12_5 | DPY19L2 | 4.94E-06 | 4.64E-05 | 1.024 | 3.186 | chr12 | 63952693 | 64062719 | - | protein_coding |
| ENSG00000139173.10_5 | TMEM117 | 0.00026937 | 0.00195261 | 1.023 | 1.983 | chr12 | 44229770 | 44783545 | + | protein_coding |
| ENSG00000170604.5_3 | IRF2BP1 | 0.00019642 | 0.00145284 | 1.023 | 5.427 | chr19 | 46386866 | 46389399 | - | protein_coding |
| ENSG00000119121.22_5 | TRPM6 | 5.62E-06 | 5.23E-05 | 1.022 | 3.468 | chr9 | 77337411 | 77503010 | - | protein_coding |
| ENSG00000226711.7_6 | FAM66C | 0.00043896 | 0.00305605 | 1.022 | 2.059 | chr12 | 8332805 | 8368747 | + | lncRNA |
| ENSG00000124615.20_7 | MOCS1 | 2.28E-05 | 0.00019558 | 1.021 | 2.799 | chr6 | 39867354 | 39902290 | - | protein_coding |
| ENSG00000080031.10_7 | PTPRH | 1.45E-05 | 0.00012724 | 1.021 | 3.102 | chr19 | 55692615 | 55720874 | - | protein_coding |
| ENSG00000089486.17_6 | CDIP1 | 2.12E-06 | 2.09E-05 | 1.021 | 3.457 | chr16 | 4560670 | 4588829 | - | protein_coding |
| ENSG00000166987.15_7 | MBD6 | 1.30E-06 | 1.32E-05 | 1.021 | 3.664 | chr12 | 57914493 | 57923931 | + | protein_coding |
| ENSG00000229848.1_6 | AC139149.1 | 0.00683221 | 0.03776848 | 1.020 | 1.191 | chr17 | 79481073 | 79494802 | + | lncRNA |
| ENSG00000118402.6_4 | ELOVL4 | 0.00047686 | 0.0032992 | 1.020 | 1.896 | chr6 | 80624531 | 80657270 | - | protein_coding |
| ENSG00000198816.7_4 | ZNF358 | 4.27E-05 | 0.00035193 | 1.019 | 2.807 | chr19 | 7580178 | 7585911 | + | protein_coding |
| ENSG00000162755.14_4 | KLHDC9 | 0.00436097 | 0.02520541 | 1.019 | 0.772 | chr1 | 161068151 | 161070136 | + | protein_coding |
| ENSG00000258056.2_7 | AC009779.3 | 0.00012766 | 0.00097344 | 1.017 | 2.295 | chr12 | 56122888 | 56124636 | + | lncRNA |
| ENSG00000006015.17_5 | REX1BD | 0.00022762 | 0.00166769 | 1.016 | 4.550 | chr19 | 18699495 | 18703146 | + | protein_coding |
| ENSG00000156966.7_3 | B3GNT7 | 7.18E-05 | 0.0005688 | 1.015 | 3.239 | chr2 | 232260421 | 232265875 | + | protein_coding |
| ENSG00000127561.15_3 | SYNGR3 | 7.80E-06 | 7.11E-05 | 1.015 | 3.459 | chr16 | 2039661 | 2044276 | + | protein_coding |
| ENSG00000183878.15_7 | UTY | 2.56E-08 | 3.25E-07 | 1.015 | 4.886 | chrY | 15360259 | 15592553 | - | protein_coding |
| ENSG00000272077.1_6 | AC124045.1 | 0.00473604 | 0.02711737 | 1.014 | 0.723 | chr3 | 44708904 | 44710856 | + | lncRNA |
| ENSG00000170949.17_4 | ZNF160 | 2.31E-08 | 2.97E-07 | 1.014 | 5.454 | chr19 | 53569859 | 53606689 | - | protein_coding |
| ENSG00000213759.10_5 | UGT2B11 | 0.02806616 | 0.13458877 | 1.013 | -0.339 | chr4 | 70065669 | 70080466 | - | protein_coding |
| ENSG00000235052.1_7 | AL021154.1 | 0.03674881 | 0.17177484 | 1.012 | -0.770 | chr1 | 23875630 | 23877406 | - | lncRNA |
| ENSG00000172830.13_6 | SSH3 | 1.05E-08 | 1.42E-07 | 1.012 | 4.863 | chr11 | 67070949 | 67080078 | + | protein_coding |
| ENSG00000282885.2_1 | AL627171.4 | 0.0354102 | 0.16621648 | 1.012 | -0.514 | chr14 | 50327894 | 50331044 | - | lncRNA |
| ENSG00000287569.1_1 | AL078587.2 | 0.02214571 | 0.10929521 | 1.012 | -0.224 | chr20 | 25990428 | 25991879 | + | lncRNA |
| ENSG00000127528.5_2 | KLF2 | 0.00036211 | 0.00256317 | 1.011 | 2.205 | chr19 | 16435628 | 16438685 | + | protein_coding |
| ENSG00000198324.13_7 | PHETA1 | 1.48E-05 | 0.00012989 | 1.010 | 3.292 | chr12 | 111798455 | 111806925 | - | protein_coding |
| ENSG00000120256.11_5 | LRP11 | 0.00079545 | 0.00526375 | 1.010 | 1.626 | chr6 | 150139893 | 150185495 | - | protein_coding |
| ENSG00000170846.17_5 | AC093323.1 | 3.92E-08 | 4.88E-07 | 1.010 | 4.565 | chr4 | 6665123 | 6678482 | + | protein_coding |
| ENSG00000123096.11_3 | SSPN | 0.01996116 | 0.09956481 | 1.007 | -0.082 | chr12 | 26274924 | 26452223 | + | protein_coding |
| ENSG00000107338.10_5 | SHB | 5.79E-07 | 6.16E-06 | 1.007 | 4.178 | chr9 | 37915895 | 38069224 | - | protein_coding |
| ENSG00000198393.8_4 | ZNF26 | 0.00401333 | 0.0233633 | 1.006 | 1.110 | chr12 | 133562951 | 133609538 | + | protein_coding |
| ENSG00000142065.14_4 | ZFP14 | 2.96E-08 | 3.74E-07 | 1.006 | 4.786 | chr19 | 36825355 | 36870103 | - | protein_coding |

| | | | | | | | | | | |
|---|---|---|---|---|---|---|---|---|---|---|
| ENSG00000114423.22_9 | CBLB | 6.24E-08 | 7.61E-07 | 1.006 | 4.425 | chr3 | 105374305 | 105588396 | - | protein_coding |
| ENSG00000105472.13_3 | CLEC11A | 0.00285557 | 0.01702512 | 1.006 | 1.789 | chr19 | 51226621 | 51228975 | + | protein_coding |
| ENSG00000197457.10_6 | STMN3 | 7.55E-06 | 6.90E-05 | 1.004 | 3.554 | chr20 | 62271058 | 62289035 | - | protein_coding |
| ENSG00000255836.1_5 | AC131206.1 | 0.00772266 | 0.04230913 | 1.004 | 0.579 | chr12 | 7591376 | 7592586 | - | processed_pseudogene |
| ENSG00000276449.1_6 | AC004076.2 | 0.03196107 | 0.15166995 | 1.003 | 0.043 | chr19 | 57961057 | 57964379 | + | lncRNA |
| ENSG00000177946.4 | CENPBD1 | 3.15E-05 | 0.00026495 | 0.999 | 3.017 | chr16 | 90036206 | 90038942 | - | protein_coding |
| ENSG00000173598.14_5 | NUDT4 | 1.14E-09 | 1.80E-08 | 0.999 | 5.435 | chr12 | 93771659 | 93801922 | + | protein_coding |
| ENSG00000152894.14_8 | PTPRK | 1.28E-10 | 2.30E-09 | 0.998 | 6.226 | chr6 | 128289924 | 128841819 | - | protein_coding |
| ENSG00000075399.14_8 | VPS9D1 | 0.00011165 | 0.00085685 | 0.998 | 3.059 | chr16 | 89773542 | 89787306 | - | protein_coding |
| ENSG00000254004.7_4 | ZNF260 | 3.89E-10 | 6.50E-09 | 0.996 | 6.342 | chr19 | 37001589 | 37019173 | - | protein_coding |
| ENSG00000116514.16_4 | RNF19B | 1.25E-07 | 1.47E-06 | 0.995 | 4.589 | chr1 | 33402046 | 33430286 | - | protein_coding |
| ENSG00000177045.10_7 | SIX5 | 0.00019577 | 0.00144866 | 0.995 | 2.966 | chr19 | 46268043 | 46272510 | - | protein_coding |
| ENSG00000168993.15_5 | CPLX1 | 0.04744482 | 0.21443708 | 0.995 | -1.015 | chr4 | 778745 | 819917 | - | protein_coding |
| ENSG00000067646.12_4 | ZFY | 5.32E-08 | 6.53E-07 | 0.994 | 4.670 | chrY | 2803322 | 2850547 | + | protein_coding |
| ENSG00000186951.16_5 | PPARA | 3.84E-06 | 3.65E-05 | 0.994 | 3.965 | chr22 | 46546424 | 46639653 | + | protein_coding |
| ENSG00000159184.7_3 | HOXB13 | 1.70E-06 | 1.70E-05 | 0.994 | 3.633 | chr17 | 46802125 | 46806540 | - | protein_coding |
| ENSG00000287356.1_1 | AL590822.3 | 0.05991237 | 0.26160311 | 0.994 | 0.540 | chr1 | 2246479 | 2254524 | + | lncRNA |
| ENSG00000283526.1_5 | PRRT1B | 0.0001662 | 0.00124645 | 0.994 | 2.521 | chr9 | 134420901 | 134434007 | + | protein_coding |
| ENSG00000259660.2_4 | DNM1P47 | 0.03565804 | 0.16724824 | 0.994 | -0.567 | chr15 | 102291235 | 102334505 | + | transcribed_unprocessed_pseudogene |
| ENSG00000259363.6_6 | AC090825.1 | 0.01581799 | 0.08071175 | 0.993 | 0.002 | chr15 | 100345998 | 100457023 | + | lncRNA |
| ENSG00000095739.11_3 | BAMBI | 1.93E-10 | 3.37E-09 | 0.993 | 6.093 | chr10 | 28966439 | 28971861 | + | protein_coding |
| ENSG00000230953.2_5 | AC099677.1 | 0.03933674 | 0.18228894 | 0.993 | -0.672 | chr1 | 53386094 | 53386268 | + | processed_pseudogene |
| ENSG00000267265.5_6 | AC011476.3 | 0.00660962 | 0.03665205 | 0.992 | 0.802 | chr19 | 55517561 | 55559454 | + | lncRNA |
| ENSG00000135052.16_6 | GOLM1 | 1.81E-11 | 3.63E-10 | 0.992 | 7.965 | chr9 | 88641061 | 88715088 | - | protein_coding |
| ENSG00000138411.13_6 | HECW2 | 6.59E-07 | 6.95E-06 | 0.992 | 4.097 | chr2 | 197053823 | 197458408 | - | protein_coding |
| ENSG00000257489.6_6 | AC010203.1 | 0.03593922 | 0.16838926 | 0.991 | -0.480 | chr12 | 100566974 | 100593561 | - | transcribed_unprocessed_pseudogene |
| ENSG00000227533.6_5 | SLC2A1-AS1 | 0.01513106 | 0.07757302 | 0.991 | 0.155 | chr1 | 43424720 | 43462485 | + | lncRNA |
| ENSG00000197959.14_5 | DNM3 | 0.00072329 | 0.0048293 | 0.991 | 1.936 | chr1 | 171810638 | 172387606 | + | protein_coding |
| ENSG00000198682.13_5 | PAPSS2 | 1.93E-10 | 3.37E-09 | 0.990 | 6.411 | chr10 | 89419370 | 89507462 | + | protein_coding |
| ENSG00000253958.2_3 | CLDN23 | 0.04507474 | 0.20513395 | 0.989 | -0.673 | chr8 | 8559447 | 8561606 | + | protein_coding |
| ENSG00000180921.7_4 | FAM83H | 2.70E-05 | 0.00022977 | 0.989 | 6.368 | chr8 | 144806103 | 144820404 | - | protein_coding |
| ENSG00000129824.16_4 | RPS4Y1 | 2.72E-11 | 5.30E-10 | 0.988 | 7.544 | chrY | 2709643 | 2800041 | + | protein_coding |
| ENSG00000007944.15_3 | MYLIP | 9.09E-07 | 9.47E-06 | 0.988 | 4.111 | chr6 | 16129317 | 16148479 | + | protein_coding |
| ENSG00000165626.17_5 | BEND7 | 1.94E-07 | 2.21E-06 | 0.987 | 4.227 | chr10 | 13480484 | 13570974 | - | protein_coding |
| ENSG00000172955.17_6 | ADH6 | 0.03234543 | 0.15316663 | 0.986 | -0.443 | chr4 | 100123795 | 100140694 | - | protein_coding |
| ENSG00000185250.16_7 | PPIL6 | 0.00159267 | 0.00993925 | 0.986 | 1.522 | chr6 | 109711418 | 109762374 | - | protein_coding |
| ENSG00000065357.20_4 | DGKA | 2.15E-08 | 2.78E-07 | 0.986 | 4.954 | chr12 | 56321103 | 56347811 | + | protein_coding |
| ENSG00000122592.8_6 | HOXA7 | 0.01847292 | 0.09295056 | 0.985 | 0.173 | chr7 | 27193335 | 27197555 | - | protein_coding |
| ENSG00000001461.17_3 | NIPAL3 | 1.26E-07 | 1.47E-06 | 0.985 | 4.464 | chr1 | 24742292 | 24799466 | + | protein_coding |
| ENSG00000197653.16_7 | DNAH10 | 0.00208635 | 0.01279097 | 0.984 | 1.558 | chr12 | 124246735 | 124420267 | + | protein_coding |
| ENSG00000213799.13_5 | ZNF845 | 1.55E-08 | 2.04E-07 | 0.984 | 5.547 | chr19 | 53837002 | 53860159 | + | protein_coding |
| ENSG00000166825.14_6 | ANPEP | 3.59E-06 | 3.44E-05 | 0.983 | 3.647 | chr15 | 90328126 | 90358633 | - | protein_coding |
| ENSG00000101049.17_7 | SGK2 | 1.63E-06 | 1.63E-05 | 0.983 | 3.760 | chr20 | 42187608 | 42216877 | + | protein_coding |
| ENSG00000112425.15_4 | EPM2A | 0.00019156 | 0.00141987 | 0.982 | 2.443 | chr6 | 145703671 | 146057159 | - | protein_coding |
| ENSG00000197483.10_4 | ZNF628 | 0.00070202 | 0.00471028 | 0.982 | 2.701 | chr19 | 55987984 | 55995854 | + | protein_coding |
| ENSG00000137331.12_4 | IER3 | 0.00154341 | 0.0096726 | 0.981 | 2.229 | chr6 | 30710976 | 30712325 | - | protein_coding |
| ENSG00000145217.14_5 | SLC26A1 | 6.38E-05 | 0.00051065 | 0.981 | 3.010 | chr4 | 972861 | 987228 | - | protein_coding |
| ENSG00000263002.8_3 | ZNF234 | 1.32E-06 | 1.34E-05 | 0.980 | 3.905 | chr19 | 44645707 | 44664466 | + | protein_coding |
| ENSG00000138771.16_5 | SHROOM3 | 1.28E-09 | 2.01E-08 | 0.980 | 5.956 | chr4 | 77356382 | 77704406 | + | protein_coding |
| ENSG00000142544.7_4 | CTU1 | 0.00062111 | 0.00420232 | 0.979 | 2.392 | chr19 | 51600863 | 51611666 | - | protein_coding |
| ENSG00000271605.1 | MILR1 | 0.02285848 | 0.11243817 | 0.979 | 0.049 | chr17 | 62461569 | 62464760 | + | protein_coding |
| ENSG00000133488.15_4 | SEC14L4 | 4.68E-06 | 4.40E-05 | 0.979 | 3.892 | chr22 | 30884889 | 30901698 | - | protein_coding |
| ENSG00000273045.6_5 | C2orf15 | 0.00320478 | 0.018921 | 0.979 | 1.243 | chr2 | 99757948 | 99767950 | + | protein_coding |
| ENSG00000182791.5_3 | CCDC87 | 0.0025147 | 0.01517105 | 0.978 | 1.600 | chr11 | 66357647 | 66360534 | - | protein_coding |
| ENSG00000146530.14_7 | VWDE | 0.0007285 | 0.00485856 | 0.978 | 2.114 | chr7 | 12370511 | 12443567 | - | protein_coding |
| ENSG00000166147.13_4 | FBN1 | 9.30E-09 | 1.27E-07 | 0.977 | 5.168 | chr15 | 48700503 | 48938046 | - | protein_coding |
| ENSG00000168528.12_4 | SERINC2 | 1.11E-05 | 9.85E-05 | 0.977 | 3.649 | chr1 | 31882412 | 31907527 | + | protein_coding |
| ENSG00000167074.15_3 | TEF | 0.0002073 | 0.00152511 | 0.976 | 2.398 | chr22 | 41763337 | 41795330 | + | protein_coding |
| ENSG00000122863.6_3 | CHST3 | 5.25E-10 | 8.61E-09 | 0.975 | 6.654 | chr10 | 73724153 | 73773316 | + | protein_coding |

| | | | | | | | | | | |
|---|---|---|---|---|---|---|---|---|---|---|
| ENSG00000279794.1_1 | AC024580.2 | 0.04458444 | 0.20321494 | 0.975 | -0.848 | chr19 | 56661186 | 56661859 | + | TEC |
| ENSG00000127720.8_4 | METTL25 | 0.00010714 | 0.00082403 | 0.975 | 2.986 | chr12 | 82752307 | 82873018 | + | protein_coding |
| ENSG00000196118.12_7 | CCDC189 | 0.0526347 | 0.23449365 | 0.975 | -0.962 | chr16 | 30768744 | 30773542 | - | protein_coding |
| ENSG00000111700.12_5 | SLCO1B3 | 0.00010267 | 0.00079174 | 0.971 | 3.691 | chr12 | 20963636 | 21069845 | + | protein_coding |
| ENSG00000186765.11_2 | FSCN2 | 0.00263854 | 0.01584819 | 0.971 | 1.766 | chr17 | 79495422 | 79504156 | + | protein_coding |
| ENSG00000101439.9_5 | CST3 | 1.28E-09 | 2.01E-08 | 0.971 | 6.778 | chr20 | 23607343 | 23619110 | - | protein_coding |
| ENSG00000251369.8_3 | ZNF550 | 5.77E-06 | 5.35E-05 | 0.971 | 3.727 | chr19 | 58046625 | 58071231 | - | protein_coding |
| ENSG00000160867.15_4 | FGFR4 | 1.49E-05 | 0.00013058 | 0.969 | 5.815 | chr5 | 176513906 | 176525145 | + | protein_coding |
| ENSG00000158828.8_4 | PINK1 | 4.59E-07 | 4.94E-06 | 0.968 | 4.243 | chr1 | 20959951 | 20978004 | + | protein_coding |
| ENSG00000185551.15_4 | NR2F2 | 1.39E-07 | 1.62E-06 | 0.968 | 4.848 | chr15 | 96869167 | 96883492 | + | protein_coding |
| ENSG00000198829.6_3 | SUCNR1 | 0.02132121 | 0.10572537 | 0.968 | 0.286 | chr3 | 151591431 | 151602407 | + | protein_coding |
| ENSG00000164056.11_5 | SPRY1 | 0.00059744 | 0.00405605 | 0.967 | 2.152 | chr4 | 124317950 | 124324915 | + | protein_coding |
| ENSG00000160932.11_5 | LY6E | 1.17E-05 | 0.00010399 | 0.967 | 6.417 | chr8 | 144099399 | 144105249 | + | protein_coding |
| ENSG00000273018.6_9 | FAM106A | 0.01779903 | 0.09004422 | 0.967 | 0.222 | chr17 | 18414535 | 18455019 | - | lncRNA |
| ENSG00000154358.21_5 | OBSCN | 3.70E-08 | 4.64E-07 | 0.967 | 6.116 | chr1 | 228395764 | 228566577 | + | protein_coding |
| ENSG00000235316.1_4 | DUSP8P5 | 0.0262059 | 0.12665945 | 0.966 | 0.152 | chr10 | 75491582 | 75493396 | + | processed_pseudogene |
| ENSG00000165269.13_5 | AQP7 | 0.01472733 | 0.07573388 | 0.966 | 0.422 | chr9 | 33383177 | 33402680 | - | protein_coding |
| ENSG00000204264.11_6 | PSMB8 | 2.50E-05 | 0.00021317 | 0.964 | 3.259 | chr6 | 32808494 | 32812456 | - | protein_coding |
| ENSG00000229619.4_6 | MBNL1-AS1 | 0.04768738 | 0.21542379 | 0.964 | -0.643 | chr3 | 151963051 | 151987346 | - | lncRNA |
| ENSG00000185215.9_6 | TNFAIP2 | 1.97E-10 | 3.42E-09 | 0.964 | 9.090 | chr14 | 103589779 | 103603776 | + | protein_coding |
| ENSG00000228801.6_6 | AC064807.1 | 0.00233582 | 0.01415916 | 0.963 | 1.676 | chr8 | 52811828 | 52862434 | + | lncRNA |
| ENSG00000264278.1_8 | ZNF236-DT | 0.0415721 | 0.19130056 | 0.962 | -0.375 | chr18 | 74506688 | 74534251 | - | lncRNA |
| ENSG00000285938.1_2 | AC072022.2 | 0.07390282 | 0.31303439 | 0.961 | -1.242 | chr3 | 187461474 | 187463816 | + | protein_coding |
| ENSG00000153071.15_6 | DAB2 | 7.76E-07 | 8.13E-06 | 0.961 | 4.302 | chr5 | 39371777 | 39462402 | - | protein_coding |
| ENSG00000186496.12_6 | ZNF396 | 0.00405542 | 0.02356541 | 0.961 | 1.190 | chr18 | 32946658 | 32957301 | - | protein_coding |
| ENSG00000103316.11_4 | CRYM | 0.00097786 | 0.00637094 | 0.960 | 1.972 | chr16 | 21250195 | 21314404 | - | protein_coding |
| ENSG00000197892.13_4 | KIF13B | 1.90E-09 | 2.89E-08 | 0.960 | 6.248 | chr8 | 28924795 | 29120641 | - | protein_coding |
| ENSG00000135100.17_2 | HNF1A | 6.96E-06 | 6.39E-05 | 0.959 | 3.566 | chr12 | 121416346 | 121440315 | + | protein_coding |
| ENSG00000131115.16_4 | ZNF227 | 7.72E-07 | 8.10E-06 | 0.958 | 4.305 | chr19 | 44711700 | 44741421 | + | protein_coding |
| ENSG00000106078.19_6 | COBL | 2.20E-06 | 2.16E-05 | 0.957 | 4.145 | chr7 | 51083909 | 51384515 | - | protein_coding |
| ENSG00000228672.4_3 | PROB1 | 0.0003184 | 0.00227371 | 0.957 | 2.964 | chr5 | 138726281 | 138730793 | - | protein_coding |
| ENSG00000099812.9_5 | MISP | 1.40E-05 | 0.00012276 | 0.957 | 7.906 | chr19 | 751112 | 764318 | + | protein_coding |
| ENSG00000176842.15_4 | IRX5 | 0.00300133 | 0.01782241 | 0.957 | 1.657 | chr16 | 54964777 | 54968397 | + | protein_coding |
| ENSG00000217801.10_6 | AL390719.1 | 0.00557818 | 0.03143275 | 0.956 | 1.033 | chr1 | 995088 | 1004735 | + | transcribed_unprocessed_pseudogene |
| ENSG00000269069.1_6 | AC007842.1 | 0.00266962 | 0.01602403 | 0.956 | 1.512 | chr19 | 40529291 | 40531409 | - | processed_pseudogene |
| ENSG00000146674.15_8 | IGFBP3 | 6.20E-10 | 1.01E-08 | 0.955 | 7.191 | chr7 | 45951844 | 45961473 | - | protein_coding |
| ENSG00000205085.11_5 | FAM71F2 | 0.02273728 | 0.11199092 | 0.955 | 0.100 | chr7 | 128312342 | 128327926 | + | protein_coding |
| ENSG00000162520.15_7 | SYNC | 0.00105368 | 0.00683488 | 0.955 | 2.080 | chr1 | 33145507 | 33169197 | - | protein_coding |
| ENSG00000258867.6_6 | LINC01146 | 0.00962754 | 0.05166187 | 0.954 | 0.724 | chr14 | 88490863 | 88564369 | + | lncRNA |
| ENSG00000130045.16_3 | NXNL2 | 0.04646106 | 0.21047263 | 0.954 | -0.802 | chr9 | 91148948 | 91199189 | + | protein_coding |
| ENSG00000104774.13_5 | MAN2B1 | 1.45E-06 | 1.46E-05 | 0.953 | 4.427 | chr19 | 12757325 | 12777556 | - | protein_coding |
| ENSG00000066185.13_4 | ZMYND12 | 0.05322751 | 0.23677846 | 0.953 | -0.701 | chr1 | 42896000 | 42921924 | - | protein_coding |
| ENSG00000132517.15_5 | SLC52A1 | 0.0020258 | 0.01245842 | 0.952 | 1.718 | chr17 | 4935895 | 4955304 | - | protein_coding |
| ENSG00000267128.2_6 | RNF157-AS1 | 0.04841495 | 0.2181558 | 0.950 | -0.756 | chr17 | 74134513 | 74150731 | + | lncRNA |
| ENSG00000230623.6_7 | AC104461.1 | 0.05393527 | 0.23944727 | 0.950 | -0.914 | chr1 | 200302321 | 200447797 | + | lncRNA |
| ENSG00000039068.19_6 | CDH1 | 2.21E-08 | 2.85E-07 | 0.950 | 5.332 | chr16 | 68771195 | 68869444 | + | protein_coding |
| ENSG00000127663.15_7 | KDM4B | 8.22E-06 | 7.47E-05 | 0.948 | 3.947 | chr19 | 4969124 | 5153609 | + | protein_coding |
| ENSG00000064547.14_5 | LPAR2 | 0.00889861 | 0.04802601 | 0.948 | 0.940 | chr19 | 19734464 | 19739739 | - | protein_coding |
| ENSG00000215068.8_6 | AC025171.2 | 1.82E-05 | 0.00015805 | 0.948 | 3.615 | chr5 | 43041677 | 43054705 | + | lncRNA |
| ENSG00000228775.8_7 | WEE2-AS1 | 0.04668793 | 0.21139258 | 0.948 | -0.477 | chr7 | 141403803 | 141438146 | - | lncRNA |
| ENSG00000110697.13_6 | PITPNM1 | 8.10E-05 | 0.00063544 | 0.947 | 6.083 | chr11 | 67259239 | 67273734 | - | protein_coding |
| ENSG00000105708.9_4 | ZNF14 | 5.03E-05 | 0.000409 | 0.947 | 3.137 | chr19 | 19821281 | 19843921 | - | protein_coding |
| ENSG00000130787.14_3 | HIP1R | 6.21E-06 | 5.74E-05 | 0.947 | 5.315 | chr12 | 123319000 | 123347508 | + | protein_coding |
| ENSG00000180998.12_5 | GPR137C | 0.05117965 | 0.22887208 | 0.947 | -0.804 | chr14 | 53019554 | 53104431 | + | protein_coding |
| ENSG00000261652.3_3 | C15orf65 | 0.00105543 | 0.00684372 | 0.946 | 1.990 | chr15 | 55700693 | 55710996 | + | protein_coding |
| ENSG00000164850.15_6 | GPER1 | 0.06465834 | 0.27964573 | 0.946 | -0.378 | chr7 | 1121844 | 1133451 | + | protein_coding |
| ENSG00000255867.1_6 | DENND5B-AS1 | 0.0229956 | 0.11298753 | 0.945 | 0.068 | chr12 | 31742857 | 31768600 | + | lncRNA |
| ENSG00000168961.17_8 | LGALS9 | 0.04270114 | 0.19568509 | 0.945 | -0.514 | chr17 | 25956824 | 25976586 | + | protein_coding |

| | | | | | | | | | | |
|---|---|---|---|---|---|---|---|---|---|---|
| ENSG00000196155.13_5 | AC040160.1 | 1.90E-08 | 2.47E-07 | 0.943 | 5.589 | chr16 | 67311413 | 67323402 | + | protein_coding |
| ENSG00000266278.1_6 | LINC01910 | 0.00462492 | 0.02654957 | 0.943 | 1.342 | chr18 | 68047376 | 68051827 | - | lncRNA |
| ENSG00000151090.19_8 | THRB | 9.26E-05 | 0.00072015 | 0.942 | 3.550 | chr3 | 24158644 | 24537247 | - | protein_coding |
| ENSG00000163644.15_5 | PPM1K | 0.00491238 | 0.02801869 | 0.942 | 1.380 | chr4 | 89178772 | 89205921 | - | protein_coding |
| ENSG00000121417.14_4 | ZNF211 | 4.82E-06 | 4.53E-05 | 0.942 | 4.084 | chr19 | 58141763 | 58155409 | + | protein_coding |
| ENSG00000233725.7_7 | NRAD1 | 0.02293756 | 0.11276471 | 0.941 | 0.174 | chr13 | 44482805 | 44604597 | + | lncRNA |
| ENSG00000137642.13_3 | SORL1 | 1.15E-07 | 1.35E-06 | 0.941 | 5.356 | chr11 | 121323023 | 121504472 | + | protein_coding |
| ENSG00000106688.12_3 | SLC1A1 | 8.25E-05 | 0.00064661 | 0.941 | 2.986 | chr9 | 4490468 | 4587469 | + | protein_coding |
| ENSG00000185261.14_6 | KIAA0825 | 0.00577034 | 0.03244353 | 0.940 | 1.525 | chr5 | 93488671 | 93954302 | - | protein_coding |
| ENSG00000171798.18_3 | KNDC1 | 0.00392611 | 0.02288555 | 0.940 | 1.946 | chr10 | 134973723 | 135039916 | + | protein_coding |
| ENSG00000089163.4_2 | SIRT4 | 0.01892633 | 0.09501672 | 0.939 | 0.300 | chr12 | 120740119 | 120751052 | + | protein_coding |
| ENSG00000076513.17_4 | ANKRD13A | 1.12E-07 | 1.32E-06 | 0.938 | 5.263 | chr12 | 110436991 | 110477568 | + | protein_coding |
| ENSG00000161328.11_3 | LRRC56 | 0.02366338 | 0.11572456 | 0.937 | 1.167 | chr11 | 537527 | 554912 | + | protein_coding |
| ENSG00000266714.9_8 | MYO15B | 5.94E-09 | 8.38E-08 | 0.936 | 6.909 | chr17 | 73584139 | 73622929 | + | protein_coding |
| ENSG00000228024.1_4 | CYP1D1P | 0.06478364 | 0.28005131 | 0.936 | -1.049 | chr9 | 75657661 | 75671444 | + | unprocessed_pseudogene |
| ENSG00000263004.2_7 | AC007114.1 | 0.01450445 | 0.07469582 | 0.936 | 0.733 | chr17 | 55149175 | 55162385 | - | lncRNA |
| ENSG00000072609.17_5 | CHFR | 1.99E-07 | 2.27E-06 | 0.936 | 5.258 | chr12 | 133398773 | 133532890 | - | protein_coding |
| ENSG00000182912.6_5 | TSPEAR-AS2 | 0.00225126 | 0.01370269 | 0.936 | 1.911 | chr21 | 45937099 | 45945835 | + | lncRNA |
| ENSG00000141497.14_5 | ZMYND15 | 0.04404669 | 0.20112504 | 0.936 | -0.565 | chr17 | 4643128 | 4649414 | + | protein_coding |
| ENSG00000106415.13_5 | GLCCI1 | 0.00297716 | 0.01769073 | 0.935 | 1.623 | chr7 | 8008427 | 8133902 | + | protein_coding |
| ENSG00000178229.8_3 | ZNF543 | 9.93E-06 | 8.90E-05 | 0.935 | 4.044 | chr19 | 57831840 | 57842138 | + | protein_coding |
| ENSG00000169891.18_3 | REPS2 | 2.68E-05 | 0.00022787 | 0.934 | 3.406 | chrX | 16964781 | 17171395 | + | protein_coding |
| ENSG00000088836.14_9 | SLC4A11 | 6.14E-05 | 0.00049289 | 0.934 | 3.496 | chr20 | 3208063 | 3220205 | - | protein_coding |
| ENSG00000139278.10_5 | GLIPR1 | 5.06E-05 | 0.00041126 | 0.934 | 3.926 | chr12 | 75874533 | 75897643 | + | protein_coding |
| ENSG00000198732.10_3 | SMOC1 | 9.44E-07 | 9.81E-06 | 0.933 | 4.517 | chr14 | 70320848 | 70499083 | + | protein_coding |
| ENSG00000128283.7_7 | CDC42EP1 | 6.30E-07 | 6.66E-06 | 0.933 | 4.969 | chr22 | 37956487 | 37965412 | + | protein_coding |
| ENSG00000238261.5 | RP11-435B5.5 | 0.01661501 | 0.08439078 | 0.932 | 0.463 | chr1 | 143377281 | 143401533 | + | lincRNA |
| ENSG00000226471.6_7 | Z93930.2 | 0.04472633 | 0.20370489 | 0.932 | -0.432 | chr22 | 29196671 | 29244547 | + | lncRNA |
| ENSG00000257869.1_6 | AL139023.1 | 0.02694238 | 0.12993628 | 0.931 | 0.405 | chr14 | 28787347 | 28887818 | + | lncRNA |
| ENSG00000142961.14_4 | MOB3C | 8.48E-05 | 0.00066266 | 0.931 | 3.176 | chr1 | 47073387 | 47082563 | - | protein_coding |
| ENSG00000111275.13_4 | ALDH2 | 6.69E-09 | 9.37E-08 | 0.930 | 6.075 | chr12 | 112204691 | 112255336 | + | protein_coding |
| ENSG00000172183.15_3 | ISG20 | 4.18E-05 | 0.00034472 | 0.930 | 3.535 | chr15 | 89179384 | 89199714 | + | protein_coding |
| ENSG00000167216.17_8 | KATNAL2 | 0.00142422 | 0.0089954 | 0.930 | 2.036 | chr18 | 44497455 | 44628614 | + | protein_coding |
| ENSG00000162552.15_4 | WNT4 | 0.00790306 | 0.04321752 | 0.930 | 0.983 | chr1 | 22443806 | 22470462 | - | protein_coding |
| ENSG00000111540.16_5 | RAB5B | 2.83E-09 | 4.21E-08 | 0.929 | 6.410 | chr12 | 56367697 | 56390467 | + | protein_coding |
| ENSG00000242147.2_8 | LINC02657 | 0.00595726 | 0.0333678 | 0.927 | 1.210 | chr10 | 5636947 | 5674398 | - | lncRNA |
| ENSG00000173281.5_4 | PPP1R3B | 3.72E-06 | 3.55E-05 | 0.927 | 4.133 | chr8 | 8993765 | 9009084 | - | protein_coding |
| ENSG00000259374.2_5 | NDUFB4P11 | 0.05252303 | 0.23411355 | 0.927 | -0.660 | chr14 | 103606012 | 103608296 | + | transcribed_processed_pseudogene |
| ENSG00000267288.2_7 | AC138150.2 | 0.00114476 | 0.00737456 | 0.926 | 2.094 | chr17 | 43246167 | 43248951 | - | lncRNA |
| ENSG00000134253.10_7 | TRIM45 | 9.86E-05 | 0.00076303 | 0.926 | 3.125 | chr1 | 117653682 | 117665209 | - | protein_coding |
| ENSG00000170581.14_6 | STAT2 | 1.59E-08 | 2.09E-07 | 0.926 | 5.770 | chr12 | 56735381 | 56753951 | - | protein_coding |
| ENSG00000198692.10_4 | EIF1AY | 1.83E-07 | 2.09E-06 | 0.925 | 5.160 | chrY | 22737662 | 22755040 | + | protein_coding |
| ENSG00000131401.11_4 | NAPSB | 0.02015165 | 0.10040223 | 0.925 | 0.367 | chr19 | 50837053 | 50848024 | - | transcribed_unprocessed_pseudogene |
| ENSG00000081386.12_3 | ZNF510 | 5.61E-07 | 5.98E-06 | 0.925 | 4.929 | chr9 | 99518147 | 99540411 | - | protein_coding |
| ENSG00000068971.14_4 | PPP2R5B | 0.00012346 | 0.00094183 | 0.924 | 3.329 | chr11 | 64685025 | 64701947 | + | protein_coding |
| ENSG00000173926.6_4 | MARCHF3 | 0.00506095 | 0.02876452 | 0.923 | 1.497 | chr5 | 126203406 | 126366250 | - | protein_coding |
| ENSG00000164845.16_5 | FAM86FP | 0.00795182 | 0.04345737 | 0.923 | 1.289 | chr12 | 8385108 | 8395544 | - | transcribed_unprocessed_pseudogene |
| ENSG00000100647.8_3 | SUSD6 | 1.39E-05 | 0.00012195 | 0.922 | 3.903 | chr14 | 70078313 | 70181861 | + | protein_coding |
| ENSG00000213015.9_6 | ZNF580 | 4.85E-05 | 0.00039666 | 0.922 | 3.767 | chr19 | 56146382 | 56154836 | + | protein_coding |
| ENSG00000254531.1_6 | AP001816.1 | 0.00207055 | 0.01271602 | 0.922 | 2.012 | chr4 | 102268937 | 102270040 | + | lncRNA |
| ENSG00000242110.8_6 | AMACR | 0.01698673 | 0.08618024 | 0.921 | 0.585 | chr5 | 33986270 | 34008209 | - | protein_coding |
| ENSG00000138617.15_4 | PARP16 | 3.38E-06 | 3.24E-05 | 0.919 | 4.210 | chr15 | 65526798 | 65592956 | - | protein_coding |
| ENSG00000237004.4_4 | ZNRF2P1 | 0.02121395 | 0.10522283 | 0.919 | 0.951 | chr7 | 32768181 | 32793356 | + | transcribed_processed_pseudogene |
| ENSG00000205436.7_5 | EXOC3L4 | 0.00152287 | 0.00956083 | 0.919 | 2.682 | chr14 | 103566481 | 103576896 | + | protein_coding |
| ENSG00000135452.10_5 | TSPAN31 | 3.70E-06 | 3.53E-05 | 0.918 | 4.334 | chr12 | 58131796 | 58144002 | + | protein_coding |
| ENSG00000171159.5_3 | C9orf16 | 0.00037765 | 0.00266364 | 0.918 | 3.954 | chr9 | 130922544 | 130926203 | + | protein_coding |
| ENSG00000160999.10_1 | SH2B2 | 0.01137043 | 0.05989278 | 0.918 | 1.096 | chr7 | 101928382 | 101962178 | + | protein_coding |

| | | | | | | | | | | |
|---|---|---|---|---|---|---|---|---|---|---|
| ENSG00000111676.15_4 | ATN1 | 5.58E-07 | 5.95E-06 | 0.917 | 4.884 | chr12 | 7033626 | 7051484 | + | protein_coding |
| ENSG00000141526.16_3 | SLC16A3 | 0.00125831 | 0.00804479 | 0.916 | 6.373 | chr17 | 80186273 | 80219005 | + | protein_coding |
| ENSG00000254739.1_6 | AC137894.1 | 0.08154567 | 0.34138123 | 0.916 | -0.999 | chr11 | 528907 | 529659 | + | lncRNA |
| ENSG00000111653.20_5 | ING4 | 1.14E-05 | 0.00010115 | 0.915 | 4.036 | chr12 | 6759467 | 6772308 | - | protein_coding |
| ENSG00000150977.10_3 | RILPL2 | 0.00016095 | 0.00121116 | 0.914 | 3.298 | chr12 | 123895230 | 123921264 | - | protein_coding |
| ENSG00000200312.1 | RN7SKP255 | 0.0382164 | 0.17760678 | 0.914 | 0.171 | chr14 | 90178855 | 90179167 | + | misc_RNA |
| ENSG00000111276.11_6 | CDKN1B | 4.56E-08 | 5.64E-07 | 0.914 | 5.576 | chr12 | 12867992 | 12875303 | + | protein_coding |
| ENSG00000183044.12_5 | ABAT | 0.0264703 | 0.12779834 | 0.914 | 0.406 | chr16 | 8768453 | 8878432 | + | protein_coding |
| ENSG00000125384.7_3 | PTGER2 | 0.06751902 | 0.29011135 | 0.913 | -0.294 | chr14 | 52781023 | 52795316 | + | protein_coding |
| ENSG00000175764.14_4 | TTLL11 | 0.01838138 | 0.09259482 | 0.913 | 0.677 | chr9 | 124584207 | 124855885 | - | protein_coding |
| ENSG00000159714.11_5 | ZDHHC1 | 0.00035246 | 0.00250246 | 0.913 | 3.175 | chr16 | 67428055 | 67450736 | - | protein_coding |
| ENSG00000183066.14_6 | WBP2NL | 0.05892171 | 0.25814877 | 0.912 | -0.774 | chr22 | 42394729 | 42454460 | + | protein_coding |
| ENSG00000226174.7_5 | TEX22 | 0.0226092 | 0.1114553 | 0.912 | 0.584 | chr14 | 105864875 | 105916443 | + | protein_coding |
| ENSG00000254615.3_6 | AC027031.2 | 0.06150469 | 0.26750401 | 0.912 | -0.832 | chr8 | 107282372 | 107285130 | + | lncRNA |
| ENSG00000167528.12_4 | ZNF641 | 1.52E-06 | 1.53E-05 | 0.911 | 4.498 | chr12 | 48730963 | 48745197 | - | protein_coding |
| ENSG00000154188.10_5 | ANGPT1 | 0.00094754 | 0.00619172 | 0.911 | 2.499 | chr8 | 108261710 | 108510283 | - | protein_coding |
| ENSG00000181588.16_5 | MEX3D | 0.00017879 | 0.00133302 | 0.910 | 3.522 | chr19 | 1554668 | 1568057 | - | protein_coding |
| ENSG00000275183.2_4 | LENG9 | 0.00055051 | 0.00376622 | 0.910 | 2.827 | chr19 | 54972912 | 54974961 | - | protein_coding |
| ENSG00000186815.12_5 | TPCN1 | 1.09E-08 | 1.46E-07 | 0.910 | 6.504 | chr12 | 113658855 | 113736390 | + | protein_coding |
| ENSG00000180353.11_4 | HCLS1 | 0.07460425 | 0.31544355 | 0.910 | -1.067 | chr3 | 121350246 | 121379774 | - | protein_coding |
| ENSG00000205464.12_3 | ATP6AP1L | 0.04233438 | 0.19425488 | 0.910 | -0.329 | chr5 | 81575281 | 81682796 | + | protein_coding |
| ENSG00000182141.10_8 | ZNF708 | 0.00025464 | 0.00185117 | 0.908 | 2.897 | chr19 | 21473962 | 21512227 | - | protein_coding |
| ENSG00000284948.1_3 | AC107959.4 | 0.05212012 | 0.2326679 | 0.908 | -0.373 | chr8 | 22941868 | 22961071 | + | lncRNA |
| ENSG00000235978.6_7 | AC018816.1 | 0.06645261 | 0.28601345 | 0.907 | -0.729 | chr3 | 4855978 | 4928977 | - | lncRNA |
| ENSG00000208892.1 | SNORA49 | 0.0030305 | 0.01798361 | 0.907 | 2.095 | chr12 | 132515769 | 132515904 | + | snoRNA |
| ENSG00000007384.15_5 | RHBDF1 | 1.48E-05 | 0.00012963 | 0.907 | 4.798 | chr16 | 108058 | 126354 | - | protein_coding |
| ENSG00000230565.1_7 | ZNF32-AS2 | 0.03565316 | 0.16724824 | 0.906 | 0.121 | chr10 | 44141390 | 44143467 | + | lncRNA |
| ENSG00000169660.16_6 | HEXD | 2.96E-06 | 2.85E-05 | 0.906 | 4.887 | chr17 | 80376194 | 80400521 | + | protein_coding |
| ENSG00000175003.15_6 | SLC22A1 | 0.07824961 | 0.32936763 | 0.906 | -1.067 | chr6 | 160542847 | 160579750 | + | protein_coding |
| ENSG00000163840.10_3 | DTX3L | 7.69E-08 | 9.26E-07 | 0.906 | 5.574 | chr3 | 122283185 | 122294050 | + | protein_coding |
| ENSG00000178951.9_5 | ZBTB7A | 4.16E-06 | 3.93E-05 | 0.906 | 4.785 | chr19 | 4043301 | 4066897 | - | protein_coding |
| ENSG00000196218.13_7 | RYR1 | 0.00024827 | 0.00180631 | 0.905 | 3.114 | chr19 | 38924331 | 39078204 | + | protein_coding |
| ENSG00000160207.9_5 | HSF2BP | 0.01754599 | 0.08884002 | 0.905 | 0.974 | chr21 | 44949066 | 45079369 | - | protein_coding |
| ENSG00000212694.8_7 | LINC01089 | 0.00053203 | 0.00364541 | 0.905 | 2.922 | chr12 | 122233173 | 122241812 | - | lncRNA |
| ENSG00000153904.20_6 | DDAH1 | 5.06E-07 | 5.43E-06 | 0.905 | 5.097 | chr1 | 85784164 | 86044046 | - | protein_coding |
| ENSG00000113369.9_4 | ARRDC3 | 3.77E-07 | 4.10E-06 | 0.904 | 5.610 | chr5 | 90664448 | 90679134 | - | protein_coding |
| ENSG00000069020.18_3 | MAST4 | 2.33E-06 | 2.28E-05 | 0.903 | 4.713 | chr5 | 65892189 | 66465423 | + | protein_coding |
| ENSG00000167701.14_4 | GPT | 0.00046665 | 0.00323484 | 0.903 | 2.807 | chr8 | 145728356 | 145732557 | + | protein_coding |
| ENSG00000132357.14_4 | CARD6 | 1.36E-05 | 0.00012024 | 0.903 | 4.236 | chr5 | 40841410 | 40860277 | + | protein_coding |
| ENSG00000239697.11_4 | TNFSF12 | 0.0101809 | 0.05440122 | 0.903 | 1.343 | chr17 | 7452375 | 7461207 | + | protein_coding |
| ENSG00000143631.11_3 | FLG | 0.01415333 | 0.07307825 | 0.901 | 0.918 | chr1 | 152274641 | 152297715 | - | protein_coding |
| ENSG00000267682.1_7 | AC016590.2 | 0.06545839 | 0.28255568 | 0.901 | -0.728 | chr19 | 37828138 | 37828645 | + | lncRNA |
| ENSG00000244503.1_5 | AC108751.5 | 0.07731794 | 0.32583205 | 0.901 | -0.946 | chr3 | 149212447 | 149213782 | + | processed_pseudogene |
| ENSG00000138759.19_4 | FRAS1 | 5.77E-08 | 7.06E-07 | 0.900 | 6.240 | chr4 | 78978477 | 79465423 | + | protein_coding |
| ENSG00000186564.5_2 | FOXD2 | 0.00804307 | 0.04386404 | 0.899 | 1.352 | chr1 | 47901689 | 47906363 | + | protein_coding |
| ENSG00000085063.16_6 | CD59 | 3.94E-09 | 5.76E-08 | 0.899 | 7.400 | chr11 | 33724556 | 33758037 | - | protein_coding |
| ENSG00000198298.13_7 | ZNF485 | 0.00100327 | 0.00652935 | 0.899 | 2.589 | chr10 | 44101867 | 44113352 | + | protein_coding |
| ENSG00000162814.11_3 | SPATA17 | 0.00126538 | 0.00808416 | 0.899 | 2.322 | chr1 | 217804666 | 218045038 | + | protein_coding |
| ENSG00000107872.12_4 | FBXL15 | 0.00113064 | 0.00729415 | 0.899 | 2.578 | chr10 | 104178946 | 104182893 | + | protein_coding |
| ENSG00000205277.9_2 | MUC12 | 0.03334784 | 0.15736813 | 0.898 | 0.225 | chr7 | 100612904 | 100662230 | + | protein_coding |
| ENSG00000279080.1_6 | AL022322.2 | 0.06039391 | 0.26325283 | 0.898 | -0.632 | chr22 | 38526223 | 38546619 | + | lncRNA |
| ENSG00000151164.18_3 | RAD9B | 0.04323148 | 0.1977584 | 0.898 | 0.003 | chr12 | 110939460 | 110969891 | + | protein_coding |
| ENSG00000119711.13_5 | ALDH6A1 | 4.20E-06 | 3.97E-05 | 0.898 | 4.670 | chr14 | 74523550 | 74551195 | - | protein_coding |
| ENSG00000135677.11_6 | GNS | 2.24E-09 | 3.38E-08 | 0.897 | 7.891 | chr12 | 65107225 | 65153211 | - | protein_coding |
| ENSG00000140682.19_5 | TGFB1I1 | 5.22E-06 | 4.88E-05 | 0.897 | 4.564 | chr16 | 31482906 | 31489281 | + | protein_coding |
| ENSG00000088881.20_4 | EBF4 | 0.00191879 | 0.01185369 | 0.896 | 2.268 | chr20 | 2673524 | 2740754 | + | protein_coding |
| ENSG00000272092.1_6 | AC087623.2 | 0.03329798 | 0.15717462 | 0.896 | 0.218 | chr8 | 38239882 | 38240979 | + | lncRNA |
| ENSG00000107902.14_3 | LHPP | 0.000309 | 0.00221285 | 0.895 | 3.064 | chr10 | 126150392 | 126306457 | + | protein_coding |
| ENSG00000118785.14_5 | SPP1 | 4.69E-09 | 6.75E-08 | 0.895 | 7.055 | chr4 | 88896802 | 88904578 | + | protein_coding |
| ENSG00000221883.3_3 | ARIH2OS | 0.0630222 | 0.27343476 | 0.895 | -0.633 | chr3 | 48955215 | 48956818 | - | protein_coding |

| | | | | | | | | | | |
|---|---|---|---|---|---|---|---|---|---|---|
| ENSG00000123933.17_6 | MXD4 | 1.57E-06 | 1.58E-05 | 0.894 | 4.870 | chr4 | 2249159 | 2263836 | - | protein_coding |
| ENSG00000186399.10_3 | GOLGA8R | 0.07758908 | 0.32681964 | 0.894 | -0.895 | chr15 | 30695943 | 30706365 | - | protein_coding |
| ENSG00000179715.13_5 | PCED1B | 1.29E-07 | 1.50E-06 | 0.893 | 5.872 | chr12 | 47473386 | 47630445 | + | protein_coding |
| ENSG00000166145.14_3 | SPINT1 | 0.02960099 | 0.14133858 | 0.893 | 0.219 | chr15 | 41136216 | 41150405 | + | protein_coding |
| ENSG00000078487.17_5 | ZCWPW1 | 0.00350911 | 0.02059673 | 0.893 | 1.938 | chr7 | 99998449 | 100026615 | - | protein_coding |
| ENSG00000163814.8_5 | CDCP1 | 2.65E-06 | 2.57E-05 | 0.891 | 4.699 | chr3 | 45123769 | 45187914 | - | protein_coding |
| ENSG00000163683.12_8 | SMIM14 | 2.54E-08 | 3.24E-07 | 0.891 | 6.373 | chr4 | 39547956 | 39640522 | - | protein_coding |
| ENSG00000179023.8_2 | KLHDC7A | 4.06E-05 | 0.00033529 | 0.890 | 4.083 | chr1 | 18807476 | 18812620 | + | protein_coding |
| ENSG00000204386.11_5 | NEU1 | 0.00077285 | 0.00513314 | 0.890 | 2.791 | chr6 | 31825436 | 31830599 | - | protein_coding |
| ENSG00000183072.10_6 | NKX2-5 | 0.00369438 | 0.02161278 | 0.889 | 2.002 | chr5 | 172659112 | 172662314 | - | protein_coding |
| ENSG00000070729.14_5 | CNGB1 | 0.07216197 | 0.30679628 | 0.889 | -0.788 | chr16 | 57916244 | 58005032 | - | protein_coding |
| ENSG00000203499.11_7 | IQANK1 | 0.00985277 | 0.05280671 | 0.887 | 1.578 | chr8 | 144816310 | 144872814 | + | protein_coding |
| ENSG00000172819.17_6 | RARG | 3.49E-06 | 3.34E-05 | 0.887 | 4.969 | chr12 | 53604351 | 53626764 | - | protein_coding |
| ENSG00000286629.1_1 | AC005803.1 | 0.077226 | 0.32559902 | 0.887 | -1.015 | chr17 | 62144979 | 62147630 | + | lncRNA |
| ENSG00000187566.6_3 | NHLRC1 | 0.00502308 | 0.02856758 | 0.887 | 1.685 | chr6 | 18120671 | 18122908 | - | protein_coding |
| ENSG00000167535.8_4 | CACNB3 | 1.54E-05 | 0.00013434 | 0.886 | 4.370 | chr12 | 49207577 | 49222724 | + | protein_coding |
| ENSG00000216915.2_4 | GPR89P | 0.07154889 | 0.3048463 | 0.885 | -0.788 | chr6 | 27704779 | 27706273 | - | processed_pseudogene |
| ENSG00000125534.10_4 | PPDPF | 0.01603048 | 0.08170212 | 0.885 | 4.310 | chr20 | 62152118 | 62153559 | + | protein_coding |
| ENSG00000136802.11_2 | LRRC8A | 2.35E-08 | 3.01E-07 | 0.885 | 6.651 | chr9 | 131644391 | 131680318 | + | protein_coding |
| ENSG00000237975.7_7 | FLG-AS1 | 0.01481175 | 0.07607989 | 0.885 | 1.107 | chr1 | 152140601 | 152417932 | + | lncRNA |
| ENSG00000108511.10_5 | HOXB6 | 0.00029297 | 0.0021074 | 0.884 | 3.351 | chr17 | 46673113 | 46682354 | - | protein_coding |
| ENSG00000277443.3_5 | MARCKS | 2.32E-08 | 2.98E-07 | 0.884 | 7.482 | chr6 | 114178524 | 114184652 | + | protein_coding |
| ENSG00000137699.17_5 | TRIM29 | 0.04476311 | 0.20382015 | 0.884 | -0.001 | chr11 | 119981994 | 120056237 | - | protein_coding |
| ENSG00000261787.2_4 | TCF24 | 0.04812327 | 0.21700664 | 0.884 | -0.202 | chr8 | 67858736 | 67874826 | - | protein_coding |
| ENSG00000214814.7_3 | FER1L6 | 0.04316069 | 0.19754968 | 0.884 | -0.036 | chr8 | 124864227 | 125132302 | + | protein_coding |
| ENSG00000196810.5_7 | CTBP1-DT | 9.54E-05 | 0.00074021 | 0.883 | 3.873 | chr4 | 1243088 | 1282079 | + | lncRNA |
| ENSG00000186318.16_3 | BACE1 | 0.00031211 | 0.00223328 | 0.883 | 3.250 | chr11 | 117155923 | 117186975 | - | protein_coding |
| ENSG00000095015.6_3 | MAP3K1 | 4.90E-06 | 4.60E-05 | 0.882 | 4.961 | chr5 | 56111376 | 56191979 | + | protein_coding |
| ENSG00000188295.14_3 | ZNF669 | 0.00119855 | 0.00770428 | 0.882 | 2.940 | chr1 | 247263264 | 247267674 | - | protein_coding |
| ENSG00000137878.17_5 | GCOM1 | 0.02189619 | 0.10821407 | 0.880 | 0.653 | chr15 | 57884106 | 58006943 | + | protein_coding |
| ENSG00000254670.1_9 | RASSF10-DT | 0.06375961 | 0.2761621 | 0.879 | -0.359 | chr11 | 13022637 | 13030706 | - | lncRNA |
| ENSG00000184939.16_3 | ZFP90 | 4.91E-07 | 5.27E-06 | 0.879 | 5.726 | chr16 | 68563993 | 68609975 | + | protein_coding |
| ENSG00000177359.20_7 | AC024940.2 | 0.08627551 | 0.35839536 | 0.879 | -0.743 | chr12 | 31264586 | 31354169 | - | transcribed_unprocessed_pseudogene |
| ENSG00000148180.19_5 | GSN | 1.08E-06 | 1.11E-05 | 0.878 | 6.080 | chr9 | 123970072 | 124095121 | + | protein_coding |
| ENSG00000103888.17_3 | CEMIP | 0.00838907 | 0.04552491 | 0.878 | 1.708 | chr15 | 81071684 | 81244117 | + | protein_coding |
| ENSG00000136542.9_3 | GALNT5 | 2.30E-06 | 2.26E-05 | 0.878 | 4.913 | chr2 | 158114217 | 158175003 | + | protein_coding |
| ENSG00000257594.4_6 | GALNT4 | 0.06006276 | 0.26200224 | 0.878 | -0.476 | chr12 | 89913189 | 89918573 | - | protein_coding |
| ENSG00000232859.9_4 | LYRM9 | 0.04985155 | 0.2237778 | 0.877 | 0.674 | chr17 | 26205340 | 26221778 | - | protein_coding |
| ENSG00000271303.1_4 | SRXN1 | 0.09016701 | 0.37238428 | 0.877 | 0.173 | chr20 | 627259 | 634014 | - | protein_coding |
| ENSG00000269867.1_8 | AC010326.4 | 0.07878664 | 0.33117548 | 0.876 | -0.910 | chr19 | 58378406 | 58379540 | - | lncRNA |
| ENSG00000113504.21_8 | SLC12A7 | 1.84E-05 | 0.00015959 | 0.875 | 5.258 | chr5 | 1050499 | 1112178 | - | protein_coding |
| ENSG00000196693.15_4 | ZNF33B | 2.49E-06 | 2.42E-05 | 0.875 | 5.197 | chr10 | 43069633 | 43134018 | - | protein_coding |
| ENSG00000128408.9_4 | RIBC2 | 0.002618 | 0.01574076 | 0.875 | 2.251 | chr22 | 45809574 | 45828389 | + | protein_coding |
| ENSG00000005073.6_3 | HOXA11 | 0.06982646 | 0.29886938 | 0.874 | -0.656 | chr7 | 27220776 | 27224851 | - | protein_coding |
| ENSG00000198855.7_3 | FICD | 0.00011637 | 0.00089039 | 0.874 | 3.726 | chr12 | 108909054 | 108919614 | + | protein_coding |
| ENSG00000236384.7_4 | LINC00479 | 0.06463742 | 0.27962332 | 0.874 | -0.352 | chr21 | 43131680 | 43135935 | - | lncRNA |
| ENSG00000073060.16_6 | SCARB1 | 1.97E-07 | 2.24E-06 | 0.873 | 6.469 | chr12 | 125261402 | 125367214 | - | protein_coding |
| ENSG00000138347.15_2 | MYPN | 0.0024002 | 0.01450486 | 0.873 | 2.748 | chr10 | 69865874 | 69971774 | + | protein_coding |
| ENSG00000196352.15_7 | CD55 | 3.29E-06 | 3.16E-05 | 0.873 | 5.821 | chr1 | 207494721 | 207560149 | + | protein_coding |
| ENSG00000176871.8_4 | WSB2 | 1.11E-07 | 1.31E-06 | 0.872 | 6.556 | chr12 | 118470499 | 118500235 | - | protein_coding |
| ENSG00000151229.13_5 | SLC2A13 | 6.39E-06 | 5.89E-05 | 0.871 | 4.721 | chr12 | 40148827 | 40499891 | - | protein_coding |
| ENSG00000272142.4_9 | AL359643.3 | 0.01369702 | 0.0708663 | 0.871 | 1.194 | chr6 | 5004008 | 5045900 | + | lncRNA |
| ENSG00000131398.14_4 | KCNC3 | 0.07094684 | 0.30278946 | 0.870 | -0.700 | chr19 | 50815194 | 50836772 | - | protein_coding |
| ENSG00000143412.10_3 | ANXA9 | 0.01068464 | 0.05671766 | 0.870 | 1.678 | chr1 | 150954725 | 150968110 | + | protein_coding |
| ENSG00000139531.13_5 | SUOX | 5.74E-06 | 5.33E-05 | 0.870 | 4.848 | chr12 | 56390964 | 56400425 | + | protein_coding |
| ENSG00000010539.11_6 | ZNF200 | 3.40E-05 | 0.00028452 | 0.869 | 4.326 | chr16 | 3272325 | 3286221 | - | protein_coding |
| ENSG00000105357.18_8 | MYH14 | 0.00331673 | 0.01951914 | 0.869 | 4.838 | chr19 | 50691443 | 50813799 | + | protein_coding |
| ENSG00000105639.19_4 | JAK3 | 0.00235624 | 0.01426836 | 0.869 | 2.420 | chr19 | 17935589 | 17958880 | - | protein_coding |
| ENSG00000106665.16_8 | CLIP2 | 6.85E-07 | 7.21E-06 | 0.869 | 5.942 | chr7 | 73703737 | 73820265 | + | protein_coding |
| ENSG00000005243.10_5 | COPZ2 | 0.00036461 | 0.00257779 | 0.869 | 3.259 | chr17 | 46103533 | 46115392 | - | protein_coding |

| | | | | | | | | | | |
|---|---|---|---|---|---|---|---|---|---|---|
| ENSG00000231107.2_7 | LINC01508 | 0.02937134 | 0.14043073 | 0.869 | 0.445 | chr9 | 93063167 | 93195848 | - | lncRNA |
| ENSG00000184640.18_6 | SEPTIN9 | 1.47E-05 | 0.00012854 | 0.868 | 8.749 | chr17 | 75276651 | 75496678 | + | protein_coding |
| ENSG00000187678.9_5 | SPRY4 | 0.0003028 | 0.00217628 | 0.868 | 3.847 | chr5 | 141689992 | 141706020 | - | protein_coding |
| ENSG00000171560.16_5 | FGA | 1.26E-06 | 1.28E-05 | 0.866 | 5.986 | chr4 | 155504280 | 155511897 | - | protein_coding |
| ENSG00000158555.15_5 | GDPD5 | 4.39E-05 | 0.00036156 | 0.866 | 4.168 | chr11 | 75145685 | 75236986 | - | protein_coding |
| ENSG00000175197.12_5 | DDIT3 | 0.0054929 | 0.03099154 | 0.866 | 2.035 | chr12 | 57910371 | 57915520 | - | protein_coding |
| ENSG00000169228.14_5 | RAB24 | 0.03305027 | 0.15612977 | 0.866 | 0.483 | chr5 | 176728199 | 176730745 | - | protein_coding |
| ENSG00000163110.15_7 | PDLIM5 | 4.06E-08 | 5.05E-07 | 0.866 | 9.536 | chr4 | 95373008 | 95589378 | + | protein_coding |
| ENSG00000261221.3_2 | ZNF865 | 0.00386448 | 0.0225485 | 0.865 | 3.050 | chr19 | 56116771 | 56128635 | + | protein_coding |
| ENSG00000268049.1_6 | AC012313.2 | 0.02761714 | 0.13272212 | 0.865 | 0.605 | chr19 | 58869365 | 58870969 | + | lncRNA |
| ENSG00000258346.1_6 | AC079384.1 | 0.07952349 | 0.33370192 | 0.865 | -0.758 | chr12 | 116806569 | 116827276 | + | lncRNA |
| ENSG00000253552.8_6 | HOXA-AS2 | 0.0029132 | 0.01733966 | 0.864 | 2.549 | chr7 | 27147396 | 27173921 | + | lncRNA |
| ENSG00000242574.9_7 | HLA-DMB | 0.01181155 | 0.06203245 | 0.864 | 1.626 | chr6 | 32902406 | 32908805 | - | protein_coding |
| ENSG00000137142.5_4 | IGFBPL1 | 6.05E-05 | 0.00048664 | 0.864 | 4.039 | chr9 | 38406525 | 38424451 | - | protein_coding |
| ENSG00000168062.10_6 | BATF2 | 0.04612725 | 0.2091204 | 0.863 | 0.377 | chr11 | 64755417 | 64764490 | - | protein_coding |
| ENSG00000109743.11_4 | BST1 | 0.00028703 | 0.00206799 | 0.862 | 3.468 | chr4 | 15704688 | 15739936 | + | protein_coding |
| ENSG00000170631.15_5 | ZNF16 | 0.00045695 | 0.00317275 | 0.862 | 3.357 | chr8 | 146155744 | 146176274 | - | protein_coding |
| ENSG00000139998.15_6 | RAB15 | 1.94E-07 | 2.21E-06 | 0.862 | 6.588 | chr14 | 65412532 | 65439944 | - | protein_coding |
| ENSG00000224231.2_5 | LINC01832 | 0.0972184 | 0.39552996 | 0.862 | -0.976 | chr2 | 138859448 | 138864748 | + | lncRNA |
| ENSG00000156273.16_6 | BACH1 | 1.58E-06 | 1.59E-05 | 0.862 | 5.532 | chr21 | 30566392 | 31003071 | + | protein_coding |
| ENSG00000182095.14_6 | TNRC18 | 0.00912979 | 0.04913934 | 0.862 | 7.150 | chr7 | 5346421 | 5465045 | - | protein_coding |
| ENSG00000147255.19_8 | IGSF1 | 0.0065498 | 0.03636571 | 0.861 | 2.013 | chrX | 130407480 | 130712873 | - | protein_coding |
| ENSG00000132170.21_7 | PPARG | 1.23E-06 | 1.26E-05 | 0.861 | 5.511 | chr3 | 12328867 | 12475855 | + | protein_coding |
| ENSG00000258343.2_7 | AC090001.1 | 0.03206881 | 0.15205937 | 0.861 | 0.756 | chr12 | 96185511 | 96252617 | - | lncRNA |
| ENSG00000279233.1_6 | AC122688.3 | 0.01339847 | 0.06950381 | 0.861 | 1.265 | chr12 | 125622791 | 125626257 | + | TEC |
| ENSG00000230061.2_7 | TRPM2-AS | 0.08151044 | 0.34131416 | 0.861 | -0.440 | chr21 | 45834471 | 45845155 | - | lncRNA |
| ENSG00000102931.8_4 | ARL2BP | 0.0896112 | 0.37034712 | 0.860 | -0.758 | chr16 | 57279171 | 57287547 | + | protein_coding |
| ENSG00000249210.1_2 | GAPDHP38 | 0.07258868 | 0.30838906 | 0.859 | -0.255 | chr19 | 47061898 | 47062899 | + | processed_pseudogene |
| ENSG00000188157.15_5 | AGRN | 0.03588371 | 0.16817356 | 0.859 | 7.943 | chr1 | 955500 | 991498 | + | protein_coding |
| ENSG00000140691.17_8 | ARMC5 | 0.0044397 | 0.02560212 | 0.858 | 3.516 | chr16 | 31469401 | 31478487 | + | protein_coding |
| ENSG00000169564.7_4 | PCBP1 | 0.07939934 | 0.33325963 | 0.858 | 0.806 | chr2 | 70314609 | 70316335 | + | protein_coding |
| ENSG00000142959.5_4 | BEST4 | 0.10270192 | 0.41470568 | 0.857 | -0.759 | chr1 | 45249257 | 45253842 | - | protein_coding |
| ENSG00000196922.10_5 | ZNF252P | 1.37E-06 | 1.39E-05 | 0.857 | 5.639 | chr8 | 146198975 | 146228281 | - | transcribed_unprocessed_pseudogene |
| ENSG00000125170.11_6 | DOK4 | 3.53E-07 | 3.85E-06 | 0.856 | 6.744 | chr16 | 57505834 | 57521239 | - | protein_coding |
| ENSG00000050165.18_4 | DKK3 | 1.26E-05 | 0.00011195 | 0.855 | 4.802 | chr11 | 11977754 | 12031316 | - | protein_coding |
| ENSG00000121039.10_3 | RDH10 | 1.46E-07 | 1.69E-06 | 0.855 | 6.711 | chr8 | 74206837 | 74237516 | + | protein_coding |
| ENSG00000051523.11_5 | CYBA | 0.01812046 | 0.09141004 | 0.854 | 6.661 | chr16 | 88709697 | 88717462 | - | protein_coding |
| ENSG00000225762.1_7 | LINC01389 | 0.06574181 | 0.28364128 | 0.852 | -0.406 | chr1 | 47846600 | 47874149 | - | lncRNA |
| ENSG00000103742.12_4 | IGDCC4 | 0.00070437 | 0.00472249 | 0.852 | 3.255 | chr15 | 65673822 | 65715285 | - | protein_coding |
| ENSG00000167930.17_6 | FAM234A | 2.18E-06 | 2.14E-05 | 0.852 | 5.783 | chr16 | 284520 | 322182 | + | protein_coding |
| ENSG00000140332.16_5 | TLE3 | 0.00012029 | 0.0009188 | 0.850 | 5.072 | chr15 | 70340129 | 70390515 | - | protein_coding |
| ENSG00000047634.15_5 | SCML1 | 0.00043712 | 0.00304562 | 0.850 | 3.554 | chrX | 17755569 | 17773108 | + | protein_coding |
| ENSG00000231298.7_7 | MANCR | 0.03169578 | 0.15061219 | 0.849 | 1.181 | chr10 | 4692350 | 4720377 | - | lncRNA |
| ENSG00000121380.12_9 | BCL2L14 | 0.06050893 | 0.26368953 | 0.849 | -0.125 | chr12 | 12202778 | 12364018 | + | protein_coding |
| ENSG00000237575.4_2 | PYY2 | 0.06166154 | 0.26812054 | 0.848 | -0.263 | chr17 | 26553589 | 26555091 | + | transcribed_unprocessed_pseudogene |
| ENSG00000131116.12_4 | ZNF428 | 0.01192925 | 0.06257659 | 0.848 | 2.993 | chr19 | 44111376 | 44123781 | - | protein_coding |
| ENSG00000168778.12_3 | TCTN2 | 4.65E-05 | 0.00038197 | 0.847 | 4.418 | chr12 | 124155660 | 124192946 | + | protein_coding |
| ENSG00000287608.1_2 | AC022137.4 | 0.01039196 | 0.05539574 | 0.847 | 2.123 | chr19 | 53937914 | 53977190 | - | lncRNA |
| ENSG00000231925.11_3 | TAPBP | 4.78E-05 | 0.00039123 | 0.846 | 4.622 | chr6 | 33267471 | 33282164 | - | protein_coding |
| ENSG00000237441.10_4 | RGL2 | 0.00080334 | 0.00530793 | 0.846 | 3.197 | chr6 | 33259431 | 33266719 | - | protein_coding |
| ENSG00000004838.14_5 | ZMYND10 | 0.07587879 | 0.32022291 | 0.846 | -0.416 | chr3 | 50378541 | 50383163 | - | protein_coding |
| ENSG00000108984.15_7 | MAP2K6 | 1.05E-06 | 1.09E-05 | 0.845 | 6.240 | chr17 | 67410838 | 67550006 | + | protein_coding |
| ENSG00000134864.10_4 | GGACT | 0.01148067 | 0.06043763 | 0.845 | 1.626 | chr13 | 101182418 | 101241782 | - | protein_coding |
| ENSG00000204054.14_7 | LINC00963 | 1.09E-06 | 1.12E-05 | 0.845 | 5.890 | chr9 | 132239225 | 132275966 | + | lncRNA |
| ENSG00000266904.6_8 | LINC00663 | 0.12544831 | 0.49028599 | 0.844 | -1.105 | chr19 | 19868175 | 19887232 | - | lncRNA |
| ENSG00000177989.13_6 | ODF3B | 0.04212063 | 0.19338759 | 0.842 | 2.138 | chr22 | 50968139 | 50971009 | - | protein_coding |
| ENSG00000165895.19_5 | ARHGAP42 | 8.46E-06 | 7.67E-05 | 0.842 | 5.446 | chr11 | 100558019 | 100864672 | + | protein_coding |
| ENSG00000196220.16_6 | SRGAP3 | 0.00237389 | 0.01436054 | 0.842 | 3.061 | chr3 | 9022275 | 9404737 | - | protein_coding |
| ENSG00000214145.7_6 | LINC00887 | 0.01559144 | 0.07971599 | 0.841 | 1.381 | chr3 | 194013980 | 194040660 | - | lncRNA |

| | | | | | | | | | | |
|---|---|---|---|---|---|---|---|---|---|---|
| ENSG00000230882.1_5 | AC005077.4 | 0.03400566 | 0.16017461 | 0.841 | 0.670 | chr7 | 75700787 | 75704281 | - | processed_pseudogene |
| ENSG00000152939.17_7 | MARVELD2 | 0.00017879 | 0.00133302 | 0.840 | 4.054 | chr5 | 68710892 | 68740157 | + | protein_coding |
| ENSG00000175866.15_6 | BAIAP2 | 0.00016341 | 0.00122811 | 0.840 | 3.999 | chr17 | 79008922 | 79091232 | + | protein_coding |
| ENSG00000132326.12_5 | PER2 | 3.96E-06 | 3.75E-05 | 0.840 | 5.693 | chr2 | 239152685 | 239198743 | - | protein_coding |
| ENSG00000115257.15_2 | PCSK4 | 0.01990751 | 0.09932512 | 0.840 | 1.686 | chr19 | 1481427 | 1490751 | - | protein_coding |
| ENSG00000172037.14_4 | LAMB2 | 2.45E-07 | 2.75E-06 | 0.840 | 7.197 | chr3 | 49158547 | 49170551 | - | protein_coding |
| ENSG00000233217.2_3 | MROH3P | 0.07122538 | 0.30368644 | 0.839 | -0.455 | chr1 | 200886588 | 200935796 | + | transcribed_unitary_pseudogene |
| ENSG00000139433.10_5 | GLTP | 3.90E-06 | 3.71E-05 | 0.839 | 5.350 | chr12 | 110288750 | 110318346 | - | protein_coding |
| ENSG00000141736.13_5 | ERBB2 | 2.47E-06 | 2.41E-05 | 0.839 | 6.787 | chr17 | 37844167 | 37886679 | + | protein_coding |
| ENSG00000181873.13_3 | IBA57 | 0.0024565 | 0.01482997 | 0.839 | 3.348 | chr1 | 228353505 | 228369958 | + | protein_coding |
| ENSG00000280138.1_7 | AC027290.3 | 0.00823427 | 0.04480803 | 0.838 | 1.923 | chr12 | 123354606 | 123373267 | - | TEC |
| ENSG00000149761.9_6 | NUDT22 | 0.00041792 | 0.00292214 | 0.837 | 3.645 | chr11 | 63993413 | 63998158 | + | protein_coding |
| ENSG00000090615.15_6 | GOLGA3 | 1.11E-06 | 1.14E-05 | 0.836 | 7.349 | chr12 | 133345500 | 133405664 | - | protein_coding |
| ENSG00000213071.11_5 | LPAL2 | 0.0119586 | 0.06271205 | 0.835 | 1.753 | chr6 | 160874460 | 160941301 | - | transcribed_unprocessed_pseudogene |
| ENSG00000254389.3_6 | RHPN1-AS1 | 0.06175898 | 0.26841288 | 0.835 | 0.203 | chr8 | 144448801 | 144450718 | - | lncRNA |
| ENSG00000204860.5_5 | FAM201A | 0.00140104 | 0.00886159 | 0.835 | 3.545 | chr9 | 38620471 | 38624987 | + | lncRNA |
| ENSG00000197385.6_3 | ZNF860 | 0.005862 | 0.03289646 | 0.835 | 2.392 | chr3 | 32023242 | 32033215 | + | protein_coding |
| ENSG00000198933.9_5 | TBKBP1 | 0.00277994 | 0.01661884 | 0.834 | 2.885 | chr17 | 45771447 | 45789416 | + | protein_coding |
| ENSG00000175970.11_4 | UNC119B | 9.41E-06 | 8.48E-05 | 0.834 | 5.316 | chr12 | 121148261 | 121161443 | + | protein_coding |
| ENSG00000230445.4_5 | LRRC37A6P | 0.0389227 | 0.18060582 | 0.834 | 0.793 | chr10 | 27535279 | 27548384 | - | transcribed_processed_pseudogene |
| ENSG00000002330.13_8 | BAD | 0.00967763 | 0.05191499 | 0.832 | 3.693 | chr11 | 64037302 | 64052176 | - | protein_coding |
| ENSG00000133739.16_3 | LRRCC1 | 0.00041541 | 0.00290692 | 0.832 | 3.657 | chr8 | 86019450 | 86058315 | + | protein_coding |
| ENSG00000149260.18_6 | CAPN5 | 0.00084696 | 0.00557949 | 0.831 | 3.670 | chr11 | 76778008 | 76837201 | + | protein_coding |
| ENSG00000108379.10_4 | WNT3 | 0.0004791 | 0.00331342 | 0.831 | 3.810 | chr17 | 44839872 | 44910520 | - | protein_coding |
| ENSG00000221491.2 | SNORA34 | 0.03541935 | 0.16621648 | 0.831 | 0.889 | chr12 | 49048165 | 49048301 | - | snoRNA |
| ENSG00000260658.6_9 | AC138305.1 | 0.03637223 | 0.17022165 | 0.830 | 0.678 | chr16 | 63348168 | 63651950 | - | lncRNA |
| ENSG00000069424.15_3 | KCNAB2 | 0.00113291 | 0.00730616 | 0.829 | 3.632 | chr1 | 6050987 | 6161253 | + | protein_coding |
| ENSG00000130307.12_6 | USHBP1 | 0.09825333 | 0.39926841 | 0.829 | -0.881 | chr19 | 17359980 | 17393595 | - | protein_coding |
| ENSG00000267107.8_7 | PCAT19 | 0.07200278 | 0.30641277 | 0.829 | -0.242 | chr19 | 41955425 | 42007133 | - | lncRNA |
| ENSG00000114861.21_7 | FOXP1 | 0.00069589 | 0.0046727 | 0.829 | 3.533 | chr3 | 71001968 | 71633144 | - | protein_coding |
| ENSG00000090581.10_4 | GNPTG | 0.0003034 | 0.00217977 | 0.828 | 4.216 | chr16 | 1401932 | 1414114 | + | protein_coding |
| ENSG00000139160.13_6 | ETFBKMT | 0.00942898 | 0.05067295 | 0.828 | 2.341 | chr12 | 31800094 | 31826048 | + | protein_coding |
| ENSG00000139644.13_8 | TMBIM6 | 2.16E-07 | 2.45E-06 | 0.828 | 9.360 | chr12 | 50101508 | 50158717 | + | protein_coding |
| ENSG00000216588.9_6 | IGSF23 | 0.08259595 | 0.3453712 | 0.827 | -0.239 | chr19 | 45116873 | 45140081 | + | protein_coding |
| ENSG00000112699.11_6 | GMDS | 0.00253264 | 0.01526891 | 0.826 | 2.954 | chr6 | 1624041 | 2245839 | - | protein_coding |
| ENSG00000152763.17_6 | WDR78 | 0.02770735 | 0.13308366 | 0.826 | 1.302 | chr1 | 67278568 | 67390539 | - | protein_coding |
| ENSG00000268516.2_8 | AC020915.3 | 0.00701776 | 0.03868587 | 0.826 | 2.204 | chr19 | 58768636 | 58790174 | - | lncRNA |
| ENSG00000231185.7_8 | SPRY4-AS1 | 0.05070227 | 0.22696579 | 0.826 | 0.524 | chr5 | 141704858 | 142051566 | + | lncRNA |
| ENSG00000146477.6_3 | SLC22A3 | 1.36E-06 | 1.38E-05 | 0.825 | 6.565 | chr6 | 160769410 | 160873609 | + | protein_coding |
| ENSG00000169131.13_6 | ZNF354A | 0.00010034 | 0.00077543 | 0.825 | 4.330 | chr5 | 178138513 | 178157660 | - | protein_coding |
| ENSG00000107960.11_6 | STN1 | 3.40E-05 | 0.00028481 | 0.825 | 5.053 | chr10 | 105637327 | 105677942 | - | protein_coding |
| ENSG00000228709.1_6 | LINC02575 | 0.02016269 | 0.10042907 | 0.824 | 1.381 | chr21 | 45905460 | 45910171 | + | lncRNA |
| ENSG00000143457.11_4 | GOLPH3L | 7.57E-06 | 6.91E-05 | 0.824 | 5.561 | chr1 | 150618706 | 150669630 | - | protein_coding |
| ENSG00000207313.1 | SNORA2B | 0.10389979 | 0.41901888 | 0.823 | -0.831 | chr12 | 49061240 | 49061376 | - | snoRNA |
| ENSG00000174989.13_7 | FBXW8 | 9.25E-05 | 0.00071937 | 0.823 | 4.798 | chr12 | 117348755 | 117468953 | + | protein_coding |
| ENSG00000112812.16_5 | PRSS16 | 0.02141618 | 0.10610742 | 0.823 | 1.736 | chr6 | 27215480 | 27224403 | + | protein_coding |
| ENSG00000166801.15_4 | FAM111A | 3.46E-06 | 3.32E-05 | 0.822 | 6.430 | chr11 | 58910221 | 58922512 | + | protein_coding |
| ENSG00000154359.13_4 | LONRF1 | 0.00059213 | 0.00402156 | 0.821 | 3.797 | chr8 | 12579415 | 12613582 | - | protein_coding |
| ENSG00000250132.6_8 | AC004803.1 | 0.0210917 | 0.10470416 | 0.821 | 1.472 | chr12 | 1083299 | 1100356 | - | lncRNA |
| ENSG00000151135.10_3 | TMEM263 | 1.41E-06 | 1.42E-05 | 0.821 | 6.476 | chr12 | 107349497 | 107372556 | + | protein_coding |
| ENSG00000006757.12_5 | PNPLA4 | 0.00126394 | 0.00807786 | 0.821 | 3.516 | chrX | 7866288 | 7895780 | - | protein_coding |
| ENSG00000230091.6_6 | TMEM254-AS1 | 0.05101896 | 0.22826843 | 0.821 | 0.416 | chr10 | 81806616 | 81838668 | - | lncRNA |
| ENSG00000267480.1_7 | AP001542.3 | 0.08980228 | 0.37105047 | 0.820 | -0.685 | chr18 | 12031177 | 12032180 | - | lncRNA |
| ENSG00000152443.13_4 | ZNF776 | 3.05E-05 | 0.00025783 | 0.820 | 4.944 | chr19 | 58258183 | 58269516 | + | protein_coding |
| ENSG00000197324.9_5 | LRP10 | 1.68E-06 | 1.68E-05 | 0.818 | 7.578 | chr14 | 23340949 | 23350922 | + | protein_coding |
| ENSG00000261455.1_5 | LINC01003 | 0.0454719 | 0.2065182 | 0.818 | 0.668 | chr7 | 152160871 | 152162634 | + | lncRNA |
| ENSG00000166689.16_6 | PLEKHA7 | 3.28E-05 | 0.00027552 | 0.818 | 5.254 | chr11 | 16798844 | 17035961 | - | protein_coding |
| ENSG00000100994.12_4 | PYGB | 1.03E-06 | 1.07E-05 | 0.818 | 8.902 | chr20 | 25228721 | 25278648 | + | protein_coding |

| | | | | | | | | | | |
|---|---|---|---|---|---|---|---|---|---|---|
| ENSG00000068903.20_6 | SIRT2 | 2.66E-05 | 0.00022677 | 0.818 | 5.369 | chr19 | 39369195 | 39390502 | - | protein_coding |
| ENSG00000214655.10_6 | ZSWIM8 | 1.86E-05 | 0.0001615 | 0.818 | 5.451 | chr10 | 75545340 | 75561555 | + | protein_coding |
| ENSG00000163820.15_4 | FYCO1 | 6.76E-05 | 0.00053833 | 0.817 | 4.794 | chr3 | 45959391 | 46037316 | - | protein_coding |
| ENSG00000099337.5_3 | KCNK6 | 0.0015287 | 0.00958714 | 0.816 | 3.643 | chr19 | 38810485 | 38822716 | + | protein_coding |
| ENSG00000135838.13_2 | NPL | 0.04804302 | 0.21678774 | 0.816 | 0.514 | chr1 | 182758428 | 182799519 | + | protein_coding |
| ENSG00000226944.1_8 | AL031847.1 | 0.09333244 | 0.38199436 | 0.815 | -0.685 | chr1 | 6264900 | 6265840 | - | lncRNA |
| ENSG00000141542.11_4 | RAB40B | 0.00048414 | 0.00334436 | 0.815 | 4.029 | chr17 | 80612849 | 80656574 | - | protein_coding |
| ENSG00000131584.19_4 | ACAP3 | 0.00152539 | 0.00957075 | 0.815 | 4.916 | chr1 | 1227770 | 1244989 | - | protein_coding |
| ENSG00000179922.6_3 | ZNF784 | 0.00558856 | 0.03147102 | 0.814 | 2.711 | chr19 | 56132107 | 56135932 | - | protein_coding |
| ENSG00000164002.11_7 | EXO5 | 0.02028724 | 0.10093628 | 0.814 | 1.671 | chr1 | 40974413 | 40982228 | + | protein_coding |
| ENSG00000104518.11_6 | GSDMD | 4.86E-05 | 0.00039739 | 0.813 | 5.811 | chr8 | 144635377 | 144645232 | + | protein_coding |
| ENSG00000117407.17_6 | ARTN | 0.10576509 | 0.42572005 | 0.812 | -0.743 | chr1 | 44398992 | 44402912 | + | protein_coding |
| ENSG00000115363.14_5 | EVA1A | 0.01783137 | 0.09018216 | 0.812 | 1.729 | chr2 | 75696428 | 75796848 | - | protein_coding |
| ENSG00000161955.16_3 | TNFSF13 | 0.08600064 | 0.35742066 | 0.811 | -0.553 | chr17 | 7461609 | 7464925 | + | protein_coding |
| ENSG00000288398.1_2 | AL109627.1 | 0.01304807 | 0.06790422 | 0.811 | 2.261 | chr1 | 16555169 | 16557830 | - | lncRNA |
| ENSG00000171703.17_4 | TCEA2 | 0.00468307 | 0.02684006 | 0.810 | 3.805 | chr20 | 62681189 | 62703700 | + | protein_coding |
| ENSG00000114854.8_5 | TNNC1 | 0.05395265 | 0.23946458 | 0.810 | 0.411 | chr3 | 52485116 | 52488057 | - | protein_coding |
| ENSG00000121895.8_6 | TMEM156 | 3.72E-05 | 0.00030952 | 0.810 | 5.157 | chr4 | 38968365 | 39034542 | - | protein_coding |
| ENSG00000164695.5_3 | CHMP4C | 0.00897447 | 0.04836195 | 0.809 | 2.634 | chr8 | 82644683 | 82671750 | + | protein_coding |
| ENSG00000132749.11_8 | TESMIN | 0.00893929 | 0.04821628 | 0.809 | 2.255 | chr11 | 68474908 | 68518988 | - | protein_coding |
| ENSG00000120215.10_3 | MLANA | 0.09326978 | 0.38182584 | 0.808 | -0.404 | chr9 | 5890889 | 5910606 | + | protein_coding |
| ENSG00000215196.5_8 | BASP1-AS1 | 0.0174516 | 0.08843771 | 0.808 | 1.824 | chr5 | 17089405 | 17217156 | - | lncRNA |
| ENSG00000205078.6_6 | SYCE1L | 0.03353332 | 0.15811743 | 0.808 | 1.469 | chr16 | 77233305 | 77247112 | + | protein_coding |
| ENSG00000229809.8_5 | ZNF688 | 0.01227389 | 0.06423278 | 0.807 | 2.518 | chr16 | 30580667 | 30584055 | - | protein_coding |
| ENSG00000134243.12_5 | SORT1 | 8.80E-06 | 7.96E-05 | 0.807 | 6.143 | chr1 | 109852190 | 109940540 | - | protein_coding |
| ENSG00000228509.5_7 | AC006460.1 | 0.12023856 | 0.47378099 | 0.807 | -0.895 | chr2 | 191541670 | 191573442 | - | lncRNA |
| ENSG00000162461.8_3 | SLC25A34 | 0.07137948 | 0.30419746 | 0.806 | -0.026 | chr1 | 16062753 | 16067887 | + | protein_coding |
| ENSG00000172780.16_4 | RAB43 | 0.07407095 | 0.31363648 | 0.806 | 0.117 | chr3 | 128806412 | 128841644 | - | protein_coding |
| ENSG00000095380.11_3 | NANS | 5.72E-05 | 0.00046153 | 0.805 | 5.202 | chr9 | 100819014 | 100845359 | + | protein_coding |
| ENSG00000157637.13_6 | SLC38A10 | 0.00927742 | 0.0498886 | 0.805 | 6.933 | chr17 | 79218611 | 79269347 | - | protein_coding |
| ENSG00000149823.9_6 | VPS51 | 0.00348247 | 0.0204539 | 0.804 | 5.540 | chr11 | 64856796 | 64879334 | + | protein_coding |
| ENSG00000165959.12_5 | CLMN | 1.65E-05 | 0.00014414 | 0.804 | 5.969 | chr14 | 95648277 | 95786245 | - | protein_coding |
| ENSG00000117461.15_5 | PIK3R3 | 0.00682383 | 0.03773386 | 0.803 | 2.545 | chr1 | 46505812 | 46598708 | - | protein_coding |
| ENSG00000125449.7_3 | ARMC7 | 0.00055001 | 0.00376574 | 0.802 | 4.293 | chr17 | 73106047 | 73126367 | + | protein_coding |
| ENSG00000274315.1_6 | AC009318.3 | 0.09262514 | 0.37971178 | 0.802 | -0.406 | chr12 | 29484367 | 29484869 | - | lncRNA |
| ENSG00000173918.15_4 | C1QTNF1 | 0.00066434 | 0.00447432 | 0.802 | 4.020 | chr17 | 77018896 | 77045870 | + | protein_coding |
| ENSG00000250539.1_4 | KRT8P33 | 0.01702226 | 0.08633588 | 0.802 | 2.239 | chr5 | 122736616 | 122738038 | - | processed_pseudogene |
| ENSG00000198133.8_6 | TMEM229B | 0.09576521 | 0.39042231 | 0.802 | -0.398 | chr14 | 67913801 | 68000456 | - | protein_coding |
| ENSG00000260018.1_6 | AC040169.1 | 0.09341912 | 0.38226105 | 0.801 | -0.360 | chr16 | 84150656 | 84151176 | + | lncRNA |
| ENSG00000160991.16_4 | ORAI2 | 3.04E-05 | 0.00025708 | 0.800 | 5.696 | chr7 | 102073553 | 102097272 | + | protein_coding |
| ENSG00000260727.1_4 | SLC7A5P1 | 0.06072587 | 0.26444031 | 0.800 | 0.586 | chr16 | 29624425 | 29624961 | - | unprocessed_pseudogene |
| ENSG00000169903.7_4 | TM4SF4 | 0.00044103 | 0.00306923 | 0.799 | 4.929 | chr3 | 149192484 | 149221181 | + | protein_coding |
| ENSG00000148204.12_3 | CRB2 | 0.09202025 | 0.37796486 | 0.799 | -0.193 | chr9 | 126118449 | 126142603 | + | protein_coding |
| ENSG00000279716.1_5 | AC006128.1 | 0.06330943 | 0.27447991 | 0.799 | 0.285 | chr19 | 15509507 | 15511167 | + | TEC |
| ENSG00000142192.21_7 | APP | 6.10E-07 | 6.47E-06 | 0.799 | 8.866 | chr21 | 27252861 | 27543446 | - | protein_coding |
| ENSG00000285976.2_3 | AL135905.2 | 0.07685017 | 0.32416834 | 0.798 | -0.033 | chr6 | 64282377 | 64293492 | + | protein_coding |
| ENSG00000279744.1_7 | AC132938.6 | 0.08595803 | 0.35732717 | 0.798 | -0.193 | chr17 | 80420477 | 80422131 | + | TEC |
| ENSG00000155974.12_7 | GRIP1 | 0.00336915 | 0.01980795 | 0.797 | 3.169 | chr12 | 66741211 | 67462942 | - | protein_coding |
| ENSG00000206127.11_7 | GOLGA8O | 0.11423843 | 0.45477798 | 0.797 | -0.848 | chr15 | 32734115 | 32747835 | - | protein_coding |
| ENSG00000149599.15_4 | DUSP15 | 0.14289848 | 0.54670053 | 0.797 | -0.961 | chr20 | 30435440 | 30458550 | - | protein_coding |
| ENSG00000167685.15_6 | ZNF444 | 0.02402856 | 0.11728445 | 0.796 | 4.347 | chr19 | 56643968 | 56672262 | + | protein_coding |
| ENSG00000179912.20_7 | R3HDM2 | 0.0005017 | 0.00345357 | 0.796 | 4.521 | chr12 | 57647545 | 57824788 | - | protein_coding |
| ENSG00000280007.1_7 | AC008079.1 | 0.10138575 | 0.40995009 | 0.795 | -0.657 | chr22 | 18593525 | 18613921 | - | lncRNA |
| ENSG00000272808.4_7 | AC015712.6 | 0.03696792 | 0.17266273 | 0.794 | 1.676 | chr15 | 101390036 | 101417041 | + | lncRNA |
| ENSG00000204946.10_5 | ZNF783 | 0.0013376 | 0.00848754 | 0.794 | 4.052 | chr7 | 148959262 | 148994393 | + | protein_coding |
| ENSG00000256294.8_5 | AC021092.2 | 0.00151828 | 0.00954204 | 0.794 | 3.713 | chr19 | 44616334 | 44638975 | + | protein_coding |
| ENSG00000103021.9_4 | CCDC113 | 0.00222737 | 0.01358055 | 0.794 | 3.406 | chr16 | 58265061 | 58317740 | + | protein_coding |
| ENSG00000139428.12_4 | MMAB | 0.00030395 | 0.0021828 | 0.794 | 4.957 | chr12 | 109991520 | 110011385 | - | protein_coding |
| ENSG00000236609.4_3 | ZNF853 | 0.00751077 | 0.04123731 | 0.793 | 2.802 | chr7 | 6655241 | 6663921 | + | protein_coding |
| ENSG00000184995.7_3 | IFNE | 0.03044731 | 0.14518449 | 0.793 | 1.375 | chr9 | 21480838 | 21482312 | - | protein_coding |

| | | | | | | | | | | |
|---|---|---|---|---|---|---|---|---|---|---|
| ENSG00000153113.23_7 | CAST | 3.35E-06 | 3.21E-05 | 0.793 | 8.607 | chr5 | 95860971 | 96115299 | + | protein_coding |
| ENSG00000124429.18_4 | POF1B | 9.08E-05 | 0.00070743 | 0.793 | 5.584 | chrX | 84532402 | 84634722 | - | protein_coding |
| ENSG00000172059.11_3 | KLF11 | 0.00252731 | 0.01524195 | 0.793 | 3.409 | chr2 | 10182976 | 10194963 | + | protein_coding |
| ENSG00000057294.14_6 | PKP2 | 0.00022262 | 0.00163374 | 0.793 | 4.806 | chr12 | 32943679 | 33049774 | - | protein_coding |
| ENSG00000080503.24_6 | SMARCA2 | 3.43E-05 | 0.00028713 | 0.793 | 6.035 | chr9 | 1980290 | 2193624 | + | protein_coding |
| ENSG00000174099.11_4 | MSRB3 | 8.67E-06 | 7.86E-05 | 0.793 | 6.257 | chr12 | 65672423 | 65885210 | + | protein_coding |
| ENSG00000213316.10_5 | LTC4S | 0.12399998 | 0.48547986 | 0.793 | -0.899 | chr5 | 179220981 | 179223648 | + | protein_coding |
| ENSG00000185186.10_6 | LINC00313 | 0.07565086 | 0.31933684 | 0.792 | -0.006 | chr21 | 44881840 | 44899414 | - | lncRNA |
| ENSG00000008853.16_2 | RHOBTB2 | 0.00041159 | 0.00288245 | 0.792 | 4.907 | chr8 | 22844930 | 22877712 | + | protein_coding |
| ENSG00000215440.12_4 | NPEPL1 | 0.00696767 | 0.03844554 | 0.792 | 3.036 | chr20 | 57264187 | 57294294 | + | protein_coding |
| ENSG00000246705.4_4 | H2AJ | 0.00046044 | 0.00319428 | 0.791 | 6.057 | chr12 | 14927317 | 14930936 | + | protein_coding |
| ENSG00000158423.17_3 | RIBC1 | 0.04391997 | 0.20059798 | 0.791 | 0.917 | chrX | 53449639 | 53458068 | + | protein_coding |
| ENSG00000142303.14_4 | ADAMTS10 | 0.00016415 | 0.00123321 | 0.791 | 5.908 | chr19 | 8645124 | 8675620 | - | protein_coding |
| ENSG00000198464.14_3 | ZNF480 | 0.00018206 | 0.00135514 | 0.791 | 5.932 | chr19 | 52800422 | 52829175 | + | protein_coding |
| ENSG00000198157.11_7 | HMGN5 | 0.07887226 | 0.33143887 | 0.790 | -0.041 | chrX | 80369198 | 80457412 | - | protein_coding |
| ENSG00000185201.16_6 | IFITM2 | 0.00169046 | 0.01052364 | 0.790 | 4.343 | chr11 | 307631 | 315272 | + | protein_coding |
| ENSG00000132024.17_5 | CC2D1A | 0.00039934 | 0.00280439 | 0.790 | 5.009 | chr19 | 14017014 | 14041692 | + | protein_coding |
| ENSG00000211451.12_3 | GNRHR2 | 0.10475655 | 0.42213874 | 0.789 | -0.658 | chr1 | 145509752 | 145516076 | - | transcribed_unitary_pseudogene |
| ENSG00000125430.9_4 | HS3ST3B1 | 0.03793847 | 0.17645361 | 0.789 | 1.166 | chr17 | 14204398 | 14252721 | + | protein_coding |
| ENSG00000164300.17_4 | SERINC5 | 5.16E-05 | 0.00041949 | 0.788 | 5.651 | chr5 | 79407474 | 79551867 | - | protein_coding |
| ENSG00000164292.13_4 | RHOBTB3 | 5.66E-05 | 0.00045662 | 0.788 | 5.572 | chr5 | 95049226 | 95160087 | + | protein_coding |
| ENSG00000107140.16_3 | TESK1 | 0.00484579 | 0.02767443 | 0.788 | 3.551 | chr9 | 35605259 | 35610038 | + | protein_coding |
| ENSG00000180881.19_8 | CAPS2 | 0.00801717 | 0.04377413 | 0.788 | 2.779 | chr12 | 75669759 | 75784708 | - | protein_coding |
| ENSG00000250159.7_7 | AC106791.1 | 0.0573102 | 0.25222449 | 0.788 | 0.764 | chr5 | 137145469 | 137225083 | - | lncRNA |
| ENSG00000148803.12_6 | FUOM | 0.01318174 | 0.06851959 | 0.788 | 2.472 | chr10 | 135168662 | 135171529 | - | protein_coding |
| ENSG00000203722.8_4 | RAET1G | 0.07323267 | 0.31060513 | 0.787 | 0.413 | chr6 | 150238014 | 150244257 | - | protein_coding |
| ENSG00000136014.12_6 | USP44 | 0.05200068 | 0.23242667 | 0.787 | 0.770 | chr12 | 95910336 | 95945252 | - | protein_coding |
| ENSG00000140853.15_6 | NLRC5 | 0.00602733 | 0.03372838 | 0.787 | 3.016 | chr16 | 57023397 | 57117443 | + | protein_coding |
| ENSG00000196498.13_8 | NCOR2 | 0.01138518 | 0.05995275 | 0.787 | 7.009 | chr12 | 124808961 | 125052135 | - | protein_coding |
| ENSG00000198001.13_5 | AC093012.1 | 0.00070833 | 0.00474366 | 0.787 | 4.395 | chr12 | 44152747 | 44183346 | + | protein_coding |
| ENSG00000139178.11_6 | C1RL | 7.38E-05 | 0.00058284 | 0.786 | 5.289 | chr12 | 7242183 | 7261834 | - | protein_coding |
| ENSG00000140548.10_4 | ZNF710 | 0.00158668 | 0.00990535 | 0.786 | 3.922 | chr15 | 90544556 | 90625438 | + | protein_coding |
| ENSG00000110042.8_3 | DTX4 | 0.01172776 | 0.0616462 | 0.786 | 3.170 | chr11 | 58938903 | 58976061 | + | protein_coding |
| ENSG00000172725.14_5 | CORO1B | 0.0019362 | 0.01194458 | 0.786 | 6.254 | chr11 | 67202981 | 67211292 | - | protein_coding |
| ENSG00000178878.12_3 | APOLD1 | 0.01955377 | 0.09777985 | 0.785 | 1.906 | chr12 | 12878851 | 12982909 | + | protein_coding |
| ENSG00000160972.9_3 | PPP1R16A | 0.04195987 | 0.19278536 | 0.785 | 4.365 | chr8 | 145703352 | 145727504 | + | protein_coding |
| ENSG00000075651.16_5 | PLD1 | 9.22E-05 | 0.00071786 | 0.784 | 5.391 | chr3 | 171318194 | 171528740 | - | protein_coding |
| ENSG00000279821.1_5 | AC145098.2 | 0.10206724 | 0.41223655 | 0.784 | -0.528 | chr5 | 176903632 | 176906657 | - | TEC |
| ENSG00000136044.12_5 | APPL2 | 2.01E-05 | 0.000174 | 0.784 | 6.154 | chr12 | 105567075 | 105629981 | - | protein_coding |
| ENSG00000175274.18_7 | TP53I11 | 7.73E-05 | 0.0006085 | 0.784 | 6.252 | chr11 | 44907454 | 44972857 | - | protein_coding |
| ENSG00000074964.17_4 | ARHGEF10L | 0.00070083 | 0.00470409 | 0.784 | 4.243 | chr1 | 17866193 | 18024369 | + | protein_coding |
| ENSG00000154914.17_6 | USP43 | 0.00158 | 0.00987058 | 0.783 | 3.894 | chr17 | 9548015 | 9633004 | + | protein_coding |
| ENSG00000267871.6_7 | ZNF460-AS1 | 0.05294288 | 0.23568932 | 0.783 | 0.718 | chr19 | 57778579 | 57791702 | - | lncRNA |
| ENSG00000287424.1_2 | AC080079.2 | 0.01979027 | 0.09879565 | 0.783 | 1.960 | chr4 | 166568788 | 166678819 | - | lncRNA |
| ENSG00000275765.5_8 | AC091982.3 | 0.04100555 | 0.18908531 | 0.783 | 1.280 | chr5 | 151149803 | 151151069 | - | lncRNA |
| ENSG00000177666.17_4 | PNPLA2 | 0.00236404 | 0.01430581 | 0.783 | 4.271 | chr11 | 818914 | 825573 | + | protein_coding |
| ENSG00000132635.17_5 | PCED1A | 0.00032244 | 0.00229701 | 0.782 | 5.484 | chr20 | 2815960 | 2821836 | - | protein_coding |
| ENSG00000135114.12_3 | OASL | 0.12394139 | 0.48535742 | 0.782 | -0.863 | chr12 | 121456914 | 121477045 | - | protein_coding |
| ENSG00000146966.13_5 | DENND2A | 0.04843487 | 0.21818382 | 0.782 | 1.130 | chr7 | 140218220 | 140373793 | - | protein_coding |
| ENSG00000110719.10_5 | TCIRG1 | 0.02526805 | 0.12279471 | 0.782 | 7.028 | chr11 | 67806492 | 67818362 | + | protein_coding |
| ENSG00000262920.5_6 | AC129507.4 | 0.10481492 | 0.42227814 | 0.781 | -0.576 | chr17 | 171183 | 172244 | + | lncRNA |
| ENSG00000176485.11_6 | PLAAT3 | 5.15E-05 | 0.00041858 | 0.781 | 5.727 | chr11 | 63340667 | 63384355 | - | protein_coding |
| ENSG00000165861.14_4 | ZFYVE1 | 0.00172446 | 0.01070614 | 0.780 | 3.940 | chr14 | 73436159 | 73493839 | - | protein_coding |
| ENSG00000177303.10_7 | CASKIN2 | 0.00065469 | 0.00441434 | 0.780 | 5.155 | chr17 | 73496342 | 73511618 | - | protein_coding |
| ENSG00000110011.13_3 | DNAJC4 | 0.00868861 | 0.04700684 | 0.779 | 2.910 | chr11 | 63997750 | 64001758 | + | protein_coding |
| ENSG00000181409.14_7 | AATK | 0.05624735 | 0.24856635 | 0.779 | 0.993 | chr17 | 79084287 | 79140021 | - | protein_coding |
| ENSG00000205746.9_6 | AC126755.1 | 0.05284019 | 0.23535013 | 0.778 | 0.925 | chr16 | 18428257 | 18446333 | - | transcribed_unprocessed_pseudogene |
| ENSG00000179163.11_3 | FUCA1 | 0.00208375 | 0.01277942 | 0.777 | 4.186 | chr1 | 24171567 | 24194784 | - | protein_coding |
| ENSG00000283538.3_7 | AC005972.3 | 0.09153759 | 0.3763824 | 0.777 | -0.018 | chr17 | 60987382 | 61045910 | - | lncRNA |
| ENSG00000168300.14_5 | PCMTD1 | 0.00064088 | 0.00432945 | 0.777 | 5.276 | chr8 | 52730135 | 52811746 | - | protein_coding |

| | | | | | | | | | | |
|---|---|---|---|---|---|---|---|---|---|---|
| ENSG00000089159.16_7 | PXN | 0.00048954 | 0.00337903 | 0.776 | 5.355 | chr12 | 120648242 | 120703574 | - | protein_coding |
| ENSG00000126106.14_4 | TMEM53 | 0.02288895 | 0.11255685 | 0.776 | 2.159 | chr1 | 45100910 | 45140153 | - | protein_coding |
| ENSG00000144677.15_4 | CTDSPL | 0.00014496 | 0.00109828 | 0.776 | 5.543 | chr3 | 37903371 | 38025960 | + | protein_coding |
| ENSG00000186111.9_4 | PIP5K1C | 0.00198267 | 0.01220585 | 0.775 | 4.449 | chr19 | 3630181 | 3700477 | - | protein_coding |
| ENSG00000286011.1_3 | AC062037.4 | 0.05462728 | 0.24233782 | 0.775 | 0.973 | chr2 | 85645627 | 85653847 | + | lncRNA |
| ENSG00000111269.3_3 | CREBL2 | 0.00016822 | 0.00126052 | 0.775 | 5.832 | chr12 | 12764761 | 12798042 | + | protein_coding |
| ENSG00000056972.20_8 | TRAF3IP2 | 0.00272619 | 0.01633049 | 0.775 | 3.950 | chr6 | 111876584 | 111928109 | - | protein_coding |
| ENSG00000259120.3_3 | SMIM6 | 0.00861157 | 0.04663267 | 0.774 | 3.157 | chr17 | 73642323 | 73644055 | + | protein_coding |
| ENSG00000232160.7_6 | RAP2C-AS1 | 0.08853298 | 0.36631704 | 0.774 | -0.044 | chrX | 131351081 | 131566890 | + | lncRNA |
| ENSG00000276814.1_6 | AC004801.6 | 0.14587545 | 0.55605429 | 0.774 | -0.963 | chr12 | 48433567 | 48434544 | - | lncRNA |
| ENSG00000117266.15_5 | CDK18 | 0.04497877 | 0.20474965 | 0.774 | 1.222 | chr1 | 205473723 | 205501921 | + | protein_coding |
| ENSG00000249599.2_7 | BMPR1B-DT | 0.07352643 | 0.31162789 | 0.773 | 0.421 | chr4 | 95664819 | 95678832 | - | lncRNA |
| ENSG00000078399.19_5 | HOXA9 | 0.12196383 | 0.47919757 | 0.773 | -0.662 | chr7 | 27202057 | 27214799 | - | protein_coding |
| ENSG00000138449.10_4 | SLC40A1 | 0.00047672 | 0.0032992 | 0.773 | 5.152 | chr2 | 190425305 | 190448484 | - | protein_coding |
| ENSG00000119139.20_8 | TJP2 | 0.09383471 | 0.38343177 | 0.772 | 0.041 | chr9 | 71736180 | 71889531 | + | protein_coding |
| ENSG00000177374.13_6 | HIC1 | 0.10397761 | 0.41918997 | 0.772 | 0.073 | chr17 | 1957448 | 1966535 | + | protein_coding |
| ENSG00000189171.14_4 | S100A13 | 0.00322176 | 0.01901065 | 0.772 | 3.643 | chr1 | 153591263 | 153603836 | - | protein_coding |
| ENSG00000133250.14_5 | ZNF414 | 0.05696853 | 0.25103173 | 0.772 | 1.409 | chr19 | 8574562 | 8579051 | - | protein_coding |
| ENSG00000104140.7_4 | RHOV | 0.04381364 | 0.20016376 | 0.771 | 1.915 | chr15 | 41164412 | 41166432 | - | protein_coding |
| ENSG00000244486.8_4 | SCARF2 | 0.12675451 | 0.49419516 | 0.771 | -0.335 | chr22 | 20779105 | 20792113 | - | protein_coding |
| ENSG00000188026.13_6 | RILPL1 | 0.00659066 | 0.03655829 | 0.770 | 3.628 | chr12 | 123954601 | 124018266 | - | protein_coding |
| ENSG00000166002.2 | SMCO4 | 0.018201 | 0.09173819 | 0.770 | 2.397 | chr11 | 93211638 | 93276674 | - | protein_coding |
| ENSG00000232533.1_7 | AC093673.1 | 0.02013509 | 0.10034785 | 0.770 | 2.204 | chr7 | 143076785 | 143077588 | - | lncRNA |
| ENSG00000197448.14_3 | GSTK1 | 5.29E-05 | 0.00042923 | 0.770 | 5.997 | chr7 | 142941186 | 142967947 | + | protein_coding |
| ENSG00000128000.16_5 | ZNF780B | 0.00027373 | 0.00198097 | 0.770 | 5.242 | chr19 | 40534167 | 40562138 | - | protein_coding |
| ENSG00000268001.1_7 | CARD8-AS1 | 0.09975035 | 0.40398095 | 0.770 | -0.244 | chr19 | 48758932 | 48761456 | + | lncRNA |
| ENSG00000125652.8_3 | ALKBH7 | 0.02870989 | 0.13745317 | 0.769 | 3.325 | chr19 | 6372805 | 6375261 | + | protein_coding |
| ENSG00000118620.13_5 | ZNF430 | 0.0059002 | 0.03308994 | 0.769 | 3.262 | chr19 | 21203440 | 21242856 | + | protein_coding |
| ENSG00000237489.5_7 | C10orf143 | 0.09530086 | 0.38879687 | 0.768 | -0.003 | chr10 | 131818289 | 131909094 | - | protein_coding |
| ENSG00000285928.1_3 | AC103591.4 | 0.11610356 | 0.46065476 | 0.767 | -0.715 | chr1 | 78488249 | 78490545 | - | lncRNA |
| ENSG00000160471.13_5 | COX6B2 | 0.09176487 | 0.37722963 | 0.767 | 0.167 | chr19 | 55860674 | 55866087 | - | protein_coding |
| ENSG00000124140.13_6 | SLC12A5 | 0.05950901 | 0.26016366 | 0.766 | 0.860 | chr20 | 44650329 | 44688791 | + | protein_coding |
| ENSG00000140479.17_5 | PCSK6 | 0.00135448 | 0.00858548 | 0.766 | 4.288 | chr15 | 101837347 | 102065405 | - | protein_coding |
| ENSG00000273145.1_6 | BX537318.1 | 0.09663862 | 0.39335128 | 0.764 | 0.167 | chr22 | 46409486 | 46411378 | + | lncRNA |
| ENSG00000276045.3_5 | ORAI1 | 0.00212654 | 0.01302387 | 0.764 | 4.507 | chr12 | 122064455 | 122080583 | + | protein_coding |
| ENSG00000111679.17_4 | PTPN6 | 0.00131935 | 0.0083867 | 0.762 | 4.644 | chr12 | 7055631 | 7070479 | + | protein_coding |
| ENSG00000170608.3_4 | FOXA3 | 0.02493006 | 0.12128489 | 0.762 | 2.323 | chr19 | 46367247 | 46377055 | + | protein_coding |
| ENSG00000231806.3_5 | PCAT7 | 0.0556763 | 0.24637629 | 0.761 | 1.615 | chr9 | 97317336 | 97366272 | + | lncRNA |
| ENSG00000223403.6_5 | MEG9 | 0.1089464 | 0.43644887 | 0.761 | -0.158 | chr14 | 101534620 | 101539274 | + | lncRNA |
| ENSG00000103528.17_4 | SYT17 | 0.00453225 | 0.0260597 | 0.761 | 3.673 | chr16 | 19179293 | 19279654 | + | protein_coding |
| ENSG00000153064.12_6 | BANK1 | 0.0181889 | 0.09170324 | 0.761 | 2.500 | chr4 | 102332443 | 102995969 | + | protein_coding |
| ENSG00000114841.17_5 | DNAH1 | 0.00548622 | 0.03097354 | 0.761 | 3.631 | chr3 | 52350335 | 52434507 | + | protein_coding |
| ENSG00000233967.7_6 | AL359715.1 | 0.14437152 | 0.5510309 | 0.760 | -0.928 | chr6 | 81151012 | 81176391 | + | lncRNA |
| ENSG00000251661.3_7 | AC136475.1 | 0.09148522 | 0.37626794 | 0.760 | 0.002 | chr11 | 318640 | 325631 | + | lncRNA |
| ENSG00000233218.1 | SNX18P16 | 0.04320135 | 0.19767151 | 0.759 | 1.412 | chr1 | 142688379 | 142688925 | - | pseudogene |
| ENSG00000133134.11_6 | BEX2 | 0.11907721 | 0.47021106 | 0.759 | -0.191 | chrX | 102564274 | 102565974 | - | protein_coding |
| ENSG00000250644.3_7 | AC068580.4 | 0.12465117 | 0.48770696 | 0.759 | -0.771 | chr11 | 1756051 | 1785184 | - | protein_coding |
| ENSG00000288096.1_1 | AC109322.2 | 0.09991209 | 0.40450514 | 0.758 | 0.091 | chr8 | 144962668 | 144975756 | + | lncRNA |
| ENSG00000185664.14_8 | PMEL | 0.06713502 | 0.28874071 | 0.758 | 0.750 | chr12 | 56347889 | 56367101 | - | protein_coding |
| ENSG00000162639.16_8 | HENMT1 | 0.00571607 | 0.03215879 | 0.758 | 3.451 | chr1 | 109190912 | 109204148 | - | protein_coding |
| ENSG00000125898.13_6 | FAM110A | 0.03222165 | 0.15262117 | 0.758 | 2.487 | chr20 | 814358 | 838106 | + | protein_coding |
| ENSG00000139718.10_4 | SETD1B | 0.00326798 | 0.01926419 | 0.757 | 4.882 | chr12 | 122242086 | 122270490 | + | protein_coding |
| ENSG00000186280.7_4 | KDM4D | 0.08479367 | 0.35306519 | 0.757 | 0.394 | chr11 | 94706873 | 94732683 | + | protein_coding |
| ENSG00000121104.8_6 | FAM117A | 0.05744965 | 0.25265047 | 0.757 | 1.518 | chr17 | 47787694 | 47866542 | - | protein_coding |
| ENSG00000225138.8_6 | SLC9A3-AS1 | 0.02395189 | 0.1169745 | 0.756 | 2.623 | chr5 | 473351 | 480999 | + | lncRNA |
| ENSG00000177738.4_8 | AC025171.1 | 0.01567763 | 0.08011058 | 0.755 | 2.816 | chr5 | 43013162 | 43067554 | - | lncRNA |
| ENSG00000070540.13_6 | WIPI1 | 0.00432281 | 0.02501739 | 0.755 | 3.872 | chr17 | 66417089 | 66453654 | - | protein_coding |
| ENSG00000103740.10_4 | ACSBG1 | 0.11089474 | 0.44345416 | 0.755 | -0.244 | chr15 | 78459810 | 78538030 | - | protein_coding |
| ENSG00000151611.16_5 | MMAA | 0.05646097 | 0.24911082 | 0.754 | 1.345 | chr4 | 146520194 | 146581185 | + | protein_coding |
| ENSG00000172640.3_2 | OR10AD1 | 0.13509777 | 0.52146036 | 0.754 | -0.928 | chr12 | 48595866 | 48597170 | - | protein_coding |
| ENSG00000165105.10_4 | RASEF | 0.00014718 | 0.00111418 | 0.753 | 6.005 | chr9 | 85594505 | 85678092 | - | protein_coding |

| | | | | | | | | | | |
|---|---|---|---|---|---|---|---|---|---|---|
| ENSG00000187624.9_6 | C17orf97 | 0.04153472 | 0.19117808 | 0.752 | 1.782 | chr17 | 260116 | 280853 | + | protein_coding |
| ENSG00000160345.13_4 | C9orf116 | 0.02921778 | 0.13977174 | 0.752 | 2.151 | chr9 | 138387027 | 138393580 | - | protein_coding |
| ENSG00000196576.15_5 | PLXNB2 | 0.00155748 | 0.00974759 | 0.752 | 8.097 | chr22 | 50713408 | 50746075 | - | protein_coding |
| ENSG00000127578.7_4 | WFIKKN1 | 0.08951554 | 0.37003782 | 0.751 | 0.467 | chr16 | 679239 | 684117 | + | protein_coding |
| ENSG00000140950.16_6 | MEAK7 | 0.00109209 | 0.00707113 | 0.751 | 4.828 | chr16 | 84509961 | 84587639 | - | protein_coding |
| ENSG00000183186.8_4 | C2CD4C | 0.1057084 | 0.42558829 | 0.750 | -0.126 | chr19 | 405445 | 409147 | - | protein_coding |
| ENSG00000204128.6_3 | C2orf72 | 0.00366103 | 0.02143179 | 0.750 | 4.126 | chr2 | 231902238 | 231914434 | + | protein_coding |
| ENSG00000185262.9_3 | UBALD2 | 0.01470061 | 0.07561834 | 0.750 | 3.304 | chr17 | 74261429 | 74267379 | + | protein_coding |
| ENSG00000176182.6_4 | MYPOP | 0.02175332 | 0.1075938 | 0.749 | 2.542 | chr19 | 46393281 | 46405871 | - | protein_coding |
| ENSG00000275793.1_4 | RIMBP3 | 0.07431486 | 0.31459426 | 0.749 | 0.711 | chr22 | 20455682 | 20461786 | - | protein_coding |
| ENSG00000136840.19_5 | ST6GALNAC4 | 0.00601224 | 0.03365451 | 0.749 | 3.881 | chr9 | 130670165 | 130679320 | - | protein_coding |
| ENSG00000197776.8_3 | KLHDC1 | 0.07878251 | 0.33117548 | 0.749 | 0.667 | chr14 | 50159823 | 50219868 | + | protein_coding |
| ENSG00000024862.17_4 | CCDC28A | 0.00910162 | 0.04901745 | 0.749 | 3.395 | chr6 | 139094646 | 139114456 | + | protein_coding |
| ENSG00000139083.11_5 | ETV6 | 0.00275406 | 0.01648079 | 0.748 | 4.838 | chr12 | 11802608 | 12048311 | + | protein_coding |
| ENSG00000036530.9_3 | CYP46A1 | 0.1508434 | 0.57094903 | 0.748 | -1.104 | chr14 | 100150635 | 100193638 | + | protein_coding |
| ENSG00000270141.3_4 | TERC | 0.06938916 | 0.29735581 | 0.748 | 5.876 | chr3 | 169482308 | 169482848 | - | lncRNA |
| ENSG00000109062.12_5 | SLC9A3R1 | 0.00047975 | 0.00331532 | 0.748 | 5.666 | chr17 | 72744767 | 72765492 | + | protein_coding |
| ENSG00000130703.16_5 | OSBPL2 | 0.00079547 | 0.00526375 | 0.747 | 5.039 | chr20 | 60806978 | 60871269 | + | protein_coding |
| ENSG00000232530.1_6 | LIF-AS1 | 0.14474693 | 0.55210792 | 0.747 | -0.631 | chr22 | 30635183 | 30636527 | + | lncRNA |
| ENSG00000099617.4_3 | EFNA2 | 0.14383749 | 0.54946477 | 0.747 | -0.930 | chr19 | 1285872 | 1301430 | + | protein_coding |
| ENSG00000259448.3_6 | LINC02352 | 0.0834523 | 0.34829846 | 0.747 | 0.673 | chr15 | 31507825 | 31516648 | + | lncRNA |
| ENSG00000134533.6_7 | RERG | 0.04250276 | 0.19487019 | 0.747 | 1.879 | chr12 | 15260717 | 15501609 | - | protein_coding |
| ENSG00000260778.5_8 | AC009065.5 | 0.06355618 | 0.27541529 | 0.746 | 1.091 | chr16 | 2318156 | 2323419 | + | lncRNA |
| ENSG00000215717.7_5 | TMEM167B | 0.00738404 | 0.04059166 | 0.746 | 3.691 | chr1 | 109633386 | 109639556 | + | protein_coding |
| ENSG00000105509.10_4 | HAS1 | 0.12238593 | 0.48032492 | 0.746 | -0.360 | chr19 | 52216365 | 52227247 | - | protein_coding |
| ENSG00000230333.6_8 | AC004160.1 | 0.10391159 | 0.41901888 | 0.745 | 0.150 | chr7 | 11220529 | 11559802 | + | lncRNA |
| ENSG00000168306.13_4 | ACOX2 | 0.01909271 | 0.09574379 | 0.745 | 2.788 | chr3 | 58490863 | 58523010 | - | protein_coding |
| ENSG00000167861.16_4 | HID1 | 0.00150565 | 0.00947271 | 0.744 | 4.697 | chr17 | 72946837 | 72969261 | - | protein_coding |
| ENSG00000167460.17_8 | TPM4 | 3.68E-05 | 0.00030622 | 0.744 | 9.230 | chr19 | 16177831 | 16213812 | + | protein_coding |
| ENSG00000114805.17_6 | PLCH1 | 0.00045715 | 0.00317275 | 0.743 | 5.503 | chr3 | 155093369 | 155462856 | - | protein_coding |
| ENSG00000118307.19_8 | CASC1 | 0.13741025 | 0.528549 | 0.743 | -0.563 | chr12 | 25261223 | 25348096 | - | protein_coding |
| ENSG00000177981.11_5 | ASB8 | 0.00196187 | 0.01209456 | 0.743 | 4.808 | chr12 | 48541572 | 48574996 | - | protein_coding |
| ENSG00000125848.10_4 | FLRT3 | 0.00055468 | 0.00379177 | 0.743 | 5.859 | chr20 | 14303631 | 14318260 | - | protein_coding |
| ENSG00000139323.14_5 | POC1B | 0.00090292 | 0.00592178 | 0.743 | 5.240 | chr12 | 89813495 | 89919824 | - | protein_coding |
| ENSG00000235823.3_5 | OLMALINC | 0.01799599 | 0.09088538 | 0.743 | 2.767 | chr10 | 102132856 | 102213800 | + | lncRNA |
| ENSG00000198915.11_7 | RASGEF1A | 0.01475875 | 0.0758296 | 0.743 | 3.447 | chr10 | 43689983 | 43762367 | - | protein_coding |
| ENSG00000120075.5_4 | HOXB5 | 0.05444831 | 0.24160416 | 0.742 | 1.535 | chr17 | 46668619 | 46671323 | - | protein_coding |
| ENSG00000251595.7_5 | ABCA11P | 0.13576201 | 0.52379667 | 0.742 | -0.879 | chr4 | 419224 | 467918 | - | transcribed_processed_pseudogene |
| ENSG00000164855.16_4 | TMEM184A | 0.00206557 | 0.01269424 | 0.742 | 5.496 | chr7 | 1581871 | 1600457 | - | protein_coding |
| ENSG00000179913.11_4 | B3GNT3 | 0.00800923 | 0.04374424 | 0.742 | 3.867 | chr19 | 17905637 | 17924385 | + | protein_coding |
| ENSG00000151376.16_7 | ME3 | 0.00192796 | 0.01189793 | 0.741 | 4.942 | chr11 | 86152150 | 86383678 | - | protein_coding |
| ENSG00000286451.1_1 | AL499602.1 | 0.08326442 | 0.34775714 | 0.741 | 0.893 | chr9 | 85678313 | 85684058 | + | lncRNA |
| ENSG00000175727.9 | MLXIP | 0.00087378 | 0.00574339 | 0.741 | 5.877 | chr12 | 122516628 | 122631894 | + | protein_coding |
| ENSG00000119514.7_4 | GALNT12 | 0.02469821 | 0.12025569 | 0.741 | 2.699 | chr9 | 101569952 | 101612363 | + | protein_coding |
| ENSG00000136881.11_3 | BAAT | 0.00650201 | 0.03612295 | 0.740 | 3.865 | chr9 | 104122699 | 104145801 | - | protein_coding |
| ENSG00000168679.18_6 | SLC16A4 | 0.00057964 | 0.00394731 | 0.740 | 5.795 | chr1 | 110905473 | 110933704 | - | protein_coding |
| ENSG00000125895.5_6 | TMEM74B | 0.11038121 | 0.44159941 | 0.740 | 0.226 | chr20 | 1161205 | 1166059 | - | protein_coding |
| ENSG00000164638.10_6 | SLC29A4 | 0.01206457 | 0.0632118 | 0.740 | 5.025 | chr7 | 5314000 | 5346501 | + | protein_coding |
| ENSG00000128833.13_4 | MYO5C | 0.00027649 | 0.00199771 | 0.740 | 6.226 | chr15 | 52484519 | 52588001 | - | protein_coding |
| ENSG00000102904.14_5 | TSNAXIP1 | 0.07038909 | 0.30091504 | 0.740 | 1.115 | chr16 | 67840668 | 67866051 | + | protein_coding |
| ENSG00000204519.11_4 | ZNF551 | 0.00255492 | 0.01538752 | 0.739 | 4.593 | chr19 | 58193337 | 58228669 | + | protein_coding |
| ENSG00000141469.18_6 | SLC14A1 | 0.13729691 | 0.52834184 | 0.739 | -0.832 | chr18 | 43266990 | 43332485 | + | protein_coding |
| ENSG00000164463.12_5 | CREBRF | 0.0112574 | 0.05939563 | 0.738 | 3.967 | chr5 | 172483355 | 172566287 | + | protein_coding |
| ENSG00000156875.14_4 | MFSD14A | 0.08496715 | 0.35362176 | 0.738 | 0.976 | chr1 | 100503651 | 100548933 | + | protein_coding |
| ENSG00000153790.12_5 | C7orf31 | 0.0502629 | 0.22533962 | 0.738 | 1.706 | chr7 | 25174311 | 25219975 | - | protein_coding |
| ENSG00000235374.2_3 | SSR4P1 | 0.09271609 | 0.37999694 | 0.737 | 0.581 | chr21 | 46490867 | 46494080 | - | transcribed_processed_pseudogene |
| ENSG00000163995.20_6 | ABLIM2 | 0.13443333 | 0.51945999 | 0.737 | -0.644 | chr4 | 7967037 | 8160559 | - | protein_coding |
| ENSG00000112245.12_5 | PTP4A1 | 0.08744667 | 0.36248253 | 0.736 | 0.692 | chr6 | 64231651 | 64293341 | + | protein_coding |

| | | | | | | | | | | |
|---|---|---|---|---|---|---|---|---|---|---|
| ENSG00000263006.6_2 | ROCK1P1 | 0.14040665 | 0.53832648 | 0.736 | -0.603 | chr18 | 109065 | 122219 | + | transcribed_unprocessed_pseudogene |
| ENSG00000110921.14_6 | MVK | 0.00273867 | 0.01639422 | 0.736 | 4.870 | chr12 | 110011060 | 110035930 | + | protein_coding |
| ENSG00000162813.18_6 | BPNT1 | 0.00433903 | 0.02509492 | 0.735 | 4.691 | chr1 | 220230824 | 220263804 | - | protein_coding |
| ENSG00000198113.3_3 | TOR4A | 0.08310764 | 0.34718398 | 0.735 | 5.602 | chr9 | 140172178 | 140177093 | + | protein_coding |
| ENSG00000130402.12_5 | ACTN4 | 0.00022414 | 0.00164424 | 0.735 | 10.564 | chr19 | 39138289 | 39222229 | + | protein_coding |
| ENSG00000112977.15_4 | DAP | 0.00128175 | 0.00817107 | 0.735 | 5.483 | chr5 | 10679342 | 10761384 | - | protein_coding |
| ENSG00000247121.7_7 | AC009126.1 | 0.12436293 | 0.48668638 | 0.735 | -0.478 | chr5 | 96149731 | 96271513 | - | lncRNA |
| ENSG00000173801.17_6 | JUP | 0.00031181 | 0.00223206 | 0.734 | 8.757 | chr17 | 39910856 | 39943183 | - | protein_coding |
| ENSG00000188316.15_5 | ENO4 | 0.12705358 | 0.49471479 | 0.734 | -0.338 | chr10 | 118609010 | 118671299 | + | protein_coding |
| ENSG00000076258.10_3 | FMO4 | 0.0838357 | 0.34981422 | 0.734 | 0.736 | chr1 | 171283322 | 171311223 | + | protein_coding |
| ENSG00000110057.8_6 | UNC93B1 | 0.02018186 | 0.10049638 | 0.733 | 6.409 | chr11 | 67758571 | 67772452 | - | protein_coding |
| ENSG00000103254.10_5 | ANTKMT | 0.02528851 | 0.12286054 | 0.733 | 4.476 | chr16 | 770581 | 772590 | + | protein_coding |
| ENSG00000117984.14_6 | CTSD | 0.02074115 | 0.10310801 | 0.733 | 6.585 | chr11 | 1773985 | 1785803 | - | protein_coding |
| ENSG00000239335.4_6 | LLPH-DT | 0.16271791 | 0.60595455 | 0.733 | -0.835 | chr12 | 66524531 | 66528229 | + | lncRNA |
| ENSG00000167588.13_6 | GPD1 | 0.15561082 | 0.58426406 | 0.733 | -0.642 | chr12 | 50497765 | 50505096 | + | protein_coding |
| ENSG00000260400.1_8 | AL513534.3 | 0.13675062 | 0.52669592 | 0.733 | -0.604 | chr10 | 70458257 | 70460551 | + | lncRNA |
| ENSG00000105877.18_4 | DNAH11 | 0.07378172 | 0.31263528 | 0.732 | 1.343 | chr7 | 21582657 | 21941457 | + | protein_coding |
| ENSG00000135506.16_6 | OS9 | 0.00021019 | 0.00154441 | 0.731 | 6.918 | chr12 | 58087738 | 58115340 | + | protein_coding |
| ENSG00000186272.12_3 | ZNF17 | 0.01893603 | 0.09503856 | 0.731 | 3.292 | chr19 | 57922531 | 57933307 | + | protein_coding |
| ENSG00000064601.18_4 | CTSA | 0.00131811 | 0.00838485 | 0.731 | 5.376 | chr20 | 44518783 | 44527459 | + | protein_coding |
| ENSG00000170558.9_5 | CDH2 | 0.00153501 | 0.00962335 | 0.731 | 5.386 | chr18 | 25530930 | 25757094 | - | protein_coding |
| ENSG00000137171.15_5 | KLC4 | 0.01306855 | 0.06799092 | 0.730 | 3.893 | chr6 | 43008515 | 43042833 | + | protein_coding |
| ENSG00000130702.15_4 | LAMA5 | 0.05116753 | 0.22887208 | 0.730 | 8.862 | chr20 | 60883011 | 60942368 | - | protein_coding |
| ENSG00000286672.1_2 | AL096712.2 | 0.16780271 | 0.62121931 | 0.729 | -1.106 | chr6 | 37268986 | 37272952 | - | lncRNA |
| ENSG00000234498.3_4 | RPL13AP20 | 0.08880764 | 0.36728245 | 0.729 | 0.782 | chr12 | 13028433 | 13029041 | + | processed_pseudogene |
| ENSG00000220201.7_4 | ZGLP1 | 0.13825108 | 0.53123778 | 0.729 | -0.172 | chr19 | 10415479 | 10420556 | - | protein_coding |
| ENSG00000141576.16_6 | RNF157 | 0.00838449 | 0.04551397 | 0.728 | 3.883 | chr17 | 74138546 | 74236574 | - | protein_coding |
| ENSG00000099204.20_7 | ABLIM1 | 0.00082463 | 0.00544451 | 0.728 | 6.638 | chr10 | 116190872 | 116527820 | - | protein_coding |
| ENSG00000133067.17_4 | LGR6 | 0.04685996 | 0.21195542 | 0.728 | 1.883 | chr1 | 202163029 | 202288909 | + | protein_coding |
| ENSG00000176399.4_3 | DMRTA1 | 0.02746621 | 0.13209538 | 0.728 | 2.948 | chr9 | 22446823 | 22455739 | + | protein_coding |
| ENSG00000159882.13_4 | ZNF230 | 0.01505258 | 0.07722762 | 0.728 | 3.420 | chr19 | 44507109 | 44518076 | + | protein_coding |
| ENSG00000238266.2_5 | LINC00707 | 0.01299096 | 0.06764663 | 0.728 | 4.024 | chr10 | 6821511 | 6921412 | + | lncRNA |
| ENSG00000149489.8_5 | ROM1 | 0.09410894 | 0.3844639 | 0.727 | 0.701 | chr11 | 62379194 | 62382592 | + | protein_coding |
| ENSG00000160271.16_7 | RALGDS | 0.04117461 | 0.18976637 | 0.727 | 2.096 | chr9 | 135973107 | 136024721 | - | protein_coding |
| ENSG00000173267.14_3 | SNCG | 0.04288588 | 0.19648105 | 0.727 | 2.116 | chr10 | 88718356 | 88723015 | + | protein_coding |
| ENSG00000161671.16_5 | EMC10 | 0.00156752 | 0.00979951 | 0.727 | 6.087 | chr19 | 50979657 | 50994127 | + | protein_coding |
| ENSG00000269473.1_6 | AC012313.8 | 0.10956711 | 0.43853994 | 0.727 | 0.289 | chr19 | 58951815 | 58957216 | + | lncRNA |
| ENSG00000129932.10_3 | DOHH | 0.06280108 | 0.27260854 | 0.726 | 3.073 | chr19 | 3490819 | 3500672 | - | protein_coding |
| ENSG00000100116.16_5 | GCAT | 0.07047592 | 0.3012138 | 0.726 | 1.669 | chr22 | 38203912 | 38213183 | + | protein_coding |
| ENSG00000151917.18_4 | BEND6 | 0.04857592 | 0.21865933 | 0.726 | 2.064 | chr6 | 56819905 | 56892144 | + | protein_coding |
| ENSG00000245571.7_8 | FAM111A-DT | 0.05487494 | 0.24325428 | 0.726 | 1.808 | chr11 | 58897606 | 58910488 | - | lncRNA |
| ENSG00000110628.16_7 | SLC22A18 | 0.01115565 | 0.05890118 | 0.726 | 4.161 | chr11 | 2920951 | 2946476 | + | protein_coding |
| ENSG00000149809.16_5 | TM7SF2 | 0.02862204 | 0.13710643 | 0.725 | 3.230 | chr11 | 64879317 | 64883880 | + | protein_coding |
| ENSG00000100075.10_4 | SLC25A1 | 0.00137046 | 0.00867745 | 0.725 | 6.360 | chr22 | 19163094 | 19166252 | - | protein_coding |
| ENSG00000170004.17_7 | CHD3 | 0.00023319 | 0.00170425 | 0.725 | 7.953 | chr17 | 7788114 | 7816078 | + | protein_coding |
| ENSG00000169962.5_3 | TAS1R3 | 0.05563509 | 0.24631662 | 0.724 | 1.777 | chr1 | 1266660 | 1270694 | + | protein_coding |
| ENSG00000181284.3_4 | TMEM102 | 0.05002917 | 0.22446166 | 0.724 | 1.872 | chr17 | 7338754 | 7340998 | + | protein_coding |
| ENSG00000070081.17_7 | NUCB2 | 0.00438332 | 0.02530163 | 0.723 | 5.448 | chr11 | 17229700 | 17371527 | + | protein_coding |
| ENSG00000196116.8_3 | TDRD7 | 0.0014552 | 0.00917154 | 0.723 | 5.441 | chr9 | 100174378 | 100258407 | + | protein_coding |
| ENSG00000173511.9_4 | VEGFB | 0.01070785 | 0.05680691 | 0.723 | 4.433 | chr11 | 64002010 | 64006265 | + | protein_coding |
| ENSG00000244926.7_7 | ALKBH3-AS1 | 0.14837937 | 0.56414643 | 0.723 | -0.742 | chr11 | 43930839 | 43942494 | - | lncRNA |
| ENSG00000186480.13_4 | INSIG1 | 0.00049281 | 0.00340029 | 0.722 | 6.476 | chr7 | 155089486 | 155101945 | + | protein_coding |
| ENSG00000277459.1_6 | AP001527.2 | 0.22400934 | 0.7772647 | 0.722 | -0.998 | chr11 | 101980558 | 101981188 | - | lncRNA |
| ENSG00000215241.4_6 | LINC02449 | 0.13082736 | 0.50740224 | 0.722 | -0.116 | chr12 | 8388011 | 8395160 | + | lncRNA |
| ENSG00000164938.14_4 | TP53INP1 | 0.00442845 | 0.02554551 | 0.722 | 4.721 | chr8 | 95938200 | 95961606 | - | protein_coding |
| ENSG00000168140.5_3 | VASN | 0.01890263 | 0.09492454 | 0.722 | 3.416 | chr16 | 4421849 | 4433539 | + | protein_coding |
| ENSG00000124564.17_5 | SLC17A3 | 0.12956948 | 0.50307277 | 0.721 | -0.273 | chr6 | 25833294 | 25882514 | - | protein_coding |
| ENSG00000198286.9_3 | CARD11 | 0.09818907 | 0.39911351 | 0.721 | 0.972 | chr7 | 2945775 | 3083579 | - | protein_coding |
| ENSG00000196544.8_3 | BORCS6 | 0.06174927 | 0.26841288 | 0.721 | 2.523 | chr17 | 8091663 | 8093498 | - | protein_coding |
| ENSG00000170430.10_5 | MGMT | 0.0271391 | 0.13070764 | 0.721 | 2.996 | chr10 | 131265454 | 131569247 | + | protein_coding |

| | | | | | | | | | | |
|---|---|---|---|---|---|---|---|---|---|---|
| ENSG00000250546.6_7 | AC079160.1 | 0.11327528 | 0.45175353 | 0.720 | 0.300 | chr4 | 84717589 | 85220342 | - | lncRNA |
| ENSG00000167470.12_4 | MIDN | 0.00740619 | 0.04070085 | 0.720 | 4.817 | chr19 | 1248552 | 1259139 | + | protein_coding |
| ENSG00000136717.15_6 | BIN1 | 0.00460409 | 0.02643856 | 0.720 | 5.773 | chr2 | 127805603 | 127864864 | - | protein_coding |
| ENSG00000225335.3_7 | AC016027.1 | 0.13995071 | 0.53692598 | 0.719 | -0.410 | chr22 | 18559293 | 18561650 | - | lncRNA |
| ENSG00000089820.15_6 | ARHGAP4 | 0.07236795 | 0.30759841 | 0.719 | 1.377 | chrX | 153172821 | 153200452 | - | protein_coding |
| ENSG00000111684.11_4 | LPCAT3 | 0.01095083 | 0.05788855 | 0.719 | 4.085 | chr12 | 7085347 | 7125781 | - | protein_coding |
| ENSG00000189180.16_3 | ZNF33A | 0.00673214 | 0.03725006 | 0.719 | 5.892 | chr10 | 38299578 | 38354016 | + | protein_coding |
| ENSG00000155530.3_3 | LRGUK | 0.12504818 | 0.4888297 | 0.718 | -0.086 | chr7 | 133812052 | 133949343 | + | protein_coding |
| ENSG00000207425.1 | Y_RNA | 0.14191122 | 0.54327442 | 0.718 | -0.308 | chr16 | 15009314 | 15009413 | - | misc_RNA |
| ENSG00000182836.10_4 | PLCXD3 | 0.00746605 | 0.0410096 | 0.718 | 4.391 | chr5 | 41307054 | 41510730 | - | protein_coding |
| ENSG00000127580.17_6 | WDR24 | 0.02133448 | 0.10576165 | 0.718 | 3.642 | chr16 | 734622 | 740444 | - | protein_coding |
| ENSG00000069493.15_7 | CLEC2D | 0.05593044 | 0.24737768 | 0.717 | 2.112 | chr12 | 9817565 | 9852149 | + | protein_coding |
| ENSG00000248409.2_5 | AP000344.2 | 0.11790225 | 0.46695705 | 0.717 | 0.594 | chr22 | 23734740 | 23736979 | + | unprocessed_pseudogene |
| ENSG00000224051.7_3 | CPTP | 0.05884682 | 0.25796483 | 0.717 | 3.908 | chr1 | 1260136 | 1264276 | + | protein_coding |
| ENSG00000240399.1_5 | AC004801.2 | 0.14877978 | 0.56506421 | 0.717 | -0.701 | chr12 | 48448596 | 48449374 | - | transcribed_processed_pseudogene |
| ENSG00000160094.15_5 | ZNF362 | 0.04053563 | 0.18716127 | 0.717 | 3.245 | chr1 | 33722093 | 33766320 | + | protein_coding |
| ENSG00000100299.18_4 | ARSA | 0.01170906 | 0.061567 | 0.716 | 4.389 | chr22 | 51061182 | 51066601 | - | protein_coding |
| ENSG00000271614.1_5 | ATP2B1-AS1 | 0.06541485 | 0.28243638 | 0.716 | 1.630 | chr12 | 90102736 | 90106367 | + | lncRNA |
| ENSG00000132359.15_6 | AC002316.1 | 0.00594825 | 0.03334252 | 0.716 | 5.130 | chr17 | 2658999 | 2941035 | + | protein_coding |
| ENSG00000239552.2_7 | HOXB-AS2 | 0.22032306 | 0.76826521 | 0.716 | -0.834 | chr17 | 46634624 | 46637695 | + | lncRNA |
| ENSG00000096696.14_4 | DSP | 0.00057841 | 0.00394039 | 0.716 | 7.689 | chr6 | 7541850 | 7586947 | + | protein_coding |
| ENSG00000131002.12_5 | TXLNGY | 0.00647663 | 0.03600447 | 0.715 | 6.287 | chrY | 21729199 | 21768160 | + | transcribed_unprocessed_pseudogene |
| ENSG00000149596.6_3 | JPH2 | 0.07874962 | 0.33117548 | 0.714 | 1.316 | chr20 | 42740335 | 42816218 | - | protein_coding |
| ENSG00000022840.16_6 | RNF10 | 0.00078918 | 0.00523184 | 0.714 | 7.312 | chr12 | 120971283 | 121015391 | + | protein_coding |
| ENSG00000238549.1 | snoU13 | 0.1814026 | 0.66207476 | 0.713 | -1.121 | chr20 | 34304393 | 34304492 | + | snoRNA |
| ENSG00000187097.12_4 | ENTPD5 | 0.00728896 | 0.04010623 | 0.713 | 4.460 | chr14 | 74424713 | 74486102 | - | protein_coding |
| ENSG00000167363.14_6 | FN3K | 0.08194657 | 0.34297873 | 0.713 | 1.568 | chr17 | 80693491 | 80709072 | + | protein_coding |
| ENSG00000237854.3_4 | LINC00674 | 0.04153196 | 0.19117808 | 0.712 | 2.929 | chr17 | 66098049 | 66111659 | + | transcribed_unprocessed_pseudogene |
| ENSG00000124507.11_3 | PACSIN1 | 0.12696428 | 0.49461564 | 0.712 | 0.162 | chr6 | 34433838 | 34503006 | + | protein_coding |
| ENSG00000073792.15_4 | IGF2BP2 | 0.01119984 | 0.05911694 | 0.712 | 4.127 | chr3 | 185361527 | 185542844 | - | protein_coding |
| ENSG00000204186.10_5 | ZDBF2 | 0.00814921 | 0.04439956 | 0.711 | 4.570 | chr2 | 207139387 | 207179151 | + | protein_coding |
| ENSG00000197915.6_4 | HRNR | 0.10439372 | 0.42077207 | 0.711 | 0.727 | chr1 | 152184552 | 152196669 | - | protein_coding |
| ENSG00000251630.1_5 | AC119751.6 | 0.16745406 | 0.62059798 | 0.711 | -1.067 | chr4 | 49563302 | 49563487 | - | processed_pseudogene |
| ENSG00000272894.5_7 | AC004982.2 | 0.14379331 | 0.54942589 | 0.711 | -0.384 | chr7 | 7592093 | 7606627 | - | lncRNA |
| ENSG00000124074.12_4 | ENKD1 | 0.04991233 | 0.22399402 | 0.711 | 2.913 | chr16 | 67696848 | 67701168 | - | protein_coding |
| ENSG00000285427.1_4 | SOD2-OT1 | 0.16842093 | 0.62275436 | 0.710 | -1.066 | chr6 | 160181290 | 160183364 | - | lncRNA |
| ENSG00000189058.9_6 | APOD | 0.17011728 | 0.6278508 | 0.710 | -1.066 | chr3 | 195295576 | 195310904 | - | protein_coding |
| ENSG00000232284.7_7 | GNG12-AS1 | 0.12226036 | 0.48004417 | 0.710 | 0.103 | chr1 | 68297986 | 68668670 | + | lncRNA |
| ENSG00000186300.12_3 | ZNF555 | 0.03687741 | 0.17228529 | 0.709 | 3.030 | chr19 | 2841435 | 2860469 | + | protein_coding |
| ENSG00000260285.1_6 | AL133367.1 | 0.06931798 | 0.29719414 | 0.709 | 1.614 | chr14 | 103991347 | 103995409 | - | lncRNA |
| ENSG00000185567.7_4 | AHNAK2 | 0.0006179 | 0.00418536 | 0.709 | 9.566 | chr14 | 105403581 | 105444711 | - | protein_coding |
| ENSG00000130202.10_4 | NECTIN2 | 0.00402724 | 0.02342892 | 0.708 | 6.575 | chr19 | 45349432 | 45392480 | + | protein_coding |
| ENSG00000154099.18_3 | DNAAF1 | 0.14435426 | 0.5510309 | 0.708 | -0.383 | chr16 | 84178892 | 84212373 | + | protein_coding |
| ENSG00000135951.16_5 | TSGA10 | 0.06215014 | 0.2699808 | 0.708 | 2.053 | chr2 | 99613724 | 99771427 | - | protein_coding |
| ENSG00000013588.8_6 | GPRC5A | 0.00042319 | 0.0029578 | 0.708 | 8.961 | chr12 | 13043716 | 13070871 | + | protein_coding |
| ENSG00000152669.9_4 | CCNO | 0.08642225 | 0.35883713 | 0.707 | 2.201 | chr5 | 54526980 | 54529436 | - | protein_coding |
| ENSG00000082438.17_8 | COBLL1 | 0.02066618 | 0.10276408 | 0.707 | 3.799 | chr2 | 165510134 | 165700189 | - | protein_coding |
| ENSG00000151176.8_3 | PLBD2 | 0.02309557 | 0.11341597 | 0.707 | 3.820 | chr12 | 113796371 | 113829434 | + | protein_coding |
| ENSG00000263105.1_7 | AC009171.2 | 0.14056748 | 0.53882685 | 0.707 | -0.243 | chr16 | 4230118 | 4233516 | - | lncRNA |
| ENSG00000171206.14_3 | TRIM8 | 0.00396441 | 0.02309366 | 0.706 | 5.349 | chr10 | 104402067 | 104420437 | + | protein_coding |
| ENSG00000132846.6_9 | ZBED3 | 0.07319924 | 0.31053746 | 0.706 | 1.984 | chr5 | 76367897 | 76383110 | - | protein_coding |
| ENSG00000135736.6_4 | CCDC102A | 0.04304678 | 0.19711654 | 0.706 | 4.013 | chr16 | 57546093 | 57570483 | - | protein_coding |
| ENSG00000185189.18_5 | NRBP2 | 0.00804351 | 0.04386404 | 0.705 | 4.763 | chr8 | 144915753 | 144923145 | - | protein_coding |
| ENSG00000188763.5_3 | FZD9 | 0.12482158 | 0.48805128 | 0.705 | 0.330 | chr7 | 72848108 | 72850450 | + | protein_coding |
| ENSG00000213445.10_4 | SIPA1 | 0.01652819 | 0.08404588 | 0.704 | 4.622 | chr11 | 65405572 | 65418389 | + | protein_coding |
| ENSG00000102189.17_6 | EEA1 | 0.00415179 | 0.02407468 | 0.704 | 5.766 | chr12 | 93164413 | 93323107 | - | protein_coding |
| ENSG00000024422.12_4 | EHD2 | 0.01474287 | 0.07579184 | 0.704 | 6.783 | chr19 | 48216679 | 48246391 | + | protein_coding |
| ENSG00000197841.15_7 | ZNF181 | 0.0086234 | 0.04668249 | 0.704 | 4.983 | chr19 | 35225060 | 35236283 | + | protein_coding |

| | | | | | | | | | | |
|---|---|---|---|---|---|---|---|---|---|---|
| ENSG00000165475.15_5 | CRYL1 | 0.05292918 | 0.2356874 | 0.704 | 2.644 | chr13 | 20977805 | 21100012 | - | protein_coding |
| ENSG00000006534.16_5 | ALDH3B1 | 0.00186153 | 0.01151078 | 0.704 | 8.055 | chr11 | 67776048 | 67796749 | + | protein_coding |
| ENSG00000099864.18_5 | PALM | 0.02338383 | 0.11460979 | 0.703 | 4.369 | chr19 | 708935 | 748329 | + | protein_coding |
| ENSG00000141232.5_5 | TOB1 | 0.00251341 | 0.01516842 | 0.703 | 7.182 | chr17 | 48939584 | 48945339 | - | protein_coding |
| ENSG00000092841.18_5 | MYL6 | 0.00128355 | 0.00817675 | 0.703 | 8.472 | chr12 | 56551945 | 56557280 | + | protein_coding |
| ENSG00000259673.6_6 | IQCH-AS1 | 0.0371537 | 0.17330263 | 0.703 | 3.056 | chr15 | 67582974 | 67814182 | - | lncRNA |
| ENSG00000111321.11_5 | LTBR | 0.00122261 | 0.0078504 | 0.703 | 7.357 | chr12 | 6484211 | 6500737 | + | protein_coding |
| ENSG00000151320.11_5 | AKAP6 | 0.04444306 | 0.20262248 | 0.702 | 2.666 | chr14 | 32798504 | 33306890 | + | protein_coding |
| ENSG00000177106.16_7 | EPS8L2 | 0.037293 | 0.17381553 | 0.702 | 6.845 | chr11 | 694438 | 727727 | + | protein_coding |
| ENSG00000111799.21_5 | COL12A1 | 0.02704159 | 0.13030863 | 0.702 | 3.669 | chr6 | 75794042 | 75915769 | - | protein_coding |
| ENSG00000287839.1_3 | AL353807.5 | 0.12117 | 0.47666366 | 0.701 | 0.573 | chr1 | 155596548 | 155607395 | - | lncRNA |
| ENSG00000072778.20_3 | ACADVL | 0.00123305 | 0.00791176 | 0.701 | 8.242 | chr17 | 7120444 | 7128585 | + | protein_coding |
| ENSG00000182796.14_5 | TMEM198B | 0.07056577 | 0.30145274 | 0.701 | 2.085 | chr12 | 56223392 | 56230030 | + | transcribed_unitary_pseudogene |
| ENSG00000185112.6_3 | FAM43A | 0.09591183 | 0.39093037 | 0.701 | 1.548 | chr3 | 194406612 | 194409766 | + | protein_coding |
| ENSG00000171603.18_4 | CLSTN1 | 0.00179805 | 0.01114652 | 0.701 | 7.018 | chr1 | 9788984 | 9884042 | - | protein_coding |
| ENSG00000158106.14_4 | RHPN1 | 0.03699166 | 0.17268281 | 0.700 | 5.594 | chr8 | 144451046 | 144466391 | + | protein_coding |
| ENSG00000026297.15_7 | RNASET2 | 0.09422851 | 0.38486391 | 0.700 | 1.709 | chr6 | 167342992 | 167370679 | - | protein_coding |
| ENSG00000189195.13_6 | BTBD8 | 0.03809275 | 0.17712481 | 0.699 | 3.344 | chr1 | 92545862 | 92650282 | + | protein_coding |
| ENSG00000257497.2_7 | AC121761.1 | 0.13799108 | 0.53043867 | 0.699 | 0.116 | chr12 | 75877234 | 75883600 | - | lncRNA |
| ENSG00000224914.4_6 | LINC00863 | 0.05050202 | 0.22635452 | 0.699 | 2.591 | chr10 | 89101442 | 89117639 | + | lncRNA |
| ENSG00000179988.14_5 | PSTK | 0.05353597 | 0.23791235 | 0.696 | 2.523 | chr10 | 124713897 | 124757029 | + | protein_coding |
| ENSG00000269054.2_7 | AC012313.6 | 0.12109755 | 0.47653161 | 0.696 | 0.684 | chr19 | 58873949 | 58878960 | + | lncRNA |
| ENSG00000135124.14_3 | P2RX4 | 0.01503893 | 0.07717988 | 0.696 | 4.360 | chr12 | 121647660 | 121671909 | + | protein_coding |
| ENSG00000204764.14_4 | RANBP17 | 0.01601613 | 0.08165238 | 0.695 | 4.593 | chr5 | 170288874 | 170727019 | + | protein_coding |
| ENSG00000188177.14_3 | ZC3H6 | 0.06056452 | 0.263803 | 0.695 | 2.721 | chr2 | 113033174 | 113097640 | + | protein_coding |
| ENSG00000123364.5_3 | HOXC13 | 0.04839407 | 0.21811705 | 0.694 | 2.833 | chr12 | 54332615 | 54340328 | + | protein_coding |
| ENSG00000179832.17_2 | MROH1 | 0.04304051 | 0.19711654 | 0.694 | 3.445 | chr8 | 145202919 | 145316843 | + | protein_coding |
| ENSG00000143409.15_7 | MINDY1 | 0.04047704 | 0.18693934 | 0.694 | 3.424 | chr1 | 150968562 | 150980851 | - | protein_coding |
| ENSG00000143850.16_8 | PLEKHA6 | 0.00364712 | 0.02135741 | 0.694 | 6.294 | chr1 | 204187981 | 204346793 | - | protein_coding |
| ENSG00000135678.12_8 | CPM | 0.03624532 | 0.16973393 | 0.693 | 3.284 | chr12 | 69235977 | 69365350 | - | protein_coding |
| ENSG00000267508.5_4 | ZNF285 | 0.06477962 | 0.28005131 | 0.693 | 2.297 | chr19 | 44886460 | 44905774 | - | protein_coding |
| ENSG00000225969.2_5 | ABHD11-AS1 | 0.15961632 | 0.59678098 | 0.692 | -0.262 | chr7 | 73149368 | 73150488 | + | transcribed_unitary_pseudogene |
| ENSG00000153029.14_2 | MR1 | 0.02434754 | 0.11873025 | 0.692 | 4.002 | chr1 | 181002561 | 181031074 | + | protein_coding |
| ENSG00000119699.7_4 | TGFB3 | 0.12616913 | 0.49256141 | 0.692 | 0.622 | chr14 | 76424442 | 76449334 | - | protein_coding |
| ENSG00000170791.18_5 | CHCHD7 | 0.02431666 | 0.1186255 | 0.691 | 4.060 | chr8 | 57124245 | 57131368 | + | protein_coding |
| ENSG00000240771.8_5 | ARHGEF25 | 0.08276958 | 0.34601582 | 0.691 | 1.844 | chr12 | 58003963 | 58011028 | + | protein_coding |
| ENSG00000201098.1 | RNY1 | 0.13547709 | 0.52281094 | 0.691 | 0.741 | chr7 | 148684228 | 148684340 | - | misc_RNA |
| ENSG00000124602.10_6 | UNC5CL | 0.09802199 | 0.39861677 | 0.691 | 1.437 | chr6 | 40994634 | 41006960 | - | protein_coding |
| ENSG00000177548.13_6 | RABEP2 | 0.03250513 | 0.15375894 | 0.691 | 4.322 | chr16 | 28915742 | 28947847 | - | protein_coding |
| ENSG00000228549.4_8 | BX284668.2 | 0.08648262 | 0.3590039 | 0.691 | 1.923 | chr1 | 17197440 | 17210154 | + | lncRNA |
| ENSG00000142875.19_5 | PRKACB | 0.00811029 | 0.04420107 | 0.690 | 5.428 | chr1 | 84543745 | 84704181 | + | protein_coding |
| ENSG00000160325.14_2 | CACFD1 | 0.05061208 | 0.22667627 | 0.690 | 3.378 | chr9 | 136325087 | 136335970 | + | protein_coding |
| ENSG00000263874.2_5 | LINC00672 | 0.14433791 | 0.5510309 | 0.689 | 0.235 | chr17 | 37081421 | 37085637 | + | protein_coding |
| ENSG00000135828.11_4 | RNASEL | 0.04097813 | 0.18900789 | 0.689 | 3.547 | chr1 | 182542769 | 182558391 | - | protein_coding |
| ENSG00000245105.4_6 | A2M-AS1 | 0.17988572 | 0.65816097 | 0.688 | 0.610 | chr12 | 9217759 | 9221285 | + | lncRNA |
| ENSG00000178209.15_7 | PLEC | 0.10589136 | 0.42603529 | 0.688 | 10.308 | chr8 | 144989315 | 145050902 | - | protein_coding |
| ENSG00000130827.6_4 | PLXNA3 | 0.03118434 | 0.14845987 | 0.687 | 3.825 | chrX | 153686621 | 153706118 | + | protein_coding |
| ENSG00000237037.9_6 | NDUFA6-DT | 0.13492941 | 0.52092371 | 0.687 | 0.452 | chr22 | 42486935 | 42533751 | + | lncRNA |
| ENSG00000179152.20_3 | TCAIM | 0.0254894 | 0.12373504 | 0.687 | 4.529 | chr3 | 44379611 | 44450943 | + | protein_coding |
| ENSG00000122861.16_4 | PLAU | 0.00559028 | 0.03147102 | 0.687 | 6.264 | chr10 | 75668935 | 75677254 | + | protein_coding |
| ENSG00000105520.10_3 | PLPPR2 | 0.04968752 | 0.22313176 | 0.686 | 3.294 | chr19 | 11466062 | 11476374 | + | protein_coding |
| ENSG00000159445.13_4 | THEM4 | 0.01319119 | 0.06852861 | 0.686 | 5.164 | chr1 | 151843342 | 151882113 | - | protein_coding |
| ENSG00000238058.2_7 | AL355574.1 | 0.07075248 | 0.30210509 | 0.686 | 2.340 | chr9 | 138799658 | 138807972 | + | lncRNA |
| ENSG00000260978.1_7 | AC126407.1 | 0.17520655 | 0.64409161 | 0.685 | -0.771 | chr15 | 23812411 | 23814804 | - | lncRNA |
| ENSG00000136807.14_6 | CDK9 | 0.00836762 | 0.04543626 | 0.685 | 5.678 | chr9 | 130547958 | 130553071 | + | protein_coding |
| ENSG00000204947.9_4 | ZNF425 | 0.11932516 | 0.47091295 | 0.685 | 1.209 | chr7 | 148799876 | 148823416 | - | protein_coding |
| ENSG00000148700.15_4 | ADD3 | 0.01228785 | 0.06428687 | 0.685 | 5.759 | chr10 | 111756126 | 111895323 | + | protein_coding |
| ENSG00000198169.9_5 | ZNF251 | 0.01615706 | 0.08225288 | 0.685 | 4.677 | chr8 | 145946292 | 145981802 | - | protein_coding |
| ENSG00000174080.11_5 | CTSF | 0.01311209 | 0.06817751 | 0.685 | 5.438 | chr11 | 66330935 | 66336312 | - | protein_coding |
| ENSG00000104522.16_5 | TSTA3 | 0.01179462 | 0.06196185 | 0.685 | 5.360 | chr8 | 144694788 | 144700218 | - | protein_coding |

| | | | | | | | | | | |
|---|---|---|---|---|---|---|---|---|---|---|
| ENSG00000234139.4_8 | AL035420.1 | 0.21474145 | 0.75297245 | 0.684 | -0.965 | chr20 | 34639413 | 34673625 | + | lncRNA |
| ENSG00000090020.11_4 | SLC9A1 | 0.04442731 | 0.20260265 | 0.684 | 3.516 | chr1 | 27425300 | 27493472 | - | protein_coding |
| ENSG00000144366.16_7 | GULP1 | 0.00567193 | 0.03192053 | 0.683 | 6.432 | chr2 | 189156396 | 189460658 | + | protein_coding |
| ENSG00000183421.12_4 | RIPK4 | 0.02778758 | 0.13339692 | 0.683 | 4.086 | chr21 | 43159529 | 43187249 | - | protein_coding |
| ENSG00000185669.6_4 | SNAI3 | 0.12118477 | 0.47666366 | 0.682 | 1.207 | chr16 | 88744096 | 88752915 | - | protein_coding |
| ENSG00000169951.9_5 | ZNF764 | 0.06323172 | 0.2742099 | 0.682 | 2.989 | chr16 | 30565085 | 30569819 | - | protein_coding |
| ENSG00000277268.2_7 | LHX1-DT | 0.14786745 | 0.56293729 | 0.682 | 0.216 | chr17 | 35218935 | 35293969 | - | lncRNA |
| ENSG00000176834.14_5 | VSIG10 | 0.01310097 | 0.06813962 | 0.682 | 5.343 | chr12 | 118501398 | 118573831 | - | protein_coding |
| ENSG00000242247.11_5 | ARFGAP3 | 0.01362549 | 0.07057846 | 0.681 | 5.145 | chr22 | 43192508 | 43254112 | - | protein_coding |
| ENSG00000150527.17_6 | MIA2 | 0.00885226 | 0.04780491 | 0.681 | 6.456 | chr14 | 39699435 | 39857717 | + | protein_coding |
| ENSG00000188643.11_4 | S100A16 | 0.00446644 | 0.02572295 | 0.680 | 6.963 | chr1 | 153579362 | 153585621 | - | protein_coding |
| ENSG00000172915.18_4 | NBEA | 0.01087652 | 0.05758137 | 0.680 | 5.638 | chr13 | 35516424 | 36247159 | + | protein_coding |
| ENSG00000111752.11_4 | PHC1 | 0.0523105 | 0.23322477 | 0.680 | 3.622 | chr12 | 9066492 | 9094063 | + | protein_coding |
| ENSG00000166925.9_7 | TSC22D4 | 0.08746342 | 0.36248253 | 0.680 | 4.933 | chr7 | 100060982 | 100076855 | - | protein_coding |
| ENSG00000162772.17_6 | ATF3 | 0.11863664 | 0.46914207 | 0.680 | 1.019 | chr1 | 212738676 | 212794119 | + | protein_coding |
| ENSG00000170214.5_5 | ADRA1B | 0.05217633 | 0.2328018 | 0.680 | 3.687 | chr5 | 159292087 | 159400019 | + | protein_coding |
| ENSG00000121900.19_4 | TMEM54 | 0.05331904 | 0.23706692 | 0.680 | 3.300 | chr1 | 33360195 | 33367039 | - | protein_coding |
| ENSG00000106686.16_5 | SPATA6L | 0.07078437 | 0.30216863 | 0.680 | 2.553 | chr9 | 4553386 | 4666674 | - | protein_coding |
| ENSG00000111110.12_4 | PPM1H | 0.01196668 | 0.0627359 | 0.680 | 5.744 | chr12 | 63037774 | 63328930 | - | protein_coding |
| ENSG00000256001.2_8 | AC079949.1 | 0.14465619 | 0.55188028 | 0.680 | 0.533 | chr12 | 127626574 | 127631077 | - | lncRNA |
| ENSG00000204344.14_3 | STK19 | 0.1213734 | 0.47725774 | 0.679 | 1.095 | chr6 | 31938868 | 31950598 | + | protein_coding |
| ENSG00000175221.15_9 | MED16 | 0.02747185 | 0.13209538 | 0.679 | 4.676 | chr19 | 867630 | 893218 | - | protein_coding |
| ENSG00000164142.16_5 | FAM160A1 | 0.16215108 | 0.60422363 | 0.678 | -0.085 | chr4 | 152330328 | 152591655 | + | protein_coding |
| ENSG00000180806.5_4 | HOXC9 | 0.07289613 | 0.30947324 | 0.678 | 2.527 | chr12 | 54388679 | 54397121 | + | protein_coding |
| ENSG00000285644.1_1 | AC108448.3 | 0.17026544 | 0.62826708 | 0.678 | -0.235 | chr11 | 3078768 | 3085937 | + | lncRNA |
| ENSG00000261764.1_2 | KRT18P18 | 0.2053646 | 0.72755989 | 0.678 | -0.925 | chr16 | 72762515 | 72763793 | + | transcribed_processed_pseudogene |
| ENSG00000006283.18_5 | CACNA1G | 0.02347969 | 0.11498448 | 0.678 | 5.495 | chr17 | 48638076 | 48704835 | + | protein_coding |
| ENSG00000112406.5_3 | HECA | 0.06336554 | 0.27465616 | 0.677 | 2.875 | chr6 | 139456217 | 139501939 | + | protein_coding |
| ENSG00000268575.1_8 | AL031282.2 | 0.12796063 | 0.49769625 | 0.677 | 1.028 | chr1 | 1634175 | 1669127 | - | lncRNA |
| ENSG00000080845.17_6 | DLGAP4 | 0.04044439 | 0.18683712 | 0.677 | 3.813 | chr20 | 34894258 | 35157040 | + | protein_coding |
| ENSG00000123989.14_5 | CHPF | 0.06967152 | 0.29834993 | 0.677 | 4.194 | chr2 | 220403670 | 220408531 | - | protein_coding |
| ENSG00000170561.13_4 | IRX2 | 0.11673878 | 0.46296828 | 0.677 | 1.215 | chr5 | 2745959 | 2751791 | - | protein_coding |
| ENSG00000230328.2_5 | AL359893.1 | 0.2080929 | 0.73529466 | 0.676 | -1.202 | chr9 | 104121214 | 104123524 | + | processed_pseudogene |
| ENSG00000169855.20_4 | ROBO1 | 0.01367648 | 0.07078068 | 0.676 | 5.511 | chr3 | 78646389 | 79817148 | - | protein_coding |
| ENSG00000160113.5_4 | NR2F6 | 0.04787433 | 0.21610346 | 0.675 | 6.023 | chr19 | 17342692 | 17356749 | - | protein_coding |
| ENSG00000178297.13_4 | TMPRSS9 | 0.16186336 | 0.6034046 | 0.675 | 0.208 | chr19 | 2360236 | 2426237 | + | protein_coding |
| ENSG00000267680.5_4 | ZNF224 | 0.03516944 | 0.16521824 | 0.675 | 4.105 | chr19 | 44598492 | 44614039 | + | protein_coding |
| ENSG00000162069.16_7 | BICDL2 | 0.19519148 | 0.69890509 | 0.675 | -1.143 | chr16 | 3077683 | 3086945 | - | protein_coding |
| ENSG00000205571.13_5 | SMN2 | 0.1227065 | 0.48127883 | 0.675 | 1.291 | chr5 | 69345439 | 69374349 | + | protein_coding |
| ENSG00000184719.12_4 | RNLS | 0.05562804 | 0.24631662 | 0.674 | 3.248 | chr10 | 90033621 | 90344287 | - | protein_coding |
| ENSG00000196739.15_3 | COL27A1 | 0.01475322 | 0.07582309 | 0.674 | 5.638 | chr9 | 116917817 | 117074791 | + | protein_coding |
| ENSG00000164211.13_4 | STARD4 | 0.01595643 | 0.08137137 | 0.673 | 5.695 | chr5 | 110831731 | 110848288 | - | protein_coding |
| ENSG00000101298.15_5 | SNPH | 0.04425277 | 0.20192315 | 0.673 | 3.807 | chr20 | 1246924 | 1289972 | + | protein_coding |
| ENSG00000254862.5_7 | LGR4-AS1 | 0.15537153 | 0.58364798 | 0.673 | 0.303 | chr11 | 27493276 | 27503980 | + | lncRNA |
| ENSG00000042317.17_8 | AL049834.1 | 0.05224692 | 0.23299974 | 0.673 | 3.705 | chr14 | 88851268 | 88936694 | + | protein_coding |
| ENSG00000101940.18_3 | WDR13 | 0.01444719 | 0.07444411 | 0.673 | 5.880 | chrX | 48448430 | 48467257 | + | protein_coding |
| ENSG00000119943.13_5 | PYROXD2 | 0.04316361 | 0.19754968 | 0.673 | 3.896 | chr10 | 100143322 | 100174939 | - | protein_coding |
| ENSG00000120832.10_7 | MTERF2 | 0.05710769 | 0.25158254 | 0.672 | 3.431 | chr12 | 107371055 | 107380938 | - | protein_coding |
| ENSG00000165312.6_2 | OTUD1 | 0.06579131 | 0.28371706 | 0.672 | 3.032 | chr10 | 23728387 | 23731319 | + | protein_coding |
| ENSG00000186715.11_4 | MST1L | 0.18196541 | 0.66368753 | 0.672 | -0.553 | chr1 | 17081405 | 17096732 | - | transcribed_unprocessed_pseudogene |
| ENSG00000117151.13_5 | CTBS | 0.09548242 | 0.38944815 | 0.671 | 2.084 | chr1 | 85015294 | 85040163 | - | protein_coding |
| ENSG00000255150.2_6 | EID3 | 0.04460839 | 0.20327192 | 0.671 | 3.957 | chr12 | 104697517 | 104698983 | + | protein_coding |
| ENSG00000176125.6_5 | UFSP1 | 0.13600236 | 0.52461004 | 0.671 | 1.036 | chr7 | 100486342 | 100487336 | - | protein_coding |
| ENSG00000163932.15_7 | PRKCD | 0.06830487 | 0.29313349 | 0.671 | 3.225 | chr3 | 53190025 | 53226733 | + | protein_coding |
| ENSG00000257704.4_3 | INAFM1 | 0.12626848 | 0.49284096 | 0.670 | 1.384 | chr19 | 47778142 | 47778980 | + | protein_coding |
| ENSG00000133639.6_5 | BTG1 | 0.01435402 | 0.07404989 | 0.670 | 5.758 | chr12 | 92534054 | 92539622 | - | protein_coding |
| ENSG00000112787.13_7 | FBRSL1 | 0.090913 | 0.3745076 | 0.670 | 4.427 | chr12 | 133066137 | 133161774 | + | protein_coding |
| ENSG00000050820.17_9 | BCAR1 | 0.03755998 | 0.17478476 | 0.670 | 6.274 | chr16 | 75262079 | 75301951 | - | protein_coding |
| ENSG00000090238.12_5 | YPEL3 | 0.0905469 | 0.3735191 | 0.669 | 2.289 | chr16 | 30103635 | 30108236 | - | protein_coding |

| | | | | | | | | | | |
|---|---|---|---|---|---|---|---|---|---|---|
| ENSG00000196247.11_3 | ZNF107 | 0.04341652 | 0.19840054 | 0.669 | 4.553 | chr7 | 64126461 | 64171960 | + | protein_coding |
| ENSG00000143061.17_5 | IGSF3 | 0.01057834 | 0.05627107 | 0.669 | 6.594 | chr1 | 117117021 | 117210377 | - | protein_coding |
| ENSG00000197472.15_5 | ZNF695 | 0.15162078 | 0.57352488 | 0.668 | 0.476 | chr1 | 247108849 | 247171395 | - | protein_coding |
| ENSG00000258359.1_4 | PCNPP1 | 0.171104 | 0.63057556 | 0.668 | -0.242 | chr12 | 112107656 | 112108166 | - | processed_pseudogene |
| ENSG00000116885.18_5 | OSCP1 | 0.09570187 | 0.39025363 | 0.668 | 2.322 | chr1 | 36881428 | 36916052 | - | protein_coding |
| ENSG00000256594.8_6 | AC010186.2 | 0.14734048 | 0.56127749 | 0.668 | 0.699 | chr12 | 9800643 | 9811008 | + | transcribed_unprocessed_pseudogene |
| ENSG00000213892.12_4 | CEACAM16 | 0.1739216 | 0.63989751 | 0.667 | -0.306 | chr19 | 45202421 | 45213990 | + | protein_coding |
| ENSG00000006837.12_6 | CDKL3 | 0.17721479 | 0.65039405 | 0.667 | -0.281 | chr5 | 133622041 | 133706738 | - | protein_coding |
| ENSG00000127328.21_3 | RAB3IP | 0.03086922 | 0.14707788 | 0.666 | 4.885 | chr12 | 70132461 | 70216984 | + | protein_coding |
| ENSG00000150672.17_10 | DLG2 | 0.18653681 | 0.67540239 | 0.666 | -0.687 | chr11 | 83166055 | 85338966 | - | protein_coding |
| ENSG00000103152.11_3 | MPG | 0.05651744 | 0.24929128 | 0.666 | 4.330 | chr16 | 127006 | 135852 | + | protein_coding |
| ENSG00000262903.1_7 | AC027796.4 | 0.10941363 | 0.43802431 | 0.666 | 2.041 | chr17 | 3558915 | 3561386 | - | lncRNA |
| ENSG00000163346.17_5 | PBXIP1 | 0.03968393 | 0.18361051 | 0.665 | 5.230 | chr1 | 154916552 | 154928599 | - | protein_coding |
| ENSG00000126870.16_4 | WDR60 | 0.04584793 | 0.20806644 | 0.665 | 4.081 | chr7 | 158649249 | 158749438 | + | protein_coding |
| ENSG00000184441.4_6 | AP001062.1 | 0.17916162 | 0.6560647 | 0.665 | -0.297 | chr21 | 45751117 | 45755734 | + | lncRNA |
| ENSG00000273369.1_7 | AC096586.2 | 0.192101 | 0.69090253 | 0.665 | -0.830 | chr4 | 44695963 | 44696403 | - | lncRNA |
| ENSG00000278129.2_4 | ZNF8 | 0.03582447 | 0.16798452 | 0.664 | 4.540 | chr19 | 58790321 | 58814157 | + | protein_coding |
| ENSG00000127334.10_7 | DYRK2 | 0.02523454 | 0.1226808 | 0.664 | 5.287 | chr12 | 68042118 | 68059186 | + | protein_coding |
| ENSG00000177721.4_4 | ANXA2R | 0.12319844 | 0.4829803 | 0.664 | 1.583 | chr5 | 43039183 | 43043272 | - | protein_coding |
| ENSG00000119537.18_6 | KDSR | 0.02772972 | 0.13315513 | 0.664 | 5.061 | chr18 | 60994959 | 61034461 | - | protein_coding |
| ENSG00000167536.14_4 | DHRS13 | 0.12689375 | 0.49461564 | 0.664 | 1.403 | chr17 | 27224799 | 27230097 | - | protein_coding |
| ENSG00000166823.5_3 | MESP1 | 0.11828778 | 0.46816153 | 0.664 | 1.784 | chr15 | 90291892 | 90294541 | - | protein_coding |
| ENSG00000173114.13_3 | LRRN3 | 0.15031572 | 0.56958698 | 0.664 | 0.762 | chr7 | 110731062 | 110765510 | + | protein_coding |
| ENSG00000165046.12_3 | LETM2 | 0.1297206 | 0.50354947 | 0.664 | 1.563 | chr8 | 38243725 | 38267045 | + | protein_coding |
| ENSG00000256092.2_7 | SBNO1-AS1 | 0.13236189 | 0.51205344 | 0.663 | 1.377 | chr12 | 123848415 | 123850660 | + | lncRNA |
| ENSG00000132437.18_4 | DDC | 0.14687273 | 0.55961565 | 0.663 | 0.864 | chr7 | 50526134 | 50633102 | - | protein_coding |
| ENSG00000148450.13_5 | MSRB2 | 0.06031483 | 0.26297264 | 0.662 | 3.710 | chr10 | 23384508 | 23410942 | + | protein_coding |
| ENSG00000088053.11_5 | GP6 | 0.20000666 | 0.71184262 | 0.662 | -0.929 | chr19 | 55525073 | 55549632 | - | protein_coding |
| ENSG00000140564.12_4 | FURIN | 0.0232888 | 0.11420705 | 0.662 | 6.224 | chr15 | 91411818 | 91426688 | + | protein_coding |
| ENSG00000186976.15_6 | EFCAB6 | 0.15179658 | 0.57396587 | 0.662 | 0.956 | chr22 | 43924624 | 44208217 | - | protein_coding |
| ENSG00000260430.2_6 | AC099518.1 | 0.17047847 | 0.62879198 | 0.662 | 0.014 | chr16 | 19098153 | 19109654 | + | lncRNA |
| ENSG00000142687.18_5 | KIAA0319L | 0.02104576 | 0.10450598 | 0.662 | 5.840 | chr1 | 35899093 | 36023551 | - | protein_coding |
| ENSG00000173068.18_7 | BNC2 | 0.05892985 | 0.25814877 | 0.662 | 3.975 | chr9 | 16409501 | 16870841 | - | protein_coding |
| ENSG00000226330.1_7 | AL606489.1 | 0.13640407 | 0.52593124 | 0.662 | 1.301 | chr1 | 204346978 | 204404974 | + | lncRNA |
| ENSG00000247081.8_9 | BAALC-AS1 | 0.18012681 | 0.65890735 | 0.661 | -0.381 | chr8 | 104165622 | 104311000 | - | lncRNA |
| ENSG00000247095.3_4 | MIR210HG | 0.12928036 | 0.50216969 | 0.661 | 1.349 | chr11 | 565660 | 568457 | - | lncRNA |
| ENSG00000196535.16_8 | MYO18A | 0.01515832 | 0.07768038 | 0.661 | 7.686 | chr17 | 27398142 | 27507430 | - | protein_coding |
| ENSG00000127129.10_4 | EDN2 | 0.19975907 | 0.71138893 | 0.661 | -0.787 | chr1 | 41944446 | 41950354 | - | protein_coding |
| ENSG00000181577.16_5 | C6orf223 | 0.15405171 | 0.58000356 | 0.661 | 0.686 | chr6 | 43968317 | 43975349 | + | protein_coding |
| ENSG00000116962.15_4 | NID1 | 0.07097227 | 0.30282526 | 0.661 | 3.403 | chr1 | 236139130 | 236228409 | - | protein_coding |
| ENSG00000287431.1_1 | AC027601.6 | 0.07515728 | 0.31755514 | 0.660 | 3.256 | chr17 | 79312983 | 79319766 | - | lncRNA |
| ENSG00000164010.15_8 | ERMAP | 0.07808973 | 0.32885049 | 0.660 | 3.307 | chr1 | 43282793 | 43310662 | + | protein_coding |
| ENSG00000172031.7_3 | EPHX4 | 0.19198703 | 0.69077229 | 0.659 | -0.831 | chr1 | 92495542 | 92529095 | + | protein_coding |
| ENSG00000108846.16_3 | ABCC3 | 0.01058491 | 0.05628918 | 0.659 | 9.348 | chr17 | 48712138 | 48769614 | + | protein_coding |
| ENSG00000064763.11_6 | FAR2 | 0.11944522 | 0.47107285 | 0.659 | 1.891 | chr12 | 29302036 | 29493913 | + | protein_coding |
| ENSG00000099330.9_3 | OCEL1 | 0.09288481 | 0.38060067 | 0.658 | 2.836 | chr19 | 17337022 | 17340028 | + | protein_coding |
| ENSG00000185482.8_5 | STAC3 | 0.19297382 | 0.69305961 | 0.658 | 0.027 | chr12 | 57637236 | 57644971 | - | protein_coding |
| ENSG00000254109.6_7 | RBPMS-AS1 | 0.15990145 | 0.59772123 | 0.658 | 0.580 | chr8 | 30239635 | 30242917 | - | lncRNA |
| ENSG00000152495.11_3 | CAMK4 | 0.17790967 | 0.65240889 | 0.658 | -0.013 | chr5 | 110559351 | 110830584 | + | protein_coding |
| ENSG00000135045.7_4 | C9orf40 | 0.06810332 | 0.29240978 | 0.658 | 3.737 | chr9 | 77561499 | 77567828 | - | protein_coding |
| ENSG00000259215.1_8 | AC027237.4 | 0.21415721 | 0.75151712 | 0.657 | -0.980 | chr15 | 69750861 | 69754145 | - | lncRNA |
| ENSG00000005206.17_4 | SPPL2B | 0.09020477 | 0.37245369 | 0.657 | 4.487 | chr19 | 2328614 | 2355093 | + | protein_coding |
| ENSG00000111452.13_5 | ADGRD1 | 0.15274206 | 0.57638688 | 0.657 | 0.945 | chr12 | 131438452 | 131626014 | + | protein_coding |
| ENSG00000129991.13_6 | TNNI3 | 0.16872833 | 0.62350173 | 0.657 | 0.523 | chr19 | 55663135 | 55669141 | - | protein_coding |
| ENSG00000179859.9_9 | RNF227 | 0.12610319 | 0.49241218 | 0.656 | 1.894 | chr17 | 7816642 | 7819594 | - | protein_coding |
| ENSG00000171466.10_5 | ZNF562 | 0.0329124 | 0.15551985 | 0.656 | 5.467 | chr19 | 9752483 | 9785776 | - | protein_coding |
| ENSG00000175536.7_5 | LIPT2 | 0.11332348 | 0.45178472 | 0.656 | 2.287 | chr11 | 74201564 | 74204769 | - | protein_coding |
| ENSG00000166401.15_5 | SERPINB8 | 0.11833812 | 0.46816153 | 0.656 | 2.152 | chr18 | 61637263 | 61687013 | + | protein_coding |
| ENSG00000117305.15_6 | HMGCL | 0.06612042 | 0.28479067 | 0.656 | 3.838 | chr1 | 24128375 | 24165110 | - | protein_coding |

| | | | | | | | | | | |
|---|---|---|---|---|---|---|---|---|---|---|
| ENSG00000255108.1_7 | AP006621.1 | 0.20038407 | 0.7127575 | 0.655 | -0.656 | chr11 | 823634 | 832883 | - | lncRNA |
| ENSG00000173846.13_3 | PLK3 | 0.07179326 | 0.30572038 | 0.655 | 3.587 | chr1 | 45266049 | 45271662 | + | protein_coding |
| ENSG00000158856.18_4 | DMTN | 0.04226341 | 0.19397935 | 0.654 | 5.207 | chr8 | 21906506 | 21940038 | + | protein_coding |
| ENSG00000101353.14_6 | MROH8 | 0.13689751 | 0.52714738 | 0.654 | 1.492 | chr20 | 35729629 | 35807991 | - | protein_coding |
| ENSG00000158769.18_5 | F11R | 0.04174088 | 0.1919281 | 0.654 | 4.848 | chr1 | 160965001 | 160991133 | - | protein_coding |
| ENSG00000240429.1_4 | LRRFIP1P1 | 0.14340616 | 0.54828863 | 0.654 | 1.300 | chr3 | 178976816 | 178979061 | - | processed_pseudogene |
| ENSG00000091622.16_5 | PITPNM3 | 0.11886673 | 0.4697298 | 0.654 | 2.118 | chr17 | 6354583 | 6459875 | - | protein_coding |
| ENSG00000172716.16_5 | SLFN11 | 0.05781042 | 0.25411128 | 0.654 | 4.229 | chr17 | 33677324 | 33700720 | - | protein_coding |
| ENSG00000089639.11_5 | GMIP | 0.11794164 | 0.46700892 | 0.653 | 2.691 | chr19 | 19740285 | 19754466 | - | protein_coding |
| ENSG00000287580.1_2 | AC073349.4 | 0.19468706 | 0.69766136 | 0.652 | -0.698 | chr7 | 64262336 | 64264495 | - | lncRNA |
| ENSG00000100197.22_6 | CYP2D6 | 0.1694528 | 0.6257884 | 0.652 | 0.452 | chr22 | 42522501 | 42526883 | - | protein_coding |
| ENSG00000131069.20_7 | ACSS2 | 0.04811091 | 0.21700597 | 0.652 | 4.844 | chr20 | 33459949 | 33515765 | + | protein_coding |
| ENSG00000107738.20_6 | VSIR | 0.03835867 | 0.17822137 | 0.651 | 5.607 | chr10 | 73507313 | 73533277 | - | protein_coding |
| ENSG00000118600.12_4 | RXYLT1 | 0.04844572 | 0.21818382 | 0.651 | 4.856 | chr12 | 64173622 | 64203342 | + | protein_coding |
| ENSG00000134531.10_5 | EMP1 | 0.05630482 | 0.24872394 | 0.651 | 4.588 | chr12 | 13349657 | 13372875 | + | protein_coding |
| ENSG00000203485.13_4 | INF2 | 0.09313114 | 0.38143408 | 0.650 | 7.673 | chr14 | 105155943 | 105188872 | + | protein_coding |
| ENSG00000111671.9_4 | SPSB2 | 0.05075147 | 0.2271288 | 0.650 | 4.979 | chr12 | 6980099 | 6998522 | - | protein_coding |
| ENSG00000185332.8_7 | AC027601.1 | 0.15272701 | 0.57638688 | 0.650 | 1.356 | chr17 | 79285070 | 79304474 | - | lncRNA |
| ENSG00000060762.19_4 | MPC1 | 0.13776266 | 0.5296752 | 0.650 | 1.891 | chr6 | 166778407 | 166796501 | - | protein_coding |
| ENSG00000167302.10_9 | TEPSIN | 0.07446779 | 0.31495369 | 0.649 | 3.967 | chr17 | 79202077 | 79212891 | - | protein_coding |
| ENSG00000181722.16_6 | ZBTB20 | 0.04623677 | 0.20956344 | 0.649 | 5.887 | chr3 | 114033348 | 114866118 | - | protein_coding |
| ENSG00000119574.13_5 | ZBTB45 | 0.08035745 | 0.33696268 | 0.649 | 4.019 | chr19 | 59024897 | 59050278 | - | protein_coding |
| ENSG00000139540.12_4 | SLC39A5 | 0.18633881 | 0.67523592 | 0.649 | -0.069 | chr12 | 56623833 | 56631630 | + | protein_coding |
| ENSG00000109089.7_3 | CDR2L | 0.04650618 | 0.21062334 | 0.649 | 6.185 | chr17 | 72983727 | 73001895 | + | protein_coding |
| ENSG00000111775.3_4 | COX6A1 | 0.05725042 | 0.25208632 | 0.649 | 5.236 | chr12 | 120875893 | 120878540 | + | protein_coding |
| ENSG00000172663.9_7 | TMEM134 | 0.08886307 | 0.36742617 | 0.648 | 3.684 | chr11 | 67229181 | 67236743 | - | protein_coding |
| ENSG00000259772.6_7 | AC012236.1 | 0.18381609 | 0.66882253 | 0.648 | 0.006 | chr15 | 31514202 | 31523041 | - | lncRNA |
| ENSG00000063169.10_6 | BICRA | 0.15310422 | 0.5771701 | 0.647 | 1.234 | chr19 | 48111453 | 48206534 | + | protein_coding |
| ENSG00000214530.9_7 | STARD10 | 0.09943717 | 0.4031721 | 0.647 | 3.523 | chr11 | 72465774 | 72504726 | - | protein_coding |
| ENSG00000149050.10_4 | ZNF214 | 0.15432336 | 0.58080261 | 0.647 | 1.367 | chr11 | 7018316 | 7041577 | - | protein_coding |
| ENSG00000163517.15_5 | HDAC11 | 0.09072627 | 0.37399853 | 0.647 | 3.482 | chr3 | 13521224 | 13547924 | + | protein_coding |
| ENSG00000135697.10_3 | BCO1 | 0.10685066 | 0.42911747 | 0.647 | 2.834 | chr16 | 81272294 | 81324747 | + | protein_coding |
| ENSG00000248905.10_8 | FMN1 | 0.06625307 | 0.28529283 | 0.647 | 4.905 | chr15 | 33057745 | 33486915 | - | protein_coding |
| ENSG00000259212.1_7 | AC103739.1 | 0.17683308 | 0.64926499 | 0.646 | 0.712 | chr15 | 91139072 | 91139679 | - | lncRNA |
| ENSG00000254681.6_3 | PKD1P5 | 0.19068948 | 0.68718323 | 0.646 | -0.100 | chr16 | 18468378 | 18495797 | - | transcribed_unprocessed_pseudogene |
| ENSG00000091073.19_3 | DTX2 | 0.06576775 | 0.28368429 | 0.646 | 4.946 | chr7 | 76090993 | 76135312 | + | protein_coding |
| ENSG00000260473.2_8 | AC068987.2 | 0.17116509 | 0.63065706 | 0.646 | 0.520 | chr12 | 52208827 | 52235890 | + | lncRNA |
| ENSG00000225680.1_7 | AL163953.1 | 0.21328803 | 0.74935504 | 0.646 | -0.995 | chr14 | 54066550 | 54080744 | - | lncRNA |
| ENSG00000235217.6_4 | TSPY26P | 0.18051503 | 0.65987608 | 0.645 | 0.117 | chr20 | 30774280 | 30778330 | - | transcribed_processed_pseudogene |
| ENSG00000076604.15_4 | TRAF4 | 0.05618641 | 0.24838617 | 0.645 | 5.665 | chr17 | 27071023 | 27077974 | + | protein_coding |
| ENSG00000127554.13_3 | GFER | 0.1172978 | 0.46487396 | 0.645 | 3.059 | chr16 | 2034194 | 2037750 | + | protein_coding |
| ENSG00000245685.7_7 | FRG1-DT | 0.20126768 | 0.71547075 | 0.645 | -0.699 | chr4 | 190685545 | 190861888 | - | lncRNA |
| ENSG00000266304.1_7 | LIVAR | 0.20833467 | 0.73573487 | 0.644 | -0.789 | chr18 | 68003165 | 68004546 | - | lncRNA |
| ENSG00000175602.4_3 | CCDC85B | 0.22313277 | 0.77520532 | 0.644 | 3.190 | chr11 | 65658144 | 65659106 | + | protein_coding |
| ENSG00000078070.13_6 | MCCC1 | 0.0672539 | 0.28911204 | 0.644 | 4.420 | chr3 | 182733006 | 182833863 | - | protein_coding |
| ENSG00000274750.2_3 | H3C6 | 0.06968905 | 0.29835311 | 0.644 | 4.387 | chr6 | 26224427 | 26227701 | + | protein_coding |
| ENSG00000179397.18_6 | CATSPERE | 0.18925873 | 0.68316943 | 0.643 | -0.126 | chr1 | 244617679 | 244804479 | + | protein_coding |
| ENSG00000153391.15_5 | INO80C | 0.10855475 | 0.43517425 | 0.643 | 2.992 | chr18 | 33032194 | 33077955 | - | protein_coding |
| ENSG00000111801.16_5 | BTN3A3 | 0.15692102 | 0.58868576 | 0.643 | 1.335 | chr6 | 26440700 | 26453643 | + | protein_coding |
| ENSG00000174123.11_4 | TLR10 | 0.20398362 | 0.72338861 | 0.643 | -0.616 | chr4 | 38773859 | 38784611 | - | protein_coding |
| ENSG00000250131.2_6 | AC078881.1 | 0.21556547 | 0.75545676 | 0.643 | -1.104 | chr4 | 178363673 | 178602535 | + | lncRNA |
| ENSG00000228812.7_6 | LAMA5-AS1 | 0.15030205 | 0.56958698 | 0.643 | 1.746 | chr20 | 60928051 | 60931536 | + | lncRNA |
| ENSG00000215790.7_6 | SLC35E2A | 0.09630026 | 0.39233357 | 0.642 | 3.539 | chr1 | 1656277 | 1677438 | - | protein_coding |
| ENSG00000204524.7_3 | ZNF805 | 0.06261332 | 0.27192638 | 0.642 | 4.748 | chr19 | 57752000 | 57774096 | + | protein_coding |
| ENSG00000073350.13_4 | LLGL2 | 0.04774882 | 0.21564462 | 0.642 | 6.844 | chr17 | 73521161 | 73571289 | + | protein_coding |
| ENSG00000140839.11_3 | CLEC18B | 0.23931168 | 0.81785891 | 0.641 | -0.881 | chr16 | 74442168 | 74455851 | - | protein_coding |
| ENSG00000187446.12_3 | CHP1 | 0.02532903 | 0.12302374 | 0.641 | 8.506 | chr15 | 41523037 | 41574085 | + | protein_coding |
| ENSG00000113269.14_6 | RNF130 | 0.05910949 | 0.25884175 | 0.641 | 5.403 | chr5 | 179338651 | 179499113 | - | protein_coding |
| ENSG00000129968.15_3 | ABHD17A | 0.1482652 | 0.563833 | 0.641 | 4.847 | chr19 | 1876809 | 1885546 | - | protein_coding |

| | | | | | | | | | | |
|---|---|---|---|---|---|---|---|---|---|---|
| ENSG00000197070.14_4 | ARRDC1 | 0.07614078 | 0.32125226 | 0.641 | 4.832 | chr9 | 140500137 | 140509812 | + | protein_coding |
| ENSG00000174326.13_5 | SLC16A11 | 0.19429627 | 0.69661673 | 0.640 | -0.157 | chr17 | 6944949 | 6947242 | - | protein_coding |
| ENSG00000184209.15_4 | SNRNP35 | 0.09633559 | 0.39238754 | 0.640 | 3.764 | chr12 | 123942635 | 123957701 | + | protein_coding |
| ENSG00000175130.7_4 | MARCKSL1 | 0.05507761 | 0.24403088 | 0.640 | 5.489 | chr1 | 32799440 | 32801834 | - | protein_coding |
| ENSG00000170345.10_4 | FOS | 0.14928272 | 0.56685318 | 0.640 | 1.622 | chr14 | 75745529 | 75748933 | + | protein_coding |
| ENSG00000106785.15_4 | TRIM14 | 0.08989612 | 0.37135186 | 0.639 | 4.249 | chr9 | 100831557 | 100881504 | - | protein_coding |
| ENSG00000185022.12_7 | MAFF | 0.14310802 | 0.54734298 | 0.639 | 2.077 | chr22 | 38596774 | 38612514 | + | protein_coding |
| ENSG00000186230.7_4 | ZNF749 | 0.08043491 | 0.33720791 | 0.639 | 4.388 | chr19 | 57946693 | 57958469 | + | protein_coding |
| ENSG00000135414.10_7 | GDF11 | 0.11307703 | 0.45116542 | 0.639 | 3.231 | chr12 | 56136906 | 56151048 | + | protein_coding |
| ENSG00000198353.8_4 | HOXC4 | 0.15443328 | 0.58094773 | 0.639 | 1.505 | chr12 | 54410715 | 54449814 | + | protein_coding |
| ENSG00000070961.15_4 | ATP2B1 | 0.03752046 | 0.1746466 | 0.639 | 8.539 | chr12 | 89981826 | 90103077 | - | protein_coding |
| ENSG00000169231.13_4 | THBS3 | 0.09202882 | 0.37796486 | 0.639 | 4.021 | chr1 | 155165379 | 155178842 | - | protein_coding |
| ENSG00000145730.20_6 | PAM | 0.06535699 | 0.28225515 | 0.638 | 4.864 | chr5 | 102089685 | 102366809 | + | protein_coding |
| ENSG00000254231.3_7 | AC103760.1 | 0.18728003 | 0.67754108 | 0.638 | 0.187 | chr8 | 87345503 | 87355658 | - | lncRNA |
| ENSG00000261819.1_6 | AC138932.4 | 0.21427418 | 0.75163066 | 0.638 | -0.531 | chr16 | 15082116 | 15084017 | - | processed_pseudogene |
| ENSG00000101213.7_4 | PTK6 | 0.15232327 | 0.57556945 | 0.637 | 1.989 | chr20 | 62159354 | 62168729 | - | protein_coding |
| ENSG00000119673.14_4 | ACOT2 | 0.08831368 | 0.36575037 | 0.637 | 4.387 | chr14 | 74034324 | 74042362 | + | protein_coding |
| ENSG00000274104.1_7 | AC020910.5 | 0.1980691 | 0.70692922 | 0.637 | -0.162 | chr19 | 35224203 | 35224742 | - | lncRNA |
| ENSG00000170653.19_7 | ATF7 | 0.11880838 | 0.46969365 | 0.636 | 3.346 | chr12 | 53901640 | 54020194 | - | protein_coding |
| ENSG00000135437.9_3 | RDH5 | 0.18079583 | 0.66040286 | 0.636 | 0.709 | chr12 | 56114151 | 56118489 | + | protein_coding |
| ENSG00000235244.4_6 | DANT2 | 0.18389247 | 0.66882651 | 0.636 | 0.727 | chrX | 114929200 | 115085463 | - | lncRNA |
| ENSG00000165698.16_6 | SPACA9 | 0.13269865 | 0.51309181 | 0.636 | 2.535 | chr9 | 135753414 | 135765588 | + | protein_coding |
| ENSG00000246898.1_5 | LINC00920 | 0.14803618 | 0.5633238 | 0.636 | 1.992 | chr16 | 66442427 | 66446038 | + | lncRNA |
| ENSG00000162522.11_3 | KIAA1522 | 0.06442051 | 0.27875286 | 0.635 | 5.675 | chr1 | 33207431 | 33240571 | + | protein_coding |
| ENSG00000149925.20_7 | ALDOA | 0.15016747 | 0.56948247 | 0.635 | 2.602 | chr16 | 30075485 | 30081778 | + | protein_coding |
| ENSG00000142784.16_4 | WDTC1 | 0.09246295 | 0.37939705 | 0.634 | 4.453 | chr1 | 27561123 | 27635127 | + | protein_coding |
| ENSG00000178913.8_5 | TAF7 | 0.06029461 | 0.26294904 | 0.633 | 6.221 | chr5 | 140639793 | 140700351 | - | protein_coding |
| ENSG00000262692.1_6 | AC116914.2 | 0.19921664 | 0.70974175 | 0.633 | 0.091 | chr17 | 3624922 | 3625782 | + | lncRNA |
| ENSG00000126005.17_7 | MMP24OS | 0.12030208 | 0.47392616 | 0.633 | 3.414 | chr20 | 33789548 | 33865934 | - | protein_coding |
| ENSG00000167183.3_4 | PRR15L | 0.19256909 | 0.6921657 | 0.633 | 0.224 | chr17 | 46029333 | 46035249 | - | protein_coding |
| ENSG00000146072.6_4 | TNFRSF21 | 0.06893722 | 0.29563302 | 0.633 | 5.608 | chr6 | 47199268 | 47277641 | - | protein_coding |
| ENSG00000196663.16_3 | TECPR2 | 0.09077653 | 0.37403214 | 0.632 | 4.773 | chr14 | 102829278 | 102968814 | + | protein_coding |
| ENSG00000173825.7_3 | TIGD3 | 0.19816985 | 0.70714676 | 0.632 | -0.053 | chr11 | 65122222 | 65125084 | + | protein_coding |
| ENSG00000130433.7_2 | CACNG6 | 0.09365864 | 0.38306471 | 0.632 | 5.229 | chr19 | 54495542 | 54515923 | + | protein_coding |
| ENSG00000244306.11_6 | AL589743.1 | 0.07898718 | 0.33176485 | 0.632 | 5.267 | chr14 | 19856543 | 19925390 | - | transcribed_processed_pseudogene |
| ENSG00000175806.15_6 | MSRA | 0.12849553 | 0.49957353 | 0.631 | 3.029 | chr8 | 9911802 | 10286401 | + | protein_coding |
| ENSG00000170425.3_2 | ADORA2B | 0.11349848 | 0.45233904 | 0.631 | 3.792 | chr17 | 15848231 | 15879060 | + | protein_coding |
| ENSG00000189042.13_5 | ZNF567 | 0.09131181 | 0.3757298 | 0.631 | 4.832 | chr19 | 37178514 | 37218603 | + | protein_coding |
| ENSG00000246859.3_7 | STARD4-AS1 | 0.16185092 | 0.6034046 | 0.631 | 2.011 | chr5 | 110846094 | 111075423 | + | lncRNA |
| ENSG00000167646.13_8 | DNAAF3 | 0.16009471 | 0.59819193 | 0.631 | 1.764 | chr19 | 55670029 | 55678090 | - | protein_coding |
| ENSG00000161914.10_6 | ZNF653 | 0.16292777 | 0.60635478 | 0.631 | 1.852 | chr19 | 11594242 | 11616654 | - | protein_coding |
| ENSG00000165516.11_3 | KLHDC2 | 0.07117028 | 0.30352434 | 0.631 | 5.395 | chr14 | 50234848 | 50253103 | + | protein_coding |
| ENSG00000247809.9_10 | NR2F2-AS1 | 0.18427359 | 0.66984046 | 0.631 | 0.687 | chr15 | 96653269 | 96870590 | - | lncRNA |
| ENSG00000280273.2_5 | AF131216.4 | 0.22219396 | 0.77330463 | 0.630 | -0.945 | chr8 | 11140990 | 11142577 | - | lncRNA |
| ENSG00000219438.8_4 | TAFA5 | 0.13913817 | 0.53392394 | 0.630 | 2.861 | chr22 | 48885272 | 49246724 | + | protein_coding |
| ENSG00000127418.15_4 | FGFRL1 | 0.09827517 | 0.39926841 | 0.629 | 4.593 | chr4 | 1003724 | 1020686 | + | protein_coding |
| ENSG00000232040.4_6 | ZBED9 | 0.23577113 | 0.80867677 | 0.629 | -0.895 | chr6 | 28538312 | 28583989 | - | protein_coding |
| ENSG00000257242.8_7 | LINC01619 | 0.21407002 | 0.75135956 | 0.629 | -0.713 | chr12 | 92378752 | 92536690 | - | lncRNA |
| ENSG00000260197.1_6 | AC010889.1 | 0.22490229 | 0.77982808 | 0.628 | -0.756 | chrY | 21853827 | 21856492 | - | lncRNA |
| ENSG00000270006.2_6 | AC010531.6 | 0.20980624 | 0.73931526 | 0.628 | -0.031 | chr16 | 87351115 | 87351649 | + | lncRNA |
| ENSG00000111252.11_3 | SH2B3 | 0.12718328 | 0.49510635 | 0.628 | 3.415 | chr12 | 111843727 | 111889427 | + | protein_coding |
| ENSG00000183283.16_3 | DAZAP2 | 0.0546766 | 0.24249605 | 0.628 | 7.706 | chr12 | 51632508 | 51665146 | + | protein_coding |
| ENSG00000139436.21_6 | GIT2 | 0.09070972 | 0.37399853 | 0.628 | 4.808 | chr12 | 110367607 | 110434194 | - | protein_coding |
| ENSG00000250742.4_7 | LINC02381 | 0.18898262 | 0.68245004 | 0.628 | 0.768 | chr12 | 54519866 | 54541269 | + | lncRNA |
| ENSG00000053501.13_4 | USE1 | 0.15732803 | 0.58983903 | 0.628 | 2.592 | chr19 | 17326155 | 17330638 | + | protein_coding |
| ENSG00000235245.1_8 | AL360181.2 | 0.19431713 | 0.69661673 | 0.628 | 0.452 | chr10 | 135108691 | 135109481 | - | lncRNA |
| ENSG00000213516.10_7 | RBMXL1 | 0.11273387 | 0.44999834 | 0.628 | 3.986 | chr1 | 89445139 | 89458643 | - | protein_coding |
| ENSG00000143507.18_4 | DUSP10 | 0.14109763 | 0.54062574 | 0.628 | 3.209 | chr1 | 221874766 | 221915431 | - | protein_coding |
| ENSG00000144645.14_5 | OSBPL10 | 0.10819561 | 0.43412631 | 0.627 | 4.740 | chr3 | 31699382 | 32119072 | - | protein_coding |
| ENSG00000136156.15_6 | ITM2B | 0.06130661 | 0.26670775 | 0.627 | 7.086 | chr13 | 48806748 | 48844493 | + | protein_coding |

| | | | | | | | | | | |
|---|---|---|---|---|---|---|---|---|---|---|
| ENSG00000099769.5_4 | IGFALS | 0.21955911 | 0.76623584 | 0.627 | -0.550 | chr16 | 1840414 | 1844972 | - | protein_coding |
| ENSG00000176894.10_4 | PXMP2 | 0.18662099 | 0.6755695 | 0.627 | 1.135 | chr12 | 133264173 | 133281571 | + | protein_coding |
| ENSG00000267221.3_9 | C17orf113 | 0.18385396 | 0.66882335 | 0.627 | 1.403 | chr17 | 40190250 | 40202619 | - | protein_coding |
| ENSG00000123739.11_5 | PLA2G12A | 0.12880336 | 0.50064522 | 0.626 | 3.643 | chr4 | 110631145 | 110651226 | - | protein_coding |
| ENSG00000138119.17_5 | MYOF | 0.05579977 | 0.24686118 | 0.626 | 9.602 | chr10 | 95066186 | 95242091 | - | protein_coding |
| ENSG00000137825.11_4 | ITPKA | 0.19071878 | 0.68718323 | 0.626 | 0.790 | chr15 | 41785591 | 41795749 | + | protein_coding |
| ENSG00000130997.16_6 | POLN | 0.20600636 | 0.72910657 | 0.625 | -0.264 | chr4 | 2073645 | 2243848 | - | protein_coding |
| ENSG00000205213.14_4 | LGR4 | 0.05809327 | 0.25510212 | 0.625 | 8.818 | chr11 | 27387508 | 27494337 | - | protein_coding |
| ENSG00000153048.11_7 | CARHSP1 | 0.09201134 | 0.37796486 | 0.625 | 5.397 | chr16 | 8946799 | 8962869 | - | protein_coding |
| ENSG00000163219.12_3 | ARHGAP25 | 0.20876049 | 0.73632765 | 0.625 | -0.037 | chr2 | 68906733 | 69053965 | + | protein_coding |
| ENSG00000011422.12_6 | PLAUR | 0.11780829 | 0.46668895 | 0.624 | 4.516 | chr19 | 44150247 | 44174699 | - | protein_coding |
| ENSG00000168264.10_3 | IRF2BP2 | 0.07255003 | 0.30829857 | 0.624 | 7.069 | chr1 | 234740015 | 234745271 | - | protein_coding |
| ENSG00000116525.14_4 | TRIM62 | 0.18064727 | 0.66006261 | 0.624 | 1.437 | chr1 | 33611000 | 33647660 | - | protein_coding |
| ENSG00000280303.2_5 | ERICD | 0.1903427 | 0.68670508 | 0.624 | 0.883 | chr8 | 141646380 | 141648382 | + | lncRNA |
| ENSG00000132122.12_7 | SPATA6 | 0.17079888 | 0.62958167 | 0.623 | 2.069 | chr1 | 48761045 | 48937880 | - | protein_coding |
| ENSG00000100218.12_4 | RSPH14 | 0.21527839 | 0.75470624 | 0.623 | -0.551 | chr22 | 23401593 | 23487208 | - | protein_coding |
| ENSG00000188747.8_4 | NOXA1 | 0.21259498 | 0.74721564 | 0.623 | 3.217 | chr9 | 140317802 | 140328858 | + | protein_coding |
| ENSG00000176022.7_7 | B3GALT6 | 0.11977225 | 0.47204826 | 0.623 | 4.582 | chr1 | 1167617 | 1170421 | + | protein_coding |
| ENSG00000148498.16_7 | PARD3 | 0.05967531 | 0.26070451 | 0.623 | 8.608 | chr10 | 34398488 | 35104253 | - | protein_coding |
| ENSG00000205517.12_6 | RGL3 | 0.17831148 | 0.65347774 | 0.623 | 1.753 | chr19 | 11495017 | 11530018 | - | protein_coding |
| ENSG00000211553.1 | AC118278.1 | 0.22337562 | 0.77574562 | 0.623 | -0.714 | chr4 | 130242 | 130349 | + | miRNA |
| ENSG00000177051.6_6 | FBXO46 | 0.16839453 | 0.62275436 | 0.623 | 3.991 | chr19 | 46213887 | 46234154 | - | protein_coding |
| ENSG00000011105.14_5 | TSPAN9 | 0.13309873 | 0.51452677 | 0.622 | 4.001 | chr12 | 3186521 | 3395730 | + | protein_coding |
| ENSG00000225975.7_6 | LINC01534 | 0.16590896 | 0.61603021 | 0.622 | 2.543 | chr19 | 37173538 | 37178351 | - | lncRNA |
| ENSG00000257512.1_5 | AC124947.2 | 0.21762788 | 0.76099086 | 0.622 | -0.671 | chr12 | 93708585 | 93709717 | + | transcribed_processed_pseudogene |
| ENSG00000109436.8_5 | TBC1D9 | 0.0912839 | 0.3757298 | 0.621 | 5.651 | chr4 | 141541936 | 141677539 | - | protein_coding |
| ENSG00000169972.12_3 | PUSL1 | 0.16053566 | 0.59933537 | 0.621 | 2.941 | chr1 | 1243977 | 1247057 | + | protein_coding |
| ENSG00000101460.13_3 | MAP1LC3A | 0.181052 | 0.66093092 | 0.621 | 1.938 | chr20 | 33134658 | 33148149 | + | protein_coding |
| ENSG00000245156.1_8 | AP001107.1 | 0.21056796 | 0.74126432 | 0.621 | -0.144 | chr11 | 66037303 | 66045996 | - | lncRNA |
| ENSG00000232973.13_8 | CYP1B1-AS1 | 0.21029486 | 0.74059638 | 0.621 | 0.072 | chr2 | 38300590 | 38458793 | + | lncRNA |
| ENSG00000176788.9_4 | BASP1 | 0.07914006 | 0.33224989 | 0.620 | 8.475 | chr5 | 17065707 | 17276952 | + | protein_coding |
| ENSG00000118507.17_5 | AKAP7 | 0.1627172 | 0.60595455 | 0.620 | 2.819 | chr6 | 131456607 | 131604675 | + | protein_coding |
| ENSG00000162729.14_6 | IGSF8 | 0.15815396 | 0.59231059 | 0.620 | 4.086 | chr1 | 160061130 | 160068733 | - | protein_coding |
| ENSG00000078808.17_5 | SDF4 | 0.11358256 | 0.45257261 | 0.620 | 6.213 | chr1 | 1152288 | 1167447 | - | protein_coding |
| ENSG00000151474.23_7 | FRMD4A | 0.15043894 | 0.56978228 | 0.619 | 3.357 | chr10 | 13685706 | 14504141 | - | protein_coding |
| ENSG00000062370.16_7 | ZNF112 | 0.14790292 | 0.56293729 | 0.619 | 3.448 | chr19 | 44830708 | 44871377 | - | protein_coding |
| ENSG00000109079.10_5 | TNFAIP1 | 0.08630102 | 0.35841752 | 0.619 | 6.557 | chr17 | 26662628 | 26674035 | + | protein_coding |
| ENSG00000267571.1_7 | AC104532.2 | 0.21993032 | 0.76722992 | 0.619 | -0.514 | chr19 | 5911589 | 5913910 | - | lncRNA |
| ENSG00000151790.9_3 | TDO2 | 0.1999984 | 0.71184262 | 0.618 | 1.349 | chr4 | 156775890 | 156841558 | + | protein_coding |
| ENSG00000173227.14_4 | SYT12 | 0.159584 | 0.59678098 | 0.618 | 3.073 | chr11 | 66774249 | 66818334 | + | protein_coding |
| ENSG00000248383.4_4 | PCDHAC1 | 0.13709256 | 0.52771593 | 0.618 | 4.063 | chr5 | 140305954 | 140391929 | + | protein_coding |
| ENSG00000153404.14_4 | PLEKHG4B | 0.1908911 | 0.68752527 | 0.617 | 1.795 | chr5 | 92266 | 190087 | + | protein_coding |
| ENSG00000155957.17_4 | TMBIM4 | 0.16836024 | 0.62275436 | 0.617 | 3.239 | chr12 | 66529626 | 66563852 | - | protein_coding |
| ENSG00000215788.10_8 | TNFRSF25 | 0.20859821 | 0.73608031 | 0.617 | 0.708 | chr1 | 6520846 | 6526255 | - | protein_coding |
| ENSG00000127125.9_4 | PPCS | 0.12220962 | 0.47995099 | 0.617 | 5.056 | chr1 | 42921788 | 42939056 | + | protein_coding |
| ENSG00000101849.17_7 | TBL1X | 0.10670359 | 0.42872064 | 0.617 | 5.669 | chrX | 9431335 | 9709077 | + | protein_coding |
| ENSG00000178093.14_6 | TSSK6 | 0.20812725 | 0.73529466 | 0.616 | 0.405 | chr19 | 19623227 | 19626357 | - | protein_coding |
| ENSG00000130479.11_6 | MAP1S | 0.18319563 | 0.66724821 | 0.616 | 3.758 | chr19 | 17830051 | 17845322 | + | protein_coding |
| ENSG00000196935.9_4 | SRGAP1 | 0.11179035 | 0.44673396 | 0.616 | 5.804 | chr12 | 64238480 | 64555997 | + | protein_coding |
| ENSG00000132825.7_3 | PPP1R3D | 0.16511569 | 0.61346937 | 0.616 | 3.271 | chr20 | 58511718 | 58515360 | - | protein_coding |
| ENSG00000133138.20_6 | TBC1D8B | 0.18920962 | 0.68313094 | 0.615 | 2.389 | chrX | 106045903 | 106119380 | + | protein_coding |
| ENSG00000257815.6_9 | CNOT2-DT | 0.19718835 | 0.7047768 | 0.615 | 1.437 | chr12 | 70295698 | 70637159 | - | lncRNA |
| ENSG00000106012.18_5 | IQCE | 0.1182135 | 0.46798113 | 0.615 | 5.495 | chr7 | 2598606 | 2654367 | + | protein_coding |
| ENSG00000049239.12_3 | H6PD | 0.11832456 | 0.46816153 | 0.615 | 5.830 | chr1 | 9294834 | 9331396 | + | protein_coding |
| ENSG00000121310.17_5 | ECHDC2 | 0.13858273 | 0.53236744 | 0.615 | 4.348 | chr1 | 53361582 | 53392884 | - | protein_coding |
| ENSG00000154642.11_6 | C21orf91 | 0.17076147 | 0.62957439 | 0.615 | 2.767 | chr21 | 19161291 | 19191703 | - | protein_coding |
| ENSG00000186376.15_4 | ZNF75D | 0.13615769 | 0.5250952 | 0.615 | 4.428 | chrX | 134382867 | 134478034 | - | protein_coding |
| ENSG00000197746.14_5 | PSAP | 0.09074899 | 0.37400542 | 0.615 | 9.416 | chr10 | 73576055 | 73611082 | - | protein_coding |
| ENSG00000198171.13_4 | DDRGK1 | 0.12202703 | 0.47933988 | 0.613 | 5.879 | chr20 | 3170996 | 3185331 | - | protein_coding |
| ENSG00000135093.13_7 | USP30 | 0.16290396 | 0.60635478 | 0.613 | 3.723 | chr12 | 109460894 | 109525828 | + | protein_coding |

| | | | | | | | | | | |
|---|---|---|---|---|---|---|---|---|---|---|
| ENSG00000255455.2_6 | AP003486.1 | 0.20152963 | 0.7161154 | 0.613 | 1.526 | chr11 | 130736149 | 130740142 | - | lncRNA |
| ENSG00000181035.14_3 | SLC25A42 | 0.19316407 | 0.69360268 | 0.613 | 1.771 | chr19 | 19174803 | 19223839 | + | protein_coding |
| ENSG00000108557.19_6 | RAI1 | 0.15538104 | 0.58364798 | 0.613 | 6.353 | chr17 | 17584772 | 17714767 | + | protein_coding |
| ENSG00000099992.15_5 | TBC1D10A | 0.19191493 | 0.69077229 | 0.612 | 1.837 | chr22 | 30687979 | 30723035 | - | protein_coding |
| ENSG00000173264.14_6 | GPR137 | 0.15157277 | 0.57346539 | 0.612 | 4.291 | chr11 | 64037534 | 64056972 | + | protein_coding |
| ENSG00000166669.13_5 | ATF7IP2 | 0.16403591 | 0.60996778 | 0.612 | 3.692 | chr16 | 10420291 | 10577495 | + | protein_coding |
| ENSG00000261189.1_7 | AL031058.1 | 0.22299754 | 0.77517691 | 0.612 | 0.037 | chr6 | 7540684 | 7541571 | - | lncRNA |
| ENSG00000187091.13_4 | PLCD1 | 0.17809984 | 0.6529715 | 0.612 | 3.136 | chr3 | 38048987 | 38071253 | - | protein_coding |
| ENSG00000272447.1_6 | AL135925.1 | 0.21013953 | 0.74019608 | 0.612 | 0.744 | chr10 | 81585658 | 81587358 | + | lncRNA |
| ENSG00000147535.17_7 | PLPP5 | 0.14004496 | 0.5371716 | 0.612 | 4.948 | chr8 | 38120648 | 38126761 | - | protein_coding |
| ENSG00000142409.6_4 | ZNF787 | 0.20011115 | 0.71207187 | 0.612 | 4.769 | chr19 | 56598732 | 56632664 | - | protein_coding |
| ENSG00000143819.12_4 | EPHX1 | 0.11560265 | 0.45948822 | 0.612 | 6.710 | chr1 | 225997794 | 226033264 | + | protein_coding |
| ENSG00000134986.14_6 | NREP | 0.2077882 | 0.73468118 | 0.611 | 0.986 | chr5 | 110998318 | 111333161 | - | protein_coding |
| ENSG00000146826.17_6 | MAP11 | 0.17779402 | 0.65211941 | 0.611 | 3.230 | chr7 | 99752043 | 99756346 | - | protein_coding |
| ENSG00000145860.11_5 | RNF145 | 0.11568809 | 0.45962215 | 0.611 | 6.920 | chr5 | 158584417 | 158637061 | - | protein_coding |
| ENSG00000086544.3_3 | ITPKC | 0.15436202 | 0.58080261 | 0.610 | 4.308 | chr19 | 41223017 | 41246765 | + | protein_coding |
| ENSG00000164733.21_5 | CTSB | 0.11549719 | 0.45917174 | 0.610 | 7.864 | chr8 | 11700033 | 11726957 | - | protein_coding |
| ENSG00000179403.12_5 | VWA1 | 0.17206783 | 0.63360189 | 0.610 | 4.432 | chr1 | 1370241 | 1378262 | + | protein_coding |
| ENSG00000181896.12_7 | ZNF101 | 0.16437339 | 0.61096695 | 0.610 | 3.829 | chr19 | 19779605 | 19794318 | + | protein_coding |
| ENSG00000184363.10_4 | PKP3 | 0.21637156 | 0.75734307 | 0.610 | 5.435 | chr11 | 392614 | 404908 | + | protein_coding |
| ENSG00000127220.6_5 | ABHD8 | 0.19207479 | 0.69090253 | 0.609 | 2.737 | chr19 | 17402940 | 17421045 | - | protein_coding |
| ENSG00000225190.11_5 | PLEKHM1 | 0.17144731 | 0.63144785 | 0.609 | 3.687 | chr17 | 43513266 | 43568115 | - | protein_coding |
| ENSG00000261170.2_7 | AC009053.4 | 0.24547104 | 0.83341945 | 0.609 | -0.300 | chr16 | 74456018 | 74475835 | + | lncRNA |
| ENSG00000041353.10_5 | RAB27B | 0.12663243 | 0.49393598 | 0.608 | 6.504 | chr18 | 52385091 | 52562747 | + | protein_coding |
| ENSG00000180263.14_6 | FGD6 | 0.12480408 | 0.48805128 | 0.608 | 7.159 | chr12 | 95470525 | 95611258 | - | protein_coding |
| ENSG00000183317.17_6 | EPHA10 | 0.19654531 | 0.70304421 | 0.608 | 2.375 | chr1 | 38179552 | 38230805 | - | protein_coding |
| ENSG00000162572.21_4 | SCNN1D | 0.20379438 | 0.72300612 | 0.608 | 2.005 | chr1 | 1215816 | 1227409 | + | protein_coding |
| ENSG00000223820.5_2 | CFL1P1 | 0.24936995 | 0.84279168 | 0.608 | -0.962 | chr10 | 89577685 | 89605369 | + | transcribed_unprocessed_pseudogene |
| ENSG00000144063.4_3 | MALL | 0.15948785 | 0.59655173 | 0.607 | 4.897 | chr2 | 110841447 | 110873599 | - | protein_coding |
| ENSG00000171428.14_3 | NAT1 | 0.18577074 | 0.67400219 | 0.607 | 3.094 | chr8 | 18027986 | 18081198 | + | protein_coding |
| ENSG00000269514.3_6 | AC024257.3 | 0.2164336 | 0.75741101 | 0.607 | 0.846 | chr12 | 48592140 | 48624058 | + | lncRNA |
| ENSG00000167632.16_7 | TRAPPC9 | 0.15545997 | 0.58382103 | 0.607 | 5.176 | chr8 | 140739968 | 141468678 | - | protein_coding |
| ENSG00000130244.12_5 | FAM98C | 0.1755545 | 0.6449704 | 0.606 | 3.886 | chr19 | 38893775 | 38899728 | + | protein_coding |
| ENSG00000181031.16_7 | RPH3AL | 0.16045852 | 0.59917331 | 0.606 | 5.292 | chr17 | 62180 | 236045 | - | protein_coding |
| ENSG00000280095.1_8 | CU639417.4 | 0.24585659 | 0.8340909 | 0.606 | -0.514 | chr21 | 44875562 | 44876954 | - | lncRNA |
| ENSG00000118804.8_3 | STBD1 | 0.22474401 | 0.77943135 | 0.606 | 0.110 | chr4 | 77227179 | 77232752 | + | protein_coding |
| ENSG00000100228.12_2 | RAB36 | 0.18358843 | 0.6682679 | 0.606 | 3.585 | chr22 | 23487513 | 23506537 | + | protein_coding |
| ENSG00000270066.3_6 | AL356488.2 | 0.16116154 | 0.60129296 | 0.605 | 5.438 | chr1 | 109642815 | 109643241 | + | lncRNA |
| ENSG00000204592.9_3 | HLA-E | 0.17543294 | 0.64479044 | 0.605 | 4.210 | chr6 | 30457286 | 30461971 | + | protein_coding |
| ENSG00000219682.4_5 | AL133268.1 | 0.26593798 | 0.88516502 | 0.605 | -1.181 | chr6 | 25140231 | 25141631 | + | processed_pseudogene |
| ENSG00000106546.14_4 | AHR | 0.14643355 | 0.55806197 | 0.604 | 6.349 | chr7 | 16955983 | 17385776 | + | protein_coding |
| ENSG00000183723.12_4 | CMTM4 | 0.14379638 | 0.54942589 | 0.604 | 6.567 | chr16 | 66648653 | 66730610 | - | protein_coding |
| ENSG00000213023.11_5 | SYT3 | 0.23762363 | 0.81330359 | 0.604 | -0.479 | chr19 | 51124564 | 51143138 | - | protein_coding |
| ENSG00000006025.12_6 | OSBPL7 | 0.1739038 | 0.63989751 | 0.604 | 4.505 | chr17 | 45884738 | 45899200 | - | protein_coding |
| ENSG00000101082.14_5 | SLA2 | 0.22672392 | 0.784766 | 0.604 | 0.245 | chr20 | 35240721 | 35274599 | - | protein_coding |
| ENSG00000120093.11_4 | HOXB3 | 0.18494798 | 0.67197768 | 0.604 | 4.026 | chr17 | 46626232 | 46682274 | - | protein_coding |
| ENSG00000236095.1_7 | AL807757.2 | 0.21343003 | 0.74970568 | 0.603 | 2.055 | chr9 | 97662711 | 97667036 | + | lncRNA |
| ENSG00000169242.12_3 | EFNA1 | 0.18304283 | 0.66696513 | 0.603 | 4.326 | chr1 | 155100352 | 155107375 | + | protein_coding |
| ENSG00000198715.13_9 | GLMP | 0.18099031 | 0.66084151 | 0.603 | 4.276 | chr1 | 156259880 | 156265480 | - | protein_coding |
| ENSG00000104883.8_5 | PEX11G | 0.21800518 | 0.76201024 | 0.602 | 1.415 | chr19 | 7541761 | 7562335 | - | protein_coding |
| ENSG00000164078.13_5 | MST1R | 0.18600605 | 0.67458043 | 0.602 | 4.060 | chr3 | 49924435 | 49941306 | - | protein_coding |
| ENSG00000257727.6_5 | CNPY2 | 0.20874962 | 0.73632765 | 0.602 | 2.537 | chr12 | 56703626 | 56709903 | - | protein_coding |
| ENSG00000160781.17_5 | PAQR6 | 0.19898707 | 0.70906605 | 0.602 | 3.399 | chr1 | 156213111 | 156217908 | - | protein_coding |
| ENSG00000178585.15_4 | CTNNBIP1 | 0.20600368 | 0.72910657 | 0.602 | 2.690 | chr1 | 9908334 | 9970394 | - | protein_coding |
| ENSG00000178531.6_4 | CTXN1 | 0.2306904 | 0.79517614 | 0.601 | 0.236 | chr19 | 7989376 | 7991020 | - | protein_coding |
| ENSG00000106460.19_7 | TMEM106B | 0.15865206 | 0.59392564 | 0.601 | 7.631 | chr7 | 12250896 | 12282993 | + | protein_coding |
| ENSG00000007255.10_3 | TRAPPC6A | 0.19665122 | 0.70328146 | 0.601 | 3.724 | chr19 | 45666186 | 45681495 | - | protein_coding |
| ENSG00000132386.11_3 | SERPINF1 | 0.21845066 | 0.76311696 | 0.601 | 2.029 | chr17 | 1665323 | 1680859 | + | protein_coding |
| ENSG00000139971.15_8 | ARMH4 | 0.27051608 | 0.89735983 | 0.600 | -1.124 | chr14 | 58466453 | 58764857 | - | protein_coding |
| ENSG00000184925.12_3 | LCN12 | 0.22086215 | 0.76942311 | 0.600 | 1.433 | chr9 | 139844003 | 139849949 | + | protein_coding |

| | | | | | | | | | | |
|---|---|---|---|---|---|---|---|---|---|---|
| ENSG00000214960.10_6 | CRPPA | 0.22114256 | 0.76994707 | 0.599 | 1.516 | chr7 | 16127150 | 16461163 | - | protein_coding |
| ENSG00000228146.8_4 | CASP16P | 0.23592959 | 0.80875188 | 0.599 | -0.156 | chr16 | 3194016 | 3199964 | + | transcribed_unitary_pseudogene |
| ENSG00000203804.4_4 | ADAMTSL4-AS1 | 0.22785281 | 0.78775264 | 0.599 | 0.668 | chr1 | 150532678 | 150547028 | - | lncRNA |
| ENSG00000196391.11_5 | ZNF774 | 0.2235395 | 0.77616303 | 0.599 | 1.234 | chr15 | 90895516 | 90912378 | + | protein_coding |
| ENSG00000157353.17_5 | FCSK | 0.19327256 | 0.69371849 | 0.599 | 4.377 | chr16 | 70488498 | 70514177 | + | protein_coding |
| ENSG00000123353.10_3 | ORMDL2 | 0.20197226 | 0.71740126 | 0.599 | 3.920 | chr12 | 56211825 | 56215663 | + | protein_coding |
| ENSG00000198520.12_8 | ARMH1 | 0.23016758 | 0.79390164 | 0.599 | 0.204 | chr1 | 45140364 | 45191263 | + | protein_coding |
| ENSG00000090863.11_4 | GLG1 | 0.16394727 | 0.60976579 | 0.598 | 7.675 | chr16 | 74481325 | 74641042 | - | protein_coding |
| ENSG00000251615.3_6 | AC104825.1 | 0.23789421 | 0.81407287 | 0.598 | -0.076 | chr4 | 8356817 | 8360065 | - | lncRNA |
| ENSG00000013364.19_4 | MVP | 0.17966451 | 0.65762243 | 0.598 | 7.743 | chr16 | 29831715 | 29859360 | + | protein_coding |
| ENSG00000075826.17_5 | SEC31B | 0.21123648 | 0.74317596 | 0.598 | 2.995 | chr10 | 102246403 | 102279621 | - | protein_coding |
| ENSG00000122986.13_6 | HVCN1 | 0.23497847 | 0.80611362 | 0.598 | -0.134 | chr12 | 111065646 | 111142755 | - | protein_coding |
| ENSG00000142552.8_5 | RCN3 | 0.22688193 | 0.78515997 | 0.598 | 1.272 | chr19 | 50031260 | 50050219 | + | protein_coding |
| ENSG00000185275.6_4 | CD24P4 | 0.17836734 | 0.65354764 | 0.598 | 6.172 | chrY | 21154353 | 21154595 | - | processed_pseudogene |
| ENSG00000110851.12_7 | PRDM4 | 0.18373658 | 0.66867019 | 0.597 | 5.729 | chr12 | 108126648 | 108155049 | - | protein_coding |
| ENSG00000265678.1_6 | AC129510.1 | 0.24689405 | 0.8364925 | 0.597 | -0.684 | chr17 | 80111306 | 80112691 | + | lncRNA |
| ENSG00000063046.17_4 | EIF4B | 0.17454745 | 0.64193429 | 0.596 | 9.261 | chr12 | 53399942 | 53435993 | + | protein_coding |
| ENSG00000135503.13_5 | ACVR1B | 0.20303137 | 0.72058696 | 0.596 | 4.835 | chr12 | 52345483 | 52390862 | + | protein_coding |
| ENSG00000168310.11_6 | IRF2 | 0.20922664 | 0.73771179 | 0.596 | 3.702 | chr4 | 185308883 | 185395704 | - | protein_coding |
| ENSG00000170473.17_5 | PYM1 | 0.2041755 | 0.72392458 | 0.596 | 4.517 | chr12 | 56295197 | 56326402 | - | protein_coding |
| ENSG00000163702.20_6 | IL17RC | 0.21616691 | 0.75677587 | 0.596 | 3.793 | chr3 | 9958758 | 9975314 | + | protein_coding |
| ENSG00000099821.14_4 | POLRMT | 0.22103202 | 0.76971305 | 0.596 | 6.128 | chr19 | 617221 | 633537 | - | protein_coding |
| ENSG00000201793.1 | RN7SKP9 | 0.2619067 | 0.8748442 | 0.595 | -0.875 | chr13 | 99857962 | 99858260 | + | misc_RNA |
| ENSG00000181191.11_6 | PJA1 | 0.2016024 | 0.71623073 | 0.595 | 4.907 | chrX | 68380694 | 68385636 | - | protein_coding |
| ENSG00000102871.16_4 | TRADD | 0.21812708 | 0.76228634 | 0.595 | 3.605 | chr16 | 67188088 | 67193812 | - | protein_coding |
| ENSG00000157837.16_5 | SPPL3 | 0.19856915 | 0.70828712 | 0.595 | 5.251 | chr12 | 121200313 | 121342161 | - | protein_coding |
| ENSG00000235437.8_7 | LINC01278 | 0.22396096 | 0.7772647 | 0.595 | 2.452 | chrX | 62442872 | 62780975 | - | lncRNA |
| ENSG00000169252.5_2 | ADRB2 | 0.26556218 | 0.88453169 | 0.595 | -0.912 | chr5 | 148204808 | 148208250 | + | protein_coding |
| ENSG00000159069.14_6 | FBXW5 | 0.23356875 | 0.80263346 | 0.595 | 6.796 | chr9 | 139834887 | 139839190 | - | protein_coding |
| ENSG00000126561.16_5 | STAT5A | 0.22214627 | 0.77329011 | 0.595 | 3.130 | chr17 | 40439565 | 40463961 | + | protein_coding |
| ENSG00000248092.8_8 | NNT-AS1 | 0.22250121 | 0.77422228 | 0.595 | 3.191 | chr5 | 43571696 | 43603332 | - | lncRNA |
| ENSG00000090661.12_4 | CERS4 | 0.22873116 | 0.79017433 | 0.594 | 1.909 | chr19 | 8271620 | 8327305 | + | protein_coding |
| ENSG00000174137.12_5 | FAM53A | 0.23625853 | 0.80956707 | 0.594 | 0.226 | chr4 | 1619642 | 1686029 | - | protein_coding |
| ENSG00000145743.16_4 | FBXL17 | 0.21255203 | 0.74721248 | 0.593 | 4.871 | chr5 | 107194736 | 107717799 | - | protein_coding |
| ENSG00000163132.8_5 | MSX1 | 0.23448272 | 0.80503467 | 0.593 | 1.103 | chr4 | 4861392 | 4865663 | + | protein_coding |
| ENSG00000203721.7_7 | LINC00862 | 0.23089267 | 0.79563033 | 0.593 | 2.029 | chr1 | 200222547 | 200369833 | - | lncRNA |
| ENSG00000065970.9_3 | FOXJ2 | 0.21566256 | 0.75545676 | 0.593 | 4.636 | chr12 | 8185312 | 8208113 | + | protein_coding |
| ENSG00000111707.12_3 | SUDS3 | 0.20645622 | 0.73040773 | 0.592 | 6.379 | chr12 | 118814360 | 118855838 | + | protein_coding |
| ENSG00000105373.19_6 | NOP53 | 0.22905312 | 0.79113272 | 0.592 | 6.375 | chr19 | 48248803 | 48260315 | + | protein_coding |
| ENSG00000150687.12_5 | PRSS23 | 0.20283804 | 0.72018856 | 0.592 | 9.161 | chr11 | 86502101 | 86663952 | + | protein_coding |
| ENSG00000205903.7_4 | ZNF316 | 0.22793798 | 0.78789378 | 0.591 | 4.148 | chr7 | 6676949 | 6697910 | + | protein_coding |
| ENSG00000157150.5_4 | TIMP4 | 0.22863898 | 0.79000949 | 0.591 | 3.582 | chr3 | 12194568 | 12200412 | - | protein_coding |
| ENSG00000071205.12_3 | ARHGAP10 | 0.22751307 | 0.78688431 | 0.590 | 4.325 | chr4 | 148653214 | 148993927 | + | protein_coding |
| ENSG00000139631.18_6 | CSAD | 0.23360261 | 0.80263346 | 0.590 | 2.883 | chr12 | 53551447 | 53574693 | - | protein_coding |
| ENSG00000138696.11_6 | BMPR1B | 0.21970894 | 0.76660815 | 0.590 | 6.253 | chr4 | 95679106 | 96079599 | + | protein_coding |
| ENSG00000136830.12_5 | NIBAN2 | 0.2272533 | 0.7861389 | 0.590 | 8.012 | chr9 | 130267618 | 130341268 | - | protein_coding |
| ENSG00000197603.14_9 | CPLANE1 | 0.22332917 | 0.77573593 | 0.590 | 6.204 | chr5 | 37106330 | 37249530 | - | protein_coding |
| ENSG00000111077.17_2 | TNS2 | 0.22072514 | 0.7690966 | 0.590 | 5.897 | chr12 | 53440753 | 53458156 | + | protein_coding |
| ENSG00000140876.11_6 | NUDT7 | 0.23988399 | 0.81930389 | 0.590 | 0.943 | chr16 | 77756389 | 77776157 | + | protein_coding |
| ENSG00000196700.9_5 | ZNF512B | 0.22619565 | 0.78324268 | 0.590 | 5.717 | chr20 | 62588057 | 62601283 | - | protein_coding |
| ENSG00000219607.3_2 | PPP1R3G | 0.24076075 | 0.82148394 | 0.590 | 0.741 | chr6 | 5084815 | 5089721 | + | protein_coding |
| ENSG00000260916.7_5 | CCPG1 | 0.2295054 | 0.79223283 | 0.590 | 4.558 | chr15 | 55632230 | 55700708 | - | protein_coding |
| ENSG00000131725.14_5 | WDR44 | 0.22539198 | 0.78112542 | 0.590 | 5.372 | chrX | 117480036 | 117583924 | + | protein_coding |
| ENSG00000229117.9_3 | RPL41 | 0.21562964 | 0.75545676 | 0.590 | 8.671 | chr12 | 56510374 | 56511751 | + | protein_coding |
| ENSG00000196923.14_6 | PDLIM7 | 0.23053209 | 0.79485031 | 0.589 | 4.798 | chr5 | 176910395 | 176924607 | - | protein_coding |
| ENSG00000163794.6_4 | UCN | 0.24277022 | 0.82661909 | 0.589 | 0.157 | chr2 | 27530268 | 27531313 | - | protein_coding |
| ENSG00000114745.13_3 | GORASP1 | 0.23101546 | 0.79589911 | 0.589 | 5.228 | chr3 | 39138150 | 39149854 | - | protein_coding |
| ENSG00000163703.18_4 | CRELD1 | 0.24264785 | 0.82651925 | 0.589 | 0.865 | chr3 | 9975500 | 9987090 | + | protein_coding |
| ENSG00000130707.18_3 | ASS1 | 0.23724913 | 0.81225454 | 0.588 | 3.545 | chr9 | 133320348 | 133376661 | + | protein_coding |

| | | | | | | | | | | |
|---|---|---|---|---|---|---|---|---|---|---|
| ENSG00000261971.8_7 | MMP25-AS1 | 0.24598512 | 0.83420838 | 0.588 | -0.318 | chr16 | 3087401 | 3109371 | - | lncRNA |
| ENSG00000230148.8_6 | HOXB-AS1 | 0.24326232 | 0.82751591 | 0.588 | 0.729 | chr17 | 46620913 | 46628612 | + | lncRNA |
| ENSG00000166848.7_4 | TERF2IP | 0.23588439 | 0.80875188 | 0.588 | 4.875 | chr16 | 75681671 | 75795770 | + | protein_coding |
| ENSG00000259172.1_6 | AC023024.1 | 0.24473995 | 0.8318911 | 0.588 | 0.145 | chr15 | 101835624 | 101845942 | + | lncRNA |
| ENSG00000254427.2_7 | LINC02696 | 0.25510234 | 0.85759758 | 0.587 | -0.462 | chr11 | 45376922 | 45389041 | + | lncRNA |
| ENSG00000106927.12_4 | AMBP | 0.24090132 | 0.8215151 | 0.587 | 3.798 | chr9 | 116822407 | 116840608 | - | protein_coding |
| ENSG00000251131.2_6 | AC025171.3 | 0.24717075 | 0.83711068 | 0.587 | -0.157 | chr5 | 43018531 | 43024780 | + | lncRNA |
| ENSG00000253320.7_8 | AZIN1-AS1 | 0.24419921 | 0.83037079 | 0.587 | 1.497 | chr8 | 103876499 | 104014652 | + | lncRNA |
| ENSG00000025039.15_4 | RRAGD | 0.24326648 | 0.82751591 | 0.587 | 2.795 | chr6 | 90074335 | 90121992 | - | protein_coding |
| ENSG00000259250.1_6 | AC018904.1 | 0.25315988 | 0.85284255 | 0.587 | -0.655 | chr15 | 58879706 | 58883875 | + | lncRNA |
| ENSG00000127419.17_8 | TMEM175 | 0.2435765 | 0.82841188 | 0.587 | 3.467 | chr4 | 926175 | 952444 | + | protein_coding |
| ENSG00000111224.14_5 | PARP11 | 0.24562946 | 0.83363871 | 0.586 | 2.367 | chr12 | 3900213 | 3982614 | - | protein_coding |
| ENSG00000104866.11_5 | PPP1R37 | 0.24507435 | 0.83270911 | 0.586 | 5.601 | chr19 | 45594654 | 45651335 | + | protein_coding |
| ENSG00000231274.5_5 | SBK3 | 0.24627723 | 0.83479393 | 0.586 | 1.618 | chr19 | 56052023 | 56056909 | - | protein_coding |
| ENSG00000088367.23_7 | EPB41L1 | 0.24524517 | 0.83297093 | 0.586 | 7.269 | chr20 | 34679426 | 34820721 | + | protein_coding |
| ENSG00000117758.14_4 | STX12 | 0.24701405 | 0.83673948 | 0.586 | 4.715 | chr1 | 28099730 | 28150954 | + | protein_coding |
| ENSG00000270419.1_3 | CAHM | 0.25122666 | 0.84761566 | 0.585 | -0.421 | chr6 | 163834097 | 163834992 | - | lncRNA |
| ENSG00000235944.8_3 | ZNF815P | 0.24779716 | 0.83875254 | 0.585 | 0.694 | chr7 | 5862791 | 5893996 | + | transcribed_unprocessed_pseudogene |
| ENSG00000140406.4_6 | TLNRD1 | 0.250298 | 0.84512438 | 0.585 | 4.248 | chr15 | 81293264 | 81298129 | + | protein_coding |
| ENSG00000226332.2_7 | AL354836.1 | 0.24970639 | 0.84376825 | 0.585 | 2.652 | chr20 | 60880488 | 60881381 | - | lncRNA |
| ENSG00000165943.5_4 | MOAP1 | 0.25226049 | 0.85013508 | 0.584 | 4.353 | chr14 | 93648544 | 93651268 | - | protein_coding |
| ENSG00000159335.16_3 | PTMS | 0.25547396 | 0.858522 | 0.584 | 7.262 | chr12 | 6874682 | 6880118 | + | protein_coding |
| ENSG00000139211.6_3 | AMIGO2 | 0.25818462 | 0.86468731 | 0.584 | 8.826 | chr12 | 47469490 | 47473734 | - | protein_coding |
| ENSG00000155016.18_3 | CYP2U1 | 0.25372041 | 0.85424492 | 0.584 | 3.522 | chr4 | 108852705 | 108874617 | + | protein_coding |
| ENSG00000073464.12_6 | CLCN4 | 0.2545801 | 0.85632804 | 0.584 | 3.847 | chrX | 10124985 | 10205700 | + | protein_coding |
| ENSG00000213676.13_7 | ATF6B | 0.25443523 | 0.85600277 | 0.584 | 4.045 | chr6 | 32083041 | 32096030 | - | protein_coding |
| ENSG00000215769.8_7 | ARHGAP27P1-BPTFP1-KPNA2P3 | 0.24990604 | 0.84428229 | 0.583 | -0.053 | chr17 | 62745781 | 62777825 | - | lncRNA |
| ENSG00000078687.17_5 | TNRC6C | 0.25659099 | 0.86097303 | 0.583 | 3.724 | chr17 | 75955322 | 76104916 | + | protein_coding |
| ENSG00000164088.18_5 | PPM1M | 0.25493114 | 0.8573465 | 0.583 | 3.090 | chr3 | 52279775 | 52284615 | + | protein_coding |
| ENSG00000198863.7_3 | RUNDC1 | 0.2610126 | 0.87234968 | 0.583 | 5.182 | chr17 | 41132582 | 41145707 | + | protein_coding |
| ENSG00000141448.10_4 | GATA6 | 0.2520689 | 0.84981179 | 0.583 | 1.525 | chr18 | 19749353 | 19782491 | + | protein_coding |
| ENSG00000169758.13_3 | TMEM266 | 0.25052969 | 0.8454247 | 0.583 | 0.537 | chr15 | 76352299 | 76521462 | + | protein_coding |
| ENSG00000205456.11_3 | TP53TG3D | 0.25617255 | 0.85989375 | 0.583 | -0.657 | chr16 | 32264040 | 32267243 | + | protein_coding |
| ENSG00000169403.12_4 | PTAFR | 0.25585446 | 0.85909282 | 0.583 | -0.615 | chr1 | 28473677 | 28520447 | - | protein_coding |
| ENSG00000254911.3_6 | SCARNA9 | 0.25190135 | 0.84940811 | 0.582 | 0.884 | chr11 | 93454679 | 93455031 | + | lncRNA |
| ENSG00000188735.13_3 | TMEM120B | 0.26308332 | 0.87794918 | 0.582 | 4.996 | chr12 | 122150658 | 122220907 | + | protein_coding |
| ENSG00000141696.13_9 | P3H4 | 0.26724579 | 0.88850224 | 0.582 | 5.753 | chr17 | 39958199 | 39968856 | - | protein_coding |
| ENSG00000106479.11_6 | ZNF862 | 0.2584421 | 0.86538649 | 0.582 | 3.001 | chr7 | 149535464 | 149564568 | + | protein_coding |
| ENSG00000101417.12_4 | PXMP4 | 0.26086249 | 0.87201203 | 0.582 | 3.639 | chr20 | 32290564 | 32308125 | - | protein_coding |
| ENSG00000161714.12_6 | PLCD3 | 0.2726262 | 0.9026742 | 0.581 | 7.109 | chr17 | 43186327 | 43210721 | - | protein_coding |
| ENSG00000188368.9_5 | PRR19 | 0.25423189 | 0.85548068 | 0.581 | 1.017 | chr19 | 42806250 | 42814973 | + | protein_coding |
| ENSG00000065320.9_3 | NTN1 | 0.25296425 | 0.85234511 | 0.581 | 0.444 | chr17 | 8924827 | 9147317 | + | protein_coding |
| ENSG00000268362.6_8 | AC092279.2 | 0.26721235 | 0.88850224 | 0.581 | 4.313 | chr19 | 24216203 | 24260179 | + | lncRNA |
| ENSG00000180891.13_6 | CUEDC1 | 0.27924252 | 0.91759139 | 0.580 | 6.974 | chr17 | 55938604 | 56032773 | - | protein_coding |
| ENSG00000085871.9_3 | MGST2 | 0.2680902 | 0.89064296 | 0.580 | 4.256 | chr4 | 140586922 | 140661899 | + | protein_coding |
| ENSG00000225529.2 | BX088645.3 | 0.25573541 | 0.85907563 | 0.580 | -0.538 | chr9 | 39722430 | 39817308 | - | pseudogene |
| ENSG00000059122.16_2 | FLYWCH1 | 0.27113845 | 0.89923746 | 0.579 | 5.125 | chr16 | 2961938 | 3001209 | + | protein_coding |
| ENSG00000105287.12_5 | PRKD2 | 0.27604798 | 0.91060983 | 0.579 | 4.695 | chr19 | 47177532 | 47220384 | - | protein_coding |
| ENSG00000185340.15_3 | GAS2L1 | 0.27185042 | 0.90094505 | 0.579 | 4.561 | chr22 | 29702572 | 29708774 | + | protein_coding |
| ENSG00000019485.13_5 | PRDM11 | 0.27763731 | 0.91466385 | 0.579 | 4.419 | chr11 | 45117357 | 45256661 | + | protein_coding |
| ENSG00000170703.15_3 | TTLL6 | 0.26915482 | 0.89351153 | 0.578 | 3.165 | chr17 | 46839597 | 46894576 | - | protein_coding |
| ENSG00000132471.12_8 | WBP2 | 0.28630503 | 0.93318318 | 0.578 | 5.600 | chr17 | 73841780 | 73852588 | - | protein_coding |
| ENSG00000164897.14_6 | TMUB1 | 0.27538364 | 0.90909348 | 0.578 | 4.379 | chr7 | 150778172 | 150780580 | - | protein_coding |
| ENSG00000165695.10_4 | AK8 | 0.25651988 | 0.86089701 | 0.578 | 0.397 | chr9 | 135600965 | 135754164 | - | protein_coding |
| ENSG00000160097.18_6 | FNDC5 | 0.26268928 | 0.87696364 | 0.578 | -0.741 | chr1 | 33327869 | 33338083 | - | protein_coding |
| ENSG00000110925.7_6 | CSRNP2 | 0.27950765 | 0.91810178 | 0.578 | 4.359 | chr12 | 51454988 | 51477447 | - | protein_coding |
| ENSG00000135722.9_4 | FBXL8 | 0.26636081 | 0.88638931 | 0.577 | 2.457 | chr16 | 67193835 | 67198473 | + | protein_coding |
| ENSG00000249641.2_7 | HOXC13-AS | 0.25665289 | 0.86101812 | 0.577 | -0.201 | chr12 | 54329112 | 54333427 | - | lncRNA |

| | | | | | | | | | | |
|---|---|---|---|---|---|---|---|---|---|---|
| ENSG00000083290.20_5 | ULK2 | 0.29631292 | 0.95632908 | 0.577 | 5.699 | chr17 | 19674142 | 19771249 | - | protein_coding |
| ENSG00000145536.15_3 | ADAMTS16 | 0.2654313 | 0.88445571 | 0.577 | 1.832 | chr5 | 5140443 | 5320417 | + | protein_coding |
| ENSG00000262136.1_7 | AC092115.4 | 0.25732485 | 0.86262089 | 0.577 | 0.051 | chr16 | 69760916 | 69776466 | + | lncRNA |
| ENSG00000174370.10_4 | C11orf45 | 0.270493 | 0.89735983 | 0.577 | 2.521 | chr11 | 128769460 | 128775964 | - | protein_coding |
| ENSG00000212443.1 | SNORA53 | 0.26430472 | 0.88152845 | 0.577 | -0.832 | chr12 | 98993413 | 98993661 | + | snoRNA |
| ENSG00000078902.16_5 | TOLLIP | 0.28822962 | 0.93802837 | 0.576 | 4.896 | chr11 | 1295601 | 1330884 | - | protein_coding |
| ENSG00000188766.12_2 | SPRED3 | 0.265497 | 0.88450876 | 0.576 | 1.650 | chr19 | 38879061 | 38890227 | + | protein_coding |
| ENSG00000187994.14_7 | RINL | 0.27118333 | 0.89923746 | 0.576 | 2.296 | chr19 | 39358470 | 39368915 | - | protein_coding |
| ENSG00000181513.14_5 | ACBD4 | 0.28936913 | 0.94040996 | 0.576 | 4.509 | chr17 | 43209967 | 43221548 | + | protein_coding |
| ENSG00000225697.13_6 | SLC26A6 | 0.30038239 | 0.96351218 | 0.576 | 5.148 | chr3 | 48663156 | 48672926 | - | protein_coding |
| ENSG00000163354.15_4 | DCST2 | 0.25923971 | 0.86756676 | 0.575 | 0.166 | chr1 | 154990996 | 155006257 | - | protein_coding |
| ENSG00000137414.6_3 | FAM8A1 | 0.29714791 | 0.95745862 | 0.575 | 4.534 | chr6 | 17600533 | 17611946 | + | protein_coding |
| ENSG00000223298.1 | RNY3P8 | 0.26024498 | 0.8704391 | 0.575 | 0.278 | chr13 | 95963084 | 95963209 | - | misc_RNA |
| ENSG00000152904.11_7 | GGPS1 | 0.28457039 | 0.9294048 | 0.575 | 3.583 | chr1 | 235490665 | 235507847 | + | protein_coding |
| ENSG00000109944.11_4 | JHY | 0.2767386 | 0.9125837 | 0.575 | 2.706 | chr11 | 122753391 | 122834570 | + | protein_coding |
| ENSG00000164023.14_3 | SGMS2 | 0.30086805 | 0.96472166 | 0.575 | 5.027 | chr4 | 108745719 | 108836203 | + | protein_coding |
| ENSG00000149474.13_5 | KAT14 | 0.29838241 | 0.95952221 | 0.575 | 4.676 | chr20 | 18118762 | 18169031 | + | protein_coding |
| ENSG00000232721.2_6 | AC239800.2 | 0.27202707 | 0.90136234 | 0.575 | -0.929 | chr1 | 149230716 | 149234156 | + | lncRNA |
| ENSG00000213420.8_4 | GPC2 | 0.26155193 | 0.87382346 | 0.575 | 0.491 | chr7 | 99767229 | 99775004 | - | protein_coding |
| ENSG00000144560.15_7 | VGLL4 | 0.2946378 | 0.95317367 | 0.574 | 4.433 | chr3 | 11597543 | 11812824 | - | protein_coding |
| ENSG00000150722.10_4 | PPP1R1C | 0.26068419 | 0.87157999 | 0.574 | -0.465 | chr2 | 182818968 | 182996125 | + | protein_coding |
| ENSG00000205707.11_8 | ETFRF1 | 0.28670418 | 0.93414141 | 0.573 | 3.496 | chr12 | 25348150 | 25362579 | + | protein_coding |
| ENSG00000163590.14_3 | PPM1L | 0.28367892 | 0.92700447 | 0.573 | 2.795 | chr3 | 160473390 | 160796690 | + | protein_coding |
| ENSG00000137842.7_4 | TMEM62 | 0.2947407 | 0.95333298 | 0.573 | 3.894 | chr15 | 43415477 | 43477342 | + | protein_coding |
| ENSG00000170439.7_5 | METTL7B | 0.28657809 | 0.93390187 | 0.573 | 3.430 | chr12 | 56075462 | 56078395 | + | protein_coding |
| ENSG00000105429.13_4 | MEGF8 | 0.30285585 | 0.96934605 | 0.573 | 7.025 | chr19 | 42829761 | 42882921 | + | protein_coding |
| ENSG00000253686.2_7 | LINC01484 | 0.26560345 | 0.88453169 | 0.573 | 0.849 | chr5 | 173134617 | 173173282 | - | lncRNA |
| ENSG00000173621.9_4 | LRFN4 | 0.27496028 | 0.90792303 | 0.573 | 4.432 | chr11 | 66624118 | 66627946 | + | protein_coding |
| ENSG00000163701.19_4 | IL17RE | 0.27209157 | 0.90140797 | 0.572 | -0.698 | chr3 | 9944296 | 9958086 | + | protein_coding |
| ENSG00000100243.21_5 | CYB5R3 | 0.32112657 | 1 | 0.572 | 5.974 | chr22 | 43013846 | 43045398 | - | protein_coding |
| ENSG00000172382.10_4 | PRSS27 | 0.27321928 | 0.90446934 | 0.572 | 1.546 | chr16 | 2762420 | 2770552 | - | protein_coding |
| ENSG00000125841.13_4 | NRSN2 | 0.33305513 | 1 | 0.572 | 5.854 | chr20 | 327426 | 340304 | + | protein_coding |
| ENSG00000166908.18_4 | PIP4K2C | 0.33082529 | 1 | 0.572 | 5.598 | chr12 | 57984957 | 57997201 | + | protein_coding |
| ENSG00000159363.18_5 | ATP13A2 | 0.30580044 | 0.97613302 | 0.572 | 4.824 | chr1 | 17312453 | 17338423 | - | protein_coding |
| ENSG00000186594.14_5 | MIR22HG | 0.3202561 | 1 | 0.571 | 4.807 | chr17 | 1614787 | 1620468 | - | lncRNA |
| ENSG00000132849.20_6 | PATJ | 0.3377308 | 1 | 0.571 | 5.850 | chr1 | 62208078 | 62644347 | + | protein_coding |
| ENSG00000167797.7_3 | CDK2AP2 | 0.31032581 | 0.98562008 | 0.571 | 4.574 | chr11 | 67273968 | 67276120 | - | protein_coding |
| ENSG00000287260.1_3 | AL357075.3 | 0.26429833 | 0.88152845 | 0.571 | -0.266 | chr6 | 154831925 | 154843441 | + | lncRNA |
| ENSG00000150782.12_5 | IL18 | 0.33649851 | 1 | 0.571 | 5.910 | chr11 | 112013976 | 112034819 | - | protein_coding |
| ENSG00000257038.1_7 | AP002761.3 | 0.27719906 | 0.91338941 | 0.571 | -0.979 | chr11 | 73018280 | 73020406 | - | lncRNA |
| ENSG00000204248.11_5 | COL11A2 | 0.26594309 | 0.88516502 | 0.571 | 0.632 | chr6 | 33130458 | 33160276 | - | protein_coding |
| ENSG00000054611.14_4 | TBC1D22A | 0.31272281 | 0.99110707 | 0.571 | 4.230 | chr22 | 47158514 | 47571342 | + | protein_coding |
| ENSG00000275703.1_7 | U47924.3 | 0.26380661 | 0.88019756 | 0.571 | -0.203 | chr12 | 7008916 | 7009376 | + | lncRNA |
| ENSG00000163362.11_9 | INAVA | 0.31800136 | 1 | 0.571 | 6.135 | chr1 | 200860176 | 200884870 | + | protein_coding |
| ENSG00000161999.12_6 | JMJD8 | 0.30960212 | 0.98423807 | 0.571 | 4.244 | chr16 | 731670 | 734528 | - | protein_coding |
| ENSG00000130844.18_6 | ZNF331 | 0.33271136 | 1 | 0.571 | 5.432 | chr19 | 54022781 | 54083523 | + | protein_coding |
| ENSG00000167671.12_5 | UBXN6 | 0.31972084 | 1 | 0.571 | 5.317 | chr19 | 4444996 | 4457791 | - | protein_coding |
| ENSG00000122970.16_5 | IFT81 | 0.34878638 | 1 | 0.570 | 6.495 | chr12 | 110562140 | 110656598 | + | protein_coding |
| ENSG00000173334.4_3 | TRIB1 | 0.33666393 | 1 | 0.570 | 5.364 | chr8 | 126442600 | 126450645 | + | protein_coding |
| ENSG00000141519.15_7 | CCDC40 | 0.30535186 | 0.97487628 | 0.570 | 3.568 | chr17 | 78010431 | 78074412 | + | protein_coding |
| ENSG00000272849.1_7 | AC084018.1 | 0.27006915 | 0.89637934 | 0.570 | 0.797 | chr12 | 122235417 | 122239878 | + | lncRNA |
| ENSG00000116133.13_5 | DHCR24 | 0.36185355 | 1 | 0.569 | 6.693 | chr1 | 55315300 | 55352868 | - | protein_coding |
| ENSG00000165886.5_3 | UBTD1 | 0.29584479 | 0.95558481 | 0.569 | 3.091 | chr10 | 99258681 | 99330963 | + | protein_coding |
| ENSG00000170542.6_4 | SERPINB9 | 0.3400231 | 1 | 0.569 | 5.146 | chr6 | 2887504 | 2903543 | - | protein_coding |
| ENSG00000085831.15_7 | TTC39A | 0.29992019 | 0.96272484 | 0.569 | 3.026 | chr1 | 51752930 | 51810788 | - | protein_coding |
| ENSG00000129657.16_8 | SEC14L1 | 0.35533381 | 1 | 0.568 | 5.640 | chr17 | 75084831 | 75213183 | + | protein_coding |
| ENSG00000184378.3_5 | ACTRT3 | 0.28746994 | 0.93594987 | 0.568 | 2.044 | chr3 | 169484709 | 169487349 | - | protein_coding |
| ENSG00000124766.7_4 | SOX4 | 0.37396581 | 1 | 0.568 | 6.485 | chr6 | 21593982 | 21598850 | + | protein_coding |
| ENSG00000131037.15_3 | EPS8L1 | 0.28908799 | 0.93966818 | 0.568 | 2.060 | chr19 | 55583388 | 55599291 | + | protein_coding |
| ENSG00000205578.4 | POM121B | 0.27426873 | 0.90700564 | 0.567 | -0.742 | chr7 | 72707497 | 72716496 | + | pseudogene |
| ENSG00000132481.7_6 | TRIM47 | 0.34857274 | 1 | 0.567 | 5.655 | chr17 | 73870245 | 73874662 | - | protein_coding |

| | | | | | | | | | | |
|---|---|---|---|---|---|---|---|---|---|---|
| ENSG00000010310.9_5 | GIPR | 0.27231693 | 0.90198638 | 0.567 | 0.568 | chr19 | 46171479 | 46186980 | + | protein_coding |
| ENSG00000177189.14_7 | RPS6KA3 | 0.38640915 | 1 | 0.567 | 7.975 | chrX | 20168029 | 20285637 | - | protein_coding |
| ENSG00000185761.11_4 | ADAMTSL5 | 0.28290064 | 0.92514185 | 0.566 | 1.320 | chr19 | 1505021 | 1513603 | - | protein_coding |
| ENSG00000110917.8_3 | MLEC | 0.39734863 | 1 | 0.566 | 8.330 | chr12 | 121124952 | 121139662 | + | protein_coding |
| ENSG00000136295.15_3 | TTYH3 | 0.29330167 | 0.95023537 | 0.566 | 5.641 | chr7 | 2671620 | 2704436 | + | protein_coding |
| ENSG00000166596.15_4 | CFAP52 | 0.28023113 | 0.91962809 | 0.566 | 1.253 | chr17 | 9479944 | 9546764 | + | protein_coding |
| ENSG00000175556.17_3 | LONRF3 | 0.32421273 | 1 | 0.565 | 3.671 | chrX | 118108577 | 118152318 | + | protein_coding |
| ENSG00000100284.21_6 | TOM1 | 0.32934759 | 1 | 0.565 | 3.739 | chr22 | 35695268 | 35743985 | + | protein_coding |
| ENSG00000135205.15_6 | CCDC146 | 0.29327427 | 0.95023537 | 0.564 | 1.983 | chr7 | 76751751 | 76958850 | + | protein_coding |
| ENSG00000286656.1_1 | AC008875.2 | 0.28254566 | 0.92415109 | 0.564 | 1.203 | chr5 | 42995236 | 42997582 | + | lncRNA |
| ENSG00000121653.11_4 | MAPK8IP1 | 0.31118957 | 0.98790027 | 0.564 | 3.215 | chr11 | 45907202 | 45928016 | + | protein_coding |
| ENSG00000112972.15_7 | HMGCS1 | 0.38470934 | 1 | 0.564 | 7.157 | chr5 | 43287572 | 43313614 | - | protein_coding |
| ENSG00000231742.6_5 | LINC01273 | 0.28115632 | 0.92147176 | 0.564 | 0.959 | chr20 | 48788346 | 48793213 | + | lncRNA |
| ENSG00000252447.1 | snoU13 | 0.27693471 | 0.91285695 | 0.563 | 0.494 | chr11 | 47749098 | 47749189 | + | snoRNA |
| ENSG00000105655.19_7 | ISYNA1 | 0.34234539 | 1 | 0.562 | 4.886 | chr19 | 18545198 | 18548977 | - | protein_coding |
| ENSG00000165985.10_4 | C1QL3 | 0.30502199 | 0.97417323 | 0.562 | 2.195 | chr10 | 16555733 | 16563878 | - | protein_coding |
| ENSG00000187535.14_5 | IFT140 | 0.37572509 | 1 | 0.562 | 5.299 | chr16 | 1560428 | 1662073 | - | protein_coding |
| ENSG00000182197.11_7 | EXT1 | 0.39051085 | 1 | 0.562 | 5.822 | chr8 | 118806729 | 119124092 | - | protein_coding |
| ENSG00000153363.13_5 | LINC00467 | 0.32168449 | 1 | 0.562 | 3.018 | chr1 | 211556078 | 211608912 | + | lncRNA |
| ENSG00000198355.5_3 | PIM3 | 0.33598385 | 1 | 0.562 | 4.363 | chr22 | 50354416 | 50357720 | + | protein_coding |
| ENSG00000135469.13_4 | COQ10A | 0.35284755 | 1 | 0.562 | 4.129 | chr12 | 56660642 | 56664750 | + | protein_coding |
| ENSG00000116035.4_6 | VAX2 | 0.32100258 | 1 | 0.562 | 1.488 | chr2 | 71127706 | 71192503 | + | protein_coding |
| ENSG00000131400.8_5 | NAPSA | 0.27841239 | 0.91580688 | 0.562 | 0.312 | chr19 | 50861734 | 50869087 | - | protein_coding |
| ENSG00000177119.16_5 | ANO6 | 0.44168155 | 1 | 0.561 | 7.821 | chr12 | 45609770 | 45834187 | + | protein_coding |
| ENSG00000262484.1_4 | CCER2 | 0.27578888 | 0.90992406 | 0.561 | 0.116 | chr19 | 39399620 | 39402798 | - | protein_coding |
| ENSG00000251179.1_7 | TMEM92-AS1 | 0.28089533 | 0.92112723 | 0.561 | -0.668 | chr17 | 48358938 | 48365216 | - | lncRNA |
| ENSG00000256967.1_7 | AC018653.3 | 0.29612565 | 0.95596622 | 0.561 | 1.661 | chr12 | 7281675 | 7283794 | - | lncRNA |
| ENSG00000244405.8_6 | ETV5 | 0.41562289 | 1 | 0.560 | 5.938 | chr3 | 185764103 | 185828107 | - | protein_coding |
| ENSG00000213213.14_4 | CCDC183 | 0.28030722 | 0.91970792 | 0.560 | 0.610 | chr9 | 139690790 | 139702193 | + | protein_coding |
| ENSG00000232386.9_7 | AC015712.1 | 0.28052906 | 0.92026585 | 0.560 | 0.375 | chr15 | 101389766 | 101401961 | + | lncRNA |
| ENSG00000128567.17_4 | PODXL | 0.42480353 | 1 | 0.559 | 6.197 | chr7 | 131185021 | 131242976 | - | protein_coding |
| ENSG00000171105.14_5 | INSR | 0.37566961 | 1 | 0.559 | 4.509 | chr19 | 7112266 | 7294425 | - | protein_coding |
| ENSG00000178338.11_4 | ZNF354B | 0.36373597 | 1 | 0.559 | 3.961 | chr5 | 178286954 | 178315123 | + | protein_coding |
| ENSG00000056998.20_5 | GYG2 | 0.35296605 | 1 | 0.558 | 3.719 | chrX | 2746863 | 2800861 | + | protein_coding |
| ENSG00000105771.14_5 | SMG9 | 0.42849448 | 1 | 0.558 | 5.706 | chr19 | 44232135 | 44259114 | - | protein_coding |
| ENSG00000104833.12_7 | TUBB4A | 0.30706979 | 0.97877808 | 0.558 | 1.777 | chr19 | 6494330 | 6502859 | - | protein_coding |
| ENSG00000013583.10_7 | HEBP1 | 0.33190386 | 1 | 0.558 | 2.736 | chr12 | 13127804 | 13153199 | - | protein_coding |
| ENSG00000213145.10_4 | CRIP1 | 0.29004797 | 0.94209914 | 0.558 | 1.033 | chr14 | 105952654 | 105955284 | + | protein_coding |
| ENSG00000253954.3_4 | HMGN1P38 | 0.28074973 | 0.92081973 | 0.557 | -0.281 | chr15 | 93255038 | 93255340 | + | processed_pseudogene |
| ENSG00000033170.17_4 | FUT8 | 0.40701373 | 1 | 0.557 | 5.203 | chr14 | 65877310 | 66210839 | + | protein_coding |
| ENSG00000213250.5_2 | RBMS2P1 | 0.28212755 | 0.9232997 | 0.557 | 0.193 | chr12 | 94817520 | 94818745 | - | processed_pseudogene |
| ENSG00000183615.6_3 | FAM167B | 0.28017802 | 0.91962365 | 0.557 | -0.183 | chr1 | 32712823 | 32714457 | + | protein_coding |
| ENSG00000157216.15_6 | SSBP3 | 0.35364072 | 1 | 0.556 | 3.261 | chr1 | 54691105 | 54879152 | - | protein_coding |
| ENSG00000101935.10_5 | AMMECR1 | 0.41321107 | 1 | 0.556 | 5.604 | chrX | 109437414 | 109683461 | - | protein_coding |
| ENSG00000064932.16_4 | SBNO2 | 0.31823867 | 1 | 0.556 | 5.906 | chr19 | 1107636 | 1174267 | - | protein_coding |
| ENSG00000089248.7_4 | ERP29 | 0.456192 | 1 | 0.555 | 6.660 | chr12 | 112451152 | 112461253 | + | protein_coding |
| ENSG00000196569.12_3 | LAMA2 | 0.28215792 | 0.9232997 | 0.555 | -0.003 | chr6 | 129204286 | 129837714 | + | protein_coding |
| ENSG00000089505.17_5 | CMTM1 | 0.29864243 | 0.96018458 | 0.555 | 1.401 | chr16 | 66600296 | 66613040 | + | protein_coding |
| ENSG00000264281.3_7 | AC016596.3 | 0.28714402 | 0.9352315 | 0.555 | -0.399 | chr5 | 55240484 | 55240561 | + | processed_pseudogene |
| ENSG00000128739.22_7 | SNRPN | 0.32710382 | 1 | 0.554 | 2.326 | chr15 | 25068784 | 25223870 | + | protein_coding |
| ENSG00000139613.12_5 | SMARCC2 | 0.47324338 | 1 | 0.554 | 6.880 | chr12 | 56556143 | 56583351 | - | protein_coding |
| ENSG00000130751.10_3 | NPAS1 | 0.29988988 | 0.96272484 | 0.554 | 1.351 | chr19 | 47523094 | 47549033 | + | protein_coding |
| ENSG00000260265.1_7 | LINC02562 | 0.30736211 | 0.9788318 | 0.554 | 0.920 | chr4 | 76006912 | 76009927 | - | lncRNA |
| ENSG00000134202.11_4 | GSTM3 | 0.35711658 | 1 | 0.553 | 3.460 | chr1 | 110276554 | 110283660 | - | protein_coding |
| ENSG00000099725.14_3 | PRKY | 0.35030392 | 1 | 0.553 | 3.203 | chrY | 7142013 | 7249589 | + | protein_coding |
| ENSG00000182158.15_4 | CREB3L2 | 0.46413833 | 1 | 0.553 | 5.792 | chr7 | 137559725 | 137686832 | - | protein_coding |
| ENSG00000161920.10_4 | MED11 | 0.37659179 | 1 | 0.553 | 3.739 | chr17 | 4634723 | 4636902 | + | protein_coding |
| ENSG00000197530.12_7 | MIB2 | 0.3335418 | 1 | 0.553 | 4.348 | chr1 | 1550795 | 1565990 | + | protein_coding |

| | | | | | | | | | | |
|---|---|---|---|---|---|---|---|---|---|---|
| ENSG00000249550.7_6 | LINC01234 | 0.44883848 | 1 | 0.552 | 5.480 | chr12 | 114021691 | 114211531 | - | lncRNA |
| ENSG00000164674.15_3 | SYTL3 | 0.29090859 | 0.94454913 | 0.552 | 0.243 | chr6 | 159071046 | 159185908 | + | protein_coding |
| ENSG00000132510.10_3 | KDM6B | 0.3283935 | 1 | 0.552 | 2.585 | chr17 | 7743222 | 7758114 | + | protein_coding |
| ENSG00000237792.3 | RP11-203I2.1 | 0.29402222 | 0.95204897 | 0.552 | 0.874 | chr9 | 65585692 | 65661972 | + | pseudogene |
| ENSG00000159167.12_4 | STC1 | 0.39225493 | 1 | 0.551 | 3.919 | chr8 | 23699442 | 23712319 | - | protein_coding |
| ENSG00000109787.13_3 | KLF3 | 0.46923281 | 1 | 0.551 | 5.813 | chr4 | 38665818 | 38703138 | + | protein_coding |
| ENSG00000287262.1_2 | AC021148.3 | 0.29377348 | 0.95141696 | 0.551 | 0.388 | chr4 | 39541015 | 39547693 | - | lncRNA |
| ENSG00000177169.10_5 | ULK1 | 0.44898457 | 1 | 0.551 | 6.074 | chr12 | 132379167 | 132407695 | + | protein_coding |
| ENSG00000267801.1_8 | AC087289.5 | 0.32994155 | 1 | 0.551 | 1.083 | chr17 | 73872453 | 73875627 | + | lncRNA |
| ENSG00000253919.1_5 | THAP12P7 | 0.30594556 | 0.97624547 | 0.551 | -0.896 | chr8 | 79672377 | 79674666 | - | processed_pseudogene |
| ENSG00000177427.13_5 | MIEF2 | 0.3862443 | 1 | 0.550 | 3.847 | chr17 | 18163911 | 18169866 | + | protein_coding |
| ENSG00000166888.12_6 | STAT6 | 0.51263403 | 1 | 0.550 | 6.951 | chr12 | 57489191 | 57525922 | - | protein_coding |
| ENSG00000262468.6_6 | LINC01569 | 0.28694043 | 0.93473974 | 0.550 | -0.442 | chr16 | 4293944 | 4303818 | - | lncRNA |
| ENSG00000162736.17_6 | NCSTN | 0.48157746 | 1 | 0.550 | 5.998 | chr1 | 160313106 | 160328742 | + | protein_coding |
| ENSG00000259318.1_5 | AL356801.1 | 0.29233348 | 0.94865551 | 0.549 | -0.629 | chr14 | 55861658 | 55861951 | - | processed_pseudogene |
| ENSG00000132915.11_5 | PDE6A | 0.28861672 | 0.93882337 | 0.549 | -0.192 | chr5 | 149237516 | 149324356 | - | protein_coding |
| ENSG00000270039.1_7 | AC025165.5 | 0.28798405 | 0.93745193 | 0.549 | -0.672 | chr12 | 58197621 | 58198198 | + | lncRNA |
| ENSG00000196961.12_2 | AP2A1 | 0.48552546 | 1 | 0.548 | 6.876 | chr19 | 50270225 | 50310370 | + | protein_coding |
| ENSG00000223979.2_4 | SMCR2 | 0.29463687 | 0.95317367 | 0.548 | -0.944 | chr17 | 17577340 | 17581002 | - | lncRNA |
| ENSG00000141503.17_5 | MINK1 | 0.50292618 | 1 | 0.548 | 7.223 | chr17 | 4736635 | 4801356 | + | protein_coding |
| ENSG00000168502.17_3 | MTCL1 | 0.50492143 | 1 | 0.548 | 7.547 | chr18 | 8705659 | 8832776 | + | protein_coding |
| ENSG00000182870.8 | GALNT9 | 0.29548047 | 0.95485658 | 0.548 | 0.324 | chr12 | 132680924 | 132905935 | - | protein_coding |
| ENSG00000170161.7_6 | AL512625.1 | 0.32319045 | 1 | 0.548 | 0.803 | chr9 | 66553192 | 66555928 | + | lncRNA |
| ENSG00000272752.6_4 | STAG3L5P-PVRIG2P-PILRB | 0.28957072 | 0.94089301 | 0.548 | -0.713 | chr7 | 99933727 | 99965454 | + | lncRNA |
| ENSG00000215493.3_7 | AC007731.2 | 0.29213041 | 0.9481697 | 0.547 | 0.004 | chr22 | 20804409 | 20806111 | + | processed_pseudogene |
| ENSG00000177576.11_5 | C18orf32 | 0.30638076 | 0.97675704 | 0.547 | 0.806 | chr18 | 47003620 | 47013622 | - | protein_coding |
| ENSG00000181027.11_6 | FKRP | 0.44773652 | 1 | 0.547 | 4.720 | chr19 | 47249303 | 47280245 | + | protein_coding |
| ENSG00000107821.14_2 | KAZALD1 | 0.38173048 | 1 | 0.547 | 3.257 | chr10 | 102821598 | 102827888 | + | protein_coding |
| ENSG00000183580.10_3 | FBXL7 | 0.33354306 | 1 | 0.547 | 1.778 | chr5 | 15500289 | 15939904 | + | protein_coding |
| ENSG00000224546.2_4 | EIF4BP3 | 0.30837928 | 0.9813675 | 0.546 | 1.025 | chr9 | 98908289 | 98910138 | + | processed_pseudogene |
| ENSG00000221821.4_3 | C6orf226 | 0.30718312 | 0.9788318 | 0.546 | 0.652 | chr6 | 42858003 | 42858559 | - | protein_coding |
| ENSG00000124104.19_3 | SNX21 | 0.38657184 | 1 | 0.546 | 3.364 | chr20 | 44462438 | 44471915 | + | protein_coding |
| ENSG00000231064.7_7 | AC234582.1 | 0.30145069 | 0.96597234 | 0.545 | 0.549 | chr1 | 155164795 | 155175286 | + | lncRNA |
| ENSG00000110721.12_4 | CHKA | 0.43940676 | 1 | 0.545 | 4.405 | chr11 | 67820326 | 67888911 | - | protein_coding |
| ENSG00000147124.12_5 | ZNF41 | 0.44617381 | 1 | 0.545 | 4.245 | chrX | 47305278 | 47342345 | - | protein_coding |
| ENSG00000244560.7_5 | AC004890.2 | 0.36022369 | 1 | 0.545 | 1.938 | chr7 | 148982372 | 148994403 | + | transcribed_unprocessed_pseudogene |
| ENSG00000272720.1_6 | AL022322.1 | 0.29710583 | 0.95745862 | 0.545 | 0.071 | chr22 | 38486134 | 38487566 | + | lncRNA |
| ENSG00000183935.5_4 | HTR7P1 | 0.35787769 | 1 | 0.545 | 2.312 | chr12 | 13153354 | 13157764 | + | transcribed_processed_pseudogene |
| ENSG00000163393.13_3 | SLC22A15 | 0.30469535 | 0.97386856 | 0.544 | 0.720 | chr1 | 116519134 | 116612675 | + | protein_coding |
| ENSG00000148358.19_3 | GPR107 | 0.55848472 | 1 | 0.544 | 6.588 | chr9 | 132815705 | 132902448 | + | protein_coding |
| ENSG00000243279.4_5 | PRAF2 | 0.32371127 | 1 | 0.544 | 1.207 | chrX | 48928818 | 48931661 | - | protein_coding |
| ENSG00000006831.10_3 | ADIPOR2 | 0.57538725 | 1 | 0.544 | 7.085 | chr12 | 1797740 | 1897840 | + | protein_coding |
| ENSG00000281691.1_5 | RBM5-AS1 | 0.29586717 | 0.95558481 | 0.544 | -0.849 | chr3 | 50137036 | 50138421 | - | lncRNA |
| ENSG00000188811.14_4 | NHLRC3 | 0.44461668 | 1 | 0.543 | 4.424 | chr13 | 39612443 | 39624246 | + | protein_coding |
| ENSG00000170629.14_4 | DPY19L2P2 | 0.39571477 | 1 | 0.543 | 3.073 | chr7 | 102815580 | 102920857 | - | transcribed_unprocessed_pseudogene |
| ENSG00000105737.9_3 | GRIK5 | 0.29522712 | 0.95450396 | 0.543 | -0.802 | chr19 | 42502473 | 42573650 | - | protein_coding |
| ENSG00000232352.1_7 | SEMA3B-AS1 | 0.29527668 | 0.95450396 | 0.543 | -0.245 | chr3 | 50304073 | 50304803 | - | lncRNA |
| ENSG00000135407.10_8 | AVIL | 0.33883494 | 1 | 0.543 | 1.775 | chr12 | 58191159 | 58212487 | - | protein_coding |
| ENSG00000100321.15_6 | SYNGR1 | 0.40210944 | 1 | 0.543 | 3.142 | chr22 | 39745930 | 39781580 | + | protein_coding |
| ENSG00000251136.9_7 | AF117829.1 | 0.39654776 | 1 | 0.543 | 3.209 | chr8 | 90598101 | 90770040 | - | lncRNA |
| ENSG00000021355.13_4 | SERPINB1 | 0.55050914 | 1 | 0.542 | 6.497 | chr6 | 2832566 | 2842193 | - | protein_coding |
| ENSG00000120889.13_4 | TNFRSF10B | 0.58879188 | 1 | 0.542 | 7.353 | chr8 | 22877646 | 22926544 | - | protein_coding |
| ENSG00000174132.9_3 | FAM174A | 0.40401919 | 1 | 0.542 | 3.176 | chr5 | 99871078 | 99922445 | + | protein_coding |
| ENSG00000176058.13_6 | TPRN | 0.39396383 | 1 | 0.542 | 3.306 | chr9 | 140086069 | 140098645 | - | protein_coding |
| ENSG00000125827.9_4 | TMX4 | 0.52990068 | 1 | 0.542 | 5.993 | chr20 | 7957993 | 8000452 | - | protein_coding |

| | | | | | | | | | | |
|---|---|---|---|---|---|---|---|---|---|---|
| ENSG00000182327.8_4 | GLTPD2 | 0.30305023 | 0.96961883 | 0.542 | 0.566 | chr17 | 4692259 | 4693884 | + | protein_coding |
| ENSG00000185044.10 | RP11-435B5.4 | 0.2991105 | 0.9615155 | 0.542 | -0.759 | chr1 | 143354963 | 143467632 | - | lincRNA |
| ENSG00000079739.17_5 | PGM1 | 0.52307823 | 1 | 0.542 | 5.635 | chr1 | 64059082 | 64125916 | + | protein_coding |
| ENSG00000257093.7_6 | DENND11 | 0.55829904 | 1 | 0.541 | 6.502 | chr7 | 141356528 | 141401966 | - | protein_coding |
| ENSG00000162458.13_5 | FBLIM1 | 0.47931821 | 1 | 0.541 | 4.691 | chr1 | 16083102 | 16113089 | + | protein_coding |
| ENSG00000021762.20_5 | OSBPL5 | 0.43782023 | 1 | 0.541 | 4.149 | chr11 | 3108337 | 3187969 | - | protein_coding |
| ENSG00000181885.18_5 | CLDN7 | 0.29815058 | 0.95937666 | 0.541 | -0.896 | chr17 | 7163222 | 7167302 | - | protein_coding |
| ENSG00000242028.6_4 | HYPK | 0.31056612 | 0.98620701 | 0.541 | 0.401 | chr15 | 44088340 | 44095241 | + | protein_coding |
| ENSG00000186364.12_4 | NUDT17 | 0.35333615 | 1 | 0.541 | 1.917 | chr1 | 145586114 | 145589438 | - | protein_coding |
| ENSG00000119632.4_5 | IFI27L2 | 0.40579371 | 1 | 0.541 | 3.152 | chr14 | 94594116 | 94596590 | - | protein_coding |
| ENSG00000183023.18_5 | SLC8A1 | 0.39663286 | 1 | 0.541 | 3.100 | chr2 | 40324410 | 40838193 | - | protein_coding |
| ENSG00000229431.1_7 | MED8-AS1 | 0.30147034 | 0.96597234 | 0.541 | -1.049 | chr1 | 43850784 | 43854826 | + | lncRNA |
| ENSG00000254837.2_5 | AP001372.2 | 0.35444374 | 1 | 0.541 | 2.111 | chr11 | 74204411 | 74209578 | + | lncRNA |
| ENSG00000122035.6_2 | RASL11A | 0.31297458 | 0.99128567 | 0.541 | 0.620 | chr13 | 27844464 | 27847827 | + | protein_coding |
| ENSG00000187098.15_8 | MITF | 0.53512183 | 1 | 0.540 | 5.633 | chr3 | 69788586 | 70017488 | + | protein_coding |
| ENSG00000179899.8_4 | PHC1P1 | 0.31374568 | 0.99325688 | 0.540 | 0.852 | chr12 | 55805511 | 55808571 | - | processed_pseudogene |
| ENSG00000268947.1_7 | AC002128.1 | 0.35557604 | 1 | 0.540 | 2.182 | chr19 | 35753749 | 35755707 | + | lncRNA |
| ENSG00000274487.2_6 | NPEPPSP1 | 0.29771358 | 0.95823861 | 0.540 | -0.488 | chr17 | 36351759 | 36413199 | - | transcribed_unprocessed_pseudogene |
| ENSG00000157110.16_5 | RBPMS | 0.49292639 | 1 | 0.540 | 4.795 | chr8 | 30242027 | 30429773 | + | protein_coding |
| ENSG00000204653.10_5 | ASPDH | 0.29929315 | 0.96158071 | 0.540 | -0.407 | chr19 | 51014857 | 51017947 | - | protein_coding |
| ENSG00000148411.8_5 | NACC2 | 0.53751492 | 1 | 0.540 | 6.782 | chr9 | 138898383 | 138987135 | - | protein_coding |
| ENSG00000150961.15_5 | SEC24D | 0.51654423 | 1 | 0.540 | 5.076 | chr4 | 119643978 | 119759838 | - | protein_coding |
| ENSG00000175294.6_4 | CATSPER1 | 0.30501709 | 0.97417323 | 0.539 | 0.248 | chr11 | 65784223 | 65793950 | - | protein_coding |
| ENSG00000005882.12_6 | PDK2 | 0.45642162 | 1 | 0.539 | 3.963 | chr17 | 48172101 | 48189516 | + | protein_coding |
| ENSG00000152292.17_6 | SH2D6 | 0.30725883 | 0.9788318 | 0.539 | 0.207 | chr2 | 85646054 | 85664152 | + | protein_coding |
| ENSG00000142798.20_8 | HSPG2 | 0.44088718 | 1 | 0.539 | 7.920 | chr1 | 22148737 | 22263803 | - | protein_coding |
| ENSG00000130313.7_4 | PGLS | 0.46272156 | 1 | 0.539 | 4.898 | chr19 | 17622445 | 17632097 | + | protein_coding |
| ENSG00000168938.6_4 | PPIC | 0.40073884 | 1 | 0.539 | 2.844 | chr5 | 122358945 | 122372420 | - | protein_coding |
| ENSG00000255774.2_8 | LINC02747 | 0.37874434 | 1 | 0.538 | 2.320 | chr11 | 69290335 | 69296313 | - | lncRNA |
| ENSG00000178722.13_3 | C5orf64 | 0.29948453 | 0.96202164 | 0.538 | -0.489 | chr5 | 60906251 | 61089228 | + | lncRNA |
| ENSG00000225830.13_5 | ERCC6 | 0.48754105 | 1 | 0.538 | 4.292 | chr10 | 50662516 | 50747584 | - | protein_coding |
| ENSG00000067606.17_5 | PRKCZ | 0.40464483 | 1 | 0.538 | 2.929 | chr1 | 1981850 | 2116834 | + | protein_coding |
| ENSG00000089060.11_5 | SLC8B1 | 0.45752739 | 1 | 0.538 | 3.888 | chr12 | 113736564 | 113797298 | - | protein_coding |
| ENSG00000130517.14_6 | PGPEP1 | 0.48152323 | 1 | 0.537 | 4.289 | chr19 | 18451408 | 18480760 | + | protein_coding |
| ENSG00000128311.14_5 | TST | 0.51624578 | 1 | 0.537 | 5.284 | chr22 | 37406900 | 37416224 | - | protein_coding |
| ENSG00000188906.16_7 | LRRK2 | 0.59149873 | 1 | 0.537 | 6.531 | chr12 | 40590546 | 40763087 | + | protein_coding |
| ENSG00000100211.10_2 | CBY1 | 0.38121177 | 1 | 0.537 | 2.419 | chr22 | 39052641 | 39069859 | + | protein_coding |
| ENSG00000102878.16_3 | HSF4 | 0.40792575 | 1 | 0.537 | 3.186 | chr16 | 67198584 | 67203848 | + | protein_coding |
| ENSG00000151746.14_6 | BICD1 | 0.45743125 | 1 | 0.537 | 3.868 | chr12 | 32259769 | 32536567 | + | protein_coding |
| ENSG00000139624.14_6 | CERS5 | 0.48224986 | 1 | 0.536 | 4.357 | chr12 | 50523072 | 50561316 | - | protein_coding |
| ENSG00000204842.18_9 | ATXN2 | 0.58277227 | 1 | 0.536 | 5.766 | chr12 | 111881289 | 112037480 | - | protein_coding |
| ENSG00000110318.15_5 | CEP126 | 0.44411095 | 1 | 0.536 | 3.866 | chr11 | 101785741 | 101871793 | + | protein_coding |
| ENSG00000090924.15_5 | PLEKHG2 | 0.57937367 | 1 | 0.536 | 6.313 | chr19 | 39903309 | 39919055 | + | protein_coding |
| ENSG00000287315.1_2 | AL670729.3 | 0.32133587 | 1 | 0.536 | 0.850 | chr1 | 228544713 | 228556255 | - | lncRNA |
| ENSG00000197256.10_8 | KANK2 | 0.57204711 | 1 | 0.536 | 6.167 | chr19 | 11274943 | 11308467 | - | protein_coding |
| ENSG00000181929.13_7 | PRKAG1 | 0.58208454 | 1 | 0.536 | 5.574 | chr12 | 49396057 | 49412590 | - | protein_coding |
| ENSG00000111647.13_8 | UHRF1BP1L | 0.60638111 | 1 | 0.536 | 6.241 | chr12 | 100422233 | 100536652 | - | protein_coding |
| ENSG00000178449.9_4 | COX14 | 0.47214003 | 1 | 0.535 | 4.062 | chr12 | 50505865 | 50514240 | + | protein_coding |
| ENSG00000283154.2_5 | IQCJ-SCHIP1 | 0.34303964 | 1 | 0.535 | 1.606 | chr3 | 158680024 | 159615155 | + | protein_coding |
| ENSG00000182400.15_5 | TRAPPC6B | 0.40135429 | 1 | 0.535 | 2.670 | chr14 | 39617015 | 39639736 | - | protein_coding |
| ENSG00000185504.17_7 | FAAP100 | 0.37834794 | 1 | 0.535 | 5.783 | chr17 | 79506911 | 79520987 | - | protein_coding |
| ENSG00000172765.17_7 | TMCC1 | 0.55550086 | 1 | 0.534 | 5.372 | chr3 | 129366635 | 129612449 | - | protein_coding |
| ENSG00000232442.1_6 | MHENCR | 0.36160035 | 1 | 0.534 | 1.925 | chr20 | 62258580 | 62260177 | + | lncRNA |
| ENSG00000226085.3_4 | UQCRFS1P1 | 0.31281958 | 0.99123694 | 0.534 | 0.781 | chr22 | 40271293 | 40272112 | + | processed_pseudogene |
| ENSG00000068137.15_4 | PLEKHH3 | 0.43061609 | 1 | 0.534 | 4.953 | chr17 | 40819932 | 40829012 | - | protein_coding |
| ENSG00000272512.1_6 | AL645608.7 | 0.30476215 | 0.97386856 | 0.534 | -0.383 | chr1 | 931346 | 933431 | - | lncRNA |
| ENSG00000083812.12_4 | ZNF324 | 0.44055069 | 1 | 0.534 | 3.879 | chr19 | 58978412 | 58986803 | + | protein_coding |

| | | | | | | | | | | |
|---|---|---|---|---|---|---|---|---|---|---|
| ENSG00000178149.17_6 | DALRD3 | 0.44365106 | 1 | 0.534 | 3.907 | chr3 | 49052921 | 49059726 | - | protein_coding |
| ENSG00000076944.16_6 | STXBP2 | 0.41747565 | 1 | 0.534 | 2.853 | chr19 | 7701767 | 7712759 | + | protein_coding |
| ENSG00000106004.5_5 | HOXA5 | 0.36645433 | 1 | 0.533 | 1.874 | chr7 | 27180671 | 27183300 | - | protein_coding |
| ENSG00000132003.9_6 | ZSWIM4 | 0.34881815 | 1 | 0.533 | 1.482 | chr19 | 13906274 | 13943044 | + | protein_coding |
| ENSG00000232811.1_6 | AL360270.1 | 0.31821906 | 1 | 0.533 | 0.427 | chr1 | 111486089 | 111495470 | + | lncRNA |
| ENSG00000075945.13_4 | KIFAP3 | 0.52647694 | 1 | 0.533 | 4.806 | chr1 | 169890467 | 170054349 | - | protein_coding |
| ENSG00000162545.6_4 | CAMK2N1 | 0.55661843 | 1 | 0.533 | 4.976 | chr1 | 20808884 | 20812703 | - | protein_coding |
| ENSG00000249242.8_6 | TMEM150C | 0.37355848 | 1 | 0.533 | 1.285 | chr4 | 83404323 | 83483510 | - | protein_coding |
| ENSG00000123144.11_7 | TRIR | 0.56423292 | 1 | 0.532 | 5.862 | chr19 | 12841454 | 12845498 | - | protein_coding |
| ENSG00000232434.2_3 | AJM1 | 0.35940551 | 1 | 0.532 | 3.264 | chr9 | 139738867 | 139743253 | + | protein_coding |
| ENSG00000184967.7_3 | NOC4L | 0.45800822 | 1 | 0.532 | 4.873 | chr12 | 132629002 | 132637018 | + | protein_coding |
| ENSG00000277775.1_3 | H3C7 | 0.56634801 | 1 | 0.532 | 5.460 | chr6 | 26250423 | 26250833 | - | protein_coding |
| ENSG00000153561.13_3 | RMND5A | 0.66560678 | 1 | 0.532 | 6.687 | chr2 | 86947414 | 87005164 | + | protein_coding |
| ENSG00000239521.8_6 | CASTOR3 | 0.54972277 | 1 | 0.532 | 4.860 | chr7 | 99798276 | 99869855 | - | protein_coding |
| ENSG00000131171.13_4 | SH3BGRL | 0.62752385 | 1 | 0.532 | 6.352 | chrX | 80457601 | 80554046 | + | protein_coding |
| ENSG00000183458.14_6 | AC138932.1 | 0.35474705 | 1 | 0.531 | 2.617 | chr16 | 15005408 | 15029565 | + | transcribed_unprocessed_pseudogene |
| ENSG00000254986.7_6 | DPP3 | 0.59390489 | 1 | 0.531 | 6.055 | chr11 | 66247484 | 66277128 | + | protein_coding |
| ENSG00000197008.9_2 | ZNF138 | 0.42144292 | 1 | 0.531 | 3.243 | chr7 | 64254766 | 64294059 | + | protein_coding |
| ENSG00000113231.13_5 | PDE8B | 0.41464273 | 1 | 0.531 | 2.802 | chr5 | 76506274 | 76725632 | + | protein_coding |
| ENSG00000248538.8_6 | AC022784.1 | 0.31026907 | 0.98562008 | 0.531 | -0.233 | chr8 | 9009205 | 9283034 | + | lncRNA |
| ENSG00000180336.18_6 | MEIOC | 0.57859442 | 1 | 0.530 | 5.098 | chr17 | 42733772 | 42767676 | + | protein_coding |
| ENSG00000214021.16_5 | TTLL3 | 0.45318073 | 1 | 0.530 | 3.197 | chr3 | 9849770 | 9896822 | + | protein_coding |
| ENSG00000141429.13_3 | GALNT1 | 0.67239501 | 1 | 0.530 | 7.439 | chr18 | 33161081 | 33291798 | + | protein_coding |
| ENSG00000178498.16_6 | DTX3 | 0.54546326 | 1 | 0.530 | 4.707 | chr12 | 57998405 | 58003587 | + | protein_coding |
| ENSG00000010017.13_4 | RANBP9 | 0.56658454 | 1 | 0.530 | 4.888 | chr6 | 13621730 | 13712067 | - | protein_coding |
| ENSG00000116663.11_3 | FBXO6 | 0.39518241 | 1 | 0.530 | 2.494 | chr1 | 11724257 | 11734411 | + | protein_coding |
| ENSG00000101421.4_3 | CHMP4B | 0.59878801 | 1 | 0.530 | 5.468 | chr20 | 32399154 | 32442172 | + | protein_coding |
| ENSG00000103490.14_6 | PYCARD | 0.46129572 | 1 | 0.530 | 3.373 | chr16 | 31212807 | 31214771 | - | protein_coding |
| ENSG00000116809.11_3 | ZBTB17 | 0.50026849 | 1 | 0.529 | 4.170 | chr1 | 16268364 | 16302627 | - | protein_coding |
| ENSG00000263753.8_5 | LINC00667 | 0.48130577 | 1 | 0.529 | 3.631 | chr18 | 5237825 | 5290607 | + | lncRNA |
| ENSG00000250295.6_7 | RDH10-AS1 | 0.334046 | 1 | 0.529 | 0.782 | chr8 | 74209946 | 74268696 | - | lncRNA |
| ENSG00000067715.14_4 | SYT1 | 0.51125336 | 1 | 0.529 | 5.135 | chr12 | 79257773 | 79845788 | + | protein_coding |
| ENSG00000130921.8_4 | C12orf65 | 0.53411562 | 1 | 0.529 | 4.553 | chr12 | 123717983 | 123742507 | + | protein_coding |
| ENSG00000196456.13_7 | ZNF775 | 0.34967257 | 1 | 0.529 | 2.088 | chr7 | 150065879 | 150095718 | + | protein_coding |
| ENSG00000235706.8_7 | DICER1-AS1 | 0.37397597 | 1 | 0.528 | 1.584 | chr14 | 95623982 | 95647812 | + | lncRNA |
| ENSG00000158292.7_4 | GPR153 | 0.34481519 | 1 | 0.528 | 1.158 | chr1 | 6307413 | 6321158 | - | protein_coding |
| ENSG00000161618.10_4 | ALDH16A1 | 0.43439264 | 1 | 0.528 | 5.007 | chr19 | 49956482 | 49974307 | + | protein_coding |
| ENSG00000112992.18_6 | NNT | 0.64324403 | 1 | 0.528 | 5.727 | chr5 | 43602794 | 43707507 | + | protein_coding |
| ENSG00000072133.11_4 | RPS6KA6 | 0.55433681 | 1 | 0.527 | 4.463 | chrX | 83313354 | 83443207 | - | protein_coding |
| ENSG00000186017.14_4 | ZNF566 | 0.60275078 | 1 | 0.527 | 5.131 | chr19 | 36936021 | 36980804 | - | protein_coding |
| ENSG00000213144.2_5 | AC084880.1 | 0.33148832 | 1 | 0.527 | -0.961 | chr12 | 119632655 | 119633199 | + | processed_pseudogene |
| ENSG00000233757.6_5 | AC092835.1 | 0.39508569 | 1 | 0.527 | 2.116 | chr2 | 95873283 | 95925522 | + | protein_coding |
| ENSG00000028203.18_4 | VEZT | 0.71196967 | 1 | 0.527 | 7.348 | chr12 | 95611522 | 95696575 | + | protein_coding |
| ENSG00000179526.17_4 | SHARPIN | 0.53319424 | 1 | 0.527 | 5.280 | chr8 | 145153536 | 145163027 | - | protein_coding |
| ENSG00000142444.7_5 | TIMM29 | 0.48392018 | 1 | 0.527 | 3.492 | chr19 | 11039487 | 11044211 | + | protein_coding |
| ENSG00000257896.1_6 | AC093012.2 | 0.31962287 | 1 | 0.526 | 0.045 | chr12 | 44130431 | 44134870 | + | transcribed_unprocessed_pseudogene |
| ENSG00000111412.6_5 | C12orf49 | 0.66779536 | 1 | 0.526 | 6.008 | chr12 | 117147976 | 117175875 | - | protein_coding |
| ENSG00000224687.2_6 | RASAL2-AS1 | 0.32574317 | 1 | 0.526 | 0.275 | chr1 | 178059812 | 178063119 | - | lncRNA |
| ENSG00000112541.15_5 | PDE10A | 0.66301804 | 1 | 0.526 | 6.167 | chr6 | 165740776 | 166401566 | - | protein_coding |
| ENSG00000198466.12_5 | ZNF587 | 0.54095732 | 1 | 0.526 | 4.284 | chr19 | 58361227 | 58376485 | + | protein_coding |
| ENSG00000280088.1_6 | AC126474.2 | 0.40593056 | 1 | 0.526 | 2.252 | chr12 | 100426103 | 100427852 | - | TEC |
| ENSG00000228363.2_7 | AC015971.1 | 0.33402258 | 1 | 0.526 | 0.605 | chr2 | 86789193 | 86845889 | + | lncRNA |
| ENSG00000140092.14_5 | FBLN5 | 0.36911684 | 1 | 0.525 | 1.348 | chr14 | 92335756 | 92414331 | - | protein_coding |
| ENSG00000178607.16_4 | ERN1 | 0.56625218 | 1 | 0.525 | 4.439 | chr17 | 62116502 | 62208179 | - | protein_coding |
| ENSG00000130669.17_4 | PAK4 | 0.54626089 | 1 | 0.525 | 6.281 | chr19 | 39616410 | 39673456 | + | protein_coding |
| ENSG00000251348.1_4 | HSPD1P11 | 0.31618485 | 0.99926427 | 0.525 | -0.296 | chr5 | 95104703 | 95106404 | + | processed_pseudogene |
| ENSG00000042062.13_8 | RIPOR3 | 0.32733876 | 1 | 0.525 | 0.203 | chr20 | 49202645 | 49308079 | - | protein_coding |
| ENSG00000130164.14_8 | LDLR | 0.69221277 | 1 | 0.525 | 6.804 | chr19 | 11200138 | 11244496 | + | protein_coding |
| ENSG00000039139.9_4 | DNAH5 | 0.47240347 | 1 | 0.525 | 3.182 | chr5 | 13690440 | 13944652 | - | protein_coding |

| | | | | | | | | | | |
|---|---|---|---|---|---|---|---|---|---|---|
| ENSG00000179588.9_3 | ZFPM1 | 0.36596891 | 1 | 0.525 | 2.404 | chr16 | 88519688 | 88603439 | + | protein_coding |
| ENSG00000151148.14_8 | UBE3B | 0.65283327 | 1 | 0.524 | 5.771 | chr12 | 109915207 | 109974510 | + | protein_coding |
| ENSG00000171503.12_5 | ETFDH | 0.51996854 | 1 | 0.524 | 3.819 | chr4 | 159593277 | 159630775 | + | protein_coding |
| ENSG00000111644.8_5 | ACRBP | 0.33361292 | 1 | 0.524 | -0.646 | chr12 | 6747241 | 6756599 | - | protein_coding |
| ENSG00000078246.17_4 | TULP3 | 0.6343663 | 1 | 0.524 | 5.221 | chr12 | 2986389 | 3050304 | + | protein_coding |
| ENSG00000234776.5_4 | C11orf94 | 0.31467286 | 0.99568883 | 0.524 | -0.728 | chr11 | 45928064 | 45928823 | - | protein_coding |
| ENSG00000275560.1_7 | AC008115.3 | 0.35512932 | 1 | 0.524 | 0.697 | chr12 | 12871907 | 12872455 | + | lncRNA |
| ENSG00000204991.11_6 | SPIRE2 | 0.54918257 | 1 | 0.524 | 4.672 | chr16 | 89884587 | 89937727 | + | protein_coding |
| ENSG00000135127.11_5 | BICDL1 | 0.44882715 | 1 | 0.523 | 3.090 | chr12 | 120427673 | 120532298 | + | protein_coding |
| ENSG00000226824.7_7 | AC006001.2 | 0.32921915 | 1 | 0.523 | -0.095 | chr7 | 66119525 | 66136510 | + | lncRNA |
| ENSG00000197329.12_4 | PELI1 | 0.42318415 | 1 | 0.523 | 2.592 | chr2 | 64319786 | 64371554 | - | protein_coding |
| ENSG00000108861.9_4 | DUSP3 | 0.7469593 | 1 | 0.523 | 7.235 | chr17 | 41843493 | 41856345 | - | protein_coding |
| ENSG00000143384.13_5 | MCL1 | 0.74580351 | 1 | 0.523 | 7.624 | chr1 | 150547027 | 150552214 | - | protein_coding |
| ENSG00000025423.11_4 | HSD17B6 | 0.3255906 | 1 | 0.523 | 0.162 | chr12 | 57145945 | 57181574 | + | protein_coding |
| ENSG00000136213.10_6 | CHST12 | 0.53038793 | 1 | 0.522 | 4.219 | chr7 | 2443223 | 2488119 | + | protein_coding |
| ENSG00000167371.21_7 | PRRT2 | 0.34250603 | 1 | 0.522 | 0.747 | chr16 | 29822703 | 29827213 | + | protein_coding |
| ENSG00000176046.8_3 | NUPR1 | 0.4364834 | 1 | 0.522 | 2.783 | chr16 | 28544029 | 28550495 | - | protein_coding |
| ENSG00000051108.15_7 | HERPUD1 | 0.65860485 | 1 | 0.522 | 5.646 | chr16 | 56966054 | 56978776 | + | protein_coding |
| ENSG00000160216.20_5 | AGPAT3 | 0.66027054 | 1 | 0.522 | 6.057 | chr21 | 45285104 | 45407473 | + | protein_coding |
| ENSG00000167657.14_5 | DAPK3 | 0.54008768 | 1 | 0.521 | 4.440 | chr19 | 3958451 | 3971121 | - | protein_coding |
| ENSG00000052795.13_4 | FNIP2 | 0.52816755 | 1 | 0.521 | 3.935 | chr4 | 159690178 | 159829202 | + | protein_coding |
| ENSG00000261253.2_6 | AC137932.2 | 0.3390816 | 1 | 0.521 | 0.485 | chr16 | 89387541 | 89391518 | + | lncRNA |
| ENSG00000269337.1 | AL591479.1 | 0.41422042 | 1 | 0.521 | 2.039 | chr9 | 70646589 | 70648956 | + | protein_coding |
| ENSG00000177951.17_3 | BET1L | 0.57188428 | 1 | 0.520 | 4.211 | chr11 | 167784 | 207428 | - | protein_coding |
| ENSG00000243176.6_6 | AC092944.1 | 0.36222984 | 1 | 0.520 | 0.990 | chr3 | 156891990 | 157099054 | + | lncRNA |
| ENSG00000102034.16_3 | ELF4 | 0.62317081 | 1 | 0.520 | 5.127 | chrX | 129198849 | 129244691 | - | protein_coding |
| ENSG00000126821.8_4 | SGPP1 | 0.69025442 | 1 | 0.520 | 5.856 | chr14 | 64150934 | 64194783 | - | protein_coding |
| ENSG00000237015.1_6 | AL031186.1 | 0.34690394 | 1 | 0.520 | 0.561 | chr22 | 29656878 | 29658026 | + | lncRNA |
| ENSG00000197933.13_6 | ZNF823 | 0.50611809 | 1 | 0.520 | 3.479 | chr19 | 11832080 | 11849824 | - | protein_coding |
| ENSG00000137691.13_5 | CFAP300 | 0.42966217 | 1 | 0.519 | 2.519 | chr11 | 101918168 | 101955285 | + | protein_coding |
| ENSG00000196705.8_5 | ZNF431 | 0.56329921 | 1 | 0.519 | 4.136 | chr19 | 21324827 | 21378856 | + | protein_coding |
| ENSG00000127774.7_5 | EMC6 | 0.32572045 | 1 | 0.519 | -0.462 | chr17 | 3572106 | 3572962 | + | protein_coding |
| ENSG00000197747.9_4 | S100A10 | 0.73359107 | 1 | 0.519 | 7.323 | chr1 | 151955391 | 151966335 | - | protein_coding |
| ENSG00000278385.1_6 | AC121338.2 | 0.32492492 | 1 | 0.519 | -0.962 | chr12 | 48298905 | 48300648 | + | lncRNA |
| ENSG00000103260.9_6 | METRN | 0.39978498 | 1 | 0.519 | 5.777 | chr16 | 765118 | 769655 | + | protein_coding |
| ENSG00000135750.15_3 | KCNK1 | 0.56747726 | 1 | 0.518 | 4.129 | chr1 | 233749852 | 233808260 | + | protein_coding |
| ENSG00000173875.14_3 | ZNF791 | 0.56752983 | 1 | 0.518 | 4.413 | chr19 | 12721732 | 12744654 | + | protein_coding |
| ENSG00000176973.8_4 | FAM89B | 0.3734637 | 1 | 0.518 | 1.160 | chr11 | 65339820 | 65341669 | + | protein_coding |
| ENSG00000153395.10_4 | LPCAT1 | 0.70248373 | 1 | 0.518 | 6.148 | chr5 | 1456595 | 1524077 | - | protein_coding |
| ENSG00000103343.13_4 | ZNF174 | 0.5197275 | 1 | 0.518 | 3.358 | chr16 | 3451215 | 3459364 | + | protein_coding |
| ENSG00000175662.17_5 | TOM1L2 | 0.68542559 | 1 | 0.518 | 5.622 | chr17 | 17746825 | 17875736 | - | protein_coding |
| ENSG00000233382.7_4 | NKAPP1 | 0.37710808 | 1 | 0.518 | 0.884 | chrX | 119254863 | 119379122 | - | transcribed_processed_pseudogene |
| ENSG00000130511.16_4 | SSBP4 | 0.40198494 | 1 | 0.517 | 4.616 | chr19 | 18529674 | 18545372 | + | protein_coding |
| ENSG00000204498.11_4 | NFKBIL1 | 0.33983049 | 1 | 0.517 | 0.174 | chr6 | 31514647 | 31526606 | + | protein_coding |
| ENSG00000105464.3_2 | GRIN2D | 0.54605283 | 1 | 0.517 | 4.338 | chr19 | 48898132 | 48948188 | + | protein_coding |
| ENSG00000136485.15_3 | DCAF7 | 0.7664131 | 1 | 0.517 | 7.020 | chr17 | 61627838 | 61671641 | + | protein_coding |
| ENSG00000183690.13_5 | EFHC2 | 0.32331736 | 1 | 0.517 | -0.576 | chrX | 44007118 | 44202918 | - | protein_coding |
| ENSG00000123411.15_4 | IKZF4 | 0.37774369 | 1 | 0.517 | 1.398 | chr12 | 56401443 | 56432219 | + | protein_coding |
| ENSG00000280195.1_6 | AC245140.2 | 0.33619199 | 1 | 0.517 | 0.036 | chrX | 153625219 | 153627157 | - | lncRNA |
| ENSG00000177700.6_4 | POLR2L | 0.65211291 | 1 | 0.517 | 6.892 | chr11 | 837356 | 842529 | - | protein_coding |
| ENSG00000104823.9_5 | ECH1 | 0.71145126 | 1 | 0.516 | 6.589 | chr19 | 39306062 | 39322481 | - | protein_coding |
| ENSG00000154265.16_5 | ABCA5 | 0.54401946 | 1 | 0.516 | 3.855 | chr17 | 67240452 | 67323385 | - | protein_coding |
| ENSG00000111786.9_4 | SRSF9 | 0.61676443 | 1 | 0.516 | 6.030 | chr12 | 120899475 | 120907551 | - | protein_coding |
| ENSG00000243433.1_7 | AC010973.1 | 0.33071811 | 1 | 0.516 | -0.865 | chr7 | 150725868 | 150726841 | - | lncRNA |
| ENSG00000179476.8_3 | C14orf28 | 0.4493635 | 1 | 0.516 | 2.474 | chr14 | 45366478 | 45376460 | + | protein_coding |
| ENSG00000123609.11_5 | NMI | 0.48470559 | 1 | 0.516 | 2.895 | chr2 | 152126984 | 152146408 | - | protein_coding |
| ENSG00000276851.1_6 | AC002401.4 | 0.34906268 | 1 | 0.516 | 0.420 | chr17 | 48171429 | 48172011 | + | lncRNA |
| ENSG00000136842.14_3 | TMOD1 | 0.36332871 | 1 | 0.516 | 0.743 | chr9 | 100263462 | 100364025 | + | protein_coding |
| ENSG00000186940.6_5 | CHCHD2P9 | 0.32749393 | 1 | 0.516 | -0.512 | chr9 | 82006219 | 82006674 | + | processed_pseudogene |
| ENSG00000138623.10_6 | SEMA7A | 0.32501713 | 1 | 0.516 | -0.538 | chr15 | 74701630 | 74726299 | - | protein_coding |

| | | | | | | | | | | |
|---|---|---|---|---|---|---|---|---|---|---|
| ENSG00000258839.3_3 | MC1R | 0.42750473 | 1 | 0.516 | 2.300 | chr16 | 89978527 | 89987385 | + | protein_coding |
| ENSG00000275180.1_7 | AC048341.2 | 0.53853126 | 1 | 0.515 | 3.987 | chr12 | 62997689 | 62998179 | + | lncRNA |
| ENSG00000120860.11_6 | WASHC3 | 0.62221612 | 1 | 0.515 | 4.669 | chr12 | 102406618 | 102455927 | - | protein_coding |
| ENSG00000074621.14_5 | SLC24A1 | 0.55712527 | 1 | 0.515 | 3.864 | chr15 | 65903704 | 65953333 | + | protein_coding |
| ENSG00000145476.16_4 | CYP4V2 | 0.54202787 | 1 | 0.515 | 3.713 | chr4 | 187112721 | 187134617 | + | protein_coding |
| ENSG00000174165.8_4 | ZDHHC24 | 0.60865861 | 1 | 0.515 | 4.494 | chr11 | 66288108 | 66313706 | - | protein_coding |
| ENSG00000197183.14_6 | NOL4L | 0.69882196 | 1 | 0.515 | 5.387 | chr20 | 31030862 | 31172876 | - | protein_coding |
| ENSG00000025708.14_6 | TYMP | 0.53867596 | 1 | 0.515 | 3.476 | chr22 | 50964181 | 50968461 | - | protein_coding |
| ENSG00000160445.11_6 | ZER1 | 0.63884005 | 1 | 0.514 | 4.833 | chr9 | 131492065 | 131534693 | - | protein_coding |
| ENSG00000055211.14_4 | GINM1 | 0.65420383 | 1 | 0.514 | 4.956 | chr6 | 149887430 | 149912884 | + | protein_coding |
| ENSG00000240038.6_3 | AMY2B | 0.32784699 | 1 | 0.514 | -0.296 | chr1 | 104096437 | 104122156 | + | protein_coding |
| ENSG00000132205.11_3 | EMILIN2 | 0.50074401 | 1 | 0.514 | 3.052 | chr18 | 2847004 | 2916001 | + | protein_coding |
| ENSG00000171298.13_5 | GAA | 0.64683649 | 1 | 0.514 | 6.451 | chr17 | 78075355 | 78093680 | + | protein_coding |
| ENSG00000186187.12_6 | ZNRF1 | 0.54168925 | 1 | 0.513 | 3.740 | chr16 | 75032922 | 75144892 | + | protein_coding |
| ENSG00000165171.11_7 | METTL27 | 0.51755632 | 1 | 0.512 | 3.192 | chr7 | 73248920 | 73256846 | - | protein_coding |
| ENSG00000105698.16_4 | USF2 | 0.73491413 | 1 | 0.512 | 6.532 | chr19 | 35759865 | 35770724 | + | protein_coding |
| ENSG00000182057.5_6 | OGFRP1 | 0.4765836 | 1 | 0.512 | 2.243 | chr22 | 42665709 | 42675540 | + | lncRNA |
| ENSG00000182612.10_3 | TSPAN10 | 0.39190478 | 1 | 0.512 | 1.549 | chr17 | 79604197 | 79615779 | + | protein_coding |
| ENSG00000125864.14_5 | BFSP1 | 0.5190865 | 1 | 0.512 | 3.125 | chr20 | 17474550 | 17549865 | - | protein_coding |
| ENSG00000134508.12_3 | CABLES1 | 0.56097284 | 1 | 0.512 | 3.576 | chr18 | 20714528 | 20840431 | + | protein_coding |
| ENSG00000170122.5_2 | FOXD4 | 0.33013892 | 1 | 0.511 | -0.539 | chr9 | 116231 | 118204 | - | protein_coding |
| ENSG00000213722.9_7 | DDAH2 | 0.5002898 | 1 | 0.511 | 3.247 | chr6 | 31694815 | 31698394 | - | protein_coding |
| ENSG00000279970.1_6 | AC023024.2 | 0.33175779 | 1 | 0.511 | -0.125 | chr15 | 101839861 | 101841853 | - | TEC |
| ENSG00000186834.3_2 | HEXIM1 | 0.69596862 | 1 | 0.511 | 5.368 | chr17 | 43225869 | 43229468 | + | protein_coding |
| ENSG00000197562.10_4 | RAB40C | 0.55721775 | 1 | 0.511 | 3.987 | chr16 | 639357 | 679272 | + | protein_coding |
| ENSG00000278970.2_7 | HEIH | 0.44980533 | 1 | 0.510 | 2.111 | chr5 | 180253871 | 180258605 | - | lncRNA |
| ENSG00000030582.18_4 | GRN | 0.78213384 | 1 | 0.510 | 7.281 | chr17 | 42422614 | 42430474 | + | protein_coding |
| ENSG00000169856.9_4 | ONECUT1 | 0.34864373 | 1 | 0.510 | 0.282 | chr15 | 53047250 | 53083275 | - | protein_coding |
| ENSG00000177000.13_7 | MTHFR | 0.62024351 | 1 | 0.510 | 4.267 | chr1 | 11845780 | 11866512 | - | protein_coding |
| ENSG00000245532.9_5 | NEAT1 | 0.51295668 | 1 | 0.510 | 9.391 | chr11 | 65190245 | 65213011 | + | lncRNA |
| ENSG00000043355.12_4 | ZIC2 | 0.49297767 | 1 | 0.510 | 2.740 | chr13 | 100634038 | 100639019 | + | protein_coding |
| ENSG00000130675.15_6 | MNX1 | 0.45315922 | 1 | 0.509 | 2.887 | chr7 | 156786745 | 156803357 | - | protein_coding |
| ENSG00000121753.12_6 | ADGRB2 | 0.53256094 | 1 | 0.509 | 3.943 | chr1 | 32192718 | 32230494 | - | protein_coding |
| ENSG00000171792.11_5 | RHNO1 | 0.75734539 | 1 | 0.509 | 5.883 | chr12 | 2985424 | 2998690 | + | protein_coding |
| ENSG00000275481.1_7 | AC025031.4 | 0.42570696 | 1 | 0.509 | 1.459 | chr12 | 46782639 | 46785909 | + | lncRNA |
| ENSG00000203644.3_7 | AC083799.1 | 0.48843159 | 1 | 0.509 | 2.629 | chr3 | 129565891 | 129566800 | - | lncRNA |
| ENSG00000203943.8_5 | SAMD13 | 0.3567446 | 1 | 0.508 | 0.256 | chr1 | 84764049 | 84855640 | + | protein_coding |
| ENSG00000214063.11_4 | TSPAN4 | 0.74925117 | 1 | 0.508 | 6.874 | chr11 | 842812 | 867116 | + | protein_coding |
| ENSG00000100239.16_6 | PPP6R2 | 0.68427573 | 1 | 0.508 | 5.131 | chr22 | 50781733 | 50883519 | + | protein_coding |
| ENSG00000133056.13_7 | PIK3C2B | 0.63422729 | 1 | 0.508 | 4.549 | chr1 | 204391756 | 204463852 | - | protein_coding |
| ENSG00000280332.1_6 | AC020917.4 | 0.3732382 | 1 | 0.508 | -0.867 | chr19 | 16467140 | 16469138 | - | TEC |
| ENSG00000214548.18_8 | MEG3 | 0.69256317 | 1 | 0.508 | 5.330 | chr14 | 101245747 | 101327368 | + | lncRNA |
| ENSG00000145247.12_7 | OCIAD2 | 0.63620713 | 1 | 0.508 | 4.254 | chr4 | 48887036 | 48908954 | - | protein_coding |
| ENSG00000112414.15_5 | ADGRG6 | 0.83269652 | 1 | 0.507 | 8.430 | chr6 | 142622991 | 142767403 | + | protein_coding |
| ENSG00000214719.13_6 | AC005562.1 | 0.34215899 | 1 | 0.507 | 0.170 | chr17 | 28903482 | 28999807 | + | lncRNA |
| ENSG00000166340.17_6 | TPP1 | 0.61713027 | 1 | 0.507 | 4.239 | chr11 | 6633999 | 6640679 | - | protein_coding |
| ENSG00000157184.7_4 | CPT2 | 0.53775078 | 1 | 0.507 | 3.224 | chr1 | 53662464 | 53679869 | + | protein_coding |
| ENSG00000063245.14_6 | EPN1 | 0.41258574 | 1 | 0.507 | 6.782 | chr19 | 56186592 | 56221224 | + | protein_coding |
| ENSG00000111596.13_6 | CNOT2 | 0.78313482 | 1 | 0.507 | 6.594 | chr12 | 70636774 | 70749037 | + | protein_coding |
| ENSG00000185033.14_5 | SEMA4B | 0.6841857 | 1 | 0.507 | 6.453 | chr15 | 90703836 | 90772911 | + | protein_coding |
| ENSG00000149115.14_7 | TNKS1BP1 | 0.67722067 | 1 | 0.506 | 6.469 | chr11 | 57067112 | 57092426 | - | protein_coding |
| ENSG00000203724.11_3 | C1orf53 | 0.37350952 | 1 | 0.506 | 0.989 | chr1 | 197871760 | 197876497 | + | protein_coding |
| ENSG00000122674.12_4 | CCZ1 | 0.56349866 | 1 | 0.506 | 3.715 | chr7 | 5938356 | 5966181 | + | protein_coding |
| ENSG00000287277.1_2 | AL392086.3 | 0.33608697 | 1 | 0.506 | -0.576 | chr10 | 6866237 | 7160431 | - | lncRNA |
| ENSG00000183255.12_5 | PTTG1IP | 0.83582883 | 1 | 0.506 | 7.129 | chr21 | 46269500 | 46293818 | - | protein_coding |
| ENSG00000228906.1_7 | AL353804.1 | 0.33619311 | 1 | 0.505 | -0.467 | chrX | 73168808 | 73169393 | + | lncRNA |
| ENSG00000211450.10_9 | SELENOH | 0.44845566 | 1 | 0.505 | 2.223 | chr11 | 57508722 | 57511026 | + | protein_coding |
| ENSG00000128872.10_4 | TMOD2 | 0.50908285 | 1 | 0.505 | 2.948 | chr15 | 52043794 | 52108560 | + | protein_coding |
| ENSG00000245112.2_7 | SMARCA5-AS1 | 0.35335255 | 1 | 0.505 | 0.168 | chr4 | 144434625 | 144435788 | - | lncRNA |
| ENSG00000272523.1_5 | LINC01023 | 0.33579326 | 1 | 0.505 | -0.657 | chr5 | 108063526 | 108063961 | - | lncRNA |

| | | | | | | | | | | |
|---|---|---|---|---|---|---|---|---|---|---|
| ENSG00000121671.12_4 | CRY2 | 0.56398903 | 1 | 0.505 | 3.358 | chr11 | 45868669 | 45904799 | + | protein_coding |
| ENSG00000181045.15_6 | SLC26A11 | 0.57456867 | 1 | 0.505 | 4.194 | chr17 | 78193498 | 78227299 | + | protein_coding |
| ENSG00000117425.14_4 | PTCH2 | 0.33608531 | 1 | 0.505 | -0.657 | chr1 | 45285516 | 45308925 | - | protein_coding |
| ENSG00000231312.7_8 | MAP4K3-DT | 0.45258464 | 1 | 0.505 | 2.343 | chr2 | 39663671 | 39892483 | + | lncRNA |
| ENSG00000054282.16_3 | SDCCAG8 | 0.61373471 | 1 | 0.505 | 4.175 | chr1 | 243419336 | 243663393 | + | protein_coding |
| ENSG00000141934.10_6 | PLPP2 | 0.7559091 | 1 | 0.504 | 6.257 | chr19 | 281040 | 291403 | - | protein_coding |
| ENSG00000133466.14_4 | C1QTNF6 | 0.68480492 | 1 | 0.504 | 4.580 | chr22 | 37576206 | 37595425 | - | protein_coding |
| ENSG00000226200.7_8 | SGMS1-AS1 | 0.47064932 | 1 | 0.504 | 2.364 | chr10 | 52383226 | 52401211 | + | lncRNA |
| ENSG00000125966.10_3 | MMP24 | 0.80507311 | 1 | 0.504 | 6.347 | chr20 | 33814493 | 33864801 | + | protein_coding |
| ENSG00000155085.15_4 | AK9 | 0.42611209 | 1 | 0.503 | 2.017 | chr6 | 109814059 | 110012420 | - | protein_coding |
| ENSG00000116604.18_6 | MEF2D | 0.72204425 | 1 | 0.503 | 5.348 | chr1 | 156433519 | 156470571 | - | protein_coding |
| ENSG00000089692.9_4 | LAG3 | 0.37328984 | 1 | 0.503 | 0.453 | chr12 | 6881678 | 6887621 | + | protein_coding |
| ENSG00000133247.14_6 | KMT5C | 0.5298926 | 1 | 0.502 | 3.527 | chr19 | 55851221 | 55859489 | + | protein_coding |
| ENSG00000129353.15_5 | SLC44A2 | 0.51622815 | 1 | 0.502 | 3.108 | chr19 | 10713133 | 10755233 | + | protein_coding |
| ENSG00000160446.19_7 | ZDHHC12 | 0.66989386 | 1 | 0.502 | 5.044 | chr9 | 131483149 | 131486406 | - | protein_coding |
| ENSG00000131779.11_4 | PEX11B | 0.63501664 | 1 | 0.502 | 4.022 | chr1 | 145516372 | 145523730 | + | protein_coding |
| ENSG00000125505.17_6 | MBOAT7 | 0.74775792 | 1 | 0.502 | 6.536 | chr19 | 54677106 | 54693733 | - | protein_coding |
| ENSG00000273032.2_6 | DGCR9 | 0.3556731 | 1 | 0.502 | 0.126 | chr22 | 18958038 | 19018755 | + | lncRNA |
| ENSG00000161082.13_3 | CELF5 | 0.35349104 | 1 | 0.502 | -0.051 | chr19 | 3224659 | 3297074 | + | protein_coding |
| ENSG00000188313.13_6 | PLSCR1 | 0.77040906 | 1 | 0.502 | 5.838 | chr3 | 146232967 | 146262643 | - | protein_coding |
| ENSG00000143570.18_6 | SLC39A1 | 0.35009494 | 1 | 0.502 | -0.049 | chr1 | 153931575 | 153940660 | - | protein_coding |
| ENSG00000248866.1_5 | USP46-AS1 | 0.38936155 | 1 | 0.502 | 0.900 | chr4 | 53525573 | 53527835 | + | lncRNA |
| ENSG00000076108.11_8 | BAZ2A | 0.81871957 | 1 | 0.501 | 6.269 | chr12 | 56989380 | 57030600 | - | protein_coding |
| ENSG00000139679.15_3 | LPAR6 | 0.52937023 | 1 | 0.501 | 2.693 | chr13 | 48963703 | 49018840 | - | protein_coding |
| ENSG00000228824.7_8 | MIR4500HG | 0.37484086 | 1 | 0.500 | 0.394 | chr13 | 88079469 | 88326490 | - | lncRNA |
| ENSG00000118557.16_7 | PMFBP1 | 0.34868507 | 1 | 0.500 | -0.154 | chr16 | 72146056 | 72210777 | - | protein_coding |
| ENSG00000179918.19_5 | SEPHS2 | 0.67471481 | 1 | 0.500 | 4.593 | chr16 | 30454952 | 30457195 | - | protein_coding |
| ENSG00000079459.13_4 | FDFT1 | 0.86541475 | 1 | 0.500 | 7.012 | chr8 | 11653082 | 11696813 | + | protein_coding |
| ENSG00000198879.12_4 | SFMBT2 | 0.73235362 | 1 | 0.500 | 4.913 | chr10 | 7200586 | 7453448 | - | protein_coding |
| ENSG00000143013.13_3 | LMO4 | 0.75352089 | 1 | 0.500 | 5.302 | chr1 | 87794563 | 87814606 | + | protein_coding |
| ENSG00000037042.9_3 | TUBG2 | 0.73176448 | 1 | 0.500 | 5.241 | chr17 | 40811302 | 40819024 | + | protein_coding |
| ENSG00000105443.15_5 | CYTH2 | 0.80673987 | 1 | 0.500 | 6.337 | chr19 | 48972465 | 48985571 | + | protein_coding |
| ENSG00000104946.13_5 | TBC1D17 | 0.73032711 | 1 | 0.500 | 4.889 | chr19 | 50380682 | 50392007 | + | protein_coding |
| ENSG00000068354.16_5 | TBC1D25 | 0.53399659 | 1 | 0.499 | 2.917 | chrX | 48398102 | 48420997 | + | protein_coding |
| ENSG00000197283.17_9 | SYNGAP1 | 0.45101025 | 1 | 0.499 | 1.610 | chr6 | 33387438 | 33421466 | + | protein_coding |
| ENSG00000137103.20_7 | TMEM8B | 0.46717074 | 1 | 0.499 | 1.844 | chr9 | 35814448 | 35865515 | + | protein_coding |
| ENSG00000227619.3_7 | AL391056.1 | 0.46900561 | 1 | 0.499 | 2.159 | chr9 | 132337396 | 132346835 | + | lncRNA |
| ENSG00000105825.14_6 | TFPI2 | 0.8643376 | 1 | 0.499 | 6.970 | chr7 | 93514708 | 93520065 | - | protein_coding |
| ENSG00000170175.11_4 | CHRNB1 | 0.44512869 | 1 | 0.499 | 1.839 | chr17 | 7348380 | 7361029 | + | protein_coding |
| ENSG00000131242.18_5 | RAB11FIP4 | 0.5424924 | 1 | 0.499 | 2.932 | chr17 | 29718693 | 29865230 | + | protein_coding |
| ENSG00000158458.20_5 | NRG2 | 0.46881563 | 1 | 0.498 | 2.206 | chr5 | 139226364 | 139422884 | - | protein_coding |
| ENSG00000115286.20_4 | NDUFS7 | 0.4418023 | 1 | 0.498 | 1.554 | chr19 | 1383526 | 1395588 | + | protein_coding |
| ENSG00000175215.11_6 | CTDSP2 | 0.82342825 | 1 | 0.498 | 6.281 | chr12 | 58213710 | 58240512 | - | protein_coding |
| ENSG00000128563.13_4 | PRKRIP1 | 0.66789957 | 1 | 0.498 | 4.481 | chr7 | 102004319 | 102067123 | + | protein_coding |
| ENSG00000126461.15_6 | SCAF1 | 0.45984111 | 1 | 0.498 | 6.291 | chr19 | 50145466 | 50161899 | + | protein_coding |
| ENSG00000125772.14_5 | GPCPD1 | 0.84436078 | 1 | 0.498 | 6.401 | chr20 | 5525085 | 5591652 | - | protein_coding |
| ENSG00000116704.8_4 | SLC35D1 | 0.70897522 | 1 | 0.498 | 4.460 | chr1 | 67465033 | 67519831 | - | protein_coding |
| ENSG00000132256.19_5 | TRIM5 | 0.74343191 | 1 | 0.497 | 4.917 | chr11 | 5684425 | 5959849 | - | protein_coding |
| ENSG00000136379.12_6 | ABHD17C | 0.73753446 | 1 | 0.497 | 4.797 | chr15 | 80972025 | 81047962 | + | protein_coding |
| ENSG00000238228.1_2 | OR7E7P | 0.34792829 | 1 | 0.497 | -0.136 | chr7 | 97576299 | 97577310 | - | unprocessed_pseudogene |
| ENSG00000213096.10_4 | AC092279.1 | 0.72769967 | 1 | 0.496 | 5.074 | chr19 | 24216207 | 24312763 | + | protein_coding |
| ENSG00000159840.16_5 | ZYX | 0.7404397 | 1 | 0.496 | 5.678 | chr7 | 143078388 | 143088204 | + | protein_coding |
| ENSG00000157578.13_3 | LCA5L | 0.41644338 | 1 | 0.496 | 1.096 | chr21 | 40777770 | 40817731 | - | protein_coding |
| ENSG00000102858.13_8 | MGRN1 | 0.71648164 | 1 | 0.496 | 5.464 | chr16 | 4666494 | 4740975 | + | protein_coding |
| ENSG00000160963.14_4 | COL26A1 | 0.35034446 | 1 | 0.496 | -0.628 | chr7 | 101006156 | 101202304 | + | protein_coding |
| ENSG00000132005.9_4 | RFX1 | 0.55490089 | 1 | 0.495 | 3.231 | chr19 | 14072342 | 14117851 | - | protein_coding |
| ENSG00000185624.15_4 | P4HB | 0.8797734 | 1 | 0.495 | 8.785 | chr17 | 79801037 | 79818500 | - | protein_coding |
| ENSG00000227038.3_6 | GTF2IP7 | 0.45208261 | 1 | 0.495 | 1.711 | chr7 | 75719749 | 75738097 | - | transcribed_unprocessed_pseudogene |
| ENSG00000175832.13_5 | ETV4 | 0.79635865 | 1 | 0.495 | 5.690 | chr17 | 41605212 | 41656988 | - | protein_coding |
| ENSG00000043143.20_3 | JADE2 | 0.83209328 | 1 | 0.495 | 6.776 | chr5 | 133860003 | 133918920 | + | protein_coding |

| | | | | | | | | | | |
|---|---|---|---|---|---|---|---|---|---|---|
| ENSG00000135519.8_4 | KCNH3 | 0.53771279 | 1 | 0.495 | 2.760 | chr12 | 49932813 | 49952120 | + | protein_coding |
| ENSG00000076067.13_4 | RBMS2 | 0.85195253 | 1 | 0.495 | 6.476 | chr12 | 56915604 | 56989977 | + | protein_coding |
| ENSG00000129566.13_7 | TEP1 | 0.7896588 | 1 | 0.495 | 5.465 | chr14 | 20833826 | 20881660 | - | protein_coding |
| ENSG00000151470.13_5 | C4orf33 | 0.53251174 | 1 | 0.494 | 3.118 | chr4 | 130014472 | 130037795 | + | protein_coding |
| ENSG00000145949.11_5 | MYLK4 | 0.35993253 | 1 | 0.494 | 0.087 | chr6 | 2663863 | 2751156 | - | protein_coding |
| ENSG00000126746.17_4 | ZNF384 | 0.79139784 | 1 | 0.494 | 5.315 | chr12 | 6775643 | 6798738 | - | protein_coding |
| ENSG00000249199.2_7 | AC026785.3 | 0.57011451 | 1 | 0.493 | 2.971 | chr5 | 17367697 | 17375762 | - | lncRNA |
| ENSG00000196150.13_4 | ZNF250 | 0.57254288 | 1 | 0.493 | 3.776 | chr8 | 146101882 | 146127553 | - | protein_coding |
| ENSG00000166326.7_3 | TRIM44 | 0.85542832 | 1 | 0.493 | 6.835 | chr11 | 35684323 | 35839557 | + | protein_coding |
| ENSG00000179981.10_6 | TSHZ1 | 0.62021376 | 1 | 0.493 | 3.460 | chr18 | 72922710 | 73001905 | + | protein_coding |
| ENSG00000103197.18_5 | TSC2 | 0.84362389 | 1 | 0.492 | 6.634 | chr16 | 2097968 | 2139492 | + | protein_coding |
| ENSG00000140939.14_5 | NOL3 | 0.66810796 | 1 | 0.492 | 4.387 | chr16 | 67204057 | 67209638 | + | protein_coding |
| ENSG00000070423.18_6 | RNF126 | 0.66827838 | 1 | 0.492 | 4.717 | chr19 | 647526 | 663233 | - | protein_coding |
| ENSG00000203799.13_4 | CCDC162P | 0.5524459 | 1 | 0.492 | 3.049 | chr6 | 109487034 | 109676266 | + | transcribed_unitary_pseudogene |
| ENSG00000105738.11_5 | SIPA1L3 | 0.81658378 | 1 | 0.491 | 8.175 | chr19 | 38397848 | 38699009 | + | protein_coding |
| ENSG00000245146.7_7 | MALINC1 | 0.45042069 | 1 | 0.491 | 1.083 | chr5 | 139450897 | 139488859 | - | lncRNA |
| ENSG00000228532.4_5 | AC005000.1 | 0.38037992 | 1 | 0.491 | 0.088 | chrX | 114953262 | 114953549 | - | processed_pseudogene |
| ENSG00000132334.16_4 | PTPRE | 0.43469024 | 1 | 0.491 | 1.242 | chr10 | 129705325 | 129884119 | + | protein_coding |
| ENSG00000262877.5_7 | AC110285.2 | 0.47374304 | 1 | 0.491 | 1.981 | chr17 | 79361926 | 79364119 | - | lncRNA |
| ENSG00000139405.16_5 | RITA1 | 0.77091923 | 1 | 0.491 | 5.050 | chr12 | 113623331 | 113630173 | + | protein_coding |
| ENSG00000204681.11_5 | GABBR1 | 0.42955367 | 1 | 0.490 | 1.180 | chr6 | 29523406 | 29601753 | - | protein_coding |
| ENSG00000083097.14_5 | DOP1A | 0.70533747 | 1 | 0.490 | 4.214 | chr6 | 83777385 | 83881069 | + | protein_coding |
| ENSG00000240230.6_5 | COX19 | 0.67208374 | 1 | 0.490 | 4.105 | chr7 | 938415 | 1015185 | - | protein_coding |
| ENSG00000140675.13_3 | SLC5A2 | 0.36132451 | 1 | 0.490 | -0.383 | chr16 | 31494323 | 31502181 | + | protein_coding |
| ENSG00000213047.13_5 | DENND1B | 0.69880804 | 1 | 0.490 | 4.369 | chr1 | 197473878 | 197744826 | - | protein_coding |
| ENSG00000111361.13_4 | EIF2B1 | 0.84304836 | 1 | 0.490 | 5.830 | chr12 | 124104953 | 124118313 | - | protein_coding |
| ENSG00000227888.4_3 | FAM66A | 0.35697416 | 1 | 0.490 | -0.978 | chr8 | 12219528 | 12245805 | + | lncRNA |
| ENSG00000257680.1_5 | AC016144.1 | 0.36225997 | 1 | 0.490 | -0.106 | chr12 | 41122613 | 41123699 | + | processed_pseudogene |
| ENSG00000170776.21_5 | AKAP13 | 0.89041665 | 1 | 0.489 | 7.562 | chr15 | 85923802 | 86292589 | + | protein_coding |
| ENSG00000141295.14_6 | SCRN2 | 0.78528719 | 1 | 0.489 | 5.211 | chr17 | 45915058 | 45918655 | - | protein_coding |
| ENSG00000110801.14_5 | PSMD9 | 0.60807785 | 1 | 0.489 | 3.092 | chr12 | 122326638 | 122356203 | + | protein_coding |
| ENSG00000172456.17_4 | FGGY | 0.46971828 | 1 | 0.488 | 1.843 | chr1 | 59762310 | 60276319 | + | protein_coding |
| ENSG00000173531.15_4 | MST1 | 0.53701469 | 1 | 0.488 | 2.360 | chr3 | 49721380 | 49726934 | - | protein_coding |
| ENSG00000251400.1_4 | ALDH7A1P1 | 0.35652384 | 1 | 0.488 | -0.350 | chr5 | 76586093 | 76587625 | + | processed_pseudogene |
| ENSG00000111737.12_5 | RAB35 | 0.85963853 | 1 | 0.488 | 6.026 | chr12 | 120532903 | 120555306 | - | protein_coding |
| ENSG00000177181.15_5 | RIMKLA | 0.59185224 | 1 | 0.488 | 3.202 | chr1 | 42846463 | 42889903 | + | protein_coding |
| ENSG00000158792.16_6 | SPATA2L | 0.55266022 | 1 | 0.488 | 2.908 | chr16 | 89762765 | 89768113 | - | protein_coding |
| ENSG00000118816.10_7 | CCNI | 0.90722231 | 1 | 0.488 | 7.052 | chr4 | 77968308 | 77997142 | - | protein_coding |
| ENSG00000279382.1_5 | AC018665.1 | 0.47840643 | 1 | 0.488 | 1.924 | chr17 | 74086389 | 74087750 | + | TEC |
| ENSG00000100350.14_4 | FOXRED2 | 0.78834055 | 1 | 0.488 | 4.878 | chr22 | 36883237 | 36903148 | - | protein_coding |
| ENSG00000160703.16_4 | NLRX1 | 0.69836063 | 1 | 0.488 | 4.628 | chr11 | 119037277 | 119054725 | + | protein_coding |
| ENSG00000126243.9_3 | LRFN3 | 0.53994769 | 1 | 0.488 | 4.035 | chr19 | 36426260 | 36437526 | + | protein_coding |
| ENSG00000168936.11_5 | TMEM129 | 0.74580693 | 1 | 0.487 | 4.982 | chr4 | 1717679 | 1723085 | - | protein_coding |
| ENSG00000254369.6_6 | HOXA-AS3 | 0.35618782 | 1 | 0.487 | -0.486 | chr7 | 27169596 | 27195547 | + | lncRNA |
| ENSG00000198911.12_5 | SREBF2 | 0.86088073 | 1 | 0.487 | 6.414 | chr22 | 42229083 | 42303311 | + | protein_coding |
| ENSG00000213463.5_5 | SYNJ2BP | 0.84945904 | 1 | 0.487 | 6.136 | chr14 | 70833216 | 70883807 | - | protein_coding |
| ENSG00000111229.16_6 | ARPC3 | 0.88200988 | 1 | 0.487 | 7.422 | chr12 | 110872628 | 110888227 | - | protein_coding |
| ENSG00000146278.11_3 | PNRC1 | 0.82309046 | 1 | 0.487 | 5.593 | chr6 | 89790470 | 89794879 | + | protein_coding |
| ENSG00000084110.11_4 | HAL | 0.35665671 | 1 | 0.487 | -0.710 | chr12 | 96366440 | 96390143 | - | protein_coding |
| ENSG00000260493.1_7 | AC011773.4 | 0.36208597 | 1 | 0.487 | -1.083 | chr8 | 86084312 | 86089297 | - | lncRNA |
| ENSG00000049769.14_6 | PPP1R3F | 0.50495166 | 1 | 0.487 | 2.170 | chrX | 49126256 | 49157940 | + | protein_coding |
| ENSG00000262119.1_7 | AL079343.1 | 0.35781438 | 1 | 0.486 | -0.630 | chr14 | 27061953 | 27063723 | - | lncRNA |
| ENSG00000213977.8_5 | TAX1BP3 | 0.53617769 | 1 | 0.486 | 2.711 | chr17 | 3566189 | 3571973 | - | protein_coding |
| ENSG00000162390.17_2 | ACOT11 | 0.493896 | 1 | 0.486 | 1.964 | chr1 | 55007930 | 55104865 | + | protein_coding |
| ENSG00000136021.19_4 | SCYL2 | 0.87075755 | 1 | 0.486 | 6.630 | chr12 | 100660918 | 100735493 | + | protein_coding |
| ENSG00000173230.15_8 | GOLGB1 | 0.8726863 | 1 | 0.486 | 7.620 | chr3 | 121382046 | 121468614 | - | protein_coding |
| ENSG00000187185.4_7 | AC092118.1 | 0.75780317 | 1 | 0.485 | 4.794 | chr16 | 57832098 | 57850850 | + | lncRNA |
| ENSG00000198363.18_5 | ASPH | 0.93612854 | 1 | 0.485 | 10.638 | chr8 | 62413115 | 62627199 | - | protein_coding |
| ENSG00000240322.2 | RN7SL481P | 0.36156336 | 1 | 0.485 | -0.882 | chr7 | 142995609 | 142995905 | + | misc_RNA |

| | | | | | | | | | | |
|---|---|---|---|---|---|---|---|---|---|---|
| ENSG00000136367.14_5 | ZFHX2 | 0.45391374 | 1 | 0.485 | 1.344 | chr14 | 23990066 | 24025401 | - | protein_coding |
| ENSG00000177432.8_5 | NAP1L5 | 0.42556331 | 1 | 0.485 | 1.028 | chr4 | 89617064 | 89618980 | - | protein_coding |
| ENSG00000160789.20_5 | LMNA | 0.80561334 | 1 | 0.485 | 8.941 | chr1 | 156052364 | 156109880 | + | protein_coding |
| ENSG00000223336.1 | RNU2-6P | 0.37722057 | 1 | 0.485 | 0.240 | chr13 | 46948536 | 46948725 | - | snRNA |
| ENSG00000167880.7_5 | EVPL | 0.6564174 | 1 | 0.485 | 6.011 | chr17 | 74000583 | 74023533 | - | protein_coding |
| ENSG00000198162.12_4 | MAN1A2 | 0.87552687 | 1 | 0.484 | 6.580 | chr1 | 117910071 | 118071494 | + | protein_coding |
| ENSG00000112379.9_3 | ARFGEF3 | 0.74641686 | 1 | 0.484 | 4.491 | chr6 | 138483076 | 138665800 | + | protein_coding |
| ENSG00000105991.9_5 | HOXA1 | 0.50424634 | 1 | 0.484 | 1.837 | chr7 | 27132932 | 27135615 | - | protein_coding |
| ENSG00000260369.2_6 | AC120024.1 | 0.37027589 | 1 | 0.484 | -0.089 | chr17 | 78427535 | 78428529 | - | lncRNA |
| ENSG00000112343.11_3 | TRIM38 | 0.71591188 | 1 | 0.484 | 4.247 | chr6 | 25963030 | 25991459 | + | protein_coding |
| ENSG00000257740.1_7 | AC073896.3 | 0.3805278 | 1 | 0.484 | 0.196 | chr12 | 56702652 | 56703233 | + | lncRNA |
| ENSG00000063854.13_4 | HAGH | 0.79466822 | 1 | 0.484 | 4.827 | chr16 | 1845621 | 1877158 | - | protein_coding |
| ENSG00000260618.1_6 | AC025917.1 | 0.36787822 | 1 | 0.484 | -1.083 | chr15 | 52870039 | 52890906 | + | lncRNA |
| ENSG00000165923.16_5 | AGBL2 | 0.48315734 | 1 | 0.484 | 1.675 | chr11 | 47681143 | 47736941 | - | protein_coding |
| ENSG00000120837.8_7 | NFYB | 0.84688069 | 1 | 0.483 | 5.713 | chr12 | 104510864 | 104532019 | - | protein_coding |
| ENSG00000156639.12_3 | ZFAND3 | 0.74022541 | 1 | 0.483 | 4.324 | chr6 | 37787503 | 38122400 | + | protein_coding |
| ENSG00000108176.15_4 | DNAJC12 | 0.75465231 | 1 | 0.483 | 4.778 | chr10 | 69556427 | 69597946 | - | protein_coding |
| ENSG00000083720.13_4 | OXCT1 | 0.57865236 | 1 | 0.483 | 2.843 | chr5 | 41730167 | 41870527 | - | protein_coding |
| ENSG00000007520.4_4 | TSR3 | 0.62212689 | 1 | 0.483 | 4.448 | chr16 | 1399241 | 1401879 | - | protein_coding |
| ENSG00000266236.1_6 | NARF-IT1 | 0.36064172 | 1 | 0.482 | -0.929 | chr17 | 80439974 | 80441264 | + | lncRNA |
| ENSG00000145198.14_2 | VWA5B2 | 0.46510575 | 1 | 0.482 | 1.418 | chr3 | 183948217 | 183960117 | + | protein_coding |
| ENSG00000160602.14_8 | NEK8 | 0.62535554 | 1 | 0.482 | 3.141 | chr17 | 27052915 | 27070473 | + | protein_coding |
| ENSG00000249129.1_4 | SUDS3P1 | 0.36701187 | 1 | 0.482 | -0.833 | chr5 | 177398317 | 177399282 | + | processed_pseudogene |
| ENSG00000102554.14_4 | KLF5 | 0.9177952 | 1 | 0.482 | 6.917 | chr13 | 73629114 | 73651679 | + | protein_coding |
| ENSG00000196782.12_6 | MAML3 | 0.88202444 | 1 | 0.482 | 6.058 | chr4 | 140637907 | 141075338 | - | protein_coding |
| ENSG00000224078.15_5 | SNHG14 | 0.92774284 | 1 | 0.481 | 7.139 | chr15 | 25223730 | 25665483 | + | lncRNA |
| ENSG00000077454.16_6 | LRCH4 | 0.51568512 | 1 | 0.481 | 2.210 | chr7 | 100171634 | 100183752 | - | protein_coding |
| ENSG00000151458.12_4 | ANKRD50 | 0.88043736 | 1 | 0.481 | 6.091 | chr4 | 125585203 | 125633887 | - | protein_coding |
| ENSG00000202058.1 | RN7SKP80 | 0.38558241 | 1 | 0.481 | 0.700 | chr22 | 42961054 | 42961336 | - | misc_RNA |
| ENSG00000166619.15_8 | BLCAP | 0.82287875 | 1 | 0.481 | 5.094 | chr20 | 36120874 | 36156333 | - | protein_coding |
| ENSG00000149257.14_6 | SERPINH1 | 0.88300057 | 1 | 0.480 | 6.367 | chr11 | 75273101 | 75283828 | + | protein_coding |
| ENSG00000110046.13_4 | ATG2A | 0.6575341 | 1 | 0.480 | 5.454 | chr11 | 64662018 | 64684683 | - | protein_coding |
| ENSG00000167306.20_5 | MYO5B | 0.68200488 | 1 | 0.480 | 3.514 | chr18 | 47349159 | 47721517 | - | protein_coding |
| ENSG00000196411.10_4 | EPHB4 | 0.81340961 | 1 | 0.480 | 6.377 | chr7 | 100400187 | 100425145 | - | protein_coding |
| ENSG00000235920.1_5 | AC073109.1 | 0.46071999 | 1 | 0.480 | 1.244 | chr7 | 11859943 | 11860920 | - | processed_pseudogene |
| ENSG00000163157.15_6 | TMOD4 | 0.36888067 | 1 | 0.480 | -0.453 | chr1 | 151142462 | 151148760 | - | protein_coding |
| ENSG00000042493.16_4 | CAPG | 0.86951596 | 1 | 0.480 | 5.987 | chr2 | 85621876 | 85645555 | - | protein_coding |
| ENSG00000286760.1_1 | AL022069.3 | 0.36467685 | 1 | 0.480 | -0.554 | chr6 | 166756120 | 166764954 | + | lncRNA |
| ENSG00000234545.8_5 | FAM133B | 0.77238888 | 1 | 0.480 | 4.281 | chr7 | 92190072 | 92219707 | - | protein_coding |
| ENSG00000204791.10_5 | SMPD5 | 0.36687659 | 1 | 0.480 | -0.202 | chr8 | 145103980 | 145106423 | + | transcribed_unitary_pseudogene |
| ENSG00000254912.2_6 | AC135983.4 | 0.36390546 | 1 | 0.480 | -0.592 | chr15 | 32786204 | 32792547 | - | unprocessed_pseudogene |
| ENSG00000065361.15_4 | ERBB3 | 0.56189214 | 1 | 0.480 | 2.620 | chr12 | 56470583 | 56497289 | + | protein_coding |
| ENSG00000120756.13_6 | PLS1 | 0.89649156 | 1 | 0.480 | 6.500 | chr3 | 142315235 | 142432506 | + | protein_coding |
| ENSG00000179277.9_2 | MEIS3P1 | 0.5910129 | 1 | 0.480 | 2.991 | chr17 | 15689932 | 15690889 | + | processed_pseudogene |
| ENSG00000251562.8_4 | MALAT1 | 0.91193952 | 1 | 0.480 | 11.022 | chr11 | 65265159 | 65273987 | + | lncRNA |
| ENSG00000258384.1_7 | AC068831.2 | 0.3724468 | 1 | 0.480 | -0.286 | chr15 | 91495469 | 91498455 | - | lncRNA |
| ENSG00000200090.1 | Y_RNA | 0.36487407 | 1 | 0.479 | -0.671 | chr11 | 47748446 | 47748544 | - | misc_RNA |
| ENSG00000166900.17_5 | STX3 | 0.82330585 | 1 | 0.479 | 4.921 | chr11 | 59480929 | 59573355 | + | protein_coding |
| ENSG00000165568.18_5 | AKR1E2 | 0.81228325 | 1 | 0.479 | 4.760 | chr10 | 4828821 | 4890254 | + | protein_coding |
| ENSG00000260751.2_8 | AC008870.3 | 0.37210297 | 1 | 0.479 | -0.138 | chr16 | 23579994 | 23581017 | + | lncRNA |
| ENSG00000160226.16_6 | CFAP410 | 0.47389937 | 1 | 0.479 | 1.355 | chr21 | 45748827 | 45759285 | - | protein_coding |
| ENSG00000248593.3_2 | DSTNP2 | 0.68765251 | 1 | 0.479 | 3.691 | chr12 | 6993846 | 6994950 | + | transcribed_unprocessed_pseudogene |
| ENSG00000103495.14_5 | MAZ | 0.69234872 | 1 | 0.479 | 4.092 | chr16 | 29817427 | 29822485 | + | protein_coding |
| ENSG00000102452.17_5 | NALCN | 0.76572199 | 1 | 0.479 | 4.231 | chr13 | 101706128 | 102068859 | - | protein_coding |
| ENSG00000138378.19_8 | STAT4 | 0.59560642 | 1 | 0.479 | 2.712 | chr2 | 191894302 | 192016322 | - | protein_coding |
| ENSG00000122386.10_2 | ZNF205 | 0.46146359 | 1 | 0.478 | 2.917 | chr16 | 3162561 | 3170518 | + | protein_coding |
| ENSG00000166405.15_6 | RIC3 | 0.59225275 | 1 | 0.478 | 2.855 | chr11 | 8127603 | 8190602 | - | protein_coding |
| ENSG00000176428.6_3 | VPS37D | 0.47188745 | 1 | 0.478 | 1.839 | chr7 | 73082161 | 73086442 | + | protein_coding |

| | | | | | | | | | | |
|---|---|---|---|---|---|---|---|---|---|---|
| ENSG00000087448.11_4 | KLHL42 | 0.86911889 | 1 | 0.478 | 5.828 | chr12 | 27932981 | 27955973 | + | protein_coding |
| ENSG00000167508.12_5 | MVD | 0.57931897 | 1 | 0.478 | 4.578 | chr16 | 88718343 | 88729569 | - | protein_coding |
| ENSG00000102312.22_5 | PORCN | 0.65297966 | 1 | 0.478 | 3.107 | chrX | 48367350 | 48379202 | + | protein_coding |
| ENSG00000137203.13_5 | TFAP2A | 0.69899818 | 1 | 0.477 | 3.546 | chr6 | 10393419 | 10419892 | - | protein_coding |
| ENSG00000123427.17_7 | EEF1AKMT3 | 0.67282108 | 1 | 0.477 | 3.496 | chr12 | 58165275 | 58176324 | + | protein_coding |
| ENSG00000203880.12_4 | PCMTD2 | 0.86438394 | 1 | 0.477 | 5.387 | chr20 | 62887048 | 62919174 | + | protein_coding |
| ENSG00000123064.12_6 | DDX54 | 0.90827722 | 1 | 0.477 | 6.644 | chr12 | 113594979 | 113623284 | - | protein_coding |
| ENSG00000132881.12_5 | CPLANE2 | 0.48564774 | 1 | 0.477 | 1.738 | chr1 | 16558187 | 16563678 | - | protein_coding |
| ENSG00000005020.13_6 | SKAP2 | 0.85355606 | 1 | 0.477 | 5.932 | chr7 | 26706687 | 27034858 | - | protein_coding |
| ENSG00000148730.7_3 | EIF4EBP2 | 0.9138928 | 1 | 0.477 | 6.210 | chr10 | 72163901 | 72188374 | + | protein_coding |
| ENSG00000233098.9_8 | CCDC144NL-AS1 | 0.36737004 | 1 | 0.477 | -0.630 | chr17 | 20771746 | 20905589 | + | lncRNA |
| ENSG00000196465.10_3 | MYL6B | 0.83034576 | 1 | 0.476 | 4.956 | chr12 | 56546040 | 56553431 | + | protein_coding |
| ENSG00000156603.18_6 | MED19 | 0.76124437 | 1 | 0.476 | 4.164 | chr11 | 57471186 | 57479795 | - | protein_coding |
| ENSG00000268438.1_4 | BNIP3P27 | 0.38106916 | 1 | 0.476 | -0.326 | chr19 | 22009315 | 22010081 | + | processed_pseudogene |
| ENSG00000256045.3_3 | MTRNR2L10 | 0.36798349 | 1 | 0.476 | -0.801 | chrX | 55207825 | 55208875 | - | protein_coding |
| ENSG00000277287.1_7 | AL109976.1 | 0.38940916 | 1 | 0.475 | 0.168 | chr20 | 3220351 | 3226028 | + | lncRNA |
| ENSG00000198909.7_6 | MAP3K3 | 0.85315846 | 1 | 0.475 | 5.015 | chr17 | 61699775 | 61773663 | + | protein_coding |
| ENSG00000285589.1_3 | AC010422.8 | 0.38422017 | 1 | 0.475 | 0.112 | chr19 | 12778885 | 12792648 | - | protein_coding |
| ENSG00000204228.4_3 | HSD17B8 | 0.60453133 | 1 | 0.475 | 2.318 | chr6 | 33172432 | 33174608 | + | protein_coding |
| ENSG00000167272.11_4 | POP5 | 0.82678071 | 1 | 0.475 | 4.923 | chr12 | 121016567 | 121019205 | - | protein_coding |
| ENSG00000131797.12_4 | CLUHP3 | 0.71523244 | 1 | 0.475 | 3.638 | chr16 | 31711911 | 31723307 | + | transcribed_unprocessed_pseudogene |
| ENSG00000267280.5_8 | TBX2-AS1 | 0.47598992 | 1 | 0.475 | 1.385 | chr17 | 59470817 | 59488916 | - | lncRNA |
| ENSG00000272668.2_8 | AL590560.3 | 0.46188051 | 1 | 0.474 | 0.926 | chr1 | 159824660 | 159837475 | + | lncRNA |
| ENSG00000066117.15_5 | SMARCD1 | 0.91181044 | 1 | 0.474 | 6.390 | chr12 | 50478983 | 50494490 | + | protein_coding |
| ENSG00000003056.3 | M6PR | 0.93688488 | 1 | 0.474 | 6.591 | chr12 | 9092959 | 9102551 | - | protein_coding |
| ENSG00000273796.1_6 | BX322562.1 | 0.37409295 | 1 | 0.473 | -0.462 | chr21 | 46823724 | 46824284 | - | lncRNA |
| ENSG00000262772.2_5 | LINC01977 | 0.46393317 | 1 | 0.473 | 1.090 | chr17 | 77792882 | 77801503 | + | lncRNA |
| ENSG00000186716.21_5 | BCR | 0.83706175 | 1 | 0.473 | 5.680 | chr22 | 23521891 | 23660224 | + | protein_coding |
| ENSG00000198270.13_7 | TMEM116 | 0.71634679 | 1 | 0.473 | 3.653 | chr12 | 112369086 | 112450989 | - | protein_coding |
| ENSG00000130304.17_6 | SLC27A1 | 0.61903299 | 1 | 0.473 | 2.871 | chr19 | 17579578 | 17616977 | + | protein_coding |
| ENSG00000124587.14_4 | PEX6 | 0.75670978 | 1 | 0.473 | 4.567 | chr6 | 42931603 | 42946919 | - | protein_coding |
| ENSG00000099364.16_6 | FBXL19 | 0.65534141 | 1 | 0.473 | 4.969 | chr16 | 30934376 | 30960104 | + | protein_coding |
| ENSG00000143624.14_4 | INTS3 | 0.91901613 | 1 | 0.472 | 6.602 | chr1 | 153700526 | 153747284 | + | protein_coding |
| ENSG00000123815.12_7 | COQ8B | 0.88817937 | 1 | 0.472 | 5.948 | chr19 | 41197435 | 41224112 | - | protein_coding |
| ENSG00000133983.16_6 | COX16 | 0.54619626 | 1 | 0.472 | 2.100 | chr14 | 70791798 | 70883701 | - | protein_coding |
| ENSG00000232268.6_3 | OR52I1 | 0.37552709 | 1 | 0.472 | -1.086 | chr11 | 4614844 | 4617804 | + | protein_coding |
| ENSG00000182544.8_4 | MFSD5 | 0.85254456 | 1 | 0.472 | 5.274 | chr12 | 53645035 | 53648189 | + | protein_coding |
| ENSG00000103855.18_3 | CD276 | 0.85512064 | 1 | 0.472 | 5.358 | chr15 | 73976307 | 74006855 | + | protein_coding |
| ENSG00000226015.2_2 | CCT8P1 | 0.48278879 | 1 | 0.472 | 0.853 | chr1 | 146674857 | 146676513 | - | processed_pseudogene |
| ENSG00000189051.5_3 | RNF222 | 0.37510651 | 1 | 0.472 | -0.329 | chr17 | 8294022 | 8301144 | - | protein_coding |
| ENSG00000139714.12_4 | MORN3 | 0.41422465 | 1 | 0.472 | 0.168 | chr12 | 122086648 | 122110537 | - | protein_coding |
| ENSG00000144655.15_4 | CSRNP1 | 0.74179028 | 1 | 0.471 | 4.220 | chr3 | 39183346 | 39196053 | - | protein_coding |
| ENSG00000141504.12_3 | SAT2 | 0.86285827 | 1 | 0.471 | 5.058 | chr17 | 7529552 | 7531194 | - | protein_coding |
| ENSG00000181789.14_3 | COPG1 | 0.93562452 | 1 | 0.471 | 7.084 | chr3 | 128968449 | 128996616 | + | protein_coding |
| ENSG00000162836.12_6 | ACP6 | 0.80077619 | 1 | 0.471 | 4.349 | chr1 | 147101453 | 147142646 | - | protein_coding |
| ENSG00000184083.12_5 | FAM120C | 0.86124974 | 1 | 0.470 | 5.144 | chrX | 54094757 | 54209714 | - | protein_coding |
| ENSG00000120725.13_6 | SIL1 | 0.80009222 | 1 | 0.470 | 4.345 | chr5 | 138282413 | 138629246 | - | protein_coding |
| ENSG00000229874.2 | RP11-312O7.2 | 0.37511631 | 1 | 0.470 | -0.593 | chr1 | 206135293 | 206137984 | - | antisense |
| ENSG00000248206.2_7 | AC074194.1 | 0.41349702 | 1 | 0.470 | 0.591 | chr4 | 184908822 | 184917833 | + | lncRNA |
| ENSG00000162585.17_5 | FAAP20 | 0.72816572 | 1 | 0.470 | 3.615 | chr1 | 2115900 | 2144159 | - | protein_coding |
| ENSG00000184900.15_3 | SUMO3 | 0.90911667 | 1 | 0.469 | 5.988 | chr21 | 46225532 | 46238694 | - | protein_coding |
| ENSG00000104885.18_3 | DOT1L | 0.73428989 | 1 | 0.469 | 4.371 | chr19 | 2163932 | 2232577 | + | protein_coding |
| ENSG00000089234.16_5 | BRAP | 0.89064387 | 1 | 0.469 | 5.406 | chr12 | 112079950 | 112123760 | - | protein_coding |
| ENSG00000166598.15_5 | HSP90B1 | 0.95463933 | 1 | 0.469 | 9.222 | chr12 | 104323885 | 104347423 | + | protein_coding |
| ENSG00000181396.13_7 | OGFOD3 | 0.83855046 | 1 | 0.469 | 5.000 | chr17 | 80347086 | 80376513 | - | protein_coding |
| ENSG00000130758.8_4 | MAP3K10 | 0.62631289 | 1 | 0.468 | 4.357 | chr19 | 40697333 | 40721482 | + | protein_coding |
| ENSG00000156510.13_3 | HKDC1 | 0.90784998 | 1 | 0.468 | 5.791 | chr10 | 70980088 | 71027308 | + | protein_coding |

| | | | | | | | | | | |
|---|---|---|---|---|---|---|---|---|---|---|
| ENSG00000234420.7_5 | ZNF37BP | 0.83611127 | 1 | 0.467 | 4.971 | chr10 | 43008958 | 43048270 | - | transcribed_processed_pseudogene |
| ENSG00000136878.13_4 | USP20 | 0.71180007 | 1 | 0.467 | 3.415 | chr9 | 132596977 | 132644117 | + | protein_coding |
| ENSG00000261732.1_6 | AL031708.1 | 0.41717404 | 1 | 0.467 | 0.120 | chr16 | 1682260 | 1736716 | + | protein_coding |
| ENSG00000167674.15_9 | HDGFL2 | 0.8424396 | 1 | 0.467 | 5.394 | chr19 | 4472294 | 4502220 | + | protein_coding |
| ENSG00000134575.13_6 | ACP2 | 0.8290738 | 1 | 0.467 | 4.709 | chr11 | 47260853 | 47270457 | - | polymorphic_pseudogene |
| ENSG00000197321.14_3 | SVIL | 0.9166338 | 1 | 0.467 | 5.939 | chr10 | 29746267 | 30025710 | - | protein_coding |
| ENSG00000139291.14_5 | TMEM19 | 0.91122985 | 1 | 0.466 | 5.708 | chr12 | 72079862 | 72098827 | + | protein_coding |
| ENSG00000160050.14_3 | CCDC28B | 0.49001476 | 1 | 0.466 | 1.107 | chr1 | 32665987 | 32670988 | + | protein_coding |
| ENSG00000100417.12_4 | PMM1 | 0.67266692 | 1 | 0.466 | 3.111 | chr22 | 41972904 | 41985875 | - | protein_coding |
| ENSG00000111725.11_4 | PRKAB1 | 0.82581625 | 1 | 0.466 | 4.746 | chr12 | 120105669 | 120119429 | + | protein_coding |
| ENSG00000127666.9_3 | TICAM1 | 0.52179095 | 1 | 0.466 | 1.808 | chr19 | 4815944 | 4831716 | - | protein_coding |
| ENSG00000189227.6_3 | C15orf61 | 0.63485016 | 1 | 0.466 | 2.283 | chr15 | 67813469 | 67822484 | + | protein_coding |
| ENSG00000111271.14_5 | ACAD10 | 0.85762096 | 1 | 0.466 | 4.910 | chr12 | 112123860 | 112194911 | + | protein_coding |
| ENSG00000136158.12_5 | SPRY2 | 0.76666789 | 1 | 0.466 | 3.849 | chr13 | 80910111 | 80915261 | - | protein_coding |
| ENSG00000272583.1_7 | AL592494.2 | 0.38035269 | 1 | 0.465 | -0.725 | chr1 | 121260164 | 121260627 | - | lncRNA |
| ENSG00000011021.23_5 | CLCN6 | 0.79981691 | 1 | 0.465 | 4.188 | chr1 | 11866153 | 11908136 | + | protein_coding |
| ENSG00000196358.11_3 | NTNG2 | 0.46761979 | 1 | 0.465 | 0.984 | chr9 | 135037063 | 135119913 | + | protein_coding |
| ENSG00000119801.13_3 | YPEL5 | 0.91528054 | 1 | 0.465 | 5.604 | chr2 | 30369807 | 30383399 | + | protein_coding |
| ENSG00000237682.2_5 | AC234775.2 | 0.40505623 | 1 | 0.465 | -1.324 | chrX | 100792472 | 100794263 | - | processed_pseudogene |
| ENSG00000105669.14_5 | COPE | 0.84475728 | 1 | 0.464 | 5.148 | chr19 | 19010323 | 19030196 | - | protein_coding |
| ENSG00000123240.17_6 | OPTN | 0.91059503 | 1 | 0.464 | 5.458 | chr10 | 13141449 | 13180308 | + | protein_coding |
| ENSG00000233429.9_5 | HOTAIRM1 | 0.5690553 | 1 | 0.464 | 2.112 | chr7 | 27135266 | 27139884 | + | lncRNA |
| ENSG00000180233.11_4 | ZNRF2 | 0.66901319 | 1 | 0.464 | 3.490 | chr7 | 30324213 | 30407305 | + | protein_coding |
| ENSG00000107262.24_10 | BAG1 | 0.94047164 | 1 | 0.463 | 6.246 | chr9 | 33247818 | 33264718 | - | protein_coding |
| ENSG00000118515.11_5 | SGK1 | 0.8257562 | 1 | 0.463 | 4.210 | chr6 | 134490384 | 134639250 | - | protein_coding |
| ENSG00000118855.21_7 | MFSD1 | 0.90703297 | 1 | 0.463 | 5.394 | chr3 | 158449987 | 158547508 | + | protein_coding |
| ENSG00000185842.15_5 | DNAH14 | 0.88161749 | 1 | 0.463 | 5.134 | chr1 | 225083964 | 225586994 | + | protein_coding |
| ENSG00000155363.18_2 | MOV10 | 0.91101616 | 1 | 0.463 | 5.485 | chr1 | 113215763 | 113243368 | + | protein_coding |
| ENSG00000184164.14_6 | CRELD2 | 0.81250846 | 1 | 0.463 | 4.303 | chr22 | 50311815 | 50321188 | + | protein_coding |
| ENSG00000254827.6_4 | SLC22A18AS | 0.43450184 | 1 | 0.462 | 0.519 | chr11 | 2908574 | 2924805 | - | protein_coding |
| ENSG00000232712.6_8 | KIZ-AS1 | 0.38603034 | 1 | 0.462 | -0.944 | chr20 | 21134664 | 21198927 | - | lncRNA |
| ENSG00000261270.1_7 | AC012181.2 | 0.38723199 | 1 | 0.462 | -0.769 | chr16 | 56974190 | 56975254 | + | lncRNA |
| ENSG00000279283.1_6 | AC131009.4 | 0.53714013 | 1 | 0.462 | 1.604 | chr12 | 132381045 | 132382784 | + | TEC |
| ENSG00000052841.15_4 | TTC17 | 0.94157819 | 1 | 0.461 | 6.423 | chr11 | 43380470 | 43516483 | + | protein_coding |
| ENSG00000196696.13_9 | AC009022.1 | 0.53967326 | 1 | 0.461 | 1.645 | chr16 | 70010200 | 70099851 | - | lncRNA |
| ENSG00000089818.18_5 | NECAP1 | 0.85800311 | 1 | 0.461 | 4.770 | chr12 | 8229535 | 8250477 | + | protein_coding |
| ENSG00000243646.10_4 | IL10RB | 0.3860545 | 1 | 0.461 | -0.537 | chr21 | 34638672 | 34682492 | + | protein_coding |
| ENSG00000120519.14_3 | SLC10A7 | 0.58597582 | 1 | 0.461 | 2.415 | chr4 | 147175127 | 147443116 | - | protein_coding |
| ENSG00000167034.10_4 | NKX3-1 | 0.59627739 | 1 | 0.461 | 2.258 | chr8 | 23536210 | 23540451 | - | protein_coding |
| ENSG00000266258.1_5 | LINC01909 | 0.47288957 | 1 | 0.460 | 0.797 | chr18 | 68002675 | 68019695 | + | lncRNA |
| ENSG00000219410.4 | RP4-761J14.8 | 0.42195671 | 1 | 0.460 | 0.046 | chr12 | 6772426 | 6781235 | + | antisense |
| ENSG00000089693.10_4 | MLF2 | 0.94877069 | 1 | 0.460 | 7.326 | chr12 | 6857162 | 6876641 | - | protein_coding |
| ENSG00000197085.11_5 | NPSR1-AS1 | 0.40155703 | 1 | 0.460 | -1.143 | chr7 | 34386124 | 34911194 | - | lncRNA |
| ENSG00000234779.1_6 | BNC2-AS1 | 0.40348332 | 1 | 0.460 | -0.103 | chr9 | 16726812 | 16727522 | + | lncRNA |
| ENSG00000197982.14_5 | C1orf122 | 0.74354895 | 1 | 0.460 | 4.008 | chr1 | 38272651 | 38275126 | + | protein_coding |
| ENSG00000130589.16_3 | HELZ2 | 0.53607939 | 1 | 0.460 | 4.316 | chr20 | 62189439 | 62205592 | - | protein_coding |
| ENSG00000179889.19_7 | PDXDC1 | 0.96255833 | 1 | 0.460 | 6.789 | chr16 | 15068448 | 15233196 | + | protein_coding |
| ENSG00000107201.10_4 | DDX58 | 0.7796549 | 1 | 0.459 | 3.976 | chr9 | 32455300 | 32526206 | - | protein_coding |
| ENSG00000265298.1_5 | AC132812.1 | 0.49209294 | 1 | 0.459 | 0.917 | chr17 | 62746538 | 62747429 | + | unprocessed_pseudogene |
| ENSG00000056736.10_5 | IL17RB | 0.6354972 | 1 | 0.459 | 2.628 | chr3 | 53880595 | 53899821 | + | protein_coding |
| ENSG00000198858.10_5 | R3HDM4 | 0.79425946 | 1 | 0.458 | 4.012 | chr19 | 896503 | 913245 | - | protein_coding |
| ENSG00000248015.7_8 | AC005329.1 | 0.388145 | 1 | 0.458 | -0.728 | chr19 | 1392169 | 1396466 | - | lncRNA |
| ENSG00000169241.19_4 | SLC50A1 | 0.86645596 | 1 | 0.458 | 4.692 | chr1 | 155107820 | 155111333 | + | protein_coding |
| ENSG00000141258.13_4 | SGSM2 | 0.92018959 | 1 | 0.458 | 6.310 | chr17 | 2240792 | 2284352 | + | protein_coding |
| ENSG00000267986.1_6 | AC130469.1 | 0.39073389 | 1 | 0.458 | -0.645 | chr19 | 8646044 | 8647599 | + | lncRNA |
| ENSG00000226976.3_2 | COX6A1P2 | 0.55647349 | 1 | 0.457 | 2.267 | chr6 | 37012636 | 37012965 | + | processed_pseudogene |
| ENSG00000227827.3_7 | AC138969.3 | 0.45170137 | 1 | 0.457 | 3.762 | chr16 | 16450081 | 16471364 | + | unprocessed_pseudogene |

| | | | | | | | | | | |
|---|---|---|---|---|---|---|---|---|---|---|
| ENSG00000213366.13_4 | GSTM2 | 0.42690202 | 1 | 0.457 | 0.222 | chr1 | 110210644 | 110252173 | + | protein_coding |
| ENSG00000120805.14_5 | ARL1 | 0.93687047 | 1 | 0.457 | 6.376 | chr12 | 101786894 | 101801550 | - | protein_coding |
| ENSG00000143126.8_3 | CELSR2 | 0.90652408 | 1 | 0.457 | 5.272 | chr1 | 109792161 | 109818373 | + | protein_coding |
| ENSG00000054793.14_6 | ATP9A | 0.95836962 | 1 | 0.456 | 6.310 | chr20 | 50213053 | 50384929 | - | protein_coding |
| ENSG00000125650.4_3 | PSPN | 0.52001988 | 1 | 0.456 | 1.884 | chr19 | 6375159 | 6379069 | - | protein_coding |
| ENSG00000175265.17_4 | GOLGA8A | 0.77906263 | 1 | 0.456 | 3.690 | chr15 | 34671269 | 34729667 | - | protein_coding |
| ENSG00000242612.7_4 | DECR2 | 0.74323975 | 1 | 0.456 | 3.745 | chr16 | 451858 | 462487 | + | protein_coding |
| ENSG00000269313.5_3 | MAGIX | 0.45364754 | 1 | 0.455 | 0.785 | chrX | 49019325 | 49024820 | + | protein_coding |
| ENSG00000272031.3_5 | ANKRD34A | 0.41846637 | 1 | 0.455 | 0.162 | chr1 | 145470515 | 145475647 | + | protein_coding |
| ENSG00000107816.17_3 | LZTS2 | 0.86942852 | 1 | 0.455 | 5.828 | chr10 | 102756375 | 102767593 | + | protein_coding |
| ENSG00000188277.10_3 | C15orf62 | 0.470839 | 1 | 0.455 | 0.822 | chr15 | 41062178 | 41064647 | + | protein_coding |
| ENSG00000063587.14_4 | ZNF275 | 0.78151925 | 1 | 0.455 | 3.621 | chrX | 152599605 | 152625568 | + | protein_coding |
| ENSG00000094916.16_6 | CBX5 | 0.98286441 | 1 | 0.455 | 8.769 | chr12 | 54624726 | 54673917 | - | protein_coding |
| ENSG00000196123.13_4 | KIAA0895L | 0.84113915 | 1 | 0.455 | 4.085 | chr16 | 67209502 | 67217943 | - | protein_coding |
| ENSG00000124201.15_6 | ZNFX1 | 0.96147573 | 1 | 0.455 | 6.413 | chr20 | 47854483 | 47894963 | - | protein_coding |
| ENSG00000168461.13_4 | RAB31 | 0.93457037 | 1 | 0.455 | 5.431 | chr18 | 9708272 | 9862548 | + | protein_coding |
| ENSG00000172831.13_6 | CES2 | 0.88823895 | 1 | 0.454 | 5.191 | chr16 | 66968347 | 66978999 | + | protein_coding |
| ENSG00000164576.12_3 | SAP30L | 0.89059 | 1 | 0.454 | 4.924 | chr5 | 153825557 | 153840613 | + | protein_coding |
| ENSG00000166407.14_4 | LMO1 | 0.39479619 | 1 | 0.454 | -0.372 | chr11 | 8245856 | 8290263 | - | protein_coding |
| ENSG00000129535.12_4 | NRL | 0.40851321 | 1 | 0.454 | 0.073 | chr14 | 24549316 | 24584223 | - | protein_coding |
| ENSG00000139218.18_7 | SCAF11 | 0.97240395 | 1 | 0.454 | 7.777 | chr12 | 46312914 | 46385903 | - | protein_coding |
| ENSG00000277639.2_7 | AC007906.2 | 0.41995986 | 1 | 0.454 | 0.379 | chr16 | 53069602 | 53086785 | - | protein_coding |
| ENSG00000176410.8_3 | DNAJC30 | 0.6362479 | 1 | 0.454 | 2.562 | chr7 | 73095248 | 73097783 | - | protein_coding |
| ENSG00000269938.1_7 | AC068790.2 | 0.41448724 | 1 | 0.454 | 0.270 | chr12 | 124452570 | 124453126 | - | lncRNA |
| ENSG00000075240.17_4 | GRAMD4 | 0.85250862 | 1 | 0.454 | 4.498 | chr22 | 46971909 | 47075687 | + | protein_coding |
| ENSG00000165879.9_3 | FRAT1 | 0.56360039 | 1 | 0.453 | 1.671 | chr10 | 99079028 | 99081672 | + | protein_coding |
| ENSG00000102265.12_3 | TIMP1 | 0.94324126 | 1 | 0.452 | 6.068 | chrX | 47441807 | 47446188 | + | protein_coding |
| ENSG00000110931.19_7 | CAMKK2 | 0.88469771 | 1 | 0.452 | 5.258 | chr12 | 121675478 | 121736111 | - | protein_coding |
| ENSG00000010219.13_5 | DYRK4 | 0.8366802 | 1 | 0.452 | 4.022 | chr12 | 4671370 | 4724468 | + | protein_coding |
| ENSG00000179058.7_6 | C9orf50 | 0.51745894 | 1 | 0.452 | 1.214 | chr9 | 132374504 | 132383055 | - | protein_coding |
| ENSG00000099622.14_7 | CIRBP | 0.95135113 | 1 | 0.452 | 5.989 | chr19 | 1259383 | 1274879 | + | protein_coding |
| ENSG00000244176.1_5 | AP003733.1 | 0.4095714 | 1 | 0.452 | -0.183 | chr11 | 61779223 | 61780074 | - | processed_pseudogene |
| ENSG00000124496.12_4 | TRERF1 | 0.85816378 | 1 | 0.452 | 4.401 | chr6 | 42192669 | 42419789 | - | protein_coding |
| ENSG00000187954.12_5 | CYHR1 | 0.9100706 | 1 | 0.452 | 5.388 | chr8 | 145674965 | 145691060 | - | protein_coding |
| ENSG00000160352.11 | ZNF714 | 0.85243399 | 1 | 0.451 | 4.532 | chr19 | 21264965 | 21308073 | + | protein_coding |
| ENSG00000159884.12_5 | CCDC107 | 0.4199379 | 1 | 0.451 | -0.191 | chr9 | 35658287 | 35661508 | + | protein_coding |
| ENSG00000175155.10_5 | YPEL2 | 0.71516671 | 1 | 0.451 | 3.608 | chr17 | 57409016 | 57479090 | + | protein_coding |
| ENSG00000078814.16_5 | MYH7B | 0.55886407 | 1 | 0.451 | 1.609 | chr20 | 33543613 | 33590240 | + | protein_coding |
| ENSG00000111331.13_5 | OAS3 | 0.96839728 | 1 | 0.450 | 6.684 | chr12 | 113376249 | 113411056 | + | protein_coding |
| ENSG00000103196.12_5 | CRISPLD2 | 0.54858754 | 1 | 0.450 | 1.506 | chr16 | 84853591 | 84954374 | + | protein_coding |
| ENSG00000156232.7_3 | WHAMM | 0.8339125 | 1 | 0.450 | 4.167 | chr15 | 83478380 | 83504860 | + | protein_coding |
| ENSG00000262001.1_6 | DLGAP1-AS2 | 0.71774287 | 1 | 0.450 | 3.290 | chr18 | 3602998 | 3608334 | + | lncRNA |
| ENSG00000120798.17_6 | NR2C1 | 0.92933288 | 1 | 0.450 | 5.346 | chr12 | 95414005 | 95467404 | - | protein_coding |
| ENSG00000272195.1_6 | AL356512.1 | 0.40169981 | 1 | 0.450 | -1.103 | chr1 | 245132652 | 245134390 | - | lncRNA |
| ENSG00000279491.1_6 | AP003733.3 | 0.4142464 | 1 | 0.450 | -0.033 | chr11 | 61735266 | 61736962 | + | TEC |
| ENSG00000144455.14_5 | SUMF1 | 0.8928614 | 1 | 0.449 | 4.742 | chr3 | 3742498 | 4508957 | - | protein_coding |
| ENSG00000136848.17_4 | DAB2IP | 0.86300258 | 1 | 0.449 | 5.155 | chr9 | 124329336 | 124547809 | + | protein_coding |
| ENSG00000146476.11_3 | ARMT1 | 0.93878986 | 1 | 0.448 | 5.277 | chr6 | 151773393 | 151791236 | + | protein_coding |
| ENSG00000008130.15_4 | NADK | 0.90179259 | 1 | 0.448 | 5.352 | chr1 | 1682671 | 1711896 | - | protein_coding |
| ENSG00000086506.3_3 | HBQ1 | 0.46367343 | 1 | 0.448 | 0.489 | chr16 | 230458 | 231178 | + | protein_coding |
| ENSG00000250280.2_7 | AC026124.1 | 0.45023909 | 1 | 0.448 | 0.197 | chr12 | 65675437 | 65680508 | + | lncRNA |
| ENSG00000180818.5_4 | HOXC10 | 0.44098785 | 1 | 0.448 | 0.180 | chr12 | 54378849 | 54384063 | + | protein_coding |
| ENSG00000132109.10_4 | TRIM21 | 0.8501251 | 1 | 0.448 | 4.041 | chr11 | 4406127 | 4414932 | - | protein_coding |
| ENSG00000100711.13_2 | ZFYVE21 | 0.83472189 | 1 | 0.448 | 3.944 | chr14 | 104182067 | 104200005 | + | protein_coding |
| ENSG00000138061.12_7 | CYP1B1 | 0.94518519 | 1 | 0.447 | 5.746 | chr2 | 38294116 | 38337044 | - | protein_coding |
| ENSG00000078061.13_3 | ARAF | 0.91161946 | 1 | 0.447 | 5.469 | chrX | 47420499 | 47431319 | + | protein_coding |
| ENSG00000174749.6_5 | FAM241A | 0.7074938 | 1 | 0.447 | 2.933 | chr4 | 113066610 | 113116412 | + | protein_coding |
| ENSG00000260604.2_6 | AL590004.3 | 0.51633556 | 1 | 0.447 | 1.249 | chr6 | 3905154 | 3912213 | - | lncRNA |
| ENSG00000274349.4_3 | ZNF658 | 0.68181056 | 1 | 0.447 | 3.348 | chr9 | 40760700 | 40836415 | - | protein_coding |
| ENSG00000068001.14_7 | HYAL2 | 0.78943491 | 1 | 0.447 | 4.258 | chr3 | 50355221 | 50360213 | - | protein_coding |

| | | | | | | | | | | |
|---|---|---|---|---|---|---|---|---|---|---|
| ENSG00000182272.12_4 | B4GALNT4 | 0.51616481 | 1 | 0.447 | 4.241 | chr11 | 369499 | 382117 | + | protein_coding |
| ENSG00000250596.2_7 | AC096751.1 | 0.44640065 | 1 | 0.447 | 0.158 | chr4 | 175444852 | 175461495 | - | transcribed_processed_pseudogene |
| ENSG00000134996.12_3 | OSTF1 | 0.76626858 | 1 | 0.446 | 3.670 | chr9 | 77703430 | 77762181 | + | protein_coding |
| ENSG00000286216.1_3 | AC104335.2 | 0.40446062 | 1 | 0.446 | -0.430 | chr1 | 247501188 | 247511378 | - | lncRNA |
| ENSG00000130813.18_7 | SHFL | 0.68068067 | 1 | 0.446 | 2.546 | chr19 | 10196798 | 10203928 | + | protein_coding |
| ENSG00000168890.14_6 | TMEM150A | 0.70971822 | 1 | 0.446 | 3.149 | chr2 | 85825670 | 85830319 | - | protein_coding |
| ENSG00000149948.13_7 | HMGA2 | 0.91191562 | 1 | 0.446 | 5.059 | chr12 | 66217911 | 66360075 | + | protein_coding |
| ENSG00000111731.12_5 | C2CD5 | 0.93550807 | 1 | 0.446 | 5.348 | chr12 | 22601517 | 22697480 | - | protein_coding |
| ENSG00000122741.16_5 | DCAF10 | 0.95195235 | 1 | 0.446 | 5.632 | chr9 | 37800551 | 37867663 | + | protein_coding |
| ENSG00000251602.7_7 | AL928654.1 | 0.61667549 | 1 | 0.446 | 1.988 | chr14 | 105883221 | 105886076 | - | lncRNA |
| ENSG00000188483.8_3 | IER5L | 0.48844891 | 1 | 0.445 | 5.535 | chr9 | 131937831 | 131940540 | - | protein_coding |
| ENSG00000085998.14_4 | POMGNT1 | 0.96475355 | 1 | 0.445 | 5.982 | chr1 | 46654354 | 46685977 | - | protein_coding |
| ENSG00000213609.3_4 | RPL7AP50 | 0.40377085 | 1 | 0.445 | -0.614 | chr10 | 65662211 | 65663005 | - | processed_pseudogene |
| ENSG00000176463.14_4 | SLCO3A1 | 0.91051874 | 1 | 0.445 | 4.682 | chr15 | 92396938 | 92715665 | + | protein_coding |
| ENSG00000152404.15_4 | CWF19L2 | 0.80478564 | 1 | 0.445 | 3.703 | chr11 | 107197071 | 107328570 | - | protein_coding |
| ENSG00000059758.8_7 | CDK17 | 0.9389631 | 1 | 0.445 | 5.584 | chr12 | 96672039 | 96794258 | - | protein_coding |
| ENSG00000198610.11_5 | AKR1C4 | 0.45018712 | 1 | 0.444 | 0.608 | chr10 | 5237425 | 5260912 | + | protein_coding |
| ENSG00000139651.11_3 | ZNF740 | 0.95145063 | 1 | 0.444 | 5.416 | chr12 | 53574488 | 53588926 | + | protein_coding |
| ENSG00000232860.8_6 | SMG7-AS1 | 0.41892241 | 1 | 0.444 | 0.017 | chr1 | 183430009 | 183441400 | - | lncRNA |
| ENSG00000223901.2_6 | AP001469.1 | 0.44000384 | 1 | 0.444 | 0.215 | chr21 | 47640183 | 47645278 | + | lncRNA |
| ENSG00000160993.4_4 | ALKBH4 | 0.72383076 | 1 | 0.444 | 3.111 | chr7 | 102096685 | 102105310 | - | protein_coding |
| ENSG00000155666.11_4 | KDM8 | 0.64227957 | 1 | 0.444 | 2.101 | chr16 | 27214816 | 27233089 | + | protein_coding |
| ENSG00000004399.12_6 | PLXND1 | 0.92830537 | 1 | 0.443 | 6.937 | chr3 | 129274018 | 129325661 | - | protein_coding |
| ENSG00000134287.10_5 | ARF3 | 0.97772976 | 1 | 0.443 | 6.807 | chr12 | 49329506 | 49351270 | - | protein_coding |
| ENSG00000268205.1_8 | AC005261.2 | 0.72934055 | 1 | 0.443 | 4.378 | chr19 | 57815673 | 57819930 | + | lncRNA |
| ENSG00000268812.3_6 | AC004264.1 | 0.50313094 | 1 | 0.443 | 0.828 | chr22 | 30642194 | 30642987 | + | lncRNA |
| ENSG00000104142.11_3 | VPS18 | 0.92486529 | 1 | 0.443 | 5.398 | chr15 | 41186648 | 41196173 | + | protein_coding |
| ENSG00000179195.16_4 | ZNF664 | 0.98031195 | 1 | 0.443 | 7.548 | chr12 | 124456392 | 124499986 | + | protein_coding |
| ENSG00000126583.11_3 | PRKCG | 0.49419614 | 1 | 0.443 | 0.874 | chr19 | 54382444 | 54410906 | + | protein_coding |
| ENSG00000179085.7_3 | DPM3 | 0.78426224 | 1 | 0.443 | 3.507 | chr1 | 155112367 | 155113071 | - | protein_coding |
| ENSG00000234264.1_6 | DEPDC1-AS1 | 0.40575492 | 1 | 0.442 | -0.816 | chr1 | 68962359 | 69004310 | + | lncRNA |
| ENSG00000166275.16_4 | BORCS7 | 0.85117994 | 1 | 0.442 | 4.417 | chr10 | 104614016 | 104624718 | + | protein_coding |
| ENSG00000276698.1_6 | AL136295.6 | 0.41845991 | 1 | 0.442 | -0.230 | chr14 | 24667639 | 24668296 | + | lncRNA |
| ENSG00000178104.19_5 | PDE4DIP | 0.82597117 | 1 | 0.442 | 3.673 | chr1 | 144836157 | 145076186 | - | protein_coding |
| ENSG00000227500.10_4 | SCAMP4 | 0.92622759 | 1 | 0.442 | 5.287 | chr19 | 1905371 | 1926012 | + | protein_coding |
| ENSG00000100802.15_5 | C14orf93 | 0.69838568 | 1 | 0.442 | 3.207 | chr14 | 23455103 | 23479375 | - | protein_coding |
| ENSG00000105404.11_5 | RABAC1 | 0.76025335 | 1 | 0.441 | 3.452 | chr19 | 42460833 | 42463473 | - | protein_coding |
| ENSG00000091656.19_8 | ZFHX4 | 0.75242192 | 1 | 0.441 | 2.997 | chr8 | 77593474 | 77779517 | + | protein_coding |
| ENSG00000196335.13_6 | STK31 | 0.62079095 | 1 | 0.441 | 1.940 | chr7 | 23749822 | 23872132 | + | protein_coding |
| ENSG00000176597.12_4 | B3GNT5 | 0.73047863 | 1 | 0.441 | 2.954 | chr3 | 182971041 | 183016292 | + | protein_coding |
| ENSG00000122477.12_3 | LRRC39 | 0.4090676 | 1 | 0.441 | -0.698 | chr1 | 100614005 | 100643829 | - | protein_coding |
| ENSG00000139263.11_7 | LRIG3 | 0.89495656 | 1 | 0.441 | 4.454 | chr12 | 59265931 | 59314303 | - | protein_coding |
| ENSG00000106683.15_3 | LIMK1 | 0.96054532 | 1 | 0.441 | 6.253 | chr7 | 73497263 | 73536855 | + | protein_coding |
| ENSG00000171777.16_7 | RASGRP4 | 0.44869379 | 1 | 0.441 | -0.040 | chr19 | 38899691 | 38916945 | - | protein_coding |
| ENSG00000257315.2_4 | ZBED6 | 0.47372882 | 1 | 0.440 | 0.452 | chr1 | 203764842 | 203824127 | + | protein_coding |
| ENSG00000103266.11_3 | STUB1 | 0.83431313 | 1 | 0.440 | 4.294 | chr16 | 730224 | 732870 | + | protein_coding |
| ENSG00000159792.10_3 | PSKH1 | 0.86157777 | 1 | 0.440 | 4.207 | chr16 | 67927157 | 67963579 | + | protein_coding |
| ENSG00000103168.16_8 | TAF1C | 0.89910916 | 1 | 0.440 | 5.081 | chr16 | 84211453 | 84220676 | - | protein_coding |
| ENSG00000260563.3_6 | AC132872.1 | 0.44576171 | 1 | 0.440 | 0.080 | chr17 | 80251592 | 80252786 | + | lncRNA |
| ENSG00000229512.1_8 | AC068580.1 | 0.46088628 | 1 | 0.440 | 0.414 | chr11 | 1784239 | 1784979 | - | lncRNA |
| ENSG00000131408.15_9 | NR1H2 | 0.90650143 | 1 | 0.440 | 6.270 | chr19 | 50832910 | 50886645 | + | protein_coding |
| ENSG00000073711.11_5 | PPP2R3A | 0.80715733 | 1 | 0.440 | 3.733 | chr3 | 135684570 | 135866736 | + | protein_coding |
| ENSG00000060138.13_7 | YBX3 | 0.98200605 | 1 | 0.440 | 6.840 | chr12 | 10851688 | 10875922 | - | protein_coding |
| ENSG00000170088.14_7 | TMEM192 | 0.91091953 | 1 | 0.440 | 4.566 | chr4 | 165991760 | 166129701 | - | protein_coding |
| ENSG00000245293.3_8 | AC096564.1 | 0.41645507 | 1 | 0.439 | -1.010 | chr4 | 108784629 | 108910835 | - | lncRNA |
| ENSG00000285881.1_3 | AC105206.3 | 0.41261348 | 1 | 0.439 | -1.105 | chr8 | 22045882 | 22050091 | - | lncRNA |
| ENSG00000170049.9_4 | KCNAB3 | 0.59756082 | 1 | 0.439 | 1.889 | chr17 | 7825177 | 7833121 | - | protein_coding |
| ENSG00000207445.1 | SNORD15B | 0.6708151 | 1 | 0.439 | 2.742 | chr11 | 75115465 | 75115610 | + | snoRNA |
| ENSG00000100271.17_4 | TTLL1 | 0.57421186 | 1 | 0.439 | 1.424 | chr22 | 43435522 | 43485425 | - | protein_coding |

| | | | | | | | | | | |
|---|---|---|---|---|---|---|---|---|---|---|
| ENSG00000258858.1_7 | AL583722.4 | 0.54963648 | 1 | 0.438 | 1.341 | chr14 | 105156428 | 105157621 | - | lncRNA |
| ENSG00000173581.8_6 | CCDC106 | 0.86518761 | 1 | 0.438 | 4.397 | chr19 | 56152428 | 56164527 | + | protein_coding |
| ENSG00000177697.19_6 | CD151 | 0.97288424 | 1 | 0.438 | 7.844 | chr11 | 832887 | 839831 | + | protein_coding |
| ENSG00000130731.16_7 | METTL26 | 0.87314903 | 1 | 0.438 | 4.744 | chr16 | 684427 | 686366 | - | protein_coding |
| ENSG00000170482.17_8 | SLC23A1 | 0.61536147 | 1 | 0.438 | 2.047 | chr5 | 138702885 | 138720242 | - | protein_coding |
| ENSG00000125734.15_8 | GPR108 | 0.93858094 | 1 | 0.438 | 5.256 | chr19 | 6729925 | 6737614 | - | protein_coding |
| ENSG00000186591.13_6 | UBE2H | 0.96849088 | 1 | 0.438 | 6.223 | chr7 | 129470572 | 129592800 | - | protein_coding |
| ENSG00000281692.2_6 | PACRG-AS1 | 0.4116885 | 1 | 0.438 | -0.817 | chr6 | 163727939 | 163745562 | - | lncRNA |
| ENSG00000182919.14_2 | C11orf54 | 0.80170251 | 1 | 0.438 | 3.763 | chr11 | 93474757 | 93497915 | + | protein_coding |
| ENSG00000141040.15_5 | ZNF287 | 0.86365896 | 1 | 0.438 | 4.172 | chr17 | 16450268 | 16472518 | - | protein_coding |
| ENSG00000114670.14_3 | NEK11 | 0.78241192 | 1 | 0.438 | 3.209 | chr3 | 130745694 | 131069309 | + | protein_coding |
| ENSG00000198750.7 | GATSL2 | 0.46753649 | 1 | 0.437 | 0.624 | chr7 | 74807499 | 74867509 | - | protein_coding |
| ENSG00000175203.15_5 | DCTN2 | 0.9795722 | 1 | 0.437 | 6.583 | chr12 | 57923885 | 57941114 | - | protein_coding |
| ENSG00000174292.12_4 | TNK1 | 0.65310665 | 1 | 0.437 | 2.219 | chr17 | 7283853 | 7293093 | + | protein_coding |
| ENSG00000269961.1_7 | AC010359.2 | 0.41289436 | 1 | 0.437 | -0.785 | chr5 | 65220457 | 65220963 | - | lncRNA |
| ENSG00000121931.16_5 | LRIF1 | 0.91857841 | 1 | 0.437 | 4.992 | chr1 | 111489812 | 111506587 | - | protein_coding |
| ENSG00000173065.13_6 | FAM222B | 0.8520623 | 1 | 0.436 | 4.272 | chr17 | 27082996 | 27182250 | - | protein_coding |
| ENSG00000104154.7_4 | SLC30A4 | 0.77141516 | 1 | 0.436 | 3.528 | chr15 | 45771804 | 45814953 | - | protein_coding |
| ENSG00000128849.10_2 | CGNL1 | 0.76839592 | 1 | 0.436 | 3.134 | chr15 | 57668165 | 57842925 | + | protein_coding |
| ENSG00000256029.6_10 | AL590560.2 | 0.42438371 | 1 | 0.436 | -0.441 | chr1 | 159804264 | 159842843 | - | protein_coding |
| ENSG00000108039.18_4 | XPNPEP1 | 0.96655991 | 1 | 0.436 | 5.705 | chr10 | 111624524 | 111683311 | - | protein_coding |
| ENSG00000137310.12_3 | TCF19 | 0.5589943 | 1 | 0.436 | 1.451 | chr6 | 31126324 | 31134936 | + | protein_coding |
| ENSG00000162194.12_7 | LBHD1 | 0.6250833 | 1 | 0.436 | 1.814 | chr11 | 62430289 | 62439727 | - | protein_coding |
| ENSG00000257511.1_6 | AC084824.2 | 0.52053732 | 1 | 0.436 | 0.613 | chr12 | 32879317 | 32880321 | - | processed_pseudogene |
| ENSG00000092820.18_5 | EZR | 0.99238541 | 1 | 0.436 | 8.683 | chr6 | 159186773 | 159240400 | - | protein_coding |
| ENSG00000125844.16_10 | RRBP1 | 0.91621907 | 1 | 0.436 | 7.948 | chr20 | 17594323 | 17662940 | - | protein_coding |
| ENSG00000166106.3_2 | ADAMTS15 | 0.76355171 | 1 | 0.436 | 3.577 | chr11 | 130318869 | 130346536 | + | protein_coding |
| ENSG00000273026.1_6 | AL358472.3 | 0.44242247 | 1 | 0.435 | 0.012 | chr1 | 153938992 | 153939406 | + | lncRNA |
| ENSG00000089220.5_3 | PEBP1 | 0.98916145 | 1 | 0.435 | 7.629 | chr12 | 118573929 | 118583389 | + | protein_coding |
| ENSG00000237877.7_7 | LINC01473 | 0.41421545 | 1 | 0.435 | -0.912 | chr2 | 186897611 | 187287159 | - | lncRNA |
| ENSG00000130529.16_5 | TRPM4 | 0.77479458 | 1 | 0.435 | 4.268 | chr19 | 49660998 | 49715093 | + | protein_coding |
| ENSG00000181523.13_6 | SGSH | 0.86469851 | 1 | 0.435 | 4.399 | chr17 | 78180515 | 78194722 | - | protein_coding |
| ENSG00000238045.9_8 | AC009133.1 | 0.56981912 | 1 | 0.435 | 2.191 | chr16 | 29819957 | 29832573 | - | lncRNA |
| ENSG00000136574.17_2 | GATA4 | 0.57469494 | 1 | 0.435 | 1.368 | chr8 | 11534468 | 11617511 | + | protein_coding |
| ENSG00000204767.4_6 | INSYN2B | 0.44286403 | 1 | 0.434 | 0.087 | chr5 | 169288307 | 169407499 | - | protein_coding |
| ENSG00000164114.19_7 | MAP9 | 0.9311546 | 1 | 0.434 | 5.340 | chr4 | 156263810 | 156298122 | - | protein_coding |
| ENSG00000116793.16_6 | PHTF1 | 0.93808605 | 1 | 0.434 | 4.828 | chr1 | 114239453 | 114302111 | - | protein_coding |
| ENSG00000259868.3_6 | AL163952.1 | 0.41733231 | 1 | 0.434 | -0.818 | chr14 | 56363161 | 56429688 | + | lncRNA |
| ENSG00000170909.13_3 | OSCAR | 0.42468057 | 1 | 0.434 | -0.325 | chr19 | 54597933 | 54606000 | - | protein_coding |
| ENSG00000103150.7_5 | MLYCD | 0.85788687 | 1 | 0.434 | 3.949 | chr16 | 83932720 | 83985050 | + | protein_coding |
| ENSG00000103326.12_4 | CAPN15 | 0.52161287 | 1 | 0.434 | 5.465 | chr16 | 577712 | 604636 | + | protein_coding |
| ENSG00000102580.15_4 | DNAJC3 | 0.97659823 | 1 | 0.433 | 6.400 | chr13 | 96329393 | 96447242 | + | protein_coding |
| ENSG00000169136.11_4 | ATF5 | 0.80360208 | 1 | 0.433 | 3.500 | chr19 | 50431959 | 50437192 | + | protein_coding |
| ENSG00000146700.9_4 | SSC4D | 0.50613741 | 1 | 0.433 | 0.671 | chr7 | 76018651 | 76039014 | - | protein_coding |
| ENSG00000197847.12_5 | SLC22A20P | 0.44328129 | 1 | 0.433 | -0.069 | chr11 | 64981311 | 65010228 | + | transcribed_unitary_pseudogene |
| ENSG00000278730.1_6 | AC005332.6 | 0.84473495 | 1 | 0.433 | 3.891 | chr17 | 66122807 | 66125727 | + | lncRNA |
| ENSG00000072062.14_4 | PRKACA | 0.94809301 | 1 | 0.433 | 4.952 | chr19 | 14202500 | 14228896 | - | protein_coding |
| ENSG00000079691.18_6 | CARMIL1 | 0.95832969 | 1 | 0.433 | 5.873 | chr6 | 25279306 | 25620758 | + | protein_coding |
| ENSG00000102226.10_4 | USP11 | 0.97630024 | 1 | 0.433 | 6.202 | chrX | 47092265 | 47107727 | + | protein_coding |
| ENSG00000164039.15_5 | BDH2 | 0.79392314 | 1 | 0.432 | 3.081 | chr4 | 103998749 | 104021027 | - | protein_coding |
| ENSG00000088256.9_3 | GNA11 | 0.86690888 | 1 | 0.432 | 5.597 | chr19 | 3094360 | 3123997 | + | protein_coding |
| ENSG00000267904.1_8 | AC024075.1 | 0.45296994 | 1 | 0.432 | 0.009 | chr19 | 16750778 | 16751479 | - | lncRNA |
| ENSG00000185585.20_4 | OLFML2A | 0.95728624 | 1 | 0.432 | 5.382 | chr9 | 127539412 | 127577164 | + | protein_coding |
| ENSG00000187778.14_4 | MCRS1 | 0.97203374 | 1 | 0.432 | 6.133 | chr12 | 49950327 | 49961928 | - | protein_coding |
| ENSG00000214650.3_6 | AC073592.1 | 0.42209413 | 1 | 0.432 | -0.418 | chr12 | 124997768 | 125001344 | + | lncRNA |
| ENSG00000273055.1_7 | AC005046.2 | 0.46999085 | 1 | 0.432 | 0.190 | chr7 | 107582561 | 107583185 | + | lncRNA |
| ENSG00000173193.15_5 | PARP14 | 0.98306 | 1 | 0.432 | 6.716 | chr3 | 122399686 | 122449687 | + | protein_coding |
| ENSG00000158467.16_4 | AHCYL2 | 0.9267064 | 1 | 0.432 | 4.591 | chr7 | 128864864 | 129070052 | + | protein_coding |
| ENSG00000162695.12_4 | SLC30A7 | 0.97163504 | 1 | 0.431 | 5.957 | chr1 | 101361632 | 101447313 | + | protein_coding |

| | | | | | | | | | | |
|---|---|---|---|---|---|---|---|---|---|---|
| ENSG00000166225.8_3 | FRS2 | 0.97046266 | 1 | 0.431 | 5.672 | chr12 | 69864129 | 69973569 | + | protein_coding |
| ENSG00000105321.14_4 | CCDC9 | 0.8077673 | 1 | 0.431 | 3.643 | chr19 | 47759237 | 47776958 | + | protein_coding |
| ENSG00000079805.16_5 | DNM2 | 0.96818347 | 1 | 0.431 | 6.560 | chr19 | 10828755 | 10944164 | + | protein_coding |
| ENSG00000167525.13_5 | PROCA1 | 0.71372324 | 1 | 0.431 | 2.451 | chr17 | 27030215 | 27038872 | - | protein_coding |
| ENSG00000106733.21_4 | NMRK1 | 0.72994038 | 1 | 0.431 | 2.709 | chr9 | 77675489 | 77703133 | - | protein_coding |
| ENSG00000079432.7_6 | CIC | 0.77031048 | 1 | 0.431 | 6.773 | chr19 | 42772689 | 42799949 | + | protein_coding |
| ENSG00000167716.18_3 | WDR81 | 0.85200339 | 1 | 0.431 | 6.037 | chr17 | 1619817 | 1641893 | + | protein_coding |
| ENSG00000058453.17_4 | CROCC | 0.67333728 | 1 | 0.430 | 4.062 | chr1 | 17066768 | 17299459 | + | protein_coding |
| ENSG00000127948.16_5 | POR | 0.97774862 | 1 | 0.430 | 6.848 | chr7 | 75528518 | 75616173 | + | protein_coding |
| ENSG00000117115.13_4 | PADI2 | 0.85684474 | 1 | 0.430 | 3.599 | chr1 | 17393256 | 17445946 | - | protein_coding |
| ENSG00000145040.4_5 | UCN2 | 0.6866523 | 1 | 0.430 | 2.146 | chr3 | 48599151 | 48601214 | - | protein_coding |
| ENSG00000160439.16_5 | RDH13 | 0.79863389 | 1 | 0.430 | 3.363 | chr19 | 55550476 | 55582659 | - | protein_coding |
| ENSG00000287009.1_1 | AC021037.1 | 0.42083553 | 1 | 0.430 | -1.013 | chr10 | 97888812 | 97890285 | - | lncRNA |
| ENSG00000100345.21_4 | MYH9 | 0.98807757 | 1 | 0.430 | 9.578 | chr22 | 36677323 | 36784112 | - | protein_coding |
| ENSG00000259065.2_7 | AC005520.4 | 0.48910329 | 1 | 0.430 | 0.489 | chr14 | 74254058 | 74270003 | + | lncRNA |
| ENSG00000173540.13_4 | GMPPB | 0.92709269 | 1 | 0.430 | 4.585 | chr3 | 49754277 | 49761384 | - | protein_coding |
| ENSG00000117616.18_7 | RSRP1 | 0.94083461 | 1 | 0.430 | 5.178 | chr1 | 25568740 | 25664704 | - | protein_coding |
| ENSG00000007516.13_5 | BAIAP3 | 0.61818789 | 1 | 0.429 | 1.644 | chr16 | 1383602 | 1399442 | + | protein_coding |
| ENSG00000246203.2_6 | AL353807.3 | 0.49374951 | 1 | 0.429 | 0.389 | chr1 | 155584517 | 155630036 | - | unprocessed_pseudogene |
| ENSG00000141449.14_5 | GREB1L | 0.60803612 | 1 | 0.429 | 1.596 | chr18 | 18822203 | 19105378 | + | protein_coding |
| ENSG00000162300.13_5 | ZFPL1 | 0.57866475 | 1 | 0.429 | 1.456 | chr11 | 64851682 | 64855870 | + | protein_coding |
| ENSG00000126247.11_8 | CAPNS1 | 0.99417391 | 1 | 0.429 | 9.218 | chr19 | 36630855 | 36641255 | + | protein_coding |
| ENSG00000197815.4_6 | AC122129.1 | 0.52754198 | 1 | 0.428 | 0.543 | chr17 | 17761541 | 17763355 | + | lncRNA |
| ENSG00000177731.16_7 | FLII | 0.98940945 | 1 | 0.428 | 7.947 | chr17 | 18148129 | 18162052 | - | protein_coding |
| ENSG00000174928.16_8 | C3orf33 | 0.49212557 | 1 | 0.428 | 0.657 | chr3 | 155480406 | 155524067 | - | protein_coding |
| ENSG00000026652.15_6 | AGPAT4 | 0.81195465 | 1 | 0.428 | 3.215 | chr6 | 161550999 | 161695093 | - | protein_coding |
| ENSG00000277715.1_6 | AC079174.2 | 0.43642575 | 1 | 0.427 | -0.172 | chr12 | 106644537 | 106646564 | + | lncRNA |
| ENSG00000166912.17_4 | MTMR10 | 0.95918287 | 1 | 0.427 | 5.300 | chr15 | 31231144 | 31283831 | - | protein_coding |
| ENSG00000261187.1_7 | AC079322.1 | 0.42302611 | 1 | 0.427 | -1.013 | chr15 | 72757469 | 72758603 | - | lncRNA |
| ENSG00000253908.1_5 | AC104115.2 | 0.42760921 | 1 | 0.427 | -0.963 | chr5 | 179684680 | 179685366 | + | processed_pseudogene |
| ENSG00000070444.15_4 | MNT | 0.85247581 | 1 | 0.427 | 4.315 | chr17 | 2287367 | 2304398 | - | protein_coding |
| ENSG00000065057.9_7 | NTHL1 | 0.95867013 | 1 | 0.427 | 5.290 | chr16 | 2089816 | 2097867 | - | protein_coding |
| ENSG00000260647.1_6 | AC127537.1 | 0.42410714 | 1 | 0.427 | -0.787 | chr17 | 18171394 | 18172142 | + | lncRNA |
| ENSG00000177465.4_2 | ACOT4 | 0.6198377 | 1 | 0.427 | 1.726 | chr14 | 74058410 | 74063200 | + | protein_coding |
| ENSG00000197961.11_3 | ZNF121 | 0.98186399 | 1 | 0.426 | 6.212 | chr19 | 9671029 | 9695209 | - | protein_coding |
| ENSG00000139437.18_4 | TCHP | 0.955872 | 1 | 0.426 | 4.983 | chr12 | 110338069 | 110421646 | + | protein_coding |
| ENSG00000207751.3_5 | AP000553.1 | 0.42392129 | 1 | 0.426 | -1.013 | chr22 | 22006559 | 22024526 | + | lncRNA |
| ENSG00000102316.17_3 | MAGED2 | 0.98681667 | 1 | 0.426 | 6.474 | chrX | 54834032 | 54842448 | + | protein_coding |
| ENSG00000250986.1_5 | LINC02600 | 0.51816153 | 1 | 0.426 | 0.894 | chr4 | 3760475 | 3765117 | - | lncRNA |
| ENSG00000197714.9_3 | ZNF460 | 0.96411295 | 1 | 0.426 | 5.490 | chr19 | 57791419 | 57805437 | + | protein_coding |
| ENSG00000130772.14_3 | MED18 | 0.81460221 | 1 | 0.425 | 3.492 | chr1 | 28655513 | 28662476 | + | protein_coding |
| ENSG00000257027.1_6 | AC010186.3 | 0.74443956 | 1 | 0.425 | 2.526 | chr12 | 9811163 | 9814681 | + | lncRNA |
| ENSG00000187601.5_3 | MAGEH1 | 0.83202527 | 1 | 0.425 | 3.622 | chrX | 55478560 | 55479999 | + | protein_coding |
| ENSG00000104368.18_6 | PLAT | 0.69237458 | 1 | 0.425 | 2.492 | chr8 | 42032236 | 42065194 | - | protein_coding |
| ENSG00000137207.12_7 | YIPF3 | 0.94921363 | 1 | 0.425 | 5.272 | chr6 | 43479570 | 43484723 | - | protein_coding |
| ENSG00000248019.3_6 | FAM13A-AS1 | 0.4477095 | 1 | 0.425 | -0.295 | chr4 | 89630449 | 89651254 | + | lncRNA |
| ENSG00000263001.6_6 | GTF2I | 0.98544813 | 1 | 0.425 | 6.045 | chr7 | 74064571 | 74175022 | + | protein_coding |
| ENSG00000111716.14_6 | LDHB | 0.95804569 | 1 | 0.425 | 5.670 | chr12 | 21788276 | 21910791 | - | polymorphic_pseudogene |
| ENSG00000061273.18_5 | HDAC7 | 0.95417188 | 1 | 0.425 | 6.136 | chr12 | 48176505 | 48226915 | - | protein_coding |
| ENSG00000213885.4_5 | RPL13AP7 | 0.43869647 | 1 | 0.425 | -0.264 | chr21 | 26734135 | 26754069 | + | transcribed_processed_pseudogene |
| ENSG00000279041.1_6 | AC102945.2 | 0.5691891 | 1 | 0.425 | 1.173 | chr8 | 30409862 | 30411283 | + | TEC |
| ENSG00000275964.1_6 | AL355001.2 | 0.47055475 | 1 | 0.425 | 0.285 | chr13 | 20437998 | 20439188 | + | lncRNA |
| ENSG00000201684.1 | RN7SKP239 | 0.43527624 | 1 | 0.425 | -0.496 | chr11 | 67129885 | 67130179 | + | misc_RNA |
| ENSG00000181830.9_6 | SLC35C1 | 0.92302361 | 1 | 0.424 | 4.544 | chr11 | 45825623 | 45834567 | + | protein_coding |
| ENSG00000215375.6_3 | MYL5 | 0.76272387 | 1 | 0.424 | 3.292 | chr4 | 667369 | 675822 | + | protein_coding |
| ENSG00000108963.17_4 | DPH1 | 0.92681554 | 1 | 0.424 | 4.562 | chr17 | 1933404 | 1946724 | + | protein_coding |
| ENSG00000175348.11_6 | TMEM9B | 0.95442565 | 1 | 0.424 | 4.860 | chr11 | 8968749 | 8986558 | - | protein_coding |
| ENSG00000174306.21_5 | ZHX3 | 0.89005211 | 1 | 0.424 | 3.946 | chr20 | 39807088 | 39946312 | - | protein_coding |
| ENSG00000251359.4_5 | WWC2-AS2 | 0.43197483 | 1 | 0.423 | -0.444 | chr4 | 184018170 | 184020352 | - | lncRNA |

| | | | | | | | | | | |
|---|---|---|---|---|---|---|---|---|---|---|
| ENSG00000269161.1_9 | AC010618.2 | 0.42872635 | 1 | 0.423 | -0.788 | chr19 | 17629139 | 17631374 | - | lncRNA |
| ENSG00000106605.11_5 | BLVRA | 0.97034439 | 1 | 0.423 | 5.465 | chr7 | 43798279 | 43846941 | + | protein_coding |
| ENSG00000175220.12_6 | ARHGAP1 | 0.96999412 | 1 | 0.423 | 5.666 | chr11 | 46698630 | 46722169 | - | protein_coding |
| ENSG00000196547.15_7 | MAN2A2 | 0.95342886 | 1 | 0.423 | 5.083 | chr15 | 91445448 | 91465814 | + | protein_coding |
| ENSG00000138606.19_5 | SHF | 0.52257641 | 1 | 0.423 | 0.697 | chr15 | 45459412 | 45493373 | - | protein_coding |
| ENSG00000204758.8_7 | AC008429.1 | 0.60254875 | 1 | 0.423 | 1.380 | chr5 | 172381789 | 172386395 | - | lncRNA |
| ENSG00000276718.1_7 | AC005840.4 | 0.42958081 | 1 | 0.423 | -0.831 | chr12 | 6575703 | 6576301 | + | lncRNA |
| ENSG00000105137.13_4 | SYDE1 | 0.88249703 | 1 | 0.423 | 4.397 | chr19 | 15218212 | 15225796 | + | protein_coding |
| ENSG00000114796.16_5 | KLHL24 | 0.92669282 | 1 | 0.423 | 4.396 | chr3 | 183353398 | 183402307 | + | protein_coding |
| ENSG00000240137.6_6 | ERICH6-AS1 | 0.59591544 | 1 | 0.422 | 1.136 | chr3 | 150421351 | 150440792 | + | lncRNA |
| ENSG00000181790.11_4 | ADGRB1 | 0.47882958 | 1 | 0.422 | 0.271 | chr8 | 143530791 | 143626370 | + | protein_coding |
| ENSG00000185344.14_4 | ATP6V0A2 | 0.96111782 | 1 | 0.422 | 5.726 | chr12 | 124196900 | 124246302 | + | protein_coding |
| ENSG00000226149.6_8 | AL356124.1 | 0.46821895 | 1 | 0.422 | 0.020 | chr6 | 129760630 | 129897192 | - | lncRNA |
| ENSG00000101670.12_6 | LIPG | 0.43022902 | 1 | 0.422 | -0.619 | chr18 | 47087069 | 47125555 | + | protein_coding |
| ENSG00000143079.15_5 | CTTNBP2NL | 0.95474433 | 1 | 0.422 | 4.847 | chr1 | 112938836 | 113006078 | + | protein_coding |
| ENSG00000184384.14_5 | MAML2 | 0.82185479 | 1 | 0.422 | 3.163 | chr11 | 95709762 | 96076359 | - | protein_coding |
| ENSG00000174483.20_5 | BBS1 | 0.51370135 | 1 | 0.422 | 0.818 | chr11 | 66278077 | 66301084 | + | protein_coding |
| ENSG00000155158.20_5 | TTC39B | 0.82649884 | 1 | 0.422 | 3.355 | chr9 | 15163620 | 15307358 | - | protein_coding |
| ENSG00000228701.2_7 | TNKS2-AS1 | 0.43307057 | 1 | 0.422 | -0.878 | chr10 | 93542592 | 93558061 | - | lncRNA |
| ENSG00000141873.11_7 | SLC39A3 | 0.79849868 | 1 | 0.421 | 3.408 | chr19 | 2732202 | 2740026 | - | protein_coding |
| ENSG00000197702.14_5 | PARVA | 0.98298713 | 1 | 0.421 | 5.817 | chr11 | 12399110 | 12556903 | + | protein_coding |
| ENSG00000125901.6_3 | MRPS26 | 0.85474629 | 1 | 0.421 | 5.909 | chr20 | 3026698 | 3028896 | + | protein_coding |
| ENSG00000224903.2_7 | AC005534.1 | 0.43776231 | 1 | 0.421 | -0.489 | chr7 | 156446879 | 156452276 | - | lncRNA |
| ENSG00000124222.22_5 | STX16 | 0.81005457 | 1 | 0.421 | 3.387 | chr20 | 57226309 | 57254582 | + | protein_coding |
| ENSG00000070404.10_7 | FSTL3 | 0.94163126 | 1 | 0.421 | 6.328 | chr19 | 676392 | 683392 | + | protein_coding |
| ENSG00000014824.14_4 | SLC30A9 | 0.98167692 | 1 | 0.421 | 6.344 | chr4 | 41992519 | 42092478 | + | protein_coding |
| ENSG00000105556.11_6 | MIER2 | 0.84255267 | 1 | 0.421 | 3.646 | chr19 | 301444 | 344815 | - | protein_coding |
| ENSG00000163956.13_6 | LRPAP1 | 0.96671937 | 1 | 0.421 | 5.995 | chr4 | 3505339 | 3534173 | - | protein_coding |
| ENSG00000196204.11_2 | RNF216P1 | 0.89021272 | 1 | 0.421 | 3.964 | chr7 | 5013619 | 5080306 | + | transcribed_unprocessed_pseudogene |
| ENSG00000054598.9_6 | FOXC1 | 0.81360432 | 1 | 0.420 | 3.079 | chr6 | 1610150 | 1614132 | + | protein_coding |
| ENSG00000070778.13_5 | PTPN21 | 0.92519805 | 1 | 0.420 | 4.399 | chr14 | 88932122 | 89021351 | - | protein_coding |
| ENSG00000106052.13_4 | TAX1BP1 | 0.99017775 | 1 | 0.420 | 7.164 | chr7 | 27778950 | 27884183 | + | protein_coding |
| ENSG00000141179.14_3 | PCTP | 0.9049409 | 1 | 0.420 | 4.394 | chr17 | 53828412 | 53920191 | + | protein_coding |
| ENSG00000217027.1_4 | TPT1P4 | 0.43560534 | 1 | 0.420 | -0.669 | chr6 | 144521584 | 144522102 | - | processed_pseudogene |
| ENSG00000116198.13_6 | CEP104 | 0.92656771 | 1 | 0.420 | 4.457 | chr1 | 3728650 | 3773778 | - | protein_coding |
| ENSG00000213889.10_2 | PPM1N | 0.63222916 | 1 | 0.420 | 1.655 | chr19 | 45992035 | 46005768 | + | protein_coding |
| ENSG00000104812.15_5 | GYS1 | 0.96612075 | 1 | 0.419 | 5.506 | chr19 | 49471387 | 49496567 | - | protein_coding |
| ENSG00000271976.1_6 | AC012467.2 | 0.58894274 | 1 | 0.419 | 1.454 | chr3 | 53893021 | 53895603 | - | lncRNA |
| ENSG00000046604.13_3 | DSG2 | 0.99218437 | 1 | 0.419 | 7.097 | chr18 | 29078140 | 29128971 | + | protein_coding |
| ENSG00000259705.1_6 | AC084757.3 | 0.45923428 | 1 | 0.419 | -0.799 | chr15 | 48938148 | 48944213 | + | lncRNA |
| ENSG00000136877.15_5 | FPGS | 0.97137291 | 1 | 0.419 | 5.609 | chr9 | 130556876 | 130576606 | + | protein_coding |
| ENSG00000198892.6_2 | SHISA4 | 0.83570942 | 1 | 0.419 | 3.270 | chr1 | 201857808 | 201861434 | + | protein_coding |
| ENSG00000229921.7_6 | KIF25-AS1 | 0.5427679 | 1 | 0.418 | 0.879 | chr6 | 168393502 | 168397790 | - | lncRNA |
| ENSG00000139182.15_6 | CLSTN3 | 0.93827088 | 1 | 0.418 | 4.628 | chr12 | 7282294 | 7311541 | + | protein_coding |
| ENSG00000188738.15_4 | FSIP2 | 0.50124141 | 1 | 0.418 | 0.358 | chr2 | 186603531 | 186698017 | + | protein_coding |
| ENSG00000201678.1 | Y_RNA | 0.43399121 | 1 | 0.418 | -0.743 | chr12 | 49321722 | 49321816 | - | misc_RNA |
| ENSG00000137166.15_5 | FOXP4 | 0.82696113 | 1 | 0.418 | 3.827 | chr6 | 41514164 | 41570122 | + | protein_coding |
| ENSG00000116171.18_5 | SCP2 | 0.98225917 | 1 | 0.418 | 6.289 | chr1 | 53392948 | 53517370 | + | protein_coding |
| ENSG00000167280.16_5 | ENGASE | 0.96438859 | 1 | 0.418 | 5.254 | chr17 | 77071021 | 77084681 | + | protein_coding |
| ENSG00000013563.14_6 | DNASE1L1 | 0.78831049 | 1 | 0.418 | 3.251 | chrX | 153629577 | 153640449 | - | protein_coding |
| ENSG00000204713.11_4 | TRIM27 | 0.92532343 | 1 | 0.418 | 4.153 | chr6 | 28870779 | 28891765 | - | protein_coding |
| ENSG00000265688.2_7 | MAFG-DT | 0.8479488 | 1 | 0.418 | 3.711 | chr17 | 79885705 | 79888629 | + | lncRNA |
| ENSG00000060491.16_5 | OGFR | 0.91210636 | 1 | 0.418 | 5.378 | chr20 | 61436187 | 61445352 | + | protein_coding |
| ENSG00000207650.1 | MIR570 | 0.43677701 | 1 | 0.418 | -0.925 | chr3 | 195426272 | 195426368 | + | miRNA |
| ENSG00000136874.11_4 | STX17 | 0.97595758 | 1 | 0.417 | 5.363 | chr9 | 102668936 | 102736816 | + | protein_coding |
| ENSG00000061936.9_3 | SFSWAP | 0.98095426 | 1 | 0.417 | 5.719 | chr12 | 132195626 | 132284282 | + | protein_coding |
| ENSG00000174938.14_7 | SEZ6L2 | 0.89670907 | 1 | 0.417 | 4.407 | chr16 | 29882480 | 29910868 | - | protein_coding |
| ENSG00000117139.18_7 | KDM5B | 0.98657843 | 1 | 0.417 | 6.176 | chr1 | 202693623 | 202777615 | - | protein_coding |
| ENSG00000110906.13_4 | KCTD10 | 0.98079521 | 1 | 0.417 | 5.655 | chr12 | 109886460 | 109915164 | - | protein_coding |
| ENSG00000055950.16_4 | MRPL43 | 0.97461552 | 1 | 0.416 | 5.345 | chr10 | 102729215 | 102747272 | - | protein_coding |

| | | | | | | | | | | |
|---|---|---|---|---|---|---|---|---|---|---|
| ENSG00000161179.14_4 | YDJC | 0.85780867 | 1 | 0.416 | 4.776 | chr22 | 21982378 | 21984353 | - | protein_coding |
| ENSG00000164535.15_5 | DAGLB | 0.92671069 | 1 | 0.416 | 4.406 | chr7 | 6448757 | 6523821 | - | protein_coding |
| ENSG00000258344.1_8 | AC078778.1 | 0.51642257 | 1 | 0.416 | 0.301 | chr12 | 54670415 | 54738867 | + | lncRNA |
| ENSG00000196214.11_7 | ZNF766 | 0.97626049 | 1 | 0.416 | 5.379 | chr19 | 52772840 | 52799299 | + | protein_coding |
| ENSG00000114127.10_7 | XRN1 | 0.97197851 | 1 | 0.416 | 5.638 | chr3 | 142025449 | 142166904 | - | protein_coding |
| ENSG00000135392.16_7 | DNAJC14 | 0.93930582 | 1 | 0.416 | 4.404 | chr12 | 56214744 | 56224608 | - | protein_coding |
| ENSG00000165548.11_7 | TMEM63C | 0.50749061 | 1 | 0.415 | 0.501 | chr14 | 77582911 | 77725838 | + | protein_coding |
| ENSG00000037757.14_4 | MRI1 | 0.85254534 | 1 | 0.415 | 3.488 | chr19 | 13875336 | 13885096 | + | protein_coding |
| ENSG00000174996.11_5 | KLC2 | 0.87060641 | 1 | 0.415 | 5.070 | chr11 | 66024765 | 66035331 | + | protein_coding |
| ENSG00000235084.3_4 | CHCHD2P6 | 0.43856381 | 1 | 0.415 | -0.879 | chr1 | 15931092 | 15931538 | + | processed_pseudogene |
| ENSG00000166140.17_3 | ZFYVE19 | 0.93514984 | 1 | 0.415 | 4.463 | chr15 | 41099284 | 41107282 | + | protein_coding |
| ENSG00000170017.12_5 | ALCAM | 0.99655542 | 1 | 0.415 | 7.920 | chr3 | 105085753 | 105295744 | + | protein_coding |
| ENSG00000160213.7_4 | CSTB | 0.9667149 | 1 | 0.415 | 5.277 | chr21 | 45192392 | 45196326 | - | protein_coding |
| ENSG00000165813.20_7 | CCDC186 | 0.95885694 | 1 | 0.414 | 5.365 | chr10 | 115880621 | 115933991 | - | protein_coding |
| ENSG00000257698.2_8 | GIHCG | 0.73329312 | 1 | 0.414 | 2.873 | chr12 | 58323898 | 58330128 | - | lncRNA |
| ENSG00000156521.14_4 | TYSND1 | 0.96270944 | 1 | 0.414 | 5.420 | chr10 | 71897737 | 71906456 | - | protein_coding |
| ENSG00000112667.12_4 | DNPH1 | 0.9067653 | 1 | 0.414 | 4.598 | chr6 | 43193367 | 43197222 | - | protein_coding |
| ENSG00000067182.8_3 | TNFRSF1A | 0.9968762 | 1 | 0.414 | 8.293 | chr12 | 6437923 | 6451280 | - | protein_coding |
| ENSG00000084234.17_5 | APLP2 | 0.99378102 | 1 | 0.414 | 7.329 | chr11 | 129938042 | 130014706 | + | protein_coding |
| ENSG00000081026.19_4 | MAGI3 | 0.9755135 | 1 | 0.414 | 5.604 | chr1 | 113933137 | 114228545 | + | protein_coding |
| ENSG00000171467.16_7 | ZNF318 | 0.93272157 | 1 | 0.414 | 4.477 | chr6 | 43274872 | 43337385 | - | protein_coding |
| ENSG00000069974.16_6 | RAB27A | 0.76313142 | 1 | 0.414 | 2.566 | chr15 | 55495164 | 55611311 | - | protein_coding |
| ENSG00000103319.12_3 | EEF2K | 0.9871639 | 1 | 0.413 | 6.045 | chr16 | 22217599 | 22300059 | + | protein_coding |
| ENSG00000270061.1_7 | AC068790.5 | 0.45787843 | 1 | 0.413 | -0.394 | chr12 | 124454537 | 124454891 | - | lncRNA |
| ENSG00000254536.2_7 | AL360181.3 | 0.53916158 | 1 | 0.413 | 0.886 | chr10 | 135193521 | 135233775 | + | protein_coding |
| ENSG00000261207.1_7 | AL031717.1 | 0.45930626 | 1 | 0.412 | -0.271 | chr16 | 1801560 | 1802263 | - | lncRNA |
| ENSG00000279369.1_5 | AC046185.3 | 0.44461765 | 1 | 0.412 | -0.672 | chr17 | 61778207 | 61780030 | + | TEC |
| ENSG00000197951.9_5 | ZNF71 | 0.90503058 | 1 | 0.412 | 4.041 | chr19 | 57106632 | 57137849 | + | protein_coding |
| ENSG00000126391.14_4 | FRMD8 | 0.97310856 | 1 | 0.412 | 5.192 | chr11 | 65154092 | 65180996 | + | protein_coding |
| ENSG00000286158.1_2 | AL034548.2 | 0.49378919 | 1 | 0.412 | 0.277 | chr20 | 292375 | 293881 | + | lncRNA |
| ENSG00000123415.15_7 | SMUG1 | 0.96411923 | 1 | 0.412 | 4.689 | chr12 | 54515061 | 54582792 | - | protein_coding |
| ENSG00000178234.13_5 | GALNT11 | 0.96950612 | 1 | 0.412 | 5.332 | chr7 | 151722759 | 151819425 | + | protein_coding |
| ENSG00000123146.20_4 | ADGRE5 | 0.93116616 | 1 | 0.412 | 4.573 | chr19 | 14491313 | 14519537 | + | protein_coding |
| ENSG00000120054.12_4 | CPN1 | 0.47843525 | 1 | 0.411 | -0.105 | chr10 | 101801950 | 101841626 | - | protein_coding |
| ENSG00000117868.17_7 | ESYT2 | 0.99694959 | 1 | 0.411 | 7.665 | chr7 | 158523686 | 158622944 | - | protein_coding |
| ENSG00000014216.15_6 | CAPN1 | 0.98431621 | 1 | 0.411 | 7.414 | chr11 | 64948037 | 64979477 | + | protein_coding |
| ENSG00000108784.10_4 | NAGLU | 0.96341305 | 1 | 0.411 | 5.249 | chr17 | 40688259 | 40696467 | + | protein_coding |
| ENSG00000149292.16_6 | TTC12 | 0.89609826 | 1 | 0.411 | 3.688 | chr11 | 113185251 | 113254266 | + | protein_coding |
| ENSG00000100300.18_4 | TSPO | 0.5786601 | 1 | 0.411 | 5.267 | chr22 | 43547553 | 43559248 | + | protein_coding |
| ENSG00000185800.12_6 | DMWD | 0.97325154 | 1 | 0.411 | 5.256 | chr19 | 46286205 | 46296103 | - | protein_coding |
| ENSG00000185753.13_4 | CXorf38 | 0.9700301 | 1 | 0.411 | 4.920 | chrX | 40486173 | 40506813 | - | protein_coding |
| ENSG00000213123.11_5 | TCTEX1D2 | 0.51856108 | 1 | 0.411 | 0.280 | chr3 | 196018090 | 196045170 | - | protein_coding |
| ENSG00000065534.18_4 | MYLK | 0.99329113 | 1 | 0.410 | 6.697 | chr3 | 123328896 | 123603178 | - | protein_coding |
| ENSG00000174353.17_3 | STAG3L3 | 0.55279863 | 1 | 0.410 | 1.049 | chr7 | 72440227 | 72476445 | - | transcribed_unprocessed_pseudogene |
| ENSG00000110066.15_8 | KMT5B | 0.98005752 | 1 | 0.410 | 5.408 | chr11 | 67922330 | 67981295 | - | protein_coding |
| ENSG00000169499.15_5 | PLEKHA2 | 0.89206571 | 1 | 0.410 | 3.843 | chr8 | 38758753 | 38831431 | + | protein_coding |
| ENSG00000169249.12_2 | ZRSR2 | 0.71914495 | 1 | 0.410 | 2.200 | chrX | 15808595 | 15841383 | + | protein_coding |
| ENSG00000222047.8_3 | C10orf55 | 0.48636666 | 1 | 0.410 | 0.046 | chr10 | 75669727 | 75682535 | - | protein_coding |
| ENSG00000054983.17_6 | GALC | 0.9702675 | 1 | 0.410 | 5.583 | chr14 | 88304164 | 88460009 | - | protein_coding |
| ENSG00000204650.14_5 | LINC02210 | 0.73894939 | 1 | 0.410 | 2.504 | chr17 | 43697694 | 43732522 | + | transcribed_unitary_pseudogene |
| ENSG00000113615.13_3 | SEC24A | 0.98579416 | 1 | 0.409 | 5.737 | chr5 | 133984475 | 134063599 | + | protein_coding |
| ENSG00000266835.6_5 | GAPLINC | 0.6460309 | 1 | 0.409 | 1.543 | chr18 | 3466248 | 3478976 | + | lncRNA |
| ENSG00000197757.8_5 | HOXC6 | 0.7050479 | 1 | 0.409 | 2.094 | chr12 | 54384408 | 54424607 | + | protein_coding |
| ENSG00000164690.8_4 | SHH | 0.54249277 | 1 | 0.408 | 0.993 | chr7 | 155592674 | 155605157 | - | protein_coding |
| ENSG00000169372.13_4 | CRADD | 0.84001878 | 1 | 0.408 | 2.994 | chr12 | 94071151 | 94288616 | + | protein_coding |
| ENSG00000198952.8_4 | SMG5 | 0.99566852 | 1 | 0.408 | 7.032 | chr1 | 156219015 | 156252616 | - | protein_coding |
| ENSG00000168453.15_5 | HR | 0.93578307 | 1 | 0.408 | 5.574 | chr8 | 21971932 | 21990897 | - | protein_coding |
| ENSG00000182372.10_6 | CLN8 | 0.9631733 | 1 | 0.408 | 4.562 | chr8 | 1703944 | 1749877 | + | protein_coding |
| ENSG00000268006.1_6 | PTOV1-AS1 | 0.61719773 | 1 | 0.408 | 1.296 | chr19 | 50341896 | 50354933 | - | lncRNA |

| | | | | | | | | | | |
|---|---|---|---|---|---|---|---|---|---|---|
| ENSG00000234155.1_7 | LINC02535 | 0.59233284 | 1 | 0.408 | 1.273 | chr6 | 86096937 | 86099904 | - | lncRNA |
| ENSG00000066923.17_5 | STAG3 | 0.87418147 | 1 | 0.408 | 3.354 | chr7 | 99775186 | 99819111 | + | protein_coding |
| ENSG00000139636.15_5 | LMBR1L | 0.91006942 | 1 | 0.408 | 3.878 | chr12 | 49490334 | 49504683 | - | protein_coding |
| ENSG00000153132.13_8 | CLGN | 0.94563606 | 1 | 0.407 | 4.517 | chr4 | 141309607 | 141348815 | - | protein_coding |
| ENSG00000132906.18_7 | CASP9 | 0.89859408 | 1 | 0.407 | 3.769 | chr1 | 15817327 | 15853029 | - | protein_coding |
| ENSG00000166886.13_5 | NAB2 | 0.88004981 | 1 | 0.407 | 3.589 | chr12 | 57482826 | 57489259 | + | protein_coding |
| ENSG00000182173.13_5 | TSEN54 | 0.9570784 | 1 | 0.407 | 5.386 | chr17 | 73512141 | 73520816 | + | protein_coding |
| ENSG00000272836.1_6 | AL022328.1 | 0.45451275 | 1 | 0.407 | -0.603 | chr22 | 50644014 | 50644491 | - | lncRNA |
| ENSG00000185100.10_4 | ADSS1 | 0.74846931 | 1 | 0.407 | 2.817 | chr14 | 105190523 | 105213662 | + | protein_coding |
| ENSG00000174516.15_4 | PELI3 | 0.84786372 | 1 | 0.407 | 3.368 | chr11 | 66233798 | 66244808 | + | protein_coding |
| ENSG00000287225.1_2 | Z84468.2 | 0.47168827 | 1 | 0.407 | -0.036 | chr22 | 48869947 | 48885136 | - | lncRNA |
| ENSG00000257167.2_7 | TMPO-AS1 | 0.65831563 | 1 | 0.406 | 2.477 | chr12 | 98906751 | 98910200 | - | lncRNA |
| ENSG00000158863.22_3 | FAM160B2 | 0.96960801 | 1 | 0.406 | 5.209 | chr8 | 21946661 | 21962422 | + | protein_coding |
| ENSG00000161533.12_6 | ACOX1 | 0.9940766 | 1 | 0.406 | 6.345 | chr17 | 73937588 | 73975258 | - | protein_coding |
| ENSG00000173614.14_4 | NMNAT1 | 0.7835878 | 1 | 0.406 | 2.492 | chr1 | 10003486 | 10045559 | + | protein_coding |
| ENSG00000286067.1_2 | AC004263.2 | 0.58218927 | 1 | 0.406 | 1.261 | chr12 | 120655873 | 120660471 | + | lncRNA |
| ENSG00000161677.11_4 | JOSD2 | 0.78225905 | 1 | 0.406 | 4.273 | chr19 | 51009255 | 51014610 | - | protein_coding |
| ENSG00000271664.1_6 | AC004890.3 | 0.48806311 | 1 | 0.406 | -0.134 | chr7 | 148972710 | 148975083 | - | unprocessed_pseudogene |
| ENSG00000261251.1_6 | Z97055.2 | 0.45079026 | 1 | 0.405 | -0.756 | chr22 | 44208336 | 44213274 | + | lncRNA |
| ENSG00000161921.15_6 | CXCL16 | 0.88076979 | 1 | 0.405 | 3.716 | chr17 | 4636828 | 4643223 | - | protein_coding |
| ENSG00000079277.21_9 | MKNK1 | 0.93852186 | 1 | 0.405 | 4.357 | chr1 | 47023080 | 47082515 | - | protein_coding |
| ENSG00000198168.9_4 | SVIP | 0.96260207 | 1 | 0.405 | 4.780 | chr11 | 22835345 | 22851845 | - | protein_coding |
| ENSG00000140391.15_5 | TSPAN3 | 0.99789977 | 1 | 0.405 | 8.194 | chr15 | 77333745 | 77376326 | - | protein_coding |
| ENSG00000162174.12_4 | ASRGL1 | 0.82278495 | 1 | 0.405 | 3.045 | chr11 | 62104920 | 62160884 | + | protein_coding |
| ENSG00000186174.12_6 | BCL9L | 0.98871589 | 1 | 0.404 | 6.262 | chr11 | 118764584 | 118796317 | - | protein_coding |
| ENSG00000205702.11_6 | CYP2D7 | 0.55112942 | 1 | 0.404 | 0.639 | chr22 | 42536214 | 42545446 | - | polymorphic_pseudogene |
| ENSG00000044090.9_3 | CUL7 | 0.97138169 | 1 | 0.404 | 4.716 | chr6 | 43005355 | 43021683 | - | protein_coding |
| ENSG00000124813.23_6 | RUNX2 | 0.49320842 | 1 | 0.404 | 0.019 | chr6 | 45295894 | 45632086 | + | protein_coding |
| ENSG00000104723.21_4 | TUSC3 | 0.9913101 | 1 | 0.404 | 6.075 | chr8 | 15274724 | 15624158 | + | protein_coding |
| ENSG00000213672.8_8 | NCKIPSD | 0.92025538 | 1 | 0.404 | 4.152 | chr3 | 48711277 | 48723797 | - | protein_coding |
| ENSG00000140104.14_6 | CLBA1 | 0.93921555 | 1 | 0.404 | 4.123 | chr14 | 105452112 | 105476819 | + | protein_coding |
| ENSG00000274756.1_5 | AC243732.1 | 0.49899312 | 1 | 0.404 | 0.106 | chr17 | 34930301 | 34931164 | - | processed_pseudogene |
| ENSG00000129636.13_5 | ITFG1 | 0.98807995 | 1 | 0.404 | 6.137 | chr16 | 47188298 | 47498060 | - | protein_coding |
| ENSG00000058799.15_4 | YIPF1 | 0.91716818 | 1 | 0.404 | 4.127 | chr1 | 54317392 | 54355471 | - | protein_coding |
| ENSG00000162068.2_3 | NTN3 | 0.59602726 | 1 | 0.404 | 1.431 | chr16 | 2521298 | 2524146 | + | protein_coding |
| ENSG00000198431.16_6 | TXNRD1 | 0.99775343 | 1 | 0.404 | 10.914 | chr12 | 104609557 | 104744085 | + | protein_coding |
| ENSG00000179630.11_4 | LACC1 | 0.72401606 | 1 | 0.403 | 2.393 | chr13 | 44453420 | 44468068 | + | protein_coding |
| ENSG00000102760.13_3 | RGCC | 0.51181885 | 1 | 0.403 | 0.208 | chr13 | 42031686 | 42045007 | + | protein_coding |
| ENSG00000141505.12_8 | ASGR1 | 0.4669526 | 1 | 0.403 | -1.124 | chr17 | 7076750 | 7082883 | - | protein_coding |
| ENSG00000138801.9_4 | PAPSS1 | 0.99031646 | 1 | 0.403 | 5.671 | chr4 | 108511433 | 108641390 | - | protein_coding |
| ENSG00000126464.14_3 | PRR12 | 0.69735576 | 1 | 0.403 | 4.798 | chr19 | 50094439 | 50129696 | + | protein_coding |
| ENSG00000188167.9_5 | TMPPE | 0.80018938 | 1 | 0.403 | 2.490 | chr3 | 33131913 | 33138638 | - | protein_coding |
| ENSG00000071553.17_5 | ATP6AP1 | 0.99016478 | 1 | 0.403 | 5.873 | chrX | 153656991 | 153664862 | + | protein_coding |
| ENSG00000272899.4_7 | ATP6V1FNB | 0.52227781 | 1 | 0.403 | 0.278 | chr7 | 128506384 | 128512101 | + | protein_coding |
| ENSG00000261474.1_7 | AC026471.4 | 0.59766599 | 1 | 0.403 | 0.975 | chr16 | 31460856 | 31464814 | - | lncRNA |
| ENSG00000164111.15_4 | ANXA5 | 0.99744326 | 1 | 0.403 | 8.148 | chr4 | 122589101 | 122618150 | - | protein_coding |
| ENSG00000114650.20_7 | SCAP | 0.91883622 | 1 | 0.402 | 6.197 | chr3 | 47455171 | 47518616 | - | protein_coding |
| ENSG00000105726.17_4 | ATP13A1 | 0.89823601 | 1 | 0.402 | 5.652 | chr19 | 19756007 | 19774485 | - | protein_coding |
| ENSG00000163141.20_5 | BNIPL | 0.49948943 | 1 | 0.402 | 0.085 | chr1 | 151008797 | 151020196 | + | protein_coding |
| ENSG00000185019.17_7 | UBOX5 | 0.89478406 | 1 | 0.402 | 3.445 | chr20 | 3088219 | 3140842 | - | protein_coding |
| ENSG00000187676.8_4 | B3GLCT | 0.94054853 | 1 | 0.402 | 4.366 | chr13 | 31774112 | 31906413 | + | protein_coding |
| ENSG00000249042.6_7 | AC008771.1 | 0.54812678 | 1 | 0.402 | 0.550 | chr5 | 79707050 | 79783914 | - | lncRNA |
| ENSG00000006282.21_7 | SPATA20 | 0.99581406 | 1 | 0.402 | 7.712 | chr17 | 48620419 | 48633213 | + | protein_coding |
| ENSG00000006459.11_4 | KDM7A | 0.81766658 | 1 | 0.401 | 3.100 | chr7 | 139784546 | 139876783 | - | protein_coding |
| ENSG00000111726.13_3 | CMAS | 0.98748179 | 1 | 0.401 | 5.353 | chr12 | 22199152 | 22218608 | + | protein_coding |
| ENSG00000243137.8_6 | PSG4 | 0.5584054 | 1 | 0.401 | 0.684 | chr19 | 43696854 | 43711451 | - | protein_coding |
| ENSG00000165406.16_5 | MARCHF8 | 0.98554267 | 1 | 0.401 | 5.594 | chr10 | 45950033 | 46090354 | - | protein_coding |
| ENSG00000116954.8_4 | RRAGC | 0.82288412 | 1 | 0.401 | 3.000 | chr1 | 39303870 | 39325444 | - | protein_coding |
| ENSG00000189114.8_6 | BLOC1S3 | 0.86940753 | 1 | 0.401 | 3.356 | chr19 | 45682042 | 45720191 | + | protein_coding |

| | | | | | | | | | | |
|---|---|---|---|---|---|---|---|---|---|---|
| ENSG00000176108.9_5 | CHMP6 | 0.91454349 | 1 | 0.401 | 4.013 | chr17 | 78965398 | 78983317 | + | protein_coding |
| ENSG00000118046.16_6 | STK11 | 0.76016065 | 1 | 0.400 | 5.481 | chr19 | 1177557 | 1228430 | + | protein_coding |
| ENSG00000249476.2_7 | AC008467.1 | 0.45585237 | 1 | 0.400 | -0.724 | chr5 | 108572821 | 108662070 | - | lncRNA |
| ENSG00000166548.16_6 | TK2 | 0.97343727 | 1 | 0.400 | 4.717 | chr16 | 66541906 | 66586447 | - | protein_coding |
| ENSG00000102984.15_9 | ZNF821 | 0.61192213 | 1 | 0.400 | 1.079 | chr16 | 71893583 | 71929239 | - | protein_coding |
| ENSG00000177030.16_4 | DEAF1 | 0.95682118 | 1 | 0.400 | 4.677 | chr11 | 644233 | 706715 | - | protein_coding |
| ENSG00000103024.7_5 | NME3 | 0.69793404 | 1 | 0.399 | 3.694 | chr16 | 1820287 | 1821731 | - | protein_coding |
| ENSG00000139197.10_8 | PEX5 | 0.98580756 | 1 | 0.399 | 5.304 | chr12 | 7341281 | 7371170 | + | protein_coding |
| ENSG00000237476.1_5 | LINC01637 | 0.53108578 | 1 | 0.399 | 0.365 | chr22 | 21311380 | 21318967 | + | lncRNA |
| ENSG00000198792.13_7 | TMEM184B | 0.98742478 | 1 | 0.399 | 5.821 | chr22 | 38615298 | 38669016 | - | protein_coding |
| ENSG00000075790.10_6 | BCAP29 | 0.46757951 | 1 | 0.399 | -0.312 | chr7 | 107220422 | 107269615 | + | protein_coding |
| ENSG00000254332.1_5 | AF201337.1 | 0.57753716 | 1 | 0.399 | 1.174 | chr8 | 70041833 | 70042580 | + | processed_pseudogene |
| ENSG00000219200.6 | RNASEK | 0.60897653 | 1 | 0.399 | 1.051 | chr17 | 6915736 | 6917851 | + | protein_coding |
| ENSG00000249784.1 | SCARNA22 | 0.6723827 | 1 | 0.399 | 1.503 | chr4 | 1976363 | 1976487 | + | snoRNA |
| ENSG00000050426.16_7 | LETMD1 | 0.97663515 | 1 | 0.399 | 5.085 | chr12 | 51441745 | 51454207 | + | protein_coding |
| ENSG00000279425.1_6 | AC092279.3 | 0.47171235 | 1 | 0.399 | -0.304 | chr19 | 24261283 | 24263471 | + | TEC |
| ENSG00000071655.18_6 | MBD3 | 0.74795768 | 1 | 0.399 | 5.376 | chr19 | 1573595 | 1592864 | - | protein_coding |
| ENSG00000065029.15_5 | ZNF76 | 0.92779198 | 1 | 0.398 | 3.913 | chr6 | 35226686 | 35263762 | + | protein_coding |
| ENSG00000260077.1_7 | AC104794.3 | 0.48290077 | 1 | 0.398 | -0.307 | chr2 | 10179219 | 10180790 | - | lncRNA |
| ENSG00000197258.5_4 | EIF4BP6 | 0.76798882 | 1 | 0.398 | 2.390 | chr7 | 104308196 | 104310023 | + | processed_pseudogene |
| ENSG00000161395.14_5 | PGAP3 | 0.87259011 | 1 | 0.398 | 3.490 | chr17 | 37827375 | 37853050 | - | protein_coding |
| ENSG00000140943.17_5 | MBTPS1 | 0.99705803 | 1 | 0.398 | 6.909 | chr16 | 84087366 | 84150547 | - | protein_coding |
| ENSG00000092010.15_4 | PSME1 | 0.99426661 | 1 | 0.398 | 6.250 | chr14 | 24605372 | 24608176 | + | protein_coding |
| ENSG00000126107.15_4 | HECTD3 | 0.97226187 | 1 | 0.398 | 4.666 | chr1 | 45468212 | 45476996 | - | protein_coding |
| ENSG00000160813.7_4 | PPP1R35 | 0.75110812 | 1 | 0.398 | 2.167 | chr7 | 100032905 | 100034120 | - | protein_coding |
| ENSG00000159733.13_3 | ZFYVE28 | 0.64091534 | 1 | 0.398 | 1.369 | chr4 | 2271309 | 2420390 | - | protein_coding |
| ENSG00000213085.10_7 | CFAP45 | 0.45793481 | 1 | 0.398 | -1.030 | chr1 | 159842154 | 159869955 | - | protein_coding |
| ENSG00000000457.14_6 | SCYL3 | 0.8963559 | 1 | 0.397 | 3.718 | chr1 | 169818772 | 169863408 | - | protein_coding |
| ENSG00000187608.10_7 | ISG15 | 0.62107487 | 1 | 0.397 | 1.070 | chr1 | 936518 | 949920 | + | protein_coding |
| ENSG00000109680.11_5 | TBC1D19 | 0.8746476 | 1 | 0.397 | 3.314 | chr4 | 26578059 | 26757845 | + | protein_coding |
| ENSG00000184047.19_7 | DIABLO | 0.76761827 | 1 | 0.397 | 2.220 | chr12 | 122692215 | 122712003 | - | protein_coding |
| ENSG00000204256.13_7 | BRD2 | 0.99297146 | 1 | 0.397 | 5.855 | chr6 | 32936371 | 32949282 | + | protein_coding |
| ENSG00000226763.5_4 | SRRM5 | 0.50995812 | 1 | 0.397 | 0.171 | chr19 | 44100769 | 44118650 | + | protein_coding |
| ENSG00000256746.5_7 | MADD-AS1 | 0.49237291 | 1 | 0.397 | -0.033 | chr11 | 47292208 | 47293661 | - | lncRNA |
| ENSG00000197555.9_4 | SIPA1L1 | 0.92517895 | 1 | 0.397 | 3.858 | chr14 | 71787166 | 72207946 | + | protein_coding |
| ENSG00000170266.16_5 | GLB1 | 0.99054927 | 1 | 0.397 | 6.061 | chr3 | 33038101 | 33138694 | - | protein_coding |
| ENSG00000110025.13_4 | SNX15 | 0.56452412 | 1 | 0.397 | 0.510 | chr11 | 64794911 | 64808044 | + | protein_coding |
| ENSG00000125968.9_4 | ID1 | 0.98213797 | 1 | 0.397 | 6.365 | chr20 | 30193086 | 30194318 | + | protein_coding |
| ENSG00000125457.14_8 | MIF4GD | 0.95808283 | 1 | 0.397 | 4.604 | chr17 | 73262309 | 73267308 | - | protein_coding |
| ENSG00000100592.15_3 | DAAM1 | 0.92787076 | 1 | 0.396 | 3.809 | chr14 | 59655364 | 59838123 | + | protein_coding |
| ENSG00000011451.20_8 | WIZ | 0.92304197 | 1 | 0.396 | 4.774 | chr19 | 15530789 | 15560767 | - | protein_coding |
| ENSG00000228274.4_8 | AL021707.2 | 0.51465336 | 1 | 0.396 | -0.067 | chr22 | 39063590 | 39077852 | - | lncRNA |
| ENSG00000179820.16_4 | MYADM | 0.99467305 | 1 | 0.396 | 6.524 | chr19 | 54368017 | 54379689 | + | protein_coding |
| ENSG00000095321.17_8 | CRAT | 0.9838235 | 1 | 0.396 | 5.498 | chr9 | 131857073 | 131873468 | - | protein_coding |
| ENSG00000262678.1_7 | AC004771.4 | 0.46730128 | 1 | 0.396 | -0.683 | chr17 | 4889761 | 4890619 | + | lncRNA |
| ENSG00000233198.3_2 | RNF224 | 0.55200252 | 1 | 0.396 | 0.652 | chr9 | 140121723 | 140124090 | + | protein_coding |
| ENSG00000185065.6_6 | AC000068.1 | 0.46252711 | 1 | 0.396 | -0.725 | chr22 | 19435416 | 19437628 | + | lncRNA |
| ENSG00000107829.15_6 | FBXW4 | 0.95063083 | 1 | 0.396 | 4.164 | chr10 | 103370421 | 103455052 | - | protein_coding |
| ENSG00000174227.16_4 | PIGG | 0.97411026 | 1 | 0.396 | 4.698 | chr4 | 492999 | 533989 | + | protein_coding |
| ENSG00000107099.15_3 | DOCK8 | 0.46149342 | 1 | 0.395 | -0.802 | chr9 | 214854 | 465259 | + | protein_coding |
| ENSG00000119333.11_3 | WDR34 | 0.92331093 | 1 | 0.395 | 7.105 | chr9 | 131395940 | 131419066 | - | protein_coding |
| ENSG00000110047.18_5 | EHD1 | 0.93997776 | 1 | 0.395 | 6.172 | chr11 | 64619114 | 64655768 | - | protein_coding |
| ENSG00000157429.16_7 | ZNF19 | 0.71915522 | 1 | 0.395 | 1.926 | chr16 | 71498458 | 71598992 | - | protein_coding |
| ENSG00000105996.7_4 | HOXA2 | 0.58633915 | 1 | 0.395 | 0.838 | chr7 | 27139973 | 27142305 | - | protein_coding |
| ENSG00000047578.13_7 | KIAA0556 | 0.98650933 | 1 | 0.395 | 5.138 | chr16 | 27561454 | 27791690 | + | protein_coding |
| ENSG00000250536.1_5 | ABHD17AP3 | 0.4627645 | 1 | 0.395 | -0.727 | chr1 | 214778813 | 214782183 | - | unprocessed_pseudogene |
| ENSG00000237753.2_8 | AC079922.2 | 0.59023301 | 1 | 0.395 | 0.905 | chr2 | 113399409 | 113403297 | - | lncRNA |
| ENSG00000170959.14_6 | DCDC1 | 0.58848632 | 1 | 0.395 | 0.587 | chr11 | 30851916 | 31391357 | - | protein_coding |
| ENSG00000261693.1_6 | AC134682.1 | 0.50240982 | 1 | 0.394 | -0.066 | chr8 | 143485013 | 143488389 | + | lncRNA |

| | | | | | | | | | | |
|---|---|---|---|---|---|---|---|---|---|---|
| ENSG00000161642.17_4 | ZNF385A | 0.64913629 | 1 | 0.394 | 1.381 | chr12 | 54762917 | 54785082 | - | protein_coding |
| ENSG00000116863.11_3 | ADPRHL2 | 0.96048886 | 1 | 0.394 | 4.259 | chr1 | 36554493 | 36559533 | + | protein_coding |
| ENSG00000212747.5_6 | RTL8B | 0.85603185 | 1 | 0.394 | 3.223 | chrX | 134154543 | 134156572 | - | protein_coding |
| ENSG00000226287.8_4 | TMEM191A | 0.5270868 | 1 | 0.394 | 0.523 | chr22 | 21055402 | 21058894 | + | transcribed_unprocessed_pseudogene |
| ENSG00000181852.18_5 | RNF41 | 0.9870164 | 1 | 0.394 | 5.403 | chr12 | 56595963 | 56615717 | - | protein_coding |
| ENSG00000166192.15_5 | SENP8 | 0.70508407 | 1 | 0.394 | 1.664 | chr15 | 72406599 | 72436033 | + | protein_coding |
| ENSG00000204588.5_5 | LINC01123 | 0.47140023 | 1 | 0.394 | -0.369 | chr2 | 110744640 | 110753717 | + | lncRNA |
| ENSG00000137275.14_3 | RIPK1 | 0.9744096 | 1 | 0.394 | 4.780 | chr6 | 3064225 | 3115421 | + | protein_coding |
| ENSG00000174106.3_4 | LEMD3 | 0.98366988 | 1 | 0.394 | 5.477 | chr12 | 65563363 | 65642135 | + | protein_coding |
| ENSG00000066230.12_7 | SLC9A3 | 0.47224513 | 1 | 0.394 | -0.315 | chr5 | 470571 | 524564 | - | protein_coding |
| ENSG00000179627.10_4 | ZBTB42 | 0.83953763 | 1 | 0.394 | 2.687 | chr14 | 105266933 | 105271049 | + | protein_coding |
| ENSG00000154222.14_4 | CC2D1B | 0.98944883 | 1 | 0.393 | 5.659 | chr1 | 52811395 | 52831865 | - | protein_coding |
| ENSG00000150456.11_5 | EEF1AKMT1 | 0.7224165 | 1 | 0.393 | 1.973 | chr13 | 21302870 | 21348100 | - | protein_coding |
| ENSG00000213930.11_5 | GALT | 0.90881267 | 1 | 0.393 | 3.602 | chr9 | 34638130 | 34651032 | + | protein_coding |
| ENSG00000133597.11_7 | ADCK2 | 0.92712738 | 1 | 0.393 | 4.338 | chr7 | 140372745 | 140396061 | + | protein_coding |
| ENSG00000174485.15_5 | DENND4A | 0.98086605 | 1 | 0.393 | 5.738 | chr15 | 65950384 | 66084631 | - | protein_coding |
| ENSG00000226284.1_4 | ARPC3P1 | 0.47344734 | 1 | 0.392 | -0.474 | chr20 | 47751017 | 47751549 | + | processed_pseudogene |
| ENSG00000161847.14_6 | RAVER1 | 0.8734833 | 1 | 0.392 | 3.846 | chr19 | 10426888 | 10444314 | - | protein_coding |
| ENSG00000159921.16_5 | GNE | 0.99742569 | 1 | 0.392 | 6.751 | chr9 | 36214438 | 36277053 | - | protein_coding |
| ENSG00000229018.5_4 | PMS2P7 | 0.46904353 | 1 | 0.392 | -0.528 | chr7 | 72476064 | 72491632 | + | unprocessed_pseudogene |
| ENSG00000213062.5_6 | Z99572.1 | 0.63383479 | 1 | 0.392 | 0.834 | chr1 | 169455314 | 169469420 | + | lncRNA |
| ENSG00000264379.1 | SNORD39 | 0.47311723 | 1 | 0.391 | -0.475 | chr9 | 21524306 | 21524381 | - | snoRNA |
| ENSG00000172375.13_3 | C2CD2L | 0.8464645 | 1 | 0.391 | 3.087 | chr11 | 118972908 | 118989254 | + | protein_coding |
| ENSG00000135899.17_6 | SP110 | 0.80777844 | 1 | 0.391 | 2.427 | chr2 | 231032009 | 231090444 | - | protein_coding |
| ENSG00000107897.19_5 | ACBD5 | 0.9973478 | 1 | 0.391 | 6.617 | chr10 | 27484143 | 27531059 | - | protein_coding |
| ENSG00000072310.17_5 | SREBF1 | 0.93840689 | 1 | 0.391 | 8.491 | chr17 | 17713713 | 17740325 | - | protein_coding |
| ENSG00000165644.11_6 | COMTD1 | 0.78433692 | 1 | 0.391 | 3.795 | chr10 | 76993399 | 76995788 | - | protein_coding |
| ENSG00000157542.11_6 | KCNJ6 | 0.64149301 | 1 | 0.391 | 1.354 | chr21 | 38979675 | 39493439 | - | protein_coding |
| ENSG00000091136.14_5 | LAMB1 | 0.99928698 | 1 | 0.391 | 8.814 | chr7 | 107564244 | 107643632 | - | protein_coding |
| ENSG00000047056.16_5 | WDR37 | 0.9371519 | 1 | 0.391 | 3.986 | chr10 | 1095478 | 1178324 | + | protein_coding |
| ENSG00000253719.4_4 | ATXN7L3B | 0.99655952 | 1 | 0.391 | 6.174 | chr12 | 74931615 | 74939210 | + | protein_coding |
| ENSG00000151364.17_7 | KCTD14 | 0.47399135 | 1 | 0.390 | -0.577 | chr11 | 77726761 | 77757237 | - | protein_coding |
| ENSG00000080371.6_3 | RAB21 | 0.99664099 | 1 | 0.390 | 6.481 | chr12 | 72148643 | 72194066 | + | protein_coding |
| ENSG00000232450.1_5 | AL133517.1 | 0.47422715 | 1 | 0.390 | -0.347 | chr1 | 114241506 | 114242253 | - | transcribed_processed_pseudogene |
| ENSG00000049130.16_6 | KITLG | 0.98178287 | 1 | 0.390 | 4.982 | chr12 | 88886570 | 88974628 | - | protein_coding |
| ENSG00000196968.11_4 | FUT11 | 0.96830822 | 1 | 0.390 | 4.395 | chr10 | 75532034 | 75540009 | + | protein_coding |
| ENSG00000130734.10_3 | ATG4D | 0.74558304 | 1 | 0.390 | 2.747 | chr19 | 10654571 | 10664094 | + | protein_coding |
| ENSG00000230798.5_7 | FOXD3-AS1 | 0.56375126 | 1 | 0.390 | 0.554 | chr1 | 63786555 | 63790112 | - | lncRNA |
| ENSG00000069399.15_5 | BCL3 | 0.96347114 | 1 | 0.390 | 4.603 | chr19 | 45250962 | 45263301 | + | protein_coding |
| ENSG00000099785.10_5 | MARCHF2 | 0.9473546 | 1 | 0.389 | 4.209 | chr19 | 8478154 | 8503901 | + | protein_coding |
| ENSG00000176933.5_4 | TOB2P1 | 0.54577635 | 1 | 0.389 | 0.497 | chr6 | 28185421 | 28186412 | - | processed_pseudogene |
| ENSG00000158156.8_3 | XKR8 | 0.82803586 | 1 | 0.389 | 2.688 | chr1 | 28286484 | 28294604 | + | protein_coding |
| ENSG00000273576.1_6 | AC009283.1 | 0.64002946 | 1 | 0.389 | 1.333 | chr17 | 37723168 | 37723812 | + | lncRNA |
| ENSG00000260630.7_6 | SNAI3-AS1 | 0.79648564 | 1 | 0.389 | 2.470 | chr16 | 88729706 | 88753686 | + | lncRNA |
| ENSG00000133195.11_4 | SLC39A11 | 0.96677984 | 1 | 0.389 | 4.764 | chr17 | 70642088 | 71088851 | - | protein_coding |
| ENSG00000083444.17_4 | PLOD1 | 0.99527278 | 1 | 0.389 | 6.408 | chr1 | 11994262 | 12035595 | + | protein_coding |
| ENSG00000103534.17_6 | TMC5 | 0.97490866 | 1 | 0.388 | 4.918 | chr16 | 19421818 | 19510435 | + | protein_coding |
| ENSG00000147799.11_4 | ARHGAP39 | 0.89831842 | 1 | 0.388 | 4.236 | chr8 | 145754563 | 145831201 | - | protein_coding |
| ENSG00000183020.13_7 | AP2A2 | 0.99783635 | 1 | 0.388 | 6.830 | chr11 | 924894 | 1012245 | + | protein_coding |
| ENSG00000111652.10_3 | COPS7A | 0.99288804 | 1 | 0.388 | 5.637 | chr12 | 6833180 | 6841041 | + | protein_coding |
| ENSG00000082641.16_6 | NFE2L1 | 0.99871598 | 1 | 0.388 | 7.534 | chr17 | 46125691 | 46138907 | + | protein_coding |
| ENSG00000186198.4_3 | SLC51B | 0.63378062 | 1 | 0.388 | 1.389 | chr15 | 65337725 | 65345735 | + | protein_coding |
| ENSG00000197217.13_6 | ENTPD4 | 0.99765624 | 1 | 0.388 | 6.485 | chr8 | 23243296 | 23315208 | - | protein_coding |
| ENSG00000130958.13_5 | SLC35D2 | 0.80163488 | 1 | 0.388 | 2.346 | chr9 | 99075726 | 99145993 | - | protein_coding |
| ENSG00000174437.17_5 | ATP2A2 | 0.99900996 | 1 | 0.387 | 8.808 | chr12 | 110718561 | 110788898 | + | protein_coding |
| ENSG00000138463.9_3 | SLC49A4 | 0.97902149 | 1 | 0.387 | 5.011 | chr3 | 122513916 | 122599986 | + | protein_coding |
| ENSG00000172086.9_5 | KRCC1 | 0.96781524 | 1 | 0.387 | 4.809 | chr2 | 88326724 | 88363771 | - | protein_coding |
| ENSG00000112511.17_2 | PHF1 | 0.8773186 | 1 | 0.387 | 3.090 | chr6 | 33378176 | 33384230 | + | protein_coding |

| | | | | | | | | | | |
|---|---|---|---|---|---|---|---|---|---|---|
| ENSG00000126524.10_4 | SBDS | 0.98900862 | 1 | 0.386 | 5.481 | chr7 | 66452667 | 66460574 | - | protein_coding |
| ENSG00000145022.4_2 | TCTA | 0.94470498 | 1 | 0.386 | 3.776 | chr3 | 49449639 | 49453908 | + | protein_coding |
| ENSG00000116209.12_7 | TMEM59 | 0.99428933 | 1 | 0.386 | 6.207 | chr1 | 54492354 | 54519177 | - | protein_coding |
| ENSG00000108582.12_7 | CPD | 0.99894178 | 1 | 0.386 | 7.870 | chr17 | 28705945 | 28797007 | + | protein_coding |
| ENSG00000064961.19_4 | HMG20B | 0.9672259 | 1 | 0.386 | 5.243 | chr19 | 3572775 | 3579086 | + | protein_coding |
| ENSG00000177663.14_3 | IL17RA | 0.97486296 | 1 | 0.386 | 4.441 | chr22 | 17565844 | 17596584 | + | protein_coding |
| ENSG00000183955.13_6 | KMT5A | 0.9928651 | 1 | 0.386 | 5.636 | chr12 | 123868320 | 123893900 | + | protein_coding |
| ENSG00000201998.1 | SNORA23 | 0.5547782 | 1 | 0.386 | 0.284 | chr11 | 9450320 | 9450501 | + | snoRNA |
| ENSG00000158717.11_4 | RNF166 | 0.93468808 | 1 | 0.386 | 3.921 | chr16 | 88762907 | 88772816 | - | protein_coding |
| ENSG00000204673.10_5 | AKT1S1 | 0.98336131 | 1 | 0.386 | 5.017 | chr19 | 50372290 | 50381716 | - | protein_coding |
| ENSG00000229222.1_2 | KRT18P4 | 0.47285997 | 1 | 0.385 | -0.943 | chr20 | 48573282 | 48574569 | + | processed_pseudogene |
| ENSG00000176678.5_5 | FOXL1 | 0.58475996 | 1 | 0.385 | 0.875 | chr16 | 86609974 | 86615766 | + | protein_coding |
| ENSG00000176927.15_5 | EFCAB5 | 0.4824624 | 1 | 0.385 | -0.578 | chr17 | 28256218 | 28435470 | + | protein_coding |
| ENSG00000185730.8_4 | ZNF696 | 0.9321517 | 1 | 0.385 | 3.669 | chr8 | 144371846 | 144382122 | + | protein_coding |
| ENSG00000155592.16_4 | ZKSCAN2 | 0.9587914 | 1 | 0.385 | 4.001 | chr16 | 25247322 | 25269166 | - | protein_coding |
| ENSG00000163602.10_5 | RYBP | 0.98289086 | 1 | 0.385 | 5.343 | chr3 | 72420976 | 72495772 | - | protein_coding |
| ENSG00000171984.15_5 | SHLD1 | 0.8341551 | 1 | 0.384 | 2.480 | chr20 | 5731039 | 5845041 | + | protein_coding |
| ENSG00000127124.16_6 | HIVEP3 | 0.72734381 | 1 | 0.384 | 1.852 | chr1 | 41972036 | 42501596 | - | protein_coding |
| ENSG00000253210.2_7 | AC040970.1 | 0.79510442 | 1 | 0.384 | 2.195 | chr8 | 142134816 | 142140163 | - | lncRNA |
| ENSG00000254635.6_7 | WAC-AS1 | 0.96893212 | 1 | 0.384 | 4.415 | chr10 | 28811581 | 28821995 | - | lncRNA |
| ENSG00000081923.14_8 | ATP8B1 | 0.99386421 | 1 | 0.384 | 5.543 | chr18 | 55313658 | 55470547 | - | protein_coding |
| ENSG00000233006.8_7 | MIR3936HG | 0.88515266 | 1 | 0.384 | 2.988 | chr5 | 131646978 | 131705862 | - | lncRNA |
| ENSG00000135709.12_3 | KIAA0513 | 0.92815312 | 1 | 0.383 | 3.594 | chr16 | 85061357 | 85127836 | + | protein_coding |
| ENSG00000259366.1_7 | AC108449.2 | 0.52195709 | 1 | 0.383 | 0.269 | chr8 | 28913452 | 28914202 | + | lncRNA |
| ENSG00000118004.18_3 | COLEC11 | 0.47841222 | 1 | 0.383 | -0.771 | chr2 | 3642422 | 3692234 | + | protein_coding |
| ENSG00000270885.2_3 | RASL10B | 0.48616434 | 1 | 0.383 | -1.086 | chr17 | 34058658 | 34070540 | + | protein_coding |
| ENSG00000278356.1_7 | AC005911.1 | 0.47700652 | 1 | 0.383 | -0.894 | chr12 | 2994985 | 2995495 | + | lncRNA |
| ENSG00000127955.17_5 | GNAI1 | 0.99663077 | 1 | 0.383 | 6.353 | chr7 | 79397344 | 79855497 | + | protein_coding |
| ENSG00000105173.14_5 | CCNE1 | 0.98192573 | 1 | 0.383 | 4.776 | chr19 | 30302898 | 30315219 | + | protein_coding |
| ENSG00000105419.18_6 | MEIS3 | 0.97356965 | 1 | 0.383 | 4.751 | chr19 | 47906381 | 47922784 | - | protein_coding |
| ENSG00000275379.1_3 | H3C11 | 0.97541614 | 1 | 0.383 | 5.193 | chr6 | 27839683 | 27840093 | - | protein_coding |
| ENSG00000100726.15_3 | TELO2 | 0.84586966 | 1 | 0.383 | 5.294 | chr16 | 1543345 | 1560458 | + | protein_coding |
| ENSG00000272087.1_6 | AC080013.4 | 0.54352759 | 1 | 0.382 | 0.047 | chr3 | 158410909 | 158411557 | - | lncRNA |
| ENSG00000249709.8_5 | ZNF564 | 0.47601163 | 1 | 0.382 | -1.085 | chr19 | 12636187 | 12662356 | - | protein_coding |
| ENSG00000110048.12_4 | OSBP | 0.99736537 | 1 | 0.382 | 6.182 | chr11 | 59341871 | 59383247 | - | protein_coding |
| ENSG00000231105.2_7 | ECE1-AS1 | 0.48947941 | 1 | 0.382 | -0.341 | chr1 | 21619783 | 21626367 | + | lncRNA |
| ENSG00000156853.13_5 | ZNF689 | 0.94794272 | 1 | 0.381 | 4.167 | chr16 | 30613879 | 30635333 | - | protein_coding |
| ENSG00000111912.20_8 | NCOA7 | 0.99824847 | 1 | 0.381 | 6.916 | chr6 | 126102307 | 126253180 | + | protein_coding |
| ENSG00000287236.1_2 | Z73420.2 | 0.48531928 | 1 | 0.381 | -0.617 | chr22 | 37429165 | 37431553 | + | lncRNA |
| ENSG00000142188.17_5 | TMEM50B | 0.96484373 | 1 | 0.381 | 4.318 | chr21 | 34804792 | 34852281 | - | protein_coding |
| ENSG00000205913.7_7 | SRRM2-AS1 | 0.80151902 | 1 | 0.381 | 2.176 | chr16 | 2787077 | 2802601 | - | lncRNA |
| ENSG00000259549.1 | RP11-115K3.1 | 0.58303345 | 1 | 0.381 | 0.714 | chr17 | 36105206 | 36108563 | + | antisense |
| ENSG00000128335.14_5 | APOL2 | 0.95235111 | 1 | 0.381 | 3.927 | chr22 | 36622255 | 36636000 | - | protein_coding |
| ENSG00000186918.14_6 | ZNF395 | 0.97979946 | 1 | 0.381 | 4.545 | chr8 | 28203107 | 28260218 | - | protein_coding |
| ENSG00000124155.18_7 | PIGT | 0.9975951 | 1 | 0.381 | 6.507 | chr20 | 44044724 | 44085574 | + | protein_coding |
| ENSG00000125503.13_5 | PPP1R12C | 0.88949998 | 1 | 0.380 | 6.212 | chr19 | 55602282 | 55629005 | - | protein_coding |
| ENSG00000105298.14_5 | CACTIN | 0.93309523 | 1 | 0.380 | 4.034 | chr19 | 3610643 | 3626813 | - | protein_coding |
| ENSG00000029534.20_8 | ANK1 | 0.96011905 | 1 | 0.380 | 4.169 | chr8 | 41510739 | 41754280 | - | protein_coding |
| ENSG00000197774.13_5 | EME2 | 0.97771176 | 1 | 0.380 | 4.846 | chr16 | 1822811 | 1831709 | + | protein_coding |
| ENSG00000159917.17_6 | ZNF235 | 0.91760879 | 1 | 0.380 | 3.456 | chr19 | 44732882 | 44809199 | - | protein_coding |
| ENSG00000287595.1_2 | AC097369.2 | 0.47979913 | 1 | 0.380 | -0.849 | chr3 | 72788069 | 72789676 | + | lncRNA |
| ENSG00000233901.6_6 | LINC01503 | 0.6393874 | 1 | 0.380 | 1.276 | chr9 | 132094579 | 132121820 | + | lncRNA |
| ENSG00000142751.15_4 | GPN2 | 0.97344423 | 1 | 0.380 | 4.392 | chr1 | 27202623 | 27216774 | - | protein_coding |
| ENSG00000053702.15_4 | NRIP2 | 0.6654583 | 1 | 0.380 | 1.386 | chr12 | 2934514 | 2944710 | - | protein_coding |
| ENSG00000167037.18_3 | SGSM1 | 0.73689902 | 1 | 0.380 | 1.757 | chr22 | 25202136 | 25323545 | + | protein_coding |
| ENSG00000079482.13_5 | OPHN1 | 0.93790304 | 1 | 0.380 | 4.061 | chrX | 67262186 | 67653755 | - | protein_coding |
| ENSG00000214176.9_6 | PLEKHM1P1 | 0.95972113 | 1 | 0.379 | 4.211 | chr17 | 62775377 | 62833272 | - | transcribed_unprocessed_pseudogene |
| ENSG00000102531.16_5 | FNDC3A | 0.99755323 | 1 | 0.379 | 6.134 | chr13 | 49550048 | 49783915 | + | protein_coding |
| ENSG00000062716.13_4 | VMP1 | 0.99856799 | 1 | 0.379 | 9.749 | chr17 | 57784553 | 57919616 | + | protein_coding |

| | | | | | | | | | | |
|---|---|---|---|---|---|---|---|---|---|---|
| ENSG00000203697.12_6 | CAPN8 | 0.50982855 | 1 | 0.379 | -0.208 | chr1 | 223711349 | 223853425 | - | protein_coding |
| ENSG00000058272.19_7 | PPP1R12A | 0.99757572 | 1 | 0.379 | 7.144 | chr12 | 80167343 | 80329240 | - | protein_coding |
| ENSG00000139289.13_4 | PHLDA1 | 0.99823068 | 1 | 0.379 | 6.494 | chr12 | 76419227 | 76427712 | - | protein_coding |
| ENSG00000122642.11_4 | FKBP9 | 0.99830983 | 1 | 0.379 | 7.004 | chr7 | 32997016 | 33046542 | + | protein_coding |
| ENSG00000119431.9_3 | HDHD3 | 0.95663992 | 1 | 0.379 | 4.176 | chr9 | 116135699 | 116139279 | - | protein_coding |
| ENSG00000188554.14_3 | NBR1 | 0.99914809 | 1 | 0.379 | 7.202 | chr17 | 41322498 | 41363708 | + | protein_coding |
| ENSG00000260329.1_6 | AC007541.1 | 0.61657858 | 1 | 0.379 | 0.775 | chr12 | 107347807 | 107349275 | - | lncRNA |
| ENSG00000159556.10_3 | ISL2 | 0.66195538 | 1 | 0.379 | 1.304 | chr15 | 76629114 | 76634816 | + | protein_coding |
| ENSG00000088766.12_5 | CRLS1 | 0.69471471 | 1 | 0.379 | 1.720 | chr20 | 5986739 | 6020699 | + | protein_coding |
| ENSG00000185670.9_8 | ZBTB3 | 0.88231409 | 1 | 0.379 | 2.913 | chr11 | 62515791 | 62521656 | - | protein_coding |
| ENSG00000171302.17_7 | CANT1 | 0.99834539 | 1 | 0.379 | 6.472 | chr17 | 76987798 | 77005949 | - | protein_coding |
| ENSG00000160188.10_4 | RSPH1 | 0.48118497 | 1 | 0.378 | -0.896 | chr21 | 43892596 | 43916356 | - | protein_coding |
| ENSG00000153443.13_4 | UBALD1 | 0.95097899 | 1 | 0.378 | 4.161 | chr16 | 4658884 | 4665028 | - | protein_coding |
| ENSG00000134899.21_6 | ERCC5 | 0.68256289 | 1 | 0.378 | 1.290 | chr13 | 103459496 | 103528351 | + | protein_coding |
| ENSG00000172602.11_6 | RND1 | 0.88892679 | 1 | 0.378 | 3.166 | chr12 | 49250928 | 49259653 | - | protein_coding |
| ENSG00000066084.13_3 | DIP2B | 0.99877528 | 1 | 0.378 | 6.561 | chr12 | 50898768 | 51142440 | + | protein_coding |
| ENSG00000266019.1 | MIR3609 | 0.85281334 | 1 | 0.378 | 3.413 | chr7 | 98479273 | 98479352 | + | miRNA |
| ENSG00000122359.18_6 | ANXA11 | 0.99863696 | 1 | 0.377 | 7.519 | chr10 | 81910645 | 81965328 | - | protein_coding |
| ENSG00000178082.6_4 | TWF1P1 | 0.7213874 | 1 | 0.377 | 1.560 | chr17 | 27530444 | 27531492 | - | processed_pseudogene |
| ENSG00000178919.9_3 | FOXE1 | 0.93012536 | 1 | 0.377 | 3.606 | chr9 | 100615508 | 100618999 | + | protein_coding |
| ENSG00000174456.14_5 | C12orf76 | 0.85823624 | 1 | 0.377 | 2.880 | chr12 | 110464833 | 110511439 | - | protein_coding |
| ENSG00000100600.15_5 | LGMN | 0.99702286 | 1 | 0.377 | 6.108 | chr14 | 93170152 | 93215024 | - | protein_coding |
| ENSG00000154447.15_5 | SH3RF1 | 0.97189957 | 1 | 0.377 | 4.758 | chr4 | 170015410 | 170192107 | - | protein_coding |
| ENSG00000259895.1_6 | AC106820.2 | 0.62765164 | 1 | 0.376 | 0.921 | chr16 | 2514951 | 2518214 | - | lncRNA |
| ENSG00000099910.17_5 | KLHL22 | 0.95610282 | 1 | 0.376 | 4.033 | chr22 | 20795806 | 20850131 | - | protein_coding |
| ENSG00000198663.16_4 | C6orf89 | 0.99740319 | 1 | 0.376 | 5.942 | chr6 | 36839646 | 36896740 | + | protein_coding |
| ENSG00000278642.1_6 | AC015813.4 | 0.48715313 | 1 | 0.376 | -0.770 | chr17 | 56033326 | 56033504 | + | lncRNA |
| ENSG00000197971.16_8 | MBP | 0.99565976 | 1 | 0.376 | 5.859 | chr18 | 74690783 | 74845639 | - | protein_coding |
| ENSG00000123384.14_4 | LRP1 | 0.99110749 | 1 | 0.376 | 5.673 | chr12 | 57522266 | 57607144 | + | protein_coding |
| ENSG00000234311.1_7 | AL451069.3 | 0.50913712 | 1 | 0.376 | -0.416 | chr10 | 134247237 | 134254988 | - | lncRNA |
| ENSG00000205356.10_5 | TECPR1 | 0.93511993 | 1 | 0.376 | 4.538 | chr7 | 97843936 | 97881544 | - | protein_coding |
| ENSG00000134278.15_4 | SPIRE1 | 0.97808986 | 1 | 0.376 | 4.516 | chr18 | 12446511 | 12658133 | - | protein_coding |
| ENSG00000213904.9_6 | LIPE-AS1 | 0.50232896 | 1 | 0.376 | -0.326 | chr19 | 42901280 | 43156507 | + | lncRNA |
| ENSG00000147144.13_6 | CCDC120 | 0.78355245 | 1 | 0.375 | 2.244 | chrX | 48911101 | 48927509 | + | protein_coding |
| ENSG00000099624.8_5 | ATP5F1D | 0.67124979 | 1 | 0.375 | 3.647 | chr19 | 1241745 | 1244824 | + | protein_coding |
| ENSG00000140545.15_5 | MFGE8 | 0.99412041 | 1 | 0.375 | 5.585 | chr15 | 89441914 | 89456612 | - | protein_coding |
| ENSG00000111678.11_7 | C12orf57 | 0.97350987 | 1 | 0.375 | 4.268 | chr12 | 7052141 | 7055166 | + | protein_coding |
| ENSG00000114646.10_6 | CSPG5 | 0.58935753 | 1 | 0.375 | 1.015 | chr3 | 47603728 | 47622282 | - | protein_coding |
| ENSG00000130382.9_5 | MLLT1 | 0.98118936 | 1 | 0.375 | 5.158 | chr19 | 6210392 | 6279986 | - | protein_coding |
| ENSG00000169239.13_4 | CA5B | 0.89521858 | 1 | 0.375 | 3.089 | chrX | 15706953 | 15806534 | + | protein_coding |
| ENSG00000256745.1_5 | AP002784.2 | 0.62481013 | 1 | 0.375 | 0.981 | chr11 | 93921615 | 93922163 | - | processed_pseudogene |
| ENSG00000197980.13_4 | LEKR1 | 0.60605239 | 1 | 0.374 | 0.580 | chr3 | 156543270 | 156763918 | + | protein_coding |
| ENSG00000181481.14_4 | RNF135 | 0.99148886 | 1 | 0.374 | 5.036 | chr17 | 29298002 | 29326929 | + | protein_coding |
| ENSG00000100558.9_5 | PLEK2 | 0.95377284 | 1 | 0.374 | 3.726 | chr14 | 67853701 | 67878884 | - | protein_coding |
| ENSG00000116406.19_7 | EDEM3 | 0.99686843 | 1 | 0.374 | 6.022 | chr1 | 184659371 | 184724041 | - | protein_coding |
| ENSG00000087470.18_7 | DNM1L | 0.99921319 | 1 | 0.374 | 7.403 | chr12 | 32832134 | 32898584 | + | protein_coding |
| ENSG00000181029.9_6 | TRAPPC5 | 0.6316437 | 1 | 0.373 | 0.623 | chr19 | 7745719 | 7752589 | + | protein_coding |
| ENSG00000169919.17_4 | GUSB | 0.99196373 | 1 | 0.373 | 5.165 | chr7 | 65425671 | 65447202 | - | protein_coding |
| ENSG00000163521.16_5 | GLB1L | 0.87841897 | 1 | 0.373 | 2.885 | chr2 | 220101320 | 220110200 | - | protein_coding |
| ENSG00000207827.1 | MIR30A | 0.52200814 | 1 | 0.373 | -0.056 | chr6 | 72113254 | 72113324 | - | miRNA |
| ENSG00000114779.19_6 | ABHD14B | 0.97709377 | 1 | 0.373 | 4.297 | chr3 | 52002526 | 52017425 | - | protein_coding |
| ENSG00000163762.7_5 | TM4SF18 | 0.99146547 | 1 | 0.373 | 6.196 | chr3 | 149036285 | 149052201 | - | protein_coding |
| ENSG00000167522.16_8 | ANKRD11 | 0.99764956 | 1 | 0.373 | 6.942 | chr16 | 89334038 | 89556969 | - | protein_coding |
| ENSG00000257337.7_7 | AC068888.1 | 0.85936656 | 1 | 0.372 | 2.840 | chr12 | 53406668 | 53448234 | - | lncRNA |
| ENSG00000130779.20_7 | CLIP1 | 0.99947162 | 1 | 0.372 | 7.692 | chr12 | 122755979 | 122907179 | - | protein_coding |
| ENSG00000261773.1_7 | AC244090.3 | 0.49007851 | 1 | 0.372 | -0.816 | chrX | 153746164 | 153746955 | - | lncRNA |
| ENSG00000183250.12_6 | LINC01547 | 0.87101593 | 1 | 0.372 | 2.898 | chr21 | 46352729 | 46359852 | - | lncRNA |
| ENSG00000248334.6_2 | WHAMMP2 | 0.86911216 | 1 | 0.372 | 2.619 | chr15 | 28982729 | 29004350 | + | transcribed_unprocessed_pseudogene |
| ENSG00000269378.1_5 | AC022149.1 | 0.9432659 | 1 | 0.372 | 4.033 | chr19 | 14732446 | 14733054 | - | processed_pseudogene |

| | | | | | | | | | | |
|---|---|---|---|---|---|---|---|---|---|---|
| ENSG00000250286.2_7 | AC021491.2 | 0.59513328 | 1 | 0.372 | 0.581 | chr17 | 48602881 | 48616122 | - | lncRNA |
| ENSG00000132824.14_4 | SERINC3 | 0.99906697 | 1 | 0.372 | 7.222 | chr20 | 43124862 | 43150726 | - | protein_coding |
| ENSG00000167778.9_4 | SPRYD3 | 0.98759356 | 1 | 0.372 | 4.753 | chr12 | 53458100 | 53473188 | - | protein_coding |
| ENSG00000186409.17_8 | CCDC30 | 0.77597966 | 1 | 0.372 | 1.519 | chr1 | 42928892 | 43122861 | + | protein_coding |
| ENSG00000050555.18_5 | LAMC3 | 0.8493636 | 1 | 0.372 | 2.621 | chr9 | 133884561 | 133969860 | + | protein_coding |
| ENSG00000102401.20_5 | ARMCX3 | 0.98985401 | 1 | 0.371 | 4.752 | chrX | 100877787 | 100882833 | + | protein_coding |
| ENSG00000205560.12_4 | CPT1B | 0.52129234 | 1 | 0.371 | 0.076 | chr22 | 51007290 | 51016894 | - | protein_coding |
| ENSG00000214309.5_4 | MBLAC1 | 0.66137961 | 1 | 0.371 | 0.837 | chr7 | 99724408 | 99726118 | + | protein_coding |
| ENSG00000159692.15_7 | CTBP1 | 0.99497214 | 1 | 0.371 | 6.071 | chr4 | 1205236 | 1243741 | - | protein_coding |
| ENSG00000175782.11_5 | SLC35E3 | 0.98774368 | 1 | 0.371 | 4.807 | chr12 | 69139905 | 69187744 | + | protein_coding |
| ENSG00000156860.16_8 | FBRS | 0.91645646 | 1 | 0.371 | 5.479 | chr16 | 30669752 | 30682131 | + | protein_coding |
| ENSG00000232527.3 | RP11-14N7.2 | 0.52216097 | 1 | 0.371 | -0.327 | chr1 | 148928322 | 148951595 | + | lincRNA |
| ENSG00000131061.14_4 | ZNF341 | 0.86889776 | 1 | 0.371 | 2.595 | chr20 | 32319463 | 32380075 | + | protein_coding |
| ENSG00000111640.15_4 | GAPDH | 0.9992343 | 1 | 0.371 | 11.037 | chr12 | 6643678 | 6647540 | + | protein_coding |
| ENSG00000165092.13_5 | ALDH1A1 | 0.99964812 | 1 | 0.371 | 13.009 | chr9 | 75515587 | 75695358 | - | protein_coding |
| ENSG00000214226.9_5 | C17orf67 | 0.67072174 | 1 | 0.370 | 2.068 | chr17 | 54869274 | 54916134 | - | protein_coding |
| ENSG00000158457.6_3 | TSPAN33 | 0.93796389 | 1 | 0.370 | 3.434 | chr7 | 128784724 | 128809540 | + | protein_coding |
| ENSG00000145214.14_6 | DGKQ | 0.96047961 | 1 | 0.370 | 4.315 | chr4 | 952675 | 980683 | - | protein_coding |
| ENSG00000197816.15_5 | CCDC180 | 0.62660995 | 1 | 0.370 | 0.542 | chr9 | 100069586 | 100141033 | + | protein_coding |
| ENSG00000275066.5_8 | SYNRG | 0.99278004 | 1 | 0.370 | 5.585 | chr17 | 35874910 | 35969520 | - | protein_coding |
| ENSG00000267317.2_6 | AC027307.2 | 0.66531212 | 1 | 0.370 | 1.024 | chr19 | 1457669 | 1458579 | - | lncRNA |
| ENSG00000171680.22_5 | PLEKHG5 | 0.87905193 | 1 | 0.369 | 2.698 | chr1 | 6526152 | 6580121 | - | protein_coding |
| ENSG00000105568.18_5 | PPP2R1A | 0.99893152 | 1 | 0.369 | 7.803 | chr19 | 52693301 | 52732771 | + | protein_coding |
| ENSG00000171443.7_3 | ZNF524 | 0.78100663 | 1 | 0.369 | 3.043 | chr19 | 56111388 | 56114504 | + | protein_coding |
| ENSG00000196507.11_4 | TCEAL3 | 0.87549164 | 1 | 0.369 | 2.229 | chrX | 102862379 | 102884618 | + | protein_coding |
| ENSG00000067836.12_7 | ROGDI | 0.79354095 | 1 | 0.369 | 2.350 | chr16 | 4846969 | 4852951 | - | protein_coding |
| ENSG00000184634.16_5 | MED12 | 0.99712312 | 1 | 0.369 | 5.640 | chrX | 70338406 | 70362304 | + | protein_coding |
| ENSG00000166532.16_5 | RIMKLB | 0.99657614 | 1 | 0.369 | 5.741 | chr12 | 8834196 | 8935691 | + | protein_coding |
| ENSG00000170456.16_5 | DENND5B | 0.99872581 | 1 | 0.369 | 6.962 | chr12 | 31535160 | 31744070 | - | protein_coding |
| ENSG00000063761.16_5 | ADCK1 | 0.87503237 | 1 | 0.369 | 2.920 | chr14 | 78266452 | 78401357 | + | protein_coding |
| ENSG00000270362.1_6 | HMGN3-AS1 | 0.58247649 | 1 | 0.368 | 0.354 | chr6 | 79943435 | 79946514 | + | lncRNA |
| ENSG00000100092.23_7 | SH3BP1 | 0.59802195 | 1 | 0.368 | 0.527 | chr22 | 38030661 | 38052126 | + | protein_coding |
| ENSG00000186301.8_2 | MST1P2 | 0.7079059 | 1 | 0.368 | 1.329 | chr1 | 16972117 | 16976784 | + | unprocessed_pseudogene |
| ENSG00000165474.8_7 | GJB2 | 0.52841193 | 1 | 0.368 | -0.153 | chr13 | 20761602 | 20767077 | - | protein_coding |
| ENSG00000163795.14_7 | ZNF513 | 0.9479279 | 1 | 0.368 | 3.655 | chr2 | 27600102 | 27603657 | - | protein_coding |
| ENSG00000221995.5_3 | TIAF1 | 0.71002805 | 1 | 0.368 | 1.490 | chr17 | 27400539 | 27405875 | - | protein_coding |
| ENSG00000151239.13_4 | TWF1 | 0.99827294 | 1 | 0.368 | 7.480 | chr12 | 44187526 | 44200178 | - | protein_coding |
| ENSG00000013503.10_3 | POLR3B | 0.96373314 | 1 | 0.368 | 3.897 | chr12 | 106751526 | 106903976 | + | protein_coding |
| ENSG00000167653.5_5 | PSCA | 0.52645492 | 1 | 0.367 | -0.358 | chr8 | 143751726 | 143764143 | + | protein_coding |
| ENSG00000159214.13_4 | CCDC24 | 0.92566859 | 1 | 0.367 | 3.252 | chr1 | 44457031 | 44462200 | + | protein_coding |
| ENSG00000232043.1_7 | AL133230.2 | 0.64101036 | 1 | 0.367 | 0.667 | chr20 | 49187512 | 49194578 | - | lncRNA |
| ENSG00000183208.12_6 | GDPGP1 | 0.74730198 | 1 | 0.367 | 2.240 | chr15 | 90777040 | 90789043 | + | protein_coding |
| ENSG00000206897.1 | SNORA9 | 0.49561713 | 1 | 0.367 | -0.945 | chr12 | 124101255 | 124101387 | + | snoRNA |
| ENSG00000213465.8_4 | ARL2 | 0.93042592 | 1 | 0.367 | 4.561 | chr11 | 64781632 | 64789656 | + | protein_coding |
| ENSG00000084444.14_4 | FAM234B | 0.92696772 | 1 | 0.367 | 3.209 | chr12 | 13197315 | 13295455 | + | protein_coding |
| ENSG00000118762.8_3 | PKD2 | 0.99871323 | 1 | 0.367 | 6.454 | chr4 | 88928787 | 88998929 | + | protein_coding |
| ENSG00000137474.22_5 | MYO7A | 0.88817813 | 1 | 0.367 | 3.078 | chr11 | 76839292 | 76926286 | + | protein_coding |
| ENSG00000178401.16_4 | DNAJC22 | 0.99807865 | 1 | 0.367 | 5.965 | chr12 | 49740671 | 49751329 | + | protein_coding |
| ENSG00000142039.4_4 | CCDC97 | 0.99065509 | 1 | 0.367 | 5.013 | chr19 | 41816077 | 41830778 | + | protein_coding |
| ENSG00000280828.1_6 | AC090114.3 | 0.52033138 | 1 | 0.367 | -0.106 | chr7 | 128173706 | 128256024 | + | unprocessed_pseudogene |
| ENSG00000166866.13_5 | MYO1A | 0.49647193 | 1 | 0.367 | -0.815 | chr12 | 57422301 | 57444982 | - | protein_coding |
| ENSG00000179909.15_3 | ZNF154 | 0.558337 | 1 | 0.367 | 0.822 | chr19 | 58208735 | 58220562 | - | protein_coding |
| ENSG00000028277.21_6 | POU2F2 | 0.51342462 | 1 | 0.367 | -0.453 | chr19 | 42590262 | 42700737 | - | protein_coding |
| ENSG00000004700.16_7 | RECQL | 0.99888808 | 1 | 0.367 | 6.576 | chr12 | 21621844 | 21654603 | - | protein_coding |
| ENSG00000245910.8_5 | SNHG6 | 0.98503397 | 1 | 0.366 | 4.543 | chr8 | 67833919 | 67838633 | - | lncRNA |
| ENSG00000167972.14_6 | ABCA3 | 0.99760021 | 1 | 0.366 | 6.216 | chr16 | 2325882 | 2390747 | - | protein_coding |
| ENSG00000125347.14_8 | IRF1 | 0.9519615 | 1 | 0.366 | 3.749 | chr5 | 131817301 | 131826469 | - | protein_coding |
| ENSG00000157379.14_4 | DHRS1 | 0.91030096 | 1 | 0.366 | 3.185 | chr14 | 24759804 | 24768986 | - | protein_coding |
| ENSG00000149328.15_3 | GLB1L2 | 0.82083009 | 1 | 0.366 | 2.128 | chr11 | 134201768 | 134248235 | + | protein_coding |
| ENSG00000111670.16_6 | GNPTAB | 0.9973949 | 1 | 0.366 | 5.952 | chr12 | 102139277 | 102224737 | - | protein_coding |

| | | | | | | | | | | |
|---|---|---|---|---|---|---|---|---|---|---|
| ENSG00000204438.11_6 | GPANK1 | 0.65355718 | 1 | 0.366 | 1.010 | chr6 | 31629005 | 31634060 | - | protein_coding |
| ENSG00000158813.18_3 | EDA | 0.49992314 | 1 | 0.365 | -0.817 | chrX | 68835911 | 69259322 | + | protein_coding |
| ENSG00000103274.11_4 | NUBP1 | 0.92095472 | 1 | 0.365 | 3.348 | chr16 | 10837643 | 10863208 | + | protein_coding |
| ENSG00000261840.3_7 | AC093249.6 | 0.55351831 | 1 | 0.365 | -0.031 | chr16 | 30709028 | 30710379 | - | lncRNA |
| ENSG00000177508.12_4 | IRX3 | 0.69691533 | 1 | 0.365 | 1.358 | chr16 | 54317216 | 54320699 | - | protein_coding |
| ENSG00000257279.1_7 | AC127164.1 | 0.54210857 | 1 | 0.365 | -0.059 | chr12 | 117415247 | 117425142 | - | lncRNA |
| ENSG00000157259.8_5 | GATAD1 | 0.9948194 | 1 | 0.364 | 5.580 | chr7 | 92076796 | 92089389 | + | protein_coding |
| ENSG00000174282.12_4 | ZBTB4 | 0.99868281 | 1 | 0.364 | 6.152 | chr17 | 7362685 | 7387582 | - | protein_coding |
| ENSG00000144747.17_6 | TMF1 | 0.9967226 | 1 | 0.364 | 5.692 | chr3 | 69068978 | 69101490 | - | protein_coding |
| ENSG00000249459.10_7 | ZNF286B | 0.70329317 | 1 | 0.364 | 1.090 | chr17 | 18561742 | 18585541 | - | transcribed_unprocessed_pseudogene |
| ENSG00000095397.15_7 | WHRN | 0.80248608 | 1 | 0.364 | 1.681 | chr9 | 117164360 | 117267753 | - | protein_coding |
| ENSG00000122965.11_6 | RBM19 | 0.98165287 | 1 | 0.364 | 5.582 | chr12 | 114254543 | 114404130 | - | protein_coding |
| ENSG00000099326.9_6 | MZF1 | 0.99362813 | 1 | 0.363 | 5.228 | chr19 | 59073298 | 59084942 | - | protein_coding |
| ENSG00000164050.13_4 | PLXNB1 | 0.9933185 | 1 | 0.363 | 5.306 | chr3 | 48445261 | 48471495 | - | protein_coding |
| ENSG00000129103.18_4 | SUMF2 | 0.99880833 | 1 | 0.363 | 6.326 | chr7 | 56131695 | 56148363 | + | protein_coding |
| ENSG00000234567.1_7 | AL024497.2 | 0.50396829 | 1 | 0.363 | -0.863 | chr6 | 133773995 | 133777743 | - | lncRNA |
| ENSG00000167384.10_3 | ZNF180 | 0.99347968 | 1 | 0.363 | 5.024 | chr19 | 44978645 | 45004576 | - | protein_coding |
| ENSG00000101400.6_3 | SNTA1 | 0.93374284 | 1 | 0.363 | 3.471 | chr20 | 31995763 | 32031569 | - | protein_coding |
| ENSG00000185245.8_3 | GP1BA | 0.67662458 | 1 | 0.363 | 1.100 | chr17 | 4835572 | 4838318 | + | protein_coding |
| ENSG00000255182.2_8 | AC084125.2 | 0.78713393 | 1 | 0.362 | 1.799 | chr8 | 145720841 | 145730827 | - | lncRNA |
| ENSG00000136100.14_5 | VPS36 | 0.99628065 | 1 | 0.362 | 5.341 | chr13 | 52986737 | 53024769 | - | protein_coding |
| ENSG00000089009.15_5 | RPL6 | 0.99980496 | 1 | 0.362 | 9.584 | chr12 | 112842994 | 112856642 | - | protein_coding |
| ENSG00000092295.12_5 | TGM1 | 0.5730843 | 1 | 0.362 | 0.381 | chr14 | 24718320 | 24733638 | - | protein_coding |
| ENSG00000139719.11_8 | VPS33A | 0.97732225 | 1 | 0.362 | 4.114 | chr12 | 122714111 | 122751041 | - | protein_coding |
| ENSG00000090971.5_3 | NAT14 | 0.96969029 | 1 | 0.362 | 4.305 | chr19 | 55996555 | 55998935 | + | protein_coding |
| ENSG00000278949.1_6 | AC127070.4 | 0.53090706 | 1 | 0.362 | -0.085 | chr12 | 133442441 | 133444544 | - | TEC |
| ENSG00000170166.6_4 | HOXD4 | 0.50712527 | 1 | 0.362 | -0.742 | chr2 | 177016278 | 177017954 | + | protein_coding |
| ENSG00000123106.10_8 | CCDC91 | 0.99264854 | 1 | 0.362 | 4.924 | chr12 | 28286182 | 28734444 | + | protein_coding |
| ENSG00000143786.8_4 | CNIH3 | 0.52981946 | 1 | 0.362 | -0.244 | chr1 | 224622362 | 224928256 | + | protein_coding |
| ENSG00000268089.3_4 | GABRQ | 0.53634595 | 1 | 0.362 | -0.211 | chrX | 151806349 | 151826007 | + | protein_coding |
| ENSG00000283930.1_5 | AL117339.5 | 0.68982198 | 1 | 0.361 | 0.922 | chr10 | 38383296 | 38649026 | + | protein_coding |
| ENSG00000279692.1_6 | AC110285.6 | 0.53615464 | 1 | 0.361 | -0.079 | chr17 | 79369409 | 79370944 | - | TEC |
| ENSG00000185686.18_8 | PRAME | 0.99960957 | 1 | 0.361 | 7.546 | chr22 | 22890123 | 22901768 | - | protein_coding |
| ENSG00000197165.11_4 | SULT1A2 | 0.51643786 | 1 | 0.361 | -0.590 | chr16 | 28603264 | 28608371 | - | protein_coding |
| ENSG00000162236.12_5 | STX5 | 0.98653041 | 1 | 0.361 | 4.484 | chr11 | 62574332 | 62599523 | - | protein_coding |
| ENSG00000133424.20_7 | LARGE1 | 0.96848433 | 1 | 0.361 | 4.247 | chr22 | 33558212 | 34318829 | - | protein_coding |
| ENSG00000151348.14_4 | EXT2 | 0.99901605 | 1 | 0.361 | 6.897 | chr11 | 44117223 | 44273531 | + | protein_coding |
| ENSG00000149428.19_8 | HYOU1 | 0.99948595 | 1 | 0.361 | 6.906 | chr11 | 118914899 | 118927938 | - | protein_coding |
| ENSG00000242498.8_5 | ARPIN | 0.9952388 | 1 | 0.361 | 5.460 | chr15 | 90438238 | 90456184 | - | protein_coding |
| ENSG00000106049.9_4 | HIBADH | 0.99837358 | 1 | 0.361 | 5.895 | chr7 | 27565061 | 27702502 | - | protein_coding |
| ENSG00000111325.16_5 | OGFOD2 | 0.70514482 | 1 | 0.360 | 1.348 | chr12 | 123459127 | 123464590 | + | protein_coding |
| ENSG00000087502.18_5 | ERGIC2 | 0.99808166 | 1 | 0.360 | 6.296 | chr12 | 29490285 | 29534122 | - | protein_coding |
| ENSG00000196628.18_8 | TCF4 | 0.97475947 | 1 | 0.360 | 3.917 | chr18 | 52889416 | 53332018 | - | protein_coding |
| ENSG00000203772.7_3 | SPRN | 0.50900615 | 1 | 0.360 | -0.742 | chr10 | 135234170 | 135238076 | - | protein_coding |
| ENSG00000138336.9_4 | TET1 | 0.91993168 | 1 | 0.360 | 3.207 | chr10 | 70320094 | 70454244 | + | protein_coding |
| ENSG00000261582.1_7 | AL121753.1 | 0.51338788 | 1 | 0.360 | -0.629 | chr20 | 33855688 | 33867846 | - | protein_coding |
| ENSG00000173848.19_4 | NET1 | 0.99950672 | 1 | 0.360 | 7.095 | chr10 | 5454520 | 5501019 | + | protein_coding |
| ENSG00000124160.12_4 | NCOA5 | 0.99260437 | 1 | 0.360 | 5.128 | chr20 | 44689630 | 44718601 | - | protein_coding |
| ENSG00000122224.18_4 | LY9 | 0.50681008 | 1 | 0.360 | -0.864 | chr1 | 160765864 | 160798051 | + | protein_coding |
| ENSG00000188559.15_7 | RALGAPA2 | 0.98050303 | 1 | 0.359 | 4.492 | chr20 | 20370174 | 20693287 | - | protein_coding |
| ENSG00000168792.5_4 | ABHD15 | 0.99648814 | 1 | 0.359 | 5.242 | chr17 | 27887565 | 27894055 | - | protein_coding |
| ENSG00000180104.16_4 | EXOC3 | 0.99760437 | 1 | 0.359 | 5.782 | chr5 | 443290 | 472052 | + | protein_coding |
| ENSG00000044115.21_7 | CTNNA1 | 0.99983508 | 1 | 0.359 | 8.778 | chr5 | 137946656 | 138270723 | + | protein_coding |
| ENSG00000164331.10_5 | ANKRA2 | 0.9702065 | 1 | 0.359 | 3.805 | chr5 | 72848015 | 72861492 | - | protein_coding |
| ENSG00000144152.13_4 | FBLN7 | 0.50881429 | 1 | 0.358 | -0.864 | chr2 | 112895962 | 112945795 | + | protein_coding |
| ENSG00000105767.3_4 | CADM4 | 0.92842213 | 1 | 0.358 | 3.462 | chr19 | 44126520 | 44144002 | - | protein_coding |
| ENSG00000213859.6_4 | KCTD11 | 0.87938467 | 1 | 0.358 | 2.838 | chr17 | 7255208 | 7258263 | + | protein_coding |
| ENSG00000255559.1_6 | ZNF252P-AS1 | 0.53514951 | 1 | 0.358 | -0.140 | chr8 | 146228197 | 146231432 | + | lncRNA |
| ENSG00000185591.10_5 | SP1 | 0.99972339 | 1 | 0.358 | 7.355 | chr12 | 53773960 | 53810230 | + | protein_coding |

| | | | | | | | | | | |
|---|---|---|---|---|---|---|---|---|---|---|
| ENSG00000196586.16_7 | MYO6 | 0.98576604 | 1 | 0.358 | 4.757 | chr6 | 76458909 | 76629254 | + | protein_coding |
| ENSG00000174136.13_4 | RGMB | 0.83142857 | 1 | 0.358 | 2.122 | chr5 | 98104354 | 98134347 | + | protein_coding |
| ENSG00000043591.6_4 | ADRB1 | 0.67432796 | 1 | 0.358 | 1.031 | chr10 | 115803625 | 115806663 | + | protein_coding |
| ENSG00000255284.2_8 | AP006621.3 | 0.89518134 | 1 | 0.358 | 2.819 | chr11 | 777578 | 784297 | + | lncRNA |
| ENSG00000106635.8_4 | BCL7B | 0.99305033 | 1 | 0.358 | 4.961 | chr7 | 72950686 | 72972020 | - | protein_coding |
| ENSG00000150867.14_7 | PIP4K2A | 0.99766454 | 1 | 0.357 | 5.499 | chr10 | 22823783 | 23003507 | - | protein_coding |
| ENSG00000143153.13_5 | ATP1B1 | 0.99978342 | 1 | 0.357 | 8.421 | chr1 | 169074935 | 169101960 | + | protein_coding |
| ENSG00000014164.7_4 | ZC3H3 | 0.98791085 | 1 | 0.357 | 4.628 | chr8 | 144519829 | 144623617 | - | protein_coding |
| ENSG00000109323.11_8 | MANBA | 0.97445066 | 1 | 0.357 | 4.186 | chr4 | 103551927 | 103682151 | - | protein_coding |
| ENSG00000143452.16_5 | HORMAD1 | 0.96281194 | 1 | 0.357 | 4.126 | chr1 | 150670536 | 150693371 | - | protein_coding |
| ENSG00000196418.12_5 | ZNF124 | 0.98173538 | 1 | 0.357 | 4.262 | chr1 | 247285277 | 247335318 | - | protein_coding |
| ENSG00000204569.10_5 | PPP1R10 | 0.99259511 | 1 | 0.357 | 5.134 | chr6 | 30568190 | 30586389 | - | protein_coding |
| ENSG00000268879.1_2 | IGFL1P1 | 0.88148048 | 1 | 0.357 | 2.566 | chr19 | 46699495 | 46700580 | + | unprocessed_pseudogene |
| ENSG00000104221.13_4 | BRF2 | 0.8331635 | 1 | 0.357 | 2.504 | chr8 | 37700786 | 37707379 | - | protein_coding |
| ENSG00000233223.3_7 | AC016876.2 | 0.79987231 | 1 | 0.356 | 1.917 | chr17 | 7485282 | 7487404 | - | lncRNA |
| ENSG00000198740.8_4 | ZNF652 | 0.99616393 | 1 | 0.356 | 5.485 | chr17 | 47366568 | 47439835 | - | protein_coding |
| ENSG00000161277.11_4 | THAP8 | 0.84045847 | 1 | 0.356 | 2.210 | chr19 | 36525886 | 36545641 | - | protein_coding |
| ENSG00000167971.16_4 | CASKIN1 | 0.63062011 | 1 | 0.356 | 0.995 | chr16 | 2227181 | 2246606 | - | protein_coding |
| ENSG00000149182.15_5 | ARFGAP2 | 0.99830896 | 1 | 0.356 | 5.676 | chr11 | 47185850 | 47198676 | - | protein_coding |
| ENSG00000123810.8_4 | B9D2 | 0.9153925 | 1 | 0.356 | 3.027 | chr19 | 41860322 | 41870070 | - | protein_coding |
| ENSG00000137478.15_6 | FCHSD2 | 0.98923516 | 1 | 0.356 | 4.593 | chr11 | 72547790 | 72853363 | - | protein_coding |
| ENSG00000237864.1_6 | LINC00322 | 0.6205702 | 1 | 0.356 | 0.689 | chr21 | 44742297 | 44751919 | - | lncRNA |
| ENSG00000052802.13_4 | MSMO1 | 0.99598823 | 1 | 0.356 | 5.762 | chr4 | 166248819 | 166264316 | + | protein_coding |
| ENSG00000173548.8_2 | SNX33 | 0.99089347 | 1 | 0.356 | 5.193 | chr15 | 75940247 | 75954642 | + | protein_coding |
| ENSG00000227855.3_5 | DPY19L2P3 | 0.76651747 | 1 | 0.356 | 1.325 | chr7 | 29689843 | 29782210 | + | transcribed_unprocessed_pseudogene |
| ENSG00000273820.2_5 | USP27X | 0.82689531 | 1 | 0.355 | 2.202 | chrX | 49644087 | 49647162 | + | protein_coding |
| ENSG00000204130.13_4 | RUFY2 | 0.99411118 | 1 | 0.355 | 5.200 | chr10 | 70100864 | 70167051 | - | protein_coding |
| ENSG00000146067.16_6 | FAM193B | 0.99300547 | 1 | 0.355 | 5.034 | chr5 | 176946790 | 176981564 | - | protein_coding |
| ENSG00000174839.13_4 | DENND6A | 0.98626692 | 1 | 0.355 | 4.683 | chr3 | 57611181 | 57678804 | - | protein_coding |
| ENSG00000287086.1_2 | AC034193.1 | 0.85631827 | 1 | 0.355 | 2.464 | chr3 | 10191670 | 10195010 | - | lncRNA |
| ENSG00000215271.9_5 | HOMEZ | 0.9608015 | 1 | 0.355 | 3.626 | chr14 | 23741631 | 23768656 | - | protein_coding |
| ENSG00000095066.11_8 | HOOK2 | 0.94661819 | 1 | 0.355 | 3.420 | chr19 | 12873817 | 12983554 | - | protein_coding |
| ENSG00000273270.1_6 | AC090114.2 | 0.89213956 | 1 | 0.355 | 2.612 | chr7 | 128164070 | 128171123 | - | lncRNA |
| ENSG00000040199.18_7 | PHLPP2 | 0.99628926 | 1 | 0.355 | 5.277 | chr16 | 71671738 | 71758604 | - | protein_coding |
| ENSG00000186666.6_4 | BCDIN3D | 0.87975389 | 1 | 0.355 | 2.155 | chr12 | 50229826 | 50236889 | - | protein_coding |
| ENSG00000180769.10_4 | WDFY3-AS2 | 0.68940763 | 1 | 0.355 | 1.253 | chr4 | 85886687 | 85932430 | + | lncRNA |
| ENSG00000104899.8_4 | AMH | 0.78831061 | 1 | 0.354 | 3.799 | chr19 | 2249308 | 2252072 | + | protein_coding |
| ENSG00000106397.12_7 | PLOD3 | 0.9946451 | 1 | 0.354 | 5.121 | chr7 | 100849258 | 100861701 | - | protein_coding |
| ENSG00000119402.17_4 | FBXW2 | 0.99920161 | 1 | 0.354 | 6.616 | chr9 | 123514256 | 123555694 | - | protein_coding |
| ENSG00000167968.13_6 | DNASE1L2 | 0.7818558 | 1 | 0.354 | 1.853 | chr16 | 2285817 | 2288712 | + | protein_coding |
| ENSG00000139370.12_5 | SLC15A4 | 0.99359238 | 1 | 0.354 | 4.893 | chr12 | 129277739 | 129308503 | - | protein_coding |
| ENSG00000126254.12_4 | RBM42 | 0.95000538 | 1 | 0.354 | 6.933 | chr19 | 36119932 | 36128588 | + | protein_coding |
| ENSG00000108306.13_9 | FBXL20 | 0.99804711 | 1 | 0.354 | 5.523 | chr17 | 37408916 | 37558776 | - | protein_coding |
| ENSG00000276691.1_6 | AC004466.3 | 0.51367491 | 1 | 0.354 | -0.999 | chr12 | 48182209 | 48182754 | + | lncRNA |
| ENSG00000129317.14_5 | PUS7L | 0.9945376 | 1 | 0.354 | 6.108 | chr12 | 44112796 | 44152620 | - | protein_coding |
| ENSG00000096060.14_3 | FKBP5 | 0.93678034 | 1 | 0.354 | 3.374 | chr6 | 35541362 | 35696360 | - | protein_coding |
| ENSG00000149743.14_5 | TRPT1 | 0.95667028 | 1 | 0.354 | 3.555 | chr11 | 63991271 | 63993726 | - | protein_coding |
| ENSG00000165169.11_4 | DYNLT3 | 0.99710889 | 1 | 0.354 | 5.426 | chrX | 37696010 | 37706824 | - | protein_coding |
| ENSG00000173482.16_5 | PTPRM | 0.9996927 | 1 | 0.354 | 7.157 | chr18 | 7566780 | 8406859 | + | protein_coding |
| ENSG00000261040.7_5 | WFDC21P | 0.54150264 | 1 | 0.353 | -0.112 | chr17 | 58160927 | 58169246 | - | transcribed_unitary_pseudogene |
| ENSG00000103249.18_5 | CLCN7 | 0.99089319 | 1 | 0.353 | 5.699 | chr16 | 1494935 | 1525085 | - | protein_coding |
| ENSG00000047621.12_5 | C12orf4 | 0.99456784 | 1 | 0.353 | 5.199 | chr12 | 4596901 | 4647674 | - | protein_coding |
| ENSG00000183826.18_6 | BTBD9 | 0.95505702 | 1 | 0.352 | 3.776 | chr6 | 38136227 | 38607924 | - | protein_coding |
| ENSG00000117114.19_7 | ADGRL2 | 0.98818472 | 1 | 0.352 | 5.385 | chr1 | 81771845 | 82458120 | + | protein_coding |
| ENSG00000220848.5_4 | RPS18P9 | 0.6323985 | 1 | 0.352 | 0.714 | chr6 | 149913823 | 149923014 | + | transcribed_processed_pseudogene |
| ENSG00000247556.6_4 | OIP5-AS1 | 0.99938434 | 1 | 0.352 | 7.013 | chr15 | 41576188 | 41601935 | + | lncRNA |
| ENSG00000176124.14_7 | DLEU1 | 0.94457017 | 1 | 0.352 | 3.488 | chr13 | 50656305 | 51480992 | + | lncRNA |
| ENSG00000124145.6_5 | SDC4 | 0.99510899 | 1 | 0.352 | 4.937 | chr20 | 43953928 | 43977064 | - | protein_coding |
| ENSG00000286699.1_3 | AC084198.4 | 0.66478424 | 1 | 0.352 | 0.850 | chr3 | 101405671 | 101419770 | + | lncRNA |

| | | | | | | | | | | |
|---|---|---|---|---|---|---|---|---|---|---|
| ENSG00000251129.2_6 | LINC02506 | 0.96067465 | 1 | 0.352 | 3.500 | chr4 | 31998998 | 32230165 | + | lncRNA |
| ENSG00000154240.17_7 | CEP112 | 0.98375218 | 1 | 0.352 | 4.314 | chr17 | 63631655 | 64188251 | - | protein_coding |
| ENSG00000075415.12_7 | SLC25A3 | 0.99982243 | 1 | 0.351 | 8.715 | chr12 | 98987369 | 99000157 | + | protein_coding |
| ENSG00000271327.1_6 | AC010201.2 | 0.55177338 | 1 | 0.351 | -0.020 | chr12 | 89761584 | 89763078 | + | lncRNA |
| ENSG00000010295.19_5 | IFFO1 | 0.67300241 | 1 | 0.351 | 0.953 | chr12 | 6647541 | 6665249 | - | protein_coding |
| ENSG00000267493.4_7 | CIRBP-AS1 | 0.79173081 | 1 | 0.350 | 1.656 | chr19 | 1267470 | 1270259 | - | lncRNA |
| ENSG00000273973.1_6 | AC025162.2 | 0.51574323 | 1 | 0.350 | -0.961 | chr12 | 56322954 | 56323513 | + | lncRNA |
| ENSG00000173401.10_4 | GLIPR1L1 | 0.52186125 | 1 | 0.350 | -0.785 | chr12 | 75728450 | 75764340 | + | protein_coding |
| ENSG00000169174.11_3 | PCSK9 | 0.98386627 | 1 | 0.350 | 4.605 | chr1 | 55505120 | 55530525 | + | protein_coding |
| ENSG00000127585.12_4 | FBXL16 | 0.75404306 | 1 | 0.350 | 1.490 | chr16 | 742498 | 755808 | - | protein_coding |
| ENSG00000179841.8_3 | AKAP5 | 0.91600473 | 1 | 0.350 | 2.988 | chr14 | 64932217 | 64941221 | + | protein_coding |
| ENSG00000139645.10_4 | ANKRD52 | 0.99964044 | 1 | 0.350 | 6.870 | chr12 | 56631591 | 56652168 | - | protein_coding |
| ENSG00000171017.11_5 | LRRC8E | 0.53339468 | 1 | 0.350 | -0.455 | chr19 | 7953390 | 7966906 | + | protein_coding |
| ENSG00000162627.17_5 | SNX7 | 0.9865942 | 1 | 0.350 | 4.961 | chr1 | 99127257 | 99226056 | + | protein_coding |
| ENSG00000245498.7_7 | AP000866.1 | 0.71616046 | 1 | 0.350 | 1.109 | chr11 | 124670320 | 124704383 | + | lncRNA |
| ENSG00000077238.14_6 | IL4R | 0.96736244 | 1 | 0.349 | 4.228 | chr16 | 27324989 | 27376099 | + | protein_coding |
| ENSG00000178537.10_4 | SLC25A20 | 0.94419772 | 1 | 0.349 | 3.397 | chr3 | 48894359 | 48936337 | - | protein_coding |
| ENSG00000167543.16_7 | TP53I13 | 0.94442503 | 1 | 0.349 | 5.667 | chr17 | 27893070 | 27900175 | + | protein_coding |
| ENSG00000004059.11_5 | ARF5 | 0.99714338 | 1 | 0.349 | 5.475 | chr7 | 127228440 | 127231754 | + | protein_coding |
| ENSG00000153814.13_5 | JAZF1 | 0.75358327 | 1 | 0.349 | 1.618 | chr7 | 27870192 | 28220414 | - | protein_coding |
| ENSG00000105953.15_5 | OGDH | 0.99930233 | 1 | 0.349 | 6.159 | chr7 | 44646171 | 44748665 | + | protein_coding |
| ENSG00000267023.5_3 | LRRC37A16P | 0.96045441 | 1 | 0.349 | 3.452 | chr17 | 66121918 | 66148609 | - | transcribed_unprocessed_pseudogene |
| ENSG00000141985.9_6 | SH3GL1 | 0.99572721 | 1 | 0.349 | 5.379 | chr19 | 4360367 | 4400544 | - | protein_coding |
| ENSG00000170633.16_4 | RNF34 | 0.99764321 | 1 | 0.348 | 5.612 | chr12 | 121837844 | 121868426 | + | protein_coding |
| ENSG00000074842.8_6 | MYDGF | 0.98586693 | 1 | 0.348 | 5.213 | chr19 | 4641386 | 4670374 | - | protein_coding |
| ENSG00000250299.6_5 | MRPS31P4 | 0.52275282 | 1 | 0.348 | -0.430 | chr13 | 53174254 | 53211581 | + | transcribed_unprocessed_pseudogene |
| ENSG00000273311.1_4 | DGCR11 | 0.63815671 | 1 | 0.348 | 0.654 | chr22 | 19033675 | 19035888 | - | lncRNA |
| ENSG00000173258.13_4 | ZNF483 | 0.99052148 | 1 | 0.348 | 4.654 | chr9 | 114287439 | 114340124 | + | protein_coding |
| ENSG00000198513.12_7 | ATL1 | 0.81820386 | 1 | 0.348 | 1.988 | chr14 | 50999227 | 51099786 | + | protein_coding |
| ENSG00000139990.18_5 | DCAF5 | 0.99716341 | 1 | 0.348 | 5.298 | chr14 | 69517598 | 69619867 | - | protein_coding |
| ENSG00000110987.8_2 | BCL7A | 0.97596956 | 1 | 0.348 | 3.813 | chr12 | 122457328 | 122499950 | + | protein_coding |
| ENSG00000061938.19_8 | TNK2 | 0.94230852 | 1 | 0.348 | 4.180 | chr3 | 195590235 | 195638816 | - | protein_coding |
| ENSG00000014914.21_4 | MTMR11 | 0.99537218 | 1 | 0.348 | 5.061 | chr1 | 149900543 | 149908772 | - | protein_coding |
| ENSG00000256050.2_6 | AL583722.1 | 0.61420178 | 1 | 0.348 | 0.361 | chr14 | 105147483 | 105151269 | + | lncRNA |
| ENSG00000111450.14_5 | STX2 | 0.9989314 | 1 | 0.347 | 5.967 | chr12 | 131274145 | 131323811 | - | protein_coding |
| ENSG00000221909.3_6 | FAM200A | 0.93829966 | 1 | 0.347 | 3.358 | chr7 | 99143923 | 99156159 | - | protein_coding |
| ENSG00000100926.15_8 | TM9SF1 | 0.89975297 | 1 | 0.347 | 2.635 | chr14 | 24658355 | 24664893 | - | protein_coding |
| ENSG00000260032.2_5 | NORAD | 0.9997616 | 1 | 0.347 | 8.234 | chr20 | 34633540 | 34638940 | - | lncRNA |
| ENSG00000067221.14_7 | STOML1 | 0.8984558 | 1 | 0.347 | 2.806 | chr15 | 74271267 | 74286963 | - | protein_coding |
| ENSG00000165972.13_6 | CCDC38 | 0.52154227 | 1 | 0.346 | -0.745 | chr12 | 96260826 | 96336413 | - | protein_coding |
| ENSG00000178467.18_4 | P4HTM | 0.99297956 | 1 | 0.346 | 4.813 | chr3 | 49027322 | 49044586 | + | protein_coding |
| ENSG00000143393.16_8 | PI4KB | 0.99803889 | 1 | 0.346 | 5.444 | chr1 | 151264273 | 151300191 | - | protein_coding |
| ENSG00000220323.4_6 | H2BC19P | 0.80932596 | 1 | 0.346 | 1.828 | chr1 | 149814442 | 149818053 | + | transcribed_unprocessed_pseudogene |
| ENSG00000105516.11_4 | DBP | 0.84071756 | 1 | 0.346 | 2.599 | chr19 | 49133287 | 49140636 | - | protein_coding |
| ENSG00000129493.15_5 | HEATR5A | 0.99796873 | 1 | 0.346 | 5.707 | chr14 | 31760994 | 31889756 | - | protein_coding |
| ENSG00000131370.16_6 | SH3BP5 | 0.75397012 | 1 | 0.346 | 1.487 | chr3 | 15295860 | 15382875 | - | protein_coding |
| ENSG00000174945.13_2 | AMZ1 | 0.53837327 | 1 | 0.345 | -0.527 | chr7 | 2719156 | 2815134 | + | protein_coding |
| ENSG00000197208.6_4 | SLC22A4 | 0.87631545 | 1 | 0.345 | 2.454 | chr5 | 131630087 | 131679883 | + | protein_coding |
| ENSG00000174276.7_4 | ZNHIT2 | 0.89982038 | 1 | 0.345 | 2.593 | chr11 | 64883875 | 64885173 | - | protein_coding |
| ENSG00000177725.5_8 | AC105206.1 | 0.61805383 | 1 | 0.345 | 0.319 | chr8 | 22026851 | 22029223 | + | lncRNA |
| ENSG00000198087.7_4 | CD2AP | 0.99943588 | 1 | 0.345 | 7.027 | chr6 | 47445525 | 47594999 | + | protein_coding |
| ENSG00000231822.1_5 | AC019097.1 | 0.54008893 | 1 | 0.345 | -0.526 | chr2 | 99718481 | 99719215 | + | processed_pseudogene |
| ENSG00000198742.9_4 | SMURF1 | 0.99868858 | 1 | 0.345 | 5.602 | chr7 | 98625061 | 98741723 | - | protein_coding |
| ENSG00000177459.11_4 | ERICH5 | 0.91901544 | 1 | 0.344 | 2.710 | chr8 | 99076750 | 99105838 | + | protein_coding |
| ENSG00000150991.15_8 | UBC | 0.99953325 | 1 | 0.344 | 9.127 | chr12 | 125396150 | 125401914 | - | protein_coding |
| ENSG00000120451.10_4 | SNX19 | 0.99922563 | 1 | 0.344 | 6.188 | chr11 | 130745331 | 130786404 | - | protein_coding |
| ENSG00000183354.12_7 | KIAA2026 | 0.99475217 | 1 | 0.344 | 5.200 | chr9 | 5881596 | 6008482 | - | protein_coding |
| ENSG00000103485.19_5 | QPRT | 0.56038786 | 1 | 0.344 | -0.251 | chr16 | 29674600 | 29710020 | + | protein_coding |
| ENSG00000154153.13_7 | RETREG1 | 0.90987791 | 1 | 0.344 | 2.641 | chr5 | 16473147 | 16617167 | - | protein_coding |

| | | | | | | | | | | |
|---|---|---|---|---|---|---|---|---|---|---|
| ENSG00000136279.20_7 | DBNL | 0.99940747 | 1 | 0.344 | 6.444 | chr7 | 44084239 | 44109055 | + | protein_coding |
| ENSG00000134698.11_3 | AGO4 | 0.96761128 | 1 | 0.344 | 3.486 | chr1 | 36273617 | 36323491 | + | protein_coding |
| ENSG00000105053.11_7 | VRK3 | 0.99672764 | 1 | 0.343 | 5.093 | chr19 | 50479725 | 50529203 | - | protein_coding |
| ENSG00000242861.1_8 | AL591895.1 | 0.54906755 | 1 | 0.343 | -0.339 | chr1 | 226028584 | 226034223 | - | lncRNA |
| ENSG00000175198.17_5 | PCCA | 0.9506054 | 1 | 0.343 | 3.478 | chr13 | 100741269 | 101182691 | + | protein_coding |
| ENSG00000116741.8_3 | RGS2 | 0.95011173 | 1 | 0.343 | 3.555 | chr1 | 192778169 | 192781405 | + | protein_coding |
| ENSG00000211455.8_4 | STK38L | 0.99526195 | 1 | 0.343 | 5.122 | chr12 | 27396901 | 27478892 | + | protein_coding |
| ENSG00000187239.17_5 | FNBP1 | 0.99443562 | 1 | 0.343 | 4.694 | chr9 | 132649466 | 132805473 | - | protein_coding |
| ENSG00000113742.14_5 | CPEB4 | 0.99947108 | 1 | 0.343 | 6.224 | chr5 | 173315352 | 173388983 | + | protein_coding |
| ENSG00000273680.1_6 | AC009318.2 | 0.56651643 | 1 | 0.342 | -0.089 | chr12 | 29485666 | 29486316 | - | lncRNA |
| ENSG00000167615.16_7 | LENG8 | 0.9991491 | 1 | 0.342 | 6.503 | chr19 | 54960065 | 54973217 | + | protein_coding |
| ENSG00000105755.8_4 | ETHE1 | 0.95913984 | 1 | 0.342 | 3.723 | chr19 | 44010871 | 44031382 | - | protein_coding |
| ENSG00000112624.12_5 | BICRAL | 0.98452359 | 1 | 0.342 | 4.078 | chr6 | 42714696 | 42836298 | + | protein_coding |
| ENSG00000213018.2_5 | AL590762.1 | 0.55051874 | 1 | 0.342 | -1.216 | chrX | 70639933 | 70640364 | - | processed_pseudogene |
| ENSG00000070371.16_5 | CLTCL1 | 0.99308969 | 1 | 0.342 | 4.570 | chr22 | 19166986 | 19279242 | - | protein_coding |
| ENSG00000124523.16_4 | SIRT5 | 0.98119456 | 1 | 0.342 | 3.813 | chr6 | 13574761 | 13615390 | + | protein_coding |
| ENSG00000164068.16_3 | RNF123 | 0.99387042 | 1 | 0.341 | 4.836 | chr3 | 49726971 | 49758962 | + | protein_coding |
| ENSG00000255200.1_4 | PGAM1P8 | 0.53869257 | 1 | 0.341 | -0.535 | chr11 | 64941588 | 64943941 | - | transcribed_processed_pseudogene |
| ENSG00000104957.14_5 | CCDC130 | 0.97658815 | 1 | 0.341 | 3.724 | chr19 | 13842574 | 13874110 | + | protein_coding |
| ENSG00000189045.14_4 | ANKDD1B | 0.57633574 | 1 | 0.341 | -0.201 | chr5 | 74907007 | 74977598 | + | protein_coding |
| ENSG00000222489.1 | SNORA79 | 0.8744654 | 1 | 0.341 | 2.190 | chr14 | 20791338 | 20791485 | - | snoRNA |
| ENSG00000232504.5_6 | ST3GAL5-AS1 | 0.55423922 | 1 | 0.340 | -0.045 | chr2 | 86116274 | 86118259 | + | lncRNA |
| ENSG00000117308.15_7 | GALE | 0.99907317 | 1 | 0.340 | 5.693 | chr1 | 24122089 | 24127271 | - | protein_coding |
| ENSG00000090565.16_7 | RAB11FIP3 | 0.99487193 | 1 | 0.340 | 5.145 | chr16 | 475649 | 573011 | + | protein_coding |
| ENSG00000105327.17_4 | BBC3 | 0.81393168 | 1 | 0.340 | 2.655 | chr19 | 47724079 | 47736023 | - | protein_coding |
| ENSG00000087274.17_6 | ADD1 | 0.99980403 | 1 | 0.340 | 7.089 | chr4 | 2845584 | 2931803 | + | protein_coding |
| ENSG00000106608.16_6 | URGCP | 0.9977579 | 1 | 0.340 | 5.186 | chr7 | 43915493 | 43966010 | - | protein_coding |
| ENSG00000156030.13_6 | ELMSAN1 | 0.99883862 | 1 | 0.340 | 5.998 | chr14 | 74181825 | 74256988 | - | protein_coding |
| ENSG00000085733.16_3 | CTTN | 0.99978375 | 1 | 0.340 | 8.126 | chr11 | 70244510 | 70282690 | + | protein_coding |
| ENSG00000117791.16_3 | MARC2 | 0.9732159 | 1 | 0.340 | 3.543 | chr1 | 220921567 | 220958157 | + | protein_coding |
| ENSG00000062485.19_8 | CS | 0.99982006 | 1 | 0.340 | 7.236 | chr12 | 56665483 | 56694175 | - | protein_coding |
| ENSG00000125826.20_3 | RBCK1 | 0.96909244 | 1 | 0.340 | 6.499 | chr20 | 388142 | 411610 | + | protein_coding |
| ENSG00000065613.15_6 | SLK | 0.99985837 | 1 | 0.340 | 7.764 | chr10 | 105726898 | 105788991 | + | protein_coding |
| ENSG00000149179.13_4 | C11orf49 | 0.99389771 | 1 | 0.339 | 4.912 | chr11 | 46958240 | 47185936 | + | protein_coding |
| ENSG00000260219.2_7 | CD2BP2-DT | 0.69466455 | 1 | 0.339 | 0.964 | chr16 | 30366762 | 30368425 | + | lncRNA |
| ENSG00000105223.20_4 | PLD3 | 0.99861482 | 1 | 0.339 | 7.244 | chr19 | 40854363 | 40886346 | + | protein_coding |
| ENSG00000171121.16_5 | KCNMB3 | 0.71706435 | 1 | 0.339 | 1.075 | chr3 | 178954479 | 178984790 | - | protein_coding |
| ENSG00000101333.16_3 | PLCB4 | 0.99778195 | 1 | 0.339 | 5.668 | chr20 | 9049410 | 9461889 | + | protein_coding |
| ENSG00000168661.14_3 | ZNF30 | 0.97709396 | 1 | 0.339 | 3.893 | chr19 | 35417807 | 35436074 | + | protein_coding |
| ENSG00000113240.13_5 | CLK4 | 0.99360692 | 1 | 0.339 | 4.651 | chr5 | 178029665 | 178057616 | - | protein_coding |
| ENSG00000181450.18_4 | ZNF678 | 0.98649806 | 1 | 0.339 | 4.508 | chr1 | 227751244 | 227865144 | + | protein_coding |
| ENSG00000160014.17_5 | CALM3 | 0.99986491 | 1 | 0.339 | 7.527 | chr19 | 47104331 | 47114039 | + | protein_coding |
| ENSG00000258311.5_8 | AC009779.4 | 0.60044208 | 1 | 0.338 | 0.113 | chr12 | 56109820 | 56118487 | + | protein_coding |
| ENSG00000168615.12_3 | ADAM9 | 0.99992279 | 1 | 0.338 | 9.106 | chr8 | 38854388 | 38962780 | + | protein_coding |
| ENSG00000179604.10_4 | CDC42EP4 | 0.99912681 | 1 | 0.338 | 6.349 | chr17 | 71279763 | 71308144 | - | protein_coding |
| ENSG00000105856.14_6 | HBP1 | 0.99694149 | 1 | 0.338 | 5.163 | chr7 | 106809406 | 106842967 | + | protein_coding |
| ENSG00000005187.12_4 | ACSM3 | 0.96333288 | 1 | 0.338 | 3.527 | chr16 | 20621565 | 20808903 | + | protein_coding |
| ENSG00000122691.13_6 | TWIST1 | 0.7543057 | 1 | 0.338 | 1.409 | chr7 | 19060614 | 19157259 | - | protein_coding |
| ENSG00000157823.17_5 | AP3S2 | 0.99339327 | 1 | 0.338 | 4.536 | chr15 | 90373831 | 90437870 | - | protein_coding |
| ENSG00000002587.10_6 | HS3ST1 | 0.87000077 | 1 | 0.338 | 2.290 | chr4 | 11394774 | 11431188 | - | protein_coding |
| ENSG00000198105.14_8 | ZNF248 | 0.98837919 | 1 | 0.338 | 4.372 | chr10 | 38065454 | 38147034 | - | protein_coding |
| ENSG00000142765.18_5 | SYTL1 | 0.65338759 | 1 | 0.338 | 0.606 | chr1 | 27668511 | 27680428 | + | protein_coding |
| ENSG00000112144.15_3 | ICK | 0.9908431 | 1 | 0.338 | 4.410 | chr6 | 52866077 | 52926600 | - | protein_coding |
| ENSG00000286909.1_2 | AC024560.4 | 0.53972515 | 1 | 0.337 | -0.755 | chr3 | 197376174 | 197378353 | - | lncRNA |
| ENSG00000083799.17_7 | CYLD | 0.9957801 | 1 | 0.337 | 4.783 | chr16 | 50775961 | 50835846 | + | protein_coding |
| ENSG00000139637.14_4 | C12orf10 | 0.99717634 | 1 | 0.337 | 4.967 | chr12 | 53693479 | 53700961 | + | protein_coding |
| ENSG00000160404.18_4 | TOR2A | 0.89307935 | 1 | 0.337 | 2.628 | chr9 | 130493803 | 130497592 | - | protein_coding |
| ENSG00000131669.10_4 | NINJ1 | 0.99826982 | 1 | 0.336 | 5.311 | chr9 | 95883778 | 95896533 | - | protein_coding |
| ENSG00000125378.16_5 | BMP4 | 0.92498261 | 1 | 0.336 | 2.915 | chr14 | 54416454 | 54425479 | - | protein_coding |

| | | | | | | | | | | |
|---|---|---|---|---|---|---|---|---|---|---|
| ENSG00000143147.14_4 | GPR161 | 0.99732394 | 1 | 0.336 | 5.088 | chr1 | 168048781 | 168106905 | - | protein_coding |
| ENSG00000110497.14_7 | AMBRA1 | 0.9987389 | 1 | 0.336 | 5.693 | chr11 | 46417964 | 46615675 | - | protein_coding |
| ENSG00000149922.10_3 | TBX6 | 0.59026875 | 1 | 0.336 | 0.159 | chr16 | 30097114 | 30103208 | - | protein_coding |
| ENSG00000213563.7_7 | C8orf82 | 0.98874829 | 1 | 0.336 | 4.539 | chr8 | 145751117 | 145754516 | - | protein_coding |
| ENSG00000223745.8_8 | CCDC18-AS1 | 0.95652235 | 1 | 0.336 | 3.430 | chr1 | 93727743 | 93811582 | - | lncRNA |
| ENSG00000070366.14_7 | SMG6 | 0.99890754 | 1 | 0.336 | 5.360 | chr17 | 1963133 | 2207079 | - | protein_coding |
| ENSG00000282977.1_4 | PCBP2-OT1 | 0.5838767 | 1 | 0.336 | -0.261 | chr12 | 53858252 | 53858841 | + | lncRNA |
| ENSG00000259431.6_4 | THTPA | 0.91579231 | 1 | 0.336 | 2.618 | chr14 | 24025197 | 24029480 | + | protein_coding |
| ENSG00000145990.11_4 | GFOD1 | 0.96067444 | 1 | 0.336 | 3.335 | chr6 | 13358062 | 13487894 | - | protein_coding |
| ENSG00000225032.5_6 | AL162586.1 | 0.82859481 | 1 | 0.336 | 1.549 | chr9 | 130578345 | 130584799 | + | lncRNA |
| ENSG00000169762.17_4 | TAPT1 | 0.9874122 | 1 | 0.336 | 4.301 | chr4 | 16162128 | 16229033 | - | protein_coding |
| ENSG00000104679.11_8 | R3HCC1 | 0.99723011 | 1 | 0.336 | 5.121 | chr8 | 23127633 | 23153792 | + | protein_coding |
| ENSG00000160785.13_2 | SLC25A44 | 0.99500011 | 1 | 0.336 | 5.385 | chr1 | 156163723 | 156182587 | + | protein_coding |
| ENSG00000235501.6_6 | AC105942.1 | 0.78462424 | 1 | 0.335 | 1.562 | chr1 | 95392917 | 95429172 | + | lncRNA |
| ENSG00000254860.6_5 | TMEM9B-AS1 | 0.90651259 | 1 | 0.335 | 2.714 | chr11 | 8986057 | 8999074 | + | lncRNA |
| ENSG00000273447.1_6 | AC004067.1 | 0.55971053 | 1 | 0.335 | -0.285 | chr4 | 110613160 | 110613859 | + | lncRNA |
| ENSG00000085644.14_3 | ZNF213 | 0.89303578 | 1 | 0.335 | 2.699 | chr16 | 3179778 | 3192805 | + | protein_coding |
| ENSG00000262884.1_7 | AC015921.1 | 0.61943428 | 1 | 0.335 | 0.439 | chr17 | 2865542 | 2869189 | - | lncRNA |
| ENSG00000117525.14_4 | F3 | 0.88553199 | 1 | 0.334 | 2.273 | chr1 | 94994729 | 95007315 | - | protein_coding |
| ENSG00000007541.16_4 | PIGQ | 0.98570579 | 1 | 0.334 | 4.772 | chr16 | 616995 | 634136 | + | protein_coding |
| ENSG00000229950.1_6 | TFAP2A-AS1 | 0.5370652 | 1 | 0.334 | -0.787 | chr6 | 10409573 | 10416679 | + | lncRNA |
| ENSG00000123689.6_3 | G0S2 | 0.57647964 | 1 | 0.334 | -0.024 | chr1 | 209848757 | 209849735 | + | protein_coding |
| ENSG00000105662.16_4 | CRTC1 | 0.92850048 | 1 | 0.334 | 3.045 | chr19 | 18794488 | 18893143 | + | protein_coding |
| ENSG00000171130.18_5 | ATP6V0E2 | 0.99652861 | 1 | 0.334 | 5.169 | chr7 | 149570057 | 149588293 | + | protein_coding |
| ENSG00000226742.4_3 | HSBP1L1 | 0.97088415 | 1 | 0.334 | 3.679 | chr18 | 77724582 | 77730822 | + | protein_coding |
| ENSG00000204116.11_6 | CHIC1 | 0.99577551 | 1 | 0.334 | 5.069 | chrX | 72783036 | 72906937 | + | protein_coding |
| ENSG00000143222.12_3 | UFC1 | 0.99846296 | 1 | 0.334 | 5.234 | chr1 | 161122566 | 161128646 | + | protein_coding |
| ENSG00000162542.13_4 | TMCO4 | 0.95541582 | 1 | 0.334 | 3.344 | chr1 | 20008706 | 20126438 | - | protein_coding |
| ENSG00000214273.4_2 | AGGF1P1 | 0.54197829 | 1 | 0.334 | -1.218 | chr4 | 190962497 | 190964569 | + | processed_pseudogene |
| ENSG00000164403.14_4 | SHROOM1 | 0.97369788 | 1 | 0.333 | 3.646 | chr5 | 132157833 | 132166590 | - | protein_coding |
| ENSG00000255857.6_6 | PXN-AS1 | 0.84814396 | 1 | 0.333 | 2.148 | chr12 | 120639094 | 120651034 | + | lncRNA |
| ENSG00000040487.13_5 | SLC66A1 | 0.9675275 | 1 | 0.333 | 3.485 | chr1 | 19638820 | 19655794 | + | protein_coding |
| ENSG00000274561.1_6 | AC005332.3 | 0.82348046 | 1 | 0.333 | 1.872 | chr17 | 66127603 | 66128048 | + | lncRNA |
| ENSG00000171161.13_5 | ZNF672 | 0.97728045 | 1 | 0.333 | 4.791 | chr1 | 249132409 | 249143716 | + | protein_coding |
| ENSG00000133027.18_4 | PEMT | 0.94536877 | 1 | 0.333 | 3.465 | chr17 | 17408877 | 17495022 | - | protein_coding |
| ENSG00000067141.17_4 | NEO1 | 0.99981155 | 1 | 0.333 | 6.871 | chr15 | 73344051 | 73597546 | + | protein_coding |
| ENSG00000238961.1 | SNORA47 | 0.55038049 | 1 | 0.333 | -0.643 | chr5 | 76376259 | 76376396 | - | snoRNA |
| ENSG00000087111.21_5 | PIGS | 0.99935631 | 1 | 0.332 | 5.983 | chr17 | 26880401 | 26898812 | - | protein_coding |
| ENSG00000185386.15_4 | MAPK11 | 0.93356207 | 1 | 0.332 | 2.901 | chr22 | 50702142 | 50709196 | - | protein_coding |
| ENSG00000082805.20_7 | ERC1 | 0.99946915 | 1 | 0.332 | 5.989 | chr12 | 1099675 | 1605099 | + | protein_coding |
| ENSG00000167186.11_5 | COQ7 | 0.99414629 | 1 | 0.332 | 4.412 | chr16 | 19078936 | 19091417 | + | protein_coding |
| ENSG00000104497.14_6 | SNX16 | 0.90794348 | 1 | 0.332 | 2.441 | chr8 | 82711816 | 82755101 | - | protein_coding |
| ENSG00000257621.8_8 | PSMA3-AS1 | 0.9953721 | 1 | 0.332 | 4.958 | chr14 | 58732083 | 58764852 | - | lncRNA |
| ENSG00000232807.2_7 | AL137186.2 | 0.53891615 | 1 | 0.332 | -1.064 | chr10 | 5976233 | 5987863 | - | lncRNA |
| ENSG00000250571.7_5 | GLI4 | 0.96353231 | 1 | 0.332 | 3.478 | chr8 | 144349603 | 144359101 | + | protein_coding |
| ENSG00000237517.9_5 | DGCR5 | 0.88250031 | 1 | 0.331 | 2.640 | chr22 | 18958027 | 18982141 | + | transcribed_unprocessed_pseudogene |
| ENSG00000256338.2_2 | RPL41P2 | 0.5604987 | 1 | 0.331 | -0.188 | chr15 | 28560797 | 28561205 | + | processed_pseudogene |
| ENSG00000166783.22_9 | MARF1 | 0.99905582 | 1 | 0.331 | 5.451 | chr16 | 15688244 | 15737011 | - | protein_coding |
| ENSG00000183617.5_3 | MRPL54 | 0.95772845 | 1 | 0.331 | 3.561 | chr19 | 3762680 | 3768573 | + | protein_coding |
| ENSG00000081177.18_3 | EXD2 | 0.99852005 | 1 | 0.331 | 5.139 | chr14 | 69658228 | 69710737 | + | protein_coding |
| ENSG00000164742.16_4 | ADCY1 | 0.82521394 | 1 | 0.331 | 1.710 | chr7 | 45613739 | 45762715 | + | protein_coding |
| ENSG00000262879.6_7 | AC005670.3 | 0.99573954 | 1 | 0.330 | 4.597 | chr17 | 45060653 | 45177689 | - | lncRNA |
| ENSG00000137266.14_5 | SLC22A23 | 0.96779008 | 1 | 0.330 | 3.665 | chr6 | 3269196 | 3457256 | - | protein_coding |
| ENSG00000167333.13_6 | TRIM68 | 0.99109905 | 1 | 0.330 | 4.334 | chr11 | 4619902 | 4629461 | - | protein_coding |
| ENSG00000107263.18_4 | RAPGEF1 | 0.99854013 | 1 | 0.330 | 6.005 | chr9 | 134452157 | 134615461 | - | protein_coding |
| ENSG00000167526.13_4 | RPL13 | 0.98469706 | 1 | 0.330 | 9.316 | chr16 | 89627065 | 89633236 | + | protein_coding |
| ENSG00000100226.16_5 | GTPBP1 | 0.99467535 | 1 | 0.330 | 4.600 | chr22 | 39101747 | 39134304 | + | protein_coding |

| | | | | | | | | | | |
|---|---|---|---|---|---|---|---|---|---|---|
| ENSG00000164187.7_4 | LMBRD2 | 0.99405867 | 1 | 0.330 | 5.081 | chr5 | 36098509 | 36151989 | - | protein_coding |
| ENSG00000047579.19_4 | DTNBP1 | 0.86003639 | 1 | 0.329 | 1.965 | chr6 | 15523032 | 15663289 | - | protein_coding |
| ENSG00000100330.15_4 | MTMR3 | 0.8057711 | 1 | 0.329 | 1.674 | chr22 | 30279144 | 30426855 | + | protein_coding |
| ENSG00000102100.16_6 | SLC35A2 | 0.99783593 | 1 | 0.329 | 5.160 | chrX | 48760457 | 48769235 | - | protein_coding |
| ENSG00000253284.2_7 | AC092828.1 | 0.66305395 | 1 | 0.329 | 0.536 | chr12 | 19300008 | 19307593 | + | lncRNA |
| ENSG00000198707.16_7 | CEP290 | 0.9973822 | 1 | 0.329 | 5.843 | chr12 | 88442791 | 88535865 | - | protein_coding |
| ENSG00000153179.13_4 | RASSF3 | 0.9993031 | 1 | 0.329 | 5.683 | chr12 | 64900781 | 65091344 | + | protein_coding |
| ENSG00000159314.11_4 | ARHGAP27 | 0.93887774 | 1 | 0.329 | 2.900 | chr17 | 43471268 | 43511787 | - | protein_coding |
| ENSG00000155366.16_6 | RHOC | 0.99961013 | 1 | 0.329 | 5.953 | chr1 | 113243728 | 113250056 | - | protein_coding |
| ENSG00000256943.1_7 | AC148477.3 | 0.56729126 | 1 | 0.329 | -0.317 | chr12 | 132851977 | 132854915 | + | lncRNA |
| ENSG00000104973.18_6 | MED25 | 0.99468036 | 1 | 0.328 | 5.253 | chr19 | 50321536 | 50343640 | + | protein_coding |
| ENSG00000156675.16_5 | RAB11FIP1 | 0.99945587 | 1 | 0.328 | 5.808 | chr8 | 37716136 | 37757015 | - | protein_coding |
| ENSG00000114573.10_4 | ATP6V1A | 0.99985348 | 1 | 0.328 | 6.828 | chr3 | 113465880 | 113530903 | + | protein_coding |
| ENSG00000178814.17_5 | OPLAH | 0.96922535 | 1 | 0.328 | 4.302 | chr8 | 145106167 | 145118868 | - | protein_coding |
| ENSG00000280287.1_6 | AC131212.3 | 0.69657702 | 1 | 0.328 | 0.850 | chr12 | 133127315 | 133131533 | + | TEC |
| ENSG00000112294.13_4 | ALDH5A1 | 0.9950863 | 1 | 0.328 | 4.540 | chr6 | 24495095 | 24537435 | + | protein_coding |
| ENSG00000229598.1_4 | PRDX3P1 | 0.54828764 | 1 | 0.328 | -0.726 | chr22 | 39118748 | 39119510 | - | processed_pseudogene |
| ENSG00000171307.19_4 | ZDHHC16 | 0.99946665 | 1 | 0.328 | 5.863 | chr10 | 99205927 | 99217127 | + | protein_coding |
| ENSG00000110422.12_5 | HIPK3 | 0.99949314 | 1 | 0.328 | 6.744 | chr11 | 33278218 | 33378569 | + | protein_coding |
| ENSG00000166860.3_4 | ZBTB39 | 0.99446253 | 1 | 0.328 | 4.436 | chr12 | 57392620 | 57400330 | - | protein_coding |
| ENSG00000100490.9_3 | CDKL1 | 0.91600462 | 1 | 0.328 | 2.709 | chr14 | 50796122 | 50883179 | - | protein_coding |
| ENSG00000130724.9_7 | CHMP2A | 0.99973578 | 1 | 0.328 | 6.352 | chr19 | 59062933 | 59066472 | - | protein_coding |
| ENSG00000286587.1_1 | AL133335.2 | 0.55236849 | 1 | 0.328 | -0.684 | chr20 | 52794953 | 52799668 | + | lncRNA |
| ENSG00000183828.15_5 | NUDT14 | 0.97396265 | 1 | 0.328 | 4.105 | chr14 | 105639275 | 105647660 | - | protein_coding |
| ENSG00000136026.14_6 | CKAP4 | 0.99992943 | 1 | 0.327 | 7.290 | chr12 | 106631659 | 106698057 | - | protein_coding |
| ENSG00000138834.13_7 | MAPK8IP3 | 0.99334726 | 1 | 0.327 | 6.002 | chr16 | 1756184 | 1820318 | + | protein_coding |
| ENSG00000111266.9_7 | DUSP16 | 0.99683705 | 1 | 0.327 | 4.923 | chr12 | 12626216 | 12715797 | - | protein_coding |
| ENSG00000173809.18_7 | TDRD12 | 0.60777447 | 1 | 0.327 | 0.012 | chr19 | 33210659 | 33320486 | + | protein_coding |
| ENSG00000162298.18_5 | SYVN1 | 0.99765682 | 1 | 0.327 | 5.023 | chr11 | 64889252 | 64902004 | - | protein_coding |
| ENSG00000139168.8_7 | ZCRB1 | 0.9991216 | 1 | 0.327 | 5.882 | chr12 | 42705888 | 42719920 | - | protein_coding |
| ENSG00000182903.16_6 | ZNF721 | 0.98908091 | 1 | 0.327 | 4.027 | chr4 | 419604 | 492945 | - | protein_coding |
| ENSG00000182318.6_3 | ZSCAN22 | 0.96937694 | 1 | 0.326 | 3.327 | chr19 | 58838360 | 58853698 | + | protein_coding |
| ENSG00000187866.10_6 | FAM122A | 0.95515231 | 1 | 0.326 | 3.103 | chr9 | 71394981 | 71400482 | + | protein_coding |
| ENSG00000104835.14_4 | SARS2 | 0.85728191 | 1 | 0.326 | 2.008 | chr19 | 39405906 | 39421536 | - | protein_coding |
| ENSG00000109458.8_6 | GAB1 | 0.94741533 | 1 | 0.326 | 3.311 | chr4 | 144257915 | 144395721 | + | protein_coding |
| ENSG00000213347.10_4 | MXD3 | 0.98209515 | 1 | 0.326 | 4.072 | chr5 | 176728462 | 176739758 | - | protein_coding |
| ENSG00000132434.9_4 | LANCL2 | 0.99274211 | 1 | 0.326 | 4.146 | chr7 | 55433141 | 55501435 | + | protein_coding |
| ENSG00000236456.1_5 | AL035458.1 | 0.55086999 | 1 | 0.326 | -0.725 | chr20 | 32833759 | 32834343 | - | processed_pseudogene |
| ENSG00000171714.11_3 | ANO5 | 0.99793923 | 1 | 0.325 | 5.144 | chr11 | 21821480 | 22304903 | + | protein_coding |
| ENSG00000232645.5_6 | LINC01431 | 0.56192095 | 1 | 0.325 | -0.263 | chr20 | 23337231 | 23338753 | - | lncRNA |
| ENSG00000225361.4_7 | PPP1R26-AS1 | 0.66741783 | 1 | 0.325 | 0.608 | chr9 | 138354573 | 138372623 | - | lncRNA |
| ENSG00000143303.12_6 | RRNAD1 | 0.99236189 | 1 | 0.325 | 4.236 | chr1 | 156698234 | 156706752 | + | protein_coding |
| ENSG00000135119.14_3 | RNFT2 | 0.99621868 | 1 | 0.325 | 4.666 | chr12 | 117175983 | 117291436 | + | protein_coding |
| ENSG00000136147.18_6 | PHF11 | 0.95582672 | 1 | 0.325 | 2.935 | chr13 | 50069746 | 50103117 | + | protein_coding |
| ENSG00000160948.14_6 | VPS28 | 0.9979115 | 1 | 0.325 | 5.358 | chr8 | 145648984 | 145653946 | - | protein_coding |
| ENSG00000154957.14_7 | ZNF18 | 0.98364245 | 1 | 0.325 | 3.841 | chr17 | 11880756 | 11900792 | - | protein_coding |
| ENSG00000110900.15_3 | TSPAN11 | 0.5482139 | 1 | 0.324 | -1.016 | chr12 | 31079362 | 31149537 | + | protein_coding |
| ENSG00000136436.14_8 | CALCOCO2 | 0.9989583 | 1 | 0.324 | 6.631 | chr17 | 46908350 | 46943884 | + | protein_coding |
| ENSG00000170265.12_4 | ZNF282 | 0.99687891 | 1 | 0.324 | 5.198 | chr7 | 148892638 | 148923329 | + | protein_coding |
| ENSG00000060069.16_2 | CTDP1 | 0.95436584 | 1 | 0.324 | 4.854 | chr18 | 77439801 | 77516623 | + | protein_coding |
| ENSG00000159720.12_5 | ATP6V0D1 | 0.99953829 | 1 | 0.324 | 5.733 | chr16 | 67471917 | 67515084 | - | protein_coding |
| ENSG00000142185.16_7 | TRPM2 | 0.95232351 | 1 | 0.324 | 2.992 | chr21 | 45770046 | 45862964 | + | protein_coding |
| ENSG00000157954.15_3 | WIPI2 | 0.99904279 | 1 | 0.324 | 5.805 | chr7 | 5229827 | 5273471 | + | protein_coding |
| ENSG00000111206.12_5 | FOXM1 | 0.99976948 | 1 | 0.324 | 6.634 | chr12 | 2966847 | 2986321 | - | protein_coding |
| ENSG00000264548.1_6 | AC132872.2 | 0.58877638 | 1 | 0.324 | -0.233 | chr17 | 80172103 | 80175228 | + | lncRNA |
| ENSG00000206634.1 | SNORA22 | 0.59624771 | 1 | 0.324 | -0.153 | chr7 | 65220513 | 65220646 | + | snoRNA |
| ENSG00000148218.16_5 | ALAD | 0.98895407 | 1 | 0.324 | 4.169 | chr9 | 116148592 | 116163570 | - | protein_coding |
| ENSG00000151240.17_4 | DIP2C | 0.99766976 | 1 | 0.323 | 4.765 | chr10 | 320130 | 735608 | - | protein_coding |
| ENSG00000110090.13_4 | CPT1A | 0.99967099 | 1 | 0.323 | 6.069 | chr11 | 68522088 | 68611878 | - | protein_coding |

| | | | | | | | | | | |
|---|---|---|---|---|---|---|---|---|---|---|
| ENSG00000101236.17_4 | RNF24 | 0.99453067 | 1 | 0.323 | 4.343 | chr20 | 3907956 | 3996205 | - | protein_coding |
| ENSG00000175894.18_8 | TSPEAR | 0.56677301 | 1 | 0.323 | -0.374 | chr21 | 45917776 | 46131487 | - | protein_coding |
| ENSG00000223612.3_5 | AC241585.1 | 0.5705954 | 1 | 0.323 | -0.288 | chr1 | 148193338 | 148193856 | + | processed_pseudogene |
| ENSG00000075391.16_6 | RASAL2 | 0.99938859 | 1 | 0.323 | 5.820 | chr1 | 178063276 | 178453282 | + | protein_coding |
| ENSG00000198205.6_2 | ZXDA | 0.9165387 | 1 | 0.323 | 2.776 | chrX | 57933142 | 57937254 | - | protein_coding |
| ENSG00000092931.11_3 | MFSD11 | 0.99301899 | 1 | 0.322 | 4.115 | chr17 | 74731947 | 74777531 | + | protein_coding |
| ENSG00000255112.3_4 | CHMP1B | 0.9975325 | 1 | 0.322 | 5.081 | chr18 | 11851412 | 11854443 | + | protein_coding |
| ENSG00000182473.22_5 | EXOC7 | 0.99986588 | 1 | 0.322 | 6.525 | chr17 | 74077097 | 74117657 | - | protein_coding |
| ENSG00000283632.2_3 | EXOC3L2 | 0.60168488 | 1 | 0.322 | -0.211 | chr19 | 45715879 | 45748689 | - | protein_coding |
| ENSG00000010803.16_6 | SCMH1 | 0.99239347 | 1 | 0.322 | 4.338 | chr1 | 41492872 | 41707826 | - | protein_coding |
| ENSG00000204316.12_4 | MRPL38 | 0.767112 | 1 | 0.322 | 1.833 | chr17 | 73894724 | 73901494 | - | protein_coding |
| ENSG00000134882.16_6 | UBAC2 | 0.99860676 | 1 | 0.322 | 5.422 | chr13 | 99853028 | 100038758 | + | protein_coding |
| ENSG00000238227.8_8 | TMEM250 | 0.95406242 | 1 | 0.321 | 5.568 | chr9 | 139006427 | 139010721 | - | protein_coding |
| ENSG00000167123.19_6 | CERCAM | 0.99497992 | 1 | 0.321 | 5.363 | chr9 | 131174030 | 131199630 | + | protein_coding |
| ENSG00000120306.11_5 | CYSTM1 | 0.99920552 | 1 | 0.321 | 5.655 | chr5 | 139554741 | 139661637 | + | protein_coding |
| ENSG00000064490.14_5 | RFXANK | 0.9869507 | 1 | 0.321 | 3.837 | chr19 | 19303038 | 19312678 | + | protein_coding |
| ENSG00000096070.19_5 | BRPF3 | 0.99449199 | 1 | 0.321 | 4.549 | chr6 | 36164521 | 36200567 | + | protein_coding |
| ENSG00000170619.10_6 | COMMD5 | 0.9963682 | 1 | 0.321 | 4.514 | chr8 | 146066427 | 146079121 | - | protein_coding |
| ENSG00000168517.10_7 | HEXIM2 | 0.92985065 | 1 | 0.321 | 2.611 | chr17 | 43238067 | 43247407 | + | protein_coding |
| ENSG00000143387.13_5 | CTSK | 0.86626218 | 1 | 0.321 | 1.579 | chr1 | 150768684 | 150780736 | - | protein_coding |
| ENSG00000049759.18_6 | NEDD4L | 0.99985701 | 1 | 0.321 | 6.292 | chr18 | 55711458 | 56068772 | + | protein_coding |
| ENSG00000079950.14_4 | STX7 | 0.99655083 | 1 | 0.321 | 4.920 | chr6 | 132767006 | 132834337 | - | protein_coding |
| ENSG00000162496.9_5 | DHRS3 | 0.99989217 | 1 | 0.321 | 7.163 | chr1 | 12627939 | 12678216 | - | protein_coding |
| ENSG00000167565.13_5 | SERTAD3 | 0.99908739 | 1 | 0.321 | 5.477 | chr19 | 40946751 | 40950242 | - | protein_coding |
| ENSG00000197136.4_3 | PCNX3 | 0.99823787 | 1 | 0.321 | 6.671 | chr11 | 65383244 | 65404910 | + | protein_coding |
| ENSG00000105379.9_4 | ETFB | 0.99026259 | 1 | 0.321 | 4.457 | chr19 | 51848423 | 51869672 | - | protein_coding |
| ENSG00000258682.1_7 | AL132989.1 | 0.61755258 | 1 | 0.320 | 0.309 | chr14 | 58731380 | 58736399 | + | lncRNA |
| ENSG00000008283.16_5 | CYB561 | 0.99901304 | 1 | 0.320 | 5.293 | chr17 | 61509665 | 61523715 | - | protein_coding |
| ENSG00000237595.5_8 | LINC01275 | 0.57579759 | 1 | 0.320 | -0.577 | chr20 | 48779302 | 48782753 | - | lncRNA |
| ENSG00000106610.15_6 | STAG3L4 | 0.98074142 | 1 | 0.320 | 3.654 | chr7 | 66767608 | 66827937 | + | transcribed_unprocessed_pseudogene |
| ENSG00000008710.20_6 | PKD1 | 0.72507856 | 1 | 0.320 | 6.024 | chr16 | 2138709 | 2185899 | - | protein_coding |
| ENSG00000163704.12_4 | PRRT3 | 0.93292334 | 1 | 0.319 | 2.578 | chr3 | 9987226 | 9994092 | - | protein_coding |
| ENSG00000089094.19_5 | KDM2B | 0.99507767 | 1 | 0.319 | 4.980 | chr12 | 121866899 | 122018928 | - | protein_coding |
| ENSG00000166272.18_5 | WBP1L | 0.99844001 | 1 | 0.319 | 4.973 | chr10 | 104503705 | 104594273 | + | protein_coding |
| ENSG00000110841.14_8 | PPFIBP1 | 0.99957461 | 1 | 0.319 | 6.170 | chr12 | 27676364 | 27848497 | + | protein_coding |
| ENSG00000074603.19_7 | DPP8 | 0.99943355 | 1 | 0.319 | 6.186 | chr15 | 65734801 | 65810042 | - | protein_coding |
| ENSG00000101473.17_5 | ACOT8 | 0.9767867 | 1 | 0.319 | 3.511 | chr20 | 44470360 | 44486044 | - | protein_coding |
| ENSG00000130717.13_4 | UCK1 | 0.99575134 | 1 | 0.319 | 4.628 | chr9 | 134399188 | 134406651 | - | protein_coding |
| ENSG00000101577.9_5 | LPIN2 | 0.99915103 | 1 | 0.319 | 5.266 | chr18 | 2916992 | 3013313 | - | protein_coding |
| ENSG00000089154.11_4 | GCN1 | 0.99996651 | 1 | 0.318 | 8.089 | chr12 | 120565006 | 120632518 | - | protein_coding |
| ENSG00000226002.1_4 | GTF2IP14 | 0.56123894 | 1 | 0.318 | -0.654 | chr7 | 64544481 | 64560610 | + | unprocessed_pseudogene |
| ENSG00000147475.17_6 | ERLIN2 | 0.99953269 | 1 | 0.318 | 6.017 | chr8 | 37594119 | 37615940 | + | protein_coding |
| ENSG00000147586.10_6 | MRPS28 | 0.8179664 | 1 | 0.318 | 1.672 | chr8 | 80830952 | 80942524 | - | protein_coding |
| ENSG00000169814.15_7 | BTD | 0.99698535 | 1 | 0.318 | 4.651 | chr3 | 15642848 | 15763818 | + | protein_coding |
| ENSG00000233421.4_6 | LINC01783 | 0.77461173 | 1 | 0.318 | 1.410 | chr1 | 16860381 | 16862667 | - | lncRNA |
| ENSG00000134897.14_5 | BIVM | 0.97258264 | 1 | 0.318 | 3.291 | chr13 | 103451469 | 103493883 | + | protein_coding |
| ENSG00000215302.8_5 | AC127502.1 | 0.80078094 | 1 | 0.318 | 1.569 | chr15 | 30762982 | 30799826 | + | transcribed_unprocessed_pseudogene |
| ENSG00000265692.1_6 | LINC01970 | 0.65257721 | 1 | 0.317 | 0.226 | chr17 | 80247922 | 80250690 | - | lncRNA |
| ENSG00000238098.9_2 | ABCA17P | 0.72551733 | 1 | 0.317 | 0.847 | chr16 | 2389151 | 2476700 | + | transcribed_unitary_pseudogene |
| ENSG00000118705.17_4 | RPN2 | 0.99997633 | 1 | 0.317 | 8.256 | chr20 | 35806813 | 35870022 | + | protein_coding |
| ENSG00000100258.18_6 | LMF2 | 0.98599507 | 1 | 0.317 | 4.685 | chr22 | 50941378 | 50946131 | - | protein_coding |
| ENSG00000167969.13_6 | ECI1 | 0.99685332 | 1 | 0.317 | 5.133 | chr16 | 2289403 | 2302301 | - | protein_coding |
| ENSG00000087074.8_4 | PPP1R15A | 0.99832473 | 1 | 0.317 | 4.962 | chr19 | 49375678 | 49379315 | + | protein_coding |
| ENSG00000186998.16_4 | EMID1 | 0.57372422 | 1 | 0.317 | -0.243 | chr22 | 29601885 | 29655586 | + | protein_coding |
| ENSG00000159459.12_5 | UBR1 | 0.99907526 | 1 | 0.317 | 5.662 | chr15 | 43235095 | 43398311 | - | protein_coding |
| ENSG00000160570.14_6 | DEDD2 | 0.99910141 | 1 | 0.316 | 5.529 | chr19 | 42702750 | 42724292 | - | protein_coding |
| ENSG00000143622.11_5 | RIT1 | 0.99688538 | 1 | 0.316 | 4.661 | chr1 | 155867599 | 155881195 | - | protein_coding |
| ENSG00000011243.18_6 | AKAP8L | 0.99585958 | 1 | 0.316 | 4.528 | chr19 | 15490861 | 15529952 | - | protein_coding |

| | | | | | | | | | | |
|---|---|---|---|---|---|---|---|---|---|---|
| ENSG00000092330.18_8 | TINF2 | 0.99937526 | 1 | 0.316 | 5.380 | chr14 | 24708849 | 24711880 | - | protein_coding |
| ENSG00000258057.6_6 | BCDIN3D-AS1 | 0.86394149 | 1 | 0.316 | 1.834 | chr12 | 50221696 | 50234926 | + | lncRNA |
| ENSG00000197587.11_4 | DMBX1 | 0.65927727 | 1 | 0.316 | 0.564 | chr1 | 46955508 | 46981888 | + | protein_coding |
| ENSG00000100246.13_5 | DNAL4 | 0.89414074 | 1 | 0.316 | 2.162 | chr22 | 39174513 | 39190203 | - | protein_coding |
| ENSG00000157933.10_3 | SKI | 0.97267992 | 1 | 0.316 | 3.611 | chr1 | 2159758 | 2241652 | + | protein_coding |
| ENSG00000102471.14_5 | NDFIP2 | 0.99916051 | 1 | 0.316 | 5.284 | chr13 | 80055259 | 80130210 | + | protein_coding |
| ENSG00000178075.20_4 | GRAMD1C | 0.6538995 | 1 | 0.315 | 0.098 | chr3 | 113547029 | 113666021 | + | protein_coding |
| ENSG00000134297.6_5 | PLEKHA8P1 | 0.88392675 | 1 | 0.315 | 2.127 | chr12 | 45566847 | 45609824 | - | transcribed_processed_pseudogene |
| ENSG00000107819.13_3 | SFXN3 | 0.99939126 | 1 | 0.315 | 5.450 | chr10 | 102790991 | 102801001 | + | protein_coding |
| ENSG00000108474.17_4 | PIGL | 0.99059098 | 1 | 0.315 | 3.879 | chr17 | 16120505 | 16255111 | + | protein_coding |
| ENSG00000116584.20_7 | ARHGEF2 | 0.99942048 | 1 | 0.315 | 5.430 | chr1 | 155916642 | 155976861 | - | protein_coding |
| ENSG00000157077.14_3 | ZFYVE9 | 0.99855696 | 1 | 0.315 | 4.804 | chr1 | 52607766 | 52812358 | + | protein_coding |
| ENSG00000196357.12_6 | ZNF565 | 0.95798449 | 1 | 0.315 | 2.896 | chr19 | 36673178 | 36737159 | - | protein_coding |
| ENSG00000105427.10_4 | CNFN | 0.78973991 | 1 | 0.314 | 1.700 | chr19 | 42891171 | 42894449 | - | protein_coding |
| ENSG00000121749.15_6 | TBC1D15 | 0.99929874 | 1 | 0.314 | 5.177 | chr12 | 72233487 | 72321028 | + | protein_coding |
| ENSG00000167107.12_3 | ACSF2 | 0.99758618 | 1 | 0.314 | 4.523 | chr17 | 48503519 | 48552206 | + | protein_coding |
| ENSG00000279518.1_1 | AC083843.4 | 0.57796479 | 1 | 0.314 | -0.352 | chr8 | 135795937 | 135798122 | - | TEC |
| ENSG00000081377.17_6 | CDC14B | 0.99569162 | 1 | 0.314 | 4.508 | chr9 | 99252523 | 99382125 | - | protein_coding |
| ENSG00000254910.1_7 | AC136475.2 | 0.62261133 | 1 | 0.314 | -0.221 | chr11 | 310139 | 311141 | - | lncRNA |
| ENSG00000100221.10_4 | JOSD1 | 0.99970075 | 1 | 0.314 | 5.724 | chr22 | 39081548 | 39097561 | - | protein_coding |
| ENSG00000079313.15_6 | REXO1 | 0.99009225 | 1 | 0.313 | 4.376 | chr19 | 1815247 | 1848482 | - | protein_coding |
| ENSG00000228150.1_7 | AL357140.2 | 0.56237414 | 1 | 0.313 | -1.123 | chr1 | 10002981 | 10010032 | + | lncRNA |
| ENSG00000114446.5_7 | IFT57 | 0.99942691 | 1 | 0.313 | 5.619 | chr3 | 107879659 | 107941282 | - | protein_coding |
| ENSG00000073614.13_5 | KDM5A | 0.99995239 | 1 | 0.313 | 7.145 | chr12 | 389223 | 498486 | - | protein_coding |
| ENSG00000180964.17_4 | TCEAL8 | 0.99848131 | 1 | 0.313 | 4.878 | chrX | 102507923 | 102510120 | - | protein_coding |
| ENSG00000173064.13_8 | HECTD4 | 0.9998375 | 1 | 0.313 | 6.200 | chr12 | 112597992 | 112820243 | - | protein_coding |
| ENSG00000108786.10_3 | HSD17B1 | 0.85979286 | 1 | 0.313 | 1.783 | chr17 | 40701232 | 40707231 | + | protein_coding |
| ENSG00000267244.5_6 | AC012615.6 | 0.71486546 | 1 | 0.313 | 0.849 | chr19 | 1822088 | 1824541 | + | lncRNA |
| ENSG00000122390.19_5 | NAA60 | 0.99610081 | 1 | 0.312 | 4.675 | chr16 | 3493649 | 3536953 | + | protein_coding |
| ENSG00000143842.15_8 | SOX13 | 0.98660582 | 1 | 0.312 | 4.242 | chr1 | 204042243 | 204096871 | + | protein_coding |
| ENSG00000143641.10_3 | GALNT2 | 0.99984469 | 1 | 0.312 | 6.533 | chr1 | 230193737 | 230417868 | + | protein_coding |
| ENSG00000070882.13_7 | OSBPL3 | 0.99931457 | 1 | 0.312 | 5.472 | chr7 | 24836159 | 25021253 | - | protein_coding |
| ENSG00000198585.11_2 | NUDT16 | 0.99774682 | 1 | 0.312 | 4.609 | chr3 | 131100515 | 131107674 | + | protein_coding |
| ENSG00000125089.17_6 | SH3TC1 | 0.98590747 | 1 | 0.312 | 4.396 | chr4 | 8183799 | 8243530 | + | protein_coding |
| ENSG00000204642.14_4 | HLA-F | 0.57499262 | 1 | 0.312 | -0.727 | chr6 | 29690552 | 29706305 | + | protein_coding |
| ENSG00000182873.5_7 | PRKCZ-AS1 | 0.6089586 | 1 | 0.312 | -0.274 | chr1 | 2113233 | 2115828 | - | lncRNA |
| ENSG00000137824.16_7 | RMDN3 | 0.9886683 | 1 | 0.312 | 3.808 | chr15 | 41028082 | 41048049 | - | protein_coding |
| ENSG00000169981.11_5 | ZNF35 | 0.97374729 | 1 | 0.312 | 3.541 | chr3 | 44690224 | 44702283 | + | protein_coding |
| ENSG00000179698.14_3 | WDR97 | 0.927345 | 1 | 0.312 | 3.066 | chr8 | 145162629 | 145173231 | + | protein_coding |
| ENSG00000140464.20_7 | PML | 0.9593574 | 1 | 0.312 | 3.218 | chr15 | 74287014 | 74340168 | + | protein_coding |
| ENSG00000139835.14_5 | GRTP1 | 0.81649964 | 1 | 0.312 | 1.466 | chr13 | 113978478 | 114018463 | - | protein_coding |
| ENSG00000114126.17_6 | TFDP2 | 0.9997144 | 1 | 0.312 | 5.758 | chr3 | 141663270 | 141868386 | - | protein_coding |
| ENSG00000066455.13_3 | GOLGA5 | 0.9998506 | 1 | 0.312 | 6.136 | chr14 | 93260650 | 93306292 | + | protein_coding |
| ENSG00000184792.16_6 | OSBP2 | 0.99825469 | 1 | 0.312 | 4.730 | chr22 | 31089769 | 31303811 | + | protein_coding |
| ENSG00000185267.10_5 | CDNF | 0.58899413 | 1 | 0.311 | -0.248 | chr10 | 14861244 | 14880574 | - | protein_coding |
| ENSG00000278619.5_3 | MRM1 | 0.99677012 | 1 | 0.311 | 4.485 | chr17 | 34958012 | 34965400 | + | protein_coding |
| ENSG00000232648.4_7 | AC107214.1 | 0.60707382 | 1 | 0.311 | -0.170 | chr4 | 184416037 | 184425647 | - | lncRNA |
| ENSG00000175182.15_4 | FAM131A | 0.94900684 | 1 | 0.310 | 2.832 | chr3 | 184053714 | 184066209 | + | protein_coding |
| ENSG00000083750.12_6 | RRAGB | 0.9630359 | 1 | 0.310 | 3.498 | chrX | 55744172 | 55785207 | + | protein_coding |
| ENSG00000156381.9_6 | ANKRD9 | 0.9959825 | 1 | 0.310 | 4.756 | chr14 | 102968104 | 102976136 | - | protein_coding |
| ENSG00000227695.6_6 | DNMBP-AS1 | 0.57766855 | 1 | 0.310 | -0.616 | chr10 | 101686767 | 101718807 | + | lncRNA |
| ENSG00000116729.14_8 | WLS | 0.98946033 | 1 | 0.310 | 3.686 | chr1 | 68564156 | 68698803 | - | protein_coding |
| ENSG00000270959.1_5 | LPP-AS2 | 0.86120135 | 1 | 0.310 | 1.821 | chr3 | 187868994 | 187871845 | - | lncRNA |
| ENSG00000287156.1_1 | AL138479.2 | 0.57116767 | 1 | 0.310 | -0.894 | chr14 | 54421389 | 54428952 | + | lncRNA |
| ENSG00000165338.16_4 | HECTD2 | 0.9963959 | 1 | 0.310 | 4.518 | chr10 | 93169037 | 93274586 | + | protein_coding |
| ENSG00000123201.14_4 | GUCY1B2 | 0.83004783 | 1 | 0.310 | 1.647 | chr13 | 51568647 | 51654998 | - | transcribed_unitary_pseudogene |
| ENSG00000169733.12_5 | RFNG | 0.98946298 | 1 | 0.310 | 4.362 | chr17 | 80005778 | 80009707 | - | protein_coding |
| ENSG00000166924.9_4 | NYAP1 | 0.75296898 | 1 | 0.310 | 1.057 | chr7 | 100081550 | 100092425 | + | protein_coding |
| ENSG00000163694.15_5 | RBM47 | 0.99995256 | 1 | 0.310 | 7.011 | chr4 | 40425284 | 40632892 | - | protein_coding |

| | | | | | | | | | | |
|---|---|---|---|---|---|---|---|---|---|---|
| ENSG00000167378.8_4 | IRGQ | 0.99985358 | 1 | 0.310 | 5.981 | chr19 | 44088521 | 44100287 | - | protein_coding |
| ENSG00000141349.8_2 | G6PC3 | 0.99975245 | 1 | 0.310 | 5.891 | chr17 | 42148103 | 42153712 | + | protein_coding |
| ENSG00000197021.9_6 | CXorf40B | 0.98286629 | 1 | 0.309 | 3.860 | chrX | 149097745 | 149106918 | - | protein_coding |
| ENSG00000270083.1_7 | AL021878.2 | 0.60194777 | 1 | 0.309 | -0.452 | chr22 | 42485634 | 42486032 | - | lncRNA |
| ENSG00000157653.11_2 | C9orf43 | 0.61873996 | 1 | 0.309 | 0.041 | chr9 | 116172334 | 116191964 | + | protein_coding |
| ENSG00000187013.4_3 | C17orf82 | 0.57399778 | 1 | 0.309 | -0.578 | chr17 | 59489112 | 59490641 | + | lncRNA |
| ENSG00000241258.7_4 | CRCP | 0.99785583 | 1 | 0.309 | 4.851 | chr7 | 65579591 | 65619555 | + | protein_coding |
| ENSG00000261654.1_6 | AL360270.2 | 0.71175891 | 1 | 0.309 | 0.963 | chr1 | 111478991 | 111484975 | - | lncRNA |
| ENSG00000072518.20_3 | MARK2 | 0.99941095 | 1 | 0.309 | 5.191 | chr11 | 63606400 | 63678491 | + | protein_coding |
| ENSG00000163171.8_3 | CDC42EP3 | 0.99994956 | 1 | 0.309 | 6.685 | chr2 | 37869025 | 37965611 | - | protein_coding |
| ENSG00000126603.8_2 | GLIS2 | 0.95912638 | 1 | 0.309 | 3.229 | chr16 | 4364762 | 4389598 | + | protein_coding |
| ENSG00000173137.12_3 | ADCK5 | 0.96742407 | 1 | 0.309 | 3.213 | chr8 | 145596790 | 145618457 | + | protein_coding |
| ENSG00000111144.10_4 | LTA4H | 0.99996147 | 1 | 0.308 | 7.238 | chr12 | 96394531 | 96437298 | - | protein_coding |
| ENSG00000169992.10_3 | NLGN2 | 0.9772544 | 1 | 0.308 | 3.954 | chr17 | 7308193 | 7323179 | + | protein_coding |
| ENSG00000257718.1_6 | CPNE8-AS1 | 0.60381749 | 1 | 0.308 | -0.192 | chr12 | 39300253 | 39303394 | + | lncRNA |
| ENSG00000173852.14_5 | DPY19L1 | 0.99997211 | 1 | 0.308 | 7.596 | chr7 | 34968488 | 35077883 | - | protein_coding |
| ENSG00000164074.15_4 | ABHD18 | 0.97870224 | 1 | 0.308 | 3.671 | chr4 | 128886461 | 128961082 | + | protein_coding |
| ENSG00000119522.16_5 | DENND1A | 0.99931351 | 1 | 0.308 | 5.054 | chr9 | 126141933 | 126692431 | - | protein_coding |
| ENSG00000011454.17_4 | RABGAP1 | 0.99962863 | 1 | 0.308 | 6.169 | chr9 | 125703112 | 125867145 | + | protein_coding |
| ENSG00000273893.1_7 | AL133520.1 | 0.59341008 | 1 | 0.308 | -0.551 | chr20 | 45310315 | 45311014 | - | lncRNA |
| ENSG00000167182.15_7 | SP2 | 0.92255108 | 1 | 0.308 | 2.561 | chr17 | 45973516 | 46006323 | + | protein_coding |
| ENSG00000179295.17_3 | PTPN11 | 0.99997353 | 1 | 0.308 | 8.121 | chr12 | 112856155 | 112947717 | + | protein_coding |
| ENSG00000130254.12_5 | SAFB2 | 0.99600063 | 1 | 0.308 | 5.015 | chr19 | 5587010 | 5624057 | - | protein_coding |
| ENSG00000196083.10_6 | IL1RAP | 0.96365742 | 1 | 0.307 | 2.839 | chr3 | 190231840 | 190377539 | + | protein_coding |
| ENSG00000078699.21_3 | CBFA2T2 | 0.99699591 | 1 | 0.307 | 4.440 | chr20 | 32077881 | 32237842 | + | protein_coding |
| ENSG00000108840.15_5 | HDAC5 | 0.99678203 | 1 | 0.307 | 4.553 | chr17 | 42154114 | 42201070 | - | protein_coding |
| ENSG00000158480.11_4 | SPATA2 | 0.98401468 | 1 | 0.307 | 3.688 | chr20 | 48519928 | 48532066 | - | protein_coding |
| ENSG00000276550.4_2 | HERC2P2 | 0.99991634 | 1 | 0.307 | 6.402 | chr15 | 23282281 | 23377526 | - | transcribed_unprocessed_pseudogene |
| ENSG00000185163.10_4 | DDX51 | 0.99762657 | 1 | 0.307 | 5.369 | chr12 | 132621139 | 132628864 | - | protein_coding |
| ENSG00000142627.13_4 | EPHA2 | 0.99962699 | 1 | 0.307 | 7.471 | chr1 | 16450832 | 16482564 | - | protein_coding |
| ENSG00000142186.17_5 | SCYL1 | 0.99976143 | 1 | 0.307 | 6.025 | chr11 | 65292548 | 65306175 | + | protein_coding |
| ENSG00000112473.18_5 | SLC39A7 | 0.99721981 | 1 | 0.307 | 4.689 | chr6 | 33168222 | 33172216 | + | protein_coding |
| ENSG00000108839.12_4 | ALOX12 | 0.75308077 | 1 | 0.307 | 1.001 | chr17 | 6899368 | 6914073 | + | protein_coding |
| ENSG00000169221.14_4 | TBC1D10B | 0.9981628 | 1 | 0.306 | 5.627 | chr16 | 30368423 | 30381815 | - | protein_coding |
| ENSG00000166734.20_5 | CASC4 | 0.99990154 | 1 | 0.306 | 6.920 | chr15 | 44580917 | 44707956 | + | protein_coding |
| ENSG00000079387.14_7 | SENP1 | 0.99962214 | 1 | 0.306 | 5.797 | chr12 | 48436680 | 48499862 | - | protein_coding |
| ENSG00000104805.16_7 | NUCB1 | 0.99967445 | 1 | 0.306 | 6.786 | chr19 | 49403569 | 49426629 | + | protein_coding |
| ENSG00000197149.5_5 | AC107956.1 | 0.85064671 | 1 | 0.306 | 1.931 | chr11 | 17249826 | 17250698 | + | processed_pseudogene |
| ENSG00000184492.6_2 | FOXD4L1 | 0.57389196 | 1 | 0.306 | -0.894 | chr2 | 114256242 | 114258732 | + | protein_coding |
| ENSG00000070413.20_6 | DGCR2 | 0.99920127 | 1 | 0.305 | 5.812 | chr22 | 19023795 | 19109967 | - | protein_coding |
| ENSG00000110717.13_5 | NDUFS8 | 0.99963775 | 1 | 0.305 | 6.657 | chr11 | 67798084 | 67804111 | + | protein_coding |
| ENSG00000174738.13_3 | NR1D2 | 0.99873734 | 1 | 0.305 | 5.285 | chr3 | 23986777 | 24022108 | + | protein_coding |
| ENSG00000161217.12_6 | PCYT1A | 0.99941956 | 1 | 0.305 | 5.535 | chr3 | 195941093 | 196014828 | - | protein_coding |
| ENSG00000067064.11_5 | IDI1 | 0.99989128 | 1 | 0.305 | 6.252 | chr10 | 1085092 | 1095110 | - | protein_coding |
| ENSG00000075407.18_3 | ZNF37A | 0.99854693 | 1 | 0.305 | 5.326 | chr10 | 38383262 | 38439221 | + | protein_coding |
| ENSG00000113580.15_7 | NR3C1 | 0.99998349 | 1 | 0.305 | 8.138 | chr5 | 142657496 | 142815077 | - | protein_coding |
| ENSG00000261242.1_7 | AL136038.5 | 0.57254528 | 1 | 0.305 | -1.031 | chr14 | 64010287 | 64011382 | + | lncRNA |
| ENSG00000110871.15_6 | COQ5 | 0.99920078 | 1 | 0.305 | 5.179 | chr12 | 120941082 | 120972237 | - | protein_coding |
| ENSG00000049656.14_4 | CLPTM1L | 0.99996319 | 1 | 0.305 | 7.391 | chr5 | 1317867 | 1345214 | - | protein_coding |
| ENSG00000196323.14_8 | ZBTB44 | 0.99976323 | 1 | 0.305 | 6.049 | chr11 | 130096572 | 130184812 | - | protein_coding |
| ENSG00000237125.10_6 | HAND2-AS1 | 0.77849344 | 1 | 0.305 | 1.070 | chr4 | 174448421 | 174580847 | + | lncRNA |
| ENSG00000163617.11_5 | CCDC191 | 0.93403884 | 1 | 0.305 | 2.633 | chr3 | 113682984 | 113775441 | - | protein_coding |
| ENSG00000137601.16_7 | NEK1 | 0.98069798 | 1 | 0.304 | 3.381 | chr4 | 170314008 | 170533780 | - | protein_coding |
| ENSG00000166171.13_3 | DPCD | 0.98039776 | 1 | 0.304 | 3.497 | chr10 | 103330317 | 103369419 | + | protein_coding |
| ENSG00000002016.18_6 | RAD52 | 0.99688334 | 1 | 0.304 | 4.569 | chr12 | 1020902 | 1099219 | - | protein_coding |
| ENSG00000119321.9_8 | FKBP15 | 0.99954866 | 1 | 0.304 | 5.535 | chr9 | 115923286 | 115983598 | - | protein_coding |
| ENSG00000143418.19_6 | CERS2 | 0.99982729 | 1 | 0.304 | 6.447 | chr1 | 150933059 | 150947480 | - | protein_coding |
| ENSG00000196372.13_4 | ASB13 | 0.99352216 | 1 | 0.304 | 4.337 | chr10 | 5680830 | 5708558 | - | protein_coding |
| ENSG00000256537.5_3 | SMIM10L1 | 0.99687314 | 1 | 0.304 | 4.537 | chr12 | 11323793 | 11328615 | + | protein_coding |
| ENSG00000120875.9_4 | DUSP4 | 0.99998962 | 1 | 0.303 | 8.684 | chr8 | 29190581 | 29208201 | - | protein_coding |

| | | | | | | | | | | |
|---|---|---|---|---|---|---|---|---|---|---|
| ENSG00000228300.14_6 | FAM174C | 0.98310054 | 1 | 0.303 | 3.839 | chr19 | 1275529 | 1279243 | + | protein_coding |
| ENSG00000020129.16_4 | NCDN | 0.99721749 | 1 | 0.303 | 4.394 | chr1 | 36023074 | 36032875 | + | protein_coding |
| ENSG00000108821.13_4 | COL1A1 | 0.92032412 | 1 | 0.303 | 1.842 | chr17 | 48260650 | 48278993 | - | protein_coding |
| ENSG00000144026.12_6 | ZNF514 | 0.9992542 | 1 | 0.303 | 5.040 | chr2 | 95810745 | 95831158 | - | protein_coding |
| ENSG00000146776.14_6 | ATXN7L1 | 0.96323533 | 1 | 0.303 | 2.832 | chr7 | 105245514 | 105517050 | - | protein_coding |
| ENSG00000048471.14_4 | SNX29 | 0.98730553 | 1 | 0.303 | 3.814 | chr16 | 12070591 | 12668144 | + | protein_coding |
| ENSG00000008382.15_5 | MPND | 0.88479317 | 1 | 0.303 | 1.929 | chr19 | 4343524 | 4360083 | + | protein_coding |
| ENSG00000185621.11_6 | LMLN | 0.95037814 | 1 | 0.303 | 2.639 | chr3 | 197687071 | 197770591 | + | protein_coding |
| ENSG00000170871.12_3 | KIAA0232 | 0.99974468 | 1 | 0.302 | 6.018 | chr4 | 6783102 | 6885897 | + | protein_coding |
| ENSG00000173546.7_4 | CSPG4 | 0.57844597 | 1 | 0.302 | -0.617 | chr15 | 75966663 | 76005189 | - | protein_coding |
| ENSG00000133243.10_6 | BTBD2 | 0.9976038 | 1 | 0.302 | 5.770 | chr19 | 1985437 | 2034880 | - | protein_coding |
| ENSG00000203778.8_3 | FAM229B | 0.90374639 | 1 | 0.302 | 2.030 | chr6 | 112408794 | 112423993 | + | protein_coding |
| ENSG00000099381.18_3 | SETD1A | 0.99603995 | 1 | 0.302 | 4.874 | chr16 | 30969075 | 30995985 | + | protein_coding |
| ENSG00000142733.17_7 | MAP3K6 | 0.99298564 | 1 | 0.302 | 4.195 | chr1 | 27680558 | 27693452 | - | protein_coding |
| ENSG00000134824.14_5 | FADS2 | 0.99997247 | 1 | 0.302 | 9.054 | chr11 | 61560452 | 61634826 | + | protein_coding |
| ENSG00000225313.5_7 | AL513327.1 | 0.74301929 | 1 | 0.302 | 1.022 | chr1 | 33772949 | 33814846 | + | lncRNA |
| ENSG00000009724.17_4 | MASP2 | 0.66966104 | 1 | 0.302 | -0.004 | chr1 | 11086580 | 11107296 | - | protein_coding |
| ENSG00000110711.10_6 | AIP | 0.99900078 | 1 | 0.302 | 5.096 | chr11 | 67250497 | 67258574 | + | protein_coding |
| ENSG00000205138.3_2 | SDHAF1 | 0.98618634 | 1 | 0.301 | 4.104 | chr19 | 36486101 | 36487217 | + | protein_coding |
| ENSG00000166484.20_4 | MAPK7 | 0.98951274 | 1 | 0.301 | 4.051 | chr17 | 19281034 | 19286857 | + | protein_coding |
| ENSG00000189077.11_5 | TMEM120A | 0.99411239 | 1 | 0.301 | 4.284 | chr7 | 75616149 | 75623974 | - | protein_coding |
| ENSG00000033100.16_4 | CHPF2 | 0.99989171 | 1 | 0.301 | 5.991 | chr7 | 150929575 | 150935913 | + | protein_coding |
| ENSG00000260558.2_6 | AC018557.2 | 0.58836411 | 1 | 0.301 | -0.658 | chr16 | 66772141 | 66773625 | + | lncRNA |
| ENSG00000107864.15_4 | CPEB3 | 0.81384506 | 1 | 0.301 | 1.004 | chr10 | 93806449 | 94050835 | - | protein_coding |
| ENSG00000174564.13_4 | IL20RB | 0.62915937 | 1 | 0.301 | -0.251 | chr3 | 136665072 | 136729927 | + | protein_coding |
| ENSG00000232850.3_4 | AL590708.1 | 0.63002125 | 1 | 0.301 | -0.101 | chr9 | 130890808 | 130892907 | + | lncRNA |
| ENSG00000182606.16_7 | TRAK1 | 0.99926702 | 1 | 0.301 | 5.401 | chr3 | 42055294 | 42267382 | + | protein_coding |
| ENSG00000136247.14_2 | ZDHHC4 | 0.99788955 | 1 | 0.300 | 4.477 | chr7 | 6617065 | 6629005 | + | protein_coding |
| ENSG00000148841.17_6 | ITPRIP | 0.98655151 | 1 | 0.300 | 3.869 | chr10 | 106069456 | 106098223 | - | protein_coding |
| ENSG00000060971.18_5 | ACAA1 | 0.99580047 | 1 | 0.300 | 4.051 | chr3 | 38144620 | 38178733 | - | protein_coding |
| ENSG00000108509.21_6 | CAMTA2 | 0.99599994 | 1 | 0.300 | 4.480 | chr17 | 4871287 | 4890970 | - | protein_coding |
| ENSG00000124299.15_5 | PEPD | 0.99973809 | 1 | 0.300 | 6.377 | chr19 | 33877856 | 34012697 | - | protein_coding |
| ENSG00000108091.11_4 | CCDC6 | 0.99995731 | 1 | 0.300 | 6.796 | chr10 | 61548505 | 61666314 | - | protein_coding |
| ENSG00000033011.12_5 | ALG1 | 0.99006505 | 1 | 0.300 | 4.176 | chr16 | 5083924 | 5137380 | + | protein_coding |
| ENSG00000145782.13_6 | ATG12 | 0.99936049 | 1 | 0.300 | 6.084 | chr5 | 115163897 | 115177534 | - | protein_coding |
| ENSG00000205885.7_6 | C1RL-AS1 | 0.99625911 | 1 | 0.300 | 4.543 | chr12 | 7260648 | 7275097 | + | lncRNA |
| ENSG00000267395.5_7 | DM1-AS | 0.74944275 | 1 | 0.300 | 0.879 | chr19 | 46271054 | 46275762 | + | lncRNA |
| ENSG00000171045.16_8 | TSNARE1 | 0.96693487 | 1 | 0.300 | 3.128 | chr8 | 143293441 | 143484543 | - | protein_coding |
| ENSG00000112659.14_5 | CUL9 | 0.99912118 | 1 | 0.300 | 4.899 | chr6 | 43149922 | 43192325 | + | protein_coding |
| ENSG00000135482.7_6 | ZC3H10 | 0.98380977 | 1 | 0.300 | 3.304 | chr12 | 56512044 | 56521298 | + | protein_coding |
| ENSG00000103507.14_6 | BCKDK | 0.99934395 | 1 | 0.300 | 5.123 | chr16 | 31117428 | 31124112 | + | protein_coding |
| ENSG00000284634.1_5 | AC092821.4 | 0.60545179 | 1 | 0.300 | -0.337 | chr12 | 9600891 | 9810988 | + | lncRNA |
| ENSG00000107745.19_7 | MICU1 | 0.99996361 | 1 | 0.299 | 6.757 | chr10 | 74127085 | 74385949 | - | protein_coding |
| ENSG00000126522.17_4 | ASL | 0.98710989 | 1 | 0.299 | 3.748 | chr7 | 65540787 | 65559684 | + | protein_coding |
| ENSG00000184436.12_4 | THAP7 | 0.99687338 | 1 | 0.299 | 4.285 | chr22 | 21353393 | 21356485 | - | protein_coding |
| ENSG00000213983.12_5 | AP1G2 | 0.9993018 | 1 | 0.299 | 5.044 | chr14 | 24028774 | 24037279 | - | protein_coding |
| ENSG00000166136.16_5 | NDUFB8 | 0.96979052 | 1 | 0.299 | 2.980 | chr10 | 102283497 | 102289757 | - | protein_coding |
| ENSG00000196998.19_8 | WDR45 | 0.99063529 | 1 | 0.299 | 3.822 | chrX | 48932092 | 48958108 | - | protein_coding |
| ENSG00000166794.4_3 | PPIB | 0.99991882 | 1 | 0.299 | 6.186 | chr15 | 64448011 | 64455404 | - | protein_coding |
| ENSG00000260233.3_6 | ZNRD2-AS1 | 0.6870781 | 1 | 0.299 | 0.384 | chr11 | 65335953 | 65337894 | - | lncRNA |
| ENSG00000074855.11_5 | ANO8 | 0.88474618 | 1 | 0.298 | 3.316 | chr19 | 17434032 | 17445664 | - | protein_coding |
| ENSG00000179632.10_3 | MAF1 | 0.99945795 | 1 | 0.298 | 6.839 | chr8 | 145159364 | 145162514 | + | protein_coding |
| ENSG00000108641.17_9 | B9D1 | 0.99889269 | 1 | 0.298 | 4.890 | chr17 | 19237621 | 19281506 | - | protein_coding |
| ENSG00000136051.14_3 | WASHC4 | 0.99996447 | 1 | 0.298 | 7.476 | chr12 | 105501102 | 105562912 | + | protein_coding |
| ENSG00000075643.6_4 | MOCOS | 0.9862793 | 1 | 0.298 | 3.741 | chr18 | 33767460 | 33852120 | + | protein_coding |
| ENSG00000146267.12_4 | FAXC | 0.95672501 | 1 | 0.298 | 2.888 | chr6 | 99719044 | 99797938 | - | protein_coding |
| ENSG00000214776.12_7 | AC092821.2 | 0.87352853 | 1 | 0.298 | 2.046 | chr12 | 9633204 | 9728871 | + | transcribed_unprocessed_pseudogene |
| ENSG00000133858.16_4 | ZFC3H1 | 0.99928822 | 1 | 0.298 | 5.565 | chr12 | 72003252 | 72061505 | - | protein_coding |
| ENSG00000223001.1 | RNU2-61P | 0.91202606 | 1 | 0.298 | 2.503 | chr6 | 89773219 | 89773409 | - | snRNA |
| ENSG00000162923.16_7 | WDR26 | 0.99993811 | 1 | 0.298 | 6.623 | chr1 | 224572845 | 224624735 | - | protein_coding |

| | | | | | | | | | | |
|---|---|---|---|---|---|---|---|---|---|---|
| ENSG00000125124.12_5 | BBS2 | 0.99885864 | 1 | 0.298 | 5.637 | chr16 | 56500748 | 56553999 | - | protein_coding |
| ENSG00000169955.7_4 | ZNF747 | 0.95267037 | 1 | 0.298 | 2.596 | chr16 | 30541688 | 30546668 | - | protein_coding |
| ENSG00000196422.11_4 | PPP1R26 | 0.99403687 | 1 | 0.297 | 5.644 | chr9 | 138370925 | 138380739 | + | protein_coding |
| ENSG00000276141.4_4 | WHAMMP3 | 0.96467635 | 1 | 0.297 | 2.826 | chr15 | 23186883 | 23208737 | - | transcribed_unprocessed_pseudogene |
| ENSG00000132522.16_5 | GPS2 | 0.72707455 | 1 | 0.297 | 0.615 | chr17 | 7214643 | 7218883 | - | protein_coding |
| ENSG00000234252.1 | AC007308.6 | 0.58269357 | 1 | 0.297 | -1.030 | chr22 | 21077335 | 21080208 | + | antisense |
| ENSG00000135617.4_4 | PRADC1 | 0.9753768 | 1 | 0.297 | 3.121 | chr2 | 73455138 | 73460367 | - | protein_coding |
| ENSG00000141582.15_7 | CBX4 | 0.99908866 | 1 | 0.297 | 5.189 | chr17 | 77806955 | 77813239 | - | protein_coding |
| ENSG00000184619.6_6 | KRBA2 | 0.71039532 | 1 | 0.297 | 0.618 | chr17 | 8260220 | 8280022 | - | protein_coding |
| ENSG00000213753.11_5 | CENPBD1P1 | 0.99647085 | 1 | 0.297 | 4.858 | chr19 | 59084870 | 59116590 | + | transcribed_processed_pseudogene |
| ENSG00000197763.17_10 | TXNRD3 | 0.96888337 | 1 | 0.297 | 3.109 | chr3 | 126325902 | 126373967 | - | protein_coding |
| ENSG00000107021.16_3 | TBC1D13 | 0.99774601 | 1 | 0.297 | 4.322 | chr9 | 131549532 | 131572709 | + | protein_coding |
| ENSG00000100151.16_4 | PICK1 | 0.98368671 | 1 | 0.296 | 3.869 | chr22 | 38452318 | 38471708 | + | protein_coding |
| ENSG00000186222.5_4 | BLOC1S4 | 0.96181627 | 1 | 0.296 | 3.268 | chr4 | 6717901 | 6719391 | + | protein_coding |
| ENSG00000078237.7_4 | TIGAR | 0.99101951 | 1 | 0.296 | 3.956 | chr12 | 4416929 | 4469194 | + | protein_coding |
| ENSG00000177692.11_3 | DNAJC28 | 0.63241004 | 1 | 0.296 | -0.125 | chr21 | 34857837 | 34864027 | - | protein_coding |
| ENSG00000277423.1_6 | AC069234.5 | 0.58424343 | 1 | 0.296 | -1.033 | chr12 | 121141670 | 121142085 | + | lncRNA |
| ENSG00000168404.13_5 | MLKL | 0.99917586 | 1 | 0.296 | 4.897 | chr16 | 74705753 | 74734858 | - | protein_coding |
| ENSG00000170876.7_3 | TMEM43 | 0.99992533 | 1 | 0.296 | 6.106 | chr3 | 14166440 | 14185179 | + | protein_coding |
| ENSG00000166685.12_6 | COG1 | 0.99898791 | 1 | 0.296 | 4.762 | chr17 | 71188771 | 71204646 | + | protein_coding |
| ENSG00000125375.15_6 | DMAC2L | 0.98140961 | 1 | 0.296 | 3.359 | chr14 | 50779042 | 50802276 | + | protein_coding |
| ENSG00000100003.18_5 | SEC14L2 | 0.96797024 | 1 | 0.296 | 3.152 | chr22 | 30792930 | 30821291 | + | protein_coding |
| ENSG00000123297.19_6 | TSFM | 0.62117752 | 1 | 0.296 | -0.428 | chr12 | 58176544 | 58201854 | + | protein_coding |
| ENSG00000127990.19_8 | SGCE | 0.99931268 | 1 | 0.296 | 5.638 | chr7 | 94153516 | 94285884 | - | protein_coding |
| ENSG00000148719.15_5 | DNAJB12 | 0.99983998 | 1 | 0.296 | 6.005 | chr10 | 74092588 | 74114907 | - | protein_coding |
| ENSG00000188897.9_9 | AC099489.1 | 0.66168011 | 1 | 0.295 | -0.023 | chr16 | 11453702 | 11621101 | - | protein_coding |
| ENSG00000169696.16_5 | ASPSCR1 | 0.99893172 | 1 | 0.295 | 5.313 | chr17 | 79934683 | 79975282 | + | protein_coding |
| ENSG00000174808.12_4 | BTC | 0.88750306 | 1 | 0.295 | 2.092 | chr4 | 75669969 | 75719733 | - | protein_coding |
| ENSG00000143537.13_3 | ADAM15 | 0.99972402 | 1 | 0.295 | 6.260 | chr1 | 155023042 | 155035251 | + | protein_coding |
| ENSG00000106009.16_4 | BRAT1 | 0.9988812 | 1 | 0.295 | 5.813 | chr7 | 2577444 | 2595328 | - | protein_coding |
| ENSG00000272763.1_7 | AC103702.2 | 0.89191452 | 1 | 0.295 | 1.922 | chr17 | 46713285 | 46724385 | - | lncRNA |
| ENSG00000126062.4_4 | TMEM115 | 0.99864116 | 1 | 0.295 | 4.811 | chr3 | 50392181 | 50396952 | - | protein_coding |
| ENSG00000127980.16_5 | PEX1 | 0.99959885 | 1 | 0.295 | 5.350 | chr7 | 92116334 | 92157834 | - | protein_coding |
| ENSG00000176383.9_4 | B3GNT4 | 0.70966641 | 1 | 0.294 | 0.792 | chr12 | 122688228 | 122693499 | + | protein_coding |
| ENSG00000148426.13_5 | PROSER2 | 0.99236455 | 1 | 0.294 | 3.853 | chr10 | 11865338 | 11914276 | + | protein_coding |
| ENSG00000260805.1 | RP11-61J19.4 | 0.59831753 | 1 | 0.294 | -0.552 | chr1 | 212731175 | 212733073 | + | lincRNA |
| ENSG00000166946.14_4 | CCNDBP1 | 0.99889705 | 1 | 0.294 | 5.041 | chr15 | 43477316 | 43489375 | + | protein_coding |
| ENSG00000029725.17_6 | RABEP1 | 0.99997175 | 1 | 0.294 | 6.577 | chr17 | 5185560 | 5289660 | + | protein_coding |
| ENSG00000228232.1_4 | GAPDHP1 | 0.9852958 | 1 | 0.294 | 3.804 | chrX | 39646386 | 39647390 | - | processed_pseudogene |
| ENSG00000158793.14_4 | NIT1 | 0.99887724 | 1 | 0.294 | 4.754 | chr1 | 161087876 | 161095235 | + | protein_coding |
| ENSG00000102362.15_3 | SYTL4 | 0.97871332 | 1 | 0.294 | 3.115 | chrX | 99929488 | 99987110 | - | protein_coding |
| ENSG00000175895.4_3 | PLEKHF2 | 0.9916844 | 1 | 0.294 | 3.903 | chr8 | 96146013 | 96168913 | + | protein_coding |
| ENSG00000146535.14_4 | GNA12 | 0.99921412 | 1 | 0.293 | 5.205 | chr7 | 2767739 | 2883942 | - | protein_coding |
| ENSG00000251169.2_5 | LINC01843 | 0.86758765 | 1 | 0.293 | 1.780 | chr5 | 133842243 | 133844920 | + | lncRNA |
| ENSG00000184752.13_5 | NDUFA12 | 0.99630259 | 1 | 0.293 | 5.594 | chr12 | 95290831 | 95397524 | - | protein_coding |
| ENSG00000135655.16_5 | USP15 | 0.99989369 | 1 | 0.293 | 6.149 | chr12 | 62654119 | 62811211 | + | protein_coding |
| ENSG00000206417.8_6 | H1FX-AS1 | 0.70128236 | 1 | 0.293 | 0.490 | chr3 | 129034235 | 129045068 | + | lncRNA |
| ENSG00000221916.4_3 | C19orf73 | 0.60349343 | 1 | 0.293 | -0.589 | chr19 | 49621659 | 49622400 | - | protein_coding |
| ENSG00000165983.14_2 | PTER | 0.99964868 | 1 | 0.293 | 5.492 | chr10 | 16478942 | 16555744 | + | protein_coding |
| ENSG00000137497.17_6 | NUMA1 | 0.99997238 | 1 | 0.293 | 7.157 | chr11 | 71713910 | 71791739 | - | protein_coding |
| ENSG00000088812.18_4 | ATRN | 0.99996359 | 1 | 0.293 | 6.782 | chr20 | 3451665 | 3631765 | + | protein_coding |
| ENSG00000236496.2_4 | GPS2P1 | 0.59900586 | 1 | 0.293 | -0.343 | chr9 | 2875442 | 2876427 | - | processed_pseudogene |
| ENSG00000013392.8_3 | RWDD2A | 0.94385472 | 1 | 0.293 | 2.619 | chr6 | 83903076 | 83908654 | + | protein_coding |
| ENSG00000157916.20_5 | RER1 | 0.99991246 | 1 | 0.293 | 6.087 | chr1 | 2323214 | 2336881 | + | protein_coding |
| ENSG00000111254.8_9 | AKAP3 | 0.62170621 | 1 | 0.292 | -0.315 | chr12 | 4724674 | 4758217 | - | protein_coding |
| ENSG00000182154.8_3 | MRPL41 | 0.94015872 | 1 | 0.292 | 4.867 | chr9 | 140446331 | 140447007 | + | protein_coding |
| ENSG00000157978.12_3 | LDLRAP1 | 0.9979589 | 1 | 0.292 | 4.285 | chr1 | 25870097 | 25895377 | + | protein_coding |
| ENSG00000162512.16_6 | SDC3 | 0.99732791 | 1 | 0.292 | 4.394 | chr1 | 31342313 | 31381605 | - | protein_coding |

| | | | | | | | | | | |
|---|---|---|---|---|---|---|---|---|---|---|
| ENSG00000152492.15_5 | CCDC50 | 0.99973934 | 1 | 0.292 | 5.774 | chr3 | 191046874 | 191116448 | + | protein_coding |
| ENSG00000281392.2_4 | AC107204.1 | 0.62175208 | 1 | 0.292 | -0.337 | chr3 | 87138279 | 87207513 | + | lncRNA |
| ENSG00000116685.16_4 | KIAA2013 | 0.99882996 | 1 | 0.292 | 5.170 | chr1 | 11979648 | 11986485 | - | protein_coding |
| ENSG00000135486.17_3 | HNRNPA1 | 0.77914504 | 1 | 0.292 | 8.020 | chr12 | 54673977 | 54680872 | + | protein_coding |
| ENSG00000051128.19_6 | HOMER3 | 0.96114038 | 1 | 0.291 | 2.739 | chr19 | 19040010 | 19052070 | - | protein_coding |
| ENSG00000100568.11_7 | VTI1B | 0.99939122 | 1 | 0.291 | 5.037 | chr14 | 68113802 | 68141537 | - | protein_coding |
| ENSG00000174007.8_5 | CEP19 | 0.96591052 | 1 | 0.291 | 2.696 | chr3 | 196433148 | 196439121 | - | protein_coding |
| ENSG00000134283.18_5 | PPHLN1 | 0.99992049 | 1 | 0.291 | 6.136 | chr12 | 42632249 | 42853517 | + | protein_coding |
| ENSG00000132394.11_3 | EEFSEC | 0.9669444 | 1 | 0.291 | 2.845 | chr3 | 127872324 | 128127489 | + | protein_coding |
| ENSG00000139354.11_8 | GAS2L3 | 0.99936377 | 1 | 0.291 | 5.221 | chr12 | 100967461 | 101022066 | + | protein_coding |
| ENSG00000164649.20_6 | CDCA7L | 0.99942817 | 1 | 0.291 | 5.104 | chr7 | 21940517 | 21985521 | - | protein_coding |
| ENSG00000167676.4_7 | PLIN4 | 0.75325789 | 1 | 0.291 | 0.782 | chr19 | 4502192 | 4518477 | - | protein_coding |
| ENSG00000278133.1_7 | AC135050.6 | 0.95695312 | 1 | 0.291 | 2.690 | chr16 | 31133556 | 31135385 | + | lncRNA |
| ENSG00000134686.18_5 | PHC2 | 0.99983815 | 1 | 0.291 | 5.565 | chr1 | 33789224 | 33896653 | - | protein_coding |
| ENSG00000138286.15_4 | FAM149B1 | 0.99796771 | 1 | 0.290 | 4.321 | chr10 | 74927877 | 75004262 | + | protein_coding |
| ENSG00000165300.7_2 | SLITRK5 | 0.98554662 | 1 | 0.290 | 3.286 | chr13 | 88324870 | 88348527 | + | protein_coding |
| ENSG00000166682.13_6 | TMPRSS5 | 0.5928459 | 1 | 0.290 | -0.943 | chr11 | 113558269 | 113577095 | - | protein_coding |
| ENSG00000204070.10_7 | SYS1 | 0.99827632 | 1 | 0.290 | 4.611 | chr20 | 43990577 | 44005438 | + | protein_coding |
| ENSG00000166394.15_4 | CYB5R2 | 0.94101534 | 1 | 0.290 | 2.410 | chr11 | 7686331 | 7698453 | - | protein_coding |
| ENSG00000165555.9_5 | NOXRED1 | 0.62601075 | 1 | 0.290 | 0.207 | chr14 | 77860364 | 77889860 | - | protein_coding |
| ENSG00000068366.20_6 | ACSL4 | 0.99992151 | 1 | 0.290 | 6.405 | chrX | 108867473 | 108976632 | - | protein_coding |
| ENSG00000160058.18_3 | BSDC1 | 0.99966711 | 1 | 0.290 | 5.308 | chr1 | 32830704 | 32860332 | - | protein_coding |
| ENSG00000047644.18_3 | WWC3 | 0.99996728 | 1 | 0.290 | 6.595 | chrX | 9983602 | 10112518 | + | protein_coding |
| ENSG00000125304.10_5 | TM9SF2 | 0.99997727 | 1 | 0.290 | 7.458 | chr13 | 100098565 | 100216302 | + | protein_coding |
| ENSG00000196371.3_3 | FUT4 | 0.87693793 | 1 | 0.290 | 2.103 | chr11 | 94277006 | 94283064 | + | protein_coding |
| ENSG00000125834.13_3 | STK35 | 0.99973617 | 1 | 0.289 | 6.092 | chr20 | 2082473 | 2157684 | + | protein_coding |
| ENSG00000201208.1 | Y_RNA | 0.59591264 | 1 | 0.289 | -0.945 | chr12 | 113616228 | 113616324 | - | misc_RNA |
| ENSG00000230487.8_6 | PSMG3-AS1 | 0.9535393 | 1 | 0.289 | 2.615 | chr7 | 1609709 | 1629262 | + | lncRNA |
| ENSG00000186812.13_6 | ZNF397 | 0.99568005 | 1 | 0.289 | 4.176 | chr18 | 32820994 | 32847097 | + | protein_coding |
| ENSG00000253671.3_8 | AC027117.1 | 0.99248567 | 1 | 0.289 | 4.167 | chr8 | 17665870 | 17679692 | + | lncRNA |
| ENSG00000179886.6_3 | TIGD5 | 0.99528485 | 1 | 0.289 | 4.180 | chr8 | 144680001 | 144685394 | + | protein_coding |
| ENSG00000105514.8_6 | RAB3D | 0.99434894 | 1 | 0.289 | 4.174 | chr19 | 11432744 | 11456946 | - | protein_coding |
| ENSG00000176595.4_4 | KBTBD11 | 0.99274833 | 1 | 0.289 | 4.094 | chr8 | 1921843 | 1955102 | + | protein_coding |
| ENSG00000100263.14_5 | RHBDD3 | 0.99401389 | 1 | 0.288 | 4.336 | chr22 | 29655861 | 29664198 | - | protein_coding |
| ENSG00000175087.10_5 | PDIK1L | 0.99298525 | 1 | 0.288 | 3.698 | chr1 | 26437656 | 26452046 | + | protein_coding |
| ENSG00000160062.15_4 | ZBTB8A | 0.91730077 | 1 | 0.288 | 2.049 | chr1 | 33005028 | 33071542 | + | protein_coding |
| ENSG00000176101.12_3 | SSNA1 | 0.99677862 | 1 | 0.288 | 5.312 | chr9 | 140083112 | 140084822 | + | protein_coding |
| ENSG00000106261.17_4 | ZKSCAN1 | 0.9999322 | 1 | 0.288 | 6.220 | chr7 | 99613195 | 99639312 | + | protein_coding |
| ENSG00000040531.15_5 | CTNS | 0.99911252 | 1 | 0.288 | 4.917 | chr17 | 3539753 | 3564836 | + | protein_coding |
| ENSG00000169410.10_6 | PTPN9 | 0.99988745 | 1 | 0.287 | 6.039 | chr15 | 75755592 | 75871656 | - | protein_coding |
| ENSG00000154710.17_4 | RABGEF1 | 0.7566456 | 1 | 0.287 | 0.897 | chr7 | 66147151 | 66276451 | + | protein_coding |
| ENSG00000170464.10_8 | DNAJC18 | 0.97721134 | 1 | 0.287 | 3.265 | chr5 | 138744277 | 138780180 | - | protein_coding |
| ENSG00000158805.12_4 | ZNF276 | 0.99281076 | 1 | 0.287 | 3.972 | chr16 | 89786808 | 89807333 | + | protein_coding |
| ENSG00000184584.13_7 | STING1 | 0.66653309 | 1 | 0.287 | -0.103 | chr5 | 138855118 | 138862520 | - | protein_coding |
| ENSG00000066739.12_4 | ATG2B | 0.99994048 | 1 | 0.287 | 6.154 | chr14 | 96745532 | 96829678 | - | protein_coding |
| ENSG00000177685.17_9 | CRACR2B | 0.9632392 | 1 | 0.287 | 3.783 | chr11 | 826144 | 831991 | + | protein_coding |
| ENSG00000273061.1_6 | CDC37L1-DT | 0.60096681 | 1 | 0.287 | -0.817 | chr9 | 4676600 | 4679502 | - | lncRNA |
| ENSG00000166167.18_4 | BTRC | 0.99951438 | 1 | 0.287 | 5.057 | chr10 | 103113790 | 103317078 | + | protein_coding |
| ENSG00000089053.13_4 | ANAPC5 | 0.99998791 | 1 | 0.287 | 7.489 | chr12 | 121746048 | 121837699 | - | protein_coding |
| ENSG00000277972.2_3 | CISD3 | 0.99682756 | 1 | 0.286 | 4.406 | chr17 | 36886594 | 36891858 | + | protein_coding |
| ENSG00000138867.16_5 | GUCD1 | 0.99994425 | 1 | 0.286 | 6.166 | chr22 | 24936391 | 24951903 | - | protein_coding |
| ENSG00000166813.15_4 | KIF7 | 0.97930741 | 1 | 0.286 | 3.600 | chr15 | 90152020 | 90198698 | - | protein_coding |
| ENSG00000238369.1 | snoU13 | 0.61239189 | 1 | 0.286 | -0.527 | chr5 | 149408912 | 149409015 | - | snoRNA |
| ENSG00000087903.13_6 | RFX2 | 0.98005523 | 1 | 0.286 | 3.067 | chr19 | 5993175 | 6199583 | - | protein_coding |
| ENSG00000279108.1_6 | AC008537.3 | 0.6414662 | 1 | 0.286 | -0.316 | chr19 | 41346064 | 41347944 | + | TEC |
| ENSG00000167705.12_5 | RILP | 0.96060851 | 1 | 0.286 | 2.597 | chr17 | 1549439 | 1553160 | - | protein_coding |
| ENSG00000056558.11_4 | TRAF1 | 0.95604932 | 1 | 0.286 | 2.428 | chr9 | 123664671 | 123691451 | - | protein_coding |
| ENSG00000130803.15_3 | ZNF317 | 0.99978771 | 1 | 0.286 | 5.317 | chr19 | 9251073 | 9274100 | + | protein_coding |
| ENSG00000276663.1_6 | AC009090.3 | 0.60262455 | 1 | 0.286 | -0.710 | chr16 | 57279744 | 57280308 | - | lncRNA |
| ENSG00000185900.10_3 | POMK | 0.99885154 | 1 | 0.286 | 4.503 | chr8 | 42948649 | 42978577 | + | protein_coding |
| ENSG00000163959.10_6 | SLC51A | 0.72890563 | 1 | 0.285 | 0.762 | chr3 | 195938358 | 195970049 | + | protein_coding |

| | | | | | | | | | | |
|---|---|---|---|---|---|---|---|---|---|---|
| ENSG00000285872.1_5 | AC007240.3 | 0.65709473 | 1 | 0.285 | -0.052 | chr2 | 10427926 | 10442611 | - | lncRNA |
| ENSG00000278594.1_6 | AL589182.2 | 0.62623101 | 1 | 0.285 | -0.501 | chr14 | 19719543 | 19720591 | - | processed_pseudogene |
| ENSG00000158122.12_7 | PRXL2C | 0.99900322 | 1 | 0.285 | 4.504 | chr9 | 99401859 | 99417599 | - | protein_coding |
| ENSG00000100744.15_6 | GSKIP | 0.99959159 | 1 | 0.285 | 5.175 | chr14 | 96829789 | 96853625 | + | protein_coding |
| ENSG00000213380.15_8 | COG8 | 0.99402124 | 1 | 0.284 | 3.707 | chr16 | 69354043 | 69373570 | - | protein_coding |
| ENSG00000012963.15_4 | UBR7 | 0.99909588 | 1 | 0.284 | 5.118 | chr14 | 93673586 | 93695561 | + | protein_coding |
| ENSG00000148384.13_5 | INPP5E | 0.99497087 | 1 | 0.284 | 4.788 | chr9 | 139323071 | 139334297 | - | protein_coding |
| ENSG00000173991.6_3 | TCAP | 0.61627964 | 1 | 0.284 | -0.630 | chr17 | 37821602 | 37822807 | + | protein_coding |
| ENSG00000241288.8_8 | LINC02614 | 0.64339844 | 1 | 0.284 | -0.202 | chr3 | 125546081 | 125635227 | - | lncRNA |
| ENSG00000133678.14_4 | TMEM254 | 0.97603826 | 1 | 0.284 | 2.921 | chr10 | 81838402 | 81852313 | + | protein_coding |
| ENSG00000177239.15_6 | MAN1B1 | 0.99983301 | 1 | 0.284 | 6.143 | chr9 | 139981437 | 140003635 | + | protein_coding |
| ENSG00000134294.14_5 | SLC38A2 | 0.99999403 | 1 | 0.283 | 9.528 | chr12 | 46751971 | 46766556 | - | protein_coding |
| ENSG00000070269.14_4 | TMEM260 | 0.99563242 | 1 | 0.283 | 4.142 | chr14 | 56955072 | 57117324 | + | protein_coding |
| ENSG00000007080.11_3 | CCDC124 | 0.98059864 | 1 | 0.283 | 4.542 | chr19 | 18043824 | 18054800 | + | protein_coding |
| ENSG00000174238.15_5 | PITPNA | 0.99961233 | 1 | 0.283 | 5.320 | chr17 | 1421012 | 1466086 | - | protein_coding |
| ENSG00000178301.4_3 | AQP11 | 0.92889222 | 1 | 0.283 | 1.795 | chr11 | 77300436 | 77321401 | + | protein_coding |
| ENSG00000269243.1_7 | AC008894.2 | 0.65304147 | 1 | 0.283 | -0.232 | chr19 | 16234471 | 16250702 | - | lncRNA |
| ENSG00000189241.7_5 | TSPYL1 | 0.99993221 | 1 | 0.283 | 5.785 | chr6 | 116588923 | 116601066 | - | protein_coding |
| ENSG00000133961.20_6 | NUMB | 0.99973293 | 1 | 0.283 | 5.241 | chr14 | 73741815 | 73925323 | - | protein_coding |
| ENSG00000117408.11_3 | IPO13 | 0.99930707 | 1 | 0.283 | 4.756 | chr1 | 44412622 | 44433694 | + | protein_coding |
| ENSG00000284747.1_5 | AL034417.5 | 0.7006293 | 1 | 0.283 | 0.285 | chr1 | 8051194 | 8065372 | + | lncRNA |
| ENSG00000138777.20_7 | PPA2 | 0.99985122 | 1 | 0.283 | 5.784 | chr4 | 106290234 | 106395224 | - | protein_coding |
| ENSG00000135314.12_5 | KHDC1 | 0.89860849 | 1 | 0.283 | 1.759 | chr6 | 73951037 | 74020088 | - | protein_coding |
| ENSG00000162241.12_3 | SLC25A45 | 0.99191305 | 1 | 0.283 | 3.841 | chr11 | 65142663 | 65151172 | - | protein_coding |
| ENSG00000124920.13_3 | MYRF | 0.99888442 | 1 | 0.283 | 6.312 | chr11 | 61520114 | 61555990 | + | protein_coding |
| ENSG00000185122.11_4 | HSF1 | 0.94486718 | 1 | 0.282 | 6.154 | chr8 | 145515273 | 145538383 | + | protein_coding |
| ENSG00000284968.1_4 | AC093827.4 | 0.63124616 | 1 | 0.282 | -0.318 | chr4 | 87845782 | 87857354 | - | lncRNA |
| ENSG00000145348.17_4 | TBCK | 0.99647078 | 1 | 0.282 | 4.234 | chr4 | 106962756 | 107237840 | - | protein_coding |
| ENSG00000184979.10_3 | USP18 | 0.97346298 | 1 | 0.282 | 3.061 | chr22 | 18632937 | 18660164 | + | protein_coding |
| ENSG00000263345.1_7 | AC006435.2 | 0.83618517 | 1 | 0.282 | 1.407 | chr17 | 2278355 | 2282600 | - | lncRNA |
| ENSG00000185164.10 | NOMO2 | 0.99864426 | 1 | 0.282 | 4.468 | chr16 | 18511182 | 18573533 | - | protein_coding |
| ENSG00000011478.12_3 | QPCTL | 0.98792114 | 1 | 0.282 | 3.462 | chr19 | 46195924 | 46207247 | + | protein_coding |
| ENSG00000215912.12_5 | TTC34 | 0.70856656 | 1 | 0.282 | 0.360 | chr1 | 2567415 | 2718282 | - | protein_coding |
| ENSG00000175066.16_5 | GK5 | 0.99980597 | 1 | 0.282 | 5.305 | chr3 | 141876369 | 141944434 | - | protein_coding |
| ENSG00000174684.7_4 | B4GAT1 | 0.99741113 | 1 | 0.282 | 4.325 | chr11 | 66112845 | 66115100 | - | protein_coding |
| ENSG00000183087.15_4 | GAS6 | 0.9402444 | 1 | 0.281 | 2.372 | chr13 | 114523522 | 114567049 | - | protein_coding |
| ENSG00000146576.13_4 | C7orf26 | 0.99640567 | 1 | 0.281 | 4.058 | chr7 | 6629652 | 6648357 | + | protein_coding |
| ENSG00000080189.15_6 | SLC35C2 | 0.99963484 | 1 | 0.281 | 5.467 | chr20 | 44974619 | 44993097 | - | protein_coding |
| ENSG00000110013.13_4 | SIAE | 0.99332386 | 1 | 0.281 | 3.816 | chr11 | 124503009 | 124565603 | - | protein_coding |
| ENSG00000099250.18_8 | NRP1 | 0.9999943 | 1 | 0.281 | 8.443 | chr10 | 33466420 | 33625190 | - | protein_coding |
| ENSG00000111860.14_7 | CEP85L | 0.98040607 | 1 | 0.281 | 3.408 | chr6 | 118781935 | 119031238 | - | protein_coding |
| ENSG00000025434.19_5 | NR1H3 | 0.97902025 | 1 | 0.281 | 3.054 | chr11 | 47269851 | 47290584 | + | protein_coding |
| ENSG00000068650.18_4 | ATP11A | 0.99853666 | 1 | 0.281 | 4.579 | chr13 | 113344643 | 113541482 | + | protein_coding |
| ENSG00000280424.1_5 | BX537318.2 | 0.68260611 | 1 | 0.281 | 0.150 | chr22 | 46402496 | 46406657 | + | TEC |
| ENSG00000100241.21_6 | SBF1 | 0.99410602 | 1 | 0.281 | 6.561 | chr22 | 50883429 | 50913464 | - | protein_coding |
| ENSG00000228981.3_6 | AC097658.2 | 0.66918292 | 1 | 0.281 | 0.218 | chr4 | 144270428 | 144271270 | - | processed_pseudogene |
| ENSG00000237149.5_5 | ZNF503-AS2 | 0.88322423 | 1 | 0.281 | 1.348 | chr10 | 77161277 | 77168740 | + | lncRNA |
| ENSG00000129625.13_5 | REEP5 | 0.99998538 | 1 | 0.281 | 6.591 | chr5 | 112212082 | 112257986 | - | protein_coding |
| ENSG00000106484.16_5 | MEST | 0.99931252 | 1 | 0.280 | 5.026 | chr7 | 130126012 | 130146306 | + | protein_coding |
| ENSG00000137216.19_4 | TMEM63B | 0.99946391 | 1 | 0.280 | 4.798 | chr6 | 44094651 | 44123256 | + | protein_coding |
| ENSG00000230291.5_5 | AC078817.1 | 0.63211158 | 1 | 0.280 | -0.102 | chr12 | 80496679 | 80497113 | - | processed_pseudogene |
| ENSG00000151743.11_6 | AMN1 | 0.96051692 | 1 | 0.280 | 2.834 | chr12 | 31824076 | 31882055 | - | protein_coding |
| ENSG00000272240.1_6 | AC004908.1 | 0.64129961 | 1 | 0.280 | -0.083 | chr8 | 184347 | 184887 | - | lncRNA |
| ENSG00000203867.8_3 | RBM20 | 0.95390638 | 1 | 0.280 | 2.590 | chr10 | 112404094 | 112599226 | + | protein_coding |
| ENSG00000185324.22_5 | CDK10 | 0.99959907 | 1 | 0.280 | 5.753 | chr16 | 89747145 | 89762762 | + | protein_coding |
| ENSG00000267858.6_7 | MZF1-AS1 | 0.97887568 | 1 | 0.280 | 3.089 | chr19 | 59070496 | 59086164 | + | lncRNA |
| ENSG00000160310.18_3 | PRMT2 | 0.99985042 | 1 | 0.280 | 5.613 | chr21 | 48055507 | 48085036 | + | protein_coding |
| ENSG00000110237.5_7 | ARHGEF17 | 0.99895573 | 1 | 0.280 | 5.029 | chr11 | 73019321 | 73080433 | + | protein_coding |
| ENSG00000106351.13_4 | AGFG2 | 0.99952222 | 1 | 0.280 | 5.375 | chr7 | 100136826 | 100165843 | + | protein_coding |

| | | | | | | | | | | |
|---|---|---|---|---|---|---|---|---|---|---|
| ENSG00000173653.8_4 | RCE1 | 0.99707562 | 1 | 0.280 | 4.301 | chr11 | 66610306 | 66614023 | + | protein_coding |
| ENSG00000099956.19_5 | SMARCB1 | 0.99969663 | 1 | 0.279 | 5.276 | chr22 | 24129118 | 24180195 | + | protein_coding |
| ENSG00000174903.16_4 | RAB1B | 0.99962488 | 1 | 0.279 | 5.273 | chr11 | 66036061 | 66044963 | + | protein_coding |
| ENSG00000270049.3_6 | AC009061.2 | 0.84328888 | 1 | 0.279 | 1.388 | chr16 | 67515217 | 67539880 | + | lncRNA |
| ENSG00000204619.8_3 | PPP1R11 | 0.89877489 | 1 | 0.279 | 2.031 | chr6 | 30034486 | 30038110 | + | protein_coding |
| ENSG00000273058.2_6 | AL359921.2 | 0.61018404 | 1 | 0.279 | -0.865 | chr1 | 236699462 | 236700004 | - | lncRNA |
| ENSG00000273033.2_5 | LINC02035 | 0.98872022 | 1 | 0.279 | 3.545 | chr3 | 122605788 | 122611263 | + | lncRNA |
| ENSG00000180488.15_6 | MIGA1 | 0.99913928 | 1 | 0.279 | 4.902 | chr1 | 78245309 | 78345224 | + | protein_coding |
| ENSG00000130150.12_5 | MOSPD2 | 0.9969437 | 1 | 0.279 | 4.688 | chrX | 14891543 | 14940449 | + | protein_coding |
| ENSG00000234664.1_3 | HMGN2P5 | 0.9413429 | 1 | 0.279 | 2.186 | chr15 | 30022917 | 30023189 | + | processed_pseudogene |
| ENSG00000206612.1 | SNORA2A | 0.60794774 | 1 | 0.278 | -0.997 | chr12 | 49050431 | 49050565 | - | snoRNA |
| ENSG00000112167.10_5 | SAYSD1 | 0.9907754 | 1 | 0.278 | 3.523 | chr6 | 39071839 | 39082962 | - | protein_coding |
| ENSG00000182584.5_4 | ACTL10 | 0.69077861 | 1 | 0.278 | 0.429 | chr20 | 32254749 | 32256331 | + | protein_coding |
| ENSG00000235852.1_7 | AC005540.1 | 0.62987016 | 1 | 0.278 | -0.439 | chr2 | 191745523 | 191746785 | - | lncRNA |
| ENSG00000023318.8_5 | ERP44 | 0.99993902 | 1 | 0.278 | 5.776 | chr9 | 102741467 | 102861282 | - | protein_coding |
| ENSG00000102096.9_4 | PIM2 | 0.99377513 | 1 | 0.278 | 3.619 | chrX | 48770459 | 48776301 | - | protein_coding |
| ENSG00000272734.1_5 | ADIRF-AS1 | 0.92238326 | 1 | 0.278 | 1.629 | chr10 | 88725102 | 88731068 | - | lncRNA |
| ENSG00000127511.9_3 | SIN3B | 0.99852786 | 1 | 0.278 | 4.727 | chr19 | 16940211 | 16991164 | + | protein_coding |
| ENSG00000143845.15_7 | ETNK2 | 0.88897896 | 1 | 0.278 | 1.841 | chr1 | 204100190 | 204121172 | - | protein_coding |
| ENSG00000118689.15_4 | FOXO3 | 0.99642328 | 1 | 0.278 | 4.281 | chr6 | 108881038 | 109005977 | + | protein_coding |
| ENSG00000255517.6_8 | AP002748.4 | 0.86312247 | 1 | 0.277 | 1.461 | chr11 | 66240961 | 66247704 | - | lncRNA |
| ENSG00000101181.17_3 | MTG2 | 0.99924208 | 1 | 0.277 | 4.636 | chr20 | 60758085 | 60778624 | + | protein_coding |
| ENSG00000089177.18_4 | KIF16B | 0.99988404 | 1 | 0.277 | 5.734 | chr20 | 16252749 | 16554079 | - | protein_coding |
| ENSG00000088970.16_7 | KIZ | 0.99168914 | 1 | 0.277 | 3.803 | chr20 | 21106624 | 21227260 | + | protein_coding |
| ENSG00000279541.1_8 | AC005261.6 | 0.90495514 | 1 | 0.277 | 1.776 | chr19 | 57804451 | 57805837 | - | TEC |
| ENSG00000164830.18_7 | OXR1 | 0.99970607 | 1 | 0.277 | 5.365 | chr8 | 107371704 | 107764922 | + | protein_coding |
| ENSG00000185986.12_5 | SDHAP3 | 0.98111647 | 1 | 0.277 | 2.898 | chr5 | 1567610 | 1594735 | - | transcribed_unprocessed_pseudogene |
| ENSG00000245958.6_6 | AC093752.1 | 0.96176209 | 1 | 0.277 | 2.560 | chr4 | 120375946 | 120473180 | + | transcribed_unprocessed_pseudogene |
| ENSG00000183864.5_5 | TOB2 | 0.99970558 | 1 | 0.277 | 5.069 | chr22 | 41829498 | 41842805 | - | protein_coding |
| ENSG00000109083.13_7 | IFT20 | 0.99921382 | 1 | 0.277 | 4.529 | chr17 | 26655351 | 26662515 | - | protein_coding |
| ENSG00000165458.14_7 | INPPL1 | 0.99995723 | 1 | 0.277 | 7.020 | chr11 | 71934745 | 71950191 | + | protein_coding |
| ENSG00000122912.15_5 | SLC25A16 | 0.99826237 | 1 | 0.277 | 4.526 | chr10 | 70237755 | 70287280 | - | protein_coding |
| ENSG00000097033.14_4 | SH3GLB1 | 0.99991078 | 1 | 0.276 | 5.939 | chr1 | 87170253 | 87213867 | + | protein_coding |
| ENSG00000049283.18_7 | EPN3 | 0.95447613 | 1 | 0.276 | 2.312 | chr17 | 48610043 | 48621111 | + | protein_coding |
| ENSG00000164307.13_6 | ERAP1 | 0.99980773 | 1 | 0.276 | 5.150 | chr5 | 96096514 | 96143803 | - | protein_coding |
| ENSG00000141522.12_4 | ARHGDIA | 0.99991684 | 1 | 0.276 | 7.159 | chr17 | 79825597 | 79829254 | - | protein_coding |
| ENSG00000223768.2_4 | LINC00205 | 0.99940286 | 1 | 0.276 | 4.706 | chr21 | 46707965 | 46717721 | + | lncRNA |
| ENSG00000163435.16_6 | ELF3 | 0.99998597 | 1 | 0.276 | 7.722 | chr1 | 201977073 | 201986311 | + | protein_coding |
| ENSG00000140632.17_5 | GLYR1 | 0.9999465 | 1 | 0.276 | 6.026 | chr16 | 4853204 | 4897289 | - | protein_coding |
| ENSG00000111203.13_6 | ITFG2 | 0.99208796 | 1 | 0.276 | 3.810 | chr12 | 2921788 | 2968957 | + | protein_coding |
| ENSG00000139131.13_7 | YARS2 | 0.9995519 | 1 | 0.276 | 4.890 | chr12 | 32880424 | 32908831 | - | protein_coding |
| ENSG00000115604.10_5 | IL18R1 | 0.78688592 | 1 | 0.276 | 1.101 | chr2 | 102927989 | 103015235 | + | protein_coding |
| ENSG00000023697.13_5 | DERA | 0.99988789 | 1 | 0.276 | 5.577 | chr12 | 16064236 | 16190315 | + | protein_coding |
| ENSG00000200488.1 | RN7SKP203 | 0.80463856 | 1 | 0.275 | 1.289 | chr2 | 76672205 | 76672536 | - | misc_RNA |
| ENSG00000257818.1_4 | C1GALT1P1 | 0.61171546 | 1 | 0.275 | -0.960 | chr12 | 69678177 | 69679265 | + | processed_pseudogene |
| ENSG00000157870.16_5 | PRXL2B | 0.9960549 | 1 | 0.275 | 3.905 | chr1 | 2517930 | 2522908 | + | protein_coding |
| ENSG00000143653.10_3 | SCCPDH | 0.99997926 | 1 | 0.275 | 6.283 | chr1 | 246887711 | 246931439 | + | protein_coding |
| ENSG00000133460.19_5 | SLC2A11 | 0.96190786 | 1 | 0.275 | 2.565 | chr22 | 24198890 | 24228496 | + | protein_coding |
| ENSG00000141959.17_2 | PFKL | 0.92394806 | 1 | 0.275 | 6.315 | chr21 | 45719934 | 45747259 | + | protein_coding |
| ENSG00000182195.9_5 | LDOC1 | 0.98957398 | 1 | 0.275 | 3.518 | chrX | 140205784 | 140271314 | - | protein_coding |
| ENSG00000063601.17_6 | MTMR1 | 0.99987256 | 1 | 0.275 | 5.386 | chrX | 149861444 | 149933581 | + | protein_coding |
| ENSG00000182648.13_7 | LINC01006 | 0.96976617 | 1 | 0.275 | 2.792 | chr7 | 156181616 | 156433348 | - | lncRNA |
| ENSG00000163946.13_5 | TASOR | 0.99999271 | 1 | 0.275 | 7.262 | chr3 | 56654161 | 56717265 | - | protein_coding |
| ENSG00000104853.16_6 | CLPTM1 | 0.99982345 | 1 | 0.275 | 6.775 | chr19 | 45457842 | 45496599 | + | protein_coding |
| ENSG00000164070.12_4 | HSPA4L | 0.99978477 | 1 | 0.275 | 5.867 | chr4 | 128702976 | 128761888 | + | protein_coding |
| ENSG00000248489.2_6 | LINC02062 | 0.6385858 | 1 | 0.274 | -0.440 | chr5 | 98264875 | 98330717 | + | lncRNA |
| ENSG00000119899.13_4 | SLC17A5 | 0.99640016 | 1 | 0.274 | 4.026 | chr6 | 74303102 | 74363715 | - | protein_coding |
| ENSG00000104904.12_6 | OAZ1 | 0.999992 | 1 | 0.274 | 7.431 | chr19 | 2269508 | 2273489 | + | protein_coding |
| ENSG00000169169.14_6 | CPT1C | 0.93312549 | 1 | 0.274 | 2.217 | chr19 | 50194155 | 50216988 | + | protein_coding |

| | | | | | | | | | | |
|---|---|---|---|---|---|---|---|---|---|---|
| ENSG00000104983.8_5 | CCDC61 | 0.93573166 | 1 | 0.274 | 2.476 | chr19 | 46498719 | 46524576 | + | protein_coding |
| ENSG00000259291.2_7 | ZNF710-AS1 | 0.81851066 | 1 | 0.274 | 1.349 | chr15 | 90617744 | 90625439 | - | lncRNA |
| ENSG00000164877.19_4 | MICALL2 | 0.99218604 | 1 | 0.274 | 4.249 | chr7 | 1468101 | 1499106 | - | protein_coding |
| ENSG00000162227.8_5 | TAF6L | 0.98969985 | 1 | 0.274 | 3.592 | chr11 | 62538829 | 62554814 | + | protein_coding |
| ENSG00000243679.1_5 | AC018638.5 | 0.86320725 | 1 | 0.274 | 1.740 | chr7 | 128294023 | 128294776 | + | processed_pseudogene |
| ENSG00000259407.1_7 | AC021739.2 | 0.61302624 | 1 | 0.274 | -1.141 | chr15 | 86287340 | 86293512 | - | lncRNA |
| ENSG00000103111.15_6 | MON1B | 0.99973516 | 1 | 0.274 | 5.479 | chr16 | 77224732 | 77236295 | + | protein_coding |
| ENSG00000251161.4_6 | AC020661.1 | 0.63809076 | 1 | 0.274 | -0.603 | chr15 | 41199009 | 41202535 | + | lncRNA |
| ENSG00000127914.16_9 | AKAP9 | 0.99959725 | 1 | 0.274 | 7.111 | chr7 | 91570181 | 91739987 | + | protein_coding |
| ENSG00000178573.7_4 | MAF | 0.61343104 | 1 | 0.273 | -1.049 | chr16 | 79619740 | 79634634 | - | protein_coding |
| ENSG00000129933.21_4 | MAU2 | 0.99956864 | 1 | 0.273 | 5.036 | chr19 | 19431638 | 19469563 | + | protein_coding |
| ENSG00000080298.15_6 | RFX3 | 0.98516958 | 1 | 0.273 | 3.229 | chr9 | 3218297 | 3526004 | - | protein_coding |
| ENSG00000109854.13_2 | HTATIP2 | 0.99998533 | 1 | 0.273 | 6.555 | chr11 | 20385231 | 20405329 | + | protein_coding |
| ENSG00000090006.17_5 | LTBP4 | 0.9688227 | 1 | 0.273 | 5.808 | chr19 | 41098789 | 41135723 | + | protein_coding |
| ENSG00000212123.4_5 | PRR22 | 0.80779049 | 1 | 0.273 | 1.165 | chr19 | 5782971 | 5784757 | - | protein_coding |
| ENSG00000079257.8_6 | LXN | 0.99946421 | 1 | 0.273 | 5.467 | chr3 | 158363611 | 158390437 | - | protein_coding |
| ENSG00000120690.16_6 | ELF1 | 0.99564081 | 1 | 0.273 | 4.900 | chr13 | 41506060 | 41635576 | - | protein_coding |
| ENSG00000105993.15_6 | DNAJB6 | 0.99996842 | 1 | 0.273 | 6.246 | chr7 | 157128075 | 157210133 | + | protein_coding |
| ENSG00000185436.12_4 | IFNLR1 | 0.92447245 | 1 | 0.273 | 2.223 | chr1 | 24480658 | 24514449 | - | protein_coding |
| ENSG00000129003.18_8 | VPS13C | 0.99874586 | 1 | 0.272 | 7.165 | chr15 | 62144588 | 62352672 | - | protein_coding |
| ENSG00000188107.15_5 | EYS | 0.84493375 | 1 | 0.272 | 0.915 | chr6 | 64429876 | 66417119 | - | protein_coding |
| ENSG00000165507.9_8 | DEPP1 | 0.6529787 | 1 | 0.272 | -0.160 | chr10 | 45466429 | 45474257 | - | protein_coding |
| ENSG00000182087.14_8 | TMEM259 | 0.99807767 | 1 | 0.272 | 5.787 | chr19 | 1009647 | 1021178 | - | protein_coding |
| ENSG00000283973.2_6 | AC099795.1 | 0.61702471 | 1 | 0.272 | -1.049 | chr1 | 43424736 | 43427535 | - | lncRNA |
| ENSG00000149485.19_5 | FADS1 | 0.99999809 | 1 | 0.272 | 9.074 | chr11 | 61567099 | 61596790 | - | protein_coding |
| ENSG00000147044.21_8 | CASK | 0.99993863 | 1 | 0.271 | 5.971 | chrX | 41374187 | 41782898 | - | protein_coding |
| ENSG00000264247.2_6 | LINC00909 | 0.9747559 | 1 | 0.271 | 2.908 | chr18 | 72257553 | 72266121 | - | lncRNA |
| ENSG00000167110.17_5 | GOLGA2 | 0.99997157 | 1 | 0.271 | 5.996 | chr9 | 131018108 | 131038274 | - | protein_coding |
| ENSG00000135390.20_9 | ATP5MC2 | 0.99997512 | 1 | 0.271 | 7.216 | chr12 | 54026510 | 54071192 | - | protein_coding |
| ENSG00000261408.7_6 | TEN1-CDK3 | 0.62522327 | 1 | 0.271 | -0.640 | chr17 | 73975312 | 74002080 | + | protein_coding |
| ENSG00000070018.9_6 | LRP6 | 0.99998674 | 1 | 0.271 | 7.272 | chr12 | 12268959 | 12419978 | - | protein_coding |
| ENSG00000163125.15_4 | RPRD2 | 0.99988763 | 1 | 0.271 | 5.564 | chr1 | 150335567 | 150449042 | + | protein_coding |
| ENSG00000101004.15_6 | NINL | 0.99981319 | 1 | 0.271 | 5.647 | chr20 | 25433333 | 25566167 | - | protein_coding |
| ENSG00000103253.18_3 | HAGHL | 0.97395722 | 1 | 0.271 | 3.442 | chr16 | 776936 | 785525 | + | protein_coding |
| ENSG00000160712.13_4 | IL6R | 0.99429864 | 1 | 0.270 | 3.826 | chr1 | 154377669 | 154441926 | + | protein_coding |
| ENSG00000238005.4_8 | AL391832.2 | 0.70790623 | 1 | 0.270 | 0.560 | chr1 | 235093089 | 235105809 | - | lncRNA |
| ENSG00000089063.15_5 | TMEM230 | 0.99999132 | 1 | 0.270 | 6.626 | chr20 | 5048878 | 5093749 | - | protein_coding |
| ENSG00000140992.19_5 | PDPK1 | 0.99886622 | 1 | 0.270 | 4.702 | chr16 | 2587980 | 2653189 | + | protein_coding |
| ENSG00000142046.15_4 | TMEM91 | 0.89952495 | 1 | 0.270 | 1.910 | chr19 | 41856816 | 41889988 | + | protein_coding |
| ENSG00000167100.14_6 | SAMD14 | 0.67643597 | 1 | 0.270 | 0.029 | chr17 | 48187404 | 48207246 | - | protein_coding |
| ENSG00000263050.1_7 | AC090617.5 | 0.70546926 | 1 | 0.270 | 0.182 | chr17 | 1946769 | 1948262 | - | lncRNA |
| ENSG00000124097.7_4 | HMGB1P1 | 0.62172864 | 1 | 0.269 | -0.642 | chr20 | 56063448 | 56064083 | - | processed_pseudogene |
| ENSG00000183735.10_4 | TBK1 | 0.99996379 | 1 | 0.269 | 6.271 | chr12 | 64845872 | 64895894 | + | protein_coding |
| ENSG00000182993.5_5 | C12orf60 | 0.84864507 | 1 | 0.269 | 1.226 | chr12 | 14956600 | 15059520 | + | protein_coding |
| ENSG00000070214.16_3 | SLC44A1 | 0.99999345 | 1 | 0.269 | 7.074 | chr9 | 108006903 | 108201452 | + | protein_coding |
| ENSG00000148300.12_4 | REXO4 | 0.9993742 | 1 | 0.269 | 4.930 | chr9 | 136271183 | 136283216 | - | protein_coding |
| ENSG00000088356.6_4 | PDRG1 | 0.98181963 | 1 | 0.269 | 3.338 | chr20 | 30532140 | 30539849 | - | protein_coding |
| ENSG00000130714.16_4 | POMT1 | 0.99978257 | 1 | 0.269 | 5.074 | chr9 | 134378289 | 134399193 | + | protein_coding |
| ENSG00000071243.16_3 | ING3 | 0.98474396 | 1 | 0.269 | 3.361 | chr7 | 120590817 | 120617270 | + | protein_coding |
| ENSG00000165175.15_3 | MID1IP1 | 0.99988217 | 1 | 0.269 | 5.886 | chrX | 38660685 | 38665790 | + | protein_coding |
| ENSG00000081087.15_5 | OSTM1 | 0.99987066 | 1 | 0.269 | 5.452 | chr6 | 108362613 | 108487058 | - | protein_coding |
| ENSG00000148225.16_5 | WDR31 | 0.95110933 | 1 | 0.268 | 2.265 | chr9 | 116075502 | 116102578 | - | protein_coding |
| ENSG00000253341.1_5 | AC115837.1 | 0.71496638 | 1 | 0.268 | 0.287 | chr8 | 75515479 | 75516479 | - | processed_pseudogene |
| ENSG00000159267.14_6 | HLCS | 0.99972193 | 1 | 0.268 | 5.115 | chr21 | 38123189 | 38362536 | - | protein_coding |
| ENSG00000197302.10_6 | ZNF720 | 0.99689257 | 1 | 0.268 | 4.230 | chr16 | 31724550 | 31806190 | + | protein_coding |
| ENSG00000125354.23_7 | SEPTIN6 | 0.99939185 | 1 | 0.268 | 4.659 | chrX | 118749687 | 118827333 | - | protein_coding |
| ENSG00000171100.14_6 | MTM1 | 0.99727878 | 1 | 0.268 | 4.253 | chrX | 149737069 | 149841795 | + | protein_coding |
| ENSG00000124209.4_3 | RAB22A | 0.99993349 | 1 | 0.268 | 5.915 | chr20 | 56884771 | 56942563 | + | protein_coding |
| ENSG00000127084.19_4 | FGD3 | 0.64174166 | 1 | 0.268 | -0.604 | chr9 | 95709805 | 95798518 | + | protein_coding |

| | | | | | | | | | | |
|---|---|---|---|---|---|---|---|---|---|---|
| ENSG00000234608.8_8 | MAPKAPK5-AS1 | 0.99907391 | 1 | 0.267 | 4.358 | chr12 | 112277568 | 112280706 | - | lncRNA |
| ENSG00000132128.16_4 | LRRC41 | 0.99993745 | 1 | 0.267 | 5.937 | chr1 | 46726868 | 46769280 | - | protein_coding |
| ENSG00000168781.22_6 | PPIP5K1 | 0.97507841 | 1 | 0.267 | 2.770 | chr15 | 43825660 | 43882451 | - | protein_coding |
| ENSG00000111674.9_5 | ENO2 | 0.99961299 | 1 | 0.267 | 4.887 | chr12 | 7022909 | 7032861 | + | protein_coding |
| ENSG00000177225.17_9 | GATD1 | 0.99990402 | 1 | 0.267 | 5.844 | chr11 | 767220 | 777488 | - | protein_coding |
| ENSG00000175048.17_5 | ZDHHC14 | 0.89646226 | 1 | 0.267 | 1.912 | chr6 | 157802165 | 158099189 | + | protein_coding |
| ENSG00000117385.16_4 | P3H1 | 0.9998668 | 1 | 0.267 | 5.250 | chr1 | 43212006 | 43232755 | - | protein_coding |
| ENSG00000258701.1_5 | LINC00638 | 0.63690576 | 1 | 0.267 | -0.415 | chr14 | 105287538 | 105290055 | + | lncRNA |
| ENSG00000167323.11_4 | STIM1 | 0.99995418 | 1 | 0.267 | 5.635 | chr11 | 3875757 | 4114440 | + | protein_coding |
| ENSG00000175224.16_6 | ATG13 | 0.99996059 | 1 | 0.267 | 5.943 | chr11 | 46639077 | 46696368 | + | protein_coding |
| ENSG00000229932.3_4 | YWHAZP3 | 0.66979692 | 1 | 0.267 | -0.092 | chr10 | 23425853 | 23426590 | + | processed_pseudogene |
| ENSG00000113812.14_6 | ACTR8 | 0.99976102 | 1 | 0.266 | 4.925 | chr3 | 53901093 | 53916179 | - | protein_coding |
| ENSG00000103194.15_5 | USP10 | 0.99999062 | 1 | 0.266 | 6.486 | chr16 | 84733584 | 84813528 | + | protein_coding |
| ENSG00000130311.11_3 | DDA1 | 0.99934191 | 1 | 0.266 | 4.761 | chr19 | 17420327 | 17434107 | + | protein_coding |
| ENSG00000204304.12_5 | PBX2 | 0.94816273 | 1 | 0.266 | 2.257 | chr6 | 32152510 | 32157979 | - | protein_coding |
| ENSG00000169359.16_7 | SLC33A1 | 0.99738764 | 1 | 0.266 | 3.830 | chr3 | 155538813 | 155572245 | - | protein_coding |
| ENSG00000160211.19_8 | G6PD | 0.99996732 | 1 | 0.266 | 10.558 | chrX | 153759606 | 153775787 | - | protein_coding |
| ENSG00000171877.21_7 | FRMD5 | 0.97118998 | 1 | 0.266 | 2.725 | chr15 | 44162959 | 44487469 | - | protein_coding |
| ENSG00000154874.15_4 | CCDC144B | 0.99084139 | 1 | 0.266 | 3.532 | chr17 | 18441114 | 18528930 | - | transcribed_unprocessed_pseudogene |
| ENSG00000060642.10_6 | PIGV | 0.99691707 | 1 | 0.266 | 3.868 | chr1 | 27113963 | 27124889 | + | protein_coding |
| ENSG00000196428.12_5 | TSC22D2 | 0.99933826 | 1 | 0.266 | 4.590 | chr3 | 150126122 | 150184218 | + | protein_coding |
| ENSG00000173402.12_7 | DAG1 | 0.99999446 | 1 | 0.266 | 7.361 | chr3 | 49506136 | 49573051 | + | protein_coding |
| ENSG00000103479.16_4 | RBL2 | 0.99994303 | 1 | 0.266 | 5.820 | chr16 | 53467889 | 53525560 | + | protein_coding |
| ENSG00000116138.13_3 | DNAJC16 | 0.99979957 | 1 | 0.266 | 4.929 | chr1 | 15853308 | 15918874 | + | protein_coding |
| ENSG00000143862.8_4 | ARL8A | 0.99914528 | 1 | 0.266 | 4.560 | chr1 | 202102532 | 202113871 | - | protein_coding |
| ENSG00000236809.2_4 | SNX25P1 | 0.74094797 | 1 | 0.266 | 0.596 | chr1 | 211590367 | 211591276 | + | unprocessed_pseudogene |
| ENSG00000228109.1_6 | MELTF-AS1 | 0.78150709 | 1 | 0.266 | 0.725 | chr3 | 196726331 | 196731615 | + | lncRNA |
| ENSG00000235169.8_6 | SMIM1 | 0.62366392 | 1 | 0.266 | -1.050 | chr1 | 3689325 | 3692546 | + | protein_coding |
| ENSG00000174915.12_3 | PTDSS2 | 0.99960964 | 1 | 0.265 | 5.039 | chr11 | 448268 | 491399 | + | protein_coding |
| ENSG00000177854.8_7 | TMEM187 | 0.94576032 | 1 | 0.265 | 2.934 | chrX | 153238205 | 153248645 | + | protein_coding |
| ENSG00000177380.14_5 | PPFIA3 | 0.98081665 | 1 | 0.265 | 2.973 | chr19 | 49622801 | 49654283 | + | protein_coding |
| ENSG00000237310.2_4 | AC008267.5 | 0.8488759 | 1 | 0.265 | 1.213 | chr7 | 65958594 | 65960745 | + | lncRNA |
| ENSG00000167967.16_6 | E4F1 | 0.97794979 | 1 | 0.265 | 5.367 | chr16 | 2273581 | 2285743 | + | protein_coding |
| ENSG00000259051.1_4 | HNRNPUP1 | 0.62524324 | 1 | 0.265 | -0.960 | chr14 | 43769466 | 43769798 | + | processed_pseudogene |
| ENSG00000069956.12_4 | MAPK6 | 0.99999217 | 1 | 0.265 | 7.365 | chr15 | 52244303 | 52359572 | + | protein_coding |
| ENSG00000165443.12_3 | PHYHIPL | 0.67307024 | 1 | 0.265 | -0.212 | chr10 | 60936403 | 61007534 | + | protein_coding |
| ENSG00000173757.10_4 | STAT5B | 0.99994723 | 1 | 0.265 | 5.771 | chr17 | 40351195 | 40428725 | - | protein_coding |
| ENSG00000197599.12_4 | CCDC154 | 0.71910579 | 1 | 0.265 | 0.292 | chr16 | 1484384 | 1494557 | - | protein_coding |
| ENSG00000176386.9_4 | CDC26 | 0.9580197 | 1 | 0.265 | 2.502 | chr9 | 116018115 | 116037852 | - | protein_coding |
| ENSG00000115935.17_5 | WIPF1 | 0.99923724 | 1 | 0.265 | 4.316 | chr2 | 175424300 | 175547644 | - | protein_coding |
| ENSG00000130340.16_4 | SNX9 | 0.99989368 | 1 | 0.265 | 5.367 | chr6 | 158121419 | 158366109 | + | protein_coding |
| ENSG00000157895.11_6 | C12orf43 | 0.999592 | 1 | 0.265 | 4.654 | chr12 | 121438289 | 121454305 | - | protein_coding |
| ENSG00000204323.5_2 | SMIM5 | 0.83585577 | 1 | 0.265 | 1.356 | chr17 | 73629514 | 73637484 | + | protein_coding |
| ENSG00000285569.2_1 | AC099344.3 | 0.6289423 | 1 | 0.264 | -1.101 | chr2 | 11545634 | 11567312 | + | lncRNA |
| ENSG00000235106.10_6 | BRD3OS | 0.99896071 | 1 | 0.264 | 4.511 | chr9 | 136890603 | 136899788 | + | protein_coding |
| ENSG00000269091.6_7 | AC010624.3 | 0.64467993 | 1 | 0.264 | -0.440 | chr19 | 50546453 | 50554319 | - | lncRNA |
| ENSG00000233184.7_7 | AC093157.1 | 0.9648068 | 1 | 0.264 | 2.570 | chr1 | 101491400 | 101556069 | + | lncRNA |
| ENSG00000234335.1_4 | RPS4XP11 | 0.65183736 | 1 | 0.264 | -0.360 | chr10 | 32391450 | 32392221 | - | processed_pseudogene |
| ENSG00000100982.12_4 | PCIF1 | 0.99787783 | 1 | 0.264 | 4.086 | chr20 | 44563322 | 44576662 | + | protein_coding |
| ENSG00000205795.4_3 | CYS1 | 0.7726161 | 1 | 0.264 | 0.853 | chr2 | 10196907 | 10221071 | - | protein_coding |
| ENSG00000196814.15_3 | MVB12B | 0.99730568 | 1 | 0.264 | 4.049 | chr9 | 129089108 | 129269320 | + | protein_coding |
| ENSG00000206195.6 | AP000525.9 | 0.99852959 | 1 | 0.264 | 4.074 | chr22 | 16147979 | 16193004 | - | processed_transcript |
| ENSG00000232656.8_8 | IDI2-AS1 | 0.88453457 | 1 | 0.264 | 1.186 | chr10 | 1068570 | 1091365 | + | lncRNA |
| ENSG00000013288.9_4 | MAN2B2 | 0.99958393 | 1 | 0.264 | 5.136 | chr4 | 6576916 | 6625089 | + | protein_coding |
| ENSG00000177042.14_4 | TMEM80 | 0.99849623 | 1 | 0.264 | 4.093 | chr11 | 695428 | 705028 | + | protein_coding |
| ENSG00000123349.14_4 | PFDN5 | 0.99998135 | 1 | 0.263 | 7.337 | chr12 | 53689075 | 53693234 | + | protein_coding |
| ENSG00000185745.10_3 | IFIT1 | 0.96113471 | 1 | 0.263 | 2.675 | chr10 | 91152303 | 91166244 | + | protein_coding |

| | | | | | | | | | | |
|---|---|---|---|---|---|---|---|---|---|---|
| ENSG00000139726.11_3 | DENR | 0.99998966 | 1 | 0.263 | 6.751 | chr12 | 123237371 | 123255611 | + | protein_coding |
| ENSG00000087008.16_4 | ACOX3 | 0.98758722 | 1 | 0.263 | 3.547 | chr4 | 8368009 | 8442450 | - | protein_coding |
| ENSG00000197956.10_4 | S100A6 | 0.99999894 | 1 | 0.263 | 10.027 | chr1 | 153507075 | 153508720 | - | protein_coding |
| ENSG00000165714.11_4 | BORCS5 | 0.97970947 | 1 | 0.263 | 3.003 | chr12 | 12510012 | 12624167 | + | protein_coding |
| ENSG00000241878.11_5 | PISD | 0.99884503 | 1 | 0.263 | 4.359 | chr22 | 32014477 | 32058418 | - | protein_coding |
| ENSG00000010404.18_5 | IDS | 0.99787131 | 1 | 0.263 | 4.101 | chrX | 148558519 | 148602635 | - | protein_coding |
| ENSG00000172500.12_5 | FIBP | 0.99993264 | 1 | 0.263 | 5.817 | chr11 | 65651212 | 65656010 | - | protein_coding |
| ENSG00000120910.14_4 | PPP3CC | 0.99592871 | 1 | 0.263 | 3.687 | chr8 | 22298332 | 22398655 | + | protein_coding |
| ENSG00000170469.10_7 | SPATA24 | 0.93809758 | 1 | 0.262 | 2.082 | chr5 | 138732252 | 138739777 | - | protein_coding |
| ENSG00000178057.15_3 | NDUFAF3 | 0.99931131 | 1 | 0.262 | 4.560 | chr3 | 49057892 | 49060928 | + | protein_coding |
| ENSG00000184227.8_3 | ACOT1 | 0.91265343 | 1 | 0.262 | 1.933 | chr14 | 74003847 | 74010500 | + | protein_coding |
| ENSG00000277938.1_7 | AL035252.4 | 0.83840427 | 1 | 0.262 | 1.202 | chr20 | 25209786 | 25212569 | + | lncRNA |
| ENSG00000124313.16_8 | IQSEC2 | 0.99965124 | 1 | 0.262 | 4.819 | chrX | 53255010 | 53350548 | - | protein_coding |
| ENSG00000100294.13_6 | MCAT | 0.99574098 | 1 | 0.262 | 3.817 | chr22 | 43528215 | 43539404 | - | protein_coding |
| ENSG00000276651.1_4 | AC007950.2 | 0.69905303 | 1 | 0.262 | -0.170 | chr15 | 63792936 | 63795282 | - | lncRNA |
| ENSG00000057757.10_6 | PITHD1 | 0.99933098 | 1 | 0.262 | 4.598 | chr1 | 24104908 | 24114722 | + | protein_coding |
| ENSG00000079337.16_7 | RAPGEF3 | 0.86915622 | 1 | 0.262 | 0.979 | chr12 | 48128146 | 48164823 | - | protein_coding |
| ENSG00000183495.14_3 | EP400 | 0.99974479 | 1 | 0.262 | 5.867 | chr12 | 132434487 | 132565005 | + | protein_coding |
| ENSG00000196510.12_7 | ANAPC7 | 0.99998279 | 1 | 0.262 | 6.059 | chr12 | 110810705 | 110841535 | - | protein_coding |
| ENSG00000173011.12_4 | TADA2B | 0.99928052 | 1 | 0.262 | 4.408 | chr4 | 7043626 | 7059679 | + | protein_coding |
| ENSG00000177628.16_5 | GBA | 0.99987446 | 1 | 0.262 | 5.222 | chr1 | 155204243 | 155214490 | - | protein_coding |
| ENSG00000148296.7_4 | SURF6 | 0.99968981 | 1 | 0.262 | 4.968 | chr9 | 136195612 | 136203025 | - | protein_coding |
| ENSG00000253368.4_5 | TRNP1 | 0.99997184 | 1 | 0.262 | 6.760 | chr1 | 27320183 | 27327377 | + | protein_coding |
| ENSG00000132670.20_4 | PTPRA | 0.99996744 | 1 | 0.261 | 5.748 | chr20 | 2844830 | 3019722 | + | protein_coding |
| ENSG00000004975.12_5 | DVL2 | 0.99995934 | 1 | 0.261 | 5.790 | chr17 | 7128661 | 7137836 | - | protein_coding |
| ENSG00000237161.4_5 | AC068446.1 | 0.64340008 | 1 | 0.261 | -0.755 | chr15 | 21936563 | 21938111 | - | processed_pseudogene |
| ENSG00000125656.11_7 | CLPP | 0.99954894 | 1 | 0.261 | 4.643 | chr19 | 6361542 | 6370253 | + | protein_coding |
| ENSG00000255435.7_8 | AP001267.3 | 0.64941944 | 1 | 0.261 | -0.643 | chr11 | 118380988 | 118401809 | - | lncRNA |
| ENSG00000185813.10_7 | PCYT2 | 0.99900105 | 1 | 0.261 | 4.895 | chr17 | 79858841 | 79869340 | - | protein_coding |
| ENSG00000197712.12_3 | FAM114A1 | 0.99997212 | 1 | 0.261 | 5.946 | chr4 | 38869298 | 38947360 | + | protein_coding |
| ENSG00000179846.9_6 | NKPD1 | 0.85424023 | 1 | 0.261 | 1.176 | chr19 | 45653008 | 45661995 | - | protein_coding |
| ENSG00000054392.13_5 | HHAT | 0.97666175 | 1 | 0.261 | 2.872 | chr1 | 210501596 | 210849638 | + | protein_coding |
| ENSG00000248559.1_6 | AC109454.2 | 0.72749502 | 1 | 0.261 | 0.255 | chr5 | 133735186 | 133737612 | + | lncRNA |
| ENSG00000147862.17_4 | NFIB | 0.99998113 | 1 | 0.261 | 6.376 | chr9 | 14081842 | 14398982 | - | protein_coding |
| ENSG00000089157.16_6 | RPLP0 | 0.99999862 | 1 | 0.261 | 10.469 | chr12 | 120634502 | 120639038 | - | protein_coding |
| ENSG00000272086.1_6 | AC025181.2 | 0.7265376 | 1 | 0.261 | 0.087 | chr5 | 32174577 | 32175378 | + | lncRNA |
| ENSG00000170545.17_6 | SMAGP | 0.98088149 | 1 | 0.261 | 2.784 | chr12 | 51638342 | 51664199 | - | protein_coding |
| ENSG00000141527.18_7 | CARD14 | 0.86477924 | 1 | 0.261 | 1.264 | chr17 | 78143791 | 78183130 | + | protein_coding |
| ENSG00000154175.17_6 | ABI3BP | 0.80581055 | 1 | 0.261 | 0.634 | chr3 | 100468000 | 100712359 | - | protein_coding |
| ENSG00000143149.12_3 | ALDH9A1 | 0.99997416 | 1 | 0.260 | 5.902 | chr1 | 165631453 | 165668100 | - | protein_coding |
| ENSG00000176438.12_5 | SYNE3 | 0.68915624 | 1 | 0.260 | -0.066 | chr14 | 95873603 | 95942173 | - | protein_coding |
| ENSG00000278376.1_7 | AP004609.5 | 0.65413482 | 1 | 0.260 | -0.540 | chr11 | 118661963 | 118663846 | + | lncRNA |
| ENSG00000168101.14_5 | NUDT16L1 | 0.9927075 | 1 | 0.260 | 3.727 | chr16 | 4743695 | 4745860 | + | protein_coding |
| ENSG00000182310.14_4 | SPACA6 | 0.99782602 | 1 | 0.260 | 4.034 | chr19 | 52192381 | 52215640 | + | protein_coding |
| ENSG00000237438.7_3 | CECR7 | 0.88138907 | 1 | 0.260 | 1.494 | chr22 | 17517460 | 17541715 | + | transcribed_unprocessed_pseudogene |
| ENSG00000100997.20_7 | ABHD12 | 0.99994095 | 1 | 0.260 | 5.976 | chr20 | 25275378 | 25371471 | - | protein_coding |
| ENSG00000285533.2_2 | RELA-DT | 0.95753302 | 1 | 0.260 | 2.251 | chr11 | 65430477 | 65439187 | + | lncRNA |
| ENSG00000020256.20_5 | ZFP64 | 0.99748423 | 1 | 0.260 | 4.339 | chr20 | 50668202 | 50820847 | - | protein_coding |
| ENSG00000258302.2_7 | AC025034.1 | 0.71479035 | 1 | 0.260 | 0.349 | chr12 | 89954906 | 89988655 | + | lncRNA |
| ENSG00000225880.5_5 | LINC00115 | 0.66723433 | 1 | 0.260 | -0.450 | chr1 | 761586 | 762902 | - | lncRNA |
| ENSG00000156052.11_6 | GNAQ | 0.99997742 | 1 | 0.260 | 6.343 | chr9 | 80331013 | 80646727 | - | protein_coding |
| ENSG00000168118.12_4 | RAB4A | 0.99972354 | 1 | 0.259 | 5.026 | chr1 | 229406809 | 229441641 | + | protein_coding |
| ENSG00000198752.11_5 | CDC42BPB | 0.99999665 | 1 | 0.259 | 7.269 | chr14 | 103398717 | 103523886 | - | protein_coding |
| ENSG00000104886.12_4 | PLEKHJ1 | 0.99740247 | 1 | 0.259 | 4.283 | chr19 | 2230083 | 2237703 | - | protein_coding |
| ENSG00000197728.11_6 | RPS26 | 0.99997763 | 1 | 0.259 | 6.603 | chr12 | 56435135 | 56438481 | + | protein_coding |
| ENSG00000122033.14_3 | MTIF3 | 0.98995936 | 1 | 0.259 | 3.610 | chr13 | 28009780 | 28024728 | - | protein_coding |
| ENSG00000157654.19_6 | PALM2AKAP2 | 0.99968894 | 1 | 0.259 | 5.059 | chr9 | 112260605 | 112934792 | + | protein_coding |
| ENSG00000090266.13_6 | NDUFB2 | 0.9998725 | 1 | 0.259 | 5.246 | chr7 | 140390577 | 140422590 | + | protein_coding |
| ENSG00000072042.13_6 | RDH11 | 0.99997924 | 1 | 0.259 | 6.717 | chr14 | 68143517 | 68162510 | - | protein_coding |

| | | | | | | | | | | |
|---|---|---|---|---|---|---|---|---|---|---|
| ENSG00000179456.10_2 | ZBTB18 | 0.99780292 | 1 | 0.259 | 4.618 | chr1 | 244212241 | 244220778 | + | protein_coding |
| ENSG00000146856.14_5 | AGBL3 | 0.94518322 | 1 | 0.259 | 2.214 | chr7 | 134671259 | 134832715 | + | protein_coding |
| ENSG00000164081.12_7 | TEX264 | 0.99990676 | 1 | 0.259 | 5.416 | chr3 | 51696709 | 51738339 | + | protein_coding |
| ENSG00000138162.19_5 | TACC2 | 0.99998396 | 1 | 0.259 | 6.426 | chr10 | 123748678 | 124014060 | + | protein_coding |
| ENSG00000183172.9_3 | SMDT1 | 0.83986134 | 1 | 0.258 | 1.949 | chr22 | 42475695 | 42480288 | + | protein_coding |
| ENSG00000167595.15_9 | PROSER3 | 0.99913348 | 1 | 0.258 | 4.384 | chr19 | 36249044 | 36261930 | + | protein_coding |
| ENSG00000117262.19_4 | GPR89A | 0.96791537 | 1 | 0.258 | 2.543 | chr1 | 145764409 | 145827075 | - | protein_coding |
| ENSG00000132950.19_5 | ZMYM5 | 0.99327659 | 1 | 0.258 | 3.737 | chr13 | 20397622 | 20437789 | - | protein_coding |
| ENSG00000267526.1_6 | AC005702.2 | 0.64488651 | 1 | 0.258 | -0.756 | chr17 | 58140365 | 58142143 | - | processed_pseudogene |
| ENSG00000197226.12_4 | TBC1D9B | 0.99999573 | 1 | 0.258 | 7.352 | chr5 | 179289066 | 179334859 | - | protein_coding |
| ENSG00000122218.16_6 | COPA | 0.99999627 | 1 | 0.258 | 7.434 | chr1 | 160258384 | 160313063 | - | protein_coding |
| ENSG00000178719.17_4 | GRINA | 0.99976517 | 1 | 0.258 | 6.364 | chr8 | 145064224 | 145067583 | + | protein_coding |
| ENSG00000111328.7_8 | CDK2AP1 | 0.99999627 | 1 | 0.258 | 6.920 | chr12 | 123734659 | 123756881 | - | protein_coding |
| ENSG00000278991.1_6 | AC090181.3 | 0.83351758 | 1 | 0.258 | 1.232 | chr15 | 77355739 | 77357252 | - | TEC |
| ENSG00000186577.14_8 | SMIM29 | 0.99123738 | 1 | 0.258 | 3.241 | chr6 | 34214158 | 34216885 | - | protein_coding |
| ENSG00000073169.14_6 | SELENOO | 0.99628107 | 1 | 0.258 | 3.959 | chr22 | 50639408 | 50656045 | + | protein_coding |
| ENSG00000185043.12_5 | CIB1 | 0.99995805 | 1 | 0.257 | 6.005 | chr15 | 90773207 | 90777279 | - | protein_coding |
| ENSG00000276148.1_7 | AC084824.6 | 0.6349347 | 1 | 0.257 | -1.010 | chr12 | 32878182 | 32878594 | - | lncRNA |
| ENSG00000198517.10_4 | MAFK | 0.95162045 | 1 | 0.257 | 6.087 | chr7 | 1570338 | 1582679 | + | protein_coding |
| ENSG00000126562.17_4 | WNK4 | 0.90244062 | 1 | 0.257 | 1.782 | chr17 | 40932628 | 40949084 | + | protein_coding |
| ENSG00000103653.16_3 | CSK | 0.99980904 | 1 | 0.257 | 5.749 | chr15 | 75074398 | 75095539 | + | protein_coding |
| ENSG00000151116.17_4 | UEVLD | 0.99997742 | 1 | 0.257 | 5.949 | chr11 | 18551156 | 18610294 | - | protein_coding |
| ENSG00000187942.12_3 | LDLRAD2 | 0.67315295 | 1 | 0.257 | -0.358 | chr1 | 22138758 | 22151718 | + | protein_coding |
| ENSG00000116396.14_4 | KCNC4 | 0.96804322 | 1 | 0.257 | 2.683 | chr1 | 110753965 | 110825722 | + | protein_coding |
| ENSG00000159658.12_5 | EFCAB14 | 0.99999182 | 1 | 0.257 | 6.648 | chr1 | 47140831 | 47184786 | - | protein_coding |
| ENSG00000125995.16_3 | ROMO1 | 0.99952539 | 1 | 0.256 | 4.952 | chr20 | 34287194 | 34288906 | + | protein_coding |
| ENSG00000143776.18_8 | CDC42BPA | 0.9999934 | 1 | 0.256 | 7.196 | chr1 | 227177566 | 227506175 | - | protein_coding |
| ENSG00000128159.12_8 | TUBGCP6 | 0.99953794 | 1 | 0.256 | 5.403 | chr22 | 50656118 | 50683452 | - | protein_coding |
| ENSG00000089022.14_4 | MAPKAPK5 | 0.9999722 | 1 | 0.256 | 5.691 | chr12 | 112280032 | 112340026 | + | protein_coding |
| ENSG00000165006.14_4 | UBAP1 | 0.99993717 | 1 | 0.256 | 5.449 | chr9 | 34179003 | 34252521 | + | protein_coding |
| ENSG00000258457.5_6 | AL132780.2 | 0.81027641 | 1 | 0.256 | 0.804 | chr14 | 23451907 | 23468287 | + | lncRNA |
| ENSG00000184508.11_5 | HDDC3 | 0.99525986 | 1 | 0.256 | 3.458 | chr15 | 91473194 | 91478426 | - | protein_coding |
| ENSG00000005339.15_8 | CREBBP | 0.99996911 | 1 | 0.256 | 5.891 | chr16 | 3775055 | 3930714 | - | protein_coding |
| ENSG00000163348.4_5 | PYGO2 | 0.99953331 | 1 | 0.256 | 4.797 | chr1 | 154929502 | 154936329 | - | protein_coding |
| ENSG00000141376.22_7 | BCAS3 | 0.99977723 | 1 | 0.256 | 5.177 | chr17 | 58754814 | 59470199 | + | protein_coding |
| ENSG00000159788.19_8 | RGS12 | 0.99929787 | 1 | 0.256 | 4.398 | chr4 | 3294755 | 3441640 | + | protein_coding |
| ENSG00000176542.10_6 | USF3 | 0.99975722 | 1 | 0.255 | 5.243 | chr3 | 113367232 | 113415493 | - | protein_coding |
| ENSG00000108602.18_5 | ALDH3A1 | 0.99999804 | 1 | 0.255 | 10.061 | chr17 | 19641297 | 19652256 | - | protein_coding |
| ENSG00000103502.14_4 | CDIPT | 0.99993833 | 1 | 0.255 | 5.444 | chr16 | 29869678 | 29874735 | - | protein_coding |
| ENSG00000172661.18_3 | WASHC2C | 0.99964853 | 1 | 0.255 | 4.664 | chr10 | 46222648 | 46288409 | + | protein_coding |
| ENSG00000128185.10_5 | DGCR6L | 0.99388216 | 1 | 0.255 | 4.276 | chr22 | 20301761 | 20307603 | - | protein_coding |
| ENSG00000133275.16_4 | CSNK1G2 | 0.97818795 | 1 | 0.255 | 5.593 | chr19 | 1941171 | 1981337 | + | protein_coding |
| ENSG00000280300.1_5 | AC117503.4 | 0.6546436 | 1 | 0.255 | -0.684 | chr12 | 124238793 | 124239341 | - | TEC |
| ENSG00000120616.16_5 | EPC1 | 0.99890932 | 1 | 0.255 | 4.909 | chr10 | 32556679 | 32667726 | - | protein_coding |
| ENSG00000212232.1 | SNORD17 | 0.99986532 | 1 | 0.255 | 6.449 | chr20 | 17943353 | 17943589 | - | snoRNA |
| ENSG00000184949.16_7 | FAM227A | 0.95784556 | 1 | 0.255 | 2.668 | chr22 | 38974125 | 39052634 | - | protein_coding |
| ENSG00000157927.17_5 | RADIL | 0.94691435 | 1 | 0.255 | 2.068 | chr7 | 4836686 | 4923347 | - | protein_coding |
| ENSG00000114544.16_5 | SLC41A3 | 0.99878349 | 1 | 0.255 | 4.133 | chr3 | 125725198 | 125820404 | - | protein_coding |
| ENSG00000100911.16_5 | PSME2 | 0.99998814 | 1 | 0.255 | 6.122 | chr14 | 24612571 | 24616779 | - | protein_coding |
| ENSG00000274897.3_6 | PANO1 | 0.64647085 | 1 | 0.255 | -0.515 | chr11 | 797511 | 799185 | + | protein_coding |
| ENSG00000165905.18_8 | LARGE2 | 0.95314444 | 1 | 0.254 | 1.671 | chr11 | 45943172 | 45950647 | + | protein_coding |
| ENSG00000277496.1_6 | AL357033.4 | 0.65820275 | 1 | 0.254 | -0.577 | chr20 | 61280313 | 61282119 | - | lncRNA |
| ENSG00000235831.7_7 | BHLHE40-AS1 | 0.67042295 | 1 | 0.254 | -0.115 | chr3 | 4938493 | 5022099 | - | lncRNA |
| ENSG00000128609.15_4 | NDUFA5 | 0.9997862 | 1 | 0.254 | 5.734 | chr7 | 123177051 | 123197958 | - | protein_coding |
| ENSG00000143420.18_4 | ENSA | 0.9999917 | 1 | 0.254 | 6.722 | chr1 | 150573327 | 150602088 | - | protein_coding |
| ENSG00000162817.7_3 | C1orf115 | 0.99891529 | 1 | 0.254 | 4.277 | chr1 | 220863705 | 220872495 | + | protein_coding |
| ENSG00000165121.11_6 | AL353743.1 | 0.9762858 | 1 | 0.254 | 2.692 | chr9 | 88420274 | 88464457 | + | transcribed_unprocessed_pseudogene |
| ENSG00000066777.9_6 | ARFGEF1 | 0.99998529 | 1 | 0.254 | 6.654 | chr8 | 68085746 | 68256016 | - | protein_coding |
| ENSG00000125843.11_4 | AP5S1 | 0.99977728 | 1 | 0.254 | 4.807 | chr20 | 3801171 | 3809485 | + | protein_coding |

| | | | | | | | | | | |
|---|---|---|---|---|---|---|---|---|---|---|
| ENSG00000176978.14_5 | DPP7 | 0.99332004 | 1 | 0.254 | 5.736 | chr9 | 140004998 | 140009629 | - | protein_coding |
| ENSG00000136935.14_5 | GOLGA1 | 0.99990735 | 1 | 0.254 | 5.427 | chr9 | 127640554 | 127710771 | - | protein_coding |
| ENSG00000100014.20_6 | SPECC1L | 0.99875172 | 1 | 0.253 | 4.116 | chr22 | 24666785 | 24813707 | + | protein_coding |
| ENSG00000187164.20_8 | SHTN1 | 0.99998113 | 1 | 0.253 | 6.589 | chr10 | 118640988 | 118886097 | - | protein_coding |
| ENSG00000104880.18_4 | ARHGEF18 | 0.99954842 | 1 | 0.253 | 4.798 | chr19 | 7459999 | 7537370 | + | protein_coding |
| ENSG00000122729.19_3 | ACO1 | 0.99998569 | 1 | 0.253 | 6.064 | chr9 | 32384601 | 32454767 | + | protein_coding |
| ENSG00000275457.1_6 | AL117332.1 | 0.72218641 | 1 | 0.253 | 0.110 | chr20 | 21283369 | 21284342 | - | lncRNA |
| ENSG00000123472.12_6 | ATPAF1 | 0.99994053 | 1 | 0.253 | 5.447 | chr1 | 47098409 | 47139539 | - | protein_coding |
| ENSG00000211448.12_7 | DIO2 | 0.99547263 | 1 | 0.253 | 3.837 | chr14 | 80663870 | 80854100 | - | protein_coding |
| ENSG00000150990.8_4 | DHX37 | 0.99998106 | 1 | 0.253 | 6.588 | chr12 | 125431371 | 125473677 | - | protein_coding |
| ENSG00000144642.22_7 | RBMS3 | 0.98351893 | 1 | 0.253 | 2.838 | chr3 | 28616282 | 30051882 | + | protein_coding |
| ENSG00000143740.14_5 | SNAP47 | 0.99988616 | 1 | 0.253 | 5.163 | chr1 | 227916240 | 227968932 | + | protein_coding |
| ENSG00000121067.19_8 | SPOP | 0.99994891 | 1 | 0.253 | 5.442 | chr17 | 47676246 | 47755525 | - | protein_coding |
| ENSG00000197579.8_4 | TOPORS | 0.99993454 | 1 | 0.253 | 5.864 | chr9 | 32540542 | 32552584 | - | protein_coding |
| ENSG00000105426.16_5 | PTPRS | 0.99778155 | 1 | 0.253 | 7.530 | chr19 | 5158506 | 5340814 | - | protein_coding |
| ENSG00000228544.1_7 | CCDC183-AS1 | 0.82457503 | 1 | 0.253 | 0.868 | chr9 | 139698379 | 139703300 | - | lncRNA |
| ENSG00000178974.10_4 | FBXO34 | 0.99993973 | 1 | 0.253 | 5.257 | chr14 | 55738139 | 55828636 | + | protein_coding |
| ENSG00000151612.18_8 | ZNF827 | 0.99256723 | 1 | 0.252 | 3.302 | chr4 | 146678779 | 146859975 | - | protein_coding |
| ENSG00000139625.13_5 | MAP3K12 | 0.99987149 | 1 | 0.252 | 5.331 | chr12 | 53873453 | 53893847 | - | protein_coding |
| ENSG00000182934.12_5 | SRPRA | 0.99999334 | 1 | 0.252 | 6.459 | chr11 | 126132833 | 126139039 | - | protein_coding |
| ENSG00000142166.13_4 | IFNAR1 | 0.99957349 | 1 | 0.252 | 4.866 | chr21 | 34696734 | 34732170 | + | protein_coding |
| ENSG00000133619.17_5 | KRBA1 | 0.98647375 | 1 | 0.252 | 3.253 | chr7 | 149411872 | 149431664 | + | protein_coding |
| ENSG00000117009.12_3 | KMO | 0.76752683 | 1 | 0.252 | 0.950 | chr1 | 241695434 | 241758944 | + | protein_coding |
| ENSG00000214022.11_9 | REPIN1 | 0.99996403 | 1 | 0.252 | 6.445 | chr7 | 150065278 | 150071133 | + | protein_coding |
| ENSG00000139800.8_3 | ZIC5 | 0.99423587 | 1 | 0.252 | 3.400 | chr13 | 100615218 | 100624163 | - | protein_coding |
| ENSG00000185808.13_3 | PIGP | 0.989813 | 1 | 0.251 | 3.322 | chr21 | 38431470 | 38445470 | - | protein_coding |
| ENSG00000121680.16_4 | PEX16 | 0.99870733 | 1 | 0.251 | 3.962 | chr11 | 45931214 | 45940363 | - | protein_coding |
| ENSG00000006042.12_5 | TMEM98 | 0.99853271 | 1 | 0.251 | 4.499 | chr17 | 31254928 | 31272124 | + | protein_coding |
| ENSG00000276390.1_7 | AC004241.3 | 0.69064064 | 1 | 0.251 | -0.070 | chr12 | 48093184 | 48093700 | - | lncRNA |
| ENSG00000161904.12_6 | LEMD2 | 0.99994656 | 1 | 0.251 | 5.641 | chr6 | 33738979 | 33756907 | - | protein_coding |
| ENSG00000259956.2_3 | RBM15B | 0.99991285 | 1 | 0.251 | 6.056 | chr3 | 51428716 | 51435339 | + | protein_coding |
| ENSG00000132669.13_4 | RIN2 | 0.9999939 | 1 | 0.251 | 6.480 | chr20 | 19738250 | 19983103 | + | protein_coding |
| ENSG00000280039.1_4 | RN7SKP23 | 0.64733038 | 1 | 0.251 | -0.845 | chr16 | 23423701 | 23428001 | + | TEC |
| ENSG00000196182.10_4 | STK40 | 0.97671976 | 1 | 0.251 | 3.133 | chr1 | 36805225 | 36851497 | - | protein_coding |
| ENSG00000101290.14_6 | CDS2 | 0.9999978 | 1 | 0.251 | 7.071 | chr20 | 5107525 | 5178533 | + | protein_coding |
| ENSG00000160877.6_5 | NACC1 | 0.99819634 | 1 | 0.251 | 6.096 | chr19 | 13227676 | 13251961 | + | protein_coding |
| ENSG00000168564.6_4 | CDKN2AIP | 0.99909224 | 1 | 0.251 | 4.187 | chr4 | 184365788 | 184370217 | + | protein_coding |
| ENSG00000174327.7_3 | SLC16A13 | 0.9792863 | 1 | 0.251 | 3.035 | chr17 | 6939334 | 6943436 | + | protein_coding |
| ENSG00000204463.12_6 | BAG6 | 0.99938707 | 1 | 0.251 | 6.226 | chr6 | 31606805 | 31620482 | - | protein_coding |
| ENSG00000180229.12_4 | HERC2P3 | 0.97465156 | 1 | 0.251 | 2.700 | chr15 | 20584748 | 20711433 | - | transcribed_unprocessed_pseudogene |
| ENSG00000091039.17_6 | OSBPL8 | 0.99999036 | 1 | 0.251 | 6.964 | chr12 | 76745577 | 76953589 | - | protein_coding |
| ENSG00000005884.18_3 | ITGA3 | 0.99999663 | 1 | 0.251 | 9.281 | chr17 | 48133332 | 48167845 | + | protein_coding |
| ENSG00000142541.17_4 | RPL13A | 0.99999893 | 1 | 0.251 | 10.224 | chr19 | 49990811 | 49995565 | + | protein_coding |
| ENSG00000175906.5_3 | ARL4D | 0.96956464 | 1 | 0.250 | 2.824 | chr17 | 41476361 | 41478505 | + | protein_coding |
| ENSG00000064666.15_6 | CNN2 | 0.99995823 | 1 | 0.250 | 5.937 | chr19 | 1026585 | 1039067 | + | protein_coding |
| ENSG00000180846.8_6 | CSNK1G2-AS1 | 0.64940628 | 1 | 0.250 | -0.757 | chr19 | 1952530 | 1954585 | - | lncRNA |
| ENSG00000142235.11_7 | LMTK3 | 0.94743969 | 1 | 0.250 | 3.728 | chr19 | 48988528 | 49017192 | - | protein_coding |
| ENSG00000119650.12_5 | IFT43 | 0.99528023 | 1 | 0.250 | 3.958 | chr14 | 76368479 | 76550928 | + | protein_coding |
| ENSG00000215114.9_3 | UBXN2B | 0.99998381 | 1 | 0.250 | 6.061 | chr8 | 59323918 | 59364060 | + | protein_coding |
| ENSG00000065923.10_6 | SLC9A7 | 0.99999336 | 1 | 0.249 | 6.508 | chrX | 46458686 | 46618553 | - | protein_coding |
| ENSG00000100445.17_7 | SDR39U1 | 0.86205334 | 1 | 0.249 | 1.539 | chr14 | 24908972 | 24912111 | - | protein_coding |
| ENSG00000250451.5_6 | HOXC-AS1 | 0.65490953 | 1 | 0.249 | -0.616 | chr12 | 54392806 | 54393794 | - | lncRNA |
| ENSG00000149577.16_4 | SIDT2 | 0.99603782 | 1 | 0.249 | 3.556 | chr11 | 117049452 | 117068161 | + | protein_coding |
| ENSG00000257605.2_9 | AC073611.1 | 0.67571125 | 1 | 0.249 | -0.487 | chr12 | 53692439 | 53694098 | - | lncRNA |
| ENSG00000039650.12_6 | PNKP | 0.99976939 | 1 | 0.249 | 5.669 | chr19 | 50363139 | 50381608 | - | protein_coding |
| ENSG00000252817.1 | AL590431.1 | 0.69226724 | 1 | 0.249 | -0.118 | chr1 | 154226921 | 154226997 | + | miRNA |
| ENSG00000118217.6_4 | ATF6 | 0.99998867 | 1 | 0.249 | 6.223 | chr1 | 161736110 | 161933860 | + | protein_coding |
| ENSG00000117475.14_3 | BLZF1 | 0.99981872 | 1 | 0.249 | 4.933 | chr1 | 169337208 | 169365778 | + | protein_coding |

| | | | | | | | | | | |
|---|---|---|---|---|---|---|---|---|---|---|
| ENSG00000073331.18_6 | ALPK1 | 0.99875467 | 1 | 0.248 | 4.126 | chr4 | 113206665 | 113363777 | + | protein_coding |
| ENSG00000120509.10_3 | PDZD11 | 0.99989141 | 1 | 0.248 | 5.046 | chrX | 69506445 | 69510364 | - | protein_coding |
| ENSG00000198551.10_5 | ZNF627 | 0.99537093 | 1 | 0.248 | 3.636 | chr19 | 11670189 | 11729976 | + | protein_coding |
| ENSG00000139620.12_4 | KANSL2 | 0.99996105 | 1 | 0.248 | 5.379 | chr12 | 49047184 | 49076021 | - | protein_coding |
| ENSG00000247077.7_4 | PGAM5 | 0.9999897 | 1 | 0.248 | 6.550 | chr12 | 133287405 | 133299320 | + | protein_coding |
| ENSG00000270504.1_6 | AL391422.4 | 0.74923794 | 1 | 0.248 | 0.057 | chr6 | 3751345 | 3754105 | + | lncRNA |
| ENSG00000186792.17_5 | HYAL3 | 0.94105437 | 1 | 0.248 | 1.985 | chr3 | 50330262 | 50336836 | - | protein_coding |
| ENSG00000111667.13_4 | USP5 | 0.99999559 | 1 | 0.248 | 7.330 | chr12 | 6961292 | 6975796 | + | protein_coding |
| ENSG00000165716.11_4 | DIPK1B | 0.95444107 | 1 | 0.248 | 3.036 | chr9 | 139607024 | 139619194 | + | protein_coding |
| ENSG00000205352.11_7 | PRR13 | 0.99997722 | 1 | 0.247 | 5.686 | chr12 | 53835462 | 53840429 | + | protein_coding |
| ENSG00000109320.13_7 | NFKB1 | 0.99991451 | 1 | 0.247 | 5.540 | chr4 | 103422488 | 103538459 | + | protein_coding |
| ENSG00000262837.1_8 | AC027801.5 | 0.64977176 | 1 | 0.247 | -0.770 | chr17 | 47911601 | 47913816 | + | lncRNA |
| ENSG00000108239.9_3 | TBC1D12 | 0.99939942 | 1 | 0.247 | 4.335 | chr10 | 96162298 | 96296089 | + | protein_coding |
| ENSG00000080815.19_6 | PSEN1 | 0.99998393 | 1 | 0.247 | 5.899 | chr14 | 73603126 | 73690399 | + | protein_coding |
| ENSG00000099940.12_4 | SNAP29 | 0.99906834 | 1 | 0.247 | 4.445 | chr22 | 21213295 | 21245502 | + | protein_coding |
| ENSG00000133422.13_6 | MORC2 | 0.99991705 | 1 | 0.247 | 5.291 | chr22 | 31321117 | 31364760 | - | protein_coding |
| ENSG00000177337.8_6 | DLGAP1-AS1 | 0.99777315 | 1 | 0.247 | 3.929 | chr18 | 3593730 | 3598361 | + | lncRNA |
| ENSG00000101605.12_5 | MYOM1 | 0.70560794 | 1 | 0.247 | -0.252 | chr18 | 3066805 | 3220106 | - | protein_coding |
| ENSG00000167785.9_4 | ZNF558 | 0.99894478 | 1 | 0.247 | 4.114 | chr19 | 8916846 | 8942990 | - | protein_coding |
| ENSG00000165424.7_4 | ZCCHC24 | 0.98241322 | 1 | 0.247 | 3.008 | chr10 | 81142081 | 81205380 | - | protein_coding |
| ENSG00000184669.9_4 | OR7E14P | 0.86979782 | 1 | 0.247 | 0.914 | chr11 | 17035545 | 17074571 | + | transcribed_unprocessed_pseudogene |
| ENSG00000184100.6_4 | BRD7P2 | 0.78339766 | 1 | 0.247 | 0.491 | chr3 | 159818637 | 159820580 | + | processed_pseudogene |
| ENSG00000066827.16_8 | ZFAT | 0.99709033 | 1 | 0.247 | 3.920 | chr8 | 135490031 | 135725292 | - | protein_coding |
| ENSG00000233762.3_5 | AC007969.1 | 0.86502754 | 1 | 0.247 | 1.333 | chr2 | 172373780 | 172374226 | - | processed_pseudogene |
| ENSG00000197620.11_8 | CXorf40A | 0.98693279 | 1 | 0.247 | 2.979 | chrX | 148621900 | 148632055 | + | protein_coding |
| ENSG00000265478.1 | CTD-2145A24.3 | 0.70773187 | 1 | 0.246 | -0.314 | chr17 | 18713269 | 18714253 | - | lincRNA |
| ENSG00000123395.14_2 | ATG101 | 0.97441402 | 1 | 0.246 | 4.344 | chr12 | 52463030 | 52471278 | + | protein_coding |
| ENSG00000258891.1_8 | AC005480.1 | 0.72241442 | 1 | 0.246 | 0.018 | chr14 | 74362867 | 74404817 | - | lncRNA |
| ENSG00000125741.4_5 | OPA3 | 0.99997683 | 1 | 0.246 | 5.697 | chr19 | 46030685 | 46105470 | - | protein_coding |
| ENSG00000136149.6_4 | RPL13AP25 | 0.75679367 | 1 | 0.246 | 0.334 | chr13 | 55014839 | 55015450 | - | processed_pseudogene |
| ENSG00000149932.17_6 | TMEM219 | 0.99984313 | 1 | 0.246 | 5.043 | chr16 | 29952206 | 29984371 | + | protein_coding |
| ENSG00000160051.11_3 | IQCC | 0.94355682 | 1 | 0.246 | 2.051 | chr1 | 32671262 | 32674288 | + | protein_coding |
| ENSG00000214279.13_3 | SCART1 | 0.7302094 | 1 | 0.246 | -0.032 | chr10 | 135267432 | 135337062 | + | protein_coding |
| ENSG00000250903.9_6 | GMDS-DT | 0.92050201 | 1 | 0.246 | 2.308 | chr6 | 2245952 | 2526210 | + | lncRNA |
| ENSG00000123352.17_7 | SPATS2 | 0.99997235 | 1 | 0.246 | 5.599 | chr12 | 49760367 | 49921207 | + | protein_coding |
| ENSG00000135083.15_7 | CCNJL | 0.99991506 | 1 | 0.246 | 5.041 | chr5 | 159676113 | 159772403 | - | protein_coding |
| ENSG00000116237.16_4 | ICMT | 0.99999806 | 1 | 0.246 | 7.124 | chr1 | 6281253 | 6296032 | - | protein_coding |
| ENSG00000111481.10_6 | COPZ1 | 0.99999822 | 1 | 0.245 | 7.560 | chr12 | 54694986 | 54745630 | + | protein_coding |
| ENSG00000146409.12_6 | SLC18B1 | 0.99737199 | 1 | 0.245 | 3.570 | chr6 | 133090509 | 133134478 | - | protein_coding |
| ENSG00000275888.1_7 | AC132872.3 | 0.65657437 | 1 | 0.245 | -0.802 | chr17 | 80202646 | 80203467 | + | lncRNA |
| ENSG00000179941.9_6 | BBS10 | 0.999081 | 1 | 0.245 | 4.274 | chr12 | 76738254 | 76742195 | - | protein_coding |
| ENSG00000108828.16_5 | VAT1 | 0.99999211 | 1 | 0.245 | 7.427 | chr17 | 41166624 | 41177140 | - | protein_coding |
| ENSG00000134504.14_6 | KCTD1 | 0.99995483 | 1 | 0.245 | 5.688 | chr18 | 24034874 | 24237365 | - | protein_coding |
| ENSG00000172922.10_4 | RNASEH2C | 0.99894105 | 1 | 0.245 | 4.382 | chr11 | 65481476 | 65488289 | - | protein_coding |
| ENSG00000213493.3_4 | ACTN4P1 | 0.65161438 | 1 | 0.245 | -0.975 | chr4 | 117519317 | 117520767 | + | processed_pseudogene |
| ENSG00000126775.9_4 | ATG14 | 0.99919231 | 1 | 0.245 | 4.585 | chr14 | 55833109 | 55878548 | - | protein_coding |
| ENSG00000165804.16_7 | ZNF219 | 0.95558565 | 1 | 0.245 | 4.541 | chr14 | 21558236 | 21572881 | - | protein_coding |
| ENSG00000275074.2_4 | NUDT18 | 0.99660046 | 1 | 0.245 | 3.613 | chr8 | 21964385 | 21966932 | - | protein_coding |
| ENSG00000196670.13_7 | ZFP62 | 0.99966769 | 1 | 0.245 | 4.801 | chr5 | 180274611 | 180288285 | - | protein_coding |
| ENSG00000197081.14_7 | IGF2R | 0.99999457 | 1 | 0.245 | 6.606 | chr6 | 160390131 | 160534539 | + | protein_coding |
| ENSG00000055118.15_4 | KCNH2 | 0.82664517 | 1 | 0.244 | 0.843 | chr7 | 150642049 | 150675409 | - | protein_coding |
| ENSG00000101152.11_3 | DNAJC5 | 0.99998722 | 1 | 0.244 | 6.315 | chr20 | 62526479 | 62567384 | + | protein_coding |
| ENSG00000275318.1_5 | AL136981.2 | 0.87810084 | 1 | 0.244 | 1.238 | chr9 | 95603237 | 95603970 | - | unprocessed_pseudogene |
| ENSG00000127452.9_8 | FBXL12 | 0.99352798 | 1 | 0.244 | 3.168 | chr19 | 9920943 | 9938492 | - | protein_coding |
| ENSG00000135090.14_5 | TAOK3 | 0.99987995 | 1 | 0.244 | 5.299 | chr12 | 118587606 | 118810712 | - | protein_coding |
| ENSG00000271425.9_3 | NBPF10 | 0.90841868 | 1 | 0.244 | 1.233 | chr1 | 145209145 | 145358886 | + | protein_coding |
| ENSG00000177542.11_8 | SLC25A22 | 0.99742124 | 1 | 0.244 | 4.570 | chr11 | 790475 | 798281 | - | protein_coding |

| | | | | | | | | | | |
|---|---|---|---|---|---|---|---|---|---|---|
| ENSG00000242802.9_6 | AP5Z1 | 0.99466784 | 1 | 0.244 | 4.749 | chr7 | 4815246 | 4834028 | + | protein_coding |
| ENSG00000279672.1_7 | AP006621.5 | 0.78566323 | 1 | 0.244 | 0.899 | chr11 | 779617 | 780755 | + | TEC |
| ENSG00000259287.2_7 | AC010809.1 | 0.66037201 | 1 | 0.244 | -0.626 | chr15 | 34150803 | 34157026 | - | lncRNA |
| ENSG00000115290.10_7 | GRB14 | 0.99933282 | 1 | 0.244 | 4.670 | chr2 | 165348927 | 165477992 | - | protein_coding |
| ENSG00000257218.6_4 | GATC | 0.9998814 | 1 | 0.243 | 5.307 | chr12 | 120884247 | 120901552 | + | protein_coding |
| ENSG00000108946.14_5 | PRKAR1A | 0.99999908 | 1 | 0.243 | 8.150 | chr17 | 66507921 | 66547460 | + | protein_coding |
| ENSG00000178718.6_2 | RPP25 | 0.99840142 | 1 | 0.243 | 5.212 | chr15 | 75246757 | 75249805 | - | protein_coding |
| ENSG00000204178.11_4 | MACO1 | 0.99963896 | 1 | 0.243 | 4.459 | chr1 | 25757349 | 25826700 | + | protein_coding |
| ENSG00000275491.1_6 | LINC01730 | 0.71584672 | 1 | 0.243 | 0.141 | chr20 | 3789004 | 3793081 | + | lncRNA |
| ENSG00000260027.5_6 | HOXB7 | 0.99945114 | 1 | 0.243 | 4.360 | chr17 | 46684594 | 46710934 | - | protein_coding |
| ENSG00000165915.14_3 | SLC39A13 | 0.99962033 | 1 | 0.243 | 4.679 | chr11 | 47428683 | 47438047 | + | protein_coding |
| ENSG00000261716.2_8 | H2BC20P | 0.99816706 | 1 | 0.243 | 3.808 | chr1 | 149816065 | 149822504 | - | transcribed_unprocessed_pseudogene |
| ENSG00000128881.17_8 | TTBK2 | 0.99798494 | 1 | 0.243 | 4.022 | chr15 | 43030928 | 43213007 | - | protein_coding |
| ENSG00000166847.10_4 | DCTN5 | 0.99999122 | 1 | 0.243 | 5.960 | chr16 | 23652787 | 23688793 | + | protein_coding |
| ENSG00000148090.11_3 | AUH | 0.99781778 | 1 | 0.243 | 3.855 | chr9 | 93976097 | 94124195 | - | protein_coding |
| ENSG00000127314.18_5 | RAP1B | 0.99881723 | 1 | 0.243 | 5.773 | chr12 | 69004635 | 69065681 | + | protein_coding |
| ENSG00000049245.13_4 | VAMP3 | 0.99999069 | 1 | 0.243 | 6.245 | chr1 | 7831356 | 7841492 | + | protein_coding |
| ENSG00000172366.20_6 | MCRIP2 | 0.99880595 | 1 | 0.243 | 4.371 | chr16 | 686817 | 698474 | + | protein_coding |
| ENSG00000143774.16_3 | GUK1 | 0.99987571 | 1 | 0.242 | 5.628 | chr1 | 228327663 | 228336685 | + | protein_coding |
| ENSG00000151502.11_3 | VPS26B | 0.99982417 | 1 | 0.242 | 5.019 | chr11 | 134094565 | 134117682 | + | protein_coding |
| ENSG00000128394.17_3 | APOBEC3F | 0.85528477 | 1 | 0.242 | 1.020 | chr22 | 39436609 | 39451977 | + | protein_coding |
| ENSG00000146066.3_4 | HIGD2A | 0.99896554 | 1 | 0.242 | 4.398 | chr5 | 175815752 | 175816762 | + | protein_coding |
| ENSG00000205581.11_5 | HMGN1 | 0.99999624 | 1 | 0.242 | 7.182 | chr21 | 40714241 | 40721573 | - | protein_coding |
| ENSG00000083828.16_5 | ZNF586 | 0.99698344 | 1 | 0.242 | 3.566 | chr19 | 58281023 | 58331307 | + | protein_coding |
| ENSG00000166189.8_3 | HPS6 | 0.99955922 | 1 | 0.242 | 5.033 | chr10 | 103825106 | 103827793 | + | protein_coding |
| ENSG00000125731.13_5 | SH2D3A | 0.99595692 | 1 | 0.242 | 3.829 | chr19 | 6752171 | 6767474 | - | protein_coding |
| ENSG00000261243.1_6 | AC022165.1 | 0.66094254 | 1 | 0.242 | -0.927 | chr16 | 84529205 | 84531101 | + | lncRNA |
| ENSG00000105402.8_6 | NAPA | 0.99962002 | 1 | 0.242 | 4.611 | chr19 | 47990894 | 48018348 | - | protein_coding |
| ENSG00000168884.15_6 | TNIP2 | 0.99971889 | 1 | 0.242 | 4.784 | chr4 | 2743375 | 2758069 | - | protein_coding |
| ENSG00000103335.22_5 | PIEZO1 | 0.99491126 | 1 | 0.242 | 6.762 | chr16 | 88781746 | 88851628 | - | protein_coding |
| ENSG00000225210.10_4 | DUXAP9 | 0.89592595 | 1 | 0.241 | 1.644 | chr14 | 19649976 | 19718857 | + | transcribed_processed_pseudogene |
| ENSG00000189283.10_5 | FHIT | 0.65812971 | 1 | 0.241 | -0.696 | chr3 | 59733003 | 61237133 | - | protein_coding |
| ENSG00000162384.14_4 | CZIB | 0.99933166 | 1 | 0.241 | 4.322 | chr1 | 53679771 | 53686306 | - | protein_coding |
| ENSG00000254812.1_8 | AC067930.2 | 0.661543 | 1 | 0.241 | -0.802 | chr8 | 144714241 | 144715926 | - | lncRNA |
| ENSG00000196976.7_4 | LAGE3 | 0.99786113 | 1 | 0.241 | 3.983 | chrX | 153706108 | 153707596 | - | protein_coding |
| ENSG00000278993.1_7 | AC002350.2 | 0.69530576 | 1 | 0.241 | -0.462 | chr12 | 110939419 | 110941246 | + | TEC |
| ENSG00000099968.18_4 | BCL2L13 | 0.99998045 | 1 | 0.241 | 5.867 | chr22 | 18111621 | 18213621 | + | protein_coding |
| ENSG00000278383.1_6 | AL031673.1 | 0.69028243 | 1 | 0.241 | -0.238 | chr20 | 25661417 | 25661882 | + | lncRNA |
| ENSG00000168297.16_5 | PXK | 0.99231393 | 1 | 0.241 | 3.460 | chr3 | 58318607 | 58411854 | + | protein_coding |
| ENSG00000183914.14_3 | DNAH2 | 0.66032935 | 1 | 0.241 | -0.814 | chr17 | 7620672 | 7737062 | + | protein_coding |
| ENSG00000168434.13_6 | COG7 | 0.99948398 | 1 | 0.241 | 4.326 | chr16 | 23399814 | 23464510 | - | protein_coding |
| ENSG00000213995.11_3 | NAXD | 0.9990532 | 1 | 0.240 | 4.275 | chr13 | 111267807 | 111292340 | + | protein_coding |
| ENSG00000283959.1_5 | AP002851.1 | 0.93163659 | 1 | 0.240 | 1.453 | chr8 | 103668692 | 103699346 | + | lncRNA |
| ENSG00000188315.8_7 | C3orf62 | 0.86223027 | 1 | 0.240 | 1.262 | chr3 | 49306029 | 49314665 | - | protein_coding |
| ENSG00000198598.6_3 | MMP17 | 0.97988978 | 1 | 0.240 | 3.405 | chr12 | 132312938 | 132336328 | + | protein_coding |
| ENSG00000076248.10_5 | UNG | 0.99999636 | 1 | 0.240 | 6.541 | chr12 | 109535379 | 109548797 | + | protein_coding |
| ENSG00000106692.14_5 | FKTN | 0.99991219 | 1 | 0.240 | 5.523 | chr9 | 108320411 | 108403399 | + | protein_coding |
| ENSG00000130021.14_8 | PUDP | 0.99828589 | 1 | 0.240 | 4.487 | chrX | 6585906 | 7066231 | - | protein_coding |
| ENSG00000100324.14_3 | TAB1 | 0.99953797 | 1 | 0.240 | 4.418 | chr22 | 39795783 | 39833065 | + | protein_coding |
| ENSG00000233654.2_6 | AC108047.1 | 0.806726 | 1 | 0.240 | 0.853 | chr2 | 191399581 | 191434502 | + | lncRNA |
| ENSG00000118564.15_7 | FBXL5 | 0.99992205 | 1 | 0.240 | 5.822 | chr4 | 15606004 | 15683302 | - | protein_coding |
| ENSG00000146426.18_8 | TIAM2 | 0.95710538 | 1 | 0.240 | 2.012 | chr6 | 155153831 | 155578857 | + | protein_coding |
| ENSG00000237594.2_6 | AP000251.1 | 0.68678296 | 1 | 0.240 | -0.338 | chr21 | 32931558 | 32932800 | + | lncRNA |
| ENSG00000120896.13_4 | SORBS3 | 0.99988069 | 1 | 0.240 | 5.320 | chr8 | 22402499 | 22433301 | + | protein_coding |
| ENSG00000129925.11_5 | PGAP6 | 0.99984099 | 1 | 0.239 | 5.202 | chr16 | 420788 | 437113 | - | protein_coding |
| ENSG00000107249.23_6 | GLIS3 | 0.99997017 | 1 | 0.239 | 5.584 | chr9 | 3824127 | 4348392 | - | protein_coding |
| ENSG00000179902.12_4 | C1orf194 | 0.65924281 | 1 | 0.239 | -0.849 | chr1 | 109648573 | 109656479 | - | protein_coding |
| ENSG00000182853.12_4 | VMO1 | 0.73349594 | 1 | 0.239 | -0.007 | chr17 | 4688580 | 4689728 | - | protein_coding |
| ENSG00000131697.18_6 | NPHP4 | 0.99685535 | 1 | 0.239 | 3.711 | chr1 | 5922871 | 6052533 | - | protein_coding |
| ENSG00000119408.16_5 | NEK6 | 0.9999216 | 1 | 0.239 | 5.193 | chr9 | 127019885 | 127115586 | + | protein_coding |

| | | | | | | | | | | |
|---|---|---|---|---|---|---|---|---|---|---|
| ENSG00000123700.5_4 | KCNJ2 | 0.92926464 | 1 | 0.239 | 1.712 | chr17 | 68164814 | 68176185 | + | protein_coding |
| ENSG00000101115.13_4 | SALL4 | 0.98518453 | 1 | 0.239 | 2.860 | chr20 | 50398870 | 50419060 | - | protein_coding |
| ENSG00000105677.12_3 | TMEM147 | 0.99996026 | 1 | 0.239 | 6.516 | chr19 | 36036502 | 36038428 | + | protein_coding |
| ENSG00000244558.6_8 | KCNK15-AS1 | 0.68453452 | 1 | 0.239 | -0.252 | chr20 | 43323533 | 43374662 | - | lncRNA |
| ENSG00000068976.14_4 | PYGM | 0.66683832 | 1 | 0.239 | -0.727 | chr11 | 64513861 | 64527446 | - | protein_coding |
| ENSG00000142279.13_3 | WTIP | 0.99963266 | 1 | 0.239 | 4.713 | chr19 | 34972663 | 35003209 | + | protein_coding |
| ENSG00000107404.20_6 | DVL1 | 0.99395612 | 1 | 0.239 | 5.569 | chr1 | 1270656 | 1284798 | - | protein_coding |
| ENSG00000140199.12_6 | SLC12A6 | 0.99957639 | 1 | 0.238 | 4.484 | chr15 | 34521985 | 34630261 | - | protein_coding |
| ENSG00000068308.13_5 | OTUD5 | 0.9999815 | 1 | 0.238 | 5.954 | chrX | 48779305 | 48815648 | - | protein_coding |
| ENSG00000135537.17_8 | AFG1L | 0.95194575 | 1 | 0.238 | 2.041 | chr6 | 108616195 | 108847999 | + | protein_coding |
| ENSG00000213684.4_4 | LDHBP2 | 0.71704438 | 1 | 0.238 | 0.109 | chrX | 75555240 | 75556244 | - | processed_pseudogene |
| ENSG00000133703.12_6 | KRAS | 0.99990902 | 1 | 0.238 | 5.467 | chr12 | 25358180 | 25403870 | - | protein_coding |
| ENSG00000259985.1_6 | AC017100.1 | 0.73911588 | 1 | 0.238 | 0.316 | chr18 | 29265618 | 29266786 | + | lncRNA |
| ENSG00000182362.14_3 | YBEY | 0.99648524 | 1 | 0.238 | 3.429 | chr21 | 47706256 | 47717665 | + | protein_coding |
| ENSG00000196453.8_4 | ZNF777 | 0.99468849 | 1 | 0.238 | 4.241 | chr7 | 149128454 | 149158153 | - | protein_coding |
| ENSG00000134072.11_4 | CAMK1 | 0.77842218 | 1 | 0.238 | 0.609 | chr3 | 9799031 | 9811676 | - | protein_coding |
| ENSG00000139890.10_3 | REM2 | 0.73821821 | 1 | 0.237 | 0.073 | chr14 | 23352431 | 23356887 | + | protein_coding |
| ENSG00000072840.13_4 | EVC | 0.99964253 | 1 | 0.237 | 5.277 | chr4 | 5712928 | 5816032 | + | protein_coding |
| ENSG00000165028.12_3 | NIPSNAP3B | 0.91969312 | 1 | 0.237 | 1.742 | chr9 | 107526410 | 107540045 | + | protein_coding |
| ENSG00000276023.5_3 | DUSP14 | 0.99989568 | 1 | 0.237 | 4.959 | chr17 | 35849997 | 35873603 | + | protein_coding |
| ENSG00000170043.12_5 | TRAPPC1 | 0.91549395 | 1 | 0.237 | 3.825 | chr17 | 7833663 | 7835441 | - | protein_coding |
| ENSG00000242445.1_2 | RPL7AP11 | 0.66745995 | 1 | 0.237 | -0.847 | chr3 | 121212957 | 121213755 | + | processed_pseudogene |
| ENSG00000261377.5_4 | PDCD6IPP2 | 0.96258156 | 1 | 0.237 | 2.135 | chr15 | 29034810 | 29104153 | + | transcribed_unprocessed_pseudogene |
| ENSG00000126456.16_6 | IRF3 | 0.99982411 | 1 | 0.237 | 5.659 | chr19 | 50162826 | 50169132 | - | protein_coding |
| ENSG00000222020.3_6 | HDAC4-AS1 | 0.70543017 | 1 | 0.237 | -0.403 | chr2 | 240323130 | 240324058 | + | lncRNA |
| ENSG00000197568.14_4 | HHLA3 | 0.73709999 | 1 | 0.237 | 0.078 | chr1 | 70820488 | 70851022 | + | protein_coding |
| ENSG00000084092.7_4 | NOA1 | 0.99838654 | 1 | 0.237 | 4.667 | chr4 | 57829516 | 57843772 | - | protein_coding |
| ENSG00000178927.18_8 | CYBC1 | 0.99993137 | 1 | 0.237 | 6.066 | chr17 | 80400462 | 80408705 | - | protein_coding |
| ENSG00000218582.2_4 | GAPDHP63 | 0.73107957 | 1 | 0.237 | 0.007 | chr6 | 80662722 | 80663728 | - | processed_pseudogene |
| ENSG00000240616.1_4 | RPS6P25 | 0.68794529 | 1 | 0.237 | -0.038 | chr19 | 13004947 | 13005694 | + | processed_pseudogene |
| ENSG00000116874.11_4 | WARS2 | 0.99554918 | 1 | 0.237 | 3.532 | chr1 | 119573839 | 119683294 | - | protein_coding |
| ENSG00000241352.3_6 | AC007688.2 | 0.66330077 | 1 | 0.237 | -0.978 | chr12 | 13003916 | 13004674 | + | processed_pseudogene |
| ENSG00000008323.15_3 | PLEKHG6 | 0.99274685 | 1 | 0.237 | 3.174 | chr12 | 6419602 | 6437672 | + | protein_coding |
| ENSG00000107862.5_4 | GBF1 | 0.99999682 | 1 | 0.237 | 6.892 | chr10 | 104005289 | 104142656 | + | protein_coding |
| ENSG00000102302.8_4 | FGD1 | 0.98582106 | 1 | 0.236 | 3.057 | chrX | 54471887 | 54522667 | - | protein_coding |
| ENSG00000145506.14_3 | NKD2 | 0.86069513 | 1 | 0.236 | 1.590 | chr5 | 1008917 | 1039058 | + | protein_coding |
| ENSG00000142208.16_8 | AKT1 | 0.99997809 | 1 | 0.236 | 6.855 | chr14 | 105235686 | 105262088 | - | protein_coding |
| ENSG00000106868.16_5 | SUSD1 | 0.99987462 | 1 | 0.236 | 4.726 | chr9 | 114803065 | 114937688 | - | protein_coding |
| ENSG00000233578.1_4 | EIF4EP1 | 0.68779149 | 1 | 0.236 | -0.429 | chr20 | 5529486 | 5530127 | + | processed_pseudogene |
| ENSG00000107554.17_6 | DNMBP | 0.99999522 | 1 | 0.236 | 6.206 | chr10 | 101635334 | 101769704 | - | protein_coding |
| ENSG00000200170.1 | Y_RNA | 0.6755824 | 1 | 0.236 | -0.552 | chr10 | 73980510 | 73980610 | + | misc_RNA |
| ENSG00000006453.14_5 | BAIAP2L1 | 0.99999405 | 1 | 0.236 | 6.314 | chr7 | 97920962 | 98030402 | - | protein_coding |
| ENSG00000173327.8_4 | MAP3K11 | 0.99984183 | 1 | 0.236 | 5.844 | chr11 | 65365227 | 65382853 | - | protein_coding |
| ENSG00000183718.5_3 | TRIM52 | 0.99700082 | 1 | 0.236 | 3.768 | chr5 | 180681417 | 180688140 | - | protein_coding |
| ENSG00000143553.11_3 | SNAPIN | 0.99943907 | 1 | 0.235 | 4.695 | chr1 | 153631179 | 153634328 | + | protein_coding |
| ENSG00000130821.16_7 | SLC6A8 | 0.98514977 | 1 | 0.235 | 2.801 | chrX | 152953381 | 152962043 | + | protein_coding |
| ENSG00000185896.11_3 | LAMP1 | 0.99999771 | 1 | 0.235 | 6.660 | chr13 | 113951554 | 113977987 | + | protein_coding |
| ENSG00000225872.4_7 | LINC01529 | 0.67179675 | 1 | 0.235 | -0.848 | chr19 | 36276599 | 36288804 | - | transcribed_unitary_pseudogene |
| ENSG00000235910.2_7 | APOA1-AS | 0.66446143 | 1 | 0.235 | -1.162 | chr11 | 116706833 | 116726734 | + | lncRNA |
| ENSG00000235865.2_7 | GSN-AS1 | 0.75549994 | 1 | 0.235 | 0.277 | chr9 | 124043046 | 124047808 | - | lncRNA |
| ENSG00000129472.15_5 | RAB2B | 0.99940048 | 1 | 0.235 | 4.615 | chr14 | 21927179 | 21945119 | - | protein_coding |
| ENSG00000006712.15_5 | PAF1 | 0.99999107 | 1 | 0.235 | 6.344 | chr19 | 39876269 | 39881794 | - | protein_coding |
| ENSG00000117569.18_2 | PTBP2 | 0.9998253 | 1 | 0.234 | 4.855 | chr1 | 97187221 | 97289294 | + | protein_coding |
| ENSG00000125388.20_6 | GRK4 | 0.80756495 | 1 | 0.234 | 0.558 | chr4 | 2965298 | 3042487 | + | protein_coding |
| ENSG00000233435.2_2 | AGGF1P2 | 0.71290085 | 1 | 0.234 | -0.182 | chr10 | 135454293 | 135456398 | + | processed_pseudogene |
| ENSG00000189339.12_4 | SLC35E2B | 0.9996113 | 1 | 0.234 | 4.610 | chr1 | 1590990 | 1624234 | - | protein_coding |

| | | | | | | | | | | |
|---|---|---|---|---|---|---|---|---|---|---|
| ENSG00000167658.16_4 | EEF2 | 0.99999073 | 1 | 0.234 | 10.371 | chr19 | 3976054 | 3985461 | - | protein_coding |
| ENSG00000135269.18_4 | TES | 0.99998441 | 1 | 0.234 | 5.788 | chr7 | 115850560 | 115898837 | + | protein_coding |
| ENSG00000166333.13_4 | ILK | 0.74686367 | 1 | 0.234 | 0.113 | chr11 | 6624938 | 6632105 | + | protein_coding |
| ENSG00000130706.13_5 | ADRM1 | 0.9996031 | 1 | 0.233 | 5.681 | chr20 | 60877149 | 60883918 | + | protein_coding |
| ENSG00000167081.18_5 | PBX3 | 0.99957137 | 1 | 0.233 | 4.346 | chr9 | 128509624 | 128729656 | + | protein_coding |
| ENSG00000141279.16_6 | NPEPPS | 0.99999779 | 1 | 0.233 | 6.793 | chr17 | 45600308 | 45700642 | + | protein_coding |
| ENSG00000123143.12_4 | PKN1 | 0.96622783 | 1 | 0.233 | 6.320 | chr19 | 14543865 | 14582679 | + | protein_coding |
| ENSG00000196187.12_6 | TMEM63A | 0.99847438 | 1 | 0.233 | 3.917 | chr1 | 226033237 | 226070080 | - | protein_coding |
| ENSG00000100280.16_4 | AP1B1 | 0.99998254 | 1 | 0.233 | 5.865 | chr22 | 29723669 | 29819168 | - | protein_coding |
| ENSG00000287170.1_1 | AC010327.8 | 0.68883646 | 1 | 0.233 | -0.406 | chr19 | 55627788 | 55634537 | + | lncRNA |
| ENSG00000111237.19_5 | VPS29 | 0.99996133 | 1 | 0.233 | 5.674 | chr12 | 110928888 | 110939916 | - | protein_coding |
| ENSG00000183655.13_4 | KLHL25 | 0.99673567 | 1 | 0.232 | 3.699 | chr15 | 86302557 | 86338156 | - | protein_coding |
| ENSG00000038002.9_4 | AGA | 0.99934557 | 1 | 0.232 | 4.303 | chr4 | 178351928 | 178363591 | - | protein_coding |
| ENSG00000272345.1_6 | AL031775.1 | 0.66860472 | 1 | 0.232 | -0.894 | chr6 | 24701135 | 24702021 | - | lncRNA |
| ENSG00000061987.16_5 | MON2 | 0.99999581 | 1 | 0.232 | 6.509 | chr12 | 62860597 | 62994256 | + | protein_coding |
| ENSG00000164631.19_4 | ZNF12 | 0.99996288 | 1 | 0.232 | 5.525 | chr7 | 6728064 | 6746578 | - | protein_coding |
| ENSG00000169223.15_6 | LMAN2 | 0.99995977 | 1 | 0.232 | 6.769 | chr5 | 176758568 | 176778669 | - | protein_coding |
| ENSG00000099256.19_4 | PRTFDC1 | 0.99908508 | 1 | 0.232 | 4.171 | chr10 | 25137543 | 25241535 | - | protein_coding |
| ENSG00000092200.12_4 | RPGRIP1 | 0.82175346 | 1 | 0.232 | 0.785 | chr14 | 21756098 | 21819460 | + | protein_coding |
| ENSG00000184986.11_3 | TMEM121 | 0.86484915 | 1 | 0.232 | 1.359 | chr14 | 105992920 | 105996539 | + | protein_coding |
| ENSG00000180884.10_4 | ZNF792 | 0.99899181 | 1 | 0.232 | 3.992 | chr19 | 35447258 | 35455133 | - | protein_coding |
| ENSG00000156873.16_6 | PHKG2 | 0.99959491 | 1 | 0.232 | 4.275 | chr16 | 30759614 | 30772497 | + | protein_coding |
| ENSG00000173040.13_4 | EVC2 | 0.99311104 | 1 | 0.232 | 3.048 | chr4 | 5544499 | 5711275 | - | protein_coding |
| ENSG00000124574.15_5 | ABCC10 | 0.99956259 | 1 | 0.232 | 4.405 | chr6 | 43395104 | 43418165 | + | protein_coding |
| ENSG00000112658.8_3 | SRF | 0.99963231 | 1 | 0.231 | 4.377 | chr6 | 43139007 | 43149244 | + | protein_coding |
| ENSG00000105732.13_5 | ZNF574 | 0.99986651 | 1 | 0.231 | 4.917 | chr19 | 42572629 | 42585704 | + | protein_coding |
| ENSG00000119636.16_6 | BBOF1 | 0.99221312 | 1 | 0.231 | 2.265 | chr14 | 74486052 | 74549566 | + | protein_coding |
| ENSG00000197818.11_3 | SLC9A8 | 0.99994722 | 1 | 0.231 | 5.086 | chr20 | 48429250 | 48508779 | + | protein_coding |
| ENSG00000125753.14_3 | VASP | 0.99999673 | 1 | 0.231 | 6.928 | chr19 | 46009837 | 46030241 | + | protein_coding |
| ENSG00000092531.10_4 | SNAP23 | 0.99139983 | 1 | 0.231 | 3.137 | chr15 | 42783431 | 42837554 | + | protein_coding |
| ENSG00000279035.1_6 | AC022211.5 | 0.73235154 | 1 | 0.231 | -0.182 | chr17 | 73126978 | 73129138 | + | TEC |
| ENSG00000064115.11_7 | TM7SF3 | 0.99999754 | 1 | 0.230 | 6.834 | chr12 | 27124512 | 27167367 | - | protein_coding |
| ENSG00000156500.15_5 | FAM122C | 0.73350967 | 1 | 0.230 | 0.287 | chrX | 133930425 | 133988865 | + | protein_coding |
| ENSG00000283709.1_5 | FAM238C | 0.67309723 | 1 | 0.230 | -0.895 | chr10 | 27220135 | 27233347 | - | transcribed_unprocessed_pseudogene |
| ENSG00000166257.9_5 | SCN3B | 0.73418188 | 1 | 0.230 | -0.039 | chr11 | 123499895 | 123525952 | - | protein_coding |
| ENSG00000237984.3_3 | PTENP1 | 0.86147447 | 1 | 0.230 | 1.142 | chr9 | 33673502 | 33677497 | - | transcribed_processed_pseudogene |
| ENSG00000255306.1_7 | AC004923.4 | 0.82481449 | 1 | 0.230 | 0.904 | chr11 | 67792277 | 67797928 | - | lncRNA |
| ENSG00000184481.17_3 | FOXO4 | 0.94356365 | 1 | 0.230 | 1.865 | chrX | 70315701 | 70323382 | + | protein_coding |
| ENSG00000126432.14_3 | PRDX5 | 0.99999689 | 1 | 0.230 | 7.140 | chr11 | 64085593 | 64089283 | + | protein_coding |
| ENSG00000166741.7_3 | NNMT | 0.99999734 | 1 | 0.230 | 6.912 | chr11 | 114128509 | 114184007 | + | protein_coding |
| ENSG00000157601.14_4 | MX1 | 0.73544986 | 1 | 0.230 | -0.015 | chr21 | 42792231 | 42831141 | + | protein_coding |
| ENSG00000269934.3_8 | AL353593.1 | 0.72248049 | 1 | 0.230 | -0.232 | chr1 | 228425942 | 228463767 | - | lncRNA |
| ENSG00000214253.9_6 | FIS1 | 0.9999092 | 1 | 0.230 | 5.190 | chr7 | 100882739 | 100895597 | - | protein_coding |
| ENSG00000176915.15_4 | ANKLE2 | 0.9999991 | 1 | 0.230 | 7.179 | chr12 | 133302089 | 133338418 | - | protein_coding |
| ENSG00000092969.12_5 | TGFB2 | 0.9999907 | 1 | 0.229 | 6.421 | chr1 | 218518678 | 218617961 | + | protein_coding |
| ENSG00000264107.1_7 | AC138207.2 | 0.68890317 | 1 | 0.229 | -0.629 | chr17 | 29417805 | 29422468 | - | lncRNA |
| ENSG00000124214.19_5 | STAU1 | 0.99999832 | 1 | 0.229 | 6.711 | chr20 | 47729876 | 47804904 | - | protein_coding |
| ENSG00000150637.9_7 | CD226 | 0.77601429 | 1 | 0.229 | 0.441 | chr18 | 67498394 | 67629039 | - | protein_coding |
| ENSG00000224596.8_8 | ZMIZ1-AS1 | 0.6781509 | 1 | 0.229 | -0.895 | chr10 | 80703085 | 80827652 | - | lncRNA |
| ENSG00000179218.13_2 | CALR | 0.99999913 | 1 | 0.229 | 8.662 | chr19 | 13049392 | 13055303 | + | protein_coding |
| ENSG00000108733.10_5 | PEX12 | 0.98784198 | 1 | 0.229 | 2.959 | chr17 | 33901814 | 33905882 | - | protein_coding |
| ENSG00000130158.14_6 | DOCK6 | 0.99879423 | 1 | 0.229 | 4.197 | chr19 | 11309971 | 11373200 | - | protein_coding |
| ENSG00000107874.11_4 | CUEDC2 | 0.99996008 | 1 | 0.229 | 5.093 | chr10 | 104183002 | 104192341 | - | protein_coding |
| ENSG00000004534.15_4 | RBM6 | 0.99999098 | 1 | 0.228 | 5.637 | chr3 | 49977440 | 50137478 | + | protein_coding |
| ENSG00000164970.15_5 | FAM219A | 0.99914612 | 1 | 0.228 | 3.922 | chr9 | 34398182 | 34458568 | - | protein_coding |
| ENSG00000184988.8_5 | TMEM106A | 0.95470276 | 1 | 0.228 | 1.906 | chr17 | 41363854 | 41372061 | + | protein_coding |
| ENSG00000143801.17_7 | PSEN2 | 0.99991975 | 1 | 0.228 | 4.865 | chr1 | 227057885 | 227083806 | + | protein_coding |
| ENSG00000248180.1_4 | GAPDHP60 | 0.7329192 | 1 | 0.228 | -0.107 | chr4 | 88128244 | 88129238 | + | processed_pseudogene |
| ENSG00000243364.8_4 | EFNA4 | 0.96166077 | 1 | 0.228 | 2.007 | chr1 | 155036213 | 155042029 | + | protein_coding |

| | | | | | | | | | | |
|---|---|---|---|---|---|---|---|---|---|---|
| ENSG00000106089.12_4 | STX1A | 0.99993767 | 1 | 0.228 | 5.284 | chr7 | 73113536 | 73134002 | - | protein_coding |
| ENSG00000273382.1_7 | AL356488.3 | 0.87457482 | 1 | 0.228 | 0.817 | chr1 | 109630593 | 109633480 | - | lncRNA |
| ENSG00000146373.16_6 | RNF217 | 0.99989013 | 1 | 0.228 | 5.058 | chr6 | 125283691 | 125413779 | + | protein_coding |
| ENSG00000112146.16_3 | FBXO9 | 0.99996975 | 1 | 0.228 | 5.252 | chr6 | 52916789 | 52965671 | + | protein_coding |
| ENSG00000220785.7_5 | MTMR9LP | 0.97608101 | 1 | 0.228 | 2.517 | chr1 | 32697259 | 32707221 | - | transcribed_unprocessed_pseudogene |
| ENSG00000233276.5_7 | GPX1 | 0.9998325 | 1 | 0.228 | 5.465 | chr3 | 49394609 | 49395791 | - | polymorphic_pseudogene |
| ENSG00000276855.1_6 | AC015922.3 | 0.99102158 | 1 | 0.228 | 2.247 | chr17 | 15692330 | 15693019 | + | lncRNA |
| ENSG00000100348.10_4 | TXN2 | 0.99937143 | 1 | 0.228 | 3.977 | chr22 | 36863093 | 36877687 | - | protein_coding |
| ENSG00000165782.10_4 | PIP4P1 | 0.99990755 | 1 | 0.228 | 4.915 | chr14 | 20925878 | 20929771 | - | protein_coding |
| ENSG00000228212.1_4 | OFD1P17 | 0.6783565 | 1 | 0.228 | -0.858 | chr5 | 37209466 | 37212483 | + | processed_pseudogene |
| ENSG00000215030.5_4 | RPL13P12 | 0.99432992 | 1 | 0.227 | 3.535 | chr17 | 17286691 | 17287326 | - | processed_pseudogene |
| ENSG00000069966.18_3 | GNB5 | 0.99991421 | 1 | 0.227 | 5.311 | chr15 | 52407302 | 52483566 | - | protein_coding |
| ENSG00000184956.16_7 | MUC6 | 0.69046423 | 1 | 0.227 | -0.699 | chr11 | 1012823 | 1036718 | - | protein_coding |
| ENSG00000127993.16_5 | RBM48 | 0.99666665 | 1 | 0.227 | 3.370 | chr7 | 92158087 | 92169795 | + | protein_coding |
| ENSG00000164169.13_4 | PRMT9 | 0.99739036 | 1 | 0.227 | 3.537 | chr4 | 148558936 | 148605314 | - | protein_coding |
| ENSG00000103811.16_4 | CTSH | 0.99894729 | 1 | 0.227 | 4.201 | chr15 | 79213400 | 79241916 | - | protein_coding |
| ENSG00000227908.3_7 | AC016596.1 | 0.67902484 | 1 | 0.227 | -0.978 | chr5 | 55290995 | 55299477 | + | lncRNA |
| ENSG00000102981.9_2 | PARD6A | 0.94831929 | 1 | 0.227 | 1.773 | chr16 | 67694849 | 67696681 | + | protein_coding |
| ENSG00000198373.12_6 | WWP2 | 0.99998988 | 1 | 0.227 | 5.682 | chr16 | 69796209 | 69975644 | + | protein_coding |
| ENSG00000272667.1_6 | AC012306.2 | 0.79764423 | 1 | 0.227 | 0.414 | chr2 | 128644130 | 128644759 | + | lncRNA |
| ENSG00000141002.20_5 | TCF25 | 0.99999175 | 1 | 0.227 | 6.110 | chr16 | 89939978 | 89980035 | + | protein_coding |
| ENSG00000172053.18_4 | QARS1 | 0.99999578 | 1 | 0.226 | 6.684 | chr3 | 49133365 | 49142563 | - | protein_coding |
| ENSG00000059378.13_5 | PARP12 | 0.99930725 | 1 | 0.226 | 3.953 | chr7 | 139723549 | 139762751 | - | protein_coding |
| ENSG00000136003.15_2 | ISCU | 0.99999434 | 1 | 0.226 | 5.964 | chr12 | 108956358 | 108963160 | + | protein_coding |
| ENSG00000073803.14_7 | MAP3K13 | 0.99946542 | 1 | 0.226 | 4.173 | chr3 | 185000729 | 185206882 | + | protein_coding |
| ENSG00000197858.11_3 | GPAA1 | 0.99904783 | 1 | 0.226 | 6.553 | chr8 | 145137493 | 145141119 | + | protein_coding |
| ENSG00000281420.1_2 | AP001052.1 | 0.7294875 | 1 | 0.226 | -0.294 | chr21 | 45168246 | 45168349 | - | pseudogene |
| ENSG00000129214.15_5 | SHBG | 0.73619342 | 1 | 0.226 | -0.114 | chr17 | 7517264 | 7536700 | + | protein_coding |
| ENSG00000152061.23_4 | RABGAP1L | 0.99987707 | 1 | 0.226 | 4.747 | chr1 | 174128548 | 174964445 | + | protein_coding |
| ENSG00000260368.1_6 | AC027373.1 | 0.70728968 | 1 | 0.226 | -0.405 | chr8 | 101925475 | 101926616 | - | lncRNA |
| ENSG00000146587.18_5 | RBAK | 0.99994762 | 1 | 0.226 | 5.215 | chr7 | 5085452 | 5109118 | + | protein_coding |
| ENSG00000144711.16_7 | IQSEC1 | 0.99975746 | 1 | 0.226 | 5.110 | chr3 | 12938542 | 13324781 | - | protein_coding |
| ENSG00000166153.16_6 | DEPDC4 | 0.93772423 | 1 | 0.225 | 1.350 | chr12 | 100597447 | 100660857 | - | protein_coding |
| ENSG00000062598.18_6 | ELMO2 | 0.99993267 | 1 | 0.225 | 5.040 | chr20 | 44994689 | 45061624 | - | protein_coding |
| ENSG00000101695.9_4 | RNF125 | 0.68400772 | 1 | 0.225 | -0.895 | chr18 | 29598788 | 29653182 | + | protein_coding |
| ENSG00000118518.15_6 | RNF146 | 0.99957968 | 1 | 0.225 | 4.248 | chr6 | 127587755 | 127609712 | + | protein_coding |
| ENSG00000270130.1_7 | AC068790.7 | 0.681598 | 1 | 0.225 | -1.068 | chr12 | 124445264 | 124445791 | - | lncRNA |
| ENSG00000159216.18_6 | RUNX1 | 0.99763875 | 1 | 0.225 | 3.458 | chr21 | 36160098 | 37376965 | - | protein_coding |
| ENSG00000247373.3_9 | TMED2-DT | 0.97557423 | 1 | 0.225 | 2.548 | chr12 | 124060438 | 124069662 | - | lncRNA |
| ENSG00000063180.9_4 | CA11 | 0.99560949 | 1 | 0.225 | 3.162 | chr19 | 49141203 | 49149444 | - | protein_coding |
| ENSG00000031003.10_4 | FAM13B | 0.99979104 | 1 | 0.225 | 5.264 | chr5 | 137273649 | 137387650 | - | protein_coding |
| ENSG00000175416.14_4 | CLTB | 0.9998236 | 1 | 0.225 | 5.207 | chr5 | 175819456 | 175843570 | - | protein_coding |
| ENSG00000110911.16_6 | SLC11A2 | 0.99998549 | 1 | 0.225 | 5.760 | chr12 | 51373184 | 51422349 | - | protein_coding |
| ENSG00000109756.9_7 | RAPGEF2 | 0.99999017 | 1 | 0.225 | 5.602 | chr4 | 160024165 | 160281326 | + | protein_coding |
| ENSG00000261744.1_7 | AC116552.1 | 0.69459111 | 1 | 0.225 | -0.406 | chr16 | 88579368 | 88597461 | - | lncRNA |
| ENSG00000239407.5_7 | Z68871.1 | 0.96312434 | 1 | 0.225 | 2.009 | chrX | 102139342 | 102161086 | + | lncRNA |
| ENSG00000137693.14_5 | YAP1 | 0.99999045 | 1 | 0.225 | 5.716 | chr11 | 101981178 | 102104155 | + | protein_coding |
| ENSG00000246130.1_6 | AC107959.2 | 0.68070146 | 1 | 0.225 | -0.849 | chr8 | 22925742 | 22941132 | + | lncRNA |
| ENSG00000227440.1_4 | ATP5MC1P4 | 0.6948734 | 1 | 0.225 | -0.525 | chr3 | 172546526 | 172546966 | - | processed_pseudogene |
| ENSG00000167113.11_4 | COQ4 | 0.99993727 | 1 | 0.224 | 5.103 | chr9 | 131084823 | 131096351 | + | protein_coding |
| ENSG00000280120.1_6 | AC073857.1 | 0.94521751 | 1 | 0.224 | 1.990 | chr12 | 123636871 | 123637924 | - | TEC |
| ENSG00000275807.1_6 | AC145285.6 | 0.71758274 | 1 | 0.224 | -0.105 | chr16 | 28833752 | 28835290 | - | lncRNA |
| ENSG00000160714.10_4 | UBE2Q1 | 0.99999568 | 1 | 0.224 | 5.959 | chr1 | 154521053 | 154531504 | - | protein_coding |
| ENSG00000281649.2_5 | EBLN3P | 0.99999551 | 1 | 0.224 | 6.206 | chr9 | 37079642 | 37090925 | + | lncRNA |
| ENSG00000172671.20_4 | ZFAND4 | 0.99158412 | 1 | 0.224 | 3.413 | chr10 | 46110948 | 46168228 | - | protein_coding |
| ENSG00000074660.16_6 | SCARF1 | 0.68240695 | 1 | 0.223 | -0.943 | chr17 | 1537152 | 1549038 | - | protein_coding |
| ENSG00000170412.18_8 | GPRC5C | 0.99967943 | 1 | 0.223 | 5.177 | chr17 | 72420990 | 72447792 | + | protein_coding |
| ENSG00000136770.11_6 | DNAJC1 | 0.99977748 | 1 | 0.223 | 4.787 | chr10 | 22045477 | 22292698 | - | protein_coding |

| | | | | | | | | | | |
|---|---|---|---|---|---|---|---|---|---|---|
| ENSG00000213139.8_5 | CRYGS | 0.81042306 | 1 | 0.223 | 0.492 | chr3 | 186256230 | 186264491 | - | protein_coding |
| ENSG00000180902.18_6 | D2HGDH | 0.9993863 | 1 | 0.223 | 4.129 | chr2 | 242674017 | 242708231 | + | protein_coding |
| ENSG00000012211.13_5 | PRICKLE3 | 0.99860762 | 1 | 0.223 | 3.705 | chrX | 49031151 | 49042981 | - | protein_coding |
| ENSG00000070831.17_8 | CDC42 | 0.99999525 | 1 | 0.223 | 7.718 | chr1 | 22352004 | 22427853 | + | protein_coding |
| ENSG00000155111.15_7 | CDK19 | 0.99988937 | 1 | 0.223 | 4.847 | chr6 | 110931181 | 111137161 | - | protein_coding |
| ENSG00000176454.14_5 | LPCAT4 | 0.99988545 | 1 | 0.223 | 5.052 | chr15 | 34650834 | 34659397 | - | protein_coding |
| ENSG00000143178.13_4 | TBX19 | 0.76569242 | 1 | 0.223 | 0.354 | chr1 | 168250115 | 168283664 | + | protein_coding |
| ENSG00000187742.15_4 | SECISBP2 | 0.99997937 | 1 | 0.223 | 5.607 | chr9 | 91933415 | 91974578 | + | protein_coding |
| ENSG00000136936.10_3 | XPA | 0.99889001 | 1 | 0.223 | 3.861 | chr9 | 100437191 | 100459639 | - | protein_coding |
| ENSG00000271966.1_6 | AC021321.1 | 0.68873454 | 1 | 0.223 | -0.772 | chr8 | 68256210 | 68257322 | + | lncRNA |
| ENSG00000183597.16_3 | TANGO2 | 0.9898266 | 1 | 0.223 | 3.172 | chr22 | 20004537 | 20054687 | + | protein_coding |
| ENSG00000099290.17_5 | WASHC2A | 0.90948261 | 1 | 0.223 | 1.288 | chr10 | 51827648 | 51893269 | + | protein_coding |
| ENSG00000110514.19_5 | MADD | 0.99999441 | 1 | 0.222 | 5.737 | chr11 | 47290712 | 47351582 | + | protein_coding |
| ENSG00000051825.15_7 | MPHOSPH9 | 0.99999654 | 1 | 0.222 | 6.745 | chr12 | 123636867 | 123728561 | - | protein_coding |
| ENSG00000148824.19_5 | MTG1 | 0.98386929 | 1 | 0.222 | 2.647 | chr10 | 135207598 | 135236024 | + | protein_coding |
| ENSG00000146834.14_3 | MEPCE | 0.99996801 | 1 | 0.222 | 5.612 | chr7 | 100026413 | 100031741 | + | protein_coding |
| ENSG00000241973.11_5 | PI4KA | 0.99998435 | 1 | 0.222 | 5.587 | chr22 | 21061979 | 21213705 | - | protein_coding |
| ENSG00000180089.6_4 | TMEM86B | 0.78737597 | 1 | 0.222 | 0.190 | chr19 | 55738006 | 55740152 | - | protein_coding |
| ENSG00000010256.11_4 | UQCRC1 | 0.99999767 | 1 | 0.222 | 6.957 | chr3 | 48636435 | 48648409 | - | protein_coding |
| ENSG00000125648.15_5 | SLC25A23 | 0.99985603 | 1 | 0.222 | 4.934 | chr19 | 6436090 | 6465214 | - | protein_coding |
| ENSG00000170260.9_4 | ZNF212 | 0.98860039 | 1 | 0.222 | 3.168 | chr7 | 148936742 | 148952697 | + | protein_coding |
| ENSG00000197006.14_4 | METTL9 | 0.99999791 | 1 | 0.222 | 6.555 | chr16 | 21608539 | 21668794 | + | protein_coding |
| ENSG00000166295.9_3 | ANAPC16 | 0.99999216 | 1 | 0.222 | 6.341 | chr10 | 73975758 | 73995618 | + | protein_coding |
| ENSG00000091542.9_3 | ALKBH5 | 0.99999747 | 1 | 0.222 | 7.094 | chr17 | 18086392 | 18113268 | + | protein_coding |
| ENSG00000058085.15_4 | LAMC2 | 0.99993124 | 1 | 0.222 | 5.420 | chr1 | 183155373 | 183214262 | + | protein_coding |
| ENSG00000121350.16_5 | PYROXD1 | 0.99766962 | 1 | 0.222 | 3.457 | chr12 | 21590549 | 21624184 | + | protein_coding |
| ENSG00000174917.8_5 | MICOS13 | 0.99788109 | 1 | 0.222 | 3.964 | chr19 | 5678432 | 5680907 | - | protein_coding |
| ENSG00000141252.20_6 | VPS53 | 0.99995613 | 1 | 0.222 | 5.241 | chr17 | 411908 | 624957 | - | protein_coding |
| ENSG00000278768.2_6 | BACE1-AS | 0.87135117 | 1 | 0.222 | 1.062 | chr11 | 117159169 | 117164287 | + | lncRNA |
| ENSG00000219747.1_5 | AL133260.1 | 0.6863691 | 1 | 0.221 | -0.894 | chr6 | 151420665 | 151421069 | + | processed_pseudogene |
| ENSG00000162599.17_6 | NFIA | 0.99999445 | 1 | 0.221 | 5.941 | chr1 | 61330931 | 61928460 | + | protein_coding |
| ENSG00000166821.9_5 | PEX11A | 0.99043583 | 1 | 0.221 | 3.140 | chr15 | 90220995 | 90234014 | - | protein_coding |
| ENSG00000140497.16_5 | SCAMP2 | 0.99995537 | 1 | 0.221 | 5.730 | chr15 | 75136071 | 75165706 | - | protein_coding |
| ENSG00000242732.4_6 | RTL5 | 0.84022531 | 1 | 0.221 | 1.030 | chrX | 71346960 | 71351751 | - | protein_coding |
| ENSG00000125247.15_4 | TMTC4 | 0.99989608 | 1 | 0.221 | 4.614 | chr13 | 101256181 | 101327347 | - | protein_coding |
| ENSG00000114353.17_4 | GNAI2 | 0.99998329 | 1 | 0.221 | 5.623 | chr3 | 50263724 | 50296794 | + | protein_coding |
| ENSG00000233396.8_6 | LINC01719 | 0.97777942 | 1 | 0.221 | 1.852 | chr1 | 145373054 | 145382434 | - | lncRNA |
| ENSG00000169609.14_4 | C15orf40 | 0.99990482 | 1 | 0.221 | 4.622 | chr15 | 83657193 | 83680393 | - | protein_coding |
| ENSG00000187187.14_4 | ZNF546 | 0.99976884 | 1 | 0.221 | 4.527 | chr19 | 40490041 | 40526945 | + | protein_coding |
| ENSG00000235681.1_5 | AC245884.3 | 0.70122688 | 1 | 0.220 | -0.437 | chr19 | 54958008 | 54958225 | + | processed_pseudogene |
| ENSG00000037637.11_5 | FBXO42 | 0.99985701 | 1 | 0.220 | 4.528 | chr1 | 16573335 | 16678975 | - | protein_coding |
| ENSG00000247982.6_5 | LINC00926 | 0.73313402 | 1 | 0.220 | -0.039 | chr15 | 57592563 | 57599967 | + | lncRNA |
| ENSG00000111816.8_4 | FRK | 0.99996127 | 1 | 0.220 | 5.462 | chr6 | 116252312 | 116382054 | - | protein_coding |
| ENSG00000075413.18_6 | MARK3 | 0.99999858 | 1 | 0.220 | 6.394 | chr14 | 103851729 | 103970168 | + | protein_coding |
| ENSG00000175482.9_5 | POLD4 | 0.98151488 | 1 | 0.220 | 3.357 | chr11 | 67118243 | 67124443 | - | protein_coding |
| ENSG00000008128.23_6 | CDK11A | 0.73486935 | 1 | 0.220 | -0.163 | chr1 | 1633818 | 1655796 | - | protein_coding |
| ENSG00000238105.7_5 | GOLGA2P5 | 0.97805983 | 1 | 0.220 | 2.782 | chr12 | 100550135 | 100567437 | - | transcribed_unprocessed_pseudogene |
| ENSG00000139372.15_6 | TDG | 0.99992801 | 1 | 0.220 | 5.153 | chr12 | 104359600 | 104382652 | + | protein_coding |
| ENSG00000124171.9_3 | PARD6B | 0.99999126 | 1 | 0.220 | 5.729 | chr20 | 49348117 | 49373332 | + | protein_coding |
| ENSG00000156976.17_6 | EIF4A2 | 0.99999876 | 1 | 0.220 | 7.319 | chr3 | 186500994 | 186507686 | + | protein_coding |
| ENSG00000103351.13_7 | CLUAP1 | 0.9989812 | 1 | 0.220 | 3.959 | chr16 | 3550976 | 3589048 | + | protein_coding |
| ENSG00000241935.9_4 | HOGA1 | 0.97354526 | 1 | 0.220 | 2.279 | chr10 | 99344080 | 99372559 | + | protein_coding |
| ENSG00000273008.1_6 | AC010864.1 | 0.7337522 | 1 | 0.220 | -0.169 | chr10 | 43632272 | 43633782 | - | lncRNA |
| ENSG00000154305.18_7 | MIA3 | 0.99999116 | 1 | 0.220 | 6.115 | chr1 | 222791439 | 222841349 | + | protein_coding |
| ENSG00000067955.15_5 | CBFB | 0.99997768 | 1 | 0.220 | 5.806 | chr16 | 67062887 | 67134961 | + | protein_coding |
| ENSG00000224032.7_3 | EPB41L4A-AS1 | 0.99627477 | 1 | 0.220 | 3.294 | chr5 | 111496223 | 111500515 | + | lncRNA |
| ENSG00000072364.13_4 | AFF4 | 0.99999826 | 1 | 0.219 | 7.539 | chr5 | 132211087 | 132299326 | - | protein_coding |
| ENSG00000181090.21_7 | EHMT1 | 0.99997533 | 1 | 0.219 | 5.692 | chr9 | 140513444 | 140764468 | + | protein_coding |

| | | | | | | | | | | |
|---|---|---|---|---|---|---|---|---|---|---|
| ENSG00000133935.7_7 | ERG28 | 0.9998569 | 1 | 0.219 | 4.617 | chr14 | 76116134 | 76127219 | - | protein_coding |
| ENSG00000211445.12_8 | GPX3 | 0.99693487 | 1 | 0.219 | 3.257 | chr5 | 150399999 | 150408553 | + | protein_coding |
| ENSG00000144824.21_7 | PHLDB2 | 0.99999729 | 1 | 0.219 | 6.115 | chr3 | 111451344 | 111695364 | + | protein_coding |
| ENSG00000274964.1_7 | AC026356.1 | 0.70331349 | 1 | 0.219 | -0.416 | chr12 | 32492302 | 32493658 | + | lncRNA |
| ENSG00000036054.13_4 | TBC1D23 | 0.99996328 | 1 | 0.219 | 5.360 | chr3 | 99979836 | 100044095 | + | protein_coding |
| ENSG00000158286.13_3 | RNF207 | 0.99995179 | 1 | 0.219 | 4.962 | chr1 | 6265535 | 6281359 | + | protein_coding |
| ENSG00000236969.2_4 | GGT8P | 0.86143105 | 1 | 0.219 | 0.668 | chr2 | 91963970 | 91969195 | + | unprocessed_pseudogene |
| ENSG00000100418.8_4 | DESI1 | 0.99997829 | 1 | 0.219 | 5.254 | chr22 | 41994032 | 42017047 | - | protein_coding |
| ENSG00000276952.1_6 | AL121772.3 | 0.84650412 | 1 | 0.219 | 0.722 | chr20 | 25265551 | 25266224 | - | lncRNA |
| ENSG00000169583.13_4 | CLIC3 | 0.70486779 | 1 | 0.218 | -0.407 | chr9 | 139889060 | 139891020 | - | protein_coding |
| ENSG00000137404.14_5 | NRM | 0.79153666 | 1 | 0.218 | 0.510 | chr6 | 30655824 | 30659197 | - | protein_coding |
| ENSG00000090975.12_4 | PITPNM2 | 0.99860386 | 1 | 0.218 | 4.041 | chr12 | 123468027 | 123634562 | - | protein_coding |
| ENSG00000075420.13_3 | FNDC3B | 0.99999961 | 1 | 0.218 | 7.424 | chr3 | 171757368 | 172119459 | + | protein_coding |
| ENSG00000103066.13_6 | PLA2G15 | 0.99638637 | 1 | 0.218 | 3.343 | chr16 | 68279207 | 68294961 | + | protein_coding |
| ENSG00000008513.16_8 | ST3GAL1 | 0.99957701 | 1 | 0.218 | 4.195 | chr8 | 134467091 | 134584183 | - | protein_coding |
| ENSG00000130023.16_7 | ERMARD | 0.99908733 | 1 | 0.218 | 3.989 | chr6 | 170151718 | 170181696 | + | protein_coding |
| ENSG00000159348.13_6 | CYB5R1 | 0.9995398 | 1 | 0.218 | 4.197 | chr1 | 202931001 | 202936403 | - | protein_coding |
| ENSG00000106211.9_3 | HSPB1 | 0.99817767 | 1 | 0.218 | 6.539 | chr7 | 75931990 | 75933612 | + | protein_coding |
| ENSG00000164904.18_6 | ALDH7A1 | 0.99999398 | 1 | 0.217 | 5.822 | chr5 | 125866892 | 125931054 | - | protein_coding |
| ENSG00000233101.10_6 | HOXB-AS3 | 0.99392172 | 1 | 0.217 | 3.376 | chr17 | 46626992 | 46683776 | + | lncRNA |
| ENSG00000186994.11_4 | KANK3 | 0.85424205 | 1 | 0.217 | 0.947 | chr19 | 8387468 | 8408146 | - | protein_coding |
| ENSG00000198677.11_5 | TTC37 | 0.99999929 | 1 | 0.217 | 7.331 | chr5 | 94797459 | 94890711 | - | protein_coding |
| ENSG00000126903.16_5 | SLC10A3 | 0.99914723 | 1 | 0.217 | 4.320 | chrX | 153715645 | 153719029 | - | protein_coding |
| ENSG00000105497.8_3 | ZNF175 | 0.99850116 | 1 | 0.217 | 3.661 | chr19 | 52074536 | 52095763 | + | protein_coding |
| ENSG00000083838.16_4 | ZNF446 | 0.99405989 | 1 | 0.217 | 3.100 | chr19 | 58985384 | 58992597 | + | protein_coding |
| ENSG00000259953.2_7 | AL138756.1 | 0.98989968 | 1 | 0.217 | 2.130 | chr9 | 114790850 | 114801423 | - | lncRNA |
| ENSG00000154781.16_3 | CCDC174 | 0.99952943 | 1 | 0.217 | 4.082 | chr3 | 14693253 | 14714166 | + | protein_coding |
| ENSG00000100485.12_4 | SOS2 | 0.99989583 | 1 | 0.217 | 5.888 | chr14 | 50583848 | 50698296 | - | protein_coding |
| ENSG00000278952.1_5 | AP003068.4 | 0.71028834 | 1 | 0.217 | -0.500 | chr11 | 64885782 | 64886583 | + | TEC |
| ENSG00000135108.14_4 | FBXO21 | 0.99999933 | 1 | 0.217 | 6.772 | chr12 | 117579793 | 117628336 | - | protein_coding |
| ENSG00000074416.14_9 | MGLL | 0.99999862 | 1 | 0.216 | 6.779 | chr3 | 127407905 | 127771033 | - | protein_coding |
| ENSG00000105701.16_8 | FKBP8 | 0.99581951 | 1 | 0.216 | 6.418 | chr19 | 18642561 | 18654887 | - | protein_coding |
| ENSG00000266670.1 | RN7SL637P | 0.69264971 | 1 | 0.216 | -1.030 | chr3 | 185699825 | 185700083 | - | misc_RNA |
| ENSG00000247011.2_7 | AC005920.1 | 0.83881935 | 1 | 0.216 | 0.908 | chr17 | 49012350 | 49022074 | - | lncRNA |
| ENSG00000131375.10_3 | CAPN7 | 0.99998362 | 1 | 0.216 | 5.920 | chr3 | 15247659 | 15294423 | + | protein_coding |
| ENSG00000171224.9_4 | FAM241B | 0.97994636 | 1 | 0.216 | 2.438 | chr10 | 71390003 | 71393352 | + | protein_coding |
| ENSG00000072818.12_6 | ACAP1 | 0.99134059 | 1 | 0.216 | 2.997 | chr17 | 7239848 | 7254796 | + | protein_coding |
| ENSG00000178802.18_5 | MPI | 0.99998734 | 1 | 0.216 | 5.546 | chr15 | 75182346 | 75194560 | + | protein_coding |
| ENSG00000231827.3_6 | AL513523.4 | 0.68968874 | 1 | 0.216 | -1.122 | chr1 | 153766423 | 153774567 | + | transcribed_unprocessed_pseudogene |
| ENSG00000169118.18_9 | CSNK1G1 | 0.99999038 | 1 | 0.216 | 5.735 | chr15 | 64457724 | 64648372 | - | protein_coding |
| ENSG00000180879.14_4 | SSR4 | 0.99999723 | 1 | 0.216 | 6.341 | chrX | 153058971 | 153063954 | + | protein_coding |
| ENSG00000196235.14_6 | SUPT5H | 0.99999921 | 1 | 0.216 | 7.874 | chr19 | 39926796 | 39967310 | + | protein_coding |
| ENSG00000279413.1_3 | AC112497.2 | 0.69337265 | 1 | 0.215 | -1.030 | chrX | 15720725 | 15721366 | - | pseudogene |
| ENSG00000202538.1 | RNU4-2 | 0.92961257 | 1 | 0.215 | 2.927 | chr12 | 120729566 | 120729706 | - | snRNA |
| ENSG00000004766.17_4 | VPS50 | 0.999971 | 1 | 0.215 | 5.091 | chr7 | 92861653 | 92990435 | + | protein_coding |
| ENSG00000238741.1 | SCARNA7 | 0.87835911 | 1 | 0.215 | 6.161 | chr3 | 160232695 | 160233024 | - | snoRNA |
| ENSG00000089327.15_5 | FXYD5 | 0.99999183 | 1 | 0.215 | 5.890 | chr19 | 35645633 | 35660784 | + | protein_coding |
| ENSG00000268218.1_6 | AC137932.3 | 0.77738941 | 1 | 0.215 | 0.883 | chr16 | 89334512 | 89339452 | + | lncRNA |
| ENSG00000157992.12_4 | KRTCAP3 | 0.72651614 | 1 | 0.215 | -0.248 | chr2 | 27665233 | 27669348 | + | protein_coding |
| ENSG00000072786.13_5 | STK10 | 0.99992674 | 1 | 0.215 | 4.809 | chr5 | 171469083 | 171615228 | - | protein_coding |
| ENSG00000108433.16_5 | GOSR2 | 0.99994082 | 1 | 0.215 | 5.123 | chr17 | 45000441 | 45052890 | + | protein_coding |
| ENSG00000131381.12_5 | RBSN | 0.99999401 | 1 | 0.215 | 5.759 | chr3 | 15111580 | 15140670 | - | protein_coding |
| ENSG00000263968.1 | RN7SL381P | 0.72426646 | 1 | 0.215 | -0.358 | chr16 | 85692984 | 85693279 | + | misc_RNA |
| ENSG00000105607.13_4 | GCDH | 0.99689508 | 1 | 0.214 | 3.346 | chr19 | 13001974 | 13025021 | + | protein_coding |
| ENSG00000149657.20_4 | LSM14B | 0.9999804 | 1 | 0.214 | 5.239 | chr20 | 60697517 | 60710430 | + | protein_coding |
| ENSG00000257830.1_7 | AC021066.2 | 0.69273379 | 1 | 0.214 | -1.032 | chr12 | 52668431 | 52672940 | - | lncRNA |
| ENSG00000075275.16_6 | CELSR1 | 0.99999202 | 1 | 0.214 | 7.146 | chr22 | 46756731 | 46933067 | - | protein_coding |
| ENSG00000167981.7_5 | ZNF597 | 0.9899155 | 1 | 0.214 | 2.796 | chr16 | 3482414 | 3493504 | - | protein_coding |
| ENSG00000241489.8_8 | AC244197.3 | 0.74761752 | 1 | 0.214 | -0.094 | chrX | 148558635 | 148622471 | - | protein_coding |
| ENSG00000247903.1_6 | AC024896.1 | 0.79585957 | 1 | 0.214 | 0.367 | chr12 | 27124519 | 27132515 | + | lncRNA |

| | | | | | | | | | | |
|---|---|---|---|---|---|---|---|---|---|---|
| ENSG00000272468.1_6 | AL021807.1 | 0.70905111 | 1 | 0.214 | -0.416 | chr6 | 27090436 | 27091000 | - | lncRNA |
| ENSG00000139546.11_8 | TARBP2 | 0.99981033 | 1 | 0.214 | 5.027 | chr12 | 53894705 | 53900215 | + | protein_coding |
| ENSG00000125991.19_5 | ERGIC3 | 0.99999948 | 1 | 0.214 | 7.007 | chr20 | 34129770 | 34145405 | + | protein_coding |
| ENSG00000123154.12_4 | WDR83 | 0.85272657 | 1 | 0.214 | 1.026 | chr19 | 12777616 | 12786646 | + | protein_coding |
| ENSG00000261799.1_6 | AC007406.5 | 0.9607639 | 1 | 0.214 | 1.564 | chr12 | 383120 | 386289 | - | lncRNA |
| ENSG00000196396.10_3 | PTPN1 | 0.99999348 | 1 | 0.214 | 5.618 | chr20 | 49126858 | 49201778 | + | protein_coding |
| ENSG00000105072.9_6 | C19orf44 | 0.9942569 | 1 | 0.214 | 2.874 | chr19 | 16607205 | 16632163 | + | protein_coding |
| ENSG00000134901.13_5 | POGLUT2 | 0.99154792 | 1 | 0.214 | 2.947 | chr13 | 103436631 | 103451326 | - | protein_coding |
| ENSG00000230734.1_2 | RPL10P3 | 0.72662551 | 1 | 0.214 | 0.798 | chr9 | 119942907 | 119943374 | - | processed_pseudogene |
| ENSG00000188993.3_3 | LRRC66 | 0.69450036 | 1 | 0.213 | -0.943 | chr4 | 52859868 | 52883786 | - | protein_coding |
| ENSG00000196338.12_5 | NLGN3 | 0.80792657 | 1 | 0.213 | 0.501 | chrX | 70364681 | 70391051 | + | protein_coding |
| ENSG00000146666.5_6 | LINC00525 | 0.75852621 | 1 | 0.213 | -0.170 | chr7 | 47801074 | 47806370 | + | lncRNA |
| ENSG00000103126.15_5 | AXIN1 | 0.99990555 | 1 | 0.213 | 5.641 | chr16 | 337440 | 402723 | - | protein_coding |
| ENSG00000081692.12_4 | JMJD4 | 0.9964438 | 1 | 0.213 | 3.288 | chr1 | 227918126 | 227923112 | - | protein_coding |
| ENSG00000287346.1_1 | AL132712.2 | 0.69840935 | 1 | 0.213 | -0.944 | chr14 | 104339485 | 104346718 | + | lncRNA |
| ENSG00000054277.14_5 | OPN3 | 0.99963714 | 1 | 0.213 | 4.141 | chr1 | 241753404 | 241840678 | - | protein_coding |
| ENSG00000135679.25_7 | MDM2 | 0.99999876 | 1 | 0.213 | 7.395 | chr12 | 69201957 | 69244466 | + | protein_coding |
| ENSG00000171451.14_5 | DSEL | 0.70401002 | 1 | 0.213 | -0.640 | chr18 | 65173824 | 65183957 | - | protein_coding |
| ENSG00000163935.14_5 | SFMBT1 | 0.99944268 | 1 | 0.213 | 3.919 | chr3 | 52937588 | 53080766 | - | protein_coding |
| ENSG00000169710.9_5 | FASN | 0.94131952 | 1 | 0.213 | 9.337 | chr17 | 80036214 | 80056170 | - | protein_coding |
| ENSG00000166313.19_7 | APBB1 | 0.99996505 | 1 | 0.213 | 4.998 | chr11 | 6416354 | 6440644 | - | protein_coding |
| ENSG00000099251.14_3 | HSD17B7P2 | 0.92688362 | 1 | 0.212 | 1.488 | chr10 | 38645308 | 38667433 | + | transcribed_unprocessed_pseudogene |
| ENSG00000179271.3_4 | GADD45GIP1 | 0.99864004 | 1 | 0.212 | 4.893 | chr19 | 13063933 | 13068037 | - | protein_coding |
| ENSG00000157224.16_5 | CLDN12 | 0.99995546 | 1 | 0.212 | 4.888 | chr7 | 90013035 | 90142716 | + | protein_coding |
| ENSG00000119523.10_4 | ALG2 | 0.9999798 | 1 | 0.212 | 5.559 | chr9 | 101978707 | 101984224 | - | protein_coding |
| ENSG00000227232.5_3 | WASH7P | 0.74202704 | 1 | 0.212 | -0.024 | chr1 | 14404 | 29570 | - | unprocessed_pseudogene |
| ENSG00000071859.15_3 | FAM50A | 0.99998466 | 1 | 0.212 | 5.744 | chrX | 153672488 | 153679002 | + | protein_coding |
| ENSG00000187984.13_5 | ANKRD19P | 0.75008711 | 1 | 0.212 | -0.220 | chr9 | 95571850 | 95600739 | + | transcribed_unprocessed_pseudogene |
| ENSG00000206530.11_8 | CFAP44 | 0.99740958 | 1 | 0.212 | 3.330 | chr3 | 113005777 | 113160457 | - | protein_coding |
| ENSG00000102225.16_9 | CDK16 | 0.99999702 | 1 | 0.212 | 6.830 | chrX | 47077259 | 47089396 | + | protein_coding |
| ENSG00000221817.10_6 | PPP3CB-AS1 | 0.97382494 | 1 | 0.212 | 2.124 | chr10 | 75255283 | 75279828 | + | lncRNA |
| ENSG00000213742.7_6 | ZNF337-AS1 | 0.99516483 | 1 | 0.212 | 3.014 | chr20 | 25604681 | 25669668 | + | lncRNA |
| ENSG00000167733.13_4 | HSD11B1L | 0.86211754 | 1 | 0.212 | 0.796 | chr19 | 5680615 | 5688534 | + | protein_coding |
| ENSG00000141756.19_7 | FKBP10 | 0.99999814 | 1 | 0.212 | 7.273 | chr17 | 39968932 | 39979465 | + | protein_coding |
| ENSG00000272977.1_7 | AL008721.2 | 0.77260649 | 1 | 0.212 | 0.079 | chr22 | 25872185 | 25875938 | + | lncRNA |
| ENSG00000279253.1_6 | AL121753.2 | 0.88677888 | 1 | 0.212 | 0.784 | chr20 | 33850530 | 33851990 | - | TEC |
| ENSG00000164054.15_5 | SHISA5 | 0.99999896 | 1 | 0.212 | 6.611 | chr3 | 48509197 | 48542259 | - | protein_coding |
| ENSG00000200087.1 | SNORA73B | 0.99517182 | 1 | 0.212 | 5.960 | chr1 | 28835071 | 28835274 | + | snoRNA |
| ENSG00000088888.18_3 | MAVS | 0.9999998 | 1 | 0.212 | 7.787 | chr20 | 3827446 | 3856770 | + | protein_coding |
| ENSG00000285184.3_4 | AC244033.2 | 0.89662803 | 1 | 0.212 | 1.191 | chr1 | 150017636 | 150039758 | - | lncRNA |
| ENSG00000139517.9_5 | LNX2 | 0.99911041 | 1 | 0.211 | 3.854 | chr13 | 28120050 | 28194666 | - | protein_coding |
| ENSG00000104960.15_2 | PTOV1 | 0.9989377 | 1 | 0.211 | 5.462 | chr19 | 50353992 | 50364001 | + | protein_coding |
| ENSG00000072195.15_2 | SPEG | 0.92941153 | 1 | 0.211 | 1.977 | chr2 | 220299565 | 220358351 | + | protein_coding |
| ENSG00000109390.12_5 | NDUFC1 | 0.99986779 | 1 | 0.211 | 4.549 | chr4 | 140188034 | 140223705 | - | protein_coding |
| ENSG00000270441.1_6 | AC135506.1 | 0.69796179 | 1 | 0.211 | -1.031 | chr3 | 49177519 | 49198284 | - | unprocessed_pseudogene |
| ENSG00000171169.9_4 | NAIF1 | 0.99436467 | 1 | 0.211 | 3.276 | chr9 | 130823512 | 130830485 | - | protein_coding |
| ENSG00000108107.14_4 | RPL28 | 0.99951227 | 1 | 0.211 | 7.847 | chr19 | 55896713 | 55914618 | + | protein_coding |
| ENSG00000130723.20_6 | PRRC2B | 0.99999672 | 1 | 0.211 | 7.509 | chr9 | 134249023 | 134375584 | + | protein_coding |
| ENSG00000123999.5_3 | INHA | 0.73612519 | 1 | 0.211 | -0.152 | chr2 | 220433884 | 220440433 | + | protein_coding |
| ENSG00000122783.16_5 | CYREN | 0.99945179 | 1 | 0.211 | 4.308 | chr7 | 134777115 | 134855547 | - | protein_coding |
| ENSG00000272872.1 | LL22NC03-N14H11.1 | 0.73549197 | 1 | 0.211 | -0.067 | chr22 | 16154073 | 16154766 | - | sense_intronic |
| ENSG00000107796.13_7 | ACTA2 | 0.95826518 | 1 | 0.211 | 1.823 | chr10 | 90694831 | 90751096 | - | protein_coding |
| ENSG00000173545.5_4 | ZNF622 | 0.99997101 | 1 | 0.211 | 4.984 | chr5 | 16451628 | 16465909 | - | protein_coding |
| ENSG00000224870.7_7 | MRPL20-AS1 | 0.99817155 | 1 | 0.211 | 3.665 | chr1 | 1334900 | 1337426 | + | lncRNA |
| ENSG00000166579.15_6 | NDEL1 | 0.99999122 | 1 | 0.211 | 5.711 | chr17 | 8316449 | 8393729 | + | protein_coding |
| ENSG00000042445.14_6 | RETSAT | 0.99999167 | 1 | 0.210 | 5.678 | chr2 | 85569078 | 85581654 | - | protein_coding |
| ENSG00000171723.15_6 | GPHN | 0.99960196 | 1 | 0.210 | 4.063 | chr14 | 66974125 | 67648520 | + | protein_coding |

| | | | | | | | | | | |
|---|---|---|---|---|---|---|---|---|---|---|
| ENSG00000262049.1_8 | AC139530.1 | 0.9350951 | 1 | 0.210 | 1.464 | chr17 | 79668354 | 79670330 | - | lncRNA |
| ENSG00000256433.3_7 | AC005840.2 | 0.97909094 | 1 | 0.210 | 2.345 | chr12 | 6503045 | 6505314 | + | lncRNA |
| ENSG00000187860.10_4 | CCDC157 | 0.8962997 | 1 | 0.210 | 1.312 | chr22 | 30752624 | 30774647 | + | protein_coding |
| ENSG00000118276.11_3 | B4GALT6 | 0.99967358 | 1 | 0.210 | 4.150 | chr18 | 29202210 | 29265799 | - | protein_coding |
| ENSG00000185418.16_6 | TARS3 | 0.99956509 | 1 | 0.210 | 4.120 | chr15 | 102193799 | 102264676 | - | protein_coding |
| ENSG00000149930.18_4 | TAOK2 | 0.99999505 | 1 | 0.210 | 6.405 | chr16 | 29985189 | 30003582 | + | protein_coding |
| ENSG00000130943.7_3 | PKDREJ | 0.70175995 | 1 | 0.210 | -0.611 | chr22 | 46651560 | 46659240 | - | protein_coding |
| ENSG00000134107.5_3 | BHLHE40 | 0.99999632 | 1 | 0.210 | 6.574 | chr3 | 5021122 | 5027008 | + | protein_coding |
| ENSG00000111961.18_4 | SASH1 | 0.99962069 | 1 | 0.210 | 4.156 | chr6 | 148593440 | 148873184 | + | protein_coding |
| ENSG00000161558.11_6 | TMEM143 | 0.98866275 | 1 | 0.210 | 2.669 | chr19 | 48835613 | 48867316 | - | protein_coding |
| ENSG00000137413.16_3 | TAF8 | 0.99991066 | 1 | 0.210 | 4.736 | chr6 | 42018251 | 42055199 | + | protein_coding |
| ENSG00000169964.8_4 | TMEM42 | 0.9955575 | 1 | 0.210 | 3.064 | chr3 | 44903396 | 44907162 | + | protein_coding |
| ENSG00000130640.13_5 | TUBGCP2 | 0.99999779 | 1 | 0.210 | 6.488 | chr10 | 135092134 | 135125841 | - | protein_coding |
| ENSG00000180190.11_3 | TDRP | 0.9997011 | 1 | 0.210 | 4.289 | chr8 | 439792 | 495781 | - | protein_coding |
| ENSG00000229980.5_5 | TOB1-AS1 | 0.77840146 | 1 | 0.210 | 0.166 | chr17 | 48944040 | 48988135 | + | lncRNA |
| ENSG00000100991.12_5 | TRPC4AP | 0.99999915 | 1 | 0.209 | 6.630 | chr20 | 33590207 | 33680610 | - | protein_coding |
| ENSG00000196177.13_3 | ACADSB | 0.9999638 | 1 | 0.209 | 4.932 | chr10 | 124768495 | 124817806 | + | protein_coding |
| ENSG00000014123.10_3 | UFL1 | 0.999906 | 1 | 0.209 | 5.116 | chr6 | 96969471 | 97003152 | + | protein_coding |
| ENSG00000235194.9_7 | PPP1R3E | 0.99703626 | 1 | 0.209 | 3.427 | chr14 | 23764861 | 23772068 | - | protein_coding |
| ENSG00000196233.13_6 | LCOR | 0.99981716 | 1 | 0.209 | 6.473 | chr10 | 98592039 | 98755716 | + | protein_coding |
| ENSG00000138794.10_4 | CASP6 | 0.99695012 | 1 | 0.209 | 3.756 | chr4 | 110609778 | 110624739 | - | protein_coding |
| ENSG00000198961.10_4 | PJA2 | 0.99999778 | 1 | 0.209 | 6.600 | chr5 | 108670414 | 108745675 | - | protein_coding |
| ENSG00000154370.16_5 | TRIM11 | 0.99997202 | 1 | 0.209 | 5.143 | chr1 | 228581374 | 228594536 | - | protein_coding |
| ENSG00000087095.13_4 | NLK | 0.99988734 | 1 | 0.209 | 4.823 | chr17 | 26368763 | 26523407 | + | protein_coding |
| ENSG00000100815.12_8 | TRIP11 | 0.9999416 | 1 | 0.208 | 6.518 | chr14 | 92432335 | 92507240 | - | protein_coding |
| ENSG00000261386.2_8 | AC027682.4 | 0.72753792 | 1 | 0.208 | -0.442 | chr16 | 67583117 | 67597861 | - | lncRNA |
| ENSG00000240370.6_4 | RPL13P5 | 0.99437059 | 1 | 0.208 | 2.884 | chr12 | 6982553 | 6993905 | + | transcribed_processed_pseudogene |
| ENSG00000100554.12_7 | ATP6V1D | 0.99999521 | 1 | 0.208 | 5.737 | chr14 | 67761088 | 67826982 | - | protein_coding |
| ENSG00000170468.8_3 | RIOX1 | 0.99966605 | 1 | 0.208 | 4.340 | chr14 | 73957638 | 73960098 | + | protein_coding |
| ENSG00000137218.10_4 | FRS3 | 0.96457578 | 1 | 0.208 | 1.545 | chr6 | 41737914 | 41754280 | - | protein_coding |
| ENSG00000105135.16_4 | ILVBL | 0.9999338 | 1 | 0.208 | 4.988 | chr19 | 15225795 | 15236597 | - | protein_coding |
| ENSG00000046651.15_5 | OFD1 | 0.9999828 | 1 | 0.208 | 5.284 | chrX | 13752864 | 13787476 | + | protein_coding |
| ENSG00000038210.14_6 | PI4K2B | 0.99990956 | 1 | 0.208 | 4.786 | chr4 | 25162285 | 25280826 | + | protein_coding |
| ENSG00000106609.16_4 | TMEM248 | 0.99999982 | 1 | 0.208 | 7.511 | chr7 | 66386212 | 66423538 | + | protein_coding |
| ENSG00000114054.14_4 | PCCB | 0.99998551 | 1 | 0.207 | 5.185 | chr3 | 135969182 | 136056738 | + | protein_coding |
| ENSG00000240024.5_5 | LINC00888 | 0.99451397 | 1 | 0.207 | 3.020 | chr3 | 183165396 | 183173801 | + | transcribed_unprocessed_pseudogene |
| ENSG00000131653.13_5 | TRAF7 | 0.99883615 | 1 | 0.207 | 6.488 | chr16 | 2205699 | 2228130 | + | protein_coding |
| ENSG00000162426.14_3 | SLC45A1 | 0.96839207 | 1 | 0.207 | 1.973 | chr1 | 8377886 | 8404227 | + | protein_coding |
| ENSG00000247345.2_6 | AC092343.1 | 0.77878982 | 1 | 0.207 | 0.138 | chr5 | 58335588 | 58359330 | + | lncRNA |
| ENSG00000168724.17_7 | DNAJC21 | 0.99999401 | 1 | 0.207 | 5.657 | chr5 | 34929664 | 34959069 | + | protein_coding |
| ENSG00000183773.15_4 | AIFM3 | 0.96567336 | 1 | 0.207 | 2.017 | chr22 | 21319396 | 21335649 | + | protein_coding |
| ENSG00000088727.12_5 | KIF9 | 0.98555226 | 1 | 0.207 | 2.489 | chr3 | 47269516 | 47324941 | - | protein_coding |
| ENSG00000114473.14_4 | IQCG | 0.99299251 | 1 | 0.207 | 3.006 | chr3 | 197615948 | 197687013 | - | protein_coding |
| ENSG00000007047.15_5 | MARK4 | 0.9958494 | 1 | 0.207 | 3.172 | chr19 | 45582546 | 45808542 | + | protein_coding |
| ENSG00000224186.8_6 | C5orf66 | 0.86841485 | 1 | 0.207 | 0.617 | chr5 | 134368970 | 134693909 | + | lncRNA |
| ENSG00000129315.11_4 | CCNT1 | 0.99999897 | 1 | 0.207 | 6.773 | chr12 | 49082241 | 49110781 | - | protein_coding |
| ENSG00000167767.14_5 | KRT80 | 0.99999945 | 1 | 0.207 | 6.641 | chr12 | 52562780 | 52585798 | - | protein_coding |
| ENSG00000257953.1_6 | AC083805.2 | 0.70216512 | 1 | 0.207 | -1.032 | chr12 | 58230875 | 58236528 | + | lncRNA |
| ENSG00000204231.10_3 | RXRB | 0.97511982 | 1 | 0.207 | 2.383 | chr6 | 33161365 | 33168465 | - | protein_coding |
| ENSG00000127527.14_5 | EPS15L1 | 0.99956977 | 1 | 0.207 | 4.025 | chr19 | 16466050 | 16582896 | - | protein_coding |
| ENSG00000164542.12_4 | KIAA0895 | 0.99981976 | 1 | 0.207 | 4.327 | chr7 | 36363830 | 36429734 | - | protein_coding |
| ENSG00000110583.13_4 | NAA40 | 0.99998951 | 1 | 0.206 | 5.405 | chr11 | 63706431 | 63724791 | + | protein_coding |
| ENSG00000164237.9_4 | CMBL | 0.99912622 | 1 | 0.206 | 3.953 | chr5 | 10275987 | 10308014 | - | protein_coding |
| ENSG00000163596.16_4 | ICA1L | 0.98669779 | 1 | 0.206 | 2.801 | chr2 | 203637873 | 203736708 | - | protein_coding |
| ENSG00000178202.13_5 | POGLUT3 | 0.99993592 | 1 | 0.206 | 5.160 | chr11 | 108342839 | 108369111 | - | protein_coding |
| ENSG00000080603.17_6 | SRCAP | 0.97015029 | 1 | 0.206 | 2.079 | chr16 | 30709530 | 30752730 | + | protein_coding |
| ENSG00000132912.12_6 | DCTN4 | 0.99999467 | 1 | 0.206 | 5.774 | chr5 | 150088002 | 150138671 | - | protein_coding |
| ENSG00000111011.18_4 | RSRC2 | 0.99999785 | 1 | 0.206 | 6.567 | chr12 | 122988001 | 123011547 | - | protein_coding |
| ENSG00000198830.11_3 | HMGN2 | 0.99999718 | 1 | 0.206 | 6.450 | chr1 | 26798931 | 26803133 | + | protein_coding |
| ENSG00000287104.1_1 | AC097382.3 | 0.7660645 | 1 | 0.206 | 0.021 | chr4 | 6987653 | 6988763 | - | lncRNA |

| | | | | | | | | | | |
|---|---|---|---|---|---|---|---|---|---|---|
| ENSG00000103226.18_5 | NOMO3 | 0.99749348 | 1 | 0.206 | 3.217 | chr16 | 16326385 | 16388668 | + | protein_coding |
| ENSG00000167645.17_5 | YIF1B | 0.99999532 | 1 | 0.205 | 5.944 | chr19 | 38794198 | 38807913 | - | protein_coding |
| ENSG00000198467.15_7 | TPM2 | 0.99999971 | 1 | 0.205 | 7.467 | chr9 | 35681989 | 35690053 | - | protein_coding |
| ENSG00000237350.1_4 | CDC42P6 | 0.88783763 | 1 | 0.205 | 1.118 | chr4 | 22728998 | 22729573 | - | processed_pseudogene |
| ENSG00000185236.12_3 | RAB11B | 0.99807304 | 1 | 0.205 | 3.746 | chr19 | 8454865 | 8469318 | + | protein_coding |
| ENSG00000214300.7_2 | SPDYE3 | 0.98973998 | 1 | 0.205 | 2.040 | chr7 | 99905325 | 99919819 | + | protein_coding |
| ENSG00000173039.19_8 | RELA | 0.99999756 | 1 | 0.205 | 6.022 | chr11 | 65421068 | 65430561 | - | protein_coding |
| ENSG00000275410.5_5 | HNF1B | 0.99993092 | 1 | 0.205 | 4.663 | chr17 | 36046434 | 36105096 | - | protein_coding |
| ENSG00000233175.2_8 | AC008105.1 | 0.86693911 | 1 | 0.205 | 0.974 | chr17 | 43315395 | 43319101 | - | lncRNA |
| ENSG00000238363.1 | SNORA13 | 0.70961923 | 1 | 0.205 | -0.864 | chr5 | 111497182 | 111497314 | + | snoRNA |
| ENSG00000185220.12_4 | PGBD2 | 0.9941091 | 1 | 0.205 | 3.284 | chr1 | 249200395 | 249214145 | + | protein_coding |
| ENSG00000141337.12_5 | ARSG | 0.92418962 | 1 | 0.205 | 1.399 | chr17 | 66255323 | 66418872 | + | protein_coding |
| ENSG00000064651.14_4 | SLC12A2 | 0.99999958 | 1 | 0.205 | 8.782 | chr5 | 127419458 | 127525369 | + | protein_coding |
| ENSG00000161021.13_5 | MAML1 | 0.99997663 | 1 | 0.205 | 4.993 | chr5 | 179159823 | 179204284 | + | protein_coding |
| ENSG00000254759.1_2 | NAP1L1P1 | 0.70619705 | 1 | 0.205 | -0.910 | chr11 | 125937434 | 125938496 | + | processed_pseudogene |
| ENSG00000168569.8_5 | TMEM223 | 0.99902127 | 1 | 0.205 | 3.858 | chr11 | 62539101 | 62559478 | - | protein_coding |
| ENSG00000104892.17_5 | KLC3 | 0.713285 | 1 | 0.205 | -0.753 | chr19 | 45836692 | 45854778 | + | protein_coding |
| ENSG00000203684.5_5 | IBA57-DT | 0.74184734 | 1 | 0.204 | -0.450 | chr1 | 228351787 | 228353213 | - | lncRNA |
| ENSG00000138814.17_6 | PPP3CA | 0.99999922 | 1 | 0.204 | 6.530 | chr4 | 101944566 | 102269435 | - | protein_coding |
| ENSG00000160799.11_6 | CCDC12 | 0.99978356 | 1 | 0.204 | 4.284 | chr3 | 46963216 | 47023500 | - | protein_coding |
| ENSG00000140854.13_4 | KATNB1 | 0.99978205 | 1 | 0.204 | 5.037 | chr16 | 57769651 | 57791156 | + | protein_coding |
| ENSG00000131668.14_4 | BARX1 | 0.80737011 | 1 | 0.204 | 0.525 | chr9 | 96713909 | 96717637 | - | protein_coding |
| ENSG00000134910.14_6 | STT3A | 0.99999902 | 1 | 0.204 | 7.055 | chr11 | 125461607 | 125495110 | + | protein_coding |
| ENSG00000235989.4_6 | MORC2-AS1 | 0.70629052 | 1 | 0.204 | -1.032 | chr22 | 31318295 | 31328436 | + | lncRNA |
| ENSG00000148337.21_7 | CIZ1 | 0.99999715 | 1 | 0.204 | 6.092 | chr9 | 130923530 | 130966662 | - | protein_coding |
| ENSG00000241549.8_3 | GUSBP2 | 0.85543994 | 1 | 0.203 | 0.745 | chr6 | 26839263 | 26924333 | - | transcribed_unprocessed_pseudogene |
| ENSG00000184489.12_5 | PTP4A3 | 0.93714869 | 1 | 0.203 | 2.062 | chr8 | 142402095 | 142442554 | + | protein_coding |
| ENSG00000127526.15_5 | SLC35E1 | 0.99996456 | 1 | 0.203 | 4.990 | chr19 | 16660642 | 16683226 | - | protein_coding |
| ENSG00000135473.15_5 | PAN2 | 0.99999718 | 1 | 0.203 | 5.731 | chr12 | 56710007 | 56727837 | - | protein_coding |
| ENSG00000110455.14_6 | ACCS | 0.99995492 | 1 | 0.203 | 4.796 | chr11 | 44087475 | 44105787 | + | protein_coding |
| ENSG00000148290.10_5 | SURF1 | 0.99931579 | 1 | 0.203 | 4.235 | chr9 | 136218613 | 136223552 | - | protein_coding |
| ENSG00000197111.15_4 | PCBP2 | 0.99999459 | 1 | 0.203 | 5.890 | chr12 | 53845886 | 53874946 | + | protein_coding |
| ENSG00000159110.19_3 | IFNAR2 | 0.73613947 | 1 | 0.203 | -0.142 | chr21 | 34602206 | 34637980 | + | protein_coding |
| ENSG00000020181.18_4 | ADGRA2 | 0.97562409 | 1 | 0.203 | 2.651 | chr8 | 37641709 | 37702414 | + | protein_coding |
| ENSG00000184787.19_4 | UBE2G2 | 0.99999361 | 1 | 0.203 | 5.806 | chr21 | 46188495 | 46221741 | - | protein_coding |
| ENSG00000267922.1_7 | AC007785.1 | 0.77093351 | 1 | 0.203 | 0.103 | chr19 | 46604380 | 46699475 | + | lncRNA |
| ENSG00000186862.20_7 | PDZD7 | 0.8024042 | 1 | 0.203 | 0.527 | chr10 | 102767436 | 102792052 | - | protein_coding |
| ENSG00000034677.13_5 | RNF19A | 0.99999943 | 1 | 0.202 | 6.972 | chr8 | 101269288 | 101348446 | - | protein_coding |
| ENSG00000180189.10_2 | HMGB1P14 | 0.74965591 | 1 | 0.202 | -0.391 | chr14 | 58751491 | 58752081 | + | processed_pseudogene |
| ENSG00000259709.1_6 | AC023906.4 | 0.71817978 | 1 | 0.202 | -0.829 | chr15 | 52416758 | 52432443 | + | lncRNA |
| ENSG00000068323.17_4 | TFE3 | 0.99999803 | 1 | 0.202 | 6.347 | chrX | 48886237 | 48900936 | - | protein_coding |
| ENSG00000106771.13_4 | TMEM245 | 0.99999979 | 1 | 0.202 | 7.548 | chr9 | 111777415 | 111882227 | - | protein_coding |
| ENSG00000266538.1_8 | AC005838.3 | 0.71297383 | 1 | 0.202 | -0.994 | chr17 | 15434087 | 15434403 | - | lncRNA |
| ENSG00000197261.11_2 | C6orf141 | 0.99986038 | 1 | 0.202 | 4.313 | chr6 | 49518359 | 49529620 | + | protein_coding |
| ENSG00000118162.14_4 | KPTN | 0.98415971 | 1 | 0.202 | 2.580 | chr19 | 47978407 | 47987522 | - | protein_coding |
| ENSG00000197536.11_7 | IRF1-AS1 | 0.81328448 | 1 | 0.202 | 0.307 | chr5 | 131746328 | 131824394 | + | lncRNA |
| ENSG00000011485.15_3 | PPP5C | 0.99999686 | 1 | 0.201 | 6.232 | chr19 | 46850344 | 46896238 | + | protein_coding |
| ENSG00000143363.17_6 | PRUNE1 | 0.99971354 | 1 | 0.201 | 4.028 | chr1 | 150980896 | 151008189 | + | protein_coding |
| ENSG00000099917.17_5 | MED15 | 0.99994302 | 1 | 0.201 | 4.615 | chr22 | 20850200 | 20941919 | + | protein_coding |
| ENSG00000265458.2_7 | AC132938.4 | 0.89658891 | 1 | 0.201 | 0.765 | chr17 | 80412016 | 80416397 | - | lncRNA |
| ENSG00000242259.8_6 | C22orf39 | 0.99920037 | 1 | 0.201 | 3.705 | chr22 | 19338891 | 19435755 | - | protein_coding |
| ENSG00000103363.14_3 | ELOB | 0.99999342 | 1 | 0.201 | 6.215 | chr16 | 2821415 | 2827298 | - | protein_coding |
| ENSG00000246640.1_7 | PICART1 | 0.85718273 | 1 | 0.201 | 0.856 | chr17 | 48127713 | 48133103 | - | lncRNA |
| ENSG00000140859.16_6 | KIFC3 | 0.99985727 | 1 | 0.201 | 8.012 | chr16 | 57792129 | 57896957 | - | protein_coding |
| ENSG00000156804.7_4 | FBXO32 | 0.98866758 | 1 | 0.201 | 2.456 | chr8 | 124510129 | 124553446 | - | protein_coding |
| ENSG00000090674.16_4 | MCOLN1 | 0.99228496 | 1 | 0.201 | 2.776 | chr19 | 7587510 | 7598895 | + | protein_coding |
| ENSG00000132740.8_2 | IGHMBP2 | 0.99956338 | 1 | 0.201 | 4.088 | chr11 | 68671310 | 68708070 | + | protein_coding |
| ENSG00000087152.15_3 | ATXN7L3 | 0.99999831 | 1 | 0.201 | 6.460 | chr17 | 42269173 | 42277481 | - | protein_coding |
| ENSG00000165632.8_3 | TAF3 | 0.9992269 | 1 | 0.201 | 3.792 | chr10 | 7860468 | 8058594 | + | protein_coding |

| | | | | | | | | | | |
|---|---|---|---|---|---|---|---|---|---|---|
| ENSG00000134109.11_3 | EDEM1 | 0.99999858 | 1 | 0.201 | 6.179 | chr3 | 5229331 | 5261643 | + | protein_coding |
| ENSG00000230667.5_3 | SETSIP | 0.71490677 | 1 | 0.201 | -0.640 | chr1 | 92540090 | 92540998 | - | protein_coding |
| ENSG00000260093.1_6 | AC034111.1 | 0.76459755 | 1 | 0.201 | -0.125 | chr8 | 9907995 | 9911764 | - | lncRNA |
| ENSG00000147119.4_3 | CHST7 | 0.9930586 | 1 | 0.201 | 3.136 | chrX | 46433200 | 46457931 | + | protein_coding |
| ENSG00000168556.7_4 | ING2 | 0.99731551 | 1 | 0.201 | 3.331 | chr4 | 184426211 | 184433582 | + | protein_coding |
| ENSG00000166398.13_5 | KIAA0355 | 0.99999623 | 1 | 0.200 | 5.562 | chr19 | 34745457 | 34846476 | + | protein_coding |
| ENSG00000136378.15_4 | ADAMTS7 | 0.97788644 | 1 | 0.200 | 2.872 | chr15 | 79051548 | 79103806 | - | protein_coding |
| ENSG00000198242.14_3 | RPL23A | 0.99999923 | 1 | 0.200 | 7.532 | chr17 | 27047003 | 27051377 | + | protein_coding |
| ENSG00000185684.15_6 | EP400P1 | 0.99514835 | 1 | 0.200 | 2.989 | chr12 | 132568828 | 132616184 | + | transcribed_unprocessed_pseudogene |
| ENSG00000163945.18_6 | UVSSA | 0.99957948 | 1 | 0.200 | 4.187 | chr4 | 1340996 | 1389777 | + | protein_coding |
| ENSG00000109016.17_5 | DHRS7B | 0.99964656 | 1 | 0.200 | 4.045 | chr17 | 21026677 | 21096578 | + | protein_coding |
| ENSG00000170275.15_4 | CRTAP | 0.99999987 | 1 | 0.200 | 7.766 | chr3 | 33155506 | 33189265 | + | protein_coding |
| ENSG00000108469.15_6 | RECQL5 | 0.99993464 | 1 | 0.200 | 5.506 | chr17 | 73622925 | 73663269 | - | protein_coding |
| ENSG00000244055.2_7 | AC007566.1 | 0.87293294 | 1 | 0.200 | 0.791 | chr7 | 92086878 | 92124915 | + | lncRNA |
| ENSG00000108523.16_4 | RNF167 | 0.99999617 | 1 | 0.199 | 5.893 | chr17 | 4843303 | 4848517 | + | protein_coding |
| ENSG00000094841.14_4 | UPRT | 0.99796217 | 1 | 0.199 | 3.486 | chrX | 74376223 | 74524720 | + | protein_coding |
| ENSG00000197019.5_4 | SERTAD1 | 0.9997437 | 1 | 0.199 | 4.059 | chr19 | 40927496 | 40931899 | - | protein_coding |
| ENSG00000275092.1_7 | AL031710.2 | 0.71422099 | 1 | 0.199 | -0.995 | chr16 | 1757253 | 1757974 | + | lncRNA |
| ENSG00000276115.1_7 | AC026356.2 | 0.72564173 | 1 | 0.199 | -0.574 | chr12 | 32505283 | 32507078 | + | lncRNA |
| ENSG00000151553.15_3 | FAM160B1 | 0.99999308 | 1 | 0.199 | 5.582 | chr10 | 116581503 | 116659591 | + | protein_coding |
| ENSG00000102910.14_4 | LONP2 | 0.99999981 | 1 | 0.199 | 7.272 | chr16 | 48278211 | 48397033 | + | protein_coding |
| ENSG00000226055.2_4 | PAICSP1 | 0.72092022 | 1 | 0.199 | -0.911 | chr9 | 37878113 | 37879490 | + | processed_pseudogene |
| ENSG00000143373.18_3 | ZNF687 | 0.99998149 | 1 | 0.199 | 5.591 | chr1 | 151254094 | 151264652 | + | protein_coding |
| ENSG00000176826.15_5 | FKBP9P1 | 0.73443636 | 1 | 0.199 | -0.392 | chr7 | 55748767 | 55780945 | - | transcribed_unprocessed_pseudogene |
| ENSG00000133872.14_9 | SARAF | 0.99970728 | 1 | 0.199 | 6.976 | chr8 | 29920519 | 29940724 | - | protein_coding |
| ENSG00000100605.17_6 | ITPK1 | 0.99999377 | 1 | 0.199 | 6.405 | chr14 | 93403259 | 93582665 | - | protein_coding |
| ENSG00000133103.17_3 | COG6 | 0.99976616 | 1 | 0.198 | 4.443 | chr13 | 40229764 | 40365802 | + | protein_coding |
| ENSG00000116871.15_7 | MAP7D1 | 0.99765635 | 1 | 0.198 | 6.388 | chr1 | 36621180 | 36646450 | + | protein_coding |
| ENSG00000158710.14_3 | TAGLN2 | 0.99999988 | 1 | 0.198 | 8.628 | chr1 | 159887897 | 159895522 | - | protein_coding |
| ENSG00000181458.10_4 | TMEM45A | 0.86884772 | 1 | 0.198 | 0.487 | chr3 | 100211463 | 100296288 | + | protein_coding |
| ENSG00000182871.16_5 | COL18A1 | 0.82405681 | 1 | 0.198 | 7.409 | chr21 | 46825080 | 46933634 | + | protein_coding |
| ENSG00000053371.12_7 | AKR7A2 | 0.9999678 | 1 | 0.198 | 4.891 | chr1 | 19630459 | 19638640 | - | protein_coding |
| ENSG00000162413.16_5 | KLHL21 | 0.9998335 | 1 | 0.198 | 4.418 | chr1 | 6650784 | 6674667 | - | protein_coding |
| ENSG00000158195.11_4 | WASF2 | 0.99997801 | 1 | 0.198 | 5.481 | chr1 | 27730734 | 27816678 | - | protein_coding |
| ENSG00000143443.10_3 | C1orf56 | 0.99563423 | 1 | 0.198 | 3.052 | chr1 | 151020227 | 151024462 | + | protein_coding |
| ENSG00000187953.10_2 | PMS2CL | 0.99731378 | 1 | 0.198 | 3.453 | chr7 | 6749759 | 6793493 | + | transcribed_unprocessed_pseudogene |
| ENSG00000265417.1_6 | AP005119.2 | 0.74799777 | 1 | 0.198 | -0.261 | chr18 | 1362446 | 1363874 | - | processed_pseudogene |
| ENSG00000073969.18_4 | NSF | 0.99998906 | 1 | 0.198 | 5.541 | chr17 | 44668035 | 44834830 | + | protein_coding |
| ENSG00000175634.15_5 | RPS6KB2 | 0.99994844 | 1 | 0.198 | 4.690 | chr11 | 67195931 | 67202872 | + | protein_coding |
| ENSG00000287576.1_1 | AC067930.8 | 0.90877917 | 1 | 0.197 | 1.222 | chr8 | 144623818 | 144627298 | + | lncRNA |
| ENSG00000150773.10_3 | PIH1D2 | 0.86393985 | 1 | 0.197 | 0.457 | chr11 | 111934734 | 111944998 | - | protein_coding |
| ENSG00000074276.10_4 | CDHR2 | 0.7880929 | 1 | 0.197 | -0.056 | chr5 | 175969512 | 176022975 | + | protein_coding |
| ENSG00000128590.5_3 | DNAJB9 | 0.99976192 | 1 | 0.197 | 4.233 | chr7 | 108210311 | 108215294 | + | protein_coding |
| ENSG00000267632.1_7 | AC067852.3 | 0.8447847 | 1 | 0.197 | 0.524 | chr17 | 40661802 | 40663537 | + | lncRNA |
| ENSG00000113384.14_4 | GOLPH3 | 0.99999796 | 1 | 0.197 | 6.178 | chr5 | 32124822 | 32174425 | - | protein_coding |
| ENSG00000011275.18_4 | RNF216 | 0.99999885 | 1 | 0.197 | 6.159 | chr7 | 5659678 | 5821370 | - | protein_coding |
| ENSG00000163374.19_5 | YY1AP1 | 0.99998491 | 1 | 0.197 | 5.010 | chr1 | 155629234 | 155658791 | - | protein_coding |
| ENSG00000179119.15_4 | SPTY2D1 | 0.99998729 | 1 | 0.197 | 5.497 | chr11 | 18627950 | 18656338 | - | protein_coding |
| ENSG00000225973.4_7 | PIGBOS1 | 0.98820179 | 1 | 0.197 | 2.521 | chr15 | 55609382 | 55611359 | - | protein_coding |
| ENSG00000255142.1_7 | AP006621.2 | 0.86786282 | 1 | 0.197 | 0.793 | chr11 | 781645 | 782105 | + | lncRNA |
| ENSG00000214198.8_4 | TTC41P | 0.83153891 | 1 | 0.197 | 0.421 | chr12 | 104237527 | 104323989 | - | transcribed_unitary_pseudogene |
| ENSG00000113141.18_4 | IK | 0.99999928 | 1 | 0.197 | 6.576 | chr5 | 140026643 | 140042065 | + | protein_coding |
| ENSG00000137312.15_6 | FLOT1 | 0.99999724 | 1 | 0.197 | 5.980 | chr6 | 30695486 | 30710509 | - | protein_coding |
| ENSG00000125246.15_4 | CLYBL | 0.82540329 | 1 | 0.197 | 0.535 | chr13 | 100258923 | 100549388 | + | protein_coding |
| ENSG00000141665.13_10 | FBXO15 | 0.90804293 | 1 | 0.196 | 1.061 | chr18 | 71740588 | 71815078 | - | protein_coding |
| ENSG00000254470.3_4 | AP5B1 | 0.99989964 | 1 | 0.196 | 4.440 | chr11 | 65541369 | 65548447 | - | protein_coding |
| ENSG00000137145.20_3 | DENND4C | 0.99999927 | 1 | 0.196 | 6.964 | chr9 | 19230433 | 19373543 | + | protein_coding |

| | | | | | | | | | | |
|---|---|---|---|---|---|---|---|---|---|---|
| ENSG00000132376.20_5 | INPP5K | 0.99987854 | 1 | 0.196 | 4.993 | chr17 | 1397871 | 1420036 | - | protein_coding |
| ENSG00000159128.14_3 | IFNGR2 | 0.99987179 | 1 | 0.196 | 4.401 | chr21 | 34775202 | 34851655 | + | protein_coding |
| ENSG00000184840.12_3 | TMED9 | 0.9999547 | 1 | 0.196 | 6.003 | chr5 | 177019204 | 177024243 | + | protein_coding |
| ENSG00000228223.3_6 | HCG11 | 0.99946154 | 1 | 0.196 | 3.797 | chr6 | 26521937 | 26527632 | + | lncRNA |
| ENSG00000149091.15_6 | DGKZ | 0.99996074 | 1 | 0.196 | 5.638 | chr11 | 46354455 | 46402104 | + | protein_coding |
| ENSG00000037897.17_7 | METTL1 | 0.99916905 | 1 | 0.196 | 3.591 | chr12 | 58162254 | 58165902 | - | protein_coding |
| ENSG00000185049.16_8 | NELFA | 0.9997908 | 1 | 0.196 | 4.083 | chr4 | 1984444 | 2043630 | - | protein_coding |
| ENSG00000089199.10_6 | CHGB | 0.99080453 | 1 | 0.196 | 3.042 | chr20 | 5892156 | 5905999 | + | protein_coding |
| ENSG00000168096.15_7 | ANKS3 | 0.99991136 | 1 | 0.196 | 4.832 | chr16 | 4746511 | 4784379 | - | protein_coding |
| ENSG00000262814.8_4 | MRPL12 | 0.76336679 | 1 | 0.196 | 2.420 | chr17 | 79670397 | 79674547 | + | protein_coding |
| ENSG00000170638.9_3 | TRABD | 0.99158539 | 1 | 0.196 | 4.636 | chr22 | 50624344 | 50638027 | + | protein_coding |
| ENSG00000154144.13_4 | TBRG1 | 0.99990886 | 1 | 0.196 | 4.525 | chr11 | 124492732 | 124505822 | + | protein_coding |
| ENSG00000102103.16_7 | PQBP1 | 0.99992337 | 1 | 0.196 | 4.522 | chrX | 48747480 | 48760420 | + | protein_coding |
| ENSG00000151208.17_5 | DLG5 | 0.99999235 | 1 | 0.196 | 6.274 | chr10 | 79550549 | 79686513 | - | protein_coding |
| ENSG00000070047.13_4 | PHRF1 | 0.99928925 | 1 | 0.195 | 6.626 | chr11 | 576470 | 612222 | + | protein_coding |
| ENSG00000107175.12_4 | CREB3 | 0.99999537 | 1 | 0.195 | 5.509 | chr9 | 35732595 | 35736996 | + | protein_coding |
| ENSG00000088899.15_4 | LZTS3 | 0.99666321 | 1 | 0.195 | 3.320 | chr20 | 3143263 | 3154238 | - | protein_coding |
| ENSG00000186298.12_5 | PPP1CC | 0.99999962 | 1 | 0.195 | 7.507 | chr12 | 111157485 | 111180744 | - | protein_coding |
| ENSG00000197905.9_8 | TEAD4 | 0.99985586 | 1 | 0.195 | 4.743 | chr12 | 3068496 | 3149842 | + | protein_coding |
| ENSG00000178950.17_6 | GAK | 0.99990979 | 1 | 0.195 | 5.765 | chr4 | 843064 | 926161 | - | protein_coding |
| ENSG00000231683.6_8 | AL033397.1 | 0.99743177 | 1 | 0.195 | 3.415 | chr6 | 53426087 | 53481969 | - | lncRNA |
| ENSG00000186787.8_6 | SPIN2B | 0.92515065 | 1 | 0.195 | 1.258 | chrX | 57144984 | 57147979 | - | protein_coding |
| ENSG00000204529.4_6 | GUCY2EP | 0.73083842 | 1 | 0.195 | -0.755 | chr11 | 76405085 | 76430845 | - | transcribed_unprocessed_pseudogene |
| ENSG00000253293.5_5 | HOXA10 | 0.93662057 | 1 | 0.195 | 1.441 | chr7 | 27210211 | 27219880 | - | protein_coding |
| ENSG00000105617.4_4 | LENG1 | 0.9978808 | 1 | 0.195 | 3.343 | chr19 | 54658899 | 54663459 | - | protein_coding |
| ENSG00000124782.20_8 | RREB1 | 0.99999423 | 1 | 0.195 | 5.518 | chr6 | 7107830 | 7252213 | + | protein_coding |
| ENSG00000187147.18_5 | RNF220 | 0.99993231 | 1 | 0.195 | 4.512 | chr1 | 44870866 | 45117396 | + | protein_coding |
| ENSG00000161040.16_7 | FBXL13 | 0.83853417 | 1 | 0.195 | 0.691 | chr7 | 102453677 | 102715290 | - | protein_coding |
| ENSG00000164062.13_4 | APEH | 0.99999208 | 1 | 0.195 | 6.295 | chr3 | 49711447 | 49721404 | + | protein_coding |
| ENSG00000221838.10_6 | AP4M1 | 0.99926461 | 1 | 0.195 | 3.709 | chr7 | 99699172 | 99707968 | + | protein_coding |
| ENSG00000276853.1_7 | AC026124.2 | 0.72267793 | 1 | 0.195 | -0.723 | chr12 | 65565042 | 65565697 | + | lncRNA |
| ENSG00000119685.20_4 | TTLL5 | 0.9999752 | 1 | 0.195 | 5.266 | chr14 | 76099968 | 76421422 | + | protein_coding |
| ENSG00000137221.14_4 | TJAP1 | 0.99974993 | 1 | 0.195 | 4.047 | chr6 | 43445261 | 43474294 | + | protein_coding |
| ENSG00000126970.16_4 | ZC4H2 | 0.91723241 | 1 | 0.195 | 1.550 | chrX | 64135682 | 64254593 | - | protein_coding |
| ENSG00000104825.17_4 | NFKBIB | 0.99953555 | 1 | 0.195 | 4.860 | chr19 | 39390340 | 39399533 | + | protein_coding |
| ENSG00000257221.4_7 | AC007569.1 | 0.76144882 | 1 | 0.194 | -0.098 | chr12 | 109022444 | 109036125 | + | lncRNA |
| ENSG00000261824.7_8 | LINC00662 | 0.999969 | 1 | 0.194 | 5.067 | chr19 | 28171980 | 28284913 | - | lncRNA |
| ENSG00000151914.20_10 | DST | 0.99796822 | 1 | 0.194 | 9.089 | chr6 | 56322785 | 56819447 | - | protein_coding |
| ENSG00000248275.2_6 | TRIM52-AS1 | 0.99585978 | 1 | 0.194 | 2.844 | chr5 | 180688170 | 180699308 | + | lncRNA |
| ENSG00000118900.15_5 | UBN1 | 0.99999721 | 1 | 0.194 | 6.160 | chr16 | 4896666 | 4932402 | + | protein_coding |
| ENSG00000244242.2_5 | IFITM10 | 0.72514526 | 1 | 0.194 | -0.787 | chr11 | 1753636 | 1771825 | - | protein_coding |
| ENSG00000188895.12_7 | MSL1 | 0.99999941 | 1 | 0.194 | 6.656 | chr17 | 38278224 | 38293170 | + | protein_coding |
| ENSG00000278570.5_3 | NR2E3 | 0.81461679 | 1 | 0.194 | 0.412 | chr15 | 72084977 | 72110600 | + | protein_coding |
| ENSG00000110888.17_6 | CAPRIN2 | 0.99998979 | 1 | 0.194 | 5.296 | chr12 | 30862486 | 30907885 | - | protein_coding |
| ENSG00000126091.20_8 | ST3GAL3 | 0.8614021 | 1 | 0.194 | 0.713 | chr1 | 44171495 | 44396837 | + | protein_coding |
| ENSG00000249485.1_4 | RBBP4P1 | 0.72935414 | 1 | 0.194 | -0.832 | chr5 | 14797234 | 14798509 | - | processed_pseudogene |
| ENSG00000085832.17_7 | EPS15 | 0.99998685 | 1 | 0.194 | 5.635 | chr1 | 51819935 | 51984938 | - | protein_coding |
| ENSG00000235173.7_3 | HGH1 | 0.94773337 | 1 | 0.194 | 1.674 | chr8 | 145192677 | 145195754 | + | protein_coding |
| ENSG00000280670.3_6 | CCDC163 | 0.93576524 | 1 | 0.194 | 1.521 | chr1 | 45959538 | 45965751 | - | protein_coding |
| ENSG00000215154.6_5 | AC141586.1 | 0.87892295 | 1 | 0.193 | 1.009 | chr16 | 2653351 | 2680495 | + | transcribed_unprocessed_pseudogene |
| ENSG00000109066.14_3 | TMEM104 | 0.99998568 | 1 | 0.193 | 5.034 | chr17 | 72772638 | 72835892 | + | protein_coding |
| ENSG00000171617.14_6 | ENC1 | 0.99999887 | 1 | 0.193 | 6.142 | chr5 | 73923231 | 73937249 | - | protein_coding |
| ENSG00000158796.17_6 | DEDD | 0.99970327 | 1 | 0.193 | 4.072 | chr1 | 161090764 | 161102478 | - | protein_coding |
| ENSG00000085872.15_6 | CHERP | 0.99834161 | 1 | 0.193 | 3.645 | chr19 | 16628705 | 16653248 | - | protein_coding |
| ENSG00000173917.10_3 | HOXB2 | 0.8880061 | 1 | 0.193 | 0.881 | chr17 | 46618256 | 46622351 | - | protein_coding |
| ENSG00000088280.19_5 | ASAP3 | 0.99847977 | 1 | 0.193 | 3.438 | chr1 | 23755056 | 23811061 | - | protein_coding |
| ENSG00000196405.13_5 | EVL | 0.99983321 | 1 | 0.193 | 4.223 | chr14 | 100437786 | 100610573 | + | protein_coding |
| ENSG00000205572.9_3 | SERF1B | 0.80677289 | 1 | 0.193 | -0.065 | chr5 | 69321074 | 69338940 | + | protein_coding |

| | | | | | | | | | | |
|---|---|---|---|---|---|---|---|---|---|---|
| ENSG00000154229.12_4 | PRKCA | 0.99999985 | 1 | 0.193 | 7.446 | chr17 | 64298731 | 64806861 | + | protein_coding |
| ENSG00000116786.13_5 | PLEKHM2 | 0.99992448 | 1 | 0.193 | 5.256 | chr1 | 16010815 | 16061264 | + | protein_coding |
| ENSG00000167925.16_7 | GHDC | 0.99991334 | 1 | 0.193 | 4.673 | chr17 | 40340817 | 40346550 | - | protein_coding |
| ENSG00000176155.19_7 | CCDC57 | 0.99998289 | 1 | 0.193 | 5.521 | chr17 | 80059336 | 80170706 | - | protein_coding |
| ENSG00000131969.15_3 | ABHD12B | 0.75871673 | 1 | 0.193 | -0.393 | chr14 | 51338771 | 51371688 | + | protein_coding |
| ENSG00000169683.8_4 | LRRC45 | 0.9475657 | 1 | 0.193 | 5.671 | chr17 | 79981181 | 79989027 | + | protein_coding |
| ENSG00000109103.12_6 | UNC119 | 0.99992965 | 1 | 0.193 | 5.004 | chr17 | 26873726 | 26879649 | - | protein_coding |
| ENSG00000100124.15_5 | ANKRD54 | 0.99983168 | 1 | 0.192 | 4.243 | chr22 | 38226862 | 38245334 | - | protein_coding |
| ENSG00000120068.7_5 | HOXB8 | 0.97320066 | 1 | 0.192 | 2.501 | chr17 | 46688739 | 46692654 | - | protein_coding |
| ENSG00000145284.12_4 | SCD5 | 0.9997877 | 1 | 0.192 | 4.096 | chr4 | 83550692 | 83719949 | - | protein_coding |
| ENSG00000166262.16_9 | FAM227B | 0.84748486 | 1 | 0.192 | 0.647 | chr15 | 49619159 | 49913126 | - | protein_coding |
| ENSG00000152380.10_4 | FAM151B | 0.93962964 | 1 | 0.192 | 1.488 | chr5 | 79783788 | 79838382 | + | protein_coding |
| ENSG00000106330.12_3 | MOSPD3 | 0.98918278 | 1 | 0.192 | 2.416 | chr7 | 100209725 | 100213007 | + | protein_coding |
| ENSG00000101844.18_4 | ATG4A | 0.99846381 | 1 | 0.192 | 3.457 | chrX | 107334898 | 107397901 | + | protein_coding |
| ENSG00000266094.8_5 | RASSF5 | 0.99197471 | 1 | 0.192 | 2.646 | chr1 | 206680864 | 206762780 | + | protein_coding |
| ENSG00000141564.15_4 | RPTOR | 0.99999736 | 1 | 0.192 | 6.199 | chr17 | 78518619 | 78940171 | + | protein_coding |
| ENSG00000032444.16_6 | PNPLA6 | 0.99522133 | 1 | 0.192 | 6.018 | chr19 | 7598890 | 7626650 | + | protein_coding |
| ENSG00000226696.6_8 | LENG8-AS1 | 0.97307004 | 1 | 0.192 | 2.163 | chr19 | 54955985 | 54960223 | - | lncRNA |
| ENSG00000167842.16_3 | MIS12 | 0.99998984 | 1 | 0.192 | 5.395 | chr17 | 5389605 | 5394134 | + | protein_coding |
| ENSG00000140398.14_4 | NEIL1 | 0.99870021 | 1 | 0.192 | 3.403 | chr15 | 75639296 | 75649456 | + | protein_coding |
| ENSG00000158604.15_6 | TMED4 | 0.99999896 | 1 | 0.192 | 6.421 | chr7 | 44617493 | 44621886 | - | protein_coding |
| ENSG00000141568.21_5 | FOXK2 | 0.99999929 | 1 | 0.192 | 6.646 | chr17 | 80477589 | 80602538 | + | protein_coding |
| ENSG00000271816.1_9 | AC022400.5 | 0.72824496 | 1 | 0.192 | -0.754 | chr10 | 75458909 | 75490245 | - | lncRNA |
| ENSG00000147789.15_2 | ZNF7 | 0.99997168 | 1 | 0.192 | 4.956 | chr8 | 146052849 | 146072894 | + | protein_coding |
| ENSG00000185127.6_2 | C6orf120 | 0.9999273 | 1 | 0.192 | 4.413 | chr6 | 170102286 | 170104952 | + | protein_coding |
| ENSG00000153406.13_3 | NMRAL1 | 0.99998698 | 1 | 0.191 | 5.366 | chr16 | 4511681 | 4545764 | - | protein_coding |
| ENSG00000082512.15_3 | TRAF5 | 0.99993426 | 1 | 0.191 | 4.544 | chr1 | 211499957 | 211548288 | + | protein_coding |
| ENSG00000131844.16_3 | MCCC2 | 0.99999954 | 1 | 0.191 | 6.619 | chr5 | 70883115 | 70954533 | + | protein_coding |
| ENSG00000267213.4_7 | AC007773.1 | 0.98003773 | 1 | 0.191 | 1.598 | chr19 | 32880956 | 32896466 | - | lncRNA |
| ENSG00000167103.12_4 | PIP5KL1 | 0.90333878 | 1 | 0.191 | 1.158 | chr9 | 130683160 | 130693064 | - | protein_coding |
| ENSG00000188191.15_6 | PRKAR1B | 0.99960815 | 1 | 0.191 | 5.078 | chr7 | 588834 | 767287 | - | protein_coding |
| ENSG00000248161.5_8 | AC098487.1 | 0.81294157 | 1 | 0.191 | 0.225 | chr4 | 103339759 | 103371167 | - | lncRNA |
| ENSG00000165730.16_4 | STOX1 | 0.98551985 | 1 | 0.191 | 2.316 | chr10 | 70587288 | 70655188 | + | protein_coding |
| ENSG00000129691.16_4 | ASH2L | 0.99997509 | 1 | 0.191 | 4.827 | chr8 | 37963011 | 38001594 | + | protein_coding |
| ENSG00000147400.10_4 | CETN2 | 0.99999394 | 1 | 0.191 | 5.830 | chrX | 151995538 | 151999301 | - | protein_coding |
| ENSG00000236499.2_4 | LINC00896 | 0.80740868 | 1 | 0.191 | 0.283 | chr22 | 20193920 | 20196047 | + | lncRNA |
| ENSG00000170634.13_4 | ACYP2 | 0.97779522 | 1 | 0.190 | 2.156 | chr2 | 54197975 | 54532437 | + | protein_coding |
| ENSG00000167799.10_5 | NUDT8 | 0.97783704 | 1 | 0.190 | 2.168 | chr11 | 67395409 | 67397408 | - | protein_coding |
| ENSG00000134250.20_5 | NOTCH2 | 0.99999985 | 1 | 0.190 | 7.169 | chr1 | 120454176 | 120643366 | - | protein_coding |
| ENSG00000268858.2_6 | AL118506.1 | 0.98975222 | 1 | 0.190 | 2.682 | chr20 | 62492565 | 62495646 | - | lncRNA |
| ENSG00000146376.11_4 | ARHGAP18 | 0.99999942 | 1 | 0.190 | 6.812 | chr6 | 129897277 | 130031322 | - | protein_coding |
| ENSG00000148158.17_3 | SNX30 | 0.99994655 | 1 | 0.190 | 4.705 | chr9 | 115513040 | 115643951 | + | protein_coding |
| ENSG00000198718.13_4 | TOGARAM1 | 0.9999881 | 1 | 0.190 | 5.514 | chr14 | 45431393 | 45543634 | + | protein_coding |
| ENSG00000204147.10_4 | ASAH2B | 0.98304917 | 1 | 0.190 | 2.354 | chr10 | 52499078 | 52576255 | + | protein_coding |
| ENSG00000111669.15_5 | TPI1 | 0.9999999 | 1 | 0.190 | 9.710 | chr12 | 6976283 | 6980112 | + | protein_coding |
| ENSG00000112079.9_5 | STK38 | 0.99999854 | 1 | 0.190 | 5.908 | chr6 | 36461669 | 36515256 | - | protein_coding |
| ENSG00000139579.13_4 | NABP2 | 0.99999731 | 1 | 0.190 | 5.710 | chr12 | 56615799 | 56623638 | + | protein_coding |
| ENSG00000159082.17_4 | SYNJ1 | 0.99203465 | 1 | 0.190 | 2.776 | chr21 | 34001069 | 34100359 | - | protein_coding |
| ENSG00000109618.12_4 | SEPSECS | 0.99912595 | 1 | 0.190 | 3.699 | chr4 | 25121636 | 25162071 | - | protein_coding |
| ENSG00000215256.4_6 | DHRS4-AS1 | 0.99839426 | 1 | 0.189 | 3.787 | chr14 | 24407428 | 24521764 | - | lncRNA |
| ENSG00000167487.12_4 | KLHL26 | 0.98274952 | 1 | 0.189 | 2.310 | chr19 | 18747835 | 18782531 | + | protein_coding |
| ENSG00000103544.15_7 | VPS35L | 0.99999876 | 1 | 0.189 | 6.134 | chr16 | 19566562 | 19718115 | + | protein_coding |
| ENSG00000110844.13_4 | PRPF40B | 0.9988945 | 1 | 0.189 | 3.404 | chr12 | 49962001 | 50038449 | + | protein_coding |
| ENSG00000125629.15_3 | INSIG2 | 0.99852943 | 1 | 0.188 | 3.312 | chr2 | 118846028 | 118868573 | + | protein_coding |
| ENSG00000117691.10_3 | NENF | 0.99999274 | 1 | 0.188 | 5.424 | chr1 | 212606262 | 212619721 | + | protein_coding |
| ENSG00000124788.18_6 | ATXN1 | 0.99980687 | 1 | 0.188 | 4.104 | chr6 | 16299343 | 16761722 | - | protein_coding |
| ENSG00000177556.12_6 | ATOX1 | 0.99976007 | 1 | 0.188 | 4.232 | chr5 | 151121877 | 151152093 | - | protein_coding |
| ENSG00000222009.8_2 | BTBD19 | 0.99397112 | 1 | 0.188 | 3.008 | chr1 | 45274154 | 45281257 | + | protein_coding |
| ENSG00000141741.12_4 | MIEN1 | 0.99990045 | 1 | 0.188 | 4.699 | chr17 | 37884749 | 37886785 | - | protein_coding |
| ENSG00000176903.5_3 | PNMA1 | 0.99999154 | 1 | 0.188 | 5.499 | chr14 | 74178486 | 74181087 | - | protein_coding |
| ENSG00000111642.15_7 | CHD4 | 0.99946218 | 1 | 0.187 | 3.717 | chr12 | 6679248 | 6723690 | - | protein_coding |

| | | | | | | | | | | |
|---|---|---|---|---|---|---|---|---|---|---|
| ENSG00000278530.4_4 | CHMP1B2P | 0.97534382 | 1 | 0.187 | 2.088 | chrX | 79483988 | 79590863 | - | transcribed_unitary_pseudogene |
| ENSG00000164889.14_6 | SLC4A2 | 0.99886775 | 1 | 0.187 | 7.548 | chr7 | 150754297 | 150773613 | + | protein_coding |
| ENSG00000070610.14_5 | GBA2 | 0.9999959 | 1 | 0.187 | 5.739 | chr9 | 35736863 | 35749225 | - | protein_coding |
| ENSG00000006007.12_4 | GDE1 | 0.99999972 | 1 | 0.187 | 6.708 | chr16 | 19513015 | 19533445 | - | protein_coding |
| ENSG00000286753.1_2 | AC091978.1 | 0.73044553 | 1 | 0.187 | -0.962 | chr5 | 5390209 | 5422229 | - | lncRNA |
| ENSG00000186019.11_7 | AC021092.1 | 0.82230937 | 1 | 0.187 | 0.470 | chr19 | 44607160 | 44617336 | - | lncRNA |
| ENSG00000175305.17_4 | CCNE2 | 0.99252461 | 1 | 0.187 | 2.870 | chr8 | 95891998 | 95908906 | - | protein_coding |
| ENSG00000189057.11_6 | FAM111B | 0.99999489 | 1 | 0.187 | 5.602 | chr11 | 58874658 | 58894885 | + | protein_coding |
| ENSG00000287737.1_1 | AC132872.5 | 0.91977911 | 1 | 0.186 | 1.367 | chr17 | 80231624 | 80237521 | + | lncRNA |
| ENSG00000147027.4_4 | TMEM47 | 0.9740146 | 1 | 0.186 | 2.086 | chrX | 34645192 | 34675402 | - | protein_coding |
| ENSG00000051009.10_4 | FAM160A2 | 0.99974784 | 1 | 0.186 | 4.518 | chr11 | 6232565 | 6255941 | - | protein_coding |
| ENSG00000124422.12_6 | USP22 | 0.99999995 | 1 | 0.186 | 8.513 | chr17 | 20902909 | 20947073 | - | protein_coding |
| ENSG00000130985.17_7 | UBA1 | 0.99999993 | 1 | 0.186 | 8.252 | chrX | 47050260 | 47074527 | + | protein_coding |
| ENSG00000068400.13_5 | GRIPAP1 | 0.99999583 | 1 | 0.186 | 5.468 | chrX | 48830131 | 48858675 | - | protein_coding |
| ENSG00000253785.1_5 | AC008429.2 | 0.75494384 | 1 | 0.186 | -0.461 | chr5 | 172402514 | 172403377 | + | processed_pseudogene |
| ENSG00000279641.1_5 | AC120057.3 | 0.74612846 | 1 | 0.186 | -0.725 | chr17 | 7145519 | 7147133 | + | TEC |
| ENSG00000022976.15_5 | ZNF839 | 0.99783715 | 1 | 0.186 | 3.449 | chr14 | 102783714 | 102809039 | + | protein_coding |
| ENSG00000140718.21_4 | FTO | 0.99999874 | 1 | 0.186 | 5.968 | chr16 | 53735604 | 54192424 | + | protein_coding |
| ENSG00000027847.14_4 | B4GALT7 | 0.99974031 | 1 | 0.186 | 4.491 | chr5 | 177027133 | 177037331 | + | protein_coding |
| ENSG00000104859.15_5 | CLASRP | 0.99994864 | 1 | 0.185 | 5.395 | chr19 | 45542303 | 45574214 | + | protein_coding |
| ENSG00000104852.15_5 | SNRNP70 | 0.99966339 | 1 | 0.185 | 7.909 | chr19 | 49588676 | 49611862 | + | protein_coding |
| ENSG00000235552.4_5 | RPL6P27 | 0.99757054 | 1 | 0.185 | 3.107 | chr18 | 6462143 | 6463014 | - | transcribed_processed_pseudogene |
| ENSG00000157657.14_4 | ZNF618 | 0.99997352 | 1 | 0.185 | 5.075 | chr9 | 116638562 | 116818871 | + | protein_coding |
| ENSG00000160087.20_7 | UBE2J2 | 0.9996438 | 1 | 0.185 | 4.649 | chr1 | 1189289 | 1209265 | - | protein_coding |
| ENSG00000090470.15_7 | PDCD7 | 0.99996433 | 1 | 0.185 | 4.800 | chr15 | 65409717 | 65426146 | - | protein_coding |
| ENSG00000257925.1_8 | AC008083.2 | 0.7327278 | 1 | 0.185 | -1.047 | chr12 | 47631517 | 47672804 | - | lncRNA |
| ENSG00000225077.3_4 | LINC00337 | 0.8272271 | 1 | 0.185 | 0.386 | chr1 | 6294752 | 6299504 | + | lncRNA |
| ENSG00000167964.13_6 | RAB26 | 0.99993139 | 1 | 0.185 | 4.495 | chr16 | 2190804 | 2204166 | + | protein_coding |
| ENSG00000133142.17_5 | TCEAL4 | 0.99999398 | 1 | 0.185 | 5.230 | chrX | 102831159 | 102842664 | + | protein_coding |
| ENSG00000110315.7_6 | RNF141 | 0.99996834 | 1 | 0.184 | 5.576 | chr11 | 10533220 | 10562777 | - | protein_coding |
| ENSG00000101856.10_3 | PGRMC1 | 0.99999988 | 1 | 0.184 | 7.384 | chrX | 118370208 | 118378429 | + | protein_coding |
| ENSG00000107882.11_2 | SUFU | 0.99409193 | 1 | 0.184 | 2.792 | chr10 | 104263744 | 104393292 | + | protein_coding |
| ENSG00000176095.12_4 | IP6K1 | 0.9999831 | 1 | 0.184 | 5.137 | chr3 | 49761727 | 49823975 | - | protein_coding |
| ENSG00000139194.8_4 | RBP5 | 0.97995645 | 1 | 0.184 | 2.059 | chr12 | 7268332 | 7281485 | - | protein_coding |
| ENSG00000197362.15_5 | ZNF786 | 0.99923138 | 1 | 0.184 | 3.559 | chr7 | 148766733 | 148787874 | - | protein_coding |
| ENSG00000196388.8_3 | INCA1 | 0.91411983 | 1 | 0.184 | 1.395 | chr17 | 4891425 | 4900905 | - | protein_coding |
| ENSG00000140688.17_5 | C16orf58 | 0.99997214 | 1 | 0.184 | 5.031 | chr16 | 31500792 | 31520630 | - | protein_coding |
| ENSG00000272140.2_10 | AC022400.6 | 0.78698801 | 1 | 0.184 | -0.209 | chr10 | 75463493 | 75473339 | - | lncRNA |
| ENSG00000076984.17_2 | MAP2K7 | 0.99931944 | 1 | 0.184 | 3.728 | chr19 | 7968728 | 7979363 | + | protein_coding |
| ENSG00000260572.1_7 | AC069224.2 | 0.73620188 | 1 | 0.183 | -0.960 | chr3 | 160471216 | 160472930 | - | lncRNA |
| ENSG00000262587.2_5 | AC133552.2 | 0.76151201 | 1 | 0.183 | -0.345 | chr16 | 25043065 | 25069430 | + | transcribed_unprocessed_pseudogene |
| ENSG00000174579.4_5 | MSL2 | 0.99995113 | 1 | 0.183 | 4.503 | chr3 | 135867759 | 135916083 | - | protein_coding |
| ENSG00000101216.11_4 | GMEB2 | 0.99979978 | 1 | 0.183 | 4.290 | chr20 | 62218955 | 62258454 | - | protein_coding |
| ENSG00000137575.12_4 | SDCBP | 0.99999602 | 1 | 0.183 | 6.761 | chr8 | 59465483 | 59495418 | + | protein_coding |
| ENSG00000133030.21_6 | MPRIP | 0.99999956 | 1 | 0.183 | 8.575 | chr17 | 16945859 | 17120993 | + | protein_coding |
| ENSG00000122574.10_2 | WIPF3 | 0.8312458 | 1 | 0.183 | 0.106 | chr7 | 29846102 | 29956682 | + | protein_coding |
| ENSG00000216624.3_3 | GAPDHP72 | 0.73535718 | 1 | 0.183 | -0.962 | chr6 | 166471993 | 166479516 | + | transcribed_processed_pseudogene |
| ENSG00000143878.10_3 | RHOB | 0.99999703 | 1 | 0.183 | 6.220 | chr2 | 20646835 | 20649201 | + | protein_coding |
| ENSG00000276293.4_6 | PIP4K2B | 0.99999986 | 1 | 0.183 | 7.029 | chr17 | 36921942 | 36956379 | - | protein_coding |
| ENSG00000089876.12_8 | DHX32 | 0.99999785 | 1 | 0.183 | 5.985 | chr10 | 127524906 | 127585005 | - | protein_coding |
| ENSG00000125744.12_5 | RTN2 | 0.93739632 | 1 | 0.183 | 1.759 | chr19 | 45988552 | 46000313 | - | protein_coding |
| ENSG00000171067.11_7 | C11orf24 | 0.99999 | 1 | 0.183 | 5.080 | chr11 | 68028806 | 68039469 | - | protein_coding |
| ENSG00000112851.14_5 | ERBIN | 0.99999988 | 1 | 0.183 | 7.618 | chr5 | 65222303 | 65378377 | + | protein_coding |
| ENSG00000240731.1_8 | AL139287.1 | 0.86874665 | 1 | 0.183 | 0.632 | chr1 | 1252961 | 1254069 | - | lncRNA |
| ENSG00000007866.21_10 | TEAD3 | 0.99016173 | 1 | 0.182 | 2.457 | chr6 | 35441374 | 35464856 | - | protein_coding |
| ENSG00000171634.18_6 | BPTF | 0.99999955 | 1 | 0.182 | 7.477 | chr17 | 65821619 | 65980494 | + | protein_coding |
| ENSG00000100461.18_6 | RBM23 | 0.99999977 | 1 | 0.182 | 6.571 | chr14 | 23362413 | 23388391 | - | protein_coding |

| | | | | | | | | | | |
|---|---|---|---|---|---|---|---|---|---|---|
| ENSG00000149480.7_4 | MTA2 | 0.9999998 | 1 | 0.182 | 7.242 | chr11 | 62360686 | 62369337 | - | protein_coding |
| ENSG00000130962.17_3 | PRRG1 | 0.99942469 | 1 | 0.182 | 3.878 | chrX | 37208528 | 37316548 | + | protein_coding |
| ENSG00000142178.9_5 | SIK1 | 0.84405544 | 1 | 0.182 | 0.382 | chr21 | 44834363 | 44847011 | - | protein_coding |
| ENSG00000111145.8_5 | ELK3 | 0.9999964 | 1 | 0.182 | 5.744 | chr12 | 96588153 | 96663602 | + | protein_coding |
| ENSG00000134644.15_5 | PUM1 | 0.99999976 | 1 | 0.182 | 6.660 | chr1 | 31404353 | 31538838 | - | protein_coding |
| ENSG00000137094.14_5 | DNAJB5 | 0.97297119 | 1 | 0.182 | 1.865 | chr9 | 34989638 | 34998897 | + | protein_coding |
| ENSG00000134285.11_6 | FKBP11 | 0.99941613 | 1 | 0.182 | 3.673 | chr12 | 49315301 | 49320257 | - | protein_coding |
| ENSG00000067082.15_4 | KLF6 | 0.99993489 | 1 | 0.182 | 4.655 | chr10 | 3818188 | 3827473 | - | protein_coding |
| ENSG00000038532.16_7 | CLEC16A | 0.9999703 | 1 | 0.182 | 4.623 | chr16 | 11038421 | 11276043 | + | protein_coding |
| ENSG00000100614.18_6 | PPM1A | 0.9999993 | 1 | 0.182 | 6.230 | chr14 | 60712470 | 60765805 | + | protein_coding |
| ENSG00000105229.7_3 | PIAS4 | 0.99984048 | 1 | 0.182 | 4.151 | chr19 | 4007734 | 4039384 | + | protein_coding |
| ENSG00000105325.14_4 | FZR1 | 0.99996816 | 1 | 0.181 | 5.078 | chr19 | 3506271 | 3538328 | + | protein_coding |
| ENSG00000134852.15_5 | CLOCK | 0.99998768 | 1 | 0.181 | 5.740 | chr4 | 56294070 | 56413076 | - | protein_coding |
| ENSG00000116254.18_5 | CHD5 | 0.8312458 | 1 | 0.181 | 0.104 | chr1 | 6161847 | 6240381 | - | protein_coding |
| ENSG00000184677.18_5 | ZBTB40 | 0.9999968 | 1 | 0.181 | 5.877 | chr1 | 22755331 | 22857650 | + | protein_coding |
| ENSG00000186104.10_6 | CYP2R1 | 0.99983372 | 1 | 0.181 | 4.185 | chr11 | 14898986 | 14913798 | - | protein_coding |
| ENSG00000279031.1_7 | AC004232.2 | 0.74118041 | 1 | 0.181 | -0.800 | chr16 | 3342879 | 3343403 | + | TEC |
| ENSG00000143793.13_4 | C1orf35 | 0.99965536 | 1 | 0.181 | 4.102 | chr1 | 228288427 | 228293112 | - | protein_coding |
| ENSG00000185359.13_5 | HGS | 0.99958338 | 1 | 0.181 | 6.744 | chr17 | 79650356 | 79670168 | + | protein_coding |
| ENSG00000047249.18_7 | ATP6V1H | 0.99999208 | 1 | 0.181 | 5.166 | chr8 | 54628103 | 54756118 | - | protein_coding |
| ENSG00000143569.19_7 | UBAP2L | 0.99999991 | 1 | 0.181 | 7.260 | chr1 | 154192655 | 154243986 | + | protein_coding |
| ENSG00000100023.19_4 | PPIL2 | 0.99999236 | 1 | 0.181 | 5.008 | chr22 | 22020298 | 22054304 | + | protein_coding |
| ENSG00000005007.12_3 | UPF1 | 0.99996354 | 1 | 0.181 | 6.330 | chr19 | 18942747 | 18979045 | + | protein_coding |
| ENSG00000189221.9_2 | MAOA | 0.99978615 | 1 | 0.181 | 3.942 | chrX | 43514155 | 43606071 | + | protein_coding |
| ENSG00000168152.13_4 | THAP9 | 0.99122311 | 1 | 0.180 | 2.778 | chr4 | 83821837 | 83841122 | + | protein_coding |
| ENSG00000132793.11_2 | LPIN3 | 0.99996238 | 1 | 0.180 | 4.744 | chr20 | 39969560 | 39989222 | + | protein_coding |
| ENSG00000231908.1_6 | IDH1-AS1 | 0.74911142 | 1 | 0.180 | -0.799 | chr2 | 209119971 | 209120905 | + | lncRNA |
| ENSG00000272288.6_6 | AL451165.2 | 0.82534966 | 1 | 0.180 | 0.228 | chr6 | 34664094 | 34665247 | + | lncRNA |
| ENSG00000228809.2_7 | AL034428.1 | 0.74093826 | 1 | 0.180 | -0.879 | chr20 | 16695489 | 16711593 | - | lncRNA |
| ENSG00000150093.19_6 | ITGB1 | 0.99999995 | 1 | 0.180 | 10.791 | chr10 | 33189246 | 33294720 | - | protein_coding |
| ENSG00000249863.2_5 | AC021106.1 | 0.75856264 | 1 | 0.180 | -0.462 | chr4 | 37869913 | 37871599 | + | processed_pseudogene |
| ENSG00000143322.21_6 | ABL2 | 0.99999921 | 1 | 0.180 | 6.072 | chr1 | 179068465 | 179198819 | - | protein_coding |
| ENSG00000166025.18_4 | AMOTL1 | 0.99999706 | 1 | 0.180 | 5.522 | chr11 | 94439597 | 94609913 | + | protein_coding |
| ENSG00000111897.7_4 | SERINC1 | 0.99999434 | 1 | 0.180 | 7.009 | chr6 | 122764496 | 122792952 | - | protein_coding |
| ENSG00000112305.15_4 | SMAP1 | 0.99655511 | 1 | 0.180 | 3.168 | chr6 | 71377479 | 71571714 | + | protein_coding |
| ENSG00000139324.12_6 | TMTC3 | 0.99999937 | 1 | 0.179 | 7.007 | chr12 | 88536073 | 88593664 | + | protein_coding |
| ENSG00000223509.8_6 | AC135983.3 | 0.99765351 | 1 | 0.179 | 3.186 | chr15 | 32812049 | 32829127 | - | transcribed_processed_pseudogene |
| ENSG00000138772.13_5 | ANXA3 | 0.99999894 | 1 | 0.179 | 6.788 | chr4 | 79472901 | 79531605 | + | protein_coding |
| ENSG00000182220.15_8 | ATP6AP2 | 0.99999985 | 1 | 0.179 | 6.844 | chrX | 40438624 | 40466100 | + | protein_coding |
| ENSG00000169057.23_8 | MECP2 | 0.99999798 | 1 | 0.179 | 6.143 | chrX | 153287024 | 153402578 | - | protein_coding |
| ENSG00000126368.6_4 | NR1D1 | 0.96529572 | 1 | 0.179 | 1.876 | chr17 | 38249046 | 38256842 | - | protein_coding |
| ENSG00000213888.3_5 | LINC01521 | 0.95428155 | 1 | 0.179 | 1.968 | chr22 | 31742763 | 31744705 | + | lncRNA |
| ENSG00000055732.13_4 | MCOLN3 | 0.99481032 | 1 | 0.179 | 2.730 | chr1 | 85483765 | 85514183 | - | protein_coding |
| ENSG00000148343.18_5 | MIGA2 | 0.99988426 | 1 | 0.179 | 4.555 | chr9 | 131798900 | 131834361 | + | protein_coding |
| ENSG00000140400.17_8 | MAN2C1 | 0.99999921 | 1 | 0.179 | 6.196 | chr15 | 75647548 | 75660953 | - | protein_coding |
| ENSG00000231563.2_6 | AL139288.1 | 0.79549964 | 1 | 0.179 | -0.040 | chr1 | 228595066 | 228598571 | + | lncRNA |
| ENSG00000099795.7_4 | NDUFB7 | 0.99960329 | 1 | 0.178 | 4.854 | chr19 | 14676890 | 14682878 | - | protein_coding |
| ENSG00000235587.2_4 | GAPDHP65 | 0.88265583 | 1 | 0.178 | 1.516 | chrX | 46299144 | 46300149 | - | processed_pseudogene |
| ENSG00000180900.20_5 | SCRIB | 0.90648764 | 1 | 0.178 | 7.163 | chr8 | 144873090 | 144897943 | - | protein_coding |
| ENSG00000227036.8_5 | LINC00511 | 0.99999382 | 1 | 0.178 | 5.451 | chr17 | 70286232 | 70636611 | - | lncRNA |
| ENSG00000183309.11_3 | ZNF623 | 0.99999837 | 1 | 0.178 | 5.665 | chr8 | 144718183 | 144738588 | + | protein_coding |
| ENSG00000189369.9_4 | GSPT2 | 0.99985926 | 1 | 0.178 | 4.103 | chrX | 51486538 | 51489328 | + | protein_coding |
| ENSG00000160685.13_2 | ZBTB7B | 0.99998147 | 1 | 0.178 | 5.319 | chr1 | 154975106 | 154990998 | + | protein_coding |
| ENSG00000144891.17_3 | AGTR1 | 0.91817852 | 1 | 0.178 | 1.073 | chr3 | 148415571 | 148460795 | + | protein_coding |
| ENSG00000130005.13_4 | GAMT | 0.99908373 | 1 | 0.178 | 3.731 | chr19 | 1397025 | 1401569 | - | protein_coding |
| ENSG00000225259.4_4 | ST13P6 | 0.82915033 | 1 | 0.178 | 0.202 | chr8 | 134420249 | 134421350 | + | processed_pseudogene |
| ENSG00000181915.5_3 | ADO | 0.99997352 | 1 | 0.178 | 5.122 | chr10 | 64564480 | 64568239 | + | protein_coding |
| ENSG00000158985.13_2 | CDC42SE2 | 0.99999949 | 1 | 0.178 | 6.351 | chr5 | 130581186 | 130734140 | + | protein_coding |
| ENSG00000073050.11_5 | XRCC1 | 0.99997397 | 1 | 0.178 | 4.746 | chr19 | 44047192 | 44084625 | - | protein_coding |

| | | | | | | | | | | |
|---|---|---|---|---|---|---|---|---|---|---|
| ENSG00000158747.14_4 | NBL1 | 0.9995786 | 1 | 0.178 | 3.698 | chr1 | 19923473 | 19984949 | + | protein_coding |
| ENSG00000158109.15_3 | TPRG1L | 0.99977414 | 1 | 0.178 | 3.914 | chr1 | 3541579 | 3546691 | + | protein_coding |
| ENSG00000131446.16_7 | MGAT1 | 0.99954617 | 1 | 0.178 | 5.904 | chr5 | 180211782 | 180242652 | - | protein_coding |
| ENSG00000280798.1_3 | LINC00294 | 0.99970705 | 1 | 0.177 | 4.040 | chr11 | 33097695 | 33101000 | + | lncRNA |
| ENSG00000142082.15_7 | SIRT3 | 0.99974817 | 1 | 0.177 | 4.147 | chr11 | 215030 | 236931 | - | protein_coding |
| ENSG00000001460.18_6 | STPG1 | 0.99858915 | 1 | 0.177 | 3.189 | chr1 | 24683489 | 24743424 | - | protein_coding |
| ENSG00000102007.11_3 | PLP2 | 0.99999886 | 1 | 0.177 | 5.834 | chrX | 49028245 | 49031584 | + | protein_coding |
| ENSG00000081791.9_7 | DELE1 | 0.99999893 | 1 | 0.177 | 5.932 | chr5 | 141303420 | 141321612 | + | protein_coding |
| ENSG00000172216.6_3 | CEBPB | 0.99992911 | 1 | 0.177 | 4.892 | chr20 | 48807367 | 48809205 | + | protein_coding |
| ENSG00000273015.2_6 | AC008124.1 | 0.99239382 | 1 | 0.177 | 2.476 | chr12 | 46111829 | 46121558 | - | lncRNA |
| ENSG00000158865.12_3 | SLC5A11 | 0.94309151 | 1 | 0.177 | 1.678 | chr16 | 24857162 | 24922949 | + | protein_coding |
| ENSG00000100139.13_6 | MICALL1 | 0.99997697 | 1 | 0.177 | 4.940 | chr22 | 38301664 | 38338829 | + | protein_coding |
| ENSG00000107771.17_5 | CCSER2 | 0.99999627 | 1 | 0.177 | 5.965 | chr10 | 86088342 | 86278277 | + | protein_coding |
| ENSG00000188659.9_6 | SAXO2 | 0.80410451 | 1 | 0.177 | 0.402 | chr15 | 82555151 | 82577271 | + | protein_coding |
| ENSG00000204954.10_6 | C12orf73 | 0.99728782 | 1 | 0.177 | 3.067 | chr12 | 104334541 | 104359486 | - | protein_coding |
| ENSG00000173020.11_6 | GRK2 | 0.99999197 | 1 | 0.177 | 5.508 | chr11 | 67033944 | 67054027 | + | protein_coding |
| ENSG00000144746.7_3 | ARL6IP5 | 0.99999794 | 1 | 0.177 | 6.088 | chr3 | 69134088 | 69155243 | + | protein_coding |
| ENSG00000234127.9_8 | TRIM26 | 0.99126214 | 1 | 0.177 | 2.596 | chr6 | 30152232 | 30181204 | - | protein_coding |
| ENSG00000168071.22_4 | CCDC88B | 0.99654936 | 1 | 0.177 | 2.938 | chr11 | 64107676 | 64125006 | + | protein_coding |
| ENSG00000215712.11_4 | TMEM242 | 0.99810486 | 1 | 0.177 | 3.363 | chr6 | 157710057 | 157744633 | - | protein_coding |
| ENSG00000131437.15_4 | KIF3A | 0.99999402 | 1 | 0.177 | 5.223 | chr5 | 132028320 | 132073330 | - | protein_coding |
| ENSG00000246596.7_6 | AC139795.1 | 0.85336084 | 1 | 0.177 | 0.557 | chr5 | 177046060 | 177099210 | - | transcribed_unprocessed_pseudogene |
| ENSG00000169989.3_4 | TIGD4 | 0.87442911 | 1 | 0.177 | 0.448 | chr4 | 153690506 | 153700882 | - | protein_coding |
| ENSG00000224216.1_7 | AC234781.1 | 0.75131687 | 1 | 0.177 | -0.656 | chrX | 154564167 | 154581271 | - | lncRNA |
| ENSG00000159147.18_8 | DONSON | 0.99991834 | 1 | 0.177 | 4.344 | chr21 | 34931848 | 34961012 | - | protein_coding |
| ENSG00000122884.12_3 | P4HA1 | 0.99999086 | 1 | 0.177 | 5.404 | chr10 | 74766975 | 74856732 | - | protein_coding |
| ENSG00000106992.19_6 | AK1 | 0.95792666 | 1 | 0.177 | 2.011 | chr9 | 130628765 | 130639954 | - | protein_coding |
| ENSG00000170921.15_4 | TANC2 | 0.99999861 | 1 | 0.177 | 5.963 | chr17 | 61086917 | 61505060 | + | protein_coding |
| ENSG00000092529.25_7 | CAPN3 | 0.7638593 | 1 | 0.177 | -0.656 | chr15 | 42651696 | 42705147 | + | protein_coding |
| ENSG00000249936.3_4 | RAC1P2 | 0.93884405 | 1 | 0.176 | 1.257 | chr4 | 46725847 | 46726425 | - | processed_pseudogene |
| ENSG00000062725.10_4 | APPBP2 | 0.99999948 | 1 | 0.176 | 6.368 | chr17 | 58520519 | 58603603 | - | protein_coding |
| ENSG00000168026.18_4 | TTC21A | 0.91296118 | 1 | 0.176 | 1.154 | chr3 | 39149195 | 39180394 | + | protein_coding |
| ENSG00000164144.16_6 | ARFIP1 | 0.99998453 | 1 | 0.176 | 5.275 | chr4 | 153701089 | 153839615 | + | protein_coding |
| ENSG00000105792.19_7 | CFAP69 | 0.98979297 | 1 | 0.176 | 2.331 | chr7 | 89874488 | 89940377 | + | protein_coding |
| ENSG00000100897.17_7 | DCAF11 | 0.99999476 | 1 | 0.176 | 5.276 | chr14 | 24583404 | 24594451 | + | protein_coding |
| ENSG00000172977.13_5 | KAT5 | 0.99878517 | 1 | 0.176 | 3.441 | chr11 | 65479467 | 65487075 | + | protein_coding |
| ENSG00000136451.9_5 | VEZF1 | 0.9999998 | 1 | 0.176 | 7.033 | chr17 | 56048908 | 56065620 | - | protein_coding |
| ENSG00000165355.7_4 | FBXO33 | 0.99991638 | 1 | 0.175 | 4.506 | chr14 | 39866873 | 39901704 | - | protein_coding |
| ENSG00000010278.14_5 | CD9 | 0.99369973 | 1 | 0.175 | 5.860 | chr12 | 6308881 | 6347437 | + | protein_coding |
| ENSG00000091640.8_4 | SPAG7 | 0.9999929 | 1 | 0.175 | 5.250 | chr17 | 4862521 | 4871112 | - | protein_coding |
| ENSG00000033800.13_2 | PIAS1 | 0.99999938 | 1 | 0.175 | 6.107 | chr15 | 68346517 | 68490941 | + | protein_coding |
| ENSG00000094975.14_4 | SUCO | 0.99999429 | 1 | 0.175 | 5.746 | chr1 | 172501489 | 172580973 | + | protein_coding |
| ENSG00000198081.11_6 | ZBTB14 | 0.99976237 | 1 | 0.175 | 4.036 | chr18 | 5289018 | 5297052 | - | protein_coding |
| ENSG00000132535.20_6 | DLG4 | 0.9965692 | 1 | 0.175 | 3.576 | chr17 | 7093209 | 7123160 | - | protein_coding |
| ENSG00000009950.16_6 | MLXIPL | 0.94965653 | 1 | 0.175 | 1.448 | chr7 | 73007524 | 73038873 | - | protein_coding |
| ENSG00000134452.19_4 | FBH1 | 0.99999913 | 1 | 0.175 | 5.776 | chr10 | 5932166 | 5979557 | + | protein_coding |
| ENSG00000067829.19_5 | IDH3G | 0.99999663 | 1 | 0.175 | 5.338 | chrX | 153051221 | 153059978 | - | protein_coding |
| ENSG00000270110.2_7 | AL353593.2 | 0.81302423 | 1 | 0.175 | -0.094 | chr1 | 228483250 | 228490703 | - | lncRNA |
| ENSG00000168612.4_3 | ZSWIM1 | 0.99950596 | 1 | 0.175 | 3.610 | chr20 | 44509866 | 44513905 | + | protein_coding |
| ENSG00000141198.16_8 | TOM1L1 | 0.99999913 | 1 | 0.175 | 5.912 | chr17 | 52976748 | 53039317 | + | protein_coding |
| ENSG00000100242.15_5 | SUN2 | 0.99978374 | 1 | 0.175 | 5.190 | chr22 | 39130730 | 39190148 | - | protein_coding |
| ENSG00000103978.16_5 | TMEM87A | 0.99991748 | 1 | 0.175 | 5.039 | chr15 | 42502645 | 42565782 | - | protein_coding |
| ENSG00000279530.2_6 | AC092881.1 | 0.94909943 | 1 | 0.175 | 1.766 | chr12 | 70715322 | 70718054 | - | TEC |
| ENSG00000147050.14_3 | KDM6A | 0.99999507 | 1 | 0.174 | 5.379 | chrX | 44732423 | 44971847 | + | protein_coding |
| ENSG00000162852.14_4 | CNST | 0.99999221 | 1 | 0.174 | 5.088 | chr1 | 246729746 | 246831897 | + | protein_coding |
| ENSG00000189079.17_5 | ARID2 | 0.99999901 | 1 | 0.174 | 6.639 | chr12 | 46123489 | 46301823 | + | protein_coding |
| ENSG00000188092.15_6 | GPR89B | 0.76239737 | 1 | 0.174 | -0.769 | chr1 | 147400506 | 147465824 | + | protein_coding |
| ENSG00000132478.10_4 | UNK | 0.99999283 | 1 | 0.174 | 5.237 | chr17 | 73780887 | 73821880 | + | protein_coding |
| ENSG00000121068.14_3 | TBX2 | 0.98007046 | 1 | 0.174 | 2.397 | chr17 | 59477204 | 59486827 | + | protein_coding |
| ENSG00000133641.18_3 | C12orf29 | 0.99998548 | 1 | 0.174 | 5.133 | chr12 | 88427623 | 88443937 | + | protein_coding |

| | | | | | | | | | | |
|---|---|---|---|---|---|---|---|---|---|---|
| ENSG00000225131.2_4 | PSME2P2 | 0.84191292 | 1 | 0.174 | 0.304 | chr13 | 49345264 | 49345963 | + | processed_pseudogene |
| ENSG00000137100.16_5 | DCTN3 | 0.9998601 | 1 | 0.174 | 4.793 | chr9 | 34613542 | 34620520 | - | protein_coding |
| ENSG00000260083.2_6 | MIR762HG | 0.75259589 | 1 | 0.173 | -0.686 | chr16 | 30886543 | 30906541 | + | lncRNA |
| ENSG00000267147.2_5 | LINC01842 | 0.7763159 | 1 | 0.173 | -0.143 | chr19 | 14444555 | 14454728 | + | lncRNA |
| ENSG00000132570.15_4 | PCBD2 | 0.99986931 | 1 | 0.173 | 4.149 | chr5 | 134240810 | 134343649 | + | protein_coding |
| ENSG00000103353.16_5 | UBFD1 | 0.99999989 | 1 | 0.173 | 6.776 | chr16 | 23569042 | 23585710 | + | protein_coding |
| ENSG00000101191.17_4 | DIDO1 | 0.99999892 | 1 | 0.173 | 6.954 | chr20 | 61509090 | 61569304 | - | protein_coding |
| ENSG00000260855.1_6 | AL591848.4 | 0.87420305 | 1 | 0.173 | 0.587 | chr1 | 246935603 | 246939074 | + | lncRNA |
| ENSG00000236627.1_6 | AL590302.2 | 0.75074493 | 1 | 0.173 | -1.302 | chr6 | 166167677 | 166189268 | + | lncRNA |
| ENSG00000104228.13_4 | TRIM35 | 0.9999651 | 1 | 0.173 | 4.832 | chr8 | 27142403 | 27168789 | - | protein_coding |
| ENSG00000164117.14_4 | FBXO8 | 0.99851811 | 1 | 0.173 | 3.445 | chr4 | 175157809 | 175204818 | - | protein_coding |
| ENSG00000260565.7_7 | ERVK13-1 | 0.99992413 | 1 | 0.173 | 4.400 | chr16 | 2710349 | 2732380 | - | lncRNA |
| ENSG00000099999.15_8 | RNF215 | 0.9999715 | 1 | 0.173 | 4.525 | chr22 | 30764800 | 30817760 | - | protein_coding |
| ENSG00000133812.15_6 | SBF2 | 0.99999918 | 1 | 0.173 | 6.064 | chr11 | 9800214 | 10315754 | - | protein_coding |
| ENSG00000120868.13_4 | APAF1 | 0.99999764 | 1 | 0.173 | 5.338 | chr12 | 99038919 | 99129211 | + | protein_coding |
| ENSG00000011009.11_3 | LYPLA2 | 0.99998552 | 1 | 0.173 | 4.869 | chr1 | 24117635 | 24122029 | + | protein_coding |
| ENSG00000132294.15_7 | EFR3A | 0.99999993 | 1 | 0.172 | 7.320 | chr8 | 132916340 | 133025889 | + | protein_coding |
| ENSG00000061794.13_4 | MRPS35 | 0.9999998 | 1 | 0.172 | 7.034 | chr12 | 27863755 | 27909228 | + | protein_coding |
| ENSG00000135446.17_7 | CDK4 | 0.99999785 | 1 | 0.172 | 5.963 | chr12 | 58141510 | 58149796 | - | protein_coding |
| ENSG00000157426.14_4 | AASDH | 0.99988762 | 1 | 0.172 | 4.082 | chr4 | 57204453 | 57253674 | - | protein_coding |
| ENSG00000135457.10_5 | TFCP2 | 0.99993694 | 1 | 0.172 | 4.283 | chr12 | 51487439 | 51566918 | - | protein_coding |
| ENSG00000198589.13_9 | LRBA | 0.9999994 | 1 | 0.172 | 6.565 | chr4 | 151185587 | 151936879 | - | protein_coding |
| ENSG00000083123.15_3 | BCKDHB | 0.99905676 | 1 | 0.172 | 3.660 | chr6 | 80816364 | 81055987 | + | protein_coding |
| ENSG00000257923.11_6 | CUX1 | 0.99999982 | 1 | 0.172 | 6.644 | chr7 | 101459184 | 101927250 | + | protein_coding |
| ENSG00000091262.15_3 | ABCC6 | 0.96280583 | 1 | 0.172 | 1.658 | chr16 | 16242785 | 16317379 | - | protein_coding |
| ENSG00000151631.8_5 | AKR1C6P | 0.75111244 | 1 | 0.172 | -1.012 | chr10 | 4914093 | 4934167 | - | unprocessed_pseudogene |
| ENSG00000150403.18_6 | TMCO3 | 0.99999897 | 1 | 0.172 | 6.119 | chr13 | 114145336 | 114208905 | + | protein_coding |
| ENSG00000108813.11_4 | DLX4 | 0.95149894 | 1 | 0.172 | 1.427 | chr17 | 48046334 | 48052323 | + | protein_coding |
| ENSG00000186073.14_7 | C15orf41 | 0.99972677 | 1 | 0.172 | 3.880 | chr15 | 36871827 | 37102449 | + | protein_coding |
| ENSG00000125122.15_6 | LRRC29 | 0.90962441 | 1 | 0.172 | 1.115 | chr16 | 67241042 | 67260951 | - | protein_coding |
| ENSG00000165527.7_4 | ARF6 | 0.99999983 | 1 | 0.171 | 6.903 | chr14 | 50359800 | 50363772 | + | protein_coding |
| ENSG00000250506.7_3 | CDK3 | 0.75213009 | 1 | 0.171 | -1.143 | chr17 | 73996987 | 74002080 | + | protein_coding |
| ENSG00000237351.1_6 | AL513366.1 | 0.76677305 | 1 | 0.171 | -0.526 | chrX | 47016037 | 47017598 | - | processed_pseudogene |
| ENSG00000109736.15_5 | MFSD10 | 0.99642315 | 1 | 0.171 | 5.344 | chr4 | 2932288 | 2936561 | - | protein_coding |
| ENSG00000230454.1_6 | U73166.1 | 0.97273509 | 1 | 0.171 | 1.851 | chr3 | 50297735 | 50300790 | + | lncRNA |
| ENSG00000160285.15_4 | LSS | 0.99997476 | 1 | 0.171 | 5.457 | chr21 | 47608055 | 47648738 | - | protein_coding |
| ENSG00000149923.14_3 | PPP4C | 0.99999556 | 1 | 0.171 | 6.077 | chr16 | 30087299 | 30096697 | + | protein_coding |
| ENSG00000161203.13_4 | AP2M1 | 0.99999981 | 1 | 0.171 | 7.607 | chr3 | 183892477 | 183901879 | + | protein_coding |
| ENSG00000161013.17_6 | MGAT4B | 0.99999788 | 1 | 0.171 | 6.953 | chr5 | 179224597 | 179233952 | - | protein_coding |
| ENSG00000100577.19_4 | GSTZ1 | 0.9999631 | 1 | 0.171 | 4.800 | chr14 | 77787339 | 77797940 | + | protein_coding |
| ENSG00000122778.9_3 | KIAA1549 | 0.99999437 | 1 | 0.171 | 5.478 | chr7 | 138516126 | 138666064 | - | protein_coding |
| ENSG00000186469.8_4 | GNG2 | 0.96953219 | 1 | 0.171 | 1.665 | chr14 | 52292913 | 52446060 | + | protein_coding |
| ENSG00000261505.1_7 | AL031714.1 | 0.98013697 | 1 | 0.171 | 1.531 | chr16 | 1367892 | 1372846 | - | lncRNA |
| ENSG00000132475.10_5 | H3-3B | 0.99999997 | 1 | 0.171 | 8.681 | chr17 | 73772515 | 73781974 | - | protein_coding |
| ENSG00000163684.11_4 | RPP14 | 0.99956875 | 1 | 0.170 | 3.584 | chr3 | 58291974 | 58310422 | + | protein_coding |
| ENSG00000171097.14_7 | KYAT1 | 0.76187998 | 1 | 0.170 | -0.592 | chr9 | 131595221 | 131644773 | - | protein_coding |
| ENSG00000148660.20_4 | CAMK2G | 0.99999818 | 1 | 0.170 | 5.458 | chr10 | 75572259 | 75634349 | - | protein_coding |
| ENSG00000213866.3_4 | YBX1P10 | 0.97538434 | 1 | 0.170 | 2.251 | chr9 | 35971341 | 35972315 | - | processed_pseudogene |
| ENSG00000145349.17_7 | CAMK2D | 0.99999992 | 1 | 0.170 | 6.796 | chr4 | 114372188 | 114683083 | - | protein_coding |
| ENSG00000089091.16_5 | DZANK1 | 0.99636217 | 1 | 0.170 | 2.790 | chr20 | 18364011 | 18447925 | - | protein_coding |
| ENSG00000071537.14_5 | SEL1L | 0.99999986 | 1 | 0.170 | 7.165 | chr14 | 81937891 | 82000197 | - | protein_coding |
| ENSG00000159346.13_5 | ADIPOR1 | 0.99999985 | 1 | 0.170 | 6.571 | chr1 | 202909954 | 202927700 | - | protein_coding |
| ENSG00000120696.9_3 | KBTBD7 | 0.99979823 | 1 | 0.170 | 4.028 | chr13 | 41763970 | 41768705 | - | protein_coding |
| ENSG00000213719.8_4 | CLIC1 | 0.99999977 | 1 | 0.170 | 7.046 | chr6 | 31698358 | 31707540 | - | protein_coding |
| ENSG00000103512.15_4 | NOMO1 | 0.99999962 | 1 | 0.170 | 6.037 | chr16 | 14927578 | 14990014 | + | protein_coding |
| ENSG00000272533.1 | SNORA28 | 0.90307287 | 1 | 0.170 | 0.805 | chr14 | 103804186 | 103804311 | + | snoRNA |
| ENSG00000172270.20_6 | BSG | 0.99999939 | 1 | 0.170 | 8.325 | chr19 | 571277 | 583493 | + | protein_coding |
| ENSG00000188997.8_7 | KCTD21 | 0.99701239 | 1 | 0.170 | 2.785 | chr11 | 77882295 | 77899672 | - | protein_coding |
| ENSG00000198018.7_4 | ENTPD7 | 0.99999023 | 1 | 0.170 | 4.989 | chr10 | 101419266 | 101470998 | + | protein_coding |

| | | | | | | | | | | |
|---|---|---|---|---|---|---|---|---|---|---|
| ENSG00000270012.1_7 | AC232271.1 | 0.95998456 | 1 | 0.170 | 1.689 | chrX | 49129517 | 49132231 | + | lncRNA |
| ENSG00000215374.6_6 | FAM66B | 0.76503589 | 1 | 0.170 | -0.562 | chr8 | 7156266 | 7212881 | - | lncRNA |
| ENSG00000141551.14_6 | CSNK1D | 0.99997942 | 1 | 0.169 | 5.589 | chr17 | 80196899 | 80231607 | - | protein_coding |
| ENSG00000131979.19_5 | GCH1 | 0.99215867 | 1 | 0.169 | 2.508 | chr14 | 55308726 | 55369544 | - | protein_coding |
| ENSG00000258279.4_7 | LINC00592 | 0.77785173 | 1 | 0.169 | -0.407 | chr12 | 52557899 | 52617597 | + | lncRNA |
| ENSG00000168575.10_5 | SLC20A2 | 0.9999579 | 1 | 0.169 | 4.556 | chr8 | 42273993 | 42397069 | - | protein_coding |
| ENSG00000086598.11_3 | TMED2 | 0.99999983 | 1 | 0.169 | 8.338 | chr12 | 124069080 | 124083129 | + | protein_coding |
| ENSG00000167987.11_5 | VPS37C | 0.98666466 | 1 | 0.169 | 3.961 | chr11 | 60897729 | 60929087 | - | protein_coding |
| ENSG00000136866.13_4 | ZFP37 | 0.99444939 | 1 | 0.169 | 2.674 | chr9 | 115800660 | 115819039 | - | protein_coding |
| ENSG00000157014.11_4 | TATDN2 | 0.99979622 | 1 | 0.169 | 4.203 | chr3 | 10289707 | 10322902 | + | protein_coding |
| ENSG00000164808.17_8 | SPIDR | 0.99999814 | 1 | 0.169 | 5.726 | chr8 | 48173470 | 48648868 | + | protein_coding |
| ENSG00000248487.9_4 | ABHD14A | 0.94472908 | 1 | 0.169 | 1.454 | chr3 | 52005442 | 52015215 | + | protein_coding |
| ENSG00000171310.11_6 | CHST11 | 0.99999466 | 1 | 0.169 | 5.330 | chr12 | 104849073 | 105155792 | + | protein_coding |
| ENSG00000247596.9_5 | TWF2 | 0.99989026 | 1 | 0.169 | 4.381 | chr3 | 52262628 | 52273174 | - | protein_coding |
| ENSG00000274471.2_6 | AC242376.2 | 0.76640819 | 1 | 0.169 | -1.061 | chr15 | 28579891 | 28583032 | + | transcribed_unprocessed_pseudogene |
| ENSG00000137098.14_6 | SPAG8 | 0.7946381 | 1 | 0.169 | -0.044 | chr9 | 35808042 | 35812269 | - | protein_coding |
| ENSG00000107959.16_7 | PITRM1 | 0.99999977 | 1 | 0.169 | 6.464 | chr10 | 3179920 | 3215033 | - | protein_coding |
| ENSG00000161202.20_5 | DVL3 | 0.99999432 | 1 | 0.169 | 5.343 | chr3 | 183873165 | 183891402 | + | protein_coding |
| ENSG00000116128.11_5 | BCL9 | 0.99953322 | 1 | 0.169 | 3.695 | chr1 | 147013271 | 147098017 | + | protein_coding |
| ENSG00000203993.6_8 | ARRDC1-AS1 | 0.99988419 | 1 | 0.169 | 4.095 | chr9 | 140509784 | 140513358 | - | lncRNA |
| ENSG00000180776.15_6 | ZDHHC20 | 0.99999943 | 1 | 0.169 | 6.492 | chr13 | 21946712 | 22033509 | - | protein_coding |
| ENSG00000133835.16_5 | HSD17B4 | 0.99999977 | 1 | 0.168 | 6.738 | chr5 | 118788168 | 118972894 | + | protein_coding |
| ENSG00000167578.18_4 | RAB4B | 0.78332062 | 1 | 0.168 | -0.351 | chr19 | 41284121 | 41302847 | + | protein_coding |
| ENSG00000181404.17_6 | WASHC1 | 0.99273874 | 1 | 0.168 | 2.637 | chr9 | 14475 | 73865 | - | protein_coding |
| ENSG00000274925.1_6 | ZKSCAN2-DT | 0.98038987 | 1 | 0.168 | 1.891 | chr16 | 25269273 | 25272387 | + | lncRNA |
| ENSG00000105656.13_5 | ELL | 0.99142826 | 1 | 0.168 | 2.929 | chr19 | 18553473 | 18632926 | - | protein_coding |
| ENSG00000164896.20_5 | FASTK | 0.99999861 | 1 | 0.168 | 5.861 | chr7 | 150773680 | 150777953 | - | protein_coding |
| ENSG00000187051.9_4 | RPS19BP1 | 0.95605412 | 1 | 0.168 | 5.377 | chr22 | 39925098 | 39928766 | - | protein_coding |
| ENSG00000175029.17_6 | CTBP2 | 0.99999981 | 1 | 0.168 | 6.603 | chr10 | 126672886 | 126849739 | - | protein_coding |
| ENSG00000285851.1_4 | AL359762.3 | 0.76032339 | 1 | 0.168 | -1.050 | chr1 | 84964033 | 85022371 | - | lncRNA |
| ENSG00000237886.1_6 | NALT1 | 0.95775462 | 1 | 0.168 | 1.906 | chr9 | 139440664 | 139444345 | + | lncRNA |
| ENSG00000125875.14_4 | TBC1D20 | 0.99999953 | 1 | 0.168 | 5.992 | chr20 | 416124 | 443187 | - | protein_coding |
| ENSG00000245888.7_7 | NSMCE1-DT | 0.78177174 | 1 | 0.168 | 0.034 | chr16 | 27279526 | 27301813 | + | lncRNA |
| ENSG00000198265.12_6 | HELZ | 0.99999152 | 1 | 0.168 | 6.368 | chr17 | 65066560 | 65242105 | - | protein_coding |
| ENSG00000215630.6_5 | GUSBP9 | 0.86110394 | 1 | 0.168 | 0.215 | chr5 | 70493473 | 70503957 | - | unprocessed_pseudogene |
| ENSG00000115268.9_4 | RPS15 | 0.99999982 | 1 | 0.167 | 7.529 | chr19 | 1438357 | 1440493 | + | protein_coding |
| ENSG00000169895.6_3 | SYAP1 | 0.99999986 | 1 | 0.167 | 6.922 | chrX | 16737735 | 16783463 | + | protein_coding |
| ENSG00000267002.4_8 | AC060780.1 | 0.99946611 | 1 | 0.167 | 3.518 | chr17 | 41300385 | 41323054 | - | lncRNA |
| ENSG00000100403.12_3 | ZC3H7B | 0.99999927 | 1 | 0.167 | 6.358 | chr22 | 41697529 | 41756151 | + | protein_coding |
| ENSG00000257803.2_5 | AC010200.1 | 0.80299671 | 1 | 0.167 | -0.272 | chr12 | 100972522 | 100973975 | + | processed_pseudogene |
| ENSG00000269335.5_6 | IKBKG | 0.99904398 | 1 | 0.167 | 3.652 | chrX | 153769414 | 153793261 | + | protein_coding |
| ENSG00000110400.11_5 | NECTIN1 | 0.99834486 | 1 | 0.167 | 3.484 | chr11 | 119494120 | 119599910 | - | protein_coding |
| ENSG00000100949.14_3 | RABGGTA | 0.99972935 | 1 | 0.167 | 3.739 | chr14 | 24734744 | 24740945 | - | protein_coding |
| ENSG00000183161.6_4 | FANCF | 0.99991526 | 1 | 0.167 | 4.201 | chr11 | 22644079 | 22647369 | - | protein_coding |
| ENSG00000111666.11_6 | CHPT1 | 0.99999927 | 1 | 0.167 | 5.786 | chr12 | 102090725 | 102137918 | + | protein_coding |
| ENSG00000166169.17_7 | POLL | 0.99994862 | 1 | 0.167 | 4.382 | chr10 | 103338639 | 103348027 | - | protein_coding |
| ENSG00000063244.12_5 | U2AF2 | 0.99999599 | 1 | 0.167 | 7.767 | chr19 | 56165512 | 56186081 | + | protein_coding |
| ENSG00000147118.11_4 | ZNF182 | 0.99984343 | 1 | 0.167 | 4.007 | chrX | 47834250 | 47863377 | - | protein_coding |
| ENSG00000131746.13_6 | TNS4 | 0.99999992 | 1 | 0.166 | 7.063 | chr17 | 38632080 | 38657875 | - | protein_coding |
| ENSG00000143971.9_4 | ETAA1 | 0.99999885 | 1 | 0.166 | 6.232 | chr2 | 67624454 | 67639221 | + | protein_coding |
| ENSG00000204227.5_3 | RING1 | 0.99745268 | 1 | 0.166 | 3.119 | chr6 | 33176277 | 33180499 | + | protein_coding |
| ENSG00000168000.14_8 | BSCL2 | 0.91200011 | 1 | 0.166 | 0.941 | chr11 | 62457747 | 62477317 | - | protein_coding |
| ENSG00000011143.17_6 | MKS1 | 0.99958474 | 1 | 0.166 | 3.688 | chr17 | 56282802 | 56296966 | - | protein_coding |
| ENSG00000214784.4_2 | AC010468.1 | 0.81069593 | 1 | 0.166 | -0.142 | chr5 | 110527924 | 110528740 | - | processed_pseudogene |
| ENSG00000102181.21_6 | CD99L2 | 0.99999989 | 1 | 0.166 | 7.390 | chrX | 149934809 | 150067289 | - | protein_coding |
| ENSG00000182552.15_5 | RWDD4 | 0.9995442 | 1 | 0.166 | 4.282 | chr4 | 184560788 | 184580356 | - | protein_coding |
| ENSG00000165650.12_4 | PDZD8 | 0.99999997 | 1 | 0.166 | 8.115 | chr10 | 119036785 | 119134951 | - | protein_coding |

| | | | | | | | | | | |
|---|---|---|---|---|---|---|---|---|---|---|
| ENSG00000181467.5_4 | RAP2B | 0.99999883 | 1 | 0.166 | 5.787 | chr3 | 152880015 | 152888416 | + | protein_coding |
| ENSG00000075303.13_4 | SLC25A40 | 0.99997307 | 1 | 0.166 | 4.637 | chr7 | 87462883 | 87505675 | - | protein_coding |
| ENSG00000127952.17_5 | STYXL1 | 0.99993769 | 1 | 0.166 | 4.336 | chr7 | 75625656 | 75677322 | - | protein_coding |
| ENSG00000157540.21_6 | DYRK1A | 0.99999983 | 1 | 0.165 | 6.406 | chr21 | 38737875 | 38898660 | + | protein_coding |
| ENSG00000269893.8_5 | SNHG8 | 0.99997489 | 1 | 0.165 | 4.553 | chr4 | 119199864 | 119206471 | + | lncRNA |
| ENSG00000112893.9_2 | MAN2A1 | 0.99999987 | 1 | 0.165 | 6.485 | chr5 | 109025067 | 109205326 | + | protein_coding |
| ENSG00000143093.15_4 | STRIP1 | 0.99999338 | 1 | 0.165 | 4.887 | chr1 | 110574199 | 110617263 | + | protein_coding |
| ENSG00000234460.2_5 | AL772337.3 | 0.96656212 | 1 | 0.165 | 1.742 | chr9 | 90479011 | 90481206 | + | transcribed_unprocessed_pseudogene |
| ENSG00000108773.11_5 | KAT2A | 0.99999861 | 1 | 0.165 | 5.949 | chr17 | 40265129 | 40273385 | - | protein_coding |
| ENSG00000180066.9_3 | C10orf91 | 0.77762575 | 1 | 0.165 | -0.628 | chr10 | 134257831 | 134262912 | + | lncRNA |
| ENSG00000076201.15_4 | PTPN23 | 0.99997426 | 1 | 0.165 | 4.582 | chr3 | 47422501 | 47454925 | + | protein_coding |
| ENSG00000103657.14_7 | HERC1 | 0.99999871 | 1 | 0.165 | 6.642 | chr15 | 63900817 | 64126147 | - | protein_coding |
| ENSG00000273654.1_6 | AC020904.2 | 0.93072018 | 1 | 0.165 | 1.200 | chr19 | 18133212 | 18142909 | + | transcribed_processed_pseudogene |
| ENSG00000124006.15_5 | OBSL1 | 0.99997434 | 1 | 0.164 | 5.237 | chr2 | 220415450 | 220436581 | - | protein_coding |
| ENSG00000133704.10_5 | IPO8 | 0.99999985 | 1 | 0.164 | 6.419 | chr12 | 30781922 | 30848803 | - | protein_coding |
| ENSG00000163686.15_4 | ABHD6 | 0.99955936 | 1 | 0.164 | 3.585 | chr3 | 58223259 | 58281420 | + | protein_coding |
| ENSG00000068028.17_4 | RASSF1 | 0.99885034 | 1 | 0.164 | 3.417 | chr3 | 50367213 | 50378411 | - | protein_coding |
| ENSG00000164008.16_5 | C1orf50 | 0.95370797 | 1 | 0.164 | 1.003 | chr1 | 43232920 | 43245162 | + | protein_coding |
| ENSG00000108785.7_2 | HSD17B1P1 | 0.83547045 | 1 | 0.164 | 0.138 | chr17 | 40698782 | 40700724 | + | unprocessed_pseudogene |
| ENSG00000108100.18_3 | CCNY | 0.99999955 | 1 | 0.164 | 6.149 | chr10 | 35535953 | 35861597 | + | protein_coding |
| ENSG00000260267.1_6 | AC026471.1 | 0.9933081 | 1 | 0.164 | 2.971 | chr16 | 31468032 | 31471057 | - | lncRNA |
| ENSG00000188158.15_5 | NHS | 0.99999615 | 1 | 0.164 | 5.185 | chrX | 17393543 | 17754114 | + | protein_coding |
| ENSG00000111530.13_6 | CAND1 | 0.99999996 | 1 | 0.164 | 8.443 | chr12 | 67663138 | 67713733 | + | protein_coding |
| ENSG00000270194.1_6 | AC097359.2 | 0.90880472 | 1 | 0.164 | 1.019 | chr3 | 37283280 | 37285668 | - | lncRNA |
| ENSG00000099889.14_5 | ARVCF | 0.97420961 | 1 | 0.164 | 1.814 | chr22 | 19957419 | 20004346 | - | protein_coding |
| ENSG00000261485.1_6 | PAN3-AS1 | 0.85135174 | 1 | 0.163 | 0.073 | chr13 | 28710980 | 28712330 | - | lncRNA |
| ENSG00000286018.1_3 | AF129075.4 | 0.83178037 | 1 | 0.163 | -0.101 | chr21 | 30435767 | 30436689 | + | lncRNA |
| ENSG00000129245.12_4 | FXR2 | 0.99999931 | 1 | 0.163 | 6.078 | chr17 | 7494548 | 7518215 | - | protein_coding |
| ENSG00000073282.13_4 | TP63 | 0.98830511 | 1 | 0.163 | 2.256 | chr3 | 189349178 | 189615065 | + | protein_coding |
| ENSG00000285669.1_2 | AC026979.4 | 0.94379723 | 1 | 0.163 | 0.938 | chr8 | 30049945 | 30058848 | + | lncRNA |
| ENSG00000120885.22_6 | CLU | 0.99999993 | 1 | 0.163 | 8.801 | chr8 | 27454434 | 27472217 | - | protein_coding |
| ENSG00000178988.11_4 | MRFAP1L1 | 0.99999895 | 1 | 0.163 | 5.494 | chr4 | 6709428 | 6711592 | - | protein_coding |
| ENSG00000142156.15_3 | COL6A1 | 0.99990126 | 1 | 0.163 | 4.407 | chr21 | 47401683 | 47424964 | + | protein_coding |
| ENSG00000272482.1_6 | AC254633.1 | 0.96346699 | 1 | 0.163 | 1.719 | chr1 | 12678906 | 12679250 | - | lncRNA |
| ENSG00000066629.17_7 | EML1 | 0.99999801 | 1 | 0.163 | 5.949 | chr14 | 100204030 | 100408397 | + | protein_coding |
| ENSG00000182670.13_3 | TTC3 | 0.99999991 | 1 | 0.163 | 7.470 | chr21 | 38445526 | 38575413 | + | protein_coding |
| ENSG00000100154.14_7 | TTC28 | 0.99997013 | 1 | 0.162 | 4.888 | chr22 | 28374002 | 29075853 | - | protein_coding |
| ENSG00000287077.1_1 | AL157893.2 | 0.76323879 | 1 | 0.162 | -1.104 | chr10 | 88963474 | 88966372 | + | lncRNA |
| ENSG00000109501.14_5 | WFS1 | 0.99993629 | 1 | 0.162 | 4.439 | chr4 | 6271576 | 6304992 | + | protein_coding |
| ENSG00000176974.20_7 | SHMT1 | 0.99999938 | 1 | 0.162 | 6.080 | chr17 | 18231174 | 18266877 | - | protein_coding |
| ENSG00000176919.13_6 | C8G | 0.76798554 | 1 | 0.162 | -0.740 | chr9 | 139839637 | 139841427 | + | protein_coding |
| ENSG00000129083.12_4 | COPB1 | 0.99999998 | 1 | 0.162 | 7.970 | chr11 | 14464986 | 14521573 | - | protein_coding |
| ENSG00000140455.17_6 | USP3 | 0.99999969 | 1 | 0.162 | 6.032 | chr15 | 63796710 | 63886839 | + | protein_coding |
| ENSG00000152767.17_5 | FARP1 | 0.99996398 | 1 | 0.162 | 4.377 | chr13 | 98794816 | 99107430 | + | protein_coding |
| ENSG00000282988.2_6 | AL031777.2 | 0.76636695 | 1 | 0.162 | -0.893 | chr6 | 26195823 | 26199521 | - | protein_coding |
| ENSG00000147364.16_4 | FBXO25 | 0.99995513 | 1 | 0.162 | 4.784 | chr8 | 356428 | 427967 | + | protein_coding |
| ENSG00000112584.13_3 | FAM120B | 0.99991589 | 1 | 0.162 | 4.413 | chr6 | 170599791 | 170716153 | + | protein_coding |
| ENSG00000261423.1_8 | TMEM202-AS1 | 0.88343703 | 1 | 0.162 | 0.799 | chr15 | 72700119 | 72767509 | - | lncRNA |
| ENSG00000160584.16_8 | SIK3 | 0.99998866 | 1 | 0.162 | 4.767 | chr11 | 116714118 | 116969153 | - | protein_coding |
| ENSG00000101665.9_4 | SMAD7 | 0.99997571 | 1 | 0.162 | 4.871 | chr18 | 46446223 | 46478422 | - | protein_coding |
| ENSG00000160469.17_4 | BRSK1 | 0.98003034 | 1 | 0.162 | 2.197 | chr19 | 55793440 | 55823930 | + | protein_coding |
| ENSG00000128578.10_4 | STRIP2 | 0.9995331 | 1 | 0.162 | 3.664 | chr7 | 129074273 | 129128240 | + | protein_coding |
| ENSG00000280739.3_5 | EIF1B-AS1 | 0.8896798 | 1 | 0.162 | 0.353 | chr3 | 40102601 | 40351189 | - | lncRNA |
| ENSG00000103415.12_5 | HMOX2 | 0.99999013 | 1 | 0.162 | 4.876 | chr16 | 4524691 | 4560348 | + | protein_coding |
| ENSG00000101294.17_5 | HM13 | 0.99999984 | 1 | 0.162 | 6.755 | chr20 | 30102213 | 30165726 | + | protein_coding |
| ENSG00000174125.8_5 | TLR1 | 0.98721735 | 1 | 0.161 | 2.097 | chr4 | 38792298 | 38858438 | - | protein_coding |
| ENSG00000122696.14_5 | SLC25A51 | 0.9998199 | 1 | 0.161 | 3.947 | chr9 | 37879397 | 37904350 | - | protein_coding |
| ENSG00000151651.16_6 | ADAM8 | 0.99144919 | 1 | 0.161 | 2.637 | chr10 | 135075924 | 135090372 | - | protein_coding |

| | | | | | | | | | | |
|---|---|---|---|---|---|---|---|---|---|---|
| ENSG00000113761.12_5 | ZNF346 | 0.99996046 | 1 | 0.161 | 4.329 | chr5 | 176449697 | 176508190 | + | protein_coding |
| ENSG00000278175.4_5 | GLIDR | 0.95379245 | 1 | 0.161 | 1.401 | chr9 | 41893532 | 41955115 | - | lncRNA |
| ENSG00000135686.13_4 | KLHL36 | 0.99999538 | 1 | 0.161 | 5.784 | chr16 | 84682117 | 84701292 | + | protein_coding |
| ENSG00000155313.15_3 | USP25 | 0.99999961 | 1 | 0.161 | 6.549 | chr21 | 17102344 | 17252388 | + | protein_coding |
| ENSG00000157881.15_7 | PANK4 | 0.99949613 | 1 | 0.161 | 3.799 | chr1 | 2439976 | 2458050 | - | protein_coding |
| ENSG00000132254.12_4 | ARFIP2 | 0.9999998 | 1 | 0.161 | 6.130 | chr11 | 6495913 | 6502709 | - | protein_coding |
| ENSG00000136908.17_3 | DPM2 | 0.99999041 | 1 | 0.161 | 5.047 | chr9 | 130697378 | 130700763 | - | protein_coding |
| ENSG00000076928.17_3 | ARHGEF1 | 0.9999729 | 1 | 0.161 | 6.131 | chr19 | 42387228 | 42434302 | + | protein_coding |
| ENSG00000102893.16_6 | PHKB | 0.99999988 | 1 | 0.161 | 6.923 | chr16 | 47495034 | 47735434 | + | protein_coding |
| ENSG00000279900.1_6 | AP001767.4 | 0.77028496 | 1 | 0.161 | -0.769 | chr11 | 82794729 | 82795180 | + | TEC |
| ENSG00000103174.13_6 | NAGPA | 0.9997542 | 1 | 0.160 | 3.963 | chr16 | 5074845 | 5084142 | - | protein_coding |
| ENSG00000148935.11_8 | GAS2 | 0.78002967 | 1 | 0.160 | -0.629 | chr11 | 22647055 | 22834601 | + | protein_coding |
| ENSG00000084764.12_6 | MAPRE3 | 0.99837129 | 1 | 0.160 | 3.000 | chr2 | 27193505 | 27250087 | + | protein_coding |
| ENSG00000096433.11_4 | ITPR3 | 0.99999902 | 1 | 0.160 | 6.813 | chr6 | 33588142 | 33664351 | + | protein_coding |
| ENSG00000256235.2_4 | SMIM3 | 0.95893307 | 1 | 0.160 | 1.667 | chr5 | 150157508 | 150176296 | + | protein_coding |
| ENSG00000137804.14_7 | NUSAP1 | 0.78177581 | 1 | 0.160 | -0.699 | chr15 | 41612992 | 41673248 | + | protein_coding |
| ENSG00000013374.16_6 | NUB1 | 0.99999869 | 1 | 0.160 | 5.897 | chr7 | 151038785 | 151075535 | + | protein_coding |
| ENSG00000188938.17_7 | FAM120AOS | 0.99999966 | 1 | 0.160 | 5.988 | chr9 | 96193723 | 96215863 | - | protein_coding |
| ENSG00000167283.8_5 | ATP5MG | 0.99877069 | 1 | 0.160 | 3.697 | chr11 | 118272321 | 118302211 | + | protein_coding |
| ENSG00000254367.5_7 | AC087269.1 | 0.92118633 | 1 | 0.160 | 1.114 | chr8 | 8581203 | 8639989 | - | lncRNA |
| ENSG00000101017.14_7 | CD40 | 0.98661061 | 1 | 0.159 | 2.208 | chr20 | 44746917 | 44758502 | + | protein_coding |
| ENSG00000276232.1_7 | AC006064.5 | 0.99996911 | 1 | 0.159 | 6.753 | chr12 | 6619441 | 6619688 | + | lncRNA |
| ENSG00000272701.3_7 | MESTIT1 | 0.84119815 | 1 | 0.159 | 0.297 | chr7 | 130125883 | 130130874 | - | lncRNA |
| ENSG00000186635.14_5 | ARAP1 | 0.9999994 | 1 | 0.159 | 5.779 | chr11 | 72396114 | 72504644 | - | protein_coding |
| ENSG00000128342.5_4 | LIF | 0.99999895 | 1 | 0.159 | 5.969 | chr22 | 30636442 | 30642748 | - | protein_coding |
| ENSG00000182446.14_5 | NPLOC4 | 0.99999992 | 1 | 0.159 | 7.838 | chr17 | 79523913 | 79615495 | - | protein_coding |
| ENSG00000198824.7_4 | CHAMP1 | 0.99962207 | 1 | 0.159 | 3.925 | chr13 | 115079957 | 115103101 | + | protein_coding |
| ENSG00000136231.14_4 | IGF2BP3 | 0.99999736 | 1 | 0.159 | 5.360 | chr7 | 23349828 | 23510110 | - | protein_coding |
| ENSG00000099625.13_5 | CBARP | 0.8098286 | 1 | 0.159 | 0.408 | chr19 | 1228286 | 1238026 | - | protein_coding |
| ENSG00000215492.6_2 | HNRNPA1P7 | 0.85978866 | 1 | 0.158 | 0.692 | chr18 | 29992177 | 29993139 | - | processed_pseudogene |
| ENSG00000232546.1_5 | AC027644.1 | 0.80795659 | 1 | 0.158 | -0.105 | chr7 | 66313483 | 66323123 | + | unprocessed_pseudogene |
| ENSG00000109929.10_4 | SC5D | 0.99995477 | 1 | 0.158 | 4.534 | chr11 | 121163390 | 121184119 | + | protein_coding |
| ENSG00000180354.16_5 | MTURN | 0.98664849 | 1 | 0.158 | 2.190 | chr7 | 30174602 | 30202378 | + | protein_coding |
| ENSG00000246582.3_6 | AC100861.1 | 0.98405966 | 1 | 0.158 | 1.442 | chr8 | 23081984 | 23088439 | + | lncRNA |
| ENSG00000102543.14_3 | CDADC1 | 0.95097903 | 1 | 0.158 | 1.351 | chr13 | 49822061 | 49867621 | + | protein_coding |
| ENSG00000167702.12_3 | KIFC2 | 0.99993138 | 1 | 0.158 | 4.889 | chr8 | 145691426 | 145699585 | + | protein_coding |
| ENSG00000258738.1_6 | AL121603.2 | 0.89175188 | 1 | 0.158 | 0.642 | chr14 | 35343549 | 35345665 | + | lncRNA |
| ENSG00000212402.1 | SNORA74B | 0.78301232 | 1 | 0.158 | -0.739 | chr5 | 172447731 | 172447931 | + | snoRNA |
| ENSG00000265413.1_7 | AP001094.2 | 0.79874344 | 1 | 0.158 | -0.249 | chr18 | 8402845 | 8405159 | + | lncRNA |
| ENSG00000198807.13_3 | PAX9 | 0.9701086 | 1 | 0.158 | 1.909 | chr14 | 37126773 | 37148567 | + | protein_coding |
| ENSG00000180035.13_5 | ZNF48 | 0.99977597 | 1 | 0.158 | 4.450 | chr16 | 30389427 | 30411429 | + | protein_coding |
| ENSG00000106400.12_3 | ZNHIT1 | 0.99998001 | 1 | 0.158 | 5.073 | chr7 | 100861446 | 100867471 | + | protein_coding |
| ENSG00000101337.16_5 | TM9SF4 | 0.99999966 | 1 | 0.157 | 6.068 | chr20 | 30697517 | 30755061 | + | protein_coding |
| ENSG00000166887.15_4 | VPS39 | 0.99999877 | 1 | 0.157 | 5.545 | chr15 | 42450899 | 42500514 | - | protein_coding |
| ENSG00000180901.11_3 | KCTD2 | 0.99999816 | 1 | 0.157 | 5.321 | chr17 | 73028670 | 73061984 | + | protein_coding |
| ENSG00000231010.1_7 | AL121672.1 | 0.85638255 | 1 | 0.157 | 0.175 | chr22 | 46451620 | 46454040 | - | lncRNA |
| ENSG00000001036.14_4 | FUCA2 | 0.99999822 | 1 | 0.157 | 5.512 | chr6 | 143815949 | 143832857 | - | protein_coding |
| ENSG00000271711.1_5 | AC069236.1 | 0.79414794 | 1 | 0.157 | -1.307 | chr3 | 150329049 | 150329664 | - | processed_pseudogene |
| ENSG00000260231.2_6 | KDM7A-DT | 0.88940504 | 1 | 0.157 | 0.296 | chr7 | 139876984 | 139879440 | + | lncRNA |
| ENSG00000160753.16_4 | RUSC1 | 0.99999642 | 1 | 0.157 | 5.175 | chr1 | 155290685 | 155300905 | + | protein_coding |
| ENSG00000152818.18_5 | UTRN | 0.99999159 | 1 | 0.157 | 6.442 | chr6 | 144606837 | 145174170 | + | protein_coding |
| ENSG00000226383.7_7 | LINC01876 | 0.86380585 | 1 | 0.157 | 0.613 | chr2 | 156868042 | 157111462 | - | lncRNA |
| ENSG00000166747.12_6 | AP1G1 | 0.99999992 | 1 | 0.157 | 7.598 | chr16 | 71762903 | 71843104 | - | protein_coding |
| ENSG00000233818.1_7 | AP000695.2 | 0.79418452 | 1 | 0.157 | -0.417 | chr21 | 37818029 | 37904706 | + | lncRNA |
| ENSG00000163788.14_3 | SNRK | 0.99988473 | 1 | 0.157 | 3.962 | chr3 | 43328004 | 43466256 | + | protein_coding |
| ENSG00000167106.12_4 | FAM102A | 0.99999908 | 1 | 0.157 | 6.551 | chr9 | 130702861 | 130743268 | - | protein_coding |
| ENSG00000141699.11_8 | RETREG3 | 0.99999976 | 1 | 0.157 | 6.719 | chr17 | 40731531 | 40762641 | - | protein_coding |
| ENSG00000272333.6_3 | KMT2B | 0.999999 | 1 | 0.157 | 6.269 | chr19 | 36208875 | 36229779 | + | protein_coding |
| ENSG00000198814.12_6 | GK | 0.99885435 | 1 | 0.156 | 2.260 | chrX | 30671476 | 30749573 | + | protein_coding |

| | | | | | | | | | | |
|---|---|---|---|---|---|---|---|---|---|---|
| ENSG00000140948.13_5 | ZCCHC14 | 0.99997124 | 1 | 0.156 | 4.495 | chr16 | 87439852 | 87526630 | - | protein_coding |
| ENSG00000124713.6_3 | GNMT | 0.77741192 | 1 | 0.156 | -0.977 | chr6 | 42928492 | 42931618 | + | protein_coding |
| ENSG00000277449.1_7 | CEBPB-AS1 | 0.80858108 | 1 | 0.156 | -0.358 | chr20 | 48801135 | 48808035 | - | lncRNA |
| ENSG00000227885.3_7 | AL590652.1 | 0.85223051 | 1 | 0.156 | 0.007 | chr6 | 53604064 | 53659702 | - | lncRNA |
| ENSG00000130811.12_6 | EIF3G | 0.99999887 | 1 | 0.156 | 6.026 | chr19 | 10225690 | 10230594 | - | protein_coding |
| ENSG00000110880.10_6 | CORO1C | 0.99999998 | 1 | 0.156 | 7.910 | chr12 | 109038885 | 109125372 | - | protein_coding |
| ENSG00000204815.10_4 | TTC25 | 0.99251694 | 1 | 0.156 | 1.728 | chr17 | 40086870 | 40118521 | + | protein_coding |
| ENSG00000136986.10_4 | DERL1 | 0.99999953 | 1 | 0.156 | 5.925 | chr8 | 124025410 | 124054542 | - | protein_coding |
| ENSG00000104331.9_6 | IMPAD1 | 0.99999997 | 1 | 0.156 | 7.591 | chr8 | 57870490 | 57906426 | - | protein_coding |
| ENSG00000214140.11_4 | PRCD | 0.78697463 | 1 | 0.156 | -0.740 | chr17 | 74523668 | 74549660 | + | protein_coding |
| ENSG00000186350.12_5 | RXRA | 0.9999992 | 1 | 0.156 | 6.181 | chr9 | 137208944 | 137332431 | + | protein_coding |
| ENSG00000110171.20_6 | TRIM3 | 0.99997501 | 1 | 0.155 | 4.346 | chr11 | 6469843 | 6495689 | - | protein_coding |
| ENSG00000104388.15_6 | RAB2A | 0.99999994 | 1 | 0.155 | 7.215 | chr8 | 61429495 | 61536203 | + | protein_coding |
| ENSG00000143390.17_5 | RFX5 | 0.99999971 | 1 | 0.155 | 6.073 | chr1 | 151313116 | 151319833 | - | protein_coding |
| ENSG00000186908.14_5 | ZDHHC17 | 0.9999986 | 1 | 0.155 | 5.642 | chr12 | 77157368 | 77247476 | + | protein_coding |
| ENSG00000137764.20_5 | MAP2K5 | 0.99937511 | 1 | 0.155 | 3.474 | chr15 | 67835041 | 68099455 | + | protein_coding |
| ENSG00000148356.13_3 | LRSAM1 | 0.99990672 | 1 | 0.155 | 4.509 | chr9 | 130213765 | 130265780 | + | protein_coding |
| ENSG00000169020.10_5 | ATP5ME | 0.98906163 | 1 | 0.155 | 2.915 | chr4 | 666225 | 668119 | - | protein_coding |
| ENSG00000119638.13_5 | NEK9 | 0.99999919 | 1 | 0.155 | 6.057 | chr14 | 75546056 | 75594047 | - | protein_coding |
| ENSG00000185000.12_3 | DGAT1 | 0.99998604 | 1 | 0.155 | 4.843 | chr8 | 145538247 | 145550573 | - | protein_coding |
| ENSG00000179943.8_4 | FIZ1 | 0.99975218 | 1 | 0.155 | 3.874 | chr19 | 56102742 | 56113336 | - | protein_coding |
| ENSG00000166575.17_5 | TMEM135 | 0.99999928 | 1 | 0.154 | 5.675 | chr11 | 86748886 | 87039866 | + | protein_coding |
| ENSG00000204856.12_3 | FAM216A | 0.99980532 | 1 | 0.154 | 3.782 | chr12 | 110906220 | 110928190 | + | protein_coding |
| ENSG00000176896.8_3 | TCEANC | 0.97576235 | 1 | 0.154 | 1.790 | chrX | 13671308 | 13700083 | + | protein_coding |
| ENSG00000121057.13_6 | AKAP1 | 0.99999972 | 1 | 0.154 | 6.539 | chr17 | 55162453 | 55198707 | + | protein_coding |
| ENSG00000177595.18_5 | PIDD1 | 0.96270944 | 1 | 0.154 | 4.142 | chr11 | 799179 | 809753 | - | protein_coding |
| ENSG00000171953.16_5 | ATPAF2 | 0.99985217 | 1 | 0.154 | 3.987 | chr17 | 17880723 | 17942523 | - | protein_coding |
| ENSG00000163975.12_6 | MELTF | 0.99971964 | 1 | 0.154 | 5.332 | chr3 | 196715492 | 196756688 | - | protein_coding |
| ENSG00000247679.3_6 | AC139795.2 | 0.98641263 | 1 | 0.154 | 1.967 | chr5 | 177038241 | 177046755 | + | lncRNA |
| ENSG00000105397.14_6 | TYK2 | 0.99999731 | 1 | 0.154 | 5.982 | chr19 | 10461205 | 10491248 | - | protein_coding |
| ENSG00000269843.1_7 | AC008537.2 | 0.8359918 | 1 | 0.153 | 0.191 | chr19 | 41337126 | 41343115 | - | lncRNA |
| ENSG00000130881.14_5 | LRP3 | 0.93634412 | 1 | 0.153 | 5.352 | chr19 | 33668509 | 33699770 | + | protein_coding |
| ENSG00000279368.1_5 | AL031123.3 | 0.78971619 | 1 | 0.153 | -0.740 | chr6 | 6685381 | 6687217 | + | TEC |
| ENSG00000100583.4_2 | SAMD15 | 0.99296857 | 1 | 0.153 | 2.471 | chr14 | 77843032 | 77857840 | + | protein_coding |
| ENSG00000236756.4_7 | DNAJC9-AS1 | 0.80349644 | 1 | 0.153 | -0.537 | chr10 | 75007118 | 75036742 | + | lncRNA |
| ENSG00000143155.13_3 | TIPRL | 0.99999958 | 1 | 0.153 | 6.017 | chr1 | 168148200 | 168171347 | + | protein_coding |
| ENSG00000204564.12_5 | C6orf136 | 0.90422697 | 1 | 0.153 | 0.912 | chr6 | 30614816 | 30620987 | + | protein_coding |
| ENSG00000204584.1_7 | AC027801.1 | 0.94739322 | 1 | 0.152 | 1.295 | chr17 | 47923272 | 47926199 | + | lncRNA |
| ENSG00000167792.13_6 | NDUFV1 | 0.99999959 | 1 | 0.152 | 6.720 | chr11 | 67373124 | 67380025 | + | protein_coding |
| ENSG00000256628.3_6 | ZBTB11-AS1 | 0.97773702 | 1 | 0.152 | 1.726 | chr3 | 101395319 | 101398061 | + | lncRNA |
| ENSG00000132613.15_5 | MTSS2 | 0.99996653 | 1 | 0.152 | 4.581 | chr16 | 70695107 | 70719956 | - | protein_coding |
| ENSG00000107185.10_3 | RGP1 | 0.99999763 | 1 | 0.152 | 5.121 | chr9 | 35749284 | 35758582 | + | protein_coding |
| ENSG00000176715.17_7 | ACSF3 | 0.99998698 | 1 | 0.152 | 4.846 | chr16 | 89154783 | 89230529 | + | protein_coding |
| ENSG00000031823.14_5 | RANBP3 | 0.99997584 | 1 | 0.152 | 5.369 | chr19 | 5916150 | 5978153 | - | protein_coding |
| ENSG00000188095.6_5 | MESP2 | 0.77749695 | 1 | 0.152 | -1.104 | chr15 | 90303822 | 90321985 | + | protein_coding |
| ENSG00000181222.17_6 | POLR2A | 0.9999975 | 1 | 0.152 | 6.133 | chr17 | 7387685 | 7417933 | + | protein_coding |
| ENSG00000198569.10_3 | SLC34A3 | 0.97229566 | 1 | 0.152 | 1.955 | chr9 | 140125209 | 140131007 | + | protein_coding |
| ENSG00000239264.9_5 | TXNDC5 | 0.82338074 | 1 | 0.152 | 0.360 | chr6 | 7881750 | 7911021 | - | protein_coding |
| ENSG00000068383.19_4 | INPP5A | 0.99996879 | 1 | 0.152 | 4.280 | chr10 | 134351291 | 134596984 | + | protein_coding |
| ENSG00000079308.19_8 | TNS1 | 0.97071031 | 1 | 0.152 | 1.916 | chr2 | 218664311 | 218898705 | - | protein_coding |
| ENSG00000149531.15_4 | FRG1BP | 0.99956656 | 1 | 0.152 | 3.897 | chr20 | 29612048 | 29633933 | + | unprocessed_pseudogene |
| ENSG00000100105.18_5 | PATZ1 | 0.99999681 | 1 | 0.152 | 5.006 | chr22 | 31721790 | 31742332 | - | protein_coding |
| ENSG00000164124.11_4 | TMEM144 | 0.9998322 | 1 | 0.151 | 4.128 | chr4 | 159122756 | 159176568 | + | protein_coding |
| ENSG00000034152.19_6 | MAP2K3 | 0.99999126 | 1 | 0.151 | 4.833 | chr17 | 21187984 | 21218544 | + | protein_coding |
| ENSG00000004866.20_6 | ST7 | 0.99922733 | 1 | 0.151 | 3.264 | chr7 | 116593292 | 116870157 | + | protein_coding |
| ENSG00000008394.13_7 | MGST1 | 0.99999981 | 1 | 0.151 | 7.651 | chr12 | 16500076 | 16762193 | + | protein_coding |
| ENSG00000141644.17_5 | MBD1 | 0.99999749 | 1 | 0.151 | 5.223 | chr18 | 47793252 | 47808144 | - | protein_coding |
| ENSG00000144566.11_4 | RAB5A | 0.99999839 | 1 | 0.151 | 5.601 | chr3 | 19988589 | 20026667 | + | protein_coding |
| ENSG00000082014.16_5 | SMARCD3 | 0.99973418 | 1 | 0.151 | 3.847 | chr7 | 150935850 | 150974982 | - | protein_coding |
| ENSG00000204219.11_5 | TCEA3 | 0.80774577 | 1 | 0.151 | -0.476 | chr1 | 23707402 | 23751241 | - | protein_coding |

| | | | | | | | | | | |
|---|---|---|---|---|---|---|---|---|---|---|
| ENSG00000215421.9_3 | ZNF407 | 0.99986735 | 1 | 0.151 | 3.912 | chr18 | 72265106 | 72777627 | + | protein_coding |
| ENSG00000184923.12_3 | NUTM2A | 0.78166385 | 1 | 0.151 | -0.848 | chr10 | 88985205 | 88996665 | + | protein_coding |
| ENSG00000197837.3_3 | H4-16 | 0.99998947 | 1 | 0.151 | 6.591 | chr12 | 14920933 | 14924065 | - | protein_coding |
| ENSG00000065457.10_8 | ADAT1 | 0.99999665 | 1 | 0.151 | 5.245 | chr16 | 75630879 | 75657198 | - | protein_coding |
| ENSG00000137656.12_5 | BUD13 | 0.99996167 | 1 | 0.151 | 4.211 | chr11 | 116618886 | 116643703 | - | protein_coding |
| ENSG00000244968.7_6 | LIFR-AS1 | 0.79083822 | 1 | 0.151 | -0.741 | chr5 | 38556867 | 38671318 | + | lncRNA |
| ENSG00000060237.17_7 | WNK1 | 0.99999987 | 1 | 0.150 | 8.150 | chr12 | 861745 | 1020618 | + | protein_coding |
| ENSG00000283148.1_7 | AC137936.1 | 0.83930238 | 1 | 0.150 | -0.132 | chr3 | 58160205 | 58164101 | - | lncRNA |
| ENSG00000102572.14_5 | STK24 | 0.99999933 | 1 | 0.150 | 5.616 | chr13 | 99097439 | 99230194 | - | protein_coding |
| ENSG00000107551.21_7 | RASSF4 | 0.99318898 | 1 | 0.150 | 2.264 | chr10 | 45454855 | 45491339 | + | protein_coding |
| ENSG00000083896.12_4 | YTHDC1 | 0.99998237 | 1 | 0.150 | 4.572 | chr4 | 69176105 | 69215807 | - | protein_coding |
| ENSG00000111291.8_5 | GPRC5D | 0.80104412 | 1 | 0.150 | -0.477 | chr12 | 13093709 | 13105081 | - | protein_coding |
| ENSG00000197168.13_5 | NEK5 | 0.79821688 | 1 | 0.150 | -0.670 | chr13 | 52607747 | 52703228 | - | protein_coding |
| ENSG00000055609.18_6 | KMT2C | 0.99999706 | 1 | 0.150 | 6.266 | chr7 | 151832007 | 152133090 | - | protein_coding |
| ENSG00000138821.13_5 | SLC39A8 | 0.99999609 | 1 | 0.150 | 5.406 | chr4 | 103172198 | 103352415 | - | protein_coding |
| ENSG00000159202.18_4 | UBE2Z | 0.99999988 | 1 | 0.150 | 7.114 | chr17 | 46985769 | 47006418 | + | protein_coding |
| ENSG00000119661.15_3 | DNAL1 | 0.99995409 | 1 | 0.150 | 4.401 | chr14 | 74111578 | 74170435 | + | protein_coding |
| ENSG00000206149.10_3 | HERC2P9 | 0.99983255 | 1 | 0.150 | 3.691 | chr15 | 28834638 | 28930410 | + | transcribed_unprocessed_pseudogene |
| ENSG00000164880.16_4 | INTS1 | 0.99250154 | 1 | 0.150 | 7.142 | chr7 | 1509913 | 1544025 | - | protein_coding |
| ENSG00000152056.17_6 | AP1S3 | 0.99970083 | 1 | 0.150 | 3.715 | chr2 | 224616403 | 224702744 | - | protein_coding |
| ENSG00000099797.15_5 | TECR | 0.99995235 | 1 | 0.150 | 5.369 | chr19 | 14627897 | 14676792 | + | protein_coding |
| ENSG00000155252.13_3 | PI4K2A | 0.999964 | 1 | 0.150 | 4.464 | chr10 | 99400443 | 99436191 | + | protein_coding |
| ENSG00000131089.17_8 | ARHGEF9 | 0.99998548 | 1 | 0.149 | 4.529 | chrX | 62854847 | 63029154 | - | protein_coding |
| ENSG00000172340.14_5 | SUCLG2 | 0.99999816 | 1 | 0.149 | 5.941 | chr3 | 67410884 | 67705038 | - | protein_coding |
| ENSG00000139537.11_5 | CCDC65 | 0.98030003 | 1 | 0.149 | 1.933 | chr12 | 49297893 | 49325623 | + | protein_coding |
| ENSG00000088833.17_5 | NSFL1C | 0.99999813 | 1 | 0.149 | 5.593 | chr20 | 1422807 | 1454487 | - | protein_coding |
| ENSG00000165219.22_6 | GAPVD1 | 0.99999985 | 1 | 0.149 | 6.512 | chr9 | 128024073 | 128129486 | + | protein_coding |
| ENSG00000141084.11_6 | RANBP10 | 0.99990227 | 1 | 0.149 | 4.218 | chr16 | 67756973 | 67840555 | - | protein_coding |
| ENSG00000152409.9_3 | JMY | 0.99999579 | 1 | 0.149 | 4.929 | chr5 | 78531954 | 78623034 | + | protein_coding |
| ENSG00000219507.4_4 | FTH1P8 | 0.84327938 | 1 | 0.149 | 0.095 | chrX | 147133753 | 147134266 | + | processed_pseudogene |
| ENSG00000215193.12_4 | PEX26 | 0.99999878 | 1 | 0.149 | 5.278 | chr22 | 18560686 | 18613905 | + | protein_coding |
| ENSG00000065000.18_7 | AP3D1 | 0.99999986 | 1 | 0.148 | 7.162 | chr19 | 2100987 | 2164467 | - | protein_coding |
| ENSG00000100522.10_6 | GNPNAT1 | 0.99999864 | 1 | 0.148 | 5.420 | chr14 | 53241911 | 53258386 | - | protein_coding |
| ENSG00000182827.9_5 | ACBD3 | 0.99999945 | 1 | 0.148 | 5.642 | chr1 | 226332380 | 226374442 | - | protein_coding |
| ENSG00000161036.13_4 | LRWD1 | 0.99991958 | 1 | 0.148 | 4.713 | chr7 | 102105403 | 102113615 | + | protein_coding |
| ENSG00000228952.1_5 | LINC02041 | 0.79484395 | 1 | 0.148 | -0.816 | chr3 | 187166633 | 187167238 | + | lncRNA |
| ENSG00000174151.14_5 | CYB561D1 | 0.99854577 | 1 | 0.148 | 2.945 | chr1 | 110036674 | 110045554 | + | protein_coding |
| ENSG00000215014.5_7 | AL645728.1 | 0.78707441 | 1 | 0.148 | -0.894 | chr1 | 1509482 | 1512455 | + | lncRNA |
| ENSG00000232208.2_5 | AL139415.1 | 0.80880568 | 1 | 0.148 | -0.337 | chr1 | 8967452 | 8967803 | - | processed_pseudogene |
| ENSG00000145014.17_6 | TMEM44 | 0.99947167 | 1 | 0.148 | 3.290 | chr3 | 194308402 | 194354418 | - | protein_coding |
| ENSG00000243710.7_5 | CFAP57 | 0.82123652 | 1 | 0.148 | -0.203 | chr1 | 43637820 | 43720029 | + | protein_coding |
| ENSG00000117602.12_5 | RCAN3 | 0.99999644 | 1 | 0.148 | 5.079 | chr1 | 24828841 | 24867530 | + | protein_coding |
| ENSG00000154889.17_7 | MPPE1 | 0.99948575 | 1 | 0.148 | 3.258 | chr18 | 11882621 | 11908365 | - | protein_coding |
| ENSG00000177733.7_4 | HNRNPA0 | 0.9999997 | 1 | 0.148 | 6.422 | chr5 | 137081340 | 137090052 | - | protein_coding |
| ENSG00000010704.18_4 | HFE | 0.99999449 | 1 | 0.148 | 4.884 | chr6 | 26087509 | 26098571 | + | protein_coding |
| ENSG00000145020.16_7 | AMT | 0.788572 | 1 | 0.148 | -0.894 | chr3 | 49454211 | 49460118 | - | protein_coding |
| ENSG00000049246.14_2 | PER3 | 0.98410563 | 1 | 0.148 | 2.076 | chr1 | 7844380 | 7905237 | + | protein_coding |
| ENSG00000058600.16_7 | POLR3E | 0.9999997 | 1 | 0.147 | 5.758 | chr16 | 22308696 | 22346422 | + | protein_coding |
| ENSG00000154529.10 | CNTNAP3B | 0.99969908 | 1 | 0.147 | 3.871 | chr9 | 43684902 | 43924049 | + | protein_coding |
| ENSG00000261442.1_7 | AC023830.1 | 0.78994939 | 1 | 0.147 | -0.815 | chr16 | 4684330 | 4690624 | - | lncRNA |
| ENSG00000099219.14_4 | ERMP1 | 0.99999997 | 1 | 0.147 | 7.082 | chr9 | 5765076 | 5833117 | - | protein_coding |
| ENSG00000168291.13_5 | PDHB | 0.99999967 | 1 | 0.147 | 5.961 | chr3 | 58413357 | 58419584 | - | protein_coding |
| ENSG00000059588.10_4 | TARBP1 | 0.99999967 | 1 | 0.147 | 5.965 | chr1 | 234527059 | 234614925 | - | protein_coding |
| ENSG00000161091.13_5 | MFSD12 | 0.99994394 | 1 | 0.147 | 5.638 | chr19 | 3538259 | 3574288 | - | protein_coding |
| ENSG00000275342.5_4 | PRAG1 | 0.99986941 | 1 | 0.147 | 5.895 | chr8 | 8175258 | 8243949 | - | protein_coding |
| ENSG00000255468.7_6 | AP001107.9 | 0.97097417 | 1 | 0.147 | 1.156 | chr11 | 66115421 | 66132275 | + | lncRNA |
| ENSG00000100979.15_5 | PLTP | 0.93894594 | 1 | 0.147 | 1.079 | chr20 | 44527260 | 44540794 | - | protein_coding |
| ENSG00000225793.3_7 | AL080250.1 | 0.79202129 | 1 | 0.147 | -0.417 | chr6 | 75994730 | 76027241 | + | lncRNA |
| ENSG00000198960.11_5 | ARMCX6 | 0.99873911 | 1 | 0.147 | 3.242 | chrX | 100870108 | 100872991 | - | protein_coding |

| | | | | | | | | | | |
|---|---|---|---|---|---|---|---|---|---|---|
| ENSG00000276488.1_6 | AC008735.4 | 0.85101751 | 1 | 0.147 | 0.029 | chr19 | 56123856 | 56124463 | - | lncRNA |
| ENSG00000176444.19_4 | CLK2 | 0.99999366 | 1 | 0.147 | 4.985 | chr1 | 155232659 | 155248282 | - | protein_coding |
| ENSG00000077254.14_5 | USP33 | 0.99999776 | 1 | 0.146 | 6.175 | chr1 | 78161672 | 78225537 | - | protein_coding |
| ENSG00000198873.12_4 | GRK5 | 0.99843796 | 1 | 0.146 | 3.229 | chr10 | 120967083 | 121219257 | + | protein_coding |
| ENSG00000168005.9_4 | SPINDOC | 0.99991043 | 1 | 0.146 | 4.832 | chr11 | 63580928 | 63595188 | + | protein_coding |
| ENSG00000280239.1_5 | AC011498.7 | 0.93400181 | 1 | 0.146 | 0.705 | chr19 | 4448807 | 4450833 | + | TEC |
| ENSG00000266338.6_2 | NBPF15 | 0.99988831 | 1 | 0.146 | 4.146 | chr1 | 148555979 | 148596267 | + | protein_coding |
| ENSG00000264070.1_4 | DND1P1 | 0.84642464 | 1 | 0.146 | -0.160 | chr17 | 43663237 | 43664295 | + | processed_pseudogene |
| ENSG00000153774.9_7 | CFDP1 | 0.9998795 | 1 | 0.146 | 4.181 | chr16 | 75327596 | 75467401 | - | protein_coding |
| ENSG00000182718.16_8 | ANXA2 | 0.99999999 | 1 | 0.146 | 11.509 | chr15 | 60639333 | 60695082 | - | protein_coding |
| ENSG00000177410.13_6 | ZFAS1 | 0.99999969 | 1 | 0.146 | 6.239 | chr20 | 47894715 | 47916137 | + | lncRNA |
| ENSG00000227372.12_7 | TP73-AS1 | 0.99992216 | 1 | 0.146 | 4.049 | chr1 | 3652075 | 3663937 | - | transcribed_unitary_pseudogene |
| ENSG00000250899.3_6 | AC125807.2 | 0.99887647 | 1 | 0.146 | 3.104 | chr12 | 3150603 | 3154116 | + | lncRNA |
| ENSG00000159403.11 | C1R | 0.99977972 | 1 | 0.146 | 3.725 | chr12 | 7187513 | 7245203 | - | protein_coding |
| ENSG00000196914.9_3 | ARHGEF12 | 0.99999995 | 1 | 0.146 | 7.340 | chr11 | 120207122 | 120360646 | + | protein_coding |
| ENSG00000196378.12_3 | ZNF34 | 0.99598769 | 1 | 0.146 | 2.816 | chr8 | 145998499 | 146012730 | - | protein_coding |
| ENSG00000205609.12_3 | EIF3CL | 0.78632227 | 1 | 0.146 | -0.980 | chr16 | 28390900 | 28415200 | - | protein_coding |
| ENSG00000188681.11_4 | TEKT4P2 | 0.86198032 | 1 | 0.146 | 0.276 | chr21 | 9907194 | 9968585 | - | transcribed_unprocessed_pseudogene |
| ENSG00000165801.10_4 | ARHGEF40 | 0.99998593 | 1 | 0.146 | 4.719 | chr14 | 21538432 | 21558407 | + | protein_coding |
| ENSG00000005486.17_3 | RHBDD2 | 0.99999676 | 1 | 0.146 | 5.246 | chr7 | 75471920 | 75518244 | + | protein_coding |
| ENSG00000160410.15_4 | SHKBP1 | 0.99999851 | 1 | 0.146 | 6.467 | chr19 | 41082759 | 41097305 | + | protein_coding |
| ENSG00000128245.15_6 | YWHAH | 0.99999874 | 1 | 0.145 | 6.579 | chr22 | 32340508 | 32353590 | + | protein_coding |
| ENSG00000186866.16_3 | POFUT2 | 0.9998337 | 1 | 0.145 | 4.110 | chr21 | 46683843 | 46707813 | - | protein_coding |
| ENSG00000171988.19_6 | JMJD1C | 0.99999995 | 1 | 0.145 | 6.997 | chr10 | 64926981 | 65281610 | - | protein_coding |
| ENSG00000163626.16_4 | COX18 | 0.99997758 | 1 | 0.145 | 4.368 | chr4 | 73918079 | 73935472 | - | protein_coding |
| ENSG00000140848.17_4 | CPNE2 | 0.99999858 | 1 | 0.145 | 5.353 | chr16 | 57126495 | 57182281 | + | protein_coding |
| ENSG00000118707.10_6 | TGIF2 | 0.99652523 | 1 | 0.145 | 2.728 | chr20 | 35201891 | 35222353 | + | protein_coding |
| ENSG00000071246.11_4 | VASH1 | 0.99184216 | 1 | 0.145 | 2.547 | chr14 | 77227811 | 77249358 | + | protein_coding |
| ENSG00000106299.8_4 | WASL | 0.99999965 | 1 | 0.145 | 6.126 | chr7 | 123321997 | 123389057 | - | protein_coding |
| ENSG00000106346.11_4 | USP42 | 0.99998419 | 1 | 0.145 | 4.408 | chr7 | 6144515 | 6201195 | + | protein_coding |
| ENSG00000274523.5_7 | RCC1L | 0.99994546 | 1 | 0.145 | 4.271 | chr7 | 74441226 | 74490064 | - | protein_coding |
| ENSG00000154917.11_4 | RAB6B | 0.94763015 | 1 | 0.145 | 1.172 | chr3 | 133543079 | 133614726 | - | protein_coding |
| ENSG00000248323.7_8 | LUCAT1 | 0.99999729 | 1 | 0.145 | 5.161 | chr5 | 90350651 | 90610364 | - | lncRNA |
| ENSG00000040341.18_8 | STAU2 | 0.99999807 | 1 | 0.144 | 5.910 | chr8 | 74332604 | 74659943 | - | protein_coding |
| ENSG00000173120.15_3 | KDM2A | 0.99999961 | 1 | 0.144 | 6.153 | chr11 | 66886734 | 67025553 | + | protein_coding |
| ENSG00000136152.15_3 | COG3 | 0.99998815 | 1 | 0.144 | 4.818 | chr13 | 46039033 | 46110836 | + | protein_coding |
| ENSG00000034053.14_7 | APBA2 | 0.99996661 | 1 | 0.144 | 5.258 | chr15 | 29129629 | 29410518 | + | protein_coding |
| ENSG00000108604.16_7 | SMARCD2 | 0.99999993 | 1 | 0.144 | 6.531 | chr17 | 61909441 | 61920425 | - | protein_coding |
| ENSG00000271971.1_6 | AC120053.1 | 0.81574903 | 1 | 0.144 | -0.450 | chr8 | 94727606 | 94728341 | - | lncRNA |
| ENSG00000172534.14_6 | HCFC1 | 0.99999873 | 1 | 0.144 | 7.504 | chrX | 153213008 | 153237269 | - | protein_coding |
| ENSG00000071889.16_5 | FAM3A | 0.9999529 | 1 | 0.144 | 4.898 | chrX | 153734490 | 153744566 | - | protein_coding |
| ENSG00000159461.15_6 | AMFR | 0.99999996 | 1 | 0.144 | 7.136 | chr16 | 56395364 | 56459457 | - | protein_coding |
| ENSG00000004478.8_3 | FKBP4 | 0.99999994 | 1 | 0.144 | 7.334 | chr12 | 2904136 | 2914589 | + | protein_coding |
| ENSG00000177105.10_5 | RHOG | 0.99994641 | 1 | 0.144 | 4.329 | chr11 | 3848208 | 3862189 | - | protein_coding |
| ENSG00000164609.10_4 | SLU7 | 0.99999974 | 1 | 0.144 | 6.053 | chr5 | 159828648 | 159848718 | - | protein_coding |
| ENSG00000110955.9_8 | ATP5F1B | 0.9997343 | 1 | 0.144 | 8.922 | chr12 | 57031959 | 57039768 | - | protein_coding |
| ENSG00000253106.1_6 | AC090198.1 | 0.97615568 | 1 | 0.144 | 1.768 | chr8 | 125500751 | 125503884 | + | lncRNA |
| ENSG00000236751.2_7 | LINC01186 | 0.96788838 | 1 | 0.143 | 1.652 | chrX | 46116194 | 46187115 | - | lncRNA |
| ENSG00000078018.20_7 | MAP2 | 0.89804803 | 1 | 0.143 | 0.538 | chr2 | 210288782 | 210598842 | + | protein_coding |
| ENSG00000215190.9_7 | LINC00680 | 0.98775478 | 1 | 0.143 | 2.000 | chr6 | 58272352 | 58287779 | - | transcribed_unprocessed_pseudogene |
| ENSG00000204469.13_3 | PRRC2A | 0.99999982 | 1 | 0.143 | 6.467 | chr6 | 31588492 | 31605548 | + | protein_coding |
| ENSG00000225518.3_6 | LINC01703 | 0.87732458 | 1 | 0.143 | 0.466 | chr1 | 226271291 | 226278166 | + | lncRNA |
| ENSG00000100412.16_3 | ACO2 | 0.99999697 | 1 | 0.143 | 5.425 | chr22 | 41865121 | 41924993 | + | protein_coding |
| ENSG00000166337.10_5 | TAF10 | 0.98688518 | 1 | 0.143 | 2.036 | chr11 | 6627524 | 6633770 | - | protein_coding |
| ENSG00000176476.9_3 | SGF29 | 0.99922159 | 1 | 0.143 | 3.212 | chr16 | 28565236 | 28603111 | + | protein_coding |
| ENSG00000172301.11_5 | COPRS | 0.99999829 | 1 | 0.143 | 5.883 | chr17 | 30178890 | 30186310 | - | protein_coding |
| ENSG00000186056.10_6 | MATN1-AS1 | 0.91298827 | 1 | 0.143 | 0.706 | chr1 | 31191351 | 31199674 | + | lncRNA |
| ENSG00000213614.9_7 | HEXA | 0.99993964 | 1 | 0.143 | 4.582 | chr15 | 72633260 | 72668817 | - | protein_coding |
| ENSG00000268087.1_7 | AC008764.2 | 0.85826906 | 1 | 0.143 | -0.020 | chr19 | 16721222 | 16747342 | + | lncRNA |

| | | | | | | | | | | |
|---|---|---|---|---|---|---|---|---|---|---|
| ENSG00000179562.3_4 | GCC1 | 0.99999565 | 1 | 0.143 | 4.911 | chr7 | 127220682 | 127233665 | - | protein_coding |
| ENSG00000173928.3_3 | SWSAP1 | 0.93279726 | 1 | 0.143 | 0.989 | chr19 | 11485342 | 11487627 | + | protein_coding |
| ENSG00000139163.16_7 | ETNK1 | 0.99999971 | 1 | 0.143 | 6.902 | chr12 | 22778009 | 22843599 | + | protein_coding |
| ENSG00000155254.13_3 | MARVELD1 | 0.99995249 | 1 | 0.143 | 5.595 | chr10 | 99472930 | 99477907 | + | protein_coding |
| ENSG00000184208.12_5 | C22orf46 | 0.99341849 | 1 | 0.142 | 2.722 | chr22 | 42084881 | 42094140 | + | transcribed_unitary_pseudogene |
| ENSG00000107223.13_5 | EDF1 | 0.99999998 | 1 | 0.142 | 7.320 | chr9 | 139756571 | 139760760 | - | protein_coding |
| ENSG00000103042.9_5 | SLC38A7 | 0.999949 | 1 | 0.142 | 4.109 | chr16 | 58699013 | 58718674 | - | protein_coding |
| ENSG00000137942.16_2 | FNBP1L | 0.99999726 | 1 | 0.142 | 5.665 | chr1 | 93913688 | 94020218 | + | protein_coding |
| ENSG00000162704.16_5 | ARPC5 | 0.9999999 | 1 | 0.142 | 7.042 | chr1 | 183589981 | 183604918 | - | protein_coding |
| ENSG00000235652.9_7 | AL356599.1 | 0.8896481 | 1 | 0.142 | 0.238 | chr6 | 146120545 | 146208566 | + | lncRNA |
| ENSG00000063241.8_7 | ISOC2 | 0.99994143 | 1 | 0.142 | 5.126 | chr19 | 55964352 | 55973710 | - | protein_coding |
| ENSG00000018280.17_3 | SLC11A1 | 0.87542056 | 1 | 0.142 | 0.081 | chr2 | 219246752 | 219261617 | + | protein_coding |
| ENSG00000151748.15_7 | SAV1 | 0.99998745 | 1 | 0.142 | 4.612 | chr14 | 51098776 | 51135024 | - | protein_coding |
| ENSG00000139233.7_5 | LLPH | 0.99995975 | 1 | 0.142 | 4.609 | chr12 | 66510335 | 66524530 | - | protein_coding |
| ENSG00000270175.1_7 | AC023509.3 | 0.88993577 | 1 | 0.142 | 0.430 | chr12 | 53893946 | 53894720 | - | lncRNA |
| ENSG00000114166.8_3 | KAT2B | 0.99999428 | 1 | 0.142 | 4.746 | chr3 | 20081938 | 20195896 | + | protein_coding |
| ENSG00000131943.17_3 | C19orf12 | 0.9999982 | 1 | 0.142 | 5.381 | chr19 | 30189793 | 30206696 | - | protein_coding |
| ENSG00000178252.18_5 | WDR6 | 0.9999999 | 1 | 0.142 | 6.826 | chr3 | 49044495 | 49053386 | + | protein_coding |
| ENSG00000278896.1_6 | AC025031.5 | 0.80125142 | 1 | 0.141 | -0.546 | chr12 | 46798427 | 46800949 | + | TEC |
| ENSG00000169570.10_5 | DTWD2 | 0.99978536 | 1 | 0.141 | 3.896 | chr5 | 118171769 | 118324242 | - | protein_coding |
| ENSG00000130332.15_6 | LSM7 | 0.99996081 | 1 | 0.141 | 4.204 | chr19 | 2321519 | 2328610 | - | protein_coding |
| ENSG00000167130.18_5 | DOLPP1 | 0.99999407 | 1 | 0.141 | 4.802 | chr9 | 131843390 | 131852717 | + | protein_coding |
| ENSG00000107789.16_3 | MINPP1 | 0.99999959 | 1 | 0.141 | 5.593 | chr10 | 89264632 | 89313218 | + | protein_coding |
| ENSG00000283453.2_5 | AC244258.1 | 0.87542056 | 1 | 0.141 | 0.082 | chr6 | 57325808 | 57514407 | + | transcribed_unprocessed_pseudogene |
| ENSG00000141452.10_4 | RMC1 | 0.99990535 | 1 | 0.141 | 4.030 | chr18 | 21083434 | 21111786 | + | protein_coding |
| ENSG00000143761.16_4 | ARF1 | 0.99999994 | 1 | 0.140 | 7.254 | chr1 | 228270361 | 228286913 | + | protein_coding |
| ENSG00000178761.15_5 | FAM219B | 0.99991876 | 1 | 0.140 | 4.089 | chr15 | 75192333 | 75199224 | - | protein_coding |
| ENSG00000282386.1_6 | AL358472.4 | 0.87542056 | 1 | 0.140 | 0.083 | chr1 | 153936837 | 153937546 | + | lncRNA |
| ENSG00000164136.17_3 | IL15 | 0.99873796 | 1 | 0.140 | 3.008 | chr4 | 142557736 | 142655140 | + | protein_coding |
| ENSG00000204311.14_7 | PJVK | 0.95241154 | 1 | 0.140 | 1.288 | chr2 | 179316073 | 179326136 | + | protein_coding |
| ENSG00000119720.18_5 | NRDE2 | 0.99999573 | 1 | 0.140 | 4.960 | chr14 | 90734204 | 90798313 | - | protein_coding |
| ENSG00000125148.7_3 | MT2A | 0.99999972 | 1 | 0.140 | 5.835 | chr16 | 56642496 | 56643409 | + | protein_coding |
| ENSG00000109111.15_4 | SUPT6H | 0.99999997 | 1 | 0.140 | 7.195 | chr17 | 26989216 | 27029697 | + | protein_coding |
| ENSG00000246273.8_6 | SBF2-AS1 | 0.99504826 | 1 | 0.140 | 2.550 | chr11 | 9779815 | 9832882 | + | lncRNA |
| ENSG00000162231.14_6 | NXF1 | 0.99999832 | 1 | 0.140 | 5.300 | chr11 | 62559595 | 62573774 | - | protein_coding |
| ENSG00000167468.17_6 | GPX4 | 0.99999963 | 1 | 0.140 | 6.663 | chr19 | 1103925 | 1106790 | + | protein_coding |
| ENSG00000197037.11_5 | ZSCAN25 | 0.99988584 | 1 | 0.140 | 4.087 | chr7 | 99214569 | 99230031 | + | protein_coding |
| ENSG00000100393.14_5 | EP300 | 0.99999724 | 1 | 0.140 | 6.365 | chr22 | 41488596 | 41576081 | + | protein_coding |
| ENSG00000105705.16_9 | SUGP1 | 0.99996478 | 1 | 0.140 | 4.261 | chr19 | 19386827 | 19431318 | - | protein_coding |
| ENSG00000157764.13_7 | BRAF | 0.99999724 | 1 | 0.140 | 5.010 | chr7 | 140419127 | 140624728 | - | protein_coding |
| ENSG00000160194.18_3 | NDUFV3 | 0.99999805 | 1 | 0.139 | 5.066 | chr21 | 44299754 | 44333414 | + | protein_coding |
| ENSG00000132155.12_5 | RAF1 | 0.99999997 | 1 | 0.139 | 6.778 | chr3 | 12625100 | 12705725 | - | protein_coding |
| ENSG00000146250.7_3 | PRSS35 | 0.79670524 | 1 | 0.139 | -0.828 | chr6 | 84222253 | 84235423 | + | protein_coding |
| ENSG00000184207.9_5 | PGP | 0.999965 | 1 | 0.139 | 5.256 | chr16 | 2261594 | 2264841 | - | protein_coding |
| ENSG00000280927.1_5 | CTBP1-AS | 0.92067391 | 1 | 0.139 | 1.013 | chr4 | 1203908 | 1212379 | + | lncRNA |
| ENSG00000215252.11_3 | GOLGA8B | 0.99999713 | 1 | 0.139 | 5.151 | chr15 | 34817408 | 34880704 | - | protein_coding |
| ENSG00000267750.6_8 | RUNDC3A-AS1 | 0.87346864 | 1 | 0.139 | 0.319 | chr17 | 42376942 | 42392683 | - | lncRNA |
| ENSG00000262580.5_7 | AC087741.1 | 0.98536177 | 1 | 0.139 | 2.052 | chr17 | 78174472 | 78179748 | - | lncRNA |
| ENSG00000183751.15_6 | TBL3 | 0.99999277 | 1 | 0.139 | 5.199 | chr16 | 2022054 | 2032930 | + | protein_coding |
| ENSG00000171475.14_7 | WIPF2 | 0.99999977 | 1 | 0.139 | 5.906 | chr17 | 38375556 | 38440388 | + | protein_coding |
| ENSG00000135441.7_3 | BLOC1S1 | 0.99148484 | 1 | 0.139 | 3.476 | chr12 | 56109821 | 56113871 | + | protein_coding |
| ENSG00000101162.4_3 | TUBB1 | 0.88945368 | 1 | 0.139 | 0.520 | chr20 | 57594484 | 57601709 | + | protein_coding |
| ENSG00000152642.10_3 | GPD1L | 0.99999642 | 1 | 0.139 | 5.000 | chr3 | 32147181 | 32210205 | + | protein_coding |
| ENSG00000104129.10_4 | DNAJC17 | 0.99821875 | 1 | 0.139 | 2.960 | chr15 | 41057359 | 41099676 | - | protein_coding |
| ENSG00000107736.21_6 | CDH23 | 0.98640439 | 1 | 0.139 | 1.973 | chr10 | 73156677 | 73575704 | + | protein_coding |
| ENSG00000184432.10_5 | COPB2 | 0.99999994 | 1 | 0.138 | 7.373 | chr3 | 139074442 | 139108522 | - | protein_coding |
| ENSG00000166681.14_5 | BEX3 | 0.99999996 | 1 | 0.138 | 6.668 | chrX | 102631323 | 102633092 | + | protein_coding |
| ENSG00000065883.16_5 | CDK13 | 0.99999995 | 1 | 0.138 | 6.528 | chr7 | 39989720 | 40139179 | + | protein_coding |
| ENSG00000198833.7_4 | UBE2J1 | 0.99999885 | 1 | 0.138 | 5.765 | chr6 | 90036344 | 90062441 | - | protein_coding |

| | | | | | | | | | | |
|---|---|---|---|---|---|---|---|---|---|---|
| ENSG00000128891.15_8 | CCDC32 | 0.95882154 | 1 | 0.138 | 1.502 | chr15 | 40820882 | 40857256 | - | protein_coding |
| ENSG00000148396.18_5 | SEC16A | 0.99999997 | 1 | 0.138 | 6.803 | chr9 | 139334548 | 139378211 | - | protein_coding |
| ENSG00000102974.16_4 | CTCF | 0.99999995 | 1 | 0.138 | 6.516 | chr16 | 67596370 | 67673080 | + | protein_coding |
| ENSG00000003249.14_3 | DBNDD1 | 0.99999729 | 1 | 0.138 | 5.574 | chr16 | 90071279 | 90086536 | - | protein_coding |
| ENSG00000183856.11_4 | IQGAP3 | 0.99999994 | 1 | 0.138 | 6.386 | chr1 | 156495197 | 156542396 | - | protein_coding |
| ENSG00000136870.10_6 | ZNF189 | 0.99998916 | 1 | 0.138 | 5.001 | chr9 | 104161155 | 104172942 | + | protein_coding |
| ENSG00000124177.15_5 | CHD6 | 0.99999979 | 1 | 0.138 | 6.460 | chr20 | 40030723 | 40247023 | - | protein_coding |
| ENSG00000204237.5_6 | OXLD1 | 0.99988821 | 1 | 0.138 | 4.647 | chr17 | 79632066 | 79633665 | - | protein_coding |
| ENSG00000125967.16_4 | NECAB3 | 0.99996721 | 1 | 0.137 | 4.372 | chr20 | 32244893 | 32262269 | - | protein_coding |
| ENSG00000215548.2_3 | FRG1JP | 0.99077455 | 1 | 0.137 | 2.352 | chr9 | 68427861 | 68455375 | - | unprocessed_pseudogene |
| ENSG00000011295.16_7 | TTC19 | 0.99999983 | 1 | 0.137 | 5.875 | chr17 | 15903098 | 15948329 | + | protein_coding |
| ENSG00000144791.10_4 | LIMD1 | 0.99999953 | 1 | 0.137 | 5.772 | chr3 | 45596886 | 45727833 | + | protein_coding |
| ENSG00000261056.2_6 | AC079416.2 | 0.88259211 | 1 | 0.137 | 0.161 | chr16 | 53332136 | 53333704 | + | transcribed_processed_pseudogene |
| ENSG00000111785.20_5 | RIC8B | 0.99998992 | 1 | 0.137 | 4.659 | chr12 | 107168399 | 107283094 | + | protein_coding |
| ENSG00000166896.9_6 | ATP23 | 0.99790707 | 1 | 0.137 | 2.921 | chr12 | 58299822 | 58352931 | + | protein_coding |
| ENSG00000204348.10_5 | DXO | 0.95718346 | 1 | 0.137 | 1.253 | chr6 | 31937587 | 31940067 | - | protein_coding |
| ENSG00000125356.7_3 | NDUFA1 | 0.9999998 | 1 | 0.137 | 6.233 | chrX | 119005795 | 119010625 | + | protein_coding |
| ENSG00000244625.7_7 | MIATNB | 0.98320867 | 1 | 0.137 | 1.808 | chr22 | 27068710 | 27176856 | + | lncRNA |
| ENSG00000089902.10_4 | RCOR1 | 0.99999988 | 1 | 0.137 | 6.033 | chr14 | 103058986 | 103196898 | + | protein_coding |
| ENSG00000106993.12_3 | CDC37L1 | 0.999975 | 1 | 0.137 | 4.427 | chr9 | 4679559 | 4708399 | + | protein_coding |
| ENSG00000125971.16_2 | DYNLRB1 | 0.99801765 | 1 | 0.136 | 2.801 | chr20 | 33104214 | 33128762 | + | protein_coding |
| ENSG00000117479.15_7 | SLC19A2 | 0.99991307 | 1 | 0.136 | 4.022 | chr1 | 169433147 | 169455182 | - | protein_coding |
| ENSG00000132004.13_4 | FBXW9 | 0.99273929 | 1 | 0.136 | 2.731 | chr19 | 12798867 | 12807445 | - | protein_coding |
| ENSG00000205485.13_5 | AC004980.1 | 0.96576583 | 1 | 0.136 | 1.439 | chr7 | 76178677 | 76257299 | + | transcribed_unprocessed_pseudogene |
| ENSG00000143437.21_5 | ARNT | 0.99999861 | 1 | 0.136 | 5.528 | chr1 | 150782189 | 150849184 | - | protein_coding |
| ENSG00000100934.15_5 | SEC23A | 0.99999997 | 1 | 0.136 | 7.969 | chr14 | 39501123 | 39578850 | - | protein_coding |
| ENSG00000101457.13_4 | DNTTIP1 | 0.99878632 | 1 | 0.136 | 3.056 | chr20 | 44420593 | 44440066 | + | protein_coding |
| ENSG00000162430.17_7 | SELENON | 0.99999817 | 1 | 0.136 | 5.570 | chr1 | 26126667 | 26144712 | + | protein_coding |
| ENSG00000165476.14_3 | REEP3 | 0.99999983 | 1 | 0.136 | 6.860 | chr10 | 65281161 | 65384888 | + | protein_coding |
| ENSG00000024048.10_3 | UBR2 | 0.9999996 | 1 | 0.136 | 5.816 | chr6 | 42531800 | 42661242 | + | protein_coding |
| ENSG00000197779.13_2 | ZNF81 | 0.99978388 | 1 | 0.136 | 3.763 | chrX | 47696301 | 47861960 | + | protein_coding |
| ENSG00000207008.1 | SNORA54 | 0.81470391 | 1 | 0.136 | 0.201 | chr11 | 2985001 | 2985123 | - | snoRNA |
| ENSG00000049449.9_4 | RCN1 | 0.99999898 | 1 | 0.136 | 5.665 | chr11 | 32112450 | 32127301 | + | protein_coding |
| ENSG00000109670.16_7 | FBXW7 | 0.99998422 | 1 | 0.136 | 4.381 | chr4 | 153241696 | 153457244 | - | protein_coding |
| ENSG00000016864.18_6 | GLT8D1 | 0.99999902 | 1 | 0.136 | 5.359 | chr3 | 52728504 | 52740048 | - | protein_coding |
| ENSG00000099203.7_5 | TMED1 | 0.99657802 | 1 | 0.135 | 2.689 | chr19 | 10942743 | 10946994 | - | protein_coding |
| ENSG00000096654.15_3 | ZNF184 | 0.99972841 | 1 | 0.135 | 3.476 | chr6 | 27418522 | 27440897 | - | protein_coding |
| ENSG00000151276.23_4 | MAGI1 | 0.9999165 | 1 | 0.135 | 3.901 | chr3 | 65339200 | 66024509 | - | protein_coding |
| ENSG00000168610.14_5 | STAT3 | 0.99999999 | 1 | 0.135 | 7.973 | chr17 | 40465342 | 40540586 | - | protein_coding |
| ENSG00000270728.1_6 | AL035413.1 | 0.82620194 | 1 | 0.135 | -0.212 | chr1 | 19623574 | 19624397 | + | processed_pseudogene |
| ENSG00000131748.16_7 | STARD3 | 0.99997914 | 1 | 0.135 | 5.385 | chr17 | 37793343 | 37820454 | + | protein_coding |
| ENSG00000184897.6_4 | H1-10 | 0.93519865 | 1 | 0.135 | 4.940 | chr3 | 129033614 | 129035129 | - | protein_coding |
| ENSG00000070476.15_4 | ZXDC | 0.99983331 | 1 | 0.135 | 3.628 | chr3 | 126156444 | 126194734 | - | protein_coding |
| ENSG00000278266.1_6 | AC079949.2 | 0.94673486 | 1 | 0.135 | 1.249 | chr12 | 127631694 | 127634626 | + | lncRNA |
| ENSG00000197062.11_5 | ZSCAN26 | 0.99804825 | 1 | 0.135 | 3.265 | chr6 | 28234788 | 28246001 | + | protein_coding |
| ENSG00000260136.6_7 | AC008915.3 | 0.90104774 | 1 | 0.134 | 0.472 | chr16 | 23464079 | 23468927 | + | lncRNA |
| ENSG00000103199.14_7 | ZNF500 | 0.99994968 | 1 | 0.134 | 4.167 | chr16 | 4798240 | 4817625 | - | protein_coding |
| ENSG00000130429.15_9 | ARPC1B | 0.98992096 | 1 | 0.134 | 2.107 | chr7 | 98971872 | 98992439 | + | protein_coding |
| ENSG00000198026.8_4 | ZNF335 | 0.99999695 | 1 | 0.134 | 5.321 | chr20 | 44577299 | 44600842 | - | protein_coding |
| ENSG00000287269.1_1 | Z82215.1 | 0.83241729 | 1 | 0.134 | -0.428 | chr22 | 36731466 | 36739575 | + | lncRNA |
| ENSG00000277203.1_2 | F8A1 | 0.9946059 | 1 | 0.134 | 2.300 | chrX | 154114624 | 154116336 | + | protein_coding |
| ENSG00000139154.16_8 | AEBP2 | 0.99999544 | 1 | 0.134 | 4.916 | chr12 | 19556979 | 19873735 | + | protein_coding |
| ENSG00000144749.13_4 | LRIG1 | 0.99975238 | 1 | 0.134 | 3.710 | chr3 | 66429221 | 66551687 | - | protein_coding |
| ENSG00000123159.16_5 | GIPC1 | 0.99872218 | 1 | 0.134 | 6.701 | chr19 | 14588572 | 14606961 | - | protein_coding |
| ENSG00000176422.14_4 | SPRYD4 | 0.99990813 | 1 | 0.133 | 3.850 | chr12 | 56862362 | 56873492 | + | protein_coding |
| ENSG00000146540.15_6 | C7orf50 | 0.9999923 | 1 | 0.133 | 5.764 | chr7 | 1036622 | 1177896 | - | protein_coding |
| ENSG00000123191.15_5 | ATP7B | 0.99929494 | 1 | 0.133 | 3.289 | chr13 | 52504572 | 52586261 | - | protein_coding |
| ENSG00000136193.17_5 | SCRN1 | 0.99999998 | 1 | 0.133 | 7.421 | chr7 | 29959720 | 30029905 | - | protein_coding |
| ENSG00000059728.11_4 | MXD1 | 0.99035338 | 1 | 0.133 | 2.326 | chr2 | 70124820 | 70170077 | + | protein_coding |

| | | | | | | | | | | |
|---|---|---|---|---|---|---|---|---|---|---|
| ENSG00000155876.6_4 | RRAGA | 0.99999956 | 1 | 0.133 | 5.770 | chr9 | 19049425 | 19051023 | + | protein_coding |
| ENSG00000140577.16_6 | CRTC3 | 0.99999912 | 1 | 0.133 | 5.302 | chr15 | 91073155 | 91188577 | + | protein_coding |
| ENSG00000198561.14_7 | CTNND1 | 0.99991929 | 1 | 0.133 | 3.849 | chr11 | 57520715 | 57587018 | + | protein_coding |
| ENSG00000101190.13_4 | TCFL5 | 0.99989808 | 1 | 0.133 | 3.790 | chr20 | 61472357 | 61493174 | - | protein_coding |
| ENSG00000106714.17_4 | CNTNAP3 | 0.99999985 | 1 | 0.133 | 5.913 | chr9 | 39072764 | 39288312 | - | protein_coding |
| ENSG00000172732.12_4 | MUS81 | 0.99999577 | 1 | 0.133 | 4.742 | chr11 | 65624597 | 65635124 | + | protein_coding |
| ENSG00000145388.15_4 | METTL14 | 0.99997711 | 1 | 0.133 | 4.455 | chr4 | 119606547 | 119636588 | + | protein_coding |
| ENSG00000137857.17_3 | DUOX1 | 0.94263066 | 1 | 0.132 | 1.128 | chr15 | 45422131 | 45457774 | + | protein_coding |
| ENSG00000162734.12_3 | PEA15 | 0.99999974 | 1 | 0.132 | 5.737 | chr1 | 160175127 | 160185166 | + | protein_coding |
| ENSG00000140153.17_6 | WDR20 | 0.99998663 | 1 | 0.132 | 4.588 | chr14 | 102605840 | 102691184 | + | protein_coding |
| ENSG00000119396.11_5 | RAB14 | 0.99999988 | 1 | 0.132 | 6.892 | chr9 | 123940411 | 123985292 | - | protein_coding |
| ENSG00000277744.1_8 | AC011462.4 | 0.90631837 | 1 | 0.132 | 0.357 | chr19 | 41879876 | 41880324 | + | lncRNA |
| ENSG00000135862.6_3 | LAMC1 | 0.99999999 | 1 | 0.132 | 8.466 | chr1 | 182992555 | 183114727 | + | protein_coding |
| ENSG00000110536.14_4 | PTPMT1 | 0.98393029 | 1 | 0.132 | 1.750 | chr11 | 47586982 | 47595013 | + | protein_coding |
| ENSG00000161204.11_4 | ABCF3 | 0.99999342 | 1 | 0.132 | 4.596 | chr3 | 183903811 | 183911800 | + | protein_coding |
| ENSG00000179965.12_3 | ZNF771 | 0.99064824 | 1 | 0.132 | 2.433 | chr16 | 30418735 | 30442429 | + | protein_coding |
| ENSG00000101558.13_5 | VAPA | 0.99999993 | 1 | 0.132 | 7.356 | chr18 | 9913999 | 9960018 | + | protein_coding |
| ENSG00000136448.12_7 | NMT1 | 0.99999999 | 1 | 0.132 | 7.317 | chr17 | 43128978 | 43186384 | + | protein_coding |
| ENSG00000285596.1_3 | AC017116.2 | 0.81653003 | 1 | 0.131 | -0.615 | chr7 | 44051481 | 44078577 | - | lncRNA |
| ENSG00000168216.13_8 | LMBRD1 | 0.99998804 | 1 | 0.131 | 4.336 | chr6 | 70382649 | 70577128 | - | protein_coding |
| ENSG00000229422.2_6 | AL512625.2 | 0.82163306 | 1 | 0.131 | -0.463 | chr9 | 66492963 | 66494126 | - | lncRNA |
| ENSG00000136504.12_5 | KAT7 | 0.99999986 | 1 | 0.131 | 6.338 | chr17 | 47866043 | 47912388 | + | protein_coding |
| ENSG00000111877.17_4 | MCM9 | 0.99999641 | 1 | 0.131 | 4.768 | chr6 | 119134605 | 119256327 | - | protein_coding |
| ENSG00000277476.1_6 | AC005332.5 | 0.96598813 | 1 | 0.131 | 1.485 | chr17 | 66129342 | 66132076 | + | lncRNA |
| ENSG00000135720.13_6 | DYNC1LI2 | 0.99999985 | 1 | 0.131 | 6.956 | chr16 | 66754796 | 66785512 | - | protein_coding |
| ENSG00000163558.13_3 | PRKCI | 0.99999988 | 1 | 0.131 | 6.576 | chr3 | 169940212 | 170023765 | + | protein_coding |
| ENSG00000102158.19_4 | MAGT1 | 0.99999991 | 1 | 0.131 | 6.839 | chrX | 77081861 | 77151090 | - | protein_coding |
| ENSG00000142867.14_7 | BCL10 | 0.99998621 | 1 | 0.131 | 4.481 | chr1 | 85731459 | 85742315 | - | protein_coding |
| ENSG00000142669.15_4 | SH3BGRL3 | 0.99957979 | 1 | 0.131 | 4.048 | chr1 | 26606577 | 26608013 | + | protein_coding |
| ENSG00000166033.13_4 | HTRA1 | 0.99999759 | 1 | 0.131 | 5.089 | chr10 | 124218067 | 124274423 | + | protein_coding |
| ENSG00000019995.6_4 | ZRANB1 | 0.99997303 | 1 | 0.131 | 4.461 | chr10 | 126630692 | 126676758 | + | protein_coding |
| ENSG00000123091.5_3 | RNF11 | 0.99999967 | 1 | 0.130 | 5.702 | chr1 | 51701945 | 51739119 | + | protein_coding |
| ENSG00000148297.16_6 | MED22 | 0.9999992 | 1 | 0.130 | 5.362 | chr9 | 136205149 | 136214986 | - | protein_coding |
| ENSG00000175137.11_4 | SH3BP5L | 0.99999535 | 1 | 0.130 | 4.809 | chr1 | 249104645 | 249120114 | - | protein_coding |
| ENSG00000166971.17_5 | AKTIP | 0.99999592 | 1 | 0.130 | 4.844 | chr16 | 53524952 | 53538323 | - | protein_coding |
| ENSG00000108654.15_6 | DDX5 | 1 | 1 | 0.130 | 9.319 | chr17 | 62494372 | 62504317 | - | protein_coding |
| ENSG00000231707.4_4 | PABPC1P1 | 0.80855932 | 1 | 0.130 | -1.121 | chr4 | 39975064 | 39975958 | + | processed_pseudogene |
| ENSG00000106829.19_6 | TLE4 | 0.99999484 | 1 | 0.130 | 5.045 | chr9 | 82186688 | 82341797 | + | protein_coding |
| ENSG00000160460.16_6 | AC020929.1 | 0.98383641 | 1 | 0.130 | 2.091 | chr19 | 40972148 | 41082370 | + | protein_coding |
| ENSG00000087460.25_7 | GNAS | 0.99999999 | 1 | 0.130 | 8.569 | chr20 | 57414773 | 57486247 | + | protein_coding |
| ENSG00000129518.9_4 | EAPP | 0.99999826 | 1 | 0.130 | 5.072 | chr14 | 34985144 | 35008917 | - | protein_coding |
| ENSG00000264290.1_7 | AC104564.3 | 0.8180923 | 1 | 0.129 | -0.317 | chr17 | 27896598 | 27897537 | + | lncRNA |
| ENSG00000141858.12_6 | SAMD1 | 0.99999753 | 1 | 0.129 | 5.220 | chr19 | 14198663 | 14201848 | - | protein_coding |
| ENSG00000165322.18_7 | ARHGAP12 | 0.99999995 | 1 | 0.129 | 6.830 | chr10 | 32094326 | 32217804 | - | protein_coding |
| ENSG00000149591.17_4 | TAGLN | 0.99139892 | 1 | 0.129 | 2.016 | chr11 | 117070086 | 117078180 | + | protein_coding |
| ENSG00000139200.13_4 | PIANP | 0.84373921 | 1 | 0.129 | -0.211 | chr12 | 6802958 | 6809966 | - | protein_coding |
| ENSG00000135535.17_6 | CD164 | 0.99999989 | 1 | 0.129 | 7.272 | chr6 | 109687717 | 109703670 | - | protein_coding |
| ENSG00000083099.11_4 | LYRM2 | 0.99999864 | 1 | 0.129 | 5.114 | chr6 | 90277863 | 90348468 | - | protein_coding |
| ENSG00000165914.15_5 | TTC7B | 0.99995703 | 1 | 0.129 | 3.960 | chr14 | 90990908 | 91282823 | - | protein_coding |
| ENSG00000165502.7_5 | RPL36AL | 0.9999996 | 1 | 0.129 | 7.152 | chr14 | 50085248 | 50087344 | - | protein_coding |
| ENSG00000156482.11_4 | RPL30 | 0.99999984 | 1 | 0.129 | 6.637 | chr8 | 99037079 | 99058697 | - | protein_coding |
| ENSG00000142347.19_8 | MYO1F | 0.90933025 | 1 | 0.129 | 0.389 | chr19 | 8585662 | 8642461 | - | protein_coding |
| ENSG00000279838.1_6 | AL356273.6 | 0.9304278 | 1 | 0.129 | 0.771 | chr1 | 185261516 | 185263504 | - | TEC |
| ENSG00000101096.20_4 | NFATC2 | 0.99792203 | 1 | 0.129 | 2.869 | chr20 | 50003494 | 50179370 | - | protein_coding |
| ENSG00000234222.6_8 | LIX1L-AS1 | 0.82872975 | 1 | 0.129 | -0.428 | chr1 | 145475909 | 145508503 | - | lncRNA |
| ENSG00000135185.12_7 | TMEM243 | 0.99988195 | 1 | 0.129 | 3.741 | chr7 | 86825476 | 86849903 | - | protein_coding |
| ENSG00000287769.1_2 | AL353748.3 | 0.9575241 | 1 | 0.129 | 1.331 | chr9 | 90999817 | 91004139 | - | lncRNA |
| ENSG00000138593.9_5 | SECISBP2L | 0.9999991 | 1 | 0.129 | 5.575 | chr15 | 49280673 | 49338644 | - | protein_coding |
| ENSG00000160613.13_5 | PCSK7 | 0.99998402 | 1 | 0.128 | 4.394 | chr11 | 117075053 | 117103241 | - | protein_coding |
| ENSG00000151366.13_6 | NDUFC2 | 0.99348939 | 1 | 0.128 | 4.425 | chr11 | 77779343 | 77790908 | - | protein_coding |

| | | | | | | | | | | |
|---|---|---|---|---|---|---|---|---|---|---|
| ENSG00000169902.15_5 | TPST1 | 0.99988527 | 1 | 0.128 | 3.614 | chr7 | 65670304 | 65885530 | + | protein_coding |
| ENSG00000120925.16_5 | RNF170 | 0.99999118 | 1 | 0.128 | 4.984 | chr8 | 42704780 | 42752433 | - | protein_coding |
| ENSG00000134291.12_4 | TMEM106C | 0.99999994 | 1 | 0.128 | 6.614 | chr12 | 48357352 | 48362661 | + | protein_coding |
| ENSG00000161638.11_5 | ITGA5 | 0.99999661 | 1 | 0.128 | 5.515 | chr12 | 54789045 | 54813050 | - | protein_coding |
| ENSG00000232630.1_4 | PRPS1P2 | 0.92378175 | 1 | 0.128 | 0.842 | chr9 | 127912932 | 127913868 | + | processed_pseudogene |
| ENSG00000230773.9_8 | AC092650.1 | 0.81246924 | 1 | 0.128 | -0.787 | chr2 | 48151320 | 48541363 | - | lncRNA |
| ENSG00000102796.11_5 | DHRS12 | 0.84757668 | 1 | 0.128 | -0.220 | chr13 | 52342129 | 52378298 | - | protein_coding |
| ENSG00000146574.15_4 | CCZ1B | 0.9995818 | 1 | 0.128 | 3.320 | chr7 | 6833765 | 6866401 | - | protein_coding |
| ENSG00000116977.18_5 | LGALS8 | 0.99999944 | 1 | 0.128 | 5.703 | chr1 | 236681300 | 236716281 | + | protein_coding |
| ENSG00000138303.18_7 | ASCC1 | 0.99999314 | 1 | 0.128 | 5.647 | chr10 | 73855790 | 73976892 | - | protein_coding |
| ENSG00000151729.11_3 | SLC25A4 | 0.99994098 | 1 | 0.128 | 4.072 | chr4 | 186064420 | 186071536 | + | protein_coding |
| ENSG00000177732.8_3 | SOX12 | 0.99996591 | 1 | 0.128 | 5.084 | chr20 | 306239 | 310868 | + | protein_coding |
| ENSG00000101193.8_3 | GID8 | 0.99999968 | 1 | 0.127 | 5.645 | chr20 | 61569499 | 61579827 | + | protein_coding |
| ENSG00000114439.19_7 | BBX | 0.99999997 | 1 | 0.127 | 7.002 | chr3 | 107241783 | 107530186 | + | protein_coding |
| ENSG00000137075.18_5 | RNF38 | 0.99999754 | 1 | 0.127 | 4.925 | chr9 | 36336393 | 36487545 | - | protein_coding |
| ENSG00000174373.16_5 | RALGAPA1 | 0.99999856 | 1 | 0.127 | 5.569 | chr14 | 36007558 | 36278510 | - | protein_coding |
| ENSG00000138614.15_6 | INTS14 | 0.99999956 | 1 | 0.127 | 5.776 | chr15 | 65871091 | 65903627 | - | protein_coding |
| ENSG00000104936.17_5 | DMPK | 0.99998602 | 1 | 0.127 | 4.643 | chr19 | 46272975 | 46285810 | - | protein_coding |
| ENSG00000051382.9_3 | PIK3CB | 0.99996728 | 1 | 0.127 | 4.450 | chr3 | 138371540 | 138553770 | - | protein_coding |
| ENSG00000167065.13_8 | DUSP18 | 0.99596381 | 1 | 0.127 | 1.713 | chr22 | 31048038 | 31063877 | - | protein_coding |
| ENSG00000189298.14_4 | ZKSCAN3 | 0.99686506 | 1 | 0.127 | 2.461 | chr6 | 28317724 | 28336949 | + | protein_coding |
| ENSG00000112238.11_2 | PRDM13 | 0.84000932 | 1 | 0.127 | -0.142 | chr6 | 100054606 | 100063454 | + | protein_coding |
| ENSG00000233927.5_3 | RPS28 | 0.99999962 | 1 | 0.126 | 5.653 | chr19 | 8386042 | 8388224 | + | protein_coding |
| ENSG00000105672.14_5 | ETV2 | 0.86962178 | 1 | 0.126 | 0.134 | chr19 | 36132647 | 36135773 | + | protein_coding |
| ENSG00000107438.9_4 | PDLIM1 | 0.93473603 | 1 | 0.126 | 5.671 | chr10 | 96997329 | 97050769 | - | protein_coding |
| ENSG00000257303.2_7 | AC073896.2 | 0.86516765 | 1 | 0.126 | 0.176 | chr12 | 56693860 | 56711776 | + | lncRNA |
| ENSG00000278861.1_6 | AC117503.2 | 0.8389243 | 1 | 0.126 | -0.349 | chr12 | 124140075 | 124140675 | + | TEC |
| ENSG00000213553.4_4 | RPLP0P6 | 0.93585091 | 1 | 0.126 | 5.170 | chr2 | 38708993 | 38709946 | + | processed_pseudogene |
| ENSG00000173818.17_4 | ENDOV | 0.99999405 | 1 | 0.126 | 4.700 | chr17 | 78388965 | 78411886 | + | protein_coding |
| ENSG00000007545.16_4 | CRAMP1 | 0.99938256 | 1 | 0.126 | 3.444 | chr16 | 1662338 | 1727909 | + | protein_coding |
| ENSG00000163832.16_6 | ELP6 | 0.99999467 | 1 | 0.126 | 4.538 | chr3 | 47537130 | 47555202 | - | protein_coding |
| ENSG00000197889.10_3 | MEIG1 | 0.81840664 | 1 | 0.126 | -0.831 | chr10 | 15001387 | 15030049 | + | protein_coding |
| ENSG00000095139.14_3 | ARCN1 | 0.99999998 | 1 | 0.126 | 7.068 | chr11 | 118443105 | 118473748 | + | protein_coding |
| ENSG00000158315.11_5 | RHBDL2 | 0.96258752 | 1 | 0.126 | 1.237 | chr1 | 39351479 | 39407502 | - | protein_coding |
| ENSG00000136045.12_3 | PWP1 | 0.99999961 | 1 | 0.126 | 6.634 | chr12 | 108079576 | 108106939 | + | protein_coding |
| ENSG00000138640.15_7 | FAM13A | 0.99998561 | 1 | 0.125 | 4.402 | chr4 | 89647106 | 90032549 | - | protein_coding |
| ENSG00000204179.6 | PTPN20A | 0.86868609 | 1 | 0.125 | -0.121 | chr10 | 46549955 | 46641045 | - | protein_coding |
| ENSG00000100441.10_5 | KHNYN | 0.99999986 | 1 | 0.125 | 5.809 | chr14 | 24898492 | 24911049 | + | protein_coding |
| ENSG00000204164.6 | BMS1P5 | 0.84362992 | 1 | 0.125 | -0.404 | chr10 | 48901102 | 48950972 | - | pseudogene |
| ENSG00000121691.7_5 | CAT | 0.99999958 | 1 | 0.125 | 5.533 | chr11 | 34460481 | 34493607 | + | protein_coding |
| ENSG00000183770.7_4 | FOXL2 | 0.9561926 | 1 | 0.125 | 1.709 | chr3 | 138663066 | 138665979 | - | protein_coding |
| ENSG00000134851.13_4 | TMEM165 | 0.99999942 | 1 | 0.125 | 5.710 | chr4 | 56262124 | 56319564 | + | protein_coding |
| ENSG00000160959.8_6 | LRRC14 | 0.99999892 | 1 | 0.125 | 5.128 | chr8 | 145743376 | 145750556 | + | protein_coding |
| ENSG00000112357.13_3 | PEX7 | 0.99251082 | 1 | 0.125 | 2.143 | chr6 | 137143702 | 137235072 | + | protein_coding |
| ENSG00000162066.15_4 | AMDHD2 | 0.99414198 | 1 | 0.125 | 3.089 | chr16 | 2570358 | 2581423 | + | protein_coding |
| ENSG00000272690.6_6 | LINC02018 | 0.83810217 | 1 | 0.124 | -0.325 | chr3 | 75484383 | 75587180 | + | lncRNA |
| ENSG00000272669.1_7 | AL021707.6 | 0.88672155 | 1 | 0.124 | 0.033 | chr22 | 39138630 | 39139120 | + | lncRNA |
| ENSG00000067365.15_4 | METTL22 | 0.99995757 | 1 | 0.124 | 4.013 | chr16 | 8715540 | 8743511 | + | protein_coding |
| ENSG00000182685.7_4 | BRICD5 | 0.97871545 | 1 | 0.124 | 2.443 | chr16 | 2259254 | 2261951 | - | protein_coding |
| ENSG00000105254.12_5 | TBCB | 0.99999998 | 1 | 0.124 | 7.201 | chr19 | 36605191 | 36616849 | + | protein_coding |
| ENSG00000166199.13_4 | ALKBH3 | 0.99999086 | 1 | 0.124 | 4.405 | chr11 | 43902361 | 43941824 | + | protein_coding |
| ENSG00000137947.12_6 | GTF2B | 0.99998503 | 1 | 0.124 | 4.259 | chr1 | 89318316 | 89357627 | - | protein_coding |
| ENSG00000126012.11_5 | KDM5C | 0.99999999 | 1 | 0.124 | 7.374 | chrX | 53220503 | 53254604 | - | protein_coding |
| ENSG00000281376.1_5 | ABALON | 0.9659744 | 1 | 0.124 | 1.434 | chr20 | 30309310 | 30311212 | + | lncRNA |
| ENSG00000178980.15_7 | SELENOW | 0.99999878 | 1 | 0.124 | 5.190 | chr19 | 48281842 | 48287943 | + | protein_coding |
| ENSG00000166529.15_4 | ZSCAN21 | 0.99765921 | 1 | 0.124 | 2.553 | chr7 | 99647397 | 99662663 | + | protein_coding |
| ENSG00000265735.1 | RN7SL5P | 0.92652781 | 1 | 0.123 | 1.349 | chr9 | 9442060 | 9442347 | + | misc_RNA |
| ENSG00000130312.6_4 | MRPL34 | 0.99976503 | 1 | 0.123 | 3.683 | chr19 | 17403418 | 17417652 | + | protein_coding |
| ENSG00000131899.11_4 | LLGL1 | 0.99999623 | 1 | 0.123 | 5.671 | chr17 | 18128949 | 18148189 | + | protein_coding |
| ENSG00000140995.16_6 | DEF8 | 0.99999955 | 1 | 0.123 | 5.883 | chr16 | 90014333 | 90034468 | + | protein_coding |

| | | | | | | | | | | |
|---|---|---|---|---|---|---|---|---|---|---|
| ENSG00000116761.12_3 | CTH | 0.96699066 | 1 | 0.123 | 1.458 | chr1 | 70876901 | 70905534 | + | protein_coding |
| ENSG00000148429.14_3 | USP6NL | 0.99999901 | 1 | 0.123 | 5.307 | chr10 | 11495945 | 11653753 | - | protein_coding |
| ENSG00000149927.18_5 | DOC2A | 0.9941186 | 1 | 0.123 | 2.400 | chr16 | 30016835 | 30034591 | - | protein_coding |
| ENSG00000173436.15_5 | MICOS10 | 0.97662747 | 1 | 0.123 | 1.764 | chr1 | 19810897 | 19956315 | + | protein_coding |
| ENSG00000135870.12_7 | RC3H1 | 0.99999981 | 1 | 0.123 | 5.974 | chr1 | 173900222 | 173991495 | - | protein_coding |
| ENSG00000157214.14_6 | STEAP2 | 0.99781309 | 1 | 0.123 | 6.012 | chr7 | 89796904 | 89867451 | + | protein_coding |
| ENSG00000147383.11_6 | NSDHL | 0.99996127 | 1 | 0.123 | 4.291 | chrX | 151999511 | 152038273 | + | protein_coding |
| ENSG00000033627.16_4 | ATP6V0A1 | 0.99999924 | 1 | 0.123 | 5.434 | chr17 | 40610862 | 40674629 | + | protein_coding |
| ENSG00000262919.8_6 | CCNQ | 0.99998808 | 1 | 0.123 | 4.474 | chrX | 152853383 | 152865500 | - | protein_coding |
| ENSG00000196313.11_2 | POM121 | 0.99990179 | 1 | 0.123 | 4.089 | chr7 | 72349936 | 72421979 | + | protein_coding |
| ENSG00000266947.1_7 | AC022916.2 | 0.86551576 | 1 | 0.122 | -0.077 | chr17 | 33558469 | 33569982 | - | lncRNA |
| ENSG00000158528.12_4 | PPP1R9A | 0.99999986 | 1 | 0.122 | 6.346 | chr7 | 94536514 | 94925727 | + | protein_coding |
| ENSG00000095906.17_5 | NUBP2 | 0.99995963 | 1 | 0.122 | 5.354 | chr16 | 1832933 | 1839187 | + | protein_coding |
| ENSG00000172888.12_3 | ZNF621 | 0.99998529 | 1 | 0.122 | 4.517 | chr3 | 40566369 | 40616176 | + | protein_coding |
| ENSG00000126581.13_5 | BECN1 | 0.9999998 | 1 | 0.122 | 5.603 | chr17 | 40962152 | 40985367 | - | protein_coding |
| ENSG00000127920.6_3 | GNG11 | 0.99999961 | 1 | 0.122 | 6.090 | chr7 | 93551047 | 93557922 | + | protein_coding |
| ENSG00000249471.8_3 | ZNF324B | 0.99942889 | 1 | 0.122 | 3.422 | chr19 | 58962978 | 58969200 | + | protein_coding |
| ENSG00000168872.17_4 | DDX19A | 0.99990275 | 1 | 0.122 | 3.868 | chr16 | 70380764 | 70407286 | + | protein_coding |
| ENSG00000141026.6_3 | MED9 | 0.99998489 | 1 | 0.122 | 4.365 | chr17 | 17380308 | 17396535 | + | protein_coding |
| ENSG00000105246.6_3 | EBI3 | 0.83015006 | 1 | 0.122 | -0.756 | chr19 | 4229520 | 4237525 | + | protein_coding |
| ENSG00000222019.7_5 | URAHP | 0.92145416 | 1 | 0.122 | 0.365 | chr16 | 90106169 | 90114181 | - | transcribed_unprocessed_pseudogene |
| ENSG00000106804.8_3 | C5 | 0.99999993 | 1 | 0.122 | 7.576 | chr9 | 123714613 | 123812553 | - | protein_coding |
| ENSG00000119723.17_4 | COQ6 | 0.99399543 | 1 | 0.122 | 2.580 | chr14 | 74416629 | 74430373 | + | protein_coding |
| ENSG00000156011.17_5 | PSD3 | 0.99999993 | 1 | 0.122 | 6.399 | chr8 | 18384813 | 18942240 | - | protein_coding |
| ENSG00000256806.6_4 | C17orf100 | 0.9742174 | 1 | 0.121 | 1.676 | chr17 | 6555082 | 6596521 | + | protein_coding |
| ENSG00000104325.7_6 | DECR1 | 0.99999677 | 1 | 0.121 | 4.736 | chr8 | 91013633 | 91065861 | + | protein_coding |
| ENSG00000258732.1_5 | AC025884.1 | 0.91207944 | 1 | 0.121 | 0.528 | chr15 | 22566922 | 22570823 | + | unprocessed_pseudogene |
| ENSG00000157152.17_8 | SYN2 | 0.89253124 | 1 | 0.121 | 0.316 | chr3 | 12045862 | 12233532 | + | protein_coding |
| ENSG00000064652.11_3 | SNX24 | 0.99996271 | 1 | 0.120 | 4.079 | chr5 | 122179134 | 122365049 | + | protein_coding |
| ENSG00000006194.10_4 | ZNF263 | 0.99999933 | 1 | 0.120 | 5.580 | chr16 | 3313800 | 3351401 | + | protein_coding |
| ENSG00000214973.3_4 | CHCHD3P3 | 0.82228274 | 1 | 0.120 | -1.084 | chr1 | 27527325 | 27527964 | - | processed_pseudogene |
| ENSG00000185104.20_5 | FAF1 | 0.99999987 | 1 | 0.120 | 5.716 | chr1 | 50902700 | 51425939 | - | protein_coding |
| ENSG00000001626.16_8 | CFTR | 0.8233071 | 1 | 0.120 | -0.875 | chr7 | 116927174 | 117356025 | + | protein_coding |
| ENSG00000132204.14_6 | LINC00470 | 0.97879763 | 1 | 0.120 | 1.603 | chr18 | 1254384 | 1408345 | - | lncRNA |
| ENSG00000206561.13_8 | COLQ | 0.83229431 | 1 | 0.120 | -0.428 | chr3 | 15491640 | 15563258 | - | protein_coding |
| ENSG00000107372.13_4 | ZFAND5 | 0.99999999 | 1 | 0.120 | 8.029 | chr9 | 74966329 | 74980151 | - | protein_coding |
| ENSG00000168286.3_3 | THAP11 | 0.99999514 | 1 | 0.120 | 4.762 | chr16 | 67876223 | 67878098 | + | protein_coding |
| ENSG00000277283.1_6 | AC004812.2 | 0.98283931 | 1 | 0.120 | 1.673 | chr12 | 120554711 | 120556804 | + | lncRNA |
| ENSG00000161016.17_6 | RPL8 | 0.99999999 | 1 | 0.120 | 9.430 | chr8 | 146015150 | 146017972 | - | protein_coding |
| ENSG00000107521.19_6 | HPS1 | 0.99998984 | 1 | 0.120 | 4.823 | chr10 | 100175955 | 100206704 | - | protein_coding |
| ENSG00000140386.13_7 | SCAPER | 0.99986156 | 1 | 0.120 | 4.036 | chr15 | 76640245 | 77197785 | - | protein_coding |
| ENSG00000111790.14_4 | FGFR1OP2 | 0.99999896 | 1 | 0.120 | 5.500 | chr12 | 27091403 | 27119583 | + | protein_coding |
| ENSG00000153721.19_7 | CNKSR3 | 0.99932485 | 1 | 0.120 | 3.038 | chr6 | 154708649 | 154831819 | - | protein_coding |
| ENSG00000269688.1_5 | AC008982.2 | 0.99653747 | 1 | 0.120 | 2.342 | chr19 | 39335369 | 39336139 | - | lncRNA |
| ENSG00000185020.5 | RP11-111G23.1 | 0.99352802 | 1 | 0.119 | 2.126 | chr9 | 40293025 | 40339528 | + | pseudogene |
| ENSG00000173599.15_6 | PC | 0.99999227 | 1 | 0.119 | 5.854 | chr11 | 66615888 | 66725857 | - | protein_coding |
| ENSG00000124659.7_3 | TBCC | 0.99989649 | 1 | 0.119 | 3.740 | chr6 | 42712236 | 42713841 | - | protein_coding |
| ENSG00000154803.13_6 | FLCN | 0.99999914 | 1 | 0.119 | 5.144 | chr17 | 17115526 | 17140502 | - | protein_coding |
| ENSG00000160326.14_5 | SLC2A6 | 0.99955157 | 1 | 0.119 | 4.805 | chr9 | 136336216 | 136344249 | - | protein_coding |
| ENSG00000139641.12_3 | ESYT1 | 0.99999998 | 1 | 0.119 | 7.065 | chr12 | 56512034 | 56538455 | + | protein_coding |
| ENSG00000117682.17_5 | DHDDS | 0.99998241 | 1 | 0.119 | 4.605 | chr1 | 26758773 | 26797797 | + | protein_coding |
| ENSG00000265100.1_6 | AC005332.1 | 0.83907719 | 1 | 0.119 | -0.656 | chr17 | 66242770 | 66244079 | - | lncRNA |
| ENSG00000125352.6_3 | RNF113A | 0.99966746 | 1 | 0.119 | 3.426 | chrX | 119004438 | 119005696 | - | protein_coding |
| ENSG00000116299.17_5 | KIAA1324 | 0.9424499 | 1 | 0.119 | 0.841 | chr1 | 109656301 | 109749403 | + | protein_coding |
| ENSG00000129028.9_5 | THAP10 | 0.99929366 | 1 | 0.119 | 3.182 | chr15 | 71173681 | 71184772 | - | protein_coding |
| ENSG00000284669.1_5 | AC092053.4 | 0.99263847 | 1 | 0.119 | 2.116 | chr3 | 39189772 | 39214443 | + | lncRNA |
| ENSG00000135315.12_5 | CEP162 | 0.99960331 | 1 | 0.118 | 3.345 | chr6 | 84833960 | 84937361 | - | protein_coding |
| ENSG00000286555.1_2 | AP003065.3 | 0.83679534 | 1 | 0.118 | -0.405 | chr11 | 72400695 | 72406866 | + | lncRNA |

| | | | | | | | | | | |
|---|---|---|---|---|---|---|---|---|---|---|
| ENSG00000276728.1_6 | AC142472.1 | 0.94187594 | 1 | 0.118 | 1.086 | chr17 | 43224097 | 43225837 | - | lncRNA |
| ENSG00000151414.15_5 | NEK7 | 0.9999999 | 1 | 0.118 | 5.915 | chr1 | 198126124 | 198291550 | + | protein_coding |
| ENSG00000283486.2_6 | FAM95C | 0.88950544 | 1 | 0.118 | 0.073 | chr9 | 38540566 | 38577204 | - | protein_coding |
| ENSG00000108854.16_7 | SMURF2 | 0.99999959 | 1 | 0.118 | 6.089 | chr17 | 62538400 | 62658425 | - | protein_coding |
| ENSG00000240849.11_5 | TMEM189 | 0.99999042 | 1 | 0.118 | 4.550 | chr20 | 48734791 | 48770271 | - | protein_coding |
| ENSG00000089057.15_6 | SLC23A2 | 0.99999993 | 1 | 0.118 | 6.074 | chr20 | 4833002 | 4990939 | - | protein_coding |
| ENSG00000185909.15_3 | KLHDC8B | 0.99929781 | 1 | 0.118 | 2.984 | chr3 | 49209031 | 49213919 | + | protein_coding |
| ENSG00000112290.13_6 | WASF1 | 0.99970653 | 1 | 0.118 | 3.681 | chr6 | 110421022 | 110501207 | - | protein_coding |
| ENSG00000034713.8_3 | GABARAPL2 | 0.99999213 | 1 | 0.118 | 4.751 | chr16 | 75600273 | 75611779 | + | protein_coding |
| ENSG00000185219.17_4 | ZNF445 | 0.99999747 | 1 | 0.118 | 5.028 | chr3 | 44473197 | 44519162 | - | protein_coding |
| ENSG00000106006.6_6 | HOXA6 | 0.84599895 | 1 | 0.118 | -0.373 | chr7 | 27185015 | 27190222 | - | protein_coding |
| ENSG00000109220.11_5 | CHIC2 | 0.99688631 | 1 | 0.117 | 2.589 | chr4 | 54875956 | 54930772 | - | protein_coding |
| ENSG00000120029.13_6 | ARMH3 | 0.99997939 | 1 | 0.117 | 4.178 | chr10 | 103605356 | 103815950 | - | protein_coding |
| ENSG00000176986.16_4 | SEC24C | 0.99999975 | 1 | 0.117 | 5.883 | chr10 | 75504130 | 75531919 | + | protein_coding |
| ENSG00000287242.1_1 | AC073593.2 | 0.83763913 | 1 | 0.117 | -0.561 | chr12 | 125271329 | 125293336 | + | lncRNA |
| ENSG00000104133.15_9 | SPG11 | 0.99999948 | 1 | 0.117 | 5.834 | chr15 | 44854894 | 44955876 | - | protein_coding |
| ENSG00000150477.15_7 | KIAA1328 | 0.99595281 | 1 | 0.117 | 2.422 | chr18 | 34409069 | 34812135 | + | protein_coding |
| ENSG00000157593.19_5 | SLC35B2 | 0.99999919 | 1 | 0.117 | 5.384 | chr6 | 44221833 | 44225627 | - | protein_coding |
| ENSG00000196821.10_7 | ILRUN | 0.99999988 | 1 | 0.117 | 5.792 | chr6 | 34555065 | 34664636 | - | protein_coding |
| ENSG00000135365.15_8 | PHF21A | 0.99996585 | 1 | 0.117 | 4.332 | chr11 | 45950874 | 46142729 | - | protein_coding |
| ENSG00000268621.6_7 | IGFL2-AS1 | 0.99999944 | 1 | 0.117 | 5.731 | chr19 | 46692286 | 46706417 | - | lncRNA |
| ENSG00000126767.18_4 | ELK1 | 0.99999853 | 1 | 0.117 | 4.992 | chrX | 47494920 | 47510003 | - | protein_coding |
| ENSG00000247934.4_8 | AC022364.1 | 0.8556959 | 1 | 0.117 | -0.349 | chr12 | 28316231 | 28343671 | - | lncRNA |
| ENSG00000272143.1_5 | FGF14-AS2 | 0.8368481 | 1 | 0.117 | -0.684 | chr13 | 103046980 | 103048053 | + | lncRNA |
| ENSG00000166130.15_5 | IKBIP | 0.99999993 | 1 | 0.117 | 6.374 | chr12 | 99007183 | 99038891 | - | protein_coding |
| ENSG00000128581.17_5 | IFT22 | 0.99997237 | 1 | 0.117 | 4.513 | chr7 | 100954195 | 100965104 | - | protein_coding |
| ENSG00000122482.21_5 | ZNF644 | 0.99999941 | 1 | 0.117 | 6.182 | chr1 | 91380855 | 91487829 | - | protein_coding |
| ENSG00000120899.18_5 | PTK2B | 0.99999371 | 1 | 0.117 | 4.738 | chr8 | 27168999 | 27316908 | + | protein_coding |
| ENSG00000267666.2_7 | AC004156.1 | 0.85651735 | 1 | 0.116 | -0.250 | chr19 | 663482 | 669500 | + | lncRNA |
| ENSG00000189266.13_6 | PNRC2 | 0.99992933 | 1 | 0.116 | 3.648 | chr1 | 24283329 | 24289952 | + | protein_coding |
| ENSG00000203827.7 | NBPF16 | 0.98771025 | 1 | 0.116 | 1.940 | chr1 | 148739443 | 148758311 | + | protein_coding |
| ENSG00000080200.10_5 | CRYBG3 | 0.99999905 | 1 | 0.116 | 6.002 | chr3 | 97540855 | 97663828 | + | protein_coding |
| ENSG00000170365.10_4 | SMAD1 | 0.99981049 | 1 | 0.116 | 3.574 | chr4 | 146402346 | 146480328 | + | protein_coding |
| ENSG00000280388.1_5 | AC006330.1 | 0.97802979 | 1 | 0.116 | 1.508 | chr7 | 75673295 | 75675281 | - | TEC |
| ENSG00000173442.13_3 | EHBP1L1 | 0.99999995 | 1 | 0.116 | 6.245 | chr11 | 65343517 | 65360121 | + | protein_coding |
| ENSG00000105127.9_5 | AKAP8 | 0.99999358 | 1 | 0.116 | 5.019 | chr19 | 15464196 | 15490609 | - | protein_coding |
| ENSG00000160294.11_6 | MCM3AP | 0.99999962 | 1 | 0.116 | 5.977 | chr21 | 47655047 | 47706211 | - | protein_coding |
| ENSG00000105576.15_8 | TNPO2 | 0.99999956 | 1 | 0.115 | 6.012 | chr19 | 12810008 | 12834825 | - | protein_coding |
| ENSG00000197879.17_8 | MYO1C | 0.99999999 | 1 | 0.115 | 8.480 | chr17 | 1367480 | 1395980 | - | protein_coding |
| ENSG00000259891.1_6 | AC107375.1 | 0.9313876 | 1 | 0.115 | 1.006 | chr8 | 141515912 | 141518142 | - | lncRNA |
| ENSG00000269392.1_7 | AC008655.2 | 0.82921415 | 1 | 0.115 | -1.082 | chr19 | 50869137 | 50872588 | - | lncRNA |
| ENSG00000255139.2_7 | AP000442.1 | 0.83443606 | 1 | 0.115 | -0.755 | chr11 | 59333396 | 59407334 | + | lncRNA |
| ENSG00000287778.1_1 | AC006230.1 | 0.98235736 | 1 | 0.115 | 1.635 | chr4 | 11428498 | 11469426 | + | lncRNA |
| ENSG00000103769.10_4 | RAB11A | 0.99999999 | 1 | 0.115 | 7.503 | chr15 | 66018392 | 66184327 | + | protein_coding |
| ENSG00000147955.17_4 | SIGMAR1 | 0.99999997 | 1 | 0.115 | 6.390 | chr9 | 34634719 | 34637806 | - | protein_coding |
| ENSG00000157350.13_5 | ST3GAL2 | 0.99999817 | 1 | 0.115 | 5.201 | chr16 | 70409880 | 70473140 | - | protein_coding |
| ENSG00000160055.19_3 | TMEM234 | 0.99799757 | 1 | 0.115 | 2.701 | chr1 | 32680073 | 32687960 | - | protein_coding |
| ENSG00000072134.15_7 | EPN2 | 0.99998573 | 1 | 0.115 | 4.599 | chr17 | 19118928 | 19240028 | + | protein_coding |
| ENSG00000008282.8_5 | SYPL1 | 0.99999985 | 1 | 0.115 | 5.779 | chr7 | 105730949 | 105753022 | - | protein_coding |
| ENSG00000174886.13_6 | NDUFA11 | 0.99833312 | 1 | 0.115 | 2.745 | chr19 | 5891287 | 5904017 | - | protein_coding |
| ENSG00000260920.2_6 | AL031985.3 | 0.94190956 | 1 | 0.115 | 0.908 | chr1 | 40929991 | 40932439 | + | lncRNA |
| ENSG00000214046.8_4 | SMIM7 | 0.99999656 | 1 | 0.115 | 4.647 | chr19 | 16741562 | 16771253 | - | protein_coding |
| ENSG00000173013.6_4 | CCDC96 | 0.84332905 | 1 | 0.115 | -0.560 | chr4 | 7042576 | 7044728 | - | protein_coding |
| ENSG00000005981.13_3 | ASB4 | 0.85363742 | 1 | 0.115 | -0.380 | chr7 | 95107756 | 95169545 | + | protein_coding |
| ENSG00000075151.20_4 | EIF4G3 | 0.99999998 | 1 | 0.115 | 6.863 | chr1 | 21132785 | 21503381 | - | protein_coding |
| ENSG00000186451.2_3 | SPATA12 | 0.8596991 | 1 | 0.115 | -0.223 | chr3 | 57094692 | 57109460 | + | protein_coding |
| ENSG00000180346.3_2 | TIGD2 | 0.99997776 | 1 | 0.115 | 4.504 | chr4 | 90032651 | 90036050 | + | protein_coding |
| ENSG00000105963.15_8 | ADAP1 | 0.83153184 | 1 | 0.115 | -0.913 | chr7 | 937537 | 995043 | - | protein_coding |
| ENSG00000170325.14_5 | PRDM10 | 0.99996324 | 1 | 0.115 | 4.029 | chr11 | 129769601 | 129872730 | - | protein_coding |
| ENSG00000114030.13_6 | KPNA1 | 0.99999966 | 1 | 0.115 | 5.724 | chr3 | 122140749 | 122233792 | - | protein_coding |
| ENSG00000164209.17_4 | SLC25A46 | 0.99999993 | 1 | 0.114 | 6.382 | chr5 | 110073837 | 110100861 | + | protein_coding |

| | | | | | | | | | | |
|---|---|---|---|---|---|---|---|---|---|---|
| ENSG00000169155.10_4 | ZBTB43 | 0.99998661 | 1 | 0.114 | 4.226 | chr9 | 129567285 | 129600489 | + | protein_coding |
| ENSG00000067113.17_5 | PLPP1 | 0.9999439 | 1 | 0.114 | 4.069 | chr5 | 54720682 | 54830797 | - | protein_coding |
| ENSG00000100596.6_4 | SPTLC2 | 0.99999998 | 1 | 0.114 | 6.556 | chr14 | 77972340 | 78083116 | - | protein_coding |
| ENSG00000104881.16_7 | PPP1R13L | 0.9978285 | 1 | 0.114 | 2.917 | chr19 | 45882896 | 45909607 | - | protein_coding |
| ENSG00000279019.1_5 | AC009090.4 | 0.85613846 | 1 | 0.114 | -0.440 | chr16 | 57174808 | 57178498 | - | TEC |
| ENSG00000151929.9_2 | BAG3 | 0.99999948 | 1 | 0.114 | 5.707 | chr10 | 121410882 | 121437331 | + | protein_coding |
| ENSG00000065559.15_4 | MAP2K4 | 0.99999995 | 1 | 0.114 | 6.332 | chr17 | 11924146 | 12047147 | + | protein_coding |
| ENSG00000198815.9_8 | FOXJ3 | 0.99999989 | 1 | 0.114 | 6.086 | chr1 | 42642210 | 42801548 | - | protein_coding |
| ENSG00000107611.16_5 | CUBN | 0.83247545 | 1 | 0.114 | -0.994 | chr10 | 16865965 | 17171810 | - | protein_coding |
| ENSG00000131127.14_4 | ZNF141 | 0.99998131 | 1 | 0.114 | 4.740 | chr4 | 331603 | 378657 | + | protein_coding |
| ENSG00000167257.11_6 | RNF214 | 0.99897104 | 1 | 0.114 | 3.349 | chr11 | 117103341 | 117157170 | + | protein_coding |
| ENSG00000187109.14_7 | NAP1L1 | 1 | 1 | 0.113 | 9.001 | chr12 | 76430365 | 76478738 | - | protein_coding |
| ENSG00000164219.10_4 | PGGT1B | 0.99999788 | 1 | 0.113 | 5.127 | chr5 | 114539709 | 114598574 | - | protein_coding |
| ENSG00000205808.7_4 | PLPP6 | 0.99976384 | 1 | 0.113 | 4.204 | chr9 | 4662294 | 4665258 | + | protein_coding |
| ENSG00000280789.2_4 | PAGR1 | 0.99987749 | 1 | 0.113 | 3.492 | chr16 | 29827473 | 29833810 | + | protein_coding |
| ENSG00000103043.15_5 | VAC14 | 0.99999725 | 1 | 0.113 | 5.500 | chr16 | 70721342 | 70835063 | - | protein_coding |
| ENSG00000114395.10_3 | CYB561D2 | 0.99918564 | 1 | 0.113 | 2.894 | chr3 | 50388126 | 50395891 | + | protein_coding |
| ENSG00000279873.2_3 | LINC01126 | 0.96127628 | 1 | 0.113 | 1.163 | chr2 | 43454349 | 43455994 | + | lncRNA |
| ENSG00000111615.14_6 | KRR1 | 0.99999918 | 1 | 0.112 | 5.802 | chr12 | 75884643 | 75905416 | - | protein_coding |
| ENSG00000129194.8_4 | SOX15 | 0.94363183 | 1 | 0.112 | 1.008 | chr17 | 7491496 | 7493412 | - | protein_coding |
| ENSG00000227959.1_6 | AL451042.2 | 0.96123576 | 1 | 0.112 | 1.255 | chr1 | 16481706 | 16483824 | + | lncRNA |
| ENSG00000182676.5_4 | PPP1R27 | 0.88139186 | 1 | 0.112 | -0.019 | chr17 | 79791368 | 79792926 | - | protein_coding |
| ENSG00000138326.21_8 | RPS24 | 1 | 1 | 0.112 | 9.719 | chr10 | 79793518 | 79816571 | + | protein_coding |
| ENSG00000105723.12_5 | GSK3A | 0.99999971 | 1 | 0.112 | 5.763 | chr19 | 42734342 | 42746777 | - | protein_coding |
| ENSG00000185085.2_5 | INTS5 | 0.99981914 | 1 | 0.112 | 4.065 | chr11 | 62414320 | 62420774 | - | protein_coding |
| ENSG00000145012.13_6 | LPP | 0.99988402 | 1 | 0.112 | 8.268 | chr3 | 187871072 | 188608460 | + | protein_coding |
| ENSG00000172354.10_6 | GNB2 | 0.99999754 | 1 | 0.112 | 6.627 | chr7 | 100271190 | 100276797 | + | protein_coding |
| ENSG00000063322.15_5 | MED29 | 0.99999997 | 1 | 0.112 | 6.522 | chr19 | 39881943 | 39891281 | + | protein_coding |
| ENSG00000163902.12_5 | RPN1 | 0.99999999 | 1 | 0.112 | 7.934 | chr3 | 128338812 | 128399918 | - | protein_coding |
| ENSG00000188021.8_3 | UBQLN2 | 0.99999943 | 1 | 0.112 | 5.138 | chrX | 56590072 | 56594301 | + | protein_coding |
| ENSG00000267530.3_5 | LINC01836 | 0.99928779 | 1 | 0.112 | 3.415 | chr19 | 782755 | 786092 | + | lncRNA |
| ENSG00000023191.17_4 | RNH1 | 0.99999912 | 1 | 0.112 | 7.230 | chr11 | 494512 | 507300 | - | protein_coding |
| ENSG00000064270.12_5 | ATP2C2 | 0.88393296 | 1 | 0.112 | 0.201 | chr16 | 84402133 | 84497793 | + | protein_coding |
| ENSG00000100028.12_4 | SNRPD3 | 0.99999849 | 1 | 0.111 | 4.858 | chr22 | 24951926 | 24978020 | + | protein_coding |
| ENSG00000158825.6_3 | CDA | 0.95531799 | 1 | 0.111 | 1.026 | chr1 | 20915579 | 20945396 | + | protein_coding |
| ENSG00000248508.8_6 | SRP14-AS1 | 0.96870049 | 1 | 0.111 | 1.339 | chr15 | 40331443 | 40368740 | + | lncRNA |
| ENSG00000185475.11_5 | TMEM179B | 0.99997153 | 1 | 0.111 | 4.095 | chr11 | 62554874 | 62557872 | + | protein_coding |
| ENSG00000147123.10_3 | NDUFB11 | 0.99999977 | 1 | 0.111 | 5.710 | chrX | 47001615 | 47004903 | - | protein_coding |
| ENSG00000109046.15_3 | WSB1 | 0.99999987 | 1 | 0.111 | 6.308 | chr17 | 25621102 | 25642952 | + | protein_coding |
| ENSG00000030110.13_4 | BAK1 | 0.99979491 | 1 | 0.111 | 3.402 | chr6 | 33540324 | 33548070 | - | protein_coding |
| ENSG00000118482.11_4 | PHF3 | 0.99999996 | 1 | 0.111 | 7.144 | chr6 | 64345725 | 64489229 | + | protein_coding |
| ENSG00000114270.17_4 | COL7A1 | 0.99964673 | 1 | 0.111 | 8.943 | chr3 | 48601506 | 48632700 | - | protein_coding |
| ENSG00000213625.9_5 | LEPROT | 0.99999995 | 1 | 0.111 | 6.181 | chr1 | 65886270 | 65901690 | + | protein_coding |
| ENSG00000142227.11_4 | EMP3 | 0.99999994 | 1 | 0.111 | 6.567 | chr19 | 48824766 | 48833810 | + | protein_coding |
| ENSG00000071189.21_4 | SNX13 | 0.99999987 | 1 | 0.111 | 6.212 | chr7 | 17830385 | 17980124 | - | protein_coding |
| ENSG00000179262.10_4 | RAD23A | 0.99999968 | 1 | 0.111 | 5.769 | chr19 | 13056669 | 13064456 | + | protein_coding |
| ENSG00000150630.4_4 | VEGFC | 0.998614 | 1 | 0.111 | 2.747 | chr4 | 177604689 | 177714076 | - | protein_coding |
| ENSG00000015532.10_3 | XYLT2 | 0.99999912 | 1 | 0.111 | 5.576 | chr17 | 48423487 | 48440499 | + | protein_coding |
| ENSG00000173992.9_4 | CCS | 0.99996783 | 1 | 0.111 | 4.333 | chr11 | 66360624 | 66373490 | + | protein_coding |
| ENSG00000260947.1_6 | AL356489.2 | 0.89665484 | 1 | 0.111 | 0.337 | chr9 | 33697457 | 33700984 | + | lncRNA |
| ENSG00000138802.11_3 | SEC24B | 0.99999955 | 1 | 0.111 | 5.577 | chr4 | 110354928 | 110462052 | + | protein_coding |
| ENSG00000276234.5_6 | TADA2A | 0.99999844 | 1 | 0.110 | 4.906 | chr17 | 35766977 | 35839830 | + | protein_coding |
| ENSG00000251034.2_7 | AC087854.1 | 0.97474782 | 1 | 0.110 | 0.984 | chr8 | 22339101 | 22410957 | - | lncRNA |
| ENSG00000172315.6_5 | TP53RK | 0.99999186 | 1 | 0.110 | 4.350 | chr20 | 45313004 | 45318083 | - | protein_coding |
| ENSG00000271826.5_7 | PLS3-AS1 | 0.86049916 | 1 | 0.110 | 0.083 | chrX | 114752490 | 114797058 | - | lncRNA |
| ENSG00000250999.2_7 | AC136604.3 | 0.86540452 | 1 | 0.110 | -0.326 | chr5 | 179084763 | 179092137 | + | lncRNA |
| ENSG00000164512.18_5 | ANKRD55 | 0.87786301 | 1 | 0.110 | -0.201 | chr5 | 55395507 | 55529157 | - | protein_coding |
| ENSG00000283297.2_7 | TEX52 | 0.83991703 | 1 | 0.110 | -0.912 | chr12 | 2958212 | 2966205 | - | protein_coding |
| ENSG00000088038.19_6 | CNOT3 | 0.9999942 | 1 | 0.110 | 4.891 | chr19 | 54641465 | 54659419 | + | protein_coding |
| ENSG00000167770.11_3 | OTUB1 | 0.99999988 | 1 | 0.110 | 6.119 | chr11 | 63753325 | 63769283 | + | protein_coding |
| ENSG00000245060.8_6 | LINC00847 | 0.97897098 | 1 | 0.110 | 1.120 | chr5 | 180257326 | 180266742 | + | lncRNA |

| | | | | | | | | | | |
|---|---|---|---|---|---|---|---|---|---|---|
| ENSG00000090520.11_3 | DNAJB11 | 0.99999804 | 1 | 0.110 | 4.816 | chr3 | 186285192 | 186303589 | + | protein_coding |
| ENSG00000238715.1 | snoU13 | 0.83657028 | 1 | 0.110 | -1.180 | chr15 | 65863311 | 65863411 | - | snoRNA |
| ENSG00000258289.8_8 | CHURC1 | 0.99999894 | 1 | 0.110 | 5.321 | chr14 | 65381079 | 65411309 | + | protein_coding |
| ENSG00000177728.17_7 | TMEM94 | 0.99999991 | 1 | 0.110 | 6.297 | chr17 | 73437240 | 73496533 | + | protein_coding |
| ENSG00000143614.10_5 | GATAD2B | 0.99999841 | 1 | 0.109 | 4.963 | chr1 | 153761506 | 153895836 | - | protein_coding |
| ENSG00000104765.16_4 | BNIP3L | 0.99999994 | 1 | 0.109 | 6.575 | chr8 | 26240570 | 26363152 | + | protein_coding |
| ENSG00000100601.10_5 | ALKBH1 | 0.99996628 | 1 | 0.109 | 4.188 | chr14 | 78138747 | 78174366 | - | protein_coding |
| ENSG00000136828.19_5 | RALGPS1 | 0.99960589 | 1 | 0.109 | 3.074 | chr9 | 129677053 | 129985445 | + | protein_coding |
| ENSG00000133874.2_4 | RNF122 | 0.98896734 | 1 | 0.109 | 1.907 | chr8 | 33405272 | 33424646 | - | protein_coding |
| ENSG00000080802.18_4 | CNOT4 | 0.99999139 | 1 | 0.109 | 4.500 | chr7 | 135046547 | 135194875 | - | protein_coding |
| ENSG00000160741.17_6 | CRTC2 | 0.99998377 | 1 | 0.109 | 4.185 | chr1 | 153920145 | 153931091 | - | protein_coding |
| ENSG00000107186.16_6 | MPDZ | 0.99999978 | 1 | 0.109 | 6.259 | chr9 | 13105703 | 13279589 | - | protein_coding |
| ENSG00000103599.20_7 | IQCH | 0.99553042 | 1 | 0.109 | 2.360 | chr15 | 67547124 | 67794598 | + | protein_coding |
| ENSG00000187049.10_4 | TMEM216 | 0.99898033 | 1 | 0.109 | 2.845 | chr11 | 61159832 | 61166335 | + | protein_coding |
| ENSG00000224063.6_6 | AC007319.1 | 0.91826628 | 1 | 0.109 | 0.891 | chr2 | 187867947 | 188421015 | + | lncRNA |
| ENSG00000261286.1_7 | ATP2C2-AS1 | 0.84486505 | 1 | 0.109 | -0.728 | chr16 | 84492865 | 84500967 | - | lncRNA |
| ENSG00000077585.14_7 | GPR137B | 0.99974524 | 1 | 0.109 | 3.267 | chr1 | 236305805 | 236385165 | + | protein_coding |
| ENSG00000040633.13_5 | PHF23 | 0.99999992 | 1 | 0.109 | 5.769 | chr17 | 7138348 | 7143041 | - | protein_coding |
| ENSG00000281706.3_4 | AL009179.1 | 0.94476948 | 1 | 0.109 | 0.893 | chr6 | 27661814 | 27681131 | + | lncRNA |
| ENSG00000105355.9_6 | PLIN3 | 0.999893 | 1 | 0.109 | 5.668 | chr19 | 4838353 | 4867706 | - | protein_coding |
| ENSG00000157021.9_4 | FAM92A1P1 | 0.91662011 | 1 | 0.109 | 0.545 | chr15 | 41455360 | 41456572 | + | unprocessed_pseudogene |
| ENSG00000232838.4_4 | PET117 | 0.91458691 | 1 | 0.109 | 0.161 | chr20 | 18118507 | 18123813 | + | protein_coding |
| ENSG00000227417.3_5 | AC114402.2 | 0.84043445 | 1 | 0.108 | -0.696 | chr1 | 203774749 | 203775391 | - | processed_pseudogene |
| ENSG00000224975.1_4 | INE1 | 0.96221139 | 1 | 0.108 | 0.734 | chrX | 47064320 | 47065264 | + | lncRNA |
| ENSG00000183150.8_6 | GPR19 | 0.95351967 | 1 | 0.108 | 0.701 | chr12 | 12813824 | 12849141 | - | protein_coding |
| ENSG00000249115.9_4 | HAUS5 | 0.99999896 | 1 | 0.108 | 5.114 | chr19 | 36103637 | 36116257 | + | protein_coding |
| ENSG00000027697.14_7 | IFNGR1 | 0.99999523 | 1 | 0.108 | 4.773 | chr6 | 137518621 | 137540586 | - | protein_coding |
| ENSG00000125952.20_7 | MAX | 0.99999798 | 1 | 0.108 | 4.849 | chr14 | 65472892 | 65569413 | - | protein_coding |
| ENSG00000132406.12_4 | TMEM128 | 0.99907562 | 1 | 0.108 | 3.379 | chr4 | 4237269 | 4249950 | - | protein_coding |
| ENSG00000143207.21_8 | COP1 | 0.99999928 | 1 | 0.108 | 5.483 | chr1 | 175913967 | 176176422 | - | protein_coding |
| ENSG00000230513.2_6 | THAP7-AS1 | 0.87289894 | 1 | 0.108 | -0.252 | chr22 | 21356175 | 21364631 | + | lncRNA |
| ENSG00000128596.17_6 | CCDC136 | 0.98714343 | 1 | 0.107 | 1.789 | chr7 | 128430811 | 128462186 | + | protein_coding |
| ENSG00000177842.12_4 | ZNF620 | 0.99968361 | 1 | 0.107 | 3.201 | chr3 | 40518622 | 40560227 | + | protein_coding |
| ENSG00000116731.22_5 | PRDM2 | 0.99997517 | 1 | 0.107 | 4.076 | chr1 | 14026693 | 14151574 | + | protein_coding |
| ENSG00000090432.7_4 | MUL1 | 0.99999218 | 1 | 0.107 | 4.742 | chr1 | 20825941 | 20834644 | - | protein_coding |
| ENSG00000253598.3_4 | SLC10A5 | 0.86034813 | 1 | 0.107 | -0.562 | chr8 | 82605866 | 82607293 | - | protein_coding |
| ENSG00000262188.2_5 | LINC01978 | 0.87256627 | 1 | 0.107 | -0.358 | chr17 | 77893156 | 77899798 | + | lncRNA |
| ENSG00000122678.17_5 | POLM | 0.99991781 | 1 | 0.107 | 3.606 | chr7 | 44111661 | 44122129 | - | protein_coding |
| ENSG00000253428.1_5 | LINC01942 | 0.93008393 | 1 | 0.107 | 0.604 | chr5 | 173116462 | 173132852 | + | lncRNA |
| ENSG00000272630.1_7 | AL731563.4 | 0.84394691 | 1 | 0.107 | -0.914 | chr10 | 74857802 | 74861055 | - | lncRNA |
| ENSG00000255121.2_8 | CCDC84-DT | 0.97304347 | 1 | 0.107 | 1.362 | chr11 | 118865534 | 118868714 | - | lncRNA |
| ENSG00000101752.11_2 | MIB1 | 0.99999997 | 1 | 0.107 | 7.019 | chr18 | 19284918 | 19450918 | + | protein_coding |
| ENSG00000226752.9_5 | CUTALP | 0.99999926 | 1 | 0.107 | 4.920 | chr9 | 123587106 | 123616663 | + | transcribed_unitary_pseudogene |
| ENSG00000182199.11_6 | SHMT2 | 0.99999998 | 1 | 0.107 | 6.615 | chr12 | 57623356 | 57628718 | + | protein_coding |
| ENSG00000146707.14_5 | POMZP3 | 0.99236998 | 1 | 0.106 | 2.276 | chr7 | 76239303 | 76256578 | - | protein_coding |
| ENSG00000211459.2 | MT-RNR1 | 1 | 1 | 0.106 | 13.629 | chrM | 648 | 1601 | + | Mt_rRNA |
| ENSG00000233860.2_7 | AC107072.2 | 0.88268891 | 1 | 0.106 | -0.151 | chr4 | 77630006 | 77723579 | - | lncRNA |
| ENSG00000130699.18_4 | TAF4 | 0.99998223 | 1 | 0.106 | 4.165 | chr20 | 60528525 | 60640866 | - | protein_coding |
| ENSG00000144339.12_4 | TMEFF2 | 0.92267219 | 1 | 0.106 | 0.307 | chr2 | 192813769 | 193059659 | - | protein_coding |
| ENSG00000183726.11_3 | TMEM50A | 0.9999999 | 1 | 0.106 | 5.876 | chr1 | 25664808 | 25688852 | + | protein_coding |
| ENSG00000090932.11_3 | DLL3 | 0.99701553 | 1 | 0.106 | 2.651 | chr19 | 39989535 | 39999121 | + | protein_coding |
| ENSG00000132356.11_3 | PRKAA1 | 0.99999999 | 1 | 0.106 | 7.133 | chr5 | 40759481 | 40798476 | - | protein_coding |
| ENSG00000172638.13_5 | EFEMP2 | 0.98789766 | 1 | 0.106 | 1.778 | chr11 | 65633912 | 65641063 | - | protein_coding |
| ENSG00000172057.10_5 | ORMDL3 | 0.99999926 | 1 | 0.106 | 5.060 | chr17 | 38077294 | 38083854 | - | protein_coding |
| ENSG00000167377.18_8 | ZNF23 | 0.94685361 | 1 | 0.106 | 0.632 | chr16 | 71481500 | 71496998 | - | protein_coding |
| ENSG00000197586.13_8 | ENTPD6 | 0.99999996 | 1 | 0.106 | 6.705 | chr20 | 25176329 | 25208711 | + | protein_coding |
| ENSG00000105341.18_7 | DMAC2 | 0.99999991 | 1 | 0.106 | 6.128 | chr19 | 41937223 | 41946622 | - | protein_coding |
| ENSG00000078369.18_4 | GNB1 | 0.99999999 | 1 | 0.105 | 7.674 | chr1 | 1716724 | 1822556 | - | protein_coding |
| ENSG00000204590.12_3 | GNL1 | 0.99993156 | 1 | 0.105 | 4.033 | chr6 | 30509154 | 30524951 | - | protein_coding |

| | | | | | | | | | | |
|---|---|---|---|---|---|---|---|---|---|---|
| ENSG00000136925.15_4 | TSTD2 | 0.99996854 | 1 | 0.105 | 3.933 | chr9 | 100362362 | 100395650 | - | protein_coding |
| ENSG00000092203.15_9 | TOX4 | 0.99999993 | 1 | 0.105 | 5.844 | chr14 | 21944756 | 21967319 | + | protein_coding |
| ENSG00000224950.2_7 | AL390066.2 | 0.84763549 | 1 | 0.105 | -0.878 | chr1 | 117035638 | 117041834 | - | lncRNA |
| ENSG00000267080.6_8 | ASB16-AS1 | 0.9991981 | 1 | 0.105 | 3.080 | chr17 | 42253336 | 42264091 | - | lncRNA |
| ENSG00000168734.14_3 | PKIG | 0.9999916 | 1 | 0.105 | 4.294 | chr20 | 43160426 | 43252888 | + | protein_coding |
| ENSG00000169105.7_2 | CHST14 | 0.99995419 | 1 | 0.105 | 3.893 | chr15 | 40763197 | 40766770 | + | protein_coding |
| ENSG00000166068.13_4 | SPRED1 | 0.99999967 | 1 | 0.105 | 5.709 | chr15 | 38545037 | 38649450 | + | protein_coding |
| ENSG00000116260.17_3 | QSOX1 | 0.99999989 | 1 | 0.105 | 5.970 | chr1 | 180124004 | 180173165 | + | protein_coding |
| ENSG00000134884.15_7 | ARGLU1 | 0.99999998 | 1 | 0.105 | 6.681 | chr13 | 107194021 | 107220485 | - | protein_coding |
| ENSG00000170242.18_4 | USP47 | 0.99999966 | 1 | 0.105 | 6.287 | chr11 | 11862970 | 11983434 | + | protein_coding |
| ENSG00000147454.14_3 | SLC25A37 | 0.99999997 | 1 | 0.105 | 6.375 | chr8 | 23386469 | 23432976 | + | protein_coding |
| ENSG00000163898.10_4 | LIPH | 0.99999823 | 1 | 0.105 | 4.871 | chr3 | 185224050 | 185270376 | - | protein_coding |
| ENSG00000222046.2_2 | DCDC2B | 0.95020886 | 1 | 0.104 | 0.896 | chr1 | 32674695 | 32681797 | + | protein_coding |
| ENSG00000112697.17_5 | TMEM30A | 0.99999998 | 1 | 0.104 | 7.217 | chr6 | 75962640 | 75994664 | - | protein_coding |
| ENSG00000148248.14_4 | SURF4 | 1 | 1 | 0.104 | 7.966 | chr9 | 136228326 | 136243037 | - | protein_coding |
| ENSG00000228502.1_4 | EEF1A1P11 | 0.92979657 | 1 | 0.104 | 0.726 | chr1 | 96912486 | 96913874 | + | processed_pseudogene |
| ENSG00000118454.13_4 | ANKRD13C | 0.99999932 | 1 | 0.104 | 5.116 | chr1 | 70724682 | 70820417 | - | protein_coding |
| ENSG00000196227.11_5 | FAM217B | 0.99995883 | 1 | 0.104 | 3.969 | chr20 | 58508819 | 58523735 | + | protein_coding |
| ENSG00000167985.6_3 | SDHAF2 | 0.99730069 | 1 | 0.104 | 2.430 | chr11 | 61197514 | 61215001 | + | protein_coding |
| ENSG00000175854.12_5 | SWI5 | 0.9999532 | 1 | 0.104 | 4.029 | chr9 | 131037658 | 131078402 | + | protein_coding |
| ENSG00000006451.8_4 | RALA | 0.99999809 | 1 | 0.104 | 4.799 | chr7 | 39663164 | 39747719 | + | protein_coding |
| ENSG00000140987.20_5 | ZSCAN32 | 0.99989611 | 1 | 0.104 | 3.729 | chr16 | 3432081 | 3451065 | - | protein_coding |
| ENSG00000099875.14_5 | MKNK2 | 0.99999406 | 1 | 0.104 | 5.599 | chr19 | 2037464 | 2051243 | - | protein_coding |
| ENSG00000120008.16_3 | WDR11 | 0.99999997 | 1 | 0.104 | 6.683 | chr10 | 122610817 | 122669036 | + | protein_coding |
| ENSG00000255062.1_7 | AP001318.2 | 0.84600016 | 1 | 0.103 | -1.047 | chr11 | 126164193 | 126174213 | - | lncRNA |
| ENSG00000137274.13_4 | BPHL | 0.99959767 | 1 | 0.103 | 3.196 | chr6 | 3118608 | 3153812 | + | protein_coding |
| ENSG00000137494.14_5 | ANKRD42 | 0.99985407 | 1 | 0.103 | 3.363 | chr11 | 82904754 | 82971736 | + | protein_coding |
| ENSG00000100612.14_5 | DHRS7 | 0.99999914 | 1 | 0.103 | 5.111 | chr14 | 60610837 | 60636574 | - | protein_coding |
| ENSG00000107798.18_5 | LIPA | 0.9999999 | 1 | 0.103 | 6.458 | chr10 | 90973326 | 91174314 | - | protein_coding |
| ENSG00000273226.1_6 | AL391834.2 | 0.9292018 | 1 | 0.103 | 0.530 | chr9 | 19375449 | 19375994 | + | lncRNA |
| ENSG00000173295.7_4 | FAM86B3P | 0.97601484 | 1 | 0.103 | 1.306 | chr8 | 8086117 | 8102387 | + | transcribed_unprocessed_pseudogene |
| ENSG00000167625.11_4 | ZNF526 | 0.99999984 | 1 | 0.103 | 5.462 | chr19 | 42724464 | 42732353 | + | protein_coding |
| ENSG00000114383.10_5 | TUSC2 | 0.99997103 | 1 | 0.103 | 3.882 | chr3 | 50357458 | 50365682 | - | protein_coding |
| ENSG00000099377.14_5 | HSD3B7 | 0.99961379 | 1 | 0.103 | 3.109 | chr16 | 30996528 | 31000468 | + | protein_coding |
| ENSG00000113734.17_5 | BNIP1 | 0.99918369 | 1 | 0.103 | 3.229 | chr5 | 172571445 | 172591390 | + | protein_coding |
| ENSG00000125386.15_5 | FAM193A | 0.999998 | 1 | 0.103 | 4.794 | chr4 | 2538358 | 2734292 | + | protein_coding |
| ENSG00000169926.11_4 | KLF13 | 0.99999999 | 1 | 0.103 | 7.294 | chr15 | 31619038 | 31727868 | + | protein_coding |
| ENSG00000007376.8_5 | RPUSD1 | 0.99808584 | 1 | 0.102 | 4.730 | chr16 | 834974 | 838397 | - | protein_coding |
| ENSG00000163364.10_7 | LINC01116 | 0.99792329 | 1 | 0.102 | 2.699 | chr2 | 177494300 | 177502659 | - | lncRNA |
| ENSG00000100473.18_5 | COCH | 0.99999314 | 1 | 0.102 | 4.379 | chr14 | 31343720 | 31364271 | + | protein_coding |
| ENSG00000183340.7_3 | JRKL | 0.99984132 | 1 | 0.102 | 3.428 | chr11 | 96123153 | 96240738 | + | protein_coding |
| ENSG00000103051.19_7 | COG4 | 0.99999998 | 1 | 0.102 | 6.296 | chr16 | 70514471 | 70557461 | - | protein_coding |
| ENSG00000171720.10_4 | HDAC3 | 0.9999991 | 1 | 0.102 | 5.002 | chr5 | 141000443 | 141016416 | - | protein_coding |
| ENSG00000172932.15_5 | ANKRD13D | 0.99999627 | 1 | 0.102 | 5.006 | chr11 | 67056771 | 67069956 | + | protein_coding |
| ENSG00000162191.13_6 | UBXN1 | 0.99999985 | 1 | 0.102 | 5.508 | chr11 | 62443970 | 62446589 | - | protein_coding |
| ENSG00000166454.10_4 | ATMIN | 0.99999999 | 1 | 0.102 | 6.901 | chr16 | 81069447 | 81080955 | + | protein_coding |
| ENSG00000148219.16_5 | ASTN2 | 0.99958583 | 1 | 0.102 | 3.136 | chr9 | 119187504 | 120177348 | - | protein_coding |
| ENSG00000265519.1_7 | AC015922.2 | 0.98862041 | 1 | 0.102 | 1.749 | chr17 | 15691101 | 15691519 | - | lncRNA |
| ENSG00000113441.16_5 | LNPEP | 0.99999985 | 1 | 0.102 | 6.811 | chr5 | 96271098 | 96373217 | + | protein_coding |
| ENSG00000168175.15_3 | MAPK1IP1L | 1 | 1 | 0.102 | 7.441 | chr14 | 55518365 | 55536912 | + | protein_coding |
| ENSG00000278876.1_5 | AC145207.9 | 0.96536236 | 1 | 0.102 | 1.095 | chr17 | 79820654 | 79820978 | + | TEC |
| ENSG00000102390.11_4 | PBDC1 | 0.99973914 | 1 | 0.102 | 3.382 | chrX | 75392875 | 75398149 | + | protein_coding |
| ENSG00000167548.15_7 | KMT2D | 0.99999855 | 1 | 0.102 | 5.884 | chr12 | 49412758 | 49453557 | - | protein_coding |
| ENSG00000182768.8_3 | NGRN | 0.99998539 | 1 | 0.102 | 5.580 | chr15 | 90808891 | 90821373 | + | protein_coding |
| ENSG00000164091.12_4 | WDR82 | 0.99999998 | 1 | 0.101 | 6.431 | chr3 | 52288450 | 52322036 | - | protein_coding |
| ENSG00000198668.12_5 | CALM1 | 0.99999999 | 1 | 0.101 | 8.294 | chr14 | 90862846 | 90874612 | + | protein_coding |
| ENSG00000170185.10_5 | USP38 | 0.99999922 | 1 | 0.101 | 5.084 | chr4 | 144106070 | 144145027 | + | protein_coding |
| ENSG00000106404.13_8 | CLDN15 | 0.99991367 | 1 | 0.101 | 3.713 | chr7 | 100875373 | 100882101 | - | protein_coding |
| ENSG00000205771.6_4 | CATSPER2P1 | 0.88119769 | 1 | 0.101 | -0.067 | chr15 | 44019116 | 44039292 | - | transcribed_unprocessed_pseudogene |

| | | | | | | | | | | |
|---|---|---|---|---|---|---|---|---|---|---|
| ENSG00000133740.11_4 | E2F5 | 0.99991455 | 1 | 0.101 | 3.789 | chr8 | 86089389 | 86129387 | + | protein_coding |
| ENSG00000272556.2_7 | GTF2IP13 | 0.91395924 | 1 | 0.101 | 0.403 | chr7 | 45808704 | 45854862 | + | transcribed_unprocessed_pseudogene |
| ENSG00000282978.1_6 | AC110994.3 | 0.88812304 | 1 | 0.101 | -0.046 | chr3 | 101176085 | 101176576 | - | processed_pseudogene |
| ENSG00000135480.16_4 | KRT7 | 0.99999999 | 1 | 0.101 | 10.649 | chr12 | 52626304 | 52645970 | + | protein_coding |
| ENSG00000152778.9_3 | IFIT5 | 0.99999807 | 1 | 0.101 | 5.140 | chr10 | 91174325 | 91180754 | + | protein_coding |
| ENSG00000141447.18_5 | OSBPL1A | 0.99998442 | 1 | 0.101 | 4.305 | chr18 | 21742009 | 21977844 | - | protein_coding |
| ENSG00000184517.12_3 | ZFP1 | 0.9999971 | 1 | 0.101 | 4.569 | chr16 | 75182392 | 75206134 | + | protein_coding |
| ENSG00000181704.12_4 | YIPF6 | 0.99999998 | 1 | 0.100 | 6.389 | chrX | 67718404 | 67757124 | + | protein_coding |
| ENSG00000176209.11_2 | SMIM19 | 0.99998149 | 1 | 0.100 | 4.075 | chr8 | 42396298 | 42410336 | + | protein_coding |
| ENSG00000004777.18_3 | ARHGAP33 | 0.99823914 | 1 | 0.100 | 2.534 | chr19 | 36265434 | 36279724 | + | protein_coding |
| ENSG00000049860.14_4 | HEXB | 0.99999998 | 1 | 0.100 | 6.536 | chr5 | 73935848 | 74018472 | + | protein_coding |
| ENSG00000124098.10_3 | FAM210B | 0.9999957 | 1 | 0.100 | 4.484 | chr20 | 54934030 | 54943719 | + | protein_coding |
| ENSG00000196199.14_3 | MPHOSPH8 | 0.9999974 | 1 | 0.100 | 4.582 | chr13 | 20207799 | 20247581 | + | protein_coding |
| ENSG00000226950.7_6 | DANCR | 0.99994099 | 1 | 0.100 | 3.898 | chr4 | 53578571 | 53586518 | + | lncRNA |
| ENSG00000169612.4_5 | RAMAC | 0.99418735 | 1 | 0.100 | 2.403 | chr15 | 83654962 | 83659809 | + | protein_coding |
| ENSG00000095951.17_4 | HIVEP1 | 0.99999551 | 1 | 0.100 | 4.839 | chr6 | 12008995 | 12165232 | + | protein_coding |
| ENSG00000126790.12_5 | L3HYPDH | 0.99999828 | 1 | 0.100 | 4.943 | chr14 | 59927081 | 59951126 | - | protein_coding |
| ENSG00000066654.14_6 | THUMPD1 | 0.99999999 | 1 | 0.100 | 6.523 | chr16 | 20714138 | 20753406 | - | protein_coding |
| ENSG00000128228.5_3 | SDF2L1 | 0.99961547 | 1 | 0.099 | 3.316 | chr22 | 21996591 | 21998588 | + | protein_coding |
| ENSG00000071626.16_4 | DAZAP1 | 0.99999957 | 1 | 0.099 | 6.623 | chr19 | 1407568 | 1435686 | + | protein_coding |
| ENSG00000273604.2_3 | EPOP | 0.99996005 | 1 | 0.099 | 4.158 | chr17 | 36827956 | 36831210 | - | protein_coding |
| ENSG00000162909.18_3 | CAPN2 | 1 | 1 | 0.099 | 9.116 | chr1 | 223889295 | 223963720 | + | protein_coding |
| ENSG00000137076.21_5 | TLN1 | 1 | 1 | 0.099 | 8.649 | chr9 | 35696945 | 35732192 | - | protein_coding |
| ENSG00000180011.7_5 | ZADH2 | 0.99999424 | 1 | 0.099 | 4.454 | chr18 | 72907068 | 72921094 | - | protein_coding |
| ENSG00000178440.5 | LINC00843 | 0.93355769 | 1 | 0.099 | 0.337 | chr10 | 51732523 | 51741813 | + | lincRNA |
| ENSG00000105323.17_5 | HNRNPUL1 | 0.99999999 | 1 | 0.099 | 8.415 | chr19 | 41768401 | 41813692 | + | protein_coding |
| ENSG00000173786.17_6 | CNP | 0.99999995 | 1 | 0.099 | 6.818 | chr17 | 40118781 | 40129758 | + | protein_coding |
| ENSG00000142102.16_9 | PGGHG | 0.99999897 | 1 | 0.099 | 5.212 | chr11 | 289126 | 296107 | + | protein_coding |
| ENSG00000010626.15_5 | LRRC23 | 0.99975246 | 1 | 0.099 | 3.353 | chr12 | 6982733 | 7023405 | + | protein_coding |
| ENSG00000183978.8_6 | COA3 | 0.9999995 | 1 | 0.099 | 5.508 | chr17 | 40947165 | 40950722 | - | protein_coding |
| ENSG00000197343.11_8 | ZNF655 | 0.99999992 | 1 | 0.099 | 6.084 | chr7 | 99156029 | 99174076 | + | protein_coding |
| ENSG00000196459.14_4 | TRAPPC2 | 0.99999565 | 1 | 0.098 | 4.314 | chrX | 13730363 | 13752754 | - | protein_coding |
| ENSG00000100109.17_5 | TFIP11 | 0.99999883 | 1 | 0.098 | 4.850 | chr22 | 26887191 | 26908471 | - | protein_coding |
| ENSG00000113532.13_4 | ST8SIA4 | 0.99999947 | 1 | 0.098 | 5.047 | chr5 | 100142637 | 100238986 | - | protein_coding |
| ENSG00000172531.15_6 | PPP1CA | 0.99999998 | 1 | 0.098 | 7.131 | chr11 | 67165654 | 67188654 | - | protein_coding |
| ENSG00000138107.13_5 | ACTR1A | 0.99999999 | 1 | 0.098 | 6.935 | chr10 | 104221638 | 104262469 | - | protein_coding |
| ENSG00000242689.3_4 | CNTF | 0.8742117 | 1 | 0.098 | 0.022 | chr11 | 58390138 | 58393206 | + | protein_coding |
| ENSG00000124067.17_7 | SLC12A4 | 0.99999981 | 1 | 0.098 | 5.585 | chr16 | 67977377 | 68003504 | - | protein_coding |
| ENSG00000250251.6_3 | PKD1P6 | 0.99814683 | 1 | 0.098 | 3.378 | chr16 | 15219099 | 15248421 | - | transcribed_unprocessed_pseudogene |
| ENSG00000240291.1_8 | AL450384.2 | 0.89439553 | 1 | 0.098 | -0.261 | chr10 | 18802044 | 18834580 | - | lncRNA |
| ENSG00000166166.13_3 | TRMT61A | 0.99998828 | 1 | 0.098 | 4.840 | chr14 | 103995533 | 104003410 | + | protein_coding |
| ENSG00000141577.14_7 | CEP131 | 0.99828107 | 1 | 0.098 | 4.523 | chr17 | 79163393 | 79196799 | - | protein_coding |
| ENSG00000183048.12_4 | SLC25A10 | 0.99961126 | 1 | 0.098 | 3.179 | chr17 | 79679266 | 79688046 | + | protein_coding |
| ENSG00000157625.15_4 | TAB3 | 0.99999701 | 1 | 0.098 | 4.616 | chrX | 30845559 | 30993201 | - | protein_coding |
| ENSG00000257267.3_2 | ZNF271P | 0.99999823 | 1 | 0.098 | 4.586 | chr18 | 32870246 | 32908354 | + | transcribed_unitary_pseudogene |
| ENSG00000160828.13 | STAG3L2 | 0.99811455 | 1 | 0.098 | 2.426 | chr7 | 74112305 | 74306731 | - | pseudogene |
| ENSG00000113161.16_4 | HMGCR | 0.99999994 | 1 | 0.097 | 6.301 | chr5 | 74632154 | 74657926 | + | protein_coding |
| ENSG00000179335.18_3 | CLK3 | 0.99999873 | 1 | 0.097 | 5.160 | chr15 | 74890841 | 74937755 | + | protein_coding |
| ENSG00000131095.13_6 | GFAP | 0.94305361 | 1 | 0.097 | 0.738 | chr17 | 42980527 | 42994305 | - | protein_coding |
| ENSG00000138767.13_5 | CNOT6L | 0.99999668 | 1 | 0.097 | 4.464 | chr4 | 78634541 | 78740769 | - | protein_coding |
| ENSG00000158220.14_3 | ESYT3 | 0.97563769 | 1 | 0.097 | 1.324 | chr3 | 138153428 | 138200528 | + | protein_coding |
| ENSG00000266490.1_6 | AC127024.5 | 0.8573573 | 1 | 0.097 | -0.769 | chr17 | 29119390 | 29119851 | + | lncRNA |
| ENSG00000067369.13_5 | TP53BP1 | 0.99999996 | 1 | 0.097 | 6.311 | chr15 | 43695259 | 43802926 | - | protein_coding |
| ENSG00000271851.1_7 | AC087501.4 | 0.98769353 | 1 | 0.097 | 1.155 | chr17 | 9456640 | 9459013 | - | lncRNA |
| ENSG00000136826.15_6 | KLF4 | 0.99964955 | 1 | 0.097 | 3.422 | chr9 | 110247133 | 110252763 | - | protein_coding |
| ENSG00000111300.10_5 | NAA25 | 0.99999996 | 1 | 0.097 | 6.304 | chr12 | 112464493 | 112546600 | - | protein_coding |
| ENSG00000111639.8_6 | MRPL51 | 0.99999998 | 1 | 0.097 | 6.485 | chr12 | 6601052 | 6603007 | - | protein_coding |
| ENSG00000241962.9_6 | AC079447.1 | 0.86245712 | 1 | 0.097 | -0.770 | chr2 | 99757948 | 99939204 | + | protein_coding |
| ENSG00000182165.18_4 | TP53TG1 | 0.99835129 | 1 | 0.097 | 2.731 | chr7 | 86952259 | 86974844 | - | lncRNA |

| | | | | | | | | | | |
|---|---|---|---|---|---|---|---|---|---|---|
| ENSG00000163527.10_3 | STT3B | 1 | 1 | 0.096 | 8.348 | chr3 | 31574130 | 31679108 | + | protein_coding |
| ENSG00000007237.18_7 | AC005747.1 | 0.95117837 | 1 | 0.096 | 0.767 | chr17 | 9813926 | 10101868 | - | protein_coding |
| ENSG00000155975.10_5 | VPS37A | 0.99999989 | 1 | 0.096 | 5.735 | chr8 | 17104440 | 17159936 | + | protein_coding |
| ENSG00000146112.12_5 | PPP1R18 | 0.97198548 | 1 | 0.096 | 1.166 | chr6 | 30644166 | 30655672 | - | protein_coding |
| ENSG00000113300.13_4 | CNOT6 | 0.99999982 | 1 | 0.096 | 5.473 | chr5 | 179921379 | 180005358 | + | protein_coding |
| ENSG00000140575.13_4 | IQGAP1 | 1 | 1 | 0.096 | 8.981 | chr15 | 90931474 | 91045471 | + | protein_coding |
| ENSG00000008405.12_5 | CRY1 | 0.99999861 | 1 | 0.096 | 4.908 | chr12 | 107385142 | 107487327 | - | protein_coding |
| ENSG00000122966.16_7 | CIT | 0.99999998 | 1 | 0.096 | 6.656 | chr12 | 120123596 | 120315124 | - | protein_coding |
| ENSG00000172817.4_5 | CYP7B1 | 0.95920445 | 1 | 0.096 | 1.260 | chr8 | 65500320 | 65711294 | - | protein_coding |
| ENSG00000264859.6_7 | DSG2-AS1 | 0.86343799 | 1 | 0.095 | -0.846 | chr18 | 29122109 | 29136911 | - | lncRNA |
| ENSG00000156345.17_3 | CDK20 | 0.99927969 | 1 | 0.095 | 2.774 | chr9 | 90581356 | 90589668 | - | protein_coding |
| ENSG00000138459.9_6 | SLC35A5 | 0.99999957 | 1 | 0.095 | 5.408 | chr3 | 112280556 | 112304426 | + | protein_coding |
| ENSG00000101413.12_3 | RPRD1B | 0.99999835 | 1 | 0.095 | 4.899 | chr20 | 36662127 | 36756182 | + | protein_coding |
| ENSG00000164073.10_6 | MFSD8 | 0.99999638 | 1 | 0.095 | 4.465 | chr4 | 128838887 | 128887189 | - | protein_coding |
| ENSG00000232637.3 | WI2-3658N16.1 | 0.96660713 | 1 | 0.095 | 1.110 | chr1 | 146032648 | 146253110 | - | pseudogene |
| ENSG00000172046.19_6 | USP19 | 0.99999988 | 1 | 0.095 | 5.535 | chr3 | 49145479 | 49158371 | - | protein_coding |
| ENSG00000100225.18_5 | FBXO7 | 0.99999998 | 1 | 0.095 | 6.530 | chr22 | 32870663 | 32894816 | + | protein_coding |
| ENSG00000167004.13_3 | PDIA3 | 1 | 1 | 0.095 | 7.955 | chr15 | 44038608 | 44065476 | + | protein_coding |
| ENSG00000122042.10_4 | UBL3 | 0.9999991 | 1 | 0.095 | 5.067 | chr13 | 30338508 | 30424754 | - | protein_coding |
| ENSG00000089289.16_3 | IGBP1 | 0.99999967 | 1 | 0.095 | 5.448 | chrX | 69353297 | 69386174 | + | protein_coding |
| ENSG00000184992.13_6 | BRI3BP | 0.99999996 | 1 | 0.095 | 6.034 | chr12 | 125478191 | 125515777 | + | polymorphic_pseudogene |
| ENSG00000254901.8_6 | BORCS8 | 0.99448556 | 1 | 0.095 | 2.030 | chr19 | 19287712 | 19303400 | - | protein_coding |
| ENSG00000104856.14_3 | RELB | 0.9989414 | 1 | 0.095 | 3.168 | chr19 | 45504707 | 45541456 | + | protein_coding |
| ENSG00000104763.19_9 | ASAH1 | 0.99999999 | 1 | 0.095 | 6.940 | chr8 | 17913501 | 17942507 | - | protein_coding |
| ENSG00000213079.9_4 | SCAF8 | 0.99999994 | 1 | 0.095 | 5.835 | chr6 | 155054459 | 155155378 | + | protein_coding |
| ENSG00000198876.13_6 | DCAF12 | 0.99999996 | 1 | 0.095 | 6.001 | chr9 | 34086385 | 34127397 | - | protein_coding |
| ENSG00000226450.2_5 | CYP2D8P | 0.89221221 | 1 | 0.095 | -0.135 | chr22 | 42545877 | 42551007 | - | unprocessed_pseudogene |
| ENSG00000244165.2_4 | P2RY11 | 0.95557416 | 1 | 0.095 | 0.940 | chr19 | 10222369 | 10226048 | + | protein_coding |
| ENSG00000247572.8_8 | CKMT2-AS1 | 0.99804786 | 1 | 0.095 | 2.354 | chr5 | 80497160 | 80597384 | - | lncRNA |
| ENSG00000230982.1_4 | DSTNP1 | 0.92122619 | 1 | 0.094 | 0.177 | chr21 | 48073470 | 48073934 | - | processed_pseudogene |
| ENSG00000164597.13_5 | COG5 | 0.99999983 | 1 | 0.094 | 5.301 | chr7 | 106842000 | 107204959 | - | protein_coding |
| ENSG00000219451.3_4 | RPL23P8 | 0.85932014 | 1 | 0.094 | -1.009 | chr7 | 20866945 | 20867372 | + | processed_pseudogene |
| ENSG00000143486.16_7 | EIF2D | 0.99999972 | 1 | 0.094 | 5.341 | chr1 | 206744620 | 206785818 | - | protein_coding |
| ENSG00000287222.1_2 | AC234917.3 | 0.9975983 | 1 | 0.094 | 2.493 | chr8 | 144911688 | 144915031 | + | lncRNA |
| ENSG00000213839.4_4 | TMX2P1 | 0.99673956 | 1 | 0.094 | 2.762 | chr9 | 37885680 | 37886387 | + | processed_pseudogene |
| ENSG00000102057.10_5 | KCND1 | 0.90940261 | 1 | 0.094 | 0.037 | chrX | 48818639 | 48828251 | - | protein_coding |
| ENSG00000229180.7_6 | AC006001.3 | 0.99953764 | 1 | 0.094 | 2.928 | chr7 | 65991075 | 66057384 | - | transcribed_unprocessed_pseudogene |
| ENSG00000131871.15_6 | SELENOS | 0.99999995 | 1 | 0.094 | 5.823 | chr15 | 101811022 | 101817705 | - | protein_coding |
| ENSG00000177889.10_4 | UBE2N | 0.99999991 | 1 | 0.094 | 6.559 | chr12 | 93799449 | 93835723 | - | protein_coding |
| ENSG00000138669.9_3 | PRKG2 | 0.99007514 | 1 | 0.094 | 1.736 | chr4 | 82008524 | 82136271 | - | protein_coding |
| ENSG00000152580.8_3 | IGSF10 | 0.99421505 | 1 | 0.094 | 2.100 | chr3 | 151143172 | 151176497 | - | protein_coding |
| ENSG00000092036.18_5 | HAUS4 | 0.99997046 | 1 | 0.094 | 3.812 | chr14 | 23415437 | 23426370 | - | protein_coding |
| ENSG00000160633.13_3 | SAFB | 0.99999983 | 1 | 0.094 | 5.509 | chr19 | 5623046 | 5668489 | + | protein_coding |
| ENSG00000150667.8_6 | FSIP1 | 0.99343859 | 1 | 0.094 | 1.414 | chr15 | 39880558 | 40075042 | - | protein_coding |
| ENSG00000074695.6_4 | LMAN1 | 0.99999999 | 1 | 0.094 | 7.375 | chr18 | 56995055 | 57026497 | - | protein_coding |
| ENSG00000113595.15_5 | TRIM23 | 0.99999332 | 1 | 0.093 | 4.486 | chr5 | 64885517 | 64921802 | - | protein_coding |
| ENSG00000182168.15_4 | UNC5C | 0.99749533 | 1 | 0.093 | 2.290 | chr4 | 96083655 | 96470357 | - | protein_coding |
| ENSG00000133773.12_5 | CCDC59 | 0.99999188 | 1 | 0.093 | 4.820 | chr12 | 82617460 | 82752584 | - | protein_coding |
| ENSG00000145494.11_2 | NDUFS6 | 0.99999983 | 1 | 0.093 | 5.477 | chr5 | 1801514 | 1816719 | + | protein_coding |
| ENSG00000141556.21_6 | TBCD | 0.99999978 | 1 | 0.093 | 6.263 | chr17 | 80709941 | 80903790 | + | protein_coding |
| ENSG00000158470.5_4 | B4GALT5 | 1 | 1 | 0.093 | 6.833 | chr20 | 48249482 | 48330415 | - | protein_coding |
| ENSG00000137269.15_3 | LRRC1 | 0.99995808 | 1 | 0.093 | 3.685 | chr6 | 53659295 | 53788923 | + | protein_coding |
| ENSG00000169047.5_3 | IRS1 | 0.99999997 | 1 | 0.093 | 7.087 | chr2 | 227596033 | 227664475 | - | protein_coding |
| ENSG00000062582.14_5 | MRPS24 | 0.86817809 | 1 | 0.093 | -0.538 | chr7 | 43906157 | 43909492 | - | protein_coding |
| ENSG00000164885.13_4 | CDK5 | 0.99998698 | 1 | 0.093 | 4.064 | chr7 | 150750902 | 150754984 | - | protein_coding |
| ENSG00000129250.12_3 | KIF1C | 0.99999998 | 1 | 0.093 | 7.705 | chr17 | 4901245 | 4931696 | + | protein_coding |
| ENSG00000198198.17_8 | SZT2 | 0.99999901 | 1 | 0.092 | 5.267 | chr1 | 43855553 | 43919918 | + | protein_coding |

| | | | | | | | | | | |
|---|---|---|---|---|---|---|---|---|---|---|
| ENSG00000176623.12_7 | RMDN1 | 0.99999947 | 1 | 0.092 | 5.214 | chr8 | 87480486 | 87526586 | - | protein_coding |
| ENSG00000116521.11_4 | SCAMP3 | 0.99999964 | 1 | 0.092 | 5.233 | chr1 | 155225770 | 155232221 | - | protein_coding |
| ENSG00000151498.11_2 | ACAD8 | 0.99996491 | 1 | 0.092 | 3.963 | chr11 | 134123389 | 134135749 | + | protein_coding |
| ENSG00000135272.10_4 | MDFIC | 0.99999325 | 1 | 0.092 | 5.483 | chr7 | 114562209 | 114659256 | + | protein_coding |
| ENSG00000074319.13_6 | TSG101 | 0.99999985 | 1 | 0.092 | 5.742 | chr11 | 18489883 | 18548498 | - | protein_coding |
| ENSG00000164402.14_9 | SEPTIN8 | 0.99999993 | 1 | 0.092 | 5.604 | chr5 | 132086511 | 132142933 | - | protein_coding |
| ENSG00000172936.14_7 | MYD88 | 0.99770976 | 1 | 0.092 | 2.313 | chr3 | 38179969 | 38184513 | + | protein_coding |
| ENSG00000214106.9_6 | PAXIP1-AS2 | 0.99888241 | 1 | 0.092 | 2.763 | chr7 | 154720170 | 154743898 | + | lncRNA |
| ENSG00000146833.15_6 | TRIM4 | 0.99999977 | 1 | 0.092 | 5.499 | chr7 | 99474581 | 99517223 | - | protein_coding |
| ENSG00000036448.10_4 | MYOM2 | 0.93205559 | 1 | 0.092 | 0.753 | chr8 | 1993161 | 2113475 | + | protein_coding |
| ENSG00000106624.11_6 | AEBP1 | 0.86480459 | 1 | 0.092 | -0.962 | chr7 | 44143944 | 44154161 | + | protein_coding |
| ENSG00000104205.15_6 | SGK3 | 0.99998411 | 1 | 0.092 | 3.993 | chr8 | 67579787 | 67774257 | + | protein_coding |
| ENSG00000184470.21_6 | TXNRD2 | 0.99999299 | 1 | 0.092 | 4.592 | chr22 | 19863040 | 19929343 | - | protein_coding |
| ENSG00000171425.10_6 | ZNF581 | 0.99962422 | 1 | 0.092 | 3.389 | chr19 | 56146825 | 56156989 | + | protein_coding |
| ENSG00000147164.12_5 | SNX12 | 0.99999995 | 1 | 0.091 | 6.187 | chrX | 70276182 | 70293276 | - | protein_coding |
| ENSG00000179532.13_6 | DNHD1 | 0.99999453 | 1 | 0.091 | 4.364 | chr11 | 6518490 | 6614988 | + | protein_coding |
| ENSG00000170348.9_4 | TMED10 | 1 | 1 | 0.091 | 7.975 | chr14 | 75598172 | 75643315 | - | protein_coding |
| ENSG00000173889.16_6 | PHC3 | 0.9999999 | 1 | 0.091 | 6.530 | chr3 | 169804520 | 169899537 | - | protein_coding |
| ENSG00000117862.13_5 | TXNDC12 | 0.98563659 | 1 | 0.091 | 2.519 | chr1 | 52485803 | 52520863 | - | protein_coding |
| ENSG00000176909.12_8 | MAMSTR | 0.94710228 | 1 | 0.091 | 0.678 | chr19 | 49215982 | 49222982 | - | protein_coding |
| ENSG00000224424.8_6 | PRKAR2A-AS1 | 0.94694325 | 1 | 0.091 | 0.664 | chr3 | 48885005 | 48889414 | + | lncRNA |
| ENSG00000157911.10_6 | PEX10 | 0.99995051 | 1 | 0.091 | 3.925 | chr1 | 2335403 | 2345236 | - | protein_coding |
| ENSG00000172943.19_8 | PHF8 | 0.99999623 | 1 | 0.090 | 4.789 | chrX | 53963109 | 54075391 | - | protein_coding |
| ENSG00000255508.7_7 | AP002990.1 | 0.92694584 | 1 | 0.090 | 0.440 | chr11 | 62327075 | 62359003 | - | protein_coding |
| ENSG00000196924.18_7 | FLNA | 0.99999617 | 1 | 0.090 | 11.248 | chrX | 153576892 | 153603006 | - | protein_coding |
| ENSG00000136238.18_4 | RAC1 | 0.99974412 | 1 | 0.090 | 7.161 | chr7 | 6414158 | 6443598 | + | protein_coding |
| ENSG00000174514.13_4 | MFSD4A | 0.92526189 | 1 | 0.090 | 0.312 | chr1 | 205538013 | 205572046 | + | protein_coding |
| ENSG00000175104.15_4 | TRAF6 | 0.9999973 | 1 | 0.090 | 4.511 | chr11 | 36505319 | 36531822 | - | protein_coding |
| ENSG00000198920.11_8 | KIAA0753 | 0.99999985 | 1 | 0.090 | 5.333 | chr17 | 6481467 | 6544031 | - | protein_coding |
| ENSG00000136457.10_4 | CHAD | 0.87512129 | 1 | 0.090 | -0.138 | chr17 | 48541857 | 48546267 | - | protein_coding |
| ENSG00000157399.16_7 | ARSL | 0.99999987 | 1 | 0.090 | 5.526 | chrX | 2852562 | 2886516 | - | protein_coding |
| ENSG00000137776.17_5 | SLTM | 0.99999999 | 1 | 0.089 | 7.155 | chr15 | 59171249 | 59225878 | - | protein_coding |
| ENSG00000152348.16_5 | ATG10 | 0.99995662 | 1 | 0.089 | 3.750 | chr5 | 81267842 | 81572676 | + | protein_coding |
| ENSG00000107537.14_6 | PHYH | 0.99999781 | 1 | 0.089 | 4.510 | chr10 | 13319796 | 13344412 | - | protein_coding |
| ENSG00000132694.18_3 | ARHGEF11 | 0.99999999 | 1 | 0.089 | 6.543 | chr1 | 156904632 | 157015162 | - | protein_coding |
| ENSG00000125945.15_4 | ZNF436 | 0.99997113 | 1 | 0.089 | 4.042 | chr1 | 23685941 | 23696329 | - | protein_coding |
| ENSG00000164902.14_3 | PHAX | 0.99999997 | 1 | 0.089 | 5.822 | chr5 | 125936617 | 125962944 | + | protein_coding |
| ENSG00000182858.14_7 | ALG12 | 0.99989963 | 1 | 0.089 | 3.378 | chr22 | 50293877 | 50312086 | - | protein_coding |
| ENSG00000125046.15_6 | SSUH2 | 0.86754051 | 1 | 0.089 | -0.926 | chr3 | 8661086 | 8786726 | - | protein_coding |
| ENSG00000180448.10_8 | AC004151.1 | 0.99676295 | 1 | 0.089 | 2.803 | chr19 | 1065922 | 1086627 | + | protein_coding |
| ENSG00000125037.12_4 | EMC3 | 0.99999996 | 1 | 0.089 | 5.760 | chr3 | 10004221 | 10052800 | - | protein_coding |
| ENSG00000041988.15_5 | THAP3 | 0.99987616 | 1 | 0.089 | 3.445 | chr1 | 6684926 | 6695646 | + | protein_coding |
| ENSG00000170677.6_6 | SOCS6 | 0.99999999 | 1 | 0.089 | 6.952 | chr18 | 67956281 | 67997435 | + | protein_coding |
| ENSG00000263528.8_3 | IKBKE | 0.99995844 | 1 | 0.089 | 3.893 | chr1 | 206643819 | 206670222 | + | protein_coding |
| ENSG00000182093.16_6 | GET1 | 0.99997561 | 1 | 0.089 | 4.183 | chr21 | 40749624 | 40800454 | + | protein_coding |
| ENSG00000244038.9_6 | DDOST | 0.99999999 | 1 | 0.089 | 7.191 | chr1 | 20978260 | 20988037 | - | protein_coding |
| ENSG00000285581.1_3 | AC091196.1 | 0.91921475 | 1 | 0.088 | 0.286 | chr11 | 1485812 | 1488862 | - | lncRNA |
| ENSG00000281501.1_6 | SEPSECS-AS1 | 0.89201271 | 1 | 0.088 | -0.338 | chr4 | 25162263 | 25203062 | + | lncRNA |
| ENSG00000101966.12_3 | XIAP | 0.99999999 | 1 | 0.088 | 6.665 | chrX | 122993574 | 123047829 | + | protein_coding |
| ENSG00000234502.2_5 | FYTTD1P1 | 0.86783011 | 1 | 0.088 | -0.803 | chr9 | 104137626 | 104138885 | - | processed_pseudogene |
| ENSG00000033122.20_7 | LRRC7 | 0.89511904 | 1 | 0.088 | -0.370 | chr1 | 70033944 | 70617628 | + | protein_coding |
| ENSG00000279453.1_5 | Z99129.4 | 0.97567197 | 1 | 0.088 | 1.280 | chr6 | 122757934 | 122760368 | - | TEC |
| ENSG00000267500.1_5 | ZNF887P | 0.8869828 | 1 | 0.088 | -0.575 | chr19 | 11759179 | 11763692 | + | transcribed_unprocessed_pseudogene |
| ENSG00000143258.16_6 | USP21 | 0.99984344 | 1 | 0.088 | 3.416 | chr1 | 161129240 | 161135513 | + | protein_coding |
| ENSG00000015171.19_3 | ZMYND11 | 1 | 1 | 0.088 | 7.019 | chr10 | 180405 | 300577 | + | protein_coding |
| ENSG00000184160.8_5 | ADRA2C | 0.99255577 | 1 | 0.088 | 2.222 | chr4 | 3768075 | 3770253 | + | protein_coding |
| ENSG00000084652.16_3 | TXLNA | 1 | 1 | 0.087 | 6.934 | chr1 | 32645276 | 32663886 | + | protein_coding |
| ENSG00000287387.1_1 | AC010883.3 | 0.94858619 | 1 | 0.087 | 0.660 | chr2 | 43446988 | 43450366 | + | lncRNA |

| | | | | | | | | | | |
|---|---|---|---|---|---|---|---|---|---|---|
| ENSG00000131626.18_5 | PPFIA1 | 0.99999996 | 1 | 0.087 | 5.919 | chr11 | 70116806 | 70231418 | + | protein_coding |
| ENSG00000117054.13_6 | ACADM | 0.99998944 | 1 | 0.087 | 5.029 | chr1 | 76190032 | 76253260 | + | protein_coding |
| ENSG00000273899.5_4 | NOL12 | 0.99018604 | 1 | 0.087 | 1.649 | chr22 | 38077680 | 38089483 | + | protein_coding |
| ENSG00000149273.15_4 | RPS3 | 1 | 1 | 0.087 | 9.171 | chr11 | 75110559 | 75133324 | + | protein_coding |
| ENSG00000028528.15_6 | SNX1 | 0.99999999 | 1 | 0.087 | 6.267 | chr15 | 64386322 | 64438289 | + | protein_coding |
| ENSG00000120915.14_8 | EPHX2 | 0.99995403 | 1 | 0.087 | 3.764 | chr8 | 27348298 | 27403081 | + | protein_coding |
| ENSG00000140264.19_5 | SERF2 | 0.99999998 | 1 | 0.087 | 6.284 | chr15 | 44069285 | 44094787 | + | protein_coding |
| ENSG00000225733.6_8 | FGD5-AS1 | 1 | 1 | 0.087 | 7.505 | chr3 | 14961854 | 14989931 | - | lncRNA |
| ENSG00000100207.18_6 | TCF20 | 0.99999999 | 1 | 0.087 | 6.339 | chr22 | 42556019 | 42739622 | - | protein_coding |
| ENSG00000124535.16_4 | WRNIP1 | 0.9999997 | 1 | 0.087 | 5.403 | chr6 | 2765627 | 2787186 | + | protein_coding |
| ENSG00000078674.17_4 | PCM1 | 1 | 1 | 0.087 | 8.173 | chr8 | 17780349 | 17887453 | + | protein_coding |
| ENSG00000269535.1_7 | AC010320.5 | 0.93628125 | 1 | 0.087 | 0.428 | chr19 | 52803946 | 52848482 | - | lncRNA |
| ENSG00000084070.12_5 | SMAP2 | 0.99988077 | 1 | 0.087 | 3.374 | chr1 | 40810522 | 40888998 | + | protein_coding |
| ENSG00000253540.5_3 | FAM86HP | 0.90014566 | 1 | 0.087 | 0.493 | chr3 | 129817935 | 129830315 | - | transcribed_unprocessed_pseudogene |
| ENSG00000132612.16_4 | VPS4A | 0.99999988 | 1 | 0.087 | 5.565 | chr16 | 69345253 | 69360842 | + | protein_coding |
| ENSG00000067208.14_4 | EVI5 | 0.99999917 | 1 | 0.087 | 4.876 | chr1 | 92974253 | 93257961 | - | protein_coding |
| ENSG00000196975.16_3 | ANXA4 | 1 | 1 | 0.087 | 7.405 | chr2 | 69871557 | 70054244 | + | protein_coding |
| ENSG00000054654.17_7 | SYNE2 | 0.99447355 | 1 | 0.086 | 7.069 | chr14 | 64228617 | 64693151 | + | protein_coding |
| ENSG00000132640.15_6 | BTBD3 | 0.99999987 | 1 | 0.086 | 5.542 | chr20 | 11871371 | 11907257 | + | protein_coding |
| ENSG00000273599.1_6 | AL731571.1 | 0.99551811 | 1 | 0.086 | 2.153 | chr10 | 126684633 | 126690060 | + | lncRNA |
| ENSG00000112715.23_10 | VEGFA | 0.99999887 | 1 | 0.086 | 4.700 | chr6 | 43737921 | 43754224 | + | polymorphic_pseudogene |
| ENSG00000079974.17_4 | RABL2B | 0.99989711 | 1 | 0.086 | 3.559 | chr22 | 51205929 | 51222091 | - | protein_coding |
| ENSG00000254034.2_6 | INTS9-AS1 | 0.87350585 | 1 | 0.086 | -0.846 | chr8 | 28655659 | 28659602 | + | lncRNA |
| ENSG00000078043.16_8 | PIAS2 | 0.99999892 | 1 | 0.086 | 4.860 | chr18 | 44383181 | 44500123 | - | protein_coding |
| ENSG00000162368.13_2 | CMPK1 | 0.99999998 | 1 | 0.086 | 7.196 | chr1 | 47799469 | 47844511 | + | protein_coding |
| ENSG00000238279.1_6 | BX470102.1 | 0.99467052 | 1 | 0.086 | 1.459 | chr1 | 153506079 | 153507591 | + | lncRNA |
| ENSG00000154114.12_5 | TBCEL | 0.99986496 | 1 | 0.086 | 3.541 | chr11 | 120894781 | 120961484 | + | protein_coding |
| ENSG00000058063.16_5 | ATP11B | 0.99999954 | 1 | 0.086 | 5.636 | chr3 | 182511291 | 182639417 | + | protein_coding |
| ENSG00000273137.1_7 | AL022328.2 | 0.86991545 | 1 | 0.086 | -1.049 | chr22 | 50646890 | 50647971 | - | lncRNA |
| ENSG00000272129.1_6 | AL359715.3 | 0.8718409 | 1 | 0.086 | -0.847 | chr6 | 81065141 | 81066576 | + | lncRNA |
| ENSG00000143294.15_3 | PRCC | 0.99999883 | 1 | 0.086 | 4.931 | chr1 | 156720402 | 156770607 | + | protein_coding |
| ENSG00000259514.1_7 | AC027243.2 | 0.87487565 | 1 | 0.086 | -0.927 | chr15 | 76631950 | 76634404 | - | lncRNA |
| ENSG00000183666.17_6 | GUSBP1 | 0.9946071 | 1 | 0.086 | 2.098 | chr5 | 21341942 | 21589481 | + | transcribed_unprocessed_pseudogene |
| ENSG00000254893.6_3 | AC113404.3 | 0.91947886 | 1 | 0.086 | 0.468 | chr5 | 75469454 | 75470008 | - | translated_processed_pseudogene |
| ENSG00000156831.8_5 | NSMCE2 | 0.99999524 | 1 | 0.086 | 4.444 | chr8 | 126103921 | 126379362 | + | protein_coding |
| ENSG00000104231.11_7 | ZFAND1 | 0.99999935 | 1 | 0.086 | 5.059 | chr8 | 82613569 | 82645138 | - | protein_coding |
| ENSG00000131788.16_4 | PIAS3 | 0.99998468 | 1 | 0.086 | 3.904 | chr1 | 145575233 | 145586546 | + | protein_coding |
| ENSG00000231028.9_4 | LINC00271 | 0.92518779 | 1 | 0.086 | 0.275 | chr6 | 135818560 | 136135128 | + | lncRNA |
| ENSG00000185596.16_2 | WASH3P | 0.99976878 | 1 | 0.086 | 3.067 | chr15 | 102501806 | 102516746 | + | transcribed_unprocessed_pseudogene |
| ENSG00000119718.11_4 | EIF2B2 | 0.99999811 | 1 | 0.085 | 4.548 | chr14 | 75469624 | 75479069 | + | protein_coding |
| ENSG00000278311.5_5 | GGNBP2 | 0.99999999 | 1 | 0.085 | 6.448 | chr17 | 34900761 | 34946278 | + | protein_coding |
| ENSG00000153487.12_2 | ING1 | 0.99888105 | 1 | 0.085 | 2.639 | chr13 | 111365083 | 111375686 | + | protein_coding |
| ENSG00000136327.7_4 | NKX2-8 | 0.87425473 | 1 | 0.085 | -0.700 | chr14 | 37049209 | 37051819 | - | protein_coding |
| ENSG00000272172.1_6 | AC138696.2 | 0.87338891 | 1 | 0.085 | -0.671 | chr8 | 144372569 | 144372791 | - | lncRNA |
| ENSG00000182325.11_5 | FBXL6 | 0.99970472 | 1 | 0.085 | 3.241 | chr8 | 145579091 | 145583036 | - | protein_coding |
| ENSG00000225206.10_6 | MIR137HG | 0.99580801 | 1 | 0.085 | 2.222 | chr1 | 98399030 | 98515419 | - | lncRNA |
| ENSG00000123268.9_5 | ATF1 | 0.99999951 | 1 | 0.085 | 4.969 | chr12 | 51157493 | 51214945 | + | protein_coding |
| ENSG00000242951.1_5 | AC007182.2 | 0.88950397 | 1 | 0.085 | -0.426 | chr14 | 76062148 | 76062549 | - | processed_pseudogene |
| ENSG00000264522.6_4 | OTUD7B | 0.99999964 | 1 | 0.084 | 5.103 | chr1 | 149909705 | 149982675 | - | protein_coding |
| ENSG00000101997.13_3 | CCDC22 | 0.99997085 | 1 | 0.084 | 4.223 | chrX | 49091930 | 49106981 | + | protein_coding |
| ENSG00000154122.14_6 | ANKH | 0.99999688 | 1 | 0.084 | 4.568 | chr5 | 14704909 | 14871887 | - | protein_coding |
| ENSG00000172466.16_5 | ZNF24 | 0.99999997 | 1 | 0.084 | 6.433 | chr18 | 32912191 | 32925446 | - | protein_coding |
| ENSG00000184007.21_7 | PTP4A2 | 1 | 1 | 0.084 | 7.959 | chr1 | 32372022 | 32410457 | - | protein_coding |
| ENSG00000087076.9_5 | HSD17B14 | 0.99840217 | 1 | 0.084 | 2.624 | chr19 | 49316275 | 49339767 | - | protein_coding |
| ENSG00000247498.10_7 | GPRC5D-AS1 | 0.97756684 | 1 | 0.084 | 1.251 | chr12 | 13080660 | 13137579 | + | lncRNA |
| ENSG00000142546.14_7 | NOSIP | 0.99999765 | 1 | 0.084 | 5.384 | chr19 | 50058725 | 50093519 | - | protein_coding |

| | | | | | | | | | | |
|---|---|---|---|---|---|---|---|---|---|---|
| ENSG00000163312.11_4 | HELQ | 0.9999352 | 1 | 0.084 | 3.457 | chr4 | 84328496 | 84377008 | - | protein_coding |
| ENSG00000135338.14_6 | LCA5 | 0.99894896 | 1 | 0.084 | 2.634 | chr6 | 80194708 | 80247175 | - | protein_coding |
| ENSG00000100325.15_6 | ASCC2 | 0.99999902 | 1 | 0.084 | 4.886 | chr22 | 30184598 | 30234293 | - | protein_coding |
| ENSG00000169548.3_5 | ZNF280A | 0.99751877 | 1 | 0.084 | 2.298 | chr22 | 22868060 | 22874613 | - | protein_coding |
| ENSG00000286388.1_1 | AC026748.3 | 0.99981444 | 1 | 0.084 | 3.117 | chr5 | 1345312 | 1350901 | + | lncRNA |
| ENSG00000078142.13_4 | PIK3C3 | 0.99999893 | 1 | 0.084 | 4.996 | chr18 | 39535199 | 39667794 | + | protein_coding |
| ENSG00000263826.1_6 | AC112907.3 | 0.98560787 | 1 | 0.083 | 1.433 | chr3 | 186499569 | 186501968 | - | lncRNA |
| ENSG00000134779.15_5 | TPGS2 | 0.99999999 | 1 | 0.083 | 6.241 | chr18 | 34357610 | 34409179 | - | protein_coding |
| ENSG00000196652.11_2 | ZKSCAN5 | 0.99999983 | 1 | 0.083 | 5.182 | chr7 | 99102274 | 99132323 | + | protein_coding |
| ENSG00000112855.17_7 | HARS2 | 0.99999991 | 1 | 0.083 | 5.667 | chr5 | 140071015 | 140078890 | + | protein_coding |
| ENSG00000164221.13_5 | CCDC112 | 0.9999512 | 1 | 0.083 | 3.733 | chr5 | 114602887 | 114632390 | - | protein_coding |
| ENSG00000243370.2 | RN7SL775P | 0.88217274 | 1 | 0.083 | -0.814 | chr17 | 17081729 | 17082020 | + | misc_RNA |
| ENSG00000217930.8_5 | PAM16 | 0.94208778 | 1 | 0.083 | 0.330 | chr16 | 4381550 | 4405608 | - | protein_coding |
| ENSG00000084463.8_8 | WBP11 | 1 | 1 | 0.083 | 6.794 | chr12 | 14937516 | 14956420 | - | protein_coding |
| ENSG00000279742.1_5 | AP000974.1 | 0.92270005 | 1 | 0.083 | -0.057 | chr11 | 85563600 | 85565986 | - | TEC |
| ENSG00000009413.16_5 | REV3L | 0.99999969 | 1 | 0.083 | 5.576 | chr6 | 111620231 | 111804918 | - | protein_coding |
| ENSG00000271643.2_7 | AC112220.2 | 0.99808442 | 1 | 0.083 | 2.496 | chr3 | 33835136 | 33840031 | - | lncRNA |
| ENSG00000163382.12_4 | NAXE | 0.99999992 | 1 | 0.083 | 6.262 | chr1 | 156561554 | 156564091 | + | protein_coding |
| ENSG00000119979.18_6 | DENND10 | 0.99999845 | 1 | 0.083 | 5.048 | chr10 | 120863598 | 120897496 | + | protein_coding |
| ENSG00000162650.16_3 | ATXN7L2 | 0.99941325 | 1 | 0.083 | 1.963 | chr1 | 110026101 | 110035426 | + | protein_coding |
| ENSG00000140265.12_3 | ZSCAN29 | 0.99999944 | 1 | 0.083 | 4.984 | chr15 | 43650370 | 43663223 | - | protein_coding |
| ENSG00000228839.5_7 | PIK3IP1-AS1 | 0.87464324 | 1 | 0.083 | -1.142 | chr22 | 31688485 | 31734007 | + | lncRNA |
| ENSG00000182208.14_4 | MOB2 | 0.99998955 | 1 | 0.082 | 4.169 | chr11 | 1490687 | 1522477 | - | protein_coding |
| ENSG00000148341.17_4 | SH3GLB2 | 0.99999923 | 1 | 0.082 | 5.632 | chr9 | 131769315 | 131790582 | - | protein_coding |
| ENSG00000172869.14_3 | DMXL1 | 0.99999786 | 1 | 0.082 | 6.227 | chr5 | 118373467 | 118584833 | + | protein_coding |
| ENSG00000225434.3_6 | LINC01504 | 0.9996921 | 1 | 0.082 | 3.244 | chr9 | 74920283 | 74971637 | + | lncRNA |
| ENSG00000128016.7_5 | ZFP36 | 0.99994664 | 1 | 0.082 | 3.817 | chr19 | 39897487 | 39900052 | + | protein_coding |
| ENSG00000069943.10_3 | PIGB | 0.99998927 | 1 | 0.082 | 4.076 | chr15 | 55611158 | 55647846 | + | protein_coding |
| ENSG00000229970.3_6 | AC007128.1 | 0.96474949 | 1 | 0.082 | 0.915 | chr7 | 8301894 | 8384146 | + | lncRNA |
| ENSG00000126934.13_5 | MAP2K2 | 0.99452866 | 1 | 0.082 | 6.524 | chr19 | 4090319 | 4124126 | - | protein_coding |
| ENSG00000120370.12_4 | GORAB | 0.99988778 | 1 | 0.082 | 3.483 | chr1 | 170501270 | 170522587 | + | protein_coding |
| ENSG00000157617.17_5 | C2CD2 | 0.99999653 | 1 | 0.082 | 4.550 | chr21 | 43305221 | 43374127 | - | protein_coding |
| ENSG00000265800.1_8 | AC022211.4 | 0.89889386 | 1 | 0.082 | -0.261 | chr17 | 73134511 | 73137445 | - | lncRNA |
| ENSG00000148737.17_7 | TCF7L2 | 0.99997557 | 1 | 0.082 | 3.930 | chr10 | 114710006 | 114927437 | + | protein_coding |
| ENSG00000167978.17_5 | SRRM2 | 0.99999999 | 1 | 0.082 | 7.770 | chr16 | 2802627 | 2822539 | + | protein_coding |
| ENSG00000161813.22_7 | LARP4 | 0.99999996 | 1 | 0.082 | 6.804 | chr12 | 50786166 | 50873787 | + | protein_coding |
| ENSG00000088448.14_5 | ANKRD10 | 0.99999973 | 1 | 0.082 | 5.876 | chr13 | 111530887 | 111567416 | - | protein_coding |
| ENSG00000158125.10_4 | XDH | 0.99840336 | 1 | 0.081 | 2.493 | chr2 | 31557187 | 31637608 | - | protein_coding |
| ENSG00000141298.19_6 | SSH2 | 0.99999978 | 1 | 0.081 | 5.227 | chr17 | 27952956 | 28257294 | - | protein_coding |
| ENSG00000142528.16_6 | ZNF473 | 0.99999941 | 1 | 0.081 | 4.797 | chr19 | 50528971 | 50556671 | + | protein_coding |
| ENSG00000231073.1_6 | AL590133.1 | 0.92187264 | 1 | 0.081 | 0.162 | chr1 | 150945599 | 150948010 | + | lncRNA |
| ENSG00000101246.20_5 | ARFRP1 | 0.99999762 | 1 | 0.081 | 5.346 | chr20 | 62329995 | 62339377 | - | protein_coding |
| ENSG00000242372.8_6 | EIF6 | 0.99999993 | 1 | 0.081 | 6.080 | chr20 | 33866714 | 33872788 | - | protein_coding |
| ENSG00000108587.15_3 | GOSR1 | 0.99999998 | 1 | 0.081 | 5.961 | chr17 | 28804380 | 28854610 | + | protein_coding |
| ENSG00000065665.21_4 | SEC61A2 | 0.9999585 | 1 | 0.081 | 3.798 | chr10 | 12171636 | 12211960 | + | protein_coding |
| ENSG00000039319.17_5 | ZFYVE16 | 0.99999998 | 1 | 0.081 | 6.460 | chr5 | 79703832 | 79779198 | + | protein_coding |
| ENSG00000163681.15_5 | SLMAP | 0.99999994 | 1 | 0.081 | 5.856 | chr3 | 57741177 | 57915730 | + | protein_coding |
| ENSG00000066135.13_5 | KDM4A | 0.99998182 | 1 | 0.081 | 3.795 | chr1 | 44115820 | 44171189 | + | protein_coding |
| ENSG00000130749.10_6 | ZC3H4 | 0.99999993 | 1 | 0.081 | 5.878 | chr19 | 47567444 | 47617033 | - | protein_coding |
| ENSG00000004779.10_4 | NDUFAB1 | 0.99999954 | 1 | 0.081 | 5.474 | chr16 | 23592335 | 23607637 | - | protein_coding |
| ENSG00000213064.10_3 | SFT2D2 | 1 | 1 | 0.081 | 7.238 | chr1 | 168195176 | 168222259 | + | protein_coding |
| ENSG00000227543.5_5 | SPAG5-AS1 | 0.94053669 | 1 | 0.080 | 0.630 | chr17 | 26925808 | 26944395 | + | lncRNA |
| ENSG00000105983.22_6 | LMBR1 | 0.99999999 | 1 | 0.080 | 6.409 | chr7 | 156461640 | 156685910 | - | protein_coding |
| ENSG00000141569.12_6 | TRIM65 | 0.99999978 | 1 | 0.080 | 5.709 | chr17 | 73876416 | 73893032 | - | protein_coding |
| ENSG00000013375.16_8 | PGM3 | 0.99999981 | 1 | 0.080 | 5.338 | chr6 | 83870869 | 83903655 | - | protein_coding |
| ENSG00000176393.11_7 | RNPEP | 0.99999948 | 1 | 0.080 | 5.619 | chr1 | 201951500 | 201975275 | + | protein_coding |
| ENSG00000184983.11_8 | NDUFA6 | 0.99999988 | 1 | 0.080 | 5.460 | chr22 | 42481530 | 42486888 | - | protein_coding |
| ENSG00000109381.20_5 | ELF2 | 0.99999975 | 1 | 0.080 | 5.106 | chr4 | 139949266 | 140098372 | - | protein_coding |
| ENSG00000172493.21_4 | AFF1 | 0.99999988 | 1 | 0.080 | 5.977 | chr4 | 87856154 | 88062206 | + | protein_coding |
| ENSG00000253878.5_7 | AC087752.3 | 0.94329474 | 1 | 0.080 | 0.499 | chr8 | 95962265 | 95963624 | + | lncRNA |
| ENSG00000174748.21_8 | RPL15 | 1 | 1 | 0.080 | 9.335 | chr3 | 23958036 | 23965865 | + | protein_coding |

| | | | | | | | | | | |
|---|---|---|---|---|---|---|---|---|---|---|
| ENSG00000204520.14_5 | MICA | 0.99846731 | 1 | 0.080 | 2.313 | chr6 | 31367561 | 31383092 | + | protein_coding |
| ENSG00000143514.17_6 | TP53BP2 | 0.99999994 | 1 | 0.079 | 5.649 | chr1 | 223967595 | 224033656 | - | protein_coding |
| ENSG00000184708.18_5 | EIF4ENIF1 | 0.99985216 | 1 | 0.079 | 3.561 | chr22 | 31832963 | 31892094 | - | protein_coding |
| ENSG00000125872.8_4 | LRRN4 | 0.8793421 | 1 | 0.079 | -1.050 | chr20 | 6021192 | 6034706 | - | protein_coding |
| ENSG00000243477.5_5 | NAA80 | 0.99636469 | 1 | 0.079 | 2.146 | chr3 | 50333833 | 50336852 | - | protein_coding |
| ENSG00000133895.15_7 | MEN1 | 0.99998797 | 1 | 0.079 | 4.572 | chr11 | 64570982 | 64578766 | - | protein_coding |
| ENSG00000130544.12_3 | ZNF557 | 0.99989611 | 1 | 0.079 | 3.646 | chr19 | 7069714 | 7087979 | + | protein_coding |
| ENSG00000164713.10_3 | BRI3 | 0.99999797 | 1 | 0.079 | 4.766 | chr7 | 97881691 | 97939753 | + | protein_coding |
| ENSG00000271918.1_6 | AC034236.2 | 0.94685644 | 1 | 0.079 | 0.587 | chr5 | 115419504 | 115421113 | - | lncRNA |
| ENSG00000204351.12_3 | SKIV2L | 0.99999785 | 1 | 0.079 | 4.913 | chr6 | 31926894 | 31937528 | + | protein_coding |
| ENSG00000218426.5_5 | AL590867.2 | 0.99952981 | 1 | 0.079 | 2.911 | chr6 | 153603422 | 153603868 | - | processed_pseudogene |
| ENSG00000170364.12_2 | SETMAR | 0.99970255 | 1 | 0.079 | 3.196 | chr3 | 4344988 | 4359251 | + | protein_coding |
| ENSG00000253669.4_6 | GASAL1 | 0.99917227 | 1 | 0.079 | 2.838 | chr8 | 103817745 | 103822267 | + | lncRNA |
| ENSG00000100056.12_7 | ESS2 | 0.99998261 | 1 | 0.079 | 3.786 | chr22 | 19117792 | 19132197 | - | protein_coding |
| ENSG00000167394.13_4 | ZNF668 | 0.99909511 | 1 | 0.079 | 2.730 | chr16 | 31072164 | 31085561 | - | protein_coding |
| ENSG00000100395.15_3 | L3MBTL2 | 0.99999372 | 1 | 0.079 | 4.466 | chr22 | 41601286 | 41627275 | + | protein_coding |
| ENSG00000105612.9_4 | DNASE2 | 0.99999636 | 1 | 0.079 | 4.381 | chr19 | 12986023 | 12992280 | - | protein_coding |
| ENSG00000121073.15_6 | SLC35B1 | 0.99999998 | 1 | 0.079 | 5.994 | chr17 | 47778296 | 47786376 | - | protein_coding |
| ENSG00000125447.17_7 | GGA3 | 0.99999981 | 1 | 0.078 | 5.588 | chr17 | 73232694 | 73258444 | - | protein_coding |
| ENSG00000215835.2_5 | AL596087.1 | 0.88818812 | 1 | 0.078 | -0.614 | chr1 | 166244866 | 166246834 | - | processed_pseudogene |
| ENSG00000179094.16_7 | PER1 | 0.99923205 | 1 | 0.078 | 2.884 | chr17 | 8043790 | 8059824 | - | protein_coding |
| ENSG00000261216.1_6 | AC007216.2 | 0.88127866 | 1 | 0.078 | -1.103 | chr16 | 12002065 | 12002773 | + | lncRNA |
| ENSG00000113758.13_7 | DBN1 | 0.99999968 | 1 | 0.078 | 6.200 | chr5 | 176883609 | 176901402 | - | protein_coding |
| ENSG00000224505.2_7 | AC138150.1 | 0.92871517 | 1 | 0.078 | 0.390 | chr17 | 43227767 | 43238877 | - | lncRNA |
| ENSG00000271335.6_8 | AL117336.2 | 0.93646831 | 1 | 0.078 | 0.330 | chr10 | 35603480 | 35625329 | - | lncRNA |
| ENSG00000263847.1_7 | AP005899.1 | 0.93485558 | 1 | 0.078 | 0.427 | chr18 | 9112402 | 9115875 | - | lncRNA |
| ENSG00000180304.14_4 | OAZ2 | 0.99999997 | 1 | 0.078 | 6.026 | chr15 | 64979772 | 64995480 | - | protein_coding |
| ENSG00000186130.5_4 | ZBTB6 | 0.9999966 | 1 | 0.078 | 4.469 | chr9 | 125670335 | 125675602 | - | protein_coding |
| ENSG00000138785.15_8 | INTS12 | 0.9984366 | 1 | 0.077 | 2.444 | chr4 | 106603784 | 106817143 | - | protein_coding |
| ENSG00000146828.18_6 | SLC12A9 | 0.99999549 | 1 | 0.077 | 4.615 | chr7 | 100424442 | 100464632 | + | protein_coding |
| ENSG00000196911.11_4 | KPNA5 | 0.99940591 | 1 | 0.077 | 3.288 | chr6 | 117002350 | 117063030 | + | protein_coding |
| ENSG00000240065.8_4 | PSMB9 | 0.98969551 | 1 | 0.077 | 1.649 | chr6 | 32811913 | 32827628 | + | protein_coding |
| ENSG00000146872.17_5 | TLK2 | 0.99999997 | 1 | 0.077 | 5.949 | chr17 | 60536019 | 60692842 | + | protein_coding |
| ENSG00000269343.8_6 | ZNF587B | 0.99999949 | 1 | 0.077 | 5.336 | chr19 | 58331089 | 58357606 | + | protein_coding |
| ENSG00000124596.17_7 | OARD1 | 0.99997713 | 1 | 0.077 | 4.073 | chr6 | 41001366 | 41065526 | - | protein_coding |
| ENSG00000108061.11_4 | SHOC2 | 0.99999605 | 1 | 0.077 | 5.559 | chr10 | 112679305 | 112773425 | + | protein_coding |
| ENSG00000143156.14_6 | NME7 | 0.99980658 | 1 | 0.077 | 3.787 | chr1 | 169101769 | 169337186 | - | protein_coding |
| ENSG00000145439.12_5 | CBR4 | 0.9999971 | 1 | 0.077 | 4.400 | chr4 | 169784921 | 169931426 | - | protein_coding |
| ENSG00000198258.10_2 | UBL5 | 0.99999982 | 1 | 0.077 | 5.406 | chr19 | 9938568 | 9940791 | + | protein_coding |
| ENSG00000162065.14_5 | TBC1D24 | 0.99999564 | 1 | 0.077 | 4.311 | chr16 | 2525052 | 2559561 | + | protein_coding |
| ENSG00000277369.1_6 | AC010654.1 | 0.92454126 | 1 | 0.077 | 0.089 | chr16 | 11891324 | 11892131 | + | lncRNA |
| ENSG00000273151.1_6 | AC073957.3 | 0.89990872 | 1 | 0.077 | -0.488 | chr7 | 919427 | 926184 | - | lncRNA |
| ENSG00000198015.13_5 | MRPL42 | 0.99999995 | 1 | 0.077 | 6.342 | chr12 | 93861264 | 93909990 | + | protein_coding |
| ENSG00000168077.14_3 | SCARA3 | 0.99999995 | 1 | 0.077 | 6.224 | chr8 | 27491385 | 27534293 | + | protein_coding |
| ENSG00000137818.12_3 | RPLP1 | 0.99946337 | 1 | 0.077 | 7.382 | chr15 | 69745153 | 69748544 | + | protein_coding |
| ENSG00000096968.13_3 | JAK2 | 0.9999469 | 1 | 0.076 | 3.908 | chr9 | 4984390 | 5128183 | + | protein_coding |
| ENSG00000124786.11_5 | SLC35B3 | 0.99996351 | 1 | 0.076 | 3.955 | chr6 | 8411696 | 8435844 | - | protein_coding |
| ENSG00000172530.20_7 | BANP | 0.99992899 | 1 | 0.076 | 3.767 | chr16 | 87982850 | 88110924 | + | protein_coding |
| ENSG00000198246.9_5 | SLC29A3 | 0.99997824 | 1 | 0.076 | 3.991 | chr10 | 73079016 | 73141180 | + | protein_coding |
| ENSG00000243660.10_5 | ZNF487 | 0.99878584 | 1 | 0.076 | 2.512 | chr10 | 43932289 | 43978629 | + | protein_coding |
| ENSG00000256973.1_8 | AC053513.2 | 0.89781911 | 1 | 0.076 | -0.403 | chr12 | 22613453 | 22616848 | - | lncRNA |
| ENSG00000205189.12_3 | ZBTB10 | 0.99999957 | 1 | 0.076 | 5.001 | chr8 | 81397854 | 81438500 | + | protein_coding |
| ENSG00000142197.12_4 | DOP1B | 0.99999688 | 1 | 0.076 | 4.270 | chr21 | 37529080 | 37666572 | + | protein_coding |
| ENSG00000138175.9_4 | ARL3 | 0.99999839 | 1 | 0.076 | 4.682 | chr10 | 104433488 | 104474154 | - | protein_coding |
| ENSG00000105227.15_4 | PRX | 0.99815557 | 1 | 0.076 | 2.453 | chr19 | 40899673 | 40920649 | - | protein_coding |
| ENSG00000099899.15_8 | TRMT2A | 0.99999295 | 1 | 0.076 | 4.404 | chr22 | 20099398 | 20104915 | - | protein_coding |
| ENSG00000162735.18_4 | PEX19 | 0.99999939 | 1 | 0.076 | 4.766 | chr1 | 160246602 | 160256138 | - | protein_coding |
| ENSG00000175309.15_6 | PHYKPL | 0.99999899 | 1 | 0.076 | 4.599 | chr5 | 177635472 | 177659803 | - | protein_coding |
| ENSG00000116251.11_6 | RPL22 | 0.99999996 | 1 | 0.076 | 5.894 | chr1 | 6245080 | 6269449 | - | protein_coding |
| ENSG00000227512.1_6 | AL592301.1 | 0.88583626 | 1 | 0.076 | -0.671 | chr9 | 139437333 | 139438287 | + | lncRNA |

| | | | | | | | | | | |
|---|---|---|---|---|---|---|---|---|---|---|
| ENSG00000250365.7_6 | AL139353.2 | 0.90832897 | 1 | 0.076 | 0.074 | chr14 | 31889492 | 31922089 | + | lncRNA |
| ENSG00000197461.13_6 | PDGFA | 0.99999919 | 1 | 0.076 | 4.679 | chr7 | 536895 | 559933 | - | protein_coding |
| ENSG00000074527.12_4 | NTN4 | 0.99999999 | 1 | 0.076 | 6.653 | chr12 | 96051583 | 96184930 | - | protein_coding |
| ENSG00000146085.8_4 | MMUT | 0.99999528 | 1 | 0.076 | 4.326 | chr6 | 49398073 | 49430966 | - | protein_coding |
| ENSG00000108528.14_4 | SLC25A11 | 0.99999997 | 1 | 0.076 | 6.011 | chr17 | 4840425 | 4843348 | - | protein_coding |
| ENSG00000232888.4_4 | RPS11P5 | 0.99125106 | 1 | 0.076 | 1.710 | chr12 | 133402287 | 133402770 | + | processed_pseudogene |
| ENSG00000130939.20_6 | UBE4B | 0.99999995 | 1 | 0.075 | 5.596 | chr1 | 10092890 | 10241297 | + | protein_coding |
| ENSG00000114520.11_7 | SNX4 | 0.99999932 | 1 | 0.075 | 5.145 | chr3 | 125165494 | 125239046 | - | protein_coding |
| ENSG00000257913.2_6 | DDN-AS1 | 0.94089658 | 1 | 0.075 | 0.387 | chr12 | 49392150 | 49413018 | + | lncRNA |
| ENSG00000123444.14_7 | KBTBD4 | 0.99968459 | 1 | 0.075 | 3.134 | chr11 | 47593749 | 47600528 | - | protein_coding |
| ENSG00000101160.14_4 | CTSZ | 1 | 1 | 0.075 | 6.681 | chr20 | 57570240 | 57582309 | - | protein_coding |
| ENSG00000227008.2_5 | AL009174.1 | 0.90950406 | 1 | 0.075 | 0.958 | chrX | 132804999 | 132805414 | + | processed_pseudogene |
| ENSG00000182742.6_5 | HOXB4 | 0.93374775 | 1 | 0.075 | 0.270 | chr17 | 46652869 | 46655712 | - | protein_coding |
| ENSG00000128191.16_4 | DGCR8 | 0.99999986 | 1 | 0.075 | 5.160 | chr22 | 20067755 | 20099400 | + | protein_coding |
| ENSG00000261663.1_7 | AC009065.9 | 0.99006884 | 1 | 0.075 | 1.643 | chr16 | 2290488 | 2291819 | + | lncRNA |
| ENSG00000187066.8_3 | TMEM262 | 0.89852868 | 1 | 0.075 | -0.642 | chr11 | 64852451 | 64856847 | - | protein_coding |
| ENSG00000128346.11_8 | C22orf23 | 0.91651572 | 1 | 0.075 | -0.102 | chr22 | 38339057 | 38349676 | - | protein_coding |
| ENSG00000176473.13_5 | WDR25 | 0.99948494 | 1 | 0.075 | 2.898 | chr14 | 100842755 | 100996640 | + | protein_coding |
| ENSG00000285437.1_2 | POLR2J3 | 0.96572375 | 1 | 0.075 | 0.968 | chr7 | 102202580 | 102213030 | - | protein_coding |
| ENSG00000187391.21_9 | MAGI2 | 0.98037942 | 1 | 0.075 | 1.252 | chr7 | 77646372 | 79082983 | - | protein_coding |
| ENSG00000187626.9_4 | ZKSCAN4 | 0.99888605 | 1 | 0.075 | 2.508 | chr6 | 28209475 | 28220047 | - | protein_coding |
| ENSG00000129187.14_8 | DCTD | 0.99999996 | 1 | 0.075 | 5.693 | chr4 | 183811213 | 183839089 | - | protein_coding |
| ENSG00000197114.11_4 | ZGPAT | 0.99858104 | 1 | 0.075 | 2.565 | chr20 | 62338817 | 62367494 | + | protein_coding |
| ENSG00000067596.12_6 | DHX8 | 1 | 1 | 0.074 | 7.136 | chr17 | 41561233 | 41687706 | + | protein_coding |
| ENSG00000065675.15_4 | PRKCQ | 0.99992854 | 1 | 0.074 | 3.384 | chr10 | 6469105 | 6622263 | - | protein_coding |
| ENSG00000185722.18_6 | ANKFY1 | 1 | 1 | 0.074 | 6.812 | chr17 | 4067115 | 4167290 | - | protein_coding |
| ENSG00000162441.12_5 | LZIC | 0.99999924 | 1 | 0.074 | 5.227 | chr1 | 9982171 | 10003465 | - | protein_coding |
| ENSG00000198646.14_6 | NCOA6 | 0.99999772 | 1 | 0.074 | 4.332 | chr20 | 33276901 | 33413454 | - | protein_coding |
| ENSG00000253250.3_4 | C8orf88 | 0.92630978 | 1 | 0.074 | 0.071 | chr8 | 91970699 | 91997466 | - | protein_coding |
| ENSG00000158691.14_3 | ZSCAN12 | 0.9998725 | 1 | 0.074 | 3.407 | chr6 | 28346732 | 28367511 | - | protein_coding |
| ENSG00000104643.10_3 | MTMR9 | 0.99999841 | 1 | 0.074 | 4.906 | chr8 | 11142325 | 11185655 | + | protein_coding |
| ENSG00000223703.1_6 | AC027612.1 | 0.91409831 | 1 | 0.074 | -0.131 | chr2 | 91924752 | 91955465 | + | unprocessed_pseudogene |
| ENSG00000267121.6_7 | AC008105.3 | 0.99860765 | 1 | 0.074 | 2.527 | chr17 | 43268298 | 43299589 | - | lncRNA |
| ENSG00000109689.16_6 | STIM2 | 0.9999983 | 1 | 0.074 | 4.730 | chr4 | 26859223 | 27027003 | + | protein_coding |
| ENSG00000088298.13_4 | EDEM2 | 0.99999995 | 1 | 0.074 | 5.865 | chr20 | 33703167 | 33735139 | - | protein_coding |
| ENSG00000139697.14_3 | SBNO1 | 0.99999999 | 1 | 0.074 | 7.643 | chr12 | 123773656 | 123849394 | - | protein_coding |
| ENSG00000277152.1_6 | AC110048.2 | 0.89289195 | 1 | 0.073 | -0.485 | chr15 | 67152641 | 67159361 | + | lncRNA |
| ENSG00000174851.16_5 | YIF1A | 0.99998522 | 1 | 0.073 | 5.703 | chr11 | 66052051 | 66056616 | - | protein_coding |
| ENSG00000070501.12_3 | POLB | 0.99996597 | 1 | 0.073 | 3.932 | chr8 | 42195972 | 42229326 | + | protein_coding |
| ENSG00000160908.15_4 | ZNF394 | 0.99999604 | 1 | 0.073 | 4.445 | chr7 | 99071500 | 99102480 | - | protein_coding |
| ENSG00000228315.12_8 | GUSBP11 | 0.97943972 | 1 | 0.073 | 1.334 | chr22 | 23980674 | 24059543 | - | lncRNA |
| ENSG00000013523.10_6 | ANGEL1 | 0.99997789 | 1 | 0.073 | 4.270 | chr14 | 77252511 | 77292589 | - | protein_coding |
| ENSG00000124593.16_9 | AL365205.1 | 0.89103323 | 1 | 0.073 | -0.644 | chr6 | 41748087 | 41757879 | + | protein_coding |
| ENSG00000222493.1 | Y_RNA | 0.88996536 | 1 | 0.073 | -1.197 | chr12 | 2969115 | 2969199 | + | misc_RNA |
| ENSG00000113811.11_6 | SELENOK | 0.99992374 | 1 | 0.073 | 3.373 | chr3 | 53918444 | 53925989 | - | protein_coding |
| ENSG00000258710.8_7 | LINC01193 | 0.96705502 | 1 | 0.073 | 0.858 | chr15 | 21145709 | 21263769 | + | lncRNA |
| ENSG00000169251.12_7 | NMD3 | 0.9999998 | 1 | 0.073 | 5.679 | chr3 | 160822484 | 160971320 | + | protein_coding |
| ENSG00000170248.15_4 | PDCD6IP | 1 | 1 | 0.072 | 7.832 | chr3 | 33840063 | 33911199 | + | protein_coding |
| ENSG00000180917.17_6 | CMTR2 | 0.99999995 | 1 | 0.072 | 5.733 | chr16 | 71315292 | 71323618 | - | protein_coding |
| ENSG00000272072.1_6 | AC004492.1 | 0.903568 | 1 | 0.072 | -0.488 | chr7 | 106833004 | 106833745 | - | lncRNA |
| ENSG00000164751.15_5 | PEX2 | 0.99999918 | 1 | 0.072 | 4.917 | chr8 | 77892494 | 77913280 | - | protein_coding |
| ENSG00000158773.14_5 | USF1 | 0.99999974 | 1 | 0.072 | 5.174 | chr1 | 161009041 | 161015767 | - | protein_coding |
| ENSG00000147526.20_5 | TACC1 | 1 | 1 | 0.072 | 7.652 | chr8 | 38585704 | 38710546 | + | protein_coding |
| ENSG00000133315.11_5 | MACROD1 | 0.99848847 | 1 | 0.072 | 2.744 | chr11 | 63766030 | 63933585 | - | protein_coding |
| ENSG00000100811.14_6 | YY1 | 1 | 1 | 0.072 | 7.130 | chr14 | 100704635 | 100749125 | + | protein_coding |
| ENSG00000011332.19_6 | DPF1 | 0.99504 | 1 | 0.071 | 1.987 | chr19 | 38701646 | 38720354 | - | protein_coding |
| ENSG00000254909.1_5 | AP003392.2 | 0.92668776 | 1 | 0.071 | 0.289 | chr11 | 118876437 | 118876644 | - | processed_pseudogene |
| ENSG00000136169.16_3 | SETDB2 | 0.99986141 | 1 | 0.071 | 3.262 | chr13 | 50018510 | 50069139 | + | protein_coding |
| ENSG00000169925.17_9 | BRD3 | 0.99999411 | 1 | 0.071 | 4.197 | chr9 | 136895427 | 136933657 | - | protein_coding |

| | | | | | | | | | | |
|---|---|---|---|---|---|---|---|---|---|---|
| ENSG00000088247.17_5 | KHSRP | 0.99999938 | 1 | 0.071 | 6.584 | chr19 | 6413359 | 6424805 | - | protein_coding |
| ENSG00000104897.10_8 | SF3A2 | 0.99999499 | 1 | 0.071 | 4.974 | chr19 | 2236823 | 2248654 | + | protein_coding |
| ENSG00000137509.11_7 | PRCP | 0.99999982 | 1 | 0.071 | 5.654 | chr11 | 82533978 | 82681626 | - | protein_coding |
| ENSG00000176783.15_7 | RUFY1 | 0.99999993 | 1 | 0.071 | 5.436 | chr5 | 178977555 | 179037013 | + | protein_coding |
| ENSG00000139977.14_6 | NAA30 | 0.99999994 | 1 | 0.071 | 5.655 | chr14 | 57857262 | 57882624 | + | protein_coding |
| ENSG00000006715.16_5 | VPS41 | 0.99999996 | 1 | 0.071 | 6.186 | chr7 | 38762574 | 38971994 | - | protein_coding |
| ENSG00000179364.13_6 | PACS2 | 0.99998285 | 1 | 0.071 | 5.872 | chr14 | 105766900 | 105864484 | + | protein_coding |
| ENSG00000112031.16_4 | MTRF1L | 0.99996193 | 1 | 0.070 | 3.481 | chr6 | 153308497 | 153323844 | - | protein_coding |
| ENSG00000198160.14_3 | MIER1 | 0.99999864 | 1 | 0.070 | 5.431 | chr1 | 67390578 | 67454302 | + | protein_coding |
| ENSG00000108799.13_4 | EZH1 | 0.99999868 | 1 | 0.070 | 5.149 | chr17 | 40852293 | 40897067 | - | protein_coding |
| ENSG00000142534.7_4 | RPS11 | 1 | 1 | 0.070 | 9.586 | chr19 | 49999622 | 50002965 | + | protein_coding |
| ENSG00000109572.14_7 | CLCN3 | 0.99999996 | 1 | 0.070 | 5.867 | chr4 | 170533784 | 170644824 | + | protein_coding |
| ENSG00000169282.17_5 | KCNAB1 | 0.99838299 | 1 | 0.070 | 1.635 | chr3 | 155755490 | 156256927 | + | protein_coding |
| ENSG00000250337.7_6 | PURPL | 0.99999977 | 1 | 0.070 | 5.295 | chr5 | 27217821 | 27497101 | + | lncRNA |
| ENSG00000183155.5_4 | RABIF | 0.99999447 | 1 | 0.070 | 4.217 | chr1 | 202847410 | 202858277 | - | protein_coding |
| ENSG00000076555.15_5 | ACACB | 0.99999961 | 1 | 0.070 | 5.129 | chr12 | 109554400 | 109706031 | + | protein_coding |
| ENSG00000116688.17_7 | MFN2 | 0.99999999 | 1 | 0.070 | 6.208 | chr1 | 12040238 | 12073571 | + | protein_coding |
| ENSG00000283103.3_1 | AC010642.2 | 0.99098111 | 1 | 0.070 | 1.684 | chr19 | 58821093 | 58838614 | - | lncRNA |
| ENSG00000048707.15_8 | VPS13D | 0.99999993 | 1 | 0.070 | 6.045 | chr1 | 12290087 | 12572099 | + | protein_coding |
| ENSG00000071894.17_7 | CPSF1 | 0.99999992 | 1 | 0.070 | 6.335 | chr8 | 145618444 | 145634719 | - | protein_coding |
| ENSG00000228451.4_5 | SDAD1P1 | 0.98977173 | 1 | 0.069 | 1.491 | chr8 | 26233429 | 26240469 | - | transcribed_processed_pseudogene |
| ENSG00000198276.16_5 | UCKL1 | 0.99995844 | 1 | 0.069 | 3.976 | chr20 | 62571182 | 62587769 | - | protein_coding |
| ENSG00000178694.10_3 | NSUN3 | 0.99999912 | 1 | 0.069 | 4.646 | chr3 | 93781824 | 93850676 | + | protein_coding |
| ENSG00000151422.13_4 | FER | 0.99999529 | 1 | 0.069 | 5.210 | chr5 | 108083542 | 108532542 | + | protein_coding |
| ENSG00000146242.8_2 | TPBG | 0.99999994 | 1 | 0.069 | 5.620 | chr6 | 83072923 | 83080545 | + | protein_coding |
| ENSG00000277156.1_4 | AL589743.6 | 0.89646957 | 1 | 0.069 | -0.896 | chr14 | 19854816 | 19855865 | + | processed_pseudogene |
| ENSG00000188921.14_4 | HACD4 | 0.99992816 | 1 | 0.069 | 3.394 | chr9 | 20999508 | 21031639 | - | protein_coding |
| ENSG00000247735.2_6 | AC120114.1 | 0.98465463 | 1 | 0.069 | 1.373 | chr16 | 29938157 | 29940254 | + | lncRNA |
| ENSG00000151117.9_3 | TMEM86A | 0.90045616 | 1 | 0.069 | -0.784 | chr11 | 18714669 | 18726332 | + | protein_coding |
| ENSG00000167395.10_6 | ZNF646 | 0.99999896 | 1 | 0.069 | 5.476 | chr16 | 31085743 | 31095517 | + | protein_coding |
| ENSG00000132424.16_6 | PNISR | 0.9999964 | 1 | 0.069 | 6.297 | chr6 | 99845926 | 99873207 | - | protein_coding |
| ENSG00000119986.7_4 | AVPI1 | 0.99999997 | 1 | 0.069 | 5.992 | chr10 | 99437181 | 99446998 | - | protein_coding |
| ENSG00000133612.19_5 | AGAP3 | 0.99999912 | 1 | 0.069 | 5.190 | chr7 | 150782918 | 150841523 | + | protein_coding |
| ENSG00000075089.9_3 | ACTR6 | 0.99999928 | 1 | 0.068 | 5.019 | chr12 | 100592900 | 100635643 | + | protein_coding |
| ENSG00000082701.15_6 | GSK3B | 1 | 1 | 0.068 | 6.407 | chr3 | 119540170 | 119813264 | - | protein_coding |
| ENSG00000270095.1_7 | AC068790.6 | 0.89558268 | 1 | 0.068 | -1.104 | chr12 | 124456004 | 124456261 | - | lncRNA |
| ENSG00000116990.11_5 | MYCL | 0.93391051 | 1 | 0.068 | 0.209 | chr1 | 40361098 | 40367928 | - | protein_coding |
| ENSG00000188013.6_4 | MEIS3P2 | 0.98116912 | 1 | 0.068 | 0.848 | chr17 | 20492606 | 20493680 | + | processed_pseudogene |
| ENSG00000151338.18_6 | MIPOL1 | 0.99999383 | 1 | 0.068 | 4.151 | chr14 | 37667118 | 38021566 | + | protein_coding |
| ENSG00000169727.12_5 | GPS1 | 0.99999999 | 1 | 0.068 | 7.026 | chr17 | 80008567 | 80015346 | + | protein_coding |
| ENSG00000198455.4_2 | ZXDB | 0.99999719 | 1 | 0.068 | 4.378 | chrX | 57618085 | 57623978 | + | protein_coding |
| ENSG00000273559.5_5 | CWC25 | 0.99999953 | 1 | 0.068 | 4.836 | chr17 | 36956694 | 36981608 | - | protein_coding |
| ENSG00000144959.10_6 | NCEH1 | 0.99999999 | 1 | 0.068 | 6.202 | chr3 | 172348039 | 172429008 | - | protein_coding |
| ENSG00000105221.18_8 | AKT2 | 1 | 1 | 0.068 | 7.714 | chr19 | 40736224 | 40791443 | - | protein_coding |
| ENSG00000173681.16_7 | BCLAF3 | 0.99998885 | 1 | 0.068 | 3.877 | chrX | 19930978 | 19988416 | - | protein_coding |
| ENSG00000136490.9_6 | LIMD2 | 0.99863316 | 1 | 0.068 | 2.716 | chr17 | 61773248 | 61778532 | - | protein_coding |
| ENSG00000116717.13_4 | GADD45A | 0.99970243 | 1 | 0.068 | 3.099 | chr1 | 68150884 | 68154017 | + | protein_coding |
| ENSG00000140983.14_4 | RHOT2 | 0.99999883 | 1 | 0.068 | 5.851 | chr16 | 718105 | 724174 | + | protein_coding |
| ENSG00000086475.15_4 | SEPHS1 | 0.99999999 | 1 | 0.068 | 6.071 | chr10 | 13359428 | 13390298 | - | protein_coding |
| ENSG00000081913.14_4 | PHLPP1 | 0.99999811 | 1 | 0.068 | 4.336 | chr18 | 60382774 | 60647666 | + | protein_coding |
| ENSG00000120318.16_4 | ARAP3 | 0.99998739 | 1 | 0.068 | 4.423 | chr5 | 141032968 | 141061797 | - | protein_coding |
| ENSG00000174405.13_3 | LIG4 | 0.99999215 | 1 | 0.068 | 4.037 | chr13 | 108859787 | 108870716 | - | protein_coding |
| ENSG00000005075.15_3 | POLR2J | 0.99999814 | 1 | 0.068 | 4.547 | chr7 | 102113565 | 102119354 | - | protein_coding |
| ENSG00000119912.17_5 | IDE | 1 | 1 | 0.068 | 7.319 | chr10 | 94211441 | 94333850 | - | protein_coding |
| ENSG00000064995.18_5 | TAF11 | 0.99996964 | 1 | 0.067 | 3.679 | chr6 | 34845239 | 34855848 | - | protein_coding |
| ENSG00000155256.17_4 | ZFYVE27 | 0.9999995 | 1 | 0.067 | 4.844 | chr10 | 99496878 | 99520664 | + | protein_coding |
| ENSG00000141380.14_6 | SS18 | 1 | 1 | 0.067 | 6.785 | chr18 | 23596217 | 23671181 | - | protein_coding |
| ENSG00000141068.14_7 | KSR1 | 0.999991 | 1 | 0.067 | 4.403 | chr17 | 25783496 | 25953464 | + | protein_coding |
| ENSG00000278989.1_5 | AP001148.1 | 0.94894599 | 1 | 0.067 | 0.176 | chr11 | 86145966 | 86148324 | - | TEC |

| | | | | | | | | | | |
|---|---|---|---|---|---|---|---|---|---|---|
| ENSG00000163625.15_5 | WDFY3 | 0.99999994 | 1 | 0.067 | 5.803 | chr4 | 85590763 | 85887544 | - | protein_coding |
| ENSG00000132275.11_4 | RRP8 | 0.99999986 | 1 | 0.067 | 5.128 | chr11 | 6616302 | 6624846 | - | protein_coding |
| ENSG00000266896.1_7 | AL354892.3 | 0.8976027 | 1 | 0.067 | -0.978 | chr6 | 170170509 | 170172138 | + | lncRNA |
| ENSG00000073008.15_5 | PVR | 1 | 1 | 0.067 | 7.044 | chr19 | 45147098 | 45169430 | + | protein_coding |
| ENSG00000118197.14_9 | DDX59 | 0.99999877 | 1 | 0.067 | 4.740 | chr1 | 200593024 | 200639035 | - | protein_coding |
| ENSG00000175175.6_4 | PPM1E | 0.91010715 | 1 | 0.067 | -0.272 | chr17 | 56833215 | 57062540 | + | protein_coding |
| ENSG00000136816.16_3 | TOR1B | 0.99999959 | 1 | 0.067 | 4.897 | chr9 | 132565436 | 132573560 | + | protein_coding |
| ENSG00000134668.12_5 | SPOCD1 | 0.94242693 | 1 | 0.067 | 0.384 | chr1 | 32256023 | 32281652 | - | protein_coding |
| ENSG00000247950.7_7 | SEC24B-AS1 | 0.9735408 | 1 | 0.067 | 1.081 | chr4 | 110268631 | 110354973 | - | lncRNA |
| ENSG00000157741.15_3 | UBN2 | 0.99999925 | 1 | 0.067 | 4.833 | chr7 | 138915102 | 138992982 | + | protein_coding |
| ENSG00000144579.7_5 | CTDSP1 | 0.9999999 | 1 | 0.066 | 5.984 | chr2 | 219262979 | 219270664 | + | protein_coding |
| ENSG00000099308.10_4 | MAST3 | 0.99936877 | 1 | 0.066 | 2.797 | chr19 | 18208603 | 18262502 | + | protein_coding |
| ENSG00000258904.1_7 | AL157871.5 | 0.90057206 | 1 | 0.066 | -0.896 | chr14 | 100730118 | 100731742 | + | lncRNA |
| ENSG00000260442.5_7 | ATP2A1-AS1 | 0.90563786 | 1 | 0.066 | -1.065 | chr16 | 28890278 | 28891241 | - | lncRNA |
| ENSG00000064199.6_2 | SPA17 | 0.99846693 | 1 | 0.066 | 2.410 | chr11 | 124543694 | 124567414 | + | protein_coding |
| ENSG00000134086.7_2 | VHL | 0.9999977 | 1 | 0.066 | 4.820 | chr3 | 10182692 | 10193904 | + | protein_coding |
| ENSG00000121060.18_7 | TRIM25 | 0.99999996 | 1 | 0.066 | 5.883 | chr17 | 54913748 | 54991441 | - | protein_coding |
| ENSG00000171992.13_3 | SYNPO | 0.99999994 | 1 | 0.066 | 5.852 | chr5 | 149980642 | 150038769 | + | protein_coding |
| ENSG00000268460.2_7 | AC006262.1 | 0.99849932 | 1 | 0.066 | 2.413 | chr19 | 46704809 | 46718095 | + | lncRNA |
| ENSG00000279495.1_6 | AL928654.4 | 0.97901371 | 1 | 0.066 | 0.723 | chr14 | 105836755 | 105839255 | + | TEC |
| ENSG00000136161.12_5 | RCBTB2 | 0.97677893 | 1 | 0.066 | 1.228 | chr13 | 49063095 | 49107392 | - | protein_coding |
| ENSG00000161791.14_6 | FMNL3 | 0.99992349 | 1 | 0.065 | 3.313 | chr12 | 50030282 | 50101948 | - | protein_coding |
| ENSG00000180694.14_5 | TMEM64 | 0.99999998 | 1 | 0.065 | 5.868 | chr8 | 91634223 | 91803860 | - | protein_coding |
| ENSG00000172269.19_5 | DPAGT1 | 0.99999885 | 1 | 0.065 | 4.815 | chr11 | 118967213 | 118979041 | - | protein_coding |
| ENSG00000023287.13_7 | RB1CC1 | 0.99999987 | 1 | 0.065 | 7.089 | chr8 | 53535018 | 53658403 | - | protein_coding |
| ENSG00000150455.13_3 | TIRAP | 0.99653529 | 1 | 0.065 | 1.977 | chr11 | 126152960 | 126168740 | + | protein_coding |
| ENSG00000160710.17_8 | ADAR | 1 | 1 | 0.065 | 8.356 | chr1 | 154554533 | 154600489 | - | protein_coding |
| ENSG00000003756.17_7 | RBM5 | 0.9999999 | 1 | 0.065 | 5.627 | chr3 | 50126352 | 50156454 | + | protein_coding |
| ENSG00000017260.19_7 | ATP2C1 | 0.99999995 | 1 | 0.065 | 5.882 | chr3 | 130569439 | 130735556 | + | protein_coding |
| ENSG00000139675.12_4 | HNRNPA1L2 | 0.92255818 | 1 | 0.065 | 0.049 | chr13 | 53216560 | 53217931 | + | protein_coding |
| ENSG00000251003.9_6 | ZFPM2-AS1 | 0.99932709 | 1 | 0.065 | 2.875 | chr8 | 106558317 | 107072752 | - | lncRNA |
| ENSG00000253982.2_7 | AC100810.1 | 0.99995947 | 1 | 0.065 | 3.662 | chr8 | 1709220 | 1712674 | - | lncRNA |
| ENSG00000188070.9_5 | C11orf95 | 0.99978342 | 1 | 0.065 | 3.943 | chr11 | 63527364 | 63536247 | - | protein_coding |
| ENSG00000279407.1_6 | AC007191.1 | 0.99678085 | 1 | 0.065 | 2.291 | chr19 | 46217645 | 46220639 | - | TEC |
| ENSG00000119392.14_2 | GLE1 | 0.99999999 | 1 | 0.064 | 6.249 | chr9 | 131266979 | 131304567 | + | protein_coding |
| ENSG00000156650.14_7 | KAT6B | 0.99999888 | 1 | 0.064 | 4.782 | chr10 | 76584685 | 76792382 | + | protein_coding |
| ENSG00000105518.14_5 | TMEM205 | 0.99998538 | 1 | 0.064 | 4.019 | chr19 | 11453452 | 11457194 | - | protein_coding |
| ENSG00000103145.10_4 | HCFC1R1 | 0.99999978 | 1 | 0.064 | 5.617 | chr16 | 3072621 | 3074287 | - | protein_coding |
| ENSG00000107077.18_4 | KDM4C | 0.99999814 | 1 | 0.064 | 4.692 | chr9 | 6720863 | 7175648 | + | protein_coding |
| ENSG00000158062.20_5 | UBXN11 | 0.9999678 | 1 | 0.064 | 3.544 | chr1 | 26607819 | 26644854 | - | protein_coding |
| ENSG00000166676.16_7 | TVP23A | 0.91545701 | 1 | 0.064 | -0.172 | chr16 | 10854776 | 10912651 | - | protein_coding |
| ENSG00000185133.14_5 | INPP5J | 0.98103135 | 1 | 0.064 | 1.232 | chr22 | 31518717 | 31530683 | + | protein_coding |
| ENSG00000086666.19_6 | ZFAND6 | 0.99999999 | 1 | 0.064 | 6.776 | chr15 | 80351910 | 80430735 | + | protein_coding |
| ENSG00000107854.6_3 | TNKS2 | 1 | 1 | 0.064 | 6.329 | chr10 | 93558183 | 93625232 | + | protein_coding |
| ENSG00000234616.9_4 | JRK | 0.99999928 | 1 | 0.064 | 4.966 | chr8 | 143738874 | 143763386 | - | protein_coding |
| ENSG00000276791.1_6 | AC092117.1 | 0.99858479 | 1 | 0.064 | 2.303 | chr16 | 2827320 | 2830569 | + | lncRNA |
| ENSG00000173933.20_4 | RBM4 | 0.9999196 | 1 | 0.064 | 3.524 | chr11 | 66406088 | 66435845 | + | protein_coding |
| ENSG00000240184.7_5 | PCDHGC3 | 0.99869386 | 1 | 0.064 | 2.648 | chr5 | 140855514 | 140892544 | + | protein_coding |
| ENSG00000165487.14_4 | MICU2 | 0.99999991 | 1 | 0.064 | 5.332 | chr13 | 22066830 | 22178320 | - | protein_coding |
| ENSG00000100354.21_7 | TNRC6B | 0.99999996 | 1 | 0.064 | 5.909 | chr22 | 40440821 | 40731812 | + | protein_coding |
| ENSG00000064999.15_6 | ANKS1A | 0.99999808 | 1 | 0.064 | 4.293 | chr6 | 34857032 | 35059183 | + | protein_coding |
| ENSG00000173465.8_5 | ZNRD2 | 0.99994381 | 1 | 0.064 | 3.795 | chr11 | 65337931 | 65341413 | + | protein_coding |
| ENSG00000180098.9_3 | TRNAU1AP | 0.9999645 | 1 | 0.064 | 3.653 | chr1 | 28879597 | 28905057 | + | protein_coding |
| ENSG00000110031.12_3 | LPXN | 0.99443461 | 1 | 0.063 | 1.886 | chr11 | 58294344 | 58345693 | - | protein_coding |
| ENSG00000164951.16_6 | PDP1 | 0.99999832 | 1 | 0.063 | 4.524 | chr8 | 94870035 | 94938296 | + | protein_coding |
| ENSG00000125520.14_4 | SLC2A4RG | 0.99983027 | 1 | 0.063 | 5.451 | chr20 | 62371129 | 62375403 | + | protein_coding |
| ENSG00000136950.13_3 | ARPC5L | 0.99999985 | 1 | 0.063 | 5.008 | chr9 | 127624409 | 127640012 | + | protein_coding |
| ENSG00000287966.1_1 | AL590062.1 | 0.90464584 | 1 | 0.063 | -0.784 | chr6 | 26986714 | 26988059 | - | lncRNA |
| ENSG00000140941.14_4 | MAP1LC3B | 0.99999589 | 1 | 0.063 | 4.268 | chr16 | 87417559 | 87438385 | + | protein_coding |
| ENSG00000228049.7_4 | POLR2J2 | 0.91284731 | 1 | 0.063 | -0.616 | chr7 | 102305815 | 102312076 | - | protein_coding |
| ENSG00000170836.12_3 | PPM1D | 0.99999974 | 1 | 0.063 | 5.029 | chr17 | 58677544 | 58743641 | + | protein_coding |

| | | | | | | | | | | |
|---|---|---|---|---|---|---|---|---|---|---|
| ENSG00000141446.11_4 | ESCO1 | 0.99999988 | 1 | 0.063 | 5.095 | chr18 | 19109245 | 19180845 | - | protein_coding |
| ENSG00000169220.18_3 | RGS14 | 0.9925232 | 1 | 0.063 | 1.732 | chr5 | 176784925 | 176799597 | + | protein_coding |
| ENSG00000168970.22_3 | JMJD7-PLA2G4B | 0.97521464 | 1 | 0.063 | 1.055 | chr15 | 42120293 | 42140353 | + | protein_coding |
| ENSG00000127445.13_3 | PIN1 | 0.99999964 | 1 | 0.063 | 5.113 | chr19 | 9945933 | 9960358 | + | protein_coding |
| ENSG00000198799.12_4 | LRIG2 | 0.99999758 | 1 | 0.063 | 4.292 | chr1 | 113615820 | 113674882 | + | protein_coding |
| ENSG00000164989.17_4 | CCDC171 | 0.99902141 | 1 | 0.063 | 2.603 | chr9 | 15553041 | 16061661 | + | protein_coding |
| ENSG00000121851.13_5 | POLR3GL | 0.99598055 | 1 | 0.062 | 2.079 | chr1 | 145456236 | 145470400 | - | protein_coding |
| ENSG00000160392.13_6 | C19orf47 | 0.99999825 | 1 | 0.062 | 4.486 | chr19 | 40825443 | 40854434 | - | protein_coding |
| ENSG00000069667.16_7 | RORA | 0.99665504 | 1 | 0.062 | 1.938 | chr15 | 60780483 | 61521501 | - | protein_coding |
| ENSG00000088876.13_5 | ZNF343 | 0.9999826 | 1 | 0.062 | 4.011 | chr20 | 2462463 | 2505348 | - | protein_coding |
| ENSG00000103245.14_7 | CIAO3 | 0.99999729 | 1 | 0.062 | 4.439 | chr16 | 779760 | 791329 | - | protein_coding |
| ENSG00000168701.19_3 | TMEM208 | 0.9999925 | 1 | 0.062 | 4.150 | chr16 | 67261006 | 67263181 | + | protein_coding |
| ENSG00000131759.18_7 | RARA | 0.9999968 | 1 | 0.062 | 4.533 | chr17 | 38465432 | 38513895 | + | protein_coding |
| ENSG00000197157.11_6 | SND1 | 1 | 1 | 0.062 | 7.697 | chr7 | 127292248 | 127732661 | + | protein_coding |
| ENSG00000083168.11_7 | KAT6A | 0.99999999 | 1 | 0.062 | 6.222 | chr8 | 41786997 | 41909512 | - | protein_coding |
| ENSG00000092108.21_8 | SCFD1 | 0.99999987 | 1 | 0.062 | 6.039 | chr14 | 31091517 | 31205056 | + | protein_coding |
| ENSG00000140829.12_5 | DHX38 | 0.99999999 | 1 | 0.062 | 6.187 | chr16 | 72127512 | 72146811 | + | protein_coding |
| ENSG00000166557.13_4 | TMED3 | 1 | 1 | 0.062 | 6.967 | chr15 | 79603454 | 79719774 | + | protein_coding |
| ENSG00000223705.9_3 | NSUN5P1 | 0.99998568 | 1 | 0.062 | 3.812 | chr7 | 75039605 | 75046066 | + | transcribed_unprocessed_pseudogene |
| ENSG00000124164.16_3 | VAPB | 0.99999998 | 1 | 0.062 | 5.711 | chr20 | 56964285 | 57026157 | + | protein_coding |
| ENSG00000237289.9_2 | CKMT1B | 0.92337339 | 1 | 0.062 | -0.438 | chr15 | 43885252 | 43897099 | + | protein_coding |
| ENSG00000137700.18_9 | SLC37A4 | 0.99996583 | 1 | 0.062 | 3.800 | chr11 | 118894824 | 118901616 | - | protein_coding |
| ENSG00000090372.15_6 | STRN4 | 0.99999928 | 1 | 0.062 | 6.705 | chr19 | 47222768 | 47250251 | - | protein_coding |
| ENSG00000055208.19_5 | TAB2 | 0.99999998 | 1 | 0.062 | 5.718 | chr6 | 149539777 | 149732749 | + | protein_coding |
| ENSG00000172890.13_5 | NADSYN1 | 0.99999999 | 1 | 0.062 | 6.295 | chr11 | 71164155 | 71235153 | + | protein_coding |
| ENSG00000169026.12_6 | SLC49A3 | 0.97939819 | 1 | 0.062 | 0.749 | chr4 | 675618 | 683230 | - | protein_coding |
| ENSG00000248714.7_7 | AC091180.3 | 0.90520269 | 1 | 0.062 | -0.978 | chr17 | 47438512 | 47447360 | + | lncRNA |
| ENSG00000086102.19_3 | NFX1 | 0.99999997 | 1 | 0.061 | 5.788 | chr9 | 33290510 | 33371155 | + | protein_coding |
| ENSG00000175886.10_4 | RPL7AP66 | 0.90673793 | 1 | 0.061 | -0.714 | chr18 | 36914836 | 36915636 | - | processed_pseudogene |
| ENSG00000123728.10_4 | RAP2C | 0.99999996 | 1 | 0.061 | 5.558 | chrX | 131337052 | 131353508 | - | protein_coding |
| ENSG00000251584.2_6 | AC096751.2 | 0.9103363 | 1 | 0.061 | -0.525 | chr4 | 175443768 | 175462198 | + | lncRNA |
| ENSG00000164603.12_7 | BMT2 | 0.9999816 | 1 | 0.061 | 3.889 | chr7 | 112459202 | 112579930 | - | protein_coding |
| ENSG00000171119.2_2 | NRTN | 0.91118668 | 1 | 0.061 | -0.786 | chr19 | 5823813 | 5828335 | + | protein_coding |
| ENSG00000198951.12_4 | NAGA | 0.99999833 | 1 | 0.061 | 4.267 | chr22 | 42454338 | 42466846 | - | protein_coding |
| ENSG00000135241.17_8 | PNPLA8 | 0.99999976 | 1 | 0.061 | 5.210 | chr7 | 108110861 | 108210110 | - | protein_coding |
| ENSG00000174804.4_4 | FZD4 | 0.99731521 | 1 | 0.061 | 2.157 | chr11 | 86656721 | 86666437 | - | protein_coding |
| ENSG00000185650.10_4 | ZFP36L1 | 1 | 1 | 0.061 | 6.965 | chr14 | 69254377 | 69262970 | - | protein_coding |
| ENSG00000034533.11_6 | ASTE1 | 0.99950515 | 1 | 0.061 | 2.722 | chr3 | 130732719 | 130746493 | - | protein_coding |
| ENSG00000255568.2 | BRWD1-IT2 | 0.91830776 | 1 | 0.061 | -0.465 | chr21 | 40685887 | 40686886 | - | sense_intronic |
| ENSG00000168159.12_5 | RNF187 | 0.99999999 | 1 | 0.061 | 6.564 | chr1 | 228675083 | 228683889 | + | protein_coding |
| ENSG00000215908.10_4 | CROCCP2 | 0.99999952 | 1 | 0.060 | 4.968 | chr1 | 16944748 | 16983727 | - | transcribed_unprocessed_pseudogene |
| ENSG00000133606.11_8 | MKRN1 | 1 | 1 | 0.060 | 6.630 | chr7 | 140152833 | 140179336 | - | protein_coding |
| ENSG00000285863.1_2 | AC009554.2 | 0.9165713 | 1 | 0.060 | -0.754 | chr15 | 62153308 | 62186363 | + | lncRNA |
| ENSG00000096872.16_5 | IFT74 | 0.9999996 | 1 | 0.060 | 5.027 | chr9 | 26947037 | 27062928 | + | protein_coding |
| ENSG00000219481.10_4 | NBPF1 | 1 | 1 | 0.060 | 6.502 | chr1 | 16888814 | 16940057 | - | protein_coding |
| ENSG00000213903.9_4 | LTB4R | 0.99999797 | 1 | 0.060 | 4.527 | chr14 | 24780656 | 24787242 | + | protein_coding |
| ENSG00000104915.15_6 | STX10 | 0.99999985 | 1 | 0.060 | 4.980 | chr19 | 13254872 | 13261197 | - | protein_coding |
| ENSG00000011007.12_3 | ELOA | 0.99999996 | 1 | 0.060 | 5.455 | chr1 | 24069645 | 24088549 | + | protein_coding |
| ENSG00000273749.5_3 | CYFIP1 | 1 | 1 | 0.060 | 7.573 | chr15 | 22892005 | 23006016 | + | protein_coding |
| ENSG00000138380.18_6 | CARF | 0.99991557 | 1 | 0.059 | 3.322 | chr2 | 203776937 | 203852986 | + | protein_coding |
| ENSG00000068097.15_5 | HEATR6 | 0.99999995 | 1 | 0.059 | 5.585 | chr17 | 58118369 | 58156283 | - | protein_coding |
| ENSG00000170832.13_5 | USP32 | 0.99999999 | 1 | 0.059 | 6.474 | chr17 | 58254688 | 58499831 | - | protein_coding |
| ENSG00000131941.8_4 | RHPN2 | 1 | 1 | 0.059 | 7.447 | chr19 | 33469498 | 33555794 | - | protein_coding |
| ENSG00000103671.10_5 | TRIP4 | 0.99999231 | 1 | 0.059 | 4.018 | chr15 | 64679947 | 64747502 | + | protein_coding |
| ENSG00000269959.1_6 | SPACA6P-AS | 0.99746844 | 1 | 0.059 | 2.255 | chr19 | 52188616 | 52196709 | - | lncRNA |
| ENSG00000118369.13_7 | USP35 | 0.99986333 | 1 | 0.059 | 3.232 | chr11 | 77899858 | 77926278 | + | protein_coding |
| ENSG00000234648.1_5 | AL162151.2 | 0.91039983 | 1 | 0.059 | -0.979 | chr14 | 99439651 | 99439808 | + | processed_pseudogene |
| ENSG00000232480.1_7 | TGFB2-AS1 | 0.91683205 | 1 | 0.059 | -0.294 | chr1 | 218517538 | 218519020 | - | lncRNA |

| | | | | | | | | | | |
|---|---|---|---|---|---|---|---|---|---|---|
| ENSG00000236682.2_7 | MAP3K2-DT | 0.97570297 | 1 | 0.059 | 0.601 | chr2 | 128145760 | 128158156 | + | lncRNA |
| ENSG00000106100.11_6 | NOD1 | 0.99984649 | 1 | 0.059 | 3.109 | chr7 | 30464143 | 30518400 | - | protein_coding |
| ENSG00000159592.11_6 | GPBP1L1 | 1 | 1 | 0.059 | 6.447 | chr1 | 46092976 | 46153785 | - | protein_coding |
| ENSG00000100916.14_4 | BRMS1L | 0.99998845 | 1 | 0.059 | 3.856 | chr14 | 36295544 | 36401531 | + | protein_coding |
| ENSG00000205758.12_7 | CRYZL1 | 0.99986269 | 1 | 0.058 | 3.271 | chr21 | 34961647 | 35016232 | - | protein_coding |
| ENSG00000247708.8_6 | STX18-AS1 | 0.97872318 | 1 | 0.058 | 1.422 | chr4 | 4543858 | 4789086 | + | lncRNA |
| ENSG00000156931.15_8 | VPS8 | 0.99999987 | 1 | 0.058 | 5.485 | chr3 | 184529931 | 184770402 | + | protein_coding |
| ENSG00000102699.6_5 | PARP4 | 1 | 1 | 0.058 | 7.086 | chr13 | 24995069 | 25086916 | - | protein_coding |
| ENSG00000173486.13_3 | FKBP2 | 0.9164498 | 1 | 0.058 | -0.683 | chr11 | 64008475 | 64011606 | + | protein_coding |
| ENSG00000226007.3_6 | BX005266.2 | 0.92184721 | 1 | 0.058 | -0.259 | chr9 | 46843638 | 46846025 | + | lncRNA |
| ENSG00000172465.14_4 | TCEAL1 | 0.99999058 | 1 | 0.058 | 3.980 | chrX | 102883632 | 102885881 | + | protein_coding |
| ENSG00000119682.17_8 | AREL1 | 0.99999999 | 1 | 0.058 | 6.102 | chr14 | 75120140 | 75179811 | - | protein_coding |
| ENSG00000196781.16_6 | TLE1 | 0.9999958 | 1 | 0.058 | 6.360 | chr9 | 84198598 | 84304462 | - | protein_coding |
| ENSG00000009335.18_4 | UBE3C | 1 | 1 | 0.058 | 7.241 | chr7 | 156931610 | 157062064 | + | protein_coding |
| ENSG00000197375.12_2 | SLC22A5 | 0.9999999 | 1 | 0.058 | 5.313 | chr5 | 131705444 | 131731306 | + | protein_coding |
| ENSG00000279520.1_6 | AC093525.8 | 0.94440852 | 1 | 0.057 | 0.263 | chr16 | 2566659 | 2568000 | + | TEC |
| ENSG00000125257.16_7 | ABCC4 | 1 | 1 | 0.057 | 7.630 | chr13 | 95672089 | 95953729 | - | protein_coding |
| ENSG00000163378.14_6 | EOGT | 0.99999431 | 1 | 0.057 | 4.225 | chr3 | 69024368 | 69062835 | - | protein_coding |
| ENSG00000116661.11_6 | FBXO2 | 0.9988952 | 1 | 0.057 | 2.288 | chr1 | 11697075 | 11715842 | - | protein_coding |
| ENSG00000181004.10_3 | BBS12 | 0.99924751 | 1 | 0.057 | 2.592 | chr4 | 123653857 | 123666097 | + | protein_coding |
| ENSG00000214026.11_3 | MRPL23 | 0.99901128 | 1 | 0.057 | 5.354 | chr11 | 1968508 | 2005752 | + | protein_coding |
| ENSG00000258643.5_5 | BCL2L2-PABPN1 | 0.91144498 | 1 | 0.057 | -1.065 | chr14 | 23776044 | 23794578 | + | protein_coding |
| ENSG00000276255.2_7 | LINC02809 | 0.98878613 | 1 | 0.057 | 0.929 | chr1 | 228261610 | 228264251 | - | lncRNA |
| ENSG00000196689.12_6 | TRPV1 | 0.9995617 | 1 | 0.057 | 2.905 | chr17 | 3468738 | 3512705 | - | protein_coding |
| ENSG00000131013.4_4 | PPIL4 | 0.99999941 | 1 | 0.057 | 4.760 | chr6 | 149825631 | 149867179 | - | protein_coding |
| ENSG00000146463.12_4 | ZMYM4 | 0.99999999 | 1 | 0.057 | 6.353 | chr1 | 35734310 | 35887659 | + | protein_coding |
| ENSG00000236914.4_7 | LINC01852 | 0.91226365 | 1 | 0.057 | -0.978 | chr15 | 38361013 | 38365191 | - | lncRNA |
| ENSG00000102897.10_3 | LYRM1 | 0.99999906 | 1 | 0.057 | 4.504 | chr16 | 20911190 | 20936328 | + | protein_coding |
| ENSG00000136153.20_9 | LMO7 | 1 | 1 | 0.057 | 6.346 | chr13 | 76194570 | 76434006 | + | protein_coding |
| ENSG00000184743.13_5 | ATL3 | 1 | 1 | 0.057 | 7.983 | chr11 | 63391559 | 63439393 | - | protein_coding |
| ENSG00000279048.1_5 | AC080080.1 | 0.92430703 | 1 | 0.057 | -0.416 | chr7 | 17980126 | 17982545 | + | TEC |
| ENSG00000166949.16_4 | SMAD3 | 1 | 1 | 0.056 | 8.258 | chr15 | 67356101 | 67487507 | + | protein_coding |
| ENSG00000265241.6_5 | RBM8A | 0.99999998 | 1 | 0.056 | 5.731 | chr1 | 145507415 | 145517375 | + | protein_coding |
| ENSG00000146063.20_3 | TRIM41 | 0.99999838 | 1 | 0.056 | 5.095 | chr5 | 180649499 | 180662808 | + | protein_coding |
| ENSG00000156535.15_7 | CD109 | 0.99999999 | 1 | 0.056 | 7.117 | chr6 | 74405508 | 74538040 | + | protein_coding |
| ENSG00000137815.14_2 | RTF1 | 0.99999996 | 1 | 0.056 | 5.406 | chr15 | 41700606 | 41775761 | + | protein_coding |
| ENSG00000095303.17_7 | PTGS1 | 0.93038208 | 1 | 0.056 | -0.042 | chr9 | 125132809 | 125157982 | + | protein_coding |
| ENSG00000106144.20_3 | CASP2 | 0.99999991 | 1 | 0.056 | 5.064 | chr7 | 142985308 | 143004789 | + | protein_coding |
| ENSG00000147439.13_5 | BIN3 | 0.99995495 | 1 | 0.056 | 3.711 | chr8 | 22477931 | 22526661 | - | protein_coding |
| ENSG00000086232.13_7 | EIF2AK1 | 1 | 1 | 0.056 | 6.875 | chr7 | 6061878 | 6098806 | - | protein_coding |
| ENSG00000173950.16_5 | XXYLT1 | 0.99999678 | 1 | 0.056 | 4.084 | chr3 | 194789013 | 194991888 | - | protein_coding |
| ENSG00000060339.14_7 | CCAR1 | 1 | 1 | 0.056 | 7.226 | chr10 | 70480769 | 70552134 | + | protein_coding |
| ENSG00000286636.1_2 | AL512356.4 | 0.91917337 | 1 | 0.056 | -0.561 | chr14 | 105561382 | 105563282 | - | lncRNA |
| ENSG00000168256.18_4 | NKIRAS2 | 0.99999986 | 1 | 0.056 | 4.952 | chr17 | 40163400 | 40177662 | + | protein_coding |
| ENSG00000287200.1_2 | AC022506.2 | 0.94708523 | 1 | 0.056 | 0.131 | chr12 | 57851379 | 57856601 | - | lncRNA |
| ENSG00000274105.1_8 | AC084824.4 | 0.91674662 | 1 | 0.055 | -0.654 | chr12 | 32881103 | 32881958 | - | lncRNA |
| ENSG00000124570.20_6 | SERPINB6 | 0.99999998 | 1 | 0.055 | 6.203 | chr6 | 2948393 | 2972399 | - | protein_coding |
| ENSG00000151332.19_5 | MBIP | 0.99994426 | 1 | 0.055 | 3.765 | chr14 | 36767770 | 36789843 | - | protein_coding |
| ENSG00000197024.9_3 | ZNF398 | 0.99999964 | 1 | 0.055 | 4.915 | chr7 | 148823508 | 148880134 | + | protein_coding |
| ENSG00000015133.19_4 | CCDC88C | 0.99999409 | 1 | 0.055 | 4.382 | chr14 | 91737667 | 91884188 | - | protein_coding |
| ENSG00000169217.9_4 | CD2BP2 | 1 | 1 | 0.055 | 6.453 | chr16 | 30362094 | 30366629 | - | protein_coding |
| ENSG00000267575.7_7 | AC006504.5 | 0.9999957 | 1 | 0.055 | 4.237 | chr19 | 28284339 | 28475892 | + | lncRNA |
| ENSG00000254505.10_8 | CHMP4A | 0.9696093 | 1 | 0.055 | 0.424 | chr14 | 24678789 | 24683075 | - | protein_coding |
| ENSG00000229833.10_7 | PET100 | 0.96266101 | 1 | 0.055 | 0.364 | chr19 | 7694679 | 7696842 | + | protein_coding |
| ENSG00000168818.10_5 | STX18 | 0.99999924 | 1 | 0.054 | 4.483 | chr4 | 4417469 | 4544073 | - | protein_coding |
| ENSG00000145337.5_5 | PYURF | 0.99999879 | 1 | 0.054 | 4.699 | chr4 | 89442149 | 89444927 | - | protein_coding |
| ENSG00000007168.13_4 | PAFAH1B1 | 1 | 1 | 0.054 | 7.838 | chr17 | 2496504 | 2588909 | + | protein_coding |
| ENSG00000137070.17_5 | IL11RA | 0.99851019 | 1 | 0.054 | 2.535 | chr9 | 34650699 | 34661889 | + | protein_coding |
| ENSG00000179168.15_7 | GGN | 0.94993917 | 1 | 0.054 | 0.312 | chr19 | 38874907 | 38878722 | - | protein_coding |
| ENSG00000285053.1_5 | TBCE | 0.97472368 | 1 | 0.054 | 0.842 | chr1 | 235491885 | 235612267 | + | protein_coding |

| | | | | | | | | | | |
|---|---|---|---|---|---|---|---|---|---|---|
| ENSG00000165675.16_6 | ENOX2 | 0.99999733 | 1 | 0.054 | 4.076 | chrX | 129757343 | 130037291 | - | protein_coding |
| ENSG00000140259.7_4 | MFAP1 | 0.99999971 | 1 | 0.054 | 5.058 | chr15 | 44096690 | 44116888 | - | protein_coding |
| ENSG00000166716.10_4 | ZNF592 | 0.99999999 | 1 | 0.054 | 6.172 | chr15 | 85291823 | 85349676 | + | protein_coding |
| ENSG00000248734.2_6 | AC008906.1 | 0.93299069 | 1 | 0.054 | -0.180 | chr5 | 96120481 | 96121703 | + | lncRNA |
| ENSG00000243335.9_3 | KCTD7 | 0.99999482 | 1 | 0.054 | 4.267 | chr7 | 66093754 | 66114054 | + | protein_coding |
| ENSG00000100150.20_8 | DEPDC5 | 0.99880723 | 1 | 0.054 | 2.693 | chr22 | 32149853 | 32304019 | + | protein_coding |
| ENSG00000205643.11_5 | CDPF1 | 0.9996491 | 1 | 0.054 | 2.694 | chr22 | 46639908 | 46646208 | - | protein_coding |
| ENSG00000103549.21_6 | RNF40 | 0.99999996 | 1 | 0.054 | 6.574 | chr16 | 30773066 | 30787628 | + | protein_coding |
| ENSG00000181788.4_4 | SIAH2 | 0.99999134 | 1 | 0.054 | 3.881 | chr3 | 150458912 | 150481264 | - | protein_coding |
| ENSG00000186260.16_4 | MRTFB | 0.99999961 | 1 | 0.053 | 5.716 | chr16 | 14165178 | 14360630 | + | protein_coding |
| ENSG00000113318.11_4 | MSH3 | 0.9999999 | 1 | 0.053 | 5.278 | chr5 | 79950471 | 80172634 | + | protein_coding |
| ENSG00000228352.2_6 | AL354989.1 | 0.92633158 | 1 | 0.053 | -0.590 | chr9 | 34084330 | 34096223 | + | lncRNA |
| ENSG00000138688.15_3 | KIAA1109 | 0.9999999 | 1 | 0.053 | 6.343 | chr4 | 123073488 | 123283913 | + | protein_coding |
| ENSG00000258526.7_7 | AL049828.1 | 0.9323923 | 1 | 0.053 | 0.051 | chr14 | 39901803 | 40859751 | + | lncRNA |
| ENSG00000269982.1_6 | AC018809.2 | 0.98895312 | 1 | 0.053 | 1.682 | chr3 | 10000401 | 10004223 | + | lncRNA |
| ENSG00000177200.17_5 | CHD9 | 1 | 1 | 0.053 | 7.125 | chr16 | 53088945 | 53363062 | + | protein_coding |
| ENSG00000167491.17_5 | GATAD2A | 0.99999996 | 1 | 0.053 | 6.051 | chr19 | 19496635 | 19619740 | + | protein_coding |
| ENSG00000153531.13_5 | ADPRHL1 | 0.99998678 | 1 | 0.053 | 3.628 | chr13 | 114053925 | 114107839 | - | protein_coding |
| ENSG00000113621.14_4 | TXNDC15 | 0.99999999 | 1 | 0.053 | 5.969 | chr5 | 134209493 | 134237215 | + | protein_coding |
| ENSG00000138674.17_6 | SEC31A | 1 | 1 | 0.053 | 7.184 | chr4 | 83739662 | 83822319 | - | protein_coding |
| ENSG00000071127.17_4 | WDR1 | 1 | 1 | 0.052 | 8.153 | chr4 | 10075963 | 10118573 | - | protein_coding |
| ENSG00000114867.20_4 | EIF4G1 | 1 | 1 | 0.052 | 8.751 | chr3 | 184032283 | 184053146 | + | protein_coding |
| ENSG00000076351.13_7 | SLC46A1 | 0.99999785 | 1 | 0.052 | 4.286 | chr17 | 26721661 | 26734215 | - | protein_coding |
| ENSG00000279982.1_5 | AL162274.3 | 0.91980637 | 1 | 0.052 | -0.863 | chr10 | 133810242 | 133813350 | + | TEC |
| ENSG00000090316.16_4 | MAEA | 0.99999953 | 1 | 0.052 | 5.326 | chr4 | 1283675 | 1333935 | + | protein_coding |
| ENSG00000063176.16_6 | SPHK2 | 0.99984572 | 1 | 0.052 | 3.568 | chr19 | 49122548 | 49133974 | + | protein_coding |
| ENSG00000163069.12_5 | SGCB | 0.99999943 | 1 | 0.052 | 4.656 | chr4 | 52886872 | 52904648 | - | protein_coding |
| ENSG00000100364.18_5 | KIAA0930 | 0.99999996 | 1 | 0.052 | 6.518 | chr22 | 45586219 | 45636650 | - | protein_coding |
| ENSG00000131165.15_4 | CHMP1A | 0.99999991 | 1 | 0.052 | 6.143 | chr16 | 89710843 | 89724129 | - | protein_coding |
| ENSG00000175395.16_4 | ZNF25 | 0.99998889 | 1 | 0.052 | 3.805 | chr10 | 38238501 | 38265575 | - | protein_coding |
| ENSG00000091527.15_3 | CDV3 | 1 | 1 | 0.052 | 7.418 | chr3 | 133292574 | 133309105 | + | protein_coding |
| ENSG00000122705.17_4 | CLTA | 1 | 1 | 0.052 | 7.287 | chr9 | 36190853 | 36304778 | + | protein_coding |
| ENSG00000226754.2_8 | AL606760.1 | 0.92580939 | 1 | 0.052 | -0.588 | chr1 | 53704222 | 53712154 | + | lncRNA |
| ENSG00000153786.12_6 | ZDHHC7 | 0.99999999 | 1 | 0.052 | 6.512 | chr16 | 85007787 | 85045141 | - | protein_coding |
| ENSG00000182179.13_5 | UBA7 | 0.92384145 | 1 | 0.052 | -0.755 | chr3 | 49842642 | 49851386 | - | protein_coding |
| ENSG00000117620.15_7 | SLC35A3 | 0.9999978 | 1 | 0.052 | 4.561 | chr1 | 100434907 | 100501190 | + | protein_coding |
| ENSG00000182979.18_7 | MTA1 | 0.99999981 | 1 | 0.052 | 5.748 | chr14 | 105886157 | 105937066 | + | protein_coding |
| ENSG00000133884.10_4 | DPF2 | 0.99999996 | 1 | 0.052 | 5.409 | chr11 | 65101323 | 65121733 | + | protein_coding |
| ENSG00000225855.7_7 | RUSC1-AS1 | 0.99931034 | 1 | 0.051 | 2.514 | chr1 | 155286654 | 155293967 | - | lncRNA |
| ENSG00000115525.18_7 | ST3GAL5 | 1 | 1 | 0.051 | 6.329 | chr2 | 86064243 | 86132322 | - | protein_coding |
| ENSG00000189067.12_4 | LITAF | 1 | 1 | 0.051 | 7.598 | chr16 | 11641578 | 11730237 | - | protein_coding |
| ENSG00000162526.7_3 | TSSK3 | 0.92804376 | 1 | 0.051 | -0.656 | chr1 | 32817122 | 32829913 | + | protein_coding |
| ENSG00000170889.14_5 | RPS9 | 1 | 1 | 0.051 | 8.100 | chr19 | 54704677 | 54752862 | + | protein_coding |
| ENSG00000188215.10_5 | DCUN1D3 | 0.9999914 | 1 | 0.051 | 4.024 | chr16 | 20866247 | 20911680 | - | protein_coding |
| ENSG00000187792.5_4 | ZNF70 | 0.99991644 | 1 | 0.051 | 3.388 | chr22 | 24080869 | 24093299 | - | protein_coding |
| ENSG00000004961.15_3 | HCCS | 0.99999986 | 1 | 0.051 | 5.200 | chrX | 11129421 | 11141206 | + | protein_coding |
| ENSG00000155034.19_6 | FBXL18 | 0.99999046 | 1 | 0.051 | 4.574 | chr7 | 5470966 | 5553440 | - | protein_coding |
| ENSG00000112983.17_4 | BRD8 | 0.99999999 | 1 | 0.051 | 6.270 | chr5 | 137475455 | 137514675 | - | protein_coding |
| ENSG00000241852.10_5 | C8orf58 | 0.99993276 | 1 | 0.051 | 3.273 | chr8 | 22457112 | 22461663 | + | protein_coding |
| ENSG00000137871.20_5 | ZNF280D | 0.9999983 | 1 | 0.051 | 4.651 | chr15 | 56922379 | 57026284 | - | protein_coding |
| ENSG00000180071.20_5 | ANKRD18A | 0.99997361 | 1 | 0.051 | 4.048 | chr9 | 38571355 | 38620593 | - | protein_coding |
| ENSG00000183722.9_7 | LHFPL6 | 0.99847117 | 1 | 0.051 | 2.151 | chr13 | 39783253 | 40177665 | - | protein_coding |
| ENSG00000105700.11_4 | KXD1 | 0.99999871 | 1 | 0.051 | 4.535 | chr19 | 18668572 | 18680197 | + | protein_coding |
| ENSG00000120211.4_3 | INSL4 | 0.99988809 | 1 | 0.050 | 4.942 | chr9 | 5231419 | 5235304 | + | protein_coding |
| ENSG00000134014.17_7 | ELP3 | 0.99999999 | 1 | 0.050 | 6.005 | chr8 | 27947190 | 28048673 | + | protein_coding |
| ENSG00000175548.9_3 | ALG10B | 0.99999954 | 1 | 0.050 | 4.750 | chr12 | 38710564 | 38723523 | + | protein_coding |
| ENSG00000188647.13_5 | PTAR1 | 0.99999998 | 1 | 0.050 | 7.093 | chr9 | 72324438 | 72374927 | - | protein_coding |
| ENSG00000154102.11_7 | C16orf74 | 0.99173954 | 1 | 0.050 | 1.406 | chr16 | 85723690 | 85784735 | - | protein_coding |
| ENSG00000212464.1 | SNORA12 | 0.99966248 | 1 | 0.050 | 3.059 | chr10 | 101996913 | 101997059 | - | snoRNA |
| ENSG00000102901.13_5 | CENPT | 0.99999906 | 1 | 0.050 | 4.503 | chr16 | 67862060 | 67881714 | - | protein_coding |
| ENSG00000127081.14_4 | ZNF484 | 0.99999835 | 1 | 0.050 | 4.554 | chr9 | 95606464 | 95640320 | - | protein_coding |

| | | | | | | | | | | |
|---|---|---|---|---|---|---|---|---|---|---|
| ENSG00000089123.16_5 | TASP1 | 0.99998191 | 1 | 0.050 | 3.738 | chr20 | 13370039 | 13619579 | - | protein_coding |
| ENSG00000165233.18_3 | CARD19 | 0.99999447 | 1 | 0.050 | 4.376 | chr9 | 95858499 | 95875565 | + | protein_coding |
| ENSG00000229261.1_7 | AL596223.1 | 0.9298481 | 1 | 0.050 | -0.475 | chr10 | 70975089 | 70992246 | - | lncRNA |
| ENSG00000104549.12_3 | SQLE | 1 | 1 | 0.050 | 6.999 | chr8 | 126010739 | 126034525 | + | protein_coding |
| ENSG00000129355.7_4 | CDKN2D | 0.98793173 | 1 | 0.050 | 1.261 | chr19 | 10677136 | 10679735 | - | protein_coding |
| ENSG00000130396.20_7 | AFDN | 0.99999998 | 1 | 0.050 | 5.890 | chr6 | 168227602 | 168372703 | + | protein_coding |
| ENSG00000079819.19_7 | EPB41L2 | 0.99999997 | 1 | 0.050 | 5.715 | chr6 | 131160487 | 131384462 | - | protein_coding |
| ENSG00000103852.13_7 | TTC23 | 0.99999953 | 1 | 0.050 | 4.600 | chr15 | 99676528 | 99791428 | - | protein_coding |
| ENSG00000153066.12_4 | TXNDC11 | 0.99999967 | 1 | 0.050 | 5.075 | chr16 | 11772936 | 11836734 | - | protein_coding |
| ENSG00000136425.14_5 | CIB2 | 0.99994844 | 1 | 0.049 | 3.510 | chr15 | 78396948 | 78423877 | - | protein_coding |
| ENSG00000126351.12_2 | THRA | 0.9999998 | 1 | 0.049 | 5.291 | chr17 | 38214543 | 38250120 | + | protein_coding |
| ENSG00000160318.6_5 | CLDND2 | 0.97739752 | 1 | 0.049 | 0.845 | chr19 | 51870352 | 51872257 | - | protein_coding |
| ENSG00000120688.9_3 | WBP4 | 0.99999382 | 1 | 0.049 | 3.829 | chr13 | 41635645 | 41658142 | + | protein_coding |
| ENSG00000130764.10_4 | LRRC47 | 0.99999972 | 1 | 0.049 | 5.001 | chr1 | 3695123 | 3713062 | - | protein_coding |
| ENSG00000223784.2_5 | LINP1 | 0.99984531 | 1 | 0.049 | 3.199 | chr10 | 6751492 | 6782494 | + | lncRNA |
| ENSG00000180758.12_5 | GPR157 | 0.99974481 | 1 | 0.049 | 3.022 | chr1 | 9160364 | 9189161 | - | protein_coding |
| ENSG00000140598.15_7 | EFL1 | 0.99999999 | 1 | 0.049 | 6.087 | chr15 | 82422574 | 82555075 | - | protein_coding |
| ENSG00000077721.16_3 | UBE2A | 1 | 1 | 0.049 | 6.612 | chrX | 118708499 | 118725046 | + | protein_coding |
| ENSG00000075142.13_4 | SRI | 0.99999997 | 1 | 0.049 | 5.423 | chr7 | 87834433 | 87856308 | - | protein_coding |
| ENSG00000168398.6_4 | BDKRB2 | 0.99998985 | 1 | 0.049 | 4.017 | chr14 | 96671016 | 96710503 | + | protein_coding |
| ENSG00000160691.18_7 | SHC1 | 1 | 1 | 0.049 | 7.225 | chr1 | 154934774 | 154946871 | - | protein_coding |
| ENSG00000158163.15_5 | DZIP1L | 0.99945342 | 1 | 0.049 | 2.726 | chr3 | 137780832 | 137834660 | - | protein_coding |
| ENSG00000165660.8_6 | ABRAXAS2 | 0.99999885 | 1 | 0.048 | 4.987 | chr10 | 126490388 | 126525236 | + | protein_coding |
| ENSG00000273521.1_6 | AL162274.1 | 0.95739828 | 1 | 0.048 | 0.032 | chr10 | 133793744 | 133794841 | + | lncRNA |
| ENSG00000105438.9_5 | KDELR1 | 1 | 1 | 0.048 | 7.082 | chr19 | 48885832 | 48894808 | - | protein_coding |
| ENSG00000185825.17_6 | BCAP31 | 1 | 1 | 0.048 | 7.342 | chrX | 152965947 | 152990020 | - | protein_coding |
| ENSG00000164924.18_9 | YWHAZ | 1 | 1 | 0.048 | 9.206 | chr8 | 101928751 | 101965616 | - | protein_coding |
| ENSG00000243302.3_5 | AC018638.4 | 0.99669079 | 1 | 0.048 | 2.220 | chr7 | 128291239 | 128292388 | + | processed_pseudogene |
| ENSG00000214087.8_6 | ARL16 | 0.99999846 | 1 | 0.048 | 4.323 | chr17 | 79648204 | 79650954 | - | protein_coding |
| ENSG00000109919.10_5 | MTCH2 | 1 | 1 | 0.048 | 6.530 | chr11 | 47638867 | 47664159 | - | protein_coding |
| ENSG00000215142.4 | RP11-292B8.1 | 0.99962414 | 1 | 0.048 | 2.646 | chr9 | 40491622 | 40633282 | - | pseudogene |
| ENSG00000226210.3_5 | WASH8P | 0.9998072 | 1 | 0.048 | 3.174 | chr12 | 73725 | 91218 | + | unprocessed_pseudogene |
| ENSG00000160360.13_5 | GPSM1 | 0.99065851 | 1 | 0.048 | 3.920 | chr9 | 139221932 | 139254057 | + | protein_coding |
| ENSG00000174669.12_4 | SLC29A2 | 0.99999125 | 1 | 0.048 | 3.713 | chr11 | 66129992 | 66139685 | - | protein_coding |
| ENSG00000196876.9 | SCN8A | 0.99998317 | 1 | 0.048 | 3.589 | chr12 | 51984050 | 52206648 | + | protein_coding |
| ENSG00000196981.4_4 | WDR5B | 0.99999509 | 1 | 0.047 | 4.021 | chr3 | 122130693 | 122134909 | - | protein_coding |
| ENSG00000238072.1_5 | AC009244.1 | 0.92483702 | 1 | 0.047 | -1.162 | chr7 | 129049954 | 129050211 | - | processed_pseudogene |
| ENSG00000140262.17_2 | TCF12 | 1 | 1 | 0.047 | 7.027 | chr15 | 57210821 | 57591479 | + | protein_coding |
| ENSG00000100888.14_7 | CHD8 | 1 | 1 | 0.047 | 7.426 | chr14 | 21853353 | 21924285 | - | protein_coding |
| ENSG00000266074.8_2 | BAHCC1 | 0.99985746 | 1 | 0.047 | 4.925 | chr17 | 79369275 | 79433358 | + | protein_coding |
| ENSG00000165060.14_7 | FXN | 0.99960472 | 1 | 0.047 | 2.887 | chr9 | 71650479 | 71715094 | + | protein_coding |
| ENSG00000145740.19_4 | SLC30A5 | 0.99999999 | 1 | 0.047 | 6.061 | chr5 | 68389776 | 68426896 | + | protein_coding |
| ENSG00000105171.10_5 | POP4 | 0.99999999 | 1 | 0.047 | 6.312 | chr19 | 30097190 | 30108144 | + | protein_coding |
| ENSG00000179978.11_6 | NAIPP2 | 0.92759618 | 1 | 0.047 | -0.800 | chr5 | 69390486 | 69424261 | - | unprocessed_pseudogene |
| ENSG00000111554.14_3 | MDM1 | 0.99999731 | 1 | 0.047 | 4.399 | chr12 | 68666223 | 68726161 | - | protein_coding |
| ENSG00000138190.17_5 | EXOC6 | 0.99999999 | 1 | 0.047 | 6.040 | chr10 | 94586588 | 94819250 | + | protein_coding |
| ENSG00000118508.5_3 | RAB32 | 0.99999754 | 1 | 0.047 | 4.425 | chr6 | 146864969 | 146876089 | + | protein_coding |
| ENSG00000164715.6_3 | LMTK2 | 0.99999999 | 1 | 0.047 | 6.231 | chr7 | 97736174 | 97838950 | + | protein_coding |
| ENSG00000259717.1_4 | LINC00677 | 0.94968836 | 1 | 0.047 | -0.140 | chr14 | 103587184 | 103589344 | - | lncRNA |
| ENSG00000229043.3_7 | AC091729.3 | 0.99989399 | 1 | 0.047 | 3.139 | chr7 | 1200010 | 1205243 | + | lncRNA |
| ENSG00000188186.10_2 | LAMTOR4 | 0.99999988 | 1 | 0.046 | 5.250 | chr7 | 99746530 | 99753567 | + | protein_coding |
| ENSG00000285667.1_4 | AC012291.3 | 0.97862671 | 1 | 0.046 | 0.950 | chr15 | 85260705 | 85290068 | - | lncRNA |
| ENSG00000109445.11_5 | ZNF330 | 0.99999426 | 1 | 0.046 | 4.361 | chr4 | 142142041 | 142155851 | + | protein_coding |
| ENSG00000213763.4_4 | ACTBP2 | 0.93377614 | 1 | 0.046 | -0.535 | chr5 | 77080705 | 77081827 | + | processed_pseudogene |
| ENSG00000159200.18_4 | RCAN1 | 0.99998069 | 1 | 0.046 | 3.622 | chr21 | 35885440 | 35987412 | - | protein_coding |
| ENSG00000088992.18_4 | TESC | 0.99999997 | 1 | 0.046 | 5.953 | chr12 | 117476728 | 117537284 | - | protein_coding |
| ENSG00000160716.6_7 | CHRNB2 | 0.99051565 | 1 | 0.046 | 0.924 | chr1 | 154540254 | 154552489 | + | protein_coding |

| | | | | | | | | | | |
|---|---|---|---|---|---|---|---|---|---|---|
| ENSG00000156599.11_4 | ZDHHC5 | 1 | 1 | 0.046 | 7.013 | chr11 | 57435219 | 57468654 | + | protein_coding |
| ENSG00000141627.13_6 | DYM | 0.99999967 | 1 | 0.046 | 4.782 | chr18 | 46567844 | 46987717 | - | protein_coding |
| ENSG00000054965.10_4 | FAM168A | 0.99999999 | 1 | 0.046 | 5.935 | chr11 | 73111532 | 73309234 | - | protein_coding |
| ENSG00000117298.16_6 | ECE1 | 0.99999991 | 1 | 0.046 | 7.313 | chr1 | 21543740 | 21672065 | - | protein_coding |
| ENSG00000116586.12_3 | LAMTOR2 | 0.99999934 | 1 | 0.046 | 4.660 | chr1 | 156024573 | 156028297 | + | protein_coding |
| ENSG00000128309.16_2 | MPST | 0.99982817 | 1 | 0.046 | 6.016 | chr22 | 37415676 | 37425863 | + | protein_coding |
| ENSG00000185298.13_4 | CCDC137 | 0.99999999 | 1 | 0.046 | 5.648 | chr17 | 79633767 | 79640934 | + | protein_coding |
| ENSG00000126749.16_6 | EMG1 | 0.99999976 | 1 | 0.045 | 5.083 | chr12 | 7080076 | 7105520 | + | protein_coding |
| ENSG00000151552.12_5 | QDPR | 0.9999798 | 1 | 0.045 | 3.685 | chr4 | 17461884 | 17513829 | - | protein_coding |
| ENSG00000100083.19_5 | GGA1 | 0.99999985 | 1 | 0.045 | 5.245 | chr22 | 38004732 | 38029571 | + | protein_coding |
| ENSG00000257341.5_10 | AL928654.3 | 0.93136156 | 1 | 0.045 | -0.727 | chr14 | 105953536 | 105958604 | + | protein_coding |
| ENSG00000205041.1_8 | AC118344.1 | 0.99111011 | 1 | 0.045 | 1.754 | chr19 | 40779396 | 40781386 | - | lncRNA |
| ENSG00000130770.18_4 | ATP5IF1 | 0.99999959 | 1 | 0.045 | 4.871 | chr1 | 28562620 | 28573417 | + | protein_coding |
| ENSG00000149476.15_3 | TKFC | 0.99999995 | 1 | 0.045 | 5.638 | chr11 | 61100682 | 61120767 | + | protein_coding |
| ENSG00000243199.1_5 | AC115223.1 | 0.99547621 | 1 | 0.045 | 1.818 | chr4 | 66439177 | 66440046 | + | processed_pseudogene |
| ENSG00000236921.1_7 | AL157937.1 | 0.99941161 | 1 | 0.045 | 2.473 | chr9 | 22203989 | 22214671 | - | lncRNA |
| ENSG00000104312.8_3 | RIPK2 | 0.99999727 | 1 | 0.045 | 4.429 | chr8 | 90770034 | 90803292 | + | protein_coding |
| ENSG00000196531.12_5 | NACA | 1 | 1 | 0.045 | 8.257 | chr12 | 57106211 | 57125412 | - | protein_coding |
| ENSG00000153982.11_4 | GDPD1 | 0.97127613 | 1 | 0.045 | 0.723 | chr17 | 57297828 | 57353331 | + | protein_coding |
| ENSG00000077549.18_6 | CAPZB | 0.99999999 | 1 | 0.045 | 6.438 | chr1 | 19665269 | 19812033 | - | protein_coding |
| ENSG00000088035.18_6 | ALG6 | 0.99996929 | 1 | 0.045 | 3.963 | chr1 | 63833246 | 63904224 | + | protein_coding |
| ENSG00000279356.1_7 | AC007610.6 | 0.96909499 | 1 | 0.045 | 0.530 | chr16 | 50106773 | 50108897 | + | TEC |
| ENSG00000177125.5_3 | ZBTB34 | 0.99999687 | 1 | 0.044 | 3.986 | chr9 | 129622944 | 129648157 | + | protein_coding |
| ENSG00000164105.4_3 | SAP30 | 0.99998326 | 1 | 0.044 | 3.569 | chr4 | 174291120 | 174298683 | + | protein_coding |
| ENSG00000273010.1_6 | AL360270.3 | 0.93187497 | 1 | 0.044 | -0.830 | chr1 | 111505924 | 111507271 | + | lncRNA |
| ENSG00000157510.14_3 | AFAP1L1 | 0.99997507 | 1 | 0.044 | 3.554 | chr5 | 148651422 | 148723200 | + | protein_coding |
| ENSG00000074370.18_5 | ATP2A3 | 0.99963403 | 1 | 0.044 | 3.326 | chr17 | 3827164 | 3867758 | - | protein_coding |
| ENSG00000260063.1_6 | AL512408.1 | 0.94012547 | 1 | 0.044 | -0.391 | chr1 | 27018623 | 27020622 | - | lncRNA |
| ENSG00000270926.1_6 | AC008013.4 | 0.93870393 | 1 | 0.044 | -0.564 | chr12 | 31258039 | 31259764 | - | unprocessed_pseudogene |
| ENSG00000272046.1_6 | AL445647.1 | 0.99397756 | 1 | 0.044 | 1.799 | chr13 | 88459889 | 88463487 | - | lncRNA |
| ENSG00000184261.5_3 | KCNK12 | 0.93421185 | 1 | 0.044 | -0.725 | chr2 | 47736429 | 47798124 | - | protein_coding |
| ENSG00000100129.18_7 | EIF3L | 1 | 1 | 0.044 | 6.714 | chr22 | 38244875 | 38285414 | + | protein_coding |
| ENSG00000127995.17_3 | CASD1 | 0.99999996 | 1 | 0.044 | 5.475 | chr7 | 94138531 | 94186331 | + | protein_coding |
| ENSG00000179862.7_3 | CITED4 | 0.99901179 | 1 | 0.043 | 3.298 | chr1 | 41326726 | 41328035 | - | protein_coding |
| ENSG00000262061.6_6 | AC129507.1 | 0.96187121 | 1 | 0.043 | 0.325 | chr17 | 180996 | 183279 | + | lncRNA |
| ENSG00000155761.14_7 | SPAG17 | 0.93226975 | 1 | 0.043 | -0.910 | chr1 | 118496213 | 118727851 | - | protein_coding |
| ENSG00000236552.2_2 | RPL13AP5 | 0.99928175 | 1 | 0.043 | 4.632 | chr10 | 98510045 | 98510656 | + | processed_pseudogene |
| ENSG00000141562.18_6 | NARF | 0.99999995 | 1 | 0.043 | 5.360 | chr17 | 80416056 | 80448413 | + | protein_coding |
| ENSG00000270589.1_6 | AL158163.1 | 0.94258178 | 1 | 0.043 | 0.007 | chr10 | 112670354 | 112672002 | + | lncRNA |
| ENSG00000174206.12_3 | C12orf66 | 0.99999939 | 1 | 0.043 | 4.445 | chr12 | 64580096 | 64616076 | - | protein_coding |
| ENSG00000088854.12_4 | C20orf194 | 0.99999898 | 1 | 0.043 | 4.491 | chr20 | 3229951 | 3388272 | - | protein_coding |
| ENSG00000119227.8_5 | PIGZ | 0.98428925 | 1 | 0.043 | 0.978 | chr3 | 196673227 | 196695931 | - | protein_coding |
| ENSG00000108510.10_5 | MED13 | 1 | 1 | 0.043 | 7.918 | chr17 | 60019966 | 60142639 | - | protein_coding |
| ENSG00000169871.13_6 | TRIM56 | 0.99999997 | 1 | 0.043 | 5.432 | chr7 | 100728762 | 100741248 | + | protein_coding |
| ENSG00000011132.12_7 | APBA3 | 0.99978733 | 1 | 0.042 | 2.985 | chr19 | 3750770 | 3761690 | - | protein_coding |
| ENSG00000186162.10_4 | CIDECP1 | 0.99966763 | 1 | 0.042 | 2.654 | chr3 | 10055922 | 10068049 | - | transcribed_unprocessed_pseudogene |
| ENSG00000135046.14_4 | ANXA1 | 1 | 1 | 0.042 | 10.020 | chr9 | 75766781 | 75785309 | + | protein_coding |
| ENSG00000105679.9_3 | GAPDHS | 0.93724468 | 1 | 0.042 | -0.697 | chr19 | 36024357 | 36036221 | + | protein_coding |
| ENSG00000100938.17_6 | GMPR2 | 0.99999999 | 1 | 0.042 | 5.925 | chr14 | 24701628 | 24708448 | + | protein_coding |
| ENSG00000253710.4_5 | ALG11 | 0.99983386 | 1 | 0.042 | 3.018 | chr13 | 52586534 | 52607736 | + | protein_coding |
| ENSG00000143224.18_6 | PPOX | 0.99998908 | 1 | 0.042 | 3.909 | chr1 | 161135846 | 161147803 | + | protein_coding |
| ENSG00000213339.9_4 | QTRT1 | 0.99999898 | 1 | 0.042 | 4.351 | chr19 | 10812106 | 10824113 | + | protein_coding |
| ENSG00000151092.17_7 | NGLY1 | 0.99999993 | 1 | 0.042 | 5.635 | chr3 | 25760435 | 25831530 | - | protein_coding |
| ENSG00000171700.14_5 | RGS19 | 0.99998746 | 1 | 0.041 | 3.631 | chr20 | 62704534 | 62711341 | - | protein_coding |
| ENSG00000137055.15_6 | PLAA | 0.99999968 | 1 | 0.041 | 5.343 | chr9 | 26903370 | 26947240 | - | protein_coding |
| ENSG00000067066.17_5 | SP100 | 0.99999993 | 1 | 0.041 | 5.039 | chr2 | 231280657 | 231410321 | + | protein_coding |
| ENSG00000274220.1_7 | AC009163.7 | 0.94982465 | 1 | 0.041 | -0.171 | chr16 | 75467734 | 75470290 | + | lncRNA |
| ENSG00000116574.6_3 | RHOU | 0.99999743 | 1 | 0.041 | 4.017 | chr1 | 228871226 | 228882411 | + | protein_coding |
| ENSG00000100650.16_4 | SRSF5 | 0.99999996 | 1 | 0.041 | 6.694 | chr14 | 70193617 | 70238722 | + | protein_coding |

| | | | | | | | | | | |
|---|---|---|---|---|---|---|---|---|---|---|
| ENSG00000134744.14_8 | TUT4 | 0.99999994 | 1 | 0.041 | 5.794 | chr1 | 52873954 | 53019159 | - | protein_coding |
| ENSG00000174365.20_5 | SNHG11 | 0.99997212 | 1 | 0.041 | 3.496 | chr20 | 37074986 | 37079583 | + | lncRNA |
| ENSG00000129646.14_7 | QRICH2 | 0.99745309 | 1 | 0.041 | 1.964 | chr17 | 74270130 | 74304079 | - | protein_coding |
| ENSG00000198785.7_6 | GRIN3A | 0.95895225 | 1 | 0.041 | 0.249 | chr9 | 104331634 | 104500929 | - | protein_coding |
| ENSG00000244952.2_6 | AC123768.2 | 0.93711823 | 1 | 0.041 | -0.666 | chr15 | 32905934 | 32907312 | + | lncRNA |
| ENSG00000163479.14_6 | SSR2 | 1 | 1 | 0.041 | 6.332 | chr1 | 155978839 | 155990742 | - | protein_coding |
| ENSG00000064393.16_5 | HIPK2 | 1 | 1 | 0.041 | 6.886 | chr7 | 139246316 | 139477797 | - | protein_coding |
| ENSG00000132623.16_4 | ANKEF1 | 0.99999996 | 1 | 0.041 | 5.330 | chr20 | 9966774 | 10038951 | + | protein_coding |
| ENSG00000123200.16_6 | ZC3H13 | 0.99999992 | 1 | 0.041 | 5.162 | chr13 | 46528600 | 46626894 | - | protein_coding |
| ENSG00000107890.16_3 | ANKRD26 | 0.99999855 | 1 | 0.041 | 5.358 | chr10 | 27280843 | 27389427 | - | protein_coding |
| ENSG00000101350.8_3 | KIF3B | 1 | 1 | 0.041 | 6.272 | chr20 | 30865454 | 30922814 | + | protein_coding |
| ENSG00000145495.16_5 | MARCHF6 | 1 | 1 | 0.041 | 7.115 | chr5 | 10353807 | 10440500 | + | protein_coding |
| ENSG00000187105.8_4 | HEATR4 | 0.95371855 | 1 | 0.041 | -0.077 | chr14 | 73945189 | 74025651 | - | protein_coding |
| ENSG00000100890.16_5 | PRORP | 0.99989734 | 1 | 0.041 | 3.117 | chr14 | 35591052 | 35746828 | + | protein_coding |
| ENSG00000214331.8_6 | AC009053.2 | 0.99963865 | 1 | 0.041 | 2.577 | chr16 | 74366300 | 74402138 | - | transcribed_unprocessed_pseudogene |
| ENSG00000185090.14_3 | MANEAL | 0.99996594 | 1 | 0.041 | 3.272 | chr1 | 38259474 | 38266809 | + | protein_coding |
| ENSG00000150787.8_4 | PTS | 0.99999763 | 1 | 0.041 | 4.160 | chr11 | 112097090 | 112140678 | + | protein_coding |
| ENSG00000229021.2_8 | AL450992.2 | 0.93734652 | 1 | 0.040 | -0.801 | chr1 | 151967007 | 152015250 | + | lncRNA |
| ENSG00000225377.6_7 | NRSN2-AS1 | 0.99991978 | 1 | 0.040 | 3.102 | chr20 | 297504 | 329134 | - | lncRNA |
| ENSG00000279765.3_7 | AC013394.1 | 0.99286445 | 1 | 0.040 | 1.626 | chr15 | 93426643 | 93492460 | + | protein_coding |
| ENSG00000151876.13_3 | FBXO4 | 0.99997919 | 1 | 0.040 | 3.470 | chr5 | 41925356 | 41941845 | + | protein_coding |
| ENSG00000154832.15_4 | CXXC1 | 0.9999991 | 1 | 0.040 | 4.700 | chr18 | 47808713 | 47814674 | - | protein_coding |
| ENSG00000165868.14_5 | HSPA12A | 0.99999996 | 1 | 0.040 | 5.207 | chr10 | 118430703 | 118609747 | - | protein_coding |
| ENSG00000160256.13_4 | FAM207A | 0.99996972 | 1 | 0.040 | 3.750 | chr21 | 46359927 | 46396904 | + | protein_coding |
| ENSG00000165288.11_4 | BRWD3 | 0.99999681 | 1 | 0.040 | 5.625 | chrX | 79925002 | 80065376 | - | protein_coding |
| ENSG00000083845.9_5 | RPS5 | 1 | 1 | 0.040 | 9.041 | chr19 | 58897767 | 58906173 | + | protein_coding |
| ENSG00000136854.21_8 | STXBP1 | 0.99992351 | 1 | 0.040 | 3.157 | chr9 | 130341649 | 130458306 | + | protein_coding |
| ENSG00000156463.18_4 | SH3RF2 | 0.99995714 | 1 | 0.040 | 3.476 | chr5 | 145316141 | 145461354 | + | protein_coding |
| ENSG00000163041.12_6 | H3-3A | 0.99999791 | 1 | 0.040 | 4.647 | chr1 | 226249552 | 226260913 | + | protein_coding |
| ENSG00000152944.9_4 | MED21 | 0.99999994 | 1 | 0.040 | 5.651 | chr12 | 27175479 | 27219276 | + | protein_coding |
| ENSG00000162144.9_5 | CYB561A3 | 0.99999991 | 1 | 0.040 | 5.080 | chr11 | 61116217 | 61129771 | - | protein_coding |
| ENSG00000185115.6_4 | NSMCE3 | 0.99999972 | 1 | 0.040 | 5.086 | chr15 | 29557193 | 29562026 | - | protein_coding |
| ENSG00000228486.10_7 | C2orf92 | 0.96214736 | 1 | 0.040 | 0.167 | chr2 | 98280680 | 98319527 | + | protein_coding |
| ENSG00000153046.18_4 | CDYL | 0.99999999 | 1 | 0.039 | 5.655 | chr6 | 4706393 | 4955785 | + | protein_coding |
| ENSG00000102125.16_7 | TAZ | 0.99999949 | 1 | 0.039 | 4.789 | chrX | 153639860 | 153650065 | + | protein_coding |
| ENSG00000168591.16_5 | TMUB2 | 0.99999983 | 1 | 0.039 | 5.057 | chr17 | 42264338 | 42269297 | + | protein_coding |
| ENSG00000164120.14_4 | HPGD | 1 | 1 | 0.039 | 7.671 | chr4 | 175411326 | 175444305 | - | protein_coding |
| ENSG00000198934.5_4 | MAGEE1 | 0.99995531 | 1 | 0.039 | 3.234 | chrX | 75648103 | 75651735 | + | protein_coding |
| ENSG00000090857.13_4 | PDPR | 0.99999993 | 1 | 0.039 | 5.142 | chr16 | 70147529 | 70196440 | + | protein_coding |
| ENSG00000181192.12_4 | DHTKD1 | 0.99999999 | 1 | 0.039 | 5.920 | chr10 | 12110953 | 12165220 | + | protein_coding |
| ENSG00000273611.5_5 | ZNHIT3 | 0.9999999 | 1 | 0.039 | 5.350 | chr17 | 34842473 | 34855154 | + | protein_coding |
| ENSG00000105939.13_5 | ZC3HAV1 | 1 | 1 | 0.039 | 7.316 | chr7 | 138728261 | 138794466 | - | protein_coding |
| ENSG00000179743.4_8 | AL450998.2 | 0.97635115 | 1 | 0.039 | 0.871 | chr1 | 16160969 | 16174642 | - | lncRNA |
| ENSG00000250067.12_5 | YJEFN3 | 0.98751873 | 1 | 0.039 | 1.623 | chr19 | 19639670 | 19648390 | + | protein_coding |
| ENSG00000113282.13_5 | CLINT1 | 1 | 1 | 0.039 | 7.030 | chr5 | 157212751 | 157286183 | - | protein_coding |
| ENSG00000178917.16_5 | ZNF852 | 0.99208979 | 1 | 0.039 | 1.435 | chr3 | 44536339 | 44552128 | - | protein_coding |
| ENSG00000107341.5_3 | UBE2R2 | 1 | 1 | 0.039 | 6.403 | chr9 | 33817158 | 33920397 | + | protein_coding |
| ENSG00000214050.8_7 | FBXO16 | 0.99318883 | 1 | 0.039 | 1.442 | chr8 | 28205804 | 28347795 | - | protein_coding |
| ENSG00000250462.8_2 | LRRC37BP1 | 0.99994518 | 1 | 0.039 | 3.248 | chr17 | 28956698 | 28964484 | + | transcribed_unprocessed_pseudogene |
| ENSG00000072864.15_5 | NDE1 | 0.99999928 | 1 | 0.039 | 4.638 | chr16 | 15737124 | 15820210 | + | protein_coding |
| ENSG00000115963.13_3 | RND3 | 1 | 1 | 0.039 | 6.523 | chr2 | 151324709 | 151395525 | - | protein_coding |
| ENSG00000126804.13_5 | ZBTB1 | 0.99999973 | 1 | 0.039 | 5.631 | chr14 | 64970430 | 65000408 | + | protein_coding |
| ENSG00000120742.11_5 | SERP1 | 1 | 1 | 0.039 | 6.907 | chr3 | 150259785 | 150321015 | - | protein_coding |
| ENSG00000144597.14_3 | EAF1 | 0.9999993 | 1 | 0.039 | 4.528 | chr3 | 15469105 | 15492142 | + | protein_coding |
| ENSG00000234494.7_8 | SP2-AS1 | 0.99931798 | 1 | 0.038 | 2.319 | chr17 | 45974696 | 46018776 | - | lncRNA |
| ENSG00000130338.13_4 | TULP4 | 0.99999999 | 1 | 0.038 | 5.558 | chr6 | 158653268 | 158932860 | + | protein_coding |
| ENSG00000176340.4_4 | COX8A | 0.99999996 | 1 | 0.038 | 5.779 | chr11 | 63742092 | 63744015 | + | protein_coding |
| ENSG00000264920.2_7 | AC018521.5 | 0.97535152 | 1 | 0.038 | 0.751 | chr17 | 45968595 | 45973178 | - | lncRNA |
| ENSG00000182923.17_6 | CEP63 | 0.99999877 | 1 | 0.038 | 4.291 | chr3 | 134204563 | 134293859 | + | protein_coding |
| ENSG00000275223.1_6 | AL121906.2 | 0.96031517 | 1 | 0.038 | -0.170 | chr20 | 32243507 | 32244229 | - | lncRNA |

| | | | | | | | | | | |
|---|---|---|---|---|---|---|---|---|---|---|
| ENSG00000258136.2_6 | AC007622.2 | 0.98844782 | 1 | 0.038 | 1.076 | chr12 | 108130332 | 108153745 | + | lncRNA |
| ENSG00000205730.6_4 | ITPRIPL2 | 0.99999999 | 1 | 0.038 | 5.706 | chr16 | 19125254 | 19132951 | + | protein_coding |
| ENSG00000100097.12_3 | LGALS1 | 1 | 1 | 0.038 | 7.383 | chr22 | 38071643 | 38075809 | + | protein_coding |
| ENSG00000131966.14_5 | ACTR10 | 0.99999979 | 1 | 0.038 | 5.214 | chr14 | 58666798 | 58702354 | + | protein_coding |
| ENSG00000267649.1_6 | AC010327.5 | 0.9386244 | 1 | 0.038 | -1.029 | chr19 | 55728028 | 55732984 | + | lncRNA |
| ENSG00000068120.15_7 | COASY | 0.99999999 | 1 | 0.038 | 6.422 | chr17 | 40713485 | 40718295 | + | protein_coding |
| ENSG00000168246.6_5 | UBTD2 | 0.99999992 | 1 | 0.038 | 5.219 | chr5 | 171636650 | 171710768 | - | protein_coding |
| ENSG00000167173.19_8 | C15orf39 | 0.99999996 | 1 | 0.038 | 5.615 | chr15 | 75487984 | 75504510 | + | protein_coding |
| ENSG00000122557.10_6 | HERPUD2 | 0.99999984 | 1 | 0.038 | 4.716 | chr7 | 35672269 | 35735181 | - | protein_coding |
| ENSG00000057252.13_3 | SOAT1 | 0.99999999 | 1 | 0.038 | 5.844 | chr1 | 179262849 | 179327815 | + | protein_coding |
| ENSG00000163870.16_7 | TPRA1 | 0.99999748 | 1 | 0.037 | 4.539 | chr3 | 127290075 | 127317110 | - | protein_coding |
| ENSG00000033327.13_7 | GAB2 | 0.99997589 | 1 | 0.037 | 3.711 | chr11 | 77926339 | 78129394 | - | protein_coding |
| ENSG00000182831.12_6 | C16orf72 | 1 | 1 | 0.037 | 6.607 | chr16 | 9185501 | 9215492 | + | protein_coding |
| ENSG00000167461.12_4 | RAB8A | 0.99999954 | 1 | 0.037 | 4.723 | chr19 | 16222699 | 16245044 | + | protein_coding |
| ENSG00000130175.9_6 | PRKCSH | 1 | 1 | 0.037 | 6.249 | chr19 | 11546109 | 11561783 | + | protein_coding |
| ENSG00000136944.18_5 | LMX1B | 0.99324211 | 1 | 0.037 | 1.435 | chr9 | 129376722 | 129463311 | + | protein_coding |
| ENSG00000126756.12_4 | UXT | 0.99999958 | 1 | 0.037 | 4.826 | chrX | 47511195 | 47518579 | - | protein_coding |
| ENSG00000227218.7_8 | AL157935.2 | 0.99233681 | 1 | 0.037 | 1.726 | chr9 | 130696782 | 130703231 | + | lncRNA |
| ENSG00000155115.7_3 | GTF3C6 | 0.99999932 | 1 | 0.037 | 5.301 | chr6 | 111279909 | 111289093 | + | protein_coding |
| ENSG00000139180.11_6 | NDUFA9 | 0.99999543 | 1 | 0.037 | 3.960 | chr12 | 4758261 | 4803483 | + | protein_coding |
| ENSG00000072110.13_6 | ACTN1 | 1 | 1 | 0.037 | 8.140 | chr14 | 69340860 | 69446157 | - | protein_coding |
| ENSG00000170919.16_5 | TPT1-AS1 | 0.99588056 | 1 | 0.037 | 1.707 | chr13 | 45915480 | 45992110 | + | lncRNA |
| ENSG00000084112.15_7 | SSH1 | 0.99999997 | 1 | 0.037 | 5.595 | chr12 | 109171967 | 109251366 | - | protein_coding |
| ENSG00000170906.16_4 | NDUFA3 | 0.99999868 | 1 | 0.037 | 4.802 | chr19 | 54606036 | 54612564 | + | protein_coding |
| ENSG00000211456.13_5 | SACM1L | 0.9999997 | 1 | 0.037 | 4.979 | chr3 | 45730548 | 45786901 | + | polymorphic_pseudogene |
| ENSG00000166441.13_4 | RPL27A | 1 | 1 | 0.036 | 8.769 | chr11 | 8704335 | 8736306 | + | protein_coding |
| ENSG00000048649.13_5 | RSF1 | 0.99999999 | 1 | 0.036 | 5.756 | chr11 | 77371041 | 77532063 | - | protein_coding |
| ENSG00000160199.15_4 | PKNOX1 | 0.99994888 | 1 | 0.036 | 3.369 | chr21 | 44394620 | 44454041 | + | protein_coding |
| ENSG00000058262.10_3 | SEC61A1 | 1 | 1 | 0.036 | 7.260 | chr3 | 127770484 | 127790526 | + | protein_coding |
| ENSG00000127870.17_5 | RNF6 | 0.99999999 | 1 | 0.036 | 5.852 | chr13 | 26706253 | 26796451 | - | protein_coding |
| ENSG00000149634.4_3 | SPATA25 | 0.9622712 | 1 | 0.036 | 0.047 | chr20 | 44515128 | 44516274 | - | protein_coding |
| ENSG00000104164.11_5 | BLOC1S6 | 0.99999997 | 1 | 0.036 | 5.626 | chr15 | 45879412 | 45908143 | + | protein_coding |
| ENSG00000157036.13_4 | EXOG | 0.99999603 | 1 | 0.036 | 3.997 | chr3 | 38537618 | 38583652 | + | protein_coding |
| ENSG00000176018.13_4 | LYSMD3 | 0.99999392 | 1 | 0.035 | 4.788 | chr5 | 89811428 | 89825401 | - | protein_coding |
| ENSG00000121236.21_5 | TRIM6 | 0.99999963 | 1 | 0.035 | 4.706 | chr11 | 5617339 | 5634188 | + | protein_coding |
| ENSG00000268061.5_6 | NAPA-AS1 | 0.96299381 | 1 | 0.035 | 0.089 | chr19 | 47987539 | 48004854 | + | lncRNA |
| ENSG00000182326.15_5 | C1S | 0.99999991 | 1 | 0.035 | 5.022 | chr12 | 7096351 | 7178336 | + | protein_coding |
| ENSG00000177830.18_7 | CHID1 | 0.99999871 | 1 | 0.035 | 5.080 | chr11 | 867859 | 915058 | - | protein_coding |
| ENSG00000181264.8_3 | TLCD5 | 0.99999281 | 1 | 0.035 | 3.681 | chr11 | 120195838 | 120204391 | + | protein_coding |
| ENSG00000116497.18_4 | S100PBP | 0.99999992 | 1 | 0.035 | 5.110 | chr1 | 33282368 | 33324476 | + | protein_coding |
| ENSG00000163155.12_4 | LYSMD1 | 0.99990419 | 1 | 0.035 | 3.194 | chr1 | 151132224 | 151138424 | - | protein_coding |
| ENSG00000229539.1_7 | AL353194.1 | 0.97279617 | 1 | 0.035 | 0.531 | chr20 | 3868886 | 3869515 | - | lncRNA |
| ENSG00000113732.9_4 | ATP6V0E1 | 1 | 1 | 0.035 | 6.607 | chr5 | 172410774 | 172462448 | + | protein_coding |
| ENSG00000108666.10_7 | C17orf75 | 0.99999982 | 1 | 0.035 | 5.175 | chr17 | 30651584 | 30677042 | - | protein_coding |
| ENSG00000165156.15_5 | ZHX1 | 0.99999996 | 1 | 0.035 | 5.386 | chr8 | 124260691 | 124287781 | - | protein_coding |
| ENSG00000068745.15_4 | IP6K2 | 0.99999998 | 1 | 0.035 | 5.595 | chr3 | 48725436 | 48777786 | - | protein_coding |
| ENSG00000100599.16_7 | AL159141.1 | 0.9999696 | 1 | 0.035 | 4.146 | chr14 | 92980125 | 93155339 | + | protein_coding |
| ENSG00000154309.8_3 | DISP1 | 0.9972485 | 1 | 0.035 | 1.790 | chr1 | 223045613 | 223179337 | + | protein_coding |
| ENSG00000237773.7_8 | AC073332.1 | 0.95331059 | 1 | 0.035 | -0.393 | chr7 | 17009944 | 17338992 | - | lncRNA |
| ENSG00000141905.19_6 | NFIC | 0.99998865 | 1 | 0.035 | 5.536 | chr19 | 3314401 | 3469215 | + | protein_coding |
| ENSG00000103257.9_4 | SLC7A5 | 1 | 1 | 0.035 | 9.892 | chr16 | 87863629 | 87903113 | - | protein_coding |
| ENSG00000137936.18_7 | BCAR3 | 0.99999998 | 1 | 0.035 | 5.577 | chr1 | 94027298 | 94312706 | - | protein_coding |
| ENSG00000111605.18_5 | CPSF6 | 1 | 1 | 0.035 | 7.197 | chr12 | 69633349 | 69668138 | + | protein_coding |
| ENSG00000132286.12_4 | TIMM10B | 0.99999843 | 1 | 0.034 | 4.582 | chr11 | 6502731 | 6505911 | + | protein_coding |
| ENSG00000207864.1 | MIR27B | 0.99736457 | 1 | 0.034 | 2.006 | chr9 | 97847727 | 97847823 | + | miRNA |
| ENSG00000260774.1_9 | AC021087.3 | 0.95581033 | 1 | 0.034 | -0.369 | chr5 | 214013 | 217394 | - | lncRNA |
| ENSG00000175582.19_3 | RAB6A | 0.99999987 | 1 | 0.034 | 6.442 | chr11 | 73386683 | 73472182 | - | protein_coding |
| ENSG00000167553.15_3 | TUBA1C | 1 | 1 | 0.034 | 7.005 | chr12 | 49582519 | 49668386 | + | protein_coding |
| ENSG00000198728.11_4 | LDB1 | 0.99999999 | 1 | 0.034 | 5.638 | chr10 | 103866246 | 103880125 | - | protein_coding |
| ENSG00000261118.1_7 | AC092123.1 | 0.98114801 | 1 | 0.034 | 0.909 | chr16 | 89558425 | 89570868 | - | lncRNA |

| | | | | | | | | | | |
|---|---|---|---|---|---|---|---|---|---|---|
| ENSG00000167614.14_5 | TTYH1 | 0.99599147 | 1 | 0.034 | 1.595 | chr19 | 54926393 | 54948084 | + | protein_coding |
| ENSG00000128534.8_4 | LSM8 | 0.99974629 | 1 | 0.033 | 5.805 | chr7 | 117824198 | 117844089 | + | protein_coding |
| ENSG00000213965.3_2 | NUDT19 | 0.99999999 | 1 | 0.033 | 5.844 | chr19 | 33182867 | 33204702 | + | protein_coding |
| ENSG00000269929.3_10 | MIRLET7A1HG | 0.99965563 | 1 | 0.033 | 2.693 | chr9 | 96928540 | 96963184 | + | lncRNA |
| ENSG00000157800.18_6 | SLC37A3 | 0.99999997 | 1 | 0.033 | 5.314 | chr7 | 139993493 | 140104233 | - | protein_coding |
| ENSG00000223459.6_5 | TCAF1P1 | 0.99328001 | 1 | 0.033 | 1.333 | chr7 | 143295132 | 143301932 | + | unprocessed_pseudogene |
| ENSG00000204576.11_2 | PRR3 | 0.97713531 | 1 | 0.033 | 0.519 | chr6 | 30524663 | 30531500 | + | protein_coding |
| ENSG00000228302.2_7 | AL512770.1 | 0.96021444 | 1 | 0.033 | -0.292 | chr10 | 12286750 | 12289844 | - | lncRNA |
| ENSG00000245213.7_7 | AC105285.1 | 0.99954346 | 1 | 0.033 | 2.382 | chr4 | 174016019 | 174090803 | - | lncRNA |
| ENSG00000133104.14_7 | SPART | 0.99999999 | 1 | 0.033 | 5.845 | chr13 | 36875775 | 36944317 | - | protein_coding |
| ENSG00000160766.14_3 | GBAP1 | 0.99968537 | 1 | 0.033 | 2.646 | chr1 | 155183612 | 155197213 | - | transcribed_unprocessed_pseudogene |
| ENSG00000235703.6_7 | LINC00894 | 0.99796474 | 1 | 0.033 | 1.955 | chrX | 149106831 | 149392815 | + | lncRNA |
| ENSG00000163159.13_4 | VPS72 | 0.99999993 | 1 | 0.033 | 5.174 | chr1 | 151148780 | 151167797 | - | protein_coding |
| ENSG00000204394.13_6 | VARS1 | 0.99994147 | 1 | 0.033 | 6.152 | chr6 | 31745295 | 31763529 | - | protein_coding |
| ENSG00000223501.9_4 | VPS52 | 0.99998873 | 1 | 0.032 | 3.749 | chr6 | 33218049 | 33239824 | - | protein_coding |
| ENSG00000220472.1_5 | AL139095.2 | 0.96069438 | 1 | 0.032 | -0.102 | chr6 | 7339072 | 7339801 | - | processed_pseudogene |
| ENSG00000196159.11_2 | FAT4 | 0.99993582 | 1 | 0.032 | 3.234 | chr4 | 126237554 | 126414087 | + | protein_coding |
| ENSG00000154485.5_6 | MMP21 | 0.96163107 | 1 | 0.032 | -0.031 | chr10 | 127442149 | 127464390 | - | protein_coding |
| ENSG00000111231.9_4 | GPN3 | 0.99999982 | 1 | 0.032 | 4.764 | chr12 | 110890291 | 110907073 | - | protein_coding |
| ENSG00000067191.16_5 | CACNB1 | 0.99969196 | 1 | 0.032 | 2.735 | chr17 | 37329706 | 37353956 | - | protein_coding |
| ENSG00000163939.18_7 | PBRM1 | 0.99999997 | 1 | 0.032 | 6.023 | chr3 | 52579368 | 52719933 | - | protein_coding |
| ENSG00000174695.10_4 | TMEM167A | 1 | 1 | 0.032 | 6.648 | chr5 | 82348665 | 82373682 | - | protein_coding |
| ENSG00000196295.12_9 | GARS-DT | 0.99999722 | 1 | 0.032 | 4.128 | chr7 | 30555925 | 30634902 | - | lncRNA |
| ENSG00000162944.11_5 | RFTN2 | 0.96108581 | 1 | 0.032 | -0.065 | chr2 | 198432948 | 198540769 | - | protein_coding |
| ENSG00000197822.11_3 | OCLN | 0.9999983 | 1 | 0.032 | 4.506 | chr5 | 68788119 | 68853931 | + | protein_coding |
| ENSG00000068912.14_4 | ERLEC1 | 1 | 1 | 0.032 | 6.552 | chr2 | 54014181 | 54045956 | + | protein_coding |
| ENSG00000137343.18_6 | ATAT1 | 0.98342633 | 1 | 0.032 | 0.806 | chr6 | 30594619 | 30614600 | + | protein_coding |
| ENSG00000198369.10_6 | SPRED2 | 0.9999998 | 1 | 0.032 | 4.970 | chr2 | 65537985 | 65659771 | - | protein_coding |
| ENSG00000156172.6_4 | C8orf37 | 0.99998853 | 1 | 0.032 | 4.163 | chr8 | 96257141 | 96281429 | - | protein_coding |
| ENSG00000113360.16_5 | DROSHA | 1 | 1 | 0.032 | 6.131 | chr5 | 31400604 | 31532303 | - | protein_coding |
| ENSG00000260000.2_6 | AL133338.1 | 0.96661663 | 1 | 0.032 | 0.005 | chr6 | 101329347 | 101330863 | + | lncRNA |
| ENSG00000000003.15_4 | TSPAN6 | 0.99999997 | 1 | 0.032 | 5.630 | chrX | 99882105 | 99894988 | - | protein_coding |
| ENSG00000119421.7_4 | NDUFA8 | 0.99999947 | 1 | 0.032 | 4.640 | chr9 | 124906337 | 124922058 | - | protein_coding |
| ENSG00000147548.17_7 | NSD3 | 1 | 1 | 0.032 | 6.702 | chr8 | 38127222 | 38239790 | - | protein_coding |
| ENSG00000197566.10_5 | ZNF624 | 0.99999756 | 1 | 0.032 | 4.071 | chr17 | 16524048 | 16557170 | - | protein_coding |
| ENSG00000188690.15_8 | UROS | 1 | 1 | 0.032 | 5.973 | chr10 | 127473549 | 127511827 | - | protein_coding |
| ENSG00000116698.21_5 | SMG7 | 0.99999997 | 1 | 0.032 | 5.388 | chr1 | 183441351 | 183567381 | + | protein_coding |
| ENSG00000138942.16_3 | RNF185 | 0.99999925 | 1 | 0.031 | 4.392 | chr22 | 31556168 | 31603005 | + | protein_coding |
| ENSG00000174943.11_9 | KCTD13 | 0.99999012 | 1 | 0.031 | 3.721 | chr16 | 29916333 | 29937557 | - | protein_coding |
| ENSG00000127054.20_4 | INTS11 | 0.99999992 | 1 | 0.031 | 5.791 | chr1 | 1246965 | 1260071 | - | protein_coding |
| ENSG00000104889.6_5 | RNASEH2A | 0.96585489 | 1 | 0.031 | 0.070 | chr19 | 12917370 | 12926015 | + | protein_coding |
| ENSG00000229719.5_7 | MIR194-2HG | 0.9998936 | 1 | 0.031 | 3.165 | chr11 | 64657032 | 64660921 | - | lncRNA |
| ENSG00000128604.20_5 | IRF5 | 0.99998694 | 1 | 0.031 | 3.651 | chr7 | 128577511 | 128590092 | + | protein_coding |
| ENSG00000186575.18_4 | NF2 | 1 | 1 | 0.031 | 6.938 | chr22 | 29999545 | 30094587 | + | protein_coding |
| ENSG00000132199.20_5 | ENOSF1 | 0.99999997 | 1 | 0.031 | 5.291 | chr18 | 670318 | 712662 | - | protein_coding |
| ENSG00000107362.13_4 | ABHD17B | 0.99999985 | 1 | 0.031 | 4.809 | chr9 | 74477368 | 74525847 | - | protein_coding |
| ENSG00000139190.16_3 | VAMP1 | 0.99990542 | 1 | 0.031 | 3.169 | chr12 | 6571403 | 6580153 | - | protein_coding |
| ENSG00000267348.3_8 | GEMIN7-AS1 | 0.96705857 | 1 | 0.031 | 0.407 | chr19 | 45579768 | 45595893 | - | lncRNA |
| ENSG00000145901.15_9 | TNIP1 | 0.99999997 | 1 | 0.031 | 6.635 | chr5 | 150409506 | 150473138 | - | protein_coding |
| ENSG00000266412.5_3 | NCOA4 | 1 | 1 | 0.031 | 7.579 | chr10 | 51565108 | 51590734 | + | protein_coding |
| ENSG00000248671.8_9 | AP003498.1 | 0.99395354 | 1 | 0.031 | 1.442 | chr11 | 71384931 | 71529284 | - | lncRNA |
| ENSG00000109339.24_9 | MAPK10 | 0.98896449 | 1 | 0.030 | 0.986 | chr4 | 86911160 | 87515778 | - | polymorphic_pseudogene |
| ENSG00000263072.8_7 | ZNF213-AS1 | 0.9999936 | 1 | 0.030 | 3.791 | chr16 | 3150697 | 3193735 | - | lncRNA |
| ENSG00000175573.7_4 | C11orf68 | 0.9999922 | 1 | 0.030 | 3.847 | chr11 | 65684281 | 65686533 | - | protein_coding |
| ENSG00000114902.14_4 | SPCS1 | 0.99999997 | 1 | 0.030 | 5.249 | chr3 | 52738971 | 52745164 | + | protein_coding |
| ENSG00000175505.11_5 | CLCF1 | 0.99999877 | 1 | 0.030 | 4.536 | chr11 | 67131639 | 67141648 | - | protein_coding |
| ENSG00000245849.8_6 | RAD51-AS1 | 0.98944107 | 1 | 0.030 | 2.123 | chr15 | 40978381 | 40987305 | - | lncRNA |
| ENSG00000137831.15_8 | UACA | 1 | 1 | 0.030 | 6.813 | chr15 | 70946893 | 71055897 | - | protein_coding |

| | | | | | | | | | | |
|---|---|---|---|---|---|---|---|---|---|---|
| ENSG00000067601.8_6 | PMS2P4 | 0.99804762 | 1 | 0.030 | 1.324 | chr7 | 66752528 | 66767406 | - | transcribed_unprocessed_pseudogene |
| ENSG00000065268.10_3 | WDR18 | 0.99397885 | 1 | 0.030 | 4.983 | chr19 | 984271 | 998437 | + | protein_coding |
| ENSG00000135424.17_5 | ITGA7 | 0.99728797 | 1 | 0.030 | 1.211 | chr12 | 56078352 | 56109827 | - | protein_coding |
| ENSG00000160408.14_4 | ST6GALNAC6 | 0.95743307 | 1 | 0.030 | -0.189 | chr9 | 130647600 | 130667687 | - | protein_coding |
| ENSG00000077147.16_7 | TM9SF3 | 1 | 1 | 0.030 | 7.789 | chr10 | 98277867 | 98347209 | - | protein_coding |
| ENSG00000145675.15_3 | PIK3R1 | 0.99999969 | 1 | 0.030 | 4.722 | chr5 | 67511584 | 67597649 | + | protein_coding |
| ENSG00000083544.15_6 | TDRD3 | 0.99999443 | 1 | 0.030 | 4.224 | chr13 | 60970591 | 61148012 | + | protein_coding |
| ENSG00000247315.4_4 | ZCCHC3 | 0.99999856 | 1 | 0.030 | 4.364 | chr20 | 278211 | 280965 | + | protein_coding |
| ENSG00000198039.11_4 | ZNF273 | 0.99992847 | 1 | 0.030 | 3.037 | chr7 | 64330550 | 64391344 | + | protein_coding |
| ENSG00000164930.12_5 | FZD6 | 0.99999999 | 1 | 0.030 | 5.978 | chr8 | 104310661 | 104345094 | + | protein_coding |
| ENSG00000181220.17_5 | ZNF746 | 0.99999993 | 1 | 0.030 | 5.139 | chr7 | 149169787 | 149194908 | - | protein_coding |
| ENSG00000147133.15_2 | TAF1 | 0.99999999 | 1 | 0.029 | 5.974 | chrX | 70586089 | 70752224 | + | protein_coding |
| ENSG00000232098.3_7 | AC012313.1 | 0.99992554 | 1 | 0.029 | 3.059 | chr19 | 58915605 | 58919851 | - | lncRNA |
| ENSG00000112335.15_4 | SNX3 | 0.99999998 | 1 | 0.029 | 6.056 | chr6 | 108532426 | 108582450 | - | protein_coding |
| ENSG00000197724.11_4 | PHF2 | 0.99999997 | 1 | 0.029 | 5.278 | chr9 | 96338866 | 96441869 | + | protein_coding |
| ENSG00000213199.8_6 | ASIC3 | 0.99940082 | 1 | 0.029 | 2.681 | chr7 | 150745379 | 150749843 | + | protein_coding |
| ENSG00000138629.16_6 | UBL7 | 0.99999966 | 1 | 0.029 | 4.592 | chr15 | 74738318 | 74753523 | - | protein_coding |
| ENSG00000181135.16_7 | ZNF707 | 0.99989565 | 1 | 0.029 | 3.403 | chr8 | 144766622 | 144796068 | + | protein_coding |
| ENSG00000100201.21_6 | DDX17 | 1 | 1 | 0.029 | 8.842 | chr22 | 38879443 | 38903665 | - | protein_coding |
| ENSG00000158669.11_7 | GPAT4 | 1 | 1 | 0.029 | 6.227 | chr8 | 41434706 | 41482520 | + | protein_coding |
| ENSG00000128928.10_5 | IVD | 0.99993225 | 1 | 0.029 | 3.392 | chr15 | 40697686 | 40728146 | + | protein_coding |
| ENSG00000168538.16_3 | TRAPPC11 | 0.99999999 | 1 | 0.029 | 5.569 | chr4 | 184580420 | 184634747 | + | protein_coding |
| ENSG00000156384.14_2 | SFR1 | 0.99999581 | 1 | 0.029 | 4.050 | chr10 | 105881816 | 105886143 | + | protein_coding |
| ENSG00000046653.15_5 | GPM6B | 0.99491643 | 1 | 0.029 | 1.084 | chrX | 13789058 | 13956757 | - | protein_coding |
| ENSG00000133059.17_4 | DSTYK | 0.99999985 | 1 | 0.029 | 4.731 | chr1 | 205111633 | 205180830 | - | protein_coding |
| ENSG00000156858.11_4 | PRR14 | 0.99999776 | 1 | 0.028 | 3.937 | chr16 | 30662038 | 30667761 | + | protein_coding |
| ENSG00000196642.19_5 | RABL6 | 0.99833945 | 1 | 0.028 | 6.554 | chr9 | 139702395 | 139735639 | + | protein_coding |
| ENSG00000087884.14_6 | AAMDC | 0.99126687 | 1 | 0.028 | 1.097 | chr11 | 77532155 | 77629478 | + | protein_coding |
| ENSG00000111846.17_5 | GCNT2 | 0.99999995 | 1 | 0.028 | 5.337 | chr6 | 10492456 | 10629601 | + | protein_coding |
| ENSG00000178381.12_4 | ZFAND2A | 0.99999822 | 1 | 0.028 | 4.032 | chr7 | 1191707 | 1200395 | - | protein_coding |
| ENSG00000184924.5_3 | PTRHD1 | 0.99998987 | 1 | 0.028 | 3.556 | chr2 | 25012603 | 25016251 | - | protein_coding |
| ENSG00000269743.3_4 | SLC25A53 | 0.99587154 | 1 | 0.028 | 1.458 | chrX | 103343898 | 103401690 | - | protein_coding |
| ENSG00000126777.18_6 | KTN1 | 1 | 1 | 0.028 | 9.081 | chr14 | 56025790 | 56168244 | + | protein_coding |
| ENSG00000131100.13_4 | ATP6V1E1 | 0.99999999 | 1 | 0.028 | 5.963 | chr22 | 18074902 | 18111515 | - | protein_coding |
| ENSG00000170145.5_3 | SIK2 | 0.99999981 | 1 | 0.028 | 4.933 | chr11 | 111473173 | 111601579 | + | protein_coding |
| ENSG00000245205.3_4 | EEF1A1P4 | 0.96727134 | 1 | 0.028 | 0.022 | chr12 | 19609178 | 19610582 | - | processed_pseudogene |
| ENSG00000133313.15_6 | CNDP2 | 0.99999948 | 1 | 0.028 | 5.081 | chr18 | 72163051 | 72190689 | + | protein_coding |
| ENSG00000245614.4_7 | DDX11-AS1 | 0.97136485 | 1 | 0.028 | 0.265 | chr12 | 31172368 | 31226781 | - | lncRNA |
| ENSG00000013561.18_7 | RNF14 | 0.99999998 | 1 | 0.027 | 5.862 | chr5 | 141337893 | 141369857 | + | protein_coding |
| ENSG00000163904.13_4 | SENP2 | 0.99999959 | 1 | 0.027 | 4.630 | chr3 | 185300284 | 185351339 | + | protein_coding |
| ENSG00000051620.11_5 | HEBP2 | 0.99999997 | 1 | 0.027 | 5.263 | chr6 | 138724668 | 138743334 | + | protein_coding |
| ENSG00000247400.4_6 | DNAJC3-DT | 0.99558614 | 1 | 0.027 | 0.988 | chr13 | 96300987 | 96329209 | - | lncRNA |
| ENSG00000211454.14_5 | AKR7L | 0.96110141 | 1 | 0.027 | -0.668 | chr1 | 19592476 | 19600688 | - | polymorphic_pseudogene |
| ENSG00000105722.10_5 | ERF | 0.99992109 | 1 | 0.027 | 6.764 | chr19 | 42751721 | 42759280 | - | protein_coding |
| ENSG00000254632.2_7 | AP003119.1 | 0.99997938 | 1 | 0.027 | 3.539 | chr11 | 76470960 | 76479666 | - | lncRNA |
| ENSG00000109576.14_5 | AADAT | 0.99991341 | 1 | 0.027 | 2.907 | chr4 | 170981373 | 171012850 | - | protein_coding |
| ENSG00000273247.5_9 | AC097376.3 | 0.99817105 | 1 | 0.027 | 2.068 | chr4 | 140333081 | 140375188 | - | lncRNA |
| ENSG00000081019.13_6 | RSBN1 | 0.99999308 | 1 | 0.027 | 4.091 | chr1 | 114304454 | 114355098 | - | protein_coding |
| ENSG00000224877.4_6 | NDUFAF8 | 0.99999914 | 1 | 0.027 | 5.012 | chr17 | 79213105 | 79215110 | + | protein_coding |
| ENSG00000226465.3_6 | AL390198.1 | 0.95767094 | 1 | 0.027 | -0.670 | chr20 | 25677639 | 25733412 | + | lncRNA |
| ENSG00000147155.11_5 | EBP | 0.99999979 | 1 | 0.027 | 4.576 | chrX | 48380187 | 48387104 | + | protein_coding |
| ENSG00000279722.1_5 | AC007342.7 | 0.98725362 | 1 | 0.027 | 0.837 | chr16 | 53521519 | 53523855 | - | TEC |
| ENSG00000166246.14_5 | C16orf71 | 0.99367438 | 1 | 0.026 | 1.345 | chr16 | 4784345 | 4799397 | + | protein_coding |
| ENSG00000091140.14_3 | DLD | 0.99999981 | 1 | 0.026 | 6.327 | chr7 | 107531607 | 107572175 | + | protein_coding |
| ENSG00000141971.13_5 | MVB12A | 0.99996198 | 1 | 0.026 | 3.969 | chr19 | 17516531 | 17544533 | + | protein_coding |
| ENSG00000143669.14_5 | LYST | 0.99903517 | 1 | 0.026 | 4.131 | chr1 | 235824341 | 236046940 | - | protein_coding |
| ENSG00000163444.12_3 | TMEM183A | 0.99999998 | 1 | 0.026 | 5.383 | chr1 | 202976502 | 202993976 | + | protein_coding |
| ENSG00000182264.9_5 | IZUMO1 | 0.96451567 | 1 | 0.026 | -0.960 | chr19 | 49244109 | 49250166 | - | protein_coding |

| | | | | | | | | | | |
|---|---|---|---|---|---|---|---|---|---|---|
| ENSG00000143458.12_3 | GABPB2 | 0.99998562 | 1 | 0.026 | 3.535 | chr1 | 151043054 | 151098018 | + | protein_coding |
| ENSG00000168522.13_6 | FNTA | 0.99999991 | 1 | 0.026 | 5.382 | chr8 | 42889337 | 42940931 | + | protein_coding |
| ENSG00000162378.13_3 | ZYG11B | 1 | 1 | 0.026 | 5.920 | chr1 | 53192125 | 53293008 | + | protein_coding |
| ENSG00000133943.20_6 | DGLUCY | 0.99999965 | 1 | 0.026 | 4.524 | chr14 | 91526677 | 91691976 | + | protein_coding |
| ENSG00000122257.20_6 | RBBP6 | 0.99999999 | 1 | 0.026 | 6.251 | chr16 | 24549014 | 24584184 | + | protein_coding |
| ENSG00000205084.11_7 | TMEM231 | 0.99987357 | 1 | 0.026 | 2.857 | chr16 | 75570639 | 75590166 | - | protein_coding |
| ENSG00000032219.19_4 | ARID4A | 0.99999861 | 1 | 0.026 | 4.521 | chr14 | 58765222 | 58840605 | + | protein_coding |
| ENSG00000073578.17_5 | SDHA | 0.99998046 | 1 | 0.026 | 3.658 | chr5 | 218356 | 257197 | + | protein_coding |
| ENSG00000255423.1_4 | EBLN2 | 0.9953886 | 1 | 0.025 | 1.469 | chr3 | 73110810 | 73112488 | + | protein_coding |
| ENSG00000181991.16_3 | MRPS11 | 0.99999996 | 1 | 0.025 | 5.087 | chr15 | 89010684 | 89024007 | + | protein_coding |
| ENSG00000106524.9_4 | ANKMY2 | 0.9999997 | 1 | 0.025 | 4.624 | chr7 | 16639404 | 16685442 | - | protein_coding |
| ENSG00000120733.14_3 | KDM3B | 1 | 1 | 0.025 | 7.310 | chr5 | 137688374 | 137772717 | + | protein_coding |
| ENSG00000165280.16_6 | VCP | 1 | 1 | 0.025 | 9.404 | chr9 | 35056061 | 35072624 | - | protein_coding |
| ENSG00000102241.12_4 | HTATSF1 | 1 | 1 | 0.025 | 6.646 | chrX | 135579238 | 135594505 | + | protein_coding |
| ENSG00000267249.1_8 | AP005482.4 | 0.9648695 | 1 | 0.025 | -0.428 | chr18 | 12670425 | 12671144 | - | lncRNA |
| ENSG00000169490.17_5 | TM2D2 | 0.99999998 | 1 | 0.025 | 5.556 | chr8 | 38846327 | 38854343 | - | protein_coding |
| ENSG00000257103.8_5 | LSM14A | 1 | 1 | 0.025 | 7.381 | chr19 | 34663409 | 34720420 | + | protein_coding |
| ENSG00000185252.19_4 | ZNF74 | 0.99994625 | 1 | 0.025 | 3.267 | chr22 | 20748405 | 20762751 | + | protein_coding |
| ENSG00000154146.13_3 | NRGN | 0.98050557 | 1 | 0.025 | 0.506 | chr11 | 124609838 | 124617106 | + | protein_coding |
| ENSG00000163399.16_4 | ATP1A1 | 0.99999989 | 1 | 0.025 | 4.697 | chr1 | 116915290 | 116952883 | + | protein_coding |
| ENSG00000279605.1_7 | AC067930.5 | 0.97402094 | 1 | 0.025 | 0.159 | chr8 | 144660528 | 144665474 | + | TEC |
| ENSG00000224597.10_5 | SVIL-AS1 | 0.99991777 | 1 | 0.025 | 4.175 | chr10 | 29698331 | 29776674 | + | transcribed_unprocessed_pseudogene |
| ENSG00000233622.3_6 | CYP2T1P | 0.99694354 | 1 | 0.025 | 1.671 | chr19 | 41314379 | 41318005 | - | transcribed_unitary_pseudogene |
| ENSG00000154027.19_4 | AK5 | 0.99996577 | 1 | 0.025 | 3.360 | chr1 | 77747704 | 78025651 | + | protein_coding |
| ENSG00000177885.15_7 | GRB2 | 1 | 1 | 0.025 | 7.127 | chr17 | 73314157 | 73401790 | - | protein_coding |
| ENSG00000137801.11_4 | THBS1 | 1 | 1 | 0.025 | 7.416 | chr15 | 39873280 | 39891667 | + | protein_coding |
| ENSG00000175756.13_3 | AURKAIP1 | 0.99986433 | 1 | 0.025 | 5.148 | chr1 | 1309110 | 1310875 | - | protein_coding |
| ENSG00000148814.18_5 | LRRC27 | 0.99994448 | 1 | 0.024 | 3.114 | chr10 | 134145658 | 134195012 | + | protein_coding |
| ENSG00000171962.18_5 | DRC3 | 0.99845181 | 1 | 0.024 | 1.915 | chr17 | 17876127 | 17920203 | + | protein_coding |
| ENSG00000198838.13_3 | RYR3 | 0.9960551 | 1 | 0.024 | 1.763 | chr15 | 33603146 | 34158322 | + | protein_coding |
| ENSG00000151690.15_5 | MFSD6 | 0.99999998 | 1 | 0.024 | 5.304 | chr2 | 191273081 | 191373931 | + | protein_coding |
| ENSG00000126773.13_6 | PCNX4 | 1 | 1 | 0.024 | 6.464 | chr14 | 60558629 | 60635851 | + | protein_coding |
| ENSG00000268366.1_6 | AC010271.1 | 0.96999205 | 1 | 0.024 | 0.000 | chr19 | 40949343 | 40949994 | + | lncRNA |
| ENSG00000177383.5_4 | MAGEF1 | 0.99999928 | 1 | 0.024 | 4.336 | chr3 | 184428152 | 184429852 | - | protein_coding |
| ENSG00000116212.15_4 | LRRC42 | 0.99999957 | 1 | 0.024 | 4.532 | chr1 | 54411758 | 54433841 | + | protein_coding |
| ENSG00000103034.14_6 | NDRG4 | 0.99999992 | 1 | 0.024 | 4.766 | chr16 | 58496750 | 58547532 | + | protein_coding |
| ENSG00000166123.14_5 | GPT2 | 0.99999895 | 1 | 0.024 | 4.325 | chr16 | 46918274 | 46965201 | + | protein_coding |
| ENSG00000232818.2_4 | RPS2P32 | 0.97622344 | 1 | 0.024 | 0.529 | chr7 | 23530092 | 23530983 | + | processed_pseudogene |
| ENSG00000107290.14_6 | SETX | 1 | 1 | 0.024 | 7.562 | chr9 | 135136743 | 135230373 | - | protein_coding |
| ENSG00000163877.11_4 | SNIP1 | 0.99999204 | 1 | 0.024 | 3.845 | chr1 | 38000050 | 38019894 | - | protein_coding |
| ENSG00000188917.15_5 | TRMT2B | 0.99999997 | 1 | 0.024 | 5.272 | chrX | 100264335 | 100307105 | - | protein_coding |
| ENSG00000152942.19_8 | RAD17 | 0.99999968 | 1 | 0.024 | 4.800 | chr5 | 68665120 | 68710628 | + | protein_coding |
| ENSG00000237813.4_7 | AC002066.1 | 0.98547234 | 1 | 0.023 | 0.800 | chr7 | 115878314 | 116139519 | - | lncRNA |
| ENSG00000139880.19_3 | CDH24 | 0.99986217 | 1 | 0.023 | 2.987 | chr14 | 23516271 | 23526747 | - | protein_coding |
| ENSG00000139117.14_6 | CPNE8 | 0.99999991 | 1 | 0.023 | 5.248 | chr12 | 39040624 | 39301232 | - | protein_coding |
| ENSG00000116791.14_5 | CRYZ | 1 | 1 | 0.023 | 6.276 | chr1 | 75171166 | 75199092 | - | protein_coding |
| ENSG00000113108.19_4 | APBB3 | 0.99981654 | 1 | 0.023 | 2.740 | chr5 | 139937853 | 139944366 | - | protein_coding |
| ENSG00000001630.17_8 | CYP51A1 | 0.99596956 | 1 | 0.023 | 2.292 | chr7 | 91741467 | 91764117 | - | protein_coding |
| ENSG00000064726.10_6 | BTBD1 | 1 | 1 | 0.023 | 6.966 | chr15 | 83685175 | 83736004 | - | protein_coding |
| ENSG00000124541.7_3 | RRP36 | 0.99999999 | 1 | 0.023 | 5.701 | chr6 | 42989361 | 43001894 | + | protein_coding |
| ENSG00000140474.14_5 | AC091230.1 | 0.99999996 | 1 | 0.023 | 5.437 | chr15 | 75128459 | 75135687 | - | protein_coding |
| ENSG00000010292.13_4 | NCAPD2 | 1 | 1 | 0.023 | 7.763 | chr12 | 6602522 | 6641121 | + | protein_coding |
| ENSG00000267185.1_4 | PTP4A2P1 | 0.96302723 | 1 | 0.023 | -0.548 | chr17 | 40685550 | 40686027 | + | processed_pseudogene |
| ENSG00000168765.17_3 | GSTM4 | 0.99975852 | 1 | 0.023 | 2.729 | chr1 | 110198721 | 110217458 | + | protein_coding |
| ENSG00000107669.17_8 | ATE1 | 1 | 1 | 0.022 | 6.477 | chr10 | 123499936 | 123688316 | - | protein_coding |
| ENSG00000115275.13_7 | MOGS | 0.99999999 | 1 | 0.022 | 5.955 | chr2 | 74688184 | 74692537 | - | protein_coding |
| ENSG00000101347.9_6 | SAMHD1 | 0.99999981 | 1 | 0.022 | 4.737 | chr20 | 35518176 | 35580304 | - | protein_coding |
| ENSG00000166428.13_3 | PLD4 | 0.97217106 | 1 | 0.022 | 0.086 | chr14 | 105391050 | 105404127 | + | protein_coding |
| ENSG00000234546.4_5 | LNCTAM34A | 0.98313462 | 1 | 0.022 | 0.540 | chr1 | 9242063 | 9256343 | + | lncRNA |

| | | | | | | | | | | |
|---|---|---|---|---|---|---|---|---|---|---|
| ENSG00000264343.6_6 | NOTCH2NLA | 0.99973226 | 1 | 0.022 | 2.625 | chr1 | 145209119 | 145291972 | + | protein_coding |
| ENSG00000114742.14_4 | WDR48 | 0.99999999 | 1 | 0.022 | 5.615 | chr3 | 39093504 | 39138162 | + | protein_coding |
| ENSG00000133026.12_8 | MYH10 | 1 | 1 | 0.022 | 6.465 | chr17 | 8377523 | 8534079 | - | protein_coding |
| ENSG00000114554.11_3 | PLXNA1 | 0.99999919 | 1 | 0.022 | 6.338 | chr3 | 126707437 | 126756235 | + | protein_coding |
| ENSG00000136371.10_4 | MTHFS | 0.99184751 | 1 | 0.022 | 0.465 | chr15 | 80125927 | 80189721 | - | protein_coding |
| ENSG00000206053.13_7 | JPT2 | 1 | 1 | 0.022 | 7.423 | chr16 | 1728257 | 1752281 | + | protein_coding |
| ENSG00000080561.13_3 | MID2 | 0.99999357 | 1 | 0.022 | 3.569 | chrX | 107068985 | 107170423 | + | protein_coding |
| ENSG00000120278.16_6 | PLEKHG1 | 0.9622498 | 1 | 0.021 | -0.731 | chr6 | 150921019 | 151164801 | + | protein_coding |
| ENSG00000072849.11_5 | DERL2 | 0.99999998 | 1 | 0.021 | 5.896 | chr17 | 5374574 | 5390131 | - | protein_coding |
| ENSG00000218283.2_4 | MORF4L1P1 | 0.99992712 | 1 | 0.021 | 3.082 | chr1 | 220426912 | 220427878 | + | processed_pseudogene |
| ENSG00000234373.1_5 | SNX18P7 | 0.98564229 | 1 | 0.021 | 0.666 | chr9 | 33576378 | 33577012 | + | processed_pseudogene |
| ENSG00000170892.12_7 | TSEN34 | 0.99999801 | 1 | 0.021 | 6.113 | chr19 | 54693789 | 54698394 | + | protein_coding |
| ENSG00000103154.9_3 | NECAB2 | 0.99888982 | 1 | 0.021 | 1.341 | chr16 | 84002237 | 84036381 | + | protein_coding |
| ENSG00000096401.8_3 | CDC5L | 1 | 1 | 0.021 | 6.419 | chr6 | 44355443 | 44418162 | + | protein_coding |
| ENSG00000188486.3_4 | H2AX | 0.99402849 | 1 | 0.021 | 6.114 | chr11 | 118964564 | 118966177 | - | protein_coding |
| ENSG00000166188.2_4 | ZNF319 | 0.99998971 | 1 | 0.021 | 4.198 | chr16 | 58028572 | 58034357 | - | protein_coding |
| ENSG00000119616.11_3 | FCF1 | 0.99999997 | 1 | 0.021 | 5.372 | chr14 | 75179847 | 75205323 | + | protein_coding |
| ENSG00000254682.1_7 | AP002387.2 | 0.9925109 | 1 | 0.021 | 1.301 | chr11 | 71159720 | 71163203 | + | lncRNA |
| ENSG00000169083.17_7 | AR | 0.99999999 | 1 | 0.021 | 6.048 | chrX | 66763863 | 66950461 | + | protein_coding |
| ENSG00000267216.1_9 | AC020915.1 | 0.99637936 | 1 | 0.021 | 1.462 | chr19 | 58790332 | 58826563 | + | lncRNA |
| ENSG00000233830.2_4 | EIF4HP1 | 0.99723583 | 1 | 0.021 | 1.712 | chr7 | 27497782 | 27498468 | - | processed_pseudogene |
| ENSG00000150712.11_4 | MTMR12 | 0.99999986 | 1 | 0.021 | 4.852 | chr5 | 32227100 | 32313093 | - | protein_coding |
| ENSG00000280219.1_6 | AC093908.1 | 0.96738651 | 1 | 0.020 | -0.453 | chr4 | 148409123 | 148417214 | + | TEC |
| ENSG00000131094.4_4 | C1QL1 | 0.99964686 | 1 | 0.020 | 2.563 | chr17 | 43037061 | 43045671 | - | protein_coding |
| ENSG00000112033.14_5 | PPARD | 0.99979357 | 1 | 0.020 | 2.529 | chr6 | 35310335 | 35395968 | + | protein_coding |
| ENSG00000119688.21_7 | ABCD4 | 0.99999987 | 1 | 0.020 | 4.657 | chr14 | 74751972 | 74769758 | - | protein_coding |
| ENSG00000075711.21_7 | DLG1 | 0.99999998 | 1 | 0.020 | 5.524 | chr3 | 196769431 | 197026201 | - | protein_coding |
| ENSG00000167962.14_6 | ZNF598 | 0.9982665 | 1 | 0.020 | 5.458 | chr16 | 2047655 | 2059822 | - | protein_coding |
| ENSG00000236780.7_7 | LINC01829 | 0.97611585 | 1 | 0.020 | -0.086 | chr2 | 67350489 | 67625112 | - | lncRNA |
| ENSG00000099804.9_3 | CDC34 | 0.99999945 | 1 | 0.020 | 5.011 | chr19 | 531760 | 542092 | + | protein_coding |
| ENSG00000278588.1_3 | H2BC10 | 1 | 1 | 0.020 | 6.237 | chr6 | 26273204 | 26273584 | + | protein_coding |
| ENSG00000099995.19_8 | SF3A1 | 0.99999999 | 1 | 0.020 | 5.476 | chr22 | 30727977 | 30752908 | - | protein_coding |
| ENSG00000074219.13_6 | TEAD2 | 0.99999451 | 1 | 0.020 | 4.147 | chr19 | 49843852 | 49865714 | - | protein_coding |
| ENSG00000206503.13_5 | HLA-A | 0.99999729 | 1 | 0.020 | 4.107 | chr6 | 29909037 | 29913661 | + | protein_coding |
| ENSG00000152582.14_6 | SPEF2 | 0.9998467 | 1 | 0.020 | 2.878 | chr5 | 35617946 | 35814713 | + | protein_coding |
| ENSG00000122512.16_5 | PMS2 | 0.99999992 | 1 | 0.020 | 4.953 | chr7 | 6010556 | 6048737 | - | protein_coding |
| ENSG00000178896.9_5 | EXOSC4 | 0.9999975 | 1 | 0.020 | 4.086 | chr8 | 145133551 | 145135551 | + | protein_coding |
| ENSG00000226049.3_4 | TLK2P1 | 0.98170069 | 1 | 0.020 | 0.507 | chr17 | 32363700 | 32366891 | - | processed_pseudogene |
| ENSG00000168802.13_7 | CHTF8 | 0.96661181 | 1 | 0.020 | -0.772 | chr16 | 69151913 | 69166481 | - | protein_coding |
| ENSG00000152620.13_4 | NADK2 | 0.9999994 | 1 | 0.020 | 4.981 | chr5 | 36192691 | 36242381 | - | protein_coding |
| ENSG00000189420.8_2 | ZFP92 | 0.99852531 | 1 | 0.019 | 1.841 | chrX | 152683780 | 152691939 | + | protein_coding |
| ENSG00000161996.19_4 | WDR90 | 0.99999911 | 1 | 0.019 | 5.284 | chr16 | 699311 | 717833 | + | protein_coding |
| ENSG00000203709.12_8 | MIR29B2CHG | 0.99997436 | 1 | 0.019 | 3.327 | chr1 | 207974863 | 208052460 | - | lncRNA |
| ENSG00000047230.15_6 | CTPS2 | 0.99999996 | 1 | 0.019 | 5.123 | chrX | 16606122 | 16731059 | - | protein_coding |
| ENSG00000160305.18_5 | DIP2A | 0.99999997 | 1 | 0.019 | 5.271 | chr21 | 47878804 | 47989928 | + | protein_coding |
| ENSG00000135040.16_3 | NAA35 | 0.99999999 | 1 | 0.019 | 5.997 | chr9 | 88556061 | 88640377 | + | protein_coding |
| ENSG00000140990.15_5 | NDUFB10 | 1 | 1 | 0.019 | 6.450 | chr16 | 2009539 | 2011976 | + | protein_coding |
| ENSG00000168172.9_5 | HOOK3 | 0.99999999 | 1 | 0.019 | 6.010 | chr8 | 42752089 | 42885678 | + | protein_coding |
| ENSG00000100359.21_4 | SGSM3 | 0.9754478 | 1 | 0.019 | -0.200 | chr22 | 40766595 | 40806293 | + | protein_coding |
| ENSG00000143479.17_5 | DYRK3 | 0.99999801 | 1 | 0.019 | 4.576 | chr1 | 206808881 | 206857764 | + | protein_coding |
| ENSG00000270882.2_3 | H4C14 | 0.9863693 | 1 | 0.019 | 0.567 | chr1 | 149804223 | 149811339 | + | protein_coding |
| ENSG00000110080.18_6 | ST3GAL4 | 0.99998189 | 1 | 0.019 | 3.506 | chr11 | 126225535 | 126310239 | + | protein_coding |
| ENSG00000272693.4_6 | AC073107.1 | 0.97026982 | 1 | 0.019 | -0.229 | chr7 | 65111924 | 65235797 | - | transcribed_unprocessed_pseudogene |
| ENSG00000272758.6_6 | AC083798.2 | 0.99387736 | 1 | 0.019 | 0.723 | chr3 | 122135054 | 122162027 | + | lncRNA |
| ENSG00000090989.18_4 | EXOC1 | 0.99999991 | 1 | 0.019 | 4.940 | chr4 | 56719814 | 56771252 | + | protein_coding |
| ENSG00000197912.15_8 | SPG7 | 1 | 1 | 0.019 | 6.912 | chr16 | 89557127 | 89624176 | + | protein_coding |
| ENSG00000134830.6_5 | C5AR2 | 0.96674369 | 1 | 0.019 | -0.814 | chr19 | 47835432 | 47850586 | + | protein_coding |

| | | | | | | | | | | |
|---|---|---|---|---|---|---|---|---|---|---|
| ENSG00000148143.13_8 | ZNF462 | 0.9999997 | 1 | 0.019 | 4.426 | chr9 | 109625447 | 109775915 | + | protein_coding |
| ENSG00000272888.7_6 | LINC01578 | 0.99999993 | 1 | 0.019 | 5.044 | chr15 | 93362770 | 93442931 | + | lncRNA |
| ENSG00000198793.13_7 | MTOR | 1 | 1 | 0.019 | 6.650 | chr1 | 11166592 | 11322608 | - | protein_coding |
| ENSG00000005175.10_5 | RPAP3 | 1 | 1 | 0.019 | 6.219 | chr12 | 48055032 | 48099813 | - | protein_coding |
| ENSG00000272631.1_7 | AC067750.1 | 0.99915878 | 1 | 0.018 | 2.129 | chr10 | 88509511 | 88516055 | - | lncRNA |
| ENSG00000117625.13_5 | RCOR3 | 0.99999888 | 1 | 0.018 | 4.715 | chr1 | 211431719 | 211489727 | + | protein_coding |
| ENSG00000129128.13_3 | SPCS3 | 1 | 1 | 0.018 | 6.656 | chr4 | 177241117 | 177253396 | + | protein_coding |
| ENSG00000138495.6_4 | COX17 | 0.99999986 | 1 | 0.018 | 4.902 | chr3 | 119373360 | 119396301 | - | protein_coding |
| ENSG00000273188.1_6 | AL022328.3 | 0.97443607 | 1 | 0.018 | -0.326 | chr22 | 50630153 | 50630831 | - | lncRNA |
| ENSG00000066427.24_8 | ATXN3 | 0.99999991 | 1 | 0.018 | 4.960 | chr14 | 92510840 | 92572965 | - | protein_coding |
| ENSG00000230612.3_7 | AC004039.1 | 0.96776167 | 1 | 0.018 | -0.574 | chr5 | 132024373 | 132059417 | + | lncRNA |
| ENSG00000186638.16_4 | KIF24 | 0.99999987 | 1 | 0.018 | 4.649 | chr9 | 34252379 | 34311369 | - | protein_coding |
| ENSG00000231652.2_7 | AL590428.1 | 0.97252246 | 1 | 0.018 | -0.406 | chr6 | 74403626 | 74405854 | - | lncRNA |
| ENSG00000187193.9_4 | MT1X | 0.99875269 | 1 | 0.018 | 2.142 | chr16 | 56716382 | 56718108 | + | protein_coding |
| ENSG00000172273.13_6 | HINFP | 0.99999566 | 1 | 0.018 | 3.672 | chr11 | 118992290 | 119006769 | + | protein_coding |
| ENSG00000179387.10_6 | ELMOD2 | 0.99999997 | 1 | 0.018 | 5.525 | chr4 | 141445322 | 141474924 | + | protein_coding |
| ENSG00000099991.18_7 | CABIN1 | 0.99999999 | 1 | 0.018 | 5.796 | chr22 | 24407642 | 24574596 | + | protein_coding |
| ENSG00000140025.16_5 | EFCAB11 | 0.99992362 | 1 | 0.018 | 3.068 | chr14 | 90261013 | 90421121 | - | protein_coding |
| ENSG00000100422.14_4 | CERK | 0.99999997 | 1 | 0.018 | 5.439 | chr22 | 47080307 | 47134149 | - | protein_coding |
| ENSG00000287507.1_2 | AP001053.1 | 0.98889907 | 1 | 0.017 | 0.324 | chr21 | 45188201 | 45191547 | + | lncRNA |
| ENSG00000169193.12_4 | CCDC126 | 0.99985804 | 1 | 0.017 | 2.637 | chr7 | 23637001 | 23684327 | + | protein_coding |
| ENSG00000157020.18_5 | SEC13 | 1 | 1 | 0.017 | 6.823 | chr3 | 10334815 | 10362796 | - | protein_coding |
| ENSG00000103264.18_5 | FBXO31 | 0.99999925 | 1 | 0.017 | 4.328 | chr16 | 87360593 | 87425748 | - | protein_coding |
| ENSG00000072201.14_6 | LNX1 | 0.97811852 | 1 | 0.017 | -0.004 | chr4 | 54325468 | 54567572 | - | protein_coding |
| ENSG00000095203.14_3 | EPB41L4B | 0.99999968 | 1 | 0.017 | 4.595 | chr9 | 111934255 | 112083244 | - | protein_coding |
| ENSG00000188243.12_3 | COMMD6 | 0.99999995 | 1 | 0.017 | 5.068 | chr13 | 76099355 | 76123575 | - | protein_coding |
| ENSG00000272106.1_6 | AL691432.2 | 0.9931643 | 1 | 0.017 | 0.619 | chr1 | 1549138 | 1551175 | - | lncRNA |
| ENSG00000164920.9_5 | OSR2 | 0.99854646 | 1 | 0.017 | 1.786 | chr8 | 99956631 | 99964332 | + | protein_coding |
| ENSG00000234585.6_2 | CCT6P3 | 0.99666411 | 1 | 0.017 | 1.701 | chr7 | 64498732 | 64535091 | + | transcribed_unprocessed_pseudogene |
| ENSG00000129292.20_4 | PHF20L1 | 1 | 1 | 0.017 | 6.417 | chr8 | 133787604 | 133861052 | + | protein_coding |
| ENSG00000234537.1_5 | AL354751.1 | 0.96973193 | 1 | 0.017 | -0.812 | chr9 | 94894680 | 94900911 | + | unprocessed_pseudogene |
| ENSG00000155660.11_7 | PDIA4 | 1 | 1 | 0.017 | 7.975 | chr7 | 148700154 | 148725754 | - | protein_coding |
| ENSG00000147065.17_4 | MSN | 1 | 1 | 0.017 | 8.664 | chrX | 64808257 | 64961793 | + | protein_coding |
| ENSG00000166226.13_3 | CCT2 | 1 | 1 | 0.017 | 8.657 | chr12 | 69979206 | 69995350 | + | protein_coding |
| ENSG00000219545.11_3 | UMAD1 | 0.99999857 | 1 | 0.017 | 4.148 | chr7 | 7680342 | 8007651 | + | protein_coding |
| ENSG00000260988.1_8 | AC090260.2 | 0.96974755 | 1 | 0.017 | -0.960 | chr15 | 78433585 | 78435515 | + | lncRNA |
| ENSG00000138738.10_5 | PRDM5 | 0.9999905 | 1 | 0.017 | 3.683 | chr4 | 121606074 | 121844025 | - | protein_coding |
| ENSG00000154277.13_4 | UCHL1 | 1 | 1 | 0.017 | 8.657 | chr4 | 41258430 | 41270472 | + | protein_coding |
| ENSG00000170502.13_4 | NUDT9 | 0.9999976 | 1 | 0.017 | 3.787 | chr4 | 88343725 | 88380607 | + | protein_coding |
| ENSG00000125434.11_4 | SLC25A35 | 0.99984065 | 1 | 0.017 | 2.738 | chr17 | 8191081 | 8198718 | - | protein_coding |
| ENSG00000216895.9_8 | AC009403.1 | 0.99858469 | 1 | 0.017 | 1.827 | chr7 | 155403925 | 155437899 | - | lncRNA |
| ENSG00000130856.16_3 | ZNF236 | 0.99999988 | 1 | 0.017 | 4.797 | chr18 | 74534563 | 74682683 | + | protein_coding |
| ENSG00000124224.17_6 | PPP4R1L | 0.99992528 | 1 | 0.016 | 2.883 | chr20 | 56803996 | 56884507 | - | transcribed_unprocessed_pseudogene |
| ENSG00000117640.18_5 | MTFR1L | 0.99999829 | 1 | 0.016 | 3.823 | chr1 | 26145131 | 26159433 | + | protein_coding |
| ENSG00000126945.9_4 | HNRNPH2 | 0.99999962 | 1 | 0.016 | 5.792 | chrX | 100663210 | 100669121 | + | protein_coding |
| ENSG00000115641.19_6 | FHL2 | 0.99999971 | 1 | 0.016 | 4.674 | chr2 | 105974169 | 106054970 | - | protein_coding |
| ENSG00000096080.11_4 | MRPS18A | 0.99996483 | 1 | 0.016 | 3.287 | chr6 | 43639040 | 43655528 | - | protein_coding |
| ENSG00000272391.6_6 | POM121C | 0.99997605 | 1 | 0.016 | 3.632 | chr7 | 75046065 | 75115576 | - | protein_coding |
| ENSG00000204839.9_7 | MROH6 | 0.99972204 | 1 | 0.016 | 3.365 | chr8 | 144648362 | 144654942 | - | protein_coding |
| ENSG00000198156.10_4 | NPIPB6 | 0.97612151 | 1 | 0.016 | -0.279 | chr16 | 28353876 | 28374829 | - | protein_coding |
| ENSG00000105063.19_7 | PPP6R1 | 0.99997357 | 1 | 0.016 | 7.160 | chr19 | 55741147 | 55770385 | - | protein_coding |
| ENSG00000005059.16_6 | MCUB | 0.99999978 | 1 | 0.016 | 4.409 | chr4 | 110481365 | 110609875 | + | protein_coding |
| ENSG00000280417.1_5 | AC096887.2 | 0.97296807 | 1 | 0.016 | -0.439 | chr3 | 53080182 | 53082138 | + | TEC |
| ENSG00000140553.18_5 | UNC45A | 0.99999998 | 1 | 0.016 | 6.085 | chr15 | 91473410 | 91497323 | + | protein_coding |
| ENSG00000198794.12_7 | SCAMP5 | 0.99999646 | 1 | 0.016 | 3.675 | chr15 | 75249560 | 75313836 | + | protein_coding |
| ENSG00000162396.6_4 | PARS2 | 0.9998525 | 1 | 0.016 | 2.706 | chr1 | 55222571 | 55230196 | - | protein_coding |
| ENSG00000176248.9_4 | ANAPC2 | 0.99999997 | 1 | 0.016 | 5.317 | chr9 | 140069236 | 140083012 | - | protein_coding |
| ENSG00000253816.3_7 | AC138866.1 | 0.97043867 | 1 | 0.016 | -1.221 | chr5 | 69515745 | 69554757 | - | unprocessed_pseudogene |
| ENSG00000177943.14_4 | MAMDC4 | 0.99950699 | 1 | 0.015 | 5.077 | chr9 | 139745395 | 139755251 | + | protein_coding |

| | | | | | | | | | | |
|---|---|---|---|---|---|---|---|---|---|---|
| ENSG00000250608.2_7 | AC010210.1 | 0.97359966 | 1 | 0.015 | -0.058 | chr3 | 131043936 | 131100500 | - | lncRNA |
| ENSG00000172081.14_6 | MOB3A | 0.9999957 | 1 | 0.015 | 3.950 | chr19 | 2071035 | 2096672 | - | protein_coding |
| ENSG00000160007.19_4 | ARHGAP35 | 1 | 1 | 0.015 | 7.641 | chr19 | 47364254 | 47508334 | + | protein_coding |
| ENSG00000164366.4_5 | CCDC127 | 0.99999986 | 1 | 0.015 | 4.652 | chr5 | 196983 | 218268 | - | protein_coding |
| ENSG00000164308.17_8 | ERAP2 | 0.99891625 | 1 | 0.015 | 2.051 | chr5 | 96211690 | 96255407 | + | protein_coding |
| ENSG00000267342.1_8 | AC087289.2 | 0.97857206 | 1 | 0.015 | 0.061 | chr17 | 73893141 | 73896229 | + | lncRNA |
| ENSG00000158850.15_4 | B4GALT3 | 0.99999907 | 1 | 0.015 | 4.238 | chr1 | 161141100 | 161147758 | - | protein_coding |
| ENSG00000198408.14_5 | OGA | 1 | 1 | 0.015 | 7.217 | chr10 | 103544200 | 103578222 | - | protein_coding |
| ENSG00000175073.8_4 | VCPIP1 | 1 | 1 | 0.015 | 5.566 | chr8 | 67540722 | 67579466 | - | protein_coding |
| ENSG00000260924.2_6 | LINC01311 | 0.99604096 | 1 | 0.015 | 0.745 | chr22 | 19158908 | 19160352 | + | lncRNA |
| ENSG00000083937.9_3 | CHMP2B | 1 | 1 | 0.015 | 7.060 | chr3 | 87276421 | 87304706 | + | protein_coding |
| ENSG00000198420.10_8 | TCAF1 | 0.99999999 | 1 | 0.015 | 5.265 | chr7 | 143548468 | 143599291 | - | protein_coding |
| ENSG00000105819.14_3 | PMPCB | 1 | 1 | 0.015 | 6.099 | chr7 | 102937882 | 102969958 | + | protein_coding |
| ENSG00000166135.14_3 | HIF1AN | 1 | 1 | 0.015 | 5.868 | chr10 | 102288829 | 102319755 | + | protein_coding |
| ENSG00000101745.17_4 | ANKRD12 | 0.99999969 | 1 | 0.015 | 5.252 | chr18 | 9136226 | 9285983 | + | protein_coding |
| ENSG00000188529.15_5 | SRSF10 | 0.99999994 | 1 | 0.015 | 4.979 | chr1 | 24290837 | 24307417 | - | protein_coding |
| ENSG00000220205.9_6 | VAMP2 | 0.99993901 | 1 | 0.015 | 2.951 | chr17 | 8062467 | 8066864 | - | protein_coding |
| ENSG00000183621.15_4 | ZNF438 | 0.9999657 | 1 | 0.015 | 3.008 | chr10 | 31109136 | 31320866 | - | protein_coding |
| ENSG00000091157.13_4 | WDR7 | 0.99999966 | 1 | 0.015 | 4.435 | chr18 | 54318574 | 54703837 | + | protein_coding |
| ENSG00000119906.13_5 | SLF2 | 1 | 1 | 0.015 | 6.279 | chr10 | 102672720 | 102724891 | + | protein_coding |
| ENSG00000106803.10_3 | SEC61B | 1 | 1 | 0.014 | 5.863 | chr9 | 101984346 | 101992897 | + | protein_coding |
| ENSG00000175061.18_5 | SNHG29 | 1 | 1 | 0.014 | 7.622 | chr17 | 16342081 | 16381992 | + | lncRNA |
| ENSG00000182149.21_7 | IST1 | 1 | 1 | 0.014 | 6.808 | chr16 | 71879899 | 71965102 | + | protein_coding |
| ENSG00000170234.13_4 | PWWP2A | 0.99999967 | 1 | 0.014 | 4.681 | chr5 | 159488808 | 159546457 | - | protein_coding |
| ENSG00000272767.1_5 | JMJD1C-AS1 | 0.97527905 | 1 | 0.014 | -1.221 | chr10 | 65224989 | 65226323 | + | lncRNA |
| ENSG00000173805.16_8 | HAP1 | 0.99973199 | 1 | 0.014 | 2.461 | chr17 | 39873994 | 39890896 | - | protein_coding |
| ENSG00000260236.1_6 | AC099778.1 | 0.98530313 | 1 | 0.014 | 0.424 | chr3 | 47420579 | 47422489 | - | lncRNA |
| ENSG00000108848.16_6 | LUC7L3 | 1 | 1 | 0.014 | 7.069 | chr17 | 48796926 | 48833580 | + | protein_coding |
| ENSG00000062194.16_3 | GPBP1 | 1 | 1 | 0.014 | 6.603 | chr5 | 56469775 | 56560506 | + | protein_coding |
| ENSG00000004139.14_5 | SARM1 | 0.99947952 | 1 | 0.014 | 2.313 | chr17 | 26691378 | 26731067 | + | protein_coding |
| ENSG00000083807.10_7 | SLC27A5 | 0.99997204 | 1 | 0.014 | 3.536 | chr19 | 58990879 | 59023780 | - | protein_coding |
| ENSG00000131507.11_3 | NDFIP1 | 1 | 1 | 0.014 | 6.504 | chr5 | 141488344 | 141534005 | + | protein_coding |
| ENSG00000233483.3_6 | AC008105.2 | 0.99076881 | 1 | 0.014 | 0.661 | chr17 | 43322553 | 43328496 | - | transcribed_unitary_pseudogene |
| ENSG00000159842.15_6 | ABR | 1 | 1 | 0.014 | 7.317 | chr17 | 906758 | 1133032 | - | protein_coding |
| ENSG00000166822.13_6 | TMEM170A | 0.99999993 | 1 | 0.014 | 4.641 | chr16 | 75476952 | 75499395 | - | protein_coding |
| ENSG00000116151.14_7 | MORN1 | 0.9999951 | 1 | 0.014 | 3.663 | chr1 | 2252692 | 2323146 | - | protein_coding |
| ENSG00000172893.15_6 | DHCR7 | 1 | 1 | 0.014 | 6.334 | chr11 | 71139239 | 71163914 | - | protein_coding |
| ENSG00000104365.15_5 | IKBKB | 1 | 1 | 0.013 | 5.896 | chr8 | 42128820 | 42190171 | + | protein_coding |
| ENSG00000092978.11_4 | GPATCH2 | 0.99999998 | 1 | 0.013 | 5.315 | chr1 | 217600334 | 217804432 | - | protein_coding |
| ENSG00000198435.4_4 | NRARP | 0.98487799 | 1 | 0.013 | 0.206 | chr9 | 140194083 | 140196723 | - | protein_coding |
| ENSG00000143324.14_4 | XPR1 | 1 | 1 | 0.013 | 6.632 | chr1 | 180601158 | 180859415 | + | protein_coding |
| ENSG00000198919.13_5 | DZIP3 | 1 | 1 | 0.013 | 5.616 | chr3 | 108308552 | 108413687 | + | protein_coding |
| ENSG00000177873.14_4 | ZNF619 | 0.99997733 | 1 | 0.013 | 3.256 | chr3 | 40518604 | 40532544 | + | protein_coding |
| ENSG00000168374.11_5 | ARF4 | 1 | 1 | 0.013 | 6.798 | chr3 | 57557090 | 57583947 | - | protein_coding |
| ENSG00000172262.11_5 | ZNF131 | 0.99999999 | 1 | 0.013 | 5.465 | chr5 | 43065278 | 43192123 | + | protein_coding |
| ENSG00000068438.15_4 | FTSJ1 | 1 | 1 | 0.013 | 5.658 | chrX | 48334409 | 48344752 | + | protein_coding |
| ENSG00000104524.14_10 | PYCR3 | 0.99999984 | 1 | 0.013 | 5.031 | chr8 | 144685380 | 144691943 | - | protein_coding |
| ENSG00000197451.12_4 | HNRNPAB | 1 | 1 | 0.013 | 6.839 | chr5 | 177631534 | 177638164 | + | protein_coding |
| ENSG00000012983.11_5 | MAP4K5 | 1 | 1 | 0.013 | 5.784 | chr14 | 50885219 | 51027844 | - | protein_coding |
| ENSG00000139116.18_5 | KIF21A | 1 | 1 | 0.013 | 6.506 | chr12 | 39687030 | 39837192 | - | protein_coding |
| ENSG00000253352.9_6 | TUG1 | 1 | 1 | 0.013 | 6.666 | chr22 | 31365231 | 31375381 | + | lncRNA |
| ENSG00000105329.10_6 | TGFB1 | 0.99999991 | 1 | 0.013 | 5.147 | chr19 | 41807492 | 41859827 | - | protein_coding |
| ENSG00000010318.21_4 | PHF7 | 0.99729444 | 1 | 0.012 | 1.501 | chr3 | 52444676 | 52457657 | + | protein_coding |
| ENSG00000073921.18_6 | PICALM | 1 | 1 | 0.012 | 6.712 | chr11 | 85668218 | 85780924 | - | protein_coding |
| ENSG00000116679.16_4 | IVNS1ABP | 1 | 1 | 0.012 | 7.127 | chr1 | 185265520 | 185286405 | - | protein_coding |
| ENSG00000074755.15_6 | ZZEF1 | 1 | 1 | 0.012 | 6.838 | chr17 | 3907739 | 4046324 | - | protein_coding |
| ENSG00000135451.13_6 | TROAP | 0.99999999 | 1 | 0.012 | 5.244 | chr12 | 49717019 | 49725514 | + | protein_coding |
| ENSG00000189319.14_5 | FAM53B | 0.99999644 | 1 | 0.012 | 4.061 | chr10 | 126307861 | 126432947 | - | protein_coding |
| ENSG00000232406.7_8 | AL121895.1 | 0.99047736 | 1 | 0.012 | 0.583 | chr20 | 34735256 | 34743682 | - | lncRNA |

| | | | | | | | | | | |
|---|---|---|---|---|---|---|---|---|---|---|
| ENSG00000153339.14_5 | TRAPPC8 | 1 | 1 | 0.012 | 5.904 | chr18 | 29409143 | 29533099 | - | protein_coding |
| ENSG00000184304.15_6 | PRKD1 | 0.99999181 | 1 | 0.012 | 3.561 | chr14 | 30045685 | 30661104 | - | protein_coding |
| ENSG00000163257.11_4 | DCAF16 | 0.99999983 | 1 | 0.012 | 4.490 | chr4 | 17802278 | 17812381 | - | protein_coding |
| ENSG00000172508.10_3 | CARNS1 | 0.99123066 | 1 | 0.012 | 0.660 | chr11 | 67182439 | 67193078 | + | protein_coding |
| ENSG00000164244.20_2 | PRRC1 | 1 | 1 | 0.011 | 6.492 | chr5 | 126853301 | 126890781 | + | protein_coding |
| ENSG00000167995.16_3 | BEST1 | 0.99826649 | 1 | 0.011 | 2.651 | chr11 | 61717535 | 61732987 | + | protein_coding |
| ENSG00000269896.2_5 | AL513477.1 | 0.99078911 | 1 | 0.011 | 0.523 | chr1 | 2281853 | 2284259 | - | transcribed_processed_pseudogene |
| ENSG00000102870.6_4 | ZNF629 | 1 | 1 | 0.011 | 5.990 | chr16 | 30789777 | 30798526 | - | protein_coding |
| ENSG00000179134.15_4 | SAMD4B | 1 | 1 | 0.011 | 6.572 | chr19 | 39833036 | 39876350 | + | protein_coding |
| ENSG00000139908.15_6 | TSSK4 | 0.99953178 | 1 | 0.011 | 2.024 | chr14 | 24674902 | 24677568 | + | protein_coding |
| ENSG00000100380.14_5 | ST13 | 1 | 1 | 0.011 | 7.527 | chr22 | 41220539 | 41252643 | - | protein_coding |
| ENSG00000109920.13_5 | FNBP4 | 1 | 1 | 0.011 | 6.477 | chr11 | 47738046 | 47788995 | - | protein_coding |
| ENSG00000271009.3_7 | AC116667.1 | 0.98326772 | 1 | 0.011 | -0.160 | chr16 | 73094144 | 73096215 | + | lncRNA |
| ENSG00000256576.2_7 | LINC02361 | 0.9827846 | 1 | 0.011 | -0.181 | chr12 | 132671280 | 132674240 | - | lncRNA |
| ENSG00000274507.1_6 | AL121772.2 | 0.98293649 | 1 | 0.011 | -0.304 | chr20 | 25231644 | 25231940 | - | lncRNA |
| ENSG00000256673.1_6 | AC141557.1 | 0.97857646 | 1 | 0.011 | -0.302 | chr12 | 9550951 | 9567447 | - | unprocessed_pseudogene |
| ENSG00000143164.15_3 | DCAF6 | 1 | 1 | 0.011 | 6.463 | chr1 | 167905021 | 168045081 | + | protein_coding |
| ENSG00000120656.11_3 | TAF12 | 0.99999886 | 1 | 0.011 | 3.878 | chr1 | 28915835 | 28969597 | - | protein_coding |
| ENSG00000147654.15_4 | EBAG9 | 0.99999997 | 1 | 0.010 | 4.859 | chr8 | 110551940 | 110578225 | + | protein_coding |
| ENSG00000268401.2_6 | AC007785.3 | 0.98265004 | 1 | 0.010 | -0.304 | chr19 | 46530906 | 46534925 | + | lncRNA |
| ENSG00000132153.15_3 | DHX30 | 1 | 1 | 0.010 | 6.687 | chr3 | 47844399 | 47891685 | + | protein_coding |
| ENSG00000086730.17_7 | LAT2 | 0.99762733 | 1 | 0.010 | 1.509 | chr7 | 73613982 | 73644164 | + | protein_coding |
| ENSG00000142655.13_3 | PEX14 | 0.99999815 | 1 | 0.010 | 3.873 | chr1 | 10532345 | 10690815 | + | protein_coding |
| ENSG00000160049.12_4 | DFFA | 1 | 1 | 0.010 | 5.822 | chr1 | 10516579 | 10532586 | - | protein_coding |
| ENSG00000270177.1_6 | AC104109.2 | 0.99333339 | 1 | 0.010 | 0.827 | chr5 | 133562101 | 133563518 | + | lncRNA |
| ENSG00000226562.4_5 | CYP4F26P | 0.98865228 | 1 | 0.010 | 0.904 | chr9 | 33580693 | 33607872 | + | lncRNA |
| ENSG00000234771.3_7 | SLC25A25-AS1 | 0.99981476 | 1 | 0.010 | 2.947 | chr9 | 130870860 | 130880972 | - | lncRNA |
| ENSG00000002919.14_2 | SNX11 | 0.99999882 | 1 | 0.010 | 4.208 | chr17 | 46180719 | 46200436 | + | protein_coding |
| ENSG00000165891.16_6 | E2F7 | 0.99999985 | 1 | 0.010 | 4.613 | chr12 | 77415028 | 77459349 | - | protein_coding |
| ENSG00000141956.13_4 | PRDM15 | 0.99999807 | 1 | 0.010 | 3.868 | chr21 | 43218385 | 43299591 | - | protein_coding |
| ENSG00000144744.17_4 | UBA3 | 0.99999999 | 1 | 0.010 | 5.160 | chr3 | 69103881 | 69129559 | - | protein_coding |
| ENSG00000229119.3_5 | AC026403.1 | 0.99898624 | 1 | 0.009 | 1.696 | chr5 | 165809310 | 165809604 | + | processed_pseudogene |
| ENSG00000166323.13_6 | C11orf65 | 0.98385891 | 1 | 0.009 | -0.381 | chr11 | 108179246 | 108338258 | - | protein_coding |
| ENSG00000099810.21_6 | MTAP | 0.99307291 | 1 | 0.009 | 0.745 | chr9 | 21802635 | 21937650 | + | protein_coding |
| ENSG00000162517.13_4 | PEF1 | 0.99999997 | 1 | 0.009 | 4.920 | chr1 | 32095467 | 32110497 | - | protein_coding |
| ENSG00000256667.6_6 | KLRA1P | 0.99610537 | 1 | 0.009 | 1.389 | chr12 | 10740662 | 10752268 | - | transcribed_unprocessed_pseudogene |
| ENSG00000095564.14_3 | BTAF1 | 1 | 1 | 0.009 | 6.610 | chr10 | 93683527 | 93790082 | + | protein_coding |
| ENSG00000285697.1_2 | AC002059.3 | 0.98063159 | 1 | 0.009 | -0.926 | chr22 | 29783898 | 29813746 | + | transcribed_unprocessed_pseudogene |
| ENSG00000224470.8_4 | ATXN1L | 1 | 1 | 0.009 | 5.648 | chr16 | 71879879 | 71919171 | + | protein_coding |
| ENSG00000129521.14_5 | EGLN3 | 0.99998866 | 1 | 0.009 | 3.233 | chr14 | 34393433 | 34931980 | - | protein_coding |
| ENSG00000156709.14_5 | AIFM1 | 1 | 1 | 0.009 | 5.964 | chrX | 129263337 | 129299861 | - | protein_coding |
| ENSG00000172432.19_6 | GTPBP2 | 0.99999996 | 1 | 0.009 | 4.856 | chr6 | 43573053 | 43597001 | - | protein_coding |
| ENSG00000178307.9_3 | TMEM11 | 0.9999996 | 1 | 0.009 | 4.251 | chr17 | 21100593 | 21117937 | - | protein_coding |
| ENSG00000121486.12_5 | TRMT1L | 0.99999999 | 1 | 0.009 | 5.451 | chr1 | 185087233 | 185126204 | - | protein_coding |
| ENSG00000259959.1_6 | AC107068.1 | 0.99577002 | 1 | 0.009 | 0.957 | chr4 | 47842139 | 47846356 | - | lncRNA |
| ENSG00000265749.6_8 | AC135178.4 | 0.99902928 | 1 | 0.009 | 1.345 | chr17 | 8268881 | 8284646 | + | lncRNA |
| ENSG00000132017.10_3 | DCAF15 | 0.99999698 | 1 | 0.009 | 3.862 | chr19 | 14063304 | 14072261 | + | protein_coding |
| ENSG00000177868.12_4 | SVBP | 0.99943536 | 1 | 0.008 | 2.020 | chr1 | 43272723 | 43283068 | - | protein_coding |
| ENSG00000108395.14_7 | TRIM37 | 1 | 1 | 0.008 | 6.370 | chr17 | 57059999 | 57184282 | - | protein_coding |
| ENSG00000178996.14_4 | SNX18 | 0.99999997 | 1 | 0.008 | 4.901 | chr5 | 53813589 | 53842416 | + | protein_coding |
| ENSG00000018610.14_4 | CXorf56 | 0.99999998 | 1 | 0.008 | 5.067 | chrX | 118672112 | 118699371 | - | protein_coding |
| ENSG00000149716.12_5 | LTO1 | 0.99999891 | 1 | 0.008 | 4.147 | chr11 | 69467844 | 69490184 | - | protein_coding |
| ENSG00000093167.18_7 | LRRFIP2 | 1 | 1 | 0.008 | 5.926 | chr3 | 37094117 | 37225180 | - | protein_coding |
| ENSG00000156642.17_5 | NPTN | 1 | 1 | 0.008 | 7.131 | chr15 | 73852355 | 73926475 | - | protein_coding |
| ENSG00000240541.3_6 | TM4SF1-AS1 | 0.98690422 | 1 | 0.008 | 0.329 | chr3 | 149095565 | 149104370 | + | lncRNA |
| ENSG00000244045.13_5 | TMEM199 | 0.99998431 | 1 | 0.008 | 3.162 | chr17 | 26684665 | 26690705 | + | protein_coding |
| ENSG00000169718.18_6 | DUS1L | 0.99999999 | 1 | 0.008 | 6.287 | chr17 | 80015382 | 80023763 | - | protein_coding |

| | | | | | | | | | | |
|---|---|---|---|---|---|---|---|---|---|---|
| ENSG00000258951.1_4 | KRT18P7 | 0.98165392 | 1 | 0.008 | -1.048 | chr14 | 71061269 | 71062487 | + | processed_pseudogene |
| ENSG00000166411.14_5 | IDH3A | 0.99999999 | 1 | 0.008 | 5.191 | chr15 | 78423840 | 78464287 | + | protein_coding |
| ENSG00000185515.14_5 | BRCC3 | 0.99999999 | 1 | 0.008 | 5.063 | chrX | 154299695 | 154351349 | + | protein_coding |
| ENSG00000125249.7_3 | RAP2A | 0.99999996 | 1 | 0.008 | 5.196 | chr13 | 98086423 | 98121382 | + | protein_coding |
| ENSG00000134153.10_7 | EMC7 | 0.99999998 | 1 | 0.008 | 5.095 | chr15 | 34376218 | 34394063 | - | protein_coding |
| ENSG00000200795.1 | RNU4-1 | 0.99913959 | 1 | 0.008 | 2.504 | chr12 | 120730900 | 120731040 | - | snRNA |
| ENSG00000065491.8_3 | TBC1D22B | 0.99997798 | 1 | 0.008 | 3.161 | chr6 | 37225548 | 37300746 | + | protein_coding |
| ENSG00000255965.1_5 | AC073916.1 | 0.98476948 | 1 | 0.008 | -0.464 | chr12 | 124867927 | 124868056 | + | processed_pseudogene |
| ENSG00000198753.12_5 | PLXNB3 | 0.99573363 | 1 | 0.008 | 1.038 | chrX | 153029651 | 153044801 | + | protein_coding |
| ENSG00000186153.17_4 | WWOX | 0.99999618 | 1 | 0.008 | 3.578 | chr16 | 78133327 | 79246564 | + | protein_coding |
| ENSG00000122203.15_5 | KIAA1191 | 1 | 1 | 0.007 | 5.930 | chr5 | 175773065 | 175788810 | - | protein_coding |
| ENSG00000182810.6_3 | DDX28 | 0.99999741 | 1 | 0.007 | 4.011 | chr16 | 68055177 | 68057345 | - | protein_coding |
| ENSG00000157450.15_5 | RNF111 | 1 | 1 | 0.007 | 6.101 | chr15 | 59157374 | 59389618 | + | protein_coding |
| ENSG00000179454.13_3 | KLHL28 | 0.99999981 | 1 | 0.007 | 4.443 | chr14 | 45393522 | 45511525 | - | protein_coding |
| ENSG00000163191.6_4 | S100A11 | 1 | 1 | 0.007 | 7.655 | chr1 | 152004982 | 152020383 | - | protein_coding |
| ENSG00000259495.3_7 | AC016705.2 | 0.99993429 | 1 | 0.007 | 2.900 | chr15 | 80637195 | 80696556 | - | lncRNA |
| ENSG00000099246.16_2 | RAB18 | 1 | 1 | 0.007 | 6.344 | chr10 | 27793103 | 27831166 | + | protein_coding |
| ENSG00000181123.8_6 | AC004832.1 | 0.98608668 | 1 | 0.007 | -0.406 | chr22 | 30871351 | 30888791 | + | lncRNA |
| ENSG00000162437.14_2 | RAVER2 | 0.99999995 | 1 | 0.007 | 4.652 | chr1 | 65210778 | 65298915 | + | protein_coding |
| ENSG00000124357.13_5 | NAGK | 0.99999998 | 1 | 0.007 | 4.905 | chr2 | 71291474 | 71306938 | + | protein_coding |
| ENSG00000184154.14_5 | LRTOMT | 0.99977099 | 1 | 0.007 | 2.419 | chr11 | 71791377 | 71821828 | + | protein_coding |
| ENSG00000188385.12_5 | JAKMIP3 | 0.99999769 | 1 | 0.007 | 3.890 | chr10 | 133849840 | 133998362 | + | protein_coding |
| ENSG00000173905.9_5 | GOLIM4 | 1 | 1 | 0.007 | 5.407 | chr3 | 167726477 | 167813712 | - | protein_coding |
| ENSG00000126500.3_4 | AP006333.1 | 0.98362576 | 1 | 0.007 | -0.847 | chr11 | 63870660 | 63886645 | + | protein_coding |
| ENSG00000165115.15_4 | KIF27 | 0.99999574 | 1 | 0.007 | 3.459 | chr9 | 86449014 | 86536380 | - | protein_coding |
| ENSG00000163818.17_5 | LZTFL1 | 0.99999951 | 1 | 0.006 | 4.316 | chr3 | 45864808 | 45957534 | - | protein_coding |
| ENSG00000100644.17_7 | HIF1A | 1 | 1 | 0.006 | 9.007 | chr14 | 62162231 | 62214977 | + | protein_coding |
| ENSG00000141698.17_4 | NT5C3B | 1 | 1 | 0.006 | 6.250 | chr17 | 39981309 | 39992512 | - | protein_coding |
| ENSG00000148484.18_5 | RSU1 | 1 | 1 | 0.006 | 5.847 | chr10 | 16632610 | 16859462 | - | protein_coding |
| ENSG00000124608.5_5 | AARS2 | 0.99999992 | 1 | 0.006 | 4.585 | chr6 | 44266468 | 44281084 | - | protein_coding |
| ENSG00000261188.1_6 | Z95115.1 | 0.99977462 | 1 | 0.006 | 2.225 | chr22 | 26908503 | 26910534 | + | lncRNA |
| ENSG00000230267.7_4 | HERC2P4 | 0.99971806 | 1 | 0.006 | 2.102 | chr16 | 32114566 | 32199428 | - | transcribed_unprocessed_pseudogene |
| ENSG00000140416.22_8 | TPM1 | 1 | 1 | 0.006 | 8.948 | chr15 | 63334831 | 63364114 | + | protein_coding |
| ENSG00000137106.18_4 | GRHPR | 1 | 1 | 0.006 | 6.772 | chr9 | 37422663 | 37436987 | + | protein_coding |
| ENSG00000149499.11_4 | EML3 | 0.99999991 | 1 | 0.006 | 4.775 | chr11 | 62369690 | 62380237 | - | protein_coding |
| ENSG00000172590.18_3 | MRPL52 | 0.99999992 | 1 | 0.006 | 5.093 | chr14 | 23299088 | 23304246 | + | protein_coding |
| ENSG00000203879.12_3 | GDI1 | 1 | 1 | 0.006 | 6.233 | chrX | 153665259 | 153671814 | + | protein_coding |
| ENSG00000286185.1_2 | AC242842.3 | 0.98471159 | 1 | 0.006 | -0.927 | chr1 | 145209145 | 145365380 | + | protein_coding |
| ENSG00000128524.5_3 | ATP6V1F | 0.99999986 | 1 | 0.006 | 4.636 | chr7 | 128502910 | 128505901 | + | protein_coding |
| ENSG00000164919.11_6 | COX6C | 1 | 1 | 0.006 | 5.921 | chr8 | 100885428 | 100905935 | - | protein_coding |
| ENSG00000185129.7_5 | PURA | 0.99999998 | 1 | 0.006 | 5.075 | chr5 | 139487362 | 139505204 | + | protein_coding |
| ENSG00000133818.14_5 | RRAS2 | 0.99999999 | 1 | 0.006 | 5.041 | chr11 | 14299468 | 14386052 | - | protein_coding |
| ENSG00000185947.15_4 | ZNF267 | 0.99999998 | 1 | 0.006 | 5.163 | chr16 | 31885128 | 31928678 | + | protein_coding |
| ENSG00000135439.11_8 | AGAP2 | 0.99660197 | 1 | 0.006 | 0.573 | chr12 | 58117544 | 58135940 | - | protein_coding |
| ENSG00000185728.17_7 | YTHDF3 | 1 | 1 | 0.005 | 6.731 | chr8 | 64081112 | 64125344 | + | protein_coding |
| ENSG00000002834.18_4 | LASP1 | 1 | 1 | 0.005 | 8.867 | chr17 | 37026112 | 37078023 | + | protein_coding |
| ENSG00000165996.14_5 | HACD1 | 0.99999969 | 1 | 0.005 | 4.300 | chr10 | 17631031 | 17659373 | - | protein_coding |
| ENSG00000104611.12_4 | SH2D4A | 1 | 1 | 0.005 | 6.065 | chr8 | 19171204 | 19253729 | + | protein_coding |
| ENSG00000093000.19_4 | NUP50 | 1 | 1 | 0.005 | 6.100 | chr22 | 45559806 | 45583898 | + | protein_coding |
| ENSG00000186501.14_4 | TMEM222 | 0.99999984 | 1 | 0.005 | 4.513 | chr1 | 27648636 | 27662891 | + | protein_coding |
| ENSG00000136111.14_8 | TBC1D4 | 1 | 1 | 0.005 | 5.916 | chr13 | 75857639 | 76056305 | - | protein_coding |
| ENSG00000170522.9_3 | ELOVL6 | 0.99999999 | 1 | 0.005 | 5.213 | chr4 | 110967002 | 111120355 | - | protein_coding |
| ENSG00000142599.19_7 | RERE | 0.99999984 | 1 | 0.005 | 4.613 | chr1 | 8412457 | 8908980 | - | protein_coding |
| ENSG00000131873.7_5 | CHSY1 | 0.99999999 | 1 | 0.005 | 5.368 | chr15 | 101715932 | 101792253 | - | protein_coding |
| ENSG00000105248.16_4 | YJU2 | 0.99999737 | 1 | 0.005 | 3.720 | chr19 | 4247077 | 4269085 | + | protein_coding |
| ENSG00000101445.10_3 | PPP1R16B | 0.99685814 | 1 | 0.005 | 0.960 | chr20 | 37434340 | 37551667 | + | protein_coding |
| ENSG00000249007.1_8 | AL691482.4 | 0.99787485 | 1 | 0.005 | 1.388 | chr1 | 201980498 | 201984785 | + | lncRNA |
| ENSG00000122378.14_4 | PRXL2A | 1 | 1 | 0.005 | 6.494 | chr10 | 82167585 | 82196871 | + | protein_coding |
| ENSG00000100983.12_7 | GSS | 1 | 1 | 0.005 | 6.161 | chr20 | 33516235 | 33543830 | - | protein_coding |

| | | | | | | | | | | |
|---|---|---|---|---|---|---|---|---|---|---|
| ENSG00000154608.14_3 | CEP170P1 | 0.99177324 | 1 | 0.004 | -0.210 | chr4 | 119388745 | 119475255 | + | transcribed_unprocessed_pseudogene |
| ENSG00000149929.16_4 | HIRIP3 | 0.99999188 | 1 | 0.004 | 3.141 | chr16 | 30003651 | 30007395 | - | protein_coding |
| ENSG00000152556.17_9 | PFKM | 1 | 1 | 0.004 | 5.896 | chr12 | 48498922 | 48540187 | + | protein_coding |
| ENSG00000174130.12_4 | TLR6 | 0.99999991 | 1 | 0.004 | 4.539 | chr4 | 38825336 | 38858438 | - | protein_coding |
| ENSG00000101310.17_8 | SEC23B | 1 | 1 | 0.004 | 6.773 | chr20 | 18488164 | 18542059 | + | protein_coding |
| ENSG00000249592.6_8 | AC139887.2 | 0.9992702 | 1 | 0.004 | 1.789 | chr4 | 753990 | 775647 | - | lncRNA |
| ENSG00000148985.20_6 | PGAP2 | 0.99998286 | 1 | 0.004 | 3.935 | chr11 | 3818954 | 3847601 | + | protein_coding |
| ENSG00000111665.11_3 | CDCA3 | 1 | 1 | 0.004 | 5.519 | chr12 | 6953957 | 6961230 | - | protein_coding |
| ENSG00000260773.1_7 | AC055855.2 | 0.98894497 | 1 | 0.004 | -0.307 | chr15 | 66607252 | 66624041 | - | lncRNA |
| ENSG00000287721.1_1 | AC061975.8 | 0.98875742 | 1 | 0.004 | -0.588 | chr17 | 26531642 | 26550761 | + | lncRNA |
| ENSG00000103365.15_7 | GGA2 | 1 | 1 | 0.004 | 5.774 | chr16 | 23474863 | 23533316 | - | protein_coding |
| ENSG00000198954.8_6 | KIFBP | 1 | 1 | 0.004 | 5.973 | chr10 | 70748477 | 70803300 | + | protein_coding |
| ENSG00000012232.9_3 | EXTL3 | 0.9999999 | 1 | 0.004 | 5.466 | chr8 | 28457986 | 28613116 | + | protein_coding |
| ENSG00000235859.5_5 | AC006978.1 | 0.99455916 | 1 | 0.004 | 0.459 | chr7 | 30411124 | 30412357 | + | processed_pseudogene |
| ENSG00000215012.9_8 | RTL10 | 0.99999892 | 1 | 0.004 | 3.793 | chr22 | 19833661 | 19842419 | - | protein_coding |
| ENSG00000104290.11_6 | FZD3 | 0.99999997 | 1 | 0.004 | 4.990 | chr8 | 28351722 | 28431784 | + | protein_coding |
| ENSG00000078967.13_4 | UBE2D4 | 0.99995798 | 1 | 0.004 | 3.045 | chr7 | 43966035 | 43995735 | + | protein_coding |
| ENSG00000168060.16_8 | NAALADL1 | 0.98825923 | 1 | 0.004 | -1.010 | chr11 | 64812290 | 64826025 | - | protein_coding |
| ENSG00000175213.3_3 | ZNF408 | 0.99999852 | 1 | 0.003 | 3.693 | chr11 | 46722580 | 46727462 | + | protein_coding |
| ENSG00000116668.13_8 | SWT1 | 0.99997033 | 1 | 0.003 | 2.919 | chr1 | 185126212 | 185260913 | + | protein_coding |
| ENSG00000257556.2_6 | LINC02298 | 0.99994861 | 1 | 0.003 | 2.685 | chr14 | 105559946 | 105566468 | + | lncRNA |
| ENSG00000268047.1_7 | AC018766.1 | 0.99005308 | 1 | 0.003 | -0.471 | chr19 | 50356144 | 50358224 | - | lncRNA |
| ENSG00000123124.14_3 | WWP1 | 1 | 1 | 0.003 | 6.046 | chr8 | 87354776 | 87490649 | + | protein_coding |
| ENSG00000186625.14_5 | KATNA1 | 0.99999515 | 1 | 0.003 | 3.268 | chr6 | 149916009 | 149970108 | - | protein_coding |
| ENSG00000113838.13_6 | TBCCD1 | 0.99999754 | 1 | 0.003 | 3.481 | chr3 | 186263856 | 186288332 | - | protein_coding |
| ENSG00000154001.14_5 | PPP2R5E | 1 | 1 | 0.003 | 6.606 | chr14 | 63838082 | 64010095 | - | protein_coding |
| ENSG00000113583.8_4 | C5orf15 | 1 | 1 | 0.003 | 6.398 | chr5 | 133291201 | 133304365 | - | protein_coding |
| ENSG00000176396.11_4 | EID2 | 0.99999997 | 1 | 0.003 | 4.688 | chr19 | 40029347 | 40030801 | - | protein_coding |
| ENSG00000185485.14_3 | SDHAP1 | 0.99998905 | 1 | 0.003 | 3.306 | chr3 | 195686619 | 195717189 | - | transcribed_unprocessed_pseudogene |
| ENSG00000267838.2_6 | AC245884.8 | 0.99444286 | 1 | 0.003 | 0.324 | chr19 | 54949847 | 54950726 | - | lncRNA |
| ENSG00000137944.18_7 | KYAT3 | 0.99999947 | 1 | 0.003 | 3.929 | chr1 | 89401456 | 89458636 | - | protein_coding |
| ENSG00000182150.16_6 | ERCC6L2 | 1 | 1 | 0.002 | 6.121 | chr9 | 98633546 | 98883436 | + | protein_coding |
| ENSG00000230658.2_6 | KLHL7-DT | 0.99482349 | 1 | 0.002 | 0.467 | chr7 | 23139833 | 23145322 | - | lncRNA |
| ENSG00000182700.5_3 | IGIP | 0.99925732 | 1 | 0.002 | 1.590 | chr5 | 139505522 | 139508977 | + | protein_coding |
| ENSG00000237039.1_5 | AC018738.1 | 0.99054049 | 1 | 0.002 | -0.653 | chr2 | 232120810 | 232121019 | + | processed_pseudogene |
| ENSG00000224790.3_8 | AP000704.2 | 0.99463649 | 1 | 0.002 | 0.314 | chr21 | 38338761 | 38347464 | + | lncRNA |
| ENSG00000186523.14_5 | FAM86B1 | 0.99716271 | 1 | 0.002 | 0.412 | chr8 | 12039605 | 12051642 | - | protein_coding |
| ENSG00000142002.17_4 | DPP9 | 1 | 1 | 0.002 | 5.946 | chr19 | 4675236 | 4724685 | - | protein_coding |
| ENSG00000104872.11_5 | PIH1D1 | 1 | 1 | 0.002 | 5.841 | chr19 | 49949555 | 49956754 | - | protein_coding |
| ENSG00000105245.9_4 | NUMBL | 0.99999997 | 1 | 0.002 | 5.099 | chr19 | 41171810 | 41196877 | - | protein_coding |
| ENSG00000103707.10_5 | MTFMT | 0.99999999 | 1 | 0.002 | 4.794 | chr15 | 65293850 | 65321977 | - | protein_coding |
| ENSG00000100603.13_3 | SNW1 | 1 | 1 | 0.002 | 6.055 | chr14 | 78183942 | 78227550 | - | protein_coding |
| ENSG00000165138.18_4 | ANKS6 | 1 | 1 | 0.001 | 5.589 | chr9 | 101493611 | 101559247 | - | protein_coding |
| ENSG00000124217.5_3 | MOCS3 | 0.99999987 | 1 | 0.001 | 4.244 | chr20 | 49575355 | 49580466 | + | protein_coding |
| ENSG00000149792.9_3 | MRPL49 | 1 | 1 | 0.001 | 5.779 | chr11 | 64889655 | 64894843 | + | protein_coding |
| ENSG00000012660.14_5 | ELOVL5 | 0.99999998 | 1 | 0.001 | 6.274 | chr6 | 53132196 | 53213977 | - | protein_coding |
| ENSG00000171552.13_6 | BCL2L1 | 1 | 1 | 0.001 | 6.726 | chr20 | 30252255 | 30311792 | - | protein_coding |
| ENSG00000131495.8_3 | NDUFA2 | 0.99999974 | 1 | 0.001 | 3.978 | chr5 | 140018325 | 140027370 | - | protein_coding |
| ENSG00000260276.2_7 | AC022167.2 | 0.99688043 | 1 | 0.001 | 0.758 | chr16 | 8941962 | 8954313 | + | lncRNA |
| ENSG00000134590.14_6 | RTL8C | 1 | 1 | 0.001 | 5.012 | chrX | 134166385 | 134167576 | + | protein_coding |
| ENSG00000185651.14_2 | UBE2L3 | 1 | 1 | 0.001 | 6.354 | chr22 | 21903736 | 21978323 | + | protein_coding |
| ENSG00000110330.8_6 | BIRC2 | 1 | 1 | 0.001 | 5.909 | chr11 | 102217942 | 102249401 | + | protein_coding |
| ENSG00000106976.20_2 | DNM1 | 1 | 1 | 0.001 | 6.876 | chr9 | 130953934 | 131017527 | + | protein_coding |
| ENSG00000109180.14_6 | OCIAD1 | 1 | 1 | 0.001 | 6.346 | chr4 | 48807229 | 48863834 | + | protein_coding |
| ENSG00000152520.14_4 | PAN3 | 0.99999999 | 1 | 0.001 | 4.659 | chr13 | 28712643 | 28869472 | + | protein_coding |
| ENSG00000182872.16_5 | RBM10 | 1 | 1 | 0.001 | 6.255 | chrX | 47004620 | 47046212 | + | protein_coding |
| ENSG00000112378.12_4 | PERP | 1 | 1 | 0.001 | 7.763 | chr6 | 138409642 | 138428556 | - | protein_coding |
| ENSG00000214160.10_4 | ALG3 | 1 | 1 | 0.000 | 4.925 | chr3 | 183960089 | 183967336 | - | protein_coding |
| ENSG00000275764.1_7 | AC092747.4 | 0.99994592 | 1 | 0.000 | 2.602 | chr12 | 27190033 | 27191893 | + | lncRNA |

| | | | | | | | | | | |
|---|---|---|---|---|---|---|---|---|---|---|
| ENSG00000145780.8_4 | FEM1C | 0.99999999 | 1 | 0.000 | 4.810 | chr5 | 114856605 | 114880472 | - | protein_coding |
| ENSG00000121774.18_3 | KHDRBS1 | 1 | 1 | 0.000 | 7.032 | chr1 | 32479469 | 32526451 | + | protein_coding |
| ENSG00000164405.11_3 | UQCRQ | 1 | 1 | 0.000 | 5.903 | chr5 | 132202322 | 132204539 | + | protein_coding |
| ENSG00000109171.15_3 | SLAIN2 | 1 | 1 | 0.000 | 5.097 | chr4 | 48343546 | 48428218 | + | protein_coding |
| ENSG00000121210.16_8 | TMEM131L | 0.99999639 | 1 | 0.000 | 3.261 | chr4 | 154387498 | 154557863 | + | protein_coding |
| ENSG00000142892.15_6 | PIGK | 1 | 1 | 0.000 | 5.699 | chr1 | 77554674 | 77685115 | - | protein_coding |
| ENSG00000166938.13_4 | DIS3L | 1 | 1 | 0.000 | 6.672 | chr15 | 66585555 | 66626236 | + | protein_coding |
| ENSG00000138660.12_5 | AP1AR | 1 | 1 | 0.000 | 5.022 | chr4 | 113152896 | 113194266 | + | protein_coding |
| ENSG00000204084.13_6 | INPP5B | 0.99999985 | 1 | 0.000 | 3.823 | chr1 | 38326369 | 38412729 | - | protein_coding |
| ENSG00000259732.1_7 | AC092757.2 | 0.99377313 | 1 | 0.000 | -0.616 | chr15 | 59413233 | 59425449 | + | lncRNA |
| ENSG00000005469.11_2 | CROT | 0.99999997 | 1 | 0.000 | 4.292 | chr7 | 86974997 | 87029111 | + | protein_coding |
| ENSG00000213918.10_5 | DNASE1 | 0.99999994 | 1 | 0.000 | 4.091 | chr16 | 3661729 | 3730144 | + | protein_coding |
| ENSG00000035403.17_3 | VCL | 1 | 1 | 0.000 | 8.183 | chr10 | 75754951 | 75881121 | + | protein_coding |
| ENSG00000134247.10_3 | PTGFRN | 1 | 1 | -0.001 | 5.159 | chr1 | 117452538 | 117532975 | + | protein_coding |
| ENSG00000009830.11_8 | POMT2 | 1 | 1 | -0.001 | 5.414 | chr14 | 77741299 | 77787227 | - | protein_coding |
| ENSG00000167112.10_5 | TRUB2 | 1 | 1 | -0.001 | 5.873 | chr9 | 131067438 | 131084726 | - | protein_coding |
| ENSG00000147140.16_7 | NONO | 1 | 1 | -0.001 | 8.609 | chrX | 70503042 | 70521018 | + | protein_coding |
| ENSG00000048828.17_4 | FAM120A | 1 | 1 | -0.001 | 7.862 | chr9 | 96213967 | 96328394 | + | protein_coding |
| ENSG00000134444.14_6 | RELCH | 1 | 1 | -0.001 | 5.702 | chr18 | 59854488 | 59977450 | + | protein_coding |
| ENSG00000268543.1_5 | AC012313.4 | 0.99844451 | 1 | -0.001 | 0.633 | chr19 | 58929927 | 58930677 | + | TEC |
| ENSG00000088682.14_4 | COQ9 | 1 | 1 | -0.001 | 5.502 | chr16 | 57481337 | 57495182 | + | protein_coding |
| ENSG00000109814.12_5 | UGDH | 1 | 1 | -0.001 | 9.140 | chr4 | 39500375 | 39529931 | - | protein_coding |
| ENSG00000266088.6_7 | AC004585.1 | 0.9999588 | 1 | -0.001 | 2.326 | chr17 | 38673144 | 38683254 | + | lncRNA |
| ENSG00000162341.18_5 | TPCN2 | 0.99999673 | 1 | -0.001 | 2.997 | chr11 | 68816400 | 68903784 | + | protein_coding |
| ENSG00000111602.12_4 | TIMELESS | 1 | 1 | -0.001 | 5.996 | chr12 | 56810147 | 56843210 | - | protein_coding |
| ENSG00000094914.14_6 | AAAS | 1 | 1 | -0.001 | 6.256 | chr12 | 53701240 | 53718648 | - | protein_coding |
| ENSG00000204138.13_4 | PHACTR4 | 0.9999996 | 1 | -0.001 | 4.876 | chr1 | 28696093 | 28826876 | + | protein_coding |
| ENSG00000102921.8_6 | N4BP1 | 1 | 1 | -0.001 | 5.037 | chr16 | 48572637 | 48654059 | - | protein_coding |
| ENSG00000007392.16_6 | LUC7L | 1 | 1 | -0.001 | 5.415 | chr16 | 238968 | 279462 | - | protein_coding |
| ENSG00000116285.13_4 | ERRFI1 | 1 | 1 | -0.002 | 6.434 | chr1 | 8064464 | 8086369 | - | protein_coding |
| ENSG00000167118.11_3 | URM1 | 1 | 1 | -0.002 | 5.402 | chr9 | 131133640 | 131154295 | + | protein_coding |
| ENSG00000167695.15_4 | TLCD3A | 1 | 1 | -0.002 | 5.071 | chr17 | 635652 | 646208 | + | protein_coding |
| ENSG00000254087.8_4 | LYN | 1 | 1 | -0.002 | 4.994 | chr8 | 56792394 | 56926728 | + | protein_coding |
| ENSG00000100425.18_4 | BRD1 | 0.99999999 | 1 | -0.002 | 4.814 | chr22 | 50166931 | 50221160 | - | protein_coding |
| ENSG00000166855.9_4 | CLPX | 1 | 1 | -0.002 | 6.408 | chr15 | 65440557 | 65477680 | - | protein_coding |
| ENSG00000162714.12_4 | ZNF496 | 1 | 1 | -0.002 | 5.495 | chr1 | 247460714 | 247495148 | - | protein_coding |
| ENSG00000084636.18_6 | COL16A1 | 0.99999962 | 1 | -0.002 | 3.337 | chr1 | 32117864 | 32169920 | - | protein_coding |
| ENSG00000105640.13_3 | RPL18A | 1 | 1 | -0.002 | 7.652 | chr19 | 17970719 | 17974962 | + | protein_coding |
| ENSG00000114316.13_6 | USP4 | 0.99999997 | 1 | -0.002 | 4.217 | chr3 | 49314577 | 49378145 | - | protein_coding |
| ENSG00000151806.14_3 | GUF1 | 0.99999999 | 1 | -0.002 | 4.927 | chr4 | 44680437 | 44702945 | + | protein_coding |
| ENSG00000132842.13_5 | AP3B1 | 1 | 1 | -0.003 | 6.573 | chr5 | 77296349 | 77590579 | - | protein_coding |
| ENSG00000164305.19_7 | CASP3 | 0.99999997 | 1 | -0.003 | 4.667 | chr4 | 185548850 | 185570663 | - | protein_coding |
| ENSG00000232134.1_4 | RPS15AP12 | 0.99662551 | 1 | -0.003 | -1.051 | chr1 | 220317306 | 220317693 | - | processed_pseudogene |
| ENSG00000198356.12_6 | GET3 | 1 | 1 | -0.003 | 5.287 | chr19 | 12847953 | 12859137 | + | protein_coding |
| ENSG00000286634.1_1 | AC146507.3 | 0.99804522 | 1 | -0.003 | -0.283 | chr5 | 176687341 | 176690910 | + | lncRNA |
| ENSG00000173171.14_4 | MTX1 | 0.99999992 | 1 | -0.003 | 4.095 | chr1 | 155178490 | 155183615 | + | protein_coding |
| ENSG00000234219.1_4 | CDCA4P4 | 0.99731702 | 1 | -0.003 | -0.782 | chr1 | 207935929 | 207936608 | + | unprocessed_pseudogene |
| ENSG00000226328.7_7 | NUP50-DT | 0.9998085 | 1 | -0.003 | 1.288 | chr22 | 45528901 | 45559662 | - | lncRNA |
| ENSG00000177963.14_5 | RIC8A | 1 | 1 | -0.003 | 6.665 | chr11 | 207511 | 215113 | + | protein_coding |
| ENSG00000161381.14_5 | PLXDC1 | 0.99812214 | 1 | -0.003 | -0.284 | chr17 | 37219566 | 37310647 | - | protein_coding |
| ENSG00000099974.8_4 | DDTL | 0.99924014 | 1 | -0.003 | 0.397 | chr22 | 24309076 | 24314745 | + | protein_coding |
| ENSG00000166377.20_6 | ATP9B | 0.99999997 | 1 | -0.003 | 4.692 | chr18 | 76829285 | 77138287 | + | protein_coding |
| ENSG00000187231.14_6 | SESTD1 | 1 | 1 | -0.003 | 5.139 | chr2 | 179966405 | 180129559 | - | protein_coding |
| ENSG00000112514.16_5 | CUTA | 0.99999998 | 1 | -0.003 | 4.849 | chr6 | 33384219 | 33386094 | - | protein_coding |
| ENSG00000114354.14_5 | TFG | 1 | 1 | -0.003 | 6.563 | chr3 | 100428175 | 100467808 | + | protein_coding |
| ENSG00000205476.9_5 | CCDC85C | 1 | 1 | -0.003 | 5.366 | chr14 | 99966527 | 100070544 | - | protein_coding |
| ENSG00000271737.1_6 | AC008608.2 | 0.99959716 | 1 | -0.003 | 0.581 | chr5 | 133340838 | 133341253 | + | lncRNA |
| ENSG00000129255.16_4 | MPDU1 | 1 | 1 | -0.004 | 6.026 | chr17 | 7486847 | 7496107 | + | protein_coding |
| ENSG00000162302.13_5 | RPS6KA4 | 0.99971228 | 1 | -0.004 | 5.713 | chr11 | 64126620 | 64139687 | + | protein_coding |
| ENSG00000143442.22_8 | POGZ | 1 | 1 | -0.004 | 5.967 | chr1 | 151375200 | 151431970 | - | protein_coding |

| | | | | | | | | | | |
|---|---|---|---|---|---|---|---|---|---|---|
| ENSG00000177370.5_3 | TIMM22 | 1 | 1 | -0.004 | 5.304 | chr17 | 900369 | 906911 | + | protein_coding |
| ENSG00000211584.14_4 | SLC48A1 | 1 | 1 | -0.004 | 5.606 | chr12 | 48147699 | 48176534 | + | protein_coding |
| ENSG00000151718.16_4 | WWC2 | 1 | 1 | -0.004 | 5.678 | chr4 | 184020410 | 184241930 | + | protein_coding |
| ENSG00000249348.1_6 | UGDH-AS1 | 0.999972 | 1 | -0.004 | 0.713 | chr4 | 39529639 | 39596327 | + | lncRNA |
| ENSG00000198736.11_6 | MSRB1 | 0.99999988 | 1 | -0.004 | 4.332 | chr16 | 1988211 | 1993327 | - | protein_coding |
| ENSG00000164164.17_7 | OTUD4 | 0.99999999 | 1 | -0.004 | 5.447 | chr4 | 146031990 | 146101741 | - | protein_coding |
| ENSG00000102048.16_4 | ASB9 | 0.99999998 | 1 | -0.004 | 4.862 | chrX | 15253410 | 15288589 | - | protein_coding |
| ENSG00000137806.9_5 | NDUFAF1 | 0.9999936 | 1 | -0.004 | 3.211 | chr15 | 41679551 | 41694717 | - | protein_coding |
| ENSG00000235159.1_7 | AL121672.2 | 0.99994109 | 1 | -0.004 | 0.157 | chr22 | 46463236 | 46465771 | - | lncRNA |
| ENSG00000255198.4_4 | SNHG9 | 0.99996969 | 1 | -0.004 | 2.146 | chr16 | 2014960 | 2015510 | + | lncRNA |
| ENSG00000169371.14_5 | SNUPN | 0.99999941 | 1 | -0.004 | 3.619 | chr15 | 75890424 | 75918810 | - | protein_coding |
| ENSG00000136783.10_3 | NIPSNAP3A | 0.99999993 | 1 | -0.004 | 4.314 | chr9 | 107509964 | 107522401 | + | protein_coding |
| ENSG00000184343.11_4 | SRPK3 | 0.9999701 | 1 | -0.004 | 2.284 | chrX | 153041867 | 153051187 | + | protein_coding |
| ENSG00000117133.11_3 | RPF1 | 0.99999972 | 1 | -0.004 | 4.727 | chr1 | 84944942 | 84964035 | + | protein_coding |
| ENSG00000160218.13_3 | TRAPPC10 | 1 | 1 | -0.005 | 5.229 | chr21 | 45432190 | 45526433 | + | protein_coding |
| ENSG00000119541.10_5 | VPS4B | 1 | 1 | -0.005 | 5.703 | chr18 | 61056423 | 61089716 | - | protein_coding |
| ENSG00000124181.14_6 | PLCG1 | 1 | 1 | -0.005 | 6.160 | chr20 | 39765600 | 39825441 | + | protein_coding |
| ENSG00000108406.10_3 | DHX40 | 1 | 1 | -0.005 | 7.044 | chr17 | 57642919 | 57685706 | + | protein_coding |
| ENSG00000158623.14_4 | COPG2 | 0.99999789 | 1 | -0.005 | 3.335 | chr7 | 130146079 | 130353588 | - | protein_coding |
| ENSG00000090097.21_8 | PCBP4 | 0.9999939 | 1 | -0.005 | 2.865 | chr3 | 51991470 | 52008032 | - | protein_coding |
| ENSG00000137628.17_5 | DDX60 | 0.99999257 | 1 | -0.005 | 3.049 | chr4 | 169137445 | 169239955 | - | protein_coding |
| ENSG00000127080.10_4 | IPPK | 0.99999101 | 1 | -0.005 | 2.851 | chr9 | 95375465 | 95432413 | - | protein_coding |
| ENSG00000228253.1_1 | MT-ATP8 | 0.99986381 | 1 | -0.005 | 4.914 | chrM | 8366 | 8572 | + | protein_coding |
| ENSG00000276832.2_2 | AL354718.3 | 0.99970698 | 1 | -0.005 | -0.713 | chr9 | 70649851 | 70663263 | + | transcribed_unprocessed_pseudogene |
| ENSG00000158526.8_3 | TSR2 | 0.99999999 | 1 | -0.005 | 4.910 | chrX | 54466837 | 54474465 | + | protein_coding |
| ENSG00000128607.14_4 | KLHDC10 | 1 | 1 | -0.005 | 5.739 | chr7 | 129710374 | 129775545 | + | protein_coding |
| ENSG00000131374.14_6 | TBC1D5 | 1 | 1 | -0.005 | 6.957 | chr3 | 17198654 | 18486309 | - | protein_coding |
| ENSG00000109466.14_5 | KLHL2 | 0.99999983 | 1 | -0.005 | 4.051 | chr4 | 166128713 | 166244308 | + | protein_coding |
| ENSG00000221930.6_6 | DENND10P1 | 0.99840978 | 1 | -0.006 | -0.293 | chrX | 129628939 | 129630562 | + | transcribed_processed_pseudogene |
| ENSG00000203865.10_6 | ATP1A1-AS1 | 0.99936863 | 1 | -0.006 | 1.079 | chr1 | 116933197 | 116961257 | - | lncRNA |
| ENSG00000138594.14_4 | TMOD3 | 1 | 1 | -0.006 | 8.121 | chr15 | 52121825 | 52239492 | + | protein_coding |
| ENSG00000258813.2_7 | AL442663.3 | 0.99853501 | 1 | -0.006 | -0.785 | chr14 | 73524633 | 73526123 | - | lncRNA |
| ENSG00000137817.17_6 | PARP6 | 1 | 1 | -0.006 | 6.164 | chr15 | 72533522 | 72565340 | - | protein_coding |
| ENSG00000141570.11_5 | CBX8 | 0.99999986 | 1 | -0.006 | 4.577 | chr17 | 77765931 | 77775482 | - | protein_coding |
| ENSG00000134905.17_5 | CARS2 | 1 | 1 | -0.006 | 5.139 | chr13 | 111293759 | 111365950 | - | protein_coding |
| ENSG00000166164.15_6 | BRD7 | 1 | 1 | -0.006 | 6.477 | chr16 | 50347398 | 50402845 | - | protein_coding |
| ENSG00000105643.10_6 | ARRDC2 | 0.99999998 | 1 | -0.006 | 4.857 | chr19 | 18111941 | 18124911 | + | protein_coding |
| ENSG00000205726.15_7 | ITSN1 | 0.99999999 | 1 | -0.006 | 5.330 | chr21 | 35014706 | 35272165 | + | protein_coding |
| ENSG00000186088.16_5 | GSAP | 0.99999634 | 1 | -0.006 | 3.699 | chr7 | 76940068 | 77045666 | - | protein_coding |
| ENSG00000187796.15_6 | CARD9 | 0.99969995 | 1 | -0.006 | 1.827 | chr9 | 139256355 | 139268133 | - | protein_coding |
| ENSG00000075785.13_3 | RAB7A | 1 | 1 | -0.006 | 6.267 | chr3 | 128445026 | 128533641 | + | protein_coding |
| ENSG00000149679.11_7 | CABLES2 | 0.99999985 | 1 | -0.006 | 4.246 | chr20 | 60963688 | 60982341 | - | protein_coding |
| ENSG00000213222.3_5 | AC093724.1 | 0.99792324 | 1 | -0.007 | 0.154 | chr2 | 132480826 | 132481852 | - | processed_pseudogene |
| ENSG00000142252.11_4 | GEMIN7 | 0.99741512 | 1 | -0.007 | 0.024 | chr19 | 45582453 | 45594776 | + | protein_coding |
| ENSG00000246339.6_6 | EXTL3-AS1 | 0.99815532 | 1 | -0.007 | 0.345 | chr8 | 28547732 | 28559547 | - | lncRNA |
| ENSG00000063015.20_6 | SEZ6 | 0.99911035 | 1 | -0.007 | 0.954 | chr17 | 27281919 | 27333458 | - | protein_coding |
| ENSG00000185973.11_4 | TMLHE | 0.99998405 | 1 | -0.007 | 3.232 | chrX | 154718672 | 154899605 | - | protein_coding |
| ENSG00000068697.7_4 | LAPTM4A | 0.99998101 | 1 | -0.007 | 5.470 | chr2 | 20232411 | 20251389 | - | protein_coding |
| ENSG00000253741.2_7 | LNCOC1 | 0.99890897 | 1 | -0.007 | 1.104 | chr8 | 143783275 | 143808434 | - | lncRNA |
| ENSG00000125817.8_3 | CENPB | 0.99999829 | 1 | -0.007 | 7.088 | chr20 | 3764498 | 3767387 | - | protein_coding |
| ENSG00000168876.9_6 | ANKRD49 | 0.99999841 | 1 | -0.007 | 3.590 | chr11 | 94227145 | 94232744 | + | protein_coding |
| ENSG00000005483.21_7 | KMT2E | 1 | 1 | -0.007 | 6.078 | chr7 | 104654637 | 104755466 | + | protein_coding |
| ENSG00000100906.10_3 | NFKBIA | 1 | 1 | -0.007 | 6.351 | chr14 | 35870717 | 35873955 | - | protein_coding |
| ENSG00000181817.6_4 | LSM10 | 0.99962906 | 1 | -0.007 | 1.986 | chr1 | 36856839 | 36863509 | - | protein_coding |
| ENSG00000146083.12_4 | RNF44 | 0.99999964 | 1 | -0.007 | 4.257 | chr5 | 175953713 | 175965026 | - | protein_coding |
| ENSG00000083093.10_6 | PALB2 | 1 | 1 | -0.007 | 5.240 | chr16 | 23614481 | 23652631 | - | protein_coding |
| ENSG00000133318.14_8 | RTN3 | 1 | 1 | -0.007 | 7.264 | chr11 | 63448918 | 63527363 | + | protein_coding |
| ENSG00000214820.3_4 | MPRIPP1 | 0.99759333 | 1 | -0.007 | 0.412 | chr3 | 44621430 | 44622518 | - | processed_pseudogene |
| ENSG00000249992.2_3 | TMEM158 | 0.99697343 | 1 | -0.007 | -0.040 | chr3 | 45265958 | 45267779 | - | protein_coding |

| | | | | | | | | | | |
|---|---|---|---|---|---|---|---|---|---|---|
| ENSG00000136932.14_3 | TRMO | 0.9999967 | 1 | -0.008 | 3.823 | chr9 | 100666771 | 100684798 | - | protein_coding |
| ENSG00000163719.19_5 | MTMR14 | 0.99999998 | 1 | -0.008 | 4.939 | chr3 | 9691117 | 9744077 | + | protein_coding |
| ENSG00000117614.10_5 | SYF2 | 0.99999998 | 1 | -0.008 | 4.890 | chr1 | 25548767 | 25558993 | - | protein_coding |
| ENSG00000114480.13_5 | GBE1 | 1 | 1 | -0.008 | 6.075 | chr3 | 81538854 | 81810796 | - | protein_coding |
| ENSG00000185513.16_6 | L3MBTL1 | 0.9997062 | 1 | -0.008 | 1.203 | chr20 | 42118082 | 42179590 | + | protein_coding |
| ENSG00000120800.5_3 | UTP20 | 1 | 1 | -0.008 | 7.039 | chr12 | 101673883 | 101780396 | + | protein_coding |
| ENSG00000140521.15_8 | POLG | 1 | 1 | -0.008 | 6.268 | chr15 | 89848429 | 89878092 | - | protein_coding |
| ENSG00000104529.17_9 | EEF1D | 1 | 1 | -0.008 | 6.919 | chr8 | 144661867 | 144681711 | - | protein_coding |
| ENSG00000287114.1_1 | AC104024.3 | 0.99889637 | 1 | -0.008 | 1.209 | chr17 | 16931674 | 16938371 | + | lncRNA |
| ENSG00000126088.14_5 | UROD | 0.9999999 | 1 | -0.008 | 4.839 | chr1 | 45476622 | 45481247 | + | protein_coding |
| ENSG00000144036.15_6 | EXOC6B | 1 | 1 | -0.008 | 5.824 | chr2 | 72403113 | 73053170 | - | protein_coding |
| ENSG00000135801.9_3 | TAF5L | 0.99999999 | 1 | -0.009 | 5.081 | chr1 | 229728858 | 229761794 | - | protein_coding |
| ENSG00000100883.12_5 | SRP54 | 1 | 1 | -0.009 | 6.533 | chr14 | 35451163 | 35498773 | + | protein_coding |
| ENSG00000109686.18_8 | SH3D19 | 1 | 1 | -0.009 | 5.942 | chr4 | 152023903 | 152246784 | - | protein_coding |
| ENSG00000102119.11_3 | EMD | 0.99999987 | 1 | -0.009 | 5.044 | chrX | 153607633 | 153609883 | + | protein_coding |
| ENSG00000130349.10_4 | MTRES1 | 0.9999254 | 1 | -0.009 | 2.731 | chr6 | 107349403 | 107372790 | + | protein_coding |
| ENSG00000151773.13_5 | CCDC122 | 0.99988172 | 1 | -0.009 | 2.284 | chr13 | 44398045 | 44453876 | - | protein_coding |
| ENSG00000147457.14_4 | CHMP7 | 1 | 1 | -0.009 | 5.769 | chr8 | 23101150 | 23119513 | + | protein_coding |
| ENSG00000177352.10_4 | CCDC71 | 0.99999791 | 1 | -0.009 | 3.717 | chr3 | 49199968 | 49203764 | - | protein_coding |
| ENSG00000125651.14_7 | GTF2F1 | 1 | 1 | -0.009 | 6.335 | chr19 | 6379583 | 6393992 | - | protein_coding |
| ENSG00000121879.5_5 | PIK3CA | 0.99999982 | 1 | -0.009 | 4.639 | chr3 | 178865902 | 178957881 | + | protein_coding |
| ENSG00000123374.11_3 | CDK2 | 0.99999999 | 1 | -0.009 | 5.254 | chr12 | 56360565 | 56366573 | + | protein_coding |
| ENSG00000169857.8_5 | AVEN | 0.99999994 | 1 | -0.010 | 4.583 | chr15 | 34158428 | 34331405 | - | protein_coding |
| ENSG00000174197.16_2 | MGA | 1 | 1 | -0.010 | 6.982 | chr15 | 41913422 | 42065279 | + | protein_coding |
| ENSG00000270587.1_5 | AC046185.2 | 0.99308473 | 1 | -0.010 | -1.101 | chr17 | 61688132 | 61688544 | - | processed_pseudogene |
| ENSG00000278765.1_6 | AC004477.3 | 0.99473395 | 1 | -0.010 | 0.028 | chr17 | 46144066 | 46144655 | - | lncRNA |
| ENSG00000132604.11_7 | TERF2 | 0.99999991 | 1 | -0.010 | 4.534 | chr16 | 69389470 | 69442474 | - | protein_coding |
| ENSG00000143183.16_4 | TMCO1 | 1 | 1 | -0.010 | 6.545 | chr1 | 165693530 | 165796992 | - | protein_coding |
| ENSG00000144674.16_6 | GOLGA4 | 1 | 1 | -0.010 | 7.602 | chr3 | 37284668 | 37408242 | + | protein_coding |
| ENSG00000111728.10_4 | ST8SIA1 | 0.99837078 | 1 | -0.010 | -0.338 | chr12 | 22216707 | 22589975 | - | protein_coding |
| ENSG00000180867.11_3 | PDIA3P1 | 0.99998439 | 1 | -0.010 | 2.968 | chr1 | 146644323 | 146651201 | + | transcribed_processed_pseudogene |
| ENSG00000221184.1 | MIR1254-1 | 0.9926709 | 1 | -0.010 | -1.102 | chr10 | 70519075 | 70519171 | + | miRNA |
| ENSG00000147533.17_3 | GOLGA7 | 0.99999593 | 1 | -0.010 | 3.384 | chr8 | 41347915 | 41368499 | + | protein_coding |
| ENSG00000026508.19_8 | CD44 | 1 | 1 | -0.010 | 8.377 | chr11 | 35160417 | 35253949 | + | protein_coding |
| ENSG00000143158.11_4 | MPC2 | 0.99999995 | 1 | -0.010 | 5.452 | chr1 | 167885913 | 167906310 | - | protein_coding |
| ENSG00000099904.15_3 | ZDHHC8 | 0.99966451 | 1 | -0.010 | 3.705 | chr22 | 20116979 | 20135530 | + | protein_coding |
| ENSG00000180357.9_3 | ZNF609 | 1 | 1 | -0.011 | 6.973 | chr15 | 64752941 | 64978267 | + | protein_coding |
| ENSG00000126653.18_6 | NSRP1 | 0.99999909 | 1 | -0.011 | 3.877 | chr17 | 28442539 | 28513493 | + | protein_coding |
| ENSG00000133112.16_3 | TPT1 | 1 | 1 | -0.011 | 8.772 | chr13 | 45907606 | 45915505 | - | protein_coding |
| ENSG00000177082.12_6 | WDR73 | 0.99999923 | 1 | -0.011 | 3.903 | chr15 | 85182512 | 85197574 | - | protein_coding |
| ENSG00000273619.1_6 | AL121832.2 | 0.99721509 | 1 | -0.011 | 0.816 | chr20 | 60961359 | 60962026 | - | lncRNA |
| ENSG00000167900.12_4 | TK1 | 0.99998854 | 1 | -0.011 | 6.918 | chr17 | 76170172 | 76183314 | - | protein_coding |
| ENSG00000268471.6_7 | MIR4453HG | 0.99951634 | 1 | -0.011 | 1.877 | chr4 | 153457416 | 153460415 | + | lncRNA |
| ENSG00000286004.1_1 | AC124944.3 | 0.99148741 | 1 | -0.011 | -0.390 | chr3 | 195627857 | 195630288 | + | lncRNA |
| ENSG00000275484.1_6 | AP003419.3 | 0.99220281 | 1 | -0.011 | -0.261 | chr11 | 67141887 | 67142403 | + | lncRNA |
| ENSG00000272825.1_6 | AL844908.1 | 0.99109594 | 1 | -0.011 | -0.754 | chr21 | 46356218 | 46356869 | + | lncRNA |
| ENSG00000124225.16_6 | PMEPA1 | 1 | 1 | -0.011 | 7.658 | chr20 | 56223448 | 56286592 | - | protein_coding |
| ENSG00000198252.12_3 | STYX | 1 | 1 | -0.011 | 5.658 | chr14 | 53196884 | 53241707 | + | protein_coding |
| ENSG00000269900.3_3 | RMRP | 0.99667585 | 1 | -0.011 | 12.060 | chr9 | 35657748 | 35658015 | - | lncRNA |
| ENSG00000127022.15_7 | CANX | 1 | 1 | -0.011 | 10.151 | chr5 | 179105629 | 179158642 | + | protein_coding |
| ENSG00000275700.5_4 | AATF | 1 | 1 | -0.011 | 6.604 | chr17 | 35306254 | 35414171 | + | protein_coding |
| ENSG00000101474.12_5 | APMAP | 1 | 1 | -0.011 | 7.052 | chr20 | 24943561 | 24973387 | - | protein_coding |
| ENSG00000137154.13_4 | RPS6 | 1 | 1 | -0.011 | 9.816 | chr9 | 19375713 | 19380234 | - | protein_coding |
| ENSG00000113272.14_3 | THG1L | 0.9999995 | 1 | -0.011 | 4.179 | chr5 | 157158428 | 157168457 | + | protein_coding |
| ENSG00000147905.17_2 | ZCCHC7 | 0.99999977 | 1 | -0.011 | 4.513 | chr9 | 37120536 | 37358146 | + | protein_coding |
| ENSG00000130741.11_3 | EIF2S3 | 1 | 1 | -0.012 | 8.641 | chrX | 24073063 | 24096927 | + | protein_coding |
| ENSG00000185158.12_6 | LRRC37B | 0.99999425 | 1 | -0.012 | 3.386 | chr17 | 30334891 | 30380523 | + | protein_coding |
| ENSG00000129354.11_4 | AP1M2 | 0.99999763 | 1 | -0.012 | 3.941 | chr19 | 10683347 | 10697991 | - | protein_coding |
| ENSG00000116221.15_2 | MRPL37 | 1 | 1 | -0.012 | 5.765 | chr1 | 54649714 | 54691137 | + | protein_coding |
| ENSG00000204574.13_5 | ABCF1 | 0.99999999 | 1 | -0.012 | 5.308 | chr6 | 30539170 | 30564956 | + | protein_coding |

| | | | | | | | | | | |
|---|---|---|---|---|---|---|---|---|---|---|
| ENSG00000170100.13_3 | ZNF778 | 0.99999506 | 1 | -0.012 | 3.757 | chr16 | 89284111 | 89303479 | + | protein_coding |
| ENSG00000137500.9_5 | CCDC90B | 0.99999998 | 1 | -0.012 | 5.302 | chr11 | 82970139 | 82997450 | - | protein_coding |
| ENSG00000175106.16_5 | TVP23C | 0.99965381 | 1 | -0.012 | 2.043 | chr17 | 15405578 | 15466909 | - | protein_coding |
| ENSG00000143162.9_5 | CREG1 | 1 | 1 | -0.012 | 6.025 | chr1 | 167498354 | 167523042 | - | protein_coding |
| ENSG00000258427.3_4 | RBM8B | 0.98897956 | 1 | -0.012 | -0.484 | chr14 | 60864475 | 60864999 | + | processed_pseudogene |
| ENSG00000172775.17_9 | PSME3IP1 | 1 | 1 | -0.012 | 6.531 | chr16 | 57186378 | 57220028 | - | protein_coding |
| ENSG00000171574.18_4 | ZNF584 | 0.99999963 | 1 | -0.012 | 4.412 | chr19 | 58912871 | 58929694 | + | protein_coding |
| ENSG00000140009.18_6 | ESR2 | 0.99062691 | 1 | -0.012 | -0.316 | chr14 | 64550950 | 64804830 | - | protein_coding |
| ENSG00000158683.8_5 | PKD1L1 | 0.99005511 | 1 | -0.013 | -0.449 | chr7 | 47779800 | 47988088 | - | protein_coding |
| ENSG00000059573.9_4 | ALDH18A1 | 1 | 1 | -0.013 | 6.588 | chr10 | 97365698 | 97416468 | - | protein_coding |
| ENSG00000119471.15_3 | HSDL2 | 0.99999999 | 1 | -0.013 | 5.783 | chr9 | 115142360 | 115234685 | + | protein_coding |
| ENSG00000004142.12_4 | POLDIP2 | 1 | 1 | -0.013 | 7.082 | chr17 | 26673659 | 26684550 | - | protein_coding |
| ENSG00000259224.2_2 | SLC35G6 | 0.99484808 | 1 | -0.013 | 0.536 | chr17 | 7384651 | 7386815 | + | protein_coding |
| ENSG00000023734.11_4 | STRAP | 1 | 1 | -0.013 | 7.329 | chr12 | 16035321 | 16056412 | + | protein_coding |
| ENSG00000158615.9_4 | PPP1R15B | 1 | 1 | -0.013 | 6.615 | chr1 | 204372509 | 204380945 | - | protein_coding |
| ENSG00000125458.7_5 | NT5C | 0.99999985 | 1 | -0.013 | 4.851 | chr17 | 73126320 | 73127852 | - | protein_coding |
| ENSG00000256282.1_6 | AC112694.1 | 0.98886441 | 1 | -0.013 | -0.314 | chr11 | 18617006 | 18621230 | - | transcribed_processed_pseudogene |
| ENSG00000076242.15_5 | MLH1 | 1 | 1 | -0.013 | 6.381 | chr3 | 37034841 | 37092337 | + | protein_coding |
| ENSG00000197885.10_3 | NKIRAS1 | 0.99999293 | 1 | -0.013 | 3.773 | chr3 | 23933151 | 23988082 | - | protein_coding |
| ENSG00000277693.2_7 | AP003900.1 | 0.99997756 | 1 | -0.013 | 3.168 | chr21 | 11169720 | 11184046 | + | lncRNA |
| ENSG00000141428.17_5 | C18orf21 | 0.99999504 | 1 | -0.013 | 3.436 | chr18 | 33552588 | 33559249 | + | protein_coding |
| ENSG00000265491.5_4 | RNF115 | 1 | 1 | -0.014 | 5.509 | chr1 | 145611018 | 145696218 | + | protein_coding |
| ENSG00000015153.14_4 | YAF2 | 0.99999989 | 1 | -0.014 | 4.684 | chr12 | 42550906 | 42632151 | - | protein_coding |
| ENSG00000110651.11_5 | CD81 | 0.99999999 | 1 | -0.014 | 6.918 | chr11 | 2397407 | 2418649 | + | protein_coding |
| ENSG00000260735.2_7 | AC139256.2 | 0.98826483 | 1 | -0.014 | -0.535 | chr16 | 15188268 | 15203054 | + | transcribed_processed_pseudogene |
| ENSG00000173272.16_5 | MZT2A | 0.99961917 | 1 | -0.014 | 5.112 | chr2 | 132222473 | 132250316 | - | protein_coding |
| ENSG00000152076.18_6 | CCDC74B | 0.99928078 | 1 | -0.014 | 1.904 | chr2 | 130896860 | 130902707 | - | protein_coding |
| ENSG00000175893.11_3 | ZDHHC21 | 0.99999997 | 1 | -0.014 | 5.094 | chr9 | 14611069 | 14693469 | - | protein_coding |
| ENSG00000248971.2_4 | KRT8P46 | 0.99876803 | 1 | -0.014 | 1.556 | chr4 | 103649903 | 103651328 | - | processed_pseudogene |
| ENSG00000174720.16_7 | LARP7 | 0.99999999 | 1 | -0.014 | 5.126 | chr4 | 113558263 | 113578748 | + | protein_coding |
| ENSG00000158941.16_7 | CCAR2 | 0.99999999 | 1 | -0.014 | 6.096 | chr8 | 22462145 | 22479027 | + | protein_coding |
| ENSG00000017797.13_4 | RALBP1 | 1 | 1 | -0.014 | 6.355 | chr18 | 9475007 | 9538112 | + | protein_coding |
| ENSG00000169957.10_5 | ZNF768 | 0.99999906 | 1 | -0.014 | 4.723 | chr16 | 30535325 | 30538142 | - | protein_coding |
| ENSG00000146433.8_2 | TMEM181 | 1 | 1 | -0.014 | 5.577 | chr6 | 158957468 | 159056460 | + | protein_coding |
| ENSG00000183426.17_5 | NPIPA1 | 0.9998689 | 1 | -0.015 | 2.646 | chr16 | 15016659 | 15045917 | + | protein_coding |
| ENSG00000160401.15_5 | CFAP157 | 0.99662559 | 1 | -0.015 | 2.000 | chr9 | 130469267 | 130478281 | + | protein_coding |
| ENSG00000128989.10_3 | ARPP19 | 1 | 1 | -0.015 | 8.630 | chr15 | 52839242 | 52862080 | - | protein_coding |
| ENSG00000112685.14_5 | EXOC2 | 0.99999999 | 1 | -0.015 | 5.343 | chr6 | 485154 | 693139 | - | protein_coding |
| ENSG00000248333.8_4 | CDK11B | 0.99982499 | 1 | -0.015 | 2.478 | chr1 | 1570603 | 1590473 | - | protein_coding |
| ENSG00000141219.15_7 | C17orf80 | 0.99999995 | 1 | -0.015 | 4.992 | chr17 | 71228372 | 71245086 | + | protein_coding |
| ENSG00000266910.2_5 | AC008507.1 | 0.98539114 | 1 | -0.015 | -0.724 | chr19 | 30528330 | 30529088 | + | processed_pseudogene |
| ENSG00000285416.1_3 | AC137630.5 | 0.99511734 | 1 | -0.015 | -0.171 | chr3 | 49034654 | 49036875 | - | unitary_pseudogene |
| ENSG00000100319.13_4 | ZMAT5 | 0.99968824 | 1 | -0.015 | 2.246 | chr22 | 30126945 | 30163000 | - | protein_coding |
| ENSG00000126883.17_6 | NUP214 | 1 | 1 | -0.015 | 5.902 | chr9 | 134000973 | 134110050 | + | protein_coding |
| ENSG00000103381.12_4 | CPPED1 | 1 | 1 | -0.015 | 6.001 | chr16 | 12753656 | 12897744 | - | protein_coding |
| ENSG00000163093.12_4 | BBS5 | 0.99773142 | 1 | -0.016 | 1.451 | chr2 | 170335990 | 170363165 | + | protein_coding |
| ENSG00000259556.2_6 | AC090971.4 | 0.9906218 | 1 | -0.016 | 0.256 | chr15 | 52211270 | 52222824 | - | unprocessed_pseudogene |
| ENSG00000254461.1_7 | AP001107.4 | 0.99499259 | 1 | -0.016 | 0.739 | chr11 | 66027038 | 66029305 | - | lncRNA |
| ENSG00000117481.10_3 | NSUN4 | 0.99999953 | 1 | -0.016 | 4.263 | chr1 | 46805849 | 46830824 | + | protein_coding |
| ENSG00000245148.2_6 | ARAP1-AS2 | 0.99247468 | 1 | -0.016 | 0.451 | chr11 | 72411519 | 72416652 | + | lncRNA |
| ENSG00000158234.12_3 | FAIM | 0.99999127 | 1 | -0.016 | 3.651 | chr3 | 138327448 | 138352218 | + | protein_coding |
| ENSG00000196632.10_5 | WNK3 | 0.98616796 | 1 | -0.016 | -0.199 | chrX | 54219256 | 54385075 | - | protein_coding |
| ENSG00000204788.3 | CR769776.1 | 0.98389913 | 1 | -0.016 | -0.829 | chr9 | 69377902 | 69380597 | - | protein_coding |
| ENSG00000101052.13_3 | IFT52 | 0.99999997 | 1 | -0.016 | 4.870 | chr20 | 42219577 | 42275939 | + | protein_coding |
| ENSG00000071564.16_8 | TCF3 | 0.99998695 | 1 | -0.016 | 5.639 | chr19 | 1609289 | 1652614 | - | protein_coding |
| ENSG00000163512.14_4 | AZI2 | 0.99999957 | 1 | -0.016 | 4.689 | chr3 | 28356494 | 28390541 | - | protein_coding |
| ENSG00000122786.20_7 | CALD1 | 1 | 1 | -0.016 | 7.194 | chr7 | 134429003 | 134655480 | + | protein_coding |
| ENSG00000156990.14_5 | RPUSD3 | 0.99999997 | 1 | -0.016 | 5.077 | chr3 | 9879533 | 9886286 | - | protein_coding |

| | | | | | | | | | | |
|---|---|---|---|---|---|---|---|---|---|---|
| ENSG00000161265.15_7 | U2AF1L4 | 0.99438977 | 1 | -0.017 | 0.771 | chr19 | 36233365 | 36236346 | - | protein_coding |
| ENSG00000284738.1_4 | AL358472.5 | 0.99029103 | 1 | -0.017 | 0.498 | chr1 | 153895813 | 153907716 | + | lncRNA |
| ENSG00000260884.1_6 | AC009120.3 | 0.9837396 | 1 | -0.017 | -0.448 | chr16 | 74347235 | 74349532 | + | lncRNA |
| ENSG00000280193.1_5 | AC132219.2 | 0.99537452 | 1 | -0.017 | 0.822 | chr8 | 82608603 | 82610929 | + | TEC |
| ENSG00000189091.13_5 | SF3B3 | 1 | 1 | -0.017 | 8.278 | chr16 | 70557694 | 70611573 | + | protein_coding |
| ENSG00000169567.12_4 | HINT1 | 1 | 1 | -0.017 | 7.586 | chr5 | 130494720 | 130507428 | - | protein_coding |
| ENSG00000187325.5_4 | TAF9B | 0.99999999 | 1 | -0.017 | 5.557 | chrX | 77385245 | 77395147 | - | protein_coding |
| ENSG00000075856.12_7 | SART3 | 1 | 1 | -0.017 | 6.247 | chr12 | 108915991 | 108955176 | - | protein_coding |
| ENSG00000163807.6_4 | KIAA1143 | 0.99999992 | 1 | -0.017 | 5.273 | chr3 | 44779153 | 44803111 | - | protein_coding |
| ENSG00000105976.15_4 | MET | 1 | 1 | -0.017 | 8.776 | chr7 | 116312250 | 116438440 | + | protein_coding |
| ENSG00000118997.14_5 | DNAH7 | 0.99973327 | 1 | -0.017 | 2.518 | chr2 | 196602427 | 196933561 | - | protein_coding |
| ENSG00000244187.8_4 | TMEM141 | 0.99996244 | 1 | -0.017 | 3.108 | chr9 | 139685796 | 139687769 | + | protein_coding |
| ENSG00000189089.5_4 | RIMKLBP1 | 0.98278923 | 1 | -0.017 | -0.861 | chr21 | 37422512 | 37423675 | + | processed_pseudogene |
| ENSG00000168394.11_4 | TAP1 | 0.99998777 | 1 | -0.017 | 3.421 | chr6 | 32812986 | 32821755 | - | protein_coding |
| ENSG00000186919.13_4 | ZACN | 0.99301296 | 1 | -0.017 | 0.644 | chr17 | 74068042 | 74079747 | + | protein_coding |
| ENSG00000280183.1_5 | AC067852.4 | 0.99203238 | 1 | -0.018 | -0.476 | chr17 | 40750661 | 40751195 | + | TEC |
| ENSG00000160888.7_3 | IER2 | 0.99993818 | 1 | -0.018 | 3.130 | chr19 | 13261225 | 13265725 | + | protein_coding |
| ENSG00000105559.12_5 | PLEKHA4 | 0.98259413 | 1 | -0.018 | -0.756 | chr19 | 49340354 | 49371874 | - | protein_coding |
| ENSG00000234684.7_7 | SDCBP2-AS1 | 0.99985121 | 1 | -0.018 | 2.506 | chr20 | 1306049 | 1359379 | + | lncRNA |
| ENSG00000175826.12_5 | CTDNEP1 | 0.99999979 | 1 | -0.018 | 4.389 | chr17 | 7146910 | 7155810 | - | protein_coding |
| ENSG00000198843.13_7 | SELENOT | 0.99999993 | 1 | -0.018 | 6.471 | chr3 | 150320662 | 150348232 | + | protein_coding |
| ENSG00000162521.19_6 | RBBP4 | 1 | 1 | -0.018 | 6.406 | chr1 | 33116743 | 33151812 | + | protein_coding |
| ENSG00000165733.8_3 | BMS1 | 1 | 1 | -0.018 | 6.845 | chr10 | 43278243 | 43330385 | + | protein_coding |
| ENSG00000119844.15_4 | AFTPH | 1 | 1 | -0.018 | 6.191 | chr2 | 64751439 | 64820139 | + | protein_coding |
| ENSG00000116771.6_4 | AGMAT | 0.9998945 | 1 | -0.018 | 3.265 | chr1 | 15898194 | 15911546 | - | protein_coding |
| ENSG00000175390.14_5 | EIF3F | 1 | 1 | -0.018 | 7.099 | chr11 | 7991798 | 8023409 | + | protein_coding |
| ENSG00000134825.16_5 | TMEM258 | 0.9999993 | 1 | -0.018 | 4.472 | chr11 | 61535973 | 61560274 | - | protein_coding |
| ENSG00000113712.18_5 | CSNK1A1 | 1 | 1 | -0.018 | 6.982 | chr5 | 148872545 | 148931034 | - | protein_coding |
| ENSG00000122483.17_5 | CCDC18 | 0.99999967 | 1 | -0.018 | 4.529 | chr1 | 93645476 | 93744287 | + | protein_coding |
| ENSG00000223546.7_7 | LINC00630 | 0.99994614 | 1 | -0.018 | 3.145 | chrX | 102024086 | 102140334 | + | lncRNA |
| ENSG00000254433.1_7 | AP001001.1 | 0.98716659 | 1 | -0.018 | 0.041 | chr11 | 105956717 | 105972702 | - | lncRNA |
| ENSG00000110442.12_4 | COMMD9 | 0.99999991 | 1 | -0.018 | 4.764 | chr11 | 36290834 | 36310999 | - | protein_coding |
| ENSG00000160695.14_2 | VPS11 | 0.9999989 | 1 | -0.018 | 4.041 | chr11 | 118938403 | 118952688 | + | protein_coding |
| ENSG00000237118.3_6 | CYP2F2P | 0.98607679 | 1 | -0.018 | 0.077 | chr19 | 41324319 | 41332677 | - | transcribed_processed_pseudogene |
| ENSG00000141027.21_7 | NCOR1 | 1 | 1 | -0.018 | 7.942 | chr17 | 15932471 | 16121499 | - | protein_coding |
| ENSG00000131149.19_5 | GSE1 | 0.99999999 | 1 | -0.018 | 5.864 | chr16 | 85203131 | 85709810 | + | protein_coding |
| ENSG00000090621.14_7 | PABPC4 | 1 | 1 | -0.018 | 5.946 | chr1 | 40026488 | 40042462 | - | protein_coding |
| ENSG00000023171.18_6 | GRAMD1B | 1 | 1 | -0.018 | 7.777 | chr11 | 123229136 | 123498482 | + | protein_coding |
| ENSG00000148925.10_5 | BTBD10 | 1 | 1 | -0.018 | 5.657 | chr11 | 13409548 | 13484844 | - | protein_coding |
| ENSG00000136240.10_5 | KDELR2 | 1 | 1 | -0.018 | 7.629 | chr7 | 6485584 | 6523821 | - | protein_coding |
| ENSG00000214114.9_5 | MYCBP | 0.99999564 | 1 | -0.019 | 3.717 | chr1 | 39328165 | 39339040 | - | protein_coding |
| ENSG00000233016.7_6 | SNHG7 | 0.99999997 | 1 | -0.019 | 5.274 | chr9 | 139615818 | 139622636 | - | lncRNA |
| ENSG00000118007.13_5 | STAG1 | 1 | 1 | -0.019 | 5.763 | chr3 | 136055078 | 136471245 | - | protein_coding |
| ENSG00000235453.10_7 | SMIM27 | 0.99705807 | 1 | -0.019 | 1.262 | chr9 | 32551142 | 32568619 | + | protein_coding |
| ENSG00000185787.15_6 | MORF4L1 | 1 | 1 | -0.019 | 7.345 | chr15 | 79102829 | 79190481 | + | protein_coding |
| ENSG00000035664.11_6 | DAPK2 | 0.99995378 | 1 | -0.019 | 2.855 | chr15 | 64199235 | 64364232 | - | protein_coding |
| ENSG00000266472.5_3 | MRPS21 | 0.99999999 | 1 | -0.019 | 5.728 | chr1 | 150266140 | 150281414 | + | protein_coding |
| ENSG00000113249.13_5 | HAVCR1 | 1 | 1 | -0.019 | 7.397 | chr5 | 156456424 | 156486130 | - | protein_coding |
| ENSG00000183530.14_7 | PRR14L | 1 | 1 | -0.019 | 5.824 | chr22 | 32072242 | 32146126 | - | protein_coding |
| ENSG00000108788.11_2 | MLX | 0.99999998 | 1 | -0.019 | 5.107 | chr17 | 40719086 | 40725257 | + | protein_coding |
| ENSG00000122873.12_3 | CISD1 | 0.99999996 | 1 | -0.019 | 4.822 | chr10 | 60028922 | 60049346 | + | protein_coding |
| ENSG00000104375.17_7 | STK3 | 0.99999984 | 1 | -0.019 | 4.954 | chr8 | 99383456 | 99955055 | - | protein_coding |
| ENSG00000172840.7_6 | PDP2 | 0.99999847 | 1 | -0.019 | 4.091 | chr16 | 66912492 | 66929657 | + | protein_coding |
| ENSG00000165113.13_5 | GKAP1 | 0.99998006 | 1 | -0.019 | 3.209 | chr9 | 86354340 | 86444431 | - | protein_coding |
| ENSG00000269439.6_9 | AC010618.3 | 0.99086572 | 1 | -0.019 | 0.606 | chr19 | 17599799 | 17622698 | - | lncRNA |
| ENSG00000231365.6_7 | WARS2-AS1 | 0.99998001 | 1 | -0.019 | 3.236 | chr1 | 119683014 | 119818596 | + | lncRNA |
| ENSG00000026950.17_5 | BTN3A1 | 0.99995888 | 1 | -0.019 | 3.101 | chr6 | 26402465 | 26415436 | + | protein_coding |
| ENSG00000154945.7_4 | ANKRD40 | 1 | 1 | -0.019 | 6.785 | chr17 | 48770559 | 48785275 | - | protein_coding |
| ENSG00000179523.5_6 | EIF3J-DT | 0.99962006 | 1 | -0.019 | 2.300 | chr15 | 44819455 | 44829244 | - | lncRNA |
| ENSG00000113391.19_6 | FAM172A | 0.99999992 | 1 | -0.019 | 4.695 | chr5 | 92953431 | 93447404 | - | protein_coding |

| | | | | | | | | | | |
|---|---|---|---|---|---|---|---|---|---|---|
| ENSG00000076053.10_4 | RBM7 | 0.99999681 | 1 | -0.019 | 4.261 | chr11 | 114270752 | 114284925 | + | protein_coding |
| ENSG00000158290.17_6 | CUL4B | 1 | 1 | -0.019 | 6.985 | chrX | 119658467 | 119709649 | - | protein_coding |
| ENSG00000099331.13_4 | MYO9B | 0.99999998 | 1 | -0.020 | 5.766 | chr19 | 17186591 | 17325346 | + | protein_coding |
| ENSG00000124733.5_6 | MEA1 | 0.99999995 | 1 | -0.020 | 4.981 | chr6 | 42978881 | 42984606 | - | protein_coding |
| ENSG00000200152.1 | RNU6-216P | 0.97921007 | 1 | -0.020 | -0.943 | chr11 | 74679234 | 74679337 | - | snRNA |
| ENSG00000079462.7_6 | PAFAH1B3 | 0.99999997 | 1 | -0.020 | 5.708 | chr19 | 42801185 | 42807698 | - | protein_coding |
| ENSG00000268573.1_8 | AC011815.1 | 0.99831153 | 1 | -0.020 | 1.577 | chr18 | 32860831 | 32870165 | - | lncRNA |
| ENSG00000149782.11_2 | PLCB3 | 0.99999998 | 1 | -0.020 | 5.844 | chr11 | 64018995 | 64036622 | + | protein_coding |
| ENSG00000116539.13_6 | ASH1L | 0.99999995 | 1 | -0.020 | 6.856 | chr1 | 155305059 | 155532598 | - | protein_coding |
| ENSG00000126070.20_3 | AGO3 | 0.99999981 | 1 | -0.020 | 4.558 | chr1 | 36396319 | 36538101 | + | protein_coding |
| ENSG00000205155.8_4 | PSENEN | 0.99999896 | 1 | -0.020 | 4.045 | chr19 | 36236501 | 36238420 | + | protein_coding |
| ENSG00000169692.13_4 | AGPAT2 | 0.99997467 | 1 | -0.020 | 5.273 | chr9 | 139567595 | 139581909 | - | protein_coding |
| ENSG00000167863.12_5 | ATP5PD | 1 | 1 | -0.020 | 6.077 | chr17 | 73034958 | 73043080 | - | protein_coding |
| ENSG00000273783.1_6 | AL136040.1 | 0.98161025 | 1 | -0.020 | -0.346 | chr14 | 81687562 | 81688804 | + | lncRNA |
| ENSG00000254612.2_5 | AP001000.1 | 0.99769796 | 1 | -0.020 | 1.814 | chr11 | 127810808 | 127811549 | - | processed_pseudogene |
| ENSG00000166415.15_5 | WDR72 | 0.99999999 | 1 | -0.021 | 5.979 | chr15 | 53805938 | 54055075 | - | protein_coding |
| ENSG00000152726.14 | FAM21B | 0.99988539 | 1 | -0.021 | 2.846 | chr10 | 47894023 | 47949412 | + | protein_coding |
| ENSG00000224543.4_4 | SNRPGP15 | 0.98983912 | 1 | -0.021 | 0.794 | chr19 | 14600200 | 14600421 | + | processed_pseudogene |
| ENSG00000240591.2_7 | AL096701.3 | 0.97933144 | 1 | -0.021 | -0.563 | chr22 | 31848867 | 31860290 | + | lncRNA |
| ENSG00000279483.2_4 | AC090498.1 | 0.98138016 | 1 | -0.021 | -0.087 | chr7 | 137198605 | 137198679 | + | processed_pseudogene |
| ENSG00000183401.12_7 | CCDC159 | 0.99979445 | 1 | -0.021 | 2.496 | chr19 | 11455360 | 11465620 | + | protein_coding |
| ENSG00000260328.1_6 | AC104024.2 | 0.98704837 | 1 | -0.021 | 0.411 | chr17 | 16915228 | 16918304 | - | lncRNA |
| ENSG00000116649.10_7 | SRM | 0.99999983 | 1 | -0.021 | 4.855 | chr1 | 11114641 | 11120077 | - | protein_coding |
| ENSG00000101544.9_4 | ADNP2 | 1 | 1 | -0.021 | 6.013 | chr18 | 77867146 | 77905406 | + | protein_coding |
| ENSG00000278784.1_7 | AL136295.7 | 0.99405109 | 1 | -0.021 | 0.944 | chr14 | 24670818 | 24672017 | - | lncRNA |
| ENSG00000244716.3_5 | BX679664.3 | 0.99274727 | 1 | -0.021 | 0.806 | chr1 | 109534904 | 109535458 | + | processed_pseudogene |
| ENSG00000131143.10_7 | COX4I1 | 1 | 1 | -0.021 | 7.207 | chr16 | 85832239 | 85840674 | + | protein_coding |
| ENSG00000188677.14_6 | PARVB | 0.99999986 | 1 | -0.021 | 4.739 | chr22 | 44395091 | 44568829 | + | protein_coding |
| ENSG00000162600.12_5 | OMA1 | 0.99999671 | 1 | -0.021 | 3.708 | chr1 | 58881056 | 59012474 | - | protein_coding |
| ENSG00000173275.13_3 | ZNF449 | 0.9999979 | 1 | -0.021 | 4.010 | chrX | 134478721 | 134497338 | + | protein_coding |
| ENSG00000181904.9_4 | C5orf24 | 0.99999999 | 1 | -0.021 | 5.964 | chr5 | 134181370 | 134195425 | + | protein_coding |
| ENSG00000149806.11_5 | FAU | 1 | 1 | -0.021 | 6.371 | chr11 | 64888102 | 64889649 | - | protein_coding |
| ENSG00000127922.9_5 | SEM1 | 0.98966601 | 1 | -0.022 | 4.932 | chr7 | 96110938 | 96339203 | - | protein_coding |
| ENSG00000248008.3_8 | NRAV | 0.99999947 | 1 | -0.022 | 4.735 | chr12 | 120925882 | 120933749 | - | lncRNA |
| ENSG00000198964.14_6 | SGMS1 | 0.99999998 | 1 | -0.022 | 5.309 | chr10 | 52065346 | 52384923 | - | protein_coding |
| ENSG00000197381.16_4 | ADARB1 | 0.9967996 | 1 | -0.022 | 1.846 | chr21 | 46493768 | 46646475 | + | protein_coding |
| ENSG00000271780.1_6 | AL118558.3 | 0.99032169 | 1 | -0.022 | 0.612 | chr14 | 102414684 | 102415762 | + | lncRNA |
| ENSG00000112651.12_4 | MRPL2 | 0.99561631 | 1 | -0.022 | 0.505 | chr6 | 43021767 | 43027176 | - | protein_coding |
| ENSG00000214013.9_3 | GANC | 0.99998832 | 1 | -0.022 | 3.412 | chr15 | 42565431 | 42649133 | + | protein_coding |
| ENSG00000161526.15_8 | SAP30BP | 1 | 1 | -0.022 | 5.907 | chr17 | 73663331 | 73704142 | + | protein_coding |
| ENSG00000167965.18_6 | MLST8 | 0.99999783 | 1 | -0.022 | 3.951 | chr16 | 2254249 | 2259454 | + | protein_coding |
| ENSG00000275465.7_7 | AL449403.3 | 0.97567913 | 1 | -0.022 | -0.798 | chr9 | 98868023 | 98878693 | - | lncRNA |
| ENSG00000117318.9_4 | ID3 | 0.99999962 | 1 | -0.022 | 6.025 | chr1 | 23884417 | 23885992 | - | protein_coding |
| ENSG00000188419.14_4 | CHM | 0.99999998 | 1 | -0.022 | 5.471 | chrX | 85116185 | 85302565 | - | protein_coding |
| ENSG00000197180.2_5 | AC244090.1 | 0.99631194 | 1 | -0.022 | 1.462 | chrX | 153652723 | 153656825 | - | lncRNA |
| ENSG00000197301.7_8 | HMGA2-AS1 | 0.97781845 | 1 | -0.022 | -0.450 | chr12 | 66245120 | 66275947 | - | lncRNA |
| ENSG00000260274.1_6 | AC068338.2 | 0.98007203 | 1 | -0.022 | -0.198 | chr15 | 75660496 | 75661925 | + | lncRNA |
| ENSG00000175691.9_4 | ZNF77 | 0.99979345 | 1 | -0.022 | 1.694 | chr19 | 2933216 | 2944969 | - | protein_coding |
| ENSG00000166762.18_6 | CATSPER2 | 0.98941671 | 1 | -0.022 | 0.114 | chr15 | 43920701 | 43960316 | - | protein_coding |
| ENSG00000196296.13_5 | ATP2A1 | 0.99468695 | 1 | -0.022 | 1.248 | chr16 | 28889726 | 28915830 | + | protein_coding |
| ENSG00000013573.17_5 | DDX11 | 1 | 1 | -0.023 | 6.302 | chr12 | 31226794 | 31257733 | + | protein_coding |
| ENSG00000271580.1_5 | AL583832.1 | 0.9769029 | 1 | -0.023 | -0.563 | chr1 | 205060291 | 205061074 | + | processed_pseudogene |
| ENSG00000100316.16_6 | RPL3 | 1 | 1 | -0.023 | 9.543 | chr22 | 39708887 | 39716394 | - | protein_coding |
| ENSG00000113638.14_5 | TTC33 | 0.99999954 | 1 | -0.023 | 4.584 | chr5 | 40512435 | 40756065 | - | protein_coding |
| ENSG00000101882.10_6 | NKAP | 0.99999963 | 1 | -0.023 | 4.376 | chrX | 119054635 | 119077714 | - | protein_coding |
| ENSG00000213853.10_5 | EMP2 | 0.99999999 | 1 | -0.023 | 5.516 | chr16 | 10622279 | 10674489 | - | protein_coding |
| ENSG00000143127.13_3 | ITGA10 | 0.99987485 | 1 | -0.023 | 1.791 | chr1 | 145524969 | 145543868 | + | protein_coding |
| ENSG00000137133.11_5 | HINT2 | 0.97742514 | 1 | -0.023 | -0.302 | chr9 | 35812957 | 35815351 | - | protein_coding |

| | | | | | | | | | | |
|---|---|---|---|---|---|---|---|---|---|---|
| ENSG00000105982.16_2 | RNF32 | 0.98713592 | 1 | -0.023 | 1.037 | chr7 | 156432975 | 156469824 | + | protein_coding |
| ENSG00000001629.10_4 | ANKIB1 | 1 | 1 | -0.023 | 6.693 | chr7 | 91875288 | 92030697 | + | protein_coding |
| ENSG00000252498.1 | RNU6-1016P | 0.9743428 | 1 | -0.023 | -0.726 | chr6 | 76353800 | 76353893 | + | snRNA |
| ENSG00000108826.16_4 | MRPL27 | 0.99999999 | 1 | -0.023 | 5.747 | chr17 | 48445218 | 48450568 | - | protein_coding |
| ENSG00000204560.10_5 | DHX16 | 0.99999981 | 1 | -0.024 | 4.863 | chr6 | 30620896 | 30640783 | - | protein_coding |
| ENSG00000272572.1_7 | AL138762.2 | 0.97816435 | 1 | -0.024 | -0.219 | chr10 | 102670860 | 102672496 | - | lncRNA |
| ENSG00000214837.8_5 | LINC01347 | 0.9887396 | 1 | -0.024 | 0.575 | chr1 | 243219609 | 243265046 | - | transcribed_unprocessed_pseudogene |
| ENSG00000136143.16_7 | SUCLA2 | 0.99999998 | 1 | -0.024 | 5.706 | chr13 | 48319871 | 48612104 | - | protein_coding |
| ENSG00000156958.15_6 | GALK2 | 0.99999969 | 1 | -0.024 | 4.430 | chr15 | 49447853 | 49660066 | + | protein_coding |
| ENSG00000138686.10_4 | BBS7 | 0.99999915 | 1 | -0.024 | 4.175 | chr4 | 122745484 | 122791642 | - | protein_coding |
| ENSG00000270605.1_6 | AL353622.1 | 0.9846534 | 1 | -0.024 | 0.447 | chr1 | 28566020 | 28567964 | - | lncRNA |
| ENSG00000242588.6_7 | AC108010.1 | 0.99980355 | 1 | -0.024 | 2.569 | chr7 | 128214805 | 128266527 | + | lncRNA |
| ENSG00000132196.14_5 | HSD17B7 | 0.9999973 | 1 | -0.024 | 3.932 | chr1 | 162760492 | 162782607 | + | protein_coding |
| ENSG00000108389.9_4 | MTMR4 | 1 | 1 | -0.024 | 6.391 | chr17 | 56566890 | 56595266 | - | protein_coding |
| ENSG00000166024.13_3 | R3HCC1L | 0.9999996 | 1 | -0.024 | 4.475 | chr10 | 99894381 | 100004654 | + | protein_coding |
| ENSG00000105968.18_4 | H2AZ2 | 1 | 1 | -0.024 | 7.350 | chr7 | 44866390 | 44887682 | - | protein_coding |
| ENSG00000205629.12_3 | LCMT1 | 0.99999932 | 1 | -0.024 | 4.303 | chr16 | 25123052 | 25189552 | + | protein_coding |
| ENSG00000084676.15_3 | NCOA1 | 0.99999999 | 1 | -0.024 | 5.670 | chr2 | 24714783 | 24993571 | + | protein_coding |
| ENSG00000127540.12_5 | UQCR11 | 0.99995623 | 1 | -0.024 | 3.570 | chr19 | 1597168 | 1605472 | - | protein_coding |
| ENSG00000184014.8_5 | DENND5A | 1 | 1 | -0.024 | 6.015 | chr11 | 9160372 | 9286897 | - | protein_coding |
| ENSG00000146729.10_7 | NIPSNAP2 | 0.99999994 | 1 | -0.024 | 6.010 | chr7 | 56019486 | 56067874 | + | protein_coding |
| ENSG00000267169.2_9 | AC022098.1 | 0.98078854 | 1 | -0.024 | -0.536 | chr19 | 14247958 | 14282079 | + | lncRNA |
| ENSG00000129219.14_4 | PLD2 | 0.99999997 | 1 | -0.025 | 5.194 | chr17 | 4710447 | 4726729 | + | protein_coding |
| ENSG00000171763.19_3 | SPATA5L1 | 0.99996529 | 1 | -0.025 | 3.399 | chr15 | 45694534 | 45713613 | + | protein_coding |
| ENSG00000147912.13_6 | FBXO10 | 0.99852572 | 1 | -0.025 | 1.930 | chr9 | 37510889 | 37576377 | - | protein_coding |
| ENSG00000117036.12_4 | ETV3 | 0.99999991 | 1 | -0.025 | 4.763 | chr1 | 157090983 | 157108266 | - | protein_coding |
| ENSG00000079999.14_5 | KEAP1 | 0.99999999 | 1 | -0.025 | 6.144 | chr19 | 10596801 | 10614234 | - | protein_coding |
| ENSG00000169032.9_2 | MAP2K1 | 0.99999996 | 1 | -0.025 | 5.039 | chr15 | 66679155 | 66784650 | + | protein_coding |
| ENSG00000138796.16_2 | HADH | 0.99999991 | 1 | -0.025 | 4.833 | chr4 | 108910870 | 108956331 | + | protein_coding |
| ENSG00000168826.16_6 | ZBTB49 | 0.99841573 | 1 | -0.025 | 1.982 | chr4 | 4291978 | 4323513 | + | protein_coding |
| ENSG00000146425.11_5 | DYNLT1 | 0.99999825 | 1 | -0.025 | 4.441 | chr6 | 159057506 | 159065775 | - | protein_coding |
| ENSG00000076321.11_4 | KLHL20 | 0.99999878 | 1 | -0.025 | 4.148 | chr1 | 173684080 | 173755830 | + | protein_coding |
| ENSG00000175115.12_6 | PACS1 | 0.99999998 | 1 | -0.025 | 5.765 | chr11 | 65837743 | 66012215 | + | protein_coding |
| ENSG00000188986.7_5 | NELFB | 0.99985183 | 1 | -0.025 | 5.929 | chr9 | 140149779 | 140167994 | + | protein_coding |
| ENSG00000008952.17_3 | SEC62 | 1 | 1 | -0.025 | 6.362 | chr3 | 169684423 | 169716161 | + | protein_coding |
| ENSG00000121390.19_6 | PSPC1 | 0.99999998 | 1 | -0.025 | 5.431 | chr13 | 20248892 | 20357159 | - | protein_coding |
| ENSG00000213398.8_8 | LCAT | 0.99986322 | 1 | -0.025 | 3.029 | chr16 | 67973653 | 67978034 | - | protein_coding |
| ENSG00000185222.10_2 | TCEAL9 | 0.99999979 | 1 | -0.025 | 4.929 | chrX | 102611417 | 102613390 | + | protein_coding |
| ENSG00000128011.4_4 | LRFN1 | 0.98332137 | 1 | -0.025 | 1.777 | chr19 | 39797208 | 39805976 | - | protein_coding |
| ENSG00000269906.1_7 | AL606834.1 | 0.98677334 | 1 | -0.025 | 0.365 | chr14 | 51129229 | 51129896 | - | lncRNA |
| ENSG00000125912.11_4 | NCLN | 0.99957871 | 1 | -0.025 | 6.216 | chr19 | 3185561 | 3209573 | + | protein_coding |
| ENSG00000153922.10_4 | CHD1 | 1 | 1 | -0.026 | 7.176 | chr5 | 98189689 | 98264661 | - | protein_coding |
| ENSG00000160223.17_4 | ICOSLG | 0.97503564 | 1 | -0.026 | -0.293 | chr21 | 45636897 | 45660849 | - | protein_coding |
| ENSG00000182534.13_5 | MXRA7 | 0.99999998 | 1 | -0.026 | 5.183 | chr17 | 74668633 | 74707098 | - | protein_coding |
| ENSG00000204272.13_5 | NBDY | 0.99999903 | 1 | -0.026 | 4.210 | chrX | 56755674 | 56845612 | + | protein_coding |
| ENSG00000188002.10_7 | AC026412.1 | 0.9999816 | 1 | -0.026 | 3.262 | chr5 | 1599035 | 1634120 | - | transcribed_unprocessed_pseudogene |
| ENSG00000105339.10_5 | DENND3 | 0.99999999 | 1 | -0.026 | 5.575 | chr8 | 142127377 | 142205907 | + | protein_coding |
| ENSG00000215414.4_4 | PSMA6P1 | 0.98032596 | 1 | -0.026 | 0.264 | chrY | 15398518 | 15399258 | + | processed_pseudogene |
| ENSG00000119977.21_7 | TCTN3 | 0.99999999 | 1 | -0.026 | 5.696 | chr10 | 97423153 | 97453900 | - | protein_coding |
| ENSG00000147872.10_5 | PLIN2 | 0.99999949 | 1 | -0.026 | 4.952 | chr9 | 19108373 | 19149288 | - | protein_coding |
| ENSG00000261801.6_7 | LOXL1-AS1 | 0.98604041 | 1 | -0.026 | 0.790 | chr15 | 74200412 | 74220589 | - | lncRNA |
| ENSG00000149532.15_7 | CPSF7 | 1 | 1 | -0.026 | 6.455 | chr11 | 61170113 | 61197503 | - | protein_coding |
| ENSG00000273828.2_7 | AL133227.1 | 0.99328369 | 1 | -0.026 | 1.133 | chr20 | 44993190 | 45026633 | + | lncRNA |
| ENSG00000125812.16_5 | GZF1 | 0.99999997 | 1 | -0.026 | 5.125 | chr20 | 23342819 | 23353699 | + | protein_coding |
| ENSG00000139187.10_4 | KLRG1 | 0.9921623 | 1 | -0.026 | 1.239 | chr12 | 9102640 | 9163356 | + | protein_coding |
| ENSG00000143434.15_5 | SEMA6C | 0.9968934 | 1 | -0.026 | 1.607 | chr1 | 151104161 | 151119140 | - | protein_coding |
| ENSG00000278709.2_7 | NKILA | 0.99904673 | 1 | -0.026 | 1.968 | chr20 | 56285212 | 56287836 | + | lncRNA |
| ENSG00000140382.15_6 | HMG20A | 0.99999993 | 1 | -0.027 | 5.396 | chr15 | 77712754 | 77777949 | + | protein_coding |
| ENSG00000095637.22_8 | SORBS1 | 0.99403236 | 1 | -0.027 | 1.381 | chr10 | 97071528 | 97321171 | - | protein_coding |

| | | | | | | | | | | |
|---|---|---|---|---|---|---|---|---|---|---|
| ENSG00000151689.13_4 | INPP1 | 0.99996245 | 1 | -0.027 | 3.251 | chr2 | 191208296 | 191236391 | + | protein_coding |
| ENSG00000100836.10_5 | PABPN1 | 0.99999995 | 1 | -0.027 | 4.984 | chr14 | 23790498 | 23795394 | + | protein_coding |
| ENSG00000004864.13_3 | SLC25A13 | 1 | 1 | -0.027 | 6.082 | chr7 | 95749532 | 95951459 | - | protein_coding |
| ENSG00000187045.18_6 | TMPRSS6 | 0.97842601 | 1 | -0.027 | -0.629 | chr22 | 37461476 | 37505753 | - | protein_coding |
| ENSG00000171530.14_5 | TBCA | 1 | 1 | -0.027 | 6.015 | chr5 | 76986991 | 77164604 | - | protein_coding |
| ENSG00000101150.17_4 | TPD52L2 | 1 | 1 | -0.027 | 6.544 | chr20 | 62496581 | 62522898 | + | protein_coding |
| ENSG00000163803.13_7 | PLB1 | 0.9720744 | 1 | -0.027 | -0.133 | chr2 | 28680012 | 28867008 | + | protein_coding |
| ENSG00000245970.2_6 | AP003352.1 | 0.98489049 | 1 | -0.027 | 0.443 | chr8 | 99053954 | 99056349 | + | lncRNA |
| ENSG00000157107.14_5 | FCHO2 | 0.99999946 | 1 | -0.027 | 4.932 | chr5 | 72251868 | 72386349 | + | protein_coding |
| ENSG00000161956.13_5 | SENP3 | 0.99999891 | 1 | -0.027 | 4.226 | chr17 | 7465236 | 7475287 | + | protein_coding |
| ENSG00000241316.8_6 | SUCLG2-AS1 | 0.99999844 | 1 | -0.027 | 4.287 | chr3 | 67705093 | 67998137 | + | lncRNA |
| ENSG00000198690.10_5 | FAN1 | 0.99999999 | 1 | -0.027 | 5.733 | chr15 | 31182762 | 31235311 | + | protein_coding |
| ENSG00000124915.10_8 | MYRF-AS1 | 0.99407842 | 1 | -0.027 | 1.133 | chr11 | 61513965 | 61525127 | - | lncRNA |
| ENSG00000163964.15_6 | PIGX | 0.99999985 | 1 | -0.027 | 4.646 | chr3 | 196366646 | 196462878 | + | protein_coding |
| ENSG00000117632.23_5 | STMN1 | 0.99999999 | 1 | -0.028 | 6.922 | chr1 | 26210672 | 26233482 | - | protein_coding |
| ENSG00000279345.1_6 | Z98885.3 | 0.96930475 | 1 | -0.028 | -0.599 | chr22 | 50199100 | 50200856 | + | lncRNA |
| ENSG00000117448.13_5 | AKR1A1 | 0.99999999 | 1 | -0.028 | 5.716 | chr1 | 46016215 | 46035721 | + | protein_coding |
| ENSG00000158545.15_5 | ZC3H18 | 0.99999999 | 1 | -0.028 | 5.698 | chr16 | 88636789 | 88698374 | + | protein_coding |
| ENSG00000166881.10_7 | NEMP1 | 0.99999999 | 1 | -0.028 | 5.925 | chr12 | 57449426 | 57481846 | - | protein_coding |
| ENSG00000095787.23_7 | WAC | 1 | 1 | -0.028 | 7.124 | chr10 | 28821422 | 28912041 | + | protein_coding |
| ENSG00000120727.13_5 | PAIP2 | 0.9999958 | 1 | -0.028 | 4.034 | chr5 | 138677276 | 138705409 | + | protein_coding |
| ENSG00000232388.4_7 | SMIM26 | 0.99999597 | 1 | -0.028 | 4.217 | chr20 | 18547991 | 18550207 | + | protein_coding |
| ENSG00000163467.11_2 | TSACC | 0.9918922 | 1 | -0.028 | 0.487 | chr1 | 156307105 | 156316786 | + | protein_coding |
| ENSG00000178028.14_5 | DMAP1 | 0.99999781 | 1 | -0.028 | 4.075 | chr1 | 44679127 | 44686353 | + | protein_coding |
| ENSG00000273156.2_8 | AC124016.3 | 0.97139439 | 1 | -0.028 | -0.137 | chr4 | 83266029 | 83267995 | - | lncRNA |
| ENSG00000213413.2_3 | PVRIG | 0.9741919 | 1 | -0.028 | 0.176 | chr7 | 99815864 | 99819113 | + | protein_coding |
| ENSG00000234925.2_4 | ATP5PDP4 | 0.96782811 | 1 | -0.028 | -0.800 | chr12 | 69036299 | 69036773 | - | processed_pseudogene |
| ENSG00000182944.17_5 | EWSR1 | 1 | 1 | -0.028 | 7.179 | chr22 | 29663998 | 29696515 | + | protein_coding |
| ENSG00000054118.15_6 | THRAP3 | 1 | 1 | -0.028 | 7.477 | chr1 | 36690033 | 36770958 | + | protein_coding |
| ENSG00000218510.8_8 | LINC00339 | 0.99002445 | 1 | -0.028 | 0.870 | chr1 | 22351051 | 22357716 | + | lncRNA |
| ENSG00000169446.5_3 | MMGT1 | 0.99999994 | 1 | -0.028 | 5.426 | chrX | 135044229 | 135056222 | - | protein_coding |
| ENSG00000107643.16_6 | MAPK8 | 0.99999997 | 1 | -0.028 | 5.185 | chr10 | 49514682 | 49647403 | + | protein_coding |
| ENSG00000224578.5_4 | HNRNPA1P48 | 0.99149565 | 1 | -0.029 | 1.609 | chr16 | 51587347 | 51681043 | + | protein_coding |
| ENSG00000156467.9_3 | UQCRB | 1 | 1 | -0.029 | 6.829 | chr8 | 97238148 | 97247862 | - | protein_coding |
| ENSG00000182054.9_3 | IDH2 | 0.99999358 | 1 | -0.029 | 5.202 | chr15 | 90626277 | 90645736 | - | protein_coding |
| ENSG00000166452.12_6 | AKIP1 | 0.99999953 | 1 | -0.029 | 4.431 | chr11 | 8932686 | 8941631 | + | protein_coding |
| ENSG00000107863.18_6 | ARHGAP21 | 1 | 1 | -0.029 | 7.343 | chr10 | 24872538 | 25012816 | - | protein_coding |
| ENSG00000117000.9_3 | RLF | 0.99999995 | 1 | -0.029 | 5.201 | chr1 | 40627059 | 40706593 | + | protein_coding |
| ENSG00000231991.4_2 | ANXA2P2 | 0.99651428 | 1 | -0.029 | 5.951 | chr9 | 33624272 | 33625291 | + | processed_pseudogene |
| ENSG00000101363.12_2 | MANBAL | 0.99999974 | 1 | -0.029 | 5.084 | chr20 | 35918041 | 35945663 | + | protein_coding |
| ENSG00000070010.19_8 | UFD1 | 0.99999997 | 1 | -0.029 | 5.189 | chr22 | 19437434 | 19466725 | - | protein_coding |
| ENSG00000147421.18_5 | HMBOX1 | 0.9999963 | 1 | -0.029 | 4.132 | chr8 | 28747912 | 28922281 | + | protein_coding |
| ENSG00000283041.1_5 | AC008038.1 | 0.99301881 | 1 | -0.029 | 4.778 | chr7 | 132719367 | 132720680 | - | processed_pseudogene |
| ENSG00000231856.2_6 | AL162377.1 | 0.96526044 | 1 | -0.029 | -0.944 | chr13 | 52377974 | 52387968 | + | lncRNA |
| ENSG00000178188.14_6 | SH2B1 | 0.99999969 | 1 | -0.029 | 5.219 | chr16 | 28857921 | 28885533 | + | protein_coding |
| ENSG00000176994.11_3 | SMCR8 | 1 | 1 | -0.029 | 5.936 | chr17 | 18218607 | 18231370 | + | protein_coding |
| ENSG00000089682.16_6 | RBM41 | 0.99999963 | 1 | -0.030 | 4.729 | chrX | 106307650 | 106362053 | - | protein_coding |
| ENSG00000184068.2_7 | SREBF2-AS1 | 0.99719327 | 1 | -0.030 | 1.621 | chr22 | 42227219 | 42230669 | - | lncRNA |
| ENSG00000113273.17_6 | ARSB | 0.99999995 | 1 | -0.030 | 4.901 | chr5 | 78073032 | 78281910 | - | protein_coding |
| ENSG00000203896.10_9 | LIME1 | 0.99576152 | 1 | -0.030 | 1.554 | chr20 | 62367635 | 62370456 | + | protein_coding |
| ENSG00000143924.19_3 | EML4 | 1 | 1 | -0.030 | 8.020 | chr2 | 42396493 | 42559688 | + | protein_coding |
| ENSG00000114904.13_4 | NEK4 | 0.99999997 | 1 | -0.030 | 5.244 | chr3 | 52742460 | 52804962 | - | protein_coding |
| ENSG00000164294.14_3 | GPX8 | 0.99999999 | 1 | -0.030 | 6.026 | chr5 | 54455995 | 54463125 | + | protein_coding |
| ENSG00000118873.16_5 | RAB3GAP2 | 0.99999999 | 1 | -0.030 | 6.159 | chr1 | 220321635 | 220445795 | - | protein_coding |
| ENSG00000272654.1_6 | AL358472.2 | 0.97064349 | 1 | -0.030 | -0.119 | chr1 | 153950219 | 153951636 | + | lncRNA |
| ENSG00000272812.1_7 | AC004908.2 | 0.98160763 | 1 | -0.030 | -0.201 | chr8 | 183119 | 183692 | + | lncRNA |
| ENSG00000097007.19_7 | ABL1 | 1 | 1 | -0.030 | 6.490 | chr9 | 133588403 | 133763062 | + | protein_coding |
| ENSG00000168067.12_9 | MAP4K2 | 0.99999958 | 1 | -0.030 | 4.356 | chr11 | 64552390 | 64570713 | - | protein_coding |

| | | | | | | | | | | |
|---|---|---|---|---|---|---|---|---|---|---|
| ENSG00000072958.8_5 | AP1M1 | 0.99976582 | 1 | -0.030 | 5.476 | chr19 | 16308389 | 16356718 | + | protein_coding |
| ENSG00000072121.16_6 | ZFYVE26 | 0.99999998 | 1 | -0.030 | 5.446 | chr14 | 68194091 | 68283307 | - | protein_coding |
| ENSG00000265808.4_3 | SEC22B | 0.99999756 | 1 | -0.030 | 4.821 | chr1 | 145096408 | 145118353 | + | protein_coding |
| ENSG00000284976.1_2 | BX255925.3 | 0.99998918 | 1 | -0.030 | 4.198 | chr9 | 140111904 | 140113813 | + | protein_coding |
| ENSG00000224189.8_6 | HAGLR | 0.99001617 | 1 | -0.030 | 0.913 | chr2 | 177028779 | 177053686 | - | lncRNA |
| ENSG00000214174.8_4 | AMZ2P1 | 0.99993606 | 1 | -0.030 | 3.082 | chr17 | 62962668 | 62971694 | - | transcribed_unprocessed_pseudogene |
| ENSG00000145817.17_4 | YIPF5 | 0.99999997 | 1 | -0.030 | 6.091 | chr5 | 143537726 | 143550278 | - | protein_coding |
| ENSG00000272455.1_7 | AL391244.2 | 0.98596416 | 1 | -0.030 | 0.877 | chr1 | 1344476 | 1345998 | + | lncRNA |
| ENSG00000115266.11_6 | APC2 | 0.99970163 | 1 | -0.031 | 2.724 | chr19 | 1446301 | 1473243 | + | protein_coding |
| ENSG00000068489.12_5 | PRR11 | 0.99999999 | 1 | -0.031 | 5.853 | chr17 | 57232860 | 57282066 | + | protein_coding |
| ENSG00000131652.14_3 | THOC6 | 0.99999926 | 1 | -0.031 | 4.689 | chr16 | 3074028 | 3077756 | + | protein_coding |
| ENSG00000150760.12_3 | DOCK1 | 1 | 1 | -0.031 | 6.687 | chr10 | 128593978 | 129250781 | + | protein_coding |
| ENSG00000104979.9_4 | C19orf53 | 0.99999995 | 1 | -0.031 | 5.253 | chr19 | 13885270 | 13889587 | + | protein_coding |
| ENSG00000087302.9_4 | RTRAF | 1 | 1 | -0.031 | 6.938 | chr14 | 52456232 | 52477412 | + | protein_coding |
| ENSG00000183506.17_5 | PI4KAP2 | 0.99903826 | 1 | -0.031 | 2.441 | chr22 | 21827289 | 21871822 | - | transcribed_unitary_pseudogene |
| ENSG00000074410.14_5 | CA12 | 1 | 1 | -0.031 | 9.451 | chr15 | 63613577 | 63674045 | - | protein_coding |
| ENSG00000181038.13_5 | METTL23 | 0.9999995 | 1 | -0.031 | 4.432 | chr17 | 74722912 | 74730018 | + | protein_coding |
| ENSG00000156304.15_5 | SCAF4 | 0.99999938 | 1 | -0.031 | 4.513 | chr21 | 33043313 | 33104431 | - | protein_coding |
| ENSG00000107581.13_4 | EIF3A | 1 | 1 | -0.032 | 8.241 | chr10 | 120793182 | 120840335 | - | protein_coding |
| ENSG00000168003.16_4 | SLC3A2 | 0.99999733 | 1 | -0.032 | 8.914 | chr11 | 62623574 | 62656347 | + | protein_coding |
| ENSG00000085382.12_5 | HACE1 | 0.99998289 | 1 | -0.032 | 3.413 | chr6 | 105175969 | 105307794 | - | protein_coding |
| ENSG00000175600.15_4 | SUGCT | 0.97310784 | 1 | -0.032 | -0.073 | chr7 | 40174576 | 40900362 | + | protein_coding |
| ENSG00000179546.4_3 | HTR1D | 0.99999616 | 1 | -0.032 | 3.972 | chr1 | 23518388 | 23521222 | - | protein_coding |
| ENSG00000082068.9_3 | WDR70 | 0.99999987 | 1 | -0.032 | 4.785 | chr5 | 37379387 | 37753537 | + | protein_coding |
| ENSG00000189343.7_4 | RPS2P46 | 0.99974564 | 1 | -0.032 | 2.925 | chr17 | 19349283 | 19350165 | - | processed_pseudogene |
| ENSG00000160298.17_7 | C21orf58 | 0.99999879 | 1 | -0.032 | 4.354 | chr21 | 47720095 | 47743789 | - | protein_coding |
| ENSG00000075624.16_7 | ACTB | 1 | 1 | -0.032 | 11.295 | chr7 | 5566779 | 5603415 | - | protein_coding |
| ENSG00000143612.20_4 | C1orf43 | 1 | 1 | -0.032 | 7.358 | chr1 | 154179172 | 154193113 | - | protein_coding |
| ENSG00000160957.13_5 | RECQL4 | 0.99999239 | 1 | -0.032 | 5.913 | chr8 | 145736671 | 145743229 | - | protein_coding |
| ENSG00000276259.1_6 | AC009118.3 | 0.96656292 | 1 | -0.032 | -0.347 | chr16 | 58556874 | 58557746 | + | lncRNA |
| ENSG00000028310.18_5 | BRD9 | 0.9999999 | 1 | -0.032 | 5.169 | chr5 | 850406 | 892916 | - | protein_coding |
| ENSG00000179115.11_4 | FARSA | 0.99999996 | 1 | -0.033 | 5.725 | chr19 | 13033293 | 13044851 | - | protein_coding |
| ENSG00000153130.17_7 | SCOC | 0.99999992 | 1 | -0.033 | 5.837 | chr4 | 141178440 | 141306880 | + | protein_coding |
| ENSG00000278041.1_6 | AL133325.3 | 0.97145512 | 1 | -0.033 | -0.159 | chr20 | 21479899 | 21483572 | - | lncRNA |
| ENSG00000267793.1_5 | AC009977.1 | 0.96079987 | 1 | -0.033 | -0.798 | chrY | 21760074 | 21760643 | + | transcribed_processed_pseudogene |
| ENSG00000172239.14_8 | PAIP1 | 1 | 1 | -0.033 | 6.695 | chr5 | 43526369 | 43557860 | - | protein_coding |
| ENSG00000125775.15_6 | SDCBP2 | 0.96777218 | 1 | -0.033 | -0.602 | chr20 | 1290553 | 1309783 | - | protein_coding |
| ENSG00000092847.12_5 | AGO1 | 0.99999999 | 1 | -0.033 | 5.565 | chr1 | 36335409 | 36396133 | + | protein_coding |
| ENSG00000169230.10_3 | PRELID1 | 0.99999997 | 1 | -0.033 | 5.361 | chr5 | 176730800 | 176733950 | + | protein_coding |
| ENSG00000168137.17_7 | SETD5 | 1 | 1 | -0.033 | 6.953 | chr3 | 9439299 | 9520924 | + | protein_coding |
| ENSG00000217555.12_4 | CKLF | 0.95934013 | 1 | -0.033 | -0.699 | chr16 | 66586466 | 66600154 | + | protein_coding |
| ENSG00000170881.4_3 | RNF139 | 0.99999993 | 1 | -0.033 | 5.681 | chr8 | 125486979 | 125500155 | + | protein_coding |
| ENSG00000147130.14_5 | ZMYM3 | 0.99999999 | 1 | -0.034 | 6.061 | chrX | 70459474 | 70474996 | - | protein_coding |
| ENSG00000099800.8_5 | TIMM13 | 0.99999962 | 1 | -0.034 | 4.804 | chr19 | 2425623 | 2427584 | - | protein_coding |
| ENSG00000149269.9_6 | PAK1 | 0.99999998 | 1 | -0.034 | 5.409 | chr11 | 77032752 | 77185680 | - | protein_coding |
| ENSG00000227671.4_6 | AL390728.4 | 0.99989875 | 1 | -0.034 | 4.175 | chr1 | 247353153 | 247374158 | - | transcribed_unprocessed_pseudogene |
| ENSG00000100722.20_7 | ZC3H14 | 1 | 1 | -0.034 | 6.971 | chr14 | 89029314 | 89093940 | + | protein_coding |
| ENSG00000149541.10_4 | B3GAT3 | 0.99999942 | 1 | -0.034 | 4.561 | chr11 | 62382768 | 62389626 | - | protein_coding |
| ENSG00000165282.14_6 | PIGO | 0.99999999 | 1 | -0.034 | 5.685 | chr9 | 35088685 | 35096598 | - | protein_coding |
| ENSG00000106245.11_3 | BUD31 | 0.99999973 | 1 | -0.034 | 4.720 | chr7 | 99006264 | 99017239 | + | protein_coding |
| ENSG00000260279.4_7 | AC137932.1 | 0.99728635 | 1 | -0.034 | 1.921 | chr16 | 89362536 | 89364725 | + | lncRNA |
| ENSG00000135476.12_4 | ESPL1 | 1 | 1 | -0.034 | 6.145 | chr12 | 53662083 | 53687422 | + | protein_coding |
| ENSG00000129055.12_3 | ANAPC13 | 0.99999965 | 1 | -0.034 | 4.874 | chr3 | 134196548 | 134205558 | - | protein_coding |
| ENSG00000186660.15_4 | ZFP91 | 1 | 1 | -0.034 | 6.625 | chr11 | 58346536 | 58389023 | + | protein_coding |
| ENSG00000108588.14_7 | CCDC47 | 0.99999997 | 1 | -0.034 | 6.711 | chr17 | 61822615 | 61853711 | - | protein_coding |
| ENSG00000170291.15_5 | ELP5 | 0.99999975 | 1 | -0.034 | 4.716 | chr17 | 7154735 | 7163259 | + | protein_coding |
| ENSG00000277258.5_5 | PCGF2 | 0.99992245 | 1 | -0.034 | 3.063 | chr17 | 36890151 | 36906070 | - | protein_coding |
| ENSG00000260081.2_5 | AF274858.1 | 0.9596271 | 1 | -0.034 | -0.639 | chrX | 152864770 | 152879392 | + | transcribed_processed_pseudogene |

| | | | | | | | | | | |
|---|---|---|---|---|---|---|---|---|---|---|
| ENSG00000274922.1_7 | AL139384.1 | 0.960059 | 1 | -0.035 | -0.405 | chr13 | 113341083 | 113344129 | - | lncRNA |
| ENSG00000287837.1_2 | AL590064.1 | 0.97939839 | 1 | -0.035 | 0.470 | chr13 | 41448821 | 41451997 | - | lncRNA |
| ENSG00000140396.13_6 | NCOA2 | 0.99999999 | 1 | -0.035 | 6.253 | chr8 | 71022017 | 71316043 | - | protein_coding |
| ENSG00000131043.12_4 | AAR2 | 0.99999986 | 1 | -0.035 | 5.255 | chr20 | 34824381 | 34858840 | + | protein_coding |
| ENSG00000205918.9_3 | PDPK2P | 0.95877725 | 1 | -0.035 | -0.910 | chr16 | 2666122 | 2693297 | - | transcribed_unprocessed_pseudogene |
| ENSG00000161057.12_5 | PSMC2 | 1 | 1 | -0.035 | 6.850 | chr7 | 102984701 | 103009842 | + | protein_coding |
| ENSG00000116213.16_4 | WRAP73 | 0.99999366 | 1 | -0.035 | 3.963 | chr1 | 3547331 | 3569325 | - | protein_coding |
| ENSG00000205302.7_3 | SNX2 | 0.99999981 | 1 | -0.035 | 5.852 | chr5 | 122110774 | 122170238 | + | protein_coding |
| ENSG00000174574.16_4 | AKIRIN1 | 0.99999999 | 1 | -0.035 | 5.843 | chr1 | 39456948 | 39471731 | + | protein_coding |
| ENSG00000152795.17_7 | HNRNPDL | 1 | 1 | -0.035 | 7.860 | chr4 | 83343717 | 83351561 | - | protein_coding |
| ENSG00000091844.8_5 | RGS17 | 0.99999968 | 1 | -0.035 | 4.540 | chr6 | 153325594 | 153452417 | - | protein_coding |
| ENSG00000269997.1_7 | AC068790.3 | 0.96491156 | 1 | -0.035 | -0.075 | chr12 | 124450624 | 124451176 | - | lncRNA |
| ENSG00000187531.14_6 | SIRT7 | 0.99999965 | 1 | -0.035 | 4.608 | chr17 | 79869815 | 79879199 | - | protein_coding |
| ENSG00000109536.12_3 | FRG1 | 0.99999958 | 1 | -0.036 | 5.112 | chr4 | 190862025 | 190884357 | + | protein_coding |
| ENSG00000196152.10_4 | ZNF79 | 0.99999434 | 1 | -0.036 | 3.750 | chr9 | 130186653 | 130207651 | + | protein_coding |
| ENSG00000225205.5_7 | AC078883.1 | 0.95920477 | 1 | -0.036 | -0.295 | chr2 | 173345568 | 173421324 | - | lncRNA |
| ENSG00000011304.20_7 | PTBP1 | 0.99999999 | 1 | -0.036 | 8.125 | chr19 | 797075 | 812327 | + | protein_coding |
| ENSG00000163875.15_4 | MEAF6 | 0.9999996 | 1 | -0.036 | 4.769 | chr1 | 37958176 | 37980375 | - | protein_coding |
| ENSG00000176953.12_4 | NFATC2IP | 0.99999996 | 1 | -0.036 | 5.219 | chr16 | 28962128 | 28978418 | + | protein_coding |
| ENSG00000074071.14_4 | MRPS34 | 0.99988428 | 1 | -0.036 | 6.132 | chr16 | 1821891 | 1823156 | - | protein_coding |
| ENSG00000022277.12_4 | RTF2 | 0.99999999 | 1 | -0.036 | 5.674 | chr20 | 55043641 | 55094505 | + | protein_coding |
| ENSG00000156398.13_5 | SFXN2 | 0.99996728 | 1 | -0.036 | 3.346 | chr10 | 104474295 | 104503249 | + | protein_coding |
| ENSG00000198315.11_3 | ZKSCAN8 | 0.99999979 | 1 | -0.036 | 5.003 | chr6 | 28109661 | 28127238 | + | protein_coding |
| ENSG00000221983.8_5 | UBA52 | 1 | 1 | -0.037 | 7.109 | chr19 | 18682540 | 18688360 | + | protein_coding |
| ENSG00000122008.15_3 | POLK | 0.9999999 | 1 | -0.037 | 5.311 | chr5 | 74807581 | 74896969 | + | protein_coding |
| ENSG00000140961.14_5 | OSGIN1 | 0.99999996 | 1 | -0.037 | 5.494 | chr16 | 83964916 | 83999937 | + | protein_coding |
| ENSG00000137200.13_3 | CMTR1 | 0.99999995 | 1 | -0.037 | 5.213 | chr6 | 37400995 | 37450603 | + | protein_coding |
| ENSG00000128602.11_5 | SMO | 0.99999981 | 1 | -0.037 | 4.937 | chr7 | 128828474 | 128853386 | + | protein_coding |
| ENSG00000198642.7_4 | KLHL9 | 0.99999996 | 1 | -0.037 | 5.643 | chr9 | 21329664 | 21335403 | - | protein_coding |
| ENSG00000280327.1_6 | Z97633.1 | 0.95780594 | 1 | -0.037 | -0.512 | chr16 | 1626717 | 1628372 | - | TEC |
| ENSG00000164061.5_4 | BSN | 0.98972689 | 1 | -0.037 | 1.253 | chr3 | 49591910 | 49708982 | + | protein_coding |
| ENSG00000225345.3_4 | SNX18P3 | 0.99949201 | 1 | -0.037 | 2.274 | chr9 | 38566257 | 38568208 | - | transcribed_unprocessed_pseudogene |
| ENSG00000184349.13_5 | EFNA5 | 0.9999998 | 1 | -0.037 | 5.233 | chr5 | 106712590 | 107006638 | - | protein_coding |
| ENSG00000197780.10_4 | TAF13 | 0.99976166 | 1 | -0.037 | 2.892 | chr1 | 109605108 | 109618625 | - | protein_coding |
| ENSG00000150316.12_4 | CWC15 | 0.99999965 | 1 | -0.037 | 4.820 | chr11 | 94695785 | 94706750 | - | protein_coding |
| ENSG00000171853.16_6 | TRAPPC12 | 0.99999998 | 1 | -0.037 | 5.677 | chr2 | 3383446 | 3488865 | + | protein_coding |
| ENSG00000196712.18_6 | NF1 | 1 | 1 | -0.037 | 7.370 | chr17 | 29421945 | 29709134 | + | protein_coding |
| ENSG00000114388.14_5 | NPRL2 | 0.99999005 | 1 | -0.037 | 3.908 | chr3 | 50384761 | 50388257 | - | protein_coding |
| ENSG00000175697.10_5 | GPR156 | 0.96867428 | 1 | -0.037 | -0.076 | chr3 | 119883492 | 120003941 | - | protein_coding |
| ENSG00000141012.13_5 | GALNS | 0.99999961 | 1 | -0.038 | 5.071 | chr16 | 88880142 | 88923378 | - | protein_coding |
| ENSG00000107281.10_5 | NPDC1 | 0.99873036 | 1 | -0.038 | 4.429 | chr9 | 139933915 | 139940631 | - | protein_coding |
| ENSG00000022556.16_7 | NLRP2 | 1 | 1 | -0.038 | 6.501 | chr19 | 55464498 | 55512510 | + | protein_coding |
| ENSG00000099341.11_6 | PSMD8 | 1 | 1 | -0.038 | 7.926 | chr19 | 38865176 | 38874464 | + | protein_coding |
| ENSG00000258949.1_7 | AL049870.3 | 0.95424378 | 1 | -0.038 | -0.799 | chr14 | 45368111 | 45381066 | - | lncRNA |
| ENSG00000187595.16_5 | ZNF385C | 0.9733211 | 1 | -0.038 | -0.095 | chr17 | 40177594 | 40250497 | - | protein_coding |
| ENSG00000163913.11_7 | IFT122 | 0.99999975 | 1 | -0.038 | 4.904 | chr3 | 129158879 | 129239350 | + | protein_coding |
| ENSG00000264364.3_3 | DYNLL2 | 0.99999999 | 1 | -0.038 | 5.720 | chr17 | 56160780 | 56172903 | + | protein_coding |
| ENSG00000153558.16_5 | FBXL2 | 0.99999895 | 1 | -0.038 | 4.218 | chr3 | 33318517 | 33445154 | + | protein_coding |
| ENSG00000184831.14_6 | APOO | 0.99996485 | 1 | -0.038 | 3.665 | chrX | 23851470 | 23926055 | - | protein_coding |
| ENSG00000171223.6_3 | JUNB | 0.99997854 | 1 | -0.038 | 4.963 | chr19 | 12902300 | 12904129 | + | protein_coding |
| ENSG00000269427.1_7 | AC024075.3 | 0.95766147 | 1 | -0.038 | -0.461 | chr19 | 16741554 | 16754753 | + | lncRNA |
| ENSG00000240682.9_4 | ISY1 | 0.9998037 | 1 | -0.038 | 2.843 | chr3 | 128846258 | 128880136 | - | protein_coding |
| ENSG00000234814.8_5 | SVIL2P | 0.95568876 | 1 | -0.038 | -0.668 | chr10 | 30944418 | 31006195 | + | transcribed_unprocessed_pseudogene |
| ENSG00000148842.18_3 | CNNM2 | 0.99999836 | 1 | -0.038 | 4.202 | chr10 | 104678051 | 104849979 | + | protein_coding |
| ENSG00000044574.8_4 | HSPA5 | 1 | 1 | -0.038 | 7.146 | chr9 | 127997132 | 128003622 | - | protein_coding |
| ENSG00000282021.1_6 | AC100810.3 | 0.99657936 | 1 | -0.038 | 1.761 | chr8 | 1706374 | 1708613 | - | lncRNA |
| ENSG00000230373.8_5 | GOLGA6L5P | 0.96777218 | 1 | -0.038 | -0.603 | chr15 | 85051116 | 85060045 | - | transcribed_unprocessed_pseudogene |
| ENSG00000112874.10_4 | NUDT12 | 0.99975346 | 1 | -0.038 | 2.766 | chr5 | 102884556 | 102898491 | - | protein_coding |
| ENSG00000155858.6_3 | LSM11 | 0.99998635 | 1 | -0.038 | 3.635 | chr5 | 157170720 | 157187717 | + | protein_coding |

| | | | | | | | | | | |
|---|---|---|---|---|---|---|---|---|---|---|
| ENSG00000225746.12_9 | MEG8 | 0.99992207 | 1 | -0.039 | 3.712 | chr14 | 101361107 | 101505196 | + | lncRNA |
| ENSG00000242262.1_5 | AC092597.1 | 0.95373692 | 1 | -0.039 | -0.809 | chr4 | 47708389 | 47709004 | + | processed_pseudogene |
| ENSG00000154079.6_3 | SDHAF4 | 0.99920966 | 1 | -0.039 | 2.292 | chr6 | 71276620 | 71299272 | + | protein_coding |
| ENSG00000164828.18_7 | SUN1 | 1 | 1 | -0.039 | 6.276 | chr7 | 856252 | 936072 | + | protein_coding |
| ENSG00000119231.11_7 | SENP5 | 0.99999997 | 1 | -0.039 | 5.601 | chr3 | 196594727 | 196661585 | + | protein_coding |
| ENSG00000071073.13_5 | MGAT4A | 0.9999705 | 1 | -0.039 | 3.535 | chr2 | 99235569 | 99347595 | - | protein_coding |
| ENSG00000150907.10_6 | FOXO1 | 0.99972677 | 1 | -0.039 | 2.561 | chr13 | 41129804 | 41240778 | - | protein_coding |
| ENSG00000167220.12_8 | HDHD2 | 0.98328122 | 1 | -0.039 | 0.753 | chr18 | 44633779 | 44676871 | - | protein_coding |
| ENSG00000236480.1_4 | PKMP1 | 0.95177999 | 1 | -0.039 | -0.494 | chr1 | 115078616 | 115080461 | + | processed_pseudogene |
| ENSG00000084093.17_4 | REST | 1 | 1 | -0.039 | 6.291 | chr4 | 57774042 | 57832844 | + | protein_coding |
| ENSG00000105372.8_6 | RPS19 | 1 | 1 | -0.039 | 10.278 | chr19 | 42364325 | 42376993 | + | protein_coding |
| ENSG00000259098.1_5 | AC025884.2 | 0.95264828 | 1 | -0.039 | -0.768 | chr15 | 22546089 | 22546799 | + | processed_pseudogene |
| ENSG00000188493.15_8 | C19orf54 | 0.99992398 | 1 | -0.039 | 2.965 | chr19 | 41246761 | 41257458 | - | protein_coding |
| ENSG00000141013.17_4 | GAS8 | 0.99999878 | 1 | -0.040 | 4.275 | chr16 | 90086037 | 90111383 | + | protein_coding |
| ENSG00000129197.14_2 | RPAIN | 0.99999997 | 1 | -0.040 | 5.400 | chr17 | 5322961 | 5336196 | + | protein_coding |
| ENSG00000276805.2_5 | AL133216.2 | 0.97780523 | 1 | -0.040 | 0.550 | chr10 | 38692117 | 38741081 | + | transcribed_unprocessed_pseudogene |
| ENSG00000174243.10_5 | DDX23 | 1 | 1 | -0.040 | 7.682 | chr12 | 49223539 | 49246625 | - | protein_coding |
| ENSG00000179477.11_6 | ALOX12B | 0.95148256 | 1 | -0.040 | -0.670 | chr17 | 7975954 | 7991034 | - | protein_coding |
| ENSG00000135924.15_5 | DNAJB2 | 0.99999985 | 1 | -0.040 | 4.811 | chr2 | 220143989 | 220151622 | + | protein_coding |
| ENSG00000164296.7_4 | TIGD6 | 0.99986829 | 1 | -0.040 | 2.973 | chr5 | 149372681 | 149380217 | - | protein_coding |
| ENSG00000092148.13_7 | HECTD1 | 1 | 1 | -0.040 | 8.194 | chr14 | 31569318 | 31677010 | - | protein_coding |
| ENSG00000137288.10_6 | UQCC2 | 0.99999945 | 1 | -0.040 | 4.708 | chr6 | 33662070 | 33679504 | - | protein_coding |
| ENSG00000066044.15_5 | ELAVL1 | 1 | 1 | -0.040 | 6.214 | chr19 | 8023457 | 8070543 | - | protein_coding |
| ENSG00000130559.19_4 | CAMSAP1 | 0.99999999 | 1 | -0.040 | 6.266 | chr9 | 138700333 | 138799392 | - | protein_coding |
| ENSG00000129696.12_6 | TTI2 | 0.99996964 | 1 | -0.041 | 3.363 | chr8 | 33330904 | 33371119 | - | protein_coding |
| ENSG00000160801.14_6 | PTH1R | 0.97025356 | 1 | -0.041 | -0.162 | chr3 | 46919211 | 46945289 | + | protein_coding |
| ENSG00000025800.14_4 | KPNA6 | 1 | 1 | -0.041 | 6.815 | chr1 | 32573657 | 32642164 | + | protein_coding |
| ENSG00000132819.17_3 | RBM38 | 0.99991161 | 1 | -0.041 | 3.682 | chr20 | 55966452 | 55984389 | + | protein_coding |
| ENSG00000120137.7_5 | PANK3 | 1 | 1 | -0.041 | 7.549 | chr5 | 167975500 | 168006373 | - | protein_coding |
| ENSG00000054148.18_4 | PHPT1 | 0.99999997 | 1 | -0.041 | 5.855 | chr9 | 139743176 | 139745479 | + | protein_coding |
| ENSG00000156508.18_6 | EEF1A1 | 1 | 1 | -0.041 | 12.195 | chr6 | 74225473 | 74233520 | - | protein_coding |
| ENSG00000166897.15_5 | ELFN2 | 0.99998913 | 1 | -0.041 | 3.805 | chr22 | 37764000 | 37823514 | - | protein_coding |
| ENSG00000005812.11_7 | FBXL3 | 0.9999999 | 1 | -0.041 | 5.005 | chr13 | 77566733 | 77601330 | - | protein_coding |
| ENSG00000029364.12_3 | SLC39A9 | 1 | 1 | -0.042 | 7.365 | chr14 | 69864732 | 69929107 | + | protein_coding |
| ENSG00000125970.12_5 | RALY | 0.99999996 | 1 | -0.042 | 6.602 | chr20 | 32581452 | 32696114 | + | protein_coding |
| ENSG00000102309.15_7 | PIN4 | 0.99999926 | 1 | -0.042 | 4.358 | chrX | 71401203 | 71522776 | + | protein_coding |
| ENSG00000109586.12_5 | GALNT7 | 0.99999999 | 1 | -0.042 | 5.936 | chr4 | 174089962 | 174245118 | + | protein_coding |
| ENSG00000144445.17_5 | KANSL1L | 0.99991956 | 1 | -0.042 | 3.531 | chr2 | 210886145 | 211036133 | - | protein_coding |
| ENSG00000135436.8_6 | FAM186B | 0.94818626 | 1 | -0.042 | -0.496 | chr12 | 49976668 | 49999422 | - | protein_coding |
| ENSG00000261490.1_7 | AC005674.2 | 0.99022452 | 1 | -0.042 | 0.657 | chr4 | 10069713 | 10074643 | - | lncRNA |
| ENSG00000189046.11_6 | ALKBH2 | 0.99999953 | 1 | -0.042 | 4.449 | chr12 | 109525993 | 109531436 | - | protein_coding |
| ENSG00000114062.20_8 | UBE3A | 1 | 1 | -0.042 | 6.610 | chr15 | 25578875 | 25684198 | - | protein_coding |
| ENSG00000044446.12_4 | PHKA2 | 0.99999924 | 1 | -0.042 | 4.828 | chrX | 18910416 | 19002232 | - | protein_coding |
| ENSG00000174628.16_4 | IQCK | 0.9995266 | 1 | -0.042 | 2.398 | chr16 | 19727778 | 19869789 | + | protein_coding |
| ENSG00000113790.11_5 | EHHADH | 0.99978927 | 1 | -0.042 | 2.903 | chr3 | 184908412 | 184999778 | - | protein_coding |
| ENSG00000182541.18_4 | LIMK2 | 0.99979126 | 1 | -0.042 | 3.240 | chr22 | 31608225 | 31676066 | + | protein_coding |
| ENSG00000189212.12_4 | DPY19L2P1 | 0.99968397 | 1 | -0.042 | 2.729 | chr7 | 35119601 | 35225653 | - | transcribed_unprocessed_pseudogene |
| ENSG00000103496.15_5 | STX4 | 0.99999929 | 1 | -0.042 | 4.392 | chr16 | 31044210 | 31054296 | + | protein_coding |
| ENSG00000056586.16_6 | RC3H2 | 1 | 1 | -0.042 | 6.709 | chr9 | 125606835 | 125667638 | - | protein_coding |
| ENSG00000138600.10_6 | SPPL2A | 1 | 1 | -0.042 | 6.991 | chr15 | 50994463 | 51057906 | - | protein_coding |
| ENSG00000286340.1_1 | AL356776.2 | 0.96623962 | 1 | -0.042 | -0.404 | chr6 | 79787558 | 79791088 | + | lncRNA |
| ENSG00000153234.14_6 | NR4A2 | 0.99777146 | 1 | -0.042 | 2.026 | chr2 | 157180949 | 157198860 | - | protein_coding |
| ENSG00000164252.13_4 | AGGF1 | 0.99999945 | 1 | -0.042 | 4.493 | chr5 | 76325076 | 76361059 | + | protein_coding |
| ENSG00000225663.8_8 | MCRIP1 | 0.99999714 | 1 | -0.042 | 4.773 | chr17 | 79780237 | 79791178 | - | protein_coding |
| ENSG00000086065.14_3 | CHMP5 | 0.99999999 | 1 | -0.043 | 6.474 | chr9 | 33264877 | 33282068 | + | protein_coding |
| ENSG00000173473.11_5 | SMARCC1 | 1 | 1 | -0.043 | 6.547 | chr3 | 47626759 | 47823596 | - | protein_coding |
| ENSG00000117335.19_4 | CD46 | 1 | 1 | -0.043 | 7.542 | chr1 | 207925402 | 207968858 | + | protein_coding |
| ENSG00000115616.3_3 | SLC9A2 | 0.99998971 | 1 | -0.043 | 3.906 | chr2 | 103236012 | 103327814 | + | protein_coding |

| | | | | | | | | | | |
|---|---|---|---|---|---|---|---|---|---|---|
| ENSG00000119401.10_3 | TRIM32 | 0.99999992 | 1 | -0.043 | 5.117 | chr9 | 119449581 | 119463579 | + | protein_coding |
| ENSG00000197323.12_5 | TRIM33 | 0.99999984 | 1 | -0.043 | 6.033 | chr1 | 114935412 | 115053824 | - | protein_coding |
| ENSG00000251127.2_7 | AC091173.1 | 0.94830385 | 1 | -0.043 | -0.603 | chr8 | 62671653 | 62678528 | - | lncRNA |
| ENSG00000168614.13 | NBPF9 | 0.99999672 | 1 | -0.043 | 4.105 | chr1 | 144811744 | 144830413 | + | protein_coding |
| ENSG00000227741.1_7 | AL121987.2 | 0.95396379 | 1 | -0.043 | -0.358 | chr1 | 160171989 | 160178659 | - | lncRNA |
| ENSG00000119559.16_7 | C19orf25 | 0.99983251 | 1 | -0.043 | 3.681 | chr19 | 1461142 | 1479555 | - | protein_coding |
| ENSG00000155827.12_3 | RNF20 | 0.99999999 | 1 | -0.043 | 6.125 | chr9 | 104296135 | 104325626 | + | protein_coding |
| ENSG00000100767.16_5 | PAPLN | 0.99999339 | 1 | -0.043 | 4.024 | chr14 | 73704205 | 73741348 | + | protein_coding |
| ENSG00000164967.10_4 | RPP25L | 0.99991222 | 1 | -0.043 | 3.847 | chr9 | 34610483 | 34612101 | - | protein_coding |
| ENSG00000164615.5_3 | CAMLG | 0.99999972 | 1 | -0.043 | 4.706 | chr5 | 134074238 | 134087847 | + | protein_coding |
| ENSG00000107679.14_3 | PLEKHA1 | 0.99999999 | 1 | -0.043 | 5.999 | chr10 | 124134212 | 124202118 | + | protein_coding |
| ENSG00000259378.1_4 | DCAF13P3 | 0.95118611 | 1 | -0.043 | -0.523 | chr15 | 51236860 | 51238193 | + | processed_pseudogene |
| ENSG00000263126.1_7 | AC040162.3 | 0.94534114 | 1 | -0.043 | -1.031 | chr16 | 67916364 | 67920270 | + | lncRNA |
| ENSG00000249279.6_6 | LINC02057 | 0.98251264 | 1 | -0.043 | 0.520 | chr5 | 60497131 | 60600638 | - | lncRNA |
| ENSG00000122497.13 | NBPF14 | 0.99448342 | 1 | -0.044 | 2.034 | chr1 | 148003642 | 148025863 | - | protein_coding |
| ENSG00000112308.13_4 | C6orf62 | 1 | 1 | -0.044 | 6.528 | chr6 | 24705089 | 24720226 | - | protein_coding |
| ENSG00000275004.4_5 | ZNF280B | 0.99996762 | 1 | -0.044 | 3.312 | chr22 | 22838767 | 22863060 | - | protein_coding |
| ENSG00000124783.14_6 | SSR1 | 0.99999999 | 1 | -0.044 | 6.724 | chr6 | 7268539 | 7347679 | - | protein_coding |
| ENSG00000235560.4_7 | AC002310.1 | 0.94588649 | 1 | -0.044 | -0.739 | chr16 | 30546073 | 30548470 | + | lncRNA |
| ENSG00000107625.13_3 | DDX50 | 0.99999997 | 1 | -0.044 | 5.787 | chr10 | 70661042 | 70706603 | + | protein_coding |
| ENSG00000131475.7_3 | VPS25 | 1 | 1 | -0.044 | 6.034 | chr17 | 40925467 | 40931617 | + | protein_coding |
| ENSG00000176148.16_5 | TCP11L1 | 0.99999999 | 1 | -0.044 | 5.636 | chr11 | 33060963 | 33127489 | + | protein_coding |
| ENSG00000011405.13_4 | PIK3C2A | 1 | 1 | -0.044 | 7.118 | chr11 | 17099277 | 17229530 | - | protein_coding |
| ENSG00000234769.7_3 | WASH4P | 0.95869087 | 1 | -0.044 | -0.382 | chr16 | 64381 | 68068 | - | unprocessed_pseudogene |
| ENSG00000166526.18_8 | ZNF3 | 0.99999995 | 1 | -0.044 | 5.183 | chr7 | 99661656 | 99680171 | - | protein_coding |
| ENSG00000261659.2_6 | Z92544.2 | 0.96623962 | 1 | -0.044 | -0.406 | chr16 | 739001 | 742554 | + | lncRNA |
| ENSG00000166997.8_4 | CNPY4 | 0.99994961 | 1 | -0.044 | 3.231 | chr7 | 99717257 | 99723131 | + | protein_coding |
| ENSG00000178425.14_4 | NT5DC1 | 0.99999983 | 1 | -0.044 | 4.840 | chr6 | 116422014 | 116570660 | + | protein_coding |
| ENSG00000094631.21_7 | HDAC6 | 0.99999998 | 1 | -0.044 | 5.844 | chrX | 48659784 | 48683392 | + | protein_coding |
| ENSG00000197948.11_4 | FCHSD1 | 0.99997527 | 1 | -0.044 | 3.877 | chr5 | 141018869 | 141030985 | - | protein_coding |
| ENSG00000120333.4_4 | MRPS14 | 0.9999993 | 1 | -0.044 | 4.407 | chr1 | 174979925 | 174992561 | - | protein_coding |
| ENSG00000091947.10_5 | TMEM101 | 0.9999998 | 1 | -0.044 | 4.950 | chr17 | 42088556 | 42101314 | - | protein_coding |
| ENSG00000136197.12_6 | C7orf25 | 0.95802915 | 1 | -0.044 | -0.012 | chr7 | 42948325 | 42951904 | - | protein_coding |
| ENSG00000137073.22_4 | UBAP2 | 0.99999999 | 1 | -0.044 | 5.685 | chr9 | 33921691 | 34048947 | - | protein_coding |
| ENSG00000275945.1_4 | EIF3FP1 | 0.99189064 | 1 | -0.044 | 1.250 | chr21 | 11180920 | 11181725 | - | processed_pseudogene |
| ENSG00000161267.12_6 | BDH1 | 0.99886164 | 1 | -0.044 | 2.132 | chr3 | 197236654 | 197300194 | - | protein_coding |
| ENSG00000175283.8_3 | DOLK | 0.99998901 | 1 | -0.044 | 4.822 | chr9 | 131707809 | 131709882 | - | protein_coding |
| ENSG00000143771.12_4 | CNIH4 | 0.99999997 | 1 | -0.045 | 5.951 | chr1 | 224544560 | 224567161 | + | protein_coding |
| ENSG00000025770.19_5 | NCAPH2 | 0.99999914 | 1 | -0.045 | 5.104 | chr22 | 50946653 | 50963209 | + | protein_coding |
| ENSG00000103510.20_6 | KAT8 | 0.99999982 | 1 | -0.045 | 5.139 | chr16 | 31125810 | 31142714 | + | protein_coding |
| ENSG00000279331.1_4 | RBM12B-AS1 | 0.98393482 | 1 | -0.045 | 0.912 | chr8 | 94752349 | 94753001 | + | TEC |
| ENSG00000214753.4_6 | HNRNPUL2 | 0.99999846 | 1 | -0.045 | 4.228 | chr11 | 62480102 | 62494929 | - | protein_coding |
| ENSG00000196366.3_3 | C9orf163 | 0.97553343 | 1 | -0.045 | 0.598 | chr9 | 139377947 | 139380519 | + | lncRNA |
| ENSG00000159023.21_8 | EPB41 | 0.99999925 | 1 | -0.045 | 4.445 | chr1 | 29213603 | 29446558 | + | protein_coding |
| ENSG00000129749.3_5 | CHRNA10 | 0.96178013 | 1 | -0.045 | 0.284 | chr11 | 3686817 | 3692614 | - | protein_coding |
| ENSG00000129422.14_6 | MTUS1 | 1 | 1 | -0.045 | 8.923 | chr8 | 17501304 | 17658426 | - | protein_coding |
| ENSG00000274211.6_6 | SOCS7 | 0.99999995 | 1 | -0.045 | 5.664 | chr17 | 36507727 | 36561846 | + | protein_coding |
| ENSG00000170684.9_5 | ZNF296 | 0.96684125 | 1 | -0.045 | 0.245 | chr19 | 45574758 | 45579845 | - | protein_coding |
| ENSG00000119862.13_5 | LGALSL | 0.99999353 | 1 | -0.045 | 3.977 | chr2 | 64681103 | 64688515 | + | protein_coding |
| ENSG00000116273.6_3 | PHF13 | 0.99999043 | 1 | -0.045 | 4.025 | chr1 | 6673791 | 6684090 | + | protein_coding |
| ENSG00000169375.16_7 | SIN3A | 1 | 1 | -0.045 | 6.630 | chr15 | 75661720 | 75748183 | - | protein_coding |
| ENSG00000280198.1_1 | AC087163.3 | 0.94734799 | 1 | -0.046 | -0.548 | chr17 | 17929386 | 17930085 | - | TEC |
| ENSG00000149600.11_4 | COMMD7 | 0.99999566 | 1 | -0.046 | 4.009 | chr20 | 31290493 | 31331803 | - | protein_coding |
| ENSG00000171456.20_8 | ASXL1 | 1 | 1 | -0.046 | 6.772 | chr20 | 30946133 | 31027122 | + | protein_coding |
| ENSG00000230715.3_5 | AC018638.2 | 0.99371981 | 1 | -0.046 | 1.732 | chr7 | 128292895 | 128293297 | + | processed_pseudogene |
| ENSG00000160783.19_6 | PMF1 | 0.99985347 | 1 | -0.046 | 3.613 | chr1 | 156182784 | 156209833 | + | protein_coding |
| ENSG00000108298.11_5 | RPL19 | 1 | 1 | -0.046 | 10.221 | chr17 | 37356536 | 37360985 | + | protein_coding |
| ENSG00000099942.13_3 | CRKL | 0.99999999 | 1 | -0.046 | 6.119 | chr22 | 21271695 | 21308035 | + | protein_coding |

| | | | | | | | | | | |
|---|---|---|---|---|---|---|---|---|---|---|
| ENSG00000225648.5_4 | SBDSP1 | 0.99999627 | 1 | -0.046 | 4.443 | chr7 | 72300004 | 72307280 | + | transcribed_unprocessed_pseudogene |
| ENSG00000048052.21_6 | HDAC9 | 0.96750178 | 1 | -0.046 | 0.461 | chr7 | 18126572 | 19042039 | + | protein_coding |
| ENSG00000116580.18_3 | GON4L | 0.99999999 | 1 | -0.046 | 5.844 | chr1 | 155719453 | 155829191 | - | protein_coding |
| ENSG00000179021.10_3 | C3orf38 | 0.99987307 | 1 | -0.046 | 5.108 | chr3 | 88199109 | 88217879 | + | protein_coding |
| ENSG00000184277.13_5 | TM2D3 | 0.99999914 | 1 | -0.046 | 5.012 | chr15 | 102161647 | 102192594 | - | protein_coding |
| ENSG00000168066.20_5 | SF1 | 1 | 1 | -0.046 | 7.311 | chr11 | 64532078 | 64546258 | - | protein_coding |
| ENSG00000186468.13_5 | RPS23 | 1 | 1 | -0.047 | 6.745 | chr5 | 81569139 | 81574215 | - | protein_coding |
| ENSG00000153006.16_6 | SREK1IP1 | 0.9999961 | 1 | -0.047 | 4.684 | chr5 | 64013975 | 64064518 | - | protein_coding |
| ENSG00000145996.11_4 | CDKAL1 | 0.99999812 | 1 | -0.047 | 4.223 | chr6 | 20534688 | 21232635 | + | protein_coding |
| ENSG00000169087.11_4 | HSPBAP1 | 0.99988181 | 1 | -0.047 | 2.924 | chr3 | 122458846 | 122512678 | - | protein_coding |
| ENSG00000135956.8_3 | TMEM127 | 0.99999999 | 1 | -0.047 | 5.878 | chr2 | 96914254 | 96931732 | - | protein_coding |
| ENSG00000171940.13_5 | ZNF217 | 1 | 1 | -0.047 | 6.393 | chr20 | 52183604 | 52226446 | - | protein_coding |
| ENSG00000171862.11_6 | PTEN | 1 | 1 | -0.047 | 7.053 | chr10 | 89623382 | 89731687 | + | protein_coding |
| ENSG00000119414.11_3 | PPP6C | 0.99999999 | 1 | -0.047 | 6.287 | chr9 | 127908852 | 127952218 | - | protein_coding |
| ENSG00000252835.1 | SCARNA21 | 0.99999994 | 1 | -0.047 | 5.135 | chr17 | 7809440 | 7809578 | + | snoRNA |
| ENSG00000168010.11_4 | ATG16L2 | 0.99990473 | 1 | -0.047 | 3.607 | chr11 | 72525451 | 72554719 | + | protein_coding |
| ENSG00000163909.8_4 | HEYL | 0.99922264 | 1 | -0.047 | 2.536 | chr1 | 40089107 | 40105315 | - | protein_coding |
| ENSG00000255031.5_7 | AP002807.1 | 0.99901129 | 1 | -0.047 | 2.427 | chr11 | 67818207 | 67821229 | + | lncRNA |
| ENSG00000257135.7_7 | ODC1-DT | 0.95872407 | 1 | -0.047 | 0.075 | chr2 | 10588815 | 10597819 | + | lncRNA |
| ENSG00000126768.12_3 | TIMM17B | 0.99999954 | 1 | -0.048 | 5.085 | chrX | 48750730 | 48755426 | - | protein_coding |
| ENSG00000134602.16_3 | STK26 | 0.99999977 | 1 | -0.048 | 4.805 | chrX | 131157330 | 131209971 | + | protein_coding |
| ENSG00000196588.18_8 | MRTFA | 0.99999975 | 1 | -0.048 | 5.531 | chr22 | 40806285 | 41032723 | - | protein_coding |
| ENSG00000189014.6 | FAM35DP | 0.94705485 | 1 | -0.048 | -0.771 | chr10 | 47379727 | 47420712 | + | pseudogene |
| ENSG00000101343.14_4 | CRNKL1 | 0.99999999 | 1 | -0.048 | 6.330 | chr20 | 20015012 | 20036690 | - | protein_coding |
| ENSG00000178741.12_4 | COX5A | 0.99999999 | 1 | -0.048 | 6.390 | chr15 | 75212132 | 75230424 | - | protein_coding |
| ENSG00000260306.1_6 | AC092375.2 | 0.94094844 | 1 | -0.048 | -0.771 | chr16 | 21805416 | 21807080 | - | lncRNA |
| ENSG00000025796.14_4 | SEC63 | 0.99999862 | 1 | -0.048 | 4.422 | chr6 | 108188960 | 108279412 | - | protein_coding |
| ENSG00000235092.6_7 | ID2-AS1 | 0.99371826 | 1 | -0.048 | 1.679 | chr2 | 8806766 | 8821994 | - | lncRNA |
| ENSG00000148331.12_4 | ASB6 | 0.99999832 | 1 | -0.048 | 4.625 | chr9 | 132396883 | 132404448 | - | protein_coding |
| ENSG00000253676.1_2 | TAGLN2P1 | 0.97166798 | 1 | -0.048 | 0.674 | chr8 | 107709655 | 107710241 | - | processed_pseudogene |
| ENSG00000168061.16_6 | SAC3D1 | 0.99999154 | 1 | -0.048 | 4.146 | chr11 | 64808373 | 64812300 | + | protein_coding |
| ENSG00000121766.15_4 | ZCCHC17 | 0.99998355 | 1 | -0.049 | 3.534 | chr1 | 31769829 | 31837800 | + | protein_coding |
| ENSG00000020633.18_3 | RUNX3 | 0.99977736 | 1 | -0.049 | 2.879 | chr1 | 25226002 | 25291612 | - | protein_coding |
| ENSG00000164077.14_6 | MON1A | 0.99965361 | 1 | -0.049 | 2.863 | chr3 | 49944593 | 49967606 | - | protein_coding |
| ENSG00000170624.13_2 | SGCD | 0.99999598 | 1 | -0.049 | 4.304 | chr5 | 155297354 | 156194799 | + | protein_coding |
| ENSG00000175324.10_4 | LSM1 | 0.99996656 | 1 | -0.049 | 3.452 | chr8 | 38020853 | 38034248 | - | protein_coding |
| ENSG00000287584.1_1 | AC004837.4 | 0.94845606 | 1 | -0.049 | -0.551 | chr7 | 39660852 | 39662800 | - | lncRNA |
| ENSG00000104884.15_9 | ERCC2 | 0.99999925 | 1 | -0.049 | 4.470 | chr19 | 45853095 | 45874176 | - | protein_coding |
| ENSG00000079332.15_7 | SAR1A | 0.99999999 | 1 | -0.049 | 6.879 | chr10 | 71907045 | 71930279 | - | protein_coding |
| ENSG00000196367.13_4 | TRRAP | 0.99994221 | 1 | -0.049 | 6.178 | chr7 | 98475556 | 98648454 | + | protein_coding |
| ENSG00000205544.4_5 | TMEM256 | 0.99974978 | 1 | -0.049 | 3.004 | chr17 | 7306294 | 7307416 | - | protein_coding |
| ENSG00000254999.4_3 | BRK1 | 0.99999996 | 1 | -0.049 | 5.823 | chr3 | 10157359 | 10168874 | + | protein_coding |
| ENSG00000259488.2_7 | AC023355.2 | 0.97019041 | 1 | -0.049 | 0.553 | chr15 | 48604550 | 48624053 | - | lncRNA |
| ENSG00000281091.3_6 | AL117327.1 | 0.9507741 | 1 | -0.049 | -0.062 | chrX | 102659705 | 102707682 | + | lncRNA |
| ENSG00000104998.4_3 | IL27RA | 0.99971728 | 1 | -0.049 | 3.012 | chr19 | 14142574 | 14164030 | + | protein_coding |
| ENSG00000140319.11_4 | SRP14 | 0.99999989 | 1 | -0.049 | 5.568 | chr15 | 40327891 | 40331382 | - | protein_coding |
| ENSG00000263990.1_6 | AC004253.1 | 0.94586977 | 1 | -0.049 | -0.263 | chr17 | 30200945 | 30213685 | + | lncRNA |
| ENSG00000226266.6_8 | AC009961.1 | 0.95249072 | 1 | -0.049 | 0.047 | chr2 | 160527219 | 160568946 | - | lncRNA |
| ENSG00000198932.13_5 | GPRASP1 | 0.98818773 | 1 | -0.049 | 1.280 | chrX | 101906020 | 101914011 | + | protein_coding |
| ENSG00000264577.1_9 | AC010761.1 | 0.99975791 | 1 | -0.049 | 2.869 | chr17 | 27048505 | 27049895 | - | lncRNA |
| ENSG00000257557.2_6 | PPP1R12A-AS1 | 0.99543734 | 1 | -0.050 | 1.202 | chr12 | 80328681 | 80336492 | + | lncRNA |
| ENSG00000159063.13_6 | ALG8 | 0.99999998 | 1 | -0.050 | 6.047 | chr11 | 77811982 | 77850699 | - | protein_coding |
| ENSG00000185880.13_6 | TRIM69 | 0.99991123 | 1 | -0.050 | 3.102 | chr15 | 45021186 | 45060027 | + | protein_coding |
| ENSG00000196792.12_6 | STRN3 | 0.99999999 | 1 | -0.050 | 6.073 | chr14 | 31363005 | 31495607 | - | protein_coding |
| ENSG00000215305.10_4 | VPS16 | 0.99999994 | 1 | -0.050 | 5.263 | chr20 | 2821349 | 2847378 | + | protein_coding |
| ENSG00000100478.15_7 | AP4S1 | 0.99999827 | 1 | -0.050 | 4.293 | chr14 | 31494312 | 31600202 | + | protein_coding |
| ENSG00000010361.14_6 | FUZ | 0.99997421 | 1 | -0.050 | 3.410 | chr19 | 50310123 | 50320633 | - | protein_coding |
| ENSG00000155545.19_4 | MIER3 | 0.99999985 | 1 | -0.050 | 5.603 | chr5 | 56215429 | 56267502 | - | protein_coding |
| ENSG00000224713.4_7 | AC025165.1 | 0.94277577 | 1 | -0.050 | -0.405 | chr12 | 58005901 | 58013421 | - | lncRNA |

| | | | | | | | | | | |
|---|---|---|---|---|---|---|---|---|---|---|
| ENSG00000236423.6_5 | LINC01134 | 0.93612676 | 1 | -0.050 | -0.771 | chr1 | 3816916 | 3833789 | + | lncRNA |
| ENSG00000108599.15_5 | AKAP10 | 0.99999999 | 1 | -0.050 | 5.873 | chr17 | 19807615 | 19881656 | - | protein_coding |
| ENSG00000154845.15_3 | PPP4R1 | 1 | 1 | -0.050 | 6.710 | chr18 | 9546789 | 9615238 | - | protein_coding |
| ENSG00000145216.16_6 | FIP1L1 | 0.99993275 | 1 | -0.050 | 3.395 | chr4 | 54243808 | 54327029 | + | protein_coding |
| ENSG00000104450.12_2 | SPAG1 | 0.99999991 | 1 | -0.050 | 5.099 | chr8 | 101170134 | 101271506 | + | protein_coding |
| ENSG00000125351.11_7 | UPF3B | 0.99997276 | 1 | -0.050 | 3.477 | chrX | 118939274 | 118986961 | - | protein_coding |
| ENSG00000241764.4_7 | AC002467.1 | 0.96750473 | 1 | -0.050 | 0.387 | chr7 | 107380444 | 107385026 | - | lncRNA |
| ENSG00000137868.19_6 | STRA6 | 0.99999999 | 1 | -0.050 | 5.971 | chr15 | 74471807 | 74504608 | - | protein_coding |
| ENSG00000171943.7 | SRGAP2C | 0.99901903 | 1 | -0.050 | 2.412 | chr1 | 121107124 | 121129949 | + | pseudogene |
| ENSG00000251417.2_6 | AC145285.2 | 0.93696286 | 1 | -0.051 | -0.641 | chr16 | 28814064 | 28829149 | + | lncRNA |
| ENSG00000065882.16_4 | TBC1D1 | 0.99999997 | 1 | -0.051 | 5.507 | chr4 | 37892705 | 38140796 | + | protein_coding |
| ENSG00000115073.8_4 | ACTR1B | 0.99999987 | 1 | -0.051 | 5.405 | chr2 | 98272402 | 98280507 | - | protein_coding |
| ENSG00000231113.2_6 | AL035587.1 | 0.99477111 | 1 | -0.051 | 1.123 | chr6 | 42908102 | 42916098 | + | lncRNA |
| ENSG00000185278.15_4 | ZBTB37 | 0.99999907 | 1 | -0.051 | 4.868 | chr1 | 173837220 | 173872687 | + | protein_coding |
| ENSG00000112679.14_4 | DUSP22 | 0.99994714 | 1 | -0.051 | 3.463 | chr6 | 291630 | 351355 | + | protein_coding |
| ENSG00000230989.7_3 | HSBP1 | 1 | 1 | -0.051 | 7.022 | chr16 | 83841583 | 83853342 | + | protein_coding |
| ENSG00000197142.10_2 | ACSL5 | 0.94773686 | 1 | -0.051 | -0.167 | chr10 | 114133776 | 114188138 | + | protein_coding |
| ENSG00000148572.16_4 | NRBF2 | 0.99999926 | 1 | -0.051 | 4.566 | chr10 | 64893007 | 64914791 | + | protein_coding |
| ENSG00000173451.7_4 | THAP2 | 0.99964884 | 1 | -0.051 | 2.831 | chr12 | 72058081 | 72074424 | + | protein_coding |
| ENSG00000128791.12_3 | TWSG1 | 0.99999995 | 1 | -0.051 | 5.464 | chr18 | 9334765 | 9402418 | + | protein_coding |
| ENSG00000136478.7_5 | TEX2 | 0.99999994 | 1 | -0.051 | 5.464 | chr17 | 62224587 | 62340661 | - | protein_coding |
| ENSG00000092871.16_6 | RFFL | 0.99999994 | 1 | -0.051 | 5.238 | chr17 | 33333009 | 33416338 | - | protein_coding |
| ENSG00000183929.7_4 | DUSP5P1 | 0.93594863 | 1 | -0.051 | -0.524 | chr1 | 228785988 | 228787126 | + | processed_pseudogene |
| ENSG00000165533.18_6 | TTC8 | 0.99999992 | 1 | -0.051 | 5.201 | chr14 | 89290497 | 89347422 | + | protein_coding |
| ENSG00000250397.2_7 | AP006623.1 | 0.99571553 | 1 | -0.051 | 1.727 | chr11 | 856880 | 859795 | - | lncRNA |
| ENSG00000287884.1_1 | AC005594.1 | 0.93976886 | 1 | -0.051 | -0.135 | chr19 | 4692033 | 4694352 | + | lncRNA |
| ENSG00000169813.16_4 | HNRNPF | 1 | 1 | -0.051 | 8.315 | chr10 | 43881065 | 43904614 | - | protein_coding |
| ENSG00000170515.14_6 | PA2G4 | 1 | 1 | -0.052 | 6.935 | chr12 | 56498321 | 56507694 | + | protein_coding |
| ENSG00000168255.20_7 | POLR2J3 | 0.9564747 | 1 | -0.052 | -0.002 | chr7 | 102178365 | 102213100 | - | protein_coding |
| ENSG00000159723.4_3 | AGRP | 0.93387844 | 1 | -0.052 | -0.803 | chr16 | 67516474 | 67517716 | - | protein_coding |
| ENSG00000114648.12_4 | KLHL18 | 0.99999326 | 1 | -0.052 | 4.147 | chr3 | 47324407 | 47388306 | + | protein_coding |
| ENSG00000223959.8_2 | AFG3L1P | 0.9999991 | 1 | -0.052 | 4.454 | chr16 | 90038994 | 90068569 | + | transcribed_unitary_pseudogene |
| ENSG00000152332.16_3 | UHMK1 | 1 | 1 | -0.052 | 7.719 | chr1 | 162467041 | 162499421 | + | protein_coding |
| ENSG00000148672.9_4 | GLUD1 | 1 | 1 | -0.052 | 7.256 | chr10 | 88809959 | 88854600 | - | protein_coding |
| ENSG00000142794.18_3 | NBPF3 | 0.99999311 | 1 | -0.053 | 3.951 | chr1 | 21766621 | 21811498 | + | protein_coding |
| ENSG00000035687.10_5 | ADSS2 | 0.99999999 | 1 | -0.053 | 6.103 | chr1 | 244571796 | 244615211 | - | protein_coding |
| ENSG00000142396.10_5 | ERVK3-1 | 0.99956173 | 1 | -0.053 | 2.610 | chr19 | 58816685 | 58827029 | + | protein_coding |
| ENSG00000105849.6_4 | TWISTNB | 0.9999999 | 1 | -0.053 | 5.904 | chr7 | 19735084 | 19748660 | - | protein_coding |
| ENSG00000109790.16_3 | KLHL5 | 1 | 1 | -0.053 | 6.817 | chr4 | 39046659 | 39128477 | + | protein_coding |
| ENSG00000173320.12_6 | STOX2 | 0.9999989 | 1 | -0.053 | 4.536 | chr4 | 184718845 | 184944679 | + | protein_coding |
| ENSG00000170903.11_5 | MSANTD4 | 0.99999416 | 1 | -0.053 | 4.476 | chr11 | 105866350 | 105893130 | - | protein_coding |
| ENSG00000225031.1_4 | EIF4BP7 | 0.99485469 | 1 | -0.053 | 2.148 | chrX | 110862905 | 110864717 | - | processed_pseudogene |
| ENSG00000044459.15_4 | CNTLN | 0.9999998 | 1 | -0.053 | 5.009 | chr9 | 17134980 | 17503921 | + | protein_coding |
| ENSG00000025156.13_6 | HSF2 | 0.99993052 | 1 | -0.053 | 3.875 | chr6 | 122720696 | 122754264 | + | protein_coding |
| ENSG00000130177.16_4 | CDC16 | 0.99999994 | 1 | -0.053 | 5.449 | chr13 | 115000362 | 115038198 | + | protein_coding |
| ENSG00000206535.8_3 | LNP1 | 0.99239 | 1 | -0.053 | 1.413 | chr3 | 100120376 | 100175163 | + | protein_coding |
| ENSG00000131373.15_6 | HACL1 | 0.99999912 | 1 | -0.053 | 4.717 | chr3 | 15602206 | 15643359 | - | protein_coding |
| ENSG00000097096.9_4 | SYDE2 | 0.99980409 | 1 | -0.053 | 3.054 | chr1 | 85622572 | 85666699 | - | protein_coding |
| ENSG00000279801.1_7 | AC111170.4 | 0.94058771 | 1 | -0.053 | -0.878 | chr17 | 75440786 | 75443249 | + | TEC |
| ENSG00000138592.14_7 | USP8 | 1 | 1 | -0.054 | 6.714 | chr15 | 50716577 | 50806618 | + | protein_coding |
| ENSG00000164961.16_5 | WASHC5 | 1 | 1 | -0.054 | 6.270 | chr8 | 126036502 | 126104061 | - | protein_coding |
| ENSG00000124198.9_3 | ARFGEF2 | 1 | 1 | -0.054 | 6.972 | chr20 | 47538248 | 47653230 | + | protein_coding |
| ENSG00000204922.5_5 | UQCC3 | 0.99952636 | 1 | -0.054 | 2.861 | chr11 | 62437745 | 62441158 | + | protein_coding |
| ENSG00000167779.9_4 | IGFBP6 | 0.99966547 | 1 | -0.054 | 5.617 | chr12 | 53491220 | 53496129 | + | protein_coding |
| ENSG00000286584.2_2 | AC023494.1 | 0.94620644 | 1 | -0.054 | -0.067 | chr3 | 119385083 | 119389535 | + | lncRNA |
| ENSG00000141646.14_6 | SMAD4 | 0.99999997 | 1 | -0.054 | 5.790 | chr18 | 48554764 | 48611415 | + | protein_coding |
| ENSG00000244005.13_5 | NFS1 | 0.99999956 | 1 | -0.054 | 4.740 | chr20 | 34255974 | 34287277 | - | protein_coding |
| ENSG00000131504.17_5 | DIAPH1 | 0.99999998 | 1 | -0.054 | 6.058 | chr5 | 140894583 | 140998622 | - | protein_coding |
| ENSG00000188786.10_4 | MTF1 | 0.99998945 | 1 | -0.054 | 4.307 | chr1 | 38275246 | 38325264 | - | protein_coding |

| | | | | | | | | | | |
|---|---|---|---|---|---|---|---|---|---|---|
| ENSG00000134470.21_10 | IL15RA | 0.93470361 | 1 | -0.054 | -0.613 | chr10 | 5985602 | 6020150 | - | protein_coding |
| ENSG00000011114.15_7 | BTBD7 | 0.99999996 | 1 | -0.054 | 5.523 | chr14 | 93703896 | 93799438 | - | protein_coding |
| ENSG00000260193.1_8 | LINC02846 | 0.96367988 | 1 | -0.054 | 0.461 | chr9 | 138999654 | 139001270 | - | lncRNA |
| ENSG00000205423.11_5 | CNEP1R1 | 0.99999953 | 1 | -0.055 | 4.600 | chr16 | 50058321 | 50070999 | + | protein_coding |
| ENSG00000147251.15_4 | DOCK11 | 0.99999778 | 1 | -0.055 | 4.399 | chrX | 117629861 | 117820126 | + | protein_coding |
| ENSG00000146859.6_3 | TMEM140 | 0.99999629 | 1 | -0.055 | 4.045 | chr7 | 134832824 | 134850967 | + | protein_coding |
| ENSG00000136643.12_4 | RPS6KC1 | 0.99999988 | 1 | -0.055 | 5.112 | chr1 | 213224575 | 213448117 | + | protein_coding |
| ENSG00000204308.8_3 | RNF5 | 0.99980939 | 1 | -0.055 | 2.852 | chr6 | 32146182 | 32148570 | + | protein_coding |
| ENSG00000242207.1_6 | HOXB-AS4 | 0.96303383 | 1 | -0.055 | 0.673 | chr17 | 46706037 | 46712294 | + | lncRNA |
| ENSG00000111344.11_4 | RASAL1 | 0.93321613 | 1 | -0.055 | -0.995 | chr12 | 113536624 | 113574044 | - | protein_coding |
| ENSG00000122126.17_6 | OCRL | 0.99999999 | 1 | -0.055 | 6.131 | chrX | 128673826 | 128726538 | + | protein_coding |
| ENSG00000287817.1_1 | AL096701.4 | 0.93957173 | 1 | -0.055 | -0.414 | chr22 | 31886136 | 31887008 | + | lncRNA |
| ENSG00000026025.16_6 | VIM | 1 | 1 | -0.055 | 8.546 | chr10 | 17270240 | 17279592 | + | protein_coding |
| ENSG00000065485.20_5 | PDIA5 | 0.99998649 | 1 | -0.055 | 3.796 | chr3 | 122785872 | 122944074 | + | protein_coding |
| ENSG00000186184.18_5 | POLR1D | 0.99999977 | 1 | -0.055 | 5.493 | chr13 | 28194879 | 28318374 | + | protein_coding |
| ENSG00000183576.13_4 | SETD3 | 0.99999999 | 1 | -0.055 | 6.324 | chr14 | 99864085 | 99947226 | - | protein_coding |
| ENSG00000159140.21_7 | SON | 1 | 1 | -0.055 | 7.059 | chr21 | 34915344 | 34949787 | + | protein_coding |
| ENSG00000003137.8_6 | CYP26B1 | 0.99997523 | 1 | -0.055 | 3.657 | chr2 | 72356367 | 72375167 | - | protein_coding |
| ENSG00000175470.14 | PPP2R2D | 0.99999992 | 1 | -0.055 | 5.095 | chr10 | 133747955 | 133773331 | + | protein_coding |
| ENSG00000140326.13_7 | CDAN1 | 0.99995544 | 1 | -0.055 | 4.012 | chr15 | 43015742 | 43029326 | - | protein_coding |
| ENSG00000141968.8_3 | VAV1 | 0.99989347 | 1 | -0.055 | 3.303 | chr19 | 6772719 | 6857377 | + | protein_coding |
| ENSG00000070159.14_4 | PTPN3 | 0.99999996 | 1 | -0.056 | 5.580 | chr9 | 112137746 | 112260593 | - | protein_coding |
| ENSG00000138829.12_7 | FBN2 | 0.99999999 | 1 | -0.056 | 6.853 | chr5 | 127593601 | 127994878 | - | protein_coding |
| ENSG00000085719.13_5 | CPNE3 | 1 | 1 | -0.056 | 7.276 | chr8 | 87526664 | 87573726 | + | protein_coding |
| ENSG00000141664.10_5 | ZCCHC2 | 0.99999743 | 1 | -0.056 | 4.391 | chr18 | 60190258 | 60254942 | + | protein_coding |
| ENSG00000197813.6_7 | AC011450.1 | 0.94407537 | 1 | -0.056 | -0.129 | chr19 | 49834782 | 49843560 | - | lncRNA |
| ENSG00000272368.2_7 | AC074032.1 | 0.95029055 | 1 | -0.056 | 0.232 | chr12 | 50505980 | 50559401 | + | lncRNA |
| ENSG00000018408.14_4 | WWTR1 | 1 | 1 | -0.056 | 7.937 | chr3 | 149235022 | 149454501 | - | protein_coding |
| ENSG00000135387.21_4 | CAPRIN1 | 1 | 1 | -0.056 | 8.740 | chr11 | 34073278 | 34124157 | + | protein_coding |
| ENSG00000162819.12_4 | BROX | 1 | 1 | -0.056 | 6.739 | chr1 | 222885895 | 222908538 | + | protein_coding |
| ENSG00000255224.1_6 | AC109322.1 | 0.97495793 | 1 | -0.056 | 0.803 | chr8 | 145132905 | 145134168 | - | lncRNA |
| ENSG00000117153.16_5 | KLHL12 | 0.99999938 | 1 | -0.056 | 5.527 | chr1 | 202860244 | 202897764 | - | protein_coding |
| ENSG00000187713.7_4 | TMEM203 | 0.99999868 | 1 | -0.056 | 4.678 | chr9 | 140098534 | 140100100 | - | protein_coding |
| ENSG00000005189.19_5 | REXO5 | 0.99887938 | 1 | -0.056 | 2.824 | chr16 | 20817751 | 20860990 | + | protein_coding |
| ENSG00000116691.11_3 | MIIP | 0.99554203 | 1 | -0.056 | 3.106 | chr1 | 12079523 | 12092102 | + | protein_coding |
| ENSG00000230366.9_4 | DSCR9 | 0.93547853 | 1 | -0.056 | -0.741 | chr21 | 38580804 | 38594037 | + | lncRNA |
| ENSG00000176358.15_3 | TAC4 | 0.95373308 | 1 | -0.056 | 0.168 | chr17 | 47915671 | 47925379 | - | protein_coding |
| ENSG00000178502.6_4 | KLHL11 | 0.99999993 | 1 | -0.056 | 5.265 | chr17 | 40004770 | 40021676 | - | protein_coding |
| ENSG00000055147.19_5 | FAM114A2 | 0.99999906 | 1 | -0.056 | 4.497 | chr5 | 153369708 | 153418496 | - | protein_coding |
| ENSG00000132002.8_6 | DNAJB1 | 1 | 1 | -0.056 | 6.883 | chr19 | 14625581 | 14640582 | - | protein_coding |
| ENSG00000257433.6_7 | AC004241.1 | 0.93231008 | 1 | -0.057 | -0.711 | chr12 | 48099866 | 48136077 | + | lncRNA |
| ENSG00000127483.19_5 | HP1BP3 | 1 | 1 | -0.057 | 7.985 | chr1 | 21066759 | 21113816 | - | protein_coding |
| ENSG00000205323.9_5 | SARNP | 0.9473676 | 1 | -0.057 | 0.055 | chr12 | 56146247 | 56211508 | - | protein_coding |
| ENSG00000164818.16_6 | DNAAF5 | 0.99999834 | 1 | -0.057 | 6.071 | chr7 | 766336 | 826112 | + | protein_coding |
| ENSG00000140463.14_4 | BBS4 | 0.99999971 | 1 | -0.057 | 4.967 | chr15 | 72978520 | 73030816 | + | protein_coding |
| ENSG00000171084.15_4 | FAM86JP | 0.99899627 | 1 | -0.057 | 2.425 | chr3 | 125635463 | 125648867 | + | transcribed_unprocessed_pseudogene |
| ENSG00000104412.8_4 | EMC2 | 0.99999876 | 1 | -0.057 | 5.043 | chr8 | 109455830 | 109501425 | + | protein_coding |
| ENSG00000225241.3 | RP11-640M9.2 | 0.99997503 | 1 | -0.057 | 3.883 | chr1 | 144593363 | 144621656 | + | pseudogene |
| ENSG00000091317.8_4 | CMTM6 | 0.99999999 | 1 | -0.057 | 6.582 | chr3 | 32522804 | 32544344 | - | protein_coding |
| ENSG00000105552.15_6 | BCAT2 | 0.99999986 | 1 | -0.057 | 5.232 | chr19 | 49298319 | 49314286 | - | protein_coding |
| ENSG00000130726.12_4 | TRIM28 | 0.99999998 | 1 | -0.057 | 9.197 | chr19 | 59055431 | 59062089 | + | protein_coding |
| ENSG00000165259.14_5 | HDX | 0.99874173 | 1 | -0.057 | 2.572 | chrX | 83572882 | 83757487 | - | protein_coding |
| ENSG00000130348.11_4 | QRSL1 | 0.99999969 | 1 | -0.057 | 4.705 | chr6 | 107077453 | 107116292 | + | protein_coding |
| ENSG00000160447.7_3 | PKN3 | 0.99998361 | 1 | -0.057 | 4.767 | chr9 | 131464782 | 131483195 | + | protein_coding |
| ENSG00000146963.18_4 | LUC7L2 | 0.99998967 | 1 | -0.057 | 4.233 | chr7 | 139025105 | 139108203 | + | protein_coding |
| ENSG00000224094.1_4 | RPS24P8 | 0.94264708 | 1 | -0.057 | -0.067 | chr3 | 45201266 | 45201667 | - | processed_pseudogene |
| ENSG00000258474.1_6 | LINC02313 | 0.95916016 | 1 | -0.057 | 0.530 | chr14 | 32414059 | 32419588 | + | lncRNA |
| ENSG00000128731.17_6 | HERC2 | 1 | 1 | -0.057 | 7.128 | chr15 | 28356186 | 28567318 | - | protein_coding |

| | | | | | | | | | | |
|---|---|---|---|---|---|---|---|---|---|---|
| ENSG00000266903.2_9 | AC243964.3 | 0.99719952 | 1 | -0.058 | 2.246 | chr19 | 45134874 | 45228481 | - | lncRNA |
| ENSG00000143578.16_5 | CREB3L4 | 0.99995907 | 1 | -0.058 | 3.401 | chr1 | 153940010 | 153946837 | + | protein_coding |
| ENSG00000111142.14_4 | METAP2 | 1 | 1 | -0.058 | 7.131 | chr12 | 95867296 | 95909615 | + | protein_coding |
| ENSG00000076043.10_6 | REXO2 | 0.99999992 | 1 | -0.058 | 5.262 | chr11 | 114310108 | 114321001 | + | protein_coding |
| ENSG00000169621.10_5 | APLF | 0.99898225 | 1 | -0.058 | 2.951 | chr2 | 68694704 | 68882994 | + | protein_coding |
| ENSG00000100307.13_6 | CBX7 | 0.97454534 | 1 | -0.058 | 1.350 | chr22 | 39516172 | 39548685 | - | protein_coding |
| ENSG00000187607.16_5 | ZNF286A | 0.99997984 | 1 | -0.058 | 4.522 | chr17 | 15602891 | 15624101 | + | protein_coding |
| ENSG00000269190.6_6 | FBXO17 | 0.9999999 | 1 | -0.058 | 5.700 | chr19 | 39432041 | 39466382 | - | protein_coding |
| ENSG00000187257.16_6 | RSBN1L | 0.99999986 | 1 | -0.058 | 5.448 | chr7 | 77325776 | 77412339 | + | protein_coding |
| ENSG00000275532.1_7 | AC006449.2 | 0.92801556 | 1 | -0.058 | -0.440 | chr17 | 36859733 | 36862514 | - | lncRNA |
| ENSG00000174788.10_5 | PCP2 | 0.92408076 | 1 | -0.058 | -0.925 | chr19 | 7696497 | 7698605 | - | protein_coding |
| ENSG00000083535.16_4 | PIBF1 | 0.99999354 | 1 | -0.058 | 4.217 | chr13 | 73356271 | 73590599 | + | protein_coding |
| ENSG00000164978.18_3 | NUDT2 | 0.9999197 | 1 | -0.058 | 3.310 | chr9 | 34329504 | 34343709 | + | protein_coding |
| ENSG00000037749.12_4 | MFAP3 | 0.99999966 | 1 | -0.058 | 4.917 | chr5 | 153418466 | 153600038 | + | protein_coding |
| ENSG00000162419.12_2 | GMEB1 | 0.99995665 | 1 | -0.058 | 3.678 | chr1 | 28995244 | 29045865 | + | protein_coding |
| ENSG00000147677.11_6 | EIF3H | 0.99999996 | 1 | -0.058 | 6.938 | chr8 | 117654369 | 117779164 | - | protein_coding |
| ENSG00000103061.13_6 | SLC7A6OS | 0.99971882 | 1 | -0.058 | 2.972 | chr16 | 68318406 | 68344849 | - | protein_coding |
| ENSG00000166133.18_4 | RPUSD2 | 0.99994856 | 1 | -0.058 | 3.595 | chr15 | 40861498 | 40867148 | + | protein_coding |
| ENSG00000067560.12_6 | RHOA | 1 | 1 | -0.059 | 8.354 | chr3 | 49396578 | 49450431 | - | protein_coding |
| ENSG00000101161.8_3 | PRPF6 | 0.99999999 | 1 | -0.059 | 6.679 | chr20 | 62612485 | 62664453 | + | protein_coding |
| ENSG00000101210.12_7 | EEF1A2 | 0.99999955 | 1 | -0.059 | 5.999 | chr20 | 62119367 | 62130538 | - | protein_coding |
| ENSG00000234518.2_5 | PTGES3P1 | 0.99699663 | 1 | -0.059 | 1.370 | chr1 | 89569968 | 89570450 | - | processed_pseudogene |
| ENSG00000240859.3_6 | AC093627.4 | 0.92374343 | 1 | -0.059 | -0.815 | chr7 | 149549 | 155465 | + | lncRNA |
| ENSG00000072506.13_4 | HSD17B10 | 0.99999994 | 1 | -0.059 | 5.919 | chrX | 53458206 | 53461317 | - | protein_coding |
| ENSG00000165671.20_4 | NSD1 | 1 | 1 | -0.059 | 7.163 | chr5 | 176560026 | 176727216 | + | protein_coding |
| ENSG00000068394.11_4 | GPKOW | 0.99999795 | 1 | -0.059 | 4.611 | chrX | 48970345 | 48980085 | - | protein_coding |
| ENSG00000143198.13_3 | MGST3 | 0.99999997 | 1 | -0.059 | 5.730 | chr1 | 165600450 | 165631033 | + | protein_coding |
| ENSG00000204381.11_6 | LAYN | 0.9676328 | 1 | -0.059 | 0.582 | chr11 | 111411005 | 111432470 | + | protein_coding |
| ENSG00000125962.14_7 | ARMCX5 | 0.99997507 | 1 | -0.059 | 3.658 | chrX | 101854096 | 101859087 | + | protein_coding |
| ENSG00000267219.1_7 | AC010504.1 | 0.92220447 | 1 | -0.060 | -0.813 | chr19 | 34839261 | 34850317 | - | lncRNA |
| ENSG00000131634.14_4 | TMEM204 | 0.92175738 | 1 | -0.060 | -0.814 | chr16 | 1578689 | 1605581 | + | protein_coding |
| ENSG00000214941.8_5 | ZSWIM7 | 0.99999871 | 1 | -0.060 | 4.336 | chr17 | 15879874 | 15903031 | - | protein_coding |
| ENSG00000101464.10_5 | PIGU | 0.99999976 | 1 | -0.060 | 4.871 | chr20 | 33148346 | 33286594 | - | protein_coding |
| ENSG00000083223.18_5 | TUT7 | 0.99999968 | 1 | -0.060 | 5.597 | chr9 | 88902648 | 88969369 | - | protein_coding |
| ENSG00000139344.8_4 | AMDHD1 | 0.99912659 | 1 | -0.060 | 2.344 | chr12 | 96337109 | 96362498 | + | protein_coding |
| ENSG00000183977.14_5 | PP2D1 | 0.93848279 | 1 | -0.060 | -0.106 | chr3 | 20021453 | 20053759 | - | protein_coding |
| ENSG00000009780.15_4 | FAM76A | 0.99995236 | 1 | -0.060 | 3.425 | chr1 | 28052490 | 28089633 | + | protein_coding |
| ENSG00000139343.10_2 | SNRPF | 0.99948711 | 1 | -0.060 | 4.664 | chr12 | 96252706 | 96297606 | + | protein_coding |
| ENSG00000115306.16_5 | SPTBN1 | 1 | 1 | -0.060 | 10.167 | chr2 | 54683454 | 54898583 | + | protein_coding |
| ENSG00000273314.1_6 | AC005229.4 | 0.949693 | 1 | -0.060 | 0.714 | chr7 | 148393559 | 148395756 | - | lncRNA |
| ENSG00000228742.11_7 | LINC02577 | 0.9681467 | 1 | -0.060 | 1.057 | chr7 | 106415401 | 106478925 | + | lncRNA |
| ENSG00000161010.15_7 | MRNIP | 0.99999494 | 1 | -0.060 | 4.827 | chr5 | 179262133 | 179289173 | - | protein_coding |
| ENSG00000245904.4_8 | AC025164.2 | 0.92800124 | 1 | -0.060 | -0.561 | chr12 | 92539349 | 92620333 | + | lncRNA |
| ENSG00000271601.4_4 | LIX1L | 0.9999997 | 1 | -0.060 | 4.973 | chr1 | 145477069 | 145501670 | + | protein_coding |
| ENSG00000151657.12_4 | KIN | 0.99998601 | 1 | -0.061 | 3.992 | chr10 | 7792925 | 7829956 | - | protein_coding |
| ENSG00000145414.9_4 | NAF1 | 0.9999578 | 1 | -0.061 | 3.388 | chr4 | 164031225 | 164088042 | - | protein_coding |
| ENSG00000100413.17_4 | POLR3H | 0.99999873 | 1 | -0.061 | 4.846 | chr22 | 41921803 | 41940610 | - | protein_coding |
| ENSG00000128272.14_3 | ATF4 | 1 | 1 | -0.061 | 7.615 | chr22 | 39915700 | 39918690 | + | protein_coding |
| ENSG00000166224.17_6 | SGPL1 | 1 | 1 | -0.061 | 6.774 | chr10 | 72575704 | 72640941 | + | protein_coding |
| ENSG00000138768.14_4 | USO1 | 0.99999999 | 1 | -0.061 | 6.918 | chr4 | 76649777 | 76735439 | + | protein_coding |
| ENSG00000132581.10_5 | SDF2 | 0.99999996 | 1 | -0.061 | 5.557 | chr17 | 26975364 | 26989207 | - | protein_coding |
| ENSG00000130299.17_5 | GTPBP3 | 0.99987581 | 1 | -0.061 | 3.870 | chr19 | 17445729 | 17453540 | + | protein_coding |
| ENSG00000272970.3_7 | AC107294.2 | 0.92752703 | 1 | -0.061 | -0.344 | chr3 | 184430033 | 184457892 | + | lncRNA |
| ENSG00000011258.16_5 | MBTD1 | 0.99999673 | 1 | -0.061 | 4.633 | chr17 | 49254786 | 49337524 | - | protein_coding |
| ENSG00000105058.12_4 | FAM32A | 0.99999999 | 1 | -0.061 | 5.919 | chr19 | 16296191 | 16302857 | + | protein_coding |
| ENSG00000147403.16_6 | RPL10 | 1 | 1 | -0.061 | 9.158 | chrX | 153618315 | 153637504 | + | protein_coding |
| ENSG00000260196.1_6 | AC124798.1 | 0.97575095 | 1 | -0.061 | 0.878 | chr11 | 17402196 | 17405078 | + | lncRNA |
| ENSG00000164543.7_4 | STK17A | 0.99999995 | 1 | -0.061 | 5.538 | chr7 | 43622357 | 43690312 | + | protein_coding |
| ENSG00000048140.18_6 | TSPAN17 | 0.99999999 | 1 | -0.062 | 5.930 | chr5 | 176074388 | 176086055 | + | protein_coding |
| ENSG00000163931.16_4 | TKT | 1 | 1 | -0.062 | 10.810 | chr3 | 53258728 | 53290068 | - | protein_coding |

| | | | | | | | | | | |
|---|---|---|---|---|---|---|---|---|---|---|
| ENSG00000107443.16_3 | CCNJ | 0.99998485 | 1 | -0.062 | 3.890 | chr10 | 97803151 | 97820627 | + | protein_coding |
| ENSG00000140365.16_4 | COMMD4 | 0.99999998 | 1 | -0.062 | 6.008 | chr15 | 75628361 | 75635565 | + | protein_coding |
| ENSG00000266934.1_7 | AC005746.1 | 0.91805897 | 1 | -0.062 | -1.083 | chr17 | 59460146 | 59462041 | + | lncRNA |
| ENSG00000198894.8_7 | CIPC | 0.9999982 | 1 | -0.062 | 4.623 | chr14 | 77564469 | 77583630 | + | protein_coding |
| ENSG00000188613.7_4 | NANOS1 | 0.98369291 | 1 | -0.062 | 1.481 | chr10 | 120789226 | 120793242 | + | protein_coding |
| ENSG00000184990.13_6 | SIVA1 | 0.99885871 | 1 | -0.062 | 5.856 | chr14 | 105219484 | 105234831 | + | protein_coding |
| ENSG00000166173.11_5 | LARP6 | 0.99999942 | 1 | -0.062 | 4.729 | chr15 | 71121469 | 71146496 | - | protein_coding |
| ENSG00000269858.6_4 | EGLN2 | 0.99999336 | 1 | -0.062 | 4.187 | chr19 | 41304901 | 41314339 | + | protein_coding |
| ENSG00000143067.5_5 | ZNF697 | 0.99338358 | 1 | -0.062 | 1.755 | chr1 | 120162000 | 120190889 | - | protein_coding |
| ENSG00000147650.11_5 | LRP12 | 0.99999914 | 1 | -0.062 | 4.760 | chr8 | 105501459 | 105601252 | - | protein_coding |
| ENSG00000111641.11_6 | NOP2 | 0.9808865 | 1 | -0.063 | 1.085 | chr12 | 6666029 | 6677857 | - | protein_coding |
| ENSG00000137038.8_7 | DMAC1 | 0.99999917 | 1 | -0.063 | 4.798 | chr9 | 7796500 | 7888380 | - | protein_coding |
| ENSG00000123595.8_4 | RAB9A | 0.999971 | 1 | -0.063 | 4.262 | chrX | 13707247 | 13728623 | + | protein_coding |
| ENSG00000006704.10_5 | GTF2IRD1 | 0.9999825 | 1 | -0.063 | 3.763 | chr7 | 73868120 | 74016930 | + | protein_coding |
| ENSG00000157181.16_5 | ODR4 | 0.99999831 | 1 | -0.063 | 5.410 | chr1 | 186344970 | 186390510 | + | protein_coding |
| ENSG00000174177.13_3 | CTU2 | 0.99994799 | 1 | -0.063 | 3.919 | chr16 | 88772891 | 88781804 | + | protein_coding |
| ENSG00000279267.1_5 | AL078621.3 | 0.93483887 | 1 | -0.063 | -0.153 | chr2 | 114363444 | 114365485 | - | TEC |
| ENSG00000143374.17_4 | TARS2 | 0.99999942 | 1 | -0.063 | 4.835 | chr1 | 150459840 | 150480085 | + | protein_coding |
| ENSG00000279862.1_1 | AC092964.1 | 0.9326436 | 1 | -0.063 | -0.616 | chr3 | 172019444 | 172020842 | - | TEC |
| ENSG00000180370.10_3 | PAK2 | 1 | 1 | -0.063 | 6.588 | chr3 | 196466728 | 196559518 | + | protein_coding |
| ENSG00000246560.2_7 | UBE2D3-AS1 | 0.92360896 | 1 | -0.063 | -0.589 | chr4 | 103749212 | 103765232 | + | lncRNA |
| ENSG00000185917.13_5 | SETD4 | 0.99999038 | 1 | -0.063 | 3.846 | chr21 | 37406839 | 37451687 | - | protein_coding |
| ENSG00000105671.12_4 | DDX49 | 0.99999668 | 1 | -0.063 | 4.976 | chr19 | 19030514 | 19039998 | + | protein_coding |
| ENSG00000165724.6_4 | ZMYND19 | 0.99999955 | 1 | -0.063 | 4.747 | chr9 | 140476533 | 140484964 | - | protein_coding |
| ENSG00000170027.7_5 | YWHAG | 1 | 1 | -0.063 | 8.447 | chr7 | 75956116 | 75988308 | - | protein_coding |
| ENSG00000166233.15_5 | ARIH1 | 1 | 1 | -0.064 | 6.718 | chr15 | 72766671 | 72895328 | + | protein_coding |
| ENSG00000182890.5_4 | GLUD2 | 0.99743424 | 1 | -0.064 | 2.794 | chrX | 120181464 | 120183948 | + | protein_coding |
| ENSG00000164647.9_5 | STEAP1 | 0.99998166 | 1 | -0.064 | 4.580 | chr7 | 89783770 | 89794143 | + | protein_coding |
| ENSG00000132361.17_6 | CLUH | 0.99999808 | 1 | -0.064 | 7.148 | chr17 | 2592680 | 2615957 | - | protein_coding |
| ENSG00000122484.9_3 | RPAP2 | 0.99999948 | 1 | -0.064 | 4.996 | chr1 | 92764616 | 92867613 | + | protein_coding |
| ENSG00000102804.15_6 | TSC22D1 | 1 | 1 | -0.064 | 6.989 | chr13 | 45006279 | 45151283 | - | protein_coding |
| ENSG00000160752.14_2 | FDPS | 0.99999996 | 1 | -0.064 | 5.472 | chr1 | 155278539 | 155290457 | + | protein_coding |
| ENSG00000137845.15_5 | ADAM10 | 1 | 1 | -0.064 | 7.795 | chr15 | 58881008 | 59041990 | - | protein_coding |
| ENSG00000147649.10_5 | MTDH | 1 | 1 | -0.064 | 8.064 | chr8 | 98656412 | 98742488 | + | protein_coding |
| ENSG00000087365.16_6 | SF3B2 | 1 | 1 | -0.064 | 7.929 | chr11 | 65818200 | 65836779 | + | protein_coding |
| ENSG00000156026.14_4 | MCU | 1 | 1 | -0.064 | 7.006 | chr10 | 74451889 | 74647452 | + | protein_coding |
| ENSG00000145868.16_4 | FBXO38 | 0.99999994 | 1 | -0.064 | 5.472 | chr5 | 147763498 | 147822399 | + | protein_coding |
| ENSG00000236308.1_6 | AL138921.2 | 0.91601538 | 1 | -0.064 | -1.084 | chr10 | 101949793 | 101950504 | + | lncRNA |
| ENSG00000102882.12_8 | MAPK3 | 0.99999757 | 1 | -0.064 | 4.563 | chr16 | 30125426 | 30134827 | - | protein_coding |
| ENSG00000116288.13_4 | PARK7 | 0.99999997 | 1 | -0.064 | 6.464 | chr1 | 8014351 | 8045565 | + | protein_coding |
| ENSG00000158882.15_4 | TOMM40L | 0.9984883 | 1 | -0.064 | 2.516 | chr1 | 161195729 | 161200536 | + | protein_coding |
| ENSG00000142676.14_6 | RPL11 | 1 | 1 | -0.064 | 8.445 | chr1 | 24018232 | 24023325 | + | protein_coding |
| ENSG00000279500.1_6 | AC108704.2 | 0.99107637 | 1 | -0.064 | 0.974 | chr12 | 129297731 | 129299295 | - | TEC |
| ENSG00000162222.14_4 | TTC9C | 0.99999687 | 1 | -0.064 | 4.319 | chr11 | 62495541 | 62507765 | + | protein_coding |
| ENSG00000165630.14_4 | PRPF18 | 0.99965802 | 1 | -0.064 | 2.810 | chr10 | 13628939 | 13672859 | + | protein_coding |
| ENSG00000205559.5_6 | CHKB-DT | 0.98301934 | 1 | -0.065 | 1.139 | chr22 | 51021455 | 51034063 | + | lncRNA |
| ENSG00000079335.20_7 | CDC14A | 0.99953197 | 1 | -0.065 | 2.768 | chr1 | 100810557 | 100985833 | + | protein_coding |
| ENSG00000147121.16_5 | KRBOX4 | 0.99993984 | 1 | -0.065 | 3.442 | chrX | 46306727 | 46356857 | + | protein_coding |
| ENSG00000170322.14_6 | NFRKB | 0.9999996 | 1 | -0.065 | 5.036 | chr11 | 129733531 | 129765485 | - | protein_coding |
| ENSG00000198089.16_6 | SFI1 | 0.99244821 | 1 | -0.065 | 3.854 | chr22 | 31884674 | 32014574 | + | protein_coding |
| ENSG00000065154.12_3 | OAT | 0.99999999 | 1 | -0.065 | 6.336 | chr10 | 126085872 | 126107545 | - | protein_coding |
| ENSG00000287126.1_1 | AC108488.3 | 0.98836346 | 1 | -0.065 | 0.930 | chr2 | 3616095 | 3622690 | - | lncRNA |
| ENSG00000117751.18_4 | PPP1R8 | 0.99999973 | 1 | -0.065 | 4.894 | chr1 | 28157293 | 28178187 | + | protein_coding |
| ENSG00000286724.1_1 | AL732509.1 | 0.96556692 | 1 | -0.065 | 0.407 | chrX | 19002409 | 19078447 | + | lncRNA |
| ENSG00000180764.13_2 | PIPSL | 0.92831076 | 1 | -0.065 | 0.028 | chr10 | 95717948 | 95721297 | - | transcribed_processed_pseudogene |
| ENSG00000169682.18_5 | SPNS1 | 0.92018756 | 1 | -0.065 | -0.286 | chr16 | 28985542 | 28995869 | + | protein_coding |
| ENSG00000025293.17_4 | PHF20 | 0.99999987 | 1 | -0.065 | 5.424 | chr20 | 34359896 | 34538292 | + | protein_coding |
| ENSG00000152223.15_8 | EPG5 | 0.99999958 | 1 | -0.065 | 5.034 | chr18 | 43427574 | 43547295 | - | protein_coding |
| ENSG00000185803.10_6 | SLC52A2 | 0.99999712 | 1 | -0.065 | 5.380 | chr8 | 145577795 | 145584932 | + | protein_coding |
| ENSG00000259523.1_6 | AC022613.2 | 0.95901017 | 1 | -0.065 | 0.636 | chr15 | 30114834 | 30116284 | + | lncRNA |

| | | | | | | | | | | |
|---|---|---|---|---|---|---|---|---|---|---|
| ENSG00000171448.9_4 | ZBTB26 | 0.99950527 | 1 | -0.065 | 2.825 | chr9 | 125677845 | 125693791 | - | protein_coding |
| ENSG00000167523.13_5 | SPATA33 | 0.99990225 | 1 | -0.065 | 3.300 | chr16 | 89724210 | 89737680 | + | protein_coding |
| ENSG00000100036.13_4 | SLC35E4 | 0.99734094 | 1 | -0.065 | 2.519 | chr22 | 31031768 | 31065003 | + | protein_coding |
| ENSG00000197930.13_5 | ERO1A | 1 | 1 | -0.065 | 6.764 | chr14 | 53106633 | 53162618 | - | protein_coding |
| ENSG00000176472.10_2 | ZNF575 | 0.9711944 | 1 | -0.065 | 0.766 | chr19 | 44029649 | 44040282 | + | protein_coding |
| ENSG00000135299.17_7 | ANKRD6 | 0.99754581 | 1 | -0.065 | 2.148 | chr6 | 90142871 | 90343553 | + | protein_coding |
| ENSG00000185009.12_3 | AP3M1 | 0.99999966 | 1 | -0.066 | 6.089 | chr10 | 75880013 | 75910821 | - | protein_coding |
| ENSG00000151233.11_4 | GXYLT1 | 0.99999725 | 1 | -0.066 | 4.569 | chr12 | 42475647 | 42538676 | - | protein_coding |
| ENSG00000173141.5_3 | MRPL57 | 0.99999976 | 1 | -0.066 | 5.015 | chr13 | 21750797 | 21753223 | + | protein_coding |
| ENSG00000169062.15_3 | UPF3A | 0.9999839 | 1 | -0.066 | 4.363 | chr13 | 115047076 | 115071292 | + | protein_coding |
| ENSG00000166359.11_3 | WDR88 | 0.99566631 | 1 | -0.066 | 1.828 | chr19 | 33622996 | 33666705 | + | protein_coding |
| ENSG00000284707.2_1 | AC079781.5 | 0.96586583 | 1 | -0.066 | 0.747 | chr7 | 97481000 | 97602297 | - | lncRNA |
| ENSG00000089916.18_6 | GPATCH2L | 0.99999994 | 1 | -0.066 | 5.651 | chr14 | 76618259 | 76720685 | + | protein_coding |
| ENSG00000274602.5_5 | PI4KAP1 | 0.98087977 | 1 | -0.066 | 1.262 | chr22 | 20383513 | 20427835 | - | transcribed_unprocessed_pseudogene |
| ENSG00000100697.14_5 | DICER1 | 0.99999999 | 1 | -0.066 | 6.850 | chr14 | 95552565 | 95624347 | - | protein_coding |
| ENSG00000172766.19_3 | NAA16 | 0.99999893 | 1 | -0.066 | 4.673 | chr13 | 41885403 | 41951166 | + | protein_coding |
| ENSG00000184281.15_7 | TSSC4 | 0.97790821 | 1 | -0.066 | 5.103 | chr11 | 2421718 | 2425108 | + | protein_coding |
| ENSG00000065243.20_5 | PKN2 | 0.99999999 | 1 | -0.066 | 6.359 | chr1 | 89149905 | 89301938 | + | protein_coding |
| ENSG00000125733.18_6 | TRIP10 | 0.99999992 | 1 | -0.067 | 5.348 | chr19 | 6737936 | 6751541 | + | protein_coding |
| ENSG00000133794.18_4 | ARNTL | 0.99999215 | 1 | -0.067 | 4.306 | chr11 | 13298199 | 13408813 | + | protein_coding |
| ENSG00000069509.6_4 | FUNDC1 | 0.9999959 | 1 | -0.067 | 4.161 | chrX | 44382885 | 44402105 | - | protein_coding |
| ENSG00000198496.12_4 | NBR2 | 0.99561776 | 1 | -0.067 | 2.169 | chr17 | 41277568 | 41305688 | + | lncRNA |
| ENSG00000010165.20_4 | EEF1AKNMT | 0.99999962 | 1 | -0.067 | 5.337 | chr1 | 171750800 | 171783163 | + | protein_coding |
| ENSG00000100379.17_5 | KCTD17 | 0.99997796 | 1 | -0.067 | 4.095 | chr22 | 37447776 | 37459430 | + | protein_coding |
| ENSG00000204634.12_5 | TBC1D8 | 1 | 1 | -0.067 | 7.194 | chr2 | 101624079 | 101869328 | - | protein_coding |
| ENSG00000141642.9_4 | ELAC1 | 0.99680621 | 1 | -0.067 | 2.094 | chr18 | 48494361 | 48514491 | + | protein_coding |
| ENSG00000232573.1_4 | RPL3P4 | 0.99970679 | 1 | -0.067 | 3.131 | chr14 | 99439216 | 99439638 | - | processed_pseudogene |
| ENSG00000120533.13_4 | ENY2 | 0.99999997 | 1 | -0.067 | 6.109 | chr8 | 110346553 | 110358183 | + | protein_coding |
| ENSG00000188612.12_5 | SUMO2 | 1 | 1 | -0.067 | 7.022 | chr17 | 73161681 | 73179054 | - | protein_coding |
| ENSG00000188152.12_3 | NUTM2G | 0.94150521 | 1 | -0.067 | 0.273 | chr9 | 99690592 | 99702535 | + | protein_coding |
| ENSG00000272655.2_4 | POLR2J4 | 0.98305229 | 1 | -0.067 | 1.257 | chr7 | 44053161 | 44058769 | - | transcribed_unprocessed_pseudogene |
| ENSG00000122515.15_5 | ZMIZ2 | 0.99999895 | 1 | -0.067 | 4.616 | chr7 | 44788180 | 44809480 | + | protein_coding |
| ENSG00000067225.18_6 | PKM | 1 | 1 | -0.067 | 11.546 | chr15 | 72491370 | 72524163 | - | protein_coding |
| ENSG00000180182.11_4 | MED14 | 0.99999996 | 1 | -0.067 | 5.989 | chrX | 40507557 | 40595110 | - | protein_coding |
| ENSG00000169946.14_6 | ZFPM2 | 0.96396111 | 1 | -0.067 | 0.785 | chr8 | 105602961 | 106816767 | + | protein_coding |
| ENSG00000142327.13_6 | RNPEPL1 | 0.99993668 | 1 | -0.068 | 5.549 | chr2 | 241505221 | 241520789 | + | protein_coding |
| ENSG00000177640.16_7 | CASC2 | 0.99068464 | 1 | -0.068 | 1.641 | chr10 | 119805790 | 119969670 | + | lncRNA |
| ENSG00000113448.19_8 | PDE4D | 1 | 1 | -0.068 | 9.177 | chr5 | 58264865 | 59817947 | - | protein_coding |
| ENSG00000159256.13_5 | MORC3 | 0.99999954 | 1 | -0.068 | 5.140 | chr21 | 37692487 | 37758446 | + | protein_coding |
| ENSG00000243964.1_4 | RPL23AP65 | 0.91896637 | 1 | -0.068 | -0.401 | chr11 | 9627689 | 9628156 | + | processed_pseudogene |
| ENSG00000152601.17_5 | MBNL1 | 0.99999999 | 1 | -0.068 | 8.021 | chr3 | 151961617 | 152183569 | + | protein_coding |
| ENSG00000156017.13_6 | CARNMT1 | 0.99999999 | 1 | -0.068 | 5.884 | chr9 | 77595706 | 77643339 | - | protein_coding |
| ENSG00000240342.3_4 | RPS2P5 | 0.99999984 | 1 | -0.068 | 5.376 | chr12 | 118683889 | 118684767 | + | processed_pseudogene |
| ENSG00000148200.17_5 | NR6A1 | 0.99942384 | 1 | -0.068 | 2.862 | chr9 | 127279554 | 127533590 | - | protein_coding |
| ENSG00000280152.1_6 | AC009078.3 | 0.9932093 | 1 | -0.068 | 1.724 | chr16 | 75279892 | 75283975 | - | TEC |
| ENSG00000181555.20_9 | SETD2 | 0.99999999 | 1 | -0.068 | 6.472 | chr3 | 47057919 | 47205457 | - | protein_coding |
| ENSG00000121064.13_6 | SCPEP1 | 0.99999984 | 1 | -0.068 | 5.062 | chr17 | 55055490 | 55084129 | + | protein_coding |
| ENSG00000126067.12_4 | PSMB2 | 0.99999999 | 1 | -0.068 | 6.354 | chr1 | 36065142 | 36107127 | - | protein_coding |
| ENSG00000116016.14_5 | EPAS1 | 1 | 1 | -0.068 | 7.104 | chr2 | 46520806 | 46613836 | + | protein_coding |
| ENSG00000185619.18_1 | PCGF3 | 0.9999988 | 1 | -0.069 | 4.458 | chr4 | 699537 | 764428 | + | protein_coding |
| ENSG00000106077.18_3 | ABHD11 | 0.99937346 | 1 | -0.069 | 3.056 | chr7 | 73150424 | 73153197 | - | protein_coding |
| ENSG00000213261.3_4 | EEF1B2P6 | 0.96131563 | 1 | -0.069 | 0.624 | chr7 | 131346711 | 131347383 | - | processed_pseudogene |
| ENSG00000170471.15_5 | RALGAPB | 0.99997116 | 1 | -0.069 | 6.204 | chr20 | 37101459 | 37207502 | + | protein_coding |
| ENSG00000267253.1_6 | WHSC1L2P | 0.91011964 | 1 | -0.069 | -0.893 | chr17 | 41795682 | 41797804 | - | processed_pseudogene |
| ENSG00000164953.16_6 | TMEM67 | 0.99998245 | 1 | -0.069 | 3.683 | chr8 | 94767072 | 94831462 | + | protein_coding |
| ENSG00000121892.15_5 | PDS5A | 1 | 1 | -0.069 | 7.620 | chr4 | 39824483 | 39979576 | - | protein_coding |
| ENSG00000106443.16_3 | PHF14 | 0.99999998 | 1 | -0.069 | 6.385 | chr7 | 11012963 | 11209257 | + | protein_coding |

| | | | | | | | | | | |
|---|---|---|---|---|---|---|---|---|---|---|
| ENSG00000151640.13_3 | DPYSL4 | 0.9999317 | 1 | -0.069 | 3.794 | chr10 | 134000452 | 134019263 | + | protein_coding |
| ENSG00000066697.14_5 | MSANTD3 | 0.99999993 | 1 | -0.069 | 5.442 | chr9 | 103189438 | 103213993 | + | protein_coding |
| ENSG00000172992.12_5 | DCAKD | 0.99999923 | 1 | -0.069 | 4.568 | chr17 | 43100708 | 43138477 | - | protein_coding |
| ENSG00000287787.1_1 | AC092275.1 | 0.92581203 | 1 | -0.069 | -0.221 | chr16 | 85260809 | 85342782 | + | lncRNA |
| ENSG00000120662.16_5 | MTRF1 | 0.99992377 | 1 | -0.069 | 3.408 | chr13 | 41790505 | 41837713 | - | protein_coding |
| ENSG00000005022.6_3 | SLC25A5 | 0.99999912 | 1 | -0.069 | 7.787 | chrX | 118602385 | 118605359 | + | protein_coding |
| ENSG00000226416.1_7 | MRPL23-AS1 | 0.91208687 | 1 | -0.069 | -0.712 | chr11 | 2004467 | 2011150 | - | lncRNA |
| ENSG00000249898.8_7 | MCPH1-AS1 | 0.91565774 | 1 | -0.069 | -0.816 | chr8 | 6474831 | 6565745 | - | lncRNA |
| ENSG00000184616.9_6 | SPDYE12P | 0.94150999 | 1 | -0.070 | -0.133 | chr7 | 74322778 | 74329393 | - | unprocessed_pseudogene |
| ENSG00000145916.19_6 | RMND5B | 0.99999972 | 1 | -0.070 | 4.993 | chr5 | 177557997 | 177577569 | + | protein_coding |
| ENSG00000100027.16_5 | YPEL1 | 0.94150999 | 1 | -0.070 | -0.133 | chr22 | 22051825 | 22090083 | - | protein_coding |
| ENSG00000286482.1_1 | AL117344.2 | 0.97429308 | 1 | -0.070 | 0.875 | chr6 | 157798094 | 157801928 | - | lncRNA |
| ENSG00000176624.11_5 | MEX3C | 0.99999997 | 1 | -0.070 | 5.665 | chr18 | 48700920 | 48744674 | - | protein_coding |
| ENSG00000163754.17_5 | GYG1 | 0.9999991 | 1 | -0.070 | 4.550 | chr3 | 148709128 | 148745455 | + | protein_coding |
| ENSG00000234869.1_6 | AL021392.1 | 0.97336794 | 1 | -0.070 | 0.843 | chr22 | 46937392 | 46944093 | + | lncRNA |
| ENSG00000177679.16_5 | SRRM3 | 0.99580146 | 1 | -0.070 | 1.961 | chr7 | 75831214 | 75916605 | + | protein_coding |
| ENSG00000260051.1_6 | AL031600.1 | 0.94589002 | 1 | -0.070 | 0.440 | chr16 | 1501761 | 1502654 | + | lncRNA |
| ENSG00000102786.15_8 | INTS6 | 0.9999999 | 1 | -0.070 | 5.601 | chr13 | 51928213 | 52028400 | - | protein_coding |
| ENSG00000144935.15_4 | TRPC1 | 0.99999943 | 1 | -0.070 | 4.787 | chr3 | 142442876 | 142526730 | + | protein_coding |
| ENSG00000104320.13_5 | NBN | 1 | 1 | -0.070 | 7.003 | chr8 | 90945564 | 91015456 | - | protein_coding |
| ENSG00000279059.1_6 | AC007485.2 | 0.99978512 | 1 | -0.071 | 2.934 | chr17 | 53034121 | 53035787 | - | TEC |
| ENSG00000179406.8_6 | LINC00174 | 0.99750071 | 1 | -0.071 | 2.057 | chr7 | 65841031 | 65958553 | - | lncRNA |
| ENSG00000023572.9_4 | GLRX2 | 0.99999995 | 1 | -0.071 | 5.440 | chr1 | 193059996 | 193075244 | - | protein_coding |
| ENSG00000239486.1_6 | AC091390.4 | 0.90571485 | 1 | -0.071 | -0.827 | chr7 | 102020912 | 102023184 | - | unprocessed_pseudogene |
| ENSG00000140694.17_6 | PARN | 0.99999999 | 1 | -0.071 | 6.460 | chr16 | 14529557 | 14726585 | - | protein_coding |
| ENSG00000093144.19_7 | ECHDC1 | 0.99999991 | 1 | -0.071 | 5.588 | chr6 | 127609857 | 127664754 | - | protein_coding |
| ENSG00000086504.17_6 | MRPL28 | 0.99999971 | 1 | -0.071 | 5.822 | chr16 | 416969 | 421289 | - | protein_coding |
| ENSG00000181315.11_6 | ZNF322 | 0.99991156 | 1 | -0.071 | 3.282 | chr6 | 26634611 | 26659980 | - | protein_coding |
| ENSG00000241839.10_4 | PLEKHO2 | 0.99439179 | 1 | -0.071 | 2.366 | chr15 | 65134082 | 65160201 | + | protein_coding |
| ENSG00000242086.8_6 | MUC20-OT1 | 0.99999069 | 1 | -0.071 | 4.366 | chr3 | 195384933 | 195466835 | + | lncRNA |
| ENSG00000117419.16_6 | ERI3 | 0.99999935 | 1 | -0.072 | 5.220 | chr1 | 44686742 | 44820932 | - | protein_coding |
| ENSG00000086758.16_7 | HUWE1 | 1 | 1 | -0.072 | 8.949 | chrX | 53559057 | 53713673 | - | protein_coding |
| ENSG00000243107.1_8 | AC000120.2 | 0.90454021 | 1 | -0.072 | -1.046 | chr7 | 91829328 | 91836171 | - | lncRNA |
| ENSG00000105618.14_4 | PRPF31 | 0.99999563 | 1 | -0.072 | 5.646 | chr19 | 54618790 | 54635150 | + | protein_coding |
| ENSG00000110944.9_3 | IL23A | 0.90644523 | 1 | -0.072 | -0.893 | chr12 | 56727958 | 56734194 | + | protein_coding |
| ENSG00000075218.19_4 | GTSE1 | 0.99999975 | 1 | -0.072 | 5.211 | chr22 | 46692767 | 46726707 | + | protein_coding |
| ENSG00000179583.19_4 | CIITA | 0.97626591 | 1 | -0.072 | 1.108 | chr16 | 10960079 | 11036878 | + | protein_coding |
| ENSG00000112062.10_3 | MAPK14 | 0.99999995 | 1 | -0.072 | 5.522 | chr6 | 35995454 | 36079013 | + | protein_coding |
| ENSG00000118690.13_6 | ARMC2 | 0.98282845 | 1 | -0.072 | 1.326 | chr6 | 109169619 | 109295679 | + | protein_coding |
| ENSG00000198856.13_3 | OSTC | 0.99999977 | 1 | -0.072 | 5.478 | chr4 | 109571741 | 109588976 | + | protein_coding |
| ENSG00000175463.12_4 | TBC1D10C | 0.9205237 | 1 | -0.072 | -0.242 | chr11 | 67171386 | 67177560 | + | protein_coding |
| ENSG00000109113.20_8 | RAB34 | 0.99999999 | 1 | -0.073 | 7.203 | chr17 | 27041299 | 27045447 | - | protein_coding |
| ENSG00000157349.16_6 | DDX19B | 0.99989705 | 1 | -0.073 | 3.474 | chr16 | 70323566 | 70369186 | + | protein_coding |
| ENSG00000108344.15_3 | PSMD3 | 1 | 1 | -0.073 | 7.420 | chr17 | 38137060 | 38154212 | + | protein_coding |
| ENSG00000207523.1 | SNORA66 | 0.90284617 | 1 | -0.073 | -1.047 | chr1 | 93306276 | 93306408 | + | snoRNA |
| ENSG00000155629.15_5 | PIK3AP1 | 0.99421943 | 1 | -0.073 | 1.847 | chr10 | 98353072 | 98480271 | - | protein_coding |
| ENSG00000109332.20_7 | UBE2D3 | 1 | 1 | -0.073 | 7.352 | chr4 | 103715540 | 103790053 | - | protein_coding |
| ENSG00000078177.14_4 | N4BP2 | 0.99999944 | 1 | -0.073 | 5.298 | chr4 | 40058470 | 40159872 | + | protein_coding |
| ENSG00000232274.2 | RP11-782C8.2 | 0.9765761 | 1 | -0.073 | 1.288 | chr1 | 143118704 | 143211070 | - | lincRNA |
| ENSG00000141580.16_5 | WDR45B | 0.99999997 | 1 | -0.073 | 6.608 | chr17 | 80572438 | 80606429 | - | protein_coding |
| ENSG00000162104.10_5 | ADCY9 | 0.99999988 | 1 | -0.073 | 5.431 | chr16 | 4003388 | 4166443 | - | protein_coding |
| ENSG00000188163.8_5 | FAM166A | 0.91565774 | 1 | -0.073 | -0.816 | chr9 | 140138036 | 140142222 | - | protein_coding |
| ENSG00000073712.15_7 | FERMT2 | 0.99999998 | 1 | -0.073 | 5.973 | chr14 | 53323986 | 53419153 | - | protein_coding |
| ENSG00000204568.12_3 | MRPS18B | 0.99999274 | 1 | -0.073 | 4.156 | chr6 | 30585617 | 30594172 | + | protein_coding |
| ENSG00000108819.11_4 | PPP1R9B | 0.99999977 | 1 | -0.073 | 5.717 | chr17 | 48211101 | 48228038 | - | protein_coding |
| ENSG00000180801.14_5 | ARSJ | 0.99999972 | 1 | -0.073 | 4.911 | chr4 | 114821440 | 114900883 | - | protein_coding |
| ENSG00000122694.16_3 | GLIPR2 | 0.99102582 | 1 | -0.073 | 1.622 | chr9 | 36136533 | 36163910 | + | protein_coding |
| ENSG00000111196.10_4 | MAGOHB | 0.99999844 | 1 | -0.073 | 4.723 | chr12 | 10756792 | 10766208 | - | protein_coding |
| ENSG00000165699.15_7 | TSC1 | 0.99999996 | 1 | -0.073 | 5.988 | chr9 | 135766735 | 135822261 | - | protein_coding |

| | | | | | | | | | | |
|---|---|---|---|---|---|---|---|---|---|---|
| ENSG00000230185.5_6 | C9orf147 | 0.9313337 | 1 | -0.073 | -0.030 | chr9 | 115161792 | 115249484 | - | lncRNA |
| ENSG00000214765.8_7 | SEPTIN7P2 | 0.99994521 | 1 | -0.073 | 3.523 | chr7 | 45763379 | 45808584 | - | transcribed_unprocessed_pseudogene |
| ENSG00000135334.9_4 | AKIRIN2 | 0.99998477 | 1 | -0.074 | 3.797 | chr6 | 88384578 | 88411951 | - | protein_coding |
| ENSG00000286786.1_1 | AC116158.3 | 0.93939287 | 1 | -0.074 | 0.172 | chr15 | 38785509 | 38792186 | + | lncRNA |
| ENSG00000087157.19_4 | PGS1 | 0.99999282 | 1 | -0.074 | 4.677 | chr17 | 76374730 | 76421195 | + | protein_coding |
| ENSG00000198382.9_5 | UVRAG | 0.9999956 | 1 | -0.074 | 4.601 | chr11 | 75526255 | 75855276 | + | protein_coding |
| ENSG00000156253.7_4 | RWDD2B | 0.99998051 | 1 | -0.074 | 3.814 | chr21 | 30376705 | 30391681 | - | protein_coding |
| ENSG00000269609.6_7 | RPARP-AS1 | 0.99953009 | 1 | -0.074 | 2.758 | chr10 | 104209573 | 104220863 | + | lncRNA |
| ENSG00000220693.2 | AC011498.1 | 0.90159729 | 1 | -0.074 | -1.085 | chr19 | 4477048 | 4479576 | + | pseudogene |
| ENSG00000069329.18_6 | VPS35 | 1 | 1 | -0.074 | 7.851 | chr16 | 46690044 | 46723430 | - | protein_coding |
| ENSG00000112941.14_7 | TENT4A | 0.99999966 | 1 | -0.074 | 4.829 | chr5 | 6713545 | 6757157 | + | protein_coding |
| ENSG00000218350.1_4 | LYPLA1P3 | 0.90523717 | 1 | -0.074 | -0.510 | chr6 | 71874779 | 71875473 | + | processed_pseudogene |
| ENSG00000210082.2 | MT-RNR2 | 0.99999991 | 1 | -0.074 | 17.608 | chrM | 1671 | 3229 | + | Mt_rRNA |
| ENSG00000212127.6_5 | TAS2R14 | 0.96044588 | 1 | -0.074 | 0.695 | chr12 | 11090007 | 11324172 | - | protein_coding |
| ENSG00000169188.5_3 | APEX2 | 0.99999991 | 1 | -0.074 | 5.391 | chrX | 55026796 | 55035490 | + | protein_coding |
| ENSG00000237950.1 | RP11-7O11.3 | 0.9069308 | 1 | -0.075 | -0.589 | chr1 | 44410042 | 44412223 | - | antisense |
| ENSG00000179950.14_5 | PUF60 | 0.99999997 | 1 | -0.075 | 7.517 | chr8 | 144898514 | 144911522 | - | protein_coding |
| ENSG00000278540.5_6 | ACACA | 1 | 1 | -0.075 | 7.922 | chr17 | 35441927 | 35766927 | - | protein_coding |
| ENSG00000123562.17_7 | MORF4L2 | 1 | 1 | -0.075 | 8.281 | chrX | 102930424 | 102943086 | - | protein_coding |
| ENSG00000166710.20_5 | B2M | 0.99999999 | 1 | -0.075 | 7.244 | chr15 | 45003685 | 45011075 | + | protein_coding |
| ENSG00000119655.10_5 | NPC2 | 0.99999912 | 1 | -0.075 | 4.729 | chr14 | 74942895 | 74960880 | - | protein_coding |
| ENSG00000126267.11_6 | COX6B1 | 0.99999996 | 1 | -0.075 | 7.765 | chr19 | 36139225 | 36149684 | + | protein_coding |
| ENSG00000238035.8_7 | AC138035.1 | 0.90468929 | 1 | -0.075 | -0.599 | chr5 | 180733503 | 180751686 | + | lncRNA |
| ENSG00000105929.16_5 | ATP6V0A4 | 0.91692015 | 1 | -0.075 | -0.251 | chr7 | 138391039 | 138484305 | - | protein_coding |
| ENSG00000184863.11_6 | RBM33 | 0.99999998 | 1 | -0.075 | 5.816 | chr7 | 155437145 | 155574179 | + | protein_coding |
| ENSG00000161011.20_5 | SQSTM1 | 0.99999998 | 1 | -0.075 | 8.806 | chr5 | 179233398 | 179265078 | + | protein_coding |
| ENSG00000184232.9_5 | OAF | 0.9994142 | 1 | -0.075 | 2.629 | chr11 | 120081740 | 120101043 | + | protein_coding |
| ENSG00000144118.14_4 | RALB | 0.99999994 | 1 | -0.075 | 5.553 | chr2 | 120997640 | 121052286 | + | protein_coding |
| ENSG00000105971.15_3 | CAV2 | 1 | 1 | -0.075 | 7.377 | chr7 | 115927434 | 116148595 | + | protein_coding |
| ENSG00000230701.2_2 | FBXW4P1 | 0.9057778 | 1 | -0.076 | -0.754 | chr22 | 23604954 | 23607192 | + | processed_pseudogene |
| ENSG00000158716.9_3 | DUSP23 | 0.99996092 | 1 | -0.076 | 3.877 | chr1 | 159750722 | 159752333 | + | protein_coding |
| ENSG00000173715.16_5 | C11orf80 | 0.99975996 | 1 | -0.076 | 3.154 | chr11 | 66511922 | 66610799 | + | protein_coding |
| ENSG00000173575.22_6 | CHD2 | 0.99999998 | 1 | -0.076 | 6.146 | chr15 | 93443419 | 93571226 | + | protein_coding |
| ENSG00000120314.18_2 | WDR55 | 0.99999983 | 1 | -0.076 | 5.395 | chr5 | 140044261 | 140053709 | + | protein_coding |
| ENSG00000108774.14_6 | RAB5C | 0.99999934 | 1 | -0.076 | 4.812 | chr17 | 40276994 | 40307062 | - | protein_coding |
| ENSG00000130305.17_4 | NSUN5 | 0.99999771 | 1 | -0.076 | 4.625 | chr7 | 72716513 | 72722823 | - | protein_coding |
| ENSG00000090013.11_5 | BLVRB | 0.99997186 | 1 | -0.076 | 6.980 | chr19 | 40953672 | 40971671 | - | protein_coding |
| ENSG00000162910.19_6 | MRPL55 | 0.99999228 | 1 | -0.076 | 4.529 | chr1 | 228294380 | 228297013 | - | protein_coding |
| ENSG00000166037.11_5 | CEP57 | 0.99999988 | 1 | -0.076 | 5.531 | chr11 | 95523129 | 95565857 | + | protein_coding |
| ENSG00000101901.12_5 | ALG13 | 0.99999897 | 1 | -0.076 | 5.118 | chrX | 110909039 | 111003877 | + | protein_coding |
| ENSG00000087301.9_5 | TXNDC16 | 0.99738786 | 1 | -0.076 | 2.228 | chr14 | 52897314 | 53019240 | - | protein_coding |
| ENSG00000179151.13_7 | EDC3 | 0.99542166 | 1 | -0.076 | 2.121 | chr15 | 74922899 | 74988633 | - | protein_coding |
| ENSG00000241468.7_5 | ATP5MF | 0.99999921 | 1 | -0.076 | 4.691 | chr7 | 99046098 | 99063954 | - | protein_coding |
| ENSG00000139793.18_4 | MBNL2 | 0.99999987 | 1 | -0.076 | 5.653 | chr13 | 97873688 | 98046374 | + | protein_coding |
| ENSG00000111911.7_3 | HINT3 | 0.99999945 | 1 | -0.076 | 4.754 | chr6 | 126277916 | 126301390 | + | protein_coding |
| ENSG00000058668.14_4 | ATP2B4 | 0.99999951 | 1 | -0.076 | 4.835 | chr1 | 203595689 | 203713209 | + | protein_coding |
| ENSG00000261173.1_6 | AC018845.3 | 0.9009003 | 1 | -0.076 | -0.561 | chr16 | 47007900 | 47012894 | + | lncRNA |
| ENSG00000245975.2_7 | AC090515.3 | 0.92405431 | 1 | -0.076 | 0.026 | chr15 | 59060271 | 59063173 | - | lncRNA |
| ENSG00000119596.18_6 | YLPM1 | 0.99999996 | 1 | -0.077 | 6.308 | chr14 | 75230019 | 75326138 | + | protein_coding |
| ENSG00000276007.1_7 | AC079414.3 | 0.90151866 | 1 | -0.077 | -0.371 | chr16 | 78157140 | 78158229 | + | lncRNA |
| ENSG00000170855.4_4 | TRIAP1 | 0.99999979 | 1 | -0.077 | 5.144 | chr12 | 120881767 | 120884187 | - | protein_coding |
| ENSG00000224420.3_5 | ADM5 | 0.98163896 | 1 | -0.077 | 1.428 | chr19 | 50191921 | 50193832 | + | protein_coding |
| ENSG00000095587.9_4 | TLL2 | 0.89799678 | 1 | -0.077 | -0.893 | chr10 | 98124365 | 98273683 | - | protein_coding |
| ENSG00000152700.14_4 | SAR1B | 0.99999993 | 1 | -0.077 | 6.226 | chr5 | 133936839 | 133984961 | - | protein_coding |
| ENSG00000169258.7_4 | GPRIN1 | 0.9991826 | 1 | -0.077 | 4.027 | chr5 | 176022803 | 176037157 | - | protein_coding |
| ENSG00000163947.11_7 | ARHGEF3 | 0.99996156 | 1 | -0.077 | 3.596 | chr3 | 56761446 | 57113357 | - | protein_coding |
| ENSG00000214999.3_8 | AC129492.1 | 0.94069507 | 1 | -0.077 | 0.259 | chr17 | 7982800 | 7984883 | + | lncRNA |
| ENSG00000137177.20_5 | KIF13A | 0.99999999 | 1 | -0.077 | 6.083 | chr6 | 17759414 | 17987866 | - | protein_coding |
| ENSG00000205238.9_2 | SPDYE2 | 0.91014786 | 1 | -0.077 | -0.395 | chr7 | 102191679 | 102202755 | + | protein_coding |

| | | | | | | | | | | |
|---|---|---|---|---|---|---|---|---|---|---|
| ENSG00000178935.5_4 | ZNF552 | 0.99957796 | 1 | -0.077 | 3.205 | chr19 | 58315209 | 58326281 | - | protein_coding |
| ENSG00000196683.10_3 | TOMM7 | 0.99999997 | 1 | -0.077 | 5.932 | chr7 | 22852247 | 22862471 | - | protein_coding |
| ENSG00000064545.15_8 | TMEM161A | 0.99966962 | 1 | -0.078 | 3.970 | chr19 | 19229978 | 19249322 | - | protein_coding |
| ENSG00000136813.14_8 | ECPAS | 1 | 1 | -0.078 | 7.707 | chr9 | 114122972 | 114247025 | - | protein_coding |
| ENSG00000204977.10_4 | TRIM13 | 0.99999812 | 1 | -0.078 | 4.466 | chr13 | 50570024 | 50594617 | + | protein_coding |
| ENSG00000198218.11_6 | QRICH1 | 0.99999999 | 1 | -0.078 | 6.259 | chr3 | 49067140 | 49131796 | - | protein_coding |
| ENSG00000048405.10_7 | ZNF800 | 0.9999998 | 1 | -0.078 | 5.250 | chr7 | 126986844 | 127071978 | - | protein_coding |
| ENSG00000143748.18_6 | NVL | 0.99999908 | 1 | -0.078 | 4.769 | chr1 | 224415036 | 224517891 | - | protein_coding |
| ENSG00000137161.17_3 | CNPY3 | 0.99999845 | 1 | -0.078 | 5.147 | chr6 | 42896930 | 42907032 | + | protein_coding |
| ENSG00000161542.17_5 | PRPSAP1 | 0.99999992 | 1 | -0.078 | 5.436 | chr17 | 74305559 | 74380602 | - | protein_coding |
| ENSG00000132182.12_4 | NUP210 | 0.99999999 | 1 | -0.078 | 7.402 | chr3 | 13357735 | 13461822 | - | protein_coding |
| ENSG00000119403.15_7 | PHF19 | 0.99999999 | 1 | -0.078 | 6.351 | chr9 | 123617929 | 123657174 | - | protein_coding |
| ENSG00000131724.11_5 | IL13RA1 | 1 | 1 | -0.078 | 7.136 | chrX | 117861096 | 117928498 | + | protein_coding |
| ENSG00000070614.15_4 | NDST1 | 0.99999999 | 1 | -0.078 | 6.630 | chr5 | 149865381 | 149937773 | + | protein_coding |
| ENSG00000271151.1_6 | AC016737.1 | 0.89702226 | 1 | -0.078 | -0.787 | chr2 | 174833079 | 174834146 | + | lncRNA |
| ENSG00000279443.1_5 | AL513497.1 | 0.98259167 | 1 | -0.078 | 1.553 | chr1 | 28870972 | 28873054 | + | TEC |
| ENSG00000114026.21_5 | OGG1 | 0.99998511 | 1 | -0.078 | 3.827 | chr3 | 9791628 | 9829903 | + | protein_coding |
| ENSG00000027001.10_5 | MIPEP | 0.99999567 | 1 | -0.078 | 4.307 | chr13 | 24304328 | 24463539 | - | protein_coding |
| ENSG00000145919.11_6 | BOD1 | 0.99999997 | 1 | -0.078 | 5.661 | chr5 | 173034148 | 173043662 | - | protein_coding |
| ENSG00000144741.17_2 | SLC25A26 | 0.99988996 | 1 | -0.078 | 3.375 | chr3 | 66119285 | 66438540 | + | protein_coding |
| ENSG00000166170.9_3 | BAG5 | 0.99999996 | 1 | -0.078 | 6.140 | chr14 | 104022881 | 104029168 | - | protein_coding |
| ENSG00000188807.13_4 | TMEM201 | 0.99996326 | 1 | -0.078 | 3.862 | chr1 | 9648969 | 9674935 | + | protein_coding |
| ENSG00000165997.5_3 | ARL5B | 0.99999926 | 1 | -0.079 | 6.239 | chr10 | 18948360 | 18970568 | + | protein_coding |
| ENSG00000167447.12_5 | SMG8 | 0.99999989 | 1 | -0.079 | 5.424 | chr17 | 57286761 | 57292608 | + | protein_coding |
| ENSG00000242622.2_6 | AC092910.3 | 0.99778777 | 1 | -0.079 | 2.238 | chr3 | 119813742 | 119855630 | + | lncRNA |
| ENSG00000279953.1_6 | AC117503.3 | 0.89550199 | 1 | -0.079 | -0.929 | chr12 | 124133615 | 124135032 | + | TEC |
| ENSG00000117899.11_8 | MESD | 0.99999999 | 1 | -0.079 | 6.419 | chr15 | 81238630 | 81282169 | - | protein_coding |
| ENSG00000100442.11_5 | FKBP3 | 0.99999949 | 1 | -0.079 | 5.158 | chr14 | 45584802 | 45604522 | - | protein_coding |
| ENSG00000103342.13_6 | GSPT1 | 0.9999998 | 1 | -0.079 | 7.694 | chr16 | 11961985 | 12009939 | - | protein_coding |
| ENSG00000287934.1_2 | AP001885.3 | 0.89733984 | 1 | -0.079 | -0.784 | chr11 | 67019807 | 67029016 | - | lncRNA |
| ENSG00000245556.3_6 | SCAMP1-AS1 | 0.9996955 | 1 | -0.079 | 2.956 | chr5 | 77638189 | 77656331 | - | lncRNA |
| ENSG00000115762.16_3 | PLEKHB2 | 1 | 1 | -0.079 | 6.667 | chr2 | 131862420 | 132111282 | + | protein_coding |
| ENSG00000261167.1_6 | AC107027.3 | 0.98694529 | 1 | -0.079 | 1.607 | chr3 | 131173970 | 131177442 | - | lncRNA |
| ENSG00000076924.12_4 | XAB2 | 0.99999868 | 1 | -0.079 | 5.106 | chr19 | 7684411 | 7694431 | - | protein_coding |
| ENSG00000105676.14_6 | ARMC6 | 0.99999812 | 1 | -0.079 | 4.795 | chr19 | 19144384 | 19171120 | + | protein_coding |
| ENSG00000161800.13_4 | RACGAP1 | 1 | 1 | -0.079 | 7.048 | chr12 | 50370706 | 50426919 | - | protein_coding |
| ENSG00000229780.1_6 | UBE2Q1-AS1 | 0.91032475 | 1 | -0.079 | -0.223 | chr1 | 154526085 | 154527493 | + | lncRNA |
| ENSG00000111832.13_7 | RWDD1 | 0.99999706 | 1 | -0.079 | 5.342 | chr6 | 116892572 | 116918838 | + | protein_coding |
| ENSG00000187630.17_5 | DHRS4L2 | 0.9997991 | 1 | -0.079 | 3.069 | chr14 | 24439083 | 24475617 | + | protein_coding |
| ENSG00000118200.14_5 | CAMSAP2 | 0.99999999 | 1 | -0.079 | 6.853 | chr1 | 200708686 | 200829832 | + | protein_coding |
| ENSG00000277969.1_7 | AC006449.7 | 0.97469768 | 1 | -0.079 | 1.024 | chr17 | 36858705 | 36861000 | + | lncRNA |
| ENSG00000143751.10_4 | SDE2 | 0.99999975 | 1 | -0.080 | 5.250 | chr1 | 226170403 | 226187044 | - | protein_coding |
| ENSG00000253738.2_8 | OTUD6B-AS1 | 0.99999619 | 1 | -0.080 | 4.515 | chr8 | 92071546 | 92082811 | - | lncRNA |
| ENSG00000184887.13_5 | BTBD6 | 0.99999938 | 1 | -0.080 | 5.058 | chr14 | 105714827 | 105717430 | + | protein_coding |
| ENSG00000102172.16_4 | SMS | 1 | 1 | -0.080 | 7.206 | chrX | 21958827 | 22012955 | + | protein_coding |
| ENSG00000108556.10_6 | CHRNE | 0.89420339 | 1 | -0.080 | -0.964 | chr17 | 4801066 | 4837733 | - | protein_coding |
| ENSG00000143815.15_6 | LBR | 0.99999999 | 1 | -0.080 | 6.894 | chr1 | 225589204 | 225616627 | - | protein_coding |
| ENSG00000110429.14_4 | FBXO3 | 0.9999999 | 1 | -0.080 | 5.386 | chr11 | 33762485 | 33796089 | - | protein_coding |
| ENSG00000109065.12_7 | NAT9 | 0.99999983 | 1 | -0.080 | 5.206 | chr17 | 72766668 | 72772506 | - | protein_coding |
| ENSG00000178026.13_4 | LRRC75B | 0.99949433 | 1 | -0.080 | 2.859 | chr22 | 24981588 | 24989175 | - | protein_coding |
| ENSG00000141867.17_7 | BRD4 | 0.99999758 | 1 | -0.080 | 4.776 | chr19 | 15346330 | 15443356 | - | protein_coding |
| ENSG00000008869.12_5 | HEATR5B | 0.99999977 | 1 | -0.080 | 5.520 | chr2 | 37195526 | 37311515 | - | protein_coding |
| ENSG00000125779.22_3 | PANK2 | 0.99999995 | 1 | -0.080 | 5.700 | chr20 | 3869486 | 3910529 | + | protein_coding |
| ENSG00000143376.14_5 | SNX27 | 0.99999999 | 1 | -0.080 | 6.270 | chr1 | 151584482 | 151671567 | + | protein_coding |
| ENSG00000111247.14_2 | RAD51AP1 | 0.99999973 | 1 | -0.080 | 5.156 | chr12 | 4647964 | 4669214 | + | protein_coding |
| ENSG00000163156.12_5 | SCNM1 | 0.99998785 | 1 | -0.081 | 4.024 | chr1 | 151129140 | 151142772 | + | protein_coding |
| ENSG00000184260.5_3 | H2AC20 | 0.99967968 | 1 | -0.081 | 7.226 | chr1 | 149858525 | 149858914 | + | protein_coding |
| ENSG00000138468.16_5 | SENP7 | 0.99999265 | 1 | -0.081 | 4.209 | chr3 | 101043049 | 101232085 | - | protein_coding |
| ENSG00000251322.9_5 | SHANK3 | 0.98769441 | 1 | -0.081 | 2.024 | chr22 | 51112843 | 51171726 | + | protein_coding |

| | | | | | | | | | | |
|---|---|---|---|---|---|---|---|---|---|---|
| ENSG00000117010.17_4 | ZNF684 | 0.99948563 | 1 | -0.081 | 1.883 | chr1 | 40997245 | 41013839 | + | protein_coding |
| ENSG00000244879.8_7 | GABPB1-AS1 | 0.99999975 | 1 | -0.081 | 5.856 | chr15 | 50647156 | 50664399 | + | lncRNA |
| ENSG00000117676.14_6 | RPS6KA1 | 0.99999958 | 1 | -0.081 | 4.948 | chr1 | 26856252 | 26901521 | + | protein_coding |
| ENSG00000038358.15_4 | EDC4 | 0.99999996 | 1 | -0.081 | 6.199 | chr16 | 67906955 | 67918402 | + | protein_coding |
| ENSG00000138231.13_4 | DBR1 | 0.99999068 | 1 | -0.081 | 4.285 | chr3 | 137879830 | 137893763 | - | protein_coding |
| ENSG00000063177.13_6 | RPL18 | 1 | 1 | -0.082 | 8.936 | chr19 | 49118585 | 49122441 | - | protein_coding |
| ENSG00000168438.15_4 | CDC40 | 0.9999987 | 1 | -0.082 | 4.777 | chr6 | 110501344 | 110575478 | + | protein_coding |
| ENSG00000244026.6_4 | FAM86DP | 0.99997244 | 1 | -0.082 | 3.795 | chr3 | 75470703 | 75484261 | - | transcribed_unprocessed_pseudogene |
| ENSG00000117543.21_6 | DPH5 | 0.9997564 | 1 | -0.082 | 3.323 | chr1 | 101455179 | 101491644 | - | protein_coding |
| ENSG00000185608.8_2 | MRPL40 | 0.99999888 | 1 | -0.082 | 4.768 | chr22 | 19419425 | 19423598 | + | protein_coding |
| ENSG00000163728.11_8 | TTC14 | 0.99999852 | 1 | -0.082 | 4.764 | chr3 | 180319951 | 180335616 | + | protein_coding |
| ENSG00000246889.2_7 | AP000487.1 | 0.97063671 | 1 | -0.083 | 0.692 | chr11 | 70218352 | 70244594 | - | lncRNA |
| ENSG00000176920.13_5 | FUT2 | 0.95128296 | 1 | -0.083 | 0.558 | chr19 | 49199228 | 49209208 | + | protein_coding |
| ENSG00000196505.11_4 | GDAP2 | 0.99999906 | 1 | -0.083 | 4.811 | chr1 | 118406107 | 118472244 | - | protein_coding |
| ENSG00000148291.10_3 | SURF2 | 0.99974891 | 1 | -0.083 | 3.014 | chr9 | 136223426 | 136228034 | + | protein_coding |
| ENSG00000168653.11_3 | NDUFS5 | 0.99999998 | 1 | -0.083 | 6.288 | chr1 | 39491990 | 39500308 | + | protein_coding |
| ENSG00000100376.12_3 | FAM118A | 0.99988417 | 1 | -0.083 | 3.365 | chr22 | 45704849 | 45737836 | + | protein_coding |
| ENSG00000100764.14_5 | PSMC1 | 0.99999911 | 1 | -0.083 | 5.195 | chr14 | 90722871 | 90741773 | + | protein_coding |
| ENSG00000049618.23_6 | ARID1B | 0.9999998 | 1 | -0.083 | 5.440 | chr6 | 157097154 | 157531913 | + | protein_coding |
| ENSG00000136940.14_4 | PDCL | 0.99999961 | 1 | -0.083 | 5.469 | chr9 | 125560668 | 125590867 | - | protein_coding |
| ENSG00000100142.15_5 | POLR2F | 0.99996224 | 1 | -0.083 | 3.768 | chr22 | 38348614 | 38437922 | + | protein_coding |
| ENSG00000230896.1_8 | AL604028.1 | 0.93479838 | 1 | -0.084 | 0.325 | chr1 | 46160356 | 46162747 | - | lncRNA |
| ENSG00000165688.12_4 | PMPCA | 0.99999997 | 1 | -0.084 | 5.843 | chr9 | 139305093 | 139318213 | + | protein_coding |
| ENSG00000261360.1_7 | AC010491.1 | 0.93685546 | 1 | -0.084 | -0.014 | chr5 | 14661917 | 14664713 | - | lncRNA |
| ENSG00000077463.15_8 | SIRT6 | 0.99977831 | 1 | -0.084 | 3.456 | chr19 | 4174106 | 4182563 | - | protein_coding |
| ENSG00000228019.1_6 | AC006480.2 | 0.88785247 | 1 | -0.084 | -1.048 | chr7 | 66772592 | 66773343 | - | unprocessed_pseudogene |
| ENSG00000166439.6_5 | RNF169 | 0.99999984 | 1 | -0.084 | 5.353 | chr11 | 74459894 | 74553458 | + | protein_coding |
| ENSG00000283761.1_8 | AC118553.2 | 0.89294106 | 1 | -0.084 | -0.600 | chr1 | 100435567 | 100548877 | + | protein_coding |
| ENSG00000101782.15_4 | RIOK3 | 0.9999995 | 1 | -0.084 | 5.241 | chr18 | 21033251 | 21066567 | + | protein_coding |
| ENSG00000113645.14_4 | WWC1 | 0.99999999 | 1 | -0.084 | 6.835 | chr5 | 167718656 | 167899308 | + | protein_coding |
| ENSG00000173914.12_5 | RBM4B | 0.99997558 | 1 | -0.084 | 4.263 | chr11 | 66432469 | 66445358 | - | protein_coding |
| ENSG00000154930.15_4 | ACSS1 | 0.99993188 | 1 | -0.084 | 3.582 | chr20 | 24986866 | 25039616 | - | protein_coding |
| ENSG00000111445.14_3 | RFC5 | 0.99999996 | 1 | -0.084 | 5.818 | chr12 | 118451393 | 118470935 | + | protein_coding |
| ENSG00000164620.9_3 | RELL2 | 0.98428621 | 1 | -0.084 | 1.408 | chr5 | 141016517 | 141020631 | + | protein_coding |
| ENSG00000271646.1_7 | AC099343.4 | 0.89184094 | 1 | -0.084 | -0.488 | chr4 | 185395956 | 185398458 | + | lncRNA |
| ENSG00000196141.14_6 | SPATS2L | 0.99999999 | 1 | -0.084 | 6.686 | chr2 | 201170604 | 201346987 | + | protein_coding |
| ENSG00000100941.9_3 | PNN | 0.99999997 | 1 | -0.084 | 5.934 | chr14 | 39644387 | 39652424 | + | protein_coding |
| ENSG00000242125.3_5 | SNHG3 | 0.99989323 | 1 | -0.084 | 6.968 | chr1 | 28832492 | 28837404 | + | lncRNA |
| ENSG00000117533.15_4 | VAMP4 | 0.99993422 | 1 | -0.085 | 3.615 | chr1 | 171669300 | 171711214 | - | protein_coding |
| ENSG00000176731.12_5 | RBIS | 0.9989655 | 1 | -0.085 | 3.186 | chr8 | 86126283 | 86132650 | - | protein_coding |
| ENSG00000026103.22_4 | FAS | 0.99998394 | 1 | -0.085 | 3.840 | chr10 | 90750288 | 90776816 | + | protein_coding |
| ENSG00000011590.14_4 | ZBTB32 | 0.99896478 | 1 | -0.085 | 2.971 | chr19 | 36195460 | 36207940 | + | protein_coding |
| ENSG00000107758.15_4 | PPP3CB | 0.99999712 | 1 | -0.085 | 4.573 | chr10 | 75196186 | 75255782 | - | protein_coding |
| ENSG00000110881.11_5 | ASIC1 | 0.99714243 | 1 | -0.085 | 2.620 | chr12 | 50451331 | 50477394 | + | protein_coding |
| ENSG00000188566.13_4 | NDOR1 | 0.99983442 | 1 | -0.085 | 4.680 | chr9 | 140100137 | 140111461 | + | protein_coding |
| ENSG00000141994.16_7 | DUS3L | 0.99972911 | 1 | -0.085 | 4.460 | chr19 | 5784843 | 5791236 | - | protein_coding |
| ENSG00000238123.1_7 | MID1IP1-AS1 | 0.92264812 | 1 | -0.085 | 0.149 | chrX | 38660821 | 38663136 | - | lncRNA |
| ENSG00000134463.15_4 | ECHDC3 | 0.99845558 | 1 | -0.085 | 2.560 | chr10 | 11784365 | 11806069 | + | protein_coding |
| ENSG00000268189.2_6 | AC005785.1 | 0.892971 | 1 | -0.085 | -0.895 | chr19 | 15489887 | 15492005 | + | lncRNA |
| ENSG00000269293.3_8 | ZSCAN16-AS1 | 0.9777603 | 1 | -0.085 | 1.195 | chr6 | 27982900 | 28105071 | - | lncRNA |
| ENSG00000162073.14_3 | PAQR4 | 0.99990175 | 1 | -0.085 | 3.616 | chr16 | 3019271 | 3023485 | + | protein_coding |
| ENSG00000168301.13_5 | KCTD6 | 0.9998197 | 1 | -0.085 | 3.728 | chr3 | 58477823 | 58488087 | + | protein_coding |
| ENSG00000122299.12_4 | ZC3H7A | 0.99999989 | 1 | -0.086 | 5.340 | chr16 | 11844442 | 11891114 | - | protein_coding |
| ENSG00000142168.14_2 | SOD1 | 1 | 1 | -0.086 | 7.274 | chr21 | 33031935 | 33041244 | + | protein_coding |
| ENSG00000155393.13_4 | HEATR3 | 0.99999963 | 1 | -0.086 | 5.270 | chr16 | 50099881 | 50141183 | + | protein_coding |
| ENSG00000100281.14_4 | HMGXB4 | 0.99998793 | 1 | -0.086 | 4.412 | chr22 | 35653445 | 35691800 | + | protein_coding |
| ENSG00000157538.14_5 | VPS26C | 0.99999959 | 1 | -0.086 | 5.076 | chr21 | 38595721 | 38640220 | - | protein_coding |
| ENSG00000222011.9_6 | FAM185A | 0.99973637 | 1 | -0.086 | 3.093 | chr7 | 102389418 | 102449672 | + | protein_coding |

| | | | | | | | | | | |
|---|---|---|---|---|---|---|---|---|---|---|
| ENSG00000270878.1_6 | AL136038.6 | 0.91324344 | 1 | -0.086 | -0.189 | chr14 | 64060749 | 64062009 | - | processed_pseudogene |
| ENSG00000203965.13_4 | EFCAB7 | 0.99965871 | 1 | -0.086 | 3.271 | chr1 | 63989043 | 64038364 | + | protein_coding |
| ENSG00000167881.15_4 | SRP68 | 0.99999985 | 1 | -0.086 | 5.367 | chr17 | 74034856 | 74068598 | - | protein_coding |
| ENSG00000171735.19_4 | CAMTA1 | 0.99997578 | 1 | -0.086 | 3.699 | chr1 | 6845514 | 7829766 | + | protein_coding |
| ENSG00000164190.19_5 | NIPBL | 0.99999992 | 1 | -0.086 | 5.858 | chr5 | 36876871 | 37066515 | + | protein_coding |
| ENSG00000129951.18_8 | PLPPR3 | 0.92976998 | 1 | -0.086 | 0.373 | chr19 | 812488 | 821977 | - | protein_coding |
| ENSG00000148399.13_5 | DPH7 | 0.99999638 | 1 | -0.086 | 4.524 | chr9 | 140448896 | 140473377 | - | protein_coding |
| ENSG00000050393.11_6 | MCUR1 | 0.99999973 | 1 | -0.087 | 5.276 | chr6 | 13786789 | 13814800 | - | protein_coding |
| ENSG00000163597.15_3 | SNHG16 | 0.99999999 | 1 | -0.087 | 6.602 | chr17 | 74553846 | 74561430 | + | lncRNA |
| ENSG00000197150.12_7 | ABCB8 | 0.9999996 | 1 | -0.087 | 5.874 | chr7 | 150725509 | 150744869 | + | protein_coding |
| ENSG00000158711.14_3 | ELK4 | 0.99999996 | 1 | -0.087 | 6.287 | chr1 | 205566684 | 205601139 | - | protein_coding |
| ENSG00000241769.7_8 | LINC00893 | 0.91431816 | 1 | -0.087 | 0.250 | chrX | 148609130 | 148622504 | - | lncRNA |
| ENSG00000270558.1_5 | AC025449.1 | 0.96543607 | 1 | -0.087 | 0.902 | chr5 | 37286551 | 37287079 | - | processed_pseudogene |
| ENSG00000103275.20_4 | UBE2I | 0.99999936 | 1 | -0.087 | 5.246 | chr16 | 1358881 | 1377019 | + | protein_coding |
| ENSG00000141552.17_5 | ANAPC11 | 0.99999996 | 1 | -0.087 | 6.006 | chr17 | 79848666 | 79858867 | + | protein_coding |
| ENSG00000106305.10_3 | AIMP2 | 0.99999847 | 1 | -0.087 | 4.518 | chr7 | 6048886 | 6063465 | + | protein_coding |
| ENSG00000136243.12 | NUPL2 | 0.99994996 | 1 | -0.087 | 3.664 | chr7 | 23221446 | 23240630 | + | protein_coding |
| ENSG00000166478.10_4 | ZNF143 | 0.9999894 | 1 | -0.087 | 4.479 | chr11 | 9481866 | 9550071 | + | protein_coding |
| ENSG00000119946.11_3 | CNNM1 | 0.99999993 | 1 | -0.087 | 5.712 | chr10 | 101089113 | 101154087 | + | protein_coding |
| ENSG00000273002.1_7 | AL355388.1 | 0.94931501 | 1 | -0.087 | 0.562 | chr1 | 155948590 | 155952777 | + | lncRNA |
| ENSG00000010322.16_5 | NISCH | 0.99999945 | 1 | -0.087 | 5.212 | chr3 | 52489134 | 52527084 | + | protein_coding |
| ENSG00000138709.19_5 | LARP1B | 0.99999643 | 1 | -0.087 | 4.448 | chr4 | 128982441 | 129144086 | + | protein_coding |
| ENSG00000100034.14_5 | PPM1F | 0.99938517 | 1 | -0.087 | 5.079 | chr22 | 22273798 | 22307220 | - | protein_coding |
| ENSG00000065060.17_3 | UHRF1BP1 | 0.99999971 | 1 | -0.087 | 5.385 | chr6 | 34759792 | 34850915 | + | protein_coding |
| ENSG00000078269.15_5 | SYNJ2 | 0.99999999 | 1 | -0.087 | 6.822 | chr6 | 158402895 | 158520208 | + | protein_coding |
| ENSG00000133624.13_5 | ZNF767P | 0.99974234 | 1 | -0.087 | 3.568 | chr7 | 149244245 | 149321843 | - | transcribed_unprocessed_pseudogene |
| ENSG00000102595.20_5 | UGGT2 | 0.99999958 | 1 | -0.088 | 5.247 | chr13 | 96453834 | 96705736 | - | protein_coding |
| ENSG00000137502.10_5 | RAB30 | 0.99984218 | 1 | -0.088 | 3.210 | chr11 | 82684175 | 82782965 | - | protein_coding |
| ENSG00000179304.16_6 | FAM156B | 0.94740492 | 1 | -0.088 | 0.986 | chrX | 52920336 | 52937587 | + | protein_coding |
| ENSG00000145107.15_5 | TM4SF19 | 0.90752381 | 1 | -0.088 | -0.282 | chr3 | 196046213 | 196065374 | - | protein_coding |
| ENSG00000065809.13_7 | FAM107B | 1 | 1 | -0.088 | 7.888 | chr10 | 14560556 | 14816896 | - | protein_coding |
| ENSG00000130193.8_3 | THEM6 | 0.99999939 | 1 | -0.088 | 4.999 | chr8 | 143808641 | 143818345 | + | protein_coding |
| ENSG00000237238.3_5 | BMS1P10 | 0.98932847 | 1 | -0.088 | 1.138 | chr9 | 67254425 | 67340089 | - | transcribed_unprocessed_pseudogene |
| ENSG00000118418.14_4 | HMGN3 | 0.99999883 | 1 | -0.088 | 4.962 | chr6 | 79910962 | 79944455 | - | protein_coding |
| ENSG00000162738.6_3 | VANGL2 | 0.98474336 | 1 | -0.088 | 1.441 | chr1 | 160370354 | 160398460 | + | protein_coding |
| ENSG00000160767.21_7 | FAM189B | 0.99929174 | 1 | -0.088 | 3.131 | chr1 | 155216996 | 155225274 | - | protein_coding |
| ENSG00000176619.13_6 | LMNB2 | 0.99965524 | 1 | -0.088 | 7.311 | chr19 | 2427636 | 2456957 | - | protein_coding |
| ENSG00000112282.18_5 | MED23 | 0.99999945 | 1 | -0.088 | 5.233 | chr6 | 131895106 | 131949382 | - | protein_coding |
| ENSG00000128829.12_3 | EIF2AK4 | 0.99999998 | 1 | -0.088 | 6.104 | chr15 | 40226316 | 40327792 | + | protein_coding |
| ENSG00000155329.12_4 | ZCCHC10 | 0.99999293 | 1 | -0.088 | 4.352 | chr5 | 132332677 | 132362296 | - | protein_coding |
| ENSG00000090060.18_5 | PAPOLA | 1 | 1 | -0.089 | 7.893 | chr14 | 96967770 | 97033448 | + | protein_coding |
| ENSG00000101391.21_6 | CDK5RAP1 | 0.9999996 | 1 | -0.089 | 5.232 | chr20 | 31946645 | 31989367 | - | protein_coding |
| ENSG00000203999.9_6 | LINC01270 | 0.99937383 | 1 | -0.089 | 3.152 | chr20 | 48909246 | 48932025 | + | lncRNA |
| ENSG00000033030.15_6 | ZCCHC8 | 0.9999999 | 1 | -0.089 | 5.484 | chr12 | 122956146 | 122985620 | - | protein_coding |
| ENSG00000086062.12_3 | B4GALT1 | 0.99999992 | 1 | -0.089 | 6.584 | chr9 | 33104080 | 33167354 | - | protein_coding |
| ENSG00000127191.18_4 | TRAF2 | 0.99999863 | 1 | -0.089 | 5.673 | chr9 | 139776364 | 139821059 | + | protein_coding |
| ENSG00000163755.9_4 | HPS3 | 0.99999995 | 1 | -0.089 | 5.723 | chr3 | 148847425 | 148891519 | + | protein_coding |
| ENSG00000115084.14_4 | SLC35F5 | 1 | 1 | -0.089 | 7.275 | chr2 | 114462588 | 114514270 | - | protein_coding |
| ENSG00000237938.5_8 | AL450998.3 | 0.89203359 | 1 | -0.089 | -0.324 | chr1 | 16046807 | 16063391 | - | lncRNA |
| ENSG00000065621.15_3 | GSTO2 | 0.96987544 | 1 | -0.089 | 1.077 | chr10 | 106028631 | 106064708 | + | protein_coding |
| ENSG00000223947.1_7 | AC016738.1 | 0.96294452 | 1 | -0.089 | 0.526 | chr2 | 101610138 | 101618706 | - | lncRNA |
| ENSG00000162775.16_5 | RBM15 | 0.99999993 | 1 | -0.089 | 5.601 | chr1 | 110881128 | 110889303 | + | protein_coding |
| ENSG00000072736.19_6 | NFATC3 | 0.99999897 | 1 | -0.089 | 5.049 | chr16 | 68118654 | 68263162 | + | protein_coding |
| ENSG00000005194.15_6 | CIAPIN1 | 0.99999993 | 1 | -0.090 | 6.003 | chr16 | 57462099 | 57481332 | - | protein_coding |
| ENSG00000142207.7_4 | URB1 | 0.99999998 | 1 | -0.090 | 6.217 | chr21 | 33683329 | 33765321 | - | protein_coding |
| ENSG00000106153.13_4 | CHCHD2 | 0.99999999 | 1 | -0.090 | 7.519 | chr7 | 56169266 | 56174169 | - | protein_coding |
| ENSG00000175334.8_3 | BANF1 | 0.99999962 | 1 | -0.090 | 5.362 | chr11 | 65769699 | 65771620 | + | protein_coding |
| ENSG00000169180.11_7 | XPO6 | 0.99999999 | 1 | -0.090 | 6.617 | chr16 | 28109300 | 28223241 | - | protein_coding |

| | | | | | | | | | | |
|---|---|---|---|---|---|---|---|---|---|---|
| ENSG00000240898.1_5 | AC132942.1 | 0.94782973 | 1 | -0.090 | 0.573 | chr17 | 3568514 | 3568996 | + | processed_pseudogene |
| ENSG00000123643.13_5 | SLC36A1 | 0.9996804 | 1 | -0.090 | 3.423 | chr5 | 150816607 | 150871940 | + | protein_coding |
| ENSG00000241560.7_7 | ZBTB20-AS1 | 0.98543848 | 1 | -0.090 | 1.461 | chr3 | 114070618 | 114107825 | + | lncRNA |
| ENSG00000166800.10_3 | LDHAL6A | 0.89978343 | 1 | -0.090 | -0.512 | chr11 | 18477371 | 18501148 | + | protein_coding |
| ENSG00000150457.9_4 | LATS2 | 0.99999833 | 1 | -0.090 | 4.818 | chr13 | 21547175 | 21635725 | - | protein_coding |
| ENSG00000143368.10_6 | SF3B4 | 0.99999977 | 1 | -0.090 | 5.494 | chr1 | 149895209 | 149899695 | - | protein_coding |
| ENSG00000114302.16_5 | PRKAR2A | 0.99999997 | 1 | -0.090 | 6.198 | chr3 | 48782030 | 48885307 | - | protein_coding |
| ENSG00000059691.12_5 | GATB | 0.99997978 | 1 | -0.090 | 3.993 | chr4 | 152591656 | 152682159 | - | protein_coding |
| ENSG00000242485.6_4 | MRPL20 | 0.99999974 | 1 | -0.090 | 5.251 | chr1 | 1337289 | 1342673 | - | protein_coding |
| ENSG00000180822.11_4 | PSMG4 | 0.99992495 | 1 | -0.090 | 3.490 | chr6 | 3231637 | 3303607 | + | protein_coding |
| ENSG00000215301.10_8 | DDX3X | 1 | 1 | -0.090 | 8.698 | chrX | 41192537 | 41223725 | + | protein_coding |
| ENSG00000001167.14_3 | NFYA | 0.99996303 | 1 | -0.090 | 3.958 | chr6 | 41040684 | 41067715 | + | protein_coding |
| ENSG00000166913.13_4 | YWHAB | 1 | 1 | -0.090 | 7.987 | chr20 | 43514343 | 43537173 | + | protein_coding |
| ENSG00000287024.1_2 | AL022162.1 | 0.88550981 | 1 | -0.091 | -0.511 | chrX | 128809865 | 128812325 | - | lncRNA |
| ENSG00000067900.8_5 | ROCK1 | 0.99999999 | 1 | -0.091 | 6.364 | chr18 | 18526867 | 18691774 | - | protein_coding |
| ENSG00000127184.13_6 | COX7C | 0.99999999 | 1 | -0.091 | 7.250 | chr5 | 85913745 | 85916779 | + | protein_coding |
| ENSG00000270903.1_4 | HNRNPA3P9 | 0.91631908 | 1 | -0.091 | 0.131 | chr11 | 32613339 | 32614317 | - | processed_pseudogene |
| ENSG00000100813.14_6 | ACIN1 | 1 | 1 | -0.091 | 7.286 | chr14 | 23527773 | 23564823 | - | protein_coding |
| ENSG00000251593.1_2 | MSNP1 | 0.88775304 | 1 | -0.091 | -0.575 | chr5 | 25909612 | 25911343 | + | processed_pseudogene |
| ENSG00000279837.1_6 | AC112694.2 | 0.88332201 | 1 | -0.091 | -0.511 | chr11 | 18623429 | 18624196 | + | TEC |
| ENSG00000148303.17_3 | RPL7A | 1 | 1 | -0.091 | 9.790 | chr9 | 136215073 | 136218281 | + | protein_coding |
| ENSG00000144659.13_7 | SLC25A38 | 0.99999819 | 1 | -0.092 | 5.077 | chr3 | 39424861 | 39438842 | + | protein_coding |
| ENSG00000248643.5_4 | RBM14-RBM4 | 0.91461803 | 1 | -0.092 | -0.103 | chr11 | 66384097 | 66413940 | + | protein_coding |
| ENSG00000116641.18_7 | DOCK7 | 0.99999991 | 1 | -0.092 | 5.780 | chr1 | 62919969 | 63154057 | - | protein_coding |
| ENSG00000155542.12_4 | SETD9 | 0.99741455 | 1 | -0.092 | 2.253 | chr5 | 56205087 | 56221359 | + | protein_coding |
| ENSG00000171928.14_4 | TVP23B | 0.99999966 | 1 | -0.092 | 5.828 | chr17 | 18684424 | 18710027 | + | protein_coding |
| ENSG00000177613.9_4 | CSTF2T | 0.99999986 | 1 | -0.092 | 5.438 | chr10 | 53455246 | 53459355 | - | protein_coding |
| ENSG00000165525.18_4 | NEMF | 0.99999969 | 1 | -0.092 | 6.113 | chr14 | 50248801 | 50319539 | - | protein_coding |
| ENSG00000075234.17_5 | TTC38 | 0.9999862 | 1 | -0.092 | 3.997 | chr22 | 46663858 | 46689905 | + | protein_coding |
| ENSG00000112763.17_5 | BTN2A1 | 0.99999685 | 1 | -0.092 | 4.512 | chr6 | 26458132 | 26476849 | + | protein_coding |
| ENSG00000008226.20_4 | DLEC1 | 0.88864145 | 1 | -0.092 | -0.514 | chr3 | 38080696 | 38165516 | + | protein_coding |
| ENSG00000227258.6_6 | SMIM2-AS1 | 0.89874397 | 1 | -0.092 | -0.382 | chr13 | 44684587 | 44814653 | + | lncRNA |
| ENSG00000160688.19_3 | FLAD1 | 0.99999985 | 1 | -0.092 | 5.272 | chr1 | 154955814 | 154965587 | + | protein_coding |
| ENSG00000267751.5_7 | AC009005.1 | 0.94354534 | 1 | -0.092 | 0.508 | chr19 | 567210 | 572228 | - | lncRNA |
| ENSG00000136827.12_5 | TOR1A | 0.99999994 | 1 | -0.092 | 5.957 | chr9 | 132575221 | 132586523 | - | protein_coding |
| ENSG00000255302.5_4 | EID1 | 1 | 1 | -0.092 | 7.077 | chr15 | 49170331 | 49172370 | + | protein_coding |
| ENSG00000099783.12_7 | HNRNPM | 0.99999983 | 1 | -0.092 | 6.298 | chr19 | 8509651 | 8553998 | + | protein_coding |
| ENSG00000177954.14_6 | RPS27 | 1 | 1 | -0.092 | 8.860 | chr1 | 153963238 | 153964631 | + | protein_coding |
| ENSG00000197782.14_4 | ZNF780A | 0.99999995 | 1 | -0.092 | 5.946 | chr19 | 40575059 | 40596845 | - | protein_coding |
| ENSG00000259562.2_5 | AC090607.2 | 0.96194314 | 1 | -0.093 | 1.256 | chr15 | 78582869 | 78583563 | - | transcribed_processed_pseudogene |
| ENSG00000119508.18_3 | NR4A3 | 0.96085555 | 1 | -0.093 | 0.905 | chr9 | 102584137 | 102629173 | + | protein_coding |
| ENSG00000170222.12_3 | ADPRM | 0.99940187 | 1 | -0.093 | 2.927 | chr17 | 10600911 | 10614875 | + | protein_coding |
| ENSG00000237214.3_6 | AL080243.4 | 0.9011035 | 1 | -0.093 | 0.069 | chr22 | 41460056 | 41460637 | + | processed_pseudogene |
| ENSG00000245694.10_5 | CRNDE | 0.99999603 | 1 | -0.093 | 4.779 | chr16 | 54879101 | 54963101 | - | lncRNA |
| ENSG00000168495.13_4 | POLR3D | 0.99999959 | 1 | -0.093 | 5.288 | chr8 | 22102646 | 22112114 | + | protein_coding |
| ENSG00000146416.19_6 | AIG1 | 0.99998205 | 1 | -0.093 | 3.995 | chr6 | 143381633 | 143662195 | + | protein_coding |
| ENSG00000125868.16_3 | DSTN | 1 | 1 | -0.093 | 9.017 | chr20 | 17550720 | 17590564 | + | protein_coding |
| ENSG00000248278.1_5 | SUMO2P17 | 0.9058271 | 1 | -0.093 | 0.073 | chr17 | 46952222 | 46986345 | - | transcribed_processed_pseudogene |
| ENSG00000254535.4_5 | PABPC4L | 0.94356558 | 1 | -0.093 | 0.538 | chr4 | 135117488 | 135122944 | - | protein_coding |
| ENSG00000090905.19_4 | TNRC6A | 0.99999998 | 1 | -0.093 | 6.725 | chr16 | 24621530 | 24838953 | + | protein_coding |
| ENSG00000101346.14_4 | POFUT1 | 0.99999999 | 1 | -0.093 | 6.621 | chr20 | 30795683 | 30826461 | + | protein_coding |
| ENSG00000258424.1_7 | AL512791.1 | 0.90926271 | 1 | -0.093 | 0.075 | chr14 | 90868867 | 90871579 | - | lncRNA |
| ENSG00000213406.3_2 | ANXA2P1 | 0.87816801 | 1 | -0.094 | -0.520 | chr4 | 154228944 | 154229869 | - | processed_pseudogene |
| ENSG00000101608.12_6 | MYL12A | 0.99999998 | 1 | -0.094 | 6.549 | chr18 | 3247479 | 3256234 | + | protein_coding |
| ENSG00000136141.15_4 | LRCH1 | 0.99999854 | 1 | -0.094 | 4.575 | chr13 | 47127303 | 47327175 | + | protein_coding |
| ENSG00000145975.15_5 | FAM217A | 0.87985771 | 1 | -0.094 | -0.547 | chr6 | 4049668 | 4087578 | - | protein_coding |

| | | | | | | | | | | |
|---|---|---|---|---|---|---|---|---|---|---|
| ENSG00000134461.16_5 | ANKRD16 | 0.9997926 | 1 | -0.094 | 3.248 | chr10 | 5903579 | 5931869 | - | protein_coding |
| ENSG00000197296.6_4 | FITM2 | 0.9999948 | 1 | -0.094 | 4.614 | chr20 | 42931480 | 42939842 | - | protein_coding |
| ENSG00000121940.15_4 | CLCC1 | 0.99999878 | 1 | -0.094 | 4.946 | chr1 | 109472130 | 109506079 | - | protein_coding |
| ENSG00000145715.15_3 | RASA1 | 0.99999877 | 1 | -0.094 | 5.128 | chr5 | 86563700 | 86687748 | + | protein_coding |
| ENSG00000140931.20_5 | CMTM3 | 0.99999985 | 1 | -0.095 | 5.323 | chr16 | 66637777 | 66647795 | + | protein_coding |
| ENSG00000164494.12_5 | PDSS2 | 0.99999435 | 1 | -0.095 | 4.334 | chr6 | 107473766 | 107780768 | - | protein_coding |
| ENSG00000189306.11_4 | RRP7A | 0.99999934 | 1 | -0.095 | 5.827 | chr22 | 42904350 | 42915802 | - | protein_coding |
| ENSG00000225264.3_5 | ZNRF2P2 | 0.87615613 | 1 | -0.095 | -0.658 | chr7 | 29638411 | 29724871 | - | transcribed_processed_pseudogene |
| ENSG00000250910.8_8 | AC097467.3 | 0.87363286 | 1 | -0.095 | -0.798 | chr4 | 156094868 | 156302846 | + | lncRNA |
| ENSG00000170382.12_4 | LRRN2 | 0.98542531 | 1 | -0.095 | 3.488 | chr1 | 204586298 | 204654866 | - | protein_coding |
| ENSG00000103148.16_5 | NPRL3 | 0.99996256 | 1 | -0.095 | 4.091 | chr16 | 134270 | 188859 | - | protein_coding |
| ENSG00000145982.12_6 | FARS2 | 0.9999251 | 1 | -0.095 | 3.854 | chr6 | 5261277 | 5829425 | + | protein_coding |
| ENSG00000186352.9_3 | ANKRD37 | 0.88362532 | 1 | -0.095 | -0.189 | chr4 | 186317175 | 186321782 | + | protein_coding |
| ENSG00000106723.17_3 | SPIN1 | 0.99999998 | 1 | -0.095 | 6.503 | chr9 | 91003345 | 91093609 | + | protein_coding |
| ENSG00000160949.17_4 | TONSL | 0.99794315 | 1 | -0.095 | 5.831 | chr8 | 145654158 | 145669823 | - | protein_coding |
| ENSG00000164168.8_3 | TMEM184C | 0.99999992 | 1 | -0.095 | 5.949 | chr4 | 148538537 | 148593195 | + | protein_coding |
| ENSG00000112592.14_5 | TBP | 0.99971267 | 1 | -0.096 | 3.463 | chr6 | 170863390 | 170881958 | + | protein_coding |
| ENSG00000125755.19_5 | SYMPK | 0.9999997 | 1 | -0.096 | 6.962 | chr19 | 46318668 | 46366452 | - | protein_coding |
| ENSG00000145623.13_7 | OSMR | 0.99999997 | 1 | -0.096 | 6.671 | chr5 | 38845960 | 38945698 | + | protein_coding |
| ENSG00000173457.11_5 | PPP1R14B | 0.99999995 | 1 | -0.096 | 6.277 | chr11 | 64011951 | 64014415 | - | protein_coding |
| ENSG00000172113.9_5 | NME6 | 0.99998914 | 1 | -0.096 | 4.251 | chr3 | 48332212 | 48343175 | - | protein_coding |
| ENSG00000281490.1_5 | CICP14 | 0.99932027 | 1 | -0.096 | 3.028 | chr7 | 128296016 | 128298845 | - | processed_pseudogene |
| ENSG00000166482.11_4 | MFAP4 | 0.9760521 | 1 | -0.096 | 1.227 | chr17 | 19286755 | 19290553 | - | protein_coding |
| ENSG00000164414.18_4 | SLC35A1 | 0.97442571 | 1 | -0.096 | 2.207 | chr6 | 88180341 | 88222054 | + | protein_coding |
| ENSG00000108021.20_8 | TASOR2 | 1 | 1 | -0.096 | 7.281 | chr10 | 5726801 | 5805703 | + | protein_coding |
| ENSG00000093010.14_5 | COMT | 0.99999948 | 1 | -0.096 | 5.328 | chr22 | 19929256 | 19957498 | + | protein_coding |
| ENSG00000230207.1_4 | RPL4P5 | 0.96991376 | 1 | -0.096 | 1.520 | chr9 | 7477045 | 7478320 | - | processed_pseudogene |
| ENSG00000102547.19_4 | CAB39L | 0.99619651 | 1 | -0.096 | 2.418 | chr13 | 49882786 | 50018200 | - | protein_coding |
| ENSG00000214113.10_5 | LYRM4 | 0.99998123 | 1 | -0.096 | 4.102 | chr6 | 5102827 | 5261172 | - | protein_coding |
| ENSG00000170899.11_5 | GSTA4 | 0.9997805 | 1 | -0.096 | 3.172 | chr6 | 52842746 | 52860102 | - | protein_coding |
| ENSG00000177888.7_3 | ZBTB41 | 0.99999918 | 1 | -0.097 | 5.887 | chr1 | 197122810 | 197169672 | - | protein_coding |
| ENSG00000005700.15_6 | IBTK | 0.99999999 | 1 | -0.097 | 7.436 | chr6 | 82879703 | 82957471 | - | protein_coding |
| ENSG00000163961.4_4 | RNF168 | 0.99996899 | 1 | -0.097 | 4.480 | chr3 | 196195654 | 196230639 | - | protein_coding |
| ENSG00000100068.13_8 | LRP5L | 0.99824432 | 1 | -0.097 | 2.455 | chr22 | 25747385 | 25801344 | - | protein_coding |
| ENSG00000250802.7_6 | ZBED3-AS1 | 0.9799174 | 1 | -0.097 | 1.789 | chr5 | 76382513 | 76462734 | + | lncRNA |
| ENSG00000162636.16_3 | FAM102B | 0.99999881 | 1 | -0.097 | 4.803 | chr1 | 109102711 | 109187522 | + | protein_coding |
| ENSG00000075223.14_4 | SEMA3C | 1 | 1 | -0.097 | 8.208 | chr7 | 80371854 | 80551675 | - | protein_coding |
| ENSG00000225573.4_4 | RPL35P5 | 0.96815489 | 1 | -0.097 | 1.014 | chr7 | 66071725 | 66072094 | - | processed_pseudogene |
| ENSG00000108518.8_3 | PFN1 | 0.99999952 | 1 | -0.097 | 8.789 | chr17 | 4848947 | 4852356 | - | protein_coding |
| ENSG00000135540.11_8 | NHSL1 | 0.99976316 | 1 | -0.097 | 3.219 | chr6 | 138743180 | 139013708 | - | protein_coding |
| ENSG00000247853.2_6 | AC006064.2 | 0.86984779 | 1 | -0.097 | -0.798 | chr12 | 6687788 | 6693905 | + | lncRNA |
| ENSG00000275017.1_5 | AL353748.2 | 0.87469118 | 1 | -0.097 | -0.631 | chr9 | 91006277 | 91006747 | - | processed_pseudogene |
| ENSG00000160209.19_3 | PDXK | 1 | 1 | -0.097 | 8.826 | chr21 | 45138975 | 45182188 | + | protein_coding |
| ENSG00000147127.8_5 | RAB41 | 0.88718895 | 1 | -0.097 | -0.311 | chrX | 69501943 | 69504852 | + | protein_coding |
| ENSG00000274528.1_7 | AC090970.2 | 0.87474911 | 1 | -0.097 | -0.698 | chr15 | 52309364 | 52310229 | - | lncRNA |
| ENSG00000270760.1_7 | AD001527.1 | 0.94054649 | 1 | -0.097 | 0.267 | chr19 | 36634983 | 36639250 | - | lncRNA |
| ENSG00000236540.7_8 | AC006547.1 | 0.8998052 | 1 | -0.097 | -0.124 | chr22 | 20045553 | 20058092 | - | lncRNA |
| ENSG00000117528.14_5 | ABCD3 | 0.99999997 | 1 | -0.098 | 6.442 | chr1 | 94883945 | 94984222 | + | protein_coding |
| ENSG00000113163.17_6 | CERT1 | 0.99999988 | 1 | -0.098 | 5.500 | chr5 | 74652170 | 74807963 | - | protein_coding |
| ENSG00000237972.1_2 | TUBG1P | 0.86897341 | 1 | -0.098 | -0.865 | chr7 | 43958296 | 43959653 | + | processed_pseudogene |
| ENSG00000198780.12_4 | FAM169A | 0.99999736 | 1 | -0.098 | 5.018 | chr5 | 74073399 | 74162791 | - | protein_coding |
| ENSG00000100580.8_4 | TMED8 | 0.99999988 | 1 | -0.098 | 5.549 | chr14 | 77801372 | 77843437 | - | protein_coding |
| ENSG00000227495.1_7 | KIF1C-AS1 | 0.88920341 | 1 | -0.098 | -0.168 | chr17 | 4922509 | 4923388 | - | lncRNA |
| ENSG00000103187.8_5 | COTL1 | 1 | 1 | -0.098 | 8.041 | chr16 | 84599202 | 84651684 | - | protein_coding |
| ENSG00000168904.15_7 | LRRC28 | 0.99998454 | 1 | -0.098 | 4.057 | chr15 | 99791567 | 99930934 | + | protein_coding |
| ENSG00000185024.17_9 | BRF1 | 0.99999776 | 1 | -0.098 | 5.493 | chr14 | 105675623 | 105781926 | - | protein_coding |
| ENSG00000215021.9_4 | PHB2 | 0.99999999 | 1 | -0.099 | 7.528 | chr12 | 7074490 | 7079916 | - | protein_coding |
| ENSG00000100731.16_5 | PCNX1 | 0.99999998 | 1 | -0.099 | 6.712 | chr14 | 71374122 | 71582099 | + | protein_coding |

| | | | | | | | | | | |
|---|---|---|---|---|---|---|---|---|---|---|
| ENSG00000179818.14_5 | PCBP1-AS1 | 0.99969949 | 1 | -0.099 | 3.373 | chr2 | 70189395 | 70330352 | - | lncRNA |
| ENSG00000156162.16_3 | DPY19L4 | 0.99999989 | 1 | -0.099 | 5.746 | chr8 | 95731931 | 95806064 | + | protein_coding |
| ENSG00000007202.15_6 | KIAA0100 | 1 | 1 | -0.099 | 8.204 | chr17 | 26941464 | 26972472 | - | protein_coding |
| ENSG00000261136.1_6 | AC023908.3 | 0.93796367 | 1 | -0.099 | 0.616 | chr15 | 40074772 | 40077818 | + | lncRNA |
| ENSG00000179933.6_3 | C14orf119 | 0.99999991 | 1 | -0.099 | 5.707 | chr14 | 23564714 | 23569665 | + | protein_coding |
| ENSG00000158483.16_5 | FAM86C1 | 0.99823756 | 1 | -0.099 | 2.779 | chr11 | 71498556 | 71512283 | + | protein_coding |
| ENSG00000135837.16_6 | CEP350 | 0.99999993 | 1 | -0.099 | 7.139 | chr1 | 179923908 | 180084010 | + | protein_coding |
| ENSG00000105220.17_7 | GPI | 0.99999999 | 1 | -0.099 | 9.394 | chr19 | 34850385 | 34893318 | + | protein_coding |
| ENSG00000273344.1_5 | PAXIP1-AS1 | 0.99793644 | 1 | -0.099 | 2.338 | chr7 | 154795158 | 154797413 | + | lncRNA |
| ENSG00000285043.1_2 | AC093512.2 | 0.96323561 | 1 | -0.099 | 0.986 | chr16 | 30064411 | 30081741 | + | protein_coding |
| ENSG00000158006.14_4 | PAFAH2 | 0.99997686 | 1 | -0.099 | 3.979 | chr1 | 26286258 | 26324608 | - | protein_coding |
| ENSG00000100347.15_4 | SAMM50 | 0.99999706 | 1 | -0.099 | 4.731 | chr22 | 44351322 | 44406411 | + | protein_coding |
| ENSG00000112599.9_5 | GUCA1B | 0.880612 | 1 | -0.099 | -0.576 | chr6 | 42151022 | 42162694 | - | protein_coding |
| ENSG00000161970.15_8 | RPL26 | 0.99796171 | 1 | -0.099 | 5.941 | chr17 | 8280834 | 8286531 | - | protein_coding |
| ENSG00000288586.1_1 | AL357874.3 | 0.87062234 | 1 | -0.099 | -0.832 | chr9 | 35603434 | 35605136 | - | lncRNA |
| ENSG00000125485.17_5 | DDX31 | 0.99999923 | 1 | -0.100 | 4.922 | chr9 | 135468384 | 135545788 | - | protein_coding |
| ENSG00000112367.10_2 | FIG4 | 0.99996259 | 1 | -0.100 | 3.765 | chr6 | 110012515 | 110146631 | + | protein_coding |
| ENSG00000198088.10_7 | NUP62CL | 0.99996609 | 1 | -0.100 | 3.797 | chrX | 106366657 | 106449663 | - | protein_coding |
| ENSG00000188827.11_4 | SLX4 | 0.99999388 | 1 | -0.100 | 4.596 | chr16 | 3631182 | 3661607 | - | protein_coding |
| ENSG00000213693.4_4 | SEC14L1P1 | 0.93038043 | 1 | -0.100 | 0.497 | chr11 | 43919006 | 43921186 | + | processed_pseudogene |
| ENSG00000141385.10_4 | AFG3L2 | 0.99999986 | 1 | -0.100 | 5.593 | chr18 | 12328943 | 12377226 | - | protein_coding |
| ENSG00000117394.23_7 | SLC2A1 | 0.99999999 | 1 | -0.100 | 7.666 | chr1 | 43391046 | 43424539 | - | protein_coding |
| ENSG00000286808.1_3 | AC092338.3 | 0.90146117 | 1 | -0.100 | -0.150 | chr16 | 22342360 | 22354259 | - | lncRNA |
| ENSG00000198231.13_6 | DDX42 | 0.99999999 | 1 | -0.100 | 7.197 | chr17 | 61850963 | 61896677 | + | protein_coding |
| ENSG00000168807.16_5 | SNTB2 | 0.99999962 | 1 | -0.100 | 6.193 | chr16 | 69221032 | 69342955 | + | protein_coding |
| ENSG00000100030.14_3 | MAPK1 | 1 | 1 | -0.100 | 7.135 | chr22 | 22108789 | 22221970 | - | protein_coding |
| ENSG00000036549.13_6 | AC118549.1 | 0.99999996 | 1 | -0.100 | 5.962 | chr1 | 78028101 | 78149104 | - | protein_coding |
| ENSG00000279738.1_7 | AL022311.1 | 0.99308475 | 1 | -0.100 | 2.060 | chr22 | 38272155 | 38291570 | + | lncRNA |
| ENSG00000162924.14_6 | REL | 0.99997669 | 1 | -0.100 | 4.049 | chr2 | 61108656 | 61158745 | + | protein_coding |
| ENSG00000148339.12_5 | SLC25A25 | 0.99996916 | 1 | -0.101 | 4.078 | chr9 | 130830480 | 130871524 | + | protein_coding |
| ENSG00000285106.2_1 | AC016831.6 | 0.99297234 | 1 | -0.101 | 2.083 | chr7 | 130476023 | 130794920 | - | lncRNA |
| ENSG00000163848.19_4 | ZNF148 | 0.99999987 | 1 | -0.101 | 6.314 | chr3 | 124944405 | 125094198 | - | protein_coding |
| ENSG00000069345.12_5 | DNAJA2 | 0.99999999 | 1 | -0.101 | 7.002 | chr16 | 46989274 | 47007585 | - | protein_coding |
| ENSG00000105281.12_5 | SLC1A5 | 0.99999999 | 1 | -0.101 | 7.167 | chr19 | 47278140 | 47291851 | - | protein_coding |
| ENSG00000127884.5_4 | ECHS1 | 0.99999999 | 1 | -0.101 | 6.590 | chr10 | 135175989 | 135186858 | - | protein_coding |
| ENSG00000102078.16_6 | SLC25A14 | 0.99986983 | 1 | -0.101 | 3.553 | chrX | 129473862 | 129507335 | + | protein_coding |
| ENSG00000205531.13_6 | NAP1L4 | 0.99999998 | 1 | -0.101 | 6.588 | chr11 | 2965661 | 3013607 | - | protein_coding |
| ENSG00000099338.23_6 | CATSPERG | 0.90782248 | 1 | -0.101 | 0.147 | chr19 | 38826415 | 38861583 | + | protein_coding |
| ENSG00000226415.1_4 | TPI1P1 | 0.99814642 | 1 | -0.101 | 3.302 | chr1 | 77165474 | 77166223 | + | processed_pseudogene |
| ENSG00000286485.1_1 | Z80897.1 | 0.86705977 | 1 | -0.101 | -0.978 | chr22 | 37162522 | 37165066 | + | lncRNA |
| ENSG00000113845.10_3 | TIMMDC1 | 0.99999993 | 1 | -0.101 | 5.748 | chr3 | 119217394 | 119243937 | + | protein_coding |
| ENSG00000125846.15_2 | ZNF133 | 0.99998926 | 1 | -0.101 | 4.048 | chr20 | 18268927 | 18297640 | + | protein_coding |
| ENSG00000271870.1_7 | AC024060.2 | 0.96449642 | 1 | -0.102 | 0.981 | chr3 | 3194626 | 3195119 | + | lncRNA |
| ENSG00000100320.23_6 | RBFOX2 | 0.99999995 | 1 | -0.102 | 6.112 | chr22 | 36134783 | 36424872 | - | protein_coding |
| ENSG00000185946.16_4 | RNPC3 | 0.99994553 | 1 | -0.102 | 3.719 | chr1 | 104068313 | 104097861 | + | protein_coding |
| ENSG00000164066.13_3 | INTU | 0.99984595 | 1 | -0.102 | 3.699 | chr4 | 128544426 | 128647892 | + | protein_coding |
| ENSG00000258366.8_4 | RTEL1 | 0.95582896 | 1 | -0.102 | 1.159 | chr20 | 62289163 | 62327606 | + | protein_coding |
| ENSG00000156787.17_5 | TBC1D31 | 0.99999772 | 1 | -0.102 | 4.707 | chr8 | 124054208 | 124164393 | + | protein_coding |
| ENSG00000176853.16_4 | FAM91A1 | 0.99999997 | 1 | -0.102 | 6.498 | chr8 | 124780679 | 124827692 | + | protein_coding |
| ENSG00000177463.15_5 | NR2C2 | 0.99999998 | 1 | -0.102 | 6.316 | chr3 | 14989091 | 15095107 | + | protein_coding |
| ENSG00000122490.19_5 | SLC66A2 | 0.99865152 | 1 | -0.102 | 3.314 | chr18 | 77662420 | 77711657 | - | protein_coding |
| ENSG00000241990.5_7 | PRR34-AS1 | 0.99011457 | 1 | -0.102 | 1.676 | chr22 | 46449585 | 46453090 | + | lncRNA |
| ENSG00000084207.18_5 | GSTP1 | 1 | 1 | -0.102 | 8.549 | chr11 | 67351213 | 67354127 | + | protein_coding |
| ENSG00000150776.18_8 | NKAPD1 | 0.99999961 | 1 | -0.102 | 5.127 | chr11 | 111944810 | 111955874 | + | protein_coding |
| ENSG00000183475.12_2 | ASB7 | 0.99999876 | 1 | -0.102 | 4.783 | chr15 | 101142739 | 101191910 | + | protein_coding |
| ENSG00000173821.19_5 | RNF213 | 0.99999999 | 1 | -0.102 | 9.493 | chr17 | 78234665 | 78372586 | + | protein_coding |
| ENSG00000145907.15_5 | G3BP1 | 0.99999999 | 1 | -0.103 | 8.374 | chr5 | 151150606 | 151192346 | + | protein_coding |
| ENSG00000168092.14_3 | PAFAH1B2 | 0.99999999 | 1 | -0.103 | 7.350 | chr11 | 117015000 | 117047610 | + | protein_coding |
| ENSG00000110060.9_6 | PUS3 | 0.99859856 | 1 | -0.103 | 2.784 | chr11 | 125763380 | 125773116 | - | protein_coding |
| ENSG00000181826.10_4 | RELL1 | 0.99999926 | 1 | -0.103 | 4.936 | chr4 | 37592422 | 37687998 | - | protein_coding |

| | | | | | | | | | | |
|---|---|---|---|---|---|---|---|---|---|---|
| ENSG00000115977.19_6 | AAK1 | 0.99999977 | 1 | -0.103 | 5.678 | chr2 | 69685129 | 69901481 | - | protein_coding |
| ENSG00000092964.18_6 | DPYSL2 | 1 | 1 | -0.103 | 8.444 | chr8 | 26371547 | 26515694 | + | protein_coding |
| ENSG00000261560.1_7 | AC007216.3 | 0.92816501 | 1 | -0.103 | 0.507 | chr16 | 11974932 | 11976426 | - | lncRNA |
| ENSG00000196365.12_5 | LONP1 | 0.99879058 | 1 | -0.103 | 6.611 | chr19 | 5691845 | 5720583 | - | protein_coding |
| ENSG00000077063.11_7 | CTTNBP2 | 0.97989488 | 1 | -0.103 | 1.567 | chr7 | 117350705 | 117514193 | - | protein_coding |
| ENSG00000104472.10_4 | CHRAC1 | 0.99999955 | 1 | -0.103 | 5.376 | chr8 | 141521410 | 141527253 | + | protein_coding |
| ENSG00000275111.5_5 | ZNF2 | 0.99946555 | 1 | -0.103 | 3.173 | chr2 | 95831177 | 95850065 | + | protein_coding |
| ENSG00000151893.15_5 | CACUL1 | 0.99999999 | 1 | -0.103 | 6.638 | chr10 | 120433679 | 120514761 | - | protein_coding |
| ENSG00000225470.8_5 | JPX | 0.99916945 | 1 | -0.103 | 2.799 | chrX | 73164017 | 73290243 | + | lncRNA |
| ENSG00000137996.12_2 | RTCA | 0.99999736 | 1 | -0.103 | 4.782 | chr1 | 100731763 | 100758325 | + | protein_coding |
| ENSG00000264538.6_4 | SUZ12P1 | 0.99998581 | 1 | -0.104 | 4.304 | chr17 | 29036317 | 29117926 | + | transcribed_unprocessed_pseudogene |
| ENSG00000164253.14_8 | WDR41 | 0.9999996 | 1 | -0.104 | 5.531 | chr5 | 76721795 | 76916436 | - | protein_coding |
| ENSG00000122140.11_7 | MRPS2 | 0.99999007 | 1 | -0.104 | 4.549 | chr9 | 138391830 | 138396519 | + | protein_coding |
| ENSG00000123130.17_7 | ACOT9 | 0.99999996 | 1 | -0.104 | 6.321 | chrX | 23719172 | 23784592 | - | protein_coding |
| ENSG00000279901.1_6 | AC092117.2 | 0.89259272 | 1 | -0.104 | -0.359 | chr16 | 2733107 | 2734765 | + | TEC |
| ENSG00000156171.14_3 | DRAM2 | 0.99998723 | 1 | -0.104 | 4.472 | chr1 | 111659955 | 111682838 | - | protein_coding |
| ENSG00000203950.6_6 | RTL8A | 0.99903075 | 1 | -0.104 | 3.424 | chrX | 134184962 | 134186226 | - | protein_coding |
| ENSG00000129084.17_6 | PSMA1 | 0.99997342 | 1 | -0.104 | 4.678 | chr11 | 14526420 | 14665181 | - | protein_coding |
| ENSG00000197694.17_8 | SPTAN1 | 0.99999999 | 1 | -0.104 | 8.664 | chr9 | 131314837 | 131395941 | + | protein_coding |
| ENSG00000225784.5 | RP11-592B15.4 | 0.99694506 | 1 | -0.104 | 2.181 | chr10 | 51287081 | 51371070 | - | pseudogene |
| ENSG00000236088.10_5 | COX10-AS1 | 0.99998089 | 1 | -0.104 | 3.928 | chr17 | 13658891 | 13972812 | - | lncRNA |
| ENSG00000234390.4_6 | USP27X-AS1 | 0.94593385 | 1 | -0.104 | 0.780 | chrX | 49641327 | 49643844 | - | lncRNA |
| ENSG00000082213.18_5 | C5orf22 | 0.99999082 | 1 | -0.104 | 4.303 | chr5 | 31532394 | 31555160 | + | protein_coding |
| ENSG00000178982.10_7 | EIF3K | 0.99999999 | 1 | -0.104 | 7.793 | chr19 | 39109722 | 39127595 | + | protein_coding |
| ENSG00000165806.20_5 | CASP7 | 0.99998978 | 1 | -0.105 | 4.843 | chr10 | 115438921 | 115490666 | + | protein_coding |
| ENSG00000255320.1_7 | AP000759.1 | 0.87344502 | 1 | -0.105 | -0.384 | chr11 | 66012311 | 66013710 | - | lncRNA |
| ENSG00000111364.16_3 | DDX55 | 0.99999995 | 1 | -0.105 | 5.832 | chr12 | 124086624 | 124105490 | + | protein_coding |
| ENSG00000267278.6_6 | MAP3K14-AS1 | 0.99985642 | 1 | -0.105 | 3.436 | chr17 | 43325283 | 43347191 | + | lncRNA |
| ENSG00000205791.3_6 | LOH12CR2 | 0.85764395 | 1 | -0.105 | -0.980 | chr12 | 12508340 | 12510001 | - | lncRNA |
| ENSG00000140320.12_5 | BAHD1 | 0.99976876 | 1 | -0.105 | 3.438 | chr15 | 40731920 | 40760435 | + | protein_coding |
| ENSG00000221886.4_4 | ZBED8 | 0.99985437 | 1 | -0.105 | 3.503 | chr5 | 159820155 | 159827061 | - | protein_coding |
| ENSG00000154723.12_5 | ATP5PF | 0.99981191 | 1 | -0.105 | 3.872 | chr21 | 27088815 | 27107984 | - | protein_coding |
| ENSG00000181274.7_3 | FRAT2 | 0.99943125 | 1 | -0.105 | 3.007 | chr10 | 99092254 | 99094486 | - | protein_coding |
| ENSG00000134815.19_3 | DHX34 | 0.99999941 | 1 | -0.105 | 5.739 | chr19 | 47852572 | 47885961 | + | protein_coding |
| ENSG00000116350.17_8 | SRSF4 | 0.99999988 | 1 | -0.105 | 5.528 | chr1 | 29474255 | 29508412 | - | protein_coding |
| ENSG00000159377.11_3 | PSMB4 | 0.99999997 | 1 | -0.105 | 7.291 | chr1 | 151372036 | 151374413 | + | protein_coding |
| ENSG00000104756.16_4 | KCTD9 | 0.99999986 | 1 | -0.105 | 6.061 | chr8 | 25285363 | 25315992 | - | protein_coding |
| ENSG00000064102.15_4 | INTS13 | 0.99999973 | 1 | -0.105 | 5.621 | chr12 | 27058114 | 27091259 | - | protein_coding |
| ENSG00000183291.17_9 | SELENOF | 0.99999971 | 1 | -0.105 | 5.927 | chr1 | 87328128 | 87380107 | - | protein_coding |
| ENSG00000139899.11_4 | CBLN3 | 0.96263668 | 1 | -0.105 | 0.635 | chr14 | 24895751 | 24900160 | - | protein_coding |
| ENSG00000176105.14_5 | YES1 | 0.99999994 | 1 | -0.106 | 6.515 | chr18 | 721588 | 812547 | - | protein_coding |
| ENSG00000267940.1_6 | AC022762.2 | 0.87286233 | 1 | -0.106 | -0.367 | chr11 | 6223131 | 6224483 | - | lncRNA |
| ENSG00000172379.21_5 | ARNT2 | 0.99999966 | 1 | -0.106 | 5.495 | chr15 | 80696692 | 80890274 | + | protein_coding |
| ENSG00000071051.14_4 | NCK2 | 0.99276148 | 1 | -0.106 | 3.761 | chr2 | 106361369 | 106510730 | + | protein_coding |
| ENSG00000185658.13_5 | BRWD1 | 0.99999991 | 1 | -0.106 | 6.490 | chr21 | 40556102 | 40693485 | - | protein_coding |
| ENSG00000118096.8_6 | IFT46 | 0.99991648 | 1 | -0.106 | 4.159 | chr11 | 118415243 | 118443685 | - | protein_coding |
| ENSG00000268670.1_9 | AC016586.1 | 0.90279841 | 1 | -0.106 | 0.233 | chr19 | 4061613 | 4062747 | - | lncRNA |
| ENSG00000134758.14_4 | RNF138 | 0.99999391 | 1 | -0.106 | 4.386 | chr18 | 29671837 | 29711524 | + | protein_coding |
| ENSG00000104219.13_4 | ZDHHC2 | 0.99999996 | 1 | -0.106 | 6.215 | chr8 | 17013991 | 17082308 | + | protein_coding |
| ENSG00000263465.4_2 | SRSF8 | 0.99999176 | 1 | -0.106 | 4.498 | chr11 | 94800361 | 94804387 | + | protein_coding |
| ENSG00000167601.12_3 | AXL | 0.99999998 | 1 | -0.106 | 8.405 | chr19 | 41725128 | 41767671 | + | protein_coding |
| ENSG00000271943.2_6 | AC098614.3 | 0.99871293 | 1 | -0.106 | 2.823 | chr3 | 27674467 | 27680602 | + | transcribed_processed_pseudogene |
| ENSG00000259083.1_7 | AL132639.3 | 0.86863548 | 1 | -0.106 | -0.525 | chr14 | 39644089 | 39645084 | - | lncRNA |
| ENSG00000269352.1_7 | PTOV1-AS2 | 0.96677203 | 1 | -0.106 | 1.783 | chr19 | 50360227 | 50362546 | - | lncRNA |
| ENSG00000139239.7_4 | RPL14P1 | 0.97365456 | 1 | -0.107 | 2.641 | chr12 | 63359105 | 63359749 | + | processed_pseudogene |
| ENSG00000112701.18_4 | SENP6 | 0.99999998 | 1 | -0.107 | 6.433 | chr6 | 76311225 | 76427997 | + | protein_coding |
| ENSG00000038382.20_4 | TRIO | 1 | 1 | -0.107 | 7.734 | chr5 | 14143451 | 14532237 | + | protein_coding |

| | | | | | | | | | | |
|---|---|---|---|---|---|---|---|---|---|---|
| ENSG00000145882.11_3 | PCYOX1L | 0.99989276 | 1 | -0.107 | 3.423 | chr5 | 148737600 | 148749216 | + | protein_coding |
| ENSG00000245848.3_3 | CEBPA | 0.95716195 | 1 | -0.107 | 1.227 | chr19 | 33790840 | 33793440 | - | protein_coding |
| ENSG00000132466.19_7 | ANKRD17 | 0.99999972 | 1 | -0.107 | 7.015 | chr4 | 73939093 | 74124515 | - | protein_coding |
| ENSG00000269973.1_6 | AC010969.2 | 0.99755128 | 1 | -0.107 | 2.697 | chr2 | 10076489 | 10079719 | + | lncRNA |
| ENSG00000205250.9_4 | E2F4 | 0.99999992 | 1 | -0.107 | 6.024 | chr16 | 67226058 | 67232821 | + | protein_coding |
| ENSG00000229994.1_4 | RPL5P4 | 0.88835408 | 1 | -0.107 | -0.090 | chr1 | 35816323 | 35817208 | + | processed_pseudogene |
| ENSG00000277142.1_5 | LINC00235 | 0.99843917 | 1 | -0.107 | 2.737 | chr16 | 575155 | 577407 | - | lncRNA |
| ENSG00000173273.16_3 | TNKS | 0.99999987 | 1 | -0.107 | 5.993 | chr8 | 9413422 | 9639856 | + | protein_coding |
| ENSG00000072803.17_4 | FBXW11 | 0.99999996 | 1 | -0.107 | 6.517 | chr5 | 171288553 | 171433877 | - | protein_coding |
| ENSG00000214293.9_8 | APTR | 0.9986989 | 1 | -0.107 | 2.602 | chr7 | 77286976 | 77325584 | - | lncRNA |
| ENSG00000075539.15_5 | FRYL | 0.99999964 | 1 | -0.107 | 5.997 | chr4 | 48499374 | 48782339 | - | protein_coding |
| ENSG00000144369.13_3 | FAM171B | 0.99999 | 1 | -0.107 | 4.658 | chr2 | 187558787 | 187630686 | + | protein_coding |
| ENSG00000087077.14_6 | TRIP6 | 0.99984274 | 1 | -0.107 | 5.270 | chr7 | 100465009 | 100471076 | + | protein_coding |
| ENSG00000089737.17_5 | DDX24 | 0.99999999 | 1 | -0.108 | 7.151 | chr14 | 94514633 | 94547558 | - | protein_coding |
| ENSG00000255823.5_4 | MTRNR2L8 | 0.99987554 | 1 | -0.108 | 3.626 | chr11 | 10529441 | 10530733 | - | protein_coding |
| ENSG00000129158.11_5 | SERGEF | 0.9994612 | 1 | -0.108 | 3.169 | chr11 | 17809595 | 18034594 | - | protein_coding |
| ENSG00000125870.11_3 | SNRPB2 | 0.99999991 | 1 | -0.108 | 6.420 | chr20 | 16710606 | 16723209 | + | protein_coding |
| ENSG00000273343.1_7 | AC007663.4 | 0.87017616 | 1 | -0.108 | -0.384 | chr22 | 20305642 | 20306272 | - | lncRNA |
| ENSG00000225921.7_3 | NOL7 | 0.99999988 | 1 | -0.108 | 5.626 | chr6 | 13615567 | 13632971 | + | protein_coding |
| ENSG00000134982.17_6 | APC | 0.99999883 | 1 | -0.108 | 6.474 | chr5 | 112043195 | 112181936 | + | protein_coding |
| ENSG00000113558.18_6 | SKP1 | 0.99999994 | 1 | -0.108 | 6.995 | chr5 | 133484626 | 133512729 | - | protein_coding |
| ENSG00000159596.7_3 | TMEM69 | 0.99997763 | 1 | -0.108 | 3.915 | chr1 | 46153853 | 46160108 | + | protein_coding |
| ENSG00000197965.12_5 | MPZL1 | 0.99999999 | 1 | -0.108 | 7.704 | chr1 | 167690429 | 167761156 | + | protein_coding |
| ENSG00000165275.10_3 | TRMT10B | 0.99990124 | 1 | -0.108 | 3.637 | chr9 | 37753800 | 37778969 | + | protein_coding |
| ENSG00000112210.12_5 | RAB23 | 0.99995287 | 1 | -0.108 | 3.777 | chr6 | 57051790 | 57087105 | - | protein_coding |
| ENSG00000108651.10_4 | UTP6 | 0.99999995 | 1 | -0.108 | 6.126 | chr17 | 30187923 | 30228727 | - | protein_coding |
| ENSG00000070756.16_7 | PABPC1 | 1 | 1 | -0.108 | 8.411 | chr8 | 101698044 | 101735037 | - | protein_coding |
| ENSG00000174606.14_5 | ANGEL2 | 0.9999999 | 1 | -0.108 | 5.875 | chr1 | 213165524 | 213189209 | - | protein_coding |
| ENSG00000104824.17_7 | HNRNPL | 1 | 1 | -0.108 | 7.412 | chr19 | 39327028 | 39342987 | - | protein_coding |
| ENSG00000031698.13_4 | SARS1 | 0.99999997 | 1 | -0.108 | 6.280 | chr1 | 109756540 | 109780804 | + | protein_coding |
| ENSG00000076650.7_3 | GPATCH1 | 0.99999773 | 1 | -0.108 | 4.859 | chr19 | 33571805 | 33621448 | + | protein_coding |
| ENSG00000127337.7_3 | YEATS4 | 0.99999926 | 1 | -0.109 | 5.248 | chr12 | 69753490 | 69784650 | + | protein_coding |
| ENSG00000155097.12_3 | ATP6V1C1 | 0.99999994 | 1 | -0.109 | 6.555 | chr8 | 104033291 | 104085279 | + | protein_coding |
| ENSG00000143858.12_4 | SYT2 | 0.87840174 | 1 | -0.109 | -0.171 | chr1 | 202559724 | 202679582 | - | protein_coding |
| ENSG00000227063.5_2 | RPL41P1 | 0.97193567 | 1 | -0.109 | 1.271 | chr20 | 21735908 | 21735988 | + | processed_pseudogene |
| ENSG00000279026.1_5 | AC005225.4 | 0.86854946 | 1 | -0.109 | -0.473 | chr14 | 74047398 | 74049622 | + | TEC |
| ENSG00000135776.5_4 | ABCB10 | 0.99999972 | 1 | -0.109 | 5.377 | chr1 | 229652329 | 229694454 | - | protein_coding |
| ENSG00000163071.11_4 | SPATA18 | 0.99493516 | 1 | -0.109 | 2.568 | chr4 | 52917470 | 52963465 | + | protein_coding |
| ENSG00000264456.1_7 | AC138207.4 | 0.9686945 | 1 | -0.109 | 1.360 | chr17 | 29298670 | 29300330 | + | lncRNA |
| ENSG00000131591.17_9 | C1orf159 | 0.96847483 | 1 | -0.109 | 3.319 | chr1 | 1017198 | 1051741 | - | protein_coding |
| ENSG00000215270.3 | LA16c-60H5.7 | 0.89596342 | 1 | -0.109 | 0.154 | chr22 | 16122720 | 16123768 | + | pseudogene |
| ENSG00000054267.22_6 | ARID4B | 0.99999994 | 1 | -0.109 | 6.050 | chr1 | 235294949 | 235491534 | - | protein_coding |
| ENSG00000232368.1_4 | FTLP2 | 0.88224375 | 1 | -0.109 | -0.009 | chrX | 30648415 | 30648942 | + | processed_pseudogene |
| ENSG00000113716.13_4 | HMGXB3 | 0.99999986 | 1 | -0.109 | 6.256 | chr5 | 149379609 | 149432705 | + | protein_coding |
| ENSG00000086300.15_4 | SNX10 | 0.99441482 | 1 | -0.109 | 2.626 | chr7 | 26331515 | 26413949 | + | protein_coding |
| ENSG00000102030.16_7 | NAA10 | 0.99999644 | 1 | -0.109 | 4.616 | chrX | 153194678 | 153200533 | - | protein_coding |
| ENSG00000137834.15_6 | SMAD6 | 0.9999989 | 1 | -0.109 | 6.744 | chr15 | 66994574 | 67075187 | + | protein_coding |
| ENSG00000272562.2_7 | AL512343.2 | 0.86538307 | 1 | -0.109 | -0.315 | chr1 | 226233262 | 226249599 | - | lncRNA |
| ENSG00000157796.18_4 | WDR19 | 0.99987937 | 1 | -0.110 | 4.048 | chr4 | 39184124 | 39287430 | + | protein_coding |
| ENSG00000237187.9_7 | NR2F1-AS1 | 0.98556117 | 1 | -0.110 | 1.884 | chr5 | 92696485 | 92921355 | - | lncRNA |
| ENSG00000287916.1_1 | AC091607.2 | 0.87510245 | 1 | -0.110 | -0.084 | chr3 | 155861359 | 156258416 | + | lncRNA |
| ENSG00000146243.14_5 | IRAK1BP1 | 0.99870168 | 1 | -0.110 | 2.692 | chr6 | 79577268 | 79656157 | + | protein_coding |
| ENSG00000130766.5_3 | SESN2 | 0.99896234 | 1 | -0.110 | 3.059 | chr1 | 28586029 | 28609002 | + | protein_coding |
| ENSG00000104381.13_3 | GDAP1 | 0.99999631 | 1 | -0.110 | 4.871 | chr8 | 75233365 | 75401107 | + | protein_coding |
| ENSG00000258515.1_6 | AL355075.2 | 0.89011882 | 1 | -0.110 | 0.042 | chr14 | 20919464 | 20920077 | + | lncRNA |
| ENSG00000198755.11_3 | RPL10A | 1 | 1 | -0.110 | 8.401 | chr6 | 35436178 | 35438562 | + | protein_coding |
| ENSG00000089847.12_6 | ANKRD24 | 0.87958644 | 1 | -0.110 | -0.222 | chr19 | 4183351 | 4224811 | + | protein_coding |
| ENSG00000153015.16_4 | CWC27 | 0.99998634 | 1 | -0.110 | 4.555 | chr5 | 64064757 | 64314577 | + | protein_coding |
| ENSG00000070759.17_4 | TESK2 | 0.88779767 | 1 | -0.110 | 0.248 | chr1 | 45809555 | 45956835 | - | protein_coding |

| | | | | | | | | | | |
|---|---|---|---|---|---|---|---|---|---|---|
| ENSG00000167986.14_4 | DDB1 | 1 | 1 | -0.110 | 8.516 | chr11 | 61066923 | 61110068 | - | protein_coding |
| ENSG00000178538.10_4 | CA8 | 0.99999583 | 1 | -0.110 | 4.560 | chr8 | 61097971 | 61193959 | - | protein_coding |
| ENSG00000121058.5_5 | COIL | 0.99999844 | 1 | -0.110 | 5.031 | chr17 | 55015560 | 55038411 | - | protein_coding |
| ENSG00000136463.8_3 | TACO1 | 0.99999831 | 1 | -0.110 | 5.045 | chr17 | 61678241 | 61685725 | + | protein_coding |
| ENSG00000110344.9_3 | UBE4A | 0.99999994 | 1 | -0.110 | 6.107 | chr11 | 118230300 | 118269926 | + | protein_coding |
| ENSG00000146830.10_4 | GIGYF1 | 0.99999428 | 1 | -0.110 | 6.222 | chr7 | 100277130 | 100291660 | - | protein_coding |
| ENSG00000119318.13_4 | RAD23B | 1 | 1 | -0.111 | 8.162 | chr9 | 110045418 | 110094473 | + | protein_coding |
| ENSG00000188636.4_7 | RTL6 | 0.99999984 | 1 | -0.111 | 5.513 | chr22 | 44888463 | 44894113 | - | protein_coding |
| ENSG00000149658.18_4 | YTHDF1 | 0.99999992 | 1 | -0.111 | 6.007 | chr20 | 61826781 | 61847491 | - | protein_coding |
| ENSG00000125686.12_5 | MED1 | 0.99999998 | 1 | -0.111 | 6.456 | chr17 | 37560538 | 37607525 | - | protein_coding |
| ENSG00000166965.12_4 | RCCD1 | 0.99999971 | 1 | -0.111 | 5.521 | chr15 | 91498100 | 91506355 | + | protein_coding |
| ENSG00000102054.17_4 | RBBP7 | 1 | 1 | -0.111 | 7.806 | chrX | 16857406 | 16888537 | - | protein_coding |
| ENSG00000101199.13_7 | ARFGAP1 | 0.99999824 | 1 | -0.111 | 6.557 | chr20 | 61904137 | 61921142 | + | protein_coding |
| ENSG00000110074.11_3 | FOXRED1 | 0.99999639 | 1 | -0.111 | 4.733 | chr11 | 126138950 | 126148026 | + | protein_coding |
| ENSG00000235655.3_5 | H3P6 | 0.99965845 | 1 | -0.111 | 3.621 | chr2 | 175584636 | 175585046 | + | processed_pseudogene |
| ENSG00000007341.19_5 | ST7L | 0.99969252 | 1 | -0.111 | 3.351 | chr1 | 113066136 | 113163447 | - | protein_coding |
| ENSG00000169592.15_6 | INO80E | 0.99999929 | 1 | -0.111 | 5.071 | chr16 | 30007036 | 30017114 | + | protein_coding |
| ENSG00000278987.1_6 | AL031009.1 | 0.97944623 | 1 | -0.111 | 1.419 | chr16 | 1738356 | 1740537 | + | TEC |
| ENSG00000111788.10_5 | AC009533.1 | 0.99994252 | 1 | -0.111 | 3.940 | chr12 | 9429831 | 9465837 | + | unprocessed_pseudogene |
| ENSG00000136636.13_5 | KCTD3 | 0.99999998 | 1 | -0.111 | 6.504 | chr1 | 215740647 | 215795149 | + | protein_coding |
| ENSG00000143398.20_5 | PIP5K1A | 0.99999991 | 1 | -0.112 | 5.737 | chr1 | 151170425 | 151222012 | + | protein_coding |
| ENSG00000186312.10_5 | CA5BP1 | 0.99639063 | 1 | -0.112 | 2.205 | chrX | 15693055 | 15721847 | + | transcribed_unprocessed_pseudogene |
| ENSG00000177646.19_4 | ACAD9 | 0.99999836 | 1 | -0.112 | 4.927 | chr3 | 128598439 | 128634910 | + | protein_coding |
| ENSG00000009307.16_8 | CSDE1 | 0.99999999 | 1 | -0.112 | 8.502 | chr1 | 115259534 | 115301297 | - | protein_coding |
| ENSG00000205765.9_3 | C5orf51 | 0.99999866 | 1 | -0.112 | 5.356 | chr5 | 41904290 | 41921738 | + | protein_coding |
| ENSG00000102024.18_4 | PLS3 | 1 | 1 | -0.112 | 9.006 | chrX | 114795501 | 114885181 | + | protein_coding |
| ENSG00000183624.14_4 | HMCES | 0.99999318 | 1 | -0.112 | 4.479 | chr3 | 128997671 | 129025029 | + | protein_coding |
| ENSG00000168806.8_4 | LCMT2 | 0.99994893 | 1 | -0.112 | 3.843 | chr15 | 43615847 | 43622780 | - | protein_coding |
| ENSG00000186583.7 | SPATC1 | 0.84866725 | 1 | -0.112 | -0.944 | chr8 | 145086582 | 145101933 | + | protein_coding |
| ENSG00000221990.4_6 | EXOC3-AS1 | 0.99531757 | 1 | -0.112 | 1.569 | chr5 | 441613 | 443275 | - | lncRNA |
| ENSG00000137267.6_4 | TUBB2A | 0.99953833 | 1 | -0.113 | 3.281 | chr6 | 3153900 | 3157778 | - | protein_coding |
| ENSG00000155749.12_7 | FLACC1 | 0.84711084 | 1 | -0.113 | -0.768 | chr2 | 202152994 | 202222121 | - | protein_coding |
| ENSG00000198832.10_6 | SELENOM | 0.99967139 | 1 | -0.113 | 3.259 | chr22 | 31500758 | 31516055 | - | protein_coding |
| ENSG00000135953.11_4 | MFSD9 | 0.99998534 | 1 | -0.113 | 4.234 | chr2 | 103331089 | 103353347 | - | protein_coding |
| ENSG00000251669.5_4 | FAM86EP | 0.99770398 | 1 | -0.113 | 2.367 | chr4 | 3943487 | 3957146 | - | transcribed_unprocessed_pseudogene |
| ENSG00000120709.11_6 | FAM53C | 0.99999772 | 1 | -0.113 | 4.955 | chr5 | 137667624 | 137685418 | + | protein_coding |
| ENSG00000109762.16_3 | SNX25 | 0.99998103 | 1 | -0.113 | 4.147 | chr4 | 186125391 | 186291339 | + | protein_coding |
| ENSG00000171148.13_3 | TADA3 | 0.99999953 | 1 | -0.113 | 5.898 | chr3 | 9821544 | 9834695 | - | protein_coding |
| ENSG00000064607.17_4 | SUGP2 | 0.99999998 | 1 | -0.113 | 6.683 | chr19 | 19101697 | 19144832 | - | protein_coding |
| ENSG00000155506.17_4 | LARP1 | 0.99999999 | 1 | -0.113 | 8.055 | chr5 | 154092403 | 154197165 | + | protein_coding |
| ENSG00000124226.11_4 | RNF114 | 0.99999987 | 1 | -0.113 | 5.590 | chr20 | 48552873 | 48570422 | + | protein_coding |
| ENSG00000153560.12_5 | UBP1 | 0.99999999 | 1 | -0.113 | 6.610 | chr3 | 33429828 | 33482863 | - | protein_coding |
| ENSG00000204356.14_6 | NELFE | 0.99999858 | 1 | -0.113 | 5.100 | chr6 | 31919864 | 31926815 | - | protein_coding |
| ENSG00000117500.13_4 | TMED5 | 0.99999937 | 1 | -0.113 | 5.841 | chr1 | 93615299 | 93646073 | - | protein_coding |
| ENSG00000130513.6_4 | GDF15 | 0.9995255 | 1 | -0.113 | 5.184 | chr19 | 18485541 | 18499986 | + | protein_coding |
| ENSG00000117713.20_6 | ARID1A | 0.99999992 | 1 | -0.113 | 6.205 | chr1 | 27019727 | 27108595 | + | protein_coding |
| ENSG00000112234.9_4 | FBXL4 | 0.99999947 | 1 | -0.113 | 5.358 | chr6 | 99316411 | 99395882 | - | protein_coding |
| ENSG00000156239.12_5 | N6AMT1 | 0.99954895 | 1 | -0.114 | 3.243 | chr21 | 30244513 | 30257693 | - | protein_coding |
| ENSG00000176142.13_5 | TMEM39A | 0.99999878 | 1 | -0.114 | 5.006 | chr3 | 119147796 | 119187677 | - | protein_coding |
| ENSG00000153933.10_3 | DGKE | 0.99996785 | 1 | -0.114 | 4.388 | chr17 | 54911468 | 54946928 | + | protein_coding |
| ENSG00000137767.14_7 | SQOR | 0.99999284 | 1 | -0.114 | 4.815 | chr15 | 45923346 | 45983479 | + | protein_coding |
| ENSG00000213782.7_6 | DDX47 | 0.96925477 | 1 | -0.114 | 1.401 | chr12 | 12966250 | 12982915 | + | protein_coding |
| ENSG00000013275.8_3 | PSMC4 | 1 | 1 | -0.114 | 8.229 | chr19 | 40477072 | 40487671 | + | protein_coding |
| ENSG00000254473.1_6 | AL354920.1 | 0.93163856 | 1 | -0.114 | 0.366 | chr9 | 86322509 | 86328293 | + | lncRNA |
| ENSG00000197798.9_5 | FAM118B | 0.99996905 | 1 | -0.114 | 4.103 | chr11 | 126081619 | 126132879 | + | protein_coding |
| ENSG00000141098.13_5 | GFOD2 | 0.99999757 | 1 | -0.114 | 4.784 | chr16 | 67708434 | 67753242 | - | protein_coding |
| ENSG00000129911.9_5 | KLF16 | 0.99928009 | 1 | -0.114 | 3.226 | chr19 | 1852398 | 1863578 | - | protein_coding |
| ENSG00000183323.13_6 | CCDC125 | 0.99996592 | 1 | -0.114 | 3.944 | chr5 | 68576002 | 68628636 | - | protein_coding |
| ENSG00000262686.1_6 | GLIS2-AS1 | 0.92391994 | 1 | -0.114 | 0.529 | chr16 | 4374668 | 4378341 | - | lncRNA |

| | | | | | | | | | | |
|---|---|---|---|---|---|---|---|---|---|---|
| ENSG00000122958.15_4 | VPS26A | 0.99999803 | 1 | -0.114 | 4.965 | chr10 | 70883268 | 70934168 | + | protein_coding |
| ENSG00000163320.11_6 | CGGBP1 | 0.99999994 | 1 | -0.114 | 6.882 | chr3 | 88101094 | 88199035 | - | protein_coding |
| ENSG00000146282.18_5 | RARS2 | 0.99999545 | 1 | -0.114 | 4.829 | chr6 | 88223656 | 88299705 | - | protein_coding |
| ENSG00000287054.1_2 | AC113410.6 | 0.86889477 | 1 | -0.114 | -0.291 | chr5 | 132361539 | 132385870 | + | lncRNA |
| ENSG00000213185.7_5 | FAM24B | 0.99040387 | 1 | -0.115 | 1.274 | chr10 | 124608594 | 124639157 | - | protein_coding |
| ENSG00000186470.14_5 | BTN3A2 | 0.99956176 | 1 | -0.115 | 3.101 | chr6 | 26365387 | 26378548 | + | protein_coding |
| ENSG00000109084.14_4 | TMEM97 | 0.99999999 | 1 | -0.115 | 7.318 | chr17 | 26646226 | 26655711 | + | protein_coding |
| ENSG00000118939.17_7 | UCHL3 | 0.86309798 | 1 | -0.115 | 0.233 | chr13 | 76123616 | 76180156 | + | protein_coding |
| ENSG00000136280.16_7 | CCM2 | 0.99999731 | 1 | -0.115 | 5.005 | chr7 | 45039074 | 45116068 | + | protein_coding |
| ENSG00000116353.16_5 | MECR | 0.99977618 | 1 | -0.115 | 3.459 | chr1 | 29519169 | 29557454 | - | protein_coding |
| ENSG00000188428.20_5 | BLOC1S5 | 0.99881939 | 1 | -0.115 | 2.927 | chr6 | 8013800 | 8064629 | - | protein_coding |
| ENSG00000177058.12_6 | SLC38A9 | 0.99999806 | 1 | -0.115 | 4.704 | chr5 | 54921673 | 55069022 | - | protein_coding |
| ENSG00000081014.11_4 | AP4E1 | 0.99999957 | 1 | -0.115 | 5.775 | chr15 | 51200869 | 51298092 | + | protein_coding |
| ENSG00000118181.11_4 | RPS25 | 0.99999999 | 1 | -0.115 | 7.336 | chr11 | 118886422 | 118889401 | - | protein_coding |
| ENSG00000088002.11_2 | SULT2B1 | 0.99241437 | 1 | -0.115 | 1.931 | chr19 | 49055332 | 49102682 | + | protein_coding |
| ENSG00000153147.6_3 | SMARCA5 | 0.99999996 | 1 | -0.115 | 7.277 | chr4 | 144434855 | 144478639 | + | protein_coding |
| ENSG00000151413.17_4 | NUBPL | 0.99998737 | 1 | -0.115 | 4.396 | chr14 | 31959162 | 32330430 | + | protein_coding |
| ENSG00000258768.2_7 | AL356019.2 | 0.85136094 | 1 | -0.115 | -0.523 | chr14 | 20728639 | 20732467 | - | lncRNA |
| ENSG00000111358.14_4 | GTF2H3 | 0.99999844 | 1 | -0.115 | 5.320 | chr12 | 124118286 | 124147151 | + | protein_coding |
| ENSG00000172380.6_4 | GNG12 | 0.99999999 | 1 | -0.116 | 8.455 | chr1 | 68167158 | 68299150 | - | protein_coding |
| ENSG00000180185.11_2 | FAHD1 | 0.99999766 | 1 | -0.116 | 4.855 | chr16 | 1876942 | 1890208 | + | protein_coding |
| ENSG00000111696.12_7 | NT5DC3 | 0.99999876 | 1 | -0.116 | 5.082 | chr12 | 104164231 | 104235012 | - | protein_coding |
| ENSG00000176371.14_5 | ZSCAN2 | 0.99925014 | 1 | -0.116 | 3.047 | chr15 | 85144217 | 85171027 | + | protein_coding |
| ENSG00000167552.14_4 | TUBA1A | 0.99999993 | 1 | -0.116 | 6.544 | chr12 | 49578578 | 49582863 | - | protein_coding |
| ENSG00000135250.17_7 | SRPK2 | 0.99999989 | 1 | -0.116 | 5.790 | chr7 | 104751151 | 105039755 | - | protein_coding |
| ENSG00000160678.11_3 | S100A1 | 0.85935548 | 1 | -0.116 | -0.269 | chr1 | 153600402 | 153604515 | + | protein_coding |
| ENSG00000122376.11_5 | SHLD2 | 0.99999991 | 1 | -0.116 | 6.020 | chr10 | 88853918 | 88951225 | + | protein_coding |
| ENSG00000137822.12_2 | TUBGCP4 | 0.99980521 | 1 | -0.116 | 3.339 | chr15 | 43661419 | 43701969 | + | protein_coding |
| ENSG00000231074.9_7 | HCG18 | 0.8991859 | 1 | -0.116 | 0.260 | chr6 | 30254467 | 30295159 | - | lncRNA |
| ENSG00000144713.13_4 | RPL32 | 0.99999999 | 1 | -0.116 | 8.779 | chr3 | 12875984 | 12883081 | - | protein_coding |
| ENSG00000152804.11_3 | HHEX | 0.99999751 | 1 | -0.116 | 4.691 | chr10 | 94449712 | 94455404 | + | protein_coding |
| ENSG00000170852.11_5 | KBTBD2 | 0.99999954 | 1 | -0.116 | 5.689 | chr7 | 32907784 | 32933743 | - | protein_coding |
| ENSG00000068885.15_8 | IFT80 | 0.99990434 | 1 | -0.116 | 4.667 | chr3 | 159974774 | 160117668 | - | protein_coding |
| ENSG00000008988.9_3 | RPS20 | 0.99999999 | 1 | -0.116 | 8.739 | chr8 | 56979854 | 56987140 | - | protein_coding |
| ENSG00000172476.4_5 | RAB40A | 0.88326421 | 1 | -0.116 | -0.139 | chrX | 102754058 | 102774417 | - | protein_coding |
| ENSG00000149084.13_5 | HSD17B12 | 0.99999979 | 1 | -0.116 | 5.516 | chr11 | 43702230 | 43878167 | + | protein_coding |
| ENSG00000213236.3_2 | YWHAZP2 | 0.90477643 | 1 | -0.116 | 0.375 | chr2 | 127315012 | 127315739 | - | processed_pseudogene |
| ENSG00000105619.13_5 | TFPT | 0.99849138 | 1 | -0.116 | 2.778 | chr19 | 54610320 | 54619055 | - | protein_coding |
| ENSG00000185697.16_5 | MYBL1 | 0.99999662 | 1 | -0.116 | 4.958 | chr8 | 67474410 | 67526482 | - | protein_coding |
| ENSG00000173113.6_4 | TRMT112 | 0.99999996 | 1 | -0.116 | 6.544 | chr11 | 64083932 | 64085556 | - | protein_coding |
| ENSG00000110958.16_4 | PTGES3 | 0.99999999 | 1 | -0.116 | 7.961 | chr12 | 57057125 | 57082192 | - | protein_coding |
| ENSG00000143543.15_5 | JTB | 0.99948526 | 1 | -0.116 | 3.506 | chr1 | 153946745 | 153950150 | - | protein_coding |
| ENSG00000277895.1_7 | AC135279.4 | 0.85087057 | 1 | -0.116 | -0.741 | chr12 | 64845371 | 64846681 | - | lncRNA |
| ENSG00000173456.5_3 | RNF26 | 0.99995619 | 1 | -0.117 | 3.940 | chr11 | 119205237 | 119208019 | + | protein_coding |
| ENSG00000196236.13_3 | XPNPEP3 | 0.99999842 | 1 | -0.117 | 4.908 | chr22 | 41253081 | 41328819 | + | protein_coding |
| ENSG00000160679.13_4 | CHTOP | 0.99999981 | 1 | -0.117 | 5.960 | chr1 | 153606458 | 153618782 | + | protein_coding |
| ENSG00000187720.14_5 | THSD4 | 0.99999999 | 1 | -0.117 | 7.388 | chr15 | 71389291 | 72075722 | + | protein_coding |
| ENSG00000217165.1_4 | ANKRD18EP | 0.99932882 | 1 | -0.117 | 2.979 | chr6 | 39078097 | 39080728 | + | processed_pseudogene |
| ENSG00000144827.9_3 | ABHD10 | 0.99999932 | 1 | -0.117 | 5.487 | chr3 | 111697857 | 111712215 | + | protein_coding |
| ENSG00000270585.1_5 | AL445435.1 | 0.8479386 | 1 | -0.117 | -0.710 | chr1 | 220380960 | 220381624 | + | processed_pseudogene |
| ENSG00000253683.1_5 | AC027309.2 | 0.91142054 | 1 | -0.117 | 0.494 | chr5 | 172083525 | 172083716 | - | processed_pseudogene |
| ENSG00000108256.9_4 | NUFIP2 | 0.99999997 | 1 | -0.117 | 7.969 | chr17 | 27582857 | 27621166 | - | protein_coding |
| ENSG00000185950.9_4 | IRS2 | 0.99996475 | 1 | -0.117 | 5.698 | chr13 | 110405042 | 110438930 | - | protein_coding |
| ENSG00000139687.15_4 | RB1 | 0.99999988 | 1 | -0.117 | 6.182 | chr13 | 48877862 | 49173572 | + | protein_coding |
| ENSG00000118894.15_5 | EEF2KMT | 0.99901925 | 1 | -0.117 | 3.468 | chr16 | 5134285 | 5147796 | - | protein_coding |
| ENSG00000132664.12_3 | POLR3F | 0.99997636 | 1 | -0.117 | 4.543 | chr20 | 18448033 | 18465290 | + | protein_coding |
| ENSG00000113851.14_6 | CRBN | 0.99999771 | 1 | -0.117 | 5.000 | chr3 | 3186312 | 3221411 | - | protein_coding |
| ENSG00000279769.1_6 | CU639417.2 | 0.84030868 | 1 | -0.118 | -1.067 | chr21 | 44922354 | 44923757 | - | TEC |
| ENSG00000177485.7_4 | ZBTB33 | 0.99999979 | 1 | -0.118 | 5.751 | chrX | 119384607 | 119392253 | + | protein_coding |

| | | | | | | | | | | |
|---|---|---|---|---|---|---|---|---|---|---|
| ENSG00000006125.18_7 | AP2B1 | 1 | 1 | -0.118 | 8.398 | chr17 | 33905065 | 34053432 | + | protein_coding |
| ENSG00000105258.9_4 | POLR2I | 0.99974741 | 1 | -0.118 | 3.308 | chr19 | 36604611 | 36606115 | - | protein_coding |
| ENSG00000286219.1_3 | NOTCH2NLC | 0.99307313 | 1 | -0.118 | 1.459 | chr1 | 145209136 | 145285911 | + | protein_coding |
| ENSG00000095574.11_3 | IKZF5 | 0.99967792 | 1 | -0.118 | 3.177 | chr10 | 124750322 | 124768333 | - | protein_coding |
| ENSG00000112237.13_5 | CCNC | 0.9999995 | 1 | -0.118 | 5.634 | chr6 | 99990263 | 100016701 | - | protein_coding |
| ENSG00000120686.12_3 | UFM1 | 0.99999165 | 1 | -0.118 | 5.571 | chr13 | 38923986 | 38937756 | + | protein_coding |
| ENSG00000171135.15_7 | JAGN1 | 0.99999163 | 1 | -0.118 | 4.708 | chr3 | 9932258 | 9936033 | + | protein_coding |
| ENSG00000145241.11_5 | CENPC | 0.99999666 | 1 | -0.118 | 4.620 | chr4 | 68334480 | 68411221 | - | protein_coding |
| ENSG00000230438.7_7 | SERPINB9P1 | 0.84157442 | 1 | -0.118 | -0.877 | chr6 | 2851464 | 2881641 | - | lncRNA |
| ENSG00000131558.15_3 | EXOC4 | 0.99999998 | 1 | -0.118 | 6.630 | chr7 | 132937829 | 133751342 | + | protein_coding |
| ENSG00000132405.19_5 | TBC1D14 | 0.99999977 | 1 | -0.118 | 5.488 | chr4 | 6910969 | 7034845 | + | protein_coding |
| ENSG00000186106.11_7 | ANKRD46 | 0.99984897 | 1 | -0.118 | 3.626 | chr8 | 101521980 | 101572012 | - | protein_coding |
| ENSG00000198837.10_6 | DENND4B | 0.99999977 | 1 | -0.118 | 5.636 | chr1 | 153901977 | 153919194 | - | protein_coding |
| ENSG00000257150.3_5 | PGAM1P5 | 0.84362884 | 1 | -0.118 | -0.585 | chr12 | 95945358 | 96067775 | + | transcribed_processed_pseudogene |
| ENSG00000136819.15_5 | C9orf78 | 0.99999963 | 1 | -0.118 | 5.268 | chr9 | 132589569 | 132598142 | - | protein_coding |
| ENSG00000277149.5_4 | TYW1B | 0.99583456 | 1 | -0.118 | 2.213 | chr7 | 72023729 | 72298779 | - | protein_coding |
| ENSG00000214575.9_7 | CPEB1 | 0.8481561 | 1 | -0.118 | -0.559 | chr15 | 83211951 | 83317612 | - | protein_coding |
| ENSG00000256061.7_9 | DNAAF4 | 0.87422641 | 1 | -0.119 | -0.272 | chr15 | 55702723 | 55800432 | - | protein_coding |
| ENSG00000198783.6_3 | ZNF830 | 0.99992522 | 1 | -0.119 | 3.745 | chr17 | 33288559 | 33290796 | + | protein_coding |
| ENSG00000149547.15_8 | EI24 | 0.99999972 | 1 | -0.119 | 6.321 | chr11 | 125439176 | 125454579 | + | protein_coding |
| ENSG00000155621.15_4 | C9orf85 | 0.99984455 | 1 | -0.119 | 3.499 | chr9 | 74526426 | 74600970 | + | protein_coding |
| ENSG00000197182.14_4 | MIRLET7BHG | 0.99999309 | 1 | -0.119 | 4.490 | chr22 | 46449749 | 46509808 | + | lncRNA |
| ENSG00000146757.13_2 | ZNF92 | 0.99998532 | 1 | -0.119 | 4.757 | chr7 | 64838712 | 64866048 | + | protein_coding |
| ENSG00000125818.18_8 | PSMF1 | 0.99999998 | 1 | -0.119 | 6.896 | chr20 | 1093883 | 1170059 | + | protein_coding |
| ENSG00000127957.18_6 | PMS2P3 | 0.99647321 | 1 | -0.119 | 2.640 | chr7 | 75137069 | 75157478 | - | transcribed_unprocessed_pseudogene |
| ENSG00000122034.15_6 | GTF3A | 0.99999997 | 1 | -0.119 | 6.276 | chr13 | 27998756 | 28009960 | + | protein_coding |
| ENSG00000204152.6 | TIMM23B | 0.85821904 | 1 | -0.119 | -0.273 | chr10 | 51371390 | 51387768 | + | protein_coding |
| ENSG00000223478.1_7 | AL441992.2 | 0.99830498 | 1 | -0.119 | 2.788 | chr9 | 131486724 | 131495473 | + | lncRNA |
| ENSG00000134371.13_7 | CDC73 | 0.99999986 | 1 | -0.119 | 5.752 | chr1 | 193091088 | 193223945 | + | protein_coding |
| ENSG00000153975.10_5 | ZUP1 | 0.99995719 | 1 | -0.119 | 3.878 | chr6 | 116956781 | 116989957 | - | protein_coding |
| ENSG00000089050.16_5 | RBBP9 | 0.99999984 | 1 | -0.120 | 5.878 | chr20 | 18467184 | 18477869 | - | protein_coding |
| ENSG00000101182.15_5 | PSMA7 | 0.99999993 | 1 | -0.120 | 7.536 | chr20 | 60711789 | 60718496 | - | protein_coding |
| ENSG00000173418.12_3 | NAA20 | 0.99999989 | 1 | -0.120 | 5.991 | chr20 | 19997954 | 20014299 | + | protein_coding |
| ENSG00000117523.16_7 | PRRC2C | 0.99999994 | 1 | -0.120 | 7.700 | chr1 | 171454690 | 171562650 | + | protein_coding |
| ENSG00000118363.12_5 | SPCS2 | 0.9999158 | 1 | -0.120 | 3.757 | chr11 | 74660292 | 74690078 | + | protein_coding |
| ENSG00000109775.11_5 | UFSP2 | 0.99989854 | 1 | -0.120 | 3.857 | chr4 | 186320691 | 186347133 | - | protein_coding |
| ENSG00000205090.9_4 | TMEM240 | 0.8412985 | 1 | -0.120 | -0.878 | chr1 | 1470158 | 1476004 | - | protein_coding |
| ENSG00000233436.7_6 | BTBD18 | 0.90845053 | 1 | -0.120 | 0.624 | chr11 | 57510986 | 57519253 | - | protein_coding |
| ENSG00000104969.10_5 | SGTA | 0.99999729 | 1 | -0.120 | 5.819 | chr19 | 2754713 | 2783280 | - | protein_coding |
| ENSG00000143545.10_5 | RAB13 | 0.99204737 | 1 | -0.120 | 1.963 | chr1 | 153954093 | 153958834 | - | protein_coding |
| ENSG00000132716.19_6 | DCAF8 | 0.99999589 | 1 | -0.120 | 4.826 | chr1 | 160185505 | 160232339 | - | protein_coding |
| ENSG00000167775.11_5 | CD320 | 0.99998181 | 1 | -0.120 | 4.933 | chr19 | 8367011 | 8373242 | - | protein_coding |
| ENSG00000089250.19_6 | NOS1 | 0.99678356 | 1 | -0.120 | 2.316 | chr12 | 117645947 | 117889975 | - | protein_coding |
| ENSG00000180543.5_5 | TSPYL5 | 0.99999799 | 1 | -0.120 | 5.187 | chr8 | 98285716 | 98290156 | - | protein_coding |
| ENSG00000272316.1_6 | AL021368.2 | 0.87656599 | 1 | -0.120 | -0.074 | chr6 | 58234838 | 58240189 | - | lncRNA |
| ENSG00000280187.1_5 | AC022107.1 | 0.97445841 | 1 | -0.121 | 1.438 | chr5 | 68482186 | 68485279 | - | TEC |
| ENSG00000229152.2_7 | ANKRD10-IT1 | 0.99246201 | 1 | -0.121 | 2.304 | chr13 | 111546986 | 111551519 | - | lncRNA |
| ENSG00000198331.10_2 | HYLS1 | 0.99999532 | 1 | -0.121 | 4.854 | chr11 | 125753509 | 125770543 | + | protein_coding |
| ENSG00000100311.17_6 | PDGFB | 0.99776095 | 1 | -0.121 | 2.725 | chr22 | 39619364 | 39640987 | - | protein_coding |
| ENSG00000140323.6_3 | DISP2 | 0.91838451 | 1 | -0.121 | 1.003 | chr15 | 40650420 | 40670822 | + | protein_coding |
| ENSG00000272602.6_5 | ZNF595 | 0.9998965 | 1 | -0.121 | 3.734 | chr4 | 53180 | 88096 | + | protein_coding |
| ENSG00000147394.18_3 | ZNF185 | 0.9999968 | 1 | -0.121 | 4.925 | chrX | 152082986 | 152142024 | + | protein_coding |
| ENSG00000197063.11_8 | MAFG | 0.99999993 | 1 | -0.121 | 6.389 | chr17 | 79876146 | 79885611 | - | protein_coding |
| ENSG00000105393.16_6 | BABAM1 | 0.99981157 | 1 | -0.121 | 3.370 | chr19 | 17378185 | 17392058 | + | protein_coding |
| ENSG00000272153.1_6 | AL365330.1 | 0.83541907 | 1 | -0.121 | -0.769 | chr1 | 3701572 | 3702102 | + | lncRNA |
| ENSG00000226549.3_2 | SCDP1 | 0.84527554 | 1 | -0.121 | -0.377 | chr17 | 20687958 | 20689038 | + | processed_pseudogene |

| | | | | | | | | | | |
|---|---|---|---|---|---|---|---|---|---|---|
| ENSG00000136930.13_4 | PSMB7 | 0.99999995 | 1 | -0.121 | 6.815 | chr9 | 127115744 | 127177723 | - | protein_coding |
| ENSG00000140157.14_6 | NIPA2 | 0.99999997 | 1 | -0.121 | 6.747 | chr15 | 23004684 | 23034427 | - | protein_coding |
| ENSG00000113597.18_5 | TRAPPC13 | 0.99999747 | 1 | -0.121 | 4.741 | chr5 | 64920831 | 64962060 | + | protein_coding |
| ENSG00000162004.17_6 | CCDC78 | 0.9996502 | 1 | -0.121 | 3.758 | chr16 | 772582 | 776954 | - | protein_coding |
| ENSG00000226688.7_8 | ENTPD1-AS1 | 0.95414953 | 1 | -0.121 | 0.952 | chr10 | 97492733 | 97850007 | - | lncRNA |
| ENSG00000165655.17_5 | ZNF503 | 0.94877605 | 1 | -0.121 | 2.201 | chr10 | 77157588 | 77161522 | - | protein_coding |
| ENSG00000243678.11_3 | NME2 | 0.94795844 | 1 | -0.121 | 0.881 | chr17 | 49242796 | 49249108 | + | protein_coding |
| ENSG00000224660.2_6 | SH3BP5-AS1 | 0.98197415 | 1 | -0.121 | 1.520 | chr3 | 15295691 | 15306022 | + | lncRNA |
| ENSG00000138433.16_5 | CIR1 | 0.99991491 | 1 | -0.121 | 3.800 | chr2 | 175212750 | 175260440 | - | protein_coding |
| ENSG00000200959.1 | SNORA74A | 0.97361195 | 1 | -0.121 | 1.473 | chr5 | 138614470 | 138614667 | + | snoRNA |
| ENSG00000228065.11_6 | LINC01515 | 0.99761627 | 1 | -0.121 | 2.541 | chr10 | 67330096 | 67640047 | + | lncRNA |
| ENSG00000168488.18_6 | ATXN2L | 0.99999996 | 1 | -0.122 | 6.262 | chr16 | 28834356 | 28848558 | + | protein_coding |
| ENSG00000158301.18_5 | GPRASP2 | 0.99977546 | 1 | -0.122 | 3.529 | chrX | 101967104 | 101972661 | + | protein_coding |
| ENSG00000108094.15_4 | CUL2 | 0.99999953 | 1 | -0.122 | 5.688 | chr10 | 35297479 | 35379570 | - | protein_coding |
| ENSG00000196668.4_3 | LINC00173 | 0.96822935 | 1 | -0.122 | 1.710 | chr12 | 116971227 | 116974323 | + | lncRNA |
| ENSG00000120963.12_5 | ZNF706 | 0.99999575 | 1 | -0.122 | 4.953 | chr8 | 102190106 | 102218421 | - | protein_coding |
| ENSG00000116489.13_3 | CAPZA1 | 0.99999997 | 1 | -0.122 | 7.161 | chr1 | 113162427 | 113214238 | + | protein_coding |
| ENSG00000198146.4_5 | ZNF770 | 0.99999988 | 1 | -0.122 | 5.706 | chr15 | 35270542 | 35280488 | - | protein_coding |
| ENSG00000204859.13_4 | ZBTB48 | 0.99986025 | 1 | -0.122 | 3.582 | chr1 | 6640054 | 6649340 | + | protein_coding |
| ENSG00000152291.14_4 | TGOLN2 | 0.99999999 | 1 | -0.122 | 7.490 | chr2 | 85545143 | 85555419 | - | protein_coding |
| ENSG00000171160.18_4 | MORN4 | 0.99762213 | 1 | -0.122 | 2.816 | chr10 | 99374310 | 99393257 | - | protein_coding |
| ENSG00000159873.10_3 | CCDC117 | 0.99999317 | 1 | -0.122 | 5.179 | chr22 | 29168662 | 29185289 | + | protein_coding |
| ENSG00000271721.1 | RP11-337C18.9 | 0.91764885 | 1 | -0.122 | 0.492 | chr1 | 146647181 | 146649319 | + | antisense |
| ENSG00000120802.13_5 | TMPO | 0.99999999 | 1 | -0.123 | 8.488 | chr12 | 98909290 | 98944157 | + | protein_coding |
| ENSG00000143319.16_3 | ISG20L2 | 0.99999609 | 1 | -0.123 | 5.057 | chr1 | 156691683 | 156698591 | - | protein_coding |
| ENSG00000100578.17_8 | KIAA0586 | 0.99999944 | 1 | -0.123 | 5.396 | chr14 | 58894103 | 59018015 | + | protein_coding |
| ENSG00000008294.21_6 | SPAG9 | 0.99999999 | 1 | -0.123 | 7.641 | chr17 | 49039535 | 49198229 | - | protein_coding |
| ENSG00000171824.14_5 | EXOSC10 | 0.99999966 | 1 | -0.123 | 5.822 | chr1 | 11126675 | 11159926 | - | protein_coding |
| ENSG00000062822.14_5 | POLD1 | 0.9966101 | 1 | -0.123 | 6.306 | chr19 | 50887461 | 50921275 | + | protein_coding |
| ENSG00000280206.1_7 | AC026401.3 | 0.994647 | 1 | -0.123 | 2.278 | chr16 | 15795094 | 15795975 | + | lncRNA |
| ENSG00000173588.15_9 | CEP83 | 0.99998101 | 1 | -0.123 | 4.349 | chr12 | 94700225 | 94853764 | - | protein_coding |
| ENSG00000144909.8_4 | OSBPL11 | 0.9999877 | 1 | -0.123 | 4.368 | chr3 | 125247702 | 125314341 | - | protein_coding |
| ENSG00000132676.16_6 | DAP3 | 0.99999995 | 1 | -0.123 | 6.345 | chr1 | 155657751 | 155708801 | + | protein_coding |
| ENSG00000287917.1_3 | AP000944.7 | 0.84928787 | 1 | -0.123 | -0.426 | chr11 | 65072877 | 65081715 | - | lncRNA |
| ENSG00000135823.14_5 | STX6 | 0.99999946 | 1 | -0.123 | 5.277 | chr1 | 180941848 | 180992257 | - | protein_coding |
| ENSG00000130165.10_6 | ELOF1 | 0.99999119 | 1 | -0.123 | 4.734 | chr19 | 11661962 | 11670051 | - | protein_coding |
| ENSG00000163349.22_7 | HIPK1 | 0.99999977 | 1 | -0.123 | 5.672 | chr1 | 114471946 | 114520491 | + | protein_coding |
| ENSG00000085433.15_7 | WDR47 | 0.99993967 | 1 | -0.124 | 3.854 | chr1 | 109512836 | 109584735 | - | protein_coding |
| ENSG00000117242.7_8 | PINK1-AS | 0.96535013 | 1 | -0.124 | 1.185 | chr1 | 20969150 | 20978686 | - | lncRNA |
| ENSG00000233559.2_5 | LINC00513 | 0.99847298 | 1 | -0.124 | 1.907 | chr7 | 130538479 | 130615441 | + | lncRNA |
| ENSG00000006744.19_6 | ELAC2 | 0.99999826 | 1 | -0.124 | 6.835 | chr17 | 12894929 | 12921382 | - | protein_coding |
| ENSG00000188599.17_3 | NPIPP1 | 0.97522454 | 1 | -0.124 | 1.590 | chr16 | 15198169 | 15217355 | - | transcribed_unprocessed_pseudogene |
| ENSG00000120685.20_6 | PROSER1 | 0.99997899 | 1 | -0.124 | 4.246 | chr13 | 39584002 | 39612226 | - | protein_coding |
| ENSG00000107036.12_3 | RIC1 | 0.99999943 | 1 | -0.124 | 5.240 | chr9 | 5629025 | 5776557 | + | protein_coding |
| ENSG00000012061.15_7 | ERCC1 | 0.99999995 | 1 | -0.124 | 6.302 | chr19 | 45910591 | 45982086 | - | protein_coding |
| ENSG00000167862.10_6 | MRPL58 | 0.99999881 | 1 | -0.124 | 5.084 | chr17 | 73008765 | 73017356 | + | protein_coding |
| ENSG00000157778.9_4 | PSMG3 | 0.99999577 | 1 | -0.124 | 4.922 | chr7 | 1606968 | 1610641 | - | protein_coding |
| ENSG00000204536.14_6 | CCHCR1 | 0.99925928 | 1 | -0.124 | 3.185 | chr6 | 31110216 | 31126015 | - | protein_coding |
| ENSG00000170759.11_4 | KIF5B | 0.99999999 | 1 | -0.125 | 8.650 | chr10 | 32297943 | 32345353 | - | protein_coding |
| ENSG00000188343.13_7 | FAM92A | 0.99997069 | 1 | -0.125 | 4.786 | chr8 | 94710789 | 94743755 | + | protein_coding |
| ENSG00000198689.11_5 | SLC9A6 | 0.9999982 | 1 | -0.125 | 4.946 | chrX | 135056000 | 135129428 | + | protein_coding |
| ENSG00000101639.18_8 | CEP192 | 0.99999987 | 1 | -0.125 | 6.173 | chr18 | 12991361 | 13125051 | + | protein_coding |
| ENSG00000103035.11_3 | PSMD7 | 0.99999844 | 1 | -0.125 | 5.746 | chr16 | 74330712 | 74340186 | + | protein_coding |
| ENSG00000120159.13_6 | CAAP1 | 0.99999443 | 1 | -0.125 | 4.980 | chr9 | 26840683 | 26892801 | - | protein_coding |
| ENSG00000122417.15_9 | ODF2L | 0.99998504 | 1 | -0.125 | 4.344 | chr1 | 86812507 | 86862025 | - | protein_coding |
| ENSG00000107614.22_4 | TRDMT1 | 0.99971413 | 1 | -0.125 | 3.314 | chr10 | 17179335 | 17244053 | - | protein_coding |
| ENSG00000134313.15_7 | KIDINS220 | 0.99999988 | 1 | -0.125 | 6.589 | chr2 | 8861211 | 8977760 | - | protein_coding |
| ENSG00000227268.4_4 | KLLN | 0.96749377 | 1 | -0.125 | 1.251 | chr10 | 89618915 | 89623194 | - | protein_coding |
| ENSG00000273542.1_3 | H4C12 | 0.98230476 | 1 | -0.125 | 2.030 | chr6 | 27798994 | 27799305 | - | protein_coding |

| | | | | | | | | | | |
|---|---|---|---|---|---|---|---|---|---|---|
| ENSG00000117281.16_5 | CD160 | 0.85614142 | 1 | -0.126 | -0.295 | chr1 | 145695798 | 145715614 | - | protein_coding |
| ENSG00000177302.15_6 | TOP3A | 0.99999981 | 1 | -0.126 | 6.104 | chr17 | 18174742 | 18218321 | - | protein_coding |
| ENSG00000136888.7_3 | ATP6V1G1 | 0.99999642 | 1 | -0.126 | 5.837 | chr9 | 117350049 | 117361159 | + | protein_coding |
| ENSG00000240445.4_4 | FOXO3B | 0.994806 | 1 | -0.126 | 2.298 | chr17 | 18570942 | 18585575 | - | protein_coding |
| ENSG00000087269.16_4 | NOP14 | 0.99999845 | 1 | -0.126 | 5.181 | chr4 | 2939660 | 2965133 | - | protein_coding |
| ENSG00000166562.9_3 | SEC11C | 0.99999456 | 1 | -0.126 | 4.862 | chr18 | 56807098 | 56826064 | + | protein_coding |
| ENSG00000055332.18_7 | EIF2AK2 | 0.99999999 | 1 | -0.126 | 6.947 | chr2 | 37326353 | 37384208 | - | protein_coding |
| ENSG00000166181.13_5 | API5 | 0.99972012 | 1 | -0.126 | 6.660 | chr11 | 43333513 | 43366079 | + | protein_coding |
| ENSG00000153485.5_4 | TMEM251 | 0.99579888 | 1 | -0.126 | 2.605 | chr14 | 93651296 | 93653434 | + | protein_coding |
| ENSG00000116560.11_4 | SFPQ | 1 | 1 | -0.126 | 8.688 | chr1 | 35641979 | 35658746 | - | protein_coding |
| ENSG00000123965.13_5 | PMS2P5 | 0.880051 | 1 | -0.126 | 0.197 | chr7 | 74310231 | 74313951 | + | unprocessed_pseudogene |
| ENSG00000113312.11_4 | TTC1 | 0.99999625 | 1 | -0.126 | 5.767 | chr5 | 159436120 | 159492550 | + | protein_coding |
| ENSG00000137040.10_5 | RANBP6 | 0.99999895 | 1 | -0.126 | 5.404 | chr9 | 6011025 | 6015625 | - | protein_coding |
| ENSG00000198839.10_3 | ZNF277 | 0.99999635 | 1 | -0.126 | 4.707 | chr7 | 111846750 | 111983989 | + | protein_coding |
| ENSG00000280254.1_6 | AC233723.2 | 0.98371679 | 1 | -0.126 | 1.928 | chr17 | 4690352 | 4692457 | + | TEC |
| ENSG00000109189.13_5 | USP46 | 0.99998016 | 1 | -0.126 | 4.189 | chr4 | 53457126 | 53525468 | - | protein_coding |
| ENSG00000250031.1_6 | AC009927.1 | 0.84727626 | 1 | -0.126 | -0.450 | chr8 | 59337147 | 59339244 | - | processed_pseudogene |
| ENSG00000172795.16_3 | DCP2 | 0.9999997 | 1 | -0.127 | 5.746 | chr5 | 112312399 | 112357892 | + | protein_coding |
| ENSG00000168228.15_4 | ZCCHC4 | 0.99765218 | 1 | -0.127 | 2.841 | chr4 | 25314396 | 25372005 | + | protein_coding |
| ENSG00000258634.3_7 | AL160006.1 | 0.98586518 | 1 | -0.127 | 1.743 | chr1 | 110600962 | 110605177 | + | lncRNA |
| ENSG00000074657.13_5 | ZNF532 | 0.99999991 | 1 | -0.127 | 6.258 | chr18 | 56529832 | 56653712 | + | protein_coding |
| ENSG00000112306.8_3 | RPS12 | 0.99999999 | 1 | -0.127 | 8.178 | chr6 | 133135708 | 133138703 | + | protein_coding |
| ENSG00000165898.14_3 | ISCA2 | 0.99994036 | 1 | -0.127 | 3.838 | chr14 | 74960459 | 74963809 | + | protein_coding |
| ENSG00000172071.14_6 | EIF2AK3 | 0.99999366 | 1 | -0.127 | 4.571 | chr2 | 88856259 | 88926982 | - | protein_coding |
| ENSG00000105865.11_4 | DUS4L | 0.99804072 | 1 | -0.127 | 2.626 | chr7 | 107203929 | 107218968 | + | protein_coding |
| ENSG00000175595.15_3 | ERCC4 | 0.99999582 | 1 | -0.127 | 4.705 | chr16 | 14014011 | 14046205 | + | protein_coding |
| ENSG00000136942.15_4 | RPL35 | 0.99999998 | 1 | -0.127 | 8.541 | chr9 | 127620159 | 127624260 | - | protein_coding |
| ENSG00000251026.2_5 | LINC02163 | 0.89792585 | 1 | -0.127 | 0.300 | chr5 | 103415548 | 103741822 | + | lncRNA |
| ENSG00000143157.11_2 | POGK | 0.99999998 | 1 | -0.128 | 6.957 | chr1 | 166808684 | 166825581 | + | protein_coding |
| ENSG00000108669.16_8 | CYTH1 | 0.99999952 | 1 | -0.128 | 5.577 | chr17 | 76670130 | 76778379 | - | protein_coding |
| ENSG00000160145.15_4 | KALRN | 0.96763224 | 1 | -0.128 | 0.907 | chr3 | 123798870 | 124445172 | + | protein_coding |
| ENSG00000133401.16_4 | PDZD2 | 0.92954528 | 1 | -0.128 | 0.700 | chr5 | 31639238 | 32111038 | + | protein_coding |
| ENSG00000041802.11_5 | LSG1 | 0.9999999 | 1 | -0.128 | 6.006 | chr3 | 194361520 | 194393192 | - | protein_coding |
| ENSG00000100429.18_5 | HDAC10 | 0.8427233 | 1 | -0.128 | -0.741 | chr22 | 50683612 | 50689834 | - | protein_coding |
| ENSG00000235027.1_6 | AC068580.3 | 0.9820799 | 1 | -0.128 | 1.830 | chr11 | 1781578 | 1783716 | + | lncRNA |
| ENSG00000135317.13_10 | SNX14 | 0.99999973 | 1 | -0.128 | 5.895 | chr6 | 86215214 | 86303874 | - | protein_coding |
| ENSG00000234405.2_6 | Z69733.1 | 0.83027627 | 1 | -0.128 | -0.615 | chrX | 102741389 | 102772847 | + | lncRNA |
| ENSG00000268764.1_7 | AC011465.1 | 0.90528292 | 1 | -0.128 | 0.563 | chr19 | 38599140 | 38600173 | - | lncRNA |
| ENSG00000111581.10_6 | NUP107 | 0.99999998 | 1 | -0.128 | 6.854 | chr12 | 69080731 | 69139589 | + | protein_coding |
| ENSG00000241547.1_4 | ACTG1P20 | 0.84242942 | 1 | -0.128 | -0.561 | chr1 | 27651820 | 27652287 | + | processed_pseudogene |
| ENSG00000114268.12_5 | PFKFB4 | 0.99855463 | 1 | -0.128 | 3.204 | chr3 | 48555117 | 48599448 | - | protein_coding |
| ENSG00000196776.16_5 | CD47 | 0.99999989 | 1 | -0.128 | 5.980 | chr3 | 107761938 | 107810709 | - | protein_coding |
| ENSG00000170085.18_5 | SIMC1 | 0.99985193 | 1 | -0.129 | 3.571 | chr5 | 175665370 | 175772994 | + | protein_coding |
| ENSG00000146350.14_7 | TBC1D32 | 0.99853483 | 1 | -0.129 | 3.167 | chr6 | 121400640 | 121655891 | - | protein_coding |
| ENSG00000105176.18_6 | URI1 | 0.99999999 | 1 | -0.129 | 7.499 | chr19 | 30414551 | 30507519 | + | protein_coding |
| ENSG00000151532.13_2 | VTI1A | 0.99999709 | 1 | -0.129 | 4.968 | chr10 | 114206756 | 114578503 | + | protein_coding |
| ENSG00000165629.20_4 | ATP5F1C | 0.99999995 | 1 | -0.129 | 7.354 | chr10 | 7830110 | 7849778 | + | protein_coding |
| ENSG00000182902.14_3 | SLC25A18 | 0.87736591 | 1 | -0.129 | 0.097 | chr22 | 18043150 | 18073761 | + | protein_coding |
| ENSG00000119383.19_9 | PTPA | 0.9999999 | 1 | -0.129 | 7.076 | chr9 | 131873229 | 131911225 | + | protein_coding |
| ENSG00000013810.19_4 | TACC3 | 0.99999909 | 1 | -0.129 | 5.894 | chr4 | 1714618 | 1746903 | + | protein_coding |
| ENSG00000102606.18_5 | ARHGEF7 | 0.99999909 | 1 | -0.129 | 5.134 | chr13 | 111766906 | 111958084 | + | protein_coding |
| ENSG00000188342.12_4 | GTF2F2 | 0.99999779 | 1 | -0.129 | 4.844 | chr13 | 45694645 | 45859028 | + | protein_coding |
| ENSG00000108592.17_6 | FTSJ3 | 0.99999996 | 1 | -0.129 | 6.534 | chr17 | 61896793 | 61907372 | - | protein_coding |
| ENSG00000214425.8_8 | LRRC37A4P | 0.99978984 | 1 | -0.129 | 3.601 | chr17 | 43584107 | 43628903 | - | transcribed_unprocessed_pseudogene |
| ENSG00000162783.11_3 | IER5 | 0.99999373 | 1 | -0.130 | 5.685 | chr1 | 181057836 | 181062036 | + | protein_coding |
| ENSG00000005893.16_4 | LAMP2 | 0.99999989 | 1 | -0.130 | 6.799 | chrX | 119560003 | 119603220 | - | protein_coding |
| ENSG00000101158.15_5 | NELFCD | 0.99999871 | 1 | -0.130 | 5.251 | chr20 | 57556263 | 57570188 | + | protein_coding |
| ENSG00000153815.16_4 | CMIP | 0.99999994 | 1 | -0.130 | 6.544 | chr16 | 81478775 | 81745367 | + | protein_coding |

| | | | | | | | | | | |
|---|---|---|---|---|---|---|---|---|---|---|
| ENSG00000164430.16_6 | CGAS | 0.99998194 | 1 | -0.130 | 4.328 | chr6 | 74123238 | 74162020 | - | protein_coding |
| ENSG00000083520.15_4 | DIS3 | 0.99999988 | 1 | -0.130 | 6.390 | chr13 | 73326307 | 73356234 | - | protein_coding |
| ENSG00000161243.9_9 | FBXO27 | 0.99246779 | 1 | -0.130 | 1.994 | chr19 | 39481354 | 39523425 | - | protein_coding |
| ENSG00000071794.16_5 | HLTF | 0.99999999 | 1 | -0.130 | 7.301 | chr3 | 148747914 | 148804341 | - | protein_coding |
| ENSG00000102805.15_6 | CLN5 | 0.99974289 | 1 | -0.130 | 3.766 | chr13 | 77564795 | 77593278 | + | protein_coding |
| ENSG00000227057.10_5 | WDR46 | 0.99999719 | 1 | -0.130 | 4.876 | chr6 | 33246885 | 33257024 | - | protein_coding |
| ENSG00000053254.15_7 | FOXN3 | 0.99995321 | 1 | -0.130 | 4.010 | chr14 | 89591215 | 90085493 | - | protein_coding |
| ENSG00000066027.12_4 | PPP2R5A | 0.99999974 | 1 | -0.130 | 5.643 | chr1 | 212458752 | 212535195 | + | protein_coding |
| ENSG00000136720.7_4 | HS6ST1 | 0.9989608 | 1 | -0.130 | 4.714 | chr2 | 128994290 | 129076442 | - | protein_coding |
| ENSG00000100335.15_7 | MIEF1 | 0.9999979 | 1 | -0.130 | 5.371 | chr22 | 39895437 | 39914137 | + | protein_coding |
| ENSG00000131828.14_4 | PDHA1 | 0.99999998 | 1 | -0.130 | 6.718 | chrX | 19362011 | 19379836 | + | protein_coding |
| ENSG00000122335.17_7 | SERAC1 | 0.99975169 | 1 | -0.130 | 3.655 | chr6 | 158530551 | 158589312 | - | protein_coding |
| ENSG00000241529.2 | RN7SL767P | 0.91296047 | 1 | -0.130 | 0.751 | chr3 | 113351551 | 113351827 | + | misc_RNA |
| ENSG00000114450.10_5 | GNB4 | 0.9999936 | 1 | -0.130 | 5.005 | chr3 | 179113876 | 179169264 | - | protein_coding |
| ENSG00000250317.8_2 | SMIM20 | 0.99986908 | 1 | -0.131 | 3.759 | chr4 | 25863452 | 25931496 | + | protein_coding |
| ENSG00000226781.1_4 | TBCAP1 | 0.82377696 | 1 | -0.131 | -0.815 | chrX | 33059512 | 33059838 | + | processed_pseudogene |
| ENSG00000167984.18_5 | NLRC3 | 0.89302604 | 1 | -0.131 | 0.611 | chr16 | 3589033 | 3627404 | - | protein_coding |
| ENSG00000197386.12_8 | HTT | 0.99999992 | 1 | -0.131 | 6.252 | chr4 | 3043149 | 3245687 | + | protein_coding |
| ENSG00000168803.15_3 | ADAL | 0.99965123 | 1 | -0.131 | 3.319 | chr15 | 43622855 | 43646753 | + | protein_coding |
| ENSG00000088826.18_4 | SMOX | 0.99999977 | 1 | -0.131 | 6.182 | chr20 | 4101627 | 4168394 | + | protein_coding |
| ENSG00000008838.20_6 | MED24 | 0.99999964 | 1 | -0.131 | 5.861 | chr17 | 38175350 | 38217468 | - | protein_coding |
| ENSG00000183479.8 | TREX2 | 0.91190912 | 1 | -0.131 | 0.472 | chrX | 152710178 | 152736045 | - | protein_coding |
| ENSG00000140988.16_7 | RPS2 | 1 | 1 | -0.131 | 10.005 | chr16 | 2012059 | 2014842 | - | protein_coding |
| ENSG00000047410.14_4 | TPR | 0.99999998 | 1 | -0.131 | 8.480 | chr1 | 186280784 | 186344825 | - | protein_coding |
| ENSG00000077235.18_6 | GTF3C1 | 0.99999999 | 1 | -0.131 | 7.769 | chr16 | 27470876 | 27561234 | - | protein_coding |
| ENSG00000245479.3_5 | LINC01585 | 0.83579943 | 1 | -0.131 | -0.683 | chr15 | 91203438 | 91208198 | + | lncRNA |
| ENSG00000134046.12_5 | MBD2 | 0.99999882 | 1 | -0.131 | 5.900 | chr18 | 51677976 | 51751039 | - | protein_coding |
| ENSG00000100796.17_6 | PPP4R3A | 0.99999993 | 1 | -0.131 | 6.763 | chr14 | 91923955 | 91976898 | - | protein_coding |
| ENSG00000114391.13_4 | RPL24 | 0.99999989 | 1 | -0.131 | 7.724 | chr3 | 101399935 | 101405562 | - | protein_coding |
| ENSG00000198791.12_4 | CNOT7 | 0.99999998 | 1 | -0.131 | 7.259 | chr8 | 17082475 | 17104387 | - | protein_coding |
| ENSG00000286092.1_3 | AC117503.5 | 0.84875626 | 1 | -0.131 | -0.110 | chr12 | 124197955 | 124207716 | - | lncRNA |
| ENSG00000188846.14_4 | RPL14 | 0.99999999 | 1 | -0.131 | 7.960 | chr3 | 40498783 | 40510078 | + | protein_coding |
| ENSG00000135341.18_4 | MAP3K7 | 0.99999926 | 1 | -0.131 | 5.901 | chr6 | 91223292 | 91296791 | - | protein_coding |
| ENSG00000089597.18_5 | GANAB | 0.99999997 | 1 | -0.131 | 8.029 | chr11 | 62392298 | 62414198 | - | protein_coding |
| ENSG00000138036.18_3 | DYNC2LI1 | 0.99997062 | 1 | -0.131 | 4.340 | chr2 | 44001178 | 44037149 | + | protein_coding |
| ENSG00000184445.12_4 | KNTC1 | 0.99999998 | 1 | -0.131 | 7.512 | chr12 | 123011793 | 123110943 | + | protein_coding |
| ENSG00000166546.14_6 | BEAN1 | 0.95592247 | 1 | -0.131 | 1.351 | chr16 | 66461198 | 66527432 | + | protein_coding |
| ENSG00000172426.16_3 | RSPH9 | 0.86508638 | 1 | -0.132 | 0.375 | chr6 | 43612773 | 43640337 | + | protein_coding |
| ENSG00000156504.16_3 | FAM122B | 0.99997834 | 1 | -0.132 | 4.462 | chrX | 133903596 | 133931262 | - | protein_coding |
| ENSG00000065518.8_3 | NDUFB4 | 0.99999967 | 1 | -0.132 | 5.974 | chr3 | 120315175 | 120321354 | + | protein_coding |
| ENSG00000136444.10_6 | RSAD1 | 0.99999948 | 1 | -0.132 | 5.894 | chr17 | 48556221 | 48563335 | + | protein_coding |
| ENSG00000219626.9_7 | FAM228B | 0.96953971 | 1 | -0.132 | 1.362 | chr2 | 24299396 | 24392509 | + | protein_coding |
| ENSG00000086061.16_3 | DNAJA1 | 0.99999998 | 1 | -0.132 | 8.170 | chr9 | 33025271 | 33039905 | + | protein_coding |
| ENSG00000120942.13_5 | UBIAD1 | 0.99991496 | 1 | -0.132 | 4.066 | chr1 | 11333263 | 11356106 | + | protein_coding |
| ENSG00000148444.16_5 | COMMD3 | 0.99856264 | 1 | -0.132 | 2.751 | chr10 | 22605315 | 22609235 | + | protein_coding |
| ENSG00000186001.14_6 | LRCH3 | 0.99999061 | 1 | -0.133 | 4.677 | chr3 | 197518097 | 197615307 | + | protein_coding |
| ENSG00000100084.14_5 | HIRA | 0.9958645 | 1 | -0.133 | 2.493 | chr22 | 19318221 | 19434973 | - | protein_coding |
| ENSG00000235919.4_6 | ASH1L-AS1 | 0.93824098 | 1 | -0.133 | 1.031 | chr1 | 155531833 | 155533735 | + | lncRNA |
| ENSG00000159079.19_7 | CFAP298 | 0.98815336 | 1 | -0.133 | 2.352 | chr21 | 33964389 | 33984913 | - | protein_coding |
| ENSG00000185128.11 | TBC1D3F | 0.8389637 | 1 | -0.133 | -0.327 | chr17 | 36283971 | 36294915 | + | protein_coding |
| ENSG00000162139.10_3 | NEU3 | 0.99987795 | 1 | -0.133 | 3.892 | chr11 | 74699324 | 74729938 | + | protein_coding |
| ENSG00000135596.18_7 | MICAL1 | 0.99998878 | 1 | -0.133 | 4.498 | chr6 | 109765265 | 109787171 | - | protein_coding |
| ENSG00000103269.13_3 | RHBDL1 | 0.93657235 | 1 | -0.133 | 0.603 | chr16 | 725666 | 728268 | + | protein_coding |
| ENSG00000049323.16_5 | LTBP1 | 0.99999773 | 1 | -0.133 | 5.023 | chr2 | 33172020 | 33624576 | + | protein_coding |
| ENSG00000167291.16_8 | TBC1D16 | 0.99999986 | 1 | -0.133 | 6.266 | chr17 | 77906142 | 78009671 | - | protein_coding |
| ENSG00000236778.8_10 | INTS6-AS1 | 0.90469228 | 1 | -0.133 | 0.463 | chr13 | 52026500 | 52128814 | + | lncRNA |
| ENSG00000134352.20_5 | IL6ST | 0.99999997 | 1 | -0.134 | 8.222 | chr5 | 55230923 | 55290850 | - | protein_coding |
| ENSG00000231770.6_7 | TMEM44-AS1 | 0.98205028 | 1 | -0.134 | 1.646 | chr3 | 194304733 | 194310989 | + | lncRNA |
| ENSG00000155463.14_6 | OXA1L | 0.99999997 | 1 | -0.134 | 6.929 | chr14 | 23235731 | 23242251 | + | protein_coding |

| | | | | | | | | | | |
|---|---|---|---|---|---|---|---|---|---|---|
| ENSG00000125746.16_9 | EML2 | 0.99957925 | 1 | -0.134 | 3.787 | chr19 | 46110252 | 46148887 | - | protein_coding |
| ENSG00000100227.18_5 | POLDIP3 | 0.9999989 | 1 | -0.134 | 5.435 | chr22 | 42979727 | 43010968 | - | protein_coding |
| ENSG00000158966.16_5 | CACHD1 | 0.99614699 | 1 | -0.134 | 2.344 | chr1 | 64935812 | 65158741 | + | protein_coding |
| ENSG00000143493.13_5 | INTS7 | 0.9999995 | 1 | -0.134 | 5.468 | chr1 | 212113741 | 212208899 | - | protein_coding |
| ENSG00000127616.18_7 | SMARCA4 | 0.9999796 | 1 | -0.134 | 4.428 | chr19 | 11071501 | 11190102 | + | protein_coding |
| ENSG00000119487.17_7 | MAPKAP1 | 0.99999993 | 1 | -0.134 | 6.367 | chr9 | 128199672 | 128469513 | - | protein_coding |
| ENSG00000103121.9_4 | CMC2 | 0.99997016 | 1 | -0.134 | 4.501 | chr16 | 81000345 | 81053875 | - | protein_coding |
| ENSG00000247624.6_6 | CPEB2-DT | 0.81537784 | 1 | -0.134 | -1.159 | chr4 | 14911585 | 15003669 | - | lncRNA |
| ENSG00000182010.11_4 | RTKN2 | 0.99914461 | 1 | -0.134 | 3.184 | chr10 | 63942794 | 64028603 | - | protein_coding |
| ENSG00000163950.13_5 | SLBP | 0.99999942 | 1 | -0.134 | 5.865 | chr4 | 1694458 | 1714071 | - | protein_coding |
| ENSG00000167699.13_4 | GLOD4 | 0.99999995 | 1 | -0.134 | 6.510 | chr17 | 660337 | 686630 | - | protein_coding |
| ENSG00000168795.5_4 | ZBTB5 | 0.99999466 | 1 | -0.135 | 4.825 | chr9 | 37438099 | 37465447 | - | protein_coding |
| ENSG00000187534.6_4 | PRR13P5 | 0.86892966 | 1 | -0.135 | 0.171 | chr19 | 40449146 | 40449587 | - | processed_pseudogene |
| ENSG00000287562.1_1 | AL109615.4 | 0.85046375 | 1 | -0.135 | 0.013 | chr6 | 44058524 | 44063322 | - | lncRNA |
| ENSG00000123119.12_3 | NECAB1 | 0.88653023 | 1 | -0.135 | 0.446 | chr8 | 91803969 | 91971621 | + | protein_coding |
| ENSG00000175110.11_5 | MRPS22 | 0.99998714 | 1 | -0.135 | 4.609 | chr3 | 138724648 | 139076065 | + | protein_coding |
| ENSG00000101126.18_7 | ADNP | 0.99999993 | 1 | -0.135 | 6.577 | chr20 | 49505453 | 49547974 | - | protein_coding |
| ENSG00000132591.12_3 | ERAL1 | 0.99999959 | 1 | -0.135 | 5.571 | chr17 | 27182028 | 27188079 | + | protein_coding |
| ENSG00000170734.12_4 | POLH | 0.99999025 | 1 | -0.135 | 4.570 | chr6 | 43543922 | 43588260 | + | protein_coding |
| ENSG00000101751.10_5 | POLI | 0.99904849 | 1 | -0.135 | 4.365 | chr18 | 51795774 | 51847636 | + | protein_coding |
| ENSG00000177425.11_6 | PAWR | 0.99999997 | 1 | -0.135 | 7.297 | chr12 | 79968759 | 80084744 | - | protein_coding |
| ENSG00000042753.11_3 | AP2S1 | 0.99999924 | 1 | -0.135 | 6.082 | chr19 | 47341393 | 47354249 | - | protein_coding |
| ENSG00000261061.1_7 | AC092718.4 | 0.99985261 | 1 | -0.135 | 3.731 | chr16 | 81064375 | 81065090 | + | lncRNA |
| ENSG00000221978.12_6 | CCNL2 | 0.99999993 | 1 | -0.135 | 6.300 | chr1 | 1321091 | 1334715 | - | protein_coding |
| ENSG00000197429.11_5 | IPP | 0.99957833 | 1 | -0.135 | 3.385 | chr1 | 46159996 | 46216325 | - | protein_coding |
| ENSG00000228060.3_8 | PABPC4-AS1 | 0.93827011 | 1 | -0.135 | 0.916 | chr1 | 40030724 | 40039532 | + | lncRNA |
| ENSG00000078319.9_6 | PMS2P1 | 0.99904446 | 1 | -0.135 | 3.350 | chr7 | 99918615 | 99939531 | - | unprocessed_pseudogene |
| ENSG00000106086.20_6 | PLEKHA8 | 0.99998306 | 1 | -0.135 | 4.379 | chr7 | 30067020 | 30170099 | + | protein_coding |
| ENSG00000108312.15_5 | UBTF | 0.9999999 | 1 | -0.135 | 6.925 | chr17 | 42282401 | 42298994 | - | protein_coding |
| ENSG00000170540.15_5 | ARL6IP1 | 0.99999979 | 1 | -0.135 | 7.906 | chr16 | 18802991 | 18812894 | - | protein_coding |
| ENSG00000146802.13_5 | TMEM168 | 0.99999128 | 1 | -0.136 | 4.848 | chr7 | 112402432 | 112430478 | - | protein_coding |
| ENSG00000163389.12_4 | POGLUT1 | 0.99990087 | 1 | -0.136 | 4.244 | chr3 | 119187810 | 119213555 | + | protein_coding |
| ENSG00000071994.11_7 | PDCD2 | 0.99999975 | 1 | -0.136 | 5.673 | chr6 | 170884383 | 170893780 | - | protein_coding |
| ENSG00000120697.9_5 | ALG5 | 0.99998053 | 1 | -0.136 | 4.277 | chr13 | 37523875 | 37574398 | - | protein_coding |
| ENSG00000253227.2_7 | AC090192.2 | 0.99463665 | 1 | -0.136 | 2.314 | chr8 | 125283908 | 125291821 | + | lncRNA |
| ENSG00000198492.16_6 | YTHDF2 | 0.99999981 | 1 | -0.136 | 6.014 | chr1 | 29063133 | 29096287 | + | protein_coding |
| ENSG00000081307.12_4 | UBA5 | 0.99999676 | 1 | -0.136 | 5.008 | chr3 | 132373290 | 132396941 | + | protein_coding |
| ENSG00000261342.1_5 | AC006538.1 | 0.88375348 | 1 | -0.136 | 0.343 | chr19 | 2727741 | 2729325 | - | lncRNA |
| ENSG00000166477.13_4 | LEO1 | 0.99999944 | 1 | -0.136 | 5.728 | chr15 | 52230222 | 52263975 | - | protein_coding |
| ENSG00000266405.3_6 | AP005061.2 | 0.81959422 | 1 | -0.136 | -0.591 | chr18 | 2652169 | 2655394 | - | transcribed_processed_pseudogene |
| ENSG00000188994.13_6 | ZNF292 | 0.99999482 | 1 | -0.137 | 5.112 | chr6 | 87862551 | 87975661 | + | protein_coding |
| ENSG00000166902.5_5 | MRPL16 | 0.99999826 | 1 | -0.137 | 4.970 | chr11 | 59573613 | 59578251 | - | protein_coding |
| ENSG00000115561.16_5 | CHMP3 | 0.999999 | 1 | -0.137 | 5.954 | chr2 | 86730553 | 86790602 | - | protein_coding |
| ENSG00000121864.10_5 | ZNF639 | 0.99999747 | 1 | -0.137 | 4.987 | chr3 | 179040819 | 179056371 | + | protein_coding |
| ENSG00000038274.17_4 | MAT2B | 0.99999913 | 1 | -0.137 | 5.802 | chr5 | 162930120 | 162946360 | + | protein_coding |
| ENSG00000137074.19_6 | APTX | 0.99999615 | 1 | -0.137 | 5.110 | chr9 | 32886599 | 33025128 | - | protein_coding |
| ENSG00000108947.5_3 | EFNB3 | 0.96088865 | 1 | -0.137 | 1.201 | chr17 | 7608520 | 7614690 | + | protein_coding |
| ENSG00000149201.10_3 | CCDC81 | 0.81280328 | 1 | -0.137 | -0.959 | chr11 | 86085778 | 86134151 | + | protein_coding |
| ENSG00000138279.16_6 | ANXA7 | 0.9999979 | 1 | -0.137 | 6.616 | chr10 | 75134859 | 75173834 | - | protein_coding |
| ENSG00000285230.2_4 | RALY-AS1 | 0.99262499 | 1 | -0.137 | 2.052 | chr20 | 32568485 | 32582163 | - | lncRNA |
| ENSG00000123836.15_6 | PFKFB2 | 0.99999879 | 1 | -0.137 | 5.193 | chr1 | 207207711 | 207254369 | + | protein_coding |
| ENSG00000124486.13_5 | USP9X | 0.99999999 | 1 | -0.137 | 8.733 | chrX | 40944698 | 41095832 | + | protein_coding |
| ENSG00000260992.1_6 | DOCK9-DT | 0.8140679 | 1 | -0.137 | -0.994 | chr13 | 99740073 | 99740879 | + | lncRNA |
| ENSG00000221051.1 | AC118344.1 | 0.81476169 | 1 | -0.137 | -0.743 | chr19 | 40710019 | 40710106 | + | miRNA |
| ENSG00000158427.15_6 | TMSB15B | 0.95087357 | 1 | -0.137 | 1.020 | chrX | 103173479 | 103221285 | + | protein_coding |
| ENSG00000146197.9_5 | SCUBE3 | 0.98628839 | 1 | -0.137 | 1.797 | chr6 | 35181733 | 35220856 | + | protein_coding |
| ENSG00000120437.9_3 | ACAT2 | 0.99999474 | 1 | -0.137 | 5.080 | chr6 | 160183077 | 160200144 | + | protein_coding |
| ENSG00000225075.1_7 | AL603832.1 | 0.81636021 | 1 | -0.137 | -0.783 | chr1 | 113236310 | 113239243 | - | lncRNA |
| ENSG00000186283.14_5 | TOR3A | 0.99995029 | 1 | -0.137 | 5.418 | chr1 | 179051205 | 179067158 | + | protein_coding |

| | | | | | | | | | | |
|---|---|---|---|---|---|---|---|---|---|---|
| ENSG00000229519.2_5 | AL591866.1 | 0.81298457 | 1 | -0.137 | -0.880 | chr1 | 6607965 | 6608679 | + | processed_pseudogene |
| ENSG00000099194.6_3 | SCD | 1 | 1 | -0.137 | 10.254 | chr10 | 102106990 | 102124583 | + | protein_coding |
| ENSG00000130309.11_7 | COLGALT1 | 0.99999873 | 1 | -0.138 | 6.125 | chr19 | 17666458 | 17693971 | + | protein_coding |
| ENSG00000084623.11_2 | EIF3I | 0.99999998 | 1 | -0.138 | 7.140 | chr1 | 32687529 | 32697205 | + | protein_coding |
| ENSG00000136381.13_4 | IREB2 | 0.99999997 | 1 | -0.138 | 7.103 | chr15 | 78729773 | 78793795 | + | protein_coding |
| ENSG00000100055.21_4 | CYTH4 | 0.84166974 | 1 | -0.138 | -0.280 | chr22 | 37678068 | 37711382 | + | protein_coding |
| ENSG00000175606.11_4 | TMEM70 | 0.99998595 | 1 | -0.138 | 4.820 | chr8 | 74884672 | 74895018 | + | protein_coding |
| ENSG00000196812.5_3 | ZSCAN16 | 0.99679067 | 1 | -0.138 | 2.480 | chr6 | 28092387 | 28097860 | + | protein_coding |
| ENSG00000260917.1_6 | AL158212.3 | 0.99246582 | 1 | -0.138 | 2.267 | chr10 | 114583249 | 114587485 | + | lncRNA |
| ENSG00000074356.17_6 | NCBP3 | 0.99999984 | 1 | -0.138 | 6.317 | chr17 | 3705452 | 3749540 | - | protein_coding |
| ENSG00000109118.14_5 | PHF12 | 0.99999964 | 1 | -0.138 | 5.747 | chr17 | 27232268 | 27278789 | - | protein_coding |
| ENSG00000163872.16_5 | YEATS2 | 0.99999812 | 1 | -0.138 | 5.199 | chr3 | 183415585 | 183530412 | + | protein_coding |
| ENSG00000175550.8_3 | DRAP1 | 0.99999865 | 1 | -0.138 | 5.638 | chr11 | 65686745 | 65689034 | + | protein_coding |
| ENSG00000181638.19_5 | ZFP41 | 0.99870135 | 1 | -0.138 | 2.932 | chr8 | 144328991 | 144344875 | + | protein_coding |
| ENSG00000086619.14_4 | ERO1B | 0.99986212 | 1 | -0.138 | 3.897 | chr1 | 236378401 | 236445258 | - | protein_coding |
| ENSG00000230069.3_5 | LRRC37A15P | 0.80975977 | 1 | -0.139 | -0.995 | chr4 | 103648431 | 103651878 | - | processed_pseudogene |
| ENSG00000225442.2_7 | MPRIP-AS1 | 0.92985481 | 1 | -0.139 | 0.820 | chr17 | 16979352 | 16981066 | - | lncRNA |
| ENSG00000131778.19_4 | CHD1L | 0.93780684 | 1 | -0.139 | 0.877 | chr1 | 146714304 | 146767446 | + | protein_coding |
| ENSG00000197958.13_5 | RPL12 | 0.99999999 | 1 | -0.139 | 8.877 | chr9 | 130209953 | 130213685 | - | protein_coding |
| ENSG00000162062.15_8 | TEDC2 | 0.99981848 | 1 | -0.139 | 4.310 | chr16 | 2510087 | 2514964 | + | protein_coding |
| ENSG00000091483.6_3 | FH | 0.99999988 | 1 | -0.139 | 6.630 | chr1 | 241660903 | 241683061 | - | protein_coding |
| ENSG00000272892.1_6 | AL133551.1 | 0.81430143 | 1 | -0.139 | -0.683 | chr10 | 69609283 | 69610504 | + | lncRNA |
| ENSG00000048392.12_3 | RRM2B | 0.99999963 | 1 | -0.139 | 6.134 | chr8 | 103216730 | 103251189 | - | protein_coding |
| ENSG00000161048.11_4 | NAPEPLD | 0.9999715 | 1 | -0.139 | 4.445 | chr7 | 102740223 | 102790007 | - | protein_coding |
| ENSG00000244230.2 | RN7SL151P | 0.81294259 | 1 | -0.139 | -0.960 | chr9 | 21699313 | 21699569 | + | misc_RNA |
| ENSG00000104714.14_6 | ERICH1 | 0.99914356 | 1 | -0.139 | 2.965 | chr8 | 564746 | 688106 | - | protein_coding |
| ENSG00000263466.1_7 | AC006441.1 | 0.88368043 | 1 | -0.139 | 0.279 | chr17 | 37059957 | 37060853 | - | lncRNA |
| ENSG00000140367.12_5 | UBE2Q2 | 0.99999977 | 1 | -0.139 | 6.416 | chr15 | 76135648 | 76193419 | + | protein_coding |
| ENSG00000095261.14_3 | PSMD5 | 0.99999978 | 1 | -0.139 | 6.293 | chr9 | 123577774 | 123605229 | - | protein_coding |
| ENSG00000128595.17_6 | CALU | 0.99999997 | 1 | -0.139 | 7.961 | chr7 | 128379346 | 128413454 | + | protein_coding |
| ENSG00000055070.17_3 | SZRD1 | 0.99999973 | 1 | -0.139 | 5.876 | chr1 | 16679070 | 16724640 | + | protein_coding |
| ENSG00000131791.8_4 | PRKAB2 | 0.99997705 | 1 | -0.139 | 4.482 | chr1 | 146626685 | 146644129 | - | protein_coding |
| ENSG00000268041.2_7 | ERFL | 0.81097152 | 1 | -0.140 | -0.959 | chr19 | 42411857 | 42432668 | - | protein_coding |
| ENSG00000198825.15_7 | INPP5F | 0.99998232 | 1 | -0.140 | 4.561 | chr10 | 121485554 | 121588659 | + | protein_coding |
| ENSG00000227014.6_6 | AC007285.1 | 0.85043667 | 1 | -0.140 | -0.044 | chr7 | 30028216 | 30067159 | + | lncRNA |
| ENSG00000087191.13_6 | PSMC5 | 0.99999771 | 1 | -0.140 | 5.424 | chr17 | 61904512 | 61909386 | + | protein_coding |
| ENSG00000250107.1_7 | CACNA1G-AS1 | 0.88601621 | 1 | -0.140 | 0.107 | chr17 | 48633568 | 48639469 | - | lncRNA |
| ENSG00000183258.12_7 | DDX41 | 0.99999891 | 1 | -0.140 | 5.621 | chr5 | 176938578 | 176943962 | - | protein_coding |
| ENSG00000280106.1_6 | AC008555.7 | 0.91622878 | 1 | -0.140 | 0.769 | chr19 | 35439942 | 35442109 | - | TEC |
| ENSG00000108468.15_5 | CBX1 | 0.9999999 | 1 | -0.140 | 6.781 | chr17 | 46147414 | 46178840 | - | protein_coding |
| ENSG00000172939.9_3 | OXSR1 | 0.99999983 | 1 | -0.140 | 5.887 | chr3 | 38206580 | 38296975 | + | protein_coding |
| ENSG00000124688.13_2 | MAD2L1BP | 0.99989733 | 1 | -0.140 | 3.982 | chr6 | 43597277 | 43608689 | + | protein_coding |
| ENSG00000183873.16_6 | SCN5A | 0.81645644 | 1 | -0.140 | -0.685 | chr3 | 38589548 | 38691178 | - | protein_coding |
| ENSG00000115756.13_4 | HPCAL1 | 0.99999934 | 1 | -0.140 | 5.908 | chr2 | 10443015 | 10567743 | + | protein_coding |
| ENSG00000163660.11_3 | CCNL1 | 0.99999929 | 1 | -0.140 | 6.377 | chr3 | 156864297 | 156878549 | - | protein_coding |
| ENSG00000232653.8_4 | GOLGA8N | 0.84080623 | 1 | -0.140 | -0.240 | chr15 | 32885657 | 32899511 | + | protein_coding |
| ENSG00000134717.18_4 | BTF3L4 | 0.99999604 | 1 | -0.140 | 5.100 | chr1 | 52521871 | 52556388 | + | protein_coding |
| ENSG00000250569.1_5 | NTAN1P2 | 0.85809333 | 1 | -0.140 | 0.057 | chr8 | 87493983 | 87495231 | - | processed_pseudogene |
| ENSG00000006530.18_7 | AGK | 0.99999942 | 1 | -0.140 | 5.585 | chr7 | 141251078 | 141355044 | + | protein_coding |
| ENSG00000184203.8_5 | PPP1R2 | 0.99999813 | 1 | -0.140 | 5.247 | chr3 | 195241224 | 195270209 | - | protein_coding |
| ENSG00000169398.19_7 | PTK2 | 0.99999998 | 1 | -0.140 | 7.018 | chr8 | 141667999 | 142012315 | - | protein_coding |
| ENSG00000063978.16_5 | RNF4 | 0.99999973 | 1 | -0.140 | 6.134 | chr4 | 2463947 | 2627047 | + | protein_coding |
| ENSG00000274605.2_8 | PCCA-DT | 0.98888631 | 1 | -0.140 | 2.174 | chr13 | 100705328 | 100741204 | - | lncRNA |
| ENSG00000238278.3_2 | ALG1L6P | 0.96212651 | 1 | -0.141 | 1.227 | chr3 | 75464221 | 75471294 | + | unprocessed_pseudogene |
| ENSG00000130159.14_8 | ECSIT | 0.99996238 | 1 | -0.141 | 4.294 | chr19 | 11616744 | 11639987 | - | protein_coding |
| ENSG00000187210.14_5 | GCNT1 | 0.98026504 | 1 | -0.141 | 1.659 | chr9 | 79034766 | 79266119 | + | protein_coding |
| ENSG00000087206.17_6 | UIMC1 | 0.99999932 | 1 | -0.141 | 5.771 | chr5 | 176332006 | 176449634 | - | protein_coding |
| ENSG00000155287.11_4 | SLC25A28 | 0.99996118 | 1 | -0.141 | 4.031 | chr10 | 101370279 | 101380366 | - | protein_coding |

| | | | | | | | | | | |
|---|---|---|---|---|---|---|---|---|---|---|
| ENSG00000112186.12_3 | CAP2 | 0.99999461 | 1 | -0.141 | 4.892 | chr6 | 17393736 | 17558021 | + | protein_coding |
| ENSG00000134900.12_6 | TPP2 | 0.99999975 | 1 | -0.141 | 6.156 | chr13 | 103249336 | 103332308 | + | protein_coding |
| ENSG00000218226.1_5 | TATDN2P2 | 0.99151387 | 1 | -0.141 | 2.080 | chr6 | 159030738 | 159042668 | - | processed_pseudogene |
| ENSG00000198931.11_4 | APRT | 0.99999317 | 1 | -0.141 | 6.691 | chr16 | 88875747 | 88878345 | - | protein_coding |
| ENSG00000268364.2_7 | SMC5-AS1 | 0.80732588 | 1 | -0.141 | -0.847 | chr9 | 72808913 | 72873782 | - | lncRNA |
| ENSG00000176749.9_4 | CDK5R1 | 0.99368774 | 1 | -0.141 | 2.281 | chr17 | 30814011 | 30818271 | + | protein_coding |
| ENSG00000156983.16_6 | BRPF1 | 0.99998845 | 1 | -0.141 | 5.198 | chr3 | 9773413 | 9789699 | + | protein_coding |
| ENSG00000165775.18_3 | FUNDC2 | 0.99999887 | 1 | -0.141 | 5.452 | chrX | 154254255 | 154288579 | + | protein_coding |
| ENSG00000181019.13_4 | NQO1 | 0.99999998 | 1 | -0.141 | 10.599 | chr16 | 69740899 | 69760571 | - | protein_coding |
| ENSG00000171806.12_5 | METTL18 | 0.99812241 | 1 | -0.141 | 2.837 | chr1 | 169761670 | 169764104 | - | protein_coding |
| ENSG00000198874.13_3 | TYW1 | 0.99999615 | 1 | -0.142 | 4.929 | chr7 | 66460160 | 66704506 | + | protein_coding |
| ENSG00000186326.4_3 | RGS9BP | 0.82692906 | 1 | -0.142 | -0.281 | chr19 | 33166754 | 33169206 | + | protein_coding |
| ENSG00000174705.13_7 | SH3PXD2B | 0.99999959 | 1 | -0.142 | 6.122 | chr5 | 171752004 | 171881529 | - | protein_coding |
| ENSG00000100100.13_6 | PIK3IP1 | 0.92231328 | 1 | -0.142 | 0.890 | chr22 | 31677580 | 31688520 | - | protein_coding |
| ENSG00000239779.7_7 | WBP1 | 0.99164936 | 1 | -0.142 | 2.319 | chr2 | 74685527 | 74688018 | + | protein_coding |
| ENSG00000198712.1_1 | MT-CO2 | 0.89448898 | 1 | -0.142 | 9.648 | chrM | 7586 | 8269 | + | protein_coding |
| ENSG00000101138.12_4 | CSTF1 | 0.99999975 | 1 | -0.142 | 5.963 | chr20 | 54967427 | 54981418 | + | protein_coding |
| ENSG00000168994.13_3 | PXDC1 | 0.99936036 | 1 | -0.142 | 3.428 | chr6 | 3722848 | 3752260 | - | protein_coding |
| ENSG00000145850.9_4 | TIMD4 | 0.90081837 | 1 | -0.142 | 1.573 | chr5 | 156346303 | 156390237 | - | protein_coding |
| ENSG00000165272.16_7 | AQP3 | 0.99843008 | 1 | -0.142 | 3.883 | chr9 | 33441154 | 33447594 | - | protein_coding |
| ENSG00000182117.5_3 | NOP10 | 0.99999975 | 1 | -0.143 | 6.441 | chr15 | 34633914 | 34635378 | - | protein_coding |
| ENSG00000012174.12_4 | MBTPS2 | 0.99999452 | 1 | -0.143 | 4.818 | chrX | 21857735 | 21903541 | + | protein_coding |
| ENSG00000261770.1_7 | AC006504.1 | 0.87134564 | 1 | -0.143 | -0.104 | chr19 | 28248092 | 28251757 | - | lncRNA |
| ENSG00000206418.4_4 | RAB12 | 0.99998729 | 1 | -0.143 | 4.582 | chr18 | 8609435 | 8639380 | + | protein_coding |
| ENSG00000164332.8_3 | UBLCP1 | 0.99931092 | 1 | -0.143 | 5.742 | chr5 | 158690298 | 158713044 | + | protein_coding |
| ENSG00000140740.11_3 | UQCRC2 | 0.99999982 | 1 | -0.143 | 7.152 | chr16 | 21964609 | 21994981 | + | protein_coding |
| ENSG00000183397.5_2 | C19orf71 | 0.93760213 | 1 | -0.143 | 1.500 | chr19 | 3539152 | 3544028 | + | protein_coding |
| ENSG00000136938.9_3 | ANP32B | 0.99935193 | 1 | -0.143 | 4.110 | chr9 | 100745623 | 100778225 | + | protein_coding |
| ENSG00000275835.5_5 | TUBGCP5 | 0.99999888 | 1 | -0.143 | 5.732 | chr15 | 22833496 | 22889876 | + | protein_coding |
| ENSG00000091164.13_5 | TXNL1 | 0.99999956 | 1 | -0.143 | 6.054 | chr18 | 54264440 | 54318831 | - | protein_coding |
| ENSG00000072954.7_4 | TMEM38A | 0.9151294 | 1 | -0.143 | 0.630 | chr19 | 16771950 | 16800834 | + | protein_coding |
| ENSG00000277879.1_6 | AL391988.1 | 0.96539788 | 1 | -0.143 | 1.313 | chr10 | 119026627 | 119028179 | - | lncRNA |
| ENSG00000259924.1_5 | AC011939.1 | 0.81027398 | 1 | -0.143 | -0.629 | chr15 | 66063514 | 66064477 | - | processed_pseudogene |
| ENSG00000225791.7_5 | TRAM2-AS1 | 0.99155617 | 1 | -0.143 | 2.439 | chr6 | 52441585 | 52507856 | + | lncRNA |
| ENSG00000145592.14_6 | RPL37 | 0.99999994 | 1 | -0.143 | 8.760 | chr5 | 40825364 | 40835324 | - | protein_coding |
| ENSG00000091732.17_5 | ZC3HC1 | 0.99997224 | 1 | -0.143 | 4.260 | chr7 | 129658127 | 129691291 | - | protein_coding |
| ENSG00000255737.2_5 | AGAP2-AS1 | 0.85416483 | 1 | -0.144 | 0.049 | chr12 | 58120054 | 58122139 | + | lncRNA |
| ENSG00000281468.1_6 | AC006504.7 | 0.9511635 | 1 | -0.144 | 1.148 | chr19 | 28293746 | 28294380 | + | lncRNA |
| ENSG00000142937.12_3 | RPS8 | 0.99999999 | 1 | -0.144 | 8.874 | chr1 | 45240923 | 45244451 | + | protein_coding |
| ENSG00000147804.9_4 | SLC39A4 | 0.99692839 | 1 | -0.144 | 2.719 | chr8 | 145635126 | 145642279 | - | protein_coding |
| ENSG00000085274.15_2 | MYNN | 0.99996856 | 1 | -0.144 | 4.348 | chr3 | 169490619 | 169507504 | + | protein_coding |
| ENSG00000069482.7_3 | GAL | 0.93038814 | 1 | -0.144 | 0.549 | chr11 | 68451247 | 68458643 | + | protein_coding |
| ENSG00000119509.13_5 | INVS | 0.99999755 | 1 | -0.144 | 5.348 | chr9 | 102861525 | 103064457 | + | protein_coding |
| ENSG00000080822.17_7 | CLDND1 | 0.99998889 | 1 | -0.144 | 4.743 | chr3 | 98216756 | 98241910 | - | protein_coding |
| ENSG00000146205.13_6 | ANO7 | 0.962799 | 1 | -0.144 | 0.929 | chr2 | 242127924 | 242164792 | + | protein_coding |
| ENSG00000138439.12_3 | FAM117B | 0.9999037 | 1 | -0.144 | 3.891 | chr2 | 203499692 | 203634480 | + | protein_coding |
| ENSG00000146109.5_3 | ABT1 | 0.99969331 | 1 | -0.144 | 4.033 | chr6 | 26597181 | 26600967 | + | protein_coding |
| ENSG00000158055.15_2 | GRHL3 | 0.97912239 | 1 | -0.144 | 1.786 | chr1 | 24645812 | 24690972 | + | protein_coding |
| ENSG00000133193.12_6 | FAM104A | 0.99999379 | 1 | -0.144 | 4.772 | chr17 | 71203492 | 71232892 | - | protein_coding |
| ENSG00000152242.11_6 | C18orf25 | 0.99999797 | 1 | -0.144 | 5.199 | chr18 | 43753519 | 43846955 | + | protein_coding |
| ENSG00000134262.13_4 | AP4B1 | 0.99882912 | 1 | -0.144 | 3.098 | chr1 | 114436816 | 114447823 | - | protein_coding |
| ENSG00000131269.19_8 | ABCB7 | 0.99999623 | 1 | -0.144 | 4.960 | chrX | 74270883 | 74376567 | - | protein_coding |
| ENSG00000152382.6_4 | TADA1 | 0.99998705 | 1 | -0.144 | 4.492 | chr1 | 166825747 | 166845501 | - | protein_coding |
| ENSG00000180329.14_6 | CCDC43 | 0.99999402 | 1 | -0.144 | 5.103 | chr17 | 42750437 | 42767147 | - | protein_coding |
| ENSG00000141424.13_6 | SLC39A6 | 0.99999987 | 1 | -0.145 | 6.428 | chr18 | 33688494 | 33709348 | - | protein_coding |
| ENSG00000125454.12_6 | SLC25A19 | 0.99845999 | 1 | -0.145 | 3.051 | chr17 | 73269062 | 73285591 | - | protein_coding |
| ENSG00000182180.14_5 | MRPS16 | 0.99999995 | 1 | -0.145 | 6.827 | chr10 | 75008601 | 75012451 | - | protein_coding |
| ENSG00000242474.3_8 | AC093627.5 | 0.99087101 | 1 | -0.145 | 2.084 | chr7 | 135853 | 149527 | - | lncRNA |
| ENSG00000134597.16_4 | RBMX2 | 0.99999254 | 1 | -0.145 | 4.761 | chrX | 129535961 | 129547630 | + | protein_coding |
| ENSG00000235138.1_6 | AL445931.1 | 0.93376494 | 1 | -0.145 | 0.966 | chr9 | 136919412 | 136923927 | + | lncRNA |

| | | | | | | | | | | |
|---|---|---|---|---|---|---|---|---|---|---|
| ENSG00000250548.7_8 | LINC01303 | 0.8170137 | 1 | -0.145 | -0.600 | chr14 | 62022991 | 62037371 | - | lncRNA |
| ENSG00000251414.1_6 | AC138956.2 | 0.81419948 | 1 | -0.145 | -0.574 | chr5 | 175781209 | 175783171 | + | lncRNA |
| ENSG00000279207.1_5 | AC015813.6 | 0.99942487 | 1 | -0.145 | 3.593 | chr17 | 56173033 | 56178840 | + | TEC |
| ENSG00000247240.8_6 | UBL7-AS1 | 0.92486049 | 1 | -0.146 | 0.768 | chr15 | 74753606 | 74805977 | + | lncRNA |
| ENSG00000165118.15_4 | C9orf64 | 0.99999763 | 1 | -0.146 | 5.082 | chr9 | 86553226 | 86571901 | - | protein_coding |
| ENSG00000065978.19_4 | YBX1 | 0.99999999 | 1 | -0.146 | 8.732 | chr1 | 43148089 | 43169476 | + | protein_coding |
| ENSG00000178409.13_3 | BEND3 | 0.99932297 | 1 | -0.146 | 3.484 | chr6 | 107386386 | 107436473 | - | protein_coding |
| ENSG00000139684.15_5 | ESD | 0.99999917 | 1 | -0.146 | 5.989 | chr13 | 47345391 | 47371555 | - | protein_coding |
| ENSG00000186205.13_5 | MARC1 | 0.9991888 | 1 | -0.146 | 3.106 | chr1 | 220960255 | 220993001 | + | protein_coding |
| ENSG00000223508.5_4 | RPL23AP53 | 0.96636742 | 1 | -0.146 | 0.961 | chr8 | 163186 | 182231 | - | transcribed_processed_pseudogene |
| ENSG00000126458.4_4 | RRAS | 0.99999468 | 1 | -0.146 | 4.919 | chr19 | 50138549 | 50143400 | - | protein_coding |
| ENSG00000171462.15_6 | DLK2 | 0.91177991 | 1 | -0.146 | 1.236 | chr6 | 43418090 | 43424370 | - | protein_coding |
| ENSG00000269737.2_6 | AL691432.1 | 0.91764747 | 1 | -0.146 | 0.711 | chr1 | 1603429 | 1605836 | + | transcribed_processed_pseudogene |
| ENSG00000133114.18_4 | GPALPP1 | 0.99998734 | 1 | -0.146 | 4.754 | chr13 | 45563664 | 45611804 | + | protein_coding |
| ENSG00000151292.17_4 | CSNK1G3 | 0.99999957 | 1 | -0.146 | 5.972 | chr5 | 122847793 | 122952739 | + | protein_coding |
| ENSG00000176171.11_3 | BNIP3 | 0.99999926 | 1 | -0.146 | 5.693 | chr10 | 133779959 | 133795517 | - | protein_coding |
| ENSG00000128463.13_4 | EMC4 | 0.99999762 | 1 | -0.146 | 5.159 | chr15 | 34517214 | 34522357 | + | protein_coding |
| ENSG00000136146.15_6 | MED4 | 0.99536608 | 1 | -0.146 | 3.408 | chr13 | 48627459 | 48669267 | - | protein_coding |
| ENSG00000150459.12_5 | SAP18 | 0.9999992 | 1 | -0.147 | 5.877 | chr13 | 21714653 | 21723223 | + | protein_coding |
| ENSG00000113575.10_5 | PPP2CA | 0.99999989 | 1 | -0.147 | 7.190 | chr5 | 133530023 | 133561764 | - | protein_coding |
| ENSG00000214016.3_4 | RPSAP61 | 0.79885186 | 1 | -0.147 | -1.083 | chrX | 44600333 | 44601210 | + | processed_pseudogene |
| ENSG00000005889.15_3 | ZFX | 0.9999999 | 1 | -0.147 | 6.466 | chrX | 24167290 | 24234372 | + | protein_coding |
| ENSG00000258317.1_8 | AC034102.6 | 0.81041421 | 1 | -0.147 | -0.656 | chr12 | 56513817 | 56523403 | - | lncRNA |
| ENSG00000109184.15_5 | DCUN1D4 | 0.99999594 | 1 | -0.147 | 5.562 | chr4 | 52709166 | 52783003 | + | protein_coding |
| ENSG00000177084.18_6 | POLE | 0.99999993 | 1 | -0.147 | 6.831 | chr12 | 133200339 | 133263962 | - | protein_coding |
| ENSG00000197122.11_3 | SRC | 0.9999997 | 1 | -0.147 | 7.690 | chr20 | 35973088 | 36034453 | + | protein_coding |
| ENSG00000100147.13_3 | CCDC134 | 0.99053254 | 1 | -0.147 | 2.202 | chr22 | 42196683 | 42222303 | + | protein_coding |
| ENSG00000198900.6_3 | TOP1 | 0.99999816 | 1 | -0.147 | 5.587 | chr20 | 39657462 | 39753127 | + | protein_coding |
| ENSG00000124243.17_6 | BCAS4 | 0.99996818 | 1 | -0.147 | 4.409 | chr20 | 49411431 | 49499213 | + | protein_coding |
| ENSG00000011426.11_7 | ANLN | 0.99999995 | 1 | -0.147 | 8.122 | chr7 | 36429430 | 36493400 | + | protein_coding |
| ENSG00000223891.6_6 | OSER1-DT | 0.97611416 | 1 | -0.147 | 1.689 | chr20 | 42839547 | 42854667 | + | lncRNA |
| ENSG00000259727.1_6 | AC103740.2 | 0.79997856 | 1 | -0.147 | -0.927 | chr15 | 63188011 | 63191742 | + | lncRNA |
| ENSG00000167670.16_3 | CHAF1A | 0.99999596 | 1 | -0.147 | 5.334 | chr19 | 4402637 | 4445015 | + | protein_coding |
| ENSG00000112996.11_4 | MRPS30 | 0.99999979 | 1 | -0.147 | 6.128 | chr5 | 44809049 | 44820530 | + | protein_coding |
| ENSG00000183648.11_5 | NDUFB1 | 0.99999656 | 1 | -0.147 | 5.575 | chr14 | 92582466 | 92588261 | - | protein_coding |
| ENSG00000256663.1_5 | AC112777.1 | 0.97582117 | 1 | -0.148 | 2.114 | chr12 | 20704524 | 20705946 | + | processed_pseudogene |
| ENSG00000230606.6 | AC159540.1 | 0.79633825 | 1 | -0.148 | -1.083 | chr2 | 98081083 | 98099107 | - | lincRNA |
| ENSG00000185963.14_4 | BICD2 | 0.99998457 | 1 | -0.148 | 6.255 | chr9 | 95473645 | 95527115 | - | protein_coding |
| ENSG00000143630.10_3 | HCN3 | 0.99663909 | 1 | -0.148 | 2.575 | chr1 | 155247254 | 155259639 | + | protein_coding |
| ENSG00000149187.18_8 | CELF1 | 0.99999998 | 1 | -0.148 | 7.239 | chr11 | 47487485 | 47587121 | - | protein_coding |
| ENSG00000232748.3_6 | AC135050.1 | 0.83849585 | 1 | -0.148 | -0.053 | chr16 | 31067781 | 31074124 | + | lncRNA |
| ENSG00000235897.1_6 | TM4SF19-AS1 | 0.80749601 | 1 | -0.148 | -0.785 | chr3 | 196045201 | 196052441 | + | lncRNA |
| ENSG00000278845.5_3 | MRPL45 | 0.99999479 | 1 | -0.148 | 5.196 | chr17 | 36452989 | 36479100 | + | protein_coding |
| ENSG00000047188.16_3 | YTHDC2 | 0.99999978 | 1 | -0.148 | 6.071 | chr5 | 112849391 | 112930982 | + | protein_coding |
| ENSG00000237523.2_5 | LINC00857 | 0.99932445 | 1 | -0.148 | 3.304 | chr10 | 81967128 | 81979413 | + | lncRNA |
| ENSG00000188130.14_5 | MAPK12 | 0.99993791 | 1 | -0.148 | 4.625 | chr22 | 50683879 | 50700145 | - | protein_coding |
| ENSG00000173153.15_5 | ESRRA | 0.99334952 | 1 | -0.148 | 4.689 | chr11 | 64072969 | 64084215 | + | protein_coding |
| ENSG00000272886.6_7 | DCP1A | 0.99997954 | 1 | -0.148 | 4.739 | chr3 | 53317449 | 53381613 | - | protein_coding |
| ENSG00000148950.11_7 | IMMP1L | 0.99419424 | 1 | -0.148 | 2.487 | chr11 | 31453948 | 31531192 | - | protein_coding |
| ENSG00000274712.1_6 | AC005332.4 | 0.82944989 | 1 | -0.148 | -0.085 | chr17 | 66201630 | 66203634 | + | lncRNA |
| ENSG00000257122.5_5 | RRN3P3 | 0.9814523 | 1 | -0.148 | 1.717 | chr16 | 22429993 | 22449036 | - | transcribed_unprocessed_pseudogene |
| ENSG00000202260.1 | RN7SKP69 | 0.79990737 | 1 | -0.148 | -0.754 | chr20 | 17556851 | 17557155 | - | misc_RNA |
| ENSG00000132341.12_3 | RAN | 0.99999999 | 1 | -0.148 | 9.129 | chr12 | 131356582 | 131362223 | + | protein_coding |
| ENSG00000168002.12_5 | POLR2G | 0.99999931 | 1 | -0.148 | 5.582 | chr11 | 62529037 | 62534182 | + | protein_coding |
| ENSG00000140650.12_4 | PMM2 | 0.9999984 | 1 | -0.148 | 5.288 | chr16 | 8882680 | 8943182 | + | protein_coding |
| ENSG00000233885.7_8 | YEATS2-AS1 | 0.8181891 | 1 | -0.148 | -0.462 | chr3 | 183524245 | 183528571 | - | lncRNA |
| ENSG00000187735.14_4 | TCEA1 | 0.99999998 | 1 | -0.149 | 7.372 | chr8 | 54879112 | 54935016 | - | protein_coding |

| | | | | | | | | | | |
|---|---|---|---|---|---|---|---|---|---|---|
| ENSG00000135775.14_4 | COG2 | 0.99999815 | 1 | -0.149 | 5.215 | chr1 | 230778227 | 230829728 | + | protein_coding |
| ENSG00000136205.17_6 | TNS3 | 0.99999998 | 1 | -0.149 | 9.687 | chr7 | 47314752 | 47622156 | - | protein_coding |
| ENSG00000002549.12_3 | LAP3 | 0.9999981 | 1 | -0.149 | 5.362 | chr4 | 17578815 | 17609595 | + | protein_coding |
| ENSG00000144567.11_7 | RETREG2 | 0.99999928 | 1 | -0.149 | 5.781 | chr2 | 220040947 | 220050197 | + | protein_coding |
| ENSG00000124702.18_3 | KLHDC3 | 0.99996823 | 1 | -0.149 | 4.533 | chr6 | 42981841 | 42989036 | + | protein_coding |
| ENSG00000272323.1_7 | AC026801.2 | 0.83570704 | 1 | -0.149 | 0.042 | chr5 | 34837654 | 34839383 | - | lncRNA |
| ENSG00000164049.14_3 | FBXW12 | 0.80140323 | 1 | -0.149 | -0.893 | chr3 | 48413709 | 48442666 | + | protein_coding |
| ENSG00000198901.14_6 | PRC1 | 0.97704063 | 1 | -0.149 | 1.779 | chr15 | 91509270 | 91538859 | - | protein_coding |
| ENSG00000117411.16_2 | B4GALT2 | 0.99999505 | 1 | -0.149 | 5.207 | chr1 | 44444615 | 44456842 | + | protein_coding |
| ENSG00000123552.17_5 | USP45 | 0.99993663 | 1 | -0.149 | 4.100 | chr6 | 99880255 | 99969604 | - | protein_coding |
| ENSG00000058673.16_6 | ZC3H11A | 0.7949576 | 1 | -0.149 | -1.122 | chr1 | 203764782 | 203824127 | + | protein_coding |
| ENSG00000124444.16_5 | ZNF576 | 0.99925928 | 1 | -0.149 | 3.111 | chr19 | 44100544 | 44105309 | + | protein_coding |
| ENSG00000085511.20_4 | MAP3K4 | 0.99999569 | 1 | -0.149 | 4.972 | chr6 | 161412759 | 161538417 | + | protein_coding |
| ENSG00000148481.14_6 | MINDY3 | 0.99998034 | 1 | -0.149 | 4.650 | chr10 | 15820169 | 15902506 | - | protein_coding |
| ENSG00000122550.18_4 | KLHL7 | 0.99998745 | 1 | -0.150 | 4.692 | chr7 | 23145377 | 23217533 | + | protein_coding |
| ENSG00000085365.18_4 | SCAMP1 | 0.99999963 | 1 | -0.150 | 6.575 | chr5 | 77656435 | 77776562 | + | protein_coding |
| ENSG00000255100.1_7 | AP003119.2 | 0.89706972 | 1 | -0.150 | 0.619 | chr11 | 76493625 | 76494106 | - | lncRNA |
| ENSG00000231043.3_2 | AC007238.1 | 0.84275594 | 1 | -0.150 | -0.091 | chr2 | 58687427 | 58689167 | - | processed_pseudogene |
| ENSG00000162063.13_6 | CCNF | 0.9999984 | 1 | -0.150 | 5.595 | chr16 | 2479395 | 2508855 | + | protein_coding |
| ENSG00000185379.20_7 | RAD51D | 0.99999121 | 1 | -0.150 | 4.625 | chr17 | 33419227 | 33448541 | - | protein_coding |
| ENSG00000175105.7_5 | ZNF654 | 0.9999623 | 1 | -0.150 | 4.263 | chr3 | 88108405 | 88193814 | + | protein_coding |
| ENSG00000102890.16_4 | ELMO3 | 0.9907595 | 1 | -0.150 | 2.208 | chr16 | 67233014 | 67237932 | + | protein_coding |
| ENSG00000203666.12_2 | EFCAB2 | 0.99464423 | 1 | -0.150 | 2.429 | chr1 | 245133007 | 245290466 | + | protein_coding |
| ENSG00000196943.14_5 | NOP9 | 0.99999433 | 1 | -0.150 | 5.235 | chr14 | 24769056 | 24778330 | + | protein_coding |
| ENSG00000167315.18_6 | ACAA2 | 0.99999819 | 1 | -0.150 | 5.369 | chr18 | 47308534 | 47340323 | - | protein_coding |
| ENSG00000186188.10_2 | FFAR4 | 0.85465493 | 1 | -0.151 | -0.152 | chr10 | 95326422 | 95364237 | + | protein_coding |
| ENSG00000160396.9_4 | HIPK4 | 0.79703034 | 1 | -0.151 | -0.860 | chr19 | 40885178 | 40896088 | - | protein_coding |
| ENSG00000172757.13_6 | CFL1 | 0.99999999 | 1 | -0.151 | 8.708 | chr11 | 65590493 | 65629497 | - | protein_coding |
| ENSG00000273568.1_7 | AC131009.3 | 0.87441057 | 1 | -0.151 | 0.572 | chr12 | 132419187 | 132419473 | + | lncRNA |
| ENSG00000228327.3_7 | AL669831.1 | 0.91186658 | 1 | -0.151 | 0.894 | chr1 | 661265 | 714006 | - | transcribed_unprocessed_pseudogene |
| ENSG00000274460.1_6 | AC092119.2 | 0.93560792 | 1 | -0.151 | 0.949 | chr16 | 21961539 | 21963029 | - | lncRNA |
| ENSG00000158864.12_3 | NDUFS2 | 0.99999963 | 1 | -0.151 | 6.034 | chr1 | 161166894 | 161184185 | + | protein_coding |
| ENSG00000122068.13_3 | FYTTD1 | 0.999999 | 1 | -0.151 | 5.731 | chr3 | 197464050 | 197514467 | + | protein_coding |
| ENSG00000153187.20_8 | HNRNPU | 0.99999999 | 1 | -0.151 | 9.058 | chr1 | 245003940 | 245027862 | - | protein_coding |
| ENSG00000137449.16_3 | CPEB2 | 0.99999847 | 1 | -0.151 | 5.799 | chr4 | 15004298 | 15071777 | + | protein_coding |
| ENSG00000277215.2_7 | SPANXA2-OT1 | 0.82839079 | 1 | -0.151 | -0.270 | chrX | 140271469 | 140738057 | + | lncRNA |
| ENSG00000108349.17_4 | CASC3 | 0.99999984 | 1 | -0.151 | 6.472 | chr17 | 38296571 | 38328424 | + | protein_coding |
| ENSG00000109063.15_4 | MYH3 | 0.974207 | 1 | -0.151 | 1.679 | chr17 | 10531843 | 10560626 | - | protein_coding |
| ENSG00000251192.7_6 | ZNF674 | 0.9941241 | 1 | -0.151 | 2.681 | chrX | 46357162 | 46404892 | - | protein_coding |
| ENSG00000172667.11_5 | ZMAT3 | 0.99996194 | 1 | -0.151 | 4.701 | chr3 | 178677909 | 178790003 | - | protein_coding |
| ENSG00000266053.2_7 | NDUFV2-AS1 | 0.87060757 | 1 | -0.151 | 0.083 | chr18 | 9121263 | 9136643 | - | lncRNA |
| ENSG00000239789.6_4 | MRPS17 | 0.98947873 | 1 | -0.151 | 2.623 | chr7 | 56019570 | 56024193 | + | protein_coding |
| ENSG00000143337.18_5 | TOR1AIP1 | 0.99999979 | 1 | -0.151 | 6.204 | chr1 | 179851177 | 179894135 | + | protein_coding |
| ENSG00000174796.12_5 | THAP6 | 0.9999363 | 1 | -0.151 | 3.987 | chr4 | 76439156 | 76475683 | + | protein_coding |
| ENSG00000131023.13_4 | LATS1 | 0.99999531 | 1 | -0.151 | 5.435 | chr6 | 149979289 | 150039241 | - | protein_coding |
| ENSG00000197943.10_4 | PLCG2 | 0.999334 | 1 | -0.151 | 3.368 | chr16 | 81772702 | 81996290 | + | protein_coding |
| ENSG00000272047.3_4 | GTF2H5 | 0.99990566 | 1 | -0.151 | 4.266 | chr6 | 158589382 | 158620376 | + | protein_coding |
| ENSG00000196502.11_6 | SULT1A1 | 0.96879544 | 1 | -0.151 | 1.659 | chr16 | 28616517 | 28634946 | - | protein_coding |
| ENSG00000105197.11_5 | TIMM50 | 0.99999974 | 1 | -0.151 | 6.956 | chr19 | 39971052 | 39984425 | + | protein_coding |
| ENSG00000265806.1 | MIR4292 | 0.8993406 | 1 | -0.151 | 0.317 | chr9 | 139725409 | 139725475 | + | miRNA |
| ENSG00000116205.14_5 | TCEANC2 | 0.9988761 | 1 | -0.151 | 3.107 | chr1 | 54519257 | 54578192 | + | protein_coding |
| ENSG00000092098.17_5 | RNF31 | 0.84887826 | 1 | -0.151 | 0.155 | chr14 | 24615892 | 24629869 | + | protein_coding |
| ENSG00000023516.9_3 | AKAP11 | 0.99999934 | 1 | -0.152 | 6.134 | chr13 | 42846288 | 42897397 | + | protein_coding |
| ENSG00000103942.13_6 | HOMER2 | 0.99996719 | 1 | -0.152 | 4.257 | chr15 | 83505698 | 83654905 | - | protein_coding |
| ENSG00000114744.9_4 | COMMD2 | 0.99999795 | 1 | -0.152 | 5.406 | chr3 | 149456259 | 149470282 | - | protein_coding |
| ENSG00000242553.1_7 | AP001432.1 | 0.81259026 | 1 | -0.152 | -0.575 | chr21 | 38593720 | 38610045 | + | lncRNA |
| ENSG00000197265.9_6 | GTF2E2 | 0.99999894 | 1 | -0.152 | 5.464 | chr8 | 30435835 | 30515753 | - | protein_coding |
| ENSG00000227354.7_6 | RBM26-AS1 | 0.88041866 | 1 | -0.152 | 0.171 | chr13 | 79980428 | 80001452 | + | lncRNA |

| | | | | | | | | | | |
|---|---|---|---|---|---|---|---|---|---|---|
| ENSG00000157326.19_3 | DHRS4 | 0.99758428 | 1 | -0.152 | 2.790 | chr14 | 24422943 | 24438488 | + | protein_coding |
| ENSG00000156170.13_4 | NDUFAF6 | 0.99991424 | 1 | -0.152 | 4.033 | chr8 | 95907995 | 96128683 | + | protein_coding |
| ENSG00000165119.21_6 | HNRNPK | 0.99999997 | 1 | -0.152 | 8.990 | chr9 | 86582998 | 86595531 | - | protein_coding |
| ENSG00000163251.4_4 | FZD5 | 0.99999599 | 1 | -0.152 | 5.283 | chr2 | 208627322 | 208634630 | - | protein_coding |
| ENSG00000182985.17_8 | CADM1 | 0.99999799 | 1 | -0.152 | 5.792 | chr11 | 115039938 | 115375675 | - | protein_coding |
| ENSG00000259767.1_8 | AC022558.1 | 0.79242454 | 1 | -0.152 | -0.788 | chr15 | 83690988 | 83693088 | - | lncRNA |
| ENSG00000167513.9_5 | CDT1 | 0.99986868 | 1 | -0.152 | 5.133 | chr16 | 88870197 | 88875666 | + | protein_coding |
| ENSG00000134874.17_3 | DZIP1 | 0.99999427 | 1 | -0.152 | 5.068 | chr13 | 96230456 | 96296957 | - | protein_coding |
| ENSG00000041880.14_4 | PARP3 | 0.99943212 | 1 | -0.152 | 3.475 | chr3 | 51976361 | 51982883 | + | protein_coding |
| ENSG00000152219.5_3 | ARL14EP | 0.9999859 | 1 | -0.153 | 4.633 | chr11 | 30344651 | 30359770 | + | protein_coding |
| ENSG00000175449.14_5 | RFESD | 0.86568541 | 1 | -0.153 | 0.302 | chr5 | 94982458 | 95020477 | + | protein_coding |
| ENSG00000109861.16_4 | CTSC | 0.99999987 | 1 | -0.153 | 6.230 | chr11 | 88026760 | 88070929 | - | protein_coding |
| ENSG00000261527.1_6 | AC026464.5 | 0.79302291 | 1 | -0.153 | -0.895 | chr16 | 69477143 | 69478193 | - | processed_pseudogene |
| ENSG00000183741.12_4 | CBX6 | 0.99998969 | 1 | -0.153 | 6.625 | chr22 | 39257427 | 39268254 | - | protein_coding |
| ENSG00000158023.10_4 | WDR66 | 0.95799886 | 1 | -0.153 | 1.272 | chr12 | 122356498 | 122441833 | + | protein_coding |
| ENSG00000261351.2_8 | AC116913.1 | 0.87636258 | 1 | -0.153 | 0.455 | chr15 | 66780996 | 66784447 | - | lncRNA |
| ENSG00000107929.15_3 | LARP4B | 0.99999993 | 1 | -0.153 | 6.472 | chr10 | 852854 | 977645 | - | protein_coding |
| ENSG00000272645.3_6 | GTF2IP20 | 0.99606479 | 1 | -0.153 | 2.655 | chr1 | 224139096 | 224180296 | - | transcribed_unprocessed_pseudogene |
| ENSG00000160563.14_5 | MED27 | 0.99960156 | 1 | -0.153 | 3.874 | chr9 | 134728315 | 134955254 | - | protein_coding |
| ENSG00000197563.11_6 | PIGN | 0.99998697 | 1 | -0.153 | 5.127 | chr18 | 59572488 | 59854351 | - | protein_coding |
| ENSG00000085788.14_8 | DDHD2 | 0.99999719 | 1 | -0.153 | 6.055 | chr8 | 38082736 | 38133076 | + | protein_coding |
| ENSG00000146676.10_6 | PURB | 0.99999548 | 1 | -0.153 | 5.910 | chr7 | 44915898 | 44925129 | - | protein_coding |
| ENSG00000164329.13_4 | TENT2 | 0.99999689 | 1 | -0.153 | 5.831 | chr5 | 78907943 | 78982471 | + | protein_coding |
| ENSG00000261868.1_4 | MFSD1P1 | 0.82517428 | 1 | -0.153 | -0.313 | chr17 | 4204667 | 4205993 | + | processed_pseudogene |
| ENSG00000217624.2_5 | AL627402.1 | 0.80886846 | 1 | -0.153 | 0.055 | chrX | 41535013 | 41535747 | - | processed_pseudogene |
| ENSG00000227775.3_5 | AL031282.1 | 0.78851135 | 1 | -0.153 | -1.123 | chr1 | 1655951 | 1668690 | + | transcribed_processed_pseudogene |
| ENSG00000148154.10_3 | UGCG | 0.99998744 | 1 | -0.154 | 5.326 | chr9 | 114659094 | 114697649 | + | protein_coding |
| ENSG00000110768.12_3 | GTF2H1 | 0.99999729 | 1 | -0.154 | 6.083 | chr11 | 18343842 | 18388592 | + | protein_coding |
| ENSG00000106263.18_3 | EIF3B | 0.99999996 | 1 | -0.154 | 8.251 | chr7 | 2393721 | 2420380 | + | protein_coding |
| ENSG00000172889.16_5 | EGFL7 | 0.99992548 | 1 | -0.154 | 5.033 | chr9 | 139553308 | 139567130 | + | protein_coding |
| ENSG00000165689.17_7 | ENTR1 | 0.99999963 | 1 | -0.154 | 5.879 | chr9 | 139296374 | 139305066 | - | protein_coding |
| ENSG00000085491.17_5 | SLC25A24 | 0.99999885 | 1 | -0.154 | 6.099 | chr1 | 108676665 | 108743471 | - | protein_coding |
| ENSG00000130119.16_3 | GNL3L | 0.99999993 | 1 | -0.154 | 7.069 | chrX | 54556644 | 54593722 | + | protein_coding |
| ENSG00000111906.18_4 | HDDC2 | 0.99999863 | 1 | -0.154 | 5.559 | chr6 | 125541108 | 125623224 | - | protein_coding |
| ENSG00000287271.1_1 | AC011455.6 | 0.85660058 | 1 | -0.154 | 0.136 | chr19 | 39465801 | 39505167 | + | lncRNA |
| ENSG00000265205.1_9 | AC010761.3 | 0.82782595 | 1 | -0.154 | 0.013 | chr17 | 26997072 | 26999822 | - | lncRNA |
| ENSG00000130560.9_4 | UBAC1 | 0.99999589 | 1 | -0.154 | 5.089 | chr9 | 138824815 | 138853219 | - | protein_coding |
| ENSG00000015676.18_4 | NUDCD3 | 0.99999974 | 1 | -0.154 | 6.095 | chr7 | 44418718 | 44530257 | - | protein_coding |
| ENSG00000047849.21_8 | MAP4 | 0.99999998 | 1 | -0.154 | 7.598 | chr3 | 47892180 | 48130762 | - | protein_coding |
| ENSG00000288065.1_1 | AC007881.4 | 0.84261875 | 1 | -0.154 | 0.100 | chr2 | 71340007 | 71344536 | + | lncRNA |
| ENSG00000235899.2_6 | LINC01564 | 0.8553196 | 1 | -0.154 | 0.237 | chr6 | 53481269 | 53496198 | + | lncRNA |
| ENSG00000162385.11_4 | MAGOH | 0.99762884 | 1 | -0.154 | 3.240 | chr1 | 53692572 | 53704190 | - | protein_coding |
| ENSG00000141560.15_3 | FN3KRP | 0.99999891 | 1 | -0.155 | 5.630 | chr17 | 80674582 | 80688204 | + | protein_coding |
| ENSG00000283341.2_6 | AC068205.2 | 0.97560845 | 1 | -0.155 | 1.645 | chr11 | 43600439 | 43861580 | + | lncRNA |
| ENSG00000213997.3_4 | PGAM1P7 | 0.81576244 | 1 | -0.155 | -0.240 | chrX | 46506029 | 46506765 | - | processed_pseudogene |
| ENSG00000183049.12_4 | CAMK1D | 0.99994141 | 1 | -0.155 | 4.077 | chr10 | 12391481 | 12877545 | + | protein_coding |
| ENSG00000198841.4_4 | KTI12 | 0.98777343 | 1 | -0.155 | 2.111 | chr1 | 52497775 | 52499482 | - | protein_coding |
| ENSG00000196678.14_6 | ERI2 | 0.9998091 | 1 | -0.155 | 4.244 | chr16 | 20791515 | 20911671 | - | protein_coding |
| ENSG00000133731.10_4 | IMPA1 | 0.99994275 | 1 | -0.155 | 5.114 | chr8 | 82569149 | 82598566 | - | protein_coding |
| ENSG00000183853.18_7 | KIRREL1 | 0.99999986 | 1 | -0.155 | 6.335 | chr1 | 157963063 | 158070052 | + | protein_coding |
| ENSG00000119004.16_4 | CYP20A1 | 0.99999817 | 1 | -0.155 | 5.257 | chr2 | 204103700 | 204170749 | + | protein_coding |
| ENSG00000165264.6 | NDUFB6 | 0.99998783 | 1 | -0.155 | 5.461 | chr9 | 32552997 | 32573160 | - | protein_coding |
| ENSG00000108309.14_4 | RUNDC3A | 0.82879355 | 1 | -0.155 | -0.112 | chr17 | 42385781 | 42396038 | + | protein_coding |
| ENSG00000283050.2_7 | GTF2IP12 | 0.96222431 | 1 | -0.155 | 1.501 | chr4 | 120326678 | 120375793 | - | transcribed_unprocessed_pseudogene |
| ENSG00000115486.12_4 | GGCX | 0.99999971 | 1 | -0.155 | 5.932 | chr2 | 85771843 | 85788670 | - | protein_coding |
| ENSG00000141367.12_5 | CLTC | 0.99999998 | 1 | -0.155 | 9.955 | chr17 | 57697050 | 57774317 | + | protein_coding |
| ENSG00000106636.8_4 | YKT6 | 0.99999861 | 1 | -0.155 | 6.292 | chr7 | 44240567 | 44253893 | + | protein_coding |

| | | | | | | | | | | |
|---|---|---|---|---|---|---|---|---|---|---|
| ENSG00000171522.6_3 | PTGER4 | 0.97056771 | 1 | -0.156 | 1.514 | chr5 | 40680017 | 40693837 | + | protein_coding |
| ENSG00000284602.1_4 | AL031432.4 | 0.896086 | 1 | -0.156 | 0.517 | chr1 | 25559077 | 25561266 | + | lncRNA |
| ENSG00000243317.8_4 | STMP1 | 0.99999199 | 1 | -0.156 | 4.838 | chr7 | 135347244 | 135378166 | + | protein_coding |
| ENSG00000139133.7_3 | ALG10 | 0.99976134 | 1 | -0.156 | 3.542 | chr12 | 34175403 | 34182629 | + | protein_coding |
| ENSG00000259321.1_6 | AL136295.2 | 0.8011343 | 1 | -0.156 | -0.696 | chr14 | 24608654 | 24609653 | + | lncRNA |
| ENSG00000213713.3_4 | PIGCP1 | 0.9948638 | 1 | -0.156 | 2.602 | chr11 | 33097112 | 33098006 | - | processed_pseudogene |
| ENSG00000174444.15_5 | RPL4 | 0.99999998 | 1 | -0.156 | 10.955 | chr15 | 66790353 | 66816870 | - | protein_coding |
| ENSG00000119684.15_4 | MLH3 | 0.99999947 | 1 | -0.156 | 5.982 | chr14 | 75480467 | 75518235 | - | protein_coding |
| ENSG00000157017.16_5 | GHRL | 0.80930524 | 1 | -0.156 | -0.463 | chr3 | 10327350 | 10334631 | - | protein_coding |
| ENSG00000066136.20_3 | NFYC | 0.99997891 | 1 | -0.156 | 4.549 | chr1 | 41157320 | 41237275 | + | protein_coding |
| ENSG00000115350.12_3 | POLE4 | 0.99998755 | 1 | -0.156 | 4.709 | chr2 | 75185770 | 75197255 | + | protein_coding |
| ENSG00000157315.5_5 | TMED6 | 0.79841891 | 1 | -0.156 | -0.487 | chr16 | 69377153 | 69385689 | - | protein_coding |
| ENSG00000120265.17_6 | PCMT1 | 0.99999127 | 1 | -0.156 | 5.131 | chr6 | 150070579 | 150132556 | + | protein_coding |
| ENSG00000137802.14_5 | MAPKBP1 | 0.99987896 | 1 | -0.156 | 3.978 | chr15 | 42066632 | 42120053 | + | protein_coding |
| ENSG00000105401.9_7 | CDC37 | 0.99999985 | 1 | -0.157 | 6.659 | chr19 | 10501809 | 10530797 | - | protein_coding |
| ENSG00000070770.9_5 | CSNK2A2 | 0.9999989 | 1 | -0.157 | 5.455 | chr16 | 58191811 | 58232010 | - | protein_coding |
| ENSG00000148308.17_4 | GTF3C5 | 0.99999916 | 1 | -0.157 | 6.086 | chr9 | 135906062 | 135933890 | + | protein_coding |
| ENSG00000086717.18_5 | PPEF1 | 0.97622689 | 1 | -0.157 | 1.636 | chrX | 18694029 | 18846039 | + | protein_coding |
| ENSG00000104691.15_7 | UBXN8 | 0.99990329 | 1 | -0.157 | 4.122 | chr8 | 30589764 | 30624522 | + | protein_coding |
| ENSG00000065308.5_4 | TRAM2 | 0.9999996 | 1 | -0.157 | 5.776 | chr6 | 52362206 | 52441858 | - | protein_coding |
| ENSG00000120647.10_9 | CCDC77 | 0.99996954 | 1 | -0.157 | 4.404 | chr12 | 498439 | 551808 | + | protein_coding |
| ENSG00000172339.10_4 | ALG14 | 0.99927919 | 1 | -0.157 | 3.264 | chr1 | 95439961 | 95538507 | - | protein_coding |
| ENSG00000117859.19_8 | OSBPL9 | 0.99999994 | 1 | -0.157 | 6.651 | chr1 | 52042851 | 52264099 | + | protein_coding |
| ENSG00000055483.19_8 | AC022966.1 | 0.99999933 | 1 | -0.157 | 5.760 | chr17 | 76783463 | 76837523 | - | protein_coding |
| ENSG00000168779.19_7 | SHOX2 | 0.96275598 | 1 | -0.157 | 1.372 | chr3 | 157813743 | 157824292 | - | protein_coding |
| ENSG00000113657.13_7 | DPYSL3 | 0.99999428 | 1 | -0.157 | 5.064 | chr5 | 146770371 | 146889631 | - | protein_coding |
| ENSG00000277791.5_3 | PSMB3 | 0.99999993 | 1 | -0.157 | 6.987 | chr17 | 36908994 | 36920478 | + | protein_coding |
| ENSG00000166263.13_5 | STXBP4 | 0.99997851 | 1 | -0.158 | 5.358 | chr17 | 53046088 | 53250993 | + | protein_coding |
| ENSG00000138757.14_3 | G3BP2 | 0.99999917 | 1 | -0.158 | 7.455 | chr4 | 76567966 | 76649709 | - | protein_coding |
| ENSG00000236772.1_6 | AL034550.1 | 0.84939507 | 1 | -0.158 | 0.256 | chr20 | 31037558 | 31041410 | + | lncRNA |
| ENSG00000134248.13_4 | LAMTOR5 | 0.99994238 | 1 | -0.158 | 4.139 | chr1 | 110943871 | 110950564 | - | protein_coding |
| ENSG00000241685.10_4 | ARPC1A | 0.99989432 | 1 | -0.158 | 3.961 | chr7 | 98923521 | 98963885 | + | protein_coding |
| ENSG00000196562.14_7 | SULF2 | 0.99999088 | 1 | -0.158 | 4.836 | chr20 | 46285092 | 46415360 | - | protein_coding |
| ENSG00000086015.22_4 | MAST2 | 0.99999809 | 1 | -0.158 | 5.550 | chr1 | 46252659 | 46501794 | + | protein_coding |
| ENSG00000134056.12_3 | MRPS36 | 0.99998964 | 1 | -0.158 | 4.631 | chr5 | 68513587 | 68525985 | + | protein_coding |
| ENSG00000274386.5_5 | TMEM269 | 0.80216121 | 1 | -0.158 | -0.497 | chr1 | 43250678 | 43282290 | + | protein_coding |
| ENSG00000165646.14_6 | SLC18A2 | 0.84865717 | 1 | -0.158 | 0.075 | chr10 | 119000604 | 119038941 | + | protein_coding |
| ENSG00000074696.13_4 | HACD3 | 0.99999997 | 1 | -0.158 | 8.377 | chr15 | 65822756 | 65870687 | + | protein_coding |
| ENSG00000141456.16_7 | PELP1 | 0.99999975 | 1 | -0.158 | 6.381 | chr17 | 4573069 | 4607632 | - | protein_coding |
| ENSG00000103404.14_5 | USP31 | 0.99999874 | 1 | -0.158 | 5.335 | chr16 | 23072727 | 23160591 | - | protein_coding |
| ENSG00000233230.1_6 | AC079807.1 | 0.84395563 | 1 | -0.158 | -0.056 | chr2 | 48132817 | 48134949 | + | lncRNA |
| ENSG00000228395.2_9 | AL356481.1 | 0.86759159 | 1 | -0.158 | 0.423 | chr9 | 131290936 | 131314689 | - | lncRNA |
| ENSG00000172409.6_4 | CLP1 | 0.99989206 | 1 | -0.158 | 4.271 | chr11 | 57416465 | 57429337 | + | protein_coding |
| ENSG00000155275.19_5 | TRMT44 | 0.99859894 | 1 | -0.158 | 3.062 | chr4 | 8437867 | 8495258 | + | protein_coding |
| ENSG00000170584.11_4 | NUDCD2 | 0.99999837 | 1 | -0.158 | 5.434 | chr5 | 162873532 | 162887108 | - | protein_coding |
| ENSG00000235513.2_7 | L3MBTL2-AS1 | 0.8058942 | 1 | -0.158 | -0.522 | chr22 | 41603596 | 41624504 | - | lncRNA |
| ENSG00000151466.12_7 | SCLT1 | 0.99984734 | 1 | -0.159 | 3.928 | chr4 | 129786076 | 130014755 | - | protein_coding |
| ENSG00000113368.12_3 | LMNB1 | 0.99999992 | 1 | -0.159 | 6.786 | chr5 | 126112315 | 126172712 | + | protein_coding |
| ENSG00000215041.10_5 | NEURL4 | 0.99999125 | 1 | -0.159 | 5.255 | chr17 | 7218947 | 7232712 | - | protein_coding |
| ENSG00000244218.2 | RN7SL81P | 0.78724473 | 1 | -0.159 | -0.895 | chr7 | 128401391 | 128401625 | + | misc_RNA |
| ENSG00000198625.13_5 | MDM4 | 0.99999912 | 1 | -0.159 | 5.903 | chr1 | 204485507 | 204527248 | + | protein_coding |
| ENSG00000270647.6_4 | TAF15 | 0.99999299 | 1 | -0.159 | 4.927 | chr17 | 34040810 | 34191619 | + | protein_coding |
| ENSG00000175745.14_5 | NR2F1 | 0.98998041 | 1 | -0.159 | 2.070 | chr5 | 92918928 | 92930317 | + | protein_coding |
| ENSG00000108106.14_4 | UBE2S | 0.99083397 | 1 | -0.159 | 6.728 | chr19 | 55911113 | 55919156 | - | protein_coding |
| ENSG00000108953.17_4 | YWHAE | 0.99999755 | 1 | -0.159 | 8.843 | chr17 | 1247569 | 1303516 | - | protein_coding |
| ENSG00000204104.12_3 | TRAF3IP1 | 0.99994962 | 1 | -0.159 | 4.419 | chr2 | 239229082 | 239309538 | + | protein_coding |
| ENSG00000115685.15_4 | PPP1R7 | 0.99998839 | 1 | -0.159 | 4.844 | chr2 | 242088991 | 242123067 | + | protein_coding |
| ENSG00000085224.22_6 | ATRX | 0.99999988 | 1 | -0.159 | 7.342 | chrX | 76760356 | 77041719 | - | protein_coding |
| ENSG00000143633.13_6 | C1orf131 | 0.99995697 | 1 | -0.160 | 4.242 | chr1 | 231359509 | 231376933 | - | protein_coding |

| | | | | | | | | | | |
|---|---|---|---|---|---|---|---|---|---|---|
| ENSG00000167904.15_7 | TMEM68 | 0.99988268 | 1 | -0.160 | 3.984 | chr8 | 56608983 | 56685966 | - | protein_coding |
| ENSG00000168476.12_5 | REEP4 | 0.99998816 | 1 | -0.160 | 4.845 | chr8 | 21995533 | 21999464 | - | protein_coding |
| ENSG00000166595.12_7 | CIAO2B | 0.99999106 | 1 | -0.160 | 5.030 | chr16 | 66965968 | 66968326 | - | protein_coding |
| ENSG00000136048.14_4 | DRAM1 | 0.99999114 | 1 | -0.160 | 4.856 | chr12 | 102271358 | 102405908 | + | protein_coding |
| ENSG00000107938.18_4 | EDRF1 | 0.99999741 | 1 | -0.160 | 5.188 | chr10 | 127408084 | 127452712 | + | protein_coding |
| ENSG00000225171.2_4 | DUTP6 | 0.78117039 | 1 | -0.160 | -0.940 | chr1 | 166837985 | 166838446 | + | processed_pseudogene |
| ENSG00000114416.18_8 | FXR1 | 0.99999977 | 1 | -0.160 | 6.704 | chr3 | 180585929 | 180700541 | + | protein_coding |
| ENSG00000172175.15_7 | MALT1 | 0.99999941 | 1 | -0.160 | 6.292 | chr18 | 56338697 | 56421709 | + | protein_coding |
| ENSG00000131473.17_7 | ACLY | 0.99999996 | 1 | -0.160 | 9.775 | chr17 | 40023170 | 40086795 | - | protein_coding |
| ENSG00000254858.10_3 | MPV17L2 | 0.99674392 | 1 | -0.160 | 2.870 | chr19 | 18304028 | 18307758 | + | protein_coding |
| ENSG00000136011.15_5 | STAB2 | 0.77921887 | 1 | -0.160 | -1.086 | chr12 | 103981051 | 104160497 | + | protein_coding |
| ENSG00000101442.10_3 | ACTR5 | 0.9997797 | 1 | -0.160 | 3.915 | chr20 | 37377103 | 37401163 | + | protein_coding |
| ENSG00000168769.13_5 | TET2 | 0.9999882 | 1 | -0.160 | 5.000 | chr4 | 106067032 | 106200973 | + | protein_coding |
| ENSG00000039523.20_7 | RIPOR1 | 0.9994212 | 1 | -0.160 | 3.524 | chr16 | 67552321 | 67580691 | + | protein_coding |
| ENSG00000169018.6_3 | FEM1B | 0.99999971 | 1 | -0.160 | 6.927 | chr15 | 68570083 | 68588200 | + | protein_coding |
| ENSG00000282164.3_6 | PEG13 | 0.78015456 | 1 | -0.160 | -1.085 | chr8 | 141104993 | 141110642 | - | lncRNA |
| ENSG00000215481.9_5 | BCRP3 | 0.78083094 | 1 | -0.161 | -0.828 | chr22 | 25028882 | 25045979 | + | lncRNA |
| ENSG00000129235.11_3 | TXNDC17 | 0.99999759 | 1 | -0.161 | 5.641 | chr17 | 6544305 | 6547861 | + | protein_coding |
| ENSG00000198727.2_1 | MT-CYB | 0.99999999 | 1 | -0.161 | 9.313 | chrM | 14747 | 15887 | + | protein_coding |
| ENSG00000179148.9_3 | ALOXE3 | 0.78519799 | 1 | -0.161 | -0.895 | chr17 | 7999218 | 8022365 | - | protein_coding |
| ENSG00000110075.14_5 | PPP6R3 | 0.99999996 | 1 | -0.161 | 7.918 | chr11 | 68228199 | 68382802 | + | protein_coding |
| ENSG00000247796.3_6 | AC008966.1 | 0.98625381 | 1 | -0.161 | 2.008 | chr5 | 52405646 | 52423503 | + | lncRNA |
| ENSG00000249850.1_4 | KRT18P31 | 0.81407498 | 1 | -0.161 | -0.416 | chr5 | 36885308 | 36886599 | + | processed_pseudogene |
| ENSG00000274403.1_6 | AC090510.2 | 0.79156733 | 1 | -0.161 | -0.698 | chr15 | 43018781 | 43019409 | + | lncRNA |
| ENSG00000213024.11_6 | NUP62 | 0.99999963 | 1 | -0.161 | 6.412 | chr19 | 50410082 | 50433020 | - | protein_coding |
| ENSG00000178904.19_4 | DPY19L3 | 0.9999994 | 1 | -0.161 | 5.774 | chr19 | 32896449 | 32976801 | + | protein_coding |
| ENSG00000027869.11_4 | SH2D2A | 0.79686457 | 1 | -0.161 | -0.393 | chr1 | 156776035 | 156786654 | - | protein_coding |
| ENSG00000285278.1_5 | AL138885.3 | 0.94415558 | 1 | -0.162 | 1.368 | chr6 | 10404735 | 10408161 | + | lncRNA |
| ENSG00000110367.13_5 | DDX6 | 0.99999994 | 1 | -0.162 | 7.434 | chr11 | 118618472 | 118661873 | - | protein_coding |
| ENSG00000182359.15_5 | KBTBD3 | 0.97253551 | 1 | -0.162 | 1.643 | chr11 | 105921825 | 105948186 | - | protein_coding |
| ENSG00000064042.18_4 | LIMCH1 | 0.99999993 | 1 | -0.162 | 7.619 | chr4 | 41361624 | 41702061 | + | protein_coding |
| ENSG00000082781.12_5 | ITGB5 | 0.99999977 | 1 | -0.162 | 8.688 | chr3 | 124480795 | 124620265 | - | protein_coding |
| ENSG00000178464.6_4 | RPL10P16 | 0.98884837 | 1 | -0.162 | 2.171 | chr19 | 12754089 | 12754733 | - | processed_pseudogene |
| ENSG00000140612.14_4 | SEC11A | 0.9999999 | 1 | -0.162 | 7.507 | chr15 | 85212769 | 85259691 | - | protein_coding |
| ENSG00000267394.1_6 | AC004596.1 | 0.80682128 | 1 | -0.162 | -0.368 | chr17 | 42276250 | 42293933 | + | lncRNA |
| ENSG00000186889.10_5 | TMEM17 | 0.99738268 | 1 | -0.162 | 2.775 | chr2 | 62727353 | 62739029 | - | protein_coding |
| ENSG00000253948.2_9 | VPS13B-DT | 0.77965445 | 1 | -0.162 | -0.828 | chr8 | 99970505 | 100025971 | - | lncRNA |
| ENSG00000143811.19_5 | PYCR2 | 0.99995047 | 1 | -0.162 | 4.359 | chr1 | 226107577 | 226112040 | - | protein_coding |
| ENSG00000223482.8_9 | NUTM2A-AS1 | 0.99990631 | 1 | -0.162 | 4.458 | chr10 | 88961404 | 89102369 | - | lncRNA |
| ENSG00000159579.14_4 | RSPRY1 | 0.99999943 | 1 | -0.163 | 6.021 | chr16 | 57220049 | 57274381 | + | protein_coding |
| ENSG00000126698.11_4 | DNAJC8 | 0.99999835 | 1 | -0.163 | 5.878 | chr1 | 28525967 | 28559540 | - | protein_coding |
| ENSG00000188266.14_6 | HYKK | 0.99923507 | 1 | -0.163 | 3.265 | chr15 | 78799906 | 78829714 | + | protein_coding |
| ENSG00000172828.13_5 | CES3 | 0.99741323 | 1 | -0.163 | 2.931 | chr16 | 66995148 | 67009052 | + | protein_coding |
| ENSG00000178623.12_6 | GPR35 | 0.78552784 | 1 | -0.163 | -0.698 | chr2 | 241544847 | 241570676 | + | protein_coding |
| ENSG00000167977.9_4 | KCTD5 | 0.99994918 | 1 | -0.163 | 4.598 | chr16 | 2732524 | 2759031 | + | protein_coding |
| ENSG00000158161.16_6 | EYA3 | 0.99998013 | 1 | -0.163 | 4.510 | chr1 | 28296855 | 28415148 | - | protein_coding |
| ENSG00000224577.3_6 | LINC01117 | 0.78836683 | 1 | -0.163 | -0.575 | chr2 | 177359983 | 177683908 | + | lncRNA |
| ENSG00000214826.5_5 | DDX12P | 0.99977514 | 1 | -0.163 | 4.046 | chr12 | 9571269 | 9600825 | - | unprocessed_pseudogene |
| ENSG00000165672.7_4 | PRDX3 | 0.99999997 | 1 | -0.163 | 8.088 | chr10 | 120927232 | 120938324 | - | protein_coding |
| ENSG00000214756.8_4 | CSKMT | 0.99212767 | 1 | -0.163 | 2.373 | chr11 | 62432781 | 62435968 | + | protein_coding |
| ENSG00000171421.13_6 | MRPL36 | 0.99995676 | 1 | -0.163 | 4.638 | chr5 | 1798499 | 1801480 | - | protein_coding |
| ENSG00000229953.1_7 | AL590666.2 | 0.90227691 | 1 | -0.163 | 0.658 | chr1 | 156616299 | 156631216 | - | lncRNA |
| ENSG00000115255.11_6 | REEP6 | 0.98378409 | 1 | -0.163 | 2.127 | chr19 | 1491165 | 1497926 | + | protein_coding |
| ENSG00000070785.17_7 | EIF2B3 | 0.99993468 | 1 | -0.164 | 4.365 | chr1 | 45316194 | 45452394 | - | protein_coding |
| ENSG00000120071.14_6 | KANSL1 | 0.99998143 | 1 | -0.164 | 4.666 | chr17 | 44107282 | 44302755 | - | protein_coding |
| ENSG00000164181.14_5 | ELOVL7 | 0.99999129 | 1 | -0.164 | 4.907 | chr5 | 60047618 | 60140101 | - | protein_coding |
| ENSG00000273131.1 | RP6-42F4.1 | 0.95982292 | 1 | -0.164 | 1.088 | chr1 | 120693320 | 120697146 | - | processed_transcript |

| | | | | | | | | | | |
|---|---|---|---|---|---|---|---|---|---|---|
| ENSG00000127946.17_5 | HIP1 | 0.99999996 | 1 | -0.164 | 7.218 | chr7 | 75162619 | 75368280 | - | protein_coding |
| ENSG00000272145.1_6 | NFYC-AS1 | 0.91475679 | 1 | -0.164 | 0.872 | chr1 | 41156052 | 41157738 | - | lncRNA |
| ENSG00000164916.11_4 | FOXK1 | 0.99999855 | 1 | -0.164 | 6.467 | chr7 | 4721926 | 4811073 | + | protein_coding |
| ENSG00000092096.16_5 | SLC22A17 | 0.98068774 | 1 | -0.164 | 2.386 | chr14 | 23815515 | 23822121 | - | protein_coding |
| ENSG00000160072.20_3 | ATAD3B | 0.83788739 | 1 | -0.164 | 0.074 | chr1 | 1407145 | 1433228 | + | protein_coding |
| ENSG00000121542.12_5 | SEC22A | 0.99964089 | 1 | -0.164 | 3.611 | chr3 | 122920774 | 122992983 | + | protein_coding |
| ENSG00000145734.19_6 | BDP1 | 0.99999954 | 1 | -0.165 | 6.937 | chr5 | 70751478 | 70863647 | + | protein_coding |
| ENSG00000146701.12_3 | MDH2 | 0.99999984 | 1 | -0.165 | 7.190 | chr7 | 75677369 | 75696826 | + | protein_coding |
| ENSG00000170445.15_8 | HARS1 | 0.99999944 | 1 | -0.165 | 5.940 | chr5 | 140053489 | 140071122 | - | protein_coding |
| ENSG00000105538.10_5 | RASIP1 | 0.81082341 | 1 | -0.165 | -0.270 | chr19 | 49223842 | 49243867 | - | protein_coding |
| ENSG00000120253.14_4 | NUP43 | 0.99999972 | 1 | -0.165 | 6.189 | chr6 | 150045451 | 150070801 | - | protein_coding |
| ENSG00000261578.1_6 | AP003119.3 | 0.99591132 | 1 | -0.165 | 3.706 | chr11 | 76511408 | 76515599 | + | lncRNA |
| ENSG00000168028.14_4 | RPSA | 0.99999988 | 1 | -0.165 | 9.203 | chr3 | 39448207 | 39454033 | + | protein_coding |
| ENSG00000014138.9_5 | POLA2 | 0.99419757 | 1 | -0.165 | 2.543 | chr11 | 65029391 | 65073060 | + | protein_coding |
| ENSG00000164347.18_6 | GFM2 | 0.99999738 | 1 | -0.165 | 5.435 | chr5 | 74017031 | 74062972 | - | protein_coding |
| ENSG00000170315.14_6 | UBB | 0.99999983 | 1 | -0.165 | 7.145 | chr17 | 16284112 | 16286059 | + | protein_coding |
| ENSG00000254373.1_5 | AC112191.2 | 0.77507588 | 1 | -0.165 | -0.828 | chr5 | 159685260 | 159685592 | + | processed_pseudogene |
| ENSG00000258947.7_7 | TUBB3 | 0.98244966 | 1 | -0.165 | 1.972 | chr16 | 89987800 | 90005169 | + | protein_coding |
| ENSG00000178695.6_3 | KCTD12 | 0.98644869 | 1 | -0.165 | 2.192 | chr13 | 77454310 | 77460540 | - | protein_coding |
| ENSG00000054116.12_4 | TRAPPC3 | 0.99999714 | 1 | -0.165 | 5.201 | chr1 | 36602171 | 36621654 | - | protein_coding |
| ENSG00000181610.13_4 | MRPS23 | 0.99999974 | 1 | -0.165 | 6.404 | chr17 | 55912142 | 55927417 | - | protein_coding |
| ENSG00000145391.13_4 | SETD7 | 0.99999991 | 1 | -0.165 | 7.382 | chr4 | 140417095 | 140527853 | - | protein_coding |
| ENSG00000230330.1_5 | HMGN2P3 | 0.81508803 | 1 | -0.165 | -0.063 | chr16 | 26043860 | 26044132 | - | processed_pseudogene |
| ENSG00000236144.8_8 | TMEM147-AS1 | 0.99994605 | 1 | -0.165 | 4.344 | chr19 | 36031640 | 36036931 | - | lncRNA |
| ENSG00000050130.18_5 | JKAMP | 0.99999957 | 1 | -0.166 | 6.362 | chr14 | 59951161 | 59972128 | + | protein_coding |
| ENSG00000119335.16_3 | SET | 0.99999998 | 1 | -0.166 | 9.430 | chr9 | 131445703 | 131458679 | + | protein_coding |
| ENSG00000114098.18_5 | ARMC8 | 0.99996849 | 1 | -0.166 | 5.074 | chr3 | 137906090 | 138017226 | + | protein_coding |
| ENSG00000263680.2_6 | AC007639.1 | 0.83295441 | 1 | -0.166 | 0.059 | chr17 | 70422080 | 70424993 | + | lncRNA |
| ENSG00000019144.19_8 | PHLDB1 | 0.99965922 | 1 | -0.166 | 3.932 | chr11 | 118477155 | 118528741 | + | protein_coding |
| ENSG00000177971.8_3 | IMP3 | 0.99995626 | 1 | -0.166 | 4.232 | chr15 | 75931426 | 75941047 | - | protein_coding |
| ENSG00000122692.9_5 | SMU1 | 0.99999993 | 1 | -0.166 | 7.048 | chr9 | 33041763 | 33076672 | - | protein_coding |
| ENSG00000136144.12_4 | RCBTB1 | 0.99999209 | 1 | -0.166 | 4.829 | chr13 | 50106082 | 50159694 | - | protein_coding |
| ENSG00000272686.1_7 | AC006333.1 | 0.9884829 | 1 | -0.167 | 2.231 | chr7 | 123389122 | 123391220 | + | lncRNA |
| ENSG00000214135.8_6 | AC132008.2 | 0.9993408 | 1 | -0.167 | 3.584 | chr3 | 197305084 | 197354777 | - | transcribed_unprocessed_pseudogene |
| ENSG00000147669.11_3 | POLR2K | 0.99998631 | 1 | -0.167 | 5.670 | chr8 | 101162851 | 101166231 | + | protein_coding |
| ENSG00000104517.13_7 | UBR5 | 0.99999993 | 1 | -0.167 | 7.351 | chr8 | 103264501 | 103424987 | - | protein_coding |
| ENSG00000197785.14_6 | ATAD3A | 0.99978735 | 1 | -0.167 | 4.468 | chr1 | 1447542 | 1470065 | + | protein_coding |
| ENSG00000260213.6_8 | AC092718.2 | 0.80574028 | 1 | -0.167 | -0.393 | chr16 | 81050397 | 81069364 | - | lncRNA |
| ENSG00000273186.1_8 | AL359091.4 | 0.77954664 | 1 | -0.167 | -0.640 | chr9 | 131193877 | 131194285 | + | lncRNA |
| ENSG00000103591.13_5 | AAGAB | 0.99999983 | 1 | -0.167 | 6.744 | chr15 | 67493005 | 67547533 | - | protein_coding |
| ENSG00000137513.10_7 | NARS2 | 0.99998283 | 1 | -0.167 | 4.851 | chr11 | 78147014 | 78285919 | - | protein_coding |
| ENSG00000115204.15_6 | MPV17 | 0.99998813 | 1 | -0.167 | 5.083 | chr2 | 27532360 | 27548547 | - | protein_coding |
| ENSG00000132463.14_5 | GRSF1 | 0.99999978 | 1 | -0.167 | 6.867 | chr4 | 71681500 | 71705614 | - | protein_coding |
| ENSG00000204160.12_5 | ZDHHC18 | 0.99991224 | 1 | -0.167 | 4.090 | chr1 | 27153179 | 27184095 | + | protein_coding |
| ENSG00000107560.12_5 | RAB11FIP2 | 0.99999424 | 1 | -0.167 | 5.412 | chr10 | 119764427 | 119806452 | - | protein_coding |
| ENSG00000136933.17_4 | RABEPK | 0.99997989 | 1 | -0.168 | 4.575 | chr9 | 127962821 | 127996440 | + | protein_coding |
| ENSG00000125166.13_4 | GOT2 | 0.99999996 | 1 | -0.168 | 7.378 | chr16 | 58741035 | 58768246 | - | protein_coding |
| ENSG00000116199.12_4 | FAM20B | 0.99999987 | 1 | -0.168 | 6.831 | chr1 | 178994939 | 179045702 | + | protein_coding |
| ENSG00000035115.22_7 | SH3YL1 | 0.99962694 | 1 | -0.168 | 3.536 | chr2 | 217730 | 266398 | - | protein_coding |
| ENSG00000239899.2 | RN7SL674P | 0.79312839 | 1 | -0.168 | -0.524 | chr2 | 11724899 | 11725176 | + | misc_RNA |
| ENSG00000180530.11_5 | NRIP1 | 0.99999996 | 1 | -0.168 | 7.832 | chr21 | 16333556 | 16438257 | - | protein_coding |
| ENSG00000183032.12_5 | SLC25A21 | 0.80666461 | 1 | -0.168 | 0.054 | chr14 | 37147126 | 37641811 | - | protein_coding |
| ENSG00000125814.17_6 | NAPB | 0.99895082 | 1 | -0.168 | 3.464 | chr20 | 23355156 | 23402156 | - | protein_coding |
| ENSG00000234602.9_4 | MCIDAS | 0.94651678 | 1 | -0.168 | 1.226 | chr5 | 54515392 | 54523143 | - | protein_coding |
| ENSG00000126858.18_5 | RHOT1 | 0.9999987 | 1 | -0.168 | 5.987 | chr17 | 30469473 | 30580393 | + | protein_coding |
| ENSG00000286299.1_1 | AL121938.1 | 0.83117673 | 1 | -0.168 | 0.057 | chr6 | 125589233 | 125595974 | + | lncRNA |
| ENSG00000130816.16_6 | DNMT1 | 0.99999973 | 1 | -0.168 | 6.847 | chr19 | 10244021 | 10341962 | - | protein_coding |
| ENSG00000171566.12_6 | PLRG1 | 0.99999927 | 1 | -0.168 | 6.144 | chr4 | 155456157 | 155471552 | - | protein_coding |

| | | | | | | | | | | |
|---|---|---|---|---|---|---|---|---|---|---|
| ENSG00000166471.11_5 | TMEM41B | 0.99999889 | 1 | -0.168 | 6.444 | chr11 | 9302201 | 9336183 | - | protein_coding |
| ENSG00000106477.19_5 | CEP41 | 0.99985285 | 1 | -0.168 | 4.357 | chr7 | 130033612 | 130082274 | - | protein_coding |
| ENSG00000237977.1_4 | EIF4HP2 | 0.77584988 | 1 | -0.168 | -0.799 | chr22 | 31298206 | 31298768 | + | processed_pseudogene |
| ENSG00000134058.12_4 | CDK7 | 0.99998704 | 1 | -0.168 | 4.918 | chr5 | 68530622 | 68573257 | + | protein_coding |
| ENSG00000088930.8_3 | XRN2 | 0.99999995 | 1 | -0.169 | 8.010 | chr20 | 21283969 | 21370463 | + | protein_coding |
| ENSG00000267397.1_6 | AC090229.1 | 0.77052018 | 1 | -0.169 | -0.726 | chr18 | 33296334 | 33297942 | + | lncRNA |
| ENSG00000261337.1 | RP11-498D10.5 | 0.82060542 | 1 | -0.169 | -0.102 | chr16 | 71952815 | 71959638 | - | antisense |
| ENSG00000126003.7_4 | PLAGL2 | 0.99982602 | 1 | -0.169 | 3.831 | chr20 | 30780307 | 30795546 | - | protein_coding |
| ENSG00000106588.11_5 | PSMA2 | 0.99575353 | 1 | -0.169 | 2.818 | chr7 | 42956460 | 42971784 | - | protein_coding |
| ENSG00000105486.14_7 | LIG1 | 0.9999995 | 1 | -0.169 | 6.245 | chr19 | 48618702 | 48673860 | - | protein_coding |
| ENSG00000135597.18_5 | REPS1 | 0.99998684 | 1 | -0.169 | 4.929 | chr6 | 139224630 | 139309398 | - | protein_coding |
| ENSG00000287218.1_1 | AL359962.3 | 0.76676895 | 1 | -0.169 | -0.974 | chr1 | 167424432 | 167427883 | + | lncRNA |
| ENSG00000228816.1_4 | AK3P5 | 0.90950319 | 1 | -0.169 | 1.040 | chr10 | 33233409 | 33233927 | + | processed_pseudogene |
| ENSG00000154978.13_6 | VOPP1 | 0.99999874 | 1 | -0.169 | 5.642 | chr7 | 55503749 | 55640681 | - | protein_coding |
| ENSG00000122507.21_7 | BBS9 | 0.99799162 | 1 | -0.169 | 3.107 | chr7 | 33149169 | 33916792 | + | protein_coding |
| ENSG00000184117.12_5 | NIPSNAP1 | 0.9999985 | 1 | -0.169 | 6.159 | chr22 | 29950797 | 29977316 | - | protein_coding |
| ENSG00000227376.1_4 | FTH1P16 | 0.77649485 | 1 | -0.169 | -0.668 | chr11 | 77445520 | 77446071 | - | processed_pseudogene |
| ENSG00000178035.12_4 | IMPDH2 | 0.99999993 | 1 | -0.169 | 7.153 | chr3 | 49061758 | 49066841 | - | protein_coding |
| ENSG00000159176.14_5 | CSRP1 | 0.99999998 | 1 | -0.169 | 8.056 | chr1 | 201452658 | 201478584 | - | protein_coding |
| ENSG00000281183.1_3 | NPTN-IT1 | 0.97637378 | 1 | -0.169 | 1.943 | chr15 | 73859353 | 73861635 | - | lncRNA |
| ENSG00000135249.8_3 | RINT1 | 0.99987078 | 1 | -0.170 | 4.045 | chr7 | 105172616 | 105208124 | + | protein_coding |
| ENSG00000158636.16_7 | EMSY | 0.99997958 | 1 | -0.170 | 4.912 | chr11 | 76155967 | 76264069 | + | protein_coding |
| ENSG00000279608.1_5 | AL353795.3 | 0.79147739 | 1 | -0.170 | -0.367 | chr9 | 35038622 | 35044323 | - | TEC |
| ENSG00000104218.14_6 | CSPP1 | 0.99996748 | 1 | -0.170 | 4.734 | chr8 | 67974661 | 68108498 | + | protein_coding |
| ENSG00000146232.17_7 | NFKBIE | 0.99679053 | 1 | -0.170 | 2.841 | chr6 | 44225903 | 44233525 | - | protein_coding |
| ENSG00000272622.2_7 | AC010735.2 | 0.77613271 | 1 | -0.170 | -0.549 | chr2 | 227664862 | 227678457 | + | lncRNA |
| ENSG00000113456.19_4 | RAD1 | 0.99999745 | 1 | -0.170 | 5.381 | chr5 | 34905365 | 34919094 | - | protein_coding |
| ENSG00000100823.12_4 | APEX1 | 0.99999867 | 1 | -0.170 | 5.679 | chr14 | 20923350 | 20925931 | + | protein_coding |
| ENSG00000100814.18_4 | CCNB1IP1 | 0.99999101 | 1 | -0.170 | 4.837 | chr14 | 20779527 | 20801471 | - | protein_coding |
| ENSG00000174912.7_4 | METTL15P1 | 0.85882544 | 1 | -0.170 | 0.340 | chr3 | 156431673 | 156432717 | - | processed_pseudogene |
| ENSG00000149609.6_3 | C20orf144 | 0.94461581 | 1 | -0.170 | 1.300 | chr20 | 32250133 | 32253425 | + | protein_coding |
| ENSG00000167193.8_4 | CRK | 0.99999994 | 1 | -0.171 | 7.409 | chr17 | 1323983 | 1366456 | - | protein_coding |
| ENSG00000182376.2_6 | AC138028.1 | 0.80262577 | 1 | -0.171 | -0.058 | chr16 | 88809175 | 88812156 | + | lncRNA |
| ENSG00000037241.7_5 | RPL26L1 | 0.99998409 | 1 | -0.171 | 4.645 | chr5 | 172385732 | 172396774 | + | protein_coding |
| ENSG00000090054.15_5 | SPTLC1 | 0.99999954 | 1 | -0.171 | 6.310 | chr9 | 94762369 | 94877695 | - | protein_coding |
| ENSG00000105926.15_5 | MPP6 | 0.99998208 | 1 | -0.171 | 4.809 | chr7 | 24612887 | 24733812 | + | protein_coding |
| ENSG00000184524.6_4 | CEND1 | 0.94371972 | 1 | -0.171 | 0.824 | chr11 | 787115 | 790113 | - | protein_coding |
| ENSG00000140836.17_9 | ZFHX3 | 0.99999606 | 1 | -0.171 | 5.560 | chr16 | 72816784 | 73925770 | - | protein_coding |
| ENSG00000138756.17_4 | BMP2K | 0.99980684 | 1 | -0.171 | 4.129 | chr4 | 79697496 | 79837526 | + | protein_coding |
| ENSG00000127603.28_9 | MACF1 | 0.99976738 | 1 | -0.171 | 7.955 | chr1 | 39546988 | 39952849 | + | protein_coding |
| ENSG00000230457.3_2 | PA2G4P4 | 0.94278537 | 1 | -0.171 | 1.530 | chr3 | 156527340 | 156528521 | - | processed_pseudogene |
| ENSG00000176387.7_3 | HSD11B2 | 0.79575396 | 1 | -0.171 | -0.474 | chr16 | 67464555 | 67471456 | + | protein_coding |
| ENSG00000227896.2_6 | AL731569.1 | 0.96907845 | 1 | -0.171 | 1.248 | chr10 | 88281702 | 88284858 | + | lncRNA |
| ENSG00000116857.17_5 | TMEM9 | 0.99999823 | 1 | -0.171 | 5.609 | chr1 | 201103900 | 201140702 | - | protein_coding |
| ENSG00000204628.12_7 | RACK1 | 0.99999996 | 1 | -0.172 | 9.892 | chr5 | 180663897 | 180675096 | - | protein_coding |
| ENSG00000140743.8_7 | CDR2 | 0.99999593 | 1 | -0.172 | 5.155 | chr16 | 22357257 | 22448486 | - | protein_coding |
| ENSG00000198060.10_4 | MARCHF5 | 0.9999948 | 1 | -0.172 | 5.165 | chr10 | 94050924 | 94113721 | + | protein_coding |
| ENSG00000107651.13_3 | SEC23IP | 0.9999998 | 1 | -0.172 | 6.698 | chr10 | 121652242 | 121704169 | + | protein_coding |
| ENSG00000154639.19_3 | CXADR | 0.9999421 | 1 | -0.172 | 4.684 | chr21 | 18885361 | 18965897 | + | protein_coding |
| ENSG00000226026.6_7 | AC092802.2 | 0.77960241 | 1 | -0.172 | -0.640 | chr1 | 95627232 | 95699538 | - | lncRNA |
| ENSG00000167258.14_6 | CDK12 | 0.99999988 | 1 | -0.172 | 7.200 | chr17 | 37617739 | 37721160 | + | protein_coding |
| ENSG00000162885.13_4 | B3GALNT2 | 0.99996615 | 1 | -0.172 | 4.679 | chr1 | 235610505 | 235667752 | - | protein_coding |
| ENSG00000246451.2_8 | AL049840.2 | 0.98845686 | 1 | -0.172 | 2.574 | chr14 | 104148699 | 104150352 | - | lncRNA |
| ENSG00000205740.2_7 | AL359878.1 | 0.93386645 | 1 | -0.172 | 1.166 | chr10 | 1017086 | 1034281 | - | lncRNA |
| ENSG00000086589.12_5 | RBM22 | 0.99999908 | 1 | -0.172 | 5.729 | chr5 | 150070354 | 150080639 | - | protein_coding |
| ENSG00000272777.1_6 | AC019131.2 | 0.80574028 | 1 | -0.172 | -0.392 | chr4 | 99988407 | 99989276 | - | lncRNA |
| ENSG00000279789.1_6 | AC120114.3 | 0.97307036 | 1 | -0.172 | 1.655 | chr16 | 29924465 | 29926658 | - | TEC |

| | | | | | | | | | | |
|---|---|---|---|---|---|---|---|---|---|---|
| ENSG00000182952.5_4 | HMGN4 | 0.99999705 | 1 | -0.172 | 5.805 | chr6 | 26538594 | 26547161 | + | protein_coding |
| ENSG00000101367.9_3 | MAPRE1 | 0.99999965 | 1 | -0.172 | 7.088 | chr20 | 31407760 | 31438211 | + | protein_coding |
| ENSG00000152234.16_7 | ATP5F1A | 0.99999993 | 1 | -0.173 | 8.471 | chr18 | 43660214 | 43684300 | - | protein_coding |
| ENSG00000126460.11_4 | PRRG2 | 0.82699644 | 1 | -0.173 | 0.038 | chr19 | 50083903 | 50094261 | + | protein_coding |
| ENSG00000258744.2_6 | AL132800.1 | 0.90832597 | 1 | -0.173 | 0.848 | chr14 | 24912222 | 24977894 | + | lncRNA |
| ENSG00000170075.9_3 | GPR37L1 | 0.8934692 | 1 | -0.173 | 0.663 | chr1 | 202092045 | 202102720 | + | protein_coding |
| ENSG00000170485.17_6 | NPAS2 | 0.99999916 | 1 | -0.173 | 5.991 | chr2 | 101436601 | 101613291 | + | protein_coding |
| ENSG00000287965.1_2 | AC018557.3 | 0.76338407 | 1 | -0.173 | -0.942 | chr16 | 66763055 | 66775319 | + | lncRNA |
| ENSG00000160293.17_6 | VAV2 | 0.99999049 | 1 | -0.173 | 5.246 | chr9 | 136627016 | 136857726 | - | protein_coding |
| ENSG00000285692.1_4 | AC023509.6 | 0.77038934 | 1 | -0.173 | -0.562 | chr12 | 53925536 | 54015002 | + | lncRNA |
| ENSG00000144867.12_5 | SRPRB | 0.99999723 | 1 | -0.173 | 5.599 | chr3 | 133502867 | 133544616 | + | protein_coding |
| ENSG00000136492.9_3 | BRIP1 | 0.99999924 | 1 | -0.173 | 6.037 | chr17 | 59756500 | 59940889 | - | protein_coding |
| ENSG00000112984.12_4 | KIF20A | 0.99999978 | 1 | -0.174 | 6.592 | chr5 | 137514408 | 137523412 | + | protein_coding |
| ENSG00000243970.3_5 | PPIEL | 0.9977595 | 1 | -0.174 | 3.022 | chr1 | 39987952 | 40025343 | - | transcribed_unprocessed_pseudogene |
| ENSG00000123684.13_4 | LPGAT1 | 0.99999993 | 1 | -0.174 | 7.688 | chr1 | 211916799 | 212004105 | - | protein_coding |
| ENSG00000152359.15_6 | POC5 | 0.99989868 | 1 | -0.174 | 4.177 | chr5 | 74969949 | 75013273 | - | protein_coding |
| ENSG00000198937.9_4 | CCDC167 | 0.99601033 | 1 | -0.174 | 3.053 | chr6 | 37450714 | 37467669 | - | protein_coding |
| ENSG00000134108.13_3 | ARL8B | 0.99999977 | 1 | -0.174 | 6.550 | chr3 | 5163930 | 5222597 | + | protein_coding |
| ENSG00000160075.12_4 | SSU72 | 0.999999 | 1 | -0.174 | 5.747 | chr1 | 1477053 | 1510243 | - | protein_coding |
| ENSG00000132589.16_4 | FLOT2 | 0.99999971 | 1 | -0.174 | 6.398 | chr17 | 27206353 | 27224751 | - | protein_coding |
| ENSG00000169139.12_5 | UBE2V2 | 0.99981758 | 1 | -0.174 | 5.688 | chr8 | 48920975 | 48977268 | + | protein_coding |
| ENSG00000269951.1_7 | AC090181.2 | 0.91038164 | 1 | -0.174 | 1.008 | chr15 | 77359996 | 77360667 | - | lncRNA |
| ENSG00000183520.12_4 | UTP11 | 0.99998292 | 1 | -0.174 | 5.032 | chr1 | 38474930 | 38490492 | + | protein_coding |
| ENSG00000181097.5_8 | AC105219.1 | 0.82881166 | 1 | -0.174 | 0.191 | chr8 | 144778324 | 144780583 | + | lncRNA |
| ENSG00000130413.15_6 | STK33 | 0.9929203 | 1 | -0.174 | 2.523 | chr11 | 8413415 | 8615836 | - | protein_coding |
| ENSG00000136631.15_8 | VPS45 | 0.9999956 | 1 | -0.174 | 5.269 | chr1 | 150039336 | 150117507 | + | protein_coding |
| ENSG00000159352.15_4 | PSMD4 | 0.99999975 | 1 | -0.175 | 6.880 | chr1 | 151227179 | 151239955 | + | protein_coding |
| ENSG00000129559.13_5 | NEDD8 | 0.99998365 | 1 | -0.175 | 5.055 | chr14 | 24686063 | 24701573 | - | protein_coding |
| ENSG00000065526.12_4 | SPEN | 0.99999859 | 1 | -0.175 | 5.598 | chr1 | 16162590 | 16266951 | + | protein_coding |
| ENSG00000144840.9_5 | RABL3 | 0.99994496 | 1 | -0.175 | 4.979 | chr3 | 120405528 | 120461840 | - | protein_coding |
| ENSG00000121741.16_5 | ZMYM2 | 0.99999982 | 1 | -0.175 | 6.697 | chr13 | 20532810 | 20665968 | + | protein_coding |
| ENSG00000131051.24_9 | RBM39 | 0.99999985 | 1 | -0.175 | 8.236 | chr20 | 34289269 | 34330234 | - | protein_coding |
| ENSG00000162688.17_4 | AGL | 0.99985757 | 1 | -0.175 | 5.055 | chr1 | 100315917 | 100389579 | + | protein_coding |
| ENSG00000150401.15_7 | DCUN1D2 | 0.99550946 | 1 | -0.175 | 2.760 | chr13 | 114110134 | 114145266 | - | protein_coding |
| ENSG00000101040.19_6 | ZMYND8 | 0.99999747 | 1 | -0.175 | 5.682 | chr20 | 45837859 | 45985633 | - | protein_coding |
| ENSG00000225891.2_7 | DHDDS-AS1 | 0.79704648 | 1 | -0.175 | -0.301 | chr1 | 26789247 | 26794505 | - | lncRNA |
| ENSG00000161653.11_3 | NAGS | 0.82954438 | 1 | -0.175 | 0.643 | chr17 | 42081990 | 42086436 | + | protein_coding |
| ENSG00000263307.1_6 | AC007216.4 | 0.75799965 | 1 | -0.175 | -0.941 | chr16 | 11945506 | 11989468 | + | lncRNA |
| ENSG00000154174.8_5 | TOMM70 | 0.99999987 | 1 | -0.175 | 7.196 | chr3 | 100082275 | 100119933 | - | protein_coding |
| ENSG00000249738.10_7 | AC008691.1 | 0.92569283 | 1 | -0.176 | 1.013 | chr5 | 158737753 | 159232543 | + | lncRNA |
| ENSG00000153037.15_6 | SRP19 | 0.95799544 | 1 | -0.176 | 3.686 | chr5 | 112196885 | 112234068 | + | protein_coding |
| ENSG00000124532.14_5 | MRS2 | 0.99994238 | 1 | -0.176 | 4.625 | chr6 | 24403136 | 24426422 | + | protein_coding |
| ENSG00000265724.1 | MIR4284 | 0.76710382 | 1 | -0.176 | -0.768 | chr7 | 73125647 | 73125727 | + | miRNA |
| ENSG00000148943.12_4 | LIN7C | 0.9999994 | 1 | -0.176 | 6.673 | chr11 | 27515965 | 27528316 | - | protein_coding |
| ENSG00000067248.10_5 | DHX29 | 0.9999997 | 1 | -0.176 | 6.656 | chr5 | 54551883 | 54603522 | - | protein_coding |
| ENSG00000168924.15_4 | LETM1 | 0.99999909 | 1 | -0.176 | 6.647 | chr4 | 1813206 | 1857883 | - | protein_coding |
| ENSG00000223396.4_6 | RPS10P7 | 0.89806954 | 1 | -0.176 | 0.788 | chr1 | 201487831 | 201553495 | + | transcribed_processed_pseudogene |
| ENSG00000176261.15_6 | ZBTB8OS | 0.99969285 | 1 | -0.176 | 4.064 | chr1 | 33065773 | 33116504 | - | protein_coding |
| ENSG00000079134.12_5 | THOC1 | 0.99999334 | 1 | -0.176 | 5.166 | chr18 | 214520 | 268050 | - | protein_coding |
| ENSG00000228606.3_7 | AL139011.1 | 0.78302438 | 1 | -0.176 | -0.275 | chr1 | 160231531 | 160233118 | + | lncRNA |
| ENSG00000164949.8_6 | GEM | 0.99960537 | 1 | -0.176 | 3.600 | chr8 | 95261481 | 95274578 | - | protein_coding |
| ENSG00000143549.21_7 | TPM3 | 0.99999988 | 1 | -0.176 | 8.590 | chr1 | 154127780 | 154167124 | - | protein_coding |
| ENSG00000170946.15_5 | DNAJC24 | 0.99953709 | 1 | -0.177 | 4.093 | chr11 | 31391387 | 31454382 | + | protein_coding |
| ENSG00000116670.15_6 | MAD2L2 | 0.99979526 | 1 | -0.177 | 4.153 | chr1 | 11734537 | 11751707 | - | protein_coding |
| ENSG00000186615.11_6 | KTN1-AS1 | 0.98847989 | 1 | -0.177 | 2.236 | chr14 | 55963504 | 56047606 | - | lncRNA |
| ENSG00000170681.7_6 | CAVIN4 | 0.84594274 | 1 | -0.177 | 0.172 | chr9 | 103340361 | 103350671 | + | protein_coding |
| ENSG00000135164.18_7 | DMTF1 | 0.99999804 | 1 | -0.177 | 5.887 | chr7 | 86781677 | 86825653 | + | protein_coding |
| ENSG00000217128.12_5 | FNIP1 | 0.99999904 | 1 | -0.177 | 5.850 | chr5 | 130977407 | 131132756 | - | protein_coding |
| ENSG00000164172.19_6 | MOCS2 | 0.99999408 | 1 | -0.177 | 5.292 | chr5 | 52391509 | 52405893 | - | protein_coding |
| ENSG00000143756.12_3 | FBXO28 | 0.99999963 | 1 | -0.177 | 6.121 | chr1 | 224301813 | 224349749 | + | protein_coding |

| | | | | | | | | | | |
|---|---|---|---|---|---|---|---|---|---|---|
| ENSG00000188818.12_4 | ZDHHC11 | 0.99816076 | 1 | -0.177 | 3.057 | chr5 | 795721 | 851101 | - | protein_coding |
| ENSG00000132388.12_3 | UBE2G1 | 0.99999987 | 1 | -0.177 | 7.142 | chr17 | 4172554 | 4269923 | - | protein_coding |
| ENSG00000260597.1_7 | AC012531.1 | 0.88202698 | 1 | -0.177 | 0.584 | chr12 | 54413694 | 54416373 | + | lncRNA |
| ENSG00000261460.1_6 | AC009690.2 | 0.75567131 | 1 | -0.177 | -1.012 | chr15 | 72571208 | 72644135 | + | lncRNA |
| ENSG00000230453.9_2 | ANKRD18B | 0.99914445 | 1 | -0.177 | 3.457 | chr9 | 33524392 | 33573007 | + | protein_coding |
| ENSG00000138185.20_6 | ENTPD1 | 0.97872909 | 1 | -0.178 | 2.183 | chr10 | 97471536 | 97637023 | + | protein_coding |
| ENSG00000205542.11_3 | TMSB4X | 0.99999105 | 1 | -0.178 | 9.405 | chrX | 12993229 | 12995346 | + | protein_coding |
| ENSG00000103160.12_5 | HSDL1 | 0.99998732 | 1 | -0.178 | 4.992 | chr16 | 84155746 | 84178797 | - | protein_coding |
| ENSG00000247137.9_8 | AP000873.2 | 0.99390081 | 1 | -0.178 | 2.566 | chr11 | 82895533 | 82904836 | - | lncRNA |
| ENSG00000182379.10_3 | NXPH4 | 0.79431993 | 1 | -0.178 | -0.252 | chr12 | 57610577 | 57620232 | + | protein_coding |
| ENSG00000171861.11_6 | MRM3 | 0.99962029 | 1 | -0.178 | 4.225 | chr17 | 685593 | 695749 | + | protein_coding |
| ENSG00000287735.1_1 | AL355075.6 | 0.87685739 | 1 | -0.178 | 0.680 | chr14 | 20786108 | 20787866 | + | lncRNA |
| ENSG00000104671.8_4 | DCTN6 | 0.99987662 | 1 | -0.178 | 4.363 | chr8 | 30013835 | 30041155 | + | protein_coding |
| ENSG00000108375.12_5 | RNF43 | 0.96234706 | 1 | -0.178 | 1.203 | chr17 | 56429861 | 56494956 | - | protein_coding |
| ENSG00000252808.1 | SCARNA4 | 0.76991616 | 1 | -0.178 | -0.614 | chr1 | 155895749 | 155895877 | - | snoRNA |
| ENSG00000147082.18_3 | CCNB3 | 0.92339822 | 1 | -0.178 | 1.246 | chrX | 49967364 | 50094913 | + | protein_coding |
| ENSG00000224892.6_4 | RPS4XP16 | 0.75409379 | 1 | -0.178 | -0.907 | chr13 | 52028300 | 52035451 | + | transcribed_processed_pseudogene |
| ENSG00000155508.13_5 | CNOT8 | 0.99999839 | 1 | -0.178 | 5.610 | chr5 | 154237113 | 154256353 | + | protein_coding |
| ENSG00000103044.11_5 | HAS3 | 0.99962558 | 1 | -0.178 | 3.970 | chr16 | 69139556 | 69152622 | + | protein_coding |
| ENSG00000180340.7_3 | FZD2 | 0.99999161 | 1 | -0.178 | 5.944 | chr17 | 42634852 | 42638630 | + | protein_coding |
| ENSG00000076685.18_6 | NT5C2 | 0.99999647 | 1 | -0.178 | 5.969 | chr10 | 104847774 | 104953063 | - | protein_coding |
| ENSG00000106244.13_4 | PDAP1 | 0.99999956 | 1 | -0.178 | 6.361 | chr7 | 98989671 | 99006220 | - | protein_coding |
| ENSG00000163930.10_4 | BAP1 | 0.99999917 | 1 | -0.178 | 5.831 | chr3 | 52435024 | 52444024 | - | protein_coding |
| ENSG00000138182.14_2 | KIF20B | 0.99999923 | 1 | -0.178 | 6.794 | chr10 | 91461367 | 91534696 | + | protein_coding |
| ENSG00000278156.2_5 | TSC22D1-AS1 | 0.76333411 | 1 | -0.178 | -0.423 | chr13 | 45150029 | 45154568 | + | lncRNA |
| ENSG00000105011.9_4 | ASF1B | 0.99997535 | 1 | -0.179 | 4.956 | chr19 | 14230324 | 14247425 | - | protein_coding |
| ENSG00000070061.15_7 | ELP1 | 0.99999979 | 1 | -0.179 | 6.724 | chr9 | 111629797 | 111696404 | - | protein_coding |
| ENSG00000115841.20_5 | RMDN2 | 0.99449295 | 1 | -0.179 | 2.681 | chr2 | 38150330 | 38294285 | + | protein_coding |
| ENSG00000115137.12_5 | DNAJC27 | 0.99493325 | 1 | -0.179 | 2.897 | chr2 | 25166505 | 25194963 | - | protein_coding |
| ENSG00000204262.14_6 | COL5A2 | 0.99999994 | 1 | -0.179 | 8.669 | chr2 | 189896624 | 190090038 | - | protein_coding |
| ENSG00000150779.12_6 | TIMM8B | 0.99995712 | 1 | -0.179 | 4.415 | chr11 | 111955524 | 111957522 | - | protein_coding |
| ENSG00000259165.1_4 | DDX18P1 | 0.75351241 | 1 | -0.179 | -0.910 | chr14 | 69550115 | 69552106 | + | processed_pseudogene |
| ENSG00000258741.4_7 | H2AZ2P1 | 0.75558954 | 1 | -0.179 | -0.908 | chr15 | 93256707 | 93277435 | - | transcribed_processed_pseudogene |
| ENSG00000173218.15_4 | VANGL1 | 0.999996 | 1 | -0.179 | 5.454 | chr1 | 116184591 | 116240845 | + | protein_coding |
| ENSG00000169598.15_3 | DFFB | 0.98059755 | 1 | -0.179 | 1.983 | chr1 | 3773831 | 3801993 | + | protein_coding |
| ENSG00000260060.1_6 | AC009088.1 | 0.77591014 | 1 | -0.179 | -0.204 | chr16 | 31207452 | 31208284 | + | lncRNA |
| ENSG00000166503.9_8 | HDGFL3 | 0.99999889 | 1 | -0.179 | 6.387 | chr15 | 83781490 | 83876575 | - | protein_coding |
| ENSG00000170296.10_5 | GABARAP | 0.77646853 | 1 | -0.179 | -0.615 | chr17 | 7143327 | 7145768 | - | protein_coding |
| ENSG00000282936.2_7 | AC004706.3 | 0.87319108 | 1 | -0.179 | 0.494 | chr17 | 6531939 | 6543776 | - | protein_coding |
| ENSG00000175793.12_4 | SFN | 0.99999651 | 1 | -0.179 | 5.576 | chr1 | 27189640 | 27190947 | + | protein_coding |
| ENSG00000139350.11_3 | NEDD1 | 0.9999992 | 1 | -0.179 | 6.213 | chr12 | 97301001 | 97347555 | + | protein_coding |
| ENSG00000164985.15_4 | PSIP1 | 0.99999978 | 1 | -0.179 | 7.027 | chr9 | 15464064 | 15510993 | - | protein_coding |
| ENSG00000109107.14_6 | ALDOC | 0.84723544 | 1 | -0.179 | 0.449 | chr17 | 26900133 | 26903966 | - | protein_coding |
| ENSG00000149357.10_4 | LAMTOR1 | 0.99999712 | 1 | -0.179 | 5.626 | chr11 | 71796941 | 71814343 | - | protein_coding |
| ENSG00000268996.3_6 | MAN1B1-DT | 0.90399642 | 1 | -0.180 | 1.105 | chr9 | 139979398 | 139981269 | - | lncRNA |
| ENSG00000132549.18_6 | VPS13B | 0.99999675 | 1 | -0.180 | 5.822 | chr8 | 100025494 | 100889808 | + | protein_coding |
| ENSG00000101557.15_4 | USP14 | 0.99999597 | 1 | -0.180 | 5.531 | chr18 | 158383 | 214629 | + | protein_coding |
| ENSG00000119707.14_4 | RBM25 | 0.99999873 | 1 | -0.180 | 6.440 | chr14 | 73525240 | 73590607 | + | protein_coding |
| ENSG00000065989.16_5 | PDE4A | 0.94357069 | 1 | -0.180 | 1.227 | chr19 | 10527449 | 10580306 | + | protein_coding |
| ENSG00000251365.4_7 | LINC02236 | 0.97697119 | 1 | -0.180 | 2.023 | chr5 | 6766004 | 6833233 | + | lncRNA |
| ENSG00000141294.10_3 | LRRC46 | 0.95438947 | 1 | -0.180 | 1.667 | chr17 | 45909000 | 45915085 | + | protein_coding |
| ENSG00000160796.18_5 | NBEAL2 | 0.99999914 | 1 | -0.180 | 6.627 | chr3 | 47021156 | 47051194 | + | protein_coding |
| ENSG00000254656.2_6 | RTL1 | 0.83914306 | 1 | -0.180 | 0.374 | chr14 | 101346090 | 101370059 | - | protein_coding |
| ENSG00000175387.16_9 | SMAD2 | 0.99999986 | 1 | -0.180 | 7.784 | chr18 | 45335328 | 45457517 | - | protein_coding |
| ENSG00000159228.13_3 | CBR1 | 0.9999995 | 1 | -0.181 | 6.159 | chr21 | 37442239 | 37445464 | + | protein_coding |
| ENSG00000105793.15_3 | GTPBP10 | 0.9999434 | 1 | -0.181 | 4.698 | chr7 | 89964537 | 90020769 | + | protein_coding |
| ENSG00000173960.14_4 | UBXN2A | 0.99999741 | 1 | -0.181 | 5.600 | chr2 | 24150155 | 24227779 | + | protein_coding |
| ENSG00000204271.13_7 | SPIN3 | 0.99966527 | 1 | -0.181 | 4.432 | chrX | 56844731 | 57022260 | - | protein_coding |

| | | | | | | | | | | |
|---|---|---|---|---|---|---|---|---|---|---|
| ENSG00000231242.1 | RP11-87H9.3 | 0.76048016 | 1 | -0.181 | -0.533 | chr9 | 69082536 | 69108793 | - | lincRNA |
| ENSG00000114739.14_3 | ACVR2B | 0.99995355 | 1 | -0.181 | 4.420 | chr3 | 38495381 | 38534633 | + | protein_coding |
| ENSG00000078304.19_6 | PPP2R5C | 0.99999959 | 1 | -0.181 | 6.774 | chr14 | 102228135 | 102394326 | + | protein_coding |
| ENSG00000088832.17_6 | FKBP1A | 0.99999961 | 1 | -0.181 | 6.455 | chr20 | 1349622 | 1373816 | - | protein_coding |
| ENSG00000165792.17_3 | METTL17 | 0.99999621 | 1 | -0.181 | 5.273 | chr14 | 21457929 | 21465194 | + | protein_coding |
| ENSG00000188229.6_3 | TUBB4B | 0.99999981 | 1 | -0.181 | 8.331 | chr9 | 140135739 | 140138159 | + | protein_coding |
| ENSG00000242715.8_6 | CCDC169 | 0.75325007 | 1 | -0.181 | -0.978 | chr13 | 36796145 | 36871977 | - | protein_coding |
| ENSG00000176732.7_5 | PFN4 | 0.90093772 | 1 | -0.181 | 0.902 | chr2 | 24337679 | 24346334 | - | protein_coding |
| ENSG00000103248.19_8 | MTHFSD | 0.99915832 | 1 | -0.182 | 3.973 | chr16 | 86563784 | 86588841 | - | protein_coding |
| ENSG00000179761.12_4 | PIPOX | 0.78108383 | 1 | -0.182 | -0.279 | chr17 | 27277531 | 27384234 | + | protein_coding |
| ENSG00000041357.16_4 | PSMA4 | 0.99999886 | 1 | -0.182 | 7.784 | chr15 | 78832747 | 78844759 | + | protein_coding |
| ENSG00000165494.11_3 | PCF11 | 0.99999922 | 1 | -0.182 | 6.258 | chr11 | 82868030 | 82898493 | + | protein_coding |
| ENSG00000268279.4_6 | AC090004.1 | 0.75036391 | 1 | -0.182 | -0.941 | chr3 | 14176717 | 14189752 | + | protein_coding |
| ENSG00000242114.6_4 | MTFP1 | 0.86450382 | 1 | -0.182 | 0.645 | chr22 | 30821611 | 30825041 | + | protein_coding |
| ENSG00000237276.8_5 | ANO7L1 | 0.75407594 | 1 | -0.182 | -0.642 | chr1 | 16542404 | 16554522 | - | transcribed_unprocessed_pseudogene |
| ENSG00000143436.11_6 | MRPL9 | 0.99984778 | 1 | -0.182 | 4.252 | chr1 | 151732123 | 151735972 | - | protein_coding |
| ENSG00000099385.11_6 | BCL7C | 0.99438218 | 1 | -0.182 | 2.703 | chr16 | 30844947 | 30906281 | - | protein_coding |
| ENSG00000103091.15_5 | WDR59 | 0.99996946 | 1 | -0.182 | 4.777 | chr16 | 74905260 | 75034071 | - | protein_coding |
| ENSG00000167397.15_5 | VKORC1 | 0.98902123 | 1 | -0.182 | 2.533 | chr16 | 31102163 | 31107301 | - | protein_coding |
| ENSG00000149503.13_3 | INCENP | 0.99999925 | 1 | -0.183 | 6.483 | chr11 | 61891470 | 61920641 | + | protein_coding |
| ENSG00000166348.18_4 | USP54 | 0.99994858 | 1 | -0.183 | 4.615 | chr10 | 75257296 | 75385711 | - | protein_coding |
| ENSG00000078403.17_5 | MLLT10 | 0.99999797 | 1 | -0.183 | 5.605 | chr10 | 21813575 | 22032559 | + | protein_coding |
| ENSG00000164241.13_4 | C5orf63 | 0.99368031 | 1 | -0.183 | 2.630 | chr5 | 126378250 | 126409184 | - | protein_coding |
| ENSG00000227946.1_6 | AC007383.2 | 0.78200327 | 1 | -0.183 | -0.426 | chr2 | 206949329 | 206951288 | + | lncRNA |
| ENSG00000271500.1_6 | AC005183.1 | 0.85328901 | 1 | -0.183 | 0.403 | chr12 | 1804996 | 1807210 | - | processed_pseudogene |
| ENSG00000008018.9_4 | PSMB1 | 0.99999967 | 1 | -0.183 | 6.720 | chr6 | 170844208 | 170862395 | - | protein_coding |
| ENSG00000106392.11_4 | C1GALT1 | 0.99999014 | 1 | -0.183 | 5.642 | chr7 | 7196565 | 7288247 | + | protein_coding |
| ENSG00000183114.8_3 | FAM43B | 0.95049006 | 1 | -0.183 | 1.480 | chr1 | 20879066 | 20881513 | + | protein_coding |
| ENSG00000128908.17_8 | INO80 | 0.99999769 | 1 | -0.184 | 5.518 | chr15 | 41271078 | 41408478 | - | protein_coding |
| ENSG00000187824.8_6 | TMEM220 | 0.96069532 | 1 | -0.184 | 1.646 | chr17 | 10602332 | 10633633 | - | protein_coding |
| ENSG00000182551.14_4 | ADI1 | 0.99999939 | 1 | -0.184 | 6.205 | chr2 | 3501137 | 3523302 | - | protein_coding |
| ENSG00000131236.17_4 | CAP1 | 0.99999987 | 1 | -0.184 | 6.972 | chr1 | 40505905 | 40538321 | + | protein_coding |
| ENSG00000163605.15_4 | PPP4R2 | 0.99999965 | 1 | -0.184 | 6.865 | chr3 | 73045954 | 73118349 | + | protein_coding |
| ENSG00000066322.15_5 | ELOVL1 | 0.99999766 | 1 | -0.184 | 5.871 | chr1 | 43829069 | 43833745 | - | protein_coding |
| ENSG00000196655.12_4 | TRAPPC4 | 0.99976904 | 1 | -0.184 | 4.202 | chr11 | 118889473 | 118896164 | + | protein_coding |
| ENSG00000105483.18_6 | CARD8 | 0.99997645 | 1 | -0.184 | 5.161 | chr19 | 48684027 | 48759203 | - | protein_coding |
| ENSG00000091409.15_5 | ITGA6 | 0.99999986 | 1 | -0.184 | 7.355 | chr2 | 173292082 | 173371187 | + | protein_coding |
| ENSG00000183011.13_6 | NAA38 | 0.99999809 | 1 | -0.184 | 5.830 | chr17 | 7760003 | 7788556 | - | protein_coding |
| ENSG00000213619.10_5 | NDUFS3 | 0.99998897 | 1 | -0.184 | 5.323 | chr11 | 47586888 | 47606114 | + | protein_coding |
| ENSG00000212125.2_4 | TAS2R15P | 0.79332392 | 1 | -0.184 | -0.072 | chr12 | 11117024 | 11117951 | - | unprocessed_pseudogene |
| ENSG00000141391.14_5 | PRELID3A | 0.98397068 | 1 | -0.184 | 2.084 | chr18 | 12407895 | 12432237 | + | protein_coding |
| ENSG00000243829.1_5 | AC011495.1 | 0.74736722 | 1 | -0.184 | -0.944 | chr19 | 50222755 | 50223338 | - | processed_pseudogene |
| ENSG00000154767.14_4 | XPC | 0.99997441 | 1 | -0.185 | 5.230 | chr3 | 14186647 | 14220283 | - | protein_coding |
| ENSG00000134077.16_6 | THUMPD3 | 0.99999893 | 1 | -0.185 | 5.829 | chr3 | 9404655 | 9428475 | + | protein_coding |
| ENSG00000198034.11_4 | RPS4X | 0.99999989 | 1 | -0.185 | 9.812 | chrX | 71475529 | 71497098 | - | protein_coding |
| ENSG00000143702.16_6 | CEP170 | 0.99999929 | 1 | -0.185 | 6.179 | chr1 | 243287730 | 243418650 | - | protein_coding |
| ENSG00000105479.15_3 | CCDC114 | 0.75116571 | 1 | -0.185 | -0.978 | chr19 | 48799714 | 48825151 | - | protein_coding |
| ENSG00000137807.15_7 | KIF23 | 0.99999976 | 1 | -0.185 | 7.592 | chr15 | 69706585 | 69740766 | + | protein_coding |
| ENSG00000196419.12_2 | XRCC6 | 0.99999988 | 1 | -0.185 | 8.141 | chr22 | 42017123 | 42060052 | + | protein_coding |
| ENSG00000145050.18_5 | MANF | 0.99992463 | 1 | -0.185 | 4.887 | chr3 | 51422722 | 51426828 | + | protein_coding |
| ENSG00000174109.5_5 | C16orf91 | 0.99896305 | 1 | -0.186 | 3.718 | chr16 | 1469753 | 1470757 | - | protein_coding |
| ENSG00000114503.11_6 | NCBP2 | 0.99999819 | 1 | -0.186 | 5.625 | chr3 | 196662273 | 196669465 | - | protein_coding |
| ENSG00000124942.14_8 | AHNAK | 0.9999967 | 1 | -0.186 | 9.188 | chr11 | 62201014 | 62323707 | - | protein_coding |
| ENSG00000133265.11_5 | HSPBP1 | 0.99999555 | 1 | -0.186 | 5.895 | chr19 | 55773591 | 55791749 | - | protein_coding |
| ENSG00000135211.6_4 | TMEM60 | 0.99953392 | 1 | -0.186 | 3.759 | chr7 | 77423045 | 77427751 | - | protein_coding |
| ENSG00000233025.1_4 | CRYZP1 | 0.8031452 | 1 | -0.186 | 0.149 | chr7 | 102729111 | 102730057 | - | processed_pseudogene |
| ENSG00000184922.14_6 | FMNL1 | 0.99998151 | 1 | -0.186 | 5.013 | chr17 | 43298811 | 43324686 | + | protein_coding |
| ENSG00000160551.11_6 | TAOK1 | 0.99999877 | 1 | -0.186 | 8.085 | chr17 | 27717482 | 27878922 | + | protein_coding |

| | | | | | | | | | | |
|---|---|---|---|---|---|---|---|---|---|---|
| ENSG00000121281.13_7 | ADCY7 | 0.9997982 | 1 | -0.186 | 4.260 | chr16 | 50280048 | 50352046 | + | protein_coding |
| ENSG00000276966.2_3 | H4C5 | 0.98437961 | 1 | -0.186 | 6.915 | chr6 | 26204780 | 26206266 | + | protein_coding |
| ENSG00000072501.18_3 | SMC1A | 0.99999951 | 1 | -0.186 | 7.374 | chrX | 53401070 | 53449677 | - | protein_coding |
| ENSG00000186654.21_6 | PRR5 | 0.99686133 | 1 | -0.186 | 2.933 | chr22 | 45064427 | 45133561 | + | protein_coding |
| ENSG00000023839.12_5 | ABCC2 | 0.99999996 | 1 | -0.186 | 9.632 | chr10 | 101542397 | 101612351 | + | protein_coding |
| ENSG00000214199.3_4 | EEF1A1P12 | 0.83396567 | 1 | -0.187 | 0.357 | chr2 | 107313787 | 107315132 | - | processed_pseudogene |
| ENSG00000236439.4_5 | AC099336.2 | 0.79335183 | 1 | -0.187 | 0.014 | chr1 | 202440992 | 202441245 | - | processed_pseudogene |
| ENSG00000176390.12_5 | CRLF3 | 0.99996296 | 1 | -0.187 | 4.716 | chr17 | 29096406 | 29151710 | - | protein_coding |
| ENSG00000116032.5_3 | GRIN3B | 0.87206031 | 1 | -0.187 | 0.863 | chr19 | 1000418 | 1009731 | + | protein_coding |
| ENSG00000179038.8_5 | AP001885.1 | 0.75553697 | 1 | -0.187 | -0.739 | chr11 | 66963040 | 66964638 | - | transcribed_processed_pseudogene |
| ENSG00000244998.1_8 | AC100803.2 | 0.74429383 | 1 | -0.187 | -0.943 | chr8 | 142400039 | 142402674 | - | lncRNA |
| ENSG00000020426.11_4 | MNAT1 | 0.99985068 | 1 | -0.187 | 4.656 | chr14 | 61201460 | 61436683 | + | protein_coding |
| ENSG00000102178.13_5 | UBL4A | 0.99999238 | 1 | -0.187 | 5.892 | chrX | 153712056 | 153714954 | - | protein_coding |
| ENSG00000278206.1_7 | AL031320.2 | 0.92050233 | 1 | -0.187 | 1.091 | chr6 | 143806116 | 143828464 | + | lncRNA |
| ENSG00000106628.10_6 | POLD2 | 0.99999978 | 1 | -0.187 | 6.718 | chr7 | 44154280 | 44163957 | - | protein_coding |
| ENSG00000224418.1_6 | STK24-AS1 | 0.78422248 | 1 | -0.187 | -0.240 | chr13 | 99229498 | 99231084 | + | lncRNA |
| ENSG00000225193.5_4 | RPS12P26 | 0.82098982 | 1 | -0.187 | 0.259 | chr15 | 90749804 | 90750199 | + | transcribed_processed_pseudogene |
| ENSG00000106603.18_6 | COA1 | 0.99999851 | 1 | -0.187 | 5.808 | chr7 | 43648055 | 43769316 | - | protein_coding |
| ENSG00000138180.16_3 | CEP55 | 0.99999987 | 1 | -0.187 | 7.390 | chr10 | 95256369 | 95288849 | + | protein_coding |
| ENSG00000156471.13_3 | PTDSS1 | 0.99999977 | 1 | -0.188 | 6.930 | chr8 | 97274130 | 97349223 | + | protein_coding |
| ENSG00000151575.14_6 | TEX9 | 0.99848843 | 1 | -0.188 | 3.297 | chr15 | 56536207 | 56738195 | + | protein_coding |
| ENSG00000113658.18_5 | SMAD5 | 0.99999963 | 1 | -0.188 | 6.790 | chr5 | 135468534 | 135524435 | + | protein_coding |
| ENSG00000130725.8_6 | UBE2M | 0.99999927 | 1 | -0.188 | 6.686 | chr19 | 59067079 | 59070321 | - | protein_coding |
| ENSG00000147853.17_5 | AK3 | 0.99999845 | 1 | -0.188 | 6.150 | chr9 | 4709556 | 4742043 | - | protein_coding |
| ENSG00000189376.12_5 | C8orf76 | 0.92878745 | 1 | -0.188 | 1.230 | chr8 | 124232207 | 124253617 | - | protein_coding |
| ENSG00000159322.17_3 | ADPGK | 0.99999958 | 1 | -0.188 | 6.451 | chr15 | 73043710 | 73078187 | - | protein_coding |
| ENSG00000152705.8_4 | CATSPER3 | 0.75615922 | 1 | -0.188 | -0.771 | chr5 | 134303597 | 134347386 | + | protein_coding |
| ENSG00000230124.8_6 | ACBD6 | 0.99999466 | 1 | -0.188 | 5.714 | chr1 | 180238788 | 180472089 | - | protein_coding |
| ENSG00000115216.14_4 | NRBP1 | 0.99999987 | 1 | -0.188 | 7.155 | chr2 | 27650657 | 27665126 | + | protein_coding |
| ENSG00000124795.17_7 | DEK | 0.99999989 | 1 | -0.188 | 7.785 | chr6 | 18224091 | 18264779 | - | protein_coding |
| ENSG00000204209.13_7 | DAXX | 0.99749103 | 1 | -0.188 | 3.254 | chr6 | 33286335 | 33290793 | - | protein_coding |
| ENSG00000179010.14_2 | MRFAP1 | 0.99999977 | 1 | -0.188 | 7.022 | chr4 | 6641818 | 6644472 | + | protein_coding |
| ENSG00000128564.7_5 | VGF | 0.78026045 | 1 | -0.188 | -0.404 | chr7 | 100805790 | 100808850 | - | protein_coding |
| ENSG00000249673.7_6 | NOP14-AS1 | 0.99909241 | 1 | -0.189 | 3.405 | chr4 | 2936609 | 2963465 | + | lncRNA |
| ENSG00000010818.10_5 | HIVEP2 | 0.99964085 | 1 | -0.189 | 3.998 | chr6 | 143072606 | 143277835 | - | protein_coding |
| ENSG00000226564.1_4 | FTH1P20 | 0.78171195 | 1 | -0.189 | -0.087 | chr2 | 181737594 | 181738141 | - | processed_pseudogene |
| ENSG00000174953.14_8 | DHX36 | 0.99999903 | 1 | -0.189 | 6.014 | chr3 | 153990335 | 154042276 | - | protein_coding |
| ENSG00000151461.20_5 | UPF2 | 0.99999809 | 1 | -0.189 | 5.915 | chr10 | 11962021 | 12085169 | - | protein_coding |
| ENSG00000198056.15_8 | PRIM1 | 0.99996281 | 1 | -0.189 | 4.597 | chr12 | 57125080 | 57146158 | - | protein_coding |
| ENSG00000171311.13_5 | EXOSC1 | 0.99995936 | 1 | -0.189 | 4.817 | chr10 | 99195666 | 99205774 | - | protein_coding |
| ENSG00000117410.14_4 | ATP6V0B | 0.99993763 | 1 | -0.189 | 5.273 | chr1 | 44440159 | 44443967 | + | protein_coding |
| ENSG00000170385.10_4 | SLC30A1 | 0.99999942 | 1 | -0.189 | 6.734 | chr1 | 211744910 | 211752503 | - | protein_coding |
| ENSG00000077097.15_8 | TOP2B | 0.99999983 | 1 | -0.189 | 7.910 | chr3 | 25639396 | 25706398 | - | protein_coding |
| ENSG00000206567.10_8 | AC022007.1 | 0.93546158 | 1 | -0.189 | 1.392 | chr3 | 10045732 | 10052893 | - | lncRNA |
| ENSG00000163322.14_7 | ABRAXAS1 | 0.98998256 | 1 | -0.189 | 2.489 | chr4 | 84380670 | 84444501 | - | protein_coding |
| ENSG00000100426.7_4 | ZBED4 | 0.99997659 | 1 | -0.189 | 4.833 | chr22 | 50247492 | 50283728 | + | protein_coding |
| ENSG00000260404.3_5 | AC110079.1 | 0.94379711 | 1 | -0.190 | 1.376 | chr4 | 119512928 | 119554884 | + | transcribed_unprocessed_pseudogene |
| ENSG00000126249.8_5 | PDCD2L | 0.99737359 | 1 | -0.190 | 2.967 | chr19 | 34895304 | 34917073 | + | protein_coding |
| ENSG00000106290.15_7 | TAF6 | 0.99997886 | 1 | -0.190 | 4.985 | chr7 | 99704693 | 99717464 | - | protein_coding |
| ENSG00000162244.12_8 | RPL29 | 0.99999968 | 1 | -0.190 | 7.653 | chr3 | 52027538 | 52029911 | - | protein_coding |
| ENSG00000185875.13_4 | THNSL1 | 0.99876303 | 1 | -0.190 | 3.878 | chr10 | 25305541 | 25315593 | + | protein_coding |
| ENSG00000273367.1_7 | AL355472.3 | 0.74134253 | 1 | -0.190 | -0.877 | chr1 | 234507932 | 234508557 | + | lncRNA |
| ENSG00000109099.15_5 | PMP22 | 0.9999964 | 1 | -0.190 | 5.849 | chr17 | 15133094 | 15168643 | - | protein_coding |
| ENSG00000152464.15_5 | RPP38 | 0.99806717 | 1 | -0.190 | 3.158 | chr10 | 15139179 | 15181817 | + | protein_coding |
| ENSG00000154124.5_3 | OTULIN | 0.99999669 | 1 | -0.190 | 5.626 | chr5 | 14664773 | 14699959 | + | protein_coding |
| ENSG00000183840.7_4 | GPR39 | 0.99851667 | 1 | -0.190 | 3.343 | chr2 | 133174378 | 133404155 | + | protein_coding |
| ENSG00000106682.14_2 | EIF4H | 0.99999978 | 1 | -0.190 | 7.913 | chr7 | 73588575 | 73611431 | + | protein_coding |
| ENSG00000260948.1_7 | AL390195.3 | 0.76977369 | 1 | -0.190 | -0.334 | chr1 | 111973668 | 111975690 | - | lncRNA |

| | | | | | | | | | | |
|---|---|---|---|---|---|---|---|---|---|---|
| ENSG00000124172.10_5 | ATP5F1E | 0.99999154 | 1 | -0.190 | 6.940 | chr20 | 57600530 | 57607400 | - | protein_coding |
| ENSG00000164096.13_5 | C4orf3 | 0.99935934 | 1 | -0.190 | 4.519 | chr4 | 120217574 | 120225600 | - | protein_coding |
| ENSG00000136754.17_6 | ABI1 | 0.99999653 | 1 | -0.190 | 5.895 | chr10 | 27035522 | 27150016 | - | protein_coding |
| ENSG00000168566.13_3 | SNRNP48 | 0.99991951 | 1 | -0.190 | 4.667 | chr6 | 7590431 | 7612200 | + | protein_coding |
| ENSG00000134697.13_7 | GNL2 | 0.99999161 | 1 | -0.190 | 5.727 | chr1 | 38032417 | 38061538 | - | protein_coding |
| ENSG00000224358.1_7 | TMCO1-AS1 | 0.90568265 | 1 | -0.191 | 0.928 | chr1 | 165738166 | 165744413 | + | lncRNA |
| ENSG00000144681.10_4 | STAC | 0.98404546 | 1 | -0.191 | 2.158 | chr3 | 36421836 | 36589499 | + | protein_coding |
| ENSG00000275734.2_7 | AC010538.1 | 0.80581682 | 1 | -0.191 | 0.130 | chr16 | 89737976 | 89751712 | + | lncRNA |
| ENSG00000145781.9_6 | COMMD10 | 0.99997854 | 1 | -0.191 | 5.168 | chr5 | 115420713 | 115748459 | + | protein_coding |
| ENSG00000147592.9_4 | LACTB2 | 0.99987599 | 1 | -0.191 | 5.606 | chr8 | 71547553 | 71581420 | - | protein_coding |
| ENSG00000066422.5_4 | ZBTB11 | 0.99999772 | 1 | -0.191 | 5.832 | chr3 | 101367733 | 101395976 | - | protein_coding |
| ENSG00000146858.8_4 | ZC3HAV1L | 0.99969288 | 1 | -0.191 | 3.948 | chr7 | 138710452 | 138720788 | - | protein_coding |
| ENSG00000132680.11_6 | KHDC4 | 0.99999784 | 1 | -0.191 | 6.162 | chr1 | 155882836 | 155904204 | - | protein_coding |
| ENSG00000168036.18_8 | CTNNB1 | 0.99999994 | 1 | -0.191 | 8.428 | chr3 | 41236232 | 41301587 | + | protein_coding |
| ENSG00000196526.10_4 | AFAP1 | 0.99997634 | 1 | -0.191 | 4.930 | chr4 | 7760441 | 7941653 | - | protein_coding |
| ENSG00000140830.9_4 | TXNL4B | 0.99942722 | 1 | -0.191 | 3.962 | chr16 | 72078188 | 72128330 | - | protein_coding |
| ENSG00000093183.14_6 | SEC22C | 0.99999717 | 1 | -0.191 | 5.690 | chr3 | 42589461 | 42642572 | - | protein_coding |
| ENSG00000074800.16_8 | ENO1 | 0.99999985 | 1 | -0.191 | 10.992 | chr1 | 8921059 | 8939249 | - | protein_coding |
| ENSG00000214353.8_7 | VAC14-AS1 | 0.90734272 | 1 | -0.192 | 1.079 | chr16 | 70788938 | 70807154 | + | lncRNA |
| ENSG00000260898.6_6 | ADPGK-AS1 | 0.78021794 | 1 | -0.192 | -0.357 | chr15 | 73075176 | 73090540 | + | lncRNA |
| ENSG00000167515.10_2 | TRAPPC2L | 0.99997901 | 1 | -0.192 | 4.914 | chr16 | 88922628 | 88929094 | + | protein_coding |
| ENSG00000043093.15_8 | DCUN1D1 | 0.99998562 | 1 | -0.192 | 5.572 | chr3 | 182655862 | 182703741 | - | protein_coding |
| ENSG00000162928.9_3 | PEX13 | 0.99999553 | 1 | -0.192 | 5.615 | chr2 | 61244360 | 61279125 | + | protein_coding |
| ENSG00000170448.12_4 | NFXL1 | 0.99398826 | 1 | -0.192 | 3.034 | chr4 | 47849250 | 47916684 | - | protein_coding |
| ENSG00000228748.2_7 | AL450306.1 | 0.78051746 | 1 | -0.193 | -0.191 | chr10 | 79542624 | 79552934 | + | lncRNA |
| ENSG00000060656.20_4 | PTPRU | 0.99992662 | 1 | -0.193 | 4.448 | chr1 | 29563028 | 29653325 | + | protein_coding |
| ENSG00000171365.17_8 | CLCN5 | 0.99967229 | 1 | -0.193 | 3.929 | chrX | 49687206 | 49863892 | + | protein_coding |
| ENSG00000213341.11_5 | CHUK | 0.99999825 | 1 | -0.193 | 6.000 | chr10 | 101948057 | 101989353 | - | protein_coding |
| ENSG00000164941.14_6 | INTS8 | 0.9999983 | 1 | -0.193 | 6.204 | chr8 | 95825539 | 95893974 | + | protein_coding |
| ENSG00000071462.11_6 | BUD23 | 0.99999656 | 1 | -0.193 | 5.762 | chr7 | 73097355 | 73119491 | + | protein_coding |
| ENSG00000126215.14_5 | XRCC3 | 0.99963666 | 1 | -0.193 | 4.554 | chr14 | 104163946 | 104181841 | - | protein_coding |
| ENSG00000270055.1_7 | AC127502.2 | 0.98142991 | 1 | -0.193 | 2.082 | chr15 | 30780166 | 30782516 | + | lncRNA |
| ENSG00000214357.9_3 | NEURL1B | 0.9955567 | 1 | -0.193 | 3.027 | chr5 | 172068266 | 172118543 | + | protein_coding |
| ENSG00000134049.6_8 | IER3IP1 | 0.99980622 | 1 | -0.193 | 4.421 | chr18 | 44679205 | 44702735 | - | protein_coding |
| ENSG00000205937.12_8 | RNPS1 | 0.99999864 | 1 | -0.193 | 6.053 | chr16 | 2303117 | 2318398 | - | protein_coding |
| ENSG00000177600.9_4 | RPLP2 | 0.99999702 | 1 | -0.193 | 8.078 | chr11 | 809965 | 812880 | + | protein_coding |
| ENSG00000088986.11_5 | DYNLL1 | 0.99999953 | 1 | -0.193 | 7.405 | chr12 | 120907653 | 120936296 | + | protein_coding |
| ENSG00000222414.1 | RNU2-59P | 0.99843578 | 1 | -0.193 | 3.494 | chr10 | 103124602 | 103124792 | - | snRNA |
| ENSG00000279467.1_1 | AP000350.7 | 0.73301372 | 1 | -0.193 | -0.944 | chr22 | 24207435 | 24215464 | - | lncRNA |
| ENSG00000147202.18_4 | DIAPH2 | 0.9999863 | 1 | -0.193 | 5.339 | chrX | 95939662 | 96859996 | + | protein_coding |
| ENSG00000174840.9_3 | PDE12 | 0.99999307 | 1 | -0.194 | 5.317 | chr3 | 57542001 | 57552575 | + | protein_coding |
| ENSG00000257403.2_5 | AC008740.1 | 0.74036973 | 1 | -0.194 | -0.742 | chr16 | 21358460 | 21363459 | + | unprocessed_pseudo gene |
| ENSG00000127838.14_4 | PNKD | 0.99994609 | 1 | -0.194 | 5.048 | chr2 | 219135115 | 219211516 | + | protein_coding |
| ENSG00000127989.14_7 | MTERF1 | 0.99878459 | 1 | -0.194 | 4.001 | chr7 | 91321323 | 91510016 | - | protein_coding |
| ENSG00000113194.13_4 | FAF2 | 0.99999974 | 1 | -0.194 | 6.848 | chr5 | 175874629 | 175937075 | + | protein_coding |
| ENSG00000184857.8_4 | TMEM186 | 0.98188611 | 1 | -0.194 | 2.067 | chr16 | 8874241 | 8891499 | - | protein_coding |
| ENSG00000198604.11_6 | BAZ1A | 0.99999812 | 1 | -0.194 | 6.378 | chr14 | 35221937 | 35344853 | - | protein_coding |
| ENSG00000116903.7_3 | EXOC8 | 0.99998896 | 1 | -0.194 | 5.223 | chr1 | 231468499 | 231473598 | - | protein_coding |
| ENSG00000175931.13_7 | UBE2O | 0.9999973 | 1 | -0.194 | 6.085 | chr17 | 74385537 | 74449234 | - | protein_coding |
| ENSG00000181392.17_8 | SYNE4 | 0.85426059 | 1 | -0.194 | 0.804 | chr19 | 36494209 | 36499715 | - | protein_coding |
| ENSG00000110274.16_6 | CEP164 | 0.99997176 | 1 | -0.194 | 5.315 | chr11 | 117185273 | 117283982 | + | protein_coding |
| ENSG00000132600.16_8 | PRMT7 | 0.99994828 | 1 | -0.194 | 5.018 | chr16 | 68344877 | 68392466 | + | protein_coding |
| ENSG00000163900.11_4 | TMEM41A | 0.99960949 | 1 | -0.195 | 3.907 | chr3 | 185194284 | 185216845 | - | protein_coding |
| ENSG00000126602.11_6 | TRAP1 | 0.99998376 | 1 | -0.195 | 5.458 | chr16 | 3701640 | 3767554 | - | protein_coding |
| ENSG00000188211.9_3 | NCR3LG1 | 0.99601004 | 1 | -0.195 | 3.269 | chr11 | 17373347 | 17398888 | + | protein_coding |
| ENSG00000279561.2_6 | AL845472.2 | 0.91460461 | 1 | -0.195 | 1.172 | chr9 | 45352939 | 45357020 | - | TEC |
| ENSG00000142453.11_7 | CARM1 | 0.99994945 | 1 | -0.195 | 4.880 | chr19 | 10982189 | 11033746 | + | protein_coding |
| ENSG00000137992.14_3 | DBT | 0.99993044 | 1 | -0.195 | 4.597 | chr1 | 100652475 | 100715390 | - | protein_coding |
| ENSG00000130255.13_6 | RPL36 | 0.99999931 | 1 | -0.195 | 7.352 | chr19 | 5674958 | 5691886 | + | protein_coding |
| ENSG00000235109.7_7 | ZSCAN31 | 0.99121645 | 1 | -0.195 | 2.689 | chr6 | 28292470 | 28324048 | - | protein_coding |

| | | | | | | | | | | |
|---|---|---|---|---|---|---|---|---|---|---|
| ENSG00000207217.1 | SNORA42 | 0.75180601 | 1 | -0.195 | -0.260 | chr7 | 6056508 | 6056642 | + | snoRNA |
| ENSG00000148057.16_4 | IDNK | 0.98983257 | 1 | -0.195 | 2.602 | chr9 | 86237964 | 86259045 | + | protein_coding |
| ENSG00000241170.3_5 | AP001992.1 | 0.78107493 | 1 | -0.195 | -0.178 | chr11 | 74587331 | 74631067 | + | transcribed_processed_pseudogene |
| ENSG00000171757.17_5 | LRRC34 | 0.75000205 | 1 | -0.195 | -0.511 | chr3 | 169510791 | 169530774 | - | protein_coding |
| ENSG00000142621.19_4 | FHAD1 | 0.80723775 | 1 | -0.196 | 0.373 | chr1 | 15573768 | 15726779 | + | protein_coding |
| ENSG00000135002.12_5 | RFK | 0.99999883 | 1 | -0.196 | 6.801 | chr9 | 79000442 | 79009433 | - | protein_coding |
| ENSG00000106246.17_5 | PTCD1 | 0.97742243 | 1 | -0.196 | 2.128 | chr7 | 99014362 | 99063786 | - | protein_coding |
| ENSG00000165669.14_4 | FAM204A | 0.99998782 | 1 | -0.196 | 5.392 | chr10 | 120057437 | 120101840 | - | protein_coding |
| ENSG00000114853.14_3 | ZBTB47 | 0.99588959 | 1 | -0.196 | 3.178 | chr3 | 42695189 | 42709072 | + | protein_coding |
| ENSG00000100304.13_5 | TTLL12 | 0.99992286 | 1 | -0.196 | 5.128 | chr22 | 43562628 | 43583140 | - | protein_coding |
| ENSG00000143995.20_6 | MEIS1 | 0.98107028 | 1 | -0.196 | 2.224 | chr2 | 66660584 | 66801001 | + | protein_coding |
| ENSG00000115507.10_3 | OTX1 | 0.87982462 | 1 | -0.196 | 1.010 | chr2 | 63277192 | 63284971 | + | protein_coding |
| ENSG00000182240.16_4 | BACE2 | 0.99995328 | 1 | -0.196 | 4.704 | chr21 | 42539728 | 42654457 | + | protein_coding |
| ENSG00000117222.14_5 | RBBP5 | 0.99999332 | 1 | -0.196 | 5.431 | chr1 | 205055270 | 205091143 | - | protein_coding |
| ENSG00000162959.13_4 | MEMO1 | 0.98407804 | 1 | -0.196 | 1.706 | chr2 | 32090129 | 32236299 | - | protein_coding |
| ENSG00000197548.12_3 | ATG7 | 0.99992928 | 1 | -0.196 | 4.798 | chr3 | 11313995 | 11599139 | + | protein_coding |
| ENSG00000106266.11_5 | SNX8 | 0.99968604 | 1 | -0.196 | 5.039 | chr7 | 2291405 | 2393953 | - | protein_coding |
| ENSG00000118058.22_7 | KMT2A | 0.99998643 | 1 | -0.197 | 5.752 | chr11 | 118307205 | 118397547 | + | protein_coding |
| ENSG00000148175.13_5 | STOM | 0.99999964 | 1 | -0.197 | 6.901 | chr9 | 124101265 | 124132582 | - | protein_coding |
| ENSG00000241553.12_3 | ARPC4 | 0.99999585 | 1 | -0.197 | 5.617 | chr3 | 9834179 | 9849410 | + | protein_coding |
| ENSG00000273230.1_6 | AC102953.2 | 0.98995727 | 1 | -0.197 | 2.420 | chr7 | 1504133 | 1507158 | - | lncRNA |
| ENSG00000116754.13_2 | SRSF11 | 0.99999956 | 1 | -0.197 | 6.739 | chr1 | 70671365 | 70718735 | + | protein_coding |
| ENSG00000118680.13_4 | MYL12B | 0.99999954 | 1 | -0.197 | 7.727 | chr18 | 3261477 | 3278429 | + | protein_coding |
| ENSG00000271267.1_5 | AL353807.4 | 0.73830418 | 1 | -0.197 | -0.845 | chr1 | 155535841 | 155536097 | - | processed_pseudogene |
| ENSG00000254602.2_7 | AP000662.1 | 0.74513999 | 1 | -0.197 | -0.654 | chr11 | 57405497 | 57420263 | + | lncRNA |
| ENSG00000285103.2_3 | AL451123.1 | 0.74082685 | 1 | -0.197 | -0.681 | chr9 | 27529083 | 27543900 | + | lncRNA |
| ENSG00000100865.15_5 | CINP | 0.99998028 | 1 | -0.197 | 4.968 | chr14 | 102807439 | 102829253 | - | protein_coding |
| ENSG00000131323.15_5 | TRAF3 | 0.9999978 | 1 | -0.197 | 5.800 | chr14 | 103243786 | 103377837 | + | protein_coding |
| ENSG00000172613.8_5 | RAD9A | 0.99967495 | 1 | -0.197 | 4.626 | chr11 | 67085342 | 67165881 | + | protein_coding |
| ENSG00000273066.5_7 | AL355987.4 | 0.82244375 | 1 | -0.197 | 0.849 | chr9 | 139693675 | 139704494 | + | lncRNA |
| ENSG00000159479.17_4 | MED8 | 0.99931306 | 1 | -0.197 | 4.353 | chr1 | 43849588 | 43855479 | - | protein_coding |
| ENSG00000137492.8_5 | THAP12 | 0.99999815 | 1 | -0.197 | 6.631 | chr11 | 76061000 | 76092176 | - | protein_coding |
| ENSG00000087053.18_5 | MTMR2 | 0.99998476 | 1 | -0.198 | 5.363 | chr11 | 95566046 | 95658479 | - | protein_coding |
| ENSG00000258334.1_7 | AC125611.4 | 0.78942618 | 1 | -0.198 | -0.002 | chr12 | 49686414 | 49718359 | - | lncRNA |
| ENSG00000124549.14_4 | BTN2A3P | 0.97126124 | 1 | -0.198 | 1.834 | chr6 | 26421619 | 26432611 | + | transcribed_unprocessed_pseudogene |
| ENSG00000105887.11_5 | MTPN | 0.99999934 | 1 | -0.198 | 7.504 | chr7 | 135611508 | 135662107 | - | protein_coding |
| ENSG00000268093.1_8 | AC022154.1 | 0.76530901 | 1 | -0.198 | 0.018 | chr19 | 49122529 | 49127389 | - | lncRNA |
| ENSG00000121274.13_4 | TENT4B | 0.99998181 | 1 | -0.198 | 5.124 | chr16 | 50186822 | 50269221 | + | protein_coding |
| ENSG00000144224.17_3 | UBXN4 | 0.9999988 | 1 | -0.198 | 6.938 | chr2 | 136499304 | 136542626 | + | protein_coding |
| ENSG00000113593.12_3 | PPWD1 | 0.98173326 | 1 | -0.198 | 4.956 | chr5 | 64859063 | 64883376 | + | protein_coding |
| ENSG00000230897.1_4 | RPS18P12 | 0.84313505 | 1 | -0.198 | 0.652 | chr17 | 14608393 | 14608851 | - | processed_pseudogene |
| ENSG00000173210.19_3 | ABLIM3 | 0.99712958 | 1 | -0.198 | 3.155 | chr5 | 148521046 | 148640105 | + | protein_coding |
| ENSG00000153310.19_6 | FAM49B | 0.99999939 | 1 | -0.198 | 7.025 | chr8 | 130851839 | 131029375 | - | protein_coding |
| ENSG00000111711.10_5 | GOLT1B | 0.99999846 | 1 | -0.198 | 6.318 | chr12 | 21654715 | 21671342 | + | protein_coding |
| ENSG00000148362.11_6 | PAXX | 0.9998916 | 1 | -0.198 | 4.553 | chr9 | 139886874 | 139888436 | + | protein_coding |
| ENSG00000274737.1_7 | AC004466.2 | 0.83419465 | 1 | -0.198 | 0.365 | chr12 | 48211234 | 48211749 | - | lncRNA |
| ENSG00000144199.11_3 | FAHD2B | 0.99282958 | 1 | -0.198 | 2.909 | chr2 | 97749320 | 97760619 | - | protein_coding |
| ENSG00000185519.9_4 | FAM131C | 0.98046172 | 1 | -0.199 | 2.273 | chr1 | 16384264 | 16400146 | - | protein_coding |
| ENSG00000135048.14_8 | CEMIP2 | 0.99999986 | 1 | -0.199 | 7.633 | chr9 | 74298282 | 74431606 | - | protein_coding |
| ENSG00000163682.16_5 | RPL9 | 0.99574172 | 1 | -0.199 | 8.002 | chr4 | 39454141 | 39460569 | - | protein_coding |
| ENSG00000180211.5_5 | FO393411.1 | 0.82553352 | 1 | -0.199 | 0.410 | chr6 | 39926153 | 39926572 | - | processed_pseudogene |
| ENSG00000176087.15_4 | SLC35A4 | 0.99999712 | 1 | -0.199 | 5.812 | chr5 | 139944413 | 139948685 | + | protein_coding |
| ENSG00000105447.13_4 | GRWD1 | 0.99992249 | 1 | -0.199 | 6.470 | chr19 | 48949098 | 48960279 | + | protein_coding |
| ENSG00000054523.17_4 | KIF1B | 0.99999881 | 1 | -0.199 | 6.085 | chr1 | 10270863 | 10441661 | + | protein_coding |
| ENSG00000035928.16_5 | RFC1 | 0.99999887 | 1 | -0.199 | 6.378 | chr4 | 39289076 | 39367995 | - | protein_coding |
| ENSG00000116747.12_5 | RO60 | 0.99999814 | 1 | -0.199 | 7.136 | chr1 | 193028552 | 193060907 | + | protein_coding |
| ENSG00000177469.13_7 | CAVIN1 | 0.99951373 | 1 | -0.199 | 8.874 | chr17 | 40554467 | 40575274 | - | protein_coding |
| ENSG00000108175.17_3 | ZMIZ1 | 0.99999581 | 1 | -0.199 | 6.135 | chr10 | 80828723 | 81076285 | + | protein_coding |

| | | | | | | | | | | |
|---|---|---|---|---|---|---|---|---|---|---|
| ENSG00000179091.5_3 | CYC1 | 0.99999697 | 1 | -0.199 | 7.006 | chr8 | 145149942 | 145152428 | + | protein_coding |
| ENSG00000157514.16_7 | TSC22D3 | 0.99789414 | 1 | -0.199 | 3.365 | chrX | 106956451 | 107020572 | - | protein_coding |
| ENSG00000234782.3_4 | TPT1P9 | 0.8442969 | 1 | -0.200 | 0.453 | chr9 | 120845147 | 120845660 | - | processed_pseudogene |
| ENSG00000224861.1_5 | YBX1P1 | 0.97182693 | 1 | -0.200 | 1.963 | chr14 | 66479548 | 66480507 | - | processed_pseudogene |
| ENSG00000164587.13_8 | RPS14 | 0.99999985 | 1 | -0.200 | 8.833 | chr5 | 149822198 | 149829302 | - | protein_coding |
| ENSG00000165055.15_5 | METTL2B | 0.99997516 | 1 | -0.200 | 5.214 | chr7 | 128116783 | 128146656 | + | protein_coding |
| ENSG00000184319.16_5 | RPL23AP82 | 0.98540087 | 1 | -0.200 | 2.482 | chr22 | 51195376 | 51239737 | + | transcribed_unprocessed_pseudogene |
| ENSG00000141965.5_3 | FEM1A | 0.94120664 | 1 | -0.200 | 1.879 | chr19 | 4791746 | 4801285 | + | protein_coding |
| ENSG00000226360.5_4 | RPL10AP6 | 0.94653131 | 1 | -0.200 | 1.524 | chr3 | 61728129 | 61728780 | - | processed_pseudogene |
| ENSG00000139269.3_3 | INHBE | 0.72370284 | 1 | -0.200 | -1.066 | chr12 | 57846106 | 57853063 | + | protein_coding |
| ENSG00000188352.12_4 | FOCAD | 0.99999944 | 1 | -0.200 | 6.583 | chr9 | 20658308 | 20995954 | + | protein_coding |
| ENSG00000173214.7_5 | MFSD4B | 0.99987276 | 1 | -0.200 | 4.279 | chr6 | 111580530 | 111766557 | + | protein_coding |
| ENSG00000053770.12_4 | AP5M1 | 0.99999134 | 1 | -0.200 | 5.740 | chr14 | 57735642 | 57765460 | + | protein_coding |
| ENSG00000060749.15_4 | QSER1 | 0.99999924 | 1 | -0.200 | 6.789 | chr11 | 32914366 | 33014862 | + | protein_coding |
| ENSG00000279148.1_5 | AC126474.1 | 0.84282556 | 1 | -0.200 | 0.231 | chr12 | 100420026 | 100421318 | - | TEC |
| ENSG00000224831.3_5 | TMEM183B | 0.84428036 | 1 | -0.200 | 0.199 | chr3 | 149699968 | 149701095 | - | processed_pseudogene |
| ENSG00000102781.14_5 | KATNAL1 | 0.99997804 | 1 | -0.200 | 4.994 | chr13 | 30776767 | 30881688 | - | protein_coding |
| ENSG00000083642.19_3 | PDS5B | 0.9999644 | 1 | -0.200 | 5.089 | chr13 | 33160589 | 33352157 | + | protein_coding |
| ENSG00000240204.3_3 | SMKR1 | 0.9645354 | 1 | -0.201 | 2.022 | chr7 | 129142372 | 129152759 | + | protein_coding |
| ENSG00000253854.1_7 | AC010834.3 | 0.77534755 | 1 | -0.201 | -0.092 | chr8 | 94731802 | 94733395 | + | lncRNA |
| ENSG00000198176.13_5 | TFDP1 | 0.99999797 | 1 | -0.201 | 6.243 | chr13 | 114239036 | 114295788 | + | protein_coding |
| ENSG00000133028.12_5 | SCO1 | 0.99999879 | 1 | -0.201 | 6.492 | chr17 | 10575791 | 10601692 | - | protein_coding |
| ENSG00000149262.17_5 | INTS4 | 0.99994837 | 1 | -0.201 | 4.794 | chr11 | 77589766 | 77705717 | - | protein_coding |
| ENSG00000104343.21_7 | UBE2W | 0.99986747 | 1 | -0.201 | 4.987 | chr8 | 74692332 | 74791145 | - | protein_coding |
| ENSG00000165684.4_4 | SNAPC4 | 0.99974528 | 1 | -0.201 | 4.577 | chr9 | 139270029 | 139294620 | - | protein_coding |
| ENSG00000143315.8_5 | PIGM | 0.99988092 | 1 | -0.201 | 4.597 | chr1 | 159994743 | 160001780 | - | protein_coding |
| ENSG00000255248.9_7 | MIR100HG | 0.99877181 | 1 | -0.201 | 3.896 | chr11 | 121899033 | 122427429 | - | lncRNA |
| ENSG00000166352.16_6 | C11orf74 | 0.99741499 | 1 | -0.201 | 3.365 | chr11 | 36616043 | 36680840 | + | protein_coding |
| ENSG00000146556.14_2 | WASH2P | 0.72868062 | 1 | -0.201 | -0.879 | chr2 | 114346127 | 114356620 | + | transcribed_unprocessed_pseudogene |
| ENSG00000196312.14_6 | MFSD14C | 0.99993157 | 1 | -0.201 | 4.787 | chr9 | 99649655 | 99775990 | - | protein_coding |
| ENSG00000226029.1_5 | LINC01772 | 0.7737093 | 1 | -0.201 | -0.159 | chr1 | 16787443 | 16794976 | + | lncRNA |
| ENSG00000066583.12_5 | ISOC1 | 0.99985702 | 1 | -0.201 | 4.210 | chr5 | 128430442 | 128449721 | + | protein_coding |
| ENSG00000164080.13_3 | RAD54L2 | 0.99997587 | 1 | -0.201 | 5.248 | chr3 | 51575160 | 51702683 | + | protein_coding |
| ENSG00000115904.13_5 | SOS1 | 0.99999626 | 1 | -0.201 | 6.000 | chr2 | 39208690 | 39351486 | - | protein_coding |
| ENSG00000160917.15_5 | CPSF4 | 0.99996549 | 1 | -0.201 | 4.827 | chr7 | 99036545 | 99054996 | + | protein_coding |
| ENSG00000229589.1_7 | ACVR2B-AS1 | 0.81746172 | 1 | -0.202 | 0.312 | chr3 | 38492518 | 38496311 | - | lncRNA |
| ENSG00000109270.13_4 | LAMTOR3 | 0.99914592 | 1 | -0.202 | 4.090 | chr4 | 100799493 | 100815585 | - | protein_coding |
| ENSG00000257596.1_8 | SCAT2 | 0.81909233 | 1 | -0.202 | 0.290 | chr12 | 54656399 | 54672847 | + | lncRNA |
| ENSG00000100099.20_5 | HPS4 | 0.99997731 | 1 | -0.202 | 4.993 | chr22 | 26839389 | 26879803 | - | protein_coding |
| ENSG00000109881.17_6 | CCDC34 | 0.99993852 | 1 | -0.202 | 5.054 | chr11 | 27352374 | 27384781 | - | protein_coding |
| ENSG00000162105.19_8 | SHANK2 | 0.99994598 | 1 | -0.202 | 5.103 | chr11 | 70313961 | 70963623 | - | protein_coding |
| ENSG00000143499.14_3 | SMYD2 | 0.99997001 | 1 | -0.202 | 5.107 | chr1 | 214454445 | 214510474 | + | protein_coding |
| ENSG00000131508.16_4 | UBE2D2 | 0.99999908 | 1 | -0.202 | 6.492 | chr5 | 138906016 | 139008019 | + | protein_coding |
| ENSG00000188312.14_3 | CENPP | 0.99231137 | 1 | -0.202 | 2.785 | chr9 | 95088235 | 95382811 | + | protein_coding |
| ENSG00000229789.1 | RP11-391M20.1 | 0.86650745 | 1 | -0.202 | 0.437 | chr9 | 68726629 | 68745997 | + | pseudogene |
| ENSG00000262420.3_6 | AC007613.1 | 0.73626915 | 1 | -0.202 | -0.628 | chr16 | 11835766 | 11838362 | + | lncRNA |
| ENSG00000144635.9_4 | DYNC1LI1 | 0.9999969 | 1 | -0.202 | 5.964 | chr3 | 32567466 | 32612350 | - | protein_coding |
| ENSG00000136319.12_5 | TTC5 | 0.99995658 | 1 | -0.202 | 4.696 | chr14 | 20724717 | 20774119 | - | protein_coding |
| ENSG00000168090.10_3 | COPS6 | 0.99997986 | 1 | -0.202 | 5.062 | chr7 | 99686592 | 99689810 | + | protein_coding |
| ENSG00000170835.15_3 | CEL | 0.75843214 | 1 | -0.202 | -0.031 | chr9 | 135937365 | 135947250 | + | protein_coding |
| ENSG00000086848.15_6 | ALG9 | 0.98858445 | 1 | -0.202 | 2.496 | chr11 | 111652919 | 111742305 | - | protein_coding |
| ENSG00000279452.1_5 | AC006277.1 | 0.82460766 | 1 | -0.202 | 0.557 | chr19 | 2949033 | 2950642 | + | TEC |
| ENSG00000270157.1_8 | AC004918.5 | 0.97430163 | 1 | -0.202 | 2.550 | chr7 | 141362722 | 141363646 | - | lncRNA |
| ENSG00000106615.10_6 | RHEB | 0.99999766 | 1 | -0.202 | 6.021 | chr7 | 151163098 | 151217206 | - | protein_coding |
| ENSG00000185453.13_5 | ZSWIM9 | 0.91142561 | 1 | -0.202 | 1.291 | chr19 | 48673937 | 48700877 | + | protein_coding |
| ENSG00000227388.3_7 | AL133410.1 | 0.95558158 | 1 | -0.203 | 1.889 | chr9 | 35772160 | 35790429 | + | lncRNA |
| ENSG00000159588.15_5 | CCDC17 | 0.87851076 | 1 | -0.203 | 0.799 | chr1 | 46085716 | 46089729 | - | protein_coding |

| | | | | | | | | | | |
|---|---|---|---|---|---|---|---|---|---|---|
| ENSG00000263089.1_6 | AC007114.2 | 0.88306116 | 1 | -0.203 | 0.819 | chr17 | 55169506 | 55173786 | - | lncRNA |
| ENSG00000204371.11_5 | EHMT2 | 0.99931996 | 1 | -0.203 | 4.407 | chr6 | 31847536 | 31865464 | - | protein_coding |
| ENSG00000188603.19_7 | CLN3 | 0.81911716 | 1 | -0.203 | 0.529 | chr16 | 28485432 | 28506896 | - | protein_coding |
| ENSG00000163933.10_7 | RFT1 | 0.99963473 | 1 | -0.203 | 4.278 | chr3 | 53122499 | 53164469 | - | protein_coding |
| ENSG00000236829.9_5 | Z97634.1 | 0.77029049 | 1 | -0.203 | -0.054 | chr16 | 432097 | 442960 | + | transcribed_processed_pseudogene |
| ENSG00000228842.3_5 | PCDH9-AS2 | 0.71991045 | 1 | -0.204 | -1.029 | chr13 | 67399301 | 67489163 | + | lncRNA |
| ENSG00000170191.5_4 | NANP | 0.9998985 | 1 | -0.204 | 4.575 | chr20 | 25593571 | 25604650 | - | protein_coding |
| ENSG00000123066.8_7 | MED13L | 0.99999973 | 1 | -0.204 | 7.952 | chr12 | 116391677 | 116715524 | - | protein_coding |
| ENSG00000087995.15_2 | METTL2A | 0.99998061 | 1 | -0.204 | 5.049 | chr17 | 60501228 | 60528183 | + | protein_coding |
| ENSG00000099284.14_4 | MACROH2A2 | 0.99987608 | 1 | -0.204 | 4.472 | chr10 | 71812602 | 71872038 | + | protein_coding |
| ENSG00000122591.12_6 | FAM126A | 0.99997942 | 1 | -0.204 | 5.152 | chr7 | 22928990 | 23053749 | - | protein_coding |
| ENSG00000182489.9_4 | XKRX | 0.72641431 | 1 | -0.205 | -0.784 | chrX | 100168434 | 100184422 | - | protein_coding |
| ENSG00000187189.11_4 | TSPYL4 | 0.99236922 | 1 | -0.205 | 2.804 | chr6 | 116571127 | 116575238 | - | protein_coding |
| ENSG00000287408.1_2 | AC027811.1 | 0.73850386 | 1 | -0.205 | -0.574 | chr15 | 79623933 | 79629619 | - | lncRNA |
| ENSG00000173542.8_3 | MOB1B | 0.99957521 | 1 | -0.205 | 4.571 | chr4 | 71768043 | 71888166 | + | protein_coding |
| ENSG00000260293.2_7 | AC106820.4 | 0.86175939 | 1 | -0.205 | 0.705 | chr16 | 2526559 | 2532174 | + | lncRNA |
| ENSG00000173209.23_5 | AHSA2P | 0.99999458 | 1 | -0.205 | 6.116 | chr2 | 61404553 | 61418338 | + | transcribed_unitary_pseudogene |
| ENSG00000135749.19_10 | PCNX2 | 0.9999867 | 1 | -0.205 | 5.159 | chr1 | 233119181 | 233431471 | - | protein_coding |
| ENSG00000096093.16_4 | EFHC1 | 0.99995585 | 1 | -0.205 | 4.712 | chr6 | 52226921 | 52394684 | + | protein_coding |
| ENSG00000145191.15_7 | EIF2B5 | 0.99999171 | 1 | -0.205 | 5.463 | chr3 | 183852826 | 183863915 | + | protein_coding |
| ENSG00000187634.12_8 | SAMD11 | 0.97640941 | 1 | -0.205 | 2.984 | chr1 | 859308 | 879961 | + | protein_coding |
| ENSG00000279806.1_5 | AC018629.1 | 0.99611481 | 1 | -0.205 | 3.318 | chr17 | 38629806 | 38631541 | - | TEC |
| ENSG00000003987.14_4 | MTMR7 | 0.85724554 | 1 | -0.205 | 1.073 | chr8 | 17154303 | 17271037 | - | protein_coding |
| ENSG00000121644.19_3 | DESI2 | 0.99999868 | 1 | -0.205 | 6.399 | chr1 | 244816405 | 244872335 | + | protein_coding |
| ENSG00000213551.7_6 | DNAJC9 | 0.99989572 | 1 | -0.205 | 4.757 | chr10 | 74943120 | 75007013 | - | protein_coding |
| ENSG00000018189.13_4 | RUFY3 | 0.99982317 | 1 | -0.205 | 4.271 | chr4 | 71569921 | 71674336 | + | protein_coding |
| ENSG00000214944.9_4 | ARHGEF28 | 0.99999945 | 1 | -0.206 | 6.789 | chr5 | 72921983 | 73237818 | + | protein_coding |
| ENSG00000103429.11_3 | BFAR | 0.9999986 | 1 | -0.206 | 6.366 | chr16 | 14726788 | 14763093 | + | protein_coding |
| ENSG00000138111.14_4 | MFSD13A | 0.98669407 | 1 | -0.206 | 2.355 | chr10 | 104221152 | 104236802 | + | protein_coding |
| ENSG00000134970.14_6 | TMED7 | 0.99999915 | 1 | -0.206 | 7.136 | chr5 | 114948907 | 114968689 | - | protein_coding |
| ENSG00000184216.14_9 | IRAK1 | 0.99936995 | 1 | -0.206 | 7.828 | chrX | 153275957 | 153285353 | - | protein_coding |
| ENSG00000161664.7_3 | ASB16 | 0.9275074 | 1 | -0.206 | 1.393 | chr17 | 42247815 | 42256452 | + | protein_coding |
| ENSG00000272604.1_6 | AC073073.2 | 0.86926707 | 1 | -0.206 | 0.994 | chr7 | 105211530 | 105214107 | + | lncRNA |
| ENSG00000123983.14_3 | ACSL3 | 0.99999986 | 1 | -0.206 | 8.478 | chr2 | 223725660 | 223809357 | + | protein_coding |
| ENSG00000141076.18_6 | UTP4 | 0.99995438 | 1 | -0.206 | 4.910 | chr16 | 69165194 | 69265033 | + | protein_coding |
| ENSG00000237190.4_4 | CDKN2AIPNL | 0.99996321 | 1 | -0.206 | 5.078 | chr5 | 133737756 | 133747572 | - | protein_coding |
| ENSG00000239569.3_8 | KMT2E-AS1 | 0.97224088 | 1 | -0.206 | 1.552 | chr7 | 104653872 | 104654768 | - | lncRNA |
| ENSG00000082996.19_6 | RNF13 | 0.99998954 | 1 | -0.206 | 5.982 | chr3 | 149530495 | 149679926 | + | protein_coding |
| ENSG00000141854.9_5 | MISP3 | 0.95333211 | 1 | -0.206 | 1.232 | chr19 | 14183348 | 14185874 | + | protein_coding |
| ENSG00000277399.4_3 | GPR179 | 0.72990042 | 1 | -0.207 | -0.486 | chr17 | 36481413 | 36499730 | - | protein_coding |
| ENSG00000131263.13_4 | RLIM | 0.99999956 | 1 | -0.207 | 7.426 | chrX | 73802811 | 73834459 | - | protein_coding |
| ENSG00000122026.10_2 | RPL21 | 0.99999646 | 1 | -0.207 | 7.791 | chr13 | 27825446 | 27830828 | + | protein_coding |
| ENSG00000164758.7_3 | MED30 | 0.98103365 | 1 | -0.207 | 2.326 | chr8 | 118532952 | 118552501 | + | protein_coding |
| ENSG00000178952.11_5 | TUFM | 0.99999925 | 1 | -0.207 | 7.313 | chr16 | 28853732 | 28857669 | - | protein_coding |
| ENSG00000135018.14_5 | UBQLN1 | 0.99999978 | 1 | -0.207 | 7.632 | chr9 | 86274883 | 86322873 | - | protein_coding |
| ENSG00000241313.3_6 | WWTR1-AS1 | 0.78212633 | 1 | -0.207 | 0.383 | chr3 | 149374786 | 149379151 | + | lncRNA |
| ENSG00000134255.14_3 | CEPT1 | 0.99944985 | 1 | -0.207 | 3.977 | chr1 | 111682249 | 111727726 | + | protein_coding |
| ENSG00000129562.11_4 | DAD1 | 0.99999104 | 1 | -0.207 | 6.444 | chr14 | 23033807 | 23058130 | - | protein_coding |
| ENSG00000113068.10_4 | PFDN1 | 0.9999879 | 1 | -0.207 | 5.293 | chr5 | 139624620 | 139682698 | - | protein_coding |
| ENSG00000148835.11_3 | TAF5 | 0.9978436 | 1 | -0.207 | 3.502 | chr10 | 105127733 | 105148822 | + | protein_coding |
| ENSG00000174307.7_4 | PHLDA3 | 0.99994931 | 1 | -0.207 | 5.106 | chr1 | 201433406 | 201438365 | - | protein_coding |
| ENSG00000145912.8_3 | NHP2 | 0.99997713 | 1 | -0.207 | 5.301 | chr5 | 177576461 | 177580968 | - | protein_coding |
| ENSG00000148606.13_5 | POLR3A | 0.99993636 | 1 | -0.207 | 5.002 | chr10 | 79729009 | 79789273 | - | protein_coding |
| ENSG00000228782.7_5 | MRPL45P2 | 0.98033791 | 1 | -0.208 | 2.274 | chr17 | 45527934 | 45569858 | - | transcribed_unprocessed_pseudogene |
| ENSG00000186806.5_4 | VSIG10L | 0.99730009 | 1 | -0.208 | 3.578 | chr19 | 51834790 | 51845378 | - | protein_coding |
| ENSG00000285410.1_3 | GABPB1-IT1 | 0.99628803 | 1 | -0.208 | 3.405 | chr15 | 50641133 | 50647058 | - | lncRNA |

| | | | | | | | | | | |
|---|---|---|---|---|---|---|---|---|---|---|
| ENSG00000285756.2_3 | BX890604.2 | 0.99968236 | 1 | -0.208 | 4.087 | chrX | 3735569 | 3855896 | - | lncRNA |
| ENSG00000004455.17_4 | AK2 | 0.9999993 | 1 | -0.208 | 6.907 | chr1 | 33473541 | 33546597 | - | protein_coding |
| ENSG00000188706.13_5 | ZDHHC9 | 0.99951859 | 1 | -0.208 | 4.333 | chrX | 128937264 | 128977885 | - | protein_coding |
| ENSG00000008256.16_5 | CYTH3 | 0.99999682 | 1 | -0.208 | 5.869 | chr7 | 6201407 | 6312275 | - | protein_coding |
| ENSG00000170917.14_4 | NUDT6 | 0.98934913 | 1 | -0.208 | 2.433 | chr4 | 123809852 | 123844123 | - | protein_coding |
| ENSG00000116478.12_5 | HDAC1 | 0.99999844 | 1 | -0.208 | 6.157 | chr1 | 32757684 | 32799236 | + | protein_coding |
| ENSG00000142661.19_4 | MYOM3 | 0.99961287 | 1 | -0.208 | 4.239 | chr1 | 24382525 | 24438625 | - | protein_coding |
| ENSG00000108424.10_5 | KPNB1 | 0.9999999 | 1 | -0.208 | 9.334 | chr17 | 45726842 | 45762871 | + | protein_coding |
| ENSG00000144848.11_4 | ATG3 | 0.99991762 | 1 | -0.208 | 5.585 | chr3 | 112251357 | 112280893 | - | protein_coding |
| ENSG00000136449.15_6 | MYCBPAP | 0.97006923 | 1 | -0.208 | 1.942 | chr17 | 48585745 | 48608862 | + | protein_coding |
| ENSG00000198624.13_4 | CCDC69 | 0.98510596 | 1 | -0.208 | 2.324 | chr5 | 150560613 | 150603654 | - | protein_coding |
| ENSG00000269044.2_7 | AC024075.2 | 0.87385024 | 1 | -0.208 | 0.941 | chr19 | 16744608 | 16746080 | - | lncRNA |
| ENSG00000180096.12_6 | SEPTIN1 | 0.9749024 | 1 | -0.208 | 2.204 | chr16 | 30389454 | 30407312 | - | protein_coding |
| ENSG00000243701.7_6 | DUBR | 0.99740439 | 1 | -0.209 | 3.148 | chr3 | 106939591 | 107067311 | + | lncRNA |
| ENSG00000136159.4_3 | NUDT15 | 0.9999323 | 1 | -0.209 | 4.604 | chr13 | 48611862 | 48621357 | + | protein_coding |
| ENSG00000155906.19_7 | RMND1 | 0.99742139 | 1 | -0.209 | 3.561 | chr6 | 151725897 | 151773293 | - | protein_coding |
| ENSG00000119042.17_4 | SATB2 | 0.99617488 | 1 | -0.209 | 3.050 | chr2 | 200134223 | 200335989 | - | protein_coding |
| ENSG00000146223.15_4 | RPL7L1 | 0.99999956 | 1 | -0.209 | 6.987 | chr6 | 42847354 | 42857663 | + | protein_coding |
| ENSG00000143799.13_4 | PARP1 | 0.99999958 | 1 | -0.209 | 7.230 | chr1 | 226548392 | 226595794 | - | protein_coding |
| ENSG00000196715.7_6 | VKORC1L1 | 0.99999231 | 1 | -0.209 | 5.743 | chr7 | 65338061 | 65424550 | + | protein_coding |
| ENSG00000185716.12_4 | MOSMO | 0.99957425 | 1 | -0.209 | 4.032 | chr16 | 22018959 | 22098855 | + | protein_coding |
| ENSG00000269556.8_10 | TMEM185A | 0.97691974 | 1 | -0.209 | 2.244 | chrX | 148678216 | 148713568 | - | protein_coding |
| ENSG00000175643.10_4 | RMI2 | 0.99996116 | 1 | -0.209 | 4.853 | chr16 | 11343476 | 11475519 | + | protein_coding |
| ENSG00000152284.5_4 | TCF7L1 | 0.76542068 | 1 | -0.209 | -0.271 | chr2 | 85360515 | 85537510 | + | protein_coding |
| ENSG00000172586.8_3 | CHCHD1 | 0.99994761 | 1 | -0.209 | 4.918 | chr10 | 75541805 | 75543410 | + | protein_coding |
| ENSG00000114626.18_5 | ABTB1 | 0.98648992 | 1 | -0.209 | 2.816 | chr3 | 127391778 | 127399769 | + | protein_coding |
| ENSG00000235499.1_6 | AC073046.1 | 0.75736819 | 1 | -0.209 | -0.249 | chr2 | 74212259 | 74213470 | + | lncRNA |
| ENSG00000185721.13_4 | DRG1 | 0.99997419 | 1 | -0.209 | 5.130 | chr22 | 31795590 | 31926620 | + | protein_coding |
| ENSG00000099949.21_6 | LZTR1 | 0.99828984 | 1 | -0.209 | 3.645 | chr22 | 21336558 | 21353321 | + | protein_coding |
| ENSG00000175567.10_8 | UCP2 | 0.99881536 | 1 | -0.210 | 3.630 | chr11 | 73685717 | 73694291 | - | protein_coding |
| ENSG00000164944.12_7 | VIRMA | 0.99999869 | 1 | -0.210 | 6.433 | chr8 | 95499917 | 95565757 | - | protein_coding |
| ENSG00000275216.2_6 | AL161431.1 | 0.98354572 | 1 | -0.210 | 2.412 | chr13 | 109921982 | 109930860 | + | lncRNA |
| ENSG00000138246.17_7 | DNAJC13 | 0.99999937 | 1 | -0.210 | 6.870 | chr3 | 132136346 | 132257876 | + | protein_coding |
| ENSG00000078747.15_5 | ITCH | 0.99999826 | 1 | -0.210 | 6.895 | chr20 | 32951047 | 33128552 | + | protein_coding |
| ENSG00000164086.10_5 | DUSP7 | 0.98263169 | 1 | -0.210 | 2.450 | chr3 | 52082935 | 52090587 | - | protein_coding |
| ENSG00000184009.12_8 | ACTG1 | 0.99999934 | 1 | -0.210 | 11.438 | chr17 | 79476997 | 79490873 | - | protein_coding |
| ENSG00000177054.14_3 | ZDHHC13 | 0.99997481 | 1 | -0.210 | 5.207 | chr11 | 19138646 | 19197969 | + | protein_coding |
| ENSG00000212124.2_4 | TAS2R19 | 0.76743245 | 1 | -0.210 | -0.005 | chr12 | 11174218 | 11175219 | - | protein_coding |
| ENSG00000197345.13_4 | MRPL21 | 0.99995763 | 1 | -0.210 | 5.121 | chr11 | 68658744 | 68671303 | - | protein_coding |
| ENSG00000184220.12_4 | CMSS1 | 0.99997322 | 1 | -0.210 | 5.026 | chr3 | 99536681 | 99900576 | + | protein_coding |
| ENSG00000198673.10_5 | TAFA2 | 0.82400632 | 1 | -0.210 | 0.389 | chr12 | 62102040 | 62672931 | - | protein_coding |
| ENSG00000256053.7_5 | COA8 | 0.99804402 | 1 | -0.210 | 3.384 | chr14 | 104029299 | 104073860 | + | protein_coding |
| ENSG00000163866.9_5 | SMIM12 | 0.99991523 | 1 | -0.210 | 4.599 | chr1 | 35178338 | 35325356 | - | protein_coding |
| ENSG00000175581.14_4 | MRPL48 | 0.9999111 | 1 | -0.211 | 4.557 | chr11 | 73498917 | 73576178 | + | protein_coding |
| ENSG00000073417.15_6 | PDE8A | 0.99999969 | 1 | -0.211 | 7.335 | chr15 | 85523671 | 85682376 | + | protein_coding |
| ENSG00000180957.18_7 | PITPNB | 0.98969163 | 1 | -0.211 | 2.629 | chr22 | 28247657 | 28316122 | - | protein_coding |
| ENSG00000077514.9_6 | POLD3 | 0.99999388 | 1 | -0.211 | 5.594 | chr11 | 74204896 | 74380162 | + | protein_coding |
| ENSG00000265666.1_7 | RARA-AS1 | 0.83408918 | 1 | -0.211 | 0.719 | chr17 | 38497119 | 38499388 | - | lncRNA |
| ENSG00000106638.17_6 | TBL2 | 0.9999754 | 1 | -0.211 | 5.526 | chr7 | 72981867 | 72993121 | - | protein_coding |
| ENSG00000196743.9_4 | GM2A | 0.99999098 | 1 | -0.211 | 5.359 | chr5 | 150591711 | 150650001 | + | protein_coding |
| ENSG00000115042.10_5 | FAHD2A | 0.99994006 | 1 | -0.211 | 4.647 | chr2 | 96068456 | 96082364 | + | protein_coding |
| ENSG00000186310.10_5 | NAP1L3 | 0.71536453 | 1 | -0.211 | -0.783 | chrX | 92925929 | 92928577 | - | protein_coding |
| ENSG00000280924.1_3 | LINC00628 | 0.7823791 | 1 | -0.211 | 0.085 | chr1 | 204337559 | 204338847 | - | lncRNA |
| ENSG00000177602.5_6 | HASPIN | 0.99989692 | 1 | -0.211 | 4.611 | chr17 | 3627197 | 3629993 | + | protein_coding |
| ENSG00000161912.18_4 | ADCY10P1 | 0.79293524 | 1 | -0.211 | 0.015 | chr6 | 41068761 | 41108573 | + | transcribed_unprocessed_pseudogene |
| ENSG00000160179.18_6 | ABCG1 | 0.99907192 | 1 | -0.211 | 3.598 | chr21 | 43619799 | 43717354 | + | protein_coding |
| ENSG00000070761.8_4 | CFAP20 | 0.99984807 | 1 | -0.211 | 4.667 | chr16 | 58147496 | 58163285 | - | protein_coding |
| ENSG00000215251.4_5 | FASTKD5 | 0.99999424 | 1 | -0.211 | 5.611 | chr20 | 3127165 | 3140511 | - | protein_coding |
| ENSG00000174775.17_4 | HRAS | 0.99982704 | 1 | -0.211 | 4.963 | chr11 | 532242 | 537287 | - | protein_coding |
| ENSG00000134759.14_3 | ELP2 | 0.99999143 | 1 | -0.211 | 5.670 | chr18 | 33709407 | 33760520 | + | protein_coding |

| | | | | | | | | | | |
|---|---|---|---|---|---|---|---|---|---|---|
| ENSG00000232119.8_4 | MCTS1 | 0.99815127 | 1 | -0.211 | 5.884 | chrX | 119727865 | 119755014 | + | protein_coding |
| ENSG00000228106.6_8 | AL392172.1 | 0.92757972 | 1 | -0.211 | 1.360 | chr1 | 222988364 | 223010726 | + | lncRNA |
| ENSG00000253716.6_7 | MINCR | 0.98991539 | 1 | -0.211 | 2.672 | chr8 | 144362331 | 144363860 | - | lncRNA |
| ENSG00000169635.10_3 | HIC2 | 0.98386669 | 1 | -0.212 | 2.587 | chr22 | 21771660 | 21805752 | + | protein_coding |
| ENSG00000101311.15_3 | FERMT1 | 0.99998787 | 1 | -0.212 | 5.498 | chr20 | 6055492 | 6104191 | - | protein_coding |
| ENSG00000107331.17_6 | ABCA2 | 0.99985069 | 1 | -0.212 | 5.418 | chr9 | 139901686 | 139923374 | - | protein_coding |
| ENSG00000162889.10_4 | MAPKAPK2 | 0.99999497 | 1 | -0.212 | 5.708 | chr1 | 206858289 | 206907628 | + | protein_coding |
| ENSG00000178460.18_3 | MCMDC2 | 0.96340701 | 1 | -0.212 | 1.762 | chr8 | 67782984 | 67834283 | + | protein_coding |
| ENSG00000128951.14_4 | DUT | 0.99999447 | 1 | -0.212 | 5.670 | chr15 | 48623208 | 48635570 | + | protein_coding |
| ENSG00000171612.7_4 | SLC25A33 | 0.98933258 | 1 | -0.212 | 2.895 | chr1 | 9599524 | 9645231 | + | protein_coding |
| ENSG00000287910.1_2 | AC104024.4 | 0.76601562 | 1 | -0.212 | -0.102 | chr17 | 16936845 | 16945619 | - | lncRNA |
| ENSG00000157045.9_5 | NTAN1 | 0.99964092 | 1 | -0.212 | 4.147 | chr16 | 15131711 | 15149936 | - | protein_coding |
| ENSG00000157191.20_3 | NECAP2 | 0.99992732 | 1 | -0.212 | 4.978 | chr1 | 16767216 | 16786573 | + | protein_coding |
| ENSG00000122644.13_6 | ARL4A | 0.99998204 | 1 | -0.212 | 5.461 | chr7 | 12726481 | 12730583 | + | protein_coding |
| ENSG00000233231.1_4 | HNRNPA1P49 | 0.73048824 | 1 | -0.212 | -0.575 | chr6 | 166748056 | 166748959 | + | processed_pseudogene |
| ENSG00000275714.1_3 | H3C1 | 0.99998395 | 1 | -0.212 | 5.497 | chr6 | 26020718 | 26021128 | + | protein_coding |
| ENSG00000152990.14_5 | ADGRA3 | 0.99998297 | 1 | -0.212 | 5.545 | chr4 | 22346694 | 22517689 | - | protein_coding |
| ENSG00000249915.8_4 | PDCD6 | 0.99744162 | 1 | -0.213 | 3.426 | chr5 | 271736 | 315089 | + | protein_coding |
| ENSG00000140526.18_4 | ABHD2 | 0.99999979 | 1 | -0.213 | 7.589 | chr15 | 89630690 | 89745586 | + | protein_coding |
| ENSG00000105974.12_5 | CAV1 | 0.99999974 | 1 | -0.213 | 8.609 | chr7 | 116165055 | 116201238 | + | protein_coding |
| ENSG00000107164.16_4 | FUBP3 | 0.99999769 | 1 | -0.213 | 6.195 | chr9 | 133454352 | 133513739 | + | protein_coding |
| ENSG00000259775.1_6 | AL138976.2 | 0.70618472 | 1 | -0.213 | -1.030 | chr14 | 103798011 | 103798704 | - | lncRNA |
| ENSG00000188522.15_5 | FAM83G | 0.9999693 | 1 | -0.213 | 5.518 | chr17 | 18872102 | 18908077 | - | protein_coding |
| ENSG00000069275.13_4 | NUCKS1 | 0.99999967 | 1 | -0.213 | 9.435 | chr1 | 205681950 | 205719310 | - | protein_coding |
| ENSG00000258568.1_2 | RHOQP1 | 0.71321814 | 1 | -0.213 | -0.755 | chr14 | 50066712 | 50067290 | + | processed_pseudogene |
| ENSG00000188760.10_3 | TMEM198 | 0.70759873 | 1 | -0.213 | -1.031 | chr2 | 220408385 | 220415317 | + | protein_coding |
| ENSG00000138386.17_6 | NAB1 | 0.99999413 | 1 | -0.213 | 5.793 | chr2 | 191511472 | 191557492 | + | protein_coding |
| ENSG00000163785.12_5 | RYK | 0.99999526 | 1 | -0.213 | 5.749 | chr3 | 133784147 | 133969586 | - | protein_coding |
| ENSG00000122406.14_5 | RPL5 | 0.99999963 | 1 | -0.213 | 8.870 | chr1 | 93297570 | 93307481 | + | protein_coding |
| ENSG00000147604.14_4 | RPL7 | 0.99999824 | 1 | -0.214 | 9.500 | chr8 | 74202477 | 74208024 | - | protein_coding |
| ENSG00000146457.16_5 | WTAP | 0.99998396 | 1 | -0.214 | 5.811 | chr6 | 160146617 | 160177351 | + | protein_coding |
| ENSG00000115310.18_6 | RTN4 | 0.99999982 | 1 | -0.214 | 8.327 | chr2 | 55199323 | 55339757 | - | protein_coding |
| ENSG00000100196.11_3 | KDELR3 | 0.99950979 | 1 | -0.214 | 4.056 | chr22 | 38864083 | 38879452 | + | protein_coding |
| ENSG00000236901.6_4 | MIR600HG | 0.96399077 | 1 | -0.214 | 1.884 | chr9 | 125871773 | 125877756 | - | lncRNA |
| ENSG00000247516.8_7 | MIR4458HG | 0.91108757 | 1 | -0.214 | 1.455 | chr5 | 8450814 | 8487043 | + | lncRNA |
| ENSG00000221843.4_2 | C2orf16 | 0.94468493 | 1 | -0.214 | 1.630 | chr2 | 27760253 | 27805588 | + | protein_coding |
| ENSG00000245937.8_7 | LINC01184 | 0.99995183 | 1 | -0.214 | 4.813 | chr5 | 127274844 | 127418864 | - | lncRNA |
| ENSG00000157483.9_5 | MYO1E | 0.99999987 | 1 | -0.214 | 8.598 | chr15 | 59424633 | 59665070 | - | protein_coding |
| ENSG00000163659.13_4 | TIPARP | 0.99999566 | 1 | -0.214 | 6.147 | chr3 | 156391024 | 156424559 | + | protein_coding |
| ENSG00000115295.20_4 | CLIP4 | 0.99999892 | 1 | -0.214 | 6.537 | chr2 | 29320571 | 29412509 | + | protein_coding |
| ENSG00000148334.15_5 | PTGES2 | 0.9738495 | 1 | -0.214 | 5.884 | chr9 | 130882972 | 130890741 | - | protein_coding |
| ENSG00000242457.1_4 | RBBP4P2 | 0.7118482 | 1 | -0.214 | -0.755 | chr3 | 93794756 | 93796019 | - | processed_pseudogene |
| ENSG00000118579.13_4 | MED28 | 0.99951407 | 1 | -0.215 | 4.377 | chr4 | 17616264 | 17635728 | + | protein_coding |
| ENSG00000131196.17_3 | NFATC1 | 0.8386618 | 1 | -0.215 | 1.785 | chr18 | 77155856 | 77289325 | + | protein_coding |
| ENSG00000213073.4_5 | AL353625.1 | 0.72793137 | 1 | -0.215 | -0.602 | chr6 | 160514114 | 160517244 | + | transcribed_processed_pseudogene |
| ENSG00000114120.14_6 | SLC25A36 | 0.99997852 | 1 | -0.215 | 5.318 | chr3 | 140660672 | 140699837 | + | protein_coding |
| ENSG00000145354.12_6 | CISD2 | 0.99983291 | 1 | -0.215 | 4.788 | chr4 | 103790131 | 103813964 | + | protein_coding |
| ENSG00000135297.15_3 | MTO1 | 0.99976767 | 1 | -0.215 | 4.147 | chr6 | 74171301 | 74218959 | + | protein_coding |
| ENSG00000110200.8_6 | ANAPC15 | 0.96813246 | 1 | -0.215 | 2.077 | chr11 | 71817424 | 71823826 | - | protein_coding |
| ENSG00000249087.7_6 | ZNF436-AS1 | 0.90020911 | 1 | -0.216 | 1.109 | chr1 | 23695490 | 23698332 | + | lncRNA |
| ENSG00000259577.2_6 | CERNA1 | 0.74927618 | 1 | -0.216 | -0.230 | chr15 | 52472196 | 52499112 | + | lncRNA |
| ENSG00000067704.10_3 | IARS2 | 0.99999977 | 1 | -0.216 | 7.760 | chr1 | 220267474 | 220321383 | + | protein_coding |
| ENSG00000198700.10_3 | IPO9 | 0.99999931 | 1 | -0.216 | 7.243 | chr1 | 201798277 | 201853419 | + | protein_coding |
| ENSG00000164975.15_2 | SNAPC3 | 0.9999356 | 1 | -0.216 | 5.547 | chr9 | 15422702 | 15465951 | + | protein_coding |
| ENSG00000253666.2_7 | AP000424.1 | 0.79206849 | 1 | -0.216 | 0.033 | chr8 | 101487895 | 101513376 | + | lncRNA |
| ENSG00000161654.10_3 | LSM12 | 0.99964768 | 1 | -0.216 | 4.316 | chr17 | 42111696 | 42144987 | - | protein_coding |
| ENSG00000184056.14_4 | VPS33B | 0.95627341 | 1 | -0.216 | 2.189 | chr15 | 91541646 | 91565833 | - | protein_coding |
| ENSG00000231793.5_3 | DOC2GP | 0.73598424 | 1 | -0.216 | -0.367 | chr11 | 67380122 | 67383728 | - | transcribed_unitary_pseudogene |

| | | | | | | | | | | |
|---|---|---|---|---|---|---|---|---|---|---|
| ENSG00000033050.9_5 | ABCF2 | 0.99990835 | 1 | -0.216 | 4.525 | chr7 | 150908570 | 150924291 | - | protein_coding |
| ENSG00000236675.1_4 | MTX1P1 | 0.91401793 | 1 | -0.216 | 1.229 | chr1 | 155200766 | 155204116 | + | unprocessed_pseudogene |
| ENSG00000230426.4_6 | ERVMER61-1 | 0.90460181 | 1 | -0.216 | 1.304 | chr1 | 187039832 | 187655760 | + | lncRNA |
| ENSG00000006695.12_4 | COX10 | 0.9998701 | 1 | -0.217 | 4.725 | chr17 | 13972807 | 14135053 | + | protein_coding |
| ENSG00000160299.17_6 | PCNT | 0.99955035 | 1 | -0.217 | 5.665 | chr21 | 47744055 | 47865682 | + | protein_coding |
| ENSG00000105185.12_3 | PDCD5 | 0.99999714 | 1 | -0.217 | 7.046 | chr19 | 33072096 | 33078359 | + | protein_coding |
| ENSG00000259153.2_6 | AC004816.1 | 0.976548 | 1 | -0.217 | 1.665 | chr14 | 71276558 | 71282711 | + | lncRNA |
| ENSG00000102898.12_4 | NUTF2 | 0.9999946 | 1 | -0.217 | 6.725 | chr16 | 67880826 | 67906470 | + | protein_coding |
| ENSG00000068796.17_7 | KIF2A | 0.99999182 | 1 | -0.217 | 5.895 | chr5 | 61601989 | 61833076 | + | protein_coding |
| ENSG00000276107.1_7 | AC037198.1 | 0.76133861 | 1 | -0.217 | -0.020 | chr15 | 39878762 | 39879494 | + | lncRNA |
| ENSG00000278112.1_7 | AC145423.3 | 0.70536021 | 1 | -0.217 | -0.757 | chr12 | 124003937 | 124004403 | - | lncRNA |
| ENSG00000116337.19_9 | AMPD2 | 0.99987781 | 1 | -0.217 | 5.007 | chr1 | 110158726 | 110174675 | + | protein_coding |
| ENSG00000162377.6_4 | COA7 | 0.99997383 | 1 | -0.217 | 5.179 | chr1 | 53150121 | 53164019 | - | protein_coding |
| ENSG00000144647.6_4 | POMGNT2 | 0.99888982 | 1 | -0.217 | 3.691 | chr3 | 43120724 | 43147571 | - | protein_coding |
| ENSG00000186532.12_5 | SMYD4 | 0.99990628 | 1 | -0.217 | 4.875 | chr17 | 1682779 | 1733928 | - | protein_coding |
| ENSG00000101811.13_4 | CSTF2 | 0.99996791 | 1 | -0.217 | 5.104 | chrX | 100075348 | 100095921 | + | protein_coding |
| ENSG00000233766.7_7 | CAVIN2-AS1 | 0.76577124 | 1 | -0.217 | 0.101 | chr2 | 192711265 | 192909251 | + | lncRNA |
| ENSG00000167747.14_6 | C19orf48 | 0.99999525 | 1 | -0.217 | 6.640 | chr19 | 51300961 | 51308186 | - | protein_coding |
| ENSG00000164031.17_8 | DNAJB14 | 0.99996709 | 1 | -0.217 | 5.782 | chr4 | 100817405 | 100867775 | - | protein_coding |
| ENSG00000145041.15_6 | DCAF1 | 0.99998051 | 1 | -0.217 | 5.233 | chr3 | 51433298 | 51534018 | - | protein_coding |
| ENSG00000136856.18_5 | SLC2A8 | 0.98874216 | 1 | -0.217 | 3.522 | chr9 | 130159417 | 130170703 | + | protein_coding |
| ENSG00000102038.15_5 | SMARCA1 | 0.99999961 | 1 | -0.217 | 7.316 | chrX | 128580478 | 128657477 | - | protein_coding |
| ENSG00000275437.1_7 | AL121832.3 | 0.87557742 | 1 | -0.217 | 0.848 | chr20 | 60977292 | 60980991 | - | lncRNA |
| ENSG00000159111.13_4 | MRPL10 | 0.99999419 | 1 | -0.217 | 5.922 | chr17 | 45900638 | 45908907 | - | protein_coding |
| ENSG00000101197.13_3 | BIRC7 | 0.88946425 | 1 | -0.218 | 1.311 | chr20 | 61867235 | 61871847 | + | protein_coding |
| ENSG00000248774.1_7 | AC097534.1 | 0.70104529 | 1 | -0.218 | -0.926 | chr4 | 174243357 | 174250845 | - | lncRNA |
| ENSG00000121775.18_3 | TMEM39B | 0.99122034 | 1 | -0.218 | 2.780 | chr1 | 32537632 | 32568467 | + | protein_coding |
| ENSG00000228008.1_6 | AC105935.1 | 0.71182502 | 1 | -0.218 | -0.667 | chr3 | 49941278 | 49954370 | + | lncRNA |
| ENSG00000213523.10_6 | SRA1 | 0.99995804 | 1 | -0.218 | 5.038 | chr5 | 139916925 | 139937837 | - | protein_coding |
| ENSG00000109133.13_3 | TMEM33 | 0.99999838 | 1 | -0.218 | 6.497 | chr4 | 41937146 | 41962820 | + | protein_coding |
| ENSG00000155189.12_5 | AGPAT5 | 0.99999437 | 1 | -0.218 | 6.340 | chr8 | 6566163 | 6619024 | + | protein_coding |
| ENSG00000117362.13_6 | APH1A | 0.9999926 | 1 | -0.218 | 6.153 | chr1 | 150237799 | 150241980 | - | protein_coding |
| ENSG00000241399.7_5 | CD302 | 0.8974778 | 1 | -0.218 | 1.312 | chr2 | 160625139 | 160654766 | - | protein_coding |
| ENSG00000254198.1_5 | AC113191.1 | 0.99191064 | 1 | -0.218 | 2.730 | chr8 | 42981138 | 42981540 | - | processed_pseudogene |
| ENSG00000279589.1_7 | AC079416.3 | 0.70165066 | 1 | -0.218 | -1.068 | chr16 | 53344709 | 53345901 | + | TEC |
| ENSG00000223496.3_4 | EXOSC6 | 0.99999241 | 1 | -0.218 | 5.782 | chr16 | 70280681 | 70285843 | - | protein_coding |
| ENSG00000198380.12_3 | GFPT1 | 0.99999943 | 1 | -0.218 | 6.914 | chr2 | 69546901 | 69614386 | - | protein_coding |
| ENSG00000230383.1_5 | AC009245.1 | 0.7189922 | 1 | -0.218 | -0.448 | chr7 | 137406731 | 137407592 | + | processed_pseudogene |
| ENSG00000244398.1_5 | AC116533.1 | 0.99428611 | 1 | -0.218 | 3.408 | chr11 | 16996240 | 16996560 | - | processed_pseudogene |
| ENSG00000182973.18_4 | CNOT10 | 0.99999075 | 1 | -0.218 | 5.518 | chr3 | 32726637 | 32815367 | + | protein_coding |
| ENSG00000185187.13_5 | SIGIRR | 0.9989131 | 1 | -0.218 | 4.064 | chr11 | 405716 | 417455 | - | protein_coding |
| ENSG00000207741.1 | MIR590 | 0.69863718 | 1 | -0.219 | -0.891 | chr7 | 73605528 | 73605624 | + | miRNA |
| ENSG00000276075.1_7 | AC027682.6 | 0.70231682 | 1 | -0.219 | -0.892 | chr16 | 67551765 | 67562578 | - | lncRNA |
| ENSG00000117592.9_3 | PRDX6 | 0.99999954 | 1 | -0.219 | 7.460 | chr1 | 173446469 | 173457954 | + | protein_coding |
| ENSG00000169189.17_6 | NSMCE1 | 0.98828109 | 1 | -0.219 | 4.533 | chr16 | 27236315 | 27280093 | - | protein_coding |
| ENSG00000123416.15_7 | TUBA1B | 0.99999918 | 1 | -0.219 | 8.123 | chr12 | 49521565 | 49525180 | - | protein_coding |
| ENSG00000151445.16_8 | VIPAS39 | 0.99990607 | 1 | -0.219 | 4.706 | chr14 | 77893018 | 77924295 | - | protein_coding |
| ENSG00000104695.13_4 | PPP2CB | 0.99999483 | 1 | -0.219 | 5.873 | chr8 | 30631973 | 30671830 | - | protein_coding |
| ENSG00000228492.2_4 | RAB11FIP1P1 | 0.76703841 | 1 | -0.219 | 0.088 | chrX | 73422669 | 73424430 | + | processed_pseudogene |
| ENSG00000168214.20_7 | RBPJ | 0.99995804 | 1 | -0.219 | 5.452 | chr4 | 26165077 | 26436753 | + | protein_coding |
| ENSG00000104907.12_5 | TRMT1 | 0.9984431 | 1 | -0.219 | 4.486 | chr19 | 13215716 | 13228381 | - | protein_coding |
| ENSG00000199568.1 | RNU5A-1 | 0.74934967 | 1 | -0.219 | -0.141 | chr15 | 65588389 | 65588504 | + | snRNA |
| ENSG00000101447.15_4 | FAM83D | 0.99998046 | 1 | -0.219 | 6.260 | chr20 | 37554955 | 37581749 | + | protein_coding |
| ENSG00000280071.4_8 | GATD3B | 0.78698248 | 1 | -0.220 | 0.206 | chr21 | 45553521 | 45602558 | + | protein_coding |
| ENSG00000138646.9_4 | HERC5 | 0.99866535 | 1 | -0.220 | 3.617 | chr4 | 89378270 | 89427314 | + | protein_coding |
| ENSG00000181827.15_7 | RFX7 | 0.99999508 | 1 | -0.220 | 6.133 | chr15 | 56379478 | 56537280 | - | protein_coding |
| ENSG00000136521.13_8 | NDUFB5 | 0.99985065 | 1 | -0.220 | 4.947 | chr3 | 179322478 | 179345435 | + | protein_coding |

| | | | | | | | | | | |
|---|---|---|---|---|---|---|---|---|---|---|
| ENSG00000171729.14_5 | TMEM51 | 0.99902899 | 1 | -0.220 | 3.724 | chr1 | 15479028 | 15546974 | + | protein_coding |
| ENSG00000168303.9_6 | MPLKIP | 0.99960612 | 1 | -0.220 | 4.036 | chr7 | 40165626 | 40174221 | - | protein_coding |
| ENSG00000164776.9_4 | PHKG1 | 0.93691008 | 1 | -0.220 | 1.636 | chr7 | 56147976 | 56160689 | - | protein_coding |
| ENSG00000054967.13_4 | RELT | 0.99119033 | 1 | -0.220 | 3.316 | chr11 | 73087444 | 73108519 | + | protein_coding |
| ENSG00000007923.16_5 | DNAJC11 | 0.99998064 | 1 | -0.220 | 5.166 | chr1 | 6694228 | 6761984 | - | protein_coding |
| ENSG00000119397.16_3 | CNTRL | 0.99997798 | 1 | -0.220 | 5.414 | chr9 | 123837141 | 123939888 | + | protein_coding |
| ENSG00000145741.15_5 | BTF3 | 0.99999959 | 1 | -0.220 | 8.207 | chr5 | 72794233 | 72801460 | + | protein_coding |
| ENSG00000125482.13_4 | TTF1 | 0.99970088 | 1 | -0.220 | 4.202 | chr9 | 135250935 | 135282238 | - | protein_coding |
| ENSG00000173517.10_4 | PEAK1 | 0.99998078 | 1 | -0.220 | 5.364 | chr15 | 77392998 | 77712486 | - | protein_coding |
| ENSG00000062650.19_6 | WAPL | 0.99998874 | 1 | -0.220 | 6.152 | chr10 | 88195013 | 88281549 | - | protein_coding |
| ENSG00000161682.15_4 | FAM171A2 | 0.96384022 | 1 | -0.220 | 2.068 | chr17 | 42430583 | 42441221 | - | protein_coding |
| ENSG00000128274.17_5 | A4GALT | 0.99613517 | 1 | -0.220 | 3.498 | chr22 | 43088127 | 43117304 | - | protein_coding |
| ENSG00000148153.14_4 | INIP | 0.99996838 | 1 | -0.221 | 5.160 | chr9 | 115446206 | 115480429 | - | protein_coding |
| ENSG00000286305.1_2 | AC004893.3 | 0.76825254 | 1 | -0.221 | 0.089 | chr7 | 98587490 | 98591401 | - | lncRNA |
| ENSG00000057608.17_7 | GDI2 | 0.99999906 | 1 | -0.221 | 8.470 | chr10 | 5807186 | 5884095 | - | protein_coding |
| ENSG00000235508.3_4 | RPS2P7 | 0.81400414 | 1 | -0.221 | 0.486 | chr20 | 44269125 | 44270009 | - | processed_pseudogene |
| ENSG00000124120.11_3 | TTPAL | 0.99991121 | 1 | -0.221 | 4.857 | chr20 | 43104514 | 43123244 | + | protein_coding |
| ENSG00000269915.1_7 | AP006621.4 | 0.69842059 | 1 | -0.221 | -0.927 | chr11 | 708564 | 727047 | - | lncRNA |
| ENSG00000130227.17_4 | XPO7 | 0.99999816 | 1 | -0.221 | 6.417 | chr8 | 21777173 | 21864096 | + | protein_coding |
| ENSG00000114346.13_5 | ECT2 | 0.99999934 | 1 | -0.221 | 8.698 | chr3 | 172468472 | 172539264 | + | protein_coding |
| ENSG00000248698.6_6 | LINC01085 | 0.7078853 | 1 | -0.221 | -0.522 | chr4 | 14113592 | 14141676 | + | transcribed_processed_pseudogene |
| ENSG00000279632.1_6 | AP003108.4 | 0.71212547 | 1 | -0.221 | -0.283 | chr11 | 61193920 | 61194797 | - | TEC |
| ENSG00000131462.8_3 | TUBG1 | 0.99999904 | 1 | -0.221 | 6.759 | chr17 | 40761701 | 40767256 | + | protein_coding |
| ENSG00000228409.6_2 | CCT6P1 | 0.99056161 | 1 | -0.222 | 2.711 | chr7 | 65216129 | 65228341 | + | transcribed_unprocessed_pseudogene |
| ENSG00000147162.14_3 | OGT | 0.99999902 | 1 | -0.222 | 7.820 | chrX | 70752954 | 70795742 | + | protein_coding |
| ENSG00000124275.15_6 | MTRR | 0.99998988 | 1 | -0.222 | 5.828 | chr5 | 7851299 | 7906138 | + | protein_coding |
| ENSG00000213221.5_5 | DNLZ | 0.74573418 | 1 | -0.222 | -0.131 | chr9 | 139253932 | 139258196 | - | protein_coding |
| ENSG00000260874.5_7 | AC106820.5 | 0.7003727 | 1 | -0.222 | -0.757 | chr16 | 2506253 | 2509980 | - | lncRNA |
| ENSG00000256862.3_6 | AC005842.1 | 0.69481134 | 1 | -0.222 | -0.996 | chr12 | 3980745 | 4019548 | + | lncRNA |
| ENSG00000198301.12_5 | SDAD1 | 0.99991979 | 1 | -0.222 | 4.916 | chr4 | 76862103 | 76912115 | - | protein_coding |
| ENSG00000135070.15_7 | ISCA1 | 0.99997941 | 1 | -0.222 | 5.498 | chr9 | 88879461 | 88898017 | - | protein_coding |
| ENSG00000278864.1_6 | AC055811.4 | 0.98441547 | 1 | -0.222 | 2.663 | chr17 | 17084818 | 17086571 | + | TEC |
| ENSG00000205659.10_5 | LIN52 | 0.99711567 | 1 | -0.222 | 3.556 | chr14 | 74551499 | 74667938 | + | protein_coding |
| ENSG00000188185.12_7 | LINC00265 | 0.92647053 | 1 | -0.222 | 1.452 | chr7 | 39773029 | 39832691 | + | lncRNA |
| ENSG00000145332.14_4 | KLHL8 | 0.9995804 | 1 | -0.222 | 4.685 | chr4 | 88081255 | 88161466 | - | protein_coding |
| ENSG00000182405.6_3 | PGBD4 | 0.99886394 | 1 | -0.222 | 3.729 | chr15 | 34394284 | 34400887 | + | protein_coding |
| ENSG00000166260.12_6 | COX11 | 0.99999297 | 1 | -0.222 | 6.215 | chr17 | 53029263 | 53046064 | - | protein_coding |
| ENSG00000152104.12_5 | PTPN14 | 0.99999321 | 1 | -0.223 | 5.868 | chr1 | 214522043 | 214725792 | - | protein_coding |
| ENSG00000233695.2_6 | GAS6-AS1 | 0.99539989 | 1 | -0.223 | 3.459 | chr13 | 114518603 | 114548717 | + | lncRNA |
| ENSG00000184178.16_5 | SCFD2 | 0.99937027 | 1 | -0.223 | 3.975 | chr4 | 53739149 | 54232233 | - | protein_coding |
| ENSG00000100104.13_4 | SRRD | 0.99716061 | 1 | -0.223 | 3.618 | chr22 | 26879843 | 26890624 | + | protein_coding |
| ENSG00000100902.11_6 | PSMA6 | 0.98779209 | 1 | -0.223 | 2.624 | chr14 | 35747839 | 35786699 | + | protein_coding |
| ENSG00000145016.16_6 | RUBCN | 0.99113691 | 1 | -0.223 | 3.163 | chr3 | 197395738 | 197476598 | - | protein_coding |
| ENSG00000264743.2_2 | DPRXP4 | 0.72657673 | 1 | -0.223 | -0.393 | chr17 | 29302405 | 29302972 | + | processed_pseudogene |
| ENSG00000159423.17_6 | ALDH4A1 | 0.99671413 | 1 | -0.223 | 3.447 | chr1 | 19197924 | 19229218 | - | protein_coding |
| ENSG00000147174.11_2 | GCNA | 0.83939874 | 1 | -0.223 | 0.787 | chrX | 70798261 | 70833433 | + | protein_coding |
| ENSG00000170854.18_8 | RIOX2 | 0.99997677 | 1 | -0.223 | 5.383 | chr3 | 97660662 | 97691301 | - | protein_coding |
| ENSG00000129351.17_5 | ILF3 | 0.99999949 | 1 | -0.223 | 8.107 | chr19 | 10764937 | 10803093 | + | protein_coding |
| ENSG00000104047.15_4 | DTWD1 | 0.99812743 | 1 | -0.223 | 3.597 | chr15 | 49913234 | 49948429 | + | protein_coding |
| ENSG00000169379.16_3 | ARL13B | 0.99993877 | 1 | -0.223 | 4.957 | chr3 | 93698983 | 93774522 | + | protein_coding |
| ENSG00000011260.14_3 | UTP18 | 0.99999791 | 1 | -0.223 | 6.745 | chr17 | 49337907 | 49375297 | + | protein_coding |
| ENSG00000100416.15_5 | TRMU | 0.99994091 | 1 | -0.223 | 5.232 | chr22 | 46726772 | 46753237 | + | protein_coding |
| ENSG00000153767.10_5 | GTF2E1 | 0.99973903 | 1 | -0.224 | 4.752 | chr3 | 120461484 | 120501916 | + | protein_coding |
| ENSG00000228474.6_4 | OST4 | 0.99998883 | 1 | -0.224 | 5.454 | chr2 | 27293340 | 27294522 | - | protein_coding |
| ENSG00000171163.15_5 | ZNF692 | 0.99955141 | 1 | -0.224 | 4.216 | chr1 | 249144205 | 249153343 | - | protein_coding |
| ENSG00000101079.21_6 | NDRG3 | 0.99998194 | 1 | -0.224 | 5.398 | chr20 | 35280169 | 35374493 | - | protein_coding |
| ENSG00000132467.4_3 | UTP3 | 0.99900302 | 1 | -0.224 | 3.884 | chr4 | 71554249 | 71556268 | + | protein_coding |
| ENSG00000090686.16_4 | USP48 | 0.99994022 | 1 | -0.224 | 5.033 | chr1 | 22004791 | 22110099 | - | protein_coding |
| ENSG00000167920.10_8 | TMEM99 | 0.99461247 | 1 | -0.224 | 3.149 | chr17 | 38975358 | 38992522 | + | lncRNA |

| | | | | | | | | | | |
|---|---|---|---|---|---|---|---|---|---|---|
| ENSG00000048544.6_4 | MRPS10 | 0.99996597 | 1 | -0.224 | 5.557 | chr6 | 42174545 | 42185599 | - | protein_coding |
| ENSG00000287656.1_1 | AL591069.1 | 0.95410269 | 1 | -0.224 | 1.935 | chr6 | 160790232 | 160807966 | - | lncRNA |
| ENSG00000104299.15_5 | INTS9 | 0.99955401 | 1 | -0.224 | 4.343 | chr8 | 28625178 | 28747759 | - | protein_coding |
| ENSG00000163412.13_5 | EIF4E3 | 0.99408657 | 1 | -0.224 | 3.104 | chr3 | 71724565 | 71803924 | - | protein_coding |
| ENSG00000178691.11_4 | SUZ12 | 0.99999854 | 1 | -0.224 | 7.273 | chr17 | 30264026 | 30328057 | + | protein_coding |
| ENSG00000251143.1_6 | AP002490.1 | 0.87116065 | 1 | -0.224 | 0.907 | chr11 | 71725337 | 71731956 | + | lncRNA |
| ENSG00000163463.11_4 | KRTCAP2 | 0.98474277 | 1 | -0.224 | 2.530 | chr1 | 155141884 | 155145951 | - | protein_coding |
| ENSG00000207721.1 | MIR186 | 0.69024894 | 1 | -0.224 | -0.893 | chr1 | 71533314 | 71533399 | - | miRNA |
| ENSG00000280129.1_6 | AL132780.5 | 0.80824782 | 1 | -0.224 | 0.153 | chr14 | 23474708 | 23476332 | - | TEC |
| ENSG00000101546.13_4 | RBFA | 0.99932986 | 1 | -0.224 | 4.048 | chr18 | 77794389 | 77810651 | + | protein_coding |
| ENSG00000265520.1 | MIR548V | 0.69635842 | 1 | -0.225 | -0.756 | chr8 | 17539087 | 17539166 | - | miRNA |
| ENSG00000168827.14_6 | GFM1 | 0.99999719 | 1 | -0.225 | 6.351 | chr3 | 158362067 | 158410364 | + | protein_coding |
| ENSG00000225439.3_6 | BOLA3-AS1 | 0.93180289 | 1 | -0.225 | 1.181 | chr2 | 74375134 | 74379079 | + | lncRNA |
| ENSG00000130024.15_5 | PHF10 | 0.99993842 | 1 | -0.225 | 4.984 | chr6 | 170103998 | 170125662 | - | protein_coding |
| ENSG00000163001.11_3 | CFAP36 | 0.99996909 | 1 | -0.225 | 5.481 | chr2 | 55746740 | 55773015 | + | protein_coding |
| ENSG00000174891.13_6 | RSRC1 | 0.9999679 | 1 | -0.225 | 5.564 | chr3 | 157823644 | 158263519 | + | protein_coding |
| ENSG00000122085.17_5 | MTERF4 | 0.99881797 | 1 | -0.225 | 3.947 | chr2 | 242011584 | 242041747 | - | protein_coding |
| ENSG00000239306.4_3 | RBM14 | 0.99999061 | 1 | -0.225 | 5.956 | chr11 | 66384053 | 66394818 | + | protein_coding |
| ENSG00000100360.15_6 | IFT27 | 0.99823506 | 1 | -0.225 | 3.762 | chr22 | 37154246 | 37172300 | - | protein_coding |
| ENSG00000163960.12_5 | UBXN7 | 0.99983003 | 1 | -0.225 | 4.817 | chr3 | 196074533 | 196159301 | - | protein_coding |
| ENSG00000224536.2_7 | AC096677.1 | 0.75787624 | 1 | -0.225 | -0.030 | chr1 | 201476369 | 201503912 | + | lncRNA |
| ENSG00000124588.20_4 | NQO2 | 0.99999295 | 1 | -0.225 | 5.709 | chr6 | 2988221 | 3029103 | + | protein_coding |
| ENSG00000260853.2_8 | AC109460.2 | 0.98804379 | 1 | -0.225 | 2.684 | chr16 | 28964137 | 28978204 | - | lncRNA |
| ENSG00000140374.16_6 | ETFA | 0.99999931 | 1 | -0.225 | 7.599 | chr15 | 76507694 | 76603813 | - | protein_coding |
| ENSG00000100038.19_6 | TOP3B | 0.70653411 | 1 | -0.225 | -0.448 | chr22 | 22311397 | 22337213 | - | protein_coding |
| ENSG00000169738.8_5 | DCXR | 0.99971659 | 1 | -0.225 | 4.287 | chr17 | 79993012 | 79995585 | - | protein_coding |
| ENSG00000138663.9_4 | COPS4 | 0.99966761 | 1 | -0.225 | 5.116 | chr4 | 83955600 | 83996971 | + | protein_coding |
| ENSG00000141034.10_4 | GID4 | 0.99990402 | 1 | -0.226 | 4.612 | chr17 | 17942722 | 17971719 | + | protein_coding |
| ENSG00000113119.12_2 | TMCO6 | 0.99858163 | 1 | -0.226 | 3.799 | chr5 | 140019012 | 140024993 | + | protein_coding |
| ENSG00000134324.11_6 | LPIN1 | 0.99999472 | 1 | -0.226 | 6.259 | chr2 | 11817721 | 11967535 | + | protein_coding |
| ENSG00000180914.10_3 | OXTR | 0.99729986 | 1 | -0.226 | 3.406 | chr3 | 8792094 | 8811314 | - | protein_coding |
| ENSG00000165271.17_5 | NOL6 | 0.99999809 | 1 | -0.226 | 6.658 | chr9 | 33461351 | 33473928 | - | protein_coding |
| ENSG00000101842.13_5 | VSIG1 | 0.69383444 | 1 | -0.226 | -0.892 | chrX | 107288200 | 107322414 | + | protein_coding |
| ENSG00000218739.10_4 | CEBPZOS | 0.99981299 | 1 | -0.226 | 4.861 | chr2 | 37423631 | 37443336 | + | protein_coding |
| ENSG00000267710.9_7 | EDDM13 | 0.86332009 | 1 | -0.226 | 0.573 | chr19 | 56784117 | 56821823 | + | protein_coding |
| ENSG00000233954.6_2 | UQCRHL | 0.98461593 | 1 | -0.226 | 2.778 | chr1 | 16133664 | 16135843 | - | protein_coding |
| ENSG00000196597.13_5 | ZNF782 | 0.99445902 | 1 | -0.226 | 3.158 | chr9 | 99578551 | 99637905 | - | protein_coding |
| ENSG00000141127.15_6 | PRPSAP2 | 0.99998322 | 1 | -0.226 | 5.481 | chr17 | 18743398 | 18834600 | + | protein_coding |
| ENSG00000163923.10_5 | RPL39L | 0.9995489 | 1 | -0.227 | 4.177 | chr3 | 186838736 | 186898696 | - | protein_coding |
| ENSG00000117748.10_4 | RPA2 | 0.9999713 | 1 | -0.227 | 5.226 | chr1 | 28218035 | 28241257 | - | protein_coding |
| ENSG00000129347.21_5 | KRI1 | 0.99926072 | 1 | -0.227 | 4.210 | chr19 | 10663761 | 10676707 | - | protein_coding |
| ENSG00000182185.18_4 | RAD51B | 0.99161665 | 1 | -0.227 | 2.997 | chr14 | 68286496 | 69196935 | + | protein_coding |
| ENSG00000108465.15_4 | CDK5RAP3 | 0.9985746 | 1 | -0.227 | 3.811 | chr17 | 46045176 | 46059147 | + | protein_coding |
| ENSG00000131503.21_4 | ANKHD1 | 0.95957529 | 1 | -0.227 | 1.510 | chr5 | 139781399 | 139919441 | + | protein_coding |
| ENSG00000009954.11_4 | BAZ1B | 0.99999955 | 1 | -0.227 | 8.086 | chr7 | 72854736 | 72936623 | - | protein_coding |
| ENSG00000246089.3_7 | AC016065.1 | 0.93712339 | 1 | -0.228 | 1.779 | chr8 | 6261072 | 6264663 | - | lncRNA |
| ENSG00000196787.3_3 | H2AC11 | 0.98144435 | 1 | -0.228 | 7.637 | chr6 | 27100821 | 27103070 | + | protein_coding |
| ENSG00000100209.10_3 | HSCB | 0.98274368 | 1 | -0.228 | 2.489 | chr22 | 29138027 | 29153503 | + | protein_coding |
| ENSG00000279159.1_7 | AC003681.1 | 0.96899431 | 1 | -0.228 | 2.372 | chr22 | 30374939 | 30424225 | - | lncRNA |
| ENSG00000081721.12_3 | DUSP12 | 0.99982175 | 1 | -0.228 | 4.841 | chr1 | 161719548 | 161727028 | + | protein_coding |
| ENSG00000187266.14_5 | EPOR | 0.95220824 | 1 | -0.228 | 2.293 | chr19 | 11487883 | 11495018 | - | protein_coding |
| ENSG00000087087.20_6 | SRRT | 0.99999427 | 1 | -0.228 | 6.639 | chr7 | 100472725 | 100486285 | + | protein_coding |
| ENSG00000180425.11_3 | C11orf71 | 0.96904546 | 1 | -0.228 | 2.375 | chr11 | 114262165 | 114271233 | - | protein_coding |
| ENSG00000270170.2_6 | NCBP2AS2 | 0.98113878 | 1 | -0.228 | 2.407 | chr3 | 196669545 | 196670414 | + | protein_coding |
| ENSG00000240376.1_5 | AC010343.1 | 0.88176025 | 1 | -0.228 | 1.036 | chr5 | 33162285 | 33162913 | + | processed_pseudogene |
| ENSG00000116752.6_4 | BCAS2 | 0.9997427 | 1 | -0.228 | 4.773 | chr1 | 115110178 | 115124250 | - | protein_coding |
| ENSG00000174744.14_7 | BRMS1 | 0.99998498 | 1 | -0.228 | 6.030 | chr11 | 66104804 | 66112596 | - | protein_coding |
| ENSG00000172009.15_7 | THOP1 | 0.99945441 | 1 | -0.228 | 5.918 | chr19 | 2785501 | 2815805 | + | protein_coding |
| ENSG00000127311.9_2 | HELB | 0.99911979 | 1 | -0.228 | 4.622 | chr12 | 66696325 | 66741425 | + | protein_coding |
| ENSG00000102317.18_3 | RBM3 | 0.9999994 | 1 | -0.229 | 7.689 | chrX | 48432837 | 48439550 | + | protein_coding |

| | | | | | | | | | | |
|---|---|---|---|---|---|---|---|---|---|---|
| ENSG00000196449.4_4 | YRDC | 0.99743239 | 1 | -0.229 | 3.425 | chr1 | 38268617 | 38273880 | - | protein_coding |
| ENSG00000102977.16_7 | ACD | 0.99963911 | 1 | -0.229 | 4.446 | chr16 | 67691415 | 67694718 | - | protein_coding |
| ENSG00000198556.14_3 | ZNF789 | 0.99474969 | 1 | -0.229 | 3.031 | chr7 | 99070513 | 99101273 | + | protein_coding |
| ENSG00000182628.13_4 | SKA2 | 0.99999027 | 1 | -0.229 | 6.656 | chr17 | 57187218 | 57232621 | - | protein_coding |
| ENSG00000154429.11_5 | CCSAP | 0.9998588 | 1 | -0.229 | 4.519 | chr1 | 229456758 | 229479041 | - | protein_coding |
| ENSG00000008083.14_3 | JARID2 | 0.99977621 | 1 | -0.229 | 4.394 | chr6 | 15246300 | 15522273 | + | protein_coding |
| ENSG00000198746.13_5 | GPATCH3 | 0.98498688 | 1 | -0.229 | 2.559 | chr1 | 27216979 | 27226958 | - | protein_coding |
| ENSG00000183207.14_4 | RUVBL2 | 0.9999433 | 1 | -0.229 | 7.260 | chr19 | 49496819 | 49519227 | + | protein_coding |
| ENSG00000196873.15_4 | CBWD3 | 0.89401856 | 1 | -0.229 | 1.185 | chr9 | 70846919 | 70914931 | + | protein_coding |
| ENSG00000117143.13_5 | UAP1 | 0.99990524 | 1 | -0.229 | 4.861 | chr1 | 162531296 | 162569632 | + | protein_coding |
| ENSG00000213390.11_7 | ARHGAP19 | 0.99664019 | 1 | -0.229 | 3.734 | chr10 | 98981930 | 99052430 | - | protein_coding |
| ENSG00000135932.11_3 | CAB39 | 0.99999362 | 1 | -0.229 | 6.293 | chr2 | 231577557 | 231685790 | + | protein_coding |
| ENSG00000111834.13_4 | RSPH4A | 0.69291176 | 1 | -0.230 | -0.697 | chr6 | 116937642 | 116954148 | + | protein_coding |
| ENSG00000177853.14_3 | ZNF518A | 0.99998822 | 1 | -0.230 | 6.170 | chr10 | 97889472 | 97965044 | + | protein_coding |
| ENSG00000155008.14_4 | APOOL | 0.99950769 | 1 | -0.230 | 4.878 | chrX | 84258883 | 84348322 | + | protein_coding |
| ENSG00000177565.17_4 | TBL1XR1 | 0.99999332 | 1 | -0.230 | 7.477 | chr3 | 176737128 | 176945788 | - | protein_coding |
| ENSG00000232448.2_7 | AL354824.2 | 0.7765067 | 1 | -0.230 | 0.271 | chr20 | 10153349 | 10167388 | + | lncRNA |
| ENSG00000140006.11_4 | WDR89 | 0.99964297 | 1 | -0.230 | 4.230 | chr14 | 64063757 | 64108579 | - | protein_coding |
| ENSG00000101972.18_3 | STAG2 | 0.99999705 | 1 | -0.230 | 7.810 | chrX | 123094062 | 123556514 | + | protein_coding |
| ENSG00000279792.1_5 | AC015909.5 | 0.89570733 | 1 | -0.230 | 1.130 | chr17 | 48440328 | 48442593 | - | TEC |
| ENSG00000110696.10_5 | C11orf58 | 0.9999986 | 1 | -0.230 | 7.323 | chr11 | 16634679 | 16779887 | + | protein_coding |
| ENSG00000112561.18_5 | TFEB | 0.950462 | 1 | -0.230 | 1.783 | chr6 | 41651716 | 41703997 | - | protein_coding |
| ENSG00000167693.17_5 | NXN | 0.99883744 | 1 | -0.230 | 4.929 | chr17 | 702550 | 883016 | - | protein_coding |
| ENSG00000259347.7_8 | AC087482.1 | 0.75895206 | 1 | -0.230 | 0.312 | chr15 | 67276441 | 67357606 | - | lncRNA |
| ENSG00000164506.14_3 | STXBP5 | 0.99990958 | 1 | -0.230 | 4.989 | chr6 | 147525561 | 147711612 | + | protein_coding |
| ENSG00000168887.11_6 | C2orf68 | 0.99998432 | 1 | -0.230 | 5.398 | chr2 | 85832377 | 85839189 | - | protein_coding |
| ENSG00000196262.14_3 | PPIA | 0.99999962 | 1 | -0.231 | 9.759 | chr7 | 44836279 | 44864163 | + | protein_coding |
| ENSG00000255642.1_4 | PABPC1P4 | 0.68103104 | 1 | -0.231 | -1.082 | chr12 | 64215801 | 64217675 | + | processed_pseudogene |
| ENSG00000275494.1_7 | AC133552.5 | 0.68616838 | 1 | -0.231 | -1.083 | chr16 | 25117890 | 25118423 | - | lncRNA |
| ENSG00000233560.2_4 | KRT8P39 | 0.81914528 | 1 | -0.231 | 0.641 | chr12 | 69099414 | 69100846 | + | processed_pseudogene |
| ENSG00000198763.3_1 | MT-ND2 | 0.99999804 | 1 | -0.231 | 7.283 | chrM | 4470 | 5511 | + | protein_coding |
| ENSG00000223573.7_5 | TINCR | 0.74603449 | 1 | -0.231 | 0.007 | chr19 | 5558178 | 5578360 | - | protein_coding |
| ENSG00000238390.1 | SNORA81 | 0.7018883 | 1 | -0.231 | -0.669 | chr21 | 33036619 | 33036795 | + | snoRNA |
| ENSG00000105879.12_4 | CBLL1 | 0.99998334 | 1 | -0.231 | 5.753 | chr7 | 107384394 | 107402112 | + | protein_coding |
| ENSG00000186166.9_3 | CCDC84 | 0.99922834 | 1 | -0.232 | 4.493 | chr11 | 118868848 | 118886503 | + | protein_coding |
| ENSG00000115966.17_5 | ATF2 | 0.99998216 | 1 | -0.232 | 5.885 | chr2 | 175936978 | 176033110 | - | protein_coding |
| ENSG00000267197.1_8 | AC011461.1 | 0.7185352 | 1 | -0.232 | -0.379 | chr19 | 10601015 | 10601676 | - | lncRNA |
| ENSG00000136104.21_5 | RNASEH2B | 0.99994736 | 1 | -0.232 | 5.174 | chr13 | 51483883 | 51598256 | + | protein_coding |
| ENSG00000149016.16_6 | TUT1 | 0.98924846 | 1 | -0.232 | 2.828 | chr11 | 62342517 | 62359109 | - | protein_coding |
| ENSG00000130956.14_4 | HABP4 | 0.99540136 | 1 | -0.232 | 3.522 | chr9 | 99212451 | 99253618 | + | protein_coding |
| ENSG00000164053.21_7 | ATRIP | 0.99922657 | 1 | -0.232 | 3.915 | chr3 | 48488114 | 48509044 | + | protein_coding |
| ENSG00000164087.8_6 | POC1A | 0.9977226 | 1 | -0.232 | 3.485 | chr3 | 52109242 | 52188706 | - | protein_coding |
| ENSG00000264918.1 | AC005229.1 | 0.68018492 | 1 | -0.232 | -0.961 | chr7 | 148427921 | 148428020 | - | miRNA |
| ENSG00000205147.3 | AC016586.1 | 0.68085378 | 1 | -0.232 | -0.860 | chr19 | 4041127 | 4043154 | - | protein_coding |
| ENSG00000186567.13_5 | CEACAM19 | 0.98308224 | 1 | -0.232 | 2.727 | chr19 | 45165545 | 45187631 | + | protein_coding |
| ENSG00000208772.1 | SNORD94 | 0.79148303 | 1 | -0.232 | 0.462 | chr2 | 86362993 | 86363129 | + | snoRNA |
| ENSG00000143344.15_3 | RGL1 | 0.99423593 | 1 | -0.232 | 3.079 | chr1 | 183605220 | 183897665 | + | protein_coding |
| ENSG00000114021.12_3 | NIT2 | 0.99999296 | 1 | -0.232 | 6.009 | chr3 | 100053583 | 100080479 | + | protein_coding |
| ENSG00000198055.11_5 | GRK6 | 0.99990377 | 1 | -0.233 | 5.382 | chr5 | 176830205 | 176869902 | + | protein_coding |
| ENSG00000138744.16_8 | NAAA | 0.99884578 | 1 | -0.233 | 4.023 | chr4 | 76834813 | 76862166 | - | protein_coding |
| ENSG00000228158.2_4 | TLE1P1 | 0.76929112 | 1 | -0.233 | 0.041 | chrX | 64627964 | 64628552 | + | processed_pseudogene |
| ENSG00000070367.16_5 | EXOC5 | 0.99999109 | 1 | -0.233 | 6.735 | chr14 | 57667225 | 57735623 | - | protein_coding |
| ENSG00000129757.13_5 | CDKN1C | 0.68492523 | 1 | -0.233 | -0.474 | chr11 | 2904443 | 2907003 | - | protein_coding |
| ENSG00000263843.1_7 | AC022211.3 | 0.89978061 | 1 | -0.233 | 1.291 | chr17 | 73267450 | 73269976 | + | lncRNA |
| ENSG00000168014.16_8 | C2CD3 | 0.99997393 | 1 | -0.233 | 5.412 | chr11 | 73723759 | 73882255 | - | protein_coding |
| ENSG00000090861.15_4 | AARS1 | 0.99999889 | 1 | -0.233 | 7.435 | chr16 | 70286198 | 70323446 | - | protein_coding |
| ENSG00000238251.2_5 | AL133477.1 | 0.67788403 | 1 | -0.233 | -1.046 | chr9 | 99169566 | 99170079 | + | processed_pseudogene |
| ENSG00000172771.12_5 | EFCAB12 | 0.82116783 | 1 | -0.233 | 0.366 | chr3 | 129120164 | 129147494 | - | protein_coding |
| ENSG00000090061.17_5 | CCNK | 0.97449605 | 1 | -0.233 | 2.619 | chr14 | 99947506 | 100001381 | + | protein_coding |

| | | | | | | | | | | |
|---|---|---|---|---|---|---|---|---|---|---|
| ENSG00000073150.14_5 | PANX2 | 0.93277762 | 1 | -0.233 | 1.770 | chr22 | 50609160 | 50618724 | + | protein_coding |
| ENSG00000228794.10_7 | LINC01128 | 0.96874572 | 1 | -0.233 | 2.636 | chr1 | 760518 | 794826 | + | lncRNA |
| ENSG00000099282.10_4 | TSPAN15 | 0.99999199 | 1 | -0.233 | 6.358 | chr10 | 71211221 | 71267422 | + | protein_coding |
| ENSG00000106459.15_5 | NRF1 | 0.99842524 | 1 | -0.233 | 3.844 | chr7 | 129251561 | 129396922 | + | protein_coding |
| ENSG00000137955.16_4 | RABGGTB | 0.99998548 | 1 | -0.233 | 5.964 | chr1 | 76251882 | 76260771 | + | protein_coding |
| ENSG00000157306.14_6 | ZFHX2-AS1 | 0.78102676 | 1 | -0.233 | 0.403 | chr14 | 23980969 | 24029987 | + | lncRNA |
| ENSG00000110108.10_3 | TMEM109 | 0.99993775 | 1 | -0.233 | 5.491 | chr11 | 60681630 | 60690915 | + | protein_coding |
| ENSG00000167114.13_4 | SLC27A4 | 0.99993653 | 1 | -0.233 | 5.509 | chr9 | 131102806 | 131123749 | + | protein_coding |
| ENSG00000145725.19_4 | PPIP5K2 | 0.99999009 | 1 | -0.233 | 6.441 | chr5 | 102455853 | 102548500 | + | protein_coding |
| ENSG00000174899.11_5 | SLC66A1L | 0.72009705 | 1 | -0.233 | 0.023 | chr3 | 157261035 | 157395538 | + | protein_coding |
| ENSG00000125107.18_5 | CNOT1 | 0.99999962 | 1 | -0.234 | 8.485 | chr16 | 58553855 | 58663789 | - | protein_coding |
| ENSG00000166289.6_3 | PLEKHF1 | 0.95562405 | 1 | -0.234 | 2.185 | chr19 | 30156366 | 30166384 | + | protein_coding |
| ENSG00000121289.18_6 | CEP89 | 0.99999544 | 1 | -0.234 | 6.066 | chr19 | 33366831 | 33462897 | - | protein_coding |
| ENSG00000131018.24_4 | SYNE1 | 0.99996249 | 1 | -0.234 | 6.988 | chr6 | 152442819 | 152958936 | - | protein_coding |
| ENSG00000229320.3_2 | KRT8P12 | 0.98475089 | 1 | -0.234 | 2.673 | chr3 | 160283235 | 160287036 | + | transcribed_processed_pseudogene |
| ENSG00000225511.7_4 | LINC00475 | 0.67776935 | 1 | -0.234 | -0.797 | chr9 | 94903580 | 94922396 | + | transcribed_unprocessed_pseudogene |
| ENSG00000273218.1_7 | AC005776.2 | 0.68217961 | 1 | -0.234 | -0.798 | chr19 | 15456879 | 15459228 | - | lncRNA |
| ENSG00000186010.19_4 | NDUFA13 | 0.83341323 | 1 | -0.235 | 0.799 | chr19 | 19626545 | 19639863 | + | protein_coding |
| ENSG00000169641.13_5 | LUZP1 | 0.99998504 | 1 | -0.235 | 5.843 | chr1 | 23410516 | 23504301 | - | protein_coding |
| ENSG00000175741.5_4 | RWDD4P2 | 0.88696017 | 1 | -0.235 | 1.107 | chr7 | 39894093 | 39894655 | - | processed_pseudogene |
| ENSG00000175591.11_3 | P2RY2 | 0.99546621 | 1 | -0.235 | 3.358 | chr11 | 72929343 | 72947397 | + | protein_coding |
| ENSG00000236397.3_3 | DDX11L2 | 0.73786147 | 1 | -0.235 | -0.111 | chr2 | 114356613 | 114358838 | - | unprocessed_pseudogene |
| ENSG00000164291.17_3 | ARSK | 0.99586982 | 1 | -0.235 | 3.645 | chr5 | 94890805 | 94940806 | + | protein_coding |
| ENSG00000176029.14_5 | C11orf16 | 0.7279476 | 1 | -0.235 | -0.151 | chr11 | 8941623 | 8954553 | - | protein_coding |
| ENSG00000275023.5_5 | MLLT6 | 0.99999235 | 1 | -0.235 | 6.351 | chr17 | 36861526 | 36886048 | + | protein_coding |
| ENSG00000137409.19_7 | MTCH1 | 0.9999949 | 1 | -0.235 | 6.763 | chr6 | 36935917 | 36954074 | - | protein_coding |
| ENSG00000141759.15_4 | TXNL4A | 0.99998144 | 1 | -0.235 | 5.668 | chr18 | 77730813 | 77793949 | - | protein_coding |
| ENSG00000160233.8_3 | LRRC3 | 0.89223939 | 1 | -0.235 | 1.193 | chr21 | 45875393 | 45882079 | + | protein_coding |
| ENSG00000186665.9_6 | C17orf58 | 0.99767146 | 1 | -0.236 | 3.509 | chr17 | 65987217 | 65992547 | - | protein_coding |
| ENSG00000182957.16_7 | SPATA13 | 0.9973393 | 1 | -0.236 | 3.692 | chr13 | 24553944 | 24881212 | + | protein_coding |
| ENSG00000136603.14_5 | SKIL | 0.9999964 | 1 | -0.236 | 7.600 | chr3 | 170075466 | 170114623 | + | protein_coding |
| ENSG00000183696.14_5 | UPP1 | 0.99974713 | 1 | -0.236 | 4.952 | chr7 | 48128225 | 48148333 | + | protein_coding |
| ENSG00000152457.18_7 | DCLRE1C | 0.99983273 | 1 | -0.236 | 4.931 | chr10 | 14939358 | 14996431 | - | protein_coding |
| ENSG00000108479.12_6 | GALK1 | 0.99748818 | 1 | -0.236 | 5.103 | chr17 | 73747675 | 73761317 | - | protein_coding |
| ENSG00000113648.16_4 | MACROH2A1 | 0.99999719 | 1 | -0.236 | 7.065 | chr5 | 134669590 | 134735604 | - | protein_coding |
| ENSG00000276043.5_6 | UHRF1 | 0.99461849 | 1 | -0.236 | 5.600 | chr19 | 4903092 | 4962165 | + | protein_coding |
| ENSG00000198399.14_6 | ITSN2 | 0.99999177 | 1 | -0.236 | 5.938 | chr2 | 24425733 | 24583583 | - | protein_coding |
| ENSG00000123009.4_2 | NME2P1 | 0.71756058 | 1 | -0.236 | -0.021 | chr12 | 120720106 | 120720519 | + | processed_pseudogene |
| ENSG00000100852.13_5 | ARHGAP5 | 0.99999676 | 1 | -0.236 | 7.781 | chr14 | 32545320 | 32628934 | + | protein_coding |
| ENSG00000122882.11_5 | ECD | 0.99998624 | 1 | -0.236 | 5.678 | chr10 | 74889913 | 74928813 | - | protein_coding |
| ENSG00000075975.16_3 | MKRN2 | 0.99993924 | 1 | -0.236 | 5.049 | chr3 | 12598586 | 12625212 | + | protein_coding |
| ENSG00000213080.3_5 | AL354714.2 | 0.83022618 | 1 | -0.237 | 0.968 | chr1 | 160864770 | 160865866 | + | processed_pseudogene |
| ENSG00000037965.6_4 | HOXC8 | 0.821866 | 1 | -0.237 | 0.732 | chr12 | 54402769 | 54406553 | + | protein_coding |
| ENSG00000180198.16_5 | RCC1 | 0.99999434 | 1 | -0.237 | 6.208 | chr1 | 28832455 | 28865812 | + | protein_coding |
| ENSG00000117597.18_5 | UTP25 | 0.99998746 | 1 | -0.237 | 5.876 | chr1 | 210001317 | 210030910 | + | protein_coding |
| ENSG00000224281.5_6 | SLC25A5-AS1 | 0.81070652 | 1 | -0.237 | 0.616 | chrX | 118599949 | 118603091 | - | lncRNA |
| ENSG00000232237.3_3 | ASCL5 | 0.69941678 | 1 | -0.237 | -0.303 | chr1 | 201083081 | 201096312 | - | protein_coding |
| ENSG00000162804.14_5 | SNED1 | 0.8128692 | 1 | -0.237 | 0.866 | chr2 | 241938035 | 242034983 | + | protein_coding |
| ENSG00000117906.14_6 | RCN2 | 0.99999526 | 1 | -0.237 | 6.831 | chr15 | 77224079 | 77246734 | + | protein_coding |
| ENSG00000124193.16_4 | SRSF6 | 0.99993957 | 1 | -0.238 | 6.663 | chr20 | 42086533 | 42094686 | + | protein_coding |
| ENSG00000014919.12_4 | COX15 | 0.99997127 | 1 | -0.238 | 5.603 | chr10 | 101471601 | 101491857 | - | protein_coding |
| ENSG00000147316.13_3 | MCPH1 | 0.99979351 | 1 | -0.238 | 4.631 | chr8 | 6264117 | 6506029 | + | protein_coding |
| ENSG00000167264.18_6 | DUS2 | 0.99875656 | 1 | -0.238 | 4.197 | chr16 | 68021649 | 68113223 | + | protein_coding |
| ENSG00000173726.11_4 | TOMM20 | 0.99999458 | 1 | -0.238 | 7.112 | chr1 | 235272656 | 235292152 | - | protein_coding |
| ENSG00000101654.17_5 | RNMT | 0.99996547 | 1 | -0.238 | 5.702 | chr18 | 13726659 | 13764557 | + | protein_coding |

| | | | | | | | | | | |
|---|---|---|---|---|---|---|---|---|---|---|
| ENSG00000160172.10_3 | FAM86C2P | 0.98613202 | 1 | -0.238 | 2.689 | chr11 | 67559119 | 67572807 | - | transcribed_unprocessed_pseudogene |
| ENSG00000110756.18_4 | HPS5 | 0.99998548 | 1 | -0.238 | 5.790 | chr11 | 18300215 | 18343745 | - | protein_coding |
| ENSG00000198743.7_6 | SLC5A3 | 0.9998962 | 1 | -0.239 | 5.519 | chr21 | 35445878 | 35478559 | + | protein_coding |
| ENSG00000152689.18_6 | RASGRP3 | 0.8975103 | 1 | -0.239 | 1.242 | chr2 | 33661391 | 33789817 | + | protein_coding |
| ENSG00000186141.9_4 | POLR3C | 0.99996403 | 1 | -0.239 | 5.252 | chr1 | 145590666 | 145611025 | - | protein_coding |
| ENSG00000081870.11_3 | HSPB11 | 0.99913143 | 1 | -0.239 | 4.323 | chr1 | 54382247 | 54411602 | - | protein_coding |
| ENSG00000149557.14_8 | FEZ1 | 0.69687149 | 1 | -0.239 | -0.451 | chr11 | 125312777 | 125462463 | - | protein_coding |
| ENSG00000225111.1_5 | AC083900.1 | 0.66989542 | 1 | -0.239 | -1.010 | chr2 | 208733306 | 208734639 | - | lncRNA |
| ENSG00000249855.1_4 | EEF1A1P19 | 0.85993906 | 1 | -0.239 | 1.136 | chr5 | 43495175 | 43496556 | - | processed_pseudogene |
| ENSG00000163874.11_4 | ZC3H12A | 0.99883556 | 1 | -0.239 | 4.079 | chr1 | 37940181 | 37949978 | + | protein_coding |
| ENSG00000013306.16_7 | SLC25A39 | 0.99999359 | 1 | -0.240 | 6.895 | chr17 | 42396993 | 42402238 | - | protein_coding |
| ENSG00000095752.7_6 | IL11 | 0.99220253 | 1 | -0.240 | 3.219 | chr19 | 55875750 | 55881831 | - | protein_coding |
| ENSG00000124228.14_7 | DDX27 | 0.99999375 | 1 | -0.240 | 6.191 | chr20 | 47835832 | 47860614 | + | protein_coding |
| ENSG00000158042.9_4 | MRPL17 | 0.99999331 | 1 | -0.240 | 6.219 | chr11 | 6701616 | 6704571 | - | protein_coding |
| ENSG00000124784.9_3 | RIOK1 | 0.99840894 | 1 | -0.240 | 3.928 | chr6 | 7390026 | 7418270 | + | protein_coding |
| ENSG00000196154.12_5 | S100A4 | 0.99999752 | 1 | -0.240 | 8.515 | chr1 | 153516089 | 153522612 | - | protein_coding |
| ENSG00000225177.6_8 | AL590617.2 | 0.78463184 | 1 | -0.240 | 0.571 | chr6 | 139013685 | 139018425 | + | lncRNA |
| ENSG00000271270.7_6 | TMCC1-AS1 | 0.99677646 | 1 | -0.240 | 3.785 | chr3 | 129612654 | 129637418 | + | lncRNA |
| ENSG00000124562.10_4 | SNRPC | 0.99988086 | 1 | -0.240 | 5.827 | chr6 | 34725282 | 34741634 | + | protein_coding |
| ENSG00000162702.8_4 | ZNF281 | 0.99999724 | 1 | -0.240 | 6.553 | chr1 | 200374068 | 200379184 | - | protein_coding |
| ENSG00000223797.6_8 | ENTPD3-AS1 | 0.88385866 | 1 | -0.240 | 0.881 | chr3 | 40355293 | 40494820 | - | lncRNA |
| ENSG00000106948.16_4 | AKNA | 0.99468052 | 1 | -0.240 | 3.368 | chr9 | 117096436 | 117156685 | - | protein_coding |
| ENSG00000134748.13_3 | PRPF38A | 0.99969434 | 1 | -0.240 | 4.438 | chr1 | 52870274 | 52886508 | + | protein_coding |
| ENSG00000170458.14_5 | CD14 | 0.96939415 | 1 | -0.240 | 2.399 | chr5 | 140011313 | 140013286 | - | protein_coding |
| ENSG00000119326.15_5 | CTNNAL1 | 0.99998982 | 1 | -0.241 | 6.633 | chr9 | 111704849 | 111775802 | - | protein_coding |
| ENSG00000214827.11_6 | MTCP1 | 0.82503561 | 1 | -0.241 | 0.709 | chrX | 154292309 | 154376212 | - | protein_coding |
| ENSG00000144815.16_4 | NXPE3 | 0.99995908 | 1 | -0.241 | 5.379 | chr3 | 101498046 | 101547075 | + | protein_coding |
| ENSG00000156313.14_6 | RPGR | 0.99161527 | 1 | -0.241 | 4.243 | chrX | 38128423 | 38186797 | - | protein_coding |
| ENSG00000158321.18_8 | AUTS2 | 0.91509463 | 1 | -0.241 | 1.944 | chr7 | 69063282 | 70258492 | + | protein_coding |
| ENSG00000146247.14_5 | PHIP | 0.99992563 | 1 | -0.241 | 6.740 | chr6 | 79644136 | 79787971 | - | protein_coding |
| ENSG00000159899.14_2 | NPR2 | 0.98529464 | 1 | -0.241 | 2.663 | chr9 | 35792151 | 35809729 | + | protein_coding |
| ENSG00000159210.10_9 | SNF8 | 0.99996037 | 1 | -0.241 | 5.517 | chr17 | 47006678 | 47022204 | - | protein_coding |
| ENSG00000125691.13_7 | RPL23 | 0.99999427 | 1 | -0.241 | 9.213 | chr17 | 37004113 | 37010017 | - | protein_coding |
| ENSG00000087263.17_3 | OGFOD1 | 0.99995426 | 1 | -0.241 | 5.820 | chr16 | 56485433 | 56513016 | + | protein_coding |
| ENSG00000229425.3_7 | AJ009632.2 | 0.93541122 | 1 | -0.241 | 1.811 | chr21 | 16741285 | 16999661 | - | lncRNA |
| ENSG00000119965.13_4 | C10orf88 | 0.994073 | 1 | -0.241 | 3.606 | chr10 | 124690417 | 124713827 | - | protein_coding |
| ENSG00000160131.13_4 | VMA21 | 0.9999977 | 1 | -0.241 | 7.347 | chrX | 150564987 | 150577836 | + | protein_coding |
| ENSG00000124279.12_5 | FASTKD3 | 0.99915891 | 1 | -0.241 | 4.352 | chr5 | 7859272 | 7869150 | - | protein_coding |
| ENSG00000231066.3_4 | NPM1P9 | 0.6717093 | 1 | -0.241 | -0.862 | chrX | 14852598 | 14853441 | - | processed_pseudogene |
| ENSG00000011376.12_8 | LARS2 | 0.99967529 | 1 | -0.241 | 4.776 | chr3 | 45430053 | 45596218 | + | protein_coding |
| ENSG00000198918.8_4 | RPL39 | 0.99999343 | 1 | -0.241 | 7.070 | chrX | 118920467 | 118925593 | - | protein_coding |
| ENSG00000122545.20_7 | SEPTIN7 | 0.99999301 | 1 | -0.241 | 7.631 | chr7 | 35840542 | 35946715 | + | protein_coding |
| ENSG00000270019.1_6 | AC110769.2 | 0.7470984 | 1 | -0.242 | 0.107 | chr2 | 114645780 | 114647327 | - | lncRNA |
| ENSG00000182841.12_5 | RRP7BP | 0.98883141 | 1 | -0.242 | 2.837 | chr22 | 42951229 | 42978044 | - | transcribed_unprocessed_pseudogene |
| ENSG00000273297.2_8 | AC009275.1 | 0.99645909 | 1 | -0.242 | 3.919 | chr7 | 134053489 | 134117286 | - | lncRNA |
| ENSG00000163655.16_3 | GMPS | 0.99999838 | 1 | -0.242 | 7.186 | chr3 | 155588439 | 155661809 | + | protein_coding |
| ENSG00000111087.10_4 | GLI1 | 0.99966106 | 1 | -0.242 | 4.425 | chr12 | 57853568 | 57866051 | + | protein_coding |
| ENSG00000254595.1_6 | AC084337.1 | 0.7160611 | 1 | -0.242 | -0.120 | chr11 | 6509416 | 6510607 | - | processed_pseudogene |
| ENSG00000158715.6_4 | SLC45A3 | 0.95121705 | 1 | -0.242 | 1.970 | chr1 | 205626979 | 205649637 | - | protein_coding |
| ENSG00000085185.15_6 | BCORL1 | 0.96772843 | 1 | -0.242 | 2.183 | chrX | 129115083 | 129192058 | + | protein_coding |
| ENSG00000165526.9_6 | RPUSD4 | 0.99935268 | 1 | -0.242 | 4.290 | chr11 | 126071991 | 126081587 | - | protein_coding |
| ENSG00000188878.19_6 | FBF1 | 0.99644288 | 1 | -0.242 | 3.697 | chr17 | 73905655 | 73937221 | - | protein_coding |
| ENSG00000196704.12_6 | AMZ2 | 0.99999101 | 1 | -0.242 | 6.701 | chr17 | 66244071 | 66253305 | + | protein_coding |
| ENSG00000025772.8_4 | TOMM34 | 0.99987546 | 1 | -0.242 | 5.033 | chr20 | 43570771 | 43589038 | - | protein_coding |
| ENSG00000106991.13_3 | ENG | 0.93686029 | 1 | -0.243 | 1.379 | chr9 | 130577291 | 130617035 | - | protein_coding |
| ENSG00000101057.16_3 | MYBL2 | 0.99996407 | 1 | -0.243 | 5.926 | chr20 | 42295659 | 42345135 | + | protein_coding |
| ENSG00000026036.22_4 | RTEL1-TNFRSF6B | 0.95115398 | 1 | -0.243 | 2.041 | chr20 | 62290653 | 62330037 | + | protein_coding |

| | | | | | | | | | | |
|---|---|---|---|---|---|---|---|---|---|---|
| ENSG00000183682.8_4 | BMP8A | 0.96385925 | 1 | -0.243 | 2.076 | chr1 | 39957308 | 39995541 | + | protein_coding |
| ENSG00000271828.1_6 | AC008937.3 | 0.75088585 | 1 | -0.243 | 0.106 | chr5 | 56223701 | 56225400 | + | lncRNA |
| ENSG00000196369.6 | SRGAP2B | 0.99579545 | 1 | -0.243 | 3.393 | chr1 | 143913615 | 144094424 | + | pseudogene |
| ENSG00000133392.18_6 | MYH11 | 0.77668777 | 1 | -0.243 | 0.496 | chr16 | 15796992 | 15950885 | - | protein_coding |
| ENSG00000212978.6_7 | AC016747.1 | 0.99940543 | 1 | -0.243 | 4.414 | chr2 | 61368727 | 61372104 | - | lncRNA |
| ENSG00000160214.13_3 | RRP1 | 0.99980434 | 1 | -0.243 | 5.001 | chr21 | 45209394 | 45225174 | + | protein_coding |
| ENSG00000241837.7_6 | ATP5PO | 0.98642998 | 1 | -0.243 | 3.633 | chr21 | 35275757 | 35288118 | - | protein_coding |
| ENSG00000156256.15_5 | USP16 | 0.99998602 | 1 | -0.243 | 6.120 | chr21 | 30396950 | 30426809 | + | protein_coding |
| ENSG00000287265.1_2 | AL365272.1 | 0.66559673 | 1 | -0.243 | -1.047 | chr6 | 285443 | 292013 | - | lncRNA |
| ENSG00000148840.11_4 | PPRC1 | 0.99999265 | 1 | -0.243 | 6.309 | chr10 | 103892751 | 103910090 | + | protein_coding |
| ENSG00000179029.15_7 | TMEM107 | 0.99890405 | 1 | -0.244 | 4.037 | chr17 | 8075775 | 8079717 | - | protein_coding |
| ENSG00000183018.9_6 | SPNS2 | 0.97788663 | 1 | -0.244 | 2.430 | chr17 | 4402176 | 4442330 | + | protein_coding |
| ENSG00000250848.1_7 | AC021087.2 | 0.67348892 | 1 | -0.244 | -0.681 | chr5 | 288948 | 290436 | - | processed_pseudogene |
| ENSG00000032389.12_9 | EIPR1 | 0.99994242 | 1 | -0.244 | 5.360 | chr2 | 3192696 | 3381653 | - | protein_coding |
| ENSG00000148110.16_4 | MFSD14B | 0.99999511 | 1 | -0.244 | 6.500 | chr9 | 97136851 | 97223324 | + | protein_coding |
| ENSG00000103067.14_8 | ESRP2 | 0.66328305 | 1 | -0.244 | -1.012 | chr16 | 68262936 | 68272005 | - | protein_coding |
| ENSG00000253190.4_7 | AC084082.1 | 0.80338018 | 1 | -0.244 | 0.522 | chr8 | 67024902 | 67038867 | + | lncRNA |
| ENSG00000143252.14_2 | SDHC | 0.9999544 | 1 | -0.244 | 5.211 | chr1 | 161284047 | 161345130 | + | protein_coding |
| ENSG00000113387.12_4 | SUB1 | 0.99999329 | 1 | -0.244 | 7.190 | chr5 | 32531739 | 32604185 | + | protein_coding |
| ENSG00000244184.1_6 | AC091153.3 | 0.67315173 | 1 | -0.244 | -0.613 | chr17 | 4607525 | 4608824 | + | lncRNA |
| ENSG00000128536.16_4 | CDHR3 | 0.96427084 | 1 | -0.244 | 2.257 | chr7 | 105517242 | 105676878 | + | protein_coding |
| ENSG00000102978.13_4 | POLR2C | 0.99999571 | 1 | -0.244 | 6.524 | chr16 | 57496572 | 57505921 | + | protein_coding |
| ENSG00000259370.2_7 | AC103740.1 | 0.71093339 | 1 | -0.245 | -0.040 | chr15 | 63119874 | 63176233 | - | lncRNA |
| ENSG00000132763.15_3 | MMACHC | 0.99580527 | 1 | -0.245 | 3.893 | chr1 | 45965972 | 45979054 | + | protein_coding |
| ENSG00000101109.12_4 | STK4 | 0.99998282 | 1 | -0.245 | 5.854 | chr20 | 43595120 | 43708662 | + | protein_coding |
| ENSG00000132692.19_6 | BCAN | 0.95666267 | 1 | -0.245 | 2.069 | chr1 | 156611182 | 156629324 | + | protein_coding |
| ENSG00000164104.12_6 | HMGB2 | 0.99999118 | 1 | -0.245 | 6.125 | chr4 | 174252527 | 174255583 | - | protein_coding |
| ENSG00000125676.20_5 | THOC2 | 0.99999655 | 1 | -0.245 | 7.252 | chrX | 122734412 | 122866906 | - | protein_coding |
| ENSG00000126001.16_5 | CEP250 | 0.99998073 | 1 | -0.245 | 6.184 | chr20 | 34042985 | 34107108 | + | protein_coding |
| ENSG00000118412.12_5 | CASP8AP2 | 0.99996957 | 1 | -0.245 | 5.831 | chr6 | 90539613 | 90584155 | + | protein_coding |
| ENSG00000251484.3_6 | AL356488.1 | 0.69398786 | 1 | -0.245 | -0.406 | chr1 | 109646157 | 109647388 | - | processed_pseudogene |
| ENSG00000088387.19_7 | DOCK9 | 0.99997447 | 1 | -0.245 | 5.632 | chr13 | 99445683 | 99738879 | - | protein_coding |
| ENSG00000242082.2_8 | SLC5A4-AS1 | 0.78545501 | 1 | -0.246 | 0.254 | chr22 | 32601102 | 32674369 | + | lncRNA |
| ENSG00000149639.15_7 | SOGA1 | 0.99992593 | 1 | -0.246 | 5.572 | chr20 | 35405850 | 35491941 | - | protein_coding |
| ENSG00000156374.16_6 | PCGF6 | 0.99866334 | 1 | -0.246 | 3.910 | chr10 | 105062553 | 105110901 | - | protein_coding |
| ENSG00000278023.6_6 | RDM1 | 0.84746788 | 1 | -0.246 | 1.589 | chr17 | 34245070 | 34257777 | - | protein_coding |
| ENSG00000072415.9_7 | MPP5 | 0.99998061 | 1 | -0.246 | 5.942 | chr14 | 67708059 | 67802778 | + | protein_coding |
| ENSG00000136868.11_3 | SLC31A1 | 0.99983258 | 1 | -0.246 | 5.367 | chr9 | 115983824 | 116026772 | + | protein_coding |
| ENSG00000184060.11_5 | ADAP2 | 0.96294437 | 1 | -0.246 | 2.179 | chr17 | 29233362 | 29286340 | + | protein_coding |
| ENSG00000101773.19_6 | RBBP8 | 0.99992324 | 1 | -0.246 | 5.840 | chr18 | 20378224 | 20606451 | + | protein_coding |
| ENSG00000187764.11_5 | SEMA4D | 0.99885816 | 1 | -0.246 | 4.029 | chr9 | 91975702 | 92113045 | - | protein_coding |
| ENSG00000182108.11_7 | DEXI | 0.96175039 | 1 | -0.246 | 2.152 | chr16 | 11022748 | 11036325 | - | protein_coding |
| ENSG00000164022.17_3 | AIMP1 | 0.99994176 | 1 | -0.246 | 5.848 | chr4 | 107236701 | 107270613 | + | protein_coding |
| ENSG00000167202.11_4 | TBC1D2B | 0.99990569 | 1 | -0.246 | 5.043 | chr15 | 78276378 | 78370066 | - | protein_coding |
| ENSG00000179833.4_5 | SERTAD2 | 0.9997672 | 1 | -0.246 | 4.703 | chr2 | 64858755 | 64978139 | - | protein_coding |
| ENSG00000278535.5_5 | DHRS11 | 0.98862608 | 1 | -0.247 | 3.067 | chr17 | 34948309 | 34957233 | + | protein_coding |
| ENSG00000203875.13_7 | SNHG5 | 0.98430046 | 1 | -0.247 | 5.902 | chr6 | 86360209 | 86388650 | - | lncRNA |
| ENSG00000127463.15_8 | EMC1 | 0.9999911 | 1 | -0.247 | 6.020 | chr1 | 19542154 | 19578021 | - | protein_coding |
| ENSG00000157193.18_8 | LRP8 | 0.99999363 | 1 | -0.247 | 6.332 | chr1 | 53708036 | 53794141 | - | protein_coding |
| ENSG00000163867.17_6 | ZMYM6 | 0.9861825 | 1 | -0.247 | 3.090 | chr1 | 35451766 | 35497546 | - | protein_coding |
| ENSG00000135631.17_4 | RAB11FIP5 | 0.99990572 | 1 | -0.247 | 5.121 | chr2 | 73300510 | 73383849 | - | protein_coding |
| ENSG00000177156.11_3 | TALDO1 | 0.99999861 | 1 | -0.247 | 7.811 | chr11 | 747415 | 765012 | + | protein_coding |
| ENSG00000082515.18_4 | MRPL22 | 0.99681823 | 1 | -0.247 | 3.642 | chr5 | 154320633 | 154348971 | + | protein_coding |
| ENSG00000165392.11_4 | WRN | 0.99982912 | 1 | -0.248 | 5.052 | chr8 | 30891304 | 31033654 | + | protein_coding |
| ENSG00000265907.2_6 | AP000919.2 | 0.66038031 | 1 | -0.248 | -0.831 | chr18 | 2948236 | 2960754 | + | lncRNA |
| ENSG00000133398.4_4 | MED10 | 0.99333725 | 1 | -0.248 | 3.320 | chr5 | 6371987 | 6378684 | - | protein_coding |
| ENSG00000104177.18_7 | MYEF2 | 0.99714094 | 1 | -0.248 | 4.664 | chr15 | 48426829 | 48470550 | - | protein_coding |
| ENSG00000035681.9_6 | NSMAF | 0.99997859 | 1 | -0.248 | 5.900 | chr8 | 59496067 | 59572412 | - | protein_coding |
| ENSG00000172007.7_5 | RAB33B | 0.99283015 | 1 | -0.248 | 3.496 | chr4 | 140374386 | 140397763 | + | protein_coding |
| ENSG00000103202.13_6 | NME4 | 0.99997515 | 1 | -0.248 | 6.718 | chr16 | 446725 | 460367 | + | protein_coding |

| | | | | | | | | | | |
|---|---|---|---|---|---|---|---|---|---|---|
| ENSG00000129460.16_6 | NGDN | 0.99987841 | 1 | -0.248 | 5.123 | chr14 | 23938898 | 23979071 | + | protein_coding |
| ENSG00000166016.6_5 | ABTB2 | 0.99873565 | 1 | -0.248 | 4.746 | chr11 | 34172534 | 34379557 | - | protein_coding |
| ENSG00000116266.11_3 | STXBP3 | 0.99975328 | 1 | -0.248 | 5.073 | chr1 | 109289296 | 109352145 | + | protein_coding |
| ENSG00000260086.3_7 | AC007611.1 | 0.71357756 | 1 | -0.248 | -0.178 | chr16 | 48657333 | 48780229 | + | lncRNA |
| ENSG00000086712.13_4 | TXLNG | 0.99995949 | 1 | -0.248 | 5.318 | chrX | 16804555 | 16862642 | + | protein_coding |
| ENSG00000230409.3_4 | TCEA1P2 | 0.86572078 | 1 | -0.248 | 2.245 | chr3 | 37317184 | 37318089 | + | processed_pseudogene |
| ENSG00000120053.12_5 | GOT1 | 0.99966129 | 1 | -0.248 | 4.856 | chr10 | 101156627 | 101190381 | - | protein_coding |
| ENSG00000260095.1_6 | AC106820.3 | 0.67717216 | 1 | -0.249 | -0.562 | chr16 | 2502582 | 2502978 | - | lncRNA |
| ENSG00000101019.22_6 | UQCC1 | 0.99986111 | 1 | -0.249 | 4.815 | chr20 | 33890369 | 33999834 | - | protein_coding |
| ENSG00000147601.14_3 | TERF1 | 0.99965996 | 1 | -0.249 | 5.027 | chr8 | 73921099 | 73960358 | + | protein_coding |
| ENSG00000012048.23_7 | BRCA1 | 0.99999604 | 1 | -0.249 | 7.066 | chr17 | 41196312 | 41322262 | - | protein_coding |
| ENSG00000213212.3_4 | NCLP1 | 0.79587385 | 1 | -0.249 | 0.513 | chr9 | 139707240 | 139709988 | - | processed_pseudogene |
| ENSG00000181381.13_4 | DDX60L | 0.99976043 | 1 | -0.249 | 5.079 | chr4 | 169277886 | 169458937 | - | protein_coding |
| ENSG00000139351.15_5 | SYCP3 | 0.75510922 | 1 | -0.249 | 0.658 | chr12 | 102122426 | 102133250 | - | protein_coding |
| ENSG00000143753.13_4 | DEGS1 | 0.99999257 | 1 | -0.249 | 6.178 | chr1 | 224363458 | 224381143 | + | protein_coding |
| ENSG00000250182.3_4 | EEF1A1P13 | 0.94206506 | 1 | -0.250 | 1.498 | chr5 | 14652050 | 14653438 | - | processed_pseudogene |
| ENSG00000227097.5_4 | RPS28P7 | 0.99730421 | 1 | -0.250 | 3.932 | chr11 | 82400601 | 82400810 | + | processed_pseudogene |
| ENSG00000183287.14_9 | CCBE1 | 0.91319026 | 1 | -0.250 | 1.587 | chr18 | 57098171 | 57364894 | - | protein_coding |
| ENSG00000168496.4_4 | FEN1 | 0.99998788 | 1 | -0.250 | 6.457 | chr11 | 61560383 | 61564710 | + | protein_coding |
| ENSG00000280407.2_7 | AC132872.4 | 0.73129817 | 1 | -0.250 | 0.376 | chr17 | 80206943 | 80209720 | + | TEC |
| ENSG00000263740.1 | RN7SL4P | 0.99375172 | 1 | -0.250 | 4.017 | chr3 | 15780022 | 15780315 | + | misc_RNA |
| ENSG00000149571.12_6 | KIRREL3 | 0.99914363 | 1 | -0.250 | 4.294 | chr11 | 126293253 | 126873355 | - | protein_coding |
| ENSG00000090487.11_4 | SPG21 | 0.99999378 | 1 | -0.250 | 6.575 | chr15 | 65255363 | 65282648 | - | protein_coding |
| ENSG00000077157.22_7 | PPP1R12B | 0.99985986 | 1 | -0.251 | 4.958 | chr1 | 202317827 | 202561834 | + | protein_coding |
| ENSG00000248231.1_5 | KRT8P4 | 0.66854271 | 1 | -0.251 | -0.739 | chr8 | 90424850 | 90427300 | - | processed_pseudogene |
| ENSG00000145685.14_6 | LHFPL2 | 0.99999408 | 1 | -0.251 | 6.381 | chr5 | 77781038 | 78065844 | - | protein_coding |
| ENSG00000147471.12_7 | PLPBP | 0.9999004 | 1 | -0.251 | 5.270 | chr8 | 37620113 | 37637286 | + | protein_coding |
| ENSG00000078795.16_4 | PKD2L2 | 0.70200298 | 1 | -0.251 | -0.242 | chr5 | 137223657 | 137278436 | + | protein_coding |
| ENSG00000240288.8_6 | GHRLOS | 0.7017765 | 1 | -0.251 | -0.269 | chr3 | 10327438 | 10336587 | + | lncRNA |
| ENSG00000151327.12_5 | FAM177A1 | 0.99997312 | 1 | -0.251 | 5.875 | chr14 | 35514113 | 35582336 | + | protein_coding |
| ENSG00000168273.8_3 | SMIM4 | 0.97362377 | 1 | -0.251 | 2.352 | chr3 | 52568029 | 52613253 | + | protein_coding |
| ENSG00000166833.21_8 | NAV2 | 0.99953533 | 1 | -0.251 | 4.539 | chr11 | 19372271 | 20143147 | + | protein_coding |
| ENSG00000236015.1_5 | AC011290.2 | 0.75003516 | 1 | -0.251 | 0.286 | chr7 | 39608975 | 39609797 | - | processed_pseudogene |
| ENSG00000262265.2_8 | AC002558.3 | 0.7854391 | 1 | -0.251 | 0.445 | chr17 | 45237764 | 45245089 | - | lncRNA |
| ENSG00000204220.11_3 | PFDN6 | 0.99887065 | 1 | -0.252 | 3.987 | chr6 | 33257079 | 33266178 | + | protein_coding |
| ENSG00000123575.9_3 | FAM199X | 0.99999532 | 1 | -0.252 | 6.704 | chrX | 103411134 | 103440583 | + | protein_coding |
| ENSG00000103356.16_4 | EARS2 | 0.99994542 | 1 | -0.252 | 5.992 | chr16 | 23533337 | 23569052 | - | protein_coding |
| ENSG00000163608.15_7 | NEPRO | 0.999978 | 1 | -0.252 | 6.093 | chr3 | 112721291 | 112738708 | - | protein_coding |
| ENSG00000166266.14_4 | CUL5 | 0.99995895 | 1 | -0.252 | 5.682 | chr11 | 107879624 | 107978488 | + | protein_coding |
| ENSG00000124191.18_5 | TOX2 | 0.88187444 | 1 | -0.252 | 1.820 | chr20 | 42543492 | 42698256 | + | protein_coding |
| ENSG00000263563.1 | UBBP4 | 0.81999673 | 1 | -0.252 | 0.799 | chr17 | 21729601 | 21731762 | + | protein_coding |
| ENSG00000104660.19_5 | LEPROTL1 | 0.99995732 | 1 | -0.252 | 5.968 | chr8 | 29952924 | 30034724 | + | protein_coding |
| ENSG00000260318.1_2 | COX6CP1 | 0.65355666 | 1 | -0.252 | -0.798 | chr16 | 11997780 | 11997994 | - | processed_pseudogene |
| ENSG00000167005.14_6 | NUDT21 | 0.99999643 | 1 | -0.252 | 7.342 | chr16 | 56463045 | 56486111 | - | protein_coding |
| ENSG00000258890.7_7 | CEP95 | 0.99988024 | 1 | -0.252 | 5.339 | chr17 | 62502983 | 62538579 | + | protein_coding |
| ENSG00000123136.15_5 | DDX39A | 0.99997603 | 1 | -0.253 | 6.003 | chr19 | 14519610 | 14530195 | - | protein_coding |
| ENSG00000240449.1_7 | AC005586.1 | 0.7652025 | 1 | -0.253 | 0.567 | chr7 | 150060866 | 150069679 | - | lncRNA |
| ENSG00000225526.4_5 | MKRN2OS | 0.70407041 | 1 | -0.253 | -0.160 | chr3 | 12556433 | 12602558 | - | protein_coding |
| ENSG00000128513.16_6 | POT1 | 0.99976117 | 1 | -0.253 | 4.880 | chr7 | 124462440 | 124570037 | - | protein_coding |
| ENSG00000100058.12_3 | CRYBB2P1 | 0.98004943 | 1 | -0.253 | 2.622 | chr22 | 25844072 | 25916821 | + | transcribed_unprocessed_pseudogene |
| ENSG00000117477.12_4 | CCDC181 | 0.72428598 | 1 | -0.253 | -0.120 | chr1 | 169364108 | 169429907 | - | protein_coding |
| ENSG00000139842.15_3 | CUL4A | 0.99999274 | 1 | -0.253 | 6.376 | chr13 | 113862507 | 113921422 | + | protein_coding |
| ENSG00000234009.1_4 | RPL5P34 | 0.73602249 | 1 | -0.253 | 0.436 | chr22 | 43172412 | 43173302 | - | processed_pseudogene |
| ENSG00000138138.13_3 | ATAD1 | 0.99997564 | 1 | -0.253 | 6.436 | chr10 | 89511269 | 89601100 | - | protein_coding |
| ENSG00000183808.11_4 | RBM12B | 0.9998465 | 1 | -0.253 | 4.952 | chr8 | 94741584 | 94753245 | - | protein_coding |
| ENSG00000177192.14_5 | PUS1 | 0.99974497 | 1 | -0.253 | 4.977 | chr12 | 132413745 | 132430441 | + | protein_coding |

| | | | | | | | | | | |
|---|---|---|---|---|---|---|---|---|---|---|
| ENSG00000149218.5_3 | ENDOD1 | 0.99933927 | 1 | -0.253 | 4.354 | chr11 | 94823010 | 94865809 | + | protein_coding |
| ENSG00000145425.10_5 | RPS3A | 0.99851071 | 1 | -0.253 | 7.599 | chr4 | 152020776 | 152025794 | + | protein_coding |
| ENSG00000232112.3_3 | TMA7 | 0.97989293 | 1 | -0.253 | 3.286 | chr3 | 48481667 | 48485616 | + | protein_coding |
| ENSG00000147416.11_6 | ATP6V1B2 | 0.99996182 | 1 | -0.254 | 5.523 | chr8 | 20054892 | 20084330 | + | protein_coding |
| ENSG00000169976.7_3 | SF3B5 | 0.99983441 | 1 | -0.254 | 5.103 | chr6 | 144416021 | 144416710 | - | protein_coding |
| ENSG00000142230.13_5 | SAE1 | 0.9999981 | 1 | -0.254 | 7.720 | chr19 | 47616531 | 47713893 | + | protein_coding |
| ENSG00000233588.1_5 | CYP51A1P2 | 0.70328396 | 1 | -0.254 | -0.206 | chr13 | 29296523 | 29298158 | - | processed_pseudogene |
| ENSG00000235957.1_2 | COX7CP1 | 0.66189365 | 1 | -0.254 | -0.769 | chr13 | 49761615 | 49761806 | - | processed_pseudogene |
| ENSG00000136738.15_3 | STAM | 0.99992635 | 1 | -0.254 | 5.140 | chr10 | 17686150 | 17758823 | + | protein_coding |
| ENSG00000043514.17_6 | TRIT1 | 0.99896323 | 1 | -0.254 | 4.337 | chr1 | 40303782 | 40349183 | - | protein_coding |
| ENSG00000271709.1_7 | AC017033.1 | 0.66427802 | 1 | -0.254 | -0.435 | chr2 | 121623953 | 121624978 | - | lncRNA |
| ENSG00000265574.1_4 | WDR45BP1 | 0.65117104 | 1 | -0.254 | -0.831 | chr17 | 30438581 | 30439609 | - | processed_pseudogene |
| ENSG00000214078.12_8 | CPNE1 | 0.99999476 | 1 | -0.254 | 6.451 | chr20 | 34213953 | 34252878 | - | protein_coding |
| ENSG00000104131.13_4 | EIF3J | 0.99992341 | 1 | -0.254 | 5.562 | chr15 | 44829323 | 44855227 | + | protein_coding |
| ENSG00000123908.12_5 | AGO2 | 0.99996847 | 1 | -0.254 | 6.135 | chr8 | 141530255 | 141645732 | - | protein_coding |
| ENSG00000070495.14_4 | JMJD6 | 0.99986015 | 1 | -0.254 | 5.125 | chr17 | 74708914 | 74722881 | - | protein_coding |
| ENSG00000108439.10_7 | PNPO | 0.99974132 | 1 | -0.254 | 4.745 | chr17 | 46018872 | 46026674 | + | protein_coding |
| ENSG00000206573.9_9 | THUMPD3-AS1 | 0.99987807 | 1 | -0.254 | 5.264 | chr3 | 9391373 | 9440263 | - | lncRNA |
| ENSG00000124356.16_4 | STAMBP | 0.99997775 | 1 | -0.255 | 5.795 | chr2 | 74056043 | 74100786 | + | protein_coding |
| ENSG00000128915.12_7 | ICE2 | 0.99998742 | 1 | -0.255 | 6.507 | chr15 | 60711808 | 60771359 | - | protein_coding |
| ENSG00000165416.14_3 | SUGT1 | 0.99989639 | 1 | -0.255 | 5.892 | chr13 | 53226844 | 53275044 | + | protein_coding |
| ENSG00000272788.1_8 | AP000864.2 | 0.72131196 | 1 | -0.255 | 0.205 | chr18 | 8801654 | 8802303 | + | lncRNA |
| ENSG00000164051.14_8 | CCDC51 | 0.99889206 | 1 | -0.255 | 4.354 | chr3 | 48473574 | 48481866 | - | protein_coding |
| ENSG00000008517.16_5 | IL32 | 0.70053815 | 1 | -0.255 | -0.314 | chr16 | 3115298 | 3132193 | + | protein_coding |
| ENSG00000250486.4_3 | FAM218A | 0.66688543 | 1 | -0.255 | -0.381 | chr4 | 165878100 | 165880274 | + | protein_coding |
| ENSG00000133627.18_3 | ACTR3B | 0.97885429 | 1 | -0.255 | 2.718 | chr7 | 152456834 | 152552463 | + | protein_coding |
| ENSG00000214413.8_6 | BBIP1 | 0.99969692 | 1 | -0.255 | 4.627 | chr10 | 112658486 | 112678959 | - | protein_coding |
| ENSG00000162601.11_5 | MYSM1 | 0.99915483 | 1 | -0.255 | 4.779 | chr1 | 59109112 | 59165749 | - | protein_coding |
| ENSG00000174446.13_7 | SNAPC5 | 0.99973909 | 1 | -0.255 | 4.848 | chr15 | 66782473 | 66790118 | - | protein_coding |
| ENSG00000115758.13_8 | ODC1 | 0.99999661 | 1 | -0.255 | 6.956 | chr2 | 10580094 | 10588453 | - | protein_coding |
| ENSG00000129480.13_5 | DTD2 | 0.99187912 | 1 | -0.255 | 3.799 | chr14 | 31915242 | 31926712 | - | protein_coding |
| ENSG00000133069.17_6 | TMCC2 | 0.91544949 | 1 | -0.255 | 1.517 | chr1 | 205197074 | 205254760 | + | protein_coding |
| ENSG00000262655.4_4 | SPON1 | 0.65979159 | 1 | -0.255 | -0.485 | chr11 | 13984270 | 14289679 | + | protein_coding |
| ENSG00000183780.13_4 | SLC35F3 | 0.99690234 | 1 | -0.255 | 3.886 | chr1 | 234040422 | 234460257 | + | protein_coding |
| ENSG00000182704.8_5 | TSKU | 0.99991352 | 1 | -0.255 | 7.788 | chr11 | 76493295 | 76509197 | + | protein_coding |
| ENSG00000121579.13_5 | NAA50 | 0.99999486 | 1 | -0.255 | 7.498 | chr3 | 113435305 | 113465147 | - | protein_coding |
| ENSG00000108950.12_4 | FAM20A | 0.9150288 | 1 | -0.256 | 1.746 | chr17 | 66531254 | 66597508 | - | protein_coding |
| ENSG00000065833.9_4 | ME1 | 0.99999524 | 1 | -0.256 | 6.828 | chr6 | 83920121 | 84140770 | - | protein_coding |
| ENSG00000149313.11_3 | AASDHPPT | 0.99996593 | 1 | -0.256 | 5.964 | chr11 | 105946228 | 105969426 | + | protein_coding |
| ENSG00000280010.1_5 | AP001350.2 | 0.68490199 | 1 | -0.256 | -0.462 | chr11 | 58394908 | 58396001 | + | TEC |
| ENSG00000084693.16_4 | AGBL5 | 0.99991552 | 1 | -0.256 | 5.432 | chr2 | 27265232 | 27293490 | + | protein_coding |
| ENSG00000139826.6_3 | ABHD13 | 0.99874208 | 1 | -0.256 | 4.053 | chr13 | 108870740 | 108886591 | + | protein_coding |
| ENSG00000147684.9_4 | NDUFB9 | 0.99983082 | 1 | -0.256 | 5.248 | chr8 | 125551342 | 125592889 | + | protein_coding |
| ENSG00000109911.19_5 | ELP4 | 0.99871448 | 1 | -0.256 | 4.290 | chr11 | 31531302 | 31811872 | + | protein_coding |
| ENSG00000121022.14_6 | COPS5 | 0.99997635 | 1 | -0.256 | 5.756 | chr8 | 67955314 | 67996018 | - | protein_coding |
| ENSG00000064419.13_4 | TNPO3 | 0.99999517 | 1 | -0.256 | 6.893 | chr7 | 128594234 | 128695227 | - | protein_coding |
| ENSG00000273203.1_6 | AC006946.2 | 0.72380455 | 1 | -0.256 | -0.162 | chr22 | 17548711 | 17551565 | + | lncRNA |
| ENSG00000175193.13_7 | PARL | 0.96851636 | 1 | -0.256 | 1.825 | chr3 | 183547059 | 183602721 | - | protein_coding |
| ENSG00000137760.14_4 | ALKBH8 | 0.99625827 | 1 | -0.256 | 3.764 | chr11 | 107373452 | 107436472 | - | protein_coding |
| ENSG00000103037.11_3 | SETD6 | 0.99740808 | 1 | -0.257 | 3.946 | chr16 | 58549383 | 58555085 | + | protein_coding |
| ENSG00000006118.14_7 | TMEM132A | 0.9877225 | 1 | -0.257 | 6.020 | chr11 | 60691935 | 60704631 | + | protein_coding |
| ENSG00000032742.17_5 | IFT88 | 0.99972487 | 1 | -0.257 | 4.945 | chr13 | 21141208 | 21265576 | + | protein_coding |
| ENSG00000271889.1_7 | AC016747.2 | 0.68290383 | 1 | -0.257 | -0.437 | chr2 | 61378568 | 61389240 | - | lncRNA |
| ENSG00000047932.14_6 | GOPC | 0.99994685 | 1 | -0.257 | 5.486 | chr6 | 117881432 | 117923705 | - | protein_coding |
| ENSG00000267264.1_5 | AC006504.3 | 0.6579803 | 1 | -0.257 | -0.313 | chr19 | 28294446 | 28294968 | + | processed_pseudogene |
| ENSG00000122643.21_7 | NT5C3A | 0.99990432 | 1 | -0.257 | 5.296 | chr7 | 33053726 | 33102409 | - | protein_coding |
| ENSG00000144485.11_6 | HES6 | 0.90100291 | 1 | -0.257 | 2.299 | chr2 | 239146908 | 239149303 | - | protein_coding |
| ENSG00000262777.1_6 | AC032044.1 | 0.66806235 | 1 | -0.257 | -0.272 | chr17 | 1327767 | 1329778 | + | lncRNA |

| | | | | | | | | | | |
|---|---|---|---|---|---|---|---|---|---|---|
| ENSG00000203761.5_5 | MSTO2P | 0.94353818 | 1 | -0.257 | 2.389 | chr1 | 155715620 | 155719928 | + | unprocessed_pseudogene |
| ENSG00000159593.15_7 | NAE1 | 0.99998341 | 1 | -0.257 | 6.622 | chr16 | 66836778 | 66907159 | - | protein_coding |
| ENSG00000226396.1_5 | AL031727.1 | 0.65120653 | 1 | -0.257 | -0.486 | chr1 | 19934608 | 19935062 | + | processed_pseudogene |
| ENSG00000231466.2_5 | AL022324.2 | 0.66154452 | 1 | -0.258 | -0.511 | chr22 | 25745510 | 25746289 | - | processed_pseudogene |
| ENSG00000175467.15_3 | SART1 | 0.99996312 | 1 | -0.258 | 6.043 | chr11 | 65729199 | 65747608 | + | protein_coding |
| ENSG00000281332.1_5 | LINC00997 | 0.82326183 | 1 | -0.258 | 0.880 | chr7 | 32799891 | 32802536 | + | lncRNA |
| ENSG00000224699.9_5 | LAMTOR5-AS1 | 0.89262118 | 1 | -0.258 | 1.453 | chr1 | 110889738 | 110986439 | + | lncRNA |
| ENSG00000239419.2 | RN7SL535P | 0.64623155 | 1 | -0.258 | -1.013 | chr7 | 142987708 | 142988004 | + | misc_RNA |
| ENSG00000181524.6_2 | RPL24P4 | 0.75462167 | 1 | -0.258 | 1.570 | chr6 | 42924083 | 42924503 | - | processed_pseudogene |
| ENSG00000141682.11_2 | PMAIP1 | 0.99494752 | 1 | -0.258 | 3.569 | chr18 | 57567180 | 57571538 | + | protein_coding |
| ENSG00000135916.16_6 | ITM2C | 0.99996034 | 1 | -0.258 | 5.799 | chr2 | 231729354 | 231743963 | + | protein_coding |
| ENSG00000198208.12_4 | RPS6KL1 | 0.86590257 | 1 | -0.258 | 1.233 | chr14 | 75370654 | 75390005 | - | protein_coding |
| ENSG00000189350.12_3 | TOGARAM2 | 0.64601273 | 1 | -0.258 | -0.945 | chr2 | 29179477 | 29284239 | + | protein_coding |
| ENSG00000107672.15_4 | NSMCE4A | 0.99996093 | 1 | -0.258 | 5.743 | chr10 | 123716606 | 123734732 | - | protein_coding |
| ENSG00000230202.1_5 | AL450405.1 | 0.94265543 | 1 | -0.259 | 2.168 | chr6 | 118320138 | 118320611 | + | processed_pseudogene |
| ENSG00000155330.10_5 | C16orf87 | 0.9952814 | 1 | -0.259 | 3.776 | chr16 | 46830515 | 46865092 | - | protein_coding |
| ENSG00000180817.12_4 | PPA1 | 0.99999468 | 1 | -0.259 | 7.246 | chr10 | 71962591 | 71993667 | - | protein_coding |
| ENSG00000156411.9_4 | ATP5MPL | 0.99991747 | 1 | -0.259 | 5.832 | chr14 | 104378625 | 104394606 | - | protein_coding |
| ENSG00000214783.9_7 | POLR2J4 | 0.65911648 | 1 | -0.259 | -0.628 | chr7 | 43980494 | 44058774 | - | lncRNA |
| ENSG00000164346.10_6 | NSA2 | 0.99982332 | 1 | -0.259 | 6.271 | chr5 | 74062816 | 74075938 | + | protein_coding |
| ENSG00000090889.12_3 | KIF4A | 0.99999189 | 1 | -0.259 | 6.322 | chrX | 69509954 | 69640736 | + | protein_coding |
| ENSG00000119446.14_4 | RBM18 | 0.99939489 | 1 | -0.259 | 4.818 | chr9 | 124999901 | 125027119 | - | protein_coding |
| ENSG00000140350.15_5 | ANP32A | 0.99998147 | 1 | -0.260 | 6.677 | chr15 | 69070874 | 69113236 | - | protein_coding |
| ENSG00000186432.9_5 | KPNA4 | 0.99999283 | 1 | -0.260 | 7.441 | chr3 | 160212795 | 160283359 | - | protein_coding |
| ENSG00000151846.8_3 | PABPC3 | 0.67357692 | 1 | -0.260 | -0.413 | chr13 | 25670006 | 25673392 | + | protein_coding |
| ENSG00000286129.1_2 | AC245060.7 | 0.64910093 | 1 | -0.260 | -0.656 | chr22 | 22652489 | 22683480 | + | lncRNA |
| ENSG00000261334.1_6 | AL353803.4 | 0.67310674 | 1 | -0.260 | -0.313 | chr9 | 132242884 | 132244817 | - | lncRNA |
| ENSG00000120129.6_4 | DUSP1 | 0.99998965 | 1 | -0.260 | 6.583 | chr5 | 172195099 | 172198198 | - | protein_coding |
| ENSG00000198898.14_5 | CAPZA2 | 0.99997598 | 1 | -0.260 | 7.414 | chr7 | 116451124 | 116562103 | + | protein_coding |
| ENSG00000146007.11_6 | ZMAT2 | 0.99998304 | 1 | -0.260 | 6.188 | chr5 | 140078265 | 140086271 | + | protein_coding |
| ENSG00000178177.15_6 | LCORL | 0.99931597 | 1 | -0.260 | 4.867 | chr4 | 17842822 | 18023499 | - | protein_coding |
| ENSG00000229677.1_5 | AC018644.1 | 0.64552669 | 1 | -0.260 | -0.978 | chr7 | 138734202 | 138734752 | - | processed_pseudogene |
| ENSG00000125877.13_3 | ITPA | 0.9998973 | 1 | -0.260 | 5.306 | chr20 | 3189514 | 3204516 | + | protein_coding |
| ENSG00000230629.2_4 | RPS23P8 | 0.96256857 | 1 | -0.260 | 2.418 | chrX | 70182814 | 70183143 | + | processed_pseudogene |
| ENSG00000171490.13_9 | RSL1D1 | 0.9999981 | 1 | -0.261 | 7.815 | chr16 | 11927707 | 11945437 | - | protein_coding |
| ENSG00000127586.17_3 | CHTF18 | 0.86077018 | 1 | -0.261 | 5.018 | chr16 | 838046 | 850737 | + | protein_coding |
| ENSG00000177479.19_2 | ARIH2 | 0.99998295 | 1 | -0.261 | 6.034 | chr3 | 48956254 | 49023815 | + | protein_coding |
| ENSG00000169684.13_5 | CHRNA5 | 0.9929956 | 1 | -0.261 | 3.391 | chr15 | 78857862 | 78887611 | + | protein_coding |
| ENSG00000168116.14_4 | KIAA1586 | 0.99527618 | 1 | -0.261 | 3.769 | chr6 | 56911330 | 56920037 | + | protein_coding |
| ENSG00000108671.11_7 | PSMD11 | 0.99999344 | 1 | -0.261 | 7.033 | chr17 | 30771397 | 30810337 | + | protein_coding |
| ENSG00000168481.9_6 | LGI3 | 0.81858637 | 1 | -0.261 | 0.784 | chr8 | 22004343 | 22014597 | - | protein_coding |
| ENSG00000048028.11_5 | USP28 | 0.99992346 | 1 | -0.261 | 5.526 | chr11 | 113668596 | 113746292 | - | protein_coding |
| ENSG00000133226.17_6 | SRRM1 | 0.99995237 | 1 | -0.262 | 5.900 | chr1 | 24958207 | 24999772 | + | protein_coding |
| ENSG00000148335.14_2 | NTMT1 | 0.99805244 | 1 | -0.262 | 4.253 | chr9 | 132371163 | 132398410 | + | protein_coding |
| ENSG00000112293.15_4 | GPLD1 | 0.949525 | 1 | -0.262 | 2.219 | chr6 | 24424793 | 24495433 | - | protein_coding |
| ENSG00000165609.13_5 | NUDT5 | 0.9999719 | 1 | -0.262 | 6.092 | chr10 | 12207329 | 12238143 | - | protein_coding |
| ENSG00000131148.9_6 | EMC8 | 0.99979206 | 1 | -0.262 | 5.314 | chr16 | 85805364 | 85833214 | - | protein_coding |
| ENSG00000180008.9_3 | SOCS4 | 0.99995048 | 1 | -0.262 | 6.136 | chr14 | 55493948 | 55516207 | + | protein_coding |
| ENSG00000119048.8_4 | UBE2B | 0.99872433 | 1 | -0.262 | 4.264 | chr5 | 133707160 | 133727799 | + | protein_coding |
| ENSG00000100652.5_4 | SLC10A1 | 0.80279778 | 1 | -0.262 | 0.528 | chr14 | 70242133 | 70263958 | - | protein_coding |
| ENSG00000066926.11_4 | FECH | 0.9999546 | 1 | -0.262 | 5.828 | chr18 | 55211621 | 55253934 | - | protein_coding |
| ENSG00000029639.11_4 | TFB1M | 0.98500651 | 1 | -0.262 | 2.846 | chr6 | 155577268 | 155635627 | - | protein_coding |
| ENSG00000078124.12_4 | ACER3 | 0.99993914 | 1 | -0.262 | 6.025 | chr11 | 76571911 | 76737841 | + | protein_coding |
| ENSG00000106689.11_6 | LHX2 | 0.82785731 | 1 | -0.262 | 1.089 | chr9 | 126763949 | 126795580 | + | protein_coding |
| ENSG00000116898.12_4 | MRPS15 | 0.99993299 | 1 | -0.262 | 5.535 | chr1 | 36921319 | 36929985 | - | protein_coding |
| ENSG00000143450.16_8 | OAZ3 | 0.66101977 | 1 | -0.263 | -0.461 | chr1 | 151735375 | 151743810 | + | protein_coding |

| | | | | | | | | | | |
|---|---|---|---|---|---|---|---|---|---|---|
| ENSG00000169021.6_4 | UQCRFS1 | 0.99999605 | 1 | -0.263 | 7.507 | chr19 | 29696227 | 29704058 | - | protein_coding |
| ENSG00000114857.18_3 | NKTR | 0.99892084 | 1 | -0.263 | 5.735 | chr3 | 42642147 | 42690227 | + | protein_coding |
| ENSG00000110713.17_6 | NUP98 | 0.99999717 | 1 | -0.263 | 7.617 | chr11 | 3692313 | 3819022 | - | protein_coding |
| ENSG00000270977.2_8 | AC015849.5 | 0.72884136 | 1 | -0.263 | 0.168 | chr17 | 34220711 | 34238655 | - | transcribed_processed_pseudogene |
| ENSG00000128917.8_4 | DLL4 | 0.66523063 | 1 | -0.263 | -0.440 | chr15 | 41221538 | 41231271 | + | protein_coding |
| ENSG00000099994.11_3 | SUSD2 | 0.98825589 | 1 | -0.263 | 3.188 | chr22 | 24577455 | 24585074 | + | protein_coding |
| ENSG00000115806.13_3 | GORASP2 | 0.99974459 | 1 | -0.263 | 5.487 | chr2 | 171784974 | 171823640 | + | protein_coding |
| ENSG00000182512.5_4 | GLRX5 | 0.99985621 | 1 | -0.263 | 5.384 | chr14 | 95999840 | 96011061 | + | protein_coding |
| ENSG00000184465.16_7 | WDR27 | 0.99690097 | 1 | -0.263 | 4.000 | chr6 | 169857307 | 170102163 | - | protein_coding |
| ENSG00000105364.14_7 | MRPL4 | 0.99645201 | 1 | -0.263 | 4.090 | chr19 | 10362577 | 10370731 | + | protein_coding |
| ENSG00000270184.1_6 | AC018695.4 | 0.70296094 | 1 | -0.263 | -0.028 | chr16 | 85817988 | 85821223 | + | lncRNA |
| ENSG00000114993.17_6 | RTKN | 0.99914602 | 1 | -0.263 | 4.482 | chr2 | 74652962 | 74669549 | - | protein_coding |
| ENSG00000134909.18_4 | ARHGAP32 | 0.99915631 | 1 | -0.263 | 4.545 | chr11 | 128834955 | 129149219 | - | protein_coding |
| ENSG00000280063.1_6 | AC012676.5 | 0.69489558 | 1 | -0.263 | -0.068 | chr16 | 4396695 | 4398649 | - | TEC |
| ENSG00000267534.4_6 | S1PR2 | 0.74820199 | 1 | -0.263 | 0.350 | chr19 | 10332109 | 10342007 | - | protein_coding |
| ENSG00000110063.10_4 | DCPS | 0.99355394 | 1 | -0.263 | 3.651 | chr11 | 126173955 | 126219900 | + | protein_coding |
| ENSG00000261777.6_7 | AC012184.3 | 0.72017159 | 1 | -0.263 | -0.028 | chr16 | 70349345 | 70380709 | - | lncRNA |
| ENSG00000105085.10_4 | MED26 | 0.92456082 | 1 | -0.264 | 2.015 | chr19 | 16685718 | 16739873 | - | protein_coding |
| ENSG00000272356.1_7 | AL080317.2 | 0.66953184 | 1 | -0.264 | -0.415 | chr6 | 111630406 | 111634720 | + | lncRNA |
| ENSG00000128487.16_4 | SPECC1 | 0.99999088 | 1 | -0.264 | 6.859 | chr17 | 19912657 | 20222339 | + | protein_coding |
| ENSG00000147419.18_4 | CCDC25 | 0.99993597 | 1 | -0.264 | 5.608 | chr8 | 27590833 | 27630170 | - | protein_coding |
| ENSG00000154118.13_3 | JPH3 | 0.7436831 | 1 | -0.264 | 0.081 | chr16 | 87635441 | 87731762 | + | protein_coding |
| ENSG00000102081.14_8 | FMR1 | 0.99991492 | 1 | -0.264 | 5.632 | chrX | 146993469 | 147032645 | + | protein_coding |
| ENSG00000158104.11_4 | HPD | 0.71525931 | 1 | -0.264 | 0.257 | chr12 | 122277433 | 122301502 | - | protein_coding |
| ENSG00000175147.13_7 | TMEM51-AS1 | 0.72183294 | 1 | -0.264 | -0.103 | chr1 | 15438311 | 15480114 | - | lncRNA |
| ENSG00000156973.14_4 | PDE6D | 0.99724512 | 1 | -0.264 | 4.244 | chr2 | 232597143 | 232650982 | - | protein_coding |
| ENSG00000174871.11_4 | CNIH2 | 0.71850641 | 1 | -0.264 | 0.136 | chr11 | 66045646 | 66052772 | + | protein_coding |
| ENSG00000260456.6_6 | C16orf95 | 0.67576149 | 1 | -0.265 | -0.130 | chr16 | 87117168 | 87351026 | - | protein_coding |
| ENSG00000135778.12_3 | NTPCR | 0.99977359 | 1 | -0.265 | 5.415 | chr1 | 233086351 | 233119628 | + | protein_coding |
| ENSG00000166004.15_7 | CEP295 | 0.99878079 | 1 | -0.265 | 5.492 | chr11 | 93394848 | 93463524 | + | protein_coding |
| ENSG00000106462.11_5 | EZH2 | 0.99982326 | 1 | -0.265 | 5.026 | chr7 | 148504475 | 148581413 | - | protein_coding |
| ENSG00000135829.17_3 | DHX9 | 0.99999814 | 1 | -0.265 | 8.581 | chr1 | 182808482 | 182857117 | + | protein_coding |
| ENSG00000067533.6_3 | RRP15 | 0.99986 | 1 | -0.265 | 5.735 | chr1 | 218458635 | 218511325 | + | protein_coding |
| ENSG00000224093.5_7 | BCAR3-AS1 | 0.65048245 | 1 | -0.265 | -0.437 | chr1 | 94057756 | 94071130 | + | lncRNA |
| ENSG00000091428.18_3 | RAPGEF4 | 0.68362131 | 1 | -0.265 | -0.251 | chr2 | 173600002 | 173917621 | + | protein_coding |
| ENSG00000244754.8_5 | N4BP2L2 | 0.99987274 | 1 | -0.265 | 5.881 | chr13 | 33006554 | 33113022 | - | protein_coding |
| ENSG00000164754.15_4 | RAD21 | 0.99999803 | 1 | -0.265 | 8.059 | chr8 | 117858173 | 117887015 | - | protein_coding |
| ENSG00000243479.3_3 | MNX1-AS1 | 0.93738117 | 1 | -0.265 | 1.894 | chr7 | 156803499 | 156809120 | + | lncRNA |
| ENSG00000140307.11_5 | GTF2A2 | 0.99987615 | 1 | -0.265 | 6.132 | chr15 | 59930261 | 59949740 | - | protein_coding |
| ENSG00000155304.6_4 | HSPA13 | 0.99980645 | 1 | -0.265 | 5.498 | chr21 | 15743436 | 15755805 | - | protein_coding |
| ENSG00000225578.1_6 | NCBP2-AS1 | 0.65071313 | 1 | -0.265 | -0.514 | chr3 | 196666748 | 196669405 | + | lncRNA |
| ENSG00000163812.15_6 | ZDHHC3 | 0.99995862 | 1 | -0.265 | 5.727 | chr3 | 44956749 | 45017677 | - | protein_coding |
| ENSG00000203668.2_7 | CHML | 0.99998751 | 1 | -0.266 | 7.000 | chr1 | 241792155 | 241803556 | - | protein_coding |
| ENSG00000143369.15_4 | ECM1 | 0.77261831 | 1 | -0.266 | 0.586 | chr1 | 150480538 | 150486265 | + | protein_coding |
| ENSG00000261229.6_7 | AC021483.2 | 0.64831809 | 1 | -0.266 | -0.655 | chr15 | 80135889 | 80136646 | - | lncRNA |
| ENSG00000221819.6_5 | GAS8-AS1 | 0.69312119 | 1 | -0.266 | 0.114 | chr16 | 90095316 | 90096309 | - | lncRNA |
| ENSG00000167723.14_4 | TRPV3 | 0.70050416 | 1 | -0.266 | -0.168 | chr17 | 3413796 | 3461289 | - | protein_coding |
| ENSG00000136861.18_4 | CDK5RAP2 | 0.99998896 | 1 | -0.266 | 6.443 | chr9 | 123151147 | 123342448 | - | protein_coding |
| ENSG00000056277.16_4 | ZNF280C | 0.99941548 | 1 | -0.266 | 4.992 | chrX | 129336681 | 129402873 | - | protein_coding |
| ENSG00000144043.12_5 | TEX261 | 0.99998606 | 1 | -0.266 | 6.320 | chr2 | 71213072 | 71222003 | - | protein_coding |
| ENSG00000080947.15_4 | CROCCP3 | 0.95931954 | 1 | -0.266 | 2.248 | chr1 | 16793931 | 16825752 | - | transcribed_unprocessed_pseudogene |
| ENSG00000151923.17_5 | TIAL1 | 0.999985 | 1 | -0.266 | 6.428 | chr10 | 121331314 | 121356541 | - | protein_coding |
| ENSG00000125637.16_6 | PSD4 | 0.99945717 | 1 | -0.266 | 4.653 | chr2 | 113914902 | 113966973 | + | protein_coding |
| ENSG00000215033.3 | AL603965.1 | 0.78456265 | 1 | -0.266 | 0.419 | chr10 | 47746962 | 47770871 | + | protein_coding |
| ENSG00000100804.19_5 | PSMB5 | 0.99998017 | 1 | -0.266 | 6.412 | chr14 | 23485752 | 23504439 | - | protein_coding |
| ENSG00000145365.11_4 | TIFA | 0.98345075 | 1 | -0.267 | 2.892 | chr4 | 113195693 | 113207060 | - | protein_coding |
| ENSG00000254676.1_7 | AP000873.4 | 0.67056998 | 1 | -0.267 | -0.189 | chr11 | 82891186 | 82895562 | + | lncRNA |
| ENSG00000158019.20_8 | BABAM2 | 0.99984468 | 1 | -0.267 | 5.226 | chr2 | 28112808 | 28561768 | + | protein_coding |
| ENSG00000091831.23_5 | ESR1 | 0.67625232 | 1 | -0.267 | -0.301 | chr6 | 151977826 | 152450754 | + | protein_coding |

| | | | | | | | | | | |
|---|---|---|---|---|---|---|---|---|---|---|
| ENSG00000184898.7_5 | RBM43 | 0.96447246 | 1 | -0.267 | 2.563 | chr2 | 152104454 | 152118377 | - | protein_coding |
| ENSG00000176401.6_5 | EID2B | 0.98617576 | 1 | -0.267 | 2.967 | chr19 | 40021627 | 40023492 | - | protein_coding |
| ENSG00000198189.11_5 | HSD17B11 | 0.99979776 | 1 | -0.267 | 5.673 | chr4 | 88257667 | 88312340 | - | protein_coding |
| ENSG00000079435.10_6 | LIPE | 0.95776036 | 1 | -0.267 | 2.474 | chr19 | 42905666 | 42931540 | - | protein_coding |
| ENSG00000010244.18_6 | ZNF207 | 0.99998462 | 1 | -0.267 | 7.734 | chr17 | 30677136 | 30708905 | + | protein_coding |
| ENSG00000075292.19_4 | ZNF638 | 0.99999128 | 1 | -0.267 | 7.711 | chr2 | 71503691 | 71662199 | + | protein_coding |
| ENSG00000234690.7_8 | EPCAM-DT | 0.7795713 | 1 | -0.267 | 0.683 | chr2 | 47419544 | 47572213 | - | lncRNA |
| ENSG00000197170.10_4 | PSMD12 | 0.99991157 | 1 | -0.267 | 6.128 | chr17 | 65334032 | 65362721 | - | protein_coding |
| ENSG00000226318.1_4 | RPS3AP38 | 0.64451825 | 1 | -0.268 | -0.655 | chr10 | 69720459 | 69721221 | - | processed_pseudogene |
| ENSG00000113048.16_4 | MRPS27 | 0.99999475 | 1 | -0.268 | 6.952 | chr5 | 71515236 | 71616473 | - | protein_coding |
| ENSG00000156735.11_3 | BAG4 | 0.99991083 | 1 | -0.268 | 5.876 | chr8 | 38034051 | 38070819 | + | protein_coding |
| ENSG00000179041.4_4 | RRS1 | 0.99395012 | 1 | -0.268 | 3.870 | chr8 | 67341249 | 67342968 | + | protein_coding |
| ENSG00000174791.11_6 | RIN1 | 0.99213003 | 1 | -0.268 | 3.588 | chr11 | 66097712 | 66104311 | - | protein_coding |
| ENSG00000052723.11_4 | SIKE1 | 0.99939709 | 1 | -0.268 | 4.857 | chr1 | 115312100 | 115323306 | - | protein_coding |
| ENSG00000223572.9_2 | CKMT1A | 0.67129983 | 1 | -0.268 | -0.208 | chr15 | 43985084 | 43991420 | + | protein_coding |
| ENSG00000246067.8_6 | RAB30-DT | 0.98774502 | 1 | -0.268 | 3.042 | chr11 | 82783094 | 82818831 | + | lncRNA |
| ENSG00000225536.2_4 | STIP1P3 | 0.6340973 | 1 | -0.268 | -0.944 | chrX | 85339720 | 85341343 | - | processed_pseudogene |
| ENSG00000197045.13_5 | GMFB | 0.9999812 | 1 | -0.268 | 7.003 | chr14 | 54941202 | 54955744 | - | protein_coding |
| ENSG00000052749.14_8 | RRP12 | 0.99996655 | 1 | -0.268 | 6.172 | chr10 | 99116115 | 99185833 | - | protein_coding |
| ENSG00000247271.7_6 | ZBED5-AS1 | 0.99186931 | 1 | -0.268 | 3.381 | chr11 | 10879726 | 10930519 | + | lncRNA |
| ENSG00000105523.3_4 | FAM83E | 0.75131323 | 1 | -0.268 | 0.378 | chr19 | 49104067 | 49118111 | - | protein_coding |
| ENSG00000127947.16_3 | PTPN12 | 0.99988555 | 1 | -0.268 | 5.741 | chr7 | 77166612 | 77269386 | + | protein_coding |
| ENSG00000196470.12_5 | SIAH1 | 0.98871262 | 1 | -0.269 | 3.076 | chr16 | 48390275 | 48482313 | - | protein_coding |
| ENSG00000151151.6_5 | IPMK | 0.99795155 | 1 | -0.269 | 4.418 | chr10 | 59951278 | 60027654 | - | protein_coding |
| ENSG00000154813.10_4 | DPH3 | 0.99967195 | 1 | -0.269 | 4.997 | chr3 | 16298568 | 16306476 | - | protein_coding |
| ENSG00000204406.13_6 | MBD5 | 0.99219004 | 1 | -0.269 | 3.689 | chr2 | 148778580 | 149274540 | + | protein_coding |
| ENSG00000046647.13_6 | GEMIN8 | 0.98132795 | 1 | -0.269 | 3.026 | chrX | 14026398 | 14048012 | - | protein_coding |
| ENSG00000163040.14_4 | CCDC74A | 0.98935497 | 1 | -0.269 | 3.266 | chr2 | 132285248 | 132291239 | + | protein_coding |
| ENSG00000230551.4_8 | AC021078.1 | 0.99756707 | 1 | -0.269 | 4.688 | chr5 | 148873877 | 148884233 | - | lncRNA |
| ENSG00000143256.5_4 | PFDN2 | 0.9911377 | 1 | -0.269 | 3.711 | chr1 | 161070346 | 161087845 | - | protein_coding |
| ENSG00000221829.9_5 | FANCG | 0.9999044 | 1 | -0.269 | 5.758 | chr9 | 35073832 | 35080013 | - | protein_coding |
| ENSG00000162402.14_4 | USP24 | 0.99998625 | 1 | -0.269 | 6.880 | chr1 | 55532032 | 55681037 | - | protein_coding |
| ENSG00000197969.14_7 | VPS13A | 0.99991802 | 1 | -0.269 | 6.951 | chr9 | 79792361 | 80036453 | + | protein_coding |
| ENSG00000251333.3_4 | RTN3P1 | 0.63171456 | 1 | -0.269 | -0.783 | chr4 | 146296695 | 146297405 | + | processed_pseudogene |
| ENSG00000165916.8_7 | PSMC3 | 0.99999375 | 1 | -0.269 | 6.993 | chr11 | 47440320 | 47448024 | - | protein_coding |
| ENSG00000108262.15_5 | GIT1 | 0.99996891 | 1 | -0.270 | 6.615 | chr17 | 27900487 | 27921072 | - | protein_coding |
| ENSG00000095319.14_4 | NUP188 | 0.99998425 | 1 | -0.270 | 6.555 | chr9 | 131709978 | 131769375 | + | protein_coding |
| ENSG00000166750.10_3 | SLFN5 | 0.99848001 | 1 | -0.270 | 4.182 | chr17 | 33570090 | 33600674 | + | protein_coding |
| ENSG00000249740.3_7 | OSMR-AS1 | 0.71920607 | 1 | -0.270 | 0.076 | chr5 | 38682683 | 38845931 | - | lncRNA |
| ENSG00000137185.12_3 | ZSCAN9 | 0.98989204 | 1 | -0.270 | 3.476 | chr6 | 28192664 | 28201260 | + | protein_coding |
| ENSG00000271119.1_7 | AC026412.3 | 0.63978914 | 1 | -0.270 | -0.602 | chr5 | 1594741 | 1611582 | + | lncRNA |
| ENSG00000168411.14_5 | RFWD3 | 0.99999064 | 1 | -0.270 | 6.730 | chr16 | 74655297 | 74700775 | - | protein_coding |
| ENSG00000227540.1_9 | AC016394.2 | 0.74305147 | 1 | -0.270 | 0.146 | chr10 | 75012549 | 75014107 | + | lncRNA |
| ENSG00000226666.1_4 | HSPA9P1 | 0.66096661 | 1 | -0.270 | -0.110 | chr2 | 222826456 | 222828484 | + | processed_pseudogene |
| ENSG00000066933.16_6 | MYO9A | 0.99993886 | 1 | -0.270 | 6.646 | chr15 | 72114632 | 72410918 | - | protein_coding |
| ENSG00000129474.16_6 | AJUBA | 0.99998453 | 1 | -0.271 | 6.834 | chr14 | 23440386 | 23451760 | - | protein_coding |
| ENSG00000156869.13_6 | FRRS1 | 0.83244543 | 1 | -0.271 | 1.453 | chr1 | 100174188 | 100232187 | - | protein_coding |
| ENSG00000258323.1_7 | AC073575.3 | 0.62765387 | 1 | -0.271 | -0.980 | chr12 | 112501713 | 112503559 | + | lncRNA |
| ENSG00000108179.14_3 | PPIF | 0.99985203 | 1 | -0.271 | 5.679 | chr10 | 81107225 | 81115090 | + | protein_coding |
| ENSG00000198862.14_6 | LTN1 | 0.9999354 | 1 | -0.271 | 6.147 | chr21 | 30300466 | 30365277 | - | protein_coding |
| ENSG00000006432.15_4 | MAP3K9 | 0.99962038 | 1 | -0.271 | 5.186 | chr14 | 71189243 | 71276251 | - | protein_coding |
| ENSG00000143190.23_8 | POU2F1 | 0.9997845 | 1 | -0.271 | 5.990 | chr1 | 167190113 | 167396582 | + | protein_coding |
| ENSG00000140795.13_4 | MYLK3 | 0.63008317 | 1 | -0.271 | -0.814 | chr16 | 46736194 | 46824319 | - | protein_coding |
| ENSG00000064703.12_3 | DDX20 | 0.99960656 | 1 | -0.271 | 4.979 | chr1 | 112297867 | 112310622 | + | protein_coding |
| ENSG00000119760.16_4 | SUPT7L | 0.99992504 | 1 | -0.271 | 6.007 | chr2 | 27873676 | 27886707 | - | protein_coding |
| ENSG00000105186.16_5 | ANKRD27 | 0.99998351 | 1 | -0.271 | 6.364 | chr19 | 33087912 | 33167503 | - | protein_coding |
| ENSG00000010270.13_3 | STARD3NL | 0.99979456 | 1 | -0.271 | 5.724 | chr7 | 38217824 | 38270272 | + | protein_coding |
| ENSG00000232485.3_8 | AC098820.2 | 0.63271385 | 1 | -0.271 | -0.630 | chr2 | 217343753 | 217363484 | - | lncRNA |
| ENSG00000173221.14_4 | GLRX | 0.99933614 | 1 | -0.271 | 4.946 | chr5 | 95087023 | 95158430 | - | protein_coding |

| | | | | | | | | | | |
|---|---|---|---|---|---|---|---|---|---|---|
| ENSG00000122566.21_6 | HNRNPA2B1 | 0.99996535 | 1 | -0.272 | 8.488 | chr7 | 26212677 | 26241149 | - | protein_coding |
| ENSG00000155368.16_4 | DBI | 0.99999207 | 1 | -0.272 | 6.929 | chr2 | 120124497 | 120130136 | + | protein_coding |
| ENSG00000100503.24_7 | NIN | 0.99998997 | 1 | -0.272 | 6.928 | chr14 | 51186481 | 51297839 | - | protein_coding |
| ENSG00000035862.12_5 | TIMP2 | 0.9999804 | 1 | -0.272 | 6.414 | chr17 | 76849059 | 76921469 | - | protein_coding |
| ENSG00000156928.5_3 | MALSU1 | 0.99900352 | 1 | -0.272 | 4.574 | chr7 | 23338358 | 23351348 | + | protein_coding |
| ENSG00000226259.10_2 | GTF2H2B | 0.68398209 | 1 | -0.272 | 0.183 | chr5 | 69711179 | 69743842 | + | transcribed_unprocessed_pseudogene |
| ENSG00000135525.18_4 | MAP7 | 0.99999258 | 1 | -0.272 | 6.827 | chr6 | 136663419 | 136871957 | - | protein_coding |
| ENSG00000101928.13_4 | MOSPD1 | 0.99908839 | 1 | -0.272 | 4.292 | chrX | 134021662 | 134049287 | - | protein_coding |
| ENSG00000142632.17_5 | ARHGEF19 | 0.97375167 | 1 | -0.272 | 2.718 | chr1 | 16524349 | 16539147 | - | protein_coding |
| ENSG00000047315.16_4 | POLR2B | 0.99999482 | 1 | -0.272 | 7.259 | chr4 | 57843888 | 57897324 | + | protein_coding |
| ENSG00000188739.15_5 | RBM34 | 0.69056673 | 1 | -0.273 | -0.035 | chr1 | 235294498 | 235324598 | - | protein_coding |
| ENSG00000120948.17_7 | TARDBP | 0.91034626 | 1 | -0.273 | 4.493 | chr1 | 11072401 | 11086477 | + | protein_coding |
| ENSG00000272911.1_6 | AC005696.3 | 0.62798423 | 1 | -0.273 | -0.877 | chr17 | 2591767 | 2592254 | + | lncRNA |
| ENSG00000259456.4_9 | ADNP-AS1 | 0.68086663 | 1 | -0.273 | 0.445 | chr20 | 49547521 | 49561685 | + | lncRNA |
| ENSG00000139146.14_8 | SINHCAF | 0.99997151 | 1 | -0.273 | 5.916 | chr12 | 31433518 | 31479992 | - | protein_coding |
| ENSG00000198546.15_5 | ZNF511 | 0.98711091 | 1 | -0.273 | 3.207 | chr10 | 135122418 | 135126666 | + | protein_coding |
| ENSG00000125319.14_3 | HROB | 0.99443647 | 1 | -0.273 | 3.742 | chr17 | 42219274 | 42239844 | + | protein_coding |
| ENSG00000100266.19_5 | PACSIN2 | 0.99995345 | 1 | -0.273 | 6.279 | chr22 | 43231418 | 43411155 | - | protein_coding |
| ENSG00000116459.11_4 | ATP5PB | 0.99998351 | 1 | -0.273 | 6.763 | chr1 | 111991486 | 112005395 | + | protein_coding |
| ENSG00000157500.12_5 | APPL1 | 0.99992763 | 1 | -0.273 | 6.138 | chr3 | 57261754 | 57312133 | + | protein_coding |
| ENSG00000226648.2_8 | PLCG1-AS1 | 0.72831153 | 1 | -0.273 | 0.788 | chr20 | 39726659 | 39766643 | - | lncRNA |
| ENSG00000105270.15_4 | CLIP3 | 0.88294208 | 1 | -0.273 | 1.624 | chr19 | 36505562 | 36524245 | - | protein_coding |
| ENSG00000269996.1 | RP11-343N15.5 | 0.98732398 | 1 | -0.273 | 3.211 | chr1 | 121133256 | 121134581 | + | lincRNA |
| ENSG00000255165.1_7 | AC134775.1 | 0.65353826 | 1 | -0.273 | -0.498 | chr11 | 44093012 | 44093953 | + | lncRNA |
| ENSG00000131876.17_5 | SNRPA1 | 0.99924124 | 1 | -0.273 | 4.461 | chr15 | 101821715 | 101835487 | - | protein_coding |
| ENSG00000226380.9_9 | AC016831.1 | 0.88346764 | 1 | -0.273 | 2.834 | chr7 | 130561568 | 130598069 | - | lncRNA |
| ENSG00000164610.9_7 | RP9 | 0.99146246 | 1 | -0.274 | 3.569 | chr7 | 33134409 | 33149016 | - | protein_coding |
| ENSG00000126822.17_8 | PLEKHG3 | 0.99986066 | 1 | -0.274 | 5.929 | chr14 | 65170820 | 65216967 | + | protein_coding |
| ENSG00000259768.6_8 | AC004943.2 | 0.96354026 | 1 | -0.274 | 2.618 | chr16 | 72699022 | 72856680 | + | lncRNA |
| ENSG00000147679.12_4 | UTP23 | 0.99949151 | 1 | -0.274 | 4.902 | chr8 | 117778744 | 117861702 | + | protein_coding |
| ENSG00000138780.15_7 | GSTCD | 0.99953247 | 1 | -0.274 | 4.944 | chr4 | 106629935 | 106768882 | + | protein_coding |
| ENSG00000110107.9_5 | PRPF19 | 0.99992637 | 1 | -0.274 | 6.766 | chr11 | 60658019 | 60674057 | - | protein_coding |
| ENSG00000213066.13_5 | FGFR1OP | 0.98527886 | 1 | -0.274 | 3.058 | chr6 | 167412805 | 167508277 | + | protein_coding |
| ENSG00000137309.20_3 | HMGA1 | 0.99998821 | 1 | -0.274 | 8.469 | chr6 | 34204650 | 34214008 | + | protein_coding |
| ENSG00000196455.8_5 | PIK3R4 | 0.99987035 | 1 | -0.274 | 5.262 | chr3 | 130397778 | 130465673 | - | protein_coding |
| ENSG00000029993.15_7 | HMGB3 | 0.9999952 | 1 | -0.274 | 7.536 | chrX | 150148982 | 150159244 | + | protein_coding |
| ENSG00000124762.13_3 | CDKN1A | 0.99994308 | 1 | -0.274 | 5.742 | chr6 | 36644237 | 36655116 | + | protein_coding |
| ENSG00000107779.13_3 | BMPR1A | 0.99996749 | 1 | -0.274 | 5.951 | chr10 | 88516358 | 88692595 | + | protein_coding |
| ENSG00000227345.4 | PARG | 0.99853181 | 1 | -0.274 | 4.450 | chr10 | 51026325 | 51130715 | - | protein_coding |
| ENSG00000278053.5_4 | DDX52 | 0.99998139 | 1 | -0.274 | 6.669 | chr17 | 35969787 | 36003475 | - | protein_coding |
| ENSG00000116005.12_6 | PCYOX1 | 0.99999381 | 1 | -0.274 | 6.930 | chr2 | 70484518 | 70508317 | + | protein_coding |
| ENSG00000136273.13_7 | HUS1 | 0.99924443 | 1 | -0.274 | 4.413 | chr7 | 48002885 | 48019212 | - | protein_coding |
| ENSG00000166920.12_3 | C15orf48 | 0.65765957 | 1 | -0.274 | -0.218 | chr15 | 45722727 | 45740959 | + | protein_coding |
| ENSG00000182899.17_7 | RPL35A | 0.99999392 | 1 | -0.274 | 7.422 | chr3 | 197677061 | 197683481 | + | protein_coding |
| ENSG00000179564.4_3 | LSMEM2 | 0.63158771 | 1 | -0.274 | -0.814 | chr3 | 50316518 | 50325545 | + | protein_coding |
| ENSG00000004897.12_6 | CDC27 | 0.99998919 | 1 | -0.274 | 7.917 | chr17 | 45195069 | 45266788 | - | protein_coding |
| ENSG00000115556.14_5 | PLCD4 | 0.97117306 | 1 | -0.274 | 2.691 | chr2 | 219472578 | 219501907 | + | protein_coding |
| ENSG00000187555.16_7 | USP7 | 0.99998879 | 1 | -0.275 | 6.869 | chr16 | 8985954 | 9069185 | - | protein_coding |
| ENSG00000235381.1_7 | AL596202.1 | 0.66131864 | 1 | -0.275 | -0.259 | chr6 | 155574273 | 155577858 | - | lncRNA |
| ENSG00000138621.12_6 | PPCDC | 0.99546513 | 1 | -0.275 | 3.847 | chr15 | 75315927 | 75409803 | + | protein_coding |
| ENSG00000100220.12_4 | RTCB | 0.9999549 | 1 | -0.275 | 6.399 | chr22 | 32783569 | 32808235 | - | protein_coding |
| ENSG00000231887.7_9 | PRH1 | 0.68103408 | 1 | -0.275 | 0.134 | chr12 | 10977559 | 11324207 | - | protein_coding |
| ENSG00000183077.15_2 | AFMID | 0.99668299 | 1 | -0.275 | 3.883 | chr17 | 76183398 | 76203782 | + | protein_coding |
| ENSG00000164151.12_8 | ICE1 | 0.99998709 | 1 | -0.275 | 7.075 | chr5 | 5420777 | 5490333 | + | protein_coding |
| ENSG00000112339.15_7 | HBS1L | 0.99991589 | 1 | -0.275 | 5.919 | chr6 | 135281516 | 135424194 | - | protein_coding |
| ENSG00000122707.12_3 | RECK | 0.98741281 | 1 | -0.275 | 3.118 | chr9 | 36036910 | 36124452 | + | protein_coding |
| ENSG00000116473.14_2 | RAP1A | 0.99488694 | 1 | -0.275 | 3.632 | chr1 | 112084840 | 112259313 | + | protein_coding |
| ENSG00000115415.19_8 | STAT1 | 0.9999919 | 1 | -0.275 | 7.129 | chr2 | 191773186 | 191885686 | - | protein_coding |

| | | | | | | | | | | |
|---|---|---|---|---|---|---|---|---|---|---|
| ENSG00000260872.1 | RP11-680G24.5 | 0.76491712 | 1 | -0.275 | 0.373 | chr16 | 15111963 | 15114345 | - | antisense |
| ENSG00000239653.1_7 | PSMD6-AS2 | 0.74664061 | 1 | -0.276 | 0.465 | chr3 | 63989698 | 63997824 | + | lncRNA |
| ENSG00000259345.7_8 | AC013652.1 | 0.6232577 | 1 | -0.276 | -0.911 | chr15 | 39157523 | 39719396 | - | lncRNA |
| ENSG00000172809.13_5 | RPL38 | 0.9939542 | 1 | -0.276 | 8.408 | chr17 | 72199721 | 72206794 | + | protein_coding |
| ENSG00000106028.11_6 | SSBP1 | 0.99995061 | 1 | -0.276 | 6.607 | chr7 | 141438134 | 141487722 | + | protein_coding |
| ENSG00000254876.5_7 | SUGT1P4-STRA6LP | 0.76466309 | 1 | -0.276 | 0.318 | chr9 | 100000731 | 100059596 | + | lncRNA |
| ENSG00000259642.2_6 | ST20-AS1 | 0.83796272 | 1 | -0.276 | 1.293 | chr15 | 80215113 | 80219335 | + | lncRNA |
| ENSG00000149346.15_3 | SLX4IP | 0.99949114 | 1 | -0.276 | 4.694 | chr20 | 10415953 | 10617477 | + | protein_coding |
| ENSG00000174013.8_3 | FBXO45 | 0.99885065 | 1 | -0.276 | 4.617 | chr3 | 196295482 | 196315930 | + | protein_coding |
| ENSG00000065325.13_4 | GLP2R | 0.99991097 | 1 | -0.276 | 5.536 | chr17 | 9725523 | 9795416 | + | protein_coding |
| ENSG00000230109.1_8 | LINC02643 | 0.70700687 | 1 | -0.276 | 0.466 | chr10 | 21629162 | 21661879 | - | lncRNA |
| ENSG00000072682.18_7 | P4HA2 | 0.99945563 | 1 | -0.276 | 4.979 | chr5 | 131527531 | 131631008 | - | protein_coding |
| ENSG00000152952.12_6 | PLOD2 | 0.99995396 | 1 | -0.276 | 7.928 | chr3 | 145787227 | 145881440 | - | protein_coding |
| ENSG00000184076.13_4 | UQCR10 | 0.99138003 | 1 | -0.276 | 3.701 | chr22 | 30163358 | 30166402 | + | protein_coding |
| ENSG00000148120.16_3 | AOPEP | 0.99999351 | 1 | -0.276 | 7.524 | chr9 | 97488983 | 97849500 | + | protein_coding |
| ENSG00000166797.11_5 | CIAO2A | 0.99994058 | 1 | -0.277 | 5.999 | chr15 | 64364764 | 64386056 | - | protein_coding |
| ENSG00000267724.1_6 | AC012254.3 | 0.62309583 | 1 | -0.277 | -0.878 | chr18 | 44632317 | 44634433 | + | lncRNA |
| ENSG00000234678.2_7 | ELF3-AS1 | 0.98146992 | 1 | -0.277 | 3.022 | chr1 | 201964824 | 201979591 | - | lncRNA |
| ENSG00000154640.14_4 | BTG3 | 0.99306611 | 1 | -0.277 | 3.496 | chr21 | 18965971 | 18985265 | - | protein_coding |
| ENSG00000196865.4_3 | NHLRC2 | 0.99991581 | 1 | -0.278 | 6.166 | chr10 | 115614420 | 115676953 | + | protein_coding |
| ENSG00000249628.2 | LINC00942 | 0.62227592 | 1 | -0.278 | -0.878 | chr12 | 1609691 | 1616484 | - | lincRNA |
| ENSG00000228887.3_2 | EEF1DP1 | 0.6621075 | 1 | -0.278 | -0.281 | chr19 | 14181154 | 14182049 | + | processed_pseudogene |
| ENSG00000245317.2_6 | AC008393.1 | 0.71967432 | 1 | -0.278 | 0.005 | chr5 | 179286013 | 179288283 | + | lncRNA |
| ENSG00000214706.10_4 | IFRD2 | 0.99965713 | 1 | -0.278 | 5.576 | chr3 | 50325163 | 50330349 | - | protein_coding |
| ENSG00000185347.17_9 | TEDC1 | 0.92761337 | 1 | -0.278 | 4.918 | chr14 | 105956192 | 105965912 | + | protein_coding |
| ENSG00000114125.14_3 | RNF7 | 0.99975485 | 1 | -0.278 | 5.077 | chr3 | 141457091 | 141466402 | + | protein_coding |
| ENSG00000260367.2_7 | AC109460.1 | 0.62371009 | 1 | -0.278 | -0.846 | chr16 | 28985283 | 28990145 | - | lncRNA |
| ENSG00000139496.16_7 | NUP58 | 0.99998252 | 1 | -0.278 | 6.528 | chr13 | 25875694 | 25923938 | + | protein_coding |
| ENSG00000282508.2_3 | LINC01002 | 0.63006434 | 1 | -0.278 | -0.753 | chr19 | 198264 | 246544 | - | lncRNA |
| ENSG00000148187.18_4 | MRRF | 0.99994622 | 1 | -0.279 | 5.870 | chr9 | 125026882 | 125093616 | + | protein_coding |
| ENSG00000273329.1_6 | AC078846.1 | 0.99540899 | 1 | -0.279 | 3.894 | chr7 | 129244389 | 129251471 | - | lncRNA |
| ENSG00000185361.9_3 | TNFAIP8L1 | 0.98179112 | 1 | -0.279 | 2.983 | chr19 | 4639528 | 4655580 | + | protein_coding |
| ENSG00000186395.8_5 | KRT10 | 0.99532307 | 1 | -0.279 | 4.240 | chr17 | 38974369 | 38978866 | - | protein_coding |
| ENSG00000275910.1_7 | AC138932.6 | 0.6471566 | 1 | -0.279 | -0.525 | chr16 | 15109685 | 15110247 | - | lncRNA |
| ENSG00000125977.7_4 | EIF2S2 | 0.99998768 | 1 | -0.279 | 7.509 | chr20 | 32676115 | 32700049 | - | protein_coding |
| ENSG00000136261.15_3 | BZW2 | 0.9999471 | 1 | -0.279 | 6.074 | chr7 | 16685756 | 16746148 | + | protein_coding |
| ENSG00000112249.14_7 | ASCC3 | 0.99997116 | 1 | -0.279 | 7.130 | chr6 | 100956070 | 101329248 | - | protein_coding |
| ENSG00000125484.12_3 | GTF3C4 | 0.99997908 | 1 | -0.279 | 6.401 | chr9 | 135545422 | 135570340 | + | protein_coding |
| ENSG00000112695.11_6 | COX7A2 | 0.99701521 | 1 | -0.279 | 5.786 | chr6 | 75947391 | 75960039 | - | protein_coding |
| ENSG00000147465.12_4 | STAR | 0.61797932 | 1 | -0.279 | -0.847 | chr8 | 38000218 | 38008510 | - | protein_coding |
| ENSG00000125703.15_3 | ATG4C | 0.98905519 | 1 | -0.279 | 3.442 | chr1 | 63249803 | 63331187 | + | protein_coding |
| ENSG00000274213.1_6 | AC015912.3 | 0.67325539 | 1 | -0.279 | -0.157 | chr17 | 54991547 | 54991894 | + | lncRNA |
| ENSG00000108551.5_4 | RASD1 | 0.65301599 | 1 | -0.279 | -0.326 | chr17 | 17397751 | 17399709 | - | protein_coding |
| ENSG00000196550.10_2 | FAM72A | 0.96116077 | 1 | -0.279 | 2.476 | chr1 | 206136916 | 206155151 | + | protein_coding |
| ENSG00000152749.8_3 | GPR180 | 0.99928055 | 1 | -0.279 | 5.103 | chr13 | 95254111 | 95286915 | + | protein_coding |
| ENSG00000148288.13_6 | GBGT1 | 0.66900917 | 1 | -0.280 | 0.006 | chr9 | 136028335 | 136039320 | - | protein_coding |
| ENSG00000136943.11_4 | CTSV | 0.99161441 | 1 | -0.280 | 3.511 | chr9 | 99791959 | 99801925 | - | protein_coding |
| ENSG00000287574.1_1 | AC004938.2 | 0.62015972 | 1 | -0.280 | -1.140 | chr7 | 44622118 | 44623275 | + | lncRNA |
| ENSG00000178971.16_7 | CTC1 | 0.99919061 | 1 | -0.280 | 4.770 | chr17 | 8128133 | 8151374 | - | protein_coding |
| ENSG00000069248.12_4 | NUP133 | 0.99996459 | 1 | -0.280 | 6.510 | chr1 | 229576006 | 229644088 | - | protein_coding |
| ENSG00000099817.12_4 | POLR2E | 0.99984559 | 1 | -0.280 | 6.055 | chr19 | 1086573 | 1095379 | - | protein_coding |
| ENSG00000153936.17_4 | HS2ST1 | 0.99955133 | 1 | -0.280 | 5.041 | chr1 | 87380318 | 87575665 | + | protein_coding |
| ENSG00000140750.17_7 | ARHGAP17 | 0.9986912 | 1 | -0.280 | 4.327 | chr16 | 24930710 | 25026987 | - | protein_coding |
| ENSG00000105829.13_5 | BET1 | 0.998098 | 1 | -0.280 | 4.754 | chr7 | 93592074 | 93633694 | - | protein_coding |
| ENSG00000090263.16_6 | MRPS33 | 0.99925131 | 1 | -0.280 | 5.307 | chr7 | 140702410 | 140715028 | - | protein_coding |
| ENSG00000092020.10_4 | PPP2R3C | 0.99265246 | 1 | -0.280 | 3.730 | chr14 | 35554673 | 35591723 | - | protein_coding |
| ENSG00000115718.17_5 | PROC | 0.94586735 | 1 | -0.280 | 2.129 | chr2 | 128176003 | 128186822 | + | protein_coding |
| ENSG00000115998.7_4 | C2orf42 | 0.98515047 | 1 | -0.280 | 3.178 | chr2 | 70377012 | 70475747 | - | protein_coding |

| | | | | | | | | | | |
|---|---|---|---|---|---|---|---|---|---|---|
| ENSG00000231500.7_3 | RPS18 | 0.99999211 | 1 | -0.281 | 8.601 | chr6 | 33239852 | 33244288 | + | protein_coding |
| ENSG00000286431.1_1 | BX842570.1 | 0.82346085 | 1 | -0.281 | 1.318 | chrX | 3737965 | 3741939 | + | lncRNA |
| ENSG00000067167.8_4 | TRAM1 | 0.99999331 | 1 | -0.281 | 9.493 | chr8 | 71485453 | 71520651 | - | protein_coding |
| ENSG00000124370.11_5 | MCEE | 0.93737026 | 1 | -0.281 | 2.601 | chr2 | 71336814 | 71357369 | - | protein_coding |
| ENSG00000250510.8_4 | GPR162 | 0.8940477 | 1 | -0.281 | 1.929 | chr12 | 6930790 | 6939136 | + | protein_coding |
| ENSG00000122481.17_4 | RWDD3 | 0.85262262 | 1 | -0.281 | 1.356 | chr1 | 95699766 | 95712781 | + | protein_coding |
| ENSG00000167608.11_4 | TMC4 | 0.8828376 | 1 | -0.281 | 1.845 | chr19 | 54663846 | 54676944 | - | protein_coding |
| ENSG00000003400.15_5 | CASP10 | 0.97018315 | 1 | -0.281 | 2.983 | chr2 | 202047621 | 202094129 | + | protein_coding |
| ENSG00000166793.12_5 | YPEL4 | 0.93149466 | 1 | -0.281 | 2.028 | chr11 | 57412560 | 57417417 | - | protein_coding |
| ENSG00000197535.14_4 | MYO5A | 0.9999796 | 1 | -0.281 | 6.738 | chr15 | 52599480 | 52821247 | - | protein_coding |
| ENSG00000276603.1_7 | AL109614.1 | 0.62175023 | 1 | -0.281 | -0.496 | chr20 | 36155044 | 36155462 | - | lncRNA |
| ENSG00000135999.12_3 | EPC2 | 0.9996412 | 1 | -0.281 | 4.942 | chr2 | 149402009 | 149545138 | + | protein_coding |
| ENSG00000128626.11_2 | MRPS12 | 0.9997008 | 1 | -0.281 | 6.438 | chr19 | 39421188 | 39423802 | + | protein_coding |
| ENSG00000104093.13_5 | DMXL2 | 0.99996498 | 1 | -0.282 | 6.902 | chr15 | 51739908 | 51915030 | - | protein_coding |
| ENSG00000187801.15_4 | ZFP69B | 0.88739921 | 1 | -0.282 | 1.700 | chr1 | 40915774 | 40929390 | + | protein_coding |
| ENSG00000268583.2_7 | AC011466.1 | 0.61481136 | 1 | -0.282 | -0.783 | chr19 | 48707328 | 48718008 | + | lncRNA |
| ENSG00000198818.10_4 | SFT2D1 | 0.98796025 | 1 | -0.282 | 3.839 | chr6 | 166733216 | 166756064 | - | protein_coding |
| ENSG00000119328.12_4 | ABITRAM | 0.99745611 | 1 | -0.282 | 4.285 | chr9 | 111696680 | 111713024 | + | protein_coding |
| ENSG00000257950.3_4 | P2RX5-TAX1BP3 | 0.76258851 | 1 | -0.282 | 0.710 | chr17 | 3566357 | 3599488 | - | protein_coding |
| ENSG00000213281.5_4 | NRAS | 0.99997059 | 1 | -0.282 | 6.507 | chr1 | 115247090 | 115259392 | - | protein_coding |
| ENSG00000038219.13_4 | BOD1L1 | 0.99993934 | 1 | -0.282 | 6.749 | chr4 | 13570362 | 13629349 | - | protein_coding |
| ENSG00000239665.8_7 | AL157392.3 | 0.98749819 | 1 | -0.282 | 3.219 | chr10 | 13673143 | 13710445 | + | lncRNA |
| ENSG00000104442.10_7 | ARMC1 | 0.99959374 | 1 | -0.282 | 5.947 | chr8 | 66514693 | 66546452 | - | protein_coding |
| ENSG00000135637.14_7 | CCDC142 | 0.99653685 | 1 | -0.282 | 4.191 | chr2 | 74699113 | 74710535 | - | protein_coding |
| ENSG00000058056.9_4 | USP13 | 0.99823662 | 1 | -0.282 | 4.425 | chr3 | 179370828 | 179507189 | + | protein_coding |
| ENSG00000206754.1 | SNORD101 | 0.69120037 | 1 | -0.282 | 0.050 | chr6 | 133136446 | 133136518 | + | snoRNA |
| ENSG00000058729.11_7 | RIOK2 | 0.99928003 | 1 | -0.283 | 4.908 | chr5 | 96496571 | 96518951 | - | protein_coding |
| ENSG00000237399.7_7 | PITRM1-AS1 | 0.75841323 | 1 | -0.283 | 0.712 | chr10 | 3183824 | 3210164 | + | lncRNA |
| ENSG00000123358.20_7 | NR4A1 | 0.99139559 | 1 | -0.283 | 3.400 | chr12 | 52416616 | 52453291 | + | protein_coding |
| ENSG00000103018.17_6 | CYB5B | 0.99999435 | 1 | -0.283 | 8.333 | chr16 | 69458522 | 69500167 | + | protein_coding |
| ENSG00000120063.10_4 | GNA13 | 0.99998705 | 1 | -0.283 | 7.355 | chr17 | 63005407 | 63052858 | - | protein_coding |
| ENSG00000033178.13_4 | UBA6 | 0.99992552 | 1 | -0.283 | 6.645 | chr4 | 68478370 | 68566873 | - | protein_coding |
| ENSG00000175575.12_3 | PAAF1 | 0.99112787 | 1 | -0.283 | 3.423 | chr11 | 73587744 | 73642169 | + | protein_coding |
| ENSG00000203667.10_3 | COX20 | 0.88172023 | 1 | -0.283 | 1.559 | chr1 | 244998918 | 245008359 | + | protein_coding |
| ENSG00000265817.3_5 | FSBP | 0.9969097 | 1 | -0.283 | 4.019 | chr8 | 95384398 | 95449180 | - | protein_coding |
| ENSG00000152240.13_4 | HAUS1 | 0.9977443 | 1 | -0.284 | 4.644 | chr18 | 43684344 | 43708299 | + | protein_coding |
| ENSG00000264270.1_7 | AC087749.1 | 0.63764111 | 1 | -0.284 | -0.424 | chr17 | 73679623 | 73680879 | - | lncRNA |
| ENSG00000132879.14_5 | FBXO44 | 0.99138166 | 1 | -0.284 | 3.882 | chr1 | 11714432 | 11723384 | + | protein_coding |
| ENSG00000271933.1_6 | AL603756.1 | 0.76636477 | 1 | -0.284 | 0.792 | chr10 | 85898176 | 85900338 | - | lncRNA |
| ENSG00000204049.1_6 | AL391421.1 | 0.6097943 | 1 | -0.284 | -1.027 | chr10 | 79626633 | 79629368 | + | lncRNA |
| ENSG00000172336.5_3 | POP7 | 0.99850442 | 1 | -0.284 | 4.933 | chr7 | 100303744 | 100305109 | + | protein_coding |
| ENSG00000233122.2_4 | CTAGE7P | 0.76906897 | 1 | -0.284 | 0.705 | chr10 | 131904310 | 131906745 | + | processed_pseudogene |
| ENSG00000106344.8_7 | RBM28 | 0.99995407 | 1 | -0.284 | 5.990 | chr7 | 127937738 | 127983962 | - | protein_coding |
| ENSG00000101224.17_4 | CDC25B | 0.99998462 | 1 | -0.284 | 7.568 | chr20 | 3767419 | 3786768 | + | protein_coding |
| ENSG00000162616.9_4 | DNAJB4 | 0.99595729 | 1 | -0.284 | 4.445 | chr1 | 78444859 | 78483648 | + | protein_coding |
| ENSG00000102710.20_7 | SUPT20H | 0.99982178 | 1 | -0.284 | 5.465 | chr13 | 37583449 | 37633850 | - | protein_coding |
| ENSG00000162642.14_4 | C1orf52 | 0.9784909 | 1 | -0.284 | 3.456 | chr1 | 85715636 | 85725355 | - | protein_coding |
| ENSG00000151131.11_6 | C12orf45 | 0.99392569 | 1 | -0.284 | 5.011 | chr12 | 105380094 | 105467975 | + | protein_coding |
| ENSG00000256356.1_4 | HSPA8P5 | 0.63859114 | 1 | -0.284 | -0.473 | chr12 | 4206617 | 4208685 | + | processed_pseudogene |
| ENSG00000182504.11_4 | CEP97 | 0.99986229 | 1 | -0.285 | 5.522 | chr3 | 101443437 | 101489406 | + | protein_coding |
| ENSG00000241322.4 | CDRT1 | 0.79116489 | 1 | -0.285 | 1.125 | chr17 | 15468798 | 15522826 | - | protein_coding |
| ENSG00000117971.12_5 | CHRNB4 | 0.67245799 | 1 | -0.285 | -0.158 | chr15 | 78916453 | 79020096 | - | protein_coding |
| ENSG00000141543.12_7 | EIF4A3 | 0.99997704 | 1 | -0.285 | 6.844 | chr17 | 78108168 | 78120950 | - | protein_coding |
| ENSG00000187288.10_3 | CIDEC | 0.60831452 | 1 | -0.285 | -0.993 | chr3 | 9908395 | 9921938 | - | protein_coding |
| ENSG00000116191.17_6 | RALGPS2 | 0.99964169 | 1 | -0.285 | 5.367 | chr1 | 178694282 | 178890976 | + | protein_coding |
| ENSG00000163864.17_6 | NMNAT3 | 0.70114744 | 1 | -0.285 | 0.450 | chr3 | 139279022 | 139396859 | - | protein_coding |
| ENSG00000213862.4_5 | AC044787.1 | 0.73546817 | 1 | -0.285 | 0.699 | chr15 | 48022341 | 48023132 | - | processed_pseudogene |
| ENSG00000162408.11_4 | NOL9 | 0.99994747 | 1 | -0.285 | 6.005 | chr1 | 6581407 | 6614573 | - | protein_coding |

| | | | | | | | | | | |
|---|---|---|---|---|---|---|---|---|---|---|
| ENSG00000136040.9_4 | PLXNC1 | 0.75720827 | 1 | -0.285 | 0.745 | chr12 | 94542353 | 94701451 | + | protein_coding |
| ENSG00000101888.11_2 | NXT2 | 0.9986422 | 1 | -0.285 | 4.406 | chrX | 108779010 | 108787919 | + | protein_coding |
| ENSG00000023330.15_3 | ALAS1 | 0.99988351 | 1 | -0.285 | 5.481 | chr3 | 52232102 | 52248343 | + | protein_coding |
| ENSG00000225151.10_5 | GOLGA2P7 | 0.88802172 | 1 | -0.285 | 1.960 | chr15 | 84868063 | 84898888 | - | transcribed_unprocessed_pseudogene |
| ENSG00000204685.7_6 | STARD7-AS1 | 0.8507794 | 1 | -0.285 | 1.030 | chr2 | 96874141 | 96909091 | + | lncRNA |
| ENSG00000126214.21_6 | KLC1 | 0.99910914 | 1 | -0.286 | 4.734 | chr14 | 104028233 | 104180586 | + | protein_coding |
| ENSG00000278212.2_7 | AC134878.2 | 0.62736347 | 1 | -0.286 | -0.291 | chrY | 13309534 | 13370702 | - | transcribed_unprocessed_pseudogene |
| ENSG00000149548.15_3 | CCDC15 | 0.90113258 | 1 | -0.286 | 1.701 | chr11 | 124824017 | 124911385 | + | protein_coding |
| ENSG00000250634.6_6 | LINC01182 | 0.86311968 | 1 | -0.286 | 1.384 | chr4 | 13655998 | 14005380 | + | lncRNA |
| ENSG00000244462.8_5 | RBM12 | 0.99996841 | 1 | -0.286 | 6.773 | chr20 | 34236847 | 34252878 | - | protein_coding |
| ENSG00000160803.8_4 | UBQLN4 | 0.99974769 | 1 | -0.286 | 5.563 | chr1 | 156005090 | 156023589 | - | protein_coding |
| ENSG00000175768.13_5 | TOMM5 | 0.90186512 | 1 | -0.286 | 1.778 | chr9 | 37582643 | 37592601 | - | protein_coding |
| ENSG00000242294.6_2 | STAG3L5P | 0.99135864 | 1 | -0.287 | 3.646 | chr7 | 99933702 | 99949523 | + | transcribed_unprocessed_pseudogene |
| ENSG00000152422.15_3 | XRCC4 | 0.99675491 | 1 | -0.287 | 3.968 | chr5 | 82373317 | 82649606 | + | protein_coding |
| ENSG00000011566.15_6 | MAP4K3 | 0.99985084 | 1 | -0.287 | 6.105 | chr2 | 39476407 | 39664442 | - | protein_coding |
| ENSG00000249492.2_8 | AC114956.2 | 0.79484674 | 1 | -0.287 | 1.088 | chr5 | 43484015 | 43509458 | + | lncRNA |
| ENSG00000140105.18_6 | WARS1 | 0.9999272 | 1 | -0.287 | 6.210 | chr14 | 100800127 | 100843142 | - | protein_coding |
| ENSG00000163486.8 | SRGAP2 | 0.99954419 | 1 | -0.287 | 5.245 | chr1 | 206516200 | 206637783 | + | protein_coding |
| ENSG00000171109.19_4 | MFN1 | 0.99955409 | 1 | -0.287 | 5.521 | chr3 | 179065497 | 179112724 | + | protein_coding |
| ENSG00000260005.6_5 | AC027601.2 | 0.80043105 | 1 | -0.288 | 1.200 | chr17 | 79202507 | 79207783 | + | lncRNA |
| ENSG00000221914.11_7 | PPP2R2A | 0.99956057 | 1 | -0.288 | 5.083 | chr8 | 26149024 | 26230196 | + | protein_coding |
| ENSG00000170271.11_7 | FAXDC2 | 0.90729187 | 1 | -0.288 | 1.955 | chr5 | 154198052 | 154238812 | - | protein_coding |
| ENSG00000167720.13_4 | SRR | 0.97355727 | 1 | -0.288 | 2.917 | chr17 | 2206677 | 2228554 | + | protein_coding |
| ENSG00000232024.2_4 | LSM12P1 | 0.80956527 | 1 | -0.288 | 1.412 | chr8 | 35382694 | 35383281 | - | processed_pseudogene |
| ENSG00000186063.13_4 | AIDA | 0.99749428 | 1 | -0.288 | 4.195 | chr1 | 222841355 | 222886552 | - | protein_coding |
| ENSG00000249565.2_4 | SERBP1P5 | 0.69757215 | 1 | -0.288 | 0.395 | chr4 | 79102020 | 79103200 | - | processed_pseudogene |
| ENSG00000134453.16_4 | RBM17 | 0.99971712 | 1 | -0.288 | 5.688 | chr10 | 6130997 | 6159420 | + | protein_coding |
| ENSG00000287454.1_2 | AC008149.2 | 0.66722781 | 1 | -0.288 | 0.085 | chr12 | 96623341 | 96629428 | - | lncRNA |
| ENSG00000198925.12_7 | ATG9A | 0.99937088 | 1 | -0.288 | 4.880 | chr2 | 220084102 | 220094439 | - | protein_coding |
| ENSG00000197622.13_5 | CDC42SE1 | 0.99995678 | 1 | -0.288 | 6.090 | chr1 | 151023447 | 151042801 | - | protein_coding |
| ENSG00000150433.9_6 | TMEM218 | 0.99485625 | 1 | -0.288 | 3.852 | chr11 | 124966441 | 124981659 | - | protein_coding |
| ENSG00000123473.15_4 | STIL | 0.99991044 | 1 | -0.288 | 5.894 | chr1 | 47715811 | 47779819 | - | protein_coding |
| ENSG00000262766.1_7 | AC135050.5 | 0.77081189 | 1 | -0.288 | 0.910 | chr16 | 31129399 | 31130068 | + | lncRNA |
| ENSG00000229989.4_4 | MIR181A1HG | 0.83457469 | 1 | -0.288 | 1.355 | chr1 | 198746990 | 198906600 | - | lncRNA |
| ENSG00000162366.8_4 | PDZK1IP1 | 0.83035739 | 1 | -0.288 | 1.186 | chr1 | 47649254 | 47656716 | - | protein_coding |
| ENSG00000075618.18_3 | FSCN1 | 0.85583972 | 1 | -0.289 | 7.100 | chr7 | 5632447 | 5646286 | + | protein_coding |
| ENSG00000117505.13_3 | DR1 | 0.99978211 | 1 | -0.289 | 5.899 | chr1 | 93811464 | 93835050 | + | protein_coding |
| ENSG00000135372.9_4 | NAT10 | 0.99996351 | 1 | -0.289 | 6.258 | chr11 | 34127164 | 34169217 | + | protein_coding |
| ENSG00000279259.1_5 | AC087741.3 | 0.73624015 | 1 | -0.289 | 0.466 | chr17 | 78121049 | 78122395 | + | TEC |
| ENSG00000183535.9_6 | COL18A1-AS1 | 0.69785179 | 1 | -0.289 | 0.645 | chr21 | 46839631 | 46844985 | - | lncRNA |
| ENSG00000101974.14_5 | ATP11C | 0.99982705 | 1 | -0.289 | 5.600 | chrX | 138808505 | 139027435 | - | protein_coding |
| ENSG00000101452.15_5 | DHX35 | 0.99576889 | 1 | -0.289 | 4.099 | chr20 | 37590942 | 37668366 | + | protein_coding |
| ENSG00000266967.7_5 | AARSD1 | 0.79506921 | 1 | -0.289 | 0.966 | chr17 | 41102543 | 41116515 | - | protein_coding |
| ENSG00000157557.13_6 | ETS2 | 0.99982996 | 1 | -0.289 | 5.574 | chr21 | 40177107 | 40196879 | + | protein_coding |
| ENSG00000130052.13_5 | STARD8 | 0.90741515 | 1 | -0.289 | 2.129 | chrX | 67867508 | 67945684 | + | protein_coding |
| ENSG00000272084.1_7 | AL137127.1 | 0.63859114 | 1 | -0.289 | -0.476 | chr1 | 19398604 | 19402005 | - | lncRNA |
| ENSG00000162623.16_4 | TYW3 | 0.99832138 | 1 | -0.289 | 4.739 | chr1 | 75198836 | 75232362 | + | protein_coding |
| ENSG00000237649.8_3 | KIFC1 | 0.99856643 | 1 | -0.289 | 4.618 | chr6 | 33359600 | 33377673 | + | protein_coding |
| ENSG00000186566.13_5 | GPATCH8 | 0.99979485 | 1 | -0.289 | 5.719 | chr17 | 42472649 | 42580798 | - | protein_coding |
| ENSG00000177683.14_5 | THAP5 | 0.99836259 | 1 | -0.289 | 5.575 | chr7 | 108194987 | 108210194 | - | protein_coding |
| ENSG00000257365.8_4 | FNTB | 0.98125801 | 1 | -0.290 | 3.218 | chr14 | 65453613 | 65529370 | + | protein_coding |
| ENSG00000269148.1_6 | AC092301.1 | 0.61042766 | 1 | -0.290 | -0.725 | chr19 | 46333422 | 46334366 | + | lncRNA |
| ENSG00000270673.1_5 | YTHDF3-AS1 | 0.6746234 | 1 | -0.290 | 0.014 | chr8 | 64080284 | 64081001 | - | lncRNA |
| ENSG00000226851.1_8 | AC004112.1 | 0.60375709 | 1 | -0.290 | -1.030 | chr7 | 111968244 | 112049678 | - | lncRNA |
| ENSG00000135587.9_4 | SMPD2 | 0.98118015 | 1 | -0.290 | 3.172 | chr6 | 109761927 | 109765122 | + | protein_coding |
| ENSG00000179611.3_4 | DGKZP1 | 0.69523188 | 1 | -0.290 | 0.165 | chr13 | 44542560 | 44545335 | + | processed_pseudogene |
| ENSG00000176563.10_4 | CNTD1 | 0.65080692 | 1 | -0.290 | -0.064 | chr17 | 40950818 | 40963605 | + | protein_coding |

| | | | | | | | | | | |
|---|---|---|---|---|---|---|---|---|---|---|
| ENSG00000108829.10_5 | LRRC59 | 0.99999337 | 1 | -0.290 | 8.088 | chr17 | 48452420 | 48474884 | - | protein_coding |
| ENSG00000171858.18_3 | RPS21 | 0.99993016 | 1 | -0.290 | 6.589 | chr20 | 60962159 | 60963576 | + | protein_coding |
| ENSG00000077312.9_5 | SNRPA | 0.99976011 | 1 | -0.291 | 5.736 | chr19 | 41256542 | 41271294 | + | protein_coding |
| ENSG00000243156.9_7 | MICAL3 | 0.9982388 | 1 | -0.291 | 4.925 | chr22 | 18270415 | 18507327 | - | protein_coding |
| ENSG00000112078.14_5 | KCTD20 | 0.99994542 | 1 | -0.291 | 6.089 | chr6 | 36410544 | 36458920 | + | protein_coding |
| ENSG00000168268.11_4 | NT5DC2 | 0.99596588 | 1 | -0.291 | 5.033 | chr3 | 52558401 | 52569070 | - | protein_coding |
| ENSG00000198720.13_5 | ANKRD13B | 0.9934859 | 1 | -0.291 | 4.361 | chr17 | 27916787 | 27941779 | + | protein_coding |
| ENSG00000139505.11_3 | MTMR6 | 0.99959525 | 1 | -0.291 | 5.591 | chr13 | 25820339 | 25862147 | - | protein_coding |
| ENSG00000233937.7_7 | AC008443.1 | 0.69424965 | 1 | -0.291 | 0.115 | chr5 | 180673507 | 180699168 | + | lncRNA |
| ENSG00000103423.14_4 | DNAJA3 | 0.99997222 | 1 | -0.291 | 6.562 | chr16 | 4475806 | 4506776 | + | protein_coding |
| ENSG00000170632.14_4 | ARMC10 | 0.99969854 | 1 | -0.291 | 5.350 | chr7 | 102715328 | 102740206 | + | protein_coding |
| ENSG00000145293.16_4 | ENOPH1 | 0.99977557 | 1 | -0.291 | 5.377 | chr4 | 83351743 | 83382330 | + | protein_coding |
| ENSG00000115325.13_2 | DOK1 | 0.86702031 | 1 | -0.291 | 1.682 | chr2 | 74776153 | 74784681 | + | protein_coding |
| ENSG00000153207.15_5 | AHCTF1 | 0.99998181 | 1 | -0.291 | 7.246 | chr1 | 247002400 | 247095269 | - | protein_coding |
| ENSG00000271013.1_4 | LRRC37A9P | 0.73682398 | 1 | -0.291 | 0.629 | chr17 | 34239639 | 34245014 | - | processed_pseudogene |
| ENSG00000242071.3_4 | RPL7AP6 | 0.92195304 | 1 | -0.291 | 3.191 | chr14 | 70352057 | 70352857 | - | processed_pseudogene |
| ENSG00000244041.7_6 | LINC01011 | 0.66999649 | 1 | -0.291 | 0.187 | chr6 | 2988201 | 2991407 | + | lncRNA |
| ENSG00000131944.10_5 | FAAP24 | 0.99611281 | 1 | -0.291 | 3.995 | chr19 | 33463115 | 33469135 | + | protein_coding |
| ENSG00000181938.13_2 | GINS3 | 0.99757523 | 1 | -0.291 | 4.582 | chr16 | 58328984 | 58440048 | + | protein_coding |
| ENSG00000226221.1_4 | RPL26P19 | 0.93585578 | 1 | -0.291 | 2.335 | chr5 | 55800462 | 55800899 | - | processed_pseudogene |
| ENSG00000165934.13_3 | CPSF2 | 0.99998279 | 1 | -0.291 | 7.422 | chr14 | 92588313 | 92638489 | + | protein_coding |
| ENSG00000225674.1_4 | IPO7P2 | 0.62809481 | 1 | -0.292 | -0.227 | chr13 | 24696204 | 24697953 | - | processed_pseudogene |
| ENSG00000162434.13_5 | JAK1 | 0.9999945 | 1 | -0.292 | 8.347 | chr1 | 65298912 | 65533437 | - | protein_coding |
| ENSG00000130520.10_7 | LSM4 | 0.99982999 | 1 | -0.292 | 6.261 | chr19 | 18417040 | 18434084 | - | protein_coding |
| ENSG00000251022.6_8 | THAP9-AS1 | 0.99672911 | 1 | -0.292 | 4.191 | chr4 | 83814162 | 83822113 | - | lncRNA |
| ENSG00000244649.5_5 | LINC02086 | 0.97697628 | 1 | -0.292 | 3.051 | chr17 | 46724285 | 46784708 | + | lncRNA |
| ENSG00000245748.1_8 | AC097382.2 | 0.6089515 | 1 | -0.292 | -0.639 | chr4 | 7032281 | 7047958 | - | lncRNA |
| ENSG00000112242.15_5 | E2F3 | 0.9996614 | 1 | -0.292 | 5.147 | chr6 | 20402110 | 20493945 | + | protein_coding |
| ENSG00000117174.11_5 | ZNHIT6 | 0.99827448 | 1 | -0.292 | 4.623 | chr1 | 86115100 | 86174116 | - | protein_coding |
| ENSG00000243927.6_4 | MRPS6 | 0.97847389 | 1 | -0.292 | 3.170 | chr21 | 35445524 | 35515334 | + | protein_coding |
| ENSG00000283078.1_6 | AL137077.2 | 0.60112746 | 1 | -0.293 | -0.785 | chr20 | 60641886 | 60643493 | + | lncRNA |
| ENSG00000104894.12_4 | CD37 | 0.84673974 | 1 | -0.293 | 1.641 | chr19 | 49838428 | 49846592 | + | protein_coding |
| ENSG00000066379.15_3 | ZNRD1 | 0.88755387 | 1 | -0.293 | 1.395 | chr6 | 30026676 | 30032686 | + | protein_coding |
| ENSG00000188725.8_4 | SMIM15 | 0.99989658 | 1 | -0.293 | 6.289 | chr5 | 60453531 | 60458295 | - | protein_coding |
| ENSG00000143590.14_4 | EFNA3 | 0.66856359 | 1 | -0.293 | 0.078 | chr1 | 155051313 | 155060014 | + | protein_coding |
| ENSG00000272941.1_6 | AC083862.2 | 0.63513648 | 1 | -0.293 | -0.358 | chr7 | 134853155 | 134854299 | + | lncRNA |
| ENSG00000267796.8_4 | LIN37 | 0.86502202 | 1 | -0.294 | 1.521 | chr19 | 36239262 | 36245420 | + | protein_coding |
| ENSG00000160953.16_4 | PWWP3A | 0.99582561 | 1 | -0.294 | 4.614 | chr19 | 1354710 | 1378430 | + | protein_coding |
| ENSG00000129667.12_7 | RHBDF2 | 0.99905644 | 1 | -0.294 | 5.011 | chr17 | 74466973 | 74497872 | - | protein_coding |
| ENSG00000133121.21_6 | STARD13 | 0.96913652 | 1 | -0.294 | 2.790 | chr13 | 33677274 | 33924767 | - | protein_coding |
| ENSG00000109475.16_2 | RPL34 | 0.99990461 | 1 | -0.294 | 6.828 | chr4 | 109541722 | 109551568 | + | protein_coding |
| ENSG00000165097.16_5 | KDM1B | 0.99442353 | 1 | -0.294 | 4.014 | chr6 | 18155560 | 18224085 | + | protein_coding |
| ENSG00000269399.1 | CTD-3222D19.7 | 0.64058696 | 1 | -0.294 | 0.039 | chr19 | 16653434 | 16655625 | - | lincRNA |
| ENSG00000172785.18_4 | CBWD1 | 0.98110182 | 1 | -0.294 | 3.455 | chr9 | 121038 | 179147 | - | protein_coding |
| ENSG00000114023.15_3 | FAM162A | 0.99662729 | 1 | -0.294 | 4.311 | chr3 | 122103023 | 122131181 | + | protein_coding |
| ENSG00000206527.10_6 | HACD2 | 0.9999488 | 1 | -0.294 | 6.655 | chr3 | 123209667 | 123303900 | - | protein_coding |
| ENSG00000147274.14_5 | RBMX | 0.99998865 | 1 | -0.294 | 8.695 | chrX | 135930163 | 135962923 | - | protein_coding |
| ENSG00000165832.6_3 | TRUB1 | 0.99980688 | 1 | -0.294 | 5.838 | chr10 | 116697954 | 116737435 | + | protein_coding |
| ENSG00000094880.11_6 | CDC23 | 0.99984112 | 1 | -0.295 | 5.888 | chr5 | 137523339 | 137549032 | - | protein_coding |
| ENSG00000288559.1_1 | AC016394.4 | 0.85906303 | 1 | -0.295 | 1.519 | chr10 | 75007100 | 75008026 | - | lncRNA |
| ENSG00000148688.13_2 | RPP30 | 0.99906979 | 1 | -0.295 | 5.144 | chr10 | 92631473 | 92668310 | + | protein_coding |
| ENSG00000129473.9_5 | BCL2L2 | 0.99725137 | 1 | -0.295 | 4.389 | chr14 | 23767999 | 23780968 | + | protein_coding |
| ENSG00000117020.18_6 | AKT3 | 0.99956768 | 1 | -0.295 | 6.309 | chr1 | 243651535 | 244014381 | - | protein_coding |
| ENSG00000255008.3_8 | LINC02739 | 0.61888383 | 1 | -0.295 | -0.614 | chr11 | 59327771 | 59333756 | - | lncRNA |
| ENSG00000203650.9_6 | LINC01285 | 0.64517929 | 1 | -0.295 | -0.142 | chrX | 117973502 | 118053632 | + | lncRNA |
| ENSG00000119689.15_5 | DLST | 0.99992338 | 1 | -0.295 | 6.262 | chr14 | 75348594 | 75370446 | + | protein_coding |
| ENSG00000101945.16_2 | SUV39H1 | 0.99602775 | 1 | -0.295 | 4.218 | chrX | 48553945 | 48567403 | + | protein_coding |
| ENSG00000281189.1_3 | GHET1 | 0.62319143 | 1 | -0.295 | -0.337 | chr7 | 148684619 | 148686524 | + | lncRNA |

| | | | | | | | | | | |
|---|---|---|---|---|---|---|---|---|---|---|
| ENSG00000163632.14_7 | C3orf49 | 0.6241223 | 1 | -0.295 | -0.428 | chr3 | 63804975 | 63834312 | + | protein_coding |
| ENSG00000257086.1_10 | AP001453.4 | 0.59653773 | 1 | -0.296 | -0.925 | chr11 | 64014411 | 64016966 | - | lncRNA |
| ENSG00000076344.16_6 | RGS11 | 0.90723831 | 1 | -0.296 | 2.409 | chr16 | 318300 | 325980 | - | protein_coding |
| ENSG00000007372.23_8 | PAX6 | 0.67694931 | 1 | -0.296 | 0.211 | chr11 | 31806327 | 31839509 | - | protein_coding |
| ENSG00000189223.15_5 | PAX8-AS1 | 0.84511498 | 1 | -0.296 | 1.377 | chr2 | 113968998 | 114034158 | + | lncRNA |
| ENSG00000160818.16_4 | GPATCH4 | 0.99933717 | 1 | -0.296 | 5.190 | chr1 | 156564279 | 156571288 | - | protein_coding |
| ENSG00000163170.12_4 | BOLA3 | 0.97926836 | 1 | -0.296 | 3.244 | chr2 | 74362527 | 74375039 | - | protein_coding |
| ENSG00000154380.17_6 | ENAH | 0.99997558 | 1 | -0.296 | 7.569 | chr1 | 225674537 | 225840844 | - | protein_coding |
| ENSG00000119772.16_4 | DNMT3A | 0.99906482 | 1 | -0.296 | 4.664 | chr2 | 25450724 | 25565459 | - | protein_coding |
| ENSG00000069702.11_6 | TGFBR3 | 0.99871254 | 1 | -0.296 | 4.522 | chr1 | 92145900 | 92371892 | - | protein_coding |
| ENSG00000285531.1_1 | Z83840.2 | 0.59461814 | 1 | -0.296 | -0.961 | chr22 | 42072749 | 42073129 | - | processed_pseudogene |
| ENSG00000152558.15_6 | TMEM123 | 0.99994702 | 1 | -0.296 | 8.098 | chr11 | 102267063 | 102341115 | - | protein_coding |
| ENSG00000138778.13_5 | CENPE | 0.99988363 | 1 | -0.297 | 6.631 | chr4 | 104026506 | 104119602 | - | protein_coding |
| ENSG00000165283.16_4 | STOML2 | 0.99993216 | 1 | -0.297 | 6.347 | chr9 | 35099773 | 35103192 | - | protein_coding |
| ENSG00000099365.11_4 | STX1B | 0.86088954 | 1 | -0.297 | 1.486 | chr16 | 31000577 | 31021959 | - | protein_coding |
| ENSG00000213741.10_6 | RPS29 | 0.9998603 | 1 | -0.297 | 6.817 | chr14 | 50037702 | 50065882 | - | protein_coding |
| ENSG00000123636.18_5 | BAZ2B | 0.99948014 | 1 | -0.297 | 5.825 | chr2 | 160175490 | 160473080 | - | protein_coding |
| ENSG00000089775.11_3 | ZBTB25 | 0.9987058 | 1 | -0.297 | 4.745 | chr14 | 64915824 | 64971931 | - | protein_coding |
| ENSG00000173276.14_5 | ZBTB21 | 0.99771114 | 1 | -0.297 | 4.620 | chr21 | 43406940 | 43430496 | - | protein_coding |
| ENSG00000188825.14_7 | LINC00910 | 0.9914476 | 1 | -0.297 | 3.700 | chr17 | 41416109 | 41466567 | - | lncRNA |
| ENSG00000089280.18_2 | FUS | 0.99945475 | 1 | -0.297 | 5.065 | chr16 | 31191431 | 31206192 | + | protein_coding |
| ENSG00000287581.1_2 | AC016549.1 | 0.59300744 | 1 | -0.297 | -0.995 | chr5 | 14356995 | 14359394 | - | lncRNA |
| ENSG00000161888.11_8 | SPC24 | 0.99943723 | 1 | -0.298 | 5.078 | chr19 | 11242196 | 11266484 | - | protein_coding |
| ENSG00000077713.19_4 | SLC25A43 | 0.99949919 | 1 | -0.298 | 5.488 | chrX | 118533299 | 118588441 | + | protein_coding |
| ENSG00000162129.14_8 | CLPB | 0.99984984 | 1 | -0.298 | 5.660 | chr11 | 71996539 | 72145724 | - | protein_coding |
| ENSG00000234975.6_4 | FTH1P2 | 0.68015589 | 1 | -0.298 | 0.165 | chr1 | 228823162 | 228823573 | + | processed_pseudogene |
| ENSG00000152684.11_4 | PELO | 0.9997075 | 1 | -0.298 | 5.440 | chr5 | 52083750 | 52099878 | + | protein_coding |
| ENSG00000005156.12_8 | LIG3 | 0.99993218 | 1 | -0.299 | 6.186 | chr17 | 33307531 | 33336762 | + | protein_coding |
| ENSG00000171951.5_6 | SCG2 | 0.70824414 | 1 | -0.299 | 0.597 | chr2 | 224461658 | 224467079 | - | protein_coding |
| ENSG00000111880.16_4 | RNGTT | 0.99962114 | 1 | -0.299 | 5.448 | chr6 | 89319616 | 89673337 | - | protein_coding |
| ENSG00000164466.13_6 | SFXN1 | 0.99995845 | 1 | -0.299 | 6.767 | chr5 | 174904065 | 174956745 | + | protein_coding |
| ENSG00000237748.1_2 | UQCRBP1 | 0.5908969 | 1 | -0.299 | -0.961 | chrX | 56763675 | 56764010 | - | processed_pseudogene |
| ENSG00000198744.5_4 | MTCO3P12 | 0.99908613 | 1 | -0.299 | 5.666 | chr1 | 569756 | 570302 | + | unprocessed_pseudogene |
| ENSG00000145248.7_4 | SLC10A4 | 0.93322972 | 1 | -0.299 | 2.232 | chr4 | 48485360 | 48491543 | + | protein_coding |
| ENSG00000140471.17_4 | LINS1 | 0.99715374 | 1 | -0.299 | 4.303 | chr15 | 101099574 | 101143435 | - | protein_coding |
| ENSG00000092199.17_8 | HNRNPC | 0.99998901 | 1 | -0.299 | 9.224 | chr14 | 21677295 | 21737653 | - | protein_coding |
| ENSG00000255585.3_6 | AL590627.1 | 0.68835478 | 1 | -0.299 | 0.214 | chr9 | 140670991 | 140683687 | - | transcribed_processed_pseudogene |
| ENSG00000111843.14_3 | TMEM14C | 0.9998739 | 1 | -0.299 | 5.992 | chr6 | 10723148 | 10731362 | + | protein_coding |
| ENSG00000215156.5_5 | AC138409.1 | 0.60685784 | 1 | -0.299 | -0.263 | chr5 | 34190161 | 34193758 | + | processed_pseudogene |
| ENSG00000080986.13_5 | NDC80 | 0.99805417 | 1 | -0.299 | 4.360 | chr18 | 2571556 | 2616634 | + | protein_coding |
| ENSG00000189308.11_4 | LIN54 | 0.99927083 | 1 | -0.300 | 5.017 | chr4 | 83831126 | 83934079 | - | protein_coding |
| ENSG00000108883.13_4 | EFTUD2 | 0.99998967 | 1 | -0.300 | 7.757 | chr17 | 42927316 | 42976813 | - | protein_coding |
| ENSG00000011198.10_8 | ABHD5 | 0.99419286 | 1 | -0.300 | 4.260 | chr3 | 43731600 | 43775863 | + | protein_coding |
| ENSG00000174010.9_3 | KLHL15 | 0.99914386 | 1 | -0.300 | 5.070 | chrX | 24001837 | 24045303 | - | protein_coding |
| ENSG00000185420.19_7 | SMYD3 | 0.99190597 | 1 | -0.300 | 3.926 | chr1 | 245912644 | 246670614 | - | protein_coding |
| ENSG00000121690.11_4 | DEPDC7 | 0.98644143 | 1 | -0.300 | 3.638 | chr11 | 33037422 | 33055128 | + | protein_coding |
| ENSG00000114331.15_7 | ACAP2 | 0.99977232 | 1 | -0.300 | 5.968 | chr3 | 194995474 | 195163773 | - | protein_coding |
| ENSG00000162613.16_4 | FUBP1 | 0.99996691 | 1 | -0.300 | 7.434 | chr1 | 78409740 | 78444794 | - | protein_coding |
| ENSG00000151779.13_5 | NBAS | 0.99992859 | 1 | -0.300 | 6.569 | chr2 | 15307038 | 15701458 | - | protein_coding |
| ENSG00000287191.1_2 | AC055807.1 | 0.65716978 | 1 | -0.300 | 0.243 | chr15 | 99382521 | 99383376 | - | lncRNA |
| ENSG00000113971.20_5 | NPHP3 | 0.9203186 | 1 | -0.300 | 2.429 | chr3 | 132399453 | 132441258 | - | protein_coding |
| ENSG00000259498.1_7 | TPM1-AS | 0.66824934 | 1 | -0.300 | 0.094 | chr15 | 63338233 | 63341586 | - | lncRNA |
| ENSG00000108352.12_6 | RAPGEFL1 | 0.90866421 | 1 | -0.301 | 2.243 | chr17 | 38333263 | 38351908 | + | protein_coding |
| ENSG00000137364.5_4 | TPMT | 0.99180334 | 1 | -0.301 | 3.904 | chr6 | 18128542 | 18155308 | - | protein_coding |
| ENSG00000162961.14_5 | DPY30 | 0.99679141 | 1 | -0.301 | 5.033 | chr2 | 32092878 | 32264874 | - | protein_coding |
| ENSG00000144134.18_2 | RABL2A | 0.84785344 | 1 | -0.301 | 2.695 | chr2 | 114384806 | 114400973 | + | protein_coding |
| ENSG00000278576.1_8 | AL162171.3 | 0.64575774 | 1 | -0.301 | -0.010 | chr14 | 89055575 | 89058752 | + | lncRNA |
| ENSG00000132423.12_5 | COQ3 | 0.93976119 | 1 | -0.301 | 2.852 | chr6 | 99817277 | 99842071 | - | protein_coding |

| | | | | | | | | | | |
|---|---|---|---|---|---|---|---|---|---|---|
| ENSG00000132823.11_4 | OSER1 | 0.9994419 | 1 | -0.301 | 5.164 | chr20 | 42824579 | 42839411 | - | protein_coding |
| ENSG00000169599.12_3 | NFU1 | 0.9988511 | 1 | -0.301 | 5.090 | chr2 | 69622882 | 69664760 | - | protein_coding |
| ENSG00000213970.4_6 | AC006122.1 | 0.59261829 | 1 | -0.301 | -0.767 | chr12 | 4432890 | 4433977 | - | transcribed_processed_pseudogene |
| ENSG00000254093.9_7 | PINX1 | 0.95879069 | 1 | -0.301 | 2.866 | chr8 | 10622471 | 10697394 | - | protein_coding |
| ENSG00000270103.3_6 | AL360012.1 | 0.69774239 | 1 | -0.301 | 0.427 | chr1 | 28975112 | 28975242 | + | lncRNA |
| ENSG00000287624.1_2 | AL590666.5 | 0.59243429 | 1 | -0.301 | -0.697 | chr1 | 156737897 | 156739025 | - | lncRNA |
| ENSG00000166228.9_4 | PCBD1 | 0.99990308 | 1 | -0.301 | 6.393 | chr10 | 72642037 | 72648322 | - | protein_coding |
| ENSG00000157693.15_5 | TMEM268 | 0.9962077 | 1 | -0.302 | 4.091 | chr9 | 117373486 | 117408702 | + | protein_coding |
| ENSG00000207547.1 | MIR25 | 0.69146514 | 1 | -0.302 | 0.467 | chr7 | 99691183 | 99691266 | - | miRNA |
| ENSG00000108797.12_3 | CNTNAP1 | 0.99221528 | 1 | -0.302 | 3.712 | chr17 | 40834549 | 40852011 | + | protein_coding |
| ENSG00000103494.14_7 | RPGRIP1L | 0.99913689 | 1 | -0.302 | 5.302 | chr16 | 53632065 | 53737850 | - | protein_coding |
| ENSG00000155096.14_5 | AZIN1 | 0.99996458 | 1 | -0.302 | 7.411 | chr8 | 103838536 | 103906092 | - | protein_coding |
| ENSG00000124635.8_3 | H2BC11 | 0.99995392 | 1 | -0.302 | 7.029 | chr6 | 27093676 | 27100529 | - | protein_coding |
| ENSG00000185798.8_5 | WDR53 | 0.92349145 | 1 | -0.302 | 2.520 | chr3 | 196281048 | 196295545 | - | protein_coding |
| ENSG00000143889.16_8 | HNRNPLL | 0.99988699 | 1 | -0.302 | 5.917 | chr2 | 38789111 | 38830728 | - | protein_coding |
| ENSG00000172404.5_4 | DNAJB7 | 0.59146355 | 1 | -0.302 | -0.797 | chr22 | 41255553 | 41258117 | - | protein_coding |
| ENSG00000175216.15_7 | CKAP5 | 0.99998909 | 1 | -0.302 | 8.264 | chr11 | 46764598 | 46867859 | - | protein_coding |
| ENSG00000151617.17_6 | EDNRA | 0.81032649 | 1 | -0.302 | 1.172 | chr4 | 148402069 | 148466106 | + | protein_coding |
| ENSG00000049541.11_5 | RFC2 | 0.9996875 | 1 | -0.302 | 5.532 | chr7 | 73645829 | 73668788 | - | protein_coding |
| ENSG00000276410.3_4 | H2BC3 | 0.99980284 | 1 | -0.302 | 5.862 | chr6 | 26043505 | 26043885 | - | protein_coding |
| ENSG00000280214.1_6 | AC027682.7 | 0.58976976 | 1 | -0.302 | -0.797 | chr16 | 67584718 | 67586838 | - | TEC |
| ENSG00000170142.12_5 | UBE2E1 | 0.99954651 | 1 | -0.302 | 6.559 | chr3 | 23847446 | 23933131 | + | protein_coding |
| ENSG00000243547.1_4 | HNRNPKP4 | 0.83193678 | 1 | -0.302 | 1.463 | chr3 | 96068398 | 96069783 | + | processed_pseudogene |
| ENSG00000198840.2_1 | MT-ND3 | 0.99982926 | 1 | -0.302 | 6.194 | chrM | 10059 | 10404 | + | protein_coding |
| ENSG00000100353.18_4 | EIF3D | 0.99997342 | 1 | -0.302 | 7.196 | chr22 | 36906902 | 36925231 | - | protein_coding |
| ENSG00000130638.17_5 | ATXN10 | 0.99987219 | 1 | -0.302 | 6.538 | chr22 | 46067678 | 46241187 | + | protein_coding |
| ENSG00000263786.1_8 | AC022211.2 | 0.71976208 | 1 | -0.302 | 0.575 | chr17 | 73141356 | 73142641 | - | lncRNA |
| ENSG00000002079.14_3 | MYH16 | 0.64310875 | 1 | -0.302 | -0.169 | chr7 | 98836452 | 98908753 | + | transcribed_unitary_pseudogene |
| ENSG00000131931.8_6 | THAP1 | 0.98569335 | 1 | -0.303 | 3.484 | chr8 | 42691817 | 42698468 | - | protein_coding |
| ENSG00000136108.15_7 | CKAP2 | 0.99989788 | 1 | -0.303 | 6.403 | chr13 | 53029564 | 53050763 | + | protein_coding |
| ENSG00000073584.20_8 | SMARCE1 | 0.99869343 | 1 | -0.303 | 4.928 | chr17 | 38781214 | 38804906 | - | protein_coding |
| ENSG00000120992.18_5 | LYPLA1 | 0.99990353 | 1 | -0.303 | 6.895 | chr8 | 54958927 | 55014577 | - | protein_coding |
| ENSG00000248473.2_6 | LINC01962 | 0.58546917 | 1 | -0.304 | -0.926 | chr5 | 180605821 | 180618852 | - | lncRNA |
| ENSG00000162994.16_5 | CLHC1 | 0.93015249 | 1 | -0.304 | 2.409 | chr2 | 55399683 | 55459699 | - | protein_coding |
| ENSG00000198225.5_3 | FKBP1C | 0.61250147 | 1 | -0.304 | -0.170 | chr6 | 63921351 | 63922929 | + | protein_coding |
| ENSG00000116984.13_5 | MTR | 0.99969724 | 1 | -0.304 | 6.038 | chr1 | 236958592 | 237084578 | + | protein_coding |
| ENSG00000168894.10_6 | RNF181 | 0.9992393 | 1 | -0.304 | 5.207 | chr2 | 85822848 | 85824831 | + | protein_coding |
| ENSG00000131467.11_5 | PSME3 | 0.99997906 | 1 | -0.304 | 7.230 | chr17 | 40976402 | 40995777 | + | protein_coding |
| ENSG00000257270.1_7 | AL928654.2 | 0.81436098 | 1 | -0.304 | 1.396 | chr14 | 105934130 | 105936954 | - | lncRNA |
| ENSG00000137770.14_3 | CTDSPL2 | 0.99923401 | 1 | -0.304 | 5.390 | chr15 | 44719820 | 44821236 | + | protein_coding |
| ENSG00000171631.14_7 | P2RY6 | 0.97843872 | 1 | -0.305 | 3.494 | chr11 | 72975550 | 73009662 | + | protein_coding |
| ENSG00000196591.12_5 | HDAC2 | 0.9999204 | 1 | -0.305 | 6.448 | chr6 | 114254192 | 114332472 | - | protein_coding |
| ENSG00000101146.13_4 | RAE1 | 0.99979963 | 1 | -0.305 | 5.733 | chr20 | 55926279 | 55954267 | + | protein_coding |
| ENSG00000115661.14_3 | STK16 | 0.99773408 | 1 | -0.305 | 4.604 | chr2 | 220110177 | 220115059 | + | protein_coding |
| ENSG00000126246.10_7 | IGFLR1 | 0.63911301 | 1 | -0.305 | -0.005 | chr19 | 36229702 | 36233354 | - | protein_coding |
| ENSG00000232593.7_4 | KANTR | 0.96064122 | 1 | -0.305 | 2.849 | chrX | 53123327 | 53200096 | + | protein_coding |
| ENSG00000055130.17_4 | CUL1 | 0.99993595 | 1 | -0.305 | 6.558 | chr7 | 148395006 | 148498202 | + | protein_coding |
| ENSG00000259972.3_8 | AC009120.2 | 0.86814199 | 1 | -0.305 | 1.836 | chr16 | 74339025 | 74369244 | - | lncRNA |
| ENSG00000167721.11_4 | TSR1 | 0.99995509 | 1 | -0.305 | 6.612 | chr17 | 2225690 | 2239951 | - | protein_coding |
| ENSG00000167992.13_4 | VWCE | 0.58290325 | 1 | -0.305 | -0.861 | chr11 | 61025758 | 61062788 | - | protein_coding |
| ENSG00000125445.11_4 | MRPS7 | 0.99990823 | 1 | -0.305 | 6.264 | chr17 | 73257755 | 73262457 | + | protein_coding |
| ENSG00000087299.12_4 | L2HGDH | 0.99238855 | 1 | -0.305 | 3.890 | chr14 | 50704281 | 50778947 | - | protein_coding |
| ENSG00000065600.12_5 | PACC1 | 0.9932586 | 1 | -0.305 | 3.886 | chr1 | 212537273 | 212588243 | - | protein_coding |
| ENSG00000097046.13_3 | CDC7 | 0.99811872 | 1 | -0.305 | 4.586 | chr1 | 91966408 | 91991321 | + | protein_coding |
| ENSG00000271127.1 | LL22NC03-N64E9.1 | 0.67772644 | 1 | -0.305 | 0.283 | chr22 | 16179617 | 16181004 | - | sense_intronic |
| ENSG00000067057.17_4 | PFKP | 0.99992882 | 1 | -0.305 | 8.332 | chr10 | 3108525 | 3179910 | + | protein_coding |
| ENSG00000261373.1_6 | VPS9D1-AS1 | 0.84486808 | 1 | -0.305 | 1.390 | chr16 | 89778264 | 89784573 | + | lncRNA |
| ENSG00000157600.12_5 | TMEM164 | 0.99975586 | 1 | -0.306 | 5.947 | chrX | 109245859 | 109425962 | + | protein_coding |

| | | | | | | | | | | |
|---|---|---|---|---|---|---|---|---|---|---|
| ENSG00000240857.2_5 | RDH14 | 0.99204158 | 1 | -0.306 | 4.367 | chr2 | 18735989 | 18741945 | - | protein_coding |
| ENSG00000168813.17_4 | ZNF507 | 0.99986479 | 1 | -0.306 | 6.033 | chr19 | 32836500 | 32878573 | + | protein_coding |
| ENSG00000162694.14_5 | EXTL2 | 0.99551403 | 1 | -0.306 | 4.174 | chr1 | 101337928 | 101360735 | - | protein_coding |
| ENSG00000254614.2_7 | AP003068.2 | 0.67168316 | 1 | -0.306 | 0.222 | chr11 | 64945077 | 64949305 | - | lncRNA |
| ENSG00000152078.10_6 | TLCD4 | 0.99981312 | 1 | -0.306 | 5.974 | chr1 | 95582911 | 95663163 | + | protein_coding |
| ENSG00000164532.11_4 | TBX20 | 0.87380232 | 1 | -0.306 | 1.683 | chr7 | 35242042 | 35293711 | - | protein_coding |
| ENSG00000108591.10_5 | DRG2 | 0.99899869 | 1 | -0.306 | 5.416 | chr17 | 17991206 | 18011284 | + | protein_coding |
| ENSG00000163281.12_6 | GNPDA2 | 0.99735629 | 1 | -0.306 | 4.510 | chr4 | 44684217 | 44728605 | - | protein_coding |
| ENSG00000198805.11_5 | PNP | 0.99925404 | 1 | -0.306 | 5.370 | chr14 | 20937113 | 20945253 | + | protein_coding |
| ENSG00000143321.19_4 | HDGF | 0.99998756 | 1 | -0.306 | 8.710 | chr1 | 156711901 | 156736717 | - | protein_coding |
| ENSG00000171497.5_4 | PPID | 0.99845027 | 1 | -0.306 | 4.608 | chr4 | 159630279 | 159644548 | - | protein_coding |
| ENSG00000101247.18_5 | NDUFAF5 | 0.99742179 | 1 | -0.306 | 4.542 | chr20 | 13765653 | 13802226 | + | protein_coding |
| ENSG00000245522.2_8 | LINC02709 | 0.78882622 | 1 | -0.306 | 1.103 | chr11 | 9776317 | 9781080 | - | lncRNA |
| ENSG00000128789.21_4 | PSMG2 | 0.99948063 | 1 | -0.306 | 5.152 | chr18 | 12658042 | 12725739 | + | protein_coding |
| ENSG00000101843.19_4 | PSMD10 | 0.99975793 | 1 | -0.307 | 5.957 | chrX | 107327437 | 107334779 | - | protein_coding |
| ENSG00000170860.4_3 | LSM3 | 0.9987304 | 1 | -0.307 | 5.623 | chr3 | 14220317 | 14242622 | + | protein_coding |
| ENSG00000124151.19_4 | NCOA3 | 0.99997488 | 1 | -0.307 | 8.605 | chr20 | 46130631 | 46285621 | + | protein_coding |
| ENSG00000200156.1 | RNU5B-1 | 0.60478829 | 1 | -0.307 | -0.536 | chr15 | 65597015 | 65597130 | + | snRNA |
| ENSG00000136758.18_4 | YME1L1 | 0.9999763 | 1 | -0.307 | 7.864 | chr10 | 27399041 | 27444195 | - | protein_coding |
| ENSG00000156671.15_8 | SAMD8 | 0.99897398 | 1 | -0.307 | 5.685 | chr10 | 76859344 | 76941881 | + | protein_coding |
| ENSG00000171867.17_4 | PRNP | 0.99994862 | 1 | -0.307 | 7.168 | chr20 | 4666996 | 4682236 | + | protein_coding |
| ENSG00000223559.1_5 | AC073136.1 | 0.58685115 | 1 | -0.307 | -0.768 | chr7 | 56355923 | 56359118 | + | processed_pseudogene |
| ENSG00000130720.13_5 | FIBCD1 | 0.8557107 | 1 | -0.307 | 1.515 | chr9 | 133777825 | 133814673 | - | protein_coding |
| ENSG00000128591.15_3 | FLNC | 0.88613364 | 1 | -0.307 | 2.016 | chr7 | 128470431 | 128499328 | + | protein_coding |
| ENSG00000204149.12_5 | AGAP6 | 0.95836458 | 1 | -0.307 | 2.774 | chr10 | 51741950 | 51770259 | + | protein_coding |
| ENSG00000133065.11_4 | SLC41A1 | 0.99919314 | 1 | -0.307 | 5.074 | chr1 | 205758222 | 205782876 | - | protein_coding |
| ENSG00000123104.12_6 | ITPR2 | 0.99788581 | 1 | -0.308 | 4.604 | chr12 | 26488285 | 26986127 | - | protein_coding |
| ENSG00000256981.1_5 | AC134349.2 | 0.62099729 | 1 | -0.308 | -0.054 | chr12 | 11311384 | 11312293 | - | unprocessed_pseudogene |
| ENSG00000171791.13_6 | BCL2 | 0.67838477 | 1 | -0.308 | 0.804 | chr18 | 60790579 | 60987361 | - | protein_coding |
| ENSG00000105290.12_6 | APLP1 | 0.99661444 | 1 | -0.308 | 4.756 | chr19 | 36358801 | 36370693 | + | protein_coding |
| ENSG00000186517.14_5 | ARHGAP30 | 0.69342337 | 1 | -0.308 | 0.578 | chr1 | 161016736 | 161039760 | - | protein_coding |
| ENSG00000104863.12_3 | LIN7B | 0.83090678 | 1 | -0.308 | 1.560 | chr19 | 49617581 | 49621717 | + | protein_coding |
| ENSG00000280486.2_7 | AC005329.3 | 0.58860889 | 1 | -0.308 | -0.738 | chr19 | 1385472 | 1389664 | - | lncRNA |
| ENSG00000172171.11_5 | TEFM | 0.98884833 | 1 | -0.308 | 3.581 | chr17 | 29224354 | 29233256 | - | protein_coding |
| ENSG00000196754.12_5 | S100A2 | 0.79159286 | 1 | -0.308 | 1.125 | chr1 | 153533584 | 153540366 | - | protein_coding |
| ENSG00000189129.14_5 | PLAC9 | 0.57866699 | 1 | -0.309 | -0.926 | chr10 | 81891438 | 81905115 | + | protein_coding |
| ENSG00000107949.17_5 | BCCIP | 0.99591705 | 1 | -0.309 | 5.538 | chr10 | 127512115 | 127542264 | + | protein_coding |
| ENSG00000286729.1_1 | AC137695.3 | 0.58316787 | 1 | -0.309 | -0.862 | chr3 | 128996596 | 128997625 | - | lncRNA |
| ENSG00000131409.13_6 | LRRC4B | 0.64360397 | 1 | -0.309 | 0.370 | chr19 | 51020149 | 51071692 | - | protein_coding |
| ENSG00000232931.6_7 | LINC00342 | 0.91642395 | 1 | -0.309 | 2.353 | chr2 | 96472800 | 96500751 | - | lncRNA |
| ENSG00000184402.15_3 | SS18L1 | 0.99500165 | 1 | -0.309 | 4.072 | chr20 | 60718825 | 60757570 | + | protein_coding |
| ENSG00000196659.10_3 | TTC30B | 0.98878037 | 1 | -0.309 | 3.691 | chr2 | 178413726 | 178417524 | - | protein_coding |
| ENSG00000183474.15_2 | GTF2H2C | 0.96658534 | 1 | -0.309 | 3.193 | chr5 | 68856035 | 68890550 | + | protein_coding |
| ENSG00000042286.15_4 | AIFM2 | 0.99953742 | 1 | -0.309 | 5.454 | chr10 | 71857979 | 71892690 | - | protein_coding |
| ENSG00000134183.12_4 | GNAT2 | 0.69555638 | 1 | -0.309 | 0.372 | chr1 | 110145876 | 110162551 | - | protein_coding |
| ENSG00000182013.18_7 | PNMA8A | 0.99825082 | 1 | -0.309 | 4.683 | chr19 | 46969748 | 46974820 | - | protein_coding |
| ENSG00000124251.11_5 | TP53TG5 | 0.68022591 | 1 | -0.309 | 0.511 | chr20 | 44001197 | 44036529 | - | protein_coding |
| ENSG00000153246.13_5 | PLA2R1 | 0.99458129 | 1 | -0.309 | 3.995 | chr2 | 160788517 | 160919126 | - | protein_coding |
| ENSG00000176014.13_4 | TUBB6 | 0.99968486 | 1 | -0.309 | 6.333 | chr18 | 12307668 | 12344319 | + | protein_coding |
| ENSG00000284024.2_4 | HSPA14 | 0.99870948 | 1 | -0.310 | 4.965 | chr10 | 14880159 | 14889017 | + | protein_coding |
| ENSG00000118298.12_4 | CA14 | 0.60224397 | 1 | -0.310 | 0.353 | chr1 | 150229646 | 150237478 | + | protein_coding |
| ENSG00000234072.1_6 | AC074117.1 | 0.94163677 | 1 | -0.310 | 2.562 | chr2 | 27579113 | 27590489 | + | lncRNA |
| ENSG00000180992.7_4 | MRPL14 | 0.99747437 | 1 | -0.310 | 4.559 | chr6 | 44081188 | 44095189 | - | protein_coding |
| ENSG00000205981.8_5 | DNAJC19 | 0.98687531 | 1 | -0.310 | 3.741 | chr3 | 180701497 | 180707562 | - | protein_coding |
| ENSG00000131351.15_4 | HAUS8 | 0.97207671 | 1 | -0.310 | 3.203 | chr19 | 17160539 | 17186435 | - | protein_coding |
| ENSG00000118655.7_6 | DCLRE1B | 0.99204445 | 1 | -0.310 | 4.027 | chr1 | 114447835 | 114456708 | + | protein_coding |
| ENSG00000131732.12_3 | ZCCHC9 | 0.99683156 | 1 | -0.310 | 4.439 | chr5 | 80597406 | 80609116 | + | protein_coding |
| ENSG00000112200.17_6 | ZNF451 | 0.99976542 | 1 | -0.310 | 6.009 | chr6 | 56951642 | 57035103 | + | protein_coding |
| ENSG00000159217.10_4 | IGF2BP1 | 0.9999563 | 1 | -0.310 | 7.101 | chr17 | 47074747 | 47133507 | + | protein_coding |
| ENSG00000166321.14_4 | NUDT13 | 0.94369908 | 1 | -0.310 | 2.594 | chr10 | 74870133 | 74891586 | + | protein_coding |

| | | | | | | | | | | |
|---|---|---|---|---|---|---|---|---|---|---|
| ENSG00000270060.1_8 | AC090589.3 | 0.65990184 | 1 | -0.310 | 0.189 | chr11 | 47189832 | 47191114 | - | lncRNA |
| ENSG00000115652.14_3 | UXS1 | 0.99470203 | 1 | -0.310 | 4.051 | chr2 | 106709759 | 106810795 | - | protein_coding |
| ENSG00000266777.1_4 | SH3GL1P1 | 0.60725744 | 1 | -0.310 | 0.030 | chr17 | 30366993 | 30369847 | + | transcribed_processed_pseudogene |
| ENSG00000125149.12_6 | C16orf70 | 0.99830821 | 1 | -0.310 | 4.768 | chr16 | 67143844 | 67182447 | + | protein_coding |
| ENSG00000163629.13_4 | PTPN13 | 0.99935428 | 1 | -0.310 | 5.703 | chr4 | 87515468 | 87736324 | + | protein_coding |
| ENSG00000168944.15_7 | CEP120 | 0.99950964 | 1 | -0.310 | 5.502 | chr5 | 122680579 | 122759286 | - | protein_coding |
| ENSG00000132661.4_4 | NXT1 | 0.99790649 | 1 | -0.311 | 4.578 | chr20 | 23331428 | 23335408 | + | protein_coding |
| ENSG00000108559.12_5 | NUP88 | 0.99992176 | 1 | -0.311 | 6.706 | chr17 | 5264258 | 5322996 | - | protein_coding |
| ENSG00000115648.14_4 | MLPH | 0.99997944 | 1 | -0.311 | 8.080 | chr2 | 238394071 | 238463965 | + | protein_coding |
| ENSG00000165410.15_4 | CFL2 | 0.99982789 | 1 | -0.311 | 6.400 | chr14 | 35178319 | 35184029 | - | protein_coding |
| ENSG00000082146.13_5 | STRADB | 0.99794833 | 1 | -0.311 | 5.230 | chr2 | 202252581 | 202345569 | + | protein_coding |
| ENSG00000196205.8_2 | EEF1A1P5 | 0.71086522 | 1 | -0.311 | 6.124 | chr9 | 135894873 | 135896261 | + | processed_pseudogene |
| ENSG00000263731.1_7 | AC145207.5 | 0.97506923 | 1 | -0.311 | 3.572 | chr17 | 79836301 | 79838982 | - | lncRNA |
| ENSG00000180667.10_3 | YOD1 | 0.99933604 | 1 | -0.311 | 5.495 | chr1 | 207217194 | 207226325 | - | protein_coding |
| ENSG00000137168.8_4 | PPIL1 | 0.99360997 | 1 | -0.311 | 4.957 | chr6 | 36822603 | 36842579 | - | protein_coding |
| ENSG00000104976.12_4 | SNAPC2 | 0.96417217 | 1 | -0.311 | 3.520 | chr19 | 7985223 | 7988135 | + | protein_coding |
| ENSG00000260644.6_3 | HERC2P5 | 0.60228813 | 1 | -0.312 | -0.462 | chr16 | 32752635 | 32787517 | + | transcribed_unprocessed_pseudogene |
| ENSG00000229358.3_6 | DPY19L1P1 | 0.95189653 | 1 | -0.312 | 2.763 | chr7 | 32620561 | 32801399 | - | unprocessed_pseudogene |
| ENSG00000138434.17_5 | ITPRID2 | 0.999956 | 1 | -0.312 | 7.565 | chr2 | 182756457 | 182795465 | + | protein_coding |
| ENSG00000056050.7_7 | HPF1 | 0.9907205 | 1 | -0.312 | 4.161 | chr4 | 170650621 | 170679095 | - | protein_coding |
| ENSG00000104613.12_4 | INTS10 | 0.99991686 | 1 | -0.312 | 6.477 | chr8 | 19674902 | 19709594 | + | protein_coding |
| ENSG00000184678.10_3 | H2BC21 | 0.99989384 | 1 | -0.312 | 8.143 | chr1 | 149856009 | 149858202 | - | protein_coding |
| ENSG00000174231.17_5 | PRPF8 | 0.99998218 | 1 | -0.312 | 9.025 | chr17 | 1553923 | 1588161 | - | protein_coding |
| ENSG00000225484.7_9 | NUTM2B-AS1 | 0.98479598 | 1 | -0.312 | 3.754 | chr10 | 81421150 | 81586350 | - | lncRNA |
| ENSG00000100749.8_4 | VRK1 | 0.99896251 | 1 | -0.312 | 5.237 | chr14 | 97263719 | 97398059 | + | protein_coding |
| ENSG00000121903.14_2 | ZSCAN20 | 0.95461316 | 1 | -0.312 | 2.848 | chr1 | 33938246 | 33962107 | + | protein_coding |
| ENSG00000119777.20_5 | TMEM214 | 0.9999058 | 1 | -0.313 | 6.409 | chr2 | 27255778 | 27264562 | + | protein_coding |
| ENSG00000184575.12_3 | XPOT | 0.99993835 | 1 | -0.313 | 7.841 | chr12 | 64798172 | 64844905 | + | protein_coding |
| ENSG00000261338.2_6 | AC021016.2 | 0.64787104 | 1 | -0.313 | 0.106 | chr2 | 219120042 | 219122089 | + | lncRNA |
| ENSG00000261098.1_6 | AP000766.1 | 0.59522685 | 1 | -0.313 | -0.173 | chr11 | 107182858 | 107186997 | - | lncRNA |
| ENSG00000240163.1_5 | AC087385.1 | 0.75897733 | 1 | -0.313 | 1.248 | chr15 | 60682570 | 60682881 | + | processed_pseudogene |
| ENSG00000182307.14_4 | C8orf33 | 0.99971528 | 1 | -0.313 | 5.790 | chr8 | 146277851 | 146292071 | + | protein_coding |
| ENSG00000162607.13_4 | USP1 | 0.99982034 | 1 | -0.313 | 6.466 | chr1 | 62901968 | 62917475 | + | protein_coding |
| ENSG00000167641.11_4 | PPP1R14A | 0.87292687 | 1 | -0.313 | 2.102 | chr19 | 38741877 | 38747172 | - | protein_coding |
| ENSG00000236801.1_4 | RPL24P8 | 0.66828142 | 1 | -0.313 | 0.107 | chr9 | 72832111 | 72832584 | + | processed_pseudogene |
| ENSG00000078618.21_4 | NRDC | 0.9999322 | 1 | -0.313 | 6.745 | chr1 | 52254863 | 52344609 | - | protein_coding |
| ENSG00000269956.2_6 | MKNK1-AS1 | 0.57378456 | 1 | -0.313 | -0.830 | chr1 | 47004283 | 47035927 | + | lncRNA |
| ENSG00000107130.10_3 | NCS1 | 0.99683907 | 1 | -0.313 | 5.543 | chr9 | 132934683 | 132999582 | + | protein_coding |
| ENSG00000275084.3_3 | SNORD91B | 0.58001686 | 1 | -0.313 | -0.830 | chr17 | 2232310 | 2232531 | - | snoRNA |
| ENSG00000166451.13_3 | CENPN | 0.99930543 | 1 | -0.314 | 5.219 | chr16 | 81040103 | 81066719 | + | protein_coding |
| ENSG00000284735.1_5 | AL139424.3 | 0.58070043 | 1 | -0.314 | -0.770 | chr1 | 10441964 | 10447207 | - | lncRNA |
| ENSG00000179051.14_5 | RCC2 | 0.99994374 | 1 | -0.314 | 6.799 | chr1 | 17733256 | 17766173 | - | protein_coding |
| ENSG00000029363.17_8 | BCLAF1 | 0.99993783 | 1 | -0.314 | 7.933 | chr6 | 136577765 | 136610989 | - | protein_coding |
| ENSG00000126216.15_6 | TUBGCP3 | 0.99632299 | 1 | -0.314 | 4.476 | chr13 | 113139325 | 113242519 | - | protein_coding |
| ENSG00000268520.1_8 | AC008750.5 | 0.57238597 | 1 | -0.314 | -0.799 | chr19 | 51864966 | 51868798 | - | lncRNA |
| ENSG00000177606.7_3 | JUN | 0.97634553 | 1 | -0.314 | 3.629 | chr1 | 59246463 | 59249719 | - | protein_coding |
| ENSG00000143643.13_5 | TTC13 | 0.99372286 | 1 | -0.314 | 4.222 | chr1 | 231041989 | 231114621 | - | protein_coding |
| ENSG00000205339.10_3 | IPO7 | 0.99994846 | 1 | -0.314 | 8.772 | chr11 | 9406199 | 9469674 | + | protein_coding |
| ENSG00000271862.1_6 | AC104118.1 | 0.72693535 | 1 | -0.315 | 0.503 | chr5 | 82345195 | 82346783 | - | lncRNA |
| ENSG00000123131.13_3 | PRDX4 | 0.99987816 | 1 | -0.315 | 6.747 | chrX | 23682379 | 23704514 | + | protein_coding |
| ENSG00000149196.16_6 | HIKESHI | 0.99786243 | 1 | -0.315 | 4.603 | chr11 | 86013253 | 86056985 | + | protein_coding |
| ENSG00000004487.16_5 | KDM1A | 0.99994585 | 1 | -0.315 | 6.935 | chr1 | 23345936 | 23410182 | + | protein_coding |
| ENSG00000241058.4_5 | NSUN6 | 0.98233839 | 1 | -0.315 | 3.487 | chr10 | 18834490 | 18948214 | - | protein_coding |
| ENSG00000219928.2_5 | AL161787.1 | 0.70213257 | 1 | -0.315 | 0.408 | chr9 | 90140217 | 90140390 | + | processed_pseudogene |
| ENSG00000265354.4_4 | TIMM23 | 0.99923363 | 1 | -0.315 | 5.494 | chr10 | 51592080 | 51623350 | - | protein_coding |
| ENSG00000159199.14_5 | ATP5MC1 | 0.9981171 | 1 | -0.315 | 6.272 | chr17 | 46970127 | 46973233 | + | protein_coding |

| | | | | | | | | | | |
|---|---|---|---|---|---|---|---|---|---|---|
| ENSG00000228889.7_7 | UBAC2-AS1 | 0.77994491 | 1 | -0.315 | 1.029 | chr13 | 99833477 | 99853011 | - | lncRNA |
| ENSG00000207563.1 | MIR23B | 0.67012249 | 1 | -0.315 | 0.172 | chr9 | 97847490 | 97847586 | + | miRNA |
| ENSG00000196639.6_3 | HRH1 | 0.93950141 | 1 | -0.315 | 2.475 | chr3 | 11178779 | 11305243 | + | protein_coding |
| ENSG00000173611.17_6 | SCAI | 0.99230032 | 1 | -0.315 | 4.180 | chr9 | 127704887 | 127905785 | - | protein_coding |
| ENSG00000119138.4_3 | KLF9 | 0.876172 | 1 | -0.316 | 1.910 | chr9 | 72999513 | 73029540 | - | protein_coding |
| ENSG00000178977.3_4 | LINC00324 | 0.79515669 | 1 | -0.316 | 1.211 | chr17 | 8123960 | 8127361 | - | lncRNA |
| ENSG00000187653.11_4 | TMSB4XP8 | 0.87791747 | 1 | -0.316 | 3.666 | chr4 | 91760054 | 91760188 | - | processed_pseudogene |
| ENSG00000080546.13_4 | SESN1 | 0.96964985 | 1 | -0.316 | 3.125 | chr6 | 109307640 | 109416022 | - | protein_coding |
| ENSG00000161960.15_4 | EIF4A1 | 0.98864215 | 1 | -0.316 | 3.810 | chr17 | 7476024 | 7482324 | + | protein_coding |
| ENSG00000224886.2_5 | AL132656.1 | 0.60977376 | 1 | -0.316 | 0.103 | chr10 | 81444250 | 81444739 | - | transcribed_processed_pseudogene |
| ENSG00000268655.2_7 | AC008687.4 | 0.72118918 | 1 | -0.316 | 1.064 | chr19 | 49520753 | 49523780 | - | protein_coding |
| ENSG00000146414.16_6 | SHPRH | 0.99599759 | 1 | -0.316 | 4.557 | chr6 | 146185381 | 146285559 | - | protein_coding |
| ENSG00000149636.15_5 | DSN1 | 0.99962784 | 1 | -0.316 | 5.676 | chr20 | 35380194 | 35402221 | - | protein_coding |
| ENSG00000149231.14_8 | CCDC82 | 0.99807992 | 1 | -0.316 | 4.856 | chr11 | 96085933 | 96123120 | - | protein_coding |
| ENSG00000286242.1_2 | AC018797.3 | 0.63152858 | 1 | -0.316 | 0.269 | chr4 | 103735409 | 103741038 | + | lncRNA |
| ENSG00000165506.14_4 | DNAAF2 | 0.99469031 | 1 | -0.316 | 4.225 | chr14 | 50091892 | 50101948 | - | protein_coding |
| ENSG00000163635.18_4 | ATXN7 | 0.7500671 | 1 | -0.316 | 0.949 | chr3 | 63884075 | 63989129 | + | protein_coding |
| ENSG00000235726.7_6 | AC010148.1 | 0.66258179 | 1 | -0.317 | 0.263 | chr2 | 235712452 | 235822028 | - | lncRNA |
| ENSG00000112304.11_3 | ACOT13 | 0.99324739 | 1 | -0.317 | 4.316 | chr6 | 24667263 | 24705293 | + | protein_coding |
| ENSG00000220842.6_4 | RPL21P16 | 0.6050779 | 1 | -0.317 | -0.415 | chr10 | 122114213 | 122114695 | - | processed_pseudogene |
| ENSG00000223528.7_8 | AL359094.1 | 0.56818985 | 1 | -0.317 | -1.044 | chr10 | 128786359 | 128824771 | - | lncRNA |
| ENSG00000128510.12_5 | CPA4 | 0.7600433 | 1 | -0.317 | 1.127 | chr7 | 129932974 | 129964020 | + | protein_coding |
| ENSG00000239213.6_7 | NCK1-DT | 0.6793422 | 1 | -0.317 | 0.372 | chr3 | 136554187 | 136581460 | - | lncRNA |
| ENSG00000262248.1_6 | AC027796.2 | 0.57992741 | 1 | -0.317 | -0.653 | chr17 | 3522550 | 3523207 | - | processed_pseudogene |
| ENSG00000253667.2_5 | AC100821.1 | 0.66881732 | 1 | -0.317 | 0.306 | chr8 | 54883791 | 54886770 | + | processed_pseudogene |
| ENSG00000112029.10_5 | FBXO5 | 0.99887568 | 1 | -0.317 | 5.127 | chr6 | 153291654 | 153304714 | - | protein_coding |
| ENSG00000231345.3_4 | BEND3P1 | 0.95627283 | 1 | -0.317 | 2.821 | chr10 | 52415727 | 52420232 | + | transcribed_processed_pseudogene |
| ENSG00000069998.12_5 | HDHD5 | 0.99732255 | 1 | -0.317 | 4.588 | chr22 | 17618401 | 17646177 | - | protein_coding |
| ENSG00000058866.15_6 | DGKG | 0.99387066 | 1 | -0.317 | 4.123 | chr3 | 185823457 | 186080023 | - | protein_coding |
| ENSG00000186275.7 | NBPF12 | 0.9072877 | 1 | -0.317 | 2.321 | chr1 | 146373546 | 146467744 | + | protein_coding |
| ENSG00000266498.1_7 | AC055811.3 | 0.61156368 | 1 | -0.317 | -0.259 | chr17 | 17138747 | 17139432 | - | lncRNA |
| ENSG00000104427.12_3 | ZC2HC1A | 0.97049917 | 1 | -0.317 | 3.279 | chr8 | 79578324 | 79632000 | + | protein_coding |
| ENSG00000232729.8_8 | AC211433.1 | 0.6473153 | 1 | -0.318 | 0.250 | chr7 | 74103198 | 74143339 | - | lncRNA |
| ENSG00000248049.7_7 | UBA6-AS1 | 0.9246909 | 1 | -0.318 | 2.405 | chr4 | 68566927 | 68946670 | + | lncRNA |
| ENSG00000170606.15_4 | HSPA4 | 0.9999705 | 1 | -0.318 | 8.519 | chr5 | 132387705 | 132442141 | + | protein_coding |
| ENSG00000261766.1_8 | AC133550.3 | 0.565502 | 1 | -0.318 | -1.082 | chr16 | 28873487 | 28874661 | - | lncRNA |
| ENSG00000157985.19_5 | AGAP1 | 0.99855132 | 1 | -0.318 | 5.130 | chr2 | 236402687 | 237040444 | + | protein_coding |
| ENSG00000132383.12_3 | RPA1 | 0.99658734 | 1 | -0.318 | 6.572 | chr17 | 1732996 | 1803376 | + | protein_coding |
| ENSG00000185627.18_4 | PSMD13 | 0.99970056 | 1 | -0.318 | 6.356 | chr11 | 236966 | 252984 | + | protein_coding |
| ENSG00000140395.9_7 | WDR61 | 0.99942042 | 1 | -0.319 | 5.705 | chr15 | 78570177 | 78592045 | - | protein_coding |
| ENSG00000243986.2_4 | ENO1P3 | 0.60422084 | 1 | -0.319 | -0.368 | chr3 | 124580941 | 124582164 | - | processed_pseudogene |
| ENSG00000188542.9_2 | DUSP28 | 0.93004211 | 1 | -0.319 | 2.677 | chr2 | 241499471 | 241503431 | + | protein_coding |
| ENSG00000181472.5_4 | ZBTB2 | 0.99693356 | 1 | -0.319 | 4.683 | chr6 | 151685250 | 151712694 | - | protein_coding |
| ENSG00000272449.2_6 | AL139246.5 | 0.56525902 | 1 | -0.319 | -0.712 | chr1 | 2461503 | 2478899 | + | lncRNA |
| ENSG00000263412.1_7 | AC004477.1 | 0.74609323 | 1 | -0.319 | 0.984 | chr17 | 46122503 | 46125435 | - | lncRNA |
| ENSG00000171094.18_8 | ALK | 0.57306613 | 1 | -0.319 | -0.771 | chr2 | 29415640 | 30144452 | - | protein_coding |
| ENSG00000246366.6_7 | LACTB2-AS1 | 0.89546206 | 1 | -0.319 | 2.031 | chr8 | 71520812 | 71575514 | + | lncRNA |
| ENSG00000263069.6_9 | RNF213-AS1 | 0.9537861 | 1 | -0.319 | 3.087 | chr17 | 78325628 | 78388968 | - | lncRNA |
| ENSG00000004660.15_5 | CAMKK1 | 0.9934257 | 1 | -0.319 | 4.944 | chr17 | 3763609 | 3798185 | - | protein_coding |
| ENSG00000081154.12_3 | PCNP | 0.99978972 | 1 | -0.320 | 6.996 | chr3 | 101293024 | 101313309 | + | protein_coding |
| ENSG00000196363.9_2 | WDR5 | 0.99892292 | 1 | -0.320 | 5.776 | chr9 | 137000487 | 137025090 | + | protein_coding |
| ENSG00000224086.6_6 | AC245452.1 | 0.96108019 | 1 | -0.320 | 2.994 | chr22 | 22292641 | 22332004 | + | lncRNA |
| ENSG00000021776.11_5 | AQR | 0.99993702 | 1 | -0.320 | 7.519 | chr15 | 35143983 | 35261943 | - | protein_coding |
| ENSG00000273014.1_7 | AC018645.3 | 0.71181524 | 1 | -0.320 | 0.655 | chr7 | 32798494 | 32798965 | + | lncRNA |
| ENSG00000142864.14_5 | SERBP1 | 0.99997271 | 1 | -0.320 | 7.793 | chr1 | 67873493 | 67896098 | - | protein_coding |
| ENSG00000112081.17_3 | SRSF3 | 0.99994255 | 1 | -0.320 | 7.938 | chr6 | 36562130 | 36573377 | + | protein_coding |
| ENSG00000184182.18_3 | UBE2F | 0.95239877 | 1 | -0.321 | 2.752 | chr2 | 238875469 | 238951423 | + | protein_coding |

| | | | | | | | | | | |
|---|---|---|---|---|---|---|---|---|---|---|
| ENSG00000136715.18_6 | SAP130 | 0.99777876 | 1 | -0.321 | 4.711 | chr2 | 128698791 | 128785694 | - | protein_coding |
| ENSG00000129534.14_5 | MIS18BP1 | 0.99966535 | 1 | -0.321 | 6.732 | chr14 | 45672393 | 45722743 | - | protein_coding |
| ENSG00000264895.1_7 | AC006141.1 | 0.70551746 | 1 | -0.321 | 0.656 | chr17 | 49326940 | 49329109 | - | lncRNA |
| ENSG00000085514.16_6 | PILRA | 0.77899431 | 1 | -0.321 | 1.224 | chr7 | 99965153 | 99997719 | + | protein_coding |
| ENSG00000154727.10_2 | GABPA | 0.99692616 | 1 | -0.321 | 4.706 | chr21 | 27106881 | 27144771 | + | protein_coding |
| ENSG00000259515.1_7 | AL117209.1 | 0.57152698 | 1 | -0.321 | -0.770 | chr14 | 103399911 | 103403514 | + | lncRNA |
| ENSG00000171314.9_3 | PGAM1 | 0.99980178 | 1 | -0.321 | 6.550 | chr10 | 99185948 | 99193201 | + | protein_coding |
| ENSG00000134419.15_5 | RPS15A | 0.99960582 | 1 | -0.321 | 6.503 | chr16 | 18792617 | 18801705 | - | protein_coding |
| ENSG00000197312.12_4 | DDI2 | 0.99310163 | 1 | -0.321 | 4.505 | chr1 | 15943953 | 15995539 | + | protein_coding |
| ENSG00000103994.17_8 | ZNF106 | 0.99952939 | 1 | -0.321 | 6.115 | chr15 | 42705021 | 42783339 | - | protein_coding |
| ENSG00000101255.11_4 | TRIB3 | 0.99534651 | 1 | -0.321 | 5.758 | chr20 | 343479 | 378203 | + | protein_coding |
| ENSG00000198804.2_1 | MT-CO1 | 0.94674932 | 1 | -0.321 | 11.534 | chrM | 5904 | 7445 | + | protein_coding |
| ENSG00000176890.16_3 | TYMS | 0.87858166 | 1 | -0.321 | 2.217 | chr18 | 657653 | 673578 | + | protein_coding |
| ENSG00000126226.22_6 | PCID2 | 0.99870343 | 1 | -0.322 | 5.161 | chr13 | 113831850 | 113863029 | - | protein_coding |
| ENSG00000279088.1_8 | AC022400.8 | 0.90051582 | 1 | -0.322 | 2.225 | chr10 | 75502753 | 75503988 | - | TEC |
| ENSG00000115145.10_4 | STAM2 | 0.99965556 | 1 | -0.322 | 5.797 | chr2 | 152973315 | 153032277 | - | protein_coding |
| ENSG00000260466.1_6 | AC126696.2 | 0.58064337 | 1 | -0.322 | -0.464 | chr16 | 87870138 | 87871269 | + | lncRNA |
| ENSG00000197989.14_7 | SNHG12 | 0.99569442 | 1 | -0.322 | 4.501 | chr1 | 28905050 | 28909644 | - | lncRNA |
| ENSG00000143575.15_7 | HAX1 | 0.99952229 | 1 | -0.322 | 5.651 | chr1 | 154245065 | 154248351 | + | protein_coding |
| ENSG00000169045.17_7 | HNRNPH1 | 0.99771716 | 1 | -0.322 | 8.810 | chr5 | 179041179 | 179061785 | - | protein_coding |
| ENSG00000267201.1_6 | LINC01775 | 0.56077932 | 1 | -0.322 | -0.896 | chr19 | 2458933 | 2462183 | - | lncRNA |
| ENSG00000138668.19_7 | HNRNPD | 0.99779819 | 1 | -0.322 | 5.018 | chr4 | 83273651 | 83295656 | - | protein_coding |
| ENSG00000136383.6_2 | ALPK3 | 0.57624007 | 1 | -0.322 | 0.064 | chr15 | 85359911 | 85416713 | + | protein_coding |
| ENSG00000150768.15_2 | DLAT | 0.9996486 | 1 | -0.322 | 6.038 | chr11 | 111895538 | 111935114 | + | protein_coding |
| ENSG00000203705.11_6 | TATDN3 | 0.97694535 | 1 | -0.322 | 3.597 | chr1 | 212965170 | 212990172 | + | protein_coding |
| ENSG00000225092.2_8 | AL355336.2 | 0.56013084 | 1 | -0.323 | -0.801 | chr6 | 7183316 | 7185520 | + | lncRNA |
| ENSG00000280173.1_5 | AC104447.1 | 0.77905954 | 1 | -0.323 | 1.122 | chr3 | 47388440 | 47390563 | + | TEC |
| ENSG00000282381.1_7 | AC104073.4 | 0.55860836 | 1 | -0.323 | -1.009 | chr7 | 64734269 | 64771165 | + | lncRNA |
| ENSG00000271147.8_6 | ARMCX5-GPRASP2 | 0.87427819 | 1 | -0.323 | 1.875 | chrX | 101854440 | 101969599 | + | lncRNA |
| ENSG00000246263.3_6 | UBR5-AS1 | 0.63121445 | 1 | -0.323 | 0.179 | chr8 | 103251622 | 103265978 | + | lncRNA |
| ENSG00000089048.14_3 | ESF1 | 0.99830185 | 1 | -0.323 | 5.636 | chr20 | 13694969 | 13765532 | - | protein_coding |
| ENSG00000259207.8_4 | ITGB3 | 0.56684957 | 1 | -0.323 | -0.710 | chr17 | 45331212 | 45391101 | + | protein_coding |
| ENSG00000151304.6_3 | SRFBP1 | 0.99728568 | 1 | -0.323 | 4.674 | chr5 | 121297670 | 121411265 | + | protein_coding |
| ENSG00000168259.16_7 | DNAJC7 | 0.99984134 | 1 | -0.323 | 6.497 | chr17 | 40128453 | 40173394 | - | protein_coding |
| ENSG00000163882.9_2 | POLR2H | 0.99711594 | 1 | -0.323 | 4.802 | chr3 | 184079506 | 184086384 | + | protein_coding |
| ENSG00000092094.11_5 | OSGEP | 0.99836306 | 1 | -0.323 | 4.853 | chr14 | 20914560 | 20923248 | - | protein_coding |
| ENSG00000166889.14_4 | PATL1 | 0.99928636 | 1 | -0.323 | 5.661 | chr11 | 59404189 | 59436510 | - | protein_coding |
| ENSG00000112039.5_4 | FANCE | 0.94745528 | 1 | -0.324 | 2.774 | chr6 | 35420115 | 35434879 | + | protein_coding |
| ENSG00000134333.14_7 | LDHA | 0.99978482 | 1 | -0.324 | 9.659 | chr11 | 18416107 | 18429972 | + | protein_coding |
| ENSG00000171204.12_3 | TMEM126B | 0.99723006 | 1 | -0.324 | 4.952 | chr11 | 85339617 | 85347583 | + | protein_coding |
| ENSG00000260290.2_6 | AC092115.2 | 0.77699566 | 1 | -0.324 | 1.148 | chr16 | 69790439 | 69791840 | - | processed_pseudogene |
| ENSG00000267160.1_7 | AC091152.2 | 0.55744329 | 1 | -0.324 | -1.010 | chr17 | 42751057 | 42753625 | - | lncRNA |
| ENSG00000175984.15_5 | DENND2C | 0.62899329 | 1 | -0.324 | 0.083 | chr1 | 115125469 | 115213043 | - | protein_coding |
| ENSG00000244513.7_7 | AC109587.1 | 0.59462488 | 1 | -0.324 | 0.146 | chr3 | 69063092 | 69105773 | + | lncRNA |
| ENSG00000100564.9_6 | PIGH | 0.97300968 | 1 | -0.324 | 3.583 | chr14 | 68048672 | 68067003 | - | protein_coding |
| ENSG00000198590.12_6 | C3orf35 | 0.72375543 | 1 | -0.324 | 0.821 | chr3 | 37422553 | 37481677 | + | lncRNA |
| ENSG00000104341.17_5 | LAPTM4B | 0.99973894 | 1 | -0.324 | 7.581 | chr8 | 98787285 | 98865241 | + | protein_coding |
| ENSG00000167604.14_7 | NFKBID | 0.94374802 | 1 | -0.324 | 2.886 | chr19 | 36378555 | 36393205 | - | protein_coding |
| ENSG00000100908.14_4 | EMC9 | 0.9826632 | 1 | -0.325 | 3.770 | chr14 | 24608168 | 24610800 | - | protein_coding |
| ENSG00000259439.2_7 | LINC01833 | 0.92135142 | 1 | -0.325 | 2.630 | chr2 | 45148216 | 45166338 | - | lncRNA |
| ENSG00000000419.12_5 | DPM1 | 0.99886648 | 1 | -0.325 | 5.223 | chr20 | 49551404 | 49575092 | - | protein_coding |
| ENSG00000116906.13_5 | GNPAT | 0.99982243 | 1 | -0.325 | 6.347 | chr1 | 231376953 | 231413719 | + | protein_coding |
| ENSG00000124767.7_4 | GLO1 | 0.99987207 | 1 | -0.325 | 7.126 | chr6 | 38643701 | 38670921 | - | protein_coding |
| ENSG00000241127.8_5 | YAE1 | 0.97933918 | 1 | -0.325 | 3.831 | chr7 | 39605984 | 39649919 | + | protein_coding |
| ENSG00000175054.16_6 | ATR | 0.99990994 | 1 | -0.325 | 6.709 | chr3 | 142167849 | 142297575 | - | protein_coding |
| ENSG00000245275.8_8 | SAP30L-AS1 | 0.9730396 | 1 | -0.325 | 3.539 | chr5 | 153705128 | 153825410 | - | lncRNA |
| ENSG00000280132.1_1 | AC026471.6 | 0.57307012 | 1 | -0.325 | -0.575 | chr16 | 31481525 | 31483429 | - | TEC |
| ENSG00000154743.18_6 | TSEN2 | 0.99643755 | 1 | -0.325 | 4.589 | chr3 | 12525931 | 12581122 | + | protein_coding |
| ENSG00000155099.8_6 | PIP4P2 | 0.93448732 | 1 | -0.325 | 2.769 | chr8 | 92006030 | 92053100 | - | protein_coding |

| | | | | | | | | | | |
|---|---|---|---|---|---|---|---|---|---|---|
| ENSG00000267855.6_5 | NDUFA7 | 0.55974212 | 1 | -0.325 | -0.713 | chr19 | 8373652 | 8386259 | - | protein_coding |
| ENSG00000065413.20_6 | ANKRD44 | 0.92931996 | 1 | -0.326 | 2.947 | chr2 | 197831741 | 198175897 | - | protein_coding |
| ENSG00000268949.1_4 | MRPS17P1 | 0.57332182 | 1 | -0.326 | -0.522 | chr1 | 45454377 | 45457506 | - | processed_pseudogene |
| ENSG00000077458.13_4 | FAM76B | 0.9921264 | 1 | -0.326 | 4.467 | chr11 | 95502117 | 95523573 | - | protein_coding |
| ENSG00000065150.21_6 | IPO5 | 0.99994488 | 1 | -0.326 | 7.866 | chr13 | 98605912 | 98676550 | + | protein_coding |
| ENSG00000169905.12_4 | TOR1AIP2 | 0.99995526 | 1 | -0.326 | 7.444 | chr1 | 179809102 | 179846938 | - | protein_coding |
| ENSG00000147010.18_5 | SH3KBP1 | 0.99949097 | 1 | -0.326 | 6.288 | chrX | 19552095 | 19905718 | - | protein_coding |
| ENSG00000225093.1_4 | RPL3P7 | 0.55431279 | 1 | -0.326 | -0.975 | chr6 | 108325561 | 108326772 | - | processed_pseudogene |
| ENSG00000174343.6_3 | CHRNA9 | 0.55394759 | 1 | -0.326 | -1.083 | chr4 | 40337350 | 40357234 | + | protein_coding |
| ENSG00000240970.1_2 | RPL23AP64 | 0.76480267 | 1 | -0.326 | 1.287 | chr11 | 118873722 | 118874156 | - | processed_pseudogene |
| ENSG00000278463.1_3 | H2AC4 | 0.99639759 | 1 | -0.326 | 6.182 | chr6 | 26033404 | 26033796 | - | protein_coding |
| ENSG00000186314.11_5 | PRELID2 | 0.97077849 | 1 | -0.327 | 3.183 | chr5 | 144851362 | 145214932 | - | protein_coding |
| ENSG00000171262.11_4 | FAM98B | 0.91686268 | 1 | -0.327 | 3.268 | chr15 | 38746328 | 38779911 | + | protein_coding |
| ENSG00000160124.9_4 | CCDC58 | 0.97857745 | 1 | -0.327 | 4.158 | chr3 | 122078438 | 122102078 | - | protein_coding |
| ENSG00000183775.11_4 | KCTD16 | 0.56735586 | 1 | -0.327 | -1.159 | chr5 | 143550437 | 143865249 | + | protein_coding |
| ENSG00000259877.2_6 | AC009113.1 | 0.7927117 | 1 | -0.327 | 1.287 | chr16 | 89281619 | 89284061 | - | lncRNA |
| ENSG00000133316.15_6 | WDR74 | 0.9969384 | 1 | -0.327 | 4.791 | chr11 | 62599814 | 62609281 | - | protein_coding |
| ENSG00000122547.11_5 | EEPD1 | 0.9928593 | 1 | -0.327 | 4.599 | chr7 | 36192863 | 36341147 | + | protein_coding |
| ENSG00000116815.16_6 | CD58 | 0.97978802 | 1 | -0.327 | 3.720 | chr1 | 117057156 | 117113661 | - | protein_coding |
| ENSG00000156968.9_5 | MPV17L | 0.97996898 | 1 | -0.327 | 3.900 | chr16 | 15489611 | 15507128 | + | protein_coding |
| ENSG00000153914.16_3 | SREK1 | 0.99967058 | 1 | -0.328 | 6.593 | chr5 | 65435799 | 65479443 | + | protein_coding |
| ENSG00000137574.11_3 | TGS1 | 0.99923232 | 1 | -0.328 | 5.615 | chr8 | 56686005 | 56739004 | + | protein_coding |
| ENSG00000011638.10_3 | TMEM159 | 0.95917433 | 1 | -0.328 | 3.320 | chr16 | 21169698 | 21191937 | + | protein_coding |
| ENSG00000185070.11_4 | FLRT2 | 0.92463604 | 1 | -0.328 | 2.722 | chr14 | 85996488 | 86120772 | + | protein_coding |
| ENSG00000114784.4_3 | EIF1B | 0.97092278 | 1 | -0.328 | 3.298 | chr3 | 40351198 | 40353915 | + | protein_coding |
| ENSG00000116819.8_4 | TFAP2E | 0.55752176 | 1 | -0.328 | -1.046 | chr1 | 36038915 | 36060929 | + | protein_coding |
| ENSG00000110218.9_4 | PANX1 | 0.99186534 | 1 | -0.328 | 4.380 | chr11 | 93862007 | 93915134 | + | protein_coding |
| ENSG00000170270.5_6 | GON7 | 0.97773297 | 1 | -0.328 | 3.580 | chr14 | 93669239 | 93673410 | - | protein_coding |
| ENSG00000135316.18_5 | SYNCRIP | 0.9997682 | 1 | -0.328 | 7.151 | chr6 | 86317503 | 86353510 | - | protein_coding |
| ENSG00000170310.15_6 | STX8 | 0.99553801 | 1 | -0.328 | 5.000 | chr17 | 9153788 | 9479908 | - | protein_coding |
| ENSG00000125863.20_5 | MKKS | 0.99629812 | 1 | -0.329 | 4.963 | chr20 | 10381657 | 10414870 | - | protein_coding |
| ENSG00000163545.10_5 | NUAK2 | 0.99892548 | 1 | -0.329 | 5.931 | chr1 | 205271191 | 205290888 | - | protein_coding |
| ENSG00000159259.8_4 | CHAF1B | 0.99693946 | 1 | -0.329 | 4.775 | chr21 | 37757690 | 37791313 | + | protein_coding |
| ENSG00000100625.9_5 | SIX4 | 0.9659573 | 1 | -0.329 | 3.233 | chr14 | 61176257 | 61191069 | - | protein_coding |
| ENSG00000248124.7_5 | RRN3P1 | 0.95586971 | 1 | -0.329 | 3.278 | chr16 | 21807427 | 21831731 | - | transcribed_unprocessed_pseudogene |
| ENSG00000189136.9_4 | UBE2Q2P1 | 0.8545792 | 1 | -0.329 | 1.846 | chr15 | 85070012 | 85114447 | - | transcribed_unprocessed_pseudogene |
| ENSG00000113552.16_7 | GNPDA1 | 0.99897385 | 1 | -0.329 | 5.635 | chr5 | 141371314 | 141392606 | - | protein_coding |
| ENSG00000165030.4_4 | NFIL3 | 0.97566933 | 1 | -0.329 | 3.608 | chr9 | 94171327 | 94186114 | - | protein_coding |
| ENSG00000270001.1_7 | AL121894.2 | 0.55017685 | 1 | -0.329 | -0.977 | chr20 | 23612463 | 23612953 | - | lncRNA |
| ENSG00000092439.16_9 | TRPM7 | 0.99981449 | 1 | -0.330 | 7.171 | chr15 | 50844670 | 50978994 | - | protein_coding |
| ENSG00000270392.2_5 | PFN1P2 | 0.59515236 | 1 | -0.330 | -0.197 | chr1 | 144610815 | 144612662 | - | transcribed_processed_pseudogene |
| ENSG00000155229.21_6 | MMS19 | 0.99983107 | 1 | -0.330 | 6.551 | chr10 | 99218081 | 99258551 | - | protein_coding |
| ENSG00000100918.13_8 | REC8 | 0.85289864 | 1 | -0.330 | 2.066 | chr14 | 24641062 | 24649463 | + | protein_coding |
| ENSG00000107104.19_4 | KANK1 | 0.99923751 | 1 | -0.330 | 5.601 | chr9 | 470291 | 746106 | + | protein_coding |
| ENSG00000228506.2_7 | AL513550.1 | 0.75855315 | 1 | -0.330 | 1.348 | chr6 | 99872798 | 99880199 | + | lncRNA |
| ENSG00000138029.14_4 | HADHB | 0.99986479 | 1 | -0.330 | 6.825 | chr2 | 26466038 | 26513333 | + | protein_coding |
| ENSG00000069764.9_3 | PLA2G10 | 0.54902877 | 1 | -0.330 | -1.084 | chr16 | 14766402 | 14788526 | - | protein_coding |
| ENSG00000213928.9_4 | IRF9 | 0.80373231 | 1 | -0.330 | 1.516 | chr14 | 24630474 | 24635774 | + | protein_coding |
| ENSG00000255495.1_6 | AC145124.1 | 0.56239834 | 1 | -0.330 | -0.523 | chr8 | 12051976 | 12053789 | + | lncRNA |
| ENSG00000158092.7_4 | NCK1 | 0.98539824 | 1 | -0.330 | 3.813 | chr3 | 136581050 | 136670448 | + | protein_coding |
| ENSG00000112096.18_6 | SOD2 | 0.99861812 | 1 | -0.330 | 5.489 | chr6 | 160090101 | 160166218 | - | protein_coding |
| ENSG00000162733.19_6 | DDR2 | 0.55612215 | 1 | -0.331 | -0.475 | chr1 | 162601163 | 162757195 | + | protein_coding |
| ENSG00000112130.17_3 | RNF8 | 0.9935119 | 1 | -0.331 | 4.296 | chr6 | 37321755 | 37362510 | + | protein_coding |
| ENSG00000068305.17_6 | MEF2A | 0.99880642 | 1 | -0.331 | 5.368 | chr15 | 100105622 | 100256671 | + | protein_coding |
| ENSG00000103056.12_5 | SMPD3 | 0.73959243 | 1 | -0.331 | 1.016 | chr16 | 68392230 | 68482411 | - | protein_coding |
| ENSG00000138071.13_4 | ACTR2 | 0.99828096 | 1 | -0.331 | 7.150 | chr2 | 65454887 | 65498387 | + | protein_coding |
| ENSG00000127564.17_6 | PKMYT1 | 0.97132071 | 1 | -0.331 | 4.724 | chr16 | 3018025 | 3030480 | - | protein_coding |

| | | | | | | | | | | |
|---|---|---|---|---|---|---|---|---|---|---|
| ENSG00000229298.1_4 | TUBB8P1 | 0.70848131 | 1 | -0.331 | 0.557 | chr9 | 21811620 | 21812346 | - | processed_pseudogene |
| ENSG00000238018.2_7 | AC093110.1 | 0.78453238 | 1 | -0.331 | 1.548 | chr2 | 54888148 | 54907182 | - | lncRNA |
| ENSG00000100664.11_4 | EIF5 | 0.99992103 | 1 | -0.331 | 8.203 | chr14 | 103799881 | 103811362 | + | protein_coding |
| ENSG00000119929.13_4 | CUTC | 0.99626521 | 1 | -0.331 | 4.608 | chr10 | 101462315 | 101515891 | + | protein_coding |
| ENSG00000107951.14_2 | MTPAP | 0.99929892 | 1 | -0.331 | 5.530 | chr10 | 30598730 | 30663377 | - | protein_coding |
| ENSG00000090447.12_4 | TFAP4 | 0.97949281 | 1 | -0.332 | 4.275 | chr16 | 4307187 | 4323076 | - | protein_coding |
| ENSG00000113966.10_6 | ARL6 | 0.97961481 | 1 | -0.332 | 3.699 | chr3 | 97483365 | 97520073 | + | protein_coding |
| ENSG00000108064.11_4 | TFAM | 0.99973805 | 1 | -0.332 | 6.648 | chr10 | 60145105 | 60158980 | + | protein_coding |
| ENSG00000169914.6_3 | OTUD3 | 0.9780901 | 1 | -0.332 | 3.536 | chr1 | 20208888 | 20239438 | + | protein_coding |
| ENSG00000271869.1_6 | AC026979.2 | 0.57831553 | 1 | -0.332 | -0.131 | chr8 | 30013346 | 30013748 | - | lncRNA |
| ENSG00000256712.1_8 | AC134349.1 | 0.5765767 | 1 | -0.332 | -0.473 | chr12 | 11318689 | 11323952 | - | lncRNA |
| ENSG00000136522.14_4 | MRPL47 | 0.99332001 | 1 | -0.332 | 4.747 | chr3 | 179306073 | 179322437 | - | protein_coding |
| ENSG00000173890.17_4 | GPR160 | 0.85465239 | 1 | -0.332 | 1.523 | chr3 | 169755783 | 169803180 | + | protein_coding |
| ENSG00000241568.2 | RN7SL338P | 0.54677335 | 1 | -0.332 | -1.161 | chr9 | 35049268 | 35049558 | + | misc_RNA |
| ENSG00000116675.16_4 | DNAJC6 | 0.97349397 | 1 | -0.332 | 3.684 | chr1 | 65713902 | 65881554 | + | protein_coding |
| ENSG00000165678.21_4 | GHITM | 0.99990798 | 1 | -0.332 | 7.662 | chr10 | 85899265 | 85913324 | + | protein_coding |
| ENSG00000165195.15_4 | PIGA | 0.99208142 | 1 | -0.332 | 4.499 | chrX | 15337573 | 15353702 | - | protein_coding |
| ENSG00000233822.4_4 | H2BC15 | 0.99773441 | 1 | -0.332 | 6.166 | chr6 | 27805538 | 27833576 | + | protein_coding |
| ENSG00000170037.14_6 | CNTROB | 0.99932213 | 1 | -0.332 | 5.697 | chr17 | 7835419 | 7853238 | + | protein_coding |
| ENSG00000126457.21_4 | PRMT1 | 0.99984444 | 1 | -0.333 | 7.623 | chr19 | 50179043 | 50192286 | + | protein_coding |
| ENSG00000100216.6_3 | TOMM22 | 0.99881843 | 1 | -0.333 | 5.902 | chr22 | 39077962 | 39081426 | + | protein_coding |
| ENSG00000092201.10_4 | SUPT16H | 0.99989005 | 1 | -0.333 | 7.909 | chr14 | 21819635 | 21852178 | - | protein_coding |
| ENSG00000244151.1_7 | AC010973.2 | 0.55099457 | 1 | -0.333 | -0.603 | chr7 | 150771829 | 150773617 | - | lncRNA |
| ENSG00000073111.14_5 | MCM2 | 0.99634839 | 1 | -0.333 | 6.687 | chr3 | 127317253 | 127341279 | + | protein_coding |
| ENSG00000015475.18_5 | BID | 0.9985454 | 1 | -0.333 | 5.356 | chr22 | 18216904 | 18257536 | - | protein_coding |
| ENSG00000285331.2_4 | AC090517.5 | 0.82481558 | 1 | -0.333 | 1.746 | chr15 | 57087658 | 57210697 | - | lncRNA |
| ENSG00000240207.7_9 | AC080013.1 | 0.88165568 | 1 | -0.333 | 2.253 | chr3 | 158450052 | 158501859 | + | lncRNA |
| ENSG00000142949.17_4 | PTPRF | 0.99973283 | 1 | -0.333 | 7.703 | chr1 | 43990858 | 44089337 | + | protein_coding |
| ENSG00000037474.15_4 | NSUN2 | 0.99952607 | 1 | -0.334 | 6.637 | chr5 | 6599352 | 6633404 | - | protein_coding |
| ENSG00000114770.17_6 | ABCC5 | 0.99474097 | 1 | -0.334 | 4.606 | chr3 | 183637722 | 183735727 | - | protein_coding |
| ENSG00000261556.10_7 | SMG1P7 | 0.91539084 | 1 | -0.334 | 2.670 | chr16 | 70253477 | 70280515 | - | transcribed_unprocessed_pseudogene |
| ENSG00000186767.7_5 | SPIN4 | 0.98487252 | 1 | -0.334 | 4.020 | chrX | 62567107 | 62571211 | - | protein_coding |
| ENSG00000114859.16_5 | CLCN2 | 0.89559068 | 1 | -0.334 | 2.388 | chr3 | 184063973 | 184079438 | - | protein_coding |
| ENSG00000171241.9_5 | SHCBP1 | 0.9994979 | 1 | -0.334 | 6.000 | chr16 | 46612503 | 46655291 | - | protein_coding |
| ENSG00000232093.1_7 | DCST1-AS1 | 0.6048873 | 1 | -0.334 | -0.120 | chr1 | 155017667 | 155018594 | - | lncRNA |
| ENSG00000056097.16_4 | ZFR | 0.99981251 | 1 | -0.334 | 7.655 | chr5 | 32354456 | 32444846 | - | protein_coding |
| ENSG00000173567.15_6 | ADGRF3 | 0.54992333 | 1 | -0.334 | -0.601 | chr2 | 26531041 | 26569685 | - | protein_coding |
| ENSG00000287750.1_2 | AL353572.4 | 0.58551055 | 1 | -0.334 | 0.265 | chr9 | 90681360 | 90691890 | - | lncRNA |
| ENSG00000042088.14_7 | TDP1 | 0.99492167 | 1 | -0.334 | 4.551 | chr14 | 90421283 | 90511108 | + | protein_coding |
| ENSG00000105048.17_7 | TNNT1 | 0.9990825 | 1 | -0.334 | 5.550 | chr19 | 55644066 | 55660574 | - | protein_coding |
| ENSG00000237973.1_4 | MTCO1P12 | 0.88407088 | 1 | -0.334 | 7.648 | chr1 | 566454 | 567996 | + | unprocessed_pseudogene |
| ENSG00000053438.11_5 | NNAT | 0.64078888 | 1 | -0.335 | 0.082 | chr20 | 36149608 | 36152092 | + | protein_coding |
| ENSG00000238142.2_7 | BX284668.5 | 0.7285308 | 1 | -0.335 | 1.019 | chr1 | 17214072 | 17216201 | - | lncRNA |
| ENSG00000261329.6_7 | AC016597.2 | 0.55461204 | 1 | -0.335 | -0.628 | chr16 | 27690261 | 27730127 | - | lncRNA |
| ENSG00000141030.13_5 | COPS3 | 0.99975498 | 1 | -0.335 | 7.027 | chr17 | 17149930 | 17184587 | - | protein_coding |
| ENSG00000096063.16_7 | SRPK1 | 0.99931592 | 1 | -0.335 | 5.769 | chr6 | 35800743 | 35889119 | - | protein_coding |
| ENSG00000137547.9_6 | MRPL15 | 0.99509083 | 1 | -0.335 | 5.110 | chr8 | 55047801 | 55061074 | + | protein_coding |
| ENSG00000171608.16_3 | PIK3CD | 0.96551859 | 1 | -0.335 | 3.424 | chr1 | 9711789 | 9789172 | + | protein_coding |
| ENSG00000136628.18_4 | EPRS1 | 0.99992814 | 1 | -0.335 | 7.876 | chr1 | 220141942 | 220219872 | - | protein_coding |
| ENSG00000103932.12_6 | RPAP1 | 0.99410626 | 1 | -0.336 | 4.412 | chr15 | 41809374 | 41836467 | - | protein_coding |
| ENSG00000263272.1_9 | AC004148.1 | 0.92127391 | 1 | -0.336 | 2.701 | chr17 | 5328459 | 5336196 | - | lncRNA |
| ENSG00000243926.2_7 | TIPARP-AS1 | 0.64721005 | 1 | -0.336 | 0.406 | chr3 | 156389651 | 156392235 | - | lncRNA |
| ENSG00000280099.1_5 | AL603750.1 | 0.54929792 | 1 | -0.336 | -0.658 | chr1 | 85618170 | 85619886 | - | TEC |
| ENSG00000139746.15_7 | RBM26 | 0.99856487 | 1 | -0.336 | 6.397 | chr13 | 79885959 | 79980612 | - | protein_coding |
| ENSG00000121481.11_3 | RNF2 | 0.9931713 | 1 | -0.336 | 4.327 | chr1 | 185014658 | 185071735 | + | protein_coding |
| ENSG00000138835.22_4 | RGS3 | 0.99711005 | 1 | -0.336 | 5.286 | chr9 | 116207011 | 116360023 | + | protein_coding |
| ENSG00000104826.14_6 | LHB | 0.5619373 | 1 | -0.336 | 0.141 | chr19 | 49519237 | 49520347 | - | protein_coding |
| ENSG00000167325.15_4 | RRM1 | 0.99986701 | 1 | -0.336 | 7.482 | chr11 | 4115937 | 4160162 | + | protein_coding |
| ENSG00000100528.12_5 | CNIH1 | 0.99952178 | 1 | -0.337 | 6.286 | chr14 | 54890279 | 54908109 | - | protein_coding |

| | | | | | | | | | | |
|---|---|---|---|---|---|---|---|---|---|---|
| ENSG00000230530.2_7 | LIMD1-AS1 | 0.69782736 | 1 | -0.337 | 0.884 | chr3 | 45717861 | 45730692 | - | lncRNA |
| ENSG00000228594.4_7 | FNDC10 | 0.80146151 | 1 | -0.337 | 1.821 | chr1 | 1533392 | 1535515 | - | protein_coding |
| ENSG00000101871.15_4 | MID1 | 0.99978537 | 1 | -0.337 | 7.852 | chrX | 10413350 | 10851773 | - | protein_coding |
| ENSG00000119820.11_4 | YIPF4 | 0.99879297 | 1 | -0.337 | 6.085 | chr2 | 32502973 | 32541663 | + | protein_coding |
| ENSG00000129673.9_2 | AANAT | 0.54219781 | 1 | -0.337 | -1.011 | chr17 | 74449433 | 74466199 | + | protein_coding |
| ENSG00000135454.14_4 | B4GALNT1 | 0.99762455 | 1 | -0.337 | 5.327 | chr12 | 58017192 | 58027022 | - | protein_coding |
| ENSG00000106123.12_5 | EPHB6 | 0.55279836 | 1 | -0.337 | -0.603 | chr7 | 142552823 | 142568847 | + | protein_coding |
| ENSG00000171621.14_3 | SPSB1 | 0.98777454 | 1 | -0.337 | 4.480 | chr1 | 9352953 | 9429591 | + | protein_coding |
| ENSG00000023445.15_4 | BIRC3 | 0.99876392 | 1 | -0.337 | 5.592 | chr11 | 102188181 | 102210134 | + | protein_coding |
| ENSG00000100483.14_5 | VCPKMT | 0.94724766 | 1 | -0.337 | 3.022 | chr14 | 50575350 | 50583318 | - | protein_coding |
| ENSG00000175376.9_5 | EIF1AD | 0.99664728 | 1 | -0.338 | 4.891 | chr11 | 65764016 | 65769647 | - | protein_coding |
| ENSG00000078140.14_4 | UBE2K | 0.99729602 | 1 | -0.338 | 5.210 | chr4 | 39699729 | 39784412 | + | protein_coding |
| ENSG00000143621.17_4 | ILF2 | 0.99989747 | 1 | -0.338 | 7.809 | chr1 | 153634264 | 153643504 | - | protein_coding |
| ENSG00000187242.6_7 | KRT12 | 0.53964882 | 1 | -0.338 | -1.013 | chr17 | 39017555 | 39023475 | - | protein_coding |
| ENSG00000166479.10_6 | TMX3 | 0.99950724 | 1 | -0.338 | 6.284 | chr18 | 66340925 | 66382345 | - | protein_coding |
| ENSG00000285399.2_3 | AC104162.2 | 0.59875242 | 1 | -0.338 | 0.342 | chr3 | 63848831 | 63891514 | + | lncRNA |
| ENSG00000266173.7_8 | STRADA | 0.92186226 | 1 | -0.338 | 2.895 | chr17 | 61759696 | 61819346 | - | protein_coding |
| ENSG00000214770.3_8 | AL161756.1 | 0.59934366 | 1 | -0.338 | -0.009 | chr14 | 64796149 | 64805357 | - | lncRNA |
| ENSG00000103550.14_8 | KNOP1 | 0.99749611 | 1 | -0.338 | 5.001 | chr16 | 19713259 | 19729557 | - | protein_coding |
| ENSG00000100575.14_4 | TIMM9 | 0.85273813 | 1 | -0.338 | 3.834 | chr14 | 58875213 | 58894249 | - | protein_coding |
| ENSG00000141425.18_7 | RPRD1A | 0.99828921 | 1 | -0.338 | 6.103 | chr18 | 33564350 | 33647539 | - | protein_coding |
| ENSG00000145723.17_4 | GIN1 | 0.96608718 | 1 | -0.338 | 3.721 | chr5 | 102421704 | 102455842 | - | protein_coding |
| ENSG00000242960.1_4 | FTH1P23 | 0.53894212 | 1 | -0.338 | -1.013 | chr3 | 72978195 | 72978744 | + | processed_pseudogene |
| ENSG00000165434.8_4 | PGM2L1 | 0.99798578 | 1 | -0.339 | 5.526 | chr11 | 74041361 | 74109478 | - | protein_coding |
| ENSG00000175166.17_4 | PSMD2 | 0.99987766 | 1 | -0.339 | 7.775 | chr3 | 184016986 | 184026838 | + | protein_coding |
| ENSG00000178852.16_6 | EFCAB13 | 0.8873904 | 1 | -0.339 | 2.283 | chr17 | 45400656 | 45518678 | + | protein_coding |
| ENSG00000133657.15_6 | ATP13A3 | 0.99974139 | 1 | -0.339 | 6.959 | chr3 | 194123401 | 194219093 | - | protein_coding |
| ENSG00000242338.6_3 | BMS1P4 | 0.61129114 | 1 | -0.339 | 0.025 | chr10 | 75475601 | 75490227 | - | transcribed_unprocessed_pseudogene |
| ENSG00000285967.1_3 | NIPBL-DT | 0.97591138 | 1 | -0.339 | 3.552 | chr5 | 36864527 | 36876802 | - | lncRNA |
| ENSG00000141510.17_7 | TP53 | 0.99966178 | 1 | -0.339 | 6.682 | chr17 | 7565097 | 7590868 | - | protein_coding |
| ENSG00000102763.18_7 | VWA8 | 0.99681948 | 1 | -0.339 | 5.065 | chr13 | 42140971 | 42535256 | - | protein_coding |
| ENSG00000144821.10_4 | MYH15 | 0.61773418 | 1 | -0.339 | 0.342 | chr3 | 108099215 | 108248169 | - | protein_coding |
| ENSG00000104980.8_5 | TIMM44 | 0.99491698 | 1 | -0.339 | 4.611 | chr19 | 7991603 | 8008552 | - | protein_coding |
| ENSG00000256591.5_7 | AP003108.2 | 0.69343078 | 1 | -0.339 | 0.884 | chr11 | 61196692 | 61253294 | + | protein_coding |
| ENSG00000163714.18_7 | U2SURP | 0.99989027 | 1 | -0.339 | 7.921 | chr3 | 142683339 | 142779567 | + | protein_coding |
| ENSG00000182518.13_4 | FAM104B | 0.95176518 | 1 | -0.339 | 3.204 | chrX | 55169535 | 55187743 | - | protein_coding |
| ENSG00000112739.17_5 | PRPF4B | 0.99960574 | 1 | -0.340 | 6.722 | chr6 | 4021501 | 4065217 | + | protein_coding |
| ENSG00000113328.19_5 | CCNG1 | 0.99966925 | 1 | -0.340 | 7.072 | chr5 | 162864575 | 162873157 | + | protein_coding |
| ENSG00000183569.18_4 | SERHL2 | 0.75262209 | 1 | -0.340 | 1.203 | chr22 | 42949623 | 42970388 | + | protein_coding |
| ENSG00000115183.15_4 | TANC1 | 0.99716599 | 1 | -0.340 | 5.224 | chr2 | 159825152 | 160089170 | + | protein_coding |
| ENSG00000075188.9_5 | NUP37 | 0.99830916 | 1 | -0.340 | 5.384 | chr12 | 102466881 | 102513898 | - | protein_coding |
| ENSG00000196208.14_3 | GREB1 | 0.79645183 | 1 | -0.340 | 1.685 | chr2 | 11622467 | 11782914 | + | protein_coding |
| ENSG00000006576.16_2 | PHTF2 | 0.99377187 | 1 | -0.340 | 5.410 | chr7 | 77428109 | 77586820 | + | protein_coding |
| ENSG00000132485.14_6 | ZRANB2 | 0.9948459 | 1 | -0.340 | 4.671 | chr1 | 71528974 | 71546972 | - | protein_coding |
| ENSG00000213186.8_5 | TRIM59 | 0.99704917 | 1 | -0.340 | 5.141 | chr3 | 160150233 | 160203561 | - | protein_coding |
| ENSG00000180574.3_7 | EIF2S3B | 0.7620809 | 1 | -0.340 | 1.925 | chr12 | 10658201 | 10675734 | + | protein_coding |
| ENSG00000106268.15_5 | NUDT1 | 0.9789979 | 1 | -0.340 | 4.303 | chr7 | 2281857 | 2290781 | + | protein_coding |
| ENSG00000237200.1_6 | ZBTB40-IT1 | 0.53841651 | 1 | -0.340 | -0.686 | chr1 | 22843967 | 22846201 | + | lncRNA |
| ENSG00000034693.15_4 | PEX3 | 0.95606328 | 1 | -0.341 | 3.379 | chr6 | 143771942 | 143811753 | + | protein_coding |
| ENSG00000233593.10_9 | LINC02609 | 0.71824603 | 1 | -0.341 | 0.747 | chr1 | 91234643 | 91317215 | - | lncRNA |
| ENSG00000273759.1_7 | AL117379.1 | 0.56791567 | 1 | -0.341 | -0.403 | chr20 | 61559973 | 61560649 | - | lncRNA |
| ENSG00000213585.11_4 | VDAC1 | 0.99993223 | 1 | -0.341 | 8.218 | chr5 | 133307562 | 133340666 | - | protein_coding |
| ENSG00000177340.5_7 | AC024940.1 | 0.53523226 | 1 | -0.341 | -0.909 | chr12 | 31477250 | 31478763 | + | lncRNA |
| ENSG00000163472.19_5 | TMEM79 | 0.91479498 | 1 | -0.341 | 2.632 | chr1 | 156252726 | 156262976 | + | protein_coding |
| ENSG00000285693.1_1 | AP002381.2 | 0.66330613 | 1 | -0.341 | 0.800 | chr11 | 72504669 | 72525099 | - | lncRNA |
| ENSG00000117280.13_4 | RAB29 | 0.97405404 | 1 | -0.342 | 4.005 | chr1 | 205737114 | 205744610 | - | protein_coding |
| ENSG00000143952.20_4 | VPS54 | 0.9983753 | 1 | -0.342 | 5.905 | chr2 | 64119280 | 64246562 | - | protein_coding |
| ENSG00000126261.13_5 | UBA2 | 0.9997812 | 1 | -0.342 | 8.486 | chr19 | 34919257 | 34962156 | + | protein_coding |
| ENSG00000028839.10_5 | TBPL1 | 0.9725321 | 1 | -0.342 | 3.622 | chr6 | 134273308 | 134311570 | + | protein_coding |
| ENSG00000132963.8_3 | POMP | 0.99858801 | 1 | -0.342 | 5.757 | chr13 | 29233241 | 29253096 | + | protein_coding |

| | | | | | | | | | | |
|---|---|---|---|---|---|---|---|---|---|---|
| ENSG00000239523.6_7 | MYLK-AS1 | 0.79153421 | 1 | -0.342 | 1.460 | chr3 | 123304147 | 123363415 | + | lncRNA |
| ENSG00000167549.18_5 | CORO6 | 0.93800289 | 1 | -0.342 | 3.031 | chr17 | 27941774 | 27949925 | - | protein_coding |
| ENSG00000198730.9_5 | CTR9 | 0.99876336 | 1 | -0.342 | 5.418 | chr11 | 10772793 | 10823172 | + | protein_coding |
| ENSG00000251288.2_6 | AC018797.2 | 0.53563827 | 1 | -0.342 | -0.874 | chr4 | 103672558 | 103673798 | + | processed_pseudogene |
| ENSG00000164182.11_5 | NDUFAF2 | 0.98474504 | 1 | -0.342 | 4.165 | chr5 | 60241032 | 60448853 | + | protein_coding |
| ENSG00000198945.8_3 | L3MBTL3 | 0.93287694 | 1 | -0.342 | 3.005 | chr6 | 130334844 | 130462594 | + | protein_coding |
| ENSG00000286522.1_2 | H3C2 | 0.99937457 | 1 | -0.342 | 8.070 | chr6 | 26031822 | 26032327 | - | protein_coding |
| ENSG00000064313.12_5 | TAF2 | 0.99943065 | 1 | -0.342 | 6.223 | chr8 | 120743014 | 120845081 | - | protein_coding |
| ENSG00000105875.14_7 | WDR91 | 0.93182008 | 1 | -0.342 | 3.091 | chr7 | 134868591 | 134896286 | - | protein_coding |
| ENSG00000132561.14_7 | MATN2 | 0.99050459 | 1 | -0.342 | 4.418 | chr8 | 98881068 | 99048952 | + | protein_coding |
| ENSG00000163539.17_5 | CLASP2 | 0.99966756 | 1 | -0.343 | 6.421 | chr3 | 33537737 | 33759848 | - | protein_coding |
| ENSG00000093217.11_5 | XYLB | 0.96019921 | 1 | -0.343 | 3.210 | chr3 | 38388251 | 38462839 | + | protein_coding |
| ENSG00000221926.12_7 | TRIM16 | 0.99960818 | 1 | -0.343 | 6.690 | chr17 | 15531274 | 15587625 | - | protein_coding |
| ENSG00000084072.17_4 | PPIE | 0.99295313 | 1 | -0.343 | 4.860 | chr1 | 40157854 | 40229586 | + | protein_coding |
| ENSG00000259295.6_4 | CSPG4P12 | 0.57554596 | 1 | -0.343 | 0.293 | chr15 | 85734669 | 85757136 | + | transcribed_unprocessed_pseudogene |
| ENSG00000140481.15_4 | CCDC33 | 0.73560182 | 1 | -0.343 | 1.215 | chr15 | 74495046 | 74628813 | + | protein_coding |
| ENSG00000042429.12_4 | MED17 | 0.91239203 | 1 | -0.343 | 2.552 | chr11 | 93517393 | 93548129 | + | protein_coding |
| ENSG00000089351.14_4 | GRAMD1A | 0.99852299 | 1 | -0.343 | 6.802 | chr19 | 35485688 | 35517375 | + | protein_coding |
| ENSG00000255837.1_3 | TAS2R20 | 0.55818379 | 1 | -0.343 | -0.427 | chr12 | 11149094 | 11150474 | - | protein_coding |
| ENSG00000245149.4_6 | RNF139-AS1 | 0.85884831 | 1 | -0.343 | 1.935 | chr8 | 125474726 | 125486823 | - | lncRNA |
| ENSG00000223984.1_4 | HNRNPRP1 | 0.53632378 | 1 | -0.343 | -0.637 | chr10 | 21913521 | 21915215 | - | processed_pseudogene |
| ENSG00000234825.3_4 | XRCC6P2 | 0.62230672 | 1 | -0.343 | 0.339 | chrX | 149399295 | 149401116 | - | processed_pseudogene |
| ENSG00000187240.16_6 | DYNC2H1 | 0.99707423 | 1 | -0.343 | 5.280 | chr11 | 102980139 | 103350591 | + | protein_coding |
| ENSG00000155090.15_4 | KLF10 | 0.99797901 | 1 | -0.343 | 5.484 | chr8 | 103661012 | 103667953 | - | protein_coding |
| ENSG00000149100.13_4 | EIF3M | 0.99970469 | 1 | -0.343 | 7.160 | chr11 | 32605344 | 32627810 | + | protein_coding |
| ENSG00000229990.3_7 | AC022400.2 | 0.53603601 | 1 | -0.343 | -0.878 | chr10 | 75601591 | 75606873 | + | lncRNA |
| ENSG00000173706.14_6 | HEG1 | 0.98676364 | 1 | -0.344 | 4.250 | chr3 | 124684554 | 124774841 | - | protein_coding |
| ENSG00000171227.7_5 | TMEM37 | 0.82966516 | 1 | -0.344 | 1.914 | chr2 | 120187477 | 120196080 | + | protein_coding |
| ENSG00000259974.3_6 | LINC00261 | 0.84453651 | 1 | -0.344 | 2.167 | chr20 | 22528309 | 22559280 | - | lncRNA |
| ENSG00000226913.2_6 | BSN-DT | 0.53211615 | 1 | -0.344 | -0.978 | chr3 | 49586739 | 49591799 | - | lncRNA |
| ENSG00000178567.8_4 | EPM2AIP1 | 0.99914844 | 1 | -0.344 | 5.821 | chr3 | 37026534 | 37034622 | - | protein_coding |
| ENSG00000146143.19_5 | PRIM2 | 0.96251646 | 1 | -0.344 | 3.484 | chr6 | 57179603 | 57513376 | + | protein_coding |
| ENSG00000186416.16_6 | NKRF | 0.98819943 | 1 | -0.344 | 4.467 | chrX | 118722300 | 118740406 | - | protein_coding |
| ENSG00000182324.7_4 | KCNJ14 | 0.69604734 | 1 | -0.344 | 0.860 | chr19 | 48958831 | 48970237 | + | protein_coding |
| ENSG00000157212.19_4 | PAXIP1 | 0.99324897 | 1 | -0.344 | 4.593 | chr7 | 154735397 | 154794834 | - | protein_coding |
| ENSG00000163319.11_5 | MRPS18C | 0.89188129 | 1 | -0.344 | 2.311 | chr4 | 84377085 | 84390888 | + | protein_coding |
| ENSG00000154059.11_5 | IMPACT | 0.9949082 | 1 | -0.344 | 4.820 | chr18 | 22006598 | 22033495 | + | protein_coding |
| ENSG00000265763.4_3 | ZNF488 | 0.86996685 | 1 | -0.344 | 2.723 | chr10 | 48355069 | 48373866 | + | protein_coding |
| ENSG00000224609.8_7 | AC093425.1 | 0.56407593 | 1 | -0.345 | -0.451 | chr1 | 59597604 | 59762163 | - | lncRNA |
| ENSG00000161981.11_4 | SNRNP25 | 0.9942554 | 1 | -0.345 | 5.069 | chr16 | 103828 | 107669 | + | protein_coding |
| ENSG00000233077.1_6 | LINC01271 | 0.58495626 | 1 | -0.345 | -0.091 | chr20 | 48927248 | 48937879 | - | lncRNA |
| ENSG00000138080.14_3 | EMILIN1 | 0.65380951 | 1 | -0.345 | 0.859 | chr2 | 27301483 | 27309271 | + | protein_coding |
| ENSG00000198000.12_8 | NOL8 | 0.99879639 | 1 | -0.345 | 5.638 | chr9 | 95059640 | 95087918 | - | protein_coding |
| ENSG00000137501.17_5 | SYTL2 | 0.99372957 | 1 | -0.345 | 4.791 | chr11 | 85405267 | 85522202 | - | protein_coding |
| ENSG00000105810.9_3 | CDK6 | 0.99819429 | 1 | -0.345 | 7.518 | chr7 | 92234235 | 92465908 | - | protein_coding |
| ENSG00000005801.18_7 | ZNF195 | 0.99873021 | 1 | -0.345 | 5.924 | chr11 | 3360491 | 3400452 | - | protein_coding |
| ENSG00000259494.2_5 | MRPL46 | 0.92020321 | 1 | -0.345 | 2.721 | chr15 | 89002708 | 89010621 | - | protein_coding |
| ENSG00000206560.11_4 | ANKRD28 | 0.9996813 | 1 | -0.346 | 7.043 | chr3 | 15708743 | 15901278 | - | protein_coding |
| ENSG00000117155.16_4 | SSX2IP | 0.99641807 | 1 | -0.346 | 5.601 | chr1 | 85109390 | 85156486 | - | protein_coding |
| ENSG00000143552.9_4 | NUP210L | 0.66822287 | 1 | -0.346 | 0.920 | chr1 | 153965161 | 154127592 | - | protein_coding |
| ENSG00000157734.14_3 | SNX22 | 0.94572669 | 1 | -0.346 | 3.095 | chr15 | 64443914 | 64449680 | + | protein_coding |
| ENSG00000120254.15_6 | MTHFD1L | 0.99399263 | 1 | -0.346 | 5.184 | chr6 | 151186685 | 151423023 | + | protein_coding |
| ENSG00000119608.13_4 | PROX2 | 0.53693768 | 1 | -0.346 | -0.784 | chr14 | 75319574 | 75342857 | - | protein_coding |
| ENSG00000128973.13_5 | CLN6 | 0.9965237 | 1 | -0.346 | 5.136 | chr15 | 68499330 | 68549549 | - | protein_coding |
| ENSG00000105193.9_4 | RPS16 | 0.99317585 | 1 | -0.347 | 4.691 | chr19 | 39923777 | 39926589 | - | protein_coding |
| ENSG00000272344.1 | SNORD114-21 | 0.5351913 | 1 | -0.347 | -0.725 | chr14 | 101448312 | 101448383 | + | snoRNA |
| ENSG00000066279.18_5 | ASPM | 0.99967352 | 1 | -0.347 | 7.918 | chr1 | 197053257 | 197115824 | - | protein_coding |
| ENSG00000005810.18_4 | MYCBP2 | 0.99671178 | 1 | -0.347 | 5.829 | chr13 | 77618792 | 77901229 | - | protein_coding |

| | | | | | | | | | | |
|---|---|---|---|---|---|---|---|---|---|---|
| ENSG00000198888.2_1 | MT-ND1 | 0.77982358 | 1 | -0.347 | 8.141 | chrM | 3307 | 4262 | + | protein_coding |
| ENSG00000261008.7_8 | LINC01572 | 0.75970368 | 1 | -0.347 | 1.462 | chr16 | 72270180 | 72698913 | - | lncRNA |
| ENSG00000183579.16_4 | ZNRF3 | 0.99292017 | 1 | -0.347 | 4.802 | chr22 | 29279560 | 29453476 | + | protein_coding |
| ENSG00000153879.9_4 | CEBPG | 0.99946796 | 1 | -0.347 | 6.424 | chr19 | 33864591 | 33873592 | + | protein_coding |
| ENSG00000236349.1_4 | SUCLG2P2 | 0.53283293 | 1 | -0.347 | -0.816 | chr12 | 94942017 | 94943312 | + | processed_pseudogene |
| ENSG00000131238.17_6 | PPT1 | 0.99893203 | 1 | -0.347 | 5.878 | chr1 | 40537133 | 40563399 | - | protein_coding |
| ENSG00000169299.14_3 | PGM2 | 0.98630723 | 1 | -0.347 | 4.614 | chr4 | 37828282 | 37864558 | + | protein_coding |
| ENSG00000205534.6_5 | SMG1P2 | 0.7339136 | 1 | -0.348 | 1.123 | chr16 | 29538889 | 29606287 | - | unprocessed_pseudogene |
| ENSG00000280347.1_6 | AC000123.3 | 0.84917487 | 1 | -0.348 | 2.082 | chr7 | 126999839 | 127004813 | - | TEC |
| ENSG00000240489.1_4 | SETP14 | 0.86030495 | 1 | -0.348 | 2.794 | chr3 | 155705093 | 155705965 | + | processed_pseudogene |
| ENSG00000278963.1_7 | AC005921.4 | 0.72916643 | 1 | -0.348 | 1.067 | chr17 | 48799435 | 48801585 | + | TEC |
| ENSG00000020922.12_8 | MRE11 | 0.99813476 | 1 | -0.348 | 5.608 | chr11 | 94148744 | 94227074 | - | protein_coding |
| ENSG00000224875.2_6 | EML4-AS1 | 0.52601874 | 1 | -0.349 | -0.979 | chr2 | 42370378 | 42397441 | - | lncRNA |
| ENSG00000258498.8_5 | DIO3OS | 0.81975108 | 1 | -0.349 | 1.781 | chr14 | 102018558 | 102026768 | - | lncRNA |
| ENSG00000231889.7_6 | TRAF3IP2-AS1 | 0.62122096 | 1 | -0.349 | 0.383 | chr6 | 111804714 | 111919505 | + | lncRNA |
| ENSG00000101189.7_3 | MRGBP | 0.99481028 | 1 | -0.349 | 5.073 | chr20 | 61427825 | 61433081 | + | protein_coding |
| ENSG00000152193.8_5 | OBI1 | 0.97428297 | 1 | -0.349 | 3.804 | chr13 | 79188424 | 79233290 | - | protein_coding |
| ENSG00000135968.21_6 | GCC2 | 0.9994276 | 1 | -0.349 | 6.433 | chr2 | 109065647 | 109125871 | + | protein_coding |
| ENSG00000110660.15_7 | SLC35F2 | 0.99807847 | 1 | -0.349 | 5.616 | chr11 | 107661717 | 107799019 | - | protein_coding |
| ENSG00000233621.2_8 | LINC01137 | 0.91241386 | 1 | -0.349 | 2.695 | chr1 | 37816535 | 37940012 | - | lncRNA |
| ENSG00000237054.9_7 | PRMT5-AS1 | 0.53845537 | 1 | -0.349 | -0.667 | chr14 | 23388156 | 23396109 | + | lncRNA |
| ENSG00000101412.13_4 | E2F1 | 0.99539136 | 1 | -0.349 | 5.666 | chr20 | 32263283 | 32274191 | - | protein_coding |
| ENSG00000261872.1_6 | AC002558.2 | 0.53316793 | 1 | -0.349 | -0.669 | chr17 | 45247192 | 45248415 | + | lncRNA |
| ENSG00000158402.20_9 | CDC25C | 0.97897008 | 1 | -0.350 | 4.114 | chr5 | 137620954 | 137674044 | - | protein_coding |
| ENSG00000169714.16_5 | CNBP | 0.99971348 | 1 | -0.350 | 7.526 | chr3 | 128888327 | 128902765 | - | protein_coding |
| ENSG00000072422.17_4 | RHOBTB1 | 0.9964513 | 1 | -0.350 | 5.226 | chr10 | 62629196 | 62761198 | - | protein_coding |
| ENSG00000068878.15_4 | PSME4 | 0.99970244 | 1 | -0.350 | 7.170 | chr2 | 54091206 | 54198130 | - | protein_coding |
| ENSG00000158169.13_8 | FANCC | 0.99612352 | 1 | -0.350 | 4.893 | chr9 | 97861336 | 98189078 | - | protein_coding |
| ENSG00000171453.20_5 | POLR1C | 0.97874561 | 1 | -0.350 | 4.058 | chr6 | 43477440 | 43530156 | + | protein_coding |
| ENSG00000233237.8_6 | LINC00472 | 0.9950459 | 1 | -0.350 | 6.109 | chr6 | 72053130 | 72130448 | - | lncRNA |
| ENSG00000110092.4_3 | CCND1 | 0.99986207 | 1 | -0.350 | 7.681 | chr11 | 69455924 | 69469242 | + | protein_coding |
| ENSG00000156697.13_6 | UTP14A | 0.9979532 | 1 | -0.350 | 5.574 | chrX | 129040097 | 129063737 | + | protein_coding |
| ENSG00000140443.15_6 | IGF1R | 0.9994458 | 1 | -0.350 | 7.060 | chr15 | 99191768 | 99507759 | + | protein_coding |
| ENSG00000285517.1_3 | AC010198.2 | 0.92595741 | 1 | -0.350 | 3.069 | chr12 | 30948392 | 31032202 | + | lncRNA |
| ENSG00000169764.15_5 | UGP2 | 0.99765492 | 1 | -0.350 | 6.168 | chr2 | 64068074 | 64118696 | + | protein_coding |
| ENSG00000258044.1_6 | AC073569.1 | 0.55918589 | 1 | -0.350 | 0.162 | chr12 | 80217558 | 80219276 | + | lncRNA |
| ENSG00000280163.1_6 | AC040160.3 | 0.62639391 | 1 | -0.350 | 0.622 | chr16 | 67239078 | 67240752 | - | TEC |
| ENSG00000166963.13_4 | MAP1A | 0.8948319 | 1 | -0.351 | 2.599 | chr15 | 43803156 | 43823818 | + | protein_coding |
| ENSG00000260257.2_6 | AL035071.1 | 0.81235276 | 1 | -0.351 | 1.973 | chr20 | 31444427 | 31446557 | + | lncRNA |
| ENSG00000213442.5_4 | RPL18AP3 | 0.90513239 | 1 | -0.351 | 2.984 | chr12 | 104659087 | 104659614 | + | processed_pseudogene |
| ENSG00000135622.13_4 | SEMA4F | 0.98316075 | 1 | -0.351 | 4.114 | chr2 | 74881355 | 74910980 | + | protein_coding |
| ENSG00000166444.19_7 | DENND2B | 0.89329568 | 1 | -0.351 | 2.741 | chr11 | 8714898 | 8932498 | - | protein_coding |
| ENSG00000180539.8_6 | C9orf139 | 0.52673351 | 1 | -0.351 | -0.756 | chr9 | 139921916 | 139932409 | + | lncRNA |
| ENSG00000183604.14_5 | SMG1P5 | 0.88875268 | 1 | -0.351 | 2.426 | chr16 | 30278874 | 30346695 | - | transcribed_unprocessed_pseudogene |
| ENSG00000234084.1_6 | AL049552.1 | 0.53456641 | 1 | -0.351 | -0.561 | chr6 | 135622706 | 135628296 | + | lncRNA |
| ENSG00000232775.2 | AP000525.10 | 0.65651491 | 1 | -0.351 | 0.632 | chr22 | 16157079 | 16172700 | + | pseudogene |
| ENSG00000135926.15_7 | TMBIM1 | 0.99919033 | 1 | -0.351 | 6.166 | chr2 | 219138920 | 219157309 | - | protein_coding |
| ENSG00000141655.17_4 | TNFRSF11A | 0.95154387 | 1 | -0.351 | 3.572 | chr18 | 59992520 | 60058521 | + | protein_coding |
| ENSG00000272263.1_6 | AC034198.2 | 0.53904801 | 1 | -0.351 | -0.641 | chr3 | 12873718 | 12874227 | - | lncRNA |
| ENSG00000153551.14_4 | CMTM7 | 0.94249607 | 1 | -0.352 | 3.081 | chr3 | 32433190 | 32524559 | + | protein_coding |
| ENSG00000174780.17_6 | SRP72 | 0.99934977 | 1 | -0.352 | 6.723 | chr4 | 57333783 | 57369847 | + | protein_coding |
| ENSG00000284726.1_5 | AL109936.8 | 0.52627636 | 1 | -0.352 | -0.753 | chr1 | 23624290 | 23629359 | + | lncRNA |
| ENSG00000234912.12_6 | SNHG20 | 0.94562068 | 1 | -0.352 | 3.270 | chr17 | 75082798 | 75095984 | + | lncRNA |
| ENSG00000100462.16_8 | PRMT5 | 0.99543683 | 1 | -0.352 | 5.078 | chr14 | 23389734 | 23398617 | - | protein_coding |
| ENSG00000119812.19_5 | FAM98A | 0.99228376 | 1 | -0.352 | 4.700 | chr2 | 33808727 | 33824449 | - | protein_coding |
| ENSG00000163344.6_4 | PMVK | 0.94618566 | 1 | -0.352 | 3.720 | chr1 | 154897216 | 154909195 | - | protein_coding |
| ENSG00000138073.14_5 | PREB | 0.99852173 | 1 | -0.352 | 5.965 | chr2 | 27353624 | 27357534 | - | protein_coding |

| | | | | | | | | | | |
|---|---|---|---|---|---|---|---|---|---|---|
| ENSG00000156502.14_4 | SUPV3L1 | 0.98271606 | 1 | -0.352 | 4.736 | chr10 | 70939990 | 70968855 | + | protein_coding |
| ENSG00000182511.12_4 | FES | 0.72147206 | 1 | -0.352 | 1.081 | chr15 | 91426925 | 91439006 | + | protein_coding |
| ENSG00000173812.11_4 | EIF1 | 0.9991545 | 1 | -0.352 | 6.032 | chr17 | 39845137 | 39848920 | + | protein_coding |
| ENSG00000276903.1_3 | H2AC16 | 0.95323468 | 1 | -0.353 | 4.170 | chr6 | 27833133 | 27833525 | + | protein_coding |
| ENSG00000112531.17_4 | QKI | 0.99951226 | 1 | -0.353 | 7.171 | chr6 | 163835032 | 163999624 | + | protein_coding |
| ENSG00000129680.16_6 | MAP7D3 | 0.99388188 | 1 | -0.353 | 4.820 | chrX | 135295379 | 135338641 | - | protein_coding |
| ENSG00000175564.13_4 | UCP3 | 0.58440556 | 1 | -0.353 | 0.463 | chr11 | 73711322 | 73720130 | - | protein_coding |
| ENSG00000106554.13_6 | CHCHD3 | 0.9989168 | 1 | -0.353 | 6.563 | chr7 | 132469631 | 132766850 | - | protein_coding |
| ENSG00000243725.7_4 | TTC4 | 0.53931456 | 1 | -0.353 | -0.461 | chr1 | 55181534 | 55208330 | + | protein_coding |
| ENSG00000130684.14_4 | ZNF337 | 0.88629596 | 1 | -0.353 | 2.728 | chr20 | 25653831 | 25677489 | - | protein_coding |
| ENSG00000205268.10_3 | PDE7A | 0.99164255 | 1 | -0.353 | 4.891 | chr8 | 66629745 | 66754557 | - | protein_coding |
| ENSG00000267199.1_6 | AP001029.2 | 0.63634105 | 1 | -0.353 | 0.513 | chr18 | 12438889 | 12448204 | + | lncRNA |
| ENSG00000178385.15_7 | PLEKHM3 | 0.95658422 | 1 | -0.353 | 3.670 | chr2 | 208686012 | 208890251 | - | protein_coding |
| ENSG00000227006.2_7 | AL136988.2 | 0.52029685 | 1 | -0.354 | -0.815 | chr1 | 230394440 | 230407846 | - | lncRNA |
| ENSG00000164048.14_3 | ZNF589 | 0.88716436 | 1 | -0.354 | 2.844 | chr3 | 48282590 | 48340743 | + | protein_coding |
| ENSG00000114107.9_7 | CEP70 | 0.98421298 | 1 | -0.354 | 4.072 | chr3 | 138213186 | 138313380 | - | protein_coding |
| ENSG00000048342.16_8 | CC2D2A | 0.99519606 | 1 | -0.354 | 4.976 | chr4 | 15471489 | 15603180 | + | protein_coding |
| ENSG00000173638.19_6 | SLC19A1 | 0.91906687 | 1 | -0.354 | 4.300 | chr21 | 46913486 | 46993279 | - | protein_coding |
| ENSG00000132792.19_4 | CTNNBL1 | 0.99278582 | 1 | -0.354 | 5.023 | chr20 | 36322357 | 36500531 | + | protein_coding |
| ENSG00000247746.4_4 | USP51 | 0.69706275 | 1 | -0.354 | 1.435 | chrX | 55511049 | 55515635 | - | protein_coding |
| ENSG00000163743.13_3 | RCHY1 | 0.93701983 | 1 | -0.354 | 3.172 | chr4 | 76404247 | 76439974 | - | protein_coding |
| ENSG00000279428.1_5 | AC087164.2 | 0.56875569 | 1 | -0.354 | -0.250 | chr17 | 18010648 | 18013777 | - | TEC |
| ENSG00000204843.12_6 | DCTN1 | 0.99977655 | 1 | -0.355 | 7.356 | chr2 | 74588281 | 74619214 | - | protein_coding |
| ENSG00000136560.14_4 | TANK | 0.99738048 | 1 | -0.355 | 5.680 | chr2 | 161993419 | 162092741 | + | protein_coding |
| ENSG00000113810.16_5 | SMC4 | 0.99883553 | 1 | -0.355 | 7.874 | chr3 | 160117062 | 160152742 | + | protein_coding |
| ENSG00000285752.1_5 | CDC42-AS1 | 0.51858321 | 1 | -0.355 | -0.785 | chr1 | 22354810 | 22379420 | - | lncRNA |
| ENSG00000237506.3_4 | RPSAP15 | 0.5846137 | 1 | -0.355 | 0.270 | chrX | 86958343 | 86959203 | - | processed_pseudogene |
| ENSG00000224888.4_6 | AC138028.2 | 0.7200459 | 1 | -0.355 | 1.334 | chr16 | 88797588 | 88807833 | + | lncRNA |
| ENSG00000134245.18_3 | WNT2B | 0.68686936 | 1 | -0.355 | 0.912 | chr1 | 113009163 | 113072787 | + | protein_coding |
| ENSG00000214485.6_4 | RPL7P1 | 0.81739386 | 1 | -0.355 | 2.222 | chr5 | 149473865 | 149474611 | + | processed_pseudogene |
| ENSG00000231841.1_4 | FAM192BP | 0.53938677 | 1 | -0.355 | -0.486 | chr4 | 153848598 | 153849359 | + | processed_pseudogene |
| ENSG00000226193.1_6 | AL049548.1 | 0.56336193 | 1 | -0.356 | 0.071 | chr6 | 152723533 | 152726101 | + | lncRNA |
| ENSG00000258667.2_7 | HIF1A-AS3 | 0.70149842 | 1 | -0.356 | 0.987 | chr14 | 62174282 | 62217817 | - | lncRNA |
| ENSG00000248098.12_4 | BCKDHA | 0.51729923 | 1 | -0.356 | -0.814 | chr19 | 41903713 | 41930907 | + | protein_coding |
| ENSG00000115548.17_4 | KDM3A | 0.99731059 | 1 | -0.356 | 5.457 | chr2 | 86667770 | 86719839 | + | protein_coding |
| ENSG00000220804.8_6 | LINC01881 | 0.54564977 | 1 | -0.356 | -0.249 | chr2 | 243030784 | 243082789 | + | transcribed_unprocessed_pseudogene |
| ENSG00000187079.18_6 | TEAD1 | 0.99977084 | 1 | -0.356 | 7.929 | chr11 | 12695968 | 12966284 | + | protein_coding |
| ENSG00000176700.20_4 | SCAND2P | 0.84981458 | 1 | -0.356 | 2.124 | chr15 | 85174682 | 85190709 | + | transcribed_unprocessed_pseudogene |
| ENSG00000259943.1_6 | AL050341.2 | 0.76493858 | 1 | -0.356 | 1.611 | chr1 | 40722099 | 40723639 | - | lncRNA |
| ENSG00000259381.2_6 | AC090695.2 | 0.53458533 | 1 | -0.356 | -0.510 | chr15 | 101185639 | 101186860 | - | processed_pseudogene |
| ENSG00000272518.1_7 | AC036214.2 | 0.52601064 | 1 | -0.357 | -0.508 | chr8 | 80868700 | 80869616 | - | lncRNA |
| ENSG00000197894.11_7 | ADH5 | 0.99903556 | 1 | -0.357 | 6.633 | chr4 | 99992129 | 100009952 | - | protein_coding |
| ENSG00000224415.1_5 | AC007683.1 | 0.57454939 | 1 | -0.357 | -0.021 | chr7 | 102781796 | 102782386 | - | processed_pseudogene |
| ENSG00000116095.11_3 | PLEKHA3 | 0.99323839 | 1 | -0.357 | 5.111 | chr2 | 179345184 | 179381190 | + | protein_coding |
| ENSG00000276931.1_8 | AC009041.3 | 0.54655862 | 1 | -0.357 | -0.294 | chr16 | 1018375 | 1019012 | - | lncRNA |
| ENSG00000218358.2_3 | RAET1K | 0.54699209 | 1 | -0.357 | -0.170 | chr6 | 150319155 | 150326293 | - | transcribed_unprocessed_pseudogene |
| ENSG00000244509.4_3 | APOBEC3C | 0.93193098 | 1 | -0.357 | 3.142 | chr22 | 39410262 | 39416357 | + | protein_coding |
| ENSG00000253570.1_4 | RNF5P1 | 0.51610377 | 1 | -0.357 | -0.891 | chr8 | 38458179 | 38458718 | - | processed_pseudogene |
| ENSG00000261609.7_6 | GAN | 0.9771857 | 1 | -0.357 | 4.212 | chr16 | 81348571 | 81413803 | + | protein_coding |
| ENSG00000286535.1_1 | AC139019.1 | 0.68921366 | 1 | -0.357 | 0.899 | chr8 | 131371033 | 131374707 | + | lncRNA |
| ENSG00000269940.1_9 | AL049840.5 | 0.51633761 | 1 | -0.358 | -1.139 | chr14 | 104160897 | 104161507 | + | lncRNA |
| ENSG00000104043.14_4 | ATP8B4 | 0.70041026 | 1 | -0.358 | 1.098 | chr15 | 50150435 | 50475014 | - | protein_coding |
| ENSG00000117519.16_4 | CNN3 | 0.99926703 | 1 | -0.358 | 6.467 | chr1 | 95362505 | 95392779 | - | protein_coding |
| ENSG00000214439.4_5 | FAM185BP | 0.54107461 | 1 | -0.358 | -0.006 | chr7 | 76713104 | 76751433 | - | transcribed_unprocessed_pseudogene |
| ENSG00000213397.10_6 | HAUS7 | 0.59133846 | 1 | -0.358 | 0.154 | chrX | 152713124 | 152760974 | - | protein_coding |
| ENSG00000278192.1_6 | AL118505.1 | 0.53065472 | 1 | -0.358 | -0.642 | chr20 | 6046612 | 6048543 | - | lncRNA |

| | | | | | | | | | | |
|---|---|---|---|---|---|---|---|---|---|---|
| ENSG00000156802.13_5 | ATAD2 | 0.99950063 | 1 | -0.358 | 7.222 | chr8 | 124332090 | 124428590 | - | protein_coding |
| ENSG00000234911.2_4 | TEX21P | 0.62674118 | 1 | -0.358 | 0.318 | chr14 | 64808391 | 64854704 | - | transcribed_unitary_pseudogene |
| ENSG00000144843.11_3 | ADPRH | 0.7764907 | 1 | -0.358 | 2.155 | chr3 | 119298115 | 119308792 | + | protein_coding |
| ENSG00000123737.12_4 | EXOSC9 | 0.99373597 | 1 | -0.358 | 5.266 | chr4 | 122722472 | 122738176 | + | protein_coding |
| ENSG00000180573.9_4 | H2AC6 | 0.99846367 | 1 | -0.359 | 7.569 | chr6 | 26124373 | 26139344 | + | protein_coding |
| ENSG00000180979.10_7 | LRRC57 | 0.932904 | 1 | -0.359 | 3.398 | chr15 | 42830018 | 42841000 | - | protein_coding |
| ENSG00000287078.1_1 | AL513218.2 | 0.5153274 | 1 | -0.359 | -1.100 | chr1 | 52831115 | 52833537 | + | lncRNA |
| ENSG00000223803.1_4 | RPS20P14 | 0.58996924 | 1 | -0.359 | 0.312 | chr3 | 186617987 | 186618346 | + | processed_pseudogene |
| ENSG00000055163.20_6 | CYFIP2 | 0.99806853 | 1 | -0.359 | 5.649 | chr5 | 156693089 | 156822603 | + | protein_coding |
| ENSG00000100410.8_4 | PHF5A | 0.99311214 | 1 | -0.359 | 4.827 | chr22 | 41855721 | 41864696 | - | protein_coding |
| ENSG00000224165.6_6 | DNAJC27-AS1 | 0.61148503 | 1 | -0.359 | 0.481 | chr2 | 25194259 | 25262585 | + | lncRNA |
| ENSG00000144504.15_6 | ANKMY1 | 0.97740229 | 1 | -0.359 | 4.084 | chr2 | 241418839 | 241508626 | - | protein_coding |
| ENSG00000070950.10_5 | RAD18 | 0.99691213 | 1 | -0.359 | 5.425 | chr3 | 8817088 | 9005457 | - | protein_coding |
| ENSG00000005302.19_6 | MSL3 | 0.99690229 | 1 | -0.359 | 5.536 | chrX | 11776278 | 11793891 | + | protein_coding |
| ENSG00000273998.1_6 | AL049794.1 | 0.51967809 | 1 | -0.359 | -0.535 | chr20 | 16556713 | 16560260 | + | lncRNA |
| ENSG00000151500.15_4 | THYN1 | 0.97060098 | 1 | -0.360 | 3.865 | chr11 | 134118173 | 134123264 | - | protein_coding |
| ENSG00000261553.5_7 | AL137782.1 | 0.66101548 | 1 | -0.360 | 0.483 | chr13 | 76123909 | 76381256 | + | lncRNA |
| ENSG00000180398.13_4 | MCFD2 | 0.9995204 | 1 | -0.360 | 7.540 | chr2 | 47129009 | 47168994 | - | protein_coding |
| ENSG00000255135.4_7 | AP002360.2 | 0.85668654 | 1 | -0.360 | 2.248 | chr11 | 76146603 | 76155731 | - | lncRNA |
| ENSG00000102043.16_5 | MTMR8 | 0.60423682 | 1 | -0.360 | 0.285 | chrX | 63487961 | 63615332 | - | protein_coding |
| ENSG00000133789.15_3 | SWAP70 | 0.99844802 | 1 | -0.360 | 6.024 | chr11 | 9685624 | 9774540 | + | protein_coding |
| ENSG00000213057.5_4 | C1orf220 | 0.56811774 | 1 | -0.360 | 0.000 | chr1 | 178511887 | 178518024 | + | lncRNA |
| ENSG00000165209.19_5 | STRBP | 0.99783769 | 1 | -0.360 | 5.602 | chr9 | 125871779 | 126030865 | - | protein_coding |
| ENSG00000205060.11_4 | SLC35B4 | 0.99945637 | 1 | -0.360 | 6.831 | chr7 | 133974084 | 134001682 | - | protein_coding |
| ENSG00000137710.16_6 | RDX | 0.99952537 | 1 | -0.360 | 7.540 | chr11 | 109735021 | 110167437 | - | protein_coding |
| ENSG00000163577.8_4 | EIF5A2 | 0.97392676 | 1 | -0.360 | 4.078 | chr3 | 170606207 | 170626433 | - | protein_coding |
| ENSG00000162851.8_4 | TFB2M | 0.93610217 | 1 | -0.360 | 4.424 | chr1 | 246703863 | 246729563 | - | protein_coding |
| ENSG00000176407.18_4 | KCMF1 | 0.99859345 | 1 | -0.361 | 6.037 | chr2 | 85198216 | 85286595 | + | protein_coding |
| ENSG00000228929.1_4 | RPS13P2 | 0.79908723 | 1 | -0.361 | 1.507 | chr1 | 53237866 | 53238320 | - | processed_pseudogene |
| ENSG00000057019.16_6 | DCBLD2 | 0.99963169 | 1 | -0.361 | 9.622 | chr3 | 98514785 | 98620539 | - | protein_coding |
| ENSG00000166582.10_4 | CENPV | 0.99639022 | 1 | -0.361 | 5.275 | chr17 | 16245848 | 16256970 | - | protein_coding |
| ENSG00000134265.13_3 | NAPG | 0.99363459 | 1 | -0.361 | 5.140 | chr18 | 10525902 | 10552761 | + | protein_coding |
| ENSG00000166986.15_6 | MARS1 | 0.99972384 | 1 | -0.361 | 7.446 | chr12 | 57869228 | 57911352 | + | protein_coding |
| ENSG00000096092.6_3 | TMEM14A | 0.98361952 | 1 | -0.361 | 4.409 | chr6 | 52535911 | 52551386 | + | protein_coding |
| ENSG00000146909.8_3 | NOM1 | 0.98857012 | 1 | -0.361 | 4.824 | chr7 | 156742406 | 156765870 | + | protein_coding |
| ENSG00000279948.1_6 | AC008895.1 | 0.5503422 | 1 | -0.361 | -0.217 | chr19 | 47449792 | 47452413 | + | TEC |
| ENSG00000072832.15_5 | CRMP1 | 0.90092853 | 1 | -0.362 | 2.806 | chr4 | 5749811 | 5894813 | - | protein_coding |
| ENSG00000254428.1_7 | AP003392.1 | 0.68660015 | 1 | -0.362 | 1.406 | chr11 | 118874452 | 118875603 | - | lncRNA |
| ENSG00000175611.12_5 | LINC00476 | 0.72442807 | 1 | -0.362 | 1.392 | chr9 | 98521513 | 98638331 | - | lncRNA |
| ENSG00000165948.11_5 | IFI27L1 | 0.8943749 | 1 | -0.362 | 2.857 | chr14 | 94547647 | 94570192 | + | protein_coding |
| ENSG00000227627.3_7 | AL080276.2 | 0.55513589 | 1 | -0.362 | -0.039 | chr6 | 153304466 | 153312457 | + | lncRNA |
| ENSG00000155850.8_3 | SLC26A2 | 0.99855429 | 1 | -0.362 | 6.749 | chr5 | 149340321 | 149373018 | + | protein_coding |
| ENSG00000110619.17_5 | CARS1 | 0.99584468 | 1 | -0.362 | 5.268 | chr11 | 3022152 | 3078843 | - | protein_coding |
| ENSG00000181924.7_6 | COA4 | 0.99793702 | 1 | -0.362 | 5.838 | chr11 | 73583712 | 73587946 | - | protein_coding |
| ENSG00000135931.17_5 | ARMC9 | 0.96960465 | 1 | -0.362 | 4.121 | chr2 | 232063260 | 232239548 | + | protein_coding |
| ENSG00000108561.8_5 | C1QBP | 0.99967813 | 1 | -0.362 | 8.160 | chr17 | 5336097 | 5352150 | - | protein_coding |
| ENSG00000177674.16_6 | AGTRAP | 0.9801753 | 1 | -0.363 | 4.708 | chr1 | 11796141 | 11814859 | + | protein_coding |
| ENSG00000284308.1_5 | C2orf81 | 0.53149709 | 1 | -0.363 | -0.438 | chr2 | 74641303 | 74648718 | - | protein_coding |
| ENSG00000250479.8_4 | CHCHD10 | 0.93757332 | 1 | -0.363 | 3.273 | chr22 | 24108021 | 24110630 | - | protein_coding |
| ENSG00000115241.11_5 | PPM1G | 0.99957039 | 1 | -0.363 | 7.528 | chr2 | 27604062 | 27632458 | - | protein_coding |
| ENSG00000165819.12_5 | METTL3 | 0.99808207 | 1 | -0.363 | 5.752 | chr14 | 21966277 | 21979484 | - | protein_coding |
| ENSG00000277144.1_6 | AC092757.3 | 0.53129215 | 1 | -0.363 | -0.487 | chr15 | 59407746 | 59408288 | - | lncRNA |
| ENSG00000198498.10_4 | TMA16 | 0.97829094 | 1 | -0.363 | 4.522 | chr4 | 164415594 | 164441691 | + | protein_coding |
| ENSG00000168395.15_5 | ING5 | 0.98898221 | 1 | -0.363 | 4.906 | chr2 | 242626500 | 242668893 | + | protein_coding |
| ENSG00000285219.2_3 | HULC | 0.50653082 | 1 | -0.363 | -1.064 | chr6 | 8435801 | 9294366 | + | lncRNA |
| ENSG00000172728.16_6 | FUT10 | 0.97272051 | 1 | -0.363 | 3.908 | chr8 | 33228342 | 33330664 | - | protein_coding |
| ENSG00000104731.14_6 | KLHDC4 | 0.98353418 | 1 | -0.363 | 5.012 | chr16 | 87730091 | 87799598 | - | protein_coding |
| ENSG00000272341.1_7 | AL137003.1 | 0.52246006 | 1 | -0.364 | -0.366 | chr6 | 16764577 | 16767114 | + | lncRNA |

| | | | | | | | | | | |
|---|---|---|---|---|---|---|---|---|---|---|
| ENSG00000140057.9_3 | AK7 | 0.88067496 | 1 | -0.364 | 2.960 | chr14 | 96858465 | 96955764 | + | protein_coding |
| ENSG00000124508.17_7 | BTN2A2 | 0.93347574 | 1 | -0.364 | 3.276 | chr6 | 26383324 | 26395102 | + | protein_coding |
| ENSG00000148773.14_5 | MKI67 | 0.9992415 | 1 | -0.364 | 8.400 | chr10 | 129894923 | 129924687 | - | protein_coding |
| ENSG00000257285.6_6 | AL132780.1 | 0.74097152 | 1 | -0.364 | 1.619 | chr14 | 23398816 | 23425583 | + | lncRNA |
| ENSG00000233783.8_7 | AP001442.1 | 0.58226202 | 1 | -0.365 | 0.033 | chr21 | 27301066 | 27324928 | + | lncRNA |
| ENSG00000100387.9_3 | RBX1 | 0.98566785 | 1 | -0.365 | 4.799 | chr22 | 41347351 | 41369313 | + | protein_coding |
| ENSG00000099901.17_7 | RANBP1 | 0.99926535 | 1 | -0.365 | 6.612 | chr22 | 20103461 | 20114878 | + | protein_coding |
| ENSG00000151093.8_5 | OXSM | 0.95789392 | 1 | -0.365 | 3.814 | chr3 | 25824408 | 25836025 | + | protein_coding |
| ENSG00000279861.1_6 | AC073548.1 | 0.53106689 | 1 | -0.365 | -0.133 | chr19 | 47990895 | 47992776 | - | TEC |
| ENSG00000234287.1_5 | AC099560.2 | 0.77067819 | 1 | -0.366 | 1.628 | chr3 | 40803036 | 40803290 | + | processed_pseudogene |
| ENSG00000159753.14_6 | CARMIL2 | 0.60943091 | 1 | -0.366 | 0.469 | chr16 | 67678891 | 67691472 | + | protein_coding |
| ENSG00000255624.3_5 | AC073585.1 | 0.87841312 | 1 | -0.366 | 2.479 | chr10 | 124639197 | 124658131 | + | transcribed_processed_pseudogene |
| ENSG00000272750.1_6 | AL592148.3 | 0.58471313 | 1 | -0.366 | 0.209 | chr1 | 222832209 | 222834854 | - | lncRNA |
| ENSG00000164548.11_4 | TRA2A | 0.99416242 | 1 | -0.366 | 5.012 | chr7 | 23544399 | 23571660 | - | protein_coding |
| ENSG00000125944.20_6 | HNRNPR | 0.99958155 | 1 | -0.366 | 7.891 | chr1 | 23630264 | 23670829 | - | protein_coding |
| ENSG00000132964.12_4 | CDK8 | 0.97678523 | 1 | -0.366 | 4.083 | chr13 | 26828241 | 26979375 | + | protein_coding |
| ENSG00000184602.6_3 | SNN | 0.9183661 | 1 | -0.366 | 3.088 | chr16 | 11762311 | 11773008 | + | protein_coding |
| ENSG00000162929.14_3 | KIAA1841 | 0.99163039 | 1 | -0.366 | 4.917 | chr2 | 61293006 | 61365169 | + | protein_coding |
| ENSG00000163636.10_4 | PSMD6 | 0.99828751 | 1 | -0.367 | 6.148 | chr3 | 63996225 | 64009686 | - | protein_coding |
| ENSG00000267060.5_5 | PTGES3L | 0.5024217 | 1 | -0.367 | -1.064 | chr17 | 41120105 | 41132450 | - | protein_coding |
| ENSG00000084073.9_3 | ZMPSTE24 | 0.99830454 | 1 | -0.367 | 6.384 | chr1 | 40723908 | 40759852 | + | protein_coding |
| ENSG00000101266.19_7 | CSNK2A1 | 0.99958021 | 1 | -0.367 | 7.822 | chr20 | 453142 | 524479 | - | protein_coding |
| ENSG00000144535.19_7 | DIS3L2 | 0.98956724 | 1 | -0.367 | 4.893 | chr2 | 232825955 | 233209060 | + | protein_coding |
| ENSG00000087088.20_5 | BAX | 0.99495108 | 1 | -0.367 | 5.988 | chr19 | 49458072 | 49465055 | + | protein_coding |
| ENSG00000175711.8_3 | B3GNTL1 | 0.89545916 | 1 | -0.367 | 2.823 | chr17 | 80900031 | 81009686 | - | protein_coding |
| ENSG00000075239.14_3 | ACAT1 | 0.99123172 | 1 | -0.367 | 5.503 | chr11 | 107987422 | 108018330 | + | protein_coding |
| ENSG00000107566.14_5 | ERLIN1 | 0.99368973 | 1 | -0.367 | 5.267 | chr10 | 101909851 | 101945790 | - | protein_coding |
| ENSG00000005100.13_7 | DHX33 | 0.99817596 | 1 | -0.367 | 5.834 | chr17 | 5344237 | 5372302 | - | protein_coding |
| ENSG00000104067.16_5 | TJP1 | 0.9995592 | 1 | -0.367 | 8.332 | chr15 | 29991571 | 30261068 | - | protein_coding |
| ENSG00000163528.13_6 | CHCHD4 | 0.98407216 | 1 | -0.368 | 4.600 | chr3 | 14153577 | 14166370 | - | protein_coding |
| ENSG00000108852.14_5 | MPP2 | 0.91428734 | 1 | -0.368 | 3.133 | chr17 | 41952725 | 41987079 | - | protein_coding |
| ENSG00000134330.19_5 | IAH1 | 0.99581262 | 1 | -0.368 | 5.561 | chr2 | 9613787 | 9636672 | + | protein_coding |
| ENSG00000259116.2_8 | AL049869.3 | 0.54075171 | 1 | -0.368 | -0.238 | chr14 | 64980670 | 65007086 | - | lncRNA |
| ENSG00000197361.7_3 | FBXL22 | 0.50117537 | 1 | -0.368 | -0.892 | chr15 | 63889552 | 63894627 | + | protein_coding |
| ENSG00000176912.5_8 | TYMSOS | 0.55397822 | 1 | -0.368 | 0.146 | chr18 | 630886 | 658340 | - | lncRNA |
| ENSG00000236184.1_4 | TCEA1P4 | 0.50091049 | 1 | -0.368 | -1.065 | chr9 | 32979558 | 32980401 | - | processed_pseudogene |
| ENSG00000278811.4_6 | LINC00624 | 0.75960309 | 1 | -0.368 | 1.544 | chr1 | 146730539 | 146989699 | - | lncRNA |
| ENSG00000115282.20_7 | TTC31 | 0.99013991 | 1 | -0.368 | 4.742 | chr2 | 74710200 | 74722013 | + | protein_coding |
| ENSG00000160606.11_5 | TLCD1 | 0.97123646 | 1 | -0.368 | 4.183 | chr17 | 27051366 | 27054953 | - | protein_coding |
| ENSG00000152953.13_3 | STK32B | 0.99062218 | 1 | -0.368 | 4.953 | chr4 | 5053207 | 5502721 | + | protein_coding |
| ENSG00000144451.19_3 | SPAG16 | 0.89274317 | 1 | -0.368 | 3.675 | chr2 | 214149103 | 215275225 | + | protein_coding |
| ENSG00000274641.1_3 | H2BC17 | 0.99792004 | 1 | -0.368 | 7.065 | chr6 | 27861241 | 27861621 | + | protein_coding |
| ENSG00000287202.1_1 | AC008119.1 | 0.65191428 | 1 | -0.368 | 0.668 | chr12 | 108925850 | 108933514 | + | lncRNA |
| ENSG00000232882.1_4 | PHKA1P1 | 0.50034676 | 1 | -0.369 | -0.958 | chr1 | 91358549 | 91359169 | - | processed_pseudogene |
| ENSG00000175879.9_5 | HOXD8 | 0.7269085 | 1 | -0.369 | 1.648 | chr2 | 176994422 | 176997423 | + | protein_coding |
| ENSG00000234062.8_7 | AL390879.1 | 0.6843896 | 1 | -0.369 | 0.817 | chrX | 135991528 | 136104302 | + | transcribed_unprocessed_pseudogene |
| ENSG00000104064.17_4 | GABPB1 | 0.99114874 | 1 | -0.369 | 5.278 | chr15 | 50567589 | 50647605 | - | protein_coding |
| ENSG00000169129.15_6 | AFAP1L2 | 0.99563037 | 1 | -0.369 | 5.602 | chr10 | 116054583 | 116164515 | - | protein_coding |
| ENSG00000116117.18_4 | PARD3B | 0.93031723 | 1 | -0.369 | 3.395 | chr2 | 205410198 | 206484886 | + | protein_coding |
| ENSG00000147459.18_4 | DOCK5 | 0.99858121 | 1 | -0.369 | 8.454 | chr8 | 25042204 | 25275598 | + | protein_coding |
| ENSG00000100591.8_5 | AHSA1 | 0.99910612 | 1 | -0.369 | 7.134 | chr14 | 77924213 | 77935815 | + | protein_coding |
| ENSG00000116830.12_3 | TTF2 | 0.99728964 | 1 | -0.369 | 5.764 | chr1 | 117602948 | 117650075 | + | protein_coding |
| ENSG00000116652.6_4 | DLEU2L | 0.54041956 | 1 | -0.370 | 0.148 | chr1 | 64012753 | 64016307 | + | lncRNA |
| ENSG00000212719.13_8 | LINC02693 | 0.97954992 | 1 | -0.370 | 4.566 | chr17 | 21331575 | 21477781 | - | lncRNA |
| ENSG00000007968.7_4 | E2F2 | 0.96865079 | 1 | -0.370 | 4.386 | chr1 | 23832930 | 23857725 | - | protein_coding |
| ENSG00000095794.19_7 | CREM | 0.98149865 | 1 | -0.370 | 4.529 | chr10 | 35415719 | 35501886 | + | protein_coding |
| ENSG00000134001.14_6 | EIF2S1 | 0.99934902 | 1 | -0.370 | 7.283 | chr14 | 67826868 | 67853233 | + | protein_coding |
| ENSG00000267430.1_5 | AC036176.2 | 0.49834067 | 1 | -0.370 | -1.065 | chr18 | 61082550 | 61083702 | - | processed_pseudogene |

| | | | | | | | | | | |
|---|---|---|---|---|---|---|---|---|---|---|
| ENSG00000238273.3_6 | AC108058.1 | 0.56109735 | 1 | -0.370 | 0.239 | chr2 | 105979495 | 105995296 | + | lncRNA |
| ENSG00000167635.11_2 | ZNF146 | 0.99949591 | 1 | -0.370 | 7.869 | chr19 | 36705504 | 36729676 | + | protein_coding |
| ENSG00000137338.5_3 | PGBD1 | 0.93626251 | 1 | -0.371 | 3.473 | chr6 | 28249349 | 28270326 | + | protein_coding |
| ENSG00000163634.12_6 | THOC7 | 0.99123235 | 1 | -0.371 | 5.158 | chr3 | 63819546 | 63849544 | - | protein_coding |
| ENSG00000154473.18_4 | BUB3 | 0.99933661 | 1 | -0.371 | 7.639 | chr10 | 124913918 | 124929983 | + | protein_coding |
| ENSG00000164707.15_4 | SLC13A4 | 0.67663387 | 1 | -0.371 | 0.713 | chr7 | 135365985 | 135414006 | - | protein_coding |
| ENSG00000010072.16_4 | SPRTN | 0.989757 | 1 | -0.371 | 4.735 | chr1 | 231472850 | 231490769 | + | protein_coding |
| ENSG00000089195.15_4 | TRMT6 | 0.9964824 | 1 | -0.371 | 5.490 | chr20 | 5917874 | 5931204 | - | protein_coding |
| ENSG00000247828.8_7 | TMEM161B-AS1 | 0.9486592 | 1 | -0.371 | 4.366 | chr5 | 87564708 | 87735154 | + | lncRNA |
| ENSG00000127804.13_5 | METTL16 | 0.99765883 | 1 | -0.371 | 6.232 | chr17 | 2308856 | 2415185 | - | protein_coding |
| ENSG00000134490.14_6 | TMEM241 | 0.92253274 | 1 | -0.372 | 3.187 | chr18 | 20777108 | 21017925 | - | protein_coding |
| ENSG00000282458.1_3 | WASH5P | 0.49678059 | 1 | -0.372 | -1.064 | chr19 | 60951 | 71626 | - | transcribed_processed_pseudogene |
| ENSG00000059769.20_4 | DNAJC25 | 0.90858719 | 1 | -0.372 | 3.121 | chr9 | 114393614 | 114416631 | + | protein_coding |
| ENSG00000285728.2_3 | AC098484.4 | 0.70159154 | 1 | -0.372 | 1.460 | chr1 | 43124358 | 43147831 | - | lncRNA |
| ENSG00000204120.14_7 | GIGYF2 | 0.99807857 | 1 | -0.372 | 6.352 | chr2 | 233562009 | 233725285 | + | protein_coding |
| ENSG00000142507.10_3 | PSMB6 | 0.99834929 | 1 | -0.372 | 6.583 | chr17 | 4699439 | 4701797 | + | protein_coding |
| ENSG00000150756.14_6 | ATPSCKMT | 0.95626277 | 1 | -0.372 | 3.929 | chr5 | 10225619 | 10250009 | - | protein_coding |
| ENSG00000262434.1_7 | AC087392.4 | 0.54076921 | 1 | -0.372 | -0.260 | chr17 | 692984 | 693765 | - | lncRNA |
| ENSG00000135541.21_6 | AHI1 | 0.9934835 | 1 | -0.372 | 5.214 | chr6 | 135604670 | 135818903 | - | protein_coding |
| ENSG00000100162.15_4 | CENPM | 0.93400475 | 1 | -0.372 | 3.857 | chr22 | 42334741 | 42343156 | - | protein_coding |
| ENSG00000142731.11_4 | PLK4 | 0.99516488 | 1 | -0.373 | 5.336 | chr4 | 128802048 | 128820379 | + | protein_coding |
| ENSG00000198938.2_1 | MT-CO3 | 0.99884189 | 1 | -0.373 | 12.117 | chrM | 9207 | 9990 | + | protein_coding |
| ENSG00000260190.1_6 | AL807752.5 | 0.49717837 | 1 | -0.373 | -0.925 | chr9 | 139831621 | 139832440 | - | lncRNA |
| ENSG00000236896.1_7 | AL354726.1 | 0.60301174 | 1 | -0.373 | 0.465 | chr9 | 100748833 | 100749938 | - | lncRNA |
| ENSG00000144228.9_4 | SPOPL | 0.98132737 | 1 | -0.373 | 5.022 | chr2 | 139259340 | 139331117 | + | protein_coding |
| ENSG00000096384.20_3 | HSP90AB1 | 0.99957695 | 1 | -0.374 | 10.342 | chr6 | 44213903 | 44221625 | + | protein_coding |
| ENSG00000179528.16_4 | LBX2 | 0.70776817 | 1 | -0.374 | 1.231 | chr2 | 74724644 | 74730443 | - | protein_coding |
| ENSG00000184428.13_4 | TOP1MT | 0.98772742 | 1 | -0.374 | 4.726 | chr8 | 144386554 | 144442149 | - | protein_coding |
| ENSG00000103876.12_4 | FAH | 0.98423779 | 1 | -0.374 | 4.541 | chr15 | 80444832 | 80479288 | + | protein_coding |
| ENSG00000100714.17_8 | MTHFD1 | 0.99753588 | 1 | -0.374 | 5.917 | chr14 | 64854749 | 64930175 | + | protein_coding |
| ENSG00000228716.7_5 | DHFR | 0.99673065 | 1 | -0.374 | 6.045 | chr5 | 79922045 | 79950802 | - | protein_coding |
| ENSG00000101365.21_5 | IDH3B | 0.9986412 | 1 | -0.374 | 6.482 | chr20 | 2639041 | 2644865 | - | protein_coding |
| ENSG00000278600.1_9 | AC015871.3 | 0.85613043 | 1 | -0.374 | 2.705 | chr15 | 80212537 | 80214797 | - | lncRNA |
| ENSG00000066557.6_4 | LRRC40 | 0.99005847 | 1 | -0.374 | 5.094 | chr1 | 70610488 | 70671262 | - | protein_coding |
| ENSG00000165661.17_6 | QSOX2 | 0.9938102 | 1 | -0.374 | 5.648 | chr9 | 139098179 | 139137658 | - | protein_coding |
| ENSG00000103047.8_4 | TANGO6 | 0.98803942 | 1 | -0.374 | 5.080 | chr16 | 68877434 | 69119085 | + | protein_coding |
| ENSG00000163508.13_5 | EOMES | 0.61245872 | 1 | -0.374 | 0.461 | chr3 | 27757440 | 27764202 | - | protein_coding |
| ENSG00000189159.16_8 | JPT1 | 0.99934179 | 1 | -0.374 | 7.379 | chr17 | 73131343 | 73164376 | - | protein_coding |
| ENSG00000136068.14_5 | FLNB | 0.99940448 | 1 | -0.374 | 10.474 | chr3 | 57994127 | 58157978 | + | protein_coding |
| ENSG00000168234.13_5 | TTC39C | 0.92809436 | 1 | -0.375 | 3.505 | chr18 | 21572737 | 21715574 | + | protein_coding |
| ENSG00000159685.11_5 | CHCHD6 | 0.91112667 | 1 | -0.375 | 2.987 | chr3 | 126423083 | 126679263 | + | protein_coding |
| ENSG00000284862.3_6 | CCDC39 | 0.49970179 | 1 | -0.375 | -0.282 | chr3 | 180320646 | 180402730 | - | protein_coding |
| ENSG00000104689.10_5 | TNFRSF10A | 0.98940726 | 1 | -0.375 | 4.913 | chr8 | 23047965 | 23082615 | - | protein_coding |
| ENSG00000101407.13_4 | TTI1 | 0.99731415 | 1 | -0.375 | 5.848 | chr20 | 36611409 | 36661863 | - | protein_coding |
| ENSG00000159086.15_4 | PAXBP1 | 0.99418569 | 1 | -0.375 | 5.602 | chr21 | 34106210 | 34144103 | - | protein_coding |
| ENSG00000272405.1_6 | AL365181.3 | 0.92658951 | 1 | -0.375 | 3.356 | chr1 | 156611458 | 156614679 | - | lncRNA |
| ENSG00000120458.11_4 | MSANTD2 | 0.89092098 | 1 | -0.375 | 3.076 | chr11 | 124636394 | 124670569 | - | protein_coding |
| ENSG00000223345.3_3 | H2BP1 | 0.65482507 | 1 | -0.375 | 0.873 | chr1 | 120906101 | 120915073 | + | transcribed_unprocessed_pseudogene |
| ENSG00000112110.10_3 | MRPL18 | 0.99592163 | 1 | -0.375 | 5.674 | chr6 | 160210844 | 160219468 | + | protein_coding |
| ENSG00000130935.10_4 | NOL11 | 0.99688645 | 1 | -0.376 | 7.057 | chr17 | 65714047 | 65740647 | + | protein_coding |
| ENSG00000105928.16_8 | GSDME | 0.96433518 | 1 | -0.376 | 4.201 | chr7 | 24737974 | 24797559 | - | protein_coding |
| ENSG00000225805.4_6 | DEFB131B | 0.49246451 | 1 | -0.376 | -0.959 | chr11 | 71589499 | 71595607 | + | protein_coding |
| ENSG00000237491.10_9 | LINC01409 | 0.5882832 | 1 | -0.376 | 0.691 | chr1 | 714127 | 745445 | + | lncRNA |
| ENSG00000213326.4_4 | RPS7P11 | 0.59698708 | 1 | -0.376 | 1.541 | chr17 | 44798948 | 44799533 | - | processed_pseudogene |
| ENSG00000198899.2_1 | MT-ATP6 | 0.99743522 | 1 | -0.376 | 8.877 | chrM | 8527 | 9207 | + | protein_coding |
| ENSG00000224574.1_7 | COL18A1-AS2 | 0.6054181 | 1 | -0.376 | 0.820 | chr21 | 46827301 | 46829980 | - | lncRNA |
| ENSG00000136811.16_5 | ODF2 | 0.99679534 | 1 | -0.376 | 5.730 | chr9 | 131217465 | 131263571 | + | protein_coding |

| | | | | | | | | | | |
|---|---|---|---|---|---|---|---|---|---|---|
| ENSG00000125871.14_4 | MGME1 | 0.99410551 | 1 | -0.377 | 5.534 | chr20 | 17949661 | 17971765 | + | protein_coding |
| ENSG00000196421.8_7 | C20orf204 | 0.73073378 | 1 | -0.377 | 1.706 | chr20 | 62665697 | 62671315 | + | protein_coding |
| ENSG00000105778.19_5 | AVL9 | 0.99819371 | 1 | -0.377 | 6.407 | chr7 | 32535038 | 32628338 | + | protein_coding |
| ENSG00000172927.3 | MYEOV | 0.9962029 | 1 | -0.377 | 6.103 | chr11 | 69061605 | 69182494 | + | protein_coding |
| ENSG00000149311.18_4 | ATM | 0.98888067 | 1 | -0.377 | 6.611 | chr11 | 108093211 | 108239829 | + | protein_coding |
| ENSG00000172197.11_4 | MBOAT1 | 0.93894042 | 1 | -0.377 | 3.616 | chr6 | 20099915 | 20212700 | - | protein_coding |
| ENSG00000229178.1_7 | AC233280.1 | 0.49243136 | 1 | -0.377 | -0.926 | chr3 | 195382436 | 195384798 | - | lncRNA |
| ENSG00000163468.15_4 | CCT3 | 0.99952222 | 1 | -0.377 | 8.924 | chr1 | 156278759 | 156337664 | - | protein_coding |
| ENSG00000130475.14_4 | FCHO1 | 0.91311603 | 1 | -0.378 | 3.829 | chr19 | 17858527 | 17899377 | + | protein_coding |
| ENSG00000253251.3_7 | SHLD3 | 0.79375951 | 1 | -0.378 | 2.021 | chr5 | 64920854 | 64926755 | + | protein_coding |
| ENSG00000109519.13_4 | GRPEL1 | 0.98715463 | 1 | -0.378 | 5.131 | chr4 | 7060622 | 7069791 | - | protein_coding |
| ENSG00000006625.18_5 | GGCT | 0.96580766 | 1 | -0.378 | 4.166 | chr7 | 30536237 | 30544457 | - | protein_coding |
| ENSG00000140043.11_3 | PTGR2 | 0.75789946 | 1 | -0.378 | 1.866 | chr14 | 74318547 | 74353530 | + | protein_coding |
| ENSG00000280213.1_5 | UCKL1-AS1 | 0.59672878 | 1 | -0.378 | 0.619 | chr20 | 62584737 | 62588338 | + | lncRNA |
| ENSG00000034510.6_3 | TMSB10 | 0.99925026 | 1 | -0.378 | 9.625 | chr2 | 85132780 | 85133795 | + | protein_coding |
| ENSG00000264769.1_6 | AC145207.8 | 0.672811 | 1 | -0.378 | 1.053 | chr17 | 79880775 | 79882387 | + | lncRNA |
| ENSG00000116991.10_3 | SIPA1L2 | 0.97439557 | 1 | -0.378 | 4.418 | chr1 | 232533711 | 232697304 | - | protein_coding |
| ENSG00000109685.18_5 | NSD2 | 0.99926366 | 1 | -0.378 | 7.328 | chr4 | 1873120 | 1983934 | + | protein_coding |
| ENSG00000151725.12_8 | CENPU | 0.98813195 | 1 | -0.378 | 5.295 | chr4 | 185615239 | 185655284 | - | protein_coding |
| ENSG00000262333.1_4 | HNRNPA1P16 | 0.58710866 | 1 | -0.378 | 0.421 | chr17 | 2210055 | 2211009 | + | processed_pseudogene |
| ENSG00000287935.1_1 | AC136297.1 | 0.54888955 | 1 | -0.378 | 0.174 | chr11 | 1354480 | 1358298 | - | lncRNA |
| ENSG00000272563.1_6 | AC016745.2 | 0.52132158 | 1 | -0.378 | -0.240 | chr2 | 114190177 | 114193619 | + | lncRNA |
| ENSG00000257702.3_5 | LBX2-AS1 | 0.58421637 | 1 | -0.378 | 0.558 | chr2 | 74729722 | 74731805 | + | lncRNA |
| ENSG00000157869.15_4 | RAB28 | 0.93424627 | 1 | -0.379 | 3.660 | chr4 | 13362978 | 13485989 | - | protein_coding |
| ENSG00000185495.10_6 | AC138393.1 | 0.71664794 | 1 | -0.379 | 1.630 | chr1 | 224180445 | 224198314 | + | transcribed_unprocessed_pseudogene |
| ENSG00000277840.1_7 | AC026368.1 | 0.49518052 | 1 | -0.379 | -0.680 | chr12 | 117440268 | 117440957 | + | lncRNA |
| ENSG00000235142.10_8 | LINC02532 | 0.63700438 | 1 | -0.379 | 1.328 | chr6 | 107153209 | 107236023 | - | lncRNA |
| ENSG00000185920.16_9 | PTCH1 | 0.81948302 | 1 | -0.379 | 2.756 | chr9 | 98205262 | 98279339 | - | protein_coding |
| ENSG00000206603.1 | SNORA22 | 0.49222056 | 1 | -0.380 | -0.601 | chr7 | 56123058 | 56123195 | + | snoRNA |
| ENSG00000113643.9_4 | RARS1 | 0.99771278 | 1 | -0.380 | 7.550 | chr5 | 167913456 | 167946306 | + | protein_coding |
| ENSG00000262712.1_7 | AC012676.1 | 0.55247201 | 1 | -0.380 | 0.109 | chr16 | 4385871 | 4387819 | - | lncRNA |
| ENSG00000130713.16_6 | EXOSC2 | 0.99673607 | 1 | -0.380 | 5.983 | chr9 | 133569108 | 133580281 | + | protein_coding |
| ENSG00000204175.5_2 | GPRIN2 | 0.96350832 | 1 | -0.380 | 5.532 | chr10 | 46994087 | 47000573 | + | protein_coding |
| ENSG00000213906.10_4 | LTB4R2 | 0.79198346 | 1 | -0.380 | 1.946 | chr14 | 24774940 | 24781259 | + | protein_coding |
| ENSG00000137337.15_6 | MDC1 | 0.78221599 | 1 | -0.380 | 2.074 | chr6 | 30667584 | 30685224 | - | protein_coding |
| ENSG00000120158.12_5 | RCL1 | 0.97642843 | 1 | -0.380 | 4.476 | chr9 | 4792944 | 4885917 | + | protein_coding |
| ENSG00000179240.12_10 | GVQW3 | 0.92560931 | 1 | -0.380 | 3.411 | chr11 | 76092347 | 76125663 | + | protein_coding |
| ENSG00000197903.7_3 | H2BC12 | 0.99887781 | 1 | -0.380 | 8.003 | chr6 | 27114197 | 27114577 | - | protein_coding |
| ENSG00000233170.4_5 | AC138356.2 | 0.50112695 | 1 | -0.380 | -0.305 | chr8 | 37604856 | 37605522 | - | processed_pseudogene |
| ENSG00000269680.1_7 | AC008760.1 | 0.5032342 | 1 | -0.380 | -0.417 | chr19 | 6748304 | 6751478 | - | lncRNA |
| ENSG00000091490.11_6 | SEL1L3 | 0.99175821 | 1 | -0.380 | 5.381 | chr4 | 25749055 | 25865382 | - | protein_coding |
| ENSG00000148834.13_6 | GSTO1 | 0.99425853 | 1 | -0.380 | 5.766 | chr10 | 105995114 | 106027217 | + | protein_coding |
| ENSG00000187492.9_4 | CDHR4 | 0.50914156 | 1 | -0.380 | -0.220 | chr3 | 49828165 | 49837306 | - | protein_coding |
| ENSG00000167703.14_4 | SLC43A2 | 0.91822936 | 1 | -0.380 | 3.274 | chr17 | 1472561 | 1532180 | - | protein_coding |
| ENSG00000235253.1_3 | AC010240.3 | 0.48770682 | 1 | -0.381 | -0.862 | chr5 | 132278624 | 132280742 | + | processed_pseudogene |
| ENSG00000184675.11_6 | AMER1 | 0.92683867 | 1 | -0.381 | 3.432 | chrX | 63404997 | 63425588 | - | protein_coding |
| ENSG00000248690.8_6 | HAS2-AS1 | 0.6415802 | 1 | -0.381 | 1.108 | chr8 | 122651533 | 123006424 | + | lncRNA |
| ENSG00000224287.2_5 | MSL3P1 | 0.48728972 | 1 | -0.381 | -0.862 | chr2 | 234774083 | 234777090 | - | transcribed_processed_pseudogene |
| ENSG00000134186.12_3 | PRPF38B | 0.97837284 | 1 | -0.381 | 5.061 | chr1 | 109234932 | 109245550 | + | protein_coding |
| ENSG00000267476.1_8 | AC104365.2 | 0.48938374 | 1 | -0.381 | -0.738 | chr18 | 56339845 | 56419962 | - | lncRNA |
| ENSG00000152042.13 | NBPF11 | 0.50723918 | 1 | -0.381 | -0.402 | chr1 | 146032647 | 146082765 | - | protein_coding |
| ENSG00000278259.4_5 | MYO19 | 0.99876763 | 1 | -0.381 | 7.184 | chr17 | 34851477 | 34899284 | - | protein_coding |
| ENSG00000180953.11_5 | ST20 | 0.81451644 | 1 | -0.381 | 2.183 | chr15 | 80191182 | 80216044 | - | protein_coding |
| ENSG00000243244.6_5 | STON1 | 0.53955308 | 1 | -0.381 | -0.178 | chr2 | 48756522 | 48825652 | + | protein_coding |
| ENSG00000132432.14_5 | SEC61G | 0.9629002 | 1 | -0.381 | 5.429 | chr7 | 54819943 | 54827667 | - | protein_coding |
| ENSG00000168397.17_5 | ATG4B | 0.98995156 | 1 | -0.381 | 5.065 | chr2 | 242576628 | 242613272 | + | protein_coding |

| | | | | | | | | | | |
|---|---|---|---|---|---|---|---|---|---|---|
| ENSG00000230280.2_4 | HNRNPA1P59 | 0.5227782 | 1 | -0.381 | -0.180 | chr1 | 202880940 | 202881857 | + | processed_pseudogene |
| ENSG00000237765.7_4 | FAM200B | 0.95198601 | 1 | -0.381 | 4.284 | chr4 | 15683285 | 15707188 | + | protein_coding |
| ENSG00000213923.13_8 | CSNK1E | 0.97958427 | 1 | -0.381 | 4.842 | chr22 | 38686697 | 38714089 | - | protein_coding |
| ENSG00000136897.8_4 | MRPL50 | 0.9920417 | 1 | -0.381 | 5.523 | chr9 | 104149915 | 104160900 | - | protein_coding |
| ENSG00000177034.16_5 | MTX3 | 0.98464027 | 1 | -0.382 | 5.077 | chr5 | 79272554 | 79287088 | - | protein_coding |
| ENSG00000196284.16_6 | SUPT3H | 0.67720198 | 1 | -0.382 | 1.420 | chr6 | 44777054 | 45345690 | - | protein_coding |
| ENSG00000135605.13_4 | TEC | 0.75057128 | 1 | -0.382 | 1.751 | chr4 | 48137800 | 48271855 | - | protein_coding |
| ENSG00000168209.5_4 | DDIT4 | 0.85709082 | 1 | -0.382 | 4.892 | chr10 | 74033682 | 74035794 | + | protein_coding |
| ENSG00000164574.16_4 | GALNT10 | 0.99871207 | 1 | -0.382 | 7.892 | chr5 | 153570290 | 153800544 | + | protein_coding |
| ENSG00000259820.1_7 | AC083843.3 | 0.8138784 | 1 | -0.382 | 2.625 | chr8 | 135804263 | 135810515 | - | lncRNA |
| ENSG00000259994.1_6 | AL353796.1 | 0.63870886 | 1 | -0.382 | 0.870 | chr10 | 30591755 | 30594995 | - | lncRNA |
| ENSG00000102409.10_3 | BEX4 | 0.49868332 | 1 | -0.382 | -0.573 | chrX | 102470036 | 102472174 | + | protein_coding |
| ENSG00000125743.10_3 | SNRPD2 | 0.99868373 | 1 | -0.382 | 7.386 | chr19 | 46190712 | 46195827 | - | protein_coding |
| ENSG00000203817.5 | FAM72C | 0.64494762 | 1 | -0.382 | 0.921 | chr1 | 149438750 | 149459549 | - | protein_coding |
| ENSG00000227518.5_7 | AL928970.1 | 0.48525358 | 1 | -0.382 | -0.894 | chr9 | 27657 | 30891 | + | lncRNA |
| ENSG00000132382.14_5 | MYBBP1A | 0.99729404 | 1 | -0.383 | 7.053 | chr17 | 4442192 | 4458926 | - | protein_coding |
| ENSG00000133997.11_4 | MED6 | 0.98278313 | 1 | -0.383 | 4.606 | chr14 | 71047974 | 71067407 | - | protein_coding |
| ENSG00000074201.9_6 | CLNS1A | 0.99784989 | 1 | -0.383 | 6.589 | chr11 | 77225981 | 77348839 | - | protein_coding |
| ENSG00000134440.12_5 | NARS1 | 0.99905011 | 1 | -0.383 | 7.656 | chr18 | 55267888 | 55289445 | - | protein_coding |
| ENSG00000105255.11_5 | FSD1 | 0.84459712 | 1 | -0.383 | 2.613 | chr19 | 4304595 | 4323840 | + | protein_coding |
| ENSG00000214796.8_6 | AC098934.1 | 0.50992504 | 1 | -0.383 | -0.240 | chr1 | 202820956 | 202830748 | - | transcribed_unprocessed_pseudogene |
| ENSG00000218175.2_5 | AC016739.1 | 0.58135247 | 1 | -0.383 | 0.494 | chr2 | 177065636 | 177065980 | - | processed_pseudogene |
| ENSG00000197417.9_5 | SHPK | 0.80051979 | 1 | -0.383 | 2.057 | chr17 | 3511534 | 3539544 | - | protein_coding |
| ENSG00000083635.8_5 | NUFIP1 | 0.94460146 | 1 | -0.383 | 3.741 | chr13 | 45513384 | 45563606 | - | protein_coding |
| ENSG00000031081.11_5 | ARHGAP31 | 0.80619503 | 1 | -0.383 | 1.854 | chr3 | 119013230 | 119139561 | + | protein_coding |
| ENSG00000125821.12_5 | DTD1 | 0.97279175 | 1 | -0.384 | 4.305 | chr20 | 18568537 | 18747288 | + | protein_coding |
| ENSG00000270871.1_7 | AC015849.3 | 0.62931755 | 1 | -0.384 | 1.100 | chr17 | 34143721 | 34157297 | - | lncRNA |
| ENSG00000184786.6_4 | TCTE3 | 0.68907515 | 1 | -0.384 | 1.368 | chr6 | 170140205 | 170151683 | - | protein_coding |
| ENSG00000279759.1_7 | AC118344.4 | 0.71528403 | 1 | -0.384 | 1.855 | chr19 | 40861749 | 40866513 | + | TEC |
| ENSG00000123992.19_6 | DNPEP | 0.98679137 | 1 | -0.384 | 4.772 | chr2 | 220238249 | 220264744 | - | protein_coding |
| ENSG00000143353.12_3 | LYPLAL1 | 0.971962 | 1 | -0.384 | 4.328 | chr1 | 219347211 | 219386207 | + | protein_coding |
| ENSG00000216937.13_7 | CCDC7 | 0.86051075 | 1 | -0.384 | 2.744 | chr10 | 32735068 | 33171802 | + | protein_coding |
| ENSG00000065427.14_7 | KARS1 | 0.9987722 | 1 | -0.384 | 7.421 | chr16 | 75661372 | 75682541 | - | protein_coding |
| ENSG00000270820.6_6 | AC016727.1 | 0.64481971 | 1 | -0.384 | 1.123 | chr2 | 61698323 | 61711265 | + | lncRNA |
| ENSG00000160746.13_7 | ANO10 | 0.97899631 | 1 | -0.385 | 4.704 | chr3 | 43396351 | 43733086 | - | protein_coding |
| ENSG00000126453.9_3 | BCL2L12 | 0.8787209 | 1 | -0.385 | 2.831 | chr19 | 50168399 | 50177173 | + | protein_coding |
| ENSG00000122952.17_6 | ZWINT | 0.99714051 | 1 | -0.385 | 6.275 | chr10 | 58116988 | 58121034 | - | protein_coding |
| ENSG00000286532.1_1 | PARTICL | 0.64979724 | 1 | -0.385 | 1.053 | chr2 | 85764585 | 85765789 | - | lncRNA |
| ENSG00000240498.9_7 | CDKN2B-AS1 | 0.8694425 | 1 | -0.385 | 3.006 | chr9 | 21994138 | 22128102 | + | lncRNA |
| ENSG00000214295.4_4 | FOXO1B | 0.52716239 | 1 | -0.385 | -0.065 | chr5 | 180526140 | 180527666 | - | processed_pseudogene |
| ENSG00000163104.17_5 | SMARCAD1 | 0.99670224 | 1 | -0.386 | 6.232 | chr4 | 95128762 | 95212443 | + | protein_coding |
| ENSG00000106125.14_6 | MINDY4 | 0.55264743 | 1 | -0.386 | 0.023 | chr7 | 30811033 | 30932002 | + | protein_coding |
| ENSG00000168763.16_3 | CNNM3 | 0.96177069 | 1 | -0.386 | 4.582 | chr2 | 97481982 | 97501119 | + | protein_coding |
| ENSG00000285725.1_3 | AC004967.2 | 0.56735838 | 1 | -0.386 | 0.218 | chr7 | 97595689 | 97599716 | - | lncRNA |
| ENSG00000115828.17_4 | QPCT | 0.98250233 | 1 | -0.386 | 4.854 | chr2 | 37569970 | 37600465 | + | protein_coding |
| ENSG00000129484.13_2 | PARP2 | 0.97627898 | 1 | -0.386 | 4.771 | chr14 | 20811741 | 20826064 | + | protein_coding |
| ENSG00000131469.14_4 | RPL27 | 0.99806447 | 1 | -0.386 | 8.172 | chr17 | 41150290 | 41154976 | + | protein_coding |
| ENSG00000217094.2_4 | PPIAP31 | 0.51274266 | 1 | -0.386 | -0.248 | chr10 | 32477251 | 32477745 | - | processed_pseudogene |
| ENSG00000179967.11_2 | PPP1R14BP3 | 0.71608414 | 1 | -0.387 | 1.350 | chr4 | 140036084 | 140036528 | - | processed_pseudogene |
| ENSG00000149150.9_7 | SLC43A1 | 0.69691668 | 1 | -0.387 | 1.784 | chr11 | 57252007 | 57283253 | - | protein_coding |
| ENSG00000077942.19_8 | FBLN1 | 0.97566502 | 1 | -0.387 | 4.552 | chr22 | 45898118 | 45997015 | + | protein_coding |
| ENSG00000134222.16_6 | PSRC1 | 0.98104932 | 1 | -0.387 | 4.684 | chr1 | 109822178 | 109825808 | - | protein_coding |
| ENSG00000168852.13_6 | TPTE2P5 | 0.47803037 | 1 | -0.387 | -0.928 | chr13 | 41386445 | 41495885 | - | transcribed_unprocessed_pseudogene |
| ENSG00000117245.12_4 | KIF17 | 0.65199973 | 1 | -0.387 | 1.082 | chr1 | 20990507 | 21044510 | - | protein_coding |
| ENSG00000130204.13_7 | TOMM40 | 0.98957512 | 1 | -0.387 | 7.362 | chr19 | 45393826 | 45406946 | + | protein_coding |
| ENSG00000232611.1_7 | AL683813.2 | 0.52209661 | 1 | -0.387 | -0.103 | chrX | 135923090 | 135929956 | - | lncRNA |

| | | | | | | | | | | |
|---|---|---|---|---|---|---|---|---|---|---|
| ENSG00000206562.11_5 | METTL6 | 0.97555763 | 1 | -0.387 | 4.490 | chr3 | 15422782 | 15482073 | - | protein_coding |
| ENSG00000065615.14_3 | CYB5R4 | 0.97308657 | 1 | -0.387 | 4.818 | chr6 | 84569375 | 84677142 | + | protein_coding |
| ENSG00000137714.3_3 | FDX1 | 0.9530486 | 1 | -0.387 | 4.149 | chr11 | 110300672 | 110335608 | + | protein_coding |
| ENSG00000147408.14_4 | CSGALNACT1 | 0.99704922 | 1 | -0.387 | 6.219 | chr8 | 19261672 | 19615540 | - | protein_coding |
| ENSG00000183562.3_6 | AC131971.1 | 0.71647017 | 1 | -0.388 | 1.587 | chr11 | 3011093 | 3012574 | + | lncRNA |
| ENSG00000229638.1_4 | RPL4P4 | 0.94580498 | 1 | -0.388 | 4.384 | chr3 | 185135283 | 185136566 | - | processed_pseudogene |
| ENSG00000187118.14_5 | CMC1 | 0.95727407 | 1 | -0.388 | 4.100 | chr3 | 28283075 | 28366633 | + | protein_coding |
| ENSG00000267698.1_6 | AC002116.2 | 0.48464303 | 1 | -0.388 | -0.576 | chr19 | 36505410 | 36536874 | + | lncRNA |
| ENSG00000096746.17_2 | HNRNPH3 | 0.93288075 | 1 | -0.389 | 3.696 | chr10 | 70090931 | 70102948 | + | protein_coding |
| ENSG00000255561.7_5 | FDXACB1 | 0.65205643 | 1 | -0.389 | 1.224 | chr11 | 111744780 | 111751967 | - | protein_coding |
| ENSG00000067177.15_5 | PHKA1 | 0.99372381 | 1 | -0.389 | 5.840 | chrX | 71798664 | 71934167 | - | protein_coding |
| ENSG00000139629.16_4 | GALNT6 | 0.88130818 | 1 | -0.389 | 3.179 | chr12 | 51745031 | 51786651 | - | protein_coding |
| ENSG00000155868.8_5 | MED7 | 0.95813751 | 1 | -0.389 | 4.101 | chr5 | 156564435 | 156586030 | - | protein_coding |
| ENSG00000100439.10_2 | ABHD4 | 0.99237983 | 1 | -0.389 | 5.612 | chr14 | 23067146 | 23082121 | + | protein_coding |
| ENSG00000237036.5_6 | ZEB1-AS1 | 0.66524345 | 1 | -0.390 | 1.254 | chr10 | 31495207 | 31609376 | - | lncRNA |
| ENSG00000151465.14_4 | CDC123 | 0.99288008 | 1 | -0.390 | 6.217 | chr10 | 12237964 | 12292588 | + | protein_coding |
| ENSG00000222460.1 | RN7SKP271 | 0.47429708 | 1 | -0.390 | -0.830 | chr20 | 34231065 | 34231384 | + | misc_RNA |
| ENSG00000260608.1_7 | AC024270.4 | 0.48258884 | 1 | -0.390 | -0.655 | chr15 | 83681413 | 83730597 | + | lncRNA |
| ENSG00000254449.1_4 | SF3A3P2 | 0.56751757 | 1 | -0.390 | 0.815 | chr11 | 123468654 | 123470160 | - | processed_pseudogene |
| ENSG00000204934.10_7 | ATP6V0E2-AS1 | 0.74951918 | 1 | -0.390 | 2.050 | chr7 | 149564786 | 149577699 | - | lncRNA |
| ENSG00000222345.1 | SNORD19 | 0.52073036 | 1 | -0.390 | -0.128 | chr3 | 52725394 | 52725469 | + | snoRNA |
| ENSG00000237357.2_6 | BX088651.4 | 0.52029646 | 1 | -0.391 | -0.031 | chr9 | 44401766 | 44404440 | + | lncRNA |
| ENSG00000253930.1_6 | TNFRSF10A-AS1 | 0.61167422 | 1 | -0.391 | 0.877 | chr8 | 23046792 | 23048188 | + | lncRNA |
| ENSG00000178700.8_9 | DHFR2 | 0.94106291 | 1 | -0.391 | 3.808 | chr3 | 93766680 | 93782233 | - | protein_coding |
| ENSG00000200169.1 | RNU5D-1 | 0.49634174 | 1 | -0.391 | -0.426 | chr1 | 45196727 | 45196842 | - | snRNA |
| ENSG00000259001.3_6 | AL355075.4 | 0.99882065 | 1 | -0.391 | 12.308 | chr14 | 20811207 | 20811844 | - | lncRNA |
| ENSG00000145491.12_3 | ROPN1L | 0.51401839 | 1 | -0.391 | -0.208 | chr5 | 10441636 | 10472141 | + | protein_coding |
| ENSG00000061656.10_7 | SPAG4 | 0.76744752 | 1 | -0.392 | 2.015 | chr20 | 34203751 | 34209016 | + | protein_coding |
| ENSG00000228192.8_7 | AL512353.1 | 0.55632271 | 1 | -0.392 | 0.073 | chr1 | 43298193 | 43312093 | - | lncRNA |
| ENSG00000234129.8_7 | AC073529.1 | 0.52949812 | 1 | -0.392 | 0.285 | chrX | 10865697 | 11129340 | - | lncRNA |
| ENSG00000138085.16_7 | ATRAID | 0.99377426 | 1 | -0.392 | 6.262 | chr2 | 27434895 | 27440046 | + | protein_coding |
| ENSG00000213453.3_4 | FTH1P3 | 0.48485722 | 1 | -0.392 | -0.601 | chr2 | 27615651 | 27616234 | - | processed_pseudogene |
| ENSG00000149089.13_4 | APIP | 0.97097577 | 1 | -0.392 | 4.636 | chr11 | 34874641 | 34937926 | - | protein_coding |
| ENSG00000074266.21_5 | EED | 0.96017061 | 1 | -0.392 | 4.300 | chr11 | 85955795 | 85989855 | + | protein_coding |
| ENSG00000153214.11_6 | TMEM87B | 0.98778019 | 1 | -0.392 | 5.455 | chr2 | 112812846 | 112876895 | + | protein_coding |
| ENSG00000279529.1_6 | AC008764.8 | 0.52808127 | 1 | -0.392 | 0.306 | chr19 | 16683435 | 16686151 | + | lncRNA |
| ENSG00000123892.12_5 | RAB38 | 0.72095766 | 1 | -0.393 | 1.679 | chr11 | 87846419 | 87908611 | - | protein_coding |
| ENSG00000215067.10_8 | ALOX12-AS1 | 0.72396442 | 1 | -0.393 | 1.709 | chr17 | 6778551 | 6915849 | - | lncRNA |
| ENSG00000204131.9_5 | NHSL2 | 0.54713998 | 1 | -0.393 | 0.246 | chrX | 71130668 | 71381600 | + | protein_coding |
| ENSG00000006327.13_2 | TNFRSF12A | 0.9820085 | 1 | -0.393 | 6.326 | chr16 | 3068446 | 3072384 | + | protein_coding |
| ENSG00000102144.15_5 | PGK1 | 0.99840045 | 1 | -0.393 | 9.106 | chrX | 77166236 | 77384792 | + | protein_coding |
| ENSG00000102221.14_6 | JADE3 | 0.97555809 | 1 | -0.393 | 4.660 | chrX | 46771711 | 46920641 | + | protein_coding |
| ENSG00000256124.6_6 | LINC01152 | 0.4719475 | 1 | -0.393 | -0.684 | chr17 | 70026432 | 70037451 | + | lncRNA |
| ENSG00000246174.9_7 | KCTD21-AS1 | 0.59969319 | 1 | -0.393 | 0.703 | chr11 | 77850817 | 77886369 | + | lncRNA |
| ENSG00000151353.15_6 | TMEM18 | 0.97905437 | 1 | -0.393 | 4.784 | chr2 | 663877 | 677406 | - | protein_coding |
| ENSG00000152455.16_8 | SUV39H2 | 0.97431703 | 1 | -0.393 | 4.732 | chr10 | 14920819 | 14946314 | + | protein_coding |
| ENSG00000142408.4_6 | CACNG8 | 0.76471387 | 1 | -0.393 | 2.373 | chr19 | 54466294 | 54493469 | + | protein_coding |
| ENSG00000143033.18_3 | MTF2 | 0.97342743 | 1 | -0.393 | 4.655 | chr1 | 93544792 | 93604636 | + | protein_coding |
| ENSG00000086189.10_5 | DIMT1 | 0.97521828 | 1 | -0.393 | 5.542 | chr5 | 61643111 | 61699766 | - | protein_coding |
| ENSG00000197670.6_6 | AL157838.1 | 0.64522509 | 1 | -0.394 | 1.255 | chr20 | 52169309 | 52192402 | + | lncRNA |
| ENSG00000164338.10_5 | UTP15 | 0.97662033 | 1 | -0.394 | 5.717 | chr5 | 72861268 | 72879205 | + | protein_coding |
| ENSG00000234176.1_4 | HSPA8P1 | 0.48093541 | 1 | -0.394 | -0.256 | chrX | 120337036 | 120338868 | - | processed_pseudogene |
| ENSG00000109917.11_5 | ZPR1 | 0.99101727 | 1 | -0.394 | 5.507 | chr11 | 116644515 | 116658755 | - | protein_coding |
| ENSG00000229419.1_5 | RALGAPA1P1 | 0.74968444 | 1 | -0.394 | 2.030 | chr9 | 108282409 | 108288640 | + | processed_pseudogene |

| | | | | | | | | | | |
|---|---|---|---|---|---|---|---|---|---|---|
| ENSG00000237522.1_4 | NONOP2 | 0.50117719 | 1 | -0.394 | -0.271 | chr2 | 61163954 | 61165184 | - | processed_pseudogene |
| ENSG00000279289.1_6 | AL136164.3 | 0.76375191 | 1 | -0.394 | 2.441 | chr6 | 72096555 | 72100532 | - | TEC |
| ENSG00000169908.12_5 | TM4SF1 | 0.99628221 | 1 | -0.394 | 8.720 | chr3 | 149086809 | 149095479 | - | protein_coding |
| ENSG00000253276.3_7 | CCDC71L | 0.85162709 | 1 | -0.394 | 3.480 | chr7 | 106294806 | 106301604 | - | protein_coding |
| ENSG00000183337.17_5 | BCOR | 0.99112159 | 1 | -0.395 | 5.886 | chrX | 39909068 | 40036582 | - | protein_coding |
| ENSG00000183684.8_4 | ALYREF | 0.9970625 | 1 | -0.395 | 6.621 | chr17 | 79845711 | 79849462 | - | protein_coding |
| ENSG00000216775.3_5 | AL109918.1 | 0.70299526 | 1 | -0.396 | 1.968 | chr6 | 52529164 | 52533953 | + | transcribed_unprocessed_pseudogene |
| ENSG00000198786.2_1 | MT-ND5 | 0.99516769 | 1 | -0.396 | 8.953 | chrM | 12337 | 14148 | + | protein_coding |
| ENSG00000180596.7_3 | H2BC4 | 0.98913334 | 1 | -0.396 | 7.833 | chr6 | 26115101 | 26124154 | - | protein_coding |
| ENSG00000184361.13_4 | SPATA32 | 0.54912145 | 1 | -0.396 | 0.576 | chr17 | 43331760 | 43339461 | - | protein_coding |
| ENSG00000088451.11_4 | TGDS | 0.90854424 | 1 | -0.397 | 3.491 | chr13 | 95226308 | 95248496 | - | protein_coding |
| ENSG00000230510.6_6 | PPP5D1 | 0.50256675 | 1 | -0.397 | -0.250 | chr19 | 46984053 | 47104457 | - | protein_coding |
| ENSG00000204620.3_7 | AC115618.1 | 0.54954768 | 1 | -0.397 | 0.341 | chrX | 48426402 | 48433248 | - | lncRNA |
| ENSG00000260877.2_6 | AP005233.2 | 0.63202399 | 1 | -0.397 | 1.074 | chr11 | 69186231 | 69187280 | + | lncRNA |
| ENSG00000155970.12_4 | MICU3 | 0.85230616 | 1 | -0.397 | 3.088 | chr8 | 16884747 | 16980151 | + | protein_coding |
| ENSG00000272787.1_7 | AC253536.6 | 0.47333778 | 1 | -0.397 | -0.500 | chr22 | 24311400 | 24312062 | + | lncRNA |
| ENSG00000177706.8 | FAM20C | 0.94140935 | 1 | -0.397 | 4.800 | chr7 | 192969 | 300711 | + | protein_coding |
| ENSG00000156886.12_4 | ITGAD | 0.46826875 | 1 | -0.398 | -0.604 | chr16 | 31404656 | 31437826 | + | protein_coding |
| ENSG00000166436.16_5 | TRIM66 | 0.87933023 | 1 | -0.398 | 3.566 | chr11 | 8633584 | 8704241 | - | protein_coding |
| ENSG00000180423.4_7 | HARBI1 | 0.71535163 | 1 | -0.398 | 1.766 | chr11 | 46624411 | 46639459 | - | protein_coding |
| ENSG00000198042.11_4 | MAK16 | 0.9826089 | 1 | -0.398 | 5.324 | chr8 | 33342700 | 33358780 | + | protein_coding |
| ENSG00000106080.11_3 | FKBP14 | 0.92705633 | 1 | -0.398 | 4.282 | chr7 | 30050203 | 30066318 | - | protein_coding |
| ENSG00000112877.8_3 | CEP72 | 0.92807765 | 1 | -0.398 | 3.779 | chr5 | 612455 | 667283 | + | protein_coding |
| ENSG00000204923.4_4 | FBXO48 | 0.71309971 | 1 | -0.398 | 1.900 | chr2 | 68686554 | 68694426 | - | protein_coding |
| ENSG00000137692.12_4 | DCUN1D5 | 0.98061946 | 1 | -0.398 | 4.947 | chr11 | 102921415 | 102962923 | - | protein_coding |
| ENSG00000270964.1_6 | AC016355.1 | 0.52066863 | 1 | -0.398 | -0.020 | chr15 | 67833410 | 67834942 | - | lncRNA |
| ENSG00000125633.11_5 | CCDC93 | 0.97929509 | 1 | -0.398 | 4.894 | chr2 | 118673054 | 118771709 | - | protein_coding |
| ENSG00000203362.2_7 | POLH-AS1 | 0.46490082 | 1 | -0.398 | -0.640 | chr6 | 43555967 | 43559099 | - | lncRNA |
| ENSG00000181016.9_4 | LSMEM1 | 0.53266137 | 1 | -0.398 | 0.256 | chr7 | 112120908 | 112131117 | + | protein_coding |
| ENSG00000115520.8_3 | COQ10B | 0.97203077 | 1 | -0.399 | 4.876 | chr2 | 198318147 | 198340032 | + | protein_coding |
| ENSG00000111859.17_6 | NEDD9 | 0.99707661 | 1 | -0.399 | 7.009 | chr6 | 11183531 | 11382581 | - | protein_coding |
| ENSG00000260261.2_7 | AC124944.2 | 0.68702764 | 1 | -0.399 | 1.597 | chr3 | 195639949 | 195681356 | - | transcribed_unprocessed_pseudogene |
| ENSG00000040275.17_7 | SPDL1 | 0.99174403 | 1 | -0.399 | 6.411 | chr5 | 169010640 | 169031782 | + | protein_coding |
| ENSG00000106591.4_4 | MRPL32 | 0.9858806 | 1 | -0.399 | 5.443 | chr7 | 42971975 | 42988557 | + | protein_coding |
| ENSG00000180376.17_5 | CCDC66 | 0.92551661 | 1 | -0.399 | 4.149 | chr3 | 56591189 | 56655865 | + | protein_coding |
| ENSG00000196951.11_8 | SCOC-AS1 | 0.57034255 | 1 | -0.399 | 0.738 | chr4 | 141204878 | 141294557 | - | lncRNA |
| ENSG00000100504.17_4 | PYGL | 0.9959709 | 1 | -0.400 | 6.455 | chr14 | 51324609 | 51411201 | - | protein_coding |
| ENSG00000114735.9_3 | HEMK1 | 0.93422773 | 1 | -0.400 | 3.925 | chr3 | 50606583 | 50633599 | + | protein_coding |
| ENSG00000184661.14_3 | CDCA2 | 0.99602621 | 1 | -0.400 | 6.571 | chr8 | 25316715 | 25365427 | + | protein_coding |
| ENSG00000267519.6_7 | AC020916.1 | 0.97068345 | 1 | -0.400 | 5.507 | chr19 | 13934694 | 13953742 | - | lncRNA |
| ENSG00000113013.15_8 | HSPA9 | 0.99835238 | 1 | -0.400 | 9.266 | chr5 | 137889445 | 137911105 | - | protein_coding |
| ENSG00000260757.1_7 | AC093520.1 | 0.6182915 | 1 | -0.400 | 1.303 | chr16 | 31413886 | 31416020 | + | lncRNA |
| ENSG00000232346.1_5 | Z74021.1 | 0.58786049 | 1 | -0.400 | 1.024 | chr22 | 32435477 | 32435883 | + | processed_pseudogene |
| ENSG00000163428.4_4 | LRRC58 | 0.99530647 | 1 | -0.400 | 6.885 | chr3 | 120043356 | 120068201 | - | protein_coding |
| ENSG00000135723.14_5 | FHOD1 | 0.98587149 | 1 | -0.400 | 5.457 | chr16 | 67263290 | 67281384 | - | protein_coding |
| ENSG00000164323.14_6 | CFAP97 | 0.99311902 | 1 | -0.400 | 6.108 | chr4 | 186080819 | 186130658 | - | protein_coding |
| ENSG00000198003.12_5 | CCDC151 | 0.46423528 | 1 | -0.401 | -0.586 | chr19 | 11531272 | 11546603 | - | protein_coding |
| ENSG00000102886.15_4 | GDPD3 | 0.4646797 | 1 | -0.401 | -0.638 | chr16 | 30116131 | 30124858 | - | protein_coding |
| ENSG00000101574.15_5 | METTL4 | 0.90368604 | 1 | -0.401 | 3.599 | chr18 | 2537524 | 2571508 | - | protein_coding |
| ENSG00000151576.10_4 | QTRT2 | 0.99193737 | 1 | -0.401 | 5.866 | chr3 | 113724680 | 113807269 | + | protein_coding |
| ENSG00000120438.12_5 | TCP1 | 0.99786469 | 1 | -0.401 | 7.918 | chr6 | 160199530 | 160210735 | - | protein_coding |
| ENSG00000256525.8_5 | POLG2 | 0.93381115 | 1 | -0.402 | 4.430 | chr17 | 62473902 | 62493172 | - | protein_coding |
| ENSG00000113407.13_4 | TARS1 | 0.99806112 | 1 | -0.402 | 8.063 | chr5 | 33440802 | 33469644 | + | protein_coding |
| ENSG00000187772.8_5 | LIN28B | 0.82220503 | 1 | -0.402 | 2.815 | chr6 | 105384491 | 105531207 | + | protein_coding |
| ENSG00000184368.16_4 | MAP7D2 | 0.95611581 | 1 | -0.402 | 4.259 | chrX | 20024831 | 20135025 | - | protein_coding |
| ENSG00000260495.1_6 | AC009148.1 | 0.5297385 | 1 | -0.402 | 0.101 | chr16 | 81344336 | 81347028 | - | lncRNA |
| ENSG00000224321.1_4 | RPL12P14 | 0.67402465 | 1 | -0.402 | 1.545 | chr1 | 16119291 | 16119780 | + | processed_pseudogene |
| ENSG00000138698.14_4 | RAP1GDS1 | 0.97613435 | 1 | -0.402 | 5.193 | chr4 | 99182535 | 99365012 | + | protein_coding |

| | | | | | | | | | | |
|---|---|---|---|---|---|---|---|---|---|---|
| ENSG00000172803.18_3 | SNX32 | 0.66969149 | 1 | -0.403 | 1.227 | chr11 | 65601305 | 65624367 | + | protein_coding |
| ENSG00000176055.10_4 | MBLAC2 | 0.86126041 | 1 | -0.403 | 3.372 | chr5 | 89754026 | 89770588 | - | protein_coding |
| ENSG00000165480.16_4 | SKA3 | 0.97082207 | 1 | -0.403 | 4.846 | chr13 | 21727734 | 21750691 | - | protein_coding |
| ENSG00000188452.14_7 | CERKL | 0.49594877 | 1 | -0.403 | -0.036 | chr2 | 182401399 | 182545392 | - | protein_coding |
| ENSG00000100401.19_4 | RANGAP1 | 0.98525651 | 1 | -0.403 | 6.531 | chr22 | 41641615 | 41682255 | - | protein_coding |
| ENSG00000268584.1_6 | AC073389.1 | 0.47439741 | 1 | -0.403 | -0.412 | chr10 | 75385754 | 75386548 | + | lncRNA |
| ENSG00000008735.14_4 | MAPK8IP2 | 0.7233344 | 1 | -0.403 | 1.937 | chr22 | 51039222 | 51052409 | + | protein_coding |
| ENSG00000258711.2_7 | AL358334.2 | 0.53076105 | 1 | -0.403 | 0.423 | chr14 | 51422977 | 51428720 | - | lncRNA |
| ENSG00000156261.13_7 | CCT8 | 0.99275462 | 1 | -0.404 | 7.559 | chr21 | 30428126 | 30446118 | - | protein_coding |
| ENSG00000170779.11_4 | CDCA4 | 0.9776504 | 1 | -0.404 | 4.870 | chr14 | 105475910 | 105487420 | - | protein_coding |
| ENSG00000225022.1_4 | UBE2D3P1 | 0.45829509 | 1 | -0.404 | -0.694 | chr20 | 5273034 | 5273477 | - | processed_pseudogene |
| ENSG00000100519.12_7 | PSMC6 | 0.99394193 | 1 | -0.404 | 6.781 | chr14 | 53173896 | 53195308 | + | protein_coding |
| ENSG00000125459.16_5 | MSTO1 | 0.92751647 | 1 | -0.404 | 3.938 | chr1 | 155579979 | 155584762 | + | protein_coding |
| ENSG00000142686.8_4 | C1orf216 | 0.94191054 | 1 | -0.404 | 4.143 | chr1 | 36179478 | 36184495 | - | protein_coding |
| ENSG00000102218.6_3 | RP2 | 0.9615167 | 1 | -0.404 | 5.108 | chrX | 46696478 | 46741793 | + | protein_coding |
| ENSG00000085377.14_5 | PREP | 0.96953576 | 1 | -0.404 | 4.681 | chr6 | 105721095 | 105901937 | - | protein_coding |
| ENSG00000085999.13_6 | RAD54L | 0.83063652 | 1 | -0.405 | 2.800 | chr1 | 46712133 | 46744152 | + | protein_coding |
| ENSG00000147854.17_6 | UHRF2 | 0.99299155 | 1 | -0.405 | 6.015 | chr9 | 6413151 | 6507054 | + | protein_coding |
| ENSG00000164654.16_3 | MIOS | 0.98363347 | 1 | -0.405 | 5.250 | chr7 | 7606506 | 7648563 | + | protein_coding |
| ENSG00000039123.16_5 | MTREX | 0.9938537 | 1 | -0.405 | 6.646 | chr5 | 54603817 | 54721407 | + | protein_coding |
| ENSG00000277581.1_7 | AL023803.3 | 0.46439708 | 1 | -0.405 | -1.197 | chr20 | 37591115 | 37591636 | + | lncRNA |
| ENSG00000277290.1_5 | AC136475.10 | 0.45741667 | 1 | -0.405 | -0.578 | chr11 | 243099 | 243483 | - | processed_pseudogene |
| ENSG00000221968.9_7 | FADS3 | 0.95047579 | 1 | -0.405 | 4.331 | chr11 | 61640991 | 61659523 | - | protein_coding |
| ENSG00000256393.1_4 | RPL41P5 | 0.4578609 | 1 | -0.406 | -1.118 | chr12 | 93477374 | 93477451 | - | processed_pseudogene |
| ENSG00000188372.15_5 | ZP3 | 0.87794232 | 1 | -0.406 | 3.385 | chr7 | 76026835 | 76071388 | + | protein_coding |
| ENSG00000235636.1_4 | NUS1P1 | 0.45530356 | 1 | -0.406 | -0.845 | chrX | 47372001 | 47372882 | - | processed_pseudogene |
| ENSG00000151247.12_6 | EIF4E | 0.97704941 | 1 | -0.406 | 5.614 | chr4 | 99792835 | 99851788 | - | protein_coding |
| ENSG00000146094.14_5 | DOK3 | 0.79662158 | 1 | -0.406 | 2.463 | chr5 | 176928908 | 176938275 | - | protein_coding |
| ENSG00000112759.19_5 | SLC29A1 | 0.97654499 | 1 | -0.406 | 5.170 | chr6 | 44187290 | 44201879 | + | protein_coding |
| ENSG00000149308.17_5 | NPAT | 0.9644834 | 1 | -0.406 | 5.090 | chr11 | 108027942 | 108093365 | - | protein_coding |
| ENSG00000136271.10_4 | DDX56 | 0.99586514 | 1 | -0.406 | 6.785 | chr7 | 44605016 | 44614650 | - | protein_coding |
| ENSG00000233476.3_4 | EEF1A1P6 | 0.61255921 | 1 | -0.406 | 2.392 | chr7 | 22550235 | 22551618 | - | processed_pseudogene |
| ENSG00000174032.17_5 | SLC25A30 | 0.86647312 | 1 | -0.406 | 3.378 | chr13 | 45967451 | 45992590 | - | protein_coding |
| ENSG00000174939.11_6 | ASPHD1 | 0.69159393 | 1 | -0.406 | 1.886 | chr16 | 29911696 | 29931185 | + | protein_coding |
| ENSG00000107937.19_3 | GTPBP4 | 0.99664343 | 1 | -0.406 | 7.018 | chr10 | 1034374 | 1065872 | + | protein_coding |
| ENSG00000134030.14_7 | CTIF | 0.95269282 | 1 | -0.406 | 4.739 | chr18 | 46065402 | 46389588 | + | protein_coding |
| ENSG00000274214.1_5 | AP005212.2 | 0.46992898 | 1 | -0.407 | -1.322 | chr18 | 14253547 | 14254074 | + | processed_pseudogene |
| ENSG00000237788.1_5 | AL162615.1 | 0.4585285 | 1 | -0.407 | -0.459 | chr20 | 48445662 | 48447622 | + | processed_pseudogene |
| ENSG00000253636.1_6 | AC022893.1 | 0.46044586 | 1 | -0.407 | -0.358 | chr8 | 73964413 | 73975296 | + | lncRNA |
| ENSG00000144320.14_6 | LNPK | 0.97839327 | 1 | -0.407 | 5.430 | chr2 | 176788610 | 176867567 | - | protein_coding |
| ENSG00000156453.13_8 | PCDH1 | 0.49126732 | 1 | -0.407 | 0.108 | chr5 | 141232676 | 141258811 | - | protein_coding |
| ENSG00000285857.1_1 | AC016727.3 | 0.57518039 | 1 | -0.407 | 0.877 | chr2 | 61754475 | 61756295 | + | lncRNA |
| ENSG00000265415.1_7 | AC099850.4 | 0.78907263 | 1 | -0.407 | 2.676 | chr17 | 57280038 | 57281190 | - | lncRNA |
| ENSG00000254554.1_7 | AC080023.1 | 0.46906943 | 1 | -0.407 | -0.294 | chr11 | 10324204 | 10325251 | - | lncRNA |
| ENSG00000172244.9_7 | C5orf34 | 0.88475206 | 1 | -0.408 | 3.408 | chr5 | 43486811 | 43515250 | - | protein_coding |
| ENSG00000267898.1_6 | AC026803.2 | 0.48531871 | 1 | -0.408 | 0.133 | chr19 | 49467232 | 49468415 | + | lncRNA |
| ENSG00000262732.1_7 | AC010401.1 | 0.4581166 | 1 | -0.408 | -0.269 | chr16 | 14024534 | 14029492 | - | lncRNA |
| ENSG00000181322.14_6 | NME9 | 0.46038139 | 1 | -0.408 | -0.534 | chr3 | 137980279 | 138048728 | - | protein_coding |
| ENSG00000162604.12_8 | TM2D1 | 0.91225539 | 1 | -0.408 | 3.826 | chr1 | 62146718 | 62191095 | - | protein_coding |
| ENSG00000181218.5_3 | H2AW | 0.51753038 | 1 | -0.408 | 3.935 | chr1 | 228644680 | 228645574 | - | protein_coding |
| ENSG00000279863.1_5 | AC069547.1 | 0.46856428 | 1 | -0.408 | -0.412 | chr10 | 52094298 | 52095883 | + | TEC |
| ENSG00000196420.7_3 | S100A5 | 0.49343188 | 1 | -0.408 | -0.075 | chr1 | 153509623 | 153514241 | - | protein_coding |
| ENSG00000213939.4_5 | AC091153.1 | 0.46399016 | 1 | -0.408 | -0.534 | chr17 | 4608111 | 4608512 | - | processed_pseudogene |
| ENSG00000285948.1_3 | AC123768.5 | 0.46989785 | 1 | -0.408 | -0.413 | chr15 | 32916307 | 32933816 | - | lncRNA |
| ENSG00000161647.18_4 | MPP3 | 0.96935951 | 1 | -0.408 | 4.885 | chr17 | 41878167 | 41910538 | - | protein_coding |
| ENSG00000151503.12_5 | NCAPD3 | 0.98905116 | 1 | -0.408 | 6.038 | chr11 | 134020014 | 134095348 | - | protein_coding |
| ENSG00000108506.12_7 | INTS2 | 0.99177428 | 1 | -0.409 | 6.260 | chr17 | 59942728 | 60005377 | - | protein_coding |

| | | | | | | | | | | |
|---|---|---|---|---|---|---|---|---|---|---|
| ENSG00000115274.15_5 | INO80B | 0.45097484 | 1 | -0.409 | -0.909 | chr2 | 74682214 | 74685071 | + | protein_coding |
| ENSG00000117593.12_6 | DARS2 | 0.99265369 | 1 | -0.409 | 6.583 | chr1 | 173793791 | 173827946 | + | protein_coding |
| ENSG00000160117.16_6 | ANKLE1 | 0.46761418 | 1 | -0.409 | -0.209 | chr19 | 17392454 | 17398455 | + | protein_coding |
| ENSG00000106780.9_4 | MEGF9 | 0.99467201 | 1 | -0.409 | 6.799 | chr9 | 123363089 | 123476748 | - | protein_coding |
| ENSG00000115271.11_6 | GCA | 0.91109099 | 1 | -0.409 | 3.847 | chr2 | 163175350 | 163228105 | + | protein_coding |
| ENSG00000250959.2_4 | GLUD1P3 | 0.51179024 | 1 | -0.410 | 0.296 | chr10 | 75490320 | 75497069 | + | transcribed_unprocessed_pseudogene |
| ENSG00000108578.15_6 | BLMH | 0.98567645 | 1 | -0.410 | 5.670 | chr17 | 28575221 | 28619074 | - | protein_coding |
| ENSG00000112640.15_5 | PPP2R5D | 0.95330218 | 1 | -0.410 | 4.716 | chr6 | 42952291 | 42980080 | + | protein_coding |
| ENSG00000070718.12_5 | AP3M2 | 0.96885881 | 1 | -0.410 | 4.836 | chr8 | 42010464 | 42029191 | + | protein_coding |
| ENSG00000131368.8_4 | MRPS25 | 0.96778508 | 1 | -0.410 | 4.832 | chr3 | 15083967 | 15106824 | - | protein_coding |
| ENSG00000261476.1_9 | AC009163.4 | 0.45016144 | 1 | -0.410 | -0.908 | chr16 | 75492150 | 75493915 | - | lncRNA |
| ENSG00000259771.1_8 | AC092756.1 | 0.49678449 | 1 | -0.410 | 0.179 | chr15 | 59558919 | 59563361 | - | lncRNA |
| ENSG00000072401.15_3 | UBE2D1 | 0.97157603 | 1 | -0.410 | 5.055 | chr10 | 60094739 | 60130511 | + | protein_coding |
| ENSG00000171174.15_4 | RBKS | 0.74309921 | 1 | -0.410 | 2.519 | chr2 | 28004246 | 28113548 | - | protein_coding |
| ENSG00000140299.12_6 | BNIP2 | 0.99470457 | 1 | -0.411 | 6.976 | chr15 | 59951345 | 59981733 | - | protein_coding |
| ENSG00000184507.16_3 | NUTM1 | 0.44874126 | 1 | -0.411 | -0.812 | chr15 | 34635516 | 34649938 | + | protein_coding |
| ENSG00000235204.1_7 | AL162724.2 | 0.45685985 | 1 | -0.411 | -0.460 | chr9 | 127021529 | 127023435 | - | lncRNA |
| ENSG00000172543.8_5 | CTSW | 0.4669719 | 1 | -0.411 | -0.412 | chr11 | 65647308 | 65651212 | + | protein_coding |
| ENSG00000181744.8_6 | DIPK2A | 0.952525 | 1 | -0.411 | 4.382 | chr3 | 143690640 | 143767561 | + | protein_coding |
| ENSG00000272842.1_6 | AL391834.1 | 0.47208471 | 1 | -0.411 | -0.279 | chr9 | 19371384 | 19371943 | - | lncRNA |
| ENSG00000271781.1_6 | AC026740.1 | 0.45654925 | 1 | -0.411 | -0.668 | chr5 | 675941 | 676731 | + | lncRNA |
| ENSG00000273888.1_5 | FRMD6-AS1 | 0.55247174 | 1 | -0.412 | 0.696 | chr14 | 52116234 | 52118462 | - | lncRNA |
| ENSG00000008324.11_2 | SS18L2 | 0.79378883 | 1 | -0.412 | 2.605 | chr3 | 42623332 | 42636606 | + | protein_coding |
| ENSG00000138443.16_5 | ABI2 | 0.98854584 | 1 | -0.412 | 6.214 | chr2 | 204192962 | 204312451 | + | protein_coding |
| ENSG00000254088.1_4 | SLC2A3P4 | 0.47258711 | 1 | -0.412 | -0.301 | chr8 | 87515820 | 87517290 | + | processed_pseudogene |
| ENSG00000205790.1_6 | DPP9-AS1 | 0.45914927 | 1 | -0.412 | -0.437 | chr19 | 4679294 | 4685960 | + | lncRNA |
| ENSG00000104408.10_5 | EIF3E | 0.99553193 | 1 | -0.412 | 8.250 | chr8 | 109213445 | 109447562 | - | protein_coding |
| ENSG00000275052.5_6 | PPP4R3B | 0.98286349 | 1 | -0.412 | 6.494 | chr2 | 55774428 | 55846015 | - | protein_coding |
| ENSG00000175354.20_6 | PTPN2 | 0.96634175 | 1 | -0.412 | 5.011 | chr18 | 12785477 | 12929642 | - | protein_coding |
| ENSG00000234753.5_7 | FOXP4-AS1 | 0.45410235 | 1 | -0.412 | -0.461 | chr6 | 41462591 | 41516359 | - | lncRNA |
| ENSG00000136270.14_8 | TBRG4 | 0.98758126 | 1 | -0.412 | 6.442 | chr7 | 45139699 | 45151646 | - | protein_coding |
| ENSG00000212237.1_1 | RNA5SP18 | 0.44756696 | 1 | -0.412 | -0.976 | chr1 | 228783659 | 228783779 | - | rRNA_pseudogene |
| ENSG00000091127.14_4 | PUS7 | 0.98566472 | 1 | -0.412 | 5.954 | chr7 | 105080108 | 105162718 | - | protein_coding |
| ENSG00000274340.1_6 | AC032011.1 | 0.45988917 | 1 | -0.412 | -0.510 | chr15 | 62679526 | 62681074 | - | lncRNA |
| ENSG00000172469.16_4 | MANEA | 0.92508125 | 1 | -0.412 | 4.299 | chr6 | 96025411 | 96057328 | + | protein_coding |
| ENSG00000092853.14_5 | CLSPN | 0.98677172 | 1 | -0.412 | 5.712 | chr1 | 36185819 | 36235579 | - | protein_coding |
| ENSG00000132436.11_9 | FIGNL1 | 0.98317301 | 1 | -0.413 | 5.818 | chr7 | 50511826 | 50610233 | - | protein_coding |
| ENSG00000050344.9_3 | NFE2L3 | 0.94824849 | 1 | -0.413 | 5.162 | chr7 | 26191818 | 26226757 | + | protein_coding |
| ENSG00000172172.8_5 | MRPL13 | 0.97000121 | 1 | -0.413 | 5.570 | chr8 | 121393000 | 121457642 | - | protein_coding |
| ENSG00000113522.14_8 | RAD50 | 0.8896721 | 1 | -0.413 | 3.561 | chr5 | 131891711 | 131982041 | + | protein_coding |
| ENSG00000105821.15_4 | DNAJC2 | 0.97966093 | 1 | -0.413 | 5.598 | chr7 | 102952736 | 102985277 | - | protein_coding |
| ENSG00000212440.1 | SNORA75 | 0.45028065 | 1 | -0.413 | -0.784 | chr12 | 9439269 | 9439418 | - | snoRNA |
| ENSG00000272068.1_6 | AL365181.2 | 0.76767822 | 1 | -0.413 | 2.593 | chr1 | 156607575 | 156610796 | - | lncRNA |
| ENSG00000197056.11_4 | ZMYM1 | 0.95690015 | 1 | -0.413 | 5.061 | chr1 | 35497773 | 35581460 | + | protein_coding |
| ENSG00000171246.6_4 | NPTX1 | 0.83845933 | 1 | -0.413 | 3.288 | chr17 | 78440634 | 78451643 | - | protein_coding |
| ENSG00000262089.1_8 | AC040977.1 | 0.45230496 | 1 | -0.413 | -0.725 | chr17 | 6897961 | 6898508 | - | lncRNA |
| ENSG00000132622.11_3 | HSPA12B | 0.65516884 | 1 | -0.413 | 1.725 | chr20 | 3713332 | 3733758 | + | protein_coding |
| ENSG00000089006.16_4 | SNX5 | 0.99627805 | 1 | -0.414 | 7.838 | chr20 | 17922241 | 17949623 | - | protein_coding |
| ENSG00000108384.15_6 | RAD51C | 0.97345362 | 1 | -0.414 | 5.226 | chr17 | 56769934 | 56812972 | + | protein_coding |
| ENSG00000215187.11_4 | FAM166B | 0.44504243 | 1 | -0.414 | -0.782 | chr9 | 35561828 | 35563896 | - | protein_coding |
| ENSG00000231050.1_6 | AL109917.1 | 0.44479133 | 1 | -0.414 | -0.846 | chr1 | 1822910 | 1824097 | + | lncRNA |
| ENSG00000112118.20_6 | MCM3 | 0.99591728 | 1 | -0.414 | 7.300 | chr6 | 52128812 | 52149679 | - | protein_coding |
| ENSG00000236409.2_5 | NRADDP | 0.48060009 | 1 | -0.414 | -0.100 | chr3 | 47052314 | 47054955 | + | transcribed_unitary_pseudogene |
| ENSG00000125798.14_5 | FOXA2 | 0.91824279 | 1 | -0.414 | 4.966 | chr20 | 22561643 | 22566093 | - | protein_coding |
| ENSG00000166435.15_7 | XRRA1 | 0.86667069 | 1 | -0.414 | 3.370 | chr11 | 74518784 | 74660245 | - | protein_coding |
| ENSG00000277232.2_4 | GTSE1-DT | 0.46699482 | 1 | -0.414 | -0.391 | chr22 | 46691040 | 46692557 | - | lncRNA |
| ENSG00000186132.15_4 | C2orf76 | 0.74308426 | 1 | -0.415 | 2.213 | chr2 | 120059801 | 120124410 | - | protein_coding |
| ENSG00000188610.8 | FAM72B | 0.75331969 | 1 | -0.415 | 2.559 | chr1 | 120837756 | 120855681 | + | protein_coding |
| ENSG00000254577.1_8 | AC087276.2 | 0.4434741 | 1 | -0.415 | -0.976 | chr11 | 43411833 | 43417045 | + | lncRNA |

| | | | | | | | | | | |
|---|---|---|---|---|---|---|---|---|---|---|
| ENSG00000151012.13_4 | SLC7A11 | 0.99338295 | 1 | -0.415 | 9.014 | chr4 | 139085251 | 139163503 | - | protein_coding |
| ENSG00000144524.17_2 | COPS7B | 0.93263512 | 1 | -0.415 | 4.254 | chr2 | 232646381 | 232673964 | + | protein_coding |
| ENSG00000286319.1_2 | AL135978.2 | 0.44938034 | 1 | -0.415 | -0.642 | chr14 | 67750466 | 67760790 | - | lncRNA |
| ENSG00000236779.1_5 | AL512306.1 | 0.44618672 | 1 | -0.415 | -0.844 | chr1 | 204497973 | 204498820 | - | processed_pseudogene |
| ENSG00000286177.1_3 | AC011462.5 | 0.64861783 | 1 | -0.415 | 1.978 | chr19 | 41813302 | 41816317 | - | lncRNA |
| ENSG00000233846.2_5 | SELENOTP1 | 0.44573717 | 1 | -0.416 | -0.724 | chr9 | 6278667 | 6279153 | + | processed_pseudogene |
| ENSG00000158201.10_7 | ABHD3 | 0.92838058 | 1 | -0.416 | 4.406 | chr18 | 19230862 | 19284741 | - | protein_coding |
| ENSG00000243224.1_6 | AC006252.1 | 0.45794759 | 1 | -0.416 | -0.414 | chr3 | 52273274 | 52275113 | + | lncRNA |
| ENSG00000287733.1_1 | AC083862.3 | 0.4460469 | 1 | -0.416 | -0.754 | chr7 | 134855568 | 134857166 | + | lncRNA |
| ENSG00000156110.14_5 | ADK | 0.92333738 | 1 | -0.416 | 4.779 | chr10 | 75910960 | 76469721 | + | protein_coding |
| ENSG00000269653.1_8 | AC011479.4 | 0.44457959 | 1 | -0.416 | -0.754 | chr19 | 38675016 | 38676905 | + | lncRNA |
| ENSG00000116922.14_6 | C1orf109 | 0.93263091 | 1 | -0.416 | 4.252 | chr1 | 38147242 | 38157921 | - | protein_coding |
| ENSG00000260423.2_6 | LINC02367 | 0.49436303 | 1 | -0.416 | 0.228 | chr12 | 9499652 | 9552983 | + | lncRNA |
| ENSG00000171155.8_4 | C1GALT1C1 | 0.96589955 | 1 | -0.416 | 5.065 | chrX | 119759529 | 119763909 | - | protein_coding |
| ENSG00000269652.1_7 | AC011510.1 | 0.46051315 | 1 | -0.417 | 0.004 | chr19 | 41727331 | 41727956 | - | lncRNA |
| ENSG00000238197.6_7 | PAXBP1-AS1 | 0.73170404 | 1 | -0.417 | 2.371 | chr21 | 34100408 | 34119376 | + | lncRNA |
| ENSG00000265681.7_7 | RPL17 | 0.7258579 | 1 | -0.417 | 2.163 | chr18 | 47014823 | 47018893 | - | protein_coding |
| ENSG00000108590.11_4 | MED31 | 0.73991049 | 1 | -0.417 | 2.380 | chr17 | 6546631 | 6554954 | - | protein_coding |
| ENSG00000184226.15_7 | PCDH9 | 0.99217257 | 1 | -0.417 | 7.246 | chr13 | 66876966 | 67804577 | - | protein_coding |
| ENSG00000223764.2_7 | LINC02593 | 0.61553377 | 1 | -0.417 | 1.367 | chr1 | 852245 | 856396 | - | lncRNA |
| ENSG00000181026.15_3 | AEN | 0.9909424 | 1 | -0.417 | 6.479 | chr15 | 89164568 | 89175512 | + | protein_coding |
| ENSG00000110435.12_3 | PDHX | 0.98114349 | 1 | -0.417 | 5.636 | chr11 | 34937376 | 35042138 | + | protein_coding |
| ENSG00000100604.13_5 | CHGA | 0.45141766 | 1 | -0.418 | -0.536 | chr14 | 93389495 | 93401630 | + | protein_coding |
| ENSG00000144021.3_3 | CIAO1 | 0.99128602 | 1 | -0.418 | 6.380 | chr2 | 96931897 | 96939911 | + | protein_coding |
| ENSG00000214248.2_6 | AC010336.1 | 0.44351985 | 1 | -0.418 | -0.753 | chr19 | 7963689 | 7968427 | - | lncRNA |
| ENSG00000231584.8_2 | FAHD2CP | 0.51520511 | 1 | -0.418 | 0.683 | chr2 | 96676299 | 96689128 | + | transcribed_unprocessed_pseudogene |
| ENSG00000187837.3_3 | H1-2 | 0.99083779 | 1 | -0.418 | 8.746 | chr6 | 26056015 | 26056656 | - | protein_coding |
| ENSG00000119280.17_4 | C1orf198 | 0.96333413 | 1 | -0.418 | 5.051 | chr1 | 230972865 | 231005335 | - | protein_coding |
| ENSG00000132773.12_3 | TOE1 | 0.84526808 | 1 | -0.418 | 3.563 | chr1 | 45805724 | 45809645 | + | protein_coding |
| ENSG00000090273.14_4 | NUDC | 0.9810107 | 1 | -0.419 | 5.793 | chr1 | 27226729 | 27273362 | + | protein_coding |
| ENSG00000264785.1_7 | AC005722.4 | 0.57949799 | 1 | -0.419 | 0.802 | chr17 | 19640995 | 19641855 | + | lncRNA |
| ENSG00000144736.14_4 | SHQ1 | 0.95968669 | 1 | -0.419 | 4.993 | chr3 | 72798428 | 72911065 | - | protein_coding |
| ENSG00000153832.12_4 | FBXO36 | 0.77968719 | 1 | -0.419 | 2.629 | chr2 | 230787018 | 230877835 | + | protein_coding |
| ENSG00000115234.11_3 | SNX17 | 0.99172658 | 1 | -0.419 | 7.005 | chr2 | 27593363 | 27600402 | + | protein_coding |
| ENSG00000259424.1_7 | IRAIN | 0.45060725 | 1 | -0.419 | -0.536 | chr15 | 99190180 | 99190600 | - | lncRNA |
| ENSG00000188153.13_5 | COL4A5 | 0.9905047 | 1 | -0.419 | 6.529 | chrX | 107683074 | 107940775 | + | protein_coding |
| ENSG00000272097.2_7 | AL024498.1 | 0.44910635 | 1 | -0.419 | -0.563 | chr6 | 10743557 | 10747896 | - | lncRNA |
| ENSG00000286964.1_1 | AL136169.1 | 0.53375871 | 1 | -0.419 | 0.697 | chrX | 134171311 | 134174385 | + | lncRNA |
| ENSG00000270022.3_6 | Z93241.1 | 0.50964906 | 1 | -0.419 | 0.474 | chr22 | 43011250 | 43011913 | + | lncRNA |
| ENSG00000149289.11_4 | ZC3H12C | 0.90620586 | 1 | -0.419 | 4.274 | chr11 | 109964118 | 110042566 | + | protein_coding |
| ENSG00000107815.10_6 | TWNK | 0.90742464 | 1 | -0.419 | 4.270 | chr10 | 102747124 | 102754160 | + | protein_coding |
| ENSG00000001497.16_3 | LAS1L | 0.9866435 | 1 | -0.420 | 6.022 | chrX | 64732462 | 64754655 | - | protein_coding |
| ENSG00000236060.2_4 | HSPB1P1 | 0.44530078 | 1 | -0.420 | -0.397 | chr9 | 74622797 | 74623052 | - | processed_pseudogene |
| ENSG00000136122.17_5 | BORA | 0.80420409 | 1 | -0.420 | 3.383 | chr13 | 73301887 | 73330336 | + | protein_coding |
| ENSG00000129071.9_3 | MBD4 | 0.96937251 | 1 | -0.420 | 5.263 | chr3 | 129149787 | 129159022 | - | protein_coding |
| ENSG00000260747.1_5 | AC022968.1 | 0.43856194 | 1 | -0.420 | -0.786 | chr18 | 65945039 | 65945480 | - | processed_pseudogene |
| ENSG00000007402.11_4 | CACNA2D2 | 0.4564674 | 1 | -0.420 | -0.242 | chr3 | 50400230 | 50541675 | - | protein_coding |
| ENSG00000277957.1_3 | SENP3-EIF4A1 | 0.43733159 | 1 | -0.420 | -0.942 | chr17 | 7466604 | 7482033 | + | protein_coding |
| ENSG00000224934.5_6 | AL391684.1 | 0.72206607 | 1 | -0.420 | 2.167 | chr10 | 101190676 | 101223016 | + | lncRNA |
| ENSG00000163781.14_7 | TOPBP1 | 0.9918036 | 1 | -0.421 | 7.179 | chr3 | 133317019 | 133381224 | - | protein_coding |
| ENSG00000256238.1_4 | SUPT16HP1 | 0.52437932 | 1 | -0.421 | 0.512 | chr12 | 16552208 | 16555340 | - | processed_pseudogene |
| ENSG00000197147.13_5 | LRRC8B | 0.92652812 | 1 | -0.421 | 4.276 | chr1 | 89990395 | 90063423 | + | protein_coding |
| ENSG00000262943.7_2 | ALOX12P2 | 0.8705031 | 1 | -0.421 | 3.715 | chr17 | 6757180 | 6857426 | + | transcribed_unprocessed_pseudogene |
| ENSG00000166402.8_4 | TUB | 0.97321064 | 1 | -0.421 | 5.192 | chr11 | 8040791 | 8127659 | + | protein_coding |
| ENSG00000148634.15_4 | HERC4 | 0.98869907 | 1 | -0.421 | 6.709 | chr10 | 69681656 | 69835105 | - | protein_coding |
| ENSG00000109805.10_3 | NCAPG | 0.98699334 | 1 | -0.421 | 6.229 | chr4 | 17812602 | 17846488 | + | protein_coding |

| | | | | | | | | | | |
|---|---|---|---|---|---|---|---|---|---|---|
| ENSG00000151014.6_3 | NOCT | 0.85328632 | 1 | -0.421 | 3.370 | chr4 | 139936935 | 139967093 | + | protein_coding |
| ENSG00000105948.13_3 | TTC26 | 0.90397692 | 1 | -0.421 | 4.156 | chr7 | 138818490 | 138876732 | + | protein_coding |
| ENSG00000212396.1_1 | RNA5SP323 | 0.43582815 | 1 | -0.421 | -0.877 | chr10 | 95270305 | 95270437 | - | rRNA_pseudogene |
| ENSG00000144306.15_5 | SCRN3 | 0.92264181 | 1 | -0.422 | 4.480 | chr2 | 175260458 | 175294303 | + | protein_coding |
| ENSG00000147324.11_4 | MFHAS1 | 0.98059022 | 1 | -0.422 | 5.876 | chr8 | 8640864 | 8751140 | - | protein_coding |
| ENSG00000250838.1_8 | AC091133.3 | 0.47253826 | 1 | -0.422 | 0.406 | chr17 | 47072628 | 47074854 | - | lncRNA |
| ENSG00000100029.18_6 | PES1 | 0.98839679 | 1 | -0.422 | 6.562 | chr22 | 30972612 | 31003070 | - | protein_coding |
| ENSG00000168283.14_5 | BMI1 | 0.958871 | 1 | -0.422 | 5.054 | chr10 | 22610028 | 22620413 | + | protein_coding |
| ENSG00000270427.1_2 | NRBF2P5 | 0.44381381 | 1 | -0.422 | -0.347 | chr10 | 5818034 | 5818821 | + | processed_pseudogene |
| ENSG00000226965.2_7 | AC073114.1 | 0.46705775 | 1 | -0.422 | -0.282 | chr7 | 109748088 | 110174814 | - | lncRNA |
| ENSG00000067334.14_5 | DNTTIP2 | 0.97504098 | 1 | -0.422 | 6.072 | chr1 | 94331840 | 94345474 | - | protein_coding |
| ENSG00000249786.7_8 | EAF1-AS1 | 0.43506478 | 1 | -0.422 | -0.784 | chr3 | 15477678 | 15497447 | - | lncRNA |
| ENSG00000113460.13_3 | BRIX1 | 0.96061949 | 1 | -0.423 | 5.475 | chr5 | 34915782 | 34926101 | + | protein_coding |
| ENSG00000164167.12_5 | LSM6 | 0.81239683 | 1 | -0.423 | 3.227 | chr4 | 147096855 | 147121152 | + | protein_coding |
| ENSG00000115464.15_5 | USP34 | 0.99182599 | 1 | -0.423 | 7.943 | chr2 | 61414598 | 61698222 | - | protein_coding |
| ENSG00000070087.14_5 | PFN2 | 0.99082701 | 1 | -0.423 | 7.159 | chr3 | 149682691 | 149768575 | - | protein_coding |
| ENSG00000236810.6_7 | ELOA-AS1 | 0.43816026 | 1 | -0.423 | -0.561 | chr1 | 24033391 | 24104786 | - | lncRNA |
| ENSG00000276166.1_6 | AC092118.2 | 0.45490503 | 1 | -0.423 | -0.260 | chr16 | 57793270 | 57793936 | + | lncRNA |
| ENSG00000113649.11_2 | TCERG1 | 0.98767589 | 1 | -0.423 | 6.926 | chr5 | 145826874 | 145891524 | + | protein_coding |
| ENSG00000119927.14_4 | GPAM | 0.95242335 | 1 | -0.423 | 4.800 | chr10 | 113909623 | 113975135 | - | protein_coding |
| ENSG00000267809.1_2 | NDUFV2P1 | 0.51394885 | 1 | -0.423 | 0.508 | chr19 | 53727226 | 53728132 | - | processed_pseudogene |
| ENSG00000222898.1 | RN7SKP97 | 0.49650399 | 1 | -0.423 | 0.358 | chr8 | 62544239 | 62544571 | - | misc_RNA |
| ENSG00000183762.12_3 | KREMEN1 | 0.82528361 | 1 | -0.423 | 3.131 | chr22 | 29469066 | 29564321 | + | protein_coding |
| ENSG00000165637.13_4 | VDAC2 | 0.99074607 | 1 | -0.423 | 6.940 | chr10 | 76969912 | 76991206 | + | protein_coding |
| ENSG00000277157.1_3 | H4C4 | 0.93296343 | 1 | -0.423 | 6.976 | chr6 | 26188993 | 26189304 | - | protein_coding |
| ENSG00000253729.8_4 | PRKDC | 0.98679721 | 1 | -0.424 | 9.105 | chr8 | 48685672 | 48872738 | - | protein_coding |
| ENSG00000130477.15_7 | UNC13A | 0.84523192 | 1 | -0.424 | 3.379 | chr19 | 17712137 | 17799174 | - | protein_coding |
| ENSG00000271888.1_6 | AL136162.1 | 0.43315154 | 1 | -0.424 | -0.911 | chr6 | 15244154 | 15245231 | - | lncRNA |
| ENSG00000227398.4_6 | KIF9-AS1 | 0.51979955 | 1 | -0.424 | 0.699 | chr3 | 47205676 | 47288091 | + | lncRNA |
| ENSG00000246250.3_7 | AC087521.2 | 0.54530435 | 1 | -0.424 | 1.052 | chr11 | 43849067 | 43902276 | - | lncRNA |
| ENSG00000170113.16_6 | NIPA1 | 0.97762689 | 1 | -0.424 | 6.158 | chr15 | 23043279 | 23100005 | - | protein_coding |
| ENSG00000153714.6_3 | LURAP1L | 0.87315369 | 1 | -0.424 | 3.752 | chr9 | 12775019 | 12823059 | + | protein_coding |
| ENSG00000127481.15_5 | UBR4 | 0.98761071 | 1 | -0.424 | 7.620 | chr1 | 19401004 | 19536760 | - | protein_coding |
| ENSG00000170915.9_4 | PAQR8 | 0.65329214 | 1 | -0.424 | 1.733 | chr6 | 52226219 | 52272575 | + | protein_coding |
| ENSG00000189043.10_4 | NDUFA4 | 0.98492385 | 1 | -0.424 | 6.474 | chr7 | 10971570 | 10979780 | - | protein_coding |
| ENSG00000136527.18_5 | TRA2B | 0.99007775 | 1 | -0.424 | 7.348 | chr3 | 185632346 | 185655891 | - | protein_coding |
| ENSG00000136891.14_5 | TEX10 | 0.98154735 | 1 | -0.424 | 5.942 | chr9 | 103064359 | 103115224 | - | protein_coding |
| ENSG00000143891.17_5 | GALM | 0.97537376 | 1 | -0.425 | 5.495 | chr2 | 38893223 | 38968379 | + | protein_coding |
| ENSG00000271576.1_6 | AL359504.2 | 0.47053377 | 1 | -0.425 | 0.050 | chr1 | 84542014 | 84543614 | - | lncRNA |
| ENSG00000262155.2_6 | LINC02175 | 0.48266848 | 1 | -0.425 | 0.358 | chr16 | 25078258 | 25099939 | + | lncRNA |
| ENSG00000253771.6_4 | TPTE2P1 | 0.43988071 | 1 | -0.425 | -0.116 | chr13 | 25498815 | 25542625 | - | transcribed_unprocessed_pseudogene |
| ENSG00000078668.14_6 | VDAC3 | 0.98462905 | 1 | -0.425 | 7.151 | chr8 | 42249142 | 42263455 | + | protein_coding |
| ENSG00000144895.12_5 | EIF2A | 0.98416554 | 1 | -0.425 | 6.712 | chr3 | 150264465 | 150303803 | + | protein_coding |
| ENSG00000134940.14_4 | ACRV1 | 0.43492348 | 1 | -0.425 | -0.470 | chr11 | 125541417 | 125550769 | - | protein_coding |
| ENSG00000087842.11_4 | PIR | 0.97973659 | 1 | -0.425 | 5.985 | chrX | 15402921 | 15511687 | - | protein_coding |
| ENSG00000166974.13_6 | MAPRE2 | 0.95191294 | 1 | -0.425 | 5.076 | chr18 | 32556892 | 32723434 | + | protein_coding |
| ENSG00000235958.5_6 | UBOX5-AS1 | 0.45273327 | 1 | -0.425 | -0.187 | chr20 | 3087559 | 3131513 | + | lncRNA |
| ENSG00000196950.14_4 | SLC39A10 | 0.99056513 | 1 | -0.425 | 6.787 | chr2 | 196440701 | 196602426 | + | protein_coding |
| ENSG00000184270.5_4 | H2AC21 | 0.96735234 | 1 | -0.425 | 6.052 | chr1 | 149859034 | 149859563 | - | protein_coding |
| ENSG00000104081.14_5 | BMF | 0.6703359 | 1 | -0.426 | 1.885 | chr15 | 40380091 | 40401093 | - | protein_coding |
| ENSG00000075914.13_4 | EXOSC7 | 0.88507791 | 1 | -0.426 | 4.191 | chr3 | 45016733 | 45077558 | + | protein_coding |
| ENSG00000136937.13_6 | NCBP1 | 0.98569552 | 1 | -0.426 | 6.745 | chr9 | 100395950 | 100436030 | + | protein_coding |
| ENSG00000183137.14_6 | CEP57L1 | 0.85569422 | 1 | -0.426 | 3.635 | chr6 | 109416313 | 109485135 | + | protein_coding |
| ENSG00000188582.9_5 | PAQR9 | 0.49552315 | 1 | -0.426 | 0.040 | chr3 | 142668006 | 142682516 | - | protein_coding |
| ENSG00000237017.1_7 | AC245052.4 | 0.60307295 | 1 | -0.426 | 1.386 | chr19 | 54622891 | 54628722 | - | lncRNA |
| ENSG00000286108.1_3 | AL359916.1 | 0.47720624 | 1 | -0.427 | 0.274 | chr20 | 2088546 | 2090637 | - | lncRNA |
| ENSG00000255966.1_7 | AC006064.3 | 0.43410757 | 1 | -0.427 | -0.496 | chr12 | 6641456 | 6642664 | - | lncRNA |
| ENSG00000275056.1_7 | AC020663.3 | 0.43196357 | 1 | -0.427 | -0.447 | chr16 | 4889245 | 4890335 | - | lncRNA |
| ENSG00000247595.4_6 | SPTY2D1OS | 0.5475081 | 1 | -0.427 | 0.955 | chr11 | 18610328 | 18631802 | + | protein_coding |

| | | | | | | | | | | |
|---|---|---|---|---|---|---|---|---|---|---|
| ENSG00000172062.16_3 | SMN1 | 0.68371879 | 1 | -0.427 | 2.347 | chr5 | 70220857 | 70249769 | + | protein_coding |
| ENSG00000095383.20_6 | TBC1D2 | 0.85931627 | 1 | -0.427 | 4.385 | chr9 | 100961293 | 101017931 | - | protein_coding |
| ENSG00000273448.1_7 | AC006480.3 | 0.45411576 | 1 | -0.427 | -0.073 | chr7 | 66798034 | 66799370 | + | lncRNA |
| ENSG00000156469.9_6 | MTERF3 | 0.87290125 | 1 | -0.428 | 4.743 | chr8 | 97251626 | 97273838 | - | protein_coding |
| ENSG00000175414.7_4 | ARL10 | 0.45858706 | 1 | -0.428 | 0.033 | chr5 | 175792490 | 175828866 | + | protein_coding |
| ENSG00000180747.15_5 | SMG1P3 | 0.75818325 | 1 | -0.428 | 2.751 | chr16 | 21458004 | 21531765 | - | transcribed_unprocessed_pseudogene |
| ENSG00000165417.12_4 | GTF2A1 | 0.97740471 | 1 | -0.428 | 5.870 | chr14 | 81641796 | 81687721 | - | protein_coding |
| ENSG00000269028.3_2 | MTRNR2L12 | 0.68332618 | 1 | -0.428 | 6.358 | chr3 | 96336032 | 96337080 | - | protein_coding |
| ENSG00000114850.6_3 | SSR3 | 0.98341275 | 1 | -0.428 | 7.414 | chr3 | 156257929 | 156272973 | - | protein_coding |
| ENSG00000133706.18_4 | LARS1 | 0.99159397 | 1 | -0.428 | 7.495 | chr5 | 145492597 | 145562213 | - | protein_coding |
| ENSG00000158435.8_3 | CNOT11 | 0.98031498 | 1 | -0.428 | 5.923 | chr2 | 101869348 | 101886778 | + | protein_coding |
| ENSG00000233913.7_4 | RPL10P9 | 0.60237007 | 1 | -0.429 | 1.401 | chr5 | 168043357 | 168044001 | + | processed_pseudogene |
| ENSG00000072756.17_3 | TRNT1 | 0.95121328 | 1 | -0.429 | 4.950 | chr3 | 3168617 | 3192563 | + | protein_coding |
| ENSG00000166851.15_4 | PLK1 | 0.96911295 | 1 | -0.429 | 5.784 | chr16 | 23688977 | 23701688 | + | protein_coding |
| ENSG00000269486.2_7 | ERVK9-11 | 0.77145111 | 1 | -0.429 | 3.280 | chr19 | 39425937 | 39429272 | - | lncRNA |
| ENSG00000151835.16_6 | SACS | 0.86303495 | 1 | -0.429 | 7.126 | chr13 | 23902965 | 24007879 | - | protein_coding |
| ENSG00000128805.14_5 | ARHGAP22 | 0.63685232 | 1 | -0.429 | 1.717 | chr10 | 49654077 | 49864310 | - | protein_coding |
| ENSG00000207643.1 | Z95704.1 | 0.44964418 | 1 | -0.429 | -0.070 | chr4 | 65017 | 65125 | + | miRNA |
| ENSG00000261123.1_8 | AC009065.6 | 0.43827923 | 1 | -0.430 | -1.201 | chr16 | 2144831 | 2147027 | - | lncRNA |
| ENSG00000187583.11_4 | PLEKHN1 | 0.52623612 | 1 | -0.430 | 0.558 | chr1 | 901862 | 911245 | + | protein_coding |
| ENSG00000148690.12_4 | FRA10AC1 | 0.86206166 | 1 | -0.430 | 3.765 | chr10 | 95427640 | 95462349 | - | protein_coding |
| ENSG00000139668.9_4 | WDFY2 | 0.81287635 | 1 | -0.430 | 3.315 | chr13 | 52158591 | 52341845 | + | protein_coding |
| ENSG00000123179.14_4 | EBPL | 0.9216162 | 1 | -0.430 | 4.642 | chr13 | 50234810 | 50265622 | - | protein_coding |
| ENSG00000183431.12_4 | SF3A3 | 0.97782233 | 1 | -0.430 | 6.424 | chr1 | 38422647 | 38455747 | - | protein_coding |
| ENSG00000260401.1_6 | AP002761.4 | 0.71732364 | 1 | -0.430 | 2.447 | chr11 | 72950020 | 72953380 | + | lncRNA |
| ENSG00000185246.18_4 | PRPF39 | 0.9477331 | 1 | -0.430 | 5.561 | chr14 | 45553310 | 45585485 | + | protein_coding |
| ENSG00000116096.6_3 | SPR | 0.97916397 | 1 | -0.430 | 6.354 | chr2 | 73114511 | 73119287 | + | protein_coding |
| ENSG00000115421.13_3 | PAPOLG | 0.94586577 | 1 | -0.430 | 5.074 | chr2 | 60983388 | 61029221 | + | protein_coding |
| ENSG00000107371.13_5 | EXOSC3 | 0.91105824 | 1 | -0.431 | 4.749 | chr9 | 37766975 | 37801434 | - | protein_coding |
| ENSG00000174799.11_3 | CEP135 | 0.91940734 | 1 | -0.431 | 4.450 | chr4 | 56815037 | 56899527 | + | protein_coding |
| ENSG00000205930.10_9 | C21orf62-AS1 | 0.42610232 | 1 | -0.431 | -0.880 | chr21 | 34144411 | 34327744 | + | lncRNA |
| ENSG00000115839.17_3 | RAB3GAP1 | 0.98110685 | 1 | -0.431 | 6.037 | chr2 | 135809835 | 135933964 | + | protein_coding |
| ENSG00000274554.1_6 | AC083806.2 | 0.42515841 | 1 | -0.431 | -0.728 | chr12 | 117386543 | 117389227 | - | lncRNA |
| ENSG00000272335.1_6 | AC093297.2 | 0.6963635 | 1 | -0.431 | 2.594 | chr5 | 44826178 | 44828694 | + | lncRNA |
| ENSG00000183386.10_4 | FHL3 | 0.64021778 | 1 | -0.431 | 1.492 | chr1 | 38462442 | 38471278 | - | protein_coding |
| ENSG00000165185.14_5 | KIAA1958 | 0.88817899 | 1 | -0.431 | 4.084 | chr9 | 115249127 | 115431677 | + | protein_coding |
| ENSG00000164306.11_4 | PRIMPOL | 0.83330263 | 1 | -0.432 | 3.407 | chr4 | 185570821 | 185616117 | + | protein_coding |
| ENSG00000144591.18_4 | GMPPA | 0.94900836 | 1 | -0.432 | 4.974 | chr2 | 220363589 | 220371711 | + | protein_coding |
| ENSG00000164024.12_2 | METAP1 | 0.97093618 | 1 | -0.432 | 6.163 | chr4 | 99916810 | 99983960 | + | protein_coding |
| ENSG00000143476.18_3 | DTL | 0.97879665 | 1 | -0.432 | 6.396 | chr1 | 212208895 | 212280742 | + | protein_coding |
| ENSG00000269318.1_6 | AC007292.2 | 0.44857989 | 1 | -0.432 | -0.039 | chr19 | 4356634 | 4358445 | - | lncRNA |
| ENSG00000172845.15_5 | SP3 | 0.98761258 | 1 | -0.432 | 7.427 | chr2 | 174745578 | 174830430 | - | protein_coding |
| ENSG00000175787.17_3 | ZNF169 | 0.66830775 | 1 | -0.432 | 2.103 | chr9 | 97021580 | 97064111 | + | protein_coding |
| ENSG00000130695.15_4 | CEP85 | 0.95226451 | 1 | -0.433 | 5.026 | chr1 | 26560691 | 26605299 | + | protein_coding |
| ENSG00000164011.18_5 | ZNF691 | 0.73122241 | 1 | -0.433 | 2.610 | chr1 | 43312244 | 43318148 | + | protein_coding |
| ENSG00000197102.12_7 | DYNC1H1 | 0.98648874 | 1 | -0.433 | 9.013 | chr14 | 102430910 | 102522780 | + | protein_coding |
| ENSG00000231952.3_4 | DPY19L1P2 | 0.46931717 | 1 | -0.433 | 0.290 | chr7 | 32852369 | 32878182 | + | unprocessed_pseudogene |
| ENSG00000175183.10_4 | CSRP2 | 0.81826648 | 1 | -0.433 | 3.249 | chr12 | 77252489 | 77272803 | - | protein_coding |
| ENSG00000163938.17_4 | GNL3 | 0.9802051 | 1 | -0.433 | 6.720 | chr3 | 52715172 | 52728513 | + | protein_coding |
| ENSG00000172428.11_7 | COPS9 | 0.93610288 | 1 | -0.433 | 4.732 | chr2 | 241065980 | 241076224 | - | protein_coding |
| ENSG00000109534.17_3 | GAR1 | 0.71422971 | 1 | -0.434 | 2.477 | chr4 | 110736666 | 110745896 | + | protein_coding |
| ENSG00000180638.17_4 | SLC47A2 | 0.49927618 | 1 | -0.434 | 0.472 | chr17 | 19581601 | 19622292 | - | protein_coding |
| ENSG00000130347.13_4 | RTN4IP1 | 0.80044799 | 1 | -0.434 | 3.343 | chr6 | 107018646 | 107077373 | - | protein_coding |
| ENSG00000250321.1_5 | AC079140.2 | 0.56102016 | 1 | -0.434 | 1.192 | chr4 | 336610 | 336751 | - | processed_pseudogene |
| ENSG00000114982.19_6 | KANSL3 | 0.98261303 | 1 | -0.434 | 6.298 | chr2 | 97258907 | 97308524 | - | protein_coding |
| ENSG00000257790.1_4 | EIF4A1P4 | 0.44779985 | 1 | -0.434 | -0.227 | chr12 | 53547056 | 53548268 | + | processed_pseudogene |
| ENSG00000086827.9_5 | ZW10 | 0.93374138 | 1 | -0.435 | 4.916 | chr11 | 113603909 | 113644457 | - | protein_coding |
| ENSG00000120334.15_3 | CENPL | 0.90284146 | 1 | -0.435 | 4.263 | chr1 | 173768688 | 173793858 | - | protein_coding |

| | | | | | | | | | | |
|---|---|---|---|---|---|---|---|---|---|---|
| ENSG00000091651.9_4 | ORC6 | 0.9407292 | 1 | -0.435 | 5.089 | chr16 | 46723555 | 46732306 | + | protein_coding |
| ENSG00000234337.4_5 | AC026462.1 | 0.41965481 | 1 | -0.435 | -0.679 | chr16 | 52689219 | 52689957 | + | processed_pseudogene |
| ENSG00000092470.12_4 | WDR76 | 0.94991535 | 1 | -0.435 | 5.453 | chr15 | 44119178 | 44160610 | + | protein_coding |
| ENSG00000057935.13_4 | MTA3 | 0.96478983 | 1 | -0.435 | 5.473 | chr2 | 42721709 | 42984087 | + | protein_coding |
| ENSG00000106348.18_8 | IMPDH1 | 0.94249797 | 1 | -0.435 | 6.337 | chr7 | 128032331 | 128050306 | - | protein_coding |
| ENSG00000167085.11_7 | PHB | 0.98793687 | 1 | -0.436 | 6.968 | chr17 | 47481411 | 47492267 | - | protein_coding |
| ENSG00000169519.21_5 | METTL15 | 0.93295625 | 1 | -0.436 | 4.916 | chr11 | 28129795 | 28548588 | + | protein_coding |
| ENSG00000242798.1_8 | AC073842.2 | 0.48143727 | 1 | -0.436 | 0.180 | chr7 | 99712837 | 99724762 | - | lncRNA |
| ENSG00000196476.12_4 | C20orf96 | 0.74374847 | 1 | -0.436 | 2.793 | chr20 | 251504 | 271419 | - | protein_coding |
| ENSG00000172878.14_3 | METAP1D | 0.71925149 | 1 | -0.436 | 2.947 | chr2 | 172864850 | 172947158 | + | protein_coding |
| ENSG00000235945.1_6 | AC002543.1 | 0.41830679 | 1 | -0.436 | -0.767 | chr7 | 116513508 | 116516430 | - | processed_pseudogene |
| ENSG00000148019.13_6 | CEP78 | 0.969007 | 1 | -0.437 | 6.226 | chr9 | 80850978 | 80894606 | + | protein_coding |
| ENSG00000227081.5_5 | AC005912.1 | 0.62748399 | 1 | -0.437 | 1.769 | chr12 | 3320829 | 3321083 | - | processed_pseudogene |
| ENSG00000214367.8_5 | HAUS3 | 0.88171139 | 1 | -0.437 | 4.106 | chr4 | 2080725 | 2244003 | - | protein_coding |
| ENSG00000143106.13_4 | PSMA5 | 0.9595576 | 1 | -0.437 | 6.302 | chr1 | 109941664 | 109969070 | - | protein_coding |
| ENSG00000100372.15_6 | SLC25A17 | 0.90135972 | 1 | -0.437 | 4.560 | chr22 | 41165634 | 41215403 | - | protein_coding |
| ENSG00000170962.13_5 | PDGFD | 0.95282401 | 1 | -0.437 | 5.335 | chr11 | 103777917 | 104035107 | - | protein_coding |
| ENSG00000235174.1_4 | RPL39P3 | 0.6556743 | 1 | -0.437 | 3.845 | chr6 | 74082831 | 74082986 | - | processed_pseudogene |
| ENSG00000198468.9_7 | FLVCR1-DT | 0.55527022 | 1 | -0.437 | 1.146 | chr1 | 213025447 | 213031468 | - | lncRNA |
| ENSG00000161980.6_4 | POLR3K | 0.77257203 | 1 | -0.437 | 3.153 | chr16 | 96407 | 103608 | - | protein_coding |
| ENSG00000212802.4_2 | RPL15P3 | 0.57574451 | 1 | -0.437 | 2.558 | chr6 | 12514342 | 12514956 | + | processed_pseudogene |
| ENSG00000105376.5_4 | ICAM5 | 0.44352604 | 1 | -0.437 | -0.073 | chr19 | 10400628 | 10407454 | + | protein_coding |
| ENSG00000160838.14_3 | LRRC71 | 0.42749 | 1 | -0.437 | -0.522 | chr1 | 156890424 | 156902886 | + | protein_coding |
| ENSG00000086289.12_5 | EPDR1 | 0.98472202 | 1 | -0.437 | 6.637 | chr7 | 37723446 | 37991538 | + | protein_coding |
| ENSG00000005448.17_5 | WDR54 | 0.95319922 | 1 | -0.438 | 5.302 | chr2 | 74648805 | 74652882 | + | protein_coding |
| ENSG00000097021.19_4 | ACOT7 | 0.89221929 | 1 | -0.438 | 4.514 | chr1 | 6324329 | 6454451 | - | protein_coding |
| ENSG00000279069.1_5 | AC015813.5 | 0.49254736 | 1 | -0.438 | 0.743 | chr17 | 56084035 | 56085548 | + | TEC |
| ENSG00000234741.8_8 | GAS5 | 0.97109344 | 1 | -0.438 | 6.898 | chr1 | 173827697 | 173838020 | - | lncRNA |
| ENSG00000100532.12_4 | CGRRF1 | 0.82777739 | 1 | -0.438 | 3.777 | chr14 | 54976530 | 55006010 | + | protein_coding |
| ENSG00000073536.18_4 | NLE1 | 0.93582099 | 1 | -0.438 | 5.121 | chr17 | 33455749 | 33469323 | - | protein_coding |
| ENSG00000272945.1_8 | AL356299.3 | 0.41623529 | 1 | -0.438 | -0.860 | chr20 | 32869438 | 32874272 | - | lncRNA |
| ENSG00000273271.1_7 | AP000254.2 | 0.44898806 | 1 | -0.438 | 0.102 | chr21 | 33039041 | 33039560 | - | lncRNA |
| ENSG00000262171.1_6 | AC130651.1 | 0.44928555 | 1 | -0.438 | -0.168 | chr16 | 15978886 | 15980015 | + | lncRNA |
| ENSG00000145736.14_7 | GTF2H2 | 0.54255613 | 1 | -0.439 | 1.079 | chr5 | 70328497 | 70363516 | - | protein_coding |
| ENSG00000066735.14_3 | KIF26A | 0.48431135 | 1 | -0.439 | 2.537 | chr14 | 104605060 | 104647231 | + | protein_coding |
| ENSG00000105675.9_5 | ATP4A | 0.42211976 | 1 | -0.439 | -0.336 | chr19 | 36040933 | 36054560 | - | protein_coding |
| ENSG00000113070.8_4 | HBEGF | 0.52149326 | 1 | -0.439 | 0.993 | chr5 | 139712428 | 139726188 | - | protein_coding |
| ENSG00000101624.11_5 | CEP76 | 0.77270947 | 1 | -0.440 | 3.669 | chr18 | 12661832 | 12702776 | - | protein_coding |
| ENSG00000138101.18_5 | DTNB | 0.84453717 | 1 | -0.440 | 3.807 | chr2 | 25600067 | 25896516 | - | protein_coding |
| ENSG00000002746.15_4 | HECW1 | 0.88030999 | 1 | -0.440 | 4.056 | chr7 | 43152228 | 43605600 | + | protein_coding |
| ENSG00000164045.12_6 | CDC25A | 0.89197499 | 1 | -0.440 | 4.288 | chr3 | 48198636 | 48229907 | - | protein_coding |
| ENSG00000225265.2_6 | TAF1A-AS1 | 0.47086149 | 1 | -0.440 | 0.234 | chr1 | 222763167 | 222767185 | + | lncRNA |
| ENSG00000158079.16_6 | PTPDC1 | 0.85604763 | 1 | -0.440 | 3.924 | chr9 | 96793076 | 96872138 | + | protein_coding |
| ENSG00000154582.16_4 | ELOC | 0.91801852 | 1 | -0.440 | 5.309 | chr8 | 74851404 | 74884522 | - | protein_coding |
| ENSG00000170727.7 | BOP1 | 0.76360247 | 1 | -0.440 | 3.635 | chr8 | 145486055 | 145515082 | - | protein_coding |
| ENSG00000205309.14_5 | NT5M | 0.74525747 | 1 | -0.440 | 2.850 | chr17 | 17206649 | 17250977 | + | protein_coding |
| ENSG00000228000.1_4 | RPL7AP65 | 0.44100178 | 1 | -0.440 | -0.073 | chr17 | 18215218 | 18215965 | + | processed_pseudogene |
| ENSG00000280033.1_6 | AC116407.4 | 0.41251706 | 1 | -0.441 | -0.892 | chr17 | 30486330 | 30487390 | + | TEC |
| ENSG00000137970.7_4 | RPL7P9 | 0.68863529 | 1 | -0.441 | 3.403 | chr1 | 97144430 | 97145176 | - | processed_pseudogene |
| ENSG00000280046.1_6 | AC104581.4 | 0.56294433 | 1 | -0.441 | 1.388 | chr17 | 7762261 | 7769401 | + | TEC |
| ENSG00000251550.1_6 | AC091133.5 | 0.5077188 | 1 | -0.442 | 0.821 | chr17 | 47114484 | 47114791 | + | processed_pseudogene |
| ENSG00000268942.2_4 | CKS1BP3 | 0.41145642 | 1 | -0.442 | -0.740 | chr5 | 61808073 | 61808309 | - | processed_pseudogene |
| ENSG00000183022.5_4 | TPM3P8 | 0.41083988 | 1 | -0.442 | -0.860 | chr2 | 231437882 | 231438524 | - | processed_pseudogene |
| ENSG00000072135.13_4 | PTPN18 | 0.89060454 | 1 | -0.442 | 4.624 | chr2 | 131113618 | 131132978 | + | protein_coding |
| ENSG00000236383.3 | LINC00854 | 0.59153772 | 1 | -0.442 | 1.755 | chr17 | 41368960 | 41383338 | - | processed_transcript |

| | | | | | | | | | | |
|---|---|---|---|---|---|---|---|---|---|---|
| ENSG00000055917.15_6 | PUM2 | 0.97154448 | 1 | -0.443 | 6.706 | chr2 | 20448452 | 20551995 | - | protein_coding |
| ENSG00000224152.1_6 | AC009506.2 | 0.42464023 | 1 | -0.443 | -0.014 | chr2 | 160471807 | 160473593 | + | lncRNA |
| ENSG00000138081.20_5 | FBXO11 | 0.93962474 | 1 | -0.443 | 5.414 | chr2 | 48016455 | 48132932 | - | protein_coding |
| ENSG00000243667.7_5 | WDR92 | 0.69835802 | 1 | -0.443 | 2.770 | chr2 | 68350068 | 68384681 | - | protein_coding |
| ENSG00000129515.20_6 | SNX6 | 0.9777822 | 1 | -0.443 | 6.883 | chr14 | 35030299 | 35099366 | - | protein_coding |
| ENSG00000101596.15_4 | SMCHD1 | 0.95085746 | 1 | -0.443 | 6.538 | chr18 | 2655737 | 2805015 | + | protein_coding |
| ENSG00000167680.16_4 | SEMA6B | 0.74226369 | 1 | -0.443 | 3.940 | chr19 | 4542605 | 4559696 | - | protein_coding |
| ENSG00000136710.10_5 | CCDC115 | 0.85362448 | 1 | -0.443 | 3.921 | chr2 | 131095506 | 131100272 | - | protein_coding |
| ENSG00000132305.20_3 | IMMT | 0.98095197 | 1 | -0.443 | 7.372 | chr2 | 86371055 | 86422893 | - | protein_coding |
| ENSG00000174173.7_4 | TRMT10C | 0.92292585 | 1 | -0.443 | 5.241 | chr3 | 101280712 | 101285290 | + | protein_coding |
| ENSG00000180628.15_3 | PCGF5 | 0.97573917 | 1 | -0.443 | 6.539 | chr10 | 92922769 | 93044094 | + | protein_coding |
| ENSG00000225507.1_5 | AC069282.1 | 0.52010714 | 1 | -0.443 | 1.154 | chr7 | 47996390 | 47996915 | + | processed_pseudogene |
| ENSG00000168439.16_2 | STIP1 | 0.98534773 | 1 | -0.443 | 8.285 | chr11 | 63952744 | 63972015 | + | protein_coding |
| ENSG00000173226.17_7 | IQCB1 | 0.88952459 | 1 | -0.443 | 4.493 | chr3 | 121488608 | 121553926 | - | protein_coding |
| ENSG00000082212.13_6 | ME2 | 0.92533112 | 1 | -0.443 | 5.858 | chr18 | 48405450 | 48480627 | + | protein_coding |
| ENSG00000240463.1_4 | RPS19P3 | 0.41971315 | 1 | -0.444 | -0.523 | chr14 | 35037895 | 35038335 | - | processed_pseudogene |
| ENSG00000140525.17_6 | FANCI | 0.98242787 | 1 | -0.444 | 7.152 | chr15 | 89787180 | 89860492 | + | protein_coding |
| ENSG00000183060.15_5 | LYSMD4 | 0.72815186 | 1 | -0.444 | 2.959 | chr15 | 100255902 | 100273766 | - | protein_coding |
| ENSG00000142657.21_4 | PGD | 0.98851121 | 1 | -0.444 | 9.733 | chr1 | 10458649 | 10480568 | + | protein_coding |
| ENSG00000286873.1_2 | AC012306.4 | 0.41185124 | 1 | -0.444 | -0.710 | chr2 | 128646403 | 128650306 | + | lncRNA |
| ENSG00000136824.19_4 | SMC2 | 0.97836035 | 1 | -0.444 | 7.142 | chr9 | 106856541 | 106903700 | + | protein_coding |
| ENSG00000231747.1_5 | AC079922.1 | 0.44270136 | 1 | -0.444 | 0.091 | chr2 | 113379386 | 113379744 | - | processed_pseudogene |
| ENSG00000239282.7_9 | CASTOR1 | 0.40845771 | 1 | -0.445 | -0.740 | chr22 | 30681106 | 30685616 | - | protein_coding |
| ENSG00000182578.13_6 | CSF1R | 0.40848639 | 1 | -0.445 | -0.800 | chr5 | 149432854 | 149492935 | - | protein_coding |
| ENSG00000151881.14_8 | TMEM267 | 0.85001634 | 1 | -0.445 | 3.980 | chr5 | 43444354 | 43483995 | - | protein_coding |
| ENSG00000240602.7_4 | AADACP1 | 0.8860379 | 1 | -0.445 | 4.422 | chr3 | 151488216 | 151502682 | + | transcribed_unprocessed_pseudogene |
| ENSG00000160961.12_3 | ZNF333 | 0.6996828 | 1 | -0.445 | 2.656 | chr19 | 14800613 | 14844558 | + | protein_coding |
| ENSG00000100138.15_6 | SNU13 | 0.94550156 | 1 | -0.445 | 5.460 | chr22 | 42069937 | 42086508 | - | protein_coding |
| ENSG00000246214.2_7 | AC024588.1 | 0.51233399 | 1 | -0.445 | 0.633 | chr5 | 16616035 | 16682014 | + | lncRNA |
| ENSG00000122435.10_3 | TRMT13 | 0.79378988 | 1 | -0.445 | 3.377 | chr1 | 100598706 | 100616054 | + | protein_coding |
| ENSG00000100523.15_5 | DDHD1 | 0.92099284 | 1 | -0.446 | 4.934 | chr14 | 53503463 | 53620041 | - | protein_coding |
| ENSG00000230590.11_7 | FTX | 0.86367505 | 1 | -0.446 | 5.372 | chrX | 73160270 | 73513409 | - | lncRNA |
| ENSG00000101104.12_2 | PABPC1L | 0.84876772 | 1 | -0.446 | 3.962 | chr20 | 43538703 | 43587676 | + | protein_coding |
| ENSG00000287045.1_2 | AC072028.1 | 0.45242312 | 1 | -0.446 | -0.093 | chr3 | 142406824 | 142442723 | - | lncRNA |
| ENSG00000146677.7_5 | AC004453.1 | 0.54556939 | 1 | -0.446 | 1.621 | chr7 | 44507496 | 44507903 | + | processed_pseudogene |
| ENSG00000257097.1_5 | CLIP1-AS1 | 0.40694354 | 1 | -0.446 | -0.680 | chr12 | 122880089 | 122885404 | + | lncRNA |
| ENSG00000154719.13_7 | MRPL39 | 0.87068706 | 1 | -0.446 | 4.417 | chr21 | 26957968 | 26979829 | - | protein_coding |
| ENSG00000126814.7_6 | TRMT5 | 0.91797516 | 1 | -0.446 | 4.809 | chr14 | 61438159 | 61447888 | - | protein_coding |
| ENSG00000205940.8_2 | HSP90AB2P | 0.40630935 | 1 | -0.446 | -0.828 | chr4 | 13335038 | 13340281 | + | transcribed_processed_pseudogene |
| ENSG00000231177.4_7 | LINC00852 | 0.41848842 | 1 | -0.446 | -0.314 | chr3 | 10326103 | 10327430 | + | lncRNA |
| ENSG00000096717.12_3 | SIRT1 | 0.94861527 | 1 | -0.446 | 5.522 | chr10 | 69644414 | 69678147 | + | protein_coding |
| ENSG00000102900.13_7 | NUP93 | 0.9714099 | 1 | -0.447 | 6.617 | chr16 | 56764030 | 56884198 | + | protein_coding |
| ENSG00000145979.18_8 | TBC1D7 | 0.82546832 | 1 | -0.447 | 3.696 | chr6 | 13266774 | 13328815 | - | protein_coding |
| ENSG00000166012.17_5 | TAF1D | 0.95314495 | 1 | -0.447 | 6.106 | chr11 | 93463114 | 93517557 | - | protein_coding |
| ENSG00000168710.18_5 | AHCYL1 | 0.96930333 | 1 | -0.447 | 6.952 | chr1 | 110527387 | 110566364 | + | protein_coding |
| ENSG00000278828.1_3 | H3C10 | 0.92802354 | 1 | -0.447 | 7.099 | chr6 | 27777842 | 27779078 | + | protein_coding |
| ENSG00000287169.1_1 | AC107050.1 | 0.42641195 | 1 | -0.447 | -1.217 | chr4 | 187622733 | 187625240 | + | lncRNA |
| ENSG00000254397.1_7 | AC132192.1 | 0.40442861 | 1 | -0.447 | -0.893 | chr11 | 9451903 | 9455033 | - | lncRNA |
| ENSG00000174547.13_6 | MRPL11 | 0.93038593 | 1 | -0.448 | 5.352 | chr11 | 66202546 | 66234209 | - | protein_coding |
| ENSG00000229388.1_6 | LINC01715 | 0.41578193 | 1 | -0.448 | -0.291 | chr1 | 28969740 | 28975093 | + | lncRNA |
| ENSG00000108219.15_3 | TSPAN14 | 0.97816097 | 1 | -0.448 | 7.257 | chr10 | 82213922 | 82292880 | + | protein_coding |
| ENSG00000221845.4_3 | C7orf65 | 0.40900198 | 1 | -0.448 | -1.029 | chr7 | 47694842 | 47701246 | + | lncRNA |
| ENSG00000260796.1_7 | AC145285.3 | 0.40776456 | 1 | -0.448 | -1.065 | chr16 | 28831891 | 28833354 | - | lncRNA |
| ENSG00000086200.16_8 | IPO11 | 0.92016769 | 1 | -0.448 | 5.692 | chr5 | 61699799 | 61924409 | + | protein_coding |
| ENSG00000269984.1_6 | AC078795.1 | 0.54607985 | 1 | -0.448 | 1.306 | chr3 | 169494980 | 169498122 | - | lncRNA |
| ENSG00000121989.15_3 | ACVR2A | 0.77352511 | 1 | -0.448 | 3.421 | chr2 | 148602086 | 148688395 | + | protein_coding |
| ENSG00000224616.3_8 | RTCA-AS1 | 0.49294981 | 1 | -0.449 | 0.875 | chr1 | 100717084 | 100731735 | - | lncRNA |
| ENSG00000268403.2_6 | AC132192.2 | 0.53121381 | 1 | -0.449 | 1.133 | chr11 | 9481103 | 9482249 | - | lncRNA |

| | | | | | | | | | | |
|---|---|---|---|---|---|---|---|---|---|---|
| ENSG00000209482.1 | SNORD83A | 0.41526189 | 1 | -0.449 | -0.356 | chr22 | 39711218 | 39711312 | - | snoRNA |
| ENSG00000239183.1 | SNORA84 | 0.40516441 | 1 | -0.449 | -0.768 | chr9 | 95054743 | 95054875 | - | snoRNA |
| ENSG00000231119.2_6 | AL031666.1 | 0.41247393 | 1 | -0.450 | -0.426 | chr20 | 45981305 | 45983377 | + | lncRNA |
| ENSG00000167550.11_6 | RHEBL1 | 0.52523576 | 1 | -0.450 | 1.277 | chr12 | 49458459 | 49463808 | - | protein_coding |
| ENSG00000083857.14_6 | FAT1 | 0.96018446 | 1 | -0.450 | 9.099 | chr4 | 187508948 | 187647876 | - | protein_coding |
| ENSG00000169330.9_4 | MINAR1 | 0.547192 | 1 | -0.450 | 1.350 | chr15 | 79724678 | 79764646 | + | protein_coding |
| ENSG00000138613.14_4 | APH1B | 0.90105843 | 1 | -0.450 | 4.619 | chr15 | 63568217 | 63601325 | + | protein_coding |
| ENSG00000102738.8_4 | MRPS31 | 0.83689581 | 1 | -0.450 | 4.018 | chr13 | 41303264 | 41345326 | - | protein_coding |
| ENSG00000100567.13_5 | PSMA3 | 0.96433871 | 1 | -0.451 | 6.620 | chr14 | 58711561 | 58738730 | + | protein_coding |
| ENSG00000263884.1_6 | AP000845.1 | 0.42427244 | 1 | -0.451 | -0.137 | chr18 | 268148 | 270278 | + | lncRNA |
| ENSG00000273702.1_6 | AC091271.1 | 0.45638085 | 1 | -0.451 | 0.420 | chr17 | 57695914 | 57697075 | - | lncRNA |
| ENSG00000183196.10_5 | CHST6 | 0.45829264 | 1 | -0.451 | 0.361 | chr16 | 75505950 | 75529343 | - | protein_coding |
| ENSG00000175063.17_3 | UBE2C | 0.95462347 | 1 | -0.451 | 5.911 | chr20 | 44441215 | 44445596 | + | protein_coding |
| ENSG00000124641.16_6 | MED20 | 0.85012368 | 1 | -0.451 | 4.291 | chr6 | 41873092 | 41888877 | - | protein_coding |
| ENSG00000275632.1_7 | AL035461.3 | 0.43382492 | 1 | -0.451 | -0.028 | chr20 | 5981064 | 5981587 | + | lncRNA |
| ENSG00000286931.1_1 | AL136419.3 | 0.42439523 | 1 | -0.451 | -0.058 | chr14 | 24476394 | 24496991 | + | lncRNA |
| ENSG00000223802.7_6 | CERS1 | 0.4055596 | 1 | -0.451 | -1.177 | chr19 | 18979354 | 19007536 | - | protein_coding |
| ENSG00000112208.11_3 | BAG2 | 0.93726151 | 1 | -0.451 | 5.409 | chr6 | 57037124 | 57054631 | + | protein_coding |
| ENSG00000100296.13_4 | THOC5 | 0.92323225 | 1 | -0.452 | 5.291 | chr22 | 29901868 | 29951205 | - | protein_coding |
| ENSG00000260589.1_6 | STAM-AS1 | 0.46431363 | 1 | -0.452 | 0.500 | chr10 | 17683283 | 17685877 | - | lncRNA |
| ENSG00000280426.1_5 | AC084876.2 | 0.39961978 | 1 | -0.452 | -0.860 | chr12 | 110435224 | 110436932 | - | TEC |
| ENSG00000145833.16_5 | DDX46 | 0.95603559 | 1 | -0.452 | 6.945 | chr5 | 134094461 | 134190823 | + | protein_coding |
| ENSG00000153317.15_5 | ASAP1 | 0.96263452 | 1 | -0.453 | 6.447 | chr8 | 131064350 | 131455920 | - | protein_coding |
| ENSG00000227782.2_7 | AC002553.1 | 0.39915112 | 1 | -0.453 | -0.710 | chr17 | 15943786 | 15944587 | + | lncRNA |
| ENSG00000243232.6_5 | PCDHAC2 | 0.82331932 | 1 | -0.453 | 3.865 | chr5 | 140346055 | 140391932 | + | protein_coding |
| ENSG00000279432.1_6 | AC015799.1 | 0.48032272 | 1 | -0.453 | 0.909 | chr17 | 2542591 | 2545712 | + | TEC |
| ENSG00000185305.11_5 | ARL15 | 0.81395291 | 1 | -0.453 | 3.871 | chr5 | 53179772 | 53606412 | - | protein_coding |
| ENSG00000164038.16_6 | SLC9B2 | 0.69074981 | 1 | -0.453 | 2.828 | chr4 | 103941025 | 104006986 | - | protein_coding |
| ENSG00000259917.1_4 | HNRNPLP2 | 0.48711821 | 1 | -0.453 | 0.847 | chr15 | 34781203 | 34782772 | - | unprocessed_pseudogene |
| ENSG00000277363.5_9 | SRCIN1 | 0.60426964 | 1 | -0.453 | 1.834 | chr17 | 36686266 | 36762205 | - | protein_coding |
| ENSG00000198912.11_4 | C1orf174 | 0.88388799 | 1 | -0.453 | 4.482 | chr1 | 3805689 | 3816857 | - | protein_coding |
| ENSG00000100739.11_4 | BDKRB1 | 0.401383 | 1 | -0.453 | -0.683 | chr14 | 96722547 | 96735304 | + | protein_coding |
| ENSG00000157890.17_4 | MEGF11 | 0.58780347 | 1 | -0.454 | 1.716 | chr15 | 66187417 | 66546085 | - | protein_coding |
| ENSG00000265511.1_6 | AC020558.2 | 0.39931597 | 1 | -0.454 | -0.960 | chr17 | 17410665 | 17411622 | + | lncRNA |
| ENSG00000188687.17_6 | SLC4A5 | 0.41797509 | 1 | -0.454 | -0.176 | chr2 | 74443369 | 74570541 | - | protein_coding |
| ENSG00000279133.1_7 | AC018628.2 | 0.59138591 | 1 | -0.454 | 2.004 | chr17 | 60081061 | 60084879 | - | TEC |
| ENSG00000138069.18_5 | RAB1A | 0.96403602 | 1 | -0.454 | 7.092 | chr2 | 65297830 | 65357465 | - | protein_coding |
| ENSG00000276214.1_5 | AC008833.1 | 0.3974379 | 1 | -0.454 | -0.629 | chr5 | 58609936 | 58610626 | + | processed_pseudogene |
| ENSG00000260852.1_5 | FBXL19-AS1 | 0.63936999 | 1 | -0.454 | 2.410 | chr16 | 30930640 | 30934590 | - | lncRNA |
| ENSG00000163126.15_5 | ANKRD23 | 0.41932488 | 1 | -0.454 | -0.196 | chr2 | 97490263 | 97523671 | - | protein_coding |
| ENSG00000197557.7_3 | TTC30A | 0.83756886 | 1 | -0.455 | 4.183 | chr2 | 178477727 | 178483470 | - | protein_coding |
| ENSG00000287643.1_2 | AC004825.3 | 0.39631009 | 1 | -0.455 | -0.960 | chr14 | 71412357 | 71425746 | - | lncRNA |
| ENSG00000130775.16_3 | THEMIS2 | 0.40326994 | 1 | -0.455 | -0.260 | chr1 | 28199054 | 28213196 | + | protein_coding |
| ENSG00000178096.9_3 | BOLA1 | 0.73724432 | 1 | -0.455 | 3.469 | chr1 | 149859440 | 149872351 | + | protein_coding |
| ENSG00000285646.2_3 | AL021155.5 | 0.39921208 | 1 | -0.455 | -0.360 | chr1 | 11967997 | 11974355 | + | lncRNA |
| ENSG00000189419.8_6 | SPATA41 | 0.42353211 | 1 | -0.455 | -0.120 | chr15 | 100884252 | 100890930 | - | lncRNA |
| ENSG00000286153.1_2 | AP000331.1 | 0.41434103 | 1 | -0.455 | -0.177 | chr21 | 36208583 | 36257179 | + | lncRNA |
| ENSG00000147166.11_3 | ITGB1BP2 | 0.48574175 | 1 | -0.455 | 0.906 | chrX | 70521600 | 70525221 | + | protein_coding |
| ENSG00000111817.18_8 | DSE | 0.86590108 | 1 | -0.455 | 4.334 | chr6 | 116575336 | 116766024 | + | protein_coding |
| ENSG00000153044.10_3 | CENPH | 0.89285708 | 1 | -0.455 | 4.814 | chr5 | 68485401 | 68506184 | + | protein_coding |
| ENSG00000166200.15_5 | COPS2 | 0.95822998 | 1 | -0.456 | 6.488 | chr15 | 49398265 | 49447858 | - | protein_coding |
| ENSG00000172345.14_5 | STARD5 | 0.44241108 | 1 | -0.456 | 0.269 | chr15 | 81601394 | 81616524 | - | protein_coding |
| ENSG00000104147.9_4 | OIP5 | 0.56635458 | 1 | -0.456 | 1.857 | chr15 | 41601471 | 41624789 | - | protein_coding |
| ENSG00000118523.6_4 | CCN2 | 0.86429819 | 1 | -0.456 | 4.550 | chr6 | 132269316 | 132272512 | - | protein_coding |
| ENSG00000167600.14_3 | CYP2S1 | 0.79518949 | 1 | -0.456 | 4.922 | chr19 | 41699115 | 41713444 | + | protein_coding |
| ENSG00000080824.18_4 | HSP90AA1 | 0.97571339 | 1 | -0.456 | 9.299 | chr14 | 102547075 | 102606036 | - | protein_coding |
| ENSG00000279196.1_5 | AC135048.4 | 0.39451173 | 1 | -0.456 | -1.101 | chr16 | 30995951 | 30999591 | - | TEC |
| ENSG00000213593.10_4 | TMX2 | 0.94934516 | 1 | -0.456 | 6.231 | chr11 | 57480065 | 57508445 | + | protein_coding |
| ENSG00000176720.6_6 | BOK | 0.6871517 | 1 | -0.456 | 3.765 | chr2 | 242490839 | 242513546 | + | protein_coding |
| ENSG00000142513.6_6 | ACP4 | 0.48645549 | 1 | -0.456 | 0.615 | chr19 | 51293672 | 51298476 | + | protein_coding |

| | | | | | | | | | | |
|---|---|---|---|---|---|---|---|---|---|---|
| ENSG00000128294.16_5 | TPST2 | 0.66290948 | 1 | -0.456 | 2.764 | chr22 | 26917962 | 26992681 | - | protein_coding |
| ENSG00000069849.11_5 | ATP1B3 | 0.95399611 | 1 | -0.456 | 6.243 | chr3 | 141594966 | 141645391 | + | protein_coding |
| ENSG00000170689.10_5 | HOXB9 | 0.73710461 | 1 | -0.457 | 3.302 | chr17 | 46698518 | 46703720 | - | protein_coding |
| ENSG00000102287.19_5 | GABRE | 0.92422676 | 1 | -0.457 | 5.661 | chrX | 151121596 | 151143152 | - | protein_coding |
| ENSG00000273841.5_6 | TAF9 | 0.91126892 | 1 | -0.457 | 5.810 | chr5 | 68660570 | 68665840 | - | protein_coding |
| ENSG00000116044.16_8 | NFE2L2 | 0.95641754 | 1 | -0.457 | 6.691 | chr2 | 178092323 | 178257425 | - | protein_coding |
| ENSG00000115459.18_3 | ELMOD3 | 0.82736454 | 1 | -0.457 | 4.155 | chr2 | 85581517 | 85618875 | + | protein_coding |
| ENSG00000273402.1_7 | AC004908.3 | 0.39301099 | 1 | -0.457 | -0.926 | chr8 | 187045 | 187669 | + | lncRNA |
| ENSG00000128923.11_6 | MINDY2 | 0.87949855 | 1 | -0.458 | 4.991 | chr15 | 59063391 | 59154099 | + | protein_coding |
| ENSG00000109971.14_8 | HSPA8 | 0.97870986 | 1 | -0.458 | 10.978 | chr11 | 122928197 | 122933938 | - | protein_coding |
| ENSG00000277462.2_5 | ZNF670 | 0.58670677 | 1 | -0.458 | 2.213 | chr1 | 247197939 | 247242113 | - | protein_coding |
| ENSG00000169288.18_4 | MRPL1 | 0.80663215 | 1 | -0.458 | 4.229 | chr4 | 78783984 | 78873939 | + | protein_coding |
| ENSG00000149136.9_5 | SSRP1 | 0.96408551 | 1 | -0.458 | 7.929 | chr11 | 57093460 | 57103366 | - | protein_coding |
| ENSG00000122565.19_3 | CBX3 | 0.96349165 | 1 | -0.458 | 7.902 | chr7 | 26240782 | 26253227 | + | protein_coding |
| ENSG00000048991.16_4 | R3HDM1 | 0.89494671 | 1 | -0.458 | 5.059 | chr2 | 136289025 | 136482840 | + | protein_coding |
| ENSG00000224468.3_7 | LAMC1-AS1 | 0.41318407 | 1 | -0.458 | -0.139 | chr1 | 183107537 | 183110417 | - | lncRNA |
| ENSG00000272121.1_6 | AC006058.3 | 0.46816271 | 1 | -0.459 | 0.673 | chr3 | 44039573 | 44040641 | - | lncRNA |
| ENSG00000269534.5_6 | AC011466.3 | 0.39573962 | 1 | -0.459 | -0.574 | chr19 | 48621689 | 48630963 | + | lncRNA |
| ENSG00000229944.3_4 | EIF4EP2 | 0.45721304 | 1 | -0.459 | 1.097 | chr17 | 47501631 | 47502284 | - | processed_pseudogene |
| ENSG00000116161.18_4 | CACYBP | 0.95617276 | 1 | -0.459 | 7.129 | chr1 | 174968300 | 174981163 | + | protein_coding |
| ENSG00000158406.4_4 | H4C8 | 0.76282386 | 1 | -0.459 | 6.390 | chr6 | 26277837 | 26285866 | - | protein_coding |
| ENSG00000162664.17_3 | ZNF326 | 0.87496055 | 1 | -0.459 | 4.937 | chr1 | 90460669 | 90501092 | + | protein_coding |
| ENSG00000115677.17_7 | HDLBP | 0.97939898 | 1 | -0.459 | 9.196 | chr2 | 242166679 | 242256476 | - | protein_coding |
| ENSG00000236200.6_7 | KDM4A-AS1 | 0.57026219 | 1 | -0.459 | 1.727 | chr1 | 44150794 | 44173809 | - | lncRNA |
| ENSG00000163798.13_4 | SLC4A1AP | 0.93153413 | 1 | -0.459 | 5.558 | chr2 | 27886338 | 27917843 | + | protein_coding |
| ENSG00000094804.12_6 | CDC6 | 0.96106493 | 1 | -0.460 | 6.973 | chr17 | 38444131 | 38460909 | + | protein_coding |
| ENSG00000254165.1_8 | AC090739.1 | 0.41991908 | 1 | -0.460 | -0.012 | chr8 | 42392672 | 42393447 | - | lncRNA |
| ENSG00000264304.1_7 | AC024267.4 | 0.39179921 | 1 | -0.460 | -0.584 | chr17 | 27219487 | 27220360 | + | lncRNA |
| ENSG00000272918.1_6 | AC005070.3 | 0.52144 | 1 | -0.460 | 1.319 | chr7 | 104743285 | 104745930 | - | lncRNA |
| ENSG00000218274.2_7 | AL121949.2 | 0.39867893 | 1 | -0.460 | -1.029 | chr6 | 64104912 | 64107011 | - | unprocessed_pseudogene |
| ENSG00000076382.17_7 | SPAG5 | 0.96518988 | 1 | -0.460 | 7.151 | chr17 | 26904583 | 26926043 | - | protein_coding |
| ENSG00000176531.10_3 | PHLDB3 | 0.64304825 | 1 | -0.460 | 2.772 | chr19 | 43979106 | 44009087 | - | protein_coding |
| ENSG00000183665.5_3 | TRMT12 | 0.7940949 | 1 | -0.460 | 3.889 | chr8 | 125463061 | 125474391 | + | protein_coding |
| ENSG00000215817.7_7 | ZC3H11B | 0.39903203 | 1 | -0.461 | -1.101 | chr1 | 219781352 | 219786487 | - | protein_coding |
| ENSG00000009844.16_4 | VTA1 | 0.87320229 | 1 | -0.461 | 5.845 | chr6 | 142468299 | 142545822 | + | protein_coding |
| ENSG00000177688.7_6 | SUMO4 | 0.39058621 | 1 | -0.461 | -0.996 | chr6 | 149721398 | 149722414 | + | protein_coding |
| ENSG00000120539.14_3 | MASTL | 0.91156735 | 1 | -0.461 | 5.998 | chr10 | 27443753 | 27475853 | + | protein_coding |
| ENSG00000078401.7_3 | EDN1 | 0.93453024 | 1 | -0.461 | 6.697 | chr6 | 12290594 | 12297427 | + | protein_coding |
| ENSG00000132603.15_6 | NIP7 | 0.89942088 | 1 | -0.461 | 5.468 | chr16 | 69371899 | 69377009 | + | protein_coding |
| ENSG00000286619.1_2 | AC119673.3 | 0.60337014 | 1 | -0.461 | 2.115 | chr1 | 205782450 | 205865210 | + | lncRNA |
| ENSG00000187961.14_5 | KLHL17 | 0.71457485 | 1 | -0.461 | 3.569 | chr1 | 895964 | 901099 | + | protein_coding |
| ENSG00000063127.16_7 | SLC6A16 | 0.49262857 | 1 | -0.461 | 1.303 | chr19 | 49792895 | 49828472 | - | protein_coding |
| ENSG00000101166.16_4 | PRELID3B | 0.94075492 | 1 | -0.462 | 6.774 | chr20 | 57608200 | 57617864 | - | protein_coding |
| ENSG00000164134.13_3 | NAA15 | 0.94322705 | 1 | -0.462 | 6.854 | chr4 | 140222659 | 140341187 | + | protein_coding |
| ENSG00000258017.2_7 | AC011603.3 | 0.38911109 | 1 | -0.462 | -0.629 | chr12 | 49521565 | 49582267 | + | lncRNA |
| ENSG00000272308.1_6 | AC104113.1 | 0.38903264 | 1 | -0.462 | -0.895 | chr5 | 60162284 | 60162762 | - | lncRNA |
| ENSG00000152683.14_4 | SLC30A6 | 0.93753426 | 1 | -0.462 | 6.155 | chr2 | 32390910 | 32449448 | + | protein_coding |
| ENSG00000256802.2_6 | AC022613.1 | 0.82743026 | 1 | -0.462 | 4.399 | chr15 | 29967194 | 29973161 | + | lncRNA |
| ENSG00000215210.3_4 | RBMXP2 | 0.43211427 | 1 | -0.462 | 0.494 | chr9 | 30689103 | 30690270 | + | processed_pseudogene |
| ENSG00000101187.16_4 | SLCO4A1 | 0.69282405 | 1 | -0.462 | 2.970 | chr20 | 61273855 | 61317137 | + | protein_coding |
| ENSG00000284523.2_7 | AC004834.1 | 0.39679462 | 1 | -0.462 | -1.178 | chr7 | 98850075 | 98923449 | - | lncRNA |
| ENSG00000229132.2_4 | EIF4A1P10 | 0.53056234 | 1 | -0.462 | 2.550 | chrX | 91368245 | 91369460 | - | processed_pseudogene |
| ENSG00000081052.13_4 | COL4A4 | 0.84243298 | 1 | -0.463 | 4.504 | chr2 | 227867430 | 228029169 | - | protein_coding |
| ENSG00000256269.10_7 | HMBS | 0.81498291 | 1 | -0.463 | 4.187 | chr11 | 118955576 | 118964259 | + | protein_coding |
| ENSG00000174672.16_6 | BRSK2 | 0.63013747 | 1 | -0.463 | 2.800 | chr11 | 1411129 | 1483919 | + | protein_coding |
| ENSG00000168275.15_3 | COA6 | 0.78226909 | 1 | -0.463 | 4.271 | chr1 | 234509202 | 234520814 | + | protein_coding |
| ENSG00000145331.14_6 | TRMT10A | 0.7038312 | 1 | -0.464 | 3.151 | chr4 | 100467866 | 100485196 | - | protein_coding |
| ENSG00000145390.11_2 | USP53 | 0.85801853 | 1 | -0.464 | 5.925 | chr4 | 120133742 | 120216672 | + | protein_coding |
| ENSG00000246922.8_3 | UBAP1L | 0.56164106 | 1 | -0.464 | 1.983 | chr15 | 65385108 | 65407535 | - | protein_coding |

| | | | | | | | | | | |
|---|---|---|---|---|---|---|---|---|---|---|
| ENSG00000172403.11_7 | SYNPO2 | 0.78779865 | 1 | -0.464 | 4.064 | chr4 | 119771843 | 119982402 | + | protein_coding |
| ENSG00000180581.7_4 | SRP9P1 | 0.45991823 | 1 | -0.464 | 0.818 | chr10 | 93566936 | 93567196 | - | processed_pseudogene |
| ENSG00000117118.10_3 | SDHB | 0.92294652 | 1 | -0.464 | 6.189 | chr1 | 17345217 | 17380527 | - | protein_coding |
| ENSG00000003509.16_4 | NDUFAF7 | 0.85928519 | 1 | -0.464 | 4.839 | chr2 | 37458774 | 37480546 | + | protein_coding |
| ENSG00000259319.1_7 | AF111167.2 | 0.45120266 | 1 | -0.465 | 0.567 | chr14 | 75890386 | 75894444 | - | lncRNA |
| ENSG00000150510.16_4 | FAM124A | 0.47612561 | 1 | -0.465 | 1.488 | chr13 | 51796470 | 51858377 | + | protein_coding |
| ENSG00000272853.1_6 | AC069544.1 | 0.40655112 | 1 | -0.465 | 0.151 | chr10 | 14919687 | 14920685 | - | lncRNA |
| ENSG00000132507.18_3 | EIF5A | 0.95915934 | 1 | -0.465 | 8.325 | chr17 | 7210318 | 7215782 | + | protein_coding |
| ENSG00000084754.12_6 | HADHA | 0.96424675 | 1 | -0.465 | 7.600 | chr2 | 26413504 | 26467540 | - | protein_coding |
| ENSG00000183598.3_3 | H3C13 | 0.49813787 | 1 | -0.465 | 6.320 | chr1 | 149784826 | 149785236 | - | protein_coding |
| ENSG00000258704.6_6 | SRP54-AS1 | 0.42526459 | 1 | -0.465 | 0.402 | chr14 | 35390064 | 35451738 | - | transcribed_unitary_pseudogene |
| ENSG00000003402.20_4 | CFLAR | 0.92748704 | 1 | -0.465 | 5.877 | chr2 | 201980877 | 202041410 | + | protein_coding |
| ENSG00000163491.16_7 | NEK10 | 0.65943239 | 1 | -0.465 | 2.847 | chr3 | 27151576 | 27410951 | - | protein_coding |
| ENSG00000104728.16_6 | ARHGEF10 | 0.49152773 | 1 | -0.466 | 1.405 | chr8 | 1772092 | 1906807 | + | protein_coding |
| ENSG00000181227.3_4 | DLSTP1 | 0.38844953 | 1 | -0.466 | -0.322 | chr1 | 76209108 | 76210461 | - | processed_pseudogene |
| ENSG00000124831.19_5 | LRRFIP1 | 0.92528612 | 1 | -0.466 | 6.013 | chr2 | 238536230 | 238722325 | + | protein_coding |
| ENSG00000176884.15_5 | GRIN1 | 0.41247526 | 1 | -0.466 | -0.180 | chr9 | 140033606 | 140063208 | + | protein_coding |
| ENSG00000173660.12_3 | UQCRH | 0.90689756 | 1 | -0.466 | 6.464 | chr1 | 46769370 | 46782448 | + | protein_coding |
| ENSG00000088325.16_3 | TPX2 | 0.96655452 | 1 | -0.466 | 7.893 | chr20 | 30327074 | 30389608 | + | protein_coding |
| ENSG00000238832.1 | snoU109 | 0.3954093 | 1 | -0.466 | -0.004 | chr7 | 107243808 | 107243952 | + | snoRNA |
| ENSG00000120705.13_5 | ETF1 | 0.95907342 | 1 | -0.467 | 7.776 | chr5 | 137841784 | 137878925 | - | protein_coding |
| ENSG00000093009.10_3 | CDC45 | 0.8412727 | 1 | -0.467 | 4.880 | chr22 | 19466980 | 19508135 | + | protein_coding |
| ENSG00000110700.7_5 | RPS13 | 0.90986193 | 1 | -0.467 | 8.080 | chr11 | 17095935 | 17099262 | - | protein_coding |
| ENSG00000225362.9_5 | CT62 | 0.44618156 | 1 | -0.467 | 0.589 | chr15 | 71402583 | 71407839 | - | protein_coding |
| ENSG00000147099.21_5 | HDAC8 | 0.70856203 | 1 | -0.467 | 3.239 | chrX | 71549366 | 71792951 | - | protein_coding |
| ENSG00000141101.13_4 | NOB1 | 0.88330732 | 1 | -0.467 | 5.146 | chr16 | 69775774 | 69788829 | - | protein_coding |
| ENSG00000198648.11_5 | STK39 | 0.88520131 | 1 | -0.467 | 5.258 | chr2 | 168810530 | 169104105 | - | protein_coding |
| ENSG00000251229.1_6 | AL645924.2 | 0.39029932 | 1 | -0.467 | -0.157 | chr4 | 2506808 | 2508006 | - | processed_pseudogene |
| ENSG00000071282.12_4 | LMCD1 | 0.89775425 | 1 | -0.467 | 5.281 | chr3 | 8543493 | 8616354 | + | protein_coding |
| ENSG00000196368.5_5 | NUDT11 | 0.50287052 | 1 | -0.467 | 1.238 | chrX | 51232863 | 51239444 | - | protein_coding |
| ENSG00000143314.12_5 | MRPL24 | 0.91637699 | 1 | -0.467 | 5.889 | chr1 | 156707095 | 156711382 | - | protein_coding |
| ENSG00000119599.17_6 | DCAF4 | 0.77794388 | 1 | -0.468 | 4.053 | chr14 | 73393085 | 73426411 | + | protein_coding |
| ENSG00000196072.12_6 | BLOC1S2 | 0.87787591 | 1 | -0.468 | 5.417 | chr10 | 102033037 | 102046437 | - | protein_coding |
| ENSG00000144029.12_5 | MRPS5 | 0.93209737 | 1 | -0.468 | 6.044 | chr2 | 95751114 | 95815179 | - | protein_coding |
| ENSG00000100288.19_4 | CHKB | 0.39027472 | 1 | -0.468 | -0.485 | chr22 | 51017378 | 51039884 | - | protein_coding |
| ENSG00000229852.4_8 | AC019205.1 | 0.54686954 | 1 | -0.468 | 1.709 | chr6 | 73972935 | 74011512 | + | lncRNA |
| ENSG00000133111.3_2 | RFXAP | 0.4331509 | 1 | -0.468 | 0.454 | chr13 | 37393361 | 37403241 | + | protein_coding |
| ENSG00000102908.22_7 | NFAT5 | 0.84763159 | 1 | -0.468 | 6.564 | chr16 | 69598997 | 69738569 | + | protein_coding |
| ENSG00000013441.16_5 | CLK1 | 0.90401933 | 1 | -0.468 | 5.899 | chr2 | 201717732 | 201729467 | - | protein_coding |
| ENSG00000183527.12_4 | PSMG1 | 0.85718533 | 1 | -0.468 | 4.989 | chr21 | 40546695 | 40555414 | - | protein_coding |
| ENSG00000158373.8_4 | H2BC5 | 0.95491044 | 1 | -0.468 | 8.093 | chr6 | 26158374 | 26171577 | + | protein_coding |
| ENSG00000267787.7_7 | AC027097.2 | 0.38803484 | 1 | -0.468 | -0.118 | chr18 | 55306687 | 55405835 | + | lncRNA |
| ENSG00000249207.2_7 | AC079921.1 | 0.41552702 | 1 | -0.469 | 0.472 | chr4 | 39068594 | 39183878 | - | lncRNA |
| ENSG00000287809.1_2 | AC005632.6 | 0.42791502 | 1 | -0.469 | 0.428 | chr16 | 21522956 | 21526452 | + | lncRNA |
| ENSG00000246877.1_7 | DNM1P35 | 0.38589127 | 1 | -0.469 | -0.864 | chr15 | 76020011 | 76030964 | - | lncRNA |
| ENSG00000243517.1_6 | AC024940.3 | 0.38168823 | 1 | -0.469 | -0.640 | chr12 | 31404926 | 31405531 | - | processed_pseudogene |
| ENSG00000103995.14_7 | CEP152 | 0.83665604 | 1 | -0.469 | 4.642 | chr15 | 49005125 | 49103343 | - | protein_coding |
| ENSG00000145088.9_4 | EAF2 | 0.49581378 | 1 | -0.469 | 1.275 | chr3 | 121554030 | 121605373 | + | protein_coding |
| ENSG00000106105.14_4 | GARS1 | 0.95296494 | 1 | -0.470 | 7.341 | chr7 | 30634454 | 30673649 | + | protein_coding |
| ENSG00000279696.1_5 | AP001273.1 | 0.48602288 | 1 | -0.470 | 1.593 | chr11 | 93459820 | 93462971 | - | TEC |
| ENSG00000229666.1_7 | MAST4-AS1 | 0.40674752 | 1 | -0.470 | 0.017 | chr5 | 66297211 | 66299781 | - | lncRNA |
| ENSG00000198721.12_7 | ECI2 | 0.7551192 | 1 | -0.470 | 3.968 | chr6 | 4115923 | 4135831 | - | protein_coding |
| ENSG00000269821.1_5 | KCNQ1OT1 | 0.78525952 | 1 | -0.470 | 4.573 | chr11 | 2629558 | 2721224 | - | lncRNA |
| ENSG00000134480.15_5 | CCNH | 0.81976873 | 1 | -0.471 | 4.770 | chr5 | 86614233 | 86708747 | - | protein_coding |
| ENSG00000163138.19_4 | PACRGL | 0.77121329 | 1 | -0.471 | 4.052 | chr4 | 20697905 | 20754530 | + | protein_coding |
| ENSG00000134070.5_4 | IRAK2 | 0.8064962 | 1 | -0.471 | 4.227 | chr3 | 10206603 | 10285429 | + | protein_coding |
| ENSG00000269487.1_7 | AC008635.2 | 0.3992781 | 1 | -0.471 | 0.055 | chr19 | 47231860 | 47235957 | + | lncRNA |
| ENSG00000249249.2_6 | AC010226.1 | 0.51261856 | 1 | -0.471 | 1.718 | chr5 | 114937754 | 114959594 | + | lncRNA |

| | | | | | | | | | | |
|---|---|---|---|---|---|---|---|---|---|---|
| ENSG00000135698.10_4 | MPHOSPH6 | 0.77482765 | 1 | -0.471 | 4.156 | chr16 | 82181403 | 82203829 | - | protein_coding |
| ENSG00000129195.16_6 | PIMREG | 0.83626006 | 1 | -0.471 | 5.278 | chr17 | 6347761 | 6354789 | + | protein_coding |
| ENSG00000169016.17_5 | E2F6 | 0.84001691 | 1 | -0.471 | 4.953 | chr2 | 11584501 | 11606303 | - | protein_coding |
| ENSG00000154839.10_4 | SKA1 | 0.8682876 | 1 | -0.471 | 4.984 | chr18 | 47901410 | 47920538 | + | protein_coding |
| ENSG00000171848.15_5 | RRM2 | 0.93719141 | 1 | -0.471 | 6.887 | chr2 | 10260825 | 10351851 | + | protein_coding |
| ENSG00000163291.14_4 | PAQR3 | 0.73656958 | 1 | -0.471 | 3.907 | chr4 | 79808281 | 79860592 | - | protein_coding |
| ENSG00000274251.1_6 | AP003419.2 | 0.38534477 | 1 | -0.472 | -0.485 | chr11 | 67121100 | 67121819 | + | lncRNA |
| ENSG00000162757.4_4 | C1orf74 | 0.59917205 | 1 | -0.472 | 2.532 | chr1 | 209952553 | 209957904 | - | protein_coding |
| ENSG00000135945.10_5 | REV1 | 0.85805001 | 1 | -0.472 | 5.034 | chr2 | 100016938 | 100106497 | - | protein_coding |
| ENSG00000177855.8_4 | CACYBPP2 | 0.46906009 | 1 | -0.472 | 0.995 | chr2 | 184472170 | 184472847 | - | processed_pseudogene |
| ENSG00000169967.16_3 | MAP3K2 | 0.92691736 | 1 | -0.472 | 6.751 | chr2 | 128056306 | 128146041 | - | protein_coding |
| ENSG00000134690.11_3 | CDCA8 | 0.86237076 | 1 | -0.472 | 5.053 | chr1 | 38158153 | 38175391 | + | protein_coding |
| ENSG00000011347.10_7 | SYT7 | 0.47076452 | 1 | -0.473 | 0.808 | chr11 | 61281186 | 61348620 | - | protein_coding |
| ENSG00000154678.18_8 | PDE1C | 0.90639587 | 1 | -0.473 | 6.205 | chr7 | 31790793 | 32467743 | - | protein_coding |
| ENSG00000227615.1_5 | AP001324.1 | 0.4819701 | 1 | -0.473 | 1.275 | chr11 | 74456761 | 74457159 | - | processed_pseudogene |
| ENSG00000154814.14_4 | OXNAD1 | 0.7902377 | 1 | -0.473 | 4.134 | chr3 | 16306667 | 16391806 | + | protein_coding |
| ENSG00000214389.2_2 | RPS3AP26 | 0.53650488 | 1 | -0.473 | 1.757 | chr7 | 98015113 | 98015896 | - | processed_pseudogene |
| ENSG00000079785.15_5 | DDX1 | 0.93642657 | 1 | -0.473 | 7.287 | chr2 | 15731302 | 15771235 | + | protein_coding |
| ENSG00000105854.13_7 | PON2 | 0.94138642 | 1 | -0.473 | 8.081 | chr7 | 95034174 | 95064641 | - | protein_coding |
| ENSG00000138398.16_4 | PPIG | 0.89400656 | 1 | -0.474 | 5.998 | chr2 | 170440854 | 170497916 | + | protein_coding |
| ENSG00000121578.13_6 | B4GALT4 | 0.9417665 | 1 | -0.474 | 6.924 | chr3 | 118930579 | 118959793 | - | protein_coding |
| ENSG00000088179.9_6 | PTPN4 | 0.86912125 | 1 | -0.474 | 5.470 | chr2 | 120517498 | 120742475 | + | protein_coding |
| ENSG00000092445.12_4 | TYRO3 | 0.73474376 | 1 | -0.474 | 3.823 | chr15 | 41849873 | 41875787 | + | protein_coding |
| ENSG00000165240.20_5 | ATP7A | 0.89310135 | 1 | -0.474 | 5.833 | chrX | 77166153 | 77305892 | + | protein_coding |
| ENSG00000272578.5_6 | AP000347.1 | 0.37371349 | 1 | -0.474 | -0.667 | chr22 | 24000281 | 24059543 | - | transcribed_unprocessed_pseudogene |
| ENSG00000023608.5_4 | SNAPC1 | 0.7016889 | 1 | -0.475 | 3.813 | chr14 | 62229138 | 62263146 | + | protein_coding |
| ENSG00000146918.20_5 | NCAPG2 | 0.93257251 | 1 | -0.475 | 6.568 | chr7 | 158423861 | 158497495 | - | protein_coding |
| ENSG00000216657.1_4 | GLRX3P2 | 0.37262352 | 1 | -0.475 | -0.783 | chr6 | 3978296 | 3979333 | + | processed_pseudogene |
| ENSG00000244313.3_5 | AC024293.1 | 0.59207682 | 1 | -0.475 | 2.956 | chr11 | 46450203 | 46450700 | - | processed_pseudogene |
| ENSG00000115944.15_6 | COX7A2L | 0.8904284 | 1 | -0.475 | 6.440 | chr2 | 42560686 | 42652228 | - | protein_coding |
| ENSG00000152484.14_4 | USP12 | 0.79646906 | 1 | -0.475 | 5.162 | chr13 | 27640293 | 27745948 | - | protein_coding |
| ENSG00000111371.16_5 | SLC38A1 | 0.94155401 | 1 | -0.476 | 8.061 | chr12 | 46576846 | 46663800 | - | protein_coding |
| ENSG00000146733.14_4 | PSPH | 0.86517711 | 1 | -0.476 | 5.228 | chr7 | 56078744 | 56119297 | - | protein_coding |
| ENSG00000104432.14_5 | IL7 | 0.38593684 | 1 | -0.476 | -1.221 | chr8 | 79587978 | 79717758 | - | protein_coding |
| ENSG00000136098.17_5 | NEK3 | 0.61704571 | 1 | -0.476 | 2.990 | chr13 | 52706775 | 52733996 | - | protein_coding |
| ENSG00000134987.11_3 | WDR36 | 0.91405219 | 1 | -0.476 | 6.667 | chr5 | 110427414 | 110466200 | + | protein_coding |
| ENSG00000101444.13_5 | AHCY | 0.93144537 | 1 | -0.476 | 7.149 | chr20 | 32868074 | 32899608 | - | protein_coding |
| ENSG00000206538.9_5 | VGLL3 | 0.90057109 | 1 | -0.476 | 5.974 | chr3 | 86925538 | 87040299 | - | protein_coding |
| ENSG00000203711.14_7 | C6orf99 | 0.4118208 | 1 | -0.476 | 0.437 | chr6 | 159290880 | 159340137 | + | lncRNA |
| ENSG00000082516.9_4 | GEMIN5 | 0.92498121 | 1 | -0.476 | 6.374 | chr5 | 154266971 | 154317771 | - | protein_coding |
| ENSG00000216866.5_4 | RPS2P55 | 0.4684181 | 1 | -0.477 | 1.482 | chrX | 40794235 | 40795117 | - | processed_pseudogene |
| ENSG00000230479.1_7 | AP000695.1 | 0.37164118 | 1 | -0.477 | -0.813 | chr21 | 37802658 | 37853368 | + | lncRNA |
| ENSG00000288538.1_1 | AP002985.1 | 0.40608162 | 1 | -0.477 | 0.205 | chr11 | 77934861 | 77936955 | + | lncRNA |
| ENSG00000140534.14_4 | TICRR | 0.8833208 | 1 | -0.477 | 5.534 | chr15 | 90118700 | 90174287 | + | protein_coding |
| ENSG00000138134.12_4 | STAMBPL1 | 0.81802695 | 1 | -0.477 | 4.782 | chr10 | 90639491 | 90734910 | + | protein_coding |
| ENSG00000064309.14_3 | CDON | 0.64400549 | 1 | -0.477 | 3.249 | chr11 | 125825691 | 125933230 | - | protein_coding |
| ENSG00000267040.6_7 | AC027097.1 | 0.38823007 | 1 | -0.477 | -0.091 | chr18 | 55297534 | 55336528 | + | lncRNA |
| ENSG00000252690.3_6 | AC105339.2 | 0.45866642 | 1 | -0.478 | 1.206 | chr15 | 83421636 | 83425960 | + | lncRNA |
| ENSG00000113569.16_4 | NUP155 | 0.92967033 | 1 | -0.478 | 7.400 | chr5 | 37288239 | 37371208 | - | protein_coding |
| ENSG00000157456.8_4 | CCNB2 | 0.88862808 | 1 | -0.478 | 6.559 | chr15 | 59397325 | 59417244 | + | protein_coding |
| ENSG00000137522.18_4 | RNF121 | 0.83939789 | 1 | -0.478 | 4.936 | chr11 | 71640064 | 71708643 | + | protein_coding |
| ENSG00000144559.10_5 | TAMM41 | 0.59433878 | 1 | -0.478 | 2.533 | chr3 | 11831916 | 11888393 | - | protein_coding |
| ENSG00000274997.1_3 | H2AC12 | 0.88798708 | 1 | -0.478 | 7.354 | chr6 | 27114908 | 27115294 | + | protein_coding |
| ENSG00000274828.1_7 | AC068473.5 | 0.46602716 | 1 | -0.479 | 1.471 | chr18 | 77437287 | 77439358 | - | lncRNA |
| ENSG00000198870.8_4 | STKLD1 | 0.44755456 | 1 | -0.479 | 1.055 | chr9 | 136243237 | 136271220 | + | protein_coding |
| ENSG00000107957.16_4 | SH3PXD2A | 0.68980612 | 1 | -0.479 | 3.984 | chr10 | 105353784 | 105615301 | - | protein_coding |
| ENSG00000234241.1_5 | AL109618.1 | 0.37373305 | 1 | -0.479 | -0.400 | chr20 | 6194346 | 6196028 | + | processed_pseudogene |

| | | | | | | | | | | |
|---|---|---|---|---|---|---|---|---|---|---|
| ENSG00000166343.10_5 | MSS51 | 0.38609976 | 1 | -0.479 | 0.130 | chr10 | 75183337 | 75193319 | - | protein_coding |
| ENSG00000160193.12_4 | WDR4 | 0.69646385 | 1 | -0.479 | 3.813 | chr21 | 44263204 | 44299678 | - | protein_coding |
| ENSG00000174977.8_5 | AC026271.1 | 0.6040808 | 1 | -0.480 | 2.698 | chr17 | 18553508 | 18554855 | + | processed_pseudogene |
| ENSG00000131470.14_3 | PSMC3IP | 0.64593686 | 1 | -0.480 | 3.124 | chr17 | 40724333 | 40729849 | - | protein_coding |
| ENSG00000273703.1_3 | H2BC14 | 0.87755716 | 1 | -0.480 | 6.215 | chr6 | 27782822 | 27783202 | + | protein_coding |
| ENSG00000162946.22_6 | DISC1 | 0.41827209 | 1 | -0.480 | 0.737 | chr1 | 231762561 | 232177018 | + | protein_coding |
| ENSG00000236526.1_7 | AL035448.1 | 0.38555447 | 1 | -0.480 | -0.007 | chr20 | 11890052 | 11899325 | - | lncRNA |
| ENSG00000168906.13_3 | MAT2A | 0.92047652 | 1 | -0.480 | 7.297 | chr2 | 85766291 | 85772404 | + | protein_coding |
| ENSG00000149554.13_5 | CHEK1 | 0.83292223 | 1 | -0.481 | 5.653 | chr11 | 125495560 | 125546150 | + | protein_coding |
| ENSG00000121716.20_6 | PILRB | 0.78217292 | 1 | -0.481 | 4.711 | chr7 | 99949799 | 99965356 | + | protein_coding |
| ENSG00000261790.2_7 | AC005606.2 | 0.38510448 | 1 | -0.481 | -0.169 | chr16 | 2034878 | 2040478 | - | lncRNA |
| ENSG00000273373.1_6 | AL355488.1 | 0.467024 | 1 | -0.481 | 1.342 | chr1 | 110912776 | 110915625 | + | lncRNA |
| ENSG00000171055.15_8 | FEZ2 | 0.89459555 | 1 | -0.481 | 6.416 | chr2 | 36758948 | 36873230 | - | protein_coding |
| ENSG00000141480.18_5 | ARRB2 | 0.8971851 | 1 | -0.481 | 6.512 | chr17 | 4613891 | 4624794 | + | protein_coding |
| ENSG00000185168.5_5 | LINC00482 | 0.4607497 | 1 | -0.481 | 1.311 | chr17 | 79277571 | 79283048 | - | lncRNA |
| ENSG00000080819.9_7 | CPOX | 0.7642125 | 1 | -0.481 | 4.573 | chr3 | 98298290 | 98312492 | - | protein_coding |
| ENSG00000249119.1_4 | MTND6P4 | 0.36524029 | 1 | -0.481 | -0.783 | chr5 | 134259813 | 134260334 | + | processed_pseudogene |
| ENSG00000236830.7_6 | CBR3-AS1 | 0.36520388 | 1 | -0.482 | -0.726 | chr21 | 37504065 | 37548113 | - | lncRNA |
| ENSG00000142945.13_5 | KIF2C | 0.87674363 | 1 | -0.482 | 5.650 | chr1 | 45205490 | 45233439 | + | protein_coding |
| ENSG00000084731.15_5 | KIF3C | 0.89595906 | 1 | -0.482 | 6.214 | chr2 | 26149467 | 26205618 | - | protein_coding |
| ENSG00000081041.9_4 | CXCL2 | 0.53135494 | 1 | -0.482 | 2.516 | chr4 | 74962757 | 74964913 | - | protein_coding |
| ENSG00000132541.11_5 | RIDA | 0.70205664 | 1 | -0.482 | 4.474 | chr8 | 99114572 | 99129399 | - | protein_coding |
| ENSG00000277534.1_7 | AC007996.1 | 0.7169812 | 1 | -0.483 | 4.054 | chr18 | 24122935 | 24125755 | - | lncRNA |
| ENSG00000181163.13_3 | NPM1 | 0.93490978 | 1 | -0.483 | 10.126 | chr5 | 170814120 | 170838141 | + | protein_coding |
| ENSG00000104687.14_6 | GSR | 0.93603374 | 1 | -0.483 | 8.595 | chr8 | 30535583 | 30585363 | - | protein_coding |
| ENSG00000165244.7_4 | ZNF367 | 0.82989259 | 1 | -0.483 | 5.210 | chr9 | 99148223 | 99180652 | - | protein_coding |
| ENSG00000262833.1_7 | AC016245.1 | 0.36653154 | 1 | -0.483 | -0.909 | chr17 | 78775440 | 78779432 | - | lncRNA |
| ENSG00000215883.10_6 | CYB5RL | 0.51360199 | 1 | -0.484 | 2.251 | chr1 | 54635324 | 54665709 | - | protein_coding |
| ENSG00000117697.15_4 | NSL1 | 0.82542374 | 1 | -0.484 | 5.630 | chr1 | 212899495 | 212965124 | - | protein_coding |
| ENSG00000271204.1_8 | AC016831.4 | 0.39419908 | 1 | -0.484 | 0.276 | chr7 | 130614968 | 130616965 | - | lncRNA |
| ENSG00000104738.18_7 | MCM4 | 0.93585687 | 1 | -0.484 | 8.165 | chr8 | 48872745 | 48890720 | + | protein_coding |
| ENSG00000115446.11_2 | UNC50 | 0.74749132 | 1 | -0.484 | 4.643 | chr2 | 99225042 | 99234978 | + | protein_coding |
| ENSG00000143797.12_6 | MBOAT2 | 0.85563198 | 1 | -0.484 | 5.644 | chr2 | 8992820 | 9143838 | - | protein_coding |
| ENSG00000198860.12_4 | TSEN15 | 0.81419957 | 1 | -0.484 | 5.266 | chr1 | 184020785 | 184093112 | + | protein_coding |
| ENSG00000135049.15_5 | AGTPBP1 | 0.85685784 | 1 | -0.485 | 5.603 | chr9 | 88161454 | 88356944 | - | protein_coding |
| ENSG00000244627.5_7 | TPTEP2 | 0.42653439 | 1 | -0.485 | 0.534 | chr22 | 38731767 | 38794934 | - | transcribed_unprocessed_pseudogene |
| ENSG00000269815.1_7 | AC010463.3 | 0.377337 | 1 | -0.485 | -0.667 | chr19 | 17407005 | 17407504 | - | lncRNA |
| ENSG00000049883.15_4 | PTCD2 | 0.76166493 | 1 | -0.485 | 4.716 | chr5 | 71616194 | 71664222 | + | protein_coding |
| ENSG00000286688.1_3 | AP002748.6 | 0.36300147 | 1 | -0.486 | -0.974 | chr11 | 66281777 | 66287191 | - | lncRNA |
| ENSG00000287080.1_2 | H3C3 | 0.87690601 | 1 | -0.486 | 6.435 | chr6 | 26045639 | 26046097 | + | protein_coding |
| ENSG00000135930.14_9 | EIF4E2 | 0.82844718 | 1 | -0.486 | 5.428 | chr2 | 233415384 | 233448354 | + | protein_coding |
| ENSG00000142330.20_4 | CAPN10 | 0.72643383 | 1 | -0.486 | 4.528 | chr2 | 241526151 | 241557122 | + | protein_coding |
| ENSG00000109606.13_4 | DHX15 | 0.92153776 | 1 | -0.486 | 8.112 | chr4 | 24519064 | 24586177 | - | protein_coding |
| ENSG00000075568.17_4 | TMEM131 | 0.8494106 | 1 | -0.486 | 5.747 | chr2 | 98372796 | 98612411 | - | protein_coding |
| ENSG00000183765.22_7 | CHEK2 | 0.67355807 | 1 | -0.486 | 3.749 | chr22 | 29083731 | 29138410 | - | protein_coding |
| ENSG00000138413.13_3 | IDH1 | 0.91020682 | 1 | -0.486 | 7.695 | chr2 | 209100951 | 209130798 | - | protein_coding |
| ENSG00000164163.11_4 | ABCE1 | 0.88172869 | 1 | -0.486 | 7.078 | chr4 | 146019440 | 146050676 | + | protein_coding |
| ENSG00000188010.14_5 | MORN2 | 0.68750764 | 1 | -0.486 | 3.768 | chr2 | 39103103 | 39156213 | + | protein_coding |
| ENSG00000064012.22_9 | CASP8 | 0.84206996 | 1 | -0.486 | 5.537 | chr2 | 202098166 | 202152434 | + | protein_coding |
| ENSG00000112312.10_4 | GMNN | 0.7975389 | 1 | -0.487 | 5.022 | chr6 | 24775159 | 24786327 | + | protein_coding |
| ENSG00000118263.15_5 | KLF7 | 0.81422352 | 1 | -0.487 | 4.978 | chr2 | 207938861 | 208031991 | - | protein_coding |
| ENSG00000138658.15_6 | ZGRF1 | 0.75635827 | 1 | -0.487 | 4.758 | chr4 | 113460489 | 113558151 | - | protein_coding |
| ENSG00000124216.4_3 | SNAI1 | 0.37775728 | 1 | -0.487 | -0.008 | chr20 | 48599517 | 48605423 | + | protein_coding |
| ENSG00000103966.11_5 | EHD4 | 0.66521348 | 1 | -0.487 | 4.302 | chr15 | 42188131 | 42264755 | - | protein_coding |
| ENSG00000287878.1_3 | AC022382.2 | 0.37693026 | 1 | -0.487 | 0.109 | chr3 | 9734866 | 9763083 | - | lncRNA |
| ENSG00000185055.11_6 | EFCAB10 | 0.41329375 | 1 | -0.487 | 0.691 | chr7 | 105205567 | 105241322 | - | protein_coding |
| ENSG00000175455.15_7 | CCDC14 | 0.85739391 | 1 | -0.488 | 6.348 | chr3 | 123616152 | 123680255 | - | protein_coding |
| ENSG00000108448.21_5 | TRIM16L | 0.91547995 | 1 | -0.488 | 7.948 | chr17 | 18601311 | 18639431 | + | protein_coding |
| ENSG00000053900.10_2 | ANAPC4 | 0.75408666 | 1 | -0.488 | 4.938 | chr4 | 25378835 | 25420120 | + | protein_coding |

| | | | | | | | | | | |
|---|---|---|---|---|---|---|---|---|---|---|
| ENSG00000169116.11_4 | PARM1 | 0.54445498 | 1 | -0.488 | 2.475 | chr4 | 75858305 | 75975325 | + | protein_coding |
| ENSG00000142549.9_2 | IGLON5 | 0.3586665 | 1 | -0.488 | -0.563 | chr19 | 51815102 | 51833608 | + | protein_coding |
| ENSG00000013016.16_4 | EHD3 | 0.75227276 | 1 | -0.488 | 4.480 | chr2 | 31457018 | 31492317 | + | protein_coding |
| ENSG00000066855.16_5 | MTFR1 | 0.79470748 | 1 | -0.489 | 5.389 | chr8 | 66556969 | 66683496 | + | protein_coding |
| ENSG00000178921.14_8 | PFAS | 0.79771151 | 1 | -0.489 | 7.225 | chr17 | 8150936 | 8173809 | + | protein_coding |
| ENSG00000141458.13_5 | NPC1 | 0.86379211 | 1 | -0.489 | 6.196 | chr18 | 21086148 | 21166470 | - | protein_coding |
| ENSG00000231789.2_7 | PIK3CD-AS2 | 0.4695944 | 1 | -0.489 | 1.444 | chr1 | 9732484 | 9747613 | - | lncRNA |
| ENSG00000151693.11_5 | ASAP2 | 0.81842293 | 1 | -0.489 | 5.216 | chr2 | 9346894 | 9545812 | + | protein_coding |
| ENSG00000244687.11_4 | UBE2V1 | 0.5481017 | 1 | -0.490 | 2.406 | chr20 | 48697661 | 48732496 | - | protein_coding |
| ENSG00000140451.13_4 | PIF1 | 0.68249299 | 1 | -0.490 | 4.384 | chr15 | 65107831 | 65117867 | - | protein_coding |
| ENSG00000287697.1_2 | Z99127.4 | 0.35680676 | 1 | -0.490 | -0.644 | chr1 | 174967490 | 174968759 | - | lncRNA |
| ENSG00000142677.4_4 | IL22RA1 | 0.48086868 | 1 | -0.490 | 1.644 | chr1 | 24446261 | 24469630 | - | protein_coding |
| ENSG00000260339.1_6 | HEXA-AS1 | 0.39667595 | 1 | -0.490 | 0.327 | chr15 | 72668454 | 72671129 | + | lncRNA |
| ENSG00000149483.12_3 | TMEM138 | 0.80159183 | 1 | -0.490 | 5.054 | chr11 | 61129816 | 61136981 | + | protein_coding |
| ENSG00000158445.10_5 | KCNB1 | 0.37398096 | 1 | -0.490 | 0.025 | chr20 | 47909931 | 48100834 | - | protein_coding |
| ENSG00000284770.2_5 | TBCE | 0.47995677 | 1 | -0.490 | 1.839 | chr1 | 235530675 | 235615756 | + | protein_coding |
| ENSG00000146670.10_5 | CDCA5 | 0.83195635 | 1 | -0.490 | 6.038 | chr11 | 64833772 | 64851636 | - | protein_coding |
| ENSG00000149177.13_4 | PTPRJ | 0.9126525 | 1 | -0.490 | 7.440 | chr11 | 48002110 | 48192393 | + | protein_coding |
| ENSG00000075702.18_5 | WDR62 | 0.87258075 | 1 | -0.490 | 6.655 | chr19 | 36545783 | 36596010 | + | protein_coding |
| ENSG00000119703.15_5 | ZC2HC1C | 0.52177645 | 1 | -0.490 | 2.208 | chr14 | 75530812 | 75586205 | + | protein_coding |
| ENSG00000218418.2_5 | AL591135.1 | 0.37140872 | 1 | -0.490 | -0.045 | chr6 | 80774003 | 80779916 | - | processed_pseudogene |
| ENSG00000230844.3_5 | ZNF674-AS1 | 0.51262882 | 1 | -0.491 | 2.372 | chrX | 46404873 | 46408276 | + | lncRNA |
| ENSG00000101220.17_3 | C20orf27 | 0.55994046 | 1 | -0.491 | 5.741 | chr20 | 3734155 | 3749034 | - | protein_coding |
| ENSG00000224817.2_7 | AC010789.1 | 0.36505765 | 1 | -0.491 | -0.080 | chr10 | 103461751 | 103489794 | - | lncRNA |
| ENSG00000162927.14_4 | PUS10 | 0.54974627 | 1 | -0.491 | 2.607 | chr2 | 61167357 | 61245394 | - | protein_coding |
| ENSG00000170802.16_5 | FOXN2 | 0.80797283 | 1 | -0.491 | 5.840 | chr2 | 48541776 | 48606434 | + | protein_coding |
| ENSG00000144554.10_2 | FANCD2 | 0.8727564 | 1 | -0.491 | 6.243 | chr3 | 10068098 | 10143614 | + | protein_coding |
| ENSG00000126231.14_3 | PROZ | 0.3536377 | 1 | -0.491 | -0.815 | chr13 | 113812962 | 113826700 | + | protein_coding |
| ENSG00000165816.13_4 | VWA2 | 0.36944578 | 1 | -0.491 | -0.138 | chr10 | 115999013 | 116054248 | + | protein_coding |
| ENSG00000276368.1_3 | H2AC14 | 0.74871507 | 1 | -0.492 | 5.589 | chr6 | 27782132 | 27782518 | - | protein_coding |
| ENSG00000260260.1_6 | SNHG19 | 0.60099903 | 1 | -0.492 | 3.268 | chr16 | 2204798 | 2205359 | - | lncRNA |
| ENSG00000136518.17_6 | ACTL6A | 0.75593597 | 1 | -0.492 | 5.233 | chr3 | 179280674 | 179306195 | + | protein_coding |
| ENSG00000085721.12_4 | RRN3 | 0.86502759 | 1 | -0.492 | 6.211 | chr16 | 15153879 | 15188174 | - | protein_coding |
| ENSG00000272338.2_6 | AC067838.1 | 0.35395072 | 1 | -0.492 | -0.644 | chr8 | 33218357 | 33218933 | - | lncRNA |
| ENSG00000154040.20_7 | CABYR | 0.60517732 | 1 | -0.492 | 3.262 | chr18 | 21718920 | 21741567 | + | protein_coding |
| ENSG00000148516.21_5 | ZEB1 | 0.7423373 | 1 | -0.492 | 5.009 | chr10 | 31607424 | 31818742 | + | protein_coding |
| ENSG00000031691.7_3 | CENPQ | 0.65001037 | 1 | -0.492 | 3.889 | chr6 | 49431083 | 49460820 | + | protein_coding |
| ENSG00000121621.7_4 | KIF18A | 0.84768786 | 1 | -0.492 | 6.561 | chr11 | 28042166 | 28129703 | - | protein_coding |
| ENSG00000131650.14_3 | KREMEN2 | 0.38981485 | 1 | -0.493 | 0.555 | chr16 | 3014217 | 3018384 | + | protein_coding |
| ENSG00000163541.12_6 | SUCLG1 | 0.85496427 | 1 | -0.493 | 6.084 | chr2 | 84650652 | 84687169 | - | protein_coding |
| ENSG00000253200.1_6 | AC037459.2 | 0.39767518 | 1 | -0.493 | 0.603 | chr8 | 22471421 | 22474170 | - | lncRNA |
| ENSG00000251795.1 | SNORA66 | 0.36843654 | 1 | -0.493 | 0.345 | chr1 | 93303575 | 93303627 | + | snoRNA |
| ENSG00000203836.7 | NBPF24 | 0.35501676 | 1 | -0.493 | -0.909 | chr1 | 147574428 | 147634886 | - | protein_coding |
| ENSG00000103540.16_4 | CCP110 | 0.77171255 | 1 | -0.493 | 5.141 | chr16 | 19535133 | 19564730 | + | protein_coding |
| ENSG00000121988.18_5 | ZRANB3 | 0.69148814 | 1 | -0.493 | 4.134 | chr2 | 135894486 | 136288788 | - | protein_coding |
| ENSG00000261572.1_6 | AC097639.1 | 0.35239367 | 1 | -0.493 | -0.520 | chr3 | 32278180 | 32280070 | - | lncRNA |
| ENSG00000163328.13_4 | GPR155 | 0.44526616 | 1 | -0.493 | 1.599 | chr2 | 175296299 | 175351822 | - | protein_coding |
| ENSG00000167524.15_7 | RSKR | 0.55960867 | 1 | -0.494 | 3.027 | chr17 | 26928845 | 26941215 | - | protein_coding |
| ENSG00000085231.14_5 | AK6 | 0.65116524 | 1 | -0.494 | 3.976 | chr5 | 68646811 | 68665840 | - | protein_coding |
| ENSG00000242299.1_5 | AC073861.1 | 0.55274078 | 1 | -0.494 | 3.257 | chr3 | 101295333 | 101295791 | - | processed_pseudogene |
| ENSG00000132953.16_4 | XPO4 | 0.82764143 | 1 | -0.494 | 5.947 | chr13 | 21351468 | 21477187 | - | protein_coding |
| ENSG00000286599.1_2 | AC110611.1 | 0.35520401 | 1 | -0.494 | -0.497 | chr4 | 56755909 | 56769343 | - | lncRNA |
| ENSG00000095485.18_5 | CWF19L1 | 0.74981884 | 1 | -0.494 | 5.033 | chr10 | 101992055 | 102027437 | - | protein_coding |
| ENSG00000196230.13_3 | TUBB | 0.83352491 | 1 | -0.495 | 6.151 | chr6 | 30688129 | 30693203 | + | protein_coding |
| ENSG00000272696.1_7 | AL359091.3 | 0.35495794 | 1 | -0.495 | -0.448 | chr9 | 131078616 | 131079188 | + | lncRNA |
| ENSG00000268034.1_5 | AC243960.2 | 0.36629019 | 1 | -0.495 | 0.101 | chr19 | 42012068 | 42012814 | + | processed_pseudogene |
| ENSG00000138587.6_4 | MNS1 | 0.57194417 | 1 | -0.495 | 2.961 | chr15 | 56713742 | 56757335 | - | protein_coding |
| ENSG00000115020.17_6 | PIKFYVE | 0.82424155 | 1 | -0.495 | 6.259 | chr2 | 209130979 | 209223470 | + | protein_coding |
| ENSG00000224080.1_4 | UBE2FP1 | 0.34927505 | 1 | -0.495 | -0.909 | chr3 | 37185003 | 37185449 | - | processed_pseudogene |

| | | | | | | | | | | |
|---|---|---|---|---|---|---|---|---|---|---|
| ENSG00000001631.15_8 | KRIT1 | 0.50190606 | 1 | -0.495 | 2.687 | chr7 | 91828283 | 91875480 | - | protein_coding |
| ENSG00000244642.2 | RN7SL396P | 0.41559502 | 1 | -0.495 | 1.019 | chr8 | 120874902 | 120875181 | + | misc_RNA |
| ENSG00000229097.1_4 | CALM2P2 | 0.45064585 | 1 | -0.496 | 1.637 | chr10 | 71923441 | 71923880 | + | processed_pseudogene |
| ENSG00000163162.9_4 | RNF149 | 0.78706077 | 1 | -0.496 | 5.345 | chr2 | 101887681 | 101925163 | - | protein_coding |
| ENSG00000178127.13_4 | NDUFV2 | 0.50637597 | 1 | -0.496 | 2.362 | chr18 | 9102628 | 9134343 | + | protein_coding |
| ENSG00000134318.14_5 | ROCK2 | 0.86603322 | 1 | -0.496 | 8.154 | chr2 | 11319885 | 11488456 | - | protein_coding |
| ENSG00000115649.16_6 | CNPPD1 | 0.66381228 | 1 | -0.496 | 4.206 | chr2 | 220036619 | 220042828 | - | protein_coding |
| ENSG00000135521.9_4 | LTV1 | 0.64224351 | 1 | -0.496 | 4.129 | chr6 | 144164475 | 144184949 | + | protein_coding |
| ENSG00000138604.10_5 | GLCE | 0.80786272 | 1 | -0.496 | 5.985 | chr15 | 69452923 | 69564556 | + | protein_coding |
| ENSG00000101003.10_3 | GINS1 | 0.81909761 | 1 | -0.496 | 5.892 | chr20 | 25388309 | 25433264 | + | protein_coding |
| ENSG00000285979.1_2 | AC009090.6 | 0.55789407 | 1 | -0.497 | 2.768 | chr16 | 57211261 | 57215302 | + | lncRNA |
| ENSG00000188976.11_4 | NOC2L | 0.78811389 | 1 | -0.497 | 6.536 | chr1 | 879583 | 894689 | - | protein_coding |
| ENSG00000125611.15_3 | CHCHD5 | 0.58157983 | 1 | -0.497 | 3.041 | chr2 | 113341817 | 113346852 | + | protein_coding |
| ENSG00000114491.14_3 | UMPS | 0.78683556 | 1 | -0.497 | 5.617 | chr3 | 124449280 | 124468120 | + | protein_coding |
| ENSG00000091009.8_4 | RBM27 | 0.75149415 | 1 | -0.497 | 5.002 | chr5 | 145583168 | 145668786 | + | protein_coding |
| ENSG00000270084.2_6 | GAS5-AS1 | 0.49143971 | 1 | -0.497 | 2.211 | chr1 | 173831611 | 173833079 | + | lncRNA |
| ENSG00000132768.14_3 | DPH2 | 0.71768942 | 1 | -0.497 | 4.590 | chr1 | 44435672 | 44439041 | + | protein_coding |
| ENSG00000125835.18_5 | SNRPB | 0.84299896 | 1 | -0.497 | 7.550 | chr20 | 2442280 | 2451499 | - | protein_coding |
| ENSG00000235734.4_4 | HMGN1P36 | 0.35110045 | 1 | -0.497 | -0.976 | chr2 | 98443711 | 98444008 | + | processed_pseudogene |
| ENSG00000124789.11_4 | NUP153 | 0.84439776 | 1 | -0.498 | 6.318 | chr6 | 17615266 | 17707065 | - | protein_coding |
| ENSG00000217241.1_4 | CBX3P9 | 0.37789023 | 1 | -0.498 | 0.460 | chr6 | 116774177 | 116774728 | - | processed_pseudogene |
| ENSG00000237651.6_4 | C2orf74 | 0.35257703 | 1 | -0.498 | -0.190 | chr2 | 61372203 | 61391964 | + | protein_coding |
| ENSG00000255458.7_9 | AC108471.3 | 0.34572186 | 1 | -0.499 | -0.626 | chr4 | 39640727 | 39668499 | + | lncRNA |
| ENSG00000128699.14_7 | ORMDL1 | 0.7678394 | 1 | -0.499 | 5.451 | chr2 | 190634993 | 190649090 | - | protein_coding |
| ENSG00000224858.5_4 | RPL29P11 | 0.35632971 | 1 | -0.499 | -0.238 | chr3 | 37058014 | 37058505 | - | processed_pseudogene |
| ENSG00000145700.10_3 | ANKRD31 | 0.38135084 | 1 | -0.499 | 0.574 | chr5 | 74364100 | 74532736 | - | protein_coding |
| ENSG00000134574.11_2 | DDB2 | 0.83699442 | 1 | -0.499 | 6.131 | chr11 | 47236016 | 47260791 | + | protein_coding |
| ENSG00000183476.12_6 | SH2D7 | 0.34999957 | 1 | -0.499 | -0.473 | chr15 | 78370150 | 78397251 | + | protein_coding |
| ENSG00000087086.15_4 | FTL | 0.83816261 | 1 | -0.499 | 12.330 | chr19 | 49468566 | 49470136 | + | protein_coding |
| ENSG00000259529.2_7 | AL136295.5 | 0.3485556 | 1 | -0.499 | -0.402 | chr14 | 24616757 | 24635661 | + | protein_coding |
| ENSG00000121957.14_6 | GPSM2 | 0.71805117 | 1 | -0.499 | 4.864 | chr1 | 109417972 | 109477167 | + | protein_coding |
| ENSG00000006634.8_3 | DBF4 | 0.78607577 | 1 | -0.500 | 5.695 | chr7 | 87505531 | 87538868 | + | protein_coding |
| ENSG00000179598.6_5 | PLD6 | 0.45805645 | 1 | -0.500 | 1.804 | chr17 | 17104309 | 17109647 | - | protein_coding |
| ENSG00000177311.11_6 | ZBTB38 | 0.87579492 | 1 | -0.500 | 8.111 | chr3 | 141043055 | 141168634 | + | protein_coding |
| ENSG00000286416.1_2 | AL442067.3 | 0.34525651 | 1 | -0.500 | -0.498 | chr13 | 98057423 | 98086248 | - | lncRNA |
| ENSG00000151726.14_5 | ACSL1 | 0.68812527 | 1 | -0.500 | 4.327 | chr4 | 185676749 | 185747972 | - | protein_coding |
| ENSG00000130810.20_5 | PPAN | 0.43601184 | 1 | -0.500 | 1.515 | chr19 | 10217038 | 10222688 | + | protein_coding |
| ENSG00000259583.3_7 | AC015712.2 | 0.5447864 | 1 | -0.501 | 2.933 | chr15 | 101432548 | 101459596 | - | lncRNA |
| ENSG00000082269.16_4 | FAM135A | 0.73058315 | 1 | -0.501 | 5.101 | chr6 | 71122644 | 71270877 | + | protein_coding |
| ENSG00000168918.14_4 | INPP5D | 0.34317487 | 1 | -0.501 | -0.679 | chr2 | 233924677 | 234116549 | + | protein_coding |
| ENSG00000235748.1_3 | SEPTIN14P12 | 0.37063438 | 1 | -0.501 | 0.165 | chr1 | 197107878 | 197108437 | - | processed_pseudogene |
| ENSG00000198618.5_4 | PPIAP22 | 0.59365399 | 1 | -0.502 | 5.277 | chr21 | 20230097 | 20230594 | + | processed_pseudogene |
| ENSG00000163214.21_7 | DHX57 | 0.78002786 | 1 | -0.502 | 5.747 | chr2 | 39024871 | 39103075 | - | protein_coding |
| ENSG00000164663.14_6 | USP49 | 0.53630223 | 1 | -0.502 | 2.699 | chr6 | 41757634 | 41863099 | - | protein_coding |
| ENSG00000229334.1_6 | AC046143.1 | 0.34281566 | 1 | -0.502 | -0.548 | chr3 | 194353652 | 194366130 | + | lncRNA |
| ENSG00000155959.12_4 | VBP1 | 0.74113112 | 1 | -0.502 | 6.160 | chrX | 154425284 | 154468122 | + | protein_coding |
| ENSG00000231154.1_6 | MORF4L2-AS1 | 0.34325113 | 1 | -0.502 | -0.277 | chrX | 102942212 | 102946700 | + | lncRNA |
| ENSG00000099834.19_3 | CDHR5 | 0.39885649 | 1 | -0.502 | 1.025 | chr11 | 616577 | 626078 | - | protein_coding |
| ENSG00000279089.1_6 | AC005839.1 | 0.47591847 | 1 | -0.502 | 2.199 | chr17 | 49260552 | 49263764 | - | TEC |
| ENSG00000166483.11_4 | WEE1 | 0.76703424 | 1 | -0.503 | 5.666 | chr11 | 9595217 | 9615004 | + | protein_coding |
| ENSG00000057663.16_6 | ATG5 | 0.74930564 | 1 | -0.503 | 5.264 | chr6 | 106493298 | 106773666 | - | protein_coding |
| ENSG00000163516.14_4 | ANKZF1 | 0.65984727 | 1 | -0.503 | 4.536 | chr2 | 220094505 | 220101401 | + | protein_coding |
| ENSG00000255050.1_7 | AC067930.4 | 0.42087085 | 1 | -0.503 | 1.267 | chr8 | 144655660 | 144659567 | + | lncRNA |
| ENSG00000143870.12_4 | PDIA6 | 0.86322668 | 1 | -0.504 | 7.790 | chr2 | 10923517 | 10978103 | - | protein_coding |
| ENSG00000261461.1_2 | UBE2MP1 | 0.35041974 | 1 | -0.504 | -0.198 | chr16 | 34404063 | 34404612 | - | processed_pseudogene |
| ENSG00000260007.3_7 | AC107871.1 | 0.45269 | 1 | -0.504 | 1.794 | chr15 | 68476370 | 68522056 | - | protein_coding |

| | | | | | | | | | | |
|---|---|---|---|---|---|---|---|---|---|---|
| ENSG00000075213.11_7 | SEMA3A | 0.74521185 | 1 | -0.504 | 5.606 | chr7 | 83585093 | 84122040 | - | protein_coding |
| ENSG00000272870.3_8 | SAP30-DT | 0.34012049 | 1 | -0.504 | -0.626 | chr4 | 174270694 | 174291872 | - | lncRNA |
| ENSG00000167088.11_4 | SNRPD1 | 0.7185713 | 1 | -0.504 | 5.497 | chr18 | 19192275 | 19213485 | + | protein_coding |
| ENSG00000287403.1_3 | AC016245.2 | 0.34239878 | 1 | -0.504 | -0.365 | chr17 | 78797970 | 78800700 | - | lncRNA |
| ENSG00000169184.6_5 | MN1 | 0.4740632 | 1 | -0.505 | 2.316 | chr22 | 28144265 | 28197744 | - | protein_coding |
| ENSG00000083457.12_4 | ITGAE | 0.66579498 | 1 | -0.505 | 4.468 | chr17 | 3617922 | 3704482 | - | protein_coding |
| ENSG00000214612.3_2 | RPS19P1 | 0.34065913 | 1 | -0.505 | -0.707 | chr20 | 18485277 | 18485710 | + | processed_pseudogene |
| ENSG00000260418.1_6 | AL023284.4 | 0.34608252 | 1 | -0.505 | 0.046 | chr6 | 136656852 | 136657225 | - | lncRNA |
| ENSG00000113719.16_4 | ERGIC1 | 0.84726101 | 1 | -0.505 | 8.137 | chr5 | 172261254 | 172379686 | + | protein_coding |
| ENSG00000279077.1_6 | AC023090.2 | 0.3396279 | 1 | -0.505 | -1.084 | chr18 | 77112458 | 77114567 | + | TEC |
| ENSG00000186687.16_3 | LYRM7 | 0.60518727 | 1 | -0.505 | 3.990 | chr5 | 130506637 | 130541121 | + | protein_coding |
| ENSG00000178726.7_3 | THBD | 0.68460948 | 1 | -0.506 | 5.155 | chr20 | 23026270 | 23030309 | - | protein_coding |
| ENSG00000219642.2_4 | BMPR1AP1 | 0.33788567 | 1 | -0.506 | -0.654 | chr6 | 129478668 | 129480265 | - | processed_pseudogene |
| ENSG00000181350.12_5 | LRRC75A | 0.58139499 | 1 | -0.506 | 3.487 | chr17 | 16344891 | 16395507 | - | protein_coding |
| ENSG00000214954.8_4 | LRRC69 | 0.35257271 | 1 | -0.506 | 0.151 | chr8 | 92114060 | 92231464 | + | protein_coding |
| ENSG00000126787.13_5 | DLGAP5 | 0.79515937 | 1 | -0.506 | 6.819 | chr14 | 55614830 | 55658326 | - | protein_coding |
| ENSG00000137812.19_7 | KNL1 | 0.79043537 | 1 | -0.506 | 6.527 | chr15 | 40886218 | 40956540 | + | protein_coding |
| ENSG00000212673.1 | AL136115.1 | 0.35480845 | 1 | -0.506 | 0.118 | chr1 | 32379174 | 32380745 | + | protein_coding |
| ENSG00000258725.2_7 | PRC1-AS1 | 0.44170019 | 1 | -0.506 | 1.748 | chr15 | 91509570 | 91531855 | + | lncRNA |
| ENSG00000286813.1_3 | AC015871.6 | 0.36179028 | 1 | -0.506 | 0.491 | chr15 | 80186844 | 80189037 | - | lncRNA |
| ENSG00000165995.20_5 | CACNB2 | 0.38577547 | 1 | -0.506 | 0.701 | chr10 | 18429606 | 18832486 | + | protein_coding |
| ENSG00000279880.1_6 | AC134407.3 | 0.33662511 | 1 | -0.507 | -0.652 | chr17 | 65970050 | 65972188 | + | TEC |
| ENSG00000228328.2_5 | AL158201.1 | 0.34165668 | 1 | -0.507 | -0.859 | chrX | 67290362 | 67291609 | - | processed_pseudogene |
| ENSG00000169826.8_3 | CSGALNACT2 | 0.54043661 | 1 | -0.507 | 3.258 | chr10 | 43633893 | 43680750 | + | protein_coding |
| ENSG00000115993.13_6 | TRAK2 | 0.73334994 | 1 | -0.507 | 5.348 | chr2 | 202241930 | 202316223 | - | protein_coding |
| ENSG00000247092.7_6 | SNHG10 | 0.49146868 | 1 | -0.508 | 2.961 | chr14 | 95988280 | 96001226 | - | lncRNA |
| ENSG00000277687.1_7 | AL139407.1 | 0.33807726 | 1 | -0.508 | -0.796 | chr10 | 120451873 | 120453047 | - | lncRNA |
| ENSG00000134146.12_5 | DPH6 | 0.5637794 | 1 | -0.508 | 3.382 | chr15 | 35509546 | 35838394 | - | protein_coding |
| ENSG00000250698.1_6 | AC131182.1 | 0.33656153 | 1 | -0.508 | -0.357 | chr4 | 141049169 | 141055719 | + | lncRNA |
| ENSG00000119285.11_5 | HEATR1 | 0.815247 | 1 | -0.508 | 7.124 | chr1 | 236712305 | 236767816 | - | protein_coding |
| ENSG00000272923.1_7 | AC092139.1 | 0.37226424 | 1 | -0.508 | 1.009 | chr16 | 81419068 | 81421165 | + | lncRNA |
| ENSG00000172731.14_5 | LRRC20 | 0.66670142 | 1 | -0.508 | 4.483 | chr10 | 72058726 | 72142406 | - | protein_coding |
| ENSG00000173898.13_5 | SPTBN2 | 0.66008116 | 1 | -0.508 | 4.921 | chr11 | 66452719 | 66496697 | - | protein_coding |
| ENSG00000177150.13_6 | FAM210A | 0.60264263 | 1 | -0.509 | 4.006 | chr18 | 13663346 | 13726662 | - | protein_coding |
| ENSG00000176102.13_5 | CSTF3 | 0.60323206 | 1 | -0.509 | 4.513 | chr11 | 33098734 | 33183917 | - | protein_coding |
| ENSG00000171960.11_5 | PPIH | 0.53667634 | 1 | -0.509 | 3.098 | chr1 | 43124094 | 43142429 | + | protein_coding |
| ENSG00000250859.1_4 | HNRNPKP1 | 0.33741433 | 1 | -0.509 | -0.826 | chr5 | 126847156 | 126848534 | - | processed_pseudogene |
| ENSG00000241741.1_4 | RPL7AP30 | 0.39474692 | 1 | -0.509 | 1.035 | chr4 | 113709239 | 113710034 | - | processed_pseudogene |
| ENSG00000165802.22_6 | NSMF | 0.43716637 | 1 | -0.509 | 4.688 | chr9 | 140342025 | 140353786 | - | protein_coding |
| ENSG00000231784.8_3 | DBIL5P | 0.34752038 | 1 | -0.509 | -0.018 | chr17 | 655900 | 658576 | + | transcribed_unitary_pseudogene |
| ENSG00000234093.3_4 | RPS15AP11 | 0.33809557 | 1 | -0.510 | -0.368 | chr1 | 45246003 | 45246387 | - | processed_pseudogene |
| ENSG00000166206.15_7 | GABRB3 | 0.82586468 | 1 | -0.510 | 7.390 | chr15 | 26788693 | 27184686 | - | protein_coding |
| ENSG00000163462.18_3 | TRIM46 | 0.4346943 | 1 | -0.510 | 1.783 | chr1 | 155146263 | 155157447 | + | protein_coding |
| ENSG00000181754.7_4 | AMIGO1 | 0.45322364 | 1 | -0.510 | 1.627 | chr1 | 110046800 | 110052360 | - | protein_coding |
| ENSG00000185989.11_5 | RASA3 | 0.55230209 | 1 | -0.510 | 3.427 | chr13 | 114747194 | 114898098 | - | protein_coding |
| ENSG00000112365.4_4 | ZBTB24 | 0.65588969 | 1 | -0.510 | 4.681 | chr6 | 109783797 | 109804440 | - | protein_coding |
| ENSG00000167196.14_6 | FBXO22 | 0.76283047 | 1 | -0.510 | 6.559 | chr15 | 76196217 | 76234852 | + | protein_coding |
| ENSG00000285973.2_3 | AC063960.2 | 0.36646441 | 1 | -0.511 | 0.787 | chr10 | 123904621 | 123914712 | - | lncRNA |
| ENSG00000156531.16_3 | PHF6 | 0.75369253 | 1 | -0.511 | 5.928 | chrX | 133507283 | 133562821 | + | protein_coding |
| ENSG00000163482.12_6 | STK36 | 0.61995003 | 1 | -0.511 | 4.497 | chr2 | 219536792 | 219567439 | + | protein_coding |
| ENSG00000143498.18_4 | TAF1A | 0.55061034 | 1 | -0.511 | 3.351 | chr1 | 222731244 | 222763275 | - | protein_coding |
| ENSG00000273139.1_6 | AC007663.3 | 0.33196221 | 1 | -0.511 | -0.600 | chr22 | 20308262 | 20308726 | + | lncRNA |
| ENSG00000129009.13_4 | ISLR | 0.34984629 | 1 | -0.511 | 0.294 | chr15 | 74466051 | 74469213 | + | protein_coding |
| ENSG00000134308.14_6 | YWHAQ | 0.81777533 | 1 | -0.511 | 9.372 | chr2 | 9724096 | 9771126 | - | protein_coding |
| ENSG00000128944.13_6 | KNSTRN | 0.72373714 | 1 | -0.512 | 5.605 | chr15 | 40674922 | 40686447 | + | protein_coding |
| ENSG00000106333.13_3 | PCOLCE | 0.40398849 | 1 | -0.512 | 1.388 | chr7 | 100199986 | 100205798 | + | protein_coding |

| | | | | | | | | | | |
|---|---|---|---|---|---|---|---|---|---|---|
| ENSG00000144426.18_3 | NBEAL1 | 0.69189995 | 1 | -0.512 | 5.756 | chr2 | 203879602 | 204091101 | + | protein_coding |
| ENSG00000181201.3_3 | H2BU2P | 0.33072026 | 1 | -0.512 | -0.654 | chr1 | 228651804 | 228652327 | - | unitary_pseudogene |
| ENSG00000134375.11_3 | TIMM17A | 0.76770861 | 1 | -0.512 | 6.117 | chr1 | 201924631 | 201939792 | + | protein_coding |
| ENSG00000135912.11_6 | TTLL4 | 0.64377987 | 1 | -0.512 | 4.450 | chr2 | 219575558 | 219620139 | + | protein_coding |
| ENSG00000267248.3_8 | AC025048.2 | 0.32979171 | 1 | -0.512 | -0.768 | chr17 | 58200003 | 58213104 | - | lncRNA |
| ENSG00000101493.11_5 | ZNF516 | 0.51016739 | 1 | -0.512 | 2.908 | chr18 | 74069637 | 74207198 | - | protein_coding |
| ENSG00000110395.7_5 | CBL | 0.66560043 | 1 | -0.512 | 5.326 | chr11 | 119077008 | 119184636 | + | protein_coding |
| ENSG00000133302.13_4 | SLF1 | 0.68248418 | 1 | -0.513 | 5.246 | chr5 | 93954374 | 94075141 | + | protein_coding |
| ENSG00000197601.13_6 | FAR1 | 0.75608873 | 1 | -0.513 | 6.766 | chr11 | 13690215 | 13753893 | + | protein_coding |
| ENSG00000231007.5_4 | CDC20P1 | 0.34609049 | 1 | -0.513 | 0.258 | chr9 | 89626567 | 89628066 | + | processed_pseudogene |
| ENSG00000217648.1_5 | AL136116.3 | 0.33094569 | 1 | -0.513 | -0.391 | chr6 | 143663383 | 143664520 | + | processed_pseudogene |
| ENSG00000280355.1_6 | AL132656.4 | 0.40200855 | 1 | -0.514 | 1.316 | chr10 | 81441729 | 81443850 | - | TEC |
| ENSG00000176208.9_3 | ATAD5 | 0.70689664 | 1 | -0.514 | 5.754 | chr17 | 29158984 | 29222887 | + | protein_coding |
| ENSG00000258727.1_6 | AL135999.1 | 0.32748248 | 1 | -0.514 | -0.768 | chr14 | 24030306 | 24037282 | + | lncRNA |
| ENSG00000141378.14_3 | PTRH2 | 0.67447144 | 1 | -0.515 | 5.170 | chr17 | 57751997 | 57784987 | - | protein_coding |
| ENSG00000128692.8_4 | EIF2S2P4 | 0.40170542 | 1 | -0.515 | 1.641 | chr2 | 171608315 | 171609298 | + | processed_pseudogene |
| ENSG00000184307.15_5 | ZDHHC23 | 0.5574037 | 1 | -0.515 | 3.729 | chr3 | 113666748 | 113684248 | + | protein_coding |
| ENSG00000202314.1 | SNORD6 | 0.34113977 | 1 | -0.515 | -0.638 | chr11 | 93464668 | 93464740 | - | snoRNA |
| ENSG00000115107.20_4 | STEAP3 | 0.66522838 | 1 | -0.516 | 5.456 | chr2 | 119981407 | 120023228 | + | protein_coding |
| ENSG00000221890.4_6 | NPTXR | 0.475984 | 1 | -0.516 | 2.722 | chr22 | 39214457 | 39240033 | - | protein_coding |
| ENSG00000138002.15_5 | IFT172 | 0.61845337 | 1 | -0.516 | 4.525 | chr2 | 27667244 | 27712656 | - | protein_coding |
| ENSG00000279342.2_5 | AP000866.6 | 0.38120181 | 1 | -0.516 | 1.256 | chr11 | 124659136 | 124662714 | - | lncRNA |
| ENSG00000236364.4_7 | AL358115.1 | 0.34970087 | 1 | -0.516 | 0.463 | chr1 | 165858962 | 165870232 | - | lncRNA |
| ENSG00000251442.6_6 | LINC01094 | 0.32714364 | 1 | -0.517 | -0.684 | chr4 | 79559934 | 79604339 | + | lncRNA |
| ENSG00000059145.18_5 | UNKL | 0.60471595 | 1 | -0.517 | 5.016 | chr16 | 1413206 | 1464752 | - | protein_coding |
| ENSG00000102753.10_5 | KPNA3 | 0.70280939 | 1 | -0.517 | 5.880 | chr13 | 50273456 | 50366818 | - | protein_coding |
| ENSG00000187741.15_7 | FANCA | 0.62413522 | 1 | -0.517 | 5.361 | chr16 | 89803957 | 89883065 | - | protein_coding |
| ENSG00000143179.16_5 | UCK2 | 0.71131553 | 1 | -0.518 | 5.733 | chr1 | 165796851 | 165880855 | + | protein_coding |
| ENSG00000130449.6_4 | ZSWIM6 | 0.64334756 | 1 | -0.518 | 5.099 | chr5 | 60628085 | 60841999 | + | protein_coding |
| ENSG00000150753.12_6 | CCT5 | 0.79228634 | 1 | -0.518 | 8.441 | chr5 | 10250041 | 10266501 | + | protein_coding |
| ENSG00000257671.1_7 | KRT7-AS | 0.61678445 | 1 | -0.518 | 5.267 | chr12 | 52638832 | 52641232 | - | lncRNA |
| ENSG00000241343.9_6 | RPL36A | 0.47071994 | 1 | -0.519 | 2.790 | chrX | 100645812 | 100651142 | + | protein_coding |
| ENSG00000221823.11_5 | PPP3R1 | 0.58940632 | 1 | -0.519 | 4.426 | chr2 | 68405989 | 68483369 | - | protein_coding |
| ENSG00000250995.1_6 | AL391280.2 | 0.32225555 | 1 | -0.519 | -0.768 | chrX | 117903759 | 117904820 | + | processed_pseudogene |
| ENSG00000147996.12 | CBWD5 | 0.47807908 | 1 | -0.519 | 2.899 | chr9 | 70432004 | 70497240 | - | protein_coding |
| ENSG00000163161.14_4 | ERCC3 | 0.69398111 | 1 | -0.519 | 5.823 | chr2 | 128014866 | 128051742 | - | protein_coding |
| ENSG00000172331.12_4 | BPGM | 0.55565284 | 1 | -0.520 | 4.130 | chr7 | 134331563 | 134364568 | + | protein_coding |
| ENSG00000142856.16_3 | ITGB3BP | 0.6089201 | 1 | -0.520 | 4.591 | chr1 | 63906441 | 64059392 | - | protein_coding |
| ENSG00000239280.1_5 | AC108693.1 | 0.32560477 | 1 | -0.520 | -0.246 | chr3 | 113459670 | 113460101 | - | processed_pseudogene |
| ENSG00000266651.1_7 | AC093484.4 | 0.43077231 | 1 | -0.520 | 2.094 | chr17 | 16343793 | 16344266 | - | lncRNA |
| ENSG00000261087.1_7 | ZNNT1 | 0.32264493 | 1 | -0.520 | -0.309 | chr8 | 102179033 | 102181857 | - | lncRNA |
| ENSG00000033867.16_5 | SLC4A7 | 0.72000661 | 1 | -0.521 | 7.187 | chr3 | 27414212 | 27525911 | - | protein_coding |
| ENSG00000248641.1_2 | HMGA1P2 | 0.34960042 | 1 | -0.521 | -0.695 | chr4 | 73964539 | 73964862 | + | processed_pseudogene |
| ENSG00000106355.10_5 | LSM5 | 0.68238087 | 1 | -0.521 | 5.819 | chr7 | 32524950 | 32534895 | - | protein_coding |
| ENSG00000148843.15_4 | PDCD11 | 0.72971882 | 1 | -0.521 | 6.509 | chr10 | 105156383 | 105206051 | + | protein_coding |
| ENSG00000102743.15_4 | SLC25A15 | 0.55850163 | 1 | -0.522 | 4.417 | chr13 | 41363548 | 41386596 | + | protein_coding |
| ENSG00000143228.13_5 | NUF2 | 0.65119615 | 1 | -0.522 | 5.295 | chr1 | 163236366 | 163325554 | + | protein_coding |
| ENSG00000132646.11_4 | PCNA | 0.75860797 | 1 | -0.522 | 8.155 | chr20 | 5095599 | 5107272 | - | protein_coding |
| ENSG00000180228.13_5 | PRKRA | 0.5755068 | 1 | -0.522 | 4.348 | chr2 | 179296141 | 179316239 | - | protein_coding |
| ENSG00000074935.14_4 | TUBE1 | 0.46527521 | 1 | -0.522 | 3.255 | chr6 | 112391866 | 112408732 | - | protein_coding |
| ENSG00000232187.1_2 | FTH1P7 | 0.3275063 | 1 | -0.522 | 0.119 | chr13 | 23270162 | 23270713 | - | processed_pseudogene |
| ENSG00000259659.1_7 | AC009996.1 | 0.32104653 | 1 | -0.523 | -0.768 | chr15 | 44850027 | 44851386 | + | lncRNA |
| ENSG00000145220.14_6 | LYAR | 0.54843626 | 1 | -0.524 | 4.010 | chr4 | 4269428 | 4291881 | - | protein_coding |
| ENSG00000257957.1_4 | QRSL1P3 | 0.3171425 | 1 | -0.524 | -0.892 | chr14 | 36235001 | 36236316 | + | processed_pseudogene |
| ENSG00000184564.11_6 | SLITRK6 | 0.42356243 | 1 | -0.524 | 2.666 | chr13 | 86366925 | 86380818 | - | protein_coding |
| ENSG00000062282.15_7 | DGAT2 | 0.38336637 | 1 | -0.524 | 1.463 | chr11 | 75470557 | 75512580 | + | protein_coding |
| ENSG00000165821.12_8 | SALL2 | 0.41836591 | 1 | -0.524 | 2.161 | chr14 | 21989231 | 22005350 | - | protein_coding |

| | | | | | | | | | | |
|---|---|---|---|---|---|---|---|---|---|---|
| ENSG00000244578.1_6 | LINC01391 | 0.31604516 | 0.99914135 | -0.525 | -0.861 | chr3 | 138654031 | 138662862 | - | lncRNA |
| ENSG00000135845.10_4 | PIGC | 0.57778615 | 1 | -0.525 | 4.661 | chr1 | 172339329 | 172413226 | - | protein_coding |
| ENSG00000213949.10_6 | ITGA1 | 0.4065011 | 1 | -0.525 | 1.965 | chr5 | 52083750 | 52255039 | + | protein_coding |
| ENSG00000277998.1_5 | AC107075.1 | 0.32218678 | 1 | -0.525 | 0.319 | chr2 | 165430251 | 165430564 | + | unprocessed_pseudogene |
| ENSG00000138160.6_3 | KIF11 | 0.70268213 | 1 | -0.525 | 7.525 | chr10 | 94352887 | 94415152 | + | protein_coding |
| ENSG00000265394.1_8 | AC104984.3 | 0.32298632 | 1 | -0.525 | -0.892 | chr17 | 28417384 | 28444515 | - | lncRNA |
| ENSG00000119969.15_4 | HELLS | 0.69104648 | 1 | -0.526 | 7.009 | chr10 | 96261191 | 96373662 | + | protein_coding |
| ENSG00000171016.12_5 | PYGO1 | 0.57928505 | 1 | -0.526 | 5.095 | chr15 | 55831088 | 55881145 | - | protein_coding |
| ENSG00000133475.17_6 | GGT2 | 0.3157216 | 0.99847379 | -0.526 | -0.416 | chr22 | 21562262 | 21581926 | - | protein_coding |
| ENSG00000206652.1 | RNU1-1 | 0.33228087 | 1 | -0.526 | 0.454 | chr1 | 16840617 | 16840780 | - | snRNA |
| ENSG00000132780.17_6 | NASP | 0.68282976 | 1 | -0.527 | 6.587 | chr1 | 46049518 | 46084576 | + | protein_coding |
| ENSG00000131080.15_5 | EDA2R | 0.54596066 | 1 | -0.527 | 4.656 | chrX | 65815479 | 65859140 | - | protein_coding |
| ENSG00000165029.16_4 | ABCA1 | 0.63173314 | 1 | -0.527 | 5.654 | chr9 | 107543287 | 107690436 | - | protein_coding |
| ENSG00000155265.11_3 | GOLGA7B | 0.35842195 | 1 | -0.527 | 1.097 | chr10 | 99609600 | 99631337 | + | protein_coding |
| ENSG00000272620.2_6 | AFAP1-AS1 | 0.35019042 | 1 | -0.528 | 1.447 | chr4 | 7755804 | 7780655 | + | lncRNA |
| ENSG00000228061.7_7 | AC131571.1 | 0.32734509 | 1 | -0.528 | -0.800 | chr11 | 31639671 | 31789501 | - | lncRNA |
| ENSG00000198886.2_1 | MT-ND4 | 0.4262734 | 1 | -0.528 | 9.756 | chrM | 10760 | 12137 | + | protein_coding |
| ENSG00000123545.6_4 | NDUFAF4 | 0.4884995 | 1 | -0.528 | 3.955 | chr6 | 97337191 | 97345767 | - | protein_coding |
| ENSG00000164258.12_4 | NDUFS4 | 0.58094827 | 1 | -0.528 | 5.373 | chr5 | 52856463 | 52979168 | + | protein_coding |
| ENSG00000258559.2_8 | AC005519.1 | 0.32396761 | 1 | -0.528 | 0.245 | chr14 | 74755830 | 74761128 | - | lncRNA |
| ENSG00000268030.1_7 | AC005253.1 | 0.31544887 | 0.99778887 | -0.528 | -0.485 | chr19 | 18668585 | 18672370 | - | lncRNA |
| ENSG00000224431.1_5 | AC063976.2 | 0.32136068 | 1 | -0.528 | -0.957 | chr5 | 131535149 | 131539180 | + | processed_pseudogene |
| ENSG00000154920.14_4 | EME1 | 0.52784152 | 1 | -0.529 | 4.081 | chr17 | 48450581 | 48458844 | + | protein_coding |
| ENSG00000230869.1_4 | AGAP10P | 0.31227749 | 0.9898723 | -0.529 | -0.861 | chr10 | 46174140 | 46195980 | + | unprocessed_pseudogene |
| ENSG00000285813.1_2 | AP000813.1 | 0.31897182 | 1 | -0.529 | -0.957 | chr11 | 105865912 | 105869940 | + | lncRNA |
| ENSG00000198130.16_5 | HIBCH | 0.52012492 | 1 | -0.529 | 4.128 | chr2 | 191054461 | 191208919 | - | protein_coding |
| ENSG00000259581.3_2 | TYRO3P | 0.3114431 | 0.98798785 | -0.529 | -0.535 | chr15 | 76551327 | 76554031 | - | processed_pseudogene |
| ENSG00000253754.1_6 | AC087620.1 | 0.31427731 | 0.99461435 | -0.529 | -0.924 | chr8 | 109238429 | 109239773 | + | lncRNA |
| ENSG00000200534.1 | SNORA33 | 0.34323821 | 1 | -0.530 | 0.819 | chr6 | 133138358 | 133138487 | + | snoRNA |
| ENSG00000214517.10_4 | PPME1 | 0.63185376 | 1 | -0.530 | 5.698 | chr11 | 73882312 | 73965748 | + | protein_coding |
| ENSG00000149781.12_4 | FERMT3 | 0.32992158 | 1 | -0.531 | 0.504 | chr11 | 63974150 | 63991358 | + | protein_coding |
| ENSG00000053372.5_4 | MRTO4 | 0.6086107 | 1 | -0.531 | 5.927 | chr1 | 19578299 | 19586622 | + | protein_coding |
| ENSG00000104356.11_4 | POP1 | 0.56962908 | 1 | -0.531 | 4.953 | chr8 | 99129521 | 99172063 | + | protein_coding |
| ENSG00000203832.6 | NBPF20 | 0.39804683 | 1 | -0.531 | 2.158 | chr1 | 148250249 | 148347506 | - | protein_coding |
| ENSG00000270091.1_7 | AC015726.1 | 0.31151528 | 0.98798785 | -0.531 | -0.894 | chr17 | 19799903 | 19800600 | - | lncRNA |
| ENSG00000214595.12_3 | EML6 | 0.3970979 | 1 | -0.531 | 2.371 | chr2 | 54950636 | 55199161 | + | protein_coding |
| ENSG00000186529.16_6 | CYP4F3 | 0.54550559 | 1 | -0.531 | 4.842 | chr19 | 15751707 | 15773635 | + | protein_coding |
| ENSG00000112742.10_4 | TTK | 0.63222104 | 1 | -0.531 | 6.205 | chr6 | 80713604 | 80752244 | + | protein_coding |
| ENSG00000128585.18_5 | MKLN1 | 0.63064473 | 1 | -0.531 | 6.914 | chr7 | 130794855 | 131181391 | + | protein_coding |
| ENSG00000101306.11_3 | MYLK2 | 0.32129622 | 1 | -0.532 | -0.770 | chr20 | 30407111 | 30422492 | + | protein_coding |
| ENSG00000110104.13_5 | CCDC86 | 0.44811942 | 1 | -0.532 | 3.460 | chr11 | 60609586 | 60618554 | + | protein_coding |
| ENSG00000072571.20_6 | HMMR | 0.6282484 | 1 | -0.532 | 6.578 | chr5 | 162887209 | 162918947 | + | protein_coding |
| ENSG00000198836.10_5 | OPA1 | 0.65596155 | 1 | -0.532 | 6.936 | chr3 | 193310933 | 193415600 | + | protein_coding |
| ENSG00000105642.15_8 | KCNN1 | 0.33017794 | 1 | -0.532 | 0.580 | chr19 | 18062102 | 18110889 | + | protein_coding |
| ENSG00000116750.13_8 | UCHL5 | 0.58154206 | 1 | -0.532 | 5.183 | chr1 | 192981380 | 193029210 | - | protein_coding |
| ENSG00000147224.12_4 | PRPS1 | 0.60803109 | 1 | -0.532 | 6.555 | chrX | 106871654 | 106894256 | + | protein_coding |
| ENSG00000236498.1_6 | AC107081.2 | 0.31132575 | 0.98791296 | -0.532 | 0.044 | chr2 | 62095567 | 62113217 | + | lncRNA |
| ENSG00000173163.10_5 | COMMD1 | 0.48098018 | 1 | -0.532 | 3.908 | chr2 | 62115859 | 62374382 | + | protein_coding |
| ENSG00000288031.1_1 | AC093838.2 | 0.31741293 | 1 | -0.532 | -1.028 | chr2 | 132449965 | 132455527 | + | transcribed_unprocessed_pseudogene |
| ENSG00000272183.1_6 | AC005041.3 | 0.30775874 | 0.97991929 | -0.532 | -0.613 | chr2 | 74728844 | 74729492 | + | lncRNA |
| ENSG00000277767.1_6 | AL442128.2 | 0.30787274 | 0.98010659 | -0.532 | -0.561 | chr13 | 111568351 | 111570174 | + | lncRNA |
| ENSG00000104626.15_3 | ERI1 | 0.61168981 | 1 | -0.532 | 5.730 | chr8 | 8859657 | 8974256 | + | protein_coding |
| ENSG00000251307.1_6 | AC026704.1 | 0.3232632 | 1 | -0.532 | -0.991 | chr5 | 54529762 | 54591029 | + | lncRNA |
| ENSG00000137876.10_4 | RSL24D1 | 0.63119815 | 1 | -0.533 | 7.203 | chr15 | 55473004 | 55489247 | - | protein_coding |
| ENSG00000170160.17_5 | CCDC144A | 0.42051942 | 1 | -0.533 | 2.954 | chr17 | 16592851 | 16681195 | + | protein_coding |
| ENSG00000287670.1_2 | AC108479.3 | 0.32431885 | 1 | -0.533 | 0.506 | chr2 | 88316299 | 88320970 | + | lncRNA |
| ENSG00000092208.17_5 | GEMIN2 | 0.47819685 | 1 | -0.533 | 4.020 | chr14 | 39583427 | 39606177 | + | protein_coding |
| ENSG00000277701.5_1 | AC159540.2 | 0.31379247 | 0.99325688 | -0.534 | 0.101 | chr2 | 97947093 | 97957603 | + | lncRNA |

| | | | | | | | | | | |
|---|---|---|---|---|---|---|---|---|---|---|
| ENSG00000180015.12_5 | AC093909.1 | 0.31733459 | 1 | -0.534 | -0.724 | chr4 | 189659527 | 189660648 | + | processed_pseudogene |
| ENSG00000189362.12_5 | NEMP2 | 0.45572021 | 1 | -0.534 | 3.746 | chr2 | 191369064 | 191399448 | - | protein_coding |
| ENSG00000132429.10_4 | POPDC3 | 0.44642004 | 1 | -0.534 | 3.453 | chr6 | 105605775 | 105627889 | - | protein_coding |
| ENSG00000149054.16_5 | ZNF215 | 0.34791707 | 1 | -0.535 | 1.376 | chr11 | 6947635 | 7022235 | + | protein_coding |
| ENSG00000229325.1_7 | ACAP2-IT1 | 0.35049727 | 1 | -0.535 | -1.027 | chr3 | 195001452 | 195003470 | - | lncRNA |
| ENSG00000279434.1_6 | AL049776.1 | 0.31012467 | 0.98550975 | -0.535 | 0.060 | chr14 | 35555241 | 35557265 | + | TEC |
| ENSG00000247151.7_6 | CSTF3-DT | 0.32868334 | 1 | -0.535 | -1.101 | chr11 | 33183203 | 33213144 | + | lncRNA |
| ENSG00000174842.17_4 | GLMN | 0.41281642 | 1 | -0.535 | 3.397 | chr1 | 92711959 | 92764544 | - | protein_coding |
| ENSG00000276744.1_7 | AC105137.2 | 0.31619653 | 0.99926427 | -0.535 | 0.332 | chr15 | 75745305 | 75746288 | - | lncRNA |
| ENSG00000258708.1_5 | SLC25A21-AS1 | 0.33169181 | 1 | -0.535 | 0.883 | chr14 | 37641093 | 37643016 | + | lncRNA |
| ENSG00000014641.20_6 | MDH1 | 0.64017188 | 1 | -0.535 | 7.652 | chr2 | 63815743 | 63834331 | + | polymorphic_pseudogene |
| ENSG00000178081.12_2 | ULK4P3 | 0.3201286 | 1 | -0.535 | 0.531 | chr15 | 30395923 | 30423960 | + | transcribed_unprocessed_pseudogene |
| ENSG00000152256.13_5 | PDK1 | 0.42865879 | 1 | -0.536 | 3.021 | chr2 | 173420101 | 173473397 | + | protein_coding |
| ENSG00000115211.15_5 | EIF2B4 | 0.54115288 | 1 | -0.536 | 5.026 | chr2 | 27587219 | 27593353 | - | protein_coding |
| ENSG00000103064.15_5 | SLC7A6 | 0.48049404 | 1 | -0.536 | 4.408 | chr16 | 68298419 | 68335726 | + | protein_coding |
| ENSG00000196353.11_6 | CPNE4 | 0.45798734 | 1 | -0.536 | 3.584 | chr3 | 131252399 | 132004254 | - | protein_coding |
| ENSG00000204217.15_5 | BMPR2 | 0.58363968 | 1 | -0.536 | 6.620 | chr2 | 203241050 | 203432474 | + | protein_coding |
| ENSG00000178105.11_5 | DDX10 | 0.53950665 | 1 | -0.536 | 5.385 | chr11 | 108535796 | 108811654 | + | protein_coding |
| ENSG00000227344.2_4 | HAUS6P1 | 0.30473766 | 0.97386856 | -0.536 | -0.469 | chr7 | 53255081 | 53256631 | + | processed_pseudogene |
| ENSG00000176170.13_5 | SPHK1 | 0.35368284 | 1 | -0.536 | 1.472 | chr17 | 74372665 | 74383941 | + | protein_coding |
| ENSG00000260035.1_8 | AC051619.7 | 0.3063319 | 0.97675704 | -0.537 | 0.224 | chr15 | 45492523 | 45492830 | - | lncRNA |
| ENSG00000147687.19_5 | TATDN1 | 0.43794813 | 1 | -0.537 | 4.441 | chr8 | 125500726 | 125551699 | - | protein_coding |
| ENSG00000257178.5_7 | AC103702.1 | 0.31287818 | 0.99124584 | -0.537 | 0.300 | chr17 | 46656992 | 46659621 | - | lncRNA |
| ENSG00000138131.4_4 | LOXL4 | 0.38725361 | 1 | -0.537 | 2.379 | chr10 | 100007447 | 100027951 | - | protein_coding |
| ENSG00000178150.10_4 | ZNF114 | 0.38746279 | 1 | -0.537 | 2.320 | chr19 | 48773338 | 48790865 | + | protein_coding |
| ENSG00000203872.7_5 | C6orf163 | 0.30974341 | 0.98447427 | -0.538 | 0.051 | chr6 | 88054531 | 88075181 | + | protein_coding |
| ENSG00000259781.1_4 | HMGB1P6 | 0.41026432 | 1 | -0.538 | 2.900 | chr15 | 71457109 | 71457754 | - | processed_pseudogene |
| ENSG00000259201.1_7 | AC090971.3 | 0.30634512 | 0.97675704 | -0.538 | -0.961 | chr15 | 52179757 | 52193320 | - | lncRNA |
| ENSG00000270195.2_8 | AC016773.2 | 0.30852959 | 0.98149423 | -0.538 | 0.134 | chr4 | 1714548 | 1717672 | + | lncRNA |
| ENSG00000278637.1_3 | H4C1 | 0.3405326 | 1 | -0.539 | 5.479 | chr6 | 26021907 | 26022218 | + | protein_coding |
| ENSG00000115539.14_5 | PDCL3 | 0.40407112 | 1 | -0.539 | 3.679 | chr2 | 101179455 | 101193201 | + | protein_coding |
| ENSG00000172572.6_3 | PDE3A | 0.44727585 | 1 | -0.539 | 4.309 | chr12 | 20522179 | 20837315 | + | protein_coding |
| ENSG00000051180.17_4 | RAD51 | 0.41770789 | 1 | -0.539 | 3.214 | chr15 | 40986972 | 41024538 | + | protein_coding |
| ENSG00000230295.1_4 | GTF2IP23 | 0.30225505 | 0.9681207 | -0.539 | -0.994 | chr7 | 66345695 | 66347968 | + | unprocessed_pseudogene |
| ENSG00000250896.1_4 | RNPS1P1 | 0.30440309 | 0.9732463 | -0.539 | 0.079 | chr4 | 11373599 | 11374516 | + | processed_pseudogene |
| ENSG00000261884.2_9 | AC040162.1 | 0.33416523 | 1 | -0.539 | 1.216 | chr16 | 67963517 | 67969920 | - | protein_coding |
| ENSG00000153827.13_7 | TRIP12 | 0.63282503 | 1 | -0.540 | 8.109 | chr2 | 230628554 | 230787955 | - | protein_coding |
| ENSG00000114656.12_7 | KIAA1257 | 0.3096137 | 0.98423807 | -0.540 | 0.325 | chr3 | 128628709 | 128721533 | - | protein_coding |
| ENSG00000115170.15_4 | ACVR1 | 0.45180913 | 1 | -0.540 | 4.100 | chr2 | 158592956 | 158732842 | - | protein_coding |
| ENSG00000272071.1_7 | AC122710.2 | 0.2998051 | 0.96272484 | -0.540 | -0.724 | chr5 | 6779571 | 6780111 | + | lncRNA |
| ENSG00000241627.3_4 | UBQLN4P1 | 0.30036038 | 0.96351218 | -0.540 | -0.275 | chr3 | 148703655 | 148705455 | - | processed_pseudogene |
| ENSG00000172215.6_4 | CXCR6 | 0.30029754 | 0.96351218 | -0.540 | -0.251 | chr3 | 45982425 | 45989843 | + | protein_coding |
| ENSG00000115540.15_4 | MOB4 | 0.44552498 | 1 | -0.540 | 4.495 | chr2 | 198380295 | 198418423 | + | protein_coding |
| ENSG00000252602.1 | U8 | 0.30624157 | 0.97675704 | -0.540 | -0.995 | chr15 | 30935128 | 30935255 | + | snoRNA |
| ENSG00000235016.1_7 | SEMA3F-AS1 | 0.29987166 | 0.96272484 | -0.540 | -0.175 | chr3 | 50153455 | 50193518 | - | lncRNA |
| ENSG00000185834.10_2 | RPL12P4 | 0.355231 | 1 | -0.540 | 1.758 | chr20 | 53691183 | 53691679 | + | processed_pseudogene |
| ENSG00000214900.11_7 | LINC01588 | 0.36829257 | 1 | -0.540 | 2.170 | chr14 | 50394289 | 50571761 | - | lncRNA |
| ENSG00000198887.9_4 | SMC5 | 0.55143565 | 1 | -0.540 | 6.384 | chr9 | 72873894 | 72969789 | + | protein_coding |
| ENSG00000152082.14_7 | MZT2B | 0.4473426 | 1 | -0.541 | 4.452 | chr2 | 130939310 | 130948302 | + | protein_coding |
| ENSG00000280248.1_8 | AC124319.3 | 0.30152995 | 0.96597234 | -0.541 | 0.124 | chr17 | 78313698 | 78315858 | + | TEC |
| ENSG00000164815.11_4 | ORC5 | 0.4934656 | 1 | -0.541 | 5.002 | chr7 | 103766788 | 103848495 | - | protein_coding |
| ENSG00000276180.1_3 | H4C9 | 0.47923615 | 1 | -0.541 | 5.530 | chr6 | 27106367 | 27107660 | + | protein_coding |
| ENSG00000205763.13_3 | RP9P | 0.39218938 | 1 | -0.541 | 2.700 | chr7 | 32956427 | 32982788 | - | transcribed_unprocessed_pseudogene |
| ENSG00000280069.1_6 | AC127024.8 | 0.32671926 | 1 | -0.541 | 1.320 | chr17 | 29065200 | 29067293 | + | TEC |
| ENSG00000273162.1_6 | AL133215.2 | 0.31840854 | 1 | -0.541 | 0.694 | chr10 | 102819786 | 102820762 | - | lncRNA |

| | | | | | | | | | | |
|---|---|---|---|---|---|---|---|---|---|---|
| ENSG00000230224.1_4 | PHBP9 | 0.3023365 | 0.96820703 | -0.541 | 0.113 | chr10 | 102008028 | 102008852 | + | processed_pseudogene |
| ENSG00000230074.3_7 | AL162231.2 | 0.323973 | 1 | -0.541 | 0.598 | chr9 | 34665604 | 34681978 | + | lncRNA |
| ENSG00000164180.13_4 | TMEM161B | 0.49102461 | 1 | -0.542 | 4.810 | chr5 | 87485450 | 87565293 | - | protein_coding |
| ENSG00000138078.15_5 | PREPL | 0.55443869 | 1 | -0.542 | 6.281 | chr2 | 44543420 | 44589001 | - | protein_coding |
| ENSG00000102098.19_6 | SCML2 | 0.46549028 | 1 | -0.542 | 4.166 | chrX | 18257433 | 18372808 | - | protein_coding |
| ENSG00000173727.12_6 | AP000769.1 | 0.38282145 | 1 | -0.542 | 2.678 | chr11 | 65222729 | 65234191 | + | transcribed_unprocessed_pseudogene |
| ENSG00000200355.1 | SNORA72 | 0.29697633 | 0.95725299 | -0.542 | -0.424 | chr3 | 172489067 | 172489195 | - | snoRNA |
| ENSG00000247473.2_6 | CARS-AS1 | 0.29661482 | 0.95677827 | -0.542 | -0.473 | chr11 | 3050239 | 3062490 | + | lncRNA |
| ENSG00000117724.13_3 | CENPF | 0.57045763 | 1 | -0.543 | 8.875 | chr1 | 214776538 | 214837914 | + | protein_coding |
| ENSG00000118960.13_6 | HS1BP3 | 0.48754838 | 1 | -0.543 | 5.065 | chr2 | 20760208 | 20850890 | - | protein_coding |
| ENSG00000277778.2_4 | PGM5P2 | 0.30733145 | 0.9788318 | -0.543 | -1.030 | chr9 | 69080228 | 69147846 | - | transcribed_unprocessed_pseudogene |
| ENSG00000143742.13_4 | SRP9 | 0.56026183 | 1 | -0.543 | 7.552 | chr1 | 225965515 | 225978170 | + | protein_coding |
| ENSG00000269958.1_9 | AL049840.6 | 0.38534496 | 1 | -0.543 | 2.667 | chr14 | 104162690 | 104163500 | + | lncRNA |
| ENSG00000270890.1_5 | AL049844.2 | 0.29817515 | 0.95937666 | -0.543 | -0.075 | chr6 | 144179199 | 144179826 | - | processed_pseudogene |
| ENSG00000276445.1_6 | AC005393.1 | 0.30292009 | 0.96937702 | -0.543 | -0.119 | chr19 | 40303612 | 40304195 | - | lncRNA |
| ENSG00000231468.2_5 | PRDX3P2 | 0.30485083 | 0.97397674 | -0.543 | -0.928 | chr1 | 28852830 | 28853739 | - | processed_pseudogene |
| ENSG00000272374.1_6 | Z97832.2 | 0.29674762 | 0.95686297 | -0.543 | -0.641 | chr6 | 35188147 | 35192407 | - | lncRNA |
| ENSG00000074590.13_4 | NUAK1 | 0.51170536 | 1 | -0.543 | 5.341 | chr12 | 106457118 | 106533811 | - | protein_coding |
| ENSG00000030066.13_5 | NUP160 | 0.56497596 | 1 | -0.543 | 6.517 | chr11 | 47799639 | 47870107 | - | protein_coding |
| ENSG00000169031.20_6 | COL4A3 | 0.40797036 | 1 | -0.544 | 3.378 | chr2 | 228029340 | 228179508 | + | protein_coding |
| ENSG00000232176.1_5 | AL161909.1 | 0.29531763 | 0.95450396 | -0.544 | -0.696 | chr9 | 19200333 | 19201044 | - | processed_pseudogene |
| ENSG00000169504.15_4 | CLIC4 | 0.53254973 | 1 | -0.544 | 6.245 | chr1 | 25071873 | 25170812 | + | protein_coding |
| ENSG00000171811.14_8 | CFAP46 | 0.29763496 | 0.95823861 | -0.544 | -0.017 | chr10 | 134621896 | 134756327 | - | protein_coding |
| ENSG00000271122.1_6 | AC018647.2 | 0.35377627 | 1 | -0.544 | 2.368 | chr7 | 35734824 | 35739023 | + | lncRNA |
| ENSG00000224635.2_7 | AL391095.2 | 0.34034868 | 1 | -0.544 | 1.741 | chr20 | 37033540 | 37045444 | - | lncRNA |
| ENSG00000006016.11_5 | CRLF1 | 0.30065827 | 0.96422302 | -0.544 | 0.335 | chr19 | 18683030 | 18718551 | - | protein_coding |
| ENSG00000225159.1_4 | NPM1P39 | 0.30736003 | 0.9788318 | -0.544 | 0.710 | chr1 | 27533421 | 27534287 | + | processed_pseudogene |
| ENSG00000137504.13_3 | CREBZF | 0.57067744 | 1 | -0.544 | 7.322 | chr11 | 85370752 | 85393951 | - | protein_coding |
| ENSG00000197153.4_4 | H3C12 | 0.53591419 | 1 | -0.545 | 6.737 | chr6 | 27858160 | 27860884 | - | protein_coding |
| ENSG00000253476.1_7 | AC091185.1 | 0.30417501 | 0.97269211 | -0.545 | 0.320 | chr8 | 25283037 | 25284096 | - | lncRNA |
| ENSG00000156876.10_5 | SASS6 | 0.43920481 | 1 | -0.545 | 4.561 | chr1 | 100549119 | 100598511 | - | protein_coding |
| ENSG00000176946.12_4 | THAP4 | 0.50070274 | 1 | -0.545 | 5.625 | chr2 | 242523820 | 242576573 | - | protein_coding |
| ENSG00000263624.1_7 | AC055811.1 | 0.31847479 | 1 | -0.545 | 1.020 | chr17 | 17071260 | 17088868 | - | lncRNA |
| ENSG00000100632.11_4 | ERH | 0.53208598 | 1 | -0.545 | 6.621 | chr14 | 69846845 | 69865016 | - | protein_coding |
| ENSG00000185527.12_6 | PDE6G | 0.2943836 | 0.95287181 | -0.545 | -0.725 | chr17 | 79617489 | 79630142 | - | protein_coding |
| ENSG00000101076.16_4 | HNF4A | 0.49066115 | 1 | -0.545 | 5.481 | chr20 | 42984340 | 43063236 | + | protein_coding |
| ENSG00000277170.1_7 | AC012676.3 | 0.29424487 | 0.95259632 | -0.545 | -0.754 | chr16 | 4476903 | 4477381 | + | lncRNA |
| ENSG00000224072.1_6 | AL139811.1 | 0.29665507 | 0.95677827 | -0.545 | 0.005 | chrX | 46313532 | 46316634 | - | processed_pseudogene |
| ENSG00000261542.1_6 | AC011978.2 | 0.29645304 | 0.95660756 | -0.546 | 0.131 | chr8 | 64128539 | 64130592 | + | lncRNA |
| ENSG00000140280.14_4 | LYSMD2 | 0.39411469 | 1 | -0.546 | 3.361 | chr15 | 52015208 | 52043782 | - | protein_coding |
| ENSG00000279347.1_6 | AC021945.1 | 0.29305823 | 0.95013972 | -0.546 | -0.400 | chr8 | 120850976 | 120852625 | - | TEC |
| ENSG00000120699.13_4 | EXOSC8 | 0.43935321 | 1 | -0.546 | 4.741 | chr13 | 37572953 | 37583751 | + | protein_coding |
| ENSG00000185480.11_5 | PARPBP | 0.47554664 | 1 | -0.546 | 5.344 | chr12 | 102513963 | 102591298 | + | protein_coding |
| ENSG00000239559.2_4 | RPL37P2 | 0.29261074 | 0.94920851 | -0.546 | -0.358 | chr11 | 67450243 | 67450529 | - | processed_pseudogene |
| ENSG00000185010.15_6 | F8 | 0.37932164 | 1 | -0.546 | 3.283 | chrX | 154064063 | 154255215 | - | protein_coding |
| ENSG00000225200.2_5 | AC246787.1 | 0.30247626 | 0.96848004 | -0.547 | 0.390 | chr14 | 106444607 | 106445233 | - | processed_pseudogene |
| ENSG00000259330.3_4 | INAFM2 | 0.33154481 | 1 | -0.547 | 1.695 | chr15 | 40615893 | 40618916 | + | protein_coding |
| ENSG00000227620.4_4 | ALG1L8P | 0.30150793 | 0.96597234 | -0.547 | -1.031 | chr11 | 67552744 | 67559806 | + | unprocessed_pseudogene |
| ENSG00000164934.14_5 | DCAF13 | 0.51882731 | 1 | -0.547 | 6.667 | chr8 | 104426942 | 104455681 | + | protein_coding |
| ENSG00000103222.19_4 | ABCC1 | 0.52843973 | 1 | -0.547 | 7.641 | chr16 | 16043434 | 16236931 | + | protein_coding |
| ENSG00000121211.8_4 | MND1 | 0.38504755 | 1 | -0.547 | 3.279 | chr4 | 154265801 | 154336270 | + | protein_coding |
| ENSG00000124529.3 | HIST1H4B | 0.54929585 | 1 | -0.547 | 7.370 | chr6 | 26027124 | 26027480 | - | protein_coding |
| ENSG00000201217.1 | Y_RNA | 0.3289206 | 1 | -0.547 | -0.961 | chr3 | 152167061 | 152167173 | + | misc_RNA |
| ENSG00000173085.14_7 | COQ2 | 0.45435828 | 1 | -0.548 | 4.666 | chr4 | 84182689 | 84206067 | - | protein_coding |
| ENSG00000262246.6_9 | CORO7 | 0.34186459 | 1 | -0.548 | 1.445 | chr16 | 4404543 | 4475706 | - | protein_coding |

| | | | | | | | | | | |
|---|---|---|---|---|---|---|---|---|---|---|
| ENSG00000285710.1_3 | AC012184.4 | 0.29317264 | 0.95023537 | -0.548 | -0.725 | chr16 | 70382988 | 70406069 | - | lncRNA |
| ENSG00000249264.1_4 | EEF1A1P9 | 0.29279593 | 0.94946678 | -0.548 | 0.157 | chr4 | 106405855 | 106407237 | + | processed_pseudogene |
| ENSG00000136731.13_3 | UGGT1 | 0.55137789 | 1 | -0.548 | 7.365 | chr2 | 128848774 | 128953251 | + | protein_coding |
| ENSG00000178999.13_6 | AURKB | 0.51518827 | 1 | -0.548 | 6.002 | chr17 | 8108051 | 8113918 | - | protein_coding |
| ENSG00000125450.11_4 | NUP85 | 0.53015287 | 1 | -0.548 | 6.582 | chr17 | 73201754 | 73231853 | + | protein_coding |
| ENSG00000185414.20_4 | MRPL30 | 0.4703199 | 1 | -0.548 | 5.197 | chr2 | 99797615 | 99816024 | + | protein_coding |
| ENSG00000150938.10_3 | CRIM1 | 0.57032632 | 1 | -0.548 | 8.808 | chr2 | 36582921 | 36778278 | + | protein_coding |
| ENSG00000196741.5_5 | LINC01560 | 0.30880835 | 0.98220515 | -0.548 | 0.518 | chrX | 47342970 | 47344222 | + | lncRNA |
| ENSG00000272402.1_6 | AL031775.2 | 0.29833395 | 0.95952221 | -0.548 | -0.671 | chr6 | 24706975 | 24707379 | + | lncRNA |
| ENSG00000260025.1_7 | CRIM1-DT | 0.33914199 | 1 | -0.548 | 1.893 | chr2 | 36581892 | 36582257 | - | lncRNA |
| ENSG00000154134.15_3 | ROBO3 | 0.32343979 | 1 | -0.549 | 1.441 | chr11 | 124735328 | 124751367 | + | protein_coding |
| ENSG00000274421.1_8 | AL162390.2 | 0.296095 | 0.95596622 | -0.549 | -1.066 | chr9 | 72925791 | 72927036 | + | lncRNA |
| ENSG00000243147.8_4 | MRPL33 | 0.44777622 | 1 | -0.549 | 5.734 | chr2 | 27994584 | 28210954 | + | protein_coding |
| ENSG00000115392.11_5 | FANCL | 0.44662378 | 1 | -0.549 | 4.703 | chr2 | 58386378 | 58468507 | - | protein_coding |
| ENSG00000118260.15_8 | CREB1 | 0.4600262 | 1 | -0.549 | 5.576 | chr2 | 208394461 | 208470712 | + | protein_coding |
| ENSG00000172824.16_6 | CES4A | 0.31319189 | 0.99170921 | -0.550 | 1.067 | chr16 | 67022492 | 67043661 | + | protein_coding |
| ENSG00000164284.15_3 | GRPEL2 | 0.43171646 | 1 | -0.550 | 5.091 | chr5 | 148724993 | 148734146 | + | protein_coding |
| ENSG00000115694.15_5 | STK25 | 0.47658096 | 1 | -0.550 | 6.120 | chr2 | 242432085 | 242449145 | - | protein_coding |
| ENSG00000023041.11_5 | ZDHHC6 | 0.47268365 | 1 | -0.551 | 5.572 | chr10 | 114184186 | 114206675 | - | protein_coding |
| ENSG00000162869.16_4 | PPP1R21 | 0.40853554 | 1 | -0.551 | 4.078 | chr2 | 48667737 | 48742530 | + | protein_coding |
| ENSG00000186628.12_5 | FSD2 | 0.30524845 | 0.97472127 | -0.551 | 0.931 | chr15 | 83424114 | 83474822 | - | protein_coding |
| ENSG00000196305.17_4 | IARS1 | 0.53875536 | 1 | -0.551 | 8.891 | chr9 | 94972489 | 95056038 | - | protein_coding |
| ENSG00000118193.12_4 | KIF14 | 0.48520208 | 1 | -0.551 | 6.755 | chr1 | 200520625 | 200589879 | - | protein_coding |
| ENSG00000163697.17_6 | APBB2 | 0.41410095 | 1 | -0.551 | 4.393 | chr4 | 40812044 | 41218731 | - | protein_coding |
| ENSG00000148229.13_4 | POLE3 | 0.49628403 | 1 | -0.551 | 6.542 | chr9 | 116169515 | 116172955 | - | protein_coding |
| ENSG00000051341.14_5 | POLQ | 0.46601771 | 1 | -0.551 | 5.976 | chr3 | 121150274 | 121265488 | - | protein_coding |
| ENSG00000060688.13_5 | SNRNP40 | 0.4766968 | 1 | -0.551 | 5.753 | chr1 | 31732415 | 31769635 | - | protein_coding |
| ENSG00000202337.1 | RNU6-8 | 0.29451219 | 0.95311442 | -0.552 | 0.599 | chr14 | 32672369 | 32672475 | - | snRNA |
| ENSG00000198826.11_5 | ARHGAP11A | 0.5033905 | 1 | -0.552 | 6.993 | chr15 | 32907345 | 32932142 | + | protein_coding |
| ENSG00000173409.14_4 | ARV1 | 0.39452236 | 1 | -0.552 | 4.292 | chr1 | 231114727 | 231136479 | + | protein_coding |
| ENSG00000189403.15_5 | HMGB1 | 0.52204289 | 1 | -0.552 | 8.052 | chr13 | 31030841 | 31191734 | - | protein_coding |
| ENSG00000051596.9_4 | THOC3 | 0.38100742 | 1 | -0.552 | 3.761 | chr5 | 175344876 | 175461683 | - | protein_coding |
| ENSG00000273302.1_6 | AC016747.3 | 0.28516787 | 0.93087522 | -0.552 | -0.496 | chr2 | 61427114 | 61427904 | + | lncRNA |
| ENSG00000185130.5_3 | H2BC13 | 0.4977978 | 1 | -0.552 | 7.064 | chr6 | 27775222 | 27775709 | - | protein_coding |
| ENSG00000254469.7_9 | AP002495.1 | 0.33704446 | 1 | -0.553 | 2.181 | chr11 | 71576555 | 71639700 | - | protein_coding |
| ENSG00000114209.15_5 | PDCD10 | 0.41211653 | 1 | -0.553 | 5.181 | chr3 | 167401086 | 167452727 | - | protein_coding |
| ENSG00000163357.10_3 | DCST1 | 0.28896956 | 0.93962704 | -0.553 | 0.060 | chr1 | 155006300 | 155023406 | + | protein_coding |
| ENSG00000156587.16_5 | UBE2L6 | 0.37644273 | 1 | -0.553 | 3.479 | chr11 | 57319129 | 57335757 | - | protein_coding |
| ENSG00000284691.1_4 | AC073111.4 | 0.29749282 | 0.95822243 | -0.553 | 0.921 | chr7 | 150097790 | 150109558 | + | protein_coding |
| ENSG00000268798.1_7 | AC027307.3 | 0.28833475 | 0.93807792 | -0.554 | 0.140 | chr19 | 1440838 | 1441937 | + | lncRNA |
| ENSG00000073605.18_5 | GSDMB | 0.33827403 | 1 | -0.554 | 2.568 | chr17 | 38060848 | 38076107 | - | protein_coding |
| ENSG00000163166.15_5 | IWS1 | 0.46098281 | 1 | -0.554 | 5.929 | chr2 | 128193783 | 128284462 | - | protein_coding |
| ENSG00000133393.13_5 | FOPNL | 0.47235056 | 1 | -0.554 | 6.378 | chr16 | 15959576 | 15982482 | - | protein_coding |
| ENSG00000059915.17_6 | PSD | 0.28388795 | 0.9274482 | -0.554 | -0.279 | chr10 | 104162374 | 104181296 | - | protein_coding |
| ENSG00000119547.6_4 | ONECUT2 | 0.33883438 | 1 | -0.554 | 2.419 | chr18 | 55102606 | 55158530 | + | protein_coding |
| ENSG00000111875.8_3 | ASF1A | 0.39527934 | 1 | -0.555 | 4.661 | chr6 | 119215316 | 119230336 | + | protein_coding |
| ENSG00000108010.12_3 | GLRX3 | 0.46543063 | 1 | -0.555 | 6.423 | chr10 | 131934655 | 131982785 | + | protein_coding |
| ENSG00000124207.17_3 | CSE1L | 0.48989369 | 1 | -0.555 | 8.281 | chr20 | 47662783 | 47713497 | + | protein_coding |
| ENSG00000157106.12 | SMG1 | 0.45721569 | 1 | -0.555 | 7.516 | chr16 | 18816175 | 18937776 | - | protein_coding |
| ENSG00000117650.13_4 | NEK2 | 0.44677368 | 1 | -0.555 | 5.913 | chr1 | 211831999 | 211848972 | - | protein_coding |
| ENSG00000254317.1_7 | AC022973.5 | 0.28184547 | 0.92305042 | -0.555 | -0.548 | chr8 | 130844547 | 130856750 | + | lncRNA |
| ENSG00000213757.3_5 | AC020898.1 | 0.28908717 | 0.93966818 | -0.555 | -0.813 | chr5 | 78579955 | 78580741 | - | processed_pseudogene |
| ENSG00000228305.2_5 | AC016734.1 | 0.28732762 | 0.93565795 | -0.555 | 0.307 | chr2 | 63849312 | 63849965 | - | processed_pseudogene |
| ENSG00000134809.9_4 | TIMM10 | 0.41712435 | 1 | -0.555 | 4.834 | chr11 | 57295937 | 57298276 | - | protein_coding |
| ENSG00000241506.1_4 | PSMC1P1 | 0.29730323 | 0.95778539 | -0.555 | 2.169 | chr3 | 68684856 | 68686175 | + | processed_pseudogene |
| ENSG00000212533.1 | SNORA75 | 0.28595314 | 0.93265109 | -0.556 | -0.696 | chr12 | 31229801 | 31229950 | - | snoRNA |
| ENSG00000128268.12_3 | MGAT3 | 0.28584224 | 0.93265109 | -0.557 | 0.109 | chr22 | 39853017 | 39888199 | + | protein_coding |
| ENSG00000065135.12_5 | GNAI3 | 0.43373229 | 1 | -0.557 | 6.653 | chr1 | 110091237 | 110142817 | + | protein_coding |
| ENSG00000264175.1 | MIR3189 | 0.28100258 | 0.92130887 | -0.557 | -0.182 | chr19 | 18497372 | 18497444 | + | miRNA |

| | | | | | | | | | | |
|---|---|---|---|---|---|---|---|---|---|---|
| ENSG00000115760.14_4 | BIRC6 | 0.4415636 | 1 | -0.557 | 8.314 | chr2 | 32582096 | 32843966 | + | protein_coding |
| ENSG00000173674.11_4 | EIF1AX | 0.45739335 | 1 | -0.557 | 6.892 | chrX | 20142643 | 20159956 | - | protein_coding |
| ENSG00000267296.2_7 | CEBPA-DT | 0.29001021 | 0.94209914 | -0.557 | 0.840 | chr19 | 33793763 | 33795960 | + | lncRNA |
| ENSG00000233337.1_4 | UBE2FP3 | 0.28125971 | 0.92147176 | -0.557 | -0.498 | chr1 | 111980136 | 111980659 | + | processed_pseudogene |
| ENSG00000286024.1_2 | AC026803.3 | 0.36319592 | 1 | -0.557 | -0.616 | chr19 | 49310578 | 49313974 | + | lncRNA |
| ENSG00000170312.16_4 | CDK1 | 0.46413483 | 1 | -0.558 | 7.103 | chr10 | 62538089 | 62554610 | + | protein_coding |
| ENSG00000174500.13_5 | GCSAM | 0.28154907 | 0.92224968 | -0.558 | 0.121 | chr3 | 111839686 | 111852117 | - | protein_coding |
| ENSG00000198134.3_6 | PTMAP9 | 0.2859579 | 0.93265109 | -0.558 | -0.642 | chr12 | 12264097 | 12264423 | + | processed_pseudogene |
| ENSG00000249353.2_4 | NPM1P27 | 0.30329504 | 0.9702274 | -0.558 | 4.552 | chr5 | 93018544 | 93019373 | - | processed_pseudogene |
| ENSG00000279170.2_6 | TSTD3 | 0.31300234 | 0.99128567 | -0.558 | 2.348 | chr6 | 99968569 | 100037775 | + | protein_coding |
| ENSG00000227973.1_5 | PIN4P1 | 0.29327916 | 0.95023537 | -0.558 | -0.940 | chr15 | 44168047 | 44168442 | - | processed_pseudogene |
| ENSG00000215158.9_7 | AC138409.2 | 0.30943651 | 0.98402694 | -0.558 | 1.817 | chr5 | 34164803 | 34244901 | - | transcribed_unprocessed_pseudogene |
| ENSG00000168758.11_5 | SEMA4C | 0.3698836 | 1 | -0.558 | 4.412 | chr2 | 97525455 | 97536494 | - | protein_coding |
| ENSG00000237943.7_5 | PRKCQ-AS1 | 0.34846694 | 1 | -0.558 | 3.388 | chr10 | 6622381 | 6658883 | + | lncRNA |
| ENSG00000198554.12_8 | WDHD1 | 0.40881542 | 1 | -0.559 | 6.053 | chr14 | 55405667 | 55493823 | - | protein_coding |
| ENSG00000121454.6_3 | LHX4 | 0.32176129 | 1 | -0.559 | 2.598 | chr1 | 180199399 | 180248119 | + | protein_coding |
| ENSG00000087338.5_3 | GMCL1 | 0.3925812 | 1 | -0.559 | 4.680 | chr2 | 70056792 | 70108516 | + | protein_coding |
| ENSG00000221539.1 | SNORD99 | 0.28125971 | 0.92147176 | -0.559 | -0.498 | chr1 | 28905261 | 28905334 | - | snoRNA |
| ENSG00000123219.13_7 | CENPK | 0.38687039 | 1 | -0.559 | 5.436 | chr5 | 64813593 | 64858995 | - | protein_coding |
| ENSG00000279660.1_6 | AC005703.6 | 0.27823952 | 0.91579874 | -0.559 | -0.301 | chr17 | 15163830 | 15165158 | + | TEC |
| ENSG00000197061.4_3 | H4C3 | 0.43052284 | 1 | -0.559 | 7.184 | chr6 | 26104104 | 26104538 | + | protein_coding |
| ENSG00000157999.6_3 | ANKRD61 | 0.2966675 | 0.95677827 | -0.559 | 1.460 | chr7 | 6071007 | 6076183 | + | protein_coding |
| ENSG00000120526.11_4 | NUDCD1 | 0.42058596 | 1 | -0.559 | 5.949 | chr8 | 110253148 | 110346347 | - | protein_coding |
| ENSG00000029153.14_6 | ARNTL2 | 0.41525905 | 1 | -0.559 | 5.722 | chr12 | 27485787 | 27578222 | + | protein_coding |
| ENSG00000189120.5_4 | SP6 | 0.2976293 | 0.95823861 | -0.559 | 0.451 | chr17 | 45922274 | 45933240 | - | protein_coding |
| ENSG00000120049.19_6 | KCNIP2 | 0.27844818 | 0.91580688 | -0.559 | -0.060 | chr10 | 103585731 | 103603677 | - | protein_coding |
| ENSG00000081181.8_3 | ARG2 | 0.28626132 | 0.93318318 | -0.559 | -0.239 | chr14 | 68086637 | 68118425 | + | protein_coding |
| ENSG00000249859.11_5 | PVT1 | 0.37233492 | 1 | -0.559 | 5.381 | chr8 | 128806772 | 129199347 | + | lncRNA |
| ENSG00000234797.5_4 | RPS3AP6 | 0.32700849 | 1 | -0.559 | 3.338 | chr15 | 60060551 | 60061345 | + | processed_pseudogene |
| ENSG00000207005.1 | RNU1-2 | 0.28122487 | 0.92147176 | -0.560 | -0.784 | chr1 | 17222475 | 17222638 | + | snRNA |
| ENSG00000180592.17_4 | SKIDA1 | 0.32697272 | 1 | -0.560 | 2.696 | chr10 | 21802404 | 21815297 | - | protein_coding |
| ENSG00000184162.14_3 | NR2C2AP | 0.33483555 | 1 | -0.560 | 3.166 | chr19 | 19312225 | 19314233 | - | protein_coding |
| ENSG00000265261.1_6 | AC027575.1 | 0.29769564 | 0.95823861 | -0.560 | -1.104 | chr18 | 74596891 | 74600680 | - | processed_pseudogene |
| ENSG00000147100.11_5 | SLC16A2 | 0.34051096 | 1 | -0.560 | 3.475 | chrX | 73641328 | 73753752 | + | protein_coding |
| ENSG00000111012.10_5 | CYP27B1 | 0.29686116 | 0.95705539 | -0.560 | 1.417 | chr12 | 58156117 | 58162769 | - | protein_coding |
| ENSG00000224046.2_7 | AC005076.2 | 0.2770811 | 0.91317008 | -0.560 | -0.199 | chr7 | 86780738 | 86781736 | - | lncRNA |
| ENSG00000041515.16_3 | MYO16 | 0.30136401 | 0.96597234 | -0.560 | -0.973 | chr13 | 109248500 | 109860353 | + | protein_coding |
| ENSG00000212907.2_1 | MT-ND4L | 0.34613943 | 1 | -0.561 | 7.021 | chrM | 10470 | 10766 | + | protein_coding |
| ENSG00000127589.4_2 | TUBBP1 | 0.29575926 | 0.9555837 | -0.561 | 1.595 | chr8 | 30209389 | 30211034 | + | transcribed_processed_pseudogene |
| ENSG00000197771.12_3 | MCMBP | 0.40833678 | 1 | -0.561 | 6.868 | chr10 | 121588916 | 121652068 | - | protein_coding |
| ENSG00000135632.12_4 | SMYD5 | 0.40288556 | 1 | -0.561 | 5.720 | chr2 | 73441350 | 73454349 | + | protein_coding |
| ENSG00000162614.18_4 | NEXN | 0.29338491 | 0.95033177 | -0.561 | 1.184 | chr1 | 78354198 | 78409580 | + | protein_coding |
| ENSG00000171316.12_4 | CHD7 | 0.39145992 | 1 | -0.561 | 5.164 | chr8 | 61591299 | 61780587 | + | protein_coding |
| ENSG00000131174.7_6 | COX7B | 0.36878013 | 1 | -0.561 | 6.446 | chrX | 77154937 | 77162873 | + | protein_coding |
| ENSG00000137814.11_5 | HAUS2 | 0.37518914 | 1 | -0.561 | 4.834 | chr15 | 42841026 | 42862192 | + | protein_coding |
| ENSG00000120694.20_5 | HSPH1 | 0.4293035 | 1 | -0.561 | 7.651 | chr13 | 31709110 | 31736525 | - | protein_coding |
| ENSG00000080608.10_4 | PUM3 | 0.40495153 | 1 | -0.562 | 6.286 | chr9 | 2720469 | 2844095 | - | protein_coding |
| ENSG00000287686.1_1 | AC062029.1 | 0.27887314 | 0.9166955 | -0.562 | 0.307 | chr2 | 88989695 | 88990982 | - | lncRNA |
| ENSG00000230140.7_8 | NPAS2-AS1 | 0.27497798 | 0.90792303 | -0.562 | -0.287 | chr2 | 101587229 | 101594228 | - | lncRNA |
| ENSG00000270894.1_6 | AC015849.4 | 0.2744455 | 0.90707462 | -0.562 | -0.369 | chr17 | 34145403 | 34150717 | + | lncRNA |
| ENSG00000231616.9_7 | AL354733.2 | 0.27783131 | 0.91513327 | -0.562 | -0.814 | chr9 | 86446501 | 86483447 | + | lncRNA |
| ENSG00000204516.10_4 | MICB | 0.28210376 | 0.9232997 | -0.562 | 0.757 | chr6 | 31462658 | 31478901 | + | protein_coding |
| ENSG00000100297.16_4 | MCM5 | 0.40893788 | 1 | -0.562 | 7.339 | chr22 | 35796127 | 35821424 | + | protein_coding |
| ENSG00000233961.1 | RP11-87H9.2 | 0.28197769 | 0.9232997 | -0.562 | 0.895 | chr9 | 69065655 | 69084989 | + | pseudogene |
| ENSG00000146386.8_3 | ABRACL | 0.34245862 | 1 | -0.563 | 3.706 | chr6 | 139349882 | 139364439 | + | protein_coding |
| ENSG00000131097.7_3 | HIGD1B | 0.27429174 | 0.90700564 | -0.563 | -0.150 | chr17 | 42923721 | 42927844 | + | protein_coding |
| ENSG00000163466.16_3 | ARPC2 | 0.41880052 | 1 | -0.563 | 7.124 | chr2 | 219081864 | 219119079 | + | protein_coding |

| | | | | | | | | | | |
|---|---|---|---|---|---|---|---|---|---|---|
| ENSG00000163808.17_5 | KIF15 | 0.37281848 | 1 | -0.563 | 5.259 | chr3 | 44803213 | 44914868 | + | protein_coding |
| ENSG00000272034.1 | SNORD14A | 0.28519165 | 0.93087522 | -0.563 | 0.947 | chr11 | 17096201 | 17096291 | - | snoRNA |
| ENSG00000153989.8_3 | NUS1 | 0.36197408 | 1 | -0.564 | 4.577 | chr6 | 117996632 | 118031890 | + | protein_coding |
| ENSG00000240395.1_4 | RPL5P23 | 0.27363681 | 0.90568278 | -0.564 | -0.411 | chr8 | 42199285 | 42200152 | - | processed_pseudogene |
| ENSG00000261326.4_6 | LINC01355 | 0.29279723 | 0.94946678 | -0.564 | 2.072 | chr1 | 23607800 | 23613981 | - | lncRNA |
| ENSG00000151687.14_3 | ANKAR | 0.29498025 | 0.95393414 | -0.564 | 1.942 | chr2 | 190539016 | 190625919 | + | protein_coding |
| ENSG00000177426.20_8 | TGIF1 | 0.40918506 | 1 | -0.564 | 7.063 | chr18 | 3411606 | 3459976 | + | protein_coding |
| ENSG00000221164.1 | SNORA11 | 0.27446271 | 0.90707462 | -0.564 | -0.161 | chr10 | 74885838 | 74885965 | - | snoRNA |
| ENSG00000154237.13_4 | LRRK1 | 0.36889701 | 1 | -0.564 | 5.350 | chr15 | 101459532 | 101618462 | + | protein_coding |
| ENSG00000249715.12_5 | FER1L5 | 0.27376365 | 0.90576513 | -0.564 | 0.024 | chr2 | 97308474 | 97370624 | + | protein_coding |
| ENSG00000106031.9_7 | HOXA13 | 0.31596216 | 0.99905673 | -0.564 | 2.849 | chr7 | 27233122 | 27239710 | - | protein_coding |
| ENSG00000142634.13_3 | EFHD2 | 0.37622971 | 1 | -0.564 | 6.491 | chr1 | 15736384 | 15756835 | + | protein_coding |
| ENSG00000130377.13_3 | ACSBG2 | 0.27446791 | 0.90707462 | -0.565 | -0.877 | chr19 | 6135258 | 6193105 | + | protein_coding |
| ENSG00000287762.1_2 | AC108050.1 | 0.27414104 | 0.90684488 | -0.565 | -0.548 | chr4 | 41610329 | 41614428 | - | lncRNA |
| ENSG00000153140.8_3 | CETN3 | 0.33227602 | 1 | -0.565 | 4.332 | chr5 | 89688078 | 89705603 | - | protein_coding |
| ENSG00000072274.13_5 | TFRC | 0.40881417 | 1 | -0.565 | 8.085 | chr3 | 195754054 | 195808967 | - | protein_coding |
| ENSG00000224409.1_7 | AC114489.2 | 0.27840123 | 0.91580688 | -0.565 | -0.877 | chr1 | 33183335 | 33190838 | - | lncRNA |
| ENSG00000178053.19_6 | MLF1 | 0.3230044 | 1 | -0.565 | 3.589 | chr3 | 158288952 | 158325041 | + | protein_coding |
| ENSG00000131747.15_5 | TOP2A | 0.40409656 | 1 | -0.565 | 10.109 | chr17 | 38544777 | 38574148 | - | protein_coding |
| ENSG00000113749.7_5 | HRH2 | 0.28358708 | 0.9269954 | -0.566 | 1.146 | chr5 | 175085033 | 175137759 | + | protein_coding |
| ENSG00000178966.16_6 | RMI1 | 0.3170587 | 1 | -0.566 | 3.501 | chr9 | 86595626 | 86618989 | + | protein_coding |
| ENSG00000146842.17_4 | TMEM209 | 0.35631453 | 1 | -0.566 | 5.216 | chr7 | 129804553 | 129847610 | - | protein_coding |
| ENSG00000233426.3_4 | EIF3FP3 | 0.28826673 | 0.93802837 | -0.566 | 1.639 | chr2 | 58478575 | 58479660 | + | processed_pseudogene |
| ENSG00000187790.11_5 | FANCM | 0.33780144 | 1 | -0.566 | 4.990 | chr14 | 45605133 | 45670093 | + | protein_coding |
| ENSG00000137193.14_4 | PIM1 | 0.34096992 | 1 | -0.566 | 4.547 | chr6 | 37137928 | 37143204 | + | protein_coding |
| ENSG00000234997.2_6 | AC016745.1 | 0.29249755 | 0.9490146 | -0.566 | 1.890 | chr2 | 114082949 | 114183125 | + | lncRNA |
| ENSG00000102384.13_2 | CENPI | 0.35986121 | 1 | -0.566 | 5.364 | chrX | 100353207 | 100418670 | + | protein_coding |
| ENSG00000261924.1_7 | AC127496.1 | 0.27554456 | 0.90945573 | -0.567 | 0.634 | chr17 | 78940039 | 78945013 | - | lncRNA |
| ENSG00000115239.23_6 | ASB3 | 0.32650492 | 1 | -0.567 | 4.480 | chr2 | 53759810 | 54087297 | - | protein_coding |
| ENSG00000152782.16_3 | PANK1 | 0.3063128 | 0.97675704 | -0.567 | 3.179 | chr10 | 91339254 | 91405329 | - | protein_coding |
| ENSG00000068784.13_4 | SRBD1 | 0.34493335 | 1 | -0.567 | 5.049 | chr2 | 45615819 | 45839304 | - | protein_coding |
| ENSG00000134775.15_2 | FHOD3 | 0.29921588 | 0.96158071 | -0.567 | 2.482 | chr18 | 33877677 | 34360018 | + | protein_coding |
| ENSG00000005436.14_7 | GCFC2 | 0.34364537 | 1 | -0.567 | 5.173 | chr2 | 75879126 | 75938111 | - | protein_coding |
| ENSG00000154328.16_3 | NEIL2 | 0.32310207 | 1 | -0.567 | 4.183 | chr8 | 11627148 | 11644854 | + | protein_coding |
| ENSG00000145386.10_4 | CCNA2 | 0.36433929 | 1 | -0.568 | 6.327 | chr4 | 122737599 | 122745088 | - | protein_coding |
| ENSG00000183814.15_3 | LIN9 | 0.33452177 | 1 | -0.568 | 4.534 | chr1 | 226418850 | 226497570 | - | protein_coding |
| ENSG00000166197.16_6 | NOLC1 | 0.37363343 | 1 | -0.568 | 7.009 | chr10 | 103911933 | 103923628 | + | protein_coding |
| ENSG00000169689.15_7 | CENPX | 0.35390161 | 1 | -0.568 | 5.631 | chr17 | 79976578 | 79981983 | - | protein_coding |
| ENSG00000119950.21_6 | MXI1 | 0.3103201 | 0.98562008 | -0.568 | 3.584 | chr10 | 111967363 | 112047123 | + | protein_coding |
| ENSG00000021574.12_6 | SPAST | 0.35307799 | 1 | -0.568 | 5.963 | chr2 | 32288616 | 32382706 | + | protein_coding |
| ENSG00000159055.4_4 | MIS18A | 0.31121055 | 0.98790027 | -0.568 | 3.538 | chr21 | 33640539 | 33651360 | - | protein_coding |
| ENSG00000226479.4_4 | TMEM185B | 0.34883967 | 1 | -0.568 | 5.566 | chr2 | 120975055 | 120980976 | - | protein_coding |
| ENSG00000166508.18_4 | MCM7 | 0.37192698 | 1 | -0.569 | 8.235 | chr7 | 99690351 | 99699563 | - | protein_coding |
| ENSG00000101868.11_4 | POLA1 | 0.36156699 | 1 | -0.569 | 6.246 | chrX | 24712026 | 25015103 | + | protein_coding |
| ENSG00000136709.12_6 | WDR33 | 0.36314204 | 1 | -0.569 | 6.408 | chr2 | 128458601 | 128568761 | - | protein_coding |
| ENSG00000246334.2_7 | PRR7-AS1 | 0.28873892 | 0.93904895 | -0.569 | 2.461 | chr5 | 176865504 | 176874700 | - | lncRNA |
| ENSG00000042781.14_5 | USH2A | 0.27136864 | 0.899684 | -0.569 | -0.909 | chr1 | 215796233 | 216596790 | - | protein_coding |
| ENSG00000114942.14_3 | EEF1B2 | 0.360022 | 1 | -0.569 | 8.267 | chr2 | 207024309 | 207027652 | + | protein_coding |
| ENSG00000284753.1_1 | EEF1AKMT4 | 0.27979095 | 0.91869262 | -0.569 | 1.652 | chr3 | 183967438 | 183977373 | + | protein_coding |
| ENSG00000115825.10_6 | PRKD3 | 0.33256171 | 1 | -0.570 | 5.637 | chr2 | 37477645 | 37551951 | - | protein_coding |
| ENSG00000288032.1_1 | AC104472.5 | 0.26855766 | 0.89169575 | -0.570 | -0.574 | chr3 | 155604404 | 155608518 | + | lncRNA |
| ENSG00000189423.13_4 | USP32P3 | 0.27809363 | 0.9156578 | -0.570 | 1.403 | chr17 | 20311624 | 20337599 | + | transcribed_unprocessed_pseudogene |
| ENSG00000163666.10_6 | HESX1 | 0.29614668 | 0.95596622 | -0.570 | -1.222 | chr3 | 57231866 | 57261634 | - | protein_coding |
| ENSG00000226803.9_9 | ZNF451-AS1 | 0.27674908 | 0.9125837 | -0.570 | 1.362 | chr6 | 56979692 | 57039034 | - | lncRNA |
| ENSG00000157570.12_5 | TSPAN18 | 0.2746578 | 0.90737172 | -0.570 | 1.270 | chr11 | 44748015 | 44953974 | + | protein_coding |
| ENSG00000279786.1_6 | AC105235.1 | 0.27253314 | 0.90253427 | -0.570 | -0.627 | chr8 | 141830698 | 141832513 | - | TEC |
| ENSG00000173250.2_2 | GPR151 | 0.31151618 | 0.98798785 | -0.571 | -1.181 | chr5 | 145892666 | 145895753 | - | protein_coding |
| ENSG00000230615.7_8 | AL139220.2 | 0.26762655 | 0.88943526 | -0.571 | 0.608 | chr1 | 44496109 | 44581585 | + | lncRNA |
| ENSG00000279662.1_5 | AC131649.2 | 0.27164924 | 0.90044629 | -0.571 | -0.346 | chr16 | 10539166 | 10543604 | - | TEC |

| | | | | | | | | | | |
|---|---|---|---|---|---|---|---|---|---|---|
| ENSG00000136206.4_3 | SPDYE1 | 0.26440028 | 0.88168162 | -0.571 | -0.392 | chr7 | 44037496 | 44049721 | + | protein_coding |
| ENSG00000153294.12_5 | ADGRF4 | 0.28218142 | 0.9232997 | -0.571 | 2.060 | chr6 | 47653600 | 47689757 | + | protein_coding |
| ENSG00000074054.18_7 | CLASP1 | 0.34184211 | 1 | -0.572 | 6.287 | chr2 | 122095352 | 122407163 | - | protein_coding |
| ENSG00000162931.12_5 | TRIM17 | 0.27042515 | 0.89735983 | -0.572 | 1.129 | chr1 | 228595636 | 228604562 | - | protein_coding |
| ENSG00000114686.9_5 | MRPL3 | 0.33682192 | 1 | -0.572 | 6.495 | chr3 | 131181056 | 131221827 | - | protein_coding |
| ENSG00000131019.11_4 | ULBP3 | 0.28357018 | 0.9269954 | -0.572 | 2.629 | chr6 | 150382189 | 150390257 | - | protein_coding |
| ENSG00000243943.10_4 | ZNF512 | 0.31193303 | 0.98901124 | -0.573 | 4.883 | chr2 | 27805836 | 27846084 | + | protein_coding |
| ENSG00000106367.15_5 | AP1S1 | 0.31629299 | 0.99939136 | -0.573 | 5.423 | chr7 | 100797737 | 100804877 | + | protein_coding |
| ENSG00000111845.5_3 | PAK1IP1 | 0.3083232 | 0.98136479 | -0.573 | 4.630 | chr6 | 10695205 | 10710015 | + | protein_coding |
| ENSG00000183010.17_7 | PYCR1 | 0.31498055 | 0.99648493 | -0.573 | 5.620 | chr17 | 79890260 | 79900288 | - | protein_coding |
| ENSG00000143919.15_4 | CAMKMT | 0.27924898 | 0.91759139 | -0.573 | 2.369 | chr2 | 44589086 | 44999731 | + | protein_coding |
| ENSG00000165501.17_4 | LRR1 | 0.2983595 | 0.95952221 | -0.574 | 4.403 | chr14 | 50065479 | 50081390 | + | protein_coding |
| ENSG00000143375.15_6 | CGN | 0.27815643 | 0.91569487 | -0.574 | 2.458 | chr1 | 151482986 | 151511168 | + | protein_coding |
| ENSG00000125885.13_8 | MCM8 | 0.303588 | 0.97098973 | -0.574 | 5.071 | chr20 | 5931298 | 5979623 | + | protein_coding |
| ENSG00000165238.16_4 | WNK2 | 0.28009133 | 0.91950899 | -0.575 | 3.031 | chr9 | 95947198 | 96082854 | + | protein_coding |
| ENSG00000225920.2_4 | RIMKLBP2 | 0.26048671 | 0.87108365 | -0.575 | 0.075 | chr1 | 219373256 | 219373909 | - | processed_pseudogene |
| ENSG00000179409.11_7 | GEMIN4 | 0.30586832 | 0.97617433 | -0.575 | 5.935 | chr17 | 647661 | 657239 | - | protein_coding |
| ENSG00000230359.5_4 | TPI1P2 | 0.26150668 | 0.87382346 | -0.575 | 0.642 | chr7 | 128695277 | 128697293 | + | transcribed_processed_pseudogene |
| ENSG00000186185.14_4 | KIF18B | 0.30844066 | 0.98138704 | -0.575 | 6.093 | chr17 | 43002077 | 43025141 | - | protein_coding |
| ENSG00000115884.11_4 | SDC1 | 0.2992613 | 0.96158071 | -0.576 | 7.795 | chr2 | 20400558 | 20425194 | - | protein_coding |
| ENSG00000182134.16_5 | TDRKH | 0.28523019 | 0.93087522 | -0.576 | 3.906 | chr1 | 151742583 | 151763892 | - | protein_coding |
| ENSG00000175792.12_7 | RUVBL1 | 0.31129152 | 0.98791296 | -0.576 | 6.504 | chr3 | 127783621 | 127872757 | - | protein_coding |
| ENSG00000184916.9_5 | JAG2 | 0.27563912 | 0.90959886 | -0.576 | 3.918 | chr14 | 105607319 | 105635161 | - | protein_coding |
| ENSG00000261762.1_6 | AC027228.2 | 0.26749709 | 0.88917134 | -0.576 | 1.820 | chr15 | 78881465 | 78883618 | - | lncRNA |
| ENSG00000148468.17_4 | FAM171A1 | 0.28602001 | 0.93265109 | -0.576 | 4.327 | chr10 | 15253642 | 15413288 | - | protein_coding |
| ENSG00000137877.10_4 | SPTBN5 | 0.26275358 | 0.8770135 | -0.576 | 1.062 | chr15 | 42140344 | 42186251 | - | protein_coding |
| ENSG00000243766.8_5 | HOTTIP | 0.26848237 | 0.89161237 | -0.576 | -0.943 | chr7 | 27238194 | 27246878 | + | lncRNA |
| ENSG00000152102.18_4 | FAM168B | 0.31195019 | 0.98901124 | -0.577 | 7.196 | chr2 | 131805449 | 131851033 | - | protein_coding |
| ENSG00000135763.10_3 | URB2 | 0.30010376 | 0.96314009 | -0.577 | 5.611 | chr1 | 229761994 | 229795947 | + | protein_coding |
| ENSG00000116985.12_5 | BMP8B | 0.27687657 | 0.91283466 | -0.577 | 3.249 | chr1 | 40222854 | 40254537 | - | protein_coding |
| ENSG00000214188.9_4 | ST7-OT4 | 0.25713654 | 0.86224172 | -0.577 | -0.269 | chr7 | 116593953 | 116738860 | + | lncRNA |
| ENSG00000132781.18_8 | MUTYH | 0.26584552 | 0.88516502 | -0.577 | 0.617 | chr1 | 45794835 | 45806565 | - | protein_coding |
| ENSG00000139921.13_3 | TMX1 | 0.30271973 | 0.96908495 | -0.577 | 6.903 | chr14 | 51706880 | 51724373 | + | protein_coding |
| ENSG00000256073.3_5 | URB1-AS1 | 0.26651487 | 0.88656988 | -0.577 | 1.965 | chr21 | 33765439 | 33766269 | + | lncRNA |
| ENSG00000116455.14_5 | WDR77 | 0.28226613 | 0.9234068 | -0.578 | 4.501 | chr1 | 111982512 | 111991878 | - | protein_coding |
| ENSG00000170961.7_4 | HAS2 | 0.2794324 | 0.91802432 | -0.578 | 4.145 | chr8 | 122624356 | 122653680 | - | protein_coding |
| ENSG00000155893.13_7 | PXYLP1 | 0.2737359 | 0.90576513 | -0.578 | 3.420 | chr3 | 140947568 | 141086595 | + | protein_coding |
| ENSG00000182963.10_5 | GJC1 | 0.2860368 | 0.93265109 | -0.578 | 5.172 | chr17 | 42875816 | 42908184 | - | protein_coding |
| ENSG00000170035.16_4 | UBE2E3 | 0.28470994 | 0.92968968 | -0.578 | 4.945 | chr2 | 181831975 | 181941312 | + | protein_coding |
| ENSG00000118922.17_4 | KLF12 | 0.27832484 | 0.91580688 | -0.578 | 4.096 | chr13 | 74260226 | 74569193 | - | protein_coding |
| ENSG00000118420.17_5 | UBE3D | 0.26708399 | 0.88829676 | -0.578 | 2.586 | chr6 | 83602109 | 83775560 | - | protein_coding |
| ENSG00000272148.1_6 | AC013403.2 | 0.25906477 | 0.86714464 | -0.578 | -0.711 | chr2 | 27285296 | 27285775 | - | lncRNA |
| ENSG00000277161.2_5 | PIGW | 0.27852131 | 0.91587785 | -0.578 | 4.390 | chr17 | 34890814 | 34895159 | + | protein_coding |
| ENSG00000278916.1_5 | CEP83-DT | 0.25808679 | 0.86468565 | -0.579 | -0.739 | chr12 | 94853779 | 94856260 | + | lncRNA |
| ENSG00000230188.1_5 | AL158156.1 | 0.25946916 | 0.86817109 | -0.579 | -0.656 | chr9 | 37490418 | 37490890 | - | processed_pseudogene |
| ENSG00000204370.12_7 | SDHD | 0.27859301 | 0.91594413 | -0.579 | 5.081 | chr11 | 111957548 | 111990739 | + | protein_coding |
| ENSG00000108830.10_3 | RND2 | 0.25558307 | 0.85872624 | -0.579 | -0.290 | chr17 | 41177248 | 41184058 | + | protein_coding |
| ENSG00000122870.12_4 | BICC1 | 0.2907776 | 0.9442964 | -0.579 | 7.864 | chr10 | 60272632 | 60591195 | + | protein_coding |
| ENSG00000203288.4_6 | TDRKH-AS1 | 0.2796193 | 0.91829871 | -0.579 | -1.119 | chr1 | 151763280 | 151766878 | + | lncRNA |
| ENSG00000285077.2_3 | ARHGAP11B | 0.25517669 | 0.85768525 | -0.579 | 0.307 | chr15 | 30916697 | 30941732 | + | protein_coding |
| ENSG00000197238.4_3 | H4C11 | 0.25968323 | 0.86872378 | -0.579 | 1.500 | chr6 | 27791886 | 27792258 | + | protein_coding |
| ENSG00000146731.11_3 | CCT6A | 0.29206903 | 0.9481437 | -0.579 | 8.627 | chr7 | 56119378 | 56131682 | + | protein_coding |
| ENSG00000111802.14_5 | TDP2 | 0.27451682 | 0.90707462 | -0.579 | 4.783 | chr6 | 24650207 | 24667158 | - | protein_coding |
| ENSG00000183444.10_3 | OR7E38P | 0.25716322 | 0.86224172 | -0.580 | -0.710 | chr7 | 97595402 | 97596386 | - | unprocessed_pseudogene |
| ENSG00000251474.6_3 | RPL32P3 | 0.26534261 | 0.88432606 | -0.580 | 3.237 | chr3 | 129101765 | 129118498 | - | transcribed_unprocessed_pseudogene |
| ENSG00000241527.1_5 | CA15P1 | 0.25507772 | 0.85759758 | -0.580 | -0.172 | chr22 | 19019077 | 19022077 | - | unitary_pseudogene |
| ENSG00000273148.1_7 | LINC00653 | 0.25361351 | 0.85420877 | -0.580 | 0.270 | chr20 | 18775173 | 18776711 | + | lncRNA |
| ENSG00000168038.11_5 | ULK4 | 0.26204565 | 0.87514382 | -0.580 | 2.700 | chr3 | 41288090 | 42003622 | - | protein_coding |

| | | | | | | | | | | |
|---|---|---|---|---|---|---|---|---|---|---|
| ENSG00000128050.8_2 | PAICS | 0.28391911 | 0.9274482 | -0.580 | 7.956 | chr4 | 57301907 | 57330745 | + | protein_coding |
| ENSG00000272462.3_6 | U91328.1 | 0.25416803 | 0.85542781 | -0.580 | 0.547 | chr6 | 25992869 | 26002003 | + | lncRNA |
| ENSG00000284642.1_4 | AL139424.2 | 0.25369805 | 0.85424492 | -0.581 | 0.621 | chr1 | 10455473 | 10457489 | + | lncRNA |
| ENSG00000155561.15_4 | NUP205 | 0.27789297 | 0.91516672 | -0.581 | 7.608 | chr7 | 135242665 | 135333505 | + | protein_coding |
| ENSG00000186462.9_4 | NAP1L2 | 0.25805012 | 0.86468565 | -0.581 | 2.246 | chrX | 72432135 | 72434687 | - | protein_coding |
| ENSG00000144712.12_7 | CAND2 | 0.25588559 | 0.85909282 | -0.582 | 1.846 | chr3 | 12837971 | 12913415 | + | protein_coding |
| ENSG00000119185.12_5 | ITGB1BP1 | 0.26775561 | 0.88969777 | -0.582 | 5.416 | chr2 | 9543604 | 9563676 | - | protein_coding |
| ENSG00000155636.15_4 | RBM45 | 0.26227596 | 0.87574835 | -0.582 | 4.185 | chr2 | 178977151 | 179003738 | + | protein_coding |
| ENSG00000229124.7_7 | VIM-AS1 | 0.25140309 | 0.84804989 | -0.582 | -0.138 | chr10 | 17256238 | 17271984 | - | lncRNA |
| ENSG00000239672.7_6 | NME1 | 0.25808169 | 0.86468565 | -0.583 | 6.270 | chr17 | 49230897 | 49239789 | + | protein_coding |
| ENSG00000136875.12_2 | PRPF4 | 0.26499576 | 0.88333578 | -0.583 | 6.283 | chr9 | 116037623 | 116055185 | + | protein_coding |
| ENSG00000197140.15_3 | ADAM32 | 0.25389487 | 0.85467035 | -0.583 | 2.091 | chr8 | 38964509 | 39142436 | + | protein_coding |
| ENSG00000126953.8_6 | TIMM8A | 0.25690734 | 0.86170901 | -0.583 | 3.394 | chrX | 100600649 | 100603730 | - | protein_coding |
| ENSG00000277130.1_7 | AC073569.3 | 0.25580864 | 0.85909282 | -0.583 | -0.486 | chr12 | 80212564 | 80213245 | - | lncRNA |
| ENSG00000273891.1_7 | AL731566.2 | 0.25223322 | 0.85013508 | -0.583 | 1.913 | chr10 | 123725279 | 123727215 | + | lncRNA |
| ENSG00000203739.4_7 | AL645568.1 | 0.25004256 | 0.84457564 | -0.584 | 1.083 | chr1 | 173386932 | 173458546 | - | lncRNA |
| ENSG00000196517.12_7 | SLC6A9 | 0.25151707 | 0.84827332 | -0.584 | 2.967 | chr1 | 44457172 | 44497139 | - | protein_coding |
| ENSG00000227347.1_4 | HNRNPKP2 | 0.24850942 | 0.84020303 | -0.584 | 0.587 | chr2 | 136956684 | 136958073 | - | processed_pseudogene |
| ENSG00000124091.9_3 | GCNT7 | 0.25039432 | 0.84512841 | -0.584 | -0.510 | chr20 | 55066548 | 55100981 | - | protein_coding |
| ENSG00000272033.1_7 | AL136984.1 | 0.25037462 | 0.84512841 | -0.585 | -0.486 | chr1 | 167348345 | 167350237 | + | lncRNA |
| ENSG00000116918.15_8 | TSNAX | 0.24835115 | 0.83989086 | -0.585 | 5.477 | chr1 | 231664287 | 231702270 | + | protein_coding |
| ENSG00000114738.11_5 | MAPKAPK3 | 0.24826734 | 0.83986401 | -0.585 | 4.712 | chr3 | 50648951 | 50686722 | + | protein_coding |
| ENSG00000279390.1_5 | AF127577.6 | 0.24726822 | 0.8372812 | -0.586 | 1.682 | chr21 | 16422510 | 16424700 | + | TEC |
| ENSG00000198924.8_4 | DCLRE1A | 0.24629874 | 0.83479393 | -0.586 | 5.377 | chr10 | 115594484 | 115614142 | - | protein_coding |
| ENSG00000023228.14_6 | NDUFS1 | 0.24597902 | 0.83420838 | -0.586 | 7.287 | chr2 | 206979541 | 207024233 | - | protein_coding |
| ENSG00000224550.1_5 | AC114491.1 | 0.27493444 | 0.90792303 | -0.586 | -1.121 | chr1 | 108047825 | 108048070 | + | processed_pseudogene |
| ENSG00000240869.2 | RN7SL128P | 0.24622098 | 0.83479393 | -0.586 | 1.344 | chr6 | 20421812 | 20422118 | - | misc_RNA |
| ENSG00000236991.6_8 | EDRF1-AS1 | 0.24796023 | 0.8390248 | -0.586 | -0.292 | chr10 | 127414203 | 127440679 | - | lncRNA |
| ENSG00000231167.3_4 | YBX1P2 | 0.24777105 | 0.83875254 | -0.586 | -0.023 | chr7 | 105222705 | 105223703 | - | processed_pseudogene |
| ENSG00000137124.8_6 | ALDH1B1 | 0.23727141 | 0.81225454 | -0.587 | 6.178 | chr9 | 38392699 | 38398658 | + | protein_coding |
| ENSG00000088305.18_4 | DNMT3B | 0.2407997 | 0.82148394 | -0.587 | 3.218 | chr20 | 31350191 | 31397162 | + | protein_coding |
| ENSG00000106617.14_7 | PRKAG2 | 0.23851855 | 0.81559327 | -0.587 | 4.565 | chr7 | 151253210 | 151574210 | - | protein_coding |
| ENSG00000175772.11_6 | LINC01106 | 0.24274488 | 0.82661909 | -0.588 | 2.037 | chr2 | 111132715 | 111142019 | - | lncRNA |
| ENSG00000165124.18_4 | SVEP1 | 0.24467431 | 0.83182711 | -0.588 | -0.053 | chr9 | 113127528 | 113342160 | - | protein_coding |
| ENSG00000186481.16_3 | ANKRD20A5P | 0.24134248 | 0.82270372 | -0.588 | 2.605 | chr18 | 14179096 | 14252139 | + | transcribed_unprocessed_pseudogene |
| ENSG00000137941.17_5 | TTLL7 | 0.23932282 | 0.81785891 | -0.588 | 3.483 | chr1 | 84330707 | 84464833 | - | protein_coding |
| ENSG00000181544.15_6 | FANCB | 0.23852213 | 0.81559327 | -0.588 | 4.104 | chrX | 14854083 | 14891191 | - | protein_coding |
| ENSG00000134569.10_5 | LRP4 | 0.236 | 0.80883719 | -0.588 | 4.290 | chr11 | 46878268 | 46940193 | - | protein_coding |
| ENSG00000171435.14_4 | KSR2 | 0.23981712 | 0.81923298 | -0.588 | 2.729 | chr12 | 117890817 | 118406795 | - | protein_coding |
| ENSG00000171202.7_3 | TMEM126A | 0.23588522 | 0.80875188 | -0.588 | 4.276 | chr11 | 85359011 | 85367591 | + | protein_coding |
| ENSG00000249252.5_8 | C1QTNF7-AS1 | 0.24288804 | 0.82684235 | -0.588 | 0.410 | chr4 | 15006566 | 15429538 | - | lncRNA |
| ENSG00000170500.13_5 | LONRF2 | 0.23295115 | 0.80101472 | -0.588 | 5.129 | chr2 | 100888337 | 100938963 | - | protein_coding |
| ENSG00000235724.10_8 | AC009299.3 | 0.2455728 | 0.83360564 | -0.588 | -0.175 | chr2 | 162079296 | 162164900 | - | lncRNA |
| ENSG00000204790.8 | CBWD6 | 0.24086338 | 0.8215151 | -0.589 | 2.109 | chr9 | 69204538 | 69269662 | - | protein_coding |
| ENSG00000260743.1_6 | AC007823.1 | 0.28362396 | 0.9269954 | -0.589 | -0.945 | chr3 | 179058110 | 179059675 | + | lncRNA |
| ENSG00000114771.14_4 | AADAC | 0.23468731 | 0.80558142 | -0.590 | 3.274 | chr3 | 151531861 | 151546276 | + | protein_coding |
| ENSG00000227799.1_6 | AC012358.3 | 0.24030432 | 0.82058176 | -0.590 | 0.508 | chr2 | 55451416 | 55453044 | + | processed_pseudogene |
| ENSG00000164327.13_6 | RICTOR | 0.22035681 | 0.76826521 | -0.590 | 6.759 | chr5 | 38938022 | 39074501 | - | protein_coding |
| ENSG00000117877.10_5 | CD3EAP | 0.23213656 | 0.79867744 | -0.590 | 2.927 | chr19 | 45909467 | 45914024 | + | protein_coding |
| ENSG00000165752.17_5 | STK32C | 0.22502924 | 0.78011599 | -0.591 | 4.439 | chr10 | 134020996 | 134145351 | - | protein_coding |
| ENSG00000163607.17_6 | GTPBP8 | 0.22388827 | 0.77722214 | -0.591 | 4.419 | chr3 | 112709831 | 112733907 | + | protein_coding |
| ENSG00000132967.9_4 | HMGB1P5 | 0.23011322 | 0.79390164 | -0.591 | 3.226 | chr3 | 22423310 | 22424420 | + | transcribed_processed_pseudogene |
| ENSG00000179314.14_6 | WSCD1 | 0.23139609 | 0.79690139 | -0.591 | 3.007 | chr17 | 5961127 | 6027747 | + | protein_coding |
| ENSG00000106799.13_4 | TGFBR1 | 0.21072835 | 0.74153508 | -0.592 | 6.620 | chr9 | 101866320 | 101916474 | + | protein_coding |
| ENSG00000226121.2_6 | AC009487.2 | 0.23436722 | 0.80479367 | -0.592 | 1.616 | chr2 | 162357396 | 162364216 | - | processed_pseudogene |

| | | | | | | | | | | |
|---|---|---|---|---|---|---|---|---|---|---|
| ENSG00000226243.1_2 | RPL37AP1 | 0.26641323 | 0.88639774 | -0.592 | -1.158 | chr20 | 43095204 | 43095482 | - | processed_pseudogene |
| ENSG00000256968.1_2 | SNRPEP2 | 0.23847826 | 0.81559327 | -0.592 | 0.060 | chr9 | 6748703 | 6748981 | - | processed_pseudogene |
| ENSG00000254996.5_5 | ANKHD1-EIF4EBP3 | 0.23690178 | 0.81161472 | -0.592 | 0.348 | chr5 | 139781493 | 139929154 | + | protein_coding |
| ENSG00000212296.1 | SNORD72 | 0.26454969 | 0.88201434 | -0.592 | -1.120 | chr5 | 40832758 | 40832837 | - | snoRNA |
| ENSG00000185839.3_5 | AL035411.1 | 0.24891013 | 0.84139771 | -0.592 | -0.364 | chr1 | 59096513 | 59097202 | - | processed_pseudogene |
| ENSG00000266402.3_5 | SNHG25 | 0.23115723 | 0.79623312 | -0.593 | 2.137 | chr17 | 62223330 | 62223836 | + | lncRNA |
| ENSG00000198853.12_4 | RUSC2 | 0.20579427 | 0.72864623 | -0.593 | 6.084 | chr9 | 35490108 | 35561895 | + | protein_coding |
| ENSG00000261616.1_6 | AC036108.3 | 0.24052237 | 0.82101076 | -0.593 | -0.313 | chr15 | 99679522 | 99685575 | + | lncRNA |
| ENSG00000253720.2_7 | AC022973.3 | 0.24124951 | 0.82254463 | -0.593 | -0.423 | chr8 | 130952090 | 130963302 | + | lncRNA |
| ENSG00000241112.1_4 | RPL29P14 | 0.24836982 | 0.83989086 | -0.593 | -0.861 | chr5 | 148924946 | 148925373 | - | processed_pseudogene |
| ENSG00000196747.4_3 | H2AC13 | 0.20293985 | 0.72040606 | -0.593 | 7.420 | chr6 | 27775977 | 27776479 | + | protein_coding |
| ENSG00000180385.8_4 | EMC3-AS1 | 0.23198517 | 0.79846581 | -0.593 | 1.765 | chr3 | 10028577 | 10048674 | + | transcribed_unprocessed_pseudogene |
| ENSG00000182481.9_4 | KPNA2 | 0.19198245 | 0.69077229 | -0.593 | 8.810 | chr17 | 66031824 | 66042970 | + | protein_coding |
| ENSG00000261094.2_6 | AC007066.2 | 0.2318001 | 0.7980013 | -0.594 | 1.354 | chr9 | 125699902 | 125702612 | + | lncRNA |
| ENSG00000119314.15_5 | PTBP3 | 0.19293648 | 0.69305961 | -0.594 | 7.954 | chr9 | 114979996 | 115095947 | - | protein_coding |
| ENSG00000124614.16_9 | RPS10 | 0.2301433 | 0.79390164 | -0.594 | 1.526 | chr6 | 34385231 | 34393846 | - | protein_coding |
| ENSG00000161281.11_5 | COX7A1 | 0.24067953 | 0.82138941 | -0.594 | -0.271 | chr19 | 36641824 | 36643351 | - | protein_coding |
| ENSG00000205622.11_9 | AP001042.1 | 0.26844871 | 0.89161237 | -0.594 | -1.121 | chr21 | 40184802 | 40328392 | - | lncRNA |
| ENSG00000155744.9_3 | FAM126B | 0.21699243 | 0.75906762 | -0.595 | 3.352 | chr2 | 201838441 | 201936394 | - | protein_coding |
| ENSG00000159131.17_6 | GART | 0.19752591 | 0.70555748 | -0.595 | 6.613 | chr21 | 34876238 | 34915797 | - | protein_coding |
| ENSG00000272769.1_6 | AC097532.2 | 0.2333631 | 0.80212077 | -0.595 | 0.854 | chr2 | 133103189 | 133104870 | - | lncRNA |
| ENSG00000138079.14_6 | SLC3A1 | 0.24490856 | 0.83230498 | -0.595 | -0.829 | chr2 | 44502597 | 44548633 | + | protein_coding |
| ENSG00000172954.13_4 | LCLAT1 | 0.19678988 | 0.70363569 | -0.595 | 6.553 | chr2 | 30670092 | 30867091 | + | protein_coding |
| ENSG00000183092.16_5 | BEGAIN | 0.23707851 | 0.81190709 | -0.595 | -0.258 | chr14 | 101003484 | 101053750 | - | protein_coding |
| ENSG00000227351.2_2 | NANOGP6 | 0.22964 | 0.79254348 | -0.595 | 1.274 | chr10 | 101548321 | 101549167 | - | processed_pseudogene |
| ENSG00000106327.13_5 | TFR2 | 0.22590994 | 0.78256139 | -0.596 | 1.656 | chr7 | 100218039 | 100240402 | - | protein_coding |
| ENSG00000235090.1_4 | RPL7L1P3 | 0.24584027 | 0.8340909 | -0.596 | -0.861 | chr7 | 75552073 | 75552761 | + | processed_pseudogene |
| ENSG00000124383.9_3 | MPHOSPH10 | 0.19478683 | 0.69787812 | -0.596 | 5.207 | chr2 | 71357762 | 71377231 | + | protein_coding |
| ENSG00000234028.3_6 | EIF2AK3-DT | 0.23433041 | 0.80479367 | -0.596 | 0.211 | chr2 | 88927057 | 88931339 | + | lncRNA |
| ENSG00000278635.1_5 | AC141557.2 | 0.2318053 | 0.7980013 | -0.596 | 0.420 | chr12 | 9568237 | 9569314 | + | unprocessed_pseudogene |
| ENSG00000164764.11_5 | SBSPON | 0.2430472 | 0.82708669 | -0.596 | -0.613 | chr8 | 73976778 | 74036323 | - | protein_coding |
| ENSG00000171320.15_4 | ESCO2 | 0.18398688 | 0.66903293 | -0.597 | 6.022 | chr8 | 27629466 | 27670157 | + | protein_coding |
| ENSG00000011523.14_4 | CEP68 | 0.19511774 | 0.69878189 | -0.597 | 4.870 | chr2 | 65283500 | 65314138 | + | protein_coding |
| ENSG00000272902.2_5 | TBC1D8-AS1 | 0.23482379 | 0.80589419 | -0.597 | -0.008 | chr2 | 101768122 | 101771874 | + | lncRNA |
| ENSG00000100629.17_7 | CEP128 | 0.20455806 | 0.72513629 | -0.597 | 4.061 | chr14 | 80943326 | 81425861 | - | protein_coding |
| ENSG00000008086.13_5 | CDKL5 | 0.1928102 | 0.69275202 | -0.597 | 5.246 | chrX | 18443703 | 18671749 | + | protein_coding |
| ENSG00000154898.15_5 | CCDC144CP | 0.22263102 | 0.77452228 | -0.597 | 1.611 | chr17 | 20224477 | 20306870 | + | transcribed_processed_pseudogene |
| ENSG00000112419.14_4 | PHACTR2 | 0.18757463 | 0.67833064 | -0.598 | 5.484 | chr6 | 143857982 | 144152322 | + | protein_coding |
| ENSG00000135940.7_3 | COX5B | 0.18054019 | 0.65987608 | -0.598 | 6.048 | chr2 | 98262525 | 98264846 | + | protein_coding |
| ENSG00000123505.18_7 | AMD1 | 0.16923318 | 0.62510727 | -0.598 | 7.304 | chr6 | 111195973 | 111220082 | + | polymorphic_pseudogene |
| ENSG00000225037.1_6 | EIF1AX-AS1 | 0.23701144 | 0.81183385 | -0.598 | -0.184 | chrX | 20158086 | 20158562 | + | lncRNA |
| ENSG00000174038.13_6 | C9orf131 | 0.24292886 | 0.82684235 | -0.598 | -0.586 | chr9 | 35041092 | 35045983 | + | protein_coding |
| ENSG00000267270.6_6 | PARD6G-AS1 | 0.23419559 | 0.80451527 | -0.599 | -0.424 | chr18 | 77905807 | 77937722 | + | lncRNA |
| ENSG00000137135.18_4 | ARHGEF39 | 0.19190595 | 0.69077229 | -0.599 | 4.318 | chr9 | 35658872 | 35675863 | - | protein_coding |
| ENSG00000119705.10_7 | SLIRP | 0.18668131 | 0.67565021 | -0.599 | 5.718 | chr14 | 78174414 | 78227447 | + | protein_coding |
| ENSG00000273007.2_7 | AC021205.3 | 0.23032758 | 0.79429933 | -0.599 | 0.026 | chr4 | 123800933 | 123806960 | - | lncRNA |
| ENSG00000259726.1_4 | CSPG4P11 | 0.21169409 | 0.74449107 | -0.599 | 2.582 | chr15 | 84855504 | 84866136 | + | unprocessed_pseudogene |
| ENSG00000219023.1_6 | AL033519.3 | 0.2330385 | 0.80116002 | -0.599 | -0.040 | chr6 | 35523650 | 35524041 | + | processed_pseudogene |
| ENSG00000271532.1_4 | BBIP1P1 | 0.24165719 | 0.8236185 | -0.599 | -0.892 | chr2 | 101684840 | 101685118 | + | processed_pseudogene |
| ENSG00000139734.19_6 | DIAPH3 | 0.18154214 | 0.66244796 | -0.599 | 5.078 | chr13 | 60239717 | 60738062 | - | protein_coding |
| ENSG00000272320.1_6 | AL445309.1 | 0.23956343 | 0.81852373 | -0.600 | -0.380 | chr6 | 3311896 | 3313884 | + | lncRNA |
| ENSG00000277782.1_6 | AC068870.2 | 0.24531158 | 0.83303724 | -0.600 | -0.861 | chr15 | 81291934 | 81292322 | - | lncRNA |

| | | | | | | | | | | |
|---|---|---|---|---|---|---|---|---|---|---|
| ENSG00000129810.15_7 | SGO1 | 0.1904542 | 0.68678705 | -0.600 | 4.371 | chr3 | 20202085 | 20227698 | - | protein_coding |
| ENSG00000101911.12_3 | PRPS2 | 0.16276809 | 0.6060144 | -0.601 | 6.260 | chrX | 12809474 | 12842341 | + | protein_coding |
| ENSG00000174371.17_4 | EXO1 | 0.16657403 | 0.61785402 | -0.601 | 5.875 | chr1 | 242011269 | 242058450 | + | protein_coding |
| ENSG00000115524.16_4 | SF3B1 | 0.14854582 | 0.56441692 | -0.601 | 8.719 | chr2 | 198253239 | 198299815 | - | protein_coding |
| ENSG00000102967.12_5 | DHODH | 0.18507393 | 0.67216038 | -0.601 | 4.330 | chr16 | 72042487 | 72061563 | + | protein_coding |
| ENSG00000077232.18_6 | DNAJC10 | 0.15642809 | 0.58708447 | -0.601 | 7.612 | chr2 | 183580768 | 183659191 | + | protein_coding |
| ENSG00000119953.13_4 | SMNDC1 | 0.1819939 | 0.66368753 | -0.601 | 4.456 | chr10 | 112050488 | 112064696 | - | protein_coding |
| ENSG00000246477.3_7 | AF131216.1 | 0.21927189 | 0.76538382 | -0.602 | 1.105 | chr8 | 11173368 | 11182938 | - | lncRNA |
| ENSG00000270108.1_9 | AL049840.7 | 0.22940557 | 0.79204212 | -0.602 | -0.109 | chr14 | 104153913 | 104154464 | + | lncRNA |
| ENSG00000227413.1_5 | AL022238.1 | 0.23242838 | 0.79952662 | -0.602 | -0.404 | chr22 | 40829140 | 40829645 | + | processed_pseudogene |
| ENSG00000178184.16_4 | PARD6G | 0.21872823 | 0.76378626 | -0.602 | 1.218 | chr18 | 77915115 | 78005397 | - | protein_coding |
| ENSG00000105202.9_8 | FBL | 0.15048597 | 0.56983893 | -0.602 | 7.733 | chr19 | 40325098 | 40337019 | - | protein_coding |
| ENSG00000124380.11_4 | SNRNP27 | 0.18777033 | 0.67890019 | -0.602 | 3.760 | chr2 | 70121088 | 70132707 | + | protein_coding |
| ENSG00000127837.9_3 | AAMP | 0.16441353 | 0.61098832 | -0.602 | 6.349 | chr2 | 219128846 | 219134980 | - | protein_coding |
| ENSG00000198586.14_7 | TLK1 | 0.16748845 | 0.62059798 | -0.603 | 5.623 | chr2 | 171847333 | 172087824 | - | protein_coding |
| ENSG00000151967.18_4 | SCHIP1 | 0.22447068 | 0.7786354 | -0.603 | 0.359 | chr3 | 159557650 | 159615149 | + | protein_coding |
| ENSG00000073905.8_4 | VDAC1P1 | 0.22298736 | 0.77517691 | -0.603 | 0.569 | chrX | 80184999 | 80185846 | + | processed_pseudogene |
| ENSG00000156970.12_4 | BUB1B | 0.15659502 | 0.58758685 | -0.603 | 6.116 | chr15 | 40453224 | 40513337 | + | protein_coding |
| ENSG00000128482.16_6 | RNF112 | 0.24520778 | 0.83297093 | -0.603 | -0.958 | chr17 | 19314438 | 19320589 | + | protein_coding |
| ENSG00000227403.2_5 | LINC01806 | 0.216715 | 0.75824645 | -0.603 | 1.113 | chr2 | 162101241 | 162105561 | + | lncRNA |
| ENSG00000258441.1_6 | LINC00641 | 0.17916508 | 0.6560647 | -0.604 | 4.091 | chr14 | 21668238 | 21675059 | - | lncRNA |
| ENSG00000170092.14_2 | SPDYE5 | 0.23208959 | 0.79867049 | -0.605 | -0.123 | chr7 | 75122945 | 75133628 | + | protein_coding |
| ENSG00000187951.11_6 | AC091057.1 | 0.1907127 | 0.68718323 | -0.605 | 3.092 | chr15 | 30950920 | 31065196 | + | lncRNA |
| ENSG00000235363.1_4 | SNRPGP10 | 0.20879263 | 0.73632765 | -0.605 | 2.183 | chr1 | 205320375 | 205320599 | + | processed_pseudogene |
| ENSG00000233871.2_5 | DLG5-AS1 | 0.22313047 | 0.77520532 | -0.605 | 0.365 | chr10 | 79687130 | 79689582 | + | lncRNA |
| ENSG00000148908.15_5 | RGS10 | 0.1881055 | 0.67997363 | -0.605 | 3.441 | chr10 | 121259329 | 121302231 | - | protein_coding |
| ENSG00000228002.2_4 | DHX9P1 | 0.25818219 | 0.86468731 | -0.605 | -1.238 | chr13 | 77559003 | 77562078 | + | processed_pseudogene |
| ENSG00000188985.6_2 | DHFRP1 | 0.21561179 | 0.75545676 | -0.606 | 0.719 | chr18 | 23750690 | 23751248 | - | processed_pseudogene |
| ENSG00000244161.1_7 | FLNB-AS1 | 0.19222041 | 0.69119207 | -0.606 | 2.878 | chr3 | 58148274 | 58156363 | - | lncRNA |
| ENSG00000235954.7_5 | TTC28-AS1 | 0.18832581 | 0.68035474 | -0.606 | 3.178 | chr22 | 28315364 | 28404569 | + | lncRNA |
| ENSG00000152527.14_6 | PLEKHH2 | 0.16812573 | 0.62205121 | -0.606 | 5.139 | chr2 | 43864399 | 43995126 | + | protein_coding |
| ENSG00000120708.17_6 | TGFBI | 0.1238893 | 0.48526039 | -0.606 | 9.048 | chr5 | 135364677 | 135399507 | + | protein_coding |
| ENSG00000162999.13_3 | DUSP19 | 0.19055582 | 0.68701404 | -0.606 | 2.692 | chr2 | 183943287 | 183964736 | + | protein_coding |
| ENSG00000237719.1_5 | Z95152.1 | 0.25008794 | 0.84457564 | -0.607 | -0.891 | chr6 | 36059768 | 36060423 | - | processed_pseudogene |
| ENSG00000230797.3_4 | YY2 | 0.22303734 | 0.77517691 | -0.607 | -0.061 | chrX | 21874105 | 21876858 | + | protein_coding |
| ENSG00000100031.19_6 | GGT1 | 0.2093168 | 0.73788327 | -0.607 | 1.419 | chr22 | 24990778 | 25024972 | + | protein_coding |
| ENSG00000260708.1_6 | AL118516.1 | 0.21283031 | 0.7478948 | -0.608 | 0.825 | chr22 | 47157791 | 47158460 | - | lncRNA |
| ENSG00000214654.8_5 | B3GNT10 | 0.19084048 | 0.68748233 | -0.608 | 2.683 | chr9 | 123554681 | 123562196 | + | protein_coding |
| ENSG00000254598.3_5 | CSNK2A3 | 0.21838337 | 0.76303193 | -0.608 | 0.217 | chr11 | 11373489 | 11374797 | - | protein_coding |
| ENSG00000213178.3_5 | RPL22P1 | 0.21352142 | 0.74987845 | -0.608 | 0.600 | chr3 | 169201459 | 169201868 | - | processed_pseudogene |
| ENSG00000163918.10_6 | RFC4 | 0.14870375 | 0.56489619 | -0.608 | 5.298 | chr3 | 186507669 | 186524847 | - | protein_coding |
| ENSG00000116062.16_7 | MSH6 | 0.11691755 | 0.46347036 | -0.608 | 7.661 | chr2 | 47922669 | 48037240 | + | protein_coding |
| ENSG00000259706.1_2 | HSP90B2P | 0.21599625 | 0.75632742 | -0.609 | 0.342 | chr15 | 99797837 | 99800220 | + | processed_pseudogene |
| ENSG00000135336.14_4 | ORC3 | 0.14171671 | 0.54264674 | -0.609 | 5.420 | chr6 | 88299785 | 88377171 | + | protein_coding |
| ENSG00000221500.1 | SNORD100 | 0.24183103 | 0.82405293 | -0.609 | -0.522 | chr6 | 133137941 | 133138016 | + | snoRNA |
| ENSG00000130054.4_2 | FAM155B | 0.19684195 | 0.70368025 | -0.609 | 1.907 | chrX | 68725084 | 68752351 | + | protein_coding |
| ENSG00000236698.1_4 | EIF1AXP1 | 0.20469293 | 0.72546962 | -0.610 | 1.387 | chr1 | 17012116 | 17012550 | - | processed_pseudogene |
| ENSG00000259349.2_6 | AC011921.1 | 0.22092173 | 0.76947977 | -0.610 | -0.131 | chr17 | 58603654 | 58628159 | + | lncRNA |
| ENSG00000272037.1_6 | AP002907.1 | 0.2307161 | 0.79517614 | -0.610 | -0.668 | chr8 | 103268620 | 103270049 | + | lncRNA |
| ENSG00000176907.5_4 | TCIM | 0.14234971 | 0.54483571 | -0.610 | 7.056 | chr8 | 40011001 | 40012829 | + | protein_coding |
| ENSG00000163481.8_6 | RNF25 | 0.14971797 | 0.56814164 | -0.610 | 4.326 | chr2 | 219528615 | 219536725 | - | protein_coding |
| ENSG00000264943.1_4 | SH3GL1P2 | 0.22017658 | 0.76793822 | -0.611 | -0.097 | chr17 | 28951431 | 28952512 | + | processed_pseudogene |
| ENSG00000186193.9_4 | SAPCD2 | 0.20556079 | 0.72796465 | -0.611 | 4.534 | chr9 | 139956579 | 139965009 | - | protein_coding |
| ENSG00000277182.1_6 | AC006449.5 | 0.2170891 | 0.75925631 | -0.611 | 0.288 | chr17 | 36905613 | 36907710 | + | lncRNA |
| ENSG00000143869.7_3 | GDF7 | 0.24252134 | 0.82624671 | -0.611 | -0.993 | chr2 | 20866904 | 20879003 | + | protein_coding |
| ENSG00000266401.2_6 | AP002478.1 | 0.19896221 | 0.70906605 | -0.612 | 1.579 | chr18 | 3653030 | 3656282 | + | lncRNA |

| | | | | | | | | | | |
|---|---|---|---|---|---|---|---|---|---|---|
| ENSG00000144468.17_7 | RHBDD1 | 0.15279507 | 0.57638688 | -0.612 | 4.048 | chr2 | 227700297 | 227863931 | + | protein_coding |
| ENSG00000232952.1_5 | AL512844.1 | 0.23871542 | 0.81609715 | -0.612 | -0.992 | chr1 | 106434361 | 106436139 | + | processed_pseudogene |
| ENSG00000049249.8_4 | TNFRSF9 | 0.22591081 | 0.78256139 | -0.612 | -0.724 | chr1 | 7975954 | 8003225 | - | protein_coding |
| ENSG00000270690.1_5 | AC105129.3 | 0.25887377 | 0.86666861 | -0.612 | -1.239 | chr15 | 65750912 | 65751872 | + | processed_pseudogene |
| ENSG00000137962.13_4 | ARHGAP29 | 0.11689708 | 0.46347036 | -0.612 | 6.221 | chr1 | 94614544 | 94740624 | - | protein_coding |
| ENSG00000183513.9_4 | COA5 | 0.15380639 | 0.57921288 | -0.612 | 4.253 | chr2 | 99215777 | 99224978 | - | protein_coding |
| ENSG00000286064.1_2 | AL035413.2 | 0.21371269 | 0.7502536 | -0.613 | 0.217 | chr1 | 19587015 | 19588801 | - | unprocessed_pseudogene |
| ENSG00000213707.2_4 | HMGB1P10 | 0.24678105 | 0.8362691 | -0.613 | -0.958 | chr22 | 26956492 | 26957054 | + | processed_pseudogene |
| ENSG00000181619.11_4 | GPR135 | 0.16743459 | 0.62059798 | -0.613 | 3.074 | chr14 | 59895740 | 59932060 | - | protein_coding |
| ENSG00000201496.1 | RN7SKP275 | 0.24050959 | 0.82101076 | -0.613 | -1.028 | chr4 | 5460308 | 5460608 | - | misc_RNA |
| ENSG00000087586.17_3 | AURKA | 0.10180171 | 0.4112576 | -0.613 | 7.004 | chr20 | 54944445 | 54967393 | - | protein_coding |
| ENSG00000181751.10_3 | C5orf30 | 0.13740573 | 0.528549 | -0.614 | 4.697 | chr5 | 102594464 | 102614361 | + | protein_coding |
| ENSG00000145604.16_4 | SKP2 | 0.13148041 | 0.50949018 | -0.615 | 4.764 | chr5 | 36152091 | 36184421 | + | protein_coding |
| ENSG00000226756.1_6 | AC007365.1 | 0.2179731 | 0.76201024 | -0.615 | -0.390 | chr2 | 64871746 | 64873832 | + | lncRNA |
| ENSG00000114988.11_5 | LMAN2L | 0.13091293 | 0.50751256 | -0.615 | 4.769 | chr2 | 97371666 | 97405801 | - | protein_coding |
| ENSG00000204253.4_4 | HNRNPCP2 | 0.16349962 | 0.60835556 | -0.615 | 3.201 | chr2 | 190788062 | 190788942 | + | processed_pseudogene |
| ENSG00000102780.16_3 | DGKH | 0.14312802 | 0.54734298 | -0.615 | 4.560 | chr13 | 42614172 | 42830714 | + | protein_coding |
| ENSG00000146281.6_3 | PM20D2 | 0.11585596 | 0.46008334 | -0.615 | 6.008 | chr6 | 89855774 | 89875284 | + | protein_coding |
| ENSG00000074211.14_4 | PPP2R2C | 0.13660487 | 0.52624866 | -0.615 | 5.228 | chr4 | 6322305 | 6565327 | - | protein_coding |
| ENSG00000267321.3_5 | SNHG30 | 0.17717825 | 0.65039405 | -0.615 | 2.495 | chr17 | 33895095 | 33901919 | + | lncRNA |
| ENSG00000064225.12_4 | ST3GAL6 | 0.18548364 | 0.67309802 | -0.615 | 1.827 | chr3 | 98451080 | 98540045 | + | protein_coding |
| ENSG00000115232.14_5 | ITGA4 | 0.19875646 | 0.70867069 | -0.615 | 1.225 | chr2 | 182321929 | 182403667 | + | protein_coding |
| ENSG00000079150.18_6 | FKBP7 | 0.16850406 | 0.62293208 | -0.615 | 3.109 | chr2 | 179328391 | 179343327 | - | protein_coding |
| ENSG00000173530.6_4 | TNFRSF10D | 0.12354967 | 0.48414358 | -0.616 | 5.044 | chr8 | 22993101 | 23021540 | - | protein_coding |
| ENSG00000232545.1_7 | AC253536.3 | 0.20852909 | 0.73598257 | -0.616 | 0.233 | chr22 | 24497652 | 24499317 | + | lncRNA |
| ENSG00000085415.16_5 | SEH1L | 0.11083877 | 0.44333013 | -0.616 | 5.446 | chr18 | 12947132 | 12987535 | + | protein_coding |
| ENSG00000171658.8_4 | NMRAL2P | 0.14823447 | 0.563833 | -0.616 | 3.892 | chr3 | 185677732 | 185698661 | + | transcribed_unprocessed_pseudogene |
| ENSG00000139618.15_4 | BRCA2 | 0.13006179 | 0.5047636 | -0.616 | 5.964 | chr13 | 32889223 | 32974403 | + | protein_coding |
| ENSG00000164898.13_6 | FMC1 | 0.22403177 | 0.7772647 | -0.616 | -0.753 | chr7 | 139024203 | 139031074 | + | protein_coding |
| ENSG00000154479.13_6 | CCDC173 | 0.22930353 | 0.7918437 | -0.616 | -0.755 | chr2 | 170501935 | 170550915 | - | protein_coding |
| ENSG00000214193.11_8 | SH3D21 | 0.19327442 | 0.69371849 | -0.617 | 1.296 | chr1 | 36771969 | 36794941 | + | protein_coding |
| ENSG00000144048.10_3 | DUSP11 | 0.13240128 | 0.51205344 | -0.617 | 4.374 | chr2 | 73989311 | 74007284 | - | protein_coding |
| ENSG00000241907.1_4 | RPS20P4 | 0.23794051 | 0.81407452 | -0.617 | -0.781 | chr5 | 149401130 | 149401484 | - | processed_pseudogene |
| ENSG00000203995.10_4 | ZYG11A | 0.15032326 | 0.56958698 | -0.617 | 3.479 | chr1 | 53308183 | 53360670 | + | protein_coding |
| ENSG00000118961.15_5 | LDAH | 0.10716465 | 0.43026288 | -0.617 | 5.534 | chr2 | 20883774 | 21022890 | - | protein_coding |
| ENSG00000119725.19_4 | ZNF410 | 0.19728631 | 0.70484332 | -0.617 | 1.207 | chr14 | 74353320 | 74399214 | + | protein_coding |
| ENSG00000235066.7_7 | AC009303.2 | 0.20788123 | 0.73481153 | -0.617 | 0.212 | chr2 | 118752973 | 118812609 | + | lncRNA |
| ENSG00000258315.5_4 | C17orf49 | 0.23491584 | 0.8060544 | -0.617 | -0.725 | chr17 | 6917814 | 6920844 | + | protein_coding |
| ENSG00000115289.14_5 | PCGF1 | 0.14951037 | 0.56747507 | -0.618 | 3.682 | chr2 | 74732170 | 74734822 | - | protein_coding |
| ENSG00000227640.3_5 | SOX21-AS1 | 0.20532517 | 0.72755989 | -0.618 | 0.444 | chr13 | 95355708 | 95455684 | + | lncRNA |
| ENSG00000162888.4_6 | C1orf147 | 0.22848891 | 0.78964452 | -0.618 | -0.644 | chr1 | 206664449 | 206671061 | - | lncRNA |
| ENSG00000198848.12_4 | CES1 | 0.13746734 | 0.52865415 | -0.618 | 4.060 | chr16 | 55836763 | 55867249 | - | protein_coding |
| ENSG00000286408.1_2 | AC113410.5 | 0.20938018 | 0.73796027 | -0.618 | 0.068 | chr5 | 132386757 | 132387585 | - | lncRNA |
| ENSG00000066651.20_8 | TRMT11 | 0.13911333 | 0.53392394 | -0.618 | 3.960 | chr6 | 126307625 | 126524963 | + | protein_coding |
| ENSG00000105866.15_5 | SP4 | 0.12751959 | 0.4961981 | -0.620 | 4.295 | chr7 | 21467661 | 21554440 | + | protein_coding |
| ENSG00000144357.17_4 | UBR3 | 0.09200753 | 0.37796486 | -0.620 | 6.120 | chr2 | 170683964 | 170940641 | + | protein_coding |
| ENSG00000256222.3_3 | MTRNR2L3 | 0.22784929 | 0.78775264 | -0.620 | -0.843 | chr20 | 55933503 | 55934554 | - | protein_coding |
| ENSG00000120913.23_5 | PDLIM2 | 0.16006374 | 0.59819193 | -0.621 | 2.679 | chr8 | 22435792 | 22455538 | + | protein_coding |
| ENSG00000108443.14_4 | RPS6KB1 | 0.08436759 | 0.35153822 | -0.621 | 6.794 | chr17 | 57970407 | 58027935 | + | protein_coding |
| ENSG00000213533.12_6 | STIMATE | 0.22643668 | 0.78392449 | -0.621 | -0.843 | chr3 | 52870235 | 52931564 | - | protein_coding |
| ENSG00000259668.6_7 | AC066613.1 | 0.1949757 | 0.69841395 | -0.622 | 0.763 | chr15 | 51747465 | 51752779 | + | lncRNA |
| ENSG00000119640.9_4 | ACYP1 | 0.1485407 | 0.56441692 | -0.622 | 3.025 | chr14 | 75519940 | 75536186 | - | protein_coding |
| ENSG00000205269.6_3 | TMEM170B | 0.13903035 | 0.53385611 | -0.622 | 3.869 | chr6 | 11537982 | 11583757 | + | protein_coding |
| ENSG00000240087.3_4 | RPSAP12 | 0.21232083 | 0.74654745 | -0.622 | -0.438 | chr12 | 68946775 | 68947662 | - | processed_pseudogene |
| ENSG00000254685.6_5 | FPGT | 0.12776733 | 0.49705323 | -0.622 | 3.855 | chr1 | 74663896 | 74699770 | + | protein_coding |
| ENSG00000236753.7_7 | MKLN1-AS | 0.16261817 | 0.60583707 | -0.622 | 2.510 | chr7 | 130994228 | 131013071 | - | lncRNA |
| ENSG00000244945.1_7 | AC136604.2 | 0.20053951 | 0.71316764 | -0.622 | 0.423 | chr5 | 179022905 | 179030742 | - | lncRNA |

| | | | | | | | | | | |
|---|---|---|---|---|---|---|---|---|---|---|
| ENSG00000187486.5_5 | KCNJ11 | 0.2027309 | 0.71995201 | -0.623 | 0.344 | chr11 | 17407406 | 17410878 | - | protein_coding |
| ENSG00000186871.7_6 | ERCC6L | 0.09346974 | 0.38238013 | -0.623 | 5.635 | chrX | 71424507 | 71458877 | - | protein_coding |
| ENSG00000229915.1_7 | AC016999.1 | 0.21360455 | 0.75002215 | -0.623 | -0.599 | chr2 | 239335718 | 239336370 | - | lncRNA |
| ENSG00000163050.17_5 | COQ8A | 0.11473085 | 0.45632932 | -0.623 | 4.502 | chr1 | 227085237 | 227175246 | + | protein_coding |
| ENSG00000258525.2_8 | AL049830.3 | 0.20767722 | 0.73443494 | -0.623 | -0.291 | chr14 | 31345385 | 31359014 | - | lncRNA |
| ENSG00000151006.7_4 | PRSS53 | 0.20790778 | 0.73481153 | -0.623 | 1.089 | chr16 | 31094746 | 31100949 | - | protein_coding |
| ENSG00000278009.1_7 | AL139099.4 | 0.22299924 | 0.77517691 | -0.624 | -0.844 | chr14 | 50067729 | 50067842 | - | unprocessed_pseudogene |
| ENSG00000197191.6_4 | CYSRT1 | 0.19196029 | 0.69077229 | -0.624 | 1.028 | chr9 | 140119087 | 140120767 | + | protein_coding |
| ENSG00000237883.2_8 | DGUOK-AS1 | 0.18828296 | 0.68033827 | -0.624 | 0.911 | chr2 | 74174449 | 74208568 | - | lncRNA |
| ENSG00000215267.8_5 | AKR1C7P | 0.16712451 | 0.61976646 | -0.624 | 2.052 | chr10 | 5317136 | 5330433 | - | transcribed_unprocessed_pseudogene |
| ENSG00000144596.13_5 | GRIP2 | 0.18536087 | 0.67292738 | -0.624 | 1.025 | chr3 | 14530615 | 14597582 | - | protein_coding |
| ENSG00000235043.3_4 | TECRP1 | 0.21910278 | 0.76494381 | -0.625 | -0.254 | chr4 | 87870821 | 87871265 | - | processed_pseudogene |
| ENSG00000173166.18_5 | RAPH1 | 0.15702235 | 0.58881723 | -0.625 | 2.641 | chr2 | 204259068 | 204400058 | - | protein_coding |
| ENSG00000125618.17_6 | PAX8 | 0.18500366 | 0.67204253 | -0.625 | 1.097 | chr2 | 113973574 | 114036498 | - | protein_coding |
| ENSG00000197860.10_4 | SGTB | 0.10075568 | 0.40768074 | -0.625 | 4.796 | chr5 | 64961755 | 65018862 | - | protein_coding |
| ENSG00000207357.1 | RNU6-2 | 0.23265563 | 0.80015341 | -0.625 | -0.961 | chr19 | 1021521 | 1021627 | + | snRNA |
| ENSG00000115317.11_2 | HTRA2 | 0.12640229 | 0.4932549 | -0.625 | 3.791 | chr2 | 74756504 | 74760475 | + | protein_coding |
| ENSG00000230082.1_6 | PRRT3-AS1 | 0.18511718 | 0.67218005 | -0.625 | 1.285 | chr3 | 9989088 | 9996471 | + | lncRNA |
| ENSG00000137713.16_5 | PPP2R1B | 0.09219998 | 0.37849285 | -0.626 | 5.017 | chr11 | 111597632 | 111637113 | - | protein_coding |
| ENSG00000180139.11_9 | ACTA2-AS1 | 0.22540834 | 0.78112542 | -0.626 | -1.028 | chr10 | 90692147 | 90700577 | + | lncRNA |
| ENSG00000112159.12_4 | MDN1 | 0.10163964 | 0.41078968 | -0.626 | 6.383 | chr6 | 90352217 | 90529513 | - | protein_coding |
| ENSG00000204392.11_4 | LSM2 | 0.14009741 | 0.53725677 | -0.627 | 2.967 | chr6 | 31765173 | 31774743 | - | protein_coding |
| ENSG00000286244.1_3 | AC073046.3 | 0.21049016 | 0.74113729 | -0.627 | -0.412 | chr2 | 74061549 | 74084092 | - | lncRNA |
| ENSG00000118503.15_7 | TNFAIP3 | 0.17475172 | 0.64255254 | -0.627 | 1.620 | chr6 | 138188351 | 138204449 | + | protein_coding |
| ENSG00000196584.3_4 | XRCC2 | 0.09121762 | 0.3755883 | -0.627 | 5.160 | chr7 | 152341861 | 152373226 | - | protein_coding |
| ENSG00000114978.18_4 | MOB1A | 0.05202165 | 0.23246194 | -0.627 | 8.201 | chr2 | 74379655 | 74406025 | - | protein_coding |
| ENSG00000215086.2_4 | NPM1P24 | 0.1934757 | 0.69416048 | -0.627 | 0.308 | chr10 | 74677399 | 74678292 | - | processed_pseudogene |
| ENSG00000071082.11_4 | RPL31 | 0.05069933 | 0.22696579 | -0.627 | 8.496 | chr2 | 101618691 | 101640494 | + | protein_coding |
| ENSG00000237382.2_4 | RPL21P121 | 0.20822557 | 0.73549576 | -0.628 | -0.508 | chr17 | 18216074 | 18216847 | + | processed_pseudogene |
| ENSG00000279636.2_4 | AL132989.2 | 0.1944375 | 0.69690761 | -0.628 | 0.450 | chr14 | 58754751 | 58755876 | + | lncRNA |
| ENSG00000286360.1_2 | AC107081.5 | 0.18889993 | 0.68229012 | -0.628 | 0.592 | chr2 | 62106075 | 62108488 | + | lncRNA |
| ENSG00000171444.18_5 | MCC | 0.18629565 | 0.67521722 | -0.628 | 0.837 | chr5 | 112357796 | 112824520 | - | protein_coding |
| ENSG00000143970.17_5 | ASXL2 | 0.06729539 | 0.28922043 | -0.629 | 5.908 | chr2 | 25956622 | 26101356 | - | protein_coding |
| ENSG00000250329.1_4 | KDELC1P1 | 0.21069969 | 0.74153508 | -0.629 | -0.626 | chr5 | 127445178 | 127446686 | + | processed_pseudogene |
| ENSG00000008438.5_4 | PGLYRP1 | 0.22071257 | 0.7690966 | -0.629 | -1.065 | chr19 | 46522411 | 46526311 | - | protein_coding |
| ENSG00000225234.1_7 | TRAPPC12-AS1 | 0.19421512 | 0.6965321 | -0.629 | 0.327 | chr2 | 3485013 | 3486180 | - | lncRNA |
| ENSG00000230010.2_7 | AL049646.1 | 0.19833845 | 0.70760628 | -0.630 | -0.197 | chr20 | 18269987 | 18339944 | - | lncRNA |
| ENSG00000103642.12_4 | LACTB | 0.06937163 | 0.29735243 | -0.630 | 5.592 | chr15 | 63414032 | 63434260 | + | protein_coding |
| ENSG00000047648.23_7 | ARHGAP6 | 0.22725292 | 0.7861389 | -0.630 | -1.065 | chrX | 11135771 | 11684040 | - | protein_coding |
| ENSG00000197498.13_3 | RPF2 | 0.08067362 | 0.33804911 | -0.630 | 5.081 | chr6 | 111303218 | 111349466 | + | protein_coding |
| ENSG00000147894.17_5 | C9orf72 | 0.11254163 | 0.44943296 | -0.630 | 3.723 | chr9 | 27535638 | 27573864 | - | protein_coding |
| ENSG00000170340.11_3 | B3GNT2 | 0.0841101 | 0.35079442 | -0.631 | 5.190 | chr2 | 62423250 | 62451866 | + | protein_coding |
| ENSG00000144040.13_4 | SFXN5 | 0.10365391 | 0.41835961 | -0.631 | 4.538 | chr2 | 73169165 | 73302747 | - | protein_coding |
| ENSG00000117360.13_3 | PRPF3 | 0.05355345 | 0.23793054 | -0.631 | 6.586 | chr1 | 150293928 | 150325709 | + | protein_coding |
| ENSG00000248909.1_4 | HMGB1P21 | 0.20151143 | 0.7161154 | -0.632 | -0.333 | chr5 | 77940088 | 77940668 | - | processed_pseudogene |
| ENSG00000224138.1_6 | AC000123.1 | 0.21003425 | 0.73997196 | -0.632 | -0.655 | chr7 | 126990182 | 126991577 | + | lncRNA |
| ENSG00000116001.16_6 | TIA1 | 0.06081178 | 0.26468469 | -0.632 | 6.231 | chr2 | 70436576 | 70475792 | - | protein_coding |
| ENSG00000071539.14_5 | TRIP13 | 0.05728998 | 0.25219797 | -0.632 | 6.123 | chr5 | 892999 | 919472 | + | protein_coding |
| ENSG00000186684.14_5 | CYP27C1 | 0.16410648 | 0.61010251 | -0.632 | 1.678 | chr2 | 127941408 | 127977889 | - | protein_coding |
| ENSG00000164823.11_5 | OSGIN2 | 0.05734065 | 0.25229603 | -0.632 | 6.602 | chr8 | 90914077 | 90940116 | + | protein_coding |
| ENSG00000235408.5_3 | SNORA71B | 0.19726763 | 0.70484332 | -0.633 | 0.109 | chr20 | 37053731 | 37054002 | - | snoRNA |
| ENSG00000156136.10_6 | DCK | 0.08777864 | 0.3637041 | -0.633 | 4.475 | chr4 | 71858255 | 71896631 | + | protein_coding |
| ENSG00000239900.12_4 | ADSL | 0.10849444 | 0.43510519 | -0.633 | 3.588 | chr22 | 40742504 | 40786467 | + | protein_coding |
| ENSG00000127423.10_4 | AUNIP | 0.1209718 | 0.4761422 | -0.633 | 3.252 | chr1 | 26158404 | 26185949 | - | protein_coding |
| ENSG00000243759.1_4 | ST13P15 | 0.22601426 | 0.78276714 | -0.633 | -1.138 | chr3 | 143235965 | 143237071 | - | processed_pseudogene |
| ENSG00000070814.21_8 | TCOF1 | 0.04867969 | 0.21896 | -0.633 | 7.179 | chr5 | 149737192 | 149779871 | + | protein_coding |

| | | | | | | | | | | |
|---|---|---|---|---|---|---|---|---|---|---|
| ENSG00000175471.19_5 | MCTP1 | 0.11923545 | 0.47066345 | -0.634 | 3.550 | chr5 | 94039446 | 94620279 | - | protein_coding |
| ENSG00000159496.14_4 | RGL4 | 0.20838977 | 0.73578321 | -0.634 | -0.601 | chr22 | 24030323 | 24041363 | + | protein_coding |
| ENSG00000176225.14_5 | RTTN | 0.06361785 | 0.27561529 | -0.634 | 5.428 | chr18 | 67670267 | 67872962 | - | protein_coding |
| ENSG00000189292.16_7 | ALKAL2 | 0.21426647 | 0.75163066 | -0.634 | -0.460 | chr2 | 279558 | 288851 | - | protein_coding |
| ENSG00000124374.9_4 | PAIP2B | 0.16580406 | 0.6157694 | -0.634 | 1.366 | chr2 | 71409868 | 71454233 | - | protein_coding |
| ENSG00000224043.8_7 | CCNT2-AS1 | 0.18651818 | 0.67540239 | -0.634 | 0.526 | chr2 | 135493034 | 135676280 | - | lncRNA |
| ENSG00000138769.11_4 | CDKL2 | 0.16203915 | 0.60393321 | -0.634 | 1.633 | chr4 | 76501706 | 76555900 | - | protein_coding |
| ENSG00000075426.12_4 | FOSL2 | 0.04685105 | 0.21195542 | -0.634 | 6.667 | chr2 | 28615315 | 28640184 | + | protein_coding |
| ENSG00000164983.7_3 | TMEM65 | 0.05972773 | 0.26086099 | -0.635 | 5.985 | chr8 | 125318430 | 125384933 | - | protein_coding |
| ENSG00000275582.1_7 | AL031670.1 | 0.18281493 | 0.66640806 | -0.635 | 0.430 | chr20 | 3901926 | 3904047 | - | lncRNA |
| ENSG00000164118.13_7 | CEP44 | 0.08334762 | 0.34802283 | -0.635 | 4.469 | chr4 | 175204881 | 175254531 | + | protein_coding |
| ENSG00000176697.19_6 | BDNF | 0.07541625 | 0.31849788 | -0.635 | 5.006 | chr11 | 27676440 | 27743605 | - | protein_coding |
| ENSG00000115808.12_4 | STRN | 0.05209854 | 0.2326679 | -0.635 | 6.227 | chr2 | 37064841 | 37193679 | - | protein_coding |
| ENSG00000065328.16_3 | MCM10 | 0.04509453 | 0.20516204 | -0.637 | 6.302 | chr10 | 13203554 | 13253104 | + | protein_coding |
| ENSG00000183763.9_4 | TRAIP | 0.09829459 | 0.39926841 | -0.637 | 3.694 | chr3 | 49866034 | 49894007 | - | protein_coding |
| ENSG00000111850.11_5 | SMIM8 | 0.13825877 | 0.53123778 | -0.637 | 2.183 | chr6 | 88032301 | 88109467 | + | protein_coding |
| ENSG00000265590.10_8 | C21orf59-TCP10L | 0.1664099 | 0.61750306 | -0.637 | 1.076 | chr21 | 33774820 | 33985175 | - | protein_coding |
| ENSG00000077150.20_6 | NFKB2 | 0.06421221 | 0.27791919 | -0.637 | 5.117 | chr10 | 104153867 | 104162281 | + | protein_coding |
| ENSG00000197406.7_4 | DIO3 | 0.18822046 | 0.68025078 | -0.637 | 0.364 | chr14 | 102027688 | 102029789 | + | protein_coding |
| ENSG00000230650.1_5 | AC140479.2 | 0.1547625 | 0.58193975 | -0.637 | 1.533 | chr2 | 111118185 | 111121314 | - | processed_pseudogene |
| ENSG00000159388.6_4 | BTG2 | 0.11428904 | 0.45487753 | -0.637 | 3.277 | chr1 | 203274619 | 203278730 | + | protein_coding |
| ENSG00000024526.17_5 | DEPDC1 | 0.05680254 | 0.25042451 | -0.637 | 6.101 | chr1 | 68939835 | 68962904 | - | protein_coding |
| ENSG00000123178.15_4 | SPRYD7 | 0.10110871 | 0.40892293 | -0.638 | 3.532 | chr13 | 50486838 | 50510626 | - | protein_coding |
| ENSG00000112137.17_3 | PHACTR1 | 0.20389072 | 0.72320353 | -0.638 | -0.438 | chr6 | 12717037 | 13290716 | + | protein_coding |
| ENSG00000234705.2_8 | HMGA1P4 | 0.19248584 | 0.69200645 | -0.638 | -0.199 | chr9 | 131425413 | 131446756 | - | lncRNA |
| ENSG00000151136.15_4 | BTBD11 | 0.04115214 | 0.189712 | -0.638 | 6.882 | chr12 | 107712198 | 108053419 | + | protein_coding |
| ENSG00000099822.3_3 | HCN2 | 0.1157278 | 0.45967714 | -0.638 | 3.504 | chr19 | 589881 | 617159 | + | protein_coding |
| ENSG00000007952.18_5 | NOX1 | 0.19267238 | 0.69239688 | -0.638 | -0.142 | chrX | 100098313 | 100129348 | - | protein_coding |
| ENSG00000170006.12_5 | TMEM154 | 0.19412316 | 0.69634287 | -0.639 | -0.074 | chr4 | 153539780 | 153601164 | - | protein_coding |
| ENSG00000234740.1_8 | AL162386.2 | 0.19564135 | 0.70023364 | -0.639 | -0.334 | chr9 | 13274509 | 13279186 | - | lncRNA |
| ENSG00000121297.8_5 | TSHZ3 | 0.13894659 | 0.53364982 | -0.639 | 2.207 | chr19 | 31640885 | 31840342 | - | protein_coding |
| ENSG00000168385.18_6 | SEPTIN2 | 0.03322305 | 0.1568626 | -0.639 | 8.647 | chr2 | 242254515 | 242293442 | + | protein_coding |
| ENSG00000180773.15_5 | SLC36A4 | 0.08504016 | 0.35384276 | -0.639 | 4.485 | chr11 | 92877340 | 92931157 | - | protein_coding |
| ENSG00000048162.20_4 | NOP16 | 0.07391114 | 0.31303439 | -0.639 | 4.521 | chr5 | 175810941 | 175815976 | - | protein_coding |
| ENSG00000232070.9_4 | TMEM253 | 0.22281736 | 0.77501879 | -0.640 | -1.139 | chr14 | 21566970 | 21571883 | + | protein_coding |
| ENSG00000280042.1_7 | AC022336.4 | 0.16974194 | 0.62659569 | -0.640 | 0.652 | chr3 | 124509966 | 124511951 | - | TEC |
| ENSG00000286116.1_3 | AL157394.3 | 0.18970212 | 0.68463082 | -0.640 | -0.219 | chr10 | 90749802 | 90754006 | - | lncRNA |
| ENSG00000287185.1_2 | AC041005.1 | 0.09148857 | 0.37626794 | -0.641 | 3.541 | chr8 | 25036628 | 25041749 | - | lncRNA |
| ENSG00000163507.14_6 | CIP2A | 0.04242103 | 0.19455199 | -0.641 | 6.187 | chr3 | 108268711 | 108308491 | - | protein_coding |
| ENSG00000095059.16_6 | DHPS | 0.15842653 | 0.59320637 | -0.642 | 1.351 | chr19 | 12786531 | 12792694 | - | protein_coding |
| ENSG00000229951.5_8 | AC104695.2 | 0.24797204 | 0.8390248 | -0.642 | -1.140 | chr2 | 28607276 | 28617539 | - | lncRNA |
| ENSG00000115355.17_5 | CCDC88A | 0.03698368 | 0.17268281 | -0.642 | 7.085 | chr2 | 55514978 | 55647031 | - | protein_coding |
| ENSG00000077348.9_4 | EXOSC5 | 0.05325509 | 0.23684187 | -0.642 | 5.241 | chr19 | 41892279 | 41903264 | - | protein_coding |
| ENSG00000177173.5_4 | NAP1L4P1 | 0.19618368 | 0.70203332 | -0.642 | -0.534 | chr1 | 117075558 | 117076714 | - | processed_pseudogene |
| ENSG00000114019.14_6 | AMOTL2 | 0.0486165 | 0.21873114 | -0.642 | 6.874 | chr3 | 134074716 | 134094321 | - | protein_coding |
| ENSG00000165732.13_4 | DDX21 | 0.03211111 | 0.15213816 | -0.642 | 8.017 | chr10 | 70715926 | 70744824 | + | protein_coding |
| ENSG00000018510.16_7 | AGPS | 0.03820498 | 0.17760019 | -0.643 | 6.847 | chr2 | 178257485 | 178424027 | + | protein_coding |
| ENSG00000226723.2_5 | AL513175.1 | 0.205465 | 0.72777046 | -0.643 | -0.683 | chr1 | 192215838 | 192216617 | - | processed_pseudogene |
| ENSG00000228852.6_9 | AC092802.3 | 0.20085354 | 0.71414144 | -0.643 | -0.560 | chr1 | 95708723 | 95744496 | - | lncRNA |
| ENSG00000173915.16_6 | ATP5MD | 0.08245235 | 0.34485192 | -0.643 | 5.291 | chr10 | 105148798 | 105156249 | - | protein_coding |
| ENSG00000183091.19_9 | NEB | 0.16114047 | 0.60129296 | -0.643 | 1.137 | chr2 | 152341850 | 152591001 | - | protein_coding |
| ENSG00000280367.1_5 | AP002364.1 | 0.1833863 | 0.66780578 | -0.643 | -0.062 | chr11 | 89956321 | 89959706 | + | TEC |
| ENSG00000240549.2_6 | THOC7-AS1 | 0.21463097 | 0.75273362 | -0.643 | -0.974 | chr3 | 63846321 | 63847495 | + | lncRNA |
| ENSG00000250614.1_7 | AC005522.1 | 0.21152568 | 0.74404609 | -0.644 | -0.879 | chr7 | 76103904 | 76108173 | - | lncRNA |
| ENSG00000119771.15_4 | KLHL29 | 0.12258097 | 0.4809841 | -0.644 | 2.348 | chr2 | 23608050 | 23931481 | + | protein_coding |
| ENSG00000226180.3_6 | AC010536.1 | 0.18464596 | 0.67101754 | -0.645 | 0.056 | chr16 | 87727143 | 87729753 | - | lncRNA |
| ENSG00000153822.13_5 | KCNJ16 | 0.17725041 | 0.65039405 | -0.645 | 0.172 | chr17 | 68049570 | 68131749 | + | protein_coding |
| ENSG00000279852.1_5 | AC083964.2 | 0.21857168 | 0.76338964 | -0.645 | -1.178 | chr8 | 436117 | 438512 | - | TEC |

| | | | | | | | | | | |
|---|---|---|---|---|---|---|---|---|---|---|
| ENSG00000243406.6_6 | MRPS31P5 | 0.18681012 | 0.67597871 | -0.645 | -0.092 | chr13 | 52741844 | 52768600 | - | transcribed_unprocessed_pseudogene |
| ENSG00000278467.1_7 | AC138393.3 | 0.18195396 | 0.66368753 | -0.645 | -0.027 | chr1 | 224181964 | 224182898 | + | lncRNA |
| ENSG00000140511.11_4 | HAPLN3 | 0.13911964 | 0.53392394 | -0.645 | 1.863 | chr15 | 89420519 | 89438857 | - | protein_coding |
| ENSG00000081059.20_4 | TCF7 | 0.11101374 | 0.4437303 | -0.646 | 2.658 | chr5 | 133450372 | 133487556 | + | protein_coding |
| ENSG00000145569.6_4 | OTULINL | 0.09131543 | 0.3757298 | -0.646 | 3.274 | chr5 | 14581901 | 14616289 | + | protein_coding |
| ENSG00000137054.16_4 | POLR1E | 0.04135404 | 0.19049449 | -0.646 | 5.413 | chr9 | 37485945 | 37503694 | + | protein_coding |
| ENSG00000259657.1_4 | PIGHP1 | 0.20731944 | 0.73331564 | -0.646 | -0.976 | chr15 | 59968956 | 59969520 | + | processed_pseudogene |
| ENSG00000136699.19_6 | SMPD4 | 0.03936001 | 0.18230168 | -0.646 | 6.122 | chr2 | 130908965 | 130940323 | - | protein_coding |
| ENSG00000264932.3_6 | AC115989.1 | 0.16951327 | 0.62588161 | -0.647 | 0.523 | chr17 | 19552686 | 19553248 | - | lncRNA |
| ENSG00000077380.15_6 | DYNC1I2 | 0.03077204 | 0.14665418 | -0.647 | 6.300 | chr2 | 172543919 | 172604930 | + | protein_coding |
| ENSG00000240401.9_9 | AC012358.4 | 0.19894095 | 0.70906605 | -0.647 | -0.446 | chr2 | 55509486 | 55573185 | + | lncRNA |
| ENSG00000164251.5_4 | F2RL1 | 0.03427865 | 0.16128938 | -0.647 | 6.139 | chr5 | 76114758 | 76131140 | + | protein_coding |
| ENSG00000185641.6_5 | AC034236.1 | 0.14565878 | 0.55534753 | -0.647 | 1.337 | chr5 | 115387614 | 115387986 | - | processed_pseudogene |
| ENSG00000250033.5_5 | SLC7A11-AS1 | 0.11000048 | 0.44017534 | -0.647 | 2.745 | chr4 | 138978618 | 139099331 | + | lncRNA |
| ENSG00000179750.16_4 | APOBEC3B | 0.18066553 | 0.66006261 | -0.648 | 0.209 | chr22 | 39378352 | 39388809 | + | protein_coding |
| ENSG00000146263.11_4 | MMS22L | 0.04533195 | 0.20609334 | -0.648 | 5.487 | chr6 | 97590037 | 97731093 | - | protein_coding |
| ENSG00000240356.6_5 | RPL23AP7 | 0.14118165 | 0.54083105 | -0.648 | 1.700 | chr2 | 114368079 | 114384667 | - | transcribed_processed_pseudogene |
| ENSG00000169750.9_3 | RAC3 | 0.10177629 | 0.41124843 | -0.649 | 3.401 | chr17 | 79989554 | 79992080 | + | protein_coding |
| ENSG00000259191.2_9 | AC027237.3 | 0.18089823 | 0.66064108 | -0.649 | -0.166 | chr15 | 69726805 | 69727459 | - | processed_pseudogene |
| ENSG00000278918.1_6 | AC080112.4 | 0.19766921 | 0.70592739 | -0.649 | -0.355 | chr17 | 38504301 | 38506791 | - | TEC |
| ENSG00000200879.1 | SNORD14E | 0.16795531 | 0.62155008 | -0.649 | 0.348 | chr11 | 122928785 | 122928869 | - | snoRNA |
| ENSG00000274943.1_8 | AC079684.2 | 0.16777915 | 0.62121931 | -0.649 | 0.348 | chr12 | 42755069 | 42755505 | + | lncRNA |
| ENSG00000137720.8_6 | C11orf1 | 0.08732193 | 0.36213531 | -0.649 | 3.542 | chr11 | 111749659 | 111756699 | + | protein_coding |
| ENSG00000249572.1_7 | AC034231.1 | 0.18751478 | 0.67825225 | -0.650 | -0.343 | chr5 | 33424131 | 33440725 | - | lncRNA |
| ENSG00000136235.16_4 | GPNMB | 0.17314607 | 0.63744017 | -0.650 | 0.668 | chr7 | 23275586 | 23314727 | + | protein_coding |
| ENSG00000256897.1_7 | AC018410.1 | 0.16607072 | 0.61650197 | -0.650 | 0.411 | chr11 | 47241769 | 47243302 | - | lncRNA |
| ENSG00000144589.22_3 | STK11IP | 0.05915814 | 0.25894623 | -0.650 | 4.685 | chr2 | 220462579 | 220481173 | + | protein_coding |
| ENSG00000184254.17_4 | ALDH1A3 | 0.08848992 | 0.36630952 | -0.650 | 3.502 | chr15 | 101417919 | 101456831 | + | protein_coding |
| ENSG00000234978.1 | RP11-423O2.5 | 0.14243598 | 0.54504847 | -0.650 | 1.285 | chr1 | 142803224 | 142816004 | - | lincRNA |
| ENSG00000187800.13_2 | PEAR1 | 0.13129592 | 0.50888627 | -0.651 | 2.034 | chr1 | 156863490 | 156886226 | + | protein_coding |
| ENSG00000068024.16_4 | HDAC4 | 0.0584303 | 0.25632869 | -0.651 | 4.666 | chr2 | 239969864 | 240323348 | - | protein_coding |
| ENSG00000079616.12_2 | KIF22 | 0.02448083 | 0.11926264 | -0.651 | 6.271 | chr16 | 29802040 | 29816706 | + | protein_coding |
| ENSG00000119782.14_4 | FKBP1B | 0.16757572 | 0.62079187 | -0.652 | 1.052 | chr2 | 24272571 | 24286551 | + | protein_coding |
| ENSG00000049167.14_7 | ERCC8 | 0.06492875 | 0.28061035 | -0.652 | 4.198 | chr5 | 60169658 | 60240900 | - | protein_coding |
| ENSG00000067840.12_4 | PDZD4 | 0.15296838 | 0.57678039 | -0.652 | 1.004 | chrX | 153067621 | 153096020 | - | protein_coding |
| ENSG00000240344.9_4 | PPIL3 | 0.07208131 | 0.30660012 | -0.652 | 3.985 | chr2 | 201735630 | 201754026 | - | protein_coding |
| ENSG00000145545.12_3 | SRD5A1 | 0.04238965 | 0.19445827 | -0.652 | 4.931 | chr5 | 6633540 | 6674499 | + | protein_coding |
| ENSG00000165704.15_3 | HPRT1 | 0.04544191 | 0.2065182 | -0.652 | 4.876 | chrX | 133594195 | 133654543 | + | protein_coding |
| ENSG00000237821.1_5 | AC083873.1 | 0.22051488 | 0.76866546 | -0.652 | -1.044 | chr7 | 132854174 | 132855272 | - | processed_pseudogene |
| ENSG00000259248.7_7 | USP3-AS1 | 0.13710456 | 0.52771593 | -0.652 | 1.684 | chr15 | 63836446 | 63893788 | - | lncRNA |
| ENSG00000240996.1_8 | AL157400.4 | 0.19335223 | 0.69385762 | -0.653 | -0.639 | chr10 | 91454052 | 91457685 | - | lncRNA |
| ENSG00000197756.10_3 | RPL37A | 0.01329686 | 0.06901702 | -0.653 | 8.762 | chr2 | 217363548 | 217443903 | + | protein_coding |
| ENSG00000228205.1_6 | AC131235.2 | 0.16110141 | 0.60129296 | -0.653 | 0.422 | chr3 | 183867492 | 183868223 | + | processed_pseudogene |
| ENSG00000261794.1_2 | GOLGA8H | 0.1573792 | 0.58990637 | -0.653 | 0.787 | chr15 | 30896329 | 30906764 | + | protein_coding |
| ENSG00000077152.11_6 | UBE2T | 0.03316579 | 0.15663387 | -0.653 | 5.356 | chr1 | 202300785 | 202311108 | - | protein_coding |
| ENSG00000118242.16_6 | MREG | 0.09930088 | 0.40271143 | -0.653 | 2.605 | chr2 | 216807307 | 216898819 | - | protein_coding |
| ENSG00000259429.5_1 | UBE2Q2P2 | 0.17119711 | 0.63065706 | -0.654 | 0.167 | chr15 | 83023847 | 83088805 | + | transcribed_unprocessed_pseudogene |
| ENSG00000279762.3_8 | AC005899.9 | 0.15247769 | 0.57590799 | -0.654 | 0.880 | chr17 | 30750990 | 30773056 | - | lncRNA |
| ENSG00000138346.15_5 | DNA2 | 0.02559815 | 0.12419507 | -0.654 | 5.804 | chr10 | 70173821 | 70231878 | - | protein_coding |
| ENSG00000124571.18_6 | XPO5 | 0.02563094 | 0.12432019 | -0.655 | 5.883 | chr6 | 43490072 | 43543775 | - | protein_coding |
| ENSG00000138685.15_6 | FGF2 | 0.02153152 | 0.10664915 | -0.655 | 6.250 | chr4 | 123747863 | 123819391 | + | protein_coding |
| ENSG00000260526.1_6 | AC109347.1 | 0.19987391 | 0.71165525 | -0.655 | -0.943 | chr4 | 113150717 | 113152752 | - | lncRNA |
| ENSG00000115109.14_3 | EPB41L5 | 0.04190717 | 0.19259304 | -0.655 | 4.711 | chr2 | 120770653 | 120936695 | + | protein_coding |
| ENSG00000213790.2_4 | OLA1P1 | 0.19549498 | 0.69985077 | -0.656 | 0.064 | chr22 | 42503769 | 42504957 | - | processed_pseudogene |

| | | | | | | | | | | |
|---|---|---|---|---|---|---|---|---|---|---|
| ENSG00000115947.13_4 | ORC4 | 0.04466979 | 0.20349954 | -0.656 | 5.059 | chr2 | 148687966 | 148779173 | - | protein_coding |
| ENSG00000260942.1_6 | CAPN10-DT | 0.12138953 | 0.47725774 | -0.657 | 1.765 | chr2 | 241522117 | 241526116 | - | lncRNA |
| ENSG00000151849.15_4 | CENPJ | 0.04134659 | 0.19049449 | -0.657 | 5.066 | chr13 | 25456417 | 25497027 | - | protein_coding |
| ENSG00000279539.1_5 | AC006486.2 | 0.12697371 | 0.49461564 | -0.657 | 1.773 | chr19 | 42746927 | 42749125 | + | TEC |
| ENSG00000129173.13_4 | E2F8 | 0.06199145 | 0.26935732 | -0.658 | 3.734 | chr11 | 19245610 | 19263167 | - | protein_coding |
| ENSG00000143401.15_8 | ANP32E | 0.03237833 | 0.15328156 | -0.658 | 6.303 | chr1 | 150190717 | 150208504 | - | protein_coding |
| ENSG00000175279.22_4 | CENPS | 0.15200901 | 0.5746264 | -0.658 | 0.791 | chr1 | 10490490 | 10502865 | + | protein_coding |
| ENSG00000002822.15_6 | MAD1L1 | 0.1765173 | 0.64823954 | -0.659 | -0.261 | chr7 | 1855429 | 2272878 | - | protein_coding |
| ENSG00000279833.1_6 | AL031846.2 | 0.18616861 | 0.67489444 | -0.659 | -0.799 | chr22 | 39529095 | 39532765 | + | lncRNA |
| ENSG00000273117.1_4 | AC144652.1 | 0.14403337 | 0.55009479 | -0.659 | 1.329 | chr7 | 155087628 | 155089251 | - | lncRNA |
| ENSG00000154548.9_4 | SRSF12 | 0.15380911 | 0.57921288 | -0.659 | 1.008 | chr6 | 89805678 | 89827790 | - | protein_coding |
| ENSG00000261526.2_6 | AC012615.1 | 0.19801442 | 0.70687609 | -0.659 | -0.976 | chr19 | 1874870 | 1876168 | - | lncRNA |
| ENSG00000259932.1_8 | AC051619.6 | 0.20846897 | 0.7359166 | -0.659 | -1.009 | chr15 | 45490715 | 45491337 | - | lncRNA |
| ENSG00000261488.1_6 | TBILA | 0.17060938 | 0.62914421 | -0.659 | -0.099 | chr3 | 111852270 | 111854206 | + | lncRNA |
| ENSG00000213412.3_3 | HNRNPA1P33 | 0.05830384 | 0.25583707 | -0.659 | 3.814 | chr10 | 47133338 | 47133898 | - | processed_pseudogene |
| ENSG00000163510.14_4 | CWC22 | 0.05689983 | 0.2507912 | -0.659 | 4.642 | chr2 | 180809603 | 180872024 | - | protein_coding |
| ENSG00000279406.1_5 | AL359183.1 | 0.156971 | 0.58874894 | -0.660 | 0.491 | chr10 | 73124000 | 73126131 | + | TEC |
| ENSG00000235298.1_6 | AL354733.3 | 0.18545668 | 0.67309802 | -0.660 | -0.799 | chr9 | 86587148 | 86590692 | + | lncRNA |
| ENSG00000158552.13_8 | ZFAND2B | 0.05820385 | 0.25546141 | -0.660 | 3.726 | chr2 | 220059959 | 220074373 | + | protein_coding |
| ENSG00000228350.1_6 | LINC02585 | 0.20346314 | 0.72197515 | -0.660 | -1.220 | chr3 | 48701209 | 48706607 | + | lncRNA |
| ENSG00000140543.14_5 | DET1 | 0.15925907 | 0.59591323 | -0.660 | 0.280 | chr15 | 89037671 | 89089906 | - | protein_coding |
| ENSG00000162643.13_4 | WDR63 | 0.19039282 | 0.68670508 | -0.660 | -0.638 | chr1 | 85464830 | 85598821 | + | protein_coding |
| ENSG00000135900.4_4 | MRPL44 | 0.02974977 | 0.14197265 | -0.661 | 5.041 | chr2 | 224822180 | 224832431 | + | protein_coding |
| ENSG00000215424.10_7 | MCM3AP-AS1 | 0.092567 | 0.37964862 | -0.661 | 2.916 | chr21 | 47649110 | 47679304 | + | lncRNA |
| ENSG00000162620.16_8 | LRRIQ3 | 0.11962229 | 0.47166651 | -0.661 | 1.639 | chr1 | 74491699 | 74663871 | - | protein_coding |
| ENSG00000068654.16_5 | POLR1A | 0.01214701 | 0.06358747 | -0.661 | 7.228 | chr2 | 86247339 | 86333278 | - | protein_coding |
| ENSG00000130414.12_7 | NDUFA10 | 0.01501868 | 0.07709819 | -0.661 | 6.294 | chr2 | 240831867 | 240964762 | - | protein_coding |
| ENSG00000130038.10_7 | CRACR2A | 0.10851304 | 0.43510519 | -0.661 | 2.033 | chr12 | 3715799 | 3873985 | - | protein_coding |
| ENSG00000257800.1_4 | FNBP1P1 | 0.16764979 | 0.6209368 | -0.662 | -0.036 | chr2 | 74347807 | 74350345 | + | processed_pseudogene |
| ENSG00000256325.1_7 | AC025423.1 | 0.19038323 | 0.68670508 | -0.662 | -0.830 | chr12 | 69221898 | 69222333 | + | lncRNA |
| ENSG00000165572.8_3 | KBTBD6 | 0.04607784 | 0.20900307 | -0.662 | 4.029 | chr13 | 41701705 | 41706938 | - | protein_coding |
| ENSG00000232233.1_6 | LINC02043 | 0.20016776 | 0.71213069 | -0.662 | -0.831 | chr3 | 186528669 | 186543310 | - | lncRNA |
| ENSG00000235677.1_4 | NPM1P26 | 0.15240813 | 0.57576763 | -0.662 | 0.466 | chr10 | 98319942 | 98320679 | + | processed_pseudogene |
| ENSG00000214535.3_2 | RPS15AP1 | 0.14998046 | 0.56889472 | -0.663 | 0.532 | chr20 | 21146846 | 21147236 | - | processed_pseudogene |
| ENSG00000281026.1_3 | N4BP2L2-IT2 | 0.13381989 | 0.5172021 | -0.663 | 1.106 | chr13 | 33078643 | 33083532 | - | lncRNA |
| ENSG00000006062.17_6 | MAP3K14 | 0.03144384 | 0.149535 | -0.663 | 5.216 | chr17 | 43340486 | 43394395 | - | protein_coding |
| ENSG00000141540.11_3 | TTYH2 | 0.14365101 | 0.54910661 | -0.663 | 0.828 | chr17 | 72209710 | 72258159 | + | protein_coding |
| ENSG00000128203.7_3 | ASPHD2 | 0.13237373 | 0.51205344 | -0.663 | 1.069 | chr22 | 26825226 | 26840981 | + | protein_coding |
| ENSG00000140749.9_5 | IGSF6 | 0.18583676 | 0.67410406 | -0.664 | -0.828 | chr16 | 21650871 | 21663929 | - | protein_coding |
| ENSG00000161692.18_3 | DBF4B | 0.03098574 | 0.14759347 | -0.664 | 4.851 | chr17 | 42785976 | 42829632 | + | protein_coding |
| ENSG00000115307.17_4 | AUP1 | 0.01148973 | 0.06046743 | -0.664 | 6.814 | chr2 | 74753772 | 74756887 | - | protein_coding |
| ENSG00000206885.1 | SNORA75 | 0.17906385 | 0.65596441 | -0.664 | -0.547 | chr2 | 232320511 | 232320647 | - | snoRNA |
| ENSG00000280381.1_6 | AC026362.2 | 0.18348464 | 0.66802695 | -0.665 | -0.829 | chr12 | 123571803 | 123577368 | - | TEC |
| ENSG00000254726.3_4 | MEX3A | 0.12180368 | 0.47867418 | -0.665 | 1.727 | chr1 | 156041804 | 156052256 | - | protein_coding |
| ENSG00000153885.14_4 | KCTD15 | 0.02334506 | 0.11445137 | -0.665 | 5.923 | chr19 | 34286838 | 34306668 | + | protein_coding |
| ENSG00000228526.7_6 | MIR34AHG | 0.05963031 | 0.26062775 | -0.665 | 3.958 | chr1 | 9208070 | 9258965 | - | lncRNA |
| ENSG00000212371.1 | SNORA46 | 0.17595587 | 0.64631135 | -0.665 | -0.346 | chr14 | 77932627 | 77932779 | + | snoRNA |
| ENSG00000128059.8_4 | PPAT | 0.02766052 | 0.13289468 | -0.666 | 5.416 | chr4 | 57259528 | 57301781 | - | protein_coding |
| ENSG00000184110.14_4 | EIF3C | 0.11883115 | 0.46969365 | -0.666 | 1.336 | chr16 | 28699879 | 28747051 | + | protein_coding |
| ENSG00000048540.15_8 | LMO3 | 0.17425137 | 0.64097807 | -0.666 | -0.521 | chr12 | 16701306 | 16763528 | - | protein_coding |
| ENSG00000223653.6_7 | AL078459.1 | 0.21576349 | 0.75566132 | -0.667 | -1.301 | chr1 | 85742071 | 85913807 | + | lncRNA |
| ENSG00000185238.13_4 | PRMT3 | 0.02410232 | 0.11761219 | -0.667 | 5.427 | chr11 | 20409104 | 20530884 | + | protein_coding |
| ENSG00000226084.5_5 | AC113935.1 | 0.1080359 | 0.43358343 | -0.667 | 1.587 | chr1 | 77594799 | 77595353 | + | processed_pseudogene |
| ENSG00000115526.11_5 | CHST10 | 0.03637821 | 0.17022165 | -0.668 | 4.173 | chr2 | 101008322 | 101034130 | - | protein_coding |
| ENSG00000183145.9_3 | RIPPLY3 | 0.19969569 | 0.71130579 | -0.668 | -0.781 | chr21 | 38378450 | 38391962 | + | protein_coding |
| ENSG00000164932.13_3 | CTHRC1 | 0.16747865 | 0.62059798 | -0.668 | -0.207 | chr8 | 104383766 | 104395217 | + | protein_coding |

| | | | | | | | | | | |
|---|---|---|---|---|---|---|---|---|---|---|
| ENSG00000231503.4_4 | PTMAP4 | 0.1864828 | 0.67540239 | -0.668 | 0.162 | chr12 | 9392582 | 9392927 | - | processed_pseudogene |
| ENSG00000213699.9_4 | SLC35F6 | 0.00802803 | 0.04381678 | -0.669 | 6.931 | chr2 | 26987152 | 27004099 | + | protein_coding |
| ENSG00000172974.11_5 | AC007318.1 | 0.11494276 | 0.45706983 | -0.669 | 1.523 | chr2 | 65432242 | 65433122 | + | processed_pseudogene |
| ENSG00000113555.5_5 | PCDH12 | 0.14157027 | 0.54220286 | -0.669 | 0.554 | chr5 | 141323150 | 141349306 | - | protein_coding |
| ENSG00000207088.1 | SNORA7B | 0.19638169 | 0.70260039 | -0.670 | -0.890 | chr3 | 129116053 | 129116191 | - | snoRNA |
| ENSG00000145362.20_8 | ANK2 | 0.07305589 | 0.31000333 | -0.670 | 3.025 | chr4 | 113739188 | 114305377 | + | protein_coding |
| ENSG00000240891.8_5 | PLCXD2 | 0.03500424 | 0.16452918 | -0.670 | 4.171 | chr3 | 111393523 | 111565294 | + | protein_coding |
| ENSG00000131153.9_5 | GINS2 | 0.01796941 | 0.09077696 | -0.670 | 5.430 | chr16 | 85709804 | 85723679 | - | protein_coding |
| ENSG00000185630.18_5 | PBX1 | 0.00825671 | 0.04490252 | -0.670 | 6.685 | chr1 | 164524821 | 164868533 | + | protein_coding |
| ENSG00000184205.14_4 | TSPYL2 | 0.03732213 | 0.17386009 | -0.671 | 3.978 | chrX | 53111549 | 53117722 | + | protein_coding |
| ENSG00000115568.15_6 | ZNF142 | 0.02782343 | 0.13353293 | -0.671 | 4.647 | chr2 | 219502639 | 219524378 | - | protein_coding |
| ENSG00000168298.6_3 | H1-4 | 0.00647337 | 0.03599764 | -0.671 | 8.621 | chr6 | 26156582 | 26157335 | + | protein_coding |
| ENSG00000157833.13_4 | GAREM2 | 0.06056467 | 0.263803 | -0.671 | 3.168 | chr2 | 26395957 | 26412532 | + | protein_coding |
| ENSG00000279278.1_7 | AC245060.6 | 0.18649488 | 0.67540239 | -0.671 | -0.372 | chr22 | 22618984 | 22627402 | + | lncRNA |
| ENSG00000213639.10_6 | PPP1CB | 0.00802991 | 0.04381678 | -0.671 | 7.875 | chr2 | 28974506 | 29025806 | + | protein_coding |
| ENSG00000263531.1_7 | AC130324.1 | 0.19775245 | 0.70608275 | -0.671 | -1.047 | chr17 | 29190939 | 29191958 | - | lncRNA |
| ENSG00000206952.2 | SNORA50 | 0.19875197 | 0.70867069 | -0.672 | -0.859 | chr16 | 58593700 | 58593835 | - | snoRNA |
| ENSG00000085982.13_5 | USP40 | 0.00669319 | 0.03708073 | -0.673 | 6.695 | chr2 | 234384166 | 234475428 | - | protein_coding |
| ENSG00000258655.2_6 | ARHGAP5-AS1 | 0.10053811 | 0.40689306 | -0.673 | 1.669 | chr14 | 32544152 | 32545999 | - | lncRNA |
| ENSG00000235440.1_5 | AC011742.3 | 0.16625657 | 0.61706299 | -0.673 | -0.189 | chr2 | 26324186 | 26325005 | - | processed_pseudogene |
| ENSG00000124787.14_5 | RPP40 | 0.0564625 | 0.24911082 | -0.673 | 3.243 | chr6 | 4994951 | 5004297 | - | protein_coding |
| ENSG00000076554.15_6 | TPD52 | 0.01625562 | 0.0827072 | -0.673 | 5.732 | chr8 | 80946980 | 81143467 | - | protein_coding |
| ENSG00000144847.13_7 | IGSF11 | 0.03482044 | 0.16370859 | -0.674 | 4.224 | chr3 | 118619404 | 118864915 | - | protein_coding |
| ENSG00000166261.11_5 | ZNF202 | 0.03883331 | 0.1802381 | -0.674 | 4.013 | chr11 | 123594622 | 123612383 | - | protein_coding |
| ENSG00000115738.10_4 | ID2 | 0.01984131 | 0.09902265 | -0.674 | 5.050 | chr2 | 8818975 | 8824591 | + | protein_coding |
| ENSG00000185838.14_5 | GNB1L | 0.10826288 | 0.43429816 | -0.676 | 2.445 | chr22 | 19770746 | 19842462 | - | protein_coding |
| ENSG00000196497.17_6 | IPO4 | 0.15221538 | 0.57528413 | -0.676 | 0.054 | chr14 | 24649425 | 24658075 | - | protein_coding |
| ENSG00000250271.2_6 | AC068647.2 | 0.0869416 | 0.36082489 | -0.676 | 2.429 | chr3 | 151514835 | 151526037 | + | transcribed_unprocessed_pseudogene |
| ENSG00000130876.11_4 | SLC7A10 | 0.18299561 | 0.66692984 | -0.676 | -0.781 | chr19 | 33699570 | 33716756 | - | protein_coding |
| ENSG00000232762.1_6 | AL355483.2 | 0.15928361 | 0.59591323 | -0.676 | -0.141 | chr1 | 53770208 | 53773134 | + | lncRNA |
| ENSG00000187605.15_2 | TET3 | 0.01878117 | 0.09439042 | -0.676 | 4.954 | chr2 | 74213531 | 74335303 | + | protein_coding |
| ENSG00000101361.17_5 | NOP56 | 0.00418355 | 0.02424302 | -0.676 | 7.727 | chr20 | 2633239 | 2639039 | + | protein_coding |
| ENSG00000171863.15_6 | RPS7 | 0.00703176 | 0.03875103 | -0.676 | 9.105 | chr2 | 3622850 | 3628510 | + | protein_coding |
| ENSG00000137210.14_6 | TMEM14B | 0.02455172 | 0.11957521 | -0.677 | 5.245 | chr6 | 10747992 | 10852986 | + | protein_coding |
| ENSG00000080839.12_5 | RBL1 | 0.01013666 | 0.05419742 | -0.677 | 6.124 | chr20 | 35624752 | 35724400 | - | protein_coding |
| ENSG00000072210.19_6 | ALDH3A2 | 0.00312926 | 0.01849551 | -0.677 | 8.495 | chr17 | 19551449 | 19589073 | + | protein_coding |
| ENSG00000164404.8_4 | GDF9 | 0.18040221 | 0.65964317 | -0.677 | -0.752 | chr5 | 132196873 | 132202576 | - | protein_coding |
| ENSG00000227321.2_4 | MTND4P15 | 0.18428436 | 0.66984046 | -0.678 | -0.924 | chr9 | 94873147 | 94874179 | + | unprocessed_pseudogene |
| ENSG00000011028.14_4 | MRC2 | 0.02800343 | 0.13432424 | -0.678 | 6.643 | chr17 | 60705031 | 60770958 | + | protein_coding |
| ENSG00000231125.2_8 | AF129075.2 | 0.15268565 | 0.57638688 | -0.678 | -0.074 | chr21 | 30430394 | 30432416 | - | lncRNA |
| ENSG00000152133.14_2 | GPATCH11 | 0.02442149 | 0.11900622 | -0.678 | 4.420 | chr2 | 37311594 | 37326387 | + | protein_coding |
| ENSG00000281128.2_5 | PTENP1-AS | 0.15180197 | 0.57396587 | -0.678 | -0.108 | chr9 | 33676783 | 33688009 | + | lncRNA |
| ENSG00000115827.14_5 | DCAF17 | 0.03337808 | 0.15746901 | -0.678 | 5.106 | chr2 | 172290727 | 172341562 | + | protein_coding |
| ENSG00000135966.13_4 | TGFBRAP1 | 0.01133618 | 0.05974782 | -0.679 | 5.579 | chr2 | 105880848 | 105946192 | - | protein_coding |
| ENSG00000244124.1_7 | ATP1B3-AS1 | 0.17962644 | 0.65761858 | -0.679 | -0.861 | chr3 | 141637094 | 141637863 | - | lncRNA |
| ENSG00000185864.16_4 | NPIPB4 | 0.15262739 | 0.57635085 | -0.679 | -0.146 | chr16 | 21845890 | 21892148 | - | protein_coding |
| ENSG00000275413.1_3 | AC002553.2 | 0.18317359 | 0.66724821 | -0.679 | -0.831 | chr17 | 15926637 | 15926967 | - | lncRNA |
| ENSG00000255529.9_7 | POLR2M | 0.00931577 | 0.05007965 | -0.681 | 5.580 | chr15 | 57998893 | 58074960 | + | protein_coding |
| ENSG00000143882.12_6 | ATP6V1C2 | 0.13045217 | 0.50605759 | -0.681 | 0.577 | chr2 | 10861226 | 10925236 | + | protein_coding |
| ENSG00000115128.7_4 | SF3B6 | 0.00704022 | 0.03878562 | -0.681 | 6.134 | chr2 | 24290456 | 24299243 | - | protein_coding |
| ENSG00000115419.13_8 | GLS | 0.00945886 | 0.05081812 | -0.682 | 7.104 | chr2 | 191745547 | 191830278 | + | protein_coding |
| ENSG00000144182.17_4 | LIPT1 | 0.11457845 | 0.45582518 | -0.682 | 1.165 | chr2 | 99771418 | 99779620 | + | protein_coding |
| ENSG00000224401.2_4 | RPL7P57 | 0.186132 | 0.67489444 | -0.682 | -0.893 | chrX | 47700362 | 47701101 | - | processed_pseudogene |
| ENSG00000280023.1_6 | AD000813.1 | 0.14991189 | 0.56875605 | -0.682 | -0.093 | chr19 | 36561141 | 36561972 | + | TEC |
| ENSG00000153157.13_6 | SYCP2L | 0.11772547 | 0.46646487 | -0.682 | 1.032 | chr6 | 10887064 | 10979553 | + | protein_coding |
| ENSG00000189337.17_6 | KAZN | 0.1193841 | 0.47103178 | -0.683 | 0.894 | chr1 | 14219287 | 15444539 | + | protein_coding |
| ENSG00000103005.12_4 | USB1 | 0.01831663 | 0.09229481 | -0.683 | 4.694 | chr16 | 58033450 | 58055522 | + | protein_coding |

| | | | | | | | | | | |
|---|---|---|---|---|---|---|---|---|---|---|
| ENSG00000160208.13_3 | RRP1B | 0.00526156 | 0.0298379 | -0.683 | 6.477 | chr21 | 45079441 | 45115960 | + | protein_coding |
| ENSG00000244932.2_5 | AL449212.1 | 0.13451121 | 0.5196479 | -0.684 | 0.257 | chr3 | 129100141 | 129112992 | + | unprocessed_pseudogene |
| ENSG00000115875.19_6 | SRSF7 | 0.00437433 | 0.02525795 | -0.684 | 7.011 | chr2 | 38970741 | 38978636 | - | protein_coding |
| ENSG00000156795.8_6 | WDYHV1 | 0.03420424 | 0.16098189 | -0.684 | 3.903 | chr8 | 124428966 | 124482268 | + | protein_coding |
| ENSG00000121897.15_4 | LIAS | 0.03644755 | 0.17050121 | -0.684 | 3.537 | chr4 | 39460624 | 39486729 | + | protein_coding |
| ENSG00000267015.1_7 | AC023090.1 | 0.15744493 | 0.59002826 | -0.685 | -0.509 | chr18 | 77100258 | 77103972 | - | lncRNA |
| ENSG00000134871.19_7 | COL4A2 | 0.00595087 | 0.03334252 | -0.685 | 7.172 | chr13 | 110958159 | 111165556 | + | protein_coding |
| ENSG00000146411.6_4 | SLC2A12 | 0.11249093 | 0.44933147 | -0.685 | 1.155 | chr6 | 134308719 | 134373762 | - | protein_coding |
| ENSG00000114956.20_3 | DGUOK | 0.01843307 | 0.09280261 | -0.685 | 4.557 | chr2 | 74153953 | 74186088 | + | protein_coding |
| ENSG00000173894.11_3 | CBX2 | 0.01248194 | 0.06526387 | -0.686 | 5.364 | chr17 | 77751947 | 77761782 | + | protein_coding |
| ENSG00000253626.3_3 | EIF5AL1 | 0.04907388 | 0.22062135 | -0.686 | 3.035 | chr10 | 81272357 | 81276196 | + | protein_coding |
| ENSG00000173207.13_3 | CKS1B | 0.00664085 | 0.03681371 | -0.686 | 5.945 | chr1 | 154947129 | 154951727 | + | protein_coding |
| ENSG00000205208.5_4 | C4orf46 | 0.01606745 | 0.08184355 | -0.686 | 4.480 | chr4 | 159587827 | 159593407 | - | protein_coding |
| ENSG00000240694.9_4 | PNMA2 | 0.0078222 | 0.04282806 | -0.686 | 5.699 | chr8 | 26362217 | 26371608 | - | protein_coding |
| ENSG00000153237.18_5 | CCDC148 | 0.17035923 | 0.62848265 | -0.687 | -0.422 | chr2 | 159027585 | 159313265 | - | protein_coding |
| ENSG00000136630.13_4 | HLX | 0.13485473 | 0.52076543 | -0.689 | 0.125 | chr1 | 221052773 | 221058401 | + | protein_coding |
| ENSG00000281903.2_9 | LINC02246 | 0.16131074 | 0.60172325 | -0.689 | -0.628 | chr21 | 16192020 | 16290873 | - | lncRNA |
| ENSG00000236849.6_6 | LINC01474 | 0.12271025 | 0.48127883 | -0.689 | 0.558 | chr9 | 75486046 | 75489051 | - | lncRNA |
| ENSG00000218537.1_6 | MIF-AS1 | 0.1535598 | 0.57851958 | -0.689 | -0.260 | chr22 | 24236613 | 24241117 | - | lncRNA |
| ENSG00000215784.6_4 | FAM72D | 0.05928184 | 0.25929604 | -0.689 | 2.593 | chr1 | 143896448 | 143913169 | - | protein_coding |
| ENSG00000108387.15_6 | SEPTIN4 | 0.1616314 | 0.60279284 | -0.689 | -0.533 | chr17 | 56597611 | 56621729 | - | protein_coding |
| ENSG00000186907.8_4 | RTN4RL2 | 0.05992895 | 0.26161121 | -0.689 | 2.962 | chr11 | 57228001 | 57245007 | + | protein_coding |
| ENSG00000141499.16_4 | WRAP53 | 0.0104691 | 0.05574005 | -0.690 | 5.149 | chr17 | 7589389 | 7606820 | + | protein_coding |
| ENSG00000255566.1_7 | AC135279.2 | 0.15527564 | 0.58362222 | -0.690 | -0.400 | chr12 | 64836290 | 64837110 | - | processed_pseudogene |
| ENSG00000233325.3_4 | MIPEPP3 | 0.15279913 | 0.57638688 | -0.690 | -0.486 | chr13 | 21872278 | 21880512 | + | transcribed_unprocessed_pseudogene |
| ENSG00000203760.8_2 | CENPW | 0.03751639 | 0.1746466 | -0.691 | 3.317 | chr6 | 126661320 | 126670021 | + | protein_coding |
| ENSG00000212452.1 | SNORD69 | 0.17730369 | 0.65045522 | -0.691 | -0.810 | chr3 | 52726752 | 52726828 | + | snoRNA |
| ENSG00000237638.2_7 | LINC02245 | 0.14943347 | 0.56730436 | -0.691 | -0.198 | chr2 | 65128974 | 65283302 | - | lncRNA |
| ENSG00000251537.4_7 | AC005324.3 | 0.14811934 | 0.56351954 | -0.691 | -0.185 | chr17 | 15474805 | 15554967 | - | protein_coding |
| ENSG00000235001.3_4 | EIF4A1P2 | 0.10878645 | 0.43590638 | -0.692 | 1.165 | chrY | 5205786 | 5207005 | - | processed_pseudogene |
| ENSG00000214194.9_8 | SMIM30 | 0.02274235 | 0.11199092 | -0.692 | 4.318 | chr7 | 112756773 | 112758615 | - | protein_coding |
| ENSG00000233270.1_2 | SNRPEP4 | 0.12697321 | 0.49461564 | -0.692 | 0.395 | chr19 | 5576671 | 5576949 | - | processed_pseudogene |
| ENSG00000171103.11_5 | TRMT61B | 0.03210049 | 0.1521284 | -0.692 | 3.513 | chr2 | 29072687 | 29093175 | - | protein_coding |
| ENSG00000196139.14_5 | AKR1C3 | 0.00124388 | 0.00796396 | -0.692 | 10.549 | chr10 | 5077546 | 5149878 | + | protein_coding |
| ENSG00000225953.3_5 | SATB2-AS1 | 0.11892326 | 0.46984868 | -0.693 | 0.769 | chr2 | 200322423 | 200341658 | + | lncRNA |
| ENSG00000101000.6_5 | PROCR | 0.00852542 | 0.04620838 | -0.693 | 5.347 | chr20 | 33759875 | 33804043 | + | protein_coding |
| ENSG00000152117.17_5 | AC073869.1 | 0.03465322 | 0.16296551 | -0.693 | 3.653 | chr2 | 132250386 | 132279146 | + | transcribed_processed_pseudogene |
| ENSG00000171865.10_5 | RNASEH1 | 0.00452035 | 0.0260065 | -0.693 | 5.835 | chr2 | 3589020 | 3605923 | - | protein_coding |
| ENSG00000171931.13_5 | FBXW10 | 0.07506819 | 0.31725415 | -0.694 | 1.935 | chr17 | 18647339 | 18682662 | + | protein_coding |
| ENSG00000213337.9_5 | ANKRD39 | 0.09063885 | 0.3738116 | -0.694 | 1.479 | chr2 | 97502348 | 97523753 | - | protein_coding |
| ENSG00000280832.1_7 | GSEC | 0.09369089 | 0.3830785 | -0.694 | 1.280 | chr11 | 126210784 | 126225482 | - | lncRNA |
| ENSG00000161513.12_5 | FDXR | 0.03175559 | 0.1508157 | -0.694 | 5.354 | chr17 | 72858619 | 72869156 | - | protein_coding |
| ENSG00000106399.11_4 | RPA3 | 0.01235707 | 0.06462998 | -0.694 | 5.470 | chr7 | 7676149 | 7758238 | - | protein_coding |
| ENSG00000106701.12_5 | FSD1L | 0.03132581 | 0.14909342 | -0.695 | 3.641 | chr9 | 108210077 | 108314714 | + | protein_coding |
| ENSG00000229739.4_7 | AL096803.2 | 0.17552865 | 0.6449704 | -0.695 | -0.994 | chr1 | 186392613 | 186490279 | + | lncRNA |
| ENSG00000143947.13_2 | RPS27A | 0.00111774 | 0.00721613 | -0.695 | 9.204 | chr2 | 55459039 | 55462989 | + | protein_coding |
| ENSG00000180616.9_5 | SSTR2 | 0.13485911 | 0.52076543 | -0.695 | -0.063 | chr17 | 71161149 | 71172772 | + | protein_coding |
| ENSG00000003393.15_7 | ALS2 | 0.00587893 | 0.03298106 | -0.695 | 5.362 | chr2 | 202564990 | 202645679 | - | protein_coding |
| ENSG00000170537.13_4 | TMC7 | 0.03961085 | 0.1833679 | -0.695 | 3.347 | chr16 | 18995256 | 19075264 | + | protein_coding |
| ENSG00000278985.1_6 | AC092718.7 | 0.15779789 | 0.59110165 | -0.695 | -0.508 | chr16 | 81015924 | 81017699 | - | TEC |
| ENSG00000279199.1_5 | AC068669.1 | 0.12073506 | 0.47531571 | -0.696 | 0.431 | chr17 | 38269468 | 38271695 | - | TEC |
| ENSG00000018699.13_5 | TTC27 | 0.00917565 | 0.04935628 | -0.696 | 5.080 | chr2 | 32853099 | 33046118 | + | protein_coding |
| ENSG00000246422.2_7 | DIAPH1-AS1 | 0.18035049 | 0.65958977 | -0.696 | -0.992 | chr5 | 140937878 | 140944830 | + | lncRNA |
| ENSG00000078900.15_7 | TP73 | 0.16380136 | 0.60935067 | -0.696 | -0.656 | chr1 | 3569080 | 3652765 | + | protein_coding |
| ENSG00000144233.9_3 | AMMECR1L | 0.00670683 | 0.03713551 | -0.697 | 5.194 | chr2 | 128619204 | 128643496 | - | protein_coding |
| ENSG00000168288.13_4 | MMADHC | 0.00374047 | 0.02186085 | -0.697 | 6.569 | chr2 | 150426151 | 150444292 | - | protein_coding |
| ENSG00000138639.18_5 | ARHGAP24 | 0.03910771 | 0.18136956 | -0.697 | 3.012 | chr4 | 86396303 | 86923821 | + | protein_coding |
| ENSG00000145555.15_6 | MYO10 | 0.00113607 | 0.00732119 | -0.698 | 7.801 | chr5 | 16662016 | 16936397 | - | protein_coding |

| | | | | | | | | | | |
|---|---|---|---|---|---|---|---|---|---|---|
| ENSG00000143994.14_4 | ABHD1 | 0.0755676 | 0.31906121 | -0.698 | 1.667 | chr2 | 27346657 | 27353680 | + | protein_coding |
| ENSG00000130558.19_6 | OLFM1 | 0.04176278 | 0.19197911 | -0.698 | 3.035 | chr9 | 137967268 | 138013025 | + | protein_coding |
| ENSG00000106070.19_6 | GRB10 | 0.00304536 | 0.01806575 | -0.698 | 5.924 | chr7 | 50657760 | 50861159 | - | protein_coding |
| ENSG00000272525.1_6 | AC099522.2 | 0.12705485 | 0.49471479 | -0.698 | 0.094 | chr5 | 72793375 | 72794118 | - | lncRNA |
| ENSG00000115091.12_4 | ACTR3 | 0.0013571 | 0.00859897 | -0.698 | 8.072 | chr2 | 114647640 | 114720173 | + | protein_coding |
| ENSG00000243819.4_5 | RN7SL832P | 0.16909979 | 0.62474441 | -0.698 | -0.845 | chr2 | 10830470 | 10832225 | + | lncRNA |
| ENSG00000235189.2_7 | AL034405.1 | 0.17984266 | 0.65813892 | -0.698 | -1.027 | chrX | 129003040 | 129091504 | - | lncRNA |
| ENSG00000269925.1_7 | Z98884.2 | 0.15039957 | 0.56975462 | -0.698 | -0.559 | chr1 | 7836443 | 7836835 | + | lncRNA |
| ENSG00000226312.7_7 | CFLAR-AS1 | 0.09961698 | 0.40353282 | -0.698 | 1.122 | chr2 | 202005001 | 202022546 | - | lncRNA |
| ENSG00000236859.7_7 | NIFK-AS1 | 0.07167206 | 0.30529789 | -0.699 | 2.061 | chr2 | 122406896 | 122486139 | + | lncRNA |
| ENSG00000149970.16_9 | CNKSR2 | 0.0160504 | 0.08178019 | -0.699 | 4.255 | chrX | 21390919 | 21672813 | + | protein_coding |
| ENSG00000147874.11_5 | HAUS6 | 0.00228009 | 0.0138592 | -0.699 | 6.650 | chr9 | 19053139 | 19102902 | - | protein_coding |
| ENSG00000174721.10_4 | FGFBP3 | 0.10599288 | 0.42625066 | -0.699 | 0.848 | chr10 | 93666341 | 93669243 | - | protein_coding |
| ENSG00000179958.9_4 | DCTPP1 | 0.01465872 | 0.0754247 | -0.699 | 4.450 | chr16 | 30434936 | 30441351 | - | protein_coding |
| ENSG00000116127.18_6 | ALMS1 | 0.00328305 | 0.01934658 | -0.699 | 6.398 | chr2 | 73612886 | 73852293 | + | protein_coding |
| ENSG00000183199.6_4 | HSP90AB3P | 0.08308097 | 0.34715419 | -0.699 | 2.321 | chr4 | 88812995 | 88815167 | + | processed_pseudogene |
| ENSG00000061676.15_5 | NCKAP1 | 0.00127274 | 0.00812313 | -0.700 | 8.315 | chr2 | 183773843 | 183903586 | - | protein_coding |
| ENSG00000069869.16_5 | NEDD4 | 0.0017246 | 0.01070614 | -0.700 | 7.791 | chr15 | 56119120 | 56285944 | - | protein_coding |
| ENSG00000152127.9_4 | MGAT5 | 0.00458412 | 0.02634089 | -0.701 | 5.558 | chr2 | 134877554 | 135212192 | + | protein_coding |
| ENSG00000134057.15_4 | CCNB1 | 0.00097534 | 0.00635686 | -0.701 | 7.634 | chr5 | 68462962 | 68474072 | + | protein_coding |
| ENSG00000168386.18_5 | FILIP1L | 0.12306586 | 0.482567 | -0.702 | 0.321 | chr3 | 99548985 | 99833357 | - | protein_coding |
| ENSG00000187775.17_5 | DNAH17 | 0.07894211 | 0.33165395 | -0.702 | 1.672 | chr17 | 76419778 | 76573478 | - | protein_coding |
| ENSG00000137496.18_4 | IL18BP | 0.03411357 | 0.16064029 | -0.702 | 3.146 | chr11 | 71709659 | 71716761 | + | protein_coding |
| ENSG00000115425.14_5 | PECR | 0.01890149 | 0.09492454 | -0.702 | 3.902 | chr2 | 216861052 | 216947678 | - | protein_coding |
| ENSG00000133216.16_4 | EPHB2 | 0.00539375 | 0.03051935 | -0.702 | 5.631 | chr1 | 23037332 | 23247993 | + | protein_coding |
| ENSG00000238450.1 | snoU13 | 0.15434662 | 0.58080261 | -0.703 | -0.845 | chr8 | 71516767 | 71516872 | + | snoRNA |
| ENSG00000215386.13_6 | MIR99AHG | 0.06873468 | 0.29484302 | -0.703 | 1.905 | chr21 | 17300616 | 18017786 | + | lncRNA |
| ENSG00000138050.15_4 | THUMPD2 | 0.01093128 | 0.05782636 | -0.703 | 4.299 | chr2 | 39963200 | 40006407 | - | protein_coding |
| ENSG00000226419.8_6 | SLC16A1-AS1 | 0.05893237 | 0.25814877 | -0.703 | 2.050 | chr1 | 113499037 | 113589677 | + | lncRNA |
| ENSG00000229853.1_6 | AL034418.1 | 0.12235473 | 0.48030855 | -0.703 | 0.278 | chr20 | 46259010 | 46260182 | - | processed_pseudogene |
| ENSG00000232995.7_9 | AL592435.1 | 0.11333396 | 0.45178472 | -0.704 | 0.398 | chr1 | 163214295 | 163291684 | - | lncRNA |
| ENSG00000269102.1_7 | AC010320.4 | 0.16520583 | 0.61367592 | -0.704 | -0.813 | chr19 | 52788033 | 52801173 | - | lncRNA |
| ENSG00000258920.1_5 | FOXN3-AS1 | 0.11595229 | 0.46026012 | -0.704 | 0.415 | chr14 | 89883698 | 89886137 | + | lncRNA |
| ENSG00000105852.11_4 | PON3 | 0.01181667 | 0.062041 | -0.704 | 4.424 | chr7 | 94989184 | 95025687 | - | protein_coding |
| ENSG00000224531.6_3 | SMIM13 | 0.00807716 | 0.04403401 | -0.704 | 4.756 | chr6 | 11094067 | 11138966 | + | protein_coding |
| ENSG00000204899.6_4 | MZT1 | 0.00892172 | 0.04813609 | -0.704 | 4.556 | chr13 | 73282505 | 73301767 | - | protein_coding |
| ENSG00000115207.13_6 | GTF3C2 | 0.00192673 | 0.01189468 | -0.705 | 6.114 | chr2 | 27548716 | 27579901 | - | protein_coding |
| ENSG00000132952.12_4 | USPL1 | 0.01077112 | 0.05710143 | -0.705 | 4.606 | chr13 | 31191830 | 31234907 | + | protein_coding |
| ENSG00000198522.14_5 | GPN1 | 0.00492053 | 0.02804993 | -0.705 | 4.997 | chr2 | 27851114 | 27874378 | + | protein_coding |
| ENSG00000181773.7_3 | GPR3 | 0.1505271 | 0.56987321 | -0.705 | -0.738 | chr1 | 27719126 | 27722318 | + | protein_coding |
| ENSG00000220749.4_4 | RPL21P28 | 0.14549286 | 0.554834 | -0.706 | -0.404 | chr1 | 212224866 | 212225348 | - | processed_pseudogene |
| ENSG00000114933.16_5 | INO80D | 0.00752572 | 0.0413066 | -0.706 | 4.857 | chr2 | 206858445 | 206951027 | - | protein_coding |
| ENSG00000224034.1_7 | LINC02561 | 0.09881576 | 0.40101865 | -0.706 | 0.944 | chr10 | 5307996 | 5313199 | - | lncRNA |
| ENSG00000162971.11_4 | TYW5 | 0.01367602 | 0.07078068 | -0.706 | 4.399 | chr2 | 200793636 | 200820459 | - | protein_coding |
| ENSG00000273319.1_6 | AC058791.1 | 0.08388802 | 0.34995031 | -0.706 | 1.533 | chr7 | 130621223 | 130624420 | + | lncRNA |
| ENSG00000040933.15_3 | INPP4A | 0.00396117 | 0.02308233 | -0.707 | 5.247 | chr2 | 99061317 | 99210853 | + | protein_coding |
| ENSG00000010319.7_5 | SEMA3G | 0.16037019 | 0.59896934 | -0.707 | -0.637 | chr3 | 52467051 | 52479119 | - | protein_coding |
| ENSG00000236519.1_6 | LINC01424 | 0.13650383 | 0.52620169 | -0.707 | -0.209 | chr21 | 46222492 | 46224632 | + | lncRNA |
| ENSG00000077616.11_6 | NAALAD2 | 0.1059457 | 0.42615741 | -0.707 | 1.209 | chr11 | 89864683 | 89926062 | + | protein_coding |
| ENSG00000132300.19_3 | PTCD3 | 0.00096612 | 0.00630143 | -0.707 | 7.063 | chr2 | 86333346 | 86369280 | + | protein_coding |
| ENSG00000237737.6_7 | DCTN1-AS1 | 0.15285079 | 0.57645934 | -0.708 | -0.876 | chr2 | 74612601 | 74621009 | + | lncRNA |
| ENSG00000237842.2_5 | AL157713.1 | 0.1546755 | 0.58173572 | -0.708 | -0.682 | chr1 | 157029024 | 157030554 | - | processed_pseudogene |
| ENSG00000228294.2 | BMS1P17 | 0.12041086 | 0.47424956 | -0.708 | 0.135 | chr14 | 19670480 | 19686002 | - | lincRNA |
| ENSG00000182263.14_4 | FIGN | 0.02400091 | 0.11718168 | -0.708 | 3.727 | chr2 | 164449906 | 164592522 | - | protein_coding |
| ENSG00000239437.2 | RN7SL752P | 0.07435892 | 0.31470578 | -0.708 | 1.550 | chr3 | 129310192 | 129310488 | + | misc_RNA |
| ENSG00000145416.13_6 | MARCHF1 | 0.05220146 | 0.23285549 | -0.708 | 2.402 | chr4 | 164445450 | 165305202 | - | protein_coding |
| ENSG00000090530.10_5 | P3H2 | 0.0069324 | 0.03827469 | -0.708 | 5.015 | chr3 | 189674517 | 189840226 | - | protein_coding |
| ENSG00000280385.1_1 | AP000648.4 | 0.02938184 | 0.14044316 | -0.709 | 4.141 | chr11 | 89926782 | 89931288 | + | TEC |

| | | | | | | | | | | |
|---|---|---|---|---|---|---|---|---|---|---|
| ENSG00000227110.7_8 | LMCD1-AS1 | 0.09951326 | 0.40329655 | -0.709 | 0.674 | chr3 | 7992950 | 8653610 | - | lncRNA |
| ENSG00000234500.1_5 | AC008267.3 | 0.17329454 | 0.6378546 | -0.709 | -0.995 | chr7 | 65976543 | 66010053 | + | unprocessed_pseudogene |
| ENSG00000142494.13_4 | SLC47A1 | 0.00495015 | 0.02819793 | -0.709 | 4.961 | chr17 | 19398698 | 19482347 | + | protein_coding |
| ENSG00000183963.18_6 | SMTN | 0.01205367 | 0.06317333 | -0.709 | 4.572 | chr22 | 31460091 | 31500743 | + | protein_coding |
| ENSG00000168883.20_5 | USP39 | 0.00171982 | 0.01068393 | -0.709 | 6.279 | chr2 | 85829979 | 85876406 | + | protein_coding |
| ENSG00000274565.1_9 | AC080038.2 | 0.13180609 | 0.5104353 | -0.710 | -0.084 | chr17 | 60703798 | 60704951 | - | lncRNA |
| ENSG00000182253.15_3 | SYNM | 0.00194192 | 0.01197572 | -0.710 | 5.826 | chr15 | 99638420 | 99675798 | + | protein_coding |
| ENSG00000251495.1_2 | PPIAP11 | 0.18239242 | 0.66500432 | -0.711 | -1.098 | chr5 | 81305421 | 81305913 | - | processed_pseudogene |
| ENSG00000117450.14_5 | PRDX1 | 0.00032086 | 0.00228763 | -0.711 | 8.874 | chr1 | 45976708 | 45988719 | - | protein_coding |
| ENSG00000076706.17_6 | MCAM | 0.02585593 | 0.12524825 | -0.711 | 3.328 | chr11 | 119179239 | 119192231 | - | protein_coding |
| ENSG00000145375.8_4 | SPATA5 | 0.00535854 | 0.0303491 | -0.711 | 4.988 | chr4 | 123844233 | 124240588 | + | protein_coding |
| ENSG00000187514.16_4 | PTMA | 0.00050515 | 0.00347195 | -0.712 | 9.514 | chr2 | 232571605 | 232578251 | + | protein_coding |
| ENSG00000121769.8_5 | FABP3 | 0.15345453 | 0.57836797 | -0.712 | -0.564 | chr1 | 31838100 | 31849697 | - | protein_coding |
| ENSG00000131187.9_3 | F12 | 0.06949987 | 0.29768663 | -0.712 | 1.973 | chr5 | 176829141 | 176836577 | - | protein_coding |
| ENSG00000155729.13_4 | KCTD18 | 0.00973173 | 0.05218946 | -0.712 | 4.405 | chr2 | 201353681 | 201384507 | - | protein_coding |
| ENSG00000163013.11_3 | FBXO41 | 0.02946003 | 0.14077899 | -0.712 | 3.198 | chr2 | 73481810 | 73511559 | - | protein_coding |
| ENSG00000231104.8_7 | AC022395.1 | 0.12668096 | 0.4940168 | -0.712 | -0.055 | chr10 | 119776998 | 119805321 | + | lncRNA |
| ENSG00000278390.5_7 | AL354696.2 | 0.1484114 | 0.56414747 | -0.712 | -0.738 | chr13 | 41707075 | 41810822 | + | lncRNA |
| ENSG00000116030.16_3 | SUMO1 | 0.00156164 | 0.00976618 | -0.712 | 6.835 | chr2 | 203070903 | 203103331 | - | protein_coding |
| ENSG00000265273.1_4 | PGDP1 | 0.13170499 | 0.51024913 | -0.712 | -0.345 | chr18 | 29542141 | 29543581 | + | processed_pseudogene |
| ENSG00000132323.9_4 | ILKAP | 0.01078438 | 0.05712762 | -0.713 | 4.105 | chr2 | 239079043 | 239112349 | - | protein_coding |
| ENSG00000260034.1_7 | LCMT1-AS2 | 0.10658626 | 0.42834607 | -0.713 | 0.459 | chr16 | 25151898 | 25160353 | - | lncRNA |
| ENSG00000153956.16_5 | CACNA2D1 | 0.00368139 | 0.02154391 | -0.713 | 5.481 | chr7 | 81575760 | 82073093 | - | protein_coding |
| ENSG00000130066.16_2 | SAT1 | 0.00110734 | 0.00715944 | -0.713 | 7.742 | chrX | 23801290 | 23804343 | + | protein_coding |
| ENSG00000115902.11_3 | SLC1A4 | 0.00434763 | 0.02513643 | -0.714 | 4.888 | chr2 | 65215611 | 65250999 | + | protein_coding |
| ENSG00000115816.15_6 | CEBPZ | 0.0014021 | 0.00886513 | -0.714 | 6.453 | chr2 | 37428755 | 37458739 | - | protein_coding |
| ENSG00000214184.3_6 | GCC2-AS1 | 0.16783096 | 0.62121931 | -0.714 | -1.100 | chr2 | 109123971 | 109150652 | - | lncRNA |
| ENSG00000275854.1_8 | AC084824.5 | 0.12059928 | 0.47488639 | -0.715 | 0.058 | chr12 | 32889864 | 32890594 | - | lncRNA |
| ENSG00000083312.17_2 | TNPO1 | 0.00049787 | 0.0034312 | -0.715 | 8.960 | chr5 | 72112139 | 72212560 | + | protein_coding |
| ENSG00000146038.12_4 | DCDC2 | 0.01158045 | 0.06090879 | -0.715 | 4.380 | chr6 | 24171983 | 24358287 | - | protein_coding |
| ENSG00000089723.10_3 | OTUB2 | 0.05358821 | 0.23802544 | -0.715 | 2.016 | chr14 | 94492686 | 94515276 | + | protein_coding |
| ENSG00000138018.18_7 | SELENOI | 0.00120231 | 0.00772564 | -0.715 | 6.733 | chr2 | 26531415 | 26618759 | + | protein_coding |
| ENSG00000224126.2_4 | UBE2SP2 | 0.15588078 | 0.58515398 | -0.716 | -0.878 | chr17 | 18580574 | 18581070 | + | processed_pseudogene |
| ENSG00000184144.12_5 | CNTN2 | 0.15515502 | 0.58329228 | -0.716 | -1.065 | chr1 | 205012065 | 205047417 | + | protein_coding |
| ENSG00000134684.10_3 | YARS1 | 0.00123917 | 0.0079367 | -0.716 | 5.946 | chr1 | 33240838 | 33283754 | - | protein_coding |
| ENSG00000235609.7_8 | AF127577.4 | 0.04969507 | 0.22313176 | -0.716 | 3.293 | chr21 | 16191164 | 16386751 | - | lncRNA |
| ENSG00000184500.16_7 | PROS1 | 0.00831244 | 0.04515043 | -0.716 | 4.472 | chr3 | 93591895 | 93698847 | - | protein_coding |
| ENSG00000155903.13_5 | RASA2 | 0.0043707 | 0.02524522 | -0.716 | 5.043 | chr3 | 141205869 | 141334186 | + | protein_coding |
| ENSG00000138381.10_5 | ASNSD1 | 0.01094064 | 0.05785187 | -0.716 | 4.261 | chr2 | 190526111 | 190535557 | + | protein_coding |
| ENSG00000188234.13_4 | AGAP4 | 0.10275914 | 0.41484246 | -0.716 | 0.442 | chr10 | 46321042 | 46349323 | - | protein_coding |
| ENSG00000266975.1_6 | FARSA-AS1 | 0.1782531 | 0.65339858 | -0.717 | -1.280 | chr19 | 13041336 | 13044110 | + | lncRNA |
| ENSG00000233966.1_2 | UBE2SP1 | 0.03727622 | 0.17381553 | -0.717 | 2.719 | chr17 | 15607546 | 15608214 | - | processed_pseudogene |
| ENSG00000065183.16_3 | WDR3 | 0.00106341 | 0.00689294 | -0.717 | 5.960 | chr1 | 118472343 | 118509166 | + | protein_coding |
| ENSG00000185483.12_4 | ROR1 | 0.00192677 | 0.01189468 | -0.717 | 5.663 | chr1 | 64239688 | 64647181 | + | protein_coding |
| ENSG00000166128.13_3 | RAB8B | 0.00277806 | 0.01661317 | -0.717 | 5.377 | chr15 | 63481759 | 63559975 | + | protein_coding |
| ENSG00000229127.2_7 | KANSL1L-AS1 | 0.12551321 | 0.49043174 | -0.717 | -0.177 | chr2 | 210895296 | 210929080 | + | lncRNA |
| ENSG00000258011.2_4 | HMGA1P3 | 0.16651396 | 0.61776017 | -0.718 | -0.794 | chr12 | 110867764 | 110868087 | - | processed_pseudogene |
| ENSG00000084733.11_3 | RAB10 | 0.00031476 | 0.0022486 | -0.718 | 8.698 | chr2 | 26256953 | 26360323 | + | protein_coding |
| ENSG00000084090.13_6 | STARD7 | 0.00031465 | 0.0022486 | -0.718 | 7.555 | chr2 | 96850597 | 96874563 | - | protein_coding |
| ENSG00000279602.1_5 | AC109326.1 | 0.08426578 | 0.35127878 | -0.718 | 1.421 | chr17 | 41437409 | 41438729 | - | TEC |
| ENSG00000273038.2_6 | AL365203.2 | 0.01259678 | 0.06582559 | -0.718 | 3.744 | chr10 | 33176183 | 33178239 | - | lncRNA |
| ENSG00000259826.1_6 | AC072061.1 | 0.12923082 | 0.50208703 | -0.718 | -0.310 | chr7 | 39987121 | 39989354 | - | lncRNA |
| ENSG00000143951.16_5 | WDPCP | 0.03288306 | 0.15544089 | -0.718 | 2.705 | chr2 | 63346694 | 64054977 | - | protein_coding |
| ENSG00000126778.12_7 | SIX1 | 0.01787781 | 0.0903656 | -0.718 | 3.271 | chr14 | 61110139 | 61124977 | - | protein_coding |
| ENSG00000171989.6_4 | LDHAL6B | 0.07912866 | 0.33224989 | -0.718 | 1.170 | chr15 | 59499042 | 59500787 | + | protein_coding |
| ENSG00000274290.2_4 | H2BC6 | 0.0012472 | 0.00798236 | -0.719 | 6.180 | chr6 | 26172287 | 26184883 | + | protein_coding |
| ENSG00000100526.20_5 | CDKN3 | 0.0044075 | 0.02543292 | -0.719 | 5.115 | chr14 | 54863567 | 54886936 | + | protein_coding |

| | | | | | | | | | | |
|---|---|---|---|---|---|---|---|---|---|---|
| ENSG00000166845.15_7 | C18orf54 | 0.01814933 | 0.09152971 | -0.719 | 3.357 | chr18 | 51884276 | 51911588 | + | protein_coding |
| ENSG00000169019.11_4 | COMMD8 | 0.01296298 | 0.06752074 | -0.720 | 3.828 | chr4 | 47452804 | 47465719 | - | protein_coding |
| ENSG00000189149.12_6 | CRYM-AS1 | 0.09047045 | 0.373293 | -0.720 | 0.856 | chr16 | 21312170 | 21350358 | + | transcribed_unitary_pseudogene |
| ENSG00000261366.1_6 | MANEA-DT | 0.12943961 | 0.50267839 | -0.720 | -0.391 | chr6 | 96023059 | 96025326 | - | lncRNA |
| ENSG00000273760.1_7 | AC245041.1 | 0.00475919 | 0.02724118 | -0.721 | 4.632 | chr10 | 47135135 | 47138586 | - | lncRNA |
| ENSG00000235776.2_5 | AC000089.1 | 0.16028973 | 0.59879469 | -0.721 | -0.911 | chr22 | 19779817 | 19780617 | + | processed_pseudogene |
| ENSG00000279649.1_6 | AC020978.9 | 0.11406136 | 0.45417483 | -0.722 | 0.036 | chr16 | 68323990 | 68326693 | - | TEC |
| ENSG00000146648.18_5 | EGFR | 0.00020703 | 0.00152373 | -0.722 | 8.621 | chr7 | 55086710 | 55279321 | + | protein_coding |
| ENSG00000256436.1_4 | TAS2R31 | 0.10587954 | 0.42603529 | -0.722 | 0.142 | chr12 | 11182986 | 11184006 | - | protein_coding |
| ENSG00000266313.2_6 | AC026254.2 | 0.09992532 | 0.40450514 | -0.722 | 0.368 | chr17 | 25660267 | 25679854 | + | lncRNA |
| ENSG00000232600.3_6 | TONSL-AS1 | 0.09866203 | 0.40065691 | -0.723 | 0.497 | chr8 | 145663058 | 145665354 | + | lncRNA |
| ENSG00000116711.10_3 | PLA2G4A | 0.00151265 | 0.00951001 | -0.723 | 5.955 | chr1 | 186798081 | 186958113 | + | protein_coding |
| ENSG00000201428.1 | RN7SKP71 | 0.15901641 | 0.59516423 | -0.723 | -0.878 | chr12 | 112704881 | 112705198 | + | misc_RNA |
| ENSG00000188933.16_4 | USP32P1 | 0.09805545 | 0.39866161 | -0.723 | 0.546 | chr17 | 16689803 | 16707769 | + | transcribed_unprocessed_pseudogene |
| ENSG00000123500.10_8 | COL10A1 | 0.1160934 | 0.46065476 | -0.723 | -0.045 | chr6 | 116440072 | 116479910 | - | protein_coding |
| ENSG00000168497.5_7 | CAVIN2 | 0.00109396 | 0.00708063 | -0.724 | 6.006 | chr2 | 192699036 | 192711814 | - | protein_coding |
| ENSG00000100479.13_4 | POLE2 | 0.00736348 | 0.04050372 | -0.724 | 4.325 | chr14 | 50110273 | 50155140 | - | protein_coding |
| ENSG00000210196.2 | MT-TP | 0.00409662 | 0.02377805 | -0.724 | 5.072 | chrM | 15956 | 16023 | - | Mt_tRNA |
| ENSG00000157470.12_4 | FAM81A | 0.0089443 | 0.04822864 | -0.725 | 4.122 | chr15 | 59664892 | 59815754 | + | protein_coding |
| ENSG00000152413.15_4 | HOMER1 | 0.01112112 | 0.0587363 | -0.725 | 4.124 | chr5 | 78668459 | 78809957 | - | protein_coding |
| ENSG00000151694.14_7 | ADAM17 | 0.00096673 | 0.00630305 | -0.725 | 6.136 | chr2 | 9628615 | 9696861 | - | protein_coding |
| ENSG00000165490.13_5 | DDIAS | 0.00268704 | 0.01611768 | -0.725 | 4.885 | chr11 | 82611017 | 82669319 | + | protein_coding |
| ENSG00000183558.5 | HIST2H2AA3 | 0.08844981 | 0.36622881 | -0.726 | 0.681 | chr1 | 149813505 | 149814478 | - | protein_coding |
| ENSG00000172986.13_4 | GXYLT2 | 0.05743355 | 0.25264223 | -0.726 | 1.616 | chr3 | 72937197 | 73047289 | + | protein_coding |
| ENSG00000198695.2_1 | MT-ND6 | 0.00112592 | 0.00726632 | -0.726 | 5.710 | chrM | 14149 | 14673 | - | protein_coding |
| ENSG00000204682.8_6 | MIR1915HG | 0.004056 | 0.02356541 | -0.727 | 5.076 | chr10 | 21781587 | 21786189 | - | lncRNA |
| ENSG00000111837.11_5 | MAK | 0.13223619 | 0.51186065 | -0.727 | -0.521 | chr6 | 10762956 | 10838788 | - | protein_coding |
| ENSG00000196504.17_8 | PRPF40A | 0.00018353 | 0.00136545 | -0.727 | 7.842 | chr2 | 153508107 | 153574511 | - | protein_coding |
| ENSG00000151287.17_4 | TEX30 | 0.03651483 | 0.17072606 | -0.727 | 2.446 | chr13 | 103418238 | 103426161 | - | protein_coding |
| ENSG00000144231.11_5 | POLR2D | 0.00200548 | 0.01234203 | -0.727 | 5.132 | chr2 | 128601127 | 128615729 | - | protein_coding |
| ENSG00000235397.1_7 | EPN2-AS1 | 0.20616849 | 0.72953506 | -0.727 | -1.324 | chr17 | 19199909 | 19209574 | - | lncRNA |
| ENSG00000188501.11_3 | LCTL | 0.1071843 | 0.43026288 | -0.728 | 0.134 | chr15 | 66839517 | 66858317 | - | protein_coding |
| ENSG00000115687.13_3 | PASK | 0.01292173 | 0.06734537 | -0.729 | 3.943 | chr2 | 242045514 | 242089679 | - | protein_coding |
| ENSG00000259802.1_7 | AC012640.4 | 0.08050928 | 0.33744007 | -0.729 | 0.743 | chr5 | 10352813 | 10353713 | - | lncRNA |
| ENSG00000035141.8_5 | FAM136A | 0.00059129 | 0.00401741 | -0.729 | 6.189 | chr2 | 70523107 | 70529222 | - | protein_coding |
| ENSG00000107020.10_5 | PLGRKT | 0.01559092 | 0.07971599 | -0.729 | 3.376 | chr9 | 5357971 | 5437925 | - | protein_coding |
| ENSG00000255717.7_4 | SNHG1 | 0.00045701 | 0.00317275 | -0.729 | 6.818 | chr11 | 62619456 | 62623425 | - | lncRNA |
| ENSG00000166922.8_2 | SCG5 | 0.08077065 | 0.33837589 | -0.729 | 0.804 | chr15 | 32933877 | 32989299 | + | protein_coding |
| ENSG00000203811.1_3 | H3C14 | 0.1478997 | 0.56293729 | -0.730 | -0.575 | chr1 | 149811110 | 149812765 | - | protein_coding |
| ENSG00000223522.2_6 | AC093690.1 | 0.15031362 | 0.56958698 | -0.730 | -0.910 | chr2 | 28529930 | 28533427 | - | lncRNA |
| ENSG00000171492.14_4 | LRRC8D | 0.00043116 | 0.00300761 | -0.730 | 6.134 | chr1 | 90286573 | 90402170 | + | protein_coding |
| ENSG00000164162.14_6 | ANAPC10 | 0.04836853 | 0.21805727 | -0.731 | 1.748 | chr4 | 145753060 | 146019693 | - | protein_coding |
| ENSG00000119778.15_7 | ATAD2B | 0.00171663 | 0.01067159 | -0.731 | 5.183 | chr2 | 23971534 | 24149993 | - | protein_coding |
| ENSG00000165389.7_5 | SPTSSA | 0.0106528 | 0.05659937 | -0.731 | 5.271 | chr14 | 34901994 | 34931446 | - | protein_coding |
| ENSG00000101955.15_6 | SRPX | 0.04682679 | 0.21195542 | -0.732 | 1.740 | chrX | 38008589 | 38080177 | - | protein_coding |
| ENSG00000147437.10_4 | GNRH1 | 0.04941585 | 0.22199023 | -0.732 | 1.858 | chr8 | 25276774 | 25282170 | - | protein_coding |
| ENSG00000284946.1_3 | AC068831.8 | 0.01574095 | 0.0803879 | -0.732 | 3.394 | chr15 | 91509420 | 91565796 | - | protein_coding |
| ENSG00000143367.16_5 | TUFT1 | 0.00272015 | 0.01629979 | -0.732 | 4.726 | chr1 | 151512781 | 151556059 | + | protein_coding |
| ENSG00000234171.3_6 | RNASEH1-AS1 | 0.01019958 | 0.05448467 | -0.732 | 3.714 | chr2 | 3606064 | 3612432 | + | lncRNA |
| ENSG00000212664.5_5 | AC064799.1 | 0.10902082 | 0.43664853 | -0.733 | 0.015 | chr15 | 71633497 | 71634051 | - | processed_pseudogene |
| ENSG00000087510.7_3 | TFAP2C | 0.00127285 | 0.00812313 | -0.733 | 5.156 | chr20 | 55204362 | 55214339 | + | protein_coding |
| ENSG00000119787.14_6 | ATL2 | 0.00043639 | 0.00304169 | -0.733 | 6.624 | chr2 | 38521096 | 38604427 | - | protein_coding |
| ENSG00000170236.16_6 | USP50 | 0.0937267 | 0.3830785 | -0.733 | 0.287 | chr15 | 50786215 | 50838905 | - | protein_coding |
| ENSG00000054690.14_5 | PLEKHH1 | 0.01537475 | 0.07872136 | -0.734 | 3.072 | chr14 | 68000007 | 68056329 | + | protein_coding |
| ENSG00000172167.8_4 | MTBP | 0.00445931 | 0.02570686 | -0.734 | 4.142 | chr8 | 121457640 | 121554373 | + | protein_coding |
| ENSG00000140044.13_5 | JDP2 | 0.00538322 | 0.03047919 | -0.735 | 4.069 | chr14 | 75894419 | 75940814 | + | protein_coding |
| ENSG00000239415.1_8 | AP001469.3 | 0.09916725 | 0.40235316 | -0.735 | 0.249 | chr21 | 47671463 | 47674047 | - | lncRNA |
| ENSG00000138092.11_4 | CENPO | 0.00114944 | 0.00739929 | -0.736 | 5.242 | chr2 | 25016005 | 25045245 | + | protein_coding |

| | | | | | | | | | | |
|---|---|---|---|---|---|---|---|---|---|---|
| ENSG00000077684.16_5 | JADE1 | 0.00020395 | 0.00150294 | -0.736 | 6.486 | chr4 | 129730855 | 129796379 | + | protein_coding |
| ENSG00000108515.17_6 | ENO3 | 0.00361673 | 0.02120041 | -0.736 | 4.666 | chr17 | 4851387 | 4860426 | + | protein_coding |
| ENSG00000112578.10_4 | BYSL | 0.00284001 | 0.01694376 | -0.736 | 4.486 | chr6 | 41889237 | 41900784 | + | protein_coding |
| ENSG00000267226.2_7 | AC104971.1 | 0.12408768 | 0.48571618 | -0.736 | -0.416 | chr18 | 56337241 | 56339109 | - | lncRNA |
| ENSG00000235105.1_5 | AL356968.2 | 0.14098393 | 0.54030663 | -0.737 | -0.750 | chr1 | 48901639 | 48902895 | + | processed_pseudogene |
| ENSG00000232472.1_4 | EEF1B2P3 | 0.03372587 | 0.15894097 | -0.737 | 2.571 | chrX | 24806509 | 24807186 | - | processed_pseudogene |
| ENSG00000139926.15_8 | FRMD6 | 0.00187643 | 0.01159599 | -0.737 | 4.840 | chr14 | 51955818 | 52197445 | + | protein_coding |
| ENSG00000285763.1_3 | AL358777.3 | 0.09522437 | 0.38857405 | -0.738 | 0.348 | chr6 | 10659731 | 10693093 | + | lncRNA |
| ENSG00000136918.7_3 | WDR38 | 0.10499292 | 0.42289938 | -0.738 | 0.028 | chr9 | 127615696 | 127620169 | + | protein_coding |
| ENSG00000132313.15_5 | MRPL35 | 0.00066096 | 0.00445498 | -0.738 | 6.087 | chr2 | 86426478 | 86440913 | + | protein_coding |
| ENSG00000181061.13_4 | HIGD1A | 0.00389452 | 0.02271635 | -0.738 | 4.525 | chr3 | 42825790 | 42846023 | - | protein_coding |
| ENSG00000250988.7_7 | SNHG21 | 0.11909438 | 0.47021106 | -0.739 | -0.322 | chr15 | 83419316 | 83425958 | + | lncRNA |
| ENSG00000127588.5_4 | GNG13 | 0.13658693 | 0.52624866 | -0.739 | -0.667 | chr16 | 848041 | 850734 | - | protein_coding |
| ENSG00000223044.1 | RNU6-130P | 0.12569183 | 0.49102171 | -0.740 | -0.624 | chr6 | 82920055 | 82920157 | - | snRNA |
| ENSG00000065802.12_3 | ASB1 | 0.00145349 | 0.00916727 | -0.740 | 5.159 | chr2 | 239335383 | 239360891 | + | protein_coding |
| ENSG00000240254.2_6 | B4GALT4-AS1 | 0.09765748 | 0.39722538 | -0.741 | 0.173 | chr3 | 118945333 | 119009513 | + | lncRNA |
| ENSG00000173692.13_5 | PSMD1 | 7.41E-05 | 0.00058451 | -0.741 | 7.739 | chr2 | 231921578 | 232037541 | + | protein_coding |
| ENSG00000215458.8_7 | AATBC | 0.10616284 | 0.42683756 | -0.741 | -0.080 | chr21 | 45225639 | 45232448 | - | lncRNA |
| ENSG00000206712.1 | RNU6-26P | 0.0964948 | 0.39294593 | -0.741 | 0.627 | chr3 | 98523822 | 98523928 | - | snRNA |
| ENSG00000256006.1_7 | AC084117.1 | 0.1065064 | 0.42812199 | -0.742 | -0.279 | chr11 | 18427156 | 18428278 | + | lncRNA |
| ENSG00000136718.10_6 | IMP4 | 0.00094756 | 0.00619172 | -0.742 | 5.689 | chr2 | 131100450 | 131105540 | + | protein_coding |
| ENSG00000183605.17_5 | SFXN4 | 0.00164455 | 0.010245 | -0.742 | 4.821 | chr10 | 120900279 | 120925226 | - | protein_coding |
| ENSG00000246851.1_7 | AL157938.3 | 0.14756205 | 0.56200105 | -0.743 | -0.976 | chr9 | 134065297 | 134069592 | - | lncRNA |
| ENSG00000287037.1_1 | AC097709.1 | 0.07850681 | 0.330372 | -0.743 | 0.639 | chr4 | 74863650 | 74873399 | + | lncRNA |
| ENSG00000119203.14_3 | CPSF3 | 0.00011783 | 0.00090119 | -0.743 | 6.840 | chr2 | 9563780 | 9613230 | + | protein_coding |
| ENSG00000260806.1_6 | AL163051.1 | 0.10376449 | 0.41861575 | -0.743 | 0.000 | chr14 | 96967147 | 96968658 | - | lncRNA |
| ENSG00000136982.6_5 | DSCC1 | 0.00497049 | 0.02829564 | -0.743 | 4.222 | chr8 | 120846216 | 120868134 | - | protein_coding |
| ENSG00000115364.14_6 | MRPL19 | 0.00023467 | 0.00171366 | -0.744 | 6.527 | chr2 | 75873909 | 75917977 | + | protein_coding |
| ENSG00000081320.11_5 | STK17B | 0.00138937 | 0.00879096 | -0.744 | 4.882 | chr2 | 196998307 | 197041227 | - | protein_coding |
| ENSG00000231548.2_4 | OR55B1P | 0.09961128 | 0.40353282 | -0.744 | -0.034 | chr11 | 4167342 | 4172334 | - | transcribed_unitary_pseudogene |
| ENSG00000170293.9_4 | CMTM8 | 0.0476467 | 0.21529473 | -0.744 | 1.562 | chr3 | 32280171 | 32411813 | + | protein_coding |
| ENSG00000115750.17_4 | TAF1B | 0.00057016 | 0.00388917 | -0.744 | 5.518 | chr2 | 9983572 | 10074545 | + | protein_coding |
| ENSG00000226608.3_4 | FTLP3 | 0.0054525 | 0.03080271 | -0.745 | 4.407 | chr20 | 4004564 | 4005091 | + | processed_pseudogene |
| ENSG00000228393.4_7 | LINC01004 | 0.07871806 | 0.33117548 | -0.746 | 0.442 | chr7 | 104590762 | 104653491 | - | lncRNA |
| ENSG00000170089.15_5 | AC106795.1 | 0.0127857 | 0.06669511 | -0.746 | 3.090 | chr5 | 177236408 | 177377733 | + | transcribed_unprocessed_pseudogene |
| ENSG00000164611.13_4 | PTTG1 | 0.00055463 | 0.00379177 | -0.747 | 6.171 | chr5 | 159848862 | 159855746 | + | protein_coding |
| ENSG00000143674.11_5 | MAP3K21 | 0.00028054 | 0.00202537 | -0.747 | 6.034 | chr1 | 233463470 | 233520894 | + | protein_coding |
| ENSG00000185477.5_4 | GPRIN3 | 0.00017197 | 0.00128484 | -0.747 | 7.059 | chr4 | 90157534 | 90228951 | - | protein_coding |
| ENSG00000183943.6_4 | PRKX | 0.00134409 | 0.00852266 | -0.749 | 4.821 | chrX | 3522381 | 3631690 | - | protein_coding |
| ENSG00000184113.9_6 | CLDN5 | 0.15778572 | 0.59110165 | -0.749 | -0.978 | chr22 | 19510547 | 19515068 | - | protein_coding |
| ENSG00000182022.18_7 | CHST15 | 0.00174715 | 0.01083477 | -0.749 | 4.422 | chr10 | 125767184 | 125853114 | - | protein_coding |
| ENSG00000224020.1_5 | MIR181A2HG | 0.11271931 | 0.44999834 | -0.749 | -0.534 | chr9 | 127420746 | 127460910 | + | lncRNA |
| ENSG00000187840.5_3 | EIF4EBP1 | 0.00070698 | 0.00473639 | -0.749 | 5.426 | chr8 | 37888052 | 37917883 | + | protein_coding |
| ENSG00000231663.1_8 | COA6-AS1 | 0.08812089 | 0.36503706 | -0.750 | 0.211 | chr1 | 234508553 | 234509339 | - | lncRNA |
| ENSG00000151692.15_3 | RNF144A | 0.00292509 | 0.01740459 | -0.750 | 4.148 | chr2 | 7057543 | 7208417 | + | protein_coding |
| ENSG00000140403.12_3 | DNAJA4 | 0.10860159 | 0.43526379 | -0.750 | -0.402 | chr15 | 78556428 | 78574538 | + | protein_coding |
| ENSG00000273391.1_6 | AC083880.1 | 0.12737174 | 0.49573138 | -0.750 | -0.755 | chr7 | 139043778 | 139044312 | - | lncRNA |
| ENSG00000275126.1_3 | H4C13 | 0.00075372 | 0.0050155 | -0.751 | 4.972 | chr6 | 27840977 | 27841288 | - | protein_coding |
| ENSG00000067992.15_7 | PDK3 | 0.00100816 | 0.00655636 | -0.751 | 5.090 | chrX | 24483338 | 24568583 | + | protein_coding |
| ENSG00000173559.13_5 | NABP1 | 0.00169114 | 0.01052389 | -0.751 | 4.491 | chr2 | 192542794 | 192561390 | + | protein_coding |
| ENSG00000226525.5_4 | RPS7P10 | 0.08615994 | 0.35799898 | -0.751 | 0.627 | chr13 | 22202552 | 22203154 | - | processed_pseudogene |
| ENSG00000109771.15_3 | LRP2BP | 0.08282863 | 0.34618121 | -0.751 | 0.444 | chr4 | 186285033 | 186317053 | - | protein_coding |
| ENSG00000158987.21_8 | RAPGEF6 | 0.00636546 | 0.03545301 | -0.752 | 3.462 | chr5 | 130759614 | 130970929 | - | protein_coding |
| ENSG00000071967.12_3 | CYBRD1 | 0.0002548 | 0.00185155 | -0.752 | 6.002 | chr2 | 172378757 | 172414639 | + | protein_coding |
| ENSG00000164171.11_3 | ITGA2 | 0.00016494 | 0.00123855 | -0.752 | 6.311 | chr5 | 52285170 | 52390609 | + | protein_coding |
| ENSG00000188322.7_6 | SBK1 | 0.12886441 | 0.50077295 | -0.753 | -0.894 | chr16 | 28270567 | 28335170 | + | protein_coding |
| ENSG00000036257.13_5 | CUL3 | 0.00010115 | 0.00078037 | -0.753 | 6.605 | chr2 | 225334867 | 225450114 | - | protein_coding |

| | | | | | | | | | | |
|---|---|---|---|---|---|---|---|---|---|---|
| ENSG00000138430.16_4 | OLA1 | 0.00011208 | 0.00085978 | -0.753 | 6.653 | chr2 | 174937175 | 175113327 | - | protein_coding |
| ENSG00000231721.7_7 | LINC-PINT | 0.01077301 | 0.05710143 | -0.754 | 3.611 | chr7 | 130623722 | 130794935 | - | lncRNA |
| ENSG00000264991.1 | AP001205.1 | 0.12606598 | 0.49237508 | -0.754 | -0.783 | chr8 | 101720106 | 101720191 | - | miRNA |
| ENSG00000115840.14_6 | SLC25A12 | 0.00260293 | 0.01566606 | -0.754 | 4.396 | chr2 | 172639915 | 172864766 | - | protein_coding |
| ENSG00000251306.1_4 | NDUFB4P2 | 0.13090644 | 0.50751256 | -0.754 | -0.681 | chr5 | 58824687 | 58825077 | + | processed_pseudogene |
| ENSG00000102879.15_3 | CORO1A | 0.06494948 | 0.28063168 | -0.755 | 0.728 | chr16 | 30194148 | 30200397 | + | protein_coding |
| ENSG00000287933.1_1 | AL357141.1 | 0.05923521 | 0.25915587 | -0.756 | 1.029 | chr6 | 116355064 | 116367916 | + | lncRNA |
| ENSG00000136114.17_5 | THSD1 | 0.01779506 | 0.09004422 | -0.756 | 2.436 | chr13 | 52951302 | 52990508 | - | protein_coding |
| ENSG00000163376.11_3 | KBTBD8 | 0.07447067 | 0.31495369 | -0.756 | 0.637 | chr3 | 67048731 | 67061634 | + | protein_coding |
| ENSG00000256618.2_2 | MTRNR2L1 | 0.11218123 | 0.44819516 | -0.756 | -0.344 | chr17 | 22022437 | 22023989 | + | protein_coding |
| ENSG00000287857.1_3 | AL592546.3 | 0.10940853 | 0.43802431 | -0.757 | -0.425 | chr10 | 115625570 | 115629373 | - | lncRNA |
| ENSG00000166780.11_6 | BMERB1 | 0.00110941 | 0.00716792 | -0.757 | 4.561 | chr16 | 15528332 | 15718885 | + | protein_coding |
| ENSG00000279227.1_5 | AC009303.4 | 0.12657121 | 0.49380559 | -0.758 | -0.925 | chr2 | 118771750 | 118773249 | + | TEC |
| ENSG00000100077.15_6 | GRK3 | 0.00402188 | 0.02340542 | -0.758 | 3.866 | chr22 | 25960642 | 26125261 | + | protein_coding |
| ENSG00000118407.15_5 | FILIP1 | 0.11973566 | 0.47200874 | -0.759 | -0.754 | chr6 | 76001575 | 76203516 | - | protein_coding |
| ENSG00000236194.3_8 | AC099811.1 | 0.10150446 | 0.41033669 | -0.759 | -0.268 | chr17 | 40422535 | 40424701 | - | lncRNA |
| ENSG00000164032.12_5 | H2AZ1 | 3.75E-05 | 0.00031174 | -0.759 | 7.615 | chr4 | 100869243 | 100871512 | - | protein_coding |
| ENSG00000241985.1_7 | WWTR1-IT1 | 0.1318105 | 0.5104353 | -0.759 | -0.924 | chr3 | 149366784 | 149367971 | - | lncRNA |
| ENSG00000168917.9_3 | SLC35G2 | 0.00594973 | 0.03334252 | -0.759 | 3.296 | chr3 | 136537489 | 136574730 | + | protein_coding |
| ENSG00000156515.23_7 | HK1 | 0.00035835 | 0.00253856 | -0.759 | 5.313 | chr10 | 71029740 | 71161640 | + | protein_coding |
| ENSG00000251806.1 | SNORD119 | 0.13236939 | 0.51205344 | -0.760 | -0.710 | chr20 | 2443608 | 2443683 | - | snoRNA |
| ENSG00000089685.15_6 | BIRC5 | 2.30E-05 | 0.00019723 | -0.760 | 7.206 | chr17 | 76210267 | 76221717 | + | protein_coding |
| ENSG00000285943.1_3 | AC112128.1 | 0.12476485 | 0.48804428 | -0.760 | -0.893 | chr3 | 113080748 | 113233998 | - | protein_coding |
| ENSG00000175352.11_5 | NRIP3 | 0.06407637 | 0.27746647 | -0.761 | 0.781 | chr11 | 9002123 | 9025596 | - | protein_coding |
| ENSG00000230979.3_2 | AC079250.1 | 0.12849952 | 0.49957353 | -0.761 | -1.082 | chr2 | 47917855 | 47918385 | - | processed_pseudogene |
| ENSG00000164465.18_2 | DCBLD1 | 0.00013319 | 0.00101299 | -0.761 | 5.615 | chr6 | 117774980 | 117891021 | + | protein_coding |
| ENSG00000125657.5_3 | TNFSF9 | 0.01746198 | 0.08846507 | -0.762 | 2.978 | chr19 | 6531037 | 6535935 | + | protein_coding |
| ENSG00000026559.14_6 | KCNG1 | 0.04693934 | 0.21226045 | -0.762 | 1.405 | chr20 | 49620193 | 49639644 | - | protein_coding |
| ENSG00000214975.4_4 | PPIAP29 | 0.0384131 | 0.17838094 | -0.762 | 1.489 | chr6 | 24976647 | 24977210 | + | processed_pseudogene |
| ENSG00000148459.16_4 | PDSS1 | 0.00620483 | 0.0346235 | -0.762 | 3.331 | chr10 | 26986630 | 27035727 | + | protein_coding |
| ENSG00000287299.1_2 | AC012459.1 | 0.03109804 | 0.14808871 | -0.762 | 1.709 | chr2 | 201255389 | 201342741 | - | lncRNA |
| ENSG00000115129.14_4 | TP53I3 | 0.00011529 | 0.00088328 | -0.763 | 6.327 | chr2 | 24300303 | 24308731 | - | protein_coding |
| ENSG00000115514.12_5 | TXNDC9 | 0.00040842 | 0.00286362 | -0.763 | 5.180 | chr2 | 99935445 | 99957165 | - | protein_coding |
| ENSG00000143727.16_3 | ACP1 | 4.76E-05 | 0.00039017 | -0.763 | 6.955 | chr2 | 264140 | 278283 | + | protein_coding |
| ENSG00000104953.20_4 | TLE6 | 0.03631772 | 0.1700282 | -0.763 | 1.412 | chr19 | 2977536 | 2995177 | + | protein_coding |
| ENSG00000288066.1_1 | AC097448.1 | 0.00820656 | 0.04468463 | -0.763 | 3.322 | chr2 | 152653202 | 152655202 | + | lncRNA |
| ENSG00000144061.12_6 | NPHP1 | 0.01025018 | 0.05470559 | -0.763 | 2.971 | chr2 | 110879888 | 110962643 | - | protein_coding |
| ENSG00000227471.9_3 | AKR1B15 | 0.00086927 | 0.00571588 | -0.763 | 4.496 | chr7 | 134233862 | 134264627 | + | protein_coding |
| ENSG00000108370.17_5 | RGS9 | 0.00436587 | 0.02522552 | -0.764 | 3.666 | chr17 | 63096930 | 63223821 | + | protein_coding |
| ENSG00000236065.2_7 | AL020995.1 | 0.14454728 | 0.55158325 | -0.764 | -1.118 | chr1 | 33452676 | 33498070 | + | lncRNA |
| ENSG00000224490.5_5 | TTC21B-AS1 | 0.09868152 | 0.40065691 | -0.764 | -0.345 | chr2 | 166790367 | 166806401 | + | lncRNA |
| ENSG00000182247.10_4 | UBE2E2 | 0.00331497 | 0.01951527 | -0.765 | 3.784 | chr3 | 23244511 | 23633285 | + | protein_coding |
| ENSG00000163689.20_6 | C3orf67 | 0.02585752 | 0.12524825 | -0.766 | 1.820 | chr3 | 58703092 | 59035810 | - | protein_coding |
| ENSG00000276248.1_7 | AL442125.1 | 0.10099492 | 0.40855572 | -0.766 | -0.389 | chr13 | 114181575 | 114184936 | + | lncRNA |
| ENSG00000115866.11_4 | DARS1 | 8.89E-05 | 0.000694 | -0.766 | 6.558 | chr2 | 136663451 | 136743670 | - | protein_coding |
| ENSG00000182004.13_3 | SNRPE | 8.39E-05 | 0.00065625 | -0.767 | 6.279 | chr1 | 203830727 | 203840280 | + | protein_coding |
| ENSG00000234936.1_6 | AC010883.1 | 0.11630362 | 0.46134546 | -0.767 | -0.709 | chr2 | 43456712 | 43460533 | + | lncRNA |
| ENSG00000164406.8_3 | LEAP2 | 0.05642498 | 0.24911082 | -0.767 | 0.883 | chr5 | 132209136 | 132210738 | + | protein_coding |
| ENSG00000015479.19_7 | MATR3 | 0.01046758 | 0.05574005 | -0.767 | 3.030 | chr5 | 138629363 | 138667366 | + | protein_coding |
| ENSG00000224837.1_4 | GCSHP5 | 0.08450865 | 0.35204342 | -0.767 | 0.219 | chr1 | 168025139 | 168025660 | - | processed_pseudogene |
| ENSG00000249502.3_8 | AC006160.1 | 0.09047108 | 0.373293 | -0.768 | -0.177 | chr4 | 17587890 | 17616208 | - | lncRNA |
| ENSG00000071054.16_7 | MAP4K4 | 3.22E-05 | 0.00027086 | -0.768 | 6.546 | chr2 | 102313312 | 102511151 | + | protein_coding |
| ENSG00000082898.17_4 | XPO1 | 9.19E-06 | 8.29E-05 | -0.768 | 8.688 | chr2 | 61704984 | 61765761 | - | protein_coding |
| ENSG00000155100.11_4 | OTUD6B | 0.0018238 | 0.01129827 | -0.769 | 4.839 | chr8 | 92082424 | 92099323 | + | protein_coding |
| ENSG00000115761.16_4 | NOL10 | 0.00012174 | 0.00092909 | -0.769 | 5.815 | chr2 | 10710880 | 10830113 | - | protein_coding |
| ENSG00000188710.3_4 | QRFP | 0.15355988 | 0.57851958 | -0.769 | -1.300 | chr9 | 133768094 | 133772199 | - | protein_coding |
| ENSG00000255317.1_6 | AP002833.2 | 0.12375902 | 0.48485702 | -0.769 | -0.783 | chr11 | 126862348 | 126865354 | + | lncRNA |
| ENSG00000174529.7_3 | TMEM81 | 0.06292054 | 0.2730604 | -0.769 | 0.717 | chr1 | 205052257 | 205053588 | - | protein_coding |
| ENSG00000236559.1_7 | BCL2L1-AS1 | 0.08856575 | 0.36636732 | -0.770 | -0.029 | chr20 | 30274019 | 30304628 | + | lncRNA |

| | | | | | | | | | | |
|---|---|---|---|---|---|---|---|---|---|---|
| ENSG00000175592.9_5 | FOSL1 | 0.00479297 | 0.02740802 | -0.770 | 3.495 | chr11 | 65659520 | 65668044 | - | protein_coding |
| ENSG00000075336.12_4 | TIMM21 | 0.00159605 | 0.00995683 | -0.770 | 4.279 | chr18 | 71815758 | 71827766 | + | protein_coding |
| ENSG00000231006.1_4 | RPL7P32 | 0.13033545 | 0.5057152 | -0.771 | -0.991 | chr7 | 108150677 | 108151420 | + | processed_pseudogene |
| ENSG00000259209.3_6 | AC004943.1 | 0.09047108 | 0.373293 | -0.771 | -0.178 | chr16 | 72839897 | 72843771 | + | lncRNA |
| ENSG00000261335.1_6 | AC005837.1 | 0.11318013 | 0.45147537 | -0.772 | -0.845 | chr17 | 74668024 | 74669740 | + | lncRNA |
| ENSG00000241634.1_6 | AC069499.2 | 0.09296681 | 0.38084882 | -0.772 | -0.100 | chr3 | 108262214 | 108262722 | - | processed_pseudogene |
| ENSG00000169627.9_5 | BOLA2B | 0.11940829 | 0.47103178 | -0.773 | -0.960 | chr16 | 30204253 | 30205627 | - | protein_coding |
| ENSG00000267100.1_5 | ILF3-DT | 0.01093256 | 0.05782636 | -0.773 | 2.567 | chr19 | 10762538 | 10764520 | - | lncRNA |
| ENSG00000128708.13_4 | HAT1 | 0.00123493 | 0.0079181 | -0.773 | 5.791 | chr2 | 172778958 | 172848600 | + | protein_coding |
| ENSG00000138385.16_3 | SSB | 0.00010486 | 0.00080756 | -0.773 | 6.513 | chr2 | 170648443 | 170668574 | + | protein_coding |
| ENSG00000168389.17_3 | MFSD2A | 0.03208953 | 0.15211703 | -0.774 | 1.632 | chr1 | 40420784 | 40435640 | + | protein_coding |
| ENSG00000138376.11_5 | BARD1 | 0.00027485 | 0.00198832 | -0.774 | 4.978 | chr2 | 215590370 | 215674407 | - | protein_coding |
| ENSG00000088538.13_3 | DOCK3 | 0.03702427 | 0.17278965 | -0.775 | 1.315 | chr3 | 50712358 | 51421629 | + | protein_coding |
| ENSG00000128654.14_4 | MTX2 | 0.00079768 | 0.0052764 | -0.776 | 4.797 | chr2 | 177134123 | 177202753 | + | protein_coding |
| ENSG00000148655.15_9 | LRMDA | 0.05814597 | 0.25527043 | -0.776 | 0.549 | chr10 | 77191382 | 78319926 | + | protein_coding |
| ENSG00000266900.1_7 | AC027514.1 | 0.132206 | 0.51185532 | -0.776 | -1.157 | chr18 | 59913043 | 59914645 | + | lncRNA |
| ENSG00000237424.1_5 | FOXD2-AS1 | 0.05967722 | 0.26070451 | -0.777 | 0.614 | chr1 | 47897805 | 47900313 | - | lncRNA |
| ENSG00000239332.6_6 | LINC01119 | 0.11100267 | 0.4437303 | -0.777 | -0.351 | chr2 | 47043836 | 47086146 | + | lncRNA |
| ENSG00000101132.10_3 | PFDN4 | 0.0003072 | 0.00220345 | -0.777 | 5.013 | chr20 | 52824626 | 52844591 | + | protein_coding |
| ENSG00000153944.11_7 | MSI2 | 4.93E-05 | 0.00040219 | -0.778 | 5.991 | chr17 | 55333212 | 55762050 | + | protein_coding |
| ENSG00000082258.13_4 | CCNT2 | 0.00017662 | 0.00131846 | -0.778 | 5.318 | chr2 | 135675805 | 135716912 | + | protein_coding |
| ENSG00000287110.1_2 | AC112484.5 | 0.07036736 | 0.30089454 | -0.779 | 0.188 | chr3 | 128591192 | 128595061 | - | lncRNA |
| ENSG00000119737.6_5 | GPR75 | 0.05260574 | 0.23442341 | -0.780 | 0.713 | chr2 | 54080049 | 54087104 | - | protein_coding |
| ENSG00000130826.18_4 | DKC1 | 6.08E-05 | 0.0004881 | -0.780 | 6.254 | chrX | 153991017 | 154005964 | + | protein_coding |
| ENSG00000085840.13_4 | ORC1 | 0.00017114 | 0.00127972 | -0.780 | 5.168 | chr1 | 52838501 | 52870095 | - | protein_coding |
| ENSG00000171208.10_5 | NETO2 | 6.04E-06 | 5.59E-05 | -0.780 | 7.328 | chr16 | 47111614 | 47177856 | - | protein_coding |
| ENSG00000197442.10_4 | MAP3K5 | 0.00398617 | 0.02321282 | -0.780 | 3.192 | chr6 | 136878184 | 137113615 | - | protein_coding |
| ENSG00000166788.10_8 | SAAL1 | 0.00018551 | 0.00137851 | -0.781 | 5.089 | chr11 | 18091482 | 18127634 | - | protein_coding |
| ENSG00000110321.17_9 | EIF4G2 | 3.87E-06 | 3.67E-05 | -0.781 | 10.039 | chr11 | 10818597 | 10830487 | - | protein_coding |
| ENSG00000249216.1_7 | AC105389.2 | 0.09256602 | 0.37964862 | -0.784 | -0.292 | chr4 | 41690875 | 41694814 | - | lncRNA |
| ENSG00000196890.4_3 | H2BU1 | 0.00162813 | 0.01014629 | -0.785 | 4.729 | chr1 | 228645808 | 228648171 | + | protein_coding |
| ENSG00000136536.15_5 | MARCHF7 | 2.71E-05 | 0.00022999 | -0.786 | 6.817 | chr2 | 160568968 | 160627538 | + | protein_coding |
| ENSG00000136682.15_5 | CBWD2 | 0.00167781 | 0.01044851 | -0.786 | 3.912 | chr2 | 114195268 | 114253781 | + | protein_coding |
| ENSG00000229447.2_5 | AC114495.2 | 0.12151568 | 0.47764804 | -0.786 | -0.995 | chr1 | 31736092 | 31736528 | - | processed_pseudogene |
| ENSG00000092140.16_6 | G2E3 | 3.83E-05 | 0.00031807 | -0.786 | 5.658 | chr14 | 31028364 | 31089270 | + | protein_coding |
| ENSG00000134962.7_3 | KLB | 0.01804897 | 0.09110118 | -0.786 | 2.120 | chr4 | 39408550 | 39453153 | + | protein_coding |
| ENSG00000143515.18_5 | ATP8B2 | 4.28E-05 | 0.0003524 | -0.786 | 5.701 | chr1 | 154298029 | 154323780 | + | protein_coding |
| ENSG00000115970.18_4 | THADA | 1.03E-05 | 9.23E-05 | -0.787 | 6.451 | chr2 | 43457975 | 43823185 | - | protein_coding |
| ENSG00000228043.5_7 | AC114763.1 | 0.08568991 | 0.35637938 | -0.787 | -0.377 | chr2 | 139175854 | 139259268 | - | lncRNA |
| ENSG00000117395.12_5 | EBNA1BP2 | 9.80E-06 | 8.79E-05 | -0.787 | 6.542 | chr1 | 43629846 | 43736607 | - | protein_coding |
| ENSG00000287600.1_2 | AC122719.3 | 0.04684332 | 0.21195542 | -0.788 | 0.691 | chr5 | 904434 | 912702 | - | lncRNA |
| ENSG00000115942.9_4 | ORC2 | 0.00010775 | 0.00082838 | -0.788 | 5.782 | chr2 | 201773700 | 201828403 | - | protein_coding |
| ENSG00000135740.17_6 | SLC9A5 | 0.01023605 | 0.05464658 | -0.788 | 2.283 | chr16 | 67271586 | 67306094 | + | protein_coding |
| ENSG00000134954.14_7 | ETS1 | 0.00023211 | 0.00169776 | -0.788 | 4.743 | chr11 | 128328656 | 128457453 | - | protein_coding |
| ENSG00000188060.7_3 | RAB42 | 0.07185887 | 0.30587364 | -0.789 | -0.052 | chr1 | 28918712 | 28921955 | + | protein_coding |
| ENSG00000230992.3_4 | FAM201B | 0.15532104 | 0.58364798 | -0.789 | -1.198 | chr2 | 133110295 | 133110891 | + | processed_pseudogene |
| ENSG00000178752.16_7 | ERFE | 0.02571783 | 0.12467359 | -0.790 | 1.624 | chr2 | 239067611 | 239077541 | + | protein_coding |
| ENSG00000264044.1_6 | AC005726.2 | 0.08578481 | 0.35669055 | -0.791 | -0.239 | chr17 | 26960224 | 26962968 | + | lncRNA |
| ENSG00000247627.2_4 | MTND4P12 | 0.11863879 | 0.46914207 | -0.791 | -0.721 | chr5 | 134262350 | 134263726 | - | processed_pseudogene |
| ENSG00000188549.12_7 | CCDC9B | 0.00132247 | 0.00840357 | -0.791 | 3.773 | chr15 | 40623653 | 40633168 | - | protein_coding |
| ENSG00000088543.15_4 | C3orf18 | 0.00618796 | 0.03455109 | -0.791 | 2.611 | chr3 | 50595456 | 50608458 | - | protein_coding |
| ENSG00000150995.19_7 | ITPR1 | 0.00119239 | 0.00766748 | -0.791 | 3.921 | chr3 | 4535029 | 4889524 | + | protein_coding |
| ENSG00000166347.19_5 | CYB5A | 0.00010011 | 0.00077398 | -0.791 | 4.948 | chr18 | 71918081 | 71959208 | - | protein_coding |
| ENSG00000178295.15_5 | GEN1 | 3.33E-05 | 0.00027894 | -0.792 | 6.540 | chr2 | 17935125 | 17970213 | + | protein_coding |
| ENSG00000000460.17_6 | C1orf112 | 0.00038718 | 0.00272548 | -0.792 | 4.613 | chr1 | 169631245 | 169823221 | + | protein_coding |
| ENSG00000287515.1_2 | AL929602.1 | 0.08458214 | 0.35226696 | -0.792 | -0.355 | chr14 | 19912365 | 19915373 | - | transcribed_unprocessed_pseudogene |
| ENSG00000157827.20_4 | FMNL2 | 2.21E-05 | 0.00018999 | -0.793 | 5.901 | chr2 | 153191688 | 153506340 | + | protein_coding |
| ENSG00000123213.23_6 | NLN | 2.47E-06 | 2.41E-05 | -0.793 | 7.455 | chr5 | 65018032 | 65167553 | + | protein_coding |

| | | | | | | | | | | |
|---|---|---|---|---|---|---|---|---|---|---|
| ENSG00000287678.1_2 | AL031778.1 | 0.09325606 | 0.38182584 | -0.795 | -0.571 | chr6 | 41048363 | 41068795 | - | lncRNA |
| ENSG00000138031.14_4 | ADCY3 | 5.27E-05 | 0.00042758 | -0.795 | 5.720 | chr2 | 25042038 | 25142708 | - | protein_coding |
| ENSG00000165304.8_4 | MELK | 3.94E-06 | 3.74E-05 | -0.795 | 6.596 | chr9 | 36572859 | 36677680 | + | protein_coding |
| ENSG00000259118.5_6 | AL139022.1 | 0.09949853 | 0.40329655 | -0.795 | -0.798 | chr14 | 65548752 | 65560930 | + | lncRNA |
| ENSG00000138678.11_3 | GPAT3 | 5.66E-05 | 0.00045662 | -0.795 | 5.283 | chr4 | 84457067 | 84527028 | + | protein_coding |
| ENSG00000198053.11_3 | SIRPA | 0.00016851 | 0.0012622 | -0.796 | 5.081 | chr20 | 1874813 | 1921238 | + | protein_coding |
| ENSG00000152270.9_3 | PDE3B | 0.00227007 | 0.01380774 | -0.796 | 3.472 | chr11 | 14665350 | 14893590 | + | protein_coding |
| ENSG00000176273.15_4 | SLC35G1 | 0.00073583 | 0.00490194 | -0.796 | 3.939 | chr10 | 95653730 | 95715819 | + | protein_coding |
| ENSG00000212607.1 | SNORA45 | 0.09661342 | 0.39333884 | -0.797 | -0.574 | chr11 | 8706986 | 8707116 | + | snoRNA |
| ENSG00000100206.10_4 | DMC1 | 0.03969504 | 0.18361411 | -0.797 | 1.039 | chr22 | 38914953 | 38966209 | - | protein_coding |
| ENSG00000253385.1_6 | AP003696.1 | 0.09206088 | 0.37800917 | -0.799 | -0.464 | chr8 | 103866683 | 103868303 | + | lncRNA |
| ENSG00000224631.4_4 | RPS27AP16 | 0.00173476 | 0.01076547 | -0.799 | 3.420 | chr16 | 61089303 | 61089868 | - | transcribed_processed_pseudogene |
| ENSG00000276672.1_7 | AL161891.1 | 0.03858167 | 0.17911693 | -0.799 | 1.168 | chr13 | 34420327 | 34424962 | + | lncRNA |
| ENSG00000135972.9_4 | MRPS9 | 0.00072033 | 0.00481311 | -0.799 | 4.043 | chr2 | 105654527 | 105716418 | + | protein_coding |
| ENSG00000074582.14_6 | BCS1L | 0.00044972 | 0.00312607 | -0.799 | 4.371 | chr2 | 219523487 | 219528166 | + | protein_coding |
| ENSG00000242265.5_4 | PEG10 | 1.10E-05 | 9.85E-05 | -0.799 | 6.691 | chr7 | 94285637 | 94299007 | + | protein_coding |
| ENSG00000283196.2_6 | AC006453.3 | 0.07496529 | 0.31689467 | -0.800 | 0.046 | chr2 | 91805186 | 91848037 | - | transcribed_unprocessed_pseudogene |
| ENSG00000168811.7_4 | IL12A | 0.09505578 | 0.38797518 | -0.800 | -0.723 | chr3 | 159706622 | 159713806 | + | protein_coding |
| ENSG00000085449.15_6 | WDFY1 | 0.00040869 | 0.00286436 | -0.800 | 5.727 | chr2 | 224720433 | 224810074 | - | protein_coding |
| ENSG00000253298.2_6 | LINC01845 | 0.09594468 | 0.39097459 | -0.800 | -0.798 | chr5 | 158875564 | 158893299 | - | lncRNA |
| ENSG00000279865.1_6 | AC006511.4 | 0.07816787 | 0.32910156 | -0.800 | -0.088 | chr12 | 8232977 | 8234759 | - | TEC |
| ENSG00000166387.13_9 | PPFIBP2 | 3.60E-05 | 0.00030033 | -0.801 | 5.278 | chr11 | 7534529 | 7678358 | + | protein_coding |
| ENSG00000234945.8_9 | GTF3C2-AS1 | 0.08243179 | 0.3448471 | -0.801 | -0.425 | chr2 | 27558387 | 27565466 | + | lncRNA |
| ENSG00000163072.15_5 | NOSTRIN | 0.00207538 | 0.01274131 | -0.802 | 3.334 | chr2 | 169643049 | 169722024 | + | protein_coding |
| ENSG00000278879.1_6 | AP000560.1 | 0.06676732 | 0.28729835 | -0.802 | 0.024 | chr11 | 74541619 | 74543555 | - | TEC |
| ENSG00000231999.7_7 | LRRC8C-DT | 0.03595468 | 0.16841731 | -0.802 | 1.133 | chr1 | 90046850 | 90098475 | - | lncRNA |
| ENSG00000167994.13_7 | RAB3IL1 | 0.01439479 | 0.07421711 | -0.802 | 2.076 | chr11 | 61664773 | 61687741 | - | protein_coding |
| ENSG00000155760.2_2 | FZD7 | 0.00316387 | 0.01869385 | -0.802 | 3.137 | chr2 | 202899310 | 202903168 | + | protein_coding |
| ENSG00000206737.1 | RNVU1-18 | 0.0586427 | 0.25719696 | -0.803 | 0.418 | chr1 | 149224058 | 149224221 | - | snRNA |
| ENSG00000273486.1_6 | AC096992.2 | 0.11440997 | 0.45525688 | -0.803 | -0.941 | chr3 | 136556180 | 136557863 | - | lncRNA |
| ENSG00000202515.1 | VTRNA1-3 | 0.00887011 | 0.04788671 | -0.804 | 2.799 | chr5 | 140105743 | 140105831 | + | misc_RNA |
| ENSG00000143429.10_7 | LSP1P4 | 0.0567996 | 0.25042451 | -0.804 | 0.692 | chr2 | 91805186 | 91847998 | - | transcribed_unprocessed_pseudogene |
| ENSG00000232956.9_4 | SNHG15 | 0.00057151 | 0.0038964 | -0.804 | 4.550 | chr7 | 45022622 | 45026560 | - | lncRNA |
| ENSG00000249684.6_7 | AC106795.2 | 0.10368235 | 0.41837937 | -0.804 | -0.943 | chr5 | 177377336 | 177390961 | + | lncRNA |
| ENSG00000236814.1_5 | AC046176.1 | 0.05012367 | 0.22477214 | -0.804 | 0.744 | chr8 | 56962597 | 56963766 | - | processed_pseudogene |
| ENSG00000258450.1_8 | AL139099.3 | 0.11292745 | 0.45066983 | -0.805 | -0.940 | chr14 | 50096009 | 50096608 | - | lncRNA |
| ENSG00000266993.3_8 | AL050343.2 | 0.07341185 | 0.31123292 | -0.805 | -0.344 | chr1 | 52259606 | 52264826 | + | lncRNA |
| ENSG00000196756.13_8 | SNHG17 | 3.08E-05 | 0.00026037 | -0.806 | 5.455 | chr20 | 37048285 | 37064052 | - | lncRNA |
| ENSG00000144724.20_4 | PTPRG | 0.0005038 | 0.00346405 | -0.806 | 4.298 | chr3 | 61547243 | 62283284 | + | protein_coding |
| ENSG00000123685.9_4 | BATF3 | 0.01033362 | 0.05510126 | -0.806 | 2.120 | chr1 | 212859759 | 212873182 | - | protein_coding |
| ENSG00000197299.12_4 | BLM | 8.26E-06 | 7.50E-05 | -0.806 | 5.917 | chr15 | 91260577 | 91359396 | + | protein_coding |
| ENSG00000135913.11_4 | USP37 | 0.00012822 | 0.00097732 | -0.806 | 5.042 | chr2 | 219314974 | 219433074 | - | protein_coding |
| ENSG00000175701.10_7 | MTLN | 0.00068283 | 0.00459541 | -0.807 | 3.832 | chr2 | 110969106 | 111002997 | - | protein_coding |
| ENSG00000182308.7_4 | DCAF4L1 | 0.11589843 | 0.46014914 | -0.807 | -1.065 | chr4 | 41983773 | 41988482 | + | protein_coding |
| ENSG00000236867.1_5 | Z98885.1 | 0.13654476 | 0.52624527 | -0.807 | -1.138 | chr22 | 50210896 | 50211360 | + | processed_pseudogene |
| ENSG00000203814.6_4 | H2BC18 | 1.46E-06 | 1.48E-05 | -0.808 | 8.074 | chr1 | 149754245 | 149783928 | - | protein_coding |
| ENSG00000263535.2_4 | AK4P1 | 0.10678361 | 0.42894515 | -0.808 | -0.942 | chr17 | 29672537 | 29673208 | + | processed_pseudogene |
| ENSG00000131067.17_4 | GGT7 | 0.01580239 | 0.08065529 | -0.808 | 1.947 | chr20 | 33432523 | 33460659 | - | protein_coding |
| ENSG00000138375.13_8 | SMARCAL1 | 0.00030855 | 0.00221137 | -0.808 | 4.091 | chr2 | 217277137 | 217347776 | + | protein_coding |
| ENSG00000145911.6_5 | N4BP3 | 0.07751314 | 0.3265772 | -0.809 | -0.343 | chr5 | 177540533 | 177553082 | + | protein_coding |
| ENSG00000201302.1 | SNORA65 | 0.04861041 | 0.21873114 | -0.809 | 0.508 | chr9 | 130210780 | 130210909 | - | snoRNA |
| ENSG00000228203.7_6 | RNF144A-AS1 | 0.04510401 | 0.20516204 | -0.810 | 0.680 | chr2 | 7051885 | 7058865 | - | lncRNA |
| ENSG00000239382.10_7 | ALKBH6 | 0.01446396 | 0.07450888 | -0.811 | 1.622 | chr19 | 36500022 | 36505141 | - | protein_coding |
| ENSG00000224411.3_4 | HSP90AA2P | 0.03681198 | 0.17202485 | -0.811 | 1.191 | chr11 | 27910385 | 27912580 | - | processed_pseudogene |
| ENSG00000137821.11_6 | LRRC49 | 0.00051546 | 0.00353597 | -0.811 | 4.168 | chr15 | 71145578 | 71345996 | + | protein_coding |
| ENSG00000260536.1_7 | AL035071.2 | 0.08852707 | 0.36631704 | -0.811 | -0.627 | chr20 | 31430934 | 31442063 | - | lncRNA |

| | | | | | | | | | | |
|---|---|---|---|---|---|---|---|---|---|---|
| ENSG00000108055.9_2 | SMC3 | 1.82E-06 | 1.81E-05 | -0.811 | 7.042 | chr10 | 112327449 | 112364394 | + | protein_coding |
| ENSG00000006607.14_6 | FARP2 | 2.09E-05 | 0.00018058 | -0.811 | 5.474 | chr2 | 242295700 | 242434256 | + | protein_coding |
| ENSG00000271936.1_6 | AC012073.1 | 0.02277601 | 0.11209458 | -0.812 | 1.707 | chr2 | 25048479 | 25049586 | + | lncRNA |
| ENSG00000119900.9_4 | OGFRL1 | 0.00038992 | 0.00274363 | -0.812 | 4.735 | chr6 | 71998514 | 72018762 | + | protein_coding |
| ENSG00000111981.5_3 | ULBP1 | 0.0230698 | 0.11332076 | -0.812 | 1.488 | chr6 | 150285079 | 150294851 | + | protein_coding |
| ENSG00000262703.1_8 | AC009121.3 | 0.07055518 | 0.30145274 | -0.813 | -0.249 | chr16 | 11442000 | 11443178 | - | lncRNA |
| ENSG00000080854.15_6 | IGSF9B | 0.04202311 | 0.19302599 | -0.813 | 0.492 | chr11 | 133766333 | 133826880 | - | protein_coding |
| ENSG00000135624.16_3 | CCT7 | 3.05E-07 | 3.37E-06 | -0.813 | 8.958 | chr2 | 73460548 | 73480149 | + | protein_coding |
| ENSG00000065548.18_4 | ZC3H15 | 7.30E-06 | 6.70E-05 | -0.813 | 6.752 | chr2 | 187350980 | 187374088 | + | protein_coding |
| ENSG00000224429.8_4 | LINC00539 | 0.08094112 | 0.3390101 | -0.814 | -0.368 | chr13 | 21877001 | 21922860 | - | lncRNA |
| ENSG00000161509.14_4 | GRIN2C | 0.0506097 | 0.22667627 | -0.815 | 0.380 | chr17 | 72838162 | 72857627 | - | protein_coding |
| ENSG00000128266.9_3 | GNAZ | 0.00719768 | 0.03962852 | -0.815 | 2.471 | chr22 | 23412705 | 23467219 | + | protein_coding |
| ENSG00000236924.1_7 | AL162411.1 | 0.07727212 | 0.32571618 | -0.815 | -0.239 | chr9 | 6645956 | 6670635 | + | lncRNA |
| ENSG00000114767.7_4 | RRP9 | 0.00010057 | 0.00077657 | -0.816 | 4.655 | chr3 | 51967445 | 51975920 | - | protein_coding |
| ENSG00000138095.19_4 | LRPPRC | 2.57E-07 | 2.87E-06 | -0.816 | 8.615 | chr2 | 44113363 | 44223128 | - | protein_coding |
| ENSG00000276517.1_6 | AL133243.3 | 0.02600546 | 0.12582763 | -0.816 | 1.164 | chr2 | 32751571 | 32754574 | + | lncRNA |
| ENSG00000255893.1_8 | AP000786.1 | 0.01975332 | 0.09863889 | -0.816 | 1.360 | chr11 | 94206074 | 94206736 | - | lncRNA |
| ENSG00000138363.15_5 | ATIC | 0.000233 | 0.00170359 | -0.817 | 4.231 | chr2 | 216176679 | 216214496 | + | protein_coding |
| ENSG00000123607.15_4 | TTC21B | 6.87E-05 | 0.00054587 | -0.818 | 4.843 | chr2 | 166713985 | 166810326 | - | protein_coding |
| ENSG00000223477.3 | LINC00842 | 4.11E-05 | 0.00034001 | -0.818 | 4.988 | chr10 | 47011753 | 47174018 | - | lincRNA |
| ENSG00000184058.14_4 | TBX1 | 0.03000335 | 0.14314433 | -0.818 | 1.044 | chr22 | 19744226 | 19771116 | + | protein_coding |
| ENSG00000175322.11_4 | ZNF519 | 0.00541826 | 0.03063852 | -0.818 | 2.333 | chr18 | 14057456 | 14132489 | - | protein_coding |
| ENSG00000162976.13_4 | SLC66A3 | 2.94E-05 | 0.00024915 | -0.819 | 5.135 | chr2 | 11295324 | 11318996 | + | protein_coding |
| ENSG00000230732.4_7 | AC016949.1 | 0.08529087 | 0.35480285 | -0.819 | -0.627 | chr3 | 196639517 | 196641450 | + | lncRNA |
| ENSG00000239388.9_6 | ASB14 | 0.06873641 | 0.29484302 | -0.820 | -0.332 | chr3 | 57302370 | 57326713 | - | protein_coding |
| ENSG00000173145.11_4 | NOC3L | 7.43E-06 | 6.79E-05 | -0.820 | 5.825 | chr10 | 96092983 | 96122716 | - | protein_coding |
| ENSG00000153201.16_5 | RANBP2 | 4.20E-06 | 3.97E-05 | -0.820 | 7.867 | chr2 | 109335938 | 109402265 | + | protein_coding |
| ENSG00000104808.8_3 | DHDH | 0.08417449 | 0.35098057 | -0.820 | -0.753 | chr19 | 49436956 | 49448226 | + | protein_coding |
| ENSG00000143379.12_5 | SETDB1 | 4.51E-05 | 0.00037099 | -0.820 | 4.902 | chr1 | 150898739 | 150937220 | + | protein_coding |
| ENSG00000180611.7_4 | MB21D2 | 0.002686 | 0.01611689 | -0.821 | 3.018 | chr3 | 192514604 | 192635645 | - | protein_coding |
| ENSG00000254634.4_5 | SMG1P6 | 0.06309567 | 0.27368671 | -0.821 | 0.043 | chr16 | 29437121 | 29458347 | - | unprocessed_pseudogene |
| ENSG00000254907.1_7 | AC087276.3 | 0.0937121 | 0.3830785 | -0.822 | -0.781 | chr11 | 43350298 | 43380846 | - | lncRNA |
| ENSG00000141384.12_3 | TAF4B | 0.00072208 | 0.00482298 | -0.822 | 3.744 | chr18 | 23805900 | 23971649 | + | protein_coding |
| ENSG00000119041.11_7 | GTF3C3 | 4.48E-06 | 4.23E-05 | -0.822 | 5.927 | chr2 | 197627759 | 197664449 | - | protein_coding |
| ENSG00000281404.2_4 | LINC01176 | 0.06591899 | 0.28419868 | -0.823 | -0.235 | chr7 | 30430501 | 30451991 | + | lncRNA |
| ENSG00000247134.7_8 | AC090204.1 | 0.00095326 | 0.00622436 | -0.823 | 3.515 | chr8 | 32785431 | 32902963 | + | lncRNA |
| ENSG00000148798.11_4 | INA | 3.54E-05 | 0.00029575 | -0.823 | 5.064 | chr10 | 105036895 | 105050103 | + | protein_coding |
| ENSG00000118496.5_4 | FBXO30 | 6.16E-05 | 0.00049372 | -0.823 | 4.785 | chr6 | 146114638 | 146135931 | - | protein_coding |
| ENSG00000165912.16_5 | PACSIN3 | 0.00022857 | 0.00167324 | -0.824 | 4.390 | chr11 | 47199073 | 47207994 | - | protein_coding |
| ENSG00000059804.16_5 | SLC2A3 | 0.0021964 | 0.01341015 | -0.824 | 3.032 | chr12 | 8071826 | 8088783 | - | protein_coding |
| ENSG00000105261.7_3 | OVOL3 | 0.04173889 | 0.1919281 | -0.824 | 0.526 | chr19 | 36602053 | 36604613 | + | protein_coding |
| ENSG00000168675.18_5 | LDLRAD4 | 0.02952388 | 0.14100905 | -0.825 | 0.899 | chr18 | 13217497 | 13652754 | + | protein_coding |
| ENSG00000162630.6_4 | B3GALT2 | 0.09683963 | 0.3940792 | -0.825 | -0.890 | chr1 | 193147860 | 193155743 | - | protein_coding |
| ENSG00000235082.2_4 | SUMO1P3 | 0.05545223 | 0.24562944 | -0.825 | -0.037 | chr1 | 160287193 | 160287496 | + | processed_pseudogene |
| ENSG00000147813.16_6 | NAPRT | 0.00488516 | 0.02788135 | -0.826 | 3.216 | chr8 | 144656955 | 144660819 | - | protein_coding |
| ENSG00000037280.16_6 | FLT4 | 0.00403985 | 0.0234946 | -0.826 | 2.554 | chr5 | 180028506 | 180076624 | - | protein_coding |
| ENSG00000253875.2_6 | AC013643.2 | 0.07113865 | 0.30346229 | -0.827 | -0.412 | chr8 | 27590432 | 27616069 | + | lncRNA |
| ENSG00000079246.16_5 | XRCC5 | 1.47E-07 | 1.71E-06 | -0.827 | 9.052 | chr2 | 216972187 | 217071026 | + | protein_coding |
| ENSG00000279145.2_6 | AC011912.1 | 0.04909726 | 0.22067064 | -0.828 | 0.303 | chr15 | 55463576 | 55468937 | - | lncRNA |
| ENSG00000138448.12_4 | ITGAV | 3.58E-07 | 3.90E-06 | -0.828 | 8.140 | chr2 | 187454783 | 187545628 | + | protein_coding |
| ENSG00000162769.13_3 | FLVCR1 | 2.19E-05 | 0.00018846 | -0.828 | 4.965 | chr1 | 213031617 | 213072705 | + | protein_coding |
| ENSG00000131480.9_3 | AOC2 | 0.0180228 | 0.09099492 | -0.828 | 1.358 | chr17 | 40996597 | 41002724 | + | protein_coding |
| ENSG00000256196.2_7 | AP003721.1 | 0.08733887 | 0.36213531 | -0.828 | -0.893 | chr11 | 60683811 | 60692869 | - | lncRNA |
| ENSG00000265791.1_8 | AC127024.4 | 0.09119772 | 0.3755883 | -0.828 | -0.629 | chr17 | 29108511 | 29109239 | - | lncRNA |
| ENSG00000171217.5_3 | CLDN20 | 0.07717967 | 0.32548095 | -0.828 | -0.498 | chr6 | 155585147 | 155597682 | + | protein_coding |
| ENSG00000144580.13_7 | CNOT9 | 1.34E-06 | 1.37E-05 | -0.829 | 6.173 | chr2 | 219433303 | 219461803 | + | protein_coding |
| ENSG00000239462.1_7 | AC091212.1 | 0.08711961 | 0.36147926 | -0.829 | -0.892 | chr3 | 98621268 | 98737406 | + | lncRNA |
| ENSG00000177076.6_3 | ACER2 | 0.0165831 | 0.08425278 | -0.829 | 1.446 | chr9 | 19409007 | 19452503 | + | protein_coding |
| ENSG00000241217.2 | RN7SL809P | 0.07180566 | 0.30572038 | -0.830 | -0.217 | chr12 | 57064236 | 57064530 | - | misc_RNA |
| ENSG00000138035.15_4 | PNPT1 | 9.75E-07 | 1.01E-05 | -0.831 | 6.369 | chr2 | 55861196 | 55920998 | - | protein_coding |

| | | | | | | | | | | |
|---|---|---|---|---|---|---|---|---|---|---|
| ENSG00000197744.5_4 | PTMAP2 | 0.00312305 | 0.01847725 | -0.831 | 2.963 | chr5 | 118309491 | 118309817 | + | processed_pseudogene |
| ENSG00000273245.1_6 | AC092653.1 | 0.1156471 | 0.45956209 | -0.832 | -1.177 | chr2 | 73977383 | 73977913 | - | lncRNA |
| ENSG00000259644.1_7 | AC022613.3 | 0.04590651 | 0.20827911 | -0.832 | 0.326 | chr15 | 30020390 | 30021735 | - | lncRNA |
| ENSG00000158417.11_4 | EIF5B | 6.25E-05 | 0.00050101 | -0.832 | 5.425 | chr2 | 99953834 | 100017789 | + | protein_coding |
| ENSG00000104213.13_3 | PDGFRL | 0.00185168 | 0.01145503 | -0.833 | 3.231 | chr8 | 17433942 | 17501580 | + | protein_coding |
| ENSG00000244363.3_4 | RPL7P23 | 0.06756369 | 0.2902331 | -0.833 | -0.206 | chr5 | 76878201 | 76878947 | + | processed_pseudogene |
| ENSG00000223404.3_6 | LINC00397 | 0.08492126 | 0.35351358 | -0.834 | -0.709 | chr13 | 88453291 | 88456063 | - | lncRNA |
| ENSG00000100167.20_5 | SEPTIN3 | 0.00995636 | 0.0533297 | -0.835 | 1.718 | chr22 | 42365479 | 42394225 | + | protein_coding |
| ENSG00000154518.9_4 | ATP5MC3 | 1.03E-06 | 1.07E-05 | -0.835 | 6.523 | chr2 | 176040986 | 176049335 | - | protein_coding |
| ENSG00000206262.9_3 | FOXL2NB | 0.00687138 | 0.03796138 | -0.835 | 2.140 | chr3 | 138666059 | 138672832 | + | protein_coding |
| ENSG00000108423.15_5 | TUBD1 | 0.00014945 | 0.00113035 | -0.835 | 4.203 | chr17 | 57936840 | 57970306 | - | protein_coding |
| ENSG00000137563.12_4 | GGH | 5.28E-06 | 4.93E-05 | -0.836 | 5.812 | chr8 | 63927638 | 63951365 | - | protein_coding |
| ENSG00000135974.10_3 | C2orf49 | 4.43E-05 | 0.00036426 | -0.836 | 4.733 | chr2 | 105953989 | 105965668 | + | protein_coding |
| ENSG00000245573.9_8 | BDNF-AS | 0.00543228 | 0.03069826 | -0.837 | 2.544 | chr11 | 27528377 | 27719778 | + | lncRNA |
| ENSG00000005844.18_6 | ITGAL | 0.06642053 | 0.28594466 | -0.838 | -0.259 | chr16 | 30484040 | 30534506 | + | protein_coding |
| ENSG00000163738.18_5 | MTHFD2L | 0.00035632 | 0.00252625 | -0.838 | 3.759 | chr4 | 74979891 | 75168816 | + | protein_coding |
| ENSG00000263266.2_2 | RPS7P1 | 0.03135562 | 0.14919532 | -0.839 | 4.017 | chr17 | 26794840 | 26795424 | + | processed_pseudogene |
| ENSG00000174442.12_4 | ZWILCH | 1.08E-06 | 1.11E-05 | -0.840 | 6.051 | chr15 | 66797297 | 66842468 | + | protein_coding |
| ENSG00000236212.2_7 | AC009262.1 | 0.05911971 | 0.25884175 | -0.840 | -0.378 | chr7 | 34249077 | 34338624 | + | lncRNA |
| ENSG00000267374.2_9 | MIR924HG | 0.01692562 | 0.08589476 | -0.840 | 1.762 | chr18 | 36786881 | 37380286 | - | lncRNA |
| ENSG00000174501.14_4 | ANKRD36C | 5.20E-05 | 0.00042238 | -0.841 | 4.412 | chr2 | 96502667 | 96657579 | - | protein_coding |
| ENSG00000259735.1_8 | AC092868.3 | 0.10564038 | 0.42541084 | -0.841 | -0.844 | chr15 | 59640847 | 59652088 | - | lncRNA |
| ENSG00000095002.15_5 | MSH2 | 2.93E-07 | 3.24E-06 | -0.841 | 6.541 | chr2 | 47630206 | 47890285 | + | protein_coding |
| ENSG00000162997.15_3 | PRORSD1P | 0.01924068 | 0.09638695 | -0.842 | 1.100 | chr2 | 55509455 | 55511658 | + | transcribed_unitary_pseudogene |
| ENSG00000278456.1_6 | AC015712.7 | 0.09919653 | 0.40238008 | -0.842 | -0.926 | chr15 | 101428677 | 101429311 | - | lncRNA |
| ENSG00000270332.2_6 | SMC2-AS1 | 0.07341572 | 0.31123292 | -0.843 | -0.498 | chr9 | 106842305 | 106855354 | - | lncRNA |
| ENSG00000164109.14_4 | MAD2L1 | 2.29E-06 | 2.25E-05 | -0.843 | 5.885 | chr4 | 120976778 | 120988013 | - | protein_coding |
| ENSG00000084774.14_5 | CAD | 3.91E-07 | 4.24E-06 | -0.843 | 7.285 | chr2 | 27440237 | 27466811 | + | protein_coding |
| ENSG00000170417.16_3 | TMEM182 | 0.00759021 | 0.04162199 | -0.843 | 1.870 | chr2 | 103353364 | 103636358 | + | protein_coding |
| ENSG00000164687.11_3 | FABP5 | 0.00196886 | 0.01212923 | -0.844 | 3.169 | chr8 | 82192771 | 82197012 | + | protein_coding |
| ENSG00000237126.8_6 | AC073254.1 | 0.07879095 | 0.33117548 | -0.844 | -0.739 | chr2 | 233445658 | 233476681 | - | lncRNA |
| ENSG00000179292.5_3 | TMEM151A | 0.03525619 | 0.16558201 | -0.844 | 0.521 | chr11 | 66059365 | 66064135 | + | protein_coding |
| ENSG00000161547.16_4 | SRSF2 | 3.88E-07 | 4.21E-06 | -0.844 | 7.549 | chr17 | 74730197 | 74733456 | - | protein_coding |
| ENSG00000163879.11_4 | DNALI1 | 0.06595162 | 0.28427038 | -0.844 | -0.438 | chr1 | 38022520 | 38032458 | + | protein_coding |
| ENSG00000152465.18_5 | NMT2 | 1.36E-05 | 0.00012022 | -0.844 | 4.827 | chr10 | 15144583 | 15210692 | - | protein_coding |
| ENSG00000231416.1_5 | AL358472.1 | 0.09449515 | 0.38586425 | -0.844 | -1.179 | chr1 | 153968108 | 153968436 | + | processed_pseudogene |
| ENSG00000214391.3_4 | TUBAP2 | 0.01848858 | 0.09300301 | -0.844 | 1.277 | chr11 | 90015728 | 90017340 | + | processed_pseudogene |
| ENSG00000186642.16_6 | PDE2A | 0.07024026 | 0.30042338 | -0.845 | -0.640 | chr11 | 72287185 | 72385635 | - | protein_coding |
| ENSG00000138758.11_8 | SEPTIN11 | 2.06E-06 | 2.04E-05 | -0.845 | 6.068 | chr4 | 77870856 | 77961537 | + | protein_coding |
| ENSG00000272054.1_7 | AC007390.2 | 0.01585097 | 0.08085678 | -0.845 | 1.731 | chr2 | 37436018 | 37439820 | + | lncRNA |
| ENSG00000235530.7_6 | AC087294.1 | 0.01969167 | 0.09837049 | -0.846 | 1.202 | chr17 | 21117635 | 21133348 | + | lncRNA |
| ENSG00000154263.17_4 | ABCA10 | 0.07128069 | 0.30384935 | -0.846 | -0.738 | chr17 | 67143355 | 67240987 | - | protein_coding |
| ENSG00000185038.14_7 | MROH2A | 0.06602873 | 0.28446473 | -0.846 | -0.355 | chr2 | 234684325 | 234742069 | + | protein_coding |
| ENSG00000228242.6_7 | AC093495.1 | 0.01341339 | 0.06956091 | -0.847 | 1.344 | chr3 | 14186137 | 14207478 | + | lncRNA |
| ENSG00000144161.13_4 | ZC3H8 | 0.00023622 | 0.00172432 | -0.848 | 3.683 | chr2 | 112969106 | 113012713 | - | protein_coding |
| ENSG00000280067.1_5 | AC023818.1 | 0.05210305 | 0.2326679 | -0.848 | 0.920 | chr16 | 48377666 | 48379969 | - | TEC |
| ENSG00000255909.1_4 | PDCD5P1 | 0.09233198 | 0.37894719 | -0.848 | -0.957 | chr12 | 19566087 | 19566395 | + | processed_pseudogene |
| ENSG00000134013.16_5 | LOXL2 | 6.84E-07 | 7.20E-06 | -0.849 | 6.618 | chr8 | 23154410 | 23282841 | - | protein_coding |
| ENSG00000211460.12_5 | TSN | 6.10E-07 | 6.47E-06 | -0.849 | 6.282 | chr2 | 122494679 | 122525429 | + | protein_coding |
| ENSG00000244556.1_4 | ODCP | 0.08729758 | 0.3621331 | -0.849 | -0.768 | chr7 | 128668943 | 128670581 | + | processed_pseudogene |
| ENSG00000213598.4_5 | AL049873.1 | 0.04251235 | 0.19487019 | -0.850 | 0.144 | chr14 | 59261372 | 59261747 | + | processed_pseudogene |
| ENSG00000207574.1 | MIR661 | 0.06947914 | 0.29766961 | -0.851 | -0.641 | chr8 | 145019359 | 145019447 | - | miRNA |
| ENSG00000280434.1_7 | AL031595.3 | 0.05793784 | 0.25454542 | -0.851 | -0.140 | chr22 | 44535245 | 44549506 | + | lncRNA |
| ENSG00000182287.15_7 | AP1S2 | 2.45E-05 | 0.00020909 | -0.851 | 4.631 | chrX | 15843929 | 15873054 | - | protein_coding |
| ENSG00000085760.15_4 | MTIF2 | 1.12E-06 | 1.15E-05 | -0.851 | 6.175 | chr2 | 55463731 | 55496483 | - | protein_coding |
| ENSG00000196636.8_3 | SDHAF3 | 0.00044648 | 0.00310476 | -0.851 | 3.684 | chr7 | 96747010 | 96811075 | + | protein_coding |

| | | | | | | | | | | |
|---|---|---|---|---|---|---|---|---|---|---|
| ENSG00000102053.12_3 | ZC3H12B | 0.02523771 | 0.1226808 | -0.851 | 0.635 | chrX | 64586518 | 64727767 | + | protein_coding |
| ENSG00000064933.17_4 | PMS1 | 1.88E-05 | 0.00016262 | -0.851 | 4.833 | chr2 | 190648811 | 190742355 | + | protein_coding |
| ENSG00000127824.14_5 | TUBA4A | 3.78E-07 | 4.12E-06 | -0.852 | 6.650 | chr2 | 220114432 | 220142892 | - | protein_coding |
| ENSG00000197223.11_3 | C1D | 0.00010696 | 0.00082299 | -0.852 | 3.808 | chr2 | 68268262 | 68338080 | - | protein_coding |
| ENSG00000036672.16_5 | USP2 | 0.07444299 | 0.31495369 | -0.852 | -0.739 | chr11 | 119225925 | 119252421 | - | protein_coding |
| ENSG00000286835.1_1 | AL133477.2 | 0.05876121 | 0.2576531 | -0.852 | -0.322 | chr9 | 99179006 | 99179659 | - | lncRNA |
| ENSG00000246731.3_8 | AC100786.1 | 0.05774568 | 0.25388953 | -0.852 | 0.073 | chr17 | 72206119 | 72209481 | - | lncRNA |
| ENSG00000077092.19_5 | RARB | 2.36E-05 | 0.00020223 | -0.852 | 4.467 | chr3 | 25215823 | 25639423 | + | protein_coding |
| ENSG00000165152.9_5 | PGAP4 | 6.06E-05 | 0.00048688 | -0.852 | 4.589 | chr9 | 104235452 | 104295819 | - | protein_coding |
| ENSG00000213881.3_4 | NPM1P6 | 0.02138505 | 0.10598276 | -0.852 | 1.124 | chr8 | 62114909 | 62115779 | + | processed_pseudogene |
| ENSG00000234694.1_7 | AL139289.1 | 0.05345449 | 0.23760967 | -0.852 | -0.060 | chr1 | 43820355 | 43824329 | - | lncRNA |
| ENSG00000277586.2_4 | NEFL | 0.05321075 | 0.23676317 | -0.853 | -0.107 | chr8 | 24808468 | 24814624 | - | protein_coding |
| ENSG00000143977.14_4 | SNRPG | 5.81E-06 | 5.39E-05 | -0.853 | 5.974 | chr2 | 70508494 | 70520872 | - | protein_coding |
| ENSG00000124575.6_3 | H1-3 | 5.77E-08 | 7.05E-07 | -0.853 | 8.077 | chr6 | 26234496 | 26235161 | - | protein_coding |
| ENSG00000197162.10_6 | ZNF785 | 0.07060722 | 0.30155732 | -0.853 | -0.640 | chr16 | 30585061 | 30597090 | - | protein_coding |
| ENSG00000132563.16_4 | REEP2 | 0.00373411 | 0.02183801 | -0.853 | 2.587 | chr5 | 137774746 | 137782658 | + | protein_coding |
| ENSG00000155380.11_8 | SLC16A1 | 4.56E-07 | 4.91E-06 | -0.854 | 6.035 | chr1 | 113454469 | 113499635 | - | protein_coding |
| ENSG00000159224.4_4 | GIP | 0.01057529 | 0.05627107 | -0.854 | 1.759 | chr17 | 47035916 | 47045958 | - | protein_coding |
| ENSG00000228634.1_6 | AL136115.1 | 0.12327833 | 0.48318688 | -0.854 | -0.742 | chr1 | 32398621 | 32399576 | + | lncRNA |
| ENSG00000115365.12_5 | LANCL1 | 1.75E-06 | 1.74E-05 | -0.855 | 5.961 | chr2 | 211295973 | 211342376 | - | protein_coding |
| ENSG00000178429.9_2 | RPS3AP5 | 0.0353716 | 0.16608015 | -0.855 | 0.358 | chr10 | 86320199 | 86321019 | + | processed_pseudogene |
| ENSG00000256188.3_3 | TAS2R30 | 0.05950949 | 0.26016366 | -0.855 | -0.236 | chr12 | 11285557 | 11287243 | - | protein_coding |
| ENSG00000115590.14_4 | IL1R2 | 0.08345284 | 0.34829846 | -0.855 | -0.696 | chr2 | 102608422 | 102645006 | + | protein_coding |
| ENSG00000144283.21_5 | PKP4 | 4.09E-07 | 4.43E-06 | -0.855 | 6.065 | chr2 | 159313476 | 159539391 | + | protein_coding |
| ENSG00000119878.6_3 | CRIPT | 3.28E-05 | 0.00027567 | -0.855 | 4.853 | chr2 | 46843555 | 46857315 | + | protein_coding |
| ENSG00000101384.12_5 | JAG1 | 5.33E-08 | 6.53E-07 | -0.856 | 7.230 | chr20 | 10618332 | 10654647 | - | protein_coding |
| ENSG00000198121.13_5 | LPAR1 | 6.76E-07 | 7.12E-06 | -0.856 | 5.808 | chr9 | 113635543 | 113800738 | - | protein_coding |
| ENSG00000135905.19_5 | DOCK10 | 1.94E-07 | 2.22E-06 | -0.857 | 6.511 | chr2 | 225629807 | 225907162 | - | protein_coding |
| ENSG00000233328.3_4 | PFN1P1 | 0.0397204 | 0.1836836 | -0.857 | 0.409 | chr1 | 171639657 | 171640079 | - | processed_pseudogene |
| ENSG00000158411.11_5 | MITD1 | 3.82E-05 | 0.0003169 | -0.857 | 4.216 | chr2 | 99777890 | 99797521 | - | protein_coding |
| ENSG00000232445.2_7 | EMSLR | 0.00538636 | 0.03048723 | -0.857 | 2.176 | chr7 | 100951551 | 100958081 | + | lncRNA |
| ENSG00000008300.17_5 | CELSR3 | 7.35E-05 | 0.0005814 | -0.857 | 4.322 | chr3 | 48673896 | 48700319 | - | protein_coding |
| ENSG00000216285.5_5 | AC078819.1 | 0.06412985 | 0.27763038 | -0.857 | -0.311 | chr12 | 104424557 | 104425321 | + | processed_pseudogene |
| ENSG00000261889.1_6 | AC108134.2 | 0.04065495 | 0.18761468 | -0.857 | 0.070 | chr16 | 3206737 | 3207484 | - | lncRNA |
| ENSG00000264229.1 | RNU4ATAC | 0.07299636 | 0.30982473 | -0.858 | -0.875 | chr2 | 122288457 | 122288583 | + | snRNA |
| ENSG00000285704.1_3 | AC004765.1 | 0.07534896 | 0.31828933 | -0.858 | -0.548 | chr12 | 974454 | 976305 | - | lncRNA |
| ENSG00000158186.13_4 | MRAS | 0.00042733 | 0.00298322 | -0.859 | 3.281 | chr3 | 138066490 | 138124376 | + | protein_coding |
| ENSG00000236581.9_6 | STARD13-AS | 0.03729147 | 0.17381553 | -0.859 | 0.223 | chr13 | 33754538 | 33855721 | + | lncRNA |
| ENSG00000163026.12_8 | WDCP | 1.08E-05 | 9.62E-05 | -0.859 | 4.887 | chr2 | 24252217 | 24272445 | - | protein_coding |
| ENSG00000225614.3_4 | ZNF469 | 0.00122611 | 0.00787004 | -0.859 | 4.879 | chr16 | 88449397 | 88507165 | + | protein_coding |
| ENSG00000172115.9_5 | CYCS | 3.05E-07 | 3.37E-06 | -0.859 | 7.093 | chr7 | 25158275 | 25164879 | - | protein_coding |
| ENSG00000227542.1_7 | AC092614.1 | 0.08432873 | 0.3514587 | -0.860 | -1.119 | chr2 | 192093891 | 192110898 | - | lncRNA |
| ENSG00000186076.5_5 | AC012085.1 | 0.04328505 | 0.19795248 | -0.860 | -0.081 | chr12 | 94034598 | 94035362 | - | processed_pseudogene |
| ENSG00000168393.13_7 | DTYMK | 9.44E-07 | 9.81E-06 | -0.861 | 6.053 | chr2 | 242615162 | 242626359 | - | protein_coding |
| ENSG00000205639.11_7 | MFSD2B | 0.03172748 | 0.15072248 | -0.861 | 0.323 | chr2 | 24232951 | 24286191 | + | protein_coding |
| ENSG00000135477.11_5 | KRT87P | 0.00041031 | 0.00287462 | -0.861 | 4.533 | chr12 | 52644250 | 52652651 | - | transcribed_unprocessed_pseudogene |
| ENSG00000212432.1 | SNORA75 | 0.0987345 | 0.40078041 | -0.862 | -1.221 | chr12 | 9597654 | 9597801 | + | snoRNA |
| ENSG00000228519.3_5 | AC097263.1 | 0.0441826 | 0.20169381 | -0.862 | -0.166 | chrX | 25663839 | 25664444 | - | processed_pseudogene |
| ENSG00000168078.10_4 | PBK | 9.22E-07 | 9.59E-06 | -0.862 | 6.487 | chr8 | 27667141 | 27695599 | - | protein_coding |
| ENSG00000075131.10_7 | TIPIN | 3.75E-06 | 3.57E-05 | -0.862 | 4.965 | chr15 | 66628529 | 66679084 | - | protein_coding |
| ENSG00000140519.14_5 | RHCG | 0.01737829 | 0.08809132 | -0.862 | 1.199 | chr15 | 90014629 | 90039820 | - | protein_coding |
| ENSG00000271410.1_5 | AC020728.1 | 0.11899442 | 0.47002533 | -0.864 | -0.938 | chr5 | 54652136 | 54653351 | - | processed_pseudogene |
| ENSG00000067798.16_5 | NAV3 | 1.36E-05 | 0.00011979 | -0.864 | 4.889 | chr12 | 77718421 | 78606790 | + | protein_coding |
| ENSG00000226609.2_8 | AL390067.1 | 0.0600561 | 0.26200224 | -0.865 | -0.532 | chr9 | 115511182 | 115512845 | - | lncRNA |
| ENSG00000179361.17_3 | ARID3B | 0.00603743 | 0.03376363 | -0.865 | 1.822 | chr15 | 74833518 | 74890472 | + | protein_coding |
| ENSG00000267006.1_6 | AC008507.2 | 0.06101981 | 0.26552509 | -0.866 | -0.599 | chr19 | 30519648 | 30520823 | - | lncRNA |
| ENSG00000287151.1_1 | C2orf27A | 0.01274569 | 0.06654502 | -0.868 | 1.221 | chr2 | 132405563 | 132524977 | + | lncRNA |

| | | | | | | | | | | |
|---|---|---|---|---|---|---|---|---|---|---|
| ENSG00000166250.12_5 | CLMP | 3.13E-05 | 0.00026376 | -0.868 | 4.056 | chr11 | 122940580 | 123065956 | - | protein_coding |
| ENSG00000285805.1_2 | AC012378.2 | 0.06764615 | 0.29051706 | -0.868 | -0.469 | chr15 | 55880864 | 55882009 | + | lncRNA |
| ENSG00000158158.12_3 | CNNM4 | 2.37E-05 | 0.00020248 | -0.869 | 4.437 | chr2 | 97426639 | 97477611 | + | protein_coding |
| ENSG00000117399.14_3 | CDC20 | 2.19E-07 | 2.48E-06 | -0.869 | 6.477 | chr1 | 43824652 | 43828874 | + | protein_coding |
| ENSG00000138382.15_6 | METTL5 | 5.14E-06 | 4.81E-05 | -0.869 | 4.674 | chr2 | 170666591 | 170681441 | - | protein_coding |
| ENSG00000228925.1_6 | AC016722.2 | 0.05214263 | 0.23270991 | -0.870 | -0.311 | chr2 | 47126414 | 47135817 | + | lncRNA |
| ENSG00000230894.2_6 | AL133284.1 | 0.01872531 | 0.09416717 | -0.870 | 1.150 | chr9 | 119329915 | 119348607 | + | lncRNA |
| ENSG00000229867.2_7 | STEAP3-AS1 | 0.0564399 | 0.24911082 | -0.870 | -0.510 | chr2 | 120001998 | 120028290 | - | lncRNA |
| ENSG00000104419.15_9 | NDRG1 | 1.54E-06 | 1.55E-05 | -0.870 | 5.311 | chr8 | 134249414 | 134314265 | - | protein_coding |
| ENSG00000270069.1_7 | MIR222HG | 0.00060982 | 0.00413376 | -0.871 | 2.945 | chrX | 45604639 | 45629677 | - | lncRNA |
| ENSG00000257222.1_6 | AC079907.1 | 0.07288346 | 0.30947324 | -0.873 | -0.908 | chr12 | 102457133 | 102468598 | + | lncRNA |
| ENSG00000272834.1_6 | AL022238.3 | 0.0822885 | 0.34432875 | -0.873 | -0.907 | chr22 | 40811007 | 40811449 | + | lncRNA |
| ENSG00000215388.3_4 | ACTG1P3 | 0.07208082 | 0.30660012 | -0.874 | -0.827 | chr20 | 1141228 | 1142341 | + | processed_pseudogene |
| ENSG00000162972.10_6 | MAIP1 | 3.38E-05 | 0.00028365 | -0.874 | 4.023 | chr2 | 200820040 | 200873263 | + | protein_coding |
| ENSG00000265018.6_6 | AGAP12P | 0.05494162 | 0.24348908 | -0.874 | -0.271 | chr10 | 49217898 | 49239658 | - | transcribed_unprocessed_pseudogene |
| ENSG00000238334.1 | snoU13 | 0.03418358 | 0.16092733 | -0.875 | 0.150 | chr5 | 68471651 | 68471754 | - | snoRNA |
| ENSG00000198074.10_3 | AKR1B10 | 4.07E-09 | 5.93E-08 | -0.875 | 11.870 | chr7 | 134212319 | 134226164 | + | protein_coding |
| ENSG00000131471.7_4 | AOC3 | 0.04532334 | 0.20609334 | -0.875 | -0.332 | chr17 | 41003201 | 41010147 | + | protein_coding |
| ENSG00000139974.15_3 | SLC38A6 | 5.92E-05 | 0.00047608 | -0.875 | 4.196 | chr14 | 61447832 | 61550451 | + | protein_coding |
| ENSG00000135976.19_7 | ANKRD36 | 3.24E-06 | 3.12E-05 | -0.876 | 4.684 | chr2 | 97778955 | 97930171 | + | protein_coding |
| ENSG00000176845.13_5 | METRNL | 0.06077371 | 0.26458382 | -0.876 | 0.396 | chr17 | 81037485 | 81052867 | + | protein_coding |
| ENSG00000177707.11_6 | NECTIN3 | 3.17E-07 | 3.49E-06 | -0.877 | 5.987 | chr3 | 110788918 | 110994410 | + | protein_coding |
| ENSG00000279412.1_5 | AC020763.3 | 0.05802033 | 0.25484478 | -0.877 | -0.347 | chr16 | 70747890 | 70750793 | + | TEC |
| ENSG00000136450.13_6 | SRSF1 | 2.05E-07 | 2.33E-06 | -0.878 | 7.126 | chr17 | 56078280 | 56084707 | - | protein_coding |
| ENSG00000136997.19_8 | MYC | 5.45E-08 | 6.67E-07 | -0.878 | 6.149 | chr8 | 128747680 | 128755197 | + | protein_coding |
| ENSG00000238683.1 | snoU13 | 0.05565665 | 0.2463507 | -0.878 | -0.332 | chr16 | 69742475 | 69742578 | - | snoRNA |
| ENSG00000202048.1 | SNORD114-20 | 0.02709037 | 0.13050829 | -0.879 | 0.671 | chr14 | 101447341 | 101447412 | + | snoRNA |
| ENSG00000223697.3_7 | AF230666.1 | 0.07261304 | 0.30841882 | -0.879 | -0.625 | chr8 | 133850362 | 133856543 | - | lncRNA |
| ENSG00000231881.2_6 | AL109615.2 | 0.05921447 | 0.25912894 | -0.879 | -0.434 | chr6 | 44026529 | 44057025 | + | lncRNA |
| ENSG00000221944.8_5 | TIGD1 | 0.00022608 | 0.00165711 | -0.879 | 3.393 | chr2 | 233412680 | 233415267 | - | protein_coding |
| ENSG00000259804.1_7 | AC027682.1 | 0.08012002 | 0.33604637 | -0.880 | -1.063 | chr16 | 67595314 | 67596212 | - | lncRNA |
| ENSG00000234160.1_6 | AL513165.1 | 0.01084437 | 0.05742827 | -0.880 | 1.199 | chr9 | 37509147 | 37510296 | + | lncRNA |
| ENSG00000197753.11_5 | LHFPL5 | 0.01844422 | 0.09283243 | -0.880 | 0.810 | chr6 | 35764983 | 35813174 | + | protein_coding |
| ENSG00000078114.18_4 | NEBL | 2.16E-06 | 2.13E-05 | -0.880 | 5.042 | chr10 | 21068902 | 21463116 | - | protein_coding |
| ENSG00000103449.12_4 | SALL1 | 0.02222779 | 0.10966985 | -0.880 | 0.613 | chr16 | 51169886 | 51185278 | - | protein_coding |
| ENSG00000188015.10_5 | S100A3 | 4.91E-06 | 4.60E-05 | -0.881 | 4.876 | chr1 | 153519805 | 153521734 | - | protein_coding |
| ENSG00000227256.1_6 | MIS18A-AS1 | 0.09261038 | 0.37971178 | -0.881 | -1.197 | chr21 | 33650174 | 33653299 | + | lncRNA |
| ENSG00000123975.5_3 | CKS2 | 1.52E-07 | 1.76E-06 | -0.881 | 6.552 | chr9 | 91926110 | 91931618 | + | protein_coding |
| ENSG00000279187.1_5 | AC027601.5 | 0.0319135 | 0.1514847 | -0.882 | 0.134 | chr17 | 79236720 | 79239201 | + | TEC |
| ENSG00000185633.10_3 | NDUFA4L2 | 0.02479938 | 0.12071525 | -0.882 | 0.342 | chr12 | 57628686 | 57634498 | - | protein_coding |
| ENSG00000196811.13_5 | CHRNG | 0.00269076 | 0.0161346 | -0.882 | 1.971 | chr2 | 233404402 | 233412825 | + | protein_coding |
| ENSG00000180422.3_3 | LINC00304 | 0.05782966 | 0.25413297 | -0.883 | -0.334 | chr16 | 89225554 | 89230653 | + | lncRNA |
| ENSG00000119013.9_3 | NDUFB3 | 1.39E-05 | 0.00012266 | -0.883 | 5.095 | chr2 | 201936156 | 201950473 | + | protein_coding |
| ENSG00000228063.2_8 | LYPLAL1-DT | 0.0721446 | 0.30679586 | -0.884 | -0.940 | chr1 | 219156365 | 219347303 | - | lncRNA |
| ENSG00000196912.12_4 | ANKRD36B | 0.00022107 | 0.00162305 | -0.885 | 3.174 | chr2 | 98109127 | 98206428 | - | protein_coding |
| ENSG00000138032.21_6 | PPM1B | 3.58E-06 | 3.43E-05 | -0.885 | 5.027 | chr2 | 44395108 | 44471523 | + | protein_coding |
| ENSG00000116678.19_4 | LEPR | 0.00024159 | 0.00175989 | -0.886 | 3.454 | chr1 | 65886335 | 66107242 | + | protein_coding |
| ENSG00000166473.17_6 | PKD1L2 | 0.00013163 | 0.00100245 | -0.886 | 3.444 | chr16 | 81134480 | 81253975 | - | polymorphic_pseudogene |
| ENSG00000169860.7_3 | P2RY1 | 0.0046379 | 0.02661344 | -0.886 | 1.605 | chr3 | 152552920 | 152559228 | + | protein_coding |
| ENSG00000169607.13_4 | CKAP2L | 2.56E-06 | 2.49E-05 | -0.887 | 4.977 | chr2 | 113493926 | 113522241 | - | protein_coding |
| ENSG00000229816.1_4 | DDX50P1 | 0.03996521 | 0.18476761 | -0.887 | 0.178 | chr2 | 32426669 | 32429068 | + | processed_pseudogene |
| ENSG00000135919.13_6 | SERPINE2 | 7.70E-08 | 9.26E-07 | -0.887 | 6.122 | chr2 | 224839762 | 224904035 | - | protein_coding |
| ENSG00000226054.2_4 | MEMO1P1 | 0.03839525 | 0.17834467 | -0.888 | 0.286 | chr21 | 37502787 | 37503674 | + | processed_pseudogene |
| ENSG00000144120.13_5 | TMEM177 | 4.12E-05 | 0.00034054 | -0.888 | 3.744 | chr2 | 120436743 | 120444083 | + | protein_coding |
| ENSG00000252010.1 | SCARNA5 | 0.00069435 | 0.00466586 | -0.888 | 2.838 | chr2 | 234184373 | 234184648 | + | snoRNA |
| ENSG00000149527.18_6 | PLCH2 | 0.01294692 | 0.06745688 | -0.889 | 1.131 | chr1 | 2357419 | 2436971 | + | protein_coding |
| ENSG00000188883.5_4 | KLRG2 | 0.02389671 | 0.11673708 | -0.889 | 0.286 | chr7 | 139137436 | 139168419 | - | protein_coding |

| | | | | | | | | | | |
|---|---|---|---|---|---|---|---|---|---|---|
| ENSG00000261786.1_6 | AC006058.1 | 5.72E-07 | 6.09E-06 | -0.889 | 5.449 | chr3 | 44158791 | 44163857 | + | lncRNA |
| ENSG00000261220.2_6 | AC103706.1 | 0.09498516 | 0.38777606 | -0.889 | -0.912 | chr8 | 134585426 | 134586104 | + | lncRNA |
| ENSG00000115233.12_5 | PSMD14 | 1.66E-08 | 2.17E-07 | -0.890 | 7.199 | chr2 | 162164936 | 162268228 | + | protein_coding |
| ENSG00000182118.8_5 | FAM89A | 0.0002038 | 0.00150246 | -0.890 | 3.226 | chr1 | 231154704 | 231176000 | - | protein_coding |
| ENSG00000258034.1_7 | AC012157.2 | 0.06696408 | 0.28807528 | -0.890 | -0.858 | chr12 | 116398992 | 116400538 | + | lncRNA |
| ENSG00000287625.1_1 | AC022210.2 | 0.00427714 | 0.02476923 | -0.891 | 1.681 | chr2 | 85153143 | 85193808 | + | lncRNA |
| ENSG00000130635.16_6 | COL5A1 | 0.00027565 | 0.00199246 | -0.891 | 5.913 | chr9 | 137533649 | 137736689 | + | protein_coding |
| ENSG00000228663.1_4 | PSMD10P1 | 0.05715433 | 0.25172559 | -0.892 | -0.611 | chr20 | 49530268 | 49530948 | + | processed_pseudogene |
| ENSG00000109321.11_4 | AREG | 0.00405751 | 0.02356648 | -0.892 | 1.693 | chr4 | 75310853 | 75320722 | + | protein_coding |
| ENSG00000112218.9_4 | GPR63 | 0.02175857 | 0.1075938 | -0.892 | 0.731 | chr6 | 97242001 | 97285353 | - | protein_coding |
| ENSG00000277383.1_6 | AC010331.1 | 0.05994534 | 0.26161848 | -0.892 | -0.859 | chr19 | 48111130 | 48111711 | - | lncRNA |
| ENSG00000039560.14_6 | RAI14 | 8.87E-09 | 1.22E-07 | -0.893 | 6.995 | chr5 | 34656433 | 34832717 | + | protein_coding |
| ENSG00000114529.12_4 | C3orf52 | 0.0005303 | 0.00363496 | -0.894 | 2.793 | chr3 | 111805182 | 111849851 | + | protein_coding |
| ENSG00000115484.15_4 | CCT4 | 2.01E-09 | 3.05E-08 | -0.894 | 8.191 | chr2 | 62095220 | 62115806 | - | protein_coding |
| ENSG00000229692.3_7 | SOS1-IT1 | 0.02155375 | 0.10672952 | -0.894 | 1.002 | chr2 | 39219420 | 39220998 | - | lncRNA |
| ENSG00000280383.1_7 | Z95331.1 | 0.00027722 | 0.00200216 | -0.895 | 3.374 | chr22 | 46052899 | 46076010 | + | lncRNA |
| ENSG00000186197.14_4 | EDARADD | 0.00132977 | 0.00844691 | -0.895 | 2.243 | chr1 | 236511557 | 236666215 | + | protein_coding |
| ENSG00000178074.6_4 | C2orf69 | 8.98E-06 | 8.11E-05 | -0.896 | 4.683 | chr2 | 200776016 | 200820658 | + | protein_coding |
| ENSG00000144136.11_7 | SLC20A1 | 1.08E-08 | 1.45E-07 | -0.896 | 6.728 | chr2 | 113403516 | 113421402 | + | protein_coding |
| ENSG00000072163.19_5 | LIMS2 | 0.02490277 | 0.12118532 | -0.896 | 0.299 | chr2 | 128395956 | 128439360 | - | protein_coding |
| ENSG00000243789.11_4 | JMJD7 | 0.04430251 | 0.20208541 | -0.897 | -0.399 | chr15 | 42120290 | 42129779 | + | protein_coding |
| ENSG00000215417.13_4 | MIR17HG | 0.00010947 | 0.00084086 | -0.897 | 3.437 | chr13 | 92000074 | 92006833 | + | lncRNA |
| ENSG00000222881.1 | Y_RNA | 0.0370363 | 0.17280042 | -0.898 | -0.109 | chr17 | 38547484 | 38547590 | - | misc_RNA |
| ENSG00000201772.1 | SNORA5C | 0.01569455 | 0.08017399 | -0.899 | 0.625 | chr7 | 45144505 | 45144641 | - | snoRNA |
| ENSG00000226124.7_5 | FTCDNL1 | 0.00408042 | 0.02369176 | -0.900 | 1.639 | chr2 | 200625267 | 200715896 | - | protein_coding |
| ENSG00000163611.11_3 | SPICE1 | 0.0037601 | 0.02196834 | -0.900 | 1.985 | chr3 | 113161565 | 113234034 | - | protein_coding |
| ENSG00000110172.12_4 | CHORDC1 | 3.40E-07 | 3.72E-06 | -0.901 | 6.649 | chr11 | 89933597 | 89956245 | - | protein_coding |
| ENSG00000139514.13_5 | SLC7A1 | 6.86E-08 | 8.32E-07 | -0.903 | 5.654 | chr13 | 30083551 | 30169825 | - | protein_coding |
| ENSG00000272189.1_7 | AL024508.1 | 0.05179289 | 0.23155617 | -0.903 | -0.638 | chr6 | 136871799 | 136873692 | + | lncRNA |
| ENSG00000274093.1_7 | AC009032.1 | 0.04545875 | 0.2065182 | -0.903 | -0.286 | chr16 | 69666044 | 69666474 | + | lncRNA |
| ENSG00000095209.12_7 | TMEM38B | 4.20E-06 | 3.97E-05 | -0.903 | 4.187 | chr9 | 108456822 | 108538910 | + | protein_coding |
| ENSG00000269246.1_7 | AC011445.2 | 0.04926172 | 0.22135381 | -0.904 | -0.524 | chr19 | 39832413 | 39832585 | - | lncRNA |
| ENSG00000111962.8_3 | UST | 0.00031931 | 0.00227934 | -0.904 | 2.712 | chr6 | 149068166 | 149398126 | + | protein_coding |
| ENSG00000146410.12_8 | MTFR2 | 0.00021559 | 0.00158348 | -0.905 | 3.207 | chr6 | 136552162 | 136571473 | - | protein_coding |
| ENSG00000230084.6_7 | AC006059.1 | 0.00285767 | 0.01703196 | -0.905 | 1.905 | chr3 | 42643455 | 42695880 | - | lncRNA |
| ENSG00000201882.1 | snoU2-30 | 0.06535573 | 0.28225515 | -0.906 | -0.923 | chrX | 20154184 | 20154253 | - | snoRNA |
| ENSG00000197275.14_5 | RAD54B | 0.00220038 | 0.01342981 | -0.907 | 1.943 | chr8 | 95384188 | 95487343 | - | protein_coding |
| ENSG00000120057.5_4 | SFRP5 | 0.05393241 | 0.23944727 | -0.907 | -0.812 | chr10 | 99526508 | 99531756 | - | protein_coding |
| ENSG00000154127.10_3 | UBASH3B | 5.72E-09 | 8.10E-08 | -0.907 | 7.070 | chr11 | 122526430 | 122685181 | + | protein_coding |
| ENSG00000120833.14_7 | SOCS2 | 0.0001869 | 0.00138707 | -0.907 | 3.284 | chr12 | 93963590 | 93977263 | + | protein_coding |
| ENSG00000222041.11_4 | CYTOR | 3.02E-05 | 0.00025571 | -0.907 | 3.653 | chr2 | 87754300 | 87936259 | + | lncRNA |
| ENSG00000171488.15_4 | LRRC8C | 5.19E-06 | 4.86E-05 | -0.908 | 4.355 | chr1 | 90098631 | 90235462 | + | protein_coding |
| ENSG00000106809.11_4 | OGN | 0.03510029 | 0.164937 | -0.908 | -0.289 | chr9 | 95145550 | 95166978 | - | protein_coding |
| ENSG00000102393.11_4 | GLA | 7.54E-09 | 1.05E-07 | -0.909 | 6.339 | chrX | 100648261 | 100663000 | - | protein_coding |
| ENSG00000242908.6_8 | AADACL2-AS1 | 0.03346531 | 0.15783863 | -0.909 | -0.236 | chr3 | 151469231 | 151645963 | - | lncRNA |
| ENSG00000203791.15_10 | EEF1AKMT2 | 0.00020113 | 0.00148526 | -0.909 | 2.916 | chr10 | 126436718 | 126480456 | - | protein_coding |
| ENSG00000180210.15_5 | F2 | 0.01261068 | 0.06585948 | -0.910 | 0.716 | chr11 | 46740746 | 46761056 | + | protein_coding |
| ENSG00000212163.5_3 | SNORD91A | 0.07009507 | 0.29987459 | -0.910 | -1.239 | chr17 | 2233474 | 2233664 | - | snoRNA |
| ENSG00000276649.1_7 | AL117335.1 | 0.01363812 | 0.07062328 | -0.910 | 0.728 | chr20 | 1868774 | 1875020 | - | lncRNA |
| ENSG00000281195.1_7 | AC007878.1 | 0.00131917 | 0.0083867 | -0.911 | 2.286 | chr2 | 71601068 | 71603450 | + | lncRNA |
| ENSG00000058804.12_4 | NDC1 | 3.99E-09 | 5.82E-08 | -0.911 | 6.710 | chr1 | 54231151 | 54304136 | - | protein_coding |
| ENSG00000131730.16_3 | CKMT2 | 0.03936 | 0.18230168 | -0.911 | -0.324 | chr5 | 80529139 | 80562217 | + | protein_coding |
| ENSG00000166803.13_6 | PCLAF | 1.95E-06 | 1.93E-05 | -0.911 | 4.856 | chr15 | 64656503 | 64679886 | - | protein_coding |
| ENSG00000263934.4_2 | SNORD3A | 0.11367326 | 0.45283248 | -0.914 | 10.460 | chr17 | 19091329 | 19092027 | + | snoRNA |
| ENSG00000284512.1_6 | AC092718.8 | 0.05210721 | 0.2326679 | -0.914 | -0.814 | chr16 | 81055921 | 81129889 | - | protein_coding |
| ENSG00000028116.17_4 | VRK2 | 4.93E-07 | 5.29E-06 | -0.915 | 4.866 | chr2 | 58134764 | 58387055 | + | protein_coding |
| ENSG00000272599.2_8 | AC016394.3 | 0.01841139 | 0.09271969 | -0.915 | 0.637 | chr10 | 74884331 | 74885290 | - | lncRNA |
| ENSG00000269570.2_6 | AP001350.1 | 0.11386072 | 0.45347755 | -0.917 | -0.292 | chr11 | 58378592 | 58380115 | - | lncRNA |

| | | | | | | | | | | |
|---|---|---|---|---|---|---|---|---|---|---|
| ENSG00000104635.14_6 | SLC39A14 | 1.73E-09 | 2.65E-08 | -0.917 | 7.210 | chr8 | 22224762 | 22291642 | + | protein_coding |
| ENSG00000213492.2_4 | NT5C3AP1 | 0.06598844 | 0.28436007 | -0.917 | -1.139 | chr4 | 118495667 | 118497329 | - | transcribed_processed_pseudogene |
| ENSG00000268412.2_5 | TRMT112P6 | 0.06716937 | 0.28881853 | -0.917 | -1.044 | chr2 | 26251077 | 26251481 | - | processed_pseudogene |
| ENSG00000271454.1_5 | AL353729.2 | 0.06816175 | 0.29258995 | -0.917 | -0.958 | chr9 | 39159068 | 39160316 | + | processed_pseudogene |
| ENSG00000278275.2_8 | AC103705.1 | 0.02378387 | 0.11628176 | -0.917 | 0.176 | chr8 | 129021797 | 129024184 | - | transcribed_processed_pseudogene |
| ENSG00000198612.11_3 | COPS8 | 1.81E-07 | 2.08E-06 | -0.918 | 5.674 | chr2 | 237994525 | 238009117 | + | protein_coding |
| ENSG00000236018.2_6 | AC004898.1 | 0.04025983 | 0.18603288 | -0.919 | -0.449 | chr7 | 156990294 | 156991205 | - | processed_pseudogene |
| ENSG00000118965.14_4 | WDR35 | 1.29E-08 | 1.72E-07 | -0.919 | 6.180 | chr2 | 20110021 | 20189892 | - | protein_coding |
| ENSG00000166979.13_4 | EVA1C | 4.80E-06 | 4.51E-05 | -0.919 | 4.429 | chr21 | 33784314 | 33887707 | + | protein_coding |
| ENSG00000183876.9_5 | ARSI | 0.00030898 | 0.00221285 | -0.920 | 2.829 | chr5 | 149675906 | 149718870 | - | protein_coding |
| ENSG00000189050.16_5 | RNFT1 | 4.86E-05 | 0.00039747 | -0.920 | 3.250 | chr17 | 58029601 | 58042122 | - | protein_coding |
| ENSG00000197355.11_3 | UAP1L1 | 8.93E-05 | 0.00069668 | -0.921 | 3.316 | chr9 | 139971969 | 139978991 | + | protein_coding |
| ENSG00000143248.13_6 | RGS5 | 0.07966791 | 0.33422899 | -0.921 | -0.770 | chr1 | 163080911 | 163291581 | - | protein_coding |
| ENSG00000163029.16_4 | SMC6 | 5.11E-09 | 7.29E-08 | -0.921 | 6.824 | chr2 | 17845079 | 17981509 | - | protein_coding |
| ENSG00000170144.20_5 | HNRNPA3 | 1.23E-08 | 1.64E-07 | -0.922 | 6.338 | chr2 | 178077291 | 178088686 | + | protein_coding |
| ENSG00000232034.1_4 | AC092168.2 | 0.00979925 | 0.05253571 | -0.923 | 0.747 | chr2 | 101439123 | 101463682 | + | lncRNA |
| ENSG00000171132.14_5 | PRKCE | 5.76E-06 | 5.35E-05 | -0.923 | 4.101 | chr2 | 45878484 | 46415129 | + | protein_coding |
| ENSG00000265683.1_5 | SYPL1P2 | 0.02116464 | 0.1050369 | -0.923 | 0.120 | chr17 | 25678884 | 25679610 | - | processed_pseudogene |
| ENSG00000275401.1_7 | AL391095.4 | 0.01414275 | 0.07304484 | -0.923 | 0.526 | chr20 | 37047130 | 37047849 | - | lncRNA |
| ENSG00000270062.1_7 | AL606834.2 | 0.0547312 | 0.24267761 | -0.924 | -0.958 | chr14 | 51190495 | 51190990 | - | lncRNA |
| ENSG00000135835.12_4 | KIAA1614 | 0.01543569 | 0.07901061 | -0.924 | 0.638 | chr1 | 180882033 | 180920750 | + | protein_coding |
| ENSG00000199377.1 | RNU5F-1 | 0.065638 | 0.28326215 | -0.924 | -1.045 | chr1 | 45187458 | 45187574 | - | snRNA |
| ENSG00000283355.1_5 | AC074194.2 | 0.04711094 | 0.21298219 | -0.924 | -0.768 | chr4 | 184935188 | 184937535 | + | TEC |
| ENSG00000124882.4_3 | EREG | 7.68E-09 | 1.06E-07 | -0.924 | 6.808 | chr4 | 75230862 | 75254466 | + | protein_coding |
| ENSG00000144214.9_3 | LYG1 | 0.02261476 | 0.1114553 | -0.924 | 0.277 | chr2 | 99900701 | 99921205 | - | protein_coding |
| ENSG00000154511.12_4 | DIPK1A | 9.86E-06 | 8.84E-05 | -0.925 | 3.800 | chr1 | 93298294 | 93427079 | - | protein_coding |
| ENSG00000091106.19_4 | NLRC4 | 0.05319573 | 0.23675566 | -0.925 | -0.843 | chr2 | 32449522 | 32490801 | - | protein_coding |
| ENSG00000164040.18_6 | PGRMC2 | 1.04E-07 | 1.24E-06 | -0.926 | 5.191 | chr4 | 129190392 | 129209984 | - | protein_coding |
| ENSG00000187522.16_6 | HSPA14 | 3.92E-07 | 4.25E-06 | -0.926 | 5.067 | chr10 | 14880305 | 14913740 | + | protein_coding |
| ENSG00000151665.13_5 | PIGF | 3.44E-05 | 0.00028776 | -0.926 | 3.403 | chr2 | 46808076 | 46844194 | - | protein_coding |
| ENSG00000215394.4 | BMS1P18 | 0.01303672 | 0.06786503 | -0.926 | 0.593 | chr14 | 19889386 | 19904890 | + | lincRNA |
| ENSG00000136810.13_4 | TXN | 1.74E-06 | 1.74E-05 | -0.927 | 8.446 | chr9 | 113006090 | 113018787 | - | protein_coding |
| ENSG00000234036.4_4 | TXNP6 | 0.03019135 | 0.14400263 | -0.927 | -0.140 | chr4 | 77880013 | 77880322 | - | processed_pseudogene |
| ENSG00000077782.20_6 | FGFR1 | 2.02E-08 | 2.61E-07 | -0.927 | 5.933 | chr8 | 38257733 | 38326352 | - | protein_coding |
| ENSG00000109674.4_3 | NEIL3 | 2.84E-06 | 2.74E-05 | -0.928 | 5.204 | chr4 | 178231028 | 178284090 | + | protein_coding |
| ENSG00000124802.12_5 | EEF1E1 | 6.15E-05 | 0.00049361 | -0.928 | 3.358 | chr6 | 8073593 | 8102792 | - | protein_coding |
| ENSG00000035499.13_4 | DEPDC1B | 9.54E-07 | 9.91E-06 | -0.928 | 4.867 | chr5 | 59892739 | 59996017 | - | protein_coding |
| ENSG00000231187.3_6 | AL356056.2 | 0.00074146 | 0.00493758 | -0.929 | 2.100 | chr10 | 46951467 | 46968909 | + | lncRNA |
| ENSG00000276662.1_6 | AL713922.2 | 0.06998616 | 0.29948077 | -0.930 | -1.281 | chr10 | 193760 | 194691 | - | unprocessed_pseudogene |
| ENSG00000271533.1_7 | Z83843.1 | 0.00017922 | 0.00133568 | -0.931 | 2.884 | chrX | 73429811 | 73433495 | - | lncRNA |
| ENSG00000114790.13_3 | ARHGEF26 | 3.54E-06 | 3.39E-05 | -0.931 | 4.056 | chr3 | 153838792 | 153975616 | + | protein_coding |
| ENSG00000250073.3_7 | AP000866.2 | 0.02389072 | 0.11673708 | -0.931 | 0.027 | chr11 | 124629025 | 124636015 | + | lncRNA |
| ENSG00000263080.1_8 | AC009121.4 | 0.04805033 | 0.21678774 | -0.933 | -0.724 | chr16 | 11435666 | 11439068 | - | lncRNA |
| ENSG00000233608.4_4 | TWIST2 | 0.0037799 | 0.02207676 | -0.933 | 1.730 | chr2 | 239756673 | 239832230 | + | protein_coding |
| ENSG00000286708.1_1 | AP003783.1 | 0.0500945 | 0.22469804 | -0.933 | -0.768 | chr11 | 70732195 | 70742410 | + | lncRNA |
| ENSG00000147536.12_4 | GINS4 | 2.34E-07 | 2.64E-06 | -0.934 | 4.729 | chr8 | 41386737 | 41402549 | + | protein_coding |
| ENSG00000271032.1_7 | AC020907.4 | 0.01275891 | 0.06657489 | -0.934 | 0.661 | chr19 | 35505865 | 35516239 | - | lncRNA |
| ENSG00000172292.14_3 | CERS6 | 8.44E-08 | 1.01E-06 | -0.934 | 5.545 | chr2 | 169312372 | 169631647 | + | protein_coding |
| ENSG00000235651.1_5 | AC064850.1 | 0.05610497 | 0.24808786 | -0.935 | -1.082 | chr2 | 136268116 | 136269513 | - | processed_pseudogene |
| ENSG00000264940.4_2 | SNORD3C | 0.02156363 | 0.1067487 | -0.935 | 2.115 | chr17 | 19092978 | 19093558 | - | snoRNA |
| ENSG00000278840.1_7 | AC009878.1 | 0.02807723 | 0.13460553 | -0.935 | -0.280 | chr15 | 26802745 | 26803084 | - | lncRNA |
| ENSG00000286614.1_2 | AL589182.3 | 0.01173106 | 0.0616462 | -0.936 | 0.583 | chr14 | 19659990 | 19662998 | + | transcribed_unprocessed_pseudogene |
| ENSG00000133805.15_4 | AMPD3 | 0.0006205 | 0.00419978 | -0.936 | 2.295 | chr11 | 10329860 | 10529126 | + | protein_coding |
| ENSG00000112319.19_5 | EYA4 | 5.08E-07 | 5.45E-06 | -0.938 | 4.813 | chr6 | 133561736 | 133853266 | + | protein_coding |
| ENSG00000287959.1_1 | AC021236.1 | 0.01969403 | 0.09837049 | -0.938 | 0.334 | chr8 | 48853986 | 48856230 | + | lncRNA |
| ENSG00000170390.16_5 | DCLK2 | 1.70E-05 | 0.00014801 | -0.939 | 3.514 | chr4 | 150999597 | 151178590 | + | protein_coding |

| | | | | | | | | | | |
|---|---|---|---|---|---|---|---|---|---|---|
| ENSG00000273723.1_6 | AL139089.1 | 0.02848205 | 0.13647265 | -0.939 | -0.069 | chr13 | 53225440 | 53226414 | - | lncRNA |
| ENSG00000259865.1_6 | AL390728.6 | 0.00580667 | 0.03262719 | -0.939 | 1.337 | chr1 | 247350583 | 247351828 | - | lncRNA |
| ENSG00000156509.14_4 | FBXO43 | 0.00035259 | 0.00250246 | -0.940 | 3.063 | chr8 | 101145579 | 101158045 | - | protein_coding |
| ENSG00000183891.6_4 | TTC32 | 2.51E-05 | 0.00021429 | -0.941 | 3.571 | chr2 | 20096392 | 20101744 | - | protein_coding |
| ENSG00000128694.12_4 | OSGEPL1 | 6.50E-05 | 0.00051925 | -0.942 | 3.200 | chr2 | 190611386 | 190627953 | - | protein_coding |
| ENSG00000176714.9_4 | CCDC121 | 0.00039678 | 0.0027886 | -0.942 | 2.684 | chr2 | 27848506 | 27851879 | - | protein_coding |
| ENSG00000239300.6_6 | AC080162.1 | 0.05059384 | 0.22667627 | -0.943 | -0.907 | chr2 | 9641595 | 9652542 | + | lncRNA |
| ENSG00000171914.16_3 | TLN2 | 1.35E-07 | 1.57E-06 | -0.943 | 4.854 | chr15 | 62682725 | 63136830 | + | protein_coding |
| ENSG00000240476.1_6 | LINC00973 | 0.00358642 | 0.02102966 | -0.945 | 1.271 | chr3 | 98699902 | 98701940 | + | lncRNA |
| ENSG00000209042.1 | SNORD12C | 0.00541448 | 0.03062688 | -0.945 | 0.970 | chr20 | 47895477 | 47895565 | + | snoRNA |
| ENSG00000214182.5_2 | PTMAP5 | 0.00092076 | 0.00602991 | -0.947 | 2.180 | chr13 | 82264046 | 82265207 | + | transcribed_processed_pseudogene |
| ENSG00000115594.12_7 | IL1R1 | 1.24E-08 | 1.65E-07 | -0.947 | 5.606 | chr2 | 102681004 | 102796334 | + | protein_coding |
| ENSG00000169756.16_5 | LIMS1 | 1.58E-09 | 2.43E-08 | -0.947 | 6.236 | chr2 | 109150811 | 109303702 | + | protein_coding |
| ENSG00000115504.14_5 | EHBP1 | 7.56E-09 | 1.05E-07 | -0.948 | 6.702 | chr2 | 62900986 | 63273622 | + | protein_coding |
| ENSG00000003989.17_6 | SLC7A2 | 2.63E-05 | 0.00022388 | -0.948 | 3.280 | chr8 | 17354597 | 17428082 | + | protein_coding |
| ENSG00000115159.16_3 | GPD2 | 5.24E-10 | 8.60E-09 | -0.950 | 6.922 | chr2 | 157291802 | 157470247 | + | protein_coding |
| ENSG00000144362.12_4 | PHOSPHO2 | 0.00450894 | 0.02595089 | -0.951 | 1.087 | chr2 | 170550964 | 170558218 | + | protein_coding |
| ENSG00000144381.17_5 | HSPD1 | 2.51E-10 | 4.30E-09 | -0.951 | 9.254 | chr2 | 198351308 | 198381461 | - | protein_coding |
| ENSG00000138760.10_6 | SCARB2 | 8.41E-10 | 1.35E-08 | -0.951 | 6.510 | chr4 | 77079890 | 77155689 | - | protein_coding |
| ENSG00000235100.4_8 | AL157400.3 | 0.02669346 | 0.12877077 | -0.952 | -0.217 | chr10 | 91426938 | 91461031 | - | lncRNA |
| ENSG00000231607.12_8 | DLEU2 | 1.35E-05 | 0.00011884 | -0.952 | 3.603 | chr13 | 50530806 | 50699856 | - | lncRNA |
| ENSG00000158482.10_5 | SNX29P1 | 0.03585177 | 0.16806816 | -0.952 | -0.586 | chr16 | 21387969 | 21397385 | + | unprocessed_pseudogene |
| ENSG00000286194.1_2 | AC138207.9 | 0.15148231 | 0.57324518 | -0.953 | -0.799 | chr17 | 29295803 | 29297325 | + | lncRNA |
| ENSG00000147180.16_3 | ZNF711 | 4.94E-05 | 0.00040267 | -0.953 | 2.968 | chrX | 84499038 | 84528368 | + | protein_coding |
| ENSG00000197713.15_4 | RPE | 1.90E-09 | 2.89E-08 | -0.953 | 6.124 | chr2 | 210867289 | 210886984 | + | protein_coding |
| ENSG00000144045.14_5 | DQX1 | 0.04059301 | 0.18737751 | -0.954 | -0.754 | chr2 | 74745258 | 74753408 | - | protein_coding |
| ENSG00000152147.11_4 | GEMIN6 | 5.33E-08 | 6.53E-07 | -0.954 | 5.231 | chr2 | 38978676 | 39012144 | + | protein_coding |
| ENSG00000127366.5_3 | TAS2R5 | 0.05625515 | 0.24856635 | -0.954 | -0.994 | chr7 | 141490017 | 141491167 | + | protein_coding |
| ENSG00000286341.1_3 | AP002762.3 | 0.05642415 | 0.24911082 | -0.955 | -1.219 | chr11 | 122550509 | 122552650 | - | lncRNA |
| ENSG00000165197.5_7 | VEGFD | 0.02638437 | 0.12741808 | -0.955 | -0.014 | chrX | 15363718 | 15402535 | - | protein_coding |
| ENSG00000273143.2_7 | AL355512.1 | 0.01806134 | 0.09113769 | -0.956 | 0.092 | chr10 | 112188598 | 112256348 | - | lncRNA |
| ENSG00000224745.1_8 | AC063965.2 | 0.03202198 | 0.15187787 | -0.957 | -0.459 | chr10 | 89638449 | 89640184 | + | lncRNA |
| ENSG00000115053.16_5 | NCL | 2.18E-10 | 3.75E-09 | -0.957 | 8.172 | chr2 | 232318242 | 232348352 | - | protein_coding |
| ENSG00000266456.1_7 | AP001178.3 | 0.02347227 | 0.11497982 | -0.958 | -0.168 | chr18 | 650229 | 652843 | + | lncRNA |
| ENSG00000092621.12_4 | PHGDH | 6.76E-08 | 8.21E-07 | -0.959 | 5.238 | chr1 | 120191034 | 120286849 | + | protein_coding |
| ENSG00000150627.15_3 | WDR17 | 2.50E-05 | 0.00021339 | -0.960 | 3.401 | chr4 | 176986985 | 177103969 | + | protein_coding |
| ENSG00000163637.13_6 | PRICKLE2 | 1.65E-06 | 1.65E-05 | -0.960 | 4.074 | chr3 | 64077912 | 64431152 | - | protein_coding |
| ENSG00000137285.10_4 | TUBB2B | 0.00036935 | 0.00260818 | -0.960 | 2.265 | chr6 | 3224511 | 3231964 | - | protein_coding |
| ENSG00000198417.7_3 | MT1F | 0.00769355 | 0.04216262 | -0.960 | 0.758 | chr16 | 56691643 | 56694610 | + | protein_coding |
| ENSG00000144028.15_6 | SNRNP200 | 2.98E-10 | 5.06E-09 | -0.961 | 7.680 | chr2 | 96940076 | 96987009 | - | protein_coding |
| ENSG00000197696.10_5 | NMB | 7.79E-05 | 0.00061274 | -0.961 | 2.698 | chr15 | 85198360 | 85201794 | - | protein_coding |
| ENSG00000163347.6_4 | CLDN1 | 1.36E-09 | 2.13E-08 | -0.961 | 5.986 | chr3 | 190023496 | 190040235 | - | protein_coding |
| ENSG00000227906.8_7 | SNAP25-AS1 | 0.03739359 | 0.17414728 | -0.961 | -0.708 | chr20 | 9987029 | 10349424 | - | lncRNA |
| ENSG00000141753.7_3 | IGFBP4 | 1.09E-05 | 9.73E-05 | -0.962 | 9.822 | chr17 | 38599702 | 38613977 | + | protein_coding |
| ENSG00000277151.1_6 | AL138820.1 | 0.04425557 | 0.20192315 | -0.962 | -0.753 | chr13 | 33357012 | 33362316 | + | lncRNA |
| ENSG00000234456.8_8 | MAGI2-AS3 | 0.00066369 | 0.00447164 | -0.963 | 1.979 | chr7 | 79082193 | 79100524 | + | lncRNA |
| ENSG00000196290.15_4 | NIF3L1 | 7.88E-09 | 1.09E-07 | -0.963 | 5.244 | chr2 | 201754050 | 201768661 | + | protein_coding |
| ENSG00000196151.10_3 | WDSUB1 | 2.90E-05 | 0.00024577 | -0.964 | 3.132 | chr2 | 160092304 | 160143310 | - | protein_coding |
| ENSG00000142149.9_4 | HUNK | 0.00065432 | 0.00441356 | -0.966 | 1.850 | chr21 | 33245333 | 33416946 | + | protein_coding |
| ENSG00000287164.1_2 | AC112254.1 | 0.03956802 | 0.18321738 | -0.967 | -0.860 | chr4 | 7800254 | 7809778 | + | lncRNA |
| ENSG00000143933.18_6 | CALM2 | 3.79E-08 | 4.73E-07 | -0.967 | 9.611 | chr2 | 47387222 | 47404060 | - | protein_coding |
| ENSG00000231982.1_5 | AC112907.2 | 0.04637866 | 0.21015292 | -0.967 | -0.755 | chr3 | 186544338 | 186545126 | + | unprocessed_pseudogene |
| ENSG00000269947.1_7 | AC135178.6 | 0.04583824 | 0.20806644 | -0.969 | -0.710 | chr17 | 8181081 | 8181754 | + | lncRNA |
| ENSG00000120217.14_3 | CD274 | 0.01548709 | 0.07925091 | -0.969 | 0.126 | chr9 | 5450503 | 5470566 | + | protein_coding |
| ENSG00000167191.12_5 | GPRC5B | 2.79E-07 | 3.10E-06 | -0.969 | 4.432 | chr16 | 19868013 | 19897489 | - | protein_coding |
| ENSG00000166664.13_6 | CHRFAM7A | 0.02052882 | 0.10210959 | -0.969 | 0.097 | chr15 | 30649969 | 30686052 | - | protein_coding |
| ENSG00000055044.11_3 | NOP58 | 1.18E-09 | 1.86E-08 | -0.970 | 5.990 | chr2 | 203130459 | 203168384 | + | protein_coding |
| ENSG00000254854.1_6 | AP003390.1 | 0.02725655 | 0.13116663 | -0.971 | -0.290 | chr11 | 119600293 | 119610333 | + | lncRNA |
| ENSG00000260128.6_2 | ULK4P2 | 0.03196977 | 0.15167074 | -0.971 | -0.640 | chr15 | 30864941 | 30892850 | + | transcribed_unprocessed_pseudogene |

| | | | | | | | | | | |
|---|---|---|---|---|---|---|---|---|---|---|
| ENSG00000237380.7_9 | HOXD-AS2 | 0.04335989 | 0.19824373 | -0.972 | -0.890 | chr2 | 176986339 | 177001826 | - | lncRNA |
| ENSG00000126259.19_4 | KIRREL2 | 0.02312393 | 0.11349249 | -0.972 | -0.030 | chr19 | 36346763 | 36358011 | + | protein_coding |
| ENSG00000118257.16_4 | NRP2 | 7.14E-11 | 1.31E-09 | -0.975 | 7.489 | chr2 | 206546714 | 206662857 | + | protein_coding |
| ENSG00000115041.13_4 | KCNIP3 | 0.02702915 | 0.13028403 | -0.976 | -0.510 | chr2 | 95963075 | 96051825 | + | protein_coding |
| ENSG00000145779.8_5 | TNFAIP8 | 2.06E-08 | 2.66E-07 | -0.977 | 5.004 | chr5 | 118604387 | 118735383 | + | protein_coding |
| ENSG00000074181.9_4 | NOTCH3 | 7.43E-06 | 6.79E-05 | -0.977 | 3.978 | chr19 | 15269849 | 15311806 | - | protein_coding |
| ENSG00000263235.1_7 | AC006111.2 | 0.02923945 | 0.13983779 | -0.977 | -0.411 | chr16 | 3700637 | 3701704 | - | lncRNA |
| ENSG00000272994.1_6 | AC012360.3 | 0.00014289 | 0.00108396 | -0.978 | 2.369 | chr2 | 105950484 | 105953932 | - | lncRNA |
| ENSG00000099860.9_6 | GADD45B | 5.47E-06 | 5.10E-05 | -0.979 | 3.744 | chr19 | 2476120 | 2478257 | + | protein_coding |
| ENSG00000287049.1_1 | AC124862.1 | 0.04000675 | 0.18491153 | -0.979 | -0.740 | chr2 | 241497506 | 241498342 | + | lncRNA |
| ENSG00000231521.1_7 | AL162385.2 | 0.04161718 | 0.1914584 | -0.979 | -0.891 | chr9 | 100396797 | 100398333 | - | lncRNA |
| ENSG00000274995.1_7 | AC013564.1 | 0.04547003 | 0.2065182 | -0.979 | -0.861 | chr15 | 67032783 | 67033489 | + | lncRNA |
| ENSG00000001084.13_9 | GCLC | 4.16E-11 | 7.87E-10 | -0.980 | 7.500 | chr6 | 53362139 | 53481768 | - | protein_coding |
| ENSG00000155755.19_5 | TMEM237 | 9.60E-08 | 1.14E-06 | -0.980 | 4.799 | chr2 | 202484907 | 202508293 | - | protein_coding |
| ENSG00000261051.1_6 | AC107021.2 | 0.00910566 | 0.04902432 | -0.981 | 0.373 | chr3 | 145777372 | 145779466 | - | lncRNA |
| ENSG00000280211.1_6 | AC106886.5 | 0.03436942 | 0.16167366 | -0.981 | -0.573 | chr16 | 30784853 | 30787354 | - | TEC |
| ENSG00000080345.18_7 | RIF1 | 7.99E-10 | 1.29E-08 | -0.981 | 7.988 | chr2 | 152266397 | 152364527 | + | protein_coding |
| ENSG00000270276.2_3 | H4C15 | 0.01907927 | 0.09570337 | -0.981 | -0.228 | chr1 | 149825607 | 149832776 | - | protein_coding |
| ENSG00000162980.17_5 | ARL5A | 2.58E-09 | 3.86E-08 | -0.981 | 5.517 | chr2 | 152645498 | 152685006 | - | protein_coding |
| ENSG00000138028.16_4 | CGREF1 | 2.43E-05 | 0.00020802 | -0.981 | 2.805 | chr2 | 27321757 | 27341996 | - | protein_coding |
| ENSG00000005421.9_4 | PON1 | 0.01464557 | 0.07537887 | -0.982 | -0.017 | chr7 | 94926988 | 94953844 | - | protein_coding |
| ENSG00000152402.10_5 | GUCY1A2 | 0.00649342 | 0.03608652 | -0.982 | 0.748 | chr11 | 106544738 | 106889250 | - | protein_coding |
| ENSG00000176490.5_7 | DIRAS1 | 0.00243119 | 0.01468614 | -0.982 | 1.121 | chr19 | 2714565 | 2721370 | - | protein_coding |
| ENSG00000197121.15_4 | PGAP1 | 2.67E-07 | 2.98E-06 | -0.982 | 4.271 | chr2 | 197697728 | 197792520 | - | protein_coding |
| ENSG00000263606.1_5 | AP000919.1 | 0.02209887 | 0.10909436 | -0.983 | -0.390 | chr18 | 2945548 | 2946888 | - | transcribed_processed_pseudogene |
| ENSG00000279348.1_5 | AC012513.3 | 4.05E-05 | 0.00033521 | -0.984 | 2.885 | chr2 | 217076127 | 217078242 | + | TEC |
| ENSG00000159399.10_4 | HK2 | 0.03244614 | 0.1535208 | -0.985 | -0.510 | chr2 | 75061254 | 75120486 | + | protein_coding |
| ENSG00000228446.2_2 | AC073052.1 | 0.02554284 | 0.12396055 | -0.985 | -0.314 | chr2 | 225364104 | 225364817 | - | processed_pseudogene |
| ENSG00000241484.9_5 | ARHGAP8 | 0.0364692 | 0.1705576 | -0.986 | -0.813 | chr22 | 45148438 | 45258668 | + | protein_coding |
| ENSG00000281344.1_4 | HELLPAR | 5.98E-05 | 0.00048096 | -0.987 | 2.891 | chr12 | 102591363 | 102796374 | + | lncRNA |
| ENSG00000127325.19_6 | BEST3 | 0.02667464 | 0.1287149 | -0.987 | -0.320 | chr12 | 70037140 | 70093256 | - | protein_coding |
| ENSG00000251432.7_7 | LINC02615 | 1.79E-05 | 0.0001556 | -0.989 | 3.007 | chr4 | 129213906 | 129440553 | + | lncRNA |
| ENSG00000124743.6_4 | KLHL31 | 0.00534602 | 0.03029755 | -0.989 | 0.781 | chr6 | 53512714 | 53530554 | - | protein_coding |
| ENSG00000255445.1_6 | AP003072.2 | 0.04850201 | 0.21838194 | -0.989 | -1.197 | chr11 | 92885241 | 92885916 | + | processed_pseudogene |
| ENSG00000284858.1_2 | AC091185.2 | 0.01650275 | 0.08394054 | -0.991 | 0.151 | chr8 | 25134461 | 25134992 | - | processed_pseudogene |
| ENSG00000280435.1_5 | AC006058.4 | 2.33E-06 | 2.28E-05 | -0.991 | 3.780 | chr3 | 44157310 | 44158979 | - | TEC |
| ENSG00000231890.8_6 | DARS-AS1 | 0.0421236 | 0.19338759 | -0.992 | -0.842 | chr2 | 136742746 | 136780163 | + | lncRNA |
| ENSG00000264350.1_4 | SNRPGP2 | 0.00076266 | 0.0050712 | -0.993 | 1.670 | chr18 | 51933259 | 51933484 | - | processed_pseudogene |
| ENSG00000241081.1_5 | RPL22P2 | 0.01519883 | 0.0778655 | -0.993 | 0.110 | chr14 | 77352720 | 77353093 | - | processed_pseudogene |
| ENSG00000275769.1_7 | AC068792.1 | 0.03925929 | 0.18202499 | -0.994 | -0.795 | chr12 | 31596726 | 31597142 | - | lncRNA |
| ENSG00000267114.1_8 | AC011481.1 | 0.04560108 | 0.207052 | -0.994 | -0.625 | chr19 | 45453301 | 45457264 | - | lncRNA |
| ENSG00000272702.1_6 | AC010913.1 | 0.02021984 | 0.10062909 | -0.994 | -0.234 | chr2 | 73340146 | 73343035 | + | lncRNA |
| ENSG00000167772.12_5 | ANGPTL4 | 2.71E-05 | 0.00022999 | -0.995 | 2.983 | chr19 | 8428173 | 8439254 | + | protein_coding |
| ENSG00000175318.12_7 | GRAMD2A | 0.04611776 | 0.2091204 | -0.995 | -1.102 | chr15 | 72452147 | 72490128 | - | protein_coding |
| ENSG00000272269.1_7 | AL138724.2 | 0.02952404 | 0.14100905 | -0.995 | -0.767 | chr6 | 17706488 | 17707575 | + | lncRNA |
| ENSG00000006652.14_7 | IFRD1 | 6.58E-09 | 9.25E-08 | -0.996 | 5.485 | chr7 | 112063023 | 112121072 | + | protein_coding |
| ENSG00000207954.1 | MIR138-1 | 0.01042239 | 0.05552464 | -0.996 | 0.118 | chr3 | 44155704 | 44155802 | + | miRNA |
| ENSG00000197977.4_4 | ELOVL2 | 2.41E-07 | 2.72E-06 | -0.996 | 4.123 | chr6 | 10980992 | 11044538 | - | protein_coding |
| ENSG00000287957.1_2 | AC007695.1 | 0.00953487 | 0.05119551 | -0.997 | 0.502 | chr12 | 107260955 | 107265283 | - | lncRNA |
| ENSG00000091436.17_5 | MAP3K20 | 4.96E-11 | 9.24E-10 | -0.998 | 6.727 | chr2 | 173940163 | 174132743 | + | protein_coding |
| ENSG00000102996.5_3 | MMP15 | 1.10E-06 | 1.13E-05 | -0.998 | 3.794 | chr16 | 58059658 | 58080805 | + | protein_coding |
| ENSG00000183496.6_4 | MEX3B | 0.00012025 | 0.0009188 | -0.998 | 2.305 | chr15 | 82334119 | 82338460 | - | protein_coding |
| ENSG00000140905.11_6 | GCSH | 0.0039152 | 0.02282947 | -0.999 | 0.830 | chr16 | 81115550 | 81130000 | - | protein_coding |
| ENSG00000114923.17_5 | SLC4A3 | 0.00038184 | 0.00268929 | -1.000 | 1.852 | chr2 | 220492116 | 220506702 | + | protein_coding |
| ENSG00000126803.9_3 | HSPA2 | 1.30E-10 | 2.32E-09 | -1.000 | 6.368 | chr14 | 65002623 | 65012891 | + | protein_coding |
| ENSG00000137561.5_4 | TTPA | 0.00823353 | 0.04480803 | -1.000 | 0.486 | chr8 | 63961112 | 63998612 | - | protein_coding |
| ENSG00000155849.15_4 | ELMO1 | 0.00083068 | 0.00548038 | -1.000 | 1.668 | chr7 | 36893966 | 37488852 | - | protein_coding |
| ENSG00000155816.20_4 | FMN2 | 9.58E-05 | 0.00074231 | -1.000 | 2.539 | chr1 | 240177648 | 240638487 | + | protein_coding |

| | | | | | | | | | | |
|---|---|---|---|---|---|---|---|---|---|---|
| ENSG00000281398.4_6 | SNHG4 | 1.62E-08 | 2.13E-07 | -1.001 | 5.223 | chr5 | 138609791 | 138620588 | + | lncRNA |
| ENSG00000144401.14_3 | METTL21A | 3.43E-07 | 3.75E-06 | -1.003 | 3.976 | chr2 | 208445355 | 208490652 | - | protein_coding |
| ENSG00000118246.14_4 | FASTKD2 | 4.70E-10 | 7.76E-09 | -1.004 | 6.201 | chr2 | 207630081 | 207660913 | + | protein_coding |
| ENSG00000226887.8_5 | ERVMER34-1 | 0.00866496 | 0.04689321 | -1.004 | 0.230 | chr4 | 53588785 | 53617753 | - | protein_coding |
| ENSG00000179241.13_4 | LDLRAD3 | 1.46E-06 | 1.48E-05 | -1.004 | 3.754 | chr11 | 35965531 | 36253686 | + | protein_coding |
| ENSG00000247626.5_4 | MARS2 | 3.76E-06 | 3.58E-05 | -1.004 | 3.178 | chr2 | 198570093 | 198573119 | + | protein_coding |
| ENSG00000255647.3_6 | AC093510.1 | 0.03928401 | 0.1820921 | -1.004 | -0.873 | chr5 | 89705817 | 89706486 | + | lncRNA |
| ENSG00000115946.8_3 | PNO1 | 2.89E-09 | 4.30E-08 | -1.005 | 5.410 | chr2 | 68385020 | 68403370 | + | protein_coding |
| ENSG00000247675.7_6 | LRP4-AS1 | 0.04776054 | 0.21564462 | -1.005 | -1.138 | chr11 | 46867963 | 46895972 | + | lncRNA |
| ENSG00000226833.6_7 | AC092164.1 | 0.00476159 | 0.02724614 | -1.005 | 0.663 | chr2 | 28930377 | 28974588 | - | lncRNA |
| ENSG00000189007.16_5 | ADAT2 | 4.97E-05 | 0.00040528 | -1.005 | 2.645 | chr6 | 143743969 | 143771832 | - | protein_coding |
| ENSG00000227051.6_3 | C14orf132 | 1.38E-07 | 1.61E-06 | -1.006 | 4.183 | chr14 | 96505661 | 96560226 | + | protein_coding |
| ENSG00000152253.9_5 | SPC25 | 2.33E-06 | 2.28E-05 | -1.007 | 3.698 | chr2 | 169690642 | 169769881 | - | protein_coding |
| ENSG00000163811.12_4 | WDR43 | 1.92E-11 | 3.82E-10 | -1.007 | 7.554 | chr2 | 29117533 | 29171085 | + | protein_coding |
| ENSG00000213606.3_4 | AKR1B10P1 | 3.18E-07 | 3.49E-06 | -1.007 | 3.838 | chr10 | 69510042 | 69510983 | + | processed_pseudogene |
| ENSG00000280055.1_4 | TMEM75 | 0.01289204 | 0.06723009 | -1.007 | 0.636 | chr8 | 128958805 | 128960969 | - | TEC |
| ENSG00000183971.8_7 | NPW | 0.01550132 | 0.07930086 | -1.009 | -0.289 | chr16 | 2059927 | 2070756 | + | protein_coding |
| ENSG00000153107.13_5 | ANAPC1 | 2.86E-11 | 5.56E-10 | -1.010 | 6.333 | chr2 | 112369216 | 112642267 | - | protein_coding |
| ENSG00000279394.1_8 | AC015871.4 | 0.0489855 | 0.22027978 | -1.010 | -1.345 | chr15 | 80168262 | 80170521 | - | TEC |
| ENSG00000132639.12_3 | SNAP25 | 1.10E-07 | 1.29E-06 | -1.011 | 4.056 | chr20 | 10199478 | 10288066 | + | protein_coding |
| ENSG00000266302.6_6 | AC098850.3 | 0.0099491 | 0.05330691 | -1.011 | 0.273 | chr17 | 16593575 | 16707767 | + | protein_coding |
| ENSG00000241478.1_4 | HSPA8P9 | 0.02579642 | 0.12502047 | -1.012 | -0.460 | chr3 | 137599137 | 137601079 | + | processed_pseudogene |
| ENSG00000082153.18_5 | BZW1 | 6.23E-12 | 1.32E-10 | -1.016 | 7.664 | chr2 | 201675317 | 201692061 | + | protein_coding |
| ENSG00000259942.1 | RP11-292F22.7 | 0.02595566 | 0.12562086 | -1.016 | -0.653 | chr10 | 47656084 | 47657233 | + | lincRNA |
| ENSG00000186522.15_7 | SEPTIN10 | 8.80E-09 | 1.21E-07 | -1.017 | 5.862 | chr2 | 110300376 | 110371720 | - | protein_coding |
| ENSG00000019186.10_4 | CYP24A1 | 9.06E-13 | 2.20E-11 | -1.017 | 10.317 | chr20 | 52769985 | 52790525 | - | protein_coding |
| ENSG00000115318.12_6 | LOXL3 | 0.00183738 | 0.01137843 | -1.018 | 1.462 | chr2 | 74759385 | 74782817 | - | protein_coding |
| ENSG00000152661.9_6 | GJA1 | 4.16E-09 | 6.05E-08 | -1.018 | 6.488 | chr6 | 121756741 | 121770873 | + | protein_coding |
| ENSG00000167080.8_2 | B4GALNT2 | 0.03731666 | 0.17386009 | -1.018 | -0.707 | chr17 | 47209822 | 47254202 | + | protein_coding |
| ENSG00000228915.3_2 | OR7E128P | 0.02744635 | 0.13204422 | -1.019 | -0.652 | chr11 | 71604456 | 71605479 | + | unprocessed_pseudogene |
| ENSG00000223551.1_2 | TMSB4XP4 | 0.03364929 | 0.15862215 | -1.021 | -0.992 | chr9 | 131104432 | 131105049 | + | processed_pseudogene |
| ENSG00000138400.13_4 | MDH1B | 0.00169339 | 0.01053079 | -1.022 | 1.034 | chr2 | 207602487 | 207630052 | - | protein_coding |
| ENSG00000127152.18_4 | BCL11B | 0.00029073 | 0.00209298 | -1.024 | 2.006 | chr14 | 99635624 | 99738534 | - | protein_coding |
| ENSG00000197043.14_4 | ANXA6 | 1.58E-09 | 2.43E-08 | -1.024 | 5.406 | chr5 | 150480267 | 150537346 | - | protein_coding |
| ENSG00000164309.15_3 | CMYA5 | 0.03168531 | 0.15060269 | -1.024 | -0.992 | chr5 | 78985659 | 79096063 | + | protein_coding |
| ENSG00000197594.13_4 | ENPP1 | 3.96E-06 | 3.75E-05 | -1.025 | 3.241 | chr6 | 132129156 | 132216295 | + | protein_coding |
| ENSG00000164621.5_7 | SMAD5-AS1 | 0.03161254 | 0.15029705 | -1.026 | -0.908 | chr5 | 135465196 | 135470579 | - | lncRNA |
| ENSG00000125630.16_3 | POLR1B | 6.82E-11 | 1.25E-09 | -1.026 | 5.974 | chr2 | 113299492 | 113337395 | + | protein_coding |
| ENSG00000185088.14_4 | RPS27L | 4.84E-11 | 9.04E-10 | -1.026 | 7.305 | chr15 | 63418071 | 63450220 | - | protein_coding |
| ENSG00000081985.11_5 | IL12RB2 | 0.03137102 | 0.14922861 | -1.027 | -0.827 | chr1 | 67773047 | 67862773 | + | protein_coding |
| ENSG00000212195.1 | U3 | 0.004466 | 0.02572295 | -1.029 | 0.581 | chr17 | 56709003 | 56709197 | - | snoRNA |
| ENSG00000224057.1_7 | EGFR-AS1 | 0.00232674 | 0.01410892 | -1.030 | 0.909 | chr7 | 55247443 | 55256627 | - | lncRNA |
| ENSG00000235837.1_7 | AC073333.1 | 0.02881474 | 0.1378989 | -1.032 | -0.781 | chr7 | 16735496 | 16759523 | - | lncRNA |
| ENSG00000119729.12_4 | RHOQ | 1.37E-10 | 2.45E-09 | -1.033 | 6.072 | chr2 | 46768945 | 46811827 | + | protein_coding |
| ENSG00000236287.8_5 | ZBED5 | 1.01E-10 | 1.82E-09 | -1.033 | 6.245 | chr11 | 10833621 | 10880343 | - | protein_coding |
| ENSG00000198142.5_4 | SOWAHC | 2.90E-08 | 3.67E-07 | -1.033 | 4.285 | chr2 | 110371941 | 110376567 | + | protein_coding |
| ENSG00000134548.11_4 | SPX | 0.00079421 | 0.00525932 | -1.033 | 1.323 | chr12 | 21679230 | 21694183 | + | protein_coding |
| ENSG00000213277.3_4 | MARCKSL1P1 | 0.02011278 | 0.10026483 | -1.033 | -0.485 | chr10 | 104935311 | 104935851 | + | processed_pseudogene |
| ENSG00000117016.10_4 | RIMS3 | 4.02E-06 | 3.81E-05 | -1.034 | 2.938 | chr1 | 41086352 | 41131354 | - | protein_coding |
| ENSG00000154217.15_5 | PITPNC1 | 5.20E-05 | 0.00042219 | -1.034 | 2.269 | chr17 | 65373397 | 65693377 | + | protein_coding |
| ENSG00000169884.14_5 | WNT10B | 0.00464211 | 0.02661385 | -1.035 | 0.450 | chr12 | 49359123 | 49365518 | - | protein_coding |
| ENSG00000286997.1_2 | AC005856.1 | 0.00895871 | 0.04829167 | -1.035 | 0.144 | chr17 | 59196456 | 59213330 | - | lncRNA |
| ENSG00000282057.1_9 | AC092807.4 | 0.01565441 | 0.08001495 | -1.036 | -0.448 | chr1 | 85947964 | 86043933 | - | lncRNA |
| ENSG00000116120.11_5 | FARSB | 5.15E-09 | 7.34E-08 | -1.038 | 6.077 | chr2 | 223431618 | 223520811 | - | protein_coding |
| ENSG00000276334.1_6 | AL133243.2 | 0.00285433 | 0.01702347 | -1.038 | 0.732 | chr2 | 32746994 | 32748614 | + | lncRNA |
| ENSG00000285730.1_3 | Z94721.3 | 0.02754931 | 0.13243196 | -1.038 | -0.940 | chr6 | 167409223 | 167412999 | - | lncRNA |

| | | | | | | | | | | |
|---|---|---|---|---|---|---|---|---|---|---|
| ENSG00000236432.8_7 | MFF-DT | 0.00216009 | 0.01321114 | -1.039 | 0.815 | chr2 | 228085768 | 228190427 | - | lncRNA |
| ENSG00000143921.9_5 | ABCG8 | 0.02441466 | 0.11900557 | -1.039 | -0.860 | chr2 | 44059081 | 44110127 | + | protein_coding |
| ENSG00000275552.1_7 | AC243965.2 | 0.00581287 | 0.03264137 | -1.043 | 0.310 | chr14 | 23064715 | 23067855 | - | lncRNA |
| ENSG00000263280.2_7 | AC003965.1 | 0.0229709 | 0.11289736 | -1.043 | -0.783 | chr16 | 2916326 | 2918252 | - | lncRNA |
| ENSG00000091128.13_9 | LAMB4 | 0.03241546 | 0.15341646 | -1.044 | -1.119 | chr7 | 107663993 | 107770806 | - | protein_coding |
| ENSG00000235489.4_4 | DBF4P1 | 0.01657619 | 0.08424173 | -1.045 | -0.378 | chr10 | 65928720 | 65930611 | - | processed_pseudogene |
| ENSG00000232564.3_6 | MRTFA-AS1 | 0.01791626 | 0.09053421 | -1.046 | -0.599 | chr22 | 40917804 | 40922711 | + | lncRNA |
| ENSG00000170837.3_3 | GPR27 | 0.00636834 | 0.03545796 | -1.047 | 1.059 | chr3 | 71803006 | 71805647 | + | protein_coding |
| ENSG00000286145.2_2 | AC068282.1 | 0.02184695 | 0.10800075 | -1.049 | -0.625 | chr2 | 128051788 | 128132651 | + | lncRNA |
| ENSG00000166448.14_5 | TMEM130 | 0.03259085 | 0.1541234 | -1.049 | -1.029 | chr7 | 98444111 | 98468394 | - | protein_coding |
| ENSG00000237991.3_4 | RPL35P1 | 0.02593573 | 0.12555861 | -1.051 | -0.892 | chr1 | 237144639 | 237145008 | + | processed_pseudogene |
| ENSG00000184347.14_5 | SLIT3 | 2.08E-08 | 2.69E-07 | -1.051 | 4.406 | chr5 | 168088738 | 168728133 | - | protein_coding |
| ENSG00000213740.2_4 | SERBP1P1 | 0.01135888 | 0.05984973 | -1.053 | -0.203 | chrX | 68003315 | 68004909 | + | processed_pseudogene |
| ENSG00000100889.12_3 | PCK2 | 7.80E-08 | 9.36E-07 | -1.053 | 4.044 | chr14 | 24563262 | 24579807 | + | protein_coding |
| ENSG00000010610.10_4 | CD4 | 0.03395689 | 0.1599873 | -1.054 | -0.892 | chr12 | 6896024 | 6929965 | + | protein_coding |
| ENSG00000148926.10_4 | ADM | 4.29E-09 | 6.22E-08 | -1.055 | 5.396 | chr11 | 10326620 | 10328944 | + | protein_coding |
| ENSG00000168646.13_6 | AXIN2 | 0.00028631 | 0.00206364 | -1.055 | 2.144 | chr17 | 63524681 | 63557766 | - | protein_coding |
| ENSG00000115368.10_3 | WDR75 | 9.04E-10 | 1.44E-08 | -1.055 | 6.220 | chr2 | 190306172 | 190340278 | + | protein_coding |
| ENSG00000230176.3_6 | LINC01433 | 0.01876397 | 0.09433489 | -1.057 | -0.694 | chr20 | 4173579 | 4176600 | + | lncRNA |
| ENSG00000287445.1_3 | AC092803.3 | 0.02716638 | 0.13080357 | -1.057 | -0.812 | chr1 | 212603611 | 212606117 | - | lncRNA |
| ENSG00000254143.1_7 | AC022733.2 | 0.00787857 | 0.04311013 | -1.057 | -0.001 | chr8 | 40018977 | 40030131 | + | lncRNA |
| ENSG00000131016.17_4 | AKAP12 | 3.78E-13 | 9.81E-12 | -1.057 | 7.597 | chr6 | 151561102 | 151679694 | + | protein_coding |
| ENSG00000005961.19_6 | ITGA2B | 0.00019547 | 0.00144704 | -1.058 | 2.106 | chr17 | 42449548 | 42467017 | - | protein_coding |
| ENSG00000076003.5_4 | MCM6 | 7.75E-12 | 1.63E-10 | -1.058 | 6.461 | chr2 | 136597196 | 136634013 | - | protein_coding |
| ENSG00000069812.11_4 | HES2 | 0.0261559 | 0.12648661 | -1.058 | -0.976 | chr1 | 6472478 | 6484730 | - | protein_coding |
| ENSG00000130147.16_4 | SH3BP4 | 4.08E-12 | 8.89E-11 | -1.059 | 6.446 | chr2 | 235860661 | 235964358 | + | protein_coding |
| ENSG00000064205.10_3 | CCN5 | 1.16E-06 | 1.19E-05 | -1.060 | 3.430 | chr20 | 43343485 | 43357150 | + | protein_coding |
| ENSG00000258376.2_7 | AC004846.1 | 0.01338276 | 0.06944262 | -1.061 | 0.056 | chr14 | 73709359 | 73712687 | - | lncRNA |
| ENSG00000099957.16_3 | P2RX6 | 0.00840631 | 0.04559061 | -1.062 | -0.126 | chr22 | 21364097 | 21383119 | + | protein_coding |
| ENSG00000213160.10_6 | KLHL23 | 1.12E-09 | 1.77E-08 | -1.063 | 4.659 | chr2 | 170550998 | 170633499 | + | protein_coding |
| ENSG00000100784.12_7 | RPS6KA5 | 9.10E-06 | 8.21E-05 | -1.063 | 2.692 | chr14 | 91314205 | 91526985 | - | protein_coding |
| ENSG00000229337.2_8 | AC079305.4 | 0.00958134 | 0.05142951 | -1.064 | -0.017 | chr2 | 177975764 | 178077390 | - | lncRNA |
| ENSG00000144730.18_4 | IL17RD | 0.00047599 | 0.00329573 | -1.064 | 1.793 | chr3 | 57124010 | 57204334 | - | protein_coding |
| ENSG00000221962.5_3 | TMEM14EP | 0.02657805 | 0.12828366 | -1.065 | -0.585 | chr3 | 152057484 | 152058779 | - | transcribed_processed_pseudogene |
| ENSG00000137434.11_5 | C6orf52 | 0.00469528 | 0.02690133 | -1.066 | 0.509 | chr6 | 10671651 | 10695030 | - | protein_coding |
| ENSG00000131389.17_5 | SLC6A6 | 3.18E-13 | 8.31E-12 | -1.066 | 7.398 | chr3 | 14444076 | 14530857 | + | protein_coding |
| ENSG00000105711.12_6 | SCN1B | 4.95E-06 | 4.64E-05 | -1.067 | 2.927 | chr19 | 35521374 | 35531353 | + | protein_coding |
| ENSG00000204380.4_7 | PKP4-AS1 | 0.00643059 | 0.0357821 | -1.069 | 0.009 | chr2 | 159514849 | 159591514 | - | lncRNA |
| ENSG00000153165.18_3 | RGPD3 | 0.01210994 | 0.06343081 | -1.069 | -0.333 | chr2 | 107007746 | 107084832 | - | protein_coding |
| ENSG00000163806.16_5 | SPDYA | 0.00229538 | 0.0139283 | -1.071 | 0.591 | chr2 | 29005383 | 29073477 | + | protein_coding |
| ENSG00000138193.15_3 | PLCE1 | 3.66E-12 | 8.07E-11 | -1.071 | 6.118 | chr10 | 95790569 | 96092580 | + | protein_coding |
| ENSG00000258702.3_7 | AL137786.1 | 0.03757519 | 0.17480974 | -1.072 | -0.719 | chr14 | 97058736 | 97072311 | + | lncRNA |
| ENSG00000108932.12_6 | SLC16A6 | 0.02102237 | 0.10444756 | -1.072 | -0.445 | chr17 | 66263167 | 66287408 | - | protein_coding |
| ENSG00000170264.13_4 | FAM161A | 4.87E-08 | 6.01E-07 | -1.073 | 3.867 | chr2 | 62051983 | 62081278 | - | protein_coding |
| ENSG00000111664.10_5 | GNB3 | 0.00162081 | 0.01010773 | -1.073 | 0.828 | chr12 | 6949118 | 6956557 | + | protein_coding |
| ENSG00000207475.1 | SNORA42 | 0.03180025 | 0.15098743 | -1.073 | -1.064 | chr1 | 155889700 | 155889836 | - | snoRNA |
| ENSG00000214185.3_2 | XPOTP1 | 0.01526413 | 0.07817749 | -1.074 | -0.653 | chr20 | 32801301 | 32803698 | - | processed_pseudogene |
| ENSG00000273687.1_7 | AC004223.4 | 0.01490693 | 0.07654664 | -1.074 | -0.585 | chr17 | 33345679 | 33346010 | + | lncRNA |
| ENSG00000206630.1 | SNORD60 | 0.01061388 | 0.05642634 | -1.074 | 0.311 | chr16 | 2205024 | 2205106 | - | snoRNA |
| ENSG00000204745.4_5 | AC083899.1 | 0.00034826 | 0.00247505 | -1.077 | 1.328 | chr2 | 87352321 | 87423752 | + | unprocessed_pseudogene |
| ENSG00000287195.1_1 | AC131568.1 | 0.01185783 | 0.06222034 | -1.077 | -0.229 | chr8 | 130965975 | 130976880 | + | lncRNA |
| ENSG00000172965.17_8 | MIR4435-2HG | 1.78E-09 | 2.72E-08 | -1.079 | 4.452 | chr2 | 111763592 | 112280953 | - | lncRNA |
| ENSG00000133119.13_5 | RFC3 | 2.98E-11 | 5.78E-10 | -1.080 | 5.721 | chr13 | 34392206 | 34540695 | + | protein_coding |
| ENSG00000113594.10_5 | LIFR | 1.78E-09 | 2.72E-08 | -1.082 | 4.475 | chr5 | 38474770 | 38608456 | - | protein_coding |
| ENSG00000280077.1_6 | AL353763.2 | 0.02326402 | 0.11411701 | -1.082 | -0.926 | chr9 | 69085460 | 69087362 | + | TEC |
| ENSG00000213963.6_9 | AC019080.1 | 0.00055846 | 0.00381474 | -1.083 | 1.069 | chr2 | 178148236 | 178257419 | - | lncRNA |
| ENSG00000115657.14_6 | ABCB6 | 6.35E-05 | 0.0005087 | -1.083 | 2.164 | chr2 | 220074494 | 220083716 | - | protein_coding |

| | | | | | | | | | | |
|---|---|---|---|---|---|---|---|---|---|---|
| ENSG00000147509.14_4 | RGS20 | 5.99E-07 | 6.35E-06 | -1.084 | 3.283 | chr8 | 54764355 | 54871863 | + | protein_coding |
| ENSG00000236255.2_6 | AC009404.1 | 0.00025234 | 0.00183518 | -1.084 | 1.417 | chr2 | 118591513 | 118599234 | + | lncRNA |
| ENSG00000260804.3_5 | LINC01963 | 3.62E-07 | 3.94E-06 | -1.084 | 3.513 | chr2 | 217081768 | 217084915 | + | lncRNA |
| ENSG00000227934.1_5 | AL732414.1 | 0.06278529 | 0.27260661 | -1.085 | -1.484 | chr1 | 231157357 | 231157929 | + | processed_pseudogene |
| ENSG00000260912.1_6 | AL158206.1 | 0.00148045 | 0.00932072 | -1.085 | 0.789 | chr9 | 19453207 | 19455171 | + | lncRNA |
| ENSG00000269887.1_7 | AL391001.1 | 0.02593105 | 0.12555861 | -1.085 | -1.101 | chr1 | 164649322 | 164650036 | + | lncRNA |
| ENSG00000110811.20_4 | P3H3 | 7.26E-08 | 8.77E-07 | -1.086 | 4.115 | chr12 | 6937572 | 6949011 | + | protein_coding |
| ENSG00000229321.2_7 | AC008269.1 | 0.0068748 | 0.03796846 | -1.086 | 0.016 | chr2 | 207663049 | 207791600 | + | lncRNA |
| ENSG00000272800.3_7 | AC021851.1 | 0.01409658 | 0.07282757 | -1.087 | -0.368 | chr2 | 184043534 | 184080142 | + | lncRNA |
| ENSG00000089558.9_7 | KCNH4 | 0.01963537 | 0.09813266 | -1.089 | -0.752 | chr17 | 40308909 | 40333160 | - | protein_coding |
| ENSG00000273284.1_8 | AP001033.4 | 0.00873762 | 0.04725763 | -1.090 | -0.321 | chr18 | 9259386 | 9260388 | + | lncRNA |
| ENSG00000264630.5_7 | PRKCA-AS1 | 0.02161733 | 0.1069847 | -1.090 | -0.507 | chr17 | 64394187 | 64412972 | - | lncRNA |
| ENSG00000250519.7_6 | AP002784.1 | 0.00695855 | 0.03840714 | -1.091 | -0.107 | chr11 | 93918605 | 94015369 | + | lncRNA |
| ENSG00000184937.15_6 | WT1 | 5.81E-05 | 0.00046815 | -1.091 | 1.796 | chr11 | 32409321 | 32457110 | - | protein_coding |
| ENSG00000143382.15_4 | ADAMTSL4 | 0.00031232 | 0.00223391 | -1.092 | 1.207 | chr1 | 150521845 | 150533413 | + | protein_coding |
| ENSG00000075461.6_3 | CACNG4 | 1.58E-06 | 1.59E-05 | -1.093 | 3.251 | chr17 | 64960823 | 65029514 | + | protein_coding |
| ENSG00000227492.2_6 | AL138921.1 | 0.01920055 | 0.09623024 | -1.093 | -0.958 | chr10 | 101989386 | 101994155 | + | lncRNA |
| ENSG00000235961.5_2 | PNMA6A | 0.02104589 | 0.10450598 | -1.093 | -0.958 | chrX | 152240865 | 152243401 | + | protein_coding |
| ENSG00000259964.7_8 | THSD4-AS1 | 0.00842457 | 0.0456757 | -1.094 | -0.137 | chr15 | 71439989 | 71481403 | - | lncRNA |
| ENSG00000273747.1_7 | AC022558.3 | 0.02191868 | 0.10829511 | -1.094 | -1.045 | chr15 | 83688867 | 83689554 | + | lncRNA |
| ENSG00000088340.16_6 | FER1L4 | 1.19E-05 | 0.00010532 | -1.096 | 2.417 | chr20 | 34146507 | 34195484 | - | transcribed_unitary_pseudogene |
| ENSG00000183153.7_3 | GJD3 | 0.00818196 | 0.04456433 | -1.096 | 0.010 | chr17 | 38516904 | 38520989 | - | protein_coding |
| ENSG00000119630.14_5 | PGF | 0.00828935 | 0.04503878 | -1.096 | -0.269 | chr14 | 75408537 | 75422329 | - | protein_coding |
| ENSG00000143320.9_4 | CRABP2 | 0.00433356 | 0.02507142 | -1.098 | 0.150 | chr1 | 156669398 | 156675608 | - | protein_coding |
| ENSG00000199753.1 | SNORD104 | 0.00151132 | 0.00950499 | -1.098 | 0.561 | chr17 | 62223443 | 62223512 | + | snoRNA |
| ENSG00000144034.16_4 | TPRKB | 1.41E-10 | 2.51E-09 | -1.099 | 5.091 | chr2 | 73956231 | 73964527 | - | protein_coding |
| ENSG00000112320.12_3 | SOBP | 0.01767085 | 0.08944671 | -1.100 | -0.076 | chr6 | 107811310 | 107982510 | + | protein_coding |
| ENSG00000202496.1 | RNVU1-20 | 0.0243515 | 0.11873025 | -1.101 | -1.138 | chr1 | 149605917 | 149606079 | + | snRNA |
| ENSG00000198125.13_5 | MB | 5.81E-05 | 0.0004678 | -1.101 | 2.037 | chr22 | 36002811 | 36033998 | - | protein_coding |
| ENSG00000104321.11_4 | TRPA1 | 4.52E-08 | 5.60E-07 | -1.102 | 3.768 | chr8 | 72932152 | 72987819 | - | protein_coding |
| ENSG00000123600.19_5 | METTL8 | 6.81E-11 | 1.25E-09 | -1.102 | 5.103 | chr2 | 172173915 | 172291312 | - | protein_coding |
| ENSG00000280184.2_4 | AL023806.3 | 0.00796315 | 0.04350591 | -1.102 | -0.268 | chr6 | 146110406 | 146113109 | - | TEC |
| ENSG00000074047.21_3 | GLI2 | 1.49E-08 | 1.97E-07 | -1.103 | 4.711 | chr2 | 121493199 | 121750229 | + | protein_coding |
| ENSG00000286207.1_3 | AC012443.2 | 0.00083557 | 0.00551059 | -1.104 | 0.894 | chr2 | 153491782 | 153497789 | - | lncRNA |
| ENSG00000136895.19_8 | GARNL3 | 9.05E-06 | 8.17E-05 | -1.105 | 2.298 | chr9 | 129986544 | 130155939 | + | protein_coding |
| ENSG00000196843.17_4 | ARID5A | 0.00028957 | 0.00208547 | -1.106 | 1.448 | chr2 | 97202480 | 97218371 | + | protein_coding |
| ENSG00000234832.1_7 | NXT1-AS1 | 0.02628055 | 0.12698572 | -1.106 | -1.047 | chr20 | 23327578 | 23332123 | - | lncRNA |
| ENSG00000245534.7_7 | RORA-AS1 | 0.00697487 | 0.03847332 | -1.106 | -0.204 | chr15 | 60771351 | 60922836 | + | lncRNA |
| ENSG00000121964.14_5 | GTDC1 | 1.04E-07 | 1.23E-06 | -1.107 | 3.464 | chr2 | 144695635 | 145090135 | - | protein_coding |
| ENSG00000102385.12_2 | DRP2 | 0.000399 | 0.00280307 | -1.107 | 1.018 | chrX | 100474758 | 100519486 | + | protein_coding |
| ENSG00000126878.13_3 | AIF1L | 8.42E-05 | 0.00065827 | -1.107 | 1.745 | chr9 | 133971863 | 133998539 | + | protein_coding |
| ENSG00000287502.1_2 | AC092418.3 | 0.02633869 | 0.12723204 | -1.108 | -1.139 | chr3 | 57541308 | 57541781 | - | lncRNA |
| ENSG00000235036.4_5 | AL035456.1 | 0.0062146 | 0.03465625 | -1.108 | -0.155 | chr20 | 10593509 | 10594877 | - | processed_pseudogene |
| ENSG00000241120.1_4 | HMGN1P8 | 0.02620587 | 0.12665945 | -1.109 | -1.046 | chr3 | 167412143 | 167412430 | - | processed_pseudogene |
| ENSG00000064655.19_6 | EYA2 | 2.84E-05 | 0.00024099 | -1.110 | 2.163 | chr20 | 45523263 | 45817492 | + | protein_coding |
| ENSG00000286588.1_2 | AC104841.1 | 0.01613256 | 0.08215165 | -1.110 | -0.710 | chr2 | 242265749 | 242266454 | - | lncRNA |
| ENSG00000153574.9_3 | RPIA | 9.81E-07 | 1.02E-05 | -1.111 | 2.861 | chr2 | 88991191 | 89050446 | + | protein_coding |
| ENSG00000236986.6_9 | AL157938.2 | 0.00523345 | 0.02968794 | -1.111 | 0.051 | chr9 | 134008239 | 134042892 | - | lncRNA |
| ENSG00000231201.1_6 | AF127577.2 | 0.02717991 | 0.13083321 | -1.113 | -1.238 | chr21 | 16343791 | 16365175 | + | lncRNA |
| ENSG00000113083.14_6 | LOX | 0.00452153 | 0.0260065 | -1.113 | -0.006 | chr5 | 121398890 | 121414055 | - | protein_coding |
| ENSG00000134317.18_3 | GRHL1 | 0.00225609 | 0.01372741 | -1.115 | 0.251 | chr2 | 10091822 | 10142405 | + | protein_coding |
| ENSG00000264829.1_7 | AC087749.2 | 0.01878564 | 0.09439042 | -1.115 | -0.858 | chr17 | 73675554 | 73676047 | - | lncRNA |
| ENSG00000144395.18_4 | CCDC150 | 3.99E-08 | 4.97E-07 | -1.116 | 3.724 | chr2 | 197504278 | 197628214 | + | protein_coding |
| ENSG00000115163.15_4 | CENPA | 2.53E-11 | 4.96E-10 | -1.118 | 5.115 | chr2 | 26987157 | 27023935 | + | protein_coding |
| ENSG00000112309.11_4 | B3GAT2 | 0.02366316 | 0.11572456 | -1.119 | -1.081 | chr6 | 71566382 | 71666763 | - | protein_coding |
| ENSG00000091879.13_3 | ANGPT2 | 0.00736953 | 0.04052445 | -1.119 | -0.345 | chr8 | 6357172 | 6420930 | - | protein_coding |
| ENSG00000213090.2_5 | AC007256.1 | 0.16862464 | 0.62324816 | -1.119 | -1.174 | chr2 | 202275267 | 202278031 | - | processed_pseudogene |
| ENSG00000269967.1_8 | AL136115.2 | 0.01669781 | 0.08476285 | -1.119 | -0.735 | chr1 | 32317514 | 32387442 | - | lncRNA |

| | | | | | | | | | | |
|---|---|---|---|---|---|---|---|---|---|---|
| ENSG00000170523.3_5 | KRT83 | 3.11E-05 | 0.00026227 | -1.121 | 3.367 | chr12 | 52708085 | 52715182 | - | protein_coding |
| ENSG00000180739.14_5 | S1PR5 | 0.00127194 | 0.00812313 | -1.121 | 0.854 | chr19 | 10623418 | 10628607 | - | protein_coding |
| ENSG00000272980.4_7 | Z94721.2 | 0.01945467 | 0.09733908 | -1.121 | -1.082 | chr6 | 167412893 | 167552629 | + | lncRNA |
| ENSG00000225137.1_2 | DYNC1I2P1 | 0.00416896 | 0.02416639 | -1.121 | 0.390 | chr10 | 52024738 | 52026575 | + | processed_pseudogene |
| ENSG00000151632.17_5 | AKR1C2 | 8.44E-15 | 2.91E-13 | -1.122 | 11.038 | chr10 | 5029592 | 5060223 | - | protein_coding |
| ENSG00000254388.1_4 | DUTP2 | 0.02568235 | 0.12453557 | -1.122 | -0.992 | chr8 | 124364421 | 124364883 | + | processed_pseudogene |
| ENSG00000233461.6_7 | AL445524.1 | 3.47E-05 | 0.00028958 | -1.122 | 1.757 | chr1 | 231656475 | 231664364 | - | lncRNA |
| ENSG00000148677.6_3 | ANKRD1 | 1.68E-05 | 0.00014638 | -1.122 | 3.049 | chr10 | 92671853 | 92681033 | - | protein_coding |
| ENSG00000235427.1_7 | AC006159.1 | 0.02117441 | 0.10505607 | -1.122 | -0.624 | chr7 | 116182772 | 116192023 | - | lncRNA |
| ENSG00000270264.1_4 | NDUFB8P2 | 0.00789345 | 0.04317822 | -1.124 | -0.299 | chr17 | 57962376 | 57962913 | + | processed_pseudogene |
| ENSG00000107833.10_3 | NPM3 | 2.78E-10 | 4.73E-09 | -1.125 | 4.586 | chr10 | 103541082 | 103543170 | - | protein_coding |
| ENSG00000128965.13_5 | CHAC1 | 0.00023794 | 0.00173543 | -1.127 | 1.471 | chr15 | 41245160 | 41248710 | + | protein_coding |
| ENSG00000188916.9_7 | INSYN2A | 0.00341343 | 0.02006169 | -1.127 | 0.071 | chr10 | 128933690 | 128994855 | - | protein_coding |
| ENSG00000266638.1 | MIR3676 | 0.00224141 | 0.01365207 | -1.128 | 0.317 | chr17 | 8090493 | 8090577 | + | miRNA |
| ENSG00000259426.6_8 | AC027237.5 | 0.00024287 | 0.00176847 | -1.129 | 1.236 | chr15 | 69689243 | 69707368 | - | lncRNA |
| ENSG00000221866.9_3 | PLXNA4 | 0.00136951 | 0.00867451 | -1.129 | 0.709 | chr7 | 131808091 | 132333447 | - | protein_coding |
| ENSG00000114999.8_3 | TTL | 6.86E-13 | 1.72E-11 | -1.130 | 6.196 | chr2 | 113239733 | 113299316 | + | protein_coding |
| ENSG00000105649.10_4 | RAB3A | 0.00420341 | 0.0243502 | -1.130 | -0.117 | chr19 | 18307594 | 18314852 | - | protein_coding |
| ENSG00000159166.15_8 | LAD1 | 0.01668664 | 0.08473036 | -1.130 | -0.721 | chr1 | 201349961 | 201369043 | - | protein_coding |
| ENSG00000068724.17_6 | TTC7A | 8.89E-09 | 1.22E-07 | -1.132 | 5.579 | chr2 | 47143296 | 47303276 | + | protein_coding |
| ENSG00000163705.13_5 | FANCD2OS | 0.02170943 | 0.1074106 | -1.132 | -0.975 | chr3 | 10123001 | 10149939 | - | protein_coding |
| ENSG00000131620.17_5 | ANO1 | 0.00072421 | 0.00483181 | -1.133 | 0.633 | chr11 | 69924408 | 70035634 | + | protein_coding |
| ENSG00000140263.14_3 | SORD | 1.05E-09 | 1.66E-08 | -1.133 | 4.611 | chr15 | 45315379 | 45369383 | + | protein_coding |
| ENSG00000284138.1_4 | ATP6V0CP4 | 0.0109142 | 0.05776365 | -1.135 | -0.651 | chr1 | 43417873 | 43418312 | - | processed_pseudogene |
| ENSG00000131378.14_8 | RFTN1 | 1.26E-05 | 0.00011143 | -1.136 | 2.139 | chr3 | 16355081 | 16555533 | - | protein_coding |
| ENSG00000075340.23_5 | ADD2 | 2.24E-11 | 4.41E-10 | -1.138 | 5.445 | chr2 | 70834750 | 70995357 | - | protein_coding |
| ENSG00000159958.6_4 | TNFRSF13C | 0.014019 | 0.07244781 | -1.139 | -0.695 | chr22 | 42318027 | 42322822 | - | protein_coding |
| ENSG00000184185.10_4 | KCNJ12 | 0.00572523 | 0.03220009 | -1.139 | -0.217 | chr17 | 21279669 | 21323182 | + | protein_coding |
| ENSG00000272275.1_7 | AC092687.3 | 0.01904352 | 0.09555104 | -1.140 | -1.119 | chr2 | 10908001 | 10910184 | - | lncRNA |
| ENSG00000138442.9_3 | WDR12 | 9.34E-12 | 1.95E-10 | -1.141 | 5.585 | chr2 | 203739505 | 203879521 | - | protein_coding |
| ENSG00000228175.3_4 | GEMIN8P4 | 0.00296989 | 0.0176534 | -1.142 | 0.519 | chr1 | 90459152 | 90459880 | - | processed_pseudogene |
| ENSG00000223960.7_8 | CHROMR | 1.34E-09 | 2.10E-08 | -1.143 | 4.183 | chr2 | 179278362 | 179304970 | + | lncRNA |
| ENSG00000261549.2_5 | AC107954.1 | 0.04142811 | 0.19078622 | -1.144 | -0.924 | chr15 | 89532750 | 89533476 | - | processed_pseudogene |
| ENSG00000105088.8_3 | OLFM2 | 0.00041656 | 0.00291379 | -1.144 | 1.189 | chr19 | 9964394 | 10047228 | - | protein_coding |
| ENSG00000252759.1 | Y_RNA | 0.02548433 | 0.12373504 | -1.146 | -1.217 | chr2 | 201727875 | 201727970 | - | misc_RNA |
| ENSG00000279544.1_2 | AL133243.4 | 0.00680644 | 0.03764944 | -1.146 | -0.376 | chr2 | 32788395 | 32791503 | + | TEC |
| ENSG00000085978.22_4 | ATG16L1 | 3.30E-13 | 8.61E-12 | -1.146 | 6.007 | chr2 | 234118697 | 234204320 | + | protein_coding |
| ENSG00000273306.1_6 | AC018690.1 | 0.00483102 | 0.02759893 | -1.146 | -0.139 | chr2 | 100021680 | 100022305 | + | lncRNA |
| ENSG00000169314.14_3 | C22orf15 | 0.02004241 | 0.09994203 | -1.146 | -1.120 | chr22 | 24105208 | 24108048 | + | protein_coding |
| ENSG00000112276.14_4 | BVES | 8.09E-08 | 9.71E-07 | -1.147 | 3.242 | chr6 | 105544697 | 105585032 | - | protein_coding |
| ENSG00000145287.10_3 | PLAC8 | 0.00018658 | 0.00138526 | -1.150 | 1.077 | chr4 | 84011201 | 84058228 | - | protein_coding |
| ENSG00000259479.6_5 | SORD2P | 0.01125933 | 0.05939563 | -1.150 | -0.469 | chr15 | 45117945 | 45176892 | - | transcribed_unprocessed_pseudogene |
| ENSG00000156113.23_7 | KCNMA1 | 4.49E-11 | 8.42E-10 | -1.153 | 4.712 | chr10 | 78629359 | 79398127 | - | protein_coding |
| ENSG00000287555.1_1 | AC023424.3 | 0.02972369 | 0.14188632 | -1.153 | -1.062 | chr4 | 153369427 | 153434940 | + | lncRNA |
| ENSG00000053747.16_4 | LAMA3 | 1.47E-13 | 4.09E-12 | -1.154 | 6.429 | chr18 | 21269407 | 21536186 | + | protein_coding |
| ENSG00000169991.11_6 | IFFO2 | 4.74E-12 | 1.02E-10 | -1.154 | 5.241 | chr1 | 19230774 | 19283170 | - | protein_coding |
| ENSG00000172575.12_4 | RASGRP1 | 0.0003258 | 0.00232 | -1.154 | 0.914 | chr15 | 38780304 | 38857776 | - | protein_coding |
| ENSG00000118513.19_6 | MYB | 0.00715772 | 0.03942073 | -1.155 | -0.376 | chr6 | 135502446 | 135540311 | + | protein_coding |
| ENSG00000243415.2_6 | AC107021.1 | 0.00494253 | 0.02816356 | -1.155 | -0.236 | chr3 | 145781829 | 145822991 | + | lncRNA |
| ENSG00000114251.14_4 | WNT5A | 0.00224458 | 0.01366668 | -1.156 | 0.086 | chr3 | 55499743 | 55524567 | - | protein_coding |
| ENSG00000022567.9_4 | SLC45A4 | 6.46E-12 | 1.37E-10 | -1.156 | 5.267 | chr8 | 142217265 | 142318404 | - | protein_coding |
| ENSG00000201592.1 | snoU2_19 | 0.00273343 | 0.01636831 | -1.162 | 0.161 | chrX | 20154424 | 20154503 | - | snoRNA |
| ENSG00000166450.13_5 | PRTG | 1.26E-10 | 2.27E-09 | -1.163 | 4.953 | chr15 | 55903742 | 56035350 | - | protein_coding |
| ENSG00000144893.12_6 | MED12L | 6.82E-05 | 0.00054298 | -1.163 | 1.839 | chr3 | 150803484 | 151154860 | + | protein_coding |
| ENSG00000285804.2_3 | AC025774.1 | 0.00457456 | 0.02629445 | -1.163 | -0.079 | chr5 | 71383609 | 71403827 | - | lncRNA |
| ENSG00000224738.1_6 | AC099850.2 | 0.00154874 | 0.00970256 | -1.164 | 0.384 | chr17 | 57183959 | 57195628 | + | lncRNA |
| ENSG00000287306.1_1 | AC016821.1 | 0.00746699 | 0.0410096 | -1.164 | -0.472 | chr10 | 71269922 | 71304013 | + | lncRNA |

| | | | | | | | | | | |
|---|---|---|---|---|---|---|---|---|---|---|
| ENSG00000121152.10_5 | NCAPH | 5.08E-13 | 1.30E-11 | -1.165 | 5.624 | chr2 | 97001504 | 97042829 | + | protein_coding |
| ENSG00000163002.13_8 | NUP35 | 4.38E-12 | 9.47E-11 | -1.167 | 5.139 | chr2 | 183982241 | 184026408 | + | protein_coding |
| ENSG00000163053.11_7 | SLC16A14 | 0.00063761 | 0.00431067 | -1.168 | 0.919 | chr2 | 230899698 | 230933709 | - | protein_coding |
| ENSG00000231731.7_8 | AC010976.1 | 6.58E-05 | 0.00052561 | -1.169 | 1.323 | chr2 | 128212970 | 128272199 | + | lncRNA |
| ENSG00000112183.15_3 | RBM24 | 1.84E-09 | 2.81E-08 | -1.171 | 3.919 | chr6 | 17281592 | 17294102 | + | protein_coding |
| ENSG00000184434.8_4 | LRRC19 | 0.00334891 | 0.01969547 | -1.171 | 0.046 | chr9 | 26993134 | 27005670 | - | protein_coding |
| ENSG00000176928.7_4 | GCNT4 | 0.01022431 | 0.05460033 | -1.172 | -0.828 | chr5 | 74321171 | 74348383 | - | protein_coding |
| ENSG00000137273.6_6 | FOXF2 | 5.26E-06 | 4.92E-05 | -1.172 | 2.418 | chr6 | 1389811 | 1395838 | + | protein_coding |
| ENSG00000135378.4_3 | PRRG4 | 0.00348441 | 0.02045855 | -1.174 | 0.040 | chr11 | 32851473 | 32879666 | + | protein_coding |
| ENSG00000287001.1_2 | AC010624.5 | 2.22E-07 | 2.51E-06 | -1.174 | 2.843 | chr19 | 50640116 | 50651487 | - | lncRNA |
| ENSG00000267432.6_7 | DNAH17-AS1 | 0.00010427 | 0.00080341 | -1.175 | 1.193 | chr17 | 76480964 | 76499138 | + | lncRNA |
| ENSG00000171126.8_6 | KCNG3 | 0.00078129 | 0.00518344 | -1.175 | 0.718 | chr2 | 42669157 | 42721122 | - | protein_coding |
| ENSG00000274667.1_7 | AC090517.2 | 0.00145204 | 0.00916135 | -1.176 | 0.221 | chr15 | 57022130 | 57022809 | - | lncRNA |
| ENSG00000287287.1_2 | AL133476.1 | 3.19E-05 | 0.0002683 | -1.176 | 1.609 | chr9 | 35491130 | 35491966 | - | lncRNA |
| ENSG00000165061.15_5 | ZMAT4 | 5.02E-05 | 0.000409 | -1.180 | 1.447 | chr8 | 40388109 | 40755352 | - | protein_coding |
| ENSG00000254960.2_5 | KIRREL3-AS2 | 0.0224732 | 0.11081914 | -1.180 | -1.100 | chr11 | 126810642 | 126814986 | + | lncRNA |
| ENSG00000205978.6_3 | NYNRIN | 0.0007239 | 0.00483153 | -1.184 | 0.597 | chr14 | 24868209 | 24888489 | + | protein_coding |
| ENSG00000150551.11_4 | LYPD1 | 0.00065154 | 0.0043965 | -1.184 | 0.644 | chr2 | 133400859 | 133429152 | - | protein_coding |
| ENSG00000115226.10_4 | FNDC4 | 0.00050157 | 0.00345357 | -1.185 | 0.508 | chr2 | 27714750 | 27718067 | - | protein_coding |
| ENSG00000250565.7_5 | ATP6V1E2 | 4.29E-10 | 7.12E-09 | -1.185 | 4.074 | chr2 | 46717889 | 46769716 | - | protein_coding |
| ENSG00000171873.8_4 | ADRA1D | 0.000862 | 0.00567021 | -1.186 | 0.699 | chr20 | 4201277 | 4229934 | - | protein_coding |
| ENSG00000280109.3_7 | PLAC4 | 0.01260441 | 0.06584609 | -1.188 | -0.664 | chr21 | 42547158 | 42558715 | - | lncRNA |
| ENSG00000239218.2_2 | RPS20P22 | 0.01275264 | 0.06656176 | -1.188 | -0.907 | chr8 | 38291865 | 38293182 | - | transcribed_processed_pseudogene |
| ENSG00000187699.10_4 | C2orf88 | 0.00087712 | 0.00576324 | -1.188 | 0.428 | chr2 | 190744335 | 191068210 | + | protein_coding |
| ENSG00000233514.1_5 | AL356653.1 | 0.01623803 | 0.08264138 | -1.188 | -0.811 | chr1 | 44587825 | 44588184 | - | processed_pseudogene |
| ENSG00000203852.3_3 | H3C15 | 0.01341908 | 0.06957012 | -1.188 | -0.811 | chr1 | 149824181 | 149825836 | + | protein_coding |
| ENSG00000149380.12_4 | P4HA3 | 3.32E-08 | 4.17E-07 | -1.189 | 3.084 | chr11 | 73946846 | 74022685 | - | protein_coding |
| ENSG00000106483.12_6 | SFRP4 | 5.45E-07 | 5.82E-06 | -1.190 | 2.804 | chr7 | 37945534 | 38065297 | - | protein_coding |
| ENSG00000020577.14_4 | SAMD4A | 2.10E-08 | 2.71E-07 | -1.191 | 3.368 | chr14 | 55032835 | 55260033 | + | protein_coding |
| ENSG00000279652.1_6 | Z82217.1 | 0.0132555 | 0.06883741 | -1.192 | -1.046 | chr22 | 36388369 | 36396517 | + | lncRNA |
| ENSG00000185818.8_5 | NAT8L | 9.21E-06 | 8.30E-05 | -1.193 | 2.275 | chr4 | 2061054 | 2070816 | + | protein_coding |
| ENSG00000171793.16_6 | CTPS1 | 5.93E-15 | 2.08E-13 | -1.193 | 6.680 | chr1 | 41445360 | 41478237 | + | protein_coding |
| ENSG00000177483.11_4 | RBM44 | 0.00215292 | 0.01317999 | -1.193 | -0.009 | chr2 | 238707032 | 238751451 | + | protein_coding |
| ENSG00000251580.3_7 | LINC02482 | 0.00174459 | 0.0108227 | -1.194 | 0.071 | chr4 | 6650422 | 6675692 | - | lncRNA |
| ENSG00000186897.5_4 | C1QL4 | 0.00015886 | 0.00119698 | -1.194 | 1.096 | chr12 | 49726192 | 49730971 | - | protein_coding |
| ENSG00000189060.5_3 | H1-0 | 1.12E-13 | 3.15E-12 | -1.195 | 6.236 | chr22 | 38201100 | 38203443 | + | protein_coding |
| ENSG00000088205.13_3 | DDX18 | 8.79E-14 | 2.56E-12 | -1.195 | 6.646 | chr2 | 118572267 | 118589953 | + | protein_coding |
| ENSG00000248483.7_4 | POU5F2 | 0.01073528 | 0.05693545 | -1.196 | -0.861 | chr5 | 93068926 | 93077306 | - | protein_coding |
| ENSG00000167771.6_4 | RCOR2 | 0.00071245 | 0.00476764 | -1.197 | 0.317 | chr11 | 63678702 | 63684636 | - | protein_coding |
| ENSG00000181798.3_5 | LINC00471 | 0.00186199 | 0.01151078 | -1.197 | 0.071 | chr2 | 232373133 | 232379077 | - | lncRNA |
| ENSG00000168243.11_5 | GNG4 | 1.06E-05 | 9.45E-05 | -1.198 | 1.875 | chr1 | 235710985 | 235814054 | - | protein_coding |
| ENSG00000023902.14_4 | PLEKHO1 | 6.77E-06 | 6.23E-05 | -1.198 | 1.737 | chr1 | 150121373 | 150136916 | + | protein_coding |
| ENSG00000266897.1_7 | AC005546.1 | 0.01104285 | 0.05834027 | -1.199 | -0.858 | chr19 | 13242385 | 13243224 | + | lncRNA |
| ENSG00000250869.2_6 | AC087359.1 | 0.01656859 | 0.08422724 | -1.200 | -1.138 | chr8 | 68045789 | 68048146 | - | unprocessed_pseudogene |
| ENSG00000213315.5_5 | AL122020.1 | 0.01942299 | 0.09720796 | -1.203 | -1.369 | chr14 | 91232875 | 91233333 | + | processed_pseudogene |
| ENSG00000106823.12_6 | ECM2 | 0.00699469 | 0.03857064 | -1.203 | -0.610 | chr9 | 95255836 | 95298937 | - | protein_coding |
| ENSG00000230699.2_7 | AL645608.2 | 0.0125104 | 0.06539343 | -1.203 | -0.765 | chr1 | 846815 | 850328 | + | lncRNA |
| ENSG00000198796.7_5 | ALPK2 | 9.03E-11 | 1.64E-09 | -1.203 | 4.349 | chr18 | 56148479 | 56296323 | - | protein_coding |
| ENSG00000138030.13_4 | KHK | 1.22E-08 | 1.62E-07 | -1.204 | 3.392 | chr2 | 27309615 | 27323630 | + | protein_coding |
| ENSG00000112149.10_3 | CD83 | 0.00110986 | 0.00716792 | -1.204 | 0.199 | chr6 | 14117487 | 14137149 | + | protein_coding |
| ENSG00000153250.20_5 | RBMS1 | 1.70E-13 | 4.65E-12 | -1.206 | 6.253 | chr2 | 161128662 | 161350318 | - | protein_coding |
| ENSG00000262202.4_6 | AC007952.4 | 0.0022811 | 0.01386055 | -1.206 | -0.100 | chr17 | 19015313 | 19015949 | - | lncRNA |
| ENSG00000105988.6_4 | NHP2P1 | 0.00998025 | 0.05344157 | -1.207 | -0.890 | chr10 | 93976089 | 93976550 | + | processed_pseudogene |
| ENSG00000264112.1_8 | AC015813.1 | 4.24E-11 | 8.00E-10 | -1.208 | 5.073 | chr17 | 56066400 | 56072211 | - | lncRNA |
| ENSG00000279358.1_6 | AP002833.3 | 0.00298803 | 0.01774934 | -1.209 | -0.288 | chr11 | 126804782 | 126807046 | - | TEC |
| ENSG00000105613.10_3 | MAST1 | 0.00058896 | 0.00400306 | -1.209 | 0.323 | chr19 | 12944765 | 12985766 | + | protein_coding |
| ENSG00000138399.18_6 | FASTKD1 | 7.04E-12 | 1.48E-10 | -1.210 | 4.790 | chr2 | 170385018 | 170430385 | - | protein_coding |

| | | | | | | | | | | |
|---|---|---|---|---|---|---|---|---|---|---|
| ENSG00000138074.15_9 | SLC5A6 | 2.62E-14 | 8.42E-13 | -1.212 | 5.849 | chr2 | 27422455 | 27435826 | - | protein_coding |
| ENSG00000169679.15_7 | BUB1 | 1.12E-14 | 3.79E-13 | -1.213 | 6.765 | chr2 | 111395105 | 111435640 | - | protein_coding |
| ENSG00000134369.15_4 | NAV1 | 1.84E-09 | 2.80E-08 | -1.213 | 3.752 | chr1 | 201592013 | 201796097 | + | protein_coding |
| ENSG00000241472.7_7 | PTPRG-AS1 | 0.00133753 | 0.00848754 | -1.216 | 0.352 | chr3 | 62206923 | 62355081 | - | lncRNA |
| ENSG00000141314.13_5 | RHBDL3 | 6.24E-07 | 6.60E-06 | -1.216 | 2.421 | chr17 | 30592851 | 30651678 | + | protein_coding |
| ENSG00000237298.10_7 | TTN-AS1 | 9.92E-05 | 0.00076727 | -1.216 | 1.018 | chr2 | 179385910 | 179644690 | + | lncRNA |
| ENSG00000169629.12_4 | RGPD8 | 1.05E-05 | 9.37E-05 | -1.217 | 1.604 | chr2 | 113125946 | 113192065 | - | protein_coding |
| ENSG00000019582.15_6 | CD74 | 0.00108813 | 0.00704805 | -1.217 | 0.334 | chr5 | 149779604 | 149792492 | - | protein_coding |
| ENSG00000173744.17_6 | AGFG1 | 3.80E-15 | 1.36E-13 | -1.217 | 7.514 | chr2 | 228336868 | 228425930 | + | protein_coding |
| ENSG00000187498.16_6 | COL4A1 | 4.78E-13 | 1.22E-11 | -1.217 | 5.711 | chr13 | 110801310 | 110959504 | - | protein_coding |
| ENSG00000115468.12_4 | EFHD1 | 0.00158209 | 0.00988013 | -1.219 | 0.053 | chr2 | 233470767 | 233547490 | + | protein_coding |
| ENSG00000246985.8_7 | SOCS2-AS1 | 0.00034036 | 0.00242079 | -1.220 | 0.697 | chr12 | 93935798 | 93965544 | - | lncRNA |
| ENSG00000163249.12_6 | CCNYL1 | 8.17E-10 | 1.31E-08 | -1.221 | 3.909 | chr2 | 208576264 | 208626563 | + | protein_coding |
| ENSG00000272056.1_7 | AC013472.2 | 0.02385503 | 0.1165976 | -1.224 | -1.177 | chr2 | 27276486 | 27277144 | - | lncRNA |
| ENSG00000222724.1 | RNU2-63P | 0.00535447 | 0.03033573 | -1.224 | -0.597 | chr2 | 88315873 | 88316066 | - | snRNA |
| ENSG00000196839.13_4 | ADA | 2.90E-10 | 4.93E-09 | -1.225 | 3.871 | chr20 | 43248163 | 43280874 | - | protein_coding |
| ENSG00000146006.8_4 | LRRTM2 | 0.01358122 | 0.07036965 | -1.225 | -0.843 | chr5 | 138204610 | 138211057 | - | protein_coding |
| ENSG00000252213.1 | SNORA74 | 1.02E-05 | 9.13E-05 | -1.227 | 1.582 | chr5 | 138611869 | 138612009 | + | snoRNA |
| ENSG00000132330.18_5 | SCLY | 7.14E-06 | 6.55E-05 | -1.227 | 1.604 | chr2 | 238969565 | 239008054 | + | protein_coding |
| ENSG00000001617.12_5 | SEMA3F | 4.78E-05 | 0.00039169 | -1.227 | 1.331 | chr3 | 50192478 | 50226508 | + | protein_coding |
| ENSG00000118777.12_7 | ABCG2 | 5.71E-16 | 2.31E-14 | -1.233 | 6.905 | chr4 | 89011302 | 89152780 | - | protein_coding |
| ENSG00000286742.1_1 | AC000065.1 | 0.00152289 | 0.00956083 | -1.233 | -0.033 | chr7 | 92277220 | 92296397 | + | lncRNA |
| ENSG00000168958.19_7 | MFF | 3.27E-15 | 1.18E-13 | -1.233 | 6.715 | chr2 | 228189867 | 228222552 | + | protein_coding |
| ENSG00000162409.11_3 | PRKAA2 | 7.49E-11 | 1.37E-09 | -1.233 | 4.855 | chr1 | 57110987 | 57181008 | + | protein_coding |
| ENSG00000186868.15_3 | MAPT | 0.00048786 | 0.00336876 | -1.234 | 0.359 | chr17 | 43971748 | 44105700 | + | protein_coding |
| ENSG00000168079.17_5 | SCARA5 | 1.93E-11 | 3.84E-10 | -1.234 | 4.469 | chr8 | 27727400 | 27850190 | - | protein_coding |
| ENSG00000159263.16_4 | SIM2 | 1.89E-11 | 3.76E-10 | -1.234 | 4.329 | chr21 | 38071415 | 38122218 | + | protein_coding |
| ENSG00000136286.16_5 | MYO1G | 0.01380217 | 0.07138955 | -1.235 | -1.179 | chr7 | 45002261 | 45018687 | - | protein_coding |
| ENSG00000166839.17_6 | ANKDD1A | 0.00014962 | 0.00113118 | -1.235 | 1.078 | chr15 | 65204101 | 65251033 | + | protein_coding |
| ENSG00000260121.1_6 | AC138028.4 | 0.00429305 | 0.02485328 | -1.236 | -0.599 | chr16 | 88785023 | 88786867 | + | lncRNA |
| ENSG00000135097.7_5 | MSI1 | 0.01078232 | 0.05712762 | -1.238 | -0.799 | chr12 | 120779133 | 120806967 | - | protein_coding |
| ENSG00000066248.15_6 | NGEF | 1.01E-08 | 1.37E-07 | -1.239 | 3.435 | chr2 | 233743411 | 233877966 | - | protein_coding |
| ENSG00000285906.2_3 | AC083855.2 | 2.32E-05 | 0.00019897 | -1.240 | 1.515 | chr3 | 11235125 | 11267563 | - | lncRNA |
| ENSG00000159433.12_4 | STARD9 | 6.02E-10 | 9.83E-09 | -1.241 | 3.754 | chr15 | 42867804 | 43013196 | + | protein_coding |
| ENSG00000231767.4_5 | AL136454.1 | 0.0001766 | 0.00131846 | -1.241 | 1.473 | chr1 | 192685262 | 192685783 | + | protein_coding |
| ENSG00000223374.1_7 | AC005104.1 | 0.01030144 | 0.05496266 | -1.241 | -0.683 | chr2 | 242290755 | 242292519 | - | lncRNA |
| ENSG00000135074.15_5 | ADAM19 | 1.77E-08 | 2.31E-07 | -1.244 | 3.112 | chr5 | 156822542 | 157002783 | - | protein_coding |
| ENSG00000265142.9_8 | MIR133A1HG | 0.00558766 | 0.03147102 | -1.244 | -0.521 | chr18 | 19405448 | 19411500 | - | lncRNA |
| ENSG00000242602.1_5 | AC008953.1 | 0.00638909 | 0.03556234 | -1.244 | -0.389 | chr5 | 72174695 | 72175012 | - | processed_pseudogene |
| ENSG00000273449.1_6 | AC093788.1 | 0.00261785 | 0.01574076 | -1.245 | -0.203 | chr4 | 164450923 | 164451849 | + | lncRNA |
| ENSG00000215559.8_5 | ANKRD20A11P | 0.00019189 | 0.00142171 | -1.246 | 0.636 | chr21 | 15281895 | 15352758 | - | transcribed_unprocessed_pseudogene |
| ENSG00000153208.17_3 | MERTK | 8.03E-07 | 8.40E-06 | -1.246 | 2.060 | chr2 | 112656184 | 112787138 | + | protein_coding |
| ENSG00000227210.2_6 | AC079145.1 | 0.00197273 | 0.01214887 | -1.246 | 0.005 | chr2 | 20189970 | 20204556 | + | lncRNA |
| ENSG00000271639.1_5 | AC019072.1 | 0.00857702 | 0.04647392 | -1.246 | -0.708 | chr2 | 234454128 | 234455678 | + | processed_pseudogene |
| ENSG00000266037.1 | RN7SL3 | 6.89E-16 | 2.75E-14 | -1.248 | 8.757 | chr14 | 50320336 | 50320632 | - | misc_RNA |
| ENSG00000155961.5_4 | RAB39B | 0.00014469 | 0.00109672 | -1.248 | 0.689 | chrX | 154487520 | 154493776 | - | protein_coding |
| ENSG00000271092.5_6 | TLCD4-RWDD3 | 0.00609706 | 0.03407563 | -1.249 | -0.828 | chr1 | 95583479 | 95712781 | + | protein_coding |
| ENSG00000155438.12_5 | NIFK | 3.88E-12 | 8.50E-11 | -1.250 | 4.740 | chr2 | 122484521 | 122494487 | - | protein_coding |
| ENSG00000128165.9_3 | ADM2 | 0.01000542 | 0.0535602 | -1.252 | -0.830 | chr22 | 50919972 | 50924869 | + | protein_coding |
| ENSG00000285159.1_3 | AL627422.2 | 0.00825055 | 0.0448829 | -1.253 | -0.956 | chr6 | 159274620 | 159285534 | + | lncRNA |
| ENSG00000159231.6_3 | CBR3 | 0.00056092 | 0.00383008 | -1.253 | 0.295 | chr21 | 37507377 | 37518860 | + | protein_coding |
| ENSG00000101198.15_4 | NKAIN4 | 2.24E-05 | 0.00019245 | -1.253 | 1.480 | chr20 | 61872136 | 61904046 | - | protein_coding |
| ENSG00000200913.1 | SNORD46 | 0.00036259 | 0.00256559 | -1.255 | 0.453 | chr1 | 45242162 | 45242265 | + | snoRNA |
| ENSG00000165795.23_6 | NDRG2 | 3.92E-05 | 0.00032436 | -1.259 | 1.033 | chr14 | 21484922 | 21539031 | - | protein_coding |
| ENSG00000104332.12_5 | SFRP1 | 2.92E-14 | 9.23E-13 | -1.259 | 5.550 | chr8 | 41119481 | 41166992 | - | protein_coding |
| ENSG00000223658.8_5 | C1GALT1C1L | 0.00015404 | 0.00116217 | -1.262 | 0.776 | chr2 | 43902290 | 43903568 | - | protein_coding |
| ENSG00000163734.4_4 | CXCL3 | 1.74E-06 | 1.73E-05 | -1.266 | 1.762 | chr4 | 74902306 | 74904524 | - | protein_coding |
| ENSG00000262691.1_8 | AC040160.2 | 0.00133557 | 0.00848072 | -1.267 | -0.135 | chr16 | 67295011 | 67297687 | + | lncRNA |

| | | | | | | | | | | |
|---|---|---|---|---|---|---|---|---|---|---|
| ENSG00000235370.6_3 | DNM1P51 | 0.00542447 | 0.03066386 | -1.269 | -0.739 | chr15 | 84953014 | 84966399 | - | transcribed_unprocessed_pseudogene |
| ENSG00000253168.1_6 | AC124069.1 | 0.00323849 | 0.01910299 | -1.270 | -0.290 | chr8 | 17561497 | 17563696 | + | lncRNA |
| ENSG00000135362.14_5 | PRR5L | 1.14E-06 | 1.17E-05 | -1.270 | 1.944 | chr11 | 36317838 | 36486754 | + | protein_coding |
| ENSG00000114405.10_2 | C3orf14 | 3.29E-09 | 4.87E-08 | -1.271 | 3.255 | chr3 | 62304648 | 62321888 | + | protein_coding |
| ENSG00000246090.7_7 | AP002026.1 | 0.01067136 | 0.05666408 | -1.273 | -1.044 | chr4 | 100009956 | 100222513 | + | lncRNA |
| ENSG00000227666.1_4 | CYCSP24 | 0.01318783 | 0.06852861 | -1.275 | -1.391 | chr9 | 96800312 | 96800627 | + | processed_pseudogene |
| ENSG00000177182.11_4 | CLVS1 | 0.00239732 | 0.01449239 | -1.275 | -0.243 | chr8 | 61969717 | 62414204 | + | protein_coding |
| ENSG00000052850.8_6 | ALX4 | 1.26E-05 | 0.00011132 | -1.276 | 1.282 | chr11 | 44281990 | 44331689 | - | protein_coding |
| ENSG00000135636.14_4 | DYSF | 0.00490849 | 0.02800549 | -1.277 | -0.767 | chr2 | 71680852 | 71913898 | + | protein_coding |
| ENSG00000163576.18_7 | EFHB | 0.00030681 | 0.00220158 | -1.279 | 0.525 | chr3 | 19920964 | 19988517 | - | protein_coding |
| ENSG00000233080.3_7 | LINC01399 | 0.01130229 | 0.05958687 | -1.281 | -1.101 | chr22 | 35515817 | 35627049 | - | transcribed_processed_pseudogene |
| ENSG00000213626.13_4 | LBH | 1.73E-16 | 7.70E-15 | -1.283 | 7.098 | chr2 | 30454400 | 30546596 | + | protein_coding |
| ENSG00000287892.1_2 | AC090907.3 | 0.00351983 | 0.02065285 | -1.283 | -0.494 | chr15 | 101598852 | 101604109 | - | lncRNA |
| ENSG00000115221.12_6 | ITGB6 | 4.58E-05 | 0.00037645 | -1.284 | 0.941 | chr2 | 160956178 | 161056824 | - | protein_coding |
| ENSG00000271192.1_6 | AC006058.2 | 0.0030884 | 0.01829056 | -1.284 | -0.573 | chr3 | 44038388 | 44038935 | + | lncRNA |
| ENSG00000164362.21_7 | TERT | 1.67E-06 | 1.67E-05 | -1.285 | 2.410 | chr5 | 1253262 | 1295183 | - | protein_coding |
| ENSG00000173083.15_4 | HPSE | 8.59E-06 | 7.79E-05 | -1.285 | 1.508 | chr4 | 84213614 | 84256306 | - | protein_coding |
| ENSG00000260586.2_7 | AC064799.2 | 0.00486444 | 0.02777202 | -1.285 | -0.721 | chr15 | 71624459 | 71644799 | + | lncRNA |
| ENSG00000185149.6_3 | NPY2R | 0.00757704 | 0.04156262 | -1.285 | -0.941 | chr4 | 156129788 | 156138230 | + | protein_coding |
| ENSG00000177917.10_2 | ARL6IP6 | 1.74E-12 | 4.08E-11 | -1.285 | 4.619 | chr2 | 153574407 | 153617767 | + | protein_coding |
| ENSG00000213430.6_4 | HSPD1P1 | 1.37E-05 | 0.00012028 | -1.286 | 1.389 | chr5 | 21882694 | 21884419 | + | processed_pseudogene |
| ENSG00000173261.9_4 | PLAC8L1 | 0.00885142 | 0.04780491 | -1.286 | -0.859 | chr5 | 145463876 | 145485140 | - | protein_coding |
| ENSG00000232745.2 | ANKRD20A14P | 0.00307295 | 0.01821724 | -1.289 | -0.680 | chr1 | 142803532 | 142827235 | + | pseudogene |
| ENSG00000227992.1_6 | AC108463.1 | 0.0062104 | 0.03464369 | -1.291 | -0.638 | chr2 | 111961541 | 111963792 | - | processed_pseudogene |
| ENSG00000077044.10_3 | DGKD | 1.21E-16 | 5.51E-15 | -1.292 | 7.126 | chr2 | 234263153 | 234380750 | + | protein_coding |
| ENSG00000181856.15_3 | SLC2A4 | 0.0106567 | 0.05660314 | -1.294 | -1.080 | chr17 | 7185037 | 7191576 | + | protein_coding |
| ENSG00000231409.3_5 | AC018868.1 | 0.00914314 | 0.04919628 | -1.294 | -1.197 | chr15 | 35235281 | 35235595 | + | processed_pseudogene |
| ENSG00000235701.1_2 | PCBP2P1 | 0.01041509 | 0.05550238 | -1.295 | -1.197 | chr21 | 40543056 | 40544032 | - | processed_pseudogene |
| ENSG00000204176.14_7 | SYT15 | 1.17E-06 | 1.19E-05 | -1.296 | 1.770 | chr10 | 46955444 | 46971400 | - | protein_coding |
| ENSG00000213513.3_4 | IMPDH1P5 | 0.00607087 | 0.03393993 | -1.296 | -0.891 | chr10 | 79540095 | 79541634 | - | processed_pseudogene |
| ENSG00000183055.5_4 | FAM133CP | 0.00269696 | 0.01616627 | -1.296 | -0.520 | chr10 | 60475314 | 60476017 | + | processed_pseudogene |
| ENSG00000132031.13_4 | MATN3 | 2.58E-11 | 5.04E-10 | -1.296 | 3.885 | chr2 | 20191813 | 20212429 | - | protein_coding |
| ENSG00000204882.4_4 | GPR20 | 0.00100891 | 0.00655882 | -1.297 | -0.143 | chr8 | 142366570 | 142377386 | - | protein_coding |
| ENSG00000227252.2_6 | AC012063.1 | 0.0028617 | 0.01705024 | -1.298 | -0.496 | chr2 | 237819440 | 237994481 | - | lncRNA |
| ENSG00000239291.1_5 | AC002558.1 | 0.00257802 | 0.01552139 | -1.300 | -0.598 | chr17 | 45236557 | 45237359 | - | processed_pseudogene |
| ENSG00000163710.9_6 | PCOLCE2 | 2.66E-10 | 4.54E-09 | -1.301 | 3.512 | chr3 | 142534764 | 142608048 | - | protein_coding |
| ENSG00000127364.3_3 | TAS2R4 | 0.00276603 | 0.0165468 | -1.301 | -0.520 | chr7 | 141478242 | 141480619 | + | protein_coding |
| ENSG00000280374.1_8 | AC019080.5 | 0.0157633 | 0.08047892 | -1.302 | -0.972 | chr2 | 178182443 | 178183199 | - | lncRNA |
| ENSG00000287707.1_2 | AC115284.4 | 0.00085592 | 0.00563435 | -1.304 | -0.032 | chr3 | 51961087 | 51966499 | - | lncRNA |
| ENSG00000165887.12_6 | ANKRD2 | 7.25E-09 | 1.01E-07 | -1.307 | 2.811 | chr10 | 99332256 | 99343641 | + | protein_coding |
| ENSG00000115457.10_6 | IGFBP2 | 1.12E-11 | 2.29E-10 | -1.307 | 4.225 | chr2 | 217497551 | 217529159 | + | protein_coding |
| ENSG00000240032.1_7 | LNCSRLR | 0.00308584 | 0.01828525 | -1.308 | -0.559 | chr3 | 145784131 | 145786972 | - | lncRNA |
| ENSG00000281912.1_3 | LINC01144 | 0.00620379 | 0.0346235 | -1.308 | -1.081 | chr1 | 45769582 | 45771291 | + | lncRNA |
| ENSG00000197852.12_7 | INKA2 | 1.63E-06 | 1.63E-05 | -1.308 | 1.751 | chr1 | 112223252 | 112298446 | - | protein_coding |
| ENSG00000142871.17_4 | CCN1 | 2.63E-12 | 5.94E-11 | -1.308 | 4.219 | chr1 | 86046444 | 86049633 | + | protein_coding |
| ENSG00000164236.12_3 | ANKRD33B | 2.44E-13 | 6.51E-12 | -1.309 | 4.839 | chr5 | 10564182 | 10657928 | + | protein_coding |
| ENSG00000272807.1_7 | AC007038.1 | 0.00255365 | 0.01538509 | -1.311 | -0.583 | chr2 | 210893141 | 210893880 | + | lncRNA |
| ENSG00000163535.18_6 | SGO2 | 1.10E-13 | 3.12E-12 | -1.311 | 5.164 | chr2 | 201374731 | 201448819 | + | protein_coding |
| ENSG00000231305.4_7 | AC112484.1 | 0.00672734 | 0.03723507 | -1.312 | -0.874 | chr3 | 128579463 | 128590383 | - | lncRNA |
| ENSG00000170442.12_7 | AC021066.1 | 4.24E-11 | 8.00E-10 | -1.316 | 4.000 | chr12 | 52643084 | 52702947 | + | protein_coding |
| ENSG00000212534.1 | SNORD70 | 0.00905673 | 0.04879048 | -1.316 | -1.030 | chr2 | 203141154 | 203141241 | + | snoRNA |
| ENSG00000179431.7_3 | FJX1 | 1.42E-10 | 2.51E-09 | -1.319 | 3.571 | chr11 | 35640008 | 35642413 | + | protein_coding |
| ENSG00000115919.15_4 | KYNU | 1.92E-17 | 1.01E-15 | -1.321 | 7.711 | chr2 | 143635226 | 143813402 | + | protein_coding |
| ENSG00000180044.5_2 | C3orf80 | 0.01383906 | 0.07155953 | -1.322 | -1.369 | chr3 | 159943283 | 159946000 | + | protein_coding |
| ENSG00000169432.17_6 | SCN9A | 5.87E-11 | 1.09E-09 | -1.323 | 3.743 | chr2 | 167051695 | 167232511 | - | protein_coding |

| | | | | | | | | | | |
|---|---|---|---|---|---|---|---|---|---|---|
| ENSG00000163584.18_4 | RPL22L1 | 1.48E-12 | 3.52E-11 | -1.323 | 5.553 | chr3 | 170582664 | 170587997 | - | protein_coding |
| ENSG00000124126.14_7 | PREX1 | 2.13E-13 | 5.71E-12 | -1.323 | 4.818 | chr20 | 47240790 | 47444536 | - | protein_coding |
| ENSG00000138166.6_3 | DUSP5 | 1.25E-14 | 4.21E-13 | -1.324 | 5.323 | chr10 | 112257665 | 112271291 | + | protein_coding |
| ENSG00000273129.1_5 | PACERR | 0.00146966 | 0.00925607 | -1.326 | -0.507 | chr1 | 186649754 | 186650578 | + | lncRNA |
| ENSG00000225216.6_7 | AC007362.1 | 0.00016965 | 0.00126966 | -1.327 | 0.589 | chr2 | 206621193 | 206628730 | - | lncRNA |
| ENSG00000073670.14_5 | ADAM11 | 1.14E-07 | 1.35E-06 | -1.329 | 2.044 | chr17 | 42836356 | 42859214 | + | protein_coding |
| ENSG00000253978.2_7 | AC026689.1 | 0.0072053 | 0.03965814 | -1.329 | -0.421 | chr5 | 167656588 | 167696420 | - | lncRNA |
| ENSG00000162148.11_4 | PPP1R32 | 0.00301394 | 0.01789129 | -1.330 | -0.694 | chr11 | 61248592 | 61258403 | + | protein_coding |
| ENSG00000170153.11_7 | RNF150 | 3.61E-05 | 0.00030096 | -1.332 | 0.745 | chr4 | 141780961 | 142134031 | - | protein_coding |
| ENSG00000240132.1_4 | ETF1P2 | 0.00184593 | 0.01142344 | -1.333 | -0.471 | chr7 | 151198964 | 151200549 | + | processed_pseudogene |
| ENSG00000198406.7_4 | BZW1P2 | 3.73E-07 | 4.07E-06 | -1.334 | 1.813 | chr3 | 116364749 | 116365574 | - | processed_pseudogene |
| ENSG00000153094.23_5 | BCL2L11 | 3.80E-09 | 5.58E-08 | -1.334 | 2.716 | chr2 | 111876955 | 111926022 | + | protein_coding |
| ENSG00000262074.6_2 | SNORD3B-2 | 0.00016563 | 0.0012427 | -1.335 | 0.466 | chr17 | 18966659 | 18967449 | - | snoRNA |
| ENSG00000125726.11_5 | CD70 | 4.15E-10 | 6.92E-09 | -1.341 | 3.256 | chr19 | 6583194 | 6604114 | - | protein_coding |
| ENSG00000265112.1 | MIR3153 | 0.00882332 | 0.04769212 | -1.342 | -1.064 | chr9 | 91927140 | 91927221 | + | miRNA |
| ENSG00000235369.1_4 | RPL36AP15 | 0.00404407 | 0.02351142 | -1.343 | -1.010 | chr2 | 48024965 | 48025230 | - | processed_pseudogene |
| ENSG00000075886.11_3 | TUBA3D | 0.00059883 | 0.00406238 | -1.345 | 0.225 | chr2 | 132233692 | 132240507 | + | protein_coding |
| ENSG00000138311.17_5 | ZNF365 | 2.30E-05 | 0.00019737 | -1.346 | 0.863 | chr10 | 64133951 | 64430061 | + | protein_coding |
| ENSG00000112139.16_6 | MDGA1 | 1.10E-08 | 1.47E-07 | -1.348 | 2.455 | chr6 | 37598455 | 37667082 | - | protein_coding |
| ENSG00000253133.2_8 | AC009630.1 | 0.0037958 | 0.02216236 | -1.348 | -0.812 | chr8 | 41367112 | 41435940 | - | lncRNA |
| ENSG00000203727.4_4 | SAMD5 | 6.75E-11 | 1.24E-09 | -1.350 | 3.570 | chr6 | 147829826 | 148058683 | + | protein_coding |
| ENSG00000163083.6_3 | INHBB | 2.24E-16 | 9.74E-15 | -1.351 | 6.436 | chr2 | 121103712 | 121109379 | + | protein_coding |
| ENSG00000115541.11_4 | HSPE1 | 0.00073336 | 0.0048873 | -1.352 | 4.622 | chr2 | 198364864 | 198368173 | + | protein_coding |
| ENSG00000085662.14_4 | AKR1B1 | 5.72E-18 | 3.20E-16 | -1.352 | 9.588 | chr7 | 134127108 | 134144036 | - | protein_coding |
| ENSG00000189410.12_7 | SH2D5 | 5.42E-12 | 1.16E-10 | -1.352 | 3.938 | chr1 | 21046224 | 21059330 | - | protein_coding |
| ENSG00000225813.1_5 | AC009299.1 | 0.0051607 | 0.029294 | -1.356 | -0.909 | chr2 | 162138413 | 162139129 | + | processed_pseudogene |
| ENSG00000095596.12_4 | CYP26A1 | 0.01129533 | 0.05956784 | -1.358 | -0.413 | chr10 | 94833232 | 94837642 | + | protein_coding |
| ENSG00000272931.1_6 | AC099568.2 | 0.00055488 | 0.00379177 | -1.359 | -0.023 | chr1 | 90285733 | 90286427 | - | lncRNA |
| ENSG00000116017.6 | ARID3A | 1.21E-09 | 1.91E-08 | -1.360 | 3.250 | chr19 | 925781 | 975939 | + | protein_coding |
| ENSG00000283156.1_5 | AC068620.3 | 0.0008393 | 0.00553315 | -1.361 | -0.225 | chr4 | 57276682 | 57297714 | + | transcribed_unprocessed_pseudogene |
| ENSG00000183798.5_4 | EMILIN3 | 0.00117494 | 0.00755797 | -1.364 | -0.483 | chr20 | 39988602 | 39995458 | - | protein_coding |
| ENSG00000266783.1_8 | AP005136.3 | 0.00825921 | 0.04490252 | -1.370 | -1.281 | chr18 | 2561171 | 2562219 | - | lncRNA |
| ENSG00000244476.3_4 | ERVFRD-1 | 0.00157922 | 0.00986916 | -1.371 | -0.445 | chr6 | 11102722 | 11111958 | - | protein_coding |
| ENSG00000214146.3_7 | LINC02026 | 0.00280746 | 0.01676082 | -1.371 | -0.843 | chr3 | 193675161 | 193721549 | - | lncRNA |
| ENSG00000135047.15_3 | CTSL | 1.35E-18 | 8.23E-17 | -1.371 | 8.128 | chr9 | 90341024 | 90346384 | + | protein_coding |
| ENSG00000287710.1_1 | AC067852.5 | 0.00028612 | 0.00206335 | -1.371 | 0.260 | chr17 | 40729843 | 40730384 | + | lncRNA |
| ENSG00000260288.4_8 | AC019294.2 | 0.00449739 | 0.02589278 | -1.374 | -1.046 | chr15 | 76030161 | 76055663 | + | lncRNA |
| ENSG00000260229.1_5 | PPIAP51 | 0.00479515 | 0.02741169 | -1.374 | -1.045 | chr16 | 81509628 | 81510124 | - | processed_pseudogene |
| ENSG00000137261.14_5 | KIAA0319 | 7.21E-10 | 1.16E-08 | -1.374 | 2.957 | chr6 | 24544332 | 24646419 | - | protein_coding |
| ENSG00000153898.13_4 | MCOLN2 | 1.01E-07 | 1.20E-06 | -1.375 | 1.918 | chr1 | 85391266 | 85462796 | - | protein_coding |
| ENSG00000279021.1_5 | AC092139.2 | 0.0003571 | 0.00253078 | -1.376 | 0.185 | chr16 | 81497497 | 81498464 | - | TEC |
| ENSG00000133131.15_7 | MORC4 | 3.88E-18 | 2.20E-16 | -1.376 | 7.297 | chrX | 106057101 | 106243442 | - | protein_coding |
| ENSG00000234193.1_7 | ACSL3-AS1 | 0.00075415 | 0.00501643 | -1.379 | -0.377 | chr2 | 223782105 | 223784081 | - | lncRNA |
| ENSG00000224251.6_6 | AL391427.1 | 2.47E-12 | 5.63E-11 | -1.382 | 4.129 | chr10 | 5037680 | 5039572 | + | lncRNA |
| ENSG00000082074.18_9 | FYB1 | 1.50E-12 | 3.55E-11 | -1.385 | 5.172 | chr5 | 39105354 | 39274630 | - | protein_coding |
| ENSG00000279841.1_6 | AC092135.3 | 0.00121102 | 0.00777884 | -1.385 | -0.432 | chr16 | 81663798 | 81665643 | + | TEC |
| ENSG00000265185.5_2 | SNORD3B-1 | 6.83E-05 | 0.00054305 | -1.386 | 1.254 | chr17 | 18965225 | 18965982 | + | snoRNA |
| ENSG00000170525.21_6 | PFKFB3 | 1.54E-16 | 6.93E-15 | -1.387 | 8.200 | chr10 | 6186897 | 6296607 | + | protein_coding |
| ENSG00000166341.8_4 | DCHS1 | 0.00215351 | 0.01317999 | -1.387 | -0.753 | chr11 | 6642561 | 6677040 | - | protein_coding |
| ENSG00000112576.12_4 | CCND3 | 1.05E-17 | 5.63E-16 | -1.389 | 7.279 | chr6 | 41902671 | 42018095 | - | protein_coding |
| ENSG00000168280.12 | KIF5C | 6.86E-05 | 0.00054575 | -1.389 | 0.412 | chr2 | 149632819 | 149883273 | + | protein_coding |
| ENSG00000257438.1_6 | AC011595.1 | 0.00138622 | 0.00877415 | -1.390 | -0.624 | chr12 | 106496941 | 106499943 | + | lncRNA |
| ENSG00000163430.12_6 | FSTL1 | 4.93E-16 | 2.01E-14 | -1.391 | 5.459 | chr3 | 120111140 | 120169840 | - | protein_coding |
| ENSG00000178773.15_6 | CPNE7 | 2.97E-14 | 9.35E-13 | -1.395 | 5.004 | chr16 | 89642166 | 89663654 | + | protein_coding |
| ENSG00000141404.16_6 | GNAL | 1.41E-10 | 2.51E-09 | -1.396 | 3.085 | chr18 | 11689263 | 11885684 | + | protein_coding |
| ENSG00000155846.17_5 | PPARGC1B | 9.15E-09 | 1.25E-07 | -1.397 | 2.567 | chr5 | 149109861 | 149234585 | + | protein_coding |
| ENSG00000073756.12_4 | PTGS2 | 9.37E-15 | 3.21E-13 | -1.399 | 5.122 | chr1 | 186640923 | 186649555 | - | protein_coding |
| ENSG00000106366.9_3 | SERPINE1 | 1.58E-18 | 9.54E-17 | -1.399 | 7.362 | chr7 | 100770385 | 100782528 | + | protein_coding |

| | | | | | | | | | | |
|---|---|---|---|---|---|---|---|---|---|---|
| ENSG00000179348.12_5 | GATA2 | 1.99E-06 | 1.97E-05 | -1.401 | 1.182 | chr3 | 128198270 | 128212044 | - | protein_coding |
| ENSG00000251791.1 | SCARNA6 | 2.50E-09 | 3.76E-08 | -1.402 | 2.498 | chr2 | 234197322 | 234197586 | + | snoRNA |
| ENSG00000213300.5_4 | HNRNPA3P6 | 0.00070557 | 0.00472875 | -1.402 | -0.352 | chr3 | 75263782 | 75264787 | + | processed_pseudogene |
| ENSG00000145087.12_4 | STXBP5L | 0.00018189 | 0.00135445 | -1.403 | 0.046 | chr3 | 120626919 | 121143608 | + | protein_coding |
| ENSG00000184557.4_4 | SOCS3 | 1.19E-12 | 2.85E-11 | -1.403 | 4.055 | chr17 | 76352859 | 76356158 | - | protein_coding |
| ENSG00000265778.3_6 | AC018413.1 | 0.00141305 | 0.00892806 | -1.404 | -0.571 | chr18 | 74203585 | 74205874 | + | lncRNA |
| ENSG00000102003.10_4 | SYP | 2.14E-06 | 2.11E-05 | -1.404 | 1.511 | chrX | 49044257 | 49056718 | - | protein_coding |
| ENSG00000168785.8_5 | TSPAN5 | 1.04E-09 | 1.65E-08 | -1.407 | 2.627 | chr4 | 99391518 | 99579762 | - | protein_coding |
| ENSG00000234281.5_6 | LANCL1-AS1 | 0.00291294 | 0.01733966 | -1.407 | -0.419 | chr2 | 211189483 | 211333970 | + | lncRNA |
| ENSG00000241211.1_7 | IQCJ-SCHIP1-AS1 | 0.00610057 | 0.03408456 | -1.407 | -1.084 | chr3 | 159483176 | 159486401 | - | lncRNA |
| ENSG00000187957.8_4 | DNER | 0.00090612 | 0.0059406 | -1.408 | -0.534 | chr2 | 230222345 | 230579271 | - | protein_coding |
| ENSG00000114737.15_3 | CISH | 0.00125384 | 0.00801904 | -1.408 | -0.570 | chr3 | 50643921 | 50649262 | - | protein_coding |
| ENSG00000286545.1_1 | AC092645.2 | 1.33E-06 | 1.35E-05 | -1.409 | 1.497 | chr2 | 112997198 | 113001952 | + | lncRNA |
| ENSG00000278090.3_7 | LUNAR1 | 0.00116148 | 0.0074741 | -1.414 | -0.422 | chr15 | 99557755 | 99574283 | + | lncRNA |
| ENSG00000233538.1_6 | AC017104.3 | 0.00145475 | 0.00917154 | -1.415 | -0.518 | chr2 | 232316906 | 232317864 | + | lncRNA |
| ENSG00000023909.10_4 | GCLM | 1.71E-18 | 1.02E-16 | -1.416 | 7.915 | chr1 | 94350755 | 94375012 | - | protein_coding |
| ENSG00000080644.16_7 | CHRNA3 | 2.76E-05 | 0.0002342 | -1.419 | 0.536 | chr15 | 78885394 | 78913637 | - | protein_coding |
| ENSG00000223749.11_8 | MIR503HG | 4.91E-12 | 1.05E-10 | -1.419 | 3.556 | chrX | 133677149 | 133680672 | - | lncRNA |
| ENSG00000145632.15_4 | PLK2 | 3.97E-19 | 2.61E-17 | -1.420 | 7.758 | chr5 | 57749809 | 57755966 | - | protein_coding |
| ENSG00000115252.18_5 | PDE1A | 5.65E-05 | 0.00045597 | -1.421 | 0.439 | chr2 | 183004763 | 183387919 | - | protein_coding |
| ENSG00000150764.14_7 | DIXDC1 | 1.58E-11 | 3.17E-10 | -1.423 | 3.729 | chr11 | 111797868 | 111893377 | + | protein_coding |
| ENSG00000105696.9_5 | TMEM59L | 0.00013667 | 0.00103814 | -1.427 | 0.064 | chr19 | 18718240 | 18731849 | + | protein_coding |
| ENSG00000249084.1_7 | AC027627.1 | 0.000374 | 0.00263895 | -1.428 | -0.156 | chr4 | 178264158 | 178278186 | - | lncRNA |
| ENSG00000143942.5_3 | CHAC2 | 1.19E-12 | 2.86E-11 | -1.429 | 3.890 | chr2 | 53994941 | 54002333 | + | protein_coding |
| ENSG00000205334.2_6 | LINC01460 | 0.00463922 | 0.02661344 | -1.432 | -1.301 | chr2 | 27928653 | 27938599 | - | lncRNA |
| ENSG00000155657.26_5 | TTN | 6.47E-11 | 1.20E-09 | -1.433 | 3.160 | chr2 | 179390716 | 179695529 | - | protein_coding |
| ENSG00000230046.1_7 | BIRC6-AS1 | 0.00373561 | 0.02183961 | -1.433 | -1.238 | chr2 | 32602699 | 32604667 | - | lncRNA |
| ENSG00000153563.15_4 | CD8A | 0.00331966 | 0.01952996 | -1.435 | -1.120 | chr2 | 87011733 | 87035519 | - | protein_coding |
| ENSG00000234964.4_4 | FABP5P7 | 0.00129344 | 0.00823383 | -1.436 | -0.722 | chr11 | 59548791 | 59549195 | - | processed_pseudogene |
| ENSG00000134516.17_6 | DOCK2 | 1.52E-06 | 1.53E-05 | -1.437 | 1.227 | chr5 | 169064272 | 169510386 | + | protein_coding |
| ENSG00000262152.7_5 | LINC00514 | 0.00281849 | 0.01682099 | -1.438 | -1.063 | chr16 | 3038257 | 3052017 | + | protein_coding |
| ENSG00000070182.21_7 | SPTB | 2.57E-10 | 4.39E-09 | -1.439 | 2.709 | chr14 | 65213001 | 65346625 | - | protein_coding |
| ENSG00000128656.14_5 | CHN1 | 5.21E-13 | 1.33E-11 | -1.444 | 3.964 | chr2 | 175664041 | 175870109 | - | protein_coding |
| ENSG00000286223.1_3 | AC064836.4 | 0.00085764 | 0.00564361 | -1.448 | -0.624 | chr2 | 203166845 | 203168337 | - | lncRNA |
| ENSG00000164379.7_4 | FOXQ1 | 1.92E-06 | 1.90E-05 | -1.449 | 1.426 | chr6 | 1312333 | 1314993 | + | protein_coding |
| ENSG00000182600.9_6 | SNORC | 1.79E-10 | 3.14E-09 | -1.449 | 2.885 | chr2 | 233721980 | 233743418 | + | protein_coding |
| ENSG00000006740.17_3 | ARHGAP44 | 3.18E-07 | 3.49E-06 | -1.451 | 1.720 | chr17 | 12692815 | 12894960 | + | protein_coding |
| ENSG00000147852.16_3 | VLDLR | 8.69E-12 | 1.82E-10 | -1.451 | 3.388 | chr9 | 2621787 | 2660056 | + | protein_coding |
| ENSG00000228649.9_7 | SNHG26 | 2.53E-12 | 5.76E-11 | -1.452 | 3.574 | chr7 | 22893745 | 22912564 | + | lncRNA |
| ENSG00000181104.7_3 | F2R | 2.93E-15 | 1.08E-13 | -1.452 | 4.939 | chr5 | 76011951 | 76031595 | + | protein_coding |
| ENSG00000171388.12_4 | APLN | 0.00043769 | 0.00304838 | -1.456 | -0.400 | chrX | 128779236 | 128788933 | - | protein_coding |
| ENSG00000138207.14_5 | RBP4 | 3.21E-05 | 0.00026966 | -1.459 | 0.916 | chr10 | 95351444 | 95361501 | - | protein_coding |
| ENSG00000101265.16_4 | RASSF2 | 5.29E-18 | 2.97E-16 | -1.459 | 6.070 | chr20 | 4760669 | 4804254 | - | protein_coding |
| ENSG00000231039.2_6 | AL355303.1 | 0.00658138 | 0.03651823 | -1.460 | -1.155 | chr10 | 5276321 | 5305371 | + | lncRNA |
| ENSG00000276564.1_7 | AC130650.2 | 0.00019722 | 0.00145815 | -1.460 | -0.134 | chr16 | 14244690 | 14247092 | + | lncRNA |
| ENSG00000135069.14_3 | PSAT1 | 6.43E-19 | 4.11E-17 | -1.461 | 7.165 | chr9 | 80912041 | 80945009 | + | protein_coding |
| ENSG00000126882.13_5 | FAM78A | 5.43E-05 | 0.00043915 | -1.462 | 0.253 | chr9 | 134133463 | 134151897 | - | protein_coding |
| ENSG00000261634.4_7 | AC026992.2 | 0.00029255 | 0.00210523 | -1.464 | -0.321 | chr15 | 69570667 | 69591134 | - | lncRNA |
| ENSG00000162894.12_6 | FCMR | 0.00310172 | 0.01836331 | -1.465 | -1.045 | chr1 | 207076662 | 207096592 | - | protein_coding |
| ENSG00000171150.9_4 | SOCS5 | 4.06E-17 | 2.05E-15 | -1.466 | 5.757 | chr2 | 46926091 | 47007384 | + | protein_coding |
| ENSG00000005249.13_3 | PRKAR2B | 9.12E-11 | 1.66E-09 | -1.467 | 3.272 | chr7 | 106685150 | 106802256 | + | protein_coding |
| ENSG00000169896.18_7 | ITGAM | 4.25E-07 | 4.58E-06 | -1.468 | 1.493 | chr16 | 31271288 | 31344213 | + | protein_coding |
| ENSG00000154065.17_4 | ANKRD29 | 1.70E-12 | 4.00E-11 | -1.469 | 3.684 | chr18 | 21178890 | 21242875 | - | protein_coding |
| ENSG00000236671.9_8 | PRKG1-AS1 | 2.73E-06 | 2.64E-05 | -1.470 | 1.195 | chr10 | 53990158 | 54074267 | - | lncRNA |
| ENSG00000183098.11_3 | GPC6 | 7.20E-13 | 1.79E-11 | -1.470 | 3.647 | chr13 | 93879060 | 95060274 | + | protein_coding |
| ENSG00000169297.8_4 | NR0B1 | 2.89E-13 | 7.60E-12 | -1.477 | 4.632 | chrX | 30322323 | 30327507 | - | protein_coding |
| ENSG00000285155.1_4 | AC092153.1 | 0.00133862 | 0.00849101 | -1.477 | -0.651 | chr2 | 160998492 | 161128399 | - | lncRNA |
| ENSG00000068615.18_7 | REEP1 | 6.36E-06 | 5.86E-05 | -1.478 | 0.738 | chr2 | 86441116 | 86565206 | - | protein_coding |

| | | | | | | | | | | |
|---|---|---|---|---|---|---|---|---|---|---|
| ENSG00000218208.1_4 | RPS27AP11 | 0.00345526 | 0.02030078 | -1.479 | -1.280 | chr6 | 113902703 | 113903149 | + | processed_pseudogene |
| ENSG00000254244.1_4 | PAICSP4 | 0.00068833 | 0.00463063 | -1.480 | -0.693 | chr8 | 4644854 | 4646106 | + | processed_pseudogene |
| ENSG00000236824.2_3 | BCYRN1 | 1.57E-09 | 2.42E-08 | -1.484 | 2.752 | chr2 | 47562454 | 47562653 | + | scRNA |
| ENSG00000279294.1_6 | AC092135.1 | 0.0008477 | 0.00558231 | -1.487 | -0.781 | chr16 | 81652110 | 81654168 | + | TEC |
| ENSG00000156097.12_2 | GPR61 | 0.00110164 | 0.00712776 | -1.492 | -0.874 | chr1 | 110082494 | 110091028 | + | protein_coding |
| ENSG00000138750.15_5 | NUP54 | 3.72E-15 | 1.34E-13 | -1.493 | 4.878 | chr4 | 77035812 | 77069597 | - | protein_coding |
| ENSG00000171408.14_6 | PDE7B | 7.84E-14 | 2.30E-12 | -1.495 | 4.141 | chr6 | 136172839 | 136516712 | + | protein_coding |
| ENSG00000103647.13_5 | CORO2B | 0.00028348 | 0.00204574 | -1.500 | -0.300 | chr15 | 68871332 | 69020145 | + | protein_coding |
| ENSG00000213856.3_4 | VDAC1P2 | 0.00236108 | 0.01429276 | -1.500 | -1.138 | chrX | 49397103 | 49397952 | - | processed_pseudogene |
| ENSG00000164330.17_4 | EBF1 | 1.68E-06 | 1.68E-05 | -1.501 | 1.147 | chr5 | 158122924 | 158526924 | - | protein_coding |
| ENSG00000084628.10_4 | NKAIN1 | 0.00280284 | 0.01674444 | -1.502 | -1.137 | chr1 | 31652592 | 31712734 | - | protein_coding |
| ENSG00000285825.2_3 | CCDC15-DT | 0.00235577 | 0.01426836 | -1.505 | -1.138 | chr11 | 124819101 | 124823940 | - | lncRNA |
| ENSG00000162746.14_3 | FCRLB | 8.20E-05 | 0.0006433 | -1.505 | 0.078 | chr1 | 161691353 | 161697933 | + | protein_coding |
| ENSG00000225376.6_7 | TMEM246-AS1 | 0.00243185 | 0.01468614 | -1.505 | -1.138 | chr9 | 104230631 | 104244481 | + | lncRNA |
| ENSG00000287141.1_2 | AC093675.2 | 0.00123417 | 0.00791603 | -1.507 | -0.857 | chr2 | 112818661 | 112821286 | - | lncRNA |
| ENSG00000203808.12_7 | BVES-AS1 | 0.00026815 | 0.0019454 | -1.510 | -0.435 | chr6 | 105584183 | 105617827 | + | lncRNA |
| ENSG00000143847.15_3 | PPFIA4 | 0.00050812 | 0.00348967 | -1.512 | -0.435 | chr1 | 202995626 | 203047868 | + | protein_coding |
| ENSG00000236478.2_5 | AC012513.2 | 0.00665843 | 0.03689968 | -1.517 | -1.199 | chr2 | 217039619 | 217040755 | - | processed_pseudogene |
| ENSG00000143867.7_4 | OSR1 | 1.99E-09 | 3.02E-08 | -1.522 | 2.403 | chr2 | 19551246 | 19558384 | - | protein_coding |
| ENSG00000257576.1_4 | HSPD1P4 | 2.09E-05 | 0.00018074 | -1.523 | 0.297 | chr12 | 56904786 | 56906487 | + | processed_pseudogene |
| ENSG00000163449.11_4 | TMEM169 | 9.54E-05 | 0.00074038 | -1.524 | 0.072 | chr2 | 216946589 | 216967506 | + | protein_coding |
| ENSG00000169604.19_3 | ANTXR1 | 3.05E-20 | 2.41E-18 | -1.528 | 7.652 | chr2 | 69240310 | 69476459 | + | protein_coding |
| ENSG00000065911.12_3 | MTHFD2 | 1.65E-19 | 1.17E-17 | -1.528 | 6.828 | chr2 | 74425737 | 74444692 | + | protein_coding |
| ENSG00000285108.1_4 | AC103718.1 | 0.00051078 | 0.00350524 | -1.531 | -0.624 | chr8 | 130697647 | 130735269 | - | lncRNA |
| ENSG00000231566.2_7 | LINC02595 | 1.60E-09 | 2.46E-08 | -1.536 | 2.119 | chrX | 45706474 | 45710971 | - | lncRNA |
| ENSG00000113356.12_6 | POLR3G | 1.63E-13 | 4.47E-12 | -1.536 | 4.026 | chr5 | 89767565 | 89810374 | + | protein_coding |
| ENSG00000173237.4_2 | C11orf86 | 1.05E-08 | 1.42E-07 | -1.537 | 2.343 | chr11 | 66742748 | 66744475 | + | protein_coding |
| ENSG00000168874.13_3 | ATOH8 | 2.63E-06 | 2.55E-05 | -1.538 | 0.903 | chr2 | 85978467 | 86018506 | + | protein_coding |
| ENSG00000110427.16_5 | KIAA1549L | 3.96E-11 | 7.56E-10 | -1.539 | 3.098 | chr11 | 33397629 | 33695648 | + | protein_coding |
| ENSG00000241494.1_5 | AL355032.1 | 0.00500053 | 0.02844845 | -1.540 | -1.235 | chr14 | 102144280 | 102144717 | + | processed_pseudogene |
| ENSG00000238840.1 | U8 | 0.00169169 | 0.01052389 | -1.542 | -1.176 | chr10 | 5004628 | 5004760 | - | snoRNA |
| ENSG00000187134.14_4 | AKR1C1 | 1.54E-20 | 1.31E-18 | -1.544 | 8.713 | chr10 | 5005445 | 5025475 | + | protein_coding |
| ENSG00000280721.1_6 | LINC01943 | 0.0022921 | 0.01391311 | -1.549 | -1.044 | chr2 | 87739042 | 87758563 | - | lncRNA |
| ENSG00000144354.14_4 | CDCA7 | 5.35E-17 | 2.67E-15 | -1.550 | 5.532 | chr2 | 174219548 | 174233725 | + | protein_coding |
| ENSG00000225783.8_7 | MIAT | 4.97E-11 | 9.26E-10 | -1.550 | 2.771 | chr22 | 27042375 | 27072438 | + | lncRNA |
| ENSG00000188064.10_7 | WNT7B | 0.00098494 | 0.00641239 | -1.554 | 6.027 | chr22 | 46316246 | 46373042 | - | protein_coding |
| ENSG00000140284.11_3 | SLC27A2 | 3.82E-17 | 1.93E-15 | -1.556 | 5.226 | chr15 | 50474393 | 50528582 | + | protein_coding |
| ENSG00000153162.9_3 | BMP6 | 4.44E-09 | 6.42E-08 | -1.559 | 2.279 | chr6 | 7726332 | 7881961 | + | protein_coding |
| ENSG00000250116.2_7 | AC018682.1 | 8.05E-05 | 0.00063235 | -1.563 | -0.094 | chr2 | 46795395 | 46807377 | - | lncRNA |
| ENSG00000279501.1_7 | CR392039.4 | 2.44E-10 | 4.19E-09 | -1.563 | 2.447 | chr21 | 9835329 | 9857503 | - | lncRNA |
| ENSG00000126353.3_5 | CCR7 | 1.94E-08 | 2.52E-07 | -1.564 | 1.596 | chr17 | 38710021 | 38721724 | - | protein_coding |
| ENSG00000144355.15_3 | DLX1 | 6.52E-05 | 0.00052112 | -1.564 | -0.033 | chr2 | 172949468 | 172954405 | + | protein_coding |
| ENSG00000056487.16_6 | PHF21B | 0.00223543 | 0.01362498 | -1.566 | -0.482 | chr22 | 45277042 | 45405886 | - | protein_coding |
| ENSG00000168505.7_4 | GBX2 | 1.48E-05 | 0.00013001 | -1.569 | 0.153 | chr2 | 237073879 | 237077029 | - | protein_coding |
| ENSG00000167619.13_4 | TMEM145 | 3.48E-05 | 0.00029106 | -1.578 | -0.007 | chr19 | 42817461 | 42829216 | + | protein_coding |
| ENSG00000224644.1_7 | AL135787.1 | 0.00068897 | 0.00463322 | -1.579 | -0.924 | chr9 | 114336472 | 114337707 | + | lncRNA |
| ENSG00000111186.13_3 | WNT5B | 0.00094793 | 0.00619184 | -1.581 | -0.768 | chr12 | 1639057 | 1756378 | + | protein_coding |
| ENSG00000134551.13_3 | PRH2 | 0.00024445 | 0.00177925 | -1.582 | -0.506 | chr12 | 11081835 | 11087444 | + | protein_coding |
| ENSG00000206190.12_6 | ATP10A | 2.37E-15 | 8.90E-14 | -1.583 | 4.163 | chr15 | 25922418 | 26110331 | - | protein_coding |
| ENSG00000207280.1 | SNORD20 | 0.00037287 | 0.00263198 | -1.587 | -0.547 | chr2 | 232321155 | 232321234 | - | snoRNA |
| ENSG00000136010.14_6 | ALDH1L2 | 2.71E-11 | 5.28E-10 | -1.591 | 2.734 | chr12 | 105413562 | 105501421 | - | protein_coding |
| ENSG00000126861.5_4 | OMG | 4.98E-05 | 0.00040552 | -1.592 | 0.055 | chr17 | 29599031 | 29624557 | - | protein_coding |
| ENSG00000197106.7_3 | SLC6A17 | 1.76E-07 | 2.02E-06 | -1.599 | 1.248 | chr1 | 110693116 | 110744824 | + | protein_coding |
| ENSG00000205426.10_4 | KRT81 | 2.90E-15 | 1.07E-13 | -1.601 | 10.692 | chr12 | 52679697 | 52685318 | - | protein_coding |
| ENSG00000129116.19_7 | PALLD | 1.61E-19 | 1.15E-17 | -1.603 | 6.715 | chr4 | 169418203 | 169849608 | + | protein_coding |
| ENSG00000137809.17_6 | ITGA11 | 8.14E-13 | 2.01E-11 | -1.604 | 3.331 | chr15 | 68588870 | 68724502 | - | protein_coding |
| ENSG00000115353.11_4 | TACR1 | 0.00228219 | 0.01386244 | -1.604 | -1.346 | chr2 | 75273590 | 75426646 | - | protein_coding |

| | | | | | | | | | | |
|---|---|---|---|---|---|---|---|---|---|---|
| ENSG00000133816.15_7 | MICAL2 | 6.71E-18 | 3.70E-16 | -1.606 | 5.329 | chr11 | 12115555 | 12380691 | + | protein_coding |
| ENSG00000233673.7_6 | ANAPC1P1 | 0.0010052 | 0.00653951 | -1.606 | -1.138 | chr2 | 87088948 | 87140101 | + | transcribed_unprocessed_pseudogene |
| ENSG00000179331.3_3 | RAB39A | 2.41E-05 | 0.00020617 | -1.607 | 0.426 | chr11 | 107799174 | 107834208 | + | protein_coding |
| ENSG00000162444.12_3 | RBP7 | 0.00059878 | 0.00406238 | -1.610 | -0.794 | chr1 | 10057264 | 10076079 | + | protein_coding |
| ENSG00000249379.2_6 | AL033397.2 | 8.10E-07 | 8.47E-06 | -1.611 | 0.747 | chr6 | 53367907 | 53372458 | + | lncRNA |
| ENSG00000213190.4_3 | MLLT11 | 3.12E-09 | 4.64E-08 | -1.611 | 4.214 | chr1 | 151032873 | 151042020 | + | protein_coding |
| ENSG00000101203.16_2 | COL20A1 | 5.61E-05 | 0.00045326 | -1.612 | -0.225 | chr20 | 61924538 | 61966203 | + | protein_coding |
| ENSG00000221955.10_4 | SLC12A8 | 0.00035269 | 0.00250246 | -1.614 | -0.796 | chr3 | 124801480 | 124931708 | - | protein_coding |
| ENSG00000070669.17_5 | ASNS | 1.10E-05 | 9.84E-05 | -1.620 | 0.237 | chr7 | 97480989 | 97501854 | - | protein_coding |
| ENSG00000242100.2_2 | RPL9P32 | 0.00103587 | 0.00672181 | -1.621 | -1.008 | chr19 | 30447276 | 30447895 | - | processed_pseudogene |
| ENSG00000165376.11_4 | CLDN2 | 1.60E-16 | 7.16E-15 | -1.623 | 4.575 | chrX | 106143394 | 106174091 | + | protein_coding |
| ENSG00000163531.15_7 | NFASC | 0.00094307 | 0.00616689 | -1.624 | -0.844 | chr1 | 204797779 | 204991950 | + | protein_coding |
| ENSG00000168913.7_4 | ENHO | 0.00227465 | 0.01383081 | -1.626 | -1.485 | chr9 | 34521041 | 34522988 | - | protein_coding |
| ENSG00000251682.1_5 | AC122718.2 | 0.00069217 | 0.00465295 | -1.629 | -1.062 | chr5 | 60642953 | 60643543 | + | processed_pseudogene |
| ENSG00000154319.16_7 | FAM167A | 1.53E-13 | 4.24E-12 | -1.632 | 3.597 | chr8 | 11278985 | 11333417 | - | protein_coding |
| ENSG00000063660.9_4 | GPC1 | 4.86E-07 | 5.23E-06 | -1.634 | 6.921 | chr2 | 241375080 | 241407493 | + | protein_coding |
| ENSG00000128512.15 | DOCK4 | 1.83E-16 | 8.05E-15 | -1.636 | 4.387 | chr7 | 111366166 | 111846466 | - | protein_coding |
| ENSG00000091986.15_7 | CCDC80 | 7.62E-19 | 4.82E-17 | -1.638 | 5.614 | chr3 | 112315641 | 112368377 | - | protein_coding |
| ENSG00000101335.10_3 | MYL9 | 1.09E-05 | 9.76E-05 | -1.640 | 0.063 | chr20 | 35169900 | 35179850 | + | protein_coding |
| ENSG00000106853.20_7 | PTGR1 | 1.16E-20 | 1.01E-18 | -1.640 | 7.031 | chr9 | 114312002 | 114362173 | - | protein_coding |
| ENSG00000269175.1 | AC093157.1 | 0.00043596 | 0.00303995 | -1.641 | -0.940 | chr1 | 101455761 | 101457653 | + | protein_coding |
| ENSG00000143355.16_7 | LHX9 | 0.00049558 | 0.00341677 | -1.644 | -0.940 | chr1 | 197881032 | 197904608 | + | protein_coding |
| ENSG00000165521.15_8 | EML5 | 4.45E-10 | 7.38E-09 | -1.648 | 2.031 | chr14 | 89078775 | 89259096 | - | protein_coding |
| ENSG00000171303.7_3 | KCNK3 | 1.26E-10 | 2.26E-09 | -1.648 | 2.144 | chr2 | 26915558 | 26956288 | + | protein_coding |
| ENSG00000157168.20_6 | NRG1 | 1.15E-20 | 1.01E-18 | -1.651 | 6.687 | chr8 | 31496738 | 32713184 | + | protein_coding |
| ENSG00000128641.19_6 | MYO1B | 1.11E-21 | 1.21E-19 | -1.652 | 8.152 | chr2 | 192109911 | 192290115 | + | protein_coding |
| ENSG00000270299.1_7 | AL121758.1 | 0.00020273 | 0.00149596 | -1.653 | -0.547 | chr20 | 627270 | 656444 | - | protein_coding |
| ENSG00000279476.1_6 | AC092139.3 | 0.00123841 | 0.00793465 | -1.660 | -0.719 | chr16 | 81551429 | 81554222 | + | TEC |
| ENSG00000269902.1_7 | AC234772.2 | 0.00024122 | 0.0017579 | -1.664 | -0.665 | chrX | 45624175 | 45624702 | - | lncRNA |
| ENSG00000207359.1 | RNU6-925P | 0.00033382 | 0.00237523 | -1.667 | -0.923 | chr8 | 90912949 | 90913052 | - | snRNA |
| ENSG00000188042.8_5 | ARL4C | 6.18E-21 | 5.61E-19 | -1.670 | 6.559 | chr2 | 235401685 | 235405725 | - | protein_coding |
| ENSG00000281207.1_5 | SLFNL1-AS1 | 2.47E-09 | 3.71E-08 | -1.677 | 1.663 | chr1 | 41480262 | 41509562 | + | lncRNA |
| ENSG00000197565.16_5 | COL4A6 | 1.27E-19 | 9.36E-18 | -1.680 | 6.053 | chrX | 107398837 | 107682727 | - | protein_coding |
| ENSG00000182580.3_3 | EPHB3 | 1.58E-07 | 1.82E-06 | -1.690 | 1.115 | chr3 | 184279573 | 184300196 | + | protein_coding |
| ENSG00000108960.9_5 | MMD | 6.83E-15 | 2.38E-13 | -1.691 | 6.481 | chr17 | 53469983 | 53499285 | - | protein_coding |
| ENSG00000183242.11_5 | WT1-AS | 4.74E-05 | 0.00038906 | -1.692 | -0.353 | chr11 | 32457064 | 32480315 | + | lncRNA |
| ENSG00000286943.1_1 | AP001992.2 | 0.00028615 | 0.00206335 | -1.692 | -0.858 | chr11 | 74642769 | 74647982 | + | lncRNA |
| ENSG00000135245.10_4 | HILPDA | 2.06E-17 | 1.07E-15 | -1.693 | 4.606 | chr7 | 128095903 | 128098472 | + | protein_coding |
| ENSG00000237892.1_7 | KLF7-IT1 | 0.00032002 | 0.00228346 | -1.697 | -0.907 | chr2 | 207985608 | 207986768 | - | lncRNA |
| ENSG00000163006.12_6 | CCDC138 | 1.19E-13 | 3.35E-12 | -1.700 | 3.736 | chr2 | 109403213 | 109501933 | + | protein_coding |
| ENSG00000207146.1 | Y_RNA | 7.84E-05 | 0.00061646 | -1.702 | -0.557 | chrX | 107920181 | 107920287 | - | misc_RNA |
| ENSG00000278719.1_8 | MCM8-AS1 | 0.00050213 | 0.00345387 | -1.709 | -0.889 | chr20 | 5971589 | 5986467 | - | lncRNA |
| ENSG00000100987.14_4 | VSX1 | 2.88E-07 | 3.20E-06 | -1.709 | 0.646 | chr20 | 25051521 | 25063001 | - | protein_coding |
| ENSG00000164509.15_4 | IL31RA | 1.86E-08 | 2.42E-07 | -1.711 | 1.396 | chr5 | 55147185 | 55218682 | + | protein_coding |
| ENSG00000143494.15_5 | VASH2 | 2.21E-11 | 4.36E-10 | -1.716 | 2.335 | chr1 | 213123862 | 213165379 | + | protein_coding |
| ENSG00000171812.13_6 | COL8A2 | 0.00022789 | 0.00166897 | -1.716 | -0.680 | chr1 | 36560840 | 36590823 | - | protein_coding |
| ENSG00000146054.18_4 | TRIM7 | 9.83E-13 | 2.37E-11 | -1.718 | 3.085 | chr5 | 180620924 | 180632293 | - | protein_coding |
| ENSG00000165323.15_2 | FAT3 | 1.40E-12 | 3.32E-11 | -1.720 | 3.462 | chr11 | 92085262 | 92629636 | + | protein_coding |
| ENSG00000275620.1_7 | AL121827.2 | 0.00057481 | 0.00391739 | -1.722 | -0.600 | chr20 | 61885330 | 61892967 | + | lncRNA |
| ENSG00000172159.16_4 | FRMD3 | 6.11E-16 | 2.45E-14 | -1.726 | 4.043 | chr9 | 85857905 | 86153461 | - | protein_coding |
| ENSG00000084710.14_3 | EFR3B | 3.38E-18 | 1.95E-16 | -1.744 | 5.077 | chr2 | 25264945 | 25382004 | + | protein_coding |
| ENSG00000112394.17_5 | SLC16A10 | 0.0007042 | 0.00472249 | -1.747 | -1.258 | chr6 | 111408706 | 111552397 | + | protein_coding |
| ENSG00000131711.15_3 | MAP1B | 1.33E-18 | 8.12E-17 | -1.749 | 8.073 | chr5 | 71403061 | 71505392 | + | protein_coding |
| ENSG00000116852.14_4 | KIF21B | 1.95E-12 | 4.53E-11 | -1.749 | 2.737 | chr1 | 200938518 | 200992828 | - | protein_coding |
| ENSG00000135914.6_4 | HTR2B | 0.00087846 | 0.00576991 | -1.750 | -1.260 | chr2 | 231972944 | 231989756 | - | protein_coding |
| ENSG00000231638.1_7 | LUARIS | 0.0002094 | 0.00153928 | -1.750 | -0.705 | chr7 | 43548327 | 43562141 | - | lncRNA |
| ENSG00000215022.7_8 | AL008729.1 | 4.75E-05 | 0.00038909 | -1.756 | -0.519 | chr6 | 13265093 | 13295818 | - | lncRNA |
| ENSG00000121743.4_4 | GJA3 | 1.81E-05 | 0.00015753 | -1.757 | -0.098 | chr13 | 20712394 | 20735191 | - | protein_coding |
| ENSG00000125245.12_5 | GPR18 | 0.00058092 | 0.00395146 | -1.757 | -1.118 | chr13 | 99906968 | 99913998 | - | protein_coding |

| | | | | | | | | | | |
|---|---|---|---|---|---|---|---|---|---|---|
| ENSG00000231609.7_7 | AC007098.1 | 1.55E-05 | 0.00013503 | -1.762 | 0.021 | chr2 | 63184461 | 63275775 | - | lncRNA |
| ENSG00000231625.4_5 | SLC47A1P2 | 0.00096225 | 0.00627848 | -1.766 | -1.322 | chr17 | 19519102 | 19537138 | + | transcribed_unprocessed_pseudogene |
| ENSG00000148704.13_4 | VAX1 | 1.80E-05 | 0.00015662 | -1.768 | -0.195 | chr10 | 118888032 | 118897812 | - | protein_coding |
| ENSG00000169429.11_4 | CXCL8 | 1.48E-06 | 1.49E-05 | -1.769 | 0.341 | chr4 | 74606258 | 74609433 | + | protein_coding |
| ENSG00000253633.2_6 | AP002852.1 | 0.00013612 | 0.00103481 | -1.771 | -0.857 | chr8 | 103540968 | 103552171 | + | lncRNA |
| ENSG00000135842.17_5 | NIBAN1 | 2.20E-07 | 2.49E-06 | -1.773 | 0.505 | chr1 | 184759858 | 184943640 | - | protein_coding |
| ENSG00000115474.7_5 | KCNJ13 | 0.00012045 | 0.00091969 | -1.774 | -0.649 | chr2 | 233630512 | 233641275 | - | protein_coding |
| ENSG00000238317.1 | SNORD11 | 5.41E-05 | 0.00043826 | -1.776 | -0.543 | chr2 | 203157772 | 203157859 | + | snoRNA |
| ENSG00000277498.1_5 | AC107081.3 | 0.00050676 | 0.00348167 | -1.789 | -1.238 | chr2 | 62085272 | 62087433 | + | unprocessed_pseudogene |
| ENSG00000162040.7_4 | HS3ST6 | 4.44E-05 | 0.00036492 | -1.795 | -0.288 | chr16 | 1961476 | 1968416 | - | protein_coding |
| ENSG00000112655.16_7 | PTK7 | 1.55E-11 | 3.13E-10 | -1.797 | 2.559 | chr6 | 43044045 | 43129457 | + | protein_coding |
| ENSG00000100234.11_2 | TIMP3 | 0.00110727 | 0.00715944 | -1.797 | -1.560 | chr22 | 33197687 | 33259030 | + | protein_coding |
| ENSG00000131773.14_6 | KHDRBS3 | 4.92E-08 | 6.06E-07 | -1.798 | 0.830 | chr8 | 136469699 | 136668965 | + | protein_coding |
| ENSG00000244332.1_7 | AL138759.1 | 8.98E-05 | 0.0006997 | -1.800 | -0.677 | chr10 | 96337196 | 96370995 | + | lncRNA |
| ENSG00000116701.14_3 | NCF2 | 1.11E-06 | 1.14E-05 | -1.801 | 0.177 | chr1 | 183524698 | 183560011 | - | protein_coding |
| ENSG00000286313.1_2 | AL360178.1 | 3.07E-07 | 3.38E-06 | -1.803 | 0.656 | chr6 | 136175191 | 136202080 | - | lncRNA |
| ENSG00000107984.10_3 | DKK1 | 1.04E-22 | 1.45E-20 | -1.805 | 7.636 | chr10 | 54074041 | 54077802 | + | protein_coding |
| ENSG00000231131.8_7 | LNCAROD | 3.41E-11 | 6.55E-10 | -1.807 | 2.112 | chr10 | 54210634 | 54515267 | - | lncRNA |
| ENSG00000125285.6_4 | SOX21 | 5.06E-08 | 6.23E-07 | -1.809 | 0.720 | chr13 | 95361876 | 95364799 | - | protein_coding |
| ENSG00000187686.4_4 | KRT18P59 | 0.00087931 | 0.00577334 | -1.810 | -1.460 | chr11 | 124983057 | 124996677 | + | transcribed_processed_pseudogene |
| ENSG00000135111.16_6 | TBX3 | 1.64E-17 | 8.64E-16 | -1.818 | 4.315 | chr12 | 115108060 | 115121980 | - | protein_coding |
| ENSG00000237263.1_4 | MAPK6P3 | 0.00038011 | 0.00267886 | -1.819 | -1.218 | chr13 | 42643068 | 42645229 | - | processed_pseudogene |
| ENSG00000180815.14_3 | MAP3K15 | 3.22E-12 | 7.17E-11 | -1.829 | 2.197 | chrX | 19378174 | 19533379 | - | protein_coding |
| ENSG00000167889.12_3 | MGAT5B | 1.06E-15 | 4.12E-14 | -1.834 | 4.316 | chr17 | 74864538 | 74946475 | + | protein_coding |
| ENSG00000162975.5_4 | KCNF1 | 1.95E-11 | 3.86E-10 | -1.836 | 2.631 | chr2 | 11052060 | 11054351 | + | protein_coding |
| ENSG00000186854.11_7 | TRABD2A | 1.14E-08 | 1.53E-07 | -1.841 | 1.501 | chr2 | 85048774 | 85134132 | - | protein_coding |
| ENSG00000107719.9_3 | PALD1 | 3.19E-12 | 7.11E-11 | -1.842 | 3.636 | chr10 | 72238523 | 72328206 | + | protein_coding |
| ENSG00000137819.14_4 | PAQR5 | 7.36E-22 | 8.28E-20 | -1.843 | 6.296 | chr15 | 69591251 | 69700119 | + | protein_coding |
| ENSG00000279713.1_8 | AC080038.4 | 3.16E-05 | 0.00026635 | -1.846 | -0.421 | chr17 | 60756764 | 60758715 | - | TEC |
| ENSG00000229644.6_4 | NAMPTP1 | 1.45E-09 | 2.26E-08 | -1.848 | 1.338 | chr10 | 36810649 | 36813162 | - | processed_pseudogene |
| ENSG00000143473.13_7 | KCNH1 | 3.76E-10 | 6.28E-09 | -1.853 | 1.456 | chr1 | 210850165 | 211307507 | - | protein_coding |
| ENSG00000127507.18_6 | ADGRE2 | 6.26E-06 | 5.77E-05 | -1.859 | -0.243 | chr19 | 14843204 | 14889372 | - | protein_coding |
| ENSG00000140807.7_4 | NKD1 | 1.74E-05 | 0.00015141 | -1.865 | -0.494 | chr16 | 50582307 | 50683160 | + | protein_coding |
| ENSG00000107968.10_3 | MAP3K8 | 2.25E-17 | 1.16E-15 | -1.871 | 4.397 | chr10 | 30722950 | 30750762 | + | protein_coding |
| ENSG00000179046.8_3 | TRIML2 | 1.18E-18 | 7.30E-17 | -1.872 | 5.021 | chr4 | 189012427 | 189030757 | - | protein_coding |
| ENSG00000119283.15_3 | TRIM67 | 1.32E-06 | 1.34E-05 | -1.874 | 0.253 | chr1 | 231297858 | 231357302 | + | protein_coding |
| ENSG00000226853.3_6 | AC010894.2 | 0.00041491 | 0.00290452 | -1.884 | -1.175 | chr2 | 175352108 | 175353114 | + | lncRNA |
| ENSG00000248890.2_7 | HHIP-AS1 | 3.38E-07 | 3.70E-06 | -1.895 | 0.247 | chr4 | 145564074 | 145582509 | - | lncRNA |
| ENSG00000135917.15_6 | SLC19A3 | 4.37E-11 | 8.24E-10 | -1.903 | 2.069 | chr2 | 228548479 | 228582744 | - | protein_coding |
| ENSG00000186603.6_3 | HPDL | 2.64E-09 | 3.95E-08 | -1.903 | 1.133 | chr1 | 45792567 | 45794351 | + | protein_coding |
| ENSG00000223770.5_7 | CACNA2D1-AS1 | 4.27E-05 | 0.00035165 | -1.904 | -0.721 | chr7 | 81638493 | 81659271 | + | lncRNA |
| ENSG00000167157.11_3 | PRRX2 | 1.50E-07 | 1.74E-06 | -1.909 | 0.555 | chr9 | 132427926 | 132484953 | + | protein_coding |
| ENSG00000251364.7_7 | AC107884.1 | 0.00012904 | 0.00098315 | -1.912 | -0.957 | chr11 | 7440057 | 7534875 | - | lncRNA |
| ENSG00000123485.12_7 | HJURP | 1.04E-23 | 1.87E-21 | -1.925 | 7.234 | chr2 | 234742062 | 234763212 | - | protein_coding |
| ENSG00000100292.17_3 | HMOX1 | 1.82E-23 | 3.05E-21 | -1.927 | 6.898 | chr22 | 35776354 | 35790200 | + | protein_coding |
| ENSG00000279384.1_6 | AC080188.2 | 0.00079243 | 0.00525146 | -1.927 | -0.706 | chr4 | 169765139 | 169765886 | + | TEC |
| ENSG00000272636.4_5 | DOC2B | 0.00123755 | 0.00793199 | -1.934 | -0.810 | chr17 | 5810 | 31441 | - | protein_coding |
| ENSG00000184985.16_2 | SORCS2 | 3.07E-11 | 5.94E-10 | -1.942 | 1.771 | chr4 | 7194265 | 7744563 | + | protein_coding |
| ENSG00000162407.9_5 | PLPP3 | 6.50E-17 | 3.14E-15 | -1.944 | 3.871 | chr1 | 56960433 | 57110974 | - | protein_coding |
| ENSG00000218416.4_6 | AC110619.1 | 3.19E-08 | 4.02E-07 | -1.955 | 0.545 | chr2 | 241388732 | 241396131 | - | lncRNA |
| ENSG00000182752.10_4 | PAPPA | 9.26E-23 | 1.33E-20 | -1.957 | 6.668 | chr9 | 118916070 | 119164600 | + | protein_coding |
| ENSG00000166086.13_4 | JAM3 | 3.70E-14 | 1.15E-12 | -1.967 | 2.804 | chr11 | 133938966 | 134021896 | + | protein_coding |
| ENSG00000160200.17_3 | CBS | 6.64E-05 | 0.00052983 | -1.992 | -0.201 | chr21 | 44473301 | 44497053 | - | protein_coding |
| ENSG00000083454.22_7 | P2RX5 | 1.03E-09 | 1.64E-08 | -1.994 | 1.226 | chr17 | 3575493 | 3599534 | - | protein_coding |
| ENSG00000176771.17_8 | NCKAP5 | 4.37E-15 | 1.55E-13 | -2.004 | 3.060 | chr2 | 133429361 | 134326034 | - | protein_coding |
| ENSG00000262454.4_4 | MIR193BHG | 1.21E-16 | 5.51E-15 | -2.005 | 3.361 | chr16 | 14395246 | 14424924 | + | lncRNA |
| ENSG00000115112.8_4 | TFCP2L1 | 3.20E-15 | 1.16E-13 | -2.010 | 3.027 | chr2 | 121974163 | 122042778 | - | protein_coding |

| | | | | | | | | | | |
|---|---|---|---|---|---|---|---|---|---|---|
| ENSG00000183688.4_4 | RFLNB | 2.01E-22 | 2.47E-20 | -2.013 | 5.853 | chr17 | 289769 | 295730 | - | protein_coding |
| ENSG00000237596.7_9 | AL138828.1 | 1.42E-06 | 1.44E-05 | -2.014 | 0.071 | chr6 | 136313074 | 136546889 | - | lncRNA |
| ENSG00000253837.1_6 | AC090197.1 | 7.54E-08 | 9.10E-07 | -2.019 | 0.385 | chr8 | 23193684 | 23223638 | + | lncRNA |
| ENSG00000119686.10_4 | FLVCR2 | 1.43E-09 | 2.23E-08 | -2.042 | 1.165 | chr14 | 76044963 | 76129557 | + | protein_coding |
| ENSG00000144278.15_4 | GALNT13 | 8.11E-19 | 5.09E-17 | -2.047 | 4.098 | chr2 | 154728435 | 155310491 | + | protein_coding |
| ENSG00000130830.15_6 | MPP1 | 8.56E-13 | 2.09E-11 | -2.054 | 2.437 | chrX | 154006959 | 154049282 | - | protein_coding |
| ENSG00000173705.9_6 | SUSD5 | 9.68E-12 | 2.01E-10 | -2.055 | 1.644 | chr3 | 33191535 | 33260302 | - | protein_coding |
| ENSG00000171790.15_4 | SLFNL1 | 2.71E-13 | 7.14E-12 | -2.060 | 2.113 | chr1 | 41481269 | 41488909 | - | protein_coding |
| ENSG00000006071.14_7 | ABCC8 | 1.94E-10 | 3.38E-09 | -2.062 | 1.124 | chr11 | 17414045 | 17498426 | - | protein_coding |
| ENSG00000143512.13_4 | HHIPL2 | 5.94E-14 | 1.77E-12 | -2.072 | 2.465 | chr1 | 222695600 | 222721446 | - | protein_coding |
| ENSG00000225422.4_4 | RBMS1P1 | 1.46E-05 | 0.00012815 | -2.075 | -0.842 | chr12 | 66627859 | 66629014 | - | processed_pseudogene |
| ENSG00000115850.10_4 | LCT | 9.45E-06 | 8.51E-05 | -2.079 | -0.594 | chr2 | 136545420 | 136594754 | - | protein_coding |
| ENSG00000130208.9_3 | APOC1 | 3.42E-14 | 1.07E-12 | -2.085 | 2.538 | chr19 | 45417504 | 45422606 | + | protein_coding |
| ENSG00000184371.14_6 | CSF1 | 2.35E-20 | 1.91E-18 | -2.092 | 4.774 | chr1 | 110452864 | 110473614 | + | protein_coding |
| ENSG00000130829.18_3 | DUSP9 | 5.30E-07 | 5.67E-06 | -2.099 | 0.034 | chrX | 152907946 | 152916781 | + | protein_coding |
| ENSG00000248222.5_6 | AC011389.1 | 0.00015073 | 0.0011391 | -2.103 | -1.120 | chr5 | 168440232 | 168465003 | + | lncRNA |
| ENSG00000169554.21_7 | ZEB2 | 6.27E-14 | 1.86E-12 | -2.108 | 2.364 | chr2 | 145121931 | 145278624 | - | protein_coding |
| ENSG00000253796.2_7 | AC104248.1 | 5.69E-06 | 5.29E-05 | -2.110 | -0.636 | chr8 | 109883357 | 110075646 | - | lncRNA |
| ENSG00000110328.6_5 | GALNT18 | 4.70E-16 | 1.92E-14 | -2.112 | 3.397 | chr11 | 11292424 | 11643552 | - | protein_coding |
| ENSG00000164616.16_4 | FBXL21P | 4.30E-09 | 6.23E-08 | -2.119 | 0.681 | chr5 | 135266006 | 135287280 | + | transcribed_unitary_pseudogene |
| ENSG00000266969.1_7 | AP002449.1 | 0.00013179 | 0.00100323 | -2.121 | -1.367 | chr18 | 12984693 | 12991172 | - | lncRNA |
| ENSG00000279673.1_6 | AC092919.3 | 8.31E-05 | 0.00065054 | -2.122 | -0.956 | chr3 | 171178401 | 171180045 | + | TEC |
| ENSG00000105835.12_5 | NAMPT | 2.71E-24 | 5.67E-22 | -2.125 | 7.219 | chr7 | 105888744 | 105926772 | - | protein_coding |
| ENSG00000108684.14_7 | ASIC2 | 5.98E-06 | 5.54E-05 | -2.133 | -0.679 | chr17 | 31340105 | 32501983 | - | protein_coding |
| ENSG00000128262.8_2 | POM121L9P | 7.56E-05 | 0.00059602 | -2.134 | -1.177 | chr22 | 24647796 | 24661493 | + | transcribed_unprocessed_pseudogene |
| ENSG00000242173.10_5 | ARHGDIG | 1.62E-05 | 0.00014131 | -2.146 | -1.008 | chr16 | 330450 | 333010 | + | protein_coding |
| ENSG00000100505.13_3 | TRIM9 | 2.71E-18 | 1.59E-16 | -2.153 | 3.707 | chr14 | 51441980 | 51562779 | - | protein_coding |
| ENSG00000186648.15_6 | CARMIL3 | 9.23E-10 | 1.47E-08 | -2.154 | 1.090 | chr14 | 24521218 | 24538938 | + | protein_coding |
| ENSG00000180287.17_4 | PLD5 | 4.32E-13 | 1.12E-11 | -2.159 | 2.133 | chr1 | 242246288 | 242687999 | - | protein_coding |
| ENSG00000230002.3_6 | ALMS1-IT1 | 4.62E-09 | 6.67E-08 | -2.160 | 0.674 | chr2 | 73683891 | 73686609 | + | lncRNA |
| ENSG00000135312.6_3 | HTR1B | 3.11E-05 | 0.00026244 | -2.164 | -1.156 | chr6 | 78171470 | 78173490 | - | protein_coding |
| ENSG00000173698.18_5 | ADGRG2 | 1.22E-15 | 4.73E-14 | -2.168 | 2.887 | chrX | 19007425 | 19140755 | - | protein_coding |
| ENSG00000238133.6_6 | MAP3K20-AS1 | 4.91E-07 | 5.27E-06 | -2.177 | -0.234 | chr2 | 174031174 | 174146764 | - | lncRNA |
| ENSG00000154310.17_6 | TNIK | 1.28E-10 | 2.28E-09 | -2.190 | 1.073 | chr3 | 170776203 | 171178197 | - | protein_coding |
| ENSG00000127863.15_4 | TNFRSF19 | 4.01E-09 | 5.84E-08 | -2.190 | 0.628 | chr13 | 24144509 | 24250243 | + | protein_coding |
| ENSG00000138356.14_6 | AOX1 | 5.90E-12 | 1.26E-10 | -2.199 | 1.445 | chr2 | 201450737 | 201541787 | + | protein_coding |
| ENSG00000221740.1 | SNORD93 | 5.47E-05 | 0.00044256 | -2.200 | -1.217 | chr7 | 22896232 | 22896305 | + | snoRNA |
| ENSG00000149564.12_4 | ESAM | 2.67E-14 | 8.54E-13 | -2.204 | 2.205 | chr11 | 124622479 | 124632186 | - | protein_coding |
| ENSG00000175874.10_4 | CREG2 | 5.66E-10 | 9.25E-09 | -2.217 | 0.875 | chr2 | 101962012 | 102003965 | - | protein_coding |
| ENSG00000138944.8_5 | SHISAL1 | 2.70E-10 | 4.61E-09 | -2.221 | 1.059 | chr22 | 44639545 | 44708831 | - | protein_coding |
| ENSG00000114948.13_4 | ADAM23 | 1.65E-12 | 3.89E-11 | -2.222 | 1.655 | chr2 | 207308256 | 207485851 | + | protein_coding |
| ENSG00000236107.10_8 | SCN1A-AS1 | 3.91E-05 | 0.00032409 | -2.232 | -1.195 | chr2 | 166813698 | 167247281 | + | lncRNA |
| ENSG00000244128.7_6 | LINC01322 | 9.63E-09 | 1.31E-07 | -2.236 | 0.861 | chr3 | 164924717 | 165564307 | + | lncRNA |
| ENSG00000288054.1_2 | AL360178.2 | 2.52E-06 | 2.46E-05 | -2.237 | -0.543 | chr6 | 136203918 | 136209271 | - | lncRNA |
| ENSG00000165757.9_7 | JCAD | 2.19E-20 | 1.79E-18 | -2.245 | 4.502 | chr10 | 30301732 | 30404423 | - | protein_coding |
| ENSG00000109265.14_8 | CRACD | 7.92E-13 | 1.96E-11 | -2.267 | 1.746 | chr4 | 56915239 | 57194791 | + | protein_coding |
| ENSG00000267279.2_9 | AC090409.1 | 1.18E-07 | 1.38E-06 | -2.282 | -0.065 | chr18 | 59252979 | 59274239 | - | lncRNA |
| ENSG00000286288.1_2 | AL109809.4 | 9.55E-05 | 0.00074054 | -2.296 | -1.459 | chr20 | 1709684 | 1798411 | - | lncRNA |
| ENSG00000104415.14_5 | CCN4 | 1.68E-09 | 2.58E-08 | -2.302 | 0.545 | chr8 | 134203282 | 134243933 | + | protein_coding |
| ENSG00000185532.19_8 | PRKG1 | 3.19E-07 | 3.51E-06 | -2.309 | -0.285 | chr10 | 52750648 | 54058183 | + | protein_coding |
| ENSG00000169213.7_4 | RAB3B | 3.79E-23 | 5.85E-21 | -2.310 | 5.563 | chr1 | 52373628 | 52456372 | - | protein_coding |
| ENSG00000172548.14_5 | NIPAL4 | 6.78E-16 | 2.71E-14 | -2.316 | 2.596 | chr5 | 156887027 | 156901725 | + | protein_coding |
| ENSG00000166073.11_4 | GPR176 | 2.25E-12 | 5.18E-11 | -2.319 | 1.549 | chr15 | 40091209 | 40212467 | - | protein_coding |
| ENSG00000273079.5_6 | GRIN2B | 5.20E-16 | 2.11E-14 | -2.321 | 2.751 | chr12 | 13590876 | 14134891 | - | protein_coding |
| ENSG00000027075.15_5 | PRKCH | 1.37E-15 | 5.26E-14 | -2.326 | 2.389 | chr14 | 61654277 | 62017694 | + | protein_coding |
| ENSG00000158089.15_4 | GALNT14 | 1.30E-21 | 1.39E-19 | -2.326 | 4.660 | chr2 | 31133333 | 31378068 | - | protein_coding |
| ENSG00000169071.15_6 | ROR2 | 5.82E-07 | 6.18E-06 | -2.334 | -0.363 | chr9 | 94325373 | 94712510 | - | protein_coding |
| ENSG00000179111.9_7 | HES7 | 2.94E-06 | 2.84E-05 | -2.340 | -0.873 | chr17 | 8023910 | 8027424 | - | protein_coding |

| | | | | | | | | | | |
|---|---|---|---|---|---|---|---|---|---|---|
| ENSG00000101463.6_3 | SYNDIG1 | 0.00011218 | 0.00086022 | -2.340 | -1.217 | chr20 | 24450265 | 24647252 | + | protein_coding |
| ENSG00000068831.19_5 | RASGRP2 | 0.00092881 | 0.00608038 | -2.342 | -1.438 | chr11 | 64494383 | 64512928 | - | protein_coding |
| ENSG00000145703.16_5 | IQGAP2 | 9.34E-16 | 3.65E-14 | -2.357 | 3.099 | chr5 | 75699110 | 76003957 | + | protein_coding |
| ENSG00000100302.7_3 | RASD2 | 1.13E-08 | 1.52E-07 | -2.369 | 0.083 | chr22 | 35936878 | 35950046 | + | protein_coding |
| ENSG00000132718.9_3 | SYT11 | 2.44E-12 | 5.57E-11 | -2.378 | 1.664 | chr1 | 155829358 | 155854990 | + | protein_coding |
| ENSG00000067445.21_8 | TRO | 1.33E-08 | 1.77E-07 | -2.421 | 0.034 | chrX | 54946895 | 54957864 | + | protein_coding |
| ENSG00000187867.10_6 | PALM3 | 1.67E-06 | 1.67E-05 | -2.455 | -0.873 | chr19 | 14164175 | 14172888 | - | protein_coding |
| ENSG00000162745.10_6 | OLFML2B | 1.13E-06 | 1.16E-05 | -2.468 | -0.706 | chr1 | 161952982 | 161993644 | - | protein_coding |
| ENSG00000120162.10_4 | MOB3B | 1.61E-14 | 5.33E-13 | -2.471 | 2.138 | chr9 | 27325207 | 27529812 | - | protein_coding |
| ENSG00000143217.9_5 | NECTIN4 | 7.07E-05 | 0.00056067 | -2.472 | -1.345 | chr1 | 161040788 | 161059348 | - | protein_coding |
| ENSG00000226604.2_8 | PAPPA-AS2 | 1.44E-06 | 1.45E-05 | -2.476 | -0.780 | chr9 | 119048108 | 119051048 | - | lncRNA |
| ENSG00000186297.12_6 | GABRA5 | 1.03E-24 | 2.44E-22 | -2.493 | 5.636 | chr15 | 27112058 | 27194355 | + | protein_coding |
| ENSG00000179242.16_6 | CDH4 | 3.63E-20 | 2.83E-18 | -2.507 | 4.749 | chr20 | 59827317 | 60515673 | + | protein_coding |
| ENSG00000286502.1_3 | AL645768.1 | 1.58E-06 | 1.59E-05 | -2.512 | -0.747 | chr9 | 137662497 | 137667517 | - | lncRNA |
| ENSG00000119865.9_3 | CNRIP1 | 1.58E-07 | 1.83E-06 | -2.520 | -0.230 | chr2 | 68511303 | 68547183 | - | protein_coding |
| ENSG00000115414.20_7 | FN1 | 1.99E-27 | 1.01E-24 | -2.536 | 8.103 | chr2 | 216225163 | 216300796 | - | protein_coding |
| ENSG00000141574.8_8 | SECTM1 | 3.96E-09 | 5.78E-08 | -2.551 | 0.729 | chr17 | 80278900 | 80291950 | - | protein_coding |
| ENSG00000118495.19_6 | PLAGL1 | 5.15E-14 | 1.56E-12 | -2.571 | 1.735 | chr6 | 144261437 | 144385736 | - | protein_coding |
| ENSG00000170745.12_4 | KCNS3 | 3.26E-16 | 1.38E-14 | -2.574 | 2.508 | chr2 | 18059114 | 18542882 | + | protein_coding |
| ENSG00000186807.9 | ANXA8L2 | 7.89E-06 | 7.18E-05 | -2.581 | -0.778 | chr10 | 47746936 | 47763041 | + | protein_coding |
| ENSG00000182389.19_9 | CACNB4 | 2.66E-07 | 2.97E-06 | -2.589 | -0.622 | chr2 | 152689282 | 152955989 | - | protein_coding |
| ENSG00000144057.15_4 | ST6GAL2 | 3.81E-27 | 1.78E-24 | -2.590 | 6.731 | chr2 | 107418056 | 107503564 | - | protein_coding |
| ENSG00000178462.12_4 | TUBAL3 | 2.60E-06 | 2.53E-05 | -2.596 | -1.043 | chr10 | 5435064 | 5446791 | - | protein_coding |
| ENSG00000163064.7_3 | EN1 | 3.74E-06 | 3.57E-05 | -2.603 | -1.043 | chr2 | 119599747 | 119605224 | - | protein_coding |
| ENSG00000286559.1_1 | AC099845.1 | 1.27E-05 | 0.00011225 | -2.632 | -1.236 | chr18 | 37235609 | 37245458 | + | lncRNA |
| ENSG00000154864.12_5 | PIEZO2 | 9.90E-14 | 2.86E-12 | -2.634 | 1.446 | chr18 | 10666480 | 11148761 | - | protein_coding |
| ENSG00000245248.8_7 | USP2-AS1 | 8.16E-09 | 1.12E-07 | -2.640 | -0.083 | chr11 | 119235069 | 119397374 | + | lncRNA |
| ENSG00000162433.15_4 | AK4 | 1.50E-16 | 6.80E-15 | -2.670 | 3.124 | chr1 | 65613232 | 65697828 | + | protein_coding |
| ENSG00000115194.11_7 | SLC30A3 | 7.96E-09 | 1.10E-07 | -2.685 | 0.009 | chr2 | 27476552 | 27498685 | - | protein_coding |
| ENSG00000213694.5_5 | S1PR3 | 7.83E-25 | 1.93E-22 | -2.700 | 5.383 | chr9 | 91605778 | 91620070 | + | protein_coding |
| ENSG00000177283.7_4 | FZD8 | 6.35E-17 | 3.09E-15 | -2.713 | 2.742 | chr10 | 35927177 | 35931206 | - | protein_coding |
| ENSG00000203930.12_4 | LINC00632 | 1.10E-13 | 3.12E-12 | -2.735 | 2.825 | chrX | 139791727 | 139875380 | + | lncRNA |
| ENSG00000178445.9_6 | GLDC | 5.46E-24 | 1.08E-21 | -2.739 | 4.857 | chr9 | 6532464 | 6645783 | - | protein_coding |
| ENSG00000150165.13 | ANXA8L1 | 3.30E-07 | 3.62E-06 | -2.740 | -0.841 | chr10 | 47157983 | 47174093 | - | protein_coding |
| ENSG00000079156.17_4 | OSBPL6 | 5.27E-15 | 1.86E-13 | -2.766 | 1.990 | chr2 | 179059208 | 179267620 | + | protein_coding |
| ENSG00000139973.16_3 | SYT16 | 2.35E-05 | 0.00020168 | -2.780 | -1.259 | chr14 | 62278692 | 62579268 | + | protein_coding |
| ENSG00000157111.13_4 | TMEM171 | 6.75E-08 | 8.21E-07 | -2.815 | -0.529 | chr5 | 72416396 | 72427636 | + | protein_coding |
| ENSG00000263460.1_4 | BOD1P1 | 1.79E-06 | 1.78E-05 | -2.816 | -0.707 | chr18 | 3415143 | 3415578 | + | transcribed_processed_pseudogene |
| ENSG00000164161.10_3 | HHIP | 1.06E-13 | 3.04E-12 | -2.832 | 1.251 | chr4 | 145567308 | 145666423 | + | protein_coding |
| ENSG00000163735.7_4 | CXCL5 | 1.03E-10 | 1.85E-09 | -2.838 | 5.934 | chr4 | 74861359 | 74864394 | - | protein_coding |
| ENSG00000246465.1_7 | AC138904.1 | 9.62E-06 | 8.65E-05 | -2.848 | -1.344 | chr16 | 28296206 | 28303385 | - | lncRNA |
| ENSG00000113319.13_3 | RASGRF2 | 1.99E-14 | 6.46E-13 | -2.853 | 1.607 | chr5 | 80256182 | 80525981 | + | protein_coding |
| ENSG00000153823.18_3 | PID1 | 1.77E-13 | 4.83E-12 | -2.868 | 1.257 | chr2 | 229715242 | 230136001 | - | protein_coding |
| ENSG00000235531.11_9 | MSC-AS1 | 1.41E-10 | 2.51E-09 | -2.870 | 1.288 | chr8 | 72740402 | 73030628 | + | lncRNA |
| ENSG00000151892.14_4 | GFRA1 | 4.53E-24 | 9.25E-22 | -2.886 | 4.564 | chr10 | 117816436 | 118032979 | - | protein_coding |
| ENSG00000154096.13_7 | THY1 | 3.53E-16 | 1.48E-14 | -2.899 | 1.952 | chr11 | 119288088 | 119295695 | - | protein_coding |
| ENSG00000244757.1_8 | AL137024.1 | 4.65E-06 | 4.38E-05 | -2.923 | -0.920 | chr9 | 119050897 | 119080968 | - | lncRNA |
| ENSG00000130294.16_6 | KIF1A | 6.83E-09 | 9.55E-08 | -2.965 | -0.262 | chr2 | 241653178 | 241760453 | - | protein_coding |
| ENSG00000103489.12_5 | XYLT1 | 7.86E-09 | 1.09E-07 | -2.997 | -0.467 | chr16 | 17195626 | 17564817 | - | protein_coding |
| ENSG00000250041.3_7 | AC069360.1 | 6.25E-07 | 6.61E-06 | -3.022 | -1.078 | chr11 | 10905705 | 10920911 | - | lncRNA |
| ENSG00000060982.15_6 | BCAT1 | 2.31E-26 | 8.73E-24 | -3.040 | 5.553 | chr12 | 24962958 | 25102035 | - | protein_coding |
| ENSG00000205362.11_4 | MT1A | 2.54E-11 | 4.98E-10 | -3.105 | 0.289 | chr16 | 56672578 | 56673999 | + | protein_coding |
| ENSG00000204869.8_5 | IGFL4 | 4.88E-12 | 1.05E-10 | -3.105 | 0.591 | chr19 | 46543006 | 46580376 | - | protein_coding |
| ENSG00000286347.1_2 | AC114802.1 | 1.66E-07 | 1.91E-06 | -3.106 | -0.889 | chr2 | 69257174 | 69260769 | - | lncRNA |
| ENSG00000113739.10_5 | STC2 | 2.48E-20 | 2.00E-18 | -3.108 | 3.227 | chr5 | 172741716 | 172756506 | - | protein_coding |
| ENSG00000108924.14_4 | HLF | 2.89E-10 | 4.91E-09 | -3.114 | 0.001 | chr17 | 53342321 | 53402548 | + | protein_coding |
| ENSG00000010810.17_8 | FYN | 5.31E-12 | 1.14E-10 | -3.138 | 0.692 | chr6 | 111981535 | 112194655 | - | protein_coding |
| ENSG00000171056.8_5 | SOX7 | 1.74E-07 | 2.00E-06 | -3.149 | -0.530 | chr8 | 10581278 | 10588021 | - | protein_coding |
| ENSG00000248590.2_2 | GLDCP1 | 4.23E-07 | 4.57E-06 | -3.154 | -1.117 | chr4 | 57459170 | 57461847 | + | transcribed_processed_pseudogene |
| ENSG00000261655.1_7 | AC100803.3 | 4.68E-09 | 6.74E-08 | -3.161 | -0.339 | chr8 | 142363503 | 142365465 | - | lncRNA |

| | | | | | | | | | | |
|---|---|---|---|---|---|---|---|---|---|---|
| ENSG00000185924.7_5 | RTN4RL1 | 2.17E-10 | 3.74E-09 | -3.178 | 0.054 | chr17 | 1837971 | 1928628 | - | protein_coding |
| ENSG00000134853.12_6 | PDGFRA | 3.83E-09 | 5.62E-08 | -3.190 | -0.504 | chr4 | 55095447 | 55164412 | + | protein_coding |
| ENSG00000121966.7_3 | CXCR4 | 9.33E-08 | 1.11E-06 | -3.191 | -0.954 | chr2 | 136871919 | 136875719 | - | protein_coding |
| ENSG00000167281.19_6 | RBFOX3 | 1.61E-13 | 4.43E-12 | -3.198 | 2.743 | chr17 | 77085427 | 77512230 | - | protein_coding |
| ENSG00000144857.14_7 | BOC | 3.42E-16 | 1.44E-14 | -3.242 | 1.600 | chr3 | 112929850 | 113006306 | + | protein_coding |
| ENSG00000152377.14_4 | SPOCK1 | 2.00E-20 | 1.66E-18 | -3.272 | 3.237 | chr5 | 136310987 | 136934068 | - | protein_coding |
| ENSG00000177494.6_4 | ZBED2 | 2.29E-06 | 2.25E-05 | -3.273 | -1.365 | chr3 | 111311747 | 111314193 | - | protein_coding |
| ENSG00000148082.10_5 | SHC3 | 3.51E-10 | 5.88E-09 | -3.303 | 0.008 | chr9 | 91620686 | 91793733 | - | protein_coding |
| ENSG00000145920.15_5 | CPLX2 | 1.55E-30 | 1.84E-27 | -3.305 | 9.672 | chr5 | 175223313 | 175311024 | + | protein_coding |
| ENSG00000140798.16_5 | ABCC12 | 1.43E-08 | 1.89E-07 | -3.328 | -0.718 | chr16 | 48114917 | 48189929 | - | protein_coding |
| ENSG00000218336.9_6 | TENM3 | 2.56E-13 | 6.79E-12 | -3.346 | 0.693 | chr4 | 183065140 | 183724177 | + | protein_coding |
| ENSG00000128655.18_7 | PDE11A | 2.54E-14 | 8.16E-13 | -3.420 | 0.944 | chr2 | 178487972 | 178937504 | - | protein_coding |
| ENSG00000151388.11_6 | ADAMTS12 | 3.49E-21 | 3.38E-19 | -3.493 | 3.165 | chr5 | 33523640 | 33892124 | - | protein_coding |
| ENSG00000128710.5_3 | HOXD10 | 2.21E-12 | 5.11E-11 | -3.504 | 0.395 | chr2 | 176973518 | 176984670 | + | protein_coding |
| ENSG00000100626.17_4 | GALNT16 | 5.34E-13 | 1.36E-11 | -3.509 | 0.625 | chr14 | 69725994 | 69823750 | + | protein_coding |
| ENSG00000250697.2_8 | AC010343.3 | 2.51E-07 | 2.81E-06 | -3.520 | -1.176 | chr5 | 32888342 | 33298172 | - | lncRNA |
| ENSG00000105605.7_2 | CACNG7 | 1.32E-17 | 7.03E-16 | -3.553 | 1.877 | chr19 | 54412589 | 54447195 | + | protein_coding |
| ENSG00000166923.12_4 | GREM1 | 2.30E-24 | 4.93E-22 | -3.558 | 4.029 | chr15 | 33010205 | 33037307 | + | protein_coding |
| ENSG00000164220.7_5 | F2RL2 | 7.78E-18 | 4.23E-16 | -3.661 | 2.129 | chr5 | 75911307 | 75919238 | - | protein_coding |
| ENSG00000248187.1_7 | AC078850.1 | 1.28E-07 | 1.50E-06 | -3.715 | -1.235 | chr4 | 129489127 | 129491686 | - | lncRNA |
| ENSG00000169783.12_6 | LINGO1 | 6.45E-11 | 1.19E-09 | -3.774 | -0.128 | chr15 | 77905369 | 78113242 | - | protein_coding |
| ENSG00000178860.8_3 | MSC | 5.75E-18 | 3.20E-16 | -3.778 | 2.544 | chr8 | 72753784 | 72756703 | - | protein_coding |
| ENSG00000155980.12_3 | KIF5A | 5.52E-17 | 2.74E-15 | -3.783 | 1.494 | chr12 | 57943827 | 57980416 | + | protein_coding |
| ENSG00000021645.19_6 | NRXN3 | 1.02E-13 | 2.94E-12 | -3.844 | 0.751 | chr14 | 78636716 | 80334633 | + | protein_coding |
| ENSG00000168955.4_5 | TM4SF20 | 4.49E-25 | 1.19E-22 | -3.851 | 4.275 | chr2 | 228226754 | 228246711 | - | protein_coding |
| ENSG00000226674.11_8 | TEX41 | 3.45E-18 | 1.98E-16 | -3.891 | 1.932 | chr2 | 145423879 | 146020556 | + | lncRNA |
| ENSG00000157851.17_3 | DPYSL5 | 7.19E-23 | 1.04E-20 | -3.975 | 3.190 | chr2 | 27070615 | 27173219 | + | protein_coding |
| ENSG00000137491.15_5 | SLCO2B1 | 1.28E-18 | 7.84E-17 | -3.992 | 1.960 | chr11 | 74811608 | 74917594 | + | protein_coding |
| ENSG00000057657.17_8 | PRDM1 | 3.20E-19 | 2.16E-17 | -4.031 | 2.727 | chr6 | 106441338 | 106557814 | + | protein_coding |
| ENSG00000101098.12_3 | RIMS4 | 1.50E-13 | 4.15E-12 | -4.153 | 0.759 | chr20 | 43380449 | 43438979 | - | protein_coding |
| ENSG00000140379.8_4 | BCL2A1 | 9.57E-13 | 2.31E-11 | -4.184 | 0.435 | chr15 | 80253234 | 80263538 | - | protein_coding |
| ENSG00000163536.12_2 | SERPINI1 | 3.29E-15 | 1.19E-13 | -4.217 | 0.812 | chr3 | 167453031 | 167543356 | + | protein_coding |
| ENSG00000226953.8_6 | NCKAP5-AS2 | 6.38E-14 | 1.90E-12 | -4.321 | 0.565 | chr2 | 134022540 | 134043017 | + | lncRNA |
| ENSG00000164619.10_6 | BMPER | 1.89E-13 | 5.10E-12 | -4.495 | 0.268 | chr7 | 33943920 | 34196039 | + | protein_coding |
| ENSG00000286215.1_2 | BMPER | 1.69E-16 | 7.53E-15 | -4.560 | 1.398 | chr6 | 126625369 | 127534754 | - | transcribed_processed_pseudogene |
| ENSG00000122862.5_3 | SRGN | 2.20E-27 | 1.08E-24 | -4.832 | 4.327 | chr10 | 70847859 | 70864561 | + | protein_coding |
| ENSG00000151468.11_7 | CCDC3 | 3.87E-12 | 8.49E-11 | -4.892 | -0.259 | chr10 | 12938625 | 13141652 | - | protein_coding |
| ENSG00000171903.16_4 | CYP4F11 | 1.38E-30 | 1.76E-27 | -5.022 | 5.307 | chr19 | 16023177 | 16045677 | - | protein_coding |
| ENSG00000116667.14_5 | C1orf21 | 2.97E-20 | 2.36E-18 | -5.079 | 2.148 | chr1 | 184356163 | 184598153 | + | protein_coding |
| ENSG00000069188.17_5 | SDK2 | 1.99E-17 | 1.04E-15 | -5.156 | 2.277 | chr17 | 71330523 | 71640584 | - | protein_coding |
| ENSG00000173557.15_4 | FAM166C | 9.53E-11 | 1.72E-09 | -5.173 | -0.620 | chr2 | 26785453 | 26802400 | + | protein_coding |
| ENSG00000134363.12_5 | FST | 5.15E-25 | 1.33E-22 | -5.233 | 3.703 | chr5 | 52776456 | 52782964 | + | protein_coding |
| ENSG00000267056.2_5 | AC005336.1 | 1.61E-22 | 2.08E-20 | -5.336 | 2.604 | chr19 | 16021392 | 16022634 | - | processed_pseudogene |
| ENSG00000077420.16_4 | APBB1IP | 1.59E-18 | 9.58E-17 | -5.669 | 1.373 | chr10 | 26727270 | 26856732 | + | protein_coding |
| ENSG00000053108.17_5 | FSTL4 | 1.57E-20 | 1.33E-18 | -5.747 | 2.903 | chr5 | 132532147 | 132948232 | - | protein_coding |
| ENSG00000146374.14_4 | RSPO3 | 1.68E-27 | 8.80E-25 | -5.826 | 5.043 | chr6 | 127439816 | 127520626 | + | protein_coding |
| ENSG00000184845.4_4 | DRD1 | 1.31E-23 | 2.29E-21 | -6.269 | 2.718 | chr5 | 174867039 | 174871185 | - | protein_coding |
| ENSG00000091129.21_9 | NRCAM | 1.45E-34 | 8.58E-31 | -6.454 | 7.208 | chr7 | 107788068 | 108097161 | - | protein_coding |
| ENSG00000198538.11_6 | ZNF28 | 5.33E-32 | 1.88E-28 | -6.566 | 5.702 | chr19 | 53300661 | 53360853 | - | protein_coding |
| ENSG00000113389.16_5 | NPR3 | 7.41E-32 | 1.88E-28 | -6.566 | 5.273 | chr5 | 32689176 | 32791830 | + | protein_coding |
| ENSG00000185760.15_5 | KCNQ5 | 2.30E-12 | 5.30E-11 | -6.753 | -0.359 | chr6 | 73331520 | 73908574 | + | protein_coding |
| ENSG00000144583.5_4 | MARCHF4 | 7.13E-18 | 3.90E-16 | -7.094 | 1.177 | chr2 | 217122588 | 217237206 | - | protein_coding |
| ENSG00000156687.11_5 | UNC5D | 8.14E-21 | 7.23E-19 | -7.125 | 2.109 | chr8 | 35092993 | 35654068 | + | protein_coding |
| ENSG00000141052.17_2 | MYOCD | 2.45E-19 | 1.69E-17 | -7.665 | 1.745 | chr17 | 12569207 | 12672266 | + | protein_coding |
| ENSG00000140538.16_4 | NTRK3 | 2.13E-25 | 6.20E-23 | -8.771 | 3.240 | chr15 | 88402982 | 88799999 | - | protein_coding |
| ENSG00000179083.6_2 | FAM133A | 1.29E-27 | 7.47E-25 | -10.937 | 3.788 | chrX | 92929012 | 92967273 | + | protein_coding |

Supplementary Table 3B, Gene ontology analysis of DEGs results, Related to Figure 2B

| GO term name | GO term ID | FDR | -log10(FDR) | GO term size | Intersection size |
|---|---|---|---|---|---|
| cell adhesion | GO:0007155 | 5.69E-26 | 25.2448 | 1135 | 257 |
| biological adhesion | GO:0022610 | 5.69E-26 | 25.2448 | 1142 | 258 |
| cell-cell adhesion | GO:0098609 | 4.20E-11 | 10.3766 | 666 | 142 |
| cell junction organization | GO:0034330 | 2.56E-10 | 9.5923 | 487 | 111 |
| synapse organization | GO:0050808 | 4.86E-10 | 9.3131 | 253 | 70 |
| cell-cell adhesion via plasma-membrane adhesion molecules | GO:0098742 | 1.55E-07 | 6.8089 | 184 | 52 |
| cell morphogenesis | GO:0000902 | 5.72E-07 | 6.2428 | 756 | 141 |
| localization of cell | GO:0051674 | 1.11122E-06 | 5.9542 | 1474 | 239 |
| cell motility | GO:0048870 | 1.11122E-06 | 5.9542 | 1474 | 239 |
| cell migration | GO:0016477 | 1.19615E-06 | 5.9222 | 1342 | 221 |
| cell morphogenesis involved in differentiation | GO:0000904 | 1.28237E-06 | 5.8920 | 530 | 106 |
| cell population proliferation | GO:0008283 | 1.28555E-06 | 5.8909 | 1434 | 233 |
| tissue development | GO:0009888 | 1.42669E-06 | 5.8457 | 1414 | 230 |
| synapse assembly | GO:0007416 | 3.33115E-06 | 5.4774 | 116 | 36 |
| connective tissue development | GO:0061448 | 4.57275E-06 | 5.3398 | 133 | 39 |
| skeletal system development | GO:0001501 | 6.24567E-06 | 5.2044 | 296 | 67 |
| cell junction assembly | GO:0034329 | 9.31379E-06 | 5.0309 | 324 | 71 |
| calcium-dependent cell-cell adhesion via plasma membrane cell adhesion molecules | GO:0016339 | 1.0102E-05 | 4.9956 | 42 | 19 |
| animal organ morphogenesis | GO:0009887 | 1.11975E-05 | 4.9509 | 624 | 116 |
| regulation of cell population proliferation | GO:0042127 | 1.21821E-05 | 4.9143 | 1275 | 206 |
| chondrocyte differentiation | GO:0002062 | 2.83072E-05 | 4.5481 | 71 | 25 |
| axonogenesis | GO:0007409 | 3.00452E-05 | 4.5222 | 335 | 71 |
| regulation of chondrocyte differentiation | GO:0032330 | 3.53443E-05 | 4.4517 | 26 | 14 |
| extracellular matrix organization | GO:0030198 | 4.19127E-05 | 4.3777 | 294 | 64 |
| cell morphogenesis involved in neuron differentiation | GO:0048667 | 4.19127E-05 | 4.3777 | 410 | 82 |
| extracellular structure organization | GO:0043062 | 4.64642E-05 | 4.3329 | 295 | 64 |
| cartilage development | GO:0051216 | 5.89912E-05 | 4.2292 | 104 | 31 |
| ion transport | GO:0006811 | 6.48065E-05 | 4.1884 | 1330 | 209 |
| regulation of cartilage development | GO:0061035 | 7.2976E-05 | 4.1368 | 35 | 16 |
| regulation of cell differentiation | GO:0045595 | 8.07445E-05 | 4.0929 | 1388 | 216 |
| axon development | GO:0061564 | 8.73696E-05 | 4.0586 | 359 | 73 |
| cell development | GO:0048468 | 9.3134E-05 | 4.0309 | 1369 | 213 |
| homophilic cell adhesion via plasma membrane adhesion molecules | GO:0007156 | 9.3134E-05 | 4.0309 | 76 | 25 |
| enzyme linked receptor protein signaling pathway | GO:0007167 | 0.000106536 | 3.9725 | 915 | 152 |
| axon guidance | GO:0007411 | 0.000123557 | 3.9081 | 211 | 49 |
| neuron projection guidance | GO:0097485 | 0.000140332 | 3.8528 | 212 | 49 |
| anion transport | GO:0006820 | 0.000172631 | 3.7629 | 474 | 89 |
| regulation of nervous system development | GO:0051960 | 0.000172631 | 3.7629 | 578 | 104 |
| cell-substrate adhesion | GO:0031589 | 0.000195457 | 3.7089 | 271 | 58 |
| regulation of cell adhesion | GO:0030155 | 0.000205113 | 3.6880 | 574 | 103 |
| gland development | GO:0048732 | 0.000213604 | 3.6704 | 139 | 36 |
| cell part morphogenesis | GO:0032990 | 0.000262201 | 3.5814 | 494 | 91 |
| regulation of locomotion | GO:0040012 | 0.000275313 | 3.5602 | 911 | 149 |

| | | | | | |
|---|---|---|---|---|---|
| regulation of cell morphogenesis involved in differentiation | GO:0010769 | 0.000323603 | 3.4900 | 207 | 47 |
| epithelial cell proliferation | GO:0050673 | 0.000426961 | 3.3696 | 311 | 63 |
| regulation of cell morphogenesis | GO:0022604 | 0.000470141 | 3.3278 | 365 | 71 |
| neuron projection morphogenesis | GO:0048812 | 0.000485461 | 3.3138 | 468 | 86 |
| cellular component morphogenesis | GO:0032989 | 0.000485461 | 3.3138 | 566 | 100 |
| cell projection morphogenesis | GO:0048858 | 0.000515564 | 3.2877 | 483 | 88 |
| transmembrane receptor protein tyrosine kinase signaling pathway | GO:0007169 | 0.00052593 | 3.2791 | 646 | 111 |
| apoptotic process | GO:0006915 | 0.000559806 | 3.2520 | 1443 | 217 |
| regulated exocytosis | GO:0045055 | 0.000559974 | 3.2518 | 705 | 119 |
| neurogenesis | GO:0022008 | 0.000577837 | 3.2382 | 1069 | 168 |
| plasma membrane bounded cell projection morphogenesis | GO:0120039 | 0.00067324 | 3.1718 | 480 | 87 |
| regulation of axonogenesis | GO:0050770 | 0.000832441 | 3.0796 | 114 | 30 |
| negative regulation of cell differentiation | GO:0045596 | 0.000843523 | 3.0739 | 483 | 87 |
| regulation of cell death | GO:0010941 | 0.000924515 | 3.0341 | 1316 | 199 |
| regulation of epithelial cell proliferation | GO:0050678 | 0.000932265 | 3.0305 | 281 | 57 |
| regulation of cell motility | GO:2000145 | 0.001007035 | 2.9970 | 885 | 142 |
| generation of neurons | GO:0048699 | 0.0010172 | 2.9926 | 1013 | 159 |
| regulation of cellular component movement | GO:0051270 | 0.001019377 | 2.9917 | 953 | 151 |
| negative regulation of cell development | GO:0010721 | 0.001047253 | 2.9799 | 187 | 42 |
| positive regulation of cell population proliferation | GO:0008284 | 0.001047253 | 2.9799 | 680 | 114 |
| endoderm formation | GO:0001706 | 0.001079486 | 2.9668 | 48 | 17 |
| cell-cell junction organization | GO:0045216 | 0.001079486 | 2.9668 | 157 | 37 |
| exocytosis | GO:0006887 | 0.001079486 | 2.9668 | 791 | 129 |
| neuron differentiation | GO:0030182 | 0.001149499 | 2.9395 | 904 | 144 |
| negative regulation of response to wounding | GO:1903035 | 0.001149499 | 2.9395 | 73 | 22 |
| organic anion transport | GO:0015711 | 0.001156331 | 2.9369 | 378 | 71 |
| cell-cell signaling | GO:0007267 | 0.001391429 | 2.8565 | 1499 | 221 |
| regulation of anatomical structure morphogenesis | GO:0022603 | 0.001397778 | 2.8546 | 938 | 148 |
| regulation of cell migration | GO:0030334 | 0.001475054 | 2.8312 | 827 | 133 |
| vascular process in circulatory system | GO:0003018 | 0.001560132 | 2.8068 | 113 | 29 |
| positive regulation of transferase activity | GO:0051347 | 0.001818455 | 2.7403 | 574 | 98 |
| endoderm development | GO:0007492 | 0.001853554 | 2.7320 | 55 | 18 |
| morphogenesis of an epithelium | GO:0002009 | 0.001886558 | 2.7243 | 323 | 62 |
| negative regulation of neurogenesis | GO:0050768 | 0.001974215 | 2.7046 | 156 | 36 |
| canonical Wnt signaling pathway | GO:0060070 | 0.001986864 | 2.7018 | 297 | 58 |
| regulation of neuron differentiation | GO:0045664 | 0.002145968 | 2.6684 | 407 | 74 |
| regulation of neurogenesis | GO:0050767 | 0.002292318 | 2.6397 | 514 | 89 |
| taxis | GO:0042330 | 0.002292318 | 2.6397 | 514 | 89 |
| chemotaxis | GO:0006935 | 0.002292318 | 2.6397 | 514 | 89 |
| positive regulation of developmental process | GO:0051094 | 0.002383229 | 2.6228 | 1034 | 159 |
| positive regulation of epithelial cell proliferation | GO:0050679 | 0.002478702 | 2.6058 | 152 | 35 |
| cell-matrix adhesion | GO:0007160 | 0.002490236 | 2.6038 | 183 | 40 |
| regulation of apoptotic process | GO:0042981 | 0.002490236 | 2.6038 | 1189 | 179 |
| export from cell | GO:0140352 | 0.002954465 | 2.5295 | 1333 | 197 |
| tissue morphogenesis | GO:0048729 | 0.002954465 | 2.5295 | 419 | 75 |
| epidermal growth factor receptor signaling pathway | GO:0007173 | 0.003015707 | 2.5206 | 112 | 28 |
| regulation of cell projection organization | GO:0031344 | 0.003234105 | 2.4902 | 484 | 84 |
| positive regulation of locomotion | GO:0040017 | 0.003585191 | 2.4455 | 500 | 86 |
| regulation of neuron projection development | GO:0010975 | 0.003937703 | 2.4048 | 326 | 61 |

| | | | | | |
|---|---|---|---|---|---|
| regulation of response to wounding | GO:1903034 | 0.004104379 | 2.3868 | 144 | 33 |
| positive regulation of cell differentiation | GO:0045597 | 0.004429314 | 2.3537 | 723 | 116 |
| regulation of cell-cell adhesion | GO:0022407 | 0.005313841 | 2.2746 | 351 | 64 |
| regulation of plasma membrane bounded cell projection organization | GO:0120035 | 0.005313841 | 2.2746 | 478 | 82 |
| cell-cell adhesion mediated by cadherin | GO:0044331 | 0.005313841 | 2.2746 | 45 | 15 |
| regulation of programmed cell death | GO:0043067 | 0.005313841 | 2.2746 | 1216 | 180 |
| adherens junction organization | GO:0034332 | 0.005313841 | 2.2746 | 55 | 17 |
| ERBB signaling pathway | GO:0038127 | 0.005313841 | 2.2746 | 134 | 31 |
| negative regulation of cell projection organization | GO:0031345 | 0.005313841 | 2.2746 | 122 | 29 |
| positive regulation of cell migration | GO:0030335 | 0.005727231 | 2.2421 | 472 | 81 |
| regulation of cell development | GO:0060284 | 0.005902506 | 2.2290 | 618 | 101 |
| negative regulation of neuron differentiation | GO:0045665 | 0.005931728 | 2.2268 | 117 | 28 |
| anatomical structure formation involved in morphogenesis | GO:0048646 | 0.0064214 | 2.1924 | 807 | 126 |
| formation of primary germ layer | GO:0001704 | 0.0064214 | 2.1924 | 83 | 22 |
| endodermal cell differentiation | GO:0035987 | 0.0064214 | 2.1924 | 41 | 14 |
| eye morphogenesis | GO:0048592 | 0.0064214 | 2.1924 | 72 | 20 |
| skeletal system morphogenesis | GO:0048705 | 0.0064214 | 2.1924 | 106 | 26 |
| regulation of phosphorylation | GO:0042325 | 0.006506267 | 2.1867 | 1380 | 200 |
| cell surface receptor signaling pathway involved in cell-cell signaling | GO:1905114 | 0.006506267 | 2.1867 | 482 | 82 |
| positive regulation of kinase activity | GO:0033674 | 0.006692471 | 2.1744 | 497 | 84 |
| regulation of blood coagulation | GO:0030193 | 0.006736756 | 2.1715 | 67 | 19 |
| regulation of hemostasis | GO:1900046 | 0.006736756 | 2.1715 | 67 | 19 |
| tube development | GO:0035295 | 0.006736756 | 2.1715 | 696 | 111 |
| Wnt signaling pathway | GO:0016055 | 0.006811826 | 2.1667 | 433 | 75 |
| positive regulation of multicellular organismal process | GO:0051240 | 0.006998894 | 2.1550 | 1375 | 199 |
| positive regulation of cell motility | GO:2000147 | 0.006998894 | 2.1550 | 491 | 83 |
| mesenchymal cell differentiation | GO:0048762 | 0.007321234 | 2.1354 | 175 | 37 |
| epithelium development | GO:0060429 | 0.007321234 | 2.1354 | 864 | 133 |
| regulation of anatomical structure size | GO:0090066 | 0.007480154 | 2.1261 | 343 | 62 |
| organic acid transport | GO:0015849 | 0.007717944 | 2.1125 | 241 | 47 |
| secretion | GO:0046903 | 0.007776942 | 2.1092 | 1347 | 195 |
| negative regulation of nervous system development | GO:0051961 | 0.008239072 | 2.0841 | 170 | 36 |
| negative regulation of response to stimulus | GO:0048585 | 0.008239072 | 2.0841 | 1404 | 202 |
| neuron migration | GO:0001764 | 0.008409824 | 2.0752 | 74 | 20 |
| cell-cell signaling by wnt | GO:0198738 | 0.008409824 | 2.0752 | 437 | 75 |
| neuron projection development | GO:0031175 | 0.008509341 | 2.0701 | 672 | 107 |
| neuron development | GO:0048666 | 0.008509341 | 2.0701 | 732 | 115 |
| transmembrane transport | GO:0055085 | 0.008784229 | 2.0563 | 1124 | 166 |
| embryonic morphogenesis | GO:0048598 | 0.0093874 | 2.0275 | 305 | 56 |
| positive regulation of cellular component movement | GO:0051272 | 0.009730934 | 2.0118 | 505 | 84 |
| negative regulation of neuron projection development | GO:0010977 | 0.009730934 | 2.0118 | 92 | 23 |
| cell adhesion mediated by integrin | GO:0033627 | 0.009730934 | 2.0118 | 64 | 18 |
| regulation of vesicle-mediated transport | GO:0060627 | 0.009730934 | 2.0118 | 404 | 70 |
| carboxylic acid transport | GO:0046942 | 0.01010142 | 1.9956 | 238 | 46 |
| secretion by cell | GO:0032940 | 0.010137258 | 1.9941 | 1285 | 186 |
| regulation of protein phosphorylation | GO:0001932 | 0.010165398 | 1.9929 | 1238 | 180 |
| negative regulation of cell population proliferation | GO:0008285 | 0.010769263 | 1.9678 | 573 | 93 |

| | | | | | |
|---|---|---|---|---|---|
| ion transmembrane transport | GO:0034220 | 0.011155748 | 1.9525 | 837 | 128 |
| stem cell differentiation | GO:0048863 | 0.011318354 | 1.9462 | 193 | 39 |
| ERK1 and ERK2 cascade | GO:0070371 | 0.01141023 | 1.9427 | 260 | 49 |
| negative regulation of blood coagulation | GO:0030195 | 0.01141023 | 1.9427 | 44 | 14 |
| negative regulation of hemostasis | GO:1900047 | 0.01141023 | 1.9427 | 44 | 14 |
| regulation of exocytosis | GO:0017157 | 0.011473612 | 1.9403 | 142 | 31 |
| negative regulation of developmental process | GO:0051093 | 0.011534565 | 1.9380 | 732 | 114 |
| integrin-mediated signaling pathway | GO:0007229 | 0.011584411 | 1.9361 | 82 | 21 |
| regulation of transferase activity | GO:0051338 | 0.011679513 | 1.9326 | 839 | 128 |
| regulation of coagulation | GO:0050818 | 0.012043113 | 1.9193 | 71 | 19 |
| homotypic cell-cell adhesion | GO:0034109 | 0.012043113 | 1.9193 | 71 | 19 |
| regulation of wound healing | GO:0061041 | 0.013707932 | 1.8630 | 125 | 28 |
| anion transmembrane transport | GO:0098656 | 0.014622544 | 1.8350 | 157 | 33 |
| circulatory system development | GO:0072359 | 0.014830142 | 1.8289 | 814 | 124 |
| response to wounding | GO:0009611 | 0.014830142 | 1.8289 | 484 | 80 |
| regulation of kinase activity | GO:0043549 | 0.014910406 | 1.8265 | 738 | 114 |
| cation transport | GO:0006812 | 0.019209426 | 1.7165 | 881 | 132 |
| regulation of blood vessel diameter | GO:0097746 | 0.020173658 | 1.6952 | 74 | 19 |
| regulation of tube size | GO:0035150 | 0.020173658 | 1.6952 | 74 | 19 |
| regulation of tube diameter | GO:0035296 | 0.020173658 | 1.6952 | 74 | 19 |
| negative regulation of wound healing | GO:0061045 | 0.021139295 | 1.6749 | 63 | 17 |
| positive regulation of cell adhesion | GO:0045785 | 0.02297624 | 1.6387 | 325 | 57 |
| reproductive system development | GO:0061458 | 0.023117688 | 1.6361 | 142 | 30 |
| reproductive structure development | GO:0048608 | 0.023117688 | 1.6361 | 142 | 30 |
| monocarboxylic acid transport | GO:0015718 | 0.023370245 | 1.6313 | 117 | 26 |
| mesenchyme development | GO:0060485 | 0.023370245 | 1.6313 | 208 | 40 |
| tube morphogenesis | GO:0035239 | 0.023370245 | 1.6313 | 634 | 99 |
| regulation of protein kinase activity | GO:0045859 | 0.023370245 | 1.6313 | 649 | 101 |
| embryo development | GO:0009790 | 0.023523054 | 1.6285 | 463 | 76 |
| positive regulation of wound healing | GO:0090303 | 0.024487771 | 1.6111 | 53 | 15 |
| platelet degranulation | GO:0002576 | 0.0251443 | 1.5996 | 124 | 27 |
| positive regulation of cell death | GO:0010942 | 0.025612158 | 1.5916 | 576 | 91 |
| negative regulation of coagulation | GO:0050819 | 0.026150256 | 1.5825 | 48 | 14 |
| retina morphogenesis in camera-type eye | GO:0060042 | 0.02660637 | 1.5750 | 28 | 10 |
| positive regulation of nervous system development | GO:0051962 | 0.027043277 | 1.5679 | 321 | 56 |
| cellular hormone metabolic process | GO:0034754 | 0.027142671 | 1.5663 | 100 | 23 |
| cell-cell junction assembly | GO:0007043 | 0.028819524 | 1.5403 | 119 | 26 |
| negative regulation of response to external stimulus | GO:0032102 | 0.030511802 | 1.5155 | 330 | 57 |
| positive regulation of protein kinase activity | GO:0045860 | 0.031305883 | 1.5044 | 432 | 71 |
| regulation of MAPK cascade | GO:0043408 | 0.033129177 | 1.4798 | 612 | 95 |
| response to lipid | GO:0033993 | 0.033324647 | 1.4772 | 522 | 83 |
| regulation of lipase activity | GO:0060191 | 0.035455326 | 1.4503 | 90 | 21 |
| ossification | GO:0001503 | 0.035774801 | 1.4464 | 241 | 44 |
| positive regulation of response to wounding | GO:1903036 | 0.036508587 | 1.4376 | 61 | 16 |
| central nervous system development | GO:0007417 | 0.036508587 | 1.4376 | 539 | 85 |
| stem cell development | GO:0048864 | 0.037072508 | 1.4309 | 50 | 14 |
| regulation of small GTPase mediated signal transduction | GO:0051056 | 0.038203379 | 1.4179 | 249 | 45 |
| sensory system development | GO:0048880 | 0.0383204 | 1.4166 | 174 | 34 |
| ameboidal-type cell migration | GO:0001667 | 0.039429057 | 1.4042 | 400 | 66 |
| positive regulation of cell-substrate adhesion | GO:0010811 | 0.039429057 | 1.4042 | 79 | 19 |
| positive regulation of epithelial to mesenchymal transition | GO:0010718 | 0.039429057 | 1.4042 | 45 | 13 |
| negative regulation of cell death | GO:0060548 | 0.039429057 | 1.4042 | 731 | 110 |

| | | | | | |
|---|---|---|---|---|---|
| visual system development | GO:0150063 | 0.04004463 | 1.3975 | 168 | 33 |
| regulation of hormone levels | GO:0010817 | 0.040758977 | 1.3898 | 357 | 60 |
| positive regulation of phosphorylation | GO:0042327 | 0.040758977 | 1.3898 | 934 | 136 |
| negative regulation of Notch signaling pathway | GO:0045746 | 0.04286883 | 1.3679 | 35 | 11 |
| regulation of canonical Wnt signaling pathway | GO:0060828 | 0.043034691 | 1.3662 | 258 | 46 |
| positive regulation of exocytosis | GO:0045921 | 0.043253386 | 1.3640 | 68 | 17 |
| circulatory system process | GO:0003013 | 0.044170117 | 1.3549 | 417 | 68 |
| regulation of Wnt signaling pathway | GO:0030111 | 0.0459328 | 1.3379 | 323 | 55 |
| camera-type eye morphogenesis | GO:0048593 | 0.046344222 | 1.3340 | 57 | 15 |
| sensory organ morphogenesis | GO:0090596 | 0.046971937 | 1.3282 | 105 | 23 |
| regulation of animal organ morphogenesis | GO:2000027 | 0.046971937 | 1.3282 | 170 | 33 |
| MAPK cascade | GO:0000165 | 0.048167208 | 1.3172 | 799 | 118 |
| regulation of ERK1 and ERK2 cascade | GO:0070372 | 0.048167208 | 1.3172 | 246 | 44 |
| negative regulation of signal transduction | GO:0009968 | 0.048521471 | 1.3141 | 1106 | 157 |
| phospholipid transport | GO:0015914 | 0.048888025 | 1.3108 | 75 | 18 |
| mesenchymal cell development | GO:0014031 | 0.049521641 | 1.3052 | 52 | 14 |
| homeostatic process | GO:0042592 | 0.049839846 | 1.3024 | 1500 | 206 |

**Supplementary Table 3C, Gene set enrichment analysis of DEGs results, Related to Figure 2C and Supplementary Figure 2D**

| Gene set name | Size | NES | FDR |
|---|---|---|---|
| HALLMARK_E2F_TARGETS | 199 | -2.3065 | 0.0000 |
| HALLMARK_MYC_TARGETS_V1 | 199 | -2.2772 | 0.0000 |
| HALLMARK_MYC_TARGETS_V2 | 57 | -2.2756 | 0.0000 |
| GO_DNA_REPLICATION_INDEPENDENT_NUCLEOSOME_ORGANIZATION | 53 | -2.2723 | 0.0001 |
| GO_CHROMATIN_REMODELING_AT_CENTROMERE | 46 | -2.3742 | 0.0001 |
| GO_CENTROMERE_COMPLEX_ASSEMBLY | 54 | -2.2043 | 0.0002 |
| GO_HOMOPHILIC_CELL_ADHESION_VIA_PLASMA_MEMBRANE_ADHESION_MOLECULES | 117 | 1.9105 | 0.0049 |
| HALLMARK_G2M_CHECKPOINT | 198 | -1.8781 | 0.0055 |
| GO_DNA_GEOMETRIC_CHANGE | 110 | -1.9712 | 0.0134 |
| GO_DNA_DEPENDENT_DNA_REPLICATION | 141 | -1.9603 | 0.0147 |
| GO_POSITIVE_REGULATION_OF_AXON_EXTENSION | 38 | -1.9506 | 0.0160 |
| GO_DNA_REPLICATION_INITIATION | 35 | -1.9383 | 0.0183 |
| GO_CALCIUM_DEPENDENT_CELL_CELL_ADHESION_VIA_PLASMA_MEMBRANE_CELL_ADHESI ON_MOLECULES | 32 | 1.8557 | 0.0212 |
| HALLMARK_INTERFERON_ALPHA_RESPONSE | 83 | 1.6677 | 0.0219 |
| HALLMARK_MTORC1_SIGNALING | 195 | -1.6669 | 0.0251 |
| GO_SPLICEOSOMAL_SNRNP_ASSEMBLY | 39 | -1.9068 | 0.0252 |
| GO_RRNA_METABOLIC_PROCESS | 213 | -1.8829 | 0.0289 |
| GO_NCRNA_EXPORT_FROM_NUCLEUS | 37 | -1.8781 | 0.0296 |
| GO_NCRNA_PROCESSING | 355 | -1.8838 | 0.0298 |
| GO_CELL_CYCLE_DNA_REPLICATION | 61 | -1.8688 | 0.0312 |
| GO_PHOSPHOLIPASE_C_ACTIVATING_G_PROTEIN_COUPLED_RECEPTOR_SIGNALING_PATH WAY | 41 | -1.8662 | 0.0312 |
| GO_DNA_REPLICATION | 253 | -1.8561 | 0.0337 |
| GO_REGULATION_OF_DNA_REPLICATION | 97 | -1.8441 | 0.0370 |
| GO_RIBONUCLEOPROTEIN_COMPLEX_BIOGENESIS | 404 | -1.8362 | 0.0391 |
| GO_RIBOSOME_BIOGENESIS | 280 | -1.8267 | 0.0416 |
| GO_MATURATION_OF_5_8S_RRNA | 26 | -1.8289 | 0.0417 |
| GO_QUINONE_METABOLIC_PROCESS | 30 | -1.8214 | 0.0421 |
| GO_REGULATION_OF_DEFENSE_RESPONSE_TO_VIRUS_BY_VIRUS | 25 | -1.8130 | 0.0452 |
| GO_RNA_EXPORT_FROM_NUCLEUS | 126 | -1.8096 | 0.0456 |
| HALLMARK_HEDGEHOG_SIGNALING | 27 | -1.5154 | 0.0504 |
| GO_CELL_CELL_ADHESION_VIA_PLASMA_MEMBRANE_ADHESION_MOLECULES | 174 | 1.8171 | 0.0515 |
| HALLMARK_WNT_BETA_CATENIN_SIGNALING | 33 | -1.4710 | 0.0528 |
| GO_RIBONUCLEOSIDE_MONOPHOSPHATE_BIOSYNTHETIC_PROCESS | 30 | -1.7854 | 0.0569 |
| GO_MRNA_EXPORT_FROM_NUCLEUS | 102 | -1.7725 | 0.0624 |
| GO_POSITIVE_REGULATION_OF_SPROUTING_ANGIOGENESIS | 31 | -1.7666 | 0.0643 |
| GO_RESPONSE_TO_ACTIVITY | 47 | -1.7432 | 0.0760 |
| HALLMARK_ALLOGRAFT_REJECTION | 108 | -1.3403 | 0.0844 |
| HALLMARK_SPERMATOGENESIS | 77 | -1.3625 | 0.0852 |
| GO_NEGATIVE_REGULATION_OF_CHROMOSOME_SEGREGATION | 41 | -1.7262 | 0.0864 |
| GO_NCRNA_3_END_PROCESSING | 35 | -1.7237 | 0.0869 |
| HALLMARK_UNFOLDED_PROTEIN_RESPONSE | 113 | -1.3629 | 0.0958 |
| GO_PROTEIN_LOCALIZATION_TO_CHROMOSOME_TELOMERIC_REGION | 29 | -1.7106 | 0.0964 |
| GO_ANOIKIS | 33 | 1.7783 | 0.0976 |

| | | | |
|---|---|---|---|
| GO_MYOBLAST_DIFFERENTIATION | 57 | -1.7031 | 0.0997 |
| GO_NEGATIVE_REGULATION_OF_BMP_SIGNALING_PATHWAY | 31 | -1.6650 | 0.1106 |
| GO_NEGATIVE_REGULATION_OF_METAPHASE_ANAPHASE_TRANSITION_OF_CELL_CYCLE | 35 | -1.6662 | 0.1110 |
| GO_REGULATION_OF_DNA_DEPENDENT_DNA_REPLICATION | 45 | -1.6879 | 0.1112 |
| GO_TRNA_METHYLATION | 38 | -1.6668 | 0.1125 |
| GO_REGULATION_OF_CHROMOSOME_SEGREGATION | 95 | -1.6813 | 0.1126 |
| GO_POSITIVE_REGULATION_OF_CELL_CYCLE_G2_M_PHASE_TRANSITION | 29 | -1.6684 | 0.1132 |
| GO_COCHLEA_DEVELOPMENT | 34 | -1.6829 | 0.1134 |
| GO_MODULATION_BY_VIRUS_OF_HOST_MORPHOLOGY_OR_PHYSIOLOGY | 30 | -1.6778 | 0.1135 |
| GO_REGULATION_OF_EXTENT_OF_CELL_GROWTH | 89 | -1.6727 | 0.1142 |
| GO_RIBONUCLEOPROTEIN_COMPLEX_SUBUNIT_ORGANIZATION | 184 | -1.6699 | 0.1143 |
| GO_RIBOSOMAL_SMALL_SUBUNIT_BIOGENESIS | 64 | -1.6744 | 0.1148 |
| GO_MITOTIC_SISTER_CHROMATID_SEGREGATION | 144 | -1.6511 | 0.1156 |
| GO_MISMATCH_REPAIR | 33 | -1.6570 | 0.1159 |
| GO_NEGATIVE_REGULATION_OF_SMOOTH_MUSCLE_CELL_PROLIFERATION | 35 | -1.6444 | 0.1164 |
| GO_TELOMERE_MAINTENANCE_VIA_SEMI_CONSERVATIVE_REPLICATION | 27 | -1.6523 | 0.1164 |
| GO_SMOOTH_MUSCLE_CONTRACTION | 71 | -1.6533 | 0.1173 |
| GO_MITOCHONDRIAL_GENE_EXPRESSION | 164 | -1.6445 | 0.1181 |
| GO_REGULATION_OF_SISTER_CHROMATID_SEGREGATION | 76 | -1.6389 | 0.1195 |
| GO_RNA_LOCALIZATION | 214 | -1.6448 | 0.1197 |
| GO_DNA_DEPENDENT_DNA_REPLICATION_MAINTENANCE_OF_FIDELITY | 42 | -1.6344 | 0.1203 |
| GO_RNA_3_END_PROCESSING | 132 | -1.6361 | 0.1204 |
| GO_DNA_CONFORMATION_CHANGE | 279 | -1.6319 | 0.1212 |
| GO_REGULATION_OF_CAMP_MEDIATED_SIGNALING | 30 | -1.6177 | 0.1289 |
| GO_RIBOSOMAL_LARGE_SUBUNIT_BIOGENESIS | 67 | -1.6192 | 0.1296 |
| GO_MODULATION_BY_SYMBIONT_OF_HOST_CELLULAR_PROCESS | 25 | -1.6213 | 0.1297 |
| GO_REPLICATION_FORK_PROCESSING | 33 | -1.6136 | 0.1298 |
| GO_MITOCHONDRIAL_TRANSLATIONAL_TERMINATION | 88 | -1.6112 | 0.1305 |
| GO_MRNA_TRANSPORT | 139 | -1.6137 | 0.1315 |
| GO_HISTONE_EXCHANGE | 55 | -1.6012 | 0.1359 |
| GO_PROTEIN_LOCALIZATION_TO_CHROMOSOME | 82 | -1.6019 | 0.1370 |
| GO_NCRNA_METABOLIC_PROCESS | 433 | -1.6027 | 0.1374 |
| GO_MITOCHONDRIAL_TRANSLATION | 136 | -1.5815 | 0.1451 |
| GO_GLUCOSE_IMPORT | 46 | -1.5835 | 0.1459 |
| GO_REGULATION_OF_CHROMOSOME_SEPARATION | 58 | -1.5822 | 0.1460 |
| GO_RRNA_TRANSCRIPTION | 33 | -1.5848 | 0.1463 |
| GO_MITOCHONDRIAL_RNA_METABOLIC_PROCESS | 47 | -1.5889 | 0.1475 |
| GO_ESTABLISHMENT_OF_RNA_LOCALIZATION | 181 | -1.5851 | 0.1478 |
| GO_REGULATION_OF_NITRIC_OXIDE_SYNTHASE_ACTIVITY | 35 | -1.5864 | 0.1484 |
| GO_CELLULAR_RESPONSE_TO_HEAT | 115 | -1.5751 | 0.1505 |
| GO_RNA_SPLICING_VIA_TRANSESTERIFICATION_REACTIONS | 318 | -1.5707 | 0.1540 |
| GO_SYNAPTIC_TRANSMISSION_GLUTAMATERGIC | 55 | -1.5651 | 0.1591 |
| GO_NUCLEOSIDE_MONOPHOSPHATE_BIOSYNTHETIC_PROCESS | 40 | -1.5575 | 0.1630 |
| GO_REGULATION_OF_CELLULAR_RESPONSE_TO_HEAT | 71 | -1.5584 | 0.1636 |
| GO_REGULATION_OF_SYNAPTIC_TRANSMISSION_GLUTAMATERGIC | 36 | -1.5590 | 0.1647 |
| GO_METAPHASE_PLATE_CONGRESSION | 55 | -1.5501 | 0.1650 |
| GO_REGULATION_OF_CELL_CYCLE_G2_M_PHASE_TRANSITION | 202 | -1.5525 | 0.1657 |
| GO_CELLULAR_RESPONSE_TO_OSMOTIC_STRESS | 35 | -1.5509 | 0.1659 |
| GO_NUCLEAR_EXPORT | 181 | -1.5538 | 0.1660 |
| GO_HISTONE_PHOSPHORYLATION | 35 | -1.5414 | 0.1737 |
| GO_ANAPHASE_PROMOTING_COMPLEX_DEPENDENT_CATABOLIC_PROCESS | 78 | -1.5324 | 0.1774 |
| GO_TRNA_METABOLIC_PROCESS | 179 | -1.5331 | 0.1781 |
| GO_METAPHASE_ANAPHASE_TRANSITION_OF_CELL_CYCLE | 54 | -1.5331 | 0.1799 |
| GO_REGULATION_OF_POSTSYNAPSE_ORGANIZATION | 76 | -1.5345 | 0.1801 |
| GO_CHROMOSOME_SEPARATION | 80 | -1.5271 | 0.1805 |

| | | | |
|---|---|---|---|
| GO_REGULATION_OF_LIPID_KINASE_ACTIVITY | 46 | -1.5287 | 0.1807 |
| GO_REGULATION_OF_SMOOTH_MUSCLE_CONTRACTION | 43 | -1.5235 | 0.1831 |
| HALLMARK_KRAS_SIGNALING_UP | 130 | 1.4481 | 0.1874 |
| HALLMARK_HYPOXIA | 174 | -1.2139 | 0.1877 |
| GO_POSITIVE_REGULATION_OF_PHOSPHATIDYLINOSITOL_3_KINASE_SIGNALING | 53 | -1.5060 | 0.1972 |
| GO_POSITIVE_REGULATION_OF_BLOOD_VESSEL_DIAMETER | 40 | -1.5063 | 0.1985 |
| GO_NEURON_MIGRATION | 117 | -1.5103 | 0.1988 |
| GO_TRANSLATIONAL_TERMINATION | 104 | -1.5034 | 0.1991 |
| GO_NEGATIVE_REGULATION_OF_NUCLEAR_DIVISION | 52 | -1.5064 | 0.2002 |
| GO_REGULATION_OF_POSTTRANSCRIPTIONAL_GENE_SILENCING | 106 | -1.5008 | 0.2011 |
| GO_MODIFICATION_BY_SYMBIONT_OF_HOST_MORPHOLOGY_OR_PHYSIOLOGY | 37 | -1.5066 | 0.2018 |
| GO_POSITIVE_REGULATION_OF_FIBROBLAST_PROLIFERATION | 42 | -1.4974 | 0.2035 |
| GO_TERMINATION_OF_RNA_POLYMERASE_I_TRANSCRIPTION | 30 | -1.4850 | 0.2114 |
| GO_ADENYLATE_CYCLASE_INHIBITING_G_PROTEIN_COUPLED_RECEPTOR_SIGNALING_PATHWAY | 36 | -1.4903 | 0.2119 |
| GO_SECRETORY_GRANULE_ORGANIZATION | 25 | -1.4873 | 0.2122 |
| GO_ENDOCRINE_HORMONE_SECRETION | 28 | -1.4856 | 0.2125 |
| GO_RNA_METHYLATION | 76 | -1.4881 | 0.2130 |
| GO_REGULATION_OF_BMP_SIGNALING_PATHWAY | 59 | -1.4806 | 0.2158 |
| GO_REGULATION_OF_NEURON_PROJECTION_REGENERATION | 25 | -1.4771 | 0.2174 |
| GO_DNA_TEMPLATED_TRANSCRIPTION_TERMINATION | 74 | -1.4775 | 0.2186 |
| GO_SNRNA_METABOLIC_PROCESS | 33 | -1.4716 | 0.2187 |
| GO_MATURATION_OF_SSU_RRNA | 45 | -1.4739 | 0.2187 |
| GO_REGULATION_OF_VASCULAR_SMOOTH_MUSCLE_CELL_PROLIFERATION | 40 | -1.4701 | 0.2191 |
| GO_MATURATION_OF_SSU_RRNA_FROM_TRICISTRONIC_RRNA_TRANSCRIPT_SSU_RRNA_5_8S_RRNA_LSU_RRNA | 33 | -1.4720 | 0.2197 |
| GO_REGULATION_OF_NEURON_PROJECTION_DEVELOPMENT | 401 | -1.4742 | 0.2201 |
| GO_INTERSTRAND_CROSS_LINK_REPAIR | 51 | -1.4677 | 0.2208 |
| HALLMARK_INTERFERON_GAMMA_RESPONSE | 155 | 1.4676 | 0.2231 |
| GO_TRNA_PROCESSING | 125 | -1.4593 | 0.2262 |
| GO_RIBONUCLEOSIDE_MONOPHOSPHATE_METABOLIC_PROCESS | 47 | -1.4611 | 0.2266 |
| GO_TRANSCRIPTION_COUPLED_NUCLEOTIDE_EXCISION_REPAIR | 72 | -1.4520 | 0.2277 |
| GO_TRNA_MODIFICATION | 84 | -1.4594 | 0.2277 |
| GO_GLYCOSYL_COMPOUND_METABOLIC_PROCESS | 107 | -1.4568 | 0.2279 |
| GO_CORONARY_VASCULATURE_DEVELOPMENT | 40 | -1.4554 | 0.2279 |
| GO_MATERNAL_PLACENTA_DEVELOPMENT | 30 | -1.4612 | 0.2283 |
| GO_RECEPTOR_LOCALIZATION_TO_SYNAPSE | 44 | -1.4526 | 0.2284 |
| GO_MRNA_PROCESSING | 457 | -1.4502 | 0.2286 |
| GO_NEGATIVE_REGULATION_OF_OSSIFICATION | 50 | -1.4531 | 0.2296 |
| GO_POSITIVE_REGULATION_OF_MUSCLE_TISSUE_DEVELOPMENT | 52 | -1.4466 | 0.2324 |
| GO_REGULATION_OF_CELL_SIZE | 145 | -1.4423 | 0.2339 |
| GO_NEURON_FATE_COMMITMENT | 30 | -1.4434 | 0.2340 |
| GO_SMOOTH_MUSCLE_CELL_PROLIFERATION | 108 | -1.4404 | 0.2348 |
| GO_AMINO_ACID_ACTIVATION | 48 | -1.4435 | 0.2355 |
| GO_TRANSLATIONAL_ELONGATION | 131 | -1.4367 | 0.2386 |
| GO_SERINE_FAMILY_AMINO_ACID_METABOLIC_PROCESS | 30 | -1.4320 | 0.2424 |
| GO_POSITIVE_CHEMOTAXIS | 35 | -1.4328 | 0.2429 |
| GO_NEURON_PROJECTION_REGENERATION | 47 | -1.4286 | 0.2463 |

### Supplementary Table 3D. Gene set variation analysis of DEGs results, related to Figure 2D

| | Lung-specific | Myc targets | Cell-cell adhesion | G2M checkpoint | Cell cycle DNA replication | ESC Wong | Proliferation Ben-Porath | NSCLC poor survival Shedden | Metastasis Ramaswamy |
|---|---|---|---|---|---|---|---|---|---|
| **Control_rep1** | -0.0438 | 0.2238 | -0.1367 | 0.3046 | 0.4245 | 0.1185 | 0.3188 | 0.3063 | 0.0238 |
| **Control_rep2** | -0.0359 | 0.1693 | -0.1772 | 0.2989 | 0.4032 | 0.1014 | 0.2819 | 0.2547 | 0.0210 |
| **Control_rep3** | -0.0236 | 0.2155 | -0.1381 | 0.3058 | 0.4164 | 0.1249 | 0.2655 | 0.3027 | 0.0181 |
| **SETDB1_LOF_rep1** | -0.0894 | -0.3290 | 0.1013 | -0.0592 | 0.0725 | -0.3069 | -0.1824 | -0.0903 | -0.1486 |
| **SETDB1_LOF_rep2** | -0.0515 | -0.3686 | 0.0903 | -0.0870 | 0.0611 | -0.3213 | -0.2457 | -0.1197 | -0.1610 |
| **SETDB1_LOF_rep3** | -0.0185 | -0.3321 | 0.1125 | -0.0679 | 0.1383 | -0.2965 | -0.2276 | -0.0683 | -0.1361 |
| **Calu3_Wyler_rep1** | 0.0140 | 0.0388 | 0.0249 | -0.1794 | 0.2003 | -0.0304 | 0.0621 | -0.1495 | -0.1538 |
| **Calu3_Wyler_rep2** | -0.0077 | 0.0559 | 0.0339 | -0.1790 | 0.2170 | -0.0245 | 0.1324 | -0.1130 | -0.2161 |
| **HBEC_Jevnikar_rep1** | 0.4402 | -0.6899 | 0.3802 | -0.6274 | -0.6448 | -0.6019 | -0.6695 | -0.6221 | -0.3537 |
| **HBEC_Jevnikar_rep2** | 0.3618 | -0.5282 | 0.1415 | -0.4989 | -0.5100 | -0.4561 | -0.5310 | -0.5803 | -0.3149 |
| **HBEC_Jevnikar_rep3** | 0.4842 | -0.6118 | 0.3425 | -0.5990 | -0.5980 | -0.5572 | -0.6517 | -0.6065 | -0.3425 |
| **HBEC_Kheirallah_rep1** | 0.1634 | -0.2464 | 0.2513 | -0.4205 | -0.5750 | -0.3009 | -0.3517 | -0.2737 | -0.1615 |
| **HBEC_Kheirallah_rep2** | 0.1849 | -0.3359 | 0.1793 | -0.4502 | -0.5734 | -0.3820 | -0.4045 | -0.3545 | -0.2163 |
| **HBEC_Kheirallah_rep3** | 0.1311 | -0.3457 | 0.2258 | -0.4465 | -0.5847 | -0.3924 | -0.3925 | -0.3352 | -0.1982 |
| **HBEC_Kheirallah_rep4** | -0.0004 | -0.4308 | 0.0691 | -0.2899 | -0.5380 | -0.3542 | -0.3371 | -0.2070 | -0.2977 |
| **HBEC_Kheirallah_rep5** | 0.0781 | -0.5134 | 0.1057 | -0.4879 | -0.6306 | -0.4752 | -0.5085 | -0.4026 | -0.3274 |
| **HBEC_Kheirallah_rep6** | 0.1067 | -0.5186 | 0.1587 | -0.4696 | -0.6091 | -0.4982 | -0.4944 | -0.4023 | -0.3128 |
| **HBEC_Wrem_rep1** | 0.3617 | -0.4232 | 0.0110 | -0.5066 | -0.4765 | -0.3643 | -0.4595 | -0.5085 | -0.1799 |
| **HBEC_Wrem_rep2** | 0.3778 | -0.3600 | 0.0601 | -0.5121 | -0.5547 | -0.2905 | -0.3316 | -0.4808 | -0.1261 |
| **HBEC_Wrem_rep3** | 0.3734 | -0.3655 | -0.0339 | -0.4837 | -0.5206 | -0.3232 | -0.3951 | -0.4626 | -0.1347 |
| **HBEC_Wrem_rep4** | 0.3956 | -0.3875 | 0.1555 | -0.4995 | -0.5385 | -0.3206 | -0.4101 | -0.4984 | -0.0908 |
| **HBE_ENCODE_rep1** | 0.1334 | -0.4814 | 0.1932 | -0.5500 | -0.5929 | -0.5184 | -0.5361 | -0.4405 | -0.3713 |
| **HBE_ENCODE_rep2** | 0.1199 | -0.2171 | 0.1241 | -0.5117 | -0.5068 | -0.3688 | -0.4840 | -0.3894 | -0.3110 |
| **HBE_Gillen_rep1** | 0.1904 | -0.6413 | 0.1833 | -0.6247 | -0.6419 | -0.6125 | -0.6048 | -0.5324 | -0.2977 |
| **HBE_Gillen_rep2** | 0.2603 | -0.6540 | 0.2046 | -0.6001 | -0.5899 | -0.5873 | -0.5615 | -0.5234 | -0.3055 |
| **HBE_Gillen_rep3** | 0.0424 | -0.3772 | -0.0657 | -0.4077 | -0.2191 | -0.3892 | -0.3928 | -0.3209 | -0.0735 |
| **HBE_Gillen_rep4** | 0.2879 | -0.6632 | 0.1392 | -0.5677 | -0.5919 | -0.5697 | -0.5258 | -0.4866 | -0.3021 |
| **HBE_Goodale_rep1** | 0.3251 | -0.4716 | -0.0162 | -0.4387 | -0.5799 | -0.4812 | -0.4417 | -0.4216 | -0.3522 |
| **HBE_Goodale_rep2** | 0.3924 | -0.5011 | -0.0940 | -0.5361 | -0.6237 | -0.4989 | -0.4761 | -0.4869 | -0.3961 |
| **HBE_Goodale_rep3** | 0.3578 | -0.6193 | 0.0221 | -0.5507 | -0.6487 | -0.5743 | -0.5535 | -0.5265 | -0.4138 |
| **HBE_Goodale_rep4** | 0.4997 | -0.6413 | 0.2503 | -0.6106 | -0.6955 | -0.6005 | -0.6353 | -0.5838 | -0.3700 |
| **HBE_Goodale_rep5** | 0.4492 | -0.6567 | 0.1283 | -0.5848 | -0.6712 | -0.6002 | -0.6114 | -0.5592 | -0.4468 |
| **HBE_Goodale_rep6** | 0.4277 | -0.6623 | 0.2103 | -0.6422 | -0.6865 | -0.6245 | -0.6692 | -0.5914 | -0.4395 |
| **HBE_Wang_rep1** | -0.0272 | 0.4486 | -0.2683 | 0.4221 | 0.3265 | 0.3475 | 0.4155 | 0.3276 | 0.1946 |
| **HBE_Wang_rep2** | -0.1149 | 0.5371 | -0.3602 | 0.3931 | 0.3269 | 0.4724 | 0.4998 | 0.3830 | 0.1083 |
| **HCC4017_Batten_rep1** | 0.0574 | 0.3325 | 0.2426 | 0.3386 | 0.3677 | 0.3084 | 0.2835 | 0.3109 | 0.1891 |
| **HCC4017_Batten_rep2** | 0.0619 | 0.3302 | 0.2377 | 0.3452 | 0.3583 | 0.3167 | 0.2827 | 0.3135 | 0.1960 |
| **HCC827_Holik_rep1** | -0.0159 | 0.4754 | 0.0131 | 0.4680 | 0.4444 | 0.4031 | 0.3709 | 0.4432 | -0.1096 |
| **HCC827_Holik_rep2** | -0.0080 | 0.4841 | 0.0011 | 0.4347 | 0.4562 | 0.4415 | 0.4027 | 0.4515 | -0.1354 |
| **NCI.H1299_Fields_rep1** | -0.3269 | -0.3915 | 0.0488 | 0.1580 | 0.2166 | -0.3061 | -0.2735 | -0.1229 | 0.1290 |
| **NCI.H1299_Fields_rep2** | -0.3560 | -0.4196 | 0.0999 | 0.1838 | 0.1964 | -0.3291 | -0.2988 | -0.1364 | 0.0854 |
| **NCI.H1299_Fields_rep3** | -0.3542 | -0.2827 | 0.0355 | 0.3403 | 0.3240 | -0.2211 | -0.0707 | 0.0713 | 0.1736 |
| **NCI.H1299_Fields_rep4** | -0.3863 | 0.3344 | -0.3415 | 0.3294 | 0.3831 | 0.2880 | 0.2533 | 0.2863 | 0.3931 |

| | | | | | | | | | |
|---|---|---|---|---|---|---|---|---|---|
| **NCI.H1299_Fields_rep5** | -0.3845 | 0.3440 | -0.3418 | 0.3824 | 0.3532 | 0.2717 | 0.3197 | 0.3027 | 0.3191 |
| **NCI.H1299_Fields_rep6** | -0.3926 | 0.3473 | -0.3328 | 0.3902 | 0.3652 | 0.2807 | 0.3331 | 0.3194 | 0.3367 |
| **NCI.H1299_Khalil_rep1** | -0.2140 | 0.2949 | -0.0991 | 0.4815 | 0.4493 | 0.2608 | 0.4173 | 0.3821 | 0.0782 |
| **NCI.H1299_Khalil_rep2** | -0.1827 | 0.2827 | -0.0344 | 0.4865 | 0.4795 | 0.2521 | 0.4072 | 0.3848 | 0.0009 |
| **NCI.H1299_Khalil_rep3** | -0.2332 | 0.3054 | -0.0749 | 0.4645 | 0.4609 | 0.2819 | 0.3811 | 0.3755 | 0.0441 |
| **NCI.H1299_Reislander_rep1** | -0.2873 | 0.2418 | -0.3245 | 0.0937 | 0.3205 | 0.2799 | 0.0713 | 0.0423 | 0.1707 |
| **NCI.H1299_Reislander_rep2** | -0.3901 | 0.6287 | -0.3837 | 0.4779 | 0.5423 | 0.5680 | 0.5310 | 0.4608 | 0.4047 |
| **NCI.H1299_Reislander_rep3** | -0.2608 | 0.3634 | -0.3344 | 0.0065 | 0.2484 | 0.3900 | 0.1639 | 0.1276 | 0.0640 |
| **NCI.H1299_Reislander_rep4** | -0.3613 | 0.6100 | -0.4050 | 0.4945 | 0.6078 | 0.5647 | 0.5116 | 0.4538 | 0.3822 |
| **NCI.H1299_Reislander_rep5** | -0.3070 | 0.3972 | -0.3312 | 0.0839 | 0.2952 | 0.4047 | 0.1973 | 0.1368 | 0.1390 |
| **NCI.H1299_Reislander_rep6** | -0.3510 | 0.6475 | -0.3810 | 0.4855 | 0.5859 | 0.5918 | 0.5240 | 0.4534 | 0.3770 |
| **NCI.H1299_Zhang_rep1** | -0.4101 | 0.3987 | -0.3946 | 0.4268 | 0.4744 | 0.3648 | 0.3847 | 0.3564 | 0.2096 |
| **NCI.H1299_Zhang_rep2** | -0.4103 | 0.2981 | -0.3392 | 0.4365 | 0.4418 | 0.2659 | 0.3833 | 0.3783 | 0.2393 |
| **NCI.H1299_Zhou_rep1** | -0.1325 | 0.2344 | 0.0485 | 0.1525 | 0.3873 | 0.3773 | 0.2763 | 0.2294 | 0.0367 |
| **NCI.H1299_Zhou_rep2** | -0.0448 | 0.1592 | 0.1634 | 0.1007 | 0.3345 | 0.3357 | 0.2379 | 0.1838 | -0.0648 |
| **NCI.H1299_Zhou_rep3** | 0.0294 | 0.0970 | 0.2436 | 0.0548 | 0.2939 | 0.3054 | 0.1482 | 0.1397 | -0.0494 |
| **NCI.H1650_Zhou_rep1** | -0.1802 | -0.0497 | -0.2500 | 0.2132 | 0.3711 | -0.0464 | 0.1026 | 0.0405 | -0.1183 |
| **NCI.H1650_Zhou_rep2** | -0.1661 | -0.0881 | -0.2264 | 0.2196 | 0.3277 | -0.0353 | 0.1122 | 0.0512 | -0.1372 |
| **NCI.H1650_Zhou_rep3** | -0.2240 | -0.0666 | -0.2495 | 0.2330 | 0.3602 | -0.0144 | 0.0852 | 0.0389 | -0.1407 |
| **NCI.H1693_Batten_rep1** | -0.2376 | 0.1353 | 0.1468 | 0.3512 | 0.3867 | 0.1674 | 0.2055 | 0.2368 | 0.0609 |
| **NCI.H1693_Batten_rep2** | -0.2699 | 0.1403 | 0.1318 | 0.3675 | 0.4059 | 0.1406 | 0.1925 | 0.2328 | 0.0314 |
| **NCI.H1819_Batten_rep1** | 0.1267 | 0.0353 | -0.0058 | 0.2177 | 0.3118 | 0.1421 | 0.0923 | 0.0575 | 0.1754 |
| **NCI.H1819_Batten_rep2** | 0.0605 | 0.0691 | -0.0161 | 0.2268 | 0.3277 | 0.1629 | 0.1303 | 0.0715 | 0.2022 |
| **NCI.H1944_Hata_rep1** | 0.1957 | 0.1548 | 0.2430 | 0.3255 | 0.3269 | 0.1335 | 0.2978 | 0.1776 | 0.3398 |
| **NCI.H1944_Hata_rep2** | 0.2853 | -0.0247 | 0.3271 | 0.1785 | 0.1757 | -0.0142 | 0.1337 | 0.0217 | 0.2901 |
| **NCI.H1975_Holik_rep1** | -0.1347 | 0.3339 | 0.0576 | 0.3705 | 0.4352 | 0.3233 | 0.3806 | 0.3445 | -0.3014 |
| **NCI.H1975_Holik_rep2** | -0.1025 | 0.3111 | 0.0177 | 0.3197 | 0.4082 | 0.2710 | 0.3523 | 0.3421 | -0.3478 |
| **NCI.H1975_Okimoto_rep1** | -0.1687 | 0.5192 | -0.1839 | 0.2545 | 0.3631 | 0.4671 | 0.4152 | 0.3102 | -0.0734 |
| **NCI.H1975_Okimoto_rep2** | -0.1912 | 0.5931 | -0.2085 | 0.2713 | 0.4226 | 0.4824 | 0.4097 | 0.3245 | -0.0731 |
| **NCI.H1993_Batten_rep1** | -0.0463 | 0.2453 | -0.2804 | 0.0100 | 0.1631 | 0.1144 | 0.2059 | 0.2028 | 0.2061 |
| **NCI.H1993_Batten_rep2** | -0.0761 | 0.1999 | -0.2657 | 0.0371 | 0.1021 | 0.1010 | 0.1897 | 0.1925 | 0.1982 |
| **NCI.H2009_Yenerall_rep1** | -0.2445 | 0.6293 | -0.0566 | 0.5912 | 0.5134 | 0.5312 | 0.5055 | 0.5205 | 0.4816 |
| **NCI.H2009_Yenerall_rep2** | -0.2448 | 0.6602 | -0.0859 | 0.5646 | 0.5236 | 0.5600 | 0.5412 | 0.5240 | 0.4298 |
| **NCI.H2086_Batten_rep1** | 0.0247 | 0.2506 | 0.1364 | 0.1735 | 0.2518 | 0.3195 | 0.3495 | 0.1418 | 0.3417 |
| **NCI.H2086_Batten_rep2** | -0.0025 | 0.2009 | 0.1758 | 0.2290 | 0.2408 | 0.2499 | 0.3205 | 0.1623 | 0.3180 |
| **NCI.H2087_Batten_rep1** | -0.0984 | 0.5243 | -0.3256 | 0.4134 | 0.4543 | 0.5116 | 0.4413 | 0.3802 | 0.0198 |
| **NCI.H2087_Batten_rep2** | -0.0903 | 0.4786 | -0.3083 | 0.4264 | 0.4268 | 0.4802 | 0.4157 | 0.3733 | 0.0701 |
| **NCI.H2228_Holik_rep1** | 0.2968 | -0.2500 | 0.2237 | 0.2587 | 0.3213 | -0.1105 | 0.2969 | 0.1238 | -0.2950 |
| **NCI.H2228_Holik_rep2** | 0.3469 | -0.2771 | 0.2412 | 0.2167 | 0.2915 | -0.1351 | 0.2872 | 0.0841 | -0.2645 |
| **NCI.H358_Hata_rep1** | 0.2670 | 0.0495 | 0.4134 | 0.2971 | 0.2867 | 0.0931 | 0.1009 | 0.0654 | 0.0852 |
| **NCI.H358_Hata_rep2** | 0.1728 | -0.1256 | 0.3731 | 0.2798 | 0.2868 | -0.0117 | 0.0001 | 0.0332 | 0.1699 |
| **NCI.H358_Hata_rep3** | 0.1960 | -0.1616 | 0.3507 | 0.1935 | 0.2724 | -0.0646 | -0.0989 | -0.0493 | 0.1147 |
| **NCI.H460_ENCODE_rep1** | 0.1072 | 0.0590 | 0.0410 | 0.1022 | -0.1710 | -0.0954 | 0.1975 | 0.1141 | -0.3351 |
| **NCI.H460_ENCODE_rep2** | 0.1531 | 0.0419 | 0.1458 | 0.0830 | -0.0791 | -0.0326 | 0.2138 | 0.1399 | -0.3544 |
| **NCI.H460_Zhou_rep1** | 0.1004 | 0.3039 | 0.2396 | 0.0946 | 0.2207 | 0.2899 | 0.1057 | 0.0789 | 0.0306 |
| **NCI.H460_Zhou_rep2** | 0.1405 | 0.3876 | 0.2491 | 0.1188 | 0.1867 | 0.3368 | 0.1528 | 0.1092 | 0.0197 |
| **NCI.H460_Zhou_rep3** | 0.0634 | 0.4160 | 0.2270 | 0.1363 | 0.2249 | 0.3571 | 0.1792 | 0.1507 | 0.0214 |
| **NCI.H596_Yenerall_rep1** | -0.1614 | 0.4733 | -0.2960 | 0.5378 | 0.6038 | 0.4704 | 0.5581 | 0.4022 | 0.3714 |
| **NCI.H596_Yenerall_rep2** | -0.1811 | 0.5747 | -0.2970 | 0.5500 | 0.6001 | 0.5365 | 0.5939 | 0.4436 | 0.4061 |
| **NCI.H838_Holik_rep1** | -0.0738 | 0.3403 | -0.3896 | 0.3701 | 0.4845 | 0.3659 | 0.4160 | 0.3933 | -0.0515 |
| **NCI.H838_Holik_rep2** | -0.0701 | 0.3366 | -0.3660 | 0.3537 | 0.4544 | 0.3474 | 0.4091 | 0.3837 | -0.1033 |
| **NHBE_Baddal_rep1** | 0.3764 | -0.3908 | 0.2168 | -0.5069 | -0.5333 | -0.4406 | -0.3669 | -0.4147 | -0.3258 |
| **NHBE_Baddal_rep2** | 0.3687 | -0.4262 | 0.2160 | -0.4824 | -0.5260 | -0.4748 | -0.4172 | -0.4302 | -0.3086 |
| **NHBE_Baddal_rep3** | 0.3636 | -0.3656 | 0.1647 | -0.4824 | -0.5127 | -0.4404 | -0.3548 | -0.3920 | -0.3677 |
| **NHBE_Laubenbacher_rep1** | 0.0891 | -0.2893 | 0.1811 | -0.4501 | -0.6060 | -0.3574 | -0.4301 | -0.2696 | 0.0990 |

| | | | | | | | | | |
|---|---|---|---|---|---|---|---|---|---|
| **NHBE_Laubenbacher_rep2** | 0.2211 | -0.1168 | 0.1882 | -0.4147 | -0.6217 | -0.2579 | -0.3913 | -0.2683 | 0.0818 |
| **NHBE_Laubenbacher_rep3** | 0.1680 | -0.3089 | 0.1718 | -0.5123 | -0.6553 | -0.3999 | -0.4657 | -0.3360 | 0.0149 |
| **NHBE_Laubenbacher_rep4** | 0.2561 | -0.1971 | 0.2342 | -0.4766 | -0.6660 | -0.3373 | -0.4706 | -0.3506 | -0.0369 |
| **NHBE_Laubenbacher_rep5** | -0.1179 | -0.1274 | -0.0078 | -0.3351 | -0.3297 | -0.2596 | -0.3561 | -0.1234 | 0.0241 |
| **NHBE_Laubenbacher_rep6** | -0.0105 | 0.1275 | 0.0494 | -0.3066 | -0.3324 | -0.1699 | -0.3243 | -0.1582 | 0.0625 |
| **NHBE_Laubenbacher_rep7** | -0.1106 | -0.1790 | 0.0374 | -0.3438 | -0.3084 | -0.2755 | -0.3736 | -0.1650 | 0.0327 |
| **NHBE_Laubenbacher_rep8** | 0.0185 | 0.0697 | 0.0614 | -0.3427 | -0.3085 | -0.1683 | -0.3967 | -0.1952 | 0.0874 |
| **NHBE_Laubenbacher_rep9** | -0.0590 | -0.2069 | 0.1354 | 0.0743 | 0.1754 | -0.0524 | -0.0495 | 0.1568 | 0.1005 |
| **NHBE_Laubenbacher_rep10** | 0.0421 | 0.0459 | 0.1765 | 0.1400 | 0.0717 | 0.1351 | -0.0442 | 0.1468 | 0.1046 |
| **NHBE_Laubenbacher_rep11** | -0.1138 | -0.1662 | 0.1619 | 0.1410 | 0.2261 | -0.0304 | -0.0108 | 0.2040 | 0.2142 |
| **NHBE_Laubenbacher_rep12** | 0.0015 | 0.0421 | 0.1992 | 0.1960 | 0.2266 | 0.1893 | -0.0367 | 0.1793 | 0.1184 |
| **NHBE_Matsuyama_rep1** | 0.4423 | -0.6654 | 0.1157 | -0.6126 | -0.6089 | -0.5818 | -0.6209 | -0.5424 | -0.4308 |
| **NHBE_Matsuyama_rep2** | 0.4470 | -0.6441 | 0.2442 | -0.6375 | -0.6549 | -0.5940 | -0.6393 | -0.5905 | -0.3980 |
| **NHBE_Matsuyama_rep3** | 0.4702 | -0.6635 | 0.3708 | -0.6636 | -0.6687 | -0.5733 | -0.6682 | -0.5975 | -0.4159 |
| **NHBE_Matsuyama_rep4** | 0.4642 | -0.6497 | 0.2914 | -0.6519 | -0.6709 | -0.5399 | -0.6576 | -0.5961 | -0.4256 |
| **NHBE_Rice_rep1** | -0.0344 | 0.1825 | -0.0555 | 0.0723 | -0.2651 | 0.1239 | 0.0783 | 0.2258 | 0.1956 |
| **NHBE_Rice_rep2** | -0.1314 | 0.1801 | -0.1762 | 0.0665 | -0.1729 | 0.1615 | 0.0962 | 0.2345 | 0.0436 |
