## Supplemental Table 4 for "SETDB1 Fuels the Lung Cancer Phenotype by Modulating Epigenome, 3D Genome Organization and Chromatin Mechanical Properties"

**Supplementary Table 4, Annotated super-enhancers by ROSE, Related to Supplementary Figure 3B**

| Control superenhancers | | | SETDB1 LOF superenhancers | | |
|---|---|---|---|---|---|
| **chrom** | **start** | **end** | **chrom** | **start** | **end** |
| chr1 | 8062189 | 8160939 | chr1 | 8062252 | 8159956 |
| chr1 | 8197642 | 8300015 | chr1 | 8213846 | 8300469 |
| chr1 | 8928487 | 8972242 | chr1 | 8916908 | 8973084 |
| chr1 | 9350040 | 9412327 | chr1 | 10433679 | 10491554 |
| chr1 | 10436234 | 1E+07 | chr1 | 12585023 | 12680437 |
| chr1 | 12585217 | 1.3E+07 | chr1 | 16084798 | 16128784 |
| chr1 | 15478598 | 1.6E+07 | chr1 | 16447117 | 16516661 |
| chr1 | 16085151 | 1.6E+07 | chr1 | 17481446 | 17537341 |
| chr1 | 16464845 | 1.7E+07 | chr1 | 23493658 | 23552834 |
| chr1 | 21156492 | 2.1E+07 | chr1 | 23872985 | 23925825 |
| chr1 | 23493856 | 2.4E+07 | chr1 | 27851926 | 27929344 |
| chr1 | 23850616 | 2.4E+07 | chr1 | 32376097 | 32422911 |
| chr1 | 27851810 | 2.8E+07 | chr1 | 33168006 | 33255189 |
| chr1 | 32375512 | 3.2E+07 | chr1 | 39623189 | 39680146 |
| chr1 | 33168018 | 3.3E+07 | chr1 | 39849861 | 39877795 |
| chr1 | 36805454 | 3.7E+07 | chr1 | 43388884 | 43434720 |
| chr1 | 37917504 | 3.8E+07 | chr1 | 45240104 | 45286625 |
| chr1 | 39623361 | 4E+07 | chr1 | 51424601 | 51451782 |
| chr1 | 41404936 | 4.1E+07 | chr1 | 65355073 | 65400898 |
| chr1 | 43388259 | 4.3E+07 | chr1 | 66722647 | 66826207 |
| chr1 | 45240043 | 4.5E+07 | chr1 | 68149927 | 68233486 |
| chr1 | 64157183 | 6.4E+07 | chr1 | 84711788 | 84770013 |
| chr1 | 65347100 | 6.5E+07 | chr1 | 85740163 | 85839333 |
| chr1 | 66696369 | 6.7E+07 | chr1 | 90350125 | 90423915 |
| chr1 | 68149927 | 6.8E+07 | chr1 | 94019962 | 94170176 |
| chr1 | 90285714 | 9E+07 | chr1 | 98490955 | 98532942 |
| chr1 | 94032282 | 9.4E+07 | chr1 | 110452755 | 110528661 |
| chr1 | 1.1E+08 | 1.1E+08 | chr1 | 150521066 | 150553190 |
| chr1 | 1.51E+08 | 1.5E+08 | chr1 | 150569056 | 150603556 |
| chr1 | 1.51E+08 | 1.5E+08 | chr1 | 150626581 | 150670983 |
| chr1 | 1.51E+08 | 1.5E+08 | chr1 | 151137394 | 151195555 |
| chr1 | 1.51E+08 | 1.5E+08 | chr1 | 151940227 | 151974305 |
| chr1 | 1.52E+08 | 1.5E+08 | chr1 | 153503932 | 153546074 |
| chr1 | 1.54E+08 | 1.5E+08 | chr1 | 153917072 | 153967025 |
| chr1 | 1.54E+08 | 1.5E+08 | chr1 | 154927887 | 155037074 |
| chr1 | 1.55E+08 | 1.6E+08 | chr1 | 155900508 | 155950592 |
| chr1 | 1.56E+08 | 1.6E+08 | chr1 | 156051845 | 156116963 |
| chr1 | 1.56E+08 | 1.6E+08 | chr1 | 156415843 | 156479435 |
| chr1 | 1.56E+08 | 1.6E+08 | chr1 | 156694876 | 156722841 |
| chr1 | 1.57E+08 | 1.6E+08 | chr1 | 157963170 | 158009037 |
| chr1 | 1.58E+08 | 1.6E+08 | chr1 | 165796171 | 165881224 |
| chr1 | 1.65E+08 | 1.6E+08 | chr1 | 169073964 | 169132966 |
| chr1 | 1.66E+08 | 1.7E+08 | chr1 | 174921607 | 174998998 |
| chr1 | 1.69E+08 | 1.7E+08 | chr1 | 179090189 | 179145390 |

| | | | | | |
|---|---|---|---|---|---|
| chr1 | 1.79E+08 | 1.8E+08 | chr1 | 181042231 | 181111697 |
| chr1 | 1.81E+08 | 1.8E+08 | chr1 | 200040121 | 200099883 |
| chr1 | 2E+08 | 2E+08 | chr1 | 200320755 | 200387121 |
| chr1 | 2.01E+08 | 2E+08 | chr1 | 200859096 | 200886602 |
| chr1 | 2.01E+08 | 2E+08 | chr1 | 201416938 | 201482156 |
| chr1 | 2.02E+08 | 2E+08 | chr1 | 201974746 | 202092352 |
| chr1 | 2.04E+08 | 2E+08 | chr1 | 204366527 | 204393494 |
| chr1 | 2.05E+08 | 2.1E+08 | chr1 | 205242369 | 205294666 |
| chr1 | 2.08E+08 | 2.1E+08 | chr1 | 207165964 | 207227223 |
| chr1 | 2.18E+08 | 2.2E+08 | chr1 | 218503630 | 218613363 |
| chr1 | 2.19E+08 | 2.2E+08 | chr1 | 218632613 | 218674507 |
| chr1 | 2.24E+08 | 2.2E+08 | chr1 | 223886112 | 223965855 |
| chr1 | 2.3E+08 | 2.3E+08 | chr1 | 230201715 | 230277159 |
| chr1 | 2.32E+08 | 2.3E+08 | chr1 | 231512030 | 231561586 |
| chr1 | 2.35E+08 | 2.3E+08 | chr1 | 233242403 | 233253235 |
| chr1 | 2.35E+08 | 2.4E+08 | chr1 | 234733610 | 234795850 |
| chr1 | 2.36E+08 | 2.4E+08 | chr1 | 235085129 | 235154121 |
| chr1 | 2.42E+08 | 2.4E+08 | chr1 | 241754096 | 241836686 |
| chr1 | 2.45E+08 | 2.5E+08 | chr1 | 244995950 | 245029907 |
| chr1 | 2.45E+08 | 2.5E+08 | chr1 | 245114629 | 245137595 |
| chr1 | 2.49E+08 | 2.5E+08 | chr10 | 3085343 | 3185242 |
| chr10 | 3085600 | 3184973 | chr10 | 3892530 | 3938747 |
| chr10 | 4699770 | 4747184 | chr10 | 4692190 | 4729921 |
| chr10 | 5022940 | 5076930 | chr10 | 5023011 | 5084986 |
| chr10 | 5110186 | 5152049 | chr10 | 5110167 | 5152135 |
| chr10 | 5312791 | 5337093 | chr10 | 5584118 | 5602393 |
| chr10 | 6313232 | 6357796 | chr10 | 6315761 | 6357514 |
| chr10 | 14563005 | 1.5E+07 | chr10 | 14548826 | 14659911 |
| chr10 | 21561786 | 2.2E+07 | chr10 | 21602888 | 21664271 |
| chr10 | 21805790 | 2.2E+07 | chr10 | 21805868 | 21856852 |
| chr10 | 30707426 | 3.1E+07 | chr10 | 27465559 | 27542413 |
| chr10 | 32201216 | 3.2E+07 | chr10 | 32201077 | 32251608 |
| chr10 | 33176590 | 3.3E+07 | chr10 | 33180144 | 33306187 |
| chr10 | 33442542 | 3.4E+07 | chr10 | 33402929 | 33630600 |
| chr10 | 34529377 | 3.5E+07 | chr10 | 34569084 | 34736828 |
| chr10 | 34970874 | 3.5E+07 | chr10 | 34984187 | 35058664 |
| chr10 | 43878798 | 4.4E+07 | chr10 | 43799635 | 43858813 |
| chr10 | 46965284 | 4.7E+07 | chr10 | 43882900 | 43917750 |
| chr10 | 60372440 | 6E+07 | chr10 | 46966153 | 47023076 |
| chr10 | 62701962 | 6.3E+07 | chr10 | 47073875 | 47151348 |
| chr10 | 64989727 | 6.5E+07 | chr10 | 64966700 | 65034307 |
| chr10 | 69539979 | 7E+07 | chr10 | 69550050 | 69615459 |
| chr10 | 69808410 | 7E+07 | chr10 | 69767482 | 69870584 |
| chr10 | 71185368 | 7.1E+07 | chr10 | 71183654 | 71268009 |
| chr10 | 73575729 | 7.4E+07 | chr10 | 73472797 | 73535002 |
| chr10 | 73928305 | 7.4E+07 | chr10 | 73577747 | 73650368 |
| chr10 | 74357753 | 7.5E+07 | chr10 | 73922645 | 74093734 |
| chr10 | 75639265 | 7.6E+07 | chr10 | 74371663 | 74515215 |
| chr10 | 75756747 | 7.6E+07 | chr10 | 75639335 | 75700856 |
| chr10 | 76944500 | 7.7E+07 | chr10 | 76944627 | 77006324 |
| chr10 | 79567673 | 8E+07 | chr10 | 80826359 | 80952857 |
| chr10 | 80826526 | 8.1E+07 | chr10 | 81907911 | 81969426 |
| chr10 | 81906856 | 8.2E+07 | chr10 | 82188064 | 82296632 |
| chr10 | 82203268 | 8.2E+07 | chr10 | 93347301 | 93377615 |
| chr10 | 88196710 | 8.8E+07 | chr10 | 94349665 | 94413966 |
| chr10 | 94350064 | 9.4E+07 | chr10 | 95046712 | 95271515 |
| chr10 | 95047393 | 9.5E+07 | chr10 | 96968789 | 97069738 |
| chr10 | 95750674 | 9.6E+07 | chr10 | 101490563 | 101611374 |
| chr10 | 96973768 | 9.7E+07 | chr10 | 101653772 | 101804592 |

| | | | | | |
|---|---|---|---|---|---|
| chr10 | 1.01E+08 | 1E+08 | chr10 | 102100744 | 102152837 |
| chr10 | 1.02E+08 | 1E+08 | chr10 | 104899599 | 104958247 |
| chr10 | 1.02E+08 | 1E+08 | chr10 | 104988975 | 105015464 |
| chr10 | 1.04E+08 | 1E+08 | chr10 | 105505196 | 105595907 |
| chr10 | 1.05E+08 | 1E+08 | chr10 | 105643785 | 105679366 |
| chr10 | 1.05E+08 | 1.1E+08 | chr10 | 112152296 | 112194078 |
| chr10 | 1.12E+08 | 1.1E+08 | chr10 | 114815967 | 114900505 |
| chr10 | 1.12E+08 | 1.1E+08 | chr10 | 119081699 | 119136313 |
| chr10 | 1.15E+08 | 1.1E+08 | chr10 | 123685073 | 123735541 |
| chr10 | 1.19E+08 | 1.2E+08 | chr10 | 123792387 | 123831089 |
| chr10 | 1.21E+08 | 1.2E+08 | chr10 | 123858879 | 123994332 |
| chr10 | 1.24E+08 | 1.2E+08 | chr10 | 126662301 | 126851500 |
| chr10 | 1.24E+08 | 1.2E+08 | chr10 | 134208955 | 134266665 |
| chr10 | 1.27E+08 | 1.3E+08 | chr11 | 795501 | 849058 |
| chr11 | 796593 | 864096 | chr11 | 1050877 | 1078511 |
| chr11 | 7621056 | 7679177 | chr11 | 2913500 | 3014550 |
| chr11 | 10876937 | 1.1E+07 | chr11 | 9585924 | 9610852 |
| chr11 | 12064947 | 1.2E+07 | chr11 | 12694811 | 12750775 |
| chr11 | 12131388 | 1.2E+07 | chr11 | 12766275 | 12893255 |
| chr11 | 12695013 | 1.3E+07 | chr11 | 13114667 | 13165428 |
| chr11 | 18378094 | 1.8E+07 | chr11 | 27368154 | 27495348 |
| chr11 | 27368885 | 2.7E+07 | chr11 | 33059917 | 33118010 |
| chr11 | 28064422 | 2.8E+07 | chr11 | 33710594 | 33759481 |
| chr11 | 33710784 | 3.4E+07 | chr11 | 34066104 | 34128865 |
| chr11 | 34168459 | 3.4E+07 | chr11 | 34653818 | 34708176 |
| chr11 | 34653445 | 3.5E+07 | chr11 | 45273751 | 45311604 |
| chr11 | 35159900 | 3.5E+07 | chr11 | 47527641 | 47601494 |
| chr11 | 46678973 | 4.7E+07 | chr11 | 47937769 | 47983300 |
| chr11 | 47502314 | 4.8E+07 | chr11 | 47995837 | 48132280 |
| chr11 | 47863882 | 4.8E+07 | chr11 | 56643840 | 56702627 |
| chr11 | 47931388 | 4.8E+07 | chr11 | 57498968 | 57569412 |
| chr11 | 47995845 | 4.8E+07 | chr11 | 59303759 | 59342787 |
| chr11 | 57508131 | 5.8E+07 | chr11 | 61447335 | 61498475 |
| chr11 | 59303725 | 5.9E+07 | chr11 | 61519604 | 61604003 |
| chr11 | 61514747 | 6.2E+07 | chr11 | 61722173 | 61747180 |
| chr11 | 61714804 | 6.2E+07 | chr11 | 62305101 | 62391599 |
| chr11 | 62305405 | 6.2E+07 | chr11 | 63375743 | 63451412 |
| chr11 | 64602690 | 6.5E+07 | chr11 | 64602837 | 64662488 |
| chr11 | 65046177 | 6.5E+07 | chr11 | 65183215 | 65278917 |
| chr11 | 65183352 | 6.5E+07 | chr11 | 65654699 | 65687867 |
| chr11 | 65237874 | 6.5E+07 | chr11 | 65867485 | 65925700 |
| chr11 | 65654740 | 6.6E+07 | chr11 | 66599903 | 66675661 |
| chr11 | 65867555 | 6.6E+07 | chr11 | 67005992 | 67086202 |
| chr11 | 66589566 | 6.7E+07 | chr11 | 67111155 | 67170044 |
| chr11 | 66713584 | 6.7E+07 | chr11 | 67763851 | 67822063 |
| chr11 | 67006663 | 6.7E+07 | chr11 | 67873245 | 67918265 |
| chr11 | 67111738 | 6.7E+07 | chr11 | 69041303 | 69070713 |
| chr11 | 67770697 | 6.8E+07 | chr11 | 71638189 | 71760568 |
| chr11 | 69048032 | 6.9E+07 | chr11 | 71774115 | 71824859 |
| chr11 | 71638266 | 7.2E+07 | chr11 | 75022685 | 75141865 |
| chr11 | 71779123 | 7.2E+07 | chr11 | 76473562 | 76510273 |
| chr11 | 73833121 | 7.4E+07 | chr11 | 86191434 | 86237142 |
| chr11 | 76455640 | 7.7E+07 | chr11 | 86278004 | 86319991 |
| chr11 | 86193615 | 8.6E+07 | chr11 | 86406638 | 86456745 |
| chr11 | 86417334 | 8.6E+07 | chr11 | 100605436 | 100663607 |
| chr11 | 1.14E+08 | 1.1E+08 | chr11 | 102175105 | 102195002 |
| chr11 | 1.19E+08 | 1.2E+08 | chr11 | 114150486 | 114189411 |
| chr11 | 1.22E+08 | 1.2E+08 | chr11 | 118777054 | 118801602 |
| chr11 | 1.22E+08 | 1.2E+08 | chr11 | 122025183 | 122073712 |
| chr11 | 1.23E+08 | 1.2E+08 | chr11 | 122499977 | 122562872 |
| chr11 | 1.28E+08 | 1.3E+08 | chr12 | 719000 | 756423 |

| | | | | | |
|---|---|---|---|---|---|
| chr12 | 718954 | 756641 | chr12 | 858332 | 938735 |
| chr12 | 858734 | 928821 | chr12 | 6304378 | 6339845 |
| chr12 | 6421623 | 6455508 | chr12 | 6386823 | 6455832 |
| chr12 | 6469686 | 6504602 | chr12 | 6469725 | 6542439 |
| chr12 | 6601342 | 6679718 | chr12 | 6601567 | 6724171 |
| chr12 | 6975982 | 7090365 | chr12 | 6975787 | 7089307 |
| chr12 | 12335670 | 1.2E+07 | chr12 | 10514208 | 10550887 |
| chr12 | 12931979 | 1.3E+07 | chr12 | 12289047 | 12385445 |
| chr12 | 13019307 | 1.3E+07 | chr12 | 12866397 | 12880189 |
| chr12 | 31576532 | 3.2E+07 | chr12 | 12931954 | 13000206 |
| chr12 | 31880753 | 3.2E+07 | chr12 | 13015954 | 13083290 |
| chr12 | 47463842 | 4.7E+07 | chr12 | 14391934 | 14437620 |
| chr12 | 47575437 | 4.8E+07 | chr12 | 14508751 | 14629463 |
| chr12 | 48146989 | 4.8E+07 | chr12 | 15835493 | 15887153 |
| chr12 | 50898175 | 5.1E+07 | chr12 | 21787004 | 21828953 |
| chr12 | 52533841 | 5.3E+07 | chr12 | 31583589 | 31681341 |
| chr12 | 53253282 | 5.3E+07 | chr12 | 31856839 | 31933500 |
| chr12 | 53811464 | 5.4E+07 | chr12 | 32108114 | 32163939 |
| chr12 | 56910033 | 5.7E+07 | chr12 | 41082176 | 41123105 |
| chr12 | 62972724 | 6.3E+07 | chr12 | 46750923 | 46785531 |
| chr12 | 63077029 | 6.3E+07 | chr12 | 46823300 | 46889621 |
| chr12 | 64236618 | 6.4E+07 | chr12 | 47463349 | 47500110 |
| chr12 | 65001248 | 6.5E+07 | chr12 | 47578067 | 47675988 |
| chr12 | 76302643 | 7.6E+07 | chr12 | 50594926 | 50721719 |
| chr12 | 95466059 | 9.6E+07 | chr12 | 50897998 | 50992076 |
| chr12 | 96176095 | 9.6E+07 | chr12 | 51155737 | 51185976 |
| chr12 | 96587328 | 9.7E+07 | chr12 | 52536157 | 52686043 |
| chr12 | 1.05E+08 | 1E+08 | chr12 | 53253080 | 53455825 |
| chr12 | 1.05E+08 | 1E+08 | chr12 | 53758559 | 53898037 |
| chr12 | 1.06E+08 | 1.1E+08 | chr12 | 53921038 | 53970820 |
| chr12 | 1.08E+08 | 1.1E+08 | chr12 | 54641974 | 54700907 |
| chr12 | 1.09E+08 | 1.1E+08 | chr12 | 56636679 | 56711624 |
| chr12 | 1.09E+08 | 1.1E+08 | chr12 | 56910847 | 56942903 |
| chr12 | 1.11E+08 | 1.1E+08 | chr12 | 57457573 | 57506935 |
| chr12 | 1.11E+08 | 1.1E+08 | chr12 | 62973804 | 63060860 |
| chr12 | 1.14E+08 | 1.1E+08 | chr12 | 63096168 | 63195151 |
| chr12 | 1.17E+08 | 1.2E+08 | chr12 | 64236575 | 64259209 |
| chr12 | 1.18E+08 | 1.2E+08 | chr12 | 64276483 | 64336511 |
| chr12 | 1.21E+08 | 1.2E+08 | chr12 | 64455250 | 64515486 |
| chr12 | 1.22E+08 | 1.2E+08 | chr12 | 65001278 | 65079959 |
| chr12 | 1.22E+08 | 1.2E+08 | chr12 | 75337000 | 75360848 |
| chr12 | 1.23E+08 | 1.2E+08 | chr12 | 76326043 | 76388895 |
| chr12 | 1.24E+08 | 1.2E+08 | chr12 | 79908021 | 79967015 |
| chr12 | 1.25E+08 | 1.2E+08 | chr12 | 80028728 | 80087613 |
| chr12 | 1.25E+08 | 1.3E+08 | chr12 | 87805678 | 87836759 |
| chr12 | 1.25E+08 | 1.3E+08 | chr12 | 89884325 | 89920794 |
| chr12 | 1.32E+08 | 1.3E+08 | chr12 | 92449650 | 92502617 |
| chr13 | 21555508 | 2.2E+07 | chr12 | 95461278 | 95730351 |
| chr13 | 67642518 | 6.8E+07 | chr12 | 95998692 | 96063505 |
| chr13 | 67757339 | 6.8E+07 | chr12 | 96178024 | 96203764 |
| chr13 | 73591805 | 7.4E+07 | chr12 | 98883763 | 98942922 |
| chr13 | 95702259 | 9.6E+07 | chr12 | 104606470 | 104618302 |
| chr14 | 23025973 | 2.3E+07 | chr12 | 104642571 | 104772290 |
| chr14 | 31417443 | 3.2E+07 | chr12 | 105338392 | 105354000 |
| chr14 | 31575111 | 3.2E+07 | chr12 | 105702855 | 105752037 |
| chr14 | 32395998 | 3.2E+07 | chr12 | 106472253 | 106534409 |
| chr14 | 32542587 | 3.3E+07 | chr12 | 107711526 | 107753658 |
| chr14 | 35789556 | 3.6E+07 | chr12 | 109053300 | 109130187 |
| chr14 | 50401200 | 5E+07 | chr12 | 109162085 | 109252253 |
| chr14 | 61978061 | 6.2E+07 | chr12 | 110462825 | 110498824 |
| chr14 | 62161345 | 6.2E+07 | chr12 | 110537659 | 110603049 |

| | | | | | |
|---|---|---|---|---|---|
| chr14 | 65170467 | 6.5E+07 | chr12 | 110630326 | 110842616 |
| chr14 | 68583046 | 6.9E+07 | chr12 | 110974019 | 111024065 |
| chr14 | 69013283 | 6.9E+07 | chr12 | 111970390 | 112039334 |
| chr14 | 69240931 | 6.9E+07 | chr12 | 112442970 | 112592297 |
| chr14 | 69347452 | 6.9E+07 | chr12 | 113611039 | 113710430 |
| chr14 | 74179054 | 7.4E+07 | chr12 | 120516938 | 120593525 |
| chr14 | 77362138 | 7.7E+07 | chr12 | 120612882 | 120715422 |
| chr14 | 77489583 | 7.8E+07 | chr12 | 122177888 | 122278747 |
| chr14 | 92968466 | 9.3E+07 | chr12 | 122476122 | 122530545 |
| chr14 | 1.02E+08 | 1E+08 | chr12 | 122580659 | 122615989 |
| chr14 | 1.03E+08 | 1E+08 | chr12 | 122843254 | 122916368 |
| chr14 | 1.04E+08 | 1E+08 | chr12 | 122971018 | 123024169 |
| chr14 | 1.05E+08 | 1.1E+08 | chr12 | 123346298 | 123383555 |
| chr14 | 1.06E+08 | 1.1E+08 | chr12 | 123826983 | 123890174 |
| chr15 | 26855524 | 2.7E+07 | chr12 | 124812002 | 124908737 |
| chr15 | 29960775 | 3E+07 | chr12 | 124955459 | 125007484 |
| chr15 | 31617129 | 3.2E+07 | chr12 | 125185140 | 125254348 |
| chr15 | 33115192 | 3.3E+07 | chr12 | 125311031 | 125394204 |
| chr15 | 41522421 | 4.2E+07 | chr12 | 132336857 | 132417008 |
| chr15 | 52121756 | 5.2E+07 | chr13 | 47169279 | 47242786 |
| chr15 | 52261836 | 5.2E+07 | chr13 | 67642581 | 67725357 |
| chr15 | 56138572 | 5.6E+07 | chr13 | 67757035 | 67779749 |
| chr15 | 58729614 | 5.9E+07 | chr13 | 73591352 | 73685869 |
| chr15 | 59396628 | 6E+07 | chr13 | 95750530 | 95954571 |
| chr15 | 59903496 | 6E+07 | chr14 | 23025463 | 23084608 |
| chr15 | 60632600 | 6.1E+07 | chr14 | 31463619 | 31561725 |
| chr15 | 63334226 | 6.3E+07 | chr14 | 31587015 | 31679071 |
| chr15 | 63614705 | 6.4E+07 | chr14 | 32543034 | 32620300 |
| chr15 | 63728626 | 6.4E+07 | chr14 | 35800718 | 35874847 |
| chr15 | 64752102 | 6.5E+07 | chr14 | 50318041 | 50387729 |
| chr15 | 66764627 | 6.7E+07 | chr14 | 50411810 | 50513097 |
| chr15 | 66990466 | 6.7E+07 | chr14 | 55490660 | 55545130 |
| chr15 | 67192111 | 6.7E+07 | chr14 | 55564101 | 55576900 |
| chr15 | 67316113 | 6.7E+07 | chr14 | 61978137 | 62053224 |
| chr15 | 69068389 | 6.9E+07 | chr14 | 62160736 | 62230440 |
| chr15 | 69590281 | 7E+07 | chr14 | 64319054 | 64391226 |
| chr15 | 71385364 | 7.1E+07 | chr14 | 64853309 | 64933043 |
| chr15 | 71476445 | 7.2E+07 | chr14 | 68582901 | 68617062 |
| chr15 | 72020752 | 7.2E+07 | chr14 | 68629880 | 68677961 |
| chr15 | 72476030 | 7.3E+07 | chr14 | 68907637 | 68937263 |
| chr15 | 75912593 | 7.6E+07 | chr14 | 68972619 | 69075959 |
| chr15 | 76578626 | 7.7E+07 | chr14 | 69240470 | 69292184 |
| chr15 | 85256733 | 8.5E+07 | chr14 | 69329380 | 69447392 |
| chr15 | 85522982 | 8.6E+07 | chr14 | 74162339 | 74262580 |
| chr15 | 89630601 | 9E+07 | chr14 | 75720905 | 75777920 |
| chr15 | 90293549 | 9E+07 | chr14 | 77407606 | 77432287 |
| chr15 | 90686588 | 9.1E+07 | chr14 | 77488170 | 77519590 |
| chr15 | 90930657 | 9.1E+07 | chr14 | 92198685 | 92247386 |
| chr15 | 91466000 | 9.2E+07 | chr14 | 95697852 | 95740338 |
| chr15 | 93340621 | 9.3E+07 | chr14 | 102720545 | 102787016 |
| chr15 | 99189570 | 9.9E+07 | chr14 | 103557547 | 103623849 |
| chr15 | 99384251 | 1E+08 | chr14 | 103697992 | 103768594 |
| chr15 | 1.01E+08 | 1E+08 | chr14 | 105131634 | 105178709 |
| chr16 | 1727723 | 1780302 | chr14 | 105423784 | 105512737 |
| chr16 | 2009096 | 2107353 | chr14 | 105912308 | 105958842 |
| chr16 | 4303093 | 4443772 | chr15 | 31616995 | 31698788 |
| chr16 | 9161816 | 9224240 | chr15 | 33110999 | 33167034 |
| chr16 | 11623277 | 1.2E+07 | chr15 | 34423960 | 34502327 |
| chr16 | 14568441 | 1.5E+07 | chr15 | 41522129 | 41557301 |
| chr16 | 15965417 | 1.6E+07 | chr15 | 52121436 | 52201158 |
| chr16 | 16013321 | 1.6E+07 | chr15 | 52261901 | 52351044 |

| | | | | | |
|---|---|---|---|---|---|
| chr16 | 19098099 | 1.9E+07 | chr15 | 52541693 | 52588933 |
| chr16 | 24795053 | 2.5E+07 | chr15 | 55650269 | 55702920 |
| chr16 | 24917329 | 2.5E+07 | chr15 | 56160343 | 56211515 |
| chr16 | 28138037 | 2.8E+07 | chr15 | 59396469 | 59681953 |
| chr16 | 30726123 | 3.1E+07 | chr15 | 59824759 | 59950480 |
| chr16 | 53075418 | 5.3E+07 | chr15 | 60635521 | 60720218 |
| chr16 | 56833217 | 5.7E+07 | chr15 | 63334044 | 63415225 |
| chr16 | 57787026 | 5.8E+07 | chr15 | 63621480 | 63674732 |
| chr16 | 67866355 | 6.8E+07 | chr15 | 63728705 | 63808936 |
| chr16 | 68288725 | 6.8E+07 | chr15 | 64894318 | 64996149 |
| chr16 | 69164900 | 6.9E+07 | chr15 | 66992768 | 67078643 |
| chr16 | 69407481 | 7E+07 | chr15 | 67315787 | 67491749 |
| chr16 | 69684792 | 7E+07 | chr15 | 70784452 | 70802923 |
| chr16 | 70408394 | 7E+07 | chr15 | 71564930 | 71593496 |
| chr16 | 70703816 | 7.1E+07 | chr15 | 71676824 | 71764935 |
| chr16 | 73017394 | 7.3E+07 | chr15 | 72471787 | 72533284 |
| chr16 | 74763090 | 7.5E+07 | chr15 | 75134796 | 75232018 |
| chr16 | 81428895 | 8.2E+07 | chr15 | 75912724 | 75965998 |
| chr16 | 84609273 | 8.5E+07 | chr15 | 85220029 | 85297993 |
| chr16 | 84732780 | 8.5E+07 | chr15 | 85522896 | 85595096 |
| chr16 | 85005800 | 8.5E+07 | chr15 | 85610136 | 85686243 |
| chr16 | 87791168 | 8.8E+07 | chr15 | 86099979 | 86163037 |
| chr16 | 89352674 | 9E+07 | chr15 | 86176810 | 86281309 |
| chr17 | 170172 | 208098 | chr15 | 89157737 | 89193308 |
| chr17 | 1326826 | 1395452 | chr15 | 89630522 | 89673592 |
| chr17 | 1613122 | 1640274 | chr15 | 90691828 | 90793164 |
| chr17 | 2282555 | 2323058 | chr15 | 90930479 | 90973880 |
| chr17 | 4577745 | 4643480 | chr15 | 93340059 | 93562582 |
| chr17 | 6478154 | 6556085 | chr15 | 99296063 | 99367158 |
| chr17 | 7107411 | 7168425 | chr15 | 99386093 | 99468449 |
| chr17 | 7451698 | 7531222 | chr16 | 1727478 | 1776583 |
| chr17 | 7737019 | 7762551 | chr16 | 2008985 | 2098860 |
| chr17 | 8021135 | 8131648 | chr16 | 4321209 | 4401320 |
| chr17 | 9720702 | 9747955 | chr16 | 9161332 | 9223255 |
| chr17 | 15884112 | 1.6E+07 | chr16 | 16038792 | 16204659 |
| chr17 | 16281168 | 1.6E+07 | chr16 | 19098020 | 19152586 |
| chr17 | 16912156 | 1.7E+07 | chr16 | 19395521 | 19458247 |
| chr17 | 17584519 | 1.8E+07 | chr16 | 52579061 | 52613779 |
| chr17 | 18084804 | 1.8E+07 | chr16 | 53111628 | 53182611 |
| chr17 | 18751744 | 1.9E+07 | chr16 | 53765785 | 53815992 |
| chr17 | 19171880 | 1.9E+07 | chr16 | 56833652 | 56903585 |
| chr17 | 19606533 | 2E+07 | chr16 | 57788104 | 57855909 |
| chr17 | 19800350 | 2E+07 | chr16 | 66535966 | 66559183 |
| chr17 | 19960386 | 2E+07 | chr16 | 69418065 | 69512237 |
| chr17 | 25620089 | 2.6E+07 | chr16 | 69687056 | 69797673 |
| chr17 | 25853506 | 2.6E+07 | chr16 | 73080256 | 73131308 |
| chr17 | 26645275 | 2.7E+07 | chr16 | 74760049 | 74809463 |
| chr17 | 27024716 | 2.7E+07 | chr16 | 81001614 | 81042625 |
| chr17 | 27423327 | 2.8E+07 | chr16 | 81462220 | 81813436 |
| chr17 | 27684016 | 2.8E+07 | chr16 | 84593406 | 84655865 |
| chr17 | 27852631 | 2.8E+07 | chr16 | 84732798 | 84772830 |
| chr17 | 33306694 | 3.3E+07 | chr16 | 85005738 | 85071403 |
| chr17 | 33894582 | 3.4E+07 | chr16 | 87859209 | 87906021 |
| chr17 | 35439561 | 3.6E+07 | chr16 | 89352493 | 89576340 |
| chr17 | 35626129 | 3.6E+07 | chr17 | 170143 | 236848 |
| chr17 | 36056777 | 3.6E+07 | chr17 | 1321334 | 1395622 |
| chr17 | 36849946 | 3.7E+07 | chr17 | 2286117 | 2323001 |
| chr17 | 37000627 | 3.7E+07 | chr17 | 2853051 | 2915051 |
| chr17 | 38254274 | 3.8E+07 | chr17 | 7107617 | 7168432 |
| chr17 | 38439702 | 3.9E+07 | chr17 | 8020753 | 8131624 |
| chr17 | 38593898 | 3.9E+07 | chr17 | 15790006 | 15825723 |

| | | | | | |
|---|---|---|---|---|---|
| chr17 | 39044001 | 3.9E+07 | chr17 | 15845209 | 15966400 |
| chr17 | 39801383 | 4E+07 | chr17 | 16912163 | 17094805 |
| chr17 | 39914999 | 4E+07 | chr17 | 17584215 | 17754471 |
| chr17 | 40020543 | 4E+07 | chr17 | 19619526 | 19660551 |
| chr17 | 40111078 | 4E+07 | chr17 | 19801049 | 19881955 |
| chr17 | 40457762 | 4.1E+07 | chr17 | 25619978 | 25681579 |
| chr17 | 40598562 | 4.1E+07 | chr17 | 25853468 | 25926646 |
| chr17 | 41131435 | 4.1E+07 | chr17 | 26638869 | 26708017 |
| chr17 | 41580365 | 4.2E+07 | chr17 | 27023834 | 27091219 |
| chr17 | 42071935 | 4.2E+07 | chr17 | 27684072 | 27787574 |
| chr17 | 42214865 | 4.2E+07 | chr17 | 33340115 | 33418025 |
| chr17 | 42607140 | 4.3E+07 | chr17 | 33894576 | 33958348 |
| chr17 | 42966370 | 4.3E+07 | chr17 | 35797520 | 35853643 |
| chr17 | 43114047 | 4.3E+07 | chr17 | 36058636 | 36126058 |
| chr17 | 43298337 | 4.3E+07 | chr17 | 37003826 | 37079592 |
| chr17 | 46313555 | 4.6E+07 | chr17 | 38465229 | 38527874 |
| chr17 | 46549695 | 4.7E+07 | chr17 | 38598567 | 38717435 |
| chr17 | 46867942 | 4.7E+07 | chr17 | 39042400 | 39079648 |
| chr17 | 47068107 | 4.7E+07 | chr17 | 39652470 | 39706532 |
| chr17 | 48103236 | 4.8E+07 | chr17 | 39801521 | 39874837 |
| chr17 | 48696555 | 4.9E+07 | chr17 | 39918788 | 39970622 |
| chr17 | 48922335 | 4.9E+07 | chr17 | 40020376 | 40087585 |
| chr17 | 53471855 | 5.4E+07 | chr17 | 40108658 | 40178185 |
| chr17 | 55926304 | 5.6E+07 | chr17 | 40470394 | 40584443 |
| chr17 | 57151836 | 5.7E+07 | chr17 | 40664271 | 40742439 |
| chr17 | 57694869 | 5.8E+07 | chr17 | 41131663 | 41177213 |
| chr17 | 61817588 | 6.2E+07 | chr17 | 42136628 | 42202121 |
| chr17 | 62615059 | 6.3E+07 | chr17 | 42611065 | 42638101 |
| chr17 | 63013429 | 6.3E+07 | chr17 | 43122534 | 43276582 |
| chr17 | 64649015 | 6.5E+07 | chr17 | 43298324 | 43385044 |
| chr17 | 70382128 | 7E+07 | chr17 | 46100231 | 46186589 |
| chr17 | 70490748 | 7.1E+07 | chr17 | 46315768 | 46347216 |
| chr17 | 71274972 | 7.1E+07 | chr17 | 46549575 | 46704686 |
| chr17 | 72977465 | 7.3E+07 | chr17 | 46867950 | 46928885 |
| chr17 | 73422264 | 7.4E+07 | chr17 | 46968593 | 47052630 |
| chr17 | 73584907 | 7.4E+07 | chr17 | 47067954 | 47106099 |
| chr17 | 73839610 | 7.4E+07 | chr17 | 48103087 | 48173349 |
| chr17 | 75276731 | 7.5E+07 | chr17 | 48702898 | 48846539 |
| chr17 | 76296118 | 7.6E+07 | chr17 | 48922427 | 49032343 |
| chr17 | 76857284 | 7.7E+07 | chr17 | 55923177 | 56038945 |
| chr17 | 78215803 | 7.8E+07 | chr17 | 57170017 | 57264507 |
| chr17 | 79816946 | 8E+07 | chr17 | 57694159 | 57950772 |
| chr17 | 80138694 | 8E+07 | chr17 | 60855231 | 60902909 |
| chr17 | 80476602 | 8.1E+07 | chr17 | 61817588 | 61870884 |
| chr18 | 2977902 | 3052848 | chr17 | 62619232 | 62705974 |
| chr18 | 3401941 | 3468835 | chr17 | 64657224 | 64730440 |
| chr18 | 3582787 | 3711671 | chr17 | 70382348 | 70473187 |
| chr18 | 7900694 | 7963936 | chr17 | 70556658 | 70613195 |
| chr18 | 8701158 | 8829485 | chr17 | 71277828 | 71309282 |
| chr18 | 9073454 | 9159059 | chr17 | 73007501 | 73209597 |
| chr18 | 9608014 | 9649339 | chr17 | 73598886 | 73785345 |
| chr18 | 24033348 | 2.4E+07 | chr17 | 73839622 | 73875609 |
| chr18 | 45386655 | 4.5E+07 | chr17 | 75276818 | 75486894 |
| chr18 | 46448227 | 4.6E+07 | chr17 | 76123719 | 76195956 |
| chr18 | 55395186 | 5.5E+07 | chr17 | 78222586 | 78269887 |
| chr18 | 56448387 | 5.6E+07 | chr17 | 79448157 | 79487399 |
| chr18 | 56516623 | 5.7E+07 | chr17 | 79779047 | 79857671 |
| chr18 | 67955173 | 6.8E+07 | chr17 | 80150499 | 80259440 |
| chr18 | 68022885 | 6.8E+07 | chr17 | 80476588 | 80561151 |
| chr18 | 68078364 | 6.8E+07 | chr18 | 3245732 | 3265593 |
| chr18 | 74765597 | 7.5E+07 | chr18 | 3441108 | 3469665 |

| | | | | | |
|---|---|---|---|---|---|
| chr19 | 749882 | 800131 | chr18 | 3581995 | 3627990 |
| chr19 | 2523348 | 2553348 | chr18 | 3647477 | 3723299 |
| chr19 | 13253371 | 1.3E+07 | chr18 | 7900497 | 7963877 |
| chr19 | 13943837 | 1.4E+07 | chr18 | 8698939 | 8831316 |
| chr19 | 16177175 | 1.6E+07 | chr18 | 9091193 | 9153768 |
| chr19 | 18384744 | 1.8E+07 | chr18 | 9594770 | 9649552 |
| chr19 | 18474004 | 1.9E+07 | chr18 | 11848736 | 11861639 |
| chr19 | 32835942 | 3.3E+07 | chr18 | 20678341 | 20740468 |
| chr19 | 33488334 | 3.4E+07 | chr18 | 24033240 | 24072093 |
| chr19 | 34650077 | 3.5E+07 | chr18 | 45393121 | 45459820 |
| chr19 | 34842202 | 3.5E+07 | chr18 | 45524139 | 45554488 |
| chr19 | 35484992 | 3.6E+07 | chr18 | 55394941 | 55488181 |
| chr19 | 36192944 | 3.6E+07 | chr18 | 56448549 | 56462902 |
| chr19 | 36526944 | 3.7E+07 | chr18 | 56515544 | 56584585 |
| chr19 | 38396712 | 3.9E+07 | chr18 | 56657740 | 56681295 |
| chr19 | 39108799 | 3.9E+07 | chr18 | 67953891 | 68054695 |
| chr19 | 39295968 | 3.9E+07 | chr18 | 68078483 | 68101964 |
| chr19 | 39601787 | 4E+07 | chr18 | 70912667 | 70964388 |
| chr19 | 39831612 | 4E+07 | chr18 | 70982643 | 71008776 |
| chr19 | 40402625 | 4E+07 | chr18 | 74765751 | 74845305 |
| chr19 | 40898611 | 4.1E+07 | chr19 | 1150963 | 1214166 |
| chr19 | 41168402 | 4.1E+07 | chr19 | 1239774 | 1276852 |
| chr19 | 41723963 | 4.2E+07 | chr19 | 13253467 | 13284482 |
| chr19 | 41793381 | 4.2E+07 | chr19 | 13943498 | 13973816 |
| chr19 | 42695418 | 4.3E+07 | chr19 | 14605532 | 14640080 |
| chr19 | 44022978 | 4.4E+07 | chr19 | 16177156 | 16223984 |
| chr19 | 44235143 | 4.4E+07 | chr19 | 18385280 | 18452201 |
| chr19 | 45127988 | 4.5E+07 | chr19 | 18473971 | 18548946 |
| chr19 | 45453791 | 4.6E+07 | chr19 | 23518964 | 23621534 |
| chr19 | 45574188 | 4.6E+07 | chr19 | 23683883 | 23755672 |
| chr19 | 45894364 | 4.6E+07 | chr19 | 23904555 | 24008573 |
| chr19 | 46044811 | 4.6E+07 | chr19 | 32834631 | 32876692 |
| chr19 | 46170780 | 4.6E+07 | chr19 | 33488391 | 33594112 |
| chr19 | 46849470 | 4.7E+07 | chr19 | 34649830 | 34725377 |
| chr19 | 47392967 | 4.7E+07 | chr19 | 34842144 | 34896177 |
| chr19 | 47571511 | 4.8E+07 | chr19 | 35476449 | 35510771 |
| chr19 | 48101331 | 4.8E+07 | chr19 | 36595578 | 36681454 |
| chr19 | 48823609 | 4.9E+07 | chr19 | 37281841 | 37342653 |
| chr19 | 48948002 | 4.9E+07 | chr19 | 37957055 | 38025237 |
| chr19 | 49042494 | 4.9E+07 | chr19 | 38117740 | 38162413 |
| chr19 | 49457287 | 5E+07 | chr19 | 38388558 | 38631298 |
| chr19 | 50083243 | 5E+07 | chr19 | 38734045 | 38811081 |
| chr19 | 54343516 | 5.4E+07 | chr19 | 39105670 | 39227409 |
| chr19 | 54480952 | 5.5E+07 | chr19 | 39295939 | 39369759 |
| chr19 | 56135734 | 5.6E+07 | chr19 | 39601292 | 39686518 |
| chr19 | 59054546 | 5.9E+07 | chr19 | 39862783 | 39984584 |
| chr2 | 9370760 | 9429450 | chr19 | 40440384 | 40488289 |
| chr2 | 9504957 | 9575172 | chr19 | 40898748 | 40972799 |
| chr2 | 9652420 | 9706894 | chr19 | 41240563 | 41285426 |
| chr2 | 10410562 | 1.1E+07 | chr19 | 41723871 | 41835366 |
| chr2 | 11510290 | 1.2E+07 | chr19 | 42363074 | 42411639 |
| chr2 | 17876850 | 1.8E+07 | chr19 | 42710724 | 42790568 |
| chr2 | 20366027 | 2E+07 | chr19 | 44143474 | 44210002 |
| chr2 | 26161352 | 2.6E+07 | chr19 | 44235307 | 44290171 |
| chr2 | 26930734 | 2.7E+07 | chr19 | 45218764 | 45312902 |
| chr2 | 27576469 | 2.8E+07 | chr19 | 45894775 | 46091644 |
| chr2 | 28612494 | 2.9E+07 | chr19 | 46170561 | 46235422 |
| chr2 | 29328174 | 2.9E+07 | chr19 | 47582796 | 47777928 |
| chr2 | 30454396 | 3.1E+07 | chr19 | 48101292 | 48182602 |
| chr2 | 36582044 | 3.7E+07 | chr19 | 48856179 | 48931409 |
| chr2 | 42313594 | 4.2E+07 | chr19 | 49046173 | 49149904 |

| | | | | | |
|---|---|---|---|---|---|
| chr2 | 43384024 | 4.3E+07 | chr19 | 49457252 | 49501731 |
| chr2 | 46858939 | 4.7E+07 | chr19 | 49946386 | 50002212 |
| chr2 | 47073555 | 4.7E+07 | chr19 | 50138554 | 50191486 |
| chr2 | 47166410 | 4.7E+07 | chr19 | 53424937 | 53450826 |
| chr2 | 47371914 | 4.7E+07 | chr19 | 54023240 | 54083313 |
| chr2 | 54750086 | 5.5E+07 | chr19 | 54341489 | 54393265 |
| chr2 | 55179791 | 5.5E+07 | chr19 | 54480934 | 54539422 |
| chr2 | 61607144 | 6.2E+07 | chr19 | 54658059 | 54715756 |
| chr2 | 65282769 | 6.5E+07 | chr19 | 56866492 | 56916403 |
| chr2 | 69960321 | 7E+07 | chr19 | 59054467 | 59093038 |
| chr2 | 70312485 | 7E+07 | chr2 | 10410512 | 10558238 |
| chr2 | 73247476 | 7.3E+07 | chr2 | 11507688 | 11558976 |
| chr2 | 85131901 | 8.5E+07 | chr2 | 17899766 | 17950216 |
| chr2 | 85636977 | 8.6E+07 | chr2 | 20365879 | 20432439 |
| chr2 | 85725522 | 8.6E+07 | chr2 | 26161399 | 26325738 |
| chr2 | 86244864 | 8.6E+07 | chr2 | 28612208 | 28673659 |
| chr2 | 1.01E+08 | 1E+08 | chr2 | 29330691 | 29358791 |
| chr2 | 1.03E+08 | 1E+08 | chr2 | 36581757 | 36808125 |
| chr2 | 1.11E+08 | 1.1E+08 | chr2 | 42310981 | 42375581 |
| chr2 | 1.14E+08 | 1.1E+08 | chr2 | 42395799 | 42525065 |
| chr2 | 1.22E+08 | 1.2E+08 | chr2 | 43202440 | 43284285 |
| chr2 | 1.28E+08 | 1.3E+08 | chr2 | 43356641 | 43428281 |
| chr2 | 1.44E+08 | 1.4E+08 | chr2 | 47166602 | 47215645 |
| chr2 | 1.44E+08 | 1.4E+08 | chr2 | 47229582 | 47317390 |
| chr2 | 1.53E+08 | 1.5E+08 | chr2 | 54750073 | 54897730 |
| chr2 | 1.61E+08 | 1.6E+08 | chr2 | 61685058 | 61766915 |
| chr2 | 1.61E+08 | 1.6E+08 | chr2 | 69952696 | 70057513 |
| chr2 | 1.78E+08 | 1.8E+08 | chr2 | 70312343 | 70371309 |
| chr2 | 1.87E+08 | 1.9E+08 | chr2 | 85131849 | 85211628 |
| chr2 | 1.88E+08 | 1.9E+08 | chr2 | 85636926 | 85700513 |
| chr2 | 1.92E+08 | 1.9E+08 | chr2 | 86104355 | 86191144 |
| chr2 | 2.01E+08 | 2E+08 | chr2 | 86916056 | 86934041 |
| chr2 | 2.07E+08 | 2.1E+08 | chr2 | 101434687 | 101542113 |
| chr2 | 2.09E+08 | 2.1E+08 | chr2 | 101618012 | 101777553 |
| chr2 | 2.28E+08 | 2.3E+08 | chr2 | 113955821 | 114000866 |
| chr2 | 2.28E+08 | 2.3E+08 | chr2 | 127810552 | 127874092 |
| chr2 | 2.32E+08 | 2.3E+08 | chr2 | 143607036 | 143644423 |
| chr2 | 2.34E+08 | 2.3E+08 | chr2 | 161060482 | 161110845 |
| chr2 | 2.34E+08 | 2.3E+08 | chr2 | 178075536 | 178141000 |
| chr2 | 2.35E+08 | 2.3E+08 | chr2 | 188390021 | 188446266 |
| chr2 | 2.35E+08 | 2.3E+08 | chr2 | 189143981 | 189189323 |
| chr2 | 2.36E+08 | 2.4E+08 | chr2 | 191702247 | 191814175 |
| chr2 | 2.38E+08 | 2.4E+08 | chr2 | 201979539 | 202023403 |
| chr2 | 2.39E+08 | 2.4E+08 | chr2 | 206540164 | 206637566 |
| chr20 | 342602 | 427037 | chr2 | 227597478 | 227666827 |
| chr20 | 3730152 | 3802600 | chr2 | 228330057 | 228382317 |
| chr20 | 4089064 | 4174404 | chr2 | 232465490 | 232513896 |
| chr20 | 4877298 | 4957688 | chr2 | 232526439 | 232552625 |
| chr20 | 5037513 | 5112453 | chr2 | 234316544 | 234350131 |
| chr20 | 5614820 | 5744571 | chr2 | 238380354 | 238450954 |
| chr20 | 5929417 | 5991295 | chr2 | 239191140 | 239209710 |
| chr20 | 10385709 | 1E+07 | chr20 | 343582 | 426275 |
| chr20 | 17535621 | 1.8E+07 | chr20 | 3747733 | 3802985 |
| chr20 | 19830019 | 2E+07 | chr20 | 4099800 | 4153641 |
| chr20 | 19944154 | 2E+07 | chr20 | 5041299 | 5114202 |
| chr20 | 22355521 | 2.2E+07 | chr20 | 5531039 | 5594415 |
| chr20 | 25175582 | 2.5E+07 | chr20 | 5622827 | 5641656 |
| chr20 | 30101474 | 3E+07 | chr20 | 5659576 | 5744551 |
| chr20 | 30248315 | 3E+07 | chr20 | 5929858 | 5989564 |
| chr20 | 31026900 | 3.1E+07 | chr20 | 17534812 | 17636965 |
| chr20 | 32943113 | 3.3E+07 | chr20 | 17917676 | 17957851 |

| | | | | | |
|---|---|---|---|---|---|
| chr20 | 33837946 | 3.4E+07 | chr20 | 19840666 | 19884581 |
| chr20 | 34667647 | 3.5E+07 | chr20 | 19944472 | 20019388 |
| chr20 | 35431028 | 3.5E+07 | chr20 | 25175533 | 25299176 |
| chr20 | 35898306 | 3.6E+07 | chr20 | 30101454 | 30222605 |
| chr20 | 36738554 | 3.7E+07 | chr20 | 30248539 | 30312383 |
| chr20 | 45881412 | 4.6E+07 | chr20 | 31031019 | 31073537 |
| chr20 | 46046226 | 4.6E+07 | chr20 | 32949364 | 33012290 |
| chr20 | 46113543 | 4.6E+07 | chr20 | 33839582 | 33873360 |
| chr20 | 48191961 | 4.8E+07 | chr20 | 34638472 | 34732679 |
| chr20 | 48719440 | 4.9E+07 | chr20 | 35963715 | 36026448 |
| chr20 | 48861792 | 4.9E+07 | chr20 | 36678045 | 36816165 |
| chr20 | 49334499 | 4.9E+07 | chr20 | 45931436 | 46032555 |
| chr20 | 49511902 | 5E+07 | chr20 | 46046513 | 46095265 |
| chr20 | 50351398 | 5E+07 | chr20 | 46121244 | 46291008 |
| chr20 | 52185888 | 5.2E+07 | chr20 | 47883574 | 47909864 |
| chr20 | 52351632 | 5.2E+07 | chr20 | 48227346 | 48345161 |
| chr20 | 52712213 | 5.3E+07 | chr20 | 48381153 | 48435797 |
| chr20 | 56230434 | 5.6E+07 | chr20 | 48702247 | 48790655 |
| chr20 | 60911301 | 6.1E+07 | chr20 | 48861732 | 48938921 |
| chr21 | 27486793 | 2.8E+07 | chr20 | 49031146 | 49068559 |
| chr21 | 33783102 | 3.4E+07 | chr20 | 49240789 | 49318619 |
| chr21 | 38251886 | 3.8E+07 | chr20 | 49337846 | 49383950 |
| chr21 | 44750301 | 4.5E+07 | chr20 | 50302818 | 50384901 |
| chr21 | 44815849 | 4.5E+07 | chr20 | 52186186 | 52242887 |
| chr21 | 45107714 | 4.5E+07 | chr20 | 52257475 | 52278624 |
| chr22 | 27136235 | 2.7E+07 | chr20 | 52351511 | 52427113 |
| chr22 | 30581739 | 3.1E+07 | chr20 | 52441878 | 52489998 |
| chr22 | 35746313 | 3.6E+07 | chr20 | 52712850 | 52794566 |
| chr22 | 36717807 | 3.7E+07 | chr20 | 55326825 | 55384077 |
| chr22 | 40849413 | 4.1E+07 | chr20 | 56229954 | 56294279 |
| chr22 | 43315154 | 4.3E+07 | chr20 | 60903810 | 60964074 |
| chr22 | 45969414 | 4.6E+07 | chr21 | 27487297 | 27543903 |
| chr22 | 46066566 | 4.6E+07 | chr21 | 30539992 | 30591090 |
| chr22 | 46298860 | 4.6E+07 | chr21 | 33783489 | 33854038 |
| chr22 | 46448015 | 4.7E+07 | chr21 | 36153331 | 36262461 |
| chr22 | 46858246 | 4.7E+07 | chr21 | 44738989 | 44766800 |
| chr3 | 5017439 | 5069349 | chr21 | 44815591 | 44941012 |
| chr3 | 8483335 | 8500814 | chr21 | 45129448 | 45210444 |
| chr3 | 8524327 | 8621237 | chr22 | 29168266 | 29220597 |
| chr3 | 9424431 | 9484552 | chr22 | 30581480 | 30659737 |
| chr3 | 10206431 | 1E+07 | chr22 | 31222128 | 31304162 |
| chr3 | 14443317 | 1.5E+07 | chr22 | 36718811 | 36795898 |
| chr3 | 15780858 | 1.6E+07 | chr22 | 37892896 | 37960821 |
| chr3 | 30636329 | 3.1E+07 | chr22 | 40867657 | 40932391 |
| chr3 | 43892509 | 4.4E+07 | chr22 | 43314757 | 43359241 |
| chr3 | 45653001 | 4.6E+07 | chr22 | 45974234 | 46042376 |
| chr3 | 46965463 | 4.7E+07 | chr22 | 46065420 | 46146542 |
| chr3 | 53249891 | 5.3E+07 | chr22 | 46447984 | 46512325 |
| chr3 | 57935309 | 5.8E+07 | chr22 | 46839832 | 46944075 |
| chr3 | 81765839 | 8.2E+07 | chr3 | 5007364 | 5069707 |
| chr3 | 98589635 | 9.9E+07 | chr3 | 8483633 | 8500272 |
| chr3 | 98679812 | 9.9E+07 | chr3 | 9424337 | 9472614 |
| chr3 | 1.02E+08 | 1E+08 | chr3 | 12327958 | 12361501 |
| chr3 | 1.19E+08 | 1.2E+08 | chr3 | 14443439 | 14515000 |
| chr3 | 1.24E+08 | 1.2E+08 | chr3 | 15779061 | 15848341 |
| chr3 | 1.27E+08 | 1.3E+08 | chr3 | 30267279 | 30306070 |
| chr3 | 1.29E+08 | 1.3E+08 | chr3 | 30360118 | 30415850 |
| chr3 | 1.41E+08 | 1.4E+08 | chr3 | 30636004 | 30739590 |
| chr3 | 1.49E+08 | 1.5E+08 | chr3 | 45634261 | 45702385 |
| chr3 | 1.49E+08 | 1.5E+08 | chr3 | 46955338 | 47021802 |
| chr3 | 1.49E+08 | 1.5E+08 | chr3 | 53250546 | 53319681 |

| | | | | | |
|---|---|---|---|---|---|
| chr3 | 1.51E+08 | 1.5E+08 | chr3 | 57935315 | 58164001 |
| chr3 | 1.52E+08 | 1.5E+08 | chr3 | 70297437 | 70333377 |
| chr3 | 1.56E+08 | 1.6E+08 | chr3 | 81765900 | 81811895 |
| chr3 | 1.56E+08 | 1.6E+08 | chr3 | 98525264 | 98644797 |
| chr3 | 1.72E+08 | 1.7E+08 | chr3 | 98680493 | 98705756 |
| chr3 | 1.72E+08 | 1.7E+08 | chr3 | 101874488 | 101915671 |
| chr3 | 1.88E+08 | 1.9E+08 | chr3 | 105082759 | 105122613 |
| chr4 | 7805084 | 7943557 | chr3 | 110650830 | 110715017 |
| chr4 | 15654589 | 1.6E+07 | chr3 | 112464808 | 112543464 |
| chr4 | 39025798 | 3.9E+07 | chr3 | 114425202 | 114494969 |
| chr4 | 39497353 | 4E+07 | chr3 | 118892324 | 118960453 |
| chr4 | 74861900 | 7.5E+07 | chr3 | 124479689 | 124620391 |
| chr4 | 74962587 | 7.5E+07 | chr3 | 127453335 | 127542399 |
| chr4 | 75573374 | 7.6E+07 | chr3 | 129305678 | 129407837 |
| chr4 | 84456364 | 8.5E+07 | chr3 | 129547421 | 129655807 |
| chr4 | 90164399 | 9E+07 | chr3 | 141110850 | 141174399 |
| chr4 | 95561227 | 9.6E+07 | chr3 | 146219041 | 146268352 |
| chr4 | 1.2E+08 | 1.2E+08 | chr3 | 149014338 | 149121630 |
| chr4 | 1.29E+08 | 1.3E+08 | chr3 | 149234945 | 149379384 |
| chr4 | 1.3E+08 | 1.3E+08 | chr3 | 151483190 | 151513664 |
| chr4 | 1.39E+08 | 1.4E+08 | chr3 | 152847784 | 152884547 |
| chr4 | 1.41E+08 | 1.4E+08 | chr3 | 156391077 | 156440400 |
| chr4 | 1.7E+08 | 1.7E+08 | chr3 | 156528464 | 156545829 |
| chr4 | 1.7E+08 | 1.7E+08 | chr3 | 160108471 | 160169403 |
| chr5 | 10549674 | 1.1E+07 | chr3 | 171509381 | 171620305 |
| chr5 | 14142249 | 1.4E+07 | chr3 | 171843601 | 171887925 |
| chr5 | 16778164 | 1.7E+07 | chr3 | 172463398 | 172532214 |
| chr5 | 17214746 | 1.7E+07 | chr3 | 185033211 | 185081564 |
| chr5 | 34655522 | 3.5E+07 | chr3 | 187869983 | 187990878 |
| chr5 | 39072804 | 3.9E+07 | chr3 | 189010602 | 189076062 |
| chr5 | 42982097 | 4.3E+07 | chr3 | 193829031 | 193860958 |
| chr5 | 52283101 | 5.2E+07 | chr4 | 7805095 | 7941473 |
| chr5 | 58262858 | 5.9E+07 | chr4 | 15001995 | 15050345 |
| chr5 | 58742892 | 5.9E+07 | chr4 | 15653186 | 15704937 |
| chr5 | 59039863 | 5.9E+07 | chr4 | 38664385 | 38698431 |
| chr5 | 60586474 | 6.1E+07 | chr4 | 39024069 | 39056564 |
| chr5 | 71402400 | 7.1E+07 | chr4 | 39497320 | 39560286 |
| chr5 | 72132747 | 7.2E+07 | chr4 | 40443262 | 40529606 |
| chr5 | 72900435 | 7.3E+07 | chr4 | 57894036 | 57985896 |
| chr5 | 77786675 | 7.8E+07 | chr4 | 74861871 | 74905521 |
| chr5 | 90568678 | 9.1E+07 | chr4 | 74962511 | 75000109 |
| chr5 | 95093895 | 9.5E+07 | chr4 | 75228456 | 75250002 |
| chr5 | 1.07E+08 | 1.1E+08 | chr4 | 75572602 | 75608972 |
| chr5 | 1.19E+08 | 1.2E+08 | chr4 | 77608902 | 77646566 |
| chr5 | 1.33E+08 | 1.3E+08 | chr4 | 79472126 | 79493255 |
| chr5 | 1.34E+08 | 1.3E+08 | chr4 | 90193063 | 90229450 |
| chr5 | 1.35E+08 | 1.4E+08 | chr4 | 95560208 | 95627092 |
| chr5 | 1.38E+08 | 1.4E+08 | chr4 | 95671133 | 95711939 |
| chr5 | 1.42E+08 | 1.4E+08 | chr4 | 95994029 | 96030808 |
| chr5 | 1.43E+08 | 1.4E+08 | chr4 | 119989168 | 120035208 |
| chr5 | 1.5E+08 | 1.5E+08 | chr4 | 129730134 | 129787918 |
| chr5 | 1.5E+08 | 1.5E+08 | chr4 | 139082326 | 139198012 |
| chr5 | 1.54E+08 | 1.5E+08 | chr4 | 140931986 | 141038671 |
| chr5 | 1.54E+08 | 1.5E+08 | chr4 | 170145730 | 170204670 |
| chr5 | 1.56E+08 | 1.6E+08 | chr4 | 187562793 | 187623075 |
| chr5 | 1.59E+08 | 1.6E+08 | chr5 | 1469573 | 1568958 |
| chr5 | 1.6E+08 | 1.6E+08 | chr5 | 14142197 | 14279065 |
| chr5 | 1.68E+08 | 1.7E+08 | chr5 | 16845747 | 16937061 |
| chr5 | 1.72E+08 | 1.7E+08 | chr5 | 17212860 | 17285736 |
| chr5 | 1.72E+08 | 1.7E+08 | chr5 | 17302757 | 17326969 |
| chr5 | 1.73E+08 | 1.7E+08 | chr5 | 39071961 | 39103727 |

| | | | | | |
|---|---|---|---|---|---|
| chr5 | 1.75E+08 | 1.8E+08 | chr5 | 42982060 | 43021416 |
| chr5 | 1.79E+08 | 1.8E+08 | chr5 | 43034676 | 43079524 |
| chr6 | 2952004 | 3025157 | chr5 | 58264047 | 58519633 |
| chr6 | 11184272 | 1.1E+07 | chr5 | 58532345 | 58629498 |
| chr6 | 12279085 | 1.2E+07 | chr5 | 58739734 | 58895163 |
| chr6 | 12449985 | 1.2E+07 | chr5 | 59038941 | 59096023 |
| chr6 | 26020177 | 2.6E+07 | chr5 | 60571235 | 60630161 |
| chr6 | 26086856 | 2.6E+07 | chr5 | 72133574 | 72204211 |
| chr6 | 26151635 | 2.6E+07 | chr5 | 72900069 | 73006756 |
| chr6 | 26272985 | 2.6E+07 | chr5 | 73090828 | 73126742 |
| chr6 | 27094365 | 2.7E+07 | chr5 | 77862037 | 77945802 |
| chr6 | 27759214 | 2.8E+07 | chr5 | 90663112 | 90680792 |
| chr6 | 34622496 | 3.5E+07 | chr5 | 95093176 | 95179819 |
| chr6 | 53357963 | 5.3E+07 | chr5 | 95827452 | 95875641 |
| chr6 | 56486609 | 5.7E+07 | chr5 | 95949215 | 96127482 |
| chr6 | 56703474 | 5.7E+07 | chr5 | 106695056 | 106737040 |
| chr6 | 64265208 | 6.4E+07 | chr5 | 106789899 | 106860487 |
| chr6 | 72093409 | 7.2E+07 | chr5 | 106874292 | 106909628 |
| chr6 | 74275264 | 7.4E+07 | chr5 | 111541389 | 111576504 |
| chr6 | 88382775 | 8.8E+07 | chr5 | 115208260 | 115277561 |
| chr6 | 1.06E+08 | 1.1E+08 | chr5 | 118590991 | 118671354 |
| chr6 | 1.18E+08 | 1.2E+08 | chr5 | 126111184 | 126183019 |
| chr6 | 1.26E+08 | 1.3E+08 | chr5 | 127243576 | 127330400 |
| chr6 | 1.27E+08 | 1.3E+08 | chr5 | 132341287 | 132423543 |
| chr6 | 1.3E+08 | 1.3E+08 | chr5 | 133292474 | 133356491 |
| chr6 | 1.34E+08 | 1.3E+08 | chr5 | 133837342 | 133872489 |
| chr6 | 1.37E+08 | 1.4E+08 | chr5 | 135328995 | 135402445 |
| chr6 | 1.37E+08 | 1.4E+08 | chr5 | 142149059 | 142260991 |
| chr6 | 1.43E+08 | 1.4E+08 | chr5 | 142711201 | 142785671 |
| chr6 | 1.51E+08 | 1.5E+08 | chr5 | 142939604 | 142980925 |
| chr6 | 1.51E+08 | 1.5E+08 | chr5 | 148851007 | 148944451 |
| chr6 | 1.58E+08 | 1.6E+08 | chr5 | 149825929 | 149903960 |
| chr6 | 1.59E+08 | 1.6E+08 | chr5 | 153643764 | 153787603 |
| chr7 | 1542567 | 1611388 | chr5 | 154133694 | 154215245 |
| chr7 | 4721317 | 4794036 | chr5 | 156457175 | 156500135 |
| chr7 | 6297552 | 6371945 | chr5 | 158584015 | 158639457 |
| chr7 | 6406741 | 6494103 | chr5 | 159324748 | 159356111 |
| chr7 | 11814410 | 1.2E+07 | chr5 | 162863352 | 162911740 |
| chr7 | 28059692 | 2.8E+07 | chr5 | 167715107 | 167883372 |
| chr7 | 31962672 | 3.2E+07 | chr5 | 172158093 | 172236667 |
| chr7 | 32602181 | 3.3E+07 | chr5 | 172317693 | 172387764 |
| chr7 | 43655242 | 4.4E+07 | chr5 | 173364793 | 173400956 |
| chr7 | 47318147 | 4.8E+07 | chr5 | 175268732 | 175299910 |
| chr7 | 47794000 | 4.8E+07 | chr5 | 179222641 | 179267357 |
| chr7 | 55086110 | 5.5E+07 | chr5 | 179659187 | 179780996 |
| chr7 | 72867383 | 7.3E+07 | chr6 | 10399675 | 10465031 |
| chr7 | 73093519 | 7.3E+07 | chr6 | 11179353 | 11234451 |
| chr7 | 73887511 | 7.4E+07 | chr6 | 12279302 | 12372902 |
| chr7 | 75267419 | 7.5E+07 | chr6 | 12460192 | 12496562 |
| chr7 | 75534589 | 7.6E+07 | chr6 | 26019853 | 26068444 |
| chr7 | 75919518 | 7.6E+07 | chr6 | 26086980 | 26129119 |
| chr7 | 80389489 | 8.1E+07 | chr6 | 26148057 | 26208228 |
| chr7 | 92346356 | 9.2E+07 | chr6 | 26272351 | 26337550 |
| chr7 | 95024530 | 9.5E+07 | chr6 | 27082561 | 27126261 |
| chr7 | 97942563 | 9.8E+07 | chr6 | 27759238 | 27808351 |
| chr7 | 1.01E+08 | 1E+08 | chr6 | 36482209 | 36585298 |
| chr7 | 1.01E+08 | 1E+08 | chr6 | 39152499 | 39197240 |
| chr7 | 1.02E+08 | 1E+08 | chr6 | 47071543 | 47138405 |
| chr7 | 1.05E+08 | 1E+08 | chr6 | 56487037 | 56598453 |
| chr7 | 1.08E+08 | 1.1E+08 | chr6 | 64268696 | 64289885 |
| chr7 | 1.08E+08 | 1.1E+08 | chr6 | 72084958 | 72131838 |

| | | | | | |
|---|---|---|---|---|---|
| chr7 | 1.08E+08 | 1.1E+08 | chr6 | 74269995 | 74310570 |
| chr7 | 1.16E+08 | 1.2E+08 | chr6 | 85443376 | 85487265 |
| chr7 | 1.16E+08 | 1.2E+08 | chr6 | 85591253 | 85621774 |
| chr7 | 1.16E+08 | 1.2E+08 | chr6 | 88374544 | 88443308 |
| chr7 | 1.31E+08 | 1.3E+08 | chr6 | 121800043 | 121844327 |
| chr7 | 1.31E+08 | 1.3E+08 | chr6 | 126209645 | 126279784 |
| chr7 | 1.31E+08 | 1.3E+08 | chr6 | 128799446 | 128842876 |
| chr7 | 1.34E+08 | 1.3E+08 | chr6 | 129956544 | 130038874 |
| chr7 | 1.34E+08 | 1.3E+08 | chr6 | 133888235 | 133969358 |
| chr7 | 1.39E+08 | 1.4E+08 | chr6 | 136708238 | 136769137 |
| chr7 | 1.51E+08 | 1.5E+08 | chr6 | 142600288 | 142710789 |
| chr7 | 1.59E+08 | 1.6E+08 | chr6 | 151325128 | 151414054 |
| chr8 | 8144823 | 8245217 | chr6 | 158395456 | 158490755 |
| chr8 | 8691352 | 8751853 | chr6 | 159184168 | 159241325 |
| chr8 | 17612333 | 1.8E+07 | chr6 | 159268140 | 159292715 |
| chr8 | 17742896 | 1.8E+07 | chr6 | 160768351 | 160828280 |
| chr8 | 19231137 | 2E+07 | chr7 | 1543152 | 1611574 |
| chr8 | 22406634 | 2.2E+07 | chr7 | 4721176 | 4791677 |
| chr8 | 23145602 | 2.3E+07 | chr7 | 5534608 | 5573105 |
| chr8 | 23376009 | 2.3E+07 | chr7 | 6258743 | 6348754 |
| chr8 | 25027512 | 2.5E+07 | chr7 | 6406811 | 6494113 |
| chr8 | 26427536 | 2.7E+07 | chr7 | 11624053 | 11711065 |
| chr8 | 27720408 | 2.8E+07 | chr7 | 11723817 | 11874265 |
| chr8 | 29188303 | 2.9E+07 | chr7 | 16830768 | 16866344 |
| chr8 | 29347993 | 2.9E+07 | chr7 | 16920268 | 16976073 |
| chr8 | 29409947 | 2.9E+07 | chr7 | 27120369 | 27224521 |
| chr8 | 29444955 | 3E+07 | chr7 | 27774201 | 27834148 |
| chr8 | 29563137 | 3E+07 | chr7 | 28068717 | 28117357 |
| chr8 | 30240017 | 3E+07 | chr7 | 36426276 | 36489460 |
| chr8 | 30358026 | 3E+07 | chr7 | 43660336 | 43715646 |
| chr8 | 30532067 | 3.1E+07 | chr7 | 47292993 | 47731428 |
| chr8 | 32160403 | 3.2E+07 | chr7 | 47797097 | 47832151 |
| chr8 | 32402709 | 3.3E+07 | chr7 | 55086047 | 55181839 |
| chr8 | 38852583 | 3.9E+07 | chr7 | 55195079 | 55275490 |
| chr8 | 40009246 | 4E+07 | chr7 | 73096846 | 73158527 |
| chr8 | 48252750 | 4.8E+07 | chr7 | 75216989 | 75322853 |
| chr8 | 48419378 | 4.8E+07 | chr7 | 75534924 | 75623892 |
| chr8 | 48718272 | 4.9E+07 | chr7 | 75919616 | 76054916 |
| chr8 | 54852850 | 5.5E+07 | chr7 | 80455166 | 80554420 |
| chr8 | 62452520 | 6.3E+07 | chr7 | 92310046 | 92466703 |
| chr8 | 67024693 | 6.7E+07 | chr7 | 93515035 | 93554104 |
| chr8 | 71018870 | 7.1E+07 | chr7 | 95024407 | 95073047 |
| chr8 | 90893625 | 9.1E+07 | chr7 | 97905061 | 98053141 |
| chr8 | 1.02E+08 | 1E+08 | chr7 | 98669821 | 98761627 |
| chr8 | 1.02E+08 | 1E+08 | chr7 | 98794151 | 98828962 |
| chr8 | 1.04E+08 | 1E+08 | chr7 | 100724240 | 100798893 |
| chr8 | 1.19E+08 | 1.2E+08 | chr7 | 101491399 | 101531893 |
| chr8 | 1.29E+08 | 1.3E+08 | chr7 | 101726230 | 101816387 |
| chr8 | 1.29E+08 | 1.3E+08 | chr7 | 104612119 | 104691010 |
| chr8 | 1.31E+08 | 1.3E+08 | chr7 | 107530897 | 107669312 |
| chr8 | 1.31E+08 | 1.3E+08 | chr7 | 111124087 | 111203295 |
| chr8 | 1.42E+08 | 1.4E+08 | chr7 | 116057234 | 116193903 |
| chr8 | 1.42E+08 | 1.4E+08 | chr7 | 116301678 | 116379078 |
| chr8 | 1.42E+08 | 1.4E+08 | chr7 | 116397873 | 116454016 |
| chr8 | 1.42E+08 | 1.4E+08 | chr7 | 130529217 | 130727967 |
| chr8 | 1.45E+08 | 1.4E+08 | chr7 | 130816587 | 130883504 |
| chr8 | 1.45E+08 | 1.5E+08 | chr7 | 130903909 | 130939507 |
| chr9 | 3502540 | 3528299 | chr7 | 131185190 | 131242238 |
| chr9 | 4114544 | 4180803 | chr7 | 134103586 | 134166145 |
| chr9 | 4258421 | 4300090 | chr7 | 134204589 | 134248913 |
| chr9 | 6645344 | 6716398 | chr7 | 137973131 | 138039618 |

| | | | | | |
|---|---|---|---|---|---|
| chr9 | 21516993 | 2.2E+07 | chr7 | 158590875 | 158623511 |
| chr9 | 21675648 | 2.2E+07 | chr8 | 8144098 | 8245317 |
| chr9 | 22123446 | 2.2E+07 | chr8 | 8455999 | 8522831 |
| chr9 | 33101330 | 3.3E+07 | chr8 | 11656602 | 11760560 |
| chr9 | 33225373 | 3.3E+07 | chr8 | 17627550 | 17679445 |
| chr9 | 35507247 | 3.6E+07 | chr8 | 17742973 | 17782981 |
| chr9 | 35707575 | 3.6E+07 | chr8 | 19242629 | 19339080 |
| chr9 | 37414529 | 3.8E+07 | chr8 | 19433194 | 19552118 |
| chr9 | 37900533 | 3.8E+07 | chr8 | 22017143 | 22086170 |
| chr9 | 74289211 | 7.4E+07 | chr8 | 22406096 | 22463734 |
| chr9 | 74906519 | 7.5E+07 | chr8 | 22831700 | 22964330 |
| chr9 | 75139381 | 7.5E+07 | chr8 | 23385352 | 23428400 |
| chr9 | 75458051 | 7.6E+07 | chr8 | 25027964 | 25349920 |
| chr9 | 75591825 | 7.6E+07 | chr8 | 27713268 | 27773698 |
| chr9 | 75636926 | 7.6E+07 | chr8 | 29187869 | 29210610 |
| chr9 | 75732497 | 7.6E+07 | chr8 | 29292313 | 29325199 |
| chr9 | 78979946 | 7.9E+07 | chr8 | 29342543 | 29389633 |
| chr9 | 84143789 | 8.4E+07 | chr8 | 29403131 | 29420826 |
| chr9 | 86534990 | 8.7E+07 | chr8 | 29446315 | 29632884 |
| chr9 | 89984804 | 9E+07 | chr8 | 30240372 | 30433718 |
| chr9 | 96212304 | 9.6E+07 | chr8 | 30538704 | 30602605 |
| chr9 | 97540773 | 9.8E+07 | chr8 | 32403411 | 32459008 |
| chr9 | 97604868 | 9.8E+07 | chr8 | 32558024 | 32594515 |
| chr9 | 97738129 | 9.8E+07 | chr8 | 37683779 | 37774308 |
| chr9 | 1.14E+08 | 1.1E+08 | chr8 | 38198187 | 38270423 |
| chr9 | 1.15E+08 | 1.1E+08 | chr8 | 38852778 | 38922916 |
| chr9 | 1.15E+08 | 1.1E+08 | chr8 | 40009922 | 40074707 |
| chr9 | 1.16E+08 | 1.2E+08 | chr8 | 48223291 | 48297980 |
| chr9 | 1.19E+08 | 1.2E+08 | chr8 | 48421550 | 48457360 |
| chr9 | 1.19E+08 | 1.2E+08 | chr8 | 48753439 | 48906086 |
| chr9 | 1.19E+08 | 1.2E+08 | chr8 | 54854006 | 54986865 |
| chr9 | 1.23E+08 | 1.2E+08 | chr8 | 62494029 | 62705226 |
| chr9 | 1.27E+08 | 1.3E+08 | chr8 | 74204149 | 74277107 |
| chr9 | 1.3E+08 | 1.3E+08 | chr8 | 90893773 | 90938824 |
| chr9 | 1.31E+08 | 1.3E+08 | chr8 | 95416010 | 95488019 |
| chr9 | 1.31E+08 | 1.3E+08 | chr8 | 101312051 | 101364739 |
| chr9 | 1.32E+08 | 1.3E+08 | chr8 | 101469509 | 101525364 |
| chr9 | 1.32E+08 | 1.3E+08 | chr8 | 101911899 | 101967475 |
| chr9 | 1.32E+08 | 1.3E+08 | chr8 | 102339440 | 102399259 |
| chr9 | 1.32E+08 | 1.3E+08 | chr8 | 102446282 | 102487789 |
| chr9 | 1.32E+08 | 1.3E+08 | chr8 | 103799768 | 103823080 |
| chr9 | 1.34E+08 | 1.3E+08 | chr8 | 126271327 | 126322657 |
| chr9 | 1.36E+08 | 1.4E+08 | chr8 | 128210581 | 128238341 |
| chr9 | 1.37E+08 | 1.4E+08 | chr8 | 128805964 | 129141748 |
| chr9 | 1.4E+08 | 1.4E+08 | chr8 | 129163992 | 129223674 |
| chrX | 10503662 | 1.1E+07 | chr8 | 130974884 | 131029768 |
| chrX | 12991858 | 1.3E+07 | chr8 | 141516983 | 141608748 |
| chrX | 23796563 | 2.4E+07 | chr8 | 145007428 | 145065503 |
| chrX | 23889199 | 2.4E+07 | chr9 | 3502350 | 3528782 |
| chrX | 24857100 | 2.5E+07 | chr9 | 4125587 | 4175639 |
| chrX | 30293778 | 3E+07 | chr9 | 4222885 | 4300773 |
| chrX | 45606755 | 4.6E+07 | chr9 | 21466240 | 21617235 |
| chrX | 46175867 | 4.6E+07 | chr9 | 21675040 | 21690822 |
| chrX | 73458691 | 7.4E+07 | chr9 | 22123380 | 22172831 |
| chrX | 1.06E+08 | 1.1E+08 | chr9 | 33116906 | 33185549 |
| chrX | 1.15E+08 | 1.1E+08 | chr9 | 33225369 | 33336245 |
| chrX | 1.34E+08 | 1.3E+08 | chr9 | 33813254 | 33889685 |
| chrX | 1.5E+08 | 1.5E+08 | chr9 | 35488707 | 35554259 |
| | | | chr9 | 36296127 | 36342688 |
| | | | chr9 | 37462880 | 37509066 |
| | | | chr9 | 37902469 | 38071262 |

| | | |
|---|---|---|
| chr9 | 74331380 | 74383874 |
| chr9 | 74906772 | 74941628 |
| chr9 | 74964115 | 74982759 |
| chr9 | 75048138 | 75107077 |
| chr9 | 75128756 | 75208441 |
| chr9 | 75440898 | 75576601 |
| chr9 | 75591811 | 75622667 |
| chr9 | 75635678 | 75710042 |
| chr9 | 75723959 | 75800047 |
| chr9 | 78979315 | 79039500 |
| chr9 | 84127261 | 84180798 |
| chr9 | 84198654 | 84306039 |
| chr9 | 89984377 | 90041003 |
| chr9 | 90111828 | 90342920 |
| chr9 | 91002064 | 91076336 |
| chr9 | 96212017 | 96264507 |
| chr9 | 97542063 | 97596707 |
| chr9 | 97609605 | 97715096 |
| chr9 | 97753049 | 97875879 |
| chr9 | 99139348 | 99184692 |
| chr9 | 111822348 | 111883037 |
| chr9 | 127013645 | 127088511 |
| chr9 | 130266457 | 130347063 |
| chr9 | 130660808 | 130797568 |
| chr9 | 130823395 | 130866894 |
| chr9 | 131375540 | 131429651 |
| chr9 | 131625489 | 131690762 |
| chr9 | 132220231 | 132270648 |
| chr9 | 132310301 | 132405270 |
| chr9 | 132489153 | 132545011 |
| chr9 | 134225422 | 134300954 |
| chr9 | 137249927 | 137305827 |
| chr9 | 139775542 | 139845789 |
| chrX | 10502952 | 10604682 |
| chrX | 12991930 | 13049376 |
| chrX | 23791493 | 23841841 |
| chrX | 23889541 | 23926884 |
| chrX | 24161197 | 24209759 |
| chrX | 24878381 | 24918964 |
| chrX | 45607201 | 45711661 |
| chrX | 46175720 | 46194270 |
| chrX | 93834020 | 93861828 |
| chrX | 114791537 | 114860761 |
| chrX | 115357993 | 115373936 |
| chrX | 115399382 | 115417652 |
| chrX | 118601851 | 118646624 |
| chrX | 123074513 | 123106242 |
| chrX | 150008756 | 150052831 |

**Supplementary Table 4, H3K27ac differential binding analysis, Related to Figure 3A, B**

| chrom | start | end | width | fold | pvalue | FDR |
|---|---|---|---|---|---|---|
| chr12 | 14508751 | 14629463 | 120713 | 3.22 | 0 | 0 |
| chr12 | 15835493 | 15887153 | 51661 | 2.02 | 0 | 0 |
| chr12 | 50594926 | 50721719 | 126794 | 2.66 | 0 | 0 |
| chr12 | 87805678 | 87836759 | 31082 | 5.25 | 0 | 0 |
| chr12 | 110974019 | 111024065 | 50047 | 3.06 | 0 | 0 |
| chr14 | 92198685 | 92247386 | 48702 | 3.62 | 0 | 0 |
| chr15 | 34423960 | 34502357 | 78398 | 3.03 | 0 | 0 |
| chr15 | 59824759 | 59950480 | 125722 | 1.25 | 0 | 0 |
| chr18 | 70912667 | 70964388 | 51722 | 3.65 | 0 | 0 |
| chr2 | 86916056 | 86934041 | 17986 | 3.34 | 0 | 0 |
| chr2 | 121515806 | 121752153 | 236348 | -1.33 | 0 | 0 |
| chr3 | 17028741 | 17058501 | 29761 | 3.66 | 0 | 0 |
| chr7 | 11624053 | 11711065 | 87013 | 2.12 | 0 | 0 |
| chr7 | 111124087 | 111203295 | 79209 | 1.72 | 0 | 0 |
| chr9 | 26240905 | 26265932 | 25028 | 3.96 | 0 | 0 |
| chr9 | 38163563 | 38205487 | 41925 | 3.02 | 0 | 0 |
| chr9 | 119026637 | 119162338 | 135702 | -3.71 | 0 | 0 |
| chr7 | 138102655 | 138168781 | 66127 | 3.85 | 9.11E-308 | 1.49E-304 |
| chr12 | 75337000 | 75360848 | 23849 | 2.5 | 2.47E-299 | 3.83E-296 |
| chr12 | 13015954 | 13083646 | 67693 | 1.24 | 1.09E-282 | 1.61E-279 |
| chr7 | 137973131 | 138039618 | 66488 | 2.65 | 2.68E-278 | 3.76E-275 |
| chr19 | 22003687 | 22038373 | 34687 | 3.11 | 8.15E-275 | 1.09E-271 |
| chr5 | 111541389 | 111576504 | 35116 | 2.11 | 1.06E-261 | 1.35E-258 |
| chr11 | 45273751 | 45311604 | 37854 | 2.32 | 1.33E-253 | 1.63E-250 |
| chr7 | 131185190 | 131242238 | 57049 | 2.15 | 8.79E-253 | 1.04E-249 |
| chr12 | 43350395 | 43362880 | 12486 | 4.04 | 2.26E-252 | 2.56E-249 |
| chr12 | 123346298 | 123383555 | 37258 | 1.93 | 2.45E-250 | 2.67E-247 |
| chr9 | 20601540 | 20623255 | 21716 | 3.01 | 1.14E-245 | 1.20E-242 |
| chr8 | 76318725 | 76340692 | 21968 | 4.35 | 2.92E-242 | 2.97E-239 |
| chr6 | 85443376 | 85487265 | 43890 | 2.01 | 2.31E-241 | 2.27E-238 |
| chr12 | 32108114 | 32163939 | 55826 | 1.93 | 2.52E-239 | 2.39E-236 |
| chr16 | 52579061 | 52613779 | 34719 | 2.73 | 9.73E-237 | 8.96E-234 |
| chr3 | 14443317 | 14532444 | 89128 | -1.33 | 2.98E-227 | 2.66E-224 |
| chr12 | 105702855 | 105752037 | 49183 | 2 | 6.31E-226 | 5.47E-223 |
| chr2 | 228189125 | 228281915 | 92791 | -1.41 | 8.60E-226 | 7.24E-223 |
| chr10 | 116263676 | 116287351 | 23676 | 2.28 | 1.92E-217 | 1.57E-214 |
| chr11 | 75012938 | 75141865 | 128928 | 1.25 | 1.99E-216 | 1.58E-213 |
| chr19 | 39105670 | 39227409 | 121740 | 0.74 | 7.16E-215 | 5.55E-212 |
| chr11 | 1041284 | 1078511 | 37228 | 1.88 | 4.30E-212 | 3.25E-209 |
| chr9 | 89984377 | 90041003 | 56627 | 1.35 | 4.91E-210 | 3.61E-207 |
| chr12 | 90517512 | 90525789 | 8278 | 2.52 | 7.83E-208 | 5.63E-205 |
| chr12 | 105338392 | 105354000 | 15609 | 2.02 | 5.43E-207 | 3.81E-204 |
| chr12 | 122177888 | 122278774 | 100887 | 1.11 | 1.25E-205 | 8.53E-203 |
| chr9 | 75635678 | 75710042 | 74365 | 1.07 | 6.33E-200 | 4.24E-197 |

| | | | | | | |
|---|---|---|---|---|---|---|
| chr7 | 11723817 | 11874265 | 150449 | 1.23 | 3.43E-197 | 2.25E-194 |
| chr12 | 15773675 | 15790463 | 16789 | 1.85 | 2.09E-193 | 1.34E-190 |
| chr5 | 32752854 | 32786163 | 33310 | -2.62 | 3.46E-193 | 2.17E-190 |
| chr2 | 46858939 | 46883570 | 24632 | -1.79 | 5.76E-190 | 3.53E-187 |
| chr12 | 84190091 | 84213491 | 23401 | 3.29 | 6.05E-190 | 3.63E-187 |
| chr1 | 177992350 | 178025835 | 33486 | 2.56 | 3.19E-187 | 1.88E-184 |
| chr12 | 31856839 | 31933500 | 76662 | 1.39 | 3.60E-186 | 2.08E-183 |
| chr3 | 193829031 | 193860958 | 31928 | 2.15 | 4.17E-186 | 2.36E-183 |
| chr22 | 49764436 | 49784567 | 20132 | 3.97 | 7.30E-186 | 4.06E-183 |
| chr10 | 112599043 | 112652204 | 53162 | 2.05 | 1.80E-182 | 9.83E-180 |
| chr8 | 24194185 | 24209208 | 15024 | 4.31 | 1.96E-182 | 1.05E-179 |
| chr4 | 106465332 | 106475633 | 10302 | 4.63 | 2.62E-182 | 1.38E-179 |
| chr1 | 84711788 | 84770013 | 58226 | 1.87 | 1.60E-181 | 8.29E-179 |
| chr12 | 95461278 | 95734708 | 273431 | 0.73 | 7.32E-181 | 3.71E-178 |
| chr12 | 47426503 | 47436147 | 9645 | 1.74 | 6.82E-179 | 3.41E-176 |
| chr11 | 16902856 | 16948724 | 45869 | 3.94 | 2.81E-178 | 1.38E-175 |
| chr10 | 30798588 | 30848949 | 50362 | -2.51 | 3.67E-176 | 1.77E-173 |
| chr12 | 62972724 | 63060860 | 88137 | 1.07 | 2.14E-172 | 1.02E-169 |
| chr3 | 161124230 | 161139758 | 15529 | 2.88 | 4.32E-170 | 2.02E-167 |
| chr8 | 128289209 | 128313245 | 24037 | 1.99 | 6.19E-169 | 2.85E-166 |
| chr12 | 47463349 | 47500110 | 36762 | 1 | 1.47E-168 | 6.68E-166 |
| chr18 | 52667076 | 52699859 | 32784 | 3.76 | 4.40E-167 | 1.97E-164 |
| chr12 | 13521574 | 13544500 | 22927 | 3.67 | 6.72E-166 | 2.95E-163 |
| chr12 | 89884325 | 89920794 | 36470 | 1.51 | 1.95E-165 | 8.44E-163 |
| chr19 | 54023240 | 54083313 | 60074 | 2.05 | 2.18E-165 | 9.32E-163 |
| chr9 | 21466240 | 21617400 | 151161 | 0.98 | 2.14E-162 | 9.01E-160 |
| chr13 | 67757035 | 67809375 | 52341 | 1.3 | 2.66E-160 | 1.10E-157 |
| chr8 | 81810279 | 81814299 | 4021 | 3.38 | 1.88E-158 | 7.68E-156 |
| chr5 | 71402400 | 71479301 | 76902 | -1.73 | 4.59E-156 | 1.85E-153 |
| chr7 | 20389558 | 20396836 | 7279 | 1.79 | 2.85E-154 | 1.13E-151 |
| chr12 | 112442970 | 112592297 | 149328 | 0.97 | 3.50E-154 | 1.38E-151 |
| chr2 | 234597302 | 234620659 | 23358 | -1.4 | 7.65E-153 | 2.96E-150 |
| chr12 | 20119493 | 20126488 | 6996 | 2.86 | 4.46E-152 | 1.70E-149 |
| chr9 | 90111828 | 90344636 | 232809 | 0.86 | 9.17E-152 | 3.46E-149 |
| chr14 | 54306291 | 54333147 | 26857 | 1.72 | 3.83E-151 | 1.43E-148 |
| chr21 | 30539992 | 30591090 | 51099 | 1.72 | 1.68E-150 | 6.18E-148 |
| chr11 | 100605436 | 100663607 | 58172 | 1.38 | 2.57E-150 | 9.34E-148 |
| chr1 | 51424601 | 51451782 | 27182 | 1.47 | 1.58E-149 | 5.67E-147 |
| chr12 | 12931954 | 13001053 | 69100 | 1.07 | 4.79E-148 | 1.70E-145 |
| chr12 | 41082176 | 41123105 | 40930 | 1.48 | 5.20E-148 | 1.82E-145 |
| chr5 | 140934829 | 140946214 | 11386 | 2.88 | 6.60E-148 | 2.29E-145 |
| chr1 | 150626581 | 150670983 | 44403 | 1.71 | 8.86E-143 | 3.04E-140 |
| chr6 | 18488566 | 18504422 | 15857 | 2.54 | 1.15E-141 | 3.90E-139 |
| chr18 | 52743713 | 52771481 | 27769 | 1.9 | 1.42E-141 | 4.74E-139 |
| chr11 | 117657004 | 117704942 | 47939 | 1.92 | 3.65E-141 | 1.21E-138 |
| chr2 | 189143981 | 189189323 | 45343 | 1.36 | 1.44E-140 | 4.70E-138 |
| chr20 | 58495136 | 58516962 | 21827 | 2.93 | 8.08E-139 | 2.62E-136 |
| chr17 | 15789958 | 15828650 | 38693 | 1.55 | 1.51E-138 | 4.84E-136 |
| chr4 | 143323658 | 143338619 | 14962 | 1.87 | 1.04E-135 | 3.30E-133 |
| chr12 | 83070310 | 83082533 | 12224 | 2.11 | 2.49E-135 | 7.79E-133 |
| chr8 | 72740518 | 72757041 | 16524 | -3.21 | 3.00E-135 | 9.30E-133 |
| chr10 | 79313754 | 79365657 | 51904 | -1.73 | 3.43E-135 | 1.05E-132 |
| chr10 | 29171864 | 29211726 | 39863 | 2.13 | 1.70E-133 | 5.18E-131 |
| chr20 | 36678045 | 36816165 | 138121 | 1.04 | 7.07E-133 | 2.12E-130 |
| chr5 | 1155098 | 1174632 | 19535 | 5 | 3.83E-132 | 1.14E-129 |
| chr5 | 175227100 | 175317570 | 90471 | -1.11 | 7.27E-132 | 2.14E-129 |
| chr12 | 79908021 | 79967015 | 58995 | 1.24 | 7.55E-132 | 2.20E-129 |
| chr7 | 137869285 | 137874604 | 5320 | 5.43 | 5.73E-131 | 1.66E-128 |
| chr17 | 73584750 | 73785345 | 200596 | 0.7 | 6.50E-131 | 1.86E-128 |

| | | | | | | |
|---|---|---|---|---|---|---|
| chr3 | 30360118 | 30415850 | 55733 | 1.24 | 9.20E-131 | 2.61E-128 |
| chr2 | 26930734 | 27014403 | 83670 | -1.21 | 1.29E-130 | 3.63E-128 |
| chr17 | 74864511 | 74897849 | 33339 | -2.85 | 2.45E-130 | 6.82E-128 |
| chr11 | 63277361 | 63313583 | 36223 | 1.67 | 2.22E-129 | 6.12E-127 |
| chr17 | 36603537 | 36624972 | 21436 | 1.89 | 7.72E-129 | 2.11E-126 |
| chr10 | 126662301 | 126851531 | 189231 | 0.71 | 5.17E-128 | 1.40E-125 |
| chrX | 106160283 | 106203160 | 42878 | -1.92 | 5.73E-128 | 1.53E-125 |
| chr2 | 47166410 | 47317390 | 150981 | -0.73 | 3.94E-127 | 1.04E-124 |
| chr8 | 30240017 | 30438997 | 198981 | 0.88 | 1.06E-126 | 2.78E-124 |
| chr12 | 12848572 | 12880189 | 31618 | 1.19 | 1.40E-126 | 3.65E-124 |
| chr15 | 62399340 | 62417884 | 18545 | 2.38 | 2.08E-126 | 5.37E-124 |
| chr3 | 146182402 | 146190665 | 8264 | 2.72 | 5.16E-126 | 1.32E-123 |
| chr3 | 114615892 | 114624516 | 8625 | 1.92 | 7.83E-126 | 1.99E-123 |
| chr2 | 109585005 | 109599718 | 14714 | 4.29 | 1.23E-125 | 3.09E-123 |
| chr2 | 30885726 | 30894555 | 8830 | 2.43 | 1.91E-125 | 4.77E-123 |
| chr12 | 86157269 | 86178247 | 20979 | 3.01 | 2.19E-125 | 5.41E-123 |
| chr7 | 94326234 | 94332894 | 6661 | 2.38 | 2.68E-123 | 6.59E-121 |
| chr12 | 130554819 | 130589725 | 34907 | 6.43 | 9.28E-123 | 2.26E-120 |
| chr20 | 47883574 | 47909864 | 26291 | 1.33 | 2.45E-122 | 5.90E-120 |
| chr10 | 26757380 | 26796886 | 39507 | -2.06 | 2.46E-122 | 5.90E-120 |
| chr10 | 95046712 | 95276079 | 229368 | 0.6 | 2.58E-122 | 6.12E-120 |
| chr5 | 29779175 | 29801629 | 22455 | 2.99 | 4.48E-121 | 1.05E-118 |
| chr8 | 23145309 | 23275449 | 130141 | -1.07 | 6.28E-121 | 1.47E-118 |
| chr5 | 129829189 | 129839097 | 9909 | 3.45 | 9.48E-120 | 2.20E-117 |
| chr14 | 38051671 | 38070326 | 18656 | 1.72 | 2.52E-119 | 5.79E-117 |
| chr14 | 56240286 | 56282183 | 41898 | 2.3 | 3.24E-119 | 7.41E-117 |
| chr5 | 37271563 | 37276903 | 5341 | 2.3 | 2.53E-117 | 5.74E-115 |
| chr5 | 42982060 | 43021416 | 39357 | 1.12 | 3.46E-117 | 7.77E-115 |
| chr5 | 142149059 | 142260991 | 111933 | 0.97 | 4.87E-117 | 1.09E-114 |
| chr10 | 93347301 | 93377689 | 30389 | 1.06 | 9.31E-117 | 2.06E-114 |
| chr16 | 66535966 | 66559654 | 23689 | 1.11 | 1.03E-115 | 2.26E-113 |
| chr1 | 207165964 | 207227223 | 61260 | 1.05 | 2.17E-115 | 4.73E-113 |
| chr1 | 228113849 | 228136837 | 22989 | 2.82 | 2.30E-115 | 4.97E-113 |
| chr5 | 66365970 | 66393237 | 27268 | 2.27 | 2.90E-115 | 6.24E-113 |
| chr1 | 200040121 | 200099883 | 59763 | 1.89 | 4.27E-115 | 9.11E-113 |
| chr21 | 41001954 | 41060430 | 58477 | 1.67 | 4.92E-115 | 1.04E-112 |
| chr12 | 53921038 | 53970820 | 49783 | 1.43 | 1.51E-114 | 3.18E-112 |
| chr12 | 95998692 | 96063505 | 64814 | 1.52 | 4.13E-114 | 8.62E-112 |
| chr6 | 128799446 | 128842876 | 43431 | 1.43 | 4.18E-114 | 8.66E-112 |
| chr12 | 78239572 | 78257626 | 18055 | 4.96 | 5.27E-114 | 1.08E-111 |
| chr1 | 80998452 | 81015755 | 17304 | 2.32 | 2.22E-113 | 4.54E-111 |
| chr2 | 161195310 | 161275877 | 80568 | -1.13 | 1.35E-111 | 2.74E-109 |
| chr13 | 73989968 | 74041289 | 51322 | 1.83 | 4.39E-111 | 8.85E-109 |
| chr20 | 60903810 | 60964836 | 61027 | 0.96 | 3.91E-110 | 7.83E-108 |
| chr3 | 146219041 | 146268352 | 49312 | 1.07 | 5.28E-110 | 1.05E-107 |
| chr3 | 78960975 | 78969011 | 8037 | 3.72 | 8.29E-110 | 1.64E-107 |
| chrX | 4579986 | 4594719 | 14734 | 2.2 | 8.42E-110 | 1.65E-107 |
| chr12 | 53253080 | 53473985 | 220906 | 0.89 | 2.76E-109 | 5.39E-107 |
| chr11 | 12131388 | 12205276 | 73889 | -1.61 | 3.33E-109 | 6.45E-107 |
| chr8 | 109953603 | 109963286 | 9684 | -2.96 | 4.78E-109 | 9.20E-107 |
| chr14 | 55561020 | 55576900 | 15881 | 1.33 | 2.15E-108 | 4.11E-106 |
| chr1 | 115000142 | 115019327 | 19186 | 2.5 | 1.24E-107 | 2.36E-105 |
| chr9 | 21675040 | 21690822 | 15783 | 0.94 | 3.71E-107 | 7.00E-105 |
| chr8 | 8931252 | 8960967 | 29716 | 1.74 | 3.76E-107 | 7.05E-105 |
| chr14 | 90966912 | 90989399 | 22488 | 2.06 | 1.17E-106 | 2.17E-104 |
| chr20 | 746455 | 750205 | 3751 | 4.24 | 2.20E-106 | 4.08E-104 |
| chr5 | 140697406 | 140735067 | 37662 | 3.08 | 1.43E-105 | 2.64E-103 |
| chr2 | 208528452 | 208616026 | 87575 | -1.32 | 4.97E-105 | 9.10E-103 |
| chrX | 35452263 | 35486094 | 33832 | 2.79 | 9.20E-105 | 1.67E-102 |
| chr4 | 77608902 | 77646566 | 37665 | 1.23 | 7.08E-104 | 1.28E-101 |
| chr12 | 100920710 | 100924524 | 3815 | 3.06 | 2.04E-102 | 3.67E-100 |

| | | | | | | |
|---|---|---|---|---|---|---|
| chr14 | 95697852 | 95740338 | 42487 | 1.35 | 3.95E-102 | 7.04E-100 |
| chr11 | 113533780 | 113540241 | 6462 | 3.47 | 3.98E-102 | 7.06E-100 |
| chr9 | 97738129 | 97883827 | 145699 | 0.63 | 1.86E-101 | 3.29E-99 |
| chr15 | 52541693 | 52588933 | 47241 | 0.99 | 3.85E-101 | 6.74E-99 |
| chr12 | 51155737 | 51185976 | 30240 | 1.19 | 6.98E-101 | 1.22E-98 |
| chr22 | 25496844 | 25534358 | 37515 | 1.55 | 1.02E-100 | 1.77E-98 |
| chr5 | 90171833 | 90237103 | 65271 | 3.77 | 1.20E-100 | 2.06E-98 |
| chr5 | 169009893 | 169028041 | 18149 | 1.41 | 1.75E-100 | 2.99E-98 |
| chr9 | 104074242 | 104091638 | 17397 | 1.61 | 3.20E-100 | 5.45E-98 |
| chr1 | 156348924 | 156375614 | 26691 | 1.94 | 1.81E-99 | 3.07E-97 |
| chr10 | 105631789 | 105679366 | 47578 | 1.08 | 1.83E-99 | 3.08E-97 |
| chr2 | 145764180 | 145794719 | 30540 | -2.08 | 9.10E-99 | 1.52E-96 |
| chr2 | 175190609 | 175209828 | 19220 | 2.82 | 2.17E-98 | 3.61E-96 |
| chr17 | 70382128 | 70478237 | 96110 | 0.6 | 3.97E-98 | 6.57E-96 |
| chr17 | 55498772 | 55549839 | 51068 | -2.2 | 8.15E-98 | 1.34E-95 |
| chr1 | 98490955 | 98532942 | 41988 | 1.23 | 1.10E-97 | 1.80E-95 |
| chr2 | 235859744 | 235965263 | 105520 | -1.26 | 3.72E-97 | 6.05E-95 |
| chr12 | 56910033 | 57040753 | 130721 | 0.81 | 1.04E-96 | 1.68E-94 |
| chr13 | 67642518 | 67725357 | 82840 | 0.77 | 1.19E-96 | 1.91E-94 |
| chr4 | 160194912 | 160220935 | 26024 | 2.88 | 1.50E-96 | 2.40E-94 |
| chr1 | 117687834 | 117710742 | 22909 | 2.13 | 1.63E-95 | 2.60E-93 |
| chr2 | 192015019 | 192045849 | 30831 | -2.48 | 4.84E-95 | 7.66E-93 |
| chr8 | 101469509 | 101525364 | 55856 | 1.11 | 7.38E-95 | 1.16E-92 |
| chr12 | 125185140 | 125254348 | 69209 | 1.53 | 7.98E-95 | 1.25E-92 |
| chr5 | 33300299 | 33314693 | 14395 | 3.77 | 1.02E-94 | 1.59E-92 |
| chr14 | 103697992 | 103768594 | 70603 | 1.61 | 1.93E-94 | 2.99E-92 |
| chrX | 107541880 | 107557590 | 15711 | -2.71 | 3.60E-94 | 5.55E-92 |
| chr12 | 110537659 | 110603049 | 65391 | 0.95 | 9.45E-94 | 1.45E-91 |
| chrX | 22114154 | 22156597 | 42444 | 1.77 | 1.84E-93 | 2.81E-91 |
| chr6 | 85683548 | 85690128 | 6581 | 2.65 | 2.50E-93 | 3.80E-91 |
| chr12 | 98883738 | 98942922 | 59185 | 1.06 | 1.09E-92 | 1.64E-90 |
| chr7 | 36426276 | 36489460 | 63185 | 1.21 | 1.26E-92 | 1.89E-90 |
| chr22 | 37888974 | 37960821 | 71848 | 1.07 | 1.29E-92 | 1.93E-90 |
| chr10 | 4385848 | 4401555 | 15708 | 1.93 | 2.28E-92 | 3.39E-90 |
| chr12 | 63077029 | 63195151 | 118123 | 0.67 | 3.57E-92 | 5.29E-90 |
| chr12 | 15339308 | 15348222 | 8915 | 5.24 | 4.02E-92 | 5.93E-90 |
| chr8 | 29115652 | 29148422 | 32771 | 1.47 | 5.24E-92 | 7.67E-90 |
| chr18 | 44447521 | 44459580 | 12060 | 3.17 | 6.49E-92 | 9.46E-90 |
| chr13 | 80283200 | 80301665 | 18466 | 1.82 | 9.04E-92 | 1.31E-89 |
| chr13 | 106092939 | 106099178 | 6240 | 2.99 | 3.16E-91 | 4.56E-89 |
| chr10 | 46965284 | 47151348 | 186065 | -0.55 | 3.84E-91 | 5.52E-89 |
| chr12 | 113270292 | 113281143 | 10852 | 1.19 | 4.34E-91 | 6.20E-89 |
| chr2 | 75464321 | 75469138 | 4818 | -1.98 | 7.03E-91 | 1.00E-88 |
| chr2 | 233699954 | 233767742 | 67789 | -1.27 | 7.28E-91 | 1.03E-88 |
| chr19 | 38268803 | 38282748 | 13946 | 4.04 | 1.06E-90 | 1.49E-88 |
| chr18 | 28972676 | 28984260 | 11585 | 2.74 | 1.59E-90 | 2.24E-88 |
| chr15 | 72020752 | 72107256 | 86505 | -0.89 | 1.85E-90 | 2.58E-88 |
| chr18 | 67953891 | 68054695 | 100805 | 0.76 | 2.16E-90 | 3.00E-88 |
| chr6 | 52976424 | 52987495 | 11072 | 3.26 | 8.77E-90 | 1.21E-87 |
| chr9 | 75723959 | 75800214 | 76256 | 0.65 | 4.85E-89 | 6.68E-87 |
| chr8 | 32402709 | 32628986 | 226278 | -0.59 | 1.38E-88 | 1.90E-86 |
| chr20 | 62819123 | 62822647 | 3525 | 4.06 | 2.36E-88 | 3.22E-86 |
| chr12 | 22776395 | 22785172 | 8778 | 1.14 | 3.53E-88 | 4.80E-86 |
| chr7 | 16920268 | 16976073 | 55806 | 1.27 | 4.97E-88 | 6.71E-86 |
| chr12 | 22729231 | 22753193 | 23963 | 1.42 | 9.63E-88 | 1.29E-85 |
| chr12 | 6469686 | 6542510 | 72825 | 0.68 | 1.20E-87 | 1.61E-85 |
| chr12 | 119613766 | 119628633 | 14868 | 2.22 | 3.56E-87 | 4.74E-85 |
| chr5 | 174803684 | 174814566 | 10883 | -3.03 | 3.80E-87 | 5.04E-85 |
| chr15 | 101458403 | 101553117 | 94715 | -0.97 | 6.32E-87 | 8.34E-85 |
| chr4 | 40181194 | 40203117 | 21924 | 1.7 | 8.81E-87 | 1.16E-84 |
| chr20 | 30248315 | 30370275 | 121961 | 0.54 | 8.97E-87 | 1.17E-84 |

| | | | | | | |
|---|---|---|---|---|---|---|
| chr6 | 2492515 | 2519343 | 26829 | 1.78 | 1.95E-86 | 2.54E-84 |
| chr4 | 56192345 | 56226045 | 33701 | 2.82 | 2.27E-86 | 2.95E-84 |
| chr7 | 11489615 | 11509773 | 20159 | 3.04 | 2.65E-86 | 3.43E-84 |
| chr12 | 86196956 | 86231333 | 34378 | 1.5 | 2.75E-86 | 3.54E-84 |
| chr12 | 78683837 | 78689273 | 5437 | 1.83 | 5.88E-86 | 7.53E-84 |
| chr2 | 134081226 | 134086963 | 5738 | -2.87 | 9.26E-86 | 1.18E-83 |
| chr1 | 61581493 | 61622688 | 41196 | 1.41 | 1.11E-85 | 1.40E-83 |
| chr10 | 80826359 | 81010399 | 184041 | -0.57 | 1.62E-85 | 2.05E-83 |
| chr19 | 58681447 | 58696317 | 14871 | 2.46 | 2.10E-85 | 2.65E-83 |
| chr14 | 32886048 | 32889426 | 3379 | 3.51 | 2.32E-85 | 2.90E-83 |
| chr1 | 199985233 | 200023807 | 38575 | 1.55 | 2.98E-85 | 3.72E-83 |
| chr8 | 128210581 | 128238341 | 27761 | 1.42 | 3.21E-85 | 3.99E-83 |
| chr2 | 174012803 | 174060562 | 47760 | -1.75 | 3.49E-85 | 4.32E-83 |
| chr7 | 106015924 | 106020948 | 5025 | -2.84 | 4.75E-85 | 5.86E-83 |
| chr9 | 130909903 | 130922353 | 12451 | 2.17 | 7.13E-85 | 8.75E-83 |
| chr1 | 224678810 | 224692228 | 13419 | 2.91 | 1.04E-84 | 1.28E-82 |
| chr3 | 70297437 | 70333377 | 35941 | 1.06 | 2.77E-84 | 3.37E-82 |
| chr11 | 108456590 | 108489262 | 32673 | 3.2 | 2.88E-84 | 3.49E-82 |
| chr10 | 115703697 | 115720989 | 17293 | 1.85 | 3.11E-84 | 3.76E-82 |
| chr12 | 1867451 | 1877412 | 9962 | 3.03 | 5.73E-84 | 6.89E-82 |
| chr12 | 110630326 | 110842616 | 212291 | 0.64 | 7.15E-84 | 8.56E-82 |
| chr15 | 77355548 | 77363990 | 8443 | 1.49 | 1.02E-83 | 1.22E-81 |
| chr9 | 75440898 | 75576601 | 135704 | 0.68 | 1.21E-83 | 1.44E-81 |
| chr15 | 69883524 | 69897063 | 13540 | 1.93 | 1.67E-83 | 1.97E-81 |
| chr9 | 75591811 | 75622667 | 30857 | 0.72 | 4.04E-83 | 4.76E-81 |
| chr3 | 160108471 | 160169403 | 60933 | 0.94 | 4.08E-83 | 4.78E-81 |
| chr17 | 26795588 | 26816652 | 21065 | 4 | 4.65E-83 | 5.43E-81 |
| chr12 | 10017422 | 10023311 | 5890 | 1.92 | 6.39E-83 | 7.43E-81 |
| chr12 | 131695226 | 131752875 | 57650 | 1.33 | 4.43E-82 | 5.14E-80 |
| chr7 | 16830768 | 16866344 | 35577 | 0.91 | 9.20E-82 | 1.06E-79 |
| chr5 | 158584015 | 158639457 | 55443 | 0.67 | 2.39E-81 | 2.75E-79 |
| chr4 | 15653186 | 15704984 | 51799 | 1 | 9.27E-81 | 1.06E-78 |
| chr15 | 37076254 | 37111874 | 35621 | 1.83 | 1.87E-80 | 2.13E-78 |
| chr5 | 115170300 | 115194466 | 24167 | 1.91 | 2.44E-80 | 2.78E-78 |
| chr12 | 74785260 | 74803103 | 17844 | 1.84 | 4.34E-80 | 4.91E-78 |
| chr19 | 7134010 | 7147663 | 13654 | 2.68 | 6.36E-80 | 7.17E-78 |
| chr16 | 48150268 | 48168794 | 18527 | -1.55 | 1.66E-79 | 1.87E-77 |
| chr15 | 69590281 | 69761814 | 171534 | -0.92 | 2.68E-79 | 3.00E-77 |
| chr9 | 131625489 | 131690762 | 65274 | 0.83 | 2.70E-79 | 3.01E-77 |
| chr6 | 4486405 | 4491423 | 5019 | 4.89 | 2.94E-79 | 3.27E-77 |
| chr7 | 16122282 | 16131860 | 9579 | 2.82 | 4.33E-79 | 4.80E-77 |
| chrX | 9116172 | 9141089 | 24918 | 1.64 | 6.39E-79 | 7.05E-77 |
| chr5 | 67690645 | 67731729 | 41085 | 1.69 | 6.84E-79 | 7.51E-77 |
| chr3 | 53511450 | 53522026 | 10577 | 2.96 | 2.56E-78 | 2.80E-76 |
| chr3 | 155369338 | 155394829 | 25492 | 1.66 | 2.72E-78 | 2.96E-76 |
| chr12 | 88195122 | 88222904 | 27783 | 1.84 | 3.60E-78 | 3.91E-76 |
| chr7 | 24858507 | 24873446 | 14940 | 1.27 | 4.00E-78 | 4.33E-76 |
| chr12 | 21786918 | 21828953 | 42036 | 1.01 | 5.26E-78 | 5.68E-76 |
| chr2 | 213765607 | 213784739 | 19133 | 1.75 | 1.06E-77 | 1.14E-75 |
| chr12 | 64236575 | 64384203 | 147629 | 0.59 | 1.38E-77 | 1.47E-75 |
| chr2 | 64331198 | 64344144 | 12947 | 2.02 | 4.56E-77 | 4.86E-75 |
| chr20 | 32365587 | 32385288 | 19702 | 1.76 | 4.87E-77 | 5.17E-75 |
| chr7 | 141729958 | 141739090 | 9133 | 2.07 | 5.59E-77 | 5.92E-75 |
| chr10 | 61882281 | 61909010 | 26730 | 2.15 | 6.47E-77 | 6.83E-75 |
| chr12 | 109007185 | 109130187 | 123003 | 0.65 | 1.50E-76 | 1.57E-74 |
| chr8 | 102768095 | 102775112 | 7018 | 3.09 | 1.80E-76 | 1.89E-74 |
| chr10 | 43799635 | 43858813 | 59179 | 1.01 | 1.87E-76 | 1.95E-74 |
| chr4 | 125347075 | 125359898 | 12824 | 1.43 | 2.29E-76 | 2.39E-74 |
| chr3 | 129541252 | 129656082 | 114831 | 1.01 | 8.17E-76 | 8.48E-74 |
| chr4 | 151582446 | 151601175 | 18730 | 2.99 | 1.50E-75 | 1.55E-73 |
| chr10 | 95750674 | 95803740 | 53067 | -1.19 | 2.12E-75 | 2.18E-73 |

| | | | | | | |
|---|---|---|---|---|---|---|
| chr15 | 55650269 | 55702920 | 52652 | 0.99 | 2.74E-75 | 2.81E-73 |
| chr8 | 101312051 | 101364739 | 52689 | 1.16 | 4.09E-75 | 4.18E-73 |
| chr15 | 93340059 | 93572415 | 232357 | 0.64 | 4.47E-75 | 4.56E-73 |
| chr2 | 235197857 | 235227530 | 29674 | -1.38 | 5.27E-75 | 5.35E-73 |
| chr3 | 194013966 | 194041853 | 27888 | 1.59 | 9.47E-75 | 9.59E-73 |
| chr12 | 110253215 | 110279614 | 26400 | 2.21 | 9.58E-75 | 9.66E-73 |
| chr10 | 79260961 | 79294204 | 33244 | -2.1 | 1.15E-74 | 1.16E-72 |
| chr2 | 46917640 | 46977578 | 59939 | -1.21 | 1.17E-74 | 1.17E-72 |
| chr9 | 78979315 | 79039500 | 60186 | 0.77 | 1.53E-74 | 1.53E-72 |
| chr17 | 53471855 | 53537665 | 65811 | -1.04 | 2.18E-74 | 2.17E-72 |
| chr14 | 75990147 | 76009538 | 19392 | -1.68 | 2.60E-74 | 2.58E-72 |
| chr18 | 29945498 | 29963791 | 18294 | 1.82 | 2.86E-74 | 2.82E-72 |
| chr14 | 70109603 | 70134479 | 24877 | 1.58 | 4.03E-74 | 3.97E-72 |
| chr5 | 174012973 | 174040631 | 27659 | 2.18 | 4.98E-74 | 4.89E-72 |
| chr12 | 122580659 | 122615989 | 35331 | 0.97 | 1.41E-73 | 1.38E-71 |
| chr10 | 71093922 | 71152919 | 58998 | -2.43 | 1.95E-73 | 1.90E-71 |
| chrX | 69696012 | 69701484 | 5473 | 2.44 | 3.77E-73 | 3.66E-71 |
| chr12 | 79449343 | 79458078 | 8736 | 1.8 | 4.72E-73 | 4.58E-71 |
| chr3 | 186459913 | 186472062 | 12150 | 1.36 | 4.97E-73 | 4.80E-71 |
| chr8 | 142237532 | 142318426 | 80895 | -1 | 1.06E-72 | 1.02E-70 |
| chr19 | 44143474 | 44210002 | 66529 | 0.94 | 2.50E-72 | 2.40E-70 |
| chr3 | 112337466 | 112375478 | 38013 | -1.49 | 2.78E-72 | 2.66E-70 |
| chr2 | 234990319 | 235001590 | 11272 | 3.01 | 4.73E-72 | 4.51E-70 |
| chr12 | 113611039 | 113710430 | 99392 | 0.93 | 4.96E-72 | 4.71E-70 |
| chr15 | 57834016 | 57854344 | 20329 | 1.4 | 4.97E-72 | 4.71E-70 |
| chr5 | 137151073 | 137174252 | 23180 | 1.81 | 5.22E-72 | 4.92E-70 |
| chr3 | 5350851 | 5392496 | 41646 | 1.87 | 5.73E-72 | 5.39E-70 |
| chr12 | 32196095 | 32222899 | 26805 | 1.58 | 8.41E-72 | 7.89E-70 |
| chr1 | 178193043 | 178238382 | 45340 | 1.2 | 9.72E-72 | 9.07E-70 |
| chr17 | 14601070 | 14605104 | 4035 | 1.71 | 9.73E-72 | 9.07E-70 |
| chr12 | 105671984 | 105689598 | 17615 | 2.19 | 1.15E-71 | 1.07E-69 |
| chr2 | 67787160 | 67802458 | 15299 | 2.06 | 1.34E-71 | 1.24E-69 |
| chr6 | 131371642 | 131385210 | 13569 | 1.75 | 1.88E-71 | 1.74E-69 |
| chr12 | 66285256 | 66297606 | 12351 | 1.29 | 2.07E-71 | 1.91E-69 |
| chr6 | 63999864 | 64018461 | 18598 | 3.55 | 3.14E-71 | 2.89E-69 |
| chr14 | 68972619 | 69075959 | 103341 | 0.89 | 3.78E-71 | 3.46E-69 |
| chr18 | 28804653 | 28857221 | 52569 | 1.55 | 6.04E-71 | 5.50E-69 |
| chr9 | 91001941 | 91076336 | 74396 | 1.04 | 1.05E-70 | 9.52E-69 |
| chr12 | 47575437 | 47676690 | 101254 | 0.58 | 1.36E-70 | 1.23E-68 |
| chr2 | 217179306 | 217184827 | 5522 | -2.5 | 1.49E-70 | 1.34E-68 |
| chr12 | 87901582 | 87913164 | 11583 | 2.37 | 2.10E-70 | 1.89E-68 |
| chr6 | 81346356 | 81350636 | 4281 | 1.54 | 2.16E-70 | 1.94E-68 |
| chr12 | 12680837 | 12717017 | 36181 | 1.24 | 2.40E-70 | 2.15E-68 |
| chr3 | 171509381 | 171620441 | 111061 | 0.74 | 2.40E-70 | 2.15E-68 |
| chr12 | 50897998 | 50992076 | 94079 | 0.89 | 3.62E-70 | 3.22E-68 |
| chr1 | 110350629 | 110372151 | 21523 | -1.97 | 5.39E-70 | 4.78E-68 |
| chr2 | 241365996 | 241407150 | 41155 | -1.66 | 7.38E-70 | 6.53E-68 |
| chr2 | 217929773 | 217932652 | 2880 | 3.91 | 7.48E-70 | 6.60E-68 |
| chr15 | 52121436 | 52201158 | 79723 | 0.53 | 9.82E-70 | 8.63E-68 |
| chr12 | 12029117 | 12045935 | 16819 | 1.75 | 1.03E-69 | 8.99E-68 |
| chr4 | 169698486 | 169787432 | 88947 | -1.08 | 1.07E-69 | 9.33E-68 |
| chr12 | 64197670 | 64210320 | 12651 | 1.71 | 1.25E-69 | 1.09E-67 |
| chr9 | 25676669 | 25679602 | 2934 | 3.75 | 1.33E-69 | 1.15E-67 |
| chr17 | 70177861 | 70195843 | 17983 | 1.73 | 1.95E-69 | 1.69E-67 |
| chr2 | 62986156 | 63054939 | 68784 | -1.4 | 2.10E-69 | 1.81E-67 |
| chr21 | 33732175 | 33766458 | 34284 | 1.38 | 3.68E-69 | 3.17E-67 |
| chr10 | 37766765 | 37784435 | 17671 | 1.63 | 4.07E-69 | 3.50E-67 |
| chr12 | 10514208 | 10550887 | 36680 | 3.15 | 5.02E-69 | 4.30E-67 |
| chr10 | 116363368 | 116410682 | 47315 | 1.41 | 7.46E-69 | 6.37E-67 |
| chr5 | 6898778 | 6907001 | 8224 | 4.78 | 8.69E-69 | 7.40E-67 |
| chr11 | 112187537 | 112193640 | 6104 | 2.59 | 9.46E-69 | 8.03E-67 |

| | | | | | | |
|---|---|---|---|---|---|---|
| chr19 | 35738274 | 35740655 | 2382 | 3.83 | 1.15E-68 | 9.76E-67 |
| chr16 | 71597779 | 71620551 | 22773 | 1.79 | 1.22E-68 | 1.03E-66 |
| chr9 | 91381357 | 91393385 | 12029 | -1.42 | 1.29E-68 | 1.09E-66 |
| chr6 | 125202234 | 125211670 | 9437 | 1.74 | 1.33E-68 | 1.12E-66 |
| chr11 | 86858518 | 86863373 | 4856 | 3.19 | 1.66E-68 | 1.39E-66 |
| chr4 | 57894036 | 57985896 | 91861 | 1.05 | 2.41E-68 | 2.01E-66 |
| chr8 | 576221 | 582429 | 6209 | 3.92 | 3.12E-68 | 2.60E-66 |
| chr2 | 154998508 | 155015331 | 16824 | -2.31 | 4.65E-68 | 3.86E-66 |
| chr10 | 13656381 | 13680727 | 24347 | 3.99 | 5.84E-68 | 4.83E-66 |
| chr1 | 168254656 | 168275403 | 20748 | 1.75 | 6.93E-68 | 5.72E-66 |
| chr3 | 100783575 | 100801587 | 18013 | 1.63 | 7.36E-68 | 6.06E-66 |
| chr2 | 73247476 | 73341088 | 93613 | -1.07 | 8.26E-68 | 6.76E-66 |
| chr14 | 107239154 | 107256589 | 17436 | 1.63 | 8.26E-68 | 6.76E-66 |
| chr1 | 152489036 | 152499339 | 10304 | 5.26 | 7.33E-67 | 5.98E-65 |
| chr16 | 55782478 | 55788656 | 6179 | 1.88 | 9.47E-67 | 7.70E-65 |
| chr6 | 141646557 | 141658967 | 12411 | 1.77 | 1.08E-66 | 8.72E-65 |
| chr17 | 76123719 | 76195956 | 72238 | 1.08 | 2.09E-66 | 1.69E-64 |
| chr14 | 64319054 | 64391226 | 72173 | 1.27 | 2.12E-66 | 1.71E-64 |
| chr19 | 53934533 | 53948981 | 14449 | 3.39 | 3.20E-66 | 2.57E-64 |
| chr12 | 9858663 | 9890444 | 31782 | 1.35 | 3.29E-66 | 2.64E-64 |
| chr7 | 16168323 | 16171529 | 3207 | 2.49 | 7.89E-66 | 6.31E-64 |
| chr12 | 118117811 | 118126614 | 8804 | 1.43 | 8.52E-66 | 6.80E-64 |
| chr19 | 606285 | 634573 | 28289 | 1.48 | 1.06E-65 | 8.47E-64 |
| chr6 | 39152499 | 39197352 | 44854 | 1.14 | 1.14E-65 | 9.07E-64 |
| chr1 | 84637915 | 84643859 | 5945 | 1.75 | 2.04E-65 | 1.62E-63 |
| chr7 | 137923226 | 137942938 | 19713 | 2.8 | 3.34E-65 | 2.63E-63 |
| chr11 | 12064712 | 12112941 | 48230 | -0.96 | 4.51E-65 | 3.55E-63 |
| chr12 | 19171476 | 19180402 | 8927 | 2.49 | 6.81E-65 | 5.35E-63 |
| chr7 | 124097906 | 124101450 | 3545 | 2.99 | 7.51E-65 | 5.89E-63 |
| chr2 | 69239718 | 69246495 | 6778 | -2.61 | 7.89E-65 | 6.17E-63 |
| chr12 | 6386823 | 6455832 | 69010 | 0.84 | 1.18E-64 | 9.23E-63 |
| chr4 | 159730713 | 159748991 | 18279 | 3.29 | 1.25E-64 | 9.71E-63 |
| chr7 | 134198960 | 134250761 | 51802 | -0.77 | 1.26E-64 | 9.73E-63 |
| chr3 | 31306655 | 31331923 | 25269 | 1.6 | 1.38E-64 | 1.07E-62 |
| chr6 | 25172881 | 25176940 | 4060 | 2.87 | 2.04E-64 | 1.58E-62 |
| chr21 | 43121523 | 43148129 | 26607 | 1.46 | 2.05E-64 | 1.58E-62 |
| chr12 | 53758559 | 53903271 | 144713 | 0.7 | 2.21E-64 | 1.70E-62 |
| chr4 | 98414433 | 98421078 | 6646 | 1.83 | 3.00E-64 | 2.29E-62 |
| chr8 | 28882388 | 28892019 | 9632 | 1.66 | 5.46E-64 | 4.17E-62 |
| chr15 | 23717608 | 23731101 | 13494 | 4.51 | 5.79E-64 | 4.40E-62 |
| chr14 | 92324438 | 92342832 | 18395 | 1.41 | 7.43E-64 | 5.64E-62 |
| chr13 | 109596261 | 109610425 | 14165 | 3.48 | 7.67E-64 | 5.81E-62 |
| chr10 | 84323393 | 84325861 | 2469 | 2.58 | 9.74E-64 | 7.36E-62 |
| chrX | 11962775 | 11995410 | 32636 | 1.04 | 1.96E-63 | 1.48E-61 |
| chr20 | 5333869 | 5346578 | 12710 | 2.8 | 2.47E-63 | 1.86E-61 |
| chr2 | 227513764 | 227538508 | 24745 | 1.22 | 2.77E-63 | 2.08E-61 |
| chr8 | 128595558 | 128599968 | 4411 | 2.32 | 2.83E-63 | 2.11E-61 |
| chr1 | 39842104 | 39877795 | 35692 | 0.94 | 2.99E-63 | 2.23E-61 |
| chr2 | 234258991 | 234406019 | 147029 | -0.69 | 3.51E-63 | 2.61E-61 |
| chrX | 12025619 | 12033013 | 7395 | 3.04 | 3.56E-63 | 2.64E-61 |
| chr4 | 158936036 | 158955519 | 19484 | 2.01 | 5.19E-63 | 3.84E-61 |
| chr1 | 172312142 | 172331756 | 19615 | 1.02 | 9.38E-63 | 6.93E-61 |
| chr3 | 87802855 | 87807654 | 4800 | 4.46 | 9.83E-63 | 7.24E-61 |
| chr3 | 174156162 | 174160807 | 4646 | 2.3 | 2.57E-62 | 1.89E-60 |
| chr3 | 114596974 | 114601774 | 4801 | 5.3 | 2.65E-62 | 1.94E-60 |
| chr4 | 148652515 | 148715330 | 62816 | 1.29 | 2.82E-62 | 2.06E-60 |
| chr10 | 90449253 | 90468211 | 18959 | 3.01 | 3.13E-62 | 2.28E-60 |
| chr18 | 3814903 | 3823562 | 8660 | 1.79 | 3.86E-62 | 2.81E-60 |
| chr12 | 42718416 | 42745656 | 27241 | 1.13 | 4.54E-62 | 3.29E-60 |
| chr12 | 65001248 | 65101354 | 100107 | 0.6 | 5.45E-62 | 3.94E-60 |
| chr17 | 72967749 | 73213860 | 246112 | 0.47 | 5.54E-62 | 4.00E-60 |

| | | | | | | |
|---|---|---|---|---|---|---|
| chr11 | 101555671 | 101578564 | 22894 | 2.76 | 2.11E-61 | 1.52E-59 |
| chr10 | 37494865 | 37516497 | 21633 | 2.6 | 4.53E-61 | 3.25E-59 |
| chr7 | 123408751 | 123438013 | 29263 | 2.2 | 5.18E-61 | 3.71E-59 |
| chr3 | 7545879 | 7551455 | 5577 | 2.93 | 7.76E-61 | 5.55E-59 |
| chr8 | 146051669 | 146079292 | 27624 | 1.67 | 1.09E-60 | 7.75E-59 |
| chr15 | 69382599 | 69387242 | 4644 | -2.04 | 1.10E-60 | 7.81E-59 |
| chr9 | 80202181 | 80210976 | 8796 | 1.59 | 1.18E-60 | 8.36E-59 |
| chr7 | 46573766 | 46609524 | 35759 | 1.72 | 1.33E-60 | 9.42E-59 |
| chr12 | 104642571 | 104772290 | 129720 | 0.65 | 1.46E-60 | 1.03E-58 |
| chr4 | 79472126 | 79493255 | 21130 | 0.94 | 1.50E-60 | 1.06E-58 |
| chr16 | 57405705 | 57428845 | 23141 | 1.43 | 1.75E-60 | 1.23E-58 |
| chr5 | 179659187 | 179780996 | 121810 | 2.89 | 2.90E-60 | 2.04E-58 |
| chr16 | 19395521 | 19458247 | 62727 | 0.87 | 3.37E-60 | 2.36E-58 |
| chr2 | 223639910 | 223662984 | 23075 | 2.13 | 4.64E-60 | 3.24E-58 |
| chr2 | 227266831 | 227278806 | 11976 | 1.38 | 5.47E-60 | 3.81E-58 |
| chr8 | 124730287 | 124763031 | 32745 | 1.35 | 7.34E-60 | 5.10E-58 |
| chr4 | 150976039 | 151002100 | 26062 | -1.95 | 7.73E-60 | 5.35E-58 |
| chr12 | 54042835 | 54070717 | 27883 | 1.17 | 8.08E-60 | 5.59E-58 |
| chrX | 117419134 | 117422195 | 3062 | 3.84 | 2.19E-59 | 1.51E-57 |
| chr1 | 89486869 | 89489253 | 2385 | 3.38 | 3.11E-59 | 2.14E-57 |
| chr15 | 66118670 | 66130530 | 11861 | 1.74 | 3.21E-59 | 2.21E-57 |
| chr12 | 14706630 | 14721269 | 14640 | 2.97 | 3.42E-59 | 2.34E-57 |
| chr9 | 117021012 | 117062287 | 41276 | -1.68 | 4.37E-59 | 2.99E-57 |
| chr1 | 196369437 | 196378430 | 8994 | 2.33 | 5.97E-59 | 4.07E-57 |
| chr6 | 47372209 | 47389357 | 17149 | 1.35 | 8.41E-59 | 5.72E-57 |
| chr12 | 65927186 | 65933321 | 6136 | 3.32 | 9.66E-59 | 6.56E-57 |
| chr11 | 126887247 | 126899011 | 11765 | 2.41 | 1.10E-58 | 7.41E-57 |
| chr12 | 38517225 | 38550651 | 33427 | 1.53 | 1.15E-58 | 7.79E-57 |
| chr8 | 52874247 | 52887062 | 12816 | 4.85 | 1.63E-58 | 1.10E-56 |
| chr13 | 83520252 | 83537138 | 16887 | 2.52 | 1.72E-58 | 1.16E-56 |
| chr2 | 228314245 | 228433813 | 119569 | -0.7 | 2.77E-58 | 1.86E-56 |
| chr4 | 151028069 | 151063958 | 35890 | -1.55 | 3.83E-58 | 2.56E-56 |
| chr12 | 88171809 | 88175225 | 3417 | 2.55 | 4.51E-58 | 3.01E-56 |
| chr1 | 75563288 | 75568084 | 4797 | 2.4 | 5.00E-58 | 3.33E-56 |
| chr4 | 143269203 | 143274759 | 5557 | 3.08 | 5.74E-58 | 3.82E-56 |
| chr1 | 221874706 | 221900014 | 25309 | 1.31 | 7.11E-58 | 4.72E-56 |
| chr7 | 92744716 | 92749109 | 4394 | 3.48 | 1.37E-57 | 9.05E-56 |
| chr12 | 41379571 | 41394416 | 14846 | 2.52 | 1.42E-57 | 9.39E-56 |
| chr2 | 234725951 | 234777560 | 51610 | -1.04 | 1.67E-57 | 1.10E-55 |
| chr3 | 71168518 | 71191945 | 23428 | 1.55 | 2.71E-57 | 1.78E-55 |
| chr12 | 20963133 | 20994235 | 31103 | 1.27 | 3.56E-57 | 2.33E-55 |
| chr12 | 85668801 | 85688652 | 19852 | 1.59 | 5.41E-57 | 3.54E-55 |
| chr3 | 115502218 | 115524569 | 22352 | 1.34 | 5.58E-57 | 3.65E-55 |
| chr10 | 33176590 | 33306187 | 129598 | 0.54 | 5.78E-57 | 3.77E-55 |
| chr12 | 66523176 | 66564614 | 41439 | 1.25 | 6.78E-57 | 4.41E-55 |
| chr2 | 47073452 | 47117040 | 43589 | -0.96 | 9.90E-57 | 6.43E-55 |
| chr19 | 16177156 | 16223984 | 46829 | 0.94 | 1.03E-56 | 6.65E-55 |
| chr7 | 121143946 | 121146263 | 2318 | 3.1 | 1.36E-56 | 8.76E-55 |
| chr11 | 61447335 | 61498475 | 51141 | 0.95 | 2.02E-56 | 1.30E-54 |
| chr7 | 134103319 | 134168615 | 65297 | -0.78 | 2.43E-56 | 1.56E-54 |
| chr8 | 96111169 | 96126836 | 15668 | 1.34 | 2.81E-56 | 1.80E-54 |
| chr20 | 10151634 | 10168421 | 16788 | 1.46 | 4.97E-56 | 3.18E-54 |
| chr15 | 79927025 | 79939243 | 12219 | 1.61 | 4.98E-56 | 3.18E-54 |
| chr14 | 32865837 | 32871244 | 5408 | 3.33 | 8.15E-56 | 5.20E-54 |
| chr12 | 76302409 | 76388956 | 86548 | 0.66 | 1.79E-55 | 1.14E-53 |
| chr11 | 67873245 | 67918265 | 45021 | 1.03 | 2.10E-55 | 1.33E-53 |
| chr5 | 72132747 | 72204211 | 71465 | -0.77 | 2.11E-55 | 1.34E-53 |
| chr1 | 234733610 | 234795850 | 62241 | 0.62 | 2.23E-55 | 1.41E-53 |
| chr7 | 155410374 | 155413415 | 3042 | 1.82 | 2.28E-55 | 1.44E-53 |
| chr5 | 127243576 | 127330400 | 86825 | 0.81 | 2.47E-55 | 1.56E-53 |
| chr12 | 113330277 | 113347479 | 17203 | 1.66 | 3.11E-55 | 1.95E-53 |

| | | | | | | |
|---|---|---|---|---|---|---|
| chr20 | 14194109 | 14216432 | 22324 | 1.6 | 3.31E-55 | 2.08E-53 |
| chr7 | 29487236 | 29491370 | 4135 | 2.14 | 3.95E-55 | 2.47E-53 |
| chr4 | 177360545 | 177366789 | 6245 | 1.83 | 4.31E-55 | 2.69E-53 |
| chr11 | 33710594 | 33759481 | 48888 | 0.85 | 4.59E-55 | 2.86E-53 |
| chr9 | 100807266 | 100849488 | 42223 | 1.26 | 4.92E-55 | 3.06E-53 |
| chr22 | 50443516 | 50447436 | 3921 | 3.54 | 1.35E-54 | 8.38E-53 |
| chr16 | 57639945 | 57682282 | 42338 | 0.96 | 2.00E-54 | 1.24E-52 |
| chr3 | 101807261 | 101849705 | 42445 | 1.98 | 2.26E-54 | 1.40E-52 |
| chr8 | 131233302 | 131282378 | 49077 | -1.21 | 2.52E-54 | 1.55E-52 |
| chr7 | 130816412 | 130883504 | 67093 | -0.71 | 3.57E-54 | 2.19E-52 |
| chr1 | 231512030 | 231561586 | 49557 | 0.72 | 3.86E-54 | 2.37E-52 |
| chr2 | 86836760 | 86877724 | 40965 | 1.26 | 4.69E-54 | 2.87E-52 |
| chr14 | 101447815 | 101461790 | 13976 | 1.3 | 4.80E-54 | 2.93E-52 |
| chr6 | 56654376 | 56676756 | 22381 | 1.17 | 5.55E-54 | 3.38E-52 |
| chr18 | 3379846 | 3387065 | 7220 | -2.78 | 8.65E-54 | 5.26E-52 |
| chr11 | 1223544 | 1247688 | 24145 | 1.48 | 9.06E-54 | 5.50E-52 |
| chr2 | 192206613 | 192261847 | 55235 | -1.4 | 1.04E-53 | 6.28E-52 |
| chr8 | 32121176 | 32129783 | 8608 | -1.53 | 1.17E-53 | 7.08E-52 |
| chr3 | 30267279 | 30306070 | 38792 | 0.88 | 1.62E-53 | 9.79E-52 |
| chr10 | 4692190 | 4749916 | 57727 | 0.7 | 1.64E-53 | 9.88E-52 |
| chr17 | 37477747 | 37515971 | 38225 | 1.19 | 2.03E-53 | 1.22E-51 |
| chr12 | 46823191 | 46890077 | 66887 | 0.77 | 3.27E-53 | 1.96E-51 |
| chr1 | 244080471 | 244114247 | 33777 | -2.48 | 4.34E-53 | 2.60E-51 |
| chr16 | 1727478 | 1780302 | 52825 | 0.79 | 6.63E-53 | 3.96E-51 |
| chr4 | 2794851 | 2814426 | 19576 | 1.45 | 7.12E-53 | 4.25E-51 |
| chr11 | 77009494 | 77012810 | 3317 | 3.24 | 8.53E-53 | 5.07E-51 |
| chr14 | 32976844 | 32984720 | 7877 | 1.52 | 9.69E-53 | 5.75E-51 |
| chr4 | 106505708 | 106523663 | 17956 | 3.81 | 1.26E-52 | 7.46E-51 |
| chr10 | 63796058 | 63809829 | 13772 | 3.17 | 1.28E-52 | 7.58E-51 |
| chr12 | 71034064 | 71054868 | 20805 | 1.44 | 1.31E-52 | 7.73E-51 |
| chr22 | 35978908 | 36001422 | 22515 | -1.45 | 1.48E-52 | 8.70E-51 |
| chr9 | 118909921 | 119014091 | 104171 | -2.08 | 2.07E-52 | 1.22E-50 |
| chr8 | 29444955 | 29633784 | 188830 | 0.52 | 2.53E-52 | 1.49E-50 |
| chr13 | 109566807 | 109573783 | 6977 | 1.62 | 2.61E-52 | 1.53E-50 |
| chr20 | 33208636 | 33228577 | 19942 | 2.08 | 2.63E-52 | 1.54E-50 |
| chr2 | 56183836 | 56194157 | 10322 | 2.34 | 2.89E-52 | 1.68E-50 |
| chr18 | 8698939 | 8831316 | 132378 | 0.62 | 6.09E-52 | 3.54E-50 |
| chr9 | 119738948 | 119752530 | 13583 | 4.13 | 6.54E-52 | 3.80E-50 |
| chr14 | 95060045 | 95079746 | 19702 | 0.96 | 7.33E-52 | 4.25E-50 |
| chr21 | 38251886 | 38302194 | 50309 | -0.78 | 7.76E-52 | 4.49E-50 |
| chr5 | 146682495 | 146691293 | 8799 | 1.76 | 9.56E-52 | 5.52E-50 |
| chr3 | 155270469 | 155290764 | 20296 | 1.92 | 1.58E-51 | 9.09E-50 |
| chr4 | 157691149 | 157714329 | 23181 | 1.81 | 2.62E-51 | 1.50E-49 |
| chr17 | 5087082 | 5096028 | 8947 | 2.37 | 2.62E-51 | 1.50E-49 |
| chr3 | 142363075 | 142387858 | 24784 | 1.14 | 2.65E-51 | 1.52E-49 |
| chr5 | 98354351 | 98373719 | 19369 | 1.79 | 2.85E-51 | 1.63E-49 |
| chr16 | 53075418 | 53182611 | 107194 | 0.64 | 3.50E-51 | 2.00E-49 |
| chr12 | 14918009 | 14929631 | 11623 | 1.06 | 3.98E-51 | 2.27E-49 |
| chr15 | 96099781 | 96103623 | 3843 | 2.54 | 4.15E-51 | 2.36E-49 |
| chr12 | 50559555 | 50565918 | 6364 | 1.74 | 5.40E-51 | 3.07E-49 |
| chr14 | 23276421 | 23307044 | 30624 | 1.66 | 6.38E-51 | 3.61E-49 |
| chr11 | 6407695 | 6412027 | 4333 | 1.92 | 6.55E-51 | 3.71E-49 |
| chr1 | 201974746 | 202092396 | 117651 | 0.61 | 8.54E-51 | 4.82E-49 |
| chr12 | 58730490 | 58736342 | 5853 | 2.17 | 1.08E-50 | 6.10E-49 |
| chr4 | 103024565 | 103028369 | 3805 | 3.66 | 1.09E-50 | 6.15E-49 |
| chr3 | 53163238 | 53180779 | 17542 | 1.61 | 1.15E-50 | 6.46E-49 |
| chr14 | 65757046 | 65782286 | 25241 | 3.16 | 1.33E-50 | 7.46E-49 |
| chr15 | 80871883 | 80902814 | 30932 | 1.4 | 2.05E-50 | 1.15E-48 |
| chr1 | 157934779 | 157939690 | 4912 | 3.08 | 2.27E-50 | 1.27E-48 |
| chr10 | 73679558 | 73690195 | 10638 | 1.64 | 2.56E-50 | 1.43E-48 |
| chr3 | 101874488 | 101915671 | 41184 | 0.97 | 2.93E-50 | 1.63E-48 |

| | | | | | | |
|---|---|---|---|---|---|---|
| chr2 | 65564332 | 65624750 | 60419 | 1.38 | 6.40E-50 | 3.55E-48 |
| chr7 | 27462877 | 27465589 | 2713 | 2.05 | 7.19E-50 | 3.98E-48 |
| chr4 | 3482493 | 3488308 | 5816 | 2.89 | 8.56E-50 | 4.73E-48 |
| chr15 | 31305855 | 31317998 | 12144 | 2.69 | 1.12E-49 | 6.18E-48 |
| chr12 | 63631392 | 63633600 | 2209 | 6.5 | 1.25E-49 | 6.86E-48 |
| chr9 | 138017897 | 138030077 | 12181 | 2.1 | 1.45E-49 | 7.97E-48 |
| chr5 | 71361616 | 71375754 | 14139 | -3.46 | 1.83E-49 | 1.00E-47 |
| chr1 | 244483104 | 244531531 | 48428 | 0.94 | 1.84E-49 | 1.01E-47 |
| chr3 | 141928056 | 141945160 | 17105 | 1.35 | 1.93E-49 | 1.05E-47 |
| chr2 | 74197608 | 74233167 | 35560 | -1.85 | 1.94E-49 | 1.06E-47 |
| chr16 | 80039968 | 80070736 | 30769 | 1 | 1.96E-49 | 1.07E-47 |
| chrX | 43719185 | 43722506 | 3322 | 1.99 | 2.11E-49 | 1.15E-47 |
| chr7 | 29601878 | 29605283 | 3406 | 2.76 | 2.97E-49 | 1.61E-47 |
| chr1 | 212643096 | 212669720 | 26625 | 1.04 | 3.46E-49 | 1.87E-47 |
| chr3 | 75184204 | 75194506 | 10303 | 2.46 | 3.72E-49 | 2.01E-47 |
| chr11 | 112020936 | 112046833 | 25898 | 0.93 | 4.17E-49 | 2.25E-47 |
| chr7 | 111692679 | 111727985 | 35307 | -1.78 | 5.32E-49 | 2.87E-47 |
| chr2 | 159332811 | 159376201 | 43391 | -1.23 | 5.64E-49 | 3.03E-47 |
| chr2 | 153257822 | 153302868 | 45047 | -1.14 | 5.75E-49 | 3.09E-47 |
| chr3 | 182970364 | 183017177 | 46814 | 1.19 | 5.90E-49 | 3.16E-47 |
| chr22 | 46447984 | 46512822 | 64839 | 0.61 | 7.99E-49 | 4.27E-47 |
| chr1 | 88353085 | 88369234 | 16150 | 2.46 | 8.40E-49 | 4.48E-47 |
| chr5 | 39071961 | 39139700 | 67740 | -0.79 | 8.60E-49 | 4.58E-47 |
| chr7 | 142975743 | 142988605 | 12863 | 1.58 | 8.76E-49 | 4.66E-47 |
| chr22 | 36717807 | 36802108 | 84302 | 0.56 | 1.04E-48 | 5.52E-47 |
| chr7 | 92543773 | 92548040 | 4268 | 8.74 | 1.44E-48 | 7.65E-47 |
| chr12 | 92345147 | 92349144 | 3998 | 2.9 | 1.57E-48 | 8.29E-47 |
| chr12 | 17449456 | 17459649 | 10194 | -1.65 | 1.91E-48 | 1.01E-46 |
| chr6 | 41984509 | 42019243 | 34735 | -1.44 | 1.98E-48 | 1.04E-46 |
| chr5 | 95949215 | 96127482 | 178268 | 0.54 | 2.85E-48 | 1.50E-46 |
| chr21 | 44141242 | 44159067 | 17826 | 1.52 | 3.20E-48 | 1.68E-46 |
| chr3 | 44089438 | 44148742 | 59305 | -1.31 | 4.64E-48 | 2.43E-46 |
| chr12 | 89247005 | 89249362 | 2358 | 2.9 | 5.40E-48 | 2.82E-46 |
| chr17 | 39042400 | 39079763 | 37364 | 0.77 | 5.56E-48 | 2.90E-46 |
| chr7 | 116253286 | 116274109 | 20824 | 1.16 | 6.15E-48 | 3.21E-46 |
| chr1 | 55969282 | 55985914 | 16633 | -3.05 | 7.31E-48 | 3.80E-46 |
| chr12 | 32715102 | 32723084 | 7983 | 2.33 | 7.32E-48 | 3.80E-46 |
| chr10 | 128109666 | 128115114 | 5449 | 4.67 | 7.52E-48 | 3.90E-46 |
| chr3 | 4781356 | 4795160 | 13805 | 1.11 | 7.85E-48 | 4.07E-46 |
| chr20 | 52256231 | 52278624 | 22394 | 0.73 | 1.02E-47 | 5.28E-46 |
| chr11 | 72758569 | 72804440 | 45872 | 1.37 | 1.15E-47 | 5.95E-46 |
| chr12 | 70379630 | 70382981 | 3352 | 2.77 | 1.28E-47 | 6.61E-46 |
| chr10 | 63511010 | 63522028 | 11019 | 2.72 | 1.42E-47 | 7.30E-46 |
| chr16 | 53765785 | 53822060 | 56276 | 0.89 | 2.29E-47 | 1.17E-45 |
| chr13 | 50639808 | 50655509 | 15702 | 1.01 | 2.55E-47 | 1.31E-45 |
| chr20 | 22355521 | 22411991 | 56471 | -0.97 | 2.73E-47 | 1.39E-45 |
| chr10 | 79146614 | 79177281 | 30668 | -2.7 | 2.83E-47 | 1.44E-45 |
| chr7 | 79901091 | 79918370 | 17280 | 0.97 | 2.97E-47 | 1.51E-45 |
| chr10 | 63836153 | 63873070 | 36918 | 1.16 | 5.51E-47 | 2.80E-45 |
| chr20 | 50519686 | 50529764 | 10079 | 1.38 | 6.70E-47 | 3.40E-45 |
| chr17 | 67461578 | 67475594 | 14017 | 1.31 | 7.05E-47 | 3.57E-45 |
| chr13 | 35390781 | 35402500 | 11720 | 3.83 | 7.29E-47 | 3.69E-45 |
| chr12 | 120516651 | 120715470 | 198820 | 0.53 | 8.70E-47 | 4.39E-45 |
| chr19 | 42975908 | 43003949 | 28042 | 1.27 | 9.56E-47 | 4.82E-45 |
| chr11 | 1568113 | 1596021 | 27909 | 2.52 | 1.42E-46 | 7.16E-45 |
| chr12 | 101444569 | 101446777 | 2209 | 3.76 | 1.85E-46 | 9.31E-45 |
| chr6 | 159150118 | 159160077 | 9960 | 3.22 | 2.20E-46 | 1.10E-44 |
| chr18 | 55394941 | 55488181 | 93241 | 0.69 | 2.37E-46 | 1.19E-44 |
| chr4 | 19776669 | 19779838 | 3170 | 5.17 | 2.46E-46 | 1.23E-44 |
| chr1 | 47777412 | 47802619 | 25208 | 1.09 | 2.68E-46 | 1.34E-44 |
| chr2 | 129037891 | 129080522 | 42632 | -1.25 | 2.76E-46 | 1.37E-44 |

| | | | | | | |
|---|---|---|---|---|---|---|
| chr17 | 55591446 | 55620155 | 28710 | -1.37 | 2.85E-46 | 1.42E-44 |
| chr3 | 172463398 | 172532214 | 68817 | 0.56 | 3.13E-46 | 1.55E-44 |
| chr9 | 20544753 | 20572220 | 27468 | 0.92 | 3.67E-46 | 1.82E-44 |
| chr2 | 240113099 | 240150802 | 37704 | -2.1 | 3.85E-46 | 1.90E-44 |
| chr10 | 95867712 | 95886532 | 18821 | -1.8 | 3.98E-46 | 1.97E-44 |
| chr10 | 91086620 | 91113227 | 26608 | 0.93 | 4.80E-46 | 2.37E-44 |
| chr4 | 139078799 | 139198265 | 119467 | 0.45 | 4.90E-46 | 2.41E-44 |
| chr15 | 89157737 | 89193308 | 35572 | 0.78 | 5.64E-46 | 2.77E-44 |
| chr12 | 48576362 | 48594596 | 18235 | 1.65 | 6.11E-46 | 3.00E-44 |
| chr11 | 78791240 | 78807373 | 16134 | -1.35 | 7.30E-46 | 3.58E-44 |
| chr6 | 64514362 | 64518519 | 4158 | 1.59 | 8.46E-46 | 4.14E-44 |
| chr4 | 111454502 | 111456368 | 1867 | 3.03 | 9.00E-46 | 4.40E-44 |
| chr10 | 24495620 | 24499876 | 4257 | 2.44 | 9.74E-46 | 4.75E-44 |
| chr4 | 76761745 | 76770328 | 8584 | -1.59 | 9.86E-46 | 4.80E-44 |
| chr6 | 155506030 | 155524222 | 18193 | 2.25 | 1.01E-45 | 4.89E-44 |
| chr8 | 103540540 | 103550734 | 10195 | -1.51 | 1.02E-45 | 4.93E-44 |
| chr14 | 64136795 | 64146910 | 10116 | 3.2 | 1.38E-45 | 6.67E-44 |
| chr3 | 59645152 | 59651568 | 6417 | -3.75 | 1.53E-45 | 7.42E-44 |
| chr13 | 113546722 | 113561114 | 14393 | 3.17 | 1.73E-45 | 8.34E-44 |
| chrX | 56146480 | 56158253 | 11774 | 2.92 | 1.94E-45 | 9.36E-44 |
| chr2 | 47371914 | 47422607 | 50694 | -0.69 | 1.98E-45 | 9.55E-44 |
| chr5 | 76441203 | 76490408 | 49206 | 1.06 | 2.19E-45 | 1.05E-43 |
| chr1 | 164557490 | 164637354 | 79865 | -1.21 | 3.28E-45 | 1.57E-43 |
| chr14 | 50317720 | 50387729 | 70010 | 0.8 | 4.27E-45 | 2.04E-43 |
| chr7 | 12246084 | 12262955 | 16872 | 1.39 | 4.33E-45 | 2.07E-43 |
| chr12 | 12502269 | 12511471 | 9203 | 1.38 | 4.83E-45 | 2.31E-43 |
| chr20 | 37497852 | 37504558 | 6707 | 2.98 | 4.92E-45 | 2.35E-43 |
| chr4 | 11489273 | 11508209 | 18937 | 2.65 | 5.45E-45 | 2.59E-43 |
| chr2 | 134021706 | 134025618 | 3913 | -6.02 | 5.55E-45 | 2.64E-43 |
| chr2 | 27926324 | 27953869 | 27546 | -0.79 | 5.68E-45 | 2.69E-43 |
| chr15 | 41522129 | 41584268 | 62140 | 0.65 | 6.27E-45 | 2.97E-43 |
| chr4 | 113003126 | 113007838 | 4713 | 1.34 | 6.51E-45 | 3.08E-43 |
| chr19 | 12399701 | 12406544 | 6844 | 3.04 | 6.77E-45 | 3.19E-43 |
| chr3 | 172277351 | 172282659 | 5309 | 2.08 | 8.04E-45 | 3.79E-43 |
| chr8 | 146010321 | 146031891 | 21571 | 1.11 | 1.04E-44 | 4.90E-43 |
| chr11 | 123300483 | 123376784 | 76302 | -0.71 | 1.09E-44 | 5.10E-43 |
| chr12 | 89318332 | 89321615 | 3284 | 2.08 | 1.13E-44 | 5.31E-43 |
| chr9 | 16504733 | 16523058 | 18326 | 2.54 | 1.24E-44 | 5.81E-43 |
| chr12 | 123209800 | 123215955 | 6156 | 3.68 | 1.57E-44 | 7.35E-43 |
| chr5 | 137086654 | 137098755 | 12102 | 1.28 | 1.74E-44 | 8.13E-43 |
| chrX | 81988973 | 82004591 | 15619 | 2.28 | 2.82E-44 | 1.31E-42 |
| chr6 | 21917258 | 21954999 | 37742 | 1.69 | 2.86E-44 | 1.33E-42 |
| chr6 | 114138829 | 114151817 | 12989 | 1.47 | 3.09E-44 | 1.44E-42 |
| chr12 | 83043052 | 83049009 | 5958 | 1.27 | 3.69E-44 | 1.71E-42 |
| chr2 | 45010 | 48111 | 3102 | 3.59 | 3.98E-44 | 1.84E-42 |
| chr9 | 16409660 | 16417913 | 8254 | 1.58 | 4.29E-44 | 1.98E-42 |
| chr3 | 101544672 | 101598496 | 53825 | 1.12 | 4.66E-44 | 2.15E-42 |
| chr3 | 71244495 | 71260321 | 15827 | 1.76 | 4.81E-44 | 2.22E-42 |
| chr12 | 130610280 | 130628186 | 17907 | 4.18 | 4.96E-44 | 2.28E-42 |
| chrX | 48332764 | 48349181 | 16418 | 1.64 | 6.46E-44 | 2.97E-42 |
| chrX | 118601851 | 118646624 | 44774 | 0.84 | 7.76E-44 | 3.56E-42 |
| chr16 | 24696240 | 24715540 | 19301 | -1.58 | 9.64E-44 | 4.42E-42 |
| chr9 | 114355042 | 114368155 | 13114 | -1.25 | 1.02E-43 | 4.67E-42 |
| chr2 | 38935482 | 38979953 | 44472 | -0.89 | 1.27E-43 | 5.81E-42 |
| chr11 | 31005218 | 31015172 | 9955 | 1.84 | 1.50E-43 | 6.86E-42 |
| chr12 | 32571682 | 32615583 | 43902 | 1.2 | 1.55E-43 | 7.06E-42 |
| chr11 | 46563402 | 46583992 | 20591 | 0.83 | 1.79E-43 | 8.13E-42 |
| chr1 | 225587550 | 225653951 | 66402 | 0.82 | 1.84E-43 | 8.33E-42 |
| chrX | 8811726 | 8818976 | 7251 | 1.64 | 1.84E-43 | 8.34E-42 |
| chr7 | 22616084 | 22624350 | 8267 | 1.67 | 1.88E-43 | 8.52E-42 |
| chr10 | 37384146 | 37406257 | 22112 | 2.02 | 2.60E-43 | 1.17E-41 |

| | | | | | | |
|---|---|---|---|---|---|---|
| chr6 | 142600288 | 142719699 | 119412 | 0.56 | 2.74E-43 | 1.24E-41 |
| chr7 | 12677499 | 12692804 | 15306 | 1.38 | 2.91E-43 | 1.31E-41 |
| chr11 | 76953579 | 76957023 | 3445 | 4.73 | 3.08E-43 | 1.38E-41 |
| chr19 | 35157627 | 35169560 | 11934 | 1.15 | 3.31E-43 | 1.49E-41 |
| chr19 | 36704472 | 36728646 | 24175 | 1.53 | 3.72E-43 | 1.67E-41 |
| chr17 | 41790888 | 41803128 | 12241 | -1.5 | 4.14E-43 | 1.85E-41 |
| chr16 | 14454148 | 14474427 | 20280 | -1.61 | 4.91E-43 | 2.19E-41 |
| chr12 | 58807440 | 58821042 | 13603 | 1.04 | 5.14E-43 | 2.29E-41 |
| chr4 | 38664385 | 38698431 | 34047 | 0.82 | 5.24E-43 | 2.34E-41 |
| chr5 | 124314359 | 124349219 | 34861 | 1.18 | 5.78E-43 | 2.57E-41 |
| chr14 | 102720545 | 102808221 | 87677 | 0.61 | 6.38E-43 | 2.84E-41 |
| chr7 | 46549642 | 46566184 | 16543 | 2.1 | 6.80E-43 | 3.02E-41 |
| chr6 | 10399239 | 10465031 | 65793 | 0.69 | 7.02E-43 | 3.11E-41 |
| chr17 | 62149863 | 62162830 | 12968 | 0.89 | 8.48E-43 | 3.75E-41 |
| chr12 | 32685150 | 32696438 | 11289 | 2.25 | 9.52E-43 | 4.20E-41 |
| chrX | 114791412 | 114860761 | 69350 | 0.68 | 9.56E-43 | 4.21E-41 |
| chr17 | 13249811 | 13252054 | 2244 | 3.71 | 1.06E-42 | 4.65E-41 |
| chr15 | 71476445 | 71765585 | 289141 | -0.38 | 1.21E-42 | 5.32E-41 |
| chr11 | 73987420 | 74000487 | 13068 | -1.33 | 1.30E-42 | 5.71E-41 |
| chr12 | 106471472 | 106534409 | 62938 | -0.73 | 1.45E-42 | 6.38E-41 |
| chr15 | 32043629 | 32050620 | 6992 | -3.53 | 2.11E-42 | 9.25E-41 |
| chr5 | 79476742 | 79490216 | 13475 | 1.28 | 2.13E-42 | 9.31E-41 |
| chr4 | 108816049 | 108819446 | 3398 | 2.01 | 2.24E-42 | 9.78E-41 |
| chr16 | 24772616 | 24867902 | 95287 | -0.58 | 2.58E-42 | 1.12E-40 |
| chr17 | 19960386 | 20014376 | 53991 | -0.64 | 2.64E-42 | 1.15E-40 |
| chr5 | 55931091 | 55945959 | 14869 | 2.94 | 2.84E-42 | 1.24E-40 |
| chr16 | 14538876 | 14545494 | 6619 | -2.34 | 3.05E-42 | 1.32E-40 |
| chr1 | 155095239 | 155113323 | 18085 | 1.01 | 3.19E-42 | 1.38E-40 |
| chr7 | 135457746 | 135460598 | 2853 | 2.28 | 3.33E-42 | 1.44E-40 |
| chr3 | 158483316 | 158500773 | 17458 | 1.24 | 3.49E-42 | 1.51E-40 |
| chr19 | 23683883 | 23755672 | 71790 | 13.89 | 3.71E-42 | 1.60E-40 |
| chr4 | 95671133 | 95711939 | 40807 | 0.65 | 3.71E-42 | 1.60E-40 |
| chr2 | 235268631 | 235285738 | 17108 | -0.9 | 3.83E-42 | 1.65E-40 |
| chr11 | 63320150 | 63357329 | 37180 | 0.85 | 5.75E-42 | 2.47E-40 |
| chr1 | 33398507 | 33431261 | 32755 | 1.06 | 6.52E-42 | 2.80E-40 |
| chr14 | 56297780 | 56301294 | 3515 | 1.63 | 6.65E-42 | 2.85E-40 |
| chr19 | 33488334 | 33610560 | 122227 | 0.43 | 7.20E-42 | 3.08E-40 |
| chr6 | 56084641 | 56107651 | 23011 | 3.03 | 7.78E-42 | 3.32E-40 |
| chr4 | 39024069 | 39056794 | 32726 | 0.61 | 8.09E-42 | 3.45E-40 |
| chr7 | 93515035 | 93554104 | 39070 | 0.72 | 8.80E-42 | 3.75E-40 |
| chr3 | 114425202 | 114494969 | 69768 | 0.93 | 1.32E-41 | 5.62E-40 |
| chr19 | 29487816 | 29493496 | 5681 | 2.44 | 1.46E-41 | 6.19E-40 |
| chr9 | 117645403 | 117673587 | 28185 | 1.24 | 1.52E-41 | 6.43E-40 |
| chr11 | 122499916 | 122613410 | 113495 | -0.63 | 1.72E-41 | 7.29E-40 |
| chr6 | 42055225 | 42111109 | 55885 | 1.01 | 1.74E-41 | 7.36E-40 |
| chr14 | 63898612 | 63907467 | 8856 | 1.49 | 1.96E-41 | 8.26E-40 |
| chr7 | 12146044 | 12151296 | 5253 | 1.56 | 2.51E-41 | 1.06E-39 |
| chr22 | 30581480 | 30659737 | 78258 | 0.56 | 2.75E-41 | 1.16E-39 |
| chr11 | 102175105 | 102195634 | 20530 | 0.66 | 2.78E-41 | 1.17E-39 |
| chr6 | 150207889 | 150219387 | 11499 | 1.45 | 2.84E-41 | 1.19E-39 |
| chr14 | 95805462 | 95808555 | 3094 | 2.88 | 3.07E-41 | 1.29E-39 |
| chr21 | 27759953 | 27764140 | 4188 | 1.49 | 3.72E-41 | 1.56E-39 |
| chrX | 15755522 | 15770864 | 15343 | 2.01 | 4.06E-41 | 1.70E-39 |
| chr10 | 35833613 | 35842235 | 8623 | 1.52 | 4.07E-41 | 1.70E-39 |
| chr2 | 42189357 | 42195753 | 6397 | 1.45 | 4.79E-41 | 1.99E-39 |
| chr3 | 105082759 | 105122613 | 39855 | 0.64 | 4.92E-41 | 2.05E-39 |
| chr3 | 185343403 | 185349043 | 5641 | 2.78 | 5.30E-41 | 2.20E-39 |
| chr7 | 17123104 | 17141231 | 18128 | 1.59 | 5.70E-41 | 2.36E-39 |
| chr12 | 74883274 | 74903817 | 20544 | 2.91 | 6.31E-41 | 2.62E-39 |
| chr15 | 76302811 | 76305492 | 2682 | 1.99 | 6.36E-41 | 2.63E-39 |
| chr12 | 31799802 | 31813774 | 13973 | 1.75 | 6.56E-41 | 2.71E-39 |

| | | | | | | |
|---|---|---|---|---|---|---|
| chr2 | 70749933 | 70762801 | 12869 | 1.15 | 8.38E-41 | 3.46E-39 |
| chr1 | 94779709 | 94801477 | 21769 | -1.38 | 9.52E-41 | 3.92E-39 |
| chr12 | 98790837 | 98813762 | 22926 | 1.35 | 1.01E-40 | 4.14E-39 |
| chr14 | 64853222 | 64933043 | 79822 | 0.65 | 1.02E-40 | 4.21E-39 |
| chr10 | 63941837 | 63945070 | 3234 | 3.41 | 1.11E-40 | 4.57E-39 |
| chr3 | 4908136 | 4940755 | 32620 | 1.79 | 1.18E-40 | 4.82E-39 |
| chr16 | 63348093 | 63350499 | 2407 | 2.07 | 1.23E-40 | 5.02E-39 |
| chr8 | 31678492 | 31692797 | 14306 | 1.32 | 1.39E-40 | 5.67E-39 |
| chr1 | 180553679 | 180570006 | 16328 | 1.31 | 1.60E-40 | 6.54E-39 |
| chr12 | 20676072 | 20684248 | 8177 | 0.98 | 1.91E-40 | 7.77E-39 |
| chr1 | 217886112 | 217897406 | 11295 | 1.58 | 2.11E-40 | 8.58E-39 |
| chr8 | 32081224 | 32106482 | 25259 | -0.79 | 2.32E-40 | 9.43E-39 |
| chr2 | 65023977 | 65035116 | 11140 | 3.16 | 2.44E-40 | 9.89E-39 |
| chr15 | 60632600 | 60744218 | 111619 | 0.44 | 2.50E-40 | 1.01E-38 |
| chr12 | 8827421 | 8862344 | 34924 | 2.22 | 2.64E-40 | 1.07E-38 |
| chr1 | 207471608 | 207496337 | 24730 | 1.14 | 2.74E-40 | 1.11E-38 |
| chr21 | 46780865 | 46807490 | 26626 | 0.98 | 2.97E-40 | 1.20E-38 |
| chr15 | 70974017 | 71015040 | 41024 | 1.1 | 3.02E-40 | 1.22E-38 |
| chr2 | 143561895 | 143571210 | 9316 | 2.8 | 3.06E-40 | 1.23E-38 |
| chr7 | 94207700 | 94214458 | 6759 | 1.8 | 3.41E-40 | 1.37E-38 |
| chr6 | 137539221 | 137563796 | 24576 | 1.1 | 3.43E-40 | 1.38E-38 |
| chrX | 153823890 | 153832988 | 9099 | 4.4 | 3.44E-40 | 1.38E-38 |
| chr4 | 39970813 | 40005938 | 35126 | 1.36 | 3.56E-40 | 1.43E-38 |
| chr4 | 157851611 | 157893803 | 42193 | 1.57 | 4.02E-40 | 1.61E-38 |
| chr1 | 95028051 | 95046622 | 18572 | 2.01 | 4.40E-40 | 1.76E-38 |
| chr2 | 219827582 | 219830080 | 2499 | -3.51 | 5.02E-40 | 2.00E-38 |
| chr8 | 48419378 | 48457360 | 37983 | 0.64 | 5.51E-40 | 2.19E-38 |
| chrX | 115067984 | 115086136 | 18153 | 2.28 | 6.34E-40 | 2.52E-38 |
| chr12 | 118485788 | 118501779 | 15992 | 1.08 | 6.56E-40 | 2.60E-38 |
| chr3 | 160385317 | 160398120 | 12804 | 1.2 | 6.68E-40 | 2.65E-38 |
| chr1 | 108211115 | 108236243 | 25129 | 0.89 | 6.77E-40 | 2.68E-38 |
| chr15 | 67315787 | 67491898 | 176112 | 0.31 | 6.82E-40 | 2.70E-38 |
| chr10 | 3892530 | 3938747 | 46218 | 0.76 | 7.23E-40 | 2.85E-38 |
| chr2 | 191331384 | 191343199 | 11816 | 1.49 | 8.24E-40 | 3.25E-38 |
| chr17 | 76296118 | 76357397 | 61280 | -0.86 | 9.60E-40 | 3.78E-38 |
| chr6 | 90925859 | 90932452 | 6594 | 1.51 | 1.09E-39 | 4.30E-38 |
| chr2 | 192370121 | 192377150 | 7030 | -1.67 | 1.10E-39 | 4.31E-38 |
| chr9 | 77417108 | 77419857 | 2750 | 2.8 | 1.27E-39 | 4.99E-38 |
| chr12 | 89766239 | 89793643 | 27405 | 1.36 | 1.33E-39 | 5.23E-38 |
| chr19 | 46570983 | 46574688 | 3706 | -2.9 | 1.42E-39 | 5.56E-38 |
| chr15 | 80234982 | 80241581 | 6600 | -3.64 | 1.43E-39 | 5.60E-38 |
| chr15 | 65693321 | 65716011 | 22691 | 1.35 | 1.84E-39 | 7.17E-38 |
| chr7 | 93809216 | 93820622 | 11407 | 0.85 | 2.09E-39 | 8.16E-38 |
| chr8 | 131305170 | 131359376 | 54207 | -1.14 | 2.10E-39 | 8.16E-38 |
| chr12 | 56359591 | 56402301 | 42711 | 1.11 | 3.07E-39 | 1.19E-37 |
| chr2 | 241307343 | 241332675 | 25333 | -0.89 | 3.49E-39 | 1.36E-37 |
| chr15 | 57769641 | 57800021 | 30381 | 1.91 | 3.73E-39 | 1.45E-37 |
| chr2 | 165752510 | 165795361 | 42852 | 1.98 | 3.74E-39 | 1.45E-37 |
| chr3 | 187869983 | 188016995 | 147013 | 0.47 | 4.49E-39 | 1.74E-37 |
| chr1 | 17448567 | 17463568 | 15002 | 1.14 | 4.74E-39 | 1.83E-37 |
| chrX | 68243474 | 68248656 | 5183 | 3.15 | 4.78E-39 | 1.84E-37 |
| chr11 | 66816448 | 66847947 | 31500 | 1.22 | 5.00E-39 | 1.93E-37 |
| chr1 | 211808937 | 211823386 | 14450 | 2.79 | 5.63E-39 | 2.16E-37 |
| chr10 | 89338147 | 89380067 | 41921 | 1.42 | 6.04E-39 | 2.32E-37 |
| chr12 | 90086910 | 90105163 | 18254 | 0.98 | 6.94E-39 | 2.66E-37 |
| chr4 | 71489067 | 71503867 | 14801 | 1.97 | 6.98E-39 | 2.67E-37 |
| chr7 | 29644836 | 29662968 | 18133 | 1.52 | 7.33E-39 | 2.80E-37 |
| chr5 | 80715531 | 80725451 | 9921 | 1.98 | 7.99E-39 | 3.05E-37 |
| chr5 | 90421190 | 90435999 | 14810 | 7.1 | 1.04E-38 | 3.96E-37 |
| chr3 | 185033211 | 185081564 | 48354 | 0.78 | 1.15E-38 | 4.38E-37 |
| chr15 | 37179596 | 37210690 | 31095 | 2.12 | 1.17E-38 | 4.45E-37 |

| | | | | | | |
|---|---|---|---|---|---|---|
| chr6 | 36482209 | 36585298 | 103090 | 0.64 | 1.46E-38 | 5.54E-37 |
| chr17 | 71788818 | 71798939 | 10122 | -1.44 | 1.67E-38 | 6.34E-37 |
| chr13 | 106425178 | 106429338 | 4161 | 2.47 | 1.68E-38 | 6.36E-37 |
| chr10 | 12634880 | 12678000 | 43121 | 1.1 | 1.75E-38 | 6.64E-37 |
| chr12 | 109162085 | 109252253 | 90169 | 0.46 | 1.77E-38 | 6.71E-37 |
| chr20 | 49031146 | 49077660 | 46515 | 0.85 | 1.78E-38 | 6.74E-37 |
| chr1 | 27851810 | 27929344 | 77535 | 0.64 | 1.79E-38 | 6.76E-37 |
| chr15 | 50138442 | 50159899 | 21458 | -1.19 | 1.80E-38 | 6.78E-37 |
| chr11 | 86278004 | 86319991 | 41988 | 0.71 | 1.84E-38 | 6.91E-37 |
| chr10 | 123792387 | 123832143 | 39757 | 0.66 | 1.85E-38 | 6.94E-37 |
| chr12 | 80192109 | 80194541 | 2433 | 3.18 | 2.05E-38 | 7.70E-37 |
| chr3 | 36949005 | 36962707 | 13703 | 2.62 | 2.78E-38 | 1.04E-36 |
| chr3 | 156845706 | 156854613 | 8908 | 1.43 | 2.82E-38 | 1.06E-36 |
| chr3 | 87346910 | 87386534 | 39625 | 0.78 | 2.86E-38 | 1.07E-36 |
| chr1 | 145434642 | 145479048 | 44407 | 1.15 | 3.26E-38 | 1.22E-36 |
| chr8 | 127569060 | 127593237 | 24178 | 1 | 3.27E-38 | 1.22E-36 |
| chr11 | 111169432 | 111181733 | 12302 | 1.45 | 3.44E-38 | 1.28E-36 |
| chr16 | 48465599 | 48513200 | 47602 | 1.34 | 3.48E-38 | 1.29E-36 |
| chr1 | 171402240 | 171411534 | 9295 | 1.69 | 3.64E-38 | 1.35E-36 |
| chr3 | 30552085 | 30596073 | 43989 | 1.28 | 3.89E-38 | 1.44E-36 |
| chr17 | 74405584 | 74462148 | 56565 | -1.05 | 4.37E-38 | 1.62E-36 |
| chr17 | 60855231 | 60902965 | 47735 | 0.7 | 4.42E-38 | 1.64E-36 |
| chr7 | 81254240 | 81269155 | 14916 | 1.81 | 4.90E-38 | 1.81E-36 |
| chr3 | 12327958 | 12361501 | 33544 | 0.66 | 5.22E-38 | 1.93E-36 |
| chr13 | 29289215 | 29293026 | 3812 | 2.62 | 5.70E-38 | 2.10E-36 |
| chr1 | 179084001 | 179145390 | 61390 | 0.73 | 6.03E-38 | 2.22E-36 |
| chr14 | 64029674 | 64040694 | 11021 | 2.82 | 6.19E-38 | 2.28E-36 |
| chr1 | 174208194 | 174209772 | 1579 | 1.67 | 6.28E-38 | 2.31E-36 |
| chr7 | 138070061 | 138081826 | 11766 | 2.1 | 6.68E-38 | 2.45E-36 |
| chrX | 130698850 | 130712872 | 14023 | 1.31 | 7.03E-38 | 2.58E-36 |
| chr7 | 158573206 | 158623720 | 50515 | 0.5 | 8.42E-38 | 3.08E-36 |
| chr8 | 118680674 | 118685179 | 4506 | 2.05 | 9.93E-38 | 3.63E-36 |
| chr8 | 101239452 | 101298428 | 58977 | 0.98 | 1.10E-37 | 4.00E-36 |
| chr12 | 87027581 | 87029926 | 2346 | 2.79 | 1.10E-37 | 4.01E-36 |
| chr1 | 33358613 | 33379170 | 20558 | 2.25 | 1.14E-37 | 4.16E-36 |
| chr12 | 28150614 | 28152496 | 1883 | 4.07 | 1.24E-37 | 4.52E-36 |
| chr9 | 123146087 | 123175906 | 29820 | -0.89 | 1.39E-37 | 5.05E-36 |
| chr1 | 113085576 | 113100979 | 15404 | 1.72 | 1.61E-37 | 5.85E-36 |
| chr10 | 112388839 | 112404861 | 16023 | 1.52 | 1.69E-37 | 6.12E-36 |
| chr9 | 100904250 | 100910539 | 6290 | 1.94 | 1.74E-37 | 6.30E-36 |
| chr6 | 82461222 | 82499328 | 38107 | 1.61 | 1.78E-37 | 6.44E-36 |
| chr5 | 162863352 | 162937756 | 74405 | 0.65 | 1.90E-37 | 6.85E-36 |
| chr9 | 132489153 | 132546156 | 57004 | 0.62 | 2.04E-37 | 7.37E-36 |
| chr20 | 47444686 | 47463898 | 19213 | -1.7 | 2.92E-37 | 1.05E-35 |
| chr7 | 107818727 | 107950135 | 131409 | -13.02 | 3.04E-37 | 1.09E-35 |
| chr18 | 2409595 | 2412329 | 2735 | 3.18 | 3.14E-37 | 1.13E-35 |
| chr19 | 16044910 | 16046869 | 1960 | -5.66 | 3.18E-37 | 1.14E-35 |
| chr4 | 11586788 | 11637982 | 51195 | 0.94 | 3.25E-37 | 1.16E-35 |
| chr6 | 72084958 | 72131838 | 46881 | 0.58 | 3.26E-37 | 1.17E-35 |
| chr19 | 23518964 | 23621534 | 102571 | 13.01 | 3.36E-37 | 1.20E-35 |
| chr17 | 28647754 | 28673847 | 26094 | 1.6 | 3.46E-37 | 1.23E-35 |
| chr4 | 38155834 | 38166652 | 10819 | 1.25 | 4.15E-37 | 1.48E-35 |
| chr12 | 96176095 | 96203764 | 27670 | 0.7 | 4.96E-37 | 1.77E-35 |
| chr10 | 60302830 | 60351713 | 48884 | -1.08 | 5.30E-37 | 1.89E-35 |
| chr12 | 32829908 | 32872248 | 42341 | 1.04 | 5.31E-37 | 1.89E-35 |
| chr3 | 45857606 | 45860939 | 3334 | 3.15 | 5.51E-37 | 1.96E-35 |
| chr9 | 118683537 | 118742897 | 59361 | -0.6 | 6.85E-37 | 2.43E-35 |
| chr21 | 43084098 | 43106660 | 22563 | 0.81 | 6.98E-37 | 2.47E-35 |
| chr3 | 32460721 | 32464373 | 3653 | 2.1 | 7.59E-37 | 2.68E-35 |
| chr7 | 27120369 | 27224521 | 104153 | 0.88 | 7.74E-37 | 2.73E-35 |
| chr10 | 81025497 | 81082226 | 56730 | -1.03 | 7.80E-37 | 2.75E-35 |

| | | | | | | |
|---|---|---|---|---|---|---|
| chr15 | 69845359 | 69855493 | 10135 | 0.81 | 8.47E-37 | 2.99E-35 |
| chr22 | 47723791 | 47734944 | 11154 | 1.12 | 9.98E-37 | 3.51E-35 |
| chr15 | 70784452 | 70802923 | 18472 | 0.64 | 1.12E-36 | 3.95E-35 |
| chr5 | 1314698 | 1326377 | 11680 | 1.6 | 1.17E-36 | 4.12E-35 |
| chr5 | 168775743 | 168782756 | 7014 | -1.83 | 1.19E-36 | 4.18E-35 |
| chr2 | 235067234 | 235121076 | 53843 | -0.9 | 1.34E-36 | 4.70E-35 |
| chr12 | 15932343 | 15943280 | 10938 | 1.56 | 1.37E-36 | 4.79E-35 |
| chr10 | 19331784 | 19338161 | 6378 | 3.14 | 1.43E-36 | 5.00E-35 |
| chr4 | 74709714 | 74737328 | 27615 | 1.8 | 1.45E-36 | 5.04E-35 |
| chr4 | 79056824 | 79101003 | 44180 | 1.7 | 1.56E-36 | 5.43E-35 |
| chr17 | 39652470 | 39706532 | 54063 | 0.78 | 1.57E-36 | 5.47E-35 |
| chr12 | 45608101 | 45630379 | 22279 | 0.69 | 1.65E-36 | 5.74E-35 |
| chr3 | 11177840 | 11189840 | 12001 | 2.43 | 1.66E-36 | 5.77E-35 |
| chr3 | 148317385 | 148325331 | 7947 | 0.93 | 1.74E-36 | 6.02E-35 |
| chr15 | 54122300 | 54129266 | 6967 | -2.72 | 1.87E-36 | 6.47E-35 |
| chr19 | 37957055 | 38025237 | 68183 | 12.87 | 1.94E-36 | 6.73E-35 |
| chr17 | 76373666 | 76423115 | 49450 | -1 | 1.97E-36 | 6.81E-35 |
| chr8 | 26005930 | 26010642 | 4713 | 1.17 | 2.04E-36 | 7.04E-35 |
| chr20 | 11666858 | 11669146 | 2289 | 3.99 | 2.05E-36 | 7.07E-35 |
| chr12 | 65361295 | 65364200 | 2906 | 1.53 | 2.15E-36 | 7.42E-35 |
| chr5 | 159324743 | 159356423 | 31681 | 0.7 | 2.16E-36 | 7.45E-35 |
| chr20 | 4527007 | 4537077 | 10071 | -5.78 | 2.25E-36 | 7.75E-35 |
| chr20 | 2280133 | 2296294 | 16162 | 1.53 | 2.45E-36 | 8.41E-35 |
| chr22 | 19246916 | 19280106 | 33191 | 1.19 | 2.52E-36 | 8.63E-35 |
| chr10 | 62516843 | 62561799 | 44957 | 1.02 | 2.52E-36 | 8.63E-35 |
| chr20 | 9047600 | 9055206 | 7607 | 0.94 | 2.70E-36 | 9.24E-35 |
| chr8 | 62942529 | 62949647 | 7119 | 1.48 | 2.77E-36 | 9.48E-35 |
| chr6 | 159184168 | 159241325 | 57158 | 0.69 | 2.88E-36 | 9.82E-35 |
| chr4 | 169981564 | 170002892 | 21329 | 1.11 | 2.92E-36 | 9.96E-35 |
| chr12 | 52345528 | 52372756 | 27229 | 1.19 | 3.07E-36 | 1.05E-34 |
| chr19 | 56866492 | 56916403 | 49912 | 12.83 | 3.38E-36 | 1.15E-34 |
| chr11 | 117572261 | 117593205 | 20945 | 1.25 | 3.58E-36 | 1.22E-34 |
| chr19 | 37697690 | 37710120 | 12431 | 1.4 | 4.12E-36 | 1.40E-34 |
| chr12 | 80028728 | 80087613 | 58886 | 0.63 | 4.33E-36 | 1.47E-34 |
| chr17 | 63623649 | 63632087 | 8439 | 1.45 | 4.40E-36 | 1.49E-34 |
| chr15 | 86099979 | 86281309 | 181331 | 0.61 | 4.60E-36 | 1.55E-34 |
| chr3 | 104670522 | 104672462 | 1941 | 4.06 | 4.90E-36 | 1.65E-34 |
| chr12 | 27412353 | 27426944 | 14592 | 2.72 | 4.93E-36 | 1.66E-34 |
| chr6 | 108321723 | 108325208 | 3486 | 1.72 | 5.15E-36 | 1.74E-34 |
| chr13 | 111553485 | 111569025 | 15541 | 1.1 | 6.38E-36 | 2.15E-34 |
| chr6 | 47071543 | 47149083 | 77541 | 0.89 | 7.10E-36 | 2.39E-34 |
| chr8 | 8455999 | 8523083 | 67085 | 0.75 | 7.82E-36 | 2.62E-34 |
| chr6 | 21021916 | 21025495 | 3580 | 7.7 | 8.02E-36 | 2.69E-34 |
| chr5 | 95533507 | 95536965 | 3459 | 2.33 | 9.70E-36 | 3.25E-34 |
| chr12 | 69194886 | 69235826 | 40941 | 0.81 | 9.97E-36 | 3.34E-34 |
| chr4 | 19739838 | 19751375 | 11538 | 1.58 | 1.12E-35 | 3.75E-34 |
| chr5 | 10497477 | 10529178 | 31702 | -1.15 | 1.17E-35 | 3.91E-34 |
| chr8 | 90837785 | 90871562 | 33778 | 1.28 | 1.19E-35 | 3.98E-34 |
| chr2 | 69364562 | 69408941 | 44380 | -1.25 | 1.33E-35 | 4.44E-34 |
| chr1 | 82195754 | 82198391 | 2638 | 4.33 | 1.35E-35 | 4.49E-34 |
| chr4 | 150325549 | 150331480 | 5932 | 2.62 | 1.39E-35 | 4.62E-34 |
| chr2 | 31348566 | 31361802 | 13237 | -3.23 | 1.46E-35 | 4.86E-34 |
| chr8 | 22017010 | 22086170 | 69161 | 0.71 | 1.47E-35 | 4.87E-34 |
| chr12 | 2983085 | 3005044 | 21960 | 0.96 | 1.53E-35 | 5.07E-34 |
| chr1 | 199743202 | 199757887 | 14686 | 2.08 | 1.63E-35 | 5.38E-34 |
| chr21 | 44815591 | 44941960 | 126370 | 0.48 | 1.72E-35 | 5.68E-34 |
| chr12 | 52533841 | 52686148 | 152308 | 0.45 | 1.72E-35 | 5.68E-34 |
| chr9 | 19157902 | 19188678 | 30777 | 0.73 | 1.85E-35 | 6.12E-34 |
| chr8 | 107218080 | 107222012 | 3933 | 2.32 | 1.90E-35 | 6.26E-34 |
| chr5 | 127207092 | 127229624 | 22533 | 1.26 | 1.98E-35 | 6.50E-34 |
| chr10 | 65458893 | 65480695 | 21803 | 1.42 | 2.25E-35 | 7.40E-34 |

| | | | | | | |
|---|---|---|---|---|---|---|
| chr9 | 124030118 | 124067168 | 37051 | 0.72 | 2.30E-35 | 7.54E-34 |
| chr8 | 82628232 | 82645402 | 17171 | 2.08 | 2.36E-35 | 7.74E-34 |
| chr3 | 182322488 | 182327828 | 5341 | 4.25 | 2.63E-35 | 8.62E-34 |
| chr1 | 246862013 | 246873758 | 11746 | 2.96 | 2.70E-35 | 8.85E-34 |
| chr18 | 42399680 | 42420836 | 21157 | 1.16 | 2.85E-35 | 9.31E-34 |
| chr10 | 8050444 | 8056282 | 5839 | 2.13 | 2.94E-35 | 9.58E-34 |
| chr7 | 12197461 | 12210953 | 13493 | 1.85 | 2.94E-35 | 9.58E-34 |
| chr9 | 88693800 | 88722235 | 28436 | 1.23 | 2.95E-35 | 9.62E-34 |
| chr17 | 36056777 | 36131047 | 74271 | 0.47 | 2.97E-35 | 9.68E-34 |
| chr1 | 81120971 | 81124717 | 3747 | 4.02 | 3.02E-35 | 9.81E-34 |
| chr11 | 86406638 | 86456745 | 50108 | 0.67 | 3.09E-35 | 1.00E-33 |
| chr21 | 40278272 | 40291753 | 13482 | 2.01 | 3.13E-35 | 1.02E-33 |
| chr12 | 10362814 | 10367689 | 4876 | 1.16 | 3.28E-35 | 1.06E-33 |
| chr3 | 16905502 | 16927953 | 22452 | 2.97 | 3.42E-35 | 1.11E-33 |
| chr12 | 7585307 | 7597456 | 12150 | 1.59 | 3.48E-35 | 1.13E-33 |
| chr20 | 36614348 | 36621222 | 6875 | 2.13 | 3.63E-35 | 1.17E-33 |
| chr19 | 52641363 | 52644120 | 2758 | 2.88 | 3.70E-35 | 1.19E-33 |
| chr2 | 99052472 | 99083173 | 30702 | -1.56 | 4.12E-35 | 1.33E-33 |
| chr7 | 95128703 | 95141842 | 13140 | -4.11 | 4.63E-35 | 1.49E-33 |
| chr3 | 186779484 | 186782755 | 3272 | 2.09 | 4.80E-35 | 1.54E-33 |
| chr11 | 16016532 | 16025259 | 8728 | 1.64 | 4.80E-35 | 1.54E-33 |
| chr5 | 169396735 | 169417577 | 20843 | 2.23 | 5.25E-35 | 1.68E-33 |
| chr2 | 150622797 | 150628129 | 5333 | 1.53 | 5.31E-35 | 1.70E-33 |
| chr4 | 139818445 | 139840301 | 21857 | 0.83 | 5.63E-35 | 1.80E-33 |
| chr6 | 13676831 | 13680416 | 3586 | 7.54 | 5.72E-35 | 1.83E-33 |
| chr4 | 165674361 | 165676611 | 2251 | 1.98 | 5.99E-35 | 1.91E-33 |
| chrX | 66814376 | 66840239 | 25864 | 0.77 | 6.78E-35 | 2.16E-33 |
| chr12 | 124812002 | 124949101 | 137100 | 0.47 | 7.01E-35 | 2.23E-33 |
| chr8 | 22831645 | 22964330 | 132686 | 0.59 | 7.31E-35 | 2.33E-33 |
| chr8 | 32160403 | 32208981 | 48579 | -0.62 | 7.33E-35 | 2.33E-33 |
| chr20 | 14959612 | 14979013 | 19402 | 0.71 | 7.57E-35 | 2.40E-33 |
| chr18 | 72656621 | 72664488 | 7868 | 3.32 | 8.56E-35 | 2.72E-33 |
| chr3 | 149011970 | 149121630 | 109661 | 0.33 | 8.94E-35 | 2.83E-33 |
| chr5 | 52283101 | 52335092 | 51992 | -0.96 | 9.25E-35 | 2.93E-33 |
| chr19 | 23904555 | 24008573 | 104019 | 12.55 | 1.05E-34 | 3.33E-33 |
| chr7 | 31962672 | 32027895 | 65224 | -0.68 | 1.09E-34 | 3.43E-33 |
| chr6 | 82723296 | 82733405 | 10110 | 2.51 | 1.19E-34 | 3.76E-33 |
| chr1 | 164704911 | 164775489 | 70579 | -0.63 | 1.23E-34 | 3.88E-33 |
| chr15 | 101888138 | 101891011 | 2874 | 2.81 | 1.31E-34 | 4.12E-33 |
| chr18 | 11938154 | 11964490 | 26337 | 1.13 | 1.44E-34 | 4.53E-33 |
| chr9 | 99405889 | 99449817 | 43929 | 1.24 | 1.45E-34 | 4.56E-33 |
| chr10 | 59449016 | 59464475 | 15460 | 0.99 | 1.68E-34 | 5.28E-33 |
| chr14 | 55798920 | 55805683 | 6764 | 1.26 | 1.73E-34 | 5.41E-33 |
| chr16 | 31884637 | 31893901 | 9265 | 1.39 | 1.80E-34 | 5.65E-33 |
| chr2 | 216690226 | 216697550 | 7325 | -1.61 | 1.83E-34 | 5.74E-33 |
| chr1 | 159249343 | 159254172 | 4830 | -2.06 | 1.89E-34 | 5.90E-33 |
| chr17 | 66372957 | 66380088 | 7132 | -1.1 | 2.15E-34 | 6.70E-33 |
| chr4 | 90164399 | 90238294 | 73896 | -0.57 | 2.15E-34 | 6.72E-33 |
| chr3 | 127608320 | 127610392 | 2073 | 4.77 | 2.23E-34 | 6.96E-33 |
| chrX | 115399382 | 115417652 | 18271 | 12.48 | 2.40E-34 | 7.46E-33 |
| chr3 | 189473428 | 189492490 | 19063 | 2.14 | 2.52E-34 | 7.85E-33 |
| chr16 | 24917324 | 25027503 | 110180 | -0.81 | 2.63E-34 | 8.17E-33 |
| chr12 | 68140794 | 68143645 | 2852 | 3.18 | 2.77E-34 | 8.61E-33 |
| chr6 | 21730605 | 21770837 | 40233 | 0.89 | 2.89E-34 | 8.97E-33 |
| chr1 | 178985553 | 179077350 | 91798 | -0.79 | 2.94E-34 | 9.10E-33 |
| chr2 | 103201522 | 103204111 | 2590 | 5 | 3.86E-34 | 1.19E-32 |
| chr14 | 51538419 | 51562881 | 24463 | -2.45 | 4.23E-34 | 1.31E-32 |
| chr15 | 66078803 | 66090674 | 11872 | 0.85 | 4.26E-34 | 1.32E-32 |
| chr9 | 104105028 | 104162218 | 57191 | 0.8 | 4.32E-34 | 1.33E-32 |
| chr2 | 231897674 | 231900230 | 2557 | 3.16 | 4.90E-34 | 1.51E-32 |
| chr12 | 32543818 | 32554155 | 10338 | 3.02 | 4.99E-34 | 1.53E-32 |

| | | | | | | |
|---|---|---|---|---|---|---|
| chr5 | 68787811 | 68817584 | 29774 | 1.06 | 5.33E-34 | 1.64E-32 |
| chr4 | 143393925 | 143407361 | 13437 | 1.62 | 5.84E-34 | 1.79E-32 |
| chr12 | 92515484 | 92555818 | 40335 | 0.86 | 5.85E-34 | 1.80E-32 |
| chr5 | 67751336 | 67755419 | 4084 | 1.98 | 5.99E-34 | 1.83E-32 |
| chr6 | 113879661 | 113895920 | 16260 | 2.71 | 5.99E-34 | 1.83E-32 |
| chr3 | 193788284 | 193813750 | 25467 | 1.39 | 6.12E-34 | 1.87E-32 |
| chr6 | 106038916 | 106059933 | 21018 | -0.69 | 6.14E-34 | 1.88E-32 |
| chr19 | 58309963 | 58341997 | 32035 | 1.13 | 6.15E-34 | 1.88E-32 |
| chr8 | 110651912 | 110658275 | 6364 | 2.69 | 6.38E-34 | 1.95E-32 |
| chr4 | 137854837 | 137858002 | 3166 | 4.26 | 7.63E-34 | 2.33E-32 |
| chr20 | 42542921 | 42588145 | 45225 | -0.92 | 8.70E-34 | 2.65E-32 |
| chr1 | 114423312 | 114431653 | 8342 | 1.98 | 8.84E-34 | 2.69E-32 |
| chr17 | 60686148 | 60730879 | 44732 | -1 | 1.02E-33 | 3.09E-32 |
| chr12 | 53609757 | 53627037 | 17281 | 1.12 | 1.14E-33 | 3.46E-32 |
| chr10 | 37872379 | 37874511 | 2133 | 4.83 | 1.15E-33 | 3.49E-32 |
| chr16 | 70703816 | 70816669 | 112854 | -0.6 | 1.22E-33 | 3.70E-32 |
| chrX | 149200557 | 149208153 | 7597 | 1.49 | 1.34E-33 | 4.04E-32 |
| chr3 | 159135371 | 159137858 | 2488 | 3.5 | 1.38E-33 | 4.16E-32 |
| chr7 | 22893488 | 22904315 | 10828 | -1.44 | 1.38E-33 | 4.18E-32 |
| chr3 | 156872030 | 156894640 | 22611 | 0.71 | 1.41E-33 | 4.24E-32 |
| chr1 | 89227406 | 89241614 | 14209 | 1.52 | 1.49E-33 | 4.48E-32 |
| chr9 | 140187468 | 140191794 | 4327 | 3.96 | 1.79E-33 | 5.38E-32 |
| chr20 | 19763336 | 19809211 | 45876 | -0.82 | 1.91E-33 | 5.73E-32 |
| chr10 | 38145171 | 38151069 | 5899 | 1.52 | 2.03E-33 | 6.09E-32 |
| chr3 | 30636004 | 30739590 | 103587 | 0.47 | 2.53E-33 | 7.60E-32 |
| chr4 | 95560208 | 95627092 | 66885 | 0.5 | 2.62E-33 | 7.84E-32 |
| chr18 | 21197007 | 21215793 | 18787 | -1.55 | 2.73E-33 | 8.16E-32 |
| chr3 | 169939184 | 169961814 | 22631 | 0.91 | 2.75E-33 | 8.24E-32 |
| chr9 | 22123380 | 22172831 | 49452 | 0.49 | 2.86E-33 | 8.54E-32 |
| chr19 | 38117740 | 38162413 | 44674 | 12.28 | 2.96E-33 | 8.83E-32 |
| chr10 | 82951860 | 82953634 | 1775 | 2.85 | 3.50E-33 | 1.04E-31 |
| chr19 | 37281841 | 37342653 | 60813 | 12.26 | 3.96E-33 | 1.18E-31 |
| chr15 | 88295052 | 88297498 | 2447 | -3.32 | 4.12E-33 | 1.22E-31 |
| chr8 | 95413315 | 95488058 | 74744 | 0.53 | 4.74E-33 | 1.41E-31 |
| chr7 | 155015535 | 155039796 | 24262 | 1.91 | 4.92E-33 | 1.46E-31 |
| chr17 | 56493584 | 56496203 | 2620 | 2.07 | 5.02E-33 | 1.49E-31 |
| chr15 | 36659323 | 36668000 | 8678 | 1.27 | 5.42E-33 | 1.61E-31 |
| chr1 | 82627279 | 82629495 | 2217 | 7.03 | 5.71E-33 | 1.69E-31 |
| chr6 | 107169142 | 107213104 | 43963 | 1.02 | 5.96E-33 | 1.76E-31 |
| chr18 | 33075936 | 33096229 | 20294 | 1.36 | 6.46E-33 | 1.91E-31 |
| chr12 | 125159910 | 125172890 | 12981 | 0.85 | 7.65E-33 | 2.26E-31 |
| chr9 | 114567444 | 114570062 | 2619 | 2.39 | 8.14E-33 | 2.40E-31 |
| chr2 | 26161352 | 26368855 | 207504 | -0.42 | 8.21E-33 | 2.42E-31 |
| chr7 | 18058363 | 18060720 | 2358 | 2.52 | 8.30E-33 | 2.44E-31 |
| chrX | 84628727 | 84635697 | 6971 | 0.9 | 9.45E-33 | 2.78E-31 |
| chr9 | 72499000 | 72514820 | 15821 | 1.5 | 9.87E-33 | 2.90E-31 |
| chr1 | 186535307 | 186548640 | 13334 | 2.24 | 1.00E-32 | 2.93E-31 |
| chr2 | 191436324 | 191445744 | 9421 | 1.2 | 1.02E-32 | 3.00E-31 |
| chr3 | 118889587 | 118960613 | 71027 | 0.39 | 1.11E-32 | 3.25E-31 |
| chr11 | 72411039 | 72464217 | 53179 | -0.92 | 1.12E-32 | 3.28E-31 |
| chr7 | 140009507 | 140017285 | 7779 | 2.16 | 1.17E-32 | 3.43E-31 |
| chr5 | 90277713 | 90281584 | 3872 | 2.16 | 1.29E-32 | 3.78E-31 |
| chr6 | 7042117 | 7059401 | 17285 | 2.58 | 1.31E-32 | 3.82E-31 |
| chr7 | 73887511 | 73959663 | 72153 | -0.71 | 1.32E-32 | 3.84E-31 |
| chr14 | 105423784 | 105512737 | 88954 | 0.55 | 1.48E-32 | 4.31E-31 |
| chr5 | 55645285 | 55655993 | 10709 | 1.82 | 1.49E-32 | 4.34E-31 |
| chr15 | 67170006 | 67178254 | 8249 | -1.37 | 1.67E-32 | 4.85E-31 |
| chr7 | 34786760 | 34796066 | 9307 | 1.47 | 1.69E-32 | 4.89E-31 |
| chr7 | 124205495 | 124208600 | 3106 | 6.38 | 1.93E-32 | 5.59E-31 |
| chr19 | 53424937 | 53450826 | 25890 | 12.11 | 2.00E-32 | 5.81E-31 |
| chr1 | 151940227 | 151974305 | 34079 | 0.48 | 2.02E-32 | 5.84E-31 |

| | | | | | | |
|---|---|---|---|---|---|---|
| chr13 | 106293949 | 106295972 | 2024 | 2.65 | 2.04E-32 | 5.89E-31 |
| chr2 | 62712552 | 62734957 | 22406 | -1.09 | 2.60E-32 | 7.50E-31 |
| chr2 | 11072078 | 11078653 | 6576 | -1.26 | 2.64E-32 | 7.61E-31 |
| chr12 | 83131162 | 83146432 | 15271 | 1.38 | 2.92E-32 | 8.42E-31 |
| chr2 | 70779909 | 70797706 | 17798 | 1.97 | 3.02E-32 | 8.69E-31 |
| chr18 | 29067336 | 29094832 | 27497 | 1.39 | 3.08E-32 | 8.87E-31 |
| chr8 | 74204149 | 74277456 | 73308 | 0.71 | 3.15E-32 | 9.05E-31 |
| chr7 | 108038936 | 108097275 | 58340 | -12.07 | 3.22E-32 | 9.25E-31 |
| chr10 | 63382926 | 63396468 | 13543 | 2.23 | 3.48E-32 | 9.97E-31 |
| chr13 | 35282398 | 35284379 | 1982 | 4.1 | 3.63E-32 | 1.04E-30 |
| chr5 | 32297323 | 32314593 | 17271 | 1.18 | 3.88E-32 | 1.11E-30 |
| chr10 | 98622356 | 98626149 | 3794 | 1.71 | 3.96E-32 | 1.13E-30 |
| chr8 | 126181303 | 126188478 | 7176 | 1.66 | 4.04E-32 | 1.15E-30 |
| chr9 | 87583096 | 87587029 | 3934 | 1.89 | 4.13E-32 | 1.18E-30 |
| chr17 | 19606533 | 19670207 | 63675 | -0.57 | 4.16E-32 | 1.19E-30 |
| chr5 | 95093176 | 95179819 | 86644 | 0.57 | 4.32E-32 | 1.23E-30 |
| chr7 | 20336139 | 20347587 | 11449 | 3.39 | 4.64E-32 | 1.32E-30 |
| chr13 | 110650197 | 110671334 | 21138 | 0.9 | 4.66E-32 | 1.33E-30 |
| chr11 | 111427610 | 111433957 | 6348 | -2.45 | 4.79E-32 | 1.36E-30 |
| chr14 | 71179996 | 71239250 | 59255 | -0.89 | 4.82E-32 | 1.37E-30 |
| chr7 | 47008397 | 47013345 | 4949 | 2.92 | 4.97E-32 | 1.41E-30 |
| chr4 | 184249100 | 184259954 | 10855 | 1.84 | 5.12E-32 | 1.45E-30 |
| chr2 | 119425028 | 119428463 | 3436 | -2.56 | 5.21E-32 | 1.48E-30 |
| chr8 | 11656602 | 11760560 | 103959 | 0.56 | 5.46E-32 | 1.54E-30 |
| chr8 | 74330629 | 74336184 | 5556 | 1.06 | 5.70E-32 | 1.61E-30 |
| chr3 | 110650830 | 110715017 | 64188 | 12.02 | 6.05E-32 | 1.71E-30 |
| chr4 | 173958923 | 173962899 | 3977 | 2.67 | 6.22E-32 | 1.75E-30 |
| chr6 | 136242596 | 136266062 | 23467 | -2.14 | 6.58E-32 | 1.85E-30 |
| chr19 | 37010788 | 37020894 | 10107 | 1.5 | 6.83E-32 | 1.92E-30 |
| chr6 | 25640337 | 25644464 | 4128 | 1.39 | 7.15E-32 | 2.01E-30 |
| chr1 | 66695795 | 66830782 | 134988 | 0.44 | 7.45E-32 | 2.09E-30 |
| chr6 | 85591253 | 85621774 | 30522 | 12 | 7.89E-32 | 2.21E-30 |
| chr4 | 25524776 | 25555405 | 30630 | 1.65 | 7.91E-32 | 2.22E-30 |
| chr5 | 10549674 | 10646751 | 97078 | -0.51 | 8.26E-32 | 2.31E-30 |
| chr7 | 101726230 | 101816387 | 90158 | 0.48 | 8.26E-32 | 2.31E-30 |
| chr6 | 56486609 | 56598453 | 111845 | 0.63 | 8.76E-32 | 2.45E-30 |
| chr16 | 3371127 | 3372965 | 1839 | 3.31 | 9.13E-32 | 2.55E-30 |
| chr9 | 116467028 | 116469486 | 2459 | 2.84 | 9.15E-32 | 2.55E-30 |
| chr1 | 109117479 | 109134830 | 17352 | 1.4 | 9.19E-32 | 2.56E-30 |
| chr6 | 127420891 | 127515320 | 94430 | -11.98 | 9.45E-32 | 2.63E-30 |
| chr2 | 138720844 | 138734022 | 13179 | 2 | 9.78E-32 | 2.72E-30 |
| chr5 | 37244125 | 37256736 | 12612 | 1.36 | 1.07E-31 | 2.97E-30 |
| chr3 | 142313392 | 142340146 | 26755 | 0.87 | 1.21E-31 | 3.36E-30 |
| chr19 | 57861934 | 57875930 | 13997 | 1.67 | 1.30E-31 | 3.61E-30 |
| chr15 | 80323632 | 80328304 | 4673 | 1.47 | 1.31E-31 | 3.64E-30 |
| chr1 | 147140822 | 147149953 | 9132 | 1.97 | 1.32E-31 | 3.66E-30 |
| chr12 | 28116733 | 28138143 | 21411 | 1.27 | 1.36E-31 | 3.75E-30 |
| chr19 | 42258287 | 42304133 | 45847 | 11.95 | 1.40E-31 | 3.87E-30 |
| chr5 | 56389973 | 56399015 | 9043 | 4.64 | 1.49E-31 | 4.12E-30 |
| chr3 | 159371508 | 159388313 | 16806 | 1.68 | 1.62E-31 | 4.46E-30 |
| chr15 | 70418833 | 70433986 | 15154 | 3.67 | 1.68E-31 | 4.62E-30 |
| chr14 | 54346897 | 54356605 | 9709 | 1.85 | 1.86E-31 | 5.11E-30 |
| chr10 | 81906856 | 81969426 | 62571 | 0.54 | 1.90E-31 | 5.21E-30 |
| chr10 | 6819129 | 6829195 | 10067 | 1.04 | 1.91E-31 | 5.23E-30 |
| chr13 | 49355180 | 49366527 | 11348 | 2.12 | 1.93E-31 | 5.30E-30 |
| chr4 | 91871221 | 91873879 | 2659 | 3.17 | 1.94E-31 | 5.31E-30 |
| chrX | 115357993 | 115373936 | 15944 | 11.92 | 1.96E-31 | 5.37E-30 |
| chr12 | 110460303 | 110498824 | 38522 | 0.82 | 2.09E-31 | 5.72E-30 |
| chr18 | 52611998 | 52643631 | 31634 | 1.27 | 2.27E-31 | 6.21E-30 |
| chr12 | 56636441 | 56711624 | 75184 | 0.6 | 2.28E-31 | 6.22E-30 |
| chr4 | 90673950 | 90675862 | 1913 | 3.34 | 2.37E-31 | 6.47E-30 |

| | | | | | | |
|---|---|---|---|---|---|---|
| chr13 | 43618255 | 43633450 | 15196 | 1.16 | 2.40E-31 | 6.53E-30 |
| chr8 | 8991414 | 8994422 | 3009 | 2.35 | 2.63E-31 | 7.18E-30 |
| chr4 | 16141583 | 16143689 | 2107 | 3.52 | 2.77E-31 | 7.54E-30 |
| chr16 | 74760049 | 74809463 | 49415 | 0.63 | 2.89E-31 | 7.85E-30 |
| chr1 | 85740163 | 85839333 | 99171 | 0.67 | 3.70E-31 | 1.01E-29 |
| chr14 | 36441880 | 36444834 | 2955 | 7.33 | 3.73E-31 | 1.01E-29 |
| chr2 | 211027011 | 211036865 | 9855 | 1.5 | 3.85E-31 | 1.04E-29 |
| chr8 | 80874515 | 80877691 | 3177 | 1.88 | 3.87E-31 | 1.05E-29 |
| chr18 | 37525120 | 37528869 | 3750 | 2.62 | 3.95E-31 | 1.07E-29 |
| chr10 | 5022940 | 5084986 | 62047 | -0.6 | 3.96E-31 | 1.07E-29 |
| chr3 | 30322788 | 30334518 | 11731 | 1.4 | 4.21E-31 | 1.14E-29 |
| chr6 | 43893506 | 43896456 | 2951 | 3.34 | 4.51E-31 | 1.22E-29 |
| chr10 | 64213373 | 64215287 | 1915 | 2.15 | 4.55E-31 | 1.23E-29 |
| chr5 | 126541237 | 126544253 | 3017 | 1.69 | 6.81E-31 | 1.84E-29 |
| chr3 | 181795404 | 181812130 | 16727 | 1.11 | 6.91E-31 | 1.86E-29 |
| chr14 | 52573114 | 52579559 | 6446 | 1.75 | 7.07E-31 | 1.90E-29 |
| chr2 | 234935904 | 234941949 | 6046 | 2.42 | 9.29E-31 | 2.50E-29 |
| chr12 | 43670703 | 43672933 | 2231 | 4.38 | 9.94E-31 | 2.67E-29 |
| chr3 | 150186114 | 150190513 | 4400 | 1.53 | 1.03E-30 | 2.77E-29 |
| chr1 | 150509901 | 150553190 | 43290 | 0.64 | 1.07E-30 | 2.87E-29 |
| chrX | 93834020 | 93861828 | 27809 | 11.77 | 1.07E-30 | 2.87E-29 |
| chrX | 100129162 | 100140683 | 11522 | 2.48 | 1.11E-30 | 2.98E-29 |
| chr2 | 175830201 | 175839453 | 9253 | -1.29 | 1.14E-30 | 3.04E-29 |
| chr14 | 90848399 | 90886436 | 38038 | 0.77 | 1.21E-30 | 3.24E-29 |
| chr11 | 114150486 | 114189411 | 38926 | 0.58 | 1.40E-30 | 3.74E-29 |
| chr14 | 81768173 | 81771101 | 2929 | 4.44 | 1.40E-30 | 3.74E-29 |
| chr18 | 70982643 | 71008776 | 26134 | 11.75 | 1.43E-30 | 3.80E-29 |
| chr7 | 64450705 | 64467770 | 17066 | 2.44 | 1.44E-30 | 3.83E-29 |
| chr9 | 123440698 | 123478211 | 37514 | -0.57 | 1.47E-30 | 3.90E-29 |
| chr5 | 115208260 | 115277561 | 69302 | 11.75 | 1.48E-30 | 3.94E-29 |
| chr14 | 32776004 | 32778426 | 2423 | 2.43 | 1.50E-30 | 3.98E-29 |
| chr10 | 5580184 | 5602393 | 22210 | 0.74 | 1.51E-30 | 4.00E-29 |
| chr8 | 102446282 | 102487789 | 41508 | 0.62 | 1.58E-30 | 4.18E-29 |
| chr1 | 205894520 | 205906067 | 11548 | 1.22 | 1.76E-30 | 4.65E-29 |
| chr7 | 17244329 | 17247813 | 3485 | 1.21 | 1.76E-30 | 4.65E-29 |
| chr12 | 122943410 | 123024169 | 80760 | 0.54 | 2.28E-30 | 6.03E-29 |
| chr12 | 123997715 | 124019443 | 21729 | 1.11 | 2.54E-30 | 6.71E-29 |
| chr1 | 207566207 | 207568442 | 2236 | 4.1 | 2.58E-30 | 6.79E-29 |
| chr21 | 30628216 | 30645082 | 16867 | 3.08 | 2.59E-30 | 6.83E-29 |
| chr19 | 57900373 | 57902356 | 1984 | 2.18 | 2.68E-30 | 7.06E-29 |
| chr5 | 131693397 | 131730828 | 37432 | 0.57 | 2.80E-30 | 7.36E-29 |
| chr3 | 57935309 | 58164001 | 228693 | 0.33 | 2.96E-30 | 7.78E-29 |
| chr5 | 43396705 | 43434501 | 37797 | 1.5 | 2.98E-30 | 7.82E-29 |
| chr5 | 95827452 | 95875641 | 48190 | 2.68 | 3.06E-30 | 8.02E-29 |
| chr9 | 119627980 | 119674963 | 46984 | 11.68 | 3.24E-30 | 8.50E-29 |
| chr17 | 39801383 | 39874837 | 73455 | 0.54 | 3.46E-30 | 9.05E-29 |
| chr1 | 16447117 | 16516816 | 69700 | 0.52 | 3.55E-30 | 9.28E-29 |
| chr12 | 89843626 | 89849153 | 5528 | 1.6 | 3.66E-30 | 9.58E-29 |
| chr16 | 59605981 | 59609311 | 3331 | -2.13 | 3.78E-30 | 9.86E-29 |
| chr11 | 16172984 | 16179235 | 6252 | 0.89 | 3.86E-30 | 1.01E-28 |
| chr8 | 126271327 | 126322657 | 51331 | 0.81 | 4.05E-30 | 1.05E-28 |
| chr10 | 70847032 | 70853592 | 6561 | -7.61 | 4.30E-30 | 1.12E-28 |
| chr1 | 88248542 | 88274106 | 25565 | 0.92 | 4.43E-30 | 1.15E-28 |
| chr3 | 30752267 | 30765083 | 12817 | 1.64 | 4.86E-30 | 1.26E-28 |
| chr3 | 191479516 | 191493355 | 13840 | 5.01 | 5.18E-30 | 1.35E-28 |
| chr7 | 63974858 | 64024198 | 49341 | 11.63 | 5.29E-30 | 1.37E-28 |
| chrX | 35357576 | 35361599 | 4024 | 2.31 | 5.46E-30 | 1.42E-28 |
| chr3 | 112453154 | 112543464 | 90311 | 0.6 | 5.47E-30 | 1.42E-28 |
| chr5 | 140367415 | 140386194 | 18780 | 0.89 | 5.69E-30 | 1.47E-28 |
| chr11 | 16192481 | 16207032 | 14552 | 3.36 | 6.05E-30 | 1.56E-28 |
| chr12 | 120150752 | 120157166 | 6415 | 1.76 | 6.88E-30 | 1.78E-28 |

| | | | | | | |
|---|---|---|---|---|---|---|
| chr9 | 33225369 | 33354986 | 129618 | 0.4 | 7.55E-30 | 1.95E-28 |
| chr10 | 30091297 | 30102558 | 11262 | 0.93 | 7.65E-30 | 1.97E-28 |
| chr2 | 231602096 | 231604573 | 2478 | 2.76 | 8.02E-30 | 2.07E-28 |
| chr5 | 110865058 | 110867366 | 2309 | 2.83 | 8.05E-30 | 2.07E-28 |
| chr3 | 16181714 | 16191114 | 9401 | -1.55 | 8.10E-30 | 2.08E-28 |
| chr3 | 148931767 | 148944080 | 12314 | 0.95 | 8.62E-30 | 2.21E-28 |
| chr15 | 52942347 | 52954064 | 11718 | 0.78 | 1.09E-29 | 2.81E-28 |
| chr10 | 127798917 | 127809241 | 10325 | 2.24 | 1.12E-29 | 2.87E-28 |
| chr6 | 108214998 | 108217469 | 2472 | 2.56 | 1.15E-29 | 2.94E-28 |
| chr12 | 57053947 | 57084782 | 30836 | 0.74 | 1.18E-29 | 3.02E-28 |
| chr2 | 201606070 | 201653077 | 47008 | -0.93 | 1.23E-29 | 3.13E-28 |
| chr2 | 167213321 | 167221976 | 8656 | -3.33 | 1.30E-29 | 3.32E-28 |
| chr4 | 88785446 | 88788764 | 3319 | 1.83 | 1.34E-29 | 3.43E-28 |
| chr2 | 228727192 | 228749738 | 22547 | -2.5 | 1.37E-29 | 3.50E-28 |
| chr3 | 69904562 | 69940813 | 36252 | 0.87 | 1.41E-29 | 3.60E-28 |
| chr7 | 38968790 | 38970696 | 1907 | 3.68 | 1.42E-29 | 3.62E-28 |
| chr2 | 216635240 | 216673862 | 38623 | -1.27 | 1.44E-29 | 3.66E-28 |
| chr5 | 10703378 | 10728453 | 25076 | 1.06 | 1.49E-29 | 3.80E-28 |
| chr7 | 116301678 | 116379078 | 77401 | 0.48 | 1.57E-29 | 4.00E-28 |
| chr3 | 129511848 | 129515919 | 4072 | 1.45 | 1.62E-29 | 4.10E-28 |
| chr19 | 23837753 | 23885187 | 47435 | 11.54 | 1.63E-29 | 4.13E-28 |
| chr16 | 27552028 | 27591296 | 39269 | -0.9 | 1.64E-29 | 4.17E-28 |
| chr1 | 60138084 | 60144807 | 6724 | 3.08 | 1.71E-29 | 4.33E-28 |
| chr4 | 169447783 | 169463738 | 15956 | -1.23 | 1.89E-29 | 4.79E-28 |
| chr20 | 16494689 | 16513321 | 18633 | 0.74 | 1.96E-29 | 4.95E-28 |
| chr9 | 38224909 | 38228799 | 3891 | 2.06 | 2.04E-29 | 5.15E-28 |
| chr3 | 13577109 | 13590462 | 13354 | 1.84 | 2.08E-29 | 5.26E-28 |
| chr12 | 40602226 | 40640764 | 38539 | 0.92 | 2.38E-29 | 6.00E-28 |
| chr9 | 85627095 | 85678849 | 51755 | 0.77 | 2.57E-29 | 6.48E-28 |
| chr1 | 210757928 | 210773417 | 15490 | -1.07 | 2.88E-29 | 7.25E-28 |
| chr19 | 53161335 | 53195437 | 34103 | 11.48 | 2.95E-29 | 7.41E-28 |
| chr10 | 74357753 | 74515230 | 157478 | 0.39 | 3.00E-29 | 7.53E-28 |
| chr12 | 72232550 | 72235248 | 2699 | 1.21 | 3.20E-29 | 8.03E-28 |
| chr14 | 24501091 | 24506130 | 5040 | -2.4 | 3.36E-29 | 8.44E-28 |
| chr11 | 10876937 | 10924235 | 47299 | -0.67 | 3.41E-29 | 8.55E-28 |
| chr4 | 86790401 | 86793797 | 3397 | 2.7 | 3.65E-29 | 9.14E-28 |
| chr14 | 54409236 | 54424082 | 14847 | 2 | 3.94E-29 | 9.87E-28 |
| chr20 | 44484849 | 44496538 | 11690 | 2.53 | 4.47E-29 | 1.12E-27 |
| chr2 | 164518089 | 164525474 | 7386 | -1.43 | 4.90E-29 | 1.22E-27 |
| chr15 | 86020936 | 86047569 | 26634 | 0.91 | 4.91E-29 | 1.23E-27 |
| chr5 | 43034676 | 43079524 | 44849 | 0.66 | 5.27E-29 | 1.32E-27 |
| chr2 | 211053594 | 211055704 | 2111 | 5.67 | 5.53E-29 | 1.38E-27 |
| chr11 | 13052355 | 13066154 | 13800 | 2.61 | 5.79E-29 | 1.44E-27 |
| chr9 | 18392258 | 18395318 | 3061 | 1.53 | 5.94E-29 | 1.48E-27 |
| chr1 | 208373800 | 208378501 | 4702 | 1.51 | 6.11E-29 | 1.52E-27 |
| chr2 | 56667381 | 56678879 | 11499 | 1.79 | 6.29E-29 | 1.56E-27 |
| chr6 | 81147599 | 81150286 | 2688 | 4.53 | 6.63E-29 | 1.64E-27 |
| chr8 | 88283832 | 88300019 | 16188 | 11.41 | 6.75E-29 | 1.67E-27 |
| chr9 | 4114544 | 4180803 | 66260 | -0.57 | 6.75E-29 | 1.67E-27 |
| chr19 | 38352923 | 38367530 | 14608 | 11.41 | 6.88E-29 | 1.70E-27 |
| chr5 | 153569075 | 153604109 | 35035 | -0.81 | 7.48E-29 | 1.85E-27 |
| chr7 | 137632000 | 137694027 | 62028 | 0.6 | 7.58E-29 | 1.87E-27 |
| chr12 | 123702926 | 123757509 | 54584 | 0.68 | 8.22E-29 | 2.03E-27 |
| chrX | 30293778 | 30327525 | 33748 | -0.71 | 8.91E-29 | 2.20E-27 |
| chr2 | 62793581 | 62839965 | 46385 | -0.59 | 9.35E-29 | 2.31E-27 |
| chr1 | 153669272 | 153682596 | 13325 | 1.44 | 9.43E-29 | 2.32E-27 |
| chr3 | 86835999 | 86844414 | 8416 | 3.38 | 9.82E-29 | 2.42E-27 |
| chr6 | 150151625 | 150190096 | 38472 | 0.63 | 1.13E-28 | 2.77E-27 |
| chr1 | 26436811 | 26454046 | 17236 | 1.4 | 1.13E-28 | 2.77E-27 |
| chr12 | 125576449 | 125587853 | 11405 | 1.53 | 1.22E-28 | 2.99E-27 |
| chr7 | 117291482 | 117324666 | 33185 | 1.28 | 1.28E-28 | 3.14E-27 |

| | | | | | | |
|---|---|---|---|---|---|---|
| chr19 | 6567528 | 6582205 | 14678 | -1.91 | 1.29E-28 | 3.16E-27 |
| chr6 | 165985749 | 165989519 | 3771 | 1.63 | 1.30E-28 | 3.19E-27 |
| chr7 | 34075130 | 34077529 | 2400 | -2.58 | 1.32E-28 | 3.24E-27 |
| chr1 | 89591171 | 89595457 | 4287 | 3.12 | 1.47E-28 | 3.60E-27 |
| chr13 | 80009304 | 80017261 | 7958 | 2.77 | 1.58E-28 | 3.87E-27 |
| chr6 | 85352768 | 85362152 | 9385 | 1.56 | 1.65E-28 | 4.02E-27 |
| chr1 | 212989538 | 213008490 | 18953 | 1.27 | 1.70E-28 | 4.15E-27 |
| chr15 | 102061404 | 102071142 | 9739 | 1.53 | 1.73E-28 | 4.22E-27 |
| chr3 | 69226878 | 69232798 | 5921 | 1.57 | 1.76E-28 | 4.29E-27 |
| chrX | 28593937 | 28605898 | 11962 | 3.53 | 1.82E-28 | 4.42E-27 |
| chr12 | 64455162 | 64515486 | 60325 | 0.62 | 1.85E-28 | 4.51E-27 |
| chr10 | 43132565 | 43138036 | 5472 | 1.18 | 1.93E-28 | 4.68E-27 |
| chr7 | 98791491 | 98833949 | 42459 | 0.61 | 2.04E-28 | 4.96E-27 |
| chr1 | 223062523 | 223072025 | 9503 | 2.81 | 2.21E-28 | 5.35E-27 |
| chr19 | 52458796 | 52494767 | 35972 | 11.3 | 2.24E-28 | 5.44E-27 |
| chr4 | 102176910 | 102221814 | 44905 | 1.22 | 2.26E-28 | 5.47E-27 |
| chr10 | 25347417 | 25352732 | 5316 | 0.91 | 2.29E-28 | 5.54E-27 |
| chr20 | 52682554 | 52699153 | 16600 | 1.4 | 2.36E-28 | 5.70E-27 |
| chr17 | 41521443 | 41547032 | 25590 | 1.42 | 2.52E-28 | 6.08E-27 |
| chr20 | 4020692 | 4023288 | 2597 | 3.32 | 2.63E-28 | 6.35E-27 |
| chr16 | 87859209 | 87906021 | 46813 | 0.57 | 2.65E-28 | 6.39E-27 |
| chr10 | 54199455 | 54215559 | 16105 | -0.84 | 2.67E-28 | 6.44E-27 |
| chr4 | 40991058 | 41012085 | 21028 | -1.19 | 2.84E-28 | 6.83E-27 |
| chr15 | 87082834 | 87101632 | 18799 | 11.28 | 2.84E-28 | 6.83E-27 |
| chr8 | 122156789 | 122178205 | 21417 | 1.16 | 2.92E-28 | 7.02E-27 |
| chr7 | 128067810 | 128102376 | 34567 | -0.73 | 3.07E-28 | 7.36E-27 |
| chr19 | 9160396 | 9191223 | 30828 | 0.85 | 3.08E-28 | 7.39E-27 |
| chr15 | 53850693 | 53853834 | 3142 | 3.41 | 3.10E-28 | 7.43E-27 |
| chr5 | 39624161 | 39626610 | 2450 | 3.13 | 3.10E-28 | 7.43E-27 |
| chr4 | 128702505 | 128706414 | 3910 | 1.7 | 3.12E-28 | 7.46E-27 |
| chr4 | 95641192 | 95654514 | 13323 | 0.74 | 3.29E-28 | 7.86E-27 |
| chr7 | 8300292 | 8302577 | 2286 | 2.4 | 3.55E-28 | 8.48E-27 |
| chr11 | 7619795 | 7679177 | 59383 | -0.73 | 3.58E-28 | 8.56E-27 |
| chr10 | 76041512 | 76054780 | 13269 | 2.04 | 4.06E-28 | 9.68E-27 |
| chr17 | 79008092 | 79031636 | 23545 | 1.38 | 4.08E-28 | 9.73E-27 |
| chr6 | 63333038 | 63334908 | 1871 | 4.48 | 4.13E-28 | 9.84E-27 |
| chr2 | 169055033 | 169078772 | 23740 | 1.39 | 4.16E-28 | 9.89E-27 |
| chr8 | 116973990 | 117008402 | 34413 | 11.25 | 4.50E-28 | 1.07E-26 |
| chr12 | 63685208 | 63692795 | 7588 | 1.22 | 4.67E-28 | 1.11E-26 |
| chr5 | 107200105 | 107206310 | 6206 | 2.62 | 4.68E-28 | 1.11E-26 |
| chr14 | 102187792 | 102199983 | 12192 | 1.58 | 4.93E-28 | 1.17E-26 |
| chr6 | 109614669 | 109618614 | 3946 | 3.68 | 5.06E-28 | 1.20E-26 |
| chr5 | 58739734 | 58895990 | 156257 | 0.3 | 5.07E-28 | 1.20E-26 |
| chr10 | 30344441 | 30348581 | 4141 | -2.55 | 5.35E-28 | 1.26E-26 |
| chr16 | 14558324 | 14686979 | 128656 | -0.49 | 5.62E-28 | 1.33E-26 |
| chr7 | 22600326 | 22603524 | 3199 | 3.5 | 5.75E-28 | 1.36E-26 |
| chr4 | 187064960 | 187071490 | 6531 | 2.33 | 5.96E-28 | 1.41E-26 |
| chr13 | 90031282 | 90033921 | 2640 | 2.63 | 5.96E-28 | 1.41E-26 |
| chr1 | 183578172 | 183605532 | 27361 | 1.19 | 6.01E-28 | 1.42E-26 |
| chr16 | 77321923 | 77333786 | 11864 | 1.2 | 6.21E-28 | 1.46E-26 |
| chr4 | 75228456 | 75264164 | 35709 | 0.68 | 6.53E-28 | 1.54E-26 |
| chr2 | 208480621 | 208500850 | 20230 | -0.93 | 6.87E-28 | 1.61E-26 |
| chr5 | 142711201 | 142785671 | 74471 | 0.45 | 7.18E-28 | 1.69E-26 |
| chr3 | 94334251 | 94336956 | 2706 | 1.75 | 7.61E-28 | 1.79E-26 |
| chr1 | 193521894 | 193528911 | 7018 | 3.12 | 7.65E-28 | 1.79E-26 |
| chr7 | 17168845 | 17183355 | 14511 | 1.32 | 7.81E-28 | 1.83E-26 |
| chr5 | 180286655 | 180301448 | 14794 | 1.97 | 8.65E-28 | 2.03E-26 |
| chr11 | 69041303 | 69089663 | 48361 | 0.49 | 8.73E-28 | 2.04E-26 |
| chr1 | 240471368 | 240478598 | 7231 | 0.93 | 8.89E-28 | 2.08E-26 |
| chr12 | 4139349 | 4142760 | 3412 | 2.2 | 1.04E-27 | 2.43E-26 |
| chr22 | 30952697 | 30970677 | 17981 | 1.32 | 1.07E-27 | 2.50E-26 |

| | | | | | | |
|---|---|---|---|---|---|---|
| chr6 | 72187839 | 72206510 | 18672 | 2.97 | 1.08E-27 | 2.52E-26 |
| chr12 | 41212745 | 41224556 | 11812 | 1.41 | 1.14E-27 | 2.66E-26 |
| chr1 | 218746471 | 218775407 | 28937 | 1.01 | 1.19E-27 | 2.76E-26 |
| chr1 | 233839821 | 233845071 | 5251 | 1.84 | 1.19E-27 | 2.77E-26 |
| chr5 | 167123466 | 167129447 | 5982 | 1.06 | 1.23E-27 | 2.85E-26 |
| chr11 | 105990954 | 105998973 | 8020 | 2.94 | 1.25E-27 | 2.90E-26 |
| chr4 | 77506500 | 77512652 | 6153 | 1.67 | 1.28E-27 | 2.98E-26 |
| chr3 | 105804544 | 105817410 | 12867 | 2.8 | 1.30E-27 | 3.02E-26 |
| chr19 | 37231586 | 37264631 | 33046 | 11.14 | 1.35E-27 | 3.13E-26 |
| chr7 | 128740083 | 128751354 | 11272 | 2.14 | 1.42E-27 | 3.29E-26 |
| chr10 | 116437565 | 116446451 | 8887 | 1.53 | 1.42E-27 | 3.29E-26 |
| chr3 | 40128283 | 40130378 | 2096 | 3.64 | 1.52E-27 | 3.51E-26 |
| chr6 | 4313116 | 4324037 | 10922 | 0.95 | 1.56E-27 | 3.60E-26 |
| chr7 | 46734961 | 46737231 | 2271 | 3.23 | 1.57E-27 | 3.61E-26 |
| chr10 | 65263995 | 65283404 | 19410 | 1.01 | 1.62E-27 | 3.74E-26 |
| chrX | 39786971 | 39794798 | 7828 | 1.43 | 1.63E-27 | 3.76E-26 |
| chr12 | 14391934 | 14437620 | 45687 | 4.36 | 1.74E-27 | 4.01E-26 |
| chr20 | 59691398 | 59696524 | 5127 | -2.04 | 1.82E-27 | 4.19E-26 |
| chr18 | 21030978 | 21041681 | 10704 | 0.88 | 1.84E-27 | 4.24E-26 |
| chr3 | 146026167 | 146033482 | 7316 | 1.49 | 1.95E-27 | 4.47E-26 |
| chr3 | 181341500 | 181343187 | 1688 | 4.34 | 1.98E-27 | 4.55E-26 |
| chr17 | 80589752 | 80647942 | 58191 | 0.71 | 1.99E-27 | 4.55E-26 |
| chr10 | 4093182 | 4110789 | 17608 | 1.6 | 2.01E-27 | 4.60E-26 |
| chr8 | 126121880 | 126143039 | 21160 | 1.43 | 2.05E-27 | 4.70E-26 |
| chr9 | 112942265 | 112969858 | 27594 | 1.23 | 2.07E-27 | 4.73E-26 |
| chr4 | 94103332 | 94106296 | 2965 | 2.47 | 2.07E-27 | 4.73E-26 |
| chr3 | 189229819 | 189232588 | 2770 | 1.96 | 2.14E-27 | 4.90E-26 |
| chr11 | 44995523 | 45005762 | 10240 | 0.83 | 2.21E-27 | 5.04E-26 |
| chr8 | 90807878 | 90817530 | 9653 | 1.17 | 2.24E-27 | 5.10E-26 |
| chr12 | 118290379 | 118292786 | 2408 | 4.28 | 2.33E-27 | 5.30E-26 |
| chr8 | 81662215 | 81680905 | 18691 | 9.13 | 2.47E-27 | 5.64E-26 |
| chr11 | 94799476 | 94823615 | 24140 | 1.4 | 2.59E-27 | 5.90E-26 |
| chr13 | 80054932 | 80067969 | 13038 | 1.02 | 2.65E-27 | 6.02E-26 |
| chr9 | 136737534 | 136763199 | 25666 | -1.09 | 2.67E-27 | 6.06E-26 |
| chr1 | 152005038 | 152022838 | 17801 | 0.65 | 2.68E-27 | 6.09E-26 |
| chr8 | 54229369 | 54237109 | 7741 | 3.82 | 2.69E-27 | 6.10E-26 |
| chr1 | 45711666 | 45713073 | 1408 | 3.15 | 2.76E-27 | 6.26E-26 |
| chr10 | 116661641 | 116678903 | 17263 | 1.97 | 3.05E-27 | 6.92E-26 |
| chr20 | 4877290 | 4990411 | 113122 | 0.53 | 3.18E-27 | 7.19E-26 |
| chr4 | 159212632 | 159243884 | 31253 | 1.01 | 3.22E-27 | 7.29E-26 |
| chr8 | 110582468 | 110634304 | 51837 | 2.22 | 3.30E-27 | 7.46E-26 |
| chr10 | 4980542 | 5008869 | 28328 | -0.75 | 3.53E-27 | 7.96E-26 |
| chrX | 9306689 | 9321560 | 14872 | 1.54 | 3.53E-27 | 7.97E-26 |
| chr7 | 105897336 | 105926996 | 29661 | -0.86 | 3.60E-27 | 8.11E-26 |
| chr12 | 54641974 | 54701003 | 59030 | 0.59 | 3.66E-27 | 8.25E-26 |
| chr20 | 31753117 | 31780293 | 27177 | 1.9 | 3.80E-27 | 8.56E-26 |
| chr12 | 104226110 | 104254776 | 28667 | 1.69 | 3.92E-27 | 8.82E-26 |
| chr6 | 149351238 | 149355323 | 4086 | -1.22 | 4.10E-27 | 9.22E-26 |
| chr12 | 41286890 | 41315476 | 28587 | 1.33 | 4.37E-27 | 9.81E-26 |
| chr17 | 76948156 | 77006686 | 58531 | 0.69 | 4.59E-27 | 1.03E-25 |
| chr5 | 125052341 | 125056264 | 3924 | 3.13 | 4.75E-27 | 1.07E-25 |
| chr12 | 79256545 | 79281449 | 24905 | 1.22 | 5.00E-27 | 1.12E-25 |
| chr4 | 95444129 | 95471174 | 27046 | 0.89 | 5.01E-27 | 1.12E-25 |
| chr18 | 3100909 | 3120699 | 19791 | 1.8 | 5.05E-27 | 1.13E-25 |
| chr17 | 74635788 | 74668804 | 33017 | 1.06 | 5.14E-27 | 1.15E-25 |
| chrX | 107638373 | 107645819 | 7447 | -1.3 | 5.15E-27 | 1.15E-25 |
| chr7 | 82665095 | 82667305 | 2211 | 1.97 | 5.58E-27 | 1.25E-25 |
| chr4 | 91416587 | 91418688 | 2102 | 2.53 | 5.64E-27 | 1.26E-25 |
| chr9 | 22200962 | 22220092 | 19131 | 0.84 | 5.71E-27 | 1.27E-25 |
| chr2 | 161060481 | 161179251 | 118771 | -0.38 | 5.79E-27 | 1.29E-25 |
| chr7 | 116057234 | 116232174 | 174941 | 0.38 | 5.85E-27 | 1.30E-25 |

| | | | | | | |
|---|---|---|---|---|---|---|
| chr6 | 24644134 | 24669890 | 25757 | -0.73 | 6.03E-27 | 1.34E-25 |
| chr2 | 129215362 | 129217878 | 2517 | 4.51 | 6.50E-27 | 1.44E-25 |
| chr19 | 58602750 | 58667038 | 64289 | 10.99 | 6.97E-27 | 1.55E-25 |
| chr17 | 38439521 | 38527874 | 88354 | -0.55 | 6.99E-27 | 1.55E-25 |
| chr6 | 7504721 | 7506824 | 2104 | 2.29 | 7.55E-27 | 1.67E-25 |
| chr8 | 112034074 | 112036200 | 2127 | 7.31 | 8.52E-27 | 1.89E-25 |
| chr16 | 88337514 | 88372399 | 34886 | -0.82 | 8.82E-27 | 1.95E-25 |
| chr12 | 69069925 | 69082439 | 12515 | 1.03 | 9.20E-27 | 2.04E-25 |
| chr5 | 82445311 | 82466489 | 21179 | 1.5 | 9.66E-27 | 2.14E-25 |
| chr8 | 54791786 | 54831648 | 39863 | -0.8 | 9.80E-27 | 2.16E-25 |
| chr10 | 12255734 | 12270709 | 14976 | 1.23 | 9.82E-27 | 2.17E-25 |
| chr8 | 144300573 | 144303275 | 2703 | 1.8 | 1.03E-26 | 2.28E-25 |
| chr2 | 7965052 | 7967151 | 2100 | 2.52 | 1.05E-26 | 2.32E-25 |
| chr12 | 6975787 | 7090365 | 114579 | 0.49 | 1.07E-26 | 2.35E-25 |
| chr5 | 127370641 | 127376803 | 6163 | 1 | 1.12E-26 | 2.46E-25 |
| chr4 | 100367199 | 100369741 | 2543 | 1.56 | 1.18E-26 | 2.59E-25 |
| chr12 | 10853420 | 10876780 | 23361 | 0.7 | 1.19E-26 | 2.61E-25 |
| chr7 | 138211232 | 138227519 | 16288 | 6.84 | 1.23E-26 | 2.71E-25 |
| chr12 | 81170264 | 81173230 | 2967 | 1.85 | 1.26E-26 | 2.76E-25 |
| chr10 | 33402929 | 33630600 | 227672 | 0.32 | 1.26E-26 | 2.77E-25 |
| chr11 | 122626653 | 122649298 | 22646 | -1.57 | 1.28E-26 | 2.81E-25 |
| chr10 | 71973119 | 71994243 | 21125 | 0.94 | 1.29E-26 | 2.82E-25 |
| chr16 | 67378861 | 67381570 | 2710 | 2.76 | 1.30E-26 | 2.85E-25 |
| chr16 | 71699369 | 71723248 | 23880 | 1.15 | 1.32E-26 | 2.88E-25 |
| chr6 | 114095633 | 114109935 | 14303 | 1.61 | 1.33E-26 | 2.91E-25 |
| chr12 | 49091765 | 49112039 | 20275 | 0.83 | 1.37E-26 | 2.99E-25 |
| chr14 | 90376103 | 90380082 | 3980 | 1.21 | 1.41E-26 | 3.07E-25 |
| chr14 | 56638927 | 56673979 | 35053 | 0.93 | 1.45E-26 | 3.16E-25 |
| chr12 | 31429591 | 31479907 | 50317 | 0.63 | 1.45E-26 | 3.16E-25 |
| chr20 | 31862501 | 31871875 | 9375 | 3.86 | 1.58E-26 | 3.44E-25 |
| chr19 | 20255574 | 20307576 | 52003 | 10.91 | 1.62E-26 | 3.51E-25 |
| chr3 | 98516547 | 98644797 | 128251 | 0.42 | 1.71E-26 | 3.72E-25 |
| chr12 | 8308557 | 8311463 | 2907 | 1.25 | 1.72E-26 | 3.73E-25 |
| chr2 | 68403519 | 68458759 | 55241 | -0.72 | 1.77E-26 | 3.84E-25 |
| chr13 | 99125665 | 99162757 | 37093 | 0.79 | 1.79E-26 | 3.88E-25 |
| chr7 | 83175724 | 83178994 | 3271 | 1.11 | 1.85E-26 | 4.02E-25 |
| chr5 | 57702178 | 57707673 | 5496 | -2.78 | 1.92E-26 | 4.17E-25 |
| chr3 | 165360242 | 165362034 | 1793 | 5.32 | 1.94E-26 | 4.20E-25 |
| chr1 | 95073027 | 95087866 | 14840 | 2.18 | 2.02E-26 | 4.37E-25 |
| chr16 | 52644435 | 52646949 | 2515 | 1.87 | 2.10E-26 | 4.53E-25 |
| chr2 | 236076371 | 236081261 | 4891 | -1.62 | 2.11E-26 | 4.56E-25 |
| chr4 | 108774163 | 108798492 | 24330 | 2.3 | 2.12E-26 | 4.57E-25 |
| chr7 | 18547646 | 18555365 | 7720 | 2.72 | 2.17E-26 | 4.67E-25 |
| chr5 | 72835778 | 72841308 | 5531 | 1.78 | 2.18E-26 | 4.70E-25 |
| chr6 | 136171950 | 136204940 | 32991 | -1.53 | 2.20E-26 | 4.73E-25 |
| chr19 | 38179559 | 38211288 | 31730 | 10.88 | 2.26E-26 | 4.85E-25 |
| chr4 | 17795304 | 17827627 | 32324 | 0.92 | 2.27E-26 | 4.87E-25 |
| chr1 | 160488922 | 160491757 | 2836 | 1.42 | 2.39E-26 | 5.13E-25 |
| chr4 | 151137582 | 151174021 | 36440 | -0.83 | 2.47E-26 | 5.31E-25 |
| chr12 | 88685040 | 88701273 | 16234 | -1.32 | 2.63E-26 | 5.64E-25 |
| chr5 | 121030261 | 121031756 | 1496 | 4.39 | 2.70E-26 | 5.78E-25 |
| chr13 | 50938906 | 50941917 | 3012 | 5.45 | 2.77E-26 | 5.93E-25 |
| chr17 | 3782596 | 3800099 | 17504 | -1.01 | 2.83E-26 | 6.05E-25 |
| chr4 | 143356697 | 143362210 | 5514 | 3.58 | 3.12E-26 | 6.67E-25 |
| chr6 | 161968191 | 161973124 | 4934 | 3.43 | 3.15E-26 | 6.72E-25 |
| chr1 | 84609014 | 84623137 | 14124 | 2.12 | 3.15E-26 | 6.72E-25 |
| chr9 | 75128756 | 75208441 | 79686 | 0.47 | 3.17E-26 | 6.77E-25 |
| chr19 | 36883725 | 36914855 | 31131 | 10.85 | 3.17E-26 | 6.77E-25 |
| chr12 | 13099606 | 13108817 | 9212 | 3.33 | 3.20E-26 | 6.81E-25 |
| chr10 | 45495823 | 45497638 | 1816 | 2.71 | 3.20E-26 | 6.82E-25 |
| chr12 | 56185696 | 56225124 | 39429 | 0.9 | 3.27E-26 | 6.95E-25 |

| | | | | | | |
|---|---|---|---|---|---|---|
| chr1 | 153699562 | 153725394 | 25833 | 0.94 | 3.38E-26 | 7.19E-25 |
| chrX | 28421274 | 28423585 | 2312 | 3.76 | 3.50E-26 | 7.44E-25 |
| chr1 | 192667294 | 192672566 | 5273 | 0.98 | 3.51E-26 | 7.46E-25 |
| chr7 | 20431906 | 20448365 | 16460 | 2.65 | 3.52E-26 | 7.46E-25 |
| chr7 | 79936504 | 79942664 | 6161 | 1.45 | 3.80E-26 | 8.07E-25 |
| chr20 | 17815530 | 17837301 | 21772 | 1.5 | 4.26E-26 | 9.02E-25 |
| chr19 | 58507478 | 58537751 | 30274 | 10.82 | 4.30E-26 | 9.11E-25 |
| chr7 | 14019802 | 14032052 | 12251 | 1.05 | 4.68E-26 | 9.90E-25 |
| chr1 | 183666459 | 183672249 | 5791 | 1.02 | 4.71E-26 | 9.96E-25 |
| chr10 | 5110167 | 5152135 | 41969 | -0.5 | 4.93E-26 | 1.04E-24 |
| chr18 | 45524139 | 45554488 | 30350 | 0.66 | 5.01E-26 | 1.06E-24 |
| chr5 | 178286327 | 178303867 | 17541 | 2.07 | 5.26E-26 | 1.11E-24 |
| chr12 | 92449650 | 92502617 | 52968 | 0.59 | 5.48E-26 | 1.15E-24 |
| chr18 | 20187956 | 20194717 | 6762 | 1.56 | 6.07E-26 | 1.28E-24 |
| chr6 | 2221808 | 2226985 | 5178 | 2.53 | 6.42E-26 | 1.35E-24 |
| chr2 | 9446693 | 9489206 | 42514 | -1.09 | 6.48E-26 | 1.36E-24 |
| chr3 | 83982057 | 84021957 | 39901 | 10.78 | 6.78E-26 | 1.43E-24 |
| chr12 | 40492664 | 40499832 | 7169 | 2.48 | 6.81E-26 | 1.43E-24 |
| chr1 | 153578654 | 153608731 | 30078 | 0.64 | 6.84E-26 | 1.44E-24 |
| chr4 | 74365375 | 74366893 | 1519 | 2.67 | 6.89E-26 | 1.45E-24 |
| chrX | 13233881 | 13253447 | 19567 | 1.3 | 6.91E-26 | 1.45E-24 |
| chr2 | 19504039 | 19505816 | 1778 | 2.12 | 7.33E-26 | 1.54E-24 |
| chr3 | 31025778 | 31027618 | 1841 | 2.75 | 7.66E-26 | 1.60E-24 |
| chr12 | 120963996 | 121001259 | 37264 | 0.71 | 7.94E-26 | 1.66E-24 |
| chr5 | 56730805 | 56733506 | 2702 | 1.76 | 8.65E-26 | 1.81E-24 |
| chr1 | 174921607 | 174998998 | 77392 | 0.76 | 8.69E-26 | 1.82E-24 |
| chr11 | 65183215 | 65278982 | 95768 | 0.36 | 8.79E-26 | 1.84E-24 |
| chr5 | 131596514 | 131603383 | 6870 | 2.16 | 8.88E-26 | 1.85E-24 |
| chr18 | 49139954 | 49144704 | 4751 | -2.51 | 9.09E-26 | 1.89E-24 |
| chr14 | 85936362 | 85953564 | 17203 | 1.37 | 9.38E-26 | 1.95E-24 |
| chr18 | 3245539 | 3265593 | 20055 | 0.69 | 9.46E-26 | 1.97E-24 |
| chr12 | 40669379 | 40695315 | 25937 | 0.77 | 9.74E-26 | 2.03E-24 |
| chr19 | 24088697 | 24107346 | 18650 | 10.76 | 9.83E-26 | 2.04E-24 |
| chr21 | 18807438 | 18840658 | 33221 | 0.99 | 9.90E-26 | 2.06E-24 |
| chr7 | 13958633 | 13961210 | 2578 | 1.73 | 1.04E-25 | 2.16E-24 |
| chr12 | 87959835 | 87961958 | 2124 | 4.08 | 1.06E-25 | 2.19E-24 |
| chr8 | 71545069 | 71582579 | 37511 | 0.74 | 1.07E-25 | 2.23E-24 |
| chr21 | 39502139 | 39515433 | 13295 | 1.05 | 1.07E-25 | 2.23E-24 |
| chr8 | 34861294 | 34889149 | 27856 | -10.73 | 1.10E-25 | 2.28E-24 |
| chr4 | 74756805 | 74759499 | 2695 | 2.89 | 1.14E-25 | 2.35E-24 |
| chr2 | 174397933 | 174416057 | 18125 | 1.86 | 1.16E-25 | 2.40E-24 |
| chr4 | 124507574 | 124522160 | 14587 | 0.92 | 1.19E-25 | 2.46E-24 |
| chr7 | 85348753 | 85363623 | 14871 | 2.47 | 1.27E-25 | 2.62E-24 |
| chr14 | 80677910 | 80684331 | 6422 | -1.34 | 1.27E-25 | 2.62E-24 |
| chr12 | 89573536 | 89587209 | 13674 | 2.32 | 1.30E-25 | 2.68E-24 |
| chr2 | 64768048 | 64776725 | 8678 | 1.66 | 1.33E-25 | 2.73E-24 |
| chr12 | 75281728 | 75291822 | 10095 | 1.3 | 1.39E-25 | 2.85E-24 |
| chr17 | 46231328 | 46241400 | 10073 | 1.81 | 1.48E-25 | 3.04E-24 |
| chr2 | 190797735 | 190803115 | 5381 | -2.37 | 1.49E-25 | 3.05E-24 |
| chr14 | 104478014 | 104481861 | 3848 | 4.34 | 1.54E-25 | 3.17E-24 |
| chr3 | 171652831 | 171686258 | 33428 | 0.75 | 1.56E-25 | 3.20E-24 |
| chr1 | 212685445 | 212692725 | 7281 | 2.32 | 1.66E-25 | 3.40E-24 |
| chr2 | 242522964 | 242525199 | 2236 | 2.21 | 1.68E-25 | 3.45E-24 |
| chr3 | 98679812 | 98705756 | 25945 | 0.49 | 1.69E-25 | 3.47E-24 |
| chr15 | 51658750 | 51662671 | 3922 | -2.94 | 1.76E-25 | 3.60E-24 |
| chr2 | 68964 | 70620 | 1657 | 2.39 | 1.88E-25 | 3.84E-24 |
| chr20 | 43882167 | 43884363 | 2197 | 2.79 | 1.90E-25 | 3.88E-24 |
| chr14 | 78470901 | 78472779 | 1879 | 4.24 | 1.93E-25 | 3.93E-24 |
| chr4 | 89055327 | 89081483 | 26157 | -1.19 | 1.98E-25 | 4.05E-24 |
| chr11 | 14911760 | 14927013 | 15254 | 6.3 | 2.04E-25 | 4.16E-24 |
| chr11 | 72947035 | 73000956 | 53922 | -0.97 | 2.05E-25 | 4.17E-24 |

| | | | | | | |
|---|---|---|---|---|---|---|
| chr2 | 44751239 | 44777573 | 26335 | 1.68 | 2.27E-25 | 4.63E-24 |
| chr1 | 47942759 | 47954041 | 11283 | 2.52 | 2.39E-25 | 4.87E-24 |
| chr12 | 105494070 | 105515618 | 21549 | 0.92 | 2.42E-25 | 4.91E-24 |
| chr11 | 59473064 | 59476066 | 3003 | -2.79 | 2.47E-25 | 5.01E-24 |
| chr17 | 52430798 | 52432094 | 1297 | 6.63 | 2.52E-25 | 5.12E-24 |
| chr4 | 48946159 | 48960701 | 14543 | 3.96 | 2.55E-25 | 5.17E-24 |
| chr2 | 75819056 | 75825148 | 6093 | 2.39 | 2.85E-25 | 5.77E-24 |
| chr11 | 86191434 | 86244822 | 53389 | 0.6 | 3.05E-25 | 6.18E-24 |
| chr14 | 21148449 | 21159290 | 10842 | 0.85 | 3.11E-25 | 6.29E-24 |
| chr10 | 32028077 | 32049964 | 21888 | 1.21 | 3.12E-25 | 6.31E-24 |
| chr3 | 111450890 | 111468485 | 17596 | 1.12 | 3.33E-25 | 6.74E-24 |
| chr1 | 171880738 | 171886806 | 6069 | 1 | 3.35E-25 | 6.77E-24 |
| chr4 | 143048327 | 143066284 | 17958 | 1.79 | 3.48E-25 | 7.04E-24 |
| chr9 | 118130349 | 118137289 | 6941 | 1.16 | 3.59E-25 | 7.25E-24 |
| chr8 | 123688589 | 123708088 | 19500 | 1.24 | 3.78E-25 | 7.62E-24 |
| chr5 | 178052392 | 178070183 | 17792 | 2.04 | 3.86E-25 | 7.78E-24 |
| chr12 | 133002704 | 133023664 | 20961 | 1.81 | 3.92E-25 | 7.90E-24 |
| chrX | 133662851 | 133713784 | 50934 | -0.68 | 3.94E-25 | 7.93E-24 |
| chr10 | 60372440 | 60432848 | 60409 | -0.56 | 4.05E-25 | 8.14E-24 |
| chr7 | 48943711 | 48945453 | 1743 | 2.38 | 4.09E-25 | 8.22E-24 |
| chr4 | 175951883 | 175954150 | 2268 | 2.72 | 4.11E-25 | 8.25E-24 |
| chr10 | 133907328 | 133912985 | 5658 | 1.1 | 4.16E-25 | 8.35E-24 |
| chr3 | 193498796 | 193501465 | 2670 | 4.49 | 4.55E-25 | 9.12E-24 |
| chr5 | 23324283 | 23326025 | 1743 | 4.45 | 4.77E-25 | 9.56E-24 |
| chr7 | 97905061 | 98053141 | 148081 | 0.34 | 4.79E-25 | 9.60E-24 |
| chr19 | 37554323 | 37571304 | 16982 | 10.59 | 4.83E-25 | 9.67E-24 |
| chr12 | 27675949 | 27679109 | 3161 | 1.18 | 5.12E-25 | 1.03E-23 |
| chr17 | 25571684 | 25597552 | 25869 | 0.91 | 5.54E-25 | 1.11E-23 |
| chrX | 133792086 | 133811380 | 19295 | -1.15 | 5.95E-25 | 1.19E-23 |
| chr7 | 90032291 | 90042360 | 10070 | 1.21 | 6.28E-25 | 1.25E-23 |
| chr13 | 93876379 | 93897055 | 20677 | -1.95 | 6.30E-25 | 1.26E-23 |
| chr2 | 43202440 | 43284285 | 81846 | 2.06 | 6.40E-25 | 1.28E-23 |
| chr12 | 6601342 | 6724171 | 122830 | 0.44 | 6.44E-25 | 1.28E-23 |
| chr3 | 105282845 | 105286740 | 3896 | 1.95 | 6.59E-25 | 1.31E-23 |
| chr17 | 27119982 | 27144243 | 24262 | 0.6 | 6.61E-25 | 1.32E-23 |
| chr19 | 20595442 | 20617024 | 21583 | 10.57 | 6.91E-25 | 1.37E-23 |
| chr22 | 50720011 | 50747768 | 27758 | 0.71 | 7.25E-25 | 1.44E-23 |
| chr2 | 119980634 | 120014478 | 33845 | -0.98 | 7.78E-25 | 1.55E-23 |
| chr13 | 73937878 | 73947991 | 10114 | 1.41 | 7.84E-25 | 1.56E-23 |
| chr1 | 234182436 | 234184194 | 1759 | 5.85 | 8.29E-25 | 1.64E-23 |
| chr3 | 85055729 | 85057403 | 1675 | 3.9 | 8.29E-25 | 1.64E-23 |
| chr8 | 101997874 | 101999944 | 2071 | 1.65 | 8.30E-25 | 1.64E-23 |
| chr19 | 10851145 | 10881147 | 30003 | 0.65 | 8.63E-25 | 1.71E-23 |
| chr21 | 40321665 | 40332066 | 10402 | 2.42 | 9.03E-25 | 1.79E-23 |
| chr3 | 78904256 | 78905692 | 1437 | 2.51 | 9.13E-25 | 1.80E-23 |
| chr11 | 10509616 | 10520550 | 10935 | -1.46 | 1.00E-24 | 1.98E-23 |
| chr12 | 88054961 | 88056698 | 1738 | 1.82 | 1.01E-24 | 1.99E-23 |
| chr4 | 169551377 | 169604848 | 53472 | -1.13 | 1.14E-24 | 2.25E-23 |
| chr19 | 6086661 | 6093219 | 6559 | 1.38 | 1.16E-24 | 2.29E-23 |
| chr8 | 58136801 | 58138999 | 2199 | 3.66 | 1.19E-24 | 2.34E-23 |
| chr3 | 143161227 | 143164320 | 3094 | 1.61 | 1.21E-24 | 2.38E-23 |
| chr19 | 53891106 | 53899170 | 8065 | 2.73 | 1.21E-24 | 2.38E-23 |
| chr15 | 38504663 | 38519944 | 15282 | 2.03 | 1.23E-24 | 2.43E-23 |
| chr5 | 141594879 | 141604045 | 9167 | 1.49 | 1.33E-24 | 2.61E-23 |
| chr19 | 44235143 | 44290171 | 55029 | 0.51 | 1.33E-24 | 2.62E-23 |
| chr4 | 145986346 | 145988393 | 2048 | 1.82 | 1.34E-24 | 2.63E-23 |
| chr10 | 24579889 | 24589165 | 9277 | 1.59 | 1.43E-24 | 2.80E-23 |
| chr20 | 2606839 | 2608873 | 2035 | 1.7 | 1.50E-24 | 2.94E-23 |
| chr19 | 46170561 | 46235422 | 64862 | 0.48 | 1.58E-24 | 3.10E-23 |
| chr6 | 82786334 | 82799731 | 13398 | 1.4 | 1.61E-24 | 3.15E-23 |
| chr5 | 55755834 | 55777813 | 21980 | 1.27 | 1.62E-24 | 3.17E-23 |

| | | | | | | |
|---|---|---|---|---|---|---|
| chr2 | 193406445 | 193411836 | 5392 | 4.63 | 1.76E-24 | 3.45E-23 |
| chr12 | 106353055 | 106355828 | 2774 | 1.48 | 1.79E-24 | 3.50E-23 |
| chr19 | 52403047 | 52440039 | 36993 | 10.48 | 1.81E-24 | 3.52E-23 |
| chr6 | 37671923 | 37686973 | 15051 | -3.31 | 1.88E-24 | 3.67E-23 |
| chr4 | 15767666 | 15782033 | 14368 | 1.03 | 1.90E-24 | 3.70E-23 |
| chr10 | 75346080 | 75362248 | 16169 | 1.32 | 1.93E-24 | 3.75E-23 |
| chr12 | 51317391 | 51329754 | 12364 | 2.77 | 1.93E-24 | 3.75E-23 |
| chr7 | 68293651 | 68297764 | 4114 | 6.56 | 1.99E-24 | 3.87E-23 |
| chr12 | 46750923 | 46785531 | 34609 | 0.58 | 2.04E-24 | 3.97E-23 |
| chr16 | 78273793 | 78278711 | 4919 | 2.7 | 2.13E-24 | 4.14E-23 |
| chr17 | 64261518 | 64273975 | 12458 | -0.99 | 2.25E-24 | 4.36E-23 |
| chr12 | 60042332 | 60047158 | 4827 | 10.45 | 2.27E-24 | 4.40E-23 |
| chr2 | 31183523 | 31193994 | 10472 | -2.76 | 2.35E-24 | 4.57E-23 |
| chr14 | 34580531 | 34583736 | 3206 | 1.43 | 2.47E-24 | 4.78E-23 |
| chr1 | 8062189 | 8160939 | 98751 | 0.51 | 2.53E-24 | 4.90E-23 |
| chr10 | 29272897 | 29298211 | 25315 | 1.41 | 2.59E-24 | 5.01E-23 |
| chr19 | 3050977 | 3071132 | 20156 | 1.11 | 2.62E-24 | 5.07E-23 |
| chr12 | 122843254 | 122916572 | 73319 | 0.38 | 2.67E-24 | 5.17E-23 |
| chrX | 13656156 | 13688252 | 32097 | 1.1 | 2.69E-24 | 5.20E-23 |
| chr15 | 72447271 | 72454632 | 7362 | -2.71 | 2.71E-24 | 5.23E-23 |
| chr13 | 30475883 | 30481184 | 5302 | 2.4 | 2.79E-24 | 5.39E-23 |
| chr5 | 60571223 | 60630161 | 58939 | 0.48 | 3.14E-24 | 6.05E-23 |
| chr10 | 21805790 | 21856852 | 51063 | 0.49 | 3.23E-24 | 6.23E-23 |
| chr15 | 53819885 | 53829264 | 9380 | 1.13 | 3.26E-24 | 6.28E-23 |
| chr7 | 121169298 | 121186579 | 17282 | 10.43 | 3.26E-24 | 6.28E-23 |
| chr1 | 100110374 | 100112822 | 2449 | 3.08 | 3.50E-24 | 6.73E-23 |
| chr4 | 78066381 | 78079907 | 13527 | 1.79 | 3.51E-24 | 6.75E-23 |
| chr15 | 70388239 | 70404301 | 16063 | 1.64 | 3.54E-24 | 6.80E-23 |
| chr4 | 74962511 | 75000283 | 37773 | 0.53 | 3.54E-24 | 6.80E-23 |
| chr12 | 107233151 | 107240114 | 6964 | 1.12 | 3.55E-24 | 6.81E-23 |
| chr17 | 74990497 | 75000352 | 9856 | -1.56 | 3.57E-24 | 6.83E-23 |
| chr10 | 29226664 | 29238936 | 12273 | 0.71 | 3.64E-24 | 6.98E-23 |
| chr11 | 56643597 | 56702723 | 59127 | 0.51 | 3.80E-24 | 7.27E-23 |
| chr8 | 124950116 | 124971766 | 21651 | 1.24 | 3.87E-24 | 7.41E-23 |
| chr12 | 13128204 | 13154816 | 26613 | 0.88 | 3.88E-24 | 7.41E-23 |
| chr8 | 42745930 | 42776424 | 30495 | 0.86 | 4.07E-24 | 7.77E-23 |
| chr8 | 128675172 | 128697207 | 22036 | 3.41 | 4.17E-24 | 7.97E-23 |
| chr12 | 120754472 | 120760647 | 6176 | 1.39 | 4.56E-24 | 8.71E-23 |
| chrX | 4969176 | 4971309 | 2134 | 2.04 | 4.82E-24 | 9.19E-23 |
| chr2 | 225542724 | 225545018 | 2295 | -2.44 | 4.87E-24 | 9.27E-23 |
| chr1 | 94423606 | 94431784 | 8179 | -1.2 | 4.95E-24 | 9.42E-23 |
| chr18 | 61344981 | 61352319 | 7339 | 10.37 | 5.06E-24 | 9.62E-23 |
| chr17 | 55441996 | 55458089 | 16094 | -0.79 | 5.18E-24 | 9.85E-23 |
| chr12 | 49873136 | 49909760 | 36625 | 0.87 | 5.23E-24 | 9.93E-23 |
| chr12 | 66024274 | 66051238 | 26965 | 1.18 | 5.23E-24 | 9.94E-23 |
| chr14 | 77488170 | 77526589 | 38420 | 0.52 | 5.42E-24 | 1.03E-22 |
| chr15 | 29948916 | 30150320 | 201405 | -0.37 | 5.55E-24 | 1.05E-22 |
| chr4 | 86396267 | 86421194 | 24928 | -0.83 | 5.63E-24 | 1.07E-22 |
| chr2 | 67621594 | 67629162 | 7569 | 0.99 | 6.13E-24 | 1.16E-22 |
| chr12 | 20361647 | 20367908 | 6262 | 0.99 | 6.32E-24 | 1.20E-22 |
| chr6 | 133887836 | 133969358 | 81523 | 0.48 | 6.35E-24 | 1.20E-22 |
| chr20 | 33263597 | 33298263 | 34667 | 0.94 | 6.42E-24 | 1.21E-22 |
| chr7 | 134039933 | 134059746 | 19814 | 0.6 | 6.44E-24 | 1.22E-22 |
| chr11 | 6339017 | 6343948 | 4932 | 10.35 | 6.49E-24 | 1.23E-22 |
| chr18 | 42259120 | 42272081 | 12962 | 2.17 | 6.53E-24 | 1.23E-22 |
| chr19 | 52227552 | 52233348 | 5797 | 6.77 | 6.55E-24 | 1.23E-22 |
| chr2 | 179922120 | 179924485 | 2366 | 3.06 | 6.84E-24 | 1.29E-22 |
| chr4 | 105410785 | 105422568 | 11784 | 1.37 | 6.94E-24 | 1.31E-22 |
| chr20 | 43223117 | 43281207 | 58091 | -0.67 | 7.06E-24 | 1.33E-22 |
| chr19 | 23416546 | 23438165 | 21620 | 10.34 | 7.28E-24 | 1.37E-22 |
| chr1 | 64312198 | 64326021 | 13824 | -4.19 | 7.32E-24 | 1.38E-22 |

| | | | | | | |
|---|---|---|---|---|---|---|
| chr7 | 106087635 | 106103593 | 15959 | -4.25 | 7.62E-24 | 1.43E-22 |
| chr12 | 87529417 | 87555613 | 26197 | 1.97 | 7.65E-24 | 1.44E-22 |
| chr14 | 68428696 | 68432680 | 3985 | 1.77 | 7.90E-24 | 1.48E-22 |
| chr12 | 56302273 | 56327511 | 25239 | 0.89 | 8.11E-24 | 1.52E-22 |
| chr3 | 99667666 | 99684320 | 16655 | 1.64 | 8.18E-24 | 1.53E-22 |
| chr3 | 17071495 | 17083435 | 11941 | 1.14 | 8.33E-24 | 1.56E-22 |
| chr5 | 140976247 | 141017114 | 40868 | 0.85 | 8.56E-24 | 1.60E-22 |
| chr12 | 48743331 | 48751048 | 7718 | 1.86 | 8.60E-24 | 1.61E-22 |
| chr12 | 90362831 | 90366715 | 3885 | 3.17 | 8.70E-24 | 1.63E-22 |
| chr7 | 10214393 | 10217524 | 3132 | 10.32 | 8.87E-24 | 1.66E-22 |
| chr1 | 212768521 | 212783677 | 15157 | 1.49 | 9.15E-24 | 1.71E-22 |
| chr13 | 23919004 | 23926242 | 7239 | 1.47 | 9.31E-24 | 1.74E-22 |
| chr11 | 126786354 | 126800281 | 13928 | -1.38 | 9.61E-24 | 1.79E-22 |
| chr7 | 51356662 | 51385263 | 28602 | 1.31 | 9.97E-24 | 1.86E-22 |
| chr2 | 53314758 | 53332485 | 17728 | 10.32 | 1.08E-23 | 2.01E-22 |
| chr5 | 57526641 | 57562827 | 36187 | -0.65 | 1.09E-23 | 2.03E-22 |
| chr7 | 17017175 | 17030522 | 13348 | 1.53 | 1.12E-23 | 2.09E-22 |
| chr12 | 106668061 | 106697863 | 29803 | 1 | 1.16E-23 | 2.16E-22 |
| chr3 | 162208814 | 162211690 | 2877 | -2.76 | 1.18E-23 | 2.19E-22 |
| chr2 | 118869937 | 118882841 | 12905 | 1.09 | 1.18E-23 | 2.19E-22 |
| chr5 | 74117372 | 74120005 | 2634 | 2.82 | 1.24E-23 | 2.30E-22 |
| chr7 | 75730873 | 75740189 | 9317 | 0.91 | 1.24E-23 | 2.30E-22 |
| chr4 | 146089625 | 146103111 | 13487 | 1.01 | 1.29E-23 | 2.38E-22 |
| chr2 | 234452179 | 234475628 | 23450 | -0.82 | 1.30E-23 | 2.40E-22 |
| chr17 | 46367226 | 46375405 | 8180 | 1.33 | 1.31E-23 | 2.42E-22 |
| chr8 | 116749829 | 116753736 | 3908 | 10.28 | 1.32E-23 | 2.43E-22 |
| chr4 | 114062362 | 114069334 | 6973 | 2.19 | 1.34E-23 | 2.48E-22 |
| chr4 | 121438613 | 121440082 | 1470 | 6.19 | 1.35E-23 | 2.49E-22 |
| chr7 | 114870224 | 114872419 | 2196 | 1.66 | 1.39E-23 | 2.57E-22 |
| chr3 | 113381707 | 113387307 | 5601 | 2.12 | 1.40E-23 | 2.59E-22 |
| chr17 | 54775122 | 54778532 | 3411 | 1.54 | 1.53E-23 | 2.83E-22 |
| chr11 | 35296112 | 35297995 | 1884 | 2.65 | 1.54E-23 | 2.83E-22 |
| chr2 | 111913044 | 111943821 | 30778 | -1.27 | 1.58E-23 | 2.92E-22 |
| chr12 | 51052495 | 51058831 | 6337 | 2.16 | 1.60E-23 | 2.94E-22 |
| chr1 | 101720901 | 101730314 | 9414 | 2.07 | 1.61E-23 | 2.96E-22 |
| chr5 | 90712621 | 90717394 | 4774 | 1.9 | 1.66E-23 | 3.04E-22 |
| chr2 | 143607036 | 143656067 | 49032 | -0.5 | 1.67E-23 | 3.08E-22 |
| chr4 | 20193780 | 20196789 | 3010 | 6.5 | 1.69E-23 | 3.10E-22 |
| chr5 | 91200799 | 91204552 | 3754 | 1.66 | 1.75E-23 | 3.21E-22 |
| chr14 | 31908733 | 31927507 | 18775 | 0.67 | 1.77E-23 | 3.25E-22 |
| chr2 | 113547800 | 113582472 | 34673 | -1.78 | 1.99E-23 | 3.65E-22 |
| chr5 | 57477150 | 57487536 | 10387 | -0.74 | 2.01E-23 | 3.69E-22 |
| chr12 | 108693961 | 108707972 | 14012 | 10.24 | 2.02E-23 | 3.69E-22 |
| chr7 | 63505632 | 63520223 | 14592 | 10.25 | 2.17E-23 | 3.97E-22 |
| chr17 | 12528465 | 12541149 | 12685 | -10.23 | 2.24E-23 | 4.10E-22 |
| chr12 | 87883254 | 87886294 | 3041 | 2.15 | 2.28E-23 | 4.16E-22 |
| chr10 | 104059759 | 104072171 | 12413 | 2.16 | 2.42E-23 | 4.42E-22 |
| chr19 | 9120897 | 9143912 | 23016 | 2.09 | 2.44E-23 | 4.44E-22 |
| chr8 | 8144098 | 8245317 | 101220 | 0.36 | 2.47E-23 | 4.50E-22 |
| chr5 | 93355581 | 93359003 | 3423 | 1.21 | 2.47E-23 | 4.50E-22 |
| chr13 | 78081238 | 78118920 | 37683 | 0.84 | 2.52E-23 | 4.59E-22 |
| chr10 | 12017654 | 12037114 | 19461 | 1.04 | 2.54E-23 | 4.62E-22 |
| chr12 | 12289047 | 12385532 | 96486 | 0.46 | 2.55E-23 | 4.63E-22 |
| chr20 | 6677855 | 6700669 | 22815 | 10.22 | 2.57E-23 | 4.66E-22 |
| chr15 | 66627051 | 66663617 | 36567 | -0.9 | 2.69E-23 | 4.88E-22 |
| chr11 | 28844261 | 28862156 | 17896 | 0.59 | 2.74E-23 | 4.97E-22 |
| chr12 | 97946265 | 97966877 | 20613 | 10.2 | 2.75E-23 | 4.99E-22 |
| chr21 | 40982421 | 40989405 | 6985 | 2.52 | 2.77E-23 | 5.02E-22 |
| chr7 | 138247497 | 138254139 | 6643 | 10.2 | 2.91E-23 | 5.27E-22 |
| chr8 | 83773324 | 83780596 | 7273 | 10.21 | 2.93E-23 | 5.31E-22 |
| chr1 | 80519412 | 80532526 | 13115 | 3.92 | 3.03E-23 | 5.49E-22 |

| | | | | | | |
|---|---|---|---|---|---|---|
| chr1 | 78001698 | 78007390 | 5693 | -1.22 | 3.10E-23 | 5.61E-22 |
| chr13 | 73591352 | 73685869 | 94518 | 0.51 | 3.11E-23 | 5.63E-22 |
| chr8 | 99737010 | 99751062 | 14053 | 1.7 | 3.20E-23 | 5.78E-22 |
| chr10 | 37823819 | 37825860 | 2042 | 2.63 | 3.24E-23 | 5.85E-22 |
| chr19 | 53029367 | 53044976 | 15610 | 10.19 | 3.27E-23 | 5.90E-22 |
| chr10 | 45434609 | 45436434 | 1826 | 4.3 | 3.46E-23 | 6.23E-22 |
| chr8 | 23737997 | 23755689 | 17693 | 1.48 | 3.49E-23 | 6.28E-22 |
| chr11 | 10414482 | 10430962 | 16481 | -1.19 | 3.54E-23 | 6.37E-22 |
| chr4 | 95915182 | 95923808 | 8627 | 1.19 | 3.54E-23 | 6.37E-22 |
| chr7 | 139404527 | 139454476 | 49950 | -1.1 | 3.56E-23 | 6.40E-22 |
| chr4 | 153405211 | 153409637 | 4427 | 1.29 | 3.70E-23 | 6.65E-22 |
| chr8 | 67419314 | 67473787 | 54474 | 0.69 | 3.73E-23 | 6.71E-22 |
| chr17 | 8953614 | 8955166 | 1553 | 2.31 | 3.78E-23 | 6.78E-22 |
| chr16 | 14143774 | 14145839 | 2066 | 1.72 | 3.92E-23 | 7.03E-22 |
| chr9 | 71792743 | 71797247 | 4505 | 1.4 | 3.93E-23 | 7.04E-22 |
| chr10 | 62041842 | 62043765 | 1924 | 3.38 | 3.96E-23 | 7.10E-22 |
| chr20 | 56821036 | 56828636 | 7601 | 1.28 | 4.27E-23 | 7.64E-22 |
| chr1 | 231683994 | 231739899 | 55906 | 0.79 | 4.42E-23 | 7.91E-22 |
| chr19 | 20632742 | 20672369 | 39628 | 10.17 | 4.52E-23 | 8.08E-22 |
| chr7 | 63766919 | 63786069 | 19151 | 10.16 | 4.61E-23 | 8.23E-22 |
| chr16 | 2008985 | 2107353 | 98369 | 0.42 | 4.86E-23 | 8.69E-22 |
| chr19 | 18025788 | 18044888 | 19101 | 1.52 | 5.04E-23 | 8.99E-22 |
| chr9 | 117540134 | 117543214 | 3081 | 1.51 | 5.24E-23 | 9.35E-22 |
| chr2 | 192079652 | 192097080 | 17429 | -0.75 | 5.31E-23 | 9.46E-22 |
| chr6 | 39383698 | 39402298 | 18601 | 1.88 | 5.39E-23 | 9.61E-22 |
| chrX | 43551866 | 43563400 | 11535 | 1.77 | 5.62E-23 | 1.00E-21 |
| chr5 | 14006852 | 14012990 | 6139 | 0.76 | 5.65E-23 | 1.01E-21 |
| chrX | 149530930 | 149536056 | 5127 | 1.19 | 5.66E-23 | 1.01E-21 |
| chr15 | 38454903 | 38457237 | 2335 | 6.18 | 5.76E-23 | 1.02E-21 |
| chr2 | 97260051 | 97304857 | 44807 | -0.78 | 5.84E-23 | 1.04E-21 |
| chr6 | 47002333 | 47015447 | 13115 | 1.34 | 5.88E-23 | 1.04E-21 |
| chr13 | 35200453 | 35203788 | 3336 | 1.66 | 5.90E-23 | 1.05E-21 |
| chr9 | 130253602 | 130347063 | 93462 | 0.42 | 5.94E-23 | 1.05E-21 |
| chr12 | 56726231 | 56728945 | 2715 | 1.02 | 6.22E-23 | 1.10E-21 |
| chr12 | 85838436 | 85840357 | 1922 | 2.34 | 6.23E-23 | 1.10E-21 |
| chr12 | 66316741 | 66348517 | 31777 | 1.08 | 6.36E-23 | 1.13E-21 |
| chr12 | 133757464 | 133770196 | 12733 | 2.59 | 6.36E-23 | 1.13E-21 |
| chr6 | 143140531 | 143168752 | 28222 | 1.12 | 6.69E-23 | 1.18E-21 |
| chr2 | 65282769 | 65358554 | 75786 | -0.58 | 6.71E-23 | 1.19E-21 |
| chr9 | 116734645 | 116772339 | 37695 | -1.04 | 6.92E-23 | 1.22E-21 |
| chr2 | 54269356 | 54272631 | 3276 | 1.31 | 7.09E-23 | 1.25E-21 |
| chr8 | 127445023 | 127448539 | 3517 | 10.12 | 7.20E-23 | 1.27E-21 |
| chr2 | 239772026 | 239797998 | 25973 | -1.21 | 7.30E-23 | 1.29E-21 |
| chr1 | 170031962 | 170044409 | 12448 | 2.32 | 7.34E-23 | 1.29E-21 |
| chr22 | 35746313 | 35827976 | 81664 | -0.63 | 7.45E-23 | 1.31E-21 |
| chr12 | 41141246 | 41154384 | 13139 | 2.2 | 7.56E-23 | 1.33E-21 |
| chr5 | 110847163 | 110849847 | 2685 | 1.7 | 7.56E-23 | 1.33E-21 |
| chr5 | 91840045 | 91842479 | 2435 | 2.88 | 7.63E-23 | 1.34E-21 |
| chr20 | 14314386 | 14326366 | 11981 | 1 | 8.40E-23 | 1.48E-21 |
| chr8 | 58633062 | 58634632 | 1571 | 2.9 | 8.63E-23 | 1.52E-21 |
| chr1 | 31572036 | 31585271 | 13236 | 1.63 | 8.75E-23 | 1.53E-21 |
| chr10 | 91460319 | 91504851 | 44533 | 0.92 | 8.75E-23 | 1.53E-21 |
| chr15 | 63334044 | 63428897 | 94854 | 0.38 | 8.78E-23 | 1.54E-21 |
| chrX | 109324444 | 109341445 | 17002 | -0.78 | 9.18E-23 | 1.61E-21 |
| chr4 | 16225717 | 16234220 | 8504 | 1.36 | 9.75E-23 | 1.71E-21 |
| chr12 | 43183596 | 43185943 | 2348 | 6.19 | 9.76E-23 | 1.71E-21 |
| chr1 | 19228458 | 19282231 | 53774 | -0.78 | 9.87E-23 | 1.72E-21 |
| chr19 | 53222119 | 53240387 | 18269 | 10.09 | 1.01E-22 | 1.76E-21 |
| chr11 | 18543062 | 18561728 | 18667 | 0.71 | 1.02E-22 | 1.78E-21 |
| chr18 | 20254075 | 20292743 | 38669 | 0.71 | 1.02E-22 | 1.78E-21 |
| chr15 | 65517314 | 65521571 | 4258 | 1.83 | 1.03E-22 | 1.80E-21 |

| | | | | | | |
|---|---|---|---|---|---|---|
| chr1 | 156147762 | 156167174 | 19413 | 1.2 | 1.03E-22 | 1.80E-21 |
| chr8 | 119559868 | 119560983 | 1116 | 3.4 | 1.04E-22 | 1.80E-21 |
| chr1 | 46435411 | 46448698 | 13288 | 1.44 | 1.04E-22 | 1.82E-21 |
| chr14 | 100658204 | 100681947 | 23744 | 1.24 | 1.12E-22 | 1.94E-21 |
| chr3 | 29217289 | 29229894 | 12606 | 3.48 | 1.14E-22 | 1.99E-21 |
| chr12 | 52416382 | 52465254 | 48873 | 0.7 | 1.16E-22 | 2.02E-21 |
| chr9 | 117867047 | 117881730 | 14684 | 2.07 | 1.17E-22 | 2.02E-21 |
| chr16 | 57309372 | 57319199 | 9828 | 1.81 | 1.20E-22 | 2.08E-21 |
| chr9 | 125621574 | 125624222 | 2649 | 2.38 | 1.21E-22 | 2.10E-21 |
| chr4 | 162498646 | 162503599 | 4954 | 4.8 | 1.21E-22 | 2.10E-21 |
| chr19 | 23455894 | 23470043 | 14150 | 10.06 | 1.22E-22 | 2.11E-21 |
| chr2 | 186700895 | 186703749 | 2855 | 1.86 | 1.23E-22 | 2.13E-21 |
| chr5 | 108741730 | 108749375 | 7646 | 0.9 | 1.25E-22 | 2.17E-21 |
| chr9 | 4657074 | 4681975 | 24902 | 0.67 | 1.29E-22 | 2.23E-21 |
| chr9 | 526459 | 575336 | 48878 | -0.71 | 1.29E-22 | 2.23E-21 |
| chr3 | 94936131 | 94948533 | 12403 | 10.04 | 1.40E-22 | 2.42E-21 |
| chr2 | 69471258 | 69482669 | 11412 | -1.04 | 1.43E-22 | 2.46E-21 |
| chr11 | 91999791 | 92003687 | 3897 | 1.02 | 1.52E-22 | 2.62E-21 |
| chr3 | 4865709 | 4874147 | 8439 | 0.91 | 1.56E-22 | 2.68E-21 |
| chr1 | 214600900 | 214623448 | 22549 | 0.84 | 1.58E-22 | 2.71E-21 |
| chr7 | 7215558 | 7223930 | 8373 | 0.8 | 1.62E-22 | 2.78E-21 |
| chr2 | 143671513 | 143757112 | 85600 | -0.67 | 1.62E-22 | 2.78E-21 |
| chr8 | 128490199 | 128492301 | 2103 | 2.2 | 1.65E-22 | 2.83E-21 |
| chr2 | 20776909 | 20797702 | 20794 | -0.87 | 1.71E-22 | 2.95E-21 |
| chr5 | 143003852 | 143018549 | 14698 | 1.1 | 1.72E-22 | 2.95E-21 |
| chr17 | 20103829 | 20132445 | 28617 | -0.94 | 1.92E-22 | 3.30E-21 |
| chr2 | 216908594 | 216931586 | 22993 | 0.83 | 1.93E-22 | 3.31E-21 |
| chr5 | 89316865 | 89318890 | 2026 | 7.51 | 1.99E-22 | 3.42E-21 |
| chr4 | 146465477 | 146468593 | 3117 | 1.79 | 2.00E-22 | 3.42E-21 |
| chr20 | 4774902 | 4776861 | 1960 | -2.22 | 2.03E-22 | 3.48E-21 |
| chr20 | 42636447 | 42648766 | 12320 | 1.05 | 2.07E-22 | 3.54E-21 |
| chr3 | 45161699 | 45187849 | 26151 | 1.22 | 2.08E-22 | 3.56E-21 |
| chr9 | 30103038 | 30122507 | 19470 | 10.01 | 2.08E-22 | 3.57E-21 |
| chr6 | 168332507 | 168367608 | 35102 | 0.93 | 2.09E-22 | 3.57E-21 |
| chr11 | 9585924 | 9610852 | 24929 | 0.53 | 2.15E-22 | 3.68E-21 |
| chr12 | 8178261 | 8192232 | 13972 | 0.79 | 2.17E-22 | 3.71E-21 |
| chr4 | 93711963 | 93714126 | 2164 | 4.7 | 2.22E-22 | 3.78E-21 |
| chr17 | 18751744 | 18836820 | 85077 | -0.52 | 2.23E-22 | 3.80E-21 |
| chr12 | 125276838 | 125394204 | 117367 | 0.46 | 2.45E-22 | 4.17E-21 |
| chr22 | 33775445 | 33788132 | 12688 | 0.87 | 2.60E-22 | 4.43E-21 |
| chr10 | 28957190 | 28975143 | 17954 | 1.56 | 2.65E-22 | 4.51E-21 |
| chr1 | 17481446 | 17537341 | 55896 | 0.71 | 2.65E-22 | 4.51E-21 |
| chr8 | 93966305 | 93979008 | 12704 | 1.51 | 2.70E-22 | 4.59E-21 |
| chr3 | 185539857 | 185556139 | 16283 | 1.32 | 2.72E-22 | 4.63E-21 |
| chr14 | 49028615 | 49036653 | 8039 | 9.98 | 2.73E-22 | 4.64E-21 |
| chr13 | 100147480 | 100159140 | 11661 | 1 | 2.77E-22 | 4.70E-21 |
| chr1 | 41404936 | 41492694 | 87759 | -0.49 | 2.89E-22 | 4.90E-21 |
| chr7 | 12583782 | 12625832 | 42051 | 9.98 | 2.94E-22 | 4.98E-21 |
| chr2 | 86244864 | 86308173 | 63310 | -0.55 | 2.98E-22 | 5.05E-21 |
| chr2 | 74640304 | 74668496 | 28193 | -0.89 | 3.00E-22 | 5.07E-21 |
| chr6 | 39136811 | 39139393 | 2583 | 4.57 | 3.05E-22 | 5.16E-21 |
| chr5 | 30098287 | 30113461 | 15175 | 1.53 | 3.13E-22 | 5.29E-21 |
| chr9 | 81477819 | 81495244 | 17426 | -0.61 | 3.18E-22 | 5.38E-21 |
| chr6 | 141872332 | 141884069 | 11738 | 2.55 | 3.28E-22 | 5.54E-21 |
| chr7 | 81333711 | 81355556 | 21846 | 1.46 | 3.33E-22 | 5.62E-21 |
| chr5 | 81652483 | 81654103 | 1621 | 3.08 | 3.45E-22 | 5.82E-21 |
| chr9 | 74744129 | 74781505 | 37377 | 9.95 | 3.57E-22 | 6.02E-21 |
| chr5 | 82847157 | 82873890 | 26734 | 9.96 | 3.59E-22 | 6.04E-21 |
| chr9 | 113672377 | 113715580 | 43204 | -0.63 | 3.67E-22 | 6.18E-21 |
| chr1 | 244459445 | 244467726 | 8282 | 1.09 | 3.70E-22 | 6.23E-21 |
| chr9 | 91605677 | 91614504 | 8828 | -6.32 | 3.77E-22 | 6.34E-21 |

| | | | | | | |
|---|---|---|---|---|---|---|
| chr1 | 221403478 | 221404987 | 1510 | 5.99 | 3.89E-22 | 6.54E-21 |
| chr12 | 69002993 | 69044184 | 41192 | 0.86 | 3.92E-22 | 6.58E-21 |
| chr19 | 21022624 | 21042790 | 20167 | 9.95 | 4.07E-22 | 6.84E-21 |
| chr1 | 8197345 | 8300469 | 103125 | 0.44 | 4.10E-22 | 6.89E-21 |
| chr1 | 239391137 | 239397677 | 6541 | 2.12 | 4.12E-22 | 6.92E-21 |
| chr10 | 124962273 | 124964495 | 2223 | 2.32 | 4.22E-22 | 7.08E-21 |
| chr7 | 63546476 | 63550990 | 4515 | 9.96 | 4.28E-22 | 7.18E-21 |
| chr9 | 123375901 | 123427177 | 51277 | -0.61 | 4.29E-22 | 7.19E-21 |
| chr9 | 134459995 | 134484731 | 24737 | -1.23 | 4.31E-22 | 7.22E-21 |
| chr6 | 47426189 | 47472840 | 46652 | 0.99 | 4.42E-22 | 7.39E-21 |
| chr13 | 106570778 | 106579331 | 8554 | 2.6 | 4.45E-22 | 7.44E-21 |
| chr10 | 122737871 | 122741851 | 3981 | 3.48 | 4.54E-22 | 7.58E-21 |
| chr2 | 43296799 | 43341500 | 44702 | 0.79 | 4.55E-22 | 7.61E-21 |
| chr12 | 18449099 | 18450104 | 1006 | 3.9 | 4.59E-22 | 7.67E-21 |
| chr3 | 193877835 | 193879812 | 1978 | 2.97 | 4.66E-22 | 7.78E-21 |
| chr8 | 21540303 | 21546524 | 6222 | -1.12 | 4.68E-22 | 7.80E-21 |
| chr10 | 101084213 | 101121338 | 37126 | 0.65 | 4.84E-22 | 8.06E-21 |
| chr12 | 62652674 | 62655943 | 3270 | 0.97 | 4.87E-22 | 8.12E-21 |
| chr7 | 79034735 | 79036760 | 2026 | 3.58 | 4.88E-22 | 8.13E-21 |
| chr16 | 64424194 | 64450908 | 26715 | 9.92 | 5.03E-22 | 8.38E-21 |
| chr19 | 52872113 | 52881841 | 9729 | 9.92 | 5.08E-22 | 8.44E-21 |
| chr8 | 117456749 | 117467231 | 10483 | 1.81 | 5.28E-22 | 8.77E-21 |
| chr5 | 90663112 | 90680792 | 17681 | 0.53 | 5.36E-22 | 8.90E-21 |
| chr17 | 57694159 | 58000044 | 305886 | 0.28 | 5.36E-22 | 8.90E-21 |
| chr4 | 148013852 | 148025315 | 11464 | 2.45 | 5.48E-22 | 9.10E-21 |
| chr12 | 21258033 | 21298591 | 40559 | 9.92 | 5.48E-22 | 9.10E-21 |
| chr10 | 99330415 | 99372469 | 42055 | -0.61 | 5.75E-22 | 9.53E-21 |
| chr15 | 80164986 | 80217291 | 52306 | 0.6 | 5.81E-22 | 9.63E-21 |
| chr15 | 33110999 | 33175680 | 64682 | -0.53 | 5.93E-22 | 9.81E-21 |
| chr6 | 167088635 | 167092778 | 4144 | 6.77 | 6.35E-22 | 1.05E-20 |
| chr10 | 92679345 | 92696215 | 16871 | 0.73 | 6.52E-22 | 1.08E-20 |
| chr7 | 158803891 | 158824130 | 20240 | 0.85 | 6.74E-22 | 1.11E-20 |
| chr4 | 129434693 | 129451175 | 16483 | -0.63 | 6.97E-22 | 1.15E-20 |
| chr12 | 69724372 | 69727234 | 2863 | 6.75 | 7.21E-22 | 1.19E-20 |
| chr8 | 66540007 | 66572641 | 32635 | 0.91 | 7.42E-22 | 1.22E-20 |
| chr1 | 204615444 | 204618250 | 2807 | -1.59 | 7.67E-22 | 1.27E-20 |
| chr7 | 3282997 | 3287337 | 4341 | 3.14 | 7.84E-22 | 1.29E-20 |
| chr10 | 53991976 | 53993298 | 1323 | 2.79 | 7.91E-22 | 1.30E-20 |
| chr7 | 143021732 | 143029086 | 7355 | 2.8 | 8.15E-22 | 1.34E-20 |
| chr12 | 107379972 | 107389654 | 9683 | 1.67 | 8.15E-22 | 1.34E-20 |
| chr2 | 113943002 | 114007358 | 64357 | -0.63 | 8.22E-22 | 1.35E-20 |
| chr4 | 69898267 | 69904812 | 6546 | -1.44 | 8.42E-22 | 1.38E-20 |
| chr1 | 231273839 | 231276853 | 3015 | -1.34 | 8.58E-22 | 1.41E-20 |
| chr11 | 85026674 | 85029396 | 2723 | 1.5 | 8.59E-22 | 1.41E-20 |
| chr4 | 16197601 | 16199161 | 1561 | 2.28 | 9.89E-22 | 1.62E-20 |
| chr12 | 113850777 | 113861576 | 10800 | 1.3 | 1.09E-21 | 1.78E-20 |
| chr1 | 7391664 | 7402497 | 10834 | 1.32 | 1.09E-21 | 1.79E-20 |
| chr6 | 55689518 | 55693336 | 3819 | 4.28 | 1.10E-21 | 1.80E-20 |
| chr2 | 102443567 | 102489701 | 46135 | -1.01 | 1.10E-21 | 1.80E-20 |
| chr17 | 31236869 | 31240282 | 3414 | 1.89 | 1.12E-21 | 1.83E-20 |
| chr2 | 46996327 | 46999976 | 3650 | -2.48 | 1.12E-21 | 1.84E-20 |
| chr19 | 13253371 | 13292969 | 39599 | 0.47 | 1.21E-21 | 1.98E-20 |
| chr14 | 55591822 | 55659238 | 67417 | 0.8 | 1.23E-21 | 2.01E-20 |
| chr6 | 6810784 | 6816473 | 5690 | 1.78 | 1.25E-21 | 2.04E-20 |
| chr5 | 148851007 | 148944451 | 93445 | 0.46 | 1.27E-21 | 2.07E-20 |
| chr12 | 56544599 | 56566895 | 22297 | 0.75 | 1.29E-21 | 2.10E-20 |
| chr19 | 37406233 | 37408521 | 2289 | 4.66 | 1.32E-21 | 2.15E-20 |
| chr5 | 109139309 | 109142959 | 3651 | 2.25 | 1.33E-21 | 2.17E-20 |
| chr1 | 53600914 | 53613555 | 12642 | 0.93 | 1.37E-21 | 2.23E-20 |
| chr12 | 93320736 | 93324318 | 3583 | 1.17 | 1.38E-21 | 2.25E-20 |
| chr6 | 143717854 | 143733065 | 15212 | -1.92 | 1.38E-21 | 2.25E-20 |

| | | | | | | |
|---|---|---|---|---|---|---|
| chr6 | 105967238 | 105971366 | 4129 | -1.83 | 1.49E-21 | 2.42E-20 |
| chr5 | 155606586 | 155629555 | 22970 | 0.74 | 1.49E-21 | 2.43E-20 |
| chr8 | 81830300 | 81854195 | 23896 | 9.81 | 1.52E-21 | 2.47E-20 |
| chr3 | 149452897 | 149455273 | 2377 | 1.83 | 1.53E-21 | 2.48E-20 |
| chr12 | 49402397 | 49426261 | 23865 | 0.78 | 1.57E-21 | 2.54E-20 |
| chr11 | 27368154 | 27495348 | 127195 | 0.31 | 1.57E-21 | 2.55E-20 |
| chr5 | 77786675 | 77945802 | 159128 | -0.35 | 1.63E-21 | 2.64E-20 |
| chr11 | 6623212 | 6641337 | 18126 | 0.72 | 1.65E-21 | 2.68E-20 |
| chr12 | 46119749 | 46127964 | 8216 | 0.74 | 1.76E-21 | 2.84E-20 |
| chr10 | 91036657 | 91044529 | 7873 | 0.95 | 1.76E-21 | 2.84E-20 |
| chr12 | 107285241 | 107287110 | 1870 | 3.72 | 1.87E-21 | 3.02E-20 |
| chr3 | 95332475 | 95336318 | 3844 | 9.8 | 1.88E-21 | 3.03E-20 |
| chr9 | 74964115 | 74982759 | 18645 | 0.57 | 1.91E-21 | 3.08E-20 |
| chr6 | 79861365 | 79870187 | 8823 | 2.15 | 1.96E-21 | 3.16E-20 |
| chr2 | 113483206 | 113522936 | 39731 | -1.21 | 1.99E-21 | 3.21E-20 |
| chr16 | 3414179 | 3416307 | 2129 | 1.95 | 2.09E-21 | 3.36E-20 |
| chr4 | 152461087 | 152463568 | 2482 | 3.07 | 2.09E-21 | 3.37E-20 |
| chr9 | 6182789 | 6184173 | 1385 | 2.2 | 2.15E-21 | 3.46E-20 |
| chr4 | 153872121 | 153892380 | 20260 | 2.57 | 2.26E-21 | 3.63E-20 |
| chr12 | 8770292 | 8772833 | 2542 | 1.68 | 2.26E-21 | 3.64E-20 |
| chr2 | 194291647 | 194294206 | 2560 | 2.2 | 2.27E-21 | 3.66E-20 |
| chr15 | 58860273 | 58867858 | 7586 | 1.93 | 2.28E-21 | 3.66E-20 |
| chr8 | 124371834 | 124409437 | 37604 | 0.72 | 2.28E-21 | 3.67E-20 |
| chr15 | 62346881 | 62356949 | 10069 | 1.55 | 2.32E-21 | 3.72E-20 |
| chr7 | 116397873 | 116454016 | 56144 | 0.52 | 2.32E-21 | 3.72E-20 |
| chr18 | 55565291 | 55577435 | 12145 | 0.92 | 2.37E-21 | 3.81E-20 |
| chr2 | 121223413 | 121229444 | 6032 | -1.89 | 2.40E-21 | 3.84E-20 |
| chr3 | 155158764 | 155160321 | 1558 | 2.9 | 2.40E-21 | 3.84E-20 |
| chr9 | 93182567 | 93205613 | 23047 | 1.07 | 2.45E-21 | 3.92E-20 |
| chr1 | 233803427 | 233825537 | 22111 | 0.81 | 2.46E-21 | 3.94E-20 |
| chr11 | 121333911 | 121339068 | 5158 | 1.2 | 2.53E-21 | 4.04E-20 |
| chr5 | 149695635 | 149701326 | 5692 | -1.11 | 2.57E-21 | 4.11E-20 |
| chr12 | 20068891 | 20076819 | 7929 | 6.28 | 2.59E-21 | 4.14E-20 |
| chr12 | 28650270 | 28652088 | 1819 | 2.68 | 2.81E-21 | 4.49E-20 |
| chr16 | 69138935 | 69257015 | 118081 | 0.5 | 2.82E-21 | 4.50E-20 |
| chr1 | 189968095 | 189970887 | 2793 | 1.51 | 2.91E-21 | 4.64E-20 |
| chr10 | 61724443 | 61737311 | 12869 | 9.75 | 2.92E-21 | 4.65E-20 |
| chr12 | 6823570 | 6878078 | 54509 | 0.59 | 2.95E-21 | 4.69E-20 |
| chr19 | 53399255 | 53409016 | 9762 | 9.75 | 3.06E-21 | 4.87E-20 |
| chr6 | 140783343 | 140785231 | 1889 | 2.66 | 3.07E-21 | 4.89E-20 |
| chr10 | 37850017 | 37856674 | 6658 | 9.73 | 3.13E-21 | 4.98E-20 |
| chr12 | 32074426 | 32087411 | 12986 | 5.05 | 3.21E-21 | 5.10E-20 |
| chr2 | 175470067 | 175489035 | 18969 | 1 | 3.23E-21 | 5.14E-20 |
| chr15 | 52261836 | 52351065 | 89230 | 0.39 | 3.33E-21 | 5.29E-20 |
| chr7 | 17638696 | 17641301 | 2606 | 2 | 3.34E-21 | 5.31E-20 |
| chr1 | 234629624 | 234661521 | 31898 | 0.71 | 3.39E-21 | 5.37E-20 |
| chr1 | 222719531 | 222722346 | 2816 | -2.18 | 3.41E-21 | 5.41E-20 |
| chr19 | 23653423 | 23666759 | 13337 | 9.73 | 3.45E-21 | 5.46E-20 |
| chr3 | 18013756 | 18016343 | 2588 | 1.7 | 3.46E-21 | 5.49E-20 |
| chr8 | 125984924 | 125986712 | 1789 | 1.68 | 3.51E-21 | 5.55E-20 |
| chr12 | 94176244 | 94181029 | 4786 | 0.69 | 3.56E-21 | 5.63E-20 |
| chr11 | 34168459 | 34303344 | 134886 | -0.46 | 3.58E-21 | 5.67E-20 |
| chr3 | 197213838 | 197250081 | 36244 | 0.73 | 3.69E-21 | 5.83E-20 |
| chr2 | 107611874 | 107649548 | 37675 | -2.58 | 3.80E-21 | 6.00E-20 |
| chr5 | 92899049 | 92900990 | 1942 | 3.48 | 4.16E-21 | 6.57E-20 |
| chr17 | 2853051 | 2915051 | 62001 | 0.67 | 4.22E-21 | 6.65E-20 |
| chr5 | 25962427 | 25970054 | 7628 | 9.71 | 4.26E-21 | 6.71E-20 |
| chr19 | 13162038 | 13172575 | 10538 | 2.16 | 4.27E-21 | 6.73E-20 |
| chr3 | 177241033 | 177246966 | 5934 | 2.28 | 4.31E-21 | 6.78E-20 |
| chrX | 9364356 | 9365931 | 1576 | 5.41 | 4.37E-21 | 6.87E-20 |
| chr14 | 80047601 | 80063896 | 16296 | -9.7 | 4.40E-21 | 6.92E-20 |

| | | | | | | |
|---|---|---|---|---|---|---|
| chr7 | 114168992 | 114174703 | 5712 | 9.7 | 4.46E-21 | 7.01E-20 |
| chr4 | 89763983 | 89768667 | 4685 | 1.83 | 4.56E-21 | 7.16E-20 |
| chr8 | 24010580 | 24015696 | 5117 | 0.8 | 4.57E-21 | 7.18E-20 |
| chr8 | 134194255 | 134208564 | 14310 | -1.13 | 4.64E-21 | 7.29E-20 |
| chr5 | 125032122 | 125037669 | 5548 | 1.28 | 4.68E-21 | 7.34E-20 |
| chr19 | 52526990 | 52532869 | 5880 | 1.34 | 4.75E-21 | 7.44E-20 |
| chr3 | 71074369 | 71114899 | 40531 | 9.74 | 4.75E-21 | 7.45E-20 |
| chr14 | 31840559 | 31851108 | 10550 | 1.09 | 4.78E-21 | 7.49E-20 |
| chr1 | 43991384 | 44053486 | 62103 | -0.76 | 4.98E-21 | 7.79E-20 |
| chr7 | 115943928 | 115958322 | 14395 | 0.66 | 5.01E-21 | 7.83E-20 |
| chr2 | 238079734 | 238111798 | 32065 | -9.68 | 5.51E-21 | 8.62E-20 |
| chr12 | 25980280 | 26006443 | 26164 | 9.67 | 5.77E-21 | 9.02E-20 |
| chr20 | 10619919 | 10656365 | 36447 | -0.64 | 5.98E-21 | 9.34E-20 |
| chr11 | 84290394 | 84305822 | 15429 | 9.67 | 6.04E-21 | 9.43E-20 |
| chr13 | 44141200 | 44148896 | 7697 | 1.49 | 6.05E-21 | 9.45E-20 |
| chr20 | 5801165 | 5830321 | 29157 | 0.92 | 6.24E-21 | 9.74E-20 |
| chr12 | 4757227 | 4763277 | 6051 | 1.25 | 6.30E-21 | 9.82E-20 |
| chr3 | 174125714 | 174128378 | 2665 | 2.53 | 6.38E-21 | 9.95E-20 |
| chr21 | 33783102 | 33854038 | 70937 | -0.47 | 6.67E-21 | 1.04E-19 |
| chr5 | 142337913 | 142345869 | 7957 | -0.78 | 7.01E-21 | 1.09E-19 |
| chr2 | 46609482 | 46616689 | 7208 | -1.99 | 7.39E-21 | 1.15E-19 |
| chr3 | 30618398 | 30622003 | 3606 | 1.81 | 7.43E-21 | 1.16E-19 |
| chr20 | 50600451 | 50604967 | 4517 | 1.31 | 7.55E-21 | 1.17E-19 |
| chr8 | 116250947 | 116270752 | 19806 | 9.64 | 7.97E-21 | 1.24E-19 |
| chr14 | 34705727 | 34707724 | 1998 | 2.28 | 8.03E-21 | 1.25E-19 |
| chr2 | 223290761 | 223306332 | 15572 | 9.71 | 8.23E-21 | 1.28E-19 |
| chr12 | 123307748 | 123331486 | 23739 | 1.27 | 8.29E-21 | 1.29E-19 |
| chr4 | 149296648 | 149298105 | 1458 | 6.17 | 8.51E-21 | 1.32E-19 |
| chr2 | 67663621 | 67692467 | 28847 | 9.63 | 8.96E-21 | 1.39E-19 |
| chr4 | 60683879 | 60685338 | 1460 | 4.4 | 9.04E-21 | 1.40E-19 |
| chr4 | 15894057 | 15923649 | 29593 | 0.72 | 9.21E-21 | 1.43E-19 |
| chr1 | 244209830 | 244224709 | 14880 | 1.35 | 9.35E-21 | 1.45E-19 |
| chr2 | 42273977 | 42289100 | 15124 | 1.08 | 9.48E-21 | 1.47E-19 |
| chrX | 24857004 | 24918964 | 61961 | 0.53 | 9.65E-21 | 1.49E-19 |
| chr4 | 91648326 | 91650587 | 2262 | 3.91 | 9.66E-21 | 1.49E-19 |
| chr14 | 57957489 | 57963148 | 5660 | 1.05 | 9.68E-21 | 1.49E-19 |
| chr6 | 2623286 | 2635479 | 12194 | 1.12 | 9.74E-21 | 1.50E-19 |
| chr1 | 199775892 | 199779264 | 3373 | 2.88 | 9.93E-21 | 1.53E-19 |
| chr3 | 186801501 | 186804182 | 2682 | 3.04 | 1.03E-20 | 1.59E-19 |
| chr1 | 212731323 | 212754048 | 22726 | 1.32 | 1.03E-20 | 1.59E-19 |
| chr7 | 11562088 | 11565095 | 3008 | 7.14 | 1.08E-20 | 1.66E-19 |
| chr9 | 32571397 | 32604883 | 33487 | 9.63 | 1.08E-20 | 1.67E-19 |
| chr3 | 183024304 | 183026050 | 1747 | 3.28 | 1.11E-20 | 1.70E-19 |
| chr17 | 5014282 | 5026409 | 12128 | 2.1 | 1.11E-20 | 1.71E-19 |
| chr15 | 45625311 | 45627796 | 2486 | 2.35 | 1.13E-20 | 1.73E-19 |
| chr1 | 191477242 | 191490813 | 13572 | 9.6 | 1.19E-20 | 1.82E-19 |
| chr12 | 57902921 | 57941313 | 38393 | 0.67 | 1.23E-20 | 1.88E-19 |
| chr16 | 74475920 | 74528245 | 52326 | 0.72 | 1.24E-20 | 1.90E-19 |
| chr1 | 227719660 | 227731205 | 11546 | 2.47 | 1.25E-20 | 1.92E-19 |
| chr6 | 125541680 | 125545940 | 4261 | 1.84 | 1.26E-20 | 1.93E-19 |
| chr11 | 28878810 | 28883936 | 5127 | 1.13 | 1.30E-20 | 2.00E-19 |
| chr5 | 45904159 | 45907437 | 3279 | 9.62 | 1.37E-20 | 2.10E-19 |
| chr4 | 176227205 | 176228794 | 1590 | 3.96 | 1.38E-20 | 2.12E-19 |
| chr12 | 121647222 | 121656067 | 8846 | 1.11 | 1.39E-20 | 2.12E-19 |
| chr8 | 38852583 | 38922916 | 70334 | 0.36 | 1.39E-20 | 2.12E-19 |
| chr14 | 37738874 | 37741070 | 2197 | 2.59 | 1.45E-20 | 2.22E-19 |
| chr4 | 95936259 | 95948746 | 12488 | -1.11 | 1.46E-20 | 2.24E-19 |
| chr18 | 52274422 | 52276367 | 1946 | 2.34 | 1.48E-20 | 2.25E-19 |
| chr4 | 129373330 | 129393300 | 19971 | -1.15 | 1.49E-20 | 2.27E-19 |
| chr3 | 192126509 | 192152396 | 25888 | 9.57 | 1.51E-20 | 2.31E-19 |
| chr2 | 197361503 | 197377367 | 15865 | 9.58 | 1.52E-20 | 2.31E-19 |

| | | | | | | |
|---|---|---|---|---|---|---|
| chr12 | 87051934 | 87058111 | 6178 | 0.99 | 1.52E-20 | 2.32E-19 |
| chr6 | 86099345 | 86117257 | 17913 | 0.73 | 1.59E-20 | 2.42E-19 |
| chr7 | 82895242 | 82897315 | 2074 | 6.14 | 1.70E-20 | 2.58E-19 |
| chr12 | 13595640 | 13597588 | 1949 | 4.49 | 1.70E-20 | 2.59E-19 |
| chr7 | 34691028 | 34696257 | 5230 | 9.59 | 1.78E-20 | 2.70E-19 |
| chr9 | 36296127 | 36342688 | 46562 | 0.49 | 1.81E-20 | 2.75E-19 |
| chr20 | 56588369 | 56595176 | 6808 | 1.01 | 1.82E-20 | 2.76E-19 |
| chr20 | 625417 | 643595 | 18179 | -0.71 | 1.83E-20 | 2.78E-19 |
| chr22 | 29168266 | 29220597 | 52332 | 0.76 | 1.85E-20 | 2.81E-19 |
| chr2 | 145695529 | 145709345 | 13817 | -1.61 | 1.87E-20 | 2.83E-19 |
| chr19 | 45574188 | 45658099 | 83912 | -0.52 | 1.91E-20 | 2.89E-19 |
| chr7 | 151777582 | 151780365 | 2784 | 3.03 | 1.92E-20 | 2.91E-19 |
| chr1 | 170531169 | 170533018 | 1850 | 2.42 | 1.97E-20 | 2.98E-19 |
| chr7 | 144531465 | 144540317 | 8853 | 1.2 | 2.00E-20 | 3.03E-19 |
| chrX | 56079554 | 56080473 | 920 | 3.84 | 2.06E-20 | 3.11E-19 |
| chr3 | 29556303 | 29568132 | 11830 | 1.87 | 2.08E-20 | 3.14E-19 |
| chr6 | 122011448 | 122017776 | 6329 | -1.4 | 2.08E-20 | 3.14E-19 |
| chr10 | 17684348 | 17716876 | 32529 | 0.7 | 2.22E-20 | 3.35E-19 |
| chr7 | 67218619 | 67242864 | 24246 | 1.27 | 2.24E-20 | 3.38E-19 |
| chr17 | 15551157 | 15564073 | 12917 | 0.69 | 2.24E-20 | 3.38E-19 |
| chr3 | 86274127 | 86281048 | 6922 | -2.32 | 2.29E-20 | 3.45E-19 |
| chr10 | 134208955 | 134266670 | 57716 | 0.55 | 2.40E-20 | 3.61E-19 |
| chr10 | 5533420 | 5538937 | 5518 | 2.16 | 2.41E-20 | 3.64E-19 |
| chr1 | 66633622 | 66650585 | 16964 | 0.85 | 2.49E-20 | 3.75E-19 |
| chr2 | 154819088 | 154821075 | 1988 | -2.29 | 2.56E-20 | 3.84E-19 |
| chr8 | 128746131 | 128780072 | 33942 | -0.5 | 2.56E-20 | 3.85E-19 |
| chr5 | 132341287 | 132423639 | 82353 | 0.5 | 2.61E-20 | 3.92E-19 |
| chr21 | 37324377 | 37327008 | 2632 | 2.13 | 2.65E-20 | 3.98E-19 |
| chr10 | 11835350 | 11941176 | 105827 | 0.47 | 2.66E-20 | 3.99E-19 |
| chr10 | 4312126 | 4324105 | 11980 | 3.59 | 2.94E-20 | 4.41E-19 |
| chr4 | 59673509 | 59677912 | 4404 | 1.93 | 3.21E-20 | 4.82E-19 |
| chr6 | 54171568 | 54175499 | 3932 | 1.83 | 3.22E-20 | 4.83E-19 |
| chr9 | 14170322 | 14181746 | 11425 | 1.36 | 3.22E-20 | 4.83E-19 |
| chr10 | 14860682 | 14865907 | 5226 | -1.23 | 3.23E-20 | 4.84E-19 |
| chr3 | 114659906 | 114674499 | 14594 | 9.5 | 3.27E-20 | 4.90E-19 |
| chr7 | 43655242 | 43715646 | 60405 | 0.45 | 3.29E-20 | 4.92E-19 |
| chr2 | 99484586 | 99487177 | 2592 | 3.2 | 3.31E-20 | 4.95E-19 |
| chr4 | 159289681 | 159306185 | 16505 | 4.13 | 3.33E-20 | 4.98E-19 |
| chr3 | 155453002 | 155484572 | 31571 | 0.61 | 3.44E-20 | 5.13E-19 |
| chr12 | 105837253 | 105840338 | 3086 | 1.87 | 3.59E-20 | 5.36E-19 |
| chr18 | 33281813 | 33284558 | 2746 | 2.16 | 3.61E-20 | 5.39E-19 |
| chr4 | 108744714 | 108748932 | 4219 | 1.29 | 3.64E-20 | 5.44E-19 |
| chr19 | 20888688 | 20901514 | 12827 | 9.48 | 3.72E-20 | 5.55E-19 |
| chr4 | 24579672 | 24607130 | 27459 | 0.61 | 3.92E-20 | 5.85E-19 |
| chr2 | 70555211 | 70557163 | 1953 | 3.81 | 4.07E-20 | 6.07E-19 |
| chr7 | 79707413 | 79717889 | 10477 | 5.65 | 4.16E-20 | 6.19E-19 |
| chr2 | 44494826 | 44523596 | 28771 | -0.7 | 4.17E-20 | 6.21E-19 |
| chr4 | 23334886 | 23336844 | 1959 | 3.75 | 4.18E-20 | 6.22E-19 |
| chr1 | 169534583 | 169542757 | 8175 | 9.47 | 4.32E-20 | 6.41E-19 |
| chr18 | 74765597 | 74845305 | 79709 | 0.43 | 4.49E-20 | 6.66E-19 |
| chr19 | 6720600 | 6745831 | 25232 | 0.78 | 4.61E-20 | 6.84E-19 |
| chr11 | 57498968 | 57570145 | 71178 | 0.41 | 4.62E-20 | 6.85E-19 |
| chr15 | 90540627 | 90548505 | 7879 | 0.97 | 4.62E-20 | 6.85E-19 |
| chr9 | 27425912 | 27428449 | 2538 | -3.22 | 4.64E-20 | 6.88E-19 |
| chr13 | 49666954 | 49688419 | 21466 | 1.45 | 4.66E-20 | 6.90E-19 |
| chr4 | 35023144 | 35030373 | 7230 | 0.86 | 4.75E-20 | 7.03E-19 |
| chr14 | 96659727 | 96692494 | 32768 | 0.85 | 4.91E-20 | 7.27E-19 |
| chr6 | 136295690 | 136318380 | 22691 | -5.17 | 5.03E-20 | 7.43E-19 |
| chr17 | 19300369 | 19302690 | 2322 | 2.02 | 5.05E-20 | 7.47E-19 |
| chr6 | 138775987 | 138792051 | 16065 | 1.31 | 5.10E-20 | 7.54E-19 |
| chr15 | 58729614 | 58768639 | 39026 | -0.54 | 5.22E-20 | 7.70E-19 |

| | | | | | | |
|---|---|---|---|---|---|---|
| chr9 | 140500082 | 140534979 | 34898 | 0.73 | 5.37E-20 | 7.92E-19 |
| chr12 | 10491469 | 10493087 | 1619 | 2.88 | 5.41E-20 | 7.99E-19 |
| chr8 | 116587969 | 116600156 | 12188 | 9.45 | 5.44E-20 | 8.02E-19 |
| chr17 | 4379790 | 4387688 | 7899 | 2.11 | 5.45E-20 | 8.04E-19 |
| chr19 | 19958331 | 19988909 | 30579 | 9.46 | 5.51E-20 | 8.11E-19 |
| chr9 | 82238381 | 82241361 | 2981 | 1.71 | 5.72E-20 | 8.43E-19 |
| chr12 | 64820526 | 64873436 | 52911 | 0.65 | 5.75E-20 | 8.47E-19 |
| chr2 | 173939941 | 173958202 | 18262 | -0.79 | 5.82E-20 | 8.56E-19 |
| chr11 | 130764000 | 130768575 | 4576 | 3.7 | 5.95E-20 | 8.75E-19 |
| chr5 | 159241690 | 159255558 | 13869 | 0.76 | 6.09E-20 | 8.95E-19 |
| chr7 | 30489552 | 30519280 | 29729 | 0.65 | 6.12E-20 | 9.00E-19 |
| chr7 | 34098791 | 34110772 | 11982 | -9.44 | 6.42E-20 | 9.43E-19 |
| chr17 | 31154255 | 31159439 | 5185 | 9.44 | 6.66E-20 | 9.78E-19 |
| chr16 | 66723285 | 66731546 | 8262 | 1.66 | 6.67E-20 | 9.79E-19 |
| chr5 | 27387144 | 27404290 | 17147 | 9.42 | 6.77E-20 | 9.92E-19 |
| chr14 | 54063520 | 54082987 | 19468 | 0.89 | 7.07E-20 | 1.04E-18 |
| chr1 | 192499297 | 192514833 | 15537 | 0.94 | 7.16E-20 | 1.05E-18 |
| chr9 | 99235208 | 99254725 | 19518 | -2.18 | 7.38E-20 | 1.08E-18 |
| chrX | 38130146 | 38139392 | 9247 | 1.48 | 7.40E-20 | 1.08E-18 |
| chr5 | 173363617 | 173400956 | 37340 | 0.67 | 7.46E-20 | 1.09E-18 |
| chr12 | 33031846 | 33055748 | 23903 | 1.32 | 7.47E-20 | 1.09E-18 |
| chr4 | 189896692 | 189911673 | 14982 | 1.45 | 7.55E-20 | 1.10E-18 |
| chr8 | 82909885 | 82915795 | 5911 | 2.28 | 7.64E-20 | 1.12E-18 |
| chr4 | 26823007 | 26829256 | 6250 | 2.13 | 7.97E-20 | 1.16E-18 |
| chr17 | 51648270 | 51649641 | 1372 | 3.28 | 8.05E-20 | 1.17E-18 |
| chr7 | 46792280 | 46804796 | 12517 | 1.9 | 8.06E-20 | 1.17E-18 |
| chr13 | 84023229 | 84025135 | 1907 | 2.26 | 8.13E-20 | 1.18E-18 |
| chr4 | 170525412 | 170544546 | 19135 | 0.92 | 8.23E-20 | 1.20E-18 |
| chr10 | 5621653 | 5674968 | 53316 | 0.8 | 8.53E-20 | 1.24E-18 |
| chr8 | 74711157 | 74736618 | 25462 | 1.21 | 8.72E-20 | 1.27E-18 |
| chr11 | 122310110 | 122312609 | 2500 | -3.92 | 8.91E-20 | 1.30E-18 |
| chr10 | 21561786 | 21664762 | 102977 | -0.33 | 8.96E-20 | 1.30E-18 |
| chr12 | 80940749 | 80947511 | 6763 | 2.15 | 8.97E-20 | 1.30E-18 |
| chr22 | 43240127 | 43270743 | 30617 | 1.04 | 9.03E-20 | 1.31E-18 |
| chr2 | 107482552 | 107504557 | 22006 | -9.39 | 9.34E-20 | 1.36E-18 |
| chr10 | 96661206 | 96665881 | 4676 | 1.42 | 9.39E-20 | 1.36E-18 |
| chr8 | 119100635 | 119125840 | 25206 | 0.66 | 9.47E-20 | 1.37E-18 |
| chr5 | 114958937 | 114970515 | 11579 | 1.15 | 9.62E-20 | 1.40E-18 |
| chr3 | 25609501 | 25646561 | 37061 | -0.81 | 9.64E-20 | 1.40E-18 |
| chr9 | 100426803 | 100478032 | 51230 | 0.77 | 9.73E-20 | 1.41E-18 |
| chr3 | 99742727 | 99768243 | 25517 | 2.74 | 9.82E-20 | 1.42E-18 |
| chr2 | 110961667 | 110970615 | 8949 | -1.35 | 9.97E-20 | 1.44E-18 |
| chr4 | 105048555 | 105050773 | 2219 | 1.9 | 1.02E-19 | 1.48E-18 |
| chr2 | 38892204 | 38921915 | 29712 | -0.7 | 1.08E-19 | 1.56E-18 |
| chr21 | 34774925 | 34818167 | 43243 | 0.56 | 1.10E-19 | 1.59E-18 |
| chr4 | 21769822 | 21782465 | 12644 | 9.39 | 1.11E-19 | 1.60E-18 |
| chr12 | 89131660 | 89143742 | 12083 | 2.55 | 1.11E-19 | 1.61E-18 |
| chr12 | 6304378 | 6339845 | 35468 | 0.5 | 1.12E-19 | 1.62E-18 |
| chr1 | 75835835 | 75841736 | 5902 | 6.98 | 1.14E-19 | 1.64E-18 |
| chr19 | 52899773 | 52902537 | 2765 | 9.37 | 1.20E-19 | 1.72E-18 |
| chr4 | 149350971 | 149364350 | 13380 | 9.36 | 1.21E-19 | 1.74E-18 |
| chr5 | 3739891 | 3744082 | 4192 | 9.37 | 1.26E-19 | 1.81E-18 |
| chr12 | 90340172 | 90348578 | 8407 | 0.55 | 1.26E-19 | 1.82E-18 |
| chr9 | 110004135 | 110012454 | 8320 | 1.22 | 1.27E-19 | 1.82E-18 |
| chr13 | 80194911 | 80197483 | 2573 | 5.18 | 1.29E-19 | 1.85E-18 |
| chr16 | 2917517 | 2977136 | 59620 | 0.69 | 1.30E-19 | 1.86E-18 |
| chr1 | 6084896 | 6087000 | 2105 | 3.91 | 1.31E-19 | 1.88E-18 |
| chr9 | 38424192 | 38444619 | 20428 | 0.86 | 1.41E-19 | 2.02E-18 |
| chr11 | 5614321 | 5646852 | 32532 | 0.62 | 1.42E-19 | 2.03E-18 |
| chr12 | 76941991 | 76954530 | 12540 | 1.2 | 1.42E-19 | 2.04E-18 |
| chr10 | 105335083 | 105339516 | 4434 | 2.96 | 1.44E-19 | 2.07E-18 |

| | | | | | | |
|---|---|---|---|---|---|---|
| chr8 | 26427536 | 26517602 | 90067 | -0.44 | 1.46E-19 | 2.09E-18 |
| chr10 | 36568057 | 36572458 | 4402 | -9.33 | 1.47E-19 | 2.10E-18 |
| chr2 | 119866723 | 119870923 | 4201 | 9.34 | 1.50E-19 | 2.14E-18 |
| chr12 | 20140426 | 20152892 | 12467 | 1.61 | 1.52E-19 | 2.18E-18 |
| chr11 | 10946008 | 10956236 | 10229 | -1.23 | 1.53E-19 | 2.18E-18 |
| chr8 | 130859863 | 130908675 | 48813 | 0.6 | 1.55E-19 | 2.22E-18 |
| chr15 | 90108787 | 90170422 | 61636 | 0.62 | 1.58E-19 | 2.26E-18 |
| chr8 | 93904229 | 93907765 | 3537 | 5.61 | 1.59E-19 | 2.26E-18 |
| chr12 | 38778546 | 38780861 | 2316 | 4.04 | 1.62E-19 | 2.31E-18 |
| chr3 | 167579150 | 167580890 | 1741 | -6.03 | 1.63E-19 | 2.33E-18 |
| chr19 | 38081519 | 38086546 | 5028 | 9.36 | 1.67E-19 | 2.39E-18 |
| chr12 | 43150192 | 43151578 | 1387 | 3.49 | 1.69E-19 | 2.41E-18 |
| chr19 | 9402581 | 9436755 | 34175 | 5.14 | 1.70E-19 | 2.43E-18 |
| chr1 | 934100 | 961476 | 27377 | 0.95 | 1.71E-19 | 2.43E-18 |
| chr17 | 2467678 | 2469996 | 2319 | 1.76 | 1.72E-19 | 2.44E-18 |
| chr19 | 20188469 | 20196436 | 7968 | 9.33 | 1.72E-19 | 2.45E-18 |
| chr7 | 27774201 | 27834148 | 59948 | 0.55 | 1.74E-19 | 2.47E-18 |
| chr9 | 77701422 | 77704638 | 3217 | 1.36 | 1.76E-19 | 2.50E-18 |
| chr1 | 54755821 | 54765740 | 9920 | 2.02 | 1.80E-19 | 2.56E-18 |
| chr15 | 96313048 | 96344193 | 31146 | 9.37 | 1.84E-19 | 2.61E-18 |
| chr4 | 59504527 | 59506554 | 2028 | 6.01 | 1.87E-19 | 2.65E-18 |
| chr6 | 66527796 | 66530438 | 2643 | -1.72 | 1.89E-19 | 2.68E-18 |
| chr4 | 13988165 | 14001481 | 13317 | -0.81 | 1.94E-19 | 2.74E-18 |
| chr2 | 24230920 | 24257114 | 26195 | -0.74 | 2.00E-19 | 2.84E-18 |
| chrX | 8834519 | 8838085 | 3567 | 1.45 | 2.05E-19 | 2.90E-18 |
| chr4 | 16030467 | 16036806 | 6340 | 3 | 2.10E-19 | 2.97E-18 |
| chr10 | 17241758 | 17281710 | 39953 | 0.54 | 2.12E-19 | 3.00E-18 |
| chr12 | 78332828 | 78342399 | 9572 | 0.7 | 2.16E-19 | 3.05E-18 |
| chr9 | 112672708 | 112675993 | 3286 | 1.49 | 2.16E-19 | 3.05E-18 |
| chr18 | 28483191 | 28496756 | 13566 | 1.93 | 2.34E-19 | 3.31E-18 |
| chr17 | 74914425 | 74949319 | 34895 | -7.47 | 2.39E-19 | 3.38E-18 |
| chr14 | 63933534 | 63972450 | 38917 | 0.7 | 2.40E-19 | 3.39E-18 |
| chr8 | 110052323 | 110066100 | 13778 | 0.66 | 2.45E-19 | 3.46E-18 |
| chr9 | 32625839 | 32628925 | 3087 | 9.29 | 2.51E-19 | 3.54E-18 |
| chr10 | 27441732 | 27448525 | 6794 | 0.74 | 2.51E-19 | 3.54E-18 |
| chr9 | 81153249 | 81156753 | 3505 | 1.54 | 2.52E-19 | 3.55E-18 |
| chr2 | 20637422 | 20650817 | 13396 | 0.63 | 2.58E-19 | 3.63E-18 |
| chrX | 43768409 | 43770283 | 1875 | 4.34 | 2.58E-19 | 3.64E-18 |
| chr14 | 36335623 | 36353514 | 17892 | 1.2 | 2.59E-19 | 3.65E-18 |
| chr1 | 46108936 | 46114079 | 5144 | 1.78 | 2.62E-19 | 3.68E-18 |
| chr9 | 112883322 | 112904299 | 20978 | 1.55 | 2.63E-19 | 3.70E-18 |
| chr2 | 240182786 | 240207616 | 24831 | -0.57 | 2.66E-19 | 3.74E-18 |
| chr17 | 9546945 | 9552912 | 5968 | 0.8 | 2.68E-19 | 3.77E-18 |
| chr7 | 139300412 | 139373365 | 72954 | -0.45 | 2.69E-19 | 3.77E-18 |
| chr11 | 44585618 | 44586944 | 1327 | 1.72 | 2.72E-19 | 3.81E-18 |
| chr14 | 31697111 | 31700241 | 3131 | 1.3 | 2.75E-19 | 3.86E-18 |
| chr21 | 43782948 | 43788101 | 5154 | 2.01 | 2.76E-19 | 3.87E-18 |
| chr9 | 4324414 | 4335463 | 11050 | 2.14 | 2.82E-19 | 3.95E-18 |
| chr6 | 41038334 | 41043036 | 4703 | 0.9 | 2.84E-19 | 3.98E-18 |
| chr7 | 81037629 | 81044443 | 6815 | 3.7 | 2.89E-19 | 4.04E-18 |
| chr17 | 4977248 | 4982767 | 5520 | 9.29 | 2.90E-19 | 4.06E-18 |
| chr5 | 74212144 | 74213692 | 1549 | 2.72 | 3.04E-19 | 4.25E-18 |
| chr10 | 13140623 | 13158990 | 18368 | 1.01 | 3.05E-19 | 4.27E-18 |
| chr12 | 121396349 | 121425122 | 28774 | 0.77 | 3.09E-19 | 4.32E-18 |
| chr2 | 1621267 | 1637679 | 16413 | 9.27 | 3.13E-19 | 4.37E-18 |
| chr6 | 84735587 | 84744096 | 8510 | 1.28 | 3.14E-19 | 4.37E-18 |
| chr18 | 12596206 | 12609075 | 12870 | 1.19 | 3.17E-19 | 4.42E-18 |
| chr4 | 173897542 | 173946214 | 48673 | 0.65 | 3.17E-19 | 4.42E-18 |
| chr10 | 98432854 | 98434237 | 1384 | 5.71 | 3.20E-19 | 4.45E-18 |
| chr15 | 52429273 | 52438046 | 8774 | 1.23 | 3.23E-19 | 4.49E-18 |
| chr12 | 127255673 | 127257871 | 2199 | 9.26 | 3.26E-19 | 4.54E-18 |

| | | | | | | |
|---|---|---|---|---|---|---|
| chr2 | 228517018 | 228523876 | 6859 | -1.34 | 3.27E-19 | 4.54E-18 |
| chr10 | 83359058 | 83363928 | 4871 | 9.26 | 3.29E-19 | 4.58E-18 |
| chr1 | 207377604 | 207392764 | 15161 | 1.26 | 3.30E-19 | 4.59E-18 |
| chr4 | 184358036 | 184367206 | 9171 | 1.08 | 3.31E-19 | 4.60E-18 |
| chr20 | 18232041 | 18244567 | 12527 | 9.26 | 3.33E-19 | 4.62E-18 |
| chr6 | 7672993 | 7700475 | 27483 | 0.69 | 3.39E-19 | 4.71E-18 |
| chr4 | 77584794 | 77588780 | 3987 | 1.72 | 3.44E-19 | 4.77E-18 |
| chr4 | 161756470 | 161771362 | 14893 | 1.82 | 3.47E-19 | 4.81E-18 |
| chr4 | 179465841 | 179480017 | 14177 | 9.26 | 3.49E-19 | 4.83E-18 |
| chr17 | 43298324 | 43385055 | 86732 | -0.33 | 3.49E-19 | 4.83E-18 |
| chr20 | 52956166 | 52964498 | 8333 | 1.12 | 3.51E-19 | 4.86E-18 |
| chr1 | 151857711 | 151860879 | 3169 | 3.72 | 3.52E-19 | 4.87E-18 |
| chr6 | 116871711 | 116894189 | 22479 | 0.97 | 3.54E-19 | 4.90E-18 |
| chr15 | 101437281 | 101443186 | 5906 | -3.88 | 3.56E-19 | 4.93E-18 |
| chr9 | 121334885 | 121341095 | 6211 | 9.25 | 3.71E-19 | 5.13E-18 |
| chr13 | 81771346 | 81773540 | 2195 | 2.64 | 3.71E-19 | 5.13E-18 |
| chrX | 135212246 | 135215449 | 3204 | 1.96 | 3.71E-19 | 5.13E-18 |
| chr2 | 83006752 | 83023468 | 16717 | 9.25 | 3.73E-19 | 5.15E-18 |
| chr1 | 23599616 | 23640350 | 40735 | 0.78 | 3.80E-19 | 5.25E-18 |
| chr13 | 43834720 | 43844729 | 10010 | 1.72 | 3.81E-19 | 5.26E-18 |
| chr13 | 48729322 | 48735722 | 6401 | 1.5 | 3.86E-19 | 5.32E-18 |
| chr4 | 72551927 | 72558554 | 6628 | 3.33 | 3.87E-19 | 5.34E-18 |
| chr12 | 78454968 | 78457900 | 2933 | 1.85 | 3.93E-19 | 5.41E-18 |
| chr19 | 24059898 | 24071888 | 11991 | 9.25 | 3.96E-19 | 5.45E-18 |
| chr17 | 33894576 | 33973521 | 78946 | 0.4 | 4.03E-19 | 5.54E-18 |
| chr8 | 116677896 | 116682376 | 4481 | 9.25 | 4.07E-19 | 5.59E-18 |
| chr7 | 46171708 | 46194301 | 22594 | -1.55 | 4.09E-19 | 5.62E-18 |
| chr2 | 47514861 | 47551223 | 36363 | -1.41 | 4.12E-19 | 5.66E-18 |
| chr2 | 24333117 | 24347769 | 14653 | -1.08 | 4.13E-19 | 5.67E-18 |
| chr13 | 104667009 | 104677882 | 10874 | 9.24 | 4.15E-19 | 5.70E-18 |
| chr10 | 88101773 | 88107598 | 5826 | 1.29 | 4.25E-19 | 5.83E-18 |
| chr12 | 125109792 | 125140974 | 31183 | 0.86 | 4.33E-19 | 5.94E-18 |
| chr12 | 80321261 | 80330760 | 9500 | 0.95 | 4.35E-19 | 5.97E-18 |
| chr3 | 95372543 | 95375592 | 3050 | 9.23 | 4.45E-19 | 6.09E-18 |
| chr6 | 55940377 | 55967319 | 26943 | 1.12 | 4.72E-19 | 6.46E-18 |
| chr9 | 77439247 | 77456251 | 17005 | 9.22 | 4.87E-19 | 6.66E-18 |
| chr5 | 80745605 | 80748937 | 3333 | 2.65 | 4.87E-19 | 6.66E-18 |
| chr1 | 173091130 | 173097939 | 6810 | 1.91 | 4.95E-19 | 6.77E-18 |
| chr14 | 85902867 | 85909693 | 6827 | 0.98 | 5.02E-19 | 6.86E-18 |
| chr14 | 76177242 | 76179519 | 2278 | 1.66 | 5.08E-19 | 6.95E-18 |
| chr10 | 122632169 | 122633812 | 1644 | 1.98 | 5.22E-19 | 7.13E-18 |
| chrX | 22077707 | 22079847 | 2141 | 2.7 | 5.24E-19 | 7.15E-18 |
| chr10 | 6313232 | 6357796 | 44565 | -0.36 | 5.37E-19 | 7.33E-18 |
| chr19 | 9684161 | 9695939 | 11779 | 1.65 | 5.42E-19 | 7.39E-18 |
| chr14 | 61978061 | 62053347 | 75287 | 0.33 | 5.51E-19 | 7.51E-18 |
| chr5 | 100471798 | 100478560 | 6763 | 4.27 | 5.53E-19 | 7.54E-18 |
| chr19 | 58379349 | 58401028 | 21680 | 9.22 | 5.60E-19 | 7.62E-18 |
| chr11 | 100551016 | 100568592 | 17577 | 1 | 5.66E-19 | 7.70E-18 |
| chr7 | 34018615 | 34028113 | 9499 | -9.19 | 5.78E-19 | 7.86E-18 |
| chr6 | 74957860 | 74967195 | 9336 | 9.2 | 5.89E-19 | 8.01E-18 |
| chr7 | 25108979 | 25113712 | 4734 | 1.83 | 5.91E-19 | 8.03E-18 |
| chr14 | 103148799 | 103158028 | 9230 | 1.86 | 5.91E-19 | 8.03E-18 |
| chr19 | 20212082 | 20221507 | 9426 | 9.2 | 5.97E-19 | 8.11E-18 |
| chr19 | 39561204 | 39573722 | 12519 | -0.86 | 6.12E-19 | 8.31E-18 |
| chr18 | 69774417 | 69783085 | 8669 | 2.6 | 6.23E-19 | 8.46E-18 |
| chr7 | 91654683 | 91678363 | 23681 | 1.03 | 6.37E-19 | 8.64E-18 |
| chr2 | 192166611 | 192185046 | 18436 | -1.03 | 6.41E-19 | 8.69E-18 |
| chr19 | 37802943 | 37839033 | 36091 | 4.95 | 6.45E-19 | 8.74E-18 |
| chr1 | 48024203 | 48044862 | 20660 | 2.13 | 6.47E-19 | 8.76E-18 |
| chr7 | 139528131 | 139531222 | 3092 | 1.16 | 6.49E-19 | 8.79E-18 |
| chrX | 107681112 | 107708777 | 27666 | -1.09 | 6.57E-19 | 8.89E-18 |

| | | | | | | |
|---|---|---|---|---|---|---|
| chr6 | 90006101 | 90009844 | 3744 | 1.2 | 6.59E-19 | 8.90E-18 |
| chr17 | 53617158 | 53619115 | 1958 | 1.71 | 6.61E-19 | 8.93E-18 |
| chr7 | 54754546 | 54756461 | 1916 | 2.62 | 6.77E-19 | 9.15E-18 |
| chr1 | 189320186 | 189327155 | 6970 | 2.82 | 6.83E-19 | 9.23E-18 |
| chr7 | 23719742 | 23726572 | 6831 | 3.19 | 7.21E-19 | 9.73E-18 |
| chr21 | 36508288 | 36510066 | 1779 | 1.99 | 7.44E-19 | 1.00E-17 |
| chr2 | 217399950 | 217405607 | 5658 | 3.3 | 7.52E-19 | 1.01E-17 |
| chr12 | 3314231 | 3315811 | 1581 | 2.47 | 7.59E-19 | 1.02E-17 |
| chr8 | 48571184 | 48574105 | 2922 | 1.1 | 7.64E-19 | 1.03E-17 |
| chr2 | 225432482 | 225451352 | 18871 | -0.96 | 7.85E-19 | 1.06E-17 |
| chr17 | 27564623 | 27574428 | 9806 | -0.88 | 7.97E-19 | 1.07E-17 |
| chr19 | 37584064 | 37598733 | 14670 | 9.16 | 7.99E-19 | 1.07E-17 |
| chr2 | 85861146 | 85890430 | 29285 | -0.97 | 8.04E-19 | 1.08E-17 |
| chr3 | 164180243 | 164217323 | 37081 | 9.18 | 8.12E-19 | 1.09E-17 |
| chr5 | 82767949 | 82773747 | 5799 | 4.01 | 8.28E-19 | 1.11E-17 |
| chr9 | 107032140 | 107033645 | 1506 | 3.57 | 8.36E-19 | 1.12E-17 |
| chr12 | 100896577 | 100899747 | 3171 | 9.17 | 8.57E-19 | 1.15E-17 |
| chr2 | 26679571 | 26691428 | 11858 | -2.19 | 8.60E-19 | 1.15E-17 |
| chr7 | 1010344 | 1018726 | 8383 | 1.14 | 9.00E-19 | 1.21E-17 |
| chr21 | 46330525 | 46360851 | 30327 | 1.01 | 9.01E-19 | 1.21E-17 |
| chr9 | 20496075 | 20515250 | 19176 | 1.07 | 9.03E-19 | 1.21E-17 |
| chr2 | 62683577 | 62695563 | 11987 | -1.28 | 9.19E-19 | 1.23E-17 |
| chr18 | 60642074 | 60645368 | 3295 | 2.29 | 9.19E-19 | 1.23E-17 |
| chr1 | 196992732 | 197013132 | 20401 | 1.99 | 9.35E-19 | 1.25E-17 |
| chr12 | 122748439 | 122764389 | 15951 | 1.01 | 9.37E-19 | 1.25E-17 |
| chr12 | 13277907 | 13288995 | 11089 | 2.07 | 9.39E-19 | 1.26E-17 |
| chr19 | 20343068 | 20355247 | 12180 | 9.15 | 9.90E-19 | 1.32E-17 |
| chr4 | 62327309 | 62334497 | 7189 | 9.14 | 1.02E-18 | 1.36E-17 |
| chr3 | 57735839 | 57743565 | 7727 | 0.92 | 1.03E-18 | 1.38E-17 |
| chr13 | 31393242 | 31407803 | 14562 | 5.04 | 1.06E-18 | 1.42E-17 |
| chr7 | 83270155 | 83278856 | 8702 | 1.72 | 1.07E-18 | 1.43E-17 |
| chr10 | 687606 | 697410 | 9805 | 1.36 | 1.11E-18 | 1.48E-17 |
| chr18 | 56722105 | 56735235 | 13131 | 1.25 | 1.12E-18 | 1.49E-17 |
| chr12 | 68040383 | 68044639 | 4257 | 0.79 | 1.14E-18 | 1.52E-17 |
| chr14 | 68582901 | 68677961 | 95061 | 0.34 | 1.14E-18 | 1.52E-17 |
| chr5 | 134580836 | 134583706 | 2871 | -1.51 | 1.14E-18 | 1.52E-17 |
| chr13 | 82575936 | 82593950 | 18015 | 9.14 | 1.15E-18 | 1.53E-17 |
| chr9 | 109677866 | 109684189 | 6324 | 1.68 | 1.17E-18 | 1.55E-17 |
| chr2 | 1711269 | 1747321 | 36053 | 9.13 | 1.21E-18 | 1.61E-17 |
| chr12 | 20170609 | 20185028 | 14420 | 9.14 | 1.22E-18 | 1.63E-17 |
| chr14 | 96774697 | 96777608 | 2912 | 1.24 | 1.26E-18 | 1.67E-17 |
| chr10 | 4004995 | 4006585 | 1591 | 2.61 | 1.26E-18 | 1.68E-17 |
| chr3 | 187720117 | 187723585 | 3469 | 1.12 | 1.30E-18 | 1.72E-17 |
| chr8 | 36967801 | 36969373 | 1573 | 4.06 | 1.32E-18 | 1.75E-17 |
| chr19 | 36817589 | 36824423 | 6835 | 9.11 | 1.33E-18 | 1.76E-17 |
| chr6 | 110110854 | 110116554 | 5701 | 1.58 | 1.34E-18 | 1.78E-17 |
| chr12 | 74763636 | 74765318 | 1683 | 2.01 | 1.35E-18 | 1.79E-17 |
| chr5 | 14032743 | 14040422 | 7680 | 0.55 | 1.38E-18 | 1.82E-17 |
| chr11 | 35639175 | 35652393 | 13219 | -2.56 | 1.40E-18 | 1.86E-17 |
| chr13 | 98943625 | 98963947 | 20323 | 2.21 | 1.54E-18 | 2.04E-17 |
| chr6 | 42243717 | 42263474 | 19758 | 0.95 | 1.55E-18 | 2.05E-17 |
| chr13 | 84703711 | 84706816 | 3106 | 9.1 | 1.56E-18 | 2.07E-17 |
| chr9 | 72358794 | 72387340 | 28547 | 0.88 | 1.57E-18 | 2.07E-17 |
| chr16 | 3354120 | 3357220 | 3101 | 1.44 | 1.57E-18 | 2.07E-17 |
| chr4 | 6675216 | 6719002 | 43787 | 0.78 | 1.60E-18 | 2.11E-17 |
| chr1 | 216773486 | 216775300 | 1815 | 1.58 | 1.64E-18 | 2.16E-17 |
| chr17 | 76697746 | 76714161 | 16416 | -1.16 | 1.64E-18 | 2.17E-17 |
| chr2 | 108977259 | 109009106 | 31848 | 0.74 | 1.67E-18 | 2.21E-17 |
| chr12 | 79309840 | 79331256 | 21417 | 2.02 | 1.67E-18 | 2.21E-17 |
| chr12 | 120105394 | 120134150 | 28757 | 0.76 | 1.73E-18 | 2.27E-17 |
| chr1 | 242107247 | 242109286 | 2040 | 1.54 | 1.74E-18 | 2.29E-17 |

| | | | | | | |
|---|---|---|---|---|---|---|
| chr12 | 42373079 | 42387881 | 14803 | 1.76 | 1.78E-18 | 2.34E-17 |
| chr17 | 32575031 | 32582993 | 7963 | 1.11 | 1.78E-18 | 2.34E-17 |
| chr19 | 24319505 | 24325146 | 5642 | 9.08 | 1.85E-18 | 2.43E-17 |
| chr12 | 45648062 | 45680480 | 32419 | 0.81 | 1.91E-18 | 2.51E-17 |
| chr6 | 106525229 | 106547301 | 22073 | -1.16 | 1.94E-18 | 2.55E-17 |
| chr9 | 123491939 | 123556616 | 64678 | 0.6 | 1.97E-18 | 2.59E-17 |
| chr7 | 95256261 | 95266097 | 9837 | 9.07 | 1.99E-18 | 2.61E-17 |
| chr3 | 96196820 | 96206665 | 9846 | 9.08 | 2.03E-18 | 2.66E-17 |
| chr7 | 145606738 | 145614302 | 7565 | 9.09 | 2.04E-18 | 2.67E-17 |
| chr17 | 48210092 | 48238875 | 28784 | -1.03 | 2.05E-18 | 2.68E-17 |
| chr19 | 24173200 | 24176584 | 3385 | 9.06 | 2.07E-18 | 2.71E-17 |
| chr19 | 52838883 | 52840426 | 1544 | 5.99 | 2.09E-18 | 2.74E-17 |
| chr11 | 73729018 | 73782833 | 53816 | -0.86 | 2.09E-18 | 2.74E-17 |
| chr18 | 24714710 | 24723222 | 8513 | 9.06 | 2.09E-18 | 2.74E-17 |
| chrX | 35574752 | 35589665 | 14914 | 2.8 | 2.19E-18 | 2.87E-17 |
| chr10 | 31290241 | 31294348 | 4108 | 2.22 | 2.20E-18 | 2.87E-17 |
| chr19 | 53546741 | 53559730 | 12990 | 9.06 | 2.22E-18 | 2.90E-17 |
| chr4 | 142460832 | 142462810 | 1979 | 3.62 | 2.26E-18 | 2.96E-17 |
| chr2 | 206540164 | 206637566 | 97403 | -0.35 | 2.27E-18 | 2.96E-17 |
| chr18 | 52488930 | 52497450 | 8521 | 0.93 | 2.28E-18 | 2.98E-17 |
| chr1 | 75599462 | 75607978 | 8517 | 9.06 | 2.30E-18 | 2.99E-17 |
| chr3 | 31570153 | 31578634 | 8482 | 0.91 | 2.32E-18 | 3.02E-17 |
| chr18 | 19860919 | 19867537 | 6619 | 1.05 | 2.32E-18 | 3.02E-17 |
| chr12 | 59158188 | 59161109 | 2922 | 1.3 | 2.34E-18 | 3.04E-17 |
| chr6 | 141803855 | 141818201 | 14347 | 1.74 | 2.37E-18 | 3.09E-17 |
| chr14 | 56690498 | 56693420 | 2923 | 1.47 | 2.38E-18 | 3.10E-17 |
| chr5 | 90458922 | 90467532 | 8611 | 7.69 | 2.41E-18 | 3.14E-17 |
| chr14 | 26862949 | 26886482 | 23534 | 9.07 | 2.44E-18 | 3.17E-17 |
| chr13 | 34727453 | 34751865 | 24413 | 0.53 | 2.49E-18 | 3.23E-17 |
| chr3 | 182002001 | 182003474 | 1474 | 4.74 | 2.49E-18 | 3.24E-17 |
| chr12 | 43085527 | 43100071 | 14545 | 9.04 | 2.51E-18 | 3.25E-17 |
| chr7 | 107992301 | 108023608 | 31308 | -9.03 | 2.57E-18 | 3.33E-17 |
| chr8 | 54256152 | 54266750 | 10599 | 1.92 | 2.61E-18 | 3.38E-17 |
| chr3 | 69248078 | 69266862 | 18785 | 1.06 | 2.73E-18 | 3.53E-17 |
| chr7 | 115316364 | 115320173 | 3810 | 9.04 | 2.75E-18 | 3.56E-17 |
| chr1 | 59604120 | 59623309 | 19190 | -0.67 | 2.77E-18 | 3.58E-17 |
| chr1 | 243407373 | 243433423 | 26051 | 0.68 | 2.79E-18 | 3.62E-17 |
| chr6 | 106241103 | 106259528 | 18426 | -1.93 | 2.87E-18 | 3.72E-17 |
| chr10 | 62462883 | 62479054 | 16172 | 9.06 | 2.88E-18 | 3.73E-17 |
| chr3 | 95927188 | 95930164 | 2977 | 9.05 | 2.91E-18 | 3.76E-17 |
| chr3 | 129439520 | 129444200 | 4681 | -1.31 | 2.93E-18 | 3.79E-17 |
| chr8 | 18809203 | 18837172 | 27970 | 0.49 | 2.95E-18 | 3.80E-17 |
| chr12 | 75772932 | 75786321 | 13390 | 0.9 | 2.96E-18 | 3.83E-17 |
| chr17 | 18863365 | 18915836 | 52472 | -0.68 | 2.99E-18 | 3.86E-17 |
| chr5 | 141735342 | 141745101 | 9760 | 0.63 | 2.99E-18 | 3.86E-17 |
| chr7 | 139612471 | 139617941 | 5471 | -1.17 | 3.04E-18 | 3.92E-17 |
| chr8 | 19231137 | 19616507 | 385371 | -0.24 | 3.10E-18 | 4.00E-17 |
| chr9 | 108080452 | 108082410 | 1959 | 2.43 | 3.16E-18 | 4.07E-17 |
| chr9 | 126679873 | 126697175 | 17303 | 0.88 | 3.21E-18 | 4.13E-17 |
| chr5 | 23630905 | 23632391 | 1487 | 7.57 | 3.23E-18 | 4.16E-17 |
| chr1 | 172213884 | 172239436 | 25553 | 1.43 | 3.24E-18 | 4.17E-17 |
| chr1 | 53571858 | 53582446 | 10589 | 0.99 | 3.26E-18 | 4.19E-17 |
| chr4 | 143456069 | 143490798 | 34730 | 3.22 | 3.35E-18 | 4.31E-17 |
| chr1 | 51338274 | 51340238 | 1965 | 1.99 | 3.42E-18 | 4.39E-17 |
| chr2 | 203050043 | 203053355 | 3313 | 3.64 | 3.43E-18 | 4.41E-17 |
| chr15 | 40021516 | 40042449 | 20934 | 0.98 | 3.45E-18 | 4.43E-17 |
| chr1 | 234095136 | 234109605 | 14470 | -2 | 3.48E-18 | 4.47E-17 |
| chr10 | 121543561 | 121547031 | 3471 | 2.24 | 3.49E-18 | 4.47E-17 |
| chr16 | 75078001 | 75100745 | 22745 | 1.13 | 3.51E-18 | 4.50E-17 |
| chr7 | 42896124 | 42909708 | 13585 | 9.02 | 3.60E-18 | 4.61E-17 |
| chr12 | 129295124 | 129309551 | 14428 | 0.93 | 3.67E-18 | 4.70E-17 |

| | | | | | | |
|---|---|---|---|---|---|---|
| chr7 | 92310046 | 92466703 | 156658 | 0.32 | 3.68E-18 | 4.72E-17 |
| chr12 | 59434384 | 59450387 | 16004 | 1.48 | 3.77E-18 | 4.82E-17 |
| chr13 | 110746567 | 110748074 | 1508 | 2.77 | 3.80E-18 | 4.86E-17 |
| chr2 | 4938439 | 4947844 | 9406 | 9 | 3.84E-18 | 4.91E-17 |
| chr1 | 156415843 | 156479435 | 63593 | 0.45 | 3.90E-18 | 4.99E-17 |
| chr5 | 52774382 | 52787204 | 12823 | -2.89 | 4.06E-18 | 5.18E-17 |
| chr7 | 81734362 | 81744221 | 9860 | -2.26 | 4.10E-18 | 5.24E-17 |
| chr18 | 32440005 | 32442019 | 2015 | 2.95 | 4.15E-18 | 5.30E-17 |
| chr4 | 38224418 | 38226203 | 1786 | 3.83 | 4.16E-18 | 5.31E-17 |
| chr21 | 31578749 | 31580375 | 1627 | -5.58 | 4.18E-18 | 5.33E-17 |
| chr12 | 15812096 | 15818045 | 5950 | 0.83 | 4.21E-18 | 5.36E-17 |
| chr12 | 51700501 | 51721733 | 21233 | 0.67 | 4.22E-18 | 5.38E-17 |
| chr3 | 57694472 | 57696093 | 1622 | 3.2 | 4.36E-18 | 5.55E-17 |
| chr9 | 29624065 | 29626456 | 2392 | 7.54 | 4.49E-18 | 5.71E-17 |
| chr3 | 121698097 | 121700058 | 1962 | 2.22 | 4.53E-18 | 5.77E-17 |
| chr16 | 10422250 | 10436996 | 14747 | 8.98 | 4.60E-18 | 5.85E-17 |
| chr6 | 160658054 | 160661206 | 3153 | 1.46 | 4.61E-18 | 5.87E-17 |
| chr12 | 47691434 | 47712148 | 20715 | 0.98 | 4.85E-18 | 6.16E-17 |
| chr1 | 196620206 | 196623633 | 3428 | 1.6 | 4.87E-18 | 6.19E-17 |
| chr2 | 37810589 | 37816804 | 6216 | 0.96 | 4.97E-18 | 6.31E-17 |
| chr19 | 15320485 | 15335170 | 14686 | -1.46 | 4.99E-18 | 6.33E-17 |
| chr4 | 89098640 | 89126650 | 28011 | -0.8 | 5.05E-18 | 6.41E-17 |
| chr12 | 75873361 | 75880453 | 7093 | 1.36 | 5.12E-18 | 6.49E-17 |
| chr3 | 150086572 | 150092326 | 5755 | 0.74 | 5.13E-18 | 6.51E-17 |
| chr3 | 15301622 | 15321794 | 20173 | -1.05 | 5.22E-18 | 6.62E-17 |
| chr11 | 28720323 | 28736692 | 16370 | 0.8 | 5.25E-18 | 6.64E-17 |
| chr7 | 5434780 | 5466685 | 31906 | 0.53 | 5.26E-18 | 6.66E-17 |
| chr17 | 48922335 | 49079638 | 157304 | 0.41 | 5.27E-18 | 6.67E-17 |
| chr17 | 43114047 | 43276582 | 162536 | 0.31 | 5.46E-18 | 6.91E-17 |
| chr7 | 63385381 | 63387598 | 2218 | 8.99 | 5.49E-18 | 6.94E-17 |
| chr19 | 48587780 | 48595819 | 8040 | 8.96 | 5.65E-18 | 7.14E-17 |
| chr18 | 52893670 | 52902480 | 8811 | 1.66 | 5.67E-18 | 7.16E-17 |
| chr12 | 70571530 | 70572663 | 1134 | 5.48 | 5.76E-18 | 7.28E-17 |
| chr8 | 10008315 | 10011083 | 2769 | 1.75 | 5.88E-18 | 7.42E-17 |
| chr19 | 20011155 | 20022829 | 11675 | 8.95 | 6.26E-18 | 7.89E-17 |
| chr7 | 100724240 | 100798893 | 74654 | -0.42 | 6.28E-18 | 7.93E-17 |
| chr11 | 17228029 | 17232447 | 4419 | 0.94 | 6.44E-18 | 8.12E-17 |
| chr10 | 6438838 | 6444900 | 6063 | -0.92 | 6.69E-18 | 8.43E-17 |
| chr20 | 5037513 | 5114202 | 76690 | 0.35 | 6.70E-18 | 8.45E-17 |
| chr16 | 66684829 | 66709499 | 24671 | -1.03 | 6.74E-18 | 8.49E-17 |
| chr7 | 35030983 | 35078396 | 47414 | 0.71 | 6.87E-18 | 8.65E-17 |
| chr19 | 57790231 | 57797207 | 6977 | 0.79 | 6.93E-18 | 8.72E-17 |
| chr5 | 54732457 | 54736834 | 4378 | 0.83 | 7.19E-18 | 9.04E-17 |
| chr8 | 17742896 | 17782981 | 40086 | 0.37 | 7.20E-18 | 9.05E-17 |
| chr13 | 76055065 | 76065112 | 10048 | 0.77 | 7.30E-18 | 9.17E-17 |
| chr14 | 29234533 | 29235780 | 1248 | 3.94 | 7.49E-18 | 9.41E-17 |
| chr12 | 59775904 | 59777323 | 1420 | 6.24 | 7.59E-18 | 9.53E-17 |
| chr6 | 146659921 | 146680716 | 20796 | 1.05 | 7.60E-18 | 9.54E-17 |
| chr15 | 88918247 | 88939994 | 21748 | -8.92 | 7.63E-18 | 9.57E-17 |
| chr1 | 117538833 | 117540711 | 1879 | 3 | 7.68E-18 | 9.63E-17 |
| chrX | 48993571 | 49048655 | 55085 | 0.69 | 8.14E-18 | 1.02E-16 |
| chr11 | 34653445 | 34708218 | 54774 | -0.4 | 8.28E-18 | 1.04E-16 |
| chr5 | 124081048 | 124085670 | 4623 | 0.95 | 8.41E-18 | 1.05E-16 |
| chr9 | 74576124 | 74584651 | 8528 | 1.71 | 8.67E-18 | 1.08E-16 |
| chr3 | 185445893 | 185447879 | 1987 | -4.23 | 8.79E-18 | 1.10E-16 |
| chr17 | 77531871 | 77546218 | 14348 | -8.9 | 9.06E-18 | 1.13E-16 |
| chr1 | 185895312 | 185904262 | 8951 | 2.61 | 9.28E-18 | 1.16E-16 |
| chr21 | 38218358 | 38234272 | 15915 | -0.86 | 9.31E-18 | 1.16E-16 |
| chr6 | 14758885 | 14779044 | 20160 | -1.22 | 1.06E-17 | 1.33E-16 |
| chr7 | 118739729 | 118745988 | 6260 | 8.89 | 1.07E-17 | 1.34E-16 |
| chr12 | 27332220 | 27345255 | 13036 | 2.04 | 1.07E-17 | 1.34E-16 |

| | | | | | | |
|---|---|---|---|---|---|---|
| chr3 | 84156058 | 84159206 | 3149 | 8.89 | 1.07E-17 | 1.34E-16 |
| chr4 | 23632604 | 23634134 | 1531 | 3.12 | 1.08E-17 | 1.35E-16 |
| chr15 | 71385328 | 71447714 | 62387 | -0.54 | 1.08E-17 | 1.35E-16 |
| chrX | 82019047 | 82022113 | 3067 | 3.32 | 1.09E-17 | 1.35E-16 |
| chr11 | 102399951 | 102402381 | 2431 | 1.87 | 1.09E-17 | 1.36E-16 |
| chr12 | 62911658 | 62928131 | 16474 | 1.29 | 1.12E-17 | 1.39E-16 |
| chr8 | 116786616 | 116792478 | 5863 | 8.88 | 1.13E-17 | 1.40E-16 |
| chrX | 106376441 | 106381087 | 4647 | 0.83 | 1.13E-17 | 1.41E-16 |
| chr2 | 8801357 | 8803543 | 2187 | -2.33 | 1.17E-17 | 1.45E-16 |
| chr5 | 49596464 | 49600857 | 4394 | 1.54 | 1.19E-17 | 1.47E-16 |
| chr3 | 120396543 | 120415375 | 18833 | 0.52 | 1.19E-17 | 1.48E-16 |
| chr5 | 57183588 | 57186531 | 2944 | -2.36 | 1.21E-17 | 1.50E-16 |
| chr6 | 168082595 | 168084847 | 2253 | 2.54 | 1.22E-17 | 1.51E-16 |
| chr17 | 31132371 | 31135880 | 3510 | 8.91 | 1.23E-17 | 1.53E-16 |
| chr9 | 88741054 | 88744273 | 3220 | 0.88 | 1.24E-17 | 1.54E-16 |
| chr8 | 141982947 | 142052116 | 69170 | -0.45 | 1.25E-17 | 1.55E-16 |
| chr3 | 175074430 | 175075828 | 1399 | 3.65 | 1.25E-17 | 1.55E-16 |
| chr5 | 125705454 | 125708045 | 2592 | 1.57 | 1.25E-17 | 1.55E-16 |
| chr16 | 85005738 | 85071403 | 65666 | 0.5 | 1.26E-17 | 1.56E-16 |
| chr2 | 99130854 | 99136059 | 5206 | -2.13 | 1.27E-17 | 1.57E-16 |
| chr16 | 66582368 | 66599728 | 17361 | 0.81 | 1.27E-17 | 1.58E-16 |
| chr5 | 30504372 | 30506850 | 2479 | 8.88 | 1.32E-17 | 1.63E-16 |
| chr12 | 115370476 | 115372776 | 2301 | -7.45 | 1.32E-17 | 1.63E-16 |
| chrX | 12991858 | 13049376 | 57519 | 0.48 | 1.32E-17 | 1.63E-16 |
| chr3 | 141564288 | 141565525 | 1238 | 2.88 | 1.35E-17 | 1.67E-16 |
| chr17 | 15280335 | 15281847 | 1513 | 5.26 | 1.35E-17 | 1.67E-16 |
| chr11 | 58328191 | 58357071 | 28881 | 0.64 | 1.41E-17 | 1.74E-16 |
| chr16 | 48532819 | 48535920 | 3102 | 1.61 | 1.43E-17 | 1.76E-16 |
| chr15 | 58813349 | 58822141 | 8793 | 1.27 | 1.45E-17 | 1.79E-16 |
| chr19 | 21106061 | 21118213 | 12153 | 8.86 | 1.48E-17 | 1.82E-16 |
| chr2 | 232045285 | 232064206 | 18922 | -1.73 | 1.48E-17 | 1.82E-16 |
| chrX | 109452489 | 109481807 | 29319 | 0.72 | 1.49E-17 | 1.83E-16 |
| chr4 | 90417766 | 90420176 | 2411 | 4.05 | 1.49E-17 | 1.83E-16 |
| chr8 | 81876721 | 81891567 | 14847 | 8.88 | 1.49E-17 | 1.83E-16 |
| chr15 | 63074222 | 63085621 | 11400 | -2.97 | 1.50E-17 | 1.84E-16 |
| chr10 | 29923312 | 29950776 | 27465 | 0.72 | 1.51E-17 | 1.86E-16 |
| chr2 | 29254535 | 29266492 | 11958 | 0.96 | 1.52E-17 | 1.86E-16 |
| chrX | 48366582 | 48399234 | 32653 | 0.86 | 1.57E-17 | 1.93E-16 |
| chr12 | 107420052 | 107422253 | 2202 | 1.63 | 1.60E-17 | 1.96E-16 |
| chr14 | 31880320 | 31890942 | 10623 | 1.05 | 1.61E-17 | 1.97E-16 |
| chr12 | 14092440 | 14108725 | 16286 | -0.61 | 1.63E-17 | 2.00E-16 |
| chr12 | 93480950 | 93485222 | 4273 | 2.27 | 1.63E-17 | 2.00E-16 |
| chr7 | 28059692 | 28117626 | 57935 | -0.45 | 1.64E-17 | 2.01E-16 |
| chr12 | 133704003 | 133708242 | 4240 | 1.42 | 1.65E-17 | 2.02E-16 |
| chr5 | 88080373 | 88082698 | 2326 | 1.87 | 1.70E-17 | 2.09E-16 |
| chr19 | 46792220 | 46832175 | 39956 | 8.85 | 1.75E-17 | 2.15E-16 |
| chr19 | 53519395 | 53532582 | 13188 | 8.83 | 1.79E-17 | 2.19E-16 |
| chr12 | 68798727 | 68800698 | 1972 | 2.42 | 1.85E-17 | 2.26E-16 |
| chr12 | 89337358 | 89349685 | 12328 | 2.11 | 1.88E-17 | 2.30E-16 |
| chr18 | 67899189 | 67914898 | 15710 | 0.67 | 1.89E-17 | 2.31E-16 |
| chr3 | 30508576 | 30519794 | 11219 | 1.26 | 1.95E-17 | 2.38E-16 |
| chr14 | 103515366 | 103542693 | 27328 | 0.61 | 1.98E-17 | 2.42E-16 |
| chr17 | 57443718 | 57460022 | 16305 | 1.75 | 2.01E-17 | 2.45E-16 |
| chr19 | 22234312 | 22260848 | 26537 | 8.87 | 2.12E-17 | 2.58E-16 |
| chr3 | 94683941 | 94686947 | 3007 | 8.82 | 2.13E-17 | 2.60E-16 |
| chr3 | 187525363 | 187528156 | 2794 | 1.59 | 2.13E-17 | 2.60E-16 |
| chr20 | 46046226 | 46098049 | 51824 | 0.43 | 2.14E-17 | 2.61E-16 |
| chr12 | 101411261 | 101416811 | 5551 | 8.82 | 2.15E-17 | 2.62E-16 |
| chr21 | 47919131 | 47932032 | 12902 | 0.85 | 2.18E-17 | 2.65E-16 |
| chr7 | 109182081 | 109192860 | 10780 | 8.83 | 2.19E-17 | 2.67E-16 |
| chr2 | 10259531 | 10296871 | 37341 | -0.71 | 2.28E-17 | 2.78E-16 |

| | | | | | | |
|---|---|---|---|---|---|---|
| chr9 | 5526096 | 5530761 | 4666 | 2.27 | 2.35E-17 | 2.86E-16 |
| chr6 | 53999585 | 54001570 | 1986 | 2.49 | 2.36E-17 | 2.86E-16 |
| chr12 | 87698030 | 87714960 | 16931 | 8.81 | 2.38E-17 | 2.89E-16 |
| chr5 | 34655522 | 34762912 | 107391 | -0.43 | 2.43E-17 | 2.95E-16 |
| chr9 | 101540411 | 101574265 | 33855 | 1.04 | 2.44E-17 | 2.96E-16 |
| chr4 | 187855911 | 187868764 | 12854 | 1 | 2.47E-17 | 2.99E-16 |
| chr8 | 82104845 | 82107299 | 2455 | 8.81 | 2.48E-17 | 3.00E-16 |
| chr19 | 53069504 | 53074499 | 4996 | 8.82 | 2.50E-17 | 3.03E-16 |
| chr4 | 72464773 | 72480933 | 16161 | 0.95 | 2.55E-17 | 3.09E-16 |
| chr11 | 13114667 | 13173290 | 58624 | 0.58 | 2.55E-17 | 3.09E-16 |
| chr4 | 74802130 | 74803507 | 1378 | 2.7 | 2.56E-17 | 3.09E-16 |
| chr12 | 103032029 | 103033392 | 1364 | 2.08 | 2.56E-17 | 3.10E-16 |
| chr14 | 104286073 | 104292986 | 6914 | 2.52 | 2.61E-17 | 3.16E-16 |
| chr7 | 137566390 | 137589275 | 22886 | 1.28 | 2.62E-17 | 3.17E-16 |
| chr14 | 88987642 | 89031469 | 43828 | 0.73 | 2.64E-17 | 3.20E-16 |
| chr2 | 46450999 | 46460317 | 9319 | 0.69 | 2.66E-17 | 3.21E-16 |
| chr4 | 69046266 | 69047802 | 1537 | 2.86 | 2.68E-17 | 3.24E-16 |
| chr10 | 105845189 | 105854747 | 9559 | 8.79 | 2.69E-17 | 3.25E-16 |
| chr14 | 53821283 | 53823421 | 2139 | 1.95 | 2.72E-17 | 3.29E-16 |
| chr2 | 210390448 | 210392211 | 1764 | 3.4 | 2.73E-17 | 3.30E-16 |
| chr1 | 232881068 | 232885220 | 4153 | 6.62 | 2.78E-17 | 3.35E-16 |
| chr5 | 166146552 | 166151955 | 5404 | 2.97 | 2.80E-17 | 3.37E-16 |
| chr20 | 5614820 | 5744571 | 129752 | 0.34 | 2.83E-17 | 3.41E-16 |
| chr12 | 79845291 | 79849752 | 4462 | 1.57 | 2.83E-17 | 3.41E-16 |
| chr12 | 3981456 | 3983104 | 1649 | 1.3 | 2.83E-17 | 3.41E-16 |
| chr12 | 44189311 | 44201098 | 11788 | 0.78 | 2.88E-17 | 3.47E-16 |
| chr11 | 47772391 | 47808858 | 36468 | 0.68 | 2.89E-17 | 3.48E-16 |
| chr1 | 199654052 | 199670262 | 16211 | 2.59 | 2.90E-17 | 3.48E-16 |
| chr14 | 59212603 | 59220637 | 8035 | 1.57 | 2.90E-17 | 3.49E-16 |
| chr18 | 8947769 | 8963420 | 15652 | 1.19 | 2.99E-17 | 3.59E-16 |
| chr6 | 22825680 | 22830040 | 4361 | 0.89 | 3.05E-17 | 3.67E-16 |
| chrX | 55736554 | 55745476 | 8923 | 1.48 | 3.13E-17 | 3.76E-16 |
| chr16 | 68669481 | 68675449 | 5969 | 1.87 | 3.19E-17 | 3.83E-16 |
| chr1 | 185613603 | 185620577 | 6975 | 1 | 3.20E-17 | 3.84E-16 |
| chrX | 125683527 | 125691965 | 8439 | 8.78 | 3.23E-17 | 3.87E-16 |
| chr2 | 191699357 | 191814237 | 114881 | -0.42 | 3.23E-17 | 3.87E-16 |
| chr5 | 113509648 | 113514231 | 4584 | 8.78 | 3.24E-17 | 3.88E-16 |
| chr20 | 57925005 | 57927686 | 2682 | -1.46 | 3.27E-17 | 3.92E-16 |
| chr12 | 124067043 | 124156399 | 89357 | 0.51 | 3.28E-17 | 3.93E-16 |
| chr3 | 150124390 | 150137961 | 13572 | 0.76 | 3.35E-17 | 4.00E-16 |
| chr1 | 181997892 | 181999738 | 1847 | 1.7 | 3.35E-17 | 4.00E-16 |
| chr5 | 130171796 | 130173678 | 1883 | 1.56 | 3.37E-17 | 4.03E-16 |
| chr11 | 103479294 | 103505409 | 26116 | -0.68 | 3.44E-17 | 4.12E-16 |
| chr11 | 75734250 | 75746592 | 12343 | 0.8 | 3.50E-17 | 4.18E-16 |
| chrX | 57312308 | 57313959 | 1652 | 4.13 | 3.52E-17 | 4.20E-16 |
| chr19 | 1239763 | 1276852 | 37090 | 0.5 | 3.53E-17 | 4.21E-16 |
| chr7 | 73223710 | 73266957 | 43248 | 0.61 | 3.54E-17 | 4.22E-16 |
| chr1 | 41227469 | 41236317 | 8849 | -2.29 | 3.55E-17 | 4.24E-16 |
| chr6 | 78001326 | 78009039 | 7714 | -1.3 | 3.65E-17 | 4.36E-16 |
| chr8 | 126225036 | 126245730 | 20695 | 0.54 | 3.66E-17 | 4.36E-16 |
| chr16 | 34832256 | 34834442 | 2187 | 8.76 | 3.68E-17 | 4.39E-16 |
| chr1 | 64157183 | 64199687 | 42505 | -0.53 | 3.69E-17 | 4.40E-16 |
| chr10 | 96749969 | 96754386 | 4418 | 8.76 | 3.72E-17 | 4.43E-16 |
| chr12 | 9216416 | 9218217 | 1802 | 1.97 | 3.86E-17 | 4.60E-16 |
| chr11 | 67417521 | 67425925 | 8405 | 1.67 | 3.92E-17 | 4.66E-16 |
| chr8 | 86868294 | 86875862 | 7569 | 1.5 | 3.94E-17 | 4.69E-16 |
| chr1 | 191259185 | 191261148 | 1964 | 8.75 | 3.99E-17 | 4.75E-16 |
| chr1 | 153503932 | 153546074 | 42143 | 0.36 | 4.03E-17 | 4.79E-16 |
| chr5 | 126590140 | 126593755 | 3616 | 8.75 | 4.08E-17 | 4.85E-16 |
| chr21 | 34732778 | 34756331 | 23554 | 0.67 | 4.20E-17 | 4.99E-16 |
| chr12 | 65995800 | 65998232 | 2433 | 1.79 | 4.31E-17 | 5.12E-16 |

| | | | | | | |
|---|---|---|---|---|---|---|
| chr12 | 89810176 | 89811896 | 1721 | 1.91 | 4.32E-17 | 5.12E-16 |
| chr2 | 129174415 | 129184380 | 9966 | 8.74 | 4.35E-17 | 5.15E-16 |
| chr3 | 9213555 | 9219582 | 6028 | 1.06 | 4.36E-17 | 5.17E-16 |
| chr12 | 43309088 | 43311021 | 1934 | 8.74 | 4.40E-17 | 5.22E-16 |
| chr18 | 2977853 | 3059093 | 81241 | 0.53 | 4.51E-17 | 5.34E-16 |
| chr2 | 19985242 | 19987229 | 1988 | -1.93 | 4.51E-17 | 5.34E-16 |
| chr13 | 101169058 | 101171541 | 2484 | 1.78 | 4.83E-17 | 5.72E-16 |
| chr12 | 70900399 | 70902584 | 2186 | 1.53 | 4.84E-17 | 5.73E-16 |
| chr15 | 41137618 | 41166875 | 29258 | 1.06 | 4.90E-17 | 5.79E-16 |
| chr7 | 37703885 | 37707376 | 3492 | -8.71 | 4.93E-17 | 5.83E-16 |
| chr1 | 85038091 | 85040959 | 2869 | 1.41 | 4.98E-17 | 5.88E-16 |
| chr10 | 58381745 | 58383319 | 1575 | 8.73 | 5.04E-17 | 5.95E-16 |
| chr13 | 86082939 | 86087644 | 4706 | -3.08 | 5.04E-17 | 5.95E-16 |
| chr19 | 44454857 | 44463457 | 8601 | 8.72 | 5.09E-17 | 6.01E-16 |
| chr16 | 81205879 | 81224771 | 18893 | -0.89 | 5.11E-17 | 6.03E-16 |
| chr3 | 112576126 | 112578314 | 2189 | 2.56 | 5.15E-17 | 6.07E-16 |
| chr9 | 121403013 | 121415113 | 12101 | 3.65 | 5.29E-17 | 6.24E-16 |
| chrX | 134544582 | 134551547 | 6966 | 8.73 | 5.32E-17 | 6.27E-16 |
| chr1 | 97736980 | 97745192 | 8213 | 1.67 | 5.36E-17 | 6.31E-16 |
| chr19 | 19931355 | 19942612 | 11258 | 8.72 | 5.44E-17 | 6.41E-16 |
| chr9 | 27117180 | 27119507 | 2328 | 4.76 | 5.52E-17 | 6.49E-16 |
| chr7 | 17483692 | 17489693 | 6002 | 1.38 | 5.53E-17 | 6.51E-16 |
| chr9 | 137216419 | 137310830 | 94412 | -0.39 | 5.70E-17 | 6.70E-16 |
| chr2 | 121141590 | 121164691 | 23102 | -0.66 | 5.78E-17 | 6.79E-16 |
| chr9 | 116861102 | 116863423 | 2322 | 2.54 | 5.79E-17 | 6.80E-16 |
| chr19 | 47948966 | 47950871 | 1906 | 2.34 | 5.80E-17 | 6.81E-16 |
| chr1 | 111458423 | 111471765 | 13343 | 1.16 | 5.81E-17 | 6.83E-16 |
| chr13 | 60181466 | 60182766 | 1301 | 4.72 | 5.92E-17 | 6.95E-16 |
| chr2 | 225382047 | 225408034 | 25988 | -1.1 | 6.09E-17 | 7.14E-16 |
| chr19 | 50311568 | 50324058 | 12491 | -0.53 | 6.11E-17 | 7.16E-16 |
| chr12 | 31947575 | 31999051 | 51477 | 2.79 | 6.15E-17 | 7.21E-16 |
| chr3 | 69280702 | 69292814 | 12113 | 2.47 | 6.29E-17 | 7.37E-16 |
| chr11 | 10041400 | 10044514 | 3115 | 2.24 | 6.30E-17 | 7.38E-16 |
| chr20 | 10536233 | 10546876 | 10644 | -6.54 | 6.35E-17 | 7.44E-16 |
| chr19 | 39831576 | 39984584 | 153009 | 0.28 | 6.39E-17 | 7.47E-16 |
| chr10 | 133468240 | 133474650 | 6411 | 8.7 | 6.55E-17 | 7.66E-16 |
| chr1 | 80324442 | 80326490 | 2049 | 5.49 | 6.75E-17 | 7.90E-16 |
| chr7 | 15435332 | 15436190 | 859 | 5.74 | 6.76E-17 | 7.90E-16 |
| chr9 | 135662794 | 135665838 | 3045 | 6.65 | 6.79E-17 | 7.94E-16 |
| chr8 | 49319396 | 49323623 | 4228 | 1.27 | 6.81E-17 | 7.96E-16 |
| chr16 | 46893222 | 46918731 | 25510 | 1.09 | 6.97E-17 | 8.14E-16 |
| chr1 | 206617831 | 206644733 | 26903 | -1.42 | 7.01E-17 | 8.18E-16 |
| chr1 | 217520819 | 217523975 | 3157 | 1.79 | 7.08E-17 | 8.26E-16 |
| chr7 | 17550917 | 17573605 | 22689 | 2.3 | 7.09E-17 | 8.27E-16 |
| chr8 | 18522393 | 18538407 | 16015 | 1.14 | 7.17E-17 | 8.35E-16 |
| chr17 | 61752722 | 61753989 | 1268 | 2.93 | 7.20E-17 | 8.39E-16 |
| chr14 | 93053470 | 93123049 | 69580 | -0.76 | 7.34E-17 | 8.55E-16 |
| chr5 | 42930315 | 42952268 | 21954 | 0.58 | 7.50E-17 | 8.73E-16 |
| chr1 | 237175648 | 237177690 | 2043 | 2.86 | 7.58E-17 | 8.82E-16 |
| chr2 | 88314885 | 88317084 | 2200 | -1.11 | 7.83E-17 | 9.11E-16 |
| chr8 | 29403131 | 29431536 | 28406 | 0.42 | 7.84E-17 | 9.12E-16 |
| chr12 | 6560267 | 6580815 | 20549 | 0.78 | 7.90E-17 | 9.18E-16 |
| chr8 | 119952481 | 119974269 | 21789 | 8.7 | 7.93E-17 | 9.21E-16 |
| chr3 | 78771354 | 78778380 | 7027 | 5.23 | 7.98E-17 | 9.27E-16 |
| chr6 | 107105708 | 107108718 | 3011 | 5.45 | 7.99E-17 | 9.27E-16 |
| chr1 | 234197211 | 234206189 | 8979 | 2.15 | 8.03E-17 | 9.31E-16 |
| chr9 | 128056898 | 128060407 | 3510 | 2.53 | 8.11E-17 | 9.41E-16 |
| chr5 | 178254997 | 178264657 | 9661 | 8.67 | 8.12E-17 | 9.41E-16 |
| chr2 | 104851785 | 104868155 | 16371 | 8.68 | 8.14E-17 | 9.43E-16 |
| chr19 | 53493949 | 53497872 | 3924 | 8.67 | 8.19E-17 | 9.49E-16 |
| chr7 | 85250453 | 85261189 | 10737 | 8.67 | 8.30E-17 | 9.61E-16 |

| | | | | | | |
|---|---|---|---|---|---|---|
| chr2 | 178026542 | 178037681 | 11140 | -0.76 | 8.32E-17 | 9.63E-16 |
| chr7 | 50800076 | 50807946 | 7871 | 1.14 | 8.43E-17 | 9.75E-16 |
| chr1 | 101178976 | 101181383 | 2408 | 2.04 | 8.44E-17 | 9.76E-16 |
| chr16 | 81001614 | 81042625 | 41012 | 0.53 | 8.46E-17 | 9.78E-16 |
| chr7 | 114756708 | 114786896 | 30189 | 8.67 | 8.50E-17 | 9.83E-16 |
| chr6 | 150066661 | 150093580 | 26920 | 1.12 | 8.59E-17 | 9.93E-16 |
| chr8 | 29082524 | 29093850 | 11327 | 0.84 | 8.79E-17 | 1.01E-15 |
| chrX | 115448573 | 115451300 | 2728 | 8.66 | 8.81E-17 | 1.02E-15 |
| chr6 | 67837516 | 67853239 | 15724 | 8.66 | 9.03E-17 | 1.04E-15 |
| chr12 | 115775076 | 115792791 | 17716 | -8.66 | 9.11E-17 | 1.05E-15 |
| chr12 | 59988907 | 59991947 | 3041 | 8.67 | 9.24E-17 | 1.07E-15 |
| chr17 | 53819881 | 53829531 | 9651 | 5.77 | 9.48E-17 | 1.09E-15 |
| chr12 | 99019920 | 99041000 | 21081 | 0.82 | 9.48E-17 | 1.09E-15 |
| chr17 | 80817358 | 80880325 | 62968 | -0.82 | 9.68E-17 | 1.12E-15 |
| chr3 | 149501025 | 149532855 | 31831 | 0.65 | 9.69E-17 | 1.12E-15 |
| chr4 | 125941107 | 125943471 | 2365 | 4.54 | 9.70E-17 | 1.12E-15 |
| chr17 | 31438920 | 31453631 | 14712 | -8.64 | 9.73E-17 | 1.12E-15 |
| chr5 | 62348428 | 62352222 | 3795 | 5.76 | 9.80E-17 | 1.13E-15 |
| chr1 | 201360967 | 201376815 | 15849 | -0.91 | 9.84E-17 | 1.13E-15 |
| chr3 | 148760124 | 148774166 | 14043 | 1.41 | 1.00E-16 | 1.15E-15 |
| chr10 | 52121428 | 52123363 | 1936 | 2.05 | 1.00E-16 | 1.15E-15 |
| chr11 | 106575964 | 106588725 | 12762 | 2.77 | 1.01E-16 | 1.16E-15 |
| chr14 | 95933936 | 95935936 | 2001 | 2.73 | 1.01E-16 | 1.16E-15 |
| chr2 | 10410512 | 10565411 | 154900 | -0.32 | 1.04E-16 | 1.19E-15 |
| chr2 | 76418846 | 76421802 | 2957 | 8.65 | 1.04E-16 | 1.19E-15 |
| chr8 | 39796882 | 39806781 | 9900 | 1.15 | 1.05E-16 | 1.20E-15 |
| chr5 | 168562724 | 168568622 | 5899 | 1.1 | 1.05E-16 | 1.21E-15 |
| chr7 | 69642858 | 69644842 | 1985 | 2.2 | 1.06E-16 | 1.21E-15 |
| chr1 | 197089337 | 197116770 | 27434 | 0.66 | 1.06E-16 | 1.21E-15 |
| chr4 | 13892078 | 13923918 | 31841 | -0.74 | 1.06E-16 | 1.21E-15 |
| chr19 | 52357515 | 52389042 | 31528 | 8.64 | 1.08E-16 | 1.23E-15 |
| chr18 | 57547873 | 57553435 | 5563 | 1.15 | 1.08E-16 | 1.24E-15 |
| chr4 | 166682930 | 166687467 | 4538 | 7.31 | 1.11E-16 | 1.27E-15 |
| chr20 | 45727843 | 45775965 | 48123 | 0.84 | 1.11E-16 | 1.27E-15 |
| chr7 | 89801469 | 89818854 | 17386 | 0.76 | 1.11E-16 | 1.27E-15 |
| chr10 | 31339795 | 31341932 | 2138 | 1.49 | 1.11E-16 | 1.27E-15 |
| chr2 | 76355696 | 76358276 | 2581 | 8.64 | 1.13E-16 | 1.29E-15 |
| chr6 | 42472239 | 42489414 | 17176 | 1.51 | 1.14E-16 | 1.30E-15 |
| chr8 | 74540225 | 74545010 | 4786 | 2.37 | 1.19E-16 | 1.36E-15 |
| chr21 | 27486793 | 27543903 | 57111 | 0.43 | 1.19E-16 | 1.36E-15 |
| chr14 | 78047041 | 78083043 | 36003 | 0.68 | 1.20E-16 | 1.37E-15 |
| chr6 | 1724010 | 1726029 | 2020 | 2.71 | 1.22E-16 | 1.39E-15 |
| chr16 | 1128069 | 1141215 | 13147 | 0.97 | 1.23E-16 | 1.40E-15 |
| chr15 | 43414534 | 43426991 | 12458 | 1.23 | 1.24E-16 | 1.41E-15 |
| chr1 | 95119014 | 95124145 | 5132 | 2.35 | 1.27E-16 | 1.44E-15 |
| chr8 | 143109573 | 143114028 | 4456 | 0.75 | 1.27E-16 | 1.44E-15 |
| chr1 | 98787784 | 98803374 | 15591 | 0.49 | 1.27E-16 | 1.45E-15 |
| chr3 | 18335831 | 18337447 | 1617 | 3.01 | 1.29E-16 | 1.47E-15 |
| chr5 | 118405174 | 118408256 | 3083 | 1.16 | 1.32E-16 | 1.50E-15 |
| chr10 | 4582951 | 4584415 | 1465 | 5.4 | 1.32E-16 | 1.50E-15 |
| chr6 | 139292152 | 139294990 | 2839 | -1.94 | 1.32E-16 | 1.50E-15 |
| chr5 | 39005780 | 39022104 | 16325 | -1.9 | 1.33E-16 | 1.50E-15 |
| chr1 | 178091099 | 178098139 | 7041 | 1.19 | 1.33E-16 | 1.51E-15 |
| chr4 | 119945674 | 119949983 | 4310 | 0.85 | 1.36E-16 | 1.54E-15 |
| chr15 | 48646258 | 48648386 | 2129 | 3.23 | 1.37E-16 | 1.55E-15 |
| chr2 | 231645958 | 231660448 | 14491 | 1.7 | 1.38E-16 | 1.56E-15 |
| chr18 | 73688824 | 73690294 | 1471 | 5.21 | 1.40E-16 | 1.58E-15 |
| chr18 | 54671168 | 54673195 | 2028 | 1.83 | 1.41E-16 | 1.60E-15 |
| chr14 | 56196814 | 56221418 | 24605 | 1.08 | 1.43E-16 | 1.62E-15 |
| chr14 | 25329490 | 25331470 | 1981 | 2.98 | 1.45E-16 | 1.64E-15 |
| chr7 | 6406741 | 6494113 | 87373 | 0.43 | 1.45E-16 | 1.64E-15 |

| | | | | | | |
|---|---|---|---|---|---|---|
| chr4 | 9997424 | 10021486 | 24063 | 1.12 | 1.45E-16 | 1.64E-15 |
| chr5 | 78906971 | 78917409 | 10439 | 0.99 | 1.46E-16 | 1.65E-15 |
| chrX | 132859486 | 132861386 | 1901 | 1.51 | 1.48E-16 | 1.67E-15 |
| chr15 | 26901171 | 26928310 | 27140 | -8.63 | 1.50E-16 | 1.69E-15 |
| chr19 | 28168446 | 28170533 | 2088 | 8.61 | 1.51E-16 | 1.71E-15 |
| chr1 | 89449777 | 89459583 | 9807 | 1.16 | 1.55E-16 | 1.75E-15 |
| chr4 | 126640732 | 126658926 | 18195 | 0.55 | 1.56E-16 | 1.76E-15 |
| chr11 | 75269586 | 75295002 | 25417 | 0.69 | 1.56E-16 | 1.76E-15 |
| chr17 | 19171278 | 19283062 | 111785 | -0.43 | 1.59E-16 | 1.79E-15 |
| chr1 | 223016507 | 223018320 | 1814 | 2.62 | 1.60E-16 | 1.81E-15 |
| chr8 | 41586737 | 41605108 | 18372 | 1.03 | 1.60E-16 | 1.81E-15 |
| chr8 | 83118782 | 83128520 | 9739 | 8.6 | 1.61E-16 | 1.81E-15 |
| chr1 | 167078034 | 167091678 | 13645 | 2.93 | 1.61E-16 | 1.81E-15 |
| chr9 | 125692688 | 125717390 | 24703 | 0.91 | 1.64E-16 | 1.84E-15 |
| chr12 | 118374378 | 118375824 | 1447 | 3.36 | 1.64E-16 | 1.85E-15 |
| chr11 | 101875803 | 101882724 | 6922 | 1.04 | 1.65E-16 | 1.86E-15 |
| chr17 | 65189123 | 65191684 | 2562 | 1.65 | 1.65E-16 | 1.86E-15 |
| chr1 | 160981774 | 160992791 | 11018 | 0.86 | 1.70E-16 | 1.91E-15 |
| chr4 | 154055675 | 154088980 | 33306 | 8.59 | 1.77E-16 | 1.99E-15 |
| chr6 | 139975985 | 139978198 | 2214 | 2.16 | 1.77E-16 | 1.99E-15 |
| chr1 | 69167582 | 69170955 | 3374 | 8.59 | 1.79E-16 | 2.01E-15 |
| chr8 | 133542345 | 133543826 | 1482 | 3.91 | 1.83E-16 | 2.05E-15 |
| chr19 | 21616953 | 21624120 | 7168 | 8.58 | 1.89E-16 | 2.12E-15 |
| chr2 | 237070201 | 237089005 | 18805 | -1.5 | 1.90E-16 | 2.13E-15 |
| chr5 | 158728582 | 158730271 | 1690 | 8.58 | 1.92E-16 | 2.15E-15 |
| chr5 | 165797865 | 165799405 | 1541 | 2.16 | 1.96E-16 | 2.19E-15 |
| chr20 | 33837946 | 33874339 | 36394 | 0.41 | 1.98E-16 | 2.22E-15 |
| chr9 | 26169337 | 26170929 | 1593 | 3 | 1.99E-16 | 2.22E-15 |
| chrX | 103674749 | 103691084 | 16336 | 8.59 | 2.00E-16 | 2.24E-15 |
| chr1 | 82135712 | 82148246 | 12535 | 3.02 | 2.03E-16 | 2.27E-15 |
| chr8 | 144135929 | 144138513 | 2585 | 8.57 | 2.04E-16 | 2.27E-15 |
| chr7 | 22706400 | 22714074 | 7675 | 1.17 | 2.05E-16 | 2.29E-15 |
| chr12 | 72163794 | 72174898 | 11105 | 1.59 | 2.10E-16 | 2.34E-15 |
| chr2 | 220070707 | 220096108 | 25402 | -0.68 | 2.10E-16 | 2.34E-15 |
| chr7 | 37010054 | 37012013 | 1960 | 2.61 | 2.10E-16 | 2.35E-15 |
| chr2 | 161417599 | 161441013 | 23415 | -1.05 | 2.11E-16 | 2.35E-15 |
| chr12 | 110300874 | 110303862 | 2989 | 1.7 | 2.11E-16 | 2.35E-15 |
| chr5 | 49950802 | 49964184 | 13383 | 8.58 | 2.12E-16 | 2.36E-15 |
| chr10 | 24620004 | 24621346 | 1343 | 3.24 | 2.13E-16 | 2.37E-15 |
| chr3 | 190250447 | 190252931 | 2485 | 1.66 | 2.16E-16 | 2.41E-15 |
| chr9 | 4220620 | 4300773 | 80154 | 0.31 | 2.17E-16 | 2.41E-15 |
| chr13 | 66875115 | 66890706 | 15592 | 0.87 | 2.17E-16 | 2.42E-15 |
| chr7 | 92205040 | 92220062 | 15023 | 1.04 | 2.18E-16 | 2.43E-15 |
| chr8 | 84038782 | 84047481 | 8700 | 8.58 | 2.20E-16 | 2.44E-15 |
| chr8 | 111847993 | 111865875 | 17883 | 8.62 | 2.21E-16 | 2.45E-15 |
| chr2 | 65056910 | 65067482 | 10573 | 0.7 | 2.21E-16 | 2.46E-15 |
| chr14 | 92295593 | 92304627 | 9035 | 8.57 | 2.23E-16 | 2.48E-15 |
| chr11 | 112066117 | 112067452 | 1336 | 4.22 | 2.27E-16 | 2.53E-15 |
| chrX | 52023597 | 52025910 | 2314 | 5.82 | 2.28E-16 | 2.53E-15 |
| chr1 | 215286794 | 215289245 | 2452 | 8.57 | 2.30E-16 | 2.55E-15 |
| chr16 | 84531991 | 84545782 | 13792 | 0.9 | 2.35E-16 | 2.60E-15 |
| chr2 | 218178340 | 218203689 | 25350 | 0.93 | 2.35E-16 | 2.61E-15 |
| chr4 | 62358022 | 62360249 | 2228 | 8.56 | 2.36E-16 | 2.62E-15 |
| chr7 | 20368361 | 20371715 | 3355 | 1.34 | 2.36E-16 | 2.62E-15 |
| chrX | 72782432 | 72801701 | 19270 | 1.9 | 2.38E-16 | 2.64E-15 |
| chr3 | 149229820 | 149380595 | 150776 | 0.22 | 2.45E-16 | 2.71E-15 |
| chr4 | 169506688 | 169511310 | 4623 | -1.89 | 2.49E-16 | 2.76E-15 |
| chrX | 12324314 | 12335698 | 11385 | 1.14 | 2.51E-16 | 2.78E-15 |
| chr10 | 7738206 | 7749836 | 11631 | 8.55 | 2.52E-16 | 2.78E-15 |
| chrX | 95707053 | 95709710 | 2658 | 1.76 | 2.55E-16 | 2.82E-15 |
| chr19 | 20149589 | 20164171 | 14583 | 8.55 | 2.55E-16 | 2.82E-15 |

| | | | | | | |
|---|---|---|---|---|---|---|
| chr8 | 95346795 | 95348955 | 2161 | 1.31 | 2.58E-16 | 2.85E-15 |
| chr4 | 32172767 | 32175554 | 2788 | 3.08 | 2.63E-16 | 2.90E-15 |
| chr5 | 158931362 | 158963935 | 32574 | 0.76 | 2.67E-16 | 2.95E-15 |
| chr2 | 62931460 | 62956307 | 24848 | -0.83 | 2.73E-16 | 3.02E-15 |
| chr12 | 122124110 | 122159019 | 34910 | 1.1 | 2.80E-16 | 3.09E-15 |
| chr8 | 134351717 | 134419481 | 67765 | 0.55 | 2.81E-16 | 3.10E-15 |
| chr13 | 104169579 | 104187866 | 18288 | 6.06 | 2.85E-16 | 3.14E-15 |
| chr7 | 22686697 | 22692169 | 5473 | 8.55 | 2.94E-16 | 3.24E-15 |
| chr14 | 85880742 | 85884309 | 3568 | 1.14 | 3.00E-16 | 3.30E-15 |
| chr6 | 138814414 | 138826546 | 12133 | 1.69 | 3.01E-16 | 3.31E-15 |
| chr15 | 26825861 | 26841628 | 15768 | -8.54 | 3.01E-16 | 3.31E-15 |
| chr2 | 122446590 | 122466349 | 19760 | 1.22 | 3.06E-16 | 3.37E-15 |
| chr12 | 88079884 | 88082396 | 2513 | 1.41 | 3.08E-16 | 3.38E-15 |
| chr5 | 143036920 | 143063886 | 26967 | 0.56 | 3.11E-16 | 3.42E-15 |
| chr11 | 130182948 | 130189361 | 6414 | 1 | 3.13E-16 | 3.44E-15 |
| chr12 | 95008205 | 95011132 | 2928 | 2.17 | 3.14E-16 | 3.45E-15 |
| chr2 | 234881705 | 234891543 | 9839 | -1.02 | 3.21E-16 | 3.52E-15 |
| chr22 | 31221640 | 31304162 | 82523 | 0.5 | 3.24E-16 | 3.56E-15 |
| chr12 | 115342329 | 115357832 | 15504 | -8.5 | 3.25E-16 | 3.56E-15 |
| chr1 | 62457930 | 62460217 | 2288 | 2.63 | 3.25E-16 | 3.56E-15 |
| chr6 | 141784720 | 141786691 | 1972 | 3.76 | 3.25E-16 | 3.57E-15 |
| chr6 | 17310869 | 17313038 | 2170 | -1.7 | 3.31E-16 | 3.63E-15 |
| chr2 | 45163368 | 45186704 | 23337 | -1.17 | 3.38E-16 | 3.71E-15 |
| chr1 | 152464161 | 152476474 | 12314 | 1.05 | 3.43E-16 | 3.76E-15 |
| chr5 | 56677634 | 56680286 | 2653 | -1.66 | 3.44E-16 | 3.77E-15 |
| chr7 | 124404346 | 124406198 | 1853 | 4.28 | 3.50E-16 | 3.83E-15 |
| chr3 | 141195680 | 141207496 | 11817 | 0.84 | 3.52E-16 | 3.85E-15 |
| chr16 | 87409082 | 87428018 | 18937 | 0.66 | 3.53E-16 | 3.86E-15 |
| chr3 | 168985110 | 168992140 | 7031 | 8.52 | 3.58E-16 | 3.92E-15 |
| chr3 | 14132326 | 14134024 | 1699 | 5.32 | 3.64E-16 | 3.98E-15 |
| chr3 | 19187857 | 19190051 | 2195 | 2.1 | 3.69E-16 | 4.03E-15 |
| chr13 | 91095201 | 91101158 | 5958 | 1.3 | 3.72E-16 | 4.07E-15 |
| chr21 | 16812132 | 16820110 | 7979 | -0.67 | 3.78E-16 | 4.13E-15 |
| chr2 | 76540321 | 76542948 | 2628 | 8.51 | 3.78E-16 | 4.13E-15 |
| chr1 | 168768570 | 168773777 | 5208 | 2.14 | 3.84E-16 | 4.19E-15 |
| chr8 | 119861508 | 119863624 | 2117 | 3.31 | 3.87E-16 | 4.22E-15 |
| chr1 | 31147856 | 31158835 | 10980 | 8.5 | 3.89E-16 | 4.24E-15 |
| chr3 | 24340228 | 24342658 | 2431 | 1.9 | 3.89E-16 | 4.24E-15 |
| chr1 | 12534413 | 12547246 | 12834 | 0.54 | 3.97E-16 | 4.33E-15 |
| chr2 | 242210689 | 242308992 | 98304 | -0.45 | 3.99E-16 | 4.35E-15 |
| chr7 | 127395117 | 127401672 | 6556 | 1.46 | 4.00E-16 | 4.35E-15 |
| chr11 | 22065165 | 22066480 | 1316 | 3.8 | 4.01E-16 | 4.36E-15 |
| chr3 | 71293337 | 71308615 | 15279 | 1.2 | 4.06E-16 | 4.42E-15 |
| chr3 | 182816273 | 182847227 | 30955 | 1.41 | 4.07E-16 | 4.42E-15 |
| chr15 | 44448690 | 44452536 | 3847 | 2.23 | 4.07E-16 | 4.42E-15 |
| chr19 | 35802285 | 35811418 | 9134 | 1.32 | 4.10E-16 | 4.46E-15 |
| chr10 | 96439199 | 96443792 | 4594 | 1.86 | 4.16E-16 | 4.52E-15 |
| chr6 | 22880126 | 22893476 | 13351 | 0.93 | 4.16E-16 | 4.52E-15 |
| chr5 | 141879332 | 141893373 | 14042 | 8.55 | 4.22E-16 | 4.58E-15 |
| chr4 | 36245040 | 36246772 | 1733 | 3.42 | 4.23E-16 | 4.59E-15 |
| chr13 | 89513750 | 89519384 | 5635 | 8.49 | 4.23E-16 | 4.59E-15 |
| chr7 | 1934674 | 1937278 | 2605 | -3.16 | 4.23E-16 | 4.59E-15 |
| chr13 | 33293282 | 33312530 | 19249 | 1.56 | 4.27E-16 | 4.63E-15 |
| chr8 | 17612333 | 17679445 | 67113 | 0.37 | 4.31E-16 | 4.67E-15 |
| chr1 | 177176944 | 177180006 | 3063 | 1.27 | 4.33E-16 | 4.69E-15 |
| chr13 | 104756935 | 104760142 | 3208 | 8.5 | 4.36E-16 | 4.72E-15 |
| chr8 | 116659861 | 116661242 | 1382 | 5.45 | 4.39E-16 | 4.75E-15 |
| chr2 | 224793782 | 224831009 | 37228 | -0.92 | 4.40E-16 | 4.76E-15 |
| chr5 | 33837929 | 33851220 | 13292 | -8.47 | 4.48E-16 | 4.85E-15 |
| chr2 | 236674071 | 236678285 | 4215 | -2.06 | 4.53E-16 | 4.90E-15 |
| chr12 | 68072030 | 68073066 | 1037 | 2.25 | 4.64E-16 | 5.02E-15 |

| | | | | | | |
|---|---|---|---|---|---|---|
| chr18 | 21264727 | 21321193 | 56467 | -0.53 | 4.77E-16 | 5.15E-15 |
| chr19 | 9894316 | 9907604 | 13289 | 1.49 | 4.78E-16 | 5.16E-15 |
| chr8 | 128046883 | 128050946 | 4064 | 8.5 | 4.83E-16 | 5.21E-15 |
| chr5 | 174770003 | 174775331 | 5329 | -8.46 | 5.02E-16 | 5.42E-15 |
| chr5 | 173958446 | 173963526 | 5081 | 0.96 | 5.04E-16 | 5.44E-15 |
| chr16 | 14394382 | 14425886 | 31505 | -0.53 | 5.13E-16 | 5.54E-15 |
| chr17 | 39367448 | 39370675 | 3228 | -1.03 | 5.19E-16 | 5.60E-15 |
| chr20 | 50672249 | 50678060 | 5812 | 2.91 | 5.21E-16 | 5.61E-15 |
| chr15 | 99860610 | 99869161 | 8552 | -0.7 | 5.24E-16 | 5.64E-15 |
| chr5 | 32851762 | 32853331 | 1570 | -3.55 | 5.27E-16 | 5.67E-15 |
| chr15 | 35988797 | 35991122 | 2326 | 1.84 | 5.28E-16 | 5.68E-15 |
| chr17 | 62488227 | 62504871 | 16645 | 0.5 | 5.29E-16 | 5.69E-15 |
| chr6 | 82406232 | 82410579 | 4348 | 3.73 | 5.33E-16 | 5.73E-15 |
| chr14 | 85985626 | 85996715 | 11090 | 1.14 | 5.36E-16 | 5.76E-15 |
| chr1 | 240944150 | 240967366 | 23217 | 3.27 | 5.40E-16 | 5.81E-15 |
| chr4 | 136427968 | 136431241 | 3274 | 8.47 | 5.41E-16 | 5.81E-15 |
| chr1 | 241091325 | 241100855 | 9531 | 8.46 | 5.50E-16 | 5.90E-15 |
| chr12 | 105582700 | 105630656 | 47957 | 0.88 | 5.51E-16 | 5.92E-15 |
| chr2 | 33238674 | 33249030 | 10357 | 1.63 | 5.70E-16 | 6.12E-15 |
| chr9 | 116216705 | 116357842 | 141138 | -0.38 | 5.82E-16 | 6.24E-15 |
| chr11 | 45110565 | 45114970 | 4406 | 1.88 | 5.87E-16 | 6.30E-15 |
| chr12 | 100304312 | 100311193 | 6882 | 0.88 | 5.96E-16 | 6.39E-15 |
| chr2 | 69709825 | 69711898 | 2074 | 2.17 | 6.02E-16 | 6.45E-15 |
| chr13 | 42353758 | 42356735 | 2978 | 1.23 | 6.12E-16 | 6.55E-15 |
| chr9 | 2014427 | 2018747 | 4321 | 1.32 | 6.25E-16 | 6.69E-15 |
| chr5 | 122632917 | 122634690 | 1774 | 8.46 | 6.26E-16 | 6.69E-15 |
| chr4 | 187562793 | 187623075 | 60283 | 0.55 | 6.34E-16 | 6.78E-15 |
| chr8 | 59664839 | 59672375 | 7537 | -1.91 | 6.39E-16 | 6.83E-15 |
| chr5 | 126365620 | 126377892 | 12273 | 8.45 | 6.48E-16 | 6.93E-15 |
| chr2 | 42647051 | 42649131 | 2081 | -1.84 | 6.49E-16 | 6.93E-15 |
| chr8 | 90768339 | 90773037 | 4699 | 0.92 | 6.55E-16 | 7.00E-15 |
| chrX | 4747379 | 4750695 | 3317 | 0.9 | 6.57E-16 | 7.02E-15 |
| chr6 | 25769542 | 25771403 | 1862 | 3.36 | 6.58E-16 | 7.03E-15 |
| chr6 | 34984575 | 34994491 | 9917 | 1.86 | 6.69E-16 | 7.14E-15 |
| chr2 | 46388123 | 46414743 | 26621 | -0.88 | 6.84E-16 | 7.30E-15 |
| chr14 | 38034729 | 38037476 | 2748 | 1.55 | 6.92E-16 | 7.38E-15 |
| chr11 | 61421507 | 61425521 | 4015 | 2.48 | 7.02E-16 | 7.49E-15 |
| chr12 | 718954 | 756641 | 37688 | 0.49 | 7.12E-16 | 7.59E-15 |
| chr9 | 100160982 | 100175487 | 14506 | 8.45 | 7.14E-16 | 7.61E-15 |
| chr12 | 95866503 | 95869928 | 3426 | 0.87 | 7.27E-16 | 7.74E-15 |
| chr20 | 25175533 | 25319727 | 144195 | 0.28 | 7.37E-16 | 7.84E-15 |
| chr1 | 197275224 | 197291352 | 16129 | 8.45 | 7.47E-16 | 7.94E-15 |
| chr11 | 45864378 | 45870008 | 5631 | 1.4 | 7.63E-16 | 8.12E-15 |
| chr6 | 3796286 | 3827551 | 31266 | 0.59 | 7.77E-16 | 8.26E-15 |
| chr19 | 15357456 | 15363445 | 5990 | 1.13 | 7.92E-16 | 8.41E-15 |
| chr18 | 68078364 | 68118011 | 39648 | 0.44 | 8.05E-16 | 8.56E-15 |
| chr1 | 181041806 | 181111704 | 69899 | -0.36 | 8.12E-16 | 8.62E-15 |
| chr17 | 38544535 | 38580298 | 35764 | 0.54 | 8.13E-16 | 8.63E-15 |
| chr22 | 23862377 | 23881950 | 19574 | 0.7 | 8.17E-16 | 8.67E-15 |
| chr1 | 178160678 | 178163453 | 2776 | 1.01 | 8.21E-16 | 8.71E-15 |
| chr13 | 109352335 | 109358754 | 6420 | 1.69 | 8.41E-16 | 8.92E-15 |
| chr5 | 33891056 | 33894863 | 3808 | -8.42 | 8.44E-16 | 8.95E-15 |
| chr9 | 137483487 | 137495528 | 12042 | -1.49 | 8.58E-16 | 9.09E-15 |
| chr16 | 16922118 | 16928015 | 5898 | -8.4 | 8.72E-16 | 9.24E-15 |
| chr16 | 16013321 | 16204659 | 191339 | -0.24 | 9.02E-16 | 9.55E-15 |
| chr19 | 37497992 | 37499647 | 1656 | 8.46 | 9.04E-16 | 9.57E-15 |
| chr8 | 131370342 | 131457926 | 87585 | -0.44 | 9.05E-16 | 9.57E-15 |
| chr22 | 40386606 | 40395390 | 8785 | 7.14 | 9.06E-16 | 9.59E-15 |
| chr4 | 120714846 | 120717022 | 2177 | 2.08 | 9.20E-16 | 9.73E-15 |
| chr5 | 178205113 | 178217023 | 11911 | 1.05 | 9.35E-16 | 9.89E-15 |
| chr2 | 153321712 | 153383157 | 61446 | -0.43 | 9.40E-16 | 9.93E-15 |

| | | | | | | |
|---|---|---|---|---|---|---|
| chrX | 9523183 | 9535222 | 12040 | 1.14 | 9.43E-16 | 9.96E-15 |
| chr3 | 35870251 | 35874864 | 4614 | -8.38 | 9.48E-16 | 1.00E-14 |
| chr7 | 137757770 | 137760689 | 2920 | 8.4 | 9.49E-16 | 1.00E-14 |
| chr17 | 41070609 | 41082472 | 11864 | -1.2 | 9.50E-16 | 1.00E-14 |
| chr1 | 121327831 | 121330157 | 2327 | 4.44 | 9.71E-16 | 1.02E-14 |
| chr4 | 48330997 | 48344687 | 13691 | 1.27 | 9.88E-16 | 1.04E-14 |
| chr5 | 168005153 | 168019663 | 14511 | 1.14 | 1.01E-15 | 1.07E-14 |
| chr12 | 9790551 | 9802346 | 11796 | 1.27 | 1.02E-15 | 1.07E-14 |
| chr5 | 88716245 | 88717965 | 1721 | 8.4 | 1.06E-15 | 1.12E-14 |
| chr12 | 56752962 | 56755197 | 2236 | 1.46 | 1.06E-15 | 1.12E-14 |
| chrX | 20983248 | 20984865 | 1618 | 4.37 | 1.07E-15 | 1.13E-14 |
| chr20 | 10263421 | 10310156 | 46736 | -1.08 | 1.08E-15 | 1.13E-14 |
| chr20 | 12957235 | 12959292 | 2058 | 2.54 | 1.08E-15 | 1.14E-14 |
| chr3 | 69953799 | 69964346 | 10548 | 1.82 | 1.08E-15 | 1.14E-14 |
| chr10 | 8405255 | 8460116 | 54862 | 0.52 | 1.09E-15 | 1.14E-14 |
| chr14 | 76759552 | 76761086 | 1535 | 5.64 | 1.09E-15 | 1.14E-14 |
| chr12 | 71551383 | 71558306 | 6924 | 8.39 | 1.09E-15 | 1.15E-14 |
| chr6 | 83390513 | 83392302 | 1790 | 2.45 | 1.10E-15 | 1.16E-14 |
| chr11 | 58909096 | 58913422 | 4327 | 1.24 | 1.12E-15 | 1.18E-14 |
| chr2 | 62079990 | 62083261 | 3272 | -1.28 | 1.14E-15 | 1.20E-14 |
| chr21 | 30443449 | 30451051 | 7603 | 0.87 | 1.14E-15 | 1.20E-14 |
| chrX | 65091529 | 65095827 | 4299 | 2.66 | 1.15E-15 | 1.21E-14 |
| chr15 | 51168311 | 51178541 | 10231 | -0.79 | 1.16E-15 | 1.21E-14 |
| chr14 | 105131634 | 105178709 | 47076 | 0.41 | 1.18E-15 | 1.24E-14 |
| chr1 | 8175444 | 8182818 | 7375 | 1.04 | 1.19E-15 | 1.25E-14 |
| chr13 | 21555508 | 21645981 | 90474 | 0.44 | 1.21E-15 | 1.27E-14 |
| chr21 | 19060883 | 19064075 | 3193 | 1.71 | 1.22E-15 | 1.27E-14 |
| chr6 | 11410996 | 11436301 | 25306 | 0.65 | 1.23E-15 | 1.28E-14 |
| chr4 | 36057021 | 36069363 | 12343 | 8.38 | 1.24E-15 | 1.30E-14 |
| chr10 | 86038998 | 86039940 | 943 | 3.38 | 1.28E-15 | 1.34E-14 |
| chr14 | 101780615 | 101783159 | 2545 | 8.37 | 1.31E-15 | 1.37E-14 |
| chr12 | 54217977 | 54223328 | 5352 | 0.94 | 1.31E-15 | 1.37E-14 |
| chr14 | 22764088 | 22765938 | 1851 | 2.46 | 1.32E-15 | 1.38E-14 |
| chr4 | 383203 | 384300 | 1098 | 6.44 | 1.34E-15 | 1.40E-14 |
| chr15 | 29810487 | 29831046 | 20560 | 8.48 | 1.34E-15 | 1.40E-14 |
| chr12 | 90005332 | 90044557 | 39226 | 0.67 | 1.36E-15 | 1.42E-14 |
| chr1 | 210591124 | 210593830 | 2707 | 1.37 | 1.36E-15 | 1.42E-14 |
| chr16 | 63647038 | 63652212 | 5175 | 1.44 | 1.37E-15 | 1.42E-14 |
| chr6 | 160673898 | 160677931 | 4034 | 1.72 | 1.38E-15 | 1.43E-14 |
| chr6 | 11794481 | 11816504 | 22024 | 1.02 | 1.38E-15 | 1.44E-14 |
| chr19 | 47808239 | 47819900 | 11662 | 1 | 1.40E-15 | 1.46E-14 |
| chr19 | 11796687 | 11805784 | 9098 | 8.37 | 1.40E-15 | 1.46E-14 |
| chr3 | 141013149 | 141056900 | 43752 | -0.73 | 1.41E-15 | 1.47E-14 |
| chr11 | 445789 | 471394 | 25606 | 0.84 | 1.42E-15 | 1.47E-14 |
| chr7 | 110681239 | 110684621 | 3383 | 1.85 | 1.43E-15 | 1.49E-14 |
| chr14 | 65436851 | 65455409 | 18559 | 1.09 | 1.44E-15 | 1.50E-14 |
| chr7 | 79133531 | 79138024 | 4494 | -1.17 | 1.46E-15 | 1.51E-14 |
| chr19 | 52302024 | 52311204 | 9181 | 8.35 | 1.49E-15 | 1.55E-14 |
| chr6 | 46950151 | 46958851 | 8701 | 1.85 | 1.50E-15 | 1.56E-14 |
| chrX | 7026379 | 7078454 | 52076 | 0.82 | 1.54E-15 | 1.59E-14 |
| chr10 | 73716084 | 73743059 | 26976 | 0.52 | 1.54E-15 | 1.60E-14 |
| chr9 | 81537693 | 81548670 | 10978 | 1.19 | 1.55E-15 | 1.61E-14 |
| chr7 | 146633551 | 146635400 | 1850 | 8.35 | 1.55E-15 | 1.61E-14 |
| chr12 | 56609880 | 56619058 | 9179 | 0.74 | 1.58E-15 | 1.64E-14 |
| chr10 | 22904694 | 22915083 | 10390 | 0.76 | 1.60E-15 | 1.66E-14 |
| chr12 | 126533880 | 126537707 | 3828 | 8.36 | 1.63E-15 | 1.69E-14 |
| chr7 | 108164285 | 108167779 | 3495 | 0.92 | 1.64E-15 | 1.69E-14 |
| chr13 | 111607189 | 111618517 | 11329 | 1.59 | 1.64E-15 | 1.70E-14 |
| chr14 | 94786071 | 94790189 | 4119 | 2.22 | 1.65E-15 | 1.70E-14 |
| chr2 | 240225520 | 240252433 | 26914 | -1.48 | 1.66E-15 | 1.71E-14 |
| chr19 | 20720541 | 20731656 | 11116 | 8.37 | 1.67E-15 | 1.72E-14 |

| | | | | | | |
|---|---|---|---|---|---|---|
| chr12 | 50281076 | 50290370 | 9295 | 1.86 | 1.69E-15 | 1.74E-14 |
| chr22 | 23620506 | 23625042 | 4537 | 1.8 | 1.69E-15 | 1.75E-14 |
| chr18 | 2569978 | 2607461 | 37484 | 0.79 | 1.72E-15 | 1.78E-14 |
| chr13 | 61503841 | 61508155 | 4315 | 8.34 | 1.74E-15 | 1.79E-14 |
| chr5 | 124743549 | 124749235 | 5687 | 1.29 | 1.74E-15 | 1.80E-14 |
| chr3 | 13880598 | 13895819 | 15222 | 8.35 | 1.75E-15 | 1.81E-14 |
| chr4 | 41134275 | 41163764 | 29490 | -1.31 | 1.78E-15 | 1.83E-14 |
| chr9 | 100879954 | 100884475 | 4522 | 1.95 | 1.78E-15 | 1.83E-14 |
| chr7 | 23502422 | 23516061 | 13640 | 0.47 | 1.80E-15 | 1.86E-14 |
| chr2 | 83282029 | 83309690 | 27662 | 8.33 | 1.82E-15 | 1.87E-14 |
| chr20 | 25364812 | 25420619 | 55808 | 0.63 | 1.82E-15 | 1.87E-14 |
| chr11 | 67763851 | 67841935 | 78085 | 0.35 | 1.82E-15 | 1.88E-14 |
| chr8 | 118409961 | 118412626 | 2666 | 1.85 | 1.85E-15 | 1.90E-14 |
| chr1 | 80665570 | 80678322 | 12753 | 8.35 | 1.85E-15 | 1.90E-14 |
| chr6 | 118886242 | 118908640 | 22399 | 1.81 | 1.86E-15 | 1.92E-14 |
| chr20 | 56702807 | 56708183 | 5377 | 1.83 | 1.86E-15 | 1.92E-14 |
| chr12 | 93516149 | 93530058 | 13910 | 8.33 | 1.93E-15 | 1.98E-14 |
| chr16 | 72399547 | 72415288 | 15742 | 8.34 | 1.93E-15 | 1.98E-14 |
| chr17 | 31202900 | 31204814 | 1915 | 2.83 | 1.96E-15 | 2.01E-14 |
| chr2 | 9370760 | 9429451 | 58692 | -0.56 | 1.98E-15 | 2.04E-14 |
| chr12 | 96791089 | 96796344 | 5256 | 0.77 | 2.04E-15 | 2.09E-14 |
| chr10 | 25605924 | 25608428 | 2505 | 1.43 | 2.04E-15 | 2.09E-14 |
| chr2 | 83220344 | 83221847 | 1504 | 5.87 | 2.04E-15 | 2.09E-14 |
| chr4 | 151870864 | 151886227 | 15364 | 1.89 | 2.05E-15 | 2.10E-14 |
| chr15 | 59976369 | 59994741 | 18373 | 0.64 | 2.06E-15 | 2.11E-14 |
| chr2 | 69345235 | 69349118 | 3884 | -3.05 | 2.06E-15 | 2.12E-14 |
| chr12 | 49657005 | 49663368 | 6364 | 0.87 | 2.07E-15 | 2.12E-14 |
| chr5 | 37662209 | 37669982 | 7774 | 0.9 | 2.08E-15 | 2.13E-14 |
| chr12 | 118558123 | 118575151 | 17029 | 0.84 | 2.09E-15 | 2.14E-14 |
| chr1 | 176172661 | 176177998 | 5338 | 0.83 | 2.10E-15 | 2.15E-14 |
| chr12 | 110903916 | 110951135 | 47220 | 0.72 | 2.10E-15 | 2.15E-14 |
| chr19 | 23257539 | 23259345 | 1807 | 8.31 | 2.17E-15 | 2.22E-14 |
| chr8 | 120790798 | 120793659 | 2862 | 1.56 | 2.19E-15 | 2.24E-14 |
| chr2 | 13822381 | 13823903 | 1523 | 5.94 | 2.21E-15 | 2.26E-14 |
| chr2 | 89032505 | 89039630 | 7126 | -1.41 | 2.24E-15 | 2.29E-14 |
| chr19 | 53684351 | 53696588 | 12238 | 8.31 | 2.28E-15 | 2.32E-14 |
| chr20 | 10486650 | 10523402 | 36753 | -0.51 | 2.29E-15 | 2.33E-14 |
| chr6 | 135043794 | 135046998 | 3205 | 7.04 | 2.29E-15 | 2.34E-14 |
| chr11 | 10063386 | 10065876 | 2491 | 6.32 | 2.34E-15 | 2.39E-14 |
| chr5 | 32805271 | 32807193 | 1923 | -1.73 | 2.36E-15 | 2.41E-14 |
| chr15 | 75862147 | 75872402 | 10256 | 0.64 | 2.40E-15 | 2.45E-14 |
| chr7 | 25131395 | 25134144 | 2750 | 1.15 | 2.43E-15 | 2.48E-14 |
| chr17 | 54862523 | 54865197 | 2675 | 2.3 | 2.44E-15 | 2.49E-14 |
| chr3 | 107316525 | 107319805 | 3281 | 1.02 | 2.49E-15 | 2.54E-14 |
| chr1 | 21941530 | 21983301 | 41772 | 0.51 | 2.53E-15 | 2.57E-14 |
| chr20 | 2310153 | 2312055 | 1903 | 6.39 | 2.57E-15 | 2.62E-14 |
| chr10 | 128063703 | 128067117 | 3415 | 8.34 | 2.61E-15 | 2.65E-14 |
| chr22 | 23052520 | 23054579 | 2060 | 2.48 | 2.64E-15 | 2.69E-14 |
| chr19 | 11241104 | 11279893 | 38790 | 0.54 | 2.65E-15 | 2.69E-14 |
| chr7 | 46460762 | 46473842 | 13081 | 1.88 | 2.65E-15 | 2.69E-14 |
| chr1 | 90017681 | 90019389 | 1709 | 1.48 | 2.69E-15 | 2.73E-14 |
| chr11 | 13894396 | 13898175 | 3780 | 8.29 | 2.73E-15 | 2.77E-14 |
| chr1 | 1092917 | 1101163 | 8247 | 1.36 | 2.76E-15 | 2.81E-14 |
| chr1 | 243447673 | 243458678 | 11006 | 2.02 | 2.77E-15 | 2.81E-14 |
| chr19 | 19716357 | 19729508 | 13152 | 1.26 | 2.78E-15 | 2.82E-14 |
| chr5 | 132039802 | 132041547 | 1746 | 2.01 | 2.79E-15 | 2.83E-14 |
| chr4 | 141555882 | 141567882 | 12001 | 1.5 | 2.81E-15 | 2.85E-14 |
| chr4 | 64502424 | 64504358 | 1935 | 8.3 | 2.83E-15 | 2.87E-14 |
| chr14 | 55486897 | 55545130 | 58234 | 0.55 | 2.87E-15 | 2.91E-14 |
| chr10 | 105455097 | 105468240 | 13144 | -2.25 | 2.88E-15 | 2.92E-14 |
| chr1 | 172938480 | 172950423 | 11944 | 0.84 | 2.89E-15 | 2.92E-14 |

| | | | | | | |
|---|---|---|---|---|---|---|
| chr19 | 54124206 | 54128433 | 4228 | 8.28 | 2.89E-15 | 2.92E-14 |
| chr6 | 80123135 | 80132287 | 9153 | 2.77 | 2.90E-15 | 2.94E-14 |
| chr2 | 166519095 | 166533189 | 14095 | 8.31 | 2.91E-15 | 2.94E-14 |
| chr5 | 43997492 | 44003691 | 6200 | 8.29 | 2.91E-15 | 2.94E-14 |
| chr1 | 169846311 | 169864200 | 17890 | 0.61 | 2.94E-15 | 2.97E-14 |
| chr9 | 35236663 | 35238747 | 2085 | 3.14 | 2.97E-15 | 3.00E-14 |
| chr2 | 153933952 | 153936887 | 2936 | 2.61 | 2.98E-15 | 3.01E-14 |
| chr5 | 66289927 | 66312114 | 22188 | 1.09 | 3.00E-15 | 3.03E-14 |
| chr2 | 102678541 | 102740179 | 61639 | -0.66 | 3.03E-15 | 3.06E-14 |
| chr13 | 85232561 | 85237124 | 4564 | 8.27 | 3.06E-15 | 3.09E-14 |
| chr6 | 14270482 | 14273260 | 2779 | 1.36 | 3.08E-15 | 3.11E-14 |
| chr12 | 79687620 | 79696569 | 8950 | 0.58 | 3.14E-15 | 3.17E-14 |
| chr4 | 120125464 | 120156248 | 30785 | 0.65 | 3.16E-15 | 3.18E-14 |
| chr12 | 133329289 | 133339961 | 10673 | 0.98 | 3.17E-15 | 3.19E-14 |
| chr8 | 119889607 | 119895172 | 5566 | 1.53 | 3.25E-15 | 3.28E-14 |
| chrX | 13211584 | 13213894 | 2311 | 1.97 | 3.26E-15 | 3.28E-14 |
| chr3 | 152847784 | 152884547 | 36764 | 0.53 | 3.26E-15 | 3.28E-14 |
| chr19 | 52987023 | 52995593 | 8571 | 8.32 | 3.26E-15 | 3.28E-14 |
| chr3 | 143198157 | 143203562 | 5406 | 2.25 | 3.27E-15 | 3.29E-14 |
| chr6 | 21219231 | 21222035 | 2805 | 1.72 | 3.27E-15 | 3.29E-14 |
| chr12 | 98986244 | 99007059 | 20816 | 0.69 | 3.28E-15 | 3.30E-14 |
| chr2 | 47435648 | 47437073 | 1426 | -3.87 | 3.35E-15 | 3.37E-14 |
| chr5 | 42846791 | 42848171 | 1381 | 3.91 | 3.36E-15 | 3.37E-14 |
| chr20 | 23603014 | 23618702 | 15689 | 0.85 | 3.38E-15 | 3.40E-14 |
| chr22 | 42931894 | 42950171 | 18278 | 1.87 | 3.39E-15 | 3.41E-14 |
| chr7 | 142817357 | 142824308 | 6952 | -1.76 | 3.41E-15 | 3.42E-14 |
| chr11 | 15986580 | 15992751 | 6172 | 0.65 | 3.42E-15 | 3.43E-14 |
| chr19 | 58445509 | 58459608 | 14100 | 8.27 | 3.44E-15 | 3.45E-14 |
| chr8 | 53705716 | 53714590 | 8875 | 8.26 | 3.52E-15 | 3.53E-14 |
| chr4 | 138675201 | 138677218 | 2018 | 3.91 | 3.56E-15 | 3.56E-14 |
| chr12 | 1053035 | 1063066 | 10032 | 0.89 | 3.57E-15 | 3.58E-14 |
| chr6 | 37018925 | 37039860 | 20936 | -0.62 | 3.59E-15 | 3.59E-14 |
| chr16 | 63397098 | 63407467 | 10370 | 8.33 | 3.60E-15 | 3.60E-14 |
| chr8 | 129661321 | 129679176 | 17856 | -1.37 | 3.65E-15 | 3.65E-14 |
| chr19 | 37742013 | 37749516 | 7504 | 8.35 | 3.67E-15 | 3.67E-14 |
| chr18 | 1269442 | 1271178 | 1737 | 3.8 | 3.68E-15 | 3.68E-14 |
| chr14 | 32460429 | 32497226 | 36798 | -0.5 | 3.73E-15 | 3.73E-14 |
| chr2 | 200914909 | 200916682 | 1774 | 4.02 | 3.81E-15 | 3.80E-14 |
| chr11 | 66078927 | 66116152 | 37226 | -0.79 | 3.81E-15 | 3.81E-14 |
| chr14 | 34504627 | 34555245 | 50619 | 0.56 | 3.87E-15 | 3.86E-14 |
| chr12 | 104322202 | 104370061 | 47860 | 0.44 | 3.87E-15 | 3.87E-14 |
| chr12 | 28763316 | 28765384 | 2069 | 2.4 | 3.91E-15 | 3.90E-14 |
| chrX | 23858657 | 23864940 | 6284 | 1.08 | 3.93E-15 | 3.92E-14 |
| chr1 | 199727821 | 199729357 | 1537 | 2.15 | 3.94E-15 | 3.93E-14 |
| chr7 | 23798553 | 23800843 | 2291 | 8.24 | 3.98E-15 | 3.97E-14 |
| chr3 | 21431453 | 21434962 | 3510 | 8.24 | 3.99E-15 | 3.98E-14 |
| chr6 | 44767009 | 44768966 | 1958 | 3.12 | 4.01E-15 | 4.00E-14 |
| chr10 | 97501583 | 97503185 | 1603 | 6.42 | 4.02E-15 | 4.01E-14 |
| chr4 | 167093411 | 167096769 | 3359 | 5.31 | 4.03E-15 | 4.01E-14 |
| chr4 | 70032371 | 70034110 | 1740 | 4.32 | 4.05E-15 | 4.04E-14 |
| chr4 | 95372262 | 95376849 | 4588 | 0.93 | 4.18E-15 | 4.16E-14 |
| chr4 | 89531950 | 89545167 | 13218 | 2.06 | 4.23E-15 | 4.21E-14 |
| chr8 | 57122472 | 57125143 | 2672 | 2.17 | 4.27E-15 | 4.25E-14 |
| chr15 | 85704557 | 85712754 | 8198 | 2.14 | 4.27E-15 | 4.25E-14 |
| chr8 | 11280636 | 11287409 | 6774 | -4.91 | 4.29E-15 | 4.26E-14 |
| chr2 | 187635371 | 187637195 | 1825 | 3.29 | 4.31E-15 | 4.28E-14 |
| chr22 | 36847091 | 36878909 | 31819 | 0.49 | 4.42E-15 | 4.39E-14 |
| chr12 | 100521434 | 100558192 | 36759 | 0.73 | 4.43E-15 | 4.40E-14 |
| chr17 | 14638942 | 14645253 | 6312 | 0.66 | 4.45E-15 | 4.42E-14 |
| chr19 | 41664494 | 41707865 | 43372 | -0.53 | 4.54E-15 | 4.50E-14 |
| chr5 | 71499013 | 71553138 | 54126 | -0.64 | 4.59E-15 | 4.55E-14 |

| | | | | | | |
|---|---|---|---|---|---|---|
| chr15 | 33065726 | 33067278 | 1553 | -2.44 | 4.81E-15 | 4.77E-14 |
| chr20 | 15042162 | 15050228 | 8067 | 2.13 | 4.83E-15 | 4.79E-14 |
| chr5 | 133983342 | 133989811 | 6470 | 0.94 | 4.84E-15 | 4.79E-14 |
| chr13 | 90052238 | 90056239 | 4002 | 1.05 | 4.86E-15 | 4.81E-14 |
| chr19 | 11998526 | 12000029 | 1504 | 8.23 | 4.95E-15 | 4.90E-14 |
| chr18 | 72959007 | 72961135 | 2129 | 1.39 | 4.96E-15 | 4.91E-14 |
| chr7 | 130523385 | 130800703 | 277319 | 0.21 | 5.02E-15 | 4.96E-14 |
| chr9 | 32693328 | 32699761 | 6434 | 8.22 | 5.14E-15 | 5.08E-14 |
| chr4 | 22970530 | 22972336 | 1807 | 2.14 | 5.19E-15 | 5.13E-14 |
| chr2 | 72303863 | 72306472 | 2610 | 8.22 | 5.29E-15 | 5.23E-14 |
| chr8 | 33056177 | 33058984 | 2808 | -2.24 | 5.32E-15 | 5.26E-14 |
| chr12 | 50502520 | 50519753 | 17234 | 0.79 | 5.34E-15 | 5.28E-14 |
| chr1 | 12439778 | 12442973 | 3196 | 1.05 | 5.35E-15 | 5.28E-14 |
| chr10 | 126919173 | 126922564 | 3392 | -0.97 | 5.41E-15 | 5.34E-14 |
| chr2 | 134178921 | 134193518 | 14598 | -8.2 | 5.43E-15 | 5.36E-14 |
| chr7 | 129545434 | 129548532 | 3099 | 1.94 | 5.51E-15 | 5.43E-14 |
| chr2 | 84524367 | 84528856 | 4490 | 8.24 | 5.55E-15 | 5.48E-14 |
| chr7 | 21656022 | 21659014 | 2993 | 0.98 | 5.60E-15 | 5.52E-14 |
| chr5 | 130151858 | 130153513 | 1656 | 2.22 | 5.63E-15 | 5.55E-14 |
| chr5 | 143292260 | 143304369 | 12110 | -1.29 | 5.74E-15 | 5.65E-14 |
| chr9 | 98135366 | 98148625 | 13260 | -1.33 | 5.78E-15 | 5.69E-14 |
| chr14 | 33366036 | 33367017 | 982 | 3.03 | 5.78E-15 | 5.69E-14 |
| chr14 | 25451214 | 25452487 | 1274 | 4.03 | 5.89E-15 | 5.80E-14 |
| chr8 | 112098202 | 112100798 | 2597 | 8.2 | 5.95E-15 | 5.85E-14 |
| chr8 | 26335746 | 26341417 | 5672 | 1.17 | 6.04E-15 | 5.94E-14 |
| chr22 | 18916715 | 18936174 | 19460 | 1.52 | 6.10E-15 | 6.00E-14 |
| chr16 | 23914338 | 23917534 | 3197 | -1.72 | 6.13E-15 | 6.03E-14 |
| chr13 | 90797228 | 90798969 | 1742 | 1.6 | 6.35E-15 | 6.24E-14 |
| chr3 | 150062294 | 150067744 | 5451 | 1.61 | 6.37E-15 | 6.26E-14 |
| chr9 | 6105705 | 6107497 | 1793 | 1.4 | 6.55E-15 | 6.43E-14 |
| chr7 | 27070995 | 27072415 | 1421 | 3.36 | 6.60E-15 | 6.48E-14 |
| chr6 | 43654771 | 43674325 | 19555 | 0.79 | 6.68E-15 | 6.55E-14 |
| chr1 | 55886256 | 55914806 | 28551 | -0.71 | 6.70E-15 | 6.58E-14 |
| chr12 | 10763984 | 10767345 | 3362 | 0.73 | 6.81E-15 | 6.68E-14 |
| chr5 | 79516020 | 79528030 | 12011 | 1.08 | 6.92E-15 | 6.78E-14 |
| chr4 | 124569377 | 124572983 | 3607 | 0.8 | 6.97E-15 | 6.83E-14 |
| chr19 | 9517191 | 9525467 | 8277 | 8.21 | 6.97E-15 | 6.83E-14 |
| chr8 | 29292313 | 29325199 | 32887 | 0.45 | 7.09E-15 | 6.94E-14 |
| chr10 | 109500324 | 109502643 | 2320 | 8.19 | 7.11E-15 | 6.95E-14 |
| chr1 | 16659855 | 16663263 | 3409 | 1.31 | 7.21E-15 | 7.06E-14 |
| chr1 | 67028803 | 67030559 | 1757 | -4.78 | 7.29E-15 | 7.13E-14 |
| chr2 | 20695760 | 20701164 | 5405 | 2.14 | 7.32E-15 | 7.16E-14 |
| chr7 | 99147942 | 99162251 | 14310 | 0.74 | 7.32E-15 | 7.16E-14 |
| chr8 | 87187105 | 87190137 | 3033 | 1.22 | 7.40E-15 | 7.23E-14 |
| chr7 | 115736797 | 115738369 | 1573 | 1.71 | 7.45E-15 | 7.28E-14 |
| chr16 | 154488 | 165223 | 10736 | -1.17 | 7.45E-15 | 7.28E-14 |
| chr11 | 40349183 | 40352391 | 3209 | 0.91 | 7.47E-15 | 7.30E-14 |
| chr3 | 46731801 | 46735647 | 3847 | 1.87 | 7.48E-15 | 7.30E-14 |
| chr10 | 115311824 | 115314327 | 2504 | 2.73 | 7.50E-15 | 7.32E-14 |
| chr11 | 85773752 | 85782147 | 8396 | 0.7 | 7.54E-15 | 7.36E-14 |
| chr7 | 114141778 | 114143508 | 1731 | 8.17 | 7.58E-15 | 7.39E-14 |
| chr21 | 16071611 | 16073321 | 1711 | 4.9 | 7.59E-15 | 7.39E-14 |
| chr6 | 127334817 | 127348918 | 14102 | 3.28 | 7.69E-15 | 7.49E-14 |
| chr3 | 167530586 | 167535138 | 4553 | -8.16 | 7.69E-15 | 7.49E-14 |
| chr15 | 57124244 | 57164757 | 40514 | 8.42 | 7.81E-15 | 7.60E-14 |
| chr18 | 61403213 | 61405618 | 2406 | 8.22 | 7.90E-15 | 7.69E-14 |
| chr11 | 130868857 | 130871295 | 2439 | 1.58 | 7.93E-15 | 7.71E-14 |
| chr19 | 44951518 | 44953119 | 1602 | 1.75 | 7.97E-15 | 7.75E-14 |
| chrX | 46425737 | 46450355 | 24619 | -0.96 | 8.06E-15 | 7.83E-14 |
| chr17 | 61042694 | 61054758 | 12065 | 1.27 | 8.06E-15 | 7.83E-14 |
| chr12 | 79801977 | 79827948 | 25972 | 0.68 | 8.41E-15 | 8.17E-14 |

| | | | | | | |
|---|---|---|---|---|---|---|
| chr14 | 71102405 | 71109510 | 7106 | 8.16 | 8.47E-15 | 8.23E-14 |
| chr2 | 64679671 | 64682449 | 2779 | 1.25 | 8.48E-15 | 8.24E-14 |
| chr8 | 29258628 | 29266647 | 8020 | 0.98 | 8.52E-15 | 8.27E-14 |
| chr10 | 13799694 | 13831081 | 31388 | 1.02 | 8.55E-15 | 8.30E-14 |
| chr17 | 46549575 | 46725755 | 176181 | 0.35 | 8.63E-15 | 8.37E-14 |
| chr1 | 95427306 | 95429022 | 1717 | 2.96 | 8.67E-15 | 8.41E-14 |
| chr1 | 69088875 | 69099472 | 10598 | 8.16 | 8.77E-15 | 8.50E-14 |
| chr19 | 40308673 | 40342233 | 33561 | -0.53 | 8.88E-15 | 8.60E-14 |
| chr8 | 55358715 | 55371484 | 12770 | 8.25 | 9.24E-15 | 8.95E-14 |
| chr9 | 4529937 | 4531357 | 1421 | 2.7 | 9.32E-15 | 9.02E-14 |
| chr19 | 53868003 | 53869568 | 1566 | 5.02 | 9.35E-15 | 9.05E-14 |
| chr3 | 98231315 | 98242318 | 11004 | 1.05 | 9.53E-15 | 9.22E-14 |
| chr4 | 16170474 | 16176892 | 6419 | 3.17 | 9.66E-15 | 9.35E-14 |
| chr3 | 179708145 | 179710329 | 2185 | -6.22 | 9.75E-15 | 9.43E-14 |
| chr5 | 124460791 | 124471521 | 10731 | 2.3 | 9.76E-15 | 9.44E-14 |
| chr16 | 56639421 | 56645736 | 6316 | 0.63 | 9.78E-15 | 9.45E-14 |
| chr15 | 35597610 | 35601778 | 4169 | 8.14 | 9.94E-15 | 9.61E-14 |
| chr6 | 160619054 | 160620734 | 1681 | 2.11 | 1.00E-14 | 9.66E-14 |
| chr18 | 9912847 | 9952737 | 39891 | 0.61 | 1.01E-14 | 9.80E-14 |
| chr8 | 53830607 | 53835526 | 4920 | 8.14 | 1.02E-14 | 9.83E-14 |
| chr9 | 113729052 | 113740024 | 10973 | -1.91 | 1.02E-14 | 9.86E-14 |
| chr6 | 151814584 | 151824357 | 9774 | 2.94 | 1.03E-14 | 9.91E-14 |
| chr12 | 75373931 | 75379300 | 5370 | 8.16 | 1.04E-14 | 1.00E-13 |
| chr3 | 47400741 | 47402366 | 1626 | 4.51 | 1.06E-14 | 1.02E-13 |
| chr16 | 71546897 | 71560890 | 13994 | 8.18 | 1.07E-14 | 1.03E-13 |
| chr2 | 20945463 | 20947536 | 2074 | 1.5 | 1.09E-14 | 1.05E-13 |
| chr5 | 20972084 | 20973517 | 1434 | 3.11 | 1.10E-14 | 1.06E-13 |
| chr4 | 102010086 | 102037835 | 27750 | 1.31 | 1.10E-14 | 1.06E-13 |
| chr3 | 138580117 | 138587637 | 7521 | 1.08 | 1.11E-14 | 1.07E-13 |
| chr14 | 37692297 | 37694061 | 1765 | 2.21 | 1.13E-14 | 1.09E-13 |
| chr16 | 24679736 | 24683581 | 3846 | -1.29 | 1.18E-14 | 1.13E-13 |
| chr4 | 127791985 | 127793059 | 1075 | 4.82 | 1.19E-14 | 1.14E-13 |
| chr18 | 54556574 | 54571258 | 14685 | 1.24 | 1.19E-14 | 1.15E-13 |
| chr14 | 24834878 | 24837176 | 2299 | 1.8 | 1.19E-14 | 1.15E-13 |
| chr22 | 45637040 | 45644098 | 7059 | 0.83 | 1.21E-14 | 1.16E-13 |
| chr15 | 73924118 | 73940708 | 16591 | 0.96 | 1.21E-14 | 1.16E-13 |
| chr10 | 27465559 | 27542413 | 76855 | 0.51 | 1.23E-14 | 1.18E-13 |
| chr2 | 232161969 | 232174689 | 12721 | 1.05 | 1.23E-14 | 1.18E-13 |
| chr15 | 68172551 | 68192811 | 20261 | 0.85 | 1.24E-14 | 1.19E-13 |
| chr19 | 23776323 | 23794083 | 17761 | 8.23 | 1.25E-14 | 1.20E-13 |
| chr2 | 180438286 | 180446279 | 7994 | 8.15 | 1.26E-14 | 1.21E-13 |
| chr4 | 23022123 | 23042381 | 20259 | 3.05 | 1.26E-14 | 1.21E-13 |
| chr16 | 66391659 | 66418305 | 26647 | 0.86 | 1.26E-14 | 1.21E-13 |
| chr9 | 115628208 | 115631130 | 2923 | 2.9 | 1.27E-14 | 1.21E-13 |
| chr9 | 126439447 | 126442879 | 3433 | 1.35 | 1.28E-14 | 1.23E-13 |
| chr2 | 128143588 | 128181344 | 37757 | -0.58 | 1.30E-14 | 1.24E-13 |
| chr3 | 96553893 | 96556456 | 2564 | 8.13 | 1.30E-14 | 1.25E-13 |
| chr12 | 16676163 | 16677891 | 1729 | 1.31 | 1.30E-14 | 1.25E-13 |
| chr2 | 34851372 | 34853207 | 1836 | 8.12 | 1.32E-14 | 1.26E-13 |
| chr7 | 95714100 | 95718879 | 4780 | 8.12 | 1.32E-14 | 1.26E-13 |
| chr6 | 22420837 | 22422276 | 1440 | 4.95 | 1.32E-14 | 1.26E-13 |
| chr2 | 56064299 | 56093110 | 28812 | 0.49 | 1.34E-14 | 1.28E-13 |
| chr2 | 102313195 | 102335890 | 22696 | -0.89 | 1.35E-14 | 1.28E-13 |
| chr7 | 90253400 | 90256249 | 2850 | 8.12 | 1.35E-14 | 1.29E-13 |
| chr19 | 11195533 | 11206998 | 11466 | 0.61 | 1.35E-14 | 1.29E-13 |
| chr2 | 28447921 | 28474011 | 26091 | -0.87 | 1.38E-14 | 1.32E-13 |
| chr15 | 69353555 | 69367691 | 14137 | -1.43 | 1.39E-14 | 1.32E-13 |
| chrX | 20532251 | 20535589 | 3339 | 1.2 | 1.41E-14 | 1.34E-13 |
| chr8 | 54852850 | 54986865 | 134016 | 0.31 | 1.41E-14 | 1.35E-13 |
| chr8 | 139842370 | 139844229 | 1860 | -1.73 | 1.43E-14 | 1.37E-13 |
| chr20 | 14692429 | 14699521 | 7093 | 1.01 | 1.44E-14 | 1.37E-13 |

| | | | | | | |
|---|---|---|---|---|---|---|
| chr5 | 23465845 | 23467855 | 2011 | 8.17 | 1.44E-14 | 1.37E-13 |
| chr19 | 20999921 | 21004646 | 4726 | 8.11 | 1.45E-14 | 1.38E-13 |
| chr16 | 78233572 | 78246761 | 13190 | 8.1 | 1.45E-14 | 1.38E-13 |
| chr19 | 53464855 | 53467100 | 2246 | 8.1 | 1.46E-14 | 1.39E-13 |
| chr8 | 53653448 | 53677820 | 24373 | 2.05 | 1.50E-14 | 1.42E-13 |
| chr1 | 21705188 | 21706389 | 1202 | 4.8 | 1.51E-14 | 1.43E-13 |
| chr12 | 32401137 | 32427135 | 25999 | 0.94 | 1.54E-14 | 1.46E-13 |
| chr7 | 97736194 | 97766238 | 30045 | 0.52 | 1.54E-14 | 1.46E-13 |
| chr15 | 70487378 | 70490442 | 3065 | -2.12 | 1.54E-14 | 1.46E-13 |
| chr9 | 14198571 | 14252320 | 53750 | 0.46 | 1.55E-14 | 1.47E-13 |
| chr12 | 76099758 | 76120157 | 20400 | 0.86 | 1.57E-14 | 1.49E-13 |
| chr13 | 50893456 | 50901690 | 8235 | 1.77 | 1.57E-14 | 1.49E-13 |
| chr1 | 33168006 | 33255189 | 87184 | 0.41 | 1.58E-14 | 1.49E-13 |
| chr7 | 111617520 | 111627894 | 10375 | -4.77 | 1.58E-14 | 1.49E-13 |
| chr10 | 112147364 | 112225875 | 78512 | -0.37 | 1.63E-14 | 1.54E-13 |
| chr14 | 38119290 | 38125589 | 6300 | 2.1 | 1.63E-14 | 1.55E-13 |
| chr8 | 119394136 | 119396714 | 2579 | 1.27 | 1.64E-14 | 1.56E-13 |
| chr14 | 29121599 | 29131083 | 9485 | 1.21 | 1.65E-14 | 1.56E-13 |
| chr1 | 145713775 | 145729614 | 15840 | 0.88 | 1.66E-14 | 1.57E-13 |
| chr4 | 98787704 | 98797358 | 9655 | 8.1 | 1.67E-14 | 1.58E-13 |
| chr9 | 113584800 | 113610491 | 25692 | -1.36 | 1.70E-14 | 1.61E-13 |
| chr18 | 63433973 | 63436625 | 2653 | -8.07 | 1.71E-14 | 1.62E-13 |
| chr17 | 26435443 | 26437825 | 2383 | 3.79 | 1.71E-14 | 1.62E-13 |
| chr8 | 116296289 | 116297797 | 1509 | 8.09 | 1.72E-14 | 1.63E-13 |
| chr9 | 113559429 | 113565795 | 6367 | -2.07 | 1.74E-14 | 1.64E-13 |
| chr18 | 70123016 | 70128966 | 5951 | 1.97 | 1.75E-14 | 1.65E-13 |
| chr16 | 52147098 | 52149422 | 2325 | 8.09 | 1.77E-14 | 1.68E-13 |
| chr14 | 50829936 | 50833093 | 3158 | 2.24 | 1.78E-14 | 1.68E-13 |
| chr20 | 10219824 | 10245035 | 25212 | -0.62 | 1.81E-14 | 1.71E-13 |
| chr1 | 3370313 | 3421567 | 51255 | 1.05 | 1.83E-14 | 1.73E-13 |
| chr8 | 79932744 | 79934907 | 2164 | 8.1 | 1.88E-14 | 1.78E-13 |
| chr12 | 57457573 | 57506935 | 49363 | 0.41 | 1.92E-14 | 1.81E-13 |
| chr20 | 61822342 | 61873189 | 50848 | -0.5 | 1.93E-14 | 1.82E-13 |
| chr6 | 82322982 | 82324814 | 1833 | 3.48 | 1.94E-14 | 1.83E-13 |
| chr3 | 7423096 | 7425996 | 2901 | 8.08 | 1.98E-14 | 1.86E-13 |
| chr8 | 71018870 | 71087596 | 68727 | -0.44 | 1.98E-14 | 1.86E-13 |
| chr6 | 11272653 | 11278810 | 6158 | 1.52 | 2.00E-14 | 1.88E-13 |
| chr3 | 16366700 | 16392301 | 25602 | -8.05 | 2.03E-14 | 1.91E-13 |
| chr5 | 156716572 | 156719499 | 2928 | 2.53 | 2.03E-14 | 1.91E-13 |
| chr13 | 91348455 | 91352202 | 3748 | 2.51 | 2.03E-14 | 1.91E-13 |
| chrX | 37954981 | 37956240 | 1260 | 2.5 | 2.04E-14 | 1.92E-13 |
| chr14 | 32512518 | 32514420 | 1903 | 2.46 | 2.05E-14 | 1.92E-13 |
| chr19 | 3699436 | 3721418 | 21983 | 1.02 | 2.08E-14 | 1.96E-13 |
| chr8 | 58767275 | 58768720 | 1446 | 6.88 | 2.15E-14 | 2.02E-13 |
| chr1 | 150185365 | 150209573 | 24209 | 0.54 | 2.18E-14 | 2.05E-13 |
| chr14 | 38165288 | 38167585 | 2298 | 2.6 | 2.19E-14 | 2.06E-13 |
| chr1 | 186121285 | 186134920 | 13636 | 8.12 | 2.21E-14 | 2.07E-13 |
| chr21 | 46286677 | 46299034 | 12358 | 0.87 | 2.21E-14 | 2.07E-13 |
| chr2 | 177259473 | 177260692 | 1220 | 3.08 | 2.22E-14 | 2.08E-13 |
| chr5 | 111109871 | 111113631 | 3761 | 4.78 | 2.23E-14 | 2.09E-13 |
| chr16 | 19175576 | 19180847 | 5272 | 1.41 | 2.25E-14 | 2.11E-13 |
| chr3 | 175244010 | 175245029 | 1020 | 8.05 | 2.25E-14 | 2.11E-13 |
| chr14 | 103557421 | 103623849 | 66429 | 0.34 | 2.25E-14 | 2.11E-13 |
| chr8 | 77338913 | 77340857 | 1945 | 6.84 | 2.25E-14 | 2.11E-13 |
| chr9 | 100551105 | 100566743 | 15639 | 1.94 | 2.27E-14 | 2.13E-13 |
| chr12 | 8229811 | 8236404 | 6594 | 0.86 | 2.29E-14 | 2.14E-13 |
| chr3 | 190283232 | 190285364 | 2133 | 1.35 | 2.35E-14 | 2.20E-13 |
| chr2 | 26620589 | 26636016 | 15428 | -1.18 | 2.35E-14 | 2.20E-13 |
| chr2 | 231914910 | 231940633 | 25724 | -0.8 | 2.36E-14 | 2.20E-13 |
| chr3 | 46955338 | 47030584 | 75247 | 0.37 | 2.38E-14 | 2.22E-13 |
| chr16 | 34739816 | 34760713 | 20898 | 8.04 | 2.39E-14 | 2.23E-13 |

| | | | | | | |
|---|---|---|---|---|---|---|
| chr19 | 20841979 | 20852365 | 10387 | 8.04 | 2.43E-14 | 2.27E-13 |
| chr5 | 95295388 | 95299088 | 3701 | 0.97 | 2.48E-14 | 2.31E-13 |
| chr9 | 76162812 | 76165409 | 2598 | 8.05 | 2.52E-14 | 2.35E-13 |
| chr3 | 27367678 | 27439530 | 71853 | 0.8 | 2.52E-14 | 2.35E-13 |
| chr8 | 33866339 | 33881372 | 15034 | 8.04 | 2.54E-14 | 2.37E-13 |
| chr9 | 22394767 | 22396835 | 2069 | 3.07 | 2.54E-14 | 2.37E-13 |
| chr2 | 69952696 | 70057709 | 105014 | -0.26 | 2.56E-14 | 2.38E-13 |
| chr2 | 76482997 | 76497572 | 14576 | 8.07 | 2.56E-14 | 2.39E-13 |
| chr2 | 221363690 | 221370714 | 7025 | 3.16 | 2.57E-14 | 2.39E-13 |
| chr7 | 144417293 | 144422747 | 5455 | 8.04 | 2.57E-14 | 2.39E-13 |
| chr8 | 105154976 | 105156653 | 1678 | 8.03 | 2.66E-14 | 2.48E-13 |
| chr5 | 32269746 | 32279030 | 9285 | 0.89 | 2.67E-14 | 2.48E-13 |
| chr17 | 19548861 | 19585149 | 36289 | -0.44 | 2.73E-14 | 2.54E-13 |
| chr13 | 104794823 | 104797682 | 2860 | 8.03 | 2.73E-14 | 2.54E-13 |
| chr13 | 107218720 | 107224029 | 5310 | 0.91 | 2.74E-14 | 2.54E-13 |
| chr19 | 16981753 | 16998012 | 16260 | 1.29 | 2.78E-14 | 2.59E-13 |
| chr6 | 52878056 | 52880200 | 2145 | 2.34 | 2.79E-14 | 2.59E-13 |
| chr11 | 554365 | 616635 | 62271 | 0.55 | 2.80E-14 | 2.60E-13 |
| chr7 | 137795571 | 137811442 | 15872 | 1.77 | 2.81E-14 | 2.61E-13 |
| chr12 | 130823885 | 130825876 | 1992 | 8.05 | 2.84E-14 | 2.63E-13 |
| chr9 | 80910835 | 80931138 | 20304 | -0.94 | 2.87E-14 | 2.66E-13 |
| chr3 | 194750931 | 194755030 | 4100 | 1.25 | 2.91E-14 | 2.70E-13 |
| chr11 | 5688650 | 5712292 | 23643 | 0.78 | 2.94E-14 | 2.73E-13 |
| chr6 | 157039311 | 157043112 | 3802 | 1.18 | 2.96E-14 | 2.74E-13 |
| chr3 | 81041970 | 81044415 | 2446 | 8.02 | 2.99E-14 | 2.77E-13 |
| chr7 | 92775880 | 92778293 | 2414 | 1.77 | 3.01E-14 | 2.79E-13 |
| chr2 | 133641064 | 133642919 | 1856 | -4.87 | 3.04E-14 | 2.82E-13 |
| chr10 | 13568478 | 13572024 | 3547 | 1.64 | 3.05E-14 | 2.82E-13 |
| chr12 | 6741606 | 6746020 | 4415 | 2.05 | 3.05E-14 | 2.82E-13 |
| chr11 | 102339267 | 102348434 | 9168 | 1.62 | 3.07E-14 | 2.84E-13 |
| chr2 | 221482706 | 221486045 | 3340 | 1.12 | 3.07E-14 | 2.84E-13 |
| chr2 | 8615997 | 8618554 | 2558 | 2.94 | 3.11E-14 | 2.87E-13 |
| chr11 | 4151055 | 4157742 | 6688 | 1.07 | 3.13E-14 | 2.89E-13 |
| chr4 | 60418711 | 60419973 | 1263 | 1.61 | 3.16E-14 | 2.92E-13 |
| chr21 | 17651730 | 17653808 | 2079 | 2.68 | 3.17E-14 | 2.93E-13 |
| chr19 | 39295939 | 39484528 | 188590 | 0.25 | 3.18E-14 | 2.93E-13 |
| chr4 | 187726362 | 187728788 | 2427 | 2.41 | 3.22E-14 | 2.97E-13 |
| chr18 | 37280862 | 37283658 | 2797 | 1.93 | 3.24E-14 | 2.99E-13 |
| chr12 | 112145309 | 112156874 | 11566 | 1.21 | 3.24E-14 | 2.99E-13 |
| chr12 | 58695677 | 58707487 | 11811 | 0.88 | 3.25E-14 | 3.00E-13 |
| chr16 | 24637356 | 24665038 | 27683 | -0.67 | 3.26E-14 | 3.00E-13 |
| chr12 | 47770614 | 47802250 | 31637 | 0.81 | 3.27E-14 | 3.02E-13 |
| chr2 | 110347968 | 110373652 | 25685 | -0.55 | 3.36E-14 | 3.10E-13 |
| chr13 | 49881798 | 49889209 | 7412 | 1.33 | 3.49E-14 | 3.21E-13 |
| chr4 | 30755916 | 30773192 | 17277 | 8 | 3.50E-14 | 3.23E-13 |
| chr2 | 133514318 | 133516103 | 1786 | -4.8 | 3.54E-14 | 3.26E-13 |
| chr2 | 218141945 | 218159848 | 17904 | 8.02 | 3.63E-14 | 3.34E-13 |
| chr3 | 22875606 | 22878610 | 3005 | 1.27 | 3.64E-14 | 3.35E-13 |
| chr6 | 117269839 | 117276794 | 6956 | 8.06 | 3.67E-14 | 3.37E-13 |
| chr20 | 56317871 | 56324814 | 6944 | -1 | 3.67E-14 | 3.38E-13 |
| chr14 | 38085094 | 38093193 | 8100 | 8.01 | 3.68E-14 | 3.38E-13 |
| chr11 | 109598148 | 109599642 | 1495 | 3.52 | 3.70E-14 | 3.40E-13 |
| chr16 | 16949975 | 16956232 | 6258 | -7.97 | 3.77E-14 | 3.46E-13 |
| chr7 | 47186483 | 47194016 | 7534 | 8 | 3.79E-14 | 3.48E-13 |
| chr17 | 52278327 | 52280205 | 1879 | 7.99 | 3.85E-14 | 3.53E-13 |
| chr2 | 56218646 | 56220668 | 2023 | 3 | 3.88E-14 | 3.56E-13 |
| chr19 | 12250421 | 12251148 | 728 | 4.75 | 3.89E-14 | 3.57E-13 |
| chr20 | 5929417 | 5991295 | 61879 | 0.32 | 3.90E-14 | 3.58E-13 |
| chr4 | 59981058 | 59986984 | 5927 | 8 | 3.93E-14 | 3.61E-13 |
| chr7 | 15593169 | 15601786 | 8618 | 0.78 | 3.97E-14 | 3.64E-13 |
| chr3 | 85209964 | 85220279 | 10316 | 7.99 | 3.98E-14 | 3.65E-13 |

| | | | | | | |
|---|---|---|---|---|---|---|
| chr19 | 39251828 | 39277591 | 25764 | 1.04 | 4.17E-14 | 3.82E-13 |
| chr17 | 4423854 | 4438620 | 14767 | 1.54 | 4.20E-14 | 3.84E-13 |
| chr12 | 4429354 | 4436998 | 7645 | 0.79 | 4.20E-14 | 3.84E-13 |
| chr5 | 13628941 | 13641835 | 12895 | 7.98 | 4.20E-14 | 3.85E-13 |
| chr5 | 95478385 | 95494945 | 16561 | 2.08 | 4.34E-14 | 3.97E-13 |
| chr1 | 68149927 | 68237474 | 87548 | 0.4 | 4.37E-14 | 4.00E-13 |
| chr9 | 127702091 | 127704988 | 2898 | 0.85 | 4.42E-14 | 4.04E-13 |
| chr1 | 180350447 | 180351940 | 1494 | 2.97 | 4.43E-14 | 4.05E-13 |
| chrX | 19715214 | 19742449 | 27236 | -1.8 | 4.45E-14 | 4.07E-13 |
| chr15 | 65450347 | 65482968 | 32622 | 0.55 | 4.47E-14 | 4.09E-13 |
| chr17 | 28032533 | 28038455 | 5923 | 0.99 | 4.55E-14 | 4.16E-13 |
| chr3 | 122282440 | 122292805 | 10366 | 1.01 | 4.60E-14 | 4.20E-13 |
| chr6 | 146864028 | 146868731 | 4704 | 1.06 | 4.61E-14 | 4.21E-13 |
| chr5 | 132575709 | 132585682 | 9974 | -0.79 | 4.63E-14 | 4.22E-13 |
| chr2 | 72970431 | 73022760 | 52330 | 0.75 | 4.75E-14 | 4.33E-13 |
| chr2 | 237620671 | 237621983 | 1313 | -3.12 | 4.76E-14 | 4.34E-13 |
| chr16 | 81428895 | 81813436 | 384542 | 0.21 | 4.80E-14 | 4.37E-13 |
| chr4 | 97904591 | 97906138 | 1548 | 6.77 | 4.81E-14 | 4.38E-13 |
| chr6 | 11392765 | 11394151 | 1387 | -2.84 | 4.84E-14 | 4.41E-13 |
| chr8 | 142082504 | 142106989 | 24486 | -0.73 | 4.87E-14 | 4.44E-13 |
| chr11 | 63375743 | 63453285 | 77543 | 0.47 | 4.92E-14 | 4.48E-13 |
| chr2 | 67516183 | 67523313 | 7131 | 7.96 | 5.02E-14 | 4.57E-13 |
| chr16 | 3069898 | 3074767 | 4870 | 0.61 | 5.11E-14 | 4.65E-13 |
| chr17 | 6267967 | 6278940 | 10974 | 1.31 | 5.13E-14 | 4.67E-13 |
| chr13 | 40688862 | 40692419 | 3558 | 0.94 | 5.14E-14 | 4.68E-13 |
| chr1 | 193939764 | 193942016 | 2253 | 1.61 | 5.20E-14 | 4.73E-13 |
| chrX | 120330736 | 120331994 | 1259 | 3.32 | 5.20E-14 | 4.73E-13 |
| chr4 | 112906077 | 112913680 | 7604 | 2.74 | 5.25E-14 | 4.77E-13 |
| chr11 | 86710290 | 86717267 | 6978 | 1.48 | 5.36E-14 | 4.86E-13 |
| chr3 | 189010602 | 189076062 | 65461 | 0.42 | 5.38E-14 | 4.89E-13 |
| chr13 | 102221785 | 102232278 | 10494 | 7.96 | 5.43E-14 | 4.93E-13 |
| chr11 | 14483636 | 14542670 | 59035 | 0.51 | 5.52E-14 | 5.01E-13 |
| chr7 | 82627205 | 82630080 | 2876 | 5.02 | 5.53E-14 | 5.02E-13 |
| chr5 | 30029889 | 30031102 | 1214 | 5.26 | 5.58E-14 | 5.06E-13 |
| chr12 | 56832972 | 56843958 | 10987 | 0.99 | 5.60E-14 | 5.08E-13 |
| chr19 | 36937591 | 36941455 | 3865 | 7.97 | 5.69E-14 | 5.16E-13 |
| chr2 | 162761146 | 162763564 | 2419 | 7.95 | 5.74E-14 | 5.20E-13 |
| chr1 | 225060780 | 225072872 | 12093 | 1.7 | 5.75E-14 | 5.20E-13 |
| chr1 | 167188487 | 167200616 | 12130 | 0.75 | 5.76E-14 | 5.21E-13 |
| chr12 | 69369815 | 69371346 | 1532 | 7.94 | 5.76E-14 | 5.21E-13 |
| chr9 | 30235109 | 30236657 | 1549 | 7.95 | 5.82E-14 | 5.26E-13 |
| chr10 | 6859615 | 6876447 | 16833 | 2.18 | 5.87E-14 | 5.31E-13 |
| chr15 | 70599818 | 70603164 | 3347 | 2.91 | 5.94E-14 | 5.37E-13 |
| chr1 | 93414742 | 93428272 | 13531 | -0.78 | 6.03E-14 | 5.45E-13 |
| chr16 | 2879376 | 2898254 | 18879 | 0.69 | 6.05E-14 | 5.47E-13 |
| chr3 | 58448422 | 58458950 | 10529 | 0.77 | 6.11E-14 | 5.52E-13 |
| chr4 | 91130359 | 91132025 | 1667 | 7.94 | 6.11E-14 | 5.52E-13 |
| chr5 | 140891686 | 140904274 | 12589 | 1.14 | 6.14E-14 | 5.54E-13 |
| chr3 | 162752349 | 162754155 | 1807 | 2.31 | 6.15E-14 | 5.55E-13 |
| chr13 | 43098261 | 43107583 | 9323 | 7.95 | 6.22E-14 | 5.61E-13 |
| chr17 | 9720702 | 9747955 | 27254 | -0.44 | 6.24E-14 | 5.63E-13 |
| chr15 | 51739332 | 51742932 | 3601 | 1.45 | 6.41E-14 | 5.78E-13 |
| chr12 | 88240058 | 88254578 | 14521 | 7.98 | 6.56E-14 | 5.92E-13 |
| chr7 | 19872699 | 19875721 | 3023 | 1.74 | 6.57E-14 | 5.92E-13 |
| chr16 | 20773910 | 20776392 | 2483 | 1.4 | 6.59E-14 | 5.93E-13 |
| chr6 | 122930717 | 122943205 | 12489 | 7.95 | 6.60E-14 | 5.94E-13 |
| chr2 | 74982492 | 74992671 | 10180 | 7.95 | 6.61E-14 | 5.95E-13 |
| chr3 | 105528089 | 105555709 | 27621 | 1.33 | 6.62E-14 | 5.96E-13 |
| chr4 | 30537663 | 30547251 | 9589 | 7.93 | 6.76E-14 | 6.08E-13 |
| chr8 | 110749768 | 110759209 | 9442 | 8.03 | 6.79E-14 | 6.11E-13 |
| chr7 | 152008913 | 152011902 | 2990 | 1.5 | 6.87E-14 | 6.18E-13 |

| | | | | | | |
|---|---|---|---|---|---|---|
| chr12 | 15087643 | 15114963 | 27321 | 1.41 | 6.96E-14 | 6.26E-13 |
| chr3 | 18140101 | 18153975 | 13875 | 7.93 | 6.97E-14 | 6.26E-13 |
| chr8 | 103612433 | 103630560 | 18128 | 0.65 | 6.99E-14 | 6.28E-13 |
| chr4 | 147852688 | 147854035 | 1348 | 2.31 | 6.99E-14 | 6.28E-13 |
| chr3 | 39301833 | 39303346 | 1514 | 3.59 | 7.03E-14 | 6.31E-13 |
| chr8 | 116849531 | 116851430 | 1900 | 7.92 | 7.04E-14 | 6.32E-13 |
| chr2 | 20188372 | 20190742 | 2371 | -1.14 | 7.09E-14 | 6.37E-13 |
| chr9 | 122724813 | 122727087 | 2275 | 1.58 | 7.13E-14 | 6.40E-13 |
| chr18 | 71915825 | 71933976 | 18152 | -0.85 | 7.19E-14 | 6.45E-13 |
| chr2 | 25712444 | 25715674 | 3231 | -2.05 | 7.20E-14 | 6.46E-13 |
| chr1 | 41798765 | 41808724 | 9960 | 7.92 | 7.22E-14 | 6.47E-13 |
| chr2 | 47136184 | 47144205 | 8022 | -0.73 | 7.24E-14 | 6.48E-13 |
| chr8 | 126556887 | 126579410 | 22524 | 0.57 | 7.44E-14 | 6.66E-13 |
| chr4 | 122705398 | 122707640 | 2243 | 1.73 | 7.50E-14 | 6.72E-13 |
| chr1 | 156389107 | 156392107 | 3001 | 2.9 | 7.54E-14 | 6.75E-13 |
| chr11 | 21675159 | 21678337 | 3179 | 6.13 | 7.59E-14 | 6.79E-13 |
| chr7 | 89839910 | 89860498 | 20589 | 0.95 | 7.78E-14 | 6.96E-13 |
| chr4 | 173990408 | 173995783 | 5376 | 6.76 | 7.79E-14 | 6.97E-13 |
| chr2 | 109186600 | 109212502 | 25903 | -0.65 | 7.82E-14 | 6.99E-13 |
| chr7 | 25888858 | 25897604 | 8747 | 1.8 | 8.04E-14 | 7.18E-13 |
| chr2 | 45685176 | 45698267 | 13092 | 0.78 | 8.11E-14 | 7.25E-13 |
| chr2 | 104659878 | 104661291 | 1414 | 7.96 | 8.14E-14 | 7.27E-13 |
| chrX | 23791493 | 23841841 | 50349 | 0.44 | 8.21E-14 | 7.34E-13 |
| chr10 | 105868530 | 105884091 | 15562 | 0.44 | 8.30E-14 | 7.41E-13 |
| chr11 | 67857744 | 67859174 | 1431 | 2.67 | 8.34E-14 | 7.45E-13 |
| chr10 | 5230310 | 5241632 | 11323 | 0.74 | 8.43E-14 | 7.52E-13 |
| chr3 | 185503785 | 185515637 | 11853 | 2.24 | 8.50E-14 | 7.58E-13 |
| chrX | 11916046 | 11918451 | 2406 | 1.39 | 8.58E-14 | 7.65E-13 |
| chr2 | 34929454 | 34931844 | 2391 | 7.9 | 8.66E-14 | 7.72E-13 |
| chr10 | 123083889 | 123086113 | 2225 | 7.92 | 8.70E-14 | 7.75E-13 |
| chr1 | 221735508 | 221750633 | 15126 | 7.9 | 8.70E-14 | 7.75E-13 |
| chr3 | 181623209 | 181627275 | 4067 | 1.22 | 8.73E-14 | 7.78E-13 |
| chr4 | 96209537 | 96226944 | 17408 | 0.97 | 8.75E-14 | 7.79E-13 |
| chr18 | 28678497 | 28682627 | 4131 | 2.03 | 8.83E-14 | 7.86E-13 |
| chr3 | 122449099 | 122451914 | 2816 | 1.04 | 9.13E-14 | 8.12E-13 |
| chr7 | 7455468 | 7456586 | 1119 | 3.44 | 9.21E-14 | 8.19E-13 |
| chr2 | 225862318 | 225892484 | 30167 | -0.69 | 9.31E-14 | 8.28E-13 |
| chr5 | 27853852 | 27856696 | 2845 | 7.92 | 9.33E-14 | 8.30E-13 |
| chr6 | 132259530 | 132280729 | 21200 | 0.86 | 9.34E-14 | 8.30E-13 |
| chr2 | 84179321 | 84181477 | 2157 | 7.9 | 9.38E-14 | 8.34E-13 |
| chr4 | 170140608 | 170204670 | 64063 | 0.34 | 9.42E-14 | 8.37E-13 |
| chr8 | 107592255 | 107594065 | 1811 | 7.93 | 9.48E-14 | 8.42E-13 |
| chr3 | 127448627 | 127542477 | 93851 | 0.32 | 9.54E-14 | 8.47E-13 |
| chr1 | 190979522 | 190986177 | 6656 | 7.89 | 9.58E-14 | 8.50E-13 |
| chr8 | 140226655 | 140229801 | 3147 | 1.2 | 9.64E-14 | 8.56E-13 |
| chr20 | 57090147 | 57100960 | 10814 | 2.66 | 9.65E-14 | 8.56E-13 |
| chr6 | 71105846 | 71136831 | 30986 | 0.83 | 9.66E-14 | 8.57E-13 |
| chr3 | 147663303 | 147664591 | 1289 | 5.09 | 9.67E-14 | 8.57E-13 |
| chrX | 120775005 | 120776914 | 1910 | 7.88 | 9.70E-14 | 8.60E-13 |
| chr8 | 107092393 | 107096788 | 4396 | 0.68 | 9.74E-14 | 8.63E-13 |
| chr3 | 190056545 | 190066436 | 9892 | 0.48 | 9.83E-14 | 8.70E-13 |
| chr8 | 33899314 | 33903510 | 4197 | 3.43 | 1.00E-13 | 8.88E-13 |
| chr4 | 91453256 | 91454743 | 1488 | 7.88 | 1.00E-13 | 8.88E-13 |
| chr11 | 68023588 | 68044177 | 20590 | 0.74 | 1.01E-13 | 8.92E-13 |
| chr20 | 5845229 | 5847205 | 1977 | 1.8 | 1.02E-13 | 9.00E-13 |
| chr7 | 12170326 | 12177388 | 7063 | 2.91 | 1.02E-13 | 9.03E-13 |
| chr8 | 63128488 | 63130600 | 2113 | 1.73 | 1.03E-13 | 9.13E-13 |
| chr10 | 70975746 | 70994741 | 18996 | 0.61 | 1.03E-13 | 9.14E-13 |
| chr4 | 120650805 | 120657094 | 6290 | 1.89 | 1.04E-13 | 9.15E-13 |
| chr19 | 9642475 | 9650313 | 7839 | 1.98 | 1.04E-13 | 9.15E-13 |
| chr8 | 39935731 | 39982512 | 46782 | -0.57 | 1.04E-13 | 9.19E-13 |

| | | | | | | |
|---|---|---|---|---|---|---|
| chr2 | 1589969 | 1598429 | 8461 | 7.88 | 1.04E-13 | 9.21E-13 |
| chr11 | 86500660 | 86528662 | 28003 | 0.48 | 1.05E-13 | 9.25E-13 |
| chr5 | 111518250 | 111519889 | 1640 | 3.03 | 1.05E-13 | 9.28E-13 |
| chr14 | 100767889 | 100785107 | 17219 | 1.64 | 1.06E-13 | 9.34E-13 |
| chr7 | 79735180 | 79736753 | 1574 | 6.06 | 1.06E-13 | 9.37E-13 |
| chr12 | 92986704 | 92999043 | 12340 | 0.71 | 1.07E-13 | 9.47E-13 |
| chr14 | 59714628 | 59746424 | 31797 | 1.11 | 1.08E-13 | 9.54E-13 |
| chr10 | 116248733 | 116250297 | 1565 | 3.55 | 1.09E-13 | 9.63E-13 |
| chr10 | 24752611 | 24756661 | 4051 | 0.96 | 1.10E-13 | 9.68E-13 |
| chr7 | 43731100 | 43737597 | 6498 | -1.22 | 1.12E-13 | 9.85E-13 |
| chr5 | 147762728 | 147781827 | 19100 | 0.75 | 1.13E-13 | 9.96E-13 |
| chr11 | 74406960 | 74408956 | 1997 | 2.8 | 1.13E-13 | 9.96E-13 |
| chr19 | 57018595 | 57020173 | 1579 | 7.88 | 1.14E-13 | 1.00E-12 |
| chr8 | 121079247 | 121081660 | 2414 | 7.87 | 1.14E-13 | 1.01E-12 |
| chr5 | 64235406 | 64237227 | 1822 | 1.73 | 1.15E-13 | 1.01E-12 |
| chr1 | 183835260 | 183851514 | 16255 | 1.8 | 1.16E-13 | 1.02E-12 |
| chr4 | 21032951 | 21034899 | 1949 | 7.86 | 1.16E-13 | 1.02E-12 |
| chrX | 28516430 | 28517853 | 1424 | 6.77 | 1.19E-13 | 1.04E-12 |
| chr3 | 124646889 | 124655599 | 8711 | 1.78 | 1.19E-13 | 1.05E-12 |
| chr2 | 230995136 | 230996322 | 1187 | 4.84 | 1.20E-13 | 1.05E-12 |
| chr2 | 63305301 | 63306986 | 1686 | 3.26 | 1.23E-13 | 1.08E-12 |
| chr7 | 72771409 | 72773942 | 2534 | 1.53 | 1.25E-13 | 1.09E-12 |
| chr11 | 94335324 | 94337226 | 1903 | 2.05 | 1.27E-13 | 1.11E-12 |
| chr10 | 134120460 | 134146596 | 26137 | -0.52 | 1.29E-13 | 1.14E-12 |
| chr15 | 61326423 | 61327847 | 1425 | 7.85 | 1.30E-13 | 1.14E-12 |
| chr19 | 44758413 | 44764848 | 6436 | 7.85 | 1.30E-13 | 1.14E-12 |
| chr5 | 40772104 | 40809166 | 37063 | 0.53 | 1.32E-13 | 1.16E-12 |
| chr2 | 173971222 | 173990319 | 19098 | -0.73 | 1.32E-13 | 1.16E-12 |
| chr12 | 2903364 | 2924837 | 21474 | 0.84 | 1.33E-13 | 1.17E-12 |
| chr11 | 73086329 | 73105060 | 18732 | -0.64 | 1.34E-13 | 1.17E-12 |
| chr14 | 92960983 | 93032440 | 71458 | -0.39 | 1.34E-13 | 1.18E-12 |
| chr4 | 65136795 | 65138648 | 1854 | 7.87 | 1.39E-13 | 1.21E-12 |
| chr7 | 81305946 | 81315354 | 9409 | 1.46 | 1.39E-13 | 1.22E-12 |
| chr8 | 71494322 | 71530237 | 35916 | 0.45 | 1.41E-13 | 1.23E-12 |
| chr20 | 56007694 | 56015384 | 7691 | 1.86 | 1.41E-13 | 1.23E-12 |
| chr13 | 74621226 | 74622977 | 1752 | 5.61 | 1.41E-13 | 1.23E-12 |
| chr12 | 16282833 | 16290209 | 7377 | 7.85 | 1.42E-13 | 1.24E-12 |
| chr4 | 93154739 | 93157093 | 2355 | 7.84 | 1.42E-13 | 1.24E-12 |
| chr2 | 85115681 | 85116691 | 1011 | 4.05 | 1.42E-13 | 1.24E-12 |
| chr4 | 177908679 | 177923678 | 15000 | 1.63 | 1.46E-13 | 1.27E-12 |
| chrX | 121009531 | 121020724 | 11194 | 7.84 | 1.47E-13 | 1.28E-12 |
| chr13 | 108805971 | 108807866 | 1896 | 7.84 | 1.47E-13 | 1.28E-12 |
| chr12 | 100731471 | 100735892 | 4422 | 2.15 | 1.51E-13 | 1.32E-12 |
| chr10 | 103317228 | 103348947 | 31720 | -1.13 | 1.53E-13 | 1.33E-12 |
| chr18 | 55929169 | 55933132 | 3964 | 1.31 | 1.55E-13 | 1.35E-12 |
| chr12 | 77156618 | 77160748 | 4131 | 0.99 | 1.56E-13 | 1.36E-12 |
| chr16 | 14699893 | 14752467 | 52575 | 0.44 | 1.59E-13 | 1.39E-12 |
| chr3 | 72771869 | 72773576 | 1708 | 2.51 | 1.61E-13 | 1.40E-12 |
| chr12 | 28173509 | 28189454 | 15946 | 0.76 | 1.62E-13 | 1.41E-12 |
| chr2 | 133131519 | 133133039 | 1521 | 5.08 | 1.62E-13 | 1.41E-12 |
| chr1 | 223886109 | 223965855 | 79747 | 0.29 | 1.63E-13 | 1.42E-12 |
| chr11 | 29140373 | 29151497 | 11125 | 0.69 | 1.65E-13 | 1.43E-12 |
| chr20 | 56229954 | 56294501 | 64548 | 0.4 | 1.65E-13 | 1.43E-12 |
| chr2 | 239067586 | 239079695 | 12110 | -1.48 | 1.66E-13 | 1.44E-12 |
| chr2 | 222381943 | 222395600 | 13658 | 7.82 | 1.70E-13 | 1.48E-12 |
| chr5 | 89842290 | 89844971 | 2682 | 7.82 | 1.77E-13 | 1.53E-12 |
| chr3 | 109070926 | 109083329 | 12404 | 1.51 | 1.79E-13 | 1.55E-12 |
| chr8 | 121821114 | 121824411 | 3298 | 1.5 | 1.83E-13 | 1.59E-12 |
| chr20 | 32651560 | 32662433 | 10874 | -1.78 | 1.83E-13 | 1.59E-12 |
| chr6 | 140805705 | 140808331 | 2627 | 1.14 | 1.83E-13 | 1.59E-12 |
| chr5 | 114678157 | 114688067 | 9911 | 1.41 | 1.83E-13 | 1.59E-12 |

| | | | | | | |
|---|---|---|---|---|---|---|
| chr10 | 36469595 | 36472166 | 2572 | -7.79 | 1.83E-13 | 1.59E-12 |
| chr16 | 90038117 | 90072050 | 33934 | 0.52 | 1.84E-13 | 1.60E-12 |
| chr7 | 67194227 | 67195547 | 1321 | 2.65 | 1.85E-13 | 1.60E-12 |
| chr9 | 32644180 | 32645792 | 1613 | 4.91 | 1.87E-13 | 1.62E-12 |
| chr2 | 197024163 | 197047803 | 23641 | 1.11 | 1.92E-13 | 1.66E-12 |
| chr5 | 43336002 | 43337746 | 1745 | 5.14 | 1.92E-13 | 1.66E-12 |
| chr16 | 9134764 | 9147586 | 12823 | -0.68 | 1.92E-13 | 1.67E-12 |
| chr3 | 64199229 | 64225928 | 26700 | -0.61 | 1.93E-13 | 1.67E-12 |
| chr14 | 56584274 | 56586871 | 2598 | 1.62 | 1.99E-13 | 1.73E-12 |
| chr11 | 46678973 | 46742153 | 63181 | -0.44 | 2.00E-13 | 1.73E-12 |
| chr4 | 113777 | 125218 | 11442 | 1.85 | 2.01E-13 | 1.73E-12 |
| chr6 | 13383361 | 13409284 | 25924 | 0.64 | 2.02E-13 | 1.75E-12 |
| chr8 | 81933833 | 81942232 | 8400 | 7.81 | 2.03E-13 | 1.76E-12 |
| chr2 | 158101971 | 158122151 | 20181 | 0.9 | 2.07E-13 | 1.79E-12 |
| chr1 | 239413343 | 239432444 | 19102 | 7.91 | 2.08E-13 | 1.80E-12 |
| chr1 | 200859096 | 200886602 | 27507 | 0.36 | 2.09E-13 | 1.80E-12 |
| chr12 | 50352774 | 50420156 | 67383 | 0.6 | 2.11E-13 | 1.82E-12 |
| chr1 | 60279647 | 60281387 | 1741 | 4.68 | 2.17E-13 | 1.87E-12 |
| chr1 | 115052031 | 115060819 | 8789 | 1.27 | 2.17E-13 | 1.87E-12 |
| chr1 | 172862481 | 172888271 | 25791 | 0.65 | 2.19E-13 | 1.88E-12 |
| chr11 | 126913433 | 126922699 | 9267 | 2.11 | 2.20E-13 | 1.89E-12 |
| chr2 | 39693929 | 39753870 | 59942 | 0.55 | 2.21E-13 | 1.90E-12 |
| chr1 | 20575520 | 20578615 | 3096 | 1.84 | 2.23E-13 | 1.92E-12 |
| chr13 | 105847819 | 105849390 | 1572 | 7.79 | 2.24E-13 | 1.93E-12 |
| chr11 | 12240248 | 12262534 | 22287 | -1.2 | 2.24E-13 | 1.93E-12 |
| chr15 | 87154514 | 87156236 | 1723 | 2.06 | 2.26E-13 | 1.95E-12 |
| chr19 | 3275171 | 3290671 | 15501 | 1.82 | 2.27E-13 | 1.95E-12 |
| chr19 | 57988193 | 58011696 | 23504 | 7.8 | 2.27E-13 | 1.95E-12 |
| chr1 | 157105161 | 157125973 | 20813 | 0.86 | 2.28E-13 | 1.96E-12 |
| chr9 | 125810324 | 125812244 | 1921 | 2.13 | 2.32E-13 | 1.99E-12 |
| chr5 | 37197617 | 37220881 | 23265 | 1.03 | 2.33E-13 | 2.00E-12 |
| chr8 | 60606349 | 60608853 | 2505 | 7.78 | 2.35E-13 | 2.02E-12 |
| chr2 | 218095766 | 218104937 | 9172 | 4.32 | 2.36E-13 | 2.02E-12 |
| chr14 | 22873519 | 22875541 | 2023 | 7.81 | 2.36E-13 | 2.02E-12 |
| chr11 | 19439879 | 19442897 | 3019 | -1.78 | 2.38E-13 | 2.04E-12 |
| chr12 | 107711526 | 107834329 | 122804 | -0.25 | 2.39E-13 | 2.05E-12 |
| chr12 | 118772315 | 118820332 | 48018 | 0.49 | 2.39E-13 | 2.05E-12 |
| chr17 | 17584215 | 17760416 | 176202 | 0.22 | 2.40E-13 | 2.06E-12 |
| chr4 | 96278751 | 96281151 | 2401 | 2.39 | 2.43E-13 | 2.08E-12 |
| chr5 | 16455496 | 16467140 | 11645 | 0.91 | 2.47E-13 | 2.12E-12 |
| chr19 | 37062986 | 37068892 | 5907 | 2.55 | 2.52E-13 | 2.16E-12 |
| chr2 | 8632614 | 8641045 | 8432 | 1.67 | 2.54E-13 | 2.18E-12 |
| chr7 | 16885069 | 16891839 | 6771 | 0.7 | 2.54E-13 | 2.18E-12 |
| chr11 | 41419270 | 41420664 | 1395 | 7.77 | 2.55E-13 | 2.18E-12 |
| chr5 | 133394278 | 133407274 | 12997 | -2.35 | 2.57E-13 | 2.20E-12 |
| chr3 | 49414460 | 49427180 | 12721 | 1.44 | 2.58E-13 | 2.20E-12 |
| chr5 | 26892708 | 26895775 | 3068 | 7.79 | 2.58E-13 | 2.21E-12 |
| chr5 | 155516249 | 155518958 | 2710 | -7.74 | 2.58E-13 | 2.21E-12 |
| chr4 | 163601779 | 163607281 | 5503 | 0.89 | 2.59E-13 | 2.22E-12 |
| chr2 | 39773792 | 39778650 | 4859 | 7.79 | 2.60E-13 | 2.22E-12 |
| chr2 | 178889458 | 178891432 | 1975 | -7.75 | 2.60E-13 | 2.22E-12 |
| chr14 | 58706219 | 58767938 | 61720 | 0.43 | 2.64E-13 | 2.25E-12 |
| chr14 | 36415884 | 36429116 | 13233 | 1.96 | 2.67E-13 | 2.28E-12 |
| chr15 | 92394728 | 92412127 | 17400 | 0.78 | 2.67E-13 | 2.28E-12 |
| chr19 | 21748721 | 21752712 | 3992 | 7.78 | 2.70E-13 | 2.30E-12 |
| chr7 | 80170982 | 80179197 | 8216 | 1.62 | 2.70E-13 | 2.30E-12 |
| chr17 | 60749165 | 60760278 | 11114 | -2.01 | 2.74E-13 | 2.34E-12 |
| chr13 | 106621581 | 106632782 | 11202 | 7.78 | 2.75E-13 | 2.34E-12 |
| chr16 | 15963487 | 16007045 | 43559 | -0.46 | 2.75E-13 | 2.35E-12 |
| chr10 | 124052753 | 124072093 | 19341 | 0.49 | 2.78E-13 | 2.37E-12 |
| chr5 | 100224813 | 100239674 | 14862 | 0.96 | 2.79E-13 | 2.37E-12 |

| | | | | | | |
|---|---|---|---|---|---|---|
| chr7 | 117950892 | 117953124 | 2233 | 7.76 | 2.80E-13 | 2.38E-12 |
| chr15 | 49085707 | 49103918 | 18212 | 0.91 | 2.83E-13 | 2.41E-12 |
| chrX | 36627289 | 36629113 | 1825 | 4.22 | 2.84E-13 | 2.42E-12 |
| chr3 | 123149636 | 123152252 | 2617 | 7.78 | 2.86E-13 | 2.43E-12 |
| chr12 | 122459338 | 122530545 | 71208 | 0.41 | 2.89E-13 | 2.45E-12 |
| chr8 | 11313803 | 11339175 | 25373 | -0.54 | 2.90E-13 | 2.47E-12 |
| chr4 | 72414838 | 72426725 | 11888 | 1.4 | 2.91E-13 | 2.48E-12 |
| chr5 | 167061336 | 167081442 | 20107 | 7.77 | 2.92E-13 | 2.48E-12 |
| chr1 | 219238686 | 219266228 | 27543 | -0.89 | 2.96E-13 | 2.52E-12 |
| chr3 | 45132612 | 45135718 | 3107 | 1.42 | 2.97E-13 | 2.52E-12 |
| chr17 | 15244409 | 15252149 | 7741 | 7.75 | 2.97E-13 | 2.53E-12 |
| chr4 | 102874340 | 102899220 | 24881 | -0.81 | 3.00E-13 | 2.55E-12 |
| chr14 | 68825615 | 68832612 | 6998 | 1.05 | 3.01E-13 | 2.56E-12 |
| chr8 | 86847883 | 86849855 | 1973 | 2.33 | 3.02E-13 | 2.56E-12 |
| chr3 | 29724074 | 29738326 | 14253 | -0.54 | 3.04E-13 | 2.57E-12 |
| chr18 | 57566035 | 57586816 | 20782 | 0.63 | 3.04E-13 | 2.58E-12 |
| chr4 | 81048515 | 81063960 | 15446 | 7.82 | 3.04E-13 | 2.58E-12 |
| chr9 | 97430735 | 97457205 | 26471 | 0.91 | 3.04E-13 | 2.58E-12 |
| chr2 | 31782788 | 31784690 | 1903 | -7.73 | 3.07E-13 | 2.60E-12 |
| chr15 | 59062027 | 59065008 | 2982 | 0.81 | 3.10E-13 | 2.63E-12 |
| chr15 | 26855058 | 26881571 | 26514 | -0.5 | 3.11E-13 | 2.64E-12 |
| chr2 | 235149338 | 235168349 | 19012 | -0.45 | 3.13E-13 | 2.65E-12 |
| chr2 | 190140281 | 190153350 | 13070 | -3.03 | 3.17E-13 | 2.68E-12 |
| chr1 | 82692788 | 82694196 | 1409 | 7.75 | 3.17E-13 | 2.68E-12 |
| chrX | 122936011 | 122970454 | 34444 | 0.54 | 3.19E-13 | 2.70E-12 |
| chr19 | 14605532 | 14640080 | 34549 | 0.49 | 3.20E-13 | 2.71E-12 |
| chr10 | 11593388 | 11595217 | 1830 | 1.77 | 3.24E-13 | 2.74E-12 |
| chr12 | 87736268 | 87742467 | 6200 | 7.82 | 3.25E-13 | 2.75E-12 |
| chr21 | 16976150 | 16991349 | 15200 | 7.78 | 3.26E-13 | 2.75E-12 |
| chr3 | 149188071 | 149198353 | 10283 | 0.47 | 3.27E-13 | 2.76E-12 |
| chr17 | 39914999 | 39993131 | 78133 | 0.32 | 3.28E-13 | 2.76E-12 |
| chr17 | 1132579 | 1137102 | 4524 | 2.61 | 3.31E-13 | 2.79E-12 |
| chr2 | 208258677 | 208274614 | 15938 | 0.8 | 3.31E-13 | 2.80E-12 |
| chr1 | 214160494 | 214162234 | 1741 | 3.41 | 3.32E-13 | 2.80E-12 |
| chr20 | 1874202 | 1910848 | 36647 | -0.81 | 3.33E-13 | 2.81E-12 |
| chr3 | 42054325 | 42122987 | 68663 | 0.47 | 3.34E-13 | 2.82E-12 |
| chr3 | 94999420 | 95000699 | 1280 | 7.74 | 3.39E-13 | 2.85E-12 |
| chr3 | 110605110 | 110611391 | 6282 | 7.75 | 3.40E-13 | 2.86E-12 |
| chr12 | 89744405 | 89749645 | 5241 | 0.51 | 3.41E-13 | 2.88E-12 |
| chr6 | 64265208 | 64289885 | 24678 | 0.39 | 3.45E-13 | 2.90E-12 |
| chr4 | 77447257 | 77448679 | 1423 | 5.59 | 3.56E-13 | 3.00E-12 |
| chr17 | 499263 | 526742 | 27480 | -0.91 | 3.58E-13 | 3.01E-12 |
| chr3 | 163981801 | 163983614 | 1814 | 7.74 | 3.58E-13 | 3.02E-12 |
| chr11 | 69140052 | 69177439 | 37388 | -0.61 | 3.61E-13 | 3.04E-12 |
| chr2 | 213804118 | 213806197 | 2080 | 5.96 | 3.70E-13 | 3.11E-12 |
| chr1 | 8539612 | 8554720 | 15109 | 1.43 | 3.72E-13 | 3.12E-12 |
| chr1 | 4728586 | 4738699 | 10114 | 4.92 | 3.73E-13 | 3.13E-12 |
| chr12 | 67661382 | 67676479 | 15098 | 0.58 | 3.76E-13 | 3.16E-12 |
| chr12 | 133403318 | 133406472 | 3155 | 1 | 3.76E-13 | 3.16E-12 |
| chr16 | 87027864 | 87034737 | 6874 | 2.13 | 3.76E-13 | 3.16E-12 |
| chr6 | 153349641 | 153361853 | 12213 | 1.14 | 3.82E-13 | 3.21E-12 |
| chr4 | 38601880 | 38604375 | 2496 | 2.86 | 3.82E-13 | 3.21E-12 |
| chr9 | 84024960 | 84033573 | 8614 | 4.69 | 3.91E-13 | 3.28E-12 |
| chr5 | 178231295 | 178233603 | 2309 | 7.72 | 3.97E-13 | 3.33E-12 |
| chr1 | 198626257 | 198639266 | 13010 | 7.72 | 3.97E-13 | 3.33E-12 |
| chr3 | 65932122 | 65942468 | 10347 | 1.16 | 4.07E-13 | 3.41E-12 |
| chr12 | 19772423 | 19773917 | 1495 | 1.41 | 4.07E-13 | 3.41E-12 |
| chr5 | 172753675 | 172757806 | 4132 | -2.45 | 4.08E-13 | 3.42E-12 |
| chr4 | 37971301 | 37980633 | 9333 | 1.58 | 4.09E-13 | 3.43E-12 |
| chr19 | 12222142 | 12224896 | 2755 | 7.72 | 4.10E-13 | 3.43E-12 |
| chr10 | 18895490 | 18916784 | 21295 | 0.52 | 4.14E-13 | 3.47E-12 |

| | | | | | | |
|---|---|---|---|---|---|---|
| chr6 | 76444218 | 76459996 | 15779 | 0.71 | 4.14E-13 | 3.47E-12 |
| chr10 | 115922767 | 115934977 | 12211 | 1.49 | 4.17E-13 | 3.49E-12 |
| chr10 | 22969601 | 22980807 | 11207 | 1.03 | 4.27E-13 | 3.57E-12 |
| chr20 | 8074771 | 8076419 | 1649 | 7.73 | 4.33E-13 | 3.62E-12 |
| chr3 | 186306477 | 186308847 | 2371 | 2.68 | 4.37E-13 | 3.65E-12 |
| chr6 | 25223154 | 25230625 | 7472 | 7.72 | 4.45E-13 | 3.72E-12 |
| chr3 | 5548636 | 5550226 | 1591 | 2.18 | 4.46E-13 | 3.73E-12 |
| chr14 | 50401200 | 50513097 | 111898 | 0.41 | 4.49E-13 | 3.75E-12 |
| chr12 | 78128298 | 78137328 | 9031 | 7.74 | 4.50E-13 | 3.76E-12 |
| chr3 | 5832913 | 5842323 | 9411 | 7.72 | 4.54E-13 | 3.79E-12 |
| chr12 | 22805832 | 22836644 | 30813 | 0.91 | 4.54E-13 | 3.79E-12 |
| chr11 | 27221885 | 27240998 | 19114 | 0.93 | 4.55E-13 | 3.79E-12 |
| chr17 | 42663239 | 42686363 | 23125 | -1.1 | 4.63E-13 | 3.86E-12 |
| chr5 | 15834324 | 15836082 | 1759 | 4.2 | 4.67E-13 | 3.89E-12 |
| chr11 | 65620730 | 65641712 | 20983 | 0.69 | 4.68E-13 | 3.90E-12 |
| chr5 | 118371508 | 118374480 | 2973 | 2.6 | 4.70E-13 | 3.91E-12 |
| chr14 | 78600734 | 78603293 | 2560 | -7.67 | 4.73E-13 | 3.94E-12 |
| chr2 | 214015599 | 214017428 | 1830 | 2.53 | 4.75E-13 | 3.96E-12 |
| chr2 | 121100660 | 121118415 | 17756 | -1.03 | 4.78E-13 | 3.98E-12 |
| chr13 | 101085755 | 101096558 | 10804 | 0.93 | 4.81E-13 | 4.01E-12 |
| chr2 | 186447354 | 186449349 | 1996 | 7.7 | 4.82E-13 | 4.01E-12 |
| chr1 | 168194333 | 168231540 | 37208 | 0.6 | 4.82E-13 | 4.01E-12 |
| chr11 | 88441356 | 88444748 | 3393 | 1.04 | 4.97E-13 | 4.13E-12 |
| chr13 | 88322681 | 88339638 | 16958 | 1.12 | 5.11E-13 | 4.25E-12 |
| chr15 | 75315171 | 75353809 | 38639 | -0.57 | 5.14E-13 | 4.27E-12 |
| chr15 | 100582791 | 100584104 | 1314 | 2.88 | 5.15E-13 | 4.27E-12 |
| chr13 | 73381865 | 73387614 | 5750 | 0.95 | 5.22E-13 | 4.33E-12 |
| chr8 | 109923354 | 109928868 | 5515 | 1.7 | 5.23E-13 | 4.34E-12 |
| chr10 | 22001818 | 22038987 | 37170 | -0.9 | 5.24E-13 | 4.35E-12 |
| chr1 | 245223643 | 245224878 | 1236 | 5.57 | 5.31E-13 | 4.40E-12 |
| chr4 | 116755534 | 116766646 | 11113 | 7.7 | 5.33E-13 | 4.42E-12 |
| chr1 | 21156492 | 21233342 | 76851 | -0.46 | 5.33E-13 | 4.42E-12 |
| chrX | 25067131 | 25069731 | 2601 | 5.58 | 5.36E-13 | 4.44E-12 |
| chr18 | 59953874 | 59956500 | 2627 | 2.54 | 5.39E-13 | 4.47E-12 |
| chr11 | 80008220 | 80009995 | 1776 | 7.69 | 5.39E-13 | 4.47E-12 |
| chr16 | 59243818 | 59259073 | 15256 | 1.27 | 5.42E-13 | 4.49E-12 |
| chr14 | 39672010 | 39704529 | 32520 | 0.88 | 5.44E-13 | 4.50E-12 |
| chr2 | 1557084 | 1558684 | 1601 | 7.7 | 5.44E-13 | 4.50E-12 |
| chr1 | 220261844 | 220296065 | 34222 | 0.74 | 5.49E-13 | 4.55E-12 |
| chr16 | 21087570 | 21112061 | 24492 | 7.68 | 5.50E-13 | 4.55E-12 |
| chr4 | 83802723 | 83824050 | 21328 | 0.48 | 5.50E-13 | 4.55E-12 |
| chr8 | 38363451 | 38407444 | 43994 | 0.56 | 5.50E-13 | 4.55E-12 |
| chr6 | 139754517 | 139758284 | 3768 | 3.81 | 5.55E-13 | 4.59E-12 |
| chr11 | 14357521 | 14380902 | 23382 | 0.88 | 5.58E-13 | 4.61E-12 |
| chr5 | 75917897 | 75919853 | 1957 | -2.67 | 5.62E-13 | 4.65E-12 |
| chr12 | 15321479 | 15325064 | 3586 | 1.72 | 5.63E-13 | 4.65E-12 |
| chr12 | 88097175 | 88099044 | 1870 | 7.75 | 5.65E-13 | 4.67E-12 |
| chr15 | 83973824 | 83975930 | 2107 | 7.75 | 5.71E-13 | 4.71E-12 |
| chr5 | 57030805 | 57033039 | 2235 | -2.2 | 5.72E-13 | 4.72E-12 |
| chr12 | 51848961 | 51855318 | 6358 | 7.71 | 5.73E-13 | 4.73E-12 |
| chr3 | 188132104 | 188158205 | 26102 | 0.81 | 5.78E-13 | 4.77E-12 |
| chrX | 4534804 | 4540991 | 6188 | 4.16 | 5.80E-13 | 4.78E-12 |
| chr7 | 21619486 | 21627047 | 7562 | 7.68 | 5.91E-13 | 4.87E-12 |
| chr1 | 186320331 | 186351666 | 31336 | 0.57 | 5.93E-13 | 4.89E-12 |
| chr2 | 231480546 | 231493680 | 13135 | 1.47 | 6.01E-13 | 4.95E-12 |
| chr9 | 114628704 | 114630240 | 1537 | 2.41 | 6.06E-13 | 5.00E-12 |
| chr1 | 110161535 | 110179663 | 18129 | -0.83 | 6.08E-13 | 5.01E-12 |
| chr10 | 129894090 | 129925480 | 31391 | 0.68 | 6.11E-13 | 5.03E-12 |
| chr3 | 111577475 | 111589600 | 12126 | 0.96 | 6.11E-13 | 5.03E-12 |
| chr20 | 4487474 | 4495168 | 7695 | 1.1 | 6.12E-13 | 5.04E-12 |
| chr18 | 19570113 | 19577063 | 6951 | 1.71 | 6.15E-13 | 5.06E-12 |

| | | | | | | |
|---|---|---|---|---|---|---|
| chr6 | 17115308 | 17117525 | 2218 | 2 | 6.26E-13 | 5.15E-12 |
| chr12 | 68630184 | 68631055 | 872 | 4.13 | 6.28E-13 | 5.17E-12 |
| chr6 | 143192766 | 143208778 | 16013 | 1.69 | 6.39E-13 | 5.25E-12 |
| chr3 | 85129950 | 85131315 | 1366 | 7.68 | 6.56E-13 | 5.39E-12 |
| chr10 | 15241327 | 15244946 | 3620 | -1.16 | 6.59E-13 | 5.42E-12 |
| chr12 | 116996579 | 116998648 | 2070 | 2.55 | 6.60E-13 | 5.42E-12 |
| chr3 | 42219728 | 42227071 | 7344 | 1.21 | 6.64E-13 | 5.46E-12 |
| chr2 | 28384007 | 28396289 | 12283 | -2.9 | 6.70E-13 | 5.50E-12 |
| chr14 | 31760091 | 31761832 | 1742 | 2.8 | 6.82E-13 | 5.60E-12 |
| chr4 | 169013336 | 169020172 | 6837 | 7.67 | 6.91E-13 | 5.67E-12 |
| chr15 | 95387534 | 95388885 | 1352 | 7.66 | 6.96E-13 | 5.71E-12 |
| chr13 | 71825894 | 71827288 | 1395 | 7.65 | 6.99E-13 | 5.73E-12 |
| chr2 | 202221395 | 202222779 | 1385 | 3.76 | 6.99E-13 | 5.73E-12 |
| chr16 | 68935927 | 68971508 | 35582 | 0.69 | 7.11E-13 | 5.83E-12 |
| chr7 | 142867725 | 142870807 | 3083 | 1.12 | 7.14E-13 | 5.85E-12 |
| chr8 | 131615818 | 131624589 | 8772 | 1.25 | 7.15E-13 | 5.85E-12 |
| chr3 | 50263919 | 50308096 | 44178 | 0.5 | 7.17E-13 | 5.87E-12 |
| chr16 | 35032419 | 35044714 | 12296 | 7.77 | 7.18E-13 | 5.88E-12 |
| chr19 | 38100830 | 38103069 | 2240 | 7.65 | 7.18E-13 | 5.88E-12 |
| chr8 | 79901334 | 79903639 | 2306 | 7.74 | 7.21E-13 | 5.90E-12 |
| chr12 | 124955459 | 125058858 | 103400 | 0.39 | 7.22E-13 | 5.90E-12 |
| chr4 | 1766814 | 1781170 | 14357 | 0.84 | 7.24E-13 | 5.92E-12 |
| chr4 | 90581839 | 90582980 | 1142 | 6.53 | 7.25E-13 | 5.93E-12 |
| chr1 | 82168207 | 82173418 | 5212 | 2.07 | 7.50E-13 | 6.13E-12 |
| chr15 | 85860888 | 85887438 | 26551 | 1.1 | 7.56E-13 | 6.17E-12 |
| chr1 | 227611479 | 227621143 | 9665 | 0.58 | 7.68E-13 | 6.27E-12 |
| chr2 | 20713943 | 20717460 | 3518 | 2.08 | 7.85E-13 | 6.41E-12 |
| chr5 | 174486757 | 174487703 | 947 | -3.92 | 8.01E-13 | 6.54E-12 |
| chr9 | 72084907 | 72092232 | 7326 | 7.67 | 8.02E-13 | 6.55E-12 |
| chr11 | 112329387 | 112336882 | 7496 | 7.64 | 8.03E-13 | 6.55E-12 |
| chr6 | 133561111 | 133578654 | 17544 | -0.55 | 8.04E-13 | 6.56E-12 |
| chr1 | 222883640 | 222893092 | 9453 | 0.76 | 8.18E-13 | 6.67E-12 |
| chr17 | 52248435 | 52251086 | 2652 | 7.66 | 8.26E-13 | 6.73E-12 |
| chr17 | 79925603 | 79996035 | 70433 | -0.85 | 8.33E-13 | 6.79E-12 |
| chr20 | 4794605 | 4804886 | 10282 | -1.33 | 8.37E-13 | 6.82E-12 |
| chr19 | 58569817 | 58573497 | 3681 | 7.64 | 8.39E-13 | 6.83E-12 |
| chr15 | 78666281 | 78668316 | 2036 | 1.38 | 8.43E-13 | 6.86E-12 |
| chr13 | 75460584 | 75467039 | 6456 | -7.64 | 8.45E-13 | 6.88E-12 |
| chr14 | 93451374 | 93482903 | 31530 | -0.86 | 8.50E-13 | 6.92E-12 |
| chr1 | 88183188 | 88184976 | 1789 | 4.62 | 8.54E-13 | 6.94E-12 |
| chr7 | 129200619 | 129235951 | 35333 | 0.68 | 8.58E-13 | 6.98E-12 |
| chr14 | 25590337 | 25591753 | 1417 | -3.91 | 8.60E-13 | 7.00E-12 |
| chr14 | 27402015 | 27403059 | 1045 | 5.2 | 8.65E-13 | 7.03E-12 |
| chr6 | 73783202 | 73790206 | 7005 | -7.61 | 8.65E-13 | 7.03E-12 |
| chr11 | 78270403 | 78286836 | 16434 | 0.74 | 8.72E-13 | 7.08E-12 |
| chr22 | 21391746 | 21400812 | 9067 | -1.51 | 8.73E-13 | 7.09E-12 |
| chr13 | 42267929 | 42275722 | 7794 | 0.87 | 8.73E-13 | 7.09E-12 |
| chr19 | 50496376 | 50554974 | 58599 | 0.54 | 8.80E-13 | 7.14E-12 |
| chr9 | 74821833 | 74832748 | 10916 | 7.64 | 8.83E-13 | 7.16E-12 |
| chr3 | 104143974 | 104167028 | 23055 | 0.86 | 8.93E-13 | 7.25E-12 |
| chr16 | 57496076 | 57520480 | 24405 | 0.5 | 8.95E-13 | 7.26E-12 |
| chr8 | 69671390 | 69673740 | 2351 | -7.6 | 8.95E-13 | 7.26E-12 |
| chr8 | 103661269 | 103675368 | 14100 | 0.55 | 8.96E-13 | 7.26E-12 |
| chr5 | 139014771 | 139048718 | 33948 | 0.64 | 9.23E-13 | 7.48E-12 |
| chr7 | 100135916 | 100165001 | 29086 | 0.42 | 9.29E-13 | 7.53E-12 |
| chr6 | 136331765 | 136339752 | 7988 | -0.66 | 9.32E-13 | 7.55E-12 |
| chr14 | 103292213 | 103352678 | 60466 | -0.87 | 9.40E-13 | 7.62E-12 |
| chr4 | 78013786 | 78015244 | 1459 | 5.93 | 9.46E-13 | 7.66E-12 |
| chr5 | 171661096 | 171664435 | 3340 | 0.92 | 9.48E-13 | 7.68E-12 |
| chr12 | 49581230 | 49629865 | 48636 | 0.59 | 9.56E-13 | 7.74E-12 |
| chr5 | 133449835 | 133486705 | 36871 | -0.77 | 9.62E-13 | 7.78E-12 |

| | | | | | | |
|---|---|---|---|---|---|---|
| chr2 | 225746243 | 225793881 | 47639 | -0.57 | 9.66E-13 | 7.81E-12 |
| chr10 | 124132748 | 124146729 | 13982 | 1.06 | 9.71E-13 | 7.85E-12 |
| chr3 | 109103798 | 109116490 | 12693 | 7.66 | 9.89E-13 | 7.99E-12 |
| chr12 | 44151830 | 44153966 | 2137 | 1.2 | 9.90E-13 | 8.00E-12 |
| chr12 | 118539537 | 118542818 | 3282 | 1.26 | 9.96E-13 | 8.05E-12 |
| chr8 | 53006125 | 53031681 | 25557 | 1.28 | 1.01E-12 | 8.17E-12 |
| chr7 | 86945171 | 86949118 | 3948 | 0.93 | 1.01E-12 | 8.17E-12 |
| chr18 | 47249779 | 47252699 | 2921 | 3.14 | 1.01E-12 | 8.19E-12 |
| chr18 | 23575600 | 23577371 | 1772 | 3.13 | 1.02E-12 | 8.20E-12 |
| chr4 | 6732274 | 6785403 | 53130 | 0.59 | 1.03E-12 | 8.29E-12 |
| chr17 | 80138694 | 80261467 | 122774 | 0.29 | 1.03E-12 | 8.29E-12 |
| chr5 | 23012809 | 23014095 | 1287 | 7.63 | 1.03E-12 | 8.30E-12 |
| chr3 | 42190769 | 42206863 | 16095 | 1 | 1.03E-12 | 8.31E-12 |
| chr7 | 152430092 | 152431424 | 1333 | 4.68 | 1.04E-12 | 8.40E-12 |
| chr6 | 58767892 | 58769710 | 1819 | 7.61 | 1.05E-12 | 8.48E-12 |
| chr17 | 53653146 | 53666129 | 12984 | 1.63 | 1.05E-12 | 8.48E-12 |
| chr5 | 142119639 | 142133500 | 13862 | 7.62 | 1.06E-12 | 8.50E-12 |
| chr7 | 114053581 | 114055899 | 2319 | 7.61 | 1.06E-12 | 8.53E-12 |
| chr1 | 78608609 | 78609782 | 1174 | 3.7 | 1.07E-12 | 8.58E-12 |
| chr1 | 151137267 | 151268143 | 130877 | 0.3 | 1.09E-12 | 8.75E-12 |
| chr21 | 17484845 | 17497442 | 12598 | 0.98 | 1.09E-12 | 8.75E-12 |
| chr13 | 65213544 | 65214817 | 1274 | -3.04 | 1.10E-12 | 8.86E-12 |
| chr1 | 197924018 | 197946167 | 22150 | 7.63 | 1.11E-12 | 8.92E-12 |
| chr19 | 24288076 | 24291570 | 3495 | 7.6 | 1.11E-12 | 8.94E-12 |
| chr5 | 178486790 | 178496270 | 9481 | 7.63 | 1.12E-12 | 9.01E-12 |
| chr3 | 8972441 | 9007584 | 35144 | 0.57 | 1.13E-12 | 9.05E-12 |
| chr12 | 17804955 | 17806449 | 1495 | 1.91 | 1.13E-12 | 9.07E-12 |
| chr15 | 85985843 | 86003393 | 17551 | 0.85 | 1.13E-12 | 9.11E-12 |
| chr12 | 110868810 | 110889445 | 20636 | 0.68 | 1.14E-12 | 9.16E-12 |
| chr11 | 26577572 | 26580092 | 2521 | 7.6 | 1.14E-12 | 9.16E-12 |
| chr4 | 144279721 | 144281226 | 1506 | 2.44 | 1.15E-12 | 9.26E-12 |
| chr2 | 4720011 | 4733172 | 13162 | 7.59 | 1.16E-12 | 9.29E-12 |
| chr7 | 115563752 | 115565563 | 1812 | 7.63 | 1.16E-12 | 9.31E-12 |
| chr6 | 75977204 | 75996285 | 19082 | 0.7 | 1.16E-12 | 9.32E-12 |
| chr1 | 191463612 | 191464582 | 971 | 5.27 | 1.16E-12 | 9.33E-12 |
| chr19 | 13840329 | 13860252 | 19924 | 0.62 | 1.17E-12 | 9.34E-12 |
| chr5 | 52961594 | 52962784 | 1191 | 2.9 | 1.17E-12 | 9.36E-12 |
| chr3 | 43811246 | 43815781 | 4536 | -1.04 | 1.18E-12 | 9.48E-12 |
| chr2 | 68821416 | 68822477 | 1062 | 5.14 | 1.18E-12 | 9.48E-12 |
| chr2 | 174218577 | 174225535 | 6959 | -1.23 | 1.19E-12 | 9.49E-12 |
| chr2 | 30367626 | 30372983 | 5358 | 1 | 1.19E-12 | 9.50E-12 |
| chr15 | 87172121 | 87173724 | 1604 | 7.59 | 1.20E-12 | 9.57E-12 |
| chr8 | 117363279 | 117365835 | 2557 | 7.59 | 1.20E-12 | 9.57E-12 |
| chr7 | 36071703 | 36078274 | 6572 | 1.28 | 1.20E-12 | 9.61E-12 |
| chr9 | 119599155 | 119601490 | 2336 | 1.93 | 1.21E-12 | 9.65E-12 |
| chrX | 81841189 | 81845479 | 4291 | 3.84 | 1.21E-12 | 9.69E-12 |
| chr1 | 206847906 | 206898314 | 50409 | -0.49 | 1.21E-12 | 9.69E-12 |
| chr14 | 53149308 | 53197872 | 48565 | 0.48 | 1.21E-12 | 9.70E-12 |
| chr3 | 49522584 | 49542838 | 20255 | 0.59 | 1.22E-12 | 9.70E-12 |
| chr8 | 117396209 | 117397323 | 1115 | 7.59 | 1.22E-12 | 9.70E-12 |
| chr7 | 105702590 | 105713170 | 10581 | 1.7 | 1.22E-12 | 9.70E-12 |
| chr3 | 114041613 | 114044441 | 2829 | 2 | 1.22E-12 | 9.75E-12 |
| chr10 | 104844186 | 104959166 | 114981 | 0.28 | 1.22E-12 | 9.75E-12 |
| chr2 | 242574853 | 242598847 | 23995 | -1.02 | 1.23E-12 | 9.78E-12 |
| chr11 | 110018124 | 110030796 | 12673 | -1.26 | 1.23E-12 | 9.81E-12 |
| chr4 | 108200249 | 108201939 | 1691 | 3.25 | 1.24E-12 | 9.89E-12 |
| chr9 | 118245941 | 118258334 | 12394 | 7.63 | 1.25E-12 | 9.95E-12 |
| chr3 | 113005199 | 113007058 | 1860 | -1.27 | 1.25E-12 | 9.97E-12 |
| chr2 | 111764520 | 111766449 | 1930 | 1.93 | 1.26E-12 | 1.01E-11 |
| chr3 | 103998386 | 104011138 | 12753 | 7.59 | 1.28E-12 | 1.02E-11 |
| chr5 | 151878561 | 151880867 | 2307 | 7.58 | 1.29E-12 | 1.02E-11 |

| | | | | | | |
|---|---|---|---|---|---|---|
| chr1 | 25188130 | 25189862 | 1733 | 2.11 | 1.30E-12 | 1.03E-11 |
| chr5 | 25558598 | 25559713 | 1116 | 5.9 | 1.30E-12 | 1.03E-11 |
| chr17 | 78666679 | 78680721 | 14043 | -1.05 | 1.31E-12 | 1.04E-11 |
| chr4 | 37431497 | 37433418 | 1922 | 7.59 | 1.32E-12 | 1.05E-11 |
| chr15 | 36448367 | 36455091 | 6725 | 1.36 | 1.32E-12 | 1.05E-11 |
| chr2 | 28803472 | 28820469 | 16998 | -0.67 | 1.32E-12 | 1.05E-11 |
| chr14 | 64219131 | 64220340 | 1210 | 3.14 | 1.33E-12 | 1.06E-11 |
| chr15 | 92691098 | 92708452 | 17355 | 1.45 | 1.33E-12 | 1.06E-11 |
| chr2 | 18479912 | 18482847 | 2936 | -1.06 | 1.33E-12 | 1.06E-11 |
| chr1 | 184561667 | 184564209 | 2543 | -7.55 | 1.36E-12 | 1.08E-11 |
| chr8 | 102618230 | 102621946 | 3717 | 0.91 | 1.38E-12 | 1.09E-11 |
| chr20 | 52185888 | 52243428 | 57541 | 0.31 | 1.38E-12 | 1.09E-11 |
| chr5 | 100641262 | 100644014 | 2753 | 1.2 | 1.40E-12 | 1.11E-11 |
| chr18 | 8115627 | 8123516 | 7890 | -1.01 | 1.41E-12 | 1.11E-11 |
| chr3 | 145936489 | 145941597 | 5109 | 0.88 | 1.42E-12 | 1.13E-11 |
| chr8 | 83907412 | 83909751 | 2340 | 7.57 | 1.45E-12 | 1.15E-11 |
| chr1 | 44996971 | 44999323 | 2353 | -1.67 | 1.46E-12 | 1.16E-11 |
| chr7 | 150062583 | 150077571 | 14989 | 0.47 | 1.47E-12 | 1.17E-11 |
| chr4 | 74461509 | 74486951 | 25443 | 0.64 | 1.49E-12 | 1.18E-11 |
| chr3 | 18372592 | 18374203 | 1612 | 7.57 | 1.49E-12 | 1.18E-11 |
| chr3 | 193560890 | 193571976 | 11087 | 1.76 | 1.51E-12 | 1.19E-11 |
| chr11 | 88511988 | 88513482 | 1495 | 4.69 | 1.51E-12 | 1.19E-11 |
| chr7 | 26852818 | 26854241 | 1424 | 5.25 | 1.51E-12 | 1.19E-11 |
| chr7 | 90868130 | 90870375 | 2246 | 7.56 | 1.51E-12 | 1.20E-11 |
| chr9 | 132447132 | 132465532 | 18401 | -2.56 | 1.52E-12 | 1.20E-11 |
| chr3 | 102698026 | 102699750 | 1725 | 7.58 | 1.53E-12 | 1.21E-11 |
| chr7 | 1542567 | 1611574 | 69008 | 0.35 | 1.56E-12 | 1.23E-11 |
| chr18 | 33014698 | 33018989 | 4292 | 1.52 | 1.56E-12 | 1.23E-11 |
| chrX | 106137547 | 106142962 | 5416 | -1.21 | 1.56E-12 | 1.23E-11 |
| chr12 | 68099136 | 68105200 | 6065 | 7.62 | 1.57E-12 | 1.24E-11 |
| chr17 | 2092650 | 2108091 | 15442 | -0.84 | 1.59E-12 | 1.25E-11 |
| chr1 | 10657723 | 10661180 | 3458 | 1.07 | 1.60E-12 | 1.26E-11 |
| chr4 | 124317437 | 124340650 | 23214 | 1.08 | 1.61E-12 | 1.27E-11 |
| chr19 | 54099983 | 54102022 | 2040 | 7.56 | 1.61E-12 | 1.27E-11 |
| chr15 | 90489633 | 90490705 | 1073 | 4.26 | 1.61E-12 | 1.27E-11 |
| chr4 | 15066297 | 15080719 | 14423 | 1.37 | 1.62E-12 | 1.28E-11 |
| chr19 | 20467969 | 20477428 | 9460 | 7.56 | 1.62E-12 | 1.28E-11 |
| chr7 | 143168825 | 143170204 | 1380 | 5.17 | 1.63E-12 | 1.28E-11 |
| chr17 | 59215738 | 59217403 | 1666 | 2.12 | 1.63E-12 | 1.29E-11 |
| chr3 | 29646794 | 29648860 | 2067 | -1.37 | 1.64E-12 | 1.29E-11 |
| chr10 | 3781730 | 3783425 | 1696 | -1.86 | 1.65E-12 | 1.29E-11 |
| chr1 | 240759396 | 240761326 | 1931 | 2.06 | 1.65E-12 | 1.30E-11 |
| chr14 | 91579747 | 91638666 | 58920 | -0.64 | 1.67E-12 | 1.31E-11 |
| chr7 | 43012003 | 43013373 | 1371 | 3.09 | 1.67E-12 | 1.32E-11 |
| chr10 | 94349665 | 94413966 | 64302 | 0.39 | 1.68E-12 | 1.32E-11 |
| chr17 | 53163785 | 53168709 | 4925 | -1.06 | 1.70E-12 | 1.33E-11 |
| chr19 | 57813127 | 57815221 | 2095 | 4.86 | 1.70E-12 | 1.33E-11 |
| chr14 | 81099954 | 81109496 | 9543 | -0.57 | 1.70E-12 | 1.34E-11 |
| chr2 | 196227520 | 196230420 | 2901 | 1.14 | 1.71E-12 | 1.34E-11 |
| chr7 | 110869576 | 110879663 | 10088 | 0.9 | 1.72E-12 | 1.35E-11 |
| chr3 | 41306444 | 41310905 | 4462 | -1.28 | 1.72E-12 | 1.35E-11 |
| chr13 | 109894785 | 109928780 | 33996 | 0.4 | 1.72E-12 | 1.35E-11 |
| chr17 | 46313555 | 46347216 | 33662 | 0.29 | 1.73E-12 | 1.35E-11 |
| chr18 | 26703263 | 26704544 | 1282 | 2.5 | 1.73E-12 | 1.35E-11 |
| chr8 | 130685732 | 130739148 | 53417 | -0.55 | 1.73E-12 | 1.35E-11 |
| chrX | 35681191 | 35684309 | 3119 | 2.63 | 1.75E-12 | 1.37E-11 |
| chr5 | 84588830 | 84589857 | 1028 | 7.56 | 1.75E-12 | 1.37E-11 |
| chr19 | 52674038 | 52675499 | 1462 | 2.17 | 1.77E-12 | 1.38E-11 |
| chr1 | 81059573 | 81060904 | 1332 | 5.17 | 1.77E-12 | 1.39E-11 |
| chr13 | 102275791 | 102281166 | 5376 | 0.8 | 1.78E-12 | 1.39E-11 |
| chr5 | 23418914 | 23421459 | 2546 | 7.55 | 1.79E-12 | 1.40E-11 |

| | | | | | | |
|---|---|---|---|---|---|---|
| chr5 | 59758717 | 59760468 | 1752 | -7.51 | 1.80E-12 | 1.41E-11 |
| chr2 | 150733845 | 150735361 | 1517 | 7.55 | 1.81E-12 | 1.41E-11 |
| chr14 | 79980692 | 79982767 | 2076 | -2.77 | 1.84E-12 | 1.44E-11 |
| chr4 | 180335797 | 180337589 | 1793 | 7.55 | 1.85E-12 | 1.44E-11 |
| chr7 | 100625819 | 100628368 | 2550 | 2.14 | 1.86E-12 | 1.45E-11 |
| chr6 | 134741045 | 134758758 | 17714 | 1.11 | 1.86E-12 | 1.46E-11 |
| chr1 | 210395195 | 210413323 | 18129 | 7.56 | 1.89E-12 | 1.48E-11 |
| chr6 | 140294776 | 140304936 | 10161 | 0.95 | 1.90E-12 | 1.48E-11 |
| chr2 | 190699426 | 190709887 | 10462 | -0.91 | 1.91E-12 | 1.49E-11 |
| chr14 | 94877255 | 94878885 | 1631 | 7.54 | 1.92E-12 | 1.50E-11 |
| chr2 | 205764273 | 205766898 | 2626 | -1.7 | 1.92E-12 | 1.50E-11 |
| chr1 | 40605208 | 40607882 | 2675 | 1.46 | 1.95E-12 | 1.52E-11 |
| chr5 | 59038941 | 59096025 | 57085 | 0.29 | 1.96E-12 | 1.53E-11 |
| chr12 | 11947877 | 11949954 | 2078 | 1.38 | 1.98E-12 | 1.54E-11 |
| chr1 | 246990203 | 246991534 | 1332 | 7.54 | 2.00E-12 | 1.56E-11 |
| chr3 | 160980752 | 160993890 | 13139 | 1.29 | 2.01E-12 | 1.57E-11 |
| chr11 | 107701008 | 107713888 | 12881 | 0.83 | 2.02E-12 | 1.58E-11 |
| chr1 | 208352749 | 208359033 | 6285 | 1.72 | 2.03E-12 | 1.58E-11 |
| chr12 | 123400576 | 123470017 | 69442 | 0.48 | 2.08E-12 | 1.62E-11 |
| chr10 | 63546011 | 63564664 | 18654 | 1.98 | 2.09E-12 | 1.63E-11 |
| chr12 | 133463409 | 133465705 | 2297 | 1.06 | 2.11E-12 | 1.64E-11 |
| chr2 | 72179616 | 72180689 | 1074 | 3.44 | 2.12E-12 | 1.65E-11 |
| chr1 | 172136757 | 172151311 | 14555 | 7.52 | 2.17E-12 | 1.69E-11 |
| chr5 | 140464242 | 140485566 | 21325 | 7.54 | 2.17E-12 | 1.69E-11 |
| chr7 | 44644939 | 44687960 | 43022 | 0.45 | 2.19E-12 | 1.70E-11 |
| chr7 | 103300250 | 103302108 | 1859 | 1.87 | 2.19E-12 | 1.70E-11 |
| chr8 | 127338318 | 127340430 | 2113 | 7.52 | 2.20E-12 | 1.71E-11 |
| chr10 | 34529356 | 34829672 | 300317 | -0.2 | 2.20E-12 | 1.71E-11 |
| chr4 | 52885733 | 52925919 | 40187 | 1.84 | 2.23E-12 | 1.73E-11 |
| chr3 | 95746016 | 95747311 | 1296 | 7.53 | 2.23E-12 | 1.74E-11 |
| chr7 | 130479018 | 130480105 | 1088 | 1.9 | 2.26E-12 | 1.75E-11 |
| chr8 | 83060003 | 83063206 | 3204 | 7.54 | 2.26E-12 | 1.76E-11 |
| chr1 | 157209235 | 157211279 | 2045 | 4.83 | 2.27E-12 | 1.76E-11 |
| chr8 | 81344557 | 81349408 | 4852 | 2.72 | 2.28E-12 | 1.77E-11 |
| chr22 | 37977112 | 38006278 | 29167 | 0.79 | 2.29E-12 | 1.78E-11 |
| chr2 | 165863510 | 165866173 | 2664 | 7.51 | 2.30E-12 | 1.78E-11 |
| chr3 | 94364626 | 94368881 | 4256 | 1.54 | 2.32E-12 | 1.80E-11 |
| chr1 | 18988215 | 18998290 | 10076 | 0.62 | 2.33E-12 | 1.81E-11 |
| chr4 | 129259333 | 129280414 | 21082 | -0.55 | 2.35E-12 | 1.82E-11 |
| chr2 | 20516356 | 20552126 | 35771 | -0.55 | 2.36E-12 | 1.83E-11 |
| chr4 | 134544388 | 134547081 | 2694 | 1.57 | 2.36E-12 | 1.83E-11 |
| chr8 | 30110078 | 30115959 | 5882 | 1.12 | 2.39E-12 | 1.85E-11 |
| chr6 | 113753065 | 113763802 | 10738 | 1.67 | 2.40E-12 | 1.86E-11 |
| chr5 | 56327162 | 56329722 | 2561 | 2.01 | 2.43E-12 | 1.88E-11 |
| chr19 | 12175519 | 12178129 | 2611 | 7.52 | 2.44E-12 | 1.89E-11 |
| chr20 | 49154222 | 49158647 | 4426 | 1.22 | 2.45E-12 | 1.89E-11 |
| chr5 | 1469573 | 1568977 | 99405 | 0.53 | 2.47E-12 | 1.91E-11 |
| chr12 | 83028191 | 83030344 | 2154 | 1.63 | 2.48E-12 | 1.92E-11 |
| chr12 | 88122616 | 88123871 | 1256 | 7.53 | 2.50E-12 | 1.93E-11 |
| chr8 | 134510822 | 134514467 | 3646 | 1.69 | 2.54E-12 | 1.96E-11 |
| chr4 | 31343076 | 31344596 | 1521 | 7.5 | 2.55E-12 | 1.97E-11 |
| chr7 | 15651838 | 15666157 | 14320 | 7.5 | 2.58E-12 | 1.99E-11 |
| chr8 | 130798751 | 130800113 | 1363 | 2.01 | 2.62E-12 | 2.02E-11 |
| chr4 | 38450481 | 38453080 | 2600 | 7.5 | 2.63E-12 | 2.03E-11 |
| chr17 | 60969474 | 60971545 | 2072 | 1.06 | 2.65E-12 | 2.04E-11 |
| chr3 | 133156155 | 133172874 | 16720 | 1.1 | 2.67E-12 | 2.06E-11 |
| chr18 | 5237015 | 5239304 | 2290 | 1.41 | 2.78E-12 | 2.15E-11 |
| chr8 | 101911851 | 101967475 | 55625 | 0.39 | 2.79E-12 | 2.15E-11 |
| chr19 | 53966147 | 53971751 | 5605 | 7.5 | 2.79E-12 | 2.15E-11 |
| chr14 | 31417443 | 31561725 | 144283 | 0.36 | 2.81E-12 | 2.16E-11 |
| chr11 | 114268780 | 114273554 | 4775 | 0.62 | 2.82E-12 | 2.17E-11 |

| | | | | | | |
|---|---|---|---|---|---|---|
| chr2 | 170929175 | 170931196 | 2022 | 7.49 | 2.84E-12 | 2.19E-11 |
| chr2 | 82122023 | 82131151 | 9129 | 7.49 | 2.88E-12 | 2.21E-11 |
| chr7 | 47210843 | 47213994 | 3152 | 7.54 | 2.90E-12 | 2.23E-11 |
| chr1 | 116324794 | 116326459 | 1666 | 2.84 | 2.93E-12 | 2.25E-11 |
| chr6 | 16575470 | 16577799 | 2330 | 1.73 | 2.94E-12 | 2.26E-11 |
| chr5 | 30849395 | 30851050 | 1656 | 7.56 | 2.95E-12 | 2.27E-11 |
| chr7 | 138779054 | 138819556 | 40503 | 0.5 | 2.95E-12 | 2.27E-11 |
| chr3 | 181556903 | 181569825 | 12923 | 1.29 | 2.95E-12 | 2.27E-11 |
| chr8 | 120075101 | 120076471 | 1371 | 7.49 | 3.00E-12 | 2.30E-11 |
| chr14 | 31077586 | 31099247 | 21662 | 0.65 | 3.00E-12 | 2.30E-11 |
| chr15 | 45540321 | 45547754 | 7434 | 1.71 | 3.00E-12 | 2.31E-11 |
| chr8 | 22406096 | 22463872 | 57777 | 0.4 | 3.01E-12 | 2.31E-11 |
| chr19 | 1150963 | 1215676 | 64714 | 0.42 | 3.01E-12 | 2.31E-11 |
| chr15 | 57410177 | 57416782 | 6606 | 1.14 | 3.03E-12 | 2.32E-11 |
| chr9 | 118021819 | 118026131 | 4313 | 0.7 | 3.05E-12 | 2.34E-11 |
| chr12 | 54260183 | 54264304 | 4122 | 0.85 | 3.05E-12 | 2.34E-11 |
| chr10 | 63730040 | 63730980 | 941 | 5.84 | 3.09E-12 | 2.37E-11 |
| chr8 | 130922923 | 131029768 | 106846 | -0.26 | 3.19E-12 | 2.44E-11 |
| chr2 | 28169144 | 28219566 | 50423 | -0.46 | 3.19E-12 | 2.45E-11 |
| chr12 | 22696112 | 22703630 | 7519 | 0.97 | 3.20E-12 | 2.45E-11 |
| chr5 | 100158323 | 100178090 | 19768 | 2.36 | 3.20E-12 | 2.45E-11 |
| chr6 | 49490410 | 49538585 | 48176 | 0.57 | 3.20E-12 | 2.45E-11 |
| chr1 | 213141387 | 213149512 | 8126 | -5.16 | 3.22E-12 | 2.46E-11 |
| chr12 | 19281390 | 19285051 | 3662 | 1.16 | 3.29E-12 | 2.52E-11 |
| chr7 | 129583330 | 129618122 | 34793 | 0.47 | 3.31E-12 | 2.53E-11 |
| chr9 | 115030341 | 115111422 | 81082 | 0.38 | 3.33E-12 | 2.55E-11 |
| chr9 | 119690178 | 119691989 | 1812 | 7.48 | 3.38E-12 | 2.58E-11 |
| chr12 | 29532369 | 29535124 | 2756 | 1.17 | 3.44E-12 | 2.63E-11 |
| chr17 | 37885481 | 37911805 | 26325 | 0.61 | 3.47E-12 | 2.65E-11 |
| chr12 | 111970390 | 112039923 | 69534 | 0.42 | 3.54E-12 | 2.71E-11 |
| chrX | 92950683 | 92952046 | 1364 | -7.45 | 3.55E-12 | 2.71E-11 |
| chr9 | 15295335 | 15307909 | 12575 | 1.22 | 3.56E-12 | 2.72E-11 |
| chr4 | 118005847 | 118017314 | 11468 | 7.54 | 3.57E-12 | 2.72E-11 |
| chr20 | 3336414 | 3338132 | 1719 | 1.59 | 3.58E-12 | 2.73E-11 |
| chr3 | 11758349 | 11763084 | 4736 | 0.78 | 3.58E-12 | 2.74E-11 |
| chr19 | 53289325 | 53291258 | 1934 | 7.46 | 3.61E-12 | 2.75E-11 |
| chr8 | 10568176 | 10569429 | 1254 | -5.74 | 3.61E-12 | 2.76E-11 |
| chr15 | 58919836 | 58921678 | 1843 | 1.15 | 3.63E-12 | 2.77E-11 |
| chr9 | 87610756 | 87611787 | 1032 | 5.11 | 3.71E-12 | 2.83E-11 |
| chr6 | 48396508 | 48400117 | 3610 | 0.85 | 3.71E-12 | 2.83E-11 |
| chr2 | 169894380 | 169908213 | 13834 | 1.57 | 3.71E-12 | 2.83E-11 |
| chr5 | 3310704 | 3311980 | 1277 | 7.46 | 3.75E-12 | 2.86E-11 |
| chr13 | 34213356 | 34224321 | 10966 | -1.15 | 3.76E-12 | 2.87E-11 |
| chr12 | 116962415 | 116978309 | 15895 | -0.65 | 3.77E-12 | 2.87E-11 |
| chr1 | 32038547 | 32043143 | 4597 | 1.27 | 3.77E-12 | 2.87E-11 |
| chr1 | 98468838 | 98476174 | 7337 | 0.99 | 3.83E-12 | 2.91E-11 |
| chr6 | 134784253 | 134790660 | 6408 | 1.01 | 3.83E-12 | 2.92E-11 |
| chr4 | 71599151 | 71606093 | 6943 | 1.34 | 3.88E-12 | 2.96E-11 |
| chr9 | 117438085 | 117454396 | 16312 | 0.78 | 3.89E-12 | 2.96E-11 |
| chr6 | 123109863 | 123120245 | 10383 | 1.5 | 3.99E-12 | 3.03E-11 |
| chr14 | 70462277 | 70515596 | 53320 | 0.51 | 4.02E-12 | 3.06E-11 |
| chr4 | 10034260 | 10048211 | 13952 | 7.45 | 4.06E-12 | 3.08E-11 |
| chr17 | 69405094 | 69421110 | 16017 | 5.8 | 4.10E-12 | 3.11E-11 |
| chr6 | 18431311 | 18435481 | 4171 | 3.59 | 4.10E-12 | 3.12E-11 |
| chr1 | 194119772 | 194120880 | 1109 | 2.7 | 4.14E-12 | 3.14E-11 |
| chr19 | 33109231 | 33167065 | 57835 | -0.44 | 4.18E-12 | 3.17E-11 |
| chr20 | 17917676 | 17957851 | 40176 | 0.37 | 4.20E-12 | 3.19E-11 |
| chr15 | 39418963 | 39446389 | 27427 | -0.49 | 4.23E-12 | 3.21E-11 |
| chr12 | 104040655 | 104047273 | 6619 | 7.45 | 4.24E-12 | 3.21E-11 |
| chr2 | 62867841 | 62901335 | 33495 | -0.62 | 4.28E-12 | 3.25E-11 |
| chr20 | 10090999 | 10092995 | 1997 | 1.82 | 4.30E-12 | 3.26E-11 |

| | | | | | | |
|---|---|---|---|---|---|---|
| chr13 | 73706971 | 73716382 | 9412 | 0.8 | 4.31E-12 | 3.27E-11 |
| chr12 | 63589374 | 63601556 | 12183 | 7.44 | 4.32E-12 | 3.27E-11 |
| chr17 | 35797520 | 35878535 | 81016 | 0.36 | 4.35E-12 | 3.30E-11 |
| chr10 | 3844774 | 3873876 | 29103 | 0.43 | 4.41E-12 | 3.34E-11 |
| chr15 | 76578626 | 76630217 | 51592 | -0.4 | 4.44E-12 | 3.36E-11 |
| chr19 | 35557112 | 35558511 | 1400 | 2.31 | 4.50E-12 | 3.40E-11 |
| chr16 | 13203431 | 13205343 | 1913 | -7.41 | 4.51E-12 | 3.41E-11 |
| chr10 | 65312220 | 65315141 | 2922 | 1.35 | 4.52E-12 | 3.42E-11 |
| chr6 | 139908447 | 139933067 | 24621 | 1.87 | 4.52E-12 | 3.42E-11 |
| chr8 | 118661986 | 118663517 | 1532 | 2.47 | 4.52E-12 | 3.42E-11 |
| chr12 | 116275646 | 116278590 | 2945 | -7.42 | 4.52E-12 | 3.42E-11 |
| chr1 | 189649684 | 189651473 | 1790 | 1.97 | 4.54E-12 | 3.43E-11 |
| chr13 | 102300950 | 102302686 | 1737 | 1.45 | 4.56E-12 | 3.45E-11 |
| chr20 | 43306537 | 43326007 | 19471 | -0.7 | 4.61E-12 | 3.48E-11 |
| chr4 | 36226422 | 36228043 | 1622 | 7.45 | 4.62E-12 | 3.49E-11 |
| chr1 | 164670346 | 164677591 | 7246 | -1.77 | 4.65E-12 | 3.51E-11 |
| chr3 | 156498356 | 156545966 | 47611 | 0.26 | 4.65E-12 | 3.51E-11 |
| chr18 | 19242688 | 19267546 | 24859 | 0.62 | 4.66E-12 | 3.52E-11 |
| chr12 | 71209142 | 71210401 | 1260 | 7.46 | 4.67E-12 | 3.52E-11 |
| chr6 | 43957413 | 43980543 | 23131 | 0.8 | 4.74E-12 | 3.58E-11 |
| chr7 | 110158484 | 110175611 | 17128 | 0.53 | 4.76E-12 | 3.59E-11 |
| chr1 | 95320130 | 95331377 | 11248 | 1.11 | 4.77E-12 | 3.59E-11 |
| chr1 | 113748476 | 113761063 | 12588 | 2.12 | 4.78E-12 | 3.61E-11 |
| chr7 | 148143733 | 148145349 | 1617 | 6.36 | 4.80E-12 | 3.62E-11 |
| chr5 | 142092268 | 142101393 | 9126 | 1.36 | 4.81E-12 | 3.62E-11 |
| chr4 | 26984444 | 26986237 | 1794 | 2.27 | 4.82E-12 | 3.63E-11 |
| chr7 | 36244125 | 36249213 | 5089 | -0.95 | 4.84E-12 | 3.64E-11 |
| chr12 | 53984591 | 54021623 | 37033 | 0.64 | 4.86E-12 | 3.66E-11 |
| chr10 | 51542546 | 51543931 | 1386 | 2.82 | 4.89E-12 | 3.68E-11 |
| chr22 | 48730730 | 48768866 | 38137 | -0.52 | 4.89E-12 | 3.68E-11 |
| chr3 | 174484939 | 174486715 | 1777 | 7.42 | 4.89E-12 | 3.68E-11 |
| chr12 | 118423761 | 118425438 | 1678 | 2.34 | 4.94E-12 | 3.71E-11 |
| chr8 | 77910804 | 77914057 | 3254 | 0.6 | 4.95E-12 | 3.72E-11 |
| chr14 | 27308861 | 27316120 | 7260 | 0.57 | 4.99E-12 | 3.75E-11 |
| chr4 | 174040305 | 174042942 | 2638 | 3.06 | 5.04E-12 | 3.79E-11 |
| chr3 | 177645442 | 177647224 | 1783 | 1.77 | 5.05E-12 | 3.79E-11 |
| chr2 | 44108894 | 44148480 | 39587 | -0.52 | 5.06E-12 | 3.80E-11 |
| chr7 | 73497394 | 73514877 | 17484 | 0.49 | 5.07E-12 | 3.80E-11 |
| chr11 | 134681431 | 134689375 | 7945 | 7.45 | 5.18E-12 | 3.89E-11 |
| chr19 | 21578446 | 21581097 | 2652 | 7.42 | 5.19E-12 | 3.89E-11 |
| chr2 | 194052431 | 194053600 | 1170 | 2.04 | 5.20E-12 | 3.90E-11 |
| chr1 | 112932850 | 112940314 | 7465 | 0.74 | 5.25E-12 | 3.94E-11 |
| chr5 | 129811539 | 129812891 | 1353 | 3.71 | 5.31E-12 | 3.98E-11 |
| chr12 | 69777451 | 69812295 | 34845 | -0.56 | 5.34E-12 | 4.00E-11 |
| chr15 | 96226868 | 96229171 | 2304 | 1.29 | 5.38E-12 | 4.03E-11 |
| chr3 | 104032799 | 104042905 | 10107 | 7.43 | 5.62E-12 | 4.21E-11 |
| chr1 | 151298603 | 151325607 | 27005 | 0.71 | 5.63E-12 | 4.22E-11 |
| chr2 | 98896013 | 98897736 | 1724 | 2.09 | 5.70E-12 | 4.27E-11 |
| chr5 | 66699241 | 66702220 | 2980 | -1.19 | 5.72E-12 | 4.28E-11 |
| chr3 | 188062501 | 188085008 | 22508 | 0.95 | 5.81E-12 | 4.35E-11 |
| chr1 | 100451595 | 100459319 | 7725 | 1.75 | 5.90E-12 | 4.41E-11 |
| chr1 | 171282497 | 171284307 | 1811 | 1.35 | 5.90E-12 | 4.41E-11 |
| chr2 | 142974995 | 142976558 | 1564 | 5.79 | 5.93E-12 | 4.43E-11 |
| chrX | 14080473 | 14092259 | 11787 | 1.6 | 5.98E-12 | 4.47E-11 |
| chr12 | 40718776 | 40736491 | 17716 | 1.62 | 6.03E-12 | 4.51E-11 |
| chr4 | 59899269 | 59900640 | 1372 | 2.09 | 6.04E-12 | 4.52E-11 |
| chr21 | 37691569 | 37712568 | 21000 | 0.56 | 6.06E-12 | 4.53E-11 |
| chr17 | 59628747 | 59660589 | 31843 | 0.36 | 6.07E-12 | 4.53E-11 |
| chr7 | 158373965 | 158375521 | 1557 | 7.4 | 6.07E-12 | 4.53E-11 |
| chr7 | 36951035 | 36953272 | 2238 | 7.41 | 6.13E-12 | 4.57E-11 |
| chr18 | 74918924 | 74921589 | 2666 | 1.87 | 6.16E-12 | 4.59E-11 |

| | | | | | | |
|---|---|---|---|---|---|---|
| chr2 | 46290294 | 46303498 | 13205 | -0.67 | 6.16E-12 | 4.60E-11 |
| chr8 | 77589765 | 77594836 | 5072 | 1.8 | 6.21E-12 | 4.63E-11 |
| chr5 | 90250586 | 90252902 | 2317 | 2.71 | 6.31E-12 | 4.71E-11 |
| chr1 | 233493174 | 233518145 | 24972 | -0.55 | 6.33E-12 | 4.72E-11 |
| chr4 | 150510268 | 150511575 | 1308 | 7.43 | 6.33E-12 | 4.72E-11 |
| chr10 | 29106463 | 29131306 | 24844 | 1.01 | 6.37E-12 | 4.74E-11 |
| chr1 | 68442627 | 68454902 | 12276 | 0.65 | 6.37E-12 | 4.75E-11 |
| chr3 | 50328403 | 50397511 | 69109 | 0.47 | 6.39E-12 | 4.76E-11 |
| chr14 | 27546082 | 27548025 | 1944 | 7.39 | 6.39E-12 | 4.76E-11 |
| chr6 | 52588753 | 52591968 | 3216 | 1.23 | 6.41E-12 | 4.77E-11 |
| chr4 | 21013746 | 21015547 | 1802 | 7.4 | 6.42E-12 | 4.78E-11 |
| chr6 | 131638632 | 131640428 | 1797 | 5.35 | 6.43E-12 | 4.78E-11 |
| chr6 | 3911175 | 3946546 | 35372 | 0.71 | 6.47E-12 | 4.81E-11 |
| chr14 | 81790067 | 81791223 | 1157 | 2.13 | 6.54E-12 | 4.87E-11 |
| chr14 | 89214985 | 89218028 | 3044 | -2.01 | 6.55E-12 | 4.87E-11 |
| chr3 | 87210201 | 87217177 | 6977 | 7.44 | 6.69E-12 | 4.98E-11 |
| chr2 | 153191138 | 153212773 | 21636 | -1.1 | 6.71E-12 | 4.98E-11 |
| chr5 | 71217709 | 71236215 | 18507 | 0.45 | 6.86E-12 | 5.10E-11 |
| chr11 | 84348106 | 84357914 | 9809 | 7.39 | 6.89E-12 | 5.12E-11 |
| chr17 | 54185748 | 54189084 | 3337 | 2.3 | 6.90E-12 | 5.12E-11 |
| chr8 | 49827233 | 49834959 | 7727 | 7.49 | 6.92E-12 | 5.14E-11 |
| chr4 | 59167026 | 59168380 | 1355 | 7.43 | 6.96E-12 | 5.17E-11 |
| chr9 | 116443711 | 116446463 | 2753 | 1.22 | 6.98E-12 | 5.18E-11 |
| chr1 | 8317825 | 8320512 | 2688 | 1.56 | 6.98E-12 | 5.18E-11 |
| chr5 | 120691664 | 120695577 | 3914 | 1.2 | 7.00E-12 | 5.19E-11 |
| chr2 | 39209504 | 39232486 | 22983 | -0.88 | 7.04E-12 | 5.22E-11 |
| chr6 | 94494244 | 94495957 | 1714 | 7.39 | 7.07E-12 | 5.24E-11 |
| chr2 | 57033212 | 57040685 | 7474 | 7.42 | 7.10E-12 | 5.26E-11 |
| chr12 | 32903987 | 32909568 | 5582 | 0.77 | 7.12E-12 | 5.27E-11 |
| chr1 | 81099815 | 81106998 | 7184 | 7.4 | 7.13E-12 | 5.28E-11 |
| chr3 | 82976093 | 82977832 | 1740 | 7.38 | 7.22E-12 | 5.35E-11 |
| chr13 | 103552241 | 103554451 | 2211 | 7.39 | 7.24E-12 | 5.36E-11 |
| chr1 | 159879230 | 159896106 | 16877 | 0.47 | 7.24E-12 | 5.36E-11 |
| chr3 | 85830069 | 85832024 | 1956 | 7.39 | 7.29E-12 | 5.39E-11 |
| chr12 | 70635321 | 70665678 | 30358 | 0.58 | 7.29E-12 | 5.39E-11 |
| chr8 | 102339428 | 102399259 | 59832 | 0.29 | 7.41E-12 | 5.48E-11 |
| chr4 | 124399421 | 124410920 | 11500 | 1.76 | 7.46E-12 | 5.52E-11 |
| chr19 | 12034865 | 12036695 | 1831 | 6.32 | 7.47E-12 | 5.53E-11 |
| chr12 | 72055184 | 72059615 | 4432 | 0.75 | 7.51E-12 | 5.55E-11 |
| chr15 | 35773400 | 35798967 | 25568 | 1.38 | 7.52E-12 | 5.56E-11 |
| chr14 | 102087931 | 102095497 | 7567 | -2.05 | 7.56E-12 | 5.59E-11 |
| chr2 | 99894834 | 99898119 | 3286 | 7.37 | 7.68E-12 | 5.67E-11 |
| chr13 | 23318568 | 23319500 | 933 | 3.25 | 7.78E-12 | 5.75E-11 |
| chr1 | 98261816 | 98263115 | 1300 | 2.55 | 7.84E-12 | 5.79E-11 |
| chr12 | 26110662 | 26113123 | 2462 | 1.24 | 7.95E-12 | 5.87E-11 |
| chr13 | 106803015 | 106805017 | 2003 | 1.66 | 8.08E-12 | 5.96E-11 |
| chr19 | 17183831 | 17219793 | 35963 | -0.8 | 8.08E-12 | 5.96E-11 |
| chr4 | 146830067 | 146831869 | 1803 | 3.24 | 8.13E-12 | 6.00E-11 |
| chr10 | 64563553 | 64577812 | 14260 | 0.7 | 8.17E-12 | 6.02E-11 |
| chr19 | 37156321 | 37159327 | 3007 | 0.82 | 8.18E-12 | 6.03E-11 |
| chr12 | 1004780 | 1030835 | 26056 | 0.66 | 8.21E-12 | 6.05E-11 |
| chr12 | 28342622 | 28356388 | 13767 | 0.94 | 8.33E-12 | 6.14E-11 |
| chr12 | 83480246 | 83482073 | 1828 | 5.06 | 8.45E-12 | 6.22E-11 |
| chr18 | 11978590 | 12006581 | 27992 | 0.54 | 8.60E-12 | 6.33E-11 |
| chr2 | 99164852 | 99179714 | 14863 | -1.77 | 8.62E-12 | 6.35E-11 |
| chr10 | 42970268 | 42972088 | 1821 | 7.38 | 8.62E-12 | 6.35E-11 |
| chr3 | 96168128 | 96169576 | 1449 | 7.37 | 8.67E-12 | 6.38E-11 |
| chr1 | 165643446 | 165668745 | 25300 | 0.74 | 8.69E-12 | 6.39E-11 |
| chr1 | 177937861 | 177939913 | 2053 | 2.29 | 8.83E-12 | 6.49E-11 |
| chr18 | 60904374 | 60906655 | 2282 | 1.68 | 8.95E-12 | 6.58E-11 |
| chr2 | 219508831 | 219537745 | 28915 | -0.79 | 9.01E-12 | 6.62E-11 |

| | | | | | | |
|---|---|---|---|---|---|---|
| chr5 | 26244912 | 26246404 | 1493 | 7.35 | 9.04E-12 | 6.64E-11 |
| chr2 | 85596568 | 85621648 | 25081 | -1.55 | 9.16E-12 | 6.73E-11 |
| chr17 | 54299234 | 54300366 | 1133 | -3.1 | 9.29E-12 | 6.82E-11 |
| chr10 | 101487227 | 101612868 | 125642 | -0.24 | 9.34E-12 | 6.86E-11 |
| chr1 | 218488618 | 218614092 | 125475 | -0.23 | 9.49E-12 | 6.97E-11 |
| chr15 | 31616995 | 31698788 | 81794 | -0.38 | 9.51E-12 | 6.98E-11 |
| chr8 | 144094535 | 144101449 | 6915 | 1.06 | 9.64E-12 | 7.08E-11 |
| chr4 | 120792403 | 120794719 | 2317 | 7.36 | 9.70E-12 | 7.12E-11 |
| chr3 | 102891806 | 102892739 | 934 | 7.34 | 9.81E-12 | 7.19E-11 |
| chr8 | 116872116 | 116885188 | 13073 | 7.37 | 9.86E-12 | 7.23E-11 |
| chr8 | 77867119 | 77876672 | 9554 | 7.41 | 9.89E-12 | 7.25E-11 |
| chr1 | 191222652 | 191233073 | 10422 | 7.5 | 9.90E-12 | 7.26E-11 |
| chr14 | 73074440 | 73076206 | 1767 | -2.6 | 9.93E-12 | 7.28E-11 |
| chr19 | 54957343 | 54979648 | 22306 | 0.45 | 9.94E-12 | 7.28E-11 |
| chr1 | 162318649 | 162320724 | 2076 | 1.33 | 9.94E-12 | 7.28E-11 |
| chr2 | 82612768 | 82613793 | 1026 | 7.33 | 1.02E-11 | 7.50E-11 |
| chr2 | 57927726 | 57929043 | 1318 | 2.71 | 1.03E-11 | 7.52E-11 |
| chr2 | 223433680 | 223442374 | 8695 | -1.21 | 1.03E-11 | 7.53E-11 |
| chr20 | 32250664 | 32275307 | 24644 | -0.77 | 1.03E-11 | 7.56E-11 |
| chr2 | 179958159 | 179984360 | 26202 | 1.38 | 1.04E-11 | 7.62E-11 |
| chr3 | 170978599 | 170980737 | 2139 | 1.37 | 1.04E-11 | 7.63E-11 |
| chr1 | 159461985 | 159463477 | 1493 | -4.02 | 1.05E-11 | 7.65E-11 |
| chr5 | 130964487 | 130973679 | 9193 | 0.61 | 1.05E-11 | 7.66E-11 |
| chr1 | 67426401 | 67451117 | 24717 | 1.02 | 1.05E-11 | 7.66E-11 |
| chr1 | 219409603 | 219411704 | 2102 | 1.33 | 1.05E-11 | 7.67E-11 |
| chr10 | 60535775 | 60581823 | 46049 | -0.74 | 1.05E-11 | 7.68E-11 |
| chr19 | 9783023 | 9786605 | 3583 | 1.05 | 1.07E-11 | 7.78E-11 |
| chr15 | 54031471 | 54040194 | 8724 | 7.35 | 1.07E-11 | 7.79E-11 |
| chr10 | 90143865 | 90147665 | 3801 | 0.96 | 1.07E-11 | 7.84E-11 |
| chr11 | 121484167 | 121485414 | 1248 | 4.74 | 1.09E-11 | 7.92E-11 |
| chr6 | 140708720 | 140710436 | 1717 | 2.2 | 1.10E-11 | 8.02E-11 |
| chr14 | 37797048 | 37813042 | 15995 | 1.09 | 1.10E-11 | 8.04E-11 |
| chr2 | 59651788 | 59653322 | 1535 | 7.32 | 1.10E-11 | 8.05E-11 |
| chr19 | 41145821 | 41153428 | 7608 | 0.47 | 1.13E-11 | 8.20E-11 |
| chr10 | 38240693 | 38246448 | 5756 | 7.33 | 1.16E-11 | 8.45E-11 |
| chr20 | 4552290 | 4556487 | 4198 | -0.61 | 1.16E-11 | 8.46E-11 |
| chr6 | 101349321 | 101350468 | 1148 | 5.09 | 1.16E-11 | 8.47E-11 |
| chr9 | 134225422 | 134353322 | 127901 | 0.37 | 1.16E-11 | 8.47E-11 |
| chr7 | 45923883 | 45929175 | 5293 | 1.1 | 1.17E-11 | 8.55E-11 |
| chr1 | 212401782 | 212403019 | 1238 | 5.7 | 1.19E-11 | 8.63E-11 |
| chr12 | 125087903 | 125094783 | 6881 | 1.53 | 1.19E-11 | 8.64E-11 |
| chr5 | 25083729 | 25091054 | 7326 | 7.47 | 1.19E-11 | 8.67E-11 |
| chr2 | 216262239 | 216285005 | 22767 | -1.2 | 1.23E-11 | 8.94E-11 |
| chr11 | 26602002 | 26603432 | 1431 | 7.32 | 1.23E-11 | 8.95E-11 |
| chrX | 103560063 | 103564592 | 4530 | 1.47 | 1.24E-11 | 9.04E-11 |
| chr12 | 58541931 | 58543042 | 1112 | 2.86 | 1.25E-11 | 9.06E-11 |
| chr3 | 94647131 | 94648498 | 1368 | 7.31 | 1.25E-11 | 9.09E-11 |
| chr15 | 33358228 | 33363016 | 4789 | 0.51 | 1.26E-11 | 9.15E-11 |
| chr8 | 122731332 | 122740989 | 9658 | -0.74 | 1.26E-11 | 9.16E-11 |
| chr11 | 71874884 | 71886062 | 11179 | 7.32 | 1.28E-11 | 9.28E-11 |
| chr4 | 574353 | 576939 | 2587 | 7.32 | 1.28E-11 | 9.28E-11 |
| chr8 | 130764231 | 130766608 | 2378 | -1.71 | 1.29E-11 | 9.36E-11 |
| chr13 | 83633804 | 83635016 | 1213 | 2.38 | 1.30E-11 | 9.42E-11 |
| chr14 | 89444794 | 89479492 | 34699 | -0.62 | 1.32E-11 | 9.54E-11 |
| chr13 | 82344951 | 82347552 | 2602 | 7.33 | 1.32E-11 | 9.56E-11 |
| chr21 | 36418710 | 36424370 | 5661 | 1.22 | 1.32E-11 | 9.58E-11 |
| chr1 | 93740754 | 93743901 | 3148 | 1.65 | 1.33E-11 | 9.64E-11 |
| chr9 | 29866185 | 29867720 | 1536 | 7.31 | 1.33E-11 | 9.65E-11 |
| chr3 | 175025253 | 175026341 | 1089 | 7.31 | 1.34E-11 | 9.73E-11 |
| chr10 | 111721216 | 111737743 | 16528 | 1.51 | 1.34E-11 | 9.74E-11 |
| chr3 | 189347315 | 189350399 | 3085 | 1.28 | 1.35E-11 | 9.77E-11 |

| | | | | | | |
|---|---|---|---|---|---|---|
| chr4 | 48604228 | 48606414 | 2187 | 2.54 | 1.35E-11 | 9.80E-11 |
| chr19 | 15946782 | 15949435 | 2654 | 0.88 | 1.36E-11 | 9.82E-11 |
| chr12 | 33071765 | 33075383 | 3619 | 0.83 | 1.37E-11 | 9.91E-11 |
| chr1 | 183106731 | 183110348 | 3618 | 1 | 1.38E-11 | 1.00E-10 |
| chr7 | 75216989 | 75374018 | 157030 | 0.26 | 1.39E-11 | 1.00E-10 |
| chr2 | 25965774 | 25970864 | 5091 | -4.68 | 1.39E-11 | 1.00E-10 |
| chr15 | 80444421 | 80463156 | 18736 | -1.05 | 1.40E-11 | 1.01E-10 |
| chr2 | 201464968 | 201484532 | 19565 | -1.45 | 1.40E-11 | 1.01E-10 |
| chr15 | 66058894 | 66060615 | 1722 | 2.17 | 1.40E-11 | 1.01E-10 |
| chr20 | 19075664 | 19078438 | 2775 | -7.27 | 1.42E-11 | 1.03E-10 |
| chr13 | 74276170 | 74277333 | 1164 | 5.08 | 1.42E-11 | 1.03E-10 |
| chr18 | 57711113 | 57722997 | 11885 | 0.79 | 1.43E-11 | 1.03E-10 |
| chr15 | 73747048 | 73749266 | 2219 | 7.32 | 1.44E-11 | 1.04E-10 |
| chr18 | 32819967 | 32822431 | 2465 | 0.89 | 1.44E-11 | 1.04E-10 |
| chr7 | 138978684 | 138984938 | 6255 | 1.42 | 1.45E-11 | 1.04E-10 |
| chrX | 122889298 | 122923145 | 33848 | 0.6 | 1.46E-11 | 1.05E-10 |
| chr3 | 155569639 | 155573501 | 3863 | 0.8 | 1.46E-11 | 1.05E-10 |
| chr19 | 22397432 | 22398542 | 1111 | 7.3 | 1.47E-11 | 1.06E-10 |
| chr3 | 10206431 | 10270898 | 64468 | -0.34 | 1.48E-11 | 1.07E-10 |
| chr6 | 7463538 | 7481270 | 17733 | 1.26 | 1.48E-11 | 1.07E-10 |
| chr15 | 56553651 | 56555601 | 1951 | 1.52 | 1.48E-11 | 1.07E-10 |
| chrX | 10502952 | 10604706 | 101755 | 0.25 | 1.49E-11 | 1.07E-10 |
| chrX | 16970326 | 16975246 | 4921 | 1.01 | 1.49E-11 | 1.07E-10 |
| chr2 | 210232412 | 210249191 | 16780 | 4.6 | 1.49E-11 | 1.07E-10 |
| chr21 | 45058755 | 45080934 | 22180 | -0.85 | 1.51E-11 | 1.09E-10 |
| chr8 | 24930424 | 24932446 | 2023 | 3.08 | 1.51E-11 | 1.09E-10 |
| chr15 | 100016444 | 100027489 | 11046 | -7.26 | 1.52E-11 | 1.09E-10 |
| chr18 | 69662125 | 69664040 | 1916 | 7.29 | 1.52E-11 | 1.09E-10 |
| chr9 | 95950876 | 95964464 | 13589 | -1.92 | 1.53E-11 | 1.10E-10 |
| chr1 | 175161002 | 175167814 | 6813 | 6.88 | 1.54E-11 | 1.10E-10 |
| chr17 | 76519353 | 76538869 | 19517 | -1.13 | 1.54E-11 | 1.11E-10 |
| chr15 | 64521707 | 64544091 | 22385 | 0.43 | 1.54E-11 | 1.11E-10 |
| chr4 | 159081909 | 159094253 | 12345 | 2.21 | 1.55E-11 | 1.11E-10 |
| chr7 | 97400179 | 97402936 | 2758 | 2.55 | 1.55E-11 | 1.11E-10 |
| chr12 | 26451486 | 26452499 | 1014 | 1.83 | 1.56E-11 | 1.12E-10 |
| chr3 | 181429836 | 181445147 | 15312 | 1.21 | 1.56E-11 | 1.12E-10 |
| chr5 | 179497955 | 179518372 | 20418 | 1.69 | 1.57E-11 | 1.13E-10 |
| chr4 | 41505766 | 41553552 | 47787 | -0.45 | 1.57E-11 | 1.13E-10 |
| chr11 | 34048808 | 34050831 | 2024 | 1.85 | 1.57E-11 | 1.13E-10 |
| chr2 | 33620177 | 33645272 | 25096 | 0.54 | 1.58E-11 | 1.13E-10 |
| chr4 | 91386315 | 91395908 | 9594 | 7.29 | 1.60E-11 | 1.15E-10 |
| chr4 | 124907054 | 124919130 | 12077 | 7.29 | 1.63E-11 | 1.17E-10 |
| chr11 | 59381134 | 59384881 | 3748 | 1.07 | 1.64E-11 | 1.18E-10 |
| chr1 | 146686471 | 146715766 | 29296 | 1.05 | 1.64E-11 | 1.18E-10 |
| chr12 | 6792981 | 6799986 | 7006 | 0.77 | 1.65E-11 | 1.18E-10 |
| chr3 | 122028845 | 122042327 | 13483 | 7.32 | 1.66E-11 | 1.19E-10 |
| chr12 | 12759556 | 12766128 | 6573 | 0.7 | 1.66E-11 | 1.19E-10 |
| chr1 | 103344534 | 103363717 | 19184 | -3.28 | 1.67E-11 | 1.19E-10 |
| chr4 | 187321623 | 187327856 | 6234 | 1.32 | 1.69E-11 | 1.21E-10 |
| chr3 | 78814708 | 78815877 | 1170 | 2.16 | 1.69E-11 | 1.21E-10 |
| chr4 | 112957714 | 112958681 | 968 | 6.27 | 1.70E-11 | 1.21E-10 |
| chr10 | 45871109 | 45872675 | 1567 | 7.29 | 1.70E-11 | 1.21E-10 |
| chr12 | 90936856 | 90937854 | 999 | 3.05 | 1.70E-11 | 1.22E-10 |
| chr1 | 65446709 | 65453819 | 7111 | 0.88 | 1.71E-11 | 1.22E-10 |
| chr2 | 166428394 | 166431313 | 2920 | 7.3 | 1.72E-11 | 1.23E-10 |
| chr2 | 238987935 | 238989411 | 1477 | 3.5 | 1.72E-11 | 1.23E-10 |
| chr13 | 89308172 | 89309930 | 1759 | 7.27 | 1.76E-11 | 1.26E-10 |
| chr14 | 56442395 | 56443989 | 1595 | 3.19 | 1.77E-11 | 1.26E-10 |
| chr14 | 103438540 | 103501208 | 62669 | 0.41 | 1.79E-11 | 1.28E-10 |
| chr4 | 101940224 | 101942810 | 2587 | 1.53 | 1.80E-11 | 1.29E-10 |
| chr11 | 111998522 | 112001063 | 2542 | 1.51 | 1.81E-11 | 1.29E-10 |

| | | | | | | |
|---|---|---|---|---|---|---|
| chr19 | 58779724 | 58791404 | 11681 | 2.55 | 1.82E-11 | 1.30E-10 |
| chr4 | 123187066 | 123189111 | 2046 | 1.2 | 1.85E-11 | 1.32E-10 |
| chr13 | 67483952 | 67490588 | 6637 | 1.48 | 1.86E-11 | 1.32E-10 |
| chr5 | 170171861 | 170187876 | 16016 | 3.98 | 1.86E-11 | 1.32E-10 |
| chr4 | 118717618 | 118718910 | 1293 | 7.27 | 1.87E-11 | 1.33E-10 |
| chr13 | 107159997 | 107165568 | 5572 | 3.94 | 1.87E-11 | 1.33E-10 |
| chr11 | 59615817 | 59634683 | 18867 | 4.88 | 1.87E-11 | 1.33E-10 |
| chr11 | 70078601 | 70080675 | 2075 | 1.41 | 1.88E-11 | 1.34E-10 |
| chr5 | 148343523 | 148359204 | 15682 | 0.86 | 1.88E-11 | 1.34E-10 |
| chr19 | 42954231 | 42960754 | 6524 | 6.39 | 1.89E-11 | 1.35E-10 |
| chr8 | 113470355 | 113472727 | 2373 | 7.3 | 1.90E-11 | 1.35E-10 |
| chr1 | 17552771 | 17616482 | 63712 | 0.42 | 1.90E-11 | 1.35E-10 |
| chr10 | 26695630 | 26697801 | 2172 | -1.66 | 1.90E-11 | 1.35E-10 |
| chr20 | 2264231 | 2266149 | 1919 | 3.01 | 1.91E-11 | 1.36E-10 |
| chr3 | 39181900 | 39223206 | 41307 | 0.51 | 1.91E-11 | 1.36E-10 |
| chr15 | 63113412 | 63114904 | 1493 | 1.85 | 1.92E-11 | 1.36E-10 |
| chr10 | 75599633 | 75621119 | 21487 | -0.95 | 1.93E-11 | 1.37E-10 |
| chr17 | 27023834 | 27091219 | 67386 | 0.32 | 1.93E-11 | 1.37E-10 |
| chr1 | 221660571 | 221668499 | 7929 | 1.75 | 1.93E-11 | 1.37E-10 |
| chr2 | 189452047 | 189464600 | 12554 | 2.21 | 1.94E-11 | 1.38E-10 |
| chr10 | 13907519 | 13909511 | 1993 | -1.34 | 1.95E-11 | 1.38E-10 |
| chr2 | 10630070 | 10638951 | 8882 | 1.2 | 1.95E-11 | 1.38E-10 |
| chr2 | 233181047 | 233205134 | 24088 | -1.45 | 1.96E-11 | 1.39E-10 |
| chr11 | 122726465 | 122728605 | 2141 | -1.2 | 1.97E-11 | 1.39E-10 |
| chr21 | 37824484 | 37851482 | 26999 | 1.16 | 1.97E-11 | 1.40E-10 |
| chr6 | 146731579 | 146732847 | 1269 | 4.78 | 1.98E-11 | 1.40E-10 |
| chr8 | 39820150 | 39823019 | 2870 | 1.46 | 1.99E-11 | 1.41E-10 |
| chr8 | 97245833 | 97251414 | 5582 | 0.88 | 1.99E-11 | 1.41E-10 |
| chr22 | 27109606 | 27118962 | 9357 | -1.27 | 1.99E-11 | 1.41E-10 |
| chr10 | 126307400 | 126377810 | 70411 | -0.61 | 1.99E-11 | 1.41E-10 |
| chr4 | 21803542 | 21813741 | 10200 | 7.25 | 2.00E-11 | 1.42E-10 |
| chr5 | 81682175 | 81702732 | 20558 | 7.25 | 2.00E-11 | 1.42E-10 |
| chr8 | 110936808 | 110938843 | 2036 | 7.25 | 2.00E-11 | 1.42E-10 |
| chr3 | 5489939 | 5490676 | 738 | 5.26 | 2.00E-11 | 1.42E-10 |
| chr16 | 86696455 | 86701261 | 4807 | 0.71 | 2.01E-11 | 1.42E-10 |
| chr5 | 94415644 | 94418537 | 2894 | 0.87 | 2.02E-11 | 1.43E-10 |
| chr8 | 47669665 | 47671039 | 1375 | 7.25 | 2.05E-11 | 1.45E-10 |
| chr14 | 38547984 | 38549479 | 1496 | 7.25 | 2.05E-11 | 1.45E-10 |
| chr19 | 37662866 | 37664308 | 1443 | 1.68 | 2.06E-11 | 1.45E-10 |
| chr9 | 139580048 | 139593955 | 13908 | 0.67 | 2.07E-11 | 1.47E-10 |
| chr17 | 79448157 | 79487399 | 39243 | 0.46 | 2.10E-11 | 1.48E-10 |
| chr6 | 106959834 | 106976656 | 16823 | 1.57 | 2.13E-11 | 1.50E-10 |
| chr6 | 123849798 | 123852132 | 2335 | -1.59 | 2.14E-11 | 1.51E-10 |
| chr17 | 64453949 | 64473119 | 19171 | -0.4 | 2.16E-11 | 1.52E-10 |
| chr3 | 148896291 | 148899679 | 3389 | 1.01 | 2.17E-11 | 1.53E-10 |
| chr5 | 52686160 | 52687784 | 1625 | 3.91 | 2.17E-11 | 1.53E-10 |
| chr12 | 87105064 | 87107233 | 2170 | 0.89 | 2.18E-11 | 1.54E-10 |
| chr4 | 71300 | 81779 | 10480 | 7.25 | 2.19E-11 | 1.54E-10 |
| chr3 | 141079357 | 141174399 | 95043 | -0.24 | 2.22E-11 | 1.56E-10 |
| chr17 | 33700143 | 33702158 | 2016 | 2.12 | 2.22E-11 | 1.56E-10 |
| chr15 | 71032563 | 71037132 | 4570 | 0.71 | 2.22E-11 | 1.57E-10 |
| chr4 | 7257288 | 7269369 | 12082 | -1.98 | 2.23E-11 | 1.57E-10 |
| chr8 | 92080571 | 92083929 | 3359 | 0.68 | 2.24E-11 | 1.58E-10 |
| chr11 | 28755538 | 28789746 | 34209 | 0.61 | 2.25E-11 | 1.58E-10 |
| chr6 | 108438811 | 108455779 | 16969 | 7.24 | 2.25E-11 | 1.59E-10 |
| chr2 | 227597478 | 227666872 | 69395 | 0.4 | 2.27E-11 | 1.60E-10 |
| chr16 | 53429670 | 53430812 | 1143 | 4.54 | 2.28E-11 | 1.61E-10 |
| chr6 | 154613166 | 154615044 | 1879 | 1.41 | 2.32E-11 | 1.63E-10 |
| chr1 | 217552035 | 217563565 | 11531 | 7.24 | 2.32E-11 | 1.63E-10 |
| chr9 | 134100918 | 134114078 | 13161 | -1.04 | 2.33E-11 | 1.64E-10 |
| chr20 | 57220042 | 57241012 | 20971 | 0.71 | 2.34E-11 | 1.65E-10 |

| | | | | | | |
|---|---|---|---|---|---|---|
| chr7 | 82467261 | 82469693 | 2433 | 2.38 | 2.37E-11 | 1.66E-10 |
| chr13 | 73763748 | 73785359 | 21612 | 1.91 | 2.37E-11 | 1.66E-10 |
| chr3 | 116995805 | 116999811 | 4007 | 0.9 | 2.37E-11 | 1.67E-10 |
| chr3 | 95985272 | 95987627 | 2356 | 7.23 | 2.37E-11 | 1.67E-10 |
| chr2 | 39101427 | 39103972 | 2546 | -1.06 | 2.38E-11 | 1.67E-10 |
| chr11 | 34908332 | 34912965 | 4634 | -2.47 | 2.39E-11 | 1.68E-10 |
| chr5 | 174622287 | 174623858 | 1572 | -3.26 | 2.39E-11 | 1.68E-10 |
| chr7 | 123386606 | 123390978 | 4373 | 0.7 | 2.41E-11 | 1.69E-10 |
| chr1 | 36948377 | 36953897 | 5521 | 1.14 | 2.44E-11 | 1.71E-10 |
| chr1 | 227937597 | 227976814 | 39218 | 0.61 | 2.46E-11 | 1.72E-10 |
| chr19 | 20119840 | 20129665 | 9826 | 7.24 | 2.46E-11 | 1.73E-10 |
| chr3 | 57566987 | 57585309 | 18323 | 0.59 | 2.48E-11 | 1.74E-10 |
| chr7 | 80222734 | 80224603 | 1870 | 1.17 | 2.50E-11 | 1.75E-10 |
| chr6 | 17905821 | 17909402 | 3582 | -0.94 | 2.51E-11 | 1.76E-10 |
| chr2 | 202644149 | 202647096 | 2948 | -0.84 | 2.51E-11 | 1.76E-10 |
| chr2 | 38123999 | 38126155 | 2157 | 7.26 | 2.52E-11 | 1.77E-10 |
| chr2 | 1679147 | 1691268 | 12122 | 7.22 | 2.54E-11 | 1.78E-10 |
| chr7 | 113724441 | 113726830 | 2390 | 7.22 | 2.56E-11 | 1.79E-10 |
| chr1 | 45563290 | 45577339 | 14050 | 1.86 | 2.58E-11 | 1.80E-10 |
| chr11 | 129715852 | 129733826 | 17975 | 0.73 | 2.58E-11 | 1.80E-10 |
| chr15 | 53744513 | 53750367 | 5855 | 1.09 | 2.58E-11 | 1.81E-10 |
| chr17 | 170143 | 236848 | 66706 | 0.31 | 2.61E-11 | 1.83E-10 |
| chr1 | 155605586 | 155622418 | 16833 | 0.88 | 2.63E-11 | 1.84E-10 |
| chr6 | 156878081 | 156879453 | 1373 | 3.4 | 2.63E-11 | 1.84E-10 |
| chr7 | 11121156 | 11125767 | 4612 | 0.79 | 2.66E-11 | 1.86E-10 |
| chr3 | 96474159 | 96475808 | 1650 | 7.26 | 2.68E-11 | 1.88E-10 |
| chr4 | 14910082 | 14922397 | 12316 | 0.5 | 2.68E-11 | 1.88E-10 |
| chr3 | 8697378 | 8701252 | 3875 | 1.5 | 2.69E-11 | 1.88E-10 |
| chr8 | 29342543 | 29389633 | 47091 | 0.33 | 2.70E-11 | 1.88E-10 |
| chr19 | 48279203 | 48282967 | 3765 | 0.7 | 2.70E-11 | 1.89E-10 |
| chr3 | 56791922 | 56793622 | 1701 | 2.08 | 2.70E-11 | 1.89E-10 |
| chr2 | 113238350 | 113252325 | 13976 | -0.91 | 2.71E-11 | 1.89E-10 |
| chr1 | 86313998 | 86316685 | 2688 | 1.02 | 2.72E-11 | 1.90E-10 |
| chr2 | 198558117 | 198571081 | 12965 | 1.92 | 2.72E-11 | 1.90E-10 |
| chr8 | 116067143 | 116069265 | 2123 | 7.22 | 2.73E-11 | 1.90E-10 |
| chr1 | 46981887 | 46988807 | 6921 | 1.29 | 2.74E-11 | 1.91E-10 |
| chr19 | 42553059 | 42553962 | 904 | 2.52 | 2.82E-11 | 1.97E-10 |
| chr18 | 47492511 | 47494010 | 1500 | 3.53 | 2.82E-11 | 1.97E-10 |
| chr1 | 55775527 | 55789882 | 14356 | -0.88 | 2.87E-11 | 2.00E-10 |
| chr7 | 3231559 | 3243764 | 12206 | 1.55 | 2.96E-11 | 2.06E-10 |
| chr18 | 3217150 | 3219103 | 1954 | 2.15 | 2.98E-11 | 2.07E-10 |
| chr1 | 66444306 | 66466822 | 22517 | 1.13 | 2.99E-11 | 2.08E-10 |
| chr13 | 23386898 | 23389139 | 2242 | 7.21 | 3.00E-11 | 2.09E-10 |
| chr1 | 231962548 | 231978057 | 15510 | -1.2 | 3.03E-11 | 2.11E-10 |
| chr2 | 228680847 | 228685698 | 4852 | -0.72 | 3.04E-11 | 2.12E-10 |
| chr18 | 60600341 | 60611232 | 10892 | 1.44 | 3.04E-11 | 2.12E-10 |
| chr13 | 32555937 | 32557310 | 1374 | 3.16 | 3.05E-11 | 2.12E-10 |
| chr21 | 36456940 | 36459273 | 2334 | 3.09 | 3.08E-11 | 2.14E-10 |
| chr12 | 16034076 | 16050633 | 16558 | 0.64 | 3.09E-11 | 2.14E-10 |
| chr19 | 21450633 | 21451991 | 1359 | 7.25 | 3.11E-11 | 2.16E-10 |
| chr4 | 140561967 | 140588615 | 26649 | 0.62 | 3.17E-11 | 2.20E-10 |
| chr4 | 105935061 | 105936198 | 1138 | 2.72 | 3.17E-11 | 2.20E-10 |
| chr12 | 76441762 | 76486271 | 44510 | 0.41 | 3.18E-11 | 2.21E-10 |
| chr1 | 42678373 | 42682648 | 4276 | 1.47 | 3.20E-11 | 2.22E-10 |
| chr16 | 77938861 | 77940058 | 1198 | 3.12 | 3.21E-11 | 2.23E-10 |
| chr19 | 31617708 | 31619069 | 1362 | -7.21 | 3.21E-11 | 2.23E-10 |
| chr5 | 95549771 | 95563738 | 13968 | 1.41 | 3.25E-11 | 2.25E-10 |
| chr20 | 24014731 | 24016760 | 2030 | -7.16 | 3.27E-11 | 2.27E-10 |
| chr19 | 23299634 | 23300913 | 1280 | 7.22 | 3.28E-11 | 2.27E-10 |
| chr8 | 132826213 | 132829909 | 3697 | 1.47 | 3.31E-11 | 2.29E-10 |
| chrX | 120888807 | 120890493 | 1687 | 7.19 | 3.32E-11 | 2.30E-10 |

| | | | | | | |
|---|---|---|---|---|---|---|
| chr3 | 30114871 | 30126993 | 12123 | 2.41 | 3.32E-11 | 2.30E-10 |
| chr20 | 3623394 | 3632584 | 9191 | -3.02 | 3.33E-11 | 2.30E-10 |
| chr2 | 179774052 | 179775428 | 1377 | 2.65 | 3.33E-11 | 2.31E-10 |
| chrX | 124422690 | 124428680 | 5991 | 2.66 | 3.33E-11 | 2.31E-10 |
| chr4 | 157919653 | 157926724 | 7072 | 1.49 | 3.36E-11 | 2.32E-10 |
| chr8 | 132915669 | 132929726 | 14058 | 0.89 | 3.36E-11 | 2.33E-10 |
| chr3 | 33617653 | 33625092 | 7440 | 0.67 | 3.37E-11 | 2.33E-10 |
| chr4 | 125866248 | 125869550 | 3303 | 1.83 | 3.38E-11 | 2.34E-10 |
| chr13 | 89253967 | 89256591 | 2625 | 7.21 | 3.38E-11 | 2.34E-10 |
| chr16 | 84429608 | 84431165 | 1558 | 1.66 | 3.39E-11 | 2.34E-10 |
| chr5 | 60742382 | 60776209 | 33828 | 0.47 | 3.41E-11 | 2.36E-10 |
| chr12 | 100353360 | 100354406 | 1047 | 7.26 | 3.42E-11 | 2.37E-10 |
| chr8 | 104455641 | 104459413 | 3773 | 0.69 | 3.44E-11 | 2.38E-10 |
| chr15 | 57475096 | 57476953 | 1858 | 1.36 | 3.45E-11 | 2.39E-10 |
| chr14 | 51123956 | 51145211 | 21256 | 0.71 | 3.47E-11 | 2.39E-10 |
| chr10 | 8810252 | 8815177 | 4926 | 7.21 | 3.47E-11 | 2.40E-10 |
| chr14 | 74459883 | 74487080 | 27198 | 0.55 | 3.48E-11 | 2.40E-10 |
| chr14 | 63202497 | 63203902 | 1406 | -7.17 | 3.53E-11 | 2.44E-10 |
| chr1 | 235454805 | 235512327 | 57523 | 0.46 | 3.54E-11 | 2.44E-10 |
| chr17 | 64297936 | 64319633 | 21698 | 0.58 | 3.54E-11 | 2.44E-10 |
| chr1 | 79506279 | 79508886 | 2608 | -2.77 | 3.56E-11 | 2.45E-10 |
| chr9 | 488216 | 491002 | 2787 | 1.11 | 3.56E-11 | 2.45E-10 |
| chr6 | 15790950 | 15792416 | 1467 | 2.59 | 3.58E-11 | 2.47E-10 |
| chr12 | 25378559 | 25404980 | 26422 | 0.7 | 3.58E-11 | 2.47E-10 |
| chr3 | 85608897 | 85610526 | 1630 | 7.22 | 3.58E-11 | 2.47E-10 |
| chr5 | 87971161 | 87976315 | 5155 | 7.18 | 3.62E-11 | 2.49E-10 |
| chr4 | 159011233 | 159012285 | 1053 | 2.61 | 3.63E-11 | 2.50E-10 |
| chr20 | 33099319 | 33129610 | 30292 | 0.51 | 3.66E-11 | 2.52E-10 |
| chr3 | 105743846 | 105749840 | 5995 | 1.68 | 3.67E-11 | 2.52E-10 |
| chr2 | 187397769 | 187478124 | 80356 | -0.35 | 3.70E-11 | 2.54E-10 |
| chr8 | 38192444 | 38270423 | 77980 | 0.39 | 3.71E-11 | 2.55E-10 |
| chr9 | 34373996 | 34387867 | 13872 | 1.24 | 3.71E-11 | 2.55E-10 |
| chr19 | 12475942 | 12477291 | 1350 | 7.17 | 3.75E-11 | 2.58E-10 |
| chr2 | 109551287 | 109553298 | 2012 | 4.35 | 3.77E-11 | 2.59E-10 |
| chr1 | 196149625 | 196151550 | 1926 | 1.24 | 3.80E-11 | 2.61E-10 |
| chr12 | 46940394 | 46957373 | 16980 | 0.71 | 3.81E-11 | 2.62E-10 |
| chr12 | 22382378 | 22383862 | 1485 | -4.18 | 3.82E-11 | 2.62E-10 |
| chr5 | 132297088 | 132300557 | 3470 | 0.71 | 3.85E-11 | 2.64E-10 |
| chr19 | 29697253 | 29704874 | 7622 | 0.53 | 3.87E-11 | 2.66E-10 |
| chr8 | 59867246 | 59868752 | 1507 | 2.04 | 3.91E-11 | 2.68E-10 |
| chr8 | 1683818 | 1735471 | 51654 | 0.39 | 3.91E-11 | 2.69E-10 |
| chr6 | 53467975 | 53493583 | 25609 | -0.47 | 3.93E-11 | 2.70E-10 |
| chr7 | 38216871 | 38219710 | 2840 | 0.82 | 3.95E-11 | 2.71E-10 |
| chr9 | 131985836 | 131999923 | 14088 | 0.93 | 3.95E-11 | 2.71E-10 |
| chr4 | 124742602 | 124743909 | 1308 | 4.77 | 3.95E-11 | 2.71E-10 |
| chr6 | 11586656 | 11614854 | 28199 | -0.7 | 4.01E-11 | 2.75E-10 |
| chr10 | 10180292 | 10181268 | 977 | 5.6 | 4.02E-11 | 2.76E-10 |
| chr21 | 32898056 | 32904170 | 6115 | 2.62 | 4.06E-11 | 2.78E-10 |
| chr19 | 37110078 | 37111540 | 1463 | 7.19 | 4.07E-11 | 2.79E-10 |
| chr1 | 240928279 | 240930559 | 2281 | 2.49 | 4.08E-11 | 2.79E-10 |
| chr6 | 121800043 | 121844530 | 44488 | 0.35 | 4.08E-11 | 2.80E-10 |
| chrX | 17753404 | 17757128 | 3725 | 1.32 | 4.11E-11 | 2.81E-10 |
| chr8 | 106635837 | 106646671 | 10835 | 7.16 | 4.14E-11 | 2.83E-10 |
| chr2 | 182819196 | 182823487 | 4292 | 1.46 | 4.15E-11 | 2.84E-10 |
| chr9 | 124113146 | 124136821 | 23676 | 0.52 | 4.17E-11 | 2.85E-10 |
| chr1 | 68638951 | 68644425 | 5475 | 3 | 4.19E-11 | 2.87E-10 |
| chr10 | 17318368 | 17319990 | 1623 | 7.16 | 4.28E-11 | 2.93E-10 |
| chr8 | 72995461 | 73021285 | 25825 | 0.6 | 4.34E-11 | 2.97E-10 |
| chr3 | 142092785 | 142093867 | 1083 | 3.25 | 4.35E-11 | 2.97E-10 |
| chr9 | 3359042 | 3360569 | 1528 | 4.39 | 4.36E-11 | 2.98E-10 |
| chr19 | 36964157 | 36965628 | 1472 | 7.18 | 4.41E-11 | 3.01E-10 |

| | | | | | | |
|---|---|---|---|---|---|---|
| chr17 | 29026721 | 29028782 | 2062 | 2.43 | 4.41E-11 | 3.01E-10 |
| chr4 | 86746707 | 86755094 | 8388 | 1.04 | 4.48E-11 | 3.06E-10 |
| chr19 | 38044078 | 38045766 | 1689 | 7.15 | 4.49E-11 | 3.06E-10 |
| chr4 | 115848815 | 115850110 | 1296 | 7.18 | 4.53E-11 | 3.09E-10 |
| chr5 | 43293556 | 43317078 | 23523 | 0.46 | 4.54E-11 | 3.10E-10 |
| chr22 | 28568366 | 28578094 | 9729 | 7.15 | 4.62E-11 | 3.15E-10 |
| chr18 | 45493262 | 45496833 | 3572 | 0.72 | 4.63E-11 | 3.16E-10 |
| chr9 | 18184056 | 18185661 | 1606 | 7.15 | 4.66E-11 | 3.18E-10 |
| chr12 | 97092340 | 97093673 | 1334 | 7.14 | 4.69E-11 | 3.20E-10 |
| chr1 | 183127770 | 183165111 | 37342 | 0.52 | 4.69E-11 | 3.20E-10 |
| chr1 | 174818105 | 174819406 | 1302 | 1.48 | 4.71E-11 | 3.21E-10 |
| chr18 | 19600159 | 19613389 | 13231 | 1.29 | 4.76E-11 | 3.24E-10 |
| chr12 | 100648477 | 100663623 | 15147 | 0.86 | 4.79E-11 | 3.26E-10 |
| chr19 | 12605701 | 12606866 | 1166 | 7.14 | 4.80E-11 | 3.27E-10 |
| chr7 | 107530897 | 107669312 | 138416 | 0.29 | 4.83E-11 | 3.29E-10 |
| chr3 | 194247498 | 194259904 | 12407 | 1.05 | 4.86E-11 | 3.31E-10 |
| chr9 | 83798098 | 83799392 | 1295 | 4.82 | 4.88E-11 | 3.32E-10 |
| chr18 | 29317962 | 29320105 | 2144 | 7.14 | 4.91E-11 | 3.34E-10 |
| chr7 | 87504012 | 87536645 | 32634 | 0.5 | 4.93E-11 | 3.35E-10 |
| chr8 | 94080208 | 94081613 | 1406 | 7.14 | 4.93E-11 | 3.35E-10 |
| chrX | 116007349 | 116008928 | 1580 | 7.14 | 4.93E-11 | 3.35E-10 |
| chr5 | 21152000 | 21153412 | 1413 | 7.15 | 4.97E-11 | 3.38E-10 |
| chr9 | 117559731 | 117569219 | 9489 | 0.69 | 4.99E-11 | 3.39E-10 |
| chr14 | 50233335 | 50264133 | 30799 | 0.49 | 4.99E-11 | 3.39E-10 |
| chr2 | 105939558 | 105955741 | 16184 | -0.78 | 5.00E-11 | 3.40E-10 |
| chr20 | 24631321 | 24637915 | 6595 | -1.36 | 5.01E-11 | 3.40E-10 |
| chr12 | 76159758 | 76179428 | 19671 | 1.25 | 5.02E-11 | 3.41E-10 |
| chr2 | 53580037 | 53581193 | 1157 | 2.8 | 5.05E-11 | 3.43E-10 |
| chr4 | 148254951 | 148256530 | 1580 | 1.82 | 5.06E-11 | 3.43E-10 |
| chr8 | 69643392 | 69646059 | 2668 | -7.13 | 5.10E-11 | 3.46E-10 |
| chr16 | 19879824 | 19896649 | 16826 | -1.46 | 5.12E-11 | 3.47E-10 |
| chr6 | 78197255 | 78203448 | 6194 | 2.25 | 5.17E-11 | 3.51E-10 |
| chr12 | 41066419 | 41067223 | 805 | 5.31 | 5.21E-11 | 3.53E-10 |
| chr3 | 124826358 | 124836196 | 9839 | -2.63 | 5.23E-11 | 3.54E-10 |
| chrX | 82276357 | 82277893 | 1537 | 5.57 | 5.24E-11 | 3.55E-10 |
| chr10 | 122999349 | 123000793 | 1445 | 4.94 | 5.26E-11 | 3.57E-10 |
| chr11 | 25086693 | 25100156 | 13464 | 7.15 | 5.27E-11 | 3.57E-10 |
| chr4 | 30946821 | 30948027 | 1207 | 7.16 | 5.28E-11 | 3.57E-10 |
| chr17 | 12413451 | 12427127 | 13677 | -7.15 | 5.28E-11 | 3.57E-10 |
| chr4 | 59963326 | 59964383 | 1058 | 7.17 | 5.29E-11 | 3.58E-10 |
| chr2 | 18446912 | 18450244 | 3333 | -7.1 | 5.29E-11 | 3.58E-10 |
| chr4 | 22720804 | 22722664 | 1861 | 7.14 | 5.31E-11 | 3.59E-10 |
| chr12 | 107348275 | 107355468 | 7194 | 0.82 | 5.37E-11 | 3.63E-10 |
| chr5 | 118590991 | 118682324 | 91334 | -0.31 | 5.41E-11 | 3.66E-10 |
| chr1 | 223161095 | 223174645 | 13551 | 7.13 | 5.46E-11 | 3.69E-10 |
| chr5 | 100415976 | 100449218 | 33243 | 0.48 | 5.46E-11 | 3.69E-10 |
| chr11 | 70545069 | 70555760 | 10692 | -2.3 | 5.47E-11 | 3.70E-10 |
| chr7 | 119184849 | 119186209 | 1361 | 7.22 | 5.48E-11 | 3.70E-10 |
| chr5 | 12815974 | 12817144 | 1171 | 4.99 | 5.53E-11 | 3.73E-10 |
| chr6 | 155406346 | 155407634 | 1289 | 5.56 | 5.57E-11 | 3.76E-10 |
| chr12 | 75619129 | 75630512 | 11384 | 7.13 | 5.59E-11 | 3.77E-10 |
| chr1 | 228961727 | 228987020 | 25294 | 0.69 | 5.63E-11 | 3.80E-10 |
| chr10 | 88406922 | 88410159 | 3238 | 2 | 5.66E-11 | 3.82E-10 |
| chrX | 123154572 | 123158967 | 4396 | 1.15 | 5.69E-11 | 3.84E-10 |
| chr13 | 42600048 | 42624268 | 24221 | 0.85 | 5.71E-11 | 3.85E-10 |
| chr15 | 95843708 | 95846086 | 2379 | 7.12 | 5.75E-11 | 3.88E-10 |
| chr19 | 35999974 | 36001815 | 1842 | 2.06 | 5.85E-11 | 3.94E-10 |
| chr2 | 237684821 | 237686688 | 1868 | -1.83 | 5.86E-11 | 3.95E-10 |
| chr2 | 69287457 | 69329752 | 42296 | -0.62 | 5.88E-11 | 3.96E-10 |
| chr15 | 92972234 | 92973617 | 1384 | -1.76 | 5.89E-11 | 3.97E-10 |
| chr2 | 208733386 | 208735999 | 2614 | -0.94 | 5.89E-11 | 3.97E-10 |

| | | | | | | |
|---|---|---|---|---|---|---|
| chr13 | 45780645 | 45817042 | 36398 | 0.82 | 5.92E-11 | 3.99E-10 |
| chr6 | 154108516 | 154112057 | 3542 | 6.12 | 5.98E-11 | 4.02E-10 |
| chr16 | 53240889 | 53260206 | 19318 | 0.51 | 6.00E-11 | 4.04E-10 |
| chr1 | 90285714 | 90424156 | 138443 | -0.25 | 6.00E-11 | 4.04E-10 |
| chr12 | 9099926 | 9104095 | 4170 | 0.68 | 6.01E-11 | 4.04E-10 |
| chrX | 13702947 | 13717887 | 14941 | 0.89 | 6.05E-11 | 4.07E-10 |
| chr14 | 36526316 | 36539556 | 13241 | 7.13 | 6.19E-11 | 4.16E-10 |
| chr19 | 52254609 | 52255876 | 1268 | 7.11 | 6.20E-11 | 4.17E-10 |
| chr12 | 74710325 | 74741077 | 30753 | 1.12 | 6.22E-11 | 4.18E-10 |
| chr8 | 131494540 | 131541519 | 46980 | -0.45 | 6.27E-11 | 4.21E-10 |
| chrX | 36406575 | 36407923 | 1349 | 4.54 | 6.28E-11 | 4.22E-10 |
| chrX | 41577553 | 41579754 | 2202 | 2.66 | 6.28E-11 | 4.22E-10 |
| chr9 | 86041605 | 86064634 | 23030 | -0.84 | 6.36E-11 | 4.27E-10 |
| chr7 | 112088553 | 112103263 | 14711 | 0.64 | 6.36E-11 | 4.27E-10 |
| chr7 | 101456738 | 101476787 | 20050 | -0.76 | 6.59E-11 | 4.42E-10 |
| chr1 | 229252998 | 229269680 | 16683 | 2.83 | 6.61E-11 | 4.43E-10 |
| chr7 | 85221278 | 85230568 | 9291 | 7.19 | 6.63E-11 | 4.45E-10 |
| chr5 | 49869982 | 49871119 | 1138 | 7.18 | 6.65E-11 | 4.46E-10 |
| chr21 | 37669918 | 37675064 | 5147 | 2.14 | 6.66E-11 | 4.46E-10 |
| chr9 | 72413188 | 72421828 | 8641 | 7.15 | 6.69E-11 | 4.48E-10 |
| chr4 | 102614436 | 102616176 | 1741 | 2.11 | 6.70E-11 | 4.49E-10 |
| chr7 | 21403494 | 21407126 | 3633 | 2.23 | 6.70E-11 | 4.49E-10 |
| chr3 | 105583506 | 105602268 | 18763 | 0.77 | 6.72E-11 | 4.50E-10 |
| chr13 | 95512821 | 95513791 | 971 | 3.08 | 6.81E-11 | 4.56E-10 |
| chr2 | 69422262 | 69424959 | 2698 | -1.88 | 6.91E-11 | 4.63E-10 |
| chr2 | 213315407 | 213317689 | 2283 | 7.11 | 6.92E-11 | 4.64E-10 |
| chr5 | 135458159 | 135470612 | 12454 | 0.98 | 6.93E-11 | 4.64E-10 |
| chr10 | 7895431 | 7900048 | 4618 | 2 | 7.11E-11 | 4.76E-10 |
| chr14 | 102000454 | 102002957 | 2504 | 1.37 | 7.12E-11 | 4.77E-10 |
| chr5 | 76785855 | 76788603 | 2749 | 0.93 | 7.13E-11 | 4.77E-10 |
| chr6 | 10517752 | 10532979 | 15228 | 0.54 | 7.20E-11 | 4.82E-10 |
| chr5 | 73044529 | 73067428 | 22900 | -0.63 | 7.24E-11 | 4.84E-10 |
| chr9 | 113770563 | 113801847 | 31285 | -0.72 | 7.25E-11 | 4.85E-10 |
| chr11 | 31530094 | 31532732 | 2639 | 1.02 | 7.25E-11 | 4.85E-10 |
| chr1 | 27780988 | 27795020 | 14033 | 0.89 | 7.32E-11 | 4.89E-10 |
| chr12 | 1097606 | 1101292 | 3687 | 0.67 | 7.34E-11 | 4.90E-10 |
| chr17 | 17379560 | 17420529 | 40970 | -0.75 | 7.35E-11 | 4.91E-10 |
| chr3 | 107148025 | 107152159 | 4135 | 0.86 | 7.39E-11 | 4.94E-10 |
| chr7 | 120953194 | 120954740 | 1547 | 7.09 | 7.40E-11 | 4.94E-10 |
| chr16 | 13011013 | 13012509 | 1497 | -3.14 | 7.41E-11 | 4.94E-10 |
| chr7 | 12981771 | 12991855 | 10085 | 7.22 | 7.46E-11 | 4.98E-10 |
| chr9 | 107662919 | 107675702 | 12784 | -2.58 | 7.46E-11 | 4.98E-10 |
| chr3 | 102765265 | 102772283 | 7019 | 7.14 | 7.46E-11 | 4.98E-10 |
| chr14 | 85683007 | 85684255 | 1249 | 4.48 | 7.51E-11 | 5.01E-10 |
| chr13 | 72021625 | 72023437 | 1813 | 7.09 | 7.52E-11 | 5.01E-10 |
| chr9 | 139413296 | 139483440 | 70145 | 0.47 | 7.53E-11 | 5.02E-10 |
| chr5 | 27825546 | 27827119 | 1574 | 7.1 | 7.53E-11 | 5.02E-10 |
| chr2 | 198435232 | 198442351 | 7120 | 1.05 | 7.73E-11 | 5.15E-10 |
| chr1 | 201785711 | 201819822 | 34112 | -0.82 | 7.74E-11 | 5.15E-10 |
| chr6 | 109549966 | 109551854 | 1889 | 2.16 | 7.76E-11 | 5.16E-10 |
| chr16 | 73685008 | 73686406 | 1399 | 4.44 | 7.77E-11 | 5.18E-10 |
| chr4 | 106878362 | 106887999 | 9638 | 7.13 | 7.78E-11 | 5.18E-10 |
| chr9 | 92931120 | 92932353 | 1234 | 7.08 | 7.80E-11 | 5.19E-10 |
| chr20 | 9095014 | 9102970 | 7957 | 1.98 | 7.85E-11 | 5.23E-10 |
| chr1 | 222816364 | 222824554 | 8191 | 0.61 | 7.87E-11 | 5.24E-10 |
| chrX | 29665624 | 29683855 | 18232 | 0.53 | 7.88E-11 | 5.24E-10 |
| chr1 | 168655348 | 168663499 | 8152 | 7.18 | 7.92E-11 | 5.26E-10 |
| chr8 | 120931435 | 120933689 | 2255 | 1.25 | 7.92E-11 | 5.27E-10 |
| chr2 | 189960077 | 189990291 | 30215 | -0.63 | 7.94E-11 | 5.27E-10 |
| chr2 | 110086738 | 110088030 | 1293 | 3.76 | 7.96E-11 | 5.29E-10 |
| chr13 | 75747395 | 75749178 | 1784 | -7.07 | 7.97E-11 | 5.29E-10 |

| | | | | | | |
|---|---|---|---|---|---|---|
| chr3 | 33831722 | 33858398 | 26677 | 0.61 | 7.99E-11 | 5.30E-10 |
| chr10 | 11757543 | 11759044 | 1502 | 5.3 | 8.02E-11 | 5.33E-10 |
| chr12 | 25053304 | 25064789 | 11486 | -7.04 | 8.05E-11 | 5.34E-10 |
| chr7 | 111640836 | 111656120 | 15285 | -1.89 | 8.08E-11 | 5.36E-10 |
| chr9 | 76257774 | 76259098 | 1325 | 7.08 | 8.09E-11 | 5.37E-10 |
| chr9 | 99327968 | 99330328 | 2361 | 1.26 | 8.11E-11 | 5.38E-10 |
| chr7 | 95223956 | 95241500 | 17545 | 3.08 | 8.12E-11 | 5.38E-10 |
| chr3 | 153317026 | 153319654 | 2629 | 7.08 | 8.12E-11 | 5.39E-10 |
| chr9 | 87561081 | 87563394 | 2314 | 2.07 | 8.15E-11 | 5.40E-10 |
| chr10 | 17103888 | 17118382 | 14495 | 1.47 | 8.15E-11 | 5.40E-10 |
| chr3 | 13373599 | 13395510 | 21912 | -0.81 | 8.18E-11 | 5.42E-10 |
| chr8 | 140190701 | 140191877 | 1177 | 4.98 | 8.26E-11 | 5.47E-10 |
| chr1 | 227001997 | 227016150 | 14154 | 2.16 | 8.29E-11 | 5.49E-10 |
| chr5 | 55345747 | 55355219 | 9473 | 0.78 | 8.38E-11 | 5.55E-10 |
| chr5 | 127684039 | 127685897 | 1859 | 1.75 | 8.45E-11 | 5.59E-10 |
| chr1 | 205446148 | 205448381 | 2234 | 1.36 | 8.49E-11 | 5.62E-10 |
| chr3 | 79986271 | 79987866 | 1596 | 7.07 | 8.52E-11 | 5.64E-10 |
| chr15 | 80770583 | 80788320 | 17738 | -0.88 | 8.53E-11 | 5.65E-10 |
| chr12 | 115206162 | 115218864 | 12703 | -7.06 | 8.61E-11 | 5.69E-10 |
| chr14 | 44784614 | 44786919 | 2306 | 1.47 | 8.65E-11 | 5.72E-10 |
| chr2 | 96856479 | 96875485 | 19007 | -0.78 | 8.67E-11 | 5.73E-10 |
| chr14 | 70215858 | 70240605 | 24748 | 0.67 | 8.74E-11 | 5.78E-10 |
| chr9 | 16442099 | 16446648 | 4550 | 1.96 | 8.78E-11 | 5.80E-10 |
| chr8 | 109455064 | 109466577 | 11514 | 0.96 | 8.82E-11 | 5.83E-10 |
| chr1 | 182150218 | 182156151 | 5934 | 1.79 | 8.85E-11 | 5.85E-10 |
| chr8 | 90791867 | 90795192 | 3326 | 1.54 | 8.86E-11 | 5.85E-10 |
| chr1 | 98371930 | 98387897 | 15968 | 1.03 | 8.86E-11 | 5.85E-10 |
| chr12 | 63285274 | 63293954 | 8681 | 1.65 | 8.87E-11 | 5.85E-10 |
| chr13 | 91489726 | 91491917 | 2192 | 1.37 | 8.88E-11 | 5.86E-10 |
| chr7 | 66385342 | 66441016 | 55675 | 0.45 | 8.90E-11 | 5.87E-10 |
| chr12 | 92171040 | 92173825 | 2786 | 1.08 | 8.90E-11 | 5.87E-10 |
| chr19 | 57049012 | 57051076 | 2065 | 1.28 | 8.92E-11 | 5.88E-10 |
| chr4 | 19808125 | 19810037 | 1913 | 7.06 | 8.95E-11 | 5.90E-10 |
| chr5 | 95194425 | 95198908 | 4484 | 1.03 | 9.02E-11 | 5.95E-10 |
| chr8 | 25027512 | 25360751 | 333240 | 0.17 | 9.07E-11 | 5.98E-10 |
| chr4 | 56049814 | 56051387 | 1574 | 4.18 | 9.09E-11 | 5.99E-10 |
| chr7 | 20646531 | 20647685 | 1155 | 3.05 | 9.10E-11 | 6.00E-10 |
| chr21 | 34574517 | 34578447 | 3931 | 0.76 | 9.24E-11 | 6.09E-10 |
| chr1 | 95257654 | 95268316 | 10663 | 1.66 | 9.31E-11 | 6.13E-10 |
| chrX | 120731882 | 120733842 | 1961 | 7.07 | 9.33E-11 | 6.14E-10 |
| chr3 | 107850393 | 107853547 | 3155 | 1.16 | 9.34E-11 | 6.15E-10 |
| chr3 | 169045081 | 169046776 | 1696 | 7.13 | 9.34E-11 | 6.15E-10 |
| chr11 | 10669943 | 10689126 | 19184 | -0.89 | 9.34E-11 | 6.15E-10 |
| chr9 | 5230295 | 5232727 | 2433 | 1.04 | 9.35E-11 | 6.15E-10 |
| chr20 | 15142772 | 15144935 | 2164 | 1.7 | 9.43E-11 | 6.20E-10 |
| chr12 | 27165580 | 27176320 | 10741 | 0.76 | 9.43E-11 | 6.20E-10 |
| chr7 | 124357620 | 124359298 | 1679 | 1.99 | 9.48E-11 | 6.23E-10 |
| chr10 | 127779082 | 127780347 | 1266 | 7.07 | 9.76E-11 | 6.41E-10 |
| chr14 | 23443036 | 23480045 | 37010 | 0.36 | 9.77E-11 | 6.42E-10 |
| chr8 | 117048956 | 117050466 | 1511 | 7.05 | 9.80E-11 | 6.44E-10 |
| chr14 | 39556122 | 39584587 | 28466 | 0.54 | 9.82E-11 | 6.45E-10 |
| chr9 | 30197133 | 30198591 | 1459 | 2.58 | 9.83E-11 | 6.45E-10 |
| chr4 | 15001977 | 15050345 | 48369 | 0.35 | 9.90E-11 | 6.50E-10 |
| chr2 | 19401410 | 19410036 | 8627 | -1.39 | 9.91E-11 | 6.51E-10 |
| chr19 | 54448127 | 54449778 | 1652 | 1.26 | 1.01E-10 | 6.62E-10 |
| chr10 | 122829130 | 122832109 | 2980 | 7.07 | 1.02E-10 | 6.70E-10 |
| chr5 | 123492122 | 123499031 | 6910 | 1.21 | 1.03E-10 | 6.75E-10 |
| chr20 | 34638440 | 34771697 | 133258 | 0.29 | 1.03E-10 | 6.78E-10 |
| chr9 | 93948708 | 93974543 | 25836 | 0.54 | 1.05E-10 | 6.88E-10 |
| chr16 | 19098020 | 19152586 | 54567 | 0.38 | 1.06E-10 | 6.95E-10 |
| chr14 | 91488868 | 91497351 | 8484 | -2.3 | 1.07E-10 | 7.01E-10 |

| | | | | | | |
|---|---|---|---|---|---|---|
| chr10 | 84387855 | 84388975 | 1121 | 7.04 | 1.08E-10 | 7.06E-10 |
| chr6 | 141199916 | 141202785 | 2870 | 1.89 | 1.08E-10 | 7.06E-10 |
| chr1 | 9825639 | 9831957 | 6319 | 1.24 | 1.08E-10 | 7.08E-10 |
| chr1 | 169209537 | 169220588 | 11052 | 1.17 | 1.09E-10 | 7.15E-10 |
| chr14 | 48890769 | 48904952 | 14184 | 7.08 | 1.09E-10 | 7.16E-10 |
| chr2 | 42051562 | 42094467 | 42906 | 0.5 | 1.10E-10 | 7.19E-10 |
| chr2 | 86362851 | 86427397 | 64547 | -0.43 | 1.11E-10 | 7.26E-10 |
| chr10 | 34923148 | 34930900 | 7753 | -1.31 | 1.11E-10 | 7.28E-10 |
| chr5 | 107715937 | 107719218 | 3282 | 0.87 | 1.12E-10 | 7.30E-10 |
| chr15 | 72630830 | 72634833 | 4004 | -0.96 | 1.12E-10 | 7.35E-10 |
| chr3 | 70049315 | 70050451 | 1137 | 6.07 | 1.13E-10 | 7.36E-10 |
| chr3 | 94131903 | 94133261 | 1359 | 7.03 | 1.13E-10 | 7.38E-10 |
| chr14 | 49972445 | 49980158 | 7714 | 7.08 | 1.14E-10 | 7.48E-10 |
| chr9 | 107502871 | 107511764 | 8894 | 0.99 | 1.15E-10 | 7.51E-10 |
| chr2 | 31546777 | 31547873 | 1097 | 2.81 | 1.15E-10 | 7.52E-10 |
| chr10 | 6898687 | 6900135 | 1449 | 2.92 | 1.16E-10 | 7.58E-10 |
| chr8 | 76465501 | 76466667 | 1167 | 7.06 | 1.16E-10 | 7.60E-10 |
| chr7 | 17336741 | 17365891 | 29151 | 0.52 | 1.18E-10 | 7.69E-10 |
| chr13 | 105790891 | 105792602 | 1712 | 1.49 | 1.18E-10 | 7.71E-10 |
| chr2 | 61921285 | 61936238 | 14954 | 1.22 | 1.18E-10 | 7.71E-10 |
| chr5 | 32400461 | 32445234 | 44774 | 0.51 | 1.20E-10 | 7.81E-10 |
| chr13 | 34391474 | 34413523 | 22050 | -0.69 | 1.20E-10 | 7.83E-10 |
| chr15 | 68229010 | 68247944 | 18935 | 0.76 | 1.20E-10 | 7.84E-10 |
| chr11 | 130546252 | 130548313 | 2062 | 2.51 | 1.21E-10 | 7.91E-10 |
| chr2 | 197096749 | 197098910 | 2162 | 1.59 | 1.22E-10 | 7.93E-10 |
| chr2 | 104905964 | 104907396 | 1433 | 7.04 | 1.22E-10 | 7.94E-10 |
| chr12 | 88277226 | 88278106 | 881 | 4.5 | 1.22E-10 | 7.97E-10 |
| chr8 | 90893625 | 90954859 | 61235 | -0.27 | 1.22E-10 | 7.97E-10 |
| chr3 | 139007318 | 139029753 | 22436 | -0.79 | 1.23E-10 | 8.01E-10 |
| chr10 | 9247156 | 9248350 | 1195 | 7.04 | 1.25E-10 | 8.11E-10 |
| chr10 | 13701956 | 13755706 | 53751 | 0.53 | 1.25E-10 | 8.14E-10 |
| chr1 | 113160484 | 113189342 | 28859 | 0.45 | 1.26E-10 | 8.23E-10 |
| chr16 | 52112053 | 52113599 | 1547 | 7.03 | 1.27E-10 | 8.27E-10 |
| chr13 | 82561032 | 82563047 | 2016 | 7.03 | 1.27E-10 | 8.28E-10 |
| chr3 | 70786457 | 70788174 | 1718 | 1.82 | 1.30E-10 | 8.43E-10 |
| chr10 | 24638051 | 24639502 | 1452 | 7.02 | 1.30E-10 | 8.45E-10 |
| chr12 | 81661533 | 81662741 | 1209 | 2.2 | 1.30E-10 | 8.45E-10 |
| chr12 | 67396065 | 67397069 | 1005 | 3.04 | 1.30E-10 | 8.46E-10 |
| chr4 | 144997 | 147119 | 2123 | 7.02 | 1.31E-10 | 8.48E-10 |
| chr13 | 80340749 | 80342010 | 1262 | 7.02 | 1.31E-10 | 8.51E-10 |
| chr19 | 7444977 | 7465770 | 20794 | 0.57 | 1.32E-10 | 8.57E-10 |
| chr19 | 9545137 | 9547426 | 2290 | 1.47 | 1.32E-10 | 8.59E-10 |
| chr10 | 79109105 | 79133674 | 24570 | -0.8 | 1.35E-10 | 8.75E-10 |
| chr4 | 111080035 | 111087404 | 7370 | -1.17 | 1.35E-10 | 8.78E-10 |
| chr13 | 45330079 | 45333587 | 3509 | -2.41 | 1.35E-10 | 8.79E-10 |
| chr6 | 153705756 | 153713940 | 8185 | 7.01 | 1.37E-10 | 8.90E-10 |
| chr16 | 21232919 | 21245385 | 12467 | 7.02 | 1.37E-10 | 8.91E-10 |
| chr14 | 70072906 | 70088602 | 15697 | 1.23 | 1.38E-10 | 8.94E-10 |
| chr3 | 47569230 | 47570491 | 1262 | 3.46 | 1.38E-10 | 8.97E-10 |
| chr1 | 101199412 | 101202653 | 3242 | 2.04 | 1.39E-10 | 8.99E-10 |
| chr6 | 74223145 | 74234481 | 11337 | 0.54 | 1.40E-10 | 9.08E-10 |
| chr10 | 119081699 | 119136354 | 54656 | 0.29 | 1.42E-10 | 9.18E-10 |
| chr5 | 142059212 | 142061802 | 2591 | 1.63 | 1.42E-10 | 9.19E-10 |
| chr4 | 120600347 | 120605474 | 5128 | 2.08 | 1.42E-10 | 9.19E-10 |
| chr1 | 75804312 | 75806106 | 1795 | 4.91 | 1.42E-10 | 9.19E-10 |
| chr11 | 16117366 | 16133108 | 15743 | 6.46 | 1.42E-10 | 9.21E-10 |
| chr11 | 26291115 | 26292933 | 1819 | 7.05 | 1.42E-10 | 9.21E-10 |
| chr12 | 25101168 | 25102620 | 1453 | -1.55 | 1.43E-10 | 9.28E-10 |
| chr5 | 50728533 | 50729881 | 1349 | 7.02 | 1.44E-10 | 9.30E-10 |
| chr4 | 59800569 | 59806573 | 6005 | 6.15 | 1.44E-10 | 9.32E-10 |
| chr9 | 91467084 | 91468687 | 1604 | -1.3 | 1.46E-10 | 9.44E-10 |

| | | | | | | |
|---|---|---|---|---|---|---|
| chr11 | 104642303 | 104644199 | 1897 | 7 | 1.48E-10 | 9.53E-10 |
| chr19 | 53103687 | 53105403 | 1717 | 7 | 1.48E-10 | 9.56E-10 |
| chr10 | 60459486 | 60465120 | 5635 | -1.54 | 1.48E-10 | 9.57E-10 |
| chr3 | 172234096 | 172241597 | 7502 | 1.77 | 1.48E-10 | 9.57E-10 |
| chr4 | 74542183 | 74550035 | 7853 | 1.18 | 1.49E-10 | 9.64E-10 |
| chr6 | 12279085 | 12385178 | 106094 | 0.25 | 1.50E-10 | 9.72E-10 |
| chr2 | 30835798 | 30837853 | 2056 | 2.86 | 1.51E-10 | 9.73E-10 |
| chr19 | 48200827 | 48261559 | 60733 | 0.34 | 1.52E-10 | 9.80E-10 |
| chr17 | 70334369 | 70336286 | 1918 | 1.39 | 1.52E-10 | 9.84E-10 |
| chr1 | 52429321 | 52443814 | 14494 | -1.58 | 1.53E-10 | 9.86E-10 |
| chr20 | 5530033 | 5594415 | 64383 | 0.33 | 1.53E-10 | 9.87E-10 |
| chr16 | 20910282 | 20926325 | 16044 | 0.48 | 1.54E-10 | 9.90E-10 |
| chr17 | 49148388 | 49211097 | 62710 | 0.4 | 1.54E-10 | 9.93E-10 |
| chr15 | 89630522 | 89693425 | 62904 | 0.32 | 1.54E-10 | 9.94E-10 |
| chr4 | 10457868 | 10459032 | 1165 | 1.91 | 1.55E-10 | 9.98E-10 |
| chr12 | 114203457 | 114212200 | 8744 | 0.86 | 1.55E-10 | 9.98E-10 |
| chr7 | 46524463 | 46527358 | 2896 | 2.14 | 1.55E-10 | 9.99E-10 |
| chr4 | 188913736 | 188943826 | 30091 | 1.34 | 1.56E-10 | 1.00E-09 |
| chr9 | 136039869 | 136044434 | 4566 | -2.97 | 1.56E-10 | 1.00E-09 |
| chr8 | 32956096 | 32964304 | 8209 | -0.75 | 1.58E-10 | 1.02E-09 |
| chr7 | 7552683 | 7569685 | 17003 | 1.72 | 1.58E-10 | 1.02E-09 |
| chr9 | 130660808 | 130806062 | 145255 | -0.23 | 1.58E-10 | 1.02E-09 |
| chr4 | 146116367 | 146117482 | 1116 | 4.46 | 1.58E-10 | 1.02E-09 |
| chr5 | 174714450 | 174716062 | 1613 | -6.96 | 1.59E-10 | 1.02E-09 |
| chr5 | 38598401 | 38608635 | 10235 | -1.35 | 1.59E-10 | 1.02E-09 |
| chr7 | 77389650 | 77399076 | 9427 | 0.96 | 1.59E-10 | 1.02E-09 |
| chr12 | 85305013 | 85307056 | 2044 | 2.53 | 1.59E-10 | 1.02E-09 |
| chr12 | 90477720 | 90495391 | 17672 | 0.59 | 1.60E-10 | 1.03E-09 |
| chr1 | 195707316 | 195709063 | 1748 | 6.99 | 1.60E-10 | 1.03E-09 |
| chr5 | 148188288 | 148189494 | 1207 | 1.82 | 1.60E-10 | 1.03E-09 |
| chr14 | 96733495 | 96742003 | 8509 | -1.78 | 1.61E-10 | 1.03E-09 |
| chr18 | 46092067 | 46094441 | 2375 | -3.02 | 1.61E-10 | 1.04E-09 |
| chr20 | 14932021 | 14944853 | 12833 | 0.77 | 1.63E-10 | 1.04E-09 |
| chr11 | 96035701 | 96077002 | 41302 | 0.52 | 1.63E-10 | 1.04E-09 |
| chr1 | 155861698 | 155882741 | 21044 | 0.61 | 1.63E-10 | 1.04E-09 |
| chr12 | 3042614 | 3086039 | 43426 | 0.59 | 1.67E-10 | 1.07E-09 |
| chr7 | 138597856 | 138667052 | 69197 | -0.55 | 1.67E-10 | 1.07E-09 |
| chr5 | 77623806 | 77625117 | 1312 | -2.96 | 1.68E-10 | 1.07E-09 |
| chr18 | 3982583 | 3996449 | 13867 | 1.42 | 1.68E-10 | 1.07E-09 |
| chr12 | 42876849 | 42878107 | 1259 | 3.06 | 1.68E-10 | 1.08E-09 |
| chr12 | 89554244 | 89560760 | 6517 | 1.17 | 1.68E-10 | 1.08E-09 |
| chr12 | 42103135 | 42104470 | 1336 | 2.5 | 1.70E-10 | 1.09E-09 |
| chr8 | 83985067 | 83986760 | 1694 | 6.99 | 1.70E-10 | 1.09E-09 |
| chrX | 86770924 | 86782025 | 11102 | 0.77 | 1.70E-10 | 1.09E-09 |
| chr3 | 109403541 | 109414325 | 10785 | 4.45 | 1.71E-10 | 1.10E-09 |
| chr9 | 110186994 | 110201634 | 14641 | 0.98 | 1.73E-10 | 1.11E-09 |
| chr6 | 51988204 | 51990851 | 2648 | 7.09 | 1.79E-10 | 1.14E-09 |
| chr18 | 33034604 | 33062756 | 28153 | 0.45 | 1.79E-10 | 1.14E-09 |
| chr1 | 200446427 | 200457968 | 11542 | 1.23 | 1.79E-10 | 1.15E-09 |
| chr4 | 154035821 | 154042853 | 7033 | 1.21 | 1.79E-10 | 1.15E-09 |
| chr7 | 13018176 | 13029351 | 11176 | 1.41 | 1.80E-10 | 1.15E-09 |
| chr2 | 222286288 | 222288813 | 2526 | 6.98 | 1.81E-10 | 1.16E-09 |
| chr14 | 100574533 | 100586566 | 12034 | -6.94 | 1.82E-10 | 1.16E-09 |
| chr3 | 165554432 | 165555211 | 780 | 4.58 | 1.85E-10 | 1.18E-09 |
| chr14 | 39624967 | 39653591 | 28625 | 0.46 | 1.88E-10 | 1.20E-09 |
| chr4 | 108117957 | 108119177 | 1221 | 4.46 | 1.90E-10 | 1.22E-09 |
| chr7 | 151029683 | 151047500 | 17818 | 1 | 1.92E-10 | 1.23E-09 |
| chr15 | 36684825 | 36700551 | 15727 | 1.32 | 1.94E-10 | 1.24E-09 |
| chr17 | 38959986 | 38976677 | 16692 | 0.6 | 1.94E-10 | 1.24E-09 |
| chr9 | 135224687 | 135232054 | 7368 | 0.78 | 1.95E-10 | 1.24E-09 |
| chr3 | 156798573 | 156808207 | 9635 | 0.81 | 1.96E-10 | 1.25E-09 |

| | | | | | | |
|---|---|---|---|---|---|---|
| chr3 | 176186602 | 176188096 | 1495 | 6.97 | 1.96E-10 | 1.25E-09 |
| chr3 | 53876394 | 53880286 | 3893 | 2.14 | 1.97E-10 | 1.26E-09 |
| chr3 | 64650970 | 64652482 | 1513 | 6.97 | 1.98E-10 | 1.26E-09 |
| chr8 | 116934472 | 116942842 | 8371 | 6.97 | 1.98E-10 | 1.26E-09 |
| chr19 | 4050734 | 4067168 | 16435 | 0.68 | 1.98E-10 | 1.26E-09 |
| chr15 | 85496262 | 85499389 | 3128 | -1.07 | 1.99E-10 | 1.27E-09 |
| chrX | 106023316 | 106024774 | 1459 | 6.96 | 1.99E-10 | 1.27E-09 |
| chr6 | 163910309 | 163935299 | 24991 | 0.82 | 1.99E-10 | 1.27E-09 |
| chr20 | 16453662 | 16459027 | 5366 | 0.83 | 2.01E-10 | 1.28E-09 |
| chr6 | 35563759 | 35571923 | 8165 | 1.23 | 2.02E-10 | 1.29E-09 |
| chr11 | 128360509 | 128394004 | 33496 | -0.33 | 2.02E-10 | 1.29E-09 |
| chr9 | 85734426 | 85754044 | 19619 | 3.96 | 2.03E-10 | 1.29E-09 |
| chr7 | 9524747 | 9526202 | 1456 | 6.96 | 2.06E-10 | 1.31E-09 |
| chr9 | 132872388 | 132892042 | 19655 | -1.08 | 2.09E-10 | 1.33E-09 |
| chr3 | 94432867 | 94434033 | 1167 | 7.03 | 2.09E-10 | 1.33E-09 |
| chr6 | 155838194 | 155850762 | 12569 | 2.36 | 2.10E-10 | 1.33E-09 |
| chr13 | 111710874 | 111720180 | 9307 | 6.96 | 2.10E-10 | 1.34E-09 |
| chr2 | 165551974 | 165573681 | 21708 | -0.59 | 2.11E-10 | 1.34E-09 |
| chr1 | 75051568 | 75052820 | 1253 | 6.97 | 2.11E-10 | 1.34E-09 |
| chr11 | 32333213 | 32335390 | 2178 | -1.47 | 2.11E-10 | 1.34E-09 |
| chr7 | 151100721 | 151107772 | 7052 | -6.92 | 2.13E-10 | 1.35E-09 |
| chr10 | 60479719 | 60481379 | 1661 | -4.15 | 2.13E-10 | 1.35E-09 |
| chr12 | 89040048 | 89052531 | 12484 | 6.97 | 2.14E-10 | 1.36E-09 |
| chr21 | 17935033 | 17944002 | 8970 | -0.98 | 2.18E-10 | 1.38E-09 |
| chr20 | 52712213 | 52794566 | 82354 | 0.2 | 2.18E-10 | 1.38E-09 |
| chr2 | 112446492 | 112464220 | 17729 | -0.72 | 2.19E-10 | 1.39E-09 |
| chr1 | 218698099 | 218732952 | 34854 | 0.5 | 2.20E-10 | 1.40E-09 |
| chr1 | 26594801 | 26633830 | 39030 | -0.69 | 2.21E-10 | 1.40E-09 |
| chr17 | 26443900 | 26453751 | 9852 | 1.16 | 2.22E-10 | 1.41E-09 |
| chr1 | 200320609 | 200387121 | 66513 | 0.35 | 2.24E-10 | 1.42E-09 |
| chr5 | 83689323 | 83690864 | 1542 | 6.98 | 2.26E-10 | 1.43E-09 |
| chr14 | 88489378 | 88492279 | 2902 | 1.81 | 2.26E-10 | 1.43E-09 |
| chr8 | 126417220 | 126447810 | 30591 | 0.39 | 2.28E-10 | 1.44E-09 |
| chr12 | 8085581 | 8134854 | 49274 | 0.59 | 2.28E-10 | 1.45E-09 |
| chr4 | 11692849 | 11701715 | 8867 | 2.39 | 2.30E-10 | 1.45E-09 |
| chr15 | 29762210 | 29764299 | 2090 | 1.97 | 2.30E-10 | 1.45E-09 |
| chr3 | 164352789 | 164353929 | 1141 | 6.96 | 2.31E-10 | 1.46E-09 |
| chr6 | 134211694 | 134213490 | 1797 | -1.49 | 2.31E-10 | 1.46E-09 |
| chr6 | 52253782 | 52255959 | 2178 | 1.55 | 2.32E-10 | 1.46E-09 |
| chr8 | 59194652 | 59198540 | 3889 | 6.98 | 2.34E-10 | 1.48E-09 |
| chr8 | 42979573 | 42981121 | 1549 | 7 | 2.35E-10 | 1.48E-09 |
| chr12 | 115437393 | 115443038 | 5646 | -6.92 | 2.35E-10 | 1.48E-09 |
| chr5 | 23776305 | 23777373 | 1069 | 7.01 | 2.36E-10 | 1.49E-09 |
| chrX | 24282635 | 24286217 | 3583 | -2.31 | 2.36E-10 | 1.49E-09 |
| chr11 | 71974157 | 71981122 | 6966 | 1.02 | 2.38E-10 | 1.50E-09 |
| chr2 | 75643627 | 75645041 | 1415 | 3.23 | 2.39E-10 | 1.51E-09 |
| chr11 | 46985117 | 47000806 | 15690 | 1.07 | 2.43E-10 | 1.53E-09 |
| chr7 | 6258743 | 6371945 | 113203 | 0.29 | 2.43E-10 | 1.53E-09 |
| chr2 | 3306629 | 3340368 | 33740 | -0.87 | 2.44E-10 | 1.54E-09 |
| chr3 | 27322877 | 27323978 | 1102 | 6.94 | 2.45E-10 | 1.55E-09 |
| chr4 | 88870666 | 88901101 | 30436 | 0.77 | 2.50E-10 | 1.58E-09 |
| chr4 | 36351017 | 36365153 | 14137 | 6.93 | 2.51E-10 | 1.58E-09 |
| chr18 | 55875321 | 55903218 | 27898 | 0.73 | 2.51E-10 | 1.58E-09 |
| chr21 | 46000334 | 46001434 | 1101 | 5.1 | 2.53E-10 | 1.60E-09 |
| chr16 | 68984807 | 69033301 | 48495 | 0.36 | 2.54E-10 | 1.60E-09 |
| chrX | 27982332 | 27985867 | 3536 | 1.07 | 2.54E-10 | 1.60E-09 |
| chr4 | 112878892 | 112882555 | 3664 | 1.45 | 2.54E-10 | 1.60E-09 |
| chr17 | 67892333 | 67894034 | 1702 | -5.03 | 2.54E-10 | 1.60E-09 |
| chr5 | 173305515 | 173348061 | 42547 | 0.46 | 2.55E-10 | 1.61E-09 |
| chr8 | 83876600 | 83877844 | 1245 | 6.94 | 2.56E-10 | 1.61E-09 |
| chr17 | 21178691 | 21202042 | 23352 | 0.49 | 2.58E-10 | 1.62E-09 |

| | | | | | | |
|---|---|---|---|---|---|---|
| chr9 | 18565500 | 18567342 | 1843 | 6.93 | 2.61E-10 | 1.64E-09 |
| chr5 | 133603086 | 133617092 | 14007 | 0.95 | 2.61E-10 | 1.64E-09 |
| chr11 | 118777054 | 118801602 | 24549 | 0.32 | 2.63E-10 | 1.66E-09 |
| chr17 | 9237709 | 9238915 | 1207 | 6.94 | 2.64E-10 | 1.66E-09 |
| chr9 | 121771169 | 121772476 | 1308 | 2.81 | 2.64E-10 | 1.66E-09 |
| chr2 | 28137029 | 28156013 | 18985 | 0.41 | 2.70E-10 | 1.70E-09 |
| chr19 | 11925106 | 11926707 | 1602 | 6.93 | 2.71E-10 | 1.70E-09 |
| chr20 | 7690346 | 7692415 | 2070 | 1.64 | 2.72E-10 | 1.71E-09 |
| chr8 | 135758252 | 135778588 | 20337 | 6.92 | 2.74E-10 | 1.72E-09 |
| chr1 | 169255834 | 169258905 | 3072 | 1.24 | 2.75E-10 | 1.73E-09 |
| chr11 | 71638189 | 71760568 | 122380 | -0.28 | 2.76E-10 | 1.74E-09 |
| chr14 | 100620708 | 100623635 | 2928 | 2.79 | 2.77E-10 | 1.74E-09 |
| chr17 | 62727711 | 62736172 | 8462 | -2.27 | 2.78E-10 | 1.75E-09 |
| chr12 | 78224196 | 78224919 | 724 | 3.56 | 2.79E-10 | 1.75E-09 |
| chr4 | 74210172 | 74212528 | 2357 | 1.68 | 2.85E-10 | 1.79E-09 |
| chr2 | 69440359 | 69441986 | 1628 | -3.5 | 2.88E-10 | 1.81E-09 |
| chr21 | 32726470 | 32739990 | 13521 | 6.91 | 2.89E-10 | 1.81E-09 |
| chr12 | 59968135 | 59969430 | 1296 | 6.93 | 2.92E-10 | 1.83E-09 |
| chr13 | 102391855 | 102400089 | 8235 | 6.92 | 2.92E-10 | 1.83E-09 |
| chr13 | 60105601 | 60106697 | 1097 | 3.29 | 2.93E-10 | 1.84E-09 |
| chr6 | 125281846 | 125336269 | 54424 | 0.53 | 2.95E-10 | 1.85E-09 |
| chr1 | 241681644 | 241695795 | 14152 | 1.27 | 2.96E-10 | 1.86E-09 |
| chr5 | 99982937 | 99984025 | 1089 | 3.62 | 2.96E-10 | 1.86E-09 |
| chr5 | 18048011 | 18049142 | 1132 | 3.64 | 2.97E-10 | 1.86E-09 |
| chr1 | 205399720 | 205419765 | 20046 | 3.68 | 2.97E-10 | 1.86E-09 |
| chr8 | 37448108 | 37471072 | 22965 | 0.53 | 2.99E-10 | 1.87E-09 |
| chr9 | 96091339 | 96093416 | 2078 | 1.82 | 3.02E-10 | 1.89E-09 |
| chr18 | 26940392 | 26942942 | 2551 | 6.91 | 3.04E-10 | 1.90E-09 |
| chr1 | 228230710 | 228231425 | 716 | 3.17 | 3.05E-10 | 1.91E-09 |
| chr22 | 25215590 | 25217208 | 1619 | 2.58 | 3.06E-10 | 1.92E-09 |
| chr17 | 48103087 | 48173349 | 70263 | 0.24 | 3.10E-10 | 1.94E-09 |
| chr7 | 65746452 | 65748705 | 2254 | -1.74 | 3.12E-10 | 1.95E-09 |
| chr5 | 134461689 | 134478817 | 17129 | 0.74 | 3.13E-10 | 1.96E-09 |
| chr10 | 35329255 | 35353239 | 23985 | 0.47 | 3.13E-10 | 1.96E-09 |
| chr2 | 74591078 | 74619850 | 28773 | -0.71 | 3.14E-10 | 1.96E-09 |
| chr7 | 63458454 | 63460440 | 1987 | 6.92 | 3.14E-10 | 1.96E-09 |
| chr5 | 124241659 | 124243186 | 1528 | 2.6 | 3.16E-10 | 1.98E-09 |
| chr2 | 38219145 | 38220419 | 1275 | 1.78 | 3.20E-10 | 2.00E-09 |
| chr18 | 11848402 | 11861785 | 13384 | 0.48 | 3.21E-10 | 2.00E-09 |
| chr5 | 87956060 | 87957738 | 1679 | 6.9 | 3.21E-10 | 2.01E-09 |
| chr13 | 28526939 | 28535815 | 8877 | 1.13 | 3.26E-10 | 2.03E-09 |
| chr8 | 115334861 | 115335747 | 887 | 6.9 | 3.26E-10 | 2.04E-09 |
| chr12 | 25536460 | 25541365 | 4906 | 0.55 | 3.29E-10 | 2.05E-09 |
| chr7 | 63708381 | 63710045 | 1665 | 6.98 | 3.32E-10 | 2.07E-09 |
| chr16 | 21637372 | 21640112 | 2741 | 1.45 | 3.34E-10 | 2.08E-09 |
| chrX | 115487635 | 115489026 | 1392 | 6.92 | 3.36E-10 | 2.10E-09 |
| chr4 | 21902380 | 21903884 | 1505 | 6.89 | 3.37E-10 | 2.10E-09 |
| chr12 | 86907358 | 86908501 | 1144 | 3.04 | 3.39E-10 | 2.11E-09 |
| chr7 | 5706986 | 5751034 | 44049 | -0.67 | 3.41E-10 | 2.13E-09 |
| chr16 | 67062075 | 67074119 | 12045 | 0.81 | 3.42E-10 | 2.13E-09 |
| chr1 | 48114720 | 48118144 | 3425 | 1.15 | 3.43E-10 | 2.14E-09 |
| chr6 | 37093237 | 37095518 | 2282 | 1.31 | 3.45E-10 | 2.15E-09 |
| chr13 | 22115167 | 22117353 | 2187 | 2.05 | 3.45E-10 | 2.15E-09 |
| chr14 | 60463952 | 60471467 | 7516 | 7.1 | 3.45E-10 | 2.15E-09 |
| chr5 | 125758401 | 125759725 | 1325 | 2.17 | 3.47E-10 | 2.16E-09 |
| chr16 | 75337523 | 75342338 | 4816 | 0.52 | 3.48E-10 | 2.17E-09 |
| chr7 | 3598874 | 3602073 | 3200 | 6.92 | 3.49E-10 | 2.17E-09 |
| chr3 | 126568485 | 126579208 | 10724 | -1.28 | 3.51E-10 | 2.19E-09 |
| chr12 | 132855848 | 132864226 | 8379 | 1.7 | 3.53E-10 | 2.19E-09 |
| chr10 | 123312644 | 123314250 | 1607 | 6.91 | 3.53E-10 | 2.19E-09 |
| chr6 | 7540698 | 7549547 | 8850 | 1.08 | 3.54E-10 | 2.20E-09 |

| | | | | | | |
|---|---|---|---|---|---|---|
| chr5 | 158521205 | 158527666 | 6462 | -0.96 | 3.58E-10 | 2.22E-09 |
| chr8 | 48223291 | 48300704 | 77414 | 0.3 | 3.58E-10 | 2.22E-09 |
| chr12 | 13712176 | 13713243 | 1068 | 3.53 | 3.59E-10 | 2.23E-09 |
| chr3 | 144852102 | 144853814 | 1713 | 6.9 | 3.59E-10 | 2.23E-09 |
| chr5 | 137126472 | 137129595 | 3124 | 1.21 | 3.60E-10 | 2.24E-09 |
| chr6 | 110849089 | 110859939 | 10851 | 1.48 | 3.61E-10 | 2.24E-09 |
| chr2 | 146971141 | 146972618 | 1478 | 6.89 | 3.64E-10 | 2.26E-09 |
| chr1 | 80363015 | 80365051 | 2037 | 6.89 | 3.69E-10 | 2.29E-09 |
| chr19 | 55717963 | 55741421 | 23459 | 0.74 | 3.70E-10 | 2.30E-09 |
| chr7 | 93468246 | 93469160 | 915 | 5.4 | 3.71E-10 | 2.30E-09 |
| chr17 | 7433058 | 7436395 | 3338 | 1.99 | 3.75E-10 | 2.32E-09 |
| chr4 | 187643639 | 187659320 | 15682 | 0.58 | 3.77E-10 | 2.34E-09 |
| chr3 | 152820176 | 152822052 | 1877 | 1.77 | 3.78E-10 | 2.34E-09 |
| chr20 | 8231027 | 8232163 | 1137 | 6.89 | 3.78E-10 | 2.34E-09 |
| chr14 | 79835488 | 79836688 | 1201 | -4.25 | 3.79E-10 | 2.35E-09 |
| chr5 | 158122937 | 158128092 | 5156 | -6.84 | 3.83E-10 | 2.37E-09 |
| chr15 | 36467756 | 36471159 | 3404 | 1.2 | 3.87E-10 | 2.40E-09 |
| chr10 | 60614355 | 60617991 | 3637 | -0.8 | 3.90E-10 | 2.41E-09 |
| chr2 | 28833311 | 28848252 | 14942 | -2.71 | 3.91E-10 | 2.42E-09 |
| chr6 | 113731141 | 113734707 | 3567 | 1.43 | 3.91E-10 | 2.42E-09 |
| chr15 | 99786020 | 99792458 | 6439 | 0.7 | 3.95E-10 | 2.45E-09 |
| chr6 | 41924806 | 41952558 | 27753 | -0.42 | 3.98E-10 | 2.46E-09 |
| chr6 | 117735474 | 117747972 | 12499 | 6.87 | 3.98E-10 | 2.46E-09 |
| chr20 | 58465638 | 58467691 | 2054 | 6.88 | 4.00E-10 | 2.47E-09 |
| chr3 | 94524950 | 94525849 | 900 | 6.9 | 4.02E-10 | 2.49E-09 |
| chr8 | 23896369 | 23897843 | 1475 | 2.36 | 4.05E-10 | 2.50E-09 |
| chr8 | 118895250 | 118896987 | 1738 | 1.68 | 4.08E-10 | 2.52E-09 |
| chr12 | 82917086 | 82918809 | 1724 | 1.24 | 4.10E-10 | 2.53E-09 |
| chr6 | 127663494 | 127669749 | 6256 | 0.77 | 4.10E-10 | 2.53E-09 |
| chr19 | 50248459 | 50297269 | 48811 | 0.5 | 4.12E-10 | 2.55E-09 |
| chr5 | 82734149 | 82747645 | 13497 | 6.9 | 4.15E-10 | 2.56E-09 |
| chr7 | 77164715 | 77170806 | 6092 | 0.57 | 4.15E-10 | 2.56E-09 |
| chr17 | 15845209 | 15966605 | 121397 | 0.27 | 4.16E-10 | 2.57E-09 |
| chr1 | 164799049 | 164801632 | 2584 | 2.34 | 4.17E-10 | 2.57E-09 |
| chr11 | 35051414 | 35084022 | 32609 | 1.64 | 4.17E-10 | 2.58E-09 |
| chr19 | 40836475 | 40877662 | 41188 | -0.5 | 4.27E-10 | 2.63E-09 |
| chr2 | 20811114 | 20851740 | 40627 | -0.55 | 4.28E-10 | 2.64E-09 |
| chr3 | 189614044 | 189633494 | 19451 | 0.87 | 4.28E-10 | 2.64E-09 |
| chr12 | 83907705 | 83908803 | 1099 | 6.89 | 4.30E-10 | 2.65E-09 |
| chr8 | 81785259 | 81787484 | 2226 | 5.95 | 4.31E-10 | 2.66E-09 |
| chr3 | 79844154 | 79845842 | 1689 | 6.92 | 4.31E-10 | 2.66E-09 |
| chr8 | 90545515 | 90570502 | 24988 | 0.87 | 4.33E-10 | 2.67E-09 |
| chr15 | 80256867 | 80263809 | 6943 | -2.71 | 4.34E-10 | 2.67E-09 |
| chr19 | 7894010 | 7902960 | 8951 | 2.49 | 4.34E-10 | 2.67E-09 |
| chr4 | 93022820 | 93023949 | 1130 | 6.86 | 4.38E-10 | 2.70E-09 |
| chr20 | 22555353 | 22568030 | 12678 | -0.56 | 4.39E-10 | 2.70E-09 |
| chr8 | 18604601 | 18612774 | 8174 | 0.85 | 4.39E-10 | 2.70E-09 |
| chr5 | 10406741 | 10428225 | 21485 | -0.84 | 4.39E-10 | 2.70E-09 |
| chr9 | 7102779 | 7105184 | 2406 | 2.31 | 4.43E-10 | 2.73E-09 |
| chr2 | 241499000 | 241527095 | 28096 | -0.5 | 4.45E-10 | 2.73E-09 |
| chr19 | 30520274 | 30527810 | 7537 | 1.65 | 4.45E-10 | 2.73E-09 |
| chr18 | 43647018 | 43648210 | 1193 | 4.6 | 4.47E-10 | 2.75E-09 |
| chr9 | 131814385 | 131815618 | 1234 | 2.16 | 4.48E-10 | 2.75E-09 |
| chr6 | 82852944 | 82855532 | 2589 | 1.24 | 4.50E-10 | 2.76E-09 |
| chr10 | 4542957 | 4544579 | 1623 | 6.87 | 4.50E-10 | 2.77E-09 |
| chr5 | 172878327 | 172883879 | 5553 | -0.82 | 4.51E-10 | 2.77E-09 |
| chr12 | 102956150 | 102962872 | 6723 | 0.68 | 4.55E-10 | 2.79E-09 |
| chr2 | 163174129 | 163176513 | 2385 | 1.14 | 4.56E-10 | 2.80E-09 |
| chr9 | 96900702 | 96903566 | 2865 | 0.88 | 4.59E-10 | 2.81E-09 |
| chr1 | 191610088 | 191611614 | 1527 | 6.85 | 4.61E-10 | 2.83E-09 |
| chr12 | 69632546 | 69661085 | 28540 | 0.53 | 4.61E-10 | 2.83E-09 |

| | | | | | | |
|---|---|---|---|---|---|---|
| chr2 | 35337999 | 35338757 | 759 | 5.03 | 4.64E-10 | 2.85E-09 |
| chr3 | 124353385 | 124356088 | 2704 | 1.57 | 4.66E-10 | 2.86E-09 |
| chr20 | 45695349 | 45704325 | 8977 | 6.95 | 4.67E-10 | 2.86E-09 |
| chr7 | 135192173 | 135206039 | 13867 | -0.64 | 4.68E-10 | 2.87E-09 |
| chr4 | 85859386 | 85889085 | 29700 | 0.75 | 4.74E-10 | 2.91E-09 |
| chr3 | 190090777 | 190105059 | 14283 | 1 | 4.75E-10 | 2.91E-09 |
| chr17 | 14574544 | 14584901 | 10358 | 6.86 | 4.80E-10 | 2.94E-09 |
| chr22 | 38613808 | 38640517 | 26710 | 0.52 | 4.80E-10 | 2.94E-09 |
| chr11 | 21573844 | 21589993 | 16150 | 7.65 | 4.83E-10 | 2.96E-09 |
| chr10 | 91132937 | 91134997 | 2061 | 1.67 | 4.84E-10 | 2.96E-09 |
| chr7 | 15170065 | 15171373 | 1309 | 5.91 | 4.85E-10 | 2.97E-09 |
| chr12 | 107854958 | 107863096 | 8139 | -1.86 | 4.87E-10 | 2.98E-09 |
| chr12 | 18318208 | 18319199 | 992 | 3.03 | 4.91E-10 | 3.00E-09 |
| chr11 | 80410103 | 80412074 | 1972 | 6.86 | 4.91E-10 | 3.00E-09 |
| chr12 | 84530174 | 84531647 | 1474 | 6.88 | 4.93E-10 | 3.01E-09 |
| chr11 | 39384761 | 39393157 | 8397 | 6.99 | 4.95E-10 | 3.03E-09 |
| chr2 | 227110450 | 227122396 | 11947 | 0.64 | 4.98E-10 | 3.05E-09 |
| chr7 | 27379269 | 27387960 | 8692 | 0.88 | 5.01E-10 | 3.06E-09 |
| chr7 | 94598688 | 94610415 | 11728 | 1.32 | 5.01E-10 | 3.06E-09 |
| chr16 | 58596121 | 58664841 | 68721 | 0.39 | 5.08E-10 | 3.11E-09 |
| chr12 | 43135935 | 43137575 | 1641 | 6.88 | 5.11E-10 | 3.12E-09 |
| chr4 | 65722397 | 65723856 | 1460 | 3.84 | 5.13E-10 | 3.13E-09 |
| chr3 | 84649544 | 84651002 | 1459 | 6.92 | 5.14E-10 | 3.14E-09 |
| chr6 | 55730527 | 55748868 | 18342 | 1.67 | 5.17E-10 | 3.16E-09 |
| chr12 | 121813997 | 121855242 | 41246 | 0.58 | 5.18E-10 | 3.16E-09 |
| chr4 | 153601508 | 153603264 | 1757 | 3.55 | 5.19E-10 | 3.17E-09 |
| chr8 | 119800122 | 119809560 | 9439 | 6.91 | 5.23E-10 | 3.19E-09 |
| chr19 | 22966402 | 22967419 | 1018 | 6.86 | 5.23E-10 | 3.19E-09 |
| chr1 | 224252947 | 224254162 | 1216 | 4.4 | 5.24E-10 | 3.20E-09 |
| chr6 | 145803522 | 145805968 | 2447 | 1.39 | 5.26E-10 | 3.20E-09 |
| chrX | 28223102 | 28225008 | 1907 | 5.9 | 5.26E-10 | 3.21E-09 |
| chr8 | 113800474 | 113801494 | 1021 | 7.01 | 5.26E-10 | 3.21E-09 |
| chr1 | 212085104 | 212103484 | 18381 | 0.46 | 5.27E-10 | 3.21E-09 |
| chr9 | 27407193 | 27408961 | 1769 | -6.8 | 5.32E-10 | 3.24E-09 |
| chr4 | 97657176 | 97658513 | 1338 | 6.85 | 5.34E-10 | 3.25E-09 |
| chr17 | 63168876 | 63181702 | 12827 | 1.14 | 5.39E-10 | 3.28E-09 |
| chr20 | 22773906 | 22792501 | 18596 | 0.62 | 5.46E-10 | 3.33E-09 |
| chr8 | 112211587 | 112224674 | 13088 | 6.94 | 5.47E-10 | 3.33E-09 |
| chr2 | 30454396 | 30525247 | 70852 | -0.35 | 5.49E-10 | 3.34E-09 |
| chr1 | 89530478 | 89531672 | 1195 | 5.07 | 5.49E-10 | 3.34E-09 |
| chr15 | 36020079 | 36026106 | 6028 | 7.01 | 5.51E-10 | 3.35E-09 |
| chr3 | 148508392 | 148512269 | 3878 | 4.47 | 5.51E-10 | 3.35E-09 |
| chr4 | 41711432 | 41713454 | 2023 | 1.61 | 5.51E-10 | 3.35E-09 |
| chr1 | 218818422 | 218850441 | 32020 | 0.4 | 5.59E-10 | 3.40E-09 |
| chr3 | 117284201 | 117285073 | 873 | 2.44 | 5.60E-10 | 3.40E-09 |
| chr4 | 159992012 | 159993750 | 1739 | 1.75 | 5.61E-10 | 3.41E-09 |
| chr4 | 30867828 | 30869483 | 1656 | 6.93 | 5.63E-10 | 3.42E-09 |
| chr3 | 104993040 | 104994785 | 1746 | 1.45 | 5.63E-10 | 3.42E-09 |
| chr12 | 60181117 | 60182043 | 927 | 1.99 | 5.67E-10 | 3.44E-09 |
| chr15 | 51520244 | 51521290 | 1047 | -2.75 | 5.68E-10 | 3.45E-09 |
| chr8 | 80007891 | 80009674 | 1784 | 6.83 | 5.69E-10 | 3.45E-09 |
| chr6 | 38372355 | 38373379 | 1025 | 3.21 | 5.71E-10 | 3.46E-09 |
| chr15 | 85522896 | 85686691 | 163796 | 0.19 | 5.71E-10 | 3.46E-09 |
| chr12 | 98334187 | 98336483 | 2297 | 6.83 | 5.72E-10 | 3.47E-09 |
| chr5 | 90338287 | 90355332 | 17046 | 1.03 | 5.84E-10 | 3.54E-09 |
| chr1 | 61787404 | 61805610 | 18207 | 0.73 | 5.85E-10 | 3.55E-09 |
| chr8 | 116112621 | 116113711 | 1091 | 6.87 | 5.88E-10 | 3.57E-09 |
| chr3 | 96531552 | 96533304 | 1753 | 6.86 | 5.89E-10 | 3.57E-09 |
| chr12 | 84015429 | 84016344 | 916 | 6.88 | 5.95E-10 | 3.61E-09 |
| chrX | 46175720 | 46194270 | 18551 | 0.29 | 5.97E-10 | 3.62E-09 |
| chr19 | 22192413 | 22193735 | 1323 | 6.83 | 5.98E-10 | 3.62E-09 |

| | | | | | | |
|---|---|---|---|---|---|---|
| chr18 | 53828928 | 53836413 | 7486 | 2 | 5.99E-10 | 3.63E-09 |
| chrX | 123074513 | 123106242 | 31730 | 0.44 | 5.99E-10 | 3.63E-09 |
| chr2 | 38405237 | 38407265 | 2029 | 1.59 | 6.00E-10 | 3.64E-09 |
| chr3 | 175536440 | 175537804 | 1365 | 6.82 | 6.01E-10 | 3.64E-09 |
| chr5 | 157315192 | 157317017 | 1826 | 6.82 | 6.02E-10 | 3.64E-09 |
| chr13 | 47105602 | 47129413 | 23812 | 0.92 | 6.07E-10 | 3.67E-09 |
| chr11 | 27900743 | 27904615 | 3873 | 1.56 | 6.08E-10 | 3.68E-09 |
| chr14 | 80891681 | 80901873 | 10193 | -6.88 | 6.10E-10 | 3.69E-09 |
| chr13 | 41110407 | 41115773 | 5367 | 0.77 | 6.11E-10 | 3.69E-09 |
| chr12 | 40396145 | 40411358 | 15214 | 3.02 | 6.13E-10 | 3.70E-09 |
| chr4 | 178228431 | 178261051 | 32621 | 0.4 | 6.14E-10 | 3.71E-09 |
| chr12 | 22464935 | 22466051 | 1117 | 4.49 | 6.26E-10 | 3.79E-09 |
| chr4 | 141647126 | 141678977 | 31852 | 0.89 | 6.36E-10 | 3.84E-09 |
| chr3 | 97661048 | 97691919 | 30872 | -0.67 | 6.38E-10 | 3.85E-09 |
| chr17 | 77805101 | 77827346 | 22246 | 0.61 | 6.46E-10 | 3.90E-09 |
| chr7 | 148487908 | 148518140 | 30233 | 0.69 | 6.48E-10 | 3.91E-09 |
| chr9 | 30642645 | 30643477 | 833 | 6.81 | 6.48E-10 | 3.91E-09 |
| chr11 | 109946660 | 109948983 | 2324 | 2.42 | 6.48E-10 | 3.91E-09 |
| chr19 | 30155663 | 30182331 | 26669 | -0.89 | 6.50E-10 | 3.92E-09 |
| chr7 | 67161459 | 67162815 | 1357 | 1.82 | 6.57E-10 | 3.96E-09 |
| chr12 | 122706010 | 122722665 | 16656 | 0.67 | 6.57E-10 | 3.96E-09 |
| chr11 | 59668371 | 59669517 | 1147 | 5.04 | 6.59E-10 | 3.97E-09 |
| chr1 | 67655506 | 67656852 | 1347 | 2.95 | 6.60E-10 | 3.98E-09 |
| chr4 | 187037488 | 187038237 | 750 | 4.19 | 6.61E-10 | 3.99E-09 |
| chr1 | 79593486 | 79605264 | 11779 | -2.19 | 6.62E-10 | 3.99E-09 |
| chr10 | 91060750 | 91063398 | 2649 | 0.78 | 6.64E-10 | 4.00E-09 |
| chr19 | 11877328 | 11879262 | 1935 | 6.81 | 6.73E-10 | 4.06E-09 |
| chr9 | 36275748 | 36277821 | 2074 | 2.79 | 6.74E-10 | 4.06E-09 |
| chr12 | 97300007 | 97313355 | 13349 | 0.87 | 6.75E-10 | 4.06E-09 |
| chr13 | 82992605 | 82994158 | 1554 | 4.67 | 6.77E-10 | 4.08E-09 |
| chr18 | 54337780 | 54338999 | 1220 | 3.72 | 6.78E-10 | 4.08E-09 |
| chr10 | 27672229 | 27673743 | 1515 | 2.48 | 6.82E-10 | 4.11E-09 |
| chr14 | 105330305 | 105333118 | 2814 | 1.2 | 6.84E-10 | 4.12E-09 |
| chr15 | 38192327 | 38195030 | 2704 | 2.12 | 6.85E-10 | 4.12E-09 |
| chr2 | 220137141 | 220145182 | 8042 | -0.67 | 6.85E-10 | 4.12E-09 |
| chr9 | 135644410 | 135646166 | 1757 | 1.38 | 6.86E-10 | 4.13E-09 |
| chr21 | 36627993 | 36638312 | 10320 | 0.87 | 6.87E-10 | 4.13E-09 |
| chr18 | 47117669 | 47118806 | 1138 | 4.31 | 6.90E-10 | 4.15E-09 |
| chr8 | 125452327 | 125464197 | 11871 | -0.91 | 6.95E-10 | 4.18E-09 |
| chr10 | 104472434 | 104490020 | 17587 | -0.87 | 7.05E-10 | 4.24E-09 |
| chr16 | 56715601 | 56736583 | 20983 | 0.78 | 7.14E-10 | 4.29E-09 |
| chr15 | 50455025 | 50475175 | 20151 | -2.37 | 7.16E-10 | 4.30E-09 |
| chr11 | 46038158 | 46039633 | 1476 | 2.07 | 7.21E-10 | 4.33E-09 |
| chr10 | 3085343 | 3185242 | 99900 | 0.3 | 7.22E-10 | 4.33E-09 |
| chr5 | 36939662 | 36941527 | 1866 | 2 | 7.23E-10 | 4.34E-09 |
| chr12 | 56109250 | 56151751 | 42502 | 0.6 | 7.23E-10 | 4.34E-09 |
| chr8 | 37159255 | 37161053 | 1799 | 2.18 | 7.27E-10 | 4.36E-09 |
| chr3 | 177001785 | 177019422 | 17638 | 0.75 | 7.30E-10 | 4.38E-09 |
| chr12 | 76411133 | 76428495 | 17363 | 0.5 | 7.38E-10 | 4.43E-09 |
| chr8 | 34713814 | 34715534 | 1721 | -6.78 | 7.38E-10 | 4.43E-09 |
| chrX | 20378071 | 20402197 | 24127 | 0.4 | 7.40E-10 | 4.44E-09 |
| chr1 | 225116865 | 225118815 | 1951 | 1.18 | 7.47E-10 | 4.48E-09 |
| chr17 | 59232643 | 59237788 | 5146 | 0.59 | 7.48E-10 | 4.48E-09 |
| chr6 | 143993461 | 144000491 | 7031 | 0.64 | 7.49E-10 | 4.49E-09 |
| chr6 | 79172097 | 79173496 | 1400 | 3.62 | 7.50E-10 | 4.49E-09 |
| chr1 | 198503461 | 198504587 | 1127 | 2.75 | 7.51E-10 | 4.50E-09 |
| chr5 | 52379541 | 52389566 | 10026 | -1.19 | 7.58E-10 | 4.54E-09 |
| chr19 | 54000684 | 54008525 | 7842 | 6.83 | 7.59E-10 | 4.55E-09 |
| chr19 | 57831004 | 57833138 | 2135 | 1.09 | 7.66E-10 | 4.58E-09 |
| chr12 | 63808357 | 63814033 | 5677 | 6.79 | 7.67E-10 | 4.59E-09 |
| chr1 | 189626842 | 189635533 | 8692 | 3.25 | 7.67E-10 | 4.59E-09 |

| | | | | | | |
|---|---|---|---|---|---|---|
| chr9 | 79070954 | 79074813 | 3860 | 5.4 | 7.74E-10 | 4.63E-09 |
| chr4 | 119989168 | 120035208 | 46041 | 0.34 | 7.74E-10 | 4.63E-09 |
| chr2 | 53108539 | 53109806 | 1268 | 6.85 | 7.78E-10 | 4.66E-09 |
| chr3 | 185254499 | 185265777 | 11279 | 0.53 | 7.81E-10 | 4.67E-09 |
| chr12 | 74930072 | 74935088 | 5017 | 0.5 | 7.84E-10 | 4.69E-09 |
| chrX | 4487846 | 4489062 | 1217 | 4.15 | 7.85E-10 | 4.69E-09 |
| chr3 | 96423707 | 96425209 | 1503 | 6.78 | 7.90E-10 | 4.72E-09 |
| chr15 | 45923404 | 45953675 | 30272 | 0.37 | 7.92E-10 | 4.73E-09 |
| chr11 | 95092362 | 95099139 | 6778 | 1.17 | 8.00E-10 | 4.78E-09 |
| chr6 | 47708674 | 47710353 | 1680 | 3.87 | 8.01E-10 | 4.78E-09 |
| chr4 | 74078377 | 74097644 | 19268 | 0.57 | 8.05E-10 | 4.80E-09 |
| chr12 | 102977856 | 102979088 | 1233 | 2.61 | 8.08E-10 | 4.82E-09 |
| chr4 | 148466664 | 148469259 | 2596 | 2.28 | 8.13E-10 | 4.85E-09 |
| chr21 | 35013034 | 35032525 | 19492 | 0.64 | 8.15E-10 | 4.86E-09 |
| chr1 | 169073964 | 169132966 | 59003 | 0.27 | 8.15E-10 | 4.86E-09 |
| chr19 | 9584977 | 9586379 | 1403 | 6.79 | 8.18E-10 | 4.88E-09 |
| chr14 | 58819258 | 58821178 | 1921 | 1.66 | 8.19E-10 | 4.88E-09 |
| chr2 | 232465490 | 232552625 | 87136 | -0.3 | 8.23E-10 | 4.91E-09 |
| chr7 | 134831751 | 134862732 | 30982 | 0.48 | 8.26E-10 | 4.93E-09 |
| chr15 | 85900511 | 85902314 | 1804 | 1.28 | 8.27E-10 | 4.93E-09 |
| chr9 | 13181019 | 13189905 | 8887 | 1.38 | 8.27E-10 | 4.93E-09 |
| chr12 | 57701821 | 57704616 | 2796 | 1 | 8.32E-10 | 4.96E-09 |
| chr5 | 65016888 | 65047200 | 30313 | -0.5 | 8.32E-10 | 4.96E-09 |
| chr4 | 138584618 | 138585989 | 1372 | 2.64 | 8.35E-10 | 4.97E-09 |
| chr12 | 27899058 | 27910650 | 11593 | 0.51 | 8.40E-10 | 5.00E-09 |
| chr1 | 223999268 | 224035295 | 36028 | 0.54 | 8.44E-10 | 5.02E-09 |
| chr2 | 111976598 | 111986136 | 9539 | -5.54 | 8.44E-10 | 5.02E-09 |
| chr6 | 20236605 | 20238153 | 1549 | 2.71 | 8.48E-10 | 5.05E-09 |
| chr3 | 105716013 | 105723653 | 7641 | 2.1 | 8.49E-10 | 5.05E-09 |
| chr12 | 85429496 | 85431031 | 1536 | 1 | 8.59E-10 | 5.11E-09 |
| chr5 | 130691553 | 130728010 | 36458 | 0.39 | 8.60E-10 | 5.11E-09 |
| chr12 | 50134342 | 50148002 | 13661 | 0.63 | 8.67E-10 | 5.15E-09 |
| chrX | 130964141 | 130965611 | 1471 | 1.94 | 8.68E-10 | 5.16E-09 |
| chr4 | 155994487 | 155996477 | 1991 | 3.76 | 8.68E-10 | 5.16E-09 |
| chr10 | 49571797 | 49575286 | 3490 | 1.23 | 8.69E-10 | 5.16E-09 |
| chr12 | 66088742 | 66090636 | 1895 | 1.59 | 8.70E-10 | 5.17E-09 |
| chr1 | 114654288 | 114657819 | 3532 | 2.08 | 8.75E-10 | 5.20E-09 |
| chr9 | 82281804 | 82283668 | 1865 | 1.7 | 8.82E-10 | 5.24E-09 |
| chr16 | 27833514 | 27834423 | 910 | 6.77 | 8.83E-10 | 5.24E-09 |
| chr13 | 99316370 | 99318751 | 2382 | 2.53 | 8.85E-10 | 5.25E-09 |
| chr14 | 98535874 | 98537222 | 1349 | 6.77 | 8.90E-10 | 5.28E-09 |
| chr2 | 238199057 | 238200969 | 1913 | -1.11 | 8.91E-10 | 5.29E-09 |
| chr6 | 129633842 | 129640439 | 6598 | 6.83 | 8.94E-10 | 5.30E-09 |
| chr12 | 63781329 | 63795219 | 13891 | 7.11 | 8.98E-10 | 5.33E-09 |
| chr14 | 22800896 | 22803613 | 2718 | 1.2 | 9.00E-10 | 5.34E-09 |
| chr7 | 150382186 | 150383533 | 1348 | 6.79 | 9.02E-10 | 5.35E-09 |
| chr4 | 122603701 | 122619105 | 15405 | 0.56 | 9.03E-10 | 5.35E-09 |
| chrX | 39811725 | 39817848 | 6124 | 1.05 | 9.07E-10 | 5.37E-09 |
| chr20 | 15014129 | 15015178 | 1050 | 6.77 | 9.07E-10 | 5.37E-09 |
| chr3 | 38417861 | 38429797 | 11937 | 2.13 | 9.07E-10 | 5.37E-09 |
| chr21 | 43178249 | 43205140 | 26892 | 0.65 | 9.11E-10 | 5.40E-09 |
| chr12 | 130537940 | 130538965 | 1026 | 3.61 | 9.13E-10 | 5.40E-09 |
| chr5 | 129761839 | 129763562 | 1724 | 3.18 | 9.13E-10 | 5.41E-09 |
| chr7 | 115849984 | 115873657 | 23674 | 0.5 | 9.14E-10 | 5.41E-09 |
| chr8 | 115724063 | 115725464 | 1402 | 6.77 | 9.14E-10 | 5.41E-09 |
| chr4 | 102784511 | 102788830 | 4320 | 0.72 | 9.16E-10 | 5.42E-09 |
| chr18 | 25773227 | 25785117 | 11891 | 2.65 | 9.23E-10 | 5.46E-09 |
| chr2 | 30101528 | 30102926 | 1399 | -5.31 | 9.23E-10 | 5.46E-09 |
| chr8 | 107669144 | 107672078 | 2935 | 0.87 | 9.25E-10 | 5.47E-09 |
| chr1 | 241754096 | 241836686 | 82591 | 0.27 | 9.28E-10 | 5.49E-09 |
| chr7 | 95278794 | 95279705 | 912 | 5.35 | 9.44E-10 | 5.58E-09 |

| | | | | | | |
|---|---|---|---|---|---|---|
| chr14 | 68744466 | 68747189 | 2724 | -1.08 | 9.45E-10 | 5.58E-09 |
| chr12 | 65808970 | 65810312 | 1343 | 4.61 | 9.47E-10 | 5.60E-09 |
| chr13 | 24316466 | 24332518 | 16053 | 0.62 | 9.53E-10 | 5.63E-09 |
| chr4 | 116592234 | 116593628 | 1395 | 6.77 | 9.56E-10 | 5.64E-09 |
| chr1 | 156497914 | 156520613 | 22700 | -0.94 | 9.61E-10 | 5.67E-09 |
| chr10 | 8091267 | 8097356 | 6090 | 1.37 | 9.61E-10 | 5.68E-09 |
| chr12 | 115731581 | 115733321 | 1741 | -6.73 | 9.62E-10 | 5.68E-09 |
| chr9 | 134916044 | 134918881 | 2838 | -1.87 | 9.70E-10 | 5.72E-09 |
| chrX | 149436753 | 149439581 | 2829 | 1.23 | 9.72E-10 | 5.74E-09 |
| chr7 | 69062413 | 69095878 | 33466 | 6.75 | 9.81E-10 | 5.79E-09 |
| chr12 | 109525001 | 109551379 | 26379 | 0.45 | 9.85E-10 | 5.81E-09 |
| chr12 | 131910847 | 131913057 | 2211 | 6.91 | 9.86E-10 | 5.81E-09 |
| chr8 | 134528943 | 134558662 | 29720 | 0.92 | 9.87E-10 | 5.82E-09 |
| chr9 | 36243555 | 36268947 | 25393 | 0.58 | 9.90E-10 | 5.83E-09 |
| chr2 | 128926168 | 128954482 | 28315 | -0.9 | 9.97E-10 | 5.88E-09 |
| chr9 | 99373898 | 99390586 | 16689 | 1.03 | 9.98E-10 | 5.88E-09 |
| chr10 | 45123899 | 45125298 | 1400 | -2.86 | 1.00E-09 | 5.89E-09 |
| chr7 | 156819874 | 156821679 | 1806 | 6.76 | 1.00E-09 | 5.92E-09 |
| chr1 | 94341868 | 94375894 | 34027 | -0.41 | 1.01E-09 | 5.93E-09 |
| chr1 | 184460287 | 184461455 | 1169 | -3.34 | 1.01E-09 | 5.93E-09 |
| chr6 | 21686696 | 21687909 | 1214 | 5.35 | 1.01E-09 | 5.95E-09 |
| chr6 | 1287004 | 1298441 | 11438 | -2.15 | 1.01E-09 | 5.95E-09 |
| chr10 | 133809932 | 133812795 | 2864 | 1.12 | 1.02E-09 | 5.98E-09 |
| chr1 | 211847996 | 211854779 | 6784 | -1.09 | 1.03E-09 | 6.04E-09 |
| chr4 | 91763674 | 91768708 | 5035 | 1.16 | 1.03E-09 | 6.04E-09 |
| chr7 | 22121852 | 22123592 | 1741 | 2.04 | 1.03E-09 | 6.05E-09 |
| chr3 | 102803804 | 102805362 | 1559 | 6.8 | 1.03E-09 | 6.05E-09 |
| chr10 | 111836095 | 111844163 | 8069 | 1.79 | 1.03E-09 | 6.06E-09 |
| chrX | 69744921 | 69747673 | 2753 | 2.15 | 1.04E-09 | 6.09E-09 |
| chr7 | 25010589 | 25021778 | 11190 | 1.12 | 1.04E-09 | 6.10E-09 |
| chr11 | 76455640 | 76517418 | 61779 | -0.28 | 1.05E-09 | 6.14E-09 |
| chr14 | 75893624 | 75938083 | 44460 | -0.5 | 1.05E-09 | 6.15E-09 |
| chr14 | 65945034 | 65945898 | 865 | 4.94 | 1.05E-09 | 6.17E-09 |
| chr21 | 44034820 | 44045247 | 10428 | 6.76 | 1.05E-09 | 6.17E-09 |
| chr20 | 23509967 | 23510621 | 655 | 6.75 | 1.06E-09 | 6.20E-09 |
| chrX | 118148529 | 118163341 | 14813 | 1.09 | 1.06E-09 | 6.22E-09 |
| chr2 | 240061819 | 240066299 | 4481 | -3.08 | 1.06E-09 | 6.22E-09 |
| chr15 | 68485148 | 68522767 | 37620 | -0.63 | 1.06E-09 | 6.24E-09 |
| chr9 | 114462679 | 114463271 | 593 | 4.77 | 1.07E-09 | 6.26E-09 |
| chr12 | 101799826 | 101802921 | 3096 | 0.83 | 1.07E-09 | 6.26E-09 |
| chr6 | 1316622 | 1336496 | 19875 | -0.95 | 1.08E-09 | 6.31E-09 |
| chr2 | 36555858 | 36567181 | 11324 | 0.65 | 1.08E-09 | 6.31E-09 |
| chr17 | 29635299 | 29641572 | 6274 | 1.39 | 1.08E-09 | 6.32E-09 |
| chr1 | 7315613 | 7339070 | 23458 | 0.64 | 1.08E-09 | 6.34E-09 |
| chr1 | 9778005 | 9780298 | 2294 | -1.29 | 1.08E-09 | 6.35E-09 |
| chr2 | 62520308 | 62521659 | 1352 | 2.16 | 1.09E-09 | 6.36E-09 |
| chr14 | 57705509 | 57715185 | 9677 | 0.96 | 1.09E-09 | 6.39E-09 |
| chr15 | 65108049 | 65150582 | 42534 | -0.71 | 1.10E-09 | 6.41E-09 |
| chr4 | 169395587 | 169412220 | 16634 | 0.47 | 1.10E-09 | 6.41E-09 |
| chr8 | 58653061 | 58658397 | 5337 | 6.78 | 1.10E-09 | 6.44E-09 |
| chr11 | 9823342 | 9845772 | 22431 | -0.95 | 1.11E-09 | 6.49E-09 |
| chr16 | 14290192 | 14297625 | 7434 | 2.53 | 1.12E-09 | 6.55E-09 |
| chr15 | 91031973 | 91034398 | 2426 | -1.96 | 1.13E-09 | 6.62E-09 |
| chr18 | 1997953 | 2002276 | 4324 | 6.74 | 1.13E-09 | 6.63E-09 |
| chr3 | 102876699 | 102878125 | 1427 | 6.74 | 1.13E-09 | 6.63E-09 |
| chr12 | 14863754 | 14864556 | 803 | 3.04 | 1.13E-09 | 6.63E-09 |
| chr8 | 86458322 | 86459980 | 1659 | 4.12 | 1.14E-09 | 6.63E-09 |
| chr9 | 74906519 | 74941628 | 35110 | 0.32 | 1.14E-09 | 6.64E-09 |
| chr13 | 83262035 | 83263341 | 1307 | 2.38 | 1.14E-09 | 6.64E-09 |
| chr2 | 40477993 | 40479001 | 1009 | 6.75 | 1.15E-09 | 6.71E-09 |
| chr14 | 85967395 | 85969788 | 2394 | 2.72 | 1.15E-09 | 6.72E-09 |

| | | | | | | |
|---|---|---|---|---|---|---|
| chr3 | 148990466 | 149001745 | 11280 | 0.76 | 1.15E-09 | 6.72E-09 |
| chr12 | 50221093 | 50237675 | 16583 | 0.77 | 1.15E-09 | 6.74E-09 |
| chr11 | 13029163 | 13035482 | 6320 | 1.2 | 1.16E-09 | 6.77E-09 |
| chr19 | 5165709 | 5178028 | 12320 | 0.48 | 1.16E-09 | 6.78E-09 |
| chr1 | 239478293 | 239479734 | 1442 | 6.73 | 1.16E-09 | 6.79E-09 |
| chr8 | 17083506 | 17084717 | 1212 | 5.91 | 1.18E-09 | 6.89E-09 |
| chrX | 115566625 | 115567878 | 1254 | 6.73 | 1.18E-09 | 6.89E-09 |
| chr14 | 53740349 | 53746727 | 6379 | -1.6 | 1.18E-09 | 6.90E-09 |
| chr5 | 126721553 | 126722843 | 1291 | 4.75 | 1.18E-09 | 6.90E-09 |
| chr12 | 97203762 | 97206186 | 2425 | 1.22 | 1.19E-09 | 6.92E-09 |
| chr1 | 212177796 | 212179126 | 1331 | 2.92 | 1.20E-09 | 6.99E-09 |
| chr20 | 49091862 | 49101288 | 9427 | 0.88 | 1.20E-09 | 6.99E-09 |
| chr13 | 80323451 | 80325061 | 1611 | 1.91 | 1.20E-09 | 6.99E-09 |
| chr5 | 90744530 | 90745682 | 1153 | 3.41 | 1.20E-09 | 7.00E-09 |
| chrX | 101185746 | 101187783 | 2038 | 1.21 | 1.21E-09 | 7.02E-09 |
| chr5 | 52702068 | 52726177 | 24110 | -0.66 | 1.21E-09 | 7.06E-09 |
| chr1 | 31296157 | 31297414 | 1258 | 1.62 | 1.21E-09 | 7.06E-09 |
| chr17 | 26383130 | 26385778 | 2649 | 2.16 | 1.21E-09 | 7.07E-09 |
| chr3 | 145967923 | 145970008 | 2086 | 1.21 | 1.22E-09 | 7.12E-09 |
| chr1 | 23425481 | 23450289 | 24809 | -0.74 | 1.23E-09 | 7.17E-09 |
| chr10 | 8657576 | 8661891 | 4316 | 1.43 | 1.24E-09 | 7.19E-09 |
| chr3 | 37934125 | 37935703 | 1579 | 2.09 | 1.24E-09 | 7.19E-09 |
| chr11 | 73238869 | 73240364 | 1496 | 2.54 | 1.24E-09 | 7.20E-09 |
| chr12 | 70131884 | 70133993 | 2110 | 1.46 | 1.24E-09 | 7.23E-09 |
| chr1 | 167516975 | 167523010 | 6036 | -1.73 | 1.26E-09 | 7.29E-09 |
| chr10 | 22604489 | 22630587 | 26099 | 0.76 | 1.26E-09 | 7.33E-09 |
| chr17 | 17478723 | 17497053 | 18331 | -0.74 | 1.28E-09 | 7.41E-09 |
| chr5 | 115337779 | 115338883 | 1105 | 6.76 | 1.28E-09 | 7.42E-09 |
| chr7 | 16245866 | 16247084 | 1219 | 6.73 | 1.28E-09 | 7.44E-09 |
| chr3 | 175389764 | 175391048 | 1285 | 6.81 | 1.28E-09 | 7.44E-09 |
| chr19 | 48394981 | 48397228 | 2248 | 5.35 | 1.29E-09 | 7.47E-09 |
| chr21 | 18883101 | 18909649 | 26549 | 0.7 | 1.29E-09 | 7.51E-09 |
| chr14 | 70720918 | 70721611 | 694 | 2.75 | 1.30E-09 | 7.54E-09 |
| chr6 | 138000935 | 138003718 | 2784 | -0.76 | 1.32E-09 | 7.65E-09 |
| chr20 | 62167903 | 62185501 | 17599 | 0.93 | 1.33E-09 | 7.70E-09 |
| chr13 | 20483913 | 20485393 | 1481 | 3.86 | 1.33E-09 | 7.70E-09 |
| chr2 | 133171882 | 133204743 | 32862 | -0.5 | 1.34E-09 | 7.78E-09 |
| chr5 | 174069418 | 174070556 | 1139 | 5.29 | 1.35E-09 | 7.81E-09 |
| chr10 | 134479501 | 134481805 | 2305 | -2.65 | 1.35E-09 | 7.83E-09 |
| chr22 | 33959583 | 33975545 | 15963 | 0.96 | 1.36E-09 | 7.87E-09 |
| chr1 | 169028152 | 169030910 | 2759 | 1.19 | 1.36E-09 | 7.89E-09 |
| chr4 | 148498362 | 148504113 | 5752 | 1.73 | 1.37E-09 | 7.92E-09 |
| chr15 | 55571504 | 55583247 | 11744 | 0.86 | 1.37E-09 | 7.93E-09 |
| chr4 | 168475886 | 168477237 | 1352 | 6.71 | 1.38E-09 | 7.96E-09 |
| chr8 | 114466728 | 114468185 | 1458 | 6.71 | 1.38E-09 | 7.96E-09 |
| chr9 | 27498908 | 27500311 | 1404 | -4.91 | 1.39E-09 | 8.07E-09 |
| chr5 | 173114276 | 173124250 | 9975 | 0.49 | 1.41E-09 | 8.14E-09 |
| chr3 | 38169127 | 38208397 | 39271 | 0.48 | 1.41E-09 | 8.18E-09 |
| chr7 | 82790885 | 82793372 | 2488 | 1.41 | 1.43E-09 | 8.26E-09 |
| chr1 | 160129253 | 160130427 | 1175 | 4.44 | 1.43E-09 | 8.28E-09 |
| chr19 | 46269803 | 46341260 | 71458 | -0.44 | 1.43E-09 | 8.28E-09 |
| chr1 | 5484376 | 5491259 | 6884 | 6.73 | 1.44E-09 | 8.31E-09 |
| chr2 | 113770116 | 113771158 | 1043 | -4.47 | 1.44E-09 | 8.31E-09 |
| chr17 | 67488265 | 67500341 | 12077 | 0.86 | 1.45E-09 | 8.36E-09 |
| chr17 | 6751889 | 6777256 | 25368 | 0.48 | 1.45E-09 | 8.38E-09 |
| chr2 | 38440391 | 38444399 | 4009 | -1.4 | 1.45E-09 | 8.39E-09 |
| chr7 | 122525804 | 122527407 | 1604 | 6.73 | 1.46E-09 | 8.40E-09 |
| chr7 | 150700313 | 150701436 | 1124 | 3.41 | 1.46E-09 | 8.42E-09 |
| chr4 | 23291744 | 23307183 | 15440 | 2.19 | 1.46E-09 | 8.42E-09 |
| chr9 | 6217717 | 6218583 | 867 | 5.85 | 1.47E-09 | 8.46E-09 |
| chr4 | 92328572 | 92329798 | 1227 | 4.33 | 1.47E-09 | 8.47E-09 |

| | | | | | | |
|---|---|---|---|---|---|---|
| chr16 | 53659909 | 53661411 | 1503 | 2.79 | 1.50E-09 | 8.63E-09 |
| chr11 | 84834372 | 84835478 | 1107 | 6.75 | 1.50E-09 | 8.64E-09 |
| chr1 | 215716145 | 215717142 | 998 | 6.73 | 1.50E-09 | 8.66E-09 |
| chr10 | 38179306 | 38180277 | 972 | 5.82 | 1.51E-09 | 8.70E-09 |
| chr13 | 32653521 | 32655181 | 1661 | 6.76 | 1.51E-09 | 8.70E-09 |
| chr2 | 38998928 | 39025598 | 26671 | -0.82 | 1.52E-09 | 8.78E-09 |
| chr4 | 8183373 | 8186470 | 3098 | 1.49 | 1.52E-09 | 8.79E-09 |
| chr16 | 11500922 | 11505550 | 4629 | -1.68 | 1.53E-09 | 8.79E-09 |
| chr8 | 83577236 | 83578822 | 1587 | 5.66 | 1.53E-09 | 8.81E-09 |
| chr12 | 84510584 | 84511600 | 1017 | 6.78 | 1.54E-09 | 8.84E-09 |
| chr9 | 8733863 | 8735545 | 1683 | 6.71 | 1.54E-09 | 8.88E-09 |
| chr12 | 79539948 | 79551066 | 11119 | 2.57 | 1.54E-09 | 8.89E-09 |
| chr19 | 5827576 | 5851130 | 23555 | 1.25 | 1.56E-09 | 8.96E-09 |
| chr19 | 35704264 | 35705613 | 1350 | 2.9 | 1.57E-09 | 9.01E-09 |
| chr6 | 13709137 | 13726024 | 16888 | 1.03 | 1.58E-09 | 9.09E-09 |
| chr6 | 20648921 | 20662948 | 14028 | 0.62 | 1.58E-09 | 9.11E-09 |
| chr20 | 33460306 | 33474458 | 14153 | 1.28 | 1.59E-09 | 9.15E-09 |
| chr2 | 25321154 | 25330505 | 9352 | -0.81 | 1.62E-09 | 9.29E-09 |
| chr6 | 50903122 | 50904646 | 1525 | 6.71 | 1.63E-09 | 9.37E-09 |
| chr6 | 101079565 | 101082202 | 2638 | 0.76 | 1.64E-09 | 9.41E-09 |
| chr5 | 148289573 | 148293916 | 4344 | 1.01 | 1.66E-09 | 9.51E-09 |
| chr9 | 127572410 | 127581637 | 9228 | 0.74 | 1.67E-09 | 9.57E-09 |
| chr5 | 95926907 | 95928434 | 1528 | 5.78 | 1.67E-09 | 9.58E-09 |
| chr8 | 22100744 | 22103914 | 3171 | 0.8 | 1.68E-09 | 9.66E-09 |
| chr1 | 191356604 | 191357907 | 1304 | 6.77 | 1.69E-09 | 9.68E-09 |
| chr8 | 35003960 | 35005513 | 1554 | -6.74 | 1.69E-09 | 9.69E-09 |
| chr5 | 167512748 | 167521236 | 8489 | 6.69 | 1.69E-09 | 9.70E-09 |
| chr18 | 9895339 | 9900210 | 4872 | 0.89 | 1.69E-09 | 9.71E-09 |
| chr7 | 142942076 | 142942918 | 843 | 5.81 | 1.69E-09 | 9.71E-09 |
| chr12 | 97988896 | 97990029 | 1134 | 6.73 | 1.70E-09 | 9.73E-09 |
| chr4 | 111574568 | 111585769 | 11202 | 6.78 | 1.72E-09 | 9.86E-09 |
| chr16 | 10597475 | 10599548 | 2074 | 4.42 | 1.73E-09 | 9.91E-09 |
| chr10 | 19447085 | 19448459 | 1375 | 6.68 | 1.73E-09 | 9.92E-09 |
| chrX | 125643398 | 125644892 | 1495 | 6.68 | 1.73E-09 | 9.92E-09 |
| chr4 | 106489557 | 106491127 | 1571 | 6.69 | 1.74E-09 | 9.95E-09 |
| chr10 | 27330014 | 27342729 | 12716 | 1.56 | 1.74E-09 | 9.99E-09 |
| chr16 | 57332697 | 57351321 | 18625 | 0.9 | 1.75E-09 | 1.00E-08 |
| chr11 | 65654699 | 65688052 | 33354 | -0.29 | 1.75E-09 | 1.00E-08 |
| chr5 | 72232999 | 72253459 | 20461 | 0.69 | 1.76E-09 | 1.01E-08 |
| chr4 | 149276426 | 149277451 | 1026 | 6.68 | 1.77E-09 | 1.01E-08 |
| chr7 | 106057456 | 106066392 | 8937 | -1.72 | 1.77E-09 | 1.01E-08 |
| chr3 | 108266936 | 108283366 | 16431 | 0.7 | 1.81E-09 | 1.03E-08 |
| chr2 | 209502765 | 209503947 | 1183 | 6.71 | 1.81E-09 | 1.03E-08 |
| chr12 | 20412214 | 20428984 | 16771 | 1.66 | 1.82E-09 | 1.04E-08 |
| chr13 | 89802247 | 89803442 | 1196 | 6.67 | 1.83E-09 | 1.05E-08 |
| chr8 | 28656862 | 28669016 | 12155 | 1.13 | 1.83E-09 | 1.05E-08 |
| chr8 | 38613608 | 38652150 | 38543 | 0.39 | 1.83E-09 | 1.05E-08 |
| chr16 | 57162764 | 57186333 | 23570 | -0.85 | 1.85E-09 | 1.05E-08 |
| chr15 | 73343223 | 73346247 | 3025 | 0.79 | 1.85E-09 | 1.05E-08 |
| chr10 | 90734140 | 90735767 | 1628 | 1.12 | 1.85E-09 | 1.06E-08 |
| chr6 | 140053551 | 140088077 | 34527 | 1.03 | 1.85E-09 | 1.06E-08 |
| chr7 | 122314596 | 122320148 | 5553 | 6.68 | 1.86E-09 | 1.06E-08 |
| chr3 | 83616601 | 83617642 | 1042 | 6.67 | 1.87E-09 | 1.07E-08 |
| chr2 | 32357701 | 32370714 | 13014 | 0.78 | 1.87E-09 | 1.07E-08 |
| chr9 | 131057385 | 131103838 | 46454 | -0.53 | 1.87E-09 | 1.07E-08 |
| chr5 | 106298146 | 106302898 | 4753 | 1.34 | 1.88E-09 | 1.07E-08 |
| chr6 | 4687629 | 4694093 | 6465 | 0.7 | 1.89E-09 | 1.08E-08 |
| chr5 | 115879976 | 115881154 | 1179 | 6.67 | 1.89E-09 | 1.08E-08 |
| chr3 | 32395295 | 32408718 | 13424 | 1.32 | 1.90E-09 | 1.08E-08 |
| chr3 | 121536606 | 121554705 | 18100 | 0.8 | 1.90E-09 | 1.08E-08 |
| chr12 | 72078973 | 72081836 | 2864 | 0.8 | 1.91E-09 | 1.09E-08 |

| | | | | | | |
|---|---|---|---|---|---|---|
| chr6 | 22025455 | 22055397 | 29943 | 1.26 | 1.91E-09 | 1.09E-08 |
| chr9 | 80637322 | 80649557 | 12236 | 0.66 | 1.92E-09 | 1.09E-08 |
| chr10 | 6394546 | 6408236 | 13691 | -1.03 | 1.93E-09 | 1.10E-08 |
| chr14 | 26749406 | 26759051 | 9646 | 0.59 | 1.96E-09 | 1.12E-08 |
| chr20 | 10349867 | 10352957 | 3091 | -0.94 | 1.96E-09 | 1.12E-08 |
| chr5 | 102161990 | 102163212 | 1223 | 2.27 | 1.96E-09 | 1.12E-08 |
| chr7 | 113687980 | 113689110 | 1131 | 2.48 | 1.96E-09 | 1.12E-08 |
| chr12 | 85470634 | 85471704 | 1071 | 6.75 | 1.96E-09 | 1.12E-08 |
| chr9 | 139775542 | 139845789 | 70248 | 0.28 | 1.96E-09 | 1.12E-08 |
| chr1 | 59964879 | 59966242 | 1364 | 2.66 | 1.96E-09 | 1.12E-08 |
| chr4 | 163134607 | 163135516 | 910 | 5.79 | 1.99E-09 | 1.13E-08 |
| chr14 | 101154991 | 101160631 | 5641 | -1.38 | 1.99E-09 | 1.13E-08 |
| chr11 | 80090102 | 80091306 | 1205 | 6.67 | 1.99E-09 | 1.13E-08 |
| chr8 | 64103312 | 64117805 | 14494 | 1.04 | 2.01E-09 | 1.14E-08 |
| chr5 | 108083107 | 108103203 | 20097 | 0.66 | 2.01E-09 | 1.14E-08 |
| chr17 | 16438306 | 16440821 | 2516 | 0.99 | 2.02E-09 | 1.15E-08 |
| chr4 | 38491473 | 38492472 | 1000 | 3.86 | 2.02E-09 | 1.15E-08 |
| chr19 | 45347262 | 45431296 | 84035 | 0.33 | 2.02E-09 | 1.15E-08 |
| chr4 | 66248916 | 66250891 | 1976 | 2.05 | 2.03E-09 | 1.15E-08 |
| chr9 | 14532444 | 14533524 | 1081 | 2.75 | 2.05E-09 | 1.16E-08 |
| chr9 | 135479693 | 135487419 | 7727 | -1.49 | 2.05E-09 | 1.16E-08 |
| chrX | 115651618 | 115652784 | 1167 | 6.66 | 2.06E-09 | 1.17E-08 |
| chr18 | 27700816 | 27708496 | 7681 | 6.66 | 2.06E-09 | 1.17E-08 |
| chr5 | 23864385 | 23865576 | 1192 | 6.66 | 2.08E-09 | 1.18E-08 |
| chr8 | 97656915 | 97658203 | 1289 | 1.63 | 2.09E-09 | 1.19E-08 |
| chr22 | 41064359 | 41065850 | 1492 | 3.02 | 2.09E-09 | 1.19E-08 |
| chr11 | 60843114 | 60844258 | 1145 | 6.66 | 2.10E-09 | 1.19E-08 |
| chr8 | 48626465 | 48652051 | 25587 | 0.48 | 2.10E-09 | 1.19E-08 |
| chr8 | 62452520 | 62705391 | 252872 | 0.16 | 2.11E-09 | 1.19E-08 |
| chr14 | 62333702 | 62334664 | 963 | 3.27 | 2.11E-09 | 1.19E-08 |
| chr4 | 11530216 | 11532508 | 2293 | 1.81 | 2.11E-09 | 1.20E-08 |
| chr9 | 85790423 | 85791468 | 1046 | 6.69 | 2.12E-09 | 1.20E-08 |
| chr1 | 10753611 | 10754460 | 850 | 3.84 | 2.13E-09 | 1.21E-08 |
| chr6 | 88717744 | 88719318 | 1575 | 1.54 | 2.15E-09 | 1.22E-08 |
| chr10 | 29057768 | 29058622 | 855 | 5.75 | 2.15E-09 | 1.22E-08 |
| chr1 | 210859895 | 210864404 | 4510 | -2.1 | 2.16E-09 | 1.22E-08 |
| chr15 | 23810208 | 23811721 | 1514 | 1.45 | 2.17E-09 | 1.23E-08 |
| chr4 | 101645829 | 101660374 | 14546 | 6.65 | 2.18E-09 | 1.24E-08 |
| chr6 | 168424557 | 168436994 | 12438 | 1.96 | 2.19E-09 | 1.24E-08 |
| chr7 | 131284721 | 131286062 | 1342 | 4.37 | 2.19E-09 | 1.24E-08 |
| chr6 | 24307065 | 24315134 | 8070 | -0.93 | 2.19E-09 | 1.24E-08 |
| chr17 | 55674790 | 55682193 | 7404 | -1.82 | 2.19E-09 | 1.24E-08 |
| chr5 | 1380409 | 1386612 | 6204 | 2.87 | 2.20E-09 | 1.25E-08 |
| chr7 | 141395118 | 141406975 | 11858 | 1.04 | 2.26E-09 | 1.28E-08 |
| chr17 | 2135971 | 2148174 | 12204 | -1.32 | 2.26E-09 | 1.28E-08 |
| chr12 | 98151304 | 98152235 | 932 | 4.42 | 2.27E-09 | 1.28E-08 |
| chr3 | 11701994 | 11722030 | 20037 | 0.9 | 2.27E-09 | 1.28E-08 |
| chr10 | 1088459 | 1103651 | 15193 | 0.62 | 2.28E-09 | 1.29E-08 |
| chr7 | 133196889 | 133199461 | 2573 | 1.28 | 2.28E-09 | 1.29E-08 |
| chr17 | 17343517 | 17351212 | 7696 | 0.81 | 2.29E-09 | 1.29E-08 |
| chr1 | 84966004 | 84973396 | 7393 | 0.77 | 2.30E-09 | 1.30E-08 |
| chr5 | 39712660 | 39717436 | 4777 | -1.25 | 2.30E-09 | 1.30E-08 |
| chr2 | 203647368 | 203648686 | 1319 | 5.25 | 2.31E-09 | 1.30E-08 |
| chr4 | 88224485 | 88239402 | 14918 | 0.93 | 2.31E-09 | 1.30E-08 |
| chr1 | 5630383 | 5631601 | 1219 | 6.69 | 2.32E-09 | 1.31E-08 |
| chr10 | 37188333 | 37190632 | 2300 | 6.64 | 2.34E-09 | 1.32E-08 |
| chr20 | 10448522 | 10457656 | 9135 | -1.2 | 2.35E-09 | 1.33E-08 |
| chr13 | 48830653 | 48833399 | 2747 | 2.22 | 2.37E-09 | 1.34E-08 |
| chr8 | 27131834 | 27144748 | 12915 | 1.48 | 2.37E-09 | 1.34E-08 |
| chr8 | 94885733 | 94893922 | 8190 | 0.9 | 2.37E-09 | 1.34E-08 |
| chr3 | 19588854 | 19590152 | 1299 | -6.6 | 2.38E-09 | 1.34E-08 |

| | | | | | | |
|---|---|---|---|---|---|---|
| chr8 | 141922153 | 141966957 | 44805 | -0.48 | 2.39E-09 | 1.35E-08 |
| chr12 | 59269669 | 59272374 | 2706 | 1.39 | 2.39E-09 | 1.35E-08 |
| chr3 | 12790895 | 12805518 | 14624 | -0.93 | 2.45E-09 | 1.38E-08 |
| chrX | 100352021 | 100360110 | 8090 | 0.91 | 2.45E-09 | 1.38E-08 |
| chr19 | 16829350 | 16831298 | 1949 | 1.45 | 2.46E-09 | 1.38E-08 |
| chr1 | 97330488 | 97332257 | 1770 | 2.11 | 2.49E-09 | 1.40E-08 |
| chr15 | 86285437 | 86338838 | 53402 | -0.52 | 2.50E-09 | 1.41E-08 |
| chr8 | 141247495 | 141249515 | 2021 | 0.99 | 2.51E-09 | 1.41E-08 |
| chr7 | 90092475 | 90093268 | 794 | 6.7 | 2.51E-09 | 1.41E-08 |
| chr7 | 90224955 | 90226468 | 1514 | 4.26 | 2.51E-09 | 1.41E-08 |
| chr5 | 98197522 | 98267519 | 69998 | 0.46 | 2.52E-09 | 1.42E-08 |
| chr2 | 72744381 | 72746392 | 2012 | 2.26 | 2.53E-09 | 1.42E-08 |
| chr4 | 23261957 | 23270519 | 8563 | 6.69 | 2.55E-09 | 1.44E-08 |
| chr4 | 113030161 | 113035894 | 5734 | 0.8 | 2.56E-09 | 1.44E-08 |
| chr14 | 103651916 | 103676098 | 24183 | 1.24 | 2.58E-09 | 1.45E-08 |
| chr12 | 55285359 | 55286542 | 1184 | 4.98 | 2.59E-09 | 1.45E-08 |
| chr17 | 28163460 | 28178043 | 14584 | -1.53 | 2.59E-09 | 1.45E-08 |
| chr12 | 74996619 | 74997627 | 1009 | 6.66 | 2.59E-09 | 1.45E-08 |
| chr3 | 103208185 | 103209135 | 951 | 6.73 | 2.60E-09 | 1.46E-08 |
| chr12 | 4108204 | 4121368 | 13165 | 6.62 | 2.61E-09 | 1.47E-08 |
| chr2 | 236452013 | 236465360 | 13348 | -1.79 | 2.64E-09 | 1.48E-08 |
| chr16 | 23081285 | 23100555 | 19271 | 0.65 | 2.64E-09 | 1.48E-08 |
| chr5 | 45793352 | 45794445 | 1094 | 6.64 | 2.66E-09 | 1.49E-08 |
| chr2 | 216001205 | 216004003 | 2799 | 1.07 | 2.69E-09 | 1.51E-08 |
| chr16 | 88824301 | 88852402 | 28102 | -0.73 | 2.72E-09 | 1.53E-08 |
| chr2 | 33515459 | 33530350 | 14892 | 1.52 | 2.73E-09 | 1.53E-08 |
| chr9 | 93893360 | 93894800 | 1441 | 2.51 | 2.74E-09 | 1.53E-08 |
| chr7 | 133253257 | 133275888 | 22632 | 0.84 | 2.74E-09 | 1.54E-08 |
| chr7 | 78910193 | 78949699 | 39507 | -0.88 | 2.76E-09 | 1.55E-08 |
| chr7 | 12442448 | 12444302 | 1855 | 1.74 | 2.76E-09 | 1.55E-08 |
| chr3 | 141593460 | 141633786 | 40327 | -0.6 | 2.77E-09 | 1.55E-08 |
| chr21 | 17030339 | 17034009 | 3671 | 0.78 | 2.79E-09 | 1.56E-08 |
| chr11 | 9858758 | 9862596 | 3839 | 1.9 | 2.82E-09 | 1.58E-08 |
| chr4 | 87477736 | 87478983 | 1248 | 2.63 | 2.82E-09 | 1.58E-08 |
| chr2 | 221137769 | 221143708 | 5940 | -0.76 | 2.83E-09 | 1.58E-08 |
| chr7 | 13056083 | 13091388 | 35306 | 0.42 | 2.83E-09 | 1.58E-08 |
| chr2 | 72425796 | 72447441 | 21646 | 0.62 | 2.85E-09 | 1.59E-08 |
| chr12 | 92378055 | 92380771 | 2717 | 1.64 | 2.85E-09 | 1.59E-08 |
| chr17 | 63013429 | 63075204 | 61776 | -0.41 | 2.85E-09 | 1.59E-08 |
| chr12 | 108078307 | 108092879 | 14573 | 0.54 | 2.87E-09 | 1.61E-08 |
| chr2 | 182390266 | 182391496 | 1231 | 6.61 | 2.88E-09 | 1.61E-08 |
| chr8 | 58714835 | 58717458 | 2624 | 6.61 | 2.88E-09 | 1.61E-08 |
| chr15 | 92611918 | 92614208 | 2291 | 0.82 | 2.89E-09 | 1.62E-08 |
| chr16 | 27704943 | 27715366 | 10424 | -2.46 | 2.90E-09 | 1.62E-08 |
| chr5 | 174870629 | 174871610 | 982 | -6.57 | 2.90E-09 | 1.62E-08 |
| chr3 | 168959082 | 168960454 | 1373 | 6.63 | 2.91E-09 | 1.63E-08 |
| chr17 | 63815976 | 63816875 | 900 | 4.46 | 2.91E-09 | 1.63E-08 |
| chr1 | 230201715 | 230294719 | 93005 | 0.36 | 2.92E-09 | 1.63E-08 |
| chr19 | 13743466 | 13744709 | 1244 | 2.41 | 2.93E-09 | 1.63E-08 |
| chr11 | 113560246 | 113562348 | 2103 | 1.24 | 2.94E-09 | 1.64E-08 |
| chr4 | 130774399 | 130775113 | 715 | 4.73 | 2.95E-09 | 1.65E-08 |
| chr13 | 110778220 | 110791661 | 13442 | 1.14 | 2.96E-09 | 1.65E-08 |
| chr8 | 122787670 | 122827421 | 39752 | 0.4 | 2.96E-09 | 1.65E-08 |
| chr20 | 4844077 | 4852386 | 8310 | 0.75 | 2.97E-09 | 1.66E-08 |
| chr11 | 16625823 | 16629942 | 4120 | 6 | 2.98E-09 | 1.66E-08 |
| chr19 | 30638688 | 30640741 | 2054 | 3.7 | 2.98E-09 | 1.66E-08 |
| chr13 | 68311024 | 68312781 | 1758 | 6.63 | 2.99E-09 | 1.67E-08 |
| chr20 | 14015768 | 14017105 | 1338 | 6.61 | 2.99E-09 | 1.67E-08 |
| chr3 | 186323630 | 186335664 | 12035 | 2.38 | 3.08E-09 | 1.71E-08 |
| chr3 | 56093493 | 56099524 | 6032 | 6.84 | 3.09E-09 | 1.72E-08 |
| chr2 | 5328840 | 5330737 | 1898 | 6.6 | 3.12E-09 | 1.74E-08 |

| | | | | | | |
|---|---|---|---|---|---|---|
| chr12 | 63748578 | 63749383 | 806 | 2.97 | 3.16E-09 | 1.76E-08 |
| chr1 | 221914349 | 221917609 | 3261 | 0.78 | 3.17E-09 | 1.77E-08 |
| chr18 | 70706631 | 70707933 | 1303 | 3.39 | 3.17E-09 | 1.77E-08 |
| chr3 | 94594954 | 94595866 | 913 | 6.68 | 3.17E-09 | 1.77E-08 |
| chr9 | 99136144 | 99184692 | 48549 | 0.35 | 3.19E-09 | 1.78E-08 |
| chr9 | 100393615 | 100406331 | 12717 | 0.45 | 3.22E-09 | 1.79E-08 |
| chr3 | 112708747 | 112720077 | 11331 | 0.73 | 3.25E-09 | 1.81E-08 |
| chr13 | 47169279 | 47242786 | 73508 | 0.35 | 3.26E-09 | 1.82E-08 |
| chr2 | 225961957 | 225985837 | 23881 | -0.39 | 3.28E-09 | 1.82E-08 |
| chr17 | 76480206 | 76501868 | 21663 | -1.62 | 3.28E-09 | 1.82E-08 |
| chr4 | 114388431 | 114390981 | 2551 | 1.56 | 3.28E-09 | 1.82E-08 |
| chr19 | 53140887 | 53141629 | 743 | 6.65 | 3.32E-09 | 1.84E-08 |
| chr10 | 112896147 | 112897518 | 1372 | 6.62 | 3.32E-09 | 1.85E-08 |
| chr8 | 102821504 | 102822865 | 1362 | 2.01 | 3.33E-09 | 1.85E-08 |
| chr13 | 45735784 | 45755464 | 19681 | 0.71 | 3.33E-09 | 1.85E-08 |
| chr13 | 82169887 | 82171129 | 1243 | 4.54 | 3.34E-09 | 1.85E-08 |
| chr9 | 128425168 | 128451527 | 26360 | 0.59 | 3.35E-09 | 1.86E-08 |
| chr19 | 52510614 | 52512508 | 1895 | 1.08 | 3.35E-09 | 1.86E-08 |
| chr13 | 101043978 | 101048933 | 4956 | 0.71 | 3.37E-09 | 1.87E-08 |
| chr12 | 30590138 | 30592147 | 2010 | 6.6 | 3.39E-09 | 1.88E-08 |
| chr1 | 160420455 | 160422257 | 1803 | 1.6 | 3.41E-09 | 1.89E-08 |
| chr12 | 3409254 | 3410060 | 807 | 1.81 | 3.41E-09 | 1.89E-08 |
| chr2 | 235401479 | 235411851 | 10373 | -0.94 | 3.42E-09 | 1.89E-08 |
| chr8 | 110503908 | 110504478 | 571 | 4.92 | 3.42E-09 | 1.89E-08 |
| chr10 | 103985220 | 104006067 | 20848 | 2.16 | 3.42E-09 | 1.89E-08 |
| chr1 | 193178526 | 193182098 | 3573 | 1.53 | 3.42E-09 | 1.90E-08 |
| chr2 | 201028062 | 201029486 | 1425 | 2.06 | 3.42E-09 | 1.90E-08 |
| chr6 | 22236251 | 22251412 | 15162 | 0.8 | 3.43E-09 | 1.90E-08 |
| chr6 | 165964454 | 165971092 | 6639 | 4.49 | 3.43E-09 | 1.90E-08 |
| chr1 | 23061475 | 23078733 | 17259 | -0.45 | 3.45E-09 | 1.91E-08 |
| chr9 | 80592249 | 80597463 | 5215 | 1.39 | 3.50E-09 | 1.94E-08 |
| chr6 | 22111972 | 22115532 | 3561 | 0.95 | 3.51E-09 | 1.94E-08 |
| chr9 | 133641781 | 133658021 | 16241 | -1.74 | 3.53E-09 | 1.96E-08 |
| chr19 | 20960930 | 20963020 | 2091 | 6.6 | 3.55E-09 | 1.96E-08 |
| chr11 | 35281590 | 35283126 | 1537 | 2.21 | 3.56E-09 | 1.97E-08 |
| chr15 | 56138572 | 56211515 | 72944 | 0.23 | 3.56E-09 | 1.97E-08 |
| chr1 | 77902013 | 77914245 | 12233 | 0.67 | 3.57E-09 | 1.97E-08 |
| chr5 | 114866982 | 114881427 | 14446 | 0.62 | 3.58E-09 | 1.98E-08 |
| chr9 | 113648691 | 113658085 | 9395 | 0.76 | 3.60E-09 | 1.99E-08 |
| chr20 | 23329440 | 23352884 | 23445 | 0.36 | 3.62E-09 | 2.00E-08 |
| chr4 | 184077212 | 184105100 | 27889 | 0.74 | 3.64E-09 | 2.01E-08 |
| chr1 | 196406611 | 196408536 | 1926 | 2.39 | 3.68E-09 | 2.03E-08 |
| chr12 | 90256365 | 90258522 | 2158 | 1.9 | 3.69E-09 | 2.04E-08 |
| chr14 | 76297093 | 76299417 | 2325 | 1.44 | 3.69E-09 | 2.04E-08 |
| chr10 | 18939185 | 18956472 | 17288 | 0.46 | 3.70E-09 | 2.04E-08 |
| chr1 | 207276484 | 207278048 | 1565 | 4.62 | 3.71E-09 | 2.05E-08 |
| chr22 | 28058819 | 28067819 | 9001 | 1.32 | 3.71E-09 | 2.05E-08 |
| chr1 | 85069249 | 85070464 | 1216 | 6.62 | 3.71E-09 | 2.05E-08 |
| chr6 | 51946085 | 51952635 | 6551 | 6.58 | 3.72E-09 | 2.05E-08 |
| chr16 | 67833662 | 67841860 | 8199 | -0.79 | 3.72E-09 | 2.05E-08 |
| chr8 | 76234542 | 76243528 | 8987 | 0.82 | 3.76E-09 | 2.07E-08 |
| chr1 | 228073203 | 228084603 | 11401 | 1.25 | 3.76E-09 | 2.08E-08 |
| chr2 | 131555037 | 131567010 | 11974 | 2.24 | 3.80E-09 | 2.09E-08 |
| chr6 | 141585270 | 141588878 | 3609 | 0.89 | 3.85E-09 | 2.12E-08 |
| chr2 | 44394434 | 44434504 | 40071 | -0.48 | 3.85E-09 | 2.12E-08 |
| chrX | 61931920 | 61933216 | 1297 | 6.57 | 3.87E-09 | 2.13E-08 |
| chr10 | 88194935 | 88299191 | 104257 | 0.38 | 3.87E-09 | 2.13E-08 |
| chr17 | 67565414 | 67586778 | 21365 | -0.53 | 3.90E-09 | 2.15E-08 |
| chr17 | 3983892 | 4091597 | 107706 | -0.3 | 3.90E-09 | 2.15E-08 |
| chr1 | 193228375 | 193230655 | 2281 | 1.65 | 3.90E-09 | 2.15E-08 |
| chr6 | 163829371 | 163842241 | 12871 | 0.57 | 3.90E-09 | 2.15E-08 |

| | | | | | | |
|---|---|---|---|---|---|---|
| chr14 | 92874679 | 92875672 | 994 | 3.04 | 3.91E-09 | 2.15E-08 |
| chr15 | 96713221 | 96729195 | 15975 | 0.56 | 3.95E-09 | 2.18E-08 |
| chr14 | 102591128 | 102615724 | 24597 | 0.67 | 3.98E-09 | 2.19E-08 |
| chr19 | 35949615 | 35951020 | 1406 | 1.56 | 4.02E-09 | 2.21E-08 |
| chr1 | 158022903 | 158055699 | 32797 | -0.63 | 4.04E-09 | 2.22E-08 |
| chr3 | 112278232 | 112282152 | 3921 | 0.83 | 4.07E-09 | 2.24E-08 |
| chr7 | 54944997 | 54956669 | 11673 | -0.74 | 4.08E-09 | 2.25E-08 |
| chr15 | 63178413 | 63193131 | 14719 | 0.34 | 4.10E-09 | 2.25E-08 |
| chr22 | 47209718 | 47228148 | 18431 | 0.46 | 4.10E-09 | 2.25E-08 |
| chr3 | 5732590 | 5733798 | 1209 | 5.81 | 4.10E-09 | 2.25E-08 |
| chr10 | 107344873 | 107346681 | 1809 | -6.53 | 4.11E-09 | 2.26E-08 |
| chr17 | 32295168 | 32296330 | 1163 | -6.53 | 4.13E-09 | 2.27E-08 |
| chr9 | 72872962 | 72875452 | 2491 | 0.8 | 4.13E-09 | 2.27E-08 |
| chr2 | 236310861 | 236326469 | 15609 | 2.37 | 4.20E-09 | 2.30E-08 |
| chr19 | 24215217 | 24238301 | 23085 | 0.74 | 4.20E-09 | 2.31E-08 |
| chr6 | 114175432 | 114200871 | 25440 | 0.62 | 4.23E-09 | 2.32E-08 |
| chr5 | 126408467 | 126413461 | 4995 | 1.39 | 4.23E-09 | 2.32E-08 |
| chr1 | 232277831 | 232286716 | 8886 | -2.62 | 4.24E-09 | 2.32E-08 |
| chr3 | 169796916 | 169803668 | 6753 | 1.65 | 4.25E-09 | 2.33E-08 |
| chr4 | 38617476 | 38624416 | 6941 | 1.43 | 4.25E-09 | 2.33E-08 |
| chr3 | 16281365 | 16289659 | 8295 | -1.98 | 4.29E-09 | 2.35E-08 |
| chr16 | 21948177 | 21949946 | 1770 | 1.27 | 4.33E-09 | 2.37E-08 |
| chr15 | 81031509 | 81045348 | 13840 | 0.59 | 4.33E-09 | 2.38E-08 |
| chr5 | 108790854 | 108791980 | 1127 | 6.57 | 4.33E-09 | 2.38E-08 |
| chr2 | 36008109 | 36009572 | 1464 | 2.42 | 4.34E-09 | 2.38E-08 |
| chr11 | 28526431 | 28527874 | 1444 | 4.84 | 4.34E-09 | 2.38E-08 |
| chr20 | 31960163 | 31990157 | 29995 | 0.87 | 4.39E-09 | 2.41E-08 |
| chr11 | 8102318 | 8114381 | 12064 | -1.89 | 4.39E-09 | 2.41E-08 |
| chr1 | 226345561 | 226375194 | 29634 | 0.56 | 4.40E-09 | 2.41E-08 |
| chr14 | 51832794 | 51834611 | 1818 | 3.3 | 4.42E-09 | 2.42E-08 |
| chr3 | 122800604 | 122809955 | 9352 | -1.58 | 4.42E-09 | 2.42E-08 |
| chr1 | 84325891 | 84326944 | 1054 | 2.27 | 4.43E-09 | 2.42E-08 |
| chr10 | 82867943 | 82869010 | 1068 | 2.47 | 4.43E-09 | 2.42E-08 |
| chrX | 67405537 | 67410543 | 5007 | 1.6 | 4.44E-09 | 2.43E-08 |
| chr4 | 7207654 | 7220119 | 12466 | -0.84 | 4.47E-09 | 2.44E-08 |
| chr20 | 9299991 | 9340438 | 40448 | 0.75 | 4.47E-09 | 2.45E-08 |
| chr4 | 102076073 | 102089042 | 12970 | 0.46 | 4.48E-09 | 2.45E-08 |
| chr6 | 55499188 | 55503825 | 4638 | 0.99 | 4.49E-09 | 2.45E-08 |
| chrX | 115336400 | 115337254 | 855 | 6.55 | 4.50E-09 | 2.46E-08 |
| chr3 | 164976939 | 164978506 | 1568 | 2.66 | 4.53E-09 | 2.48E-08 |
| chr4 | 159130116 | 159132993 | 2878 | 0.9 | 4.53E-09 | 2.48E-08 |
| chr20 | 15476259 | 15478062 | 1804 | 4.08 | 4.54E-09 | 2.48E-08 |
| chr1 | 68252910 | 68269705 | 16796 | 0.4 | 4.55E-09 | 2.49E-08 |
| chr3 | 148963092 | 148972125 | 9034 | 1.14 | 4.56E-09 | 2.49E-08 |
| chr9 | 95673385 | 95675306 | 1922 | 2.11 | 4.56E-09 | 2.49E-08 |
| chrX | 74493216 | 74499385 | 6170 | 1.21 | 4.57E-09 | 2.50E-08 |
| chr15 | 66160915 | 66178471 | 17557 | 0.51 | 4.61E-09 | 2.52E-08 |
| chr6 | 157301611 | 157316601 | 14991 | 0.98 | 4.62E-09 | 2.52E-08 |
| chr19 | 39086362 | 39087745 | 1384 | 2.85 | 4.62E-09 | 2.52E-08 |
| chr2 | 219575192 | 219614025 | 38834 | -0.65 | 4.62E-09 | 2.52E-08 |
| chr5 | 60455810 | 60461706 | 5897 | 0.5 | 4.64E-09 | 2.53E-08 |
| chr17 | 27581941 | 27622351 | 40411 | 0.38 | 4.65E-09 | 2.54E-08 |
| chr9 | 4053333 | 4078168 | 24836 | -1.73 | 4.66E-09 | 2.54E-08 |
| chr2 | 196977723 | 196978675 | 953 | 2.05 | 4.67E-09 | 2.54E-08 |
| chr19 | 12443348 | 12444454 | 1107 | 6.55 | 4.68E-09 | 2.55E-08 |
| chr5 | 114755215 | 114760452 | 5238 | 1.32 | 4.68E-09 | 2.55E-08 |
| chr9 | 15389913 | 15391282 | 1370 | 5.16 | 4.69E-09 | 2.55E-08 |
| chr7 | 16646395 | 16651762 | 5368 | 1.1 | 4.70E-09 | 2.56E-08 |
| chr10 | 89394646 | 89422761 | 28116 | 0.76 | 4.72E-09 | 2.57E-08 |
| chr13 | 76957581 | 76958885 | 1305 | 4.36 | 4.72E-09 | 2.57E-08 |
| chr1 | 169355245 | 169356657 | 1413 | 2.3 | 4.75E-09 | 2.59E-08 |

| | | | | | | |
|---|---|---|---|---|---|---|
| chr12 | 54717911 | 54729248 | 11338 | 0.6 | 4.76E-09 | 2.59E-08 |
| chr3 | 175263725 | 175265017 | 1293 | 6.6 | 4.77E-09 | 2.60E-08 |
| chr2 | 3828135 | 3833300 | 5166 | 6.54 | 4.78E-09 | 2.60E-08 |
| chr18 | 42350713 | 42353606 | 2894 | 1.54 | 4.80E-09 | 2.61E-08 |
| chr20 | 11382104 | 11383133 | 1030 | 6.67 | 4.82E-09 | 2.62E-08 |
| chr9 | 29077456 | 29078909 | 1454 | 6.54 | 4.84E-09 | 2.63E-08 |
| chr16 | 85423410 | 85424932 | 1523 | 2.99 | 4.87E-09 | 2.65E-08 |
| chr5 | 111742108 | 111756124 | 14017 | 4.22 | 4.88E-09 | 2.65E-08 |
| chr3 | 103157054 | 103158558 | 1505 | 6.56 | 4.88E-09 | 2.65E-08 |
| chr5 | 82970338 | 82971379 | 1042 | 3.18 | 4.93E-09 | 2.68E-08 |
| chr3 | 12221104 | 12240469 | 19366 | 0.61 | 4.94E-09 | 2.68E-08 |
| chr9 | 35707575 | 35775627 | 68053 | 0.34 | 4.96E-09 | 2.69E-08 |
| chr1 | 214638855 | 214648584 | 9730 | 0.66 | 4.96E-09 | 2.69E-08 |
| chr4 | 166661810 | 166665137 | 3328 | 1.1 | 4.96E-09 | 2.70E-08 |
| chr16 | 72947108 | 72988504 | 41397 | -0.81 | 4.98E-09 | 2.70E-08 |
| chr12 | 112122065 | 112124735 | 2671 | 0.86 | 4.98E-09 | 2.70E-08 |
| chr5 | 176797942 | 176816899 | 18958 | -0.89 | 4.98E-09 | 2.70E-08 |
| chr3 | 7381838 | 7382983 | 1146 | 6.6 | 5.00E-09 | 2.71E-08 |
| chr2 | 197123986 | 197125514 | 1529 | -3.05 | 5.03E-09 | 2.73E-08 |
| chr8 | 131739009 | 131740891 | 1883 | -0.98 | 5.04E-09 | 2.73E-08 |
| chr1 | 999180 | 1015983 | 16804 | 0.98 | 5.08E-09 | 2.75E-08 |
| chr5 | 135302392 | 135314291 | 11900 | -4.75 | 5.08E-09 | 2.75E-08 |
| chr2 | 209460160 | 209474130 | 13971 | 6.53 | 5.10E-09 | 2.76E-08 |
| chr2 | 40698018 | 40700295 | 2278 | 1.3 | 5.10E-09 | 2.76E-08 |
| chr7 | 50108404 | 50109498 | 1095 | 6.53 | 5.12E-09 | 2.77E-08 |
| chr4 | 160724899 | 160725639 | 741 | 6.59 | 5.13E-09 | 2.78E-08 |
| chr5 | 45385307 | 45386541 | 1235 | 6.54 | 5.14E-09 | 2.79E-08 |
| chr19 | 21688380 | 21689296 | 917 | 6.53 | 5.16E-09 | 2.79E-08 |
| chr8 | 49665434 | 49666719 | 1286 | 6.53 | 5.16E-09 | 2.79E-08 |
| chr10 | 124608003 | 124640225 | 32223 | -0.42 | 5.17E-09 | 2.80E-08 |
| chr20 | 49233396 | 49318877 | 85482 | 0.39 | 5.19E-09 | 2.81E-08 |
| chr3 | 138392731 | 138413847 | 21117 | 0.79 | 5.20E-09 | 2.81E-08 |
| chrX | 42597989 | 42599655 | 1667 | 2.68 | 5.21E-09 | 2.82E-08 |
| chr17 | 61248479 | 61249911 | 1433 | 3.22 | 5.23E-09 | 2.83E-08 |
| chr2 | 82705229 | 82706253 | 1025 | 6.56 | 5.23E-09 | 2.83E-08 |
| chr2 | 235360244 | 235373299 | 13056 | -1.06 | 5.23E-09 | 2.83E-08 |
| chr14 | 59479816 | 59481030 | 1215 | 6.57 | 5.24E-09 | 2.83E-08 |
| chr17 | 35274747 | 35322014 | 47268 | 0.41 | 5.25E-09 | 2.84E-08 |
| chr2 | 113298797 | 113323937 | 25141 | -0.72 | 5.26E-09 | 2.84E-08 |
| chr8 | 117207774 | 117208794 | 1021 | 6.55 | 5.27E-09 | 2.85E-08 |
| chr22 | 49895267 | 49906134 | 10868 | 2.12 | 5.28E-09 | 2.85E-08 |
| chr8 | 41569057 | 41571197 | 2141 | 1.88 | 5.28E-09 | 2.85E-08 |
| chr12 | 116929595 | 116947060 | 17466 | 0.66 | 5.29E-09 | 2.86E-08 |
| chr15 | 75248111 | 75258027 | 9917 | 0.65 | 5.31E-09 | 2.86E-08 |
| chr2 | 159996447 | 160004991 | 8545 | -2.21 | 5.31E-09 | 2.86E-08 |
| chr2 | 171004750 | 171014288 | 9539 | -1.81 | 5.36E-09 | 2.89E-08 |
| chr20 | 13223941 | 13225135 | 1195 | 2.76 | 5.41E-09 | 2.92E-08 |
| chr6 | 76077061 | 76086736 | 9676 | -0.78 | 5.42E-09 | 2.93E-08 |
| chr1 | 79997865 | 79998774 | 910 | 4.28 | 5.42E-09 | 2.93E-08 |
| chr12 | 25347082 | 25366670 | 19589 | 0.54 | 5.44E-09 | 2.94E-08 |
| chr8 | 102550177 | 102557767 | 7591 | 0.97 | 5.45E-09 | 2.94E-08 |
| chr12 | 25964064 | 25965546 | 1483 | 1.86 | 5.45E-09 | 2.94E-08 |
| chr15 | 33082361 | 33094545 | 12185 | -6.5 | 5.46E-09 | 2.94E-08 |
| chr3 | 130681683 | 130683838 | 2156 | 1.73 | 5.47E-09 | 2.95E-08 |
| chr2 | 67953159 | 67954193 | 1035 | 4.56 | 5.49E-09 | 2.96E-08 |
| chr4 | 169806685 | 169809523 | 2839 | -2.35 | 5.51E-09 | 2.97E-08 |
| chr3 | 82151978 | 82152589 | 612 | 4.15 | 5.52E-09 | 2.97E-08 |
| chr18 | 70881692 | 70883207 | 1516 | 6.53 | 5.54E-09 | 2.98E-08 |
| chr2 | 8369572 | 8370450 | 879 | -2.47 | 5.56E-09 | 2.99E-08 |
| chr1 | 178971441 | 178972483 | 1043 | -3.14 | 5.57E-09 | 3.00E-08 |
| chr6 | 136354011 | 136373705 | 19695 | -3.25 | 5.62E-09 | 3.02E-08 |

| | | | | | | |
|---|---|---|---|---|---|---|
| chr11 | 22850220 | 22852195 | 1976 | 1.29 | 5.62E-09 | 3.03E-08 |
| chr3 | 153740274 | 153741847 | 1574 | 1.75 | 5.67E-09 | 3.05E-08 |
| chr11 | 27677712 | 27679413 | 1702 | 3.05 | 5.73E-09 | 3.09E-08 |
| chr8 | 106511098 | 106512270 | 1173 | 6.51 | 5.77E-09 | 3.10E-08 |
| chr2 | 46042592 | 46049668 | 7077 | -2.75 | 5.77E-09 | 3.10E-08 |
| chr11 | 92149832 | 92151327 | 1496 | 2.16 | 5.77E-09 | 3.10E-08 |
| chr1 | 191143043 | 191144218 | 1176 | 6.51 | 5.77E-09 | 3.10E-08 |
| chr14 | 74162339 | 74262580 | 100242 | 0.23 | 5.79E-09 | 3.11E-08 |
| chr11 | 94963243 | 94976086 | 12844 | 6.62 | 5.86E-09 | 3.15E-08 |
| chr11 | 46258456 | 46270794 | 12339 | -0.94 | 5.88E-09 | 3.16E-08 |
| chr13 | 81165888 | 81166627 | 740 | 4.18 | 5.90E-09 | 3.17E-08 |
| chr6 | 138406533 | 138430633 | 24101 | 0.42 | 5.91E-09 | 3.18E-08 |
| chr15 | 51912813 | 51916818 | 4006 | 0.69 | 5.93E-09 | 3.18E-08 |
| chr1 | 151509190 | 151522533 | 13344 | 0.4 | 5.94E-09 | 3.19E-08 |
| chr6 | 131029070 | 131030300 | 1231 | 6.53 | 5.94E-09 | 3.19E-08 |
| chr14 | 99974766 | 100005384 | 30619 | -0.74 | 6.00E-09 | 3.22E-08 |
| chr12 | 52077131 | 52078514 | 1384 | 1.59 | 6.01E-09 | 3.23E-08 |
| chr13 | 111198805 | 111209186 | 10382 | 1.84 | 6.04E-09 | 3.24E-08 |
| chr4 | 71569060 | 71575922 | 6863 | 0.82 | 6.11E-09 | 3.28E-08 |
| chr4 | 2491400 | 2498446 | 7047 | 1.16 | 6.16E-09 | 3.30E-08 |
| chr19 | 20747642 | 20748588 | 947 | 5.14 | 6.17E-09 | 3.31E-08 |
| chr14 | 59259372 | 59263671 | 4300 | 1.57 | 6.17E-09 | 3.31E-08 |
| chr3 | 49936369 | 49944803 | 8435 | 0.65 | 6.21E-09 | 3.33E-08 |
| chr7 | 16354519 | 16355673 | 1155 | 2.42 | 6.25E-09 | 3.35E-08 |
| chr5 | 96268254 | 96272983 | 4730 | 0.59 | 6.33E-09 | 3.39E-08 |
| chr3 | 83182028 | 83182745 | 718 | 6.51 | 6.35E-09 | 3.40E-08 |
| chr4 | 72322156 | 72326851 | 4696 | 1.85 | 6.42E-09 | 3.44E-08 |
| chr2 | 33700318 | 33702646 | 2329 | 1.04 | 6.45E-09 | 3.46E-08 |
| chr10 | 63659417 | 63664502 | 5086 | 0.77 | 6.46E-09 | 3.46E-08 |
| chr17 | 28071047 | 28072547 | 1501 | 2.04 | 6.49E-09 | 3.47E-08 |
| chr1 | 224480112 | 224529425 | 49314 | 0.44 | 6.51E-09 | 3.48E-08 |
| chr13 | 107239854 | 107241136 | 1283 | 5.18 | 6.51E-09 | 3.48E-08 |
| chr8 | 145978410 | 145996428 | 18019 | 0.6 | 6.53E-09 | 3.49E-08 |
| chr3 | 132150893 | 132165175 | 14283 | 1.32 | 6.53E-09 | 3.49E-08 |
| chr12 | 1456629 | 1460689 | 4061 | 1.53 | 6.55E-09 | 3.50E-08 |
| chr10 | 27143975 | 27149984 | 6010 | 0.82 | 6.66E-09 | 3.56E-08 |
| chr21 | 30753936 | 30765268 | 11333 | 3.01 | 6.67E-09 | 3.56E-08 |
| chr10 | 83440749 | 83451769 | 11021 | 6.54 | 6.68E-09 | 3.57E-08 |
| chr8 | 31945310 | 31946492 | 1183 | 2.9 | 6.68E-09 | 3.57E-08 |
| chr3 | 140105323 | 140106231 | 909 | -3.83 | 6.69E-09 | 3.58E-08 |
| chr1 | 3447200 | 3460629 | 13430 | 1.89 | 6.71E-09 | 3.59E-08 |
| chrX | 107265938 | 107282283 | 16346 | -0.58 | 6.71E-09 | 3.59E-08 |
| chr12 | 27483633 | 27498257 | 14625 | -0.85 | 6.72E-09 | 3.59E-08 |
| chr5 | 65210127 | 65226749 | 16623 | 0.62 | 6.81E-09 | 3.64E-08 |
| chr8 | 143742545 | 143764364 | 21820 | -0.51 | 6.82E-09 | 3.64E-08 |
| chr17 | 60211009 | 60257639 | 46631 | 0.69 | 6.85E-09 | 3.65E-08 |
| chr1 | 61906325 | 61921831 | 15507 | 0.43 | 6.97E-09 | 3.72E-08 |
| chr12 | 114369686 | 114373963 | 4278 | 1.34 | 6.98E-09 | 3.73E-08 |
| chr1 | 242766485 | 242767715 | 1231 | -4.01 | 7.08E-09 | 3.78E-08 |
| chr9 | 128666275 | 128667283 | 1009 | 4.01 | 7.12E-09 | 3.80E-08 |
| chr5 | 27315656 | 27316472 | 817 | 5.85 | 7.22E-09 | 3.85E-08 |
| chr7 | 109333045 | 109334167 | 1123 | 5.68 | 7.25E-09 | 3.86E-08 |
| chr10 | 74853667 | 74857538 | 3872 | 0.62 | 7.28E-09 | 3.88E-08 |
| chr1 | 67389051 | 67417213 | 28163 | 0.33 | 7.28E-09 | 3.88E-08 |
| chr12 | 82751008 | 82753946 | 2939 | 0.66 | 7.31E-09 | 3.90E-08 |
| chr19 | 19475118 | 19518307 | 43190 | 0.55 | 7.33E-09 | 3.91E-08 |
| chr4 | 75859060 | 75873645 | 14586 | 1.83 | 7.36E-09 | 3.92E-08 |
| chr1 | 11844870 | 11867665 | 22796 | 0.67 | 7.37E-09 | 3.92E-08 |
| chr4 | 127989541 | 127990402 | 862 | 2.82 | 7.37E-09 | 3.93E-08 |
| chr1 | 155970644 | 155990897 | 20254 | 0.57 | 7.42E-09 | 3.95E-08 |
| chr19 | 33685581 | 33689649 | 4069 | -1.94 | 7.44E-09 | 3.96E-08 |

| | | | | | | |
|---|---|---|---|---|---|---|
| chr1 | 182758525 | 182765682 | 7158 | 2.05 | 7.45E-09 | 3.96E-08 |
| chr19 | 9608702 | 9609712 | 1011 | 6.48 | 7.45E-09 | 3.96E-08 |
| chr2 | 217204287 | 217205419 | 1133 | -4.54 | 7.49E-09 | 3.98E-08 |
| chr1 | 154323697 | 154324923 | 1227 | 2.91 | 7.49E-09 | 3.98E-08 |
| chr5 | 55068126 | 55069814 | 1689 | 1.28 | 7.50E-09 | 3.99E-08 |
| chr1 | 68849442 | 68851735 | 2294 | 1.89 | 7.54E-09 | 4.01E-08 |
| chr2 | 26725732 | 26727192 | 1461 | -1.3 | 7.55E-09 | 4.02E-08 |
| chr11 | 103991507 | 104005786 | 14280 | 1.16 | 7.58E-09 | 4.03E-08 |
| chr5 | 83017729 | 83020333 | 2605 | 6.52 | 7.59E-09 | 4.03E-08 |
| chr6 | 5439219 | 5440355 | 1137 | 1.72 | 7.62E-09 | 4.05E-08 |
| chr20 | 50703692 | 50704661 | 970 | 6.48 | 7.63E-09 | 4.05E-08 |
| chr1 | 178836278 | 178842072 | 5795 | 1.06 | 7.67E-09 | 4.08E-08 |
| chr2 | 29214865 | 29217348 | 2484 | 2.12 | 7.68E-09 | 4.08E-08 |
| chr3 | 160086405 | 160093233 | 6829 | 1.87 | 7.72E-09 | 4.10E-08 |
| chr8 | 8125780 | 8131481 | 5702 | 0.85 | 7.74E-09 | 4.11E-08 |
| chr2 | 61152731 | 61154615 | 1885 | 2.19 | 7.74E-09 | 4.11E-08 |
| chr5 | 85860244 | 85867648 | 7405 | 0.84 | 7.75E-09 | 4.11E-08 |
| chr20 | 48191961 | 48345161 | 153201 | 0.23 | 7.76E-09 | 4.12E-08 |
| chr5 | 13745300 | 13746366 | 1067 | 3.67 | 7.80E-09 | 4.14E-08 |
| chr3 | 67685252 | 67705784 | 20533 | 0.49 | 7.81E-09 | 4.14E-08 |
| chr1 | 33500830 | 33529175 | 28346 | 0.97 | 7.88E-09 | 4.18E-08 |
| chr21 | 17054361 | 17056006 | 1646 | 2.49 | 7.90E-09 | 4.19E-08 |
| chr11 | 91526092 | 91531004 | 4913 | 1.5 | 7.91E-09 | 4.19E-08 |
| chr12 | 56497132 | 56523632 | 26501 | 0.54 | 7.96E-09 | 4.22E-08 |
| chr12 | 78286791 | 78288937 | 2147 | 2.26 | 7.99E-09 | 4.24E-08 |
| chr9 | 16307993 | 16322225 | 14233 | 2 | 8.07E-09 | 4.27E-08 |
| chr1 | 69347383 | 69348223 | 841 | 6.47 | 8.08E-09 | 4.28E-08 |
| chr4 | 93519630 | 93520869 | 1240 | 2.73 | 8.09E-09 | 4.29E-08 |
| chr13 | 71953148 | 71953973 | 826 | 6.49 | 8.11E-09 | 4.29E-08 |
| chr12 | 26266363 | 26280003 | 13641 | 1.32 | 8.11E-09 | 4.29E-08 |
| chr14 | 37934633 | 37936480 | 1848 | 2.82 | 8.15E-09 | 4.31E-08 |
| chr11 | 640767 | 666390 | 25624 | -0.98 | 8.20E-09 | 4.34E-08 |
| chr16 | 83835669 | 83848604 | 12936 | 0.83 | 8.33E-09 | 4.41E-08 |
| chr4 | 101816579 | 101829449 | 12871 | 6.47 | 8.34E-09 | 4.41E-08 |
| chr14 | 37550476 | 37552311 | 1836 | 2.78 | 8.35E-09 | 4.42E-08 |
| chr16 | 89876712 | 89905552 | 28841 | 0.56 | 8.42E-09 | 4.46E-08 |
| chr3 | 101394159 | 101406953 | 12795 | 0.53 | 8.49E-09 | 4.49E-08 |
| chr3 | 77873918 | 77875463 | 1546 | 6.48 | 8.52E-09 | 4.50E-08 |
| chr1 | 246598398 | 246600037 | 1640 | -2.23 | 8.63E-09 | 4.56E-08 |
| chr12 | 50783868 | 50852390 | 68523 | 0.33 | 8.64E-09 | 4.57E-08 |
| chr4 | 138653517 | 138654710 | 1194 | 2.18 | 8.70E-09 | 4.60E-08 |
| chr5 | 111821339 | 111822177 | 839 | 4.43 | 8.71E-09 | 4.60E-08 |
| chr3 | 193646295 | 193647474 | 1180 | 4.59 | 8.83E-09 | 4.67E-08 |
| chr6 | 53588482 | 53601158 | 12677 | 0.87 | 8.89E-09 | 4.69E-08 |
| chr13 | 73898453 | 73913112 | 14660 | 0.49 | 8.90E-09 | 4.70E-08 |
| chr4 | 123072439 | 123074564 | 2126 | 1.07 | 8.97E-09 | 4.73E-08 |
| chr5 | 45277978 | 45279841 | 1864 | 6.49 | 9.04E-09 | 4.77E-08 |
| chr18 | 61219888 | 61221229 | 1342 | 6.46 | 9.04E-09 | 4.77E-08 |
| chr2 | 70913412 | 70915261 | 1850 | -1.26 | 9.08E-09 | 4.79E-08 |
| chr12 | 12734720 | 12738531 | 3812 | 1.36 | 9.15E-09 | 4.83E-08 |
| chr11 | 16283394 | 16284768 | 1375 | 4.1 | 9.22E-09 | 4.86E-08 |
| chr9 | 134608225 | 134623200 | 14976 | 0.8 | 9.27E-09 | 4.89E-08 |
| chr1 | 31944663 | 31956466 | 11804 | 2.34 | 9.30E-09 | 4.90E-08 |
| chr2 | 97320803 | 97332091 | 11289 | 1.54 | 9.31E-09 | 4.91E-08 |
| chr6 | 18154534 | 18156837 | 2304 | 1.45 | 9.35E-09 | 4.93E-08 |
| chr5 | 23527896 | 23528800 | 905 | 6.45 | 9.38E-09 | 4.94E-08 |
| chr19 | 12098486 | 12099602 | 1117 | 6.46 | 9.42E-09 | 4.97E-08 |
| chr6 | 108999078 | 109010824 | 11747 | 1.79 | 9.48E-09 | 5.00E-08 |
| chr13 | 45427773 | 45429388 | 1616 | 2.34 | 9.50E-09 | 5.01E-08 |
| chr17 | 5419200 | 5430744 | 11545 | -6.41 | 9.54E-09 | 5.02E-08 |
| chr16 | 14091231 | 14092972 | 1742 | 1.58 | 9.64E-09 | 5.08E-08 |

| | | | | | | |
|---|---|---|---|---|---|---|
| chrX | 35220549 | 35222281 | 1733 | 6.44 | 9.65E-09 | 5.08E-08 |
| chr1 | 117031722 | 117032724 | 1003 | 5.14 | 9.68E-09 | 5.10E-08 |
| chr19 | 23325703 | 23327308 | 1606 | 6.44 | 9.68E-09 | 5.10E-08 |
| chr20 | 54923166 | 54981426 | 58261 | 0.43 | 9.69E-09 | 5.10E-08 |
| chr5 | 19314022 | 19323529 | 9508 | 6.53 | 9.71E-09 | 5.11E-08 |
| chr10 | 71528491 | 71529820 | 1330 | -1.47 | 9.78E-09 | 5.14E-08 |
| chr13 | 86268052 | 86269355 | 1304 | -4.82 | 9.79E-09 | 5.15E-08 |
| chr9 | 98268122 | 98274142 | 6021 | -0.71 | 9.80E-09 | 5.15E-08 |
| chr8 | 122706858 | 122721800 | 14943 | 0.45 | 9.82E-09 | 5.17E-08 |
| chr2 | 27291548 | 27309468 | 17921 | -0.94 | 9.86E-09 | 5.19E-08 |
| chr10 | 84364197 | 84365468 | 1272 | 6.5 | 9.91E-09 | 5.21E-08 |
| chr6 | 141347461 | 141349109 | 1649 | 6.45 | 9.93E-09 | 5.22E-08 |
| chr5 | 133026506 | 133027908 | 1403 | -5.56 | 1.00E-08 | 5.26E-08 |
| chr18 | 30310211 | 30311055 | 845 | 6.44 | 1.00E-08 | 5.27E-08 |
| chr1 | 154698862 | 154704232 | 5371 | 0.98 | 1.01E-08 | 5.28E-08 |
| chr8 | 129638075 | 129652633 | 14559 | 1.97 | 1.02E-08 | 5.37E-08 |
| chr9 | 115773937 | 115775016 | 1080 | 4.42 | 1.03E-08 | 5.38E-08 |
| chr1 | 106207417 | 106209170 | 1754 | 6.44 | 1.03E-08 | 5.41E-08 |
| chr2 | 18413976 | 18415348 | 1373 | -6.43 | 1.03E-08 | 5.42E-08 |
| chr2 | 204191892 | 204221447 | 29556 | -0.42 | 1.04E-08 | 5.43E-08 |
| chr11 | 18033906 | 18035943 | 2038 | 1.31 | 1.04E-08 | 5.47E-08 |
| chr4 | 84127686 | 84139311 | 11626 | -0.77 | 1.04E-08 | 5.47E-08 |
| chr18 | 7616454 | 7638142 | 21689 | -1.23 | 1.05E-08 | 5.49E-08 |
| chr22 | 30191111 | 30218504 | 27394 | -0.44 | 1.05E-08 | 5.50E-08 |
| chr9 | 118377174 | 118379464 | 2291 | -1.44 | 1.06E-08 | 5.58E-08 |
| chr6 | 43137828 | 43150710 | 12883 | 0.62 | 1.07E-08 | 5.63E-08 |
| chr7 | 129007657 | 129008688 | 1032 | 6.45 | 1.07E-08 | 5.63E-08 |
| chr7 | 30981243 | 30982328 | 1086 | 6.52 | 1.08E-08 | 5.65E-08 |
| chr12 | 75256018 | 75257389 | 1372 | 2.02 | 1.08E-08 | 5.66E-08 |
| chr1 | 172412183 | 172453361 | 41179 | 0.67 | 1.09E-08 | 5.69E-08 |
| chr8 | 84078974 | 84080505 | 1532 | 6.46 | 1.09E-08 | 5.70E-08 |
| chr4 | 187702327 | 187703521 | 1195 | 2.58 | 1.09E-08 | 5.72E-08 |
| chr3 | 32001609 | 32023930 | 22322 | 0.72 | 1.09E-08 | 5.73E-08 |
| chr9 | 112010013 | 112017349 | 7337 | 0.61 | 1.10E-08 | 5.74E-08 |
| chr3 | 24639722 | 24641129 | 1408 | 2 | 1.10E-08 | 5.75E-08 |
| chr10 | 19209398 | 19210298 | 901 | 6.43 | 1.10E-08 | 5.75E-08 |
| chr5 | 178322286 | 178323488 | 1203 | 2.19 | 1.10E-08 | 5.78E-08 |
| chr18 | 71381959 | 71385746 | 3788 | 2.14 | 1.12E-08 | 5.85E-08 |
| chr10 | 44118505 | 44119213 | 709 | 4.51 | 1.12E-08 | 5.86E-08 |
| chr16 | 88448042 | 88491659 | 43618 | -0.39 | 1.13E-08 | 5.89E-08 |
| chr11 | 9912465 | 9932868 | 20404 | 0.35 | 1.13E-08 | 5.91E-08 |
| chr5 | 36874796 | 36880765 | 5970 | 0.6 | 1.13E-08 | 5.93E-08 |
| chr5 | 102593563 | 102609841 | 16279 | -0.61 | 1.13E-08 | 5.93E-08 |
| chrX | 82068332 | 82069916 | 1585 | 1.86 | 1.13E-08 | 5.93E-08 |
| chr5 | 38640151 | 38673383 | 33233 | -4.59 | 1.14E-08 | 5.95E-08 |
| chr7 | 131961153 | 131962622 | 1470 | -6.4 | 1.14E-08 | 5.97E-08 |
| chr11 | 120937218 | 120943289 | 6072 | 1.19 | 1.14E-08 | 5.97E-08 |
| chr1 | 202168094 | 202170486 | 2393 | 1.33 | 1.15E-08 | 5.98E-08 |
| chr12 | 56274296 | 56275558 | 1263 | 4.43 | 1.15E-08 | 5.99E-08 |
| chr6 | 56142287 | 56146038 | 3752 | 0.77 | 1.15E-08 | 6.00E-08 |
| chr11 | 68137754 | 68151339 | 13586 | 0.5 | 1.15E-08 | 6.03E-08 |
| chr1 | 217996424 | 217997466 | 1043 | 6.44 | 1.16E-08 | 6.07E-08 |
| chr1 | 27744231 | 27765519 | 21289 | 0.5 | 1.17E-08 | 6.08E-08 |
| chr20 | 30101454 | 30222760 | 121307 | 0.25 | 1.17E-08 | 6.08E-08 |
| chr12 | 57760412 | 57793897 | 33486 | 0.48 | 1.17E-08 | 6.08E-08 |
| chr16 | 1346457 | 1362322 | 15866 | 0.61 | 1.17E-08 | 6.08E-08 |
| chr4 | 77532284 | 77533633 | 1350 | 1.83 | 1.17E-08 | 6.10E-08 |
| chr10 | 31606633 | 31610400 | 3768 | 0.79 | 1.17E-08 | 6.12E-08 |
| chrX | 124338606 | 124340195 | 1590 | 4.05 | 1.17E-08 | 6.12E-08 |
| chr4 | 15452549 | 15454450 | 1902 | -2.66 | 1.17E-08 | 6.12E-08 |
| chr22 | 20187949 | 20207364 | 19416 | 0.75 | 1.17E-08 | 6.12E-08 |

| | | | | | | |
|---|---|---|---|---|---|---|
| chr18 | 28778884 | 28788420 | 9537 | 1.64 | 1.18E-08 | 6.12E-08 |
| chr17 | 46405553 | 46413372 | 7820 | 1.3 | 1.18E-08 | 6.16E-08 |
| chr5 | 136805632 | 136807136 | 1505 | -6.38 | 1.19E-08 | 6.18E-08 |
| chr4 | 148282826 | 148285360 | 2535 | 1.63 | 1.20E-08 | 6.24E-08 |
| chr2 | 139031439 | 139035416 | 3978 | 0.95 | 1.20E-08 | 6.26E-08 |
| chr1 | 155162015 | 155180370 | 18356 | -0.49 | 1.20E-08 | 6.26E-08 |
| chr1 | 235085129 | 235154672 | 69544 | 0.26 | 1.21E-08 | 6.27E-08 |
| chr1 | 193056345 | 193098010 | 41666 | 0.5 | 1.21E-08 | 6.28E-08 |
| chr4 | 84046708 | 84062046 | 15339 | 1.82 | 1.22E-08 | 6.32E-08 |
| chr5 | 174672744 | 174674106 | 1363 | -6.37 | 1.22E-08 | 6.36E-08 |
| chr12 | 110200329 | 110201946 | 1618 | 2.31 | 1.23E-08 | 6.39E-08 |
| chr4 | 111531814 | 111533054 | 1241 | 2.36 | 1.24E-08 | 6.42E-08 |
| chr6 | 80234857 | 80256878 | 22022 | 0.76 | 1.24E-08 | 6.42E-08 |
| chr2 | 11586519 | 11607182 | 20664 | -0.61 | 1.25E-08 | 6.50E-08 |
| chr9 | 18538678 | 18540059 | 1382 | 4.53 | 1.25E-08 | 6.50E-08 |
| chr8 | 131822831 | 131824042 | 1212 | -4.19 | 1.25E-08 | 6.52E-08 |
| chr7 | 33276758 | 33277533 | 776 | 3.02 | 1.26E-08 | 6.54E-08 |
| chr18 | 42207037 | 42218374 | 11338 | 6.41 | 1.26E-08 | 6.54E-08 |
| chr2 | 170549827 | 170552572 | 2746 | -0.77 | 1.26E-08 | 6.57E-08 |
| chr8 | 29659565 | 29663141 | 3577 | 1.29 | 1.26E-08 | 6.57E-08 |
| chr5 | 81930594 | 81932302 | 1709 | 1.59 | 1.27E-08 | 6.58E-08 |
| chr10 | 115084908 | 115087275 | 2368 | -1.14 | 1.27E-08 | 6.60E-08 |
| chr19 | 9878262 | 9880522 | 2261 | 1.31 | 1.27E-08 | 6.60E-08 |
| chr3 | 64671740 | 64672690 | 951 | 5.58 | 1.27E-08 | 6.61E-08 |
| chr10 | 64966700 | 65035418 | 68719 | 0.31 | 1.28E-08 | 6.62E-08 |
| chr3 | 29321499 | 29328102 | 6604 | 0.62 | 1.28E-08 | 6.63E-08 |
| chr5 | 54102982 | 54104411 | 1430 | 1.82 | 1.28E-08 | 6.64E-08 |
| chr1 | 92907022 | 92908286 | 1265 | 1.51 | 1.29E-08 | 6.70E-08 |
| chr19 | 35976068 | 35976599 | 532 | 5.57 | 1.29E-08 | 6.71E-08 |
| chr15 | 98502796 | 98504783 | 1988 | 2.29 | 1.30E-08 | 6.74E-08 |
| chr17 | 9810719 | 9812169 | 1451 | -1.71 | 1.31E-08 | 6.77E-08 |
| chr11 | 4567542 | 4568869 | 1328 | 3.26 | 1.31E-08 | 6.78E-08 |
| chr9 | 89951377 | 89953863 | 2487 | 1.16 | 1.32E-08 | 6.82E-08 |
| chr4 | 89015197 | 89036261 | 21065 | -0.39 | 1.32E-08 | 6.82E-08 |
| chr10 | 34020846 | 34025519 | 4674 | 1.48 | 1.32E-08 | 6.83E-08 |
| chr15 | 40293058 | 40299559 | 6502 | 0.93 | 1.33E-08 | 6.87E-08 |
| chr14 | 56002900 | 56008666 | 5767 | 2.27 | 1.33E-08 | 6.88E-08 |
| chr12 | 58565518 | 58566140 | 623 | 4.4 | 1.33E-08 | 6.88E-08 |
| chr6 | 112524188 | 112538955 | 14768 | 1.22 | 1.33E-08 | 6.89E-08 |
| chr5 | 90779981 | 90782831 | 2851 | 2.8 | 1.34E-08 | 6.92E-08 |
| chr5 | 28954232 | 28955455 | 1224 | 6.45 | 1.34E-08 | 6.95E-08 |
| chr8 | 53852870 | 53854951 | 2082 | 1.9 | 1.35E-08 | 6.96E-08 |
| chrX | 28005498 | 28006844 | 1347 | 4.01 | 1.35E-08 | 6.96E-08 |
| chr5 | 95444889 | 95461697 | 16809 | 6.46 | 1.35E-08 | 6.98E-08 |
| chr5 | 95402826 | 95431174 | 28349 | 0.54 | 1.35E-08 | 6.98E-08 |
| chr19 | 3450356 | 3460783 | 10428 | 0.74 | 1.36E-08 | 7.01E-08 |
| chr1 | 247390985 | 247391596 | 612 | 6.49 | 1.36E-08 | 7.01E-08 |
| chr12 | 51417885 | 51423062 | 5178 | 0.89 | 1.36E-08 | 7.02E-08 |
| chr17 | 41320453 | 41366331 | 45879 | 0.33 | 1.36E-08 | 7.02E-08 |
| chr13 | 43687403 | 43688648 | 1246 | 3.2 | 1.36E-08 | 7.03E-08 |
| chr4 | 64802879 | 64804263 | 1385 | 6.41 | 1.36E-08 | 7.05E-08 |
| chr6 | 11631186 | 11638153 | 6968 | 0.77 | 1.37E-08 | 7.07E-08 |
| chr10 | 52609306 | 52610389 | 1084 | 2.67 | 1.37E-08 | 7.07E-08 |
| chr16 | 11768782 | 11820467 | 51686 | -0.52 | 1.38E-08 | 7.12E-08 |
| chr3 | 65868714 | 65875602 | 6889 | 1.35 | 1.38E-08 | 7.14E-08 |
| chr12 | 69927602 | 69929547 | 1946 | 1.98 | 1.40E-08 | 7.23E-08 |
| chr12 | 27089507 | 27092849 | 3343 | 0.65 | 1.40E-08 | 7.23E-08 |
| chr18 | 29175789 | 29176939 | 1151 | 6.44 | 1.40E-08 | 7.24E-08 |
| chr12 | 115832372 | 115833060 | 689 | -6.35 | 1.41E-08 | 7.24E-08 |
| chr4 | 142599052 | 142613609 | 14558 | 0.55 | 1.41E-08 | 7.25E-08 |
| chr20 | 10828984 | 10845605 | 16622 | -0.72 | 1.41E-08 | 7.25E-08 |

| | | | | | | |
|---|---|---|---|---|---|---|
| chr4 | 118320788 | 118321976 | 1189 | 6.43 | 1.41E-08 | 7.26E-08 |
| chr7 | 137351867 | 137353314 | 1448 | 6.4 | 1.41E-08 | 7.27E-08 |
| chr7 | 14535096 | 14536269 | 1174 | 6.41 | 1.44E-08 | 7.44E-08 |
| chr1 | 161358896 | 161370752 | 11857 | 0.48 | 1.45E-08 | 7.46E-08 |
| chr11 | 35772986 | 35773919 | 934 | 3.51 | 1.45E-08 | 7.47E-08 |
| chr22 | 32473974 | 32481486 | 7513 | 1.2 | 1.45E-08 | 7.48E-08 |
| chr1 | 239332264 | 239333385 | 1122 | 6.39 | 1.46E-08 | 7.50E-08 |
| chr13 | 49548817 | 49552887 | 4071 | 0.72 | 1.46E-08 | 7.50E-08 |
| chr3 | 130968827 | 130970297 | 1471 | 6.45 | 1.46E-08 | 7.52E-08 |
| chr2 | 104756631 | 104757601 | 971 | 6.39 | 1.47E-08 | 7.56E-08 |
| chr15 | 22369464 | 22370416 | 953 | 6.4 | 1.48E-08 | 7.60E-08 |
| chr10 | 53458047 | 53460521 | 2475 | 0.62 | 1.48E-08 | 7.61E-08 |
| chr13 | 99624481 | 99625912 | 1432 | 4.12 | 1.48E-08 | 7.62E-08 |
| chr9 | 76129886 | 76130807 | 922 | 6.46 | 1.49E-08 | 7.64E-08 |
| chr2 | 221051823 | 221064629 | 12807 | 0.93 | 1.49E-08 | 7.67E-08 |
| chr13 | 71864578 | 71865967 | 1390 | 2.48 | 1.49E-08 | 7.68E-08 |
| chr3 | 151022522 | 151023930 | 1409 | 1.38 | 1.50E-08 | 7.69E-08 |
| chr8 | 48382885 | 48395645 | 12761 | 0.53 | 1.50E-08 | 7.69E-08 |
| chr9 | 78794688 | 78801970 | 7283 | 0.77 | 1.50E-08 | 7.69E-08 |
| chr12 | 102269973 | 102289707 | 19735 | 0.45 | 1.50E-08 | 7.72E-08 |
| chr1 | 233678929 | 233680299 | 1371 | 2.55 | 1.51E-08 | 7.73E-08 |
| chr2 | 13698281 | 13699290 | 1010 | 6.39 | 1.51E-08 | 7.73E-08 |
| chr10 | 16478289 | 16484044 | 5756 | 0.73 | 1.51E-08 | 7.77E-08 |
| chr6 | 74354895 | 74376629 | 21735 | 0.72 | 1.52E-08 | 7.80E-08 |
| chr15 | 33533098 | 33537049 | 3952 | 1.41 | 1.52E-08 | 7.81E-08 |
| chr2 | 83532945 | 83533780 | 836 | 4.52 | 1.52E-08 | 7.82E-08 |
| chr4 | 30398091 | 30399645 | 1555 | 6.47 | 1.52E-08 | 7.82E-08 |
| chr1 | 220052959 | 220053738 | 780 | 6.38 | 1.53E-08 | 7.85E-08 |
| chr16 | 22036129 | 22043970 | 7842 | 0.59 | 1.54E-08 | 7.91E-08 |
| chr2 | 11040605 | 11054729 | 14125 | -1.45 | 1.54E-08 | 7.92E-08 |
| chr17 | 67797911 | 67808803 | 10893 | -6.35 | 1.55E-08 | 7.92E-08 |
| chr8 | 32283431 | 32291431 | 8001 | -1.24 | 1.55E-08 | 7.95E-08 |
| chr3 | 48631058 | 48673876 | 42819 | 0.47 | 1.55E-08 | 7.95E-08 |
| chr3 | 114565100 | 114566322 | 1223 | 4.23 | 1.56E-08 | 7.98E-08 |
| chr17 | 26968642 | 26991254 | 22613 | 0.42 | 1.56E-08 | 7.98E-08 |
| chr11 | 39289021 | 39290098 | 1078 | 6.39 | 1.56E-08 | 7.99E-08 |
| chr4 | 67404794 | 67406127 | 1334 | 1.49 | 1.56E-08 | 7.99E-08 |
| chr7 | 17444775 | 17449500 | 4726 | 0.62 | 1.56E-08 | 8.01E-08 |
| chr11 | 92949480 | 92963172 | 13693 | 1.72 | 1.58E-08 | 8.07E-08 |
| chr12 | 13323882 | 13376811 | 52930 | 0.55 | 1.58E-08 | 8.08E-08 |
| chr19 | 3965788 | 3971135 | 5348 | 0.7 | 1.58E-08 | 8.09E-08 |
| chr10 | 112532963 | 112535713 | 2751 | 3.65 | 1.58E-08 | 8.10E-08 |
| chr6 | 116375837 | 116395598 | 19762 | 0.46 | 1.58E-08 | 8.10E-08 |
| chr10 | 72163183 | 72166641 | 3459 | 0.83 | 1.59E-08 | 8.12E-08 |
| chr6 | 140263561 | 140265723 | 2163 | 1.99 | 1.59E-08 | 8.13E-08 |
| chr7 | 79063770 | 79065025 | 1256 | 4.26 | 1.59E-08 | 8.13E-08 |
| chrX | 70584749 | 70588305 | 3557 | 0.73 | 1.59E-08 | 8.15E-08 |
| chr3 | 178136754 | 178139784 | 3031 | 2.9 | 1.60E-08 | 8.17E-08 |
| chr21 | 37559456 | 37562028 | 2573 | 1.59 | 1.60E-08 | 8.18E-08 |
| chr11 | 94181283 | 94191021 | 9739 | 1.37 | 1.60E-08 | 8.19E-08 |
| chr7 | 141348291 | 141377164 | 28874 | 0.38 | 1.61E-08 | 8.20E-08 |
| chr2 | 108249985 | 108251635 | 1651 | 2.43 | 1.61E-08 | 8.24E-08 |
| chr3 | 193587054 | 193604806 | 17753 | 1.8 | 1.62E-08 | 8.28E-08 |
| chr6 | 63232901 | 63234373 | 1473 | 6.4 | 1.63E-08 | 8.30E-08 |
| chr10 | 26727605 | 26728855 | 1251 | -4.46 | 1.63E-08 | 8.31E-08 |
| chr11 | 82829573 | 82836778 | 7206 | 1.6 | 1.63E-08 | 8.34E-08 |
| chr19 | 44711150 | 44717931 | 6782 | 0.83 | 1.64E-08 | 8.35E-08 |
| chr3 | 164285484 | 164286118 | 635 | 6.37 | 1.64E-08 | 8.37E-08 |
| chr7 | 45147637 | 45152564 | 4928 | 0.51 | 1.65E-08 | 8.40E-08 |
| chr10 | 52229705 | 52231554 | 1850 | 1.68 | 1.65E-08 | 8.41E-08 |
| chr11 | 59412971 | 59438000 | 25030 | 0.53 | 1.65E-08 | 8.42E-08 |

| | | | | | | |
|---|---|---|---|---|---|---|
| chr2 | 224283014 | 224283853 | 840 | 6.39 | 1.65E-08 | 8.42E-08 |
| chr10 | 9153681 | 9154681 | 1001 | 6.39 | 1.66E-08 | 8.44E-08 |
| chr2 | 194006506 | 194008500 | 1995 | 6.39 | 1.66E-08 | 8.44E-08 |
| chr4 | 41386844 | 41451994 | 65151 | 0.33 | 1.66E-08 | 8.45E-08 |
| chr20 | 3527378 | 3540904 | 13527 | 0.74 | 1.66E-08 | 8.47E-08 |
| chr17 | 17920525 | 17944576 | 24052 | -1.02 | 1.67E-08 | 8.49E-08 |
| chr10 | 127321801 | 127322939 | 1139 | 6.37 | 1.67E-08 | 8.50E-08 |
| chr12 | 52277868 | 52280684 | 2817 | 1.7 | 1.67E-08 | 8.53E-08 |
| chr15 | 91282936 | 91287429 | 4494 | -6.36 | 1.68E-08 | 8.57E-08 |
| chr9 | 447085 | 448644 | 1560 | 2.56 | 1.69E-08 | 8.59E-08 |
| chr4 | 174529949 | 174531472 | 1524 | 1.35 | 1.69E-08 | 8.62E-08 |
| chr3 | 81765839 | 81811986 | 46148 | -0.32 | 1.69E-08 | 8.62E-08 |
| chr19 | 47477793 | 47509514 | 31722 | -0.78 | 1.70E-08 | 8.64E-08 |
| chr6 | 8556345 | 8557505 | 1161 | 6.44 | 1.72E-08 | 8.75E-08 |
| chr4 | 124466487 | 124477862 | 11376 | 0.77 | 1.72E-08 | 8.76E-08 |
| chr2 | 196271325 | 196279517 | 8193 | -1.38 | 1.72E-08 | 8.76E-08 |
| chr12 | 1702655 | 1716172 | 13518 | 0.69 | 1.72E-08 | 8.77E-08 |
| chr11 | 47693637 | 47694599 | 963 | -3.39 | 1.73E-08 | 8.78E-08 |
| chr8 | 133724054 | 133725012 | 959 | 6.39 | 1.73E-08 | 8.78E-08 |
| chr8 | 6766546 | 6768459 | 1914 | 1.79 | 1.73E-08 | 8.80E-08 |
| chr1 | 9864611 | 9888250 | 23640 | 0.65 | 1.73E-08 | 8.80E-08 |
| chr9 | 95852842 | 95897258 | 44417 | -0.38 | 1.74E-08 | 8.82E-08 |
| chr7 | 31446552 | 31450463 | 3912 | 0.57 | 1.74E-08 | 8.84E-08 |
| chr14 | 68123594 | 68168130 | 44537 | 0.42 | 1.75E-08 | 8.88E-08 |
| chr17 | 14255887 | 14265912 | 10026 | 6.36 | 1.75E-08 | 8.89E-08 |
| chr4 | 181054938 | 181055892 | 955 | 6.36 | 1.75E-08 | 8.90E-08 |
| chr22 | 18255988 | 18265237 | 9250 | -1.4 | 1.77E-08 | 8.96E-08 |
| chr19 | 16372708 | 16374494 | 1787 | -2.79 | 1.77E-08 | 8.98E-08 |
| chrX | 140996444 | 140997536 | 1093 | 6.36 | 1.78E-08 | 9.01E-08 |
| chr12 | 133284513 | 133301034 | 16522 | 1.13 | 1.78E-08 | 9.01E-08 |
| chr18 | 42368927 | 42371039 | 2113 | 1.88 | 1.78E-08 | 9.05E-08 |
| chr4 | 65488636 | 65489784 | 1149 | 6.37 | 1.79E-08 | 9.06E-08 |
| chr5 | 68041057 | 68042808 | 1752 | 6.36 | 1.79E-08 | 9.07E-08 |
| chr16 | 55762473 | 55763633 | 1161 | 2.79 | 1.79E-08 | 9.09E-08 |
| chr11 | 39242750 | 39243907 | 1158 | 6.36 | 1.80E-08 | 9.12E-08 |
| chr12 | 49503108 | 49505487 | 2380 | 1.13 | 1.81E-08 | 9.16E-08 |
| chr17 | 12306260 | 12307332 | 1073 | -6.32 | 1.81E-08 | 9.19E-08 |
| chr8 | 56831615 | 56851666 | 20052 | 3.28 | 1.81E-08 | 9.19E-08 |
| chr4 | 145500510 | 145513496 | 12987 | 0.49 | 1.84E-08 | 9.33E-08 |
| chr11 | 106030875 | 106031861 | 987 | 3.15 | 1.85E-08 | 9.38E-08 |
| chr11 | 78075262 | 78077741 | 2480 | 4.38 | 1.85E-08 | 9.38E-08 |
| chr2 | 200580313 | 200582961 | 2649 | 0.7 | 1.85E-08 | 9.39E-08 |
| chr9 | 16688717 | 16708147 | 19431 | 0.44 | 1.86E-08 | 9.40E-08 |
| chr16 | 10740111 | 10748594 | 8484 | 1.19 | 1.87E-08 | 9.47E-08 |
| chr20 | 5757864 | 5788787 | 30924 | 0.41 | 1.88E-08 | 9.50E-08 |
| chr9 | 112831685 | 112852461 | 20777 | 6.63 | 1.88E-08 | 9.51E-08 |
| chr5 | 130232106 | 130233558 | 1453 | 1.59 | 1.88E-08 | 9.53E-08 |
| chr7 | 81068921 | 81077519 | 8599 | 6.38 | 1.89E-08 | 9.56E-08 |
| chr12 | 50468732 | 50479840 | 11109 | 0.65 | 1.90E-08 | 9.59E-08 |
| chr7 | 101491399 | 101668736 | 177338 | -0.24 | 1.90E-08 | 9.60E-08 |
| chr1 | 86458033 | 86459370 | 1338 | 1.86 | 1.90E-08 | 9.60E-08 |
| chr3 | 185814827 | 185828591 | 13765 | 0.59 | 1.90E-08 | 9.60E-08 |
| chr8 | 118949293 | 119044281 | 94989 | -0.32 | 1.90E-08 | 9.62E-08 |
| chr10 | 1016213 | 1050707 | 34495 | 0.85 | 1.91E-08 | 9.63E-08 |
| chr5 | 45577857 | 45578983 | 1127 | 6.35 | 1.91E-08 | 9.66E-08 |
| chr8 | 124068539 | 124071666 | 3128 | 0.91 | 1.92E-08 | 9.69E-08 |
| chr12 | 83967265 | 83968129 | 865 | 4.49 | 1.92E-08 | 9.70E-08 |
| chr14 | 26994201 | 26995210 | 1010 | 6.41 | 1.93E-08 | 9.74E-08 |
| chr1 | 78289227 | 78301625 | 12399 | 1.05 | 1.94E-08 | 9.79E-08 |
| chr4 | 150882798 | 150883967 | 1170 | 4.13 | 1.96E-08 | 9.88E-08 |
| chr8 | 120831475 | 120846500 | 15026 | 0.54 | 1.97E-08 | 9.94E-08 |

| | | | | | | |
|---|---|---|---|---|---|---|
| chr1 | 189988088 | 189990377 | 2290 | 0.95 | 1.97E-08 | 9.94E-08 |
| chr19 | 19078082 | 19082455 | 4374 | 0.77 | 1.99E-08 | 1.00E-07 |
| chr9 | 29790055 | 29791370 | 1316 | 6.35 | 2.00E-08 | 1.01E-07 |
| chrX | 42614574 | 42615740 | 1167 | 5.1 | 2.00E-08 | 1.01E-07 |
| chr7 | 141694766 | 141696631 | 1866 | 3.37 | 2.00E-08 | 1.01E-07 |
| chr21 | 40898931 | 40901167 | 2237 | 3.06 | 2.02E-08 | 1.02E-07 |
| chr3 | 182343938 | 182345498 | 1561 | 6.38 | 2.02E-08 | 1.02E-07 |
| chr3 | 187120767 | 187129995 | 9229 | 1.51 | 2.03E-08 | 1.02E-07 |
| chr17 | 31180949 | 31182176 | 1228 | 6.36 | 2.03E-08 | 1.02E-07 |
| chr19 | 37197712 | 37199102 | 1391 | 6.34 | 2.04E-08 | 1.03E-07 |
| chr19 | 30432693 | 30455265 | 22573 | 0.57 | 2.05E-08 | 1.03E-07 |
| chr2 | 64994364 | 64997269 | 2906 | 0.99 | 2.05E-08 | 1.03E-07 |
| chr16 | 19593546 | 19621597 | 28052 | -0.73 | 2.05E-08 | 1.03E-07 |
| chr3 | 123499243 | 123528163 | 28921 | -0.76 | 2.05E-08 | 1.03E-07 |
| chr6 | 12973257 | 12974412 | 1156 | 6.44 | 2.05E-08 | 1.03E-07 |
| chr19 | 12202490 | 12203920 | 1431 | 6.37 | 2.07E-08 | 1.04E-07 |
| chr9 | 114336775 | 114340707 | 3933 | -1.78 | 2.08E-08 | 1.05E-07 |
| chr5 | 17000069 | 17025442 | 25374 | -0.99 | 2.08E-08 | 1.05E-07 |
| chr4 | 156260227 | 156261570 | 1344 | -2.46 | 2.08E-08 | 1.05E-07 |
| chr10 | 63157196 | 63157965 | 770 | 2.71 | 2.09E-08 | 1.05E-07 |
| chr1 | 162350528 | 162352562 | 2035 | 1.47 | 2.09E-08 | 1.05E-07 |
| chr18 | 12730469 | 12742786 | 12318 | -1.07 | 2.09E-08 | 1.05E-07 |
| chr2 | 19010363 | 19011687 | 1325 | 2.7 | 2.09E-08 | 1.05E-07 |
| chr7 | 156319805 | 156321218 | 1414 | 4.27 | 2.09E-08 | 1.05E-07 |
| chr2 | 174074509 | 174096625 | 22117 | -0.68 | 2.10E-08 | 1.05E-07 |
| chr19 | 35410748 | 35418510 | 7763 | 6.67 | 2.11E-08 | 1.06E-07 |
| chrX | 67472214 | 67476148 | 3935 | 1.29 | 2.12E-08 | 1.07E-07 |
| chr1 | 160282257 | 160296330 | 14074 | 0.87 | 2.12E-08 | 1.07E-07 |
| chr2 | 25001340 | 25065158 | 63819 | -0.38 | 2.13E-08 | 1.07E-07 |
| chr7 | 63955203 | 63956532 | 1330 | 6.36 | 2.14E-08 | 1.07E-07 |
| chr1 | 245114629 | 245137595 | 22967 | 0.31 | 2.16E-08 | 1.08E-07 |
| chr12 | 64425489 | 64434703 | 9215 | 1.02 | 2.17E-08 | 1.09E-07 |
| chr2 | 3706953 | 3709203 | 2251 | 2.26 | 2.18E-08 | 1.09E-07 |
| chr4 | 106830548 | 106831742 | 1195 | 3.37 | 2.18E-08 | 1.09E-07 |
| chr14 | 56468189 | 56473210 | 5022 | 1 | 2.18E-08 | 1.09E-07 |
| chr6 | 160768351 | 160828970 | 60620 | 0.27 | 2.18E-08 | 1.10E-07 |
| chr8 | 29718453 | 29719974 | 1522 | 2.03 | 2.19E-08 | 1.10E-07 |
| chr10 | 111674240 | 111683761 | 9522 | 0.99 | 2.21E-08 | 1.11E-07 |
| chr4 | 8103316 | 8115675 | 12360 | 6.34 | 2.22E-08 | 1.12E-07 |
| chr18 | 43353680 | 43383894 | 30215 | 0.62 | 2.24E-08 | 1.12E-07 |
| chr20 | 4089064 | 4174404 | 85341 | -0.22 | 2.26E-08 | 1.13E-07 |
| chr9 | 128585771 | 128587400 | 1630 | 1.55 | 2.26E-08 | 1.13E-07 |
| chr13 | 100740346 | 100765190 | 24845 | 0.95 | 2.28E-08 | 1.14E-07 |
| chr8 | 131053663 | 131177485 | 123823 | -0.29 | 2.29E-08 | 1.15E-07 |
| chr3 | 55363984 | 55365425 | 1442 | 2.21 | 2.30E-08 | 1.15E-07 |
| chr1 | 2506919 | 2518032 | 11114 | 1.72 | 2.31E-08 | 1.15E-07 |
| chr20 | 37232702 | 37235179 | 2478 | 1.14 | 2.31E-08 | 1.16E-07 |
| chr11 | 126242515 | 126256459 | 13945 | 1.45 | 2.32E-08 | 1.16E-07 |
| chr12 | 46659113 | 46663905 | 4793 | 0.51 | 2.33E-08 | 1.17E-07 |
| chr7 | 2662781 | 2674142 | 11362 | 0.77 | 2.35E-08 | 1.18E-07 |
| chr4 | 4992640 | 4993681 | 1042 | -2.32 | 2.36E-08 | 1.18E-07 |
| chr22 | 19323277 | 19324534 | 1258 | 3.2 | 2.36E-08 | 1.18E-07 |
| chr4 | 95994012 | 96030808 | 36797 | 0.3 | 2.36E-08 | 1.18E-07 |
| chr8 | 16573804 | 16576542 | 2739 | -1.04 | 2.37E-08 | 1.19E-07 |
| chr1 | 216704328 | 216705493 | 1166 | 2.06 | 2.39E-08 | 1.20E-07 |
| chr17 | 4577547 | 4643816 | 66270 | -0.32 | 2.39E-08 | 1.20E-07 |
| chr7 | 158220945 | 158227452 | 6508 | 0.52 | 2.41E-08 | 1.21E-07 |
| chr17 | 38593898 | 38718809 | 124912 | -0.22 | 2.42E-08 | 1.21E-07 |
| chr21 | 35894308 | 35899960 | 5653 | 0.69 | 2.42E-08 | 1.21E-07 |
| chr1 | 81894203 | 81895287 | 1085 | 6.32 | 2.43E-08 | 1.22E-07 |
| chr17 | 70146526 | 70155899 | 9374 | 0.54 | 2.44E-08 | 1.22E-07 |

| | | | | | | |
|---|---|---|---|---|---|---|
| chr7 | 151540928 | 151542482 | 1555 | 2.42 | 2.45E-08 | 1.23E-07 |
| chr4 | 190057162 | 190065432 | 8271 | 1.39 | 2.45E-08 | 1.23E-07 |
| chr5 | 59788382 | 59789485 | 1104 | -6.29 | 2.46E-08 | 1.23E-07 |
| chr3 | 3167594 | 3192683 | 25090 | 0.48 | 2.46E-08 | 1.23E-07 |
| chr1 | 105248739 | 105249488 | 750 | 3.33 | 2.47E-08 | 1.23E-07 |
| chr14 | 95776713 | 95789663 | 12951 | 0.96 | 2.47E-08 | 1.23E-07 |
| chr1 | 28761554 | 28776909 | 15356 | 0.91 | 2.47E-08 | 1.23E-07 |
| chr10 | 84170876 | 84171976 | 1101 | 6.31 | 2.48E-08 | 1.24E-07 |
| chr3 | 17771483 | 17785642 | 14160 | 0.35 | 2.49E-08 | 1.24E-07 |
| chr4 | 36089230 | 36091106 | 1877 | 6.32 | 2.49E-08 | 1.24E-07 |
| chr5 | 21016558 | 21017483 | 926 | 6.31 | 2.51E-08 | 1.25E-07 |
| chr19 | 53360985 | 53361933 | 949 | 6.39 | 2.51E-08 | 1.25E-07 |
| chr3 | 116869077 | 116870496 | 1420 | 2.15 | 2.52E-08 | 1.26E-07 |
| chr2 | 8529231 | 8530429 | 1199 | 2.23 | 2.53E-08 | 1.26E-07 |
| chr1 | 219728337 | 219729977 | 1641 | -4.25 | 2.53E-08 | 1.26E-07 |
| chr1 | 158969771 | 158986021 | 16251 | 6.37 | 2.56E-08 | 1.28E-07 |
| chr1 | 198898190 | 198907437 | 9248 | 0.51 | 2.58E-08 | 1.28E-07 |
| chr15 | 83586629 | 83589442 | 2814 | 6.52 | 2.58E-08 | 1.29E-07 |
| chr13 | 100618112 | 100636905 | 18794 | 0.59 | 2.59E-08 | 1.29E-07 |
| chr12 | 19635327 | 19683575 | 48249 | 0.35 | 2.60E-08 | 1.29E-07 |
| chr3 | 52311172 | 52323057 | 11886 | 0.46 | 2.62E-08 | 1.30E-07 |
| chr5 | 23506639 | 23508028 | 1390 | 2.68 | 2.62E-08 | 1.30E-07 |
| chr11 | 1283235 | 1297005 | 13771 | -1.3 | 2.62E-08 | 1.31E-07 |
| chr12 | 115860787 | 115861886 | 1100 | -3.56 | 2.63E-08 | 1.31E-07 |
| chr1 | 190648272 | 190649517 | 1246 | 6.31 | 2.64E-08 | 1.31E-07 |
| chr11 | 28160792 | 28168241 | 7450 | -1.37 | 2.64E-08 | 1.31E-07 |
| chr7 | 134276031 | 134299703 | 23673 | -0.29 | 2.64E-08 | 1.31E-07 |
| chr14 | 22733076 | 22734377 | 1302 | 6.33 | 2.65E-08 | 1.31E-07 |
| chr1 | 244898324 | 244899363 | 1040 | -3.83 | 2.65E-08 | 1.32E-07 |
| chr2 | 83911213 | 83912608 | 1396 | 6.3 | 2.66E-08 | 1.32E-07 |
| chr13 | 30156769 | 30170414 | 13646 | -0.66 | 2.67E-08 | 1.33E-07 |
| chr2 | 173557173 | 173563595 | 6423 | -1.43 | 2.68E-08 | 1.33E-07 |
| chr2 | 32182628 | 32218992 | 36365 | -0.81 | 2.68E-08 | 1.33E-07 |
| chrX | 36030907 | 36037677 | 6771 | 6.32 | 2.69E-08 | 1.33E-07 |
| chr16 | 10475836 | 10481354 | 5519 | 1.2 | 2.69E-08 | 1.33E-07 |
| chr2 | 53254955 | 53256065 | 1111 | 6.3 | 2.69E-08 | 1.33E-07 |
| chr7 | 46750644 | 46754188 | 3545 | 6.3 | 2.69E-08 | 1.33E-07 |
| chr20 | 22319329 | 22322448 | 3120 | -5 | 2.70E-08 | 1.34E-07 |
| chr21 | 34833782 | 34834912 | 1131 | 3.41 | 2.72E-08 | 1.35E-07 |
| chr8 | 26148163 | 26169237 | 21075 | 0.74 | 2.72E-08 | 1.35E-07 |
| chr3 | 104128710 | 104131397 | 2688 | -2.43 | 2.72E-08 | 1.35E-07 |
| chr7 | 103072970 | 103073804 | 835 | 6.32 | 2.72E-08 | 1.35E-07 |
| chr1 | 149980687 | 149983711 | 3025 | 0.75 | 2.73E-08 | 1.36E-07 |
| chr2 | 182902999 | 182904288 | 1290 | 3.74 | 2.75E-08 | 1.36E-07 |
| chr5 | 58220261 | 58230109 | 9849 | 0.6 | 2.76E-08 | 1.37E-07 |
| chr9 | 123238010 | 123249357 | 11348 | -0.94 | 2.78E-08 | 1.38E-07 |
| chr19 | 49890566 | 49891931 | 1366 | 2.6 | 2.80E-08 | 1.39E-07 |
| chrX | 14981416 | 14982603 | 1188 | 4.06 | 2.80E-08 | 1.39E-07 |
| chr10 | 79442346 | 79443495 | 1150 | 4.73 | 2.81E-08 | 1.39E-07 |
| chr18 | 9073327 | 9159059 | 85733 | 0.28 | 2.81E-08 | 1.39E-07 |
| chr1 | 171710555 | 171713258 | 2704 | 0.74 | 2.82E-08 | 1.39E-07 |
| chr2 | 154727407 | 154729208 | 1802 | -2.06 | 2.82E-08 | 1.39E-07 |
| chr4 | 82073707 | 82086008 | 12302 | 1.51 | 2.82E-08 | 1.40E-07 |
| chr15 | 36789502 | 36793928 | 4427 | 1.27 | 2.82E-08 | 1.40E-07 |
| chr4 | 59626413 | 59627993 | 1581 | 6.31 | 2.83E-08 | 1.40E-07 |
| chr12 | 59790039 | 59791292 | 1254 | 6.36 | 2.84E-08 | 1.40E-07 |
| chr6 | 41589808 | 41590535 | 728 | 5.46 | 2.84E-08 | 1.40E-07 |
| chr1 | 87857660 | 87859378 | 1719 | 1.64 | 2.84E-08 | 1.40E-07 |
| chr5 | 67871339 | 67872496 | 1158 | 6.29 | 2.85E-08 | 1.41E-07 |
| chrX | 16788836 | 16790974 | 2139 | -0.83 | 2.85E-08 | 1.41E-07 |
| chr2 | 25886894 | 25896362 | 9469 | -1.52 | 2.86E-08 | 1.41E-07 |

| | | | | | | |
|---|---|---|---|---|---|---|
| chr2 | 240551121 | 240553598 | 2478 | 1.3 | 2.87E-08 | 1.42E-07 |
| chr17 | 10632786 | 10642080 | 9295 | 0.92 | 2.88E-08 | 1.42E-07 |
| chr4 | 91468298 | 91469134 | 837 | 5.03 | 2.88E-08 | 1.42E-07 |
| chr7 | 141284119 | 141293655 | 9537 | -0.63 | 2.91E-08 | 1.44E-07 |
| chr17 | 36472377 | 36525323 | 52947 | -0.62 | 2.91E-08 | 1.44E-07 |
| chr4 | 17031803 | 17033074 | 1272 | -6.25 | 2.92E-08 | 1.44E-07 |
| chr12 | 96335560 | 96337681 | 2122 | 0.8 | 2.93E-08 | 1.45E-07 |
| chr13 | 94615390 | 94616956 | 1567 | 1.75 | 2.93E-08 | 1.45E-07 |
| chr18 | 27674005 | 27674937 | 933 | 6.32 | 2.95E-08 | 1.46E-07 |
| chr5 | 10441191 | 10482359 | 41169 | -0.73 | 2.96E-08 | 1.46E-07 |
| chr2 | 46256258 | 46270097 | 13840 | 1.63 | 2.96E-08 | 1.46E-07 |
| chr1 | 99126406 | 99129244 | 2839 | 0.95 | 2.96E-08 | 1.46E-07 |
| chr8 | 80672908 | 80682277 | 9370 | 1.38 | 2.97E-08 | 1.46E-07 |
| chr7 | 82163515 | 82164937 | 1423 | 2.51 | 2.97E-08 | 1.46E-07 |
| chr2 | 173841891 | 173870334 | 28444 | 0.38 | 2.97E-08 | 1.46E-07 |
| chr1 | 86441277 | 86442331 | 1055 | 6.3 | 3.00E-08 | 1.48E-07 |
| chr16 | 74557825 | 74559060 | 1236 | 6.45 | 3.00E-08 | 1.48E-07 |
| chr5 | 126253064 | 126255101 | 2038 | 1.8 | 3.02E-08 | 1.49E-07 |
| chr21 | 45107714 | 45210823 | 103110 | 0.24 | 3.03E-08 | 1.49E-07 |
| chr9 | 81569152 | 81597995 | 28844 | 1.1 | 3.05E-08 | 1.50E-07 |
| chr11 | 27955403 | 27957415 | 2013 | 0.89 | 3.08E-08 | 1.52E-07 |
| chr3 | 193271951 | 193273233 | 1283 | 6.29 | 3.08E-08 | 1.52E-07 |
| chr12 | 32258979 | 32270589 | 11611 | 0.49 | 3.09E-08 | 1.52E-07 |
| chr5 | 91090187 | 91091379 | 1193 | -4.26 | 3.11E-08 | 1.53E-07 |
| chr7 | 151145216 | 151165980 | 20765 | 0.7 | 3.11E-08 | 1.53E-07 |
| chr3 | 30788718 | 30795209 | 6492 | 1.79 | 3.11E-08 | 1.53E-07 |
| chr1 | 1688810 | 1715161 | 26352 | 0.53 | 3.11E-08 | 1.53E-07 |
| chr14 | 88509829 | 88512868 | 3040 | 0.86 | 3.11E-08 | 1.53E-07 |
| chr9 | 114756648 | 114757782 | 1135 | 2.67 | 3.12E-08 | 1.53E-07 |
| chr9 | 94741140 | 94749358 | 8219 | -6.25 | 3.16E-08 | 1.56E-07 |
| chr1 | 95478837 | 95480149 | 1313 | 2.36 | 3.17E-08 | 1.56E-07 |
| chr1 | 51527704 | 51530128 | 2425 | 1.02 | 3.21E-08 | 1.58E-07 |
| chr9 | 127414484 | 127428813 | 14330 | 0.4 | 3.22E-08 | 1.58E-07 |
| chrX | 122646970 | 122648329 | 1360 | 1.38 | 3.22E-08 | 1.58E-07 |
| chr6 | 65211801 | 65215217 | 3417 | 6.28 | 3.22E-08 | 1.58E-07 |
| chr5 | 74805627 | 74809052 | 3426 | 0.72 | 3.24E-08 | 1.59E-07 |
| chr2 | 46523658 | 46565256 | 41599 | 0.35 | 3.25E-08 | 1.60E-07 |
| chr1 | 231327882 | 231347758 | 19877 | -0.62 | 3.27E-08 | 1.61E-07 |
| chr14 | 33125603 | 33127089 | 1487 | -1.57 | 3.28E-08 | 1.61E-07 |
| chr1 | 198849829 | 198873924 | 24096 | 0.93 | 3.28E-08 | 1.61E-07 |
| chr10 | 67553816 | 67555554 | 1739 | 1.91 | 3.29E-08 | 1.61E-07 |
| chr15 | 64734079 | 64996158 | 262080 | 0.21 | 3.29E-08 | 1.61E-07 |
| chr8 | 102513155 | 102533993 | 20839 | 0.77 | 3.30E-08 | 1.62E-07 |
| chr5 | 156557761 | 156571073 | 13313 | 0.69 | 3.30E-08 | 1.62E-07 |
| chr16 | 69595391 | 69674182 | 78792 | 0.34 | 3.32E-08 | 1.63E-07 |
| chr10 | 99037882 | 99058486 | 20605 | -0.52 | 3.33E-08 | 1.63E-07 |
| chr11 | 70852736 | 70864995 | 12260 | -1.64 | 3.34E-08 | 1.64E-07 |
| chr4 | 166826319 | 166827495 | 1177 | 2.31 | 3.34E-08 | 1.64E-07 |
| chr6 | 42160953 | 42228976 | 68024 | 0.4 | 3.36E-08 | 1.65E-07 |
| chr9 | 111341115 | 111343888 | 2774 | 1.08 | 3.39E-08 | 1.66E-07 |
| chr12 | 213088 | 213580 | 493 | 4.74 | 3.41E-08 | 1.67E-07 |
| chrX | 81670234 | 81671245 | 1012 | 4.74 | 3.41E-08 | 1.67E-07 |
| chr13 | 52531825 | 52533057 | 1233 | 4.11 | 3.42E-08 | 1.68E-07 |
| chr7 | 93632328 | 93634608 | 2281 | 0.94 | 3.42E-08 | 1.68E-07 |
| chr22 | 39077471 | 39115697 | 38227 | 0.4 | 3.43E-08 | 1.68E-07 |
| chr16 | 31264195 | 31278000 | 13806 | -1.22 | 3.44E-08 | 1.68E-07 |
| chr9 | 126311354 | 126319721 | 8368 | -1.34 | 3.45E-08 | 1.69E-07 |
| chr4 | 92119659 | 92120743 | 1085 | 3.94 | 3.46E-08 | 1.69E-07 |
| chr5 | 92956207 | 92958520 | 2314 | 0.9 | 3.48E-08 | 1.70E-07 |
| chr16 | 75018244 | 75064580 | 46337 | 0.53 | 3.48E-08 | 1.71E-07 |
| chr6 | 91077731 | 91079817 | 2087 | 1.18 | 3.49E-08 | 1.71E-07 |

| | | | | | | |
|---|---|---|---|---|---|---|
| chr2 | 109394482 | 109404709 | 10228 | -0.69 | 3.54E-08 | 1.73E-07 |
| chr1 | 151809316 | 151810300 | 985 | 6.26 | 3.54E-08 | 1.73E-07 |
| chr1 | 209437769 | 209450340 | 12572 | 0.84 | 3.55E-08 | 1.73E-07 |
| chr20 | 43342876 | 43376246 | 33371 | -0.5 | 3.55E-08 | 1.74E-07 |
| chr4 | 72521483 | 72523031 | 1549 | 4.2 | 3.56E-08 | 1.74E-07 |
| chr4 | 85503357 | 85505126 | 1770 | 3.27 | 3.58E-08 | 1.75E-07 |
| chr19 | 12424967 | 12425987 | 1021 | 6.27 | 3.59E-08 | 1.75E-07 |
| chr7 | 80780199 | 80805853 | 25655 | -0.5 | 3.59E-08 | 1.75E-07 |
| chr3 | 152678606 | 152692128 | 13523 | 0.51 | 3.59E-08 | 1.75E-07 |
| chr2 | 227338769 | 227349672 | 10904 | 0.45 | 3.61E-08 | 1.76E-07 |
| chr5 | 66253418 | 66260004 | 6587 | 0.63 | 3.61E-08 | 1.76E-07 |
| chr3 | 52148935 | 52163077 | 14143 | -1.31 | 3.64E-08 | 1.78E-07 |
| chr12 | 78792060 | 78809735 | 17676 | 5.97 | 3.65E-08 | 1.78E-07 |
| chr12 | 94945977 | 94947811 | 1835 | 2.1 | 3.66E-08 | 1.78E-07 |
| chr10 | 99298222 | 99313858 | 15637 | -0.61 | 3.66E-08 | 1.79E-07 |
| chr10 | 100163201 | 100175797 | 12597 | 0.88 | 3.67E-08 | 1.79E-07 |
| chr8 | 10530068 | 10549426 | 19359 | -0.85 | 3.68E-08 | 1.79E-07 |
| chr3 | 126722727 | 126730519 | 7793 | -1.54 | 3.68E-08 | 1.80E-07 |
| chr3 | 94831111 | 94832006 | 896 | 6.26 | 3.69E-08 | 1.80E-07 |
| chr1 | 229300337 | 229301556 | 1220 | 3.24 | 3.69E-08 | 1.80E-07 |
| chr6 | 140639874 | 140641910 | 2037 | 1.56 | 3.71E-08 | 1.81E-07 |
| chr19 | 9730987 | 9734069 | 3083 | 0.89 | 3.71E-08 | 1.81E-07 |
| chr15 | 71070175 | 71095345 | 25171 | -0.38 | 3.72E-08 | 1.81E-07 |
| chr1 | 215766807 | 215789436 | 22630 | 0.97 | 3.72E-08 | 1.81E-07 |
| chr8 | 115823202 | 115824019 | 818 | 6.3 | 3.73E-08 | 1.82E-07 |
| chr9 | 89656015 | 89657031 | 1017 | 1.47 | 3.75E-08 | 1.83E-07 |
| chr16 | 67951332 | 68003316 | 51985 | -0.48 | 3.78E-08 | 1.84E-07 |
| chr12 | 78482006 | 78482824 | 819 | 1.79 | 3.78E-08 | 1.84E-07 |
| chr19 | 23814951 | 23816221 | 1271 | 6.25 | 3.78E-08 | 1.84E-07 |
| chr4 | 159033753 | 159034762 | 1010 | 3.95 | 3.80E-08 | 1.85E-07 |
| chr8 | 32996590 | 32998989 | 2400 | -1.71 | 3.80E-08 | 1.85E-07 |
| chr12 | 47851858 | 47853281 | 1424 | 6.32 | 3.80E-08 | 1.85E-07 |
| chr2 | 236256071 | 236257099 | 1029 | -3.05 | 3.82E-08 | 1.86E-07 |
| chr6 | 168701991 | 168728296 | 26306 | 1.97 | 3.87E-08 | 1.88E-07 |
| chr1 | 8760740 | 8778003 | 17264 | 0.39 | 3.87E-08 | 1.88E-07 |
| chr4 | 98353089 | 98354259 | 1171 | 6.25 | 3.88E-08 | 1.89E-07 |
| chr17 | 57151836 | 57267669 | 115834 | 0.24 | 3.90E-08 | 1.89E-07 |
| chr18 | 12490547 | 12491236 | 690 | 4.02 | 3.91E-08 | 1.90E-07 |
| chr3 | 94873177 | 94874793 | 1617 | 6.26 | 3.91E-08 | 1.90E-07 |
| chr6 | 47197495 | 47211772 | 14278 | 0.63 | 3.92E-08 | 1.91E-07 |
| chr3 | 105464961 | 105467163 | 2203 | 1.28 | 3.92E-08 | 1.91E-07 |
| chr22 | 39679062 | 39717031 | 37970 | 0.38 | 3.93E-08 | 1.91E-07 |
| chr10 | 28622602 | 28624345 | 1744 | 3.62 | 3.94E-08 | 1.91E-07 |
| chr17 | 57068560 | 57082389 | 13830 | 0.66 | 3.95E-08 | 1.92E-07 |
| chr2 | 207986290 | 208001705 | 15416 | -0.79 | 3.99E-08 | 1.94E-07 |
| chr16 | 77223894 | 77226564 | 2671 | 0.56 | 3.99E-08 | 1.94E-07 |
| chr5 | 40333094 | 40334158 | 1065 | 4.47 | 4.01E-08 | 1.95E-07 |
| chr22 | 43314757 | 43371160 | 56404 | 0.3 | 4.01E-08 | 1.95E-07 |
| chr4 | 175169355 | 175182792 | 13438 | 0.84 | 4.02E-08 | 1.95E-07 |
| chr2 | 70584462 | 70588122 | 3661 | 2.03 | 4.03E-08 | 1.96E-07 |
| chr4 | 155797245 | 155799058 | 1814 | 1.02 | 4.03E-08 | 1.96E-07 |
| chr8 | 102647177 | 102654939 | 7763 | 1.17 | 4.04E-08 | 1.96E-07 |
| chr1 | 80933495 | 80934553 | 1059 | 3.64 | 4.04E-08 | 1.96E-07 |
| chr11 | 63551153 | 63552352 | 1200 | 2.14 | 4.09E-08 | 1.98E-07 |
| chr5 | 79760780 | 79764092 | 3313 | 1 | 4.10E-08 | 1.99E-07 |
| chr7 | 5127966 | 5128977 | 1012 | 3.7 | 4.11E-08 | 1.99E-07 |
| chr17 | 73839610 | 73909058 | 69449 | 0.33 | 4.12E-08 | 2.00E-07 |
| chr12 | 25648702 | 25649468 | 767 | 6.27 | 4.13E-08 | 2.00E-07 |
| chr6 | 39247378 | 39248617 | 1240 | 1.5 | 4.14E-08 | 2.00E-07 |
| chr11 | 36055998 | 36057080 | 1083 | -3.39 | 4.14E-08 | 2.01E-07 |
| chr11 | 86381044 | 86384504 | 3461 | 0.64 | 4.14E-08 | 2.01E-07 |

| | | | | | | |
|---|---|---|---|---|---|---|
| chr12 | 14480145 | 14481175 | 1031 | 6.31 | 4.15E-08 | 2.01E-07 |
| chr3 | 187840106 | 187842195 | 2090 | 1.36 | 4.16E-08 | 2.01E-07 |
| chr9 | 120465821 | 120467067 | 1247 | 6.24 | 4.18E-08 | 2.02E-07 |
| chr7 | 111800022 | 111804368 | 4347 | -1.23 | 4.18E-08 | 2.03E-07 |
| chr11 | 66510473 | 66513640 | 3168 | 1.04 | 4.20E-08 | 2.03E-07 |
| chr3 | 99594481 | 99650642 | 56162 | 0.9 | 4.21E-08 | 2.04E-07 |
| chr1 | 154297337 | 154319946 | 22610 | -0.89 | 4.21E-08 | 2.04E-07 |
| chr1 | 65151818 | 65154424 | 2607 | 1.04 | 4.21E-08 | 2.04E-07 |
| chr10 | 59387467 | 59390301 | 2835 | 0.81 | 4.25E-08 | 2.05E-07 |
| chr2 | 104673923 | 104674831 | 909 | 6.26 | 4.26E-08 | 2.06E-07 |
| chr2 | 149476763 | 149478228 | 1466 | -2.68 | 4.26E-08 | 2.06E-07 |
| chr9 | 138969210 | 139012458 | 43249 | -0.44 | 4.29E-08 | 2.07E-07 |
| chr2 | 194381904 | 194383487 | 1584 | 3.2 | 4.29E-08 | 2.07E-07 |
| chr6 | 63372365 | 63383023 | 10659 | 6.28 | 4.31E-08 | 2.08E-07 |
| chr17 | 68682635 | 68684999 | 2365 | 4.39 | 4.32E-08 | 2.09E-07 |
| chr1 | 59049089 | 59058933 | 9845 | 1.4 | 4.33E-08 | 2.09E-07 |
| chr16 | 3550811 | 3581160 | 30350 | -0.68 | 4.34E-08 | 2.10E-07 |
| chr10 | 26844447 | 26846806 | 2360 | -2.66 | 4.34E-08 | 2.10E-07 |
| chr1 | 248893994 | 248894899 | 906 | 5.42 | 4.35E-08 | 2.10E-07 |
| chr2 | 206665016 | 206671367 | 6352 | 6.24 | 4.36E-08 | 2.10E-07 |
| chr11 | 61714804 | 61749308 | 34505 | -0.27 | 4.36E-08 | 2.11E-07 |
| chr11 | 9405412 | 9464125 | 58714 | 0.34 | 4.37E-08 | 2.11E-07 |
| chr17 | 55923177 | 56093015 | 169839 | 0.16 | 4.37E-08 | 2.11E-07 |
| chr4 | 73072689 | 73073638 | 950 | 6.23 | 4.38E-08 | 2.11E-07 |
| chr12 | 30814081 | 30814981 | 901 | 2.07 | 4.41E-08 | 2.13E-07 |
| chr2 | 178416934 | 178418536 | 1603 | -0.93 | 4.41E-08 | 2.13E-07 |
| chr4 | 120983565 | 120989642 | 6078 | 0.78 | 4.42E-08 | 2.13E-07 |
| chr2 | 152583357 | 152586244 | 2888 | 1.02 | 4.44E-08 | 2.14E-07 |
| chr10 | 30061265 | 30074996 | 13732 | 0.58 | 4.45E-08 | 2.15E-07 |
| chr4 | 159689563 | 159696648 | 7086 | 1.58 | 4.48E-08 | 2.16E-07 |
| chr8 | 134440682 | 134442093 | 1412 | 2.1 | 4.48E-08 | 2.16E-07 |
| chr7 | 140857811 | 140863883 | 6073 | 3.78 | 4.53E-08 | 2.18E-07 |
| chr1 | 178693678 | 178697848 | 4171 | 0.71 | 4.55E-08 | 2.19E-07 |
| chr20 | 11432444 | 11440646 | 8203 | -0.44 | 4.57E-08 | 2.20E-07 |
| chr21 | 40419459 | 40420852 | 1394 | 2.8 | 4.58E-08 | 2.21E-07 |
| chr11 | 74081867 | 74083534 | 1668 | 1.92 | 4.61E-08 | 2.22E-07 |
| chr7 | 16385251 | 16386936 | 1686 | 2.59 | 4.61E-08 | 2.22E-07 |
| chr1 | 105629755 | 105630919 | 1165 | 6.22 | 4.63E-08 | 2.23E-07 |
| chr10 | 9110807 | 9111812 | 1006 | 6.22 | 4.63E-08 | 2.23E-07 |
| chr20 | 47777460 | 47805436 | 27977 | 0.56 | 4.64E-08 | 2.23E-07 |
| chr2 | 209111742 | 209132681 | 20940 | -0.65 | 4.65E-08 | 2.24E-07 |
| chr17 | 60063946 | 60090432 | 26487 | 0.7 | 4.65E-08 | 2.24E-07 |
| chr16 | 47163209 | 47178610 | 15402 | -0.63 | 4.65E-08 | 2.24E-07 |
| chr9 | 2466401 | 2479468 | 13068 | 1.91 | 4.66E-08 | 2.24E-07 |
| chr1 | 35729421 | 35735959 | 6539 | 0.61 | 4.69E-08 | 2.26E-07 |
| chr7 | 83874861 | 83875819 | 959 | 3.27 | 4.70E-08 | 2.26E-07 |
| chr8 | 134653992 | 134665060 | 11069 | 1.79 | 4.71E-08 | 2.26E-07 |
| chr2 | 162164007 | 162176730 | 12724 | -0.51 | 4.73E-08 | 2.27E-07 |
| chr3 | 175507812 | 175508752 | 941 | 6.22 | 4.79E-08 | 2.30E-07 |
| chr18 | 2395197 | 2396194 | 998 | 3.29 | 4.82E-08 | 2.32E-07 |
| chr14 | 95120531 | 95121924 | 1394 | 6.24 | 4.84E-08 | 2.32E-07 |
| chr10 | 102100744 | 102155751 | 55008 | 0.32 | 4.85E-08 | 2.33E-07 |
| chr5 | 82812389 | 82814401 | 2013 | 3.9 | 4.86E-08 | 2.33E-07 |
| chr9 | 17268975 | 17270273 | 1299 | 2.01 | 4.87E-08 | 2.34E-07 |
| chr10 | 95307621 | 95337551 | 29931 | -0.9 | 4.89E-08 | 2.35E-07 |
| chr4 | 170578298 | 170583127 | 4830 | 0.5 | 4.90E-08 | 2.35E-07 |
| chr9 | 22446563 | 22449114 | 2552 | 1.19 | 4.90E-08 | 2.35E-07 |
| chr3 | 83081901 | 83082915 | 1015 | 6.29 | 4.90E-08 | 2.35E-07 |
| chr1 | 26216150 | 26232906 | 16757 | 0.64 | 4.91E-08 | 2.35E-07 |
| chr6 | 155634386 | 155650670 | 16285 | 0.9 | 4.92E-08 | 2.36E-07 |
| chr8 | 34940054 | 34941191 | 1138 | -6.17 | 4.92E-08 | 2.36E-07 |

| | | | | | | |
|---|---|---|---|---|---|---|
| chr5 | 92274730 | 92282969 | 8240 | 0.7 | 4.94E-08 | 2.37E-07 |
| chr9 | 36397186 | 36401912 | 4727 | 0.67 | 4.96E-08 | 2.38E-07 |
| chr5 | 126912959 | 126913649 | 691 | 4.17 | 4.96E-08 | 2.38E-07 |
| chr3 | 85530793 | 85531771 | 979 | 6.26 | 4.96E-08 | 2.38E-07 |
| chr8 | 144534191 | 144641552 | 107362 | -0.49 | 4.97E-08 | 2.38E-07 |
| chr12 | 58119389 | 58177580 | 58192 | 0.37 | 4.97E-08 | 2.38E-07 |
| chr6 | 139728258 | 139729077 | 820 | 4.67 | 4.97E-08 | 2.38E-07 |
| chr15 | 31280249 | 31285152 | 4904 | 0.66 | 4.97E-08 | 2.38E-07 |
| chr4 | 16124757 | 16125758 | 1002 | 4.67 | 5.01E-08 | 2.40E-07 |
| chr10 | 128810970 | 128817524 | 6555 | 1.12 | 5.02E-08 | 2.40E-07 |
| chr8 | 67521020 | 67526865 | 5846 | 0.89 | 5.02E-08 | 2.40E-07 |
| chr4 | 106954001 | 106959736 | 5736 | 4.11 | 5.05E-08 | 2.42E-07 |
| chr13 | 86302850 | 86304071 | 1222 | -4.54 | 5.07E-08 | 2.43E-07 |
| chr2 | 96986634 | 97008509 | 21876 | -0.76 | 5.15E-08 | 2.46E-07 |
| chr12 | 14459980 | 14461072 | 1093 | 6.2 | 5.16E-08 | 2.47E-07 |
| chr3 | 174666629 | 174674964 | 8336 | 1.06 | 5.17E-08 | 2.47E-07 |
| chr12 | 4646801 | 4649332 | 2532 | 0.77 | 5.18E-08 | 2.48E-07 |
| chr18 | 67454729 | 67455740 | 1012 | -3.64 | 5.19E-08 | 2.48E-07 |
| chr20 | 23117299 | 23137301 | 20003 | -0.93 | 5.19E-08 | 2.48E-07 |
| chr19 | 51436127 | 51437926 | 1800 | -1.45 | 5.20E-08 | 2.49E-07 |
| chr12 | 69962421 | 69964729 | 2309 | 0.71 | 5.22E-08 | 2.49E-07 |
| chr3 | 96711671 | 96712906 | 1236 | 6.2 | 5.25E-08 | 2.51E-07 |
| chrX | 40502440 | 40512519 | 10080 | 0.77 | 5.27E-08 | 2.52E-07 |
| chr13 | 41805007 | 41806320 | 1314 | 2.26 | 5.30E-08 | 2.53E-07 |
| chrX | 42584098 | 42585114 | 1017 | 6.23 | 5.31E-08 | 2.54E-07 |
| chr1 | 169454100 | 169456388 | 2289 | 0.98 | 5.32E-08 | 2.54E-07 |
| chr3 | 160464969 | 160474127 | 9159 | 1.97 | 5.32E-08 | 2.54E-07 |
| chr5 | 155447320 | 155448379 | 1060 | 3.21 | 5.34E-08 | 2.55E-07 |
| chr4 | 159434115 | 159435179 | 1065 | 5.48 | 5.36E-08 | 2.56E-07 |
| chr18 | 20557629 | 20559188 | 1560 | 1.27 | 5.37E-08 | 2.56E-07 |
| chr14 | 73602587 | 73604869 | 2283 | 0.74 | 5.38E-08 | 2.57E-07 |
| chr2 | 188390021 | 188446506 | 56486 | 0.21 | 5.39E-08 | 2.57E-07 |
| chr13 | 77565527 | 77567362 | 1836 | 1.01 | 5.39E-08 | 2.57E-07 |
| chr7 | 91388119 | 91389219 | 1101 | 2.8 | 5.39E-08 | 2.57E-07 |
| chr8 | 87424093 | 87442980 | 18888 | 0.66 | 5.39E-08 | 2.57E-07 |
| chr8 | 99952796 | 99957918 | 5123 | -0.74 | 5.43E-08 | 2.59E-07 |
| chr11 | 16416874 | 16419585 | 2712 | 5.95 | 5.43E-08 | 2.59E-07 |
| chr12 | 57809911 | 57826298 | 16388 | 0.53 | 5.44E-08 | 2.59E-07 |
| chr15 | 57249291 | 57282611 | 33321 | 0.49 | 5.45E-08 | 2.60E-07 |
| chr5 | 18091101 | 18092205 | 1105 | 4.63 | 5.45E-08 | 2.60E-07 |
| chr14 | 44919164 | 44920310 | 1147 | 6.24 | 5.46E-08 | 2.60E-07 |
| chr3 | 160938119 | 160962539 | 24421 | 0.49 | 5.46E-08 | 2.60E-07 |
| chr13 | 81966977 | 81981407 | 14431 | 5.53 | 5.49E-08 | 2.61E-07 |
| chr20 | 46613565 | 46618825 | 5261 | 6.24 | 5.51E-08 | 2.62E-07 |
| chr6 | 100793152 | 100795234 | 2083 | 1.19 | 5.51E-08 | 2.63E-07 |
| chr5 | 115312482 | 115313565 | 1084 | 6.25 | 5.52E-08 | 2.63E-07 |
| chr6 | 143247328 | 143249290 | 1963 | 1.13 | 5.53E-08 | 2.63E-07 |
| chr1 | 168452201 | 168465247 | 13047 | 6.23 | 5.54E-08 | 2.64E-07 |
| chr2 | 191141825 | 191142450 | 626 | 6.21 | 5.55E-08 | 2.64E-07 |
| chr7 | 127615622 | 127621374 | 5753 | 3.81 | 5.55E-08 | 2.64E-07 |
| chr1 | 222234988 | 222236152 | 1165 | 4 | 5.57E-08 | 2.65E-07 |
| chr8 | 108433200 | 108448617 | 15418 | 6.43 | 5.74E-08 | 2.73E-07 |
| chr7 | 155652980 | 155655940 | 2961 | 0.88 | 5.75E-08 | 2.74E-07 |
| chr2 | 88898654 | 88900592 | 1939 | 2.07 | 5.76E-08 | 2.74E-07 |
| chr10 | 37722940 | 37723754 | 815 | 4.28 | 5.77E-08 | 2.74E-07 |
| chr6 | 110974666 | 110986724 | 12059 | 2.11 | 5.77E-08 | 2.74E-07 |
| chr14 | 25435091 | 25436281 | 1191 | 3.02 | 5.78E-08 | 2.74E-07 |
| chr13 | 46928219 | 46930118 | 1900 | 1.26 | 5.78E-08 | 2.75E-07 |
| chr10 | 26653761 | 26655259 | 1499 | -6.18 | 5.79E-08 | 2.75E-07 |
| chr21 | 38112596 | 38130353 | 17758 | -1.01 | 5.79E-08 | 2.75E-07 |
| chr7 | 151383006 | 151452775 | 69770 | -0.41 | 5.81E-08 | 2.76E-07 |

| | | | | | | |
|---|---|---|---|---|---|---|
| chr9 | 124481960 | 124483412 | 1453 | 1.44 | 5.81E-08 | 2.76E-07 |
| chr12 | 114266403 | 114279183 | 12781 | -1.11 | 5.84E-08 | 2.77E-07 |
| chr7 | 17078909 | 17083819 | 4911 | 0.88 | 5.86E-08 | 2.78E-07 |
| chr16 | 79646347 | 79647204 | 858 | 2.76 | 5.86E-08 | 2.78E-07 |
| chr5 | 105551360 | 105552531 | 1172 | 3.19 | 5.87E-08 | 2.79E-07 |
| chr1 | 151907408 | 151926860 | 19453 | 0.38 | 5.89E-08 | 2.79E-07 |
| chr19 | 36526944 | 36681454 | 154511 | 0.23 | 5.90E-08 | 2.80E-07 |
| chr7 | 91186093 | 91195485 | 9393 | 1.09 | 5.93E-08 | 2.81E-07 |
| chr18 | 59995920 | 60007399 | 11480 | -1.16 | 5.95E-08 | 2.82E-07 |
| chr6 | 4793871 | 4807302 | 13432 | 0.9 | 5.96E-08 | 2.83E-07 |
| chr11 | 93300246 | 93304487 | 4242 | 1.93 | 5.99E-08 | 2.84E-07 |
| chr7 | 12657674 | 12659502 | 1829 | 3.07 | 5.99E-08 | 2.84E-07 |
| chr18 | 19749498 | 19750968 | 1471 | 1.98 | 6.00E-08 | 2.84E-07 |
| chr9 | 3477843 | 3480116 | 2274 | 1.47 | 6.03E-08 | 2.86E-07 |
| chr20 | 50617544 | 50619345 | 1802 | 1.16 | 6.03E-08 | 2.86E-07 |
| chr15 | 41623159 | 41635420 | 12262 | 0.71 | 6.07E-08 | 2.88E-07 |
| chr5 | 94981300 | 94984783 | 3484 | 0.99 | 6.13E-08 | 2.90E-07 |
| chr16 | 48642462 | 48666744 | 24283 | 0.56 | 6.14E-08 | 2.91E-07 |
| chr8 | 24061749 | 24086971 | 25223 | 0.5 | 6.14E-08 | 2.91E-07 |
| chr5 | 152930973 | 152932001 | 1029 | -6.14 | 6.14E-08 | 2.91E-07 |
| chr1 | 160311240 | 160314152 | 2913 | 0.64 | 6.17E-08 | 2.92E-07 |
| chr5 | 14528567 | 14529952 | 1386 | 2.16 | 6.25E-08 | 2.96E-07 |
| chr10 | 70407303 | 70408511 | 1209 | 2.31 | 6.27E-08 | 2.97E-07 |
| chr19 | 48101292 | 48182602 | 81311 | 0.3 | 6.31E-08 | 2.98E-07 |
| chr5 | 44874288 | 44886307 | 12020 | 7.3 | 6.33E-08 | 2.99E-07 |
| chr9 | 111453655 | 111454893 | 1239 | 6.17 | 6.40E-08 | 3.03E-07 |
| chr8 | 140207960 | 140210303 | 2344 | 1.89 | 6.45E-08 | 3.05E-07 |
| chr12 | 6193332 | 6203646 | 10315 | 0.92 | 6.47E-08 | 3.06E-07 |
| chr4 | 164316769 | 164326170 | 9402 | 6.31 | 6.51E-08 | 3.08E-07 |
| chr10 | 116123385 | 116132322 | 8938 | -0.81 | 6.51E-08 | 3.08E-07 |
| chr5 | 129727119 | 129736848 | 9730 | 1.88 | 6.53E-08 | 3.08E-07 |
| chr11 | 11003755 | 11013861 | 10107 | -0.93 | 6.58E-08 | 3.11E-07 |
| chrX | 13751622 | 13754477 | 2856 | 0.94 | 6.61E-08 | 3.12E-07 |
| chr6 | 900976 | 908307 | 7332 | -6.16 | 6.62E-08 | 3.13E-07 |
| chr14 | 85699640 | 85701814 | 2175 | 1.35 | 6.67E-08 | 3.15E-07 |
| chr1 | 86200168 | 86201306 | 1139 | 1.77 | 6.76E-08 | 3.19E-07 |
| chr3 | 81319905 | 81320787 | 883 | 6.23 | 6.78E-08 | 3.20E-07 |
| chr5 | 81982848 | 81983932 | 1085 | 6.17 | 6.79E-08 | 3.21E-07 |
| chr2 | 76654088 | 76655433 | 1346 | 6.18 | 6.82E-08 | 3.22E-07 |
| chr3 | 38244902 | 38246292 | 1391 | -4.54 | 6.82E-08 | 3.22E-07 |
| chr11 | 26487014 | 26495805 | 8792 | 6.18 | 6.84E-08 | 3.23E-07 |
| chr6 | 85787429 | 85788528 | 1100 | 6.18 | 6.84E-08 | 3.23E-07 |
| chr14 | 91715156 | 91720370 | 5215 | 1.03 | 6.87E-08 | 3.24E-07 |
| chr13 | 106457982 | 106469430 | 11449 | 6.24 | 6.87E-08 | 3.24E-07 |
| chr4 | 123030691 | 123031456 | 766 | 1.95 | 6.91E-08 | 3.26E-07 |
| chr13 | 43881609 | 43891535 | 9927 | 6.16 | 6.98E-08 | 3.29E-07 |
| chr3 | 71060360 | 71061502 | 1143 | 3.76 | 7.02E-08 | 3.31E-07 |
| chr13 | 44614195 | 44643844 | 29650 | 0.68 | 7.06E-08 | 3.33E-07 |
| chr12 | 43562433 | 43563298 | 866 | 6.21 | 7.07E-08 | 3.33E-07 |
| chr3 | 33396902 | 33406022 | 9121 | 1 | 7.10E-08 | 3.34E-07 |
| chr2 | 157908894 | 157914051 | 5158 | 1.17 | 7.10E-08 | 3.34E-07 |
| chr7 | 103450305 | 103451262 | 958 | 6.22 | 7.11E-08 | 3.35E-07 |
| chr2 | 146517699 | 146519696 | 1998 | 1.45 | 7.11E-08 | 3.35E-07 |
| chr8 | 15559162 | 15571787 | 12626 | 6.27 | 7.16E-08 | 3.37E-07 |
| chr1 | 114470685 | 114474909 | 4225 | 0.77 | 7.19E-08 | 3.38E-07 |
| chr14 | 49261469 | 49262456 | 988 | 5.57 | 7.20E-08 | 3.39E-07 |
| chr2 | 72490171 | 72503670 | 13500 | 0.45 | 7.31E-08 | 3.44E-07 |
| chr15 | 78304351 | 78331657 | 27307 | -0.95 | 7.32E-08 | 3.44E-07 |
| chr2 | 236549948 | 236551622 | 1675 | -2.91 | 7.32E-08 | 3.44E-07 |
| chr19 | 20431748 | 20432652 | 905 | 6.15 | 7.35E-08 | 3.46E-07 |
| chr2 | 40506002 | 40517733 | 11732 | 6.19 | 7.37E-08 | 3.46E-07 |

| | | | | | | |
|---|---|---|---|---|---|---|
| chr11 | 2632435 | 2635601 | 3167 | 1.58 | 7.38E-08 | 3.47E-07 |
| chr19 | 50803288 | 50819871 | 16584 | 1.45 | 7.38E-08 | 3.47E-07 |
| chr1 | 188058138 | 188060295 | 2158 | 2 | 7.39E-08 | 3.47E-07 |
| chr19 | 55516733 | 55518398 | 1666 | 0.97 | 7.42E-08 | 3.49E-07 |
| chr1 | 106174615 | 106175666 | 1052 | 6.15 | 7.44E-08 | 3.49E-07 |
| chr9 | 102580973 | 102586606 | 5634 | 0.72 | 7.46E-08 | 3.50E-07 |
| chr22 | 42466227 | 42487825 | 21599 | 0.82 | 7.46E-08 | 3.50E-07 |
| chr13 | 81519454 | 81520416 | 963 | 6.21 | 7.46E-08 | 3.50E-07 |
| chr12 | 68914855 | 68916013 | 1159 | 1.4 | 7.52E-08 | 3.53E-07 |
| chr6 | 74269995 | 74320310 | 50316 | 0.32 | 7.52E-08 | 3.53E-07 |
| chr6 | 835094 | 837401 | 2308 | -1.76 | 7.54E-08 | 3.54E-07 |
| chr7 | 123283676 | 123284766 | 1091 | 6.15 | 7.55E-08 | 3.54E-07 |
| chr7 | 148958333 | 148961223 | 2891 | 1.05 | 7.58E-08 | 3.56E-07 |
| chrX | 2816387 | 2820522 | 4136 | 2.2 | 7.63E-08 | 3.58E-07 |
| chr3 | 102433271 | 102434244 | 974 | 6.16 | 7.66E-08 | 3.59E-07 |
| chr5 | 139697044 | 139707941 | 10898 | 0.9 | 7.69E-08 | 3.60E-07 |
| chr1 | 117635285 | 117637224 | 1940 | -1.42 | 7.69E-08 | 3.61E-07 |
| chr10 | 38053614 | 38054690 | 1077 | 6.17 | 7.71E-08 | 3.62E-07 |
| chr1 | 210516253 | 210517528 | 1276 | 4.37 | 7.72E-08 | 3.62E-07 |
| chr15 | 79271483 | 79297935 | 26453 | 0.72 | 7.73E-08 | 3.62E-07 |
| chr6 | 131720620 | 131722442 | 1823 | 2.33 | 7.74E-08 | 3.63E-07 |
| chr4 | 169479225 | 169491099 | 11875 | -1.41 | 7.80E-08 | 3.65E-07 |
| chr18 | 21629953 | 21632928 | 2976 | -1.45 | 7.81E-08 | 3.66E-07 |
| chr5 | 21048602 | 21049525 | 924 | 6.2 | 7.82E-08 | 3.67E-07 |
| chr13 | 73860579 | 73875467 | 14889 | 1.99 | 7.85E-08 | 3.68E-07 |
| chr4 | 137883950 | 137889018 | 5069 | 2.45 | 7.86E-08 | 3.68E-07 |
| chr9 | 2157288 | 2161027 | 3740 | 0.62 | 7.86E-08 | 3.68E-07 |
| chr2 | 84474210 | 84475323 | 1114 | 6.14 | 7.87E-08 | 3.68E-07 |
| chr13 | 88777135 | 88778141 | 1007 | 6.14 | 7.87E-08 | 3.68E-07 |
| chr4 | 140931986 | 141077063 | 145078 | 0.18 | 7.97E-08 | 3.73E-07 |
| chr3 | 40493457 | 40519340 | 25884 | 0.55 | 8.00E-08 | 3.74E-07 |
| chr1 | 234492215 | 234510357 | 18143 | -0.45 | 8.05E-08 | 3.77E-07 |
| chr4 | 38414833 | 38426747 | 11915 | 6.32 | 8.05E-08 | 3.77E-07 |
| chr3 | 27548383 | 27549682 | 1300 | 2.15 | 8.06E-08 | 3.77E-07 |
| chr10 | 59512543 | 59513560 | 1018 | 5.02 | 8.06E-08 | 3.77E-07 |
| chr15 | 99741968 | 99743096 | 1129 | 3.41 | 8.09E-08 | 3.78E-07 |
| chr15 | 50526764 | 50529109 | 2346 | -2.07 | 8.10E-08 | 3.79E-07 |
| chr10 | 89839130 | 89840624 | 1495 | 1.83 | 8.11E-08 | 3.79E-07 |
| chr1 | 59917160 | 59918172 | 1013 | 2.33 | 8.16E-08 | 3.81E-07 |
| chr12 | 18535697 | 18536805 | 1109 | 4.87 | 8.20E-08 | 3.83E-07 |
| chr5 | 49736048 | 49737815 | 1768 | 1.67 | 8.23E-08 | 3.84E-07 |
| chr17 | 34883494 | 34912288 | 28795 | 0.31 | 8.29E-08 | 3.87E-07 |
| chr13 | 102424929 | 102425990 | 1062 | 6.17 | 8.33E-08 | 3.89E-07 |
| chr5 | 68747295 | 68750630 | 3336 | 1.09 | 8.33E-08 | 3.89E-07 |
| chr1 | 96828074 | 96836574 | 8501 | 6.24 | 8.35E-08 | 3.90E-07 |
| chr3 | 190529855 | 190530711 | 857 | 3.48 | 8.41E-08 | 3.93E-07 |
| chr1 | 151639184 | 151655927 | 16744 | 0.71 | 8.43E-08 | 3.93E-07 |
| chr2 | 202121361 | 202137194 | 15834 | -0.54 | 8.50E-08 | 3.96E-07 |
| chr5 | 137223670 | 137231017 | 7348 | 1.27 | 8.51E-08 | 3.97E-07 |
| chr7 | 16150193 | 16150943 | 751 | 6.13 | 8.55E-08 | 3.99E-07 |
| chr2 | 164202027 | 164207350 | 5324 | -0.6 | 8.58E-08 | 4.00E-07 |
| chr11 | 1105780 | 1107038 | 1259 | 2.39 | 8.60E-08 | 4.01E-07 |
| chr13 | 113364647 | 113380484 | 15838 | 0.65 | 8.61E-08 | 4.02E-07 |
| chr3 | 134027106 | 134056879 | 29774 | -0.31 | 8.62E-08 | 4.02E-07 |
| chrX | 19620931 | 19622347 | 1417 | -1.38 | 8.63E-08 | 4.02E-07 |
| chr1 | 19747301 | 19778225 | 30925 | -0.62 | 8.64E-08 | 4.03E-07 |
| chr14 | 76485754 | 76486820 | 1067 | 6.14 | 8.65E-08 | 4.03E-07 |
| chr17 | 37201815 | 37213844 | 12030 | -2.07 | 8.65E-08 | 4.03E-07 |
| chr19 | 33863743 | 33897117 | 33375 | -0.66 | 8.65E-08 | 4.03E-07 |
| chr10 | 102729106 | 102733235 | 4130 | 1.48 | 8.79E-08 | 4.09E-07 |
| chr4 | 103939767 | 103941616 | 1850 | 0.82 | 8.81E-08 | 4.10E-07 |

| | | | | | | |
|---|---|---|---|---|---|---|
| chr6 | 18237034 | 18265864 | 28831 | 0.52 | 8.82E-08 | 4.11E-07 |
| chr15 | 86379727 | 86383831 | 4105 | 1.4 | 8.82E-08 | 4.11E-07 |
| chr8 | 21946033 | 22000084 | 54052 | -0.4 | 8.85E-08 | 4.12E-07 |
| chr11 | 64426625 | 64427694 | 1070 | -2.92 | 8.85E-08 | 4.12E-07 |
| chr7 | 30936608 | 30961739 | 25132 | 0.86 | 8.88E-08 | 4.13E-07 |
| chr1 | 64808127 | 64810946 | 2820 | 0.78 | 8.88E-08 | 4.13E-07 |
| chrX | 109404001 | 109436148 | 32148 | -0.74 | 8.89E-08 | 4.14E-07 |
| chr5 | 150283722 | 150285494 | 1773 | 0.69 | 8.93E-08 | 4.15E-07 |
| chr12 | 49520970 | 49526863 | 5894 | 0.67 | 9.08E-08 | 4.22E-07 |
| chr12 | 109628165 | 109645837 | 17673 | -0.56 | 9.11E-08 | 4.24E-07 |
| chr10 | 6244138 | 6290166 | 46029 | -0.37 | 9.12E-08 | 4.24E-07 |
| chr7 | 17044847 | 17045559 | 713 | 4.92 | 9.25E-08 | 4.30E-07 |
| chr8 | 24223765 | 24225373 | 1609 | 6.13 | 9.25E-08 | 4.30E-07 |
| chr2 | 235996332 | 236026017 | 29686 | -0.63 | 9.26E-08 | 4.30E-07 |
| chr16 | 11989395 | 12012151 | 22757 | 0.44 | 9.34E-08 | 4.34E-07 |
| chr16 | 59202216 | 59214171 | 11956 | 6.37 | 9.35E-08 | 4.34E-07 |
| chr11 | 61765628 | 61816424 | 50797 | 0.38 | 9.35E-08 | 4.34E-07 |
| chr5 | 93702069 | 93703639 | 1571 | 2.52 | 9.39E-08 | 4.36E-07 |
| chr14 | 54181193 | 54183078 | 1886 | 2.14 | 9.42E-08 | 4.37E-07 |
| chrX | 20270510 | 20285954 | 15445 | 0.65 | 9.43E-08 | 4.38E-07 |
| chr14 | 67427043 | 67428102 | 1060 | 6.2 | 9.45E-08 | 4.39E-07 |
| chr8 | 53252340 | 53256502 | 4163 | 1.72 | 9.46E-08 | 4.39E-07 |
| chr2 | 102132710 | 102137419 | 4710 | -0.67 | 9.48E-08 | 4.40E-07 |
| chr12 | 86623807 | 86625257 | 1451 | 1.38 | 9.48E-08 | 4.40E-07 |
| chr2 | 181772340 | 181774226 | 1887 | 1.78 | 9.48E-08 | 4.40E-07 |
| chr22 | 18393512 | 18406595 | 13084 | -1.25 | 9.50E-08 | 4.41E-07 |
| chr5 | 73703230 | 73712297 | 9068 | 1.15 | 9.54E-08 | 4.43E-07 |
| chr9 | 71394041 | 71396306 | 2266 | 0.92 | 9.58E-08 | 4.44E-07 |
| chr7 | 36820444 | 36821990 | 1547 | 6.17 | 9.59E-08 | 4.45E-07 |
| chr5 | 31023095 | 31024081 | 987 | 6.25 | 9.60E-08 | 4.45E-07 |
| chr5 | 45160188 | 45172364 | 12177 | 6.16 | 9.61E-08 | 4.46E-07 |
| chr3 | 15418715 | 15419893 | 1179 | -1.08 | 9.65E-08 | 4.47E-07 |
| chr2 | 64289898 | 64291855 | 1958 | 1.75 | 9.65E-08 | 4.47E-07 |
| chr3 | 30481137 | 30490397 | 9261 | 1.02 | 9.68E-08 | 4.48E-07 |
| chr14 | 21682268 | 21738997 | 56730 | 0.33 | 9.70E-08 | 4.49E-07 |
| chr14 | 35389391 | 35401160 | 11770 | 1.56 | 9.77E-08 | 4.52E-07 |
| chr5 | 13859666 | 13866283 | 6618 | 6.11 | 9.81E-08 | 4.54E-07 |
| chr3 | 187630470 | 187633270 | 2801 | 0.94 | 9.81E-08 | 4.54E-07 |
| chr4 | 23583826 | 23586328 | 2503 | 2.4 | 9.83E-08 | 4.55E-07 |
| chr3 | 190303098 | 190305409 | 2312 | 1.05 | 9.85E-08 | 4.56E-07 |
| chr5 | 40826555 | 40842786 | 16232 | 0.46 | 9.85E-08 | 4.56E-07 |
| chr10 | 116006408 | 116007667 | 1260 | 6.13 | 9.87E-08 | 4.56E-07 |
| chr18 | 72617785 | 72618747 | 963 | 6.1 | 9.90E-08 | 4.58E-07 |
| chr8 | 64079744 | 64085608 | 5865 | 0.56 | 9.91E-08 | 4.58E-07 |
| chr12 | 647421 | 649130 | 1710 | 2.37 | 9.92E-08 | 4.59E-07 |
| chr14 | 93637362 | 93652614 | 15253 | 0.66 | 9.96E-08 | 4.60E-07 |
| chr9 | 132934956 | 132950253 | 15298 | -0.75 | 9.97E-08 | 4.61E-07 |
| chr1 | 221946945 | 221949339 | 2395 | 0.99 | 9.99E-08 | 4.62E-07 |
| chr10 | 127935434 | 127937131 | 1698 | 1.46 | 1.00E-07 | 4.64E-07 |
| chr12 | 50166746 | 50192976 | 26231 | -1.1 | 1.00E-07 | 4.64E-07 |
| chr9 | 91794195 | 91796683 | 2489 | -1.14 | 1.01E-07 | 4.64E-07 |
| chr13 | 114523573 | 114524604 | 1032 | -2.72 | 1.01E-07 | 4.65E-07 |
| chr22 | 21160207 | 21162706 | 2500 | -1.44 | 1.01E-07 | 4.66E-07 |
| chr2 | 103156898 | 103158100 | 1203 | 4.91 | 1.01E-07 | 4.67E-07 |
| chr14 | 59823575 | 59831733 | 8159 | 0.52 | 1.02E-07 | 4.71E-07 |
| chr14 | 37666011 | 37668012 | 2002 | 0.78 | 1.03E-07 | 4.74E-07 |
| chr6 | 153321897 | 153325890 | 3994 | 0.69 | 1.03E-07 | 4.75E-07 |
| chr5 | 169372556 | 169374058 | 1503 | 2.17 | 1.03E-07 | 4.77E-07 |
| chr1 | 204695024 | 204698520 | 3497 | 1.36 | 1.05E-07 | 4.82E-07 |
| chr10 | 8388519 | 8389179 | 661 | 4.28 | 1.05E-07 | 4.82E-07 |
| chr6 | 45578204 | 45579593 | 1390 | 6.1 | 1.05E-07 | 4.83E-07 |

| | | | | | | |
|---|---|---|---|---|---|---|
| chr16 | 72834299 | 72854182 | 19884 | -0.92 | 1.05E-07 | 4.83E-07 |
| chr13 | 75399120 | 75400608 | 1489 | -6.06 | 1.06E-07 | 4.86E-07 |
| chr1 | 98549709 | 98556890 | 7182 | 0.58 | 1.07E-07 | 4.92E-07 |
| chr12 | 68334470 | 68335487 | 1018 | 6.16 | 1.07E-07 | 4.93E-07 |
| chr13 | 66979933 | 66980936 | 1004 | 6.09 | 1.07E-07 | 4.94E-07 |
| chr8 | 60630126 | 60630885 | 760 | 6.09 | 1.07E-07 | 4.94E-07 |
| chr2 | 151070565 | 151072037 | 1473 | 1.36 | 1.07E-07 | 4.94E-07 |
| chr12 | 96880785 | 96897022 | 16238 | 0.54 | 1.08E-07 | 4.97E-07 |
| chr2 | 28561258 | 28590084 | 28827 | -0.76 | 1.08E-07 | 4.97E-07 |
| chr1 | 202974930 | 202978146 | 3217 | 0.71 | 1.08E-07 | 4.97E-07 |
| chr9 | 101982399 | 101986505 | 4107 | 0.56 | 1.09E-07 | 5.00E-07 |
| chr10 | 5559899 | 5561322 | 1424 | 1.7 | 1.09E-07 | 5.01E-07 |
| chr12 | 51440782 | 51443442 | 2661 | 1.03 | 1.09E-07 | 5.04E-07 |
| chr1 | 82265418 | 82279869 | 14452 | -0.67 | 1.10E-07 | 5.04E-07 |
| chr1 | 227486167 | 227488171 | 2005 | 0.92 | 1.10E-07 | 5.06E-07 |
| chr12 | 75836138 | 75837253 | 1116 | 4.17 | 1.10E-07 | 5.07E-07 |
| chr4 | 161899280 | 161900441 | 1162 | 6.13 | 1.12E-07 | 5.13E-07 |
| chr19 | 53635446 | 53636178 | 733 | 6.09 | 1.12E-07 | 5.14E-07 |
| chr8 | 135559628 | 135567234 | 7607 | 0.86 | 1.12E-07 | 5.15E-07 |
| chr5 | 133945773 | 133969321 | 23549 | 0.66 | 1.12E-07 | 5.15E-07 |
| chr7 | 18317479 | 18318958 | 1480 | 2.24 | 1.12E-07 | 5.17E-07 |
| chr5 | 29262440 | 29263902 | 1463 | 2.23 | 1.13E-07 | 5.18E-07 |
| chr3 | 43892509 | 43936406 | 43898 | -0.29 | 1.13E-07 | 5.19E-07 |
| chr4 | 84810406 | 84811704 | 1299 | 1.35 | 1.13E-07 | 5.21E-07 |
| chr12 | 14954874 | 14957649 | 2776 | 0.56 | 1.13E-07 | 5.21E-07 |
| chr10 | 97168112 | 97169598 | 1487 | 3.13 | 1.14E-07 | 5.22E-07 |
| chr17 | 67709390 | 67710673 | 1284 | -4.28 | 1.14E-07 | 5.22E-07 |
| chr3 | 466079 | 469175 | 3097 | 5.98 | 1.14E-07 | 5.23E-07 |
| chr5 | 77175212 | 77190868 | 15657 | -1.13 | 1.14E-07 | 5.23E-07 |
| chr11 | 6494663 | 6504854 | 10192 | 0.53 | 1.15E-07 | 5.26E-07 |
| chr12 | 92060931 | 92062153 | 1223 | 2.68 | 1.15E-07 | 5.28E-07 |
| chr9 | 86018516 | 86019513 | 998 | -3.23 | 1.15E-07 | 5.30E-07 |
| chr1 | 209820240 | 209828318 | 8079 | 0.81 | 1.15E-07 | 5.30E-07 |
| chr15 | 95739303 | 95740225 | 923 | 6.08 | 1.16E-07 | 5.33E-07 |
| chr10 | 32734256 | 32737094 | 2839 | 0.56 | 1.16E-07 | 5.34E-07 |
| chr11 | 34066091 | 34140339 | 74249 | 0.31 | 1.17E-07 | 5.37E-07 |
| chr7 | 124561834 | 124570453 | 8620 | 0.78 | 1.17E-07 | 5.39E-07 |
| chr2 | 18594610 | 18595810 | 1201 | -2.12 | 1.19E-07 | 5.45E-07 |
| chr18 | 10139747 | 10141722 | 1976 | -2.11 | 1.19E-07 | 5.45E-07 |
| chr12 | 25909767 | 25910603 | 837 | 4.5 | 1.19E-07 | 5.46E-07 |
| chr19 | 21786930 | 21787840 | 911 | 6.08 | 1.20E-07 | 5.48E-07 |
| chr9 | 132220231 | 132270648 | 50418 | 0.21 | 1.20E-07 | 5.49E-07 |
| chr4 | 20420245 | 20421067 | 823 | 5.12 | 1.20E-07 | 5.50E-07 |
| chr14 | 102529856 | 102554819 | 24964 | 0.55 | 1.20E-07 | 5.50E-07 |
| chr1 | 233748832 | 233751420 | 2589 | 1.1 | 1.20E-07 | 5.51E-07 |
| chr22 | 18559082 | 18575790 | 16709 | 0.65 | 1.21E-07 | 5.52E-07 |
| chr10 | 89564645 | 89579278 | 14634 | 0.54 | 1.22E-07 | 5.59E-07 |
| chr10 | 65052205 | 65061260 | 9056 | -1.25 | 1.22E-07 | 5.60E-07 |
| chr6 | 3712757 | 3714322 | 1566 | 1.52 | 1.23E-07 | 5.64E-07 |
| chr1 | 97657544 | 97659043 | 1500 | 2.17 | 1.23E-07 | 5.64E-07 |
| chr4 | 124593870 | 124595436 | 1567 | 2.22 | 1.24E-07 | 5.65E-07 |
| chr7 | 121035384 | 121039135 | 3752 | 1.09 | 1.24E-07 | 5.68E-07 |
| chr6 | 106671510 | 106675378 | 3869 | -1.72 | 1.24E-07 | 5.69E-07 |
| chr11 | 60137947 | 60139091 | 1145 | 6.07 | 1.25E-07 | 5.70E-07 |
| chr1 | 92197779 | 92204797 | 7019 | 0.59 | 1.25E-07 | 5.71E-07 |
| chr6 | 81405803 | 81406836 | 1034 | 6.07 | 1.25E-07 | 5.73E-07 |
| chr9 | 18237293 | 18239815 | 2523 | 1.65 | 1.25E-07 | 5.73E-07 |
| chr1 | 60061407 | 60062648 | 1242 | 2.77 | 1.26E-07 | 5.74E-07 |
| chrX | 20234767 | 20253817 | 19051 | 0.45 | 1.26E-07 | 5.74E-07 |
| chr6 | 113951925 | 113976303 | 24379 | 0.62 | 1.26E-07 | 5.74E-07 |
| chr13 | 111063935 | 111072568 | 8634 | -0.96 | 1.26E-07 | 5.75E-07 |

| | | | | | | |
|---|---|---|---|---|---|---|
| chr13 | 102042526 | 102043441 | 916 | 3.96 | 1.26E-07 | 5.75E-07 |
| chr4 | 130012872 | 130018296 | 5425 | 0.7 | 1.26E-07 | 5.76E-07 |
| chr4 | 167307482 | 167308235 | 754 | 6.07 | 1.26E-07 | 5.77E-07 |
| chr4 | 141868254 | 141869186 | 933 | 6.07 | 1.27E-07 | 5.78E-07 |
| chr12 | 104135801 | 104137190 | 1390 | 1.87 | 1.27E-07 | 5.79E-07 |
| chr4 | 99180683 | 99183960 | 3278 | 0.71 | 1.27E-07 | 5.80E-07 |
| chr14 | 30111874 | 30124620 | 12747 | -1.64 | 1.27E-07 | 5.80E-07 |
| chr8 | 41158027 | 41160995 | 2969 | -2.57 | 1.27E-07 | 5.82E-07 |
| chr12 | 65778209 | 65779275 | 1067 | 4.25 | 1.28E-07 | 5.84E-07 |
| chr8 | 9006925 | 9009752 | 2828 | 1 | 1.28E-07 | 5.85E-07 |
| chr5 | 611801 | 616862 | 5062 | 1.13 | 1.28E-07 | 5.86E-07 |
| chr1 | 247217891 | 247218841 | 951 | 4.57 | 1.29E-07 | 5.89E-07 |
| chrX | 143076685 | 143077459 | 775 | 6.08 | 1.29E-07 | 5.90E-07 |
| chr7 | 474916 | 476795 | 1880 | 1.51 | 1.30E-07 | 5.92E-07 |
| chr5 | 137849969 | 137912894 | 62926 | 0.27 | 1.30E-07 | 5.93E-07 |
| chr2 | 192326650 | 192327716 | 1067 | 6.07 | 1.31E-07 | 5.95E-07 |
| chr4 | 126147043 | 126148215 | 1173 | 6.07 | 1.31E-07 | 5.95E-07 |
| chr17 | 63693920 | 63695363 | 1444 | 1.5 | 1.31E-07 | 5.96E-07 |
| chr9 | 101610030 | 101611466 | 1437 | 1.84 | 1.31E-07 | 5.99E-07 |
| chr20 | 5130544 | 5157167 | 26624 | -0.52 | 1.32E-07 | 6.01E-07 |
| chr17 | 64370119 | 64393779 | 23661 | 0.58 | 1.33E-07 | 6.04E-07 |
| chr17 | 74476792 | 74498451 | 21660 | -0.95 | 1.33E-07 | 6.04E-07 |
| chr18 | 11907528 | 11910152 | 2625 | 0.96 | 1.33E-07 | 6.04E-07 |
| chr7 | 99290945 | 99296821 | 5877 | 1.31 | 1.33E-07 | 6.08E-07 |
| chr4 | 63805979 | 63814420 | 8442 | 6.17 | 1.34E-07 | 6.10E-07 |
| chrX | 117349387 | 117351105 | 1719 | 6.17 | 1.34E-07 | 6.10E-07 |
| chr16 | 4062229 | 4071825 | 9597 | 0.66 | 1.35E-07 | 6.12E-07 |
| chr17 | 5163008 | 5163857 | 850 | 4.66 | 1.35E-07 | 6.12E-07 |
| chr2 | 158950621 | 158951996 | 1376 | 2.29 | 1.35E-07 | 6.13E-07 |
| chr2 | 238583498 | 238616705 | 33208 | -0.45 | 1.35E-07 | 6.13E-07 |
| chr4 | 150452455 | 150453600 | 1146 | 6.06 | 1.35E-07 | 6.13E-07 |
| chr10 | 5902330 | 5904610 | 2281 | 1.58 | 1.35E-07 | 6.15E-07 |
| chr7 | 13681133 | 13682079 | 947 | 6.06 | 1.36E-07 | 6.18E-07 |
| chr1 | 172362558 | 172369534 | 6977 | 0.85 | 1.36E-07 | 6.19E-07 |
| chr8 | 77645015 | 77645812 | 798 | 2.44 | 1.36E-07 | 6.20E-07 |
| chr1 | 205587981 | 205602147 | 14167 | 0.52 | 1.37E-07 | 6.22E-07 |
| chr8 | 18651012 | 18653187 | 2176 | 1.83 | 1.37E-07 | 6.23E-07 |
| chr3 | 81106373 | 81107086 | 714 | 6.1 | 1.38E-07 | 6.25E-07 |
| chr4 | 116936952 | 116938069 | 1118 | 6.1 | 1.38E-07 | 6.25E-07 |
| chr6 | 68071035 | 68071745 | 711 | 6.06 | 1.38E-07 | 6.26E-07 |
| chr4 | 38780076 | 38791423 | 11348 | 0.72 | 1.38E-07 | 6.26E-07 |
| chr3 | 18465846 | 18466943 | 1098 | 4.55 | 1.38E-07 | 6.26E-07 |
| chr15 | 25093517 | 25093989 | 473 | 6.09 | 1.38E-07 | 6.27E-07 |
| chr4 | 180045958 | 180046797 | 840 | 6.11 | 1.38E-07 | 6.27E-07 |
| chr3 | 133242282 | 133267676 | 25395 | 0.8 | 1.38E-07 | 6.28E-07 |
| chr5 | 20275613 | 20287326 | 11714 | -0.79 | 1.39E-07 | 6.30E-07 |
| chr1 | 214754690 | 214755996 | 1307 | -4.79 | 1.39E-07 | 6.31E-07 |
| chr10 | 35974084 | 35974941 | 858 | 2.7 | 1.39E-07 | 6.32E-07 |
| chr7 | 41187835 | 41189145 | 1311 | 3.36 | 1.40E-07 | 6.35E-07 |
| chr11 | 122289913 | 122294925 | 5013 | -2.77 | 1.41E-07 | 6.37E-07 |
| chr17 | 40948885 | 40999728 | 50844 | 0.34 | 1.41E-07 | 6.38E-07 |
| chr14 | 91192697 | 91194036 | 1340 | 1.98 | 1.41E-07 | 6.40E-07 |
| chr11 | 90896992 | 90897870 | 879 | -6.01 | 1.42E-07 | 6.42E-07 |
| chr8 | 116382319 | 116383045 | 727 | 4.11 | 1.43E-07 | 6.46E-07 |
| chr2 | 180406054 | 180408149 | 2096 | 6.07 | 1.43E-07 | 6.48E-07 |
| chr17 | 27989571 | 28019935 | 30365 | -0.84 | 1.43E-07 | 6.48E-07 |
| chr7 | 20681506 | 20682259 | 754 | 6.06 | 1.43E-07 | 6.50E-07 |
| chr15 | 33486581 | 33495598 | 9018 | 0.54 | 1.44E-07 | 6.53E-07 |
| chr3 | 42169780 | 42173070 | 3291 | 0.96 | 1.45E-07 | 6.54E-07 |
| chr11 | 93356065 | 93357002 | 938 | 4.35 | 1.45E-07 | 6.56E-07 |
| chr3 | 37901427 | 37906223 | 4797 | 1.36 | 1.45E-07 | 6.57E-07 |

| | | | | | | |
|---|---|---|---|---|---|---|
| chr10 | 92922098 | 92924418 | 2321 | 1.03 | 1.45E-07 | 6.58E-07 |
| chr20 | 32579788 | 32584104 | 4317 | 0.56 | 1.46E-07 | 6.62E-07 |
| chr5 | 43513362 | 43516272 | 2911 | 0.6 | 1.46E-07 | 6.62E-07 |
| chr2 | 232316473 | 232330574 | 14102 | -0.71 | 1.47E-07 | 6.64E-07 |
| chr2 | 104793133 | 104794385 | 1253 | 6.11 | 1.47E-07 | 6.65E-07 |
| chr1 | 213087931 | 213090529 | 2599 | -1.12 | 1.47E-07 | 6.66E-07 |
| chr5 | 10301437 | 10311570 | 10134 | 0.75 | 1.48E-07 | 6.68E-07 |
| chr8 | 78030842 | 78031818 | 977 | 6.05 | 1.48E-07 | 6.68E-07 |
| chr14 | 93724404 | 93738160 | 13757 | -0.53 | 1.48E-07 | 6.69E-07 |
| chr10 | 80721447 | 80738337 | 16891 | 0.34 | 1.48E-07 | 6.70E-07 |
| chr9 | 118090330 | 118091261 | 932 | 5.29 | 1.48E-07 | 6.70E-07 |
| chr12 | 56434169 | 56440993 | 6825 | 0.66 | 1.48E-07 | 6.71E-07 |
| chr17 | 73965945 | 73977053 | 11109 | 0.43 | 1.49E-07 | 6.74E-07 |
| chr12 | 68221264 | 68222604 | 1341 | 1.35 | 1.50E-07 | 6.75E-07 |
| chr10 | 106078819 | 106099795 | 20977 | 0.38 | 1.50E-07 | 6.75E-07 |
| chr5 | 108234565 | 108235947 | 1383 | 3.62 | 1.50E-07 | 6.76E-07 |
| chr1 | 78055575 | 78057936 | 2362 | -0.97 | 1.50E-07 | 6.77E-07 |
| chr7 | 75534589 | 75623892 | 89304 | 0.24 | 1.50E-07 | 6.78E-07 |
| chr2 | 25831439 | 25842504 | 11066 | 0.95 | 1.50E-07 | 6.78E-07 |
| chr2 | 47030702 | 47032279 | 1578 | 2.36 | 1.50E-07 | 6.78E-07 |
| chr10 | 5401140 | 5403856 | 2717 | 0.83 | 1.51E-07 | 6.80E-07 |
| chr2 | 74076353 | 74079707 | 3355 | -1.92 | 1.51E-07 | 6.80E-07 |
| chr8 | 90992794 | 90997193 | 4400 | 0.54 | 1.51E-07 | 6.81E-07 |
| chr19 | 7247108 | 7247855 | 748 | 4.29 | 1.52E-07 | 6.87E-07 |
| chr6 | 22140659 | 22143091 | 2433 | 1.22 | 1.53E-07 | 6.90E-07 |
| chr7 | 99095490 | 99103944 | 8455 | 0.6 | 1.54E-07 | 6.94E-07 |
| chr15 | 69068389 | 69114964 | 46576 | -0.3 | 1.54E-07 | 6.94E-07 |
| chr14 | 24549642 | 24564850 | 15209 | -0.89 | 1.54E-07 | 6.95E-07 |
| chr12 | 106415903 | 106426323 | 10421 | -0.89 | 1.56E-07 | 7.05E-07 |
| chr11 | 19999689 | 20071116 | 71428 | 0.4 | 1.56E-07 | 7.05E-07 |
| chr1 | 217202233 | 217203679 | 1447 | 2.13 | 1.58E-07 | 7.12E-07 |
| chr12 | 94651787 | 94656738 | 4952 | 2.06 | 1.59E-07 | 7.15E-07 |
| chr10 | 52381339 | 52386611 | 5273 | 0.53 | 1.59E-07 | 7.15E-07 |
| chr12 | 132568341 | 132581134 | 12794 | 1.04 | 1.59E-07 | 7.16E-07 |
| chr5 | 150325763 | 150326614 | 852 | 6.04 | 1.59E-07 | 7.17E-07 |
| chr1 | 165599359 | 165621660 | 22302 | 0.41 | 1.60E-07 | 7.18E-07 |
| chr2 | 113276657 | 113279104 | 2448 | -1.91 | 1.60E-07 | 7.19E-07 |
| chr19 | 35926919 | 35928993 | 2075 | 0.79 | 1.60E-07 | 7.22E-07 |
| chr2 | 133829971 | 133831184 | 1214 | -6 | 1.61E-07 | 7.23E-07 |
| chr14 | 32839625 | 32845175 | 5551 | 2.58 | 1.61E-07 | 7.24E-07 |
| chr4 | 23098605 | 23099488 | 884 | 3.2 | 1.61E-07 | 7.24E-07 |
| chr5 | 140201224 | 140214337 | 13114 | 6.17 | 1.62E-07 | 7.27E-07 |
| chr5 | 107333323 | 107339482 | 6160 | 6.09 | 1.62E-07 | 7.29E-07 |
| chr8 | 111906538 | 111915195 | 8658 | 6.07 | 1.63E-07 | 7.31E-07 |
| chr14 | 22460251 | 22462022 | 1772 | 1.95 | 1.63E-07 | 7.32E-07 |
| chr17 | 68603943 | 68605295 | 1353 | 2.49 | 1.64E-07 | 7.35E-07 |
| chr18 | 9670306 | 9684323 | 14018 | 0.45 | 1.64E-07 | 7.36E-07 |
| chr7 | 157599510 | 157600662 | 1153 | 6.04 | 1.64E-07 | 7.38E-07 |
| chr3 | 110578624 | 110579498 | 875 | 6.06 | 1.64E-07 | 7.38E-07 |
| chr9 | 104207302 | 104211963 | 4662 | 1.23 | 1.64E-07 | 7.38E-07 |
| chr7 | 81537237 | 81538908 | 1672 | 2.63 | 1.64E-07 | 7.38E-07 |
| chr3 | 191414408 | 191415305 | 898 | 6.04 | 1.67E-07 | 7.50E-07 |
| chr17 | 27403408 | 27409619 | 6212 | -1.52 | 1.67E-07 | 7.51E-07 |
| chr2 | 29116680 | 29176076 | 59397 | -0.38 | 1.67E-07 | 7.51E-07 |
| chr1 | 45139160 | 45143530 | 4371 | 0.7 | 1.67E-07 | 7.51E-07 |
| chr9 | 15328183 | 15334455 | 6273 | 1 | 1.68E-07 | 7.52E-07 |
| chr22 | 28860150 | 28861295 | 1146 | 2.22 | 1.69E-07 | 7.57E-07 |
| chr10 | 95530451 | 95536288 | 5838 | 5.88 | 1.70E-07 | 7.60E-07 |
| chr10 | 44069022 | 44070556 | 1535 | 2.12 | 1.70E-07 | 7.61E-07 |
| chr5 | 64777367 | 64779506 | 2140 | 1.68 | 1.70E-07 | 7.64E-07 |
| chr5 | 71882872 | 71897608 | 14737 | -0.81 | 1.72E-07 | 7.69E-07 |

| | | | | | | |
|---|---|---|---|---|---|---|
| chr2 | 180422555 | 180423389 | 835 | 6.06 | 1.72E-07 | 7.71E-07 |
| chr1 | 81347796 | 81358875 | 11080 | 6.12 | 1.72E-07 | 7.71E-07 |
| chr2 | 9794470 | 9849703 | 55234 | -0.34 | 1.72E-07 | 7.71E-07 |
| chr6 | 26473405 | 26476864 | 3460 | 0.85 | 1.72E-07 | 7.71E-07 |
| chr12 | 90417178 | 90419352 | 2175 | 1.45 | 1.72E-07 | 7.71E-07 |
| chr7 | 114687424 | 114688591 | 1168 | 4.59 | 1.73E-07 | 7.73E-07 |
| chr14 | 71609780 | 71610960 | 1181 | 2.66 | 1.73E-07 | 7.75E-07 |
| chr2 | 10724031 | 10726185 | 2155 | 0.71 | 1.73E-07 | 7.76E-07 |
| chr5 | 83074094 | 83076131 | 2038 | 1.63 | 1.74E-07 | 7.77E-07 |
| chr14 | 52897038 | 52898010 | 973 | -3.07 | 1.74E-07 | 7.77E-07 |
| chr9 | 28824366 | 28826676 | 2311 | 1.83 | 1.74E-07 | 7.78E-07 |
| chr5 | 155739645 | 155755284 | 15640 | 1.18 | 1.74E-07 | 7.79E-07 |
| chr14 | 85762296 | 85763256 | 961 | 6.05 | 1.74E-07 | 7.80E-07 |
| chrX | 95728350 | 95730677 | 2328 | 1.48 | 1.75E-07 | 7.82E-07 |
| chr13 | 96393111 | 96394086 | 976 | 1.6 | 1.75E-07 | 7.84E-07 |
| chr7 | 148821896 | 148825714 | 3819 | 0.6 | 1.76E-07 | 7.87E-07 |
| chr3 | 182637739 | 182639235 | 1497 | -1.22 | 1.77E-07 | 7.89E-07 |
| chr2 | 19580198 | 19580811 | 614 | -3.83 | 1.77E-07 | 7.92E-07 |
| chr15 | 99384251 | 99501110 | 116860 | -0.14 | 1.77E-07 | 7.92E-07 |
| chr5 | 50040514 | 50041634 | 1121 | 6.15 | 1.77E-07 | 7.92E-07 |
| chr1 | 227575929 | 227577085 | 1157 | 1.97 | 1.77E-07 | 7.93E-07 |
| chr10 | 122755845 | 122756842 | 998 | 4.29 | 1.78E-07 | 7.94E-07 |
| chr3 | 71630393 | 71634682 | 4290 | 0.88 | 1.78E-07 | 7.95E-07 |
| chr12 | 20014926 | 20016449 | 1524 | 1.37 | 1.78E-07 | 7.96E-07 |
| chr11 | 12560833 | 12563345 | 2513 | -2.09 | 1.79E-07 | 7.98E-07 |
| chr3 | 197023687 | 197026774 | 3088 | 1.07 | 1.79E-07 | 7.98E-07 |
| chr14 | 61644483 | 61656498 | 12016 | -0.96 | 1.80E-07 | 8.03E-07 |
| chr14 | 61927862 | 61960413 | 32552 | -0.39 | 1.80E-07 | 8.03E-07 |
| chr4 | 48779186 | 48783637 | 4452 | 0.87 | 1.80E-07 | 8.05E-07 |
| chr5 | 166720166 | 166724694 | 4529 | 0.44 | 1.81E-07 | 8.06E-07 |
| chr11 | 82803291 | 82804643 | 1353 | 2.3 | 1.81E-07 | 8.07E-07 |
| chr10 | 125689382 | 125690651 | 1270 | 2.99 | 1.81E-07 | 8.07E-07 |
| chr1 | 78430491 | 78450966 | 20476 | 0.48 | 1.83E-07 | 8.14E-07 |
| chr5 | 150419430 | 150475692 | 56263 | -0.33 | 1.83E-07 | 8.14E-07 |
| chr16 | 72867022 | 72885363 | 18342 | -0.44 | 1.83E-07 | 8.18E-07 |
| chr5 | 124222827 | 124225170 | 2344 | 0.98 | 1.83E-07 | 8.18E-07 |
| chr7 | 134670481 | 134681167 | 10687 | 0.53 | 1.84E-07 | 8.19E-07 |
| chr22 | 37562734 | 37596953 | 34220 | 0.41 | 1.84E-07 | 8.20E-07 |
| chr3 | 135684230 | 135685930 | 1701 | 1.78 | 1.84E-07 | 8.22E-07 |
| chr2 | 156877583 | 156878918 | 1336 | 2.63 | 1.85E-07 | 8.24E-07 |
| chrX | 65152789 | 65153381 | 593 | 6.01 | 1.85E-07 | 8.25E-07 |
| chr8 | 34782750 | 34784029 | 1280 | -5.97 | 1.87E-07 | 8.31E-07 |
| chr8 | 116346175 | 116346993 | 819 | 6.01 | 1.87E-07 | 8.31E-07 |
| chr10 | 25621529 | 25622755 | 1227 | 6.01 | 1.87E-07 | 8.32E-07 |
| chr8 | 106530153 | 106531265 | 1113 | 6.01 | 1.87E-07 | 8.32E-07 |
| chr5 | 39398808 | 39460506 | 61699 | 2.16 | 1.87E-07 | 8.32E-07 |
| chr2 | 237779901 | 237780714 | 814 | -3.4 | 1.87E-07 | 8.34E-07 |
| chr7 | 28724631 | 28726456 | 1826 | 1.28 | 1.87E-07 | 8.34E-07 |
| chr11 | 72849609 | 72855078 | 5470 | 0.8 | 1.88E-07 | 8.34E-07 |
| chr19 | 50036438 | 50069365 | 32928 | -1.17 | 1.88E-07 | 8.38E-07 |
| chr11 | 60330682 | 60331616 | 935 | -5.18 | 1.89E-07 | 8.39E-07 |
| chr2 | 224874126 | 224888925 | 14800 | -0.86 | 1.89E-07 | 8.40E-07 |
| chr13 | 90084984 | 90085589 | 606 | 6.01 | 1.89E-07 | 8.40E-07 |
| chrX | 27416406 | 27417136 | 731 | 6.01 | 1.89E-07 | 8.40E-07 |
| chr8 | 106728721 | 106729623 | 903 | 6.01 | 1.89E-07 | 8.41E-07 |
| chr9 | 30628769 | 30629804 | 1036 | 6.01 | 1.89E-07 | 8.41E-07 |
| chr10 | 99118467 | 99142119 | 23653 | -0.81 | 1.89E-07 | 8.42E-07 |
| chr4 | 15149989 | 15164419 | 14431 | -0.66 | 1.90E-07 | 8.43E-07 |
| chr11 | 101453420 | 101455191 | 1772 | 2.49 | 1.90E-07 | 8.43E-07 |
| chr17 | 45142942 | 45148853 | 5912 | 0.59 | 1.90E-07 | 8.46E-07 |
| chr9 | 134532972 | 134554702 | 21731 | -0.8 | 1.91E-07 | 8.47E-07 |

| | | | | | | |
|---|---|---|---|---|---|---|
| chr5 | 73611388 | 73624071 | 12684 | -1.21 | 1.92E-07 | 8.52E-07 |
| chr10 | 127272437 | 127274751 | 2315 | 6.17 | 1.92E-07 | 8.53E-07 |
| chr13 | 44804536 | 44807754 | 3219 | 0.75 | 1.92E-07 | 8.54E-07 |
| chr16 | 83170868 | 83171507 | 640 | 3.69 | 1.93E-07 | 8.55E-07 |
| chr4 | 142270400 | 142275420 | 5021 | 1.05 | 1.93E-07 | 8.55E-07 |
| chr18 | 32750883 | 32751978 | 1096 | 6.01 | 1.93E-07 | 8.55E-07 |
| chr16 | 368948 | 403819 | 34872 | 0.37 | 1.93E-07 | 8.56E-07 |
| chr17 | 25619978 | 25711771 | 91794 | 0.31 | 1.94E-07 | 8.59E-07 |
| chr1 | 75194912 | 75200171 | 5260 | 0.62 | 1.94E-07 | 8.59E-07 |
| chr7 | 32572287 | 32659043 | 86757 | -0.25 | 1.95E-07 | 8.66E-07 |
| chr4 | 159456182 | 159457401 | 1220 | 2.38 | 1.96E-07 | 8.71E-07 |
| chr4 | 44918284 | 44919050 | 767 | 6.01 | 1.97E-07 | 8.73E-07 |
| chr5 | 14442744 | 14481700 | 38957 | 0.34 | 1.97E-07 | 8.74E-07 |
| chr7 | 73093519 | 73158527 | 65009 | 0.31 | 1.97E-07 | 8.75E-07 |
| chr19 | 46354984 | 46391313 | 36330 | 0.37 | 1.98E-07 | 8.76E-07 |
| chr13 | 23488959 | 23489849 | 891 | 3.17 | 1.98E-07 | 8.78E-07 |
| chr7 | 104897826 | 104918541 | 20716 | 0.61 | 2.00E-07 | 8.84E-07 |
| chr9 | 6565950 | 6567971 | 2022 | -1.19 | 2.00E-07 | 8.85E-07 |
| chr2 | 39053530 | 39064566 | 11037 | -0.57 | 2.01E-07 | 8.88E-07 |
| chr8 | 125754978 | 125764875 | 9898 | 2.16 | 2.01E-07 | 8.90E-07 |
| chr1 | 7530020 | 7552518 | 22499 | 0.63 | 2.01E-07 | 8.90E-07 |
| chr5 | 24487769 | 24488544 | 776 | 6.04 | 2.02E-07 | 8.94E-07 |
| chr17 | 67602542 | 67605686 | 3145 | -0.96 | 2.02E-07 | 8.96E-07 |
| chr3 | 23726945 | 23744204 | 17260 | -1.23 | 2.03E-07 | 8.96E-07 |
| chr7 | 17300374 | 17301456 | 1083 | 2.97 | 2.03E-07 | 8.97E-07 |
| chrX | 47905171 | 47906262 | 1092 | 6 | 2.04E-07 | 9.01E-07 |
| chr4 | 88476868 | 88488721 | 11854 | 1.57 | 2.05E-07 | 9.06E-07 |
| chr11 | 128076888 | 128081679 | 4792 | 0.55 | 2.06E-07 | 9.12E-07 |
| chr1 | 47496063 | 47497507 | 1445 | 6 | 2.07E-07 | 9.14E-07 |
| chr14 | 55763697 | 55771561 | 7865 | 1.06 | 2.08E-07 | 9.18E-07 |
| chr5 | 55683641 | 55684869 | 1229 | 3.04 | 2.08E-07 | 9.19E-07 |
| chr8 | 36999875 | 37002340 | 2466 | 1.8 | 2.08E-07 | 9.20E-07 |
| chr16 | 8617973 | 8622264 | 4292 | 1.48 | 2.09E-07 | 9.25E-07 |
| chr12 | 92951804 | 92952788 | 985 | 1.75 | 2.09E-07 | 9.26E-07 |
| chr5 | 145404756 | 145408278 | 3523 | 1.53 | 2.10E-07 | 9.28E-07 |
| chr12 | 109898323 | 109916659 | 18337 | 0.44 | 2.11E-07 | 9.32E-07 |
| chr12 | 43732257 | 43732983 | 727 | 6 | 2.11E-07 | 9.33E-07 |
| chr17 | 53684154 | 53686095 | 1942 | 1.46 | 2.12E-07 | 9.36E-07 |
| chr4 | 128496672 | 128497837 | 1166 | 5.24 | 2.13E-07 | 9.39E-07 |
| chr4 | 30913438 | 30914448 | 1011 | 3.89 | 2.15E-07 | 9.47E-07 |
| chr4 | 166756154 | 166757434 | 1281 | 4.19 | 2.15E-07 | 9.49E-07 |
| chr10 | 120826063 | 120841669 | 15607 | 0.52 | 2.15E-07 | 9.51E-07 |
| chr2 | 19687818 | 19688730 | 913 | -3.71 | 2.16E-07 | 9.54E-07 |
| chr6 | 159126832 | 159129387 | 2556 | 1.29 | 2.16E-07 | 9.55E-07 |
| chr8 | 117242277 | 117243245 | 969 | 6 | 2.17E-07 | 9.56E-07 |
| chr14 | 21943670 | 21947274 | 3605 | 0.82 | 2.17E-07 | 9.58E-07 |
| chr5 | 140535440 | 140536265 | 826 | 5.99 | 2.18E-07 | 9.60E-07 |
| chr8 | 116206349 | 116207681 | 1333 | 6.21 | 2.18E-07 | 9.62E-07 |
| chr2 | 177446493 | 177449789 | 3297 | -0.71 | 2.19E-07 | 9.64E-07 |
| chr3 | 192634324 | 192645615 | 11292 | 0.82 | 2.20E-07 | 9.68E-07 |
| chr12 | 62859775 | 62862197 | 2423 | 0.67 | 2.20E-07 | 9.69E-07 |
| chr1 | 99368605 | 99369765 | 1161 | 5.98 | 2.22E-07 | 9.79E-07 |
| chr1 | 6216943 | 6218878 | 1936 | 1.86 | 2.22E-07 | 9.79E-07 |
| chr19 | 24023979 | 24024887 | 909 | 5.99 | 2.22E-07 | 9.80E-07 |
| chr4 | 84538555 | 84552404 | 13850 | 1.31 | 2.23E-07 | 9.81E-07 |
| chr7 | 90892768 | 90894684 | 1917 | 1.48 | 2.23E-07 | 9.83E-07 |
| chr7 | 36155078 | 36169741 | 14664 | 0.63 | 2.23E-07 | 9.84E-07 |
| chr7 | 133535570 | 133548835 | 13266 | 1.26 | 2.24E-07 | 9.85E-07 |
| chr6 | 106207249 | 106208869 | 1621 | -2.11 | 2.25E-07 | 9.89E-07 |
| chr4 | 90493113 | 90494166 | 1054 | 6.05 | 2.25E-07 | 9.92E-07 |
| chr15 | 69509617 | 69510297 | 681 | 4 | 2.27E-07 | 9.99E-07 |

| | | | | | | |
|---|---|---|---|---|---|---|
| chr2 | 189904681 | 189915665 | 10985 | -1.3 | 2.28E-07 | 1.00E-06 |
| chr10 | 135070631 | 135075462 | 4832 | -0.75 | 2.28E-07 | 1.00E-06 |
| chr1 | 236259603 | 236269844 | 10242 | 1.65 | 2.29E-07 | 1.01E-06 |
| chr2 | 242613203 | 242642730 | 29528 | -0.53 | 2.30E-07 | 1.01E-06 |
| chr5 | 92023537 | 92025369 | 1833 | 3.29 | 2.31E-07 | 1.02E-06 |
| chr12 | 40586247 | 40587054 | 808 | 6 | 2.31E-07 | 1.02E-06 |
| chr11 | 20091525 | 20092613 | 1089 | 2.75 | 2.31E-07 | 1.02E-06 |
| chr5 | 23285658 | 23286689 | 1032 | 5.99 | 2.32E-07 | 1.02E-06 |
| chr1 | 188658389 | 188659366 | 978 | 2.64 | 2.33E-07 | 1.02E-06 |
| chrX | 43448653 | 43450031 | 1379 | 2.09 | 2.33E-07 | 1.03E-06 |
| chr8 | 2074902 | 2076809 | 1908 | 1.07 | 2.34E-07 | 1.03E-06 |
| chr15 | 76115814 | 76116946 | 1133 | 6.11 | 2.35E-07 | 1.03E-06 |
| chr8 | 114117069 | 114118117 | 1049 | 6.11 | 2.35E-07 | 1.03E-06 |
| chr6 | 11296790 | 11298465 | 1676 | 1.79 | 2.36E-07 | 1.04E-06 |
| chr3 | 78920560 | 78923405 | 2846 | 1.05 | 2.36E-07 | 1.04E-06 |
| chr13 | 39855954 | 39858278 | 2325 | 2.37 | 2.37E-07 | 1.04E-06 |
| chr2 | 119936223 | 119937711 | 1489 | 2.37 | 2.37E-07 | 1.04E-06 |
| chr17 | 66452030 | 66453635 | 1606 | 1.33 | 2.37E-07 | 1.04E-06 |
| chr3 | 114528745 | 114535721 | 6977 | 3.97 | 2.38E-07 | 1.04E-06 |
| chr5 | 149643088 | 149650957 | 7870 | -1.33 | 2.38E-07 | 1.04E-06 |
| chr1 | 24103835 | 24128155 | 24321 | 0.5 | 2.38E-07 | 1.04E-06 |
| chr1 | 219346566 | 219363962 | 17397 | 0.64 | 2.38E-07 | 1.05E-06 |
| chr19 | 43752165 | 43752790 | 626 | 5.29 | 2.38E-07 | 1.05E-06 |
| chr16 | 66764571 | 66786240 | 21670 | 0.43 | 2.39E-07 | 1.05E-06 |
| chr9 | 16571590 | 16576449 | 4860 | 1.59 | 2.39E-07 | 1.05E-06 |
| chr8 | 109363183 | 109364526 | 1344 | -2.94 | 2.40E-07 | 1.05E-06 |
| chr12 | 111162352 | 111183753 | 21402 | 0.51 | 2.40E-07 | 1.05E-06 |
| chr3 | 4991747 | 4993895 | 2149 | 1.18 | 2.41E-07 | 1.06E-06 |
| chr16 | 55504222 | 55506405 | 2184 | 1.79 | 2.41E-07 | 1.06E-06 |
| chr2 | 11507688 | 11558976 | 51289 | 0.29 | 2.41E-07 | 1.06E-06 |
| chr1 | 182556854 | 182558948 | 2095 | 0.8 | 2.42E-07 | 1.06E-06 |
| chr1 | 150480035 | 150489634 | 9600 | -1.05 | 2.42E-07 | 1.06E-06 |
| chr11 | 66535188 | 66536573 | 1386 | -6.04 | 2.44E-07 | 1.07E-06 |
| chr10 | 53592359 | 53593356 | 998 | -6.02 | 2.45E-07 | 1.07E-06 |
| chr6 | 160693553 | 160695964 | 2412 | 1.73 | 2.46E-07 | 1.08E-06 |
| chr15 | 59196117 | 59228087 | 31971 | 0.34 | 2.46E-07 | 1.08E-06 |
| chr4 | 142476930 | 142493903 | 16974 | 0.58 | 2.47E-07 | 1.08E-06 |
| chr4 | 1713044 | 1731550 | 18507 | 0.5 | 2.48E-07 | 1.09E-06 |
| chr12 | 89862436 | 89869525 | 7090 | 1.06 | 2.48E-07 | 1.09E-06 |
| chr9 | 75048138 | 75107100 | 58963 | 0.28 | 2.48E-07 | 1.09E-06 |
| chrX | 16729405 | 16744914 | 15510 | 0.46 | 2.48E-07 | 1.09E-06 |
| chr4 | 131561781 | 131581599 | 19819 | 2.01 | 2.49E-07 | 1.09E-06 |
| chr11 | 72686623 | 72687669 | 1047 | 4.07 | 2.49E-07 | 1.09E-06 |
| chr21 | 39312589 | 39313609 | 1021 | 4.04 | 2.50E-07 | 1.09E-06 |
| chr2 | 82347376 | 82348051 | 676 | 6.06 | 2.51E-07 | 1.10E-06 |
| chr8 | 83602914 | 83610967 | 8054 | 5.97 | 2.55E-07 | 1.11E-06 |
| chr3 | 80238942 | 80240005 | 1064 | 6.16 | 2.55E-07 | 1.12E-06 |
| chr3 | 103010153 | 103010745 | 593 | 5.96 | 2.56E-07 | 1.12E-06 |
| chr4 | 154209186 | 154210112 | 927 | 5.96 | 2.56E-07 | 1.12E-06 |
| chr1 | 64344271 | 64401759 | 57489 | -0.4 | 2.56E-07 | 1.12E-06 |
| chr7 | 43792896 | 43844997 | 52102 | 0.37 | 2.58E-07 | 1.13E-06 |
| chr19 | 45894364 | 46150179 | 255816 | 0.19 | 2.59E-07 | 1.13E-06 |
| chr1 | 111019587 | 111021127 | 1541 | -1.99 | 2.61E-07 | 1.14E-06 |
| chr21 | 34599220 | 34604332 | 5113 | 0.97 | 2.61E-07 | 1.14E-06 |
| chr2 | 27346292 | 27358736 | 12445 | -1.06 | 2.62E-07 | 1.14E-06 |
| chr4 | 94763695 | 94764650 | 956 | 5.96 | 2.62E-07 | 1.14E-06 |
| chr7 | 46046322 | 46048494 | 2173 | 1.25 | 2.62E-07 | 1.15E-06 |
| chr16 | 12736628 | 12741739 | 5112 | -0.82 | 2.63E-07 | 1.15E-06 |
| chr8 | 49532018 | 49542747 | 10730 | 3.54 | 2.64E-07 | 1.15E-06 |
| chr4 | 187741404 | 187767051 | 25648 | 0.62 | 2.64E-07 | 1.15E-06 |
| chr2 | 190028140 | 190054040 | 25901 | -0.47 | 2.65E-07 | 1.16E-06 |

| | | | | | | |
|---|---|---|---|---|---|---|
| chr9 | 19080103 | 19128601 | 48499 | 0.48 | 2.66E-07 | 1.16E-06 |
| chr12 | 46375187 | 46386948 | 11762 | 0.58 | 2.66E-07 | 1.16E-06 |
| chr2 | 160142348 | 160147903 | 5556 | -1.18 | 2.66E-07 | 1.16E-06 |
| chr3 | 97757740 | 97758579 | 840 | 5.96 | 2.67E-07 | 1.16E-06 |
| chr3 | 133224397 | 133225782 | 1386 | 2.42 | 2.67E-07 | 1.16E-06 |
| chr10 | 3407565 | 3408653 | 1089 | 3.04 | 2.67E-07 | 1.17E-06 |
| chr6 | 148449201 | 148463678 | 14478 | 0.47 | 2.68E-07 | 1.17E-06 |
| chr6 | 151766830 | 151778811 | 11982 | 0.78 | 2.68E-07 | 1.17E-06 |
| chr11 | 64035148 | 64054073 | 18926 | 0.43 | 2.71E-07 | 1.18E-06 |
| chr3 | 37030727 | 37043724 | 12998 | 0.45 | 2.72E-07 | 1.18E-06 |
| chr6 | 25121621 | 25124407 | 2787 | 6.35 | 2.72E-07 | 1.19E-06 |
| chr11 | 62305101 | 62391599 | 86499 | 0.22 | 2.73E-07 | 1.19E-06 |
| chr6 | 67769006 | 67770358 | 1353 | 5.96 | 2.73E-07 | 1.19E-06 |
| chr17 | 70886759 | 70900874 | 14116 | -2.1 | 2.74E-07 | 1.19E-06 |
| chrX | 37771202 | 37772953 | 1752 | 5.32 | 2.74E-07 | 1.19E-06 |
| chr12 | 93808760 | 93837129 | 28370 | 0.53 | 2.75E-07 | 1.19E-06 |
| chr2 | 180127361 | 180138339 | 10979 | 0.82 | 2.75E-07 | 1.20E-06 |
| chr2 | 9652420 | 9707146 | 54727 | -0.25 | 2.76E-07 | 1.20E-06 |
| chr20 | 22742835 | 22758025 | 15191 | 0.38 | 2.77E-07 | 1.21E-06 |
| chr2 | 45836824 | 45845593 | 8770 | -0.59 | 2.77E-07 | 1.21E-06 |
| chr12 | 43847372 | 43848003 | 632 | 5.95 | 2.77E-07 | 1.21E-06 |
| chr17 | 73922696 | 73938064 | 15369 | -0.77 | 2.78E-07 | 1.21E-06 |
| chr4 | 147161501 | 147163960 | 2460 | 1.2 | 2.78E-07 | 1.21E-06 |
| chr1 | 16157828 | 16185285 | 27458 | 0.37 | 2.79E-07 | 1.21E-06 |
| chr11 | 28363295 | 28368224 | 4930 | 0.48 | 2.79E-07 | 1.21E-06 |
| chr15 | 83406566 | 83421677 | 15112 | 0.63 | 2.79E-07 | 1.21E-06 |
| chr4 | 27876512 | 27877254 | 743 | 5.99 | 2.80E-07 | 1.22E-06 |
| chr1 | 233705257 | 233706188 | 932 | 5.98 | 2.81E-07 | 1.22E-06 |
| chr1 | 118104451 | 118105868 | 1418 | 4.01 | 2.81E-07 | 1.22E-06 |
| chr7 | 12515142 | 12515910 | 769 | 3.16 | 2.82E-07 | 1.22E-06 |
| chr15 | 96117366 | 96118159 | 794 | 5.95 | 2.82E-07 | 1.22E-06 |
| chr8 | 55294065 | 55296077 | 2013 | 2.23 | 2.83E-07 | 1.23E-06 |
| chr2 | 58318848 | 58349143 | 30296 | 0.33 | 2.84E-07 | 1.23E-06 |
| chr8 | 9522607 | 9523879 | 1273 | -2.99 | 2.84E-07 | 1.23E-06 |
| chr13 | 111024050 | 111045494 | 21445 | -1.58 | 2.84E-07 | 1.23E-06 |
| chr4 | 73035502 | 73036215 | 714 | 5.95 | 2.87E-07 | 1.24E-06 |
| chr16 | 54690892 | 54691915 | 1024 | 3.53 | 2.88E-07 | 1.25E-06 |
| chr16 | 71804904 | 71807683 | 2780 | 1.5 | 2.88E-07 | 1.25E-06 |
| chr4 | 86848685 | 86859554 | 10870 | 0.57 | 2.88E-07 | 1.25E-06 |
| chr20 | 8203890 | 8205019 | 1130 | 2.86 | 2.89E-07 | 1.25E-06 |
| chr13 | 49757152 | 49773400 | 16249 | 0.94 | 2.89E-07 | 1.25E-06 |
| chr11 | 103552513 | 103554030 | 1518 | -4.78 | 2.91E-07 | 1.26E-06 |
| chr8 | 52806860 | 52813243 | 6384 | 0.52 | 2.92E-07 | 1.27E-06 |
| chr11 | 62472027 | 62499417 | 27391 | 0.34 | 2.92E-07 | 1.27E-06 |
| chr4 | 39899580 | 39936495 | 36916 | 0.55 | 2.93E-07 | 1.27E-06 |
| chr4 | 128915423 | 128919437 | 4015 | -0.85 | 2.97E-07 | 1.29E-06 |
| chr1 | 188386087 | 188387032 | 946 | 5.17 | 3.00E-07 | 1.30E-06 |
| chr5 | 42886771 | 42888015 | 1245 | 5.93 | 3.01E-07 | 1.30E-06 |
| chr20 | 57603423 | 57618622 | 15200 | 0.58 | 3.01E-07 | 1.30E-06 |
| chr2 | 83931787 | 83932846 | 1060 | 5.93 | 3.02E-07 | 1.31E-06 |
| chr2 | 231191467 | 231192993 | 1527 | 1.03 | 3.02E-07 | 1.31E-06 |
| chr2 | 122344102 | 122354954 | 10853 | -1.07 | 3.04E-07 | 1.32E-06 |
| chr1 | 186409058 | 186412459 | 3402 | 1.27 | 3.04E-07 | 1.32E-06 |
| chr6 | 83215419 | 83220466 | 5048 | 1.21 | 3.05E-07 | 1.32E-06 |
| chr16 | 20816165 | 20819368 | 3204 | 0.8 | 3.06E-07 | 1.32E-06 |
| chr20 | 22696743 | 22710116 | 13374 | 0.55 | 3.06E-07 | 1.32E-06 |
| chrX | 40579841 | 40598957 | 19117 | 0.57 | 3.06E-07 | 1.32E-06 |
| chr11 | 76726039 | 76738902 | 12864 | 0.68 | 3.06E-07 | 1.32E-06 |
| chr4 | 11652112 | 11653327 | 1216 | 1.63 | 3.07E-07 | 1.33E-06 |
| chr3 | 189537352 | 189543030 | 5679 | 1.22 | 3.08E-07 | 1.33E-06 |
| chr12 | 48415284 | 48416445 | 1162 | 5.94 | 3.09E-07 | 1.33E-06 |

| | | | | | | |
|---|---|---|---|---|---|---|
| chr10 | 76742490 | 76743801 | 1312 | -2.46 | 3.10E-07 | 1.34E-06 |
| chr7 | 92582678 | 92583691 | 1014 | 4.47 | 3.10E-07 | 1.34E-06 |
| chr17 | 79334042 | 79340116 | 6075 | 1.96 | 3.11E-07 | 1.34E-06 |
| chr20 | 31306566 | 31366333 | 59768 | -0.45 | 3.11E-07 | 1.34E-06 |
| chr12 | 89938060 | 89939687 | 1628 | 1.27 | 3.11E-07 | 1.34E-06 |
| chr3 | 113254371 | 113257280 | 2910 | 1.24 | 3.13E-07 | 1.35E-06 |
| chr12 | 123920458 | 123957152 | 36695 | 0.45 | 3.13E-07 | 1.35E-06 |
| chr1 | 68662729 | 68664268 | 1540 | 2.41 | 3.14E-07 | 1.36E-06 |
| chr10 | 80014146 | 80017143 | 2998 | -0.89 | 3.20E-07 | 1.38E-06 |
| chr4 | 3462717 | 3463807 | 1091 | 5.17 | 3.20E-07 | 1.38E-06 |
| chr10 | 89499747 | 89500788 | 1042 | 3.04 | 3.21E-07 | 1.38E-06 |
| chr9 | 8907655 | 8908678 | 1024 | 2.88 | 3.21E-07 | 1.39E-06 |
| chr1 | 221262532 | 221275308 | 12777 | 0.63 | 3.22E-07 | 1.39E-06 |
| chr10 | 73363103 | 73373562 | 10460 | 1.58 | 3.22E-07 | 1.39E-06 |
| chr3 | 94721065 | 94722004 | 940 | 6.08 | 3.23E-07 | 1.39E-06 |
| chr18 | 71515415 | 71516700 | 1286 | 3.29 | 3.23E-07 | 1.39E-06 |
| chr2 | 236504862 | 236506147 | 1286 | -5.14 | 3.28E-07 | 1.41E-06 |
| chr4 | 158839638 | 158840649 | 1012 | 4.47 | 3.28E-07 | 1.41E-06 |
| chr13 | 45284594 | 45290785 | 6192 | -0.93 | 3.28E-07 | 1.42E-06 |
| chr2 | 3033065 | 3034415 | 1351 | 2.15 | 3.29E-07 | 1.42E-06 |
| chr11 | 38992379 | 38993158 | 780 | 5.92 | 3.29E-07 | 1.42E-06 |
| chr7 | 121113832 | 121120740 | 6909 | 5.92 | 3.29E-07 | 1.42E-06 |
| chr2 | 69741082 | 69748022 | 6941 | -1.94 | 3.29E-07 | 1.42E-06 |
| chr5 | 120440982 | 120442787 | 1806 | 1.3 | 3.30E-07 | 1.42E-06 |
| chr6 | 25192613 | 25194785 | 2173 | 1.28 | 3.31E-07 | 1.43E-06 |
| chr6 | 152346685 | 152347545 | 861 | 5.94 | 3.32E-07 | 1.43E-06 |
| chr7 | 102512374 | 102529698 | 17325 | 0.47 | 3.33E-07 | 1.43E-06 |
| chr2 | 28490831 | 28500798 | 9968 | -0.38 | 3.36E-07 | 1.45E-06 |
| chr5 | 29067843 | 29068708 | 866 | 5.92 | 3.38E-07 | 1.46E-06 |
| chr21 | 30805742 | 30807060 | 1319 | 3.59 | 3.39E-07 | 1.46E-06 |
| chr12 | 1238729 | 1240535 | 1807 | 2.03 | 3.39E-07 | 1.46E-06 |
| chr12 | 106736022 | 106770741 | 34720 | 0.56 | 3.39E-07 | 1.46E-06 |
| chr1 | 53420223 | 53422297 | 2075 | 1.29 | 3.44E-07 | 1.48E-06 |
| chr16 | 22412379 | 22424571 | 12193 | 0.7 | 3.45E-07 | 1.49E-06 |
| chr5 | 138939958 | 138998404 | 58447 | 0.38 | 3.46E-07 | 1.49E-06 |
| chr4 | 78459597 | 78460994 | 1398 | 2.41 | 3.47E-07 | 1.49E-06 |
| chr17 | 42871643 | 42873943 | 2301 | -0.95 | 3.47E-07 | 1.49E-06 |
| chr4 | 75071809 | 75073281 | 1473 | 2.02 | 3.47E-07 | 1.49E-06 |
| chr18 | 56028203 | 56031940 | 3738 | -0.86 | 3.47E-07 | 1.49E-06 |
| chr11 | 116801691 | 116802972 | 1282 | 3.09 | 3.50E-07 | 1.51E-06 |
| chr3 | 101643719 | 101646742 | 3024 | 3.51 | 3.51E-07 | 1.51E-06 |
| chr10 | 44143477 | 44144923 | 1447 | 1.25 | 3.51E-07 | 1.51E-06 |
| chr6 | 155544640 | 155545929 | 1290 | 2.24 | 3.53E-07 | 1.52E-06 |
| chr3 | 191676542 | 191677388 | 847 | 5.91 | 3.56E-07 | 1.53E-06 |
| chr3 | 95015933 | 95016824 | 892 | 5.92 | 3.56E-07 | 1.53E-06 |
| chr19 | 44377011 | 44377840 | 830 | 5.91 | 3.57E-07 | 1.53E-06 |
| chr1 | 47719083 | 47738546 | 19464 | 1.02 | 3.57E-07 | 1.53E-06 |
| chr5 | 111846926 | 111847841 | 916 | 6 | 3.58E-07 | 1.54E-06 |
| chr6 | 105835383 | 105837244 | 1862 | 2.03 | 3.58E-07 | 1.54E-06 |
| chr17 | 26638869 | 26708038 | 69170 | 0.28 | 3.58E-07 | 1.54E-06 |
| chr1 | 199703033 | 199703971 | 939 | 3.42 | 3.59E-07 | 1.54E-06 |
| chr18 | 32239099 | 32240341 | 1243 | 2.83 | 3.60E-07 | 1.54E-06 |
| chr16 | 81172266 | 81176044 | 3779 | -0.95 | 3.60E-07 | 1.55E-06 |
| chr19 | 3336266 | 3406254 | 69989 | -0.36 | 3.62E-07 | 1.55E-06 |
| chr1 | 80906579 | 80911217 | 4639 | 5.99 | 3.62E-07 | 1.55E-06 |
| chr7 | 104856867 | 104866535 | 9669 | 0.97 | 3.65E-07 | 1.57E-06 |
| chr2 | 198074060 | 198075504 | 1445 | 1.88 | 3.66E-07 | 1.57E-06 |
| chr2 | 96969949 | 96973141 | 3193 | -0.65 | 3.67E-07 | 1.57E-06 |
| chrX | 18811145 | 18812685 | 1541 | -2.24 | 3.67E-07 | 1.57E-06 |
| chr2 | 43356641 | 43479000 | 122360 | 0.2 | 3.67E-07 | 1.57E-06 |
| chr19 | 37177099 | 37180867 | 3769 | 0.75 | 3.67E-07 | 1.57E-06 |

| | | | | | | |
|---|---|---|---|---|---|---|
| chr13 | 109798110 | 109810373 | 12264 | -0.4 | 3.67E-07 | 1.57E-06 |
| chr14 | 45714995 | 45724184 | 9190 | 0.54 | 3.69E-07 | 1.58E-06 |
| chr5 | 73871555 | 73878182 | 6628 | 0.92 | 3.73E-07 | 1.60E-06 |
| chr3 | 154475053 | 154476383 | 1331 | 1.61 | 3.73E-07 | 1.60E-06 |
| chr12 | 100753265 | 100754316 | 1052 | 6.06 | 3.75E-07 | 1.60E-06 |
| chr3 | 97540022 | 97542049 | 2028 | 1.02 | 3.75E-07 | 1.60E-06 |
| chr2 | 173291414 | 173320275 | 28862 | 0.35 | 3.75E-07 | 1.60E-06 |
| chr1 | 7177511 | 7178420 | 910 | 3.81 | 3.75E-07 | 1.61E-06 |
| chrX | 47400127 | 47401561 | 1435 | 1.1 | 3.76E-07 | 1.61E-06 |
| chrX | 71601143 | 71602398 | 1256 | 3.92 | 3.76E-07 | 1.61E-06 |
| chrX | 54747360 | 54748678 | 1319 | 2.14 | 3.77E-07 | 1.61E-06 |
| chr19 | 36095344 | 36104446 | 9103 | 0.99 | 3.78E-07 | 1.62E-06 |
| chr18 | 7838658 | 7884674 | 46017 | 0.31 | 3.80E-07 | 1.63E-06 |
| chr8 | 115550183 | 115551213 | 1031 | 5.98 | 3.81E-07 | 1.63E-06 |
| chr19 | 6765226 | 6768775 | 3550 | 1.01 | 3.81E-07 | 1.63E-06 |
| chr22 | 18110232 | 18123614 | 13383 | 0.43 | 3.81E-07 | 1.63E-06 |
| chr4 | 65548352 | 65549265 | 914 | 5.91 | 3.81E-07 | 1.63E-06 |
| chr7 | 2438699 | 2453204 | 14506 | 0.81 | 3.84E-07 | 1.64E-06 |
| chr14 | 57734251 | 57747176 | 12926 | 0.55 | 3.85E-07 | 1.65E-06 |
| chr20 | 50240684 | 50279839 | 39156 | -0.6 | 3.88E-07 | 1.66E-06 |
| chr13 | 71883981 | 71884695 | 715 | 5.9 | 3.89E-07 | 1.66E-06 |
| chr11 | 87835838 | 87839001 | 3164 | 0.74 | 3.89E-07 | 1.66E-06 |
| chr2 | 182700972 | 182701763 | 792 | 4.42 | 3.92E-07 | 1.67E-06 |
| chr2 | 107433194 | 107434388 | 1195 | -4.7 | 3.92E-07 | 1.67E-06 |
| chr1 | 85992549 | 85993632 | 1084 | 5.94 | 3.92E-07 | 1.67E-06 |
| chr12 | 10578077 | 10588470 | 10394 | 5.94 | 3.92E-07 | 1.67E-06 |
| chrX | 28157351 | 28161511 | 4161 | 5.62 | 3.92E-07 | 1.67E-06 |
| chr2 | 74373761 | 74406768 | 33008 | -0.5 | 3.93E-07 | 1.68E-06 |
| chr9 | 86361851 | 86363482 | 1632 | 2.01 | 3.94E-07 | 1.68E-06 |
| chr12 | 85572403 | 85578645 | 6243 | 2.53 | 3.94E-07 | 1.68E-06 |
| chr19 | 12145945 | 12146552 | 608 | 5.93 | 3.94E-07 | 1.68E-06 |
| chr19 | 58125417 | 58127639 | 2223 | 1.34 | 3.98E-07 | 1.70E-06 |
| chr2 | 201312637 | 201351968 | 39332 | -0.45 | 3.99E-07 | 1.70E-06 |
| chr7 | 90269297 | 90270017 | 721 | 5.9 | 4.00E-07 | 1.71E-06 |
| chr15 | 41078696 | 41100444 | 21749 | 0.45 | 4.00E-07 | 1.71E-06 |
| chr4 | 142868828 | 142869659 | 832 | 3.59 | 4.01E-07 | 1.71E-06 |
| chr15 | 85922719 | 85958512 | 35794 | 0.42 | 4.01E-07 | 1.71E-06 |
| chr3 | 5007364 | 5069707 | 62344 | 0.25 | 4.03E-07 | 1.72E-06 |
| chr15 | 50547950 | 50550206 | 2257 | -1.54 | 4.05E-07 | 1.72E-06 |
| chr3 | 36933918 | 36935303 | 1386 | 1.89 | 4.05E-07 | 1.72E-06 |
| chr1 | 170500143 | 170502794 | 2652 | 0.59 | 4.07E-07 | 1.74E-06 |
| chr4 | 92280422 | 92280893 | 472 | 5.89 | 4.10E-07 | 1.75E-06 |
| chr10 | 94958192 | 94959913 | 1722 | 0.98 | 4.11E-07 | 1.75E-06 |
| chr3 | 64336402 | 64341916 | 5515 | 1.76 | 4.12E-07 | 1.76E-06 |
| chr2 | 181276008 | 181277272 | 1265 | 1.92 | 4.15E-07 | 1.77E-06 |
| chr12 | 132663079 | 132675330 | 12252 | 1.98 | 4.18E-07 | 1.78E-06 |
| chr19 | 20386925 | 20387781 | 857 | 5.94 | 4.18E-07 | 1.78E-06 |
| chr2 | 165648914 | 165650151 | 1238 | 5.92 | 4.18E-07 | 1.78E-06 |
| chr2 | 106259439 | 106268117 | 8679 | 0.95 | 4.19E-07 | 1.78E-06 |
| chr3 | 105212368 | 105223256 | 10889 | 0.86 | 4.20E-07 | 1.79E-06 |
| chr1 | 48407051 | 48408304 | 1254 | 1.88 | 4.21E-07 | 1.79E-06 |
| chr1 | 200304945 | 200307651 | 2707 | 0.71 | 4.21E-07 | 1.79E-06 |
| chr9 | 77521183 | 77522523 | 1341 | 5.89 | 4.21E-07 | 1.79E-06 |
| chr5 | 67580797 | 67585916 | 5120 | 1.4 | 4.22E-07 | 1.79E-06 |
| chr16 | 52485455 | 52486247 | 793 | 5.88 | 4.22E-07 | 1.79E-06 |
| chr3 | 146091838 | 146105505 | 13668 | -3.66 | 4.25E-07 | 1.81E-06 |
| chr3 | 83276672 | 83283008 | 6337 | 5.88 | 4.26E-07 | 1.81E-06 |
| chr3 | 56139427 | 56141120 | 1694 | 1.59 | 4.26E-07 | 1.81E-06 |
| chr6 | 169691083 | 169694380 | 3298 | 0.62 | 4.26E-07 | 1.81E-06 |
| chr12 | 92320020 | 92325699 | 5680 | 1.03 | 4.31E-07 | 1.83E-06 |
| chr6 | 134051731 | 134073844 | 22114 | 0.41 | 4.32E-07 | 1.83E-06 |

| | | | | | | |
|---|---|---|---|---|---|---|
| chr9 | 99800942 | 99809347 | 8406 | 1.68 | 4.32E-07 | 1.83E-06 |
| chr10 | 115784669 | 115788226 | 3558 | 0.74 | 4.33E-07 | 1.84E-06 |
| chr4 | 89580110 | 89581833 | 1724 | 2.1 | 4.34E-07 | 1.84E-06 |
| chr7 | 89400984 | 89402302 | 1319 | 5.9 | 4.34E-07 | 1.84E-06 |
| chr3 | 165401779 | 165402452 | 674 | 5.99 | 4.36E-07 | 1.85E-06 |
| chr10 | 35282854 | 35297338 | 14485 | -1 | 4.37E-07 | 1.85E-06 |
| chr17 | 2282555 | 2323058 | 40504 | 0.28 | 4.38E-07 | 1.86E-06 |
| chr6 | 33792171 | 33810894 | 18724 | -0.53 | 4.40E-07 | 1.87E-06 |
| chr11 | 18584157 | 18612582 | 28426 | 0.35 | 4.40E-07 | 1.87E-06 |
| chr3 | 172184606 | 172195404 | 10799 | 1.13 | 4.41E-07 | 1.87E-06 |
| chr15 | 96773595 | 96774875 | 1281 | 1.73 | 4.41E-07 | 1.87E-06 |
| chr2 | 14236206 | 14236893 | 688 | 5.91 | 4.43E-07 | 1.88E-06 |
| chr6 | 141742330 | 141743871 | 1542 | 1.55 | 4.43E-07 | 1.88E-06 |
| chr15 | 66990466 | 67079056 | 88591 | 0.19 | 4.43E-07 | 1.88E-06 |
| chr12 | 76144050 | 76146165 | 2116 | 1.1 | 4.46E-07 | 1.89E-06 |
| chr12 | 20639147 | 20662638 | 23492 | 0.43 | 4.47E-07 | 1.89E-06 |
| chr15 | 76478249 | 76510567 | 32319 | -0.4 | 4.48E-07 | 1.90E-06 |
| chr1 | 92074161 | 92081153 | 6993 | 0.46 | 4.48E-07 | 1.90E-06 |
| chr4 | 79359525 | 79362594 | 3070 | 1.84 | 4.50E-07 | 1.91E-06 |
| chr18 | 32491052 | 32494720 | 3669 | -1 | 4.50E-07 | 1.91E-06 |
| chr5 | 142430473 | 142467717 | 37245 | -0.62 | 4.50E-07 | 1.91E-06 |
| chr2 | 38528594 | 38530600 | 2007 | -1.76 | 4.51E-07 | 1.91E-06 |
| chr10 | 37356347 | 37357282 | 936 | 5.88 | 4.51E-07 | 1.91E-06 |
| chr5 | 82197996 | 82198871 | 876 | 5.95 | 4.53E-07 | 1.92E-06 |
| chr7 | 30834564 | 30837500 | 2937 | 1.46 | 4.53E-07 | 1.92E-06 |
| chr12 | 34371096 | 34371509 | 414 | 3.42 | 4.56E-07 | 1.93E-06 |
| chr3 | 53914844 | 53926994 | 12151 | 0.62 | 4.64E-07 | 1.96E-06 |
| chr8 | 129595386 | 129598562 | 3177 | -1.17 | 4.64E-07 | 1.96E-06 |
| chr10 | 25136948 | 25139992 | 3045 | 0.61 | 4.65E-07 | 1.96E-06 |
| chr12 | 7119373 | 7126893 | 7521 | 0.76 | 4.65E-07 | 1.97E-06 |
| chr11 | 118065354 | 118066650 | 1297 | 2.34 | 4.66E-07 | 1.97E-06 |
| chr16 | 55701585 | 55703183 | 1599 | 3.45 | 4.68E-07 | 1.98E-06 |
| chr5 | 82353758 | 82374709 | 20952 | -0.51 | 4.69E-07 | 1.98E-06 |
| chr1 | 193377745 | 193401514 | 23770 | 3.99 | 4.70E-07 | 1.99E-06 |
| chr4 | 175285752 | 175299518 | 13767 | -1.35 | 4.71E-07 | 1.99E-06 |
| chr16 | 27324490 | 27339593 | 15104 | 0.56 | 4.72E-07 | 1.99E-06 |
| chrX | 56115852 | 56116607 | 756 | 4.5 | 4.73E-07 | 2.00E-06 |
| chr7 | 44717820 | 44720439 | 2620 | -1.36 | 4.73E-07 | 2.00E-06 |
| chr12 | 116475379 | 116476594 | 1216 | 2.05 | 4.74E-07 | 2.00E-06 |
| chr12 | 83758607 | 83759556 | 950 | 5.87 | 4.75E-07 | 2.01E-06 |
| chr15 | 96378360 | 96379178 | 819 | 5.87 | 4.75E-07 | 2.01E-06 |
| chr1 | 178498516 | 178512929 | 14414 | 0.58 | 4.78E-07 | 2.02E-06 |
| chr5 | 75767315 | 75768553 | 1239 | -5.82 | 4.78E-07 | 2.02E-06 |
| chr12 | 117780149 | 117785989 | 5841 | 1.17 | 4.80E-07 | 2.03E-06 |
| chr4 | 140215516 | 140246966 | 31451 | 0.47 | 4.81E-07 | 2.03E-06 |
| chr10 | 89600828 | 89606991 | 6164 | 0.44 | 4.82E-07 | 2.03E-06 |
| chr11 | 21493563 | 21494381 | 819 | 3.91 | 4.83E-07 | 2.04E-06 |
| chr12 | 79596152 | 79603616 | 7465 | 0.73 | 4.85E-07 | 2.05E-06 |
| chr11 | 32969361 | 32970814 | 1454 | 3.54 | 4.86E-07 | 2.05E-06 |
| chr2 | 83075097 | 83075929 | 833 | 5.87 | 4.86E-07 | 2.05E-06 |
| chr3 | 182303784 | 182304191 | 408 | 5.87 | 4.86E-07 | 2.05E-06 |
| chr10 | 96505855 | 96508307 | 2453 | 1.98 | 4.92E-07 | 2.08E-06 |
| chr8 | 122611119 | 122613251 | 2133 | 1.11 | 4.94E-07 | 2.08E-06 |
| chr5 | 173172715 | 173230971 | 58257 | -0.35 | 4.95E-07 | 2.09E-06 |
| chr2 | 186994473 | 186995560 | 1088 | 4.43 | 4.96E-07 | 2.09E-06 |
| chr14 | 24577605 | 24585123 | 7519 | 0.64 | 4.97E-07 | 2.09E-06 |
| chr3 | 80782634 | 80783597 | 964 | 5.86 | 5.00E-07 | 2.11E-06 |
| chr3 | 111792903 | 111816704 | 23802 | 0.6 | 5.00E-07 | 2.11E-06 |
| chr5 | 31197047 | 31198038 | 992 | 5.85 | 5.01E-07 | 2.11E-06 |
| chr5 | 32192614 | 32201366 | 8753 | 1.49 | 5.02E-07 | 2.11E-06 |
| chr4 | 30790926 | 30791612 | 687 | 5.96 | 5.03E-07 | 2.12E-06 |

| | | | | | | |
|---|---|---|---|---|---|---|
| chr4 | 84173502 | 84207153 | 33652 | -0.45 | 5.03E-07 | 2.12E-06 |
| chr4 | 47486679 | 47488679 | 2001 | 1.19 | 5.05E-07 | 2.12E-06 |
| chr9 | 84331953 | 84354422 | 22470 | 0.4 | 5.05E-07 | 2.13E-06 |
| chr3 | 84224266 | 84225139 | 874 | 5.85 | 5.05E-07 | 2.13E-06 |
| chr14 | 47040704 | 47041704 | 1001 | 6.04 | 5.06E-07 | 2.13E-06 |
| chr1 | 80548811 | 80550633 | 1823 | 1.57 | 5.08E-07 | 2.14E-06 |
| chr5 | 49897725 | 49898237 | 513 | 5.11 | 5.08E-07 | 2.14E-06 |
| chr12 | 29793197 | 29794425 | 1229 | 2.58 | 5.10E-07 | 2.14E-06 |
| chr6 | 2838215 | 2878523 | 40309 | 0.39 | 5.12E-07 | 2.15E-06 |
| chr7 | 140320854 | 140341139 | 20286 | -0.71 | 5.14E-07 | 2.16E-06 |
| chr4 | 7540768 | 7541415 | 648 | 2.38 | 5.17E-07 | 2.18E-06 |
| chr5 | 146110398 | 146112809 | 2412 | 1.12 | 5.22E-07 | 2.19E-06 |
| chr1 | 247553637 | 247565900 | 12264 | 6.02 | 5.23E-07 | 2.20E-06 |
| chr1 | 215731492 | 215745100 | 13609 | 0.7 | 5.23E-07 | 2.20E-06 |
| chr12 | 18394519 | 18396692 | 2174 | 0.83 | 5.26E-07 | 2.21E-06 |
| chr16 | 3312933 | 3334984 | 22052 | 0.65 | 5.27E-07 | 2.21E-06 |
| chr2 | 107586578 | 107588160 | 1583 | -1.64 | 5.28E-07 | 2.22E-06 |
| chr20 | 42470901 | 42479803 | 8903 | -0.81 | 5.33E-07 | 2.24E-06 |
| chr2 | 216444060 | 216445401 | 1342 | -1.76 | 5.33E-07 | 2.24E-06 |
| chr9 | 92864118 | 92870340 | 6223 | 2.54 | 5.34E-07 | 2.24E-06 |
| chr13 | 44680105 | 44686587 | 6483 | 1.02 | 5.34E-07 | 2.24E-06 |
| chr4 | 3193225 | 3194496 | 1272 | -2 | 5.34E-07 | 2.24E-06 |
| chr3 | 144899820 | 144900781 | 962 | 5.86 | 5.36E-07 | 2.25E-06 |
| chr4 | 114305323 | 114306384 | 1062 | 5.87 | 5.36E-07 | 2.25E-06 |
| chr9 | 419172 | 420307 | 1136 | 3 | 5.37E-07 | 2.25E-06 |
| chr16 | 25970029 | 25970687 | 659 | 5.85 | 5.37E-07 | 2.25E-06 |
| chr8 | 10509730 | 10512764 | 3035 | -1.55 | 5.39E-07 | 2.26E-06 |
| chr12 | 308604 | 309580 | 977 | 4.7 | 5.39E-07 | 2.26E-06 |
| chr15 | 91095451 | 91141835 | 46385 | -0.47 | 5.40E-07 | 2.26E-06 |
| chr8 | 102015312 | 102016595 | 1284 | 1.89 | 5.40E-07 | 2.26E-06 |
| chr10 | 70479865 | 70495703 | 15839 | 0.52 | 5.40E-07 | 2.27E-06 |
| chr12 | 32347864 | 32350827 | 2964 | 4.22 | 5.41E-07 | 2.27E-06 |
| chr20 | 47826133 | 47850442 | 24310 | 0.49 | 5.43E-07 | 2.27E-06 |
| chr6 | 89789279 | 89792949 | 3671 | 0.71 | 5.43E-07 | 2.28E-06 |
| chr7 | 105287179 | 105291513 | 4335 | -0.7 | 5.44E-07 | 2.28E-06 |
| chr11 | 128317898 | 128319043 | 1146 | 2.86 | 5.44E-07 | 2.28E-06 |
| chr1 | 90868186 | 90869190 | 1005 | 5.84 | 5.44E-07 | 2.28E-06 |
| chr13 | 22614740 | 22615670 | 931 | 5.84 | 5.44E-07 | 2.28E-06 |
| chr4 | 93681261 | 93682259 | 999 | 4.13 | 5.45E-07 | 2.28E-06 |
| chr12 | 48498551 | 48514023 | 15473 | 0.6 | 5.45E-07 | 2.28E-06 |
| chr20 | 9925925 | 9926982 | 1058 | 5.84 | 5.47E-07 | 2.29E-06 |
| chr22 | 43380376 | 43411843 | 31468 | -0.33 | 5.49E-07 | 2.30E-06 |
| chr14 | 95830918 | 95832601 | 1684 | 2.03 | 5.50E-07 | 2.30E-06 |
| chr6 | 52367628 | 52388301 | 20674 | -0.55 | 5.55E-07 | 2.32E-06 |
| chr15 | 32219377 | 32219987 | 611 | 5.16 | 5.56E-07 | 2.33E-06 |
| chrX | 142936663 | 142937646 | 984 | 4.08 | 5.57E-07 | 2.33E-06 |
| chr6 | 162794235 | 162795159 | 925 | 5.89 | 5.57E-07 | 2.33E-06 |
| chr4 | 185383667 | 185396943 | 13277 | 0.78 | 5.58E-07 | 2.33E-06 |
| chr9 | 119326304 | 119337294 | 10991 | -0.43 | 5.59E-07 | 2.34E-06 |
| chr8 | 22501602 | 22504449 | 2848 | -2.28 | 5.62E-07 | 2.35E-06 |
| chr3 | 102827976 | 102828998 | 1023 | 5.84 | 5.63E-07 | 2.35E-06 |
| chr10 | 89292000 | 89304957 | 12958 | 1.03 | 5.66E-07 | 2.36E-06 |
| chr4 | 87132388 | 87142070 | 9683 | 0.93 | 5.68E-07 | 2.37E-06 |
| chr10 | 113240887 | 113241748 | 862 | -5.79 | 5.68E-07 | 2.37E-06 |
| chr13 | 106115296 | 106116532 | 1237 | 4.26 | 5.70E-07 | 2.38E-06 |
| chr2 | 11432322 | 11434189 | 1868 | 1.12 | 5.71E-07 | 2.39E-06 |
| chr7 | 91569057 | 91579519 | 10463 | 0.61 | 5.71E-07 | 2.39E-06 |
| chr14 | 55914164 | 55926293 | 12130 | 0.71 | 5.73E-07 | 2.39E-06 |
| chr3 | 130810939 | 130811417 | 479 | 5.15 | 5.75E-07 | 2.40E-06 |
| chr20 | 1841748 | 1842973 | 1226 | 2.53 | 5.78E-07 | 2.41E-06 |
| chr14 | 68205222 | 68206927 | 1706 | 0.73 | 5.79E-07 | 2.41E-06 |

| | | | | | | |
|---|---|---|---|---|---|---|
| chr20 | 8570432 | 8583724 | 13293 | 6.25 | 5.79E-07 | 2.42E-06 |
| chr5 | 29624828 | 29625601 | 774 | 5.93 | 5.83E-07 | 2.43E-06 |
| chr11 | 77062213 | 77080143 | 17931 | 2.32 | 5.83E-07 | 2.43E-06 |
| chr5 | 28300969 | 28314190 | 13222 | 5.9 | 5.85E-07 | 2.44E-06 |
| chr3 | 122841404 | 122889353 | 47950 | -0.51 | 5.85E-07 | 2.44E-06 |
| chr11 | 95882748 | 95902610 | 19863 | -0.7 | 5.85E-07 | 2.44E-06 |
| chr20 | 1350639 | 1374471 | 23833 | -0.67 | 5.85E-07 | 2.44E-06 |
| chr4 | 142249454 | 142251451 | 1998 | 1.34 | 5.86E-07 | 2.44E-06 |
| chr5 | 173247962 | 173257161 | 9200 | -0.51 | 5.86E-07 | 2.44E-06 |
| chr20 | 32399013 | 32420648 | 21636 | 0.52 | 5.87E-07 | 2.45E-06 |
| chr7 | 112119655 | 112126211 | 6557 | 0.62 | 5.88E-07 | 2.45E-06 |
| chr4 | 31147704 | 31148787 | 1084 | 3.97 | 5.89E-07 | 2.46E-06 |
| chr14 | 37442288 | 37453214 | 10927 | 0.43 | 5.90E-07 | 2.46E-06 |
| chr8 | 33936745 | 33937697 | 953 | 5.84 | 5.93E-07 | 2.47E-06 |
| chr12 | 46008370 | 46009669 | 1300 | 5.83 | 5.93E-07 | 2.47E-06 |
| chr4 | 429995 | 430824 | 830 | 5.83 | 5.93E-07 | 2.47E-06 |
| chrX | 81066986 | 81067480 | 495 | 3.52 | 5.93E-07 | 2.47E-06 |
| chr4 | 72872088 | 72872946 | 859 | 5.89 | 5.94E-07 | 2.47E-06 |
| chr3 | 70520300 | 70521584 | 1285 | 5.86 | 5.97E-07 | 2.49E-06 |
| chr5 | 122774072 | 122775534 | 1463 | 1.98 | 5.98E-07 | 2.49E-06 |
| chr11 | 56723183 | 56724162 | 980 | 2.56 | 6.01E-07 | 2.50E-06 |
| chr9 | 100273482 | 100292470 | 18989 | 0.9 | 6.02E-07 | 2.51E-06 |
| chr8 | 107921142 | 107928800 | 7659 | 5.99 | 6.04E-07 | 2.51E-06 |
| chr9 | 130547015 | 130574558 | 27544 | 0.43 | 6.05E-07 | 2.52E-06 |
| chr14 | 20746420 | 20748488 | 2069 | 2.65 | 6.07E-07 | 2.52E-06 |
| chr3 | 14377278 | 14378607 | 1330 | 5.83 | 6.08E-07 | 2.53E-06 |
| chr1 | 53755434 | 53794267 | 38834 | -0.38 | 6.09E-07 | 2.53E-06 |
| chr14 | 64265982 | 64276468 | 10487 | 4.37 | 6.09E-07 | 2.53E-06 |
| chr5 | 100112002 | 100126506 | 14505 | 1.08 | 6.09E-07 | 2.53E-06 |
| chr20 | 35354193 | 35414957 | 60765 | -0.41 | 6.12E-07 | 2.54E-06 |
| chr7 | 106038354 | 106039194 | 841 | -5.79 | 6.13E-07 | 2.55E-06 |
| chr11 | 36167442 | 36172700 | 5259 | -0.65 | 6.14E-07 | 2.55E-06 |
| chr1 | 186917443 | 186920577 | 3135 | 1.82 | 6.19E-07 | 2.57E-06 |
| chr14 | 54215414 | 54216332 | 919 | 2.01 | 6.22E-07 | 2.58E-06 |
| chr6 | 52647637 | 52653516 | 5880 | 2.29 | 6.24E-07 | 2.59E-06 |
| chr8 | 128415762 | 128416976 | 1215 | 2.35 | 6.26E-07 | 2.60E-06 |
| chr13 | 113840113 | 113843328 | 3216 | 0.75 | 6.28E-07 | 2.61E-06 |
| chr7 | 16523722 | 16525771 | 2050 | 0.97 | 6.29E-07 | 2.61E-06 |
| chr3 | 176624458 | 176625034 | 577 | 3.27 | 6.29E-07 | 2.61E-06 |
| chr7 | 22757266 | 22766899 | 9634 | 1.41 | 6.30E-07 | 2.62E-06 |
| chr11 | 102138388 | 102161536 | 23149 | 1.02 | 6.33E-07 | 2.63E-06 |
| chrX | 21456683 | 21457453 | 771 | 5.12 | 6.34E-07 | 2.63E-06 |
| chr13 | 93458232 | 93459122 | 891 | 4.46 | 6.36E-07 | 2.64E-06 |
| chr13 | 97816928 | 97819825 | 2898 | 1.49 | 6.36E-07 | 2.64E-06 |
| chr7 | 79835116 | 79859899 | 24784 | 0.96 | 6.36E-07 | 2.64E-06 |
| chr15 | 88521012 | 88521747 | 736 | -5.77 | 6.41E-07 | 2.66E-06 |
| chr4 | 143535142 | 143537180 | 2039 | 1.23 | 6.42E-07 | 2.66E-06 |
| chr6 | 152448149 | 152458157 | 10009 | -1.28 | 6.44E-07 | 2.67E-06 |
| chr4 | 150954993 | 150955861 | 869 | 5.18 | 6.45E-07 | 2.67E-06 |
| chr12 | 27925153 | 27944122 | 18970 | 0.61 | 6.45E-07 | 2.67E-06 |
| chr4 | 147441835 | 147444156 | 2322 | 1.01 | 6.45E-07 | 2.67E-06 |
| chr4 | 108469431 | 108470239 | 809 | 2.35 | 6.46E-07 | 2.68E-06 |
| chr2 | 186404703 | 186405253 | 551 | 5.83 | 6.57E-07 | 2.72E-06 |
| chr11 | 56608294 | 56622950 | 14657 | 0.45 | 6.58E-07 | 2.72E-06 |
| chr4 | 64593603 | 64594553 | 951 | 5.81 | 6.58E-07 | 2.73E-06 |
| chr11 | 46293847 | 46303486 | 9640 | 3.33 | 6.59E-07 | 2.73E-06 |
| chr14 | 70777699 | 70778333 | 635 | 5.82 | 6.61E-07 | 2.74E-06 |
| chr12 | 27735951 | 27773245 | 37295 | 0.8 | 6.61E-07 | 2.74E-06 |
| chr6 | 134870921 | 134872956 | 2036 | 1.86 | 6.63E-07 | 2.75E-06 |
| chr12 | 63241416 | 63262789 | 21374 | 0.4 | 6.66E-07 | 2.76E-06 |
| chr6 | 141701919 | 141703297 | 1379 | 1.2 | 6.66E-07 | 2.76E-06 |

| | | | | | | |
|---|---|---|---|---|---|---|
| chr9 | 96278994 | 96303092 | 24099 | -0.65 | 6.66E-07 | 2.76E-06 |
| chr1 | 169497970 | 169499045 | 1076 | 3.53 | 6.68E-07 | 2.76E-06 |
| chr1 | 25050554 | 25052489 | 1936 | 1.2 | 6.68E-07 | 2.76E-06 |
| chr13 | 101385883 | 101387037 | 1155 | 3.93 | 6.70E-07 | 2.77E-06 |
| chr6 | 81383723 | 81384681 | 959 | 5.87 | 6.71E-07 | 2.78E-06 |
| chrX | 39872058 | 39874622 | 2565 | 0.81 | 6.73E-07 | 2.78E-06 |
| chr2 | 200602711 | 200604257 | 1547 | 2.02 | 6.76E-07 | 2.80E-06 |
| chr7 | 148753474 | 148770017 | 16544 | 0.74 | 6.76E-07 | 2.80E-06 |
| chr17 | 586143 | 620407 | 34265 | -0.41 | 6.80E-07 | 2.81E-06 |
| chr17 | 36031514 | 36032895 | 1382 | -2.21 | 6.82E-07 | 2.82E-06 |
| chr1 | 205141066 | 205146329 | 5264 | -0.55 | 6.82E-07 | 2.82E-06 |
| chr9 | 75885623 | 75886696 | 1074 | 4.81 | 6.82E-07 | 2.82E-06 |
| chrX | 9346955 | 9348905 | 1951 | 1.97 | 6.85E-07 | 2.83E-06 |
| chr10 | 73472797 | 73535002 | 62206 | 0.24 | 6.85E-07 | 2.83E-06 |
| chr16 | 53535870 | 53546153 | 10284 | 1.1 | 6.85E-07 | 2.83E-06 |
| chr14 | 37582363 | 37587015 | 4653 | 1.57 | 6.85E-07 | 2.83E-06 |
| chr12 | 4714002 | 4714957 | 956 | 1.32 | 6.86E-07 | 2.83E-06 |
| chrX | 19082110 | 19083091 | 982 | -5.82 | 6.93E-07 | 2.86E-06 |
| chr16 | 12407935 | 12410735 | 2801 | 1.11 | 6.95E-07 | 2.87E-06 |
| chr8 | 42573505 | 42597575 | 24071 | 0.86 | 7.00E-07 | 2.89E-06 |
| chr5 | 45491447 | 45492383 | 937 | 5.84 | 7.01E-07 | 2.89E-06 |
| chr2 | 163588428 | 163589257 | 830 | 4.44 | 7.02E-07 | 2.90E-06 |
| chr6 | 46892843 | 46895600 | 2758 | 0.95 | 7.02E-07 | 2.90E-06 |
| chr4 | 151934740 | 151938459 | 3720 | 0.74 | 7.03E-07 | 2.90E-06 |
| chr6 | 158178025 | 158201762 | 23738 | 0.51 | 7.03E-07 | 2.90E-06 |
| chr3 | 158881336 | 158893234 | 11899 | 0.64 | 7.03E-07 | 2.90E-06 |
| chr5 | 39670897 | 39672236 | 1340 | 2.59 | 7.04E-07 | 2.90E-06 |
| chr2 | 205186557 | 205191236 | 4680 | -1.87 | 7.05E-07 | 2.90E-06 |
| chr4 | 19456799 | 19457738 | 940 | 5.8 | 7.06E-07 | 2.91E-06 |
| chr2 | 208766627 | 208772351 | 5725 | 1.88 | 7.10E-07 | 2.92E-06 |
| chr5 | 158689074 | 158691338 | 2265 | 0.6 | 7.10E-07 | 2.93E-06 |
| chr11 | 102618079 | 102618784 | 706 | 2.1 | 7.11E-07 | 2.93E-06 |
| chr2 | 235242398 | 235244555 | 2158 | -1.88 | 7.12E-07 | 2.93E-06 |
| chr12 | 123232830 | 123242755 | 9926 | 0.59 | 7.14E-07 | 2.94E-06 |
| chr1 | 241720906 | 241723135 | 2230 | 2.26 | 7.16E-07 | 2.95E-06 |
| chr1 | 80181749 | 80182945 | 1197 | 3.76 | 7.17E-07 | 2.95E-06 |
| chr12 | 125608500 | 125611269 | 2770 | 1.61 | 7.18E-07 | 2.95E-06 |
| chr11 | 66589566 | 66689454 | 99889 | -0.22 | 7.19E-07 | 2.96E-06 |
| chr20 | 56565367 | 56575617 | 10251 | 1.48 | 7.19E-07 | 2.96E-06 |
| chr3 | 53321965 | 53326427 | 4463 | -2.04 | 7.21E-07 | 2.97E-06 |
| chr17 | 44945863 | 44948907 | 3045 | 2.04 | 7.21E-07 | 2.97E-06 |
| chr5 | 98329786 | 98330845 | 1060 | 5.91 | 7.23E-07 | 2.97E-06 |
| chr13 | 89561006 | 89562046 | 1041 | 5.8 | 7.25E-07 | 2.98E-06 |
| chr13 | 104823051 | 104824171 | 1121 | 5.8 | 7.25E-07 | 2.98E-06 |
| chr4 | 110598317 | 110599129 | 813 | 3.8 | 7.27E-07 | 2.99E-06 |
| chr16 | 71917175 | 71937083 | 19909 | 0.51 | 7.28E-07 | 2.99E-06 |
| chr5 | 36192510 | 36194068 | 1559 | 2.2 | 7.29E-07 | 3.00E-06 |
| chr1 | 214175514 | 214176615 | 1102 | 5.82 | 7.30E-07 | 3.00E-06 |
| chr3 | 53539496 | 53555039 | 15544 | 0.68 | 7.36E-07 | 3.03E-06 |
| chr1 | 234568459 | 234573860 | 5402 | -1.47 | 7.36E-07 | 3.03E-06 |
| chr19 | 6055892 | 6071741 | 15850 | 0.42 | 7.37E-07 | 3.03E-06 |
| chr2 | 28019513 | 28024212 | 4700 | -0.58 | 7.40E-07 | 3.04E-06 |
| chr3 | 11266195 | 11268623 | 2429 | -0.65 | 7.43E-07 | 3.05E-06 |
| chr20 | 24357001 | 24357909 | 909 | -3 | 7.44E-07 | 3.06E-06 |
| chr17 | 19428870 | 19450774 | 21905 | -0.82 | 7.45E-07 | 3.06E-06 |
| chr12 | 117348182 | 117358201 | 10020 | 0.68 | 7.48E-07 | 3.07E-06 |
| chr1 | 19392420 | 19416994 | 24575 | -0.52 | 7.48E-07 | 3.07E-06 |
| chr8 | 91682513 | 91684143 | 1631 | 0.85 | 7.53E-07 | 3.09E-06 |
| chr3 | 121515132 | 121522897 | 7766 | 1.48 | 7.54E-07 | 3.09E-06 |
| chr3 | 83238508 | 83239083 | 576 | 5.82 | 7.54E-07 | 3.09E-06 |
| chr10 | 25582536 | 25583907 | 1372 | 1.81 | 7.54E-07 | 3.09E-06 |

| | | | | | | |
|---|---|---|---|---|---|---|
| chr10 | 62821685 | 62827555 | 5871 | -0.93 | 7.54E-07 | 3.09E-06 |
| chr6 | 85772901 | 85773897 | 997 | 4.16 | 7.57E-07 | 3.10E-06 |
| chr10 | 54357186 | 54357955 | 770 | 2.42 | 7.57E-07 | 3.10E-06 |
| chr18 | 26977337 | 26978593 | 1257 | 5.9 | 7.57E-07 | 3.10E-06 |
| chr6 | 107472937 | 107475224 | 2288 | 1.55 | 7.60E-07 | 3.11E-06 |
| chr19 | 35476449 | 35521266 | 44818 | -0.24 | 7.60E-07 | 3.11E-06 |
| chr2 | 171034682 | 171035811 | 1130 | 4.01 | 7.61E-07 | 3.12E-06 |
| chr16 | 34979326 | 34979798 | 473 | 5.8 | 7.63E-07 | 3.13E-06 |
| chr2 | 216611437 | 216616564 | 5128 | -1.27 | 7.63E-07 | 3.13E-06 |
| chr6 | 118789929 | 118790987 | 1059 | 5.93 | 7.63E-07 | 3.13E-06 |
| chrX | 115944871 | 115945752 | 882 | 5.81 | 7.66E-07 | 3.14E-06 |
| chr16 | 56404075 | 56418525 | 14451 | 0.95 | 7.67E-07 | 3.14E-06 |
| chr22 | 46065420 | 46204275 | 138856 | -0.18 | 7.67E-07 | 3.14E-06 |
| chr10 | 129075984 | 129078247 | 2264 | 1.7 | 7.67E-07 | 3.14E-06 |
| chr6 | 141238222 | 141239669 | 1448 | 2.3 | 7.68E-07 | 3.14E-06 |
| chrX | 71237989 | 71251017 | 13029 | 1.85 | 7.70E-07 | 3.15E-06 |
| chr1 | 22740428 | 22741657 | 1230 | 1.74 | 7.71E-07 | 3.16E-06 |
| chr4 | 147978123 | 147979306 | 1184 | 2.21 | 7.75E-07 | 3.17E-06 |
| chr16 | 53467631 | 53473150 | 5520 | 0.72 | 7.76E-07 | 3.18E-06 |
| chr13 | 35096477 | 35098559 | 2083 | 1.51 | 7.77E-07 | 3.18E-06 |
| chr21 | 45844761 | 45849470 | 4710 | 1.2 | 7.77E-07 | 3.18E-06 |
| chr7 | 30024910 | 30030499 | 5590 | 0.47 | 7.78E-07 | 3.18E-06 |
| chr11 | 129058672 | 129060958 | 2287 | 0.82 | 7.82E-07 | 3.20E-06 |
| chr21 | 43893677 | 43895460 | 1784 | 1.32 | 7.82E-07 | 3.20E-06 |
| chr11 | 91341567 | 91342870 | 1304 | -2.27 | 7.84E-07 | 3.21E-06 |
| chr7 | 86362729 | 86363754 | 1026 | -2.99 | 7.85E-07 | 3.21E-06 |
| chr1 | 44489893 | 44516288 | 26396 | -0.79 | 7.85E-07 | 3.21E-06 |
| chr6 | 161779797 | 161792604 | 12808 | 4.84 | 7.85E-07 | 3.21E-06 |
| chr7 | 80369673 | 80554420 | 184748 | 0.18 | 7.86E-07 | 3.21E-06 |
| chr5 | 77655043 | 77658578 | 3536 | 0.58 | 7.91E-07 | 3.23E-06 |
| chr4 | 26855059 | 26864039 | 8981 | 0.88 | 7.92E-07 | 3.24E-06 |
| chr14 | 75117803 | 75130555 | 12753 | -1.13 | 7.94E-07 | 3.24E-06 |
| chr7 | 151614623 | 151615880 | 1258 | 3.68 | 7.97E-07 | 3.26E-06 |
| chr12 | 110396024 | 110412729 | 16706 | 0.77 | 8.00E-07 | 3.26E-06 |
| chr9 | 74491207 | 74492651 | 1445 | 5.81 | 8.04E-07 | 3.28E-06 |
| chr12 | 15680002 | 15681148 | 1147 | 2.49 | 8.06E-07 | 3.29E-06 |
| chr16 | 8942385 | 8973072 | 30688 | 0.42 | 8.07E-07 | 3.29E-06 |
| chr6 | 14621593 | 14622855 | 1263 | -2.59 | 8.07E-07 | 3.29E-06 |
| chr7 | 111289143 | 111291860 | 2718 | 0.81 | 8.07E-07 | 3.29E-06 |
| chr2 | 230029280 | 230030239 | 960 | -5.75 | 8.08E-07 | 3.30E-06 |
| chr10 | 97848655 | 97850745 | 2091 | 0.72 | 8.09E-07 | 3.30E-06 |
| chr10 | 12062572 | 12086168 | 23597 | 0.42 | 8.10E-07 | 3.30E-06 |
| chr1 | 59279262 | 59291701 | 12440 | 0.56 | 8.16E-07 | 3.33E-06 |
| chr13 | 98918148 | 98919330 | 1183 | 3.13 | 8.16E-07 | 3.33E-06 |
| chr12 | 84340045 | 84340948 | 904 | 3.62 | 8.17E-07 | 3.33E-06 |
| chr2 | 14108605 | 14109531 | 927 | 5.79 | 8.20E-07 | 3.34E-06 |
| chr9 | 6068698 | 6081178 | 12481 | 6.18 | 8.29E-07 | 3.38E-06 |
| chr6 | 12007806 | 12014530 | 6725 | 0.49 | 8.31E-07 | 3.39E-06 |
| chr14 | 69783895 | 69784931 | 1037 | -5.74 | 8.32E-07 | 3.39E-06 |
| chr2 | 10586141 | 10594044 | 7904 | 0.5 | 8.32E-07 | 3.39E-06 |
| chr12 | 19952544 | 19954086 | 1543 | 5.44 | 8.35E-07 | 3.40E-06 |
| chr2 | 97480916 | 97536380 | 55465 | -0.39 | 8.36E-07 | 3.41E-06 |
| chr15 | 68205492 | 68211480 | 5989 | 0.41 | 8.37E-07 | 3.41E-06 |
| chr4 | 40046272 | 40060426 | 14155 | 0.41 | 8.39E-07 | 3.41E-06 |
| chr3 | 165601752 | 165602315 | 564 | 5.06 | 8.39E-07 | 3.42E-06 |
| chr5 | 139989493 | 139990806 | 1314 | 3.71 | 8.41E-07 | 3.42E-06 |
| chr4 | 72902752 | 72903798 | 1047 | 5.77 | 8.42E-07 | 3.43E-06 |
| chr10 | 29319620 | 29320717 | 1098 | 5.79 | 8.42E-07 | 3.43E-06 |
| chr6 | 86457316 | 86457965 | 650 | 5.13 | 8.43E-07 | 3.43E-06 |
| chr2 | 194211857 | 194213300 | 1444 | 1.97 | 8.46E-07 | 3.44E-06 |
| chr2 | 133720444 | 133721443 | 1000 | -3.61 | 8.46E-07 | 3.44E-06 |

| | | | | | | |
|---|---|---|---|---|---|---|
| chr1 | 87150782 | 87183472 | 32691 | 0.76 | 8.47E-07 | 3.44E-06 |
| chr14 | 96552304 | 96634525 | 82222 | 0.35 | 8.47E-07 | 3.45E-06 |
| chr5 | 17951062 | 17952721 | 1660 | 1.25 | 8.47E-07 | 3.45E-06 |
| chr4 | 103605219 | 103609431 | 4213 | -1.9 | 8.47E-07 | 3.45E-06 |
| chr3 | 153988168 | 153999051 | 10884 | 0.93 | 8.47E-07 | 3.45E-06 |
| chr4 | 62309212 | 62310202 | 991 | 5.78 | 8.48E-07 | 3.45E-06 |
| chr1 | 68283306 | 68299766 | 16461 | 0.46 | 8.50E-07 | 3.46E-06 |
| chr4 | 74641307 | 74642782 | 1476 | 1.84 | 8.53E-07 | 3.47E-06 |
| chr17 | 10113149 | 10114272 | 1124 | 5.77 | 8.54E-07 | 3.47E-06 |
| chr9 | 126180785 | 126181837 | 1053 | -3.65 | 8.55E-07 | 3.47E-06 |
| chr13 | 76110852 | 76140856 | 30005 | 0.61 | 8.55E-07 | 3.47E-06 |
| chr7 | 120496971 | 120498856 | 1886 | 1.61 | 8.57E-07 | 3.48E-06 |
| chr3 | 136713436 | 136714760 | 1325 | 2.2 | 8.61E-07 | 3.49E-06 |
| chr10 | 13202737 | 13214024 | 11288 | -0.9 | 8.61E-07 | 3.49E-06 |
| chr12 | 65672046 | 65673895 | 1850 | 0.83 | 8.65E-07 | 3.51E-06 |
| chr17 | 4736700 | 4778900 | 42201 | 0.42 | 8.71E-07 | 3.53E-06 |
| chr19 | 16254228 | 16255114 | 887 | 3.17 | 8.71E-07 | 3.53E-06 |
| chr13 | 24756076 | 24759424 | 3349 | 1.83 | 8.77E-07 | 3.56E-06 |
| chr17 | 258952 | 262995 | 4044 | -0.9 | 8.85E-07 | 3.59E-06 |
| chr14 | 38511359 | 38512134 | 776 | 4.13 | 8.88E-07 | 3.60E-06 |
| chr4 | 103788094 | 103801404 | 13311 | 0.48 | 8.92E-07 | 3.62E-06 |
| chr10 | 84484203 | 84485101 | 899 | 5.8 | 8.93E-07 | 3.62E-06 |
| chr9 | 74639893 | 74641572 | 1680 | 1.86 | 8.93E-07 | 3.62E-06 |
| chr21 | 45005005 | 45040127 | 35123 | 0.27 | 9.00E-07 | 3.65E-06 |
| chr5 | 79434784 | 79436197 | 1414 | 2.03 | 9.05E-07 | 3.67E-06 |
| chr15 | 22999984 | 23045926 | 45943 | 0.36 | 9.05E-07 | 3.67E-06 |
| chrX | 106231356 | 106243398 | 12043 | -1.26 | 9.05E-07 | 3.67E-06 |
| chr3 | 153001474 | 153011739 | 10266 | 0.72 | 9.06E-07 | 3.67E-06 |
| chr1 | 108135977 | 108136959 | 983 | 4.37 | 9.07E-07 | 3.67E-06 |
| chr7 | 81288905 | 81289939 | 1035 | 5.77 | 9.07E-07 | 3.67E-06 |
| chr18 | 20131238 | 20140623 | 9386 | 1.02 | 9.09E-07 | 3.68E-06 |
| chr3 | 182472359 | 182473615 | 1257 | 4.04 | 9.09E-07 | 3.68E-06 |
| chr7 | 124047030 | 124047700 | 671 | 5.79 | 9.10E-07 | 3.68E-06 |
| chr1 | 172175204 | 172176025 | 822 | 3.14 | 9.17E-07 | 3.71E-06 |
| chr5 | 112848804 | 112868731 | 19928 | 0.5 | 9.17E-07 | 3.71E-06 |
| chr4 | 7805084 | 7943557 | 138474 | 0.2 | 9.19E-07 | 3.72E-06 |
| chr7 | 54822838 | 54828144 | 5307 | 0.57 | 9.19E-07 | 3.72E-06 |
| chr10 | 115468272 | 115470200 | 1929 | -2.09 | 9.21E-07 | 3.73E-06 |
| chr19 | 19824890 | 19825819 | 930 | 5.75 | 9.22E-07 | 3.73E-06 |
| chr2 | 26784903 | 26786121 | 1219 | -1.8 | 9.26E-07 | 3.75E-06 |
| chr2 | 5383894 | 5384703 | 810 | 5.76 | 9.27E-07 | 3.75E-06 |
| chr10 | 37263354 | 37264059 | 706 | 5.75 | 9.28E-07 | 3.75E-06 |
| chr7 | 41026229 | 41027085 | 857 | 5.75 | 9.28E-07 | 3.75E-06 |
| chr20 | 22469553 | 22472794 | 3242 | 0.62 | 9.32E-07 | 3.77E-06 |
| chr8 | 80979716 | 81019765 | 40050 | 0.35 | 9.32E-07 | 3.77E-06 |
| chr1 | 51086455 | 51087685 | 1231 | 2.11 | 9.33E-07 | 3.77E-06 |
| chr12 | 112845169 | 112862516 | 17348 | 0.42 | 9.34E-07 | 3.77E-06 |
| chr1 | 60720803 | 60721947 | 1145 | -2.82 | 9.35E-07 | 3.78E-06 |
| chr15 | 99090934 | 99092518 | 1585 | 1.73 | 9.39E-07 | 3.79E-06 |
| chr2 | 225581393 | 225582364 | 972 | 3.86 | 9.41E-07 | 3.80E-06 |
| chr10 | 73821783 | 73870157 | 48375 | 1.91 | 9.43E-07 | 3.81E-06 |
| chr13 | 93901100 | 93902496 | 1397 | 1.94 | 9.45E-07 | 3.82E-06 |
| chr1 | 229160038 | 229161510 | 1473 | -1.45 | 9.46E-07 | 3.82E-06 |
| chr15 | 37136475 | 37137644 | 1170 | 2.26 | 9.49E-07 | 3.83E-06 |
| chr2 | 166325746 | 166327097 | 1352 | 5.94 | 9.49E-07 | 3.83E-06 |
| chr2 | 208017996 | 208032674 | 14679 | -0.36 | 9.50E-07 | 3.84E-06 |
| chr8 | 110240377 | 110242773 | 2397 | 0.88 | 9.53E-07 | 3.85E-06 |
| chr2 | 53089837 | 53090803 | 967 | 5.76 | 9.55E-07 | 3.85E-06 |
| chr20 | 48598948 | 48617656 | 18709 | -0.99 | 9.56E-07 | 3.86E-06 |
| chr6 | 63475593 | 63476766 | 1174 | 3.26 | 9.56E-07 | 3.86E-06 |
| chr2 | 193450053 | 193450849 | 797 | 5.86 | 9.57E-07 | 3.86E-06 |

| | | | | | | |
|---|---|---|---|---|---|---|
| chr7 | 121878218 | 121878653 | 436 | 4.63 | 9.61E-07 | 3.88E-06 |
| chr14 | 96712375 | 96715877 | 3503 | -2.64 | 9.61E-07 | 3.88E-06 |
| chr19 | 4559613 | 4577540 | 17928 | -0.71 | 9.62E-07 | 3.88E-06 |
| chr2 | 210654588 | 210655377 | 790 | 5.02 | 9.65E-07 | 3.89E-06 |
| chr1 | 160740676 | 160741844 | 1169 | 3.33 | 9.66E-07 | 3.90E-06 |
| chr7 | 38026987 | 38028138 | 1152 | 2.17 | 9.69E-07 | 3.91E-06 |
| chr18 | 46358743 | 46385765 | 27023 | 0.47 | 9.72E-07 | 3.92E-06 |
| chr12 | 12585049 | 12603376 | 18328 | -1.54 | 9.73E-07 | 3.92E-06 |
| chr4 | 1107170 | 1107749 | 580 | 5.75 | 9.83E-07 | 3.96E-06 |
| chr1 | 150569056 | 150603556 | 34501 | 0.3 | 9.87E-07 | 3.98E-06 |
| chr7 | 109532847 | 109533992 | 1146 | 1.81 | 9.98E-07 | 4.02E-06 |
| chr3 | 24169612 | 24170981 | 1370 | 3.88 | 1.00E-06 | 4.04E-06 |
| chr16 | 61951753 | 61952248 | 496 | 5.74 | 1.01E-06 | 4.07E-06 |
| chr10 | 88841499 | 88858048 | 16550 | 0.52 | 1.01E-06 | 4.09E-06 |
| chr4 | 180556925 | 180557698 | 774 | 5.78 | 1.02E-06 | 4.09E-06 |
| chr1 | 81703468 | 81704055 | 588 | 5.77 | 1.02E-06 | 4.10E-06 |
| chr15 | 65766743 | 65794627 | 27885 | 0.56 | 1.02E-06 | 4.10E-06 |
| chr8 | 26296853 | 26319628 | 22776 | 0.36 | 1.02E-06 | 4.12E-06 |
| chr12 | 123826983 | 123890174 | 63192 | 0.3 | 1.02E-06 | 4.12E-06 |
| chr13 | 94212170 | 94213553 | 1384 | 2.23 | 1.03E-06 | 4.13E-06 |
| chr6 | 12570246 | 12604555 | 34310 | -0.38 | 1.03E-06 | 4.15E-06 |
| chr7 | 73473624 | 73477707 | 4084 | 0.67 | 1.03E-06 | 4.16E-06 |
| chr20 | 16709928 | 16712051 | 2124 | 0.55 | 1.03E-06 | 4.16E-06 |
| chr16 | 70403579 | 70489201 | 85623 | -0.28 | 1.03E-06 | 4.16E-06 |
| chr1 | 60322170 | 60323160 | 991 | 4.14 | 1.03E-06 | 4.16E-06 |
| chr1 | 214223827 | 214224874 | 1048 | 5.74 | 1.04E-06 | 4.17E-06 |
| chr1 | 77463971 | 77465186 | 1216 | 2.62 | 1.04E-06 | 4.19E-06 |
| chr12 | 115093172 | 115123377 | 30206 | -0.64 | 1.05E-06 | 4.21E-06 |
| chr19 | 37037163 | 37037890 | 728 | 5.75 | 1.05E-06 | 4.23E-06 |
| chr2 | 233308667 | 233309783 | 1117 | -1.87 | 1.06E-06 | 4.24E-06 |
| chr8 | 18067550 | 18068240 | 691 | 1.52 | 1.06E-06 | 4.24E-06 |
| chr9 | 116381816 | 116386283 | 4468 | 0.66 | 1.06E-06 | 4.24E-06 |
| chr14 | 39895556 | 39903334 | 7779 | 0.54 | 1.06E-06 | 4.24E-06 |
| chr10 | 68875389 | 68876350 | 962 | 5.74 | 1.06E-06 | 4.24E-06 |
| chr12 | 47358147 | 47358882 | 736 | 5.74 | 1.06E-06 | 4.24E-06 |
| chr5 | 13730994 | 13731889 | 896 | 5.74 | 1.06E-06 | 4.24E-06 |
| chr18 | 29567299 | 29567994 | 696 | 3.57 | 1.06E-06 | 4.25E-06 |
| chr10 | 71032067 | 71032940 | 874 | 4.62 | 1.06E-06 | 4.26E-06 |
| chr17 | 12494546 | 12503059 | 8514 | -5.83 | 1.07E-06 | 4.27E-06 |
| chrX | 99890841 | 99904234 | 13394 | 0.76 | 1.07E-06 | 4.28E-06 |
| chr1 | 215237850 | 215238896 | 1047 | 5.78 | 1.08E-06 | 4.32E-06 |
| chr5 | 140749524 | 140763240 | 13717 | 5.78 | 1.08E-06 | 4.32E-06 |
| chr4 | 34180771 | 34186015 | 5245 | -0.48 | 1.08E-06 | 4.33E-06 |
| chr12 | 32442749 | 32458209 | 15461 | 0.58 | 1.08E-06 | 4.33E-06 |
| chr17 | 48863975 | 48865000 | 1026 | -3.22 | 1.08E-06 | 4.34E-06 |
| chr8 | 61861576 | 61863354 | 1779 | 2.4 | 1.09E-06 | 4.36E-06 |
| chr16 | 50330328 | 50349119 | 18792 | -0.83 | 1.09E-06 | 4.36E-06 |
| chr4 | 112559736 | 112561916 | 2181 | 1.53 | 1.09E-06 | 4.36E-06 |
| chrX | 35177521 | 35178458 | 938 | 4.69 | 1.09E-06 | 4.37E-06 |
| chr20 | 12395382 | 12396230 | 849 | -5.75 | 1.09E-06 | 4.38E-06 |
| chr13 | 51655000 | 51655585 | 586 | 2.98 | 1.10E-06 | 4.39E-06 |
| chr2 | 71637194 | 71654736 | 17543 | 0.52 | 1.11E-06 | 4.44E-06 |
| chr2 | 20226795 | 20273078 | 46284 | 0.34 | 1.11E-06 | 4.46E-06 |
| chr12 | 41407468 | 41408780 | 1313 | 3.9 | 1.12E-06 | 4.49E-06 |
| chr1 | 160903787 | 160919979 | 16193 | 1.38 | 1.12E-06 | 4.50E-06 |
| chr12 | 91807028 | 91807845 | 818 | 4.4 | 1.12E-06 | 4.50E-06 |
| chr10 | 59777071 | 59785231 | 8161 | 0.59 | 1.13E-06 | 4.51E-06 |
| chrX | 133737847 | 133739411 | 1565 | -2.23 | 1.13E-06 | 4.51E-06 |
| chr10 | 129322817 | 129324347 | 1531 | 3.87 | 1.13E-06 | 4.52E-06 |
| chr2 | 160082295 | 160094002 | 11708 | -1.18 | 1.14E-06 | 4.54E-06 |
| chr15 | 62181565 | 62182943 | 1379 | 2.39 | 1.14E-06 | 4.55E-06 |

| | | | | | | |
|---|---|---|---|---|---|---|
| chr12 | 100645345 | 100646320 | 976 | 2.49 | 1.15E-06 | 4.59E-06 |
| chr5 | 56659981 | 56664207 | 4227 | -0.6 | 1.15E-06 | 4.60E-06 |
| chr8 | 101539082 | 101540737 | 1656 | 2.43 | 1.15E-06 | 4.60E-06 |
| chr17 | 39214250 | 39216029 | 1780 | 1.26 | 1.15E-06 | 4.60E-06 |
| chr7 | 111675518 | 111676525 | 1008 | -4.06 | 1.15E-06 | 4.60E-06 |
| chr3 | 97712535 | 97713822 | 1288 | 3.21 | 1.15E-06 | 4.62E-06 |
| chr18 | 29929625 | 29930773 | 1149 | 2.08 | 1.16E-06 | 4.62E-06 |
| chr6 | 50958430 | 50959410 | 981 | 5.75 | 1.16E-06 | 4.62E-06 |
| chr6 | 114685019 | 114686180 | 1162 | 5.74 | 1.17E-06 | 4.67E-06 |
| chr14 | 69240470 | 69292184 | 51715 | 0.27 | 1.17E-06 | 4.67E-06 |
| chr4 | 130964067 | 130965389 | 1323 | 2.04 | 1.17E-06 | 4.67E-06 |
| chr4 | 38386452 | 38388703 | 2252 | 1.79 | 1.18E-06 | 4.70E-06 |
| chr8 | 80581895 | 80582699 | 805 | -4.03 | 1.18E-06 | 4.70E-06 |
| chr20 | 14430132 | 14430751 | 620 | 5.74 | 1.18E-06 | 4.71E-06 |
| chr20 | 2113717 | 2132114 | 18398 | -0.74 | 1.18E-06 | 4.71E-06 |
| chr2 | 243017165 | 243018085 | 921 | 5.8 | 1.18E-06 | 4.72E-06 |
| chr4 | 74921244 | 74924094 | 2851 | -1.04 | 1.19E-06 | 4.74E-06 |
| chr5 | 168722967 | 168728696 | 5730 | -2.36 | 1.20E-06 | 4.78E-06 |
| chr5 | 4019202 | 4020042 | 841 | -5.67 | 1.20E-06 | 4.80E-06 |
| chr19 | 8244500 | 8250879 | 6380 | 1.86 | 1.20E-06 | 4.80E-06 |
| chr18 | 61636345 | 61647393 | 11049 | 0.64 | 1.21E-06 | 4.82E-06 |
| chr18 | 41268677 | 41269371 | 695 | 5.71 | 1.21E-06 | 4.83E-06 |
| chr16 | 4662761 | 4676552 | 13792 | 0.44 | 1.21E-06 | 4.83E-06 |
| chr11 | 126570798 | 126571826 | 1029 | -4.07 | 1.21E-06 | 4.84E-06 |
| chr10 | 35476391 | 35486340 | 9950 | 0.43 | 1.21E-06 | 4.84E-06 |
| chr12 | 96487778 | 96488422 | 645 | 5.81 | 1.22E-06 | 4.85E-06 |
| chr14 | 48368542 | 48370489 | 1948 | 5.72 | 1.22E-06 | 4.85E-06 |
| chr17 | 14346368 | 14347097 | 730 | 5.72 | 1.22E-06 | 4.85E-06 |
| chr6 | 117709733 | 117711580 | 1848 | 5.72 | 1.22E-06 | 4.85E-06 |
| chr14 | 56163838 | 56175011 | 11174 | 0.7 | 1.23E-06 | 4.89E-06 |
| chr13 | 82312130 | 82313283 | 1154 | 5.74 | 1.24E-06 | 4.93E-06 |
| chr11 | 91126246 | 91137952 | 11707 | -0.32 | 1.24E-06 | 4.93E-06 |
| chrX | 56058873 | 56060166 | 1294 | 5.71 | 1.24E-06 | 4.94E-06 |
| chr12 | 112195062 | 112230820 | 35759 | 0.37 | 1.24E-06 | 4.95E-06 |
| chr15 | 80711487 | 80712350 | 864 | -4.95 | 1.24E-06 | 4.95E-06 |
| chr1 | 191103953 | 191104628 | 676 | 5.71 | 1.25E-06 | 4.96E-06 |
| chr8 | 128575392 | 128580898 | 5507 | 1.32 | 1.25E-06 | 4.97E-06 |
| chr1 | 117908269 | 117916343 | 8075 | 0.6 | 1.25E-06 | 4.98E-06 |
| chr5 | 149736523 | 149774172 | 37650 | -0.5 | 1.25E-06 | 4.98E-06 |
| chr13 | 46670954 | 46676962 | 6009 | 1.63 | 1.25E-06 | 4.99E-06 |
| chr18 | 12946129 | 12949424 | 3296 | 0.69 | 1.26E-06 | 5.02E-06 |
| chr18 | 12639943 | 12641832 | 1890 | 1.17 | 1.26E-06 | 5.03E-06 |
| chr2 | 83058657 | 83059527 | 871 | 5.72 | 1.27E-06 | 5.03E-06 |
| chr8 | 123792360 | 123807895 | 15536 | 0.82 | 1.27E-06 | 5.05E-06 |
| chr17 | 55727012 | 55743250 | 16239 | -0.94 | 1.27E-06 | 5.06E-06 |
| chr10 | 122851925 | 122852707 | 783 | 5.71 | 1.27E-06 | 5.07E-06 |
| chrX | 27618364 | 27619353 | 990 | 3.81 | 1.27E-06 | 5.07E-06 |
| chr11 | 2003774 | 2011473 | 7700 | 0.81 | 1.28E-06 | 5.07E-06 |
| chr8 | 99977829 | 99979323 | 1495 | 1.2 | 1.28E-06 | 5.07E-06 |
| chr20 | 56793877 | 56794909 | 1033 | 5.74 | 1.28E-06 | 5.10E-06 |
| chr3 | 70246446 | 70247237 | 792 | 5.74 | 1.28E-06 | 5.10E-06 |
| chr9 | 80109448 | 80112586 | 3139 | 0.68 | 1.29E-06 | 5.14E-06 |
| chr15 | 36719091 | 36721915 | 2825 | -0.76 | 1.30E-06 | 5.15E-06 |
| chr3 | 108301117 | 108309742 | 8626 | 0.44 | 1.30E-06 | 5.15E-06 |
| chr4 | 32270412 | 32271231 | 820 | 5.12 | 1.30E-06 | 5.18E-06 |
| chr2 | 110852725 | 110874770 | 22046 | -0.33 | 1.30E-06 | 5.18E-06 |
| chr10 | 134553312 | 134554794 | 1483 | 2.43 | 1.31E-06 | 5.19E-06 |
| chr6 | 168407359 | 168409426 | 2068 | 0.87 | 1.31E-06 | 5.20E-06 |
| chr20 | 50054279 | 50075288 | 21010 | -0.63 | 1.31E-06 | 5.20E-06 |
| chr1 | 191670933 | 191672425 | 1493 | 1.16 | 1.32E-06 | 5.23E-06 |
| chr8 | 31354802 | 31355170 | 369 | 3.14 | 1.32E-06 | 5.23E-06 |

| | | | | | | |
|---|---|---|---|---|---|---|
| chr2 | 144215819 | 144225111 | 9293 | -5.66 | 1.32E-06 | 5.23E-06 |
| chr5 | 109023812 | 109027464 | 3653 | 0.58 | 1.32E-06 | 5.24E-06 |
| chr6 | 117148594 | 117149736 | 1143 | 5.69 | 1.32E-06 | 5.25E-06 |
| chr11 | 73289353 | 73290025 | 673 | 4.44 | 1.32E-06 | 5.25E-06 |
| chr2 | 203878383 | 203883165 | 4783 | 0.58 | 1.33E-06 | 5.26E-06 |
| chr1 | 110319665 | 110337960 | 18296 | -0.55 | 1.34E-06 | 5.30E-06 |
| chr17 | 78037231 | 78038745 | 1515 | -1.17 | 1.34E-06 | 5.30E-06 |
| chr12 | 111371511 | 111373107 | 1597 | 2 | 1.34E-06 | 5.31E-06 |
| chr5 | 54888929 | 54900995 | 12067 | 0.39 | 1.35E-06 | 5.33E-06 |
| chr4 | 101414753 | 101416054 | 1302 | 2.28 | 1.35E-06 | 5.34E-06 |
| chr11 | 23737351 | 23738208 | 858 | 5.7 | 1.35E-06 | 5.34E-06 |
| chr2 | 159009975 | 159011412 | 1438 | -3.44 | 1.35E-06 | 5.35E-06 |
| chr5 | 66173140 | 66200650 | 27511 | 0.33 | 1.35E-06 | 5.35E-06 |
| chr17 | 52450760 | 52451549 | 790 | 5.69 | 1.35E-06 | 5.35E-06 |
| chr19 | 20795345 | 20796048 | 704 | 5.69 | 1.35E-06 | 5.35E-06 |
| chr11 | 12523519 | 12525503 | 1985 | -1.33 | 1.35E-06 | 5.36E-06 |
| chr17 | 59693385 | 59694253 | 869 | 3.75 | 1.36E-06 | 5.37E-06 |
| chr6 | 79785247 | 79790016 | 4770 | 0.59 | 1.36E-06 | 5.38E-06 |
| chr4 | 56914879 | 56916278 | 1400 | -2.33 | 1.36E-06 | 5.39E-06 |
| chr11 | 2846626 | 2848400 | 1775 | 2.87 | 1.36E-06 | 5.39E-06 |
| chr17 | 36208906 | 36209620 | 715 | 3.12 | 1.37E-06 | 5.42E-06 |
| chr2 | 200774885 | 200797379 | 22495 | -0.56 | 1.37E-06 | 5.43E-06 |
| chr1 | 230442721 | 230444483 | 1763 | 2.77 | 1.38E-06 | 5.46E-06 |
| chr5 | 105692254 | 105693205 | 952 | 1.95 | 1.38E-06 | 5.47E-06 |
| chr12 | 123519170 | 123528923 | 9754 | -1.79 | 1.39E-06 | 5.48E-06 |
| chr5 | 32041834 | 32043179 | 1346 | 2.75 | 1.39E-06 | 5.49E-06 |
| chr2 | 75743035 | 75755963 | 12929 | 5.88 | 1.39E-06 | 5.49E-06 |
| chr4 | 101749991 | 101750744 | 754 | 4.59 | 1.39E-06 | 5.50E-06 |
| chr6 | 168595868 | 168597709 | 1842 | 2.17 | 1.40E-06 | 5.54E-06 |
| chr5 | 150136263 | 150142986 | 6724 | 0.72 | 1.40E-06 | 5.54E-06 |
| chr14 | 21775360 | 21789092 | 13733 | 0.65 | 1.40E-06 | 5.55E-06 |
| chr9 | 4092230 | 4094394 | 2165 | -1.62 | 1.40E-06 | 5.55E-06 |
| chr7 | 30323123 | 30330643 | 7521 | 0.63 | 1.40E-06 | 5.55E-06 |
| chr3 | 169826216 | 169899928 | 73713 | 0.38 | 1.41E-06 | 5.55E-06 |
| chr2 | 40716194 | 40717106 | 913 | 5.7 | 1.41E-06 | 5.56E-06 |
| chr6 | 115390843 | 115392120 | 1278 | 2.82 | 1.42E-06 | 5.60E-06 |
| chr4 | 72576597 | 72577492 | 896 | 4.12 | 1.42E-06 | 5.61E-06 |
| chr22 | 19861346 | 19879618 | 18273 | -0.46 | 1.42E-06 | 5.61E-06 |
| chr3 | 164078358 | 164079073 | 716 | 4.74 | 1.43E-06 | 5.66E-06 |
| chr2 | 61094860 | 61127522 | 32663 | 0.41 | 1.44E-06 | 5.67E-06 |
| chr15 | 71053327 | 71057221 | 3895 | 0.65 | 1.44E-06 | 5.68E-06 |
| chr3 | 95641909 | 95642430 | 522 | 5.75 | 1.44E-06 | 5.69E-06 |
| chr19 | 54480934 | 54539902 | 58969 | -0.22 | 1.44E-06 | 5.69E-06 |
| chr1 | 26676744 | 26686158 | 9415 | 1.11 | 1.45E-06 | 5.71E-06 |
| chr3 | 88103330 | 88109720 | 6391 | 0.69 | 1.45E-06 | 5.73E-06 |
| chr4 | 97275066 | 97275754 | 689 | 5.68 | 1.46E-06 | 5.74E-06 |
| chr10 | 9139579 | 9140443 | 865 | 5.91 | 1.46E-06 | 5.74E-06 |
| chr2 | 9504957 | 9575172 | 70216 | -0.3 | 1.46E-06 | 5.76E-06 |
| chr20 | 46438004 | 46439354 | 1351 | 2.17 | 1.46E-06 | 5.77E-06 |
| chr9 | 80559263 | 80568143 | 8881 | 1.13 | 1.46E-06 | 5.77E-06 |
| chr8 | 116644841 | 116645985 | 1145 | 5.82 | 1.47E-06 | 5.78E-06 |
| chr5 | 29587867 | 29588948 | 1082 | 5.68 | 1.47E-06 | 5.80E-06 |
| chr4 | 90390919 | 90391952 | 1034 | 5.72 | 1.47E-06 | 5.80E-06 |
| chr4 | 75532809 | 75559617 | 26809 | -0.28 | 1.48E-06 | 5.83E-06 |
| chr14 | 37404755 | 37406262 | 1508 | 2 | 1.48E-06 | 5.83E-06 |
| chr22 | 37700184 | 37734577 | 34394 | -0.41 | 1.49E-06 | 5.86E-06 |
| chr4 | 44018212 | 44018698 | 487 | 3.55 | 1.49E-06 | 5.86E-06 |
| chr3 | 85458742 | 85464513 | 5772 | 5.68 | 1.49E-06 | 5.86E-06 |
| chr11 | 27817502 | 27818534 | 1033 | 2.25 | 1.49E-06 | 5.86E-06 |
| chrX | 43410703 | 43411704 | 1002 | 3.26 | 1.49E-06 | 5.87E-06 |
| chr2 | 40559055 | 40560005 | 951 | 4.11 | 1.50E-06 | 5.89E-06 |

| | | | | | | |
|---|---|---|---|---|---|---|
| chr1 | 175945497 | 175955461 | 9965 | 0.95 | 1.50E-06 | 5.90E-06 |
| chr2 | 9613865 | 9632637 | 18773 | -0.44 | 1.50E-06 | 5.91E-06 |
| chr2 | 38318983 | 38337949 | 18967 | 0.44 | 1.51E-06 | 5.92E-06 |
| chr14 | 68907637 | 68937263 | 29627 | 0.24 | 1.51E-06 | 5.93E-06 |
| chr18 | 72162608 | 72167480 | 4873 | 0.81 | 1.51E-06 | 5.95E-06 |
| chr9 | 15421807 | 15447808 | 26002 | 0.51 | 1.52E-06 | 5.96E-06 |
| chr18 | 12287586 | 12288907 | 1322 | 2.52 | 1.52E-06 | 5.98E-06 |
| chr7 | 23636721 | 23638071 | 1351 | 1.38 | 1.52E-06 | 5.99E-06 |
| chr18 | 39763588 | 39764264 | 677 | -5.63 | 1.52E-06 | 5.99E-06 |
| chrX | 56614400 | 56615395 | 996 | -4.24 | 1.53E-06 | 6.01E-06 |
| chr10 | 43632582 | 43636665 | 4084 | 0.65 | 1.53E-06 | 6.01E-06 |
| chr13 | 106322587 | 106323384 | 798 | 4.57 | 1.53E-06 | 6.01E-06 |
| chr8 | 37547835 | 37554639 | 6805 | 0.58 | 1.53E-06 | 6.01E-06 |
| chr3 | 110061607 | 110065665 | 4059 | 0.5 | 1.53E-06 | 6.02E-06 |
| chr11 | 59866278 | 59866916 | 639 | 5.73 | 1.54E-06 | 6.03E-06 |
| chr13 | 80443228 | 80444749 | 1522 | 2.19 | 1.54E-06 | 6.05E-06 |
| chr1 | 169910942 | 169912581 | 1640 | 1.62 | 1.54E-06 | 6.06E-06 |
| chr17 | 12718001 | 12718599 | 599 | -5.68 | 1.54E-06 | 6.06E-06 |
| chr14 | 66298720 | 66299494 | 775 | 4.57 | 1.54E-06 | 6.06E-06 |
| chr8 | 30644646 | 30670688 | 26043 | 0.33 | 1.55E-06 | 6.08E-06 |
| chr12 | 92932172 | 92934033 | 1862 | 2.19 | 1.55E-06 | 6.08E-06 |
| chr1 | 81453255 | 81454287 | 1033 | 5.76 | 1.55E-06 | 6.09E-06 |
| chr12 | 10653184 | 10654069 | 886 | 5.76 | 1.55E-06 | 6.09E-06 |
| chr8 | 135539442 | 135541494 | 2053 | 1.66 | 1.55E-06 | 6.10E-06 |
| chr19 | 28266839 | 28267961 | 1123 | 2.95 | 1.56E-06 | 6.10E-06 |
| chr7 | 98669821 | 98761627 | 91807 | 0.37 | 1.56E-06 | 6.10E-06 |
| chr5 | 92694506 | 92696163 | 1658 | 1.26 | 1.56E-06 | 6.12E-06 |
| chrX | 64862291 | 64865708 | 3418 | 0.87 | 1.56E-06 | 6.12E-06 |
| chr7 | 105750813 | 105753857 | 3045 | 0.63 | 1.56E-06 | 6.12E-06 |
| chr8 | 32725714 | 32726874 | 1161 | 1.12 | 1.56E-06 | 6.13E-06 |
| chr19 | 53661750 | 53662285 | 536 | 5.68 | 1.57E-06 | 6.14E-06 |
| chr12 | 103827507 | 103828352 | 846 | 2.16 | 1.57E-06 | 6.14E-06 |
| chr7 | 39987448 | 39992786 | 5339 | 0.45 | 1.57E-06 | 6.16E-06 |
| chr15 | 29474819 | 29475746 | 928 | 5.7 | 1.57E-06 | 6.17E-06 |
| chr15 | 35219259 | 35226326 | 7068 | 0.78 | 1.58E-06 | 6.17E-06 |
| chr3 | 126499844 | 126503680 | 3837 | 0.66 | 1.59E-06 | 6.22E-06 |
| chrX | 22967896 | 22971188 | 3293 | 1.78 | 1.60E-06 | 6.25E-06 |
| chr4 | 59140231 | 59141055 | 825 | 5.66 | 1.60E-06 | 6.27E-06 |
| chr7 | 51276586 | 51277717 | 1132 | 2.02 | 1.60E-06 | 6.27E-06 |
| chr3 | 42514214 | 42549836 | 35623 | 6.56 | 1.60E-06 | 6.27E-06 |
| chr2 | 45395660 | 45397363 | 1704 | -1.33 | 1.60E-06 | 6.27E-06 |
| chr17 | 56160054 | 56170705 | 10652 | -0.92 | 1.61E-06 | 6.29E-06 |
| chr5 | 122370836 | 122373654 | 2819 | 0.89 | 1.61E-06 | 6.32E-06 |
| chr15 | 75063613 | 75092948 | 29336 | -0.56 | 1.62E-06 | 6.33E-06 |
| chr2 | 58253837 | 58254756 | 920 | -3.22 | 1.62E-06 | 6.33E-06 |
| chr17 | 70669154 | 70715317 | 46164 | 0.26 | 1.62E-06 | 6.35E-06 |
| chr12 | 86658778 | 86659411 | 634 | 4.99 | 1.63E-06 | 6.36E-06 |
| chr4 | 33888903 | 33889920 | 1018 | 2.25 | 1.63E-06 | 6.37E-06 |
| chrX | 86895887 | 86897401 | 1515 | 1.93 | 1.63E-06 | 6.37E-06 |
| chr7 | 156724169 | 156725974 | 1806 | 1.13 | 1.63E-06 | 6.37E-06 |
| chr6 | 80309900 | 80319126 | 9227 | 0.83 | 1.63E-06 | 6.37E-06 |
| chr4 | 175303952 | 175325885 | 21934 | 0.33 | 1.63E-06 | 6.37E-06 |
| chr10 | 130402986 | 130404302 | 1317 | 3.62 | 1.63E-06 | 6.38E-06 |
| chr7 | 133859247 | 133860219 | 973 | 3.62 | 1.63E-06 | 6.38E-06 |
| chr19 | 9275649 | 9276771 | 1123 | 5.76 | 1.64E-06 | 6.39E-06 |
| chr1 | 197254580 | 197255379 | 800 | 3.68 | 1.64E-06 | 6.40E-06 |
| chr8 | 67574857 | 67626038 | 51182 | 0.36 | 1.64E-06 | 6.41E-06 |
| chr5 | 144910609 | 144911814 | 1206 | 2.09 | 1.64E-06 | 6.42E-06 |
| chr2 | 115554280 | 115556705 | 2426 | -5.64 | 1.64E-06 | 6.42E-06 |
| chr5 | 68772988 | 68773749 | 762 | 4.11 | 1.65E-06 | 6.42E-06 |
| chrX | 122992218 | 123015982 | 23765 | 0.42 | 1.65E-06 | 6.43E-06 |

| | | | | | | |
|---|---|---|---|---|---|---|
| chr2 | 166052694 | 166053796 | 1103 | 3.3 | 1.65E-06 | 6.43E-06 |
| chr8 | 83169804 | 83170465 | 662 | 5.67 | 1.65E-06 | 6.44E-06 |
| chr4 | 29845586 | 29846326 | 741 | 5.7 | 1.65E-06 | 6.45E-06 |
| chr6 | 116531955 | 116532893 | 939 | 4.31 | 1.66E-06 | 6.48E-06 |
| chr4 | 100324451 | 100335650 | 11200 | 5.66 | 1.66E-06 | 6.48E-06 |
| chr2 | 121297786 | 121298807 | 1022 | -5.66 | 1.67E-06 | 6.50E-06 |
| chr7 | 106665451 | 106666785 | 1335 | -5.66 | 1.67E-06 | 6.50E-06 |
| chr7 | 137773754 | 137775217 | 1464 | 1.79 | 1.67E-06 | 6.51E-06 |
| chr8 | 130157191 | 130158920 | 1730 | -1 | 1.67E-06 | 6.51E-06 |
| chr10 | 86073081 | 86089582 | 16502 | 0.91 | 1.67E-06 | 6.51E-06 |
| chr8 | 118910221 | 118933496 | 23276 | 0.52 | 1.68E-06 | 6.53E-06 |
| chr2 | 207665465 | 207666577 | 1113 | -3.95 | 1.69E-06 | 6.57E-06 |
| chr6 | 27832336 | 27842869 | 10534 | 0.53 | 1.69E-06 | 6.58E-06 |
| chr15 | 101816083 | 101836291 | 20209 | 0.47 | 1.69E-06 | 6.59E-06 |
| chr5 | 23368097 | 23368712 | 616 | 5.73 | 1.69E-06 | 6.59E-06 |
| chr7 | 16338879 | 16340065 | 1187 | 1.88 | 1.69E-06 | 6.60E-06 |
| chr4 | 97598291 | 97599046 | 756 | 5.67 | 1.69E-06 | 6.60E-06 |
| chr12 | 11322771 | 11325779 | 3009 | 0.54 | 1.69E-06 | 6.60E-06 |
| chrX | 117478068 | 117481935 | 3868 | 0.65 | 1.70E-06 | 6.63E-06 |
| chr11 | 11862332 | 11865944 | 3613 | 0.52 | 1.70E-06 | 6.64E-06 |
| chr4 | 64127678 | 64128554 | 877 | 5.66 | 1.71E-06 | 6.65E-06 |
| chr8 | 87519310 | 87527498 | 8189 | 0.63 | 1.71E-06 | 6.66E-06 |
| chr3 | 78872482 | 78874023 | 1542 | 1.55 | 1.72E-06 | 6.68E-06 |
| chr19 | 5869707 | 5881233 | 11527 | 2.22 | 1.72E-06 | 6.68E-06 |
| chr19 | 46974192 | 46975155 | 964 | -2.14 | 1.72E-06 | 6.68E-06 |
| chr14 | 72117615 | 72118772 | 1158 | 3.74 | 1.72E-06 | 6.69E-06 |
| chr8 | 27107396 | 27108366 | 971 | -3.17 | 1.72E-06 | 6.69E-06 |
| chr17 | 66394762 | 66395299 | 538 | -4.47 | 1.72E-06 | 6.70E-06 |
| chr8 | 71115526 | 71139695 | 24170 | 0.54 | 1.73E-06 | 6.72E-06 |
| chr2 | 133378668 | 133389853 | 11186 | -1.75 | 1.73E-06 | 6.73E-06 |
| chr1 | 227161633 | 227168264 | 6632 | -1.93 | 1.73E-06 | 6.74E-06 |
| chr6 | 160197504 | 160217633 | 20130 | 0.53 | 1.74E-06 | 6.77E-06 |
| chr22 | 40362995 | 40363748 | 754 | 5.67 | 1.75E-06 | 6.79E-06 |
| chrX | 74966311 | 74967522 | 1212 | 2.42 | 1.75E-06 | 6.79E-06 |
| chr15 | 92482834 | 92529170 | 46337 | 0.34 | 1.75E-06 | 6.80E-06 |
| chr5 | 16774939 | 16937061 | 162123 | -0.19 | 1.75E-06 | 6.81E-06 |
| chr8 | 125332820 | 125349486 | 16667 | -0.93 | 1.75E-06 | 6.82E-06 |
| chr3 | 31198185 | 31212875 | 14691 | 5.61 | 1.76E-06 | 6.84E-06 |
| chr7 | 155985280 | 155986097 | 818 | -4.13 | 1.77E-06 | 6.89E-06 |
| chr11 | 22645146 | 22648207 | 3062 | 0.65 | 1.78E-06 | 6.89E-06 |
| chr5 | 134664726 | 134693158 | 28433 | -0.57 | 1.78E-06 | 6.91E-06 |
| chr13 | 90551623 | 90552309 | 687 | 3.8 | 1.78E-06 | 6.91E-06 |
| chr2 | 146791524 | 146792254 | 731 | 3.07 | 1.78E-06 | 6.92E-06 |
| chr2 | 54300220 | 54302542 | 2323 | 1.54 | 1.78E-06 | 6.92E-06 |
| chr5 | 17110501 | 17146572 | 36072 | 0.35 | 1.78E-06 | 6.92E-06 |
| chr6 | 160146961 | 160184064 | 37104 | 0.37 | 1.78E-06 | 6.92E-06 |
| chr14 | 65170467 | 65231852 | 61386 | -0.25 | 1.78E-06 | 6.92E-06 |
| chr4 | 103728210 | 103751332 | 23123 | 0.35 | 1.80E-06 | 6.99E-06 |
| chr3 | 186758166 | 186762605 | 4440 | 1.43 | 1.81E-06 | 7.01E-06 |
| chr10 | 33319905 | 33334881 | 14977 | 0.62 | 1.82E-06 | 7.06E-06 |
| chr13 | 50319489 | 50321304 | 1816 | 1.72 | 1.85E-06 | 7.18E-06 |
| chr6 | 149369968 | 149383340 | 13373 | -0.95 | 1.85E-06 | 7.18E-06 |
| chr9 | 87839371 | 87840343 | 973 | 5.7 | 1.85E-06 | 7.18E-06 |
| chr21 | 16854262 | 16855603 | 1342 | -2.24 | 1.85E-06 | 7.18E-06 |
| chr13 | 49402538 | 49414121 | 11584 | 0.92 | 1.85E-06 | 7.18E-06 |
| chr6 | 149967885 | 149970311 | 2427 | 0.95 | 1.86E-06 | 7.22E-06 |
| chr20 | 3176503 | 3223483 | 46981 | 0.35 | 1.87E-06 | 7.24E-06 |
| chr10 | 45469549 | 45479240 | 9692 | 0.54 | 1.87E-06 | 7.25E-06 |
| chr15 | 27005048 | 27019169 | 14122 | -2.6 | 1.87E-06 | 7.25E-06 |
| chr20 | 61424984 | 61455056 | 30073 | -0.6 | 1.88E-06 | 7.26E-06 |
| chr1 | 193016549 | 193030667 | 14119 | 0.47 | 1.88E-06 | 7.28E-06 |

| | | | | | | |
|---|---|---|---|---|---|---|
| chr3 | 58202407 | 58209107 | 6701 | 1.44 | 1.88E-06 | 7.29E-06 |
| chr5 | 71169954 | 71183696 | 13743 | 1.16 | 1.88E-06 | 7.29E-06 |
| chrX | 68371931 | 68385803 | 13873 | 1.41 | 1.89E-06 | 7.30E-06 |
| chr2 | 157888051 | 157889604 | 1554 | 1.38 | 1.89E-06 | 7.33E-06 |
| chr3 | 16090457 | 16114985 | 24529 | -0.34 | 1.89E-06 | 7.33E-06 |
| chr4 | 104763851 | 104764922 | 1072 | 5.67 | 1.90E-06 | 7.35E-06 |
| chr8 | 19064898 | 19083648 | 18751 | 1.1 | 1.90E-06 | 7.36E-06 |
| chr12 | 14987969 | 14995934 | 7966 | -1.01 | 1.91E-06 | 7.39E-06 |
| chr12 | 27048468 | 27072863 | 24396 | 0.82 | 1.91E-06 | 7.39E-06 |
| chr2 | 122277477 | 122290411 | 12935 | -0.59 | 1.91E-06 | 7.40E-06 |
| chr16 | 51082186 | 51083026 | 841 | -3.69 | 1.91E-06 | 7.40E-06 |
| chr7 | 139467574 | 139478835 | 11262 | -0.55 | 1.91E-06 | 7.40E-06 |
| chr3 | 12261815 | 12277825 | 16011 | 6.27 | 1.92E-06 | 7.41E-06 |
| chr8 | 28207883 | 28221923 | 14041 | 0.77 | 1.92E-06 | 7.43E-06 |
| chr1 | 96685765 | 96686844 | 1080 | 5.66 | 1.93E-06 | 7.45E-06 |
| chr2 | 105888949 | 105904906 | 15958 | -1.04 | 1.93E-06 | 7.47E-06 |
| chr2 | 39632066 | 39675833 | 43768 | -0.45 | 1.94E-06 | 7.50E-06 |
| chr3 | 44399210 | 44406448 | 7239 | 1.07 | 1.95E-06 | 7.54E-06 |
| chr16 | 57048305 | 57048911 | 607 | 5.63 | 1.95E-06 | 7.54E-06 |
| chr16 | 87298214 | 87298909 | 696 | 5.63 | 1.95E-06 | 7.54E-06 |
| chr3 | 155768172 | 155769634 | 1463 | -1.57 | 1.96E-06 | 7.56E-06 |
| chr3 | 69128245 | 69143947 | 15703 | 0.53 | 1.96E-06 | 7.58E-06 |
| chr2 | 73460005 | 73482960 | 22956 | -0.69 | 1.97E-06 | 7.59E-06 |
| chr17 | 30129135 | 30131781 | 2647 | 1.24 | 1.97E-06 | 7.61E-06 |
| chr1 | 192795592 | 192796934 | 1343 | 2.21 | 1.97E-06 | 7.61E-06 |
| chr5 | 111015802 | 111017927 | 2126 | 1.07 | 1.98E-06 | 7.63E-06 |
| chr7 | 80095560 | 80111350 | 15791 | 2.41 | 1.98E-06 | 7.63E-06 |
| chr4 | 144480177 | 144480823 | 647 | 2.24 | 1.98E-06 | 7.63E-06 |
| chr10 | 43662055 | 43662960 | 906 | 2.2 | 1.99E-06 | 7.66E-06 |
| chr15 | 63712136 | 63713648 | 1513 | -1.29 | 1.99E-06 | 7.69E-06 |
| chr5 | 36240329 | 36243423 | 3095 | 0.62 | 1.99E-06 | 7.69E-06 |
| chr8 | 88241208 | 88241953 | 746 | 5.67 | 2.00E-06 | 7.70E-06 |
| chr5 | 159844551 | 159850103 | 5553 | 0.6 | 2.00E-06 | 7.72E-06 |
| chr2 | 72778821 | 72799629 | 20809 | -1.51 | 2.00E-06 | 7.72E-06 |
| chr2 | 208708490 | 208713684 | 5195 | 0.59 | 2.00E-06 | 7.72E-06 |
| chr16 | 23689034 | 23723880 | 34847 | 0.56 | 2.01E-06 | 7.73E-06 |
| chr12 | 51818629 | 51824889 | 6261 | 5.63 | 2.01E-06 | 7.73E-06 |
| chr16 | 30362795 | 30390853 | 28059 | 0.32 | 2.01E-06 | 7.73E-06 |
| chr18 | 13672013 | 13679246 | 7234 | 1.16 | 2.01E-06 | 7.75E-06 |
| chr1 | 62900857 | 62911989 | 11133 | 0.45 | 2.01E-06 | 7.75E-06 |
| chr16 | 64214610 | 64215260 | 651 | 5.67 | 2.02E-06 | 7.78E-06 |
| chr15 | 57097770 | 57098934 | 1165 | 1.74 | 2.02E-06 | 7.79E-06 |
| chr7 | 14272959 | 14273832 | 874 | 5.69 | 2.02E-06 | 7.80E-06 |
| chr2 | 141398324 | 141418517 | 20194 | 0.61 | 2.03E-06 | 7.80E-06 |
| chr11 | 110247662 | 110248207 | 546 | 3.96 | 2.03E-06 | 7.81E-06 |
| chr21 | 46961111 | 46976414 | 15304 | -0.85 | 2.03E-06 | 7.82E-06 |
| chr10 | 15385936 | 15387355 | 1420 | 2.39 | 2.03E-06 | 7.82E-06 |
| chrX | 37683423 | 37684605 | 1183 | 5.63 | 2.04E-06 | 7.86E-06 |
| chr7 | 150911110 | 150949732 | 38623 | -0.47 | 2.05E-06 | 7.89E-06 |
| chr16 | 87791168 | 87841551 | 50384 | -0.34 | 2.05E-06 | 7.90E-06 |
| chr11 | 36659938 | 36660788 | 851 | 3.14 | 2.06E-06 | 7.91E-06 |
| chr19 | 9099154 | 9100333 | 1180 | 1.75 | 2.06E-06 | 7.93E-06 |
| chr5 | 122789535 | 122796940 | 7406 | 1.36 | 2.06E-06 | 7.93E-06 |
| chr6 | 75029311 | 75030716 | 1406 | 2.03 | 2.06E-06 | 7.93E-06 |
| chr8 | 135584048 | 135595069 | 11022 | 0.68 | 2.07E-06 | 7.95E-06 |
| chr19 | 9238533 | 9251991 | 13459 | 1.26 | 2.07E-06 | 7.96E-06 |
| chr18 | 54754743 | 54755789 | 1047 | 5.64 | 2.07E-06 | 7.96E-06 |
| chr13 | 24252508 | 24253965 | 1458 | -1.93 | 2.07E-06 | 7.97E-06 |
| chr2 | 65800228 | 65805341 | 5114 | 0.6 | 2.08E-06 | 8.00E-06 |
| chr3 | 197547193 | 197552030 | 4838 | -1.26 | 2.08E-06 | 8.00E-06 |
| chr15 | 53880858 | 53882506 | 1649 | -2.68 | 2.08E-06 | 8.00E-06 |

| | | | | | | |
|---|---|---|---|---|---|---|
| chr3 | 36748132 | 36749909 | 1778 | -1.23 | 2.09E-06 | 8.02E-06 |
| chr15 | 68834942 | 68836650 | 1709 | -1.67 | 2.10E-06 | 8.05E-06 |
| chr1 | 101108391 | 101110081 | 1691 | 1.37 | 2.10E-06 | 8.08E-06 |
| chr3 | 84141908 | 84142433 | 526 | 5.62 | 2.11E-06 | 8.09E-06 |
| chr9 | 98823106 | 98824246 | 1141 | -3.79 | 2.11E-06 | 8.10E-06 |
| chr3 | 110625207 | 110626990 | 1784 | 2.74 | 2.11E-06 | 8.10E-06 |
| chr22 | 42218242 | 42259508 | 41267 | 0.39 | 2.12E-06 | 8.13E-06 |
| chr12 | 122325730 | 122344656 | 18927 | 0.52 | 2.13E-06 | 8.16E-06 |
| chr1 | 199240923 | 199242578 | 1656 | 2.61 | 2.13E-06 | 8.16E-06 |
| chr8 | 50035333 | 50035764 | 432 | 5.65 | 2.14E-06 | 8.21E-06 |
| chr7 | 115805643 | 115810622 | 4980 | 1.39 | 2.14E-06 | 8.21E-06 |
| chr15 | 38233961 | 38234791 | 831 | 5.61 | 2.14E-06 | 8.21E-06 |
| chr7 | 13889491 | 13890526 | 1036 | 1.37 | 2.14E-06 | 8.21E-06 |
| chr14 | 31575111 | 31679071 | 103961 | 0.23 | 2.15E-06 | 8.24E-06 |
| chr6 | 22442994 | 22450191 | 7198 | 0.92 | 2.16E-06 | 8.27E-06 |
| chr18 | 64266272 | 64267152 | 881 | 5.67 | 2.16E-06 | 8.28E-06 |
| chr5 | 39604093 | 39608013 | 3921 | 0.8 | 2.16E-06 | 8.29E-06 |
| chr4 | 71449854 | 71450859 | 1006 | 5.62 | 2.17E-06 | 8.30E-06 |
| chr5 | 92298153 | 92307876 | 9724 | 0.69 | 2.17E-06 | 8.30E-06 |
| chr12 | 88490333 | 88508260 | 17928 | 0.79 | 2.17E-06 | 8.32E-06 |
| chr1 | 153095190 | 153095958 | 769 | 3.94 | 2.17E-06 | 8.33E-06 |
| chr15 | 31373001 | 31385792 | 12792 | 1.06 | 2.17E-06 | 8.33E-06 |
| chr6 | 125510138 | 125526253 | 16116 | 0.36 | 2.18E-06 | 8.35E-06 |
| chr3 | 13520716 | 13527375 | 6660 | 0.87 | 2.18E-06 | 8.36E-06 |
| chr10 | 77747280 | 77748334 | 1055 | -2.39 | 2.18E-06 | 8.36E-06 |
| chr14 | 103117972 | 103122794 | 4823 | 1.23 | 2.18E-06 | 8.36E-06 |
| chr3 | 30811530 | 30812723 | 1194 | 4.03 | 2.19E-06 | 8.37E-06 |
| chr6 | 46843594 | 46844736 | 1143 | 3.63 | 2.19E-06 | 8.38E-06 |
| chr6 | 157270164 | 157277403 | 7240 | 0.54 | 2.21E-06 | 8.46E-06 |
| chr15 | 25610612 | 25611731 | 1120 | 5.62 | 2.22E-06 | 8.48E-06 |
| chr3 | 175056509 | 175057670 | 1162 | 2.95 | 2.22E-06 | 8.51E-06 |
| chr8 | 82968331 | 82969145 | 815 | 4.06 | 2.23E-06 | 8.54E-06 |
| chr9 | 27565513 | 27574818 | 9306 | 0.86 | 2.23E-06 | 8.54E-06 |
| chr14 | 96043757 | 96044826 | 1070 | 5.61 | 2.24E-06 | 8.55E-06 |
| chr5 | 140165744 | 140166160 | 417 | 5.69 | 2.25E-06 | 8.59E-06 |
| chr6 | 85175553 | 85178071 | 2519 | 1.02 | 2.25E-06 | 8.59E-06 |
| chr3 | 123102924 | 123132768 | 29845 | 4.13 | 2.26E-06 | 8.65E-06 |
| chr3 | 3707659 | 3708630 | 972 | 3.65 | 2.27E-06 | 8.68E-06 |
| chr19 | 16599324 | 16608279 | 8956 | 0.67 | 2.27E-06 | 8.68E-06 |
| chr14 | 65677344 | 65725916 | 48573 | 0.54 | 2.27E-06 | 8.68E-06 |
| chr3 | 72709695 | 72713306 | 3612 | 1.46 | 2.27E-06 | 8.68E-06 |
| chr6 | 106837795 | 106838551 | 757 | 4.96 | 2.28E-06 | 8.70E-06 |
| chr8 | 110013219 | 110014385 | 1167 | 2.09 | 2.29E-06 | 8.75E-06 |
| chr6 | 2476096 | 2477597 | 1502 | 2.83 | 2.30E-06 | 8.80E-06 |
| chr12 | 39858470 | 39859553 | 1084 | 1.99 | 2.31E-06 | 8.81E-06 |
| chr17 | 14367632 | 14375168 | 7537 | 0.85 | 2.31E-06 | 8.82E-06 |
| chr11 | 47327350 | 47328682 | 1333 | -2.69 | 2.32E-06 | 8.84E-06 |
| chr11 | 86599068 | 86599813 | 746 | 5.6 | 2.32E-06 | 8.85E-06 |
| chr5 | 86678518 | 86709461 | 30944 | 0.48 | 2.32E-06 | 8.86E-06 |
| chr2 | 8020690 | 8022237 | 1548 | 1.3 | 2.32E-06 | 8.87E-06 |
| chr12 | 858332 | 938735 | 80404 | 0.29 | 2.33E-06 | 8.89E-06 |
| chr1 | 200987995 | 200992565 | 4571 | -1.13 | 2.34E-06 | 8.94E-06 |
| chr8 | 59464953 | 59467834 | 2882 | 0.53 | 2.36E-06 | 9.00E-06 |
| chr3 | 163044593 | 163045438 | 846 | 5.6 | 2.36E-06 | 9.01E-06 |
| chr2 | 128127072 | 128128180 | 1109 | -3.52 | 2.36E-06 | 9.01E-06 |
| chr2 | 165244516 | 165245725 | 1210 | 1.6 | 2.36E-06 | 9.02E-06 |
| chr5 | 126111098 | 126184364 | 73267 | 0.25 | 2.37E-06 | 9.02E-06 |
| chr19 | 11724199 | 11724707 | 509 | 4.91 | 2.37E-06 | 9.03E-06 |
| chr14 | 27432677 | 27438521 | 5845 | 5.28 | 2.38E-06 | 9.07E-06 |
| chr3 | 31959446 | 31965316 | 5871 | 1.72 | 2.38E-06 | 9.07E-06 |
| chr1 | 204419624 | 204421195 | 1572 | 1.74 | 2.38E-06 | 9.08E-06 |

| | | | | | | |
|---|---|---|---|---|---|---|
| chr2 | 75252906 | 75253727 | 822 | -5.56 | 2.39E-06 | 9.11E-06 |
| chr2 | 141353724 | 141354569 | 846 | 3.53 | 2.39E-06 | 9.12E-06 |
| chr1 | 111503291 | 111507858 | 4568 | 0.53 | 2.40E-06 | 9.15E-06 |
| chr3 | 72787678 | 72803328 | 15651 | 1.79 | 2.41E-06 | 9.16E-06 |
| chr1 | 109814211 | 109826839 | 12629 | 0.42 | 2.41E-06 | 9.17E-06 |
| chr1 | 70670476 | 70700287 | 29812 | 0.49 | 2.41E-06 | 9.17E-06 |
| chr1 | 7598353 | 7611684 | 13332 | 0.65 | 2.41E-06 | 9.18E-06 |
| chr6 | 135107856 | 135109118 | 1263 | 3.65 | 2.42E-06 | 9.20E-06 |
| chr6 | 137222084 | 137222917 | 834 | -3.68 | 2.42E-06 | 9.21E-06 |
| chr2 | 67486033 | 67488784 | 2752 | 1.35 | 2.44E-06 | 9.27E-06 |
| chr19 | 10146588 | 10149096 | 2509 | 1.39 | 2.44E-06 | 9.28E-06 |
| chr8 | 18868858 | 18872383 | 3526 | 0.72 | 2.45E-06 | 9.34E-06 |
| chr10 | 116607004 | 116617410 | 10407 | -0.82 | 2.45E-06 | 9.34E-06 |
| chr12 | 60334211 | 60335169 | 959 | 4.15 | 2.46E-06 | 9.36E-06 |
| chr17 | 66789207 | 66791754 | 2548 | 1.01 | 2.47E-06 | 9.38E-06 |
| chr1 | 197061099 | 197069322 | 8224 | 0.9 | 2.48E-06 | 9.42E-06 |
| chr8 | 129560905 | 129575829 | 14925 | -0.24 | 2.48E-06 | 9.42E-06 |
| chr5 | 103415896 | 103417143 | 1248 | 1.34 | 2.48E-06 | 9.42E-06 |
| chr6 | 87425148 | 87434047 | 8900 | 1.69 | 2.48E-06 | 9.42E-06 |
| chr18 | 21231036 | 21243092 | 12057 | -1.55 | 2.48E-06 | 9.43E-06 |
| chr11 | 19183925 | 19185650 | 1726 | 2.16 | 2.49E-06 | 9.47E-06 |
| chr6 | 157956639 | 157957711 | 1073 | 1.85 | 2.50E-06 | 9.52E-06 |
| chr6 | 5866538 | 5867669 | 1132 | 3 | 2.51E-06 | 9.53E-06 |
| chr2 | 197971659 | 197974364 | 2706 | -1.17 | 2.51E-06 | 9.53E-06 |
| chr7 | 46948032 | 46948865 | 834 | 4.47 | 2.51E-06 | 9.54E-06 |
| chr2 | 75071783 | 75095720 | 23938 | 1.58 | 2.51E-06 | 9.54E-06 |
| chr1 | 27471850 | 27490364 | 18515 | 0.62 | 2.51E-06 | 9.55E-06 |
| chr4 | 177543480 | 177560903 | 17424 | 0.55 | 2.52E-06 | 9.56E-06 |
| chr15 | 42564320 | 42568747 | 4428 | 0.48 | 2.52E-06 | 9.58E-06 |
| chr9 | 133699523 | 133739477 | 39955 | -0.37 | 2.52E-06 | 9.58E-06 |
| chr3 | 160280709 | 160285220 | 4512 | 0.47 | 2.53E-06 | 9.60E-06 |
| chrX | 70401234 | 70403008 | 1775 | 0.84 | 2.53E-06 | 9.60E-06 |
| chr4 | 169828322 | 169833747 | 5426 | -1.4 | 2.54E-06 | 9.66E-06 |
| chr3 | 174809183 | 174810006 | 824 | 5.58 | 2.55E-06 | 9.67E-06 |
| chr2 | 114563362 | 114589488 | 26127 | 0.58 | 2.55E-06 | 9.67E-06 |
| chr18 | 69827678 | 69828528 | 851 | 5.6 | 2.55E-06 | 9.69E-06 |
| chr16 | 55535954 | 55544093 | 8140 | 1.18 | 2.56E-06 | 9.69E-06 |
| chr4 | 65064873 | 65065607 | 735 | 5.58 | 2.56E-06 | 9.71E-06 |
| chr2 | 176962001 | 176981331 | 19331 | -5.55 | 2.56E-06 | 9.71E-06 |
| chr13 | 101862907 | 101864718 | 1812 | 1.03 | 2.57E-06 | 9.73E-06 |
| chr15 | 43801809 | 43804187 | 2379 | -0.88 | 2.57E-06 | 9.73E-06 |
| chr7 | 69377251 | 69378686 | 1436 | 1.41 | 2.57E-06 | 9.75E-06 |
| chr3 | 58512031 | 58523360 | 11330 | -1.19 | 2.58E-06 | 9.77E-06 |
| chr17 | 45381695 | 45384014 | 2320 | 0.9 | 2.58E-06 | 9.79E-06 |
| chr2 | 154449319 | 154450356 | 1038 | -2.19 | 2.59E-06 | 9.80E-06 |
| chr1 | 54410462 | 54431920 | 21459 | 0.55 | 2.59E-06 | 9.80E-06 |
| chr2 | 216545302 | 216578991 | 33690 | -0.52 | 2.59E-06 | 9.82E-06 |
| chr2 | 76808349 | 76809112 | 764 | 5.79 | 2.60E-06 | 9.83E-06 |
| chr12 | 76051684 | 76054531 | 2848 | 1.29 | 2.60E-06 | 9.86E-06 |
| chr10 | 70882696 | 70885908 | 3213 | 0.68 | 2.60E-06 | 9.86E-06 |
| chr1 | 55750672 | 55751881 | 1210 | -2.29 | 2.61E-06 | 9.87E-06 |
| chr19 | 2144737 | 2172417 | 27681 | 0.42 | 2.61E-06 | 9.89E-06 |
| chr1 | 21644694 | 21666958 | 22265 | 0.34 | 2.61E-06 | 9.89E-06 |
| chr14 | 28777734 | 28792446 | 14713 | 0.55 | 2.61E-06 | 9.89E-06 |
| chr8 | 142126475 | 142220511 | 94037 | 0.18 | 2.62E-06 | 9.90E-06 |
| chr4 | 104393815 | 104394370 | 556 | 5.58 | 2.62E-06 | 9.92E-06 |
| chr4 | 156699172 | 156700163 | 992 | 5.58 | 2.62E-06 | 9.92E-06 |
| chr8 | 91012373 | 91014624 | 2252 | 0.82 | 2.62E-06 | 9.92E-06 |
| chr21 | 36153331 | 36262461 | 109131 | 0.22 | 2.62E-06 | 9.93E-06 |
| chr9 | 90772197 | 90773133 | 937 | 4.6 | 2.63E-06 | 9.93E-06 |
| chr11 | 57045007 | 57046955 | 1949 | 0.71 | 2.64E-06 | 9.99E-06 |

| | | | | | | |
|---|---|---|---|---|---|---|
| chr5 | 16671446 | 16760997 | 89552 | -0.32 | 2.65E-06 | 1.00E-05 |
| chr21 | 43933416 | 43939081 | 5666 | 1.4 | 2.65E-06 | 1.00E-05 |
| chr13 | 50760480 | 50761118 | 639 | 2.84 | 2.65E-06 | 1.00E-05 |
| chr14 | 64100337 | 64110298 | 9962 | 0.51 | 2.66E-06 | 1.01E-05 |
| chr12 | 96809687 | 96821571 | 11885 | -1.21 | 2.67E-06 | 1.01E-05 |
| chr4 | 70434474 | 70435388 | 915 | 3.91 | 2.68E-06 | 1.01E-05 |
| chr11 | 130480894 | 130483670 | 2777 | 0.84 | 2.69E-06 | 1.02E-05 |
| chr2 | 102079892 | 102081151 | 1260 | -5.58 | 2.70E-06 | 1.02E-05 |
| chr3 | 170459372 | 170470717 | 11346 | -1.17 | 2.71E-06 | 1.02E-05 |
| chr11 | 6254776 | 6257379 | 2604 | 0.53 | 2.71E-06 | 1.02E-05 |
| chr5 | 172114684 | 172145368 | 30685 | 0.38 | 2.71E-06 | 1.02E-05 |
| chr5 | 86370826 | 86371739 | 914 | 2.65 | 2.71E-06 | 1.03E-05 |
| chr5 | 173090064 | 173092160 | 2097 | -2.26 | 2.72E-06 | 1.03E-05 |
| chr7 | 33290670 | 33291779 | 1110 | 1.94 | 2.73E-06 | 1.03E-05 |
| chr15 | 81270754 | 81294811 | 24058 | 0.41 | 2.74E-06 | 1.03E-05 |
| chr12 | 112080869 | 112096266 | 15398 | 0.78 | 2.74E-06 | 1.03E-05 |
| chrX | 108970836 | 108984263 | 13428 | 1.32 | 2.75E-06 | 1.04E-05 |
| chr11 | 2912734 | 3025519 | 112786 | 0.25 | 2.76E-06 | 1.04E-05 |
| chr12 | 133442430 | 133443934 | 1505 | 2.25 | 2.77E-06 | 1.04E-05 |
| chr17 | 35533415 | 35556894 | 23480 | -0.57 | 2.78E-06 | 1.05E-05 |
| chr5 | 756827 | 763151 | 6325 | 1.1 | 2.79E-06 | 1.05E-05 |
| chr7 | 142711025 | 142711865 | 841 | -5.53 | 2.79E-06 | 1.05E-05 |
| chr1 | 12405137 | 12417944 | 12808 | 0.47 | 2.79E-06 | 1.05E-05 |
| chr17 | 16184207 | 16202029 | 17823 | 0.24 | 2.80E-06 | 1.06E-05 |
| chr8 | 116626881 | 116627418 | 538 | 5.57 | 2.81E-06 | 1.06E-05 |
| chr5 | 148552666 | 148567205 | 14540 | 1.91 | 2.81E-06 | 1.06E-05 |
| chr1 | 45389718 | 45391378 | 1661 | 1.36 | 2.81E-06 | 1.06E-05 |
| chr1 | 62752119 | 62753045 | 927 | 5.64 | 2.82E-06 | 1.06E-05 |
| chr15 | 37423934 | 37424695 | 762 | 5.57 | 2.82E-06 | 1.06E-05 |
| chr6 | 43766084 | 43772733 | 6650 | 1.12 | 2.82E-06 | 1.06E-05 |
| chr16 | 58686198 | 58687243 | 1046 | 4.86 | 2.83E-06 | 1.07E-05 |
| chr3 | 189899409 | 189919791 | 20383 | -0.49 | 2.83E-06 | 1.07E-05 |
| chr5 | 93262591 | 93263408 | 818 | 1.85 | 2.85E-06 | 1.07E-05 |
| chr9 | 94986190 | 95038073 | 51884 | -0.49 | 2.86E-06 | 1.08E-05 |
| chr10 | 26624235 | 26625164 | 930 | -5.54 | 2.87E-06 | 1.08E-05 |
| chr4 | 117632210 | 117632710 | 501 | 5.59 | 2.87E-06 | 1.08E-05 |
| chr1 | 25899513 | 25901233 | 1721 | 2.17 | 2.87E-06 | 1.08E-05 |
| chr4 | 153856400 | 153857303 | 904 | 5.56 | 2.88E-06 | 1.08E-05 |
| chr1 | 45599706 | 45601363 | 1658 | 2.22 | 2.90E-06 | 1.09E-05 |
| chr7 | 106807282 | 106827459 | 20178 | 0.52 | 2.92E-06 | 1.10E-05 |
| chr6 | 155812476 | 155816924 | 4449 | 0.53 | 2.92E-06 | 1.10E-05 |
| chr16 | 24739803 | 24759726 | 19924 | -0.45 | 2.92E-06 | 1.10E-05 |
| chr8 | 99159043 | 99160177 | 1135 | -2.71 | 2.93E-06 | 1.10E-05 |
| chr2 | 158336196 | 158337440 | 1245 | -2.66 | 2.93E-06 | 1.10E-05 |
| chr8 | 81203579 | 81204446 | 868 | 4.86 | 2.93E-06 | 1.10E-05 |
| chr1 | 244435758 | 244444929 | 9172 | -0.52 | 2.93E-06 | 1.10E-05 |
| chr10 | 9051288 | 9052206 | 919 | 3.68 | 2.95E-06 | 1.11E-05 |
| chr4 | 125552088 | 125564579 | 12492 | 0.69 | 2.97E-06 | 1.12E-05 |
| chr11 | 35159900 | 35250679 | 90780 | -0.36 | 2.97E-06 | 1.12E-05 |
| chr3 | 27454223 | 27492481 | 38259 | -0.39 | 2.98E-06 | 1.12E-05 |
| chr19 | 13189264 | 13191518 | 2255 | 1.01 | 2.98E-06 | 1.12E-05 |
| chr8 | 17575850 | 17592941 | 17092 | -0.87 | 2.98E-06 | 1.12E-05 |
| chr12 | 104065876 | 104067802 | 1927 | 1.21 | 3.01E-06 | 1.13E-05 |
| chr18 | 67419855 | 67420863 | 1009 | -5.52 | 3.02E-06 | 1.13E-05 |
| chr11 | 46612278 | 46616733 | 4456 | 0.69 | 3.02E-06 | 1.13E-05 |
| chr5 | 89813838 | 89826528 | 12691 | 0.69 | 3.03E-06 | 1.14E-05 |
| chr17 | 6478154 | 6556085 | 77932 | 0.19 | 3.03E-06 | 1.14E-05 |
| chr2 | 217237174 | 217238174 | 1001 | -5.55 | 3.04E-06 | 1.14E-05 |
| chr6 | 139645916 | 139647091 | 1176 | 2.43 | 3.04E-06 | 1.14E-05 |
| chr10 | 15164881 | 15180281 | 15401 | -0.93 | 3.05E-06 | 1.14E-05 |
| chr5 | 33400461 | 33402453 | 1993 | 1.33 | 3.06E-06 | 1.15E-05 |

| | | | | | | |
|---|---|---|---|---|---|---|
| chr11 | 73539983 | 73550439 | 10457 | -2.17 | 3.06E-06 | 1.15E-05 |
| chr2 | 160456932 | 160474637 | 17706 | -0.33 | 3.07E-06 | 1.15E-05 |
| chr11 | 9737581 | 9810694 | 73114 | -0.28 | 3.07E-06 | 1.15E-05 |
| chr12 | 91251794 | 91252545 | 752 | 3.72 | 3.09E-06 | 1.16E-05 |
| chr6 | 80327200 | 80341653 | 14454 | 5.55 | 3.09E-06 | 1.16E-05 |
| chr2 | 64429174 | 64442380 | 13207 | 2.1 | 3.09E-06 | 1.16E-05 |
| chr9 | 91923677 | 91935823 | 12147 | 0.32 | 3.09E-06 | 1.16E-05 |
| chr1 | 229356365 | 229375148 | 18784 | 0.52 | 3.10E-06 | 1.16E-05 |
| chr6 | 18081360 | 18082238 | 879 | -4.4 | 3.10E-06 | 1.16E-05 |
| chr11 | 90836993 | 90838543 | 1551 | -2.26 | 3.12E-06 | 1.17E-05 |
| chr8 | 72503209 | 72504043 | 835 | -5.5 | 3.13E-06 | 1.17E-05 |
| chrX | 132810112 | 132811265 | 1154 | 5.55 | 3.13E-06 | 1.17E-05 |
| chr3 | 189766456 | 189795723 | 29268 | -0.58 | 3.13E-06 | 1.17E-05 |
| chr7 | 131033908 | 131089502 | 55595 | -0.44 | 3.13E-06 | 1.17E-05 |
| chr7 | 81658155 | 81659558 | 1404 | 1.84 | 3.14E-06 | 1.17E-05 |
| chr7 | 11475760 | 11476582 | 823 | 5.55 | 3.14E-06 | 1.18E-05 |
| chr11 | 67295793 | 67296515 | 723 | -5.5 | 3.15E-06 | 1.18E-05 |
| chr7 | 36802214 | 36803279 | 1066 | -2.19 | 3.16E-06 | 1.18E-05 |
| chr1 | 193197812 | 193199358 | 1547 | 1.35 | 3.17E-06 | 1.19E-05 |
| chr1 | 117752252 | 117780999 | 28748 | 6.79 | 3.18E-06 | 1.19E-05 |
| chr2 | 5197630 | 5211633 | 14004 | 6.08 | 3.18E-06 | 1.19E-05 |
| chr11 | 9984311 | 10013015 | 28705 | -0.36 | 3.18E-06 | 1.19E-05 |
| chr2 | 175862820 | 175871063 | 8244 | -0.95 | 3.19E-06 | 1.19E-05 |
| chr6 | 83627241 | 83628946 | 1706 | 1.74 | 3.19E-06 | 1.19E-05 |
| chr4 | 185025228 | 185027490 | 2263 | 1.27 | 3.19E-06 | 1.19E-05 |
| chrX | 35515188 | 35516319 | 1132 | 2.13 | 3.20E-06 | 1.20E-05 |
| chr2 | 109491194 | 109492376 | 1183 | 2.19 | 3.20E-06 | 1.20E-05 |
| chr1 | 198094838 | 198102168 | 7331 | 2.95 | 3.21E-06 | 1.20E-05 |
| chr16 | 49682615 | 49683453 | 839 | 5.58 | 3.22E-06 | 1.20E-05 |
| chr1 | 218793973 | 218798172 | 4200 | 1.16 | 3.22E-06 | 1.21E-05 |
| chrX | 15317698 | 15329447 | 11750 | 1.26 | 3.25E-06 | 1.21E-05 |
| chr16 | 60009614 | 60010743 | 1130 | 1.26 | 3.26E-06 | 1.22E-05 |
| chr12 | 15305548 | 15306515 | 968 | 2.91 | 3.27E-06 | 1.22E-05 |
| chr8 | 52781745 | 52784630 | 2886 | 1.15 | 3.27E-06 | 1.22E-05 |
| chr7 | 154996510 | 155001445 | 4936 | 0.74 | 3.28E-06 | 1.22E-05 |
| chr12 | 46972462 | 46980359 | 7898 | 1.59 | 3.28E-06 | 1.22E-05 |
| chr2 | 214103347 | 214117577 | 14231 | -0.74 | 3.28E-06 | 1.23E-05 |
| chr18 | 28880492 | 28881977 | 1486 | 3.56 | 3.28E-06 | 1.23E-05 |
| chr13 | 32539220 | 32539974 | 755 | 5.56 | 3.29E-06 | 1.23E-05 |
| chr4 | 39402677 | 39415965 | 13289 | -1.07 | 3.30E-06 | 1.23E-05 |
| chr19 | 34797090 | 34809409 | 12320 | -1.55 | 3.30E-06 | 1.23E-05 |
| chr4 | 66808009 | 66809369 | 1361 | 2.63 | 3.31E-06 | 1.23E-05 |
| chr9 | 124737561 | 124738944 | 1384 | 2.23 | 3.32E-06 | 1.24E-05 |
| chr10 | 90599820 | 90612040 | 12221 | 5.59 | 3.33E-06 | 1.24E-05 |
| chr2 | 5159019 | 5160015 | 997 | 3.68 | 3.33E-06 | 1.24E-05 |
| chr12 | 65549943 | 65564401 | 14459 | 1.25 | 3.35E-06 | 1.25E-05 |
| chr7 | 83816320 | 83831041 | 14722 | -0.51 | 3.36E-06 | 1.25E-05 |
| chr1 | 26184908 | 26202717 | 17810 | 0.73 | 3.37E-06 | 1.26E-05 |
| chr16 | 50653491 | 50654470 | 980 | -3.3 | 3.37E-06 | 1.26E-05 |
| chr9 | 95491081 | 95528466 | 37386 | -0.52 | 3.37E-06 | 1.26E-05 |
| chr1 | 88334683 | 88335885 | 1203 | 1.78 | 3.37E-06 | 1.26E-05 |
| chr8 | 61368038 | 61380932 | 12895 | -5.54 | 3.38E-06 | 1.26E-05 |
| chr18 | 67704213 | 67716019 | 11807 | -1.28 | 3.38E-06 | 1.26E-05 |
| chrX | 71838561 | 71844201 | 5641 | 0.52 | 3.39E-06 | 1.26E-05 |
| chr4 | 11429743 | 11432061 | 2319 | 1.03 | 3.39E-06 | 1.26E-05 |
| chr1 | 186586862 | 186588939 | 2078 | 0.72 | 3.40E-06 | 1.27E-05 |
| chr5 | 79541591 | 79553665 | 12075 | 0.62 | 3.40E-06 | 1.27E-05 |
| chr3 | 115778220 | 115779445 | 1226 | 1.97 | 3.40E-06 | 1.27E-05 |
| chr17 | 59910212 | 59914983 | 4772 | 0.94 | 3.41E-06 | 1.27E-05 |
| chr10 | 9279821 | 9280785 | 965 | 5.53 | 3.41E-06 | 1.27E-05 |
| chr13 | 91795180 | 91795720 | 541 | 5.53 | 3.41E-06 | 1.27E-05 |

| | | | | | | |
|---|---|---|---|---|---|---|
| chr6 | 85558515 | 85559197 | 683 | 5.53 | 3.41E-06 | 1.27E-05 |
| chr1 | 97514590 | 97515441 | 852 | 5.58 | 3.41E-06 | 1.27E-05 |
| chr13 | 96216634 | 96217839 | 1206 | 3.59 | 3.42E-06 | 1.27E-05 |
| chr14 | 50283376 | 50298602 | 15227 | 0.88 | 3.43E-06 | 1.28E-05 |
| chr3 | 70187435 | 70188679 | 1245 | 3.26 | 3.44E-06 | 1.28E-05 |
| chr1 | 105454265 | 105455123 | 859 | 5.53 | 3.44E-06 | 1.28E-05 |
| chr2 | 6221629 | 6222227 | 599 | 5.53 | 3.44E-06 | 1.28E-05 |
| chr20 | 5473414 | 5474505 | 1092 | -4.89 | 3.45E-06 | 1.28E-05 |
| chr2 | 109061368 | 109069616 | 8249 | -0.5 | 3.47E-06 | 1.29E-05 |
| chr1 | 110452708 | 110548450 | 95743 | -0.23 | 3.47E-06 | 1.29E-05 |
| chr6 | 84902118 | 84903909 | 1792 | 2.47 | 3.48E-06 | 1.30E-05 |
| chr18 | 28567380 | 28568196 | 817 | 5.55 | 3.49E-06 | 1.30E-05 |
| chr19 | 47791113 | 47794913 | 3801 | 0.55 | 3.49E-06 | 1.30E-05 |
| chr8 | 66893830 | 66895068 | 1239 | 2.43 | 3.49E-06 | 1.30E-05 |
| chr5 | 139621021 | 139642337 | 21317 | 0.64 | 3.51E-06 | 1.31E-05 |
| chr3 | 163078552 | 163079939 | 1388 | 1.38 | 3.51E-06 | 1.31E-05 |
| chr1 | 31000403 | 31010904 | 10502 | -5.51 | 3.52E-06 | 1.31E-05 |
| chr3 | 154528988 | 154530046 | 1059 | -5.51 | 3.52E-06 | 1.31E-05 |
| chr3 | 111757199 | 111758717 | 1519 | 1.85 | 3.53E-06 | 1.31E-05 |
| chr11 | 41398944 | 41399770 | 827 | 5.61 | 3.54E-06 | 1.31E-05 |
| chr6 | 149455140 | 149456268 | 1129 | 5.61 | 3.54E-06 | 1.31E-05 |
| chr7 | 12105270 | 12107113 | 1844 | 2.14 | 3.54E-06 | 1.31E-05 |
| chr13 | 99216502 | 99230578 | 14077 | 0.5 | 3.55E-06 | 1.32E-05 |
| chr7 | 849953 | 876474 | 26522 | 0.46 | 3.55E-06 | 1.32E-05 |
| chr22 | 39319142 | 39323937 | 4796 | 0.93 | 3.55E-06 | 1.32E-05 |
| chr4 | 168557423 | 168558407 | 985 | 2.57 | 3.56E-06 | 1.32E-05 |
| chr10 | 30129496 | 30132132 | 2637 | 1 | 3.57E-06 | 1.32E-05 |
| chr7 | 56516283 | 56516850 | 568 | 5.57 | 3.57E-06 | 1.32E-05 |
| chr3 | 33463379 | 33484193 | 20815 | 0.4 | 3.57E-06 | 1.33E-05 |
| chr14 | 77768921 | 77789150 | 20230 | 0.35 | 3.58E-06 | 1.33E-05 |
| chr1 | 171341942 | 171342899 | 958 | 1.29 | 3.59E-06 | 1.33E-05 |
| chrX | 27393397 | 27393903 | 507 | 4.86 | 3.59E-06 | 1.33E-05 |
| chr3 | 149676910 | 149690280 | 13371 | -0.34 | 3.59E-06 | 1.33E-05 |
| chr10 | 29429169 | 29449983 | 20815 | -0.87 | 3.59E-06 | 1.33E-05 |
| chr2 | 43839863 | 43840337 | 475 | -4.09 | 3.60E-06 | 1.33E-05 |
| chr3 | 50654592 | 50665719 | 11128 | -1.33 | 3.61E-06 | 1.34E-05 |
| chr3 | 123185600 | 123186483 | 884 | 4.87 | 3.62E-06 | 1.34E-05 |
| chrX | 44606796 | 44607657 | 862 | 4.83 | 3.62E-06 | 1.34E-05 |
| chr13 | 34270088 | 34272739 | 2652 | -1.17 | 3.63E-06 | 1.34E-05 |
| chr20 | 39619011 | 39637655 | 18645 | 0.57 | 3.63E-06 | 1.34E-05 |
| chr4 | 102332870 | 102334113 | 1244 | 1.28 | 3.63E-06 | 1.34E-05 |
| chr12 | 120728012 | 120741095 | 13084 | 0.4 | 3.64E-06 | 1.35E-05 |
| chrX | 153378808 | 153380343 | 1536 | 1.93 | 3.64E-06 | 1.35E-05 |
| chr7 | 117033563 | 117034109 | 547 | 5.53 | 3.64E-06 | 1.35E-05 |
| chr4 | 115099074 | 115099866 | 793 | 5.55 | 3.65E-06 | 1.35E-05 |
| chr18 | 21655500 | 21656946 | 1447 | -2.42 | 3.66E-06 | 1.35E-05 |
| chr10 | 38077631 | 38079937 | 2307 | 5.82 | 3.66E-06 | 1.36E-05 |
| chr2 | 70160523 | 70175004 | 14482 | -0.73 | 3.67E-06 | 1.36E-05 |
| chr8 | 120981571 | 121014689 | 33119 | 0.69 | 3.68E-06 | 1.36E-05 |
| chr19 | 51977791 | 51979032 | 1242 | 5.53 | 3.69E-06 | 1.37E-05 |
| chr5 | 32709648 | 32715655 | 6008 | -5.5 | 3.70E-06 | 1.37E-05 |
| chr10 | 97256276 | 97256993 | 718 | -5.48 | 3.71E-06 | 1.37E-05 |
| chr8 | 120706384 | 120707482 | 1099 | 2.84 | 3.72E-06 | 1.37E-05 |
| chr2 | 104577352 | 104578211 | 860 | 5.54 | 3.72E-06 | 1.38E-05 |
| chr9 | 89968209 | 89969518 | 1310 | 2.07 | 3.73E-06 | 1.38E-05 |
| chr7 | 131579629 | 131582127 | 2499 | -1.33 | 3.74E-06 | 1.38E-05 |
| chr6 | 83774769 | 83781364 | 6596 | 1.13 | 3.74E-06 | 1.38E-05 |
| chr11 | 119056203 | 119056761 | 559 | 4.58 | 3.76E-06 | 1.39E-05 |
| chr1 | 153894323 | 153897083 | 2761 | 0.81 | 3.76E-06 | 1.39E-05 |
| chr19 | 53798070 | 53799044 | 975 | -5.5 | 3.77E-06 | 1.39E-05 |
| chr8 | 86131688 | 86133827 | 2140 | 1.09 | 3.78E-06 | 1.40E-05 |

| | | | | | | |
|---|---|---|---|---|---|---|
| chr6 | 38019736 | 38020711 | 976 | 5.57 | 3.79E-06 | 1.40E-05 |
| chr10 | 131329233 | 131330944 | 1712 | 1.95 | 3.79E-06 | 1.40E-05 |
| chr5 | 56467582 | 56547774 | 80193 | 0.29 | 3.80E-06 | 1.40E-05 |
| chr4 | 92788583 | 92789283 | 701 | 3.99 | 3.80E-06 | 1.40E-05 |
| chr2 | 68144491 | 68145462 | 972 | -1.96 | 3.80E-06 | 1.40E-05 |
| chr21 | 16272500 | 16273202 | 703 | -5.47 | 3.80E-06 | 1.40E-05 |
| chr10 | 13236383 | 13249872 | 13490 | -0.91 | 3.81E-06 | 1.41E-05 |
| chr2 | 31155500 | 31156565 | 1066 | -2.87 | 3.82E-06 | 1.41E-05 |
| chr2 | 165416946 | 165426254 | 9309 | 0.69 | 3.82E-06 | 1.41E-05 |
| chr1 | 242382488 | 242407492 | 25005 | 0.35 | 3.83E-06 | 1.41E-05 |
| chr2 | 172777899 | 172783306 | 5408 | -0.52 | 3.84E-06 | 1.42E-05 |
| chr3 | 87038696 | 87046130 | 7435 | 0.4 | 3.85E-06 | 1.42E-05 |
| chr6 | 26019853 | 26071887 | 52035 | -0.23 | 3.85E-06 | 1.42E-05 |
| chr7 | 62246443 | 62247368 | 926 | 4.42 | 3.86E-06 | 1.42E-05 |
| chr1 | 161695585 | 161697782 | 2198 | -1.02 | 3.86E-06 | 1.42E-05 |
| chr5 | 150529513 | 150537926 | 8414 | -1.47 | 3.86E-06 | 1.42E-05 |
| chr1 | 211548376 | 211557393 | 9018 | 0.52 | 3.87E-06 | 1.43E-05 |
| chr8 | 56979916 | 56987805 | 7890 | 0.56 | 3.88E-06 | 1.43E-05 |
| chr12 | 133069634 | 133083526 | 13893 | -1.18 | 3.89E-06 | 1.44E-05 |
| chr10 | 3817519 | 3829218 | 11700 | 0.31 | 3.91E-06 | 1.44E-05 |
| chr17 | 3376534 | 3378962 | 2429 | 1.14 | 3.91E-06 | 1.44E-05 |
| chr1 | 209161548 | 209162656 | 1109 | 1.96 | 3.93E-06 | 1.45E-05 |
| chr9 | 11100408 | 11101448 | 1041 | -3.43 | 3.93E-06 | 1.45E-05 |
| chr18 | 23651255 | 23654990 | 3736 | 1.45 | 3.95E-06 | 1.46E-05 |
| chr13 | 38230951 | 38231505 | 555 | -5.5 | 3.95E-06 | 1.46E-05 |
| chrX | 46219645 | 46220229 | 585 | 3.08 | 3.96E-06 | 1.46E-05 |
| chr19 | 13358846 | 13359412 | 567 | 2.22 | 3.97E-06 | 1.46E-05 |
| chr10 | 104722734 | 104726077 | 3344 | -1.44 | 3.97E-06 | 1.46E-05 |
| chr5 | 93325118 | 93326836 | 1719 | 2.49 | 3.98E-06 | 1.46E-05 |
| chr18 | 29252670 | 29266658 | 13989 | 0.6 | 4.02E-06 | 1.48E-05 |
| chr4 | 178648797 | 178650156 | 1360 | 2.18 | 4.03E-06 | 1.48E-05 |
| chr13 | 89714142 | 89714875 | 734 | 4.22 | 4.03E-06 | 1.48E-05 |
| chr6 | 119117055 | 119117847 | 793 | 3.01 | 4.05E-06 | 1.49E-05 |
| chr2 | 52208827 | 52209833 | 1007 | -2.61 | 4.06E-06 | 1.49E-05 |
| chr22 | 31685872 | 31687517 | 1646 | 1.51 | 4.08E-06 | 1.50E-05 |
| chr5 | 79702596 | 79706462 | 3867 | 0.52 | 4.09E-06 | 1.50E-05 |
| chr16 | 1427080 | 1430425 | 3346 | -0.66 | 4.10E-06 | 1.51E-05 |
| chr8 | 54999837 | 55015231 | 15395 | 0.64 | 4.10E-06 | 1.51E-05 |
| chr4 | 3437807 | 3442071 | 4265 | -0.75 | 4.11E-06 | 1.51E-05 |
| chr4 | 162325191 | 162326133 | 943 | 5.61 | 4.11E-06 | 1.51E-05 |
| chr5 | 64113834 | 64134916 | 21083 | 1.5 | 4.12E-06 | 1.52E-05 |
| chr8 | 28585949 | 28586772 | 824 | -4.12 | 4.12E-06 | 1.52E-05 |
| chr1 | 196526988 | 196528631 | 1644 | 1.16 | 4.15E-06 | 1.52E-05 |
| chr5 | 83821663 | 83822057 | 395 | 5.5 | 4.15E-06 | 1.52E-05 |
| chr13 | 83439968 | 83440522 | 555 | 5.51 | 4.15E-06 | 1.52E-05 |
| chr4 | 149318251 | 149319162 | 912 | 5.5 | 4.16E-06 | 1.53E-05 |
| chr2 | 118900510 | 118901147 | 638 | 2.16 | 4.16E-06 | 1.53E-05 |
| chr17 | 35439561 | 35531784 | 92224 | -0.28 | 4.17E-06 | 1.53E-05 |
| chr2 | 169561642 | 169563451 | 1810 | 1.64 | 4.17E-06 | 1.53E-05 |
| chr17 | 68657338 | 68658038 | 701 | 5.5 | 4.17E-06 | 1.53E-05 |
| chr3 | 83364242 | 83365109 | 868 | 5.5 | 4.17E-06 | 1.53E-05 |
| chr1 | 224564996 | 224624015 | 59020 | 0.34 | 4.18E-06 | 1.53E-05 |
| chr8 | 123256169 | 123256699 | 531 | 3.77 | 4.19E-06 | 1.54E-05 |
| chr4 | 42413966 | 42414950 | 985 | 5.5 | 4.19E-06 | 1.54E-05 |
| chr21 | 27335095 | 27358841 | 23747 | 0.55 | 4.22E-06 | 1.55E-05 |
| chr8 | 39888777 | 39919967 | 31191 | -0.3 | 4.22E-06 | 1.55E-05 |
| chr6 | 74018428 | 74020363 | 1936 | 0.77 | 4.22E-06 | 1.55E-05 |
| chr6 | 86320249 | 86355623 | 35375 | 0.36 | 4.24E-06 | 1.55E-05 |
| chr4 | 145483198 | 145486086 | 2889 | 3.68 | 4.24E-06 | 1.56E-05 |
| chr8 | 96145262 | 96158460 | 13199 | 1.11 | 4.25E-06 | 1.56E-05 |
| chr4 | 64944924 | 64945857 | 934 | 5.52 | 4.25E-06 | 1.56E-05 |

| | | | | | | |
|---|---|---|---|---|---|---|
| chr10 | 16420359 | 16425201 | 4843 | 0.6 | 4.25E-06 | 1.56E-05 |
| chr8 | 56882650 | 56883424 | 775 | 3.97 | 4.26E-06 | 1.56E-05 |
| chr12 | 100937620 | 100949149 | 11530 | 4.47 | 4.27E-06 | 1.56E-05 |
| chr2 | 233582331 | 233584288 | 1958 | -1.68 | 4.27E-06 | 1.57E-05 |
| chr5 | 171774022 | 171787725 | 13704 | -1.1 | 4.28E-06 | 1.57E-05 |
| chr17 | 49823323 | 49823951 | 629 | -5.46 | 4.28E-06 | 1.57E-05 |
| chr10 | 93504611 | 93509609 | 4999 | 1.22 | 4.29E-06 | 1.57E-05 |
| chr1 | 25493575 | 25495554 | 1980 | 1.99 | 4.29E-06 | 1.57E-05 |
| chr14 | 51386918 | 51432084 | 45167 | -0.35 | 4.30E-06 | 1.58E-05 |
| chr14 | 80175344 | 80175951 | 608 | -5.46 | 4.31E-06 | 1.58E-05 |
| chr15 | 96014308 | 96015041 | 734 | 3.84 | 4.32E-06 | 1.58E-05 |
| chr2 | 154842144 | 154844197 | 2054 | -1.01 | 4.34E-06 | 1.59E-05 |
| chr6 | 21799939 | 21808994 | 9056 | 0.55 | 4.35E-06 | 1.59E-05 |
| chr4 | 153352044 | 153353442 | 1399 | 1.4 | 4.35E-06 | 1.59E-05 |
| chr19 | 36118517 | 36164798 | 46282 | 0.37 | 4.35E-06 | 1.59E-05 |
| chr8 | 26047366 | 26062924 | 15559 | 5.67 | 4.35E-06 | 1.59E-05 |
| chr12 | 93707421 | 93710020 | 2600 | 0.62 | 4.36E-06 | 1.59E-05 |
| chr13 | 30359559 | 30361189 | 1631 | 2.23 | 4.36E-06 | 1.59E-05 |
| chr13 | 99697914 | 99721678 | 23765 | -0.55 | 4.36E-06 | 1.60E-05 |
| chrX | 3180593 | 3182325 | 1733 | 2.16 | 4.37E-06 | 1.60E-05 |
| chr14 | 27335906 | 27337255 | 1350 | 1.54 | 4.38E-06 | 1.60E-05 |
| chr7 | 17315441 | 17316890 | 1450 | 1.4 | 4.38E-06 | 1.60E-05 |
| chr1 | 229038589 | 229039676 | 1088 | 2.53 | 4.41E-06 | 1.61E-05 |
| chr9 | 137691135 | 137692225 | 1091 | -3.81 | 4.41E-06 | 1.61E-05 |
| chr19 | 39034228 | 39036546 | 2319 | 0.63 | 4.44E-06 | 1.62E-05 |
| chr3 | 39359355 | 39359843 | 489 | 5.5 | 4.44E-06 | 1.62E-05 |
| chr11 | 8739023 | 8741536 | 2514 | -1.49 | 4.44E-06 | 1.62E-05 |
| chr5 | 89703782 | 89707114 | 3333 | 0.55 | 4.45E-06 | 1.62E-05 |
| chr6 | 85845067 | 85846054 | 988 | 5.52 | 4.45E-06 | 1.63E-05 |
| chr13 | 53224709 | 53240302 | 15594 | 0.46 | 4.47E-06 | 1.63E-05 |
| chr11 | 32861952 | 32862566 | 615 | -5.47 | 4.47E-06 | 1.63E-05 |
| chr11 | 29189848 | 29196721 | 6874 | -1.7 | 4.47E-06 | 1.63E-05 |
| chr6 | 99938362 | 99939531 | 1170 | 2.46 | 4.50E-06 | 1.64E-05 |
| chr1 | 186051088 | 186051767 | 680 | 5.49 | 4.50E-06 | 1.64E-05 |
| chr8 | 112365038 | 112365839 | 802 | 5.49 | 4.50E-06 | 1.64E-05 |
| chr8 | 57232334 | 57236228 | 3895 | 1.28 | 4.53E-06 | 1.65E-05 |
| chr14 | 21457106 | 21459480 | 2375 | 0.72 | 4.53E-06 | 1.65E-05 |
| chr12 | 65152546 | 65154580 | 2035 | 1.04 | 4.53E-06 | 1.65E-05 |
| chr5 | 159480958 | 159492775 | 11818 | -1.01 | 4.53E-06 | 1.65E-05 |
| chr5 | 176512673 | 176522150 | 9478 | 0.88 | 4.54E-06 | 1.66E-05 |
| chr15 | 44083496 | 44120813 | 37318 | 0.35 | 4.54E-06 | 1.66E-05 |
| chr5 | 158302171 | 158302953 | 783 | 2.26 | 4.55E-06 | 1.66E-05 |
| chr11 | 94269339 | 94278090 | 8752 | 0.83 | 4.56E-06 | 1.66E-05 |
| chr4 | 103496212 | 103497754 | 1543 | 1.33 | 4.56E-06 | 1.66E-05 |
| chr6 | 168250257 | 168251439 | 1183 | 1.57 | 4.57E-06 | 1.67E-05 |
| chr10 | 95972768 | 95975307 | 2540 | -3.57 | 4.57E-06 | 1.67E-05 |
| chr20 | 62465645 | 62497825 | 32181 | 0.49 | 4.58E-06 | 1.67E-05 |
| chr3 | 186925698 | 186926720 | 1023 | 5.48 | 4.58E-06 | 1.67E-05 |
| chr8 | 72899566 | 72900469 | 904 | -2.26 | 4.60E-06 | 1.68E-05 |
| chr1 | 51701169 | 51704233 | 3065 | 0.94 | 4.60E-06 | 1.68E-05 |
| chr17 | 49092397 | 49108309 | 15913 | 0.64 | 4.60E-06 | 1.68E-05 |
| chr12 | 70245517 | 70246126 | 610 | 5.48 | 4.61E-06 | 1.68E-05 |
| chr15 | 99189570 | 99367158 | 177589 | 0.16 | 4.62E-06 | 1.68E-05 |
| chr6 | 28456611 | 28458750 | 2140 | 0.82 | 4.63E-06 | 1.69E-05 |
| chr11 | 4627517 | 4630193 | 2677 | 1 | 4.64E-06 | 1.69E-05 |
| chr5 | 166336817 | 166337668 | 852 | 3.99 | 4.64E-06 | 1.69E-05 |
| chr2 | 73113711 | 73121628 | 7918 | -0.98 | 4.64E-06 | 1.69E-05 |
| chr16 | 15521512 | 15614360 | 92849 | -0.26 | 4.65E-06 | 1.69E-05 |
| chr20 | 52441878 | 52490076 | 48199 | 0.23 | 4.67E-06 | 1.70E-05 |
| chrX | 37904222 | 37905406 | 1185 | 2.68 | 4.69E-06 | 1.71E-05 |
| chr6 | 46916949 | 46918378 | 1430 | 3.56 | 4.71E-06 | 1.71E-05 |

| | | | | | | |
|---|---|---|---|---|---|---|
| chr10 | 35506841 | 35507830 | 990 | -3.56 | 4.71E-06 | 1.72E-05 |
| chr15 | 58520261 | 58521339 | 1079 | -2.58 | 4.73E-06 | 1.72E-05 |
| chrX | 48931133 | 48938653 | 7521 | 0.61 | 4.74E-06 | 1.72E-05 |
| chr16 | 52529723 | 52532267 | 2545 | 5.41 | 4.74E-06 | 1.73E-05 |
| chr19 | 1094387 | 1133395 | 39009 | 0.34 | 4.75E-06 | 1.73E-05 |
| chr5 | 9775106 | 9775899 | 794 | -5.45 | 4.76E-06 | 1.73E-05 |
| chr3 | 64635849 | 64636452 | 604 | 5.54 | 4.78E-06 | 1.74E-05 |
| chr10 | 96303258 | 96356254 | 52997 | 0.35 | 4.78E-06 | 1.74E-05 |
| chr4 | 76787650 | 76789307 | 1658 | 1.46 | 4.80E-06 | 1.74E-05 |
| chr21 | 43346055 | 43352857 | 6803 | 1.07 | 4.81E-06 | 1.75E-05 |
| chr12 | 93591611 | 93592468 | 858 | 3.71 | 4.81E-06 | 1.75E-05 |
| chr17 | 64560186 | 64612034 | 51849 | 0.37 | 4.81E-06 | 1.75E-05 |
| chr4 | 125042234 | 125043074 | 841 | 5.48 | 4.84E-06 | 1.76E-05 |
| chr1 | 200113948 | 200123819 | 9872 | 0.75 | 4.84E-06 | 1.76E-05 |
| chr7 | 26902896 | 26905447 | 2552 | 0.68 | 4.84E-06 | 1.76E-05 |
| chr1 | 88393159 | 88422864 | 29706 | 0.73 | 4.85E-06 | 1.76E-05 |
| chr7 | 44457033 | 44471901 | 14869 | -1.36 | 4.86E-06 | 1.76E-05 |
| chr8 | 54754680 | 54756961 | 2282 | 0.85 | 4.87E-06 | 1.77E-05 |
| chr12 | 65824366 | 65825562 | 1197 | 1.56 | 4.88E-06 | 1.77E-05 |
| chr8 | 109218617 | 109261819 | 43203 | 0.31 | 4.91E-06 | 1.78E-05 |
| chr3 | 118143755 | 118144496 | 742 | -3.77 | 4.91E-06 | 1.78E-05 |
| chr1 | 171992001 | 171993306 | 1306 | 2.25 | 4.92E-06 | 1.78E-05 |
| chr15 | 63829260 | 63904471 | 75212 | -0.32 | 4.92E-06 | 1.78E-05 |
| chr5 | 157222004 | 157231149 | 9146 | 1.18 | 4.94E-06 | 1.79E-05 |
| chr3 | 122511998 | 122514905 | 2908 | 0.65 | 4.96E-06 | 1.80E-05 |
| chr3 | 81580217 | 81581262 | 1046 | -3.45 | 4.96E-06 | 1.80E-05 |
| chr4 | 21658056 | 21659139 | 1084 | 5.52 | 4.96E-06 | 1.80E-05 |
| chr5 | 65266360 | 65280406 | 14047 | -0.81 | 4.97E-06 | 1.80E-05 |
| chrX | 62412048 | 62412958 | 911 | 4.38 | 4.97E-06 | 1.80E-05 |
| chr15 | 81097121 | 81098189 | 1069 | 2.78 | 4.99E-06 | 1.81E-05 |
| chr5 | 17444404 | 17445978 | 1575 | 2.02 | 5.02E-06 | 1.82E-05 |
| chr2 | 44356987 | 44363573 | 6587 | -0.77 | 5.02E-06 | 1.82E-05 |
| chr9 | 131375540 | 131471264 | 95725 | 0.2 | 5.02E-06 | 1.82E-05 |
| chr1 | 115710279 | 115734275 | 23997 | -0.63 | 5.03E-06 | 1.82E-05 |
| chr7 | 114619288 | 114650059 | 30772 | 0.5 | 5.04E-06 | 1.83E-05 |
| chr2 | 236979872 | 236998086 | 18215 | -1.19 | 5.04E-06 | 1.83E-05 |
| chr5 | 124510625 | 124512618 | 1994 | 1.69 | 5.04E-06 | 1.83E-05 |
| chr15 | 50115387 | 50125411 | 10025 | 0.73 | 5.05E-06 | 1.83E-05 |
| chr12 | 58922505 | 58927779 | 5275 | 1.31 | 5.06E-06 | 1.83E-05 |
| chr14 | 43944754 | 43945402 | 649 | 5.46 | 5.07E-06 | 1.83E-05 |
| chr9 | 98841851 | 98850873 | 9023 | -0.81 | 5.07E-06 | 1.84E-05 |
| chr3 | 116576941 | 116577604 | 664 | 5.48 | 5.07E-06 | 1.84E-05 |
| chr2 | 216480225 | 216486137 | 5913 | -0.71 | 5.10E-06 | 1.85E-05 |
| chr5 | 133070617 | 133074295 | 3679 | -1.25 | 5.10E-06 | 1.85E-05 |
| chr15 | 62379734 | 62380581 | 848 | 3.8 | 5.11E-06 | 1.85E-05 |
| chr1 | 227993427 | 227994156 | 730 | 5.51 | 5.11E-06 | 1.85E-05 |
| chr12 | 19374254 | 19376478 | 2225 | 1.22 | 5.15E-06 | 1.86E-05 |
| chr7 | 101831273 | 101852757 | 21485 | -0.61 | 5.18E-06 | 1.87E-05 |
| chr5 | 142922422 | 142980925 | 58504 | 0.27 | 5.20E-06 | 1.88E-05 |
| chr9 | 94262959 | 94267066 | 4108 | -1.74 | 5.22E-06 | 1.89E-05 |
| chr14 | 34889794 | 34890939 | 1146 | 3.34 | 5.23E-06 | 1.89E-05 |
| chr12 | 70082002 | 70083743 | 1742 | 0.63 | 5.23E-06 | 1.89E-05 |
| chr10 | 24672806 | 24673408 | 603 | 1.63 | 5.23E-06 | 1.89E-05 |
| chr1 | 42419959 | 42422010 | 2052 | 1.19 | 5.24E-06 | 1.89E-05 |
| chr1 | 232714037 | 232715936 | 1900 | 1.66 | 5.27E-06 | 1.90E-05 |
| chr16 | 22625413 | 22626318 | 906 | 5.46 | 5.28E-06 | 1.91E-05 |
| chr20 | 9597990 | 9607893 | 9904 | 5.56 | 5.28E-06 | 1.91E-05 |
| chr11 | 105946674 | 105962916 | 16243 | 0.56 | 5.30E-06 | 1.91E-05 |
| chr21 | 35050278 | 35058335 | 8058 | 0.97 | 5.30E-06 | 1.92E-05 |
| chrX | 19352648 | 19363374 | 10727 | 0.96 | 5.31E-06 | 1.92E-05 |
| chr5 | 31148768 | 31162623 | 13856 | 5.82 | 5.33E-06 | 1.92E-05 |

| | | | | | | |
|---|---|---|---|---|---|---|
| chr1 | 15478598 | 15547488 | 68891 | -0.24 | 5.33E-06 | 1.92E-05 |
| chr12 | 1798943 | 1816997 | 18055 | 0.58 | 5.35E-06 | 1.93E-05 |
| chr16 | 57125359 | 57146617 | 21259 | -0.79 | 5.36E-06 | 1.93E-05 |
| chr2 | 3469251 | 3479195 | 9945 | -1.19 | 5.37E-06 | 1.94E-05 |
| chr3 | 113774092 | 113791007 | 16916 | -0.59 | 5.37E-06 | 1.94E-05 |
| chr15 | 59700144 | 59705559 | 5416 | 0.48 | 5.38E-06 | 1.94E-05 |
| chr4 | 96329797 | 96337301 | 7505 | 1.45 | 5.39E-06 | 1.94E-05 |
| chr9 | 35646204 | 35673834 | 27631 | 0.37 | 5.39E-06 | 1.94E-05 |
| chr19 | 33752152 | 33794478 | 42327 | -0.44 | 5.40E-06 | 1.95E-05 |
| chr12 | 87444647 | 87445852 | 1206 | 3.54 | 5.40E-06 | 1.95E-05 |
| chr1 | 244593925 | 244626096 | 32172 | 0.4 | 5.40E-06 | 1.95E-05 |
| chr17 | 64534923 | 64546939 | 12017 | 0.31 | 5.40E-06 | 1.95E-05 |
| chr5 | 133837332 | 133929171 | 91840 | 0.2 | 5.41E-06 | 1.95E-05 |
| chr1 | 86283617 | 86297209 | 13593 | 1.36 | 5.42E-06 | 1.95E-05 |
| chr14 | 50908071 | 50909336 | 1266 | 1.95 | 5.43E-06 | 1.96E-05 |
| chr20 | 33369374 | 33386640 | 17267 | -1.09 | 5.43E-06 | 1.96E-05 |
| chr17 | 42516193 | 42525376 | 9184 | -0.79 | 5.47E-06 | 1.97E-05 |
| chr8 | 23608142 | 23619922 | 11781 | -1.34 | 5.47E-06 | 1.97E-05 |
| chr19 | 41337104 | 41338635 | 1532 | 1.77 | 5.48E-06 | 1.98E-05 |
| chr5 | 55903407 | 55912454 | 9048 | 5.49 | 5.49E-06 | 1.98E-05 |
| chr10 | 43038946 | 43048968 | 10023 | 1.51 | 5.49E-06 | 1.98E-05 |
| chr12 | 10826216 | 10827337 | 1122 | 2.54 | 5.50E-06 | 1.98E-05 |
| chr14 | 58181003 | 58181879 | 877 | 5.46 | 5.51E-06 | 1.99E-05 |
| chr14 | 21552456 | 21575729 | 23274 | -0.45 | 5.52E-06 | 1.99E-05 |
| chr15 | 43731354 | 43735371 | 4018 | -0.59 | 5.53E-06 | 1.99E-05 |
| chr18 | 20678341 | 20740468 | 62128 | 0.27 | 5.54E-06 | 1.99E-05 |
| chr14 | 36193226 | 36198341 | 5116 | 1.03 | 5.55E-06 | 2.00E-05 |
| chr14 | 81684752 | 81688769 | 4018 | 0.4 | 5.56E-06 | 2.00E-05 |
| chr2 | 4067426 | 4068452 | 1027 | -2.26 | 5.56E-06 | 2.00E-05 |
| chr16 | 71517392 | 71526484 | 9093 | 0.77 | 5.58E-06 | 2.01E-05 |
| chr6 | 7416938 | 7421042 | 4105 | -0.73 | 5.60E-06 | 2.01E-05 |
| chr2 | 155051060 | 155052598 | 1539 | -2.61 | 5.60E-06 | 2.02E-05 |
| chr14 | 43353291 | 43354258 | 968 | 1.87 | 5.60E-06 | 2.02E-05 |
| chr5 | 46097044 | 46097785 | 742 | 5.44 | 5.61E-06 | 2.02E-05 |
| chr5 | 92406314 | 92415706 | 9393 | 0.74 | 5.62E-06 | 2.02E-05 |
| chr12 | 25610754 | 25611948 | 1195 | 1.78 | 5.62E-06 | 2.02E-05 |
| chr7 | 66092660 | 66120378 | 27719 | 0.44 | 5.65E-06 | 2.03E-05 |
| chr2 | 32036565 | 32038687 | 2123 | -0.91 | 5.65E-06 | 2.03E-05 |
| chr2 | 175098570 | 175114182 | 15613 | -0.65 | 5.65E-06 | 2.03E-05 |
| chr6 | 134311414 | 134318815 | 7402 | 0.73 | 5.66E-06 | 2.03E-05 |
| chr4 | 184384185 | 184392108 | 7924 | 1.06 | 5.66E-06 | 2.03E-05 |
| chr1 | 191175648 | 191176343 | 696 | 5.44 | 5.67E-06 | 2.04E-05 |
| chr4 | 125329694 | 125330959 | 1266 | 1.4 | 5.68E-06 | 2.04E-05 |
| chr16 | 71641331 | 71643219 | 1889 | 2.41 | 5.71E-06 | 2.05E-05 |
| chr20 | 38289997 | 38290922 | 926 | -3.77 | 5.71E-06 | 2.05E-05 |
| chr14 | 105671927 | 105680387 | 8461 | -1.28 | 5.72E-06 | 2.05E-05 |
| chr10 | 124165140 | 124182271 | 17132 | 0.74 | 5.72E-06 | 2.06E-05 |
| chr2 | 189278463 | 189283708 | 5246 | -0.62 | 5.73E-06 | 2.06E-05 |
| chr4 | 168609249 | 168610090 | 842 | 3.67 | 5.73E-06 | 2.06E-05 |
| chr1 | 49381395 | 49398893 | 17499 | -0.74 | 5.75E-06 | 2.07E-05 |
| chr1 | 150699320 | 150703059 | 3740 | 1.06 | 5.76E-06 | 2.07E-05 |
| chr1 | 185510867 | 185512117 | 1251 | 1.49 | 5.78E-06 | 2.07E-05 |
| chr17 | 17107869 | 17142133 | 34265 | -0.35 | 5.78E-06 | 2.07E-05 |
| chr8 | 86018722 | 86020071 | 1350 | 1.39 | 5.79E-06 | 2.08E-05 |
| chr2 | 33435241 | 33468343 | 33103 | -0.51 | 5.79E-06 | 2.08E-05 |
| chr10 | 5293594 | 5337240 | 43647 | -0.3 | 5.80E-06 | 2.08E-05 |
| chr5 | 45766462 | 45767256 | 795 | 5.44 | 5.83E-06 | 2.09E-05 |
| chr7 | 17273247 | 17280109 | 6863 | 0.59 | 5.84E-06 | 2.09E-05 |
| chr8 | 122131367 | 122132029 | 663 | 3.87 | 5.84E-06 | 2.09E-05 |
| chr17 | 12568735 | 12569472 | 738 | -5.42 | 5.85E-06 | 2.10E-05 |
| chr20 | 24612440 | 24616721 | 4282 | -0.51 | 5.85E-06 | 2.10E-05 |

| | | | | | | |
|---|---|---|---|---|---|---|
| chr4 | 79300966 | 79302274 | 1309 | 2.48 | 5.86E-06 | 2.10E-05 |
| chr12 | 86070280 | 86084864 | 14585 | 4.95 | 5.86E-06 | 2.10E-05 |
| chr1 | 64094238 | 64107889 | 13652 | 1.48 | 5.87E-06 | 2.10E-05 |
| chr2 | 150504840 | 150519083 | 14244 | 1.54 | 5.90E-06 | 2.11E-05 |
| chr7 | 56085748 | 56147872 | 62125 | 0.27 | 5.91E-06 | 2.12E-05 |
| chr11 | 44322315 | 44324308 | 1994 | -1.72 | 5.92E-06 | 2.12E-05 |
| chr5 | 60476513 | 60477528 | 1016 | 2.17 | 5.93E-06 | 2.12E-05 |
| chr11 | 125491489 | 125509858 | 18370 | 0.43 | 5.94E-06 | 2.13E-05 |
| chr3 | 123401661 | 123408335 | 6675 | -1.09 | 5.95E-06 | 2.13E-05 |
| chr5 | 91688371 | 91690281 | 1911 | 1.84 | 5.96E-06 | 2.13E-05 |
| chr12 | 96427637 | 96442636 | 15000 | 0.66 | 5.98E-06 | 2.14E-05 |
| chr3 | 95719529 | 95732075 | 12547 | 5.6 | 5.99E-06 | 2.14E-05 |
| chr2 | 218251316 | 218268550 | 17235 | 0.63 | 5.99E-06 | 2.15E-05 |
| chr1 | 93543830 | 93559756 | 15927 | 0.54 | 6.00E-06 | 2.15E-05 |
| chr10 | 42862922 | 42863433 | 512 | 5.44 | 6.00E-06 | 2.15E-05 |
| chr11 | 57200825 | 57228746 | 27922 | -0.37 | 6.01E-06 | 2.15E-05 |
| chr15 | 93198358 | 93211788 | 13431 | 0.94 | 6.01E-06 | 2.15E-05 |
| chr3 | 119810369 | 119817426 | 7058 | 0.4 | 6.02E-06 | 2.15E-05 |
| chr8 | 80695343 | 80712055 | 16713 | 0.8 | 6.04E-06 | 2.16E-05 |
| chr2 | 233253664 | 233261172 | 7509 | -2.03 | 6.06E-06 | 2.17E-05 |
| chr3 | 153339572 | 153340324 | 753 | 5.52 | 6.09E-06 | 2.18E-05 |
| chr7 | 74127889 | 74135324 | 7436 | 1.14 | 6.10E-06 | 2.18E-05 |
| chr11 | 46637306 | 46661835 | 24530 | 0.37 | 6.11E-06 | 2.18E-05 |
| chr11 | 100697484 | 100734846 | 37363 | 1.6 | 6.14E-06 | 2.20E-05 |
| chr11 | 62225496 | 62248850 | 23355 | -0.78 | 6.15E-06 | 2.20E-05 |
| chr17 | 40243973 | 40244997 | 1025 | -4.37 | 6.15E-06 | 2.20E-05 |
| chr12 | 65622984 | 65623629 | 646 | 5.43 | 6.20E-06 | 2.22E-05 |
| chr4 | 123703336 | 123706085 | 2750 | -0.84 | 6.24E-06 | 2.23E-05 |
| chr7 | 46009590 | 46024673 | 15084 | -0.29 | 6.28E-06 | 2.24E-05 |
| chr8 | 1757154 | 1758173 | 1020 | 3.63 | 6.28E-06 | 2.24E-05 |
| chr2 | 83191524 | 83192037 | 514 | 5.43 | 6.33E-06 | 2.26E-05 |
| chr10 | 52363037 | 52366970 | 3934 | 1.03 | 6.35E-06 | 2.27E-05 |
| chr15 | 49053243 | 49065824 | 12582 | 1.04 | 6.37E-06 | 2.27E-05 |
| chr3 | 100914473 | 100915108 | 636 | 1.86 | 6.37E-06 | 2.28E-05 |
| chr16 | 63612703 | 63613541 | 839 | 4.77 | 6.38E-06 | 2.28E-05 |
| chr4 | 116412563 | 116413347 | 785 | 5.42 | 6.38E-06 | 2.28E-05 |
| chr1 | 40973799 | 40975151 | 1353 | 1.4 | 6.41E-06 | 2.29E-05 |
| chr4 | 84375567 | 84378233 | 2667 | 0.66 | 6.43E-06 | 2.30E-05 |
| chr1 | 194356856 | 194357536 | 681 | 3.85 | 6.45E-06 | 2.30E-05 |
| chr22 | 41900411 | 41920712 | 20302 | -0.57 | 6.48E-06 | 2.31E-05 |
| chrX | 37865223 | 37866232 | 1010 | 5.49 | 6.48E-06 | 2.31E-05 |
| chr11 | 8955845 | 8957150 | 1306 | 2.31 | 6.49E-06 | 2.31E-05 |
| chr11 | 76673589 | 76674713 | 1125 | 3.04 | 6.49E-06 | 2.31E-05 |
| chr3 | 171911257 | 171922805 | 11549 | -0.61 | 6.50E-06 | 2.32E-05 |
| chr11 | 115341983 | 115342813 | 831 | -4.31 | 6.51E-06 | 2.32E-05 |
| chr12 | 133052055 | 133052573 | 519 | 5.44 | 6.51E-06 | 2.32E-05 |
| chr4 | 122721679 | 122745778 | 24100 | 0.51 | 6.54E-06 | 2.33E-05 |
| chr1 | 191808701 | 191809299 | 599 | 5.47 | 6.55E-06 | 2.33E-05 |
| chr5 | 132201350 | 132235808 | 34459 | 0.39 | 6.55E-06 | 2.34E-05 |
| chr3 | 162011869 | 162014768 | 2900 | -0.63 | 6.56E-06 | 2.34E-05 |
| chrX | 96312174 | 96313296 | 1123 | 3.12 | 6.58E-06 | 2.34E-05 |
| chr8 | 143780487 | 143782011 | 1525 | -1.11 | 6.60E-06 | 2.35E-05 |
| chrX | 24134886 | 24136206 | 1321 | -1.31 | 6.60E-06 | 2.35E-05 |
| chr2 | 226377062 | 226378477 | 1416 | -2.33 | 6.61E-06 | 2.36E-05 |
| chr19 | 19906602 | 19907144 | 543 | 4.35 | 6.62E-06 | 2.36E-05 |
| chr12 | 123546809 | 123651288 | 104480 | -0.32 | 6.63E-06 | 2.36E-05 |
| chr22 | 23637591 | 23639212 | 1622 | 1.68 | 6.63E-06 | 2.36E-05 |
| chr11 | 62093009 | 62105799 | 12791 | 5.43 | 6.66E-06 | 2.37E-05 |
| chr9 | 120256955 | 120257604 | 650 | 5.43 | 6.66E-06 | 2.37E-05 |
| chr14 | 36788309 | 36790401 | 2093 | 0.93 | 6.67E-06 | 2.37E-05 |
| chr17 | 26480929 | 26481866 | 938 | 3.57 | 6.68E-06 | 2.38E-05 |

| | | | | | | |
|---|---|---|---|---|---|---|
| chr6 | 150712985 | 150714469 | 1485 | 1.13 | 6.70E-06 | 2.38E-05 |
| chr4 | 6910441 | 6920779 | 10339 | 0.64 | 6.71E-06 | 2.39E-05 |
| chr10 | 5454122 | 5510244 | 56123 | 0.31 | 6.76E-06 | 2.40E-05 |
| chr17 | 21064515 | 21066568 | 2054 | 4.72 | 6.80E-06 | 2.42E-05 |
| chrX | 47930437 | 47931536 | 1100 | 1.48 | 6.86E-06 | 2.44E-05 |
| chr11 | 12307119 | 12310837 | 3719 | -0.69 | 6.86E-06 | 2.44E-05 |
| chr9 | 34325883 | 34346239 | 20357 | -0.4 | 6.88E-06 | 2.44E-05 |
| chr13 | 27743290 | 27770428 | 27139 | -0.55 | 6.88E-06 | 2.45E-05 |
| chr3 | 101702973 | 101704047 | 1075 | 5.41 | 6.88E-06 | 2.45E-05 |
| chr7 | 109818949 | 109819757 | 809 | 2.58 | 6.89E-06 | 2.45E-05 |
| chr4 | 53682227 | 53682701 | 475 | 3.2 | 6.89E-06 | 2.45E-05 |
| chr8 | 100694207 | 100699140 | 4934 | -1.18 | 6.91E-06 | 2.45E-05 |
| chr4 | 114089993 | 114091084 | 1092 | 4.35 | 6.92E-06 | 2.46E-05 |
| chr11 | 34606623 | 34635401 | 28779 | -0.3 | 6.92E-06 | 2.46E-05 |
| chr15 | 97433191 | 97433719 | 529 | 3.88 | 6.93E-06 | 2.46E-05 |
| chr22 | 45072155 | 45081363 | 9209 | -0.9 | 6.94E-06 | 2.47E-05 |
| chr10 | 15503076 | 15504108 | 1033 | -5.39 | 6.95E-06 | 2.47E-05 |
| chr19 | 45453791 | 45563740 | 109950 | -0.25 | 6.96E-06 | 2.47E-05 |
| chr13 | 30968571 | 30997242 | 28672 | 0.59 | 6.98E-06 | 2.48E-05 |
| chr8 | 32775499 | 32786390 | 10892 | -1.12 | 6.99E-06 | 2.48E-05 |
| chr16 | 49994756 | 49995416 | 661 | 4.75 | 7.04E-06 | 2.50E-05 |
| chr9 | 111203399 | 111204828 | 1430 | 1.73 | 7.06E-06 | 2.51E-05 |
| chr12 | 21643098 | 21656170 | 13073 | 0.52 | 7.07E-06 | 2.51E-05 |
| chr3 | 61612661 | 61623698 | 11038 | -1.34 | 7.08E-06 | 2.51E-05 |
| chr18 | 11689295 | 11690382 | 1088 | -3.4 | 7.10E-06 | 2.52E-05 |
| chr2 | 113445456 | 113451972 | 6517 | -0.5 | 7.11E-06 | 2.52E-05 |
| chr21 | 16581326 | 16597394 | 16069 | -0.62 | 7.13E-06 | 2.53E-05 |
| chr20 | 14839986 | 14842526 | 2541 | 0.85 | 7.15E-06 | 2.54E-05 |
| chr17 | 69434523 | 69438025 | 3503 | 0.89 | 7.22E-06 | 2.56E-05 |
| chr5 | 34904702 | 34931465 | 26764 | 0.35 | 7.25E-06 | 2.57E-05 |
| chr2 | 243002227 | 243002738 | 512 | 5.47 | 7.25E-06 | 2.57E-05 |
| chr11 | 8980247 | 8987740 | 7494 | 0.55 | 7.28E-06 | 2.58E-05 |
| chr2 | 42795049 | 42805785 | 10737 | -0.7 | 7.32E-06 | 2.59E-05 |
| chr13 | 65248185 | 65263936 | 15752 | -2.57 | 7.33E-06 | 2.60E-05 |
| chr4 | 177991449 | 177992262 | 814 | -5.35 | 7.34E-06 | 2.60E-05 |
| chr1 | 200264970 | 200288885 | 23916 | 0.9 | 7.34E-06 | 2.60E-05 |
| chr19 | 31704738 | 31705508 | 771 | -3.46 | 7.36E-06 | 2.61E-05 |
| chr12 | 112272911 | 112286174 | 13264 | 0.47 | 7.38E-06 | 2.61E-05 |
| chr11 | 3663413 | 3666656 | 3244 | 1.56 | 7.38E-06 | 2.61E-05 |
| chr9 | 125658025 | 125676939 | 18915 | 0.45 | 7.39E-06 | 2.62E-05 |
| chr2 | 188288525 | 188289867 | 1343 | 1.67 | 7.43E-06 | 2.63E-05 |
| chr5 | 66563654 | 66564932 | 1279 | -2.24 | 7.45E-06 | 2.64E-05 |
| chr2 | 212008610 | 212009529 | 920 | -5.37 | 7.45E-06 | 2.64E-05 |
| chr16 | 86340887 | 86342372 | 1486 | -2.06 | 7.45E-06 | 2.64E-05 |
| chr13 | 22009462 | 22011664 | 2203 | 1.39 | 7.45E-06 | 2.64E-05 |
| chr6 | 132405467 | 132406710 | 1244 | 1.8 | 7.45E-06 | 2.64E-05 |
| chr2 | 18074153 | 18074934 | 782 | -2.63 | 7.47E-06 | 2.64E-05 |
| chr6 | 2244816 | 2260077 | 15262 | 0.8 | 7.51E-06 | 2.66E-05 |
| chr18 | 2222496 | 2223861 | 1366 | 1.81 | 7.51E-06 | 2.66E-05 |
| chr1 | 40504405 | 40526916 | 22512 | 0.39 | 7.52E-06 | 2.66E-05 |
| chr9 | 118432713 | 118457619 | 24907 | -0.4 | 7.55E-06 | 2.67E-05 |
| chr19 | 2576126 | 2628017 | 51892 | 0.49 | 7.55E-06 | 2.67E-05 |
| chr11 | 72496703 | 72543668 | 46966 | -0.33 | 7.57E-06 | 2.68E-05 |
| chr20 | 47300689 | 47301681 | 993 | -5.35 | 7.57E-06 | 2.68E-05 |
| chr4 | 145932778 | 145935457 | 2680 | 1.04 | 7.58E-06 | 2.68E-05 |
| chr13 | 113648296 | 113649124 | 829 | 4.33 | 7.59E-06 | 2.68E-05 |
| chr7 | 81475261 | 81489000 | 13740 | 5.8 | 7.60E-06 | 2.68E-05 |
| chr13 | 103244457 | 103261724 | 17268 | 0.55 | 7.60E-06 | 2.68E-05 |
| chr5 | 14570500 | 14589992 | 19493 | 3.53 | 7.60E-06 | 2.69E-05 |
| chr7 | 55560757 | 55572968 | 12212 | -1.87 | 7.60E-06 | 2.69E-05 |
| chr4 | 66516442 | 66517287 | 846 | 5.39 | 7.63E-06 | 2.70E-05 |

| | | | | | | |
|---|---|---|---|---|---|---|
| chr15 | 65269385 | 65283339 | 13955 | 0.34 | 7.66E-06 | 2.71E-05 |
| chr14 | 60431635 | 60436847 | 5213 | 3.05 | 7.67E-06 | 2.71E-05 |
| chr12 | 69293182 | 69293880 | 699 | 2.89 | 7.67E-06 | 2.71E-05 |
| chr15 | 45002913 | 45005914 | 3002 | 0.78 | 7.68E-06 | 2.71E-05 |
| chr3 | 125029116 | 125030078 | 963 | 4 | 7.70E-06 | 2.72E-05 |
| chr4 | 179772383 | 179773172 | 790 | 5.39 | 7.71E-06 | 2.72E-05 |
| chr9 | 129838177 | 129838914 | 738 | 4.33 | 7.71E-06 | 2.72E-05 |
| chr11 | 68593121 | 68609252 | 16132 | 0.53 | 7.71E-06 | 2.72E-05 |
| chr5 | 82279897 | 82287956 | 8060 | 1.7 | 7.73E-06 | 2.73E-05 |
| chr14 | 92363491 | 92378675 | 15185 | 1.32 | 7.74E-06 | 2.73E-05 |
| chr8 | 135904508 | 135905632 | 1125 | 1.72 | 7.75E-06 | 2.74E-05 |
| chr16 | 27879206 | 27879699 | 494 | 5.39 | 7.77E-06 | 2.74E-05 |
| chr10 | 73004036 | 73007552 | 3517 | 1.49 | 7.77E-06 | 2.74E-05 |
| chr10 | 120485140 | 120490911 | 5772 | 1.13 | 7.79E-06 | 2.75E-05 |
| chr2 | 33670908 | 33671453 | 546 | 3.62 | 7.82E-06 | 2.76E-05 |
| chr1 | 184482596 | 184483695 | 1100 | -5.52 | 7.82E-06 | 2.76E-05 |
| chr14 | 93174478 | 93215454 | 40977 | 0.51 | 7.84E-06 | 2.76E-05 |
| chr15 | 66909242 | 66912620 | 3379 | -1.08 | 7.84E-06 | 2.76E-05 |
| chr4 | 101111013 | 101112472 | 1460 | 2.38 | 7.85E-06 | 2.77E-05 |
| chr16 | 575282 | 590077 | 14796 | -0.56 | 7.86E-06 | 2.77E-05 |
| chr1 | 155275399 | 155298345 | 22947 | 0.39 | 7.87E-06 | 2.77E-05 |
| chr20 | 35616734 | 35618729 | 1996 | -2.86 | 7.87E-06 | 2.77E-05 |
| chr6 | 56802954 | 56821167 | 18214 | -0.64 | 7.89E-06 | 2.78E-05 |
| chr6 | 5892572 | 5893267 | 696 | 1.54 | 7.93E-06 | 2.79E-05 |
| chr21 | 38089966 | 38097630 | 7665 | -1.22 | 7.94E-06 | 2.80E-05 |
| chr17 | 37287847 | 37289241 | 1395 | -2.41 | 7.97E-06 | 2.80E-05 |
| chr20 | 61399340 | 61400174 | 835 | 5.4 | 8.02E-06 | 2.82E-05 |
| chr8 | 26120133 | 26124206 | 4074 | -1.55 | 8.03E-06 | 2.83E-05 |
| chr8 | 146104147 | 146104950 | 804 | 5.61 | 8.06E-06 | 2.84E-05 |
| chr2 | 172287330 | 172292226 | 4897 | -0.57 | 8.08E-06 | 2.84E-05 |
| chr5 | 154276699 | 154286278 | 9580 | -1.1 | 8.08E-06 | 2.84E-05 |
| chr9 | 15372290 | 15374360 | 2071 | 1.45 | 8.09E-06 | 2.85E-05 |
| chr5 | 64919104 | 64922639 | 3536 | 0.64 | 8.10E-06 | 2.85E-05 |
| chr14 | 68508580 | 68515059 | 6480 | 1 | 8.11E-06 | 2.85E-05 |
| chr5 | 102349239 | 102350348 | 1110 | 2.65 | 8.12E-06 | 2.86E-05 |
| chr2 | 173204892 | 173206577 | 1686 | 2.41 | 8.15E-06 | 2.87E-05 |
| chr10 | 25155185 | 25162780 | 7596 | 0.91 | 8.16E-06 | 2.87E-05 |
| chr5 | 60699228 | 60728897 | 29670 | 0.29 | 8.17E-06 | 2.87E-05 |
| chr12 | 115393153 | 115393890 | 738 | -5.33 | 8.17E-06 | 2.87E-05 |
| chr8 | 96000065 | 96019626 | 19562 | 0.62 | 8.18E-06 | 2.88E-05 |
| chr9 | 97410602 | 97412822 | 2221 | 1.38 | 8.19E-06 | 2.88E-05 |
| chr1 | 93907146 | 93915177 | 8032 | 0.78 | 8.19E-06 | 2.88E-05 |
| chr16 | 50295162 | 50300803 | 5642 | -1.12 | 8.21E-06 | 2.88E-05 |
| chr21 | 32881318 | 32883093 | 1776 | 1.53 | 8.24E-06 | 2.90E-05 |
| chr10 | 120528253 | 120529370 | 1118 | 3.5 | 8.29E-06 | 2.91E-05 |
| chr3 | 153451009 | 153454698 | 3690 | 7.15 | 8.30E-06 | 2.92E-05 |
| chr5 | 130598489 | 130660055 | 61567 | 0.34 | 8.31E-06 | 2.92E-05 |
| chr7 | 46071404 | 46073646 | 2243 | -1.95 | 8.33E-06 | 2.92E-05 |
| chr3 | 32085947 | 32087904 | 1958 | 0.79 | 8.33E-06 | 2.93E-05 |
| chr19 | 21324841 | 21334993 | 10153 | 1.49 | 8.35E-06 | 2.93E-05 |
| chr12 | 20751857 | 20765471 | 13615 | 0.82 | 8.38E-06 | 2.94E-05 |
| chr10 | 12164641 | 12172707 | 8067 | 0.72 | 8.42E-06 | 2.96E-05 |
| chr2 | 38501255 | 38506230 | 4976 | 1.23 | 8.43E-06 | 2.96E-05 |
| chr3 | 133286854 | 133310328 | 23475 | 0.37 | 8.45E-06 | 2.96E-05 |
| chr13 | 82079747 | 82080285 | 539 | 5.4 | 8.45E-06 | 2.97E-05 |
| chr4 | 162592848 | 162602553 | 9706 | 5.37 | 8.47E-06 | 2.97E-05 |
| chr6 | 50919630 | 50920379 | 750 | 5.37 | 8.47E-06 | 2.97E-05 |
| chr9 | 121433038 | 121433923 | 886 | 5.37 | 8.47E-06 | 2.97E-05 |
| chr4 | 99831267 | 99852165 | 20899 | 0.45 | 8.48E-06 | 2.97E-05 |
| chr13 | 52206830 | 52207561 | 732 | -3.37 | 8.54E-06 | 3.00E-05 |
| chr9 | 27460517 | 27461116 | 600 | -4.68 | 8.58E-06 | 3.01E-05 |

| | | | | | | |
|---|---|---|---|---|---|---|
| chr15 | 71944247 | 71953046 | 8800 | -1.37 | 8.60E-06 | 3.01E-05 |
| chr17 | 6810226 | 6811715 | 1490 | 1.8 | 8.60E-06 | 3.02E-05 |
| chr16 | 85455379 | 85457056 | 1678 | 2.26 | 8.61E-06 | 3.02E-05 |
| chr8 | 110547200 | 110553863 | 6664 | 0.59 | 8.61E-06 | 3.02E-05 |
| chr11 | 69793977 | 69816724 | 22748 | -0.67 | 8.62E-06 | 3.02E-05 |
| chr7 | 93578628 | 93579354 | 727 | 2.93 | 8.64E-06 | 3.03E-05 |
| chr12 | 11732574 | 11733964 | 1391 | 1.87 | 8.64E-06 | 3.03E-05 |
| chr2 | 49068077 | 49068676 | 600 | -5.32 | 8.70E-06 | 3.05E-05 |
| chr9 | 19748363 | 19749019 | 657 | -5.32 | 8.70E-06 | 3.05E-05 |
| chr11 | 117389764 | 117391075 | 1312 | 2.29 | 8.72E-06 | 3.05E-05 |
| chr5 | 73637427 | 73640410 | 2984 | 0.89 | 8.73E-06 | 3.06E-05 |
| chr4 | 151223261 | 151227983 | 4723 | 1.44 | 8.74E-06 | 3.06E-05 |
| chr14 | 27418173 | 27418646 | 474 | 3.79 | 8.75E-06 | 3.06E-05 |
| chr19 | 28466330 | 28466793 | 464 | -5.32 | 8.75E-06 | 3.06E-05 |
| chr16 | 81270732 | 81273711 | 2980 | 0.82 | 8.77E-06 | 3.07E-05 |
| chr19 | 44487571 | 44489178 | 1608 | 1.3 | 8.81E-06 | 3.08E-05 |
| chr5 | 102200986 | 102202537 | 1552 | 1.3 | 8.84E-06 | 3.09E-05 |
| chr20 | 15988394 | 15988918 | 525 | -5.32 | 8.86E-06 | 3.10E-05 |
| chr13 | 81690646 | 81691174 | 529 | 3.04 | 8.88E-06 | 3.11E-05 |
| chr2 | 55745260 | 55753868 | 8609 | -0.48 | 8.88E-06 | 3.11E-05 |
| chr12 | 19863801 | 19865173 | 1373 | 1.69 | 8.89E-06 | 3.11E-05 |
| chr6 | 109028313 | 109029312 | 1000 | 3.5 | 8.89E-06 | 3.11E-05 |
| chr16 | 27858170 | 27859074 | 905 | 3.14 | 8.95E-06 | 3.13E-05 |
| chr15 | 101359029 | 101359975 | 947 | 2.63 | 8.97E-06 | 3.14E-05 |
| chr8 | 33218140 | 33219002 | 863 | 3.9 | 8.97E-06 | 3.14E-05 |
| chr1 | 158114780 | 158120035 | 5256 | 1.77 | 8.97E-06 | 3.14E-05 |
| chr20 | 8136271 | 8138147 | 1877 | 1.35 | 8.98E-06 | 3.14E-05 |
| chr1 | 16692776 | 16725352 | 32577 | -0.39 | 8.99E-06 | 3.14E-05 |
| chr11 | 44264265 | 44270016 | 5752 | -1.49 | 9.01E-06 | 3.15E-05 |
| chr17 | 62560614 | 62585092 | 24479 | 0.54 | 9.01E-06 | 3.15E-05 |
| chr17 | 73996164 | 74002163 | 6000 | -1.15 | 9.05E-06 | 3.16E-05 |
| chr19 | 40898611 | 40972799 | 74189 | 0.21 | 9.05E-06 | 3.16E-05 |
| chr1 | 29044194 | 29046499 | 2306 | -1.47 | 9.06E-06 | 3.16E-05 |
| chr3 | 154040193 | 154043027 | 2835 | 0.53 | 9.08E-06 | 3.17E-05 |
| chr2 | 210430367 | 210431321 | 955 | 5.37 | 9.09E-06 | 3.17E-05 |
| chr1 | 240193273 | 240195618 | 2346 | -1.13 | 9.10E-06 | 3.18E-05 |
| chr10 | 71883820 | 71931350 | 47531 | -0.27 | 9.12E-06 | 3.18E-05 |
| chr14 | 99890837 | 99903688 | 12852 | -1.76 | 9.12E-06 | 3.18E-05 |
| chr1 | 64058566 | 64060916 | 2351 | 1.21 | 9.13E-06 | 3.19E-05 |
| chr5 | 155693530 | 155694808 | 1279 | 2.35 | 9.13E-06 | 3.19E-05 |
| chr1 | 46146677 | 46155107 | 8431 | 0.5 | 9.13E-06 | 3.19E-05 |
| chr4 | 63747411 | 63748162 | 752 | 5.38 | 9.14E-06 | 3.19E-05 |
| chr11 | 122024009 | 122073712 | 49704 | 0.27 | 9.17E-06 | 3.20E-05 |
| chr1 | 173159095 | 173160718 | 1624 | 1.63 | 9.19E-06 | 3.21E-05 |
| chr6 | 79356006 | 79356812 | 807 | 1.82 | 9.19E-06 | 3.21E-05 |
| chr7 | 27913544 | 27914395 | 852 | 2.6 | 9.21E-06 | 3.21E-05 |
| chr1 | 48472303 | 48473346 | 1044 | -3.94 | 9.21E-06 | 3.21E-05 |
| chr5 | 6158571 | 6160856 | 2286 | -0.92 | 9.22E-06 | 3.21E-05 |
| chr18 | 48555277 | 48558098 | 2822 | 0.74 | 9.22E-06 | 3.21E-05 |
| chr6 | 40682135 | 40687909 | 5775 | 1.53 | 9.23E-06 | 3.22E-05 |
| chr4 | 14635186 | 14635956 | 771 | 5.37 | 9.26E-06 | 3.23E-05 |
| chr4 | 163057968 | 163058875 | 908 | 5.37 | 9.26E-06 | 3.23E-05 |
| chr1 | 168677552 | 168680802 | 3251 | -0.65 | 9.33E-06 | 3.25E-05 |
| chr5 | 74755638 | 74770432 | 14795 | 1.17 | 9.34E-06 | 3.26E-05 |
| chr12 | 69118919 | 69140948 | 22030 | 0.56 | 9.35E-06 | 3.26E-05 |
| chr9 | 6928891 | 6929666 | 776 | 3 | 9.36E-06 | 3.26E-05 |
| chr9 | 106996163 | 106996687 | 525 | 5.39 | 9.38E-06 | 3.27E-05 |
| chr2 | 157285732 | 157324519 | 38788 | -0.35 | 9.43E-06 | 3.28E-05 |
| chr11 | 50426747 | 50427275 | 529 | 5.35 | 9.44E-06 | 3.29E-05 |
| chr4 | 138530909 | 138531566 | 658 | 5.47 | 9.45E-06 | 3.29E-05 |
| chr1 | 116518258 | 116519990 | 1733 | 1.32 | 9.49E-06 | 3.30E-05 |

| | | | | | | |
|---|---|---|---|---|---|---|
| chr18 | 21424311 | 21430142 | 5832 | 1.07 | 9.50E-06 | 3.31E-05 |
| chr11 | 19463478 | 19464572 | 1095 | -1.66 | 9.53E-06 | 3.32E-05 |
| chr3 | 177306714 | 177338171 | 31458 | 0.39 | 9.53E-06 | 3.32E-05 |
| chr10 | 63421851 | 63423616 | 1766 | 1.02 | 9.60E-06 | 3.34E-05 |
| chr3 | 101280046 | 101295126 | 15081 | 0.37 | 9.61E-06 | 3.35E-05 |
| chr11 | 108367861 | 108380529 | 12669 | 1.13 | 9.63E-06 | 3.35E-05 |
| chr9 | 21334091 | 21336031 | 1941 | 0.52 | 9.63E-06 | 3.35E-05 |
| chr14 | 103240498 | 103276944 | 36447 | -0.51 | 9.66E-06 | 3.36E-05 |
| chr7 | 150481441 | 150482051 | 611 | 5.35 | 9.67E-06 | 3.36E-05 |
| chr6 | 122779945 | 122793646 | 13702 | 0.53 | 9.68E-06 | 3.37E-05 |
| chr15 | 77711154 | 77714448 | 3295 | 0.55 | 9.72E-06 | 3.38E-05 |
| chr19 | 49402480 | 49431069 | 28590 | -0.43 | 9.74E-06 | 3.39E-05 |
| chr2 | 216299126 | 216301864 | 2739 | -1.84 | 9.75E-06 | 3.39E-05 |
| chr1 | 222134315 | 222139439 | 5125 | 0.78 | 9.76E-06 | 3.39E-05 |
| chr16 | 57787026 | 57933973 | 146948 | 0.16 | 9.76E-06 | 3.39E-05 |
| chr9 | 36419059 | 36421482 | 2424 | 0.78 | 9.76E-06 | 3.39E-05 |
| chr13 | 27556332 | 27558434 | 2103 | -1.01 | 9.82E-06 | 3.41E-05 |
| chr15 | 69335573 | 69345443 | 9871 | -1.44 | 9.92E-06 | 3.45E-05 |
| chr12 | 87010230 | 87010760 | 531 | 3.88 | 9.93E-06 | 3.45E-05 |
| chr3 | 29471649 | 29473486 | 1838 | 1.23 | 9.94E-06 | 3.45E-05 |
| chr9 | 26745425 | 26747835 | 2411 | 1.11 | 9.97E-06 | 3.46E-05 |
| chr7 | 101874516 | 101880822 | 6307 | -0.94 | 9.98E-06 | 3.47E-05 |
| chr3 | 35973802 | 35998486 | 24685 | 2.58 | 1.00E-05 | 3.48E-05 |
| chr2 | 72342029 | 72344047 | 2019 | 0.9 | 1.00E-05 | 3.48E-05 |
| chr1 | 43388259 | 43435387 | 47129 | 0.21 | 1.00E-05 | 3.48E-05 |
| chr4 | 140720693 | 140722298 | 1606 | 2.2 | 1.00E-05 | 3.48E-05 |
| chr11 | 3636722 | 3638126 | 1405 | -2.05 | 1.01E-05 | 3.49E-05 |
| chr2 | 170425049 | 170442511 | 17463 | -0.58 | 1.01E-05 | 3.49E-05 |
| chr18 | 20494624 | 20495165 | 542 | 4.1 | 1.01E-05 | 3.49E-05 |
| chr4 | 87926799 | 87947883 | 21085 | 0.38 | 1.01E-05 | 3.50E-05 |
| chr6 | 113912965 | 113914450 | 1486 | 3.42 | 1.01E-05 | 3.50E-05 |
| chr4 | 39847097 | 39875782 | 28686 | 0.56 | 1.01E-05 | 3.51E-05 |
| chr6 | 82646888 | 82649598 | 2711 | 1.29 | 1.01E-05 | 3.51E-05 |
| chr5 | 41903495 | 41905830 | 2336 | 0.76 | 1.01E-05 | 3.51E-05 |
| chr4 | 63467574 | 63468018 | 445 | 5.37 | 1.01E-05 | 3.52E-05 |
| chr10 | 97618872 | 97619694 | 823 | -2 | 1.02E-05 | 3.53E-05 |
| chr1 | 94510160 | 94545738 | 35579 | 0.35 | 1.02E-05 | 3.54E-05 |
| chr3 | 37972247 | 38040490 | 68244 | 1.01 | 1.02E-05 | 3.54E-05 |
| chr11 | 74991498 | 74992840 | 1343 | 2.21 | 1.02E-05 | 3.54E-05 |
| chr18 | 72263578 | 72266530 | 2953 | 0.76 | 1.02E-05 | 3.55E-05 |
| chr11 | 85826866 | 85835792 | 8927 | -1.46 | 1.03E-05 | 3.56E-05 |
| chr14 | 96828057 | 96843176 | 15120 | 0.74 | 1.03E-05 | 3.56E-05 |
| chr8 | 102586272 | 102587030 | 759 | 2.84 | 1.03E-05 | 3.56E-05 |
| chrX | 10086731 | 10091575 | 4845 | 1.48 | 1.03E-05 | 3.56E-05 |
| chr15 | 77799354 | 77800990 | 1637 | 1.74 | 1.03E-05 | 3.56E-05 |
| chr6 | 107920901 | 107925165 | 4265 | 1.19 | 1.03E-05 | 3.57E-05 |
| chr2 | 192541848 | 192554840 | 12993 | 0.49 | 1.03E-05 | 3.57E-05 |
| chr4 | 82766096 | 82766787 | 692 | -5.31 | 1.03E-05 | 3.57E-05 |
| chr17 | 42889377 | 42908272 | 18896 | -0.53 | 1.03E-05 | 3.58E-05 |
| chr16 | 51914073 | 51915502 | 1430 | 2.1 | 1.03E-05 | 3.58E-05 |
| chr10 | 103112986 | 103128226 | 15241 | 0.61 | 1.04E-05 | 3.59E-05 |
| chr9 | 28882775 | 28928262 | 45488 | 1.01 | 1.04E-05 | 3.59E-05 |
| chr14 | 71331786 | 71332831 | 1046 | -5.29 | 1.04E-05 | 3.59E-05 |
| chrX | 4827106 | 4827820 | 715 | 2.43 | 1.04E-05 | 3.60E-05 |
| chr4 | 41899004 | 41900514 | 1511 | 2.38 | 1.04E-05 | 3.60E-05 |
| chr19 | 55788212 | 55814062 | 25851 | -0.76 | 1.04E-05 | 3.61E-05 |
| chr3 | 116093359 | 116094141 | 783 | 4.14 | 1.04E-05 | 3.61E-05 |
| chr9 | 97210303 | 97218386 | 8084 | -0.87 | 1.05E-05 | 3.62E-05 |
| chr14 | 34606404 | 34609574 | 3171 | 1.03 | 1.05E-05 | 3.63E-05 |
| chrX | 115519487 | 115520290 | 804 | 5.35 | 1.05E-05 | 3.63E-05 |
| chr14 | 81004144 | 81006061 | 1918 | 0.98 | 1.05E-05 | 3.64E-05 |

| | | | | | | |
|---|---|---|---|---|---|---|
| chr3 | 42692846 | 42697267 | 4422 | -0.69 | 1.05E-05 | 3.64E-05 |
| chr4 | 113205857 | 113219633 | 13777 | 0.71 | 1.06E-05 | 3.65E-05 |
| chr6 | 105933015 | 105933726 | 712 | -5.29 | 1.06E-05 | 3.66E-05 |
| chr12 | 76713029 | 76713631 | 603 | 2.24 | 1.06E-05 | 3.66E-05 |
| chr10 | 14992830 | 15003295 | 10466 | 0.39 | 1.06E-05 | 3.67E-05 |
| chr4 | 187686023 | 187687784 | 1762 | 1.62 | 1.07E-05 | 3.68E-05 |
| chr9 | 77501449 | 77502899 | 1451 | 1.37 | 1.07E-05 | 3.69E-05 |
| chr8 | 124624609 | 124625331 | 723 | -4.09 | 1.07E-05 | 3.69E-05 |
| chr3 | 189508004 | 189508949 | 946 | 2.99 | 1.07E-05 | 3.69E-05 |
| chr3 | 155252380 | 155253647 | 1268 | 1.73 | 1.07E-05 | 3.69E-05 |
| chr8 | 124868634 | 124871496 | 2863 | 1.94 | 1.07E-05 | 3.70E-05 |
| chr21 | 45595794 | 45596616 | 823 | 4.51 | 1.07E-05 | 3.70E-05 |
| chr4 | 186279127 | 186281112 | 1986 | -1.24 | 1.07E-05 | 3.70E-05 |
| chr3 | 25534301 | 25536168 | 1868 | 2.14 | 1.07E-05 | 3.71E-05 |
| chr2 | 226600298 | 226601497 | 1200 | -2.35 | 1.08E-05 | 3.71E-05 |
| chr5 | 137798788 | 137802901 | 4114 | 0.5 | 1.08E-05 | 3.71E-05 |
| chr13 | 29231907 | 29235308 | 3402 | 0.54 | 1.08E-05 | 3.73E-05 |
| chr4 | 160356634 | 160357273 | 640 | 3.8 | 1.08E-05 | 3.74E-05 |
| chr19 | 15386895 | 15416769 | 29875 | -0.38 | 1.08E-05 | 3.74E-05 |
| chr7 | 123197113 | 123198679 | 1567 | 0.75 | 1.09E-05 | 3.75E-05 |
| chr3 | 160700049 | 160701035 | 987 | 2.85 | 1.09E-05 | 3.75E-05 |
| chr14 | 101052481 | 101053688 | 1208 | -1.8 | 1.09E-05 | 3.75E-05 |
| chr7 | 127478349 | 127478831 | 483 | 3.53 | 1.09E-05 | 3.76E-05 |
| chr7 | 1170479 | 1178961 | 8483 | -0.7 | 1.09E-05 | 3.77E-05 |
| chr2 | 234190308 | 234203114 | 12807 | -0.92 | 1.10E-05 | 3.80E-05 |
| chr5 | 144843442 | 144844402 | 961 | 5.33 | 1.10E-05 | 3.80E-05 |
| chr6 | 8743450 | 8743977 | 528 | 5.33 | 1.10E-05 | 3.80E-05 |
| chrX | 119762587 | 119765119 | 2533 | 0.78 | 1.10E-05 | 3.81E-05 |
| chr8 | 142578187 | 142579404 | 1218 | -2.56 | 1.11E-05 | 3.81E-05 |
| chr19 | 41361230 | 41362354 | 1125 | 1.93 | 1.11E-05 | 3.81E-05 |
| chr3 | 17519689 | 17521704 | 2016 | -1.29 | 1.11E-05 | 3.82E-05 |
| chr20 | 18774550 | 18775393 | 844 | 0.88 | 1.11E-05 | 3.83E-05 |
| chr1 | 234844872 | 234863621 | 18750 | 0.38 | 1.12E-05 | 3.84E-05 |
| chr9 | 91505403 | 91506223 | 821 | -3.42 | 1.12E-05 | 3.85E-05 |
| chr6 | 18022772 | 18023922 | 1151 | 1.99 | 1.12E-05 | 3.86E-05 |
| chr4 | 119839098 | 119840178 | 1081 | 3.66 | 1.12E-05 | 3.86E-05 |
| chr7 | 65203941 | 65217376 | 13436 | 0.44 | 1.12E-05 | 3.86E-05 |
| chr17 | 69177509 | 69177979 | 471 | 5.33 | 1.12E-05 | 3.87E-05 |
| chr11 | 122953734 | 122955611 | 1878 | 1.06 | 1.13E-05 | 3.88E-05 |
| chr3 | 37328880 | 37341800 | 12921 | -0.6 | 1.13E-05 | 3.88E-05 |
| chr20 | 35268989 | 35276915 | 7927 | 0.65 | 1.13E-05 | 3.89E-05 |
| chr13 | 100881347 | 100888230 | 6884 | 1.05 | 1.13E-05 | 3.90E-05 |
| chr6 | 33538721 | 33591022 | 52302 | 0.39 | 1.14E-05 | 3.91E-05 |
| chrX | 35898081 | 35898705 | 625 | 4.08 | 1.14E-05 | 3.91E-05 |
| chr12 | 49063338 | 49076960 | 13623 | 0.57 | 1.14E-05 | 3.91E-05 |
| chr7 | 86780570 | 86784519 | 3950 | 0.49 | 1.14E-05 | 3.91E-05 |
| chr9 | 114490548 | 114505732 | 15185 | 2.52 | 1.14E-05 | 3.91E-05 |
| chr2 | 41962260 | 41963158 | 899 | -2.16 | 1.14E-05 | 3.92E-05 |
| chr10 | 90299915 | 90301040 | 1126 | 3.64 | 1.15E-05 | 3.94E-05 |
| chr5 | 55724613 | 55738042 | 13430 | 1.03 | 1.15E-05 | 3.96E-05 |
| chr22 | 40439705 | 40442119 | 2415 | 1 | 1.15E-05 | 3.96E-05 |
| chr8 | 126203252 | 126205496 | 2245 | 0.78 | 1.15E-05 | 3.96E-05 |
| chr7 | 50696759 | 50700729 | 3971 | -1.66 | 1.15E-05 | 3.96E-05 |
| chr15 | 99650842 | 99668963 | 18122 | -0.31 | 1.15E-05 | 3.97E-05 |
| chr12 | 19584220 | 19607430 | 23211 | 0.68 | 1.16E-05 | 3.97E-05 |
| chr4 | 26370153 | 26371823 | 1671 | 1.69 | 1.16E-05 | 3.97E-05 |
| chr5 | 81266299 | 81269257 | 2959 | 0.46 | 1.16E-05 | 3.98E-05 |
| chr22 | 25288879 | 25297682 | 8804 | 0.49 | 1.16E-05 | 3.98E-05 |
| chr3 | 185405709 | 185420950 | 15242 | 0.85 | 1.16E-05 | 3.98E-05 |
| chr20 | 58406386 | 58407074 | 689 | 4.27 | 1.16E-05 | 3.99E-05 |
| chr16 | 75119822 | 75128061 | 8240 | 0.94 | 1.16E-05 | 3.99E-05 |

| | | | | | | |
|---|---|---|---|---|---|---|
| chr13 | 73732395 | 73746641 | 14247 | 1.03 | 1.16E-05 | 3.99E-05 |
| chr2 | 114548030 | 114550576 | 2547 | 0.77 | 1.16E-05 | 4.00E-05 |
| chr12 | 18298834 | 18299479 | 646 | 3.99 | 1.17E-05 | 4.00E-05 |
| chr6 | 64136608 | 64171432 | 34825 | 0.28 | 1.17E-05 | 4.01E-05 |
| chr20 | 48381153 | 48476834 | 95682 | 0.23 | 1.17E-05 | 4.01E-05 |
| chr19 | 3868486 | 3868996 | 511 | 5.43 | 1.17E-05 | 4.02E-05 |
| chr7 | 87079523 | 87080587 | 1065 | 2.61 | 1.17E-05 | 4.02E-05 |
| chr19 | 24154197 | 24155282 | 1086 | 5.31 | 1.17E-05 | 4.02E-05 |
| chr4 | 67879813 | 67880316 | 504 | 5.31 | 1.17E-05 | 4.02E-05 |
| chr13 | 83795900 | 83796967 | 1068 | 2.32 | 1.18E-05 | 4.04E-05 |
| chr13 | 34797147 | 34799124 | 1978 | 1.01 | 1.18E-05 | 4.04E-05 |
| chr12 | 124574303 | 124575454 | 1152 | 2.61 | 1.18E-05 | 4.05E-05 |
| chr1 | 242742093 | 242746621 | 4529 | -0.69 | 1.18E-05 | 4.05E-05 |
| chr4 | 173883209 | 173884584 | 1376 | 1.02 | 1.18E-05 | 4.05E-05 |
| chr2 | 120821876 | 120826219 | 4344 | -0.5 | 1.18E-05 | 4.05E-05 |
| chr18 | 25643023 | 25672311 | 29289 | 0.5 | 1.18E-05 | 4.05E-05 |
| chr19 | 11039487 | 11072793 | 33307 | 0.46 | 1.18E-05 | 4.05E-05 |
| chr17 | 15155327 | 15194561 | 39235 | 0.27 | 1.18E-05 | 4.06E-05 |
| chr3 | 86078261 | 86079635 | 1375 | 1.81 | 1.19E-05 | 4.06E-05 |
| chr8 | 118858975 | 118861629 | 2655 | 1.39 | 1.19E-05 | 4.07E-05 |
| chr15 | 75466151 | 75475784 | 9634 | 0.4 | 1.19E-05 | 4.08E-05 |
| chr1 | 99568490 | 99568983 | 494 | 5.38 | 1.19E-05 | 4.09E-05 |
| chr7 | 103127879 | 103128662 | 784 | 5.38 | 1.19E-05 | 4.09E-05 |
| chr16 | 30568535 | 30584291 | 15757 | 0.43 | 1.20E-05 | 4.09E-05 |
| chr17 | 60102353 | 60144055 | 41703 | 0.3 | 1.20E-05 | 4.10E-05 |
| chr8 | 82935049 | 82936367 | 1319 | 3 | 1.20E-05 | 4.10E-05 |
| chr4 | 104343276 | 104347294 | 4019 | 5.73 | 1.20E-05 | 4.10E-05 |
| chr2 | 223723896 | 223791169 | 67274 | -0.29 | 1.20E-05 | 4.10E-05 |
| chr10 | 105505196 | 105596286 | 91091 | 0.24 | 1.20E-05 | 4.11E-05 |
| chr19 | 12273883 | 12274937 | 1055 | 1.42 | 1.20E-05 | 4.12E-05 |
| chr10 | 29907317 | 29910767 | 3451 | 1.37 | 1.21E-05 | 4.13E-05 |
| chr5 | 59005113 | 59006440 | 1328 | 1.63 | 1.21E-05 | 4.14E-05 |
| chr12 | 85763204 | 85764136 | 933 | 2.77 | 1.21E-05 | 4.14E-05 |
| chr11 | 30343935 | 30346315 | 2381 | 0.74 | 1.21E-05 | 4.15E-05 |
| chr15 | 57612669 | 57616436 | 3768 | 0.77 | 1.22E-05 | 4.16E-05 |
| chr7 | 12796142 | 12817742 | 21601 | -1.43 | 1.22E-05 | 4.17E-05 |
| chr8 | 79577388 | 79579600 | 2213 | 0.76 | 1.22E-05 | 4.18E-05 |
| chr8 | 95907015 | 95918293 | 11279 | -0.92 | 1.23E-05 | 4.19E-05 |
| chr20 | 46808527 | 46810148 | 1622 | 1.4 | 1.23E-05 | 4.21E-05 |
| chr3 | 27234403 | 27244695 | 10293 | 0.4 | 1.23E-05 | 4.21E-05 |
| chr10 | 133438493 | 133439345 | 853 | 5.35 | 1.23E-05 | 4.21E-05 |
| chr7 | 117337300 | 117355904 | 18605 | 5.14 | 1.23E-05 | 4.21E-05 |
| chr1 | 204677012 | 204678439 | 1428 | 1.63 | 1.24E-05 | 4.23E-05 |
| chr7 | 37349384 | 37350446 | 1063 | -2.69 | 1.24E-05 | 4.24E-05 |
| chr1 | 242153624 | 242163112 | 9489 | 0.94 | 1.24E-05 | 4.25E-05 |
| chr1 | 20189887 | 20191027 | 1141 | 5.42 | 1.25E-05 | 4.26E-05 |
| chr7 | 17596834 | 17600230 | 3397 | 0.64 | 1.25E-05 | 4.27E-05 |
| chr8 | 131198875 | 131210509 | 11635 | -0.66 | 1.25E-05 | 4.28E-05 |
| chr4 | 7703498 | 7704563 | 1066 | -2.33 | 1.25E-05 | 4.28E-05 |
| chr15 | 85383384 | 85389459 | 6076 | 0.78 | 1.25E-05 | 4.28E-05 |
| chr1 | 201851465 | 201865645 | 14181 | -0.9 | 1.26E-05 | 4.28E-05 |
| chr5 | 149177863 | 149187118 | 9256 | -0.49 | 1.26E-05 | 4.29E-05 |
| chr1 | 77684299 | 77696800 | 12502 | 0.75 | 1.26E-05 | 4.30E-05 |
| chr9 | 107949257 | 107955244 | 5988 | 0.57 | 1.26E-05 | 4.30E-05 |
| chr18 | 25755815 | 25758978 | 3164 | 1.39 | 1.26E-05 | 4.30E-05 |
| chr15 | 54201986 | 54202797 | 812 | -4.63 | 1.26E-05 | 4.31E-05 |
| chr6 | 21586917 | 21598558 | 11642 | 0.55 | 1.27E-05 | 4.31E-05 |
| chr1 | 109189115 | 109190320 | 1206 | 2.14 | 1.27E-05 | 4.31E-05 |
| chr2 | 178195944 | 178204100 | 8157 | -1.3 | 1.27E-05 | 4.32E-05 |
| chrX | 119682133 | 119702429 | 20297 | 0.37 | 1.27E-05 | 4.33E-05 |
| chr4 | 48018199 | 48019442 | 1244 | 1.86 | 1.28E-05 | 4.37E-05 |

| | | | | | | |
|---|---|---|---|---|---|---|
| chr20 | 48632358 | 48661145 | 28788 | -0.37 | 1.28E-05 | 4.38E-05 |
| chr10 | 97204659 | 97206312 | 1654 | -1.92 | 1.28E-05 | 4.38E-05 |
| chr15 | 101707746 | 101720775 | 13030 | 0.82 | 1.29E-05 | 4.38E-05 |
| chr2 | 202695029 | 202696064 | 1036 | 3.56 | 1.29E-05 | 4.40E-05 |
| chr19 | 21841630 | 21842540 | 911 | 5.29 | 1.29E-05 | 4.40E-05 |
| chr4 | 155320880 | 155321366 | 487 | 5.29 | 1.29E-05 | 4.40E-05 |
| chr8 | 96202433 | 96211305 | 8873 | 1.51 | 1.29E-05 | 4.40E-05 |
| chr14 | 75696016 | 75706439 | 10424 | 0.63 | 1.29E-05 | 4.40E-05 |
| chr13 | 94382740 | 94383607 | 868 | 2.69 | 1.29E-05 | 4.41E-05 |
| chr14 | 22888457 | 22903071 | 14615 | 5.8 | 1.30E-05 | 4.42E-05 |
| chr13 | 31189075 | 31193609 | 4535 | 0.44 | 1.30E-05 | 4.42E-05 |
| chr1 | 94583265 | 94592168 | 8904 | 1.36 | 1.30E-05 | 4.42E-05 |
| chr3 | 50628053 | 50640051 | 11999 | -0.97 | 1.30E-05 | 4.43E-05 |
| chr4 | 83987629 | 83997427 | 9799 | -0.78 | 1.31E-05 | 4.44E-05 |
| chr12 | 10902241 | 10902837 | 597 | 1.71 | 1.31E-05 | 4.45E-05 |
| chr8 | 42395730 | 42399427 | 3698 | 0.46 | 1.31E-05 | 4.46E-05 |
| chr5 | 91633147 | 91633984 | 838 | 3.52 | 1.32E-05 | 4.47E-05 |
| chr6 | 34622496 | 34691948 | 69453 | -0.29 | 1.32E-05 | 4.47E-05 |
| chr6 | 106772310 | 106774539 | 2230 | 0.65 | 1.32E-05 | 4.47E-05 |
| chr8 | 86157331 | 86158398 | 1068 | 1.57 | 1.32E-05 | 4.47E-05 |
| chr19 | 35223821 | 35226166 | 2346 | 0.49 | 1.32E-05 | 4.48E-05 |
| chr13 | 35251699 | 35253209 | 1511 | 2 | 1.32E-05 | 4.48E-05 |
| chr7 | 16782590 | 16794289 | 11700 | 1.08 | 1.32E-05 | 4.49E-05 |
| chr4 | 171001412 | 171012154 | 10743 | -0.88 | 1.33E-05 | 4.51E-05 |
| chr7 | 57508841 | 57509441 | 601 | 4.08 | 1.33E-05 | 4.51E-05 |
| chr19 | 18384744 | 18461211 | 76468 | 0.22 | 1.33E-05 | 4.52E-05 |
| chr12 | 126357178 | 126357623 | 446 | 5.29 | 1.33E-05 | 4.52E-05 |
| chr3 | 104307042 | 104307580 | 539 | 5.29 | 1.33E-05 | 4.52E-05 |
| chr16 | 18797799 | 18832449 | 34651 | 0.34 | 1.33E-05 | 4.53E-05 |
| chr5 | 83543743 | 83544422 | 680 | 5.31 | 1.33E-05 | 4.53E-05 |
| chr12 | 32992354 | 32997158 | 4805 | 1.09 | 1.34E-05 | 4.54E-05 |
| chr7 | 30466968 | 30475557 | 8590 | 0.66 | 1.34E-05 | 4.54E-05 |
| chr11 | 2798659 | 2801305 | 2647 | 1.08 | 1.34E-05 | 4.55E-05 |
| chr22 | 44843681 | 44846036 | 2356 | -1.44 | 1.34E-05 | 4.55E-05 |
| chr1 | 8471935 | 8485241 | 13307 | 0.78 | 1.35E-05 | 4.57E-05 |
| chr17 | 66831299 | 66831839 | 541 | 5.29 | 1.35E-05 | 4.58E-05 |
| chr2 | 236401965 | 236432298 | 30334 | -0.47 | 1.35E-05 | 4.59E-05 |
| chr17 | 34110281 | 34114406 | 4126 | 2.07 | 1.35E-05 | 4.59E-05 |
| chr7 | 48178577 | 48184773 | 6197 | 1.1 | 1.35E-05 | 4.59E-05 |
| chr10 | 94731211 | 94732758 | 1548 | 1.03 | 1.35E-05 | 4.59E-05 |
| chr10 | 115966410 | 115967161 | 752 | 4.27 | 1.36E-05 | 4.60E-05 |
| chr3 | 58183977 | 58184986 | 1010 | 3.42 | 1.36E-05 | 4.60E-05 |
| chr12 | 120050841 | 120051754 | 914 | 3.14 | 1.36E-05 | 4.60E-05 |
| chr6 | 152782510 | 152793237 | 10728 | -2.11 | 1.36E-05 | 4.61E-05 |
| chr12 | 104874592 | 104881624 | 7033 | -1.67 | 1.37E-05 | 4.64E-05 |
| chr2 | 121831784 | 121833808 | 2025 | -1.75 | 1.37E-05 | 4.64E-05 |
| chr6 | 35692580 | 35700795 | 8216 | 0.62 | 1.37E-05 | 4.64E-05 |
| chr5 | 122454736 | 122456025 | 1290 | 0.95 | 1.37E-05 | 4.64E-05 |
| chrX | 36273492 | 36274261 | 770 | 3.2 | 1.37E-05 | 4.64E-05 |
| chr1 | 29502861 | 29509770 | 6910 | 0.56 | 1.37E-05 | 4.65E-05 |
| chr20 | 45637233 | 45638637 | 1405 | 1.79 | 1.37E-05 | 4.65E-05 |
| chr9 | 123291181 | 123310338 | 19158 | 0.42 | 1.38E-05 | 4.66E-05 |
| chr19 | 6603458 | 6612063 | 8606 | -1.44 | 1.38E-05 | 4.67E-05 |
| chr3 | 9833283 | 9852798 | 19516 | 0.29 | 1.38E-05 | 4.69E-05 |
| chr2 | 203129165 | 203153078 | 23914 | -0.47 | 1.39E-05 | 4.69E-05 |
| chr11 | 9479731 | 9502162 | 22432 | 0.28 | 1.39E-05 | 4.70E-05 |
| chr11 | 28064316 | 28145235 | 80920 | 0.22 | 1.39E-05 | 4.72E-05 |
| chr10 | 60752680 | 60767511 | 14832 | 0.81 | 1.40E-05 | 4.72E-05 |
| chr1 | 157963103 | 158009037 | 45935 | -0.2 | 1.40E-05 | 4.73E-05 |
| chr13 | 84128440 | 84129216 | 777 | 4.69 | 1.40E-05 | 4.75E-05 |
| chr7 | 12961061 | 12961580 | 520 | 2.41 | 1.40E-05 | 4.75E-05 |

| | | | | | | |
|---|---|---|---|---|---|---|
| chr17 | 8888622 | 8889188 | 567 | 5.55 | 1.40E-05 | 4.75E-05 |
| chr2 | 181438287 | 181438811 | 525 | 5.3 | 1.41E-05 | 4.76E-05 |
| chr10 | 121943132 | 121943738 | 607 | -4.63 | 1.41E-05 | 4.76E-05 |
| chr17 | 36664596 | 36669158 | 4563 | 1.17 | 1.41E-05 | 4.76E-05 |
| chr1 | 6885914 | 6886991 | 1078 | 2.16 | 1.41E-05 | 4.77E-05 |
| chr8 | 49166496 | 49167066 | 571 | 3.77 | 1.42E-05 | 4.79E-05 |
| chr20 | 35200911 | 35204248 | 3338 | 1.45 | 1.42E-05 | 4.79E-05 |
| chr9 | 116789944 | 116844888 | 54945 | -0.41 | 1.42E-05 | 4.79E-05 |
| chr12 | 51374068 | 51379408 | 5341 | 1.39 | 1.42E-05 | 4.80E-05 |
| chr3 | 40565367 | 40567474 | 2108 | 0.6 | 1.42E-05 | 4.81E-05 |
| chr2 | 158986316 | 158989006 | 2691 | 0.9 | 1.42E-05 | 4.81E-05 |
| chr12 | 46736541 | 46737753 | 1213 | -1.98 | 1.43E-05 | 4.83E-05 |
| chr15 | 25164596 | 25165262 | 667 | 5.38 | 1.43E-05 | 4.83E-05 |
| chr10 | 128264577 | 128265053 | 477 | 5.27 | 1.44E-05 | 4.85E-05 |
| chr7 | 145767842 | 145768234 | 393 | 5.27 | 1.44E-05 | 4.85E-05 |
| chr3 | 164668537 | 164669593 | 1057 | 1.84 | 1.44E-05 | 4.85E-05 |
| chr10 | 70281915 | 70290604 | 8690 | 0.62 | 1.44E-05 | 4.85E-05 |
| chrX | 120654785 | 120655564 | 780 | 5.27 | 1.44E-05 | 4.86E-05 |
| chr12 | 111877964 | 111944296 | 66333 | 0.29 | 1.44E-05 | 4.87E-05 |
| chr16 | 66440302 | 66443796 | 3495 | 0.49 | 1.44E-05 | 4.87E-05 |
| chr9 | 7027032 | 7027740 | 709 | -4 | 1.45E-05 | 4.89E-05 |
| chr9 | 136324123 | 136326892 | 2770 | 0.79 | 1.45E-05 | 4.89E-05 |
| chr3 | 182156030 | 182156635 | 606 | 4.36 | 1.45E-05 | 4.89E-05 |
| chr3 | 113993223 | 113994557 | 1335 | -1.38 | 1.45E-05 | 4.89E-05 |
| chr6 | 132832305 | 132834943 | 2639 | 0.92 | 1.45E-05 | 4.90E-05 |
| chr11 | 27332338 | 27341378 | 9041 | 0.68 | 1.46E-05 | 4.93E-05 |
| chr13 | 74122447 | 74123307 | 861 | 5.27 | 1.47E-05 | 4.95E-05 |
| chrX | 140434570 | 140437122 | 2553 | -0.62 | 1.47E-05 | 4.96E-05 |
| chr2 | 238844241 | 238849094 | 4854 | -2.85 | 1.47E-05 | 4.96E-05 |
| chr3 | 96369270 | 96369876 | 607 | 4.32 | 1.47E-05 | 4.97E-05 |
| chr2 | 239221976 | 239230447 | 8472 | -0.89 | 1.47E-05 | 4.97E-05 |
| chr1 | 52470786 | 52474431 | 3646 | 0.92 | 1.48E-05 | 4.97E-05 |
| chr14 | 25476618 | 25479697 | 3080 | 1.86 | 1.48E-05 | 4.98E-05 |
| chr2 | 134599096 | 134600384 | 1289 | -3.46 | 1.48E-05 | 5.00E-05 |
| chr4 | 3586948 | 3587443 | 496 | 5.27 | 1.48E-05 | 5.00E-05 |
| chr11 | 329044 | 329919 | 876 | 3.45 | 1.49E-05 | 5.01E-05 |
| chr20 | 58829811 | 58831053 | 1243 | 3.48 | 1.49E-05 | 5.02E-05 |
| chr7 | 4721176 | 4794036 | 72861 | 0.19 | 1.49E-05 | 5.03E-05 |
| chr11 | 91364391 | 91364895 | 505 | -5.22 | 1.49E-05 | 5.03E-05 |
| chr1 | 198172620 | 198173462 | 843 | -2.45 | 1.49E-05 | 5.03E-05 |
| chr17 | 79431304 | 79438656 | 7353 | -1.43 | 1.50E-05 | 5.04E-05 |
| chr3 | 112389686 | 112409059 | 19374 | -0.42 | 1.50E-05 | 5.06E-05 |
| chr5 | 158547261 | 158548084 | 824 | 2.36 | 1.50E-05 | 5.07E-05 |
| chr12 | 67462090 | 67464304 | 2215 | 1.18 | 1.51E-05 | 5.07E-05 |
| chr6 | 107584335 | 107586494 | 2160 | 0.94 | 1.51E-05 | 5.07E-05 |
| chrX | 95690646 | 95691661 | 1016 | 2.83 | 1.51E-05 | 5.08E-05 |
| chr11 | 112160477 | 112173919 | 13443 | 0.92 | 1.51E-05 | 5.09E-05 |
| chr5 | 120608669 | 120610923 | 2255 | 0.48 | 1.52E-05 | 5.11E-05 |
| chr16 | 48191617 | 48200545 | 8929 | -0.76 | 1.52E-05 | 5.11E-05 |
| chr9 | 79678500 | 79679441 | 942 | -5.22 | 1.52E-05 | 5.12E-05 |
| chr12 | 100792313 | 100797950 | 5638 | 4.37 | 1.52E-05 | 5.12E-05 |
| chr16 | 4587876 | 4589390 | 1515 | 2.1 | 1.52E-05 | 5.12E-05 |
| chr17 | 19474029 | 19474981 | 953 | -2.38 | 1.52E-05 | 5.13E-05 |
| chr17 | 4113001 | 4152039 | 39039 | -0.38 | 1.53E-05 | 5.15E-05 |
| chr15 | 87037802 | 87038855 | 1054 | 2.17 | 1.53E-05 | 5.16E-05 |
| chr5 | 106184654 | 106185623 | 970 | 2.24 | 1.54E-05 | 5.16E-05 |
| chr14 | 54937101 | 54937933 | 833 | 2.17 | 1.54E-05 | 5.17E-05 |
| chr13 | 48876567 | 48879307 | 2741 | 0.54 | 1.55E-05 | 5.21E-05 |
| chr4 | 134109215 | 134109736 | 522 | 5.27 | 1.55E-05 | 5.22E-05 |
| chr9 | 95147827 | 95148439 | 613 | 4.24 | 1.56E-05 | 5.25E-05 |
| chr20 | 15183665 | 15184397 | 733 | 3.36 | 1.56E-05 | 5.26E-05 |

| | | | | | | |
|---|---|---|---|---|---|---|
| chr15 | 71184025 | 71186056 | 2032 | -0.69 | 1.57E-05 | 5.26E-05 |
| chr13 | 66559515 | 66560750 | 1236 | 3.43 | 1.57E-05 | 5.28E-05 |
| chr11 | 130716554 | 130718071 | 1518 | 1.91 | 1.57E-05 | 5.28E-05 |
| chr4 | 187693693 | 187694422 | 730 | -3.96 | 1.58E-05 | 5.29E-05 |
| chr10 | 105032747 | 105033706 | 960 | -1.79 | 1.59E-05 | 5.33E-05 |
| chr2 | 57072983 | 57073330 | 348 | 5.27 | 1.59E-05 | 5.33E-05 |
| chr14 | 62065822 | 62070195 | 4374 | 1.73 | 1.59E-05 | 5.34E-05 |
| chr5 | 169599807 | 169601132 | 1326 | -1.44 | 1.59E-05 | 5.35E-05 |
| chr8 | 9638626 | 9640672 | 2047 | 1.01 | 1.60E-05 | 5.36E-05 |
| chr15 | 52488221 | 52528079 | 39859 | 0.51 | 1.60E-05 | 5.36E-05 |
| chr2 | 207629118 | 207631416 | 2299 | -0.71 | 1.60E-05 | 5.37E-05 |
| chr2 | 172885670 | 172887632 | 1963 | 0.94 | 1.61E-05 | 5.39E-05 |
| chr6 | 82949880 | 82971555 | 21676 | 0.3 | 1.61E-05 | 5.39E-05 |
| chr10 | 87380924 | 87382859 | 1936 | 1.04 | 1.61E-05 | 5.40E-05 |
| chr2 | 33500187 | 33502796 | 2610 | 0.94 | 1.61E-05 | 5.41E-05 |
| chr8 | 42696323 | 42699858 | 3536 | 0.63 | 1.62E-05 | 5.42E-05 |
| chr3 | 14918068 | 14919513 | 1446 | 1.32 | 1.62E-05 | 5.42E-05 |
| chr12 | 85862228 | 85864018 | 1791 | 1.15 | 1.62E-05 | 5.43E-05 |
| chr5 | 57735988 | 57762864 | 26877 | -0.38 | 1.62E-05 | 5.44E-05 |
| chr2 | 15922060 | 15922888 | 829 | -4.57 | 1.63E-05 | 5.47E-05 |
| chr17 | 66015197 | 66042353 | 27157 | 0.43 | 1.63E-05 | 5.47E-05 |
| chr2 | 7889170 | 7889816 | 647 | 4.65 | 1.63E-05 | 5.48E-05 |
| chr3 | 95425015 | 95425489 | 475 | 5.25 | 1.64E-05 | 5.49E-05 |
| chr17 | 20969506 | 20970182 | 677 | -5.3 | 1.64E-05 | 5.50E-05 |
| chr5 | 39517533 | 39527675 | 10143 | 1.15 | 1.64E-05 | 5.51E-05 |
| chr16 | 51441672 | 51443717 | 2046 | 1.06 | 1.65E-05 | 5.52E-05 |
| chr2 | 224556760 | 224566704 | 9945 | 0.51 | 1.66E-05 | 5.55E-05 |
| chr17 | 76656689 | 76679310 | 22622 | 0.38 | 1.66E-05 | 5.57E-05 |
| chr2 | 203101219 | 203114844 | 13626 | -0.72 | 1.67E-05 | 5.59E-05 |
| chr4 | 7587802 | 7588742 | 941 | 3.03 | 1.67E-05 | 5.61E-05 |
| chr3 | 180319309 | 180321405 | 2097 | 0.86 | 1.68E-05 | 5.62E-05 |
| chr5 | 156452078 | 156500135 | 48058 | 0.2 | 1.68E-05 | 5.63E-05 |
| chr15 | 70144432 | 70147749 | 3318 | 1.32 | 1.69E-05 | 5.65E-05 |
| chr7 | 906789 | 917070 | 10282 | 0.68 | 1.69E-05 | 5.66E-05 |
| chr14 | 93752682 | 93800693 | 48012 | -0.27 | 1.69E-05 | 5.67E-05 |
| chr8 | 129238636 | 129243250 | 4615 | 1.12 | 1.69E-05 | 5.67E-05 |
| chr11 | 101619096 | 101632759 | 13664 | 3.65 | 1.69E-05 | 5.67E-05 |
| chr17 | 31074563 | 31075520 | 958 | 3.24 | 1.69E-05 | 5.67E-05 |
| chr20 | 18464993 | 18492507 | 27515 | 0.46 | 1.70E-05 | 5.68E-05 |
| chr2 | 95965697 | 95973434 | 7738 | 5.98 | 1.70E-05 | 5.68E-05 |
| chr12 | 58281947 | 58299232 | 17286 | 0.55 | 1.70E-05 | 5.68E-05 |
| chr1 | 43918343 | 43948322 | 29980 | 0.31 | 1.70E-05 | 5.68E-05 |
| chr3 | 95075389 | 95075962 | 574 | 5.25 | 1.70E-05 | 5.69E-05 |
| chr10 | 5956720 | 5957651 | 932 | -3.78 | 1.70E-05 | 5.70E-05 |
| chr5 | 71799767 | 71800633 | 867 | -1.82 | 1.70E-05 | 5.70E-05 |
| chr13 | 24368197 | 24370013 | 1817 | 1.04 | 1.71E-05 | 5.70E-05 |
| chr3 | 130826497 | 130828610 | 2114 | 1.21 | 1.71E-05 | 5.70E-05 |
| chr12 | 112380126 | 112381162 | 1037 | 2.42 | 1.71E-05 | 5.71E-05 |
| chr13 | 60969545 | 60976961 | 7417 | 0.52 | 1.71E-05 | 5.73E-05 |
| chr1 | 10143429 | 10168984 | 25556 | -0.81 | 1.72E-05 | 5.74E-05 |
| chr7 | 87198358 | 87199225 | 868 | 5.3 | 1.72E-05 | 5.74E-05 |
| chr5 | 157272667 | 157299387 | 26721 | 0.4 | 1.72E-05 | 5.74E-05 |
| chr14 | 47059789 | 47061571 | 1783 | 1.04 | 1.72E-05 | 5.74E-05 |
| chr19 | 16434280 | 16462807 | 28528 | 0.43 | 1.72E-05 | 5.74E-05 |
| chr2 | 67536114 | 67536922 | 809 | 5.28 | 1.73E-05 | 5.76E-05 |
| chr18 | 47073657 | 47075621 | 1965 | 1.17 | 1.73E-05 | 5.77E-05 |
| chr21 | 44232143 | 44232675 | 533 | 4.72 | 1.73E-05 | 5.78E-05 |
| chr5 | 13605778 | 13607191 | 1414 | 1.62 | 1.74E-05 | 5.79E-05 |
| chr1 | 105340513 | 105341201 | 689 | 5.24 | 1.74E-05 | 5.79E-05 |
| chr1 | 172640420 | 172641431 | 1012 | 5.24 | 1.74E-05 | 5.79E-05 |
| chr4 | 19695535 | 19696102 | 568 | 5.24 | 1.74E-05 | 5.79E-05 |

| | | | | | | |
|---|---|---|---|---|---|---|
| chr7 | 124121552 | 124130921 | 9370 | 6.39 | 1.74E-05 | 5.79E-05 |
| chr7 | 92075933 | 92078782 | 2850 | 0.64 | 1.74E-05 | 5.80E-05 |
| chr10 | 102901574 | 102902148 | 575 | -2.33 | 1.74E-05 | 5.82E-05 |
| chr7 | 134463613 | 134465062 | 1450 | 1.31 | 1.74E-05 | 5.82E-05 |
| chr3 | 122479317 | 122481165 | 1849 | 1.75 | 1.75E-05 | 5.85E-05 |
| chr7 | 114293706 | 114294454 | 749 | 4.22 | 1.75E-05 | 5.85E-05 |
| chr14 | 105863696 | 105866170 | 2475 | 0.68 | 1.76E-05 | 5.86E-05 |
| chr3 | 134346136 | 134347438 | 1303 | 1.42 | 1.76E-05 | 5.87E-05 |
| chr11 | 68337926 | 68365178 | 27253 | -0.54 | 1.76E-05 | 5.87E-05 |
| chr20 | 55326825 | 55384077 | 57253 | 0.21 | 1.76E-05 | 5.88E-05 |
| chr22 | 42573093 | 42585474 | 12382 | 1.03 | 1.76E-05 | 5.88E-05 |
| chr16 | 72908707 | 72924596 | 15890 | -0.49 | 1.77E-05 | 5.89E-05 |
| chr6 | 7899254 | 7911551 | 12298 | -0.74 | 1.77E-05 | 5.90E-05 |
| chr8 | 121983505 | 121984006 | 502 | 4.59 | 1.77E-05 | 5.91E-05 |
| chr1 | 231194109 | 231195935 | 1827 | -1.32 | 1.77E-05 | 5.91E-05 |
| chr2 | 192282434 | 192283621 | 1188 | -2.39 | 1.78E-05 | 5.93E-05 |
| chr15 | 70271020 | 70271771 | 752 | -5.19 | 1.78E-05 | 5.93E-05 |
| chr2 | 236286391 | 236287600 | 1210 | 1.67 | 1.78E-05 | 5.94E-05 |
| chr6 | 105880315 | 105881413 | 1099 | -2.92 | 1.78E-05 | 5.94E-05 |
| chr8 | 128259430 | 128266510 | 7081 | 1.09 | 1.79E-05 | 5.97E-05 |
| chr11 | 34447888 | 34461195 | 13308 | -0.94 | 1.80E-05 | 5.99E-05 |
| chr14 | 60296288 | 60297103 | 816 | -3.46 | 1.80E-05 | 5.99E-05 |
| chr8 | 27965000 | 27977446 | 12447 | 0.5 | 1.81E-05 | 6.01E-05 |
| chr6 | 83717469 | 83719938 | 2470 | 1.17 | 1.81E-05 | 6.03E-05 |
| chr16 | 70185960 | 70196904 | 10945 | -0.97 | 1.81E-05 | 6.04E-05 |
| chr5 | 156226260 | 156226935 | 676 | 3.76 | 1.82E-05 | 6.05E-05 |
| chr12 | 116570700 | 116610534 | 39835 | 0.43 | 1.82E-05 | 6.06E-05 |
| chr11 | 69026378 | 69027197 | 820 | 3.72 | 1.82E-05 | 6.07E-05 |
| chr1 | 183812247 | 183813360 | 1114 | 2.59 | 1.83E-05 | 6.10E-05 |
| chr4 | 119888940 | 119889714 | 775 | 2.91 | 1.83E-05 | 6.10E-05 |
| chr6 | 133008010 | 133009233 | 1224 | 1.93 | 1.84E-05 | 6.13E-05 |
| chr3 | 141835701 | 141837157 | 1457 | 1.75 | 1.84E-05 | 6.13E-05 |
| chr10 | 46075804 | 46076492 | 689 | 5.22 | 1.85E-05 | 6.14E-05 |
| chr3 | 151482797 | 151537111 | 54315 | -0.17 | 1.85E-05 | 6.16E-05 |
| chr10 | 3022886 | 3028475 | 5590 | -1.9 | 1.86E-05 | 6.17E-05 |
| chr4 | 138544243 | 138544782 | 540 | 3.96 | 1.86E-05 | 6.19E-05 |
| chr13 | 32500993 | 32505643 | 4651 | 1.93 | 1.87E-05 | 6.21E-05 |
| chr1 | 228247505 | 228248588 | 1084 | 1.47 | 1.87E-05 | 6.22E-05 |
| chr14 | 90797027 | 90799166 | 2140 | 0.62 | 1.87E-05 | 6.23E-05 |
| chr15 | 26806748 | 26807977 | 1230 | -2.49 | 1.88E-05 | 6.23E-05 |
| chr11 | 60920632 | 60930059 | 9428 | 0.64 | 1.88E-05 | 6.23E-05 |
| chr19 | 48823609 | 48931419 | 107811 | 0.2 | 1.89E-05 | 6.27E-05 |
| chr17 | 52977632 | 52981408 | 3777 | 0.57 | 1.89E-05 | 6.27E-05 |
| chr11 | 106268711 | 106271041 | 2331 | -1.28 | 1.89E-05 | 6.28E-05 |
| chr14 | 74299489 | 74301489 | 2001 | 1.57 | 1.89E-05 | 6.29E-05 |
| chr5 | 92358098 | 92360021 | 1924 | 2.07 | 1.89E-05 | 6.29E-05 |
| chr17 | 33774599 | 33776238 | 1640 | 1.44 | 1.89E-05 | 6.29E-05 |
| chr5 | 23703897 | 23704409 | 513 | 3.89 | 1.90E-05 | 6.29E-05 |
| chr20 | 37610723 | 37614844 | 4122 | 1.42 | 1.90E-05 | 6.29E-05 |
| chr14 | 73869128 | 73899565 | 30438 | 0.41 | 1.90E-05 | 6.30E-05 |
| chr17 | 13323669 | 13324317 | 649 | 2.83 | 1.90E-05 | 6.31E-05 |
| chr2 | 177177998 | 177192143 | 14146 | 0.61 | 1.90E-05 | 6.32E-05 |
| chr19 | 56763860 | 56764304 | 445 | 4 | 1.91E-05 | 6.32E-05 |
| chr19 | 1847065 | 1906426 | 59362 | -0.32 | 1.91E-05 | 6.33E-05 |
| chr12 | 79514902 | 79519940 | 5039 | 1.16 | 1.91E-05 | 6.33E-05 |
| chr7 | 116764953 | 116773698 | 8746 | 0.72 | 1.91E-05 | 6.35E-05 |
| chr8 | 12610476 | 12614002 | 3527 | 0.7 | 1.93E-05 | 6.40E-05 |
| chr19 | 51160490 | 51162851 | 2362 | 0.57 | 1.93E-05 | 6.40E-05 |
| chr1 | 193665985 | 193679429 | 13445 | 0.38 | 1.94E-05 | 6.43E-05 |
| chr17 | 64408929 | 64437972 | 29044 | 0.43 | 1.94E-05 | 6.43E-05 |
| chr9 | 88551581 | 88589669 | 38089 | 0.37 | 1.94E-05 | 6.43E-05 |

| | | | | | | |
|---|---|---|---|---|---|---|
| chr11 | 102028295 | 102051001 | 22707 | 0.63 | 1.94E-05 | 6.43E-05 |
| chr6 | 100010806 | 100017456 | 6651 | 0.81 | 1.94E-05 | 6.43E-05 |
| chr4 | 148213548 | 148220842 | 7295 | 1.98 | 1.94E-05 | 6.43E-05 |
| chr14 | 96967362 | 97023796 | 56435 | 0.26 | 1.95E-05 | 6.45E-05 |
| chrX | 149935335 | 149945237 | 9903 | -0.83 | 1.95E-05 | 6.45E-05 |
| chr9 | 86750364 | 86762849 | 12486 | 4.02 | 1.95E-05 | 6.47E-05 |
| chr10 | 43158628 | 43160862 | 2235 | 5.84 | 1.96E-05 | 6.49E-05 |
| chr9 | 112810835 | 112812536 | 1702 | 1.52 | 1.96E-05 | 6.50E-05 |
| chr3 | 120360192 | 120362352 | 2161 | -1.72 | 1.96E-05 | 6.51E-05 |
| chr3 | 44666034 | 44667167 | 1134 | 1.32 | 1.97E-05 | 6.51E-05 |
| chr14 | 80561462 | 80562114 | 653 | -5.16 | 1.98E-05 | 6.54E-05 |
| chr12 | 27396349 | 27398392 | 2044 | 0.77 | 1.98E-05 | 6.55E-05 |
| chr8 | 126643357 | 126677510 | 34154 | 0.48 | 1.98E-05 | 6.55E-05 |
| chr11 | 72712454 | 72714241 | 1788 | 2.15 | 1.98E-05 | 6.57E-05 |
| chr1 | 22394469 | 22416785 | 22317 | 0.64 | 1.99E-05 | 6.59E-05 |
| chr4 | 158854975 | 158855922 | 948 | 3.56 | 1.99E-05 | 6.59E-05 |
| chr5 | 73400550 | 73401459 | 910 | 3.66 | 1.99E-05 | 6.59E-05 |
| chr17 | 45365405 | 45366555 | 1151 | -2.39 | 1.99E-05 | 6.59E-05 |
| chr15 | 60756945 | 60772344 | 15400 | 0.24 | 2.01E-05 | 6.63E-05 |
| chr1 | 105786099 | 105786587 | 489 | 5.2 | 2.01E-05 | 6.64E-05 |
| chr1 | 151870145 | 151882773 | 12629 | 0.59 | 2.01E-05 | 6.64E-05 |
| chr12 | 126308247 | 126308902 | 656 | 5.23 | 2.01E-05 | 6.66E-05 |
| chr4 | 128544111 | 128555093 | 10983 | -0.99 | 2.02E-05 | 6.67E-05 |
| chr3 | 24699257 | 24700233 | 977 | 5.21 | 2.02E-05 | 6.67E-05 |
| chr3 | 161067072 | 161068263 | 1192 | 3.43 | 2.02E-05 | 6.67E-05 |
| chr19 | 40561226 | 40562886 | 1661 | 0.69 | 2.03E-05 | 6.71E-05 |
| chr3 | 141738589 | 141748732 | 10144 | 1.06 | 2.03E-05 | 6.71E-05 |
| chr6 | 6843515 | 6851916 | 8402 | 2.75 | 2.03E-05 | 6.71E-05 |
| chr7 | 110932046 | 110932576 | 531 | 4.56 | 2.03E-05 | 6.71E-05 |
| chr6 | 111910160 | 111937530 | 27371 | 0.34 | 2.04E-05 | 6.73E-05 |
| chrX | 24248719 | 24264287 | 15569 | -1.56 | 2.04E-05 | 6.73E-05 |
| chr7 | 7603621 | 7607875 | 4255 | 0.78 | 2.04E-05 | 6.74E-05 |
| chr7 | 62311949 | 62312585 | 637 | 5.2 | 2.04E-05 | 6.74E-05 |
| chr7 | 95174522 | 95179874 | 5353 | 1.18 | 2.04E-05 | 6.74E-05 |
| chr2 | 23824622 | 23826132 | 1511 | -2.36 | 2.04E-05 | 6.75E-05 |
| chr6 | 152779574 | 152780128 | 555 | 3.81 | 2.05E-05 | 6.75E-05 |
| chr8 | 116507817 | 116508331 | 515 | 5.28 | 2.05E-05 | 6.75E-05 |
| chr5 | 36442137 | 36443360 | 1224 | -2.34 | 2.05E-05 | 6.76E-05 |
| chr15 | 91002758 | 91020175 | 17418 | -0.63 | 2.05E-05 | 6.77E-05 |
| chr10 | 25971242 | 25971659 | 418 | 5.3 | 2.05E-05 | 6.78E-05 |
| chr17 | 59278207 | 59286366 | 8160 | -0.86 | 2.06E-05 | 6.78E-05 |
| chr1 | 9649146 | 9653825 | 4680 | -1.6 | 2.06E-05 | 6.78E-05 |
| chr18 | 7728331 | 7736531 | 8201 | -0.99 | 2.06E-05 | 6.78E-05 |
| chr1 | 246760256 | 246773538 | 13283 | 0.59 | 2.06E-05 | 6.79E-05 |
| chr17 | 42541335 | 42550913 | 9579 | -1.04 | 2.06E-05 | 6.80E-05 |
| chr22 | 32865526 | 32872409 | 6884 | 0.55 | 2.07E-05 | 6.83E-05 |
| chr12 | 71148027 | 71148479 | 453 | 5.22 | 2.08E-05 | 6.84E-05 |
| chr13 | 86239164 | 86239848 | 685 | -2.58 | 2.08E-05 | 6.84E-05 |
| chrX | 12378035 | 12380005 | 1971 | -1.14 | 2.08E-05 | 6.85E-05 |
| chr5 | 178820915 | 178821794 | 880 | 5.32 | 2.09E-05 | 6.88E-05 |
| chrX | 115586865 | 115587581 | 717 | 5.32 | 2.09E-05 | 6.88E-05 |
| chr5 | 73021614 | 73026733 | 5120 | 0.64 | 2.09E-05 | 6.88E-05 |
| chrX | 46119483 | 46132125 | 12643 | 0.77 | 2.09E-05 | 6.90E-05 |
| chr11 | 70144570 | 70181593 | 37024 | -0.52 | 2.10E-05 | 6.91E-05 |
| chr21 | 17078920 | 17079736 | 817 | 2.53 | 2.11E-05 | 6.94E-05 |
| chr8 | 37683677 | 37774308 | 90632 | 0.24 | 2.12E-05 | 6.97E-05 |
| chr18 | 60407868 | 60409664 | 1797 | 1.41 | 2.12E-05 | 6.99E-05 |
| chr11 | 3174736 | 3187068 | 12333 | -0.96 | 2.13E-05 | 7.01E-05 |
| chr3 | 33509150 | 33510195 | 1046 | -3.01 | 2.13E-05 | 7.01E-05 |
| chr4 | 38315428 | 38337264 | 21837 | 0.54 | 2.13E-05 | 7.02E-05 |
| chr3 | 169481687 | 169492857 | 11171 | 0.42 | 2.14E-05 | 7.03E-05 |

| | | | | | | |
|---|---|---|---|---|---|---|
| chr12 | 85818893 | 85819234 | 342 | 2.11 | 2.14E-05 | 7.04E-05 |
| chr17 | 6346408 | 6361126 | 14719 | -0.72 | 2.14E-05 | 7.05E-05 |
| chrX | 37313464 | 37315213 | 1750 | 2.05 | 2.15E-05 | 7.06E-05 |
| chr20 | 42983314 | 43031875 | 48562 | -0.32 | 2.15E-05 | 7.06E-05 |
| chr18 | 7900497 | 7963936 | 63440 | 0.19 | 2.15E-05 | 7.08E-05 |
| chr15 | 49015045 | 49021144 | 6100 | 1.45 | 2.16E-05 | 7.11E-05 |
| chr3 | 66659587 | 66661159 | 1573 | -1.75 | 2.17E-05 | 7.14E-05 |
| chr1 | 108740909 | 108744146 | 3238 | 0.6 | 2.17E-05 | 7.14E-05 |
| chr6 | 161649757 | 161650773 | 1017 | 2.5 | 2.17E-05 | 7.14E-05 |
| chr14 | 73762297 | 73779148 | 16852 | 0.63 | 2.17E-05 | 7.14E-05 |
| chr3 | 105680635 | 105682377 | 1743 | 1.7 | 2.17E-05 | 7.15E-05 |
| chr5 | 37301676 | 37318056 | 16381 | 0.82 | 2.18E-05 | 7.18E-05 |
| chr21 | 34657553 | 34674241 | 16689 | 0.4 | 2.19E-05 | 7.19E-05 |
| chr15 | 67522215 | 67523670 | 1456 | -2.21 | 2.20E-05 | 7.22E-05 |
| chr2 | 234120071 | 234120597 | 527 | 5.21 | 2.20E-05 | 7.24E-05 |
| chr20 | 58620894 | 58621877 | 984 | 2.71 | 2.21E-05 | 7.25E-05 |
| chr11 | 112096474 | 112098652 | 2179 | 0.6 | 2.22E-05 | 7.28E-05 |
| chr22 | 39599132 | 39611608 | 12477 | 0.64 | 2.22E-05 | 7.30E-05 |
| chr11 | 124350864 | 124352998 | 2135 | 1.51 | 2.22E-05 | 7.30E-05 |
| chr8 | 145925305 | 145925941 | 637 | 5.18 | 2.23E-05 | 7.31E-05 |
| chr10 | 79092804 | 79095329 | 2526 | -0.89 | 2.23E-05 | 7.33E-05 |
| chr7 | 12462256 | 12464143 | 1888 | 1.84 | 2.24E-05 | 7.34E-05 |
| chr22 | 29916372 | 29917775 | 1404 | -1.35 | 2.24E-05 | 7.35E-05 |
| chr2 | 61426870 | 61448456 | 21587 | 0.68 | 2.24E-05 | 7.35E-05 |
| chr16 | 9081016 | 9085991 | 4976 | 1.52 | 2.24E-05 | 7.36E-05 |
| chr4 | 187487235 | 187487771 | 537 | -3.94 | 2.25E-05 | 7.38E-05 |
| chr7 | 11538767 | 11539829 | 1063 | 3.75 | 2.25E-05 | 7.40E-05 |
| chr2 | 109465398 | 109466544 | 1147 | -1.97 | 2.26E-05 | 7.41E-05 |
| chr9 | 26310397 | 26310897 | 501 | 5.18 | 2.26E-05 | 7.42E-05 |
| chr7 | 117013248 | 117013831 | 584 | 5.2 | 2.27E-05 | 7.43E-05 |
| chr1 | 242987511 | 242988474 | 964 | -3.74 | 2.27E-05 | 7.44E-05 |
| chr12 | 19999352 | 20000306 | 955 | 1.81 | 2.27E-05 | 7.45E-05 |
| chr10 | 22517240 | 22520208 | 2969 | 0.71 | 2.28E-05 | 7.46E-05 |
| chr16 | 4186000 | 4191687 | 5688 | 1.11 | 2.28E-05 | 7.47E-05 |
| chr12 | 111500826 | 111502140 | 1315 | -3.47 | 2.28E-05 | 7.47E-05 |
| chr8 | 94928212 | 94930250 | 2039 | 0.97 | 2.29E-05 | 7.51E-05 |
| chr2 | 74493992 | 74495028 | 1037 | 3.03 | 2.29E-05 | 7.52E-05 |
| chr15 | 43396717 | 43399290 | 2574 | 0.67 | 2.30E-05 | 7.53E-05 |
| chr14 | 37721519 | 37723926 | 2408 | 2.37 | 2.30E-05 | 7.53E-05 |
| chr11 | 16301634 | 16302290 | 657 | 5.18 | 2.31E-05 | 7.57E-05 |
| chr7 | 87920654 | 87921152 | 499 | 5.18 | 2.31E-05 | 7.57E-05 |
| chr17 | 26723493 | 26733681 | 10189 | -0.79 | 2.31E-05 | 7.58E-05 |
| chr6 | 83756522 | 83758243 | 1722 | 1.36 | 2.32E-05 | 7.59E-05 |
| chr3 | 131310019 | 131317037 | 7019 | 1.86 | 2.32E-05 | 7.59E-05 |
| chr6 | 154875496 | 154877114 | 1619 | 1.23 | 2.32E-05 | 7.59E-05 |
| chr15 | 29252471 | 29254054 | 1584 | 1.06 | 2.32E-05 | 7.60E-05 |
| chr1 | 46296484 | 46306972 | 10489 | -1.58 | 2.33E-05 | 7.62E-05 |
| chr1 | 42896526 | 42897615 | 1090 | 3.88 | 2.33E-05 | 7.64E-05 |
| chr7 | 64254029 | 64255893 | 1865 | 0.82 | 2.33E-05 | 7.64E-05 |
| chr16 | 52668229 | 52668652 | 424 | 5.19 | 2.34E-05 | 7.64E-05 |
| chr2 | 20290857 | 20294338 | 3482 | -1.23 | 2.34E-05 | 7.64E-05 |
| chr7 | 25163435 | 25166228 | 2794 | 0.51 | 2.34E-05 | 7.65E-05 |
| chr8 | 17397760 | 17398266 | 507 | 5.22 | 2.34E-05 | 7.65E-05 |
| chr19 | 43967711 | 43980613 | 12903 | 0.71 | 2.34E-05 | 7.67E-05 |
| chr4 | 77295060 | 77295858 | 799 | 4.21 | 2.35E-05 | 7.67E-05 |
| chr2 | 218882454 | 218883816 | 1363 | -1.67 | 2.35E-05 | 7.69E-05 |
| chr12 | 18876593 | 18877379 | 787 | 3.45 | 2.35E-05 | 7.70E-05 |
| chr1 | 76054622 | 76055219 | 598 | 3.44 | 2.36E-05 | 7.72E-05 |
| chr2 | 128511865 | 128531800 | 19936 | -0.64 | 2.37E-05 | 7.76E-05 |
| chr21 | 45735542 | 45736031 | 490 | -3.5 | 2.37E-05 | 7.76E-05 |
| chr19 | 36192725 | 36250409 | 57685 | -0.23 | 2.37E-05 | 7.76E-05 |

| | | | | | | |
|---|---|---|---|---|---|---|
| chrX | 47414912 | 47421824 | 6913 | 0.55 | 2.38E-05 | 7.78E-05 |
| chr2 | 131811296 | 131814987 | 3692 | -1.48 | 2.38E-05 | 7.78E-05 |
| chr6 | 117760582 | 117771173 | 10592 | 0.56 | 2.39E-05 | 7.80E-05 |
| chr10 | 98590058 | 98601434 | 11377 | 0.6 | 2.39E-05 | 7.80E-05 |
| chr3 | 85014035 | 85014896 | 862 | 5.2 | 2.39E-05 | 7.80E-05 |
| chr12 | 104606470 | 104618302 | 11833 | 0.24 | 2.39E-05 | 7.80E-05 |
| chr2 | 234427639 | 234429690 | 2052 | -1.45 | 2.39E-05 | 7.80E-05 |
| chr18 | 71128425 | 71129545 | 1121 | 2.86 | 2.39E-05 | 7.80E-05 |
| chr4 | 76051520 | 76052101 | 582 | 2.23 | 2.39E-05 | 7.81E-05 |
| chr3 | 58549049 | 58549755 | 707 | -4.14 | 2.40E-05 | 7.84E-05 |
| chr9 | 85957914 | 85959133 | 1220 | -3.52 | 2.40E-05 | 7.84E-05 |
| chr1 | 55416104 | 55422048 | 5945 | 0.97 | 2.40E-05 | 7.85E-05 |
| chr4 | 102266099 | 102279387 | 13289 | 0.46 | 2.41E-05 | 7.87E-05 |
| chr13 | 102528045 | 102528482 | 438 | 5.19 | 2.42E-05 | 7.89E-05 |
| chrX | 80456369 | 80459189 | 2821 | 0.66 | 2.42E-05 | 7.89E-05 |
| chr11 | 78954072 | 78956492 | 2421 | 0.95 | 2.42E-05 | 7.90E-05 |
| chr11 | 126001390 | 126003150 | 1761 | 1.26 | 2.43E-05 | 7.92E-05 |
| chr10 | 4765289 | 4776487 | 11199 | 0.53 | 2.43E-05 | 7.92E-05 |
| chr14 | 35619409 | 35631575 | 12167 | -0.64 | 2.43E-05 | 7.92E-05 |
| chr1 | 68948438 | 68963355 | 14918 | 0.6 | 2.44E-05 | 7.96E-05 |
| chr15 | 64338064 | 64365824 | 27761 | -1.07 | 2.46E-05 | 8.03E-05 |
| chr5 | 17796429 | 17806189 | 9761 | 0.96 | 2.47E-05 | 8.05E-05 |
| chr9 | 99955632 | 99956190 | 559 | 3.54 | 2.47E-05 | 8.05E-05 |
| chr2 | 133951017 | 133959575 | 8559 | -3.98 | 2.47E-05 | 8.06E-05 |
| chr9 | 26137634 | 26138341 | 708 | 3.8 | 2.47E-05 | 8.06E-05 |
| chr18 | 55843335 | 55855184 | 11850 | 0.57 | 2.48E-05 | 8.09E-05 |
| chr5 | 87563446 | 87566598 | 3153 | 0.76 | 2.48E-05 | 8.09E-05 |
| chr8 | 134686615 | 134688038 | 1424 | 1.85 | 2.49E-05 | 8.11E-05 |
| chr2 | 7931999 | 7932680 | 682 | 5.16 | 2.49E-05 | 8.12E-05 |
| chr4 | 105648414 | 105650794 | 2381 | 1.06 | 2.50E-05 | 8.13E-05 |
| chr6 | 86131704 | 86132415 | 712 | -3.88 | 2.50E-05 | 8.13E-05 |
| chr18 | 2258786 | 2259426 | 641 | 3.36 | 2.50E-05 | 8.14E-05 |
| chr2 | 27711362 | 27713354 | 1993 | -0.85 | 2.50E-05 | 8.14E-05 |
| chr5 | 151063060 | 151083121 | 20062 | 1.67 | 2.50E-05 | 8.15E-05 |
| chr11 | 65028619 | 65159966 | 131348 | 0.2 | 2.50E-05 | 8.15E-05 |
| chr4 | 178935661 | 178936207 | 547 | 5.16 | 2.51E-05 | 8.17E-05 |
| chr7 | 39453411 | 39454232 | 822 | 5.24 | 2.52E-05 | 8.19E-05 |
| chr11 | 118185954 | 118186443 | 490 | 5.19 | 2.52E-05 | 8.21E-05 |
| chr1 | 212937365 | 212950697 | 13333 | -1.46 | 2.52E-05 | 8.21E-05 |
| chrX | 4896908 | 4897786 | 879 | 4.55 | 2.53E-05 | 8.22E-05 |
| chr10 | 5089909 | 5091537 | 1629 | -1.99 | 2.54E-05 | 8.26E-05 |
| chr7 | 19443404 | 19446058 | 2655 | 0.7 | 2.54E-05 | 8.27E-05 |
| chr5 | 133770890 | 133775763 | 4874 | 0.61 | 2.54E-05 | 8.27E-05 |
| chr11 | 128017057 | 128017825 | 769 | 5.16 | 2.55E-05 | 8.31E-05 |
| chr11 | 13918087 | 13918901 | 815 | 3.19 | 2.55E-05 | 8.31E-05 |
| chr11 | 120174940 | 120175886 | 947 | 2.58 | 2.56E-05 | 8.31E-05 |
| chr17 | 26851494 | 26852885 | 1392 | 1.17 | 2.56E-05 | 8.33E-05 |
| chr6 | 100705157 | 100706897 | 1741 | -0.81 | 2.56E-05 | 8.33E-05 |
| chr5 | 337617 | 344486 | 6870 | 2.29 | 2.57E-05 | 8.35E-05 |
| chr11 | 88287628 | 88289201 | 1574 | 1.37 | 2.57E-05 | 8.36E-05 |
| chr1 | 110923281 | 110951614 | 28334 | 0.34 | 2.57E-05 | 8.37E-05 |
| chr10 | 104283013 | 104283850 | 838 | -2.97 | 2.57E-05 | 8.37E-05 |
| chr2 | 82086381 | 82087013 | 633 | 5.16 | 2.58E-05 | 8.37E-05 |
| chr4 | 158102056 | 158102589 | 534 | 5.16 | 2.58E-05 | 8.37E-05 |
| chr5 | 34523517 | 34526349 | 2833 | -2.09 | 2.58E-05 | 8.38E-05 |
| chr16 | 25952278 | 25954476 | 2199 | 1.19 | 2.58E-05 | 8.38E-05 |
| chr7 | 112578368 | 112580963 | 2596 | 0.89 | 2.58E-05 | 8.38E-05 |
| chrX | 53482833 | 53484125 | 1293 | -1.71 | 2.59E-05 | 8.42E-05 |
| chr2 | 159047579 | 159048197 | 619 | 2.71 | 2.60E-05 | 8.43E-05 |
| chr2 | 224719668 | 224753204 | 33537 | -0.32 | 2.60E-05 | 8.43E-05 |
| chr1 | 61764788 | 61768325 | 3538 | 0.86 | 2.60E-05 | 8.44E-05 |

| | | | | | | |
|---|---|---|---|---|---|---|
| chr2 | 120243984 | 120250358 | 6375 | 0.84 | 2.61E-05 | 8.46E-05 |
| chr7 | 29533198 | 29535138 | 1941 | 1 | 2.62E-05 | 8.50E-05 |
| chr14 | 77810567 | 77811167 | 601 | -5.11 | 2.62E-05 | 8.50E-05 |
| chr11 | 13949936 | 13950429 | 494 | 5.16 | 2.62E-05 | 8.51E-05 |
| chr15 | 63688018 | 63689155 | 1138 | 1.4 | 2.63E-05 | 8.55E-05 |
| chr1 | 236331994 | 236341006 | 9013 | -0.87 | 2.63E-05 | 8.55E-05 |
| chr8 | 63625027 | 63625542 | 516 | -5.11 | 2.64E-05 | 8.56E-05 |
| chr10 | 93884136 | 93885223 | 1088 | 1.95 | 2.64E-05 | 8.56E-05 |
| chr12 | 107932230 | 107934521 | 2292 | -2.13 | 2.64E-05 | 8.57E-05 |
| chr8 | 41648461 | 41656811 | 8351 | 0.44 | 2.64E-05 | 8.58E-05 |
| chr5 | 81454802 | 81461970 | 7169 | 0.54 | 2.65E-05 | 8.58E-05 |
| chr7 | 39661978 | 39668682 | 6705 | 0.51 | 2.65E-05 | 8.60E-05 |
| chr2 | 208103735 | 208104522 | 788 | -3.15 | 2.65E-05 | 8.61E-05 |
| chr4 | 160691420 | 160692177 | 758 | 5.17 | 2.66E-05 | 8.61E-05 |
| chr21 | 42954813 | 42955262 | 450 | 4.19 | 2.66E-05 | 8.62E-05 |
| chr6 | 3472683 | 3473690 | 1008 | -1.53 | 2.66E-05 | 8.63E-05 |
| chr20 | 56935966 | 56948652 | 12687 | 1.38 | 2.66E-05 | 8.63E-05 |
| chr12 | 120868647 | 120908519 | 39873 | 0.44 | 2.67E-05 | 8.64E-05 |
| chr11 | 21468334 | 21469723 | 1390 | 1.7 | 2.67E-05 | 8.65E-05 |
| chr12 | 93637332 | 93638675 | 1344 | 1.94 | 2.67E-05 | 8.67E-05 |
| chr16 | 47897916 | 47899815 | 1900 | 2.08 | 2.68E-05 | 8.68E-05 |
| chr2 | 88837870 | 88838747 | 878 | -5.1 | 2.68E-05 | 8.69E-05 |
| chr18 | 7668263 | 7669120 | 858 | -2.51 | 2.70E-05 | 8.74E-05 |
| chr8 | 128658396 | 128658938 | 543 | 3.57 | 2.70E-05 | 8.75E-05 |
| chr17 | 38209941 | 38232338 | 22398 | -0.46 | 2.71E-05 | 8.76E-05 |
| chr4 | 120051476 | 120061452 | 9977 | 1.5 | 2.71E-05 | 8.78E-05 |
| chr3 | 149858497 | 149859570 | 1074 | -2.09 | 2.72E-05 | 8.79E-05 |
| chr2 | 170747552 | 170752813 | 5262 | -0.91 | 2.72E-05 | 8.80E-05 |
| chrX | 27501385 | 27502001 | 617 | 5.16 | 2.72E-05 | 8.80E-05 |
| chr3 | 181744762 | 181746122 | 1361 | 2.64 | 2.73E-05 | 8.84E-05 |
| chr11 | 13461136 | 13468642 | 7507 | 1.02 | 2.73E-05 | 8.84E-05 |
| chr3 | 112930134 | 112931771 | 1638 | -2.88 | 2.74E-05 | 8.85E-05 |
| chr5 | 148441366 | 148443013 | 1648 | 1.26 | 2.74E-05 | 8.87E-05 |
| chr5 | 74935403 | 74936869 | 1467 | 2.2 | 2.75E-05 | 8.88E-05 |
| chr10 | 103957160 | 103957881 | 722 | 3.21 | 2.75E-05 | 8.90E-05 |
| chr7 | 17735189 | 17737314 | 2126 | -0.68 | 2.76E-05 | 8.92E-05 |
| chr1 | 171917506 | 171918103 | 598 | 5.17 | 2.76E-05 | 8.93E-05 |
| chr4 | 28094609 | 28095181 | 573 | 5.17 | 2.76E-05 | 8.93E-05 |
| chr17 | 37461286 | 37463707 | 2422 | 1.15 | 2.76E-05 | 8.93E-05 |
| chr12 | 51595206 | 51596204 | 999 | 3.67 | 2.76E-05 | 8.93E-05 |
| chr6 | 11179353 | 11234451 | 55099 | 0.17 | 2.77E-05 | 8.95E-05 |
| chr5 | 72568792 | 72570546 | 1755 | 1.75 | 2.77E-05 | 8.96E-05 |
| chr5 | 95666258 | 95667518 | 1261 | 1.49 | 2.78E-05 | 8.98E-05 |
| chr1 | 30180659 | 30183882 | 3224 | -0.61 | 2.78E-05 | 8.98E-05 |
| chr2 | 25993221 | 25995411 | 2191 | -1.13 | 2.78E-05 | 8.99E-05 |
| chr6 | 8392383 | 8393455 | 1073 | 3.34 | 2.79E-05 | 9.00E-05 |
| chr2 | 67099500 | 67100339 | 840 | 5.14 | 2.79E-05 | 9.01E-05 |
| chr7 | 132755329 | 132767608 | 12280 | -0.74 | 2.79E-05 | 9.02E-05 |
| chr12 | 10169293 | 10170325 | 1033 | 1.28 | 2.79E-05 | 9.02E-05 |
| chr12 | 105290177 | 105290773 | 597 | 3.12 | 2.80E-05 | 9.03E-05 |
| chr9 | 137394127 | 137395961 | 1835 | -0.93 | 2.80E-05 | 9.05E-05 |
| chr12 | 127567007 | 127569113 | 2107 | 0.57 | 2.81E-05 | 9.06E-05 |
| chr19 | 44575554 | 44576856 | 1303 | 0.98 | 2.81E-05 | 9.07E-05 |
| chr6 | 157402199 | 157430893 | 28695 | 0.71 | 2.81E-05 | 9.08E-05 |
| chr8 | 60556645 | 60557174 | 530 | 5.19 | 2.81E-05 | 9.08E-05 |
| chr18 | 19019200 | 19020259 | 1060 | 2.29 | 2.81E-05 | 9.08E-05 |
| chr10 | 64271553 | 64272641 | 1089 | 1.78 | 2.82E-05 | 9.09E-05 |
| chr4 | 65330188 | 65330851 | 664 | 5.26 | 2.82E-05 | 9.10E-05 |
| chr13 | 28180818 | 28208729 | 27912 | 0.66 | 2.82E-05 | 9.11E-05 |
| chr17 | 32946908 | 32947649 | 742 | -4.47 | 2.82E-05 | 9.11E-05 |
| chr5 | 176328704 | 176337694 | 8991 | 0.52 | 2.82E-05 | 9.11E-05 |

| | | | | | | |
|---|---|---|---|---|---|---|
| chr4 | 143010815 | 143018232 | 7418 | 0.4 | 2.83E-05 | 9.12E-05 |
| chr17 | 67965882 | 67966348 | 467 | -4.07 | 2.83E-05 | 9.12E-05 |
| chr10 | 21687526 | 21691851 | 4326 | -1.34 | 2.83E-05 | 9.13E-05 |
| chr6 | 53657403 | 53660363 | 2961 | 0.85 | 2.84E-05 | 9.14E-05 |
| chr3 | 175650211 | 175651227 | 1017 | 2.86 | 2.84E-05 | 9.15E-05 |
| chr13 | 66922629 | 66928594 | 5966 | 1.74 | 2.84E-05 | 9.16E-05 |
| chr16 | 89352493 | 89576340 | 223848 | -0.14 | 2.84E-05 | 9.16E-05 |
| chr1 | 185373783 | 185374152 | 370 | 5.14 | 2.85E-05 | 9.19E-05 |
| chr9 | 71735383 | 71737710 | 2328 | 1.28 | 2.86E-05 | 9.22E-05 |
| chr9 | 140549023 | 140585305 | 36283 | -0.53 | 2.86E-05 | 9.22E-05 |
| chr2 | 8737654 | 8746756 | 9103 | -0.95 | 2.86E-05 | 9.22E-05 |
| chr10 | 88159074 | 88175705 | 16632 | 1.6 | 2.87E-05 | 9.25E-05 |
| chrX | 36450255 | 36450876 | 622 | 4.19 | 2.87E-05 | 9.25E-05 |
| chr12 | 90168140 | 90177915 | 9776 | 0.85 | 2.87E-05 | 9.26E-05 |
| chr12 | 49161832 | 49184209 | 22378 | 0.74 | 2.89E-05 | 9.30E-05 |
| chr6 | 159268140 | 159292715 | 24576 | 0.24 | 2.89E-05 | 9.31E-05 |
| chr1 | 220440543 | 220446542 | 6000 | 0.59 | 2.89E-05 | 9.32E-05 |
| chr7 | 40938554 | 40939432 | 879 | 3.57 | 2.91E-05 | 9.35E-05 |
| chr4 | 110532049 | 110532968 | 920 | 2.19 | 2.91E-05 | 9.38E-05 |
| chr5 | 32222495 | 32235909 | 13415 | 0.53 | 2.92E-05 | 9.38E-05 |
| chr4 | 180147259 | 180147905 | 647 | 3.12 | 2.92E-05 | 9.40E-05 |
| chr19 | 41723871 | 41917491 | 193621 | -0.13 | 2.93E-05 | 9.42E-05 |
| chr1 | 41326593 | 41344656 | 18064 | 0.32 | 2.93E-05 | 9.42E-05 |
| chr4 | 115675866 | 115676335 | 470 | 5.14 | 2.93E-05 | 9.42E-05 |
| chr10 | 99078204 | 99095393 | 17190 | -0.46 | 2.93E-05 | 9.43E-05 |
| chr12 | 21589811 | 21598100 | 8290 | 0.41 | 2.94E-05 | 9.47E-05 |
| chr20 | 9564750 | 9577328 | 12579 | 0.61 | 2.95E-05 | 9.48E-05 |
| chr8 | 9911172 | 9913353 | 2182 | 0.72 | 2.95E-05 | 9.49E-05 |
| chr1 | 186631671 | 186632601 | 931 | 1.51 | 2.96E-05 | 9.51E-05 |
| chr4 | 104917087 | 104918589 | 1503 | 1.3 | 2.96E-05 | 9.51E-05 |
| chr3 | 16224587 | 16226146 | 1560 | -3.41 | 2.96E-05 | 9.52E-05 |
| chr5 | 88044605 | 88045408 | 804 | 3.17 | 2.97E-05 | 9.53E-05 |
| chr7 | 143596008 | 143599618 | 3611 | 0.56 | 2.97E-05 | 9.56E-05 |
| chr4 | 174563465 | 174564203 | 739 | 4.56 | 2.98E-05 | 9.59E-05 |
| chr3 | 127789146 | 127843635 | 54490 | -0.39 | 2.98E-05 | 9.59E-05 |
| chr2 | 210336783 | 210339466 | 2684 | -0.62 | 2.99E-05 | 9.59E-05 |
| chr5 | 126564494 | 126565750 | 1257 | 1.38 | 2.99E-05 | 9.59E-05 |
| chr1 | 172901422 | 172906201 | 4780 | 0.37 | 2.99E-05 | 9.60E-05 |
| chr1 | 226027261 | 226034048 | 6788 | -1.21 | 2.99E-05 | 9.62E-05 |
| chr7 | 116797861 | 116810760 | 12900 | 1.42 | 3.00E-05 | 9.64E-05 |
| chr9 | 102690440 | 102692963 | 2524 | 1.69 | 3.00E-05 | 9.64E-05 |
| chr11 | 123431823 | 123433144 | 1322 | -3.01 | 3.01E-05 | 9.65E-05 |
| chr6 | 132341282 | 132343774 | 2493 | 1.76 | 3.01E-05 | 9.65E-05 |
| chr2 | 112416288 | 112418111 | 1824 | 0.98 | 3.01E-05 | 9.65E-05 |
| chr11 | 82986034 | 82998446 | 12413 | 0.56 | 3.01E-05 | 9.65E-05 |
| chr2 | 191858819 | 191862280 | 3462 | -2.74 | 3.01E-05 | 9.66E-05 |
| chr13 | 43941571 | 43942941 | 1371 | 3.41 | 3.01E-05 | 9.66E-05 |
| chr20 | 25439928 | 25490684 | 50757 | -0.34 | 3.02E-05 | 9.69E-05 |
| chr12 | 56861423 | 56871718 | 10296 | 0.54 | 3.02E-05 | 9.69E-05 |
| chr5 | 85615589 | 85617179 | 1591 | 1.34 | 3.02E-05 | 9.69E-05 |
| chr21 | 43049660 | 43063276 | 13617 | 0.58 | 3.02E-05 | 9.70E-05 |
| chr17 | 27965591 | 27976585 | 10995 | -0.57 | 3.03E-05 | 9.71E-05 |
| chr3 | 85401530 | 85402155 | 626 | 5.14 | 3.03E-05 | 9.72E-05 |
| chr15 | 74609667 | 74610635 | 969 | -2.86 | 3.03E-05 | 9.72E-05 |
| chr10 | 74926887 | 74929635 | 2749 | 0.59 | 3.04E-05 | 9.73E-05 |
| chr1 | 151019641 | 151045259 | 25619 | -0.28 | 3.04E-05 | 9.73E-05 |
| chr2 | 164427131 | 164429440 | 2310 | 0.93 | 3.04E-05 | 9.73E-05 |
| chr5 | 167988758 | 167990409 | 1652 | 1.19 | 3.04E-05 | 9.76E-05 |
| chr7 | 99275872 | 99277671 | 1800 | 0.98 | 3.05E-05 | 9.76E-05 |
| chr11 | 33002693 | 33003758 | 1066 | -1.77 | 3.05E-05 | 9.77E-05 |
| chr4 | 189120064 | 189121350 | 1287 | -1.27 | 3.05E-05 | 9.78E-05 |

| | | | | | | |
|---|---|---|---|---|---|---|
| chr3 | 100859773 | 100860278 | 506 | 5.17 | 3.06E-05 | 9.80E-05 |
| chr5 | 174117013 | 174121719 | 4707 | 5.44 | 3.06E-05 | 9.81E-05 |
| chr8 | 57085838 | 57087157 | 1320 | 2.01 | 3.06E-05 | 9.81E-05 |
| chr15 | 64383517 | 64390770 | 7254 | 0.41 | 3.06E-05 | 9.81E-05 |
| chr2 | 236920470 | 236922492 | 2023 | -1.76 | 3.07E-05 | 9.83E-05 |
| chr11 | 108336798 | 108338916 | 2119 | 0.62 | 3.08E-05 | 9.86E-05 |
| chr11 | 101435693 | 101436344 | 652 | 4.2 | 3.09E-05 | 9.89E-05 |
| chr8 | 122335824 | 122337803 | 1980 | 1.28 | 3.09E-05 | 9.89E-05 |
| chr7 | 110996963 | 111005040 | 8078 | 1.23 | 3.09E-05 | 9.89E-05 |
| chr2 | 225711855 | 225726626 | 14772 | -1.23 | 3.09E-05 | 9.90E-05 |
| chr12 | 132335754 | 132417008 | 81255 | 0.19 | 3.10E-05 | 9.91E-05 |
| chr21 | 44629670 | 44631119 | 1450 | -1.24 | 3.10E-05 | 9.93E-05 |
| chr9 | 3502350 | 3528782 | 26433 | 0.21 | 3.10E-05 | 9.93E-05 |
| chr9 | 99533128 | 99541221 | 8094 | 2.7 | 3.11E-05 | 9.93E-05 |
| chr11 | 44179036 | 44179749 | 714 | -3.37 | 3.11E-05 | 9.94E-05 |
| chr17 | 51792075 | 51799707 | 7633 | 5.5 | 3.11E-05 | 9.95E-05 |
| chr3 | 191067949 | 191075308 | 7360 | 1.73 | 3.11E-05 | 9.95E-05 |
| chr2 | 209516770 | 209517649 | 880 | 5.14 | 3.12E-05 | 9.96E-05 |
| chr15 | 93160083 | 93162054 | 1972 | 1.32 | 3.13E-05 | 9.99E-05 |
| chr4 | 96652836 | 96653427 | 592 | 5.12 | 3.13E-05 | 1.00E-04 |
| chr10 | 124090351 | 124098483 | 8133 | 0.86 | 3.13E-05 | 1.00E-04 |
| chr8 | 40849762 | 40851851 | 2090 | -1.86 | 3.13E-05 | 1.00E-04 |
| chr20 | 19668606 | 19673268 | 4663 | -1.83 | 3.14E-05 | 1.00E-04 |
| chr5 | 32574351 | 32596891 | 22541 | 0.41 | 3.15E-05 | 0.000101 |
| chr12 | 678712 | 688738 | 10027 | 0.39 | 3.16E-05 | 0.000101 |
| chr2 | 27233249 | 27257176 | 23928 | -0.66 | 3.16E-05 | 0.000101 |
| chr14 | 55218170 | 55219150 | 981 | -5.08 | 3.17E-05 | 0.000101 |
| chr12 | 62946222 | 62948524 | 2303 | 0.93 | 3.17E-05 | 0.000101 |
| chr4 | 153384773 | 153386270 | 1498 | 2.1 | 3.17E-05 | 0.000101 |
| chrX | 41239803 | 41249096 | 9294 | 0.41 | 3.18E-05 | 0.000101 |
| chr9 | 29185582 | 29185995 | 414 | 4.52 | 3.18E-05 | 0.000102 |
| chr14 | 80989009 | 80990759 | 1751 | 1.75 | 3.19E-05 | 0.000102 |
| chr15 | 56756679 | 56758000 | 1322 | -0.85 | 3.19E-05 | 0.000102 |
| chr6 | 134495320 | 134504599 | 9280 | 0.84 | 3.20E-05 | 0.000102 |
| chr15 | 36870607 | 36894424 | 23818 | 0.67 | 3.20E-05 | 0.000102 |
| chr20 | 32076736 | 32078902 | 2167 | 0.63 | 3.22E-05 | 0.000103 |
| chr3 | 125328426 | 125329721 | 1296 | 1.8 | 3.22E-05 | 0.000103 |
| chr13 | 32622209 | 32623693 | 1485 | 2.09 | 3.23E-05 | 0.000103 |
| chr5 | 50006802 | 50007344 | 543 | 5.12 | 3.23E-05 | 0.000103 |
| chr20 | 14469912 | 14475204 | 5293 | 3.98 | 3.24E-05 | 0.000103 |
| chr5 | 149257768 | 149267898 | 10131 | -1.72 | 3.25E-05 | 0.000104 |
| chr6 | 132654052 | 132656356 | 2305 | 0.97 | 3.25E-05 | 0.000104 |
| chr1 | 209957141 | 209958775 | 1635 | 0.69 | 3.25E-05 | 0.000104 |
| chr11 | 94500835 | 94510569 | 9735 | 0.95 | 3.25E-05 | 0.000104 |
| chr1 | 98675088 | 98677148 | 2061 | -1.04 | 3.26E-05 | 0.000104 |
| chrX | 48885735 | 48911692 | 25958 | 0.43 | 3.27E-05 | 0.000104 |
| chr8 | 127353497 | 127353957 | 461 | 4.53 | 3.28E-05 | 0.000105 |
| chr17 | 70653085 | 70654597 | 1513 | -1.7 | 3.29E-05 | 0.000105 |
| chr4 | 85569550 | 85571015 | 1466 | 2.9 | 3.29E-05 | 0.000105 |
| chr16 | 48012399 | 48013289 | 891 | -2.25 | 3.29E-05 | 0.000105 |
| chr2 | 210993682 | 210994250 | 569 | 5.11 | 3.29E-05 | 0.000105 |
| chr17 | 20058690 | 20060464 | 1775 | 0.99 | 3.30E-05 | 0.000105 |
| chr6 | 4864610 | 4865356 | 747 | 1.96 | 3.30E-05 | 0.000105 |
| chr11 | 95744262 | 95747860 | 3599 | 0.9 | 3.33E-05 | 0.000106 |
| chr11 | 45129716 | 45151479 | 21764 | 5.01 | 3.34E-05 | 0.000106 |
| chr3 | 13692252 | 13692889 | 638 | 3.44 | 3.34E-05 | 0.000106 |
| chr3 | 93877316 | 93878899 | 1584 | 1.04 | 3.35E-05 | 0.000107 |
| chr8 | 109989982 | 109991205 | 1224 | 1.67 | 3.35E-05 | 0.000107 |
| chr20 | 43964024 | 43977683 | 13660 | 0.35 | 3.35E-05 | 0.000107 |
| chr3 | 37950466 | 37951658 | 1193 | 1.98 | 3.36E-05 | 0.000107 |
| chr4 | 144104419 | 144110554 | 6136 | 0.56 | 3.37E-05 | 0.000107 |

| | | | | | | |
|---|---|---|---|---|---|---|
| chr10 | 99429221 | 99479601 | 50381 | 0.24 | 3.37E-05 | 0.000107 |
| chrX | 39524810 | 39531054 | 6245 | 1.37 | 3.37E-05 | 0.000107 |
| chr7 | 100224415 | 100231698 | 7284 | -1.04 | 3.38E-05 | 0.000107 |
| chr17 | 2536268 | 2592158 | 55891 | 0.3 | 3.38E-05 | 0.000108 |
| chr1 | 204639384 | 204640359 | 976 | -2.23 | 3.38E-05 | 0.000108 |
| chr5 | 140514754 | 140515546 | 793 | 2.08 | 3.39E-05 | 0.000108 |
| chr3 | 36155550 | 36156831 | 1282 | -2.02 | 3.40E-05 | 0.000108 |
| chr2 | 162242813 | 162244012 | 1200 | -1.97 | 3.40E-05 | 0.000108 |
| chr8 | 117483610 | 117485333 | 1724 | 1.86 | 3.40E-05 | 0.000108 |
| chr15 | 57178387 | 57180925 | 2539 | 0.56 | 3.40E-05 | 0.000108 |
| chr2 | 100089473 | 100125007 | 35535 | 0.67 | 3.41E-05 | 0.000108 |
| chr2 | 120565901 | 120567812 | 1912 | -1.18 | 3.41E-05 | 0.000109 |
| chr8 | 99837068 | 99849151 | 12084 | 0.7 | 3.43E-05 | 0.000109 |
| chrX | 67526761 | 67528493 | 1733 | 1.49 | 3.43E-05 | 0.000109 |
| chr9 | 100972460 | 101019760 | 47301 | -0.25 | 3.43E-05 | 0.000109 |
| chr8 | 49282405 | 49284471 | 2067 | 1.14 | 3.44E-05 | 0.000109 |
| chr17 | 7816478 | 7820579 | 4102 | 0.83 | 3.44E-05 | 0.000109 |
| chr6 | 76159625 | 76169678 | 10054 | -1.5 | 3.46E-05 | 0.00011 |
| chr3 | 406913 | 407535 | 623 | 5.12 | 3.47E-05 | 0.00011 |
| chr12 | 113415879 | 113416666 | 788 | 2.17 | 3.47E-05 | 0.00011 |
| chr4 | 154883797 | 154884553 | 757 | 3.57 | 3.48E-05 | 0.000111 |
| chr12 | 121497459 | 121498891 | 1433 | 1.24 | 3.48E-05 | 0.000111 |
| chr1 | 167682611 | 167722842 | 40232 | -0.33 | 3.49E-05 | 0.000111 |
| chr6 | 6876235 | 6880189 | 3955 | 1.57 | 3.50E-05 | 0.000111 |
| chr11 | 44600438 | 44606669 | 6232 | 1.94 | 3.50E-05 | 0.000111 |
| chr1 | 61332966 | 61333516 | 551 | 5.09 | 3.51E-05 | 0.000112 |
| chrX | 27566646 | 27567430 | 785 | 5.09 | 3.51E-05 | 0.000112 |
| chr12 | 28263187 | 28284586 | 21400 | 5.91 | 3.52E-05 | 0.000112 |
| chr4 | 77491264 | 77492921 | 1658 | 1.65 | 3.52E-05 | 0.000112 |
| chr13 | 102333889 | 102334904 | 1016 | 3.51 | 3.53E-05 | 0.000112 |
| chr4 | 64787476 | 64788100 | 625 | 5.09 | 3.53E-05 | 0.000112 |
| chr11 | 70933009 | 70934062 | 1054 | -3.29 | 3.53E-05 | 0.000112 |
| chr7 | 6198919 | 6243600 | 44682 | -0.43 | 3.54E-05 | 0.000112 |
| chr5 | 122846774 | 122849365 | 2592 | 0.52 | 3.54E-05 | 0.000112 |
| chr18 | 72873336 | 72874156 | 821 | 3.32 | 3.54E-05 | 0.000112 |
| chr13 | 86103064 | 86103639 | 576 | -4.51 | 3.54E-05 | 0.000112 |
| chr3 | 146342543 | 146343561 | 1019 | 5.2 | 3.55E-05 | 0.000112 |
| chrX | 130668958 | 130670087 | 1130 | 2.17 | 3.55E-05 | 0.000112 |
| chr7 | 23116566 | 23117149 | 584 | 5.09 | 3.56E-05 | 0.000113 |
| chr3 | 97620106 | 97630199 | 10094 | 1.17 | 3.57E-05 | 0.000113 |
| chr11 | 95867101 | 95868701 | 1601 | 1.7 | 3.57E-05 | 0.000113 |
| chr2 | 119144524 | 119146170 | 1647 | -2.56 | 3.57E-05 | 0.000113 |
| chr12 | 66269241 | 66271363 | 2123 | 1.94 | 3.57E-05 | 0.000113 |
| chr4 | 111005921 | 111013422 | 7502 | 1.04 | 3.57E-05 | 0.000113 |
| chr20 | 50647796 | 50649386 | 1591 | -0.97 | 3.58E-05 | 0.000113 |
| chr11 | 77184122 | 77186009 | 1888 | 1.12 | 3.58E-05 | 0.000113 |
| chr19 | 21541788 | 21543078 | 1291 | 1.65 | 3.59E-05 | 0.000114 |
| chr19 | 22443029 | 22444117 | 1089 | 3.53 | 3.60E-05 | 0.000114 |
| chr9 | 7798510 | 7800258 | 1749 | 0.76 | 3.60E-05 | 0.000114 |
| chr16 | 3439305 | 3452229 | 12925 | 0.79 | 3.60E-05 | 0.000114 |
| chr20 | 14756033 | 14757051 | 1019 | -3.24 | 3.61E-05 | 0.000114 |
| chr2 | 20160299 | 20161322 | 1024 | -3.46 | 3.62E-05 | 0.000115 |
| chr5 | 150603138 | 150614881 | 11744 | -5.38 | 3.63E-05 | 0.000115 |
| chr1 | 68344655 | 68385739 | 41085 | 0.32 | 3.63E-05 | 0.000115 |
| chr2 | 223170383 | 223171335 | 953 | 5.09 | 3.64E-05 | 0.000115 |
| chr3 | 62430847 | 62431590 | 744 | 5.12 | 3.64E-05 | 0.000115 |
| chr22 | 27503874 | 27543281 | 39408 | -0.38 | 3.65E-05 | 0.000115 |
| chr2 | 69272632 | 69275606 | 2975 | 1.22 | 3.66E-05 | 0.000116 |
| chr7 | 94571325 | 94571931 | 607 | 4.12 | 3.67E-05 | 0.000116 |
| chr6 | 136930093 | 136935418 | 5326 | 0.65 | 3.67E-05 | 0.000116 |
| chr2 | 84329986 | 84332129 | 2144 | 5.57 | 3.67E-05 | 0.000116 |

| | | | | | | |
|---|---|---|---|---|---|---|
| chr13 | 40138760 | 40140710 | 1951 | 1.05 | 3.68E-05 | 0.000116 |
| chr10 | 12305673 | 12314485 | 8813 | 0.47 | 3.68E-05 | 0.000116 |
| chr22 | 34141835 | 34143300 | 1466 | 2.21 | 3.69E-05 | 0.000117 |
| chr9 | 71939524 | 71940051 | 528 | 5.32 | 3.69E-05 | 0.000117 |
| chr20 | 6070680 | 6085872 | 15193 | -0.97 | 3.69E-05 | 0.000117 |
| chr3 | 101230442 | 101233065 | 2624 | 0.56 | 3.70E-05 | 0.000117 |
| chr15 | 55951098 | 55954223 | 3126 | -1.5 | 3.71E-05 | 0.000117 |
| chr11 | 92128826 | 92129976 | 1151 | 2.3 | 3.71E-05 | 0.000117 |
| chr19 | 36978829 | 36981539 | 2711 | 0.43 | 3.72E-05 | 0.000117 |
| chr2 | 145592299 | 145593569 | 1271 | -1.33 | 3.72E-05 | 0.000118 |
| chr14 | 76025896 | 76026527 | 632 | -5.06 | 3.73E-05 | 0.000118 |
| chr8 | 63075266 | 63079001 | 3736 | 0.42 | 3.73E-05 | 0.000118 |
| chr15 | 37269585 | 37270587 | 1003 | 3.2 | 3.73E-05 | 0.000118 |
| chr20 | 8608055 | 8608746 | 692 | 5.18 | 3.74E-05 | 0.000118 |
| chr14 | 74352308 | 74364029 | 11722 | 0.42 | 3.75E-05 | 0.000118 |
| chr7 | 37939100 | 37991082 | 51983 | -0.29 | 3.75E-05 | 0.000118 |
| chr6 | 38476390 | 38478070 | 1681 | -2.41 | 3.75E-05 | 0.000119 |
| chr4 | 136957953 | 136958672 | 720 | 3.15 | 3.76E-05 | 0.000119 |
| chr6 | 101318342 | 101330287 | 11946 | 0.71 | 3.76E-05 | 0.000119 |
| chr8 | 19164249 | 19217605 | 53357 | -0.26 | 3.76E-05 | 0.000119 |
| chr5 | 148398233 | 148399529 | 1297 | 2.53 | 3.78E-05 | 0.000119 |
| chr16 | 74829790 | 74835875 | 6086 | 5.85 | 3.79E-05 | 0.000119 |
| chr12 | 73915953 | 73916548 | 596 | 5.11 | 3.79E-05 | 0.000119 |
| chr8 | 103739754 | 103751696 | 11943 | -0.48 | 3.79E-05 | 0.00012 |
| chr3 | 61630127 | 61630788 | 662 | 3.61 | 3.79E-05 | 0.00012 |
| chr1 | 227531267 | 227547184 | 15918 | 0.36 | 3.80E-05 | 0.00012 |
| chr14 | 74315640 | 74320502 | 4863 | 0.44 | 3.80E-05 | 0.00012 |
| chr12 | 12466855 | 12480872 | 14018 | -1.73 | 3.80E-05 | 0.00012 |
| chr3 | 11588213 | 11624653 | 36441 | -0.45 | 3.81E-05 | 0.00012 |
| chr3 | 181763682 | 181764663 | 982 | 2.93 | 3.81E-05 | 0.00012 |
| chr2 | 238036348 | 238037568 | 1221 | -1.44 | 3.82E-05 | 0.00012 |
| chr13 | 50421937 | 50435168 | 13232 | -0.39 | 3.83E-05 | 0.000121 |
| chr2 | 25585048 | 25600559 | 15512 | -1.05 | 3.84E-05 | 0.000121 |
| chr14 | 21077404 | 21101733 | 24330 | 0.39 | 3.84E-05 | 0.000121 |
| chr11 | 73489530 | 73507515 | 17986 | 0.38 | 3.84E-05 | 0.000121 |
| chr18 | 61121556 | 61122508 | 953 | 2.63 | 3.85E-05 | 0.000121 |
| chr10 | 70820474 | 70822254 | 1781 | 0.75 | 3.86E-05 | 0.000122 |
| chr16 | 13052191 | 13052920 | 730 | -3.48 | 3.86E-05 | 0.000122 |
| chr17 | 11639775 | 11640510 | 736 | -5.03 | 3.87E-05 | 0.000122 |
| chr9 | 135281007 | 135285884 | 4878 | -0.67 | 3.88E-05 | 0.000122 |
| chr4 | 175238645 | 175239518 | 874 | 3.5 | 3.88E-05 | 0.000122 |
| chr2 | 189540206 | 189540899 | 694 | -3.59 | 3.88E-05 | 0.000122 |
| chr2 | 58766576 | 58768221 | 1646 | 1.61 | 3.89E-05 | 0.000122 |
| chr3 | 54915753 | 54926753 | 11001 | 0.8 | 3.89E-05 | 0.000123 |
| chr2 | 226978937 | 227025502 | 46566 | 0.31 | 3.89E-05 | 0.000123 |
| chr12 | 105949453 | 105957861 | 8409 | 0.44 | 3.90E-05 | 0.000123 |
| chr15 | 71345940 | 71348814 | 2875 | 0.55 | 3.91E-05 | 0.000123 |
| chr11 | 61514747 | 61639446 | 124700 | 0.18 | 3.92E-05 | 0.000123 |
| chr12 | 56580949 | 56587509 | 6561 | 0.65 | 3.92E-05 | 0.000123 |
| chr20 | 21163467 | 21164452 | 986 | 2.71 | 3.93E-05 | 0.000124 |
| chr3 | 83922480 | 83923033 | 554 | 5.09 | 3.93E-05 | 0.000124 |
| chr16 | 27775845 | 27777132 | 1288 | 3.38 | 3.93E-05 | 0.000124 |
| chr19 | 30500308 | 30502456 | 2149 | 0.72 | 3.93E-05 | 0.000124 |
| chrX | 118467169 | 118467595 | 427 | 5.07 | 3.93E-05 | 0.000124 |
| chr12 | 57851429 | 57856659 | 5231 | 0.5 | 3.94E-05 | 0.000124 |
| chr8 | 67867146 | 67867846 | 701 | 2.18 | 3.94E-05 | 0.000124 |
| chr1 | 145540916 | 145544281 | 3366 | 0.59 | 3.95E-05 | 0.000124 |
| chr9 | 94064855 | 94066019 | 1165 | 0.9 | 3.95E-05 | 0.000124 |
| chr4 | 100077777 | 100079010 | 1234 | 2.18 | 3.95E-05 | 0.000124 |
| chr17 | 7785667 | 7801971 | 16305 | 0.39 | 3.96E-05 | 0.000124 |
| chr8 | 123501423 | 123502655 | 1233 | 3.4 | 3.96E-05 | 0.000124 |

| | | | | | | |
|---|---|---|---|---|---|---|
| chr2 | 35305100 | 35317652 | 12553 | 1.3 | 3.96E-05 | 0.000124 |
| chr10 | 19169689 | 19170197 | 509 | 5.07 | 3.97E-05 | 0.000125 |
| chr11 | 87315030 | 87315611 | 582 | -5.06 | 3.97E-05 | 0.000125 |
| chr1 | 153479066 | 153489279 | 10214 | 0.7 | 3.99E-05 | 0.000125 |
| chr16 | 75655572 | 75658009 | 2438 | 0.72 | 3.99E-05 | 0.000125 |
| chr3 | 69730928 | 69741231 | 10304 | 0.95 | 3.99E-05 | 0.000125 |
| chr19 | 3175915 | 3187023 | 11109 | 0.59 | 4.00E-05 | 0.000126 |
| chr6 | 28179680 | 28193818 | 14139 | 0.78 | 4.01E-05 | 0.000126 |
| chr11 | 102096092 | 102097959 | 1868 | -1.99 | 4.01E-05 | 0.000126 |
| chr6 | 42938107 | 42952991 | 14885 | -0.5 | 4.02E-05 | 0.000126 |
| chr10 | 102849072 | 102852183 | 3112 | -1.2 | 4.02E-05 | 0.000126 |
| chr20 | 35233024 | 35253898 | 20875 | 0.42 | 4.02E-05 | 0.000126 |
| chr2 | 113626520 | 113627722 | 1203 | 2.32 | 4.03E-05 | 0.000126 |
| chr3 | 165523172 | 165525009 | 1838 | 3.42 | 4.04E-05 | 0.000127 |
| chr4 | 160148441 | 160151433 | 2993 | 4.76 | 4.04E-05 | 0.000127 |
| chr8 | 116480059 | 116481038 | 980 | 5.07 | 4.04E-05 | 0.000127 |
| chr9 | 91768844 | 91770406 | 1563 | -1.32 | 4.05E-05 | 0.000127 |
| chr2 | 68546431 | 68547209 | 779 | -2.84 | 4.06E-05 | 0.000127 |
| chr6 | 21987086 | 21988344 | 1259 | 1.86 | 4.06E-05 | 0.000127 |
| chr19 | 33807198 | 33808294 | 1097 | 3.39 | 4.07E-05 | 0.000128 |
| chr10 | 91390677 | 91405870 | 15194 | -1.15 | 4.07E-05 | 0.000128 |
| chr1 | 21471382 | 21504163 | 32782 | 0.33 | 4.07E-05 | 0.000128 |
| chr4 | 117240580 | 117241244 | 665 | 5.07 | 4.08E-05 | 0.000128 |
| chr4 | 98325783 | 98338727 | 12945 | 5.15 | 4.08E-05 | 0.000128 |
| chr1 | 186671954 | 186688547 | 16594 | 0.31 | 4.08E-05 | 0.000128 |
| chr5 | 105508565 | 105509586 | 1022 | -2.12 | 4.09E-05 | 0.000128 |
| chr12 | 14369089 | 14369857 | 769 | 2.12 | 4.09E-05 | 0.000128 |
| chr4 | 143299703 | 143301623 | 1921 | 1.44 | 4.10E-05 | 0.000128 |
| chr18 | 25501333 | 25502572 | 1240 | 1.81 | 4.11E-05 | 0.000129 |
| chr2 | 38545061 | 38553506 | 8446 | -1.13 | 4.13E-05 | 0.000129 |
| chr11 | 9333677 | 9341783 | 8107 | 0.47 | 4.13E-05 | 0.000129 |
| chr14 | 70825073 | 70827269 | 2197 | 0.64 | 4.14E-05 | 0.00013 |
| chr20 | 42793981 | 42816945 | 22965 | 0.63 | 4.16E-05 | 0.00013 |
| chr19 | 6508947 | 6539290 | 30344 | 0.49 | 4.16E-05 | 0.00013 |
| chr1 | 36348837 | 36360609 | 11773 | -1.13 | 4.17E-05 | 0.00013 |
| chr19 | 40402625 | 40488289 | 85665 | 0.15 | 4.17E-05 | 0.000131 |
| chr8 | 146227570 | 146229010 | 1441 | 0.85 | 4.18E-05 | 0.000131 |
| chr3 | 114864108 | 114868189 | 4082 | 0.38 | 4.19E-05 | 0.000131 |
| chr17 | 53024760 | 53047705 | 22946 | 0.4 | 4.20E-05 | 0.000132 |
| chr2 | 8765194 | 8777696 | 12503 | -0.52 | 4.20E-05 | 0.000132 |
| chr16 | 759837 | 791754 | 31918 | -0.47 | 4.21E-05 | 0.000132 |
| chr18 | 29691262 | 29702548 | 11287 | 0.78 | 4.22E-05 | 0.000132 |
| chr6 | 37995383 | 37995958 | 576 | -5.04 | 4.22E-05 | 0.000132 |
| chr14 | 50691903 | 50699368 | 7466 | 0.57 | 4.24E-05 | 0.000133 |
| chr16 | 14106993 | 14108420 | 1428 | 1.56 | 4.26E-05 | 0.000133 |
| chr12 | 80543870 | 80544438 | 569 | 2.81 | 4.28E-05 | 0.000134 |
| chr12 | 18075684 | 18083163 | 7480 | 1.32 | 4.28E-05 | 0.000134 |
| chr4 | 38729436 | 38731312 | 1877 | 1.1 | 4.29E-05 | 0.000134 |
| chr12 | 15910125 | 15911704 | 1580 | 1.26 | 4.29E-05 | 0.000134 |
| chr15 | 39871278 | 39889444 | 18167 | 0.42 | 4.29E-05 | 0.000134 |
| chr18 | 5293896 | 5297513 | 3618 | 0.76 | 4.30E-05 | 0.000134 |
| chr2 | 69170576 | 69171553 | 978 | -1.85 | 4.32E-05 | 0.000135 |
| chr12 | 110317451 | 110351148 | 33698 | 0.56 | 4.33E-05 | 0.000135 |
| chr3 | 45882735 | 45884077 | 1343 | -0.79 | 4.33E-05 | 0.000135 |
| chr2 | 214536788 | 214537229 | 442 | 5.11 | 4.33E-05 | 0.000135 |
| chr14 | 32542587 | 32620310 | 77724 | 0.22 | 4.34E-05 | 0.000135 |
| chr8 | 25365708 | 25372634 | 6927 | 0.97 | 4.35E-05 | 0.000136 |
| chr7 | 788893 | 789982 | 1090 | -5.02 | 4.35E-05 | 0.000136 |
| chr1 | 167012954 | 167013875 | 922 | 2.98 | 4.35E-05 | 0.000136 |
| chr2 | 173345836 | 173346727 | 892 | 2.89 | 4.36E-05 | 0.000136 |
| chr1 | 172008822 | 172010338 | 1517 | 1.41 | 4.37E-05 | 0.000136 |

| | | | | | | |
|---|---|---|---|---|---|---|
| chr2 | 183811341 | 183818838 | 7498 | -0.67 | 4.37E-05 | 0.000137 |
| chr1 | 145497995 | 145525264 | 27270 | 0.44 | 4.38E-05 | 0.000137 |
| chr1 | 46915286 | 46916361 | 1076 | -2.36 | 4.39E-05 | 0.000137 |
| chr9 | 4205266 | 4205986 | 721 | -3.65 | 4.39E-05 | 0.000137 |
| chr4 | 40720568 | 40734487 | 13920 | 0.81 | 4.39E-05 | 0.000137 |
| chr12 | 96587118 | 96636778 | 49661 | 0.2 | 4.40E-05 | 0.000137 |
| chr19 | 53605469 | 53607399 | 1931 | 0.66 | 4.40E-05 | 0.000137 |
| chr7 | 48577535 | 48578101 | 567 | 3.71 | 4.40E-05 | 0.000137 |
| chrX | 15681990 | 15695397 | 13408 | 0.62 | 4.41E-05 | 0.000138 |
| chr12 | 72033255 | 72039788 | 6534 | 0.91 | 4.41E-05 | 0.000138 |
| chr4 | 90348715 | 90349847 | 1133 | 2.27 | 4.42E-05 | 0.000138 |
| chr5 | 141628186 | 141629069 | 884 | 3.43 | 4.42E-05 | 0.000138 |
| chr2 | 201751950 | 201774907 | 22958 | -0.34 | 4.42E-05 | 0.000138 |
| chr14 | 65637171 | 65651622 | 14452 | 1.38 | 4.43E-05 | 0.000138 |
| chr7 | 155596317 | 155602384 | 6068 | 1.16 | 4.43E-05 | 0.000138 |
| chrX | 109098410 | 109099248 | 839 | 3.48 | 4.44E-05 | 0.000138 |
| chr2 | 195415179 | 195415846 | 668 | -4.04 | 4.44E-05 | 0.000138 |
| chr3 | 24403941 | 24418174 | 14234 | 0.61 | 4.44E-05 | 0.000138 |
| chr17 | 42784730 | 42788555 | 3826 | -0.47 | 4.46E-05 | 0.000139 |
| chr2 | 118613006 | 118615152 | 2147 | -1.06 | 4.47E-05 | 0.000139 |
| chr8 | 83232624 | 83233004 | 381 | 5.04 | 4.50E-05 | 0.00014 |
| chr9 | 93317425 | 93318479 | 1055 | 2.5 | 4.50E-05 | 0.00014 |
| chr8 | 79600293 | 79600983 | 691 | 3.93 | 4.50E-05 | 0.00014 |
| chr1 | 81139674 | 81140361 | 688 | 4.42 | 4.51E-05 | 0.00014 |
| chr8 | 28242170 | 28270377 | 28208 | 0.51 | 4.51E-05 | 0.00014 |
| chr7 | 5534211 | 5573105 | 38895 | 0.29 | 4.52E-05 | 0.000141 |
| chr17 | 19800350 | 19881985 | 81636 | 0.24 | 4.52E-05 | 0.000141 |
| chr1 | 178439261 | 178441533 | 2273 | 2.22 | 4.54E-05 | 0.000141 |
| chr8 | 91656698 | 91662812 | 6115 | 0.82 | 4.54E-05 | 0.000141 |
| chr9 | 121370785 | 121373011 | 2227 | 0.64 | 4.55E-05 | 0.000141 |
| chr7 | 158648167 | 158662554 | 14388 | 0.45 | 4.55E-05 | 0.000141 |
| chr11 | 11989937 | 12031426 | 41490 | 0.33 | 4.56E-05 | 0.000142 |
| chr21 | 35747121 | 35757027 | 9907 | 0.51 | 4.57E-05 | 0.000142 |
| chr12 | 39725354 | 39726405 | 1052 | 1.34 | 4.57E-05 | 0.000142 |
| chr12 | 49794670 | 49823405 | 28736 | -0.64 | 4.62E-05 | 0.000144 |
| chr12 | 123779165 | 123791009 | 11845 | 0.79 | 4.62E-05 | 0.000144 |
| chr9 | 90476504 | 90495366 | 18863 | 0.35 | 4.62E-05 | 0.000144 |
| chr7 | 82434138 | 82435651 | 1514 | 0.96 | 4.63E-05 | 0.000144 |
| chr2 | 18537718 | 18538302 | 585 | -3.65 | 4.64E-05 | 0.000144 |
| chr12 | 20098414 | 20100591 | 2178 | 1.1 | 4.64E-05 | 0.000144 |
| chr5 | 80848643 | 80849276 | 634 | 3.48 | 4.64E-05 | 0.000144 |
| chr3 | 64486623 | 64488018 | 1396 | 1.65 | 4.65E-05 | 0.000144 |
| chr16 | 3980493 | 3981184 | 692 | 5.05 | 4.65E-05 | 0.000144 |
| chr8 | 40333218 | 40345275 | 12058 | -1.01 | 4.66E-05 | 0.000145 |
| chr1 | 94882165 | 94892554 | 10390 | 0.53 | 4.68E-05 | 0.000145 |
| chr7 | 47749142 | 47774494 | 25353 | -1.18 | 4.68E-05 | 0.000145 |
| chr1 | 180026185 | 180028279 | 2095 | -1.23 | 4.69E-05 | 0.000146 |
| chr9 | 4197380 | 4198796 | 1417 | 2.37 | 4.69E-05 | 0.000146 |
| chr1 | 193874464 | 193875433 | 970 | 2.46 | 4.69E-05 | 0.000146 |
| chr4 | 146539589 | 146545853 | 6265 | 0.52 | 4.69E-05 | 0.000146 |
| chr2 | 54750073 | 54901085 | 151013 | 0.15 | 4.70E-05 | 0.000146 |
| chr12 | 121036605 | 121041055 | 4451 | 1.55 | 4.70E-05 | 0.000146 |
| chr2 | 96911569 | 96932851 | 21283 | -0.43 | 4.70E-05 | 0.000146 |
| chr17 | 48189453 | 48189996 | 544 | -4.99 | 4.71E-05 | 0.000146 |
| chr2 | 72290535 | 72291099 | 565 | 3.67 | 4.71E-05 | 0.000146 |
| chr5 | 60655623 | 60656501 | 879 | -4.99 | 4.71E-05 | 0.000146 |
| chr4 | 24857076 | 24858947 | 1872 | 2.87 | 4.73E-05 | 0.000147 |
| chr1 | 147188562 | 147191831 | 3270 | 1.09 | 4.73E-05 | 0.000147 |
| chr1 | 190443809 | 190444932 | 1124 | 2.64 | 4.73E-05 | 0.000147 |
| chrX | 149352325 | 149353420 | 1096 | 2.49 | 4.75E-05 | 0.000147 |
| chr7 | 83613467 | 83614186 | 720 | -4.99 | 4.76E-05 | 0.000147 |

| | | | | | | |
|---|---|---|---|---|---|---|
| chr16 | 46842836 | 46845247 | 2412 | 1.17 | 4.76E-05 | 0.000147 |
| chr4 | 491913 | 494232 | 2320 | 0.66 | 4.77E-05 | 0.000148 |
| chr5 | 122469650 | 122470027 | 378 | 4.47 | 4.77E-05 | 0.000148 |
| chr11 | 70460108 | 70461240 | 1133 | -2.31 | 4.77E-05 | 0.000148 |
| chr17 | 7712411 | 7713658 | 1248 | 2.73 | 4.78E-05 | 0.000148 |
| chr11 | 1781017 | 1804973 | 23957 | -0.43 | 4.78E-05 | 0.000148 |
| chr2 | 44846921 | 44848659 | 1739 | 1.82 | 4.78E-05 | 0.000148 |
| chr7 | 95755299 | 95765688 | 10390 | 1.09 | 4.79E-05 | 0.000149 |
| chr8 | 66266118 | 66268476 | 2359 | -0.92 | 4.79E-05 | 0.000149 |
| chr2 | 198044491 | 198045239 | 749 | -3.27 | 4.80E-05 | 0.000149 |
| chrX | 153624638 | 153641135 | 16498 | 0.33 | 4.81E-05 | 0.000149 |
| chr6 | 149320414 | 149321681 | 1268 | 1.81 | 4.81E-05 | 0.000149 |
| chr9 | 74396703 | 74422043 | 25341 | 0.43 | 4.81E-05 | 0.000149 |
| chr5 | 73090538 | 73127607 | 37070 | 0.28 | 4.85E-05 | 0.00015 |
| chr2 | 204675076 | 204677569 | 2494 | 1.03 | 4.85E-05 | 0.00015 |
| chr7 | 154719504 | 154721538 | 2035 | 1.03 | 4.87E-05 | 0.000151 |
| chr2 | 240030463 | 240035746 | 5284 | -1.42 | 4.89E-05 | 0.000151 |
| chr8 | 38322304 | 38326500 | 4197 | -0.98 | 4.90E-05 | 0.000152 |
| chr5 | 152191419 | 152192037 | 619 | -3.54 | 4.91E-05 | 0.000152 |
| chr2 | 181230371 | 181233398 | 3028 | 0.92 | 4.91E-05 | 0.000152 |
| chr12 | 94070330 | 94072514 | 2185 | 0.42 | 4.92E-05 | 0.000152 |
| chr11 | 82718733 | 82719545 | 813 | -2.11 | 4.92E-05 | 0.000152 |
| chr13 | 29267849 | 29268634 | 786 | 1.86 | 4.92E-05 | 0.000152 |
| chr2 | 133561565 | 133562366 | 802 | -3.23 | 4.92E-05 | 0.000152 |
| chr3 | 170025359 | 170028848 | 3490 | -1.32 | 4.93E-05 | 0.000152 |
| chr8 | 122861923 | 122873941 | 12019 | 5.38 | 4.93E-05 | 0.000152 |
| chr5 | 139417786 | 139423672 | 5887 | 0.88 | 4.95E-05 | 0.000153 |
| chr1 | 42798805 | 42807539 | 8735 | 0.62 | 4.95E-05 | 0.000153 |
| chr6 | 107231101 | 107253046 | 21946 | 0.51 | 4.95E-05 | 0.000153 |
| chr9 | 135142554 | 135169077 | 26524 | 0.4 | 4.96E-05 | 0.000153 |
| chr2 | 105535916 | 105536365 | 450 | 5.02 | 4.97E-05 | 0.000154 |
| chr18 | 69884678 | 69885397 | 720 | 5.04 | 4.98E-05 | 0.000154 |
| chr19 | 21419950 | 21420478 | 529 | 5.04 | 4.98E-05 | 0.000154 |
| chr17 | 7252455 | 7256981 | 4527 | 0.44 | 4.98E-05 | 0.000154 |
| chr6 | 143223616 | 143224569 | 954 | 1.84 | 4.99E-05 | 0.000154 |
| chr3 | 8910332 | 8912710 | 2379 | 0.68 | 4.99E-05 | 0.000154 |
| chr7 | 27955616 | 27961387 | 5772 | 0.8 | 5.00E-05 | 0.000154 |
| chr14 | 101614274 | 101618234 | 3961 | 1.25 | 5.01E-05 | 0.000155 |
| chr5 | 77715904 | 77717495 | 1592 | 1.62 | 5.01E-05 | 0.000155 |
| chr4 | 38890226 | 38892543 | 2318 | 1.39 | 5.01E-05 | 0.000155 |
| chr1 | 81474044 | 81474569 | 526 | 4.45 | 5.01E-05 | 0.000155 |
| chr12 | 100966825 | 100968732 | 1908 | 0.85 | 5.02E-05 | 0.000155 |
| chr4 | 129186179 | 129210538 | 24360 | -0.47 | 5.02E-05 | 0.000155 |
| chr20 | 18409037 | 18410341 | 1305 | 1.25 | 5.03E-05 | 0.000155 |
| chr8 | 32829080 | 32837975 | 8896 | -2.02 | 5.03E-05 | 0.000155 |
| chr3 | 110847457 | 110848244 | 788 | -2.31 | 5.03E-05 | 0.000155 |
| chr18 | 24147988 | 24161711 | 13724 | 0.29 | 5.04E-05 | 0.000156 |
| chr8 | 127544214 | 127544582 | 369 | 5.02 | 5.04E-05 | 0.000156 |
| chr17 | 4210712 | 4271372 | 60661 | -0.27 | 5.04E-05 | 0.000156 |
| chr11 | 77226275 | 77226653 | 379 | 5.05 | 5.05E-05 | 0.000156 |
| chr5 | 41122274 | 41123128 | 855 | -3.36 | 5.05E-05 | 0.000156 |
| chr1 | 200529234 | 200541019 | 11786 | 0.7 | 5.06E-05 | 0.000156 |
| chr16 | 47960999 | 47961828 | 830 | 2.84 | 5.06E-05 | 0.000156 |
| chr15 | 57899196 | 57904609 | 5414 | 1.02 | 5.07E-05 | 0.000156 |
| chr5 | 56365574 | 56366053 | 480 | 5.17 | 5.08E-05 | 0.000156 |
| chr7 | 20663361 | 20666034 | 2674 | 5.67 | 5.08E-05 | 0.000157 |
| chrX | 82342762 | 82343319 | 558 | 5.09 | 5.08E-05 | 0.000157 |
| chr9 | 77566103 | 77570265 | 4163 | 0.51 | 5.09E-05 | 0.000157 |
| chr20 | 14801534 | 14802383 | 850 | 3.13 | 5.09E-05 | 0.000157 |
| chr15 | 101667527 | 101668539 | 1013 | 3.32 | 5.09E-05 | 0.000157 |
| chr4 | 92154140 | 92154637 | 498 | 4.43 | 5.10E-05 | 0.000157 |

| | | | | | | |
|---|---|---|---|---|---|---|
| chr19 | 24269063 | 24271163 | 2101 | 0.58 | 5.10E-05 | 0.000157 |
| chr3 | 9640546 | 9642785 | 2240 | 2.68 | 5.10E-05 | 0.000157 |
| chr9 | 135934689 | 135937055 | 2367 | -1.57 | 5.11E-05 | 0.000157 |
| chr8 | 37887297 | 37895691 | 8395 | -0.54 | 5.11E-05 | 0.000157 |
| chr3 | 36359392 | 36360908 | 1517 | 1.7 | 5.12E-05 | 0.000158 |
| chr4 | 162793325 | 162794018 | 694 | 2.42 | 5.14E-05 | 0.000158 |
| chrX | 66911826 | 66921172 | 9347 | 0.78 | 5.14E-05 | 0.000158 |
| chr1 | 207261267 | 207262941 | 1675 | 1.63 | 5.15E-05 | 0.000159 |
| chrX | 73832426 | 73835711 | 3286 | 0.35 | 5.15E-05 | 0.000159 |
| chr1 | 76189340 | 76202639 | 13300 | 0.62 | 5.18E-05 | 0.00016 |
| chr20 | 40161552 | 40163034 | 1483 | 1.51 | 5.18E-05 | 0.00016 |
| chr7 | 151318660 | 151329662 | 11003 | -0.85 | 5.19E-05 | 0.00016 |
| chr9 | 128275620 | 128302960 | 27341 | -0.75 | 5.19E-05 | 0.00016 |
| chr6 | 80852641 | 80853436 | 796 | 2.21 | 5.19E-05 | 0.00016 |
| chr17 | 9783068 | 9785135 | 2068 | 1.14 | 5.20E-05 | 0.00016 |
| chr3 | 45239686 | 45242915 | 3230 | 0.61 | 5.20E-05 | 0.00016 |
| chr17 | 1884716 | 1906479 | 21764 | -0.49 | 5.23E-05 | 0.000161 |
| chr6 | 3610072 | 3613851 | 3780 | 0.98 | 5.23E-05 | 0.000161 |
| chr2 | 25759951 | 25761199 | 1249 | -1.28 | 5.23E-05 | 0.000161 |
| chrX | 131156722 | 131158540 | 1819 | 1.23 | 5.24E-05 | 0.000161 |
| chr2 | 239332241 | 239358130 | 25890 | -0.51 | 5.24E-05 | 0.000161 |
| chr17 | 59467086 | 59505125 | 38040 | -0.47 | 5.25E-05 | 0.000161 |
| chr19 | 11909412 | 11909951 | 540 | 5.04 | 5.25E-05 | 0.000161 |
| chrX | 117670480 | 117679642 | 9163 | -5.06 | 5.26E-05 | 0.000162 |
| chr3 | 152661420 | 152662426 | 1007 | 3.55 | 5.26E-05 | 0.000162 |
| chr15 | 63525370 | 63527273 | 1904 | -1.26 | 5.26E-05 | 0.000162 |
| chr6 | 151325072 | 151414054 | 88983 | 0.19 | 5.27E-05 | 0.000162 |
| chr7 | 105374692 | 105375362 | 671 | 5.02 | 5.27E-05 | 0.000162 |
| chr22 | 50324801 | 50364490 | 39690 | 0.32 | 5.27E-05 | 0.000162 |
| chr12 | 47392283 | 47392831 | 549 | 3.78 | 5.27E-05 | 0.000162 |
| chr22 | 39756644 | 39757716 | 1073 | 1.97 | 5.29E-05 | 0.000162 |
| chr8 | 11141248 | 11144750 | 3503 | 0.6 | 5.29E-05 | 0.000162 |
| chr20 | 52351511 | 52427651 | 76141 | -0.17 | 5.29E-05 | 0.000162 |
| chr4 | 92830872 | 92831510 | 639 | 3.94 | 5.30E-05 | 0.000163 |
| chr1 | 233544796 | 233546204 | 1409 | 1.97 | 5.30E-05 | 0.000163 |
| chr10 | 75488826 | 75505432 | 16607 | 0.38 | 5.30E-05 | 0.000163 |
| chr13 | 47369918 | 47372004 | 2087 | 0.76 | 5.30E-05 | 0.000163 |
| chr1 | 33800267 | 33816519 | 16253 | 0.36 | 5.31E-05 | 0.000163 |
| chr10 | 116030424 | 116031299 | 876 | 2.28 | 5.32E-05 | 0.000163 |
| chr9 | 84127261 | 84306039 | 178779 | 0.15 | 5.33E-05 | 0.000164 |
| chr1 | 9350040 | 9412327 | 62288 | -0.27 | 5.34E-05 | 0.000164 |
| chr14 | 95102864 | 95103521 | 658 | 2.46 | 5.34E-05 | 0.000164 |
| chr4 | 151973490 | 151974548 | 1059 | 2.26 | 5.35E-05 | 0.000164 |
| chr15 | 56034053 | 56036168 | 2116 | -0.7 | 5.35E-05 | 0.000164 |
| chr9 | 115140608 | 115143266 | 2659 | 0.56 | 5.35E-05 | 0.000164 |
| chr4 | 48081549 | 48083397 | 1849 | 1.1 | 5.36E-05 | 0.000164 |
| chr7 | 127020537 | 127033722 | 13186 | 0.48 | 5.38E-05 | 0.000165 |
| chr2 | 153429530 | 153437789 | 8260 | -1.16 | 5.38E-05 | 0.000165 |
| chr14 | 39843632 | 39847215 | 3584 | 0.46 | 5.39E-05 | 0.000165 |
| chr6 | 134151171 | 134159600 | 8430 | 0.72 | 5.40E-05 | 0.000165 |
| chr17 | 12905908 | 12922606 | 16699 | 0.63 | 5.41E-05 | 0.000166 |
| chr20 | 15290843 | 15291403 | 561 | 3.5 | 5.41E-05 | 0.000166 |
| chr1 | 116087659 | 116089650 | 1992 | 0.86 | 5.41E-05 | 0.000166 |
| chr7 | 27700828 | 27720029 | 19202 | 0.47 | 5.41E-05 | 0.000166 |
| chr4 | 57331016 | 57335142 | 4127 | 0.54 | 5.41E-05 | 0.000166 |
| chr12 | 54519832 | 54530051 | 10220 | 1.31 | 5.42E-05 | 0.000166 |
| chr10 | 128769863 | 128772378 | 2516 | 0.68 | 5.42E-05 | 0.000166 |
| chr14 | 31720291 | 31734373 | 14083 | 1.02 | 5.43E-05 | 0.000166 |
| chr2 | 55382543 | 55391785 | 9243 | 0.77 | 5.43E-05 | 0.000166 |
| chr12 | 92896901 | 92898253 | 1353 | 2.18 | 5.44E-05 | 0.000167 |
| chr12 | 69741872 | 69760598 | 18727 | 0.47 | 5.46E-05 | 0.000167 |

| | | | | | | |
|---|---|---|---|---|---|---|
| chr1 | 173717986 | 173728592 | 10607 | -1.31 | 5.47E-05 | 0.000167 |
| chr15 | 72329398 | 72330396 | 999 | 2.64 | 5.47E-05 | 0.000167 |
| chr21 | 38821902 | 38848026 | 26125 | -0.47 | 5.47E-05 | 0.000167 |
| chr10 | 22716057 | 22726671 | 10615 | 0.73 | 5.48E-05 | 0.000168 |
| chrX | 73136420 | 73138290 | 1871 | 0.98 | 5.48E-05 | 0.000168 |
| chr5 | 145304288 | 145326001 | 21714 | 0.54 | 5.49E-05 | 0.000168 |
| chr5 | 91377570 | 91378421 | 852 | 2.59 | 5.51E-05 | 0.000169 |
| chr6 | 49636214 | 49636711 | 498 | 5.17 | 5.53E-05 | 0.000169 |
| chr16 | 69065322 | 69124075 | 58754 | 0.22 | 5.53E-05 | 0.000169 |
| chr10 | 30926864 | 30928099 | 1236 | -2.77 | 5.53E-05 | 0.000169 |
| chr2 | 74152312 | 74184563 | 32252 | -0.32 | 5.53E-05 | 0.000169 |
| chr6 | 2683928 | 2685171 | 1244 | 1.74 | 5.54E-05 | 0.00017 |
| chr1 | 27517909 | 27520193 | 2285 | 0.96 | 5.55E-05 | 0.00017 |
| chr7 | 32915795 | 32945535 | 29741 | 0.26 | 5.56E-05 | 0.00017 |
| chr3 | 114957828 | 114959619 | 1792 | 1.08 | 5.57E-05 | 0.00017 |
| chr7 | 118003527 | 118004396 | 870 | 3.48 | 5.57E-05 | 0.00017 |
| chr12 | 96090553 | 96127753 | 37201 | 0.47 | 5.57E-05 | 0.00017 |
| chr13 | 74181681 | 74193641 | 11961 | 1.13 | 5.58E-05 | 0.00017 |
| chr12 | 100838856 | 100839401 | 546 | 2.15 | 5.58E-05 | 0.00017 |
| chr15 | 45570618 | 45572326 | 1709 | 0.96 | 5.58E-05 | 0.000171 |
| chr8 | 129746414 | 129747197 | 784 | -2.72 | 5.59E-05 | 0.000171 |
| chr11 | 102500370 | 102500985 | 616 | 3.85 | 5.59E-05 | 0.000171 |
| chr4 | 63260553 | 63261065 | 513 | 5.04 | 5.59E-05 | 0.000171 |
| chr7 | 15055405 | 15055759 | 355 | 5.04 | 5.59E-05 | 0.000171 |
| chr5 | 23352102 | 23352957 | 856 | 2.94 | 5.60E-05 | 0.000171 |
| chr1 | 80502571 | 80503016 | 446 | 5 | 5.60E-05 | 0.000171 |
| chrX | 47640882 | 47641258 | 377 | 5 | 5.60E-05 | 0.000171 |
| chr10 | 118752084 | 118767968 | 15885 | 0.34 | 5.61E-05 | 0.000171 |
| chr7 | 123363432 | 123368714 | 5283 | 0.65 | 5.62E-05 | 0.000172 |
| chr7 | 137615074 | 137618772 | 3699 | 0.67 | 5.62E-05 | 0.000172 |
| chr1 | 244553948 | 244559651 | 5704 | 0.66 | 5.63E-05 | 0.000172 |
| chr17 | 76268490 | 76269468 | 979 | 5 | 5.63E-05 | 0.000172 |
| chr3 | 56656214 | 56657104 | 891 | 2.88 | 5.64E-05 | 0.000172 |
| chr10 | 8313294 | 8314677 | 1384 | 1.67 | 5.64E-05 | 0.000172 |
| chr4 | 131965277 | 131966347 | 1071 | 1.99 | 5.64E-05 | 0.000172 |
| chr2 | 201577382 | 201578903 | 1522 | -0.98 | 5.65E-05 | 0.000172 |
| chr22 | 27869832 | 27885817 | 15986 | 0.75 | 5.66E-05 | 0.000173 |
| chr12 | 117255435 | 117258427 | 2993 | 0.59 | 5.67E-05 | 0.000173 |
| chr1 | 66003491 | 66004092 | 602 | 5.05 | 5.67E-05 | 0.000173 |
| chr8 | 37818195 | 37820313 | 2119 | 2.24 | 5.68E-05 | 0.000173 |
| chr11 | 95709539 | 95725692 | 16154 | 0.4 | 5.68E-05 | 0.000173 |
| chr1 | 203104882 | 203105507 | 626 | 3.71 | 5.68E-05 | 0.000173 |
| chr7 | 32995748 | 32998636 | 2889 | 0.47 | 5.69E-05 | 0.000173 |
| chr19 | 17610289 | 17623589 | 13301 | 0.54 | 5.69E-05 | 0.000173 |
| chr21 | 15914010 | 15914793 | 784 | 3.61 | 5.69E-05 | 0.000173 |
| chr7 | 2143603 | 2144127 | 525 | -4.97 | 5.69E-05 | 0.000174 |
| chr4 | 181370829 | 181371320 | 492 | 5 | 5.70E-05 | 0.000174 |
| chr8 | 57904310 | 57907308 | 2999 | 0.78 | 5.70E-05 | 0.000174 |
| chr20 | 30575498 | 30576664 | 1167 | -2.29 | 5.70E-05 | 0.000174 |
| chr6 | 157096885 | 157104296 | 7412 | 0.44 | 5.71E-05 | 0.000174 |
| chr8 | 75703978 | 75704646 | 669 | -2.32 | 5.74E-05 | 0.000175 |
| chr3 | 111387545 | 111396633 | 9089 | 0.44 | 5.75E-05 | 0.000175 |
| chr13 | 88454510 | 88463954 | 9445 | 0.62 | 5.75E-05 | 0.000175 |
| chr19 | 37446979 | 37447352 | 374 | 5.02 | 5.76E-05 | 0.000175 |
| chr2 | 48448591 | 48449773 | 1183 | 2.01 | 5.77E-05 | 0.000176 |
| chr5 | 179103819 | 179134007 | 30189 | 0.34 | 5.77E-05 | 0.000176 |
| chr1 | 163290676 | 163294250 | 3575 | 0.57 | 5.77E-05 | 0.000176 |
| chr3 | 52080232 | 52091289 | 11058 | -0.63 | 5.78E-05 | 0.000176 |
| chr12 | 44229409 | 44230847 | 1439 | 0.86 | 5.79E-05 | 0.000176 |
| chr16 | 84025350 | 84029095 | 3746 | -1.28 | 5.79E-05 | 0.000176 |
| chr10 | 98328666 | 98348232 | 19567 | 0.33 | 5.80E-05 | 0.000176 |

| | | | | | | |
|---|---|---|---|---|---|---|
| chr12 | 63359870 | 63361534 | 1665 | 4.74 | 5.81E-05 | 0.000177 |
| chr4 | 48164110 | 48178020 | 13911 | -0.72 | 5.81E-05 | 0.000177 |
| chr3 | 189749053 | 189750879 | 1827 | -0.62 | 5.82E-05 | 0.000177 |
| chr1 | 97558523 | 97560770 | 2248 | 1.16 | 5.82E-05 | 0.000177 |
| chr8 | 27361955 | 27362420 | 466 | -4.38 | 5.82E-05 | 0.000177 |
| chr12 | 105424409 | 105431825 | 7417 | 1.25 | 5.83E-05 | 0.000177 |
| chr6 | 132709468 | 132710089 | 622 | 5 | 5.84E-05 | 0.000178 |
| chr17 | 41889870 | 41923756 | 33887 | -0.28 | 5.85E-05 | 0.000178 |
| chr7 | 133572567 | 133573632 | 1066 | 1.87 | 5.85E-05 | 0.000178 |
| chr22 | 35552594 | 35553281 | 688 | 1.64 | 5.86E-05 | 0.000178 |
| chr10 | 75003241 | 75014019 | 10779 | 0.33 | 5.86E-05 | 0.000178 |
| chr5 | 167715107 | 167883372 | 168266 | 0.13 | 5.87E-05 | 0.000178 |
| chr5 | 125861497 | 125874923 | 13427 | 3.47 | 5.87E-05 | 0.000178 |
| chr3 | 20492052 | 20492590 | 539 | -4.95 | 5.88E-05 | 0.000179 |
| chr20 | 322044 | 329883 | 7840 | 0.61 | 5.88E-05 | 0.000179 |
| chr11 | 60638919 | 60640417 | 1499 | -1.87 | 5.90E-05 | 0.000179 |
| chr16 | 21186441 | 21200411 | 13971 | -1.39 | 5.92E-05 | 0.00018 |
| chr7 | 107975989 | 107976692 | 704 | -4.94 | 5.92E-05 | 0.00018 |
| chr4 | 56750176 | 56761008 | 10833 | 1 | 5.92E-05 | 0.00018 |
| chr7 | 65703427 | 65703864 | 438 | 3.42 | 5.97E-05 | 0.000181 |
| chr6 | 166060150 | 166063146 | 2997 | 2.37 | 5.98E-05 | 0.000181 |
| chr4 | 91238867 | 91239910 | 1044 | 2.39 | 5.98E-05 | 0.000182 |
| chr3 | 172313660 | 172314572 | 913 | 3.08 | 5.99E-05 | 0.000182 |
| chr10 | 117705988 | 117706823 | 836 | -2.2 | 6.00E-05 | 0.000182 |
| chr4 | 103655688 | 103659224 | 3537 | 1.47 | 6.00E-05 | 0.000182 |
| chr5 | 95066226 | 95068533 | 2308 | 1.12 | 6.01E-05 | 0.000182 |
| chr9 | 72120594 | 72121087 | 494 | 3.89 | 6.01E-05 | 0.000182 |
| chr3 | 121150589 | 121151914 | 1326 | 1.84 | 6.01E-05 | 0.000182 |
| chr10 | 7274805 | 7276716 | 1912 | 1.75 | 6.01E-05 | 0.000182 |
| chr17 | 58467984 | 58471440 | 3457 | 0.4 | 6.02E-05 | 0.000182 |
| chr1 | 151058203 | 151059429 | 1227 | -1.1 | 6.03E-05 | 0.000183 |
| chr5 | 33049161 | 33050588 | 1428 | -1.58 | 6.03E-05 | 0.000183 |
| chr8 | 145007428 | 145065503 | 58076 | 0.21 | 6.06E-05 | 0.000184 |
| chr4 | 41257813 | 41262810 | 4998 | -0.5 | 6.06E-05 | 0.000184 |
| chr6 | 96968860 | 96971610 | 2751 | 0.59 | 6.07E-05 | 0.000184 |
| chr15 | 101584291 | 101586127 | 1837 | -1.34 | 6.09E-05 | 0.000185 |
| chr14 | 54465403 | 54468617 | 3215 | 1.03 | 6.10E-05 | 0.000185 |
| chrX | 3731524 | 3735156 | 3633 | 0.66 | 6.11E-05 | 0.000185 |
| chr5 | 10688855 | 10690490 | 1636 | -1.32 | 6.13E-05 | 0.000186 |
| chr3 | 78888155 | 78889113 | 959 | 3.44 | 6.13E-05 | 0.000186 |
| chr15 | 70353042 | 70359506 | 6465 | -1.11 | 6.13E-05 | 0.000186 |
| chr19 | 42363074 | 42411885 | 48812 | 0.22 | 6.13E-05 | 0.000186 |
| chr16 | 29626830 | 29627369 | 540 | 1.92 | 6.13E-05 | 0.000186 |
| chr8 | 38590906 | 38594440 | 3535 | 0.55 | 6.14E-05 | 0.000186 |
| chr12 | 112740525 | 112749998 | 9474 | -0.88 | 6.16E-05 | 0.000187 |
| chr2 | 81708519 | 81709262 | 744 | 3.79 | 6.17E-05 | 0.000187 |
| chr6 | 109517259 | 109518770 | 1512 | 3.24 | 6.17E-05 | 0.000187 |
| chr1 | 109718608 | 109741700 | 23093 | 0.48 | 6.19E-05 | 0.000187 |
| chr3 | 25704372 | 25707553 | 3182 | 0.57 | 6.19E-05 | 0.000187 |
| chr6 | 15948788 | 15956686 | 7899 | 3.22 | 6.19E-05 | 0.000187 |
| chr18 | 74275915 | 74286042 | 10128 | 0.96 | 6.20E-05 | 0.000188 |
| chr7 | 129359425 | 129362719 | 3295 | -0.83 | 6.22E-05 | 0.000188 |
| chr6 | 22470803 | 22472077 | 1275 | 1.36 | 6.23E-05 | 0.000189 |
| chr12 | 22187695 | 22200808 | 13114 | 1.03 | 6.24E-05 | 0.000189 |
| chr15 | 25585282 | 25585916 | 635 | 3.79 | 6.24E-05 | 0.000189 |
| chr14 | 54862906 | 54866409 | 3504 | -0.68 | 6.24E-05 | 0.000189 |
| chr19 | 32778917 | 32779382 | 466 | -4.03 | 6.25E-05 | 0.000189 |
| chr6 | 41882214 | 41912016 | 29803 | -0.3 | 6.31E-05 | 0.000191 |
| chr5 | 46112083 | 46112542 | 460 | 4.97 | 6.31E-05 | 0.000191 |
| chr10 | 32561991 | 32591277 | 29287 | 0.59 | 6.32E-05 | 0.000191 |
| chr12 | 72914445 | 72914905 | 461 | 4.97 | 6.32E-05 | 0.000191 |

| | | | | | | |
|---|---|---|---|---|---|---|
| chr2 | 190271936 | 190277019 | 5084 | -1.12 | 6.34E-05 | 0.000192 |
| chr1 | 19325187 | 19340613 | 15427 | 0.34 | 6.34E-05 | 0.000192 |
| chr22 | 31884883 | 31893354 | 8472 | 0.56 | 6.34E-05 | 0.000192 |
| chr5 | 96303277 | 96304659 | 1383 | 1.66 | 6.35E-05 | 0.000192 |
| chr4 | 113151336 | 113154574 | 3239 | 0.69 | 6.37E-05 | 0.000193 |
| chr13 | 50697087 | 50701063 | 3977 | 0.62 | 6.37E-05 | 0.000193 |
| chr10 | 21776837 | 21786893 | 10057 | -0.41 | 6.37E-05 | 0.000193 |
| chr16 | 87614541 | 87616928 | 2388 | 0.63 | 6.38E-05 | 0.000193 |
| chr11 | 21696594 | 21697154 | 561 | 3.77 | 6.39E-05 | 0.000193 |
| chr2 | 24581898 | 24588983 | 7086 | 0.53 | 6.39E-05 | 0.000193 |
| chrX | 133756017 | 133756786 | 770 | -3.2 | 6.39E-05 | 0.000193 |
| chr1 | 87458471 | 87460158 | 1688 | -1.14 | 6.39E-05 | 0.000193 |
| chr12 | 90592561 | 90593241 | 681 | 2.87 | 6.39E-05 | 0.000193 |
| chr1 | 9448471 | 9474219 | 25749 | -1.01 | 6.40E-05 | 0.000193 |
| chr1 | 195621637 | 195622073 | 437 | 4.97 | 6.40E-05 | 0.000193 |
| chr2 | 81635562 | 81636044 | 483 | 4.97 | 6.40E-05 | 0.000193 |
| chr2 | 99682200 | 99683240 | 1041 | 2.26 | 6.40E-05 | 0.000193 |
| chr3 | 24276259 | 24295444 | 19186 | 2.5 | 6.41E-05 | 0.000193 |
| chr12 | 131309480 | 131324807 | 15328 | 0.55 | 6.41E-05 | 0.000194 |
| chr6 | 154751166 | 154752631 | 1466 | 1.67 | 6.42E-05 | 0.000194 |
| chr22 | 48171951 | 48172482 | 532 | 5 | 6.42E-05 | 0.000194 |
| chr20 | 32307539 | 32339792 | 32254 | 0.32 | 6.42E-05 | 0.000194 |
| chr14 | 95289546 | 95291217 | 1672 | 1.41 | 6.43E-05 | 0.000194 |
| chr2 | 27576469 | 27637413 | 60945 | -0.27 | 6.43E-05 | 0.000194 |
| chr15 | 41766177 | 41769702 | 3526 | 1.03 | 6.45E-05 | 0.000194 |
| chr22 | 30439994 | 30440830 | 837 | 4.04 | 6.45E-05 | 0.000195 |
| chr2 | 127974793 | 127985262 | 10470 | -0.63 | 6.46E-05 | 0.000195 |
| chr2 | 133274545 | 133275477 | 933 | -2.02 | 6.47E-05 | 0.000195 |
| chr3 | 152217532 | 152218256 | 725 | 2.47 | 6.49E-05 | 0.000196 |
| chr1 | 36395393 | 36397971 | 2579 | 0.54 | 6.50E-05 | 0.000196 |
| chr12 | 43390782 | 43391280 | 499 | 4.97 | 6.50E-05 | 0.000196 |
| chr21 | 32931918 | 32932692 | 775 | 2.72 | 6.51E-05 | 0.000196 |
| chr7 | 69985105 | 69986332 | 1228 | 3.18 | 6.53E-05 | 0.000197 |
| chr4 | 168700779 | 168701503 | 725 | 2.24 | 6.57E-05 | 0.000198 |
| chr15 | 57024042 | 57030990 | 6949 | 0.53 | 6.57E-05 | 0.000198 |
| chr7 | 105542143 | 105543363 | 1221 | -2.22 | 6.58E-05 | 0.000198 |
| chr13 | 34508775 | 34509729 | 955 | 2.24 | 6.61E-05 | 0.000199 |
| chr2 | 150976706 | 151045795 | 69090 | 0.33 | 6.62E-05 | 0.000199 |
| chr2 | 197503644 | 197505383 | 1740 | -1.01 | 6.64E-05 | 2.00E-04 |
| chr9 | 78935428 | 78939060 | 3633 | 0.99 | 6.65E-05 | 2.00E-04 |
| chr8 | 24108641 | 24110115 | 1475 | 2.64 | 6.66E-05 | 0.000201 |
| chr4 | 114374015 | 114375384 | 1370 | 1.59 | 6.66E-05 | 0.000201 |
| chr13 | 33398164 | 33399954 | 1791 | 1.18 | 6.68E-05 | 0.000201 |
| chrX | 46039078 | 46040637 | 1560 | -1.27 | 6.68E-05 | 0.000201 |
| chr16 | 67693223 | 67716316 | 23094 | -0.56 | 6.69E-05 | 0.000201 |
| chr12 | 17137488 | 17138390 | 903 | 2.25 | 6.70E-05 | 0.000201 |
| chr2 | 193302563 | 193303111 | 549 | 4.97 | 6.70E-05 | 0.000202 |
| chrX | 62654175 | 62654775 | 601 | 4.97 | 6.70E-05 | 0.000202 |
| chr2 | 37227228 | 37228567 | 1340 | -1.66 | 6.71E-05 | 0.000202 |
| chr1 | 153912331 | 153967025 | 54695 | 0.22 | 6.73E-05 | 0.000202 |
| chr4 | 130466628 | 130467863 | 1236 | -2.32 | 6.74E-05 | 0.000203 |
| chr10 | 99155699 | 99167623 | 11925 | -0.33 | 6.75E-05 | 0.000203 |
| chr3 | 59795080 | 59805135 | 10056 | -1.22 | 6.77E-05 | 0.000204 |
| chr12 | 80168799 | 80179559 | 10761 | 0.6 | 6.78E-05 | 0.000204 |
| chr6 | 1110949 | 1111722 | 774 | 2.31 | 6.78E-05 | 0.000204 |
| chr14 | 77452089 | 77452839 | 751 | -2.65 | 6.79E-05 | 0.000204 |
| chr14 | 55368436 | 55370249 | 1814 | 1 | 6.83E-05 | 0.000205 |
| chr1 | 229388611 | 229390130 | 1520 | 1.27 | 6.83E-05 | 0.000205 |
| chr3 | 141294832 | 141295498 | 667 | 2.44 | 6.85E-05 | 0.000206 |
| chr8 | 134259824 | 134281693 | 21870 | -2.11 | 6.85E-05 | 0.000206 |
| chr2 | 225084405 | 225087888 | 3484 | -0.47 | 6.86E-05 | 0.000206 |

| | | | | | | |
|---|---|---|---|---|---|---|
| chr5 | 138087979 | 138106656 | 18678 | 0.43 | 6.87E-05 | 0.000206 |
| chr18 | 48362659 | 48363020 | 362 | 4.44 | 6.87E-05 | 0.000206 |
| chr3 | 83484158 | 83484667 | 510 | 3.88 | 6.87E-05 | 0.000206 |
| chr5 | 131924709 | 131934665 | 9957 | -0.97 | 6.88E-05 | 0.000207 |
| chr12 | 64610040 | 64617024 | 6985 | 0.52 | 6.88E-05 | 0.000207 |
| chr17 | 81066276 | 81068419 | 2144 | 0.69 | 6.89E-05 | 0.000207 |
| chr4 | 175203258 | 175206976 | 3719 | 0.5 | 6.90E-05 | 0.000207 |
| chr12 | 93557612 | 93561176 | 3565 | 4.68 | 6.90E-05 | 0.000207 |
| chr1 | 84991708 | 84992220 | 513 | 3.32 | 6.91E-05 | 0.000207 |
| chrX | 43930205 | 43932602 | 2398 | 1.69 | 6.92E-05 | 0.000208 |
| chr5 | 43601658 | 43608758 | 7101 | 0.75 | 6.93E-05 | 0.000208 |
| chr20 | 10111603 | 10113400 | 1798 | 0.99 | 6.93E-05 | 0.000208 |
| chr2 | 191500721 | 191516257 | 15537 | 0.54 | 6.93E-05 | 0.000208 |
| chr2 | 217121931 | 217122482 | 552 | -4.98 | 6.94E-05 | 0.000208 |
| chr1 | 36472823 | 36475363 | 2541 | 1.07 | 6.95E-05 | 0.000209 |
| chr3 | 23709065 | 23711235 | 2171 | -0.65 | 6.97E-05 | 0.000209 |
| chr1 | 3503681 | 3504294 | 614 | -2.78 | 6.98E-05 | 0.000209 |
| chr11 | 70782739 | 70783591 | 853 | -4.93 | 6.98E-05 | 0.000209 |
| chr8 | 31611198 | 31614925 | 3728 | 0.39 | 6.99E-05 | 0.00021 |
| chrX | 18635098 | 18636867 | 1770 | 1.3 | 6.99E-05 | 0.00021 |
| chr8 | 134142714 | 134143609 | 896 | -2.01 | 6.99E-05 | 0.00021 |
| chr6 | 37068795 | 37073480 | 4686 | 0.44 | 7.00E-05 | 0.00021 |
| chr3 | 197436766 | 197478611 | 41846 | 0.3 | 7.02E-05 | 0.00021 |
| chr4 | 169525853 | 169534310 | 8458 | -1.57 | 7.03E-05 | 0.000211 |
| chr10 | 43455328 | 43457003 | 1676 | 2.39 | 7.03E-05 | 0.000211 |
| chr2 | 48666200 | 48680308 | 14109 | -0.41 | 7.05E-05 | 0.000211 |
| chr4 | 108640125 | 108642480 | 2356 | 1.23 | 7.07E-05 | 0.000212 |
| chr17 | 37122672 | 37132188 | 9517 | 0.62 | 7.08E-05 | 0.000212 |
| chr6 | 168648069 | 168675478 | 27410 | 0.55 | 7.09E-05 | 0.000212 |
| chr19 | 3499781 | 3507287 | 7507 | -0.49 | 7.09E-05 | 0.000212 |
| chr7 | 109298934 | 109299882 | 949 | 3.28 | 7.09E-05 | 0.000212 |
| chr5 | 7905746 | 7906762 | 1017 | 1.86 | 7.10E-05 | 0.000213 |
| chr1 | 15665638 | 15694870 | 29233 | -0.28 | 7.10E-05 | 0.000213 |
| chrX | 27768134 | 27768555 | 422 | 4.35 | 7.11E-05 | 0.000213 |
| chr5 | 129861695 | 129868860 | 7166 | 0.66 | 7.13E-05 | 0.000214 |
| chr9 | 131754491 | 131799486 | 44996 | -0.43 | 7.13E-05 | 0.000214 |
| chr4 | 187149021 | 187149718 | 698 | 4.99 | 7.14E-05 | 0.000214 |
| chr19 | 35911023 | 35912607 | 1585 | 1.32 | 7.14E-05 | 0.000214 |
| chr3 | 42615301 | 42658971 | 43671 | 0.34 | 7.15E-05 | 0.000214 |
| chr8 | 84440455 | 84440822 | 368 | 4.95 | 7.17E-05 | 0.000214 |
| chr17 | 78933300 | 78936021 | 2722 | -1.9 | 7.18E-05 | 0.000215 |
| chr12 | 66171777 | 66172951 | 1175 | 1.43 | 7.18E-05 | 0.000215 |
| chr1 | 95285110 | 95287913 | 2804 | 1.47 | 7.18E-05 | 0.000215 |
| chr11 | 9186991 | 9203524 | 16534 | -0.83 | 7.19E-05 | 0.000215 |
| chr10 | 27377156 | 27423746 | 46591 | 0.39 | 7.20E-05 | 0.000215 |
| chr12 | 83002823 | 83003475 | 653 | 3.35 | 7.21E-05 | 0.000216 |
| chr13 | 74245566 | 74253317 | 7752 | 3.79 | 7.21E-05 | 0.000216 |
| chr17 | 25553272 | 25554263 | 992 | 2.51 | 7.21E-05 | 0.000216 |
| chr13 | 20419497 | 20451852 | 32356 | 0.41 | 7.22E-05 | 0.000216 |
| chr19 | 47267547 | 47268943 | 1397 | 1.52 | 7.22E-05 | 0.000216 |
| chr4 | 148752116 | 148754184 | 2069 | 0.71 | 7.23E-05 | 0.000216 |
| chr14 | 69698072 | 69698617 | 546 | 4.95 | 7.25E-05 | 0.000217 |
| chr2 | 63947762 | 63948528 | 767 | 4.95 | 7.25E-05 | 0.000217 |
| chr1 | 244402328 | 244403351 | 1024 | -1.89 | 7.26E-05 | 0.000217 |
| chr5 | 162584731 | 162585269 | 539 | -4.94 | 7.26E-05 | 0.000217 |
| chr2 | 128560525 | 128569430 | 8906 | -0.54 | 7.26E-05 | 0.000217 |
| chr1 | 28639762 | 28640899 | 1138 | -1.6 | 7.27E-05 | 0.000217 |
| chr8 | 58423896 | 58427545 | 3650 | 0.8 | 7.28E-05 | 0.000217 |
| chr15 | 45679756 | 45680208 | 453 | 3.61 | 7.29E-05 | 0.000218 |
| chr9 | 106560838 | 106567441 | 6604 | 4.97 | 7.30E-05 | 0.000218 |
| chr14 | 100600340 | 100612055 | 11716 | -0.95 | 7.30E-05 | 0.000218 |

| | | | | | | |
|---|---|---|---|---|---|---|
| chr21 | 45663382 | 45663885 | 504 | 4.95 | 7.30E-05 | 0.000218 |
| chr9 | 102883534 | 102894641 | 11108 | 1.49 | 7.31E-05 | 0.000218 |
| chr3 | 112789376 | 112798993 | 9618 | -4.99 | 7.31E-05 | 0.000218 |
| chr2 | 69533243 | 69538031 | 4789 | -0.57 | 7.33E-05 | 0.000219 |
| chrX | 20047086 | 20049121 | 2036 | 1.3 | 7.33E-05 | 0.000219 |
| chr3 | 179687255 | 179687971 | 717 | -3.46 | 7.33E-05 | 0.000219 |
| chr2 | 44160741 | 44176181 | 15441 | -0.61 | 7.35E-05 | 0.000219 |
| chrX | 2845050 | 2848283 | 3234 | 0.52 | 7.36E-05 | 0.000219 |
| chr8 | 49405885 | 49428220 | 22336 | 0.57 | 7.36E-05 | 0.00022 |
| chr4 | 7430834 | 7433707 | 2874 | 1.03 | 7.38E-05 | 0.00022 |
| chr9 | 71114982 | 71116653 | 1672 | 1.33 | 7.38E-05 | 0.00022 |
| chr11 | 84861098 | 84861663 | 566 | 3.8 | 7.39E-05 | 0.00022 |
| chr3 | 188984706 | 188995143 | 10438 | 0.77 | 7.39E-05 | 0.00022 |
| chr2 | 102775668 | 102785080 | 9413 | -1.52 | 7.39E-05 | 0.00022 |
| chr6 | 110650849 | 110651632 | 784 | 3.65 | 7.41E-05 | 0.000221 |
| chr8 | 29831587 | 29834412 | 2826 | 0.68 | 7.42E-05 | 0.000221 |
| chr10 | 61750621 | 61756257 | 5637 | 6.36 | 7.42E-05 | 0.000221 |
| chr22 | 30681485 | 30724359 | 42875 | -0.44 | 7.42E-05 | 0.000221 |
| chrX | 118108135 | 118111594 | 3460 | 0.74 | 7.42E-05 | 0.000221 |
| chr3 | 138048125 | 138049015 | 891 | -1.48 | 7.43E-05 | 0.000221 |
| chr1 | 9169656 | 9173846 | 4191 | 0.89 | 7.43E-05 | 0.000221 |
| chr1 | 188675249 | 188677006 | 1758 | 1.48 | 7.43E-05 | 0.000221 |
| chr2 | 42706495 | 42707550 | 1056 | -4.92 | 7.43E-05 | 0.000221 |
| chr22 | 28809713 | 28810776 | 1064 | 3.23 | 7.44E-05 | 0.000222 |
| chr1 | 46644421 | 46665072 | 20652 | 0.46 | 7.44E-05 | 0.000222 |
| chr5 | 149921844 | 149930304 | 8461 | -1.14 | 7.44E-05 | 0.000222 |
| chr9 | 31346920 | 31347419 | 500 | 4.94 | 7.44E-05 | 0.000222 |
| chr2 | 57334289 | 57335324 | 1036 | 3.2 | 7.46E-05 | 0.000222 |
| chr12 | 32887871 | 32889943 | 2073 | 1.36 | 7.48E-05 | 0.000223 |
| chr2 | 61885856 | 61900412 | 14557 | 0.38 | 7.50E-05 | 0.000223 |
| chr9 | 139964910 | 139982157 | 17248 | -0.63 | 7.51E-05 | 0.000224 |
| chr15 | 83349280 | 83349853 | 574 | 2.51 | 7.52E-05 | 0.000224 |
| chr19 | 57077834 | 57079919 | 2086 | 0.95 | 7.56E-05 | 0.000225 |
| chr2 | 217017924 | 217029994 | 12071 | -0.79 | 7.56E-05 | 0.000225 |
| chr6 | 43394712 | 43405118 | 10407 | 0.75 | 7.57E-05 | 0.000225 |
| chr2 | 231772841 | 231774025 | 1185 | -1.97 | 7.58E-05 | 0.000225 |
| chr14 | 60580549 | 60581897 | 1349 | 1.98 | 7.58E-05 | 0.000226 |
| chr4 | 23375190 | 23376066 | 877 | 3.51 | 7.59E-05 | 0.000226 |
| chr2 | 170584414 | 170591342 | 6929 | 1.49 | 7.60E-05 | 0.000226 |
| chr17 | 33867738 | 33868977 | 1240 | 1.37 | 7.60E-05 | 0.000226 |
| chr1 | 162530279 | 162541138 | 10860 | 0.6 | 7.61E-05 | 0.000226 |
| chr6 | 141484869 | 141486702 | 1834 | 4.66 | 7.62E-05 | 0.000226 |
| chr7 | 26977038 | 26978334 | 1297 | 1.9 | 7.64E-05 | 0.000227 |
| chr2 | 128631658 | 128645514 | 13857 | -0.98 | 7.65E-05 | 0.000228 |
| chr17 | 58350331 | 58350848 | 518 | -3.75 | 7.67E-05 | 0.000228 |
| chr3 | 130611929 | 130625156 | 13228 | 0.57 | 7.69E-05 | 0.000229 |
| chr4 | 4249083 | 4250887 | 1805 | 0.71 | 7.69E-05 | 0.000229 |
| chr2 | 158006538 | 158009227 | 2690 | 1.42 | 7.70E-05 | 0.000229 |
| chr9 | 134693574 | 134694313 | 740 | 2.96 | 7.71E-05 | 0.000229 |
| chr2 | 25500968 | 25501734 | 767 | -3.31 | 7.71E-05 | 0.000229 |
| chr8 | 41997176 | 42011778 | 14603 | -0.86 | 7.72E-05 | 0.000229 |
| chr14 | 65472034 | 65487272 | 15239 | -0.7 | 7.74E-05 | 0.00023 |
| chr8 | 131646747 | 131664004 | 17258 | 0.75 | 7.76E-05 | 0.000231 |
| chr7 | 127713548 | 127729224 | 15677 | -0.68 | 7.76E-05 | 0.000231 |
| chr15 | 78964069 | 78977854 | 13786 | -1.37 | 7.76E-05 | 0.000231 |
| chr19 | 40653266 | 40662151 | 8886 | -1.49 | 7.76E-05 | 0.000231 |
| chr5 | 140253899 | 140256131 | 2233 | 0.75 | 7.77E-05 | 0.000231 |
| chr17 | 48440653 | 48475371 | 34719 | -0.35 | 7.81E-05 | 0.000232 |
| chr7 | 155087941 | 155092950 | 5010 | 0.57 | 7.81E-05 | 0.000232 |
| chr2 | 19064946 | 19065591 | 646 | -3.18 | 7.83E-05 | 0.000232 |
| chr3 | 105831755 | 105850295 | 18541 | 0.46 | 7.87E-05 | 0.000234 |

| | | | | | | |
|---|---|---|---|---|---|---|
| chr1 | 209252922 | 209254637 | 1716 | 1.15 | 7.92E-05 | 0.000235 |
| chr4 | 187522928 | 187524191 | 1264 | 2.62 | 7.92E-05 | 0.000235 |
| chr2 | 153537219 | 153576387 | 39169 | -0.38 | 7.93E-05 | 0.000235 |
| chr5 | 65504217 | 65506192 | 1976 | 1.14 | 7.96E-05 | 0.000236 |
| chr9 | 107572621 | 107574032 | 1412 | 1.02 | 7.97E-05 | 0.000236 |
| chr11 | 66349620 | 66423505 | 73886 | 0.28 | 7.97E-05 | 0.000236 |
| chr16 | 16219664 | 16244181 | 24518 | -0.4 | 7.97E-05 | 0.000237 |
| chr3 | 13031702 | 13037449 | 5748 | 1.23 | 7.99E-05 | 0.000237 |
| chr19 | 36335987 | 36337837 | 1851 | 1.71 | 7.99E-05 | 0.000237 |
| chr4 | 137240709 | 137242468 | 1760 | 0.89 | 8.00E-05 | 0.000237 |
| chr13 | 106499442 | 106500214 | 773 | 4.92 | 8.04E-05 | 0.000238 |
| chr4 | 41288174 | 41292980 | 4807 | 1.13 | 8.07E-05 | 0.000239 |
| chr17 | 59612250 | 59612808 | 559 | -2.61 | 8.07E-05 | 0.000239 |
| chr19 | 14134091 | 14154570 | 20480 | -0.67 | 8.08E-05 | 0.00024 |
| chr3 | 38571841 | 38572748 | 908 | -3.26 | 8.08E-05 | 0.00024 |
| chr18 | 21578835 | 21579470 | 636 | 4.94 | 8.09E-05 | 0.00024 |
| chr22 | 32640643 | 32641127 | 485 | 4.94 | 8.09E-05 | 0.00024 |
| chr5 | 139486314 | 139494903 | 8590 | 0.47 | 8.11E-05 | 0.00024 |
| chr1 | 198411623 | 198419188 | 7566 | 4.7 | 8.13E-05 | 0.000241 |
| chr6 | 134981147 | 134981612 | 466 | 3.76 | 8.16E-05 | 0.000242 |
| chr16 | 34815329 | 34815844 | 516 | 4.92 | 8.16E-05 | 0.000242 |
| chr5 | 28524327 | 28524863 | 537 | 4.92 | 8.16E-05 | 0.000242 |
| chr15 | 32968530 | 32981330 | 12801 | 0.65 | 8.20E-05 | 0.000243 |
| chrX | 45482080 | 45483407 | 1328 | 1.3 | 8.20E-05 | 0.000243 |
| chr1 | 146642826 | 146645480 | 2655 | 0.56 | 8.21E-05 | 0.000243 |
| chr4 | 11472043 | 11472847 | 805 | 2.57 | 8.23E-05 | 0.000244 |
| chr12 | 31829930 | 31836240 | 6311 | 1.62 | 8.24E-05 | 0.000244 |
| chr21 | 39543030 | 39544427 | 1398 | 1.95 | 8.24E-05 | 0.000244 |
| chr5 | 63461357 | 63462225 | 869 | 2.7 | 8.26E-05 | 0.000244 |
| chr11 | 63605162 | 63665019 | 59858 | -0.27 | 8.27E-05 | 0.000245 |
| chr15 | 52968298 | 52972375 | 4078 | 0.56 | 8.29E-05 | 0.000245 |
| chr1 | 234819582 | 234820933 | 1352 | 1.89 | 8.29E-05 | 0.000245 |
| chr10 | 112802346 | 112802975 | 630 | 4.92 | 8.30E-05 | 0.000245 |
| chr11 | 73197847 | 73198477 | 631 | 4.92 | 8.30E-05 | 0.000245 |
| chr2 | 230649633 | 230739093 | 89461 | -0.32 | 8.31E-05 | 0.000246 |
| chr18 | 19177908 | 19193647 | 15740 | 0.34 | 8.31E-05 | 0.000246 |
| chr10 | 19927251 | 19928148 | 898 | 2.93 | 8.32E-05 | 0.000246 |
| chr3 | 178864842 | 178867868 | 3027 | 0.65 | 8.32E-05 | 0.000246 |
| chr19 | 38893070 | 38909383 | 16314 | 0.87 | 8.34E-05 | 0.000246 |
| chr13 | 68759567 | 68760248 | 682 | 3.2 | 8.34E-05 | 0.000246 |
| chr12 | 100377972 | 100379805 | 1834 | 1.03 | 8.35E-05 | 0.000247 |
| chr11 | 36030082 | 36031973 | 1892 | -1.46 | 8.38E-05 | 0.000248 |
| chr10 | 32871418 | 32872982 | 1565 | -1.05 | 8.39E-05 | 0.000248 |
| chr21 | 39292872 | 39294355 | 1484 | 1.56 | 8.39E-05 | 0.000248 |
| chr18 | 20861321 | 20864942 | 3622 | 0.89 | 8.41E-05 | 0.000248 |
| chr21 | 33888747 | 33897062 | 8316 | 1.16 | 8.41E-05 | 0.000248 |
| chr2 | 24044718 | 24052946 | 8229 | -0.79 | 8.43E-05 | 0.000249 |
| chr6 | 123976424 | 123977253 | 830 | 2.6 | 8.44E-05 | 0.000249 |
| chr3 | 85988039 | 85988579 | 541 | 5.01 | 8.44E-05 | 0.000249 |
| chr3 | 61880287 | 61881523 | 1237 | 1.75 | 8.45E-05 | 0.000249 |
| chr13 | 50229291 | 50230245 | 955 | -1.94 | 8.47E-05 | 0.00025 |
| chr7 | 43516991 | 43529222 | 12232 | 0.42 | 8.47E-05 | 0.00025 |
| chr5 | 141571067 | 141573076 | 2010 | 2.26 | 8.47E-05 | 0.00025 |
| chr15 | 53599457 | 53601328 | 1872 | -1.13 | 8.48E-05 | 0.00025 |
| chr8 | 83807014 | 83807511 | 498 | 4.98 | 8.48E-05 | 0.00025 |
| chr3 | 12917491 | 12918236 | 746 | -3.33 | 8.51E-05 | 0.000251 |
| chr7 | 44923552 | 44926695 | 3144 | 0.42 | 8.53E-05 | 0.000252 |
| chr8 | 41981082 | 41984582 | 3501 | 0.78 | 8.53E-05 | 0.000252 |
| chr3 | 166171086 | 166172016 | 931 | 2.42 | 8.53E-05 | 0.000252 |
| chrX | 18466718 | 18467911 | 1194 | -2.02 | 8.53E-05 | 0.000252 |
| chr15 | 72845886 | 72848233 | 2348 | 0.8 | 8.54E-05 | 0.000252 |

| | | | | | | |
|---|---|---|---|---|---|---|
| chr11 | 68846679 | 68848700 | 2022 | -0.97 | 8.57E-05 | 0.000253 |
| chr6 | 134837815 | 134852115 | 14301 | 0.65 | 8.58E-05 | 0.000253 |
| chr5 | 100613221 | 100614927 | 1707 | 1.37 | 8.59E-05 | 0.000253 |
| chr20 | 14560948 | 14561677 | 730 | 2.29 | 8.59E-05 | 0.000253 |
| chr2 | 74424681 | 74437552 | 12872 | -0.35 | 8.63E-05 | 0.000255 |
| chr8 | 28798486 | 28811266 | 12781 | 1.01 | 8.63E-05 | 0.000255 |
| chr3 | 112950229 | 112951236 | 1008 | -2.1 | 8.65E-05 | 0.000255 |
| chr13 | 49481963 | 49483685 | 1723 | 1.51 | 8.67E-05 | 0.000256 |
| chr5 | 96517048 | 96529613 | 12566 | 0.52 | 8.69E-05 | 0.000256 |
| chr12 | 63326921 | 63329564 | 2644 | 1 | 8.69E-05 | 0.000256 |
| chr5 | 68483551 | 68486886 | 3336 | -0.44 | 8.73E-05 | 0.000257 |
| chr4 | 166299678 | 166300590 | 913 | 2.81 | 8.73E-05 | 0.000257 |
| chr7 | 94829312 | 94829789 | 478 | -4.86 | 8.74E-05 | 0.000257 |
| chr12 | 66205867 | 66221206 | 15340 | 0.61 | 8.74E-05 | 0.000257 |
| chr3 | 194590698 | 194594259 | 3562 | 0.91 | 8.74E-05 | 0.000258 |
| chr8 | 56900972 | 56902400 | 1429 | 1.79 | 8.75E-05 | 0.000258 |
| chr7 | 43873789 | 43875157 | 1369 | 1.83 | 8.76E-05 | 0.000258 |
| chr9 | 81216876 | 81217692 | 817 | -2.33 | 8.76E-05 | 0.000258 |
| chr2 | 193966943 | 193969112 | 2170 | 1.27 | 8.79E-05 | 0.000259 |
| chr9 | 134131145 | 134132305 | 1161 | -1.86 | 8.81E-05 | 0.000259 |
| chr13 | 110958973 | 110973486 | 14514 | -0.63 | 8.83E-05 | 0.00026 |
| chr4 | 166354588 | 166355029 | 442 | 4.02 | 8.84E-05 | 0.00026 |
| chr2 | 84510334 | 84510835 | 502 | 4.35 | 8.85E-05 | 0.00026 |
| chr1 | 57034828 | 57045895 | 11068 | -0.91 | 8.86E-05 | 0.000261 |
| chr11 | 96835675 | 96836367 | 693 | -3.47 | 8.87E-05 | 0.000261 |
| chr5 | 10746624 | 10762230 | 15607 | 0.5 | 8.87E-05 | 0.000261 |
| chr15 | 68935692 | 68936896 | 1205 | -1.85 | 8.88E-05 | 0.000261 |
| chr10 | 38023718 | 38024850 | 1133 | 3.48 | 8.88E-05 | 0.000261 |
| chr3 | 72494837 | 72496951 | 2115 | 0.98 | 8.88E-05 | 0.000261 |
| chr10 | 63752216 | 63753394 | 1179 | 1.96 | 8.89E-05 | 0.000261 |
| chr2 | 120516536 | 120520355 | 3820 | -0.54 | 8.91E-05 | 0.000262 |
| chr18 | 61137792 | 61145717 | 7926 | 3.17 | 8.91E-05 | 0.000262 |
| chr9 | 92805899 | 92808176 | 2278 | 0.73 | 8.91E-05 | 0.000262 |
| chr5 | 90302932 | 90304883 | 1952 | 1.32 | 8.93E-05 | 0.000262 |
| chr15 | 70086997 | 70087815 | 819 | 2.32 | 8.94E-05 | 0.000263 |
| chr9 | 28997686 | 29000390 | 2705 | 1.34 | 8.95E-05 | 0.000263 |
| chr12 | 21489354 | 21490175 | 822 | 2.23 | 9.00E-05 | 0.000264 |
| chr9 | 119308851 | 119313759 | 4909 | -0.44 | 9.00E-05 | 0.000265 |
| chrX | 43664639 | 43665191 | 553 | 3.32 | 9.01E-05 | 0.000265 |
| chr6 | 170892110 | 170894533 | 2424 | 0.88 | 9.01E-05 | 0.000265 |
| chr17 | 55161195 | 55199067 | 37873 | 0.34 | 9.03E-05 | 0.000265 |
| chr6 | 152415484 | 152416906 | 1423 | 1.12 | 9.04E-05 | 0.000266 |
| chr16 | 55869272 | 55876833 | 7562 | 0.92 | 9.05E-05 | 0.000266 |
| chr13 | 84557855 | 84558398 | 544 | 3.98 | 9.07E-05 | 0.000266 |
| chr20 | 8095449 | 8095861 | 413 | 4.89 | 9.09E-05 | 0.000267 |
| chr7 | 35838849 | 35846017 | 7169 | 0.4 | 9.09E-05 | 0.000267 |
| chr11 | 86149366 | 86150362 | 997 | 1.78 | 9.09E-05 | 0.000267 |
| chr6 | 149948727 | 149951597 | 2871 | 1.17 | 9.10E-05 | 0.000267 |
| chr14 | 23754259 | 23792323 | 38065 | 0.31 | 9.11E-05 | 0.000267 |
| chr15 | 34539833 | 34545288 | 5456 | -1.05 | 9.12E-05 | 0.000268 |
| chr2 | 192791782 | 192792951 | 1170 | 1.57 | 9.13E-05 | 0.000268 |
| chr1 | 235314952 | 235325542 | 10591 | 0.49 | 9.13E-05 | 0.000268 |
| chr14 | 102637379 | 102658443 | 21065 | -0.47 | 9.14E-05 | 0.000268 |
| chr3 | 148725459 | 148726248 | 790 | -3.07 | 9.14E-05 | 0.000268 |
| chr7 | 131103180 | 131109143 | 5964 | -0.97 | 9.15E-05 | 0.000268 |
| chr15 | 90858695 | 90864081 | 5387 | 1.1 | 9.16E-05 | 0.000269 |
| chr5 | 157856623 | 157857163 | 541 | 4.9 | 9.16E-05 | 0.000269 |
| chr3 | 81298013 | 81299594 | 1582 | 5.29 | 9.17E-05 | 0.000269 |
| chr18 | 48760932 | 48763181 | 2250 | -1.49 | 9.18E-05 | 0.000269 |
| chr22 | 40672354 | 40673450 | 1097 | 1.93 | 9.20E-05 | 0.00027 |
| chr3 | 196501139 | 196501686 | 548 | -4.85 | 9.21E-05 | 0.00027 |

| | | | | | | |
|---|---|---|---|---|---|---|
| chr6 | 106748591 | 106749238 | 648 | -4.85 | 9.21E-05 | 0.00027 |
| chr7 | 95098329 | 95108790 | 10462 | -0.35 | 9.21E-05 | 0.00027 |
| chr1 | 89166121 | 89167521 | 1401 | 1.84 | 9.22E-05 | 0.00027 |
| chr2 | 161037593 | 161038198 | 606 | -2.95 | 9.24E-05 | 0.000271 |
| chr5 | 153417309 | 153419414 | 2106 | 0.67 | 9.24E-05 | 0.000271 |
| chrX | 42656595 | 42657638 | 1044 | 3.39 | 9.24E-05 | 0.000271 |
| chr17 | 29570694 | 29602447 | 31754 | -0.4 | 9.25E-05 | 0.000271 |
| chr4 | 10657405 | 10657905 | 501 | 4.89 | 9.27E-05 | 0.000272 |
| chr8 | 33994265 | 33994617 | 353 | 4.89 | 9.27E-05 | 0.000272 |
| chr4 | 26569499 | 26571247 | 1749 | 0.98 | 9.30E-05 | 0.000272 |
| chr16 | 23651166 | 23654381 | 3216 | 0.64 | 9.30E-05 | 0.000272 |
| chr3 | 24535184 | 24538874 | 3691 | 0.54 | 9.30E-05 | 0.000272 |
| chr17 | 65712890 | 65756216 | 43327 | 0.32 | 9.31E-05 | 0.000273 |
| chr9 | 35253826 | 35254640 | 815 | 3.28 | 9.33E-05 | 0.000273 |
| chr16 | 85800721 | 85802716 | 1996 | -0.83 | 9.33E-05 | 0.000273 |
| chr12 | 11882086 | 11885208 | 3123 | 0.99 | 9.34E-05 | 0.000273 |
| chr5 | 111665003 | 111675796 | 10794 | 2.81 | 9.36E-05 | 0.000274 |
| chr6 | 144661484 | 144697348 | 35865 | 0.36 | 9.37E-05 | 0.000274 |
| chr10 | 67154212 | 67155483 | 1272 | -1.38 | 9.37E-05 | 0.000274 |
| chr3 | 142442603 | 142445108 | 2506 | 0.54 | 9.38E-05 | 0.000275 |
| chrX | 36549235 | 36549791 | 557 | 2.62 | 9.39E-05 | 0.000275 |
| chr5 | 141976826 | 141983028 | 6203 | -0.54 | 9.39E-05 | 0.000275 |
| chr4 | 166570070 | 166570763 | 694 | 3.53 | 9.41E-05 | 0.000275 |
| chr17 | 19750221 | 19750814 | 594 | 4.92 | 9.42E-05 | 0.000275 |
| chr7 | 69118845 | 69120174 | 1330 | 1.58 | 9.42E-05 | 0.000275 |
| chrX | 150564658 | 150567908 | 3251 | 0.62 | 9.44E-05 | 0.000276 |
| chr10 | 62144036 | 62145521 | 1486 | 1.33 | 9.47E-05 | 0.000277 |
| chr3 | 185477013 | 185477438 | 426 | 4.03 | 9.47E-05 | 0.000277 |
| chr9 | 14626938 | 14641130 | 14193 | 1.7 | 9.48E-05 | 0.000277 |
| chr5 | 134565644 | 134567374 | 1731 | -1.55 | 9.48E-05 | 0.000277 |
| chr12 | 51457090 | 51477805 | 20716 | 0.56 | 9.48E-05 | 0.000277 |
| chr2 | 97452233 | 97467040 | 14808 | -0.82 | 9.52E-05 | 0.000278 |
| chr11 | 88116047 | 88118185 | 2139 | 0.83 | 9.52E-05 | 0.000278 |
| chr19 | 22601564 | 22602110 | 547 | 4.89 | 9.52E-05 | 0.000278 |
| chr4 | 45648891 | 45649327 | 437 | 4.93 | 9.54E-05 | 0.000279 |
| chr18 | 54534379 | 54535152 | 774 | -3.41 | 9.55E-05 | 0.000279 |
| chr2 | 239171430 | 239209710 | 38281 | -0.2 | 9.56E-05 | 0.000279 |
| chr3 | 188396985 | 188398495 | 1511 | 1.74 | 9.59E-05 | 0.00028 |
| chr16 | 72324117 | 72326643 | 2527 | 0.67 | 9.60E-05 | 0.00028 |
| chr2 | 40590939 | 40592079 | 1141 | 2.02 | 9.65E-05 | 0.000282 |
| chr1 | 37917504 | 37965876 | 48373 | 0.28 | 9.67E-05 | 0.000282 |
| chr5 | 57130130 | 57139151 | 9022 | 0.28 | 9.69E-05 | 0.000283 |
| chr1 | 2425857 | 2431928 | 6072 | -1.73 | 9.71E-05 | 0.000283 |
| chr15 | 42963273 | 42964478 | 1206 | -2.27 | 9.71E-05 | 0.000284 |
| chr1 | 212298464 | 212299217 | 754 | -4.06 | 9.72E-05 | 0.000284 |
| chr3 | 4631459 | 4631986 | 528 | -4.87 | 9.74E-05 | 0.000284 |
| chr2 | 53160497 | 53164140 | 3644 | 5.69 | 9.75E-05 | 0.000284 |
| chr1 | 185284720 | 185288029 | 3310 | 0.51 | 9.76E-05 | 0.000285 |
| chr3 | 187753514 | 187756189 | 2676 | 0.78 | 9.79E-05 | 0.000286 |
| chr17 | 79345788 | 79346656 | 869 | -3.23 | 9.81E-05 | 0.000286 |
| chr4 | 81727285 | 81727728 | 444 | 4.3 | 9.84E-05 | 0.000287 |
| chr7 | 105986357 | 105988030 | 1674 | -1.75 | 9.87E-05 | 0.000288 |
| chr1 | 209608323 | 209609097 | 775 | -3.54 | 9.87E-05 | 0.000288 |
| chr9 | 79920420 | 79924015 | 3596 | -1.37 | 9.87E-05 | 0.000288 |
| chr17 | 76731042 | 76738585 | 7544 | 0.71 | 9.89E-05 | 0.000288 |
| chr3 | 58472543 | 58489183 | 16641 | 0.74 | 9.89E-05 | 0.000288 |
| chr15 | 39370550 | 39377901 | 7352 | 1.19 | 9.90E-05 | 0.000289 |
| chr3 | 181588281 | 181598331 | 10051 | 1.28 | 9.91E-05 | 0.000289 |
| chr1 | 113241906 | 113250877 | 8972 | 0.47 | 9.93E-05 | 0.000289 |
| chr15 | 64295697 | 64320559 | 24863 | 0.51 | 9.94E-05 | 0.00029 |
| chr10 | 71624027 | 71629860 | 5834 | 1.45 | 9.97E-05 | 0.00029 |

| | | | | | | |
|---|---|---|---|---|---|---|
| chr6 | 142333723 | 142336875 | 3153 | 0.7 | 9.97E-05 | 0.000291 |
| chr5 | 156549161 | 156549537 | 377 | 4.95 | 9.99E-05 | 0.000291 |
| chr7 | 64154307 | 64154845 | 539 | 4.9 | 9.99E-05 | 0.000291 |
| chr3 | 24491405 | 24494197 | 2793 | 1 | 1.00E-04 | 0.000292 |
| chr1 | 184760688 | 184762842 | 2155 | 0.86 | 1.00E-04 | 0.000292 |
| chr8 | 115934366 | 115934851 | 486 | 4.91 | 1.00E-04 | 0.000292 |
| chr8 | 83681658 | 83682239 | 582 | 4.98 | 1.00E-04 | 0.000292 |
| chr4 | 113281951 | 113282649 | 699 | 3 | 0.000101 | 0.000293 |
| chr6 | 133051189 | 133052415 | 1227 | 1.88 | 0.000101 | 0.000293 |
| chr6 | 136546752 | 136552610 | 5859 | -0.82 | 0.000101 | 0.000293 |
| chr6 | 15543528 | 15543859 | 332 | 3.78 | 0.000101 | 0.000294 |
| chr17 | 79702327 | 79703056 | 730 | -2.09 | 0.000101 | 0.000294 |
| chr1 | 10269332 | 10278785 | 9454 | 0.54 | 0.000101 | 0.000295 |
| chr21 | 35855536 | 35856348 | 813 | 2.75 | 0.000102 | 0.000296 |
| chr10 | 128831780 | 128832819 | 1040 | -1.85 | 0.000102 | 0.000297 |
| chr2 | 189864478 | 189879989 | 15512 | 0.91 | 0.000102 | 0.000298 |
| chr16 | 53638766 | 53639574 | 809 | -3.05 | 0.000102 | 0.000298 |
| chr5 | 55380287 | 55439460 | 59174 | -0.25 | 0.000103 | 0.000298 |
| chr4 | 137124649 | 137125453 | 805 | 2.47 | 0.000103 | 0.000299 |
| chr9 | 35178996 | 35185612 | 6617 | 2.62 | 0.000103 | 0.000299 |
| chr11 | 119351555 | 119352013 | 459 | 4.87 | 0.000103 | 0.000299 |
| chr1 | 217689301 | 217701975 | 12675 | 1.37 | 0.000103 | 3.00E-04 |
| chr11 | 109665547 | 109665963 | 417 | 4.87 | 0.000103 | 3.00E-04 |
| chr10 | 126552282 | 126554224 | 1943 | 1.44 | 0.000103 | 3.00E-04 |
| chr3 | 129295786 | 129409077 | 113292 | 0.16 | 0.000103 | 3.00E-04 |
| chrX | 64886658 | 64941974 | 55317 | -0.26 | 0.000103 | 3.00E-04 |
| chr12 | 38569027 | 38570100 | 1074 | 3.34 | 0.000103 | 3.00E-04 |
| chr1 | 60119334 | 60120022 | 689 | 2.14 | 0.000103 | 3.00E-04 |
| chr11 | 86343132 | 86349497 | 6366 | -0.78 | 0.000103 | 3.00E-04 |
| chr1 | 30324147 | 30324638 | 492 | 3.53 | 0.000103 | 3.00E-04 |
| chr1 | 205859119 | 205860002 | 884 | -1.87 | 0.000103 | 3.00E-04 |
| chr10 | 76727775 | 76728784 | 1010 | 3.24 | 0.000104 | 0.000301 |
| chr19 | 4624478 | 4645133 | 20656 | -0.85 | 0.000104 | 0.000301 |
| chr8 | 93939566 | 93940568 | 1003 | 2.39 | 0.000104 | 0.000301 |
| chr10 | 45406244 | 45406640 | 397 | 4.87 | 0.000104 | 0.000302 |
| chr16 | 66008679 | 66010059 | 1381 | -1.3 | 0.000104 | 0.000302 |
| chr10 | 27792488 | 27825816 | 33329 | 0.35 | 0.000104 | 0.000303 |
| chr13 | 105636650 | 105637182 | 533 | 3.97 | 0.000104 | 0.000303 |
| chr8 | 116463948 | 116464584 | 637 | 4.87 | 0.000104 | 0.000303 |
| chr2 | 165370620 | 165400087 | 29468 | 0.6 | 0.000105 | 0.000304 |
| chr2 | 183923621 | 183925293 | 1673 | -0.79 | 0.000105 | 0.000304 |
| chr3 | 174576029 | 174577926 | 1898 | 5.07 | 0.000105 | 0.000305 |
| chr4 | 130263410 | 130264596 | 1187 | 2.67 | 0.000105 | 0.000305 |
| chr2 | 217910483 | 217913191 | 2709 | 0.98 | 0.000105 | 0.000306 |
| chr5 | 38806655 | 38810886 | 4232 | -1.24 | 0.000105 | 0.000306 |
| chr8 | 95209769 | 95232452 | 22684 | 0.64 | 0.000106 | 0.000306 |
| chr1 | 10516857 | 10535936 | 19080 | 0.43 | 0.000106 | 0.000306 |
| chr15 | 55609641 | 55612479 | 2839 | 0.4 | 0.000106 | 0.000306 |
| chrX | 12228044 | 12245132 | 17089 | 2.67 | 0.000106 | 0.000306 |
| chr1 | 42137626 | 42150235 | 12610 | -1.45 | 0.000106 | 0.000307 |
| chr11 | 68284878 | 68318410 | 33533 | -0.37 | 0.000106 | 0.000307 |
| chr16 | 30996493 | 31010563 | 14071 | -0.37 | 0.000106 | 0.000307 |
| chr12 | 94152986 | 94154077 | 1092 | 1.33 | 0.000106 | 0.000307 |
| chr2 | 227994766 | 227995348 | 583 | -4.84 | 0.000106 | 0.000307 |
| chr8 | 8339567 | 8341806 | 2240 | 0.87 | 0.000106 | 0.000308 |
| chr12 | 59685266 | 59685630 | 365 | 4.87 | 0.000106 | 0.000308 |
| chr4 | 40252245 | 40264050 | 11806 | 0.42 | 0.000107 | 0.000309 |
| chr10 | 114498158 | 114504059 | 5902 | 0.6 | 0.000107 | 0.000309 |
| chr5 | 140602919 | 140603483 | 565 | 4.87 | 0.000107 | 0.00031 |
| chr8 | 73391219 | 73391754 | 536 | 4.87 | 0.000107 | 0.00031 |
| chr9 | 115660900 | 115661515 | 616 | 4.87 | 0.000107 | 0.00031 |

| | | | | | | |
|---|---|---|---|---|---|---|
| chr4 | 123964732 | 123965508 | 777 | 3.97 | 0.000107 | 0.00031 |
| chr10 | 7342922 | 7362002 | 19081 | 0.45 | 0.000107 | 0.00031 |
| chr3 | 64536836 | 64538581 | 1746 | 4.44 | 0.000107 | 0.000311 |
| chr3 | 129101842 | 129160507 | 58666 | -0.25 | 0.000107 | 0.000311 |
| chr10 | 93314974 | 93317281 | 2308 | 2.04 | 0.000108 | 0.000312 |
| chr3 | 139258368 | 139258989 | 622 | 3.31 | 0.000108 | 0.000313 |
| chr2 | 128612459 | 128616359 | 3901 | -0.8 | 0.000108 | 0.000313 |
| chr7 | 30059170 | 30069457 | 10288 | 0.54 | 0.000108 | 0.000313 |
| chr17 | 29923006 | 29926676 | 3671 | -0.85 | 0.000108 | 0.000313 |
| chr2 | 137084522 | 137085669 | 1148 | 1.87 | 0.000109 | 0.000315 |
| chr11 | 68585488 | 68586035 | 548 | -3.7 | 0.000109 | 0.000315 |
| chr3 | 34097261 | 34099362 | 2102 | 0.53 | 0.000109 | 0.000315 |
| chr10 | 38442075 | 38442691 | 617 | 4.39 | 0.000109 | 0.000316 |
| chr5 | 133538920 | 133588596 | 49677 | 0.29 | 0.000109 | 0.000316 |
| chr14 | 65271331 | 65275437 | 4107 | 0.37 | 0.000109 | 0.000317 |
| chr10 | 49614020 | 49623028 | 9009 | -1.66 | 0.00011 | 0.000317 |
| chr5 | 97645342 | 97645758 | 417 | -4.82 | 0.00011 | 0.000317 |
| chr1 | 234679336 | 234681894 | 2559 | 1.43 | 0.00011 | 0.000317 |
| chr2 | 108930970 | 108932513 | 1544 | 1.57 | 0.00011 | 0.000318 |
| chr3 | 46155494 | 46156506 | 1013 | 1.94 | 0.00011 | 0.000318 |
| chr1 | 110870680 | 110899938 | 29259 | 0.34 | 0.00011 | 0.000319 |
| chr20 | 3287785 | 3296748 | 8964 | -0.48 | 0.00011 | 0.000319 |
| chr4 | 14179856 | 14183840 | 3985 | -0.76 | 0.000111 | 0.000319 |
| chr9 | 82553980 | 82555562 | 1583 | 1.23 | 0.000111 | 0.00032 |
| chr16 | 59585909 | 59586677 | 769 | -2.58 | 0.000111 | 0.00032 |
| chr2 | 24207733 | 24217670 | 9938 | -0.8 | 0.000111 | 0.00032 |
| chr9 | 118641676 | 118648216 | 6541 | -2.4 | 0.000111 | 0.00032 |
| chr2 | 223383101 | 223383625 | 525 | 4.87 | 0.000111 | 0.00032 |
| chr1 | 88065902 | 88067144 | 1243 | 1.7 | 0.000111 | 0.000321 |
| chr6 | 128764193 | 128765996 | 1804 | 1.02 | 0.000111 | 0.000321 |
| chr7 | 2158214 | 2158863 | 650 | -4.85 | 0.000111 | 0.000321 |
| chr8 | 87331828 | 87334273 | 2446 | -0.84 | 0.000111 | 0.000321 |
| chr15 | 83803546 | 83804085 | 540 | -3.28 | 0.000111 | 0.000321 |
| chr12 | 92080735 | 92081931 | 1197 | 1.51 | 0.000112 | 0.000322 |
| chr9 | 114110107 | 114120031 | 9925 | 0.59 | 0.000112 | 0.000322 |
| chr10 | 43114952 | 43115927 | 976 | 2.3 | 0.000112 | 0.000322 |
| chr15 | 76176864 | 76179309 | 2446 | 1.02 | 0.000112 | 0.000322 |
| chr10 | 13341083 | 13345391 | 4309 | 0.42 | 0.000112 | 0.000324 |
| chr1 | 211453576 | 211464155 | 10580 | -0.65 | 0.000112 | 0.000324 |
| chr4 | 101128998 | 101130139 | 1142 | 1.57 | 0.000112 | 0.000324 |
| chr12 | 11094440 | 11095781 | 1342 | 0.97 | 0.000113 | 0.000326 |
| chr12 | 76843060 | 76849635 | 6576 | 0.59 | 0.000113 | 0.000326 |
| chr12 | 4252082 | 4252665 | 584 | 3.64 | 0.000113 | 0.000326 |
| chr19 | 59054467 | 59093038 | 38572 | -0.2 | 0.000113 | 0.000327 |
| chr10 | 62173513 | 62187581 | 14069 | 2.04 | 0.000113 | 0.000327 |
| chr19 | 17402632 | 17433607 | 30976 | 0.35 | 0.000113 | 0.000327 |
| chrX | 46988921 | 46989740 | 820 | 3.35 | 0.000114 | 0.000327 |
| chr3 | 172258739 | 172260243 | 1505 | 1.44 | 0.000114 | 0.000328 |
| chr11 | 55919736 | 55920400 | 665 | -4.81 | 0.000114 | 0.000328 |
| chr3 | 36403584 | 36404888 | 1305 | 1.59 | 0.000114 | 0.000328 |
| chr3 | 138528566 | 138555074 | 26509 | 0.37 | 0.000114 | 0.000329 |
| chr5 | 143154534 | 143155814 | 1281 | -1.53 | 0.000114 | 0.000329 |
| chr10 | 31531164 | 31532228 | 1065 | -1.29 | 0.000114 | 0.00033 |
| chr17 | 21029630 | 21031553 | 1924 | 0.67 | 0.000114 | 0.00033 |
| chr7 | 127517759 | 127524954 | 7196 | -1.17 | 0.000114 | 0.00033 |
| chr17 | 72565125 | 72566580 | 1456 | -1.35 | 0.000114 | 0.00033 |
| chr10 | 6760049 | 6785961 | 25913 | 0.28 | 0.000115 | 0.00033 |
| chr6 | 63593821 | 63596009 | 2189 | 0.9 | 0.000115 | 0.00033 |
| chr14 | 92499912 | 92507815 | 7904 | 0.45 | 0.000115 | 0.000331 |
| chr12 | 67948858 | 67950077 | 1220 | 2.06 | 0.000115 | 0.000331 |
| chr12 | 39821467 | 39839208 | 17742 | 0.42 | 0.000115 | 0.000332 |

| | | | | | | |
|---|---|---|---|---|---|---|
| chr16 | 68822742 | 68834314 | 11573 | -0.57 | 0.000115 | 0.000332 |
| chr5 | 76031624 | 76036701 | 5078 | -0.49 | 0.000115 | 0.000332 |
| chr14 | 50968122 | 50969486 | 1365 | 1.84 | 0.000115 | 0.000332 |
| chr1 | 234699629 | 234701369 | 1741 | 1.13 | 0.000115 | 0.000332 |
| chr16 | 53645394 | 53646522 | 1129 | 3.32 | 0.000116 | 0.000333 |
| chr11 | 129657971 | 129658672 | 702 | 4.87 | 0.000116 | 0.000334 |
| chr4 | 76743431 | 76745812 | 2382 | -2.88 | 0.000116 | 0.000335 |
| chr1 | 84784665 | 84786021 | 1357 | 1.48 | 0.000116 | 0.000335 |
| chr8 | 102091426 | 102122738 | 31313 | -0.58 | 0.000116 | 0.000335 |
| chr7 | 86991549 | 87005893 | 14345 | -0.8 | 0.000116 | 0.000335 |
| chr1 | 185335389 | 185336665 | 1277 | 2.14 | 0.000117 | 0.000335 |
| chr10 | 42836716 | 42837004 | 289 | 4.84 | 0.000117 | 0.000335 |
| chr1 | 247610980 | 247611967 | 988 | 1.02 | 0.000117 | 0.000336 |
| chr14 | 95756545 | 95762240 | 5696 | 1.3 | 0.000117 | 0.000336 |
| chr2 | 131774201 | 131775895 | 1695 | 1.32 | 0.000117 | 0.000336 |
| chr9 | 96918430 | 96942086 | 23657 | 0.31 | 0.000117 | 0.000336 |
| chr1 | 24371960 | 24374501 | 2542 | -0.98 | 0.000117 | 0.000336 |
| chr14 | 37853916 | 37855328 | 1413 | 1.58 | 0.000117 | 0.000337 |
| chr2 | 179292850 | 179317248 | 24399 | -0.38 | 0.000117 | 0.000337 |
| chr13 | 53383833 | 53384468 | 636 | 3.94 | 0.000117 | 0.000337 |
| chr2 | 134430654 | 134433133 | 2480 | -1.18 | 0.000117 | 0.000337 |
| chr19 | 54162222 | 54162582 | 361 | 4.84 | 0.000117 | 0.000338 |
| chr3 | 168935826 | 168936240 | 415 | 4.94 | 0.000118 | 0.000338 |
| chr3 | 84190178 | 84190623 | 446 | 4.89 | 0.000118 | 0.000338 |
| chr3 | 31842691 | 31843349 | 659 | -4.81 | 0.000118 | 0.000339 |
| chr17 | 30245320 | 30246232 | 913 | 2.19 | 0.000118 | 0.000339 |
| chr5 | 118323080 | 118325132 | 2053 | 0.71 | 0.000118 | 0.00034 |
| chr3 | 117611967 | 117612694 | 728 | 2.74 | 0.000118 | 0.00034 |
| chr12 | 89511241 | 89511783 | 543 | 2.02 | 0.000119 | 0.000341 |
| chr11 | 109811093 | 109817957 | 6865 | -0.58 | 0.000119 | 0.000341 |
| chr17 | 462676 | 476839 | 14164 | -1.24 | 0.000119 | 0.000341 |
| chr4 | 42331043 | 42334802 | 3760 | 4.94 | 0.000119 | 0.000341 |
| chr5 | 153617122 | 153796732 | 179611 | -0.14 | 0.000119 | 0.000341 |
| chr1 | 225695170 | 225708488 | 13319 | 0.45 | 0.000119 | 0.000341 |
| chr1 | 26962889 | 26963849 | 961 | 0.75 | 0.000119 | 0.000341 |
| chr15 | 36613250 | 36613986 | 737 | 2.95 | 0.000119 | 0.000341 |
| chr13 | 113737666 | 113738212 | 547 | 4.84 | 0.000119 | 0.000342 |
| chr4 | 47717920 | 47718502 | 583 | 4.84 | 0.000119 | 0.000342 |
| chr4 | 130916149 | 130916622 | 474 | 4.84 | 0.000119 | 0.000342 |
| chr9 | 23825658 | 23826288 | 631 | 4.9 | 0.000119 | 0.000342 |
| chr14 | 20584950 | 20585335 | 386 | -4.79 | 0.000119 | 0.000342 |
| chr15 | 51339052 | 51339773 | 722 | 1.96 | 0.00012 | 0.000344 |
| chr22 | 32338828 | 32350557 | 11730 | 0.39 | 0.00012 | 0.000345 |
| chr13 | 75017984 | 75018420 | 437 | -4.82 | 0.00012 | 0.000345 |
| chr8 | 48325714 | 48355826 | 30113 | -0.5 | 0.000121 | 0.000346 |
| chr6 | 151606943 | 151608107 | 1165 | 1.65 | 0.000121 | 0.000346 |
| chr4 | 151489097 | 151492772 | 3676 | 0.83 | 0.000121 | 0.000347 |
| chr2 | 172949255 | 172964566 | 15312 | -0.85 | 0.000121 | 0.000347 |
| chr14 | 92118787 | 92119625 | 839 | 3.39 | 0.000121 | 0.000347 |
| chrX | 9810265 | 9815824 | 5560 | 1.23 | 0.000121 | 0.000348 |
| chr18 | 8609472 | 8632007 | 22536 | -0.67 | 0.000121 | 0.000348 |
| chr5 | 35479005 | 35479914 | 910 | 1.74 | 0.000122 | 0.000348 |
| chr16 | 58482718 | 58493813 | 11096 | 0.67 | 0.000122 | 0.000348 |
| chrX | 43828789 | 43830023 | 1235 | 1.83 | 0.000122 | 0.00035 |
| chr2 | 128880964 | 128883225 | 2262 | -1.36 | 0.000122 | 0.00035 |
| chr12 | 32630774 | 32639832 | 9059 | 3.29 | 0.000122 | 0.00035 |
| chr1 | 100125594 | 100126115 | 522 | 4.05 | 0.000122 | 0.000351 |
| chr12 | 15743005 | 15743998 | 994 | 1.17 | 0.000123 | 0.000351 |
| chr17 | 53364003 | 53365281 | 1279 | -2.15 | 0.000123 | 0.000351 |
| chr12 | 42983718 | 42984579 | 862 | 3.2 | 0.000123 | 0.000352 |
| chr17 | 76776002 | 76797972 | 21971 | -0.48 | 0.000123 | 0.000352 |

| | | | | | | |
|---|---|---|---|---|---|---|
| chr22 | 28424768 | 28426144 | 1377 | 1.39 | 0.000123 | 0.000353 |
| chr7 | 35733189 | 35735814 | 2626 | 0.45 | 0.000123 | 0.000353 |
| chr9 | 38257299 | 38258102 | 804 | 2.96 | 0.000123 | 0.000353 |
| chr11 | 77169879 | 77170545 | 667 | 4.24 | 0.000124 | 0.000354 |
| chr2 | 88925768 | 88928670 | 2903 | 0.41 | 0.000124 | 0.000355 |
| chr9 | 132309277 | 132405312 | 96036 | -0.23 | 0.000124 | 0.000355 |
| chr5 | 177715154 | 177717024 | 1871 | 4.05 | 0.000124 | 0.000356 |
| chr10 | 126604321 | 126608065 | 3745 | 0.54 | 0.000125 | 0.000357 |
| chr8 | 140906899 | 140907581 | 683 | -3.33 | 0.000125 | 0.000358 |
| chr11 | 88167326 | 88168638 | 1313 | -1.82 | 0.000125 | 0.000358 |
| chr9 | 83928857 | 83935222 | 6366 | 0.94 | 0.000125 | 0.000358 |
| chr19 | 5572309 | 5573382 | 1074 | 2.48 | 0.000125 | 0.000358 |
| chr20 | 14597342 | 14598289 | 948 | 1.99 | 0.000125 | 0.000358 |
| chr7 | 139024137 | 139027996 | 3860 | 0.34 | 0.000125 | 0.000359 |
| chr1 | 98046095 | 98048624 | 2530 | 0.89 | 0.000126 | 0.000359 |
| chr15 | 88020796 | 88021777 | 982 | -1.73 | 0.000126 | 0.000359 |
| chr6 | 7127137 | 7153466 | 26330 | 0.33 | 0.000126 | 0.000361 |
| chr20 | 32141867 | 32155292 | 13426 | 0.77 | 0.000127 | 0.000362 |
| chrX | 47049911 | 47094715 | 44805 | 0.24 | 0.000127 | 0.000362 |
| chr19 | 16557818 | 16583613 | 25796 | -0.62 | 0.000127 | 0.000362 |
| chr20 | 13800144 | 13800906 | 763 | 2.47 | 0.000127 | 0.000363 |
| chr16 | 1204138 | 1205792 | 1655 | 1.1 | 0.000127 | 0.000363 |
| chr11 | 123088601 | 123090425 | 1825 | -1.4 | 0.000127 | 0.000363 |
| chr6 | 55622787 | 55624467 | 1681 | 1.46 | 0.000128 | 0.000367 |
| chr19 | 20818427 | 20819018 | 592 | 4.84 | 0.000128 | 0.000367 |
| chr3 | 96584563 | 96585046 | 484 | 4.84 | 0.000128 | 0.000367 |
| chr7 | 72300275 | 72301970 | 1696 | 1.65 | 0.000129 | 0.000367 |
| chr1 | 8657996 | 8659186 | 1191 | -2.16 | 0.000129 | 0.000367 |
| chr7 | 149118289 | 149120312 | 2024 | 0.73 | 0.000129 | 0.000367 |
| chr13 | 98219571 | 98221615 | 2045 | 1.09 | 0.000129 | 0.000368 |
| chr22 | 26142428 | 26157123 | 14696 | 0.86 | 0.000129 | 0.000368 |
| chr8 | 91411571 | 91412246 | 676 | 2.24 | 0.000129 | 0.000368 |
| chr16 | 14492397 | 14494520 | 2124 | -1.12 | 0.000129 | 0.000368 |
| chr6 | 18387119 | 18407091 | 19973 | 5.7 | 0.000129 | 0.000369 |
| chr12 | 121063575 | 121064355 | 781 | 1.26 | 0.000129 | 0.000369 |
| chr9 | 102668285 | 102670497 | 2213 | 0.65 | 0.000129 | 0.000369 |
| chr3 | 69713656 | 69715235 | 1580 | 0.82 | 0.000129 | 0.000369 |
| chr2 | 28934661 | 28939850 | 5190 | -1.03 | 0.000129 | 0.000369 |
| chr4 | 175630514 | 175635623 | 5110 | -0.74 | 0.00013 | 0.000369 |
| chr4 | 1513762 | 1515712 | 1951 | 1.68 | 0.00013 | 0.00037 |
| chr13 | 29318011 | 29329762 | 11752 | 0.43 | 0.00013 | 0.00037 |
| chr1 | 113930644 | 113948721 | 18078 | 0.52 | 0.00013 | 0.00037 |
| chr7 | 47794000 | 47859600 | 65601 | 0.15 | 0.00013 | 0.00037 |
| chr11 | 125742322 | 125773057 | 30736 | 0.39 | 0.00013 | 0.000371 |
| chr10 | 118896122 | 118898084 | 1963 | -1.38 | 0.00013 | 0.000371 |
| chr6 | 152823311 | 152823879 | 569 | -4.8 | 0.00013 | 0.000371 |
| chrX | 25031851 | 25038859 | 7009 | 2.37 | 0.000131 | 0.000372 |
| chrX | 35936738 | 35938394 | 1657 | 1.97 | 0.000131 | 0.000373 |
| chr6 | 83072815 | 83075523 | 2709 | 0.6 | 0.000131 | 0.000374 |
| chr10 | 61664831 | 61668998 | 4168 | 0.58 | 0.000131 | 0.000374 |
| chr9 | 95372356 | 95381014 | 8659 | -0.86 | 0.000131 | 0.000374 |
| chr12 | 113362486 | 113377289 | 14804 | 0.68 | 0.000132 | 0.000375 |
| chr5 | 172570924 | 172581744 | 10821 | 0.6 | 0.000132 | 0.000376 |
| chr15 | 55484286 | 55490190 | 5905 | 0.48 | 0.000132 | 0.000376 |
| chr9 | 117994597 | 117997246 | 2650 | 0.92 | 0.000132 | 0.000376 |
| chr5 | 162799932 | 162807493 | 7562 | 0.96 | 0.000132 | 0.000376 |
| chr9 | 133453403 | 133456780 | 3378 | 0.58 | 0.000132 | 0.000377 |
| chr4 | 138291715 | 138292206 | 492 | 4.81 | 0.000132 | 0.000377 |
| chrX | 37176630 | 37177095 | 466 | 4.81 | 0.000132 | 0.000377 |
| chr17 | 38350617 | 38351188 | 572 | -3.48 | 0.000133 | 0.000377 |
| chr3 | 158519145 | 158520655 | 1511 | 1.09 | 0.000133 | 0.000377 |

| | | | | | | |
|---|---|---|---|---|---|---|
| chr4 | 115536696 | 115537197 | 502 | 4.81 | 0.000133 | 0.000377 |
| chr2 | 160056833 | 160057594 | 762 | -2.76 | 0.000133 | 0.000378 |
| chrX | 46357763 | 46358364 | 602 | 3.62 | 0.000133 | 0.000378 |
| chrX | 96643802 | 96657125 | 13324 | 0.49 | 0.000133 | 0.000378 |
| chr1 | 182354388 | 182361662 | 7275 | 0.83 | 0.000133 | 0.000378 |
| chr15 | 77453216 | 77453654 | 439 | 3.67 | 0.000133 | 0.000378 |
| chr4 | 2143647 | 2144546 | 900 | 3.67 | 0.000133 | 0.000378 |
| chr1 | 32375512 | 32429825 | 54314 | 0.19 | 0.000133 | 0.000379 |
| chr5 | 124394126 | 124395367 | 1242 | 1.93 | 0.000133 | 0.000379 |
| chr10 | 82049499 | 82049828 | 330 | 4.27 | 0.000134 | 0.00038 |
| chr4 | 24520031 | 24522626 | 2596 | 1.58 | 0.000134 | 0.00038 |
| chr10 | 127095482 | 127096609 | 1128 | 2.19 | 0.000134 | 0.00038 |
| chr16 | 47921310 | 47922816 | 1507 | -2.86 | 0.000134 | 0.000381 |
| chr4 | 170840025 | 170841942 | 1918 | 4.63 | 0.000134 | 0.000382 |
| chr2 | 240850246 | 240867959 | 17714 | 0.47 | 0.000135 | 0.000383 |
| chr8 | 102199943 | 102220141 | 20199 | 0.34 | 0.000135 | 0.000384 |
| chr13 | 78676404 | 78677134 | 731 | -3.38 | 0.000135 | 0.000384 |
| chr17 | 41105096 | 41118291 | 13196 | 0.41 | 0.000136 | 0.000385 |
| chr2 | 232092273 | 232093566 | 1294 | -1.78 | 0.000136 | 0.000385 |
| chr4 | 139251284 | 139251864 | 581 | -4.77 | 0.000136 | 0.000385 |
| chr11 | 124705827 | 124707482 | 1656 | 1.09 | 0.000136 | 0.000386 |
| chr2 | 147390955 | 147393361 | 2407 | 0.83 | 0.000136 | 0.000386 |
| chr7 | 18575582 | 18594259 | 18678 | 0.52 | 0.000136 | 0.000386 |
| chr3 | 142281155 | 142298964 | 17810 | 0.41 | 0.000136 | 0.000387 |
| chr1 | 47753908 | 47756909 | 3002 | 1.02 | 0.000137 | 0.000388 |
| chr18 | 1011051 | 1011992 | 942 | -3.08 | 0.000137 | 0.000388 |
| chr7 | 32201115 | 32204142 | 3028 | -0.66 | 0.000137 | 0.000389 |
| chr5 | 27701818 | 27703102 | 1285 | 3.32 | 0.000137 | 0.000389 |
| chr4 | 184579390 | 184593936 | 14547 | 0.56 | 0.000137 | 0.000389 |
| chr2 | 35519740 | 35520331 | 592 | -3.21 | 0.000137 | 0.00039 |
| chr13 | 80703889 | 80708255 | 4367 | 0.47 | 0.000138 | 0.00039 |
| chr8 | 43446523 | 43447188 | 666 | 4.81 | 0.000138 | 0.000391 |
| chr10 | 98783315 | 98784843 | 1529 | -1.17 | 0.000138 | 0.000391 |
| chr1 | 109617385 | 109660207 | 42823 | 0.36 | 0.000138 | 0.000391 |
| chr22 | 38127119 | 38159384 | 32266 | 0.25 | 0.000138 | 0.000392 |
| chr5 | 72623656 | 72624284 | 629 | -3.37 | 0.000138 | 0.000392 |
| chr2 | 173891882 | 173893128 | 1247 | -1.62 | 0.000138 | 0.000392 |
| chr16 | 27185690 | 27187687 | 1998 | 1.96 | 0.000139 | 0.000393 |
| chr5 | 176722767 | 176779485 | 56719 | 0.3 | 0.000139 | 0.000394 |
| chr14 | 33033754 | 33034326 | 573 | 4.05 | 0.000139 | 0.000395 |
| chr17 | 4869271 | 4872789 | 3519 | 0.54 | 0.000139 | 0.000395 |
| chr11 | 77486682 | 77487283 | 602 | -3.39 | 0.00014 | 0.000396 |
| chr4 | 113066625 | 113068220 | 1596 | 1.26 | 0.00014 | 0.000396 |
| chr12 | 95097865 | 95099170 | 1306 | 0.84 | 0.00014 | 0.000396 |
| chr6 | 20129694 | 20132175 | 2482 | 0.9 | 0.00014 | 0.000397 |
| chr4 | 95719452 | 95720291 | 840 | -2.49 | 0.00014 | 0.000397 |
| chr11 | 78048872 | 78049203 | 332 | 3.66 | 0.00014 | 0.000397 |
| chr5 | 168526900 | 168528153 | 1254 | -1.88 | 0.00014 | 0.000397 |
| chr20 | 10573108 | 10586443 | 13336 | 0.44 | 0.00014 | 0.000398 |
| chr3 | 49156517 | 49172177 | 15661 | 0.46 | 0.000141 | 0.000398 |
| chr7 | 79805088 | 79813075 | 7988 | 1.23 | 0.000141 | 0.000399 |
| chr13 | 100866991 | 100867819 | 829 | -1.92 | 0.000141 | 4.00E-04 |
| chr22 | 29105553 | 29109473 | 3921 | -0.67 | 0.000142 | 0.000401 |
| chr12 | 60014080 | 60014555 | 476 | 3.41 | 0.000142 | 0.000402 |
| chr12 | 14034816 | 14053162 | 18347 | -4.59 | 0.000142 | 0.000402 |
| chr7 | 46623009 | 46644653 | 21645 | 3.83 | 0.000142 | 0.000402 |
| chr12 | 117956884 | 117957319 | 436 | 4.21 | 0.000142 | 0.000402 |
| chr4 | 32142236 | 32143244 | 1009 | 2.42 | 0.000142 | 0.000402 |
| chr12 | 96223148 | 96254226 | 31079 | 0.45 | 0.000142 | 0.000402 |
| chr5 | 127148137 | 127177017 | 28881 | 1.69 | 0.000142 | 0.000402 |
| chr20 | 40040656 | 40042925 | 2270 | 1.03 | 0.000142 | 0.000402 |

| | | | | | | |
|---|---|---|---|---|---|---|
| chrX | 23889199 | 23926884 | 37686 | 0.2 | 0.000142 | 0.000402 |
| chr11 | 45920149 | 45922241 | 2093 | -0.89 | 0.000142 | 0.000403 |
| chr10 | 91620106 | 91621027 | 922 | 3.1 | 0.000142 | 0.000403 |
| chr1 | 173475259 | 173478410 | 3152 | 0.68 | 0.000143 | 0.000403 |
| chr22 | 42418232 | 42419152 | 921 | 2.21 | 0.000143 | 0.000403 |
| chr3 | 41722561 | 41741098 | 18538 | 0.75 | 0.000143 | 0.000405 |
| chr22 | 21956368 | 21963020 | 6653 | -0.71 | 0.000143 | 0.000405 |
| chr1 | 21416766 | 21426207 | 9442 | -0.88 | 0.000143 | 0.000405 |
| chr11 | 10756911 | 10758504 | 1594 | -1 | 0.000144 | 0.000406 |
| chr4 | 98832662 | 98833493 | 832 | 2.68 | 0.000144 | 0.000407 |
| chr14 | 45365438 | 45367735 | 2298 | 0.52 | 0.000144 | 0.000408 |
| chr1 | 226184573 | 226188161 | 3589 | 0.5 | 0.000144 | 0.000408 |
| chr11 | 101979756 | 101993201 | 13446 | 0.28 | 0.000145 | 0.000408 |
| chr20 | 47961172 | 47964464 | 3293 | 0.8 | 0.000145 | 0.000409 |
| chr2 | 197450196 | 197473410 | 23215 | 0.67 | 0.000145 | 0.000409 |
| chr2 | 152630838 | 152631669 | 832 | 2.27 | 0.000145 | 0.00041 |
| chr9 | 138446369 | 138447644 | 1276 | 2.91 | 0.000145 | 0.00041 |
| chr11 | 116818824 | 116820291 | 1468 | 1.55 | 0.000145 | 0.000411 |
| chr6 | 3354918 | 3411632 | 56715 | 0.24 | 0.000146 | 0.000411 |
| chr3 | 24186541 | 24199353 | 12813 | -0.51 | 0.000146 | 0.000411 |
| chr22 | 37169397 | 37172152 | 2756 | -0.98 | 0.000146 | 0.000411 |
| chr10 | 131613345 | 131613851 | 507 | 4.24 | 0.000146 | 0.000412 |
| chr10 | 73123667 | 73137580 | 13914 | -0.72 | 0.000146 | 0.000413 |
| chr1 | 32572838 | 32609465 | 36628 | -0.34 | 0.000147 | 0.000414 |
| chr9 | 110099056 | 110099796 | 741 | -3.64 | 0.000147 | 0.000414 |
| chr7 | 7801197 | 7805807 | 4611 | 1.33 | 0.000147 | 0.000414 |
| chr7 | 91762062 | 91765070 | 3009 | 0.51 | 0.000147 | 0.000415 |
| chr5 | 52856033 | 52873135 | 17103 | -0.8 | 0.000147 | 0.000415 |
| chrX | 129473044 | 129474691 | 1648 | -0.78 | 0.000147 | 0.000416 |
| chr16 | 18944055 | 18958325 | 14271 | 0.68 | 0.000147 | 0.000416 |
| chr1 | 109925662 | 109941331 | 15670 | 0.95 | 0.000148 | 0.000417 |
| chr10 | 92075769 | 92080081 | 4313 | 1.09 | 0.000148 | 0.000417 |
| chr9 | 100954188 | 100956043 | 1856 | 1.59 | 0.000148 | 0.000418 |
| chr10 | 111626899 | 111644814 | 17916 | -0.71 | 0.000149 | 0.000419 |
| chr18 | 57620206 | 57622513 | 2308 | 1.51 | 0.000149 | 0.000419 |
| chrX | 69242599 | 69243791 | 1193 | 2.4 | 0.000149 | 0.000421 |
| chr12 | 115250668 | 115251304 | 637 | -4.77 | 0.00015 | 0.000422 |
| chr11 | 96219921 | 96221143 | 1223 | 1.66 | 0.00015 | 0.000422 |
| chr12 | 105534388 | 105538868 | 4481 | 0.97 | 0.00015 | 0.000422 |
| chr5 | 176353623 | 176363879 | 10257 | -0.94 | 0.00015 | 0.000423 |
| chr1 | 199900691 | 199901912 | 1222 | 1.39 | 0.00015 | 0.000423 |
| chr14 | 97369315 | 97371150 | 1836 | 1.74 | 0.00015 | 0.000423 |
| chr6 | 85895963 | 85902947 | 6985 | 4.95 | 0.00015 | 0.000423 |
| chr14 | 58494267 | 58494763 | 497 | 4.78 | 0.00015 | 0.000424 |
| chr13 | 39610402 | 39613975 | 3574 | 0.57 | 0.000151 | 0.000424 |
| chr2 | 177418181 | 177428633 | 10453 | -0.95 | 0.000151 | 0.000424 |
| chr11 | 110064089 | 110079810 | 15722 | 0.54 | 0.000151 | 0.000425 |
| chr7 | 81410616 | 81411269 | 654 | 3.06 | 0.000151 | 0.000425 |
| chr14 | 61568251 | 61595587 | 27337 | -0.31 | 0.000151 | 0.000425 |
| chr18 | 43901119 | 43903275 | 2157 | 1.54 | 0.000151 | 0.000425 |
| chr11 | 26620403 | 26621059 | 657 | 4.78 | 0.000151 | 0.000425 |
| chr3 | 49498462 | 49509690 | 11229 | 0.49 | 0.000151 | 0.000425 |
| chr2 | 25160450 | 25161177 | 728 | 2.84 | 0.000152 | 0.000427 |
| chr1 | 162375360 | 162375809 | 450 | 2.51 | 0.000152 | 0.000428 |
| chr21 | 17613567 | 17615093 | 1527 | 1.32 | 0.000152 | 0.000428 |
| chr12 | 28625931 | 28626532 | 602 | 1.97 | 0.000153 | 0.000429 |
| chr6 | 134237954 | 134240044 | 2091 | -1.24 | 0.000153 | 0.000429 |
| chr7 | 10978106 | 10981083 | 2978 | 0.46 | 0.000153 | 0.00043 |
| chrX | 149367948 | 149370015 | 2068 | 1.55 | 0.000153 | 0.000431 |
| chr1 | 43281142 | 43290638 | 9497 | 0.48 | 0.000154 | 0.000433 |
| chr1 | 26946058 | 26948789 | 2732 | 0.46 | 0.000154 | 0.000434 |

| | | | | | | |
|---|---|---|---|---|---|---|
| chr8 | 54089219 | 54089753 | 535 | 4.79 | 0.000154 | 0.000434 |
| chr2 | 205809906 | 205813746 | 3841 | 0.92 | 0.000154 | 0.000434 |
| chrX | 96515580 | 96516195 | 616 | 2.99 | 0.000154 | 0.000434 |
| chr1 | 244762753 | 244763924 | 1172 | 1.52 | 0.000155 | 0.000435 |
| chrX | 24159379 | 24223780 | 64402 | 0.22 | 0.000155 | 0.000436 |
| chr19 | 1016432 | 1045142 | 28711 | 0.36 | 0.000155 | 0.000436 |
| chr16 | 56833217 | 56903601 | 70385 | 0.23 | 0.000155 | 0.000436 |
| chr2 | 112986959 | 112994740 | 7782 | -0.82 | 0.000156 | 0.000438 |
| chr14 | 100702559 | 100729084 | 26526 | 0.36 | 0.000156 | 0.000438 |
| chr13 | 40976094 | 40981873 | 5780 | 0.64 | 0.000156 | 0.000438 |
| chr5 | 70916546 | 70921944 | 5399 | -1.2 | 0.000156 | 0.000438 |
| chr10 | 101441134 | 101442775 | 1642 | 1.48 | 0.000156 | 0.000439 |
| chr1 | 2381451 | 2384900 | 3450 | -1 | 0.000156 | 0.000439 |
| chr2 | 31272126 | 31272628 | 503 | -3.61 | 0.000156 | 0.000439 |
| chr7 | 13564350 | 13564851 | 502 | 4.29 | 0.000156 | 0.000439 |
| chr1 | 192455151 | 192459669 | 4519 | 1.72 | 0.000157 | 0.000441 |
| chr10 | 73078458 | 73109304 | 30847 | -0.32 | 0.000157 | 0.000441 |
| chr17 | 48696555 | 48846539 | 149985 | 0.13 | 0.000157 | 0.000441 |
| chr2 | 11269694 | 11274490 | 4797 | -0.78 | 0.000157 | 0.000441 |
| chr2 | 64166295 | 64167212 | 918 | -2.35 | 0.000157 | 0.000442 |
| chr1 | 211904577 | 211912692 | 8116 | -0.45 | 0.000157 | 0.000442 |
| chr10 | 17429931 | 17430694 | 764 | 2.36 | 0.000158 | 0.000442 |
| chr4 | 59737624 | 59738456 | 833 | 1.9 | 0.000158 | 0.000442 |
| chr8 | 126605383 | 126619339 | 13957 | 0.73 | 0.000158 | 0.000443 |
| chr8 | 102702131 | 102703812 | 1682 | 1.07 | 0.000158 | 0.000443 |
| chr17 | 49717226 | 49718428 | 1203 | -1.74 | 0.000158 | 0.000444 |
| chr9 | 137787148 | 137789071 | 1924 | -1.52 | 0.000158 | 0.000444 |
| chr1 | 27625538 | 27626426 | 889 | 2.92 | 0.000158 | 0.000444 |
| chr14 | 20770029 | 20774215 | 4187 | -0.72 | 0.000159 | 0.000445 |
| chr18 | 33160552 | 33163148 | 2597 | 1.01 | 0.000159 | 0.000446 |
| chr5 | 15992668 | 15993826 | 1159 | 1.37 | 0.000159 | 0.000446 |
| chr12 | 131354454 | 131360109 | 5656 | 0.41 | 0.00016 | 0.000448 |
| chr4 | 57623414 | 57625336 | 1923 | 1.14 | 0.00016 | 0.000448 |
| chr18 | 7979680 | 7988232 | 8553 | -0.82 | 0.00016 | 0.000448 |
| chr13 | 52432542 | 52433525 | 984 | 1.73 | 0.00016 | 0.000448 |
| chr1 | 150798426 | 150809511 | 11086 | 1.16 | 0.00016 | 0.000448 |
| chr7 | 100025458 | 100049159 | 23702 | 0.29 | 0.00016 | 0.000448 |
| chr3 | 9553931 | 9554717 | 787 | -2.81 | 0.000161 | 0.00045 |
| chr2 | 238057709 | 238058826 | 1118 | 2.78 | 0.000161 | 0.000452 |
| chr6 | 126209645 | 126279784 | 70140 | 0.17 | 0.000161 | 0.000452 |
| chr14 | 77362138 | 77432287 | 70150 | 0.18 | 0.000161 | 0.000452 |
| chr19 | 30321792 | 30337380 | 15589 | 0.59 | 0.000161 | 0.000452 |
| chr2 | 71214948 | 71228655 | 13708 | -0.71 | 0.000162 | 0.000452 |
| chrX | 48812385 | 48817026 | 4642 | 0.5 | 0.000162 | 0.000453 |
| chr10 | 127822177 | 127823961 | 1785 | 1.49 | 0.000162 | 0.000453 |
| chr1 | 117657318 | 117672215 | 14898 | 0.66 | 0.000162 | 0.000453 |
| chr1 | 210501433 | 210503223 | 1791 | 0.74 | 0.000162 | 0.000453 |
| chr12 | 96688574 | 96689906 | 1333 | 1.3 | 0.000162 | 0.000453 |
| chr5 | 146858985 | 146864936 | 5952 | 0.98 | 0.000162 | 0.000454 |
| chr1 | 212058374 | 212071219 | 12846 | 0.55 | 0.000162 | 0.000454 |
| chr6 | 27435606 | 27447894 | 12289 | 0.53 | 0.000162 | 0.000454 |
| chr2 | 153165536 | 153167385 | 1850 | -0.83 | 0.000163 | 0.000455 |
| chr16 | 53706108 | 53706818 | 711 | 1.91 | 0.000163 | 0.000455 |
| chr1 | 207063429 | 207083506 | 20078 | -0.4 | 0.000163 | 0.000455 |
| chr20 | 21325004 | 21344311 | 19308 | 0.51 | 0.000163 | 0.000455 |
| chr12 | 54120455 | 54122008 | 1554 | 1.2 | 0.000163 | 0.000456 |
| chr18 | 67784009 | 67784730 | 722 | 2.43 | 0.000163 | 0.000457 |
| chr18 | 8882363 | 8892595 | 10233 | 0.42 | 0.000163 | 0.000457 |
| chr5 | 31404039 | 31414178 | 10140 | -0.89 | 0.000163 | 0.000457 |
| chr1 | 149814738 | 149822201 | 7464 | -0.43 | 0.000163 | 0.000457 |
| chr10 | 121651213 | 121669910 | 18698 | 0.48 | 0.000164 | 0.000457 |

| | | | | | | |
|---|---|---|---|---|---|---|
| chr8 | 50967962 | 50970701 | 2740 | -3.17 | 0.000164 | 0.000459 |
| chr11 | 117000737 | 117016326 | 15590 | 0.59 | 0.000164 | 0.000459 |
| chr4 | 147256727 | 147258276 | 1550 | 2.14 | 0.000164 | 0.000459 |
| chr2 | 32552958 | 32561270 | 8313 | 0.93 | 0.000164 | 0.000459 |
| chr11 | 69311265 | 69312618 | 1354 | 1.07 | 0.000164 | 0.000459 |
| chr17 | 79509383 | 79545635 | 36253 | -0.27 | 0.000164 | 0.000459 |
| chr8 | 59742265 | 59771465 | 29201 | 0.27 | 0.000164 | 0.000459 |
| chrX | 46404112 | 46407945 | 3834 | 0.57 | 0.000165 | 0.00046 |
| chr19 | 35263383 | 35264140 | 758 | 1.27 | 0.000165 | 0.000461 |
| chr5 | 114795941 | 114798166 | 2226 | -1.52 | 0.000165 | 0.000461 |
| chr11 | 93458164 | 93479972 | 21809 | 0.28 | 0.000165 | 0.000461 |
| chr13 | 76333692 | 76377790 | 44099 | 0.25 | 0.000165 | 0.000461 |
| chr6 | 142049271 | 142050337 | 1067 | 1.54 | 0.000166 | 0.000462 |
| chr13 | 24432836 | 24434304 | 1469 | 2.12 | 0.000166 | 0.000463 |
| chr2 | 228625586 | 228637323 | 11738 | -0.69 | 0.000166 | 0.000463 |
| chr5 | 127534448 | 127535799 | 1352 | 1.88 | 0.000166 | 0.000463 |
| chr5 | 40734149 | 40756552 | 22404 | 0.55 | 0.000166 | 0.000465 |
| chr6 | 127618007 | 127623974 | 5968 | 1.15 | 0.000167 | 0.000466 |
| chr21 | 40718374 | 40722666 | 4293 | 0.59 | 0.000167 | 0.000466 |
| chr3 | 196289612 | 196304080 | 14469 | 0.66 | 0.000167 | 0.000466 |
| chr3 | 24729461 | 24731447 | 1987 | 5.49 | 0.000167 | 0.000466 |
| chr10 | 13121525 | 13122061 | 537 | 4.81 | 0.000167 | 0.000467 |
| chr2 | 16052012 | 16053043 | 1032 | -3.01 | 0.000167 | 0.000467 |
| chr13 | 45986587 | 45993055 | 6469 | 0.65 | 0.000167 | 0.000467 |
| chr1 | 15415408 | 15417913 | 2506 | 4.32 | 0.000168 | 0.000467 |
| chrX | 37233972 | 37234647 | 676 | 2.6 | 0.000168 | 0.000467 |
| chr3 | 105404981 | 105408487 | 3507 | 0.77 | 0.000168 | 0.000468 |
| chr1 | 172500751 | 172504063 | 3313 | 0.48 | 0.000168 | 0.000468 |
| chr7 | 46689222 | 46700935 | 11714 | 4.09 | 0.000168 | 0.000468 |
| chr15 | 81148638 | 81150047 | 1410 | 1.31 | 0.000168 | 0.000468 |
| chr8 | 100475955 | 100476937 | 983 | 2.29 | 0.000168 | 0.000469 |
| chr4 | 129083408 | 129085502 | 2095 | 1.04 | 0.000168 | 0.000469 |
| chr4 | 78738041 | 78742220 | 4180 | 0.66 | 0.000168 | 0.000469 |
| chr9 | 77641198 | 77644174 | 2977 | 0.61 | 0.000168 | 0.000469 |
| chr2 | 75274407 | 75275033 | 627 | -3.34 | 0.000168 | 0.000469 |
| chr9 | 37061560 | 37062269 | 710 | -3.84 | 0.000169 | 0.00047 |
| chr11 | 122238206 | 122238881 | 676 | -3.25 | 0.000169 | 0.00047 |
| chr11 | 86666071 | 86667741 | 1671 | 0.87 | 0.000169 | 0.00047 |
| chr20 | 56883775 | 56886276 | 2502 | 0.65 | 0.000169 | 0.000471 |
| chr10 | 38264016 | 38266450 | 2435 | 0.51 | 0.000169 | 0.000472 |
| chr11 | 70489711 | 70490722 | 1012 | -3.07 | 0.000169 | 0.000472 |
| chr20 | 11871042 | 11887960 | 16919 | 0.43 | 0.00017 | 0.000472 |
| chr6 | 126164373 | 126195527 | 31155 | 0.39 | 0.00017 | 0.000472 |
| chr12 | 131627056 | 131628763 | 1708 | 0.91 | 0.00017 | 0.000472 |
| chr17 | 12251327 | 12252578 | 1252 | -4.84 | 0.00017 | 0.000473 |
| chr10 | 127849901 | 127850923 | 1023 | 2.75 | 0.00017 | 0.000473 |
| chr14 | 50045550 | 50066717 | 21168 | 0.34 | 0.00017 | 0.000474 |
| chr13 | 100808769 | 100809887 | 1119 | 1.9 | 0.000171 | 0.000475 |
| chr2 | 178498768 | 178499564 | 797 | 4.91 | 0.000171 | 0.000476 |
| chr5 | 149416037 | 149433480 | 17444 | -0.6 | 0.000171 | 0.000476 |
| chr7 | 5110515 | 5111165 | 651 | 4.75 | 0.000171 | 0.000476 |
| chrX | 30697224 | 30698145 | 922 | 2.42 | 0.000171 | 0.000477 |
| chr4 | 67306990 | 67307507 | 518 | 4.76 | 0.000171 | 0.000477 |
| chr7 | 152351554 | 152374221 | 22668 | 0.34 | 0.000172 | 0.000477 |
| chr20 | 43103907 | 43126425 | 22519 | -0.48 | 0.000172 | 0.000478 |
| chr4 | 85218987 | 85220076 | 1090 | 1.2 | 0.000172 | 0.000479 |
| chr7 | 129073562 | 129079624 | 6063 | 1.64 | 0.000172 | 0.000479 |
| chr1 | 59001669 | 59012997 | 11329 | 0.61 | 0.000172 | 0.000479 |
| chr2 | 48540164 | 48584881 | 44718 | 0.22 | 0.000172 | 0.00048 |
| chr5 | 29759196 | 29759806 | 611 | 2.35 | 0.000173 | 0.00048 |
| chr9 | 5900507 | 5901286 | 780 | 3.08 | 0.000173 | 0.000481 |

| | | | | | | |
|---|---|---|---|---|---|---|
| chr3 | 146322762 | 146323305 | 544 | 3.39 | 0.000173 | 0.000481 |
| chr2 | 31208141 | 31208626 | 486 | -4.8 | 0.000173 | 0.000481 |
| chr1 | 112462608 | 112463460 | 853 | -2.81 | 0.000173 | 0.000481 |
| chr9 | 114860292 | 114939644 | 79353 | -0.26 | 0.000173 | 0.000482 |
| chr7 | 144166728 | 144167251 | 524 | 4.78 | 0.000173 | 0.000482 |
| chr13 | 82538561 | 82538947 | 387 | 4.76 | 0.000174 | 0.000482 |
| chr9 | 116917766 | 116940049 | 22284 | -0.89 | 0.000174 | 0.000484 |
| chr10 | 79047243 | 79052611 | 5369 | -3.86 | 0.000174 | 0.000485 |
| chr6 | 154830140 | 154833725 | 3586 | 0.71 | 0.000175 | 0.000485 |
| chr3 | 30176618 | 30177946 | 1329 | 1.38 | 0.000175 | 0.000485 |
| chr12 | 3185221 | 3187438 | 2218 | 0.87 | 0.000175 | 0.000485 |
| chr4 | 90317739 | 90319211 | 1473 | 3.14 | 0.000175 | 0.000486 |
| chr4 | 137724345 | 137725177 | 833 | 2.9 | 0.000175 | 0.000486 |
| chr5 | 134889520 | 134890046 | 527 | -3.12 | 0.000175 | 0.000486 |
| chr1 | 14074996 | 14077310 | 2315 | 0.9 | 0.000176 | 0.000487 |
| chr1 | 116586668 | 116587052 | 385 | -4.71 | 0.000176 | 0.000489 |
| chr2 | 55130803 | 55131838 | 1036 | -1.48 | 0.000176 | 0.000489 |
| chr3 | 36718667 | 36721623 | 2957 | 1.08 | 0.000176 | 0.000489 |
| chr10 | 44408124 | 44408638 | 515 | -4.14 | 0.000176 | 0.000489 |
| chr15 | 58838024 | 58841422 | 3399 | -0.55 | 0.000176 | 0.000489 |
| chr18 | 9016278 | 9018741 | 2464 | -0.49 | 0.000177 | 0.00049 |
| chr5 | 17301640 | 17326988 | 25349 | 0.21 | 0.000177 | 0.00049 |
| chr6 | 17472718 | 17473655 | 938 | 2.13 | 0.000177 | 0.000491 |
| chr11 | 125838034 | 125838782 | 749 | 2.94 | 0.000177 | 0.000491 |
| chr4 | 90561756 | 90562307 | 552 | 3.54 | 0.000177 | 0.000491 |
| chr2 | 152329342 | 152339124 | 9783 | -0.88 | 0.000177 | 0.000492 |
| chr14 | 73618136 | 73618495 | 360 | 4.19 | 0.000177 | 0.000492 |
| chr5 | 68658173 | 68685360 | 27188 | 0.33 | 0.000177 | 0.000492 |
| chr2 | 1235280 | 1236210 | 931 | 3.12 | 0.000178 | 0.000492 |
| chr2 | 67727006 | 67727547 | 542 | 4.76 | 0.000178 | 0.000494 |
| chr15 | 60419778 | 60420509 | 732 | 1.58 | 0.000178 | 0.000494 |
| chr11 | 10025683 | 10027499 | 1817 | 1.6 | 0.000178 | 0.000494 |
| chr8 | 133036757 | 133038169 | 1413 | -1.78 | 0.000179 | 0.000496 |
| chr12 | 32027573 | 32029030 | 1458 | 0.9 | 0.000179 | 0.000497 |
| chr2 | 46768086 | 46780496 | 12411 | -0.52 | 0.00018 | 0.000498 |
| chr15 | 64715318 | 64719916 | 4599 | 0.91 | 0.00018 | 0.000498 |
| chr13 | 96327341 | 96330626 | 3286 | 0.66 | 0.00018 | 0.000499 |
| chr11 | 104916669 | 104919087 | 2419 | 0.75 | 0.00018 | 0.000499 |
| chr10 | 97312417 | 97321173 | 8757 | 1.68 | 0.000181 | 0.000501 |
| chr2 | 9139579 | 9145266 | 5688 | 0.98 | 0.000181 | 0.000501 |
| chr3 | 151643672 | 151651611 | 7940 | -0.74 | 0.000181 | 0.000501 |
| chrX | 146992692 | 146994911 | 2220 | 0.74 | 0.000181 | 0.000502 |
| chr16 | 70591829 | 70624508 | 32680 | -0.46 | 0.000182 | 0.000503 |
| chr2 | 206919384 | 206920967 | 1584 | 1.72 | 0.000182 | 0.000504 |
| chr12 | 12407526 | 12426429 | 18904 | 0.37 | 0.000182 | 0.000504 |
| chr19 | 58891283 | 58921685 | 30403 | -0.24 | 0.000182 | 0.000504 |
| chr3 | 136732986 | 136733550 | 565 | -2.62 | 0.000182 | 0.000505 |
| chr8 | 42109540 | 42110235 | 696 | 3.14 | 0.000183 | 0.000506 |
| chr9 | 94298249 | 94299529 | 1281 | -2.07 | 0.000183 | 0.000506 |
| chr17 | 35618112 | 35730590 | 112479 | -0.22 | 0.000183 | 0.000506 |
| chr3 | 52250564 | 52251819 | 1256 | 1.85 | 0.000183 | 0.000506 |
| chr11 | 6588536 | 6590392 | 1857 | -1.86 | 0.000183 | 0.000507 |
| chrX | 82234767 | 82235140 | 374 | 4.75 | 0.000184 | 0.000508 |
| chr2 | 11878841 | 11911442 | 32602 | -0.32 | 0.000184 | 0.000508 |
| chr9 | 34087310 | 34096598 | 9289 | -0.86 | 0.000184 | 0.000509 |
| chr19 | 56014643 | 56015763 | 1121 | 1.81 | 0.000184 | 0.000509 |
| chr16 | 48595246 | 48596918 | 1673 | 1.85 | 0.000184 | 0.000509 |
| chr7 | 100659351 | 100662511 | 3161 | -0.59 | 0.000184 | 0.000509 |
| chr8 | 66679686 | 66680381 | 696 | -4.69 | 0.000184 | 0.00051 |
| chr4 | 72995848 | 72996434 | 587 | 4.78 | 0.000184 | 0.00051 |
| chr3 | 123167971 | 123169203 | 1233 | 2.63 | 0.000185 | 0.000511 |

| | | | | | | |
|---|---|---|---|---|---|---|
| chr20 | 50845865 | 50848777 | 2913 | 2.83 | 0.000185 | 0.000512 |
| chr12 | 59463122 | 59464853 | 1732 | 0.98 | 0.000186 | 0.000513 |
| chr3 | 65839894 | 65841237 | 1344 | 1.51 | 0.000186 | 0.000513 |
| chr5 | 72319880 | 72335814 | 15935 | 0.7 | 0.000186 | 0.000513 |
| chr2 | 178437960 | 178438432 | 473 | -3.53 | 0.000186 | 0.000514 |
| chr4 | 38648740 | 38649321 | 582 | 3.65 | 0.000186 | 0.000514 |
| chr9 | 118848327 | 118852018 | 3692 | 0.44 | 0.000186 | 0.000515 |
| chr5 | 82928580 | 82929513 | 934 | 1.89 | 0.000186 | 0.000515 |
| chr8 | 53735850 | 53740751 | 4902 | 0.96 | 0.000187 | 0.000517 |
| chr12 | 58428032 | 58428745 | 714 | 3.12 | 0.000187 | 0.000517 |
| chr13 | 44595200 | 44597200 | 2001 | 0.96 | 0.000188 | 0.000518 |
| chr4 | 74121718 | 74126532 | 4815 | 0.47 | 0.000188 | 0.000518 |
| chr8 | 48677325 | 48906176 | 228852 | 0.14 | 0.000188 | 0.00052 |
| chr9 | 77613645 | 77621339 | 7695 | 1.26 | 0.000188 | 0.00052 |
| chr7 | 94137469 | 94144694 | 7226 | 0.43 | 0.000189 | 0.00052 |
| chr1 | 221290242 | 221292779 | 2538 | 0.82 | 0.000189 | 0.000521 |
| chr22 | 25348195 | 25405122 | 56928 | 0.3 | 0.000189 | 0.000522 |
| chr3 | 116970426 | 116981217 | 10792 | 1.4 | 0.000189 | 0.000522 |
| chr5 | 102293678 | 102295604 | 1927 | 0.95 | 0.00019 | 0.000524 |
| chr7 | 105094217 | 105105241 | 11025 | -0.79 | 0.00019 | 0.000525 |
| chr5 | 175157122 | 175157842 | 721 | -2.76 | 0.00019 | 0.000525 |
| chr11 | 119438375 | 119439640 | 1266 | -1.51 | 0.00019 | 0.000525 |
| chr16 | 89306344 | 89313787 | 7444 | 0.6 | 0.000191 | 0.000526 |
| chr3 | 71030889 | 71033113 | 2225 | 0.59 | 0.000191 | 0.000526 |
| chr15 | 49966212 | 49968045 | 1834 | 1.11 | 0.000191 | 0.000528 |
| chr5 | 114631341 | 114632315 | 975 | 1.18 | 0.000192 | 0.000529 |
| chr8 | 63146257 | 63162081 | 15825 | -0.86 | 0.000192 | 0.00053 |
| chr14 | 61223547 | 61225397 | 1851 | -2.83 | 0.000192 | 0.00053 |
| chr13 | 91775963 | 91776652 | 690 | 4.75 | 0.000192 | 0.00053 |
| chr2 | 53428889 | 53429378 | 490 | 4.75 | 0.000192 | 0.00053 |
| chr10 | 45915566 | 45922252 | 6687 | 4.68 | 0.000192 | 0.000531 |
| chr2 | 105050724 | 105051284 | 561 | 3.35 | 0.000193 | 0.000531 |
| chr2 | 128032141 | 128052846 | 20706 | -0.45 | 0.000193 | 0.000531 |
| chr18 | 61033120 | 61046907 | 13788 | 0.91 | 0.000193 | 0.000532 |
| chr1 | 232901938 | 232904065 | 2128 | 1.22 | 0.000193 | 0.000532 |
| chr6 | 141835019 | 141835570 | 552 | 3.19 | 0.000193 | 0.000532 |
| chr4 | 152065533 | 152068717 | 3185 | -1.45 | 0.000193 | 0.000532 |
| chr1 | 45637537 | 45645793 | 8257 | 3.39 | 0.000194 | 0.000533 |
| chr11 | 14566154 | 14566557 | 404 | -4.69 | 0.000194 | 0.000533 |
| chr12 | 108152152 | 108155814 | 3663 | 0.6 | 0.000194 | 0.000533 |
| chr6 | 679982 | 693524 | 13543 | -0.65 | 0.000194 | 0.000534 |
| chr8 | 121409001 | 121417868 | 8868 | 0.71 | 0.000194 | 0.000534 |
| chrX | 130912765 | 130951517 | 38753 | -0.44 | 0.000194 | 0.000534 |
| chr16 | 57277787 | 57280875 | 3089 | 0.64 | 0.000194 | 0.000535 |
| chrX | 48978890 | 48980970 | 2081 | 0.76 | 0.000195 | 0.000536 |
| chr16 | 14204694 | 14212132 | 7439 | 0.77 | 0.000195 | 0.000536 |
| chr9 | 999055 | 999656 | 602 | 3.19 | 0.000195 | 0.000536 |
| chr3 | 191046211 | 191050192 | 3982 | 0.66 | 0.000195 | 0.000537 |
| chr17 | 42607140 | 42640125 | 32986 | -0.18 | 0.000195 | 0.000537 |
| chr8 | 17940017 | 17943083 | 3067 | 0.48 | 0.000195 | 0.000538 |
| chr3 | 194908190 | 194919539 | 11350 | -1.26 | 0.000196 | 0.000539 |
| chr2 | 13978058 | 13978679 | 622 | 4.73 | 0.000196 | 0.00054 |
| chr20 | 36535546 | 36536007 | 462 | 3.48 | 0.000197 | 0.00054 |
| chr5 | 37637580 | 37638160 | 581 | -2.33 | 0.000197 | 0.000541 |
| chr18 | 9808473 | 9839281 | 30809 | 0.29 | 0.000197 | 0.000542 |
| chr11 | 70908227 | 70909791 | 1565 | -1.51 | 0.000197 | 0.000542 |
| chr4 | 56539263 | 56540279 | 1017 | -1.73 | 0.000197 | 0.000542 |
| chr6 | 4577622 | 4583192 | 5571 | 1.93 | 0.000197 | 0.000542 |
| chr1 | 219038567 | 219052508 | 13942 | 0.93 | 0.000197 | 0.000543 |
| chr2 | 151323533 | 151353370 | 29838 | 0.26 | 0.000197 | 0.000543 |
| chr6 | 166365776 | 166402591 | 36816 | 0.34 | 0.000198 | 0.000543 |

| | | | | | | |
|---|---|---|---|---|---|---|
| chrX | 47509559 | 47519361 | 9803 | 0.51 | 0.000198 | 0.000544 |
| chr7 | 80263490 | 80264493 | 1004 | 1.86 | 0.000198 | 0.000544 |
| chr16 | 1963130 | 1993124 | 29995 | -0.55 | 0.000198 | 0.000544 |
| chr14 | 58876502 | 58877745 | 1244 | -1.68 | 0.000198 | 0.000545 |
| chr8 | 59049101 | 59050067 | 967 | 4.78 | 0.000198 | 0.000545 |
| chr4 | 10090510 | 10119557 | 29048 | 0.36 | 0.000199 | 0.000545 |
| chr17 | 61112630 | 61118437 | 5808 | 0.85 | 0.000199 | 0.000546 |
| chr4 | 26320089 | 26323558 | 3470 | 0.53 | 0.000199 | 0.000546 |
| chr2 | 131819099 | 131879424 | 60326 | -0.26 | 0.000199 | 0.000547 |
| chrX | 19904856 | 19925137 | 20282 | 0.47 | 2.00E-04 | 0.000549 |
| chr14 | 60617517 | 60632826 | 15310 | 0.47 | 2.00E-04 | 0.000549 |
| chr1 | 184349452 | 184350038 | 587 | -4.73 | 2.00E-04 | 0.000549 |
| chr3 | 114690583 | 114691266 | 684 | 3.54 | 2.00E-04 | 0.000549 |
| chr5 | 54590585 | 54605740 | 15156 | 0.36 | 2.00E-04 | 0.000549 |
| chr21 | 19032522 | 19032831 | 310 | 4.79 | 2.00E-04 | 0.000549 |
| chr17 | 66770673 | 66771850 | 1178 | 1.55 | 2.00E-04 | 0.00055 |
| chr5 | 108135957 | 108156301 | 20345 | 0.79 | 0.000201 | 0.00055 |
| chrX | 108779577 | 108781469 | 1893 | 0.67 | 0.000201 | 0.000551 |
| chr3 | 99565102 | 99566055 | 954 | 2.11 | 0.000201 | 0.000552 |
| chr14 | 58217984 | 58218916 | 933 | 3.12 | 0.000202 | 0.000554 |
| chr5 | 78011833 | 78024720 | 12888 | -0.5 | 0.000202 | 0.000554 |
| chr7 | 81383185 | 81384015 | 831 | 2.24 | 0.000202 | 0.000554 |
| chr12 | 67986942 | 67989114 | 2173 | -1.73 | 0.000202 | 0.000555 |
| chr9 | 75008384 | 75012553 | 4170 | 0.53 | 0.000203 | 0.000556 |
| chr1 | 95178681 | 95180188 | 1508 | 1.49 | 0.000203 | 0.000556 |
| chr22 | 29898475 | 29901440 | 2966 | -0.86 | 0.000203 | 0.000556 |
| chr4 | 142557017 | 142564936 | 7920 | 0.61 | 0.000203 | 0.000556 |
| chr1 | 151762536 | 151764253 | 1718 | 1.17 | 0.000204 | 0.00056 |
| chr11 | 32719709 | 32729718 | 10010 | 0.48 | 0.000205 | 0.000561 |
| chr8 | 57856732 | 57857204 | 473 | 4.72 | 0.000205 | 0.000562 |
| chr8 | 103779785 | 103823766 | 43982 | -0.17 | 0.000206 | 0.000565 |
| chr18 | 64350066 | 64350503 | 438 | 4.73 | 0.000206 | 0.000566 |
| chr2 | 85540582 | 85556591 | 16010 | -0.44 | 0.000207 | 0.000567 |
| chr4 | 34273275 | 34273983 | 709 | 2.61 | 0.000207 | 0.000567 |
| chr6 | 1480817 | 1486641 | 5825 | -1.32 | 0.000207 | 0.000568 |
| chr7 | 115670462 | 115671464 | 1003 | 1.52 | 0.000208 | 0.000568 |
| chr8 | 131807840 | 131809018 | 1179 | -1.95 | 0.000208 | 0.000569 |
| chr19 | 44506414 | 44508304 | 1891 | 0.78 | 0.000208 | 0.000569 |
| chr4 | 143640664 | 143641416 | 753 | 3.16 | 0.000208 | 0.000569 |
| chr11 | 102214840 | 102220175 | 5336 | 0.42 | 0.000208 | 0.00057 |
| chr11 | 689520 | 728886 | 39367 | 0.31 | 0.000208 | 0.00057 |
| chr1 | 51975184 | 51984920 | 9737 | 0.72 | 0.000208 | 0.00057 |
| chr2 | 196382324 | 196410852 | 28529 | -0.37 | 0.000209 | 0.000571 |
| chr2 | 153398082 | 153403744 | 5663 | 0.62 | 0.000209 | 0.000572 |
| chr15 | 32926806 | 32928140 | 1335 | 1.91 | 0.000209 | 0.000572 |
| chr10 | 74318472 | 74328195 | 9724 | -0.89 | 0.000209 | 0.000572 |
| chr3 | 119878464 | 119879444 | 981 | -3.06 | 0.000209 | 0.000573 |
| chr1 | 35496184 | 35498200 | 2017 | 0.86 | 0.000209 | 0.000573 |
| chr6 | 108908276 | 108913387 | 5112 | 0.57 | 0.00021 | 0.000573 |
| chr5 | 149208233 | 149211441 | 3209 | -1.83 | 0.00021 | 0.000574 |
| chr1 | 171484414 | 171506702 | 22289 | 0.66 | 0.00021 | 0.000574 |
| chr21 | 30331007 | 30332863 | 1857 | 0.9 | 0.00021 | 0.000574 |
| chr9 | 122262279 | 122263196 | 918 | -2.02 | 0.00021 | 0.000574 |
| chr4 | 180979003 | 180979759 | 757 | 3.46 | 0.00021 | 0.000574 |
| chr9 | 35400884 | 35401296 | 413 | -4.67 | 0.00021 | 0.000575 |
| chr16 | 15707622 | 15745621 | 38000 | 0.33 | 0.00021 | 0.000575 |
| chr4 | 15119686 | 15134210 | 14525 | 1.16 | 0.000211 | 0.000575 |
| chr18 | 41064522 | 41065303 | 782 | -2.17 | 0.000211 | 0.000576 |
| chr15 | 52470729 | 52473047 | 2319 | 0.53 | 0.000211 | 0.000578 |
| chr1 | 207880353 | 207880895 | 543 | -3.61 | 0.000212 | 0.000578 |
| chr16 | 1795079 | 1797463 | 2385 | 1.05 | 0.000212 | 0.000578 |

| | | | | | | |
|---|---|---|---|---|---|---|
| chr8 | 60358941 | 60359933 | 993 | -2.11 | 0.000212 | 0.00058 |
| chr7 | 92860929 | 92862415 | 1487 | 1.12 | 0.000213 | 0.000581 |
| chrX | 48722384 | 48727091 | 4708 | 1.23 | 0.000213 | 0.000581 |
| chr14 | 70408000 | 70409266 | 1267 | 1.54 | 0.000213 | 0.000582 |
| chr15 | 57359771 | 57360381 | 611 | -2.63 | 0.000213 | 0.000582 |
| chr2 | 233801848 | 233802713 | 866 | -1.79 | 0.000214 | 0.000585 |
| chr17 | 16891331 | 16892049 | 719 | -3.3 | 0.000215 | 0.000586 |
| chr5 | 115366522 | 115368875 | 2354 | 5.39 | 0.000215 | 0.000586 |
| chr12 | 67709355 | 67710318 | 964 | 1.95 | 0.000215 | 0.000587 |
| chr17 | 30600895 | 30614403 | 13509 | -3.81 | 0.000215 | 0.000587 |
| chrX | 131349997 | 131365573 | 15577 | 0.55 | 0.000215 | 0.000587 |
| chr9 | 2241411 | 2249640 | 8230 | 1.24 | 0.000215 | 0.000587 |
| chr11 | 18511553 | 18515677 | 4125 | 0.98 | 0.000215 | 0.000588 |
| chr2 | 196520915 | 196528049 | 7135 | -0.4 | 0.000215 | 0.000588 |
| chr1 | 178289847 | 178315550 | 25704 | 0.91 | 0.000216 | 0.000589 |
| chr14 | 59889680 | 59895622 | 5943 | -1.44 | 0.000216 | 0.000589 |
| chr2 | 85032443 | 85035208 | 2766 | 0.83 | 0.000216 | 0.000589 |
| chr6 | 139854240 | 139862304 | 8065 | 0.54 | 0.000216 | 0.00059 |
| chr10 | 113291742 | 113292438 | 697 | -4.82 | 0.000216 | 0.00059 |
| chr10 | 7396313 | 7402458 | 6146 | -1.03 | 0.000216 | 0.00059 |
| chr16 | 86587745 | 86625006 | 37262 | 0.45 | 0.000216 | 0.00059 |
| chr8 | 24857248 | 24861754 | 4507 | -0.76 | 0.000216 | 0.00059 |
| chr10 | 91151510 | 91158596 | 7087 | 0.91 | 0.000217 | 0.000591 |
| chr9 | 126581782 | 126582982 | 1201 | -1.58 | 0.000217 | 0.000591 |
| chr12 | 120492712 | 120494227 | 1516 | 0.99 | 0.000217 | 0.000593 |
| chr1 | 81623979 | 81624535 | 557 | 4.79 | 0.000218 | 0.000594 |
| chr12 | 6959719 | 6963026 | 3308 | 0.76 | 0.000218 | 0.000595 |
| chr11 | 795501 | 867088 | 71588 | 0.19 | 0.000218 | 0.000595 |
| chr7 | 40866795 | 40867268 | 474 | 4.75 | 0.000218 | 0.000595 |
| chr18 | 56189042 | 56221129 | 32088 | 0.24 | 0.000219 | 0.000596 |
| chr8 | 114323561 | 114324052 | 492 | 4.82 | 0.000219 | 0.000598 |
| chr3 | 86157562 | 86169578 | 12017 | 4.89 | 0.000219 | 0.000598 |
| chr1 | 231469725 | 231494376 | 24652 | 0.39 | 0.000219 | 0.000598 |
| chr11 | 28829226 | 28830647 | 1422 | 1.15 | 0.00022 | 0.000599 |
| chr12 | 54366933 | 54427502 | 60570 | 0.29 | 0.00022 | 6.00E-04 |
| chr10 | 101467638 | 101469921 | 2284 | -1 | 0.000221 | 0.000601 |
| chr19 | 44668234 | 44670043 | 1810 | 0.72 | 0.000221 | 0.000601 |
| chr21 | 38710157 | 38712413 | 2257 | -0.69 | 0.000221 | 0.000602 |
| chrX | 44730590 | 44734934 | 4345 | 0.43 | 0.000221 | 0.000602 |
| chr1 | 173816530 | 173844159 | 27630 | 0.32 | 0.000222 | 0.000604 |
| chr16 | 82030305 | 82031885 | 1581 | 1.29 | 0.000222 | 0.000605 |
| chr17 | 8263101 | 8264303 | 1203 | 2.71 | 0.000222 | 0.000605 |
| chr10 | 87399650 | 87406299 | 6650 | 0.21 | 0.000223 | 0.000607 |
| chr4 | 14857208 | 14858824 | 1617 | 1.3 | 0.000223 | 0.000607 |
| chr6 | 136707828 | 136769823 | 61996 | 0.19 | 0.000223 | 0.000608 |
| chr1 | 24542789 | 24549979 | 7191 | 1.03 | 0.000223 | 0.000608 |
| chr10 | 85898056 | 85901073 | 3018 | 0.41 | 0.000223 | 0.000608 |
| chr2 | 198151231 | 198153070 | 1840 | -0.91 | 0.000224 | 0.000608 |
| chr2 | 64500621 | 64504947 | 4327 | -0.89 | 0.000224 | 0.000609 |
| chr1 | 27691316 | 27694407 | 3092 | 0.59 | 0.000224 | 0.000609 |
| chr6 | 128671606 | 128672743 | 1138 | 1.79 | 0.000224 | 0.000609 |
| chr6 | 109217321 | 109218764 | 1444 | -2.11 | 0.000224 | 0.000609 |
| chr5 | 76067482 | 76135324 | 67843 | -0.25 | 0.000224 | 0.00061 |
| chr2 | 187776262 | 187776693 | 432 | -4.7 | 0.000225 | 0.000611 |
| chr1 | 230797557 | 230798054 | 498 | -3.26 | 0.000225 | 0.000612 |
| chr9 | 124079977 | 124094961 | 14985 | -0.39 | 0.000225 | 0.000613 |
| chr4 | 40443262 | 40534018 | 90757 | 0.17 | 0.000226 | 0.000613 |
| chr20 | 8499693 | 8500524 | 832 | 3.09 | 0.000226 | 0.000614 |
| chr18 | 45659470 | 45660421 | 952 | 3.21 | 0.000226 | 0.000614 |
| chr16 | 80984531 | 80985504 | 974 | 2.31 | 0.000226 | 0.000614 |
| chr2 | 227828098 | 227828950 | 853 | -1.65 | 0.000226 | 0.000615 |

| | | | | | | |
|---|---|---|---|---|---|---|
| chr13 | 105204259 | 105204697 | 439 | 3.62 | 0.000226 | 0.000615 |
| chr2 | 9953731 | 9954387 | 657 | -3.75 | 0.000226 | 0.000615 |
| chr2 | 161789043 | 161790514 | 1472 | -1.25 | 0.000227 | 0.000615 |
| chr12 | 114298282 | 114314425 | 16144 | -1.02 | 0.000227 | 0.000616 |
| chr7 | 79578824 | 79579462 | 639 | 4.73 | 0.000227 | 0.000618 |
| chr12 | 27699828 | 27727489 | 27662 | 0.47 | 0.000227 | 0.000618 |
| chr1 | 191340706 | 191341381 | 676 | 4.91 | 0.000228 | 0.00062 |
| chr2 | 27192279 | 27194293 | 2015 | -0.93 | 0.000229 | 0.00062 |
| chr12 | 16588404 | 16589993 | 1590 | 1.73 | 0.000229 | 0.000623 |
| chr4 | 140372113 | 140376927 | 4815 | 0.55 | 0.00023 | 0.000623 |
| chr5 | 134420623 | 134422036 | 1414 | 1.65 | 0.00023 | 0.000624 |
| chr2 | 23603170 | 23607990 | 4821 | 1.07 | 0.00023 | 0.000624 |
| chr6 | 99460946 | 99461546 | 601 | -4.64 | 0.00023 | 0.000624 |
| chr17 | 57642497 | 57650312 | 7816 | -0.55 | 0.00023 | 0.000625 |
| chr3 | 152004886 | 152050843 | 45958 | 0.21 | 0.00023 | 0.000625 |
| chr5 | 14105435 | 14114267 | 8833 | 0.75 | 0.000231 | 0.000626 |
| chr7 | 82326697 | 82330205 | 3509 | 3.45 | 0.000231 | 0.000627 |
| chr16 | 4836943 | 4838960 | 2018 | -1.62 | 0.000231 | 0.000627 |
| chr10 | 61408175 | 61408793 | 619 | -3.33 | 0.000231 | 0.000628 |
| chr10 | 37434977 | 37435408 | 432 | 4.7 | 0.000232 | 0.000628 |
| chr4 | 91514388 | 91514857 | 470 | 4.7 | 0.000232 | 0.000628 |
| chr5 | 55257052 | 55325861 | 68810 | 0.22 | 0.000232 | 0.000629 |
| chr6 | 109186055 | 109187617 | 1563 | -1.5 | 0.000232 | 0.00063 |
| chr3 | 143183823 | 143185010 | 1188 | 3.06 | 0.000232 | 0.00063 |
| chr16 | 67396254 | 67399513 | 3260 | 3.63 | 0.000232 | 0.00063 |
| chr5 | 83583381 | 83601566 | 18186 | 5.44 | 0.000233 | 0.000631 |
| chr2 | 17622499 | 17622905 | 407 | -4.64 | 0.000233 | 0.000632 |
| chr8 | 9044623 | 9046660 | 2038 | 0.85 | 0.000233 | 0.000632 |
| chr1 | 212894434 | 212899526 | 5093 | 0.83 | 0.000234 | 0.000633 |
| chr14 | 29211533 | 29212463 | 931 | 2.02 | 0.000234 | 0.000634 |
| chr2 | 151381675 | 151387444 | 5770 | 0.37 | 0.000234 | 0.000634 |
| chr7 | 86961654 | 86975557 | 13904 | 0.81 | 0.000235 | 0.000636 |
| chr3 | 305741 | 307098 | 1358 | 1.89 | 0.000235 | 0.000636 |
| chr16 | 67359690 | 67364291 | 4602 | 1.28 | 0.000235 | 0.000636 |
| chr5 | 146783522 | 146784542 | 1021 | -1.84 | 0.000235 | 0.000636 |
| chr3 | 124690749 | 124706163 | 15415 | -0.74 | 0.000236 | 0.000638 |
| chr2 | 67826403 | 67828861 | 2459 | 1.23 | 0.000236 | 0.000639 |
| chr22 | 39567139 | 39574562 | 7424 | -0.37 | 0.000236 | 0.000639 |
| chr6 | 157800572 | 157803516 | 2945 | 0.9 | 0.000237 | 0.000641 |
| chr1 | 155817778 | 155829131 | 11354 | 0.52 | 0.000237 | 0.000642 |
| chr6 | 37934082 | 37934978 | 897 | 2.46 | 0.000237 | 0.000642 |
| chr18 | 34324833 | 34333750 | 8918 | -4.73 | 0.000237 | 0.000642 |
| chr11 | 27517058 | 27529309 | 12252 | 0.47 | 0.000238 | 0.000644 |
| chr18 | 56074564 | 56085696 | 11133 | -1.59 | 0.000238 | 0.000644 |
| chr5 | 139776954 | 139789366 | 12413 | 0.41 | 0.000238 | 0.000644 |
| chr2 | 95786258 | 95788025 | 1768 | -0.6 | 0.000238 | 0.000645 |
| chr12 | 7340906 | 7343091 | 2186 | 0.52 | 0.000239 | 0.000648 |
| chr12 | 30906311 | 30908833 | 2523 | 0.79 | 0.000239 | 0.000648 |
| chr12 | 17214737 | 17236657 | 21921 | 0.39 | 0.00024 | 0.00065 |
| chr7 | 57411140 | 57411823 | 684 | 4.7 | 0.000241 | 0.000651 |
| chr4 | 56719822 | 56733108 | 13287 | 0.8 | 0.000241 | 0.000651 |
| chr17 | 3598179 | 3599874 | 1696 | -1.91 | 0.000241 | 0.000651 |
| chr2 | 102351084 | 102386693 | 35610 | -0.31 | 0.000241 | 0.000652 |
| chr8 | 30618890 | 30621048 | 2159 | -1.56 | 0.000242 | 0.000655 |
| chr6 | 20200498 | 20214207 | 13710 | 0.33 | 0.000242 | 0.000655 |
| chr4 | 84519971 | 84520972 | 1002 | 2.56 | 0.000242 | 0.000656 |
| chr20 | 34356735 | 34390539 | 33805 | -0.42 | 0.000242 | 0.000656 |
| chr14 | 60557552 | 60561520 | 3969 | 0.36 | 0.000243 | 0.000656 |
| chr5 | 76439924 | 76440746 | 823 | -1.82 | 0.000243 | 0.000656 |
| chr20 | 56212772 | 56214516 | 1745 | 1.2 | 0.000243 | 0.000657 |
| chr13 | 114302128 | 114302552 | 425 | 3.64 | 0.000243 | 0.000658 |

| | | | | | | |
|---|---|---|---|---|---|---|
| chr8 | 146125163 | 146128260 | 3098 | 0.58 | 0.000244 | 0.000658 |
| chr12 | 20700679 | 20702173 | 1495 | 1.32 | 0.000244 | 0.000659 |
| chr6 | 16747125 | 16771096 | 23972 | 0.45 | 0.000245 | 0.000662 |
| chr5 | 113655335 | 113655914 | 580 | 3.39 | 0.000245 | 0.000663 |
| chr7 | 93207318 | 93209673 | 2356 | 2.16 | 0.000246 | 0.000664 |
| chr8 | 117876538 | 117887439 | 10902 | 0.39 | 0.000246 | 0.000664 |
| chr2 | 69566118 | 69615605 | 49488 | -0.25 | 0.000247 | 0.000666 |
| chr2 | 160372109 | 160373811 | 1703 | 0.65 | 0.000247 | 0.000668 |
| chr1 | 79788369 | 79789952 | 1584 | 0.99 | 0.000248 | 0.000669 |
| chr1 | 172715207 | 172716108 | 902 | 2.23 | 0.000248 | 0.000669 |
| chr11 | 44140310 | 44141522 | 1213 | -1.5 | 0.000249 | 0.000671 |
| chr12 | 91971391 | 91971825 | 435 | 1.39 | 0.000249 | 0.000671 |
| chr5 | 127008873 | 127009576 | 704 | -2.36 | 0.000249 | 0.000672 |
| chr6 | 136807197 | 136871653 | 64457 | -0.22 | 0.000249 | 0.000673 |
| chr17 | 16232363 | 16257443 | 25081 | -0.43 | 0.00025 | 0.000675 |
| chr9 | 139111446 | 139137626 | 26181 | -0.65 | 0.00025 | 0.000675 |
| chr12 | 65211910 | 65212868 | 959 | 2.64 | 0.00025 | 0.000675 |
| chr17 | 4416931 | 4417784 | 854 | -2.3 | 0.00025 | 0.000675 |
| chr9 | 14371593 | 14373818 | 2226 | 1.05 | 0.000251 | 0.000676 |
| chr21 | 28058394 | 28058716 | 323 | 4.69 | 0.000251 | 0.000677 |
| chr2 | 62752258 | 62768407 | 16150 | 0.77 | 0.000251 | 0.000678 |
| chr7 | 149190547 | 149195394 | 4848 | -0.62 | 0.000251 | 0.000678 |
| chr17 | 46834004 | 46847210 | 13207 | 0.96 | 0.000252 | 0.00068 |
| chr1 | 227379963 | 227380561 | 599 | -3.2 | 0.000253 | 0.000681 |
| chr2 | 148765434 | 148780495 | 15062 | 0.55 | 0.000253 | 0.000682 |
| chrX | 47208350 | 47226126 | 17777 | 0.64 | 0.000253 | 0.000683 |
| chr6 | 88182118 | 88184229 | 2112 | 0.8 | 0.000253 | 0.000683 |
| chr19 | 4693345 | 4725125 | 31781 | 0.34 | 0.000253 | 0.000683 |
| chr3 | 57644045 | 57648258 | 4214 | -0.84 | 0.000254 | 0.000685 |
| chr3 | 43625473 | 43626157 | 685 | -1.88 | 0.000254 | 0.000685 |
| chr4 | 92009738 | 92010701 | 964 | 1.61 | 0.000255 | 0.000687 |
| chr19 | 22177840 | 22178473 | 634 | 4.66 | 0.000255 | 0.000687 |
| chr1 | 90528859 | 90529202 | 344 | 4.66 | 0.000255 | 0.000688 |
| chr10 | 101355300 | 101356198 | 899 | -1.72 | 0.000255 | 0.000688 |
| chr4 | 83701233 | 83708070 | 6838 | 0.68 | 0.000255 | 0.000688 |
| chr2 | 29235423 | 29236662 | 1240 | -1.88 | 0.000257 | 0.000691 |
| chr11 | 10771395 | 10773804 | 2410 | 0.59 | 0.000257 | 0.000693 |
| chr21 | 39591267 | 39592158 | 892 | 1.63 | 0.000258 | 0.000695 |
| chr12 | 125623991 | 125626225 | 2235 | 0.89 | 0.000259 | 0.000696 |
| chr4 | 103350090 | 103361812 | 11723 | 0.36 | 0.000259 | 0.000697 |
| chr18 | 56159680 | 56168862 | 9183 | -1.01 | 0.000259 | 0.000697 |
| chr14 | 107281104 | 107281823 | 720 | 2.57 | 0.000259 | 0.000698 |
| chrX | 29552640 | 29561181 | 8542 | -0.6 | 0.000259 | 0.000698 |
| chr5 | 17743797 | 17749044 | 5248 | 1.44 | 0.00026 | 0.000698 |
| chr1 | 173963715 | 173993874 | 30160 | 0.31 | 0.00026 | 7.00E-04 |
| chr18 | 21450710 | 21464727 | 14018 | -0.75 | 0.00026 | 7.00E-04 |
| chr2 | 30337854 | 30338705 | 852 | 1.82 | 0.000261 | 0.000701 |
| chr10 | 80382868 | 80384604 | 1737 | 1.46 | 0.000261 | 0.000702 |
| chr9 | 96074207 | 96076591 | 2385 | 0.47 | 0.000261 | 0.000702 |
| chr12 | 95942235 | 95943311 | 1077 | 1.45 | 0.000261 | 0.000703 |
| chr15 | 64436607 | 64456416 | 19810 | -0.41 | 0.000262 | 0.000704 |
| chr15 | 62898013 | 62899501 | 1489 | 1.67 | 0.000262 | 0.000705 |
| chr3 | 72067391 | 72068247 | 857 | -2.22 | 0.000263 | 0.000707 |
| chr19 | 14031849 | 14032429 | 581 | 1.98 | 0.000263 | 0.000707 |
| chr5 | 71658281 | 71662642 | 4362 | -0.97 | 0.000264 | 0.000709 |
| chr16 | 22288229 | 22289970 | 1742 | -1.66 | 0.000264 | 0.00071 |
| chr10 | 103872216 | 103896639 | 24424 | -0.37 | 0.000264 | 0.00071 |
| chr17 | 5388172 | 5391979 | 3808 | 0.44 | 0.000267 | 0.000716 |
| chr2 | 167086055 | 167087179 | 1125 | 3.13 | 0.000267 | 0.000716 |
| chr1 | 28098615 | 28100687 | 2073 | 0.98 | 0.000267 | 0.000716 |
| chr14 | 102173847 | 102174878 | 1032 | -1.69 | 0.000267 | 0.000717 |

| | | | | | | |
|---|---|---|---|---|---|---|
| chr20 | 34604065 | 34607001 | 2937 | 0.43 | 0.000267 | 0.000718 |
| chr7 | 37258115 | 37259090 | 976 | 1.86 | 0.000268 | 0.000719 |
| chr5 | 178570045 | 178570547 | 503 | 4.66 | 0.000268 | 0.000719 |
| chr16 | 83868678 | 83870388 | 1711 | -1.26 | 0.000268 | 0.000719 |
| chr7 | 19480142 | 19481039 | 898 | 2.47 | 0.000268 | 0.00072 |
| chr3 | 118976344 | 118977593 | 1250 | 2.2 | 0.000269 | 0.000723 |
| chr11 | 35408505 | 35413809 | 5305 | 1 | 0.00027 | 0.000724 |
| chr11 | 76220985 | 76224087 | 3103 | 0.96 | 0.00027 | 0.000725 |
| chr5 | 95580689 | 95596008 | 15320 | 0.32 | 0.00027 | 0.000725 |
| chr12 | 104457454 | 104459008 | 1555 | 1.24 | 0.00027 | 0.000726 |
| chr6 | 11253497 | 11254453 | 957 | 2.44 | 0.00027 | 0.000726 |
| chr1 | 108179550 | 108186737 | 7188 | 2.17 | 0.000271 | 0.000726 |
| chr12 | 63889440 | 63890787 | 1348 | 1.67 | 0.000271 | 0.000726 |
| chr1 | 229405538 | 229411403 | 5866 | 0.68 | 0.000271 | 0.000727 |
| chr1 | 95669201 | 95670452 | 1252 | 2.18 | 0.000271 | 0.000727 |
| chr11 | 9519628 | 9553206 | 33579 | 0.38 | 0.000271 | 0.000727 |
| chr8 | 29067101 | 29067992 | 892 | -3.3 | 0.000271 | 0.000727 |
| chr7 | 83306153 | 83306960 | 808 | 2.31 | 0.000271 | 0.000727 |
| chr3 | 8780892 | 8785940 | 5049 | 0.44 | 0.000271 | 0.000727 |
| chr20 | 45881412 | 46032555 | 151144 | -0.15 | 0.000271 | 0.000727 |
| chr5 | 169645716 | 169647657 | 1942 | 0.89 | 0.000271 | 0.000728 |
| chr5 | 81115008 | 81116962 | 1955 | 0.88 | 0.000271 | 0.000728 |
| chr19 | 58738630 | 58751988 | 13359 | 3.62 | 0.000272 | 0.000729 |
| chr13 | 109961710 | 109963295 | 1586 | 1.36 | 0.000272 | 0.00073 |
| chr13 | 80908006 | 80917764 | 9759 | 0.34 | 0.000272 | 0.00073 |
| chr14 | 23321904 | 23357642 | 35739 | 0.31 | 0.000273 | 0.000733 |
| chr17 | 72927455 | 72928224 | 770 | -3.02 | 0.000273 | 0.000733 |
| chr6 | 22717233 | 22717736 | 504 | 3.89 | 0.000274 | 0.000734 |
| chr2 | 44587361 | 44590175 | 2815 | -0.43 | 0.000274 | 0.000735 |
| chr20 | 42218282 | 42225163 | 6882 | 0.51 | 0.000275 | 0.000736 |
| chr20 | 46964564 | 46968804 | 4241 | -5.01 | 0.000275 | 0.000737 |
| chr3 | 119233371 | 119234633 | 1263 | 1.18 | 0.000275 | 0.000737 |
| chr13 | 31019607 | 31020474 | 868 | 2.51 | 0.000275 | 0.000737 |
| chr17 | 79301957 | 79320977 | 19021 | 0.27 | 0.000275 | 0.000737 |
| chr5 | 160365457 | 160366091 | 635 | -3.19 | 0.000276 | 0.000738 |
| chrX | 15352342 | 15354886 | 2545 | 0.55 | 0.000276 | 0.000739 |
| chr1 | 89266487 | 89268947 | 2461 | 1.55 | 0.000277 | 0.000741 |
| chr2 | 103129913 | 103137806 | 7894 | 4.85 | 0.000277 | 0.000741 |
| chr7 | 151493143 | 151506133 | 12991 | -3.13 | 0.000277 | 0.000741 |
| chr4 | 94440151 | 94440825 | 675 | -3.57 | 0.000277 | 0.000741 |
| chr10 | 79388557 | 79391146 | 2590 | 0.8 | 0.000277 | 0.000742 |
| chr8 | 143084023 | 143089555 | 5533 | -1.12 | 0.000277 | 0.000742 |
| chr13 | 45716135 | 45718121 | 1987 | 0.81 | 0.000277 | 0.000742 |
| chr2 | 164589007 | 164594044 | 5038 | -0.51 | 0.000277 | 0.000743 |
| chr14 | 105820115 | 105821106 | 992 | -3.09 | 0.000278 | 0.000743 |
| chr12 | 27378851 | 27380198 | 1348 | 1.21 | 0.000278 | 0.000743 |
| chr6 | 129810475 | 129822810 | 12336 | 0.44 | 0.000278 | 0.000744 |
| chr1 | 192716314 | 192719055 | 2742 | 1.12 | 0.000278 | 0.000745 |
| chr18 | 46064947 | 46075350 | 10404 | -0.79 | 0.000279 | 0.000746 |
| chr2 | 28359999 | 28360545 | 547 | 3.41 | 0.000279 | 0.000747 |
| chr6 | 133033150 | 133034129 | 980 | 1.86 | 0.00028 | 0.000748 |
| chr11 | 43701579 | 43713939 | 12361 | 0.36 | 0.00028 | 0.000748 |
| chr1 | 41362244 | 41385855 | 23612 | 0.55 | 0.00028 | 0.000749 |
| chr19 | 54145778 | 54146262 | 485 | 4.67 | 0.000282 | 0.000753 |
| chr2 | 158029226 | 158041376 | 12151 | 0.36 | 0.000282 | 0.000754 |
| chr11 | 35683339 | 35685796 | 2458 | 0.49 | 0.000282 | 0.000754 |
| chr1 | 25344151 | 25359816 | 15666 | 0.52 | 0.000283 | 0.000756 |
| chr12 | 68752351 | 68760064 | 7714 | 0.64 | 0.000283 | 0.000756 |
| chr4 | 188468422 | 188469058 | 637 | 1.86 | 0.000283 | 0.000757 |
| chr17 | 3506867 | 3512756 | 5890 | -1.26 | 0.000284 | 0.000758 |
| chr9 | 14084816 | 14104662 | 19847 | 0.43 | 0.000284 | 0.00076 |

| | | | | | | |
|---|---|---|---|---|---|---|
| chr8 | 128341799 | 128347906 | 6108 | 1.17 | 0.000284 | 0.00076 |
| chr5 | 82391250 | 82392493 | 1244 | -3.01 | 0.000285 | 0.000763 |
| chr12 | 114164937 | 114165412 | 476 | -3.5 | 0.000286 | 0.000763 |
| chr5 | 169351027 | 169351935 | 909 | 3.15 | 0.000286 | 0.000764 |
| chr8 | 108151348 | 108172705 | 21358 | 0.47 | 0.000287 | 0.000766 |
| chrX | 102086542 | 102088668 | 2127 | -0.64 | 0.000287 | 0.000767 |
| chr4 | 186989536 | 186990839 | 1304 | 1.48 | 0.000288 | 0.000768 |
| chr16 | 81109705 | 81112083 | 2379 | 0.43 | 0.000289 | 0.000771 |
| chr12 | 39630456 | 39631262 | 807 | 3.03 | 0.000289 | 0.000771 |
| chr16 | 67477096 | 67499806 | 22711 | -0.36 | 0.000289 | 0.000772 |
| chr2 | 198285202 | 198301232 | 16031 | -0.38 | 0.00029 | 0.000774 |
| chr2 | 155303244 | 155307973 | 4730 | 0.74 | 0.00029 | 0.000774 |
| chr4 | 56170189 | 56170613 | 425 | 4.63 | 0.00029 | 0.000774 |
| chr7 | 119494545 | 119494988 | 444 | 4.63 | 0.00029 | 0.000774 |
| chr9 | 71712667 | 71720767 | 8101 | 3.35 | 0.00029 | 0.000774 |
| chr12 | 131886872 | 131888252 | 1381 | 1.64 | 0.00029 | 0.000775 |
| chr12 | 6276852 | 6290033 | 13182 | -4.59 | 0.000291 | 0.000777 |
| chr7 | 127876473 | 127877751 | 1279 | -1.68 | 0.000292 | 0.000778 |
| chr13 | 109115193 | 109116687 | 1495 | 1.4 | 0.000292 | 0.000779 |
| chr10 | 35701458 | 35750672 | 49215 | 0.2 | 0.000292 | 0.000779 |
| chr16 | 87462749 | 87463621 | 873 | -2.82 | 0.000292 | 0.00078 |
| chr10 | 24738165 | 24738684 | 520 | 3.76 | 0.000292 | 0.00078 |
| chr5 | 32531310 | 32538305 | 6996 | -0.91 | 0.000293 | 0.00078 |
| chr17 | 72673810 | 72674380 | 571 | -2.98 | 0.000293 | 0.000783 |
| chr4 | 74568559 | 74574934 | 6376 | -0.44 | 0.000294 | 0.000783 |
| chr1 | 81032536 | 81032892 | 357 | 4.63 | 0.000294 | 0.000783 |
| chr5 | 107586869 | 107587291 | 423 | 4.63 | 0.000294 | 0.000783 |
| chr3 | 15671439 | 15690017 | 18579 | -0.24 | 0.000294 | 0.000785 |
| chr7 | 42144499 | 42145048 | 550 | 4.63 | 0.000294 | 0.000785 |
| chr14 | 104333491 | 104348744 | 15254 | 0.51 | 0.000295 | 0.000785 |
| chr2 | 230764097 | 230793788 | 29692 | -0.38 | 0.000295 | 0.000786 |
| chr9 | 132969316 | 132977636 | 8321 | -1.56 | 0.000295 | 0.000787 |
| chr6 | 83040714 | 83041786 | 1073 | 1.57 | 0.000295 | 0.000787 |
| chrX | 66999991 | 67001266 | 1276 | 1.5 | 0.000295 | 0.000787 |
| chr2 | 203498728 | 203501938 | 3211 | 0.54 | 0.000296 | 0.000788 |
| chr1 | 195732313 | 195733444 | 1132 | 1.79 | 0.000296 | 0.000788 |
| chr1 | 192608278 | 192610197 | 1920 | 1.55 | 0.000296 | 0.000788 |
| chr2 | 3148128 | 3155368 | 7241 | -1.18 | 0.000297 | 0.000791 |
| chr1 | 35788667 | 35790709 | 2043 | 1.52 | 0.000297 | 0.000791 |
| chr2 | 16237035 | 16245242 | 8208 | 1.12 | 0.000298 | 0.000793 |
| chr6 | 122973643 | 122974846 | 1204 | 2.14 | 0.000298 | 0.000793 |
| chr5 | 105857089 | 105860245 | 3157 | 0.67 | 0.000298 | 0.000793 |
| chr12 | 65173911 | 65175254 | 1344 | 2.03 | 0.000298 | 0.000793 |
| chr15 | 52368706 | 52369913 | 1208 | -1.98 | 0.000298 | 0.000794 |
| chr16 | 27733705 | 27736473 | 2769 | -1.7 | 0.000298 | 0.000794 |
| chr5 | 156119696 | 156121049 | 1354 | 1.28 | 0.000299 | 0.000795 |
| chr5 | 111066592 | 111067381 | 790 | 3.06 | 0.000299 | 0.000796 |
| chr10 | 22375437 | 22376597 | 1161 | 1.82 | 3.00E-04 | 0.000799 |
| chr2 | 121194166 | 121195451 | 1286 | -2.52 | 3.00E-04 | 0.000799 |
| chr2 | 118715188 | 118716552 | 1365 | 1.58 | 0.000302 | 0.000803 |
| chr3 | 71544975 | 71554506 | 9532 | -0.65 | 0.000302 | 0.000803 |
| chr5 | 130810256 | 130832014 | 21759 | -0.45 | 0.000302 | 0.000804 |
| chr12 | 121561506 | 121567454 | 5949 | 1.14 | 0.000303 | 0.000806 |
| chr1 | 173423116 | 173423488 | 373 | -2.68 | 0.000304 | 0.000808 |
| chr6 | 4647378 | 4648619 | 1242 | 1.01 | 0.000304 | 0.000808 |
| chr13 | 113339914 | 113346338 | 6425 | 0.97 | 0.000304 | 0.000809 |
| chr5 | 37518562 | 37518907 | 346 | 4.1 | 0.000305 | 0.00081 |
| chr2 | 183222665 | 183238762 | 16098 | -0.28 | 0.000305 | 0.00081 |
| chr4 | 125120750 | 125134383 | 13634 | 2.82 | 0.000305 | 0.000811 |
| chr2 | 207138853 | 207140648 | 1796 | 0.88 | 0.000306 | 0.000813 |
| chr2 | 227048488 | 227056038 | 7551 | 0.36 | 0.000306 | 0.000813 |

| | | | | | | |
|---|---|---|---|---|---|---|
| chr2 | 11722478 | 11724257 | 1780 | -1.66 | 0.000306 | 0.000813 |
| chr12 | 90727135 | 90727855 | 721 | 1.7 | 0.000306 | 0.000814 |
| chr18 | 29009375 | 29010377 | 1003 | 2.37 | 0.000306 | 0.000814 |
| chrX | 40471231 | 40472327 | 1097 | 1.43 | 0.000307 | 0.000815 |
| chr4 | 152085925 | 152088268 | 2344 | 1.03 | 0.000307 | 0.000815 |
| chr7 | 27086081 | 27093123 | 7043 | 0.95 | 0.000307 | 0.000817 |
| chr14 | 21175422 | 21176003 | 582 | 3.35 | 0.000308 | 0.000817 |
| chr2 | 174516392 | 174523714 | 7323 | -0.58 | 0.000308 | 0.000817 |
| chr9 | 125341699 | 125343749 | 2051 | 0.65 | 0.000308 | 0.000818 |
| chr2 | 186719205 | 186729380 | 10176 | 4.25 | 0.000309 | 0.000819 |
| chr13 | 95200270 | 95202772 | 2503 | 0.49 | 0.000309 | 0.00082 |
| chr14 | 75029345 | 75051798 | 22454 | 0.27 | 0.000309 | 0.00082 |
| chr14 | 93251852 | 93262256 | 10405 | 0.51 | 0.000309 | 0.000822 |
| chr7 | 45072712 | 45080562 | 7851 | -1.02 | 0.00031 | 0.000822 |
| chr3 | 151838650 | 151839684 | 1035 | -2.45 | 0.00031 | 0.000822 |
| chr10 | 129970786 | 129973099 | 2314 | -0.7 | 0.00031 | 0.000822 |
| chr1 | 101241653 | 101242166 | 514 | 3.57 | 0.00031 | 0.000823 |
| chr5 | 55700770 | 55711874 | 11105 | 2.21 | 0.00031 | 0.000824 |
| chr13 | 24791163 | 24796830 | 5668 | -0.74 | 0.000311 | 0.000824 |
| chr4 | 91824870 | 91825672 | 803 | 2.62 | 0.000311 | 0.000824 |
| chr17 | 26003334 | 26004075 | 742 | -3.26 | 0.000311 | 0.000825 |
| chr4 | 2712295 | 2713059 | 765 | 2.18 | 0.000312 | 0.000827 |
| chr11 | 107963737 | 107965777 | 2041 | 0.8 | 0.000312 | 0.000827 |
| chr10 | 8743695 | 8745801 | 2107 | 0.55 | 0.000312 | 0.000828 |
| chr2 | 159394213 | 159397190 | 2978 | 0.84 | 0.000312 | 0.000828 |
| chr2 | 196312179 | 196313092 | 914 | 2 | 0.000313 | 0.000829 |
| chr10 | 87527733 | 87528377 | 645 | 4.63 | 0.000313 | 0.00083 |
| chr9 | 130610829 | 130611161 | 333 | 4.63 | 0.000313 | 0.00083 |
| chr10 | 127033243 | 127034048 | 806 | 2.34 | 0.000314 | 0.000832 |
| chr9 | 6271779 | 6272385 | 607 | 3.05 | 0.000314 | 0.000832 |
| chr8 | 31732141 | 31732904 | 764 | 3.1 | 0.000314 | 0.000833 |
| chr14 | 24082652 | 24083672 | 1021 | 3.11 | 0.000315 | 0.000834 |
| chr7 | 2592812 | 2613388 | 20577 | 0.4 | 0.000315 | 0.000835 |
| chr3 | 186499671 | 186509595 | 9925 | 0.43 | 0.000315 | 0.000835 |
| chr7 | 63360709 | 63361033 | 325 | 4.66 | 0.000316 | 0.000837 |
| chr2 | 106126817 | 106128447 | 1631 | 1.49 | 0.000317 | 0.000838 |
| chr10 | 74969088 | 74970212 | 1125 | 1.88 | 0.000317 | 0.000839 |
| chr11 | 102906997 | 102907418 | 422 | 3.21 | 0.000317 | 0.000839 |
| chr22 | 38239488 | 38247155 | 7668 | 0.48 | 0.000317 | 0.000841 |
| chr10 | 90041121 | 90042175 | 1055 | 1.77 | 0.000318 | 0.000841 |
| chr7 | 128043636 | 128051296 | 7661 | -0.72 | 0.000318 | 0.000842 |
| chr2 | 193332687 | 193333869 | 1183 | 1.57 | 0.000319 | 0.000845 |
| chr11 | 74203145 | 74217133 | 13989 | -0.59 | 0.000319 | 0.000845 |
| chr21 | 43516877 | 43518907 | 2031 | 0.68 | 0.000319 | 0.000845 |
| chr17 | 68234578 | 68235510 | 933 | 2.33 | 0.00032 | 0.000846 |
| chr4 | 137451659 | 137452130 | 472 | 3.03 | 0.00032 | 0.000847 |
| chr16 | 11349369 | 11351154 | 1786 | 0.95 | 0.00032 | 0.000847 |
| chr11 | 111402111 | 111402947 | 837 | -3.19 | 0.00032 | 0.000848 |
| chrX | 21943607 | 22016299 | 72693 | 0.23 | 0.000321 | 0.000848 |
| chr16 | 47367963 | 47368356 | 394 | -4.6 | 0.000321 | 0.000848 |
| chr1 | 227750691 | 227753150 | 2460 | 0.6 | 0.000321 | 0.000849 |
| chr11 | 6375881 | 6376763 | 883 | 2 | 0.000321 | 0.00085 |
| chr11 | 70650852 | 70651749 | 898 | -2.25 | 0.000322 | 0.000851 |
| chr3 | 108190013 | 108192156 | 2144 | -0.98 | 0.000322 | 0.000851 |
| chr19 | 42854038 | 42868862 | 14825 | -0.73 | 0.000322 | 0.000852 |
| chr2 | 20365879 | 20432439 | 66561 | -0.19 | 0.000322 | 0.000852 |
| chr9 | 87515093 | 87515445 | 353 | 4.06 | 0.000322 | 0.000852 |
| chr11 | 35358053 | 35359414 | 1362 | 1.75 | 0.000322 | 0.000852 |
| chr2 | 207941452 | 207942093 | 642 | -4.56 | 0.000324 | 0.000856 |
| chr2 | 119495986 | 119496868 | 883 | -1.47 | 0.000324 | 0.000856 |
| chr10 | 106568953 | 106569370 | 418 | -4.56 | 0.000325 | 0.000858 |

| | | | | | | |
|---|---|---|---|---|---|---|
| chr5 | 105536449 | 105537052 | 604 | 3.09 | 0.000325 | 0.000858 |
| chrX | 54664893 | 54667070 | 2178 | 0.56 | 0.000325 | 0.000859 |
| chrX | 149252290 | 149264180 | 11891 | 1.11 | 0.000325 | 0.00086 |
| chr9 | 15999563 | 16000087 | 525 | -4.56 | 0.000326 | 0.000861 |
| chr6 | 145118789 | 145120166 | 1378 | 1.62 | 0.000329 | 0.000868 |
| chr2 | 208119159 | 208124767 | 5609 | 0.51 | 0.000329 | 0.000869 |
| chr11 | 115358959 | 115376233 | 17275 | -0.6 | 0.000329 | 0.000869 |
| chr12 | 56171901 | 56173130 | 1230 | 2.19 | 0.000329 | 0.000869 |
| chr2 | 24925131 | 24926440 | 1310 | 1.19 | 0.000329 | 0.000869 |
| chr3 | 52483008 | 52490332 | 7325 | 0.61 | 0.000329 | 0.000869 |
| chr8 | 142378706 | 142386955 | 8250 | -1.06 | 0.00033 | 0.000871 |
| chr4 | 57609712 | 57610379 | 668 | 3.83 | 0.00033 | 0.000871 |
| chr8 | 53375277 | 53380762 | 5486 | 2.51 | 0.00033 | 0.000871 |
| chr1 | 68680031 | 68681003 | 973 | 2.52 | 0.00033 | 0.000872 |
| chr15 | 66420939 | 66423220 | 2282 | 4.8 | 0.000331 | 0.000875 |
| chr10 | 16935759 | 16936255 | 497 | 4.6 | 0.000332 | 0.000876 |
| chr11 | 27302899 | 27305031 | 2133 | 0.63 | 0.000332 | 0.000877 |
| chr8 | 126521934 | 126541592 | 19659 | -0.47 | 0.000332 | 0.000877 |
| chr12 | 60603553 | 60604150 | 598 | 1.76 | 0.000333 | 0.000878 |
| chrX | 4628107 | 4632077 | 3971 | -1.06 | 0.000333 | 0.000879 |
| chr14 | 92080202 | 92081202 | 1001 | -1.19 | 0.000333 | 0.000879 |
| chr18 | 805563 | 813654 | 8092 | 0.52 | 0.000334 | 0.000882 |
| chr18 | 7566125 | 7578075 | 11951 | -0.43 | 0.000334 | 0.000882 |
| chr2 | 173912279 | 173925276 | 12998 | -0.4 | 0.000335 | 0.000883 |
| chr8 | 145718870 | 145755033 | 36164 | -0.33 | 0.000335 | 0.000884 |
| chr11 | 65404410 | 65432027 | 27618 | 0.31 | 0.000335 | 0.000884 |
| chr2 | 45346606 | 45347711 | 1106 | 3.14 | 0.000335 | 0.000884 |
| chr4 | 13626853 | 13644748 | 17896 | 0.49 | 0.000336 | 0.000885 |
| chr6 | 27247300 | 27265147 | 17848 | -0.52 | 0.000336 | 0.000885 |
| chr2 | 62458754 | 62459091 | 338 | -4.56 | 0.000337 | 0.000889 |
| chr16 | 31517956 | 31520931 | 2976 | 0.43 | 0.000337 | 0.000889 |
| chr10 | 99571714 | 99573532 | 1819 | -0.94 | 0.000338 | 0.00089 |
| chr12 | 28317427 | 28319073 | 1647 | 1.11 | 0.000338 | 0.00089 |
| chr13 | 45827347 | 45828321 | 975 | -2.41 | 0.000339 | 0.000892 |
| chr11 | 17025789 | 17037577 | 11789 | 0.46 | 0.000339 | 0.000893 |
| chr11 | 62725656 | 62726684 | 1029 | 2.18 | 0.000339 | 0.000893 |
| chr9 | 129746323 | 129749154 | 2832 | 1.15 | 0.00034 | 0.000895 |
| chr10 | 21704714 | 21717703 | 12990 | -1.05 | 0.00034 | 0.000896 |
| chr1 | 24985328 | 24985735 | 408 | -4.57 | 0.00034 | 0.000896 |
| chr9 | 117110770 | 117112225 | 1456 | -1.1 | 0.000341 | 0.000897 |
| chr16 | 30850362 | 30851286 | 925 | 3.32 | 0.000341 | 0.000899 |
| chr1 | 233654333 | 233656992 | 2660 | 1.59 | 0.000341 | 0.000899 |
| chr12 | 74434418 | 74438553 | 4136 | 0.31 | 0.000342 | 0.000899 |
| chr2 | 215547877 | 215548675 | 799 | 1.6 | 0.000342 | 9.00E-04 |
| chr4 | 59642354 | 59643263 | 910 | 1.98 | 0.000342 | 9.00E-04 |
| chr5 | 85335329 | 85335909 | 581 | 2.81 | 0.000342 | 0.000901 |
| chr8 | 126103047 | 126105269 | 2223 | 0.54 | 0.000343 | 0.000902 |
| chr7 | 110659168 | 110661638 | 2471 | 1.35 | 0.000343 | 0.000902 |
| chr4 | 176210811 | 176212176 | 1366 | 2.99 | 0.000343 | 0.000902 |
| chr15 | 76109494 | 76109943 | 450 | -2.83 | 0.000343 | 0.000903 |
| chr7 | 137843184 | 137855228 | 12045 | 5.22 | 0.000343 | 0.000903 |
| chr19 | 52189344 | 52209405 | 20062 | -0.27 | 0.000344 | 0.000905 |
| chr12 | 40014083 | 40014444 | 362 | 3.09 | 0.000344 | 0.000906 |
| chr20 | 42718503 | 42726704 | 8202 | -4.91 | 0.000344 | 0.000906 |
| chrX | 17602091 | 17633053 | 30963 | 1.89 | 0.000344 | 0.000906 |
| chr9 | 79917492 | 79918470 | 979 | 1.94 | 0.000344 | 0.000906 |
| chr8 | 104409061 | 104432641 | 23581 | 0.3 | 0.000344 | 0.000906 |
| chr2 | 197264191 | 197265656 | 1466 | -1.44 | 0.000346 | 0.000909 |
| chr1 | 234242697 | 234260007 | 17311 | -0.67 | 0.000347 | 0.000911 |
| chr1 | 99197207 | 99199631 | 2425 | 1 | 0.000348 | 0.000914 |
| chr15 | 92644812 | 92646173 | 1362 | 1.95 | 0.000348 | 0.000914 |

| | | | | | | |
|---|---|---|---|---|---|---|
| chr18 | 71230573 | 71233173 | 2601 | 0.93 | 0.000348 | 0.000915 |
| chr2 | 153703565 | 153704059 | 495 | -3.48 | 0.000348 | 0.000915 |
| chr9 | 2790096 | 2845693 | 55598 | -0.22 | 0.000349 | 0.000917 |
| chr12 | 104516806 | 104532831 | 16026 | 0.46 | 0.000349 | 0.000917 |
| chr22 | 34267341 | 34268322 | 982 | 2.34 | 0.000349 | 0.000917 |
| chr2 | 215393946 | 215395051 | 1106 | -2.43 | 0.000349 | 0.000917 |
| chr12 | 29301445 | 29302570 | 1126 | 1.88 | 0.000349 | 0.000918 |
| chr13 | 76582753 | 76589460 | 6708 | 0.76 | 0.00035 | 0.000918 |
| chr6 | 134356313 | 134357584 | 1272 | 2.09 | 0.00035 | 0.000919 |
| chr2 | 242400483 | 242401994 | 1512 | 1.57 | 0.000351 | 0.000921 |
| chr7 | 28338231 | 28339093 | 863 | -3.06 | 0.000351 | 0.000922 |
| chr6 | 21109126 | 21112555 | 3430 | 0.66 | 0.000351 | 0.000922 |
| chr21 | 30390017 | 30407661 | 17645 | 0.4 | 0.000352 | 0.000924 |
| chr3 | 149886973 | 149888110 | 1138 | -1.59 | 0.000352 | 0.000924 |
| chr3 | 57436353 | 57437655 | 1303 | -1.77 | 0.000353 | 0.000926 |
| chr7 | 142956024 | 142961450 | 5427 | 0.83 | 0.000353 | 0.000926 |
| chr1 | 161084578 | 161104565 | 19988 | 0.37 | 0.000353 | 0.000926 |
| chr15 | 99880349 | 99881621 | 1273 | 1.38 | 0.000353 | 0.000928 |
| chr3 | 105485887 | 105486443 | 557 | 3.22 | 0.000354 | 0.000928 |
| chr3 | 185000127 | 185001907 | 1781 | 0.73 | 0.000355 | 0.000931 |
| chr7 | 105070485 | 105080457 | 9973 | 1.55 | 0.000355 | 0.000931 |
| chr3 | 143689186 | 143693363 | 4178 | 0.44 | 0.000355 | 0.000932 |
| chr14 | 24860768 | 24868552 | 7785 | 0.97 | 0.000356 | 0.000933 |
| chr1 | 194013895 | 194015235 | 1341 | 1.71 | 0.000356 | 0.000934 |
| chr1 | 85929425 | 85931669 | 2245 | 1.09 | 0.000356 | 0.000934 |
| chr11 | 17372924 | 17376003 | 3080 | 0.69 | 0.000356 | 0.000934 |
| chr7 | 117256004 | 117258504 | 2501 | 0.93 | 0.000356 | 0.000935 |
| chr5 | 129885712 | 129887668 | 1957 | 1.41 | 0.000357 | 0.000935 |
| chr4 | 6034040 | 6035198 | 1159 | -1.82 | 0.000357 | 0.000935 |
| chr3 | 136673276 | 136680922 | 7647 | 0.52 | 0.000357 | 0.000936 |
| chr22 | 19678803 | 19680327 | 1525 | 0.66 | 0.000357 | 0.000937 |
| chr18 | 47792054 | 47814996 | 22943 | 0.38 | 0.000357 | 0.000937 |
| chrX | 15736006 | 15738234 | 2229 | 1.02 | 0.000358 | 0.000937 |
| chr5 | 132097733 | 132099388 | 1656 | -2.36 | 0.000358 | 0.000938 |
| chrX | 102911035 | 102912495 | 1461 | -1.11 | 0.000358 | 0.000939 |
| chr17 | 57286198 | 57298690 | 12493 | 0.48 | 0.000359 | 0.000941 |
| chr12 | 46167188 | 46168289 | 1102 | 1.81 | 0.00036 | 0.000943 |
| chr17 | 29616174 | 29617813 | 1640 | 1.43 | 0.00036 | 0.000944 |
| chr14 | 103415195 | 103419578 | 4384 | -1.02 | 0.000361 | 0.000945 |
| chr12 | 122429718 | 122434892 | 5175 | 1.14 | 0.000361 | 0.000946 |
| chr19 | 19773644 | 19781584 | 7941 | 0.56 | 0.000361 | 0.000947 |
| chr12 | 124672698 | 124673807 | 1110 | 2.08 | 0.000362 | 0.000948 |
| chr1 | 156694876 | 156760245 | 65370 | 0.19 | 0.000362 | 0.000948 |
| chr9 | 91143820 | 91159788 | 15969 | 0.37 | 0.000362 | 0.000949 |
| chr1 | 246195076 | 246195992 | 917 | 2 | 0.000363 | 0.000949 |
| chr2 | 182644232 | 182657229 | 12998 | 0.35 | 0.000363 | 0.000952 |
| chr2 | 172127567 | 172129674 | 2108 | 1.17 | 0.000364 | 0.000952 |
| chr6 | 158632128 | 158654286 | 22159 | 0.62 | 0.000364 | 0.000953 |
| chr22 | 41797240 | 41814626 | 17387 | 0.39 | 0.000364 | 0.000954 |
| chr4 | 151770539 | 151771039 | 501 | 3.75 | 0.000365 | 0.000954 |
| chr4 | 133449691 | 133450298 | 608 | 1.23 | 0.000365 | 0.000955 |
| chr3 | 37166412 | 37219350 | 52939 | 0.24 | 0.000366 | 0.000956 |
| chr14 | 63622225 | 63622602 | 378 | -3.48 | 0.000366 | 0.000957 |
| chr11 | 91246235 | 91247232 | 998 | -2.56 | 0.000366 | 0.000958 |
| chr11 | 126391713 | 126392924 | 1212 | -1.17 | 0.000367 | 0.000961 |
| chr5 | 80595935 | 80599150 | 3216 | 0.49 | 0.000368 | 0.000963 |
| chr11 | 86883994 | 86886065 | 2072 | 0.62 | 0.00037 | 0.000967 |
| chr12 | 130644942 | 130645588 | 647 | 3.54 | 0.000371 | 0.000971 |
| chr2 | 14616163 | 14616788 | 626 | -2.75 | 0.000371 | 0.000971 |
| chr22 | 36072453 | 36073082 | 630 | 3.06 | 0.000372 | 0.000972 |
| chr2 | 219051243 | 219051871 | 629 | -1.31 | 0.000372 | 0.000973 |

| | | | | | | |
|---|---|---|---|---|---|---|
| chr3 | 143549816 | 143550670 | 855 | 2.28 | 0.000373 | 0.000974 |
| chr5 | 179604086 | 179605017 | 932 | 2.34 | 0.000373 | 0.000975 |
| chr8 | 120547344 | 120548193 | 850 | 2.79 | 0.000373 | 0.000975 |
| chr2 | 170675395 | 170684720 | 9326 | -0.55 | 0.000373 | 0.000975 |
| chr11 | 126208684 | 126227317 | 18634 | -0.3 | 0.000373 | 0.000975 |
| chr3 | 194276729 | 194277854 | 1126 | -1.08 | 0.000373 | 0.000976 |
| chr7 | 100323134 | 100323977 | 844 | 3.14 | 0.000374 | 0.000976 |
| chr21 | 16738618 | 16739022 | 405 | 4.65 | 0.000374 | 0.000977 |
| chr8 | 61157585 | 61158545 | 961 | 1.77 | 0.000374 | 0.000977 |
| chr3 | 127390590 | 127428812 | 38223 | -0.48 | 0.000374 | 0.000977 |
| chr4 | 138045700 | 138046858 | 1159 | 2.31 | 0.000374 | 0.000977 |
| chr2 | 217046371 | 217057417 | 11047 | -0.8 | 0.000374 | 0.000977 |
| chrX | 153706655 | 153719892 | 13238 | 0.48 | 0.000374 | 0.000977 |
| chr5 | 78052708 | 78053837 | 1130 | -2.19 | 0.000374 | 0.000978 |
| chr14 | 74003238 | 74005764 | 2527 | 1.05 | 0.000375 | 0.000978 |
| chr4 | 54231312 | 54245136 | 13825 | 0.88 | 0.000375 | 0.000978 |
| chr1 | 228352283 | 228355407 | 3125 | 0.49 | 0.000375 | 0.000979 |
| chr21 | 36399305 | 36400794 | 1490 | 1.77 | 0.000375 | 0.00098 |
| chr7 | 68428117 | 68429669 | 1553 | 1.18 | 0.000376 | 0.00098 |
| chrX | 7894624 | 7896287 | 1664 | 0.72 | 0.000376 | 0.00098 |
| chr2 | 82510016 | 82510500 | 485 | -2.47 | 0.000376 | 0.00098 |
| chr11 | 111793856 | 111816248 | 22393 | -0.69 | 0.000376 | 0.00098 |
| chr17 | 62184394 | 62185957 | 1564 | 1.45 | 0.000376 | 0.000981 |
| chr10 | 91229237 | 91242539 | 13303 | 1.32 | 0.000376 | 0.000982 |
| chr11 | 73690754 | 73695999 | 5246 | 0.45 | 0.000376 | 0.000982 |
| chr15 | 90198092 | 90209398 | 11307 | 5.82 | 0.000377 | 0.000984 |
| chr10 | 62607489 | 62608600 | 1112 | 1.67 | 0.000378 | 0.000986 |
| chr20 | 8028374 | 8029227 | 854 | 3.05 | 0.000379 | 0.000987 |
| chr18 | 2301107 | 2305711 | 4605 | 0.42 | 0.000379 | 0.000988 |
| chr2 | 43056780 | 43064773 | 7994 | 0.82 | 0.000379 | 0.000988 |
| chr7 | 72867279 | 72972797 | 105519 | 0.19 | 0.000379 | 0.000988 |
| chr4 | 7104654 | 7106159 | 1506 | 1.28 | 0.000379 | 0.000988 |
| chr3 | 128078552 | 128081096 | 2545 | 0.93 | 0.00038 | 0.000991 |
| chr22 | 24435873 | 24436889 | 1017 | -2.96 | 0.000381 | 0.000992 |
| chr5 | 154133594 | 154218624 | 85031 | -0.18 | 0.000381 | 0.000992 |
| chr13 | 104535818 | 104536186 | 369 | 4.57 | 0.000381 | 0.000992 |
| chr7 | 55600343 | 55641082 | 40740 | -0.31 | 0.000381 | 0.000993 |
| chr14 | 52171001 | 52184009 | 13009 | -1.16 | 0.000382 | 0.000995 |
| chr1 | 18806017 | 18808766 | 2750 | 0.74 | 0.000382 | 0.000995 |
| chr2 | 55179558 | 55249970 | 70413 | -0.23 | 0.000383 | 0.000998 |
| chr11 | 40409023 | 40409936 | 914 | 3.07 | 0.000385 | 0.001 |
| chr11 | 44961450 | 44977727 | 16278 | 0.38 | 0.000385 | 0.001 |
| chr5 | 164557385 | 164558105 | 721 | -3.69 | 0.000385 | 0.001 |
| chr4 | 130949890 | 130950286 | 397 | 4.56 | 0.000386 | 0.001 |
| chr1 | 32154256 | 32181400 | 27145 | -0.32 | 0.000386 | 0.001 |
| chr1 | 145641855 | 145669780 | 27926 | -0.37 | 0.000386 | 0.00101 |
| chr11 | 13228612 | 13239128 | 10517 | 0.65 | 0.000387 | 0.00101 |
| chr7 | 139650834 | 139651399 | 566 | 2.87 | 0.000388 | 0.00101 |
| chr13 | 46716952 | 46717747 | 796 | 2.84 | 0.000388 | 0.00101 |
| chr3 | 44753683 | 44754890 | 1208 | 1.28 | 0.000389 | 0.00101 |
| chr20 | 44844301 | 44845078 | 778 | -2.93 | 0.00039 | 0.00101 |
| chr5 | 134160835 | 134191156 | 30322 | 0.32 | 0.000391 | 0.00102 |
| chr3 | 182271799 | 182272666 | 868 | 2.52 | 0.000391 | 0.00102 |
| chr6 | 1797413 | 1798411 | 999 | 2.04 | 0.000391 | 0.00102 |
| chr5 | 10352587 | 10378678 | 26092 | 0.3 | 0.000392 | 0.00102 |
| chr2 | 108832092 | 108836383 | 4292 | -1.21 | 0.000394 | 0.00102 |
| chr10 | 101988013 | 101990701 | 2689 | 0.52 | 0.000394 | 0.00103 |
| chr5 | 54455586 | 54457805 | 2220 | 0.47 | 0.000395 | 0.00103 |
| chr1 | 249199629 | 249201705 | 2077 | 0.56 | 0.000395 | 0.00103 |
| chr2 | 159886981 | 159887950 | 970 | -1.36 | 0.000396 | 0.00103 |
| chr1 | 62676263 | 62676717 | 455 | 4.57 | 0.000397 | 0.00103 |

| | | | | | | |
|---|---|---|---|---|---|---|
| chr4 | 22031826 | 22032227 | 402 | 4.57 | 0.000397 | 0.00103 |
| chr2 | 55314513 | 55346349 | 31837 | 0.27 | 0.000398 | 0.00103 |
| chr17 | 40335858 | 40359199 | 23342 | -0.63 | 0.000398 | 0.00104 |
| chr8 | 26239627 | 26242140 | 2514 | 0.71 | 0.000399 | 0.00104 |
| chr9 | 140600028 | 140603561 | 3534 | -1.98 | 0.000399 | 0.00104 |
| chr9 | 36867551 | 36868103 | 553 | 3.16 | 4.00E-04 | 0.00104 |
| chrX | 109072994 | 109074931 | 1938 | 1 | 4.00E-04 | 0.00104 |
| chr10 | 30518930 | 30519369 | 440 | -4.52 | 4.00E-04 | 0.00104 |
| chr2 | 55709140 | 55727994 | 18855 | -0.85 | 0.000401 | 0.00104 |
| chr1 | 46267873 | 46270653 | 2781 | 0.85 | 0.000401 | 0.00104 |
| chr18 | 65085111 | 65086614 | 1504 | 1 | 0.000402 | 0.00104 |
| chr16 | 4117384 | 4117923 | 540 | -3.22 | 0.000402 | 0.00104 |
| chr14 | 53247570 | 53259711 | 12142 | 0.48 | 0.000402 | 0.00104 |
| chrX | 133930708 | 133942205 | 11498 | 0.61 | 0.000403 | 0.00105 |
| chr16 | 75680579 | 75689376 | 8798 | 0.57 | 0.000403 | 0.00105 |
| chr10 | 60502306 | 60522897 | 20592 | 0.35 | 0.000404 | 0.00105 |
| chr2 | 9892308 | 9923061 | 30754 | -0.48 | 0.000404 | 0.00105 |
| chr19 | 58238292 | 58239563 | 1272 | 1.48 | 0.000404 | 0.00105 |
| chr1 | 55400010 | 55401195 | 1186 | 1.71 | 0.000404 | 0.00105 |
| chr2 | 191177430 | 191185805 | 8376 | -0.6 | 0.000405 | 0.00105 |
| chr1 | 168147300 | 168157033 | 9734 | 0.51 | 0.000405 | 0.00105 |
| chr3 | 142165116 | 142168221 | 3106 | 0.44 | 0.000405 | 0.00105 |
| chr14 | 21904221 | 21925777 | 21557 | 0.3 | 0.000405 | 0.00105 |
| chr4 | 168273952 | 168274385 | 434 | 3.49 | 0.000406 | 0.00105 |
| chr2 | 239313593 | 239316382 | 2790 | -1.73 | 0.000407 | 0.00105 |
| chr7 | 27674939 | 27687442 | 12504 | 0.68 | 0.000407 | 0.00105 |
| chr16 | 3788348 | 3797069 | 8722 | -1.11 | 0.000407 | 0.00105 |
| chr8 | 41053653 | 41060275 | 6623 | -0.78 | 0.000408 | 0.00106 |
| chr6 | 48969304 | 48971696 | 2393 | 4.38 | 0.000408 | 0.00106 |
| chr7 | 140612532 | 140625749 | 13218 | 0.49 | 0.000408 | 0.00106 |
| chr2 | 137133023 | 137133723 | 701 | -3.08 | 0.000409 | 0.00106 |
| chr4 | 123985316 | 123986551 | 1236 | 2.23 | 0.000409 | 0.00106 |
| chr8 | 99096896 | 99098161 | 1266 | 1.31 | 0.00041 | 0.00106 |
| chr12 | 59144570 | 59144949 | 380 | -4.5 | 0.00041 | 0.00106 |
| chr14 | 23181709 | 23183139 | 1431 | 1.05 | 0.00041 | 0.00106 |
| chr21 | 17904587 | 17910928 | 6342 | -0.41 | 0.00041 | 0.00106 |
| chr9 | 123341085 | 123343339 | 2255 | -0.6 | 0.000411 | 0.00107 |
| chr9 | 120545079 | 120546057 | 979 | 2.61 | 0.000412 | 0.00107 |
| chr7 | 148982384 | 148982866 | 483 | 3.05 | 0.000412 | 0.00107 |
| chr3 | 143833428 | 143834271 | 844 | -1.92 | 0.000412 | 0.00107 |
| chr1 | 149898768 | 149919792 | 21025 | 0.32 | 0.000412 | 0.00107 |
| chr17 | 37436915 | 37438577 | 1663 | 1.36 | 0.000412 | 0.00107 |
| chr22 | 18183140 | 18218050 | 34911 | 0.26 | 0.000412 | 0.00107 |
| chr4 | 8538843 | 8539297 | 455 | 3.72 | 0.000413 | 0.00107 |
| chr3 | 40940097 | 40940815 | 719 | -3.46 | 0.000413 | 0.00107 |
| chr19 | 8452068 | 8463116 | 11049 | 0.35 | 0.000413 | 0.00107 |
| chr18 | 53965606 | 53967183 | 1578 | 1.02 | 0.000414 | 0.00107 |
| chr8 | 99913705 | 99925593 | 11889 | 5.05 | 0.000414 | 0.00107 |
| chr6 | 161906982 | 161908442 | 1461 | -1.12 | 0.000414 | 0.00107 |
| chr10 | 115547182 | 115549742 | 2561 | -0.51 | 0.000415 | 0.00107 |
| chr22 | 46959419 | 46976908 | 17490 | -0.69 | 0.000415 | 0.00107 |
| chr19 | 14444131 | 14446209 | 2079 | 0.97 | 0.000415 | 0.00107 |
| chr14 | 61822485 | 61823251 | 767 | -3 | 0.000415 | 0.00107 |
| chr2 | 118752419 | 118752789 | 371 | -4.5 | 0.000416 | 0.00108 |
| chr1 | 9615426 | 9616455 | 1030 | -2.35 | 0.000416 | 0.00108 |
| chr11 | 12577413 | 12578861 | 1449 | -1.26 | 0.000416 | 0.00108 |
| chr13 | 76867066 | 76868466 | 1401 | 2.13 | 0.000417 | 0.00108 |
| chr15 | 90435995 | 90438625 | 2631 | 0.43 | 0.000417 | 0.00108 |
| chrX | 13503149 | 13507949 | 4801 | 0.68 | 0.000418 | 0.00108 |
| chr12 | 108876004 | 108877248 | 1245 | 1.14 | 0.000418 | 0.00108 |
| chr2 | 57623362 | 57624346 | 985 | -2.05 | 0.000418 | 0.00108 |

| | | | | | | |
|---|---|---|---|---|---|---|
| chr1 | 44286808 | 44288858 | 2051 | 0.69 | 0.000418 | 0.00108 |
| chr9 | 87667573 | 87668730 | 1158 | 1.31 | 0.000418 | 0.00108 |
| chr19 | 5773357 | 5773828 | 472 | -4.51 | 0.000418 | 0.00108 |
| chr12 | 127630174 | 127631474 | 1301 | 0.82 | 0.000418 | 0.00108 |
| chr7 | 41146826 | 41148149 | 1324 | 1.35 | 0.000419 | 0.00108 |
| chr7 | 17408515 | 17412638 | 4124 | 0.34 | 0.00042 | 0.00108 |
| chr16 | 67260133 | 67283098 | 22966 | -0.27 | 0.00042 | 0.00108 |
| chr6 | 12449985 | 12496774 | 46790 | -0.21 | 0.00042 | 0.00109 |
| chr2 | 151496039 | 151501583 | 5545 | 1.64 | 0.00042 | 0.00109 |
| chr2 | 236199561 | 236200896 | 1336 | 1.91 | 0.00042 | 0.00109 |
| chr4 | 57013375 | 57014305 | 931 | -1.67 | 0.000421 | 0.00109 |
| chrX | 153192295 | 153200405 | 8111 | 0.5 | 0.000421 | 0.00109 |
| chr17 | 29814341 | 29815836 | 1496 | 1.34 | 0.000422 | 0.00109 |
| chrX | 114606184 | 114607893 | 1710 | 0.71 | 0.000423 | 0.00109 |
| chr2 | 69894800 | 69923692 | 28893 | 0.54 | 0.000423 | 0.00109 |
| chr4 | 160169445 | 160170667 | 1223 | 2.17 | 0.000423 | 0.00109 |
| chr2 | 236352896 | 236353834 | 939 | 1.77 | 0.000424 | 0.00109 |
| chr6 | 168789230 | 168789946 | 717 | 2.51 | 0.000424 | 0.00109 |
| chr2 | 111420034 | 111436733 | 16700 | -0.42 | 0.000424 | 0.00109 |
| chr11 | 17002826 | 17009934 | 7109 | 0.73 | 0.000424 | 0.00109 |
| chr15 | 100889886 | 100892270 | 2385 | -0.46 | 0.000424 | 0.00109 |
| chr17 | 39108531 | 39114995 | 6465 | 0.63 | 0.000424 | 0.0011 |
| chr18 | 60188907 | 60195506 | 6600 | 0.51 | 0.000425 | 0.0011 |
| chr10 | 54230606 | 54249281 | 18676 | -0.39 | 0.000425 | 0.0011 |
| chr7 | 107382815 | 107387681 | 4867 | 0.41 | 0.000426 | 0.0011 |
| chr10 | 90749869 | 90752200 | 2332 | 0.75 | 0.000428 | 0.0011 |
| chr8 | 72946844 | 72948375 | 1532 | -0.85 | 0.000428 | 0.0011 |
| chr15 | 81134786 | 81135332 | 547 | 3.36 | 0.000428 | 0.00111 |
| chr17 | 77712792 | 77752855 | 40064 | -0.41 | 0.000429 | 0.00111 |
| chr6 | 27759214 | 27808433 | 49220 | -0.2 | 0.000429 | 0.00111 |
| chr1 | 33446035 | 33463972 | 17938 | 0.31 | 0.00043 | 0.00111 |
| chr2 | 106069798 | 106073304 | 3507 | 0.36 | 0.000431 | 0.00111 |
| chr1 | 36102420 | 36108682 | 6263 | 0.41 | 0.000431 | 0.00111 |
| chr15 | 28378057 | 28379096 | 1040 | 2.41 | 0.000431 | 0.00111 |
| chr22 | 28955908 | 28956885 | 978 | -3.19 | 0.000431 | 0.00111 |
| chrX | 141046249 | 141047198 | 950 | 2.21 | 0.000432 | 0.00111 |
| chr2 | 42812470 | 42815402 | 2933 | 1.24 | 0.000433 | 0.00112 |
| chr9 | 19048116 | 19051428 | 3313 | 0.58 | 0.000433 | 0.00112 |
| chr12 | 83360542 | 83361051 | 510 | 3 | 0.000433 | 0.00112 |
| chr12 | 45342211 | 45342560 | 350 | 3.13 | 0.000434 | 0.00112 |
| chr1 | 173683443 | 173685915 | 2473 | 0.63 | 0.000434 | 0.00112 |
| chr13 | 105223615 | 105223965 | 351 | 4.56 | 0.000435 | 0.00112 |
| chr17 | 7567912 | 7568286 | 375 | 4.56 | 0.000435 | 0.00112 |
| chr20 | 46550719 | 46551767 | 1049 | 2.54 | 0.000435 | 0.00112 |
| chr21 | 16421810 | 16439105 | 17296 | 0.22 | 0.000436 | 0.00112 |
| chr9 | 98030851 | 98032845 | 1995 | 1.75 | 0.000436 | 0.00112 |
| chr11 | 40466640 | 40467053 | 414 | 4.53 | 0.000437 | 0.00112 |
| chr18 | 61517637 | 61517986 | 350 | 4.53 | 0.000437 | 0.00112 |
| chr13 | 34252832 | 34254697 | 1866 | -0.65 | 0.000437 | 0.00113 |
| chr13 | 37571835 | 37576717 | 4883 | 0.39 | 0.000437 | 0.00113 |
| chr7 | 106958492 | 106959716 | 1225 | 1.3 | 0.000437 | 0.00113 |
| chr4 | 106780095 | 106781049 | 955 | 2.77 | 0.000438 | 0.00113 |
| chr5 | 124612521 | 124614066 | 1546 | 1.41 | 0.000438 | 0.00113 |
| chr6 | 148851434 | 148852433 | 1000 | -1.32 | 0.000439 | 0.00113 |
| chr22 | 40654122 | 40655106 | 985 | 1.81 | 0.000439 | 0.00113 |
| chrX | 128622152 | 128623061 | 910 | -2.9 | 0.000439 | 0.00113 |
| chr2 | 111877655 | 111880775 | 3121 | -1.03 | 0.000439 | 0.00113 |
| chr12 | 48337235 | 48338028 | 794 | 3.67 | 0.000441 | 0.00113 |
| chr2 | 55535905 | 55537629 | 1725 | -1.59 | 0.000441 | 0.00114 |
| chr17 | 62221730 | 62228398 | 6669 | 0.39 | 0.000441 | 0.00114 |
| chr2 | 15689993 | 15702453 | 12461 | -0.5 | 0.000443 | 0.00114 |

| | | | | | | |
|---|---|---|---|---|---|---|
| chr1 | 161044250 | 161053984 | 9735 | -0.96 | 0.000443 | 0.00114 |
| chr10 | 60796008 | 60796746 | 739 | 3.02 | 0.000444 | 0.00114 |
| chr8 | 110343800 | 110347901 | 4102 | 0.41 | 0.000444 | 0.00114 |
| chr22 | 29304965 | 29314302 | 9338 | -0.6 | 0.000444 | 0.00114 |
| chr1 | 68024872 | 68037549 | 12678 | -1.17 | 0.000445 | 0.00114 |
| chr3 | 20065073 | 20065919 | 847 | 1.86 | 0.000445 | 0.00115 |
| chr17 | 9827650 | 9832752 | 5103 | -1.51 | 0.000446 | 0.00115 |
| chr6 | 150489243 | 150490313 | 1071 | 1.14 | 0.000446 | 0.00115 |
| chr7 | 131909371 | 131910609 | 1239 | -1.63 | 0.000447 | 0.00115 |
| chr6 | 76232465 | 76235996 | 3532 | -0.71 | 0.000448 | 0.00115 |
| chr15 | 78348372 | 78375325 | 26954 | -0.49 | 0.000449 | 0.00115 |
| chr9 | 18460238 | 18460553 | 316 | 4.53 | 0.000449 | 0.00116 |
| chr1 | 100787810 | 100789529 | 1720 | 0.98 | 0.00045 | 0.00116 |
| chr7 | 140290739 | 140291599 | 861 | 2.04 | 0.00045 | 0.00116 |
| chr5 | 32686852 | 32688115 | 1264 | 1.4 | 0.000451 | 0.00116 |
| chr12 | 77024431 | 77025149 | 719 | 2.57 | 0.000451 | 0.00116 |
| chr5 | 130497583 | 130507554 | 9972 | 0.37 | 0.000452 | 0.00116 |
| chr7 | 66459002 | 66460344 | 1343 | 1.19 | 0.000453 | 0.00116 |
| chr5 | 134238853 | 134247730 | 8878 | 0.35 | 0.000454 | 0.00117 |
| chr8 | 124529134 | 124561344 | 32211 | 0.61 | 0.000454 | 0.00117 |
| chr9 | 71073559 | 71074707 | 1149 | 1.83 | 0.000454 | 0.00117 |
| chr8 | 56813329 | 56815051 | 1723 | -0.83 | 0.000455 | 0.00117 |
| chr17 | 39002983 | 39016542 | 13560 | 0.29 | 0.000455 | 0.00117 |
| chr10 | 96696253 | 96696938 | 686 | 2.36 | 0.000455 | 0.00117 |
| chr13 | 114102660 | 114104800 | 2141 | 0.62 | 0.000456 | 0.00117 |
| chr10 | 105725792 | 105743819 | 18028 | 0.36 | 0.000456 | 0.00117 |
| chr3 | 63895883 | 63904450 | 8568 | 0.48 | 0.000456 | 0.00117 |
| chr1 | 198124721 | 198129691 | 4971 | 0.49 | 0.000457 | 0.00117 |
| chr22 | 42335189 | 42343722 | 8534 | 0.54 | 0.000457 | 0.00117 |
| chr20 | 10713817 | 10714309 | 493 | -2.96 | 0.000458 | 0.00118 |
| chr20 | 22979783 | 22994943 | 15161 | 0.4 | 0.000458 | 0.00118 |
| chr20 | 55622048 | 55623424 | 1377 | -1.04 | 0.000459 | 0.00118 |
| chr11 | 66138267 | 66140677 | 2411 | 0.7 | 0.000459 | 0.00118 |
| chr3 | 9164074 | 9176587 | 12514 | 2.36 | 0.000461 | 0.00118 |
| chr9 | 136374404 | 136375234 | 831 | 2.88 | 0.000461 | 0.00118 |
| chr14 | 65597584 | 65598878 | 1295 | 2.2 | 0.000461 | 0.00118 |
| chr11 | 102421488 | 102447612 | 26125 | 0.3 | 0.000462 | 0.00118 |
| chr3 | 11812164 | 11812966 | 803 | 1.93 | 0.000462 | 0.00119 |
| chr5 | 95033722 | 95034683 | 962 | 1.68 | 0.000462 | 0.00119 |
| chr8 | 33073206 | 33074309 | 1104 | -2 | 0.000464 | 0.00119 |
| chr2 | 81885746 | 81886158 | 413 | 4.53 | 0.000464 | 0.00119 |
| chr5 | 27266417 | 27266789 | 373 | 4.54 | 0.000465 | 0.00119 |
| chr3 | 57763697 | 57791826 | 28130 | -0.35 | 0.000466 | 0.00119 |
| chr20 | 24472585 | 24473033 | 449 | -4.48 | 0.000466 | 0.00119 |
| chr2 | 190494220 | 190494621 | 402 | -3.44 | 0.000467 | 0.0012 |
| chr17 | 34127059 | 34185433 | 58375 | 0.23 | 0.000468 | 0.0012 |
| chr11 | 61061484 | 61062732 | 1249 | -1.97 | 0.000469 | 0.0012 |
| chr21 | 17551798 | 17569614 | 17817 | 0.26 | 0.000469 | 0.0012 |
| chr17 | 40398545 | 40440979 | 42435 | -0.37 | 0.00047 | 0.0012 |
| chr11 | 69450687 | 69471062 | 20376 | 0.26 | 0.000471 | 0.00121 |
| chr19 | 53836235 | 53837826 | 1592 | 1.19 | 0.000471 | 0.00121 |
| chr6 | 3118012 | 3140192 | 22181 | 0.65 | 0.000472 | 0.00121 |
| chr6 | 22928986 | 22929646 | 661 | 2.85 | 0.000473 | 0.00121 |
| chr17 | 69334522 | 69335194 | 673 | 2.13 | 0.000473 | 0.00121 |
| chr2 | 63244957 | 63246468 | 1512 | -1.44 | 0.000473 | 0.00121 |
| chr12 | 40306228 | 40306777 | 550 | 2.19 | 0.000473 | 0.00121 |
| chr1 | 113652223 | 113653646 | 1424 | 2.23 | 0.000473 | 0.00121 |
| chr5 | 95898582 | 95899184 | 603 | 3.25 | 0.000474 | 0.00121 |
| chrX | 53621372 | 53623018 | 1647 | 1.48 | 0.000476 | 0.00122 |
| chr9 | 71993845 | 71995227 | 1383 | 4.29 | 0.000476 | 0.00122 |
| chr15 | 83834636 | 83835095 | 460 | 3.11 | 0.000477 | 0.00122 |

| | | | | | | |
|---|---|---|---|---|---|---|
| chr10 | 14778325 | 14791986 | 13662 | 0.52 | 0.000477 | 0.00122 |
| chr9 | 4849434 | 4854602 | 5169 | -1.04 | 0.000478 | 0.00122 |
| chr7 | 18977386 | 18978933 | 1548 | 1.12 | 0.000479 | 0.00122 |
| chr7 | 23552865 | 23555221 | 2357 | 1.24 | 0.000479 | 0.00122 |
| chr5 | 861763 | 862618 | 856 | -2.89 | 0.000479 | 0.00123 |
| chr6 | 145214503 | 145219715 | 5213 | 1.25 | 0.000479 | 0.00123 |
| chr15 | 31901291 | 31901862 | 572 | -3.06 | 0.00048 | 0.00123 |
| chr5 | 61630718 | 61631183 | 466 | -3.3 | 0.00048 | 0.00123 |
| chr14 | 91111695 | 91113214 | 1520 | 1.35 | 0.00048 | 0.00123 |
| chr9 | 93087904 | 93107239 | 19336 | -0.65 | 0.00048 | 0.00123 |
| chr3 | 185958719 | 185961281 | 2563 | -0.53 | 0.000481 | 0.00123 |
| chr5 | 14943444 | 14960874 | 17431 | 0.64 | 0.000481 | 0.00123 |
| chr15 | 98418232 | 98419185 | 954 | 1.7 | 0.000481 | 0.00123 |
| chr2 | 129199008 | 129200938 | 1931 | 1.81 | 0.000482 | 0.00123 |
| chrX | 107727798 | 107729285 | 1488 | -1.99 | 0.000482 | 0.00123 |
| chr17 | 8367261 | 8375793 | 8533 | -0.75 | 0.000482 | 0.00123 |
| chr15 | 57317674 | 57318784 | 1111 | -1.86 | 0.000482 | 0.00123 |
| chr8 | 81397175 | 81400212 | 3038 | 0.68 | 0.000482 | 0.00123 |
| chr1 | 196392590 | 196393323 | 734 | 3.09 | 0.000484 | 0.00124 |
| chr1 | 185758856 | 185759842 | 987 | 2.94 | 0.000486 | 0.00124 |
| chr11 | 75807128 | 75808463 | 1336 | -1.87 | 0.000486 | 0.00124 |
| chr15 | 68691238 | 68697001 | 5764 | 0.35 | 0.000487 | 0.00124 |
| chr7 | 115898694 | 115929223 | 30530 | 0.23 | 0.000487 | 0.00124 |
| chr5 | 41788548 | 41795489 | 6942 | 0.67 | 0.000488 | 0.00125 |
| chr12 | 26306581 | 26307065 | 485 | 2.67 | 0.000488 | 0.00125 |
| chr6 | 35655213 | 35657452 | 2240 | 1.28 | 0.000489 | 0.00125 |
| chr19 | 4909356 | 4938867 | 29512 | 0.35 | 0.00049 | 0.00125 |
| chr12 | 122062750 | 122080120 | 17371 | 0.31 | 0.00049 | 0.00125 |
| chrX | 40011836 | 40036700 | 24865 | -0.4 | 0.00049 | 0.00125 |
| chr6 | 135027456 | 135028574 | 1119 | 2.51 | 0.000491 | 0.00125 |
| chr20 | 10685819 | 10698991 | 13173 | -0.24 | 0.000491 | 0.00125 |
| chr1 | 220905761 | 220922695 | 16935 | 0.47 | 0.000492 | 0.00126 |
| chr4 | 26788964 | 26790626 | 1663 | 1.03 | 0.000493 | 0.00126 |
| chr6 | 109169143 | 109170613 | 1471 | -0.93 | 0.000493 | 0.00126 |
| chr5 | 143229551 | 143230364 | 814 | 1.84 | 0.000494 | 0.00126 |
| chr16 | 84113896 | 84115978 | 2083 | -1.39 | 0.000495 | 0.00126 |
| chr11 | 35384990 | 35395930 | 10941 | 0.83 | 0.000495 | 0.00126 |
| chr16 | 47697797 | 47699486 | 1690 | 1.11 | 0.000496 | 0.00126 |
| chr12 | 111354930 | 111356143 | 1214 | 1.14 | 0.000496 | 0.00126 |
| chr15 | 59104535 | 59105223 | 689 | -1.98 | 0.000496 | 0.00127 |
| chr13 | 110317957 | 110318601 | 645 | -3.32 | 0.000496 | 0.00127 |
| chr8 | 9304995 | 9307009 | 2015 | 0.9 | 0.000497 | 0.00127 |
| chr9 | 80781981 | 80782327 | 347 | 4.59 | 0.000498 | 0.00127 |
| chr11 | 62408463 | 62458329 | 49867 | 0.23 | 0.000499 | 0.00127 |
| chr4 | 66864800 | 66867273 | 2474 | 0.65 | 0.000499 | 0.00127 |
| chr4 | 53578584 | 53589239 | 10656 | 0.59 | 0.000499 | 0.00127 |
| chr4 | 170360134 | 170368676 | 8543 | 1.38 | 5.00E-04 | 0.00127 |
| chr3 | 45505574 | 45507050 | 1477 | -1.7 | 0.000501 | 0.00128 |
| chr2 | 155097771 | 155098272 | 502 | -3.44 | 0.000501 | 0.00128 |
| chr2 | 145273547 | 145275606 | 2060 | -0.98 | 0.000501 | 0.00128 |
| chr18 | 54876933 | 54891271 | 14339 | 0.75 | 0.000502 | 0.00128 |
| chr14 | 97342074 | 97343925 | 1852 | 1.52 | 0.000502 | 0.00128 |
| chr7 | 134000541 | 134014912 | 14372 | 0.92 | 0.000502 | 0.00128 |
| chr19 | 57259436 | 57259891 | 456 | 4.5 | 0.000503 | 0.00128 |
| chr3 | 148285969 | 148287073 | 1105 | 3.02 | 0.000505 | 0.00129 |
| chr10 | 75756484 | 75883605 | 127122 | 0.16 | 0.000506 | 0.00129 |
| chr4 | 176691365 | 176692970 | 1606 | 0.91 | 0.000507 | 0.00129 |
| chr1 | 40009911 | 40010758 | 848 | 1.85 | 0.000509 | 0.0013 |
| chr18 | 20980891 | 20987622 | 6732 | 0.72 | 0.000509 | 0.0013 |
| chr7 | 87640766 | 87641206 | 441 | 4.5 | 0.00051 | 0.0013 |
| chr17 | 46755627 | 46756774 | 1148 | -2.33 | 0.00051 | 0.0013 |

| | | | | | | |
|---|---|---|---|---|---|---|
| chr12 | 19525474 | 19536097 | 10624 | -0.64 | 0.00051 | 0.0013 |
| chr12 | 93115741 | 93116120 | 380 | 4.5 | 0.000511 | 0.0013 |
| chr14 | 26799826 | 26800281 | 456 | 4.5 | 0.000511 | 0.0013 |
| chr2 | 208781165 | 208782045 | 881 | -2.13 | 0.000512 | 0.0013 |
| chr1 | 70780134 | 70787453 | 7320 | 0.68 | 0.000512 | 0.0013 |
| chr11 | 9878473 | 9890690 | 12218 | -0.39 | 0.000512 | 0.0013 |
| chr1 | 218382411 | 218383789 | 1379 | -1.44 | 0.000512 | 0.0013 |
| chr17 | 17802738 | 17827527 | 24790 | -0.24 | 0.000512 | 0.0013 |
| chr1 | 214432746 | 214433285 | 540 | -3.03 | 0.000513 | 0.0013 |
| chr6 | 35359254 | 35359867 | 614 | -3.03 | 0.000513 | 0.0013 |
| chr12 | 132432104 | 132442619 | 10516 | 0.47 | 0.000513 | 0.0013 |
| chr3 | 123966987 | 123982832 | 15846 | -1.24 | 0.000514 | 0.00131 |
| chr11 | 82744316 | 82785249 | 40934 | 0.31 | 0.000515 | 0.00131 |
| chr10 | 129788456 | 129790520 | 2065 | 1.07 | 0.000515 | 0.00131 |
| chr1 | 10821056 | 10822395 | 1340 | 1.48 | 0.000516 | 0.00131 |
| chr8 | 98305771 | 98306829 | 1059 | -1.81 | 0.000516 | 0.00131 |
| chr1 | 28261198 | 28262066 | 869 | 1.6 | 0.000517 | 0.00132 |
| chr6 | 17932404 | 17933447 | 1044 | 1.72 | 0.000517 | 0.00132 |
| chr19 | 15468582 | 15471446 | 2865 | -0.77 | 0.000518 | 0.00132 |
| chr1 | 247170736 | 247172113 | 1378 | 1.8 | 0.000518 | 0.00132 |
| chr11 | 113072601 | 113073678 | 1078 | -2.96 | 0.000519 | 0.00132 |
| chr1 | 42629978 | 42632183 | 2206 | 1.33 | 0.000519 | 0.00132 |
| chr5 | 149674221 | 149682751 | 8531 | -1.24 | 0.00052 | 0.00132 |
| chr18 | 32508117 | 32528520 | 20404 | -0.43 | 0.000521 | 0.00132 |
| chr22 | 33888337 | 33888928 | 592 | -2.45 | 0.000522 | 0.00132 |
| chr3 | 189207539 | 189209682 | 2144 | 1.04 | 0.000522 | 0.00132 |
| chr11 | 11878959 | 11879332 | 374 | 3.36 | 0.000522 | 0.00133 |
| chr8 | 133519461 | 133520608 | 1148 | 1.95 | 0.000523 | 0.00133 |
| chr17 | 67738535 | 67739949 | 1415 | -1.73 | 0.000523 | 0.00133 |
| chr4 | 31045155 | 31045685 | 531 | 3.69 | 0.000525 | 0.00133 |
| chr10 | 127383100 | 127409533 | 26434 | 0.66 | 0.000525 | 0.00133 |
| chr6 | 14499517 | 14500478 | 962 | -1.89 | 0.000526 | 0.00133 |
| chr8 | 49700690 | 49702100 | 1411 | 1.03 | 0.000527 | 0.00134 |
| chr10 | 33145351 | 33146081 | 731 | -2.08 | 0.000527 | 0.00134 |
| chr18 | 12543407 | 12544156 | 750 | 2.31 | 0.000527 | 0.00134 |
| chr8 | 116088679 | 116089062 | 384 | 2.46 | 0.000527 | 0.00134 |
| chr21 | 16511876 | 16515002 | 3127 | 0.64 | 0.000528 | 0.00134 |
| chr14 | 32395702 | 32439551 | 43850 | -0.19 | 0.000528 | 0.00134 |
| chr3 | 33237668 | 33238089 | 422 | 3.23 | 0.000528 | 0.00134 |
| chr2 | 31605124 | 31609701 | 4578 | 0.35 | 0.000529 | 0.00134 |
| chr5 | 56110184 | 56113250 | 3067 | 0.71 | 0.000529 | 0.00134 |
| chr1 | 182830306 | 182839204 | 8899 | 0.62 | 0.00053 | 0.00134 |
| chr6 | 116707642 | 116719924 | 12283 | 1.4 | 0.00053 | 0.00134 |
| chr4 | 110650446 | 110651990 | 1545 | 1.19 | 0.000531 | 0.00135 |
| chr18 | 52403702 | 52418326 | 14625 | 0.61 | 0.000532 | 0.00135 |
| chr8 | 140887346 | 140892300 | 4955 | -1.79 | 0.000533 | 0.00135 |
| chr2 | 136426011 | 136426605 | 595 | -3.9 | 0.000535 | 0.00136 |
| chr3 | 9610763 | 9611200 | 438 | -4.46 | 0.000535 | 0.00136 |
| chr6 | 84131146 | 84144918 | 13773 | 0.36 | 0.000536 | 0.00136 |
| chr12 | 40448128 | 40449406 | 1279 | 1.08 | 0.000539 | 0.00137 |
| chr8 | 53886209 | 53886900 | 692 | 1.75 | 0.000539 | 0.00137 |
| chr10 | 116063943 | 116066733 | 2791 | -1.78 | 0.000539 | 0.00137 |
| chr1 | 55259872 | 55262714 | 2843 | 1.92 | 0.000541 | 0.00137 |
| chr8 | 65669183 | 65669946 | 764 | -2.24 | 0.000542 | 0.00137 |
| chr3 | 138312240 | 138314210 | 1971 | 0.66 | 0.000543 | 0.00138 |
| chr12 | 91299551 | 91300018 | 468 | -2.77 | 0.000543 | 0.00138 |
| chr14 | 102947604 | 102949041 | 1438 | 1.35 | 0.000544 | 0.00138 |
| chr18 | 48571156 | 48580542 | 9387 | 0.95 | 0.000545 | 0.00138 |
| chr3 | 181951707 | 181952442 | 736 | 2.6 | 0.000545 | 0.00138 |
| chr19 | 35635632 | 35654464 | 18833 | -0.71 | 0.000546 | 0.00138 |
| chr7 | 20146803 | 20148056 | 1254 | 1.24 | 0.000546 | 0.00138 |

| | | | | | | |
|---|---|---|---|---|---|---|
| chr11 | 113915014 | 113915445 | 432 | 4.49 | 0.000546 | 0.00138 |
| chrX | 118532489 | 118562107 | 29619 | -0.33 | 0.000546 | 0.00138 |
| chr1 | 160509820 | 160513298 | 3479 | 0.72 | 0.000547 | 0.00139 |
| chr20 | 9165594 | 9187396 | 21803 | 0.32 | 0.000548 | 0.00139 |
| chr1 | 52650360 | 52651397 | 1038 | 1.58 | 0.000548 | 0.00139 |
| chr12 | 34757339 | 34757878 | 540 | 4.5 | 0.000548 | 0.00139 |
| chr2 | 57181908 | 57182205 | 298 | 4.5 | 0.000548 | 0.00139 |
| chr14 | 71163580 | 71166021 | 2442 | 0.72 | 0.000549 | 0.00139 |
| chr1 | 247266581 | 247275247 | 8667 | 0.84 | 0.00055 | 0.00139 |
| chr2 | 29328174 | 29396571 | 68398 | -0.13 | 0.000551 | 0.00139 |
| chr1 | 88793050 | 88793956 | 907 | 2.77 | 0.000551 | 0.00139 |
| chr15 | 56497083 | 56498086 | 1004 | -1.31 | 0.000553 | 0.0014 |
| chr1 | 52194621 | 52215367 | 20747 | 0.39 | 0.000553 | 0.0014 |
| chr15 | 81003737 | 81004981 | 1245 | 1.84 | 0.000553 | 0.0014 |
| chr18 | 21811323 | 21812609 | 1287 | 1.36 | 0.000554 | 0.0014 |
| chr3 | 152545651 | 152552987 | 7337 | 4.52 | 0.000555 | 0.0014 |
| chr15 | 77097171 | 77098334 | 1164 | -1.36 | 0.000555 | 0.0014 |
| chr17 | 45192918 | 45201453 | 8536 | 0.69 | 0.000555 | 0.0014 |
| chr3 | 15779061 | 15858679 | 79619 | -0.17 | 0.000556 | 0.0014 |
| chr8 | 92051572 | 92054155 | 2584 | 0.72 | 0.000557 | 0.00141 |
| chr5 | 81047218 | 81065911 | 18694 | 0.52 | 0.000557 | 0.00141 |
| chr9 | 86940340 | 86941045 | 706 | 2.96 | 0.000557 | 0.00141 |
| chr3 | 42844373 | 42846585 | 2213 | 0.73 | 0.000558 | 0.00141 |
| chr7 | 50879786 | 50906410 | 26625 | -0.59 | 0.000559 | 0.00141 |
| chr2 | 38573678 | 38605261 | 31584 | -0.39 | 0.000559 | 0.00141 |
| chr3 | 189825800 | 189839574 | 13775 | -0.52 | 0.000559 | 0.00141 |
| chr4 | 1856489 | 1898360 | 41872 | 0.34 | 0.000559 | 0.00141 |
| chr21 | 45611968 | 45614178 | 2211 | 1.4 | 0.000559 | 0.00141 |
| chr3 | 28060409 | 28065088 | 4680 | 0.74 | 0.00056 | 0.00141 |
| chr18 | 65026821 | 65028820 | 2000 | 0.76 | 0.000561 | 0.00142 |
| chr1 | 236552017 | 236564131 | 12115 | -0.48 | 0.000561 | 0.00142 |
| chr1 | 235871711 | 235872757 | 1047 | -1.78 | 0.000561 | 0.00142 |
| chr7 | 65303294 | 65306470 | 3177 | -1.12 | 0.000561 | 0.00142 |
| chr19 | 7933753 | 7936930 | 3178 | 1.31 | 0.000563 | 0.00142 |
| chr3 | 113751902 | 113752999 | 1098 | -1.92 | 0.000563 | 0.00142 |
| chr5 | 41438621 | 41439542 | 922 | 2.49 | 0.000563 | 0.00142 |
| chr2 | 207448998 | 207449552 | 555 | -2.47 | 0.000563 | 0.00142 |
| chr21 | 17958473 | 17961380 | 2908 | 0.51 | 0.000563 | 0.00142 |
| chr15 | 100167853 | 100169514 | 1662 | 0.71 | 0.000563 | 0.00142 |
| chr4 | 143182231 | 143187632 | 5402 | 1 | 0.000563 | 0.00142 |
| chr3 | 14987757 | 14991628 | 3872 | 0.4 | 0.000563 | 0.00142 |
| chr16 | 78495161 | 78495925 | 765 | 2.5 | 0.000564 | 0.00142 |
| chr2 | 225666911 | 225684899 | 17989 | -0.51 | 0.000565 | 0.00143 |
| chr9 | 140352528 | 140354978 | 2451 | -0.83 | 0.000565 | 0.00143 |
| chrX | 135577907 | 135581215 | 3309 | 0.46 | 0.000566 | 0.00143 |
| chr10 | 23726994 | 23729611 | 2618 | 0.76 | 0.000567 | 0.00143 |
| chr16 | 56389936 | 56391634 | 1699 | -1.11 | 0.000567 | 0.00143 |
| chr2 | 170567983 | 170574388 | 6406 | -1.39 | 0.000567 | 0.00143 |
| chr6 | 3438200 | 3439708 | 1509 | -0.86 | 0.000568 | 0.00143 |
| chr1 | 235665901 | 235670038 | 4138 | 0.41 | 0.000568 | 0.00143 |
| chr10 | 104613553 | 104629843 | 16291 | -0.51 | 0.000569 | 0.00143 |
| chr16 | 1569341 | 1570181 | 841 | -2.53 | 0.000569 | 0.00143 |
| chr17 | 72406165 | 72451122 | 44958 | 0.29 | 0.00057 | 0.00144 |
| chr8 | 8274341 | 8287596 | 13256 | 0.39 | 0.000571 | 0.00144 |
| chr10 | 90106144 | 90106944 | 801 | 2.98 | 0.000571 | 0.00144 |
| chr15 | 31213873 | 31254489 | 40617 | 0.25 | 0.000571 | 0.00144 |
| chr11 | 117615080 | 117615619 | 540 | -3.27 | 0.000571 | 0.00144 |
| chr2 | 238630131 | 238666536 | 36406 | -0.36 | 0.000571 | 0.00144 |
| chr15 | 49821878 | 49823158 | 1281 | 1.63 | 0.000571 | 0.00144 |
| chr1 | 202894846 | 202897196 | 2351 | 0.57 | 0.000572 | 0.00144 |
| chr11 | 88406037 | 88406723 | 687 | 2.1 | 0.000572 | 0.00144 |

| | | | | | | |
|---|---|---|---|---|---|---|
| chr22 | 45969414 | 46044957 | 75544 | 0.16 | 0.000573 | 0.00144 |
| chr9 | 84946325 | 84948287 | 1963 | 0.84 | 0.000573 | 0.00144 |
| chr11 | 7025434 | 7026149 | 716 | 2.21 | 0.000573 | 0.00144 |
| chr4 | 129672247 | 129673192 | 946 | -1.68 | 0.000573 | 0.00144 |
| chr3 | 196842948 | 196845270 | 2323 | -0.87 | 0.000574 | 0.00145 |
| chr3 | 99721271 | 99722367 | 1097 | 1.99 | 0.000575 | 0.00145 |
| chr15 | 57064947 | 57066606 | 1660 | 4.36 | 0.000575 | 0.00145 |
| chr3 | 20222319 | 20228787 | 6469 | 0.56 | 0.000575 | 0.00145 |
| chr18 | 43771922 | 43781712 | 9791 | -0.67 | 0.000576 | 0.00145 |
| chr10 | 30002900 | 30004232 | 1333 | -1.62 | 0.000577 | 0.00145 |
| chr16 | 50840333 | 50840583 | 251 | 4.46 | 0.000578 | 0.00145 |
| chr4 | 96535385 | 96535840 | 456 | 4.46 | 0.000578 | 0.00145 |
| chr5 | 86562923 | 86565513 | 2591 | 0.55 | 0.000578 | 0.00145 |
| chr18 | 11120302 | 11121492 | 1191 | -2.1 | 0.000579 | 0.00146 |
| chr9 | 38671822 | 38690839 | 19018 | 0.35 | 0.00058 | 0.00146 |
| chr2 | 113541732 | 113542480 | 749 | 3.15 | 0.00058 | 0.00146 |
| chr1 | 33592011 | 33593957 | 1947 | 0.77 | 0.00058 | 0.00146 |
| chr1 | 111991080 | 111993323 | 2244 | 0.88 | 0.000581 | 0.00146 |
| chr10 | 121484659 | 121498792 | 14134 | 0.62 | 0.000582 | 0.00146 |
| chr1 | 190071132 | 190073135 | 2004 | 4.71 | 0.000583 | 0.00147 |
| chr3 | 20043441 | 20043848 | 408 | -3.2 | 0.000583 | 0.00147 |
| chr20 | 32526536 | 32527887 | 1352 | 2.55 | 0.000583 | 0.00147 |
| chr3 | 122988073 | 122992659 | 4587 | 0.71 | 0.000584 | 0.00147 |
| chr9 | 133607429 | 133609377 | 1949 | 0.88 | 0.000584 | 0.00147 |
| chr16 | 19650932 | 19651311 | 380 | -4.42 | 0.000585 | 0.00147 |
| chr1 | 211532975 | 211534325 | 1351 | -1.79 | 0.000586 | 0.00147 |
| chr15 | 65423337 | 65426964 | 3628 | 0.54 | 0.000586 | 0.00147 |
| chr5 | 57104724 | 57105575 | 852 | 2.75 | 0.000586 | 0.00147 |
| chr7 | 105305054 | 105318895 | 13842 | -0.86 | 0.000587 | 0.00147 |
| chr19 | 16389691 | 16405742 | 16052 | 0.38 | 0.000587 | 0.00147 |
| chr1 | 201276475 | 201281345 | 4871 | 1.4 | 0.000588 | 0.00148 |
| chr5 | 106694897 | 106955801 | 260905 | 0.09 | 0.000588 | 0.00148 |
| chr15 | 74066701 | 74067926 | 1226 | -1.67 | 0.000588 | 0.00148 |
| chr14 | 24680793 | 24713252 | 32460 | 0.23 | 0.00059 | 0.00148 |
| chr11 | 3116620 | 3117662 | 1043 | 2.79 | 0.000591 | 0.00149 |
| chr1 | 165706252 | 165718406 | 12155 | -0.7 | 0.000592 | 0.00149 |
| chr3 | 145995942 | 145996271 | 330 | 3.68 | 0.000592 | 0.00149 |
| chr11 | 36682827 | 36683321 | 495 | -2.7 | 0.000593 | 0.00149 |
| chr6 | 710697 | 711634 | 938 | -2.22 | 0.000593 | 0.00149 |
| chr10 | 93847769 | 93848261 | 493 | -3.89 | 0.000594 | 0.00149 |
| chr1 | 28928018 | 28929441 | 1424 | -1.7 | 0.000595 | 0.00149 |
| chr3 | 148360033 | 148366190 | 6158 | -0.43 | 0.000595 | 0.00149 |
| chr8 | 104310302 | 104312074 | 1773 | 0.63 | 0.000595 | 0.00149 |
| chr4 | 130249657 | 130250057 | 401 | 4.46 | 0.000596 | 0.00149 |
| chr6 | 63517411 | 63519196 | 1786 | 4.62 | 0.000596 | 0.0015 |
| chr6 | 8939349 | 8940407 | 1059 | 1.5 | 0.000596 | 0.0015 |
| chr12 | 105037093 | 105038673 | 1581 | -1.74 | 0.000596 | 0.0015 |
| chr12 | 97113525 | 97114645 | 1121 | 1.27 | 0.000598 | 0.0015 |
| chr2 | 220209208 | 220209936 | 729 | 2.02 | 0.000598 | 0.0015 |
| chr14 | 60646036 | 60651203 | 5168 | 4.31 | 0.000598 | 0.0015 |
| chr3 | 105656870 | 105657598 | 729 | 2.96 | 0.000599 | 0.0015 |
| chr8 | 61563278 | 61565449 | 2172 | 0.48 | 0.000599 | 0.0015 |
| chr7 | 155435779 | 155459524 | 23746 | 0.36 | 6.00E-04 | 0.0015 |
| chr16 | 27528157 | 27534507 | 6351 | -0.84 | 6.00E-04 | 0.0015 |
| chr3 | 44036050 | 44074348 | 38299 | -0.32 | 0.000601 | 0.00151 |
| chr1 | 155409132 | 155412753 | 3622 | -1.02 | 0.000601 | 0.00151 |
| chr17 | 39289686 | 39297653 | 7968 | 1.46 | 0.000603 | 0.00151 |
| chr19 | 1542185 | 1542688 | 504 | 2.35 | 0.000603 | 0.00151 |
| chr3 | 45988526 | 45989586 | 1061 | -1.99 | 0.000603 | 0.00151 |
| chr12 | 121471445 | 121477329 | 5885 | 1.16 | 0.000603 | 0.00151 |
| chr2 | 15441564 | 15442697 | 1134 | -1.81 | 0.000604 | 0.00151 |

| | | | | | | |
|---|---|---|---|---|---|---|
| chr9 | 139668399 | 139668870 | 472 | 4.58 | 0.000606 | 0.00152 |
| chr22 | 20939657 | 20940633 | 977 | -2.02 | 0.000606 | 0.00152 |
| chr18 | 21794900 | 21796412 | 1513 | 1.9 | 0.000607 | 0.00152 |
| chr19 | 33366958 | 33380886 | 13929 | -0.68 | 0.000607 | 0.00152 |
| chr16 | 68765460 | 68795586 | 30127 | 0.27 | 0.000607 | 0.00152 |
| chr5 | 124439240 | 124439553 | 314 | 3.61 | 0.000608 | 0.00152 |
| chr5 | 126339454 | 126339886 | 433 | 3.61 | 0.000608 | 0.00152 |
| chr14 | 62160736 | 62230961 | 70226 | 0.19 | 0.000608 | 0.00152 |
| chr11 | 71855736 | 71856768 | 1033 | 2.02 | 0.000608 | 0.00152 |
| chr2 | 232981639 | 232982011 | 373 | -3.89 | 0.000609 | 0.00152 |
| chr12 | 26077306 | 26078168 | 863 | 3 | 0.000609 | 0.00153 |
| chr12 | 120213390 | 120222707 | 9318 | 0.77 | 0.000611 | 0.00153 |
| chr7 | 5588857 | 5610489 | 21633 | -0.3 | 0.000612 | 0.00153 |
| chr12 | 110433298 | 110441618 | 8321 | 0.48 | 0.000612 | 0.00153 |
| chr10 | 76235066 | 76237138 | 2073 | -0.98 | 0.000612 | 0.00153 |
| chr2 | 239756059 | 239757741 | 1683 | -1.03 | 0.000613 | 0.00153 |
| chr2 | 18568343 | 18569081 | 739 | -1.65 | 0.000613 | 0.00154 |
| chr16 | 20861408 | 20894925 | 33518 | -0.28 | 0.000614 | 0.00154 |
| chr14 | 24453741 | 24454520 | 780 | -3.24 | 0.000615 | 0.00154 |
| chr14 | 85779617 | 85780178 | 562 | 2.33 | 0.000616 | 0.00154 |
| chr1 | 85084447 | 85099883 | 15437 | 0.56 | 0.000616 | 0.00154 |
| chr10 | 59752098 | 59753313 | 1216 | 1.22 | 0.000617 | 0.00154 |
| chr5 | 884095 | 919778 | 35684 | -0.27 | 0.000619 | 0.00155 |
| chr14 | 68282123 | 68287251 | 5129 | 0.54 | 0.000619 | 0.00155 |
| chr3 | 195836818 | 195838179 | 1362 | 1.15 | 0.000619 | 0.00155 |
| chr10 | 87610550 | 87612283 | 1734 | 4.3 | 0.000619 | 0.00155 |
| chr6 | 9851632 | 9852339 | 708 | 2.55 | 0.00062 | 0.00155 |
| chr11 | 11310804 | 11311313 | 510 | -3.25 | 0.00062 | 0.00155 |
| chr7 | 95515532 | 95516621 | 1090 | 2.19 | 0.000622 | 0.00155 |
| chr15 | 55542128 | 55543052 | 925 | 3.07 | 0.000622 | 0.00156 |
| chr15 | 89009018 | 89017239 | 8222 | 0.36 | 0.000623 | 0.00156 |
| chr10 | 111802481 | 111803320 | 840 | 2.13 | 0.000623 | 0.00156 |
| chr17 | 58732470 | 58756513 | 24044 | 0.52 | 0.000623 | 0.00156 |
| chr7 | 54841249 | 54841990 | 742 | 0.98 | 0.000624 | 0.00156 |
| chr2 | 190476932 | 190478884 | 1953 | -0.63 | 0.000624 | 0.00156 |
| chr1 | 186797290 | 186823999 | 26710 | 0.29 | 0.000624 | 0.00156 |
| chr16 | 89118654 | 89120135 | 1482 | 0.93 | 0.000625 | 0.00156 |
| chr13 | 38922699 | 38925177 | 2479 | 0.39 | 0.000627 | 0.00157 |
| chr4 | 180386152 | 180387369 | 1218 | 1.04 | 0.000627 | 0.00157 |
| chr11 | 92681552 | 92682474 | 923 | 1.87 | 0.000628 | 0.00157 |
| chr20 | 14048002 | 14049207 | 1206 | 1.87 | 0.000629 | 0.00157 |
| chr9 | 34247616 | 34257761 | 10146 | -0.48 | 0.000629 | 0.00157 |
| chr13 | 33413057 | 33413862 | 806 | 1.97 | 0.00063 | 0.00157 |
| chr6 | 157467901 | 157471883 | 3983 | 0.53 | 0.000632 | 0.00158 |
| chr17 | 46005589 | 46058517 | 52929 | -0.32 | 0.000632 | 0.00158 |
| chr22 | 41105403 | 41129258 | 23856 | -0.5 | 0.000632 | 0.00158 |
| chr11 | 47652862 | 47665067 | 12206 | 0.48 | 0.000633 | 0.00158 |
| chr1 | 174764579 | 174768820 | 4242 | 1.34 | 0.000634 | 0.00158 |
| chr1 | 162185306 | 162186263 | 958 | 2.73 | 0.000634 | 0.00158 |
| chr8 | 59010903 | 59011391 | 489 | 2.22 | 0.000634 | 0.00158 |
| chr3 | 110825389 | 110827202 | 1814 | -1.22 | 0.000635 | 0.00158 |
| chr13 | 26794124 | 26797252 | 3129 | 0.4 | 0.000635 | 0.00158 |
| chr10 | 124201259 | 124202264 | 1006 | 1.98 | 0.000636 | 0.00159 |
| chr1 | 244995950 | 245029921 | 33972 | 0.17 | 0.000636 | 0.00159 |
| chr10 | 15901664 | 15903102 | 1439 | 0.76 | 0.000637 | 0.00159 |
| chr8 | 130547768 | 130548697 | 930 | -1.98 | 0.000638 | 0.00159 |
| chr8 | 47829271 | 47830045 | 775 | 2.51 | 0.000639 | 0.00159 |
| chr12 | 53644477 | 53647392 | 2916 | 0.68 | 0.000639 | 0.00159 |
| chrX | 46233804 | 46236581 | 2778 | -0.74 | 0.00064 | 0.0016 |
| chr17 | 10585638 | 10602210 | 16573 | 0.39 | 0.000641 | 0.0016 |
| chr3 | 187596848 | 187597208 | 361 | 3.31 | 0.000641 | 0.0016 |

| | | | | | | |
|---|---|---|---|---|---|---|
| chr3 | 135984692 | 135986561 | 1870 | -1.23 | 0.000641 | 0.0016 |
| chr8 | 28741299 | 28749401 | 8103 | 0.39 | 0.000641 | 0.0016 |
| chr2 | 149001664 | 149017764 | 16101 | -1.1 | 0.000641 | 0.0016 |
| chr9 | 4489596 | 4491475 | 1880 | 1.75 | 0.000642 | 0.0016 |
| chr5 | 39274343 | 39275470 | 1128 | 1.7 | 0.000643 | 0.0016 |
| chrX | 19191728 | 19193739 | 2012 | 1.01 | 0.000644 | 0.0016 |
| chr16 | 12322154 | 12322714 | 561 | -3.61 | 0.000644 | 0.0016 |
| chr17 | 61817588 | 61927135 | 109548 | 0.15 | 0.000644 | 0.0016 |
| chr1 | 191870739 | 191871558 | 820 | -2.91 | 0.000645 | 0.00161 |
| chr1 | 43499586 | 43500966 | 1381 | 1.3 | 0.000645 | 0.00161 |
| chr3 | 189716368 | 189736477 | 20110 | 0.28 | 0.000645 | 0.00161 |
| chr7 | 30894206 | 30899981 | 5776 | -1.56 | 0.000646 | 0.00161 |
| chr4 | 120580467 | 120581515 | 1049 | 1.31 | 0.000646 | 0.00161 |
| chr13 | 35564430 | 35566212 | 1783 | 4.34 | 0.000647 | 0.00161 |
| chr10 | 37550516 | 37559864 | 9349 | 5.16 | 0.000648 | 0.00161 |
| chr16 | 31918627 | 31918979 | 353 | 4.47 | 0.000648 | 0.00161 |
| chr1 | 117183790 | 117184310 | 521 | 3.08 | 0.000648 | 0.00161 |
| chr16 | 14164354 | 14169304 | 4951 | 0.71 | 0.000648 | 0.00161 |
| chr6 | 71376664 | 71379245 | 2582 | 0.65 | 0.000649 | 0.00161 |
| chr20 | 45343375 | 45345141 | 1767 | 1.05 | 0.00065 | 0.00162 |
| chr1 | 154120918 | 154159795 | 38878 | 0.33 | 0.00065 | 0.00162 |
| chr6 | 11494859 | 11497271 | 2413 | -0.99 | 0.00065 | 0.00162 |
| chr3 | 106236452 | 106238514 | 2063 | 0.69 | 0.000652 | 0.00162 |
| chr12 | 20779635 | 20784003 | 4369 | 0.42 | 0.000654 | 0.00163 |
| chr15 | 44485923 | 44487024 | 1102 | 2.1 | 0.000655 | 0.00163 |
| chr2 | 157038398 | 157039299 | 902 | 2.12 | 0.000656 | 0.00163 |
| chr7 | 99056206 | 99077134 | 20929 | 0.36 | 0.000656 | 0.00163 |
| chr10 | 94216230 | 94217911 | 1682 | -1.68 | 0.000657 | 0.00163 |
| chr9 | 21801340 | 21832429 | 31090 | 0.25 | 0.000657 | 0.00163 |
| chr2 | 179106343 | 179107601 | 1259 | -1.08 | 0.000658 | 0.00163 |
| chr2 | 74307990 | 74308407 | 418 | -3.87 | 0.000659 | 0.00164 |
| chr15 | 80471120 | 80472084 | 965 | 1.98 | 0.00066 | 0.00164 |
| chr10 | 96161424 | 96164854 | 3431 | 0.63 | 0.000661 | 0.00164 |
| chr1 | 15735082 | 15760367 | 25286 | 0.23 | 0.000661 | 0.00164 |
| chr22 | 48795655 | 48804335 | 8681 | 0.28 | 0.000661 | 0.00164 |
| chr5 | 106593423 | 106594082 | 660 | -2.75 | 0.000663 | 0.00165 |
| chr8 | 34743893 | 34744298 | 406 | -4.49 | 0.000663 | 0.00165 |
| chr11 | 111204691 | 111215400 | 10710 | 1.39 | 0.000663 | 0.00165 |
| chr16 | 14046942 | 14055564 | 8623 | 0.29 | 0.000664 | 0.00165 |
| chr15 | 35563793 | 35564116 | 324 | 3.41 | 0.000664 | 0.00165 |
| chr6 | 64372392 | 64397854 | 25463 | 0.61 | 0.000665 | 0.00165 |
| chr6 | 54711012 | 54711918 | 907 | 1.8 | 0.000667 | 0.00166 |
| chr6 | 13302055 | 13329665 | 27611 | 0.25 | 0.000667 | 0.00166 |
| chr2 | 205706487 | 205708498 | 2012 | -0.75 | 0.000668 | 0.00166 |
| chr2 | 74275975 | 74276963 | 989 | -2.89 | 0.000668 | 0.00166 |
| chr12 | 69417558 | 69419630 | 2073 | 2.72 | 0.000669 | 0.00166 |
| chr14 | 99660501 | 99661255 | 755 | -1.77 | 0.00067 | 0.00166 |
| chr11 | 12433259 | 12440024 | 6766 | -1.3 | 0.000671 | 0.00167 |
| chr13 | 52025067 | 52028929 | 3863 | 0.6 | 0.000673 | 0.00167 |
| chr6 | 44001657 | 44011363 | 9707 | 0.39 | 0.000673 | 0.00167 |
| chr17 | 58432918 | 58436316 | 3399 | 0.96 | 0.000673 | 0.00167 |
| chr1 | 24861592 | 24895451 | 33860 | 0.38 | 0.000674 | 0.00167 |
| chr4 | 30718149 | 30718497 | 349 | 3.62 | 0.000674 | 0.00167 |
| chrX | 117858602 | 117874967 | 16366 | 0.37 | 0.000674 | 0.00167 |
| chr20 | 6054736 | 6057320 | 2585 | -0.97 | 0.000674 | 0.00167 |
| chr2 | 236791968 | 236793102 | 1135 | -1.67 | 0.000675 | 0.00168 |
| chr2 | 240087461 | 240088136 | 676 | -3.06 | 0.000676 | 0.00168 |
| chr6 | 135095362 | 135095974 | 613 | -2.6 | 0.000676 | 0.00168 |
| chr15 | 58959069 | 58973274 | 14206 | -0.81 | 0.000676 | 0.00168 |
| chr8 | 29890695 | 29891416 | 722 | 2.89 | 0.000677 | 0.00168 |
| chr3 | 191460175 | 191460696 | 522 | 3.54 | 0.000678 | 0.00168 |

| | | | | | | |
|---|---|---|---|---|---|---|
| chr6 | 78175963 | 78176303 | 341 | 3.61 | 0.000678 | 0.00168 |
| chr4 | 75955641 | 75956564 | 924 | 2.47 | 0.000678 | 0.00168 |
| chr14 | 57046159 | 57047723 | 1565 | 0.57 | 0.00068 | 0.00169 |
| chr14 | 67824733 | 67854296 | 29564 | 0.29 | 0.000681 | 0.00169 |
| chr4 | 89742928 | 89745865 | 2938 | 0.48 | 0.000681 | 0.00169 |
| chr6 | 165929092 | 165930389 | 1298 | 1.4 | 0.000682 | 0.00169 |
| chr12 | 117232574 | 117233271 | 698 | 2.57 | 0.000683 | 0.00169 |
| chr9 | 33619029 | 33620516 | 1488 | -1.63 | 0.000686 | 0.0017 |
| chr2 | 27839036 | 27853427 | 14392 | -0.6 | 0.000687 | 0.0017 |
| chr1 | 37998935 | 38001743 | 2809 | 0.66 | 0.000688 | 0.0017 |
| chr4 | 39305586 | 39307815 | 2230 | -0.77 | 0.000688 | 0.0017 |
| chr7 | 128685329 | 128696154 | 10826 | 0.34 | 0.000688 | 0.0017 |
| chr3 | 159681407 | 159682950 | 1544 | 1.46 | 0.000688 | 0.0017 |
| chr8 | 104032444 | 104034085 | 1642 | 0.78 | 0.000689 | 0.00171 |
| chr1 | 154357845 | 154359602 | 1758 | 1.62 | 0.00069 | 0.00171 |
| chr11 | 103255303 | 103256336 | 1034 | 1.06 | 0.00069 | 0.00171 |
| chr1 | 79297544 | 79299967 | 2424 | 1.28 | 0.00069 | 0.00171 |
| chr6 | 111521038 | 111522675 | 1638 | 0.86 | 0.00069 | 0.00171 |
| chr3 | 29105987 | 29108911 | 2925 | 0.68 | 0.000691 | 0.00171 |
| chr2 | 169657918 | 169660763 | 2846 | 0.71 | 0.000691 | 0.00171 |
| chr5 | 39754813 | 39762126 | 7314 | -0.44 | 0.000692 | 0.00171 |
| chr2 | 113032184 | 113034432 | 2249 | -0.46 | 0.000692 | 0.00171 |
| chr6 | 34949441 | 34950664 | 1224 | -1.56 | 0.000694 | 0.00172 |
| chr2 | 189006546 | 189008328 | 1783 | 1.22 | 0.000694 | 0.00172 |
| chr19 | 17665943 | 17678207 | 12265 | 0.45 | 0.000694 | 0.00172 |
| chr4 | 34418779 | 34419131 | 353 | -4.39 | 0.000694 | 0.00172 |
| chr8 | 29276125 | 29276499 | 375 | -4.39 | 0.000694 | 0.00172 |
| chr3 | 25823719 | 25832525 | 8807 | 0.43 | 0.000696 | 0.00172 |
| chr4 | 109039212 | 109039592 | 381 | 4.42 | 0.000697 | 0.00172 |
| chr14 | 54517540 | 54518480 | 941 | 3.01 | 7.00E-04 | 0.00173 |
| chrX | 129379422 | 129380082 | 661 | -2.08 | 7.00E-04 | 0.00173 |
| chr6 | 5190255 | 5191671 | 1417 | 1.95 | 0.000701 | 0.00173 |
| chr13 | 109295455 | 109296927 | 1473 | 1.49 | 0.000703 | 0.00174 |
| chr8 | 124780030 | 124781912 | 1883 | 0.56 | 0.000703 | 0.00174 |
| chr9 | 125023641 | 125038740 | 15100 | 0.29 | 0.000706 | 0.00175 |
| chr12 | 118391203 | 118392150 | 948 | 2.9 | 0.000706 | 0.00175 |
| chr17 | 64899925 | 64902870 | 2946 | -0.46 | 0.000706 | 0.00175 |
| chr1 | 53479852 | 53481540 | 1689 | 0.94 | 0.000708 | 0.00175 |
| chr11 | 126080962 | 126085777 | 4816 | 0.79 | 0.000709 | 0.00175 |
| chrX | 10157350 | 10160010 | 2661 | -0.61 | 0.00071 | 0.00176 |
| chr4 | 74861871 | 74905734 | 43864 | 0.19 | 0.00071 | 0.00176 |
| chr8 | 94947166 | 94950035 | 2870 | 0.49 | 0.000711 | 0.00176 |
| chr17 | 17845989 | 17895465 | 49477 | -0.25 | 0.000712 | 0.00176 |
| chr20 | 31142185 | 31143086 | 902 | -1.98 | 0.000713 | 0.00176 |
| chrX | 76883612 | 76892726 | 9115 | 0.99 | 0.000713 | 0.00176 |
| chr5 | 123964025 | 124000103 | 36079 | 0.26 | 0.000713 | 0.00176 |
| chr12 | 11071061 | 11072830 | 1770 | 1.16 | 0.000713 | 0.00176 |
| chr1 | 180499277 | 180500328 | 1052 | 1.68 | 0.000714 | 0.00176 |
| chr5 | 123032135 | 123044551 | 12417 | 0.51 | 0.000714 | 0.00176 |
| chr8 | 108314117 | 108316340 | 2224 | 1.34 | 0.000714 | 0.00176 |
| chr9 | 20730855 | 20735905 | 5051 | -0.58 | 0.000714 | 0.00176 |
| chr1 | 9061406 | 9062001 | 596 | 1.95 | 0.000715 | 0.00176 |
| chr6 | 132678145 | 132679030 | 886 | 1.57 | 0.000716 | 0.00177 |
| chr12 | 120425839 | 120427376 | 1538 | 1.09 | 0.000717 | 0.00177 |
| chr5 | 159402602 | 159403097 | 496 | -3.58 | 0.000717 | 0.00177 |
| chr10 | 93390758 | 93394002 | 3245 | 0.71 | 0.000718 | 0.00177 |
| chr15 | 90058030 | 90058607 | 578 | -3.26 | 0.000718 | 0.00177 |
| chr9 | 7184879 | 7187193 | 2315 | 1.15 | 0.000719 | 0.00177 |
| chr17 | 45400506 | 45402372 | 1867 | 0.44 | 0.000721 | 0.00178 |
| chr7 | 139141398 | 139142444 | 1047 | -2.99 | 0.000721 | 0.00178 |
| chr6 | 37785871 | 37789101 | 3231 | 0.41 | 0.000722 | 0.00178 |

| | | | | | | |
|---|---|---|---|---|---|---|
| chr2 | 48694787 | 48709664 | 14878 | 0.46 | 0.000723 | 0.00178 |
| chr4 | 96044012 | 96052503 | 8492 | 0.37 | 0.000724 | 0.00179 |
| chr12 | 11167994 | 11169320 | 1327 | 1.53 | 0.000725 | 0.00179 |
| chr17 | 17990473 | 18012742 | 22270 | -0.4 | 0.000725 | 0.00179 |
| chr10 | 43277351 | 43279202 | 1852 | 0.46 | 0.000725 | 0.00179 |
| chr2 | 155956549 | 155957137 | 589 | -3.23 | 0.000725 | 0.00179 |
| chr12 | 95419002 | 95435228 | 16227 | 0.61 | 0.000727 | 0.00179 |
| chr1 | 53685063 | 53687094 | 2032 | 0.75 | 0.000727 | 0.00179 |
| chr1 | 76620087 | 76620919 | 833 | 2.61 | 0.000727 | 0.00179 |
| chr15 | 26093886 | 26109072 | 15187 | -0.47 | 0.000728 | 0.00179 |
| chr8 | 117670448 | 117675622 | 5175 | -1.24 | 0.000729 | 0.0018 |
| chr2 | 106805344 | 106811443 | 6100 | -0.86 | 0.000729 | 0.0018 |
| chr11 | 47863882 | 47908310 | 44429 | -0.19 | 0.00073 | 0.0018 |
| chr10 | 29392614 | 29410790 | 18177 | 0.34 | 0.00073 | 0.0018 |
| chr8 | 18428666 | 18429579 | 914 | 1.73 | 0.000731 | 0.0018 |
| chr4 | 125013327 | 125014555 | 1229 | 2.87 | 0.000732 | 0.0018 |
| chr9 | 108319628 | 108321545 | 1918 | 0.8 | 0.000733 | 0.00181 |
| chr7 | 133373998 | 133374547 | 550 | 2.99 | 0.000733 | 0.00181 |
| chr6 | 80713503 | 80730822 | 17320 | 0.44 | 0.000734 | 0.00181 |
| chr22 | 40634002 | 40636176 | 2175 | 0.79 | 0.000734 | 0.00181 |
| chr6 | 15434884 | 15435831 | 948 | 1.93 | 0.000735 | 0.00181 |
| chr3 | 22785131 | 22785999 | 869 | 2.91 | 0.000735 | 0.00181 |
| chr2 | 63661981 | 63662886 | 906 | 1.97 | 0.000736 | 0.00181 |
| chr16 | 46721418 | 46730134 | 8717 | 0.42 | 0.000736 | 0.00181 |
| chr12 | 10321502 | 10323361 | 1860 | 1.76 | 0.000736 | 0.00181 |
| chr10 | 105126733 | 105128380 | 1648 | 0.93 | 0.000739 | 0.00182 |
| chr12 | 70359823 | 70360901 | 1079 | 1.7 | 0.00074 | 0.00182 |
| chr8 | 52753723 | 52760991 | 7269 | 1.22 | 0.00074 | 0.00182 |
| chr20 | 10329076 | 10337138 | 8063 | 0.21 | 0.00074 | 0.00182 |
| chr22 | 48818926 | 48829210 | 10285 | -0.82 | 0.000741 | 0.00182 |
| chr1 | 200936918 | 200941759 | 4842 | -1.55 | 0.000741 | 0.00182 |
| chr7 | 47114189 | 47114926 | 738 | 2.92 | 0.000744 | 0.00183 |
| chr2 | 110401953 | 110403114 | 1162 | 1.47 | 0.000745 | 0.00183 |
| chr8 | 48589248 | 48607071 | 17824 | 0.51 | 0.000747 | 0.00184 |
| chr10 | 12621708 | 12622959 | 1252 | 1.36 | 0.000748 | 0.00184 |
| chrX | 38659797 | 38676851 | 17055 | 0.32 | 0.000749 | 0.00184 |
| chr1 | 194225632 | 194226042 | 411 | 3.89 | 0.000751 | 0.00185 |
| chr19 | 18840579 | 18841723 | 1145 | -2.84 | 0.000752 | 0.00185 |
| chr15 | 44191854 | 44193432 | 1579 | -1.2 | 0.000752 | 0.00185 |
| chr11 | 125170947 | 125171557 | 611 | 3.13 | 0.000753 | 0.00185 |
| chr2 | 192496804 | 192503615 | 6812 | -0.86 | 0.000753 | 0.00185 |
| chr1 | 74662937 | 74664986 | 2050 | 0.48 | 0.000753 | 0.00185 |
| chr4 | 125630486 | 125635533 | 5048 | 0.47 | 0.000753 | 0.00185 |
| chr12 | 31576532 | 31702105 | 125574 | 0.12 | 0.000754 | 0.00185 |
| chr2 | 147106710 | 147107133 | 424 | 2.37 | 0.000754 | 0.00185 |
| chr8 | 107738327 | 107739064 | 738 | 1.05 | 0.000755 | 0.00186 |
| chr21 | 39812073 | 39813473 | 1401 | 1.22 | 0.000756 | 0.00186 |
| chr13 | 52921751 | 52923713 | 1963 | 1.69 | 0.000757 | 0.00186 |
| chr1 | 111965375 | 111966316 | 942 | 2.99 | 0.000757 | 0.00186 |
| chr10 | 18969936 | 18971996 | 2061 | 0.82 | 0.000758 | 0.00186 |
| chr2 | 190744977 | 190745896 | 920 | -2.83 | 0.000759 | 0.00186 |
| chr1 | 230984635 | 231005309 | 20675 | -0.59 | 0.000759 | 0.00186 |
| chr5 | 173042356 | 173057245 | 14890 | -0.54 | 0.000759 | 0.00187 |
| chr2 | 223572683 | 223575062 | 2380 | 0.87 | 0.000761 | 0.00187 |
| chr20 | 1246168 | 1247272 | 1105 | 1.17 | 0.000761 | 0.00187 |
| chr2 | 192758868 | 192764991 | 6124 | -0.8 | 0.000762 | 0.00187 |
| chr11 | 108798250 | 108810299 | 12050 | -0.33 | 0.000762 | 0.00187 |
| chr7 | 103846190 | 103849318 | 3129 | 0.54 | 0.000762 | 0.00187 |
| chr2 | 239007356 | 239008381 | 1026 | 1.76 | 0.000763 | 0.00187 |
| chrX | 154027106 | 154028496 | 1391 | -1.52 | 0.000764 | 0.00188 |
| chr1 | 146987462 | 146988067 | 606 | 2.01 | 0.000765 | 0.00188 |

| | | | | | | |
|---|---|---|---|---|---|---|
| chr7 | 87563010 | 87564048 | 1039 | 2.07 | 0.000765 | 0.00188 |
| chrX | 21545983 | 21546980 | 998 | 1.51 | 0.000766 | 0.00188 |
| chr11 | 126350338 | 126354765 | 4428 | -1.38 | 0.000766 | 0.00188 |
| chr10 | 29697860 | 29699762 | 1903 | 0.86 | 0.000767 | 0.00188 |
| chr14 | 32920257 | 32920719 | 463 | 3.1 | 0.00077 | 0.00189 |
| chr9 | 99482862 | 99483422 | 561 | 4.39 | 0.00077 | 0.00189 |
| chr10 | 34338191 | 34339715 | 1525 | -1.41 | 0.000772 | 0.00189 |
| chr14 | 36216008 | 36217766 | 1759 | 0.92 | 0.000774 | 0.0019 |
| chr17 | 75155838 | 75157801 | 1964 | 1.41 | 0.000774 | 0.0019 |
| chr1 | 95534660 | 95539183 | 4524 | 0.88 | 0.000774 | 0.0019 |
| chr7 | 98539629 | 98550669 | 11041 | -0.58 | 0.000774 | 0.0019 |
| chr8 | 133786952 | 133800469 | 13518 | 0.57 | 0.000777 | 0.00191 |
| chr12 | 79718133 | 79722165 | 4033 | -0.56 | 0.000777 | 0.00191 |
| chr5 | 168186481 | 168187366 | 886 | -1.64 | 0.00078 | 0.00191 |
| chr6 | 131334518 | 131351701 | 17184 | 0.63 | 0.000781 | 0.00191 |
| chr5 | 179974711 | 179985948 | 11238 | 0.36 | 0.000782 | 0.00192 |
| chr17 | 5135066 | 5136555 | 1490 | 1.29 | 0.000782 | 0.00192 |
| chr3 | 104914348 | 104915304 | 957 | 4.2 | 0.000784 | 0.00192 |
| chr21 | 32671278 | 32672071 | 794 | 3 | 0.000784 | 0.00192 |
| chr9 | 135412747 | 135425289 | 12543 | 4.73 | 0.000784 | 0.00192 |
| chr6 | 27856314 | 27871524 | 15211 | -0.24 | 0.000784 | 0.00192 |
| chr4 | 117407444 | 117408102 | 659 | 2.2 | 0.000785 | 0.00192 |
| chr2 | 46578172 | 46580519 | 2348 | -1.61 | 0.000787 | 0.00193 |
| chr1 | 97954546 | 97955769 | 1224 | 1.58 | 0.000788 | 0.00193 |
| chr1 | 179174632 | 179204407 | 29776 | 0.34 | 0.000788 | 0.00193 |
| chr17 | 65240244 | 65243165 | 2922 | 0.91 | 0.000789 | 0.00193 |
| chr6 | 157212158 | 157212713 | 556 | 2.52 | 0.000789 | 0.00193 |
| chrX | 10851189 | 10852253 | 1065 | 1.91 | 0.00079 | 0.00194 |
| chr1 | 207318575 | 207319311 | 737 | 2.14 | 0.000791 | 0.00194 |
| chr2 | 217362209 | 217373002 | 10794 | -0.38 | 0.000791 | 0.00194 |
| chr12 | 82854955 | 82856167 | 1213 | 1.18 | 0.000792 | 0.00194 |
| chr1 | 47069156 | 47083255 | 14100 | 0.52 | 0.000792 | 0.00194 |
| chr8 | 87249700 | 87250211 | 512 | 4.38 | 0.000793 | 0.00194 |
| chr7 | 91873261 | 91876646 | 3386 | 0.38 | 0.000796 | 0.00195 |
| chr7 | 94518142 | 94518650 | 509 | 2.36 | 0.000797 | 0.00195 |
| chr2 | 190067152 | 190098749 | 31598 | -0.19 | 0.000799 | 0.00196 |
| chr5 | 176407614 | 176434756 | 27143 | 0.29 | 0.000801 | 0.00196 |
| chr21 | 42738622 | 42739227 | 606 | -2.3 | 0.000803 | 0.00196 |
| chr2 | 121461195 | 121461674 | 480 | -3.2 | 0.000803 | 0.00196 |
| chr19 | 44860426 | 44861329 | 904 | 1.16 | 0.000804 | 0.00197 |
| chr1 | 168943181 | 168944326 | 1146 | -2.9 | 0.000808 | 0.00198 |
| chr5 | 178450327 | 178451599 | 1273 | 1.34 | 0.00081 | 0.00198 |
| chr10 | 82031853 | 82033409 | 1557 | -1.19 | 0.00081 | 0.00198 |
| chr16 | 74465117 | 74466021 | 905 | -2.86 | 0.000811 | 0.00198 |
| chr3 | 128496995 | 128507118 | 10124 | -1.35 | 0.000811 | 0.00198 |
| chr20 | 24653678 | 24654628 | 951 | -1.96 | 0.000811 | 0.00198 |
| chr6 | 63561979 | 63562556 | 578 | 4.45 | 0.000811 | 0.00198 |
| chr3 | 24358053 | 24360215 | 2163 | 1.72 | 0.000812 | 0.00198 |
| chr5 | 154093729 | 154094789 | 1061 | 2.44 | 0.000812 | 0.00199 |
| chr19 | 569925 | 589419 | 19495 | -0.37 | 0.000812 | 0.00199 |
| chr12 | 87479702 | 87480658 | 957 | 1.87 | 0.000814 | 0.00199 |
| chr5 | 170059916 | 170061202 | 1287 | 1.28 | 0.000815 | 0.00199 |
| chr18 | 51794962 | 51807542 | 12581 | 0.38 | 0.000815 | 0.00199 |
| chr12 | 79097179 | 79098716 | 1538 | 0.83 | 0.000816 | 0.00199 |
| chr1 | 152161283 | 152162409 | 1127 | 0.9 | 0.000816 | 0.00199 |
| chr1 | 227430177 | 227432957 | 2781 | 0.67 | 0.000816 | 0.00199 |
| chr10 | 102671778 | 102686346 | 14569 | 0.47 | 0.000817 | 0.002 |
| chr8 | 6263045 | 6265487 | 2443 | 0.48 | 0.000818 | 0.002 |
| chr14 | 94856469 | 94857511 | 1043 | 1.55 | 0.000818 | 0.002 |
| chr7 | 36298177 | 36355903 | 57727 | 0.21 | 0.000818 | 0.002 |
| chr1 | 213026613 | 213044212 | 17600 | 0.39 | 0.000818 | 0.002 |

| | | | | | | |
|---|---|---|---|---|---|---|
| chr18 | 59273031 | 59274987 | 1957 | -0.88 | 0.00082 | 0.002 |
| chr16 | 47443928 | 47445103 | 1176 | -1.83 | 0.000825 | 0.00201 |
| chr8 | 50022571 | 50023591 | 1021 | -2.95 | 0.000826 | 0.00202 |
| chr1 | 231108335 | 231134462 | 26128 | -0.42 | 0.000826 | 0.00202 |
| chr1 | 110305526 | 110307090 | 1565 | 0.95 | 0.000828 | 0.00202 |
| chrX | 118921689 | 118927375 | 5687 | 0.45 | 0.000829 | 0.00202 |
| chr3 | 27663480 | 27675117 | 11638 | -0.32 | 0.00083 | 0.00203 |
| chr10 | 16874456 | 16875217 | 762 | 3.26 | 0.000831 | 0.00203 |
| chr2 | 171077856 | 171079050 | 1195 | -1.8 | 0.000832 | 0.00203 |
| chrX | 44879946 | 44882115 | 2170 | -1.53 | 0.000832 | 0.00203 |
| chr4 | 124425765 | 124430140 | 4376 | 0.57 | 0.000833 | 0.00203 |
| chr1 | 246468633 | 246471542 | 2910 | -0.54 | 0.000835 | 0.00204 |
| chr1 | 197963321 | 197965667 | 2347 | 0.64 | 0.000836 | 0.00204 |
| chr15 | 100781416 | 100782332 | 917 | -1.35 | 0.000836 | 0.00204 |
| chr8 | 101407698 | 101408770 | 1073 | -1.51 | 0.000837 | 0.00204 |
| chr17 | 55388599 | 55390037 | 1439 | -1.5 | 0.000837 | 0.00204 |
| chrX | 14870901 | 14871941 | 1041 | 1.77 | 0.00084 | 0.00205 |
| chr6 | 138548203 | 138553702 | 5500 | 1.01 | 0.000841 | 0.00205 |
| chr22 | 21462450 | 21463884 | 1435 | -1.09 | 0.000841 | 0.00205 |
| chr13 | 80789289 | 80810584 | 21296 | 0.61 | 0.000842 | 0.00205 |
| chr3 | 47951658 | 47953855 | 2198 | -1.18 | 0.000842 | 0.00205 |
| chr4 | 38181510 | 38191372 | 9863 | 1.13 | 0.000843 | 0.00206 |
| chr2 | 43567927 | 43568770 | 844 | -2.05 | 0.000844 | 0.00206 |
| chrX | 107862411 | 107862917 | 507 | -2.76 | 0.000845 | 0.00206 |
| chr22 | 30331750 | 30332984 | 1235 | 1.3 | 0.000845 | 0.00206 |
| chr5 | 140418534 | 140420794 | 2261 | 0.45 | 0.000845 | 0.00206 |
| chr18 | 12678432 | 12680256 | 1825 | 0.89 | 0.000845 | 0.00206 |
| chr9 | 94860038 | 94860537 | 500 | -4.32 | 0.000847 | 0.00206 |
| chr22 | 44207332 | 44220316 | 12985 | 0.61 | 0.000847 | 0.00206 |
| chr6 | 25602014 | 25602749 | 736 | 2.15 | 0.000848 | 0.00206 |
| chr5 | 71310496 | 71311450 | 955 | 2.23 | 0.000848 | 0.00207 |
| chr17 | 48670814 | 48683710 | 12897 | -4.6 | 0.00085 | 0.00207 |
| chr3 | 10286133 | 10291561 | 5429 | 0.69 | 0.00085 | 0.00207 |
| chr10 | 64372786 | 64373922 | 1137 | -1.79 | 0.00085 | 0.00207 |
| chr15 | 100104801 | 100107431 | 2631 | 0.66 | 0.000851 | 0.00207 |
| chr2 | 158276992 | 158278010 | 1019 | -1.8 | 0.000851 | 0.00207 |
| chr12 | 120275295 | 120275685 | 391 | -3.3 | 0.000852 | 0.00207 |
| chr9 | 27371158 | 27377725 | 6568 | -2.09 | 0.000852 | 0.00208 |
| chr1 | 209568657 | 209586057 | 17401 | -2.59 | 0.000853 | 0.00208 |
| chr2 | 102641479 | 102651429 | 9951 | -0.55 | 0.000854 | 0.00208 |
| chr13 | 45935552 | 45943154 | 7603 | 0.86 | 0.000854 | 0.00208 |
| chr5 | 58181663 | 58199154 | 17492 | -0.26 | 0.000854 | 0.00208 |
| chr8 | 49748221 | 49748656 | 436 | 3.11 | 0.000855 | 0.00208 |
| chr14 | 39764730 | 39765620 | 891 | 1.67 | 0.000855 | 0.00208 |
| chr6 | 169612611 | 169614019 | 1409 | 1.06 | 0.000857 | 0.00208 |
| chr11 | 33653775 | 33658664 | 4890 | -0.52 | 0.000858 | 0.00209 |
| chr1 | 64411534 | 64411957 | 424 | -3.19 | 0.000858 | 0.00209 |
| chrX | 117914426 | 117915254 | 829 | 2.26 | 0.000859 | 0.00209 |
| chr15 | 101389198 | 101391758 | 2561 | 0.72 | 0.00086 | 0.00209 |
| chr8 | 119451007 | 119456953 | 5947 | 0.7 | 0.000861 | 0.00209 |
| chr17 | 78351120 | 78369575 | 18456 | -0.42 | 0.000862 | 0.0021 |
| chr3 | 58983834 | 58985257 | 1424 | -2 | 0.000862 | 0.0021 |
| chrX | 51545779 | 51547194 | 1416 | 0.98 | 0.000865 | 0.0021 |
| chr17 | 71823649 | 71824598 | 950 | -1.45 | 0.000865 | 0.0021 |
| chr2 | 150588063 | 150603381 | 15319 | 0.41 | 0.000867 | 0.00211 |
| chr1 | 43670406 | 43674003 | 3598 | -0.4 | 0.000867 | 0.00211 |
| chr5 | 118569395 | 118570132 | 738 | -2.75 | 0.000867 | 0.00211 |
| chr15 | 83654192 | 83656526 | 2335 | 0.69 | 0.000868 | 0.00211 |
| chr15 | 44067263 | 44070608 | 3346 | 0.8 | 0.000868 | 0.00211 |
| chr18 | 13796666 | 13797829 | 1164 | 1.94 | 0.000871 | 0.00212 |
| chrX | 67808855 | 67809417 | 563 | -2.51 | 0.000871 | 0.00212 |

| | | | | | | |
|---|---|---|---|---|---|---|
| chr10 | 8719015 | 8719707 | 693 | 2.28 | 0.000872 | 0.00212 |
| chr12 | 50339106 | 50339965 | 860 | 1.68 | 0.000872 | 0.00212 |
| chr14 | 26839472 | 26840077 | 606 | 3.23 | 0.000872 | 0.00212 |
| chr5 | 129239457 | 129239935 | 479 | 3.14 | 0.000873 | 0.00212 |
| chr5 | 149789567 | 149790814 | 1248 | 1.04 | 0.000873 | 0.00212 |
| chr15 | 45123223 | 45123889 | 667 | 2.07 | 0.000873 | 0.00212 |
| chr5 | 171422859 | 171435057 | 12199 | 0.37 | 0.000873 | 0.00212 |
| chr15 | 45489655 | 45493790 | 4136 | 0.47 | 0.000874 | 0.00212 |
| chr8 | 124169105 | 124170843 | 1739 | 1.8 | 0.000875 | 0.00212 |
| chr8 | 22507832 | 22509297 | 1466 | 1.29 | 0.000875 | 0.00212 |
| chr4 | 110623209 | 110625209 | 2001 | 0.83 | 0.000875 | 0.00212 |
| chr8 | 143896129 | 143898147 | 2019 | -0.68 | 0.000876 | 0.00213 |
| chr12 | 76255743 | 76271763 | 16021 | -0.57 | 0.000877 | 0.00213 |
| chr3 | 128216083 | 128216972 | 890 | -1.29 | 0.000877 | 0.00213 |
| chr4 | 100775346 | 100775868 | 523 | 3.15 | 0.000878 | 0.00213 |
| chr1 | 224430951 | 224434735 | 3785 | 0.66 | 0.00088 | 0.00214 |
| chr3 | 122822790 | 122823974 | 1185 | -1.8 | 0.000882 | 0.00214 |
| chr8 | 134228364 | 134230727 | 2364 | 0.86 | 0.000883 | 0.00214 |
| chr2 | 235300858 | 235301496 | 639 | -2.74 | 0.000884 | 0.00214 |
| chr6 | 142863923 | 142864535 | 613 | -2.4 | 0.000884 | 0.00214 |
| chr8 | 123587177 | 123600833 | 13657 | 0.84 | 0.000884 | 0.00215 |
| chr3 | 12411186 | 12413288 | 2103 | 1.05 | 0.000885 | 0.00215 |
| chr5 | 148381230 | 148382635 | 1406 | 1.5 | 0.000886 | 0.00215 |
| chr16 | 75174362 | 75183430 | 9069 | 1.2 | 0.000886 | 0.00215 |
| chr8 | 107281849 | 107283420 | 1572 | 1.22 | 0.000887 | 0.00215 |
| chr4 | 56244569 | 56246664 | 2096 | 2.13 | 0.000887 | 0.00215 |
| chr3 | 113592545 | 113593492 | 948 | 2.93 | 0.000888 | 0.00215 |
| chr3 | 131068408 | 131081436 | 13029 | 0.57 | 0.000889 | 0.00215 |
| chr19 | 2523257 | 2553370 | 30114 | -0.25 | 0.000889 | 0.00215 |
| chr7 | 128577434 | 128599164 | 21731 | -0.64 | 0.00089 | 0.00216 |
| chr10 | 72237686 | 72239704 | 2019 | -1.16 | 0.00089 | 0.00216 |
| chr5 | 74244273 | 74245977 | 1705 | 1.88 | 0.00089 | 0.00216 |
| chr12 | 110123859 | 110124822 | 964 | 2.36 | 0.000891 | 0.00216 |
| chr18 | 46116294 | 46117511 | 1218 | -1.9 | 0.000892 | 0.00216 |
| chr19 | 22816427 | 22816849 | 423 | 3.68 | 0.000893 | 0.00216 |
| chr2 | 214967926 | 214968842 | 917 | 1.93 | 0.000895 | 0.00217 |
| chr7 | 45730373 | 45730943 | 571 | 3 | 0.000896 | 0.00217 |
| chr20 | 60756869 | 60759376 | 2508 | 0.47 | 0.000896 | 0.00217 |
| chr7 | 139948117 | 139953065 | 4949 | 0.78 | 0.000897 | 0.00217 |
| chr12 | 125464240 | 125479660 | 15421 | 0.45 | 0.000898 | 0.00217 |
| chr13 | 91475913 | 91476907 | 995 | 4.32 | 0.000898 | 0.00217 |
| chr4 | 40655401 | 40677830 | 22430 | -0.25 | 0.000899 | 0.00218 |
| chr1 | 94445577 | 94446104 | 528 | -3.17 | 0.000901 | 0.00218 |
| chr12 | 102503952 | 102520704 | 16753 | 0.37 | 0.000902 | 0.00218 |
| chr22 | 42011648 | 42018405 | 6758 | 0.54 | 0.000904 | 0.00219 |
| chr12 | 57718954 | 57719673 | 720 | -2.62 | 0.000904 | 0.00219 |
| chr22 | 45264851 | 45267082 | 2232 | -0.75 | 0.000905 | 0.00219 |
| chr5 | 71126284 | 71127293 | 1010 | 1.59 | 0.000906 | 0.00219 |
| chr3 | 151688242 | 151705174 | 16933 | -0.6 | 0.000908 | 0.0022 |
| chr3 | 134070674 | 134095321 | 24648 | 0.19 | 0.000908 | 0.0022 |
| chr9 | 134573176 | 134590435 | 17260 | 0.43 | 0.000908 | 0.0022 |
| chrX | 66873963 | 66874560 | 598 | 2.1 | 0.000909 | 0.0022 |
| chr13 | 72740224 | 72740594 | 371 | -3.8 | 0.000911 | 0.0022 |
| chr3 | 26479490 | 26481186 | 1697 | 1.32 | 0.000912 | 0.00221 |
| chr3 | 31134841 | 31136208 | 1368 | 1.4 | 0.000912 | 0.00221 |
| chr10 | 133729038 | 133762630 | 33593 | -0.43 | 0.000913 | 0.00221 |
| chr12 | 26149963 | 26151447 | 1485 | 1.22 | 0.000914 | 0.00221 |
| chr5 | 61817905 | 61818897 | 993 | -1.74 | 0.000914 | 0.00221 |
| chr2 | 145964255 | 145964693 | 439 | -3 | 0.000914 | 0.00221 |
| chr3 | 111482520 | 111487720 | 5201 | -1.03 | 0.000914 | 0.00221 |
| chr10 | 120670237 | 120671623 | 1387 | -1.63 | 0.000915 | 0.00221 |

| | | | | | | |
|---|---|---|---|---|---|---|
| chr7 | 22963216 | 22963827 | 612 | 2.19 | 0.000915 | 0.00221 |
| chr3 | 183088128 | 183089772 | 1645 | 1.17 | 0.000916 | 0.00221 |
| chr20 | 40222842 | 40229132 | 6291 | 4.41 | 0.000916 | 0.00222 |
| chr1 | 89148290 | 89164464 | 16175 | 0.46 | 0.000917 | 0.00222 |
| chr10 | 72117105 | 72147433 | 30329 | -0.5 | 0.000917 | 0.00222 |
| chr14 | 36276428 | 36279478 | 3051 | 0.62 | 0.000919 | 0.00222 |
| chr1 | 76524674 | 76535519 | 10846 | 4.5 | 0.000921 | 0.00222 |
| chrX | 48659013 | 48662005 | 2993 | 0.47 | 0.000923 | 0.00223 |
| chr12 | 75212044 | 75220694 | 8651 | 0.96 | 0.000923 | 0.00223 |
| chr13 | 114828944 | 114834885 | 5942 | -0.32 | 0.000923 | 0.00223 |
| chr2 | 99502137 | 99502554 | 418 | 4.38 | 0.000923 | 0.00223 |
| chr12 | 117923173 | 117934175 | 11003 | -0.42 | 0.000925 | 0.00223 |
| chr16 | 75375570 | 75377553 | 1984 | 1.54 | 0.000926 | 0.00224 |
| chr10 | 131754454 | 131755137 | 684 | 1.99 | 0.000927 | 0.00224 |
| chr15 | 64149238 | 64150149 | 912 | 1.48 | 0.000927 | 0.00224 |
| chr5 | 60969014 | 60969658 | 645 | 3.14 | 0.000927 | 0.00224 |
| chr17 | 33306306 | 33418025 | 111720 | 0.11 | 0.00093 | 0.00225 |
| chr2 | 163438496 | 163439814 | 1319 | -1.8 | 0.000933 | 0.00225 |
| chr12 | 10183547 | 10185339 | 1793 | 0.93 | 0.000935 | 0.00226 |
| chr2 | 9247458 | 9252976 | 5519 | -1.15 | 0.000936 | 0.00226 |
| chr2 | 63112877 | 63123376 | 10500 | -2.39 | 0.000937 | 0.00226 |
| chr5 | 149821929 | 149904432 | 82504 | -0.2 | 0.000937 | 0.00226 |
| chrX | 18718534 | 18740913 | 22380 | -0.31 | 0.000938 | 0.00226 |
| chr7 | 93685219 | 93698131 | 12913 | 3.27 | 0.000938 | 0.00226 |
| chr14 | 78558577 | 78558905 | 329 | -4.36 | 0.000939 | 0.00226 |
| chr6 | 8063637 | 8065612 | 1976 | 0.57 | 0.000939 | 0.00226 |
| chr22 | 30742988 | 30754126 | 11139 | -0.41 | 0.000939 | 0.00227 |
| chr4 | 107503619 | 107506576 | 2958 | 3.99 | 0.00094 | 0.00227 |
| chr18 | 25693517 | 25709745 | 16229 | 1.29 | 0.000941 | 0.00227 |
| chr13 | 103525556 | 103526430 | 875 | 2.38 | 0.000944 | 0.00228 |
| chr3 | 48561151 | 48562549 | 1399 | -1.69 | 0.000944 | 0.00228 |
| chr14 | 31159834 | 31161080 | 1247 | 0.62 | 0.000946 | 0.00228 |
| chr3 | 40340519 | 40352761 | 12243 | 0.53 | 0.000946 | 0.00228 |
| chr2 | 200941642 | 200943079 | 1438 | 1.16 | 0.000947 | 0.00228 |
| chr17 | 36172410 | 36189027 | 16618 | -0.33 | 0.000951 | 0.00229 |
| chr21 | 33710904 | 33714057 | 3154 | 0.78 | 0.000952 | 0.00229 |
| chr7 | 47140212 | 47141034 | 823 | 1.87 | 0.000953 | 0.0023 |
| chr17 | 78008306 | 78011433 | 3128 | -0.43 | 0.000955 | 0.0023 |
| chr18 | 20578676 | 20583635 | 4960 | 1.19 | 0.000956 | 0.0023 |
| chr9 | 6756301 | 6759911 | 3611 | 0.36 | 0.000956 | 0.0023 |
| chr2 | 172143765 | 172144234 | 470 | -2.9 | 0.000958 | 0.00231 |
| chr15 | 40630208 | 40636136 | 5929 | -0.54 | 0.000958 | 0.00231 |
| chr4 | 166603950 | 166604858 | 909 | 2.27 | 0.000959 | 0.00231 |
| chr7 | 120767069 | 120767386 | 318 | 3.35 | 0.000961 | 0.00232 |
| chr1 | 24433580 | 24447130 | 13551 | -0.52 | 0.000966 | 0.00233 |
| chr2 | 28999485 | 29040056 | 40572 | -0.25 | 0.000966 | 0.00233 |
| chr8 | 50313623 | 50314481 | 859 | 1.28 | 0.000968 | 0.00233 |
| chr3 | 33699557 | 33701929 | 2373 | 0.71 | 0.000968 | 0.00233 |
| chr1 | 7460572 | 7478312 | 17741 | 0.36 | 0.000969 | 0.00233 |
| chr8 | 117651685 | 117656411 | 4727 | 0.9 | 0.000969 | 0.00233 |
| chr8 | 110274397 | 110276090 | 1694 | 1.72 | 0.00097 | 0.00234 |
| chr5 | 156057466 | 156058606 | 1141 | 2.84 | 0.000971 | 0.00234 |
| chr17 | 19896337 | 19896963 | 627 | -1.17 | 0.000972 | 0.00234 |
| chr2 | 219420455 | 219447628 | 27174 | -0.31 | 0.000972 | 0.00234 |
| chr13 | 96576546 | 96577616 | 1071 | 1.69 | 0.000972 | 0.00234 |
| chr14 | 74077481 | 74080556 | 3076 | 1.3 | 0.000973 | 0.00234 |
| chr2 | 61491309 | 61511117 | 19809 | -0.45 | 0.000978 | 0.00235 |
| chrX | 139846727 | 139850477 | 3751 | -3.12 | 0.000979 | 0.00236 |
| chr2 | 239046960 | 239051541 | 4582 | -1.16 | 0.000981 | 0.00236 |
| chr10 | 32702213 | 32702650 | 438 | -3.18 | 0.000984 | 0.00237 |
| chr6 | 109572226 | 109574582 | 2357 | 0.78 | 0.000985 | 0.00237 |

| | | | | | | |
|---|---|---|---|---|---|---|
| chr3 | 114367810 | 114380001 | 12192 | 1.07 | 0.000985 | 0.00237 |
| chr6 | 146054887 | 146071473 | 16587 | 0.38 | 0.000985 | 0.00237 |
| chr11 | 67230511 | 67251078 | 20568 | 0.43 | 0.000985 | 0.00237 |
| chr9 | 106876276 | 106877023 | 748 | 2.29 | 0.000986 | 0.00237 |
| chr16 | 71741377 | 71758440 | 17064 | 0.45 | 0.000986 | 0.00237 |
| chr10 | 96365921 | 96367055 | 1135 | -1.88 | 0.000988 | 0.00238 |
| chr7 | 89873909 | 89875346 | 1438 | -0.69 | 0.000989 | 0.00238 |
| chr2 | 181154102 | 181157617 | 3516 | -0.78 | 0.000989 | 0.00238 |
| chr2 | 205556301 | 205557308 | 1008 | -2.01 | 0.000989 | 0.00238 |
| chr3 | 41358984 | 41360173 | 1190 | 1.94 | 0.00099 | 0.00238 |
| chr5 | 33996947 | 34009501 | 12555 | 0.46 | 0.000991 | 0.00238 |
| chr10 | 88677968 | 88678247 | 280 | -4.29 | 0.000992 | 0.00238 |
| chr2 | 169478565 | 169480002 | 1438 | 1.34 | 0.000992 | 0.00238 |
| chr6 | 34597284 | 34598851 | 1568 | 1.14 | 0.000992 | 0.00238 |
| chr8 | 27458277 | 27495201 | 36925 | -0.27 | 0.000992 | 0.00238 |
| chr10 | 11636848 | 11654648 | 17801 | 0.49 | 0.000993 | 0.00239 |
| chrX | 149105766 | 149108669 | 2904 | 0.6 | 0.000996 | 0.00239 |
| chr11 | 64999218 | 65001063 | 1846 | 1.55 | 0.000998 | 0.0024 |
| chr15 | 66942567 | 66943374 | 808 | -2.43 | 0.001 | 0.0024 |
| chr20 | 4065821 | 4072638 | 6818 | -0.71 | 0.001 | 0.0024 |
| chr20 | 39765404 | 39782138 | 16735 | -0.56 | 0.001 | 0.00241 |
| chr1 | 194143808 | 194154738 | 10931 | 4.22 | 0.001 | 0.00241 |
| chr17 | 42733065 | 42735019 | 1955 | 0.98 | 0.001 | 0.00241 |
| chr1 | 167051393 | 167052829 | 1437 | 1.16 | 0.001 | 0.00241 |
| chr17 | 46391020 | 46392231 | 1212 | 1.52 | 0.001 | 0.00241 |
| chr14 | 96890315 | 96890819 | 505 | 1.87 | 0.00101 | 0.00242 |
| chr11 | 16247365 | 16248016 | 652 | 2.23 | 0.00101 | 0.00242 |
| chr1 | 101436455 | 101437046 | 592 | 2.83 | 0.00101 | 0.00242 |
| chr6 | 136784506 | 136791636 | 7131 | -0.55 | 0.00101 | 0.00242 |
| chr20 | 25505730 | 25566779 | 61050 | -0.22 | 0.00101 | 0.00243 |
| chr10 | 79014576 | 79015306 | 731 | -1.92 | 0.00101 | 0.00243 |
| chr3 | 185774926 | 185776330 | 1405 | -1.48 | 0.00101 | 0.00243 |
| chr2 | 5660560 | 5661435 | 876 | 2.87 | 0.00102 | 0.00244 |
| chr2 | 237993605 | 237996382 | 2778 | -0.5 | 0.00102 | 0.00245 |
| chr13 | 63251835 | 63252789 | 955 | 1.54 | 0.00102 | 0.00245 |
| chr3 | 12446182 | 12448334 | 2153 | 0.74 | 0.00102 | 0.00245 |
| chr17 | 28844711 | 28849371 | 4661 | -0.72 | 0.00102 | 0.00245 |
| chr2 | 235328903 | 235338155 | 9253 | 0.37 | 0.00102 | 0.00245 |
| chr4 | 123866811 | 123867365 | 555 | 2.95 | 0.00102 | 0.00246 |
| chr15 | 83477241 | 83481591 | 4351 | 0.46 | 0.00103 | 0.00246 |
| chr22 | 19362104 | 19362606 | 503 | -4.29 | 0.00103 | 0.00246 |
| chr6 | 113989426 | 113991383 | 1958 | 0.8 | 0.00103 | 0.00247 |
| chr7 | 56160490 | 56174935 | 14446 | 0.38 | 0.00103 | 0.00247 |
| chr11 | 44875669 | 44885294 | 9626 | 0.42 | 0.00103 | 0.00247 |
| chr8 | 105677029 | 105678496 | 1468 | -0.93 | 0.00103 | 0.00247 |
| chr10 | 64133549 | 64140164 | 6616 | -1.41 | 0.00103 | 0.00247 |
| chr15 | 77574546 | 77575299 | 754 | -2.1 | 0.00103 | 0.00247 |
| chr6 | 46620033 | 46621233 | 1201 | 1.74 | 0.00103 | 0.00248 |
| chr5 | 390687 | 396006 | 5320 | 1.11 | 0.00104 | 0.00248 |
| chr18 | 61455476 | 61455964 | 489 | 4.36 | 0.00104 | 0.00248 |
| chr5 | 52460274 | 52460788 | 515 | 2.98 | 0.00104 | 0.00248 |
| chrX | 69617169 | 69619378 | 2210 | 1.15 | 0.00104 | 0.00249 |
| chr7 | 70201681 | 70202520 | 840 | 2.88 | 0.00104 | 0.00249 |
| chr2 | 46343145 | 46350566 | 7422 | -0.64 | 0.00104 | 0.00249 |
| chr11 | 2872741 | 2874115 | 1375 | -1.46 | 0.00104 | 0.00249 |
| chr1 | 25557940 | 25575130 | 17191 | 0.32 | 0.00104 | 0.00249 |
| chr6 | 34723487 | 34726229 | 2743 | 0.51 | 0.00104 | 0.00249 |
| chr7 | 108207284 | 108212337 | 5054 | 0.38 | 0.00104 | 0.0025 |
| chr15 | 26967556 | 26982745 | 15190 | -0.97 | 0.00104 | 0.0025 |
| chr6 | 22081889 | 22082950 | 1062 | 2.34 | 0.00105 | 0.0025 |
| chr2 | 43616640 | 43630102 | 13463 | -1 | 0.00105 | 0.0025 |

| | | | | | | |
|---|---|---|---|---|---|---|
| chr10 | 34087872 | 34091326 | 3455 | 1.5 | 0.00105 | 0.00251 |
| chr2 | 220334076 | 220334690 | 615 | -2.62 | 0.00105 | 0.00251 |
| chr17 | 79390677 | 79404959 | 14283 | -0.21 | 0.00105 | 0.00251 |
| chr4 | 150744288 | 150745486 | 1199 | 1.37 | 0.00105 | 0.00251 |
| chr8 | 42797034 | 42797355 | 322 | 3.86 | 0.00105 | 0.00251 |
| chr12 | 110010319 | 110012099 | 1781 | 0.63 | 0.00105 | 0.00252 |
| chr7 | 23762340 | 23763086 | 747 | 1.38 | 0.00105 | 0.00252 |
| chr9 | 136343319 | 136351600 | 8282 | 0.73 | 0.00105 | 0.00252 |
| chr11 | 21928168 | 21929177 | 1010 | 1.6 | 0.00105 | 0.00252 |
| chr6 | 128688471 | 128693980 | 5510 | 2.07 | 0.00105 | 0.00252 |
| chr7 | 50965886 | 50966626 | 741 | 2.38 | 0.00105 | 0.00252 |
| chr6 | 154974344 | 154975706 | 1363 | 1.92 | 0.00105 | 0.00252 |
| chr3 | 94242891 | 94244195 | 1305 | 1.37 | 0.00105 | 0.00252 |
| chr6 | 105403463 | 105407791 | 4329 | 0.51 | 0.00106 | 0.00252 |
| chr8 | 143423315 | 143424025 | 711 | -1.91 | 0.00106 | 0.00252 |
| chr12 | 69861080 | 69891972 | 30893 | 0.36 | 0.00106 | 0.00253 |
| chr1 | 10115401 | 10125807 | 10407 | -0.81 | 0.00106 | 0.00253 |
| chr2 | 26853557 | 26853909 | 353 | -3.06 | 0.00106 | 0.00253 |
| chr16 | 60027012 | 60027512 | 501 | 2.44 | 0.00106 | 0.00253 |
| chr5 | 6711679 | 6770425 | 58747 | 0.22 | 0.00106 | 0.00254 |
| chr9 | 21057149 | 21079316 | 22168 | 0.98 | 0.00106 | 0.00254 |
| chr12 | 98868215 | 98869751 | 1537 | 0.62 | 0.00106 | 0.00254 |
| chr2 | 241159679 | 241172743 | 13065 | 2.91 | 0.00106 | 0.00254 |
| chr19 | 58554631 | 58554993 | 363 | 4.31 | 0.00107 | 0.00254 |
| chr4 | 201447 | 201901 | 455 | 4.31 | 0.00107 | 0.00254 |
| chr2 | 230270496 | 230275480 | 4985 | 0.26 | 0.00107 | 0.00255 |
| chr10 | 29352443 | 29354498 | 2056 | 0.76 | 0.00107 | 0.00255 |
| chr5 | 174150824 | 174152557 | 1734 | 1.27 | 0.00107 | 0.00256 |
| chr6 | 106127847 | 106128583 | 737 | -2.18 | 0.00107 | 0.00256 |
| chr2 | 45417179 | 45418100 | 922 | -2.21 | 0.00107 | 0.00256 |
| chr9 | 137643844 | 137644391 | 548 | -4.3 | 0.00107 | 0.00256 |
| chr3 | 87612783 | 87613988 | 1206 | -1.52 | 0.00107 | 0.00256 |
| chr3 | 156252398 | 156254067 | 1670 | 1.31 | 0.00107 | 0.00257 |
| chr12 | 120306803 | 120316088 | 9286 | 0.56 | 0.00108 | 0.00257 |
| chr19 | 14491482 | 14496588 | 5107 | -0.33 | 0.00108 | 0.00257 |
| chr3 | 167450457 | 167453882 | 3426 | 0.48 | 0.00108 | 0.00257 |
| chr6 | 6686474 | 6725338 | 38865 | 2.14 | 0.00108 | 0.00257 |
| chr5 | 141347662 | 141351933 | 4272 | 0.39 | 0.00108 | 0.00258 |
| chr11 | 56991672 | 56992525 | 854 | -2.29 | 0.00108 | 0.00258 |
| chr20 | 14123655 | 14124476 | 822 | 1.84 | 0.00108 | 0.00258 |
| chr4 | 140807284 | 140808757 | 1474 | 1.5 | 0.00108 | 0.00258 |
| chr5 | 102897070 | 102898793 | 1724 | 0.95 | 0.00108 | 0.00258 |
| chr7 | 112429160 | 112430947 | 1788 | 0.49 | 0.00108 | 0.00258 |
| chr2 | 178482938 | 178484322 | 1385 | -0.61 | 0.00108 | 0.00258 |
| chr3 | 51532443 | 51535338 | 2896 | 0.53 | 0.00108 | 0.00258 |
| chr9 | 7117770 | 7122686 | 4917 | 0.57 | 0.00109 | 0.00259 |
| chr6 | 90527890 | 90540679 | 12790 | 0.41 | 0.00109 | 0.00259 |
| chr2 | 83891302 | 83894061 | 2760 | 5.2 | 0.00109 | 0.00259 |
| chr5 | 72665902 | 72667598 | 1697 | -0.91 | 0.00109 | 0.0026 |
| chr2 | 27543314 | 27547374 | 4061 | 0.35 | 0.00109 | 0.0026 |
| chr19 | 14098155 | 14118823 | 20669 | -0.34 | 0.00109 | 0.0026 |
| chr15 | 51933651 | 51934199 | 549 | 2.95 | 0.00109 | 0.0026 |
| chr1 | 241852067 | 241852804 | 738 | 2.2 | 0.00109 | 0.00261 |
| chr9 | 133003160 | 133006019 | 2860 | -1.38 | 0.00109 | 0.00261 |
| chrX | 133851460 | 133862394 | 10935 | -0.51 | 0.0011 | 0.00262 |
| chr20 | 13976138 | 13977192 | 1055 | 1.74 | 0.0011 | 0.00263 |
| chr6 | 66635769 | 66644062 | 8294 | 1.04 | 0.0011 | 0.00263 |
| chr18 | 1509103 | 1509517 | 415 | 4.31 | 0.00111 | 0.00264 |
| chr13 | 90634633 | 90635419 | 787 | 1.64 | 0.00111 | 0.00264 |
| chr3 | 188740874 | 188741574 | 701 | 1.55 | 0.00111 | 0.00264 |
| chr3 | 16522696 | 16523705 | 1010 | -2.05 | 0.00111 | 0.00264 |

| | | | | | | |
|---|---|---|---|---|---|---|
| chr2 | 31623503 | 31653464 | 29962 | -0.41 | 0.00111 | 0.00264 |
| chr12 | 67023007 | 67024267 | 1261 | 1.95 | 0.00111 | 0.00265 |
| chr5 | 66455261 | 66463021 | 7761 | 0.46 | 0.00111 | 0.00265 |
| chr2 | 174887288 | 174919859 | 32572 | 0.21 | 0.00111 | 0.00265 |
| chr1 | 31916996 | 31918630 | 1635 | 0.82 | 0.00112 | 0.00266 |
| chr5 | 112130919 | 112134374 | 3456 | -0.66 | 0.00112 | 0.00266 |
| chr13 | 103021289 | 103022683 | 1395 | -1.78 | 0.00112 | 0.00266 |
| chr6 | 149724709 | 149727756 | 3048 | -1.44 | 0.00112 | 0.00266 |
| chr17 | 81005878 | 81010441 | 4564 | -0.56 | 0.00112 | 0.00266 |
| chr5 | 149045534 | 149046280 | 747 | -3.06 | 0.00112 | 0.00266 |
| chr19 | 56135734 | 56209328 | 73595 | -0.17 | 0.00112 | 0.00266 |
| chr7 | 34214974 | 34216439 | 1466 | 0.83 | 0.00112 | 0.00267 |
| chr10 | 105306335 | 105307436 | 1102 | -1.89 | 0.00112 | 0.00267 |
| chr15 | 96212552 | 96213009 | 458 | 2.17 | 0.00112 | 0.00267 |
| chr5 | 73233678 | 73237347 | 3670 | -0.64 | 0.00113 | 0.00268 |
| chr2 | 102012015 | 102013174 | 1160 | -1.22 | 0.00113 | 0.00268 |
| chr4 | 76577252 | 76599873 | 22622 | 0.33 | 0.00113 | 0.00268 |
| chr17 | 70839544 | 70843221 | 3678 | 2.05 | 0.00113 | 0.00269 |
| chr5 | 78787011 | 78811284 | 24274 | 0.51 | 0.00113 | 0.00269 |
| chr10 | 76308095 | 76309351 | 1257 | 1.82 | 0.00113 | 0.00269 |
| chr20 | 35898306 | 36028220 | 129915 | -0.13 | 0.00113 | 0.00269 |
| chr6 | 14713417 | 14713995 | 579 | -3.25 | 0.00113 | 0.00269 |
| chr11 | 91214794 | 91215784 | 991 | 1.26 | 0.00113 | 0.00269 |
| chr20 | 45581963 | 45600064 | 18102 | -0.29 | 0.00113 | 0.00269 |
| chr13 | 49465442 | 49467118 | 1677 | 0.83 | 0.00113 | 0.00269 |
| chr6 | 162477516 | 162484994 | 7479 | 0.74 | 0.00113 | 0.0027 |
| chr12 | 13970169 | 13970700 | 532 | -2.93 | 0.00114 | 0.0027 |
| chr7 | 12745224 | 12762676 | 17453 | 0.23 | 0.00114 | 0.0027 |
| chr18 | 30208933 | 30209698 | 766 | -2.63 | 0.00114 | 0.00271 |
| chr12 | 11800981 | 11809796 | 8816 | 0.36 | 0.00114 | 0.00271 |
| chr1 | 230009888 | 230010837 | 950 | -1.47 | 0.00114 | 0.00272 |
| chr18 | 37033500 | 37034412 | 913 | 2.08 | 0.00115 | 0.00272 |
| chr1 | 230808725 | 230818355 | 9631 | 0.84 | 0.00115 | 0.00272 |
| chr5 | 58156591 | 58157252 | 662 | -2.17 | 0.00115 | 0.00272 |
| chr16 | 75142118 | 75156770 | 14653 | 0.68 | 0.00115 | 0.00273 |
| chr15 | 75493361 | 75508133 | 14773 | -0.41 | 0.00115 | 0.00273 |
| chr2 | 162015011 | 162034815 | 19805 | -0.36 | 0.00115 | 0.00273 |
| chr2 | 219207705 | 219233341 | 25637 | -0.45 | 0.00115 | 0.00273 |
| chr13 | 35427336 | 35442312 | 14977 | -0.53 | 0.00115 | 0.00273 |
| chr6 | 37597815 | 37607260 | 9446 | -0.76 | 0.00115 | 0.00273 |
| chr1 | 109262427 | 109263885 | 1459 | -1.45 | 0.00115 | 0.00274 |
| chr16 | 64404100 | 64404947 | 848 | 2.85 | 0.00115 | 0.00274 |
| chr1 | 86172509 | 86175482 | 2974 | 0.55 | 0.00115 | 0.00274 |
| chr4 | 128629696 | 128630828 | 1133 | 1.33 | 0.00115 | 0.00274 |
| chr11 | 68871087 | 68906668 | 35582 | 0.3 | 0.00115 | 0.00274 |
| chr9 | 95985022 | 95986354 | 1333 | -1.07 | 0.00116 | 0.00274 |
| chr1 | 101359575 | 101363198 | 3624 | 0.37 | 0.00116 | 0.00274 |
| chr15 | 73435043 | 73435900 | 858 | 1.28 | 0.00116 | 0.00274 |
| chr5 | 74952142 | 74984703 | 32562 | 0.55 | 0.00116 | 0.00274 |
| chr17 | 41264777 | 41282355 | 17579 | -0.32 | 0.00116 | 0.00274 |
| chr15 | 40128011 | 40128573 | 563 | -2.95 | 0.00116 | 0.00275 |
| chr22 | 47702603 | 47703748 | 1146 | -1.4 | 0.00116 | 0.00275 |
| chr17 | 28590624 | 28619744 | 29121 | -0.57 | 0.00116 | 0.00276 |
| chr5 | 176829698 | 176900642 | 70945 | -0.28 | 0.00116 | 0.00276 |
| chr2 | 187502319 | 187507527 | 5209 | -0.84 | 0.00116 | 0.00276 |
| chr19 | 42006662 | 42008208 | 1547 | 0.64 | 0.00117 | 0.00276 |
| chr8 | 27621113 | 27645098 | 23986 | -0.38 | 0.00117 | 0.00277 |
| chr8 | 80928303 | 80943160 | 14858 | 0.62 | 0.00117 | 0.00277 |
| chr1 | 80858209 | 80858608 | 400 | 4.32 | 0.00117 | 0.00277 |
| chr4 | 151689504 | 151691494 | 1991 | 0.74 | 0.00117 | 0.00277 |
| chr4 | 2865891 | 2875943 | 10053 | 0.67 | 0.00117 | 0.00277 |

| | | | | | | |
|---|---|---|---|---|---|---|
| chr5 | 150502895 | 150503841 | 947 | -1.79 | 0.00117 | 0.00277 |
| chr11 | 130686001 | 130697795 | 11795 | 2.87 | 0.00117 | 0.00278 |
| chr10 | 126210874 | 126212659 | 1786 | 1.3 | 0.00117 | 0.00278 |
| chr17 | 58154954 | 58160900 | 5947 | -0.41 | 0.00118 | 0.00279 |
| chr20 | 11963506 | 11964107 | 602 | 2.85 | 0.00118 | 0.00279 |
| chr19 | 55573533 | 55592855 | 19323 | 1.59 | 0.00118 | 0.00279 |
| chr18 | 3905880 | 3906875 | 996 | 2.37 | 0.00118 | 0.00279 |
| chr20 | 17854219 | 17870007 | 15789 | 2.34 | 0.00118 | 0.00279 |
| chrX | 64728527 | 64729236 | 710 | -3 | 0.00118 | 0.00279 |
| chr2 | 193365238 | 193366466 | 1229 | 2.09 | 0.00118 | 0.00279 |
| chr10 | 25781937 | 25783447 | 1511 | 1.23 | 0.00118 | 0.00279 |
| chr4 | 126187299 | 126189801 | 2503 | 0.95 | 0.00118 | 0.0028 |
| chr6 | 159463098 | 159463717 | 620 | 2.85 | 0.00119 | 0.00281 |
| chr3 | 11740316 | 11741348 | 1033 | -1.82 | 0.00119 | 0.00281 |
| chr7 | 50576166 | 50578038 | 1873 | 1.42 | 0.00119 | 0.00281 |
| chr19 | 4246388 | 4248290 | 1903 | 0.65 | 0.00119 | 0.00281 |
| chr1 | 154897855 | 155037074 | 139220 | 0.14 | 0.00119 | 0.00281 |
| chr9 | 82473046 | 82473549 | 504 | 2.81 | 0.00119 | 0.00282 |
| chr10 | 99893609 | 99895657 | 2049 | 0.43 | 0.00119 | 0.00282 |
| chr9 | 114286906 | 114288553 | 1648 | 1.48 | 0.00119 | 0.00282 |
| chr17 | 78723507 | 78779391 | 55885 | -0.24 | 0.00119 | 0.00282 |
| chr4 | 57251744 | 57278031 | 26288 | 0.28 | 0.00119 | 0.00283 |
| chr5 | 55596591 | 55598539 | 1949 | 1.23 | 0.00119 | 0.00283 |
| chr20 | 49334499 | 49486038 | 151540 | 0.12 | 0.0012 | 0.00283 |
| chr20 | 3564653 | 3565003 | 351 | -2.83 | 0.0012 | 0.00283 |
| chr8 | 96237505 | 96238415 | 911 | -1.41 | 0.0012 | 0.00283 |
| chr10 | 86183577 | 86185384 | 1808 | 0.79 | 0.0012 | 0.00283 |
| chr4 | 153248956 | 153250772 | 1817 | 0.93 | 0.0012 | 0.00284 |
| chr1 | 98961773 | 98962733 | 961 | 2.27 | 0.0012 | 0.00284 |
| chr22 | 35957962 | 35958574 | 613 | -2.74 | 0.0012 | 0.00284 |
| chr11 | 84382653 | 84383008 | 356 | 3.12 | 0.0012 | 0.00285 |
| chr4 | 143378586 | 143380585 | 2000 | 1.29 | 0.00121 | 0.00286 |
| chr1 | 212963204 | 212966361 | 3158 | 0.5 | 0.00121 | 0.00286 |
| chr10 | 20971569 | 20972380 | 812 | -2.88 | 0.00121 | 0.00286 |
| chr22 | 24697660 | 24701409 | 3750 | 0.86 | 0.00121 | 0.00287 |
| chrX | 82487235 | 82487737 | 503 | 2.91 | 0.00122 | 0.00287 |
| chr1 | 164033946 | 164034345 | 400 | -3.53 | 0.00122 | 0.00288 |
| chr3 | 196466561 | 196491263 | 24703 | 0.34 | 0.00122 | 0.00288 |
| chr11 | 15528843 | 15529983 | 1141 | 2.37 | 0.00122 | 0.00288 |
| chr2 | 219773073 | 219774873 | 1801 | -0.69 | 0.00122 | 0.00288 |
| chr2 | 6343278 | 6343695 | 418 | 4.27 | 0.00122 | 0.00288 |
| chr6 | 145562017 | 145563125 | 1109 | -1.27 | 0.00122 | 0.00289 |
| chr3 | 44379019 | 44381275 | 2257 | 0.63 | 0.00122 | 0.00289 |
| chr5 | 77510371 | 77518411 | 8041 | 0.78 | 0.00122 | 0.00289 |
| chr7 | 33944165 | 33944600 | 436 | -3.34 | 0.00122 | 0.00289 |
| chr16 | 84047101 | 84047519 | 419 | 3.13 | 0.00122 | 0.00289 |
| chr2 | 160932626 | 160933390 | 765 | 2.04 | 0.00123 | 0.00289 |
| chr17 | 64134944 | 64135615 | 672 | 2.34 | 0.00123 | 0.00289 |
| chr6 | 44077984 | 44102289 | 24306 | -0.41 | 0.00123 | 0.00289 |
| chr6 | 80087023 | 80095966 | 8944 | 1.3 | 0.00123 | 0.0029 |
| chr5 | 127059012 | 127060009 | 998 | 1.5 | 0.00123 | 0.0029 |
| chr14 | 42609636 | 42610369 | 734 | -1.69 | 0.00123 | 0.0029 |
| chr2 | 131581362 | 131595015 | 13654 | 0.92 | 0.00123 | 0.0029 |
| chr2 | 175960933 | 175966470 | 5538 | 1.04 | 0.00123 | 0.0029 |
| chr10 | 134498181 | 134499308 | 1128 | -1.78 | 0.00123 | 0.00291 |
| chr5 | 124884587 | 124885820 | 1234 | 1.35 | 0.00123 | 0.00291 |
| chr17 | 7338770 | 7340918 | 2149 | 0.79 | 0.00123 | 0.00291 |
| chr3 | 97408600 | 97409549 | 950 | -1.88 | 0.00123 | 0.00291 |
| chr2 | 173796042 | 173796559 | 518 | -3.04 | 0.00123 | 0.00291 |
| chr19 | 11484727 | 11494213 | 9487 | 0.36 | 0.00123 | 0.00291 |
| chr6 | 136228106 | 136228604 | 499 | -3.13 | 0.00123 | 0.00291 |

| | | | | | | |
|---|---|---|---|---|---|---|
| chr14 | 27234309 | 27245721 | 11413 | 0.73 | 0.00123 | 0.00291 |
| chr7 | 8029246 | 8029886 | 641 | 2.49 | 0.00124 | 0.00291 |
| chr1 | 196755880 | 196756385 | 506 | 1.72 | 0.00124 | 0.00292 |
| chr2 | 238131653 | 238137081 | 5429 | -4.64 | 0.00124 | 0.00292 |
| chr5 | 133292474 | 133356491 | 64018 | 0.17 | 0.00124 | 0.00292 |
| chr19 | 5952577 | 5954592 | 2016 | -1.22 | 0.00124 | 0.00292 |
| chr2 | 70994017 | 71005154 | 11138 | -2.7 | 0.00124 | 0.00292 |
| chr16 | 56798049 | 56798821 | 773 | -2.58 | 0.00124 | 0.00293 |
| chr2 | 72830143 | 72831240 | 1098 | -1.27 | 0.00124 | 0.00293 |
| chrX | 119004099 | 119007490 | 3392 | 0.48 | 0.00124 | 0.00293 |
| chr6 | 128259537 | 128261397 | 1861 | -0.74 | 0.00124 | 0.00293 |
| chr14 | 27064562 | 27068287 | 3726 | 0.58 | 0.00125 | 0.00293 |
| chr9 | 20683156 | 20686944 | 3789 | 0.5 | 0.00125 | 0.00294 |
| chr18 | 34435966 | 34437326 | 1361 | 2.29 | 0.00125 | 0.00294 |
| chr14 | 22817028 | 22817431 | 404 | 4.3 | 0.00125 | 0.00294 |
| chr4 | 173644628 | 173648925 | 4298 | 0.41 | 0.00125 | 0.00294 |
| chr11 | 1394432 | 1410890 | 16459 | 0.68 | 0.00125 | 0.00294 |
| chr12 | 9065864 | 9067952 | 2089 | 0.53 | 0.00125 | 0.00294 |
| chr20 | 22725154 | 22726137 | 984 | 2.83 | 0.00125 | 0.00295 |
| chr12 | 57328719 | 57336502 | 7784 | -0.53 | 0.00125 | 0.00295 |
| chrX | 53409643 | 53411610 | 1968 | -1.37 | 0.00125 | 0.00295 |
| chr1 | 204131269 | 204131593 | 325 | 3.04 | 0.00126 | 0.00296 |
| chr1 | 240994892 | 240995373 | 482 | 3.04 | 0.00126 | 0.00296 |
| chr4 | 5365737 | 5388371 | 22635 | -0.26 | 0.00126 | 0.00296 |
| chr7 | 158484389 | 158501162 | 16774 | 0.42 | 0.00126 | 0.00296 |
| chr6 | 140945404 | 140946571 | 1168 | 1.76 | 0.00126 | 0.00296 |
| chr8 | 42910026 | 42914396 | 4371 | 0.41 | 0.00126 | 0.00296 |
| chr15 | 84214745 | 84215665 | 921 | -2 | 0.00126 | 0.00296 |
| chr15 | 72471787 | 72535823 | 64037 | -0.14 | 0.00126 | 0.00296 |
| chr2 | 178618005 | 178619811 | 1807 | 1.18 | 0.00126 | 0.00297 |
| chr1 | 62146218 | 62191847 | 45630 | 0.26 | 0.00126 | 0.00297 |
| chr2 | 74729171 | 74735526 | 6356 | -0.49 | 0.00126 | 0.00297 |
| chr1 | 115800508 | 115801095 | 588 | 2.51 | 0.00126 | 0.00297 |
| chr1 | 93955809 | 93956802 | 994 | 1.55 | 0.00126 | 0.00297 |
| chr6 | 42747379 | 42752577 | 5199 | 0.29 | 0.00126 | 0.00297 |
| chr4 | 93319838 | 93320597 | 760 | 1.86 | 0.00126 | 0.00297 |
| chr2 | 147335948 | 147337859 | 1912 | -0.96 | 0.00127 | 0.00298 |
| chr3 | 51646089 | 51655784 | 9696 | -1.01 | 0.00127 | 0.00298 |
| chr8 | 37911469 | 37916869 | 5401 | -0.86 | 0.00127 | 0.00298 |
| chr17 | 67410248 | 67412511 | 2264 | 0.83 | 0.00127 | 0.00298 |
| chr1 | 174009628 | 174010184 | 557 | 3.19 | 0.00127 | 0.00298 |
| chr6 | 70505471 | 70507757 | 2287 | 0.67 | 0.00127 | 0.00299 |
| chr12 | 110517238 | 110518248 | 1011 | 1.38 | 0.00127 | 0.00299 |
| chr3 | 101172828 | 101173393 | 566 | -2.95 | 0.00127 | 0.00299 |
| chr2 | 112640220 | 112655907 | 15688 | -0.74 | 0.00127 | 0.003 |
| chr12 | 49759541 | 49769821 | 10281 | 0.4 | 0.00128 | 0.003 |
| chr17 | 74618932 | 74620184 | 1253 | 1.46 | 0.00128 | 0.003 |
| chr10 | 101639760 | 101806364 | 166605 | 0.11 | 0.00128 | 0.003 |
| chr5 | 39892660 | 39893194 | 535 | 2.8 | 0.00128 | 0.003 |
| chr6 | 91617323 | 91618200 | 878 | -2.73 | 0.00128 | 0.003 |
| chr6 | 25302347 | 25313913 | 11567 | 0.53 | 0.00128 | 0.003 |
| chr9 | 130073558 | 130074420 | 863 | 1.22 | 0.00128 | 0.00301 |
| chr5 | 72018969 | 72020267 | 1299 | 1.37 | 0.00128 | 0.00301 |
| chr17 | 38194673 | 38196010 | 1338 | -1.12 | 0.00129 | 0.00302 |
| chr8 | 23074167 | 23088920 | 14754 | 0.32 | 0.00129 | 0.00303 |
| chr1 | 85312517 | 85313838 | 1322 | 1.77 | 0.00129 | 0.00303 |
| chr10 | 49683929 | 49685026 | 1098 | -2.05 | 0.00129 | 0.00303 |
| chr4 | 69922801 | 69924423 | 1623 | 1.12 | 0.00129 | 0.00303 |
| chr2 | 159515360 | 159525269 | 9910 | -0.87 | 0.00129 | 0.00303 |
| chr2 | 3443775 | 3444875 | 1101 | 1.89 | 0.00129 | 0.00303 |
| chr12 | 109685557 | 109699193 | 13637 | -0.84 | 0.00129 | 0.00303 |

| | | | | | | |
|---|---|---|---|---|---|---|
| chr17 | 80403637 | 80417317 | 13681 | 0.42 | 0.0013 | 0.00304 |
| chr2 | 188850324 | 188851653 | 1330 | -1.08 | 0.0013 | 0.00304 |
| chr20 | 44406825 | 44421765 | 14941 | 0.42 | 0.0013 | 0.00305 |
| chr3 | 105625756 | 105626895 | 1140 | 2.62 | 0.0013 | 0.00305 |
| chr20 | 23167824 | 23169020 | 1197 | 2.26 | 0.0013 | 0.00305 |
| chr13 | 95619430 | 95621253 | 1824 | -0.76 | 0.0013 | 0.00305 |
| chrX | 18771567 | 18773051 | 1485 | 0.9 | 0.0013 | 0.00305 |
| chr1 | 169295619 | 169308415 | 12797 | -0.91 | 0.0013 | 0.00306 |
| chr10 | 11524637 | 11525330 | 694 | 1.89 | 0.0013 | 0.00306 |
| chr1 | 192637776 | 192638251 | 476 | 2.03 | 0.0013 | 0.00306 |
| chr16 | 57546689 | 57571021 | 24333 | -0.45 | 0.00131 | 0.00306 |
| chr11 | 9682407 | 9686989 | 4583 | 0.44 | 0.00131 | 0.00307 |
| chr6 | 33605921 | 33607183 | 1263 | 1.82 | 0.00131 | 0.00307 |
| chr13 | 64007823 | 64011056 | 3234 | 4.63 | 0.00131 | 0.00307 |
| chr16 | 74729956 | 74744281 | 14326 | 0.41 | 0.00131 | 0.00307 |
| chr5 | 10232799 | 10235924 | 3126 | -1.1 | 0.00131 | 0.00307 |
| chr2 | 36581757 | 36808125 | 226369 | -0.08 | 0.00131 | 0.00307 |
| chr4 | 119199009 | 119203023 | 4015 | 0.47 | 0.00131 | 0.00308 |
| chr5 | 73795075 | 73796579 | 1505 | 0.87 | 0.00131 | 0.00308 |
| chr4 | 25812841 | 25817218 | 4378 | 0.53 | 0.00132 | 0.00308 |
| chr14 | 64243219 | 64244473 | 1255 | 2.86 | 0.00132 | 0.00308 |
| chr4 | 3359560 | 3360425 | 866 | 2.29 | 0.00132 | 0.00308 |
| chr6 | 27144099 | 27165323 | 21225 | 0.6 | 0.00132 | 0.00309 |
| chr20 | 47339106 | 47340411 | 1306 | -1.21 | 0.00132 | 0.00309 |
| chr1 | 227053436 | 227072577 | 19142 | 0.5 | 0.00132 | 0.00309 |
| chr2 | 217782078 | 217782775 | 698 | -2.69 | 0.00132 | 0.0031 |
| chr3 | 14319954 | 14320467 | 514 | -4.23 | 0.00132 | 0.0031 |
| chr15 | 36638983 | 36640592 | 1610 | 2.15 | 0.00132 | 0.0031 |
| chr12 | 28577712 | 28578623 | 912 | 2.08 | 0.00133 | 0.00311 |
| chr6 | 169970968 | 169972186 | 1219 | 1.4 | 0.00133 | 0.00311 |
| chr17 | 1090181 | 1090574 | 394 | -3.69 | 0.00133 | 0.00311 |
| chr4 | 157502717 | 157503305 | 589 | 2.96 | 0.00133 | 0.00311 |
| chr2 | 153516868 | 153517370 | 503 | 3.09 | 0.00133 | 0.00311 |
| chr20 | 16536645 | 16557108 | 20464 | 0.45 | 0.00133 | 0.00311 |
| chr3 | 47420282 | 47423593 | 3312 | 0.42 | 0.00133 | 0.00312 |
| chr19 | 55538913 | 55543944 | 5032 | 1.16 | 0.00133 | 0.00312 |
| chr2 | 84713049 | 84714291 | 1243 | 1.26 | 0.00133 | 0.00312 |
| chr13 | 34110806 | 34118606 | 7801 | 0.31 | 0.00134 | 0.00313 |
| chr4 | 168203320 | 168204412 | 1093 | 1.2 | 0.00134 | 0.00314 |
| chr10 | 32661388 | 32676273 | 14886 | 0.39 | 0.00134 | 0.00315 |
| chr3 | 190230889 | 190233162 | 2274 | 0.81 | 0.00135 | 0.00315 |
| chr4 | 184201933 | 184210823 | 8891 | 1.21 | 0.00135 | 0.00315 |
| chr15 | 44607144 | 44608768 | 1625 | 1.14 | 0.00135 | 0.00315 |
| chrX | 150994014 | 150995342 | 1329 | -1.12 | 0.00135 | 0.00315 |
| chr7 | 80568281 | 80574353 | 6073 | 0.26 | 0.00135 | 0.00315 |
| chr2 | 172563975 | 172581768 | 17794 | -0.85 | 0.00135 | 0.00316 |
| chr3 | 121255149 | 121268322 | 13174 | 0.4 | 0.00135 | 0.00316 |
| chr12 | 39274325 | 39301218 | 26894 | 0.34 | 0.00135 | 0.00316 |
| chr3 | 186284290 | 186289767 | 5478 | 0.51 | 0.00135 | 0.00317 |
| chr3 | 57093313 | 57113941 | 20629 | 0.35 | 0.00136 | 0.00317 |
| chr2 | 98972836 | 98973836 | 1001 | 2.89 | 0.00136 | 0.00317 |
| chr12 | 70698766 | 70709614 | 10849 | 2.18 | 0.00136 | 0.00317 |
| chr5 | 17890646 | 17891627 | 982 | -1.66 | 0.00136 | 0.00318 |
| chr2 | 107464763 | 107465163 | 401 | -3.15 | 0.00136 | 0.00318 |
| chr1 | 243801533 | 243801926 | 394 | 2.73 | 0.00136 | 0.00318 |
| chr1 | 214739705 | 214742030 | 2326 | -1.11 | 0.00136 | 0.00318 |
| chr10 | 125753899 | 125764124 | 10226 | -0.68 | 0.00136 | 0.00319 |
| chr6 | 147522690 | 147526344 | 3655 | 0.51 | 0.00137 | 0.00319 |
| chr9 | 36502585 | 36503996 | 1412 | -1.54 | 0.00137 | 0.0032 |
| chr6 | 56349453 | 56350182 | 730 | -2.22 | 0.00137 | 0.0032 |
| chr12 | 776156 | 777210 | 1055 | -1.31 | 0.00137 | 0.0032 |

| | | | | | | |
|---|---|---|---|---|---|---|
| chr8 | 131693901 | 131704494 | 10594 | -0.99 | 0.00137 | 0.0032 |
| chr19 | 43768649 | 43770092 | 1444 | 1.19 | 0.00137 | 0.00321 |
| chr4 | 108620366 | 108621871 | 1506 | 1.81 | 0.00137 | 0.00321 |
| chr18 | 46219565 | 46220661 | 1097 | -1.54 | 0.00137 | 0.00321 |
| chr7 | 158300590 | 158301160 | 571 | 1.89 | 0.00137 | 0.00321 |
| chr4 | 80993340 | 80994815 | 1476 | 1.21 | 0.00137 | 0.00321 |
| chr11 | 73918290 | 73919328 | 1039 | -1.77 | 0.00138 | 0.00322 |
| chr12 | 96148463 | 96158320 | 9858 | 0.67 | 0.00138 | 0.00322 |
| chrX | 151070953 | 151072707 | 1755 | 0.88 | 0.00138 | 0.00322 |
| chr11 | 101312322 | 101313002 | 681 | 1.83 | 0.00138 | 0.00322 |
| chr12 | 34488387 | 34516783 | 28397 | 0.69 | 0.00138 | 0.00322 |
| chr20 | 42285107 | 42311083 | 25977 | -0.34 | 0.00138 | 0.00323 |
| chr12 | 112779945 | 112781075 | 1131 | -1.63 | 0.00138 | 0.00323 |
| chr18 | 24084919 | 24102026 | 17108 | -0.37 | 0.00138 | 0.00323 |
| chr1 | 114976868 | 114978315 | 1448 | -1.86 | 0.00139 | 0.00323 |
| chr6 | 140858133 | 140861140 | 3008 | 0.54 | 0.00139 | 0.00325 |
| chr3 | 183462377 | 183484072 | 21696 | 0.44 | 0.00139 | 0.00325 |
| chr11 | 3391858 | 3401106 | 9249 | 0.66 | 0.00139 | 0.00325 |
| chr10 | 11991012 | 12004898 | 13887 | 0.48 | 0.00139 | 0.00325 |
| chr8 | 28982511 | 28985238 | 2728 | 2.42 | 0.0014 | 0.00326 |
| chr16 | 19221549 | 19222414 | 866 | 2.09 | 0.0014 | 0.00326 |
| chr21 | 30312245 | 30313270 | 1026 | 2 | 0.0014 | 0.00326 |
| chr2 | 40973617 | 40973941 | 325 | 3.25 | 0.0014 | 0.00327 |
| chr14 | 56317919 | 56318329 | 411 | 4.28 | 0.0014 | 0.00327 |
| chr5 | 123916481 | 123936769 | 20289 | -0.28 | 0.0014 | 0.00327 |
| chr11 | 116843749 | 116844407 | 659 | 3.31 | 0.00141 | 0.00328 |
| chr14 | 77842273 | 77868522 | 26250 | -0.47 | 0.00141 | 0.00328 |
| chr19 | 5469772 | 5471157 | 1386 | 1.52 | 0.00141 | 0.00328 |
| chrX | 35639769 | 35642684 | 2916 | 0.52 | 0.00141 | 0.00329 |
| chr1 | 29276353 | 29277373 | 1021 | -1.45 | 0.00141 | 0.00329 |
| chr14 | 103363256 | 103370656 | 7401 | -0.7 | 0.00141 | 0.00329 |
| chr13 | 61145050 | 61147675 | 2626 | 0.81 | 0.00141 | 0.00329 |
| chr5 | 130108907 | 130110706 | 1800 | 4.46 | 0.00141 | 0.00329 |
| chrX | 40939763 | 40950439 | 10677 | 0.29 | 0.00141 | 0.0033 |
| chr1 | 228945183 | 228946583 | 1401 | 1.25 | 0.00141 | 0.0033 |
| chr8 | 105660863 | 105661702 | 840 | -2.77 | 0.00142 | 0.0033 |
| chr3 | 197098629 | 197099055 | 427 | 3.44 | 0.00142 | 0.0033 |
| chr12 | 71791369 | 71791930 | 562 | 2.36 | 0.00142 | 0.0033 |
| chr11 | 70997951 | 71014390 | 16440 | -0.43 | 0.00142 | 0.00331 |
| chr3 | 176638435 | 176639278 | 844 | 1.82 | 0.00142 | 0.00331 |
| chr17 | 58537747 | 58557824 | 20078 | -0.26 | 0.00142 | 0.00331 |
| chr11 | 2097464 | 2099459 | 1996 | 0.66 | 0.00142 | 0.00331 |
| chrX | 96818211 | 96818999 | 789 | 1.51 | 0.00142 | 0.00331 |
| chr8 | 126463690 | 126475789 | 12100 | 0.81 | 0.00142 | 0.00331 |
| chr2 | 218225115 | 218232732 | 7618 | 2.86 | 0.00143 | 0.00332 |
| chr10 | 104780642 | 104782774 | 2133 | 0.85 | 0.00143 | 0.00332 |
| chr1 | 36886633 | 36887192 | 560 | -2.26 | 0.00143 | 0.00332 |
| chr9 | 33101330 | 33185549 | 84220 | 0.14 | 0.00143 | 0.00332 |
| chr6 | 131578143 | 131581165 | 3023 | 3.52 | 0.00143 | 0.00333 |
| chr9 | 3223290 | 3224487 | 1198 | 1.95 | 0.00143 | 0.00333 |
| chr10 | 104252335 | 104264599 | 12265 | 0.56 | 0.00143 | 0.00333 |
| chr4 | 184942009 | 184942476 | 468 | 3.03 | 0.00143 | 0.00333 |
| chr4 | 1580772 | 1582266 | 1495 | 0.94 | 0.00143 | 0.00333 |
| chr3 | 160880647 | 160882015 | 1369 | 4.38 | 0.00143 | 0.00333 |
| chr3 | 127871976 | 127873082 | 1107 | 1.13 | 0.00144 | 0.00334 |
| chr6 | 57106848 | 57108579 | 1732 | 1.06 | 0.00144 | 0.00334 |
| chr2 | 236645337 | 236647082 | 1746 | -1.62 | 0.00144 | 0.00334 |
| chr7 | 115715307 | 115715907 | 601 | -2.43 | 0.00144 | 0.00334 |
| chr1 | 27985999 | 28009265 | 23267 | 0.93 | 0.00144 | 0.00335 |
| chr17 | 64487822 | 64507501 | 19680 | -0.2 | 0.00144 | 0.00335 |
| chr14 | 61153608 | 61154729 | 1122 | 1.14 | 0.00144 | 0.00336 |

| | | | | | | |
|---|---|---|---|---|---|---|
| chr9 | 78504812 | 78506094 | 1283 | 1.68 | 0.00144 | 0.00336 |
| chr9 | 107616652 | 107649388 | 32737 | -0.37 | 0.00145 | 0.00336 |
| chr10 | 60868129 | 60869934 | 1806 | 0.95 | 0.00145 | 0.00337 |
| chr9 | 125938339 | 125939042 | 704 | 2.09 | 0.00145 | 0.00337 |
| chr5 | 78978028 | 78978431 | 404 | 3.74 | 0.00145 | 0.00337 |
| chr16 | 12967534 | 12976063 | 8530 | -3.91 | 0.00145 | 0.00338 |
| chr1 | 86070797 | 86079558 | 8762 | -0.32 | 0.00146 | 0.00338 |
| chr18 | 51699525 | 51700319 | 795 | -2.48 | 0.00146 | 0.00339 |
| chr1 | 8875973 | 8883696 | 7724 | 0.57 | 0.00146 | 0.00339 |
| chr10 | 52261120 | 52273171 | 12052 | 0.64 | 0.00146 | 0.00339 |
| chr13 | 110394937 | 110443920 | 48984 | 0.26 | 0.00146 | 0.00339 |
| chr3 | 37389189 | 37390460 | 1272 | -1.27 | 0.00146 | 0.00339 |
| chr7 | 141796793 | 141810754 | 13962 | 0.29 | 0.00146 | 0.00339 |
| chr1 | 18046338 | 18046947 | 610 | -1.85 | 0.00146 | 0.00339 |
| chr19 | 12661100 | 12663893 | 2794 | 0.64 | 0.00146 | 0.00339 |
| chr4 | 105817688 | 105818473 | 786 | 2.04 | 0.00147 | 0.0034 |
| chr5 | 43482308 | 43486614 | 4307 | 0.39 | 0.00147 | 0.0034 |
| chr2 | 47647413 | 47671235 | 23823 | -0.51 | 0.00147 | 0.00341 |
| chr8 | 61620438 | 61622174 | 1737 | 0.74 | 0.00147 | 0.00341 |
| chr11 | 13482896 | 13485832 | 2937 | 0.46 | 0.00147 | 0.00341 |
| chr7 | 106931796 | 106932473 | 678 | 2.33 | 0.00147 | 0.00341 |
| chr7 | 100539551 | 100540754 | 1204 | -1.51 | 0.00147 | 0.00341 |
| chrX | 41386065 | 41387032 | 968 | -1.87 | 0.00147 | 0.00341 |
| chr15 | 34610294 | 34611499 | 1206 | 0.78 | 0.00147 | 0.00342 |
| chr16 | 9563708 | 9564055 | 348 | 3.03 | 0.00147 | 0.00342 |
| chr15 | 101995465 | 101996953 | 1489 | -1.43 | 0.00147 | 0.00342 |
| chr2 | 18497990 | 18498886 | 897 | -1.59 | 0.00148 | 0.00343 |
| chr1 | 28994315 | 28996553 | 2239 | 0.45 | 0.00148 | 0.00343 |
| chr2 | 70387623 | 70388546 | 924 | 1.86 | 0.00148 | 0.00343 |
| chr2 | 153487152 | 153487592 | 441 | -3.19 | 0.00148 | 0.00343 |
| chr17 | 71331300 | 71340277 | 8978 | -0.36 | 0.00148 | 0.00343 |
| chr2 | 204261306 | 204293722 | 32417 | -0.32 | 0.00148 | 0.00343 |
| chr8 | 56675941 | 56688154 | 12214 | 0.33 | 0.00148 | 0.00344 |
| chr11 | 47931388 | 47983300 | 51913 | -0.14 | 0.00149 | 0.00345 |
| chr18 | 2654791 | 2659619 | 4829 | 0.41 | 0.00149 | 0.00345 |
| chr4 | 88340356 | 88345031 | 4676 | 0.46 | 0.00149 | 0.00345 |
| chr8 | 60911290 | 60912044 | 755 | -2.77 | 0.00149 | 0.00346 |
| chr2 | 120613641 | 120620412 | 6772 | 0.55 | 0.00149 | 0.00346 |
| chr3 | 86963617 | 86963956 | 340 | -2.83 | 0.00149 | 0.00346 |
| chr5 | 43208322 | 43209328 | 1007 | 2.44 | 0.0015 | 0.00347 |
| chr1 | 40722494 | 40724854 | 2361 | 0.54 | 0.0015 | 0.00347 |
| chr6 | 35915393 | 35916499 | 1107 | -1.43 | 0.0015 | 0.00347 |
| chr3 | 196392465 | 196393608 | 1144 | 1.37 | 0.0015 | 0.00347 |
| chr3 | 49043840 | 49067419 | 23580 | 0.29 | 0.0015 | 0.00347 |
| chr4 | 147224807 | 147225256 | 450 | 4.22 | 0.0015 | 0.00348 |
| chrX | 51580144 | 51580915 | 772 | 2.09 | 0.0015 | 0.00348 |
| chr2 | 62100093 | 62116811 | 16719 | -0.32 | 0.0015 | 0.00348 |
| chr1 | 85872044 | 85874472 | 2429 | 0.47 | 0.0015 | 0.00348 |
| chr19 | 4759302 | 4760497 | 1196 | -1.64 | 0.0015 | 0.00348 |
| chr19 | 49457252 | 49503506 | 46255 | -0.24 | 0.00151 | 0.00349 |
| chr4 | 69132603 | 69134208 | 1606 | 3.29 | 0.00151 | 0.00349 |
| chr7 | 111894689 | 111897730 | 3042 | 1.05 | 0.00151 | 0.00349 |
| chr13 | 113969639 | 113970482 | 844 | 2.06 | 0.00151 | 0.0035 |
| chr2 | 162946340 | 162949860 | 3521 | 0.86 | 0.00151 | 0.0035 |
| chr18 | 55112316 | 55114120 | 1805 | -0.89 | 0.00151 | 0.0035 |
| chr21 | 34696337 | 34698411 | 2075 | 0.52 | 0.00152 | 0.00351 |
| chr11 | 67005992 | 67086202 | 80211 | -0.18 | 0.00152 | 0.00351 |
| chr17 | 830854 | 837731 | 6878 | -0.45 | 0.00152 | 0.00351 |
| chr3 | 106064121 | 106065021 | 901 | -1.96 | 0.00152 | 0.00351 |
| chrX | 153595830 | 153609290 | 13461 | 0.29 | 0.00152 | 0.00351 |
| chr2 | 69197295 | 69200454 | 3160 | -2.73 | 0.00152 | 0.00352 |

| | | | | | | |
|---|---|---|---|---|---|---|
| chr1 | 153986502 | 153987236 | 735 | 2.5 | 0.00152 | 0.00352 |
| chr9 | 88286687 | 88288203 | 1517 | -1.44 | 0.00152 | 0.00352 |
| chr19 | 10627449 | 10628483 | 1035 | -2.08 | 0.00152 | 0.00352 |
| chr6 | 96023947 | 96026671 | 2725 | 0.51 | 0.00152 | 0.00352 |
| chr5 | 90485291 | 90486447 | 1157 | 1.17 | 0.00152 | 0.00352 |
| chr6 | 21633855 | 21645117 | 11263 | 0.86 | 0.00153 | 0.00353 |
| chr2 | 63997072 | 63997528 | 457 | -2.72 | 0.00153 | 0.00353 |
| chr11 | 126859782 | 126873347 | 13566 | -0.34 | 0.00153 | 0.00354 |
| chr1 | 226084600 | 226086349 | 1750 | 1.26 | 0.00153 | 0.00354 |
| chr11 | 28279660 | 28280765 | 1106 | -2.26 | 0.00153 | 0.00355 |
| chr12 | 113795468 | 113797325 | 1858 | 0.83 | 0.00153 | 0.00355 |
| chrX | 19643244 | 19649930 | 6687 | -0.92 | 0.00154 | 0.00355 |
| chr19 | 14223453 | 14230839 | 7387 | 0.73 | 0.00154 | 0.00356 |
| chr18 | 65183425 | 65193269 | 9845 | 4.4 | 0.00154 | 0.00356 |
| chr4 | 183159345 | 183162686 | 3342 | -4.58 | 0.00154 | 0.00356 |
| chr1 | 10783803 | 10791869 | 8067 | 1.08 | 0.00154 | 0.00357 |
| chr11 | 126301604 | 126302331 | 728 | -2.86 | 0.00154 | 0.00357 |
| chr14 | 54778763 | 54779791 | 1029 | 2 | 0.00155 | 0.00357 |
| chr3 | 15351560 | 15353319 | 1760 | 1.02 | 0.00155 | 0.00358 |
| chr3 | 177174441 | 177175043 | 603 | 2.93 | 0.00155 | 0.00358 |
| chr12 | 41170022 | 41171453 | 1432 | 0.99 | 0.00155 | 0.00358 |
| chr11 | 96121709 | 96124216 | 2508 | 0.5 | 0.00155 | 0.00358 |
| chr15 | 90630298 | 90631091 | 794 | 2.19 | 0.00155 | 0.00358 |
| chr13 | 53017458 | 53038429 | 20972 | 0.36 | 0.00155 | 0.00358 |
| chr1 | 11795559 | 11797494 | 1936 | -0.93 | 0.00155 | 0.00358 |
| chr10 | 133453371 | 133455278 | 1908 | 4.71 | 0.00155 | 0.00359 |
| chr5 | 137641125 | 137641958 | 834 | -2.7 | 0.00155 | 0.00359 |
| chr15 | 33305696 | 33315216 | 9521 | 1.15 | 0.00155 | 0.00359 |
| chr8 | 41540351 | 41543500 | 3150 | -0.8 | 0.00156 | 0.0036 |
| chr15 | 52729588 | 52730820 | 1233 | -1.15 | 0.00156 | 0.0036 |
| chr9 | 116412182 | 116417823 | 5642 | -1.05 | 0.00156 | 0.0036 |
| chr7 | 68794308 | 68794989 | 682 | 2.88 | 0.00156 | 0.0036 |
| chr5 | 141906479 | 141910130 | 3652 | 1.89 | 0.00156 | 0.0036 |
| chr1 | 98433151 | 98433919 | 769 | 1.75 | 0.00156 | 0.0036 |
| chrX | 134185029 | 134187043 | 2015 | 0.61 | 0.00156 | 0.0036 |
| chr15 | 42782584 | 42788725 | 6142 | 0.41 | 0.00156 | 0.00361 |
| chr1 | 171453556 | 171471614 | 18059 | 0.37 | 0.00157 | 0.00361 |
| chr9 | 5493320 | 5494405 | 1086 | -1.44 | 0.00157 | 0.00362 |
| chr17 | 10658189 | 10659725 | 1537 | -1.89 | 0.00157 | 0.00362 |
| chr13 | 90159836 | 90168276 | 8441 | 0.31 | 0.00157 | 0.00363 |
| chr12 | 116673872 | 116726219 | 52348 | 0.34 | 0.00157 | 0.00363 |
| chr12 | 82800285 | 82806507 | 6223 | 2.71 | 0.00158 | 0.00364 |
| chr3 | 24127715 | 24143541 | 15827 | 0.74 | 0.00158 | 0.00364 |
| chr2 | 170654730 | 170657183 | 2454 | -0.69 | 0.00158 | 0.00365 |
| chr17 | 78846641 | 78848279 | 1639 | -1.06 | 0.00158 | 0.00365 |
| chr1 | 193583615 | 193584735 | 1121 | -1.45 | 0.00159 | 0.00366 |
| chr2 | 11924850 | 11925726 | 877 | -3 | 0.00159 | 0.00366 |
| chr13 | 24132869 | 24134201 | 1333 | 0.98 | 0.00159 | 0.00366 |
| chr17 | 40756319 | 40777078 | 20760 | 0.4 | 0.00159 | 0.00366 |
| chr15 | 23064027 | 23071772 | 7746 | -1.1 | 0.00159 | 0.00367 |
| chr9 | 34218429 | 34223263 | 4835 | -0.75 | 0.00159 | 0.00367 |
| chrX | 129280650 | 129305700 | 25051 | -0.48 | 0.00159 | 0.00367 |
| chr1 | 28672392 | 28681432 | 9041 | 0.43 | 0.00159 | 0.00367 |
| chr6 | 119082458 | 119083669 | 1212 | 1.31 | 0.00159 | 0.00367 |
| chr14 | 102413869 | 102470072 | 56204 | 0.19 | 0.0016 | 0.00368 |
| chr4 | 165877546 | 165878803 | 1258 | 0.82 | 0.0016 | 0.00369 |
| chr9 | 117348827 | 117355160 | 6334 | 0.37 | 0.0016 | 0.00369 |
| chr5 | 74887291 | 74888136 | 846 | 1.81 | 0.0016 | 0.00369 |
| chr12 | 99990383 | 99991170 | 788 | -1.77 | 0.0016 | 0.00369 |
| chrX | 41189201 | 41196318 | 7118 | 0.22 | 0.0016 | 0.00369 |
| chr7 | 73989443 | 74027192 | 37750 | -0.28 | 0.00161 | 0.0037 |

| | | | | | | |
|---|---|---|---|---|---|---|
| chr12 | 41040852 | 41042166 | 1315 | 0.97 | 0.00161 | 0.0037 |
| chr6 | 26148057 | 26208228 | 60172 | 0.17 | 0.00161 | 0.0037 |
| chr19 | 58192640 | 58194423 | 1784 | 0.9 | 0.00161 | 0.00371 |
| chr8 | 109479761 | 109493181 | 13421 | 0.51 | 0.00161 | 0.00371 |
| chr11 | 117778193 | 117778867 | 675 | 1.77 | 0.00161 | 0.00371 |
| chr16 | 14343616 | 14355923 | 12308 | -1.07 | 0.00161 | 0.00371 |
| chr8 | 74903006 | 74904137 | 1132 | 1.42 | 0.00161 | 0.00371 |
| chr10 | 70066106 | 70069241 | 3136 | 0.99 | 0.00161 | 0.00371 |
| chr1 | 6709675 | 6710678 | 1004 | -1.88 | 0.00162 | 0.00372 |
| chr19 | 51667686 | 51668032 | 347 | -4.15 | 0.00162 | 0.00372 |
| chr14 | 54717440 | 54720933 | 3494 | 2.92 | 0.00162 | 0.00373 |
| chr19 | 18473971 | 18548946 | 74976 | -0.19 | 0.00162 | 0.00373 |
| chr14 | 52326987 | 52341091 | 14105 | 0.63 | 0.00162 | 0.00373 |
| chr10 | 22853264 | 22853872 | 609 | -2.97 | 0.00162 | 0.00373 |
| chr5 | 64398397 | 64399991 | 1595 | 0.6 | 0.00162 | 0.00373 |
| chr1 | 189282435 | 189282851 | 417 | 2.43 | 0.00162 | 0.00373 |
| chr6 | 160450845 | 160452279 | 1435 | 1.58 | 0.00162 | 0.00373 |
| chr5 | 158212307 | 158213680 | 1374 | -1.5 | 0.00162 | 0.00373 |
| chr10 | 90342267 | 90343628 | 1362 | 0.64 | 0.00162 | 0.00373 |
| chr7 | 1082817 | 1084646 | 1830 | 0.6 | 0.00163 | 0.00374 |
| chr17 | 31610808 | 31611833 | 1026 | 1.55 | 0.00163 | 0.00374 |
| chr17 | 11895176 | 11901698 | 6523 | 0.45 | 0.00163 | 0.00374 |
| chr20 | 48861732 | 48938981 | 77250 | -0.17 | 0.00163 | 0.00375 |
| chr11 | 76412747 | 76413650 | 904 | -2.08 | 0.00163 | 0.00375 |
| chr22 | 29698813 | 29707749 | 8937 | -0.51 | 0.00163 | 0.00375 |
| chr17 | 66096758 | 66098507 | 1750 | 0.73 | 0.00163 | 0.00375 |
| chr7 | 83338038 | 83338942 | 905 | -2.01 | 0.00164 | 0.00376 |
| chr4 | 153818820 | 153819977 | 1158 | -1.23 | 0.00164 | 0.00376 |
| chr16 | 72041876 | 72060273 | 18398 | -0.42 | 0.00164 | 0.00377 |
| chr4 | 189048391 | 189052172 | 3782 | 0.79 | 0.00164 | 0.00377 |
| chr14 | 77624023 | 77625774 | 1752 | -1.04 | 0.00164 | 0.00377 |
| chr19 | 40354950 | 40355292 | 343 | -2.59 | 0.00164 | 0.00377 |
| chr20 | 20528804 | 20529835 | 1032 | 1.64 | 0.00164 | 0.00377 |
| chr11 | 98235317 | 98237503 | 2187 | 0.88 | 0.00164 | 0.00377 |
| chr4 | 183786230 | 183787157 | 928 | 1.87 | 0.00164 | 0.00377 |
| chr7 | 40259808 | 40260718 | 911 | 2.82 | 0.00164 | 0.00377 |
| chr3 | 11432020 | 11433268 | 1249 | 2.08 | 0.00164 | 0.00377 |
| chr1 | 220510011 | 220511300 | 1290 | 1.71 | 0.00165 | 0.00378 |
| chr22 | 50825236 | 50835818 | 10583 | 1.28 | 0.00165 | 0.00379 |
| chr1 | 109202978 | 109204532 | 1555 | 0.59 | 0.00165 | 0.00379 |
| chr11 | 77704754 | 77707229 | 2476 | 0.52 | 0.00165 | 0.00379 |
| chr2 | 233432320 | 233471488 | 39169 | -0.57 | 0.00165 | 0.0038 |
| chr3 | 13258912 | 13260622 | 1711 | -1.43 | 0.00166 | 0.0038 |
| chr10 | 123358293 | 123358596 | 304 | 4.18 | 0.00166 | 0.0038 |
| chr4 | 122790625 | 122792218 | 1594 | 0.77 | 0.00166 | 0.0038 |
| chr18 | 3321298 | 3322027 | 730 | -2.23 | 0.00166 | 0.0038 |
| chr1 | 212186477 | 212187769 | 1293 | -1.75 | 0.00166 | 0.0038 |
| chr15 | 44380245 | 44380799 | 555 | -2.88 | 0.00166 | 0.00381 |
| chr16 | 46645937 | 46664032 | 18096 | -0.57 | 0.00166 | 0.00381 |
| chr9 | 116880270 | 116881780 | 1511 | 4.2 | 0.00166 | 0.00381 |
| chr2 | 181593149 | 181608324 | 15176 | -0.66 | 0.00166 | 0.00381 |
| chr13 | 34922293 | 34951320 | 29028 | 0.27 | 0.00166 | 0.00381 |
| chr13 | 30392293 | 30392947 | 655 | 3.01 | 0.00166 | 0.00381 |
| chrX | 9971214 | 10010463 | 39250 | -0.22 | 0.00166 | 0.00381 |
| chr11 | 85497222 | 85507296 | 10075 | 0.3 | 0.00167 | 0.00382 |
| chr19 | 48064151 | 48065838 | 1688 | 1.08 | 0.00167 | 0.00382 |
| chr1 | 113683390 | 113685106 | 1717 | -1.43 | 0.00167 | 0.00383 |
| chr9 | 78883694 | 78886681 | 2988 | 0.56 | 0.00167 | 0.00383 |
| chr14 | 68085515 | 68087154 | 1640 | 0.69 | 0.00167 | 0.00384 |
| chr10 | 75715619 | 75728436 | 12818 | 0.26 | 0.00167 | 0.00384 |
| chr1 | 39812168 | 39813392 | 1225 | -1.33 | 0.00168 | 0.00384 |

| | | | | | | |
|---|---|---|---|---|---|---|
| chr3 | 114806128 | 114820111 | 13984 | 2.07 | 0.00168 | 0.00384 |
| chr11 | 108070045 | 108071543 | 1499 | -1.05 | 0.00168 | 0.00384 |
| chr15 | 96794946 | 96812782 | 17837 | 1.88 | 0.00168 | 0.00384 |
| chr3 | 124669443 | 124670973 | 1531 | -1.27 | 0.00168 | 0.00384 |
| chr3 | 43779802 | 43780557 | 756 | -1.87 | 0.00168 | 0.00384 |
| chr9 | 91361753 | 91362915 | 1163 | -2.14 | 0.00168 | 0.00385 |
| chr6 | 80145523 | 80146053 | 531 | 2.74 | 0.00168 | 0.00386 |
| chr12 | 69977883 | 69992397 | 14515 | 0.36 | 0.00169 | 0.00386 |
| chr17 | 2338061 | 2368418 | 30358 | -0.51 | 0.00169 | 0.00386 |
| chr16 | 68738003 | 68742745 | 4743 | 0.55 | 0.00169 | 0.00386 |
| chr6 | 147475053 | 147476364 | 1312 | 2.76 | 0.00169 | 0.00387 |
| chr6 | 151308276 | 151309175 | 900 | -2.09 | 0.00169 | 0.00387 |
| chr8 | 122753507 | 122762427 | 8921 | -0.35 | 0.00169 | 0.00387 |
| chr20 | 10764237 | 10765577 | 1341 | -1.29 | 0.00169 | 0.00387 |
| chr4 | 26387048 | 26387589 | 542 | -3.06 | 0.00169 | 0.00388 |
| chr11 | 45063835 | 45068738 | 4904 | 0.59 | 0.00169 | 0.00388 |
| chr8 | 18244667 | 18245044 | 378 | 2.38 | 0.0017 | 0.00388 |
| chr2 | 133235387 | 133237759 | 2373 | 4.58 | 0.0017 | 0.00388 |
| chr19 | 10596470 | 10614892 | 18423 | -0.37 | 0.0017 | 0.00389 |
| chr1 | 24513238 | 24529830 | 16593 | -0.5 | 0.0017 | 0.00389 |
| chr9 | 77385210 | 77386250 | 1041 | 1.13 | 0.0017 | 0.0039 |
| chr18 | 19207430 | 19220108 | 12679 | 0.7 | 0.0017 | 0.0039 |
| chr16 | 82686926 | 82688071 | 1146 | -1.51 | 0.00171 | 0.0039 |
| chr7 | 214315 | 227172 | 12858 | -0.58 | 0.00171 | 0.0039 |
| chr4 | 68564920 | 68580286 | 15367 | 0.36 | 0.00171 | 0.0039 |
| chr2 | 28107128 | 28115460 | 8333 | 0.26 | 0.00171 | 0.0039 |
| chr5 | 171530003 | 171544593 | 14591 | -0.72 | 0.00171 | 0.00391 |
| chr2 | 43650915 | 43714174 | 63260 | -0.17 | 0.00171 | 0.00391 |
| chr1 | 114352119 | 114355871 | 3753 | 0.62 | 0.00171 | 0.00391 |
| chr6 | 165591834 | 165595990 | 4157 | 2.77 | 0.00171 | 0.00391 |
| chr2 | 54681283 | 54716513 | 35231 | -0.35 | 0.00171 | 0.00391 |
| chr19 | 36689259 | 36689855 | 597 | -2.1 | 0.00171 | 0.00392 |
| chr8 | 224163 | 225054 | 892 | 2.76 | 0.00171 | 0.00392 |
| chr13 | 25874929 | 25893120 | 18192 | 0.36 | 0.00172 | 0.00392 |
| chr1 | 61718810 | 61719888 | 1079 | 1.25 | 0.00172 | 0.00393 |
| chr17 | 75228787 | 75252681 | 23895 | 0.36 | 0.00172 | 0.00393 |
| chr8 | 23309782 | 23315732 | 5951 | 0.71 | 0.00172 | 0.00393 |
| chr22 | 39493100 | 39493776 | 677 | -1.54 | 0.00172 | 0.00393 |
| chr9 | 139949550 | 139950643 | 1094 | 1.63 | 0.00172 | 0.00393 |
| chr12 | 79757406 | 79758902 | 1497 | -1.1 | 0.00172 | 0.00394 |
| chr11 | 1357349 | 1364742 | 7394 | 1.87 | 0.00173 | 0.00394 |
| chr11 | 18654633 | 18657015 | 2383 | 0.39 | 0.00173 | 0.00395 |
| chr22 | 22429170 | 22429561 | 392 | -2.68 | 0.00173 | 0.00395 |
| chr3 | 148342209 | 148344115 | 1907 | 1.01 | 0.00173 | 0.00395 |
| chr2 | 26829111 | 26838799 | 9689 | -4.76 | 0.00173 | 0.00395 |
| chr11 | 56550679 | 56551056 | 378 | 2.8 | 0.00174 | 0.00397 |
| chr14 | 80735956 | 80736378 | 423 | -2.35 | 0.00174 | 0.00397 |
| chr12 | 76034196 | 76035798 | 1603 | 0.88 | 0.00174 | 0.00397 |
| chr14 | 95649896 | 95650957 | 1062 | 1.49 | 0.00174 | 0.00398 |
| chr22 | 50699012 | 50701668 | 2657 | -0.98 | 0.00175 | 0.00399 |
| chr3 | 11408557 | 11409500 | 944 | 1.11 | 0.00175 | 0.00399 |
| chr2 | 24122988 | 24171045 | 48058 | -0.25 | 0.00175 | 0.00399 |
| chr7 | 127224647 | 127229446 | 4800 | 0.46 | 0.00175 | 0.004 |
| chr7 | 38885434 | 38904469 | 19036 | 5.27 | 0.00175 | 0.004 |
| chr5 | 38834371 | 38855471 | 21101 | 0.38 | 0.00175 | 0.004 |
| chr15 | 59279199 | 59288915 | 9717 | 0.42 | 0.00175 | 0.004 |
| chr13 | 45905590 | 45916812 | 11223 | 0.42 | 0.00175 | 0.004 |
| chr17 | 45675545 | 45701709 | 26165 | 0.22 | 0.00176 | 0.004 |
| chr10 | 94865607 | 94867451 | 1845 | 1.2 | 0.00176 | 0.004 |
| chr7 | 12071067 | 12071644 | 578 | 1.92 | 0.00176 | 0.00401 |
| chr19 | 47986500 | 48019703 | 33204 | -0.3 | 0.00176 | 0.00401 |

| | | | | | | |
|---|---|---|---|---|---|---|
| chr13 | 114808449 | 114810991 | 2543 | -0.94 | 0.00176 | 0.00401 |
| chr2 | 197755882 | 197756225 | 344 | -2.84 | 0.00176 | 0.00401 |
| chr19 | 41929998 | 41946341 | 16344 | -0.49 | 0.00176 | 0.00401 |
| chr15 | 85433052 | 85433851 | 800 | -1.8 | 0.00176 | 0.00401 |
| chr11 | 125982852 | 125987164 | 4313 | -0.39 | 0.00176 | 0.00402 |
| chr10 | 72734094 | 72734610 | 517 | -3.08 | 0.00176 | 0.00402 |
| chr16 | 11262430 | 11276442 | 14013 | -1.24 | 0.00176 | 0.00402 |
| chr11 | 102380911 | 102382030 | 1120 | 1.99 | 0.00176 | 0.00402 |
| chr2 | 153025293 | 153033338 | 8046 | -0.24 | 0.00177 | 0.00403 |
| chr8 | 95071050 | 95071694 | 645 | 2.21 | 0.00177 | 0.00403 |
| chr1 | 194202280 | 194203147 | 868 | 1.55 | 0.00177 | 0.00403 |
| chr2 | 128255666 | 128268565 | 12900 | -0.41 | 0.00177 | 0.00403 |
| chr1 | 240025571 | 240026056 | 486 | 2.54 | 0.00177 | 0.00403 |
| chr16 | 89665884 | 89676287 | 10404 | -1.13 | 0.00177 | 0.00403 |
| chrX | 18604179 | 18604926 | 748 | -2.88 | 0.00177 | 0.00403 |
| chr1 | 41947698 | 41964808 | 17111 | -0.45 | 0.00177 | 0.00404 |
| chr6 | 138169977 | 138170468 | 492 | 2.74 | 0.00178 | 0.00405 |
| chr4 | 186346112 | 186348015 | 1904 | -0.49 | 0.00178 | 0.00405 |
| chr12 | 57359646 | 57359958 | 313 | 3.67 | 0.00178 | 0.00405 |
| chr7 | 38409041 | 38409541 | 501 | 2.84 | 0.00178 | 0.00406 |
| chr11 | 76534324 | 76548102 | 13779 | -0.47 | 0.00178 | 0.00406 |
| chr13 | 115078683 | 115080871 | 2189 | 0.62 | 0.00178 | 0.00406 |
| chr7 | 32078198 | 32111446 | 33249 | 0.21 | 0.00178 | 0.00406 |
| chr19 | 41402832 | 41403939 | 1108 | -2.06 | 0.00178 | 0.00406 |
| chr1 | 209979646 | 209980003 | 358 | 2.57 | 0.00179 | 0.00406 |
| chr5 | 100596064 | 100597490 | 1427 | 1.04 | 0.00179 | 0.00407 |
| chrX | 41977883 | 41979625 | 1743 | 1.13 | 0.00179 | 0.00407 |
| chr10 | 114148834 | 114154948 | 6115 | 4.71 | 0.00179 | 0.00407 |
| chrX | 40633254 | 40633683 | 430 | 2.2 | 0.00179 | 0.00407 |
| chr7 | 99213504 | 99216073 | 2570 | 0.52 | 0.00179 | 0.00408 |
| chr10 | 22534354 | 22541727 | 7374 | -0.31 | 0.00179 | 0.00408 |
| chr1 | 222761919 | 222774376 | 12458 | 0.58 | 0.0018 | 0.00409 |
| chr12 | 78725687 | 78726383 | 697 | 2.22 | 0.0018 | 0.00409 |
| chr10 | 12740529 | 12763038 | 22510 | -0.33 | 0.0018 | 0.00409 |
| chr7 | 7154122 | 7155083 | 962 | 1.82 | 0.0018 | 0.00409 |
| chr9 | 124307704 | 124320694 | 12991 | -1.12 | 0.0018 | 0.00409 |
| chr15 | 66147292 | 66147776 | 485 | 3.21 | 0.0018 | 0.0041 |
| chr4 | 77751527 | 77753650 | 2124 | -0.48 | 0.0018 | 0.0041 |
| chr13 | 25860359 | 25862036 | 1678 | 0.69 | 0.00181 | 0.00411 |
| chr7 | 80001628 | 80015788 | 14161 | 4.63 | 0.00181 | 0.00412 |
| chr1 | 209268608 | 209280126 | 11519 | 1.65 | 0.00181 | 0.00412 |
| chr14 | 70179201 | 70180506 | 1306 | -1.66 | 0.00182 | 0.00413 |
| chr5 | 149509273 | 149523390 | 14118 | -0.75 | 0.00182 | 0.00413 |
| chr2 | 235582756 | 235612703 | 29948 | -0.28 | 0.00182 | 0.00413 |
| chr22 | 27281848 | 27292364 | 10517 | 0.39 | 0.00182 | 0.00414 |
| chr6 | 113843096 | 113843893 | 798 | 1.76 | 0.00182 | 0.00414 |
| chr1 | 187460323 | 187461446 | 1124 | -1.42 | 0.00182 | 0.00414 |
| chr6 | 45412743 | 45415046 | 2304 | 0.99 | 0.00182 | 0.00414 |
| chr17 | 8020753 | 8131648 | 110896 | 0.11 | 0.00182 | 0.00414 |
| chr10 | 89698380 | 89721973 | 23594 | -0.3 | 0.00183 | 0.00415 |
| chr7 | 40026692 | 40056229 | 29538 | 0.36 | 0.00183 | 0.00415 |
| chr2 | 121954913 | 121955509 | 597 | -2.18 | 0.00183 | 0.00415 |
| chr11 | 126455239 | 126456623 | 1385 | 1.21 | 0.00183 | 0.00416 |
| chr12 | 85447114 | 85447486 | 373 | -3.04 | 0.00183 | 0.00416 |
| chr10 | 112676694 | 112684826 | 8133 | 0.27 | 0.00183 | 0.00416 |
| chr1 | 217262739 | 217263852 | 1114 | 1.22 | 0.00183 | 0.00416 |
| chr9 | 36922972 | 36925743 | 2772 | 0.78 | 0.00184 | 0.00417 |
| chr2 | 109148674 | 109154487 | 5814 | -0.4 | 0.00184 | 0.00417 |
| chr4 | 105341486 | 105342446 | 961 | 2.21 | 0.00184 | 0.00418 |
| chr6 | 46702272 | 46703076 | 805 | 2.51 | 0.00184 | 0.00418 |
| chr1 | 232972590 | 232973808 | 1219 | 1.23 | 0.00184 | 0.00419 |

| | | | | | | |
|---|---|---|---|---|---|---|
| chr2 | 139153495 | 139154146 | 652 | -2.45 | 0.00184 | 0.00419 |
| chr2 | 32322026 | 32323238 | 1213 | -1.42 | 0.00185 | 0.00419 |
| chr9 | 94876650 | 94878567 | 1918 | 0.4 | 0.00185 | 0.00421 |
| chr8 | 63950752 | 63952940 | 2189 | -0.48 | 0.00186 | 0.00421 |
| chr10 | 134036610 | 134044316 | 7707 | -0.95 | 0.00186 | 0.00422 |
| chr11 | 77379734 | 77383910 | 4177 | 0.68 | 0.00186 | 0.00422 |
| chr2 | 55554712 | 55555340 | 629 | 2.24 | 0.00186 | 0.00422 |
| chr21 | 46587013 | 46588258 | 1246 | -1.55 | 0.00186 | 0.00423 |
| chr12 | 113573764 | 113598519 | 24756 | 0.81 | 0.00186 | 0.00423 |
| chr4 | 185733644 | 185735555 | 1912 | -1.74 | 0.00186 | 0.00423 |
| chr8 | 31029592 | 31030422 | 831 | -1.71 | 0.00187 | 0.00423 |
| chr4 | 177514900 | 177515845 | 946 | 2.92 | 0.00187 | 0.00424 |
| chr16 | 55726069 | 55727591 | 1523 | 1.17 | 0.00187 | 0.00424 |
| chr6 | 37123056 | 37140783 | 17728 | 0.37 | 0.00187 | 0.00425 |
| chr9 | 108209486 | 108221835 | 12350 | -0.44 | 0.00187 | 0.00425 |
| chr22 | 30882398 | 30883426 | 1029 | 1.76 | 0.00188 | 0.00425 |
| chr21 | 48054396 | 48056693 | 2298 | 0.57 | 0.00188 | 0.00425 |
| chr9 | 135777540 | 135786904 | 9365 | -1.06 | 0.00188 | 0.00426 |
| chr4 | 111332457 | 111333153 | 697 | 1.48 | 0.00188 | 0.00426 |
| chr21 | 39695140 | 39696409 | 1270 | 1.42 | 0.00188 | 0.00426 |
| chr13 | 33845150 | 33860657 | 15508 | 0.27 | 0.00188 | 0.00427 |
| chr13 | 100307272 | 100311394 | 4123 | 0.39 | 0.00189 | 0.00427 |
| chrX | 73807374 | 73808350 | 977 | 1.96 | 0.00189 | 0.00428 |
| chr10 | 103441774 | 103455562 | 13789 | 0.34 | 0.00189 | 0.00428 |
| chr1 | 22581643 | 22586744 | 5102 | 0.81 | 0.00189 | 0.00428 |
| chr8 | 81077302 | 81084954 | 7653 | -0.41 | 0.00189 | 0.00429 |
| chr6 | 148677799 | 148678798 | 1000 | 1.84 | 0.00189 | 0.00429 |
| chr7 | 48039203 | 48040609 | 1407 | 1.69 | 0.00189 | 0.00429 |
| chrX | 4158568 | 4159582 | 1015 | 2.06 | 0.00189 | 0.00429 |
| chr12 | 58328951 | 58336074 | 7124 | 0.34 | 0.0019 | 0.0043 |
| chr8 | 29171846 | 29173647 | 1802 | 0.78 | 0.0019 | 0.0043 |
| chr11 | 73308647 | 73310525 | 1879 | 0.77 | 0.0019 | 0.00431 |
| chr7 | 46969509 | 46970681 | 1173 | 1.24 | 0.0019 | 0.00431 |
| chr6 | 21557403 | 21558107 | 705 | 2.63 | 0.0019 | 0.00431 |
| chr11 | 69294107 | 69296338 | 2232 | 0.6 | 0.0019 | 0.00431 |
| chr19 | 58815825 | 58818100 | 2276 | 0.48 | 0.0019 | 0.00431 |
| chr5 | 36690129 | 36690913 | 785 | 1.4 | 0.00191 | 0.00432 |
| chr9 | 114061151 | 114061738 | 588 | -1.82 | 0.00191 | 0.00432 |
| chr18 | 76915946 | 76917961 | 2016 | 1.28 | 0.00191 | 0.00433 |
| chr15 | 86085722 | 86087133 | 1412 | 1.13 | 0.00191 | 0.00433 |
| chr16 | 53927316 | 53928709 | 1394 | 1.36 | 0.00191 | 0.00433 |
| chr1 | 193158992 | 193160242 | 1251 | -1.37 | 0.00191 | 0.00433 |
| chr6 | 141167839 | 141169041 | 1203 | 1.51 | 0.00191 | 0.00433 |
| chr4 | 166033053 | 166035185 | 2133 | 0.74 | 0.00192 | 0.00434 |
| chr15 | 44827960 | 44839056 | 11097 | 0.46 | 0.00192 | 0.00434 |
| chr12 | 13860817 | 13861230 | 414 | -3.33 | 0.00192 | 0.00434 |
| chr2 | 71727356 | 71731560 | 4205 | -1.46 | 0.00192 | 0.00435 |
| chr10 | 79192046 | 79192437 | 392 | -4.1 | 0.00192 | 0.00435 |
| chr6 | 159051911 | 159066742 | 14832 | 0.33 | 0.00193 | 0.00436 |
| chr1 | 218457854 | 218459419 | 1566 | 0.58 | 0.00193 | 0.00436 |
| chr6 | 137315909 | 137323094 | 7186 | 2.73 | 0.00193 | 0.00437 |
| chr1 | 63078978 | 63089583 | 10606 | 0.58 | 0.00193 | 0.00437 |
| chr10 | 96723813 | 96724182 | 370 | 3.21 | 0.00193 | 0.00437 |
| chr13 | 34838692 | 34841762 | 3071 | 0.84 | 0.00193 | 0.00437 |
| chr19 | 53589841 | 53590714 | 874 | 2.81 | 0.00194 | 0.00438 |
| chr15 | 57737165 | 57738925 | 1761 | -0.7 | 0.00194 | 0.00438 |
| chr1 | 147305332 | 147305909 | 578 | 3.01 | 0.00194 | 0.00438 |
| chr3 | 5291939 | 5293381 | 1443 | 4.02 | 0.00194 | 0.00439 |
| chr4 | 77221858 | 77228802 | 6945 | 0.48 | 0.00194 | 0.00439 |
| chr6 | 75340576 | 75341989 | 1414 | -1.93 | 0.00195 | 0.0044 |
| chr9 | 33366888 | 33372188 | 5301 | -0.96 | 0.00195 | 0.0044 |

| | | | | | | |
|---|---|---|---|---|---|---|
| chr11 | 106353616 | 106354093 | 478 | 2.67 | 0.00195 | 0.0044 |
| chr15 | 29560290 | 29562796 | 2507 | 0.46 | 0.00196 | 0.00442 |
| chr2 | 169102577 | 169109131 | 6555 | 0.64 | 0.00196 | 0.00444 |
| chr21 | 32475757 | 32476134 | 378 | 4.14 | 0.00197 | 0.00444 |
| chr20 | 20615475 | 20615981 | 507 | -2.51 | 0.00197 | 0.00444 |
| chr6 | 22954596 | 22956024 | 1429 | 1.01 | 0.00197 | 0.00444 |
| chr16 | 3283936 | 3286810 | 2875 | 0.68 | 0.00197 | 0.00445 |
| chr11 | 117959088 | 117959960 | 873 | 2.79 | 0.00197 | 0.00445 |
| chr7 | 149146645 | 149158739 | 12095 | -0.84 | 0.00197 | 0.00446 |
| chr16 | 1375736 | 1382882 | 7147 | -0.51 | 0.00198 | 0.00446 |
| chr19 | 50681902 | 50682699 | 798 | 1.57 | 0.00198 | 0.00446 |
| chr2 | 20062958 | 20063790 | 833 | -2.23 | 0.00198 | 0.00447 |
| chr7 | 75496034 | 75516579 | 20546 | -0.46 | 0.00199 | 0.00449 |
| chr11 | 3062710 | 3080250 | 17541 | -0.3 | 0.00199 | 0.00449 |
| chr4 | 128981322 | 128984322 | 3001 | 0.54 | 0.00199 | 0.0045 |
| chr16 | 75539840 | 75551659 | 11820 | 0.69 | 0.00199 | 0.0045 |
| chr7 | 81097508 | 81130559 | 33052 | -0.47 | 0.002 | 0.00451 |
| chr4 | 99402999 | 99405751 | 2753 | -0.58 | 0.002 | 0.00451 |
| chr5 | 53996083 | 53999731 | 3649 | 4.8 | 0.002 | 0.00452 |
| chr1 | 36200759 | 36201592 | 834 | -2.96 | 0.00201 | 0.00453 |
| chr21 | 35132765 | 35136295 | 3531 | 0.33 | 0.00201 | 0.00453 |
| chrX | 119737236 | 119739909 | 2674 | 0.65 | 0.00201 | 0.00453 |
| chr1 | 150736178 | 150738928 | 2751 | 0.37 | 0.00201 | 0.00453 |
| chrX | 63141778 | 63142228 | 451 | -3.02 | 0.00201 | 0.00453 |
| chr17 | 47177878 | 47184749 | 6872 | -0.64 | 0.00201 | 0.00453 |
| chr3 | 33790945 | 33791503 | 559 | -2.64 | 0.00201 | 0.00453 |
| chr13 | 46625735 | 46636996 | 11262 | 0.63 | 0.00201 | 0.00454 |
| chrX | 13445415 | 13451337 | 5923 | -0.45 | 0.00202 | 0.00455 |
| chr22 | 45098259 | 45104430 | 6172 | -0.96 | 0.00202 | 0.00455 |
| chr5 | 180050237 | 180050869 | 633 | -1.85 | 0.00202 | 0.00456 |
| chr2 | 38735746 | 38737091 | 1346 | -1.21 | 0.00202 | 0.00456 |
| chr20 | 25057245 | 25058939 | 1695 | -1.11 | 0.00202 | 0.00456 |
| chrX | 109559386 | 109562477 | 3092 | 0.55 | 0.00202 | 0.00456 |
| chr7 | 129419046 | 129421190 | 2145 | 0.68 | 0.00202 | 0.00456 |
| chr15 | 55919069 | 55921259 | 2191 | 0.55 | 0.00202 | 0.00456 |
| chr1 | 115258065 | 115260325 | 2261 | 0.57 | 0.00203 | 0.00457 |
| chr17 | 7209040 | 7234330 | 25291 | 0.24 | 0.00203 | 0.00458 |
| chr16 | 27214261 | 27224358 | 10098 | 0.4 | 0.00204 | 0.00459 |
| chr20 | 883467 | 884174 | 708 | -2.77 | 0.00204 | 0.00459 |
| chr15 | 75706848 | 75766669 | 59822 | 0.2 | 0.00204 | 0.0046 |
| chr12 | 64388268 | 64404995 | 16728 | -0.53 | 0.00204 | 0.0046 |
| chr15 | 35742196 | 35743105 | 910 | 1.74 | 0.00204 | 0.0046 |
| chr11 | 6240084 | 6241354 | 1271 | 1.52 | 0.00204 | 0.0046 |
| chr10 | 24919324 | 24920557 | 1234 | -1.15 | 0.00205 | 0.00461 |
| chr18 | 56272133 | 56279527 | 7395 | 0.93 | 0.00205 | 0.00461 |
| chr17 | 42808108 | 42815782 | 7675 | -0.83 | 0.00205 | 0.00461 |
| chr1 | 2242649 | 2246573 | 3925 | -0.9 | 0.00205 | 0.00462 |
| chr7 | 152160233 | 152162483 | 2251 | 0.66 | 0.00205 | 0.00462 |
| chr18 | 70835383 | 70836785 | 1403 | 1.67 | 0.00206 | 0.00463 |
| chr2 | 232403301 | 232403679 | 379 | 4.22 | 0.00206 | 0.00463 |
| chr16 | 68692092 | 68693234 | 1143 | 1.06 | 0.00206 | 0.00463 |
| chr9 | 124454282 | 124455544 | 1263 | 1.47 | 0.00206 | 0.00464 |
| chr15 | 52653081 | 52654299 | 1219 | -1.32 | 0.00206 | 0.00464 |
| chr1 | 27240578 | 27250181 | 9604 | -0.52 | 0.00206 | 0.00464 |
| chr7 | 5512043 | 5512671 | 629 | -2.06 | 0.00206 | 0.00464 |
| chr14 | 69657452 | 69659267 | 1816 | 0.6 | 0.00207 | 0.00465 |
| chr1 | 52034550 | 52036428 | 1879 | 0.75 | 0.00207 | 0.00465 |
| chr13 | 32604880 | 32606542 | 1663 | 3.98 | 0.00207 | 0.00465 |
| chr6 | 13422711 | 13423504 | 794 | 1.92 | 0.00207 | 0.00465 |
| chr4 | 141588024 | 141589332 | 1309 | 1.29 | 0.00207 | 0.00466 |
| chr3 | 189427447 | 189440394 | 12948 | 0.84 | 0.00207 | 0.00466 |

| | | | | | | |
|---|---|---|---|---|---|---|
| chr11 | 66005346 | 66008880 | 3535 | -1.41 | 0.00207 | 0.00466 |
| chr4 | 14668762 | 14670050 | 1289 | 1.31 | 0.00207 | 0.00466 |
| chr11 | 130785456 | 130786745 | 1290 | 0.95 | 0.00207 | 0.00466 |
| chr2 | 109104566 | 109109905 | 5340 | -0.74 | 0.00208 | 0.00467 |
| chr13 | 39880829 | 39881243 | 415 | 3.42 | 0.00208 | 0.00467 |
| chr1 | 8720967 | 8730940 | 9974 | -1.07 | 0.00208 | 0.00467 |
| chr3 | 9904523 | 9941462 | 36940 | 0.21 | 0.00208 | 0.00468 |
| chr3 | 132440105 | 132442562 | 2458 | 0.66 | 0.00208 | 0.00468 |
| chr1 | 98981520 | 98982458 | 939 | -2.71 | 0.00208 | 0.00468 |
| chr1 | 9598561 | 9600892 | 2332 | 0.67 | 0.00209 | 0.00469 |
| chr7 | 111369456 | 111370768 | 1313 | 1.79 | 0.00209 | 0.00469 |
| chr19 | 42695281 | 42807756 | 112476 | -0.13 | 0.00209 | 0.0047 |
| chr12 | 104917064 | 104918022 | 959 | 1.32 | 0.00209 | 0.0047 |
| chr5 | 40677039 | 40690586 | 13548 | 0.45 | 0.00209 | 0.0047 |
| chr1 | 173445221 | 173449891 | 4671 | 0.41 | 0.00209 | 0.0047 |
| chr10 | 90791481 | 90792846 | 1366 | 1.64 | 0.00209 | 0.0047 |
| chr6 | 113699566 | 113713436 | 13871 | 1.43 | 0.00209 | 0.0047 |
| chr3 | 187704151 | 187704975 | 825 | -1.87 | 0.0021 | 0.00471 |
| chr3 | 13202101 | 13202774 | 674 | -2.88 | 0.0021 | 0.00471 |
| chr3 | 120314536 | 120318475 | 3940 | 0.49 | 0.0021 | 0.00471 |
| chr18 | 47314449 | 47315999 | 1551 | 0.7 | 0.0021 | 0.00471 |
| chr10 | 8494724 | 8497538 | 2815 | 0.97 | 0.0021 | 0.00471 |
| chr7 | 31034712 | 31035988 | 1277 | -1.55 | 0.0021 | 0.00471 |
| chr3 | 31982091 | 31983478 | 1388 | 1.02 | 0.0021 | 0.00472 |
| chr12 | 66134241 | 66135933 | 1693 | 0.78 | 0.0021 | 0.00472 |
| chr15 | 73562720 | 73563643 | 924 | -1.39 | 0.0021 | 0.00472 |
| chr2 | 208337458 | 208338477 | 1020 | -1.52 | 0.0021 | 0.00472 |
| chr12 | 33187918 | 33188410 | 493 | -2.84 | 0.0021 | 0.00472 |
| chr19 | 32912416 | 32913200 | 785 | -1.51 | 0.00211 | 0.00473 |
| chr10 | 133587404 | 133588482 | 1079 | 1.87 | 0.00211 | 0.00473 |
| chr7 | 141062493 | 141063368 | 876 | -2.73 | 0.00211 | 0.00474 |
| chr11 | 14013911 | 14015146 | 1236 | 1.92 | 0.00211 | 0.00474 |
| chr13 | 109309534 | 109315480 | 5947 | 0.91 | 0.00211 | 0.00474 |
| chr4 | 67429248 | 67444101 | 14854 | -0.25 | 0.00211 | 0.00474 |
| chr1 | 64546032 | 64546973 | 942 | -1.64 | 0.00211 | 0.00474 |
| chr15 | 82424822 | 82425469 | 648 | -2.69 | 0.00212 | 0.00474 |
| chr9 | 130937279 | 130981502 | 44224 | 0.17 | 0.00212 | 0.00475 |
| chr9 | 140444267 | 140451667 | 7401 | 0.65 | 0.00212 | 0.00476 |
| chr3 | 31082617 | 31083828 | 1212 | 3.63 | 0.00212 | 0.00476 |
| chr5 | 133701072 | 133714631 | 13560 | 0.29 | 0.00212 | 0.00476 |
| chr2 | 9345758 | 9352563 | 6806 | -0.46 | 0.00212 | 0.00476 |
| chr2 | 114030274 | 114042668 | 12395 | -0.3 | 0.00213 | 0.00478 |
| chr12 | 18428358 | 18429041 | 684 | 2.01 | 0.00213 | 0.00478 |
| chr6 | 1142621 | 1143038 | 418 | 2.94 | 0.00213 | 0.00478 |
| chr12 | 94852164 | 94854747 | 2584 | 0.37 | 0.00214 | 0.00479 |
| chr4 | 146402261 | 146404487 | 2227 | 0.55 | 0.00214 | 0.00479 |
| chr11 | 70371030 | 70372121 | 1092 | -1.7 | 0.00214 | 0.00479 |
| chr17 | 2062009 | 2078729 | 16721 | -0.53 | 0.00214 | 0.00479 |
| chr4 | 170864009 | 170865814 | 1806 | 4.36 | 0.00214 | 0.00479 |
| chr15 | 42766746 | 42767788 | 1043 | -1.51 | 0.00214 | 0.00479 |
| chr19 | 40993486 | 40994302 | 817 | -1.15 | 0.00214 | 0.0048 |
| chr4 | 72363468 | 72365074 | 1607 | 4.03 | 0.00214 | 0.0048 |
| chr7 | 139703612 | 139704697 | 1086 | -2.85 | 0.00215 | 0.00481 |
| chr2 | 180870871 | 180872562 | 1692 | -0.44 | 0.00216 | 0.00483 |
| chr13 | 73544256 | 73545200 | 945 | 1.32 | 0.00216 | 0.00484 |
| chr15 | 72402247 | 72411661 | 9415 | 0.47 | 0.00216 | 0.00485 |
| chr10 | 46089084 | 46090909 | 1826 | 0.67 | 0.00216 | 0.00485 |
| chr9 | 17195336 | 17196496 | 1161 | 1.64 | 0.00217 | 0.00486 |
| chr7 | 79081067 | 79092335 | 11269 | -0.77 | 0.00218 | 0.00487 |
| chr5 | 102453553 | 102457349 | 3797 | 0.33 | 0.00218 | 0.00488 |
| chr3 | 126325691 | 126326762 | 1072 | -1.62 | 0.00218 | 0.00488 |

| | | | | | | |
|---|---|---|---|---|---|---|
| chr4 | 65270900 | 65271260 | 361 | 3.65 | 0.00218 | 0.00488 |
| chr16 | 24596703 | 24603982 | 7280 | -4.16 | 0.00218 | 0.00489 |
| chr6 | 90389549 | 90401164 | 11616 | -0.66 | 0.00218 | 0.00489 |
| chr15 | 57959252 | 57960038 | 787 | 2.1 | 0.00218 | 0.00489 |
| chr14 | 65877778 | 65880612 | 2835 | 0.49 | 0.00219 | 0.00489 |
| chr17 | 47067954 | 47134241 | 66288 | 0.12 | 0.00219 | 0.0049 |
| chr6 | 25138205 | 25139704 | 1500 | 0.89 | 0.00219 | 0.00491 |
| chr8 | 74862537 | 74890216 | 27680 | 0.3 | 0.00219 | 0.00491 |
| chr11 | 122661941 | 122663608 | 1668 | -1.22 | 0.0022 | 0.00491 |
| chr1 | 92291576 | 92318006 | 26431 | 0.23 | 0.0022 | 0.00492 |
| chr18 | 46925996 | 46927931 | 1936 | 1.13 | 0.0022 | 0.00492 |
| chr6 | 137582245 | 137583490 | 1246 | 1.48 | 0.00221 | 0.00493 |
| chrX | 24750026 | 24751450 | 1425 | 1.52 | 0.00221 | 0.00493 |
| chr1 | 240515631 | 240517337 | 1707 | 0.94 | 0.00221 | 0.00493 |
| chr4 | 6096762 | 6098053 | 1292 | -1.39 | 0.00221 | 0.00495 |
| chr3 | 183208240 | 183209271 | 1032 | 1.8 | 0.00221 | 0.00495 |
| chr22 | 44453188 | 44457846 | 4659 | -0.46 | 0.00221 | 0.00495 |
| chr1 | 239941133 | 239942149 | 1017 | -1.64 | 0.00222 | 0.00495 |
| chr2 | 68382914 | 68386225 | 3312 | -0.34 | 0.00222 | 0.00495 |
| chr3 | 105136414 | 105160199 | 23786 | 0.47 | 0.00222 | 0.00496 |
| chr19 | 57751368 | 57753153 | 1786 | 0.44 | 0.00222 | 0.00496 |
| chr16 | 80231449 | 80233468 | 2020 | -0.51 | 0.00222 | 0.00496 |
| chr4 | 154313792 | 154315004 | 1213 | 1.76 | 0.00222 | 0.00497 |
| chr15 | 58712549 | 58713655 | 1107 | -1.8 | 0.00222 | 0.00497 |
| chr3 | 70422976 | 70424082 | 1107 | 2.12 | 0.00222 | 0.00497 |
| chr7 | 100952160 | 100966017 | 13858 | -0.44 | 0.00223 | 0.00497 |
| chr11 | 74270629 | 74271140 | 512 | 2.22 | 0.00223 | 0.00498 |
| chr3 | 113275855 | 113277688 | 1834 | 0.77 | 0.00223 | 0.00499 |
| chr2 | 242088263 | 242090609 | 2347 | -0.89 | 0.00223 | 0.00499 |
| chr10 | 14967984 | 14969466 | 1483 | 1.4 | 0.00224 | 0.00499 |
| chr3 | 25452755 | 25500668 | 47914 | -0.2 | 0.00224 | 0.005 |
| chr11 | 92664763 | 92665948 | 1186 | 1.7 | 0.00224 | 0.005 |
| chr1 | 39767998 | 39772119 | 4122 | 0.58 | 0.00224 | 0.005 |
| chr7 | 65877975 | 65879171 | 1197 | 1.11 | 0.00224 | 0.00501 |
| chr4 | 108611029 | 108611996 | 968 | -2.07 | 0.00224 | 0.00501 |
| chr11 | 12460896 | 12461733 | 838 | 2.25 | 0.00224 | 0.00501 |
| chr15 | 59729540 | 59739683 | 10144 | -1.19 | 0.00224 | 0.00501 |
| chr8 | 135805779 | 135806905 | 1127 | 1.52 | 0.00224 | 0.00501 |
| chr1 | 85961504 | 85961977 | 474 | 2.59 | 0.00224 | 0.00501 |
| chr11 | 87009127 | 87012631 | 3505 | 1.04 | 0.00225 | 0.00501 |
| chr3 | 16592524 | 16593196 | 673 | -2.25 | 0.00225 | 0.00501 |
| chr1 | 45956274 | 45992827 | 36554 | -0.21 | 0.00225 | 0.00502 |
| chr1 | 94682802 | 94683954 | 1153 | -1.69 | 0.00225 | 0.00503 |
| chr6 | 14001211 | 14002538 | 1328 | 1.11 | 0.00226 | 0.00504 |
| chr1 | 233242403 | 233253235 | 10833 | 0.16 | 0.00226 | 0.00504 |
| chr7 | 117822781 | 117826609 | 3829 | 0.33 | 0.00226 | 0.00504 |
| chr12 | 100866965 | 100868261 | 1297 | 4.36 | 0.00226 | 0.00505 |
| chr9 | 32663666 | 32675067 | 11402 | 5.55 | 0.00226 | 0.00505 |
| chr15 | 53081927 | 53083049 | 1123 | -1.42 | 0.00226 | 0.00505 |
| chr2 | 73440963 | 73442716 | 1754 | -0.58 | 0.00226 | 0.00505 |
| chr2 | 101396608 | 101403713 | 7106 | 0.81 | 0.00226 | 0.00505 |
| chr12 | 41269536 | 41270359 | 824 | 1.78 | 0.00227 | 0.00505 |
| chr6 | 54800004 | 54801002 | 999 | 1.65 | 0.00227 | 0.00505 |
| chr15 | 77875081 | 77875788 | 708 | 2.11 | 0.00227 | 0.00507 |
| chr2 | 242692452 | 242693629 | 1178 | -1.5 | 0.00227 | 0.00507 |
| chr3 | 141804774 | 141809107 | 4334 | 2.13 | 0.00227 | 0.00507 |
| chr11 | 206041 | 217614 | 11574 | 0.3 | 0.00228 | 0.00507 |
| chr15 | 99826852 | 99828425 | 1574 | -1.99 | 0.00228 | 0.00508 |
| chr1 | 112135195 | 112136202 | 1008 | 1.89 | 0.00228 | 0.00508 |
| chr18 | 412951 | 414072 | 1122 | -1.32 | 0.00228 | 0.00508 |
| chr1 | 41156775 | 41159265 | 2491 | -0.51 | 0.00228 | 0.00508 |

| | | | | | | |
|---|---|---|---|---|---|---|
| chrX | 70445008 | 70446306 | 1299 | 1.57 | 0.00228 | 0.00508 |
| chr10 | 13169320 | 13171092 | 1773 | -1.55 | 0.00228 | 0.00508 |
| chr21 | 48086937 | 48088230 | 1294 | -1.52 | 0.00228 | 0.00508 |
| chr12 | 64173355 | 64175322 | 1968 | 0.55 | 0.00228 | 0.00508 |
| chr1 | 232799415 | 232814902 | 15488 | 5.46 | 0.00229 | 0.0051 |
| chr6 | 53146700 | 53147605 | 906 | -1.97 | 0.00229 | 0.0051 |
| chr3 | 69488366 | 69489002 | 637 | 1.19 | 0.00229 | 0.0051 |
| chr6 | 132301658 | 132303241 | 1584 | 1.34 | 0.0023 | 0.00511 |
| chr18 | 20354289 | 20354885 | 597 | 2.98 | 0.00231 | 0.00514 |
| chr5 | 57315418 | 57318895 | 3478 | -1.02 | 0.00231 | 0.00514 |
| chr12 | 28692404 | 28704771 | 12368 | 0.66 | 0.00231 | 0.00514 |
| chr2 | 228017206 | 228030146 | 12941 | 0.7 | 0.00231 | 0.00514 |
| chr10 | 96084644 | 96085131 | 488 | -3.13 | 0.00231 | 0.00514 |
| chr12 | 69326018 | 69327492 | 1475 | 1.37 | 0.00231 | 0.00514 |
| chr2 | 148887900 | 148890890 | 2991 | 0.86 | 0.00231 | 0.00514 |
| chr11 | 18785170 | 18786604 | 1435 | -1.32 | 0.00231 | 0.00514 |
| chr10 | 90638970 | 90658238 | 19269 | -0.22 | 0.00231 | 0.00514 |
| chr9 | 116968763 | 116970220 | 1458 | -1.75 | 0.00232 | 0.00515 |
| chr4 | 154153923 | 154168658 | 14736 | 3.31 | 0.00232 | 0.00515 |
| chr3 | 131171891 | 131173433 | 1543 | -1.52 | 0.00232 | 0.00515 |
| chr3 | 142606867 | 142608384 | 1518 | -0.97 | 0.00232 | 0.00515 |
| chr10 | 112240994 | 112296416 | 55423 | -0.15 | 0.00232 | 0.00516 |
| chr20 | 44034424 | 44049311 | 14888 | 0.36 | 0.00233 | 0.00517 |
| chr8 | 29876840 | 29878036 | 1197 | 1.38 | 0.00233 | 0.00517 |
| chr10 | 95363795 | 95364539 | 745 | 2.27 | 0.00233 | 0.00517 |
| chr10 | 103638969 | 103640621 | 1653 | -1.1 | 0.00233 | 0.00518 |
| chr1 | 51939072 | 51940206 | 1135 | 1.65 | 0.00233 | 0.00519 |
| chr1 | 207970176 | 208043090 | 72915 | 0.16 | 0.00234 | 0.0052 |
| chr1 | 185703349 | 185703827 | 479 | 2.62 | 0.00234 | 0.0052 |
| chr1 | 159566435 | 159569232 | 2798 | -0.4 | 0.00234 | 0.00521 |
| chr1 | 42326432 | 42329768 | 3337 | -0.59 | 0.00235 | 0.00522 |
| chr8 | 54605316 | 54606082 | 767 | -1.32 | 0.00235 | 0.00522 |
| chr16 | 9161332 | 9224240 | 62909 | 0.17 | 0.00235 | 0.00523 |
| chrX | 36233314 | 36233815 | 502 | 2.56 | 0.00235 | 0.00523 |
| chr4 | 134632778 | 134633627 | 850 | 1.83 | 0.00236 | 0.00525 |
| chr18 | 66380477 | 66383222 | 2746 | 0.62 | 0.00236 | 0.00525 |
| chr1 | 63782554 | 63789490 | 6937 | 0.73 | 0.00236 | 0.00525 |
| chr6 | 40904478 | 40941196 | 36719 | 0.39 | 0.00236 | 0.00525 |
| chr21 | 33503887 | 33505403 | 1517 | -1.13 | 0.00236 | 0.00525 |
| chr8 | 33316135 | 33372070 | 55936 | 0.17 | 0.00236 | 0.00525 |
| chr9 | 33906661 | 33954208 | 47548 | -0.23 | 0.00237 | 0.00526 |
| chr5 | 55053592 | 55054883 | 1292 | 1.59 | 0.00237 | 0.00527 |
| chr15 | 66678147 | 66681041 | 2895 | 0.59 | 0.00237 | 0.00527 |
| chr5 | 76011336 | 76013042 | 1707 | -1.1 | 0.00237 | 0.00527 |
| chr2 | 37330638 | 37346700 | 16063 | -0.46 | 0.00238 | 0.00527 |
| chr5 | 178156812 | 178157660 | 849 | 1.26 | 0.00238 | 0.00527 |
| chr1 | 12008935 | 12009781 | 847 | 2.74 | 0.00238 | 0.00528 |
| chr14 | 81130412 | 81131467 | 1056 | -1.38 | 0.00238 | 0.00528 |
| chr8 | 80031366 | 80033441 | 2076 | 4.36 | 0.00238 | 0.00528 |
| chr2 | 183771133 | 183795144 | 24012 | -0.46 | 0.00239 | 0.0053 |
| chr2 | 28884128 | 28885473 | 1346 | 1.69 | 0.00239 | 0.0053 |
| chr4 | 3343563 | 3346011 | 2449 | 0.49 | 0.0024 | 0.00532 |
| chr18 | 9333800 | 9349531 | 15732 | 0.73 | 0.0024 | 0.00532 |
| chr3 | 99535854 | 99537522 | 1669 | 0.58 | 0.0024 | 0.00533 |
| chr6 | 421872 | 424053 | 2182 | 0.44 | 0.0024 | 0.00533 |
| chr4 | 38116095 | 38135673 | 19579 | 0.58 | 0.0024 | 0.00533 |
| chr14 | 50777761 | 50788739 | 10979 | 0.45 | 0.0024 | 0.00533 |
| chr7 | 73754946 | 73778988 | 24043 | -0.4 | 0.0024 | 0.00533 |
| chr11 | 67329584 | 67332160 | 2577 | -0.9 | 0.0024 | 0.00533 |
| chr8 | 87120284 | 87121902 | 1619 | 1.13 | 0.00241 | 0.00534 |
| chr4 | 124299924 | 124301121 | 1198 | 2.42 | 0.00242 | 0.00536 |

| | | | | | | |
|---|---|---|---|---|---|---|
| chr22 | 51020305 | 51021957 | 1653 | 0.79 | 0.00242 | 0.00536 |
| chr12 | 43025264 | 43026187 | 924 | 2.72 | 0.00242 | 0.00536 |
| chr5 | 82325624 | 82326350 | 727 | -1.83 | 0.00242 | 0.00537 |
| chr8 | 26206905 | 26208542 | 1638 | -1.23 | 0.00242 | 0.00537 |
| chr3 | 17225004 | 17226396 | 1393 | -1.02 | 0.00242 | 0.00537 |
| chr6 | 10372910 | 10384558 | 11649 | 0.56 | 0.00242 | 0.00537 |
| chr16 | 75419002 | 75419564 | 563 | -1.45 | 0.00242 | 0.00537 |
| chr7 | 7678831 | 7681921 | 3091 | 0.41 | 0.00242 | 0.00537 |
| chr1 | 46215376 | 46217340 | 1965 | 0.46 | 0.00242 | 0.00537 |
| chr6 | 17579807 | 17601804 | 21998 | 0.6 | 0.00243 | 0.00538 |
| chr3 | 111518509 | 111530734 | 12226 | -0.29 | 0.00243 | 0.00538 |
| chr22 | 18008372 | 18010026 | 1655 | 0.95 | 0.00243 | 0.00538 |
| chr7 | 11012174 | 11015569 | 3396 | 0.38 | 0.00244 | 0.0054 |
| chr1 | 228782828 | 228785804 | 2977 | 0.4 | 0.00244 | 0.0054 |
| chr1 | 233266646 | 233271985 | 5340 | -1.02 | 0.00244 | 0.00541 |
| chr3 | 122130923 | 122147708 | 16786 | 0.33 | 0.00244 | 0.00541 |
| chr5 | 154108491 | 154109463 | 973 | 2.19 | 0.00245 | 0.00542 |
| chr10 | 95461031 | 95463362 | 2332 | 0.45 | 0.00245 | 0.00542 |
| chr16 | 82044510 | 82045121 | 612 | 1.95 | 0.00245 | 0.00542 |
| chr2 | 37788087 | 37790115 | 2029 | 0.67 | 0.00245 | 0.00542 |
| chr8 | 86107496 | 86108121 | 626 | -2.06 | 0.00245 | 0.00543 |
| chr11 | 63974533 | 63975993 | 1461 | -1.74 | 0.00245 | 0.00543 |
| chr18 | 46516489 | 46548347 | 31859 | -0.24 | 0.00246 | 0.00544 |
| chr2 | 20725011 | 20735978 | 10968 | -1.36 | 0.00246 | 0.00544 |
| chr17 | 37773697 | 37775454 | 1758 | -0.69 | 0.00246 | 0.00544 |
| chr2 | 234505717 | 234506398 | 682 | -2.82 | 0.00246 | 0.00544 |
| chr3 | 100520582 | 100521829 | 1248 | 4.32 | 0.00246 | 0.00544 |
| chr8 | 101554129 | 101555857 | 1729 | -0.81 | 0.00246 | 0.00545 |
| chr11 | 28587252 | 28589514 | 2263 | -0.45 | 0.00246 | 0.00545 |
| chr17 | 47372989 | 47375739 | 2751 | 0.85 | 0.00247 | 0.00546 |
| chr2 | 75549006 | 75549546 | 541 | -2.7 | 0.00247 | 0.00547 |
| chr4 | 15939330 | 15940812 | 1483 | 2.03 | 0.00247 | 0.00547 |
| chr4 | 31558975 | 31559780 | 806 | 1.6 | 0.00248 | 0.00548 |
| chr11 | 95940495 | 95977696 | 37202 | 0.34 | 0.00248 | 0.00548 |
| chr1 | 228639647 | 228647859 | 8213 | -0.51 | 0.00248 | 0.00548 |
| chr15 | 31780024 | 31782348 | 2325 | -1.16 | 0.00248 | 0.00548 |
| chrX | 67652700 | 67683550 | 30851 | 0.41 | 0.00248 | 0.00548 |
| chr1 | 204448055 | 204495322 | 47268 | 0.22 | 0.00248 | 0.00548 |
| chr12 | 87610912 | 87611930 | 1019 | 4.04 | 0.00248 | 0.00548 |
| chr4 | 124981347 | 124982290 | 944 | 2.67 | 0.00248 | 0.00549 |
| chr1 | 169336155 | 169338592 | 2438 | 0.57 | 0.00248 | 0.00549 |
| chr19 | 58858331 | 58859909 | 1579 | 1.16 | 0.00249 | 0.0055 |
| chr18 | 3966016 | 3968781 | 2766 | -0.66 | 0.00249 | 0.0055 |
| chr4 | 1075640 | 1076734 | 1095 | -1.54 | 0.00249 | 0.00551 |
| chr10 | 111965388 | 111970944 | 5557 | -0.86 | 0.00249 | 0.00551 |
| chr10 | 75935302 | 75952607 | 17306 | 0.32 | 0.0025 | 0.00552 |
| chr15 | 75890683 | 75894079 | 3397 | -1.44 | 0.0025 | 0.00552 |
| chr22 | 46398615 | 46433690 | 35076 | 0.25 | 0.00251 | 0.00554 |
| chr2 | 3621604 | 3633770 | 12167 | -0.35 | 0.00251 | 0.00554 |
| chr4 | 2668950 | 2669505 | 556 | 2.86 | 0.00251 | 0.00555 |
| chr8 | 103248330 | 103252998 | 4669 | 0.35 | 0.00251 | 0.00555 |
| chr5 | 61910121 | 61912592 | 2472 | 0.77 | 0.00251 | 0.00555 |
| chr18 | 13452220 | 13452632 | 413 | -2.72 | 0.00251 | 0.00555 |
| chr1 | 70874368 | 70877747 | 3380 | 0.42 | 0.00251 | 0.00555 |
| chr10 | 120706508 | 120720006 | 13499 | -0.46 | 0.00251 | 0.00555 |
| chr7 | 83638409 | 83656796 | 18388 | -0.46 | 0.00251 | 0.00555 |
| chr13 | 65045855 | 65047205 | 1351 | -1.07 | 0.00252 | 0.00556 |
| chr21 | 35916566 | 35918916 | 2351 | 0.82 | 0.00252 | 0.00557 |
| chr10 | 72620609 | 72621007 | 399 | 3.65 | 0.00252 | 0.00557 |
| chr9 | 33505888 | 33535662 | 29775 | 0.32 | 0.00252 | 0.00557 |
| chr20 | 37050208 | 37102475 | 52268 | 0.26 | 0.00252 | 0.00557 |

| | | | | | | |
|---|---|---|---|---|---|---|
| chr4 | 48377157 | 48379771 | 2615 | 0.75 | 0.00253 | 0.00558 |
| chr15 | 51322140 | 51323842 | 1703 | -0.7 | 0.00253 | 0.00558 |
| chr20 | 18538854 | 18549782 | 10929 | 0.37 | 0.00253 | 0.00558 |
| chr11 | 119991749 | 119993655 | 1907 | 1.45 | 0.00253 | 0.00558 |
| chr8 | 30012847 | 30032488 | 19642 | 0.33 | 0.00253 | 0.00558 |
| chr2 | 3270834 | 3293821 | 22988 | -1.04 | 0.00253 | 0.00558 |
| chr9 | 139885541 | 139891687 | 6147 | 0.55 | 0.00253 | 0.00558 |
| chr14 | 91525693 | 91533689 | 7997 | -0.51 | 0.00253 | 0.00558 |
| chr2 | 205409304 | 205411086 | 1783 | -0.7 | 0.00253 | 0.00559 |
| chr1 | 160398201 | 160399369 | 1169 | 1.73 | 0.00253 | 0.00559 |
| chr3 | 106176014 | 106177095 | 1082 | -1.3 | 0.00253 | 0.00559 |
| chr20 | 15087910 | 15088606 | 697 | 1.67 | 0.00254 | 0.00559 |
| chr14 | 22750511 | 22751334 | 824 | 1.7 | 0.00254 | 0.00559 |
| chr15 | 65404239 | 65406386 | 2148 | -1.37 | 0.00254 | 0.0056 |
| chr18 | 47360415 | 47386332 | 25918 | 0.31 | 0.00254 | 0.0056 |
| chr20 | 464484 | 525192 | 60709 | 0.2 | 0.00254 | 0.0056 |
| chr15 | 99887383 | 99888177 | 795 | -1.71 | 0.00254 | 0.00561 |
| chr8 | 22209400 | 22259358 | 49959 | 0.22 | 0.00254 | 0.00561 |
| chr3 | 150037453 | 150038509 | 1057 | 1.99 | 0.00254 | 0.00561 |
| chr12 | 93919965 | 93936491 | 16527 | 0.97 | 0.00255 | 0.00562 |
| chr1 | 113849619 | 113869184 | 19566 | 0.39 | 0.00255 | 0.00563 |
| chr3 | 193719963 | 193723703 | 3741 | 0.47 | 0.00255 | 0.00563 |
| chr1 | 31434673 | 31461858 | 27186 | 0.38 | 0.00255 | 0.00563 |
| chr1 | 111746355 | 111747400 | 1046 | 1.77 | 0.00255 | 0.00563 |
| chr4 | 25160944 | 25163966 | 3023 | 0.51 | 0.00256 | 0.00563 |
| chr12 | 96501754 | 96502518 | 765 | 1.9 | 0.00256 | 0.00564 |
| chr2 | 159312498 | 159316231 | 3734 | -0.51 | 0.00256 | 0.00564 |
| chr1 | 205669612 | 205720272 | 50661 | 0.22 | 0.00256 | 0.00565 |
| chrX | 51635747 | 51638556 | 2810 | 0.75 | 0.00256 | 0.00565 |
| chr3 | 161607466 | 161609066 | 1601 | 1.08 | 0.00257 | 0.00566 |
| chr17 | 11924021 | 11945330 | 21310 | 0.41 | 0.00257 | 0.00566 |
| chr2 | 47496363 | 47500510 | 4148 | -0.44 | 0.00257 | 0.00566 |
| chr16 | 74328967 | 74334977 | 6011 | 0.27 | 0.00257 | 0.00567 |
| chr5 | 16539146 | 16539989 | 844 | 2 | 0.00257 | 0.00567 |
| chr1 | 8013528 | 8023250 | 9723 | 0.39 | 0.00257 | 0.00567 |
| chr6 | 119029001 | 119031406 | 2406 | 0.48 | 0.00258 | 0.00567 |
| chrX | 151038698 | 151041039 | 2342 | -0.94 | 0.00258 | 0.00568 |
| chr14 | 99419334 | 99420507 | 1174 | -1.75 | 0.00259 | 0.0057 |
| chr7 | 36565518 | 36568126 | 2609 | 0.49 | 0.00259 | 0.0057 |
| chr2 | 69766071 | 69766512 | 442 | -2.03 | 0.00259 | 0.0057 |
| chr19 | 12720954 | 12723124 | 2171 | 0.55 | 0.00259 | 0.0057 |
| chr3 | 108880355 | 108880858 | 504 | 3.05 | 0.00259 | 0.0057 |
| chr16 | 3888713 | 3900606 | 11894 | -0.71 | 0.00259 | 0.0057 |
| chr22 | 32136147 | 32150700 | 14554 | 0.34 | 0.00259 | 0.00571 |
| chr9 | 100144048 | 100145757 | 1710 | 1.27 | 0.00259 | 0.00571 |
| chr2 | 160121883 | 160122338 | 456 | -2.38 | 0.00259 | 0.00571 |
| chr7 | 81802233 | 81803193 | 961 | -1.83 | 0.00259 | 0.00571 |
| chr17 | 37334027 | 37336588 | 2562 | 0.83 | 0.00259 | 0.00571 |
| chr9 | 91956185 | 91974246 | 18062 | -0.52 | 0.0026 | 0.00571 |
| chr1 | 165796171 | 165881224 | 85054 | -0.15 | 0.0026 | 0.00571 |
| chr6 | 17349523 | 17356400 | 6878 | -0.67 | 0.0026 | 0.00572 |
| chr7 | 107724331 | 107744930 | 20600 | 0.5 | 0.0026 | 0.00572 |
| chr3 | 177390887 | 177395311 | 4425 | -1.04 | 0.0026 | 0.00573 |
| chrX | 19665434 | 19666737 | 1304 | -1.74 | 0.0026 | 0.00573 |
| chr1 | 10838710 | 10840191 | 1482 | 1.05 | 0.00261 | 0.00573 |
| chr5 | 77961374 | 77962513 | 1140 | -0.96 | 0.00261 | 0.00573 |
| chr6 | 157935525 | 157940455 | 4931 | 1.68 | 0.00261 | 0.00574 |
| chr7 | 40756607 | 40758105 | 1499 | 0.84 | 0.00261 | 0.00574 |
| chr1 | 200707270 | 200710364 | 3095 | 0.41 | 0.00261 | 0.00574 |
| chrX | 77347690 | 77365264 | 17575 | 0.37 | 0.00261 | 0.00575 |
| chr6 | 42895948 | 42898764 | 2817 | 0.42 | 0.00262 | 0.00575 |

| | | | | | | |
|---|---|---|---|---|---|---|
| chr18 | 74236891 | 74237919 | 1029 | 1.46 | 0.00262 | 0.00575 |
| chr8 | 18770835 | 18771596 | 762 | -2.75 | 0.00262 | 0.00576 |
| chr16 | 84593406 | 84655886 | 62481 | 0.17 | 0.00262 | 0.00577 |
| chr1 | 117882017 | 117883352 | 1336 | 1.87 | 0.00263 | 0.00578 |
| chr19 | 34558414 | 34558984 | 571 | 2.06 | 0.00264 | 0.00579 |
| chr1 | 188990719 | 188991820 | 1102 | 1.76 | 0.00264 | 0.00579 |
| chr4 | 126763713 | 126764677 | 965 | 1.74 | 0.00264 | 0.0058 |
| chr9 | 16921659 | 16922149 | 491 | 2.77 | 0.00264 | 0.0058 |
| chr2 | 45240781 | 45253102 | 12322 | -0.73 | 0.00264 | 0.00581 |
| chr2 | 183291171 | 183291949 | 779 | -1.37 | 0.00265 | 0.00582 |
| chr3 | 172000092 | 172008043 | 7952 | -0.7 | 0.00265 | 0.00582 |
| chr7 | 46825464 | 46826079 | 616 | 1.76 | 0.00265 | 0.00582 |
| chr2 | 209676005 | 209676374 | 370 | 2.34 | 0.00266 | 0.00583 |
| chr20 | 40205859 | 40206974 | 1116 | 2.15 | 0.00266 | 0.00584 |
| chr7 | 45433419 | 45435052 | 1634 | -1.24 | 0.00266 | 0.00585 |
| chr11 | 85008379 | 85011694 | 3316 | 0.39 | 0.00266 | 0.00585 |
| chrX | 152987833 | 152993598 | 5766 | 0.56 | 0.00267 | 0.00585 |
| chr12 | 70231224 | 70231884 | 661 | 2.67 | 0.00267 | 0.00586 |
| chr11 | 69233052 | 69258921 | 25870 | -0.47 | 0.00267 | 0.00586 |
| chr9 | 138694546 | 138695162 | 617 | -2.9 | 0.00267 | 0.00586 |
| chr11 | 122451104 | 122452584 | 1481 | 0.76 | 0.00267 | 0.00586 |
| chrX | 44686808 | 44687798 | 991 | 2.71 | 0.00268 | 0.00587 |
| chr5 | 39194768 | 39198926 | 4159 | -0.39 | 0.00268 | 0.00587 |
| chr4 | 137784927 | 137785591 | 665 | 1.73 | 0.00268 | 0.00588 |
| chr3 | 141240113 | 141260026 | 19914 | 0.38 | 0.00268 | 0.00588 |
| chr5 | 53734725 | 53735626 | 902 | -2.68 | 0.00269 | 0.00589 |
| chr11 | 2398735 | 2423720 | 24986 | 0.24 | 0.00269 | 0.00589 |
| chr4 | 114678886 | 114685107 | 6222 | 0.35 | 0.00269 | 0.00589 |
| chr13 | 91137733 | 91138422 | 690 | 2.25 | 0.00269 | 0.00591 |
| chr12 | 51554516 | 51567826 | 13311 | 0.36 | 0.0027 | 0.00592 |
| chr7 | 87757072 | 87771014 | 13943 | 4.04 | 0.00271 | 0.00593 |
| chr3 | 23897724 | 23908898 | 11175 | 0.45 | 0.00271 | 0.00593 |
| chr7 | 123485969 | 123487257 | 1289 | 1.1 | 0.00271 | 0.00594 |
| chr16 | 15905476 | 15906692 | 1217 | 2.77 | 0.00271 | 0.00594 |
| chr1 | 98224015 | 98225022 | 1008 | 2.36 | 0.00271 | 0.00594 |
| chr11 | 63684222 | 63686392 | 2171 | -1.3 | 0.00271 | 0.00594 |
| chr1 | 89343306 | 89359619 | 16314 | 0.38 | 0.00271 | 0.00594 |
| chr3 | 5228331 | 5241149 | 12819 | 0.31 | 0.00271 | 0.00594 |
| chr11 | 10079472 | 10080425 | 954 | 1.9 | 0.00271 | 0.00595 |
| chr8 | 411106 | 412773 | 1668 | 0.98 | 0.00271 | 0.00595 |
| chr12 | 9752591 | 9753170 | 580 | 4.06 | 0.00272 | 0.00595 |
| chr6 | 9481408 | 9483252 | 1845 | 1.04 | 0.00272 | 0.00596 |
| chr4 | 15563875 | 15564607 | 733 | -2.11 | 0.00272 | 0.00597 |
| chr1 | 45240043 | 45286625 | 46583 | 0.16 | 0.00273 | 0.00597 |
| chr1 | 155677105 | 155678420 | 1316 | 2.37 | 0.00273 | 0.00597 |
| chr14 | 103051484 | 103061863 | 10380 | 0.34 | 0.00273 | 0.00597 |
| chr17 | 70863650 | 70864935 | 1286 | 1.26 | 0.00273 | 0.00598 |
| chr1 | 166458686 | 166459893 | 1208 | 1.16 | 0.00273 | 0.00598 |
| chr11 | 56747913 | 56748846 | 934 | 1.11 | 0.00273 | 0.00599 |
| chr1 | 240739152 | 240740164 | 1013 | 1.29 | 0.00273 | 0.00599 |
| chr12 | 85922802 | 85925209 | 2408 | 4.95 | 0.00274 | 0.00599 |
| chr12 | 16498901 | 16515299 | 16399 | 0.21 | 0.00274 | 0.00599 |
| chr16 | 88665376 | 88665851 | 476 | -3.24 | 0.00274 | 0.00599 |
| chr15 | 39747371 | 39748324 | 954 | -1.52 | 0.00274 | 0.006 |
| chr1 | 172828174 | 172829053 | 880 | 1.81 | 0.00274 | 0.006 |
| chr6 | 41423758 | 41425390 | 1633 | 1.34 | 0.00274 | 0.006 |
| chr8 | 129511618 | 129539218 | 27601 | 0.47 | 0.00274 | 0.006 |
| chr13 | 52337574 | 52339404 | 1831 | 0.89 | 0.00275 | 0.00601 |
| chr1 | 152432365 | 152432884 | 520 | 2.3 | 0.00275 | 0.00601 |
| chr10 | 101378819 | 101382495 | 3677 | 0.33 | 0.00275 | 0.00602 |
| chr10 | 32146670 | 32187089 | 40420 | 0.19 | 0.00275 | 0.00602 |

| | | | | | | |
|---|---|---|---|---|---|---|
| chr5 | 87404292 | 87405669 | 1378 | -0.64 | 0.00275 | 0.00602 |
| chr1 | 242309054 | 242311540 | 2487 | 0.48 | 0.00275 | 0.00603 |
| chr2 | 106328591 | 106330177 | 1587 | 1.33 | 0.00276 | 0.00603 |
| chr3 | 66522756 | 66528080 | 5325 | -0.68 | 0.00276 | 0.00603 |
| chr6 | 53984688 | 53985567 | 880 | 1.92 | 0.00276 | 0.00604 |
| chr11 | 116966812 | 116970449 | 3638 | -0.47 | 0.00276 | 0.00604 |
| chrX | 150000624 | 150068360 | 67737 | -0.14 | 0.00277 | 0.00605 |
| chr12 | 16063162 | 16065774 | 2613 | 0.44 | 0.00277 | 0.00607 |
| chr7 | 87848202 | 87857226 | 9025 | 0.41 | 0.00277 | 0.00607 |
| chr19 | 42461328 | 42464323 | 2996 | 0.45 | 0.00278 | 0.00608 |
| chr11 | 100588638 | 100591906 | 3269 | 0.31 | 0.00278 | 0.00609 |
| chr18 | 33551635 | 33553417 | 1783 | 0.74 | 0.00279 | 0.0061 |
| chr4 | 124281735 | 124283947 | 2213 | 0.93 | 0.00279 | 0.0061 |
| chr18 | 29664805 | 29677510 | 12706 | 0.34 | 0.00279 | 0.00611 |
| chr8 | 29754717 | 29756000 | 1284 | 1.75 | 0.0028 | 0.00611 |
| chr10 | 96938025 | 96944440 | 6416 | 0.51 | 0.0028 | 0.00611 |
| chr5 | 155107524 | 155107940 | 417 | -4.02 | 0.0028 | 0.00612 |
| chr10 | 34944560 | 34955671 | 11112 | -0.66 | 0.00281 | 0.00613 |
| chr5 | 142290584 | 142309961 | 19378 | -0.51 | 0.00281 | 0.00615 |
| chrX | 17420973 | 17432177 | 11205 | 0.57 | 0.00282 | 0.00616 |
| chr1 | 52337549 | 52345828 | 8280 | -0.29 | 0.00282 | 0.00616 |
| chr15 | 53646893 | 53647935 | 1043 | 1.44 | 0.00282 | 0.00617 |
| chr7 | 115274921 | 115275295 | 375 | 2.83 | 0.00283 | 0.00618 |
| chr15 | 59144830 | 59170496 | 25667 | 0.56 | 0.00283 | 0.00618 |
| chr1 | 216394737 | 216395166 | 430 | 1.72 | 0.00283 | 0.00618 |
| chr18 | 12271199 | 12272114 | 916 | 0.86 | 0.00283 | 0.00619 |
| chr1 | 31819708 | 31820176 | 469 | -2.59 | 0.00283 | 0.00619 |
| chr1 | 161283471 | 161285212 | 1742 | 0.44 | 0.00283 | 0.00619 |
| chr6 | 107028807 | 107029505 | 699 | -2.08 | 0.00284 | 0.00619 |
| chr1 | 2125859 | 2170229 | 44371 | -0.23 | 0.00284 | 0.0062 |
| chr9 | 123833295 | 123838097 | 4803 | -0.45 | 0.00284 | 0.0062 |
| chr5 | 170813555 | 170846827 | 33273 | -0.32 | 0.00284 | 0.0062 |
| chr10 | 70957939 | 70960495 | 2557 | -1.53 | 0.00284 | 0.0062 |
| chr1 | 95006667 | 95008415 | 1749 | 0.84 | 0.00284 | 0.0062 |
| chr18 | 23985301 | 23986298 | 998 | -2.02 | 0.00284 | 0.0062 |
| chr1 | 182612831 | 182613395 | 565 | -2.23 | 0.00285 | 0.00621 |
| chr9 | 136690395 | 136691525 | 1131 | -2.61 | 0.00285 | 0.00621 |
| chr17 | 55237925 | 55238689 | 765 | -2.37 | 0.00285 | 0.00621 |
| chr4 | 162474907 | 162477302 | 2396 | 0.43 | 0.00286 | 0.00624 |
| chr17 | 70266904 | 70267297 | 394 | 2.92 | 0.00286 | 0.00624 |
| chr10 | 99184942 | 99190440 | 5499 | 0.44 | 0.00286 | 0.00624 |
| chr11 | 7998490 | 8009969 | 11480 | -0.91 | 0.00286 | 0.00625 |
| chr7 | 43469640 | 43471290 | 1651 | -1.02 | 0.00286 | 0.00625 |
| chr1 | 179843030 | 179853611 | 10582 | 0.25 | 0.00288 | 0.00629 |
| chr6 | 133420079 | 133421650 | 1572 | 0.87 | 0.00289 | 0.00629 |
| chr1 | 64276565 | 64290020 | 13456 | -0.5 | 0.00289 | 0.0063 |
| chr4 | 122098387 | 122104303 | 5917 | 4.51 | 0.00289 | 0.0063 |
| chr5 | 14767698 | 14775242 | 7545 | 1 | 0.00289 | 0.0063 |
| chr4 | 6987466 | 7046350 | 58885 | -0.2 | 0.00289 | 0.00631 |
| chr14 | 71757443 | 71758626 | 1184 | 0.99 | 0.00289 | 0.00631 |
| chr12 | 20328338 | 20329362 | 1025 | 1.5 | 0.0029 | 0.00631 |
| chr3 | 125044383 | 125051396 | 7014 | 1.17 | 0.0029 | 0.00632 |
| chr8 | 122350756 | 122361691 | 10936 | -2.84 | 0.0029 | 0.00632 |
| chr3 | 152313926 | 152314418 | 493 | 2.02 | 0.0029 | 0.00633 |
| chr6 | 24803199 | 24803657 | 459 | 2.84 | 0.00292 | 0.00636 |
| chr13 | 111356923 | 111369273 | 12351 | 0.29 | 0.00292 | 0.00636 |
| chr12 | 123976874 | 123977257 | 384 | 2.67 | 0.00292 | 0.00636 |
| chrX | 153338862 | 153349234 | 10373 | -0.56 | 0.00292 | 0.00636 |
| chr8 | 131755726 | 131756740 | 1015 | -1.74 | 0.00292 | 0.00636 |
| chr9 | 136843336 | 136858718 | 15383 | -0.46 | 0.00293 | 0.00637 |
| chr12 | 69517121 | 69519082 | 1962 | 4.34 | 0.00293 | 0.00637 |

| | | | | | | |
|---|---|---|---|---|---|---|
| chr10 | 100092281 | 100093211 | 931 | 1.38 | 0.00293 | 0.00638 |
| chr7 | 27317045 | 27318118 | 1074 | 1.19 | 0.00293 | 0.00638 |
| chr7 | 158988556 | 158989336 | 781 | 2.27 | 0.00293 | 0.00638 |
| chr13 | 40725976 | 40726364 | 389 | 2.96 | 0.00293 | 0.00638 |
| chr22 | 43283420 | 43288590 | 5171 | -0.98 | 0.00293 | 0.00639 |
| chr3 | 193950864 | 193951495 | 632 | -2.75 | 0.00293 | 0.00639 |
| chr6 | 111059598 | 111060807 | 1210 | 0.7 | 0.00294 | 0.0064 |
| chr7 | 44601227 | 44622998 | 21772 | 0.29 | 0.00294 | 0.0064 |
| chr10 | 93168606 | 93171187 | 2582 | 0.36 | 0.00294 | 0.00641 |
| chrX | 103173277 | 103174211 | 935 | 1.45 | 0.00295 | 0.00641 |
| chr2 | 102758604 | 102772678 | 14075 | -0.76 | 0.00295 | 0.00642 |
| chr19 | 58258182 | 58259490 | 1309 | 0.66 | 0.00295 | 0.00642 |
| chr1 | 32774688 | 32801689 | 27002 | 0.27 | 0.00296 | 0.00643 |
| chr9 | 73027606 | 73037045 | 9440 | -0.29 | 0.00296 | 0.00643 |
| chr2 | 114696025 | 114696595 | 571 | 3.09 | 0.00296 | 0.00643 |
| chr11 | 68172559 | 68175351 | 2793 | 1.04 | 0.00296 | 0.00643 |
| chr18 | 43835700 | 43845196 | 9497 | 0.73 | 0.00296 | 0.00644 |
| chr14 | 102274108 | 102306483 | 32376 | 0.25 | 0.00296 | 0.00644 |
| chr10 | 120068862 | 120082617 | 13756 | 0.64 | 0.00296 | 0.00645 |
| chr2 | 33719115 | 33720986 | 1872 | 1.23 | 0.00297 | 0.00645 |
| chr5 | 94806837 | 94807927 | 1091 | 1.93 | 0.00297 | 0.00645 |
| chr6 | 36062447 | 36065116 | 2670 | 0.79 | 0.00297 | 0.00647 |
| chr4 | 102806487 | 102807682 | 1196 | 1.53 | 0.00298 | 0.00647 |
| chr6 | 127891801 | 127914542 | 22742 | 1.13 | 0.00298 | 0.00648 |
| chr7 | 18215211 | 18219336 | 4126 | -0.46 | 0.00298 | 0.00649 |
| chr13 | 80376310 | 80385928 | 9619 | 0.59 | 0.00299 | 0.00649 |
| chrX | 129010772 | 129011840 | 1069 | 1.05 | 0.003 | 0.00651 |
| chr8 | 32642179 | 32643035 | 857 | -1.54 | 0.003 | 0.00652 |
| chr3 | 52639138 | 52643870 | 4733 | 0.45 | 0.00301 | 0.00654 |
| chr4 | 105982343 | 105983108 | 766 | 2.18 | 0.00301 | 0.00654 |
| chr10 | 97878120 | 97891194 | 13075 | 0.29 | 0.00301 | 0.00654 |
| chr6 | 166969901 | 166971049 | 1149 | 2.04 | 0.00301 | 0.00654 |
| chr17 | 3465928 | 3466857 | 930 | -1.74 | 0.00301 | 0.00654 |
| chr2 | 70453800 | 70485953 | 32154 | -0.21 | 0.00302 | 0.00655 |
| chr22 | 44319686 | 44320986 | 1301 | 1.42 | 0.00302 | 0.00656 |
| chr3 | 33523594 | 33524186 | 593 | 2.08 | 0.00302 | 0.00656 |
| chr9 | 80366254 | 80379143 | 12890 | -0.97 | 0.00302 | 0.00656 |
| chr12 | 107162610 | 107174493 | 11884 | 0.34 | 0.00302 | 0.00657 |
| chr9 | 134024806 | 134026421 | 1616 | -1.03 | 0.00303 | 0.00658 |
| chr9 | 112748305 | 112749804 | 1500 | 4.16 | 0.00303 | 0.00659 |
| chr16 | 77755672 | 77757243 | 1572 | 0.69 | 0.00304 | 0.00659 |
| chr9 | 4738862 | 4742528 | 3667 | 0.56 | 0.00304 | 0.00659 |
| chr13 | 85508953 | 85509513 | 561 | 2.11 | 0.00304 | 0.00659 |
| chr17 | 8338219 | 8347130 | 8912 | 0.31 | 0.00304 | 0.00659 |
| chr4 | 38968394 | 38969600 | 1207 | -1.33 | 0.00304 | 0.00659 |
| chr7 | 110737647 | 110738819 | 1173 | 1.18 | 0.00304 | 0.0066 |
| chr19 | 33732342 | 33732708 | 367 | -3 | 0.00304 | 0.0066 |
| chr21 | 39029700 | 39044159 | 14460 | 0.53 | 0.00304 | 0.0066 |
| chr2 | 11751705 | 11753011 | 1307 | -1.51 | 0.00304 | 0.0066 |
| chr4 | 75022133 | 75025249 | 3117 | 0.33 | 0.00305 | 0.00661 |
| chr17 | 17318953 | 17320669 | 1717 | -0.6 | 0.00306 | 0.00663 |
| chr3 | 25332814 | 25334042 | 1229 | 1.54 | 0.00306 | 0.00664 |
| chr1 | 247346116 | 247346832 | 717 | 2.71 | 0.00306 | 0.00664 |
| chr21 | 17101378 | 17135000 | 33623 | 0.18 | 0.00306 | 0.00664 |
| chr15 | 40526759 | 40531717 | 4959 | -1.23 | 0.00306 | 0.00664 |
| chr5 | 120712990 | 120714317 | 1328 | 1.5 | 0.00306 | 0.00665 |
| chr12 | 70683275 | 70684849 | 1575 | 1.38 | 0.00307 | 0.00666 |
| chr6 | 138018155 | 138019529 | 1375 | 0.91 | 0.00307 | 0.00666 |
| chr11 | 99127435 | 99128808 | 1374 | 1.07 | 0.00308 | 0.00667 |
| chr1 | 154552264 | 154556829 | 4566 | -0.89 | 0.00308 | 0.00668 |
| chr7 | 99552701 | 99554119 | 1419 | 1.41 | 0.00309 | 0.00669 |

| | | | | | | |
|---|---|---|---|---|---|---|
| chr11 | 68811223 | 68820697 | 9475 | -0.56 | 0.00309 | 0.00669 |
| chr17 | 41236850 | 41237801 | 952 | 1.3 | 0.00309 | 0.00669 |
| chrX | 77907543 | 77909966 | 2424 | 0.59 | 0.0031 | 0.00672 |
| chr3 | 110246461 | 110246830 | 370 | -4 | 0.0031 | 0.00672 |
| chr1 | 180226227 | 180282086 | 55860 | 0.18 | 0.00311 | 0.00674 |
| chr12 | 30788534 | 30789292 | 759 | 1.75 | 0.00311 | 0.00674 |
| chr1 | 91748053 | 91752397 | 4345 | -0.62 | 0.00311 | 0.00675 |
| chr16 | 89753027 | 89789133 | 36107 | -0.31 | 0.00311 | 0.00675 |
| chr5 | 159199750 | 159201227 | 1478 | 0.66 | 0.00312 | 0.00676 |
| chr11 | 75977619 | 75978415 | 797 | 1.54 | 0.00313 | 0.00678 |
| chr2 | 64202098 | 64225139 | 23042 | 0.41 | 0.00313 | 0.00679 |
| chr4 | 128800660 | 128812043 | 11384 | 0.45 | 0.00314 | 0.00679 |
| chr13 | 22176674 | 22191592 | 14919 | 0.73 | 0.00314 | 0.00681 |
| chr1 | 65298094 | 65299324 | 1231 | -1.27 | 0.00315 | 0.00681 |
| chr3 | 148838327 | 148849359 | 11033 | 0.42 | 0.00315 | 0.00682 |
| chr15 | 26075770 | 26077332 | 1563 | -0.97 | 0.00315 | 0.00682 |
| chr19 | 43942412 | 43944343 | 1932 | 0.7 | 0.00315 | 0.00682 |
| chr18 | 57686033 | 57690464 | 4432 | -0.88 | 0.00315 | 0.00682 |
| chr3 | 119216070 | 119218583 | 2514 | 0.45 | 0.00315 | 0.00682 |
| chr4 | 113554219 | 113560167 | 5949 | 0.36 | 0.00315 | 0.00682 |
| chr22 | 42394127 | 42395402 | 1276 | 1.28 | 0.00315 | 0.00682 |
| chr4 | 138231534 | 138232729 | 1196 | 1.55 | 0.00315 | 0.00682 |
| chr5 | 172940158 | 172941411 | 1254 | 1.37 | 0.00315 | 0.00683 |
| chr9 | 139916850 | 139929811 | 12962 | 0.97 | 0.00316 | 0.00683 |
| chr10 | 59977145 | 59978269 | 1125 | 1.89 | 0.00316 | 0.00684 |
| chr19 | 43662464 | 43668743 | 6280 | 0.86 | 0.00316 | 0.00684 |
| chr7 | 92156580 | 92159161 | 2582 | 0.43 | 0.00316 | 0.00684 |
| chr10 | 60833055 | 60840081 | 7027 | 1.04 | 0.00317 | 0.00685 |
| chr20 | 13745374 | 13746608 | 1235 | 1.22 | 0.00318 | 0.00687 |
| chr9 | 13139013 | 13140996 | 1984 | 0.86 | 0.00319 | 0.00691 |
| chr11 | 14664669 | 14666666 | 1998 | -1.02 | 0.0032 | 0.00692 |
| chr3 | 159478428 | 159517358 | 38931 | 0.21 | 0.0032 | 0.00692 |
| chr15 | 39565296 | 39566715 | 1420 | 0.97 | 0.0032 | 0.00692 |
| chr8 | 67024693 | 67045990 | 21298 | -0.17 | 0.0032 | 0.00692 |
| chr19 | 43032279 | 43033555 | 1277 | 0.68 | 0.0032 | 0.00692 |
| chrX | 47375571 | 47376565 | 995 | 2.63 | 0.0032 | 0.00692 |
| chr8 | 83959523 | 83961712 | 2190 | 4.72 | 0.00321 | 0.00694 |
| chr6 | 4007721 | 4024192 | 16472 | 0.29 | 0.00321 | 0.00694 |
| chr5 | 73359255 | 73364075 | 4821 | 4.3 | 0.00321 | 0.00694 |
| chr2 | 218550196 | 218550709 | 514 | -2.19 | 0.00322 | 0.00696 |
| chr1 | 63893421 | 63895038 | 1618 | 1.53 | 0.00322 | 0.00696 |
| chr1 | 117939111 | 117939905 | 795 | 2.04 | 0.00323 | 0.00698 |
| chr1 | 209212338 | 209213480 | 1143 | 1.23 | 0.00323 | 0.00698 |
| chr4 | 38063847 | 38081467 | 17621 | 0.56 | 0.00323 | 0.00698 |
| chr19 | 4972732 | 4976959 | 4228 | -0.99 | 0.00323 | 0.00698 |
| chr17 | 79056543 | 79072283 | 15741 | 0.37 | 0.00323 | 0.00698 |
| chr7 | 155675089 | 155690328 | 15240 | -3.14 | 0.00324 | 0.00699 |
| chr6 | 149699707 | 149700521 | 815 | -2.01 | 0.00324 | 0.00699 |
| chr8 | 102801929 | 102804083 | 2155 | 0.69 | 0.00324 | 0.00699 |
| chr6 | 47275190 | 47277663 | 2474 | 0.75 | 0.00324 | 0.00699 |
| chr6 | 80891474 | 80892465 | 992 | 1.54 | 0.00324 | 0.00701 |
| chr8 | 100559428 | 100560180 | 753 | 1.64 | 0.00324 | 0.00701 |
| chr12 | 77078725 | 77079516 | 792 | 1.47 | 0.00325 | 0.00701 |
| chr2 | 101368470 | 101375109 | 6640 | 1.01 | 0.00325 | 0.00701 |
| chr4 | 114935444 | 114941448 | 6005 | 0.62 | 0.00325 | 0.00701 |
| chr3 | 150283360 | 150284607 | 1248 | -1.68 | 0.00325 | 0.00702 |
| chr20 | 39945403 | 39970366 | 24964 | 0.58 | 0.00325 | 0.00702 |
| chr2 | 138816379 | 138831633 | 15255 | 0.49 | 0.00325 | 0.00702 |
| chr16 | 12499850 | 12502419 | 2570 | -0.69 | 0.00325 | 0.00702 |
| chr18 | 24008817 | 24017230 | 8414 | -0.68 | 0.00325 | 0.00702 |
| chr3 | 71438677 | 71440060 | 1384 | 0.83 | 0.00325 | 0.00702 |

| | | | | | | |
|---|---|---|---|---|---|---|
| chr20 | 30738105 | 30748488 | 10384 | -0.88 | 0.00327 | 0.00705 |
| chr12 | 51656125 | 51666042 | 9918 | 0.53 | 0.00327 | 0.00705 |
| chr2 | 142888149 | 142889605 | 1457 | 3.95 | 0.00328 | 0.00707 |
| chr14 | 37961090 | 37961986 | 897 | 1.65 | 0.00328 | 0.00708 |
| chr4 | 84455960 | 84504457 | 48498 | -0.15 | 0.00329 | 0.00709 |
| chr7 | 127806870 | 127808078 | 1209 | 1.9 | 0.00329 | 0.0071 |
| chr19 | 55657669 | 55703328 | 45660 | 0.33 | 0.00329 | 0.0071 |
| chr6 | 74483103 | 74493051 | 9949 | 0.6 | 0.00329 | 0.0071 |
| chr11 | 115079501 | 115080033 | 533 | -1.98 | 0.00329 | 0.0071 |
| chr5 | 142485541 | 142498317 | 12777 | 0.71 | 0.0033 | 0.00711 |
| chr7 | 150125339 | 150126897 | 1559 | 1.01 | 0.0033 | 0.00711 |
| chr11 | 11643782 | 11644182 | 401 | -1.51 | 0.0033 | 0.00711 |
| chr9 | 114188823 | 114190803 | 1981 | -0.88 | 0.0033 | 0.00712 |
| chr13 | 97792738 | 97795518 | 2781 | 0.51 | 0.00331 | 0.00713 |
| chr7 | 91693580 | 91694267 | 688 | 2.62 | 0.00331 | 0.00713 |
| chr5 | 54625509 | 54677002 | 51494 | 0.29 | 0.00331 | 0.00714 |
| chr5 | 30043821 | 30044969 | 1149 | 1.45 | 0.00331 | 0.00714 |
| chr7 | 80683207 | 80683592 | 386 | -2.37 | 0.00331 | 0.00714 |
| chr10 | 79221047 | 79221765 | 719 | -2.56 | 0.00332 | 0.00715 |
| chr4 | 135122489 | 135123173 | 685 | -1.65 | 0.00332 | 0.00715 |
| chr8 | 32926996 | 32927763 | 768 | -2.1 | 0.00332 | 0.00715 |
| chr12 | 74162996 | 74163627 | 632 | 2.02 | 0.00332 | 0.00715 |
| chr18 | 37354989 | 37355980 | 992 | 1.57 | 0.00332 | 0.00716 |
| chr7 | 99435134 | 99435769 | 636 | 1.6 | 0.00332 | 0.00716 |
| chrX | 123197692 | 123200420 | 2729 | 1.12 | 0.00333 | 0.00717 |
| chr17 | 62476832 | 62477339 | 508 | -2.98 | 0.00333 | 0.00717 |
| chr22 | 39051211 | 39053926 | 2716 | -0.38 | 0.00333 | 0.00717 |
| chr12 | 288059 | 288966 | 908 | -1.98 | 0.00333 | 0.00718 |
| chr11 | 99563604 | 99565871 | 2268 | 0.66 | 0.00334 | 0.00719 |
| chr9 | 6774944 | 6796222 | 21279 | -0.37 | 0.00334 | 0.0072 |
| chr3 | 87841367 | 87842542 | 1176 | -1.24 | 0.00334 | 0.0072 |
| chr13 | 48796361 | 48808794 | 12434 | 0.39 | 0.00334 | 0.0072 |
| chr17 | 29798636 | 29799546 | 911 | -1.66 | 0.00335 | 0.00721 |
| chr7 | 17719305 | 17721173 | 1869 | -1.21 | 0.00335 | 0.00722 |
| chr9 | 80322040 | 80328361 | 6322 | 1.39 | 0.00335 | 0.00722 |
| chr7 | 33756637 | 33762825 | 6189 | -1.69 | 0.00336 | 0.00723 |
| chr5 | 132655294 | 132656147 | 854 | -2.67 | 0.00336 | 0.00724 |
| chr6 | 27630573 | 27662662 | 32090 | 0.31 | 0.00336 | 0.00724 |
| chr2 | 134574581 | 134576891 | 2311 | -0.68 | 0.00336 | 0.00724 |
| chr3 | 23616854 | 23622544 | 5691 | 0.48 | 0.00337 | 0.00724 |
| chr4 | 184121223 | 184122242 | 1020 | 1.28 | 0.00337 | 0.00726 |
| chr15 | 64257902 | 64258707 | 806 | -2.3 | 0.00337 | 0.00726 |
| chr11 | 35588950 | 35602550 | 13601 | -0.41 | 0.00339 | 0.00729 |
| chr5 | 168541424 | 168541912 | 489 | -2.71 | 0.00339 | 0.0073 |
| chr6 | 106439828 | 106443304 | 3477 | -1.03 | 0.0034 | 0.00731 |
| chr20 | 36632910 | 36634136 | 1227 | -1.27 | 0.0034 | 0.00731 |
| chr13 | 22050762 | 22051282 | 521 | 1.69 | 0.0034 | 0.00731 |
| chr17 | 79360233 | 79376101 | 15869 | -0.28 | 0.00341 | 0.00732 |
| chr2 | 234837660 | 234838328 | 669 | -2.34 | 0.00341 | 0.00732 |
| chr6 | 131314102 | 131315372 | 1271 | 1.44 | 0.00341 | 0.00733 |
| chr11 | 43332982 | 43343751 | 10770 | 0.43 | 0.00341 | 0.00734 |
| chr5 | 131429901 | 131440040 | 10140 | -0.66 | 0.00341 | 0.00734 |
| chr7 | 139544517 | 139545076 | 560 | 2.94 | 0.00341 | 0.00734 |
| chr14 | 103894257 | 103950316 | 56060 | 0.32 | 0.00342 | 0.00734 |
| chr7 | 5274583 | 5275096 | 514 | 2.69 | 0.00342 | 0.00734 |
| chr4 | 54643219 | 54649245 | 6027 | -0.47 | 0.00342 | 0.00736 |
| chr10 | 65332654 | 65333598 | 945 | 1.75 | 0.00343 | 0.00737 |
| chr4 | 174249764 | 174257855 | 8092 | 0.37 | 0.00343 | 0.00737 |
| chr15 | 67192111 | 67257210 | 65100 | -0.2 | 0.00343 | 0.00738 |
| chr2 | 219248566 | 219272621 | 24056 | -0.23 | 0.00344 | 0.00739 |
| chr17 | 63836863 | 63838188 | 1326 | 0.93 | 0.00345 | 0.0074 |

| | | | | | | |
|---|---|---|---|---|---|---|
| chr13 | 25826696 | 25828773 | 2078 | 0.93 | 0.00345 | 0.00741 |
| chr5 | 134706811 | 134735830 | 29020 | 0.2 | 0.00346 | 0.00743 |
| chr5 | 72383917 | 72384446 | 530 | -2.02 | 0.00346 | 0.00743 |
| chr11 | 111588858 | 111608301 | 19444 | 0.32 | 0.00346 | 0.00744 |
| chr3 | 32822063 | 32823722 | 1660 | 1.12 | 0.00347 | 0.00745 |
| chr2 | 63208684 | 63227900 | 19217 | -0.43 | 0.00347 | 0.00745 |
| chr14 | 32287632 | 32288869 | 1238 | 1.12 | 0.00347 | 0.00745 |
| chr6 | 81233334 | 81246589 | 13256 | 0.77 | 0.00348 | 0.00747 |
| chr5 | 70856466 | 70857579 | 1114 | -1.23 | 0.00349 | 0.00749 |
| chr2 | 203037125 | 203038101 | 977 | -1.67 | 0.00349 | 0.00749 |
| chr17 | 76583427 | 76601249 | 17823 | -0.19 | 0.00349 | 0.0075 |
| chr2 | 236302439 | 236303281 | 843 | -1.74 | 0.0035 | 0.00751 |
| chr11 | 57424183 | 57449518 | 25336 | 0.24 | 0.0035 | 0.00752 |
| chr15 | 96596073 | 96608776 | 12704 | 4.69 | 0.00351 | 0.00752 |
| chr13 | 83675970 | 83684113 | 8144 | 1.64 | 0.00351 | 0.00753 |
| chr12 | 69501332 | 69502333 | 1002 | 1.54 | 0.00351 | 0.00753 |
| chr18 | 3786812 | 3800368 | 13557 | 4.51 | 0.00351 | 0.00753 |
| chr13 | 20748491 | 20749952 | 1462 | 0.85 | 0.00351 | 0.00753 |
| chr5 | 32639913 | 32643652 | 3740 | 0.53 | 0.00351 | 0.00753 |
| chr6 | 82618733 | 82619411 | 679 | 2.17 | 0.00351 | 0.00754 |
| chrX | 21122534 | 21127066 | 4533 | 2.68 | 0.00351 | 0.00754 |
| chr4 | 79433361 | 79434288 | 928 | 2.09 | 0.00352 | 0.00754 |
| chr2 | 169439782 | 169441562 | 1781 | -0.82 | 0.00352 | 0.00755 |
| chr16 | 31711921 | 31733837 | 21917 | 0.62 | 0.00352 | 0.00755 |
| chr12 | 19198032 | 19198986 | 955 | 2.68 | 0.00352 | 0.00756 |
| chr1 | 154177994 | 154209147 | 31154 | 0.24 | 0.00352 | 0.00756 |
| chr14 | 72265318 | 72271469 | 6152 | 4.27 | 0.00353 | 0.00757 |
| chr3 | 70400944 | 70402177 | 1234 | -1.29 | 0.00354 | 0.00758 |
| chr7 | 137714124 | 137716236 | 2113 | 0.97 | 0.00354 | 0.00758 |
| chr5 | 53454801 | 53455678 | 878 | -1.47 | 0.00354 | 0.0076 |
| chr2 | 193010607 | 193011339 | 733 | -2.56 | 0.00355 | 0.00761 |
| chr1 | 98758313 | 98759583 | 1271 | 1.11 | 0.00355 | 0.00761 |
| chr22 | 48231062 | 48231612 | 551 | 1.94 | 0.00355 | 0.00762 |
| chr12 | 56795052 | 56795380 | 329 | 2.96 | 0.00356 | 0.00762 |
| chr15 | 48622232 | 48625346 | 3115 | 0.43 | 0.00356 | 0.00763 |
| chr14 | 89126136 | 89127283 | 1148 | 1.59 | 0.00356 | 0.00763 |
| chr9 | 29211788 | 29213574 | 1787 | 1.58 | 0.00356 | 0.00763 |
| chr19 | 4668799 | 4671887 | 3089 | 0.61 | 0.00356 | 0.00764 |
| chr11 | 59770701 | 59771521 | 821 | -1.8 | 0.00357 | 0.00764 |
| chr19 | 43401511 | 43405367 | 3857 | 0.57 | 0.00357 | 0.00765 |
| chr3 | 72359556 | 72368548 | 8993 | -4.21 | 0.00357 | 0.00765 |
| chr13 | 103482780 | 103500740 | 17961 | 0.35 | 0.00357 | 0.00766 |
| chr20 | 23735712 | 23737393 | 1682 | 1.6 | 0.00358 | 0.00766 |
| chr3 | 44902434 | 44904415 | 1982 | 0.48 | 0.00358 | 0.00767 |
| chr2 | 208370074 | 208371060 | 987 | 1.76 | 0.00358 | 0.00767 |
| chr4 | 132297819 | 132301206 | 3388 | 1.56 | 0.00359 | 0.00768 |
| chr4 | 13210425 | 13210859 | 435 | -2.57 | 0.00359 | 0.00769 |
| chr17 | 75116933 | 75142663 | 25731 | 0.27 | 0.00359 | 0.00769 |
| chr15 | 41325966 | 41353361 | 27396 | 0.44 | 0.0036 | 0.0077 |
| chr15 | 90807556 | 90816899 | 9344 | 0.34 | 0.0036 | 0.00771 |
| chr8 | 29187869 | 29210610 | 22742 | 0.21 | 0.00361 | 0.00772 |
| chr9 | 100000079 | 100001610 | 1532 | 0.62 | 0.00361 | 0.00772 |
| chrX | 10124341 | 10126216 | 1876 | 0.79 | 0.00361 | 0.00772 |
| chr2 | 225346309 | 225362607 | 16299 | -0.36 | 0.00361 | 0.00772 |
| chr19 | 40388782 | 40389226 | 445 | 1.91 | 0.00361 | 0.00772 |
| chr3 | 190580613 | 190581059 | 447 | 1.64 | 0.00361 | 0.00773 |
| chr6 | 82575425 | 82576117 | 693 | 1.87 | 0.00361 | 0.00773 |
| chr17 | 70490748 | 70614576 | 123829 | 0.14 | 0.00361 | 0.00773 |
| chr5 | 173930482 | 173931238 | 757 | 2.05 | 0.00361 | 0.00773 |
| chr15 | 68676152 | 68677772 | 1621 | -1.02 | 0.00361 | 0.00773 |
| chr5 | 112042420 | 112044490 | 2071 | 0.68 | 0.00362 | 0.00775 |

| | | | | | | |
|---|---|---|---|---|---|---|
| chr19 | 37852194 | 37862978 | 10785 | 5.11 | 0.00362 | 0.00775 |
| chr3 | 149590091 | 149591509 | 1419 | -1.16 | 0.00362 | 0.00775 |
| chr18 | 60152827 | 60154017 | 1191 | 1.23 | 0.00362 | 0.00775 |
| chrX | 132677477 | 132679250 | 1774 | 4.32 | 0.00363 | 0.00776 |
| chr8 | 142464601 | 142465501 | 901 | -1.56 | 0.00363 | 0.00777 |
| chr5 | 133507157 | 133514447 | 7291 | 0.28 | 0.00363 | 0.00777 |
| chr4 | 163621156 | 163622103 | 948 | -1.79 | 0.00364 | 0.00778 |
| chr19 | 4539567 | 4540747 | 1181 | 1.21 | 0.00364 | 0.00778 |
| chr4 | 5206479 | 5208876 | 2398 | -1.39 | 0.00364 | 0.00779 |
| chr21 | 27777311 | 27779226 | 1916 | -0.84 | 0.00365 | 0.0078 |
| chr17 | 71006363 | 71010580 | 4218 | 0.89 | 0.00365 | 0.0078 |
| chr13 | 73828785 | 73829540 | 756 | 2.61 | 0.00365 | 0.0078 |
| chr18 | 73201237 | 73202472 | 1236 | -1.19 | 0.00365 | 0.00781 |
| chr4 | 142957572 | 142959649 | 2078 | 1.08 | 0.00366 | 0.00782 |
| chr3 | 8664849 | 8674838 | 9990 | 0.31 | 0.00366 | 0.00782 |
| chr5 | 54929249 | 54976793 | 47545 | -0.34 | 0.00366 | 0.00782 |
| chr7 | 105188550 | 105190115 | 1566 | -2.13 | 0.00366 | 0.00782 |
| chr17 | 74720613 | 74735187 | 14575 | 0.27 | 0.00366 | 0.00782 |
| chr21 | 35319066 | 35321638 | 2573 | 0.55 | 0.00367 | 0.00783 |
| chr5 | 73189415 | 73213887 | 24473 | 0.46 | 0.00367 | 0.00784 |
| chr13 | 108921817 | 108922856 | 1040 | 1.83 | 0.00368 | 0.00785 |
| chr1 | 173894244 | 173895637 | 1394 | 1.22 | 0.00368 | 0.00785 |
| chr12 | 101025324 | 101026915 | 1592 | 1.31 | 0.00368 | 0.00786 |
| chr5 | 90514339 | 90516318 | 1980 | 0.91 | 0.00368 | 0.00786 |
| chr7 | 40131651 | 40132928 | 1278 | 1.96 | 0.00368 | 0.00786 |
| chr1 | 214719964 | 214727098 | 7135 | 0.33 | 0.00368 | 0.00786 |
| chr7 | 138423463 | 138438217 | 14755 | 0.72 | 0.00368 | 0.00786 |
| chr17 | 41471088 | 41507551 | 36464 | -0.32 | 0.00369 | 0.00787 |
| chr12 | 92814800 | 92815652 | 853 | 1.54 | 0.00369 | 0.00788 |
| chr16 | 59390417 | 59392153 | 1737 | 1.08 | 0.00369 | 0.00788 |
| chrX | 102708916 | 102709407 | 492 | -2.89 | 0.00369 | 0.00788 |
| chr5 | 172409947 | 172457324 | 47378 | -0.21 | 0.00369 | 0.00788 |
| chr9 | 140375847 | 140377679 | 1833 | -0.59 | 0.0037 | 0.00789 |
| chr11 | 123045103 | 123066269 | 21167 | -0.68 | 0.0037 | 0.00789 |
| chr17 | 18281228 | 18281890 | 663 | -1.82 | 0.0037 | 0.00789 |
| chr18 | 46147466 | 46159040 | 11575 | -0.36 | 0.0037 | 0.0079 |
| chr12 | 41915332 | 41916450 | 1119 | 4.24 | 0.00371 | 0.00791 |
| chr5 | 17882757 | 17883843 | 1087 | 1.74 | 0.00371 | 0.00791 |
| chr9 | 37146048 | 37147116 | 1069 | 0.79 | 0.00371 | 0.00791 |
| chr15 | 84540577 | 84541145 | 569 | -2.68 | 0.00371 | 0.00791 |
| chr22 | 49479781 | 49480599 | 819 | -1.89 | 0.00371 | 0.00791 |
| chr1 | 23574781 | 23576512 | 1732 | 1.22 | 0.00371 | 0.00792 |
| chr14 | 50848376 | 50870973 | 22598 | 0.55 | 0.00372 | 0.00793 |
| chr2 | 138864656 | 138871029 | 6374 | 0.72 | 0.00372 | 0.00794 |
| chr6 | 56020860 | 56022328 | 1469 | 1.4 | 0.00372 | 0.00794 |
| chr1 | 53162804 | 53174897 | 12094 | 0.5 | 0.00373 | 0.00794 |
| chr15 | 69417966 | 69419203 | 1238 | 1.12 | 0.00373 | 0.00795 |
| chr8 | 145846185 | 145847871 | 1687 | -1.47 | 0.00373 | 0.00796 |
| chr18 | 29032796 | 29035138 | 2343 | 0.67 | 0.00373 | 0.00796 |
| chr7 | 135660365 | 135667710 | 7346 | 0.55 | 0.00374 | 0.00797 |
| chr14 | 103827607 | 103871749 | 44143 | 0.27 | 0.00375 | 0.00798 |
| chr4 | 107264655 | 107266576 | 1922 | -1.24 | 0.00376 | 0.00801 |
| chr14 | 34794468 | 34802144 | 7677 | 0.71 | 0.00376 | 0.00801 |
| chr22 | 40991353 | 41006748 | 15396 | -0.62 | 0.00376 | 0.00801 |
| chr6 | 127805048 | 127806293 | 1246 | 1.04 | 0.00376 | 0.00801 |
| chr2 | 159569341 | 159584372 | 15032 | -0.32 | 0.00376 | 0.00802 |
| chr20 | 3351726 | 3355329 | 3604 | 0.8 | 0.00377 | 0.00802 |
| chr15 | 102263478 | 102264672 | 1195 | 0.86 | 0.00377 | 0.00803 |
| chr13 | 45008536 | 45012238 | 3703 | 0.37 | 0.00377 | 0.00803 |
| chr6 | 139889415 | 139890694 | 1280 | 1.46 | 0.00377 | 0.00803 |
| chr12 | 6771051 | 6773649 | 2599 | 0.44 | 0.00377 | 0.00803 |

| | | | | | | |
|---|---|---|---|---|---|---|
| chr7 | 104603821 | 104604794 | 974 | -1.34 | 0.00377 | 0.00804 |
| chr17 | 71274972 | 71309297 | 34326 | -0.14 | 0.00378 | 0.00805 |
| chr2 | 11317302 | 11318711 | 1410 | -1.59 | 0.00378 | 0.00805 |
| chr15 | 80351122 | 80366541 | 15420 | 0.26 | 0.00379 | 0.00806 |
| chr15 | 35718913 | 35720082 | 1170 | 1.6 | 0.00379 | 0.00806 |
| chr20 | 2240653 | 2242610 | 1958 | 3.53 | 0.00379 | 0.00808 |
| chr18 | 3554839 | 3568368 | 13530 | -0.66 | 0.00379 | 0.00808 |
| chr4 | 87013567 | 87014580 | 1014 | -1.28 | 0.0038 | 0.00808 |
| chr10 | 7962781 | 7970429 | 7649 | 3.47 | 0.0038 | 0.00808 |
| chr1 | 52362776 | 52364564 | 1789 | -4.1 | 0.0038 | 0.00809 |
| chr21 | 43268503 | 43270646 | 2144 | 1.09 | 0.0038 | 0.00809 |
| chr8 | 126340660 | 126355576 | 14917 | 0.38 | 0.0038 | 0.00809 |
| chr11 | 14308254 | 14344443 | 36190 | 0.35 | 0.00381 | 0.0081 |
| chr8 | 142350838 | 142363072 | 12235 | -3.4 | 0.00381 | 0.0081 |
| chr12 | 77271898 | 77274153 | 2256 | 0.68 | 0.00381 | 0.00811 |
| chr8 | 28822975 | 28823978 | 1004 | -1.28 | 0.00382 | 0.00813 |
| chr10 | 100300672 | 100301162 | 491 | 2.34 | 0.00382 | 0.00813 |
| chr3 | 72951669 | 72952659 | 991 | -1.95 | 0.00382 | 0.00813 |
| chr19 | 19843197 | 19844829 | 1633 | 0.97 | 0.00382 | 0.00813 |
| chr7 | 13257169 | 13258318 | 1150 | 1.62 | 0.00382 | 0.00813 |
| chr3 | 53194483 | 53195795 | 1313 | 1.23 | 0.00383 | 0.00813 |
| chr9 | 37079116 | 37080837 | 1722 | 0.66 | 0.00383 | 0.00813 |
| chr6 | 99345408 | 99346389 | 982 | 1.69 | 0.00383 | 0.00814 |
| chr4 | 140834513 | 140856148 | 21636 | 0.49 | 0.00383 | 0.00814 |
| chr1 | 220999499 | 221001713 | 2215 | 1.27 | 0.00383 | 0.00815 |
| chr1 | 233085335 | 233089140 | 3806 | -0.6 | 0.00384 | 0.00816 |
| chr20 | 33329968 | 33342624 | 12657 | -0.66 | 0.00384 | 0.00816 |
| chr8 | 62788820 | 62790698 | 1879 | -0.81 | 0.00384 | 0.00816 |
| chr2 | 88875586 | 88876081 | 496 | -2.75 | 0.00384 | 0.00817 |
| chr12 | 2112445 | 2114896 | 2452 | 0.47 | 0.00384 | 0.00817 |
| chr9 | 123728429 | 123746210 | 17782 | -0.79 | 0.00385 | 0.00817 |
| chr13 | 60488942 | 60489648 | 707 | 1.67 | 0.00385 | 0.00817 |
| chr5 | 13566683 | 13568305 | 1623 | 0.9 | 0.00385 | 0.00818 |
| chr20 | 4572666 | 4573734 | 1069 | -1.55 | 0.00385 | 0.00818 |
| chr20 | 17268456 | 17268984 | 529 | -2.65 | 0.00385 | 0.00819 |
| chr8 | 8381651 | 8415173 | 33523 | -0.35 | 0.00386 | 0.00819 |
| chr2 | 221014789 | 221015327 | 539 | 1.8 | 0.00386 | 0.0082 |
| chr15 | 92078598 | 92082236 | 3639 | -0.38 | 0.00386 | 0.0082 |
| chr2 | 166947495 | 166948637 | 1143 | -1.34 | 0.00387 | 0.00822 |
| chr4 | 28314765 | 28317499 | 2735 | 0.65 | 0.00387 | 0.00822 |
| chr7 | 139155284 | 139158535 | 3252 | -0.36 | 0.00387 | 0.00822 |
| chr2 | 37382707 | 37395793 | 13087 | 0.4 | 0.00388 | 0.00824 |
| chr7 | 69790732 | 69791243 | 512 | -2.23 | 0.00388 | 0.00824 |
| chr8 | 98655106 | 98660207 | 5102 | 0.37 | 0.00388 | 0.00824 |
| chr1 | 150846900 | 150850372 | 3473 | 0.45 | 0.00389 | 0.00825 |
| chr18 | 2932463 | 2949317 | 16855 | -0.46 | 0.00389 | 0.00825 |
| chr6 | 83669268 | 83678606 | 9339 | -1.21 | 0.00389 | 0.00825 |
| chr3 | 59034920 | 59036542 | 1623 | -0.53 | 0.00389 | 0.00825 |
| chr3 | 39092735 | 39094270 | 1536 | 0.43 | 0.00389 | 0.00825 |
| chr4 | 37827171 | 37829079 | 1909 | 0.56 | 0.00389 | 0.00825 |
| chr5 | 14142197 | 14367032 | 224836 | 0.09 | 0.00389 | 0.00826 |
| chr3 | 157077670 | 157088076 | 10407 | -2.54 | 0.0039 | 0.00827 |
| chr4 | 185455866 | 185459631 | 3766 | 0.42 | 0.00391 | 0.00829 |
| chr14 | 100738926 | 100764025 | 25100 | -0.49 | 0.00391 | 0.0083 |
| chr16 | 25565194 | 25565722 | 529 | -2.81 | 0.00392 | 0.00831 |
| chr17 | 27423327 | 27508375 | 85049 | -0.14 | 0.00392 | 0.00831 |
| chr2 | 181542029 | 181542787 | 759 | 2.4 | 0.00392 | 0.00832 |
| chr11 | 392845 | 407116 | 14272 | -0.3 | 0.00392 | 0.00832 |
| chr2 | 162270805 | 162271532 | 728 | -1.7 | 0.00392 | 0.00832 |
| chr3 | 100753846 | 100756266 | 2421 | 0.35 | 0.00393 | 0.00833 |
| chr16 | 10613757 | 10628617 | 14861 | -0.59 | 0.00393 | 0.00833 |

| | | | | | | |
|---|---|---|---|---|---|---|
| chr6 | 117922225 | 117924578 | 2354 | 0.67 | 0.00393 | 0.00833 |
| chr12 | 9502896 | 9503243 | 348 | 2.82 | 0.00393 | 0.00833 |
| chr5 | 145380811 | 145381755 | 945 | -0.94 | 0.00393 | 0.00833 |
| chr7 | 69173675 | 69174581 | 907 | 2.61 | 0.00394 | 0.00834 |
| chrX | 55934979 | 55935549 | 571 | 1.71 | 0.00394 | 0.00836 |
| chr17 | 8278424 | 8289655 | 11232 | 0.31 | 0.00394 | 0.00836 |
| chr5 | 124378934 | 124381362 | 2429 | 0.66 | 0.00394 | 0.00836 |
| chr1 | 150898143 | 150982086 | 83944 | -0.18 | 0.00395 | 0.00836 |
| chr22 | 43139229 | 43157649 | 18421 | -0.32 | 0.00395 | 0.00836 |
| chr17 | 78864588 | 78876277 | 11690 | -0.51 | 0.00395 | 0.00836 |
| chr14 | 91685273 | 91686074 | 802 | -2.74 | 0.00395 | 0.00837 |
| chr8 | 138167913 | 138169130 | 1218 | -1.05 | 0.00395 | 0.00837 |
| chr8 | 98556858 | 98557665 | 808 | -1.7 | 0.00395 | 0.00837 |
| chr7 | 91714008 | 91714389 | 382 | 2.82 | 0.00395 | 0.00837 |
| chr8 | 16683627 | 16684387 | 761 | 0.9 | 0.00396 | 0.00839 |
| chr6 | 49293325 | 49294857 | 1533 | 4.23 | 0.00396 | 0.00839 |
| chr19 | 30296731 | 30315956 | 19226 | -0.58 | 0.00397 | 0.0084 |
| chr2 | 73510805 | 73512059 | 1255 | -1.12 | 0.00397 | 0.0084 |
| chr6 | 100986014 | 100986892 | 879 | 1.91 | 0.00397 | 0.0084 |
| chr11 | 76571191 | 76572976 | 1786 | 0.75 | 0.00399 | 0.00844 |
| chr1 | 155895546 | 155953677 | 58132 | 0.15 | 0.00399 | 0.00844 |
| chr7 | 150833784 | 150834510 | 727 | -2.61 | 0.00399 | 0.00844 |
| chr5 | 141765809 | 141770397 | 4589 | 0.56 | 0.00399 | 0.00845 |
| chr10 | 119412038 | 119423578 | 11541 | 4.11 | 0.00399 | 0.00845 |
| chr19 | 16933404 | 16940962 | 7559 | 0.69 | 0.00399 | 0.00845 |
| chr2 | 29087859 | 29094393 | 6535 | -0.44 | 0.00399 | 0.00845 |
| chrX | 16886015 | 16911390 | 25376 | 0.4 | 0.004 | 0.00846 |
| chr1 | 45671014 | 45672828 | 1815 | 1.28 | 0.004 | 0.00847 |
| chr11 | 128774799 | 128778875 | 4077 | 0.77 | 0.00401 | 0.00848 |
| chr11 | 59521708 | 59525054 | 3347 | 0.52 | 0.00401 | 0.00848 |
| chr4 | 151805529 | 151806913 | 1385 | 0.75 | 0.00401 | 0.00848 |
| chr8 | 121789688 | 121790866 | 1179 | 3.86 | 0.00401 | 0.00848 |
| chr17 | 26878919 | 26904595 | 25677 | -0.54 | 0.00401 | 0.00849 |
| chr6 | 44272740 | 44281109 | 8370 | -0.64 | 0.00401 | 0.00849 |
| chr10 | 27099317 | 27100128 | 812 | -1.7 | 0.00401 | 0.00849 |
| chr4 | 124445824 | 124447234 | 1411 | 1.05 | 0.00401 | 0.00849 |
| chr21 | 40378561 | 40379184 | 624 | 2.41 | 0.00402 | 0.00849 |
| chr4 | 34047424 | 34049721 | 2298 | 0.71 | 0.00402 | 0.0085 |
| chr15 | 36289984 | 36290538 | 555 | 2.69 | 0.00402 | 0.00851 |
| chrX | 150081514 | 150082748 | 1235 | -1.67 | 0.00403 | 0.00851 |
| chr2 | 202041527 | 202048470 | 6944 | -0.58 | 0.00403 | 0.00852 |
| chr20 | 13472627 | 13473461 | 835 | 2.61 | 0.00403 | 0.00853 |
| chr3 | 119895558 | 119897575 | 2018 | -0.55 | 0.00404 | 0.00854 |
| chr3 | 190334813 | 190335923 | 1111 | 1.65 | 0.00404 | 0.00854 |
| chr15 | 74950279 | 74956826 | 6548 | -1.13 | 0.00404 | 0.00855 |
| chr4 | 156775222 | 156776755 | 1534 | 0.56 | 0.00404 | 0.00855 |
| chr7 | 43287813 | 43301633 | 13821 | -0.33 | 0.00404 | 0.00855 |
| chr22 | 21290297 | 21313147 | 22851 | -0.33 | 0.00404 | 0.00855 |
| chr8 | 21905772 | 21924855 | 19084 | -0.22 | 0.00405 | 0.00855 |
| chr17 | 77960786 | 77980263 | 19478 | 0.35 | 0.00405 | 0.00855 |
| chr4 | 83270427 | 83297093 | 26667 | 0.26 | 0.00405 | 0.00856 |
| chr15 | 25682305 | 25685090 | 2786 | 0.42 | 0.00405 | 0.00856 |
| chr9 | 93753946 | 93754989 | 1044 | -1.35 | 0.00405 | 0.00856 |
| chr13 | 28146299 | 28158234 | 11936 | 3.6 | 0.00406 | 0.00858 |
| chr4 | 17550912 | 17551921 | 1010 | 1.14 | 0.00406 | 0.00858 |
| chr11 | 118271054 | 118287393 | 16340 | 0.36 | 0.00407 | 0.0086 |
| chr7 | 81675171 | 81683049 | 7879 | -0.25 | 0.00407 | 0.0086 |
| chr15 | 41939093 | 41964700 | 25608 | 0.55 | 0.00407 | 0.0086 |
| chr10 | 52305011 | 52305928 | 918 | -1.74 | 0.00407 | 0.0086 |
| chr4 | 109541225 | 109543688 | 2464 | 0.44 | 0.00407 | 0.0086 |
| chr19 | 8219055 | 8226763 | 7709 | 4.55 | 0.00407 | 0.0086 |

| | | | | | | |
|---|---|---|---|---|---|---|
| chr2 | 209193821 | 209199469 | 5649 | -0.83 | 0.00408 | 0.00861 |
| chr2 | 9210771 | 9211357 | 587 | 1.45 | 0.00409 | 0.00863 |
| chr8 | 19673845 | 19693534 | 19690 | 0.3 | 0.0041 | 0.00865 |
| chr6 | 139456839 | 139457692 | 854 | 1.35 | 0.0041 | 0.00866 |
| chr6 | 44728041 | 44728817 | 777 | 1.12 | 0.0041 | 0.00866 |
| chr7 | 46921179 | 46922013 | 835 | 1.41 | 0.00411 | 0.00867 |
| chr1 | 196543064 | 196555409 | 12346 | 4.65 | 0.00411 | 0.00868 |
| chr3 | 110036878 | 110037655 | 778 | 1.5 | 0.00411 | 0.00868 |
| chr11 | 111741184 | 111751066 | 9883 | 0.38 | 0.00412 | 0.0087 |
| chr7 | 67553702 | 67554445 | 744 | -2.15 | 0.00413 | 0.00872 |
| chr3 | 30234936 | 30241587 | 6652 | 1.04 | 0.00413 | 0.00872 |
| chr4 | 113091111 | 113092643 | 1533 | -0.84 | 0.00414 | 0.00873 |
| chr20 | 10015371 | 10028642 | 13272 | -0.4 | 0.00415 | 0.00875 |
| chr5 | 40641296 | 40647068 | 5773 | 3.9 | 0.00415 | 0.00875 |
| chrX | 10918553 | 10919851 | 1299 | 1 | 0.00415 | 0.00875 |
| chr21 | 42606572 | 42608033 | 1462 | 4.05 | 0.00415 | 0.00876 |
| chr4 | 37584845 | 37586408 | 1564 | 3.84 | 0.00415 | 0.00876 |
| chr11 | 125461049 | 125463776 | 2728 | 0.35 | 0.00416 | 0.00877 |
| chr17 | 26219680 | 26220763 | 1084 | 0.92 | 0.00416 | 0.00877 |
| chr1 | 246665806 | 246678184 | 12379 | 0.68 | 0.00416 | 0.00877 |
| chr17 | 27009373 | 27010760 | 1388 | -1.73 | 0.00417 | 0.00878 |
| chr8 | 27612525 | 27613458 | 934 | 1.65 | 0.00417 | 0.00879 |
| chr7 | 16183270 | 16184869 | 1600 | -1.07 | 0.00417 | 0.00879 |
| chr3 | 49275478 | 49283373 | 7896 | 0.68 | 0.00417 | 0.00879 |
| chr11 | 45985955 | 45987052 | 1098 | 1.32 | 0.00417 | 0.0088 |
| chr7 | 80951565 | 80952375 | 811 | 1.74 | 0.00419 | 0.00883 |
| chr9 | 107017925 | 107018474 | 550 | 2.2 | 0.0042 | 0.00885 |
| chr17 | 65205759 | 65221043 | 15285 | 0.59 | 0.0042 | 0.00885 |
| chr3 | 151911151 | 151922013 | 10863 | 0.47 | 0.0042 | 0.00885 |
| chr15 | 74752720 | 74754342 | 1623 | 0.73 | 0.0042 | 0.00885 |
| chr16 | 90148558 | 90149556 | 999 | 0.81 | 0.0042 | 0.00886 |
| chr19 | 50030606 | 50031152 | 547 | 1.71 | 0.00421 | 0.00886 |
| chr11 | 85586187 | 85587280 | 1094 | 1.12 | 0.00421 | 0.00887 |
| chr22 | 31926260 | 31927522 | 1263 | 0.74 | 0.00421 | 0.00887 |
| chr4 | 88740364 | 88741078 | 715 | -1.53 | 0.00421 | 0.00887 |
| chr11 | 121162889 | 121164296 | 1408 | 1.04 | 0.00422 | 0.00888 |
| chr13 | 83321772 | 83322528 | 757 | 1.72 | 0.00422 | 0.00889 |
| chr22 | 42753647 | 42754183 | 537 | -2.71 | 0.00422 | 0.00889 |
| chr2 | 208877635 | 208891063 | 13429 | -0.51 | 0.00423 | 0.0089 |
| chr9 | 109984137 | 109984989 | 853 | 1.52 | 0.00423 | 0.0089 |
| chr17 | 59412308 | 59412673 | 366 | -2.76 | 0.00423 | 0.0089 |
| chr9 | 83747298 | 83747628 | 331 | -2.65 | 0.00423 | 0.00891 |
| chr17 | 46867942 | 47052921 | 184980 | -0.12 | 0.00423 | 0.00891 |
| chr18 | 47337905 | 47341120 | 3216 | 0.42 | 0.00423 | 0.00891 |
| chr6 | 109414588 | 109417762 | 3175 | 0.5 | 0.00424 | 0.00893 |
| chr7 | 11579385 | 11580838 | 1454 | 1.29 | 0.00426 | 0.00896 |
| chr8 | 18924927 | 18925957 | 1031 | -1.49 | 0.00426 | 0.00896 |
| chr2 | 139002274 | 139003091 | 818 | 2.5 | 0.00426 | 0.00896 |
| chr6 | 104973353 | 104973682 | 330 | 2.92 | 0.00426 | 0.00897 |
| chr7 | 139874905 | 139877762 | 2858 | 0.72 | 0.00427 | 0.00897 |
| chr1 | 230707694 | 230709094 | 1401 | 4.17 | 0.00428 | 0.009 |
| chr1 | 31701718 | 31702133 | 416 | -2.35 | 0.00429 | 0.00902 |
| chr12 | 91015958 | 91016779 | 822 | -2.18 | 0.00429 | 0.00902 |
| chr3 | 195161508 | 195164630 | 3123 | 0.54 | 0.00429 | 0.00902 |
| chr3 | 119316016 | 119317048 | 1033 | 1.31 | 0.00429 | 0.00902 |
| chr14 | 95622368 | 95625574 | 3207 | 0.4 | 0.00429 | 0.00902 |
| chr7 | 112757676 | 112759347 | 1672 | 0.8 | 0.0043 | 0.00904 |
| chrX | 46771283 | 46773741 | 2459 | 0.89 | 0.00431 | 0.00906 |
| chr11 | 70422409 | 70423517 | 1109 | -1.69 | 0.00431 | 0.00907 |
| chr2 | 20967035 | 20967876 | 842 | -1.69 | 0.00432 | 0.00908 |
| chr3 | 123220368 | 123221066 | 699 | -1.88 | 0.00432 | 0.00908 |

| | | | | | | |
|---|---|---|---|---|---|---|
| chr3 | 177351202 | 177352099 | 898 | 1.2 | 0.00432 | 0.00908 |
| chr7 | 107219526 | 107221836 | 2311 | 0.37 | 0.00432 | 0.00909 |
| chr10 | 35177588 | 35179069 | 1482 | 1.33 | 0.00433 | 0.00911 |
| chr18 | 56234537 | 56258581 | 24045 | -0.27 | 0.00433 | 0.00911 |
| chr11 | 128050300 | 128051348 | 1049 | -1.44 | 0.00435 | 0.00913 |
| chrX | 130574924 | 130576089 | 1166 | 1.85 | 0.00435 | 0.00913 |
| chr12 | 70936853 | 70937972 | 1120 | 1.14 | 0.00435 | 0.00914 |
| chr17 | 7080551 | 7093226 | 12676 | -1.14 | 0.00435 | 0.00914 |
| chr9 | 32525176 | 32527347 | 2172 | 1.02 | 0.00435 | 0.00915 |
| chr4 | 110480356 | 110496130 | 15775 | 0.46 | 0.00436 | 0.00915 |
| chr1 | 44435688 | 44458503 | 22816 | -0.38 | 0.00436 | 0.00915 |
| chr4 | 91049569 | 91049972 | 404 | 3.9 | 0.00437 | 0.00917 |
| chr5 | 68710462 | 68711633 | 1172 | 1.08 | 0.00437 | 0.00917 |
| chr10 | 28698749 | 28699423 | 675 | 1.85 | 0.00437 | 0.00918 |
| chr13 | 110352154 | 110358982 | 6829 | 0.51 | 0.00437 | 0.00918 |
| chr15 | 50682753 | 50685378 | 2626 | -1 | 0.00438 | 0.00919 |
| chr6 | 144328874 | 144329421 | 548 | -1.89 | 0.00438 | 0.00919 |
| chr2 | 152144331 | 152147578 | 3248 | 0.45 | 0.00438 | 0.0092 |
| chr17 | 26554089 | 26555627 | 1539 | 0.59 | 0.00438 | 0.0092 |
| chr18 | 13709990 | 13711405 | 1416 | 1.72 | 0.00438 | 0.0092 |
| chrX | 39680168 | 39690291 | 10124 | 1.02 | 0.00439 | 0.00921 |
| chr17 | 20157611 | 20171344 | 13734 | -0.57 | 0.00439 | 0.00921 |
| chr5 | 137259234 | 137259703 | 470 | 2.17 | 0.0044 | 0.00924 |
| chr11 | 34991753 | 35036170 | 44418 | 0.18 | 0.00441 | 0.00925 |
| chrX | 62780019 | 62781747 | 1729 | 0.77 | 0.00442 | 0.00926 |
| chr2 | 180323020 | 180325879 | 2860 | 4.28 | 0.00442 | 0.00927 |
| chr1 | 193455557 | 193456662 | 1106 | 0.86 | 0.00442 | 0.00927 |
| chr1 | 224752241 | 224753834 | 1594 | -1.31 | 0.00443 | 0.00928 |
| chr2 | 85475678 | 85476886 | 1209 | 1.25 | 0.00443 | 0.00929 |
| chr4 | 105669733 | 105670760 | 1028 | 1.41 | 0.00443 | 0.00929 |
| chr14 | 58564792 | 58565659 | 868 | 1.8 | 0.00443 | 0.0093 |
| chr6 | 38512758 | 38513239 | 482 | -1.97 | 0.00444 | 0.00931 |
| chr1 | 202681717 | 202683061 | 1345 | 1.26 | 0.00444 | 0.00931 |
| chr10 | 13424789 | 13425766 | 978 | 2.58 | 0.00444 | 0.00931 |
| chr6 | 48792549 | 48796561 | 4013 | 0.23 | 0.00444 | 0.00931 |
| chr5 | 30289352 | 30289755 | 404 | 3.16 | 0.00444 | 0.00931 |
| chr6 | 11039395 | 11040712 | 1318 | -1.16 | 0.00444 | 0.00931 |
| chr1 | 65614136 | 65625720 | 11585 | -0.78 | 0.00445 | 0.00933 |
| chrX | 24982067 | 24982750 | 684 | 1.79 | 0.00445 | 0.00933 |
| chr5 | 104245480 | 104248008 | 2529 | 4.63 | 0.00446 | 0.00935 |
| chr4 | 190013975 | 190014520 | 546 | 2.19 | 0.00446 | 0.00936 |
| chr1 | 9309550 | 9310222 | 673 | -2.13 | 0.00447 | 0.00937 |
| chr2 | 138686186 | 138701381 | 15196 | 1.11 | 0.00447 | 0.00937 |
| chr1 | 232597737 | 232598584 | 848 | -2.28 | 0.00447 | 0.00938 |
| chr1 | 32644440 | 32648303 | 3864 | 0.51 | 0.00448 | 0.00938 |
| chr4 | 103421293 | 103434061 | 12769 | 0.33 | 0.00448 | 0.00938 |
| chr7 | 129338348 | 129339220 | 873 | -1.75 | 0.00449 | 0.0094 |
| chr2 | 206987069 | 206994757 | 7689 | -0.7 | 0.00449 | 0.0094 |
| chr14 | 60714028 | 60718962 | 4935 | 0.32 | 0.0045 | 0.00942 |
| chr16 | 49696713 | 49698940 | 2228 | 4.3 | 0.0045 | 0.00943 |
| chr14 | 23008644 | 23009275 | 632 | -1.83 | 0.0045 | 0.00944 |
| chr6 | 88481661 | 88484007 | 2347 | 0.65 | 0.00451 | 0.00945 |
| chr9 | 124975818 | 124976294 | 477 | 2.15 | 0.00452 | 0.00946 |
| chr6 | 69773578 | 69774125 | 548 | -2.39 | 0.00452 | 0.00946 |
| chr7 | 139042307 | 139082540 | 40234 | 0.2 | 0.00453 | 0.00948 |
| chr17 | 61773769 | 61783399 | 9631 | -0.88 | 0.00454 | 0.0095 |
| chr4 | 25862498 | 25864380 | 1883 | -0.87 | 0.00454 | 0.00951 |
| chr6 | 114071391 | 114072066 | 676 | 2.43 | 0.00454 | 0.00951 |
| chr11 | 4208009 | 4208998 | 990 | 0.74 | 0.00455 | 0.00953 |
| chr5 | 100682640 | 100683496 | 857 | 2.24 | 0.00455 | 0.00953 |
| chr1 | 71545496 | 71547749 | 2254 | 0.37 | 0.00455 | 0.00953 |

| | | | | | | |
|---|---|---|---|---|---|---|
| chr6 | 148662734 | 148664922 | 2189 | 0.52 | 0.00456 | 0.00955 |
| chr2 | 232419227 | 232420297 | 1071 | 1.65 | 0.00456 | 0.00955 |
| chr12 | 99837106 | 99850240 | 13135 | -1.14 | 0.00457 | 0.00955 |
| chrX | 21856829 | 21862117 | 5289 | 0.37 | 0.00457 | 0.00955 |
| chr1 | 153625116 | 153646156 | 21041 | 0.29 | 0.00457 | 0.00955 |
| chr1 | 233136038 | 233137008 | 971 | -1.44 | 0.00457 | 0.00956 |
| chr6 | 89371480 | 89372721 | 1242 | 1.41 | 0.00457 | 0.00957 |
| chr5 | 55199370 | 55231531 | 32162 | -0.33 | 0.00457 | 0.00957 |
| chr7 | 21590271 | 21590804 | 534 | 2.58 | 0.00458 | 0.00958 |
| chr21 | 18511232 | 18512823 | 1592 | 0.77 | 0.00458 | 0.00958 |
| chr1 | 27018035 | 27044170 | 26136 | -0.2 | 0.00458 | 0.00958 |
| chr3 | 193460654 | 193462416 | 1763 | 1.12 | 0.00458 | 0.00958 |
| chr2 | 71114365 | 71129526 | 15162 | -0.63 | 0.00458 | 0.00958 |
| chr2 | 204623690 | 204624927 | 1238 | -1.07 | 0.00459 | 0.0096 |
| chr14 | 53415672 | 53418996 | 3325 | 0.37 | 0.0046 | 0.00961 |
| chr3 | 131049812 | 131050506 | 695 | 0.93 | 0.00461 | 0.00964 |
| chr8 | 16883708 | 16886056 | 2349 | 0.48 | 0.00461 | 0.00964 |
| chr1 | 95698084 | 95700506 | 2423 | 0.5 | 0.00461 | 0.00964 |
| chr1 | 86146168 | 86158359 | 12192 | -0.62 | 0.00461 | 0.00965 |
| chr4 | 71553157 | 71555605 | 2449 | 0.46 | 0.00462 | 0.00965 |
| chr6 | 20740629 | 20741378 | 750 | 2.07 | 0.00462 | 0.00965 |
| chrX | 8789414 | 8789756 | 343 | 2.28 | 0.00462 | 0.00965 |
| chr13 | 113951021 | 113952527 | 1507 | 0.93 | 0.00462 | 0.00966 |
| chr8 | 103401938 | 103425882 | 23945 | 0.31 | 0.00462 | 0.00966 |
| chr1 | 167885356 | 167920990 | 35635 | 0.17 | 0.00462 | 0.00966 |
| chr7 | 150049036 | 150049767 | 732 | 1.95 | 0.00463 | 0.00967 |
| chr7 | 77107666 | 77112850 | 5185 | 0.3 | 0.00463 | 0.00967 |
| chr1 | 30556029 | 30557499 | 1471 | -0.76 | 0.00463 | 0.00967 |
| chr19 | 11372336 | 11380121 | 7786 | 0.67 | 0.00463 | 0.00968 |
| chr17 | 17460356 | 17464824 | 4469 | -2.85 | 0.00463 | 0.00968 |
| chr20 | 18117960 | 18126460 | 8501 | 0.4 | 0.00464 | 0.00969 |
| chr3 | 149824026 | 149825831 | 1806 | 0.85 | 0.00465 | 0.0097 |
| chr8 | 102903311 | 102903956 | 646 | 1.85 | 0.00465 | 0.00971 |
| chrX | 15807829 | 15809488 | 1660 | 0.78 | 0.00465 | 0.00971 |
| chrX | 15470107 | 15471796 | 1690 | -1.35 | 0.00465 | 0.00971 |
| chr2 | 37157892 | 37194870 | 36979 | -0.37 | 0.00466 | 0.00972 |
| chr5 | 28468003 | 28468605 | 603 | 1.72 | 0.00466 | 0.00972 |
| chr15 | 25199701 | 25202499 | 2799 | 0.55 | 0.00466 | 0.00972 |
| chr13 | 114145352 | 114148720 | 3369 | 0.66 | 0.00466 | 0.00973 |
| chr3 | 176853433 | 176854767 | 1335 | 1.71 | 0.00467 | 0.00975 |
| chr2 | 233561076 | 233565722 | 4647 | -0.58 | 0.00467 | 0.00975 |
| chr16 | 73017394 | 73131308 | 113915 | -0.12 | 0.00467 | 0.00975 |
| chr8 | 23848571 | 23849499 | 929 | 1.55 | 0.00468 | 0.00977 |
| chr13 | 34071074 | 34071844 | 771 | -2.06 | 0.00468 | 0.00977 |
| chr22 | 24167615 | 24169211 | 1597 | -0.97 | 0.00469 | 0.00978 |
| chr4 | 106105363 | 106106520 | 1158 | 1.28 | 0.00469 | 0.00978 |
| chr14 | 73646355 | 73660636 | 14282 | 0.45 | 0.00469 | 0.00979 |
| chr2 | 151554152 | 151555161 | 1010 | 1.35 | 0.0047 | 0.0098 |
| chr9 | 126074874 | 126075558 | 685 | -2.24 | 0.0047 | 0.00981 |
| chr11 | 102363470 | 102366205 | 2736 | 0.68 | 0.00472 | 0.00983 |
| chr2 | 218295978 | 218297275 | 1298 | -1.37 | 0.00473 | 0.00986 |
| chr8 | 38087830 | 38090401 | 2572 | 0.41 | 0.00473 | 0.00986 |
| chr5 | 167105466 | 167106134 | 669 | 2.69 | 0.00473 | 0.00987 |
| chr5 | 37043414 | 37056169 | 12756 | 0.76 | 0.00474 | 0.00988 |
| chr11 | 1989061 | 1990831 | 1771 | 1.11 | 0.00475 | 0.0099 |
| chr18 | 54481745 | 54484728 | 2984 | 0.45 | 0.00475 | 0.00991 |
| chr17 | 2739067 | 2758552 | 19486 | 0.38 | 0.00476 | 0.00992 |
| chr1 | 76413823 | 76431218 | 17396 | 3.15 | 0.00476 | 0.00992 |
| chr15 | 66833350 | 66834636 | 1287 | -0.89 | 0.00476 | 0.00992 |
| chr2 | 63135965 | 63147962 | 11998 | -1.95 | 0.00476 | 0.00992 |
| chr2 | 160567241 | 160631595 | 64355 | -0.17 | 0.00476 | 0.00992 |

| | | | | | | |
|---|---|---|---|---|---|---|
| chr9 | 37900533 | 38071262 | 170730 | -0.09 | 0.00477 | 0.00993 |
| chr4 | 188399918 | 188400621 | 704 | -2.2 | 0.00477 | 0.00993 |
| chr3 | 182399926 | 182401442 | 1517 | 1.6 | 0.00477 | 0.00993 |
| chr20 | 44623822 | 44624659 | 838 | 2.03 | 0.00477 | 0.00994 |
| chr5 | 87733530 | 87734868 | 1339 | -2.5 | 0.00478 | 0.00995 |
| chr7 | 129287098 | 129288106 | 1009 | 1.35 | 0.00478 | 0.00996 |
| chr9 | 95268473 | 95269993 | 1521 | -1.42 | 0.00479 | 0.00999 |
| chr4 | 56261122 | 56265367 | 4246 | 0.37 | 0.0048 | 0.00999 |
| chr11 | 11592725 | 11607134 | 14410 | -4.53 | 0.0048 | 0.01 |
| chr9 | 109645466 | 109656920 | 11455 | 0.37 | 0.0048 | 0.01 |
| chrX | 40652930 | 40654220 | 1291 | -0.99 | 0.0048 | 0.01 |
| chr2 | 171992858 | 171994324 | 1467 | -0.83 | 0.00481 | 0.01 |
| chr2 | 25097136 | 25109869 | 12734 | -3.28 | 0.00481 | 0.01 |
| chr11 | 8692491 | 8722220 | 29730 | 0.24 | 0.00482 | 0.01 |
| chr4 | 77693365 | 77704086 | 10722 | -0.83 | 0.00482 | 0.01 |
| chr4 | 79213607 | 79215729 | 2123 | 1.09 | 0.00483 | 0.01 |
| chr3 | 187453707 | 187469585 | 15879 | 0.27 | 0.00483 | 0.0101 |
| chr16 | 89283117 | 89284829 | 1713 | 0.62 | 0.00484 | 0.0101 |
| chr8 | 68285642 | 68286581 | 940 | 1.72 | 0.00484 | 0.0101 |
| chr12 | 58087903 | 58088682 | 780 | 0.91 | 0.00487 | 0.0101 |
| chr2 | 33807778 | 33825243 | 17466 | -0.4 | 0.00487 | 0.0101 |
| chr3 | 193692359 | 193693821 | 1463 | 1.41 | 0.00488 | 0.0102 |
| chr4 | 16084053 | 16084577 | 525 | 2.67 | 0.00489 | 0.0102 |
| chr15 | 93277039 | 93278124 | 1086 | 0.82 | 0.0049 | 0.0102 |
| chr8 | 27949240 | 27952384 | 3145 | 0.37 | 0.00491 | 0.0102 |
| chr2 | 239264981 | 239266727 | 1747 | 0.99 | 0.00492 | 0.0102 |
| chr14 | 90764103 | 90777217 | 13115 | -0.74 | 0.00492 | 0.0102 |
| chr6 | 138983184 | 138985538 | 2355 | -0.88 | 0.00492 | 0.0102 |
| chr20 | 18636382 | 18637357 | 976 | 1.84 | 0.00492 | 0.0102 |
| chr1 | 32816286 | 32818264 | 1979 | 0.64 | 0.00493 | 0.0103 |
| chr2 | 72594627 | 72595147 | 521 | -2 | 0.00493 | 0.0103 |
| chr3 | 64069100 | 64080850 | 11751 | -0.36 | 0.00494 | 0.0103 |
| chr21 | 30670540 | 30673509 | 2970 | 0.59 | 0.00494 | 0.0103 |
| chr1 | 15849524 | 15859935 | 10412 | 0.3 | 0.00494 | 0.0103 |
| chr6 | 56703474 | 56757557 | 54084 | -0.17 | 0.00494 | 0.0103 |
| chr7 | 43767875 | 43770304 | 2430 | 0.49 | 0.00494 | 0.0103 |
| chrX | 74374983 | 74376748 | 1766 | 0.57 | 0.00495 | 0.0103 |
| chr3 | 194840126 | 194841729 | 1604 | -1.13 | 0.00495 | 0.0103 |
| chr14 | 50083043 | 50127832 | 44790 | 0.18 | 0.00495 | 0.0103 |
| chr12 | 66695651 | 66697584 | 1934 | 0.56 | 0.00496 | 0.0103 |
| chr2 | 97425824 | 97430245 | 4422 | -0.73 | 0.00496 | 0.0103 |
| chr6 | 119468005 | 119469156 | 1152 | 1.78 | 0.00497 | 0.0103 |
| chr3 | 11866074 | 11867474 | 1401 | 1.46 | 0.00497 | 0.0103 |
| chr19 | 56090860 | 56122206 | 31347 | 0.17 | 0.00498 | 0.0103 |
| chr13 | 32914595 | 32922829 | 8235 | 0.85 | 0.00499 | 0.0104 |
| chr17 | 7834468 | 7837069 | 2602 | 0.53 | 0.00499 | 0.0104 |
| chrX | 47383046 | 47383577 | 532 | -2.09 | 0.00499 | 0.0104 |
| chr2 | 206836746 | 206855548 | 18803 | 0.26 | 0.005 | 0.0104 |
| chr13 | 64210587 | 64211110 | 524 | 1.82 | 0.005 | 0.0104 |
| chr13 | 36906917 | 36920997 | 14081 | 0.33 | 0.005 | 0.0104 |
| chr17 | 58449919 | 58451181 | 1263 | -1.51 | 0.005 | 0.0104 |
| chr2 | 25345948 | 25346972 | 1025 | -1.58 | 0.005 | 0.0104 |
| chrX | 107018520 | 107019949 | 1430 | 1.2 | 0.005 | 0.0104 |
| chr11 | 74302427 | 74305116 | 2690 | -0.43 | 0.005 | 0.0104 |
| chr5 | 110829956 | 110831133 | 1178 | -1.08 | 0.005 | 0.0104 |
| chr2 | 190663719 | 190665310 | 1592 | -1.45 | 0.00501 | 0.0104 |
| chr1 | 149215064 | 149231499 | 16436 | 0.28 | 0.00501 | 0.0104 |
| chr4 | 128953736 | 128954627 | 892 | 2.07 | 0.00501 | 0.0104 |
| chr3 | 32581643 | 32586800 | 5158 | 0.59 | 0.00502 | 0.0104 |
| chr5 | 68587131 | 68587927 | 797 | 1.96 | 0.00502 | 0.0104 |
| chr9 | 80747383 | 80748005 | 623 | 1.54 | 0.00502 | 0.0104 |

| | | | | | | |
|---|---|---|---|---|---|---|
| chr1 | 173638495 | 173639052 | 558 | -2.29 | 0.00502 | 0.0104 |
| chr9 | 4983636 | 4986715 | 3080 | 0.41 | 0.00502 | 0.0104 |
| chr17 | 16487799 | 16492733 | 4935 | 4.49 | 0.00503 | 0.0104 |
| chr18 | 9473957 | 9477053 | 3097 | 0.54 | 0.00503 | 0.0104 |
| chr7 | 105128070 | 105138038 | 9969 | -0.41 | 0.00503 | 0.0104 |
| chr7 | 114937178 | 114940947 | 3770 | 0.82 | 0.00504 | 0.0104 |
| chr2 | 241074591 | 241076320 | 1730 | -0.84 | 0.00504 | 0.0104 |
| chr9 | 34177699 | 34205765 | 28067 | 0.2 | 0.00504 | 0.0105 |
| chr6 | 64196925 | 64248428 | 51504 | 0.19 | 0.00504 | 0.0105 |
| chr15 | 64124802 | 64127024 | 2223 | 0.48 | 0.00505 | 0.0105 |
| chr15 | 83679085 | 83705668 | 26584 | 0.29 | 0.00505 | 0.0105 |
| chr17 | 37604313 | 37647947 | 43635 | 0.18 | 0.00505 | 0.0105 |
| chr3 | 12744131 | 12745104 | 974 | -2.09 | 0.00505 | 0.0105 |
| chrX | 57617326 | 57623662 | 6337 | 0.42 | 0.00505 | 0.0105 |
| chr5 | 179892574 | 179922688 | 30115 | 0.29 | 0.00505 | 0.0105 |
| chr16 | 54959982 | 54971381 | 11400 | 0.32 | 0.00506 | 0.0105 |
| chrX | 39714680 | 39726146 | 11467 | 0.71 | 0.00506 | 0.0105 |
| chr20 | 30538339 | 30539939 | 1601 | 0.56 | 0.00506 | 0.0105 |
| chrX | 135228779 | 135229280 | 502 | 2.13 | 0.00506 | 0.0105 |
| chr11 | 21910243 | 21910940 | 698 | 1.9 | 0.00506 | 0.0105 |
| chr7 | 50732089 | 50745250 | 13162 | -2.11 | 0.00507 | 0.0105 |
| chr15 | 68563616 | 68581814 | 18199 | 0.24 | 0.00507 | 0.0105 |
| chr15 | 52626399 | 52633315 | 6917 | -0.46 | 0.00507 | 0.0105 |
| chr6 | 89848724 | 89858120 | 9397 | 0.33 | 0.00508 | 0.0105 |
| chr3 | 27559087 | 27579284 | 20198 | -0.21 | 0.00508 | 0.0105 |
| chr2 | 38688987 | 38689973 | 987 | 1.42 | 0.00508 | 0.0105 |
| chr7 | 12131689 | 12132617 | 929 | -1.46 | 0.00509 | 0.0105 |
| chr11 | 11414409 | 11415178 | 770 | -2.09 | 0.00509 | 0.0105 |
| chr1 | 151701742 | 151702668 | 927 | 1.5 | 0.0051 | 0.0106 |
| chrX | 154254012 | 154257607 | 3596 | 0.37 | 0.00511 | 0.0106 |
| chr8 | 96067010 | 96087765 | 20756 | 0.51 | 0.00512 | 0.0106 |
| chr6 | 25831837 | 25832851 | 1015 | 1.47 | 0.00512 | 0.0106 |
| chr7 | 96746133 | 96748059 | 1927 | 0.6 | 0.00513 | 0.0106 |
| chr15 | 99525135 | 99527577 | 2443 | -0.7 | 0.00514 | 0.0106 |
| chr16 | 19316035 | 19316771 | 737 | -1.47 | 0.00514 | 0.0106 |
| chr5 | 32151215 | 32175801 | 24587 | 0.22 | 0.00515 | 0.0106 |
| chr4 | 107490513 | 107490980 | 468 | 2.46 | 0.00515 | 0.0107 |
| chr9 | 86853597 | 86854536 | 940 | -1.19 | 0.00516 | 0.0107 |
| chr20 | 37176449 | 37177721 | 1273 | 1.22 | 0.00516 | 0.0107 |
| chr20 | 33542500 | 33544442 | 1943 | 0.47 | 0.00517 | 0.0107 |
| chr15 | 39908618 | 39909367 | 750 | 1.57 | 0.00518 | 0.0107 |
| chr5 | 80801990 | 80802826 | 837 | 1.72 | 0.00519 | 0.0107 |
| chr13 | 90484322 | 90485938 | 1617 | 0.86 | 0.00519 | 0.0107 |
| chr9 | 588835 | 590254 | 1420 | -1.31 | 0.00519 | 0.0107 |
| chr5 | 56790096 | 56795418 | 5323 | -3.97 | 0.0052 | 0.0107 |
| chr4 | 87884008 | 87904367 | 20360 | -0.41 | 0.0052 | 0.0108 |
| chr17 | 4529051 | 4529833 | 783 | -2.04 | 0.00521 | 0.0108 |
| chr18 | 56515544 | 56681295 | 165752 | 0.1 | 0.00523 | 0.0108 |
| chr5 | 157246865 | 157261572 | 14708 | -0.48 | 0.00524 | 0.0108 |
| chr6 | 108395077 | 108402172 | 7096 | 0.64 | 0.00524 | 0.0108 |
| chr4 | 26023442 | 26024624 | 1183 | 1.51 | 0.00526 | 0.0109 |
| chr10 | 27608438 | 27609133 | 696 | 1.31 | 0.00527 | 0.0109 |
| chr7 | 77127902 | 77128566 | 665 | -2.58 | 0.00527 | 0.0109 |
| chrX | 3608009 | 3608982 | 974 | -1.54 | 0.00527 | 0.0109 |
| chr3 | 170444331 | 170445660 | 1330 | -1.08 | 0.00528 | 0.0109 |
| chr5 | 65106221 | 65121222 | 15002 | 0.34 | 0.00528 | 0.0109 |
| chr17 | 1670021 | 1750869 | 80849 | 0.18 | 0.00528 | 0.0109 |
| chr6 | 20866179 | 20880590 | 14412 | 0.33 | 0.00528 | 0.0109 |
| chrX | 106045159 | 106046525 | 1367 | 0.93 | 0.00528 | 0.0109 |
| chr8 | 135842543 | 135852234 | 9692 | 0.34 | 0.00529 | 0.0109 |
| chr19 | 44597945 | 44600204 | 2260 | 0.62 | 0.00529 | 0.0109 |

| | | | | | | |
|---|---|---|---|---|---|---|
| chr2 | 203829707 | 203831528 | 1822 | -0.44 | 0.00529 | 0.0109 |
| chr14 | 50571132 | 50584231 | 13100 | 0.34 | 0.00529 | 0.0109 |
| chr4 | 155497868 | 155519836 | 21969 | 0.36 | 0.0053 | 0.0109 |
| chr14 | 81051039 | 81052225 | 1187 | 1.1 | 0.0053 | 0.0109 |
| chr1 | 239352520 | 239353572 | 1053 | 1.87 | 0.0053 | 0.0109 |
| chr15 | 48460042 | 48471051 | 11010 | 0.36 | 0.00531 | 0.011 |
| chr3 | 189403617 | 189404334 | 718 | 1.44 | 0.00531 | 0.011 |
| chr2 | 203808721 | 203810062 | 1342 | -1.2 | 0.00532 | 0.011 |
| chr4 | 77068469 | 77070372 | 1904 | -0.7 | 0.00532 | 0.011 |
| chr10 | 103003627 | 103005134 | 1508 | -0.94 | 0.00532 | 0.011 |
| chr2 | 236171003 | 236177870 | 6868 | -0.34 | 0.00532 | 0.011 |
| chr2 | 182435574 | 182435903 | 330 | 2.79 | 0.00532 | 0.011 |
| chr5 | 37319857 | 37320601 | 745 | -1.91 | 0.00532 | 0.011 |
| chr12 | 26522523 | 26522989 | 467 | -2.55 | 0.00532 | 0.011 |
| chr9 | 130823395 | 130866894 | 43500 | 0.15 | 0.00533 | 0.011 |
| chr20 | 35722876 | 35725188 | 2313 | -0.4 | 0.00533 | 0.011 |
| chr20 | 22672415 | 22673139 | 725 | -1.02 | 0.00533 | 0.011 |
| chr7 | 140064435 | 140098735 | 34301 | -0.4 | 0.00534 | 0.011 |
| chr6 | 22363053 | 22364549 | 1497 | 1.34 | 0.00534 | 0.011 |
| chrX | 34974491 | 34975438 | 948 | 1.06 | 0.00534 | 0.011 |
| chr10 | 112062326 | 112096739 | 34414 | -0.32 | 0.00534 | 0.011 |
| chr4 | 30210076 | 30211130 | 1055 | 2.1 | 0.00535 | 0.011 |
| chr2 | 172174936 | 172175386 | 451 | -2.55 | 0.00535 | 0.011 |
| chr11 | 3109316 | 3110327 | 1012 | -2.18 | 0.00535 | 0.011 |
| chr9 | 100683355 | 100686094 | 2740 | 0.34 | 0.00535 | 0.011 |
| chr1 | 150292988 | 150299663 | 6676 | 0.33 | 0.00537 | 0.0111 |
| chr6 | 151038152 | 151038655 | 504 | 2.64 | 0.00537 | 0.0111 |
| chr18 | 46428027 | 46429433 | 1407 | -0.67 | 0.00538 | 0.0111 |
| chr16 | 86798516 | 86800417 | 1902 | 4.31 | 0.00538 | 0.0111 |
| chr4 | 34677635 | 34678758 | 1124 | 1.27 | 0.00538 | 0.0111 |
| chr6 | 119000198 | 119005743 | 5546 | 0.89 | 0.00539 | 0.0111 |
| chr7 | 16274169 | 16275068 | 900 | 1.56 | 0.00539 | 0.0111 |
| chr15 | 39267575 | 39268678 | 1104 | -1.38 | 0.0054 | 0.0111 |
| chr2 | 9318709 | 9320511 | 1803 | 1.05 | 0.0054 | 0.0111 |
| chr11 | 501786 | 518698 | 16913 | 0.28 | 0.0054 | 0.0111 |
| chr9 | 15509577 | 15512254 | 2678 | -0.39 | 0.0054 | 0.0111 |
| chr18 | 21524138 | 21524841 | 704 | -1.54 | 0.0054 | 0.0111 |
| chr21 | 34055170 | 34055639 | 470 | 1.88 | 0.0054 | 0.0111 |
| chr4 | 111436954 | 111437630 | 677 | 1.61 | 0.00541 | 0.0111 |
| chr2 | 46235396 | 46236217 | 822 | -2.5 | 0.00541 | 0.0111 |
| chr5 | 108062985 | 108064846 | 1862 | 0.42 | 0.00541 | 0.0111 |
| chr1 | 35544357 | 35551590 | 7234 | 0.54 | 0.00542 | 0.0112 |
| chr14 | 30692485 | 30693281 | 797 | 2.39 | 0.00543 | 0.0112 |
| chr4 | 169929808 | 169932354 | 2547 | 0.39 | 0.00543 | 0.0112 |
| chr13 | 31143305 | 31147607 | 4303 | 0.6 | 0.00543 | 0.0112 |
| chr15 | 39463957 | 39506853 | 42897 | 0.77 | 0.00544 | 0.0112 |
| chr8 | 67367347 | 67384907 | 17561 | 0.39 | 0.00544 | 0.0112 |
| chr2 | 24065462 | 24066799 | 1338 | -1.14 | 0.00544 | 0.0112 |
| chr14 | 50987408 | 51000197 | 12790 | 0.38 | 0.00544 | 0.0112 |
| chr16 | 28101375 | 28122153 | 20779 | -0.48 | 0.00545 | 0.0112 |
| chr6 | 2166679 | 2167788 | 1110 | 1.54 | 0.00545 | 0.0112 |
| chr1 | 110834770 | 110835175 | 406 | 2.47 | 0.00546 | 0.0112 |
| chr2 | 11460241 | 11461091 | 851 | -1.69 | 0.00546 | 0.0112 |
| chr18 | 19399074 | 19399443 | 370 | -2.86 | 0.00547 | 0.0113 |
| chr7 | 75919518 | 76055933 | 136416 | 0.1 | 0.00547 | 0.0113 |
| chr19 | 57702288 | 57704179 | 1892 | 0.64 | 0.00549 | 0.0113 |
| chr4 | 22502383 | 22518427 | 16045 | -0.51 | 0.00549 | 0.0113 |
| chr4 | 7957746 | 7980529 | 22784 | 0.22 | 0.00549 | 0.0113 |
| chr12 | 47549065 | 47558745 | 9681 | 0.49 | 0.00549 | 0.0113 |
| chr20 | 57198200 | 57199833 | 1634 | -0.67 | 0.0055 | 0.0113 |
| chr5 | 66156094 | 66156870 | 777 | 1.94 | 0.0055 | 0.0113 |

| | | | | | | |
|---|---|---|---|---|---|---|
| chr6 | 14877157 | 14891330 | 14174 | 0.4 | 0.0055 | 0.0113 |
| chr7 | 73831533 | 73850020 | 18488 | -0.69 | 0.00551 | 0.0113 |
| chr10 | 49797527 | 49798579 | 1053 | -1.87 | 0.00551 | 0.0113 |
| chr11 | 47395503 | 47396607 | 1105 | -1.82 | 0.00551 | 0.0113 |
| chr2 | 74346933 | 74348070 | 1138 | -0.74 | 0.00552 | 0.0113 |
| chr10 | 80122392 | 80122959 | 568 | -1.89 | 0.00552 | 0.0113 |
| chr11 | 88131261 | 88136435 | 5175 | 0.5 | 0.00552 | 0.0113 |
| chr20 | 47625429 | 47626218 | 790 | 2.22 | 0.00552 | 0.0113 |
| chr1 | 223841741 | 223842983 | 1243 | -1.31 | 0.00553 | 0.0114 |
| chr1 | 35227963 | 35230058 | 2096 | 0.8 | 0.00553 | 0.0114 |
| chr12 | 101182959 | 101183421 | 463 | -2.65 | 0.00553 | 0.0114 |
| chr17 | 64186638 | 64201094 | 14457 | -0.42 | 0.00553 | 0.0114 |
| chr1 | 160695939 | 160697121 | 1183 | -1 | 0.00553 | 0.0114 |
| chr20 | 47513217 | 47514568 | 1352 | -1.24 | 0.00553 | 0.0114 |
| chr5 | 130075200 | 130076620 | 1421 | -3.38 | 0.00554 | 0.0114 |
| chr5 | 71747228 | 71749552 | 2325 | 0.44 | 0.00554 | 0.0114 |
| chr10 | 74626571 | 74635030 | 8460 | 0.72 | 0.00554 | 0.0114 |
| chr1 | 235631710 | 235633184 | 1475 | 1.22 | 0.00556 | 0.0114 |
| chr10 | 4925083 | 4926072 | 990 | -1.53 | 0.00557 | 0.0114 |
| chr1 | 173298634 | 173301535 | 2902 | -0.65 | 0.00558 | 0.0115 |
| chr9 | 463477 | 471489 | 8013 | 1.44 | 0.00558 | 0.0115 |
| chr10 | 61560691 | 61561383 | 693 | 1.96 | 0.00558 | 0.0115 |
| chr20 | 18394074 | 18394447 | 374 | -2.55 | 0.00558 | 0.0115 |
| chr8 | 1915221 | 1922659 | 7439 | 0.64 | 0.00559 | 0.0115 |
| chr4 | 110353276 | 110371194 | 17919 | 0.35 | 0.00559 | 0.0115 |
| chr3 | 157036137 | 157037335 | 1199 | 1.46 | 0.00559 | 0.0115 |
| chrX | 95661450 | 95663905 | 2456 | 0.55 | 0.0056 | 0.0115 |
| chr5 | 134644460 | 134645512 | 1053 | 1.69 | 0.0056 | 0.0115 |
| chr1 | 26099664 | 26102605 | 2942 | 0.61 | 0.0056 | 0.0115 |
| chr11 | 88268677 | 88272090 | 3414 | 0.93 | 0.00561 | 0.0115 |
| chr8 | 140963394 | 140964043 | 650 | 2.58 | 0.00561 | 0.0115 |
| chr7 | 100099992 | 100113079 | 13088 | -0.6 | 0.00562 | 0.0115 |
| chr6 | 138328190 | 138328660 | 471 | -2.36 | 0.00562 | 0.0115 |
| chr10 | 94189877 | 94198561 | 8685 | -1.2 | 0.00563 | 0.0116 |
| chr8 | 142065251 | 142066535 | 1285 | -1.57 | 0.00563 | 0.0116 |
| chr17 | 62762295 | 62770669 | 8375 | 0.67 | 0.00563 | 0.0116 |
| chr12 | 86020094 | 86020927 | 834 | 2.02 | 0.00563 | 0.0116 |
| chr2 | 27505373 | 27526445 | 21073 | -0.82 | 0.00563 | 0.0116 |
| chr5 | 43120063 | 43144072 | 24010 | 0.25 | 0.00564 | 0.0116 |
| chr10 | 97226757 | 97227606 | 850 | 2.39 | 0.00564 | 0.0116 |
| chr15 | 75986740 | 75988119 | 1380 | -2.46 | 0.00564 | 0.0116 |
| chr6 | 89440141 | 89441045 | 905 | 1.34 | 0.00565 | 0.0116 |
| chr10 | 94818415 | 94819212 | 798 | 2.54 | 0.00565 | 0.0116 |
| chr15 | 74279270 | 74288232 | 8963 | 0.4 | 0.00566 | 0.0116 |
| chr12 | 90210896 | 90220682 | 9787 | 0.26 | 0.00566 | 0.0116 |
| chr19 | 41426148 | 41428536 | 2389 | -0.43 | 0.00566 | 0.0116 |
| chr1 | 225727241 | 225736461 | 9221 | -1.06 | 0.00566 | 0.0116 |
| chr7 | 79526526 | 79527492 | 967 | 1.76 | 0.00567 | 0.0116 |
| chr20 | 43689378 | 43689841 | 464 | -2.66 | 0.00568 | 0.0116 |
| chr3 | 196354105 | 196360152 | 6048 | -0.55 | 0.00569 | 0.0117 |
| chr1 | 92021312 | 92022348 | 1037 | -1.97 | 0.0057 | 0.0117 |
| chr4 | 149755686 | 149756859 | 1174 | 1.54 | 0.0057 | 0.0117 |
| chr3 | 150261684 | 150268872 | 7189 | 0.32 | 0.0057 | 0.0117 |
| chr20 | 3725146 | 3802985 | 77840 | 0.13 | 0.00571 | 0.0117 |
| chr4 | 105496549 | 105497473 | 925 | 2.63 | 0.00572 | 0.0117 |
| chr12 | 57109828 | 57122715 | 12888 | 0.29 | 0.00572 | 0.0117 |
| chrX | 37705735 | 37707840 | 2106 | 0.67 | 0.00573 | 0.0117 |
| chr8 | 79617916 | 79618819 | 904 | -1.38 | 0.00573 | 0.0117 |
| chr2 | 19768537 | 19780925 | 12389 | 0.42 | 0.00573 | 0.0117 |
| chr12 | 18947060 | 18952840 | 5781 | 4.73 | 0.00574 | 0.0118 |
| chr11 | 62264106 | 62280813 | 16708 | -0.27 | 0.00576 | 0.0118 |

| | | | | | | |
|---|---|---|---|---|---|---|
| chr11 | 46360364 | 46385590 | 25227 | -0.46 | 0.00576 | 0.0118 |
| chr2 | 190211448 | 190213681 | 2234 | -0.68 | 0.00576 | 0.0118 |
| chr14 | 34407127 | 34421367 | 14241 | 0.49 | 0.00577 | 0.0118 |
| chr4 | 56813759 | 56817187 | 3429 | 0.43 | 0.00577 | 0.0118 |
| chr4 | 15553429 | 15554394 | 966 | 2 | 0.00578 | 0.0118 |
| chr6 | 123075075 | 123075615 | 541 | 2.27 | 0.00578 | 0.0118 |
| chr10 | 17156877 | 17158101 | 1225 | -1.18 | 0.00578 | 0.0118 |
| chr20 | 42602636 | 42614745 | 12110 | -2.2 | 0.00578 | 0.0118 |
| chr7 | 24796136 | 24797994 | 1859 | 0.56 | 0.00579 | 0.0119 |
| chrX | 37543763 | 37545683 | 1921 | 1.09 | 0.0058 | 0.0119 |
| chr16 | 11588164 | 11602850 | 14687 | -0.29 | 0.0058 | 0.0119 |
| chrX | 62973879 | 62976043 | 2165 | 0.52 | 0.0058 | 0.0119 |
| chr5 | 53328226 | 53346803 | 18578 | 0.22 | 0.0058 | 0.0119 |
| chr13 | 21384957 | 21389219 | 4263 | -0.81 | 0.00581 | 0.0119 |
| chr19 | 16796341 | 16796843 | 503 | 2.16 | 0.00582 | 0.0119 |
| chr2 | 71294619 | 71300019 | 5401 | -0.43 | 0.00582 | 0.0119 |
| chr14 | 56333843 | 56334616 | 774 | 2.51 | 0.00583 | 0.0119 |
| chr1 | 23036836 | 23044410 | 7575 | -0.8 | 0.00583 | 0.0119 |
| chr2 | 11622752 | 11623401 | 650 | -2.08 | 0.00584 | 0.0119 |
| chr4 | 1241002 | 1249314 | 8313 | 0.6 | 0.00585 | 0.012 |
| chr22 | 31794386 | 31796857 | 2472 | 0.56 | 0.00585 | 0.012 |
| chr11 | 9231283 | 9237218 | 5936 | -0.98 | 0.00586 | 0.012 |
| chr17 | 74153412 | 74154291 | 880 | -1.75 | 0.00586 | 0.012 |
| chr17 | 28245872 | 28259137 | 13266 | 0.35 | 0.00586 | 0.012 |
| chr7 | 68249774 | 68250967 | 1194 | 1.31 | 0.00587 | 0.012 |
| chr9 | 137589610 | 137590943 | 1334 | -1.27 | 0.00587 | 0.012 |
| chr7 | 114561508 | 114564540 | 3033 | 0.39 | 0.00588 | 0.012 |
| chr3 | 15766988 | 15768106 | 1119 | -1.29 | 0.00589 | 0.012 |
| chr9 | 32549480 | 32553755 | 4276 | 0.39 | 0.00589 | 0.012 |
| chr4 | 146855649 | 146860472 | 4824 | 0.46 | 0.0059 | 0.0121 |
| chr5 | 127510641 | 127511510 | 870 | 1.91 | 0.00591 | 0.0121 |
| chr2 | 15528443 | 15530352 | 1910 | -0.88 | 0.00592 | 0.0121 |
| chr13 | 68652096 | 68653940 | 1845 | 0.86 | 0.00592 | 0.0121 |
| chr3 | 196594001 | 196596191 | 2191 | 0.55 | 0.00592 | 0.0121 |
| chr19 | 58977737 | 58993133 | 15397 | -0.39 | 0.00592 | 0.0121 |
| chr6 | 83604198 | 83605047 | 850 | -1.56 | 0.00593 | 0.0121 |
| chr2 | 230844650 | 230845863 | 1214 | -1.58 | 0.00593 | 0.0121 |
| chr14 | 69310452 | 69310795 | 344 | -2.1 | 0.00593 | 0.0121 |
| chr2 | 61011669 | 61012252 | 584 | 2.01 | 0.00593 | 0.0121 |
| chr2 | 17535675 | 17538021 | 2347 | 0.32 | 0.00594 | 0.0121 |
| chr4 | 75929021 | 75929976 | 956 | -1.48 | 0.00594 | 0.0121 |
| chr1 | 161854107 | 161855302 | 1196 | 1.29 | 0.00594 | 0.0121 |
| chr4 | 40899038 | 40911357 | 12320 | 0.66 | 0.00595 | 0.0122 |
| chr13 | 89237970 | 89238906 | 937 | 1.6 | 0.00596 | 0.0122 |
| chr7 | 131800504 | 131803623 | 3120 | -0.59 | 0.00596 | 0.0122 |
| chr1 | 198202154 | 198204330 | 2177 | 0.58 | 0.00596 | 0.0122 |
| chr14 | 70160855 | 70161909 | 1055 | 1.11 | 0.00596 | 0.0122 |
| chr14 | 101910196 | 101910885 | 690 | -1.94 | 0.00596 | 0.0122 |
| chr11 | 107940346 | 107942154 | 1809 | 1.12 | 0.00596 | 0.0122 |
| chr3 | 49705289 | 49712528 | 7240 | 0.37 | 0.00598 | 0.0122 |
| chr14 | 72206142 | 72206895 | 754 | 1.54 | 0.00598 | 0.0122 |
| chr17 | 57103918 | 57106461 | 2544 | -0.49 | 0.00599 | 0.0122 |
| chr6 | 10114437 | 10115217 | 781 | 1.94 | 0.006 | 0.0122 |
| chr2 | 122229314 | 122230552 | 1239 | -1.46 | 0.006 | 0.0122 |
| chr11 | 62597734 | 62600670 | 2937 | 0.47 | 0.006 | 0.0122 |
| chr6 | 100731368 | 100749379 | 18012 | -0.22 | 0.006 | 0.0122 |
| chr15 | 41693899 | 41695550 | 1652 | 0.63 | 0.006 | 0.0123 |
| chr18 | 29521888 | 29524321 | 2434 | 0.51 | 0.00601 | 0.0123 |
| chr2 | 242443336 | 242448934 | 5599 | -0.73 | 0.00601 | 0.0123 |
| chr7 | 11605117 | 11611316 | 6200 | 2.87 | 0.00602 | 0.0123 |
| chr13 | 104387078 | 104387616 | 539 | -2.58 | 0.00603 | 0.0123 |

| | | | | | | |
|---|---|---|---|---|---|---|
| chr10 | 12207955 | 12239719 | 31765 | 0.24 | 0.00603 | 0.0123 |
| chr19 | 47571511 | 47777928 | 206418 | 0.09 | 0.00603 | 0.0123 |
| chr6 | 149846575 | 149847949 | 1375 | -1.74 | 0.00603 | 0.0123 |
| chr17 | 77985997 | 77987057 | 1061 | -1.46 | 0.00604 | 0.0123 |
| chr3 | 172448771 | 172449289 | 519 | -2.45 | 0.00604 | 0.0123 |
| chr3 | 49365400 | 49400729 | 35330 | 0.27 | 0.00604 | 0.0123 |
| chr20 | 36637233 | 36638211 | 979 | 1.31 | 0.00604 | 0.0123 |
| chr11 | 32455745 | 32457486 | 1742 | -1.14 | 0.00605 | 0.0123 |
| chr10 | 64892279 | 64899823 | 7545 | 0.33 | 0.00605 | 0.0123 |
| chr4 | 151430889 | 151440160 | 9272 | 0.79 | 0.00606 | 0.0123 |
| chr21 | 28904712 | 28904999 | 288 | 2.81 | 0.00606 | 0.0123 |
| chr1 | 52071470 | 52135566 | 64097 | 0.19 | 0.00606 | 0.0123 |
| chr11 | 70813477 | 70814368 | 892 | -1.64 | 0.00606 | 0.0124 |
| chr17 | 55098676 | 55099580 | 905 | 1.18 | 0.00606 | 0.0124 |
| chr4 | 41030520 | 41031489 | 970 | 1.68 | 0.00606 | 0.0124 |
| chr18 | 57978759 | 57979780 | 1022 | 3.68 | 0.00606 | 0.0124 |
| chr7 | 20056277 | 20056775 | 499 | -2.65 | 0.00606 | 0.0124 |
| chr22 | 38349069 | 38350714 | 1646 | -0.61 | 0.00606 | 0.0124 |
| chr2 | 135878657 | 135879964 | 1308 | 1.3 | 0.00607 | 0.0124 |
| chr15 | 82431946 | 82440392 | 8447 | 0.59 | 0.00607 | 0.0124 |
| chr11 | 67271119 | 67277214 | 6096 | 0.32 | 0.00607 | 0.0124 |
| chr1 | 86009874 | 86011353 | 1480 | 1.28 | 0.00607 | 0.0124 |
| chr3 | 37247761 | 37248214 | 454 | -2.2 | 0.00608 | 0.0124 |
| chr13 | 96705241 | 96706736 | 1496 | 0.58 | 0.00608 | 0.0124 |
| chr3 | 189947290 | 189950836 | 3547 | 0.53 | 0.00608 | 0.0124 |
| chr5 | 118873174 | 118874125 | 952 | 1.36 | 0.00608 | 0.0124 |
| chr11 | 20381839 | 20410985 | 29147 | 0.17 | 0.00609 | 0.0124 |
| chr15 | 27111431 | 27112872 | 1442 | -3.84 | 0.00609 | 0.0124 |
| chr1 | 35656662 | 35663800 | 7139 | 0.33 | 0.0061 | 0.0124 |
| chr13 | 31773796 | 31775487 | 1692 | 0.7 | 0.0061 | 0.0124 |
| chr1 | 114446199 | 114448989 | 2791 | 0.57 | 0.0061 | 0.0124 |
| chr5 | 81315559 | 81316125 | 567 | -2.48 | 0.00611 | 0.0124 |
| chr1 | 211686728 | 211691754 | 5027 | 0.35 | 0.00611 | 0.0124 |
| chr7 | 8343563 | 8344637 | 1075 | 0.77 | 0.00611 | 0.0124 |
| chrX | 36900257 | 36901381 | 1125 | 1.49 | 0.00612 | 0.0125 |
| chr13 | 21121360 | 21121765 | 406 | 2.61 | 0.00612 | 0.0125 |
| chr7 | 156839483 | 156840383 | 901 | 2.52 | 0.00612 | 0.0125 |
| chr3 | 119731893 | 119737126 | 5234 | 0.83 | 0.00613 | 0.0125 |
| chr17 | 33546025 | 33546555 | 531 | 2.27 | 0.00613 | 0.0125 |
| chr19 | 11313880 | 11314449 | 570 | -1.96 | 0.00616 | 0.0125 |
| chr22 | 36481990 | 36483144 | 1155 | 1.69 | 0.00616 | 0.0125 |
| chr1 | 110414449 | 110421261 | 6813 | -0.37 | 0.00616 | 0.0125 |
| chr5 | 159509571 | 159511198 | 1628 | -0.97 | 0.00616 | 0.0125 |
| chr11 | 32604421 | 32621447 | 17027 | 0.3 | 0.00617 | 0.0126 |
| chr7 | 80078856 | 80080694 | 1839 | 0.91 | 0.00618 | 0.0126 |
| chr3 | 71136198 | 71136699 | 502 | -2.79 | 0.00619 | 0.0126 |
| chr1 | 95349179 | 95350341 | 1163 | 1.21 | 0.0062 | 0.0126 |
| chr11 | 86262181 | 86264094 | 1914 | 0.8 | 0.0062 | 0.0126 |
| chr7 | 43098309 | 43098894 | 586 | 1.6 | 0.00621 | 0.0126 |
| chr1 | 167389185 | 167390868 | 1684 | -0.75 | 0.00622 | 0.0126 |
| chr3 | 13128610 | 13129618 | 1009 | -2.54 | 0.00624 | 0.0127 |
| chr6 | 3582472 | 3593457 | 10986 | 0.44 | 0.00624 | 0.0127 |
| chr6 | 54757617 | 54758032 | 416 | -2.09 | 0.00624 | 0.0127 |
| chr5 | 180196340 | 180197793 | 1454 | 1.54 | 0.00625 | 0.0127 |
| chr16 | 53849905 | 53857329 | 7425 | -0.46 | 0.00626 | 0.0127 |
| chr7 | 34299478 | 34300516 | 1039 | -2.43 | 0.00627 | 0.0127 |
| chr4 | 141263738 | 141268270 | 4533 | 2.99 | 0.00627 | 0.0127 |
| chr9 | 119245114 | 119263342 | 18229 | -3.43 | 0.00628 | 0.0128 |
| chr2 | 189932881 | 189933932 | 1052 | -1.71 | 0.00629 | 0.0128 |
| chr17 | 69055770 | 69068701 | 12932 | 4.56 | 0.00629 | 0.0128 |
| chr2 | 239832163 | 239832729 | 567 | -2.11 | 0.00629 | 0.0128 |

| | | | | | | |
|---|---|---|---|---|---|---|
| chr4 | 25487576 | 25488026 | 451 | 1.82 | 0.00629 | 0.0128 |
| chr3 | 112816353 | 112818082 | 1730 | -0.68 | 0.0063 | 0.0128 |
| chr21 | 33002282 | 33003119 | 838 | 1.31 | 0.0063 | 0.0128 |
| chr8 | 141901552 | 141908617 | 7066 | -1.04 | 0.0063 | 0.0128 |
| chr6 | 106993805 | 106995714 | 1910 | 0.71 | 0.00631 | 0.0128 |
| chr19 | 18574935 | 18633940 | 59006 | -0.27 | 0.00631 | 0.0128 |
| chr2 | 182257946 | 182258999 | 1054 | 0.94 | 0.00631 | 0.0128 |
| chrX | 48595097 | 48596783 | 1687 | 0.62 | 0.00632 | 0.0128 |
| chr15 | 26007121 | 26033691 | 26571 | -0.29 | 0.00632 | 0.0128 |
| chr22 | 23483394 | 23488869 | 5476 | -0.36 | 0.00632 | 0.0128 |
| chr4 | 146018141 | 146022016 | 3876 | 0.38 | 0.00632 | 0.0128 |
| chr3 | 161033130 | 161043045 | 9916 | 4.88 | 0.00632 | 0.0128 |
| chr1 | 91556993 | 91558198 | 1206 | 1.18 | 0.00633 | 0.0128 |
| chr12 | 93095679 | 93097548 | 1870 | 4.44 | 0.00633 | 0.0128 |
| chr11 | 56930724 | 56931593 | 870 | -2.48 | 0.00633 | 0.0128 |
| chr13 | 60841785 | 60842246 | 462 | 2.48 | 0.00633 | 0.0128 |
| chr14 | 104207139 | 104208046 | 908 | -2.25 | 0.00634 | 0.0129 |
| chr4 | 89241362 | 89242445 | 1084 | 0.97 | 0.00634 | 0.0129 |
| chr12 | 94135218 | 94137263 | 2046 | 0.54 | 0.00634 | 0.0129 |
| chr14 | 32029506 | 32031521 | 2016 | -0.43 | 0.00635 | 0.0129 |
| chr4 | 125824714 | 125826608 | 1895 | 0.72 | 0.00635 | 0.0129 |
| chr17 | 74022344 | 74050324 | 27981 | -0.36 | 0.00635 | 0.0129 |
| chr20 | 33890825 | 33917491 | 26667 | -0.16 | 0.00635 | 0.0129 |
| chr2 | 179146589 | 179147701 | 1113 | -1.26 | 0.00636 | 0.0129 |
| chr16 | 87490432 | 87499408 | 8977 | 0.54 | 0.00636 | 0.0129 |
| chr20 | 25158612 | 25159969 | 1358 | -1.57 | 0.00636 | 0.0129 |
| chr22 | 29372616 | 29374777 | 2162 | 0.87 | 0.00636 | 0.0129 |
| chr2 | 239505234 | 239507036 | 1803 | -0.57 | 0.00637 | 0.0129 |
| chr11 | 117101012 | 117110561 | 9550 | 0.6 | 0.00637 | 0.0129 |
| chr6 | 141105654 | 141106790 | 1137 | 1.17 | 0.00637 | 0.0129 |
| chr14 | 73921190 | 73933357 | 12168 | 0.38 | 0.00637 | 0.0129 |
| chr8 | 53606551 | 53609511 | 2961 | 0.67 | 0.00638 | 0.0129 |
| chr1 | 169476612 | 169485102 | 8491 | 3.97 | 0.00638 | 0.0129 |
| chr13 | 74095164 | 74096835 | 1672 | 1.26 | 0.00638 | 0.0129 |
| chr19 | 18770499 | 18771772 | 1274 | -0.82 | 0.00638 | 0.0129 |
| chr3 | 23636142 | 23642454 | 6313 | -0.93 | 0.00639 | 0.0129 |
| chr1 | 179884456 | 179886187 | 1732 | 1.45 | 0.00639 | 0.0129 |
| chr3 | 114214571 | 114220127 | 5557 | 0.37 | 0.00639 | 0.0129 |
| chr5 | 138861137 | 138873063 | 11927 | -0.9 | 0.00639 | 0.0129 |
| chr17 | 6376018 | 6376564 | 547 | -2.06 | 0.00641 | 0.013 |
| chr18 | 60541622 | 60549866 | 8245 | 0.62 | 0.00641 | 0.013 |
| chr1 | 174127715 | 174129815 | 2101 | 0.62 | 0.00642 | 0.013 |
| chr9 | 131524517 | 131561510 | 36994 | -0.28 | 0.00642 | 0.013 |
| chr1 | 194278249 | 194279153 | 905 | 1.31 | 0.00642 | 0.013 |
| chr15 | 59012050 | 59043137 | 31088 | -0.28 | 0.00643 | 0.013 |
| chr5 | 166309415 | 166311168 | 1754 | 0.79 | 0.00643 | 0.013 |
| chr6 | 162166755 | 162172616 | 5862 | 1.26 | 0.00644 | 0.013 |
| chr2 | 19946647 | 19947426 | 780 | -1.93 | 0.00645 | 0.0131 |
| chr15 | 100468761 | 100469374 | 614 | 1.62 | 0.00645 | 0.0131 |
| chr1 | 249111478 | 249168952 | 57475 | 0.17 | 0.00646 | 0.0131 |
| chr10 | 95482112 | 95483014 | 903 | -1.81 | 0.00647 | 0.0131 |
| chr11 | 84274149 | 84276665 | 2517 | 0.32 | 0.00648 | 0.0131 |
| chr11 | 130449793 | 130451276 | 1484 | 1.14 | 0.00648 | 0.0131 |
| chr3 | 180706451 | 180708191 | 1741 | 0.74 | 0.00648 | 0.0131 |
| chr20 | 24725551 | 24728399 | 2849 | 0.45 | 0.00649 | 0.0131 |
| chr2 | 165824211 | 165825056 | 846 | 2.47 | 0.0065 | 0.0131 |
| chr16 | 74907697 | 74909636 | 1940 | -0.79 | 0.0065 | 0.0132 |
| chr5 | 72792705 | 72797511 | 4807 | 0.39 | 0.0065 | 0.0132 |
| chr16 | 27374529 | 27377594 | 3066 | 0.54 | 0.00651 | 0.0132 |
| chr1 | 246746112 | 246747552 | 1441 | 0.88 | 0.00651 | 0.0132 |
| chr19 | 30096066 | 30103618 | 7553 | 0.23 | 0.00652 | 0.0132 |

| | | | | | | |
|---|---|---|---|---|---|---|
| chr12 | 87220778 | 87221710 | 933 | 1.28 | 0.00652 | 0.0132 |
| chr9 | 12858916 | 12861831 | 2916 | 0.43 | 0.00652 | 0.0132 |
| chr2 | 63814484 | 63832805 | 18322 | -0.35 | 0.00653 | 0.0132 |
| chr17 | 77396825 | 77462501 | 65677 | -3.62 | 0.00653 | 0.0132 |
| chr20 | 35431028 | 35493522 | 62495 | -0.22 | 0.00653 | 0.0132 |
| chr4 | 34966133 | 34966668 | 536 | 1.92 | 0.00654 | 0.0132 |
| chr8 | 24799754 | 24800231 | 478 | -1.95 | 0.00655 | 0.0132 |
| chr3 | 183530274 | 183531400 | 1127 | -1.36 | 0.00655 | 0.0132 |
| chr2 | 27667777 | 27669531 | 1755 | 1.15 | 0.00655 | 0.0132 |
| chr20 | 40297902 | 40299006 | 1105 | -1.2 | 0.00655 | 0.0132 |
| chr18 | 43546324 | 43547288 | 965 | 0.91 | 0.00655 | 0.0132 |
| chr14 | 57132731 | 57133530 | 800 | -1.72 | 0.00655 | 0.0132 |
| chr18 | 69928620 | 69928904 | 285 | 2.81 | 0.00655 | 0.0132 |
| chr10 | 125795082 | 125797403 | 2322 | -0.9 | 0.00656 | 0.0132 |
| chr4 | 28450941 | 28451318 | 378 | 2.66 | 0.00656 | 0.0132 |
| chr22 | 42552057 | 42559497 | 7441 | -0.49 | 0.00656 | 0.0132 |
| chr2 | 189244872 | 189247967 | 3096 | 0.85 | 0.00656 | 0.0133 |
| chr22 | 42362494 | 42363245 | 752 | 2.11 | 0.00656 | 0.0133 |
| chr19 | 17493229 | 17501870 | 8642 | 0.94 | 0.00657 | 0.0133 |
| chr5 | 657267 | 674937 | 17671 | 0.35 | 0.00657 | 0.0133 |
| chr2 | 113374439 | 113422840 | 48402 | -0.22 | 0.00658 | 0.0133 |
| chr17 | 66470131 | 66471457 | 1327 | 3.66 | 0.00658 | 0.0133 |
| chr4 | 37009494 | 37010631 | 1138 | -1.16 | 0.00659 | 0.0133 |
| chr11 | 118436272 | 118444597 | 8326 | 0.46 | 0.0066 | 0.0133 |
| chr7 | 102613270 | 102635041 | 21772 | 0.35 | 0.00661 | 0.0133 |
| chr16 | 22009082 | 22020505 | 11424 | -0.54 | 0.00661 | 0.0133 |
| chr9 | 114773470 | 114837417 | 63948 | -0.15 | 0.00662 | 0.0134 |
| chr3 | 67643880 | 67670843 | 26964 | -0.32 | 0.00662 | 0.0134 |
| chr1 | 42921072 | 42930072 | 9001 | -0.32 | 0.00662 | 0.0134 |
| chr12 | 57521148 | 57522136 | 989 | 0.94 | 0.00662 | 0.0134 |
| chr6 | 143264405 | 143269497 | 5093 | 0.43 | 0.00663 | 0.0134 |
| chr14 | 35449523 | 35465839 | 16317 | 0.23 | 0.00663 | 0.0134 |
| chr8 | 37980438 | 38036845 | 56408 | -0.23 | 0.00664 | 0.0134 |
| chr17 | 63088064 | 63103350 | 15287 | -0.44 | 0.00664 | 0.0134 |
| chr13 | 40101534 | 40102895 | 1362 | -1.06 | 0.00665 | 0.0134 |
| chr3 | 129486723 | 129497013 | 10291 | -0.79 | 0.00665 | 0.0134 |
| chrX | 71864776 | 71865626 | 851 | -1.71 | 0.00665 | 0.0134 |
| chr20 | 35855956 | 35857229 | 1274 | -1.36 | 0.00665 | 0.0134 |
| chr12 | 130671056 | 130673014 | 1959 | 4.35 | 0.00666 | 0.0134 |
| chr7 | 157000439 | 157004922 | 4484 | -1.02 | 0.00667 | 0.0135 |
| chr1 | 76138073 | 76139488 | 1416 | -0.96 | 0.00667 | 0.0135 |
| chr5 | 112196175 | 112198811 | 2637 | 0.42 | 0.00667 | 0.0135 |
| chr9 | 130205834 | 130215287 | 9454 | 0.39 | 0.00668 | 0.0135 |
| chr13 | 74707551 | 74711073 | 3523 | 0.41 | 0.00668 | 0.0135 |
| chr1 | 203763804 | 203766254 | 2451 | 0.4 | 0.00668 | 0.0135 |
| chr11 | 92787376 | 92789690 | 2315 | 0.39 | 0.00668 | 0.0135 |
| chr20 | 2903489 | 2908464 | 4976 | 0.85 | 0.00669 | 0.0135 |
| chrX | 21788359 | 21789565 | 1207 | -3.75 | 0.00669 | 0.0135 |
| chr6 | 42531652 | 42538790 | 7139 | 0.44 | 0.0067 | 0.0135 |
| chr8 | 120733824 | 120734771 | 948 | 2.48 | 0.0067 | 0.0135 |
| chr9 | 88998342 | 88999053 | 712 | 1.37 | 0.0067 | 0.0135 |
| chr13 | 107085289 | 107085923 | 635 | 2.68 | 0.00671 | 0.0135 |
| chr7 | 24441826 | 24442238 | 413 | 2.53 | 0.00671 | 0.0135 |
| chr1 | 224759121 | 224760661 | 1541 | 1.34 | 0.00672 | 0.0135 |
| chr18 | 28765778 | 28766115 | 338 | 2.62 | 0.00672 | 0.0135 |
| chr1 | 12482872 | 12519158 | 36287 | 0.22 | 0.00672 | 0.0135 |
| chr8 | 75262383 | 75263351 | 969 | 0.81 | 0.00673 | 0.0136 |
| chr10 | 59996195 | 60030739 | 34545 | 0.25 | 0.00673 | 0.0136 |
| chr10 | 92980427 | 92981816 | 1390 | 0.67 | 0.00673 | 0.0136 |
| chr16 | 11422088 | 11423722 | 1635 | -1.03 | 0.00673 | 0.0136 |
| chr4 | 144256713 | 144259189 | 2477 | 0.62 | 0.00673 | 0.0136 |

| | | | | | | |
|---|---|---|---|---|---|---|
| chr15 | 69318515 | 69319643 | 1129 | -3.67 | 0.00674 | 0.0136 |
| chr9 | 107525576 | 107527230 | 1655 | 0.57 | 0.00674 | 0.0136 |
| chr4 | 88274514 | 88279030 | 4517 | 0.57 | 0.00674 | 0.0136 |
| chr10 | 100220274 | 100228153 | 7880 | 3.88 | 0.00674 | 0.0136 |
| chrX | 39478733 | 39479547 | 815 | 2.1 | 0.00674 | 0.0136 |
| chr5 | 138758238 | 138758689 | 452 | -2.03 | 0.00675 | 0.0136 |
| chr18 | 55952322 | 55958165 | 5844 | 0.5 | 0.00675 | 0.0136 |
| chr7 | 94695129 | 94699797 | 4669 | -0.48 | 0.00675 | 0.0136 |
| chr5 | 34827158 | 34840451 | 13294 | -0.38 | 0.00675 | 0.0136 |
| chr16 | 57700781 | 57712347 | 11567 | -0.76 | 0.00676 | 0.0136 |
| chr5 | 169283154 | 169284322 | 1169 | -1.03 | 0.00676 | 0.0136 |
| chrX | 119020185 | 119022348 | 2164 | 1.29 | 0.00676 | 0.0136 |
| chr15 | 63270128 | 63271453 | 1326 | -1.33 | 0.00677 | 0.0136 |
| chr10 | 67650832 | 67652328 | 1497 | 0.63 | 0.00677 | 0.0136 |
| chr7 | 46331018 | 46331914 | 897 | 1.67 | 0.00677 | 0.0136 |
| chr15 | 51413444 | 51425240 | 11797 | 4.33 | 0.00677 | 0.0136 |
| chr5 | 150079164 | 150095409 | 16246 | 0.39 | 0.00678 | 0.0136 |
| chr3 | 52347092 | 52348510 | 1419 | -1.18 | 0.00678 | 0.0136 |
| chr9 | 123927174 | 123976113 | 48940 | 0.24 | 0.00678 | 0.0136 |
| chr2 | 191398183 | 191408245 | 10063 | 0.44 | 0.00678 | 0.0136 |
| chr1 | 22777062 | 22779670 | 2609 | 0.59 | 0.00678 | 0.0136 |
| chr4 | 187542969 | 187544651 | 1683 | -1.12 | 0.00679 | 0.0137 |
| chr2 | 61825917 | 61826781 | 865 | 1.26 | 0.00679 | 0.0137 |
| chr2 | 219471939 | 219473007 | 1069 | 1.48 | 0.0068 | 0.0137 |
| chr2 | 44271271 | 44271974 | 704 | -0.9 | 0.00681 | 0.0137 |
| chr1 | 236664172 | 236695806 | 31635 | 0.21 | 0.00681 | 0.0137 |
| chr2 | 240904902 | 240906158 | 1257 | -1.35 | 0.00681 | 0.0137 |
| chr3 | 41289600 | 41290754 | 1155 | 0.99 | 0.00682 | 0.0137 |
| chr4 | 3108656 | 3120708 | 12053 | -1.26 | 0.00682 | 0.0137 |
| chr7 | 35869564 | 35881120 | 11557 | -0.68 | 0.00682 | 0.0137 |
| chr15 | 33570765 | 33572404 | 1640 | -0.79 | 0.00683 | 0.0137 |
| chr3 | 128444188 | 128457239 | 13052 | -0.39 | 0.00683 | 0.0137 |
| chr8 | 130831958 | 130832485 | 528 | 1.09 | 0.00683 | 0.0137 |
| chr1 | 113585043 | 113585625 | 583 | -2.5 | 0.00683 | 0.0137 |
| chr4 | 174088136 | 174114046 | 25911 | 0.28 | 0.00684 | 0.0137 |
| chr7 | 151347628 | 151349025 | 1398 | -1.38 | 0.00684 | 0.0137 |
| chr11 | 63570118 | 63592760 | 22643 | -0.44 | 0.00684 | 0.0137 |
| chr9 | 125164236 | 125177378 | 13143 | 0.66 | 0.00684 | 0.0137 |
| chr1 | 193323515 | 193325193 | 1679 | 2.61 | 0.00685 | 0.0138 |
| chr7 | 23144530 | 23146493 | 1964 | 0.49 | 0.00685 | 0.0138 |
| chr12 | 262604 | 262996 | 393 | 2.72 | 0.00685 | 0.0138 |
| chr9 | 83642836 | 83643859 | 1024 | 1.49 | 0.00685 | 0.0138 |
| chr20 | 22901596 | 22903301 | 1706 | 0.85 | 0.00685 | 0.0138 |
| chr2 | 393913 | 394853 | 941 | 1.33 | 0.00686 | 0.0138 |
| chr2 | 53979617 | 53980382 | 766 | -1.78 | 0.00686 | 0.0138 |
| chr4 | 137034907 | 137035564 | 658 | 2.55 | 0.00687 | 0.0138 |
| chr12 | 123258931 | 123260130 | 1200 | 1.03 | 0.00687 | 0.0138 |
| chr17 | 6409539 | 6411170 | 1632 | 0.82 | 0.00687 | 0.0138 |
| chr5 | 82789359 | 82790901 | 1543 | 4.38 | 0.00689 | 0.0138 |
| chr6 | 21371274 | 21385323 | 14050 | 0.49 | 0.0069 | 0.0138 |
| chr16 | 28596548 | 28621169 | 24622 | 0.47 | 0.0069 | 0.0139 |
| chr7 | 148786692 | 148789265 | 2574 | 0.38 | 0.0069 | 0.0139 |
| chr14 | 54006387 | 54010215 | 3829 | 0.88 | 0.00691 | 0.0139 |
| chr11 | 68530994 | 68531796 | 803 | 1.76 | 0.00691 | 0.0139 |
| chr6 | 158872180 | 158872684 | 505 | -2.57 | 0.00691 | 0.0139 |
| chr11 | 17352677 | 17353430 | 754 | 2.6 | 0.00692 | 0.0139 |
| chr5 | 43104839 | 43106473 | 1635 | 1.02 | 0.00692 | 0.0139 |
| chr10 | 21318827 | 21319808 | 982 | -1.2 | 0.00695 | 0.0139 |
| chr6 | 89524436 | 89534287 | 9852 | 0.89 | 0.00695 | 0.0139 |
| chr4 | 86640602 | 86647382 | 6781 | -0.47 | 0.00695 | 0.0139 |
| chr11 | 44006868 | 44007251 | 384 | 2.24 | 0.00696 | 0.014 |

| | | | | | | |
|---|---|---|---|---|---|---|
| chr8 | 37031806 | 37032068 | 263 | 2.72 | 0.00696 | 0.014 |
| chr1 | 185441236 | 185458707 | 17472 | 0.48 | 0.00697 | 0.014 |
| chr17 | 42060852 | 42061824 | 973 | 1.53 | 0.00697 | 0.014 |
| chr2 | 149371683 | 149373026 | 1344 | 1.07 | 0.00697 | 0.014 |
| chr2 | 158289721 | 158290075 | 355 | 2.62 | 0.00697 | 0.014 |
| chr4 | 136397115 | 136397814 | 700 | 2.29 | 0.00697 | 0.014 |
| chr16 | 21609922 | 21612213 | 2292 | 0.54 | 0.00698 | 0.014 |
| chr6 | 138482607 | 138483720 | 1114 | 1.34 | 0.00699 | 0.014 |
| chr11 | 2311489 | 2312142 | 654 | 1.84 | 0.00699 | 0.014 |
| chr1 | 64504468 | 64518077 | 13610 | -0.68 | 0.007 | 0.014 |
| chr17 | 17957767 | 17958033 | 267 | -3.73 | 0.007 | 0.014 |
| chr11 | 34752535 | 34752843 | 309 | -2.65 | 0.007 | 0.014 |
| chr6 | 100623556 | 100624803 | 1248 | 1.05 | 0.00701 | 0.014 |
| chr1 | 178386594 | 178387457 | 864 | 1.38 | 0.00702 | 0.0141 |
| chr10 | 43750237 | 43751435 | 1199 | 1.21 | 0.00702 | 0.0141 |
| chr11 | 77988273 | 77989208 | 936 | 1.59 | 0.00702 | 0.0141 |
| chr2 | 146663705 | 146664288 | 584 | 2.56 | 0.00703 | 0.0141 |
| chr1 | 109288695 | 109292075 | 3381 | 0.43 | 0.00703 | 0.0141 |
| chr4 | 15471084 | 15482221 | 11138 | -0.67 | 0.00703 | 0.0141 |
| chr10 | 43878798 | 43917750 | 38953 | 0.16 | 0.00703 | 0.0141 |
| chr17 | 9989078 | 9990257 | 1180 | 1.33 | 0.00703 | 0.0141 |
| chr3 | 142718112 | 142723679 | 5568 | 0.29 | 0.00704 | 0.0141 |
| chr10 | 61469170 | 61469785 | 616 | 1.7 | 0.00704 | 0.0141 |
| chr1 | 226558106 | 226568036 | 9931 | -0.65 | 0.00704 | 0.0141 |
| chr18 | 20045434 | 20051218 | 5785 | 0.28 | 0.00705 | 0.0141 |
| chr2 | 27272300 | 27275550 | 3251 | -0.49 | 0.00706 | 0.0141 |
| chr7 | 25607787 | 25608625 | 839 | 1.13 | 0.00706 | 0.0141 |
| chr22 | 33723730 | 33734972 | 11243 | 0.33 | 0.00707 | 0.0142 |
| chr2 | 242881287 | 242881844 | 558 | -1.79 | 0.00708 | 0.0142 |
| chr6 | 7235869 | 7244261 | 8393 | 0.45 | 0.00708 | 0.0142 |
| chr5 | 52650710 | 52665424 | 14715 | -0.41 | 0.00708 | 0.0142 |
| chr5 | 88039013 | 88040004 | 992 | -1.13 | 0.0071 | 0.0142 |
| chr15 | 28464158 | 28464933 | 776 | 1.73 | 0.0071 | 0.0142 |
| chr1 | 40308945 | 40312339 | 3395 | -1.22 | 0.0071 | 0.0142 |
| chr15 | 70766452 | 70771546 | 5095 | 0.34 | 0.0071 | 0.0142 |
| chr7 | 34876894 | 34893414 | 16521 | -0.26 | 0.0071 | 0.0142 |
| chr13 | 90661250 | 90662021 | 772 | 1.72 | 0.0071 | 0.0142 |
| chr14 | 37130601 | 37132482 | 1882 | 1.1 | 0.00711 | 0.0142 |
| chr8 | 42660163 | 42661635 | 1473 | -0.91 | 0.00711 | 0.0142 |
| chr9 | 127215266 | 127216080 | 815 | 1.85 | 0.00711 | 0.0142 |
| chr7 | 37069675 | 37074163 | 4489 | 0.3 | 0.00711 | 0.0142 |
| chr13 | 74171890 | 74173216 | 1327 | -1.49 | 0.00712 | 0.0142 |
| chr20 | 20874510 | 20876123 | 1614 | -0.93 | 0.00712 | 0.0142 |
| chr7 | 55052527 | 55065320 | 12794 | 0.22 | 0.00712 | 0.0142 |
| chr17 | 7184018 | 7184796 | 779 | -0.92 | 0.00713 | 0.0143 |
| chr1 | 11967349 | 11979538 | 12190 | -0.28 | 0.00713 | 0.0143 |
| chr7 | 6116190 | 6122026 | 5837 | -0.38 | 0.00714 | 0.0143 |
| chr20 | 34441177 | 34442814 | 1638 | -1.32 | 0.00714 | 0.0143 |
| chr11 | 93275402 | 93277531 | 2130 | 0.77 | 0.00715 | 0.0143 |
| chr12 | 42803843 | 42821192 | 17350 | 0.66 | 0.00715 | 0.0143 |
| chr15 | 91160303 | 91162231 | 1929 | -1.11 | 0.00716 | 0.0143 |
| chr7 | 129647753 | 129650800 | 3048 | -0.38 | 0.00717 | 0.0143 |
| chr1 | 43585986 | 43587491 | 1506 | 0.95 | 0.00717 | 0.0143 |
| chr10 | 94321747 | 94335664 | 13918 | -0.29 | 0.00718 | 0.0143 |
| chr8 | 129270121 | 129295946 | 25826 | -2.92 | 0.00718 | 0.0143 |
| chr3 | 177062309 | 177079033 | 16725 | -0.22 | 0.00719 | 0.0144 |
| chr5 | 43532770 | 43558113 | 25344 | 0.31 | 0.00719 | 0.0144 |
| chrX | 39611584 | 39620860 | 9277 | 0.23 | 0.00719 | 0.0144 |
| chr17 | 45887077 | 45941063 | 53987 | -0.22 | 0.0072 | 0.0144 |
| chr18 | 60087027 | 60128797 | 41771 | 0.24 | 0.0072 | 0.0144 |
| chr7 | 30246054 | 30246844 | 791 | 1.81 | 0.0072 | 0.0144 |

| | | | | | | |
|---|---|---|---|---|---|---|
| chr16 | 56763094 | 56784514 | 21421 | -0.32 | 0.0072 | 0.0144 |
| chr10 | 75530594 | 75546534 | 15941 | 0.31 | 0.00721 | 0.0144 |
| chr1 | 12465605 | 12468356 | 2752 | 0.73 | 0.00722 | 0.0144 |
| chr6 | 117802927 | 117877116 | 74190 | -0.14 | 0.00723 | 0.0144 |
| chr2 | 192814774 | 192816227 | 1454 | -0.71 | 0.00723 | 0.0144 |
| chr12 | 124455929 | 124477450 | 21522 | 0.31 | 0.00724 | 0.0145 |
| chr13 | 26963960 | 26964913 | 954 | 1.77 | 0.00724 | 0.0145 |
| chr13 | 99562169 | 99580921 | 18753 | 0.87 | 0.00724 | 0.0145 |
| chr1 | 186865351 | 186866521 | 1171 | -1.16 | 0.00725 | 0.0145 |
| chr11 | 102866445 | 102867475 | 1031 | 1.53 | 0.00725 | 0.0145 |
| chr18 | 70168108 | 70169621 | 1514 | 3.32 | 0.00725 | 0.0145 |
| chr18 | 20452853 | 20453609 | 757 | 1.56 | 0.00726 | 0.0145 |
| chr6 | 2028804 | 2029817 | 1014 | 1.56 | 0.00726 | 0.0145 |
| chr3 | 10156712 | 10158991 | 2280 | 0.46 | 0.00727 | 0.0145 |
| chr7 | 43908370 | 43922212 | 13843 | -0.58 | 0.00728 | 0.0145 |
| chr20 | 5494199 | 5495500 | 1302 | 0.78 | 0.00729 | 0.0145 |
| chr2 | 183203109 | 183203958 | 850 | 1.53 | 0.0073 | 0.0146 |
| chr17 | 80008746 | 80024530 | 15785 | -0.43 | 0.0073 | 0.0146 |
| chr6 | 108615041 | 108617238 | 2198 | 0.36 | 0.0073 | 0.0146 |
| chr8 | 40573128 | 40603111 | 29984 | -0.57 | 0.0073 | 0.0146 |
| chr12 | 88636125 | 88641594 | 5470 | -0.57 | 0.0073 | 0.0146 |
| chr4 | 90899086 | 90899981 | 896 | 2.45 | 0.0073 | 0.0146 |
| chr6 | 4665578 | 4666309 | 732 | -2.25 | 0.00732 | 0.0146 |
| chr5 | 139889122 | 139889747 | 626 | 2.36 | 0.00733 | 0.0146 |
| chr19 | 33950669 | 33951120 | 452 | -1.57 | 0.00733 | 0.0146 |
| chr4 | 124042028 | 124043493 | 1466 | -0.98 | 0.00734 | 0.0146 |
| chr1 | 247367158 | 247374627 | 7470 | 0.53 | 0.00734 | 0.0146 |
| chr7 | 110026602 | 110028106 | 1505 | 0.97 | 0.00735 | 0.0147 |
| chr15 | 67807270 | 67815600 | 8331 | 0.33 | 0.00736 | 0.0147 |
| chr14 | 67694919 | 67715147 | 20229 | 0.26 | 0.00736 | 0.0147 |
| chr21 | 27960158 | 27960904 | 747 | 1.58 | 0.00736 | 0.0147 |
| chr11 | 45578834 | 45579397 | 564 | -3 | 0.00736 | 0.0147 |
| chr19 | 54315950 | 54317536 | 1587 | 1.36 | 0.00736 | 0.0147 |
| chr16 | 10694913 | 10696447 | 1535 | 1.22 | 0.00737 | 0.0147 |
| chr3 | 121240328 | 121241004 | 677 | 1.93 | 0.00737 | 0.0147 |
| chr4 | 47527244 | 47534054 | 6811 | 1.02 | 0.00738 | 0.0147 |
| chr10 | 63926188 | 63927351 | 1164 | 0.83 | 0.00739 | 0.0147 |
| chr2 | 164695333 | 164695987 | 655 | 2.36 | 0.00739 | 0.0147 |
| chr1 | 235752035 | 235753081 | 1047 | -1.02 | 0.00739 | 0.0147 |
| chr3 | 181853753 | 181854984 | 1232 | 1.15 | 0.0074 | 0.0147 |
| chr2 | 99207777 | 99226676 | 18900 | -0.33 | 0.0074 | 0.0147 |
| chr12 | 46613057 | 46616088 | 3032 | -0.79 | 0.00741 | 0.0148 |
| chr18 | 51674816 | 51676882 | 2067 | 0.69 | 0.00741 | 0.0148 |
| chr18 | 42231172 | 42232091 | 920 | 1.62 | 0.00742 | 0.0148 |
| chr17 | 16555924 | 16557207 | 1284 | 0.53 | 0.00742 | 0.0148 |
| chr2 | 191623716 | 191625227 | 1512 | 0.96 | 0.00742 | 0.0148 |
| chr21 | 37506130 | 37514640 | 8511 | -0.64 | 0.00742 | 0.0148 |
| chr20 | 34096466 | 34102475 | 6010 | -0.76 | 0.00743 | 0.0148 |
| chr12 | 101004334 | 101005133 | 800 | 1.36 | 0.00743 | 0.0148 |
| chr4 | 88578959 | 88581213 | 2255 | 0.76 | 0.00743 | 0.0148 |
| chr2 | 158686382 | 158715505 | 29124 | 0.26 | 0.00744 | 0.0148 |
| chr10 | 48524062 | 48525095 | 1034 | -1.46 | 0.00745 | 0.0148 |
| chr2 | 144708342 | 144709167 | 826 | 1.9 | 0.00745 | 0.0148 |
| chr15 | 57295551 | 57296325 | 775 | -1.55 | 0.00745 | 0.0148 |
| chr2 | 209144970 | 209146599 | 1630 | -1.34 | 0.00746 | 0.0149 |
| chr4 | 48129034 | 48130517 | 1484 | 1.13 | 0.00746 | 0.0149 |
| chr13 | 96548187 | 96549571 | 1385 | 0.85 | 0.00746 | 0.0149 |
| chr7 | 77648342 | 77649949 | 1608 | 0.68 | 0.00747 | 0.0149 |
| chr2 | 28731003 | 28731700 | 698 | -1.69 | 0.00747 | 0.0149 |
| chr4 | 74681690 | 74683693 | 2004 | 0.67 | 0.00747 | 0.0149 |
| chr5 | 176646173 | 176646824 | 652 | -2.04 | 0.00747 | 0.0149 |

| | | | | | | |
|---|---|---|---|---|---|---|
| chr2 | 23980423 | 23981547 | 1125 | -1.28 | 0.00749 | 0.0149 |
| chr2 | 190304900 | 190309180 | 4281 | -0.36 | 0.0075 | 0.0149 |
| chr8 | 133333272 | 133334352 | 1081 | 0.84 | 0.0075 | 0.0149 |
| chr11 | 63360811 | 63361364 | 554 | -2.09 | 0.0075 | 0.0149 |
| chrX | 41006665 | 41013003 | 6339 | 0.52 | 0.00751 | 0.0149 |
| chr7 | 19417406 | 19418230 | 825 | 1.79 | 0.00751 | 0.0149 |
| chr5 | 79919125 | 79928377 | 9253 | 0.49 | 0.00752 | 0.0149 |
| chr11 | 9142238 | 9142658 | 421 | 1.82 | 0.00752 | 0.015 |
| chr4 | 25235675 | 25237278 | 1604 | 0.89 | 0.00752 | 0.015 |
| chr4 | 164414106 | 164418223 | 4118 | 0.3 | 0.00752 | 0.015 |
| chr15 | 78891507 | 78892015 | 509 | -1.71 | 0.00753 | 0.015 |
| chr18 | 52432301 | 52446040 | 13740 | 0.28 | 0.00753 | 0.015 |
| chr15 | 83991510 | 83993623 | 2114 | 0.65 | 0.00754 | 0.015 |
| chr12 | 108813400 | 108814558 | 1159 | -1.09 | 0.00754 | 0.015 |
| chr4 | 184718332 | 184719751 | 1420 | 1.49 | 0.00755 | 0.015 |
| chr4 | 129900743 | 129902529 | 1787 | 0.81 | 0.00755 | 0.015 |
| chr18 | 48493609 | 48495633 | 2025 | 0.45 | 0.00756 | 0.015 |
| chr10 | 77792797 | 77794501 | 1705 | 0.59 | 0.00757 | 0.015 |
| chr12 | 132969586 | 132973620 | 4035 | -0.49 | 0.00758 | 0.0151 |
| chr7 | 138939528 | 138940963 | 1436 | -1.72 | 0.00758 | 0.0151 |
| chr8 | 67088231 | 67089814 | 1584 | -0.88 | 0.00758 | 0.0151 |
| chr12 | 57195740 | 57196282 | 543 | 1.94 | 0.00759 | 0.0151 |
| chr9 | 28866718 | 28867559 | 842 | 1.73 | 0.00759 | 0.0151 |
| chr2 | 202330219 | 202331182 | 964 | -1.49 | 0.00759 | 0.0151 |
| chr2 | 238318904 | 238341996 | 23093 | -4.49 | 0.0076 | 0.0151 |
| chr4 | 186738318 | 186739186 | 869 | 1.23 | 0.00761 | 0.0151 |
| chr17 | 37000627 | 37079637 | 79011 | -0.13 | 0.00763 | 0.0151 |
| chr2 | 98523099 | 98523529 | 431 | -2.63 | 0.00763 | 0.0152 |
| chr5 | 133262418 | 133263786 | 1369 | 1.14 | 0.00763 | 0.0152 |
| chr3 | 186440178 | 186440794 | 617 | 2.56 | 0.00764 | 0.0152 |
| chr9 | 33390825 | 33392274 | 1450 | 1.25 | 0.00764 | 0.0152 |
| chr16 | 68027346 | 68028003 | 658 | -1.29 | 0.00764 | 0.0152 |
| chr1 | 32057318 | 32058039 | 722 | 2.55 | 0.00764 | 0.0152 |
| chr2 | 75005355 | 75006342 | 988 | -1.88 | 0.00764 | 0.0152 |
| chr5 | 112293313 | 112342070 | 48758 | 0.26 | 0.00765 | 0.0152 |
| chr20 | 39747210 | 39749925 | 2716 | 0.81 | 0.00767 | 0.0152 |
| chr11 | 93063571 | 93064141 | 571 | 2.28 | 0.00768 | 0.0152 |
| chr13 | 91519933 | 91520610 | 678 | 1.8 | 0.00768 | 0.0152 |
| chr13 | 78272517 | 78273167 | 651 | 2.2 | 0.00768 | 0.0152 |
| chr6 | 109760281 | 109763004 | 2724 | -0.43 | 0.00768 | 0.0152 |
| chr10 | 118706240 | 118731505 | 25266 | 0.27 | 0.00769 | 0.0152 |
| chr3 | 101725830 | 101726547 | 718 | 1.37 | 0.0077 | 0.0153 |
| chr16 | 10549082 | 10549595 | 514 | 1.91 | 0.0077 | 0.0153 |
| chr15 | 99558002 | 99559087 | 1086 | 0.6 | 0.0077 | 0.0153 |
| chr4 | 114840668 | 114841341 | 674 | -1.84 | 0.0077 | 0.0153 |
| chr10 | 863023 | 865502 | 2480 | -1.01 | 0.0077 | 0.0153 |
| chr4 | 28169250 | 28180051 | 10802 | -0.63 | 0.00771 | 0.0153 |
| chr3 | 151876546 | 151877853 | 1308 | 1.03 | 0.00771 | 0.0153 |
| chr8 | 145776760 | 145777394 | 635 | -2.72 | 0.00772 | 0.0153 |
| chr4 | 132371572 | 132372159 | 588 | 2.17 | 0.00772 | 0.0153 |
| chr15 | 41230638 | 41311231 | 80594 | 0.21 | 0.00773 | 0.0153 |
| chr12 | 102455045 | 102479300 | 24256 | 0.4 | 0.00774 | 0.0153 |
| chrX | 69728109 | 69730044 | 1936 | 1.45 | 0.00774 | 0.0153 |
| chr12 | 75853874 | 75855032 | 1159 | 3.72 | 0.00774 | 0.0154 |
| chr6 | 44354229 | 44356418 | 2190 | 0.54 | 0.00775 | 0.0154 |
| chrX | 3478113 | 3478600 | 488 | 1.86 | 0.00775 | 0.0154 |
| chr4 | 176986212 | 176987712 | 1501 | -0.74 | 0.00775 | 0.0154 |
| chr2 | 232825534 | 232827704 | 2171 | -0.45 | 0.00776 | 0.0154 |
| chr9 | 16725851 | 16728027 | 2177 | 0.47 | 0.00777 | 0.0154 |
| chr12 | 106812163 | 106813222 | 1060 | 1.31 | 0.00777 | 0.0154 |
| chr7 | 91508819 | 91510662 | 1844 | 0.48 | 0.00778 | 0.0154 |

| | | | | | | |
|---|---|---|---|---|---|---|
| chr5 | 156182635 | 156183183 | 549 | -2.59 | 0.0078 | 0.0154 |
| chr20 | 3371140 | 3372059 | 920 | 1.82 | 0.0078 | 0.0155 |
| chr22 | 32450797 | 32451714 | 918 | 1.49 | 0.0078 | 0.0155 |
| chr8 | 145079239 | 145079979 | 741 | -2.46 | 0.00781 | 0.0155 |
| chr2 | 33328958 | 33330392 | 1435 | -0.76 | 0.00781 | 0.0155 |
| chr16 | 23400801 | 23401509 | 709 | -2.18 | 0.00782 | 0.0155 |
| chrX | 106897623 | 106898672 | 1050 | 1.57 | 0.00782 | 0.0155 |
| chr6 | 56929413 | 56930468 | 1056 | -1.63 | 0.00782 | 0.0155 |
| chr22 | 28313804 | 28317179 | 3376 | 0.48 | 0.00783 | 0.0155 |
| chr3 | 170047893 | 170108018 | 60126 | 0.16 | 0.00783 | 0.0155 |
| chr22 | 27233930 | 27234573 | 644 | -1.98 | 0.00784 | 0.0155 |
| chr2 | 42310981 | 42375595 | 64615 | 0.13 | 0.00786 | 0.0156 |
| chr6 | 3282381 | 3282835 | 455 | -2.66 | 0.00786 | 0.0156 |
| chr6 | 35148996 | 35153983 | 4988 | 0.66 | 0.00786 | 0.0156 |
| chr17 | 28286225 | 28286924 | 700 | -2.32 | 0.00787 | 0.0156 |
| chr20 | 61273251 | 61285189 | 11939 | 0.44 | 0.00787 | 0.0156 |
| chr19 | 34285210 | 34311538 | 26329 | -0.69 | 0.00788 | 0.0156 |
| chrX | 48431823 | 48457002 | 25180 | 0.21 | 0.00788 | 0.0156 |
| chr16 | 89216271 | 89217339 | 1069 | -1.59 | 0.00788 | 0.0156 |
| chr15 | 76868869 | 76870466 | 1598 | 1 | 0.00789 | 0.0156 |
| chr10 | 6130881 | 6167550 | 36670 | -0.28 | 0.00789 | 0.0156 |
| chr3 | 17541961 | 17542615 | 655 | 1.98 | 0.0079 | 0.0156 |
| chr15 | 68033584 | 68033957 | 374 | -2.11 | 0.0079 | 0.0156 |
| chr13 | 34823708 | 34824328 | 621 | 1.88 | 0.0079 | 0.0156 |
| chr3 | 87486279 | 87487390 | 1112 | 1.46 | 0.00791 | 0.0156 |
| chr19 | 11848050 | 11850895 | 2846 | 0.48 | 0.00791 | 0.0156 |
| chr10 | 64191737 | 64194270 | 2534 | -0.54 | 0.00791 | 0.0157 |
| chr7 | 65957603 | 65959167 | 1565 | 0.52 | 0.00793 | 0.0157 |
| chr3 | 107085470 | 107087342 | 1873 | 0.74 | 0.00794 | 0.0157 |
| chr1 | 218632342 | 218674832 | 42491 | -0.13 | 0.00795 | 0.0157 |
| chr3 | 107383996 | 107384818 | 823 | -1.3 | 0.00795 | 0.0157 |
| chr1 | 183178601 | 183184005 | 5405 | 1.14 | 0.00795 | 0.0157 |
| chr9 | 131002863 | 131003847 | 985 | -1.72 | 0.00796 | 0.0157 |
| chr2 | 239591068 | 239595706 | 4639 | -0.32 | 0.00797 | 0.0158 |
| chr16 | 78218576 | 78219102 | 527 | 2.13 | 0.00797 | 0.0158 |
| chrX | 126842117 | 126842674 | 558 | -1.98 | 0.00797 | 0.0158 |
| chr2 | 218792892 | 218793491 | 600 | -1.7 | 0.00798 | 0.0158 |
| chr5 | 90568467 | 90629242 | 60776 | -0.13 | 0.008 | 0.0158 |
| chr16 | 28526549 | 28566107 | 39559 | 0.33 | 0.008 | 0.0158 |
| chr14 | 58879828 | 58934016 | 54189 | 0.21 | 0.008 | 0.0158 |
| chr9 | 98899523 | 98902264 | 2742 | -1.17 | 0.008 | 0.0158 |
| chr1 | 233332841 | 233347786 | 14946 | -0.45 | 0.008 | 0.0158 |
| chr19 | 36034479 | 36037830 | 3352 | -0.25 | 0.00801 | 0.0158 |
| chr4 | 90133843 | 90135321 | 1479 | 1 | 0.00801 | 0.0158 |
| chr19 | 49042494 | 49149908 | 107415 | -0.12 | 0.00802 | 0.0158 |
| chr12 | 63439177 | 63439727 | 551 | 2.32 | 0.00802 | 0.0158 |
| chr11 | 4414226 | 4415116 | 891 | 0.94 | 0.00803 | 0.0159 |
| chr18 | 24033240 | 24072093 | 38854 | 0.12 | 0.00803 | 0.0159 |
| chr17 | 73802897 | 73824000 | 21104 | -0.33 | 0.00804 | 0.0159 |
| chr3 | 15467862 | 15470633 | 2772 | 0.39 | 0.00804 | 0.0159 |
| chr12 | 79195850 | 79196934 | 1085 | 1.15 | 0.00804 | 0.0159 |
| chr7 | 151769038 | 151769984 | 947 | -1.6 | 0.00804 | 0.0159 |
| chr14 | 81931006 | 81947791 | 16786 | -0.69 | 0.00806 | 0.0159 |
| chrX | 45735481 | 45736802 | 1322 | 1.11 | 0.00807 | 0.0159 |
| chr3 | 128383954 | 128384756 | 803 | -1.14 | 0.00807 | 0.0159 |
| chr8 | 129440270 | 129440796 | 527 | -2.16 | 0.00807 | 0.0159 |
| chr2 | 41942280 | 41943427 | 1148 | 0.93 | 0.00807 | 0.0159 |
| chr15 | 78774736 | 78800930 | 26195 | 0.24 | 0.00807 | 0.0159 |
| chr19 | 7199373 | 7200434 | 1062 | 2.47 | 0.00809 | 0.016 |
| chr6 | 10480655 | 10493244 | 12590 | 4.42 | 0.0081 | 0.016 |
| chr4 | 143765804 | 143768383 | 2580 | 3.96 | 0.0081 | 0.016 |

| | | | | | | |
|---|---|---|---|---|---|---|
| chr3 | 157502583 | 157504886 | 2304 | -1.02 | 0.00811 | 0.016 |
| chr3 | 189654324 | 189666937 | 12614 | 0.56 | 0.00812 | 0.016 |
| chr12 | 90452087 | 90453088 | 1002 | 1.56 | 0.00812 | 0.016 |
| chr9 | 90680426 | 90692359 | 11934 | 0.54 | 0.00813 | 0.016 |
| chr4 | 159573812 | 159574490 | 679 | 1.63 | 0.00813 | 0.016 |
| chr8 | 9490918 | 9491572 | 655 | -1.58 | 0.00815 | 0.0161 |
| chr15 | 31555532 | 31596111 | 40580 | -0.22 | 0.00817 | 0.0161 |
| chr11 | 93378098 | 93378720 | 623 | 2.51 | 0.00817 | 0.0161 |
| chr16 | 73266672 | 73267196 | 525 | 1.65 | 0.00817 | 0.0161 |
| chr14 | 102486965 | 102512623 | 25659 | 0.37 | 0.00818 | 0.0161 |
| chr5 | 73833591 | 73841217 | 7627 | 0.28 | 0.00818 | 0.0161 |
| chr12 | 101673643 | 101674796 | 1154 | 0.65 | 0.00819 | 0.0162 |
| chr20 | 22942723 | 22945304 | 2582 | -0.45 | 0.00819 | 0.0162 |
| chr14 | 64008507 | 64011333 | 2827 | 0.61 | 0.0082 | 0.0162 |
| chr2 | 112895792 | 112896226 | 435 | -2.62 | 0.0082 | 0.0162 |
| chr1 | 113733858 | 113735255 | 1398 | -0.91 | 0.00821 | 0.0162 |
| chr3 | 81641113 | 81642201 | 1089 | 1.54 | 0.00821 | 0.0162 |
| chr1 | 179922198 | 179925907 | 3710 | 0.5 | 0.00821 | 0.0162 |
| chr1 | 201114370 | 201124118 | 9749 | -0.87 | 0.00822 | 0.0162 |
| chr16 | 11438342 | 11444006 | 5665 | -0.56 | 0.00822 | 0.0162 |
| chr9 | 138780060 | 138781836 | 1777 | -1.54 | 0.00822 | 0.0162 |
| chr17 | 11997950 | 11998440 | 491 | -1.89 | 0.00822 | 0.0162 |
| chr13 | 110220613 | 110246413 | 25801 | -0.2 | 0.00823 | 0.0162 |
| chr8 | 66841108 | 66846437 | 5330 | 0.77 | 0.00823 | 0.0162 |
| chr1 | 244325867 | 244328572 | 2706 | 0.32 | 0.00824 | 0.0162 |
| chr6 | 127587191 | 127589998 | 2808 | 0.53 | 0.00825 | 0.0163 |
| chr12 | 29499467 | 29500176 | 710 | 1.67 | 0.00826 | 0.0163 |
| chr4 | 155581306 | 155581889 | 584 | 2.47 | 0.00826 | 0.0163 |
| chr6 | 9204095 | 9205024 | 930 | 2.05 | 0.00826 | 0.0163 |
| chr14 | 73267753 | 73268146 | 394 | -2.31 | 0.00827 | 0.0163 |
| chr6 | 53693861 | 53695113 | 1253 | 1.94 | 0.00827 | 0.0163 |
| chr4 | 77119072 | 77136044 | 16973 | 0.37 | 0.00827 | 0.0163 |
| chr2 | 71589453 | 71606857 | 17405 | -0.55 | 0.00829 | 0.0163 |
| chr6 | 150036983 | 150040082 | 3100 | 0.4 | 0.00829 | 0.0163 |
| chr7 | 91855643 | 91856337 | 695 | 2.46 | 0.00829 | 0.0163 |
| chr21 | 37484852 | 37486372 | 1521 | 1.09 | 0.0083 | 0.0163 |
| chr4 | 72051925 | 72055766 | 3842 | 0.67 | 0.00832 | 0.0164 |
| chr10 | 29871771 | 29872565 | 795 | -2.46 | 0.00833 | 0.0164 |
| chr11 | 76287861 | 76289429 | 1569 | 0.97 | 0.00834 | 0.0164 |
| chr9 | 103140902 | 103143066 | 2165 | -0.57 | 0.00835 | 0.0164 |
| chr7 | 24826342 | 24827172 | 831 | -1.26 | 0.00835 | 0.0164 |
| chr2 | 11479652 | 11486738 | 7087 | 0.21 | 0.00836 | 0.0165 |
| chr10 | 71605536 | 71606378 | 843 | 1.7 | 0.00837 | 0.0165 |
| chr17 | 49229281 | 49248990 | 19710 | 0.18 | 0.00837 | 0.0165 |
| chr16 | 12703779 | 12723588 | 19810 | 0.19 | 0.00837 | 0.0165 |
| chr1 | 15609179 | 15617022 | 7844 | -0.38 | 0.00837 | 0.0165 |
| chr15 | 52750942 | 52751465 | 524 | 1.42 | 0.00838 | 0.0165 |
| chr12 | 58024707 | 58027493 | 2787 | -0.6 | 0.00838 | 0.0165 |
| chr3 | 196755118 | 196757019 | 1902 | -0.98 | 0.00838 | 0.0165 |
| chr6 | 100952306 | 100953457 | 1152 | 1.46 | 0.00839 | 0.0165 |
| chr12 | 116784220 | 116895860 | 111641 | -0.14 | 0.00839 | 0.0165 |
| chr22 | 44419487 | 44434474 | 14988 | -0.44 | 0.0084 | 0.0165 |
| chr2 | 133300217 | 133302388 | 2172 | -0.89 | 0.0084 | 0.0165 |
| chr14 | 37889968 | 37906965 | 16998 | 2.86 | 0.0084 | 0.0165 |
| chr17 | 30810289 | 30814685 | 4397 | -0.72 | 0.00841 | 0.0165 |
| chr3 | 99978897 | 99981156 | 2260 | 0.37 | 0.00841 | 0.0165 |
| chr2 | 122182740 | 122195418 | 12679 | -0.67 | 0.00841 | 0.0165 |
| chr3 | 69099571 | 69103177 | 3607 | 0.29 | 0.00842 | 0.0166 |
| chr8 | 117766703 | 117780459 | 13757 | 0.32 | 0.00843 | 0.0166 |
| chr19 | 10945723 | 10948005 | 2283 | 0.48 | 0.00843 | 0.0166 |
| chr20 | 9427501 | 9428423 | 923 | -1.31 | 0.00844 | 0.0166 |

| | | | | | | |
|---|---|---|---|---|---|---|
| chr2 | 65638199 | 65665026 | 26828 | 0.21 | 0.00844 | 0.0166 |
| chr8 | 16460745 | 16462418 | 1674 | 0.51 | 0.00844 | 0.0166 |
| chr10 | 126149653 | 126151230 | 1578 | 0.63 | 0.00845 | 0.0166 |
| chr3 | 3220471 | 3221745 | 1275 | 0.79 | 0.00845 | 0.0166 |
| chr12 | 60197461 | 60198219 | 759 | 2.42 | 0.00845 | 0.0166 |
| chr5 | 134824294 | 134830405 | 6112 | -1.07 | 0.00845 | 0.0166 |
| chr3 | 171429500 | 171430685 | 1186 | -1.46 | 0.00845 | 0.0166 |
| chr1 | 161883144 | 161884068 | 925 | 1.23 | 0.00846 | 0.0166 |
| chr3 | 28388722 | 28391686 | 2965 | 0.31 | 0.00847 | 0.0166 |
| chr4 | 13484492 | 13487282 | 2791 | 0.38 | 0.00847 | 0.0166 |
| chr13 | 46244375 | 46244997 | 623 | 1.52 | 0.00848 | 0.0167 |
| chr3 | 188598222 | 188600389 | 2168 | -0.78 | 0.00849 | 0.0167 |
| chr7 | 82605077 | 82605682 | 606 | 1.92 | 0.00849 | 0.0167 |
| chr7 | 143076334 | 143089387 | 13054 | 0.34 | 0.0085 | 0.0167 |
| chr3 | 42297106 | 42298430 | 1325 | 1.35 | 0.00851 | 0.0167 |
| chr11 | 63518844 | 63521751 | 2908 | -1.26 | 0.00851 | 0.0167 |
| chr20 | 49598237 | 49604294 | 6058 | 3.78 | 0.00852 | 0.0167 |
| chr16 | 19764590 | 19767368 | 2779 | 0.41 | 0.00853 | 0.0167 |
| chr20 | 60812756 | 60814569 | 1814 | 0.86 | 0.00855 | 0.0168 |
| chr2 | 191370015 | 191372452 | 2438 | 1.1 | 0.00855 | 0.0168 |
| chr15 | 67833400 | 67836201 | 2802 | 0.38 | 0.00855 | 0.0168 |
| chr3 | 106447391 | 106447831 | 441 | -1.96 | 0.00856 | 0.0168 |
| chr5 | 75664241 | 75670047 | 5807 | -0.38 | 0.00856 | 0.0168 |
| chr18 | 29881713 | 29898085 | 16373 | 0.8 | 0.00858 | 0.0168 |
| chr10 | 76073963 | 76074543 | 581 | 2.43 | 0.00859 | 0.0169 |
| chr7 | 72991994 | 72994237 | 2244 | 0.51 | 0.0086 | 0.0169 |
| chr3 | 57676560 | 57679662 | 3103 | 0.47 | 0.0086 | 0.0169 |
| chr12 | 111834579 | 111850853 | 16275 | 0.45 | 0.00862 | 0.0169 |
| chr5 | 156770893 | 156771286 | 394 | -2.08 | 0.00862 | 0.0169 |
| chr9 | 132020836 | 132021422 | 587 | -1.98 | 0.00862 | 0.0169 |
| chr8 | 122101163 | 122110232 | 9070 | 3.57 | 0.00863 | 0.0169 |
| chr5 | 174905042 | 174922718 | 17677 | 0.35 | 0.00864 | 0.0169 |
| chr16 | 14240403 | 14241633 | 1231 | 1.46 | 0.00864 | 0.0169 |
| chr2 | 172864381 | 172865830 | 1450 | -0.67 | 0.00866 | 0.017 |
| chr14 | 34671634 | 34673423 | 1790 | 3.99 | 0.00867 | 0.017 |
| chrX | 40439436 | 40441311 | 1876 | 0.74 | 0.00868 | 0.017 |
| chr5 | 91313655 | 91315998 | 2344 | 0.87 | 0.00868 | 0.017 |
| chr14 | 92571484 | 92574012 | 2529 | 0.5 | 0.00869 | 0.017 |
| chr5 | 153810034 | 153810367 | 334 | -3.62 | 0.00869 | 0.017 |
| chr3 | 128564309 | 128565267 | 959 | -1.2 | 0.00869 | 0.017 |
| chr7 | 86845896 | 86850246 | 4351 | 0.48 | 0.0087 | 0.017 |
| chr4 | 158915847 | 158916635 | 789 | 1.3 | 0.0087 | 0.017 |
| chr1 | 36897987 | 36898858 | 872 | 2.56 | 0.00872 | 0.0171 |
| chr5 | 54686931 | 54687849 | 919 | 1.82 | 0.00873 | 0.0171 |
| chr3 | 12757849 | 12762825 | 4977 | -0.67 | 0.00873 | 0.0171 |
| chr8 | 12582591 | 12585498 | 2908 | 0.38 | 0.00873 | 0.0171 |
| chr5 | 123737197 | 123738593 | 1397 | 1.18 | 0.00873 | 0.0171 |
| chr11 | 19733420 | 19734279 | 860 | 0.68 | 0.00874 | 0.0171 |
| chr15 | 76825139 | 76826328 | 1190 | 1.48 | 0.00875 | 0.0171 |
| chr12 | 92601382 | 92603327 | 1946 | 0.58 | 0.00875 | 0.0171 |
| chr10 | 7323673 | 7325214 | 1542 | 1 | 0.00875 | 0.0171 |
| chr9 | 125109155 | 125109893 | 739 | 1.38 | 0.00876 | 0.0172 |
| chr1 | 30205299 | 30205797 | 499 | -2.54 | 0.00877 | 0.0172 |
| chr11 | 85752101 | 85753393 | 1293 | -1.27 | 0.00878 | 0.0172 |
| chr8 | 23376009 | 23428400 | 52392 | 0.14 | 0.00878 | 0.0172 |
| chr3 | 15642167 | 15650161 | 7995 | 0.36 | 0.00878 | 0.0172 |
| chr4 | 102711088 | 102712771 | 1684 | 0.43 | 0.00879 | 0.0172 |
| chrX | 41523659 | 41525197 | 1539 | 1.48 | 0.00879 | 0.0172 |
| chrX | 43886009 | 43886409 | 401 | 2.25 | 0.0088 | 0.0172 |
| chr6 | 36461297 | 36462379 | 1083 | 1.41 | 0.00881 | 0.0172 |
| chr7 | 106410163 | 106417241 | 7079 | 0.3 | 0.00881 | 0.0172 |

| | | | | | | |
|---|---|---|---|---|---|---|
| chr1 | 159823627 | 159837896 | 14270 | 0.55 | 0.00881 | 0.0172 |
| chr1 | 183773801 | 183775084 | 1284 | 0.89 | 0.00882 | 0.0173 |
| chr1 | 32468944 | 32489608 | 20665 | 0.33 | 0.00882 | 0.0173 |
| chrX | 27641173 | 27641527 | 355 | 2.56 | 0.00882 | 0.0173 |
| chr8 | 59723523 | 59724952 | 1430 | 0.96 | 0.00883 | 0.0173 |
| chr11 | 68648900 | 68650141 | 1242 | -1.3 | 0.00884 | 0.0173 |
| chr11 | 66487306 | 66495479 | 8174 | -1.32 | 0.00884 | 0.0173 |
| chr5 | 79782434 | 79785368 | 2935 | 0.59 | 0.00885 | 0.0173 |
| chr9 | 102728726 | 102730247 | 1522 | 1.14 | 0.00885 | 0.0173 |
| chr1 | 245208380 | 245209800 | 1421 | 1.59 | 0.00885 | 0.0173 |
| chr5 | 33439287 | 33445458 | 6172 | 0.25 | 0.00887 | 0.0173 |
| chr7 | 68067339 | 68067670 | 332 | -2.69 | 0.00888 | 0.0174 |
| chr14 | 53368059 | 53387483 | 19425 | -0.43 | 0.00888 | 0.0174 |
| chr5 | 32351784 | 32353365 | 1582 | -1.38 | 0.00888 | 0.0174 |
| chr3 | 195603185 | 195604685 | 1501 | -1.22 | 0.00888 | 0.0174 |
| chr6 | 134571186 | 134571775 | 590 | 2.51 | 0.00889 | 0.0174 |
| chr4 | 145678574 | 145679822 | 1249 | -1.41 | 0.00889 | 0.0174 |
| chr7 | 99646752 | 99648410 | 1659 | 0.49 | 0.0089 | 0.0174 |
| chrX | 45606755 | 45711661 | 104907 | 0.12 | 0.0089 | 0.0174 |
| chr13 | 99972928 | 99973910 | 983 | 1.61 | 0.0089 | 0.0174 |
| chr3 | 108348818 | 108354106 | 5289 | 0.93 | 0.0089 | 0.0174 |
| chr16 | 57094629 | 57095968 | 1340 | 1.32 | 0.00892 | 0.0174 |
| chr7 | 102983408 | 102989459 | 6052 | 0.26 | 0.00892 | 0.0174 |
| chr11 | 69189499 | 69196693 | 7195 | -0.73 | 0.00893 | 0.0174 |
| chr4 | 6799212 | 6809786 | 10575 | -0.68 | 0.00893 | 0.0174 |
| chr9 | 74219655 | 74220073 | 419 | -2.56 | 0.00893 | 0.0175 |
| chr22 | 48879453 | 48889146 | 9694 | -0.39 | 0.00894 | 0.0175 |
| chr15 | 50426385 | 50430497 | 4113 | -0.38 | 0.00895 | 0.0175 |
| chr8 | 74789592 | 74792032 | 2441 | 0.51 | 0.00896 | 0.0175 |
| chr12 | 76741114 | 76743183 | 2070 | 0.47 | 0.00896 | 0.0175 |
| chr10 | 31025216 | 31026774 | 1559 | -0.73 | 0.00897 | 0.0175 |
| chr11 | 110182135 | 110183763 | 1629 | -0.73 | 0.00898 | 0.0175 |
| chrX | 149961512 | 149962397 | 886 | -1.43 | 0.00899 | 0.0176 |
| chrX | 20497843 | 20498761 | 919 | 1.31 | 0.00899 | 0.0176 |
| chr7 | 6543395 | 6544645 | 1251 | 1.03 | 0.009 | 0.0176 |
| chr5 | 39219278 | 39225052 | 5775 | 3.79 | 0.009 | 0.0176 |
| chr13 | 111179513 | 111180222 | 710 | 2.44 | 0.009 | 0.0176 |
| chr8 | 82584723 | 82599651 | 14929 | 0.44 | 0.009 | 0.0176 |
| chr1 | 35971163 | 35972838 | 1676 | 1.3 | 0.00901 | 0.0176 |
| chr6 | 85725050 | 85725912 | 863 | 1.44 | 0.00901 | 0.0176 |
| chr1 | 52728460 | 52729752 | 1293 | 1.41 | 0.00902 | 0.0176 |
| chr5 | 148205262 | 148206585 | 1324 | 1.27 | 0.00903 | 0.0176 |
| chr5 | 56203930 | 56207163 | 3234 | 0.31 | 0.00903 | 0.0176 |
| chr9 | 94047852 | 94048763 | 912 | 1.55 | 0.00904 | 0.0176 |
| chr10 | 98047157 | 98047670 | 514 | 2.55 | 0.00905 | 0.0177 |
| chr13 | 48545057 | 48546641 | 1585 | 0.84 | 0.00905 | 0.0177 |
| chr19 | 52550593 | 52603802 | 53210 | 5.55 | 0.00906 | 0.0177 |
| chr6 | 160503697 | 160504033 | 337 | -2.44 | 0.00907 | 0.0177 |
| chr12 | 26471320 | 26471827 | 508 | 1.85 | 0.00908 | 0.0177 |
| chr4 | 150419206 | 150419939 | 734 | 1.63 | 0.00908 | 0.0177 |
| chr14 | 66480785 | 66481354 | 570 | -2.14 | 0.00909 | 0.0177 |
| chr6 | 126852399 | 126853403 | 1005 | 1.43 | 0.00911 | 0.0178 |
| chr14 | 24297862 | 24300344 | 2483 | 0.52 | 0.00911 | 0.0178 |
| chr2 | 10826073 | 10831272 | 5200 | -0.4 | 0.00916 | 0.0179 |
| chr5 | 137328135 | 137372458 | 44324 | 0.16 | 0.00918 | 0.0179 |
| chr4 | 173822257 | 173822711 | 455 | 2.31 | 0.00918 | 0.0179 |
| chr11 | 115114379 | 115115829 | 1451 | 1.14 | 0.00918 | 0.0179 |
| chr13 | 42162161 | 42164886 | 2726 | -0.57 | 0.00918 | 0.0179 |
| chr12 | 58225116 | 58260149 | 35034 | 0.27 | 0.0092 | 0.0179 |
| chr1 | 93569270 | 93569769 | 500 | 2.65 | 0.0092 | 0.0179 |
| chr11 | 22174470 | 22174856 | 387 | 2.65 | 0.0092 | 0.0179 |

| | | | | | | |
|---|---|---|---|---|---|---|
| chr4 | 109570818 | 109573208 | 2391 | 0.36 | 0.00922 | 0.018 |
| chr4 | 84091862 | 84092765 | 904 | -1.13 | 0.00922 | 0.018 |
| chr11 | 116881171 | 116898106 | 16936 | -0.45 | 0.00923 | 0.018 |
| chr18 | 74171343 | 74172666 | 1324 | -0.98 | 0.00923 | 0.018 |
| chr3 | 149469124 | 149471218 | 2095 | 0.54 | 0.00924 | 0.018 |
| chr9 | 111758634 | 111759753 | 1120 | 1.11 | 0.00925 | 0.018 |
| chr9 | 128468191 | 128470200 | 2010 | 0.44 | 0.00926 | 0.018 |
| chr13 | 24440503 | 24443247 | 2745 | -0.87 | 0.00926 | 0.018 |
| chr13 | 52367198 | 52379029 | 11832 | -0.35 | 0.00927 | 0.0181 |
| chr3 | 119388805 | 119396929 | 8125 | 0.53 | 0.00927 | 0.0181 |
| chr12 | 14647612 | 14653321 | 5710 | 1.07 | 0.00931 | 0.0181 |
| chr1 | 172289582 | 172296299 | 6718 | -0.21 | 0.00931 | 0.0181 |
| chr3 | 67926275 | 67927009 | 735 | -1.47 | 0.00931 | 0.0181 |
| chr2 | 46500053 | 46501487 | 1435 | -1.19 | 0.00932 | 0.0182 |
| chr1 | 17959104 | 17959680 | 577 | -1.59 | 0.00933 | 0.0182 |
| chr2 | 136332881 | 136333240 | 360 | -2.49 | 0.00933 | 0.0182 |
| chr9 | 88224773 | 88226131 | 1359 | 1.06 | 0.00934 | 0.0182 |
| chr12 | 91164461 | 91165552 | 1092 | 0.87 | 0.00935 | 0.0182 |
| chr2 | 151442761 | 151459364 | 16604 | -0.39 | 0.00936 | 0.0182 |
| chr10 | 37999725 | 38001025 | 1301 | 1.35 | 0.00936 | 0.0182 |
| chr14 | 29072585 | 29083797 | 11213 | 3.84 | 0.00936 | 0.0182 |
| chr4 | 124670806 | 124671835 | 1030 | 2 | 0.00936 | 0.0182 |
| chr5 | 18093532 | 18094153 | 622 | -1.66 | 0.00936 | 0.0182 |
| chr21 | 34851175 | 34864595 | 13421 | 0.68 | 0.00937 | 0.0182 |
| chr1 | 9845514 | 9846267 | 754 | -2.47 | 0.00937 | 0.0182 |
| chr6 | 128889871 | 128892176 | 2306 | 0.6 | 0.00938 | 0.0182 |
| chr13 | 106781260 | 106782181 | 922 | 1.76 | 0.00938 | 0.0183 |
| chr4 | 186887513 | 186888333 | 821 | 1.64 | 0.00938 | 0.0183 |
| chr4 | 58629010 | 58629895 | 886 | -1.7 | 0.00939 | 0.0183 |
| chr13 | 36173226 | 36173952 | 727 | -2.16 | 0.00939 | 0.0183 |
| chr14 | 43477578 | 43506032 | 28455 | 2.2 | 0.0094 | 0.0183 |
| chr3 | 12486060 | 12487254 | 1195 | 1.18 | 0.0094 | 0.0183 |
| chr6 | 110011272 | 110013101 | 1830 | 0.55 | 0.00942 | 0.0183 |
| chr11 | 34323613 | 34325003 | 1391 | -1.76 | 0.00943 | 0.0183 |
| chr8 | 67227408 | 67229043 | 1636 | -0.99 | 0.00944 | 0.0184 |
| chr5 | 62037614 | 62048409 | 10796 | 4.32 | 0.00944 | 0.0184 |
| chr1 | 212457380 | 212465343 | 7964 | 0.28 | 0.00944 | 0.0184 |
| chr4 | 66657082 | 66658474 | 1393 | 1.13 | 0.00945 | 0.0184 |
| chr11 | 100802606 | 100803317 | 712 | 2.43 | 0.00945 | 0.0184 |
| chr15 | 22975392 | 22985934 | 10543 | 0.36 | 0.00945 | 0.0184 |
| chr12 | 48098850 | 48112227 | 13378 | 0.41 | 0.00945 | 0.0184 |
| chr17 | 4841061 | 4855597 | 14537 | 0.18 | 0.00946 | 0.0184 |
| chr11 | 108091222 | 108104556 | 13335 | 0.31 | 0.00947 | 0.0184 |
| chr16 | 2496092 | 2496530 | 439 | -2.65 | 0.00949 | 0.0184 |
| chr3 | 61545921 | 61547753 | 1833 | -0.71 | 0.00949 | 0.0185 |
| chr7 | 156625522 | 156626248 | 727 | -1.48 | 0.00951 | 0.0185 |
| chr1 | 226864656 | 226865404 | 749 | 1.53 | 0.00951 | 0.0185 |
| chr6 | 36034252 | 36035020 | 769 | -1.77 | 0.00951 | 0.0185 |
| chr10 | 5803201 | 5856956 | 53756 | 0.18 | 0.00952 | 0.0185 |
| chr12 | 102223550 | 102225296 | 1747 | 0.72 | 0.00952 | 0.0185 |
| chr17 | 44835405 | 44860743 | 25339 | 0.31 | 0.00953 | 0.0185 |
| chr10 | 120859669 | 120905508 | 45840 | 0.2 | 0.00954 | 0.0185 |
| chr18 | 44675841 | 44678071 | 2231 | 0.5 | 0.00955 | 0.0186 |
| chr9 | 28955489 | 28957096 | 1608 | 0.84 | 0.00956 | 0.0186 |
| chr1 | 28516276 | 28518234 | 1959 | -0.5 | 0.00956 | 0.0186 |
| chr18 | 8384871 | 8385853 | 983 | -1.51 | 0.00957 | 0.0186 |
| chr13 | 52702340 | 52703685 | 1346 | -1.15 | 0.00957 | 0.0186 |
| chr6 | 84936707 | 84937796 | 1090 | -0.8 | 0.00957 | 0.0186 |
| chr1 | 52814451 | 52814931 | 481 | -2.07 | 0.00958 | 0.0186 |
| chr15 | 34629515 | 34635954 | 6440 | 0.36 | 0.0096 | 0.0186 |
| chr2 | 217082207 | 217082795 | 589 | -1.75 | 0.0096 | 0.0186 |

| | | | | | | |
|---|---|---|---|---|---|---|
| chr1 | 206784811 | 206786709 | 1899 | 0.46 | 0.00961 | 0.0187 |
| chr1 | 201235793 | 201247055 | 11263 | 0.46 | 0.00961 | 0.0187 |
| chr7 | 50936059 | 50938953 | 2895 | 0.53 | 0.00962 | 0.0187 |
| chr1 | 147070550 | 147085528 | 14979 | 0.43 | 0.00962 | 0.0187 |
| chr11 | 72895764 | 72930148 | 34385 | -0.3 | 0.00962 | 0.0187 |
| chr1 | 28305428 | 28306242 | 815 | -1.21 | 0.00963 | 0.0187 |
| chr6 | 166416482 | 166424427 | 7946 | 0.25 | 0.00963 | 0.0187 |
| chr4 | 99979874 | 99983704 | 3831 | 0.39 | 0.00964 | 0.0187 |
| chr8 | 61590349 | 61593753 | 3405 | 0.45 | 0.00965 | 0.0187 |
| chr17 | 76564074 | 76564903 | 830 | -2.41 | 0.00965 | 0.0187 |
| chr2 | 43495084 | 43539127 | 44044 | -0.25 | 0.00966 | 0.0187 |
| chr9 | 93155682 | 93156407 | 726 | 1.31 | 0.00966 | 0.0187 |
| chr12 | 42840150 | 42841335 | 1186 | 1.34 | 0.00966 | 0.0187 |
| chr3 | 5430271 | 5431931 | 1661 | 3.64 | 0.00967 | 0.0188 |
| chr2 | 66998233 | 66999579 | 1347 | -1.61 | 0.00967 | 0.0188 |
| chr5 | 154316767 | 154321610 | 4844 | 0.4 | 0.00967 | 0.0188 |
| chr17 | 20704685 | 20705609 | 925 | 0.75 | 0.00968 | 0.0188 |
| chr6 | 114014768 | 114016373 | 1606 | 0.99 | 0.00968 | 0.0188 |
| chr5 | 142612559 | 142629861 | 17303 | 0.24 | 0.00968 | 0.0188 |
| chr3 | 45585954 | 45590284 | 4331 | 0.62 | 0.00969 | 0.0188 |
| chr8 | 101426889 | 101430242 | 3354 | 0.57 | 0.00969 | 0.0188 |
| chr11 | 6703293 | 6705382 | 2090 | 0.58 | 0.00969 | 0.0188 |
| chr10 | 69425549 | 69426925 | 1377 | -1.05 | 0.00969 | 0.0188 |
| chr3 | 157216741 | 157221969 | 5229 | 4.49 | 0.0097 | 0.0188 |
| chr4 | 174173778 | 174174532 | 755 | 1.88 | 0.0097 | 0.0188 |
| chr1 | 109676974 | 109678030 | 1057 | 1.56 | 0.00971 | 0.0188 |
| chr17 | 21001869 | 21003188 | 1320 | -0.77 | 0.00971 | 0.0188 |
| chr3 | 194124935 | 194125530 | 596 | 1.05 | 0.00973 | 0.0189 |
| chr5 | 143535954 | 143550970 | 15017 | 0.42 | 0.00974 | 0.0189 |
| chr15 | 42211902 | 42265727 | 53826 | -0.29 | 0.00974 | 0.0189 |
| chrX | 101853690 | 101855442 | 1753 | 0.49 | 0.00975 | 0.0189 |
| chr8 | 100739807 | 100741033 | 1227 | -1.09 | 0.00975 | 0.0189 |
| chr8 | 71286673 | 71288217 | 1545 | 0.79 | 0.00976 | 0.0189 |
| chr7 | 133812100 | 133812682 | 583 | -1.01 | 0.00976 | 0.0189 |
| chr5 | 136645623 | 136647035 | 1413 | 1.02 | 0.00976 | 0.0189 |
| chrX | 133506534 | 133509062 | 2529 | 0.52 | 0.00978 | 0.0189 |
| chr10 | 72084420 | 72092420 | 8001 | -0.6 | 0.00978 | 0.0189 |
| chr12 | 28932321 | 28933101 | 781 | 1.48 | 0.00979 | 0.019 |
| chr7 | 77495771 | 77497319 | 1549 | 0.88 | 0.0098 | 0.019 |
| chr20 | 3817362 | 3914175 | 96814 | -0.13 | 0.00981 | 0.019 |
| chr9 | 108673070 | 108675980 | 2911 | -0.54 | 0.00982 | 0.019 |
| chr13 | 86160517 | 86161965 | 1449 | -1.45 | 0.00983 | 0.019 |
| chr4 | 153022759 | 153028577 | 5819 | 0.38 | 0.00985 | 0.0191 |
| chr6 | 42624204 | 42624795 | 592 | 1.94 | 0.00986 | 0.0191 |
| chr5 | 125890586 | 125891124 | 539 | -2.48 | 0.00986 | 0.0191 |
| chr2 | 239552277 | 239554099 | 1823 | -0.67 | 0.00986 | 0.0191 |
| chr12 | 75393044 | 75393791 | 748 | 3.8 | 0.00986 | 0.0191 |
| chr14 | 74100182 | 74142596 | 42415 | 0.24 | 0.00987 | 0.0191 |
| chr9 | 95397211 | 95397734 | 524 | -1.98 | 0.00988 | 0.0191 |
| chr3 | 57041203 | 57051727 | 10525 | -0.63 | 0.00988 | 0.0191 |
| chr8 | 95979533 | 95981083 | 1551 | 0.94 | 0.00988 | 0.0191 |
| chr11 | 352866 | 364580 | 11715 | 0.33 | 0.00988 | 0.0191 |
| chr12 | 110175481 | 110176213 | 733 | 1.5 | 0.00988 | 0.0191 |
| chr22 | 25550084 | 25551019 | 936 | 1.56 | 0.00989 | 0.0191 |
| chr14 | 102874665 | 102875588 | 924 | 1.27 | 0.0099 | 0.0191 |
| chr7 | 54856649 | 54857096 | 448 | 2.11 | 0.0099 | 0.0191 |
| chr3 | 104838083 | 104838423 | 341 | 1.53 | 0.0099 | 0.0192 |
| chr6 | 38325865 | 38326341 | 477 | -2.22 | 0.00991 | 0.0192 |
| chr12 | 57559091 | 57561664 | 2574 | -0.54 | 0.00992 | 0.0192 |
| chr4 | 124778066 | 124778794 | 729 | 1.16 | 0.00992 | 0.0192 |
| chr7 | 29310702 | 29311292 | 591 | 2.43 | 0.00993 | 0.0192 |

| | | | | | | |
|---|---|---|---|---|---|---|
| chr1 | 38390944 | 38413428 | 22485 | 0.38 | 0.00993 | 0.0192 |
| chr10 | 64384874 | 64385669 | 796 | 1.17 | 0.00993 | 0.0192 |
| chr9 | 33000095 | 33003179 | 3085 | 0.41 | 0.00993 | 0.0192 |
| chr1 | 110777999 | 110779428 | 1430 | 0.92 | 0.00993 | 0.0192 |
| chr1 | 241970058 | 241971310 | 1253 | -0.91 | 0.00994 | 0.0192 |
| chr11 | 94658630 | 94661090 | 2461 | 4.05 | 0.00995 | 0.0192 |
| chr9 | 112045225 | 112045825 | 601 | 2.47 | 0.00995 | 0.0192 |
| chr2 | 227860418 | 227861024 | 607 | -1.98 | 0.00995 | 0.0192 |
| chr3 | 141456233 | 141461154 | 4922 | 0.33 | 0.00997 | 0.0193 |
| chr11 | 18299475 | 18299972 | 498 | -2.46 | 0.00998 | 0.0193 |
| chr13 | 44715104 | 44718485 | 3382 | 0.43 | 0.00998 | 0.0193 |
| chr4 | 35359191 | 35360445 | 1255 | 4.08 | 0.00998 | 0.0193 |
| chr14 | 73524127 | 73573095 | 48969 | 0.18 | 0.00999 | 0.0193 |
| chr19 | 908399 | 949481 | 41083 | -0.28 | 0.01 | 0.0193 |
| chr11 | 29355380 | 29355918 | 539 | -1.69 | 0.01 | 0.0193 |
| chrX | 13104907 | 13142692 | 37786 | 0.19 | 0.01 | 0.0193 |
| chr1 | 160000604 | 160002668 | 2065 | 0.72 | 0.01 | 0.0194 |
| chr21 | 47703934 | 47707570 | 3637 | 0.49 | 0.01 | 0.0194 |
| chr12 | 99146265 | 99148944 | 2680 | 0.46 | 0.01 | 0.0194 |
| chr7 | 29762433 | 29764426 | 1994 | 3.9 | 0.01 | 0.0194 |
| chr1 | 100502359 | 100541654 | 39296 | 0.24 | 0.0101 | 0.0194 |
| chr7 | 7138358 | 7139912 | 1555 | 1.52 | 0.0101 | 0.0194 |
| chr8 | 95021768 | 95022715 | 948 | 1.09 | 0.0101 | 0.0194 |
| chr11 | 110122055 | 110124314 | 2260 | 0.56 | 0.0101 | 0.0194 |
| chr20 | 33487529 | 33511454 | 23926 | 0.35 | 0.0101 | 0.0195 |
| chr2 | 108882189 | 108889258 | 7070 | -0.28 | 0.0101 | 0.0195 |
| chr17 | 2023737 | 2027445 | 3709 | -0.84 | 0.0101 | 0.0195 |
| chr7 | 36636388 | 36637591 | 1204 | -0.9 | 0.0101 | 0.0195 |
| chr19 | 39728145 | 39729412 | 1268 | 1 | 0.0101 | 0.0196 |
| chr3 | 170407042 | 170408088 | 1047 | -1.82 | 0.0101 | 0.0196 |
| chr17 | 19911759 | 19915852 | 4094 | -0.36 | 0.0102 | 0.0196 |
| chr20 | 9273399 | 9292679 | 19281 | 0.29 | 0.0102 | 0.0196 |
| chr12 | 115878105 | 115889946 | 11842 | -4.33 | 0.0102 | 0.0196 |
| chr2 | 97379516 | 97406619 | 27104 | -0.3 | 0.0102 | 0.0196 |
| chr14 | 74444268 | 74446165 | 1898 | 1.01 | 0.0102 | 0.0196 |
| chr8 | 119427792 | 119428531 | 740 | 1.5 | 0.0102 | 0.0196 |
| chr6 | 153409646 | 153453721 | 44076 | 0.18 | 0.0102 | 0.0197 |
| chr1 | 46334193 | 46362957 | 28765 | 0.22 | 0.0102 | 0.0197 |
| chr9 | 79291915 | 79292978 | 1064 | 1.1 | 0.0102 | 0.0198 |
| chr8 | 95557698 | 95567112 | 9415 | 0.37 | 0.0103 | 0.0198 |
| chr12 | 49960796 | 49963079 | 2284 | 0.43 | 0.0103 | 0.0198 |
| chr6 | 170850175 | 170864671 | 14497 | 0.27 | 0.0103 | 0.0198 |
| chr7 | 156684542 | 156687197 | 2656 | 0.44 | 0.0103 | 0.0198 |
| chr1 | 70814712 | 70828443 | 13732 | 0.36 | 0.0103 | 0.0198 |
| chr1 | 178061446 | 178064992 | 3547 | 0.45 | 0.0103 | 0.0198 |
| chr5 | 80564417 | 80565734 | 1318 | 1.11 | 0.0103 | 0.0198 |
| chr5 | 179176457 | 179196772 | 20316 | -0.45 | 0.0103 | 0.0198 |
| chr17 | 1480265 | 1502273 | 22009 | 0.35 | 0.0103 | 0.0198 |
| chr3 | 10321490 | 10364106 | 42617 | -0.26 | 0.0103 | 0.0199 |
| chr8 | 138053137 | 138054274 | 1138 | 0.79 | 0.0103 | 0.0199 |
| chr9 | 127661674 | 127664698 | 3025 | -0.87 | 0.0103 | 0.0199 |
| chr11 | 22213893 | 22215762 | 1870 | 0.61 | 0.0103 | 0.0199 |
| chr17 | 59327828 | 59328674 | 847 | -1.13 | 0.0103 | 0.0199 |
| chr17 | 62934599 | 62935734 | 1136 | -1.39 | 0.0103 | 0.0199 |
| chr10 | 133971174 | 133976215 | 5042 | 0.86 | 0.0103 | 0.0199 |
| chr2 | 240321339 | 240324316 | 2978 | -0.47 | 0.0104 | 0.0199 |
| chr2 | 223508675 | 223521900 | 13226 | -0.34 | 0.0104 | 0.0199 |
| chr12 | 80605430 | 80605852 | 423 | 1.51 | 0.0104 | 0.02 |
| chr11 | 118560066 | 118561589 | 1524 | 0.73 | 0.0104 | 0.02 |
| chr15 | 31506574 | 31531652 | 25079 | 0.26 | 0.0104 | 0.02 |
| chr2 | 102417996 | 102424266 | 6271 | -0.36 | 0.0104 | 0.02 |

| | | | | | | |
|---|---|---|---|---|---|---|
| chr13 | 23947623 | 23949654 | 2032 | 0.69 | 0.0104 | 0.0201 |
| chr3 | 107241280 | 107244932 | 3653 | 0.38 | 0.0104 | 0.0201 |
| chr1 | 93810448 | 93819670 | 9223 | 0.42 | 0.0104 | 0.0201 |
| chr2 | 162213290 | 162214048 | 759 | 1.76 | 0.0104 | 0.0201 |
| chr2 | 200818581 | 200821461 | 2881 | -0.38 | 0.0104 | 0.0201 |
| chr17 | 4165876 | 4170199 | 4324 | -0.44 | 0.0104 | 0.0201 |
| chr11 | 65939485 | 65990246 | 50762 | -0.42 | 0.0104 | 0.0201 |
| chr9 | 22237411 | 22245382 | 7972 | 0.4 | 0.0104 | 0.0201 |
| chr16 | 68101951 | 68126237 | 24287 | -0.32 | 0.0104 | 0.0201 |
| chr1 | 21687741 | 21688311 | 571 | -1.68 | 0.0104 | 0.0201 |
| chr22 | 44184937 | 44186366 | 1430 | 0.88 | 0.0104 | 0.0201 |
| chr4 | 103533557 | 103549614 | 16058 | 0.33 | 0.0105 | 0.0201 |
| chr1 | 97186583 | 97189095 | 2513 | 0.49 | 0.0105 | 0.0201 |
| chrX | 48956907 | 48961006 | 4100 | 0.66 | 0.0105 | 0.0202 |
| chr3 | 177801549 | 177804074 | 2526 | 0.52 | 0.0105 | 0.0202 |
| chr1 | 226890429 | 226891903 | 1475 | 0.58 | 0.0105 | 0.0202 |
| chr16 | 52503315 | 52506578 | 3264 | 1.1 | 0.0105 | 0.0202 |
| chr7 | 128863652 | 128865841 | 2190 | 0.49 | 0.0105 | 0.0202 |
| chr2 | 164733724 | 164734365 | 642 | -1.76 | 0.0105 | 0.0202 |
| chr1 | 244931356 | 244932057 | 702 | -1.59 | 0.0105 | 0.0202 |
| chr1 | 169748543 | 169749666 | 1124 | -1.13 | 0.0105 | 0.0202 |
| chr7 | 140177158 | 140179895 | 2738 | 0.4 | 0.0105 | 0.0202 |
| chr7 | 109846178 | 109846659 | 482 | 2.37 | 0.0105 | 0.0202 |
| chr2 | 165454229 | 165457679 | 3451 | -0.69 | 0.0105 | 0.0202 |
| chr1 | 32858752 | 32867175 | 8424 | 0.53 | 0.0105 | 0.0202 |
| chr9 | 6006228 | 6016261 | 10034 | 0.35 | 0.0105 | 0.0202 |
| chr19 | 43685439 | 43686472 | 1034 | 1.08 | 0.0105 | 0.0202 |
| chr13 | 80658268 | 80661008 | 2741 | 0.77 | 0.0105 | 0.0202 |
| chr3 | 192596406 | 192618774 | 22369 | -0.19 | 0.0105 | 0.0203 |
| chr4 | 186480356 | 186495407 | 15052 | 0.66 | 0.0105 | 0.0203 |
| chr5 | 68564532 | 68573579 | 9048 | 0.49 | 0.0106 | 0.0203 |
| chr5 | 126765638 | 126766925 | 1288 | 0.97 | 0.0106 | 0.0203 |
| chr2 | 207479187 | 207481595 | 2409 | -0.76 | 0.0106 | 0.0203 |
| chr12 | 1325972 | 1329920 | 3949 | 2.73 | 0.0106 | 0.0203 |
| chr2 | 210359436 | 210373264 | 13829 | -0.37 | 0.0106 | 0.0203 |
| chr9 | 28808046 | 28810790 | 2745 | 1.12 | 0.0106 | 0.0203 |
| chr4 | 95726826 | 95727726 | 901 | 1.37 | 0.0106 | 0.0203 |
| chr3 | 57530182 | 57543690 | 13509 | 0.39 | 0.0106 | 0.0203 |
| chr11 | 115419713 | 115420576 | 864 | 1.78 | 0.0106 | 0.0204 |
| chr8 | 42927129 | 42927884 | 756 | 2.07 | 0.0106 | 0.0204 |
| chr9 | 97021047 | 97043817 | 22771 | -0.3 | 0.0106 | 0.0204 |
| chr4 | 119963358 | 119970395 | 7038 | 0.67 | 0.0106 | 0.0204 |
| chr12 | 83290296 | 83291437 | 1142 | 1.23 | 0.0106 | 0.0204 |
| chr10 | 84109734 | 84110752 | 1019 | 3.95 | 0.0106 | 0.0204 |
| chr16 | 57999888 | 58044899 | 45012 | -0.35 | 0.0106 | 0.0204 |
| chr4 | 102674425 | 102675422 | 998 | 1.43 | 0.0106 | 0.0204 |
| chr1 | 78085541 | 78086672 | 1132 | 1.05 | 0.0107 | 0.0205 |
| chr12 | 70446352 | 70447465 | 1114 | 1.03 | 0.0107 | 0.0205 |
| chr2 | 197851933 | 197853632 | 1700 | 0.49 | 0.0107 | 0.0205 |
| chr1 | 178124618 | 178125380 | 763 | 1.65 | 0.0107 | 0.0205 |
| chr15 | 90879568 | 90880227 | 660 | -1.48 | 0.0107 | 0.0205 |
| chr5 | 124692764 | 124694150 | 1387 | 1.23 | 0.0107 | 0.0205 |
| chr3 | 151553353 | 151563767 | 10415 | -0.28 | 0.0107 | 0.0205 |
| chr3 | 131621983 | 131623507 | 1525 | -1.79 | 0.0107 | 0.0205 |
| chr8 | 17012895 | 17015321 | 2427 | 0.63 | 0.0107 | 0.0206 |
| chr1 | 39735114 | 39736016 | 903 | -1.49 | 0.0107 | 0.0206 |
| chr11 | 102956500 | 102963891 | 7392 | 0.43 | 0.0108 | 0.0206 |
| chr9 | 98636046 | 98670922 | 34877 | 0.22 | 0.0108 | 0.0206 |
| chr3 | 139107370 | 139110185 | 2816 | 0.29 | 0.0108 | 0.0207 |
| chr3 | 105242224 | 105243135 | 912 | -1.29 | 0.0108 | 0.0207 |
| chr2 | 62011987 | 62013877 | 1891 | 0.81 | 0.0108 | 0.0207 |

| | | | | | | |
|---|---|---|---|---|---|---|
| chr2 | 234517219 | 234517551 | 333 | 2.59 | 0.0108 | 0.0207 |
| chr1 | 202452784 | 202453346 | 563 | 1.68 | 0.0108 | 0.0207 |
| chr14 | 100941789 | 100944294 | 2506 | 0.45 | 0.0108 | 0.0207 |
| chr1 | 236227374 | 236228752 | 1379 | 1.33 | 0.0108 | 0.0207 |
| chr15 | 39236711 | 39237413 | 703 | 1.84 | 0.0108 | 0.0208 |
| chr8 | 9575459 | 9575933 | 475 | -1.75 | 0.0108 | 0.0208 |
| chr11 | 18633891 | 18636025 | 2135 | -1.2 | 0.0108 | 0.0208 |
| chr3 | 137831772 | 137835151 | 3380 | 0.39 | 0.0108 | 0.0208 |
| chr1 | 54481262 | 54483107 | 1846 | 0.89 | 0.0108 | 0.0208 |
| chr12 | 122101387 | 122111708 | 10322 | 0.44 | 0.0108 | 0.0208 |
| chr16 | 1463679 | 1479035 | 15357 | 0.24 | 0.0109 | 0.0208 |
| chr11 | 45201103 | 45203871 | 2769 | 0.3 | 0.0109 | 0.0208 |
| chr11 | 5300143 | 5303735 | 3593 | -0.38 | 0.0109 | 0.0208 |
| chr4 | 100866271 | 100872488 | 6218 | 0.35 | 0.0109 | 0.0208 |
| chr6 | 13573531 | 13576393 | 2863 | 0.43 | 0.0109 | 0.0208 |
| chr2 | 10054400 | 10055274 | 875 | -1.27 | 0.0109 | 0.0208 |
| chr15 | 73831328 | 73833232 | 1905 | 0.83 | 0.0109 | 0.0208 |
| chr2 | 224949604 | 224952031 | 2428 | -0.64 | 0.0109 | 0.0208 |
| chr11 | 119485878 | 119487161 | 1284 | 0.83 | 0.0109 | 0.0209 |
| chr13 | 28051129 | 28052132 | 1004 | 1.2 | 0.0109 | 0.0209 |
| chr10 | 103697542 | 103699838 | 2297 | -1.04 | 0.0109 | 0.0209 |
| chr1 | 237983182 | 237984636 | 1455 | 1.16 | 0.0109 | 0.0209 |
| chr20 | 22526699 | 22527543 | 845 | 1.09 | 0.0109 | 0.0209 |
| chr8 | 18671926 | 18672725 | 800 | -1.71 | 0.0109 | 0.0209 |
| chr9 | 15937036 | 15937546 | 511 | 1.61 | 0.0109 | 0.021 |
| chr5 | 6614660 | 6615533 | 874 | -1.26 | 0.011 | 0.021 |
| chr19 | 47281347 | 47380158 | 98812 | -0.14 | 0.011 | 0.021 |
| chrX | 20333048 | 20333768 | 721 | 1.46 | 0.011 | 0.021 |
| chr7 | 93852109 | 93853549 | 1441 | 1.03 | 0.011 | 0.021 |
| chr11 | 74810903 | 74816575 | 5673 | -3.91 | 0.011 | 0.021 |
| chr20 | 44533330 | 44543910 | 10581 | 0.58 | 0.011 | 0.021 |
| chr1 | 160253795 | 160255373 | 1579 | 0.53 | 0.011 | 0.021 |
| chr2 | 86033028 | 86033488 | 461 | 1.72 | 0.011 | 0.0211 |
| chr2 | 235043854 | 235056990 | 13137 | -1.79 | 0.011 | 0.0211 |
| chr1 | 197138966 | 197172021 | 33056 | 0.18 | 0.011 | 0.0211 |
| chr17 | 1321334 | 1395622 | 74289 | 0.15 | 0.011 | 0.0211 |
| chr3 | 30443159 | 30444061 | 903 | 1.9 | 0.011 | 0.0211 |
| chr18 | 47439420 | 47440030 | 611 | -2.43 | 0.011 | 0.0211 |
| chr2 | 46162874 | 46167419 | 4546 | -0.32 | 0.0111 | 0.0212 |
| chr22 | 41939546 | 41941348 | 1803 | -0.9 | 0.0111 | 0.0212 |
| chr20 | 3136444 | 3155456 | 19013 | 0.31 | 0.0111 | 0.0212 |
| chr11 | 66024239 | 66057750 | 33512 | -0.28 | 0.0111 | 0.0212 |
| chr11 | 62553147 | 62573461 | 20315 | 0.24 | 0.0111 | 0.0212 |
| chr5 | 441959 | 444481 | 2523 | 0.52 | 0.0111 | 0.0212 |
| chr8 | 123029145 | 123030331 | 1187 | 1.13 | 0.0111 | 0.0212 |
| chr20 | 30423104 | 30430946 | 7843 | -0.57 | 0.0111 | 0.0212 |
| chr5 | 132009882 | 132013680 | 3799 | 0.76 | 0.0111 | 0.0212 |
| chr4 | 100551795 | 100553218 | 1424 | 1.4 | 0.0111 | 0.0212 |
| chr11 | 13094734 | 13095670 | 937 | -1.1 | 0.0111 | 0.0213 |
| chr6 | 28091720 | 28110980 | 19261 | 0.34 | 0.0111 | 0.0213 |
| chr1 | 173011930 | 173022518 | 10589 | 0.87 | 0.0111 | 0.0213 |
| chr5 | 93446333 | 93448178 | 1846 | 0.35 | 0.0111 | 0.0213 |
| chr1 | 246302160 | 246303779 | 1620 | 0.84 | 0.0111 | 0.0213 |
| chr3 | 33758263 | 33760732 | 2470 | 0.36 | 0.0111 | 0.0213 |
| chr2 | 53795672 | 53796258 | 587 | 1.6 | 0.0112 | 0.0213 |
| chr1 | 155737565 | 155743608 | 6044 | 0.44 | 0.0112 | 0.0213 |
| chr7 | 39605207 | 39606739 | 1533 | 0.53 | 0.0112 | 0.0214 |
| chr17 | 36849946 | 36937868 | 87923 | -0.14 | 0.0112 | 0.0214 |
| chr2 | 152265579 | 152316639 | 51061 | -0.21 | 0.0112 | 0.0214 |
| chr6 | 158702924 | 158707650 | 4727 | 0.25 | 0.0112 | 0.0214 |
| chr10 | 111987103 | 111987810 | 708 | 2 | 0.0112 | 0.0214 |

| | | | | | | |
|---|---|---|---|---|---|---|
| chr12 | 42137203 | 42138034 | 832 | 2.42 | 0.0112 | 0.0214 |
| chr21 | 37632407 | 37633199 | 793 | 0.88 | 0.0112 | 0.0214 |
| chr1 | 196576929 | 196578555 | 1627 | 0.93 | 0.0112 | 0.0214 |
| chr1 | 62206796 | 62213906 | 7111 | 0.3 | 0.0112 | 0.0214 |
| chr17 | 40020376 | 40087916 | 67541 | -0.14 | 0.0112 | 0.0214 |
| chr6 | 163950495 | 163969892 | 19398 | 0.31 | 0.0112 | 0.0215 |
| chr12 | 19425781 | 19428018 | 2238 | 0.51 | 0.0112 | 0.0215 |
| chr15 | 50190324 | 50190962 | 639 | 1.3 | 0.0112 | 0.0215 |
| chr11 | 74380979 | 74385990 | 5012 | -0.29 | 0.0112 | 0.0215 |
| chrX | 16761637 | 16762449 | 813 | -1.4 | 0.0112 | 0.0215 |
| chr7 | 126987658 | 126988418 | 761 | 1.58 | 0.0113 | 0.0215 |
| chr2 | 25438594 | 25439969 | 1376 | -1.1 | 0.0113 | 0.0215 |
| chr2 | 113340132 | 113343363 | 3232 | -0.28 | 0.0113 | 0.0215 |
| chr10 | 7416512 | 7421471 | 4960 | -0.56 | 0.0113 | 0.0216 |
| chr11 | 17158453 | 17158865 | 413 | 1.97 | 0.0113 | 0.0216 |
| chr20 | 43200881 | 43202237 | 1357 | -0.89 | 0.0113 | 0.0216 |
| chr1 | 64127334 | 64141091 | 13758 | -0.57 | 0.0113 | 0.0216 |
| chr16 | 1542156 | 1544945 | 2790 | 0.44 | 0.0113 | 0.0216 |
| chr5 | 176560019 | 176574499 | 14481 | 0.32 | 0.0113 | 0.0216 |
| chr15 | 81058134 | 81059065 | 932 | -1.74 | 0.0113 | 0.0216 |
| chr22 | 31159950 | 31160999 | 1050 | 1.06 | 0.0113 | 0.0217 |
| chr16 | 85355419 | 85356215 | 797 | 1.3 | 0.0113 | 0.0217 |
| chr2 | 201085241 | 201086556 | 1316 | 1.16 | 0.0113 | 0.0217 |
| chr8 | 101224612 | 101226532 | 1921 | 0.59 | 0.0114 | 0.0217 |
| chr6 | 136903911 | 136905600 | 1690 | -0.98 | 0.0114 | 0.0217 |
| chr7 | 111571254 | 111571900 | 647 | 1.54 | 0.0114 | 0.0217 |
| chr7 | 83742118 | 83743284 | 1167 | -1.43 | 0.0114 | 0.0217 |
| chr1 | 40839402 | 40851034 | 11633 | 0.59 | 0.0114 | 0.0217 |
| chr2 | 110913760 | 110914762 | 1003 | -1.55 | 0.0114 | 0.0217 |
| chr7 | 41229036 | 41230366 | 1331 | 1.1 | 0.0114 | 0.0217 |
| chr2 | 64863918 | 64882620 | 18703 | -0.17 | 0.0114 | 0.0218 |
| chr14 | 57856503 | 57869805 | 13303 | 0.35 | 0.0114 | 0.0218 |
| chr6 | 89472781 | 89473671 | 891 | -1.48 | 0.0114 | 0.0218 |
| chr18 | 48776413 | 48780666 | 4254 | -0.68 | 0.0114 | 0.0218 |
| chr1 | 211499399 | 211517700 | 18302 | 0.36 | 0.0114 | 0.0218 |
| chr1 | 196039870 | 196048750 | 8881 | 0.72 | 0.0114 | 0.0218 |
| chr5 | 77641364 | 77642053 | 690 | -2.37 | 0.0114 | 0.0218 |
| chr9 | 127001988 | 127120592 | 118605 | 0.1 | 0.0114 | 0.0218 |
| chr8 | 27218770 | 27220251 | 1482 | 0.57 | 0.0114 | 0.0218 |
| chr19 | 2840462 | 2841270 | 809 | 0.93 | 0.0115 | 0.0219 |
| chr10 | 101155284 | 101156709 | 1426 | 1.14 | 0.0115 | 0.0219 |
| chr1 | 59192536 | 59194101 | 1566 | 0.93 | 0.0115 | 0.0219 |
| chr5 | 55481260 | 55482228 | 969 | -1.46 | 0.0115 | 0.0219 |
| chr6 | 16277384 | 16278562 | 1179 | 1.25 | 0.0115 | 0.0219 |
| chr7 | 94952030 | 94960541 | 8512 | 0.82 | 0.0115 | 0.0219 |
| chr2 | 45500594 | 45501334 | 741 | 1.95 | 0.0115 | 0.0219 |
| chr1 | 236579390 | 236585652 | 6263 | 0.64 | 0.0115 | 0.0219 |
| chr14 | 31027645 | 31058668 | 31024 | 0.33 | 0.0115 | 0.022 |
| chr7 | 121078138 | 121078933 | 796 | 1.47 | 0.0115 | 0.022 |
| chr8 | 99305303 | 99307070 | 1768 | 0.46 | 0.0115 | 0.022 |
| chr19 | 44437507 | 44439982 | 2476 | 0.56 | 0.0116 | 0.022 |
| chr6 | 138708475 | 138709866 | 1392 | 1.4 | 0.0116 | 0.0221 |
| chr2 | 118572235 | 118583673 | 11439 | -0.42 | 0.0116 | 0.0221 |
| chr10 | 127567043 | 127585544 | 18502 | 0.45 | 0.0116 | 0.0221 |
| chr15 | 63139862 | 63161838 | 21977 | 0.23 | 0.0116 | 0.0221 |
| chr2 | 201794071 | 201833945 | 39875 | -0.19 | 0.0116 | 0.0221 |
| chr12 | 109956953 | 109957259 | 307 | -2.87 | 0.0116 | 0.0222 |
| chr2 | 27870375 | 27871372 | 998 | -1.32 | 0.0117 | 0.0222 |
| chr7 | 940462 | 941142 | 681 | 1.62 | 0.0117 | 0.0222 |
| chr2 | 190999933 | 191002403 | 2471 | 0.63 | 0.0117 | 0.0222 |
| chr2 | 40101960 | 40103502 | 1543 | -3.81 | 0.0117 | 0.0222 |

| | | | | | | |
|---|---|---|---|---|---|---|
| chr2 | 172271687 | 172272716 | 1030 | 1.23 | 0.0117 | 0.0222 |
| chr2 | 208435639 | 208445876 | 10238 | -0.61 | 0.0117 | 0.0222 |
| chr1 | 47104132 | 47120057 | 15926 | 0.49 | 0.0117 | 0.0222 |
| chrX | 35375538 | 35376477 | 940 | 1.13 | 0.0117 | 0.0222 |
| chr13 | 36871307 | 36872414 | 1108 | 0.87 | 0.0117 | 0.0223 |
| chr15 | 75395545 | 75418797 | 23253 | 0.21 | 0.0117 | 0.0223 |
| chr7 | 155554600 | 155564147 | 9548 | -0.7 | 0.0117 | 0.0223 |
| chr3 | 64251377 | 64252982 | 1606 | -0.81 | 0.0117 | 0.0223 |
| chr11 | 134081206 | 134094403 | 13198 | 0.65 | 0.0117 | 0.0223 |
| chr10 | 133561069 | 133563270 | 2202 | 1.21 | 0.0117 | 0.0223 |
| chr6 | 161231138 | 161231503 | 366 | 2.58 | 0.0118 | 0.0224 |
| chr16 | 10782634 | 10783345 | 712 | 2.38 | 0.0118 | 0.0224 |
| chr10 | 24771547 | 24774074 | 2528 | 0.59 | 0.0118 | 0.0224 |
| chr1 | 39512672 | 39513882 | 1211 | -1.46 | 0.0118 | 0.0224 |
| chr5 | 34466921 | 34488040 | 21120 | 1.78 | 0.0118 | 0.0224 |
| chr14 | 51207268 | 51209287 | 2020 | -0.47 | 0.0118 | 0.0224 |
| chr6 | 28350032 | 28351787 | 1756 | 0.72 | 0.0118 | 0.0224 |
| chr12 | 132239778 | 132240513 | 736 | 1.75 | 0.0118 | 0.0225 |
| chr10 | 105434977 | 105437391 | 2415 | 0.9 | 0.0118 | 0.0225 |
| chr17 | 40457762 | 40584443 | 126682 | -0.11 | 0.0118 | 0.0225 |
| chr1 | 94019962 | 94170451 | 150490 | 0.1 | 0.0118 | 0.0225 |
| chr7 | 141824810 | 141845883 | 21074 | 0.41 | 0.0118 | 0.0225 |
| chr9 | 126773074 | 126774335 | 1262 | -1.04 | 0.0118 | 0.0225 |
| chr9 | 135198442 | 135200099 | 1658 | 1.02 | 0.0118 | 0.0225 |
| chr21 | 39355760 | 39367231 | 11472 | 0.57 | 0.0118 | 0.0225 |
| chr16 | 10406277 | 10407275 | 999 | 1.41 | 0.0119 | 0.0226 |
| chr15 | 50502125 | 50503571 | 1447 | 0.75 | 0.0119 | 0.0226 |
| chr9 | 92039849 | 92041123 | 1275 | -1.7 | 0.0119 | 0.0226 |
| chr10 | 124913783 | 124923348 | 9566 | 0.39 | 0.0119 | 0.0226 |
| chr2 | 85724877 | 85789548 | 64672 | -0.13 | 0.0119 | 0.0226 |
| chr22 | 47589208 | 47589766 | 559 | 1.95 | 0.0119 | 0.0226 |
| chr7 | 101945214 | 101960417 | 15204 | -0.63 | 0.0119 | 0.0227 |
| chr4 | 155547868 | 155548496 | 629 | 2.42 | 0.0119 | 0.0227 |
| chr2 | 182675280 | 182677945 | 2666 | 0.37 | 0.0119 | 0.0227 |
| chr15 | 39592890 | 39593902 | 1013 | 1.27 | 0.0119 | 0.0227 |
| chr8 | 27881746 | 27882178 | 433 | -2.58 | 0.0119 | 0.0227 |
| chr22 | 30812272 | 30822846 | 10575 | 0.25 | 0.0119 | 0.0227 |
| chr14 | 63013331 | 63013999 | 669 | -1.82 | 0.012 | 0.0227 |
| chr3 | 119513491 | 119514922 | 1432 | -1.05 | 0.012 | 0.0227 |
| chr12 | 114348573 | 114349325 | 753 | -1.75 | 0.012 | 0.0227 |
| chr5 | 67977990 | 67987754 | 9765 | 3.44 | 0.012 | 0.0227 |
| chr10 | 79567673 | 79709970 | 142298 | -0.13 | 0.012 | 0.0227 |
| chr13 | 81877584 | 81879084 | 1501 | 1.19 | 0.012 | 0.0227 |
| chr6 | 127744699 | 127745650 | 952 | 1.4 | 0.012 | 0.0227 |
| chr14 | 90085242 | 90085656 | 415 | -1.81 | 0.012 | 0.0227 |
| chr7 | 66204267 | 66207311 | 3045 | 0.44 | 0.012 | 0.0227 |
| chr2 | 55999048 | 56010781 | 11734 | 0.25 | 0.012 | 0.0228 |
| chr5 | 110073870 | 110075794 | 1925 | 0.51 | 0.012 | 0.0228 |
| chr1 | 31624779 | 31644556 | 19778 | 0.3 | 0.012 | 0.0228 |
| chr6 | 160409934 | 160410689 | 756 | -1.19 | 0.012 | 0.0228 |
| chr7 | 101194289 | 101194801 | 513 | -2.28 | 0.012 | 0.0228 |
| chr9 | 137323882 | 137349008 | 25127 | -0.47 | 0.012 | 0.0228 |
| chrX | 46590476 | 46590835 | 360 | 1.83 | 0.012 | 0.0228 |
| chr19 | 51059156 | 51060450 | 1295 | -1.14 | 0.0121 | 0.0229 |
| chr7 | 14778932 | 14779504 | 573 | 1.55 | 0.0121 | 0.0229 |
| chr2 | 43821348 | 43824327 | 2980 | -0.31 | 0.0121 | 0.0229 |
| chr16 | 1659432 | 1710650 | 51219 | -0.24 | 0.0121 | 0.0229 |
| chr1 | 10346870 | 10347955 | 1086 | -1.64 | 0.0121 | 0.0229 |
| chr15 | 63219056 | 63233628 | 14573 | -0.64 | 0.0121 | 0.0229 |
| chr2 | 159496573 | 159497430 | 858 | -1.85 | 0.0121 | 0.0229 |
| chr1 | 203296069 | 203297858 | 1790 | 0.57 | 0.0121 | 0.0229 |

| | | | | | | |
|---|---|---|---|---|---|---|
| chr12 | 90970464 | 90971152 | 689 | -2.48 | 0.0121 | 0.0229 |
| chr8 | 101454882 | 101456032 | 1151 | 1.16 | 0.0121 | 0.0229 |
| chr1 | 198242671 | 198244001 | 1331 | 1.04 | 0.0121 | 0.0229 |
| chr4 | 6344764 | 6346490 | 1727 | -0.87 | 0.0121 | 0.0229 |
| chr15 | 65093888 | 65095012 | 1125 | 1.37 | 0.0121 | 0.023 |
| chr16 | 8890658 | 8894773 | 4116 | 0.29 | 0.0121 | 0.023 |
| chr1 | 29172272 | 29172975 | 704 | -1.27 | 0.0121 | 0.023 |
| chr5 | 72084067 | 72085715 | 1649 | 1.02 | 0.0121 | 0.023 |
| chr8 | 41009354 | 41019927 | 10574 | 0.93 | 0.0121 | 0.023 |
| chr16 | 86888069 | 86889524 | 1456 | 0.64 | 0.0121 | 0.023 |
| chr6 | 170090433 | 170104016 | 13584 | 0.46 | 0.0121 | 0.023 |
| chr17 | 58308803 | 58309820 | 1018 | -1.22 | 0.0122 | 0.023 |
| chr2 | 25116091 | 25143802 | 27712 | -0.54 | 0.0122 | 0.023 |
| chr4 | 54458311 | 54459758 | 1448 | 0.62 | 0.0122 | 0.0231 |
| chr6 | 149609474 | 149610325 | 852 | -1.84 | 0.0122 | 0.0231 |
| chr2 | 190526192 | 190540334 | 14143 | -0.39 | 0.0122 | 0.0231 |
| chr4 | 146074020 | 146074499 | 480 | 1.96 | 0.0122 | 0.0231 |
| chr3 | 182798921 | 182800577 | 1657 | -0.87 | 0.0122 | 0.0231 |
| chr13 | 35125449 | 35125838 | 390 | 2.4 | 0.0122 | 0.0231 |
| chr6 | 155332845 | 155333628 | 784 | 1.61 | 0.0122 | 0.0231 |
| chr1 | 186723487 | 186733674 | 10188 | 0.37 | 0.0122 | 0.0231 |
| chr2 | 204315478 | 204316635 | 1158 | -1.13 | 0.0123 | 0.0232 |
| chr20 | 18205512 | 18206643 | 1132 | 0.81 | 0.0123 | 0.0233 |
| chr16 | 28830979 | 28837235 | 6257 | 0.32 | 0.0123 | 0.0233 |
| chr2 | 26044942 | 26076388 | 31447 | -0.28 | 0.0123 | 0.0233 |
| chr1 | 84830911 | 84832268 | 1358 | 0.85 | 0.0123 | 0.0233 |
| chr19 | 49284264 | 49323747 | 39484 | -0.27 | 0.0123 | 0.0233 |
| chr12 | 46367033 | 46367793 | 761 | 1.51 | 0.0123 | 0.0233 |
| chr1 | 19529327 | 19538482 | 9156 | 0.3 | 0.0123 | 0.0233 |
| chr8 | 92034963 | 92035674 | 712 | -1.47 | 0.0123 | 0.0233 |
| chr8 | 144392975 | 144394036 | 1062 | 1.54 | 0.0123 | 0.0233 |
| chr1 | 178816119 | 178817266 | 1148 | 1.2 | 0.0123 | 0.0233 |
| chr6 | 84569259 | 84578983 | 9725 | -0.32 | 0.0123 | 0.0233 |
| chr13 | 109988967 | 109997808 | 8842 | -0.59 | 0.0123 | 0.0233 |
| chr1 | 18944829 | 18958480 | 13652 | 1.25 | 0.0123 | 0.0233 |
| chr22 | 37850414 | 37853444 | 3031 | -0.48 | 0.0123 | 0.0233 |
| chr2 | 217505200 | 217505755 | 556 | -2.43 | 0.0123 | 0.0233 |
| chr9 | 109975794 | 109976835 | 1042 | -1.14 | 0.0123 | 0.0234 |
| chr6 | 44185406 | 44233839 | 48434 | 0.18 | 0.0124 | 0.0234 |
| chr6 | 134351147 | 134351765 | 619 | -1.66 | 0.0124 | 0.0234 |
| chr14 | 91065124 | 91066820 | 1697 | -0.66 | 0.0124 | 0.0234 |
| chr14 | 91263796 | 91264604 | 809 | -1.62 | 0.0124 | 0.0234 |
| chr18 | 77889853 | 77890767 | 915 | -1.23 | 0.0124 | 0.0234 |
| chr5 | 155559897 | 155583690 | 23794 | 0.18 | 0.0124 | 0.0234 |
| chr6 | 7388823 | 7402075 | 13253 | 0.39 | 0.0124 | 0.0234 |
| chr22 | 35587060 | 35588247 | 1188 | 1.51 | 0.0124 | 0.0234 |
| chr2 | 70823368 | 70824635 | 1268 | -1.1 | 0.0124 | 0.0234 |
| chr1 | 52725769 | 52726365 | 597 | -2 | 0.0124 | 0.0235 |
| chr2 | 216973291 | 216987544 | 14254 | -0.25 | 0.0124 | 0.0235 |
| chr12 | 77243640 | 77245048 | 1409 | -1.54 | 0.0124 | 0.0235 |
| chr14 | 45660436 | 45662809 | 2374 | -1.02 | 0.0124 | 0.0235 |
| chr17 | 16206174 | 16210812 | 4639 | -1.09 | 0.0125 | 0.0235 |
| chr14 | 85631686 | 85632077 | 392 | 2.44 | 0.0125 | 0.0235 |
| chr11 | 73516666 | 73517239 | 574 | -2.16 | 0.0125 | 0.0236 |
| chr12 | 123505550 | 123506414 | 865 | -1.58 | 0.0125 | 0.0236 |
| chr15 | 101741783 | 101742228 | 446 | 1.54 | 0.0125 | 0.0236 |
| chr3 | 136648229 | 136657830 | 9602 | 0.43 | 0.0125 | 0.0236 |
| chr6 | 128784814 | 128786300 | 1487 | 1.84 | 0.0125 | 0.0236 |
| chr6 | 41487149 | 41487948 | 800 | 1.06 | 0.0125 | 0.0236 |
| chr1 | 6595270 | 6615104 | 19835 | 0.54 | 0.0125 | 0.0237 |
| chr7 | 29962894 | 29993880 | 30987 | -0.27 | 0.0125 | 0.0237 |

| | | | | | | |
|---|---|---|---|---|---|---|
| chr14 | 23502870 | 23505254 | 2385 | -0.29 | 0.0125 | 0.0237 |
| chr14 | 94453105 | 94454023 | 919 | -1.29 | 0.0125 | 0.0237 |
| chr5 | 131825357 | 131839018 | 13662 | 0.18 | 0.0126 | 0.0237 |
| chr10 | 38298745 | 38301130 | 2386 | 0.37 | 0.0126 | 0.0237 |
| chr12 | 124613494 | 124616833 | 3340 | 0.45 | 0.0126 | 0.0237 |
| chr2 | 209218451 | 209218868 | 418 | -1.83 | 0.0126 | 0.0237 |
| chr18 | 10057515 | 10058429 | 915 | -1.54 | 0.0126 | 0.0238 |
| chr5 | 33088278 | 33089034 | 757 | -1.92 | 0.0126 | 0.0238 |
| chr5 | 135538225 | 135538796 | 572 | 1.92 | 0.0126 | 0.0238 |
| chr1 | 15067926 | 15070065 | 2140 | -0.69 | 0.0126 | 0.0238 |
| chr3 | 127307768 | 127327257 | 19490 | -0.27 | 0.0126 | 0.0238 |
| chr7 | 12866770 | 12867276 | 507 | 2.05 | 0.0126 | 0.0238 |
| chr9 | 129182787 | 129183583 | 797 | 1.56 | 0.0126 | 0.0238 |
| chr6 | 25911256 | 25911621 | 366 | 2.56 | 0.0126 | 0.0238 |
| chr13 | 50017329 | 50019805 | 2477 | 0.58 | 0.0126 | 0.0239 |
| chr11 | 108756598 | 108778537 | 21940 | -0.35 | 0.0127 | 0.0239 |
| chr22 | 41883225 | 41883667 | 443 | -2.08 | 0.0127 | 0.0239 |
| chr2 | 19168456 | 19173806 | 5351 | -0.36 | 0.0127 | 0.0239 |
| chr2 | 146557375 | 146558795 | 1421 | 0.94 | 0.0127 | 0.0239 |
| chr4 | 87259804 | 87260686 | 883 | -1.52 | 0.0127 | 0.0239 |
| chr10 | 104448685 | 104449495 | 811 | 1.77 | 0.0127 | 0.0239 |
| chr4 | 148604406 | 148615313 | 10908 | 0.4 | 0.0127 | 0.0239 |
| chr20 | 1396447 | 1404997 | 8551 | -1.04 | 0.0127 | 0.0239 |
| chr1 | 110434556 | 110438861 | 4306 | -0.66 | 0.0127 | 0.0239 |
| chr7 | 20854567 | 20855548 | 982 | 1.25 | 0.0127 | 0.0239 |
| chr1 | 200623252 | 200640793 | 17542 | 0.2 | 0.0127 | 0.0239 |
| chr17 | 18084804 | 18197132 | 112329 | -0.13 | 0.0127 | 0.024 |
| chr3 | 12944887 | 12946652 | 1766 | -1.24 | 0.0127 | 0.024 |
| chr19 | 36359222 | 36361466 | 2245 | -0.67 | 0.0127 | 0.024 |
| chrX | 103400577 | 103412511 | 11935 | 0.36 | 0.0127 | 0.024 |
| chr18 | 21753684 | 21754737 | 1054 | -1.29 | 0.0127 | 0.024 |
| chr17 | 79667544 | 79689683 | 22140 | -0.29 | 0.0127 | 0.024 |
| chr18 | 51780505 | 51781523 | 1019 | 1.58 | 0.0127 | 0.024 |
| chr16 | 31453685 | 31496741 | 43057 | -0.24 | 0.0128 | 0.024 |
| chrX | 100871852 | 100878904 | 7053 | 0.51 | 0.0128 | 0.024 |
| chr9 | 20811343 | 20812713 | 1371 | -0.99 | 0.0128 | 0.024 |
| chr13 | 80880054 | 80880758 | 705 | 1.89 | 0.0128 | 0.0241 |
| chr4 | 177240504 | 177242976 | 2473 | 0.54 | 0.0128 | 0.0241 |
| chr6 | 9710166 | 9712059 | 1894 | -0.7 | 0.0128 | 0.0242 |
| chr8 | 56756328 | 56761207 | 4880 | 0.48 | 0.0128 | 0.0242 |
| chr10 | 90187650 | 90189285 | 1636 | 1.05 | 0.0128 | 0.0242 |
| chr7 | 67502425 | 67511841 | 9417 | -0.24 | 0.0128 | 0.0242 |
| chr5 | 33457734 | 33473445 | 15712 | -0.36 | 0.0128 | 0.0242 |
| chr14 | 43320952 | 43321411 | 460 | 2.4 | 0.0129 | 0.0242 |
| chr4 | 14720304 | 14729279 | 8976 | 0.42 | 0.0129 | 0.0242 |
| chr20 | 9146624 | 9147453 | 830 | 0.96 | 0.0129 | 0.0242 |
| chr6 | 20445937 | 20447789 | 1853 | -1.16 | 0.0129 | 0.0242 |
| chr17 | 60023671 | 60025169 | 1499 | 1.22 | 0.0129 | 0.0242 |
| chr3 | 188035594 | 188036065 | 472 | -2.16 | 0.0129 | 0.0242 |
| chr9 | 36550348 | 36551547 | 1200 | -1.26 | 0.0129 | 0.0243 |
| chr3 | 15056248 | 15107243 | 50996 | -0.24 | 0.0129 | 0.0243 |
| chr11 | 108829492 | 108830289 | 798 | -1.39 | 0.0129 | 0.0243 |
| chr1 | 183430238 | 183463337 | 33100 | 0.2 | 0.0129 | 0.0243 |
| chr12 | 71003485 | 71004384 | 900 | 1.01 | 0.0129 | 0.0243 |
| chrX | 49759414 | 49761001 | 1588 | 1.12 | 0.0129 | 0.0243 |
| chrX | 46617317 | 46653438 | 36122 | 0.22 | 0.0129 | 0.0243 |
| chr11 | 10476357 | 10478311 | 1955 | -0.56 | 0.0129 | 0.0243 |
| chr8 | 145132004 | 145160593 | 28590 | 0.21 | 0.0129 | 0.0244 |
| chr15 | 99751573 | 99752442 | 870 | -1.59 | 0.0129 | 0.0244 |
| chr18 | 8048155 | 8052303 | 4149 | 0.46 | 0.013 | 0.0244 |
| chr5 | 148982516 | 148989111 | 6596 | 0.42 | 0.013 | 0.0244 |

| | | | | | | |
|---|---|---|---|---|---|---|
| chr11 | 36452974 | 36454295 | 1322 | -0.94 | 0.013 | 0.0244 |
| chr11 | 77731235 | 77773233 | 41999 | 0.23 | 0.013 | 0.0244 |
| chr14 | 81975911 | 82001113 | 25203 | 0.26 | 0.013 | 0.0245 |
| chr10 | 35414746 | 35421789 | 7044 | 0.25 | 0.013 | 0.0245 |
| chrX | 79590242 | 79591337 | 1096 | 0.79 | 0.013 | 0.0245 |
| chr18 | 2746041 | 2746703 | 663 | -1.74 | 0.013 | 0.0245 |
| chr1 | 117209525 | 117211232 | 1708 | 0.7 | 0.0131 | 0.0246 |
| chr6 | 781347 | 781996 | 650 | -2.13 | 0.0131 | 0.0246 |
| chr17 | 17154941 | 17156533 | 1593 | -1.03 | 0.0131 | 0.0246 |
| chr4 | 120549502 | 120550638 | 1137 | 1.49 | 0.0131 | 0.0246 |
| chr11 | 56213513 | 56214064 | 552 | -2.17 | 0.0131 | 0.0246 |
| chr1 | 214550905 | 214551802 | 898 | -1.55 | 0.0131 | 0.0246 |
| chr2 | 137003401 | 137004235 | 835 | -1.46 | 0.0131 | 0.0246 |
| chr3 | 4818205 | 4819600 | 1396 | 1.33 | 0.0131 | 0.0246 |
| chr15 | 84048034 | 84049577 | 1544 | 0.8 | 0.0131 | 0.0246 |
| chr4 | 152172893 | 152180049 | 7157 | 0.72 | 0.0131 | 0.0246 |
| chr13 | 67597231 | 67598542 | 1312 | 0.94 | 0.0131 | 0.0247 |
| chr2 | 63192706 | 63193635 | 930 | -2.25 | 0.0131 | 0.0247 |
| chr14 | 28937166 | 28938965 | 1800 | -0.55 | 0.0131 | 0.0247 |
| chrX | 106953360 | 106980806 | 27447 | 0.22 | 0.0131 | 0.0247 |
| chr1 | 204592388 | 204592743 | 356 | -2.57 | 0.0132 | 0.0247 |
| chr5 | 67614552 | 67616830 | 2279 | 0.49 | 0.0132 | 0.0247 |
| chr10 | 31435634 | 31436582 | 949 | -1.49 | 0.0132 | 0.0247 |
| chr2 | 218766804 | 218768322 | 1519 | 1.04 | 0.0132 | 0.0247 |
| chr3 | 160964948 | 160967654 | 2707 | 0.93 | 0.0132 | 0.0248 |
| chr21 | 35573897 | 35574665 | 769 | -1.61 | 0.0132 | 0.0248 |
| chr7 | 75714398 | 75717227 | 2830 | 0.7 | 0.0132 | 0.0248 |
| chr6 | 119255475 | 119256845 | 1371 | 0.78 | 0.0132 | 0.0248 |
| chr11 | 121526415 | 121527353 | 939 | 0.88 | 0.0132 | 0.0248 |
| chr6 | 107348923 | 107350215 | 1293 | 0.99 | 0.0132 | 0.0248 |
| chr5 | 139598773 | 139599884 | 1112 | 2.04 | 0.0132 | 0.0248 |
| chr6 | 27196862 | 27220323 | 23462 | 0.34 | 0.0132 | 0.0248 |
| chr6 | 24329421 | 24343272 | 13852 | -0.33 | 0.0132 | 0.0248 |
| chr3 | 123540640 | 123541906 | 1267 | -1.37 | 0.0132 | 0.0248 |
| chr8 | 121068659 | 121069729 | 1071 | -1.11 | 0.0132 | 0.0249 |
| chr15 | 25653172 | 25654454 | 1283 | 1.67 | 0.0133 | 0.0249 |
| chr1 | 44114095 | 44117031 | 2937 | 0.51 | 0.0133 | 0.0249 |
| chr8 | 63014986 | 63015793 | 808 | 1.54 | 0.0133 | 0.0249 |
| chr2 | 166477626 | 166478627 | 1002 | 0.89 | 0.0133 | 0.0249 |
| chr21 | 38935284 | 38938647 | 3364 | -0.29 | 0.0133 | 0.0249 |
| chr6 | 149678684 | 149679679 | 996 | -1.03 | 0.0133 | 0.0249 |
| chr1 | 178269325 | 178269798 | 474 | 2.36 | 0.0133 | 0.0249 |
| chr21 | 36280212 | 36281200 | 989 | 1.53 | 0.0133 | 0.0249 |
| chr2 | 110311847 | 110322681 | 10835 | -0.26 | 0.0133 | 0.0249 |
| chr3 | 136265840 | 136266510 | 671 | 2.36 | 0.0133 | 0.0249 |
| chr3 | 100710797 | 100711842 | 1046 | 1.77 | 0.0133 | 0.0249 |
| chr7 | 154788779 | 154796420 | 7642 | 0.36 | 0.0133 | 0.025 |
| chr19 | 14543752 | 14546595 | 2844 | 0.68 | 0.0133 | 0.025 |
| chr3 | 4382110 | 4382791 | 682 | -1.32 | 0.0133 | 0.025 |
| chr1 | 234528447 | 234549137 | 20691 | -0.43 | 0.0133 | 0.025 |
| chr7 | 6143141 | 6147033 | 3893 | 0.32 | 0.0133 | 0.025 |
| chr8 | 48960246 | 48967313 | 7068 | 0.66 | 0.0133 | 0.025 |
| chr11 | 105114940 | 105115657 | 718 | -1.93 | 0.0133 | 0.025 |
| chr5 | 71642083 | 71644840 | 2758 | -0.56 | 0.0133 | 0.025 |
| chr8 | 99053852 | 99059279 | 5428 | 0.34 | 0.0133 | 0.025 |
| chr10 | 94579701 | 94584041 | 4341 | 0.21 | 0.0134 | 0.025 |
| chr1 | 172598609 | 172624538 | 25930 | 0.38 | 0.0134 | 0.025 |
| chr8 | 22525194 | 22531591 | 6398 | 0.27 | 0.0134 | 0.0251 |
| chr2 | 203358449 | 203359962 | 1514 | 0.82 | 0.0134 | 0.0251 |
| chr6 | 107266942 | 107269540 | 2599 | -0.63 | 0.0134 | 0.0251 |
| chr2 | 170979037 | 170980849 | 1813 | 0.85 | 0.0134 | 0.0251 |

| | | | | | | |
|---|---|---|---|---|---|---|
| chr6 | 107139393 | 107151063 | 11671 | 4.03 | 0.0134 | 0.0251 |
| chr17 | 78565757 | 78595072 | 29316 | -0.31 | 0.0134 | 0.0251 |
| chr2 | 63961204 | 63968617 | 7414 | -0.63 | 0.0135 | 0.0252 |
| chr10 | 22291109 | 22293406 | 2298 | 0.41 | 0.0135 | 0.0252 |
| chr18 | 32869090 | 32871210 | 2121 | 0.58 | 0.0135 | 0.0252 |
| chr11 | 31677075 | 31677933 | 859 | -1.32 | 0.0135 | 0.0252 |
| chr8 | 95869504 | 95876041 | 6538 | 0.52 | 0.0135 | 0.0253 |
| chr10 | 50737105 | 50748148 | 11044 | 0.49 | 0.0135 | 0.0253 |
| chr20 | 53710931 | 53711374 | 444 | -2.37 | 0.0135 | 0.0253 |
| chr13 | 27595473 | 27619688 | 24216 | -0.26 | 0.0135 | 0.0253 |
| chr16 | 84913292 | 84916311 | 3020 | 0.68 | 0.0135 | 0.0254 |
| chr12 | 74285692 | 74296788 | 11097 | 2.04 | 0.0136 | 0.0254 |
| chr5 | 64087697 | 64088755 | 1059 | 1.25 | 0.0136 | 0.0255 |
| chr2 | 214148166 | 214150514 | 2349 | -0.37 | 0.0136 | 0.0255 |
| chr17 | 67322384 | 67324126 | 1743 | 0.45 | 0.0136 | 0.0255 |
| chr1 | 234389892 | 234391121 | 1230 | 1.92 | 0.0136 | 0.0255 |
| chr12 | 106638225 | 106643123 | 4899 | 0.38 | 0.0136 | 0.0255 |
| chr17 | 19726236 | 19727122 | 887 | -1.08 | 0.0136 | 0.0255 |
| chr1 | 100816032 | 100819016 | 2985 | -0.37 | 0.0137 | 0.0256 |
| chr3 | 38870703 | 38871172 | 470 | -2.36 | 0.0137 | 0.0256 |
| chr1 | 231156298 | 231156895 | 598 | -1.62 | 0.0137 | 0.0256 |
| chr1 | 22429487 | 22429996 | 510 | 1.92 | 0.0137 | 0.0256 |
| chr3 | 195105201 | 195106553 | 1353 | 1.06 | 0.0137 | 0.0256 |
| chr6 | 82543356 | 82549053 | 5698 | 0.46 | 0.0137 | 0.0256 |
| chr18 | 6414642 | 6415151 | 510 | 2.2 | 0.0137 | 0.0256 |
| chr5 | 141162547 | 141163098 | 552 | -2.32 | 0.0137 | 0.0256 |
| chr6 | 144978328 | 144983720 | 5393 | 0.58 | 0.0137 | 0.0256 |
| chr12 | 54321652 | 54343122 | 21471 | 4.11 | 0.0137 | 0.0257 |
| chr14 | 65557938 | 65570907 | 12970 | 0.38 | 0.0137 | 0.0257 |
| chr19 | 8738805 | 8751630 | 12826 | -0.23 | 0.0137 | 0.0257 |
| chr1 | 233376294 | 233431859 | 55566 | -0.17 | 0.0137 | 0.0257 |
| chr1 | 27808711 | 27817826 | 9116 | 0.31 | 0.0138 | 0.0257 |
| chr16 | 20837158 | 20838705 | 1548 | -0.98 | 0.0138 | 0.0257 |
| chr9 | 80408722 | 80456197 | 47476 | -0.24 | 0.0138 | 0.0257 |
| chr10 | 35077489 | 35105056 | 27568 | -0.26 | 0.0138 | 0.0257 |
| chr6 | 119213830 | 119225163 | 11334 | 0.21 | 0.0138 | 0.0257 |
| chr1 | 232101234 | 232101690 | 457 | -2.31 | 0.0138 | 0.0258 |
| chr16 | 72588858 | 72589804 | 947 | -1.75 | 0.0138 | 0.0258 |
| chr14 | 35654608 | 35685137 | 30530 | -0.33 | 0.0138 | 0.0258 |
| chr18 | 69264685 | 69265155 | 471 | 1.77 | 0.0138 | 0.0258 |
| chr13 | 46592201 | 46593351 | 1151 | 1.23 | 0.0138 | 0.0258 |
| chr5 | 38782638 | 38783714 | 1077 | 1.2 | 0.0138 | 0.0259 |
| chr10 | 64323934 | 64324703 | 770 | -1.18 | 0.0139 | 0.0259 |
| chr5 | 89768512 | 89776864 | 8353 | -0.43 | 0.0139 | 0.0259 |
| chr1 | 163316664 | 163317593 | 930 | 1.65 | 0.0139 | 0.0259 |
| chr5 | 146130255 | 146130754 | 500 | 2.13 | 0.0139 | 0.0259 |
| chr17 | 58867345 | 58869074 | 1730 | 3.77 | 0.0139 | 0.0259 |
| chr4 | 4444827 | 4446381 | 1555 | -0.7 | 0.0139 | 0.026 |
| chr13 | 80509345 | 80512541 | 3197 | 0.28 | 0.0139 | 0.026 |
| chr4 | 178362138 | 178364721 | 2584 | 0.43 | 0.0139 | 0.026 |
| chr2 | 187304029 | 187305852 | 1824 | 0.76 | 0.0139 | 0.026 |
| chr6 | 125566391 | 125568198 | 1808 | 1.48 | 0.0139 | 0.026 |
| chr2 | 239095716 | 239097102 | 1387 | -1.39 | 0.014 | 0.0261 |
| chr4 | 95283945 | 95284961 | 1017 | 1.4 | 0.014 | 0.0261 |
| chr9 | 136908167 | 136932883 | 24717 | -0.29 | 0.014 | 0.0261 |
| chr6 | 23424847 | 23425348 | 502 | -2.37 | 0.014 | 0.0261 |
| chr1 | 22755140 | 22756041 | 902 | 1.11 | 0.014 | 0.0261 |
| chr9 | 90947864 | 90948951 | 1088 | 1.6 | 0.014 | 0.0262 |
| chr2 | 128012825 | 128013529 | 705 | -1.66 | 0.014 | 0.0262 |
| chr4 | 129300881 | 129309369 | 8489 | -0.33 | 0.014 | 0.0262 |
| chr6 | 157006751 | 157009408 | 2658 | 0.41 | 0.014 | 0.0262 |

| | | | | | | |
|---|---|---|---|---|---|---|
| chr6 | 161693833 | 161695936 | 2104 | 0.39 | 0.014 | 0.0262 |
| chr2 | 30669113 | 30688779 | 19667 | 0.24 | 0.014 | 0.0262 |
| chr21 | 38885991 | 38887363 | 1373 | 0.82 | 0.014 | 0.0262 |
| chr10 | 102183933 | 102189468 | 5536 | 0.75 | 0.014 | 0.0262 |
| chr15 | 49911865 | 49920202 | 8338 | 0.33 | 0.0141 | 0.0263 |
| chr10 | 24969478 | 24970426 | 949 | 1.1 | 0.0141 | 0.0263 |
| chr8 | 55466765 | 55467419 | 655 | 1.46 | 0.0141 | 0.0263 |
| chr7 | 150095731 | 150104486 | 8756 | -0.49 | 0.0141 | 0.0264 |
| chrX | 6966959 | 6981102 | 14144 | 0.56 | 0.0141 | 0.0264 |
| chr12 | 46320051 | 46334864 | 14814 | 0.41 | 0.0141 | 0.0264 |
| chr3 | 28337153 | 28338125 | 973 | 1.43 | 0.0142 | 0.0264 |
| chr4 | 143738677 | 143739443 | 767 | 1.83 | 0.0142 | 0.0264 |
| chr22 | 44756319 | 44760763 | 4445 | -0.42 | 0.0142 | 0.0264 |
| chr3 | 43662743 | 43688386 | 25644 | -0.47 | 0.0142 | 0.0264 |
| chr14 | 20880227 | 20884833 | 4607 | 0.38 | 0.0142 | 0.0265 |
| chr2 | 135674668 | 135702510 | 27843 | -0.18 | 0.0142 | 0.0265 |
| chr7 | 50598518 | 50604350 | 5833 | 0.24 | 0.0142 | 0.0265 |
| chr1 | 32253340 | 32254898 | 1559 | -0.49 | 0.0142 | 0.0265 |
| chr9 | 97986944 | 97988441 | 1498 | 0.98 | 0.0143 | 0.0266 |
| chr22 | 24869860 | 24870487 | 628 | -1.33 | 0.0143 | 0.0266 |
| chr5 | 140069429 | 140106669 | 37241 | 0.17 | 0.0143 | 0.0266 |
| chrX | 72433938 | 72435394 | 1457 | 0.87 | 0.0143 | 0.0266 |
| chr8 | 21887665 | 21889655 | 1991 | 0.61 | 0.0143 | 0.0266 |
| chr17 | 6135242 | 6136137 | 896 | -1.58 | 0.0143 | 0.0266 |
| chr6 | 75952522 | 75954542 | 2021 | 0.35 | 0.0143 | 0.0266 |
| chr21 | 40554274 | 40556764 | 2491 | 0.42 | 0.0143 | 0.0266 |
| chr14 | 64155965 | 64169474 | 13510 | 0.5 | 0.0143 | 0.0266 |
| chr15 | 101406525 | 101420337 | 13813 | -3.95 | 0.0143 | 0.0266 |
| chr15 | 91313533 | 91314271 | 739 | 1.16 | 0.0143 | 0.0266 |
| chr3 | 23985815 | 24005904 | 20090 | 0.27 | 0.0143 | 0.0267 |
| chr4 | 148538008 | 148548634 | 10627 | 0.47 | 0.0143 | 0.0267 |
| chr5 | 74631258 | 74634321 | 3064 | 0.35 | 0.0143 | 0.0267 |
| chr10 | 114657846 | 114677360 | 19515 | 0.2 | 0.0143 | 0.0267 |
| chr14 | 105912308 | 105967433 | 55126 | -0.14 | 0.0144 | 0.0268 |
| chr9 | 37280704 | 37294334 | 13631 | 0.25 | 0.0144 | 0.0268 |
| chr6 | 43543184 | 43545697 | 2514 | 0.66 | 0.0144 | 0.0268 |
| chr12 | 95155615 | 95156725 | 1111 | 1.76 | 0.0144 | 0.0268 |
| chr21 | 19190731 | 19192672 | 1942 | 0.56 | 0.0144 | 0.0268 |
| chr17 | 80691989 | 80694147 | 2159 | 0.65 | 0.0144 | 0.0268 |
| chr20 | 25095074 | 25101327 | 6254 | 0.3 | 0.0144 | 0.0268 |
| chr16 | 68157704 | 68181806 | 24103 | -0.36 | 0.0144 | 0.0268 |
| chr14 | 74034463 | 74037689 | 3227 | 0.37 | 0.0144 | 0.0268 |
| chr8 | 54171323 | 54180337 | 9015 | 4.19 | 0.0144 | 0.0269 |
| chr1 | 24151154 | 24152583 | 1430 | 0.66 | 0.0145 | 0.0269 |
| chr1 | 29556811 | 29566517 | 9707 | 0.54 | 0.0145 | 0.0269 |
| chr8 | 144513098 | 144513958 | 861 | 0.95 | 0.0145 | 0.0269 |
| chr7 | 116501890 | 116504543 | 2654 | 0.41 | 0.0145 | 0.0269 |
| chr11 | 28347645 | 28348187 | 543 | 1.74 | 0.0145 | 0.027 |
| chr17 | 79045257 | 79046917 | 1661 | 1.04 | 0.0145 | 0.027 |
| chr11 | 73467232 | 73473117 | 5886 | 0.41 | 0.0145 | 0.027 |
| chr14 | 91120782 | 91121746 | 965 | -1.39 | 0.0145 | 0.027 |
| chr2 | 242673486 | 242675274 | 1789 | -0.71 | 0.0145 | 0.027 |
| chr8 | 17501538 | 17534435 | 32898 | 0.26 | 0.0145 | 0.027 |
| chr9 | 5436787 | 5451042 | 14256 | 0.39 | 0.0145 | 0.027 |
| chr6 | 24774718 | 24780514 | 5797 | 0.53 | 0.0145 | 0.027 |
| chr15 | 51717200 | 51718221 | 1022 | 1.99 | 0.0145 | 0.027 |
| chr12 | 42518217 | 42519130 | 914 | 1.47 | 0.0145 | 0.0271 |
| chr2 | 61394183 | 61409630 | 15448 | -0.3 | 0.0145 | 0.0271 |
| chr16 | 30456058 | 30457852 | 1795 | 0.52 | 0.0146 | 0.0271 |
| chr6 | 107306034 | 107307419 | 1386 | 1.12 | 0.0146 | 0.0271 |
| chr6 | 86157421 | 86175315 | 17895 | 0.22 | 0.0146 | 0.0271 |

| | | | | | | |
|---|---|---|---|---|---|---|
| chr19 | 10735771 | 10736704 | 934 | 1.2 | 0.0146 | 0.0271 |
| chr17 | 79092397 | 79094316 | 1920 | 0.71 | 0.0146 | 0.0271 |
| chr18 | 39534636 | 39536039 | 1404 | 0.55 | 0.0146 | 0.0271 |
| chr1 | 220473536 | 220474519 | 984 | 1.28 | 0.0146 | 0.0272 |
| chr9 | 19361751 | 19381448 | 19698 | 0.24 | 0.0146 | 0.0272 |
| chr13 | 21957085 | 21958274 | 1190 | 1.38 | 0.0146 | 0.0272 |
| chr7 | 16748206 | 16749417 | 1212 | -1.03 | 0.0146 | 0.0272 |
| chr7 | 148460328 | 148471053 | 10726 | 0.48 | 0.0146 | 0.0272 |
| chr8 | 144449787 | 144452096 | 2310 | -0.66 | 0.0146 | 0.0272 |
| chr15 | 40508939 | 40509521 | 583 | 2.3 | 0.0147 | 0.0272 |
| chr5 | 76925810 | 76927201 | 1392 | -1.08 | 0.0147 | 0.0273 |
| chr2 | 73214801 | 73227929 | 13129 | -4.12 | 0.0147 | 0.0273 |
| chr11 | 88090104 | 88092037 | 1934 | 0.45 | 0.0147 | 0.0273 |
| chr10 | 93999787 | 94001194 | 1408 | 0.88 | 0.0147 | 0.0273 |
| chr22 | 18326590 | 18327318 | 729 | -1.56 | 0.0147 | 0.0273 |
| chr7 | 91639203 | 91640105 | 903 | 1.55 | 0.0147 | 0.0273 |
| chr18 | 18910016 | 18910978 | 963 | 1.21 | 0.0147 | 0.0273 |
| chr14 | 61747218 | 61748013 | 796 | -1.2 | 0.0147 | 0.0273 |
| chr18 | 76302504 | 76303270 | 767 | 1.17 | 0.0147 | 0.0273 |
| chr4 | 7330357 | 7331355 | 999 | 0.91 | 0.0147 | 0.0273 |
| chr3 | 185893697 | 185894749 | 1053 | 1.68 | 0.0147 | 0.0274 |
| chr17 | 27810012 | 27921466 | 111455 | -0.13 | 0.0147 | 0.0274 |
| chr4 | 178284402 | 178285708 | 1307 | -1.39 | 0.0147 | 0.0274 |
| chr6 | 12957504 | 12958012 | 509 | 2.43 | 0.0147 | 0.0274 |
| chr10 | 66490988 | 66492588 | 1601 | -0.59 | 0.0148 | 0.0274 |
| chr1 | 5956146 | 5957070 | 925 | -1.1 | 0.0148 | 0.0274 |
| chr6 | 105490082 | 105490789 | 708 | 1.63 | 0.0148 | 0.0274 |
| chr15 | 69545167 | 69546499 | 1333 | -0.75 | 0.0148 | 0.0274 |
| chr15 | 22868174 | 22868865 | 692 | -1.9 | 0.0148 | 0.0274 |
| chr10 | 73890255 | 73895593 | 5339 | -0.86 | 0.0148 | 0.0275 |
| chr1 | 211055736 | 211056824 | 1089 | -1.02 | 0.0148 | 0.0275 |
| chr3 | 128734340 | 128746490 | 12151 | -1.29 | 0.0148 | 0.0275 |
| chr3 | 48954779 | 48964854 | 10076 | 0.34 | 0.0148 | 0.0275 |
| chr8 | 67501664 | 67503394 | 1731 | -0.95 | 0.0148 | 0.0275 |
| chr14 | 101292071 | 101293589 | 1519 | 0.95 | 0.0148 | 0.0275 |
| chr18 | 55671482 | 55684814 | 13333 | 0.32 | 0.0148 | 0.0275 |
| chr3 | 160822571 | 160823183 | 613 | 2.19 | 0.0148 | 0.0275 |
| chr3 | 37135919 | 37147018 | 11100 | 0.27 | 0.0148 | 0.0275 |
| chr21 | 18376039 | 18376675 | 637 | 2 | 0.0148 | 0.0275 |
| chr19 | 4181811 | 4182611 | 801 | 0.97 | 0.0149 | 0.0276 |
| chr2 | 220435165 | 220436603 | 1439 | -0.75 | 0.0149 | 0.0276 |
| chrX | 80376053 | 80377166 | 1114 | 1.28 | 0.0149 | 0.0276 |
| chr17 | 25853468 | 25926813 | 73346 | -0.13 | 0.0149 | 0.0277 |
| chr4 | 55147131 | 55148312 | 1182 | 3.42 | 0.0149 | 0.0277 |
| chr5 | 167912948 | 167914952 | 2005 | 0.58 | 0.0149 | 0.0277 |
| chr3 | 177195481 | 177197508 | 2028 | 0.58 | 0.0149 | 0.0277 |
| chr10 | 115841416 | 115843489 | 2074 | 0.88 | 0.0149 | 0.0277 |
| chr13 | 20692137 | 20702588 | 10452 | 0.4 | 0.0149 | 0.0277 |
| chr2 | 55919612 | 55921776 | 2165 | -0.44 | 0.015 | 0.0277 |
| chr1 | 113215567 | 113218629 | 3063 | 0.44 | 0.015 | 0.0277 |
| chr2 | 187213865 | 187214554 | 690 | 1.92 | 0.015 | 0.0277 |
| chr7 | 157069187 | 157075278 | 6092 | 1.13 | 0.015 | 0.0278 |
| chr20 | 13316151 | 13321041 | 4891 | -0.36 | 0.015 | 0.0278 |
| chr3 | 180629204 | 180632104 | 2901 | 0.34 | 0.015 | 0.0278 |
| chrX | 100431090 | 100431556 | 467 | 2.36 | 0.015 | 0.0278 |
| chr17 | 57367062 | 57368712 | 1651 | 3.72 | 0.015 | 0.0278 |
| chr4 | 186716971 | 186717505 | 535 | -1.75 | 0.015 | 0.0278 |
| chr6 | 86225452 | 86239562 | 14111 | 0.56 | 0.015 | 0.0279 |
| chr16 | 71322185 | 71324185 | 2001 | 0.51 | 0.015 | 0.0279 |
| chr14 | 54954240 | 54956841 | 2602 | 0.31 | 0.015 | 0.0279 |
| chr22 | 46691751 | 46698774 | 7024 | 0.6 | 0.0151 | 0.0279 |

| | | | | | | |
|---|---|---|---|---|---|---|
| chr20 | 36845567 | 36849428 | 3862 | 0.63 | 0.0151 | 0.0279 |
| chr6 | 151177708 | 151287632 | 109925 | -0.12 | 0.0151 | 0.0279 |
| chr9 | 97518950 | 97519906 | 957 | -1.52 | 0.0151 | 0.0279 |
| chr8 | 28939289 | 28944574 | 5286 | 0.66 | 0.0151 | 0.0279 |
| chr20 | 34623393 | 34623913 | 521 | -1.81 | 0.0151 | 0.0279 |
| chr12 | 13224498 | 13225345 | 848 | -1.33 | 0.0151 | 0.0279 |
| chr6 | 114316288 | 114317760 | 1473 | 1.12 | 0.0151 | 0.028 |
| chr9 | 98926090 | 98940198 | 14109 | -0.2 | 0.0151 | 0.028 |
| chr14 | 21963108 | 21980330 | 17223 | 0.21 | 0.0151 | 0.028 |
| chr16 | 28289256 | 28289710 | 455 | -1.22 | 0.0151 | 0.028 |
| chrX | 24781749 | 24788527 | 6779 | -0.72 | 0.0151 | 0.028 |
| chr1 | 230680539 | 230681049 | 511 | -1.56 | 0.0152 | 0.028 |
| chr2 | 214052359 | 214053498 | 1140 | 1.02 | 0.0152 | 0.028 |
| chr1 | 162583254 | 162584714 | 1461 | 1.23 | 0.0152 | 0.0281 |
| chr7 | 138526463 | 138527513 | 1051 | -1.04 | 0.0152 | 0.0281 |
| chr8 | 26890301 | 26891793 | 1493 | 0.87 | 0.0152 | 0.0281 |
| chr10 | 104125789 | 104139263 | 13475 | 0.32 | 0.0152 | 0.0281 |
| chr6 | 11114905 | 11127766 | 12862 | -0.94 | 0.0152 | 0.0281 |
| chr15 | 65676933 | 65678041 | 1109 | -1.18 | 0.0152 | 0.0281 |
| chr17 | 65417218 | 65438255 | 21038 | 0.27 | 0.0152 | 0.0281 |
| chr6 | 97344475 | 97347159 | 2685 | 0.41 | 0.0152 | 0.0281 |
| chrX | 67791224 | 67791949 | 726 | -2.27 | 0.0152 | 0.0281 |
| chr16 | 9053705 | 9061693 | 7989 | -0.64 | 0.0152 | 0.0281 |
| chr8 | 101859069 | 101860541 | 1473 | -0.64 | 0.0152 | 0.0282 |
| chr11 | 10202844 | 10203443 | 600 | 1.45 | 0.0152 | 0.0282 |
| chr8 | 134741966 | 134742875 | 910 | 1.19 | 0.0152 | 0.0282 |
| chr20 | 32879539 | 32903089 | 23551 | 0.28 | 0.0152 | 0.0282 |
| chr11 | 129938821 | 129940956 | 2136 | 0.6 | 0.0152 | 0.0282 |
| chr2 | 145194271 | 145196361 | 2091 | 3.74 | 0.0153 | 0.0282 |
| chr10 | 79963561 | 79964635 | 1075 | 1.18 | 0.0153 | 0.0282 |
| chr17 | 9603912 | 9605759 | 1848 | -1.01 | 0.0153 | 0.0282 |
| chr9 | 112075656 | 112092120 | 16465 | 0.44 | 0.0153 | 0.0282 |
| chr1 | 111652775 | 111653206 | 432 | 1.89 | 0.0153 | 0.0282 |
| chr4 | 72597702 | 72598590 | 889 | -1.17 | 0.0153 | 0.0282 |
| chr8 | 31109951 | 31110598 | 648 | 2.3 | 0.0153 | 0.0283 |
| chr17 | 59938819 | 59941802 | 2984 | 0.35 | 0.0153 | 0.0283 |
| chr4 | 41592133 | 41621142 | 29010 | -0.25 | 0.0153 | 0.0283 |
| chr5 | 14695417 | 14715742 | 20326 | -0.38 | 0.0153 | 0.0283 |
| chr2 | 98484834 | 98488724 | 3891 | -0.35 | 0.0153 | 0.0284 |
| chr10 | 112112687 | 112124902 | 12216 | -0.61 | 0.0153 | 0.0284 |
| chr20 | 43181287 | 43183041 | 1755 | -0.59 | 0.0153 | 0.0284 |
| chr1 | 33098329 | 33118943 | 20615 | 0.27 | 0.0153 | 0.0284 |
| chr19 | 47102974 | 47110519 | 7546 | 0.21 | 0.0154 | 0.0284 |
| chr7 | 43965034 | 43967136 | 2103 | 0.44 | 0.0154 | 0.0284 |
| chr1 | 151466684 | 151486941 | 20258 | 2.53 | 0.0154 | 0.0284 |
| chr2 | 217460915 | 217461584 | 670 | 2.33 | 0.0154 | 0.0284 |
| chr19 | 33974626 | 33990825 | 16200 | -1.94 | 0.0154 | 0.0284 |
| chr1 | 100231737 | 100232490 | 754 | 0.91 | 0.0154 | 0.0284 |
| chr4 | 111112270 | 111120791 | 8522 | 0.25 | 0.0154 | 0.0284 |
| chr12 | 132267238 | 132288955 | 21718 | 0.25 | 0.0154 | 0.0284 |
| chr19 | 52691836 | 52695976 | 4141 | 0.31 | 0.0154 | 0.0284 |
| chr6 | 139057858 | 139058335 | 478 | -1.5 | 0.0154 | 0.0284 |
| chr6 | 88207432 | 88208377 | 946 | -1.28 | 0.0154 | 0.0285 |
| chr6 | 138334061 | 138334545 | 485 | 2.17 | 0.0154 | 0.0285 |
| chr1 | 242688093 | 242688710 | 618 | -1.65 | 0.0154 | 0.0285 |
| chr1 | 176117870 | 176131351 | 13482 | 1.71 | 0.0155 | 0.0285 |
| chr9 | 83978527 | 83979355 | 829 | -1.28 | 0.0155 | 0.0285 |
| chr1 | 22323827 | 22324347 | 521 | 1.71 | 0.0155 | 0.0285 |
| chr20 | 45618749 | 45619641 | 893 | 1.64 | 0.0155 | 0.0285 |
| chr7 | 44582316 | 44583076 | 761 | 1.2 | 0.0155 | 0.0285 |
| chr17 | 3881724 | 3887540 | 5817 | 1.37 | 0.0155 | 0.0285 |

| | | | | | | |
|---|---|---|---|---|---|---|
| chr2 | 170363796 | 170364971 | 1176 | 1.26 | 0.0155 | 0.0286 |
| chr14 | 58660529 | 58668773 | 8245 | 0.38 | 0.0155 | 0.0286 |
| chr2 | 236367053 | 236378294 | 11242 | 0.91 | 0.0155 | 0.0286 |
| chr7 | 44740782 | 44741998 | 1217 | -1.18 | 0.0155 | 0.0286 |
| chr6 | 161411881 | 161413832 | 1952 | 0.68 | 0.0155 | 0.0286 |
| chr4 | 75507847 | 75509924 | 2078 | 0.74 | 0.0155 | 0.0286 |
| chr17 | 41773362 | 41774361 | 1000 | -1.71 | 0.0155 | 0.0287 |
| chrX | 140314000 | 140316746 | 2747 | -0.51 | 0.0156 | 0.0287 |
| chr13 | 45478247 | 45480139 | 1893 | 0.67 | 0.0156 | 0.0287 |
| chr1 | 233462458 | 233479479 | 17022 | -0.39 | 0.0156 | 0.0287 |
| chr2 | 183483002 | 183484094 | 1093 | -1.07 | 0.0156 | 0.0287 |
| chr22 | 39189200 | 39190830 | 1631 | -0.61 | 0.0156 | 0.0288 |
| chr13 | 34350411 | 34351282 | 872 | 2.31 | 0.0156 | 0.0288 |
| chr5 | 95629122 | 95631215 | 2094 | -1.08 | 0.0156 | 0.0288 |
| chr1 | 29292731 | 29300966 | 8236 | 0.62 | 0.0156 | 0.0288 |
| chr5 | 124782879 | 124783600 | 722 | 2.27 | 0.0156 | 0.0288 |
| chr4 | 38990234 | 38991060 | 827 | 1.48 | 0.0156 | 0.0288 |
| chr8 | 106896018 | 106896648 | 631 | -1.78 | 0.0156 | 0.0288 |
| chr14 | 86400814 | 86401568 | 755 | 0.84 | 0.0157 | 0.0288 |
| chr2 | 32877720 | 32878522 | 803 | -1.97 | 0.0157 | 0.0289 |
| chr12 | 117458367 | 117537606 | 79240 | -0.16 | 0.0157 | 0.0289 |
| chr13 | 100084714 | 100085645 | 932 | -1.18 | 0.0157 | 0.0289 |
| chr16 | 48569785 | 48571243 | 1459 | -1.37 | 0.0157 | 0.0289 |
| chr13 | 79955021 | 79960456 | 5436 | 0.89 | 0.0157 | 0.0289 |
| chr11 | 3971244 | 3972872 | 1629 | -0.71 | 0.0157 | 0.029 |
| chr6 | 166754700 | 166757009 | 2310 | 0.34 | 0.0157 | 0.029 |
| chr2 | 233164596 | 233166989 | 2394 | 0.58 | 0.0158 | 0.029 |
| chr12 | 126467343 | 126468915 | 1573 | 3.77 | 0.0158 | 0.0291 |
| chr1 | 46527958 | 46529465 | 1508 | 1.08 | 0.0158 | 0.0291 |
| chr3 | 167791839 | 167792733 | 895 | 1.44 | 0.0158 | 0.0291 |
| chr16 | 1401230 | 1414038 | 12809 | -0.53 | 0.0158 | 0.0291 |
| chr12 | 50015887 | 50037387 | 21501 | -0.43 | 0.0158 | 0.0291 |
| chr17 | 62050877 | 62052533 | 1657 | -0.86 | 0.0158 | 0.0291 |
| chr9 | 136709486 | 136711443 | 1958 | -1.05 | 0.0158 | 0.0291 |
| chr4 | 107207694 | 107209779 | 2086 | 0.51 | 0.0158 | 0.0291 |
| chr2 | 85386236 | 85387092 | 857 | -1.91 | 0.0158 | 0.0291 |
| chr1 | 212359990 | 212360618 | 629 | 1.5 | 0.0158 | 0.0292 |
| chr2 | 72364823 | 72373184 | 8362 | 0.57 | 0.0158 | 0.0292 |
| chr20 | 8736429 | 8738357 | 1929 | -0.96 | 0.0158 | 0.0292 |
| chrX | 153671843 | 153673345 | 1503 | 0.9 | 0.0158 | 0.0292 |
| chr2 | 169951945 | 169952507 | 563 | -1.62 | 0.0159 | 0.0292 |
| chr7 | 111879035 | 111879713 | 679 | -1.12 | 0.0159 | 0.0292 |
| chr5 | 39794530 | 39795059 | 530 | 1.65 | 0.0159 | 0.0292 |
| chr7 | 35710500 | 35711054 | 555 | -1.27 | 0.0159 | 0.0292 |
| chr7 | 83847952 | 83849306 | 1355 | 0.99 | 0.0159 | 0.0292 |
| chr9 | 125788730 | 125789241 | 512 | 1.64 | 0.0159 | 0.0292 |
| chr7 | 94640480 | 94641688 | 1209 | -1.61 | 0.0159 | 0.0292 |
| chr8 | 124835377 | 124836420 | 1044 | -1.31 | 0.0159 | 0.0292 |
| chr21 | 35171038 | 35175088 | 4051 | 0.62 | 0.0159 | 0.0292 |
| chr11 | 9267689 | 9268312 | 624 | -1.62 | 0.0159 | 0.0292 |
| chr21 | 18914264 | 18915745 | 1482 | 1.23 | 0.0159 | 0.0292 |
| chr6 | 135775745 | 135777153 | 1409 | -1.19 | 0.0159 | 0.0293 |
| chr12 | 51610701 | 51617482 | 6782 | 0.53 | 0.0159 | 0.0293 |
| chr11 | 17134139 | 17134799 | 661 | -1.46 | 0.0159 | 0.0293 |
| chr20 | 46680864 | 46681283 | 420 | 2.06 | 0.0159 | 0.0293 |
| chr1 | 38324198 | 38329000 | 4803 | -0.35 | 0.0159 | 0.0293 |
| chr10 | 25034987 | 25035716 | 730 | 1.82 | 0.0159 | 0.0293 |
| chr1 | 29208949 | 29214645 | 5697 | -0.36 | 0.0159 | 0.0293 |
| chr2 | 219341008 | 219344070 | 3063 | -0.82 | 0.0159 | 0.0293 |
| chr22 | 36697571 | 36701707 | 4137 | 0.62 | 0.016 | 0.0293 |
| chr10 | 75639265 | 75701300 | 62036 | 0.15 | 0.016 | 0.0294 |

| | | | | | | |
|---|---|---|---|---|---|---|
| chr10 | 59732921 | 59733414 | 494 | 2.05 | 0.016 | 0.0294 |
| chr20 | 33349412 | 33354969 | 5558 | 0.69 | 0.016 | 0.0294 |
| chr14 | 89290266 | 89297265 | 7000 | -0.34 | 0.016 | 0.0295 |
| chr1 | 235180399 | 235184661 | 4263 | -0.67 | 0.0161 | 0.0295 |
| chr6 | 163867560 | 163883530 | 15971 | 0.36 | 0.0161 | 0.0295 |
| chr16 | 810142 | 811690 | 1549 | 1.03 | 0.0161 | 0.0295 |
| chr15 | 42840128 | 42842266 | 2139 | 0.42 | 0.0161 | 0.0295 |
| chr19 | 8046085 | 8071313 | 25229 | 0.35 | 0.0161 | 0.0295 |
| chr3 | 87745451 | 87746226 | 776 | -1.16 | 0.0161 | 0.0296 |
| chr1 | 90137346 | 90138224 | 879 | -1.61 | 0.0161 | 0.0296 |
| chr1 | 155785766 | 155786589 | 824 | 1.5 | 0.0161 | 0.0296 |
| chr3 | 31245847 | 31246623 | 777 | 1.09 | 0.0161 | 0.0296 |
| chr14 | 75585730 | 75644408 | 58679 | -0.14 | 0.0162 | 0.0297 |
| chr11 | 46431318 | 46436694 | 5377 | -0.53 | 0.0162 | 0.0297 |
| chr22 | 22535580 | 22536316 | 737 | 1.66 | 0.0162 | 0.0297 |
| chr2 | 191877414 | 191916532 | 39119 | -0.2 | 0.0162 | 0.0297 |
| chr22 | 49001463 | 49002039 | 577 | -1.3 | 0.0162 | 0.0297 |
| chr3 | 56714574 | 56725946 | 11373 | 0.36 | 0.0162 | 0.0298 |
| chr3 | 36565340 | 36568111 | 2772 | -0.6 | 0.0162 | 0.0298 |
| chr13 | 31103328 | 31104000 | 673 | -2.21 | 0.0163 | 0.0298 |
| chr8 | 144074580 | 144075886 | 1307 | 1.09 | 0.0163 | 0.0298 |
| chr4 | 178828930 | 178829559 | 630 | 1.82 | 0.0163 | 0.0298 |
| chr16 | 81327153 | 81328686 | 1534 | 1.05 | 0.0163 | 0.0299 |
| chr12 | 60649867 | 60661293 | 11427 | 0.35 | 0.0163 | 0.0299 |
| chr15 | 78590781 | 78596613 | 5833 | 0.31 | 0.0163 | 0.0299 |
| chr4 | 77467800 | 77468510 | 711 | 1.53 | 0.0163 | 0.0299 |
| chr12 | 51764625 | 51766303 | 1679 | 0.5 | 0.0163 | 0.0299 |
| chr2 | 161978757 | 161996265 | 17509 | -0.18 | 0.0163 | 0.0299 |
| chr3 | 141685946 | 141686980 | 1035 | 2.3 | 0.0163 | 0.0299 |
| chrX | 100074406 | 100076090 | 1685 | -0.45 | 0.0163 | 0.0299 |
| chr3 | 132135659 | 132138330 | 2672 | 0.55 | 0.0163 | 0.0299 |
| chr5 | 75986852 | 75987819 | 968 | 1.4 | 0.0163 | 0.0299 |
| chr22 | 31062887 | 31064765 | 1879 | -0.48 | 0.0163 | 0.0299 |
| chr17 | 5300699 | 5301353 | 655 | 1.54 | 0.0163 | 0.0299 |
| chr7 | 98664655 | 98668005 | 3351 | -0.76 | 0.0163 | 0.0299 |
| chr20 | 32803649 | 32804497 | 849 | -1.11 | 0.0163 | 0.0299 |
| chr10 | 30007653 | 30008731 | 1079 | 1.73 | 0.0163 | 0.03 |
| chr20 | 25037315 | 25039533 | 2219 | 0.41 | 0.0163 | 0.03 |
| chr8 | 91704625 | 91705729 | 1105 | 3.53 | 0.0164 | 0.03 |
| chr2 | 178284140 | 178284718 | 579 | -1.76 | 0.0164 | 0.03 |
| chr4 | 102997594 | 102998531 | 938 | 1.36 | 0.0164 | 0.0301 |
| chr2 | 106048915 | 106049571 | 657 | 1.68 | 0.0164 | 0.0301 |
| chr21 | 47861939 | 47864219 | 2281 | -0.71 | 0.0164 | 0.0301 |
| chr15 | 101297934 | 101301071 | 3138 | -0.3 | 0.0164 | 0.0301 |
| chr3 | 100321099 | 100322785 | 1687 | 0.71 | 0.0164 | 0.0301 |
| chr3 | 39447234 | 39457490 | 10257 | 0.24 | 0.0165 | 0.0302 |
| chr3 | 122397919 | 122400704 | 2786 | 0.37 | 0.0165 | 0.0302 |
| chr11 | 33312270 | 33314013 | 1744 | 0.91 | 0.0165 | 0.0302 |
| chr15 | 64403725 | 64405801 | 2077 | -0.71 | 0.0165 | 0.0302 |
| chr16 | 18866012 | 18867361 | 1350 | -1.19 | 0.0165 | 0.0302 |
| chr4 | 115238869 | 115240246 | 1378 | 3.44 | 0.0165 | 0.0302 |
| chr6 | 8434308 | 8436379 | 2072 | 0.37 | 0.0165 | 0.0302 |
| chr10 | 65370684 | 65398584 | 27901 | 0.2 | 0.0165 | 0.0302 |
| chr11 | 73795684 | 73797498 | 1815 | -1.01 | 0.0165 | 0.0302 |
| chr6 | 26440042 | 26441339 | 1298 | 0.81 | 0.0165 | 0.0302 |
| chr9 | 32970567 | 32971164 | 598 | -2 | 0.0165 | 0.0303 |
| chr11 | 73612541 | 73614238 | 1698 | -0.92 | 0.0165 | 0.0303 |
| chr4 | 141149861 | 141150841 | 981 | -0.92 | 0.0166 | 0.0303 |
| chr3 | 162108671 | 162109901 | 1231 | -0.99 | 0.0166 | 0.0303 |
| chr11 | 46320524 | 46321344 | 821 | -1.4 | 0.0166 | 0.0303 |
| chr1 | 186943748 | 186944684 | 937 | 1.16 | 0.0166 | 0.0304 |

| | | | | | | |
|---|---|---|---|---|---|---|
| chr3 | 196664083 | 196670684 | 6602 | 0.35 | 0.0166 | 0.0304 |
| chr19 | 39601292 | 39686518 | 85227 | -0.13 | 0.0166 | 0.0304 |
| chr7 | 5665235 | 5694021 | 28787 | -0.38 | 0.0166 | 0.0304 |
| chr8 | 54426485 | 54437463 | 10979 | -2.67 | 0.0166 | 0.0304 |
| chr13 | 34573693 | 34575033 | 1341 | 1.36 | 0.0166 | 0.0304 |
| chr2 | 232770324 | 232770891 | 568 | -2.24 | 0.0166 | 0.0304 |
| chr12 | 53190670 | 53221908 | 31239 | 0.17 | 0.0166 | 0.0304 |
| chr21 | 38519264 | 38520670 | 1407 | -1.31 | 0.0166 | 0.0305 |
| chr2 | 206137978 | 206140394 | 2417 | 1.29 | 0.0166 | 0.0305 |
| chr4 | 6420369 | 6421703 | 1335 | -1.24 | 0.0167 | 0.0305 |
| chr8 | 61698534 | 61699129 | 596 | -2.29 | 0.0167 | 0.0305 |
| chr18 | 9362168 | 9363452 | 1285 | 1.2 | 0.0167 | 0.0305 |
| chr5 | 53034898 | 53035276 | 379 | -2.29 | 0.0167 | 0.0305 |
| chr1 | 174470116 | 174470799 | 684 | -1.78 | 0.0167 | 0.0305 |
| chr6 | 80745651 | 80746216 | 566 | 1.89 | 0.0167 | 0.0305 |
| chr4 | 107235415 | 107246888 | 11474 | 0.23 | 0.0167 | 0.0305 |
| chr22 | 35863510 | 35877588 | 14079 | 0.45 | 0.0167 | 0.0305 |
| chr4 | 5673644 | 5674251 | 608 | -1.57 | 0.0167 | 0.0305 |
| chr9 | 2040808 | 2041460 | 653 | -2.1 | 0.0167 | 0.0306 |
| chr16 | 75267523 | 75303849 | 36327 | 0.14 | 0.0167 | 0.0306 |
| chr7 | 66056128 | 66058514 | 2387 | 0.4 | 0.0167 | 0.0306 |
| chr1 | 95193141 | 95194110 | 970 | 1.12 | 0.0167 | 0.0306 |
| chr16 | 85695765 | 85708326 | 12562 | -0.53 | 0.0167 | 0.0306 |
| chr1 | 48275019 | 48275802 | 784 | 1.28 | 0.0167 | 0.0306 |
| chr1 | 66848600 | 66849499 | 900 | -1.58 | 0.0168 | 0.0306 |
| chr3 | 48836733 | 48837151 | 419 | 2.07 | 0.0168 | 0.0306 |
| chr11 | 124543028 | 124544927 | 1900 | -0.36 | 0.0168 | 0.0306 |
| chr17 | 80289610 | 80297829 | 8220 | -0.25 | 0.0168 | 0.0307 |
| chr9 | 36486060 | 36488777 | 2718 | 0.46 | 0.0168 | 0.0307 |
| chr14 | 61492205 | 61492938 | 734 | -1.51 | 0.0168 | 0.0307 |
| chr2 | 200319930 | 200328673 | 8744 | -0.31 | 0.0168 | 0.0307 |
| chr6 | 122719939 | 122721842 | 1904 | 0.63 | 0.0168 | 0.0307 |
| chr1 | 44773644 | 44774100 | 457 | -2.32 | 0.0168 | 0.0307 |
| chr13 | 102067790 | 102070045 | 2256 | 0.54 | 0.0168 | 0.0307 |
| chr18 | 24499676 | 24500479 | 804 | 1.47 | 0.0168 | 0.0307 |
| chr1 | 214578044 | 214578637 | 594 | -2.24 | 0.0168 | 0.0307 |
| chr12 | 58889508 | 58899606 | 10099 | 4.38 | 0.0168 | 0.0308 |
| chr19 | 51777017 | 51778249 | 1233 | -3.39 | 0.0169 | 0.0308 |
| chr18 | 11361736 | 11362661 | 926 | -1.33 | 0.0169 | 0.0308 |
| chr17 | 60170334 | 60171927 | 1594 | -0.67 | 0.0169 | 0.0308 |
| chr12 | 120378238 | 120380039 | 1802 | 0.74 | 0.0169 | 0.0308 |
| chr11 | 120081030 | 120081825 | 796 | -1.38 | 0.0169 | 0.0308 |
| chrX | 149817445 | 149819057 | 1613 | 0.8 | 0.0169 | 0.0309 |
| chr12 | 7244496 | 7262486 | 17991 | 0.26 | 0.0169 | 0.0309 |
| chr6 | 133661514 | 133663641 | 2128 | -0.77 | 0.0169 | 0.0309 |
| chr8 | 48496242 | 48497130 | 889 | -1.51 | 0.0169 | 0.0309 |
| chr10 | 27057926 | 27071064 | 13139 | 0.61 | 0.017 | 0.031 |
| chr2 | 9982759 | 9984894 | 2136 | -0.43 | 0.017 | 0.031 |
| chr11 | 130051594 | 130060641 | 9048 | 0.82 | 0.017 | 0.031 |
| chr2 | 74775679 | 74786141 | 10463 | -0.5 | 0.017 | 0.031 |
| chr9 | 139615238 | 139623884 | 8647 | -0.42 | 0.017 | 0.031 |
| chr22 | 45927370 | 45928294 | 925 | -1.4 | 0.017 | 0.031 |
| chr6 | 75883352 | 75883850 | 499 | -1.68 | 0.017 | 0.031 |
| chr12 | 93956661 | 93966080 | 9420 | 0.35 | 0.017 | 0.031 |
| chr9 | 134210119 | 134210513 | 395 | -2.23 | 0.017 | 0.031 |
| chr11 | 62149426 | 62163263 | 13838 | 0.53 | 0.017 | 0.031 |
| chr13 | 46038183 | 46040585 | 2403 | 0.5 | 0.0171 | 0.0311 |
| chr13 | 35235536 | 35236320 | 785 | 1.51 | 0.0171 | 0.0311 |
| chr19 | 4122629 | 4150470 | 27842 | -0.22 | 0.0171 | 0.0311 |
| chr7 | 155711574 | 155712984 | 1411 | -0.74 | 0.0171 | 0.0311 |
| chr18 | 61011107 | 61012318 | 1212 | 1.27 | 0.0171 | 0.0312 |

| | | | | | | |
|---|---|---|---|---|---|---|
| chr1 | 159849661 | 159852927 | 3267 | 0.49 | 0.0171 | 0.0312 |
| chrX | 63614816 | 63615304 | 489 | 1.43 | 0.0171 | 0.0312 |
| chr13 | 21542749 | 21543447 | 699 | 1.53 | 0.0171 | 0.0313 |
| chr20 | 60837669 | 60839409 | 1741 | -0.91 | 0.0171 | 0.0313 |
| chr20 | 47255074 | 47255947 | 874 | -1.22 | 0.0171 | 0.0313 |
| chr14 | 34648923 | 34650731 | 1809 | 0.99 | 0.0172 | 0.0313 |
| chr3 | 24851938 | 24852641 | 704 | 1.43 | 0.0172 | 0.0313 |
| chr5 | 37389550 | 37390360 | 811 | 1.27 | 0.0172 | 0.0313 |
| chr14 | 76044264 | 76044711 | 448 | -1.66 | 0.0172 | 0.0313 |
| chr3 | 113439765 | 113467219 | 27455 | 0.19 | 0.0172 | 0.0314 |
| chr7 | 97802211 | 97808767 | 6557 | 0.77 | 0.0172 | 0.0314 |
| chr2 | 98384624 | 98385371 | 748 | -1.45 | 0.0172 | 0.0314 |
| chr7 | 156985590 | 156988250 | 2661 | 1.1 | 0.0173 | 0.0315 |
| chr2 | 54557153 | 54558912 | 1760 | -0.49 | 0.0173 | 0.0315 |
| chr17 | 47437348 | 47440356 | 3009 | 0.47 | 0.0173 | 0.0315 |
| chr22 | 45596512 | 45609707 | 13196 | -0.33 | 0.0173 | 0.0315 |
| chr2 | 42395787 | 42525065 | 129279 | 0.11 | 0.0173 | 0.0315 |
| chr16 | 23328376 | 23348514 | 20139 | 0.4 | 0.0173 | 0.0316 |
| chr18 | 28622305 | 28623334 | 1030 | 1.28 | 0.0173 | 0.0316 |
| chr19 | 40807725 | 40808849 | 1125 | -0.99 | 0.0173 | 0.0316 |
| chr1 | 87191736 | 87192950 | 1215 | 1.4 | 0.0173 | 0.0316 |
| chr10 | 35820693 | 35821806 | 1114 | -1.47 | 0.0173 | 0.0316 |
| chr4 | 41634006 | 41634887 | 882 | -1.49 | 0.0173 | 0.0316 |
| chrX | 107314707 | 107315355 | 649 | -1.25 | 0.0173 | 0.0316 |
| chr1 | 112297127 | 112301289 | 4163 | 0.41 | 0.0174 | 0.0316 |
| chr8 | 21613997 | 21615122 | 1126 | -1.31 | 0.0174 | 0.0317 |
| chr19 | 4342152 | 4343923 | 1772 | 0.75 | 0.0174 | 0.0317 |
| chr5 | 137280811 | 137297733 | 16923 | 0.34 | 0.0174 | 0.0317 |
| chr3 | 29306723 | 29307661 | 939 | -1.83 | 0.0174 | 0.0317 |
| chr17 | 66287088 | 66315017 | 27930 | -0.48 | 0.0174 | 0.0317 |
| chr20 | 10056399 | 10058653 | 2255 | 0.64 | 0.0174 | 0.0317 |
| chr4 | 121613579 | 121614529 | 951 | 1.21 | 0.0174 | 0.0317 |
| chr17 | 66754527 | 66756576 | 2050 | -0.28 | 0.0175 | 0.0318 |
| chr7 | 2019326 | 2020151 | 826 | -1.63 | 0.0175 | 0.0318 |
| chr18 | 72919386 | 72924233 | 4848 | 0.39 | 0.0175 | 0.0318 |
| chr6 | 56235028 | 56247873 | 12846 | 3.85 | 0.0175 | 0.0318 |
| chr2 | 135779233 | 135779725 | 493 | -2 | 0.0175 | 0.0318 |
| chr2 | 128239244 | 128240211 | 968 | -1.47 | 0.0175 | 0.0318 |
| chr10 | 29593636 | 29608735 | 15100 | 0.48 | 0.0175 | 0.0319 |
| chr20 | 6012853 | 6013385 | 533 | -1.76 | 0.0175 | 0.0319 |
| chr16 | 87441343 | 87441757 | 415 | 2.53 | 0.0175 | 0.0319 |
| chr19 | 1478696 | 1513983 | 35288 | 0.48 | 0.0175 | 0.0319 |
| chr2 | 37762825 | 37763521 | 697 | 0.77 | 0.0175 | 0.0319 |
| chr1 | 166136315 | 166136719 | 405 | 2.09 | 0.0175 | 0.0319 |
| chr11 | 61690759 | 61691429 | 671 | 1.78 | 0.0175 | 0.0319 |
| chr8 | 23059393 | 23060252 | 860 | -1.27 | 0.0175 | 0.0319 |
| chr4 | 186695782 | 186697338 | 1557 | 0.72 | 0.0175 | 0.0319 |
| chr1 | 188458492 | 188459097 | 606 | 1.6 | 0.0175 | 0.0319 |
| chr14 | 30374280 | 30376681 | 2402 | 0.57 | 0.0175 | 0.0319 |
| chr2 | 24012218 | 24014742 | 2525 | -0.46 | 0.0175 | 0.0319 |
| chr18 | 74934922 | 74935737 | 816 | 2.28 | 0.0175 | 0.0319 |
| chr4 | 39948769 | 39951272 | 2504 | -0.84 | 0.0176 | 0.032 |
| chr8 | 38123685 | 38137001 | 13317 | 0.26 | 0.0176 | 0.032 |
| chr15 | 28340521 | 28342277 | 1757 | 0.9 | 0.0176 | 0.032 |
| chr1 | 147735283 | 147753116 | 17834 | -0.32 | 0.0176 | 0.032 |
| chr17 | 39440680 | 39444371 | 3692 | -0.57 | 0.0176 | 0.032 |
| chr18 | 72782150 | 72783633 | 1484 | 0.77 | 0.0176 | 0.032 |
| chr12 | 9903573 | 9904021 | 449 | 1.97 | 0.0176 | 0.0321 |
| chr3 | 13343006 | 13343896 | 891 | -1.12 | 0.0177 | 0.0321 |
| chr19 | 35659444 | 35660387 | 944 | -1.13 | 0.0177 | 0.0321 |
| chr16 | 71878879 | 71893919 | 15041 | 0.32 | 0.0177 | 0.0321 |

| | | | | | | |
|---|---|---|---|---|---|---|
| chr16 | 4302942 | 4443772 | 140831 | -0.14 | 0.0177 | 0.0321 |
| chr16 | 80965895 | 80966799 | 905 | 1.95 | 0.0177 | 0.0322 |
| chr1 | 26405577 | 26407225 | 1649 | -0.87 | 0.0177 | 0.0322 |
| chr4 | 159587185 | 159594632 | 7448 | 0.47 | 0.0177 | 0.0322 |
| chr11 | 1467635 | 1468895 | 1261 | -1.54 | 0.0177 | 0.0322 |
| chr16 | 5121149 | 5122670 | 1522 | 0.63 | 0.0177 | 0.0322 |
| chr14 | 29104336 | 29104739 | 404 | 2.31 | 0.0177 | 0.0322 |
| chr19 | 56601948 | 56655098 | 53151 | 0.16 | 0.0177 | 0.0322 |
| chrX | 140879461 | 140880063 | 603 | -1.91 | 0.0177 | 0.0322 |
| chr8 | 128431494 | 128432974 | 1481 | 4.07 | 0.0177 | 0.0322 |
| chr20 | 18268077 | 18270243 | 2167 | 0.35 | 0.0177 | 0.0322 |
| chr13 | 25575570 | 25577560 | 1991 | 0.71 | 0.0178 | 0.0322 |
| chr7 | 26330993 | 26332069 | 1077 | 1.05 | 0.0178 | 0.0322 |
| chr3 | 33340888 | 33360627 | 19740 | -0.34 | 0.0178 | 0.0322 |
| chr15 | 40073901 | 40077136 | 3236 | -0.51 | 0.0178 | 0.0323 |
| chr15 | 91999758 | 92000299 | 542 | 1.73 | 0.0178 | 0.0323 |
| chr1 | 221603898 | 221614834 | 10937 | 0.4 | 0.0178 | 0.0323 |
| chr22 | 40551795 | 40552770 | 976 | 1.03 | 0.0178 | 0.0324 |
| chr10 | 12109221 | 12124794 | 15574 | 0.4 | 0.0178 | 0.0324 |
| chr9 | 29332431 | 29333801 | 1371 | 1.13 | 0.0178 | 0.0324 |
| chr22 | 42062245 | 42070280 | 8036 | 0.26 | 0.0178 | 0.0324 |
| chr4 | 39286649 | 39289710 | 3062 | -0.84 | 0.0179 | 0.0324 |
| chr4 | 169967226 | 169967615 | 390 | 2.24 | 0.0179 | 0.0324 |
| chr7 | 132448389 | 132449537 | 1149 | 0.98 | 0.0179 | 0.0324 |
| chr8 | 37494610 | 37495389 | 780 | 1.54 | 0.0179 | 0.0324 |
| chr4 | 41659135 | 41660285 | 1151 | -1.2 | 0.0179 | 0.0325 |
| chr5 | 54829710 | 54831650 | 1941 | 0.51 | 0.0179 | 0.0325 |
| chr8 | 66711892 | 66713203 | 1312 | -1.19 | 0.0179 | 0.0325 |
| chr22 | 25457279 | 25458317 | 1039 | -1.05 | 0.0179 | 0.0325 |
| chr6 | 88344119 | 88344871 | 753 | 1.56 | 0.0179 | 0.0325 |
| chr3 | 151952541 | 151954860 | 2320 | -0.41 | 0.0179 | 0.0325 |
| chr8 | 34032727 | 34033222 | 496 | -1.55 | 0.0179 | 0.0325 |
| chr15 | 52244867 | 52245861 | 995 | -1.19 | 0.0179 | 0.0325 |
| chr17 | 37417025 | 37417878 | 854 | -1.35 | 0.018 | 0.0326 |
| chr20 | 25115863 | 25119431 | 3569 | -0.25 | 0.018 | 0.0326 |
| chr7 | 38834956 | 38835749 | 794 | -2.26 | 0.018 | 0.0326 |
| chr15 | 68131206 | 68156550 | 25345 | -0.37 | 0.018 | 0.0326 |
| chr14 | 86871032 | 86871573 | 542 | -2.38 | 0.018 | 0.0326 |
| chr6 | 168100030 | 168100682 | 653 | 2.35 | 0.018 | 0.0326 |
| chrX | 11885788 | 11886967 | 1180 | -1.07 | 0.018 | 0.0327 |
| chr5 | 179149078 | 179161307 | 12230 | 0.37 | 0.018 | 0.0327 |
| chr10 | 121591892 | 121610891 | 19000 | 0.28 | 0.018 | 0.0327 |
| chr1 | 161170821 | 161173659 | 2839 | 0.32 | 0.018 | 0.0327 |
| chr5 | 61873923 | 61889894 | 15972 | -0.49 | 0.018 | 0.0327 |
| chr8 | 95095539 | 95102487 | 6949 | 0.81 | 0.018 | 0.0327 |
| chrX | 14890210 | 14892565 | 2356 | 0.51 | 0.018 | 0.0327 |
| chr13 | 51503591 | 51504252 | 662 | -1.6 | 0.0181 | 0.0327 |
| chr22 | 30097396 | 30098520 | 1125 | 1.23 | 0.0181 | 0.0328 |
| chr11 | 124492110 | 124493847 | 1738 | 0.38 | 0.0181 | 0.0328 |
| chr8 | 124889248 | 124890972 | 1725 | -0.88 | 0.0181 | 0.0328 |
| chr15 | 41116607 | 41118919 | 2313 | 1.03 | 0.0181 | 0.0329 |
| chr16 | 84732780 | 84789088 | 56309 | 0.14 | 0.0182 | 0.0329 |
| chr17 | 59996262 | 60018457 | 22196 | 0.28 | 0.0182 | 0.0329 |
| chr1 | 118277593 | 118278397 | 805 | 1.13 | 0.0182 | 0.0329 |
| chr8 | 23363949 | 23365299 | 1351 | 1.31 | 0.0182 | 0.0329 |
| chr9 | 97959939 | 97960665 | 727 | -2.21 | 0.0182 | 0.0329 |
| chr8 | 74572990 | 74574563 | 1574 | -1.33 | 0.0182 | 0.033 |
| chr8 | 27713268 | 27806856 | 93589 | 0.1 | 0.0182 | 0.033 |
| chr10 | 102649981 | 102657345 | 7365 | 0.46 | 0.0182 | 0.033 |
| chr2 | 216175758 | 216194867 | 19110 | -0.24 | 0.0182 | 0.033 |
| chr2 | 224648882 | 224702902 | 54021 | 0.19 | 0.0182 | 0.033 |

| | | | | | | |
|---|---|---|---|---|---|---|
| chr19 | 44555283 | 44556913 | 1631 | 0.56 | 0.0183 | 0.0331 |
| chr7 | 80146113 | 80146868 | 756 | 1.74 | 0.0183 | 0.0331 |
| chr17 | 71183020 | 71230202 | 47183 | -0.21 | 0.0183 | 0.0331 |
| chr3 | 49687035 | 49687708 | 674 | -1.57 | 0.0183 | 0.0331 |
| chr5 | 81568960 | 81575385 | 6426 | 0.27 | 0.0183 | 0.0331 |
| chr1 | 232940580 | 232941997 | 1418 | 0.56 | 0.0183 | 0.0332 |
| chr3 | 151001397 | 151002308 | 912 | 2.24 | 0.0183 | 0.0332 |
| chr1 | 165737297 | 165743082 | 5786 | 0.34 | 0.0183 | 0.0332 |
| chr10 | 99002881 | 99003433 | 553 | 2.39 | 0.0183 | 0.0332 |
| chr11 | 57266813 | 57267780 | 968 | -1.29 | 0.0183 | 0.0332 |
| chr3 | 150442806 | 150443483 | 678 | 1.29 | 0.0184 | 0.0332 |
| chr4 | 41309184 | 41367303 | 58120 | -0.17 | 0.0184 | 0.0333 |
| chr2 | 216134069 | 216134693 | 625 | -1.31 | 0.0184 | 0.0333 |
| chr2 | 86054722 | 86055279 | 558 | -1.37 | 0.0184 | 0.0333 |
| chr17 | 19764542 | 19771722 | 7181 | 0.43 | 0.0184 | 0.0333 |
| chr15 | 56491900 | 56493098 | 1199 | 1.47 | 0.0184 | 0.0333 |
| chr16 | 12953036 | 12953744 | 709 | 1.26 | 0.0184 | 0.0334 |
| chr1 | 113497066 | 113500872 | 3807 | 0.39 | 0.0185 | 0.0334 |
| chr3 | 141985704 | 141986933 | 1230 | 3.75 | 0.0185 | 0.0334 |
| chr1 | 210465528 | 210466286 | 759 | 1.33 | 0.0185 | 0.0335 |
| chr7 | 31803942 | 31804673 | 732 | -2.23 | 0.0185 | 0.0335 |
| chr1 | 168830480 | 168834216 | 3737 | -0.49 | 0.0185 | 0.0335 |
| chr2 | 10951339 | 10953665 | 2327 | -0.45 | 0.0185 | 0.0335 |
| chr14 | 52104352 | 52122487 | 18136 | 0.27 | 0.0185 | 0.0335 |
| chr16 | 87594295 | 87594623 | 329 | 1.88 | 0.0186 | 0.0335 |
| chr10 | 122610195 | 122612032 | 1838 | 0.41 | 0.0186 | 0.0335 |
| chr2 | 236050984 | 236051855 | 872 | 1.92 | 0.0186 | 0.0335 |
| chr9 | 79839504 | 79840465 | 962 | -1.42 | 0.0186 | 0.0335 |
| chr15 | 93257487 | 93258343 | 857 | 0.96 | 0.0186 | 0.0336 |
| chr3 | 136305131 | 136305867 | 737 | 1.6 | 0.0186 | 0.0336 |
| chr19 | 46455743 | 46457187 | 1445 | -0.82 | 0.0186 | 0.0336 |
| chr15 | 53872670 | 53873284 | 615 | 2.25 | 0.0186 | 0.0336 |
| chr12 | 30975013 | 30976261 | 1249 | 0.75 | 0.0186 | 0.0336 |
| chr19 | 38852687 | 38879811 | 27125 | 0.22 | 0.0186 | 0.0336 |
| chr4 | 157468276 | 157468771 | 496 | 1.74 | 0.0186 | 0.0337 |
| chr13 | 73300261 | 73304966 | 4706 | 0.39 | 0.0187 | 0.0337 |
| chr4 | 48832537 | 48834709 | 2173 | 0.32 | 0.0187 | 0.0338 |
| chr5 | 114802785 | 114803581 | 797 | 1.17 | 0.0187 | 0.0338 |
| chr2 | 158731506 | 158733726 | 2221 | -0.5 | 0.0187 | 0.0338 |
| chr19 | 7693547 | 7702948 | 9402 | 0.42 | 0.0187 | 0.0338 |
| chr19 | 48705499 | 48709987 | 4489 | -0.3 | 0.0187 | 0.0338 |
| chr3 | 190955824 | 190956516 | 693 | -1.52 | 0.0187 | 0.0338 |
| chr18 | 45346735 | 45370568 | 23834 | 0.31 | 0.0187 | 0.0338 |
| chr20 | 18503294 | 18514627 | 11334 | -0.51 | 0.0188 | 0.0339 |
| chr4 | 87751749 | 87753493 | 1745 | 0.82 | 0.0188 | 0.0339 |
| chr2 | 203672023 | 203673610 | 1588 | -1.17 | 0.0188 | 0.0339 |
| chr8 | 80893868 | 80894716 | 849 | 0.89 | 0.0188 | 0.0339 |
| chr22 | 17638888 | 17653411 | 14524 | 0.38 | 0.0188 | 0.0339 |
| chr3 | 14692173 | 14695913 | 3741 | 0.29 | 0.0188 | 0.034 |
| chr8 | 122273372 | 122274352 | 981 | -1 | 0.0188 | 0.034 |
| chr2 | 162879028 | 162879607 | 580 | 2.36 | 0.0188 | 0.034 |
| chr3 | 110212013 | 110214656 | 2644 | 1 | 0.0188 | 0.034 |
| chr8 | 48485724 | 48487101 | 1378 | 1.36 | 0.0189 | 0.034 |
| chr6 | 24355369 | 24361824 | 6456 | 0.33 | 0.0189 | 0.0341 |
| chr1 | 151627579 | 151628149 | 571 | -2.25 | 0.0189 | 0.0341 |
| chr3 | 121449983 | 121450564 | 582 | -1.86 | 0.0189 | 0.0341 |
| chr6 | 13952794 | 13954207 | 1414 | 0.83 | 0.0189 | 0.0341 |
| chr5 | 92389197 | 92390200 | 1004 | 1.22 | 0.0189 | 0.0341 |
| chr2 | 171829433 | 171830961 | 1529 | -0.68 | 0.019 | 0.0342 |
| chrX | 123051047 | 123052886 | 1840 | 0.66 | 0.019 | 0.0342 |
| chr2 | 64097194 | 64132506 | 35313 | 0.22 | 0.019 | 0.0343 |

| | | | | | | |
|---|---|---|---|---|---|---|
| chr5 | 133746029 | 133756700 | 10672 | 0.37 | 0.019 | 0.0343 |
| chr4 | 41953405 | 41954931 | 1527 | -0.82 | 0.019 | 0.0343 |
| chrX | 20847036 | 20849576 | 2541 | 0.55 | 0.019 | 0.0343 |
| chr18 | 55780253 | 55786238 | 5986 | 0.81 | 0.019 | 0.0343 |
| chr19 | 6460410 | 6464890 | 4481 | 1.2 | 0.0191 | 0.0344 |
| chr8 | 21856175 | 21868699 | 12525 | -0.84 | 0.0191 | 0.0344 |
| chr1 | 145743505 | 145744745 | 1241 | 1.18 | 0.0191 | 0.0344 |
| chr20 | 3995611 | 3999656 | 4046 | -0.58 | 0.0191 | 0.0344 |
| chr20 | 11812955 | 11813617 | 663 | -1.89 | 0.0191 | 0.0344 |
| chr4 | 174927341 | 174927965 | 625 | 2.24 | 0.0191 | 0.0344 |
| chr19 | 54600047 | 54620054 | 20008 | -0.28 | 0.0191 | 0.0344 |
| chr2 | 37086006 | 37086627 | 622 | 1.24 | 0.0191 | 0.0345 |
| chr15 | 42082797 | 42084082 | 1286 | -1.11 | 0.0191 | 0.0345 |
| chr10 | 33992932 | 33993725 | 794 | -2.22 | 0.0191 | 0.0345 |
| chr11 | 33795269 | 33796655 | 1387 | 0.79 | 0.0191 | 0.0345 |
| chr2 | 206903221 | 206903742 | 522 | -1.92 | 0.0191 | 0.0345 |
| chr15 | 82454979 | 82455541 | 563 | -1.48 | 0.0192 | 0.0345 |
| chr2 | 178992637 | 178993340 | 704 | -1.8 | 0.0192 | 0.0345 |
| chr3 | 186418113 | 186418683 | 571 | -1.5 | 0.0192 | 0.0345 |
| chr11 | 66971514 | 66973778 | 2265 | -0.55 | 0.0192 | 0.0345 |
| chr4 | 86664111 | 86665122 | 1012 | -1.45 | 0.0192 | 0.0345 |
| chr7 | 2197002 | 2216174 | 19173 | 0.49 | 0.0192 | 0.0346 |
| chr12 | 10306001 | 10307011 | 1011 | 0.67 | 0.0192 | 0.0346 |
| chr2 | 239987038 | 239988135 | 1098 | -1.46 | 0.0192 | 0.0346 |
| chr4 | 110551093 | 110553595 | 2503 | 0.49 | 0.0192 | 0.0347 |
| chr1 | 28051387 | 28053997 | 2611 | -0.45 | 0.0192 | 0.0347 |
| chr2 | 234665064 | 234666056 | 993 | -1.2 | 0.0192 | 0.0347 |
| chr2 | 170895652 | 170896595 | 944 | 1.26 | 0.0193 | 0.0347 |
| chr12 | 13249905 | 13256328 | 6424 | 0.18 | 0.0193 | 0.0347 |
| chr8 | 74498146 | 74498677 | 532 | 1.44 | 0.0193 | 0.0347 |
| chr21 | 17705285 | 17706092 | 808 | 1.65 | 0.0193 | 0.0347 |
| chr9 | 27539324 | 27540585 | 1262 | -1.14 | 0.0193 | 0.0347 |
| chr18 | 215283 | 216172 | 890 | 1.19 | 0.0193 | 0.0347 |
| chr5 | 61625674 | 61630006 | 4333 | 1.48 | 0.0193 | 0.0347 |
| chr7 | 81917518 | 81919360 | 1843 | 0.86 | 0.0193 | 0.0348 |
| chr2 | 85998932 | 86000550 | 1619 | -0.68 | 0.0193 | 0.0348 |
| chr11 | 106831461 | 106833064 | 1604 | -0.95 | 0.0193 | 0.0348 |
| chr11 | 89929916 | 89957657 | 27742 | 0.23 | 0.0193 | 0.0348 |
| chr5 | 39775503 | 39776479 | 977 | 1.68 | 0.0193 | 0.0348 |
| chr18 | 60291925 | 60293357 | 1433 | 1.78 | 0.0193 | 0.0348 |
| chr4 | 189775923 | 189776510 | 588 | 1.78 | 0.0194 | 0.0348 |
| chr14 | 93135715 | 93138135 | 2421 | -0.64 | 0.0194 | 0.0348 |
| chr14 | 73465093 | 73465507 | 415 | 2.08 | 0.0194 | 0.0348 |
| chr15 | 63614705 | 63674732 | 60028 | -0.09 | 0.0194 | 0.0349 |
| chr4 | 41642164 | 41643554 | 1391 | 1.22 | 0.0194 | 0.0349 |
| chr2 | 8079923 | 8080494 | 572 | 1.7 | 0.0194 | 0.0349 |
| chr2 | 20338761 | 20342003 | 3243 | -0.59 | 0.0194 | 0.035 |
| chr8 | 133995973 | 133996438 | 466 | -1.81 | 0.0194 | 0.035 |
| chr12 | 57144271 | 57151367 | 7097 | 0.41 | 0.0195 | 0.035 |
| chr12 | 1945603 | 1946193 | 591 | 2.25 | 0.0195 | 0.035 |
| chr1 | 115968504 | 115973490 | 4987 | -0.97 | 0.0195 | 0.035 |
| chr4 | 141293544 | 141295857 | 2314 | 0.5 | 0.0195 | 0.0351 |
| chr11 | 129116553 | 129120111 | 3559 | -0.59 | 0.0195 | 0.0351 |
| chr8 | 135516367 | 135517023 | 657 | -1.43 | 0.0195 | 0.0351 |
| chr21 | 30486735 | 30487622 | 888 | 1.67 | 0.0195 | 0.0351 |
| chr6 | 9463824 | 9468714 | 4891 | 0.9 | 0.0195 | 0.0351 |
| chr1 | 222909477 | 222910915 | 1439 | 0.95 | 0.0196 | 0.0351 |
| chr12 | 131512368 | 131513625 | 1258 | -0.99 | 0.0196 | 0.0352 |
| chr1 | 210611243 | 210614021 | 2779 | -0.64 | 0.0196 | 0.0352 |
| chr6 | 139093969 | 139095771 | 1803 | 0.55 | 0.0196 | 0.0353 |
| chr3 | 112736742 | 112739254 | 2513 | 0.36 | 0.0196 | 0.0353 |

| | | | | | | |
|---|---|---|---|---|---|---|
| chr1 | 184985934 | 184987274 | 1341 | -1.06 | 0.0196 | 0.0353 |
| chr12 | 59360478 | 59361269 | 792 | -1.26 | 0.0196 | 0.0353 |
| chr2 | 223486471 | 223488857 | 2387 | -0.78 | 0.0196 | 0.0353 |
| chr1 | 86860847 | 86862714 | 1868 | 0.44 | 0.0196 | 0.0353 |
| chr9 | 115298238 | 115298602 | 365 | -1.81 | 0.0197 | 0.0353 |
| chr4 | 83930209 | 83935582 | 5374 | 0.31 | 0.0197 | 0.0354 |
| chr15 | 64188009 | 64190002 | 1994 | -0.55 | 0.0197 | 0.0354 |
| chr2 | 175075581 | 175076279 | 699 | 1.6 | 0.0197 | 0.0354 |
| chr1 | 60196182 | 60197476 | 1295 | -0.7 | 0.0197 | 0.0354 |
| chr3 | 177137567 | 177139799 | 2233 | 0.85 | 0.0197 | 0.0354 |
| chr7 | 64532102 | 64532452 | 351 | 2.4 | 0.0197 | 0.0354 |
| chr8 | 87505070 | 87506035 | 966 | 1.61 | 0.0197 | 0.0355 |
| chr6 | 42134412 | 42134772 | 361 | 1.22 | 0.0198 | 0.0355 |
| chr12 | 32050547 | 32051368 | 822 | 1.33 | 0.0198 | 0.0355 |
| chr3 | 126678213 | 126679673 | 1461 | -0.8 | 0.0198 | 0.0355 |
| chr5 | 5334548 | 5351946 | 17399 | 0.3 | 0.0198 | 0.0355 |
| chr11 | 116937017 | 116945091 | 8075 | -0.28 | 0.0198 | 0.0355 |
| chr4 | 37649347 | 37672423 | 23077 | 0.31 | 0.0198 | 0.0355 |
| chr20 | 18521654 | 18522689 | 1036 | 1.45 | 0.0198 | 0.0355 |
| chr8 | 145514200 | 145520201 | 6002 | 0.31 | 0.0198 | 0.0355 |
| chr20 | 39192233 | 39192664 | 432 | 1.58 | 0.0198 | 0.0355 |
| chr9 | 102195583 | 102196706 | 1124 | 1 | 0.0198 | 0.0355 |
| chr18 | 74531966 | 74562260 | 30295 | 0.18 | 0.0198 | 0.0355 |
| chrX | 71791758 | 71793620 | 1863 | 0.58 | 0.0198 | 0.0356 |
| chr10 | 530324 | 531394 | 1071 | -0.97 | 0.0198 | 0.0356 |
| chr1 | 187694044 | 187694986 | 943 | -1.4 | 0.0198 | 0.0356 |
| chr2 | 205204018 | 205205549 | 1532 | -0.6 | 0.0198 | 0.0356 |
| chr13 | 103424990 | 103426679 | 1690 | 0.6 | 0.0198 | 0.0356 |
| chr2 | 232347113 | 232348342 | 1230 | -0.8 | 0.0199 | 0.0356 |
| chr1 | 204366527 | 204393494 | 26968 | 0.12 | 0.0199 | 0.0356 |
| chr11 | 14443129 | 14444783 | 1655 | -0.76 | 0.0199 | 0.0357 |
| chr15 | 73588018 | 73588814 | 797 | -1.39 | 0.0199 | 0.0357 |
| chr17 | 35987793 | 36004715 | 16923 | 0.27 | 0.0199 | 0.0357 |
| chr15 | 53975151 | 53976595 | 1445 | 0.78 | 0.0199 | 0.0357 |
| chrX | 86664783 | 86665111 | 329 | 2.53 | 0.0199 | 0.0357 |
| chr1 | 54447583 | 54448681 | 1099 | 1.68 | 0.0199 | 0.0358 |
| chr17 | 4800682 | 4813961 | 13280 | 0.97 | 0.02 | 0.0358 |
| chr2 | 28303348 | 28308842 | 5495 | 1.03 | 0.02 | 0.0358 |
| chr8 | 9370099 | 9370941 | 843 | 1.31 | 0.02 | 0.0358 |
| chr6 | 125420217 | 125424729 | 4513 | 0.53 | 0.02 | 0.0358 |
| chr4 | 124260931 | 124262360 | 1430 | 0.59 | 0.02 | 0.0359 |
| chr17 | 48618092 | 48652484 | 34393 | -0.23 | 0.02 | 0.0359 |
| chr8 | 54570007 | 54571536 | 1530 | -0.98 | 0.02 | 0.0359 |
| chr2 | 75873000 | 75875235 | 2236 | -0.37 | 0.0201 | 0.0359 |
| chr12 | 104151804 | 104201849 | 50046 | 0.18 | 0.0201 | 0.036 |
| chr4 | 39697256 | 39733444 | 36189 | 0.19 | 0.0201 | 0.036 |
| chr11 | 62651465 | 62657938 | 6474 | -0.56 | 0.0201 | 0.036 |
| chr11 | 117033039 | 117033764 | 726 | -1.58 | 0.0201 | 0.036 |
| chr15 | 28507014 | 28508124 | 1111 | -1.18 | 0.0201 | 0.036 |
| chr5 | 149540919 | 149541658 | 740 | -1.91 | 0.0201 | 0.036 |
| chr1 | 247016011 | 247035410 | 19400 | 0.36 | 0.0201 | 0.036 |
| chr9 | 138947167 | 138949350 | 2184 | -0.63 | 0.0201 | 0.0361 |
| chr15 | 78530467 | 78531379 | 913 | 1.25 | 0.0201 | 0.0361 |
| chr15 | 100243390 | 100249540 | 6151 | 0.63 | 0.0201 | 0.0361 |
| chr10 | 97802656 | 97804757 | 2102 | 0.42 | 0.0201 | 0.0361 |
| chr14 | 71630040 | 71630406 | 367 | 2.17 | 0.0202 | 0.0361 |
| chr22 | 23562153 | 23569440 | 7288 | 0.41 | 0.0202 | 0.0361 |
| chr20 | 43149519 | 43165501 | 15983 | 0.26 | 0.0202 | 0.0361 |
| chr7 | 139933247 | 139934092 | 846 | 1.45 | 0.0202 | 0.0361 |
| chr17 | 6872166 | 6873203 | 1038 | 1.21 | 0.0202 | 0.0362 |
| chr12 | 62893276 | 62895338 | 2063 | 0.83 | 0.0202 | 0.0362 |

| | | | | | | |
|---|---|---|---|---|---|---|
| chr10 | 127689898 | 127703028 | 13131 | 0.9 | 0.0202 | 0.0362 |
| chr9 | 131353965 | 131366450 | 12486 | -0.44 | 0.0202 | 0.0362 |
| chr3 | 188437930 | 188439031 | 1102 | -1.3 | 0.0203 | 0.0363 |
| chr12 | 121546082 | 121546836 | 755 | 2.23 | 0.0203 | 0.0363 |
| chr8 | 33265791 | 33266231 | 441 | -2.29 | 0.0203 | 0.0363 |
| chr2 | 37367013 | 37368370 | 1358 | -1.08 | 0.0203 | 0.0363 |
| chr6 | 72153583 | 72153984 | 402 | 1.83 | 0.0203 | 0.0364 |
| chr1 | 244255189 | 244268268 | 13080 | 2.48 | 0.0203 | 0.0364 |
| chr10 | 35642493 | 35658683 | 16191 | 0.45 | 0.0203 | 0.0364 |
| chr1 | 234962927 | 234963485 | 559 | -1.76 | 0.0203 | 0.0364 |
| chr12 | 95884058 | 95899081 | 15024 | 0.57 | 0.0204 | 0.0364 |
| chr5 | 124646618 | 124653584 | 6967 | -0.69 | 0.0204 | 0.0365 |
| chr1 | 62706789 | 62707800 | 1012 | 0.83 | 0.0204 | 0.0365 |
| chr17 | 38082719 | 38115906 | 33188 | -0.32 | 0.0204 | 0.0365 |
| chr1 | 232839556 | 232854914 | 15359 | 3.29 | 0.0204 | 0.0365 |
| chr5 | 159817848 | 159828190 | 10343 | 0.31 | 0.0204 | 0.0365 |
| chr2 | 19273073 | 19274137 | 1065 | -0.96 | 0.0204 | 0.0365 |
| chr5 | 13821477 | 13822058 | 582 | -1.79 | 0.0204 | 0.0365 |
| chr12 | 122353675 | 122366127 | 12453 | -0.36 | 0.0204 | 0.0365 |
| chr9 | 94157679 | 94158254 | 576 | -1.94 | 0.0204 | 0.0366 |
| chr1 | 154226440 | 154228325 | 1886 | 1.31 | 0.0204 | 0.0366 |
| chr3 | 119498420 | 119499187 | 768 | -1.13 | 0.0204 | 0.0366 |
| chr8 | 145596873 | 145598570 | 1698 | 0.72 | 0.0205 | 0.0366 |
| chr6 | 152622299 | 152652747 | 30449 | -0.3 | 0.0205 | 0.0366 |
| chr14 | 42654556 | 42664617 | 10062 | 0.22 | 0.0205 | 0.0366 |
| chr10 | 101445353 | 101445735 | 383 | -2.15 | 0.0205 | 0.0367 |
| chr1 | 197632195 | 197632996 | 802 | 1.39 | 0.0205 | 0.0367 |
| chr9 | 37309090 | 37310313 | 1224 | -1.13 | 0.0205 | 0.0367 |
| chr13 | 42376497 | 42377494 | 998 | -1.65 | 0.0205 | 0.0367 |
| chr9 | 101142089 | 101143363 | 1275 | 0.96 | 0.0206 | 0.0367 |
| chr16 | 1918927 | 1922959 | 4033 | 0.26 | 0.0206 | 0.0367 |
| chr2 | 46218349 | 46221239 | 2891 | -0.32 | 0.0206 | 0.0368 |
| chr13 | 103046141 | 103047524 | 1384 | -0.58 | 0.0206 | 0.0368 |
| chr1 | 50489269 | 50489838 | 570 | -1.85 | 0.0206 | 0.0368 |
| chr19 | 19701654 | 19702774 | 1121 | 0.85 | 0.0206 | 0.0369 |
| chr15 | 66231444 | 66233800 | 2357 | 0.62 | 0.0206 | 0.0369 |
| chr18 | 3281486 | 3297987 | 16502 | -0.51 | 0.0206 | 0.0369 |
| chr14 | 20795132 | 20827741 | 32610 | 0.17 | 0.0206 | 0.0369 |
| chr11 | 13353008 | 13364375 | 11368 | -0.17 | 0.0207 | 0.0369 |
| chr2 | 56967862 | 56970727 | 2866 | -0.29 | 0.0207 | 0.0369 |
| chr12 | 80838084 | 80840033 | 1950 | 0.74 | 0.0207 | 0.037 |
| chr10 | 34092051 | 34092663 | 613 | -1.51 | 0.0207 | 0.037 |
| chr14 | 36652248 | 36653185 | 938 | -1.84 | 0.0207 | 0.037 |
| chr11 | 67311431 | 67312981 | 1551 | 1.14 | 0.0207 | 0.037 |
| chr11 | 59303725 | 59348475 | 44751 | 0.12 | 0.0207 | 0.037 |
| chr12 | 69180579 | 69182131 | 1553 | 0.51 | 0.0208 | 0.037 |
| chr16 | 84129789 | 84151745 | 21957 | 0.25 | 0.0208 | 0.0371 |
| chr2 | 86073668 | 86086971 | 13304 | -0.37 | 0.0208 | 0.0371 |
| chr3 | 61235974 | 61237680 | 1707 | 0.87 | 0.0208 | 0.0371 |
| chr11 | 118954981 | 118978734 | 23754 | -0.19 | 0.0208 | 0.0371 |
| chr2 | 120436456 | 120437853 | 1398 | -0.78 | 0.0208 | 0.0371 |
| chr5 | 78208006 | 78208567 | 562 | 1.98 | 0.0208 | 0.0371 |
| chr13 | 41050404 | 41051397 | 994 | 1.58 | 0.0208 | 0.0371 |
| chr4 | 140096860 | 140110725 | 13866 | 0.46 | 0.0208 | 0.0372 |
| chr3 | 45223950 | 45225666 | 1717 | 0.61 | 0.0208 | 0.0372 |
| chr3 | 17425450 | 17427338 | 1889 | 0.76 | 0.0209 | 0.0372 |
| chr21 | 47563474 | 47564042 | 569 | 0.96 | 0.0209 | 0.0372 |
| chr2 | 42537827 | 42596889 | 59063 | -0.24 | 0.0209 | 0.0372 |
| chr2 | 201526919 | 201544941 | 18023 | 2.15 | 0.0209 | 0.0373 |
| chr11 | 21971119 | 21972392 | 1274 | 1.38 | 0.0209 | 0.0373 |
| chr4 | 189135678 | 189136831 | 1154 | -1.32 | 0.0209 | 0.0373 |

| | | | | | | |
|---|---|---|---|---|---|---|
| chr17 | 62008654 | 62009268 | 615 | -2.3 | 0.0209 | 0.0373 |
| chr13 | 84973102 | 84974157 | 1056 | 0.86 | 0.0209 | 0.0373 |
| chr7 | 304081 | 304896 | 816 | 1.35 | 0.0209 | 0.0373 |
| chr2 | 672615 | 678814 | 6200 | -0.26 | 0.021 | 0.0374 |
| chr5 | 121464506 | 121465829 | 1324 | 1.34 | 0.021 | 0.0374 |
| chr2 | 69782103 | 69810728 | 28626 | -0.87 | 0.021 | 0.0374 |
| chr14 | 54391207 | 54391929 | 723 | 1.39 | 0.021 | 0.0374 |
| chr5 | 37369916 | 37372177 | 2262 | 0.34 | 0.021 | 0.0374 |
| chr3 | 58291495 | 58293866 | 2372 | 0.49 | 0.021 | 0.0375 |
| chr15 | 79222955 | 79237685 | 14731 | 0.97 | 0.021 | 0.0375 |
| chr9 | 124793580 | 124795005 | 1426 | -0.98 | 0.0211 | 0.0375 |
| chr6 | 80815608 | 80816998 | 1391 | 0.86 | 0.0211 | 0.0376 |
| chr9 | 36371880 | 36372781 | 902 | 1.3 | 0.0211 | 0.0376 |
| chr22 | 36448410 | 36462924 | 14515 | -0.53 | 0.0211 | 0.0376 |
| chr6 | 8157913 | 8158559 | 647 | 2.24 | 0.0211 | 0.0376 |
| chr17 | 16423979 | 16424761 | 783 | -0.53 | 0.0211 | 0.0377 |
| chr2 | 225640287 | 225640947 | 661 | -1.64 | 0.0211 | 0.0377 |
| chr19 | 2385302 | 2386035 | 734 | 1.04 | 0.0211 | 0.0377 |
| chr5 | 34793386 | 34799427 | 6042 | -0.77 | 0.0212 | 0.0377 |
| chr8 | 119068081 | 119082056 | 13976 | -0.67 | 0.0212 | 0.0377 |
| chr10 | 62404129 | 62423972 | 19844 | 4.02 | 0.0212 | 0.0378 |
| chr18 | 42478421 | 42479136 | 716 | 1.77 | 0.0212 | 0.0378 |
| chr5 | 172898599 | 172900394 | 1796 | 0.83 | 0.0212 | 0.0378 |
| chrX | 53840036 | 53840743 | 708 | -1.47 | 0.0213 | 0.0379 |
| chr4 | 39353871 | 39368632 | 14762 | 0.28 | 0.0213 | 0.0379 |
| chr9 | 5720785 | 5722317 | 1533 | 0.81 | 0.0213 | 0.0379 |
| chr6 | 140494983 | 140495720 | 738 | -1.14 | 0.0213 | 0.0379 |
| chr22 | 19897694 | 19898063 | 370 | -2.43 | 0.0213 | 0.0379 |
| chr22 | 31733333 | 31744111 | 10779 | -0.35 | 0.0213 | 0.0379 |
| chr2 | 29415139 | 29416141 | 1003 | -1.13 | 0.0213 | 0.038 |
| chr8 | 53755232 | 53758306 | 3075 | 0.45 | 0.0213 | 0.038 |
| chr1 | 28413840 | 28415721 | 1882 | 0.57 | 0.0214 | 0.038 |
| chr3 | 25675633 | 25680487 | 4855 | -0.49 | 0.0214 | 0.038 |
| chr9 | 126382366 | 126383105 | 740 | -1.07 | 0.0214 | 0.038 |
| chr3 | 187778909 | 187779534 | 626 | 1.59 | 0.0214 | 0.0381 |
| chr6 | 134373231 | 134382082 | 8852 | -0.95 | 0.0214 | 0.0381 |
| chr4 | 28288613 | 28290519 | 1907 | -0.59 | 0.0214 | 0.0381 |
| chr9 | 2761064 | 2771553 | 10490 | -0.45 | 0.0214 | 0.0381 |
| chr7 | 2347551 | 2354041 | 6491 | 0.22 | 0.0214 | 0.0381 |
| chrX | 38739036 | 38740161 | 1126 | -1.23 | 0.0214 | 0.0381 |
| chr2 | 50371204 | 50371567 | 364 | -2.21 | 0.0214 | 0.0381 |
| chr16 | 53736575 | 53738740 | 2166 | -0.5 | 0.0214 | 0.0381 |
| chr8 | 30891266 | 30892696 | 1431 | 0.6 | 0.0214 | 0.0381 |
| chr4 | 143437406 | 143438426 | 1021 | 3.32 | 0.0214 | 0.0382 |
| chr7 | 48075045 | 48076020 | 976 | -1.02 | 0.0215 | 0.0382 |
| chr20 | 47599571 | 47612769 | 13199 | -0.34 | 0.0215 | 0.0382 |
| chr3 | 93780715 | 93783875 | 3161 | 0.24 | 0.0215 | 0.0382 |
| chr8 | 117740578 | 117754068 | 13491 | 0.46 | 0.0215 | 0.0382 |
| chr4 | 25909839 | 25922186 | 12348 | 0.44 | 0.0215 | 0.0382 |
| chr10 | 227946 | 228639 | 694 | 1.26 | 0.0215 | 0.0383 |
| chr15 | 35276728 | 35293188 | 16461 | 0.28 | 0.0215 | 0.0383 |
| chr5 | 168489373 | 168508600 | 19228 | -0.48 | 0.0215 | 0.0383 |
| chr11 | 73577278 | 73592672 | 15395 | 0.27 | 0.0215 | 0.0383 |
| chr16 | 120941 | 140905 | 19965 | 0.26 | 0.0215 | 0.0383 |
| chr1 | 11137552 | 11138361 | 810 | -1.15 | 0.0216 | 0.0383 |
| chr5 | 80167256 | 80167972 | 717 | -1.53 | 0.0216 | 0.0383 |
| chr1 | 57151349 | 57155482 | 4134 | -0.38 | 0.0216 | 0.0383 |
| chr18 | 47833824 | 47835247 | 1424 | 0.88 | 0.0216 | 0.0383 |
| chr5 | 42811419 | 42812727 | 1309 | 3.73 | 0.0216 | 0.0383 |
| chr3 | 35424923 | 35426267 | 1345 | -3.52 | 0.0216 | 0.0383 |
| chr5 | 37717358 | 37718474 | 1117 | 1.53 | 0.0216 | 0.0384 |

| | | | | | | |
|---|---|---|---|---|---|---|
| chr7 | 38350548 | 38358498 | 7951 | 1.06 | 0.0216 | 0.0384 |
| chr2 | 31760430 | 31761558 | 1129 | 1.3 | 0.0216 | 0.0384 |
| chr6 | 42280527 | 42307621 | 27095 | 0.24 | 0.0216 | 0.0384 |
| chr13 | 97773976 | 97775751 | 1776 | 0.9 | 0.0216 | 0.0384 |
| chr8 | 29982049 | 30000308 | 18260 | 0.26 | 0.0216 | 0.0384 |
| chr22 | 29260486 | 29261551 | 1066 | -1.78 | 0.0216 | 0.0384 |
| chr4 | 111152635 | 111153939 | 1305 | 1.35 | 0.0216 | 0.0384 |
| chr5 | 112093043 | 112102748 | 9706 | -0.7 | 0.0216 | 0.0384 |
| chr10 | 74870109 | 74899609 | 29501 | 0.2 | 0.0216 | 0.0384 |
| chr13 | 21871565 | 21884222 | 12658 | -0.6 | 0.0216 | 0.0384 |
| chr4 | 83903902 | 83911463 | 7562 | 0.29 | 0.0216 | 0.0384 |
| chr12 | 46149140 | 46149594 | 455 | 1.62 | 0.0217 | 0.0385 |
| chr19 | 55753360 | 55771862 | 18503 | 0.2 | 0.0217 | 0.0385 |
| chr4 | 6641424 | 6643734 | 2311 | 0.5 | 0.0217 | 0.0385 |
| chr5 | 99870161 | 99872577 | 2417 | 0.35 | 0.0217 | 0.0385 |
| chr17 | 65465309 | 65465926 | 618 | -1.56 | 0.0217 | 0.0385 |
| chr3 | 41011891 | 41014678 | 2788 | 0.35 | 0.0217 | 0.0385 |
| chr6 | 155567982 | 155575523 | 7542 | 0.51 | 0.0217 | 0.0385 |
| chr3 | 29632913 | 29633836 | 924 | 1.13 | 0.0217 | 0.0385 |
| chr15 | 73517417 | 73518805 | 1389 | -0.92 | 0.0217 | 0.0385 |
| chr3 | 119040822 | 119043079 | 2258 | 0.31 | 0.0217 | 0.0385 |
| chr19 | 58089308 | 58090838 | 1531 | 0.64 | 0.0217 | 0.0385 |
| chr2 | 25219165 | 25220603 | 1439 | -0.56 | 0.0217 | 0.0385 |
| chr1 | 202856599 | 202872489 | 15891 | 0.34 | 0.0217 | 0.0386 |
| chr17 | 42853337 | 42857438 | 4102 | -0.67 | 0.0217 | 0.0386 |
| chr22 | 39636222 | 39651875 | 15654 | -0.44 | 0.0217 | 0.0386 |
| chr5 | 67510177 | 67513181 | 3005 | 0.41 | 0.0217 | 0.0386 |
| chr2 | 231966104 | 231967034 | 931 | -1.29 | 0.0218 | 0.0386 |
| chr15 | 58987232 | 58998965 | 11734 | -0.55 | 0.0218 | 0.0386 |
| chr4 | 168138887 | 168139653 | 767 | 1.16 | 0.0218 | 0.0386 |
| chr16 | 30124312 | 30135194 | 10883 | -0.57 | 0.0218 | 0.0386 |
| chr16 | 80004303 | 80005359 | 1057 | -1.23 | 0.0218 | 0.0387 |
| chr7 | 95848388 | 95849334 | 947 | 1.33 | 0.0218 | 0.0387 |
| chr11 | 120677837 | 120679800 | 1964 | -0.77 | 0.0218 | 0.0387 |
| chr3 | 12595212 | 12626055 | 30844 | 0.23 | 0.0219 | 0.0388 |
| chr19 | 36868305 | 36870709 | 2405 | 0.46 | 0.0219 | 0.0388 |
| chr11 | 47026045 | 47026858 | 814 | 1.55 | 0.0219 | 0.0388 |
| chr13 | 24606564 | 24607552 | 989 | 1.21 | 0.0219 | 0.0388 |
| chr22 | 40029988 | 40031798 | 1811 | 0.72 | 0.0219 | 0.0388 |
| chr10 | 52498970 | 52500348 | 1379 | 0.63 | 0.0219 | 0.0388 |
| chr2 | 65144972 | 65146258 | 1287 | -0.86 | 0.0219 | 0.0388 |
| chr10 | 43539005 | 43540799 | 1795 | 0.7 | 0.0219 | 0.0388 |
| chr2 | 211286679 | 211295163 | 8485 | -1.91 | 0.0219 | 0.0388 |
| chr15 | 53903293 | 53904331 | 1039 | 1.03 | 0.0219 | 0.0389 |
| chr12 | 104513871 | 104516067 | 2197 | -0.91 | 0.0219 | 0.0389 |
| chr15 | 52023725 | 52030546 | 6822 | -0.65 | 0.022 | 0.0389 |
| chr8 | 90623052 | 90625087 | 2036 | 0.38 | 0.022 | 0.0389 |
| chr4 | 90032685 | 90051577 | 18893 | 0.2 | 0.022 | 0.0389 |
| chr9 | 3809493 | 3810454 | 962 | 1.3 | 0.022 | 0.039 |
| chr9 | 99867174 | 99867699 | 526 | 1.51 | 0.022 | 0.039 |
| chr14 | 35161967 | 35192739 | 30773 | 0.15 | 0.022 | 0.039 |
| chr12 | 90628237 | 90628855 | 619 | -1.53 | 0.022 | 0.039 |
| chr1 | 206674385 | 206674910 | 526 | -2.04 | 0.022 | 0.039 |
| chr8 | 48172692 | 48174285 | 1594 | 0.42 | 0.022 | 0.0391 |
| chr3 | 56165304 | 56167298 | 1995 | 0.87 | 0.0221 | 0.0391 |
| chr4 | 177698924 | 177716168 | 17245 | 0.33 | 0.0221 | 0.0391 |
| chr1 | 22160146 | 22160638 | 493 | 1.87 | 0.0221 | 0.0391 |
| chr15 | 45018464 | 45022459 | 3996 | 0.29 | 0.0221 | 0.0392 |
| chr2 | 72946618 | 72951465 | 4848 | 0.79 | 0.0221 | 0.0392 |
| chr17 | 49327932 | 49348392 | 20461 | 0.2 | 0.0222 | 0.0393 |
| chr2 | 223701923 | 223709607 | 7685 | 0.3 | 0.0222 | 0.0393 |

| | | | | | | |
|---|---|---|---|---|---|---|
| chr8 | 101800660 | 101801905 | 1246 | -0.73 | 0.0222 | 0.0393 |
| chr7 | 129825151 | 129830069 | 4919 | -0.59 | 0.0223 | 0.0394 |
| chr1 | 94269594 | 94294343 | 24750 | -0.52 | 0.0223 | 0.0394 |
| chr17 | 4934462 | 4939105 | 4644 | 0.72 | 0.0223 | 0.0394 |
| chr3 | 15719637 | 15733904 | 14268 | -0.39 | 0.0223 | 0.0395 |
| chr3 | 187085989 | 187086406 | 418 | 1.73 | 0.0223 | 0.0395 |
| chr16 | 48269227 | 48285106 | 15880 | 0.29 | 0.0223 | 0.0395 |
| chr7 | 83671387 | 83672094 | 708 | 1.29 | 0.0223 | 0.0395 |
| chr3 | 184276483 | 184277898 | 1416 | 0.82 | 0.0223 | 0.0395 |
| chr2 | 131094897 | 131114252 | 19356 | -0.36 | 0.0224 | 0.0396 |
| chr16 | 4937356 | 4947963 | 10608 | -0.51 | 0.0224 | 0.0396 |
| chr7 | 156609918 | 156611402 | 1485 | -0.78 | 0.0224 | 0.0396 |
| chr5 | 138179905 | 138181589 | 1685 | 0.83 | 0.0224 | 0.0396 |
| chr15 | 58681316 | 58682341 | 1026 | 1.24 | 0.0224 | 0.0397 |
| chr9 | 133900255 | 133901294 | 1040 | -1.33 | 0.0224 | 0.0397 |
| chr2 | 135987636 | 135988298 | 663 | -1.38 | 0.0225 | 0.0397 |
| chr18 | 12552905 | 12553632 | 728 | -1.92 | 0.0225 | 0.0398 |
| chr10 | 82422664 | 82423272 | 609 | 1.4 | 0.0225 | 0.0398 |
| chr14 | 68533468 | 68561707 | 28240 | 0.24 | 0.0225 | 0.0398 |
| chr3 | 111414458 | 111415212 | 755 | -1.24 | 0.0225 | 0.0398 |
| chr5 | 74861242 | 74875182 | 13941 | 0.24 | 0.0225 | 0.0398 |
| chr2 | 219366218 | 219367680 | 1463 | -0.95 | 0.0225 | 0.0399 |
| chr17 | 3658470 | 3658943 | 474 | 1.89 | 0.0226 | 0.0399 |
| chr7 | 73294803 | 73296156 | 1354 | 1.42 | 0.0226 | 0.0399 |
| chr2 | 4961705 | 4963166 | 1462 | 3.56 | 0.0226 | 0.0399 |
| chr10 | 74774512 | 74775984 | 1473 | -0.9 | 0.0226 | 0.0399 |
| chr9 | 124361505 | 124375574 | 14070 | -0.78 | 0.0226 | 0.0399 |
| chr18 | 12656281 | 12658962 | 2682 | 0.55 | 0.0226 | 0.0399 |
| chrX | 13185424 | 13186287 | 864 | 1.44 | 0.0226 | 0.0399 |
| chr3 | 181929993 | 181931418 | 1426 | 0.82 | 0.0226 | 0.0399 |
| chr11 | 64836695 | 64864984 | 28290 | -0.26 | 0.0226 | 0.04 |
| chr2 | 3208530 | 3209364 | 835 | -1.44 | 0.0226 | 0.04 |
| chr8 | 58907128 | 58907666 | 539 | 2.26 | 0.0227 | 0.04 |
| chr9 | 82424600 | 82427987 | 3388 | 2.34 | 0.0227 | 0.04 |
| chr11 | 73017652 | 73046915 | 29264 | -0.2 | 0.0227 | 0.04 |
| chr9 | 114986820 | 114987491 | 672 | -1.1 | 0.0227 | 0.04 |
| chr4 | 78782698 | 78785259 | 2562 | 0.31 | 0.0227 | 0.0401 |
| chr14 | 69617789 | 69620911 | 3123 | 0.42 | 0.0227 | 0.0401 |
| chr7 | 20777037 | 20778675 | 1639 | 0.77 | 0.0227 | 0.0401 |
| chr10 | 43301211 | 43302060 | 850 | 1.56 | 0.0227 | 0.0401 |
| chr8 | 21441136 | 21441524 | 389 | -1.9 | 0.0227 | 0.0401 |
| chrX | 139013628 | 139016582 | 2955 | 0.41 | 0.0227 | 0.0401 |
| chr6 | 36610793 | 36611627 | 835 | -1.26 | 0.0227 | 0.0401 |
| chr4 | 145958512 | 145959314 | 803 | 2.18 | 0.0227 | 0.0401 |
| chr4 | 186758907 | 186769591 | 10685 | 0.33 | 0.0227 | 0.0401 |
| chr8 | 52929426 | 52930049 | 624 | -1.7 | 0.0227 | 0.0402 |
| chr2 | 128101879 | 128102689 | 811 | -2.19 | 0.0228 | 0.0402 |
| chr11 | 45684390 | 45685001 | 612 | -1.59 | 0.0228 | 0.0402 |
| chr15 | 78440419 | 78455233 | 14815 | -0.34 | 0.0228 | 0.0402 |
| chr17 | 28760318 | 28761206 | 889 | -1.02 | 0.0228 | 0.0402 |
| chr17 | 80476588 | 80568256 | 91669 | -0.12 | 0.0228 | 0.0402 |
| chr10 | 116639229 | 116639916 | 688 | 1.89 | 0.0228 | 0.0402 |
| chr2 | 10790156 | 10790671 | 516 | -1.57 | 0.0228 | 0.0402 |
| chr18 | 26743157 | 26746207 | 3051 | -0.52 | 0.0228 | 0.0402 |
| chr1 | 179783464 | 179784973 | 1510 | 0.83 | 0.0228 | 0.0403 |
| chr2 | 73496703 | 73497087 | 385 | 1.86 | 0.0228 | 0.0403 |
| chr2 | 183942846 | 183944263 | 1418 | 0.75 | 0.0228 | 0.0403 |
| chr15 | 45878476 | 45881202 | 2727 | 0.28 | 0.0228 | 0.0403 |
| chr19 | 50832912 | 50837731 | 4820 | 0.52 | 0.0229 | 0.0403 |
| chr1 | 178280054 | 178280375 | 322 | -1.55 | 0.0229 | 0.0403 |
| chr20 | 11523684 | 11525337 | 1654 | -0.64 | 0.0229 | 0.0404 |

| | | | | | | |
|---|---|---|---|---|---|---|
| chr1 | 200147692 | 200153995 | 6304 | -0.4 | 0.0229 | 0.0404 |
| chr11 | 12646900 | 12647870 | 971 | 1.29 | 0.0229 | 0.0404 |
| chr15 | 70877326 | 70890305 | 12980 | 3.82 | 0.0229 | 0.0404 |
| chr16 | 75411206 | 75412510 | 1305 | 1.01 | 0.0229 | 0.0404 |
| chrX | 54834409 | 54835994 | 1586 | 0.82 | 0.0229 | 0.0404 |
| chr10 | 79730485 | 79743753 | 13269 | -0.59 | 0.0229 | 0.0404 |
| chr12 | 118643571 | 118657162 | 13592 | -0.59 | 0.0229 | 0.0404 |
| chr2 | 174435586 | 174440060 | 4475 | 0.39 | 0.0229 | 0.0404 |
| chr18 | 29449079 | 29449629 | 551 | 1.52 | 0.0229 | 0.0405 |
| chrX | 133645943 | 133647238 | 1296 | -1.07 | 0.023 | 0.0405 |
| chr3 | 123343444 | 123369177 | 25734 | -0.43 | 0.023 | 0.0405 |
| chr16 | 4048844 | 4049771 | 928 | -2.13 | 0.023 | 0.0405 |
| chr10 | 86070153 | 86071128 | 976 | -1.11 | 0.023 | 0.0405 |
| chr2 | 113011715 | 113013235 | 1521 | -0.61 | 0.023 | 0.0405 |
| chr5 | 156886662 | 156892136 | 5475 | -0.86 | 0.023 | 0.0405 |
| chr18 | 10213194 | 10214139 | 946 | 2.18 | 0.023 | 0.0405 |
| chr15 | 56923618 | 56924921 | 1304 | 0.84 | 0.023 | 0.0406 |
| chr8 | 66976335 | 66976953 | 619 | -3.21 | 0.023 | 0.0406 |
| chr12 | 12486944 | 12487417 | 474 | 1.43 | 0.023 | 0.0406 |
| chr8 | 82192297 | 82194763 | 2467 | -0.61 | 0.023 | 0.0406 |
| chr2 | 121968119 | 121969489 | 1371 | -1.06 | 0.0231 | 0.0406 |
| chr4 | 103119944 | 103120222 | 279 | 2.18 | 0.0231 | 0.0406 |
| chr12 | 25307616 | 25308772 | 1157 | 0.89 | 0.0231 | 0.0407 |
| chr5 | 111092192 | 111094737 | 2546 | 0.38 | 0.0231 | 0.0407 |
| chr9 | 83498300 | 83502757 | 4458 | 3.8 | 0.0231 | 0.0407 |
| chr20 | 2476822 | 2506420 | 29599 | -0.42 | 0.0231 | 0.0407 |
| chrX | 118369755 | 118373429 | 3675 | -0.44 | 0.0231 | 0.0407 |
| chr18 | 48593927 | 48595489 | 1563 | 0.79 | 0.0231 | 0.0408 |
| chr11 | 2762371 | 2763900 | 1530 | -0.76 | 0.0231 | 0.0408 |
| chr16 | 74611325 | 74673239 | 61915 | 0.18 | 0.0232 | 0.0408 |
| chr20 | 14858105 | 14862559 | 4455 | -0.2 | 0.0232 | 0.0408 |
| chr17 | 16108246 | 16121597 | 13352 | 0.19 | 0.0232 | 0.0409 |
| chr10 | 104236144 | 104237148 | 1005 | -1.15 | 0.0232 | 0.0409 |
| chr13 | 109510581 | 109513564 | 2984 | 0.38 | 0.0232 | 0.0409 |
| chr6 | 41604323 | 41605206 | 884 | 1.63 | 0.0232 | 0.0409 |
| chr20 | 45033632 | 45035598 | 1967 | 0.59 | 0.0233 | 0.0409 |
| chr16 | 58884027 | 58884293 | 267 | -1.96 | 0.0233 | 0.0409 |
| chr9 | 130150170 | 130169775 | 19606 | -0.58 | 0.0233 | 0.041 |
| chr1 | 202250498 | 202251670 | 1173 | -1.35 | 0.0233 | 0.041 |
| chr14 | 81406788 | 81409094 | 2307 | -0.61 | 0.0233 | 0.041 |
| chr11 | 64529540 | 64547851 | 18312 | 0.28 | 0.0233 | 0.0411 |
| chr19 | 39018663 | 39019289 | 627 | 1.23 | 0.0233 | 0.0411 |
| chr3 | 113213211 | 113235475 | 22265 | -0.27 | 0.0233 | 0.0411 |
| chr7 | 134330265 | 134332617 | 2353 | 0.39 | 0.0233 | 0.0411 |
| chr12 | 39538824 | 39539781 | 958 | 1.08 | 0.0234 | 0.0411 |
| chr12 | 56472324 | 56474074 | 1751 | 0.56 | 0.0234 | 0.0411 |
| chr3 | 142677521 | 142697371 | 19851 | 0.25 | 0.0234 | 0.0411 |
| chr5 | 106637655 | 106638330 | 676 | -1.11 | 0.0234 | 0.0412 |
| chr2 | 128211698 | 128213712 | 2015 | 0.69 | 0.0234 | 0.0412 |
| chr6 | 168555556 | 168556065 | 510 | -1.93 | 0.0234 | 0.0412 |
| chr6 | 154724833 | 154736666 | 11834 | 0.27 | 0.0234 | 0.0412 |
| chr4 | 127770749 | 127772651 | 1903 | 0.84 | 0.0234 | 0.0412 |
| chr3 | 28966164 | 28967065 | 902 | -1.85 | 0.0235 | 0.0412 |
| chr19 | 10024568 | 10026112 | 1545 | -0.55 | 0.0235 | 0.0413 |
| chr12 | 42628916 | 42633158 | 4243 | 0.32 | 0.0235 | 0.0413 |
| chr15 | 30356255 | 30356700 | 446 | -1.45 | 0.0235 | 0.0413 |
| chr6 | 11650636 | 11656401 | 5766 | -0.73 | 0.0235 | 0.0413 |
| chr4 | 186316213 | 186318378 | 2166 | 0.49 | 0.0235 | 0.0414 |
| chr6 | 74139145 | 74139591 | 447 | 2.39 | 0.0235 | 0.0414 |
| chr17 | 68118773 | 68119906 | 1134 | -3.22 | 0.0236 | 0.0414 |
| chr2 | 187380120 | 187381554 | 1435 | -0.95 | 0.0236 | 0.0415 |

| | | | | | | |
|---|---|---|---|---|---|---|
| chr4 | 139930332 | 139987544 | 57213 | 0.16 | 0.0236 | 0.0415 |
| chr16 | 19786220 | 19787483 | 1264 | -1.26 | 0.0236 | 0.0415 |
| chr19 | 6674731 | 6675485 | 755 | -1.12 | 0.0236 | 0.0415 |
| chr6 | 79942761 | 79944947 | 2187 | 0.62 | 0.0236 | 0.0416 |
| chr20 | 4270420 | 4271123 | 704 | 2.04 | 0.0237 | 0.0416 |
| chr11 | 104838557 | 104839943 | 1387 | 0.52 | 0.0237 | 0.0416 |
| chr15 | 76552681 | 76555713 | 3033 | -0.77 | 0.0237 | 0.0416 |
| chr1 | 218877077 | 218886273 | 9197 | 0.17 | 0.0237 | 0.0416 |
| chr22 | 36972936 | 36983952 | 11017 | -1.09 | 0.0237 | 0.0417 |
| chr9 | 93062276 | 93065176 | 2901 | 0.34 | 0.0237 | 0.0417 |
| chr4 | 111556979 | 111558584 | 1606 | 0.87 | 0.0238 | 0.0417 |
| chr20 | 441935 | 443795 | 1861 | 0.56 | 0.0238 | 0.0417 |
| chr17 | 21219412 | 21226406 | 6995 | -3.31 | 0.0238 | 0.0417 |
| chr11 | 77548973 | 77550479 | 1507 | -0.88 | 0.0238 | 0.0417 |
| chr11 | 3851354 | 3882800 | 31447 | 0.21 | 0.0238 | 0.0417 |
| chr6 | 137978394 | 137979962 | 1569 | 0.67 | 0.0238 | 0.0418 |
| chr3 | 52187315 | 52189512 | 2198 | 0.54 | 0.0238 | 0.0419 |
| chr7 | 2371502 | 2372878 | 1377 | -1.42 | 0.0239 | 0.0419 |
| chr16 | 18938418 | 18939138 | 721 | 1.29 | 0.0239 | 0.0419 |
| chr11 | 85745222 | 85745894 | 673 | 1.58 | 0.0239 | 0.0419 |
| chr8 | 103209188 | 103209734 | 547 | 1.62 | 0.0239 | 0.0419 |
| chr9 | 123883333 | 123885065 | 1733 | 0.51 | 0.0239 | 0.042 |
| chr2 | 150467565 | 150467986 | 422 | -2.35 | 0.0239 | 0.042 |
| chr4 | 41991639 | 41993929 | 2291 | 0.49 | 0.0239 | 0.042 |
| chr4 | 24913687 | 24915090 | 1404 | 0.56 | 0.0239 | 0.042 |
| chr1 | 6803613 | 6804159 | 547 | 1.31 | 0.0239 | 0.042 |
| chr11 | 77406267 | 77422313 | 16047 | 0.49 | 0.0239 | 0.042 |
| chr12 | 52207757 | 52222285 | 14529 | 0.32 | 0.024 | 0.0421 |
| chr10 | 21419743 | 21425185 | 5443 | 0.35 | 0.024 | 0.0421 |
| chr18 | 12882626 | 12885030 | 2405 | 0.43 | 0.0241 | 0.0423 |
| chr9 | 110918523 | 110926558 | 8036 | -3.27 | 0.0241 | 0.0423 |
| chr18 | 43329629 | 43330953 | 1325 | 0.8 | 0.0241 | 0.0423 |
| chr4 | 153561692 | 153562518 | 827 | 1.21 | 0.0241 | 0.0423 |
| chr3 | 105510771 | 105512635 | 1865 | 0.61 | 0.0241 | 0.0424 |
| chr10 | 44795900 | 44797752 | 1853 | 0.69 | 0.0242 | 0.0424 |
| chr7 | 30292525 | 30293062 | 538 | 2.18 | 0.0242 | 0.0424 |
| chr2 | 63272272 | 63287702 | 15431 | -0.46 | 0.0242 | 0.0424 |
| chr12 | 21035863 | 21036862 | 1000 | 0.95 | 0.0242 | 0.0425 |
| chr1 | 153852967 | 153853847 | 881 | 2.18 | 0.0242 | 0.0425 |
| chr5 | 1343098 | 1346193 | 3096 | 0.43 | 0.0242 | 0.0425 |
| chr9 | 115473166 | 115481267 | 8102 | 0.38 | 0.0242 | 0.0425 |
| chrX | 101967494 | 101976133 | 8640 | 0.61 | 0.0243 | 0.0425 |
| chr1 | 116652212 | 116653352 | 1141 | 0.89 | 0.0243 | 0.0425 |
| chr6 | 162338620 | 162346094 | 7475 | 0.38 | 0.0243 | 0.0426 |
| chr4 | 134486262 | 134487363 | 1102 | 3.3 | 0.0243 | 0.0426 |
| chr3 | 176912729 | 176931166 | 18438 | 0.22 | 0.0243 | 0.0427 |
| chr10 | 32383588 | 32384289 | 702 | 1.67 | 0.0243 | 0.0427 |
| chr20 | 13737233 | 13740789 | 3557 | -0.7 | 0.0244 | 0.0427 |
| chr7 | 11101460 | 11102024 | 565 | 1.66 | 0.0244 | 0.0427 |
| chr10 | 92944647 | 92945753 | 1107 | -1.19 | 0.0244 | 0.0427 |
| chr1 | 237034527 | 237035281 | 755 | -1.29 | 0.0244 | 0.0427 |
| chrX | 107429917 | 107430830 | 914 | -1.97 | 0.0244 | 0.0427 |
| chr10 | 35595465 | 35596449 | 985 | 1.07 | 0.0244 | 0.0428 |
| chr18 | 7749056 | 7762251 | 13196 | -0.39 | 0.0244 | 0.0428 |
| chr8 | 27837825 | 27862756 | 24932 | -0.35 | 0.0244 | 0.0428 |
| chr12 | 72014335 | 72015500 | 1166 | 1.07 | 0.0244 | 0.0428 |
| chrX | 37208588 | 37210205 | 1618 | 0.66 | 0.0245 | 0.0428 |
| chr2 | 166809442 | 166811294 | 1853 | -0.52 | 0.0245 | 0.0428 |
| chr16 | 15951917 | 15952447 | 531 | -1.7 | 0.0245 | 0.0429 |
| chr2 | 233603486 | 233604138 | 653 | -2.12 | 0.0245 | 0.0429 |
| chr6 | 100676432 | 100677442 | 1011 | 0.96 | 0.0245 | 0.0429 |

| | | | | | | |
|---|---|---|---|---|---|---|
| chr16 | 11834950 | 11852399 | 17450 | 0.23 | 0.0245 | 0.043 |
| chrX | 17565092 | 17586086 | 20995 | -0.34 | 0.0245 | 0.043 |
| chr9 | 4814639 | 4815983 | 1345 | 0.81 | 0.0246 | 0.043 |
| chr1 | 204041979 | 204100826 | 58848 | -0.16 | 0.0246 | 0.0431 |
| chr9 | 6152376 | 6154172 | 1797 | 3.71 | 0.0246 | 0.0431 |
| chr16 | 30405381 | 30442218 | 36838 | -0.23 | 0.0246 | 0.0431 |
| chr1 | 20125457 | 20127400 | 1944 | 0.39 | 0.0246 | 0.0431 |
| chr2 | 19807436 | 19813344 | 5909 | -0.46 | 0.0246 | 0.0431 |
| chr4 | 31997499 | 32006439 | 8941 | 0.25 | 0.0246 | 0.0431 |
| chr10 | 26669397 | 26670483 | 1087 | -3.25 | 0.0246 | 0.0431 |
| chr2 | 205846765 | 205847544 | 780 | -1.27 | 0.0246 | 0.0431 |
| chr2 | 121244205 | 121245005 | 801 | -1.99 | 0.0247 | 0.0432 |
| chr16 | 87239923 | 87241706 | 1784 | 0.67 | 0.0247 | 0.0432 |
| chr3 | 94159414 | 94160960 | 1547 | 3.6 | 0.0247 | 0.0432 |
| chr1 | 21298741 | 21299235 | 495 | -1.75 | 0.0247 | 0.0432 |
| chrX | 103587457 | 103588400 | 944 | 1.06 | 0.0247 | 0.0432 |
| chr3 | 47322475 | 47325758 | 3284 | -0.23 | 0.0247 | 0.0432 |
| chr3 | 64163607 | 64177660 | 14054 | 0.19 | 0.0247 | 0.0432 |
| chr15 | 51794771 | 51795778 | 1008 | -0.88 | 0.0247 | 0.0432 |
| chr12 | 112392283 | 112393364 | 1082 | -1.57 | 0.0247 | 0.0433 |
| chr19 | 8007652 | 8009545 | 1894 | 0.42 | 0.0247 | 0.0433 |
| chr11 | 64739169 | 64740157 | 989 | 1.17 | 0.0248 | 0.0433 |
| chr1 | 164527323 | 164533563 | 6241 | 0.44 | 0.0248 | 0.0433 |
| chr7 | 129663874 | 129669171 | 5298 | 0.24 | 0.0248 | 0.0434 |
| chr3 | 101659248 | 101659834 | 587 | 1.64 | 0.0248 | 0.0434 |
| chr2 | 238968515 | 238970971 | 2457 | -0.51 | 0.0248 | 0.0434 |
| chr14 | 103285428 | 103286459 | 1032 | 1.12 | 0.0248 | 0.0434 |
| chr7 | 73810849 | 73812098 | 1250 | -1.12 | 0.0248 | 0.0434 |
| chr18 | 55981001 | 55989361 | 8361 | -0.6 | 0.0248 | 0.0434 |
| chr14 | 96854015 | 96859451 | 5437 | -0.39 | 0.0248 | 0.0434 |
| chr4 | 146600441 | 146601936 | 1496 | 0.91 | 0.0249 | 0.0435 |
| chr9 | 133837541 | 133838741 | 1201 | 1.1 | 0.0249 | 0.0435 |
| chr6 | 9141062 | 9142008 | 947 | 0.96 | 0.0249 | 0.0435 |
| chr19 | 33071143 | 33094569 | 23427 | 0.25 | 0.0249 | 0.0435 |
| chr15 | 74526177 | 74535450 | 9274 | -0.58 | 0.0249 | 0.0435 |
| chr12 | 38710576 | 38711241 | 666 | 0.76 | 0.0249 | 0.0435 |
| chrX | 64815540 | 64822089 | 6550 | 0.89 | 0.0249 | 0.0436 |
| chrX | 17366234 | 17367227 | 994 | 0.85 | 0.025 | 0.0436 |
| chr14 | 97646534 | 97647296 | 763 | -2.21 | 0.025 | 0.0436 |
| chr11 | 19261672 | 19263606 | 1935 | -0.64 | 0.025 | 0.0436 |
| chr12 | 125517351 | 125518403 | 1053 | -1.32 | 0.025 | 0.0437 |
| chr10 | 65071786 | 65072654 | 869 | 1.09 | 0.025 | 0.0437 |
| chr17 | 74117452 | 74138012 | 20561 | -0.42 | 0.025 | 0.0437 |
| chr22 | 24862428 | 24863503 | 1076 | 1.15 | 0.025 | 0.0438 |
| chr2 | 61231170 | 61253216 | 22047 | 0.21 | 0.0251 | 0.0438 |
| chr4 | 175075197 | 175076950 | 1754 | 0.48 | 0.0251 | 0.0438 |
| chr8 | 10676332 | 10677089 | 758 | -1.19 | 0.0251 | 0.0438 |
| chr14 | 34360699 | 34361969 | 1271 | 3.36 | 0.0251 | 0.0438 |
| chr15 | 48104065 | 48105973 | 1909 | 0.88 | 0.0251 | 0.0438 |
| chr17 | 1106862 | 1107480 | 619 | 1.05 | 0.0251 | 0.0439 |
| chr1 | 25664825 | 25666734 | 1910 | 0.54 | 0.0251 | 0.0439 |
| chr22 | 30840568 | 30841320 | 753 | -1.11 | 0.0252 | 0.0439 |
| chr12 | 52976397 | 52977242 | 846 | -1.42 | 0.0252 | 0.0439 |
| chr20 | 47489097 | 47489907 | 811 | -1.21 | 0.0252 | 0.044 |
| chr18 | 2791405 | 2800241 | 8837 | 0.66 | 0.0252 | 0.044 |
| chr18 | 45987122 | 45988248 | 1127 | 1.46 | 0.0252 | 0.0441 |
| chr11 | 63741624 | 63755792 | 14169 | -0.21 | 0.0253 | 0.0441 |
| chr13 | 107569744 | 107570694 | 951 | 0.72 | 0.0253 | 0.0441 |
| chr1 | 246727914 | 246733258 | 5345 | 0.31 | 0.0253 | 0.0441 |
| chr1 | 203805636 | 203832748 | 27113 | 0.25 | 0.0253 | 0.0441 |
| chr12 | 107450251 | 107452162 | 1912 | 3.89 | 0.0253 | 0.0442 |

| | | | | | | |
|---|---|---|---|---|---|---|
| chr11 | 235538 | 238440 | 2903 | 0.32 | 0.0253 | 0.0442 |
| chr8 | 23820390 | 23821655 | 1266 | 1.13 | 0.0253 | 0.0442 |
| chr12 | 80304485 | 80306686 | 2202 | -0.7 | 0.0254 | 0.0443 |
| chr8 | 145829861 | 145830901 | 1041 | -1.25 | 0.0254 | 0.0443 |
| chr22 | 28197735 | 28198605 | 871 | -0.98 | 0.0255 | 0.0444 |
| chr1 | 222839353 | 222839979 | 627 | 1.34 | 0.0255 | 0.0444 |
| chr12 | 123154376 | 123167463 | 13088 | 0.31 | 0.0255 | 0.0444 |
| chr18 | 55747605 | 55748259 | 655 | 1.4 | 0.0255 | 0.0444 |
| chr16 | 69344417 | 69380450 | 36034 | 0.2 | 0.0255 | 0.0444 |
| chr6 | 17838414 | 17839556 | 1143 | -1.94 | 0.0255 | 0.0444 |
| chr4 | 20701228 | 20703108 | 1881 | -0.41 | 0.0255 | 0.0444 |
| chr14 | 34766827 | 34767637 | 811 | -1.29 | 0.0255 | 0.0445 |
| chr6 | 83962691 | 83963131 | 441 | -2.15 | 0.0255 | 0.0445 |
| chr1 | 166879919 | 166880394 | 476 | 1.48 | 0.0255 | 0.0445 |
| chr22 | 23412763 | 23413226 | 464 | -1.8 | 0.0255 | 0.0445 |
| chr17 | 1415106 | 1467103 | 51998 | -0.18 | 0.0256 | 0.0446 |
| chr7 | 138706454 | 138745958 | 39505 | 0.23 | 0.0256 | 0.0446 |
| chr3 | 127432146 | 127433456 | 1311 | 1.1 | 0.0256 | 0.0446 |
| chr5 | 54994251 | 54994807 | 557 | -2.17 | 0.0256 | 0.0447 |
| chr20 | 55530657 | 55532774 | 2118 | 1.24 | 0.0256 | 0.0447 |
| chr12 | 26843981 | 26858766 | 14786 | -1.06 | 0.0256 | 0.0447 |
| chr7 | 66145576 | 66147878 | 2303 | 0.36 | 0.0256 | 0.0447 |
| chr20 | 36919327 | 36919948 | 622 | 1.11 | 0.0256 | 0.0447 |
| chr8 | 68150042 | 68155271 | 5230 | 0.76 | 0.0257 | 0.0447 |
| chr19 | 15488266 | 15493162 | 4897 | -0.33 | 0.0257 | 0.0447 |
| chrX | 53009086 | 53010854 | 1769 | -0.86 | 0.0257 | 0.0447 |
| chr18 | 28590932 | 28592084 | 1153 | 3.27 | 0.0257 | 0.0447 |
| chr6 | 86279109 | 86304755 | 25647 | 0.31 | 0.0257 | 0.0448 |
| chr13 | 78554581 | 78555337 | 757 | -1.1 | 0.0257 | 0.0448 |
| chr12 | 17033676 | 17034682 | 1007 | 0.77 | 0.0257 | 0.0448 |
| chr4 | 159367165 | 159367759 | 595 | -2.01 | 0.0257 | 0.0448 |
| chr13 | 96032690 | 96033556 | 867 | -1.25 | 0.0258 | 0.0449 |
| chr20 | 35082975 | 35122731 | 39757 | 0.16 | 0.0258 | 0.0449 |
| chrX | 57936941 | 57937595 | 655 | 1.16 | 0.0258 | 0.0449 |
| chr9 | 128693675 | 128694654 | 980 | -1.29 | 0.0258 | 0.0449 |
| chr22 | 24230130 | 24238454 | 8325 | -0.31 | 0.0258 | 0.0449 |
| chr2 | 70073701 | 70075787 | 2087 | -0.91 | 0.0258 | 0.045 |
| chr11 | 85532337 | 85566789 | 34453 | 0.32 | 0.0258 | 0.045 |
| chr2 | 30830914 | 30831893 | 980 | -1.01 | 0.0258 | 0.045 |
| chr18 | 55709916 | 55722497 | 12582 | 0.33 | 0.0259 | 0.045 |
| chr3 | 118656928 | 118658639 | 1712 | 0.57 | 0.0259 | 0.0451 |
| chr4 | 183996103 | 183996554 | 452 | 2.26 | 0.0259 | 0.0451 |
| chr3 | 11990355 | 11990831 | 477 | -1.55 | 0.0259 | 0.0451 |
| chr5 | 151119035 | 151121734 | 2700 | -0.73 | 0.0259 | 0.0451 |
| chr4 | 111027068 | 111029327 | 2260 | 0.39 | 0.0259 | 0.0451 |
| chr15 | 48936934 | 48938607 | 1674 | 1.52 | 0.0259 | 0.0451 |
| chr2 | 221698947 | 221700330 | 1384 | 1.1 | 0.026 | 0.0452 |
| chrX | 96412646 | 96419312 | 6667 | 0.25 | 0.026 | 0.0452 |
| chr3 | 17812089 | 17813347 | 1259 | -0.93 | 0.026 | 0.0452 |
| chr6 | 3053619 | 3089514 | 35896 | 0.25 | 0.026 | 0.0453 |
| chr7 | 41744417 | 41745435 | 1019 | 0.97 | 0.026 | 0.0453 |
| chr11 | 3801861 | 3830366 | 28506 | 0.21 | 0.026 | 0.0453 |
| chr17 | 47885831 | 47929233 | 43403 | 0.19 | 0.026 | 0.0453 |
| chr4 | 75718923 | 75720245 | 1323 | 0.43 | 0.026 | 0.0453 |
| chr21 | 30364076 | 30376058 | 11983 | 0.34 | 0.026 | 0.0453 |
| chr8 | 1795182 | 1795770 | 589 | -2.13 | 0.026 | 0.0453 |
| chr5 | 138692304 | 138740562 | 48259 | 0.14 | 0.0261 | 0.0453 |
| chr11 | 16876070 | 16877287 | 1218 | 1.2 | 0.0261 | 0.0453 |
| chr13 | 21713593 | 21724880 | 11288 | 0.28 | 0.0261 | 0.0453 |
| chr12 | 32495956 | 32498029 | 2074 | -0.7 | 0.0261 | 0.0453 |
| chr14 | 54446563 | 54447577 | 1015 | 1.08 | 0.0261 | 0.0454 |

| | | | | | | |
|---|---|---|---|---|---|---|
| chr19 | 58951018 | 58963535 | 12518 | 2.74 | 0.0261 | 0.0454 |
| chr5 | 73890740 | 73895780 | 5041 | 0.95 | 0.0261 | 0.0454 |
| chr9 | 108005925 | 108020897 | 14973 | 0.39 | 0.0261 | 0.0454 |
| chr4 | 2747162 | 2758055 | 10894 | -0.57 | 0.0261 | 0.0454 |
| chr11 | 86625909 | 86630762 | 4854 | 0.34 | 0.0261 | 0.0454 |
| chr2 | 237591223 | 237592123 | 901 | -2.16 | 0.0261 | 0.0454 |
| chr12 | 808990 | 822226 | 13237 | 1.48 | 0.0261 | 0.0454 |
| chr21 | 42791934 | 42792987 | 1054 | 0.76 | 0.0261 | 0.0454 |
| chrX | 41350548 | 41351491 | 944 | 1.11 | 0.0261 | 0.0454 |
| chr6 | 54738583 | 54752231 | 13649 | 4.04 | 0.0262 | 0.0454 |
| chr22 | 18427899 | 18485060 | 57162 | -0.16 | 0.0262 | 0.0455 |
| chr11 | 8190170 | 8190962 | 793 | 1.44 | 0.0262 | 0.0455 |
| chr10 | 134923069 | 134924543 | 1475 | 2.59 | 0.0262 | 0.0455 |
| chr9 | 97540773 | 97596707 | 55935 | -0.11 | 0.0262 | 0.0455 |
| chr4 | 88816961 | 88818119 | 1159 | 3.06 | 0.0262 | 0.0455 |
| chr3 | 9290548 | 9291977 | 1430 | 0.92 | 0.0262 | 0.0455 |
| chr2 | 203409162 | 203412271 | 3110 | -0.86 | 0.0262 | 0.0455 |
| chr20 | 60579845 | 60580890 | 1046 | -1.36 | 0.0262 | 0.0456 |
| chr9 | 136809443 | 136810221 | 779 | -1.5 | 0.0262 | 0.0456 |
| chr1 | 200743393 | 200744810 | 1418 | -0.99 | 0.0262 | 0.0456 |
| chr18 | 9771261 | 9784269 | 13009 | 0.49 | 0.0263 | 0.0456 |
| chr6 | 16391652 | 16392157 | 506 | -1.78 | 0.0263 | 0.0457 |
| chr3 | 125312248 | 125315790 | 3543 | 0.34 | 0.0263 | 0.0457 |
| chr1 | 213187589 | 213189839 | 2251 | 0.39 | 0.0263 | 0.0457 |
| chr8 | 48510215 | 48546468 | 36254 | -0.22 | 0.0264 | 0.0458 |
| chr4 | 157997003 | 157997937 | 935 | 0.97 | 0.0264 | 0.0458 |
| chr1 | 10587487 | 10589468 | 1982 | 0.55 | 0.0264 | 0.0459 |
| chr11 | 108205693 | 108206618 | 926 | -1.23 | 0.0264 | 0.0459 |
| chr15 | 83447562 | 83448084 | 523 | -1.55 | 0.0264 | 0.0459 |
| chr11 | 13508888 | 13509323 | 436 | 1.41 | 0.0265 | 0.0459 |
| chr11 | 9960962 | 9966016 | 5055 | 0.54 | 0.0265 | 0.0459 |
| chr22 | 38863575 | 38864479 | 905 | -1.09 | 0.0265 | 0.046 |
| chr17 | 41045802 | 41047977 | 2176 | -0.67 | 0.0265 | 0.046 |
| chr9 | 34456894 | 34459391 | 2498 | -0.45 | 0.0265 | 0.046 |
| chr1 | 246444393 | 246445806 | 1414 | -0.95 | 0.0265 | 0.046 |
| chr4 | 6471600 | 6473888 | 2289 | -0.43 | 0.0265 | 0.046 |
| chr22 | 28082007 | 28083459 | 1453 | 0.64 | 0.0265 | 0.046 |
| chr7 | 143670198 | 143677613 | 7416 | 0.3 | 0.0266 | 0.0461 |
| chr2 | 100023965 | 100039346 | 15382 | -0.39 | 0.0266 | 0.0461 |
| chr8 | 110986354 | 110986749 | 396 | 2.21 | 0.0266 | 0.0461 |
| chr17 | 61326707 | 61327309 | 603 | 1.63 | 0.0266 | 0.0461 |
| chr11 | 63948579 | 63961959 | 13381 | 0.31 | 0.0266 | 0.0462 |
| chr14 | 75148542 | 75159440 | 10899 | -0.44 | 0.0266 | 0.0462 |
| chr10 | 75253387 | 75256847 | 3461 | 0.31 | 0.0266 | 0.0462 |
| chr3 | 36421404 | 36422562 | 1159 | -1 | 0.0266 | 0.0462 |
| chr2 | 121479440 | 121500086 | 20647 | 0.19 | 0.0266 | 0.0462 |
| chr5 | 171577275 | 171578016 | 742 | -1.3 | 0.0267 | 0.0462 |
| chr4 | 77995907 | 77998298 | 2392 | 0.39 | 0.0267 | 0.0462 |
| chr12 | 71987840 | 71988491 | 652 | 1.35 | 0.0267 | 0.0462 |
| chr13 | 101317604 | 101327809 | 10206 | 0.47 | 0.0267 | 0.0463 |
| chr1 | 205633162 | 205633986 | 825 | 1.57 | 0.0267 | 0.0463 |
| chr12 | 124758028 | 124758778 | 751 | -1.3 | 0.0267 | 0.0463 |
| chr7 | 65669633 | 65671111 | 1479 | 0.68 | 0.0267 | 0.0463 |
| chr16 | 83932086 | 83933572 | 1487 | 0.88 | 0.0267 | 0.0463 |
| chr9 | 132044298 | 132045284 | 987 | 1.22 | 0.0267 | 0.0464 |
| chr20 | 34816368 | 34825430 | 9063 | 0.35 | 0.0267 | 0.0464 |
| chr1 | 186152790 | 186158223 | 5434 | 0.8 | 0.0267 | 0.0464 |
| chr9 | 101904295 | 101904688 | 394 | 1.63 | 0.0268 | 0.0464 |
| chr19 | 54411675 | 54413091 | 1417 | -1.49 | 0.0268 | 0.0464 |
| chr13 | 34593458 | 34611465 | 18008 | 4.16 | 0.0268 | 0.0464 |
| chr22 | 33914270 | 33921643 | 7374 | 0.39 | 0.0268 | 0.0464 |

| | | | | | | |
|---|---|---|---|---|---|---|
| chr3 | 87252897 | 87259191 | 6295 | 0.26 | 0.0268 | 0.0464 |
| chr10 | 4812278 | 4814313 | 2036 | 0.38 | 0.0268 | 0.0464 |
| chr9 | 80997490 | 80998673 | 1184 | -0.95 | 0.0268 | 0.0465 |
| chr12 | 3861935 | 3862643 | 709 | -1.37 | 0.0269 | 0.0465 |
| chr14 | 101537365 | 101541217 | 3853 | -0.4 | 0.0269 | 0.0465 |
| chr16 | 54227451 | 54228140 | 690 | -2.14 | 0.0269 | 0.0465 |
| chr2 | 161958906 | 161960096 | 1191 | 1.04 | 0.0269 | 0.0465 |
| chr4 | 186239175 | 186240835 | 1661 | 0.57 | 0.0269 | 0.0466 |
| chr5 | 159413865 | 159414501 | 637 | 1.63 | 0.0269 | 0.0466 |
| chr11 | 46763141 | 46766129 | 2989 | -1.06 | 0.0269 | 0.0466 |
| chr2 | 85636926 | 85704260 | 67335 | 0.12 | 0.0269 | 0.0466 |
| chr20 | 39656441 | 39716686 | 60246 | -0.19 | 0.0269 | 0.0466 |
| chr1 | 59313428 | 59316471 | 3044 | -0.4 | 0.0269 | 0.0466 |
| chr4 | 179319260 | 179319835 | 576 | -1.55 | 0.0269 | 0.0466 |
| chr7 | 100463670 | 100488676 | 25007 | 0.18 | 0.027 | 0.0468 |
| chr10 | 63030558 | 63044197 | 13640 | -0.58 | 0.027 | 0.0468 |
| chr8 | 74843110 | 74848287 | 5178 | -0.35 | 0.0271 | 0.0468 |
| chr9 | 139683079 | 139686736 | 3658 | 0.53 | 0.0271 | 0.0468 |
| chr10 | 128209492 | 128210581 | 1090 | 1.15 | 0.0271 | 0.0469 |
| chr3 | 194298282 | 194320802 | 22521 | -0.24 | 0.0271 | 0.0469 |
| chr15 | 91564492 | 91566794 | 2303 | 0.31 | 0.0271 | 0.0469 |
| chr1 | 74058517 | 74059756 | 1240 | 1.36 | 0.0271 | 0.0469 |
| chr8 | 132352192 | 132353939 | 1748 | 0.61 | 0.0271 | 0.0469 |
| chr5 | 142381535 | 142383430 | 1896 | -0.83 | 0.0271 | 0.0469 |
| chr4 | 69960427 | 69962939 | 2513 | 0.48 | 0.0271 | 0.0469 |
| chr1 | 112049908 | 112051844 | 1937 | 0.5 | 0.0271 | 0.047 |
| chr21 | 18979472 | 18985489 | 6018 | 0.34 | 0.0272 | 0.047 |
| chr7 | 23372222 | 23398284 | 26063 | 0.46 | 0.0272 | 0.0471 |
| chrX | 63424486 | 63427087 | 2602 | 0.35 | 0.0273 | 0.0471 |
| chr11 | 925345 | 982224 | 56880 | 0.16 | 0.0273 | 0.0471 |
| chr7 | 77082953 | 77089774 | 6822 | -0.6 | 0.0273 | 0.0472 |
| chr5 | 16763800 | 16765037 | 1238 | -1.27 | 0.0273 | 0.0472 |
| chr2 | 44074876 | 44076285 | 1410 | 1.71 | 0.0273 | 0.0472 |
| chr5 | 32559310 | 32559829 | 520 | 1.55 | 0.0273 | 0.0472 |
| chr5 | 106016066 | 106016823 | 758 | 1.31 | 0.0273 | 0.0472 |
| chr11 | 78773846 | 78774994 | 1149 | -1.08 | 0.0273 | 0.0473 |
| chr16 | 19294366 | 19303330 | 8965 | -0.67 | 0.0273 | 0.0473 |
| chr3 | 9381479 | 9405959 | 24481 | 0.23 | 0.0274 | 0.0473 |
| chr12 | 8878171 | 8878822 | 652 | -1.49 | 0.0274 | 0.0473 |
| chr1 | 169792998 | 169793854 | 857 | -1.45 | 0.0274 | 0.0473 |
| chr4 | 177800089 | 177805875 | 5787 | -0.46 | 0.0274 | 0.0473 |
| chr2 | 64975902 | 64979634 | 3733 | 0.39 | 0.0274 | 0.0474 |
| chr14 | 61549335 | 61550091 | 757 | -1.82 | 0.0274 | 0.0474 |
| chr1 | 147051123 | 147052376 | 1254 | 3.18 | 0.0274 | 0.0474 |
| chr2 | 231084216 | 231085536 | 1321 | 0.51 | 0.0274 | 0.0474 |
| chr9 | 106855047 | 106871013 | 15967 | 0.24 | 0.0274 | 0.0474 |
| chr3 | 172340129 | 172353860 | 13732 | 0.27 | 0.0274 | 0.0474 |
| chr9 | 16262311 | 16268225 | 5915 | 0.81 | 0.0274 | 0.0474 |
| chr8 | 122572715 | 122573817 | 1103 | 1.02 | 0.0274 | 0.0474 |
| chr2 | 44564900 | 44565293 | 394 | -1.68 | 0.0274 | 0.0474 |
| chr11 | 32985321 | 32999315 | 13995 | 0.46 | 0.0275 | 0.0475 |
| chr20 | 10965317 | 10966206 | 890 | 1.07 | 0.0275 | 0.0475 |
| chr19 | 46662922 | 46663767 | 846 | 1.52 | 0.0275 | 0.0475 |
| chr10 | 99628668 | 99658842 | 30175 | 0.26 | 0.0275 | 0.0475 |
| chr15 | 40986214 | 40999039 | 12826 | 0.33 | 0.0276 | 0.0476 |
| chr16 | 21963112 | 21965893 | 2782 | 0.27 | 0.0276 | 0.0476 |
| chr9 | 96438231 | 96438782 | 552 | -1.95 | 0.0276 | 0.0476 |
| chr2 | 143794848 | 143807254 | 12407 | -0.56 | 0.0276 | 0.0476 |
| chr4 | 189292975 | 189293964 | 990 | 1.64 | 0.0276 | 0.0476 |
| chr4 | 154110198 | 154110798 | 601 | 1.29 | 0.0276 | 0.0476 |
| chr17 | 67426074 | 67426990 | 917 | -1.15 | 0.0276 | 0.0476 |

| | | | | | | |
|---|---|---|---|---|---|---|
| chr4 | 145708455 | 145709630 | 1176 | -1.04 | 0.0276 | 0.0477 |
| chr9 | 139331765 | 139337306 | 5542 | -0.59 | 0.0276 | 0.0477 |
| chr3 | 123302917 | 123326141 | 23225 | -0.15 | 0.0276 | 0.0477 |
| chr3 | 122785481 | 122787925 | 2445 | -0.42 | 0.0276 | 0.0477 |
| chr2 | 231982977 | 232000393 | 17417 | -0.23 | 0.0276 | 0.0477 |
| chr6 | 20984252 | 20984671 | 420 | -1.85 | 0.0276 | 0.0477 |
| chr16 | 81900550 | 81901275 | 726 | 1.43 | 0.0277 | 0.0477 |
| chr20 | 3235372 | 3235956 | 585 | -1.54 | 0.0277 | 0.0478 |
| chr5 | 33556208 | 33557154 | 947 | -1.14 | 0.0277 | 0.0478 |
| chr4 | 156873576 | 156875587 | 2012 | 0.38 | 0.0277 | 0.0478 |
| chr2 | 151767907 | 151768841 | 935 | -1.33 | 0.0277 | 0.0478 |
| chr14 | 103798242 | 103805773 | 7532 | 0.27 | 0.0278 | 0.0479 |
| chr19 | 7869468 | 7870383 | 916 | 2.12 | 0.0278 | 0.048 |
| chr6 | 27181761 | 27182957 | 1197 | 0.81 | 0.0278 | 0.048 |
| chr6 | 15401527 | 15402860 | 1334 | 0.84 | 0.0278 | 0.048 |
| chr22 | 27054356 | 27080912 | 26557 | -0.21 | 0.0278 | 0.048 |
| chr21 | 34912574 | 34918525 | 5952 | 0.2 | 0.0278 | 0.048 |
| chr6 | 134812416 | 134813169 | 754 | 2.12 | 0.0279 | 0.0481 |
| chr20 | 32445667 | 32447848 | 2182 | -0.76 | 0.0279 | 0.0482 |
| chr20 | 50300103 | 50384907 | 84805 | 0.09 | 0.0279 | 0.0482 |
| chr7 | 33146972 | 33149985 | 3014 | 0.5 | 0.0279 | 0.0482 |
| chr7 | 65817028 | 65817779 | 752 | 1.31 | 0.028 | 0.0482 |
| chr14 | 69925864 | 69928003 | 2140 | -0.57 | 0.028 | 0.0482 |
| chr1 | 171220660 | 171220970 | 311 | 1.93 | 0.028 | 0.0482 |
| chr14 | 64239901 | 64240590 | 690 | -1.24 | 0.028 | 0.0482 |
| chr7 | 82069912 | 82074150 | 4239 | -0.3 | 0.028 | 0.0482 |
| chr16 | 19812363 | 19828913 | 16551 | -0.16 | 0.028 | 0.0482 |
| chr12 | 8043399 | 8044430 | 1032 | 0.7 | 0.028 | 0.0482 |
| chr2 | 223455831 | 223458949 | 3119 | -1.09 | 0.028 | 0.0483 |
| chrX | 128710985 | 128712704 | 1720 | -0.74 | 0.028 | 0.0483 |
| chr10 | 17658158 | 17660367 | 2210 | 0.34 | 0.0281 | 0.0483 |
| chr1 | 31406427 | 31408196 | 1770 | -0.77 | 0.0281 | 0.0483 |
| chr15 | 67094171 | 67098433 | 4263 | -0.22 | 0.0281 | 0.0484 |
| chr1 | 196946323 | 196946907 | 585 | 1.84 | 0.0281 | 0.0484 |
| chr6 | 149553589 | 149556833 | 3245 | 0.36 | 0.0281 | 0.0484 |
| chr3 | 171178442 | 171179292 | 851 | -1.36 | 0.0281 | 0.0484 |
| chr1 | 109101781 | 109104407 | 2627 | 0.63 | 0.0281 | 0.0485 |
| chr20 | 3019415 | 3032700 | 13286 | -0.33 | 0.0281 | 0.0485 |
| chr5 | 64848491 | 64861631 | 13141 | 0.24 | 0.0281 | 0.0485 |
| chr18 | 55331864 | 55332652 | 789 | -1.28 | 0.0282 | 0.0485 |
| chr7 | 105856497 | 105858216 | 1720 | -0.53 | 0.0282 | 0.0485 |
| chr5 | 75698194 | 75699142 | 949 | -1.24 | 0.0282 | 0.0485 |
| chr9 | 117415547 | 117415916 | 370 | -2.16 | 0.0282 | 0.0485 |
| chr3 | 45314962 | 45315725 | 764 | -1.77 | 0.0282 | 0.0485 |
| chr22 | 31974998 | 31983051 | 8054 | 0.61 | 0.0282 | 0.0485 |
| chr2 | 235703685 | 235704851 | 1167 | -0.97 | 0.0282 | 0.0486 |
| chr7 | 131618645 | 131638584 | 19940 | -0.26 | 0.0282 | 0.0486 |
| chr16 | 21658213 | 21659100 | 888 | -1.3 | 0.0282 | 0.0486 |
| chr6 | 20693540 | 20708671 | 15132 | 0.55 | 0.0283 | 0.0487 |
| chr11 | 32913241 | 32926315 | 13075 | 0.21 | 0.0283 | 0.0487 |
| chr10 | 35806521 | 35807091 | 571 | -1.84 | 0.0283 | 0.0488 |
| chr2 | 225006808 | 225011858 | 5051 | -0.59 | 0.0283 | 0.0488 |
| chr15 | 44578970 | 44590811 | 11842 | 0.22 | 0.0283 | 0.0488 |
| chr4 | 7068298 | 7070537 | 2240 | 0.62 | 0.0283 | 0.0488 |
| chr7 | 55432333 | 55435369 | 3037 | 0.26 | 0.0284 | 0.0488 |
| chr16 | 29953005 | 29957893 | 4889 | 0.19 | 0.0284 | 0.0488 |
| chr10 | 25179022 | 25192626 | 13605 | -0.26 | 0.0284 | 0.0488 |
| chrX | 77394398 | 77395671 | 1274 | 0.56 | 0.0284 | 0.0488 |
| chr2 | 208343801 | 208344824 | 1024 | 1.36 | 0.0284 | 0.0488 |
| chr3 | 185140505 | 185141608 | 1104 | -0.96 | 0.0284 | 0.0488 |
| chr1 | 205781412 | 205783244 | 1833 | -0.62 | 0.0284 | 0.0489 |

| | | | | | | |
|---|---|---|---|---|---|---|
| chrX | 46670217 | 46674816 | 4600 | -0.5 | 0.0284 | 0.0489 |
| chr14 | 65538557 | 65540996 | 2440 | -1.11 | 0.0285 | 0.049 |
| chr1 | 247493679 | 247506778 | 13100 | -0.28 | 0.0285 | 0.049 |
| chr9 | 80540864 | 80541922 | 1059 | 0.73 | 0.0285 | 0.049 |
| chr6 | 116988887 | 117003848 | 14962 | 0.53 | 0.0285 | 0.049 |
| chr3 | 5163441 | 5180056 | 16616 | -0.28 | 0.0285 | 0.049 |
| chr8 | 87267220 | 87268584 | 1365 | 3.47 | 0.0285 | 0.049 |
| chr14 | 93496703 | 93532264 | 35562 | 0.13 | 0.0285 | 0.0491 |
| chr18 | 74398475 | 74399731 | 1257 | 1.99 | 0.0285 | 0.0491 |
| chr9 | 126905690 | 126907506 | 1817 | -0.8 | 0.0286 | 0.0491 |
| chr9 | 91635422 | 91636925 | 1504 | -0.83 | 0.0286 | 0.0491 |
| chr13 | 113164563 | 113165621 | 1059 | -1.26 | 0.0286 | 0.0492 |
| chr2 | 214933493 | 214934641 | 1149 | 0.99 | 0.0286 | 0.0492 |
| chr8 | 128805964 | 129223734 | 417771 | 0.04 | 0.0286 | 0.0492 |
| chr5 | 156104856 | 156106524 | 1669 | 3.52 | 0.0286 | 0.0492 |
| chr3 | 194979967 | 194992950 | 12984 | -0.42 | 0.0286 | 0.0492 |
| chr1 | 118363630 | 118369929 | 6300 | 0.42 | 0.0287 | 0.0492 |
| chr2 | 194709734 | 194710709 | 976 | -1.32 | 0.0287 | 0.0493 |
| chr7 | 2550683 | 2570963 | 20281 | -0.25 | 0.0287 | 0.0493 |
| chr2 | 58929122 | 58929641 | 520 | 2.18 | 0.0287 | 0.0493 |
| chr17 | 8191278 | 8204042 | 12765 | 0.27 | 0.0287 | 0.0493 |
| chr9 | 5796612 | 5800173 | 3562 | -0.61 | 0.0287 | 0.0493 |
| chr10 | 120924270 | 120939258 | 14989 | 0.24 | 0.0287 | 0.0493 |
| chr2 | 106762325 | 106764656 | 2332 | -0.62 | 0.0287 | 0.0494 |
| chr10 | 71077142 | 71079350 | 2209 | -0.72 | 0.0287 | 0.0494 |
| chr7 | 25219215 | 25220591 | 1377 | 0.55 | 0.0287 | 0.0494 |
| chr12 | 107619533 | 107620072 | 540 | 1.37 | 0.0288 | 0.0494 |
| chr3 | 97601734 | 97602678 | 945 | 1.67 | 0.0288 | 0.0494 |
| chr6 | 4125742 | 4137024 | 11283 | -0.45 | 0.0288 | 0.0494 |
| chr1 | 243504076 | 243504910 | 835 | 1.25 | 0.0288 | 0.0494 |
| chr10 | 134328370 | 134332178 | 3809 | 0.33 | 0.0288 | 0.0495 |
| chr12 | 105776507 | 105777253 | 747 | 1.14 | 0.0289 | 0.0496 |
| chr2 | 191835730 | 191846257 | 10528 | -0.46 | 0.0289 | 0.0496 |
| chr4 | 2562353 | 2563033 | 681 | -1.52 | 0.0289 | 0.0496 |
| chr10 | 133487392 | 133488269 | 878 | 2.27 | 0.0289 | 0.0497 |
| chr2 | 175018781 | 175019389 | 609 | -1.66 | 0.0289 | 0.0497 |
| chr17 | 38177321 | 38180765 | 3445 | -0.77 | 0.029 | 0.0497 |
| chr22 | 36924030 | 36943213 | 19184 | -0.35 | 0.029 | 0.0498 |
| chr15 | 49427084 | 49448793 | 21710 | 0.32 | 0.029 | 0.0498 |
| chr17 | 9671611 | 9672039 | 429 | -2.33 | 0.029 | 0.0498 |
| chr3 | 159645204 | 159647974 | 2771 | 0.45 | 0.029 | 0.0498 |
| chr17 | 48502901 | 48505880 | 2980 | 0.39 | 0.0291 | 0.0498 |
| chr4 | 156313647 | 156314584 | 938 | 1.04 | 0.0291 | 0.0499 |
| chr2 | 161459969 | 161460601 | 633 | 1.19 | 0.0292 | 0.05 |
| chr3 | 131099947 | 131101544 | 1598 | 0.58 | 0.0292 | 0.0501 |
| chr17 | 48422777 | 48426591 | 3815 | -0.62 | 0.0292 | 0.0501 |
| chr2 | 164066906 | 164071094 | 4189 | -0.41 | 0.0292 | 0.0501 |
| chr10 | 93193825 | 93194589 | 765 | 1.55 | 0.0292 | 0.0501 |
| chr1 | 227501977 | 227507068 | 5092 | 0.29 | 0.0292 | 0.0501 |
| chrX | 43871547 | 43872629 | 1083 | 1.04 | 0.0292 | 0.0502 |
| chr8 | 56918955 | 56920266 | 1312 | 0.99 | 0.0293 | 0.0502 |
| chr1 | 214688898 | 214690074 | 1177 | 0.83 | 0.0293 | 0.0502 |
| chr11 | 63992616 | 64020210 | 27595 | 0.24 | 0.0293 | 0.0502 |
| chr1 | 156861672 | 156877792 | 16121 | -0.26 | 0.0293 | 0.0502 |
| chr4 | 99063781 | 99065304 | 1524 | 0.46 | 0.0293 | 0.0502 |
| chr3 | 196063052 | 196075383 | 12332 | 0.2 | 0.0293 | 0.0502 |
| chr20 | 14899738 | 14909424 | 9687 | 0.42 | 0.0293 | 0.0502 |
| chr6 | 99871753 | 99875154 | 3402 | 0.3 | 0.0293 | 0.0503 |
| chr17 | 36944803 | 36982698 | 37896 | -0.16 | 0.0293 | 0.0503 |
| chr2 | 217416243 | 217417015 | 773 | -1.47 | 0.0294 | 0.0503 |
| chr3 | 100421245 | 100456996 | 35752 | 0.21 | 0.0294 | 0.0503 |

| | | | | | | |
|---|---|---|---|---|---|---|
| chr6 | 42371557 | 42373468 | 1912 | 0.42 | 0.0294 | 0.0504 |
| chr3 | 119357664 | 119358101 | 438 | -1.91 | 0.0294 | 0.0504 |
| chr21 | 46237841 | 46238756 | 916 | 0.79 | 0.0294 | 0.0505 |
| chr10 | 14541063 | 14542736 | 1674 | -0.89 | 0.0295 | 0.0505 |
| chr11 | 60809276 | 60810107 | 832 | 1.19 | 0.0295 | 0.0505 |
| chr8 | 644073 | 654831 | 10759 | -0.69 | 0.0295 | 0.0505 |
| chr8 | 37419516 | 37419999 | 484 | 2.1 | 0.0295 | 0.0506 |
| chr17 | 51703471 | 51705472 | 2002 | 0.68 | 0.0296 | 0.0506 |
| chrX | 19505201 | 19507287 | 2087 | 0.54 | 0.0296 | 0.0507 |
| chr1 | 23654574 | 23672709 | 18136 | 0.22 | 0.0296 | 0.0507 |
| chr16 | 84839621 | 84840696 | 1076 | 0.87 | 0.0296 | 0.0507 |
| chr21 | 33058698 | 33068186 | 9489 | 0.67 | 0.0296 | 0.0507 |
| chr16 | 58155105 | 58164247 | 9143 | -0.26 | 0.0296 | 0.0508 |
| chr5 | 111496383 | 111503320 | 6938 | 0.33 | 0.0296 | 0.0508 |
| chr14 | 31809681 | 31810182 | 502 | 1.71 | 0.0296 | 0.0508 |
| chr10 | 99392155 | 99417253 | 25099 | -0.18 | 0.0297 | 0.0509 |
| chr15 | 60095179 | 60096004 | 826 | 3.19 | 0.0297 | 0.0509 |
| chr4 | 100007519 | 100011180 | 3662 | 0.33 | 0.0297 | 0.0509 |
| chr2 | 190117953 | 190119674 | 1722 | -2.98 | 0.0297 | 0.0509 |
| chr5 | 150631654 | 150633438 | 1785 | 0.6 | 0.0297 | 0.0509 |
| chr5 | 6675696 | 6676489 | 794 | -1.94 | 0.0298 | 0.051 |
| chr10 | 106013892 | 106029710 | 15819 | 0.2 | 0.0298 | 0.051 |
| chr8 | 107947170 | 107952274 | 5105 | 0.63 | 0.0298 | 0.051 |
| chrX | 153058696 | 153061292 | 2597 | 0.36 | 0.0298 | 0.051 |
| chr8 | 58615178 | 58616206 | 1029 | 1.21 | 0.0298 | 0.051 |
| chr2 | 196425318 | 196427772 | 2455 | -0.71 | 0.0298 | 0.051 |
| chr9 | 22341979 | 22355449 | 13471 | 3.8 | 0.0298 | 0.051 |
| chr7 | 133043523 | 133044211 | 689 | -1.55 | 0.0298 | 0.051 |
| chr12 | 102931507 | 102942885 | 11379 | -0.4 | 0.0298 | 0.0511 |
| chr16 | 75588490 | 75603960 | 15471 | -0.35 | 0.0298 | 0.0511 |
| chr20 | 30392855 | 30395331 | 2477 | -0.92 | 0.0298 | 0.0511 |
| chr5 | 176963914 | 176986663 | 22750 | 0.39 | 0.0298 | 0.0511 |
| chr1 | 9969505 | 9971751 | 2247 | 0.57 | 0.0298 | 0.0511 |
| chr12 | 132932741 | 132934548 | 1808 | 0.87 | 0.0299 | 0.0511 |
| chr5 | 95516034 | 95517222 | 1189 | 1.03 | 0.0299 | 0.0511 |
| chr11 | 1884057 | 1885170 | 1114 | -1.18 | 0.0299 | 0.0512 |
| chrX | 41297208 | 41309264 | 12057 | 0.36 | 0.03 | 0.0512 |
| chr14 | 39960536 | 39961755 | 1220 | -1.1 | 0.03 | 0.0512 |
| chr8 | 125700739 | 125701245 | 507 | 1.36 | 0.03 | 0.0513 |
| chr15 | 52246462 | 52247977 | 1516 | 1.3 | 0.03 | 0.0513 |
| chr10 | 76944500 | 77006494 | 61995 | 0.12 | 0.03 | 0.0513 |
| chr7 | 43317223 | 43342190 | 24968 | -2.97 | 0.0301 | 0.0514 |
| chr7 | 157091631 | 157111578 | 19948 | -0.31 | 0.0301 | 0.0514 |
| chr3 | 12464176 | 12466909 | 2734 | -0.44 | 0.0301 | 0.0514 |
| chr4 | 144433794 | 144453766 | 19973 | 0.25 | 0.0301 | 0.0514 |
| chr18 | 20538429 | 20539429 | 1001 | -1.48 | 0.0301 | 0.0514 |
| chr1 | 23850616 | 23925900 | 75285 | 0.11 | 0.0301 | 0.0514 |
| chr22 | 43522375 | 43552279 | 29905 | 0.25 | 0.0301 | 0.0514 |
| chr6 | 28302684 | 28325413 | 22730 | 0.23 | 0.0301 | 0.0514 |
| chr10 | 32620168 | 32637013 | 16846 | -0.21 | 0.0301 | 0.0515 |
| chr2 | 86975415 | 86982531 | 7117 | 0.72 | 0.0301 | 0.0515 |
| chr13 | 68630699 | 68633073 | 2375 | 0.37 | 0.0301 | 0.0515 |
| chr15 | 63307047 | 63312400 | 5354 | -0.52 | 0.0301 | 0.0515 |
| chrX | 85247605 | 85248231 | 627 | 1.54 | 0.0301 | 0.0515 |
| chr9 | 131872426 | 131941234 | 68809 | 0.1 | 0.0302 | 0.0515 |
| chr20 | 60695816 | 60719899 | 24084 | 0.2 | 0.0302 | 0.0516 |
| chr6 | 131254540 | 131260296 | 5757 | 0.55 | 0.0302 | 0.0516 |
| chr4 | 69940588 | 69941497 | 910 | 1.4 | 0.0302 | 0.0516 |
| chr1 | 55841114 | 55845359 | 4246 | -0.88 | 0.0302 | 0.0516 |
| chr1 | 59394685 | 59409729 | 15045 | 0.29 | 0.0302 | 0.0516 |
| chr2 | 182755362 | 182783033 | 27672 | 0.21 | 0.0302 | 0.0516 |

| | | | | | | |
|---|---|---|---|---|---|---|
| chr1 | 56750408 | 56752124 | 1717 | -0.76 | 0.0302 | 0.0516 |
| chr9 | 20355883 | 20357430 | 1548 | 1.11 | 0.0302 | 0.0516 |
| chr1 | 31885794 | 31899250 | 13457 | 0.61 | 0.0302 | 0.0516 |
| chr7 | 33492603 | 33492995 | 393 | -1.75 | 0.0303 | 0.0517 |
| chr5 | 76506002 | 76507085 | 1084 | 0.99 | 0.0303 | 0.0517 |
| chr6 | 140382462 | 140384393 | 1932 | 0.75 | 0.0303 | 0.0518 |
| chr4 | 89888450 | 89889945 | 1496 | 0.73 | 0.0303 | 0.0518 |
| chr2 | 174778996 | 174790773 | 11778 | -0.56 | 0.0304 | 0.0518 |
| chr14 | 28526667 | 28527204 | 538 | 1.28 | 0.0304 | 0.0519 |
| chr22 | 44350265 | 44363289 | 13025 | -0.47 | 0.0304 | 0.0519 |
| chr13 | 49106114 | 49107858 | 1745 | 0.4 | 0.0304 | 0.0519 |
| chr17 | 5256394 | 5270083 | 13690 | -0.48 | 0.0304 | 0.052 |
| chr4 | 140164015 | 140167863 | 3849 | -0.54 | 0.0304 | 0.052 |
| chr1 | 186761740 | 186769614 | 7875 | 3.88 | 0.0305 | 0.052 |
| chr4 | 178494396 | 178495502 | 1107 | -1.26 | 0.0305 | 0.052 |
| chr5 | 57012347 | 57014798 | 2452 | -0.67 | 0.0305 | 0.052 |
| chr6 | 90121185 | 90122539 | 1355 | 1.12 | 0.0305 | 0.0521 |
| chr8 | 58449204 | 58457351 | 8148 | 1.02 | 0.0306 | 0.0522 |
| chr7 | 42222810 | 42224072 | 1263 | -0.84 | 0.0306 | 0.0522 |
| chr8 | 49647471 | 49648208 | 738 | 1.28 | 0.0306 | 0.0522 |
| chr18 | 53522487 | 53524795 | 2309 | 0.44 | 0.0306 | 0.0522 |
| chr9 | 20980714 | 20982421 | 1708 | 0.74 | 0.0306 | 0.0523 |
| chr1 | 2454534 | 2458062 | 3529 | -0.73 | 0.0306 | 0.0523 |
| chr1 | 67870908 | 67897510 | 26603 | 0.27 | 0.0306 | 0.0523 |
| chr10 | 116079612 | 116088656 | 9045 | 0.43 | 0.0307 | 0.0523 |
| chr4 | 143088966 | 143097155 | 8190 | 3.35 | 0.0307 | 0.0523 |
| chr2 | 157332710 | 157333472 | 763 | -1.22 | 0.0307 | 0.0523 |
| chr6 | 52319243 | 52320044 | 802 | 1.37 | 0.0307 | 0.0524 |
| chr5 | 92497890 | 92499379 | 1490 | 0.81 | 0.0307 | 0.0524 |
| chr16 | 69934549 | 69935142 | 594 | -1.7 | 0.0307 | 0.0524 |
| chr2 | 32283058 | 32290010 | 6953 | -0.36 | 0.0308 | 0.0525 |
| chr1 | 208409029 | 208418585 | 9557 | 3.87 | 0.0308 | 0.0525 |
| chr7 | 5490127 | 5490479 | 353 | -2.15 | 0.0308 | 0.0525 |
| chr9 | 90400767 | 90409398 | 8632 | 0.3 | 0.0308 | 0.0525 |
| chr2 | 74755476 | 74757893 | 2418 | -0.35 | 0.0308 | 0.0525 |
| chr11 | 18628910 | 18629278 | 369 | 2.27 | 0.0308 | 0.0525 |
| chr11 | 126137446 | 126154094 | 16649 | 0.27 | 0.0308 | 0.0526 |
| chr15 | 66734999 | 66736156 | 1158 | -1.16 | 0.0309 | 0.0526 |
| chr3 | 142754142 | 142756139 | 1998 | 0.68 | 0.0309 | 0.0526 |
| chr5 | 145214248 | 145215816 | 1569 | 0.52 | 0.0309 | 0.0526 |
| chrX | 14954488 | 14955410 | 923 | 0.89 | 0.0309 | 0.0526 |
| chr4 | 104020455 | 104021719 | 1265 | 0.9 | 0.0309 | 0.0526 |
| chr9 | 92755914 | 92763107 | 7194 | 0.35 | 0.0309 | 0.0527 |
| chr17 | 58964545 | 58965813 | 1269 | 0.32 | 0.0309 | 0.0527 |
| chr8 | 68254065 | 68257002 | 2938 | 0.33 | 0.0309 | 0.0527 |
| chr19 | 55477884 | 55478962 | 1079 | 1 | 0.0309 | 0.0527 |
| chr10 | 98134776 | 98135233 | 458 | -1.23 | 0.0309 | 0.0527 |
| chr9 | 127614560 | 127635989 | 21430 | -0.21 | 0.031 | 0.0527 |
| chr3 | 167809260 | 167814873 | 5614 | 0.37 | 0.031 | 0.0528 |
| chr5 | 56630428 | 56635210 | 4783 | -0.58 | 0.031 | 0.0528 |
| chr11 | 92190168 | 92190579 | 412 | -1.66 | 0.031 | 0.0528 |
| chr13 | 78051933 | 78052959 | 1027 | -0.74 | 0.031 | 0.0528 |
| chr20 | 42184036 | 42194455 | 10420 | 0.24 | 0.031 | 0.0528 |
| chr17 | 9579037 | 9581243 | 2207 | 0.53 | 0.031 | 0.0528 |
| chr2 | 198215148 | 198216076 | 929 | 1.09 | 0.031 | 0.0528 |
| chr7 | 38989890 | 38990701 | 812 | -1.31 | 0.0311 | 0.0529 |
| chr6 | 159314135 | 159316968 | 2834 | 0.36 | 0.0311 | 0.0529 |
| chr6 | 150325981 | 150327148 | 1168 | 0.86 | 0.0311 | 0.0529 |
| chr2 | 121124937 | 121125861 | 925 | 1.2 | 0.0311 | 0.0529 |
| chr11 | 35257055 | 35258420 | 1366 | 0.97 | 0.0311 | 0.053 |
| chr3 | 146524721 | 146525603 | 883 | 0.92 | 0.0311 | 0.053 |

| | | | | | | |
|---|---|---|---|---|---|---|
| chr9 | 124910981 | 124922964 | 11984 | -0.41 | 0.0311 | 0.053 |
| chr18 | 67668333 | 67668743 | 411 | -1.73 | 0.0312 | 0.053 |
| chr11 | 102678463 | 102678789 | 327 | 1.58 | 0.0312 | 0.0531 |
| chr18 | 13435396 | 13436327 | 932 | 1.65 | 0.0312 | 0.0531 |
| chr3 | 171241647 | 171242547 | 901 | -1.31 | 0.0312 | 0.0532 |
| chr12 | 121011129 | 121022899 | 11771 | 0.53 | 0.0312 | 0.0532 |
| chr16 | 86417126 | 86418570 | 1445 | 0.73 | 0.0313 | 0.0532 |
| chr11 | 109870314 | 109871736 | 1423 | 0.91 | 0.0313 | 0.0533 |
| chr2 | 196351134 | 196353034 | 1901 | -0.61 | 0.0313 | 0.0533 |
| chr20 | 10981707 | 10983864 | 2158 | 0.36 | 0.0313 | 0.0533 |
| chr5 | 148513646 | 148534473 | 20828 | 0.35 | 0.0314 | 0.0533 |
| chr17 | 61742948 | 61743633 | 686 | -2.09 | 0.0314 | 0.0533 |
| chr1 | 19922442 | 19943547 | 21106 | -0.3 | 0.0314 | 0.0533 |
| chr7 | 89974674 | 89977660 | 2987 | 0.27 | 0.0314 | 0.0534 |
| chr2 | 233414383 | 233416840 | 2458 | -0.36 | 0.0314 | 0.0534 |
| chr1 | 163244741 | 163245525 | 785 | -1.19 | 0.0314 | 0.0534 |
| chr4 | 148141794 | 148142603 | 810 | 1.13 | 0.0314 | 0.0534 |
| chr11 | 106877041 | 106877683 | 643 | 1.54 | 0.0314 | 0.0534 |
| chr13 | 20832497 | 20833032 | 536 | 1.22 | 0.0314 | 0.0534 |
| chr21 | 44074116 | 44074749 | 634 | 2.11 | 0.0314 | 0.0534 |
| chr9 | 133556211 | 133589366 | 33156 | 0.2 | 0.0314 | 0.0534 |
| chr3 | 160759279 | 160760669 | 1391 | 3.17 | 0.0315 | 0.0535 |
| chr6 | 131689236 | 131689931 | 696 | 3.02 | 0.0315 | 0.0535 |
| chr20 | 342602 | 427037 | 84436 | -0.11 | 0.0315 | 0.0535 |
| chr16 | 19937645 | 19938912 | 1268 | -1.08 | 0.0315 | 0.0535 |
| chr7 | 132966738 | 132967600 | 863 | -1.39 | 0.0315 | 0.0535 |
| chr10 | 75217039 | 75217483 | 445 | -1.32 | 0.0315 | 0.0535 |
| chr3 | 43327357 | 43329026 | 1670 | 0.75 | 0.0315 | 0.0535 |
| chr10 | 21452632 | 21498304 | 45673 | 0.32 | 0.0315 | 0.0535 |
| chr12 | 17108046 | 17108938 | 893 | 1.01 | 0.0315 | 0.0535 |
| chr20 | 47537106 | 47539687 | 2582 | 0.34 | 0.0315 | 0.0536 |
| chr3 | 127680365 | 127681385 | 1021 | 1.03 | 0.0316 | 0.0536 |
| chr4 | 86116816 | 86118681 | 1866 | -0.62 | 0.0316 | 0.0537 |
| chr19 | 4360954 | 4379583 | 18630 | -0.21 | 0.0316 | 0.0537 |
| chr2 | 202315216 | 202317564 | 2349 | -0.44 | 0.0316 | 0.0537 |
| chr11 | 44805414 | 44805836 | 423 | -1.82 | 0.0316 | 0.0537 |
| chr14 | 74959191 | 74961503 | 2313 | 0.56 | 0.0316 | 0.0537 |
| chr17 | 7570630 | 7593773 | 23144 | 0.13 | 0.0316 | 0.0537 |
| chr17 | 26609111 | 26610433 | 1323 | -1.05 | 0.0317 | 0.0537 |
| chr1 | 21577204 | 21622510 | 45307 | -0.15 | 0.0317 | 0.0538 |
| chr5 | 100725619 | 100726137 | 519 | -1.16 | 0.0317 | 0.0538 |
| chr5 | 71269866 | 71270429 | 564 | -1.23 | 0.0317 | 0.0538 |
| chr16 | 4505048 | 4545611 | 40564 | 0.2 | 0.0317 | 0.0538 |
| chr17 | 79211940 | 79227241 | 15302 | -0.33 | 0.0317 | 0.0539 |
| chr8 | 25991078 | 25992195 | 1118 | 0.93 | 0.0317 | 0.0539 |
| chr9 | 108242350 | 108246688 | 4339 | -0.64 | 0.0317 | 0.0539 |
| chr1 | 218990487 | 218997496 | 7010 | -0.92 | 0.0318 | 0.0539 |
| chr5 | 150051669 | 150052445 | 777 | -1.36 | 0.0318 | 0.0539 |
| chr9 | 88963309 | 88970100 | 6792 | 0.29 | 0.0318 | 0.0539 |
| chr17 | 35412230 | 35425958 | 13729 | -0.5 | 0.0318 | 0.054 |
| chr4 | 113533846 | 113540989 | 7144 | 0.57 | 0.0318 | 0.054 |
| chr7 | 114913695 | 114914591 | 897 | -1.11 | 0.0318 | 0.054 |
| chr3 | 169995275 | 169996447 | 1173 | 0.91 | 0.0318 | 0.054 |
| chr5 | 115419759 | 115421822 | 2064 | 0.31 | 0.0319 | 0.0541 |
| chr1 | 6051727 | 6052584 | 858 | -0.59 | 0.0319 | 0.0541 |
| chr6 | 152234822 | 152236311 | 1490 | 0.97 | 0.0319 | 0.0541 |
| chr1 | 172649434 | 172650235 | 802 | -1.03 | 0.0319 | 0.0541 |
| chr7 | 91989480 | 91990359 | 880 | -0.92 | 0.0319 | 0.0541 |
| chr10 | 63990279 | 64004482 | 14204 | -0.3 | 0.0319 | 0.0542 |
| chr21 | 46392434 | 46393097 | 664 | -1.62 | 0.0319 | 0.0542 |
| chr14 | 97543446 | 97544063 | 618 | -1.34 | 0.0319 | 0.0542 |

| | | | | | | |
|---|---|---|---|---|---|---|
| chr10 | 74549429 | 74607712 | 58284 | -0.16 | 0.032 | 0.0542 |
| chr11 | 124615896 | 124617349 | 1454 | 0.81 | 0.032 | 0.0543 |
| chr1 | 45173015 | 45209496 | 36482 | -0.22 | 0.032 | 0.0543 |
| chr1 | 185124421 | 185127317 | 2897 | 0.28 | 0.032 | 0.0543 |
| chr10 | 97147471 | 97154799 | 7329 | 1.28 | 0.032 | 0.0543 |
| chr13 | 98866317 | 98869038 | 2722 | -1.29 | 0.032 | 0.0543 |
| chr1 | 150038224 | 150040906 | 2683 | 0.35 | 0.0321 | 0.0544 |
| chr22 | 44298453 | 44298854 | 402 | -1.73 | 0.0321 | 0.0544 |
| chr15 | 74934920 | 74935933 | 1014 | -1.03 | 0.0321 | 0.0544 |
| chr6 | 20160392 | 20161470 | 1079 | 1.22 | 0.0321 | 0.0544 |
| chr20 | 52813596 | 52825985 | 12390 | -0.29 | 0.0321 | 0.0545 |
| chr1 | 189743568 | 189745003 | 1436 | 0.66 | 0.0321 | 0.0545 |
| chr9 | 137029239 | 137031199 | 1961 | -0.33 | 0.0322 | 0.0545 |
| chr13 | 67565730 | 67571572 | 5843 | 0.26 | 0.0322 | 0.0545 |
| chr18 | 13036465 | 13037269 | 805 | 1.13 | 0.0322 | 0.0545 |
| chr1 | 119703360 | 119703983 | 624 | 1.7 | 0.0322 | 0.0545 |
| chr4 | 103172444 | 103172918 | 475 | 2.14 | 0.0322 | 0.0545 |
| chr10 | 104986600 | 105015778 | 29179 | 0.12 | 0.0322 | 0.0546 |
| chrX | 129219069 | 129266155 | 47087 | 0.19 | 0.0323 | 0.0546 |
| chr7 | 156741827 | 156748219 | 6393 | -0.44 | 0.0323 | 0.0547 |
| chr19 | 58070348 | 58071853 | 1506 | 0.73 | 0.0323 | 0.0547 |
| chr2 | 134876991 | 134879194 | 2204 | -0.88 | 0.0323 | 0.0547 |
| chr12 | 88148902 | 88150821 | 1920 | 0.76 | 0.0323 | 0.0547 |
| chr17 | 15139316 | 15140283 | 968 | -1.58 | 0.0323 | 0.0547 |
| chr3 | 159336977 | 159356279 | 19303 | 0.24 | 0.0323 | 0.0548 |
| chr2 | 86440044 | 86440969 | 926 | 1.09 | 0.0324 | 0.0548 |
| chr16 | 4986666 | 5007982 | 21317 | 0.26 | 0.0324 | 0.0548 |
| chr20 | 45443165 | 45444341 | 1177 | -0.69 | 0.0324 | 0.0548 |
| chr8 | 74760583 | 74764703 | 4121 | -0.81 | 0.0324 | 0.0548 |
| chr12 | 15368708 | 15369908 | 1201 | 3.54 | 0.0324 | 0.0549 |
| chr17 | 45016017 | 45019357 | 3341 | -1.06 | 0.0324 | 0.0549 |
| chr7 | 99663640 | 99664098 | 459 | 2.21 | 0.0324 | 0.0549 |
| chr11 | 46401756 | 46402841 | 1086 | 0.91 | 0.0325 | 0.055 |
| chr20 | 24880765 | 24881350 | 586 | 1.55 | 0.0325 | 0.055 |
| chr4 | 167921131 | 167922114 | 984 | 1.73 | 0.0325 | 0.055 |
| chr8 | 63442294 | 63443119 | 826 | 1.48 | 0.0325 | 0.055 |
| chr14 | 23937937 | 23939687 | 1751 | 0.45 | 0.0325 | 0.055 |
| chr1 | 65317095 | 65331625 | 14531 | -0.37 | 0.0326 | 0.0551 |
| chr2 | 45607459 | 45617110 | 9652 | 0.72 | 0.0326 | 0.0551 |
| chr3 | 173655835 | 173656761 | 927 | -1.36 | 0.0326 | 0.0551 |
| chr12 | 93291455 | 93295432 | 3978 | 0.21 | 0.0326 | 0.0552 |
| chr4 | 73933630 | 73936556 | 2927 | 0.31 | 0.0326 | 0.0552 |
| chr2 | 196206207 | 196208187 | 1981 | 0.51 | 0.0326 | 0.0552 |
| chr11 | 33151323 | 33188812 | 37490 | 0.17 | 0.0326 | 0.0552 |
| chr4 | 128885843 | 128897345 | 11503 | 0.35 | 0.0326 | 0.0552 |
| chr3 | 162583427 | 162583812 | 386 | 1.76 | 0.0326 | 0.0552 |
| chr16 | 67142862 | 67151275 | 8414 | -0.42 | 0.0327 | 0.0552 |
| chr19 | 33181679 | 33210269 | 28591 | 0.2 | 0.0327 | 0.0553 |
| chr5 | 55822961 | 55839936 | 16976 | 3.51 | 0.0327 | 0.0553 |
| chr14 | 45694148 | 45696030 | 1883 | 0.82 | 0.0327 | 0.0553 |
| chr1 | 64262720 | 64263399 | 680 | -1.27 | 0.0328 | 0.0554 |
| chr15 | 71804350 | 71822774 | 18425 | 0.54 | 0.0328 | 0.0554 |
| chr1 | 226424026 | 226434761 | 10736 | -0.74 | 0.0328 | 0.0554 |
| chr5 | 33195526 | 33196753 | 1228 | -1.03 | 0.0328 | 0.0555 |
| chr8 | 146175672 | 146176709 | 1038 | 0.54 | 0.0328 | 0.0555 |
| chr8 | 614094 | 614938 | 845 | 1.17 | 0.0328 | 0.0555 |
| chr12 | 132221262 | 132222582 | 1321 | 0.7 | 0.0329 | 0.0555 |
| chr2 | 39173661 | 39174962 | 1302 | -0.7 | 0.0329 | 0.0556 |
| chr7 | 133389649 | 133390993 | 1345 | 0.9 | 0.0329 | 0.0556 |
| chr6 | 105158587 | 105159918 | 1332 | 1.07 | 0.0329 | 0.0556 |
| chr10 | 86284851 | 86285270 | 420 | -2.18 | 0.0329 | 0.0557 |

| | | | | | | |
|---|---|---|---|---|---|---|
| chr10 | 96025276 | 96026436 | 1161 | -1.55 | 0.033 | 0.0558 |
| chr7 | 35292568 | 35294407 | 1840 | 0.43 | 0.033 | 0.0558 |
| chr10 | 97435250 | 97436108 | 859 | -1.33 | 0.033 | 0.0558 |
| chr11 | 29039081 | 29043228 | 4148 | 0.25 | 0.033 | 0.0558 |
| chr2 | 192677147 | 192677782 | 636 | -1.92 | 0.033 | 0.0558 |
| chr1 | 115123153 | 115125119 | 1967 | 0.37 | 0.0331 | 0.0559 |
| chr5 | 159276498 | 159277949 | 1452 | -1.08 | 0.0331 | 0.0559 |
| chr8 | 134299531 | 134311125 | 11595 | -0.2 | 0.0331 | 0.0559 |
| chr4 | 89443580 | 89446022 | 2443 | 0.37 | 0.0331 | 0.0559 |
| chr7 | 156470032 | 156470909 | 878 | -1.49 | 0.0331 | 0.056 |
| chr8 | 133358244 | 133359434 | 1191 | -0.64 | 0.0332 | 0.056 |
| chr1 | 169883168 | 169884338 | 1171 | 3.26 | 0.0332 | 0.056 |
| chr10 | 69643425 | 69656581 | 13157 | 0.31 | 0.0332 | 0.056 |
| chr12 | 1771985 | 1776473 | 4489 | 0.87 | 0.0332 | 0.0561 |
| chrX | 65844627 | 65859984 | 15358 | -0.33 | 0.0332 | 0.0561 |
| chr6 | 41513142 | 41519286 | 6145 | -0.55 | 0.0332 | 0.0561 |
| chr3 | 111042703 | 111043566 | 864 | 1.04 | 0.0333 | 0.0562 |
| chr9 | 98948775 | 98949287 | 513 | 2.06 | 0.0333 | 0.0562 |
| chr18 | 18624083 | 18630263 | 6181 | 0.65 | 0.0333 | 0.0563 |
| chr6 | 34854584 | 34857865 | 3282 | 0.28 | 0.0333 | 0.0563 |
| chr17 | 437230 | 438276 | 1047 | -0.94 | 0.0334 | 0.0563 |
| chr1 | 118147899 | 118155991 | 8093 | -1.95 | 0.0334 | 0.0563 |
| chr2 | 105653968 | 105655689 | 1722 | -0.43 | 0.0334 | 0.0563 |
| chr2 | 32010771 | 32012250 | 1480 | 0.77 | 0.0334 | 0.0564 |
| chr6 | 56613876 | 56638020 | 24145 | -0.31 | 0.0334 | 0.0564 |
| chr5 | 997098 | 1005503 | 8406 | -1.23 | 0.0335 | 0.0564 |
| chr9 | 107894354 | 107895268 | 915 | -2.03 | 0.0335 | 0.0565 |
| chr15 | 88781897 | 88782814 | 918 | -1.09 | 0.0335 | 0.0565 |
| chr1 | 79933935 | 79934980 | 1046 | 1.36 | 0.0335 | 0.0565 |
| chr6 | 28047162 | 28049874 | 2713 | 0.29 | 0.0335 | 0.0565 |
| chr13 | 51305618 | 51306035 | 418 | -1.74 | 0.0335 | 0.0565 |
| chr2 | 212886379 | 212887474 | 1096 | 1.13 | 0.0335 | 0.0566 |
| chr2 | 37871385 | 37917442 | 46058 | 0.17 | 0.0336 | 0.0566 |
| chr1 | 235358343 | 235359953 | 1611 | 1.09 | 0.0336 | 0.0566 |
| chrX | 70473705 | 70475924 | 2220 | -0.84 | 0.0336 | 0.0567 |
| chr3 | 66643589 | 66645988 | 2400 | -0.49 | 0.0336 | 0.0567 |
| chr3 | 197316972 | 197317897 | 926 | 1.14 | 0.0336 | 0.0567 |
| chr2 | 224623226 | 224629147 | 5922 | 0.3 | 0.0336 | 0.0567 |
| chr8 | 124283737 | 124288135 | 4399 | 0.35 | 0.0337 | 0.0568 |
| chr4 | 5787207 | 5805576 | 18370 | -0.41 | 0.0337 | 0.0568 |
| chr11 | 64602690 | 64662488 | 59799 | -0.11 | 0.0337 | 0.0568 |
| chr14 | 93427466 | 93431848 | 4383 | -0.45 | 0.0337 | 0.0568 |
| chr4 | 148078766 | 148080374 | 1609 | 0.58 | 0.0337 | 0.0569 |
| chr11 | 31390780 | 31392159 | 1380 | 0.5 | 0.0337 | 0.0569 |
| chr7 | 141436599 | 141439533 | 2935 | 0.24 | 0.0337 | 0.0569 |
| chrX | 15422683 | 15429579 | 6897 | -0.58 | 0.0338 | 0.0569 |
| chr1 | 203273872 | 203275851 | 1980 | 0.45 | 0.0338 | 0.0569 |
| chr4 | 95780501 | 95800680 | 20180 | -0.36 | 0.0338 | 0.057 |
| chr10 | 125011825 | 125012456 | 632 | 1.28 | 0.0338 | 0.057 |
| chr8 | 32146046 | 32146640 | 595 | -1.28 | 0.0339 | 0.0571 |
| chr17 | 5233256 | 5234152 | 897 | -1.09 | 0.0339 | 0.0571 |
| chr6 | 7311707 | 7314301 | 2595 | 0.42 | 0.0339 | 0.0571 |
| chr6 | 13924112 | 13925921 | 1810 | 1.14 | 0.0339 | 0.0571 |
| chr4 | 37686738 | 37689113 | 2376 | 0.53 | 0.0339 | 0.0572 |
| chr17 | 45175781 | 45179126 | 3346 | 0.4 | 0.0339 | 0.0572 |
| chr2 | 216399734 | 216401659 | 1926 | 3.31 | 0.034 | 0.0572 |
| chr7 | 107203610 | 107205860 | 2251 | 0.34 | 0.034 | 0.0573 |
| chr1 | 95552114 | 95554289 | 2176 | 0.38 | 0.034 | 0.0573 |
| chr3 | 130660262 | 130661605 | 1344 | -1.01 | 0.0341 | 0.0574 |
| chr12 | 53107679 | 53108585 | 907 | -1.09 | 0.0341 | 0.0575 |
| chr2 | 131147908 | 131149933 | 2026 | 0.67 | 0.0341 | 0.0575 |

| | | | | | | |
|---|---|---|---|---|---|---|
| chr1 | 47183211 | 47185749 | 2539 | 0.46 | 0.0341 | 0.0575 |
| chr3 | 45207183 | 45210745 | 3563 | 0.21 | 0.0341 | 0.0575 |
| chr7 | 155887058 | 155887564 | 507 | -1.43 | 0.0341 | 0.0575 |
| chr7 | 132632456 | 132632962 | 507 | -1.45 | 0.0341 | 0.0575 |
| chr1 | 28654736 | 28656888 | 2153 | 0.34 | 0.0342 | 0.0575 |
| chr2 | 134933654 | 134934179 | 526 | -1.6 | 0.0342 | 0.0576 |
| chr6 | 52746891 | 52747654 | 764 | 0.85 | 0.0342 | 0.0576 |
| chr1 | 145396577 | 145400469 | 3893 | 0.26 | 0.0342 | 0.0576 |
| chr2 | 198315964 | 198326705 | 10742 | 0.2 | 0.0342 | 0.0576 |
| chr1 | 9621076 | 9622159 | 1084 | 1.08 | 0.0342 | 0.0576 |
| chr4 | 79450700 | 79456058 | 5359 | 0.89 | 0.0342 | 0.0576 |
| chr1 | 167265418 | 167275665 | 10248 | 0.3 | 0.0343 | 0.0578 |
| chr5 | 66501646 | 66508456 | 6811 | 0.38 | 0.0343 | 0.0578 |
| chr14 | 51026598 | 51028431 | 1834 | 0.69 | 0.0343 | 0.0578 |
| chr15 | 39049876 | 39050722 | 847 | 1.61 | 0.0343 | 0.0578 |
| chr5 | 159530921 | 159546506 | 15586 | -0.39 | 0.0343 | 0.0578 |
| chr14 | 57928278 | 57929359 | 1082 | 0.9 | 0.0344 | 0.0578 |
| chr19 | 44331464 | 44332439 | 976 | 0.73 | 0.0344 | 0.0578 |
| chr9 | 138370690 | 138373033 | 2344 | 0.67 | 0.0344 | 0.0579 |
| chr6 | 158895401 | 158903727 | 8327 | -0.47 | 0.0344 | 0.0579 |
| chr5 | 95319871 | 95320968 | 1098 | 1.44 | 0.0344 | 0.0579 |
| chr9 | 81804008 | 81805424 | 1417 | -0.83 | 0.0345 | 0.058 |
| chr10 | 11975660 | 11979141 | 3482 | -0.68 | 0.0345 | 0.058 |
| chr8 | 82695605 | 82697165 | 1561 | -1.36 | 0.0345 | 0.058 |
| chr18 | 71983483 | 71984904 | 1422 | -0.82 | 0.0345 | 0.058 |
| chr7 | 32528887 | 32556406 | 27520 | 0.17 | 0.0345 | 0.058 |
| chr20 | 57465686 | 57483929 | 18244 | 0.28 | 0.0345 | 0.0581 |
| chr17 | 43047396 | 43061391 | 13996 | 0.47 | 0.0346 | 0.0581 |
| chr2 | 200715010 | 200716454 | 1445 | -0.57 | 0.0346 | 0.0581 |
| chr1 | 32225765 | 32227298 | 1534 | -0.77 | 0.0346 | 0.0582 |
| chr16 | 67866355 | 67928601 | 62247 | 0.12 | 0.0346 | 0.0582 |
| chr15 | 65059111 | 65068493 | 9383 | -3.37 | 0.0346 | 0.0582 |
| chr1 | 79518536 | 79519110 | 575 | 1.46 | 0.0346 | 0.0582 |
| chr4 | 147353042 | 147353797 | 756 | -1.13 | 0.0346 | 0.0582 |
| chr4 | 2964079 | 2965108 | 1030 | 0.75 | 0.0347 | 0.0582 |
| chr10 | 75982950 | 75989883 | 6934 | -0.33 | 0.0347 | 0.0583 |
| chr17 | 36570706 | 36585497 | 14792 | -1.51 | 0.0347 | 0.0583 |
| chr2 | 202614697 | 202615371 | 675 | -1.23 | 0.0347 | 0.0583 |
| chr15 | 30209725 | 30227240 | 17516 | -0.19 | 0.0347 | 0.0584 |
| chr5 | 167932317 | 167932986 | 670 | -1.71 | 0.0347 | 0.0584 |
| chr13 | 45196007 | 45197346 | 1340 | -0.81 | 0.0348 | 0.0584 |
| chr17 | 78828977 | 78833639 | 4663 | -0.27 | 0.0348 | 0.0585 |
| chr4 | 187111619 | 187113774 | 2156 | 0.47 | 0.0348 | 0.0585 |
| chr4 | 129472726 | 129524370 | 51645 | -0.13 | 0.0348 | 0.0585 |
| chr10 | 64027930 | 64029154 | 1225 | 0.77 | 0.0349 | 0.0586 |
| chr10 | 14026881 | 14027328 | 448 | -1.5 | 0.0349 | 0.0586 |
| chr6 | 74159579 | 74172400 | 12822 | 0.23 | 0.0349 | 0.0586 |
| chr17 | 41977457 | 41979337 | 1881 | -0.6 | 0.0349 | 0.0586 |
| chr19 | 47833572 | 47835801 | 2230 | 0.84 | 0.0349 | 0.0587 |
| chr21 | 34939685 | 34962484 | 22800 | 0.23 | 0.0349 | 0.0587 |
| chr22 | 44893021 | 44894954 | 1934 | -0.41 | 0.0349 | 0.0587 |
| chr8 | 28558012 | 28571660 | 13649 | -0.47 | 0.035 | 0.0587 |
| chr4 | 87979496 | 87993087 | 13592 | -0.62 | 0.035 | 0.0587 |
| chr17 | 8149432 | 8157802 | 8371 | 0.16 | 0.035 | 0.0587 |
| chr19 | 49836610 | 49837905 | 1296 | -0.84 | 0.035 | 0.0588 |
| chr13 | 73526608 | 73527570 | 963 | -2 | 0.035 | 0.0588 |
| chr2 | 183896607 | 183904249 | 7643 | -0.25 | 0.035 | 0.0588 |
| chr9 | 20379608 | 20406971 | 27364 | 3.2 | 0.035 | 0.0588 |
| chrX | 53587698 | 53588172 | 475 | -1.69 | 0.035 | 0.0588 |
| chr8 | 6692057 | 6692675 | 619 | -1.78 | 0.0351 | 0.0588 |
| chr22 | 36954232 | 36960011 | 5780 | -0.65 | 0.0351 | 0.0589 |

| | | | | | | |
|---|---|---|---|---|---|---|
| chr6 | 105657427 | 105658382 | 956 | -1.05 | 0.0351 | 0.0589 |
| chr9 | 139293378 | 139306156 | 12779 | -0.3 | 0.0351 | 0.0589 |
| chr8 | 30532067 | 30602605 | 70539 | -0.14 | 0.0351 | 0.0589 |
| chr1 | 204182310 | 204183980 | 1671 | 0.71 | 0.0351 | 0.0589 |
| chr11 | 56838741 | 56851931 | 13191 | 0.25 | 0.0351 | 0.0589 |
| chr6 | 4907252 | 4910799 | 3548 | -0.7 | 0.0351 | 0.0589 |
| chr1 | 241887764 | 241889023 | 1260 | -1.1 | 0.0351 | 0.059 |
| chr16 | 66304111 | 66304457 | 347 | 1.91 | 0.0352 | 0.059 |
| chr12 | 28722154 | 28723753 | 1600 | 0.74 | 0.0352 | 0.059 |
| chr13 | 45854309 | 45855931 | 1623 | -1.12 | 0.0352 | 0.059 |
| chr4 | 170074794 | 170075859 | 1066 | -1.26 | 0.0352 | 0.059 |
| chr2 | 101792188 | 101797248 | 5061 | 2.83 | 0.0352 | 0.059 |
| chr20 | 10903245 | 10904134 | 890 | -0.91 | 0.0353 | 0.0592 |
| chr5 | 95252993 | 95260845 | 7853 | 0.71 | 0.0353 | 0.0592 |
| chr7 | 5228918 | 5231734 | 2817 | 0.42 | 0.0353 | 0.0592 |
| chr4 | 81123278 | 81124108 | 831 | 1.32 | 0.0353 | 0.0592 |
| chr2 | 203903360 | 203903870 | 511 | -1.4 | 0.0354 | 0.0593 |
| chr7 | 43944614 | 43946806 | 2193 | 0.53 | 0.0354 | 0.0593 |
| chr1 | 197742758 | 197745635 | 2878 | 0.33 | 0.0354 | 0.0594 |
| chr22 | 48649758 | 48650934 | 1177 | 1.26 | 0.0354 | 0.0594 |
| chr12 | 90287522 | 90288963 | 1442 | -0.68 | 0.0354 | 0.0594 |
| chr9 | 6851080 | 6878571 | 27492 | -0.32 | 0.0355 | 0.0594 |
| chr8 | 8537210 | 8543757 | 6548 | 1.19 | 0.0355 | 0.0594 |
| chr11 | 46065731 | 46066679 | 949 | 1.48 | 0.0355 | 0.0594 |
| chr4 | 143431071 | 143431437 | 367 | -1.81 | 0.0355 | 0.0594 |
| chr13 | 20251597 | 20260235 | 8639 | 0.56 | 0.0355 | 0.0595 |
| chr1 | 54276312 | 54277549 | 1238 | -1.05 | 0.0355 | 0.0595 |
| chr6 | 27027698 | 27028376 | 679 | -2.07 | 0.0355 | 0.0595 |
| chr5 | 77411258 | 77413829 | 2572 | 0.53 | 0.0355 | 0.0595 |
| chr7 | 79049520 | 79050545 | 1026 | 0.95 | 0.0356 | 0.0596 |
| chr8 | 38696967 | 38697400 | 434 | -1.63 | 0.0356 | 0.0596 |
| chr4 | 177116204 | 177117521 | 1318 | 0.5 | 0.0356 | 0.0596 |
| chr18 | 267096 | 269416 | 2321 | 0.33 | 0.0357 | 0.0597 |
| chr16 | 23372943 | 23377934 | 4992 | 0.38 | 0.0357 | 0.0597 |
| chr22 | 22750960 | 22752424 | 1465 | -0.73 | 0.0357 | 0.0598 |
| chr20 | 45316651 | 45319161 | 2511 | 0.36 | 0.0357 | 0.0598 |
| chr8 | 18918350 | 18919377 | 1028 | 1.11 | 0.0357 | 0.0598 |
| chr18 | 53500374 | 53501499 | 1126 | -1.14 | 0.0357 | 0.0598 |
| chr17 | 28087652 | 28088894 | 1243 | 0.58 | 0.0357 | 0.0598 |
| chr4 | 101453791 | 101454295 | 505 | 1.53 | 0.0358 | 0.0599 |
| chr8 | 38153257 | 38161652 | 8396 | 0.54 | 0.0358 | 0.0599 |
| chr15 | 44394397 | 44395699 | 1303 | 0.98 | 0.0358 | 0.06 |
| chr1 | 161122690 | 161148218 | 25529 | -0.15 | 0.0358 | 0.06 |
| chr8 | 119908792 | 119910273 | 1482 | 3.56 | 0.0359 | 0.0601 |
| chr1 | 213243977 | 213244458 | 482 | 1.76 | 0.0359 | 0.0601 |
| chr1 | 246116961 | 246117929 | 969 | 0.73 | 0.0359 | 0.0601 |
| chr11 | 58213672 | 58214205 | 534 | -2.05 | 0.0359 | 0.0601 |
| chr10 | 16686258 | 16686962 | 705 | -1.21 | 0.0359 | 0.0601 |
| chr15 | 76030453 | 76031907 | 1455 | -0.75 | 0.0359 | 0.0601 |
| chr1 | 54328088 | 54328821 | 734 | -1.48 | 0.0359 | 0.0601 |
| chr22 | 31089700 | 31090586 | 887 | 0.7 | 0.036 | 0.0602 |
| chr19 | 487539 | 489526 | 1988 | 0.55 | 0.036 | 0.0602 |
| chr3 | 38494802 | 38497531 | 2730 | -0.38 | 0.036 | 0.0603 |
| chr2 | 128394929 | 128396016 | 1088 | -0.8 | 0.036 | 0.0603 |
| chr16 | 10520735 | 10521674 | 940 | 1.21 | 0.0361 | 0.0603 |
| chr11 | 120894259 | 120895894 | 1636 | -0.56 | 0.0361 | 0.0603 |
| chr10 | 17071701 | 17072105 | 405 | -1.42 | 0.0361 | 0.0604 |
| chr13 | 60265463 | 60266068 | 606 | 1.41 | 0.0361 | 0.0604 |
| chr14 | 69139472 | 69172051 | 32580 | -0.21 | 0.0361 | 0.0604 |
| chr8 | 16948665 | 16949582 | 918 | -1.33 | 0.0361 | 0.0604 |
| chr17 | 30573908 | 30574830 | 923 | -0.67 | 0.0361 | 0.0604 |

| | | | | | | |
|---|---|---|---|---|---|---|
| chr6 | 43264861 | 43265198 | 338 | -1.71 | 0.0361 | 0.0604 |
| chr22 | 46839795 | 46944075 | 104281 | -0.11 | 0.0361 | 0.0604 |
| chr11 | 66911427 | 66915078 | 3652 | 0.5 | 0.0361 | 0.0604 |
| chr6 | 90061284 | 90063527 | 2244 | 0.31 | 0.0361 | 0.0604 |
| chr7 | 149172420 | 149175785 | 3366 | 0.79 | 0.0361 | 0.0604 |
| chr9 | 93220180 | 93220845 | 666 | 1.41 | 0.0361 | 0.0604 |
| chr10 | 5183815 | 5194329 | 10515 | 0.14 | 0.0362 | 0.0605 |
| chr11 | 72144734 | 72146303 | 1570 | -0.43 | 0.0362 | 0.0605 |
| chr10 | 24876462 | 24892809 | 16348 | 0.47 | 0.0362 | 0.0605 |
| chr1 | 64238155 | 64242175 | 4021 | -0.53 | 0.0362 | 0.0605 |
| chr16 | 29275444 | 29275719 | 276 | -2.28 | 0.0362 | 0.0605 |
| chr7 | 37382427 | 37382835 | 409 | -1.94 | 0.0362 | 0.0605 |
| chr1 | 192775258 | 192779156 | 3899 | 0.27 | 0.0362 | 0.0605 |
| chr9 | 113539219 | 113539745 | 527 | 1.36 | 0.0362 | 0.0606 |
| chr6 | 135948299 | 135949110 | 812 | 1.29 | 0.0363 | 0.0606 |
| chr11 | 102468590 | 102492383 | 23794 | -0.18 | 0.0363 | 0.0606 |
| chr12 | 113233500 | 113234649 | 1150 | -0.85 | 0.0363 | 0.0606 |
| chr7 | 72722285 | 72723543 | 1259 | 0.76 | 0.0363 | 0.0606 |
| chr1 | 226456214 | 226460308 | 4095 | -0.85 | 0.0363 | 0.0606 |
| chr2 | 201674348 | 201684832 | 10485 | -0.23 | 0.0363 | 0.0606 |
| chr1 | 25071227 | 25073423 | 2197 | 0.6 | 0.0363 | 0.0606 |
| chr15 | 41912590 | 41915210 | 2621 | 0.4 | 0.0363 | 0.0607 |
| chrX | 37741981 | 37743118 | 1138 | 1.02 | 0.0363 | 0.0607 |
| chr16 | 90014370 | 90023836 | 9467 | -0.48 | 0.0364 | 0.0607 |
| chr8 | 19976857 | 19977737 | 881 | -1.27 | 0.0364 | 0.0608 |
| chr12 | 78604385 | 78605629 | 1245 | 0.76 | 0.0365 | 0.0609 |
| chr3 | 127935985 | 127937057 | 1073 | -2 | 0.0365 | 0.0609 |
| chr4 | 3782501 | 3782956 | 456 | 1.73 | 0.0365 | 0.0609 |
| chr4 | 48249162 | 48249740 | 579 | -1.43 | 0.0365 | 0.0609 |
| chr22 | 41229889 | 41254469 | 24581 | 0.21 | 0.0365 | 0.061 |
| chr7 | 30632558 | 30638480 | 5923 | 0.18 | 0.0365 | 0.061 |
| chr5 | 149309470 | 149321459 | 11990 | -0.18 | 0.0365 | 0.061 |
| chr1 | 161070446 | 161070966 | 521 | -1.52 | 0.0365 | 0.061 |
| chr3 | 81492176 | 81493878 | 1703 | -0.53 | 0.0365 | 0.061 |
| chr10 | 728840 | 740484 | 11645 | 0.26 | 0.0365 | 0.061 |
| chr17 | 33759779 | 33761336 | 1558 | 0.66 | 0.0365 | 0.061 |
| chr14 | 55737023 | 55740081 | 3059 | 0.32 | 0.0365 | 0.061 |
| chr15 | 43621783 | 43623440 | 1658 | 0.46 | 0.0366 | 0.0611 |
| chr20 | 33972265 | 33972775 | 511 | -2.05 | 0.0367 | 0.0612 |
| chr16 | 84565101 | 84567235 | 2135 | 0.49 | 0.0367 | 0.0612 |
| chr3 | 42673122 | 42677197 | 4076 | 0.61 | 0.0367 | 0.0613 |
| chr2 | 136144722 | 136145545 | 824 | 1.88 | 0.0367 | 0.0613 |
| chr2 | 32891313 | 32892209 | 897 | -1.16 | 0.0368 | 0.0613 |
| chr3 | 66101173 | 66102497 | 1325 | 1.21 | 0.0368 | 0.0614 |
| chr9 | 110399650 | 110401091 | 1442 | 0.62 | 0.0368 | 0.0614 |
| chr10 | 97451971 | 97453771 | 1801 | -0.32 | 0.0368 | 0.0614 |
| chr2 | 233844206 | 233855061 | 10856 | -0.26 | 0.0368 | 0.0614 |
| chr21 | 38736981 | 38753862 | 16882 | 0.18 | 0.0368 | 0.0614 |
| chr13 | 33089136 | 33090102 | 967 | -1.34 | 0.0368 | 0.0615 |
| chr4 | 188703570 | 188704389 | 820 | 1.22 | 0.0369 | 0.0615 |
| chr10 | 112662174 | 112663371 | 1198 | -1.05 | 0.0369 | 0.0616 |
| chr1 | 2203092 | 2208539 | 5448 | -0.76 | 0.0369 | 0.0616 |
| chr13 | 51563308 | 51563996 | 689 | 1.01 | 0.037 | 0.0617 |
| chr7 | 91605431 | 91606624 | 1194 | -1.12 | 0.037 | 0.0617 |
| chr14 | 94512463 | 94514338 | 1876 | -0.87 | 0.037 | 0.0617 |
| chr3 | 79066782 | 79069404 | 2623 | 0.45 | 0.0371 | 0.0619 |
| chr10 | 125685765 | 125686647 | 883 | -1.98 | 0.0371 | 0.0619 |
| chrX | 102508925 | 102510336 | 1412 | 0.85 | 0.0372 | 0.0619 |
| chr2 | 203774728 | 203777209 | 2482 | -0.39 | 0.0372 | 0.062 |
| chr14 | 78206048 | 78207530 | 1483 | 0.96 | 0.0372 | 0.062 |
| chr13 | 95247423 | 95255566 | 8144 | 0.41 | 0.0372 | 0.062 |

| | | | | | | |
|---|---|---|---|---|---|---|
| chr3 | 137892265 | 137907496 | 15232 | 0.29 | 0.0372 | 0.0621 |
| chr2 | 188268108 | 188269245 | 1138 | -1.02 | 0.0373 | 0.0621 |
| chr5 | 17400897 | 17414012 | 13116 | 0.5 | 0.0373 | 0.0621 |
| chr11 | 64139959 | 64141207 | 1249 | 0.84 | 0.0373 | 0.0621 |
| chr13 | 111267240 | 111269546 | 2307 | 0.44 | 0.0373 | 0.0622 |
| chr7 | 93929184 | 93944351 | 15168 | 0.34 | 0.0373 | 0.0622 |
| chr1 | 24468992 | 24477389 | 8398 | -0.7 | 0.0373 | 0.0622 |
| chr6 | 11702316 | 11703394 | 1079 | -1.13 | 0.0373 | 0.0622 |
| chr17 | 62262194 | 62263575 | 1382 | -0.98 | 0.0374 | 0.0623 |
| chr12 | 102642539 | 102644568 | 2030 | 3.81 | 0.0374 | 0.0623 |
| chr7 | 80928053 | 80929637 | 1585 | 0.6 | 0.0374 | 0.0623 |
| chr1 | 175990079 | 175991122 | 1044 | -1.88 | 0.0374 | 0.0623 |
| chr6 | 7178705 | 7179649 | 945 | 1.24 | 0.0374 | 0.0623 |
| chr22 | 36815159 | 36832043 | 16885 | -0.35 | 0.0374 | 0.0623 |
| chr14 | 65377153 | 65422377 | 45225 | 0.12 | 0.0374 | 0.0623 |
| chr1 | 120459537 | 120460650 | 1114 | -1.14 | 0.0375 | 0.0625 |
| chrX | 2766239 | 2768162 | 1924 | 0.59 | 0.0375 | 0.0625 |
| chr19 | 58361083 | 58363150 | 2068 | 0.43 | 0.0375 | 0.0625 |
| chr1 | 9187886 | 9190187 | 2302 | 0.79 | 0.0376 | 0.0626 |
| chr13 | 49821299 | 49823159 | 1861 | 0.51 | 0.0376 | 0.0626 |
| chr6 | 147271679 | 147272069 | 391 | 2.11 | 0.0376 | 0.0627 |
| chr7 | 75059893 | 75060500 | 608 | 1.55 | 0.0377 | 0.0627 |
| chr2 | 188679518 | 188680215 | 698 | -1.35 | 0.0377 | 0.0627 |
| chr12 | 52024942 | 52025976 | 1035 | -0.84 | 0.0377 | 0.0628 |
| chr20 | 4044378 | 4046002 | 1625 | 0.6 | 0.0377 | 0.0628 |
| chr6 | 111194778 | 111208783 | 14006 | 0.23 | 0.0378 | 0.0628 |
| chr15 | 68091947 | 68092475 | 529 | -1.15 | 0.0378 | 0.0628 |
| chr1 | 21139699 | 21141107 | 1409 | -0.63 | 0.0378 | 0.0628 |
| chr1 | 31534880 | 31539628 | 4749 | 0.26 | 0.0378 | 0.0629 |
| chr1 | 155352131 | 155394925 | 42795 | 0.2 | 0.0378 | 0.0629 |
| chr20 | 13375570 | 13376752 | 1183 | -0.76 | 0.0378 | 0.0629 |
| chr1 | 156009320 | 156025918 | 16599 | -0.28 | 0.0378 | 0.0629 |
| chr4 | 177822299 | 177822784 | 486 | -1.25 | 0.0378 | 0.0629 |
| chr2 | 45139547 | 45140539 | 993 | 1.41 | 0.0379 | 0.063 |
| chr1 | 53637942 | 53642888 | 4947 | 0.52 | 0.038 | 0.0631 |
| chr11 | 12378623 | 12379049 | 427 | 1.48 | 0.038 | 0.0632 |
| chr3 | 188258965 | 188277199 | 18235 | 0.26 | 0.038 | 0.0632 |
| chr5 | 60139092 | 60140964 | 1873 | 0.65 | 0.038 | 0.0632 |
| chr22 | 38070361 | 38083247 | 12887 | 0.34 | 0.038 | 0.0633 |
| chr3 | 152626577 | 152628317 | 1741 | 0.48 | 0.038 | 0.0633 |
| chr2 | 120890979 | 120892709 | 1731 | 0.73 | 0.0381 | 0.0633 |
| chr10 | 69251779 | 69252990 | 1212 | 1.51 | 0.0381 | 0.0633 |
| chr6 | 109126779 | 109127694 | 916 | 1.55 | 0.0381 | 0.0633 |
| chr9 | 118157807 | 118158797 | 991 | 2.87 | 0.0381 | 0.0633 |
| chr4 | 132919217 | 132919672 | 456 | 1.66 | 0.0382 | 0.0634 |
| chr6 | 166796690 | 166797845 | 1156 | 0.92 | 0.0382 | 0.0634 |
| chr6 | 160720616 | 160721807 | 1192 | 0.96 | 0.0382 | 0.0634 |
| chr1 | 89196920 | 89197959 | 1040 | 0.97 | 0.0382 | 0.0634 |
| chr17 | 30313745 | 30319384 | 5640 | 0.57 | 0.0382 | 0.0634 |
| chr12 | 57657316 | 57675182 | 17867 | 0.31 | 0.0382 | 0.0635 |
| chr3 | 41240136 | 41243352 | 3217 | 0.37 | 0.0382 | 0.0635 |
| chr9 | 75814344 | 75815709 | 1366 | 0.93 | 0.0382 | 0.0635 |
| chr20 | 35552173 | 35553065 | 893 | -1.02 | 0.0382 | 0.0635 |
| chr3 | 160242259 | 160250434 | 8176 | 0.72 | 0.0382 | 0.0635 |
| chr21 | 33087771 | 33089022 | 1252 | -0.89 | 0.0383 | 0.0636 |
| chr19 | 19029020 | 19031341 | 2322 | 0.52 | 0.0383 | 0.0636 |
| chr7 | 30651363 | 30654122 | 2760 | -0.48 | 0.0383 | 0.0636 |
| chr4 | 99351258 | 99352724 | 1467 | -0.61 | 0.0383 | 0.0636 |
| chr3 | 132185774 | 132208507 | 22734 | -0.31 | 0.0383 | 0.0636 |
| chr5 | 78277298 | 78282483 | 5186 | 0.27 | 0.0383 | 0.0637 |
| chr10 | 89822161 | 89823236 | 1076 | 0.56 | 0.0383 | 0.0637 |

| | | | | | | |
|---|---|---|---|---|---|---|
| chr18 | 71749838 | 71750496 | 659 | 1.13 | 0.0383 | 0.0637 |
| chr2 | 236610871 | 236619246 | 8376 | -0.41 | 0.0384 | 0.0638 |
| chr3 | 171804426 | 171823405 | 18980 | -0.31 | 0.0385 | 0.0639 |
| chr15 | 48706417 | 48708079 | 1663 | 0.77 | 0.0385 | 0.0639 |
| chr15 | 37390662 | 37395532 | 4871 | 0.28 | 0.0385 | 0.0639 |
| chr5 | 159657685 | 159658536 | 852 | -1.98 | 0.0385 | 0.0639 |
| chr4 | 162930862 | 162932993 | 2132 | 0.6 | 0.0385 | 0.0639 |
| chr4 | 2625177 | 2627044 | 1868 | -1.07 | 0.0385 | 0.064 |
| chr4 | 56410596 | 56414626 | 4031 | 0.31 | 0.0386 | 0.064 |
| chr3 | 23846864 | 23853637 | 6774 | -0.26 | 0.0386 | 0.064 |
| chr4 | 100805820 | 100816868 | 11049 | 0.39 | 0.0386 | 0.0641 |
| chr10 | 21386033 | 21387452 | 1420 | -0.77 | 0.0386 | 0.0641 |
| chr4 | 78557391 | 78558839 | 1449 | 0.57 | 0.0387 | 0.0642 |
| chr19 | 59009932 | 59033628 | 23697 | 0.19 | 0.0387 | 0.0642 |
| chr2 | 195890933 | 195891558 | 626 | -2 | 0.0387 | 0.0643 |
| chr6 | 136434521 | 136435628 | 1108 | -1.19 | 0.0387 | 0.0643 |
| chr12 | 17257546 | 17258043 | 498 | 1.63 | 0.0387 | 0.0643 |
| chrX | 64753535 | 64757859 | 4325 | 0.57 | 0.0387 | 0.0643 |
| chr1 | 222695142 | 222696350 | 1209 | 0.81 | 0.0387 | 0.0643 |
| chr1 | 220297785 | 220324131 | 26347 | 0.28 | 0.0387 | 0.0643 |
| chr10 | 27361058 | 27362874 | 1817 | -0.77 | 0.0388 | 0.0644 |
| chr4 | 154427031 | 154428360 | 1330 | 1.67 | 0.0388 | 0.0644 |
| chr14 | 51808061 | 51809591 | 1531 | 0.78 | 0.0388 | 0.0644 |
| chr16 | 50268069 | 50281091 | 13023 | 0.43 | 0.0389 | 0.0645 |
| chr4 | 177758382 | 177760926 | 2545 | 0.63 | 0.0389 | 0.0645 |
| chr2 | 42148568 | 42165848 | 17281 | 0.2 | 0.0389 | 0.0645 |
| chr7 | 98922524 | 99020383 | 97860 | 0.12 | 0.0389 | 0.0645 |
| chr2 | 63082287 | 63101766 | 19480 | -0.28 | 0.0389 | 0.0645 |
| chr1 | 160959806 | 160967991 | 8186 | 0.55 | 0.039 | 0.0646 |
| chr1 | 150441776 | 150461061 | 19286 | 0.3 | 0.039 | 0.0647 |
| chr4 | 40418838 | 40426684 | 7847 | 0.23 | 0.039 | 0.0647 |
| chr10 | 23384460 | 23385586 | 1127 | 0.49 | 0.039 | 0.0647 |
| chr10 | 62492713 | 62493469 | 757 | 1.14 | 0.0391 | 0.0647 |
| chr11 | 82553833 | 82554903 | 1071 | 0.99 | 0.0391 | 0.0648 |
| chr13 | 52979674 | 53005711 | 26038 | -0.25 | 0.0391 | 0.0648 |
| chr20 | 37117376 | 37118531 | 1156 | -1.39 | 0.0391 | 0.0648 |
| chr3 | 128522144 | 128528059 | 5916 | -0.47 | 0.0391 | 0.0648 |
| chr19 | 4516783 | 4517226 | 444 | -1.56 | 0.0391 | 0.0648 |
| chr2 | 145107076 | 145117091 | 10016 | -0.34 | 0.0391 | 0.0649 |
| chr1 | 152139445 | 152141459 | 2015 | -0.73 | 0.0391 | 0.0649 |
| chr11 | 47081604 | 47083713 | 2110 | -0.69 | 0.0392 | 0.0649 |
| chr2 | 72107734 | 72110715 | 2982 | -3.29 | 0.0392 | 0.065 |
| chr2 | 210307607 | 210308446 | 840 | 2.02 | 0.0392 | 0.065 |
| chr1 | 200811381 | 200812385 | 1005 | -1 | 0.0392 | 0.065 |
| chr9 | 107688223 | 107690875 | 2653 | -0.42 | 0.0392 | 0.065 |
| chr12 | 54582252 | 54583457 | 1206 | 0.69 | 0.0392 | 0.065 |
| chr8 | 118628202 | 118629427 | 1226 | 0.9 | 0.0393 | 0.0651 |
| chr3 | 19961384 | 20022370 | 60987 | 0.18 | 0.0393 | 0.0651 |
| chr22 | 32953272 | 32955302 | 2031 | -0.9 | 0.0393 | 0.0652 |
| chr3 | 28309705 | 28310707 | 1003 | -1.4 | 0.0394 | 0.0652 |
| chr12 | 17359572 | 17359981 | 410 | 1.7 | 0.0394 | 0.0652 |
| chr13 | 45495223 | 45507583 | 12361 | 0.32 | 0.0394 | 0.0653 |
| chr10 | 114707849 | 114902514 | 194666 | -0.07 | 0.0395 | 0.0654 |
| chr6 | 154923390 | 154937412 | 14023 | 0.89 | 0.0395 | 0.0654 |
| chr17 | 80091145 | 80092370 | 1226 | -1.1 | 0.0395 | 0.0654 |
| chr14 | 38014983 | 38017474 | 2492 | 0.3 | 0.0395 | 0.0654 |
| chr4 | 161238310 | 161242882 | 4573 | -0.19 | 0.0395 | 0.0654 |
| chr9 | 79707496 | 79708158 | 663 | -1.26 | 0.0395 | 0.0654 |
| chr5 | 127913911 | 127915870 | 1960 | 0.53 | 0.0395 | 0.0654 |
| chr5 | 141227579 | 141230761 | 3183 | 0.3 | 0.0395 | 0.0655 |
| chr3 | 184031723 | 184064167 | 32445 | 0.23 | 0.0395 | 0.0655 |

| | | | | | | |
|---|---|---|---|---|---|---|
| chr7 | 21983911 | 22012304 | 28394 | 4.13 | 0.0395 | 0.0655 |
| chr21 | 35340430 | 35349606 | 9177 | 0.43 | 0.0396 | 0.0655 |
| chr22 | 37186678 | 37187196 | 519 | -1.77 | 0.0396 | 0.0655 |
| chr2 | 118691354 | 118692128 | 775 | -0.91 | 0.0396 | 0.0656 |
| chr8 | 125208056 | 125216725 | 8670 | 0.25 | 0.0396 | 0.0656 |
| chr7 | 46672764 | 46673417 | 654 | -1.14 | 0.0396 | 0.0656 |
| chr1 | 90204581 | 90205197 | 617 | -1.84 | 0.0397 | 0.0656 |
| chr14 | 76617694 | 76619718 | 2025 | 0.45 | 0.0397 | 0.0657 |
| chr11 | 119075905 | 119078737 | 2833 | 0.5 | 0.0397 | 0.0657 |
| chr2 | 3172047 | 3176891 | 4845 | -0.99 | 0.0398 | 0.0658 |
| chr11 | 47481987 | 47489495 | 7509 | 0.39 | 0.0398 | 0.0659 |
| chr4 | 154330220 | 154331539 | 1320 | 0.86 | 0.0399 | 0.0659 |
| chrX | 23511663 | 23523262 | 11600 | 3.26 | 0.0399 | 0.066 |
| chr1 | 185527561 | 185555619 | 28059 | 3.1 | 0.0399 | 0.066 |
| chr6 | 2103613 | 2104651 | 1039 | 0.96 | 0.0399 | 0.066 |
| chr5 | 93303152 | 93304138 | 987 | 1.08 | 0.0399 | 0.066 |
| chr3 | 123452550 | 123478240 | 25691 | -0.14 | 0.04 | 0.0661 |
| chr6 | 125245294 | 125246124 | 831 | 2.03 | 0.04 | 0.0661 |
| chr1 | 47132943 | 47148023 | 15081 | 0.45 | 0.04 | 0.0661 |
| chr2 | 149401854 | 149404351 | 2498 | -0.34 | 0.04 | 0.0661 |
| chr1 | 53662033 | 53663342 | 1310 | 0.48 | 0.04 | 0.0662 |
| chr4 | 123679346 | 123687543 | 8198 | -0.34 | 0.04 | 0.0662 |
| chr20 | 12916433 | 12918064 | 1632 | 0.77 | 0.04 | 0.0662 |
| chr3 | 123678845 | 123680527 | 1683 | 0.4 | 0.04 | 0.0662 |
| chr1 | 70712954 | 70714314 | 1361 | 1.51 | 0.0401 | 0.0663 |
| chr3 | 32799594 | 32800095 | 502 | 1.5 | 0.0401 | 0.0663 |
| chr1 | 95459261 | 95460626 | 1366 | 0.61 | 0.0401 | 0.0663 |
| chr2 | 157339739 | 157340966 | 1228 | 0.87 | 0.0401 | 0.0663 |
| chr17 | 12011290 | 12011980 | 691 | 0.98 | 0.0401 | 0.0663 |
| chr4 | 167952690 | 167953506 | 817 | 1 | 0.0402 | 0.0665 |
| chr12 | 90908724 | 90910274 | 1551 | 0.54 | 0.0403 | 0.0665 |
| chr2 | 238480235 | 238481730 | 1496 | -1.44 | 0.0403 | 0.0665 |
| chr13 | 83472575 | 83473577 | 1003 | 1.01 | 0.0403 | 0.0666 |
| chr15 | 80295204 | 80296628 | 1425 | -1.36 | 0.0403 | 0.0666 |
| chr15 | 90619004 | 90628905 | 9902 | -1.34 | 0.0403 | 0.0666 |
| chr1 | 78120782 | 78121591 | 810 | -1.58 | 0.0403 | 0.0666 |
| chr1 | 86092686 | 86096896 | 4211 | 0.2 | 0.0403 | 0.0666 |
| chr2 | 201870935 | 201874305 | 3371 | 0.54 | 0.0404 | 0.0667 |
| chrX | 13304784 | 13306019 | 1236 | 1.01 | 0.0404 | 0.0667 |
| chr13 | 31037163 | 31041278 | 4116 | 0.28 | 0.0404 | 0.0667 |
| chr1 | 232299480 | 232301110 | 1631 | 0.64 | 0.0404 | 0.0668 |
| chr9 | 127483161 | 127498154 | 14994 | 0.49 | 0.0404 | 0.0668 |
| chr10 | 120965269 | 120967021 | 1753 | 0.42 | 0.0404 | 0.0668 |
| chr9 | 75401625 | 75410693 | 9069 | 0.38 | 0.0404 | 0.0668 |
| chr7 | 140395200 | 140397977 | 2778 | 0.31 | 0.0404 | 0.0668 |
| chr2 | 128847533 | 128861184 | 13652 | -0.24 | 0.0404 | 0.0668 |
| chr19 | 2944599 | 2950723 | 6125 | 0.4 | 0.0404 | 0.0668 |
| chr2 | 45901624 | 45903100 | 1477 | -0.74 | 0.0404 | 0.0668 |
| chr1 | 149997719 | 149998163 | 445 | 2.03 | 0.0405 | 0.0668 |
| chr15 | 50594596 | 50597336 | 2741 | -0.95 | 0.0405 | 0.0668 |
| chr18 | 7645561 | 7646192 | 632 | -2.04 | 0.0405 | 0.0668 |
| chr6 | 41813651 | 41815641 | 1991 | -0.62 | 0.0405 | 0.0669 |
| chr17 | 39165190 | 39181267 | 16078 | 1.89 | 0.0405 | 0.0669 |
| chr7 | 120589516 | 120593083 | 3568 | 0.35 | 0.0405 | 0.0669 |
| chrX | 30906510 | 30908359 | 1850 | 0.49 | 0.0405 | 0.0669 |
| chr10 | 63596689 | 63597881 | 1193 | 0.75 | 0.0405 | 0.0669 |
| chr16 | 2723735 | 2733111 | 9377 | 0.35 | 0.0406 | 0.0669 |
| chr8 | 17102260 | 17114681 | 12422 | -0.19 | 0.0406 | 0.0669 |
| chr1 | 203594605 | 203597378 | 2774 | -0.34 | 0.0406 | 0.067 |
| chr2 | 161825770 | 161827345 | 1576 | -0.31 | 0.0406 | 0.067 |
| chr1 | 180165820 | 180166894 | 1075 | -1.96 | 0.0407 | 0.0671 |

| | | | | | | |
|---|---|---|---|---|---|---|
| chr14 | 95651661 | 95652334 | 674 | -1.31 | 0.0407 | 0.0671 |
| chr21 | 16664783 | 16666731 | 1949 | 0.51 | 0.0407 | 0.0671 |
| chr5 | 141700528 | 141722350 | 21823 | 0.21 | 0.0407 | 0.0671 |
| chr16 | 4630509 | 4631106 | 598 | 1.48 | 0.0407 | 0.0672 |
| chr14 | 103017770 | 103019870 | 2101 | 0.62 | 0.0407 | 0.0672 |
| chr7 | 73416414 | 73417745 | 1332 | 0.51 | 0.0407 | 0.0672 |
| chr22 | 24535857 | 24558243 | 22387 | 0.22 | 0.0408 | 0.0672 |
| chr12 | 60729281 | 60730117 | 837 | 1.02 | 0.0408 | 0.0672 |
| chr1 | 155192946 | 155198212 | 5267 | 0.33 | 0.0408 | 0.0672 |
| chr6 | 88541868 | 88542404 | 537 | 0.99 | 0.0408 | 0.0673 |
| chrX | 128640316 | 128641426 | 1111 | -0.87 | 0.0408 | 0.0673 |
| chr5 | 6376930 | 6379688 | 2759 | 0.32 | 0.0408 | 0.0673 |
| chr5 | 169949043 | 169950030 | 988 | 1.99 | 0.0409 | 0.0674 |
| chr4 | 147096316 | 147098182 | 1867 | 0.4 | 0.0409 | 0.0674 |
| chr1 | 77496857 | 77497746 | 890 | 1.25 | 0.0409 | 0.0674 |
| chr2 | 8274478 | 8275054 | 577 | 2.23 | 0.0409 | 0.0674 |
| chr2 | 152683800 | 152685638 | 1839 | -0.49 | 0.0409 | 0.0675 |
| chr18 | 45386655 | 45459820 | 73166 | 0.11 | 0.041 | 0.0675 |
| chrX | 19001549 | 19016480 | 14932 | -0.29 | 0.041 | 0.0675 |
| chr3 | 53333822 | 53334222 | 401 | 1.38 | 0.041 | 0.0676 |
| chr16 | 53499385 | 53500267 | 883 | -1.28 | 0.041 | 0.0676 |
| chr14 | 75506617 | 75537057 | 30441 | 0.18 | 0.041 | 0.0676 |
| chr10 | 5922371 | 5933530 | 11160 | -0.35 | 0.041 | 0.0676 |
| chrX | 154841235 | 154843113 | 1879 | 0.41 | 0.041 | 0.0676 |
| chr18 | 778644 | 779241 | 598 | 1.86 | 0.041 | 0.0676 |
| chr14 | 71543025 | 71543605 | 581 | -1.29 | 0.0411 | 0.0677 |
| chr16 | 69562249 | 69569312 | 7064 | 0.17 | 0.0411 | 0.0677 |
| chr6 | 88497765 | 88505178 | 7414 | 0.2 | 0.0411 | 0.0677 |
| chr7 | 2882745 | 2884884 | 2140 | 0.64 | 0.0411 | 0.0677 |
| chr1 | 32528016 | 32538796 | 10781 | -0.64 | 0.0411 | 0.0677 |
| chr12 | 79985685 | 79992606 | 6922 | 0.41 | 0.0411 | 0.0677 |
| chr4 | 89204674 | 89205716 | 1043 | 0.97 | 0.0411 | 0.0677 |
| chr15 | 66584954 | 66591660 | 6707 | -0.31 | 0.0411 | 0.0677 |
| chr4 | 152019846 | 152029161 | 9316 | 0.29 | 0.0411 | 0.0677 |
| chr5 | 156419843 | 156420359 | 517 | -1.67 | 0.0412 | 0.0678 |
| chr5 | 130522111 | 130523018 | 908 | -1.22 | 0.0412 | 0.0678 |
| chr8 | 66885993 | 66886704 | 712 | -1.31 | 0.0412 | 0.0678 |
| chrX | 54069427 | 54072829 | 3403 | 0.29 | 0.0412 | 0.0679 |
| chr18 | 46448227 | 46487919 | 39693 | -0.14 | 0.0412 | 0.0679 |
| chr3 | 182041166 | 182042089 | 924 | 1.44 | 0.0413 | 0.0679 |
| chr8 | 102740344 | 102749012 | 8669 | 3.55 | 0.0413 | 0.0679 |
| chr9 | 16625863 | 16627356 | 1494 | 0.54 | 0.0413 | 0.068 |
| chr2 | 27650391 | 27660329 | 9939 | -0.29 | 0.0413 | 0.068 |
| chr17 | 29457930 | 29458637 | 708 | -1.3 | 0.0413 | 0.068 |
| chr7 | 77419606 | 77429375 | 9770 | 0.32 | 0.0413 | 0.068 |
| chr14 | 53684105 | 53685874 | 1770 | 0.48 | 0.0413 | 0.068 |
| chr11 | 123451255 | 123472667 | 21413 | 0.17 | 0.0414 | 0.0681 |
| chr7 | 6745421 | 6747226 | 1806 | 0.39 | 0.0414 | 0.0681 |
| chr15 | 33437434 | 33438315 | 882 | -1.28 | 0.0414 | 0.0681 |
| chr6 | 10721848 | 10763616 | 41769 | -0.18 | 0.0415 | 0.0682 |
| chr1 | 173389230 | 173399885 | 10656 | 0.77 | 0.0415 | 0.0682 |
| chr17 | 29231958 | 29234559 | 2602 | 0.34 | 0.0415 | 0.0683 |
| chr1 | 203661369 | 203662570 | 1202 | 0.71 | 0.0416 | 0.0684 |
| chr12 | 105811666 | 105812050 | 385 | 1.35 | 0.0416 | 0.0685 |
| chr1 | 168105648 | 168107174 | 1527 | 0.91 | 0.0416 | 0.0685 |
| chr3 | 169683924 | 169698386 | 14463 | 0.3 | 0.0416 | 0.0685 |
| chr15 | 68440381 | 68441413 | 1033 | -1.34 | 0.0416 | 0.0685 |
| chr2 | 201084081 | 201084859 | 779 | -1.15 | 0.0417 | 0.0685 |
| chr16 | 31043283 | 31045690 | 2408 | 0.37 | 0.0417 | 0.0685 |
| chr15 | 91300418 | 91301421 | 1004 | -0.99 | 0.0417 | 0.0686 |
| chr19 | 36389218 | 36398084 | 8867 | 0.24 | 0.0417 | 0.0686 |

| | | | | | | |
|---|---|---|---|---|---|---|
| chr5 | 179942603 | 179960632 | 18030 | 0.32 | 0.0418 | 0.0686 |
| chr10 | 102790569 | 102828394 | 37826 | 0.14 | 0.0418 | 0.0687 |
| chr15 | 79151244 | 79177922 | 26679 | 0.18 | 0.0418 | 0.0687 |
| chr7 | 98519625 | 98521396 | 1772 | 0.71 | 0.0418 | 0.0687 |
| chr9 | 96857189 | 96857838 | 650 | 1.06 | 0.0418 | 0.0687 |
| chr14 | 103162697 | 103189706 | 27010 | -0.34 | 0.0418 | 0.0688 |
| chr1 | 178251761 | 178253885 | 2125 | 0.59 | 0.0419 | 0.0688 |
| chr21 | 35464821 | 35465305 | 485 | 1.48 | 0.042 | 0.0691 |
| chr2 | 32454724 | 32455466 | 743 | -1.61 | 0.0421 | 0.0691 |
| chr17 | 30114781 | 30115838 | 1058 | -0.95 | 0.0421 | 0.0691 |
| chr14 | 36769352 | 36770060 | 709 | 1.21 | 0.0421 | 0.0691 |
| chr5 | 115074162 | 115085229 | 11068 | 3.44 | 0.0421 | 0.0692 |
| chr10 | 89452621 | 89472600 | 19980 | 0.26 | 0.0421 | 0.0692 |
| chr1 | 36087640 | 36088103 | 464 | -1.84 | 0.0421 | 0.0692 |
| chr2 | 182740696 | 182742643 | 1948 | 0.39 | 0.0422 | 0.0693 |
| chrX | 107333110 | 107335835 | 2726 | 0.32 | 0.0422 | 0.0694 |
| chr9 | 125007817 | 125008673 | 857 | -1.58 | 0.0422 | 0.0694 |
| chr2 | 3508993 | 3524055 | 15063 | 0.29 | 0.0423 | 0.0694 |
| chr10 | 123855671 | 124029255 | 173585 | 0.08 | 0.0423 | 0.0694 |
| chr5 | 35617002 | 35619251 | 2250 | -0.36 | 0.0423 | 0.0695 |
| chr22 | 50219915 | 50228297 | 8383 | 0.56 | 0.0423 | 0.0695 |
| chr10 | 88514510 | 88519268 | 4759 | 0.24 | 0.0424 | 0.0695 |
| chr7 | 110562689 | 110563812 | 1124 | 0.81 | 0.0424 | 0.0696 |
| chr19 | 47128522 | 47137502 | 8981 | -0.67 | 0.0424 | 0.0696 |
| chr12 | 27862398 | 27870105 | 7708 | 0.25 | 0.0424 | 0.0696 |
| chr2 | 182450072 | 182452132 | 2061 | 0.51 | 0.0424 | 0.0696 |
| chr17 | 64072100 | 64072868 | 769 | -1.04 | 0.0424 | 0.0696 |
| chr12 | 113772183 | 113773570 | 1388 | 0.73 | 0.0424 | 0.0696 |
| chr9 | 119854368 | 119855868 | 1501 | -0.55 | 0.0424 | 0.0696 |
| chr2 | 54732400 | 54732877 | 478 | -1.68 | 0.0424 | 0.0697 |
| chrX | 122764584 | 122765917 | 1334 | -0.82 | 0.0425 | 0.0697 |
| chr12 | 14075368 | 14075853 | 486 | -1.99 | 0.0425 | 0.0697 |
| chr2 | 152623823 | 152624363 | 541 | -1.34 | 0.0425 | 0.0697 |
| chr1 | 33709337 | 33723923 | 14587 | 0.41 | 0.0426 | 0.0699 |
| chr3 | 133362594 | 133366062 | 3469 | 0.46 | 0.0426 | 0.0699 |
| chr1 | 78353570 | 78355849 | 2280 | -0.57 | 0.0426 | 0.0699 |
| chr9 | 96319308 | 96352843 | 33536 | -0.23 | 0.0426 | 0.07 |
| chr7 | 105435182 | 105447743 | 12562 | -0.54 | 0.0427 | 0.0701 |
| chr22 | 40811476 | 40945632 | 134157 | -0.09 | 0.0427 | 0.0701 |
| chr8 | 81142388 | 81143640 | 1253 | -0.65 | 0.0428 | 0.0701 |
| chr3 | 122171706 | 122188808 | 17103 | -0.3 | 0.0428 | 0.0702 |
| chr5 | 6868909 | 6869350 | 442 | 1.61 | 0.0428 | 0.0702 |
| chr9 | 38119178 | 38120652 | 1475 | 3.1 | 0.0428 | 0.0702 |
| chr15 | 34148393 | 34148742 | 350 | -1.5 | 0.0428 | 0.0702 |
| chr17 | 19350166 | 19351157 | 992 | -0.85 | 0.0429 | 0.0703 |
| chr14 | 27460500 | 27463044 | 2545 | 2.21 | 0.0429 | 0.0703 |
| chr9 | 129155412 | 129156290 | 879 | -1.21 | 0.0429 | 0.0703 |
| chr8 | 122020219 | 122020768 | 550 | 1.2 | 0.0429 | 0.0703 |
| chr5 | 146889042 | 146889568 | 527 | -1.4 | 0.0429 | 0.0703 |
| chr12 | 109659336 | 109663659 | 4324 | -0.53 | 0.0429 | 0.0703 |
| chr17 | 54901544 | 54912487 | 10944 | 0.37 | 0.0429 | 0.0703 |
| chr5 | 5135659 | 5146868 | 11210 | 0.61 | 0.0429 | 0.0704 |
| chr3 | 127374162 | 127375174 | 1013 | -1.26 | 0.043 | 0.0704 |
| chr11 | 94883774 | 94885300 | 1527 | 0.62 | 0.043 | 0.0705 |
| chr11 | 46938356 | 46970771 | 32416 | 0.15 | 0.043 | 0.0705 |
| chr4 | 23244228 | 23245170 | 943 | 1.04 | 0.043 | 0.0705 |
| chr4 | 40614590 | 40634535 | 19946 | 0.16 | 0.043 | 0.0705 |
| chr10 | 73453133 | 73454005 | 873 | 1.2 | 0.043 | 0.0705 |
| chr15 | 99707304 | 99718795 | 11492 | -0.35 | 0.0431 | 0.0705 |
| chr13 | 60439136 | 60439596 | 461 | 1.83 | 0.0431 | 0.0705 |
| chr1 | 23120899 | 23121733 | 835 | -1.3 | 0.0431 | 0.0706 |

| | | | | | | |
|---|---|---|---|---|---|---|
| chr15 | 56331961 | 56345541 | 13581 | 0.3 | 0.0431 | 0.0707 |
| chr6 | 91227455 | 91228638 | 1184 | 1.14 | 0.0431 | 0.0707 |
| chr12 | 50866387 | 50867096 | 710 | 1.3 | 0.0432 | 0.0707 |
| chr8 | 42127282 | 42196672 | 69391 | -0.13 | 0.0432 | 0.0707 |
| chr8 | 40500425 | 40501879 | 1455 | -0.8 | 0.0432 | 0.0707 |
| chr1 | 22107856 | 22111080 | 3225 | 0.33 | 0.0432 | 0.0707 |
| chr11 | 92027649 | 92040975 | 13327 | 0.38 | 0.0432 | 0.0707 |
| chr2 | 43859968 | 43865371 | 5404 | -0.3 | 0.0432 | 0.0708 |
| chr3 | 138633625 | 138641314 | 7690 | 0.43 | 0.0432 | 0.0708 |
| chr4 | 185609003 | 185610454 | 1452 | 0.68 | 0.0433 | 0.0708 |
| chr14 | 101971508 | 101985769 | 14262 | -1.95 | 0.0433 | 0.0709 |
| chr16 | 89698421 | 89700223 | 1803 | -0.56 | 0.0433 | 0.0709 |
| chr20 | 35305198 | 35306433 | 1236 | -1.18 | 0.0433 | 0.0709 |
| chr11 | 129899061 | 129899474 | 414 | 1.49 | 0.0434 | 0.071 |
| chr12 | 117112254 | 117113273 | 1020 | -0.81 | 0.0434 | 0.071 |
| chr9 | 98783463 | 98804006 | 20544 | -0.35 | 0.0434 | 0.071 |
| chr20 | 47644846 | 47647891 | 3046 | -0.86 | 0.0434 | 0.071 |
| chr22 | 42608157 | 42608910 | 754 | -1.25 | 0.0434 | 0.071 |
| chr14 | 72301441 | 72301775 | 335 | 2.16 | 0.0434 | 0.0711 |
| chr20 | 13702279 | 13703527 | 1249 | -0.91 | 0.0434 | 0.0711 |
| chr12 | 56152647 | 56153318 | 672 | 1.33 | 0.0434 | 0.0711 |
| chr12 | 105477127 | 105478288 | 1162 | -0.72 | 0.0434 | 0.0711 |
| chr6 | 43784514 | 43785237 | 724 | -1.12 | 0.0435 | 0.0711 |
| chr6 | 36934866 | 36939920 | 5055 | -0.6 | 0.0435 | 0.0712 |
| chr4 | 8303698 | 8318516 | 14819 | -0.31 | 0.0435 | 0.0712 |
| chr1 | 163259731 | 163260227 | 497 | 1.33 | 0.0436 | 0.0713 |
| chr8 | 48920056 | 48947052 | 26997 | 0.21 | 0.0436 | 0.0713 |
| chr22 | 32224645 | 32226541 | 1897 | -0.59 | 0.0436 | 0.0713 |
| chr22 | 46298860 | 46375306 | 76447 | -0.12 | 0.0436 | 0.0713 |
| chr3 | 156391077 | 156440400 | 49324 | 0.11 | 0.0436 | 0.0714 |
| chr12 | 91890080 | 91890472 | 393 | 1.76 | 0.0436 | 0.0714 |
| chr5 | 61690532 | 61719850 | 29319 | -0.16 | 0.0437 | 0.0714 |
| chr7 | 111024382 | 111025441 | 1060 | 0.81 | 0.0437 | 0.0714 |
| chr16 | 12518506 | 12520363 | 1858 | -0.57 | 0.0437 | 0.0715 |
| chr2 | 740889 | 741366 | 478 | 1.01 | 0.0437 | 0.0715 |
| chr1 | 55938375 | 55942555 | 4181 | -0.64 | 0.0437 | 0.0715 |
| chr14 | 38260608 | 38261080 | 473 | 1.54 | 0.0438 | 0.0716 |
| chr3 | 183966023 | 183968198 | 2176 | -0.4 | 0.0438 | 0.0716 |
| chr15 | 73856484 | 73864319 | 7836 | -0.5 | 0.0438 | 0.0716 |
| chr17 | 1655421 | 1656823 | 1403 | 0.39 | 0.0438 | 0.0716 |
| chr1 | 162106359 | 162107573 | 1215 | -0.81 | 0.0438 | 0.0716 |
| chr21 | 40211050 | 40212422 | 1373 | -0.88 | 0.0438 | 0.0717 |
| chr6 | 14074865 | 14089621 | 14757 | 3.47 | 0.0439 | 0.0717 |
| chr9 | 17005265 | 17005704 | 440 | 1.98 | 0.0439 | 0.0717 |
| chr1 | 94488078 | 94488601 | 524 | -1.38 | 0.0439 | 0.0718 |
| chr10 | 46046053 | 46047195 | 1143 | 0.82 | 0.0439 | 0.0718 |
| chr6 | 80837930 | 80838547 | 618 | 0.95 | 0.044 | 0.0718 |
| chr6 | 81319875 | 81333420 | 13546 | 3.49 | 0.044 | 0.0718 |
| chr1 | 87258468 | 87259344 | 877 | 0.76 | 0.044 | 0.0719 |
| chr8 | 28636293 | 28636996 | 704 | -1.48 | 0.044 | 0.0719 |
| chr5 | 55146627 | 55147888 | 1262 | -0.74 | 0.044 | 0.0719 |
| chr10 | 105190765 | 105212542 | 21778 | -0.35 | 0.044 | 0.072 |
| chr3 | 66270753 | 66272214 | 1462 | 0.55 | 0.0441 | 0.072 |
| chr16 | 12632448 | 12632929 | 482 | -1.64 | 0.0441 | 0.072 |
| chr20 | 34403243 | 34420995 | 17753 | -0.37 | 0.0441 | 0.072 |
| chr4 | 95541437 | 95546959 | 5523 | 0.33 | 0.0441 | 0.072 |
| chr1 | 156560469 | 156586692 | 26224 | -0.21 | 0.0441 | 0.0721 |
| chr9 | 82104118 | 82105177 | 1060 | -0.85 | 0.0441 | 0.0721 |
| chr11 | 7533921 | 7535733 | 1813 | 0.74 | 0.0442 | 0.0722 |
| chr6 | 116449009 | 116449998 | 990 | 1.28 | 0.0442 | 0.0722 |
| chr2 | 28886380 | 28887343 | 964 | -1.24 | 0.0442 | 0.0722 |

| | | | | | | |
|---|---|---|---|---|---|---|
| chr9 | 93340142 | 93340579 | 438 | 1.71 | 0.0442 | 0.0722 |
| chrX | 47862454 | 47863960 | 1507 | 0.49 | 0.0442 | 0.0722 |
| chr3 | 122081253 | 122103645 | 22393 | -0.22 | 0.0442 | 0.0722 |
| chr2 | 151476186 | 151478169 | 1984 | 0.73 | 0.0443 | 0.0722 |
| chr16 | 63099743 | 63101333 | 1591 | 0.68 | 0.0443 | 0.0723 |
| chr8 | 124682924 | 124683911 | 988 | -0.94 | 0.0443 | 0.0724 |
| chrX | 40966934 | 40981468 | 14535 | -0.37 | 0.0443 | 0.0724 |
| chrX | 150102377 | 150103527 | 1151 | 0.96 | 0.0443 | 0.0724 |
| chr2 | 47567123 | 47596928 | 29806 | 0.16 | 0.0445 | 0.0726 |
| chr20 | 59828159 | 59828789 | 631 | -1.24 | 0.0445 | 0.0726 |
| chr8 | 87486413 | 87488590 | 2178 | 0.74 | 0.0445 | 0.0726 |
| chr5 | 131971120 | 131972167 | 1048 | -0.97 | 0.0445 | 0.0726 |
| chr20 | 5170045 | 5177653 | 7609 | -0.59 | 0.0445 | 0.0726 |
| chr14 | 67652754 | 67654558 | 1805 | -0.61 | 0.0445 | 0.0726 |
| chr12 | 12911303 | 12912616 | 1314 | 0.81 | 0.0445 | 0.0726 |
| chrX | 40781157 | 40805684 | 24528 | 0.27 | 0.0445 | 0.0726 |
| chr2 | 10654203 | 10655698 | 1496 | 2.71 | 0.0445 | 0.0726 |
| chr7 | 82759601 | 82760758 | 1158 | 0.98 | 0.0445 | 0.0726 |
| chr17 | 17175860 | 17185861 | 10002 | -0.24 | 0.0446 | 0.0727 |
| chr20 | 32011042 | 32011541 | 500 | -1.41 | 0.0446 | 0.0727 |
| chr13 | 41494638 | 41496398 | 1761 | 0.47 | 0.0446 | 0.0727 |
| chr16 | 30639415 | 30647049 | 7635 | 0.27 | 0.0446 | 0.0728 |
| chrX | 68329164 | 68330470 | 1307 | -0.71 | 0.0446 | 0.0728 |
| chrX | 41637341 | 41639107 | 1767 | 0.5 | 0.0447 | 0.0728 |
| chr5 | 176211373 | 176212320 | 948 | -0.93 | 0.0447 | 0.0728 |
| chr17 | 74067299 | 74100765 | 33467 | -0.25 | 0.0447 | 0.0729 |
| chr8 | 41385960 | 41387857 | 1898 | -0.45 | 0.0447 | 0.073 |
| chr17 | 53006004 | 53009437 | 3434 | 0.79 | 0.0448 | 0.073 |
| chr18 | 8981888 | 8983515 | 1628 | 0.8 | 0.0448 | 0.073 |
| chr20 | 34244958 | 34253705 | 8748 | 0.19 | 0.0448 | 0.0731 |
| chr1 | 97249813 | 97250648 | 836 | 1.4 | 0.0449 | 0.0731 |
| chr11 | 84880483 | 84883136 | 2654 | 0.48 | 0.0449 | 0.0731 |
| chrX | 49124849 | 49127462 | 2614 | 0.31 | 0.0449 | 0.0731 |
| chr13 | 46056384 | 46057511 | 1128 | -1.05 | 0.0449 | 0.0732 |
| chr7 | 149551561 | 149552119 | 559 | 1.72 | 0.0449 | 0.0733 |
| chr12 | 86260537 | 86261026 | 490 | 1.47 | 0.0449 | 0.0733 |
| chrX | 55168222 | 55171190 | 2969 | 0.48 | 0.045 | 0.0733 |
| chr14 | 63156289 | 63160310 | 4022 | 0.19 | 0.045 | 0.0733 |
| chr4 | 39010458 | 39011120 | 663 | 1.45 | 0.045 | 0.0733 |
| chr8 | 108110968 | 108132035 | 21068 | -0.17 | 0.045 | 0.0733 |
| chr8 | 74473019 | 74474007 | 989 | -0.89 | 0.045 | 0.0734 |
| chr1 | 20878432 | 20879436 | 1005 | -1.13 | 0.045 | 0.0734 |
| chr1 | 112161894 | 112163909 | 2016 | 0.57 | 0.045 | 0.0734 |
| chr9 | 33021093 | 33053344 | 32252 | 0.16 | 0.0451 | 0.0734 |
| chr21 | 27105966 | 27109499 | 3534 | 0.26 | 0.0451 | 0.0734 |
| chr8 | 125152208 | 125160963 | 8756 | -2.61 | 0.0451 | 0.0734 |
| chr16 | 57017225 | 57023941 | 6717 | 0.4 | 0.0451 | 0.0734 |
| chr1 | 160331530 | 160332753 | 1224 | -1.18 | 0.0451 | 0.0735 |
| chr7 | 139761751 | 139763431 | 1681 | 0.62 | 0.0451 | 0.0735 |
| chr19 | 41075964 | 41122155 | 46192 | 0.12 | 0.0451 | 0.0735 |
| chr2 | 24491601 | 24493077 | 1477 | -0.54 | 0.0452 | 0.0735 |
| chr15 | 68334032 | 68347943 | 13912 | 0.3 | 0.0452 | 0.0735 |
| chr8 | 11593443 | 11594033 | 591 | 0.83 | 0.0452 | 0.0736 |
| chr7 | 158527715 | 158535527 | 7813 | 0.31 | 0.0452 | 0.0736 |
| chr22 | 33757618 | 33758572 | 955 | -1.09 | 0.0452 | 0.0736 |
| chr5 | 148699395 | 148707057 | 7663 | 0.25 | 0.0452 | 0.0736 |
| chr4 | 73840342 | 73841707 | 1366 | 0.48 | 0.0453 | 0.0737 |
| chr8 | 23539514 | 23540701 | 1188 | 0.84 | 0.0453 | 0.0737 |
| chr5 | 72511370 | 72512569 | 1200 | -0.81 | 0.0453 | 0.0737 |
| chr14 | 54801819 | 54802887 | 1069 | -0.81 | 0.0453 | 0.0737 |
| chr1 | 7830701 | 7844979 | 14279 | 0.38 | 0.0453 | 0.0738 |

| | | | | | | |
|---|---|---|---|---|---|---|
| chr18 | 31733677 | 31734431 | 755 | 1.42 | 0.0454 | 0.0738 |
| chr12 | 49982042 | 49983101 | 1060 | -1.19 | 0.0454 | 0.0738 |
| chr7 | 83243048 | 83244408 | 1361 | 1.19 | 0.0454 | 0.0739 |
| chr5 | 13591495 | 13591909 | 415 | 1.23 | 0.0454 | 0.0739 |
| chr7 | 33063902 | 33103541 | 39640 | 0.19 | 0.0454 | 0.0739 |
| chr11 | 16759078 | 16766802 | 7725 | 0.2 | 0.0455 | 0.074 |
| chr13 | 109485815 | 109486902 | 1088 | 1.62 | 0.0455 | 0.0741 |
| chr7 | 99774542 | 99774953 | 412 | 1.92 | 0.0456 | 0.0741 |
| chr6 | 151676814 | 151678761 | 1948 | -0.68 | 0.0456 | 0.0741 |
| chr6 | 158588446 | 158590272 | 1827 | 0.56 | 0.0456 | 0.0741 |
| chr7 | 102035640 | 102081716 | 46077 | 0.16 | 0.0456 | 0.0742 |
| chr2 | 36875998 | 36877964 | 1967 | -0.44 | 0.0456 | 0.0742 |
| chr18 | 42305091 | 42323307 | 18217 | 1.27 | 0.0457 | 0.0742 |
| chr2 | 165923962 | 165925349 | 1388 | -0.48 | 0.0457 | 0.0742 |
| chrX | 149335749 | 149336802 | 1054 | 0.92 | 0.0457 | 0.0743 |
| chr1 | 230402055 | 230404815 | 2761 | -0.84 | 0.0457 | 0.0743 |
| chr9 | 6433086 | 6433885 | 800 | -1.33 | 0.0458 | 0.0744 |
| chr11 | 36508755 | 36514937 | 6183 | 0.76 | 0.0458 | 0.0745 |
| chrX | 47880141 | 47881495 | 1355 | 0.35 | 0.0458 | 0.0745 |
| chr7 | 84152641 | 84168528 | 15888 | -0.14 | 0.0458 | 0.0745 |
| chr6 | 89672588 | 89674231 | 1644 | 0.51 | 0.0458 | 0.0745 |
| chr7 | 105967429 | 105968955 | 1527 | -3.14 | 0.0458 | 0.0745 |
| chr11 | 94371515 | 94372161 | 647 | 1.51 | 0.0458 | 0.0745 |
| chr2 | 102813827 | 102814442 | 616 | -1.36 | 0.0459 | 0.0746 |
| chr18 | 32620019 | 32630153 | 10135 | -0.35 | 0.0459 | 0.0746 |
| chr11 | 82866431 | 82906578 | 40148 | 0.15 | 0.0459 | 0.0746 |
| chr11 | 119460767 | 119462216 | 1450 | 3.01 | 0.0459 | 0.0746 |
| chr1 | 10092025 | 10094621 | 2597 | 0.35 | 0.0459 | 0.0746 |
| chr5 | 96211477 | 96216813 | 5337 | 0.29 | 0.0459 | 0.0746 |
| chr5 | 124819056 | 124823521 | 4466 | 0.19 | 0.0459 | 0.0746 |
| chr3 | 136467105 | 136477445 | 10341 | 0.18 | 0.0459 | 0.0746 |
| chr13 | 27936057 | 27936701 | 645 | -1.36 | 0.0459 | 0.0746 |
| chr10 | 128961472 | 128962707 | 1236 | 0.93 | 0.046 | 0.0748 |
| chr11 | 126272532 | 126273515 | 984 | 1.07 | 0.046 | 0.0748 |
| chr13 | 27946702 | 27947131 | 430 | 1.78 | 0.046 | 0.0748 |
| chr9 | 35161387 | 35162936 | 1550 | 0.44 | 0.0461 | 0.0748 |
| chr8 | 1875334 | 1876163 | 830 | -1.31 | 0.0461 | 0.0749 |
| chr5 | 7850840 | 7890580 | 39741 | 0.18 | 0.0461 | 0.0749 |
| chr2 | 151660125 | 151660594 | 470 | -1.47 | 0.0461 | 0.0749 |
| chr11 | 129985329 | 129985701 | 373 | 1.4 | 0.0461 | 0.0749 |
| chr2 | 217276149 | 217278105 | 1957 | -0.4 | 0.0462 | 0.0749 |
| chr14 | 75955254 | 75956246 | 993 | -0.69 | 0.0462 | 0.0749 |
| chr5 | 77390502 | 77391666 | 1165 | 1.1 | 0.0462 | 0.075 |
| chr8 | 134124155 | 134124916 | 762 | -1.19 | 0.0462 | 0.075 |
| chr5 | 94236820 | 94237787 | 968 | 1.27 | 0.0462 | 0.075 |
| chr19 | 32933713 | 32950908 | 17196 | -0.43 | 0.0462 | 0.075 |
| chr10 | 70938571 | 70941441 | 2871 | 0.23 | 0.0463 | 0.0752 |
| chr22 | 37663052 | 37664679 | 1628 | -0.53 | 0.0463 | 0.0752 |
| chr14 | 27896213 | 27897921 | 1709 | 3.27 | 0.0464 | 0.0753 |
| chr19 | 49710485 | 49711138 | 654 | 1.42 | 0.0464 | 0.0753 |
| chr7 | 81252360 | 81253033 | 674 | -1.34 | 0.0464 | 0.0753 |
| chr8 | 38299695 | 38308392 | 8698 | -0.4 | 0.0465 | 0.0754 |
| chr22 | 18631843 | 18633759 | 1917 | 0.43 | 0.0465 | 0.0754 |
| chr9 | 131709352 | 131711309 | 1958 | 0.46 | 0.0465 | 0.0755 |
| chr7 | 65596176 | 65636440 | 40265 | 0.24 | 0.0465 | 0.0755 |
| chr2 | 242556239 | 242557740 | 1502 | -0.96 | 0.0466 | 0.0755 |
| chr10 | 64528960 | 64530172 | 1213 | -0.67 | 0.0466 | 0.0756 |
| chr1 | 23206341 | 23206976 | 636 | -1.32 | 0.0466 | 0.0756 |
| chr2 | 61334015 | 61334853 | 839 | -1.47 | 0.0466 | 0.0756 |
| chr5 | 141851705 | 141852834 | 1130 | -1.08 | 0.0466 | 0.0756 |
| chr8 | 54643573 | 54644413 | 841 | 1.03 | 0.0466 | 0.0756 |

| | | | | | | |
|---|---|---|---|---|---|---|
| chr16 | 50776003 | 50776641 | 639 | 0.91 | 0.0466 | 0.0756 |
| chr4 | 152205350 | 152206407 | 1058 | 1.13 | 0.0467 | 0.0757 |
| chr15 | 57598318 | 57599548 | 1231 | -0.84 | 0.0467 | 0.0757 |
| chr5 | 142519507 | 142520869 | 1363 | 0.98 | 0.0468 | 0.0758 |
| chr11 | 102979461 | 102981861 | 2401 | -0.31 | 0.0468 | 0.0758 |
| chr2 | 109172019 | 109172376 | 358 | -2 | 0.0468 | 0.0758 |
| chr17 | 48074167 | 48076342 | 2176 | -0.8 | 0.0468 | 0.0758 |
| chr12 | 52712597 | 52740213 | 27617 | 0.42 | 0.0468 | 0.0759 |
| chr4 | 84030209 | 84033162 | 2954 | -0.32 | 0.0468 | 0.0759 |
| chr20 | 37257136 | 37258194 | 1059 | 0.85 | 0.0468 | 0.0759 |
| chrX | 24745844 | 24746248 | 405 | -1.52 | 0.0469 | 0.076 |
| chr14 | 45601657 | 45605825 | 4169 | 0.28 | 0.0469 | 0.076 |
| chr13 | 80123046 | 80123816 | 771 | 1.36 | 0.0469 | 0.076 |
| chr6 | 90458269 | 90458955 | 687 | -1.32 | 0.0469 | 0.076 |
| chr1 | 41702874 | 41708759 | 5886 | 0.3 | 0.047 | 0.0761 |
| chr10 | 76344863 | 76350861 | 5999 | 3.91 | 0.047 | 0.0761 |
| chr20 | 54996676 | 54998209 | 1534 | 1.07 | 0.047 | 0.0762 |
| chr12 | 86531425 | 86531960 | 536 | 1.05 | 0.047 | 0.0762 |
| chr5 | 130989413 | 131014461 | 25049 | -0.22 | 0.047 | 0.0762 |
| chr13 | 35032954 | 35044458 | 11505 | 0.53 | 0.047 | 0.0762 |
| chr15 | 102180822 | 102193824 | 13003 | 0.2 | 0.047 | 0.0762 |
| chr17 | 26585080 | 26585727 | 648 | 1.5 | 0.0471 | 0.0762 |
| chr20 | 14773153 | 14773747 | 595 | 1.1 | 0.0471 | 0.0763 |
| chr21 | 16235401 | 16261493 | 26093 | 0.24 | 0.0471 | 0.0763 |
| chr19 | 6827814 | 6828900 | 1087 | 0.75 | 0.0471 | 0.0763 |
| chr6 | 1390026 | 1391160 | 1135 | -1.27 | 0.0471 | 0.0763 |
| chr6 | 2429395 | 2430420 | 1026 | 1.03 | 0.0471 | 0.0764 |
| chr9 | 4000398 | 4001000 | 603 | 2.06 | 0.0472 | 0.0764 |
| chr5 | 158281397 | 158284108 | 2712 | -0.43 | 0.0472 | 0.0764 |
| chr5 | 68423991 | 68426772 | 2782 | -0.6 | 0.0472 | 0.0764 |
| chr17 | 40260119 | 40307916 | 47798 | 0.12 | 0.0472 | 0.0764 |
| chr15 | 52384662 | 52396621 | 11960 | 0.19 | 0.0472 | 0.0764 |
| chr20 | 1808650 | 1809718 | 1069 | 1.29 | 0.0472 | 0.0765 |
| chr4 | 125311823 | 125312375 | 553 | 1.37 | 0.0472 | 0.0765 |
| chr12 | 62690553 | 62691444 | 892 | -1.51 | 0.0472 | 0.0765 |
| chr17 | 38863948 | 38864720 | 773 | -0.95 | 0.0473 | 0.0765 |
| chr16 | 73204236 | 73204740 | 505 | -1.62 | 0.0473 | 0.0765 |
| chr18 | 47002463 | 47024291 | 21829 | 0.19 | 0.0473 | 0.0765 |
| chr16 | 68918382 | 68919585 | 1204 | 1.07 | 0.0473 | 0.0765 |
| chr7 | 21466189 | 21468764 | 2576 | 0.3 | 0.0473 | 0.0766 |
| chr15 | 34874559 | 34881198 | 6640 | 0.52 | 0.0473 | 0.0766 |
| chr15 | 70289086 | 70291501 | 2416 | 3.51 | 0.0474 | 0.0766 |
| chr15 | 90455243 | 90456825 | 1583 | -0.46 | 0.0474 | 0.0767 |
| chr17 | 37271387 | 37271890 | 504 | -1.94 | 0.0474 | 0.0767 |
| chr13 | 67077718 | 67078990 | 1273 | 0.54 | 0.0474 | 0.0767 |
| chr19 | 48947849 | 49017836 | 69988 | -0.16 | 0.0476 | 0.077 |
| chr9 | 91564908 | 91566454 | 1547 | -0.65 | 0.0477 | 0.0772 |
| chrX | 135055109 | 135068393 | 13285 | 0.54 | 0.0477 | 0.0772 |
| chr16 | 58283295 | 58284569 | 1275 | 0.69 | 0.0477 | 0.0772 |
| chr3 | 128278119 | 128278811 | 693 | 1.17 | 0.0478 | 0.0772 |
| chr5 | 79968910 | 79969425 | 516 | -1.33 | 0.0478 | 0.0774 |
| chr3 | 156270856 | 156274082 | 3227 | 0.3 | 0.0478 | 0.0774 |
| chr20 | 3960258 | 3962328 | 2071 | 0.3 | 0.0478 | 0.0774 |
| chr22 | 24108902 | 24110716 | 1815 | -0.62 | 0.0479 | 0.0774 |
| chr18 | 59924558 | 59925553 | 996 | 0.96 | 0.0479 | 0.0774 |
| chr1 | 205113552 | 205116969 | 3418 | -0.73 | 0.0479 | 0.0774 |
| chr2 | 172378007 | 172380987 | 2981 | -0.35 | 0.0479 | 0.0774 |
| chr5 | 1799013 | 1809069 | 10057 | 0.29 | 0.0479 | 0.0774 |
| chr10 | 64939395 | 64951265 | 11871 | 0.43 | 0.0479 | 0.0774 |
| chr1 | 231406713 | 231408765 | 2053 | 0.88 | 0.0479 | 0.0775 |
| chr2 | 181971742 | 181972146 | 405 | -1.12 | 0.0479 | 0.0775 |

| | | | | | | |
|---|---|---|---|---|---|---|
| chr12 | 74190916 | 74192886 | 1971 | 0.65 | 0.0479 | 0.0775 |
| chr4 | 84771529 | 84777722 | 6194 | 0.22 | 0.048 | 0.0775 |
| chr8 | 49726719 | 49728283 | 1565 | 3.03 | 0.048 | 0.0776 |
| chr15 | 52088560 | 52089938 | 1379 | 0.74 | 0.048 | 0.0776 |
| chr2 | 143353944 | 143358584 | 4641 | 0.53 | 0.048 | 0.0776 |
| chr4 | 69597900 | 69598527 | 628 | -2.93 | 0.048 | 0.0776 |
| chr12 | 11992124 | 11992993 | 870 | 1.31 | 0.048 | 0.0776 |
| chr1 | 41831834 | 41832699 | 866 | 1.35 | 0.048 | 0.0776 |
| chr12 | 16084205 | 16085570 | 1366 | -0.63 | 0.048 | 0.0776 |
| chrX | 19533091 | 19534305 | 1215 | -0.85 | 0.0481 | 0.0777 |
| chr17 | 61258646 | 61260019 | 1374 | -3.08 | 0.0481 | 0.0777 |
| chr3 | 169713757 | 169714634 | 878 | -1.02 | 0.0481 | 0.0777 |
| chr6 | 53675556 | 53676665 | 1110 | 1.07 | 0.0481 | 0.0777 |
| chr5 | 108325486 | 108326588 | 1103 | 1.01 | 0.0481 | 0.0778 |
| chr1 | 212574737 | 212588861 | 14125 | -0.46 | 0.0481 | 0.0778 |
| chr14 | 71838951 | 71839757 | 807 | 1.07 | 0.0481 | 0.0778 |
| chr2 | 177052736 | 177053851 | 1116 | 1.03 | 0.0482 | 0.0778 |
| chr7 | 69197484 | 69207536 | 10053 | 3.81 | 0.0482 | 0.0779 |
| chr20 | 22443058 | 22452580 | 9523 | -0.22 | 0.0483 | 0.0779 |
| chr2 | 146534390 | 146535755 | 1366 | 0.84 | 0.0483 | 0.078 |
| chr18 | 12038562 | 12039103 | 542 | -1.24 | 0.0483 | 0.078 |
| chr7 | 38921775 | 38922295 | 521 | -1.7 | 0.0483 | 0.078 |
| chr5 | 68338238 | 68343571 | 5334 | -0.37 | 0.0483 | 0.078 |
| chr1 | 244045181 | 244046332 | 1152 | -0.84 | 0.0484 | 0.0781 |
| chr19 | 4790667 | 4791184 | 518 | -1.17 | 0.0484 | 0.0781 |
| chr18 | 53478897 | 53480498 | 1602 | 2.77 | 0.0484 | 0.0781 |
| chr20 | 25602595 | 25606098 | 3504 | 0.19 | 0.0484 | 0.0781 |
| chr17 | 78074457 | 78076211 | 1755 | 0.47 | 0.0484 | 0.0781 |
| chr10 | 60726835 | 60727819 | 985 | -0.81 | 0.0485 | 0.0782 |
| chr2 | 85932455 | 85933049 | 595 | -1.3 | 0.0485 | 0.0783 |
| chr1 | 228510511 | 228510835 | 325 | 2.11 | 0.0485 | 0.0783 |
| chr19 | 38388558 | 38811081 | 422524 | 0.05 | 0.0485 | 0.0783 |
| chr14 | 34691144 | 34692099 | 956 | 2.95 | 0.0486 | 0.0784 |
| chr20 | 46414819 | 46416014 | 1196 | 1.04 | 0.0486 | 0.0785 |
| chr7 | 23333091 | 23340278 | 7188 | 0.28 | 0.0487 | 0.0785 |
| chr8 | 102036789 | 102076312 | 39524 | 0.11 | 0.0487 | 0.0786 |
| chr1 | 157014244 | 157016526 | 2283 | 0.34 | 0.0488 | 0.0787 |
| chr5 | 128429775 | 128431590 | 1816 | 0.6 | 0.0488 | 0.0788 |
| chr2 | 24432707 | 24442800 | 10094 | -0.56 | 0.0488 | 0.0788 |
| chr1 | 205087222 | 205092423 | 5202 | 0.22 | 0.0488 | 0.0788 |
| chr14 | 91943450 | 91978274 | 34825 | 0.14 | 0.0489 | 0.0788 |
| chr10 | 88825553 | 88827149 | 1597 | -0.76 | 0.0489 | 0.0788 |
| chr1 | 231913649 | 231918264 | 4616 | 0.64 | 0.0489 | 0.0788 |
| chr10 | 135148263 | 135149831 | 1569 | -0.7 | 0.0489 | 0.0788 |
| chr12 | 48477557 | 48482156 | 4600 | 0.8 | 0.0489 | 0.0788 |
| chr3 | 79000657 | 79000995 | 339 | 1.99 | 0.0489 | 0.0788 |
| chr1 | 6550152 | 6552449 | 2298 | 0.43 | 0.0489 | 0.0788 |
| chr4 | 170091679 | 170127492 | 35814 | 0.18 | 0.0489 | 0.0789 |
| chr13 | 50114195 | 50115324 | 1130 | -1.01 | 0.049 | 0.0789 |
| chr14 | 51229737 | 51230702 | 966 | 0.98 | 0.049 | 0.0789 |
| chr9 | 107703390 | 107704101 | 712 | -1.26 | 0.049 | 0.079 |
| chr11 | 129870894 | 129874727 | 3834 | 0.2 | 0.049 | 0.079 |
| chr9 | 75281847 | 75282835 | 989 | 1.23 | 0.049 | 0.079 |
| chr16 | 69287765 | 69288200 | 436 | 1.51 | 0.049 | 0.079 |
| chr8 | 99772392 | 99773058 | 667 | 1.21 | 0.049 | 0.079 |
| chr2 | 192352744 | 192353572 | 829 | -0.79 | 0.0491 | 0.0792 |
| chrX | 35882468 | 35882914 | 447 | 2.09 | 0.0492 | 0.0792 |
| chr20 | 8834488 | 8835382 | 895 | -1.19 | 0.0492 | 0.0792 |
| chr4 | 142141576 | 142143294 | 1719 | 0.48 | 0.0492 | 0.0793 |
| chr2 | 11294213 | 11296566 | 2354 | -0.28 | 0.0492 | 0.0793 |
| chr15 | 89730910 | 89732771 | 1862 | -0.93 | 0.0492 | 0.0793 |

| | | | | | | |
|---|---|---|---|---|---|---|
| chr12 | 91236350 | 91236793 | 444 | 1.4 | 0.0492 | 0.0793 |
| chr3 | 48342018 | 48344030 | 2013 | 0.33 | 0.0493 | 0.0793 |
| chr3 | 148537630 | 148539464 | 1835 | 3.1 | 0.0493 | 0.0795 |
| chr2 | 112730518 | 112733650 | 3133 | 0.6 | 0.0494 | 0.0795 |
| chr16 | 51589380 | 51603879 | 14500 | 0.32 | 0.0494 | 0.0796 |
| chr3 | 184934006 | 184934458 | 453 | 1.26 | 0.0494 | 0.0796 |
| chr5 | 121296609 | 121318890 | 22282 | 0.21 | 0.0494 | 0.0796 |
| chr10 | 134346553 | 134352012 | 5460 | 0.5 | 0.0494 | 0.0796 |
| chrX | 41765476 | 41766870 | 1395 | 2.98 | 0.0495 | 0.0798 |
| chr3 | 72987591 | 72988127 | 537 | -1.9 | 0.0496 | 0.0798 |
| chr15 | 92464176 | 92465324 | 1149 | 0.8 | 0.0496 | 0.0798 |
| chr20 | 42930552 | 42955821 | 25270 | 0.3 | 0.0496 | 0.0798 |
| chr5 | 73546637 | 73547296 | 660 | 1.38 | 0.0496 | 0.0798 |
| chr3 | 47820624 | 47846634 | 26011 | 0.18 | 0.0496 | 0.0798 |
| chr10 | 14548826 | 14659920 | 111095 | 0.1 | 0.0496 | 0.0799 |
| chr8 | 122466681 | 122467499 | 819 | -1.34 | 0.0496 | 0.0799 |
| chrX | 49643270 | 49649631 | 6362 | 0.59 | 0.0497 | 0.08 |
| chr9 | 130636748 | 130640576 | 3829 | -0.51 | 0.0497 | 0.08 |
| chr12 | 1121500 | 1123884 | 2385 | 0.39 | 0.0497 | 0.08 |
| chr2 | 239957201 | 239957517 | 317 | -2.06 | 0.0497 | 0.08 |
| chr10 | 135206958 | 135209163 | 2206 | 0.39 | 0.0497 | 0.08 |
| chr1 | 117229807 | 117231001 | 1195 | -1 | 0.0497 | 0.08 |
| chr1 | 8402946 | 8404524 | 1579 | 0.49 | 0.0497 | 0.08 |
| chr3 | 37261133 | 37299226 | 38094 | 0.15 | 0.0497 | 0.08 |
| chr18 | 19880590 | 19881455 | 866 | 1.27 | 0.0498 | 0.0801 |
| chr16 | 87982568 | 87990637 | 8070 | 0.21 | 0.0498 | 0.0801 |
| chr4 | 105314889 | 105315490 | 602 | 1.37 | 0.0498 | 0.0801 |
| chr12 | 53679586 | 53719813 | 40228 | 0.14 | 0.0498 | 0.0801 |
| chr10 | 12442328 | 12448605 | 6278 | -0.3 | 0.0498 | 0.0801 |
| chr12 | 68877202 | 68877956 | 755 | -1.18 | 0.0498 | 0.0801 |
| chr4 | 54948625 | 54949095 | 471 | 1.52 | 0.0498 | 0.0801 |
| chr21 | 38780055 | 38780744 | 690 | -1.13 | 0.0499 | 0.0802 |
| chr8 | 38686871 | 38688283 | 1413 | 0.82 | 0.0499 | 0.0803 |
| chr19 | 15560630 | 15561375 | 746 | -1.05 | 0.0499 | 0.0803 |
| chr6 | 36634326 | 36652205 | 17880 | -0.27 | 0.0499 | 0.0803 |
| chr12 | 13492828 | 13493525 | 698 | 1.21 | 0.0499 | 0.0803 |
| chr11 | 63770091 | 63779535 | 9445 | -0.32 | 0.05 | 0.0805 |
| chr17 | 42071935 | 42202217 | 130283 | -0.1 | 0.0501 | 0.0806 |
| chr13 | 35923730 | 35924425 | 696 | 1.16 | 0.0501 | 0.0806 |
| chr2 | 70093524 | 70095465 | 1942 | -2.85 | 0.0502 | 0.0807 |
| chr16 | 50185659 | 50202047 | 16389 | -0.34 | 0.0502 | 0.0807 |
| chr6 | 14558917 | 14567527 | 8611 | 0.29 | 0.0502 | 0.0807 |
| chr9 | 29044 | 29394 | 351 | -1.37 | 0.0502 | 0.0807 |
| chr8 | 17433856 | 17435368 | 1513 | -0.73 | 0.0502 | 0.0807 |
| chr5 | 143344235 | 143345301 | 1067 | -1.18 | 0.0502 | 0.0807 |
| chr2 | 176097987 | 176099071 | 1085 | 1.07 | 0.0502 | 0.0807 |
| chr12 | 100623637 | 100624162 | 526 | 1.39 | 0.0503 | 0.0808 |
| chr20 | 50416843 | 50418494 | 1652 | -0.49 | 0.0504 | 0.0809 |
| chr16 | 21271569 | 21275997 | 4429 | -0.31 | 0.0504 | 0.0809 |
| chr8 | 132868505 | 132871379 | 2875 | 0.36 | 0.0504 | 0.081 |
| chr9 | 72915089 | 72915631 | 543 | 1.26 | 0.0505 | 0.0811 |
| chr10 | 60174266 | 60175079 | 814 | -1 | 0.0505 | 0.0811 |
| chr11 | 57071542 | 57128963 | 57422 | 0.14 | 0.0505 | 0.0812 |
| chr3 | 56957259 | 56958776 | 1518 | 0.53 | 0.0505 | 0.0812 |
| chr9 | 737988 | 745139 | 7152 | -0.5 | 0.0506 | 0.0813 |
| chr3 | 17646144 | 17647214 | 1071 | -0.99 | 0.0506 | 0.0813 |
| chr10 | 35930232 | 35931174 | 943 | -1.12 | 0.0506 | 0.0813 |
| chr17 | 35382022 | 35383315 | 1294 | 1.04 | 0.0506 | 0.0813 |
| chr12 | 104570504 | 104575682 | 5179 | 0.41 | 0.0507 | 0.0813 |
| chr10 | 65224410 | 65236824 | 12415 | 0.28 | 0.0507 | 0.0813 |
| chr9 | 36640114 | 36666562 | 26449 | 0.25 | 0.0507 | 0.0814 |

| | | | | | | |
|---|---|---|---|---|---|---|
| chr11 | 73906767 | 73907415 | 649 | 1.03 | 0.0507 | 0.0814 |
| chr20 | 33145613 | 33180457 | 34845 | -0.26 | 0.0508 | 0.0815 |
| chr8 | 124933672 | 124935346 | 1675 | -0.44 | 0.0508 | 0.0815 |
| chr10 | 74798700 | 74812360 | 13661 | 0.31 | 0.0508 | 0.0815 |
| chr2 | 234912528 | 234913243 | 716 | -0.83 | 0.0508 | 0.0815 |
| chr8 | 64130602 | 64133833 | 3232 | 0.34 | 0.0508 | 0.0815 |
| chrX | 41729225 | 41749609 | 20385 | 0.71 | 0.0508 | 0.0816 |
| chr16 | 3730870 | 3768497 | 37628 | -0.18 | 0.0508 | 0.0816 |
| chr9 | 129226969 | 129227861 | 893 | -1.14 | 0.0508 | 0.0816 |
| chr2 | 37980180 | 38031196 | 51017 | 0.15 | 0.0509 | 0.0816 |
| chr9 | 80048357 | 80048973 | 617 | -1.29 | 0.0509 | 0.0817 |
| chr3 | 131800397 | 131805270 | 4874 | 0.24 | 0.0509 | 0.0817 |
| chr3 | 184561001 | 184562214 | 1214 | -0.74 | 0.0509 | 0.0817 |
| chr3 | 195793507 | 195794881 | 1375 | 1.39 | 0.051 | 0.0818 |
| chr18 | 55504889 | 55522234 | 17346 | -0.59 | 0.051 | 0.0818 |
| chr11 | 33364146 | 33364760 | 615 | -1.23 | 0.051 | 0.0818 |
| chr1 | 98569730 | 98570339 | 610 | 1.31 | 0.051 | 0.0818 |
| chr1 | 35246591 | 35247467 | 877 | 1.08 | 0.051 | 0.0818 |
| chr9 | 36368747 | 36371363 | 2617 | -1.26 | 0.051 | 0.0818 |
| chr1 | 104068321 | 104070858 | 2538 | 0.31 | 0.051 | 0.0819 |
| chr10 | 126043197 | 126043990 | 794 | -1.07 | 0.051 | 0.0819 |
| chr3 | 49966859 | 49988843 | 21985 | 0.34 | 0.0511 | 0.0819 |
| chr7 | 30452996 | 30453912 | 917 | 1.14 | 0.0512 | 0.082 |
| chr7 | 111078157 | 111079396 | 1240 | 0.9 | 0.0512 | 0.0821 |
| chr14 | 52083292 | 52084320 | 1029 | 0.84 | 0.0512 | 0.0821 |
| chr6 | 34807232 | 34808168 | 937 | -0.66 | 0.0513 | 0.0822 |
| chr17 | 41838048 | 41869734 | 31687 | 0.18 | 0.0513 | 0.0822 |
| chr16 | 67426457 | 67450921 | 24465 | -0.3 | 0.0513 | 0.0822 |
| chr22 | 18538468 | 18544858 | 6391 | 0.34 | 0.0513 | 0.0822 |
| chr5 | 102529925 | 102530369 | 445 | -1.63 | 0.0513 | 0.0822 |
| chr5 | 114597593 | 114599143 | 1551 | 0.3 | 0.0513 | 0.0822 |
| chr2 | 230309046 | 230320507 | 11462 | -0.28 | 0.0513 | 0.0823 |
| chr18 | 60051234 | 60067450 | 16217 | -0.77 | 0.0513 | 0.0823 |
| chr10 | 16586813 | 16587194 | 382 | 1.58 | 0.0513 | 0.0823 |
| chr15 | 71863076 | 71865845 | 2770 | -0.56 | 0.0514 | 0.0823 |
| chr8 | 91479338 | 91480654 | 1317 | 0.57 | 0.0514 | 0.0823 |
| chr9 | 103090423 | 103091213 | 791 | -1.27 | 0.0514 | 0.0823 |
| chr14 | 71488116 | 71488797 | 682 | 1.75 | 0.0514 | 0.0823 |
| chr3 | 23558810 | 23559746 | 937 | 1.26 | 0.0514 | 0.0823 |
| chr2 | 55360754 | 55369823 | 9070 | 0.41 | 0.0514 | 0.0823 |
| chr3 | 33073604 | 33075753 | 2150 | 0.38 | 0.0514 | 0.0824 |
| chr16 | 30194219 | 30199335 | 5117 | -0.56 | 0.0514 | 0.0824 |
| chr1 | 186606156 | 186606643 | 488 | 1.79 | 0.0515 | 0.0825 |
| chr6 | 41965011 | 41965607 | 597 | -1.22 | 0.0515 | 0.0825 |
| chr8 | 42623793 | 42624443 | 651 | -1.45 | 0.0515 | 0.0826 |
| chr4 | 14959551 | 14960964 | 1414 | 2.27 | 0.0516 | 0.0826 |
| chr10 | 119173011 | 119174658 | 1648 | 0.68 | 0.0516 | 0.0826 |
| chr12 | 46547653 | 46548177 | 525 | -1.26 | 0.0516 | 0.0826 |
| chr16 | 28138037 | 28224214 | 86178 | -0.16 | 0.0516 | 0.0826 |
| chr7 | 18706974 | 18707819 | 846 | 0.87 | 0.0516 | 0.0826 |
| chr3 | 101203375 | 101204353 | 979 | -1.12 | 0.0517 | 0.0828 |
| chr19 | 18107953 | 18133781 | 25829 | 0.16 | 0.0517 | 0.0828 |
| chr22 | 26948771 | 26949648 | 878 | -0.88 | 0.0517 | 0.0828 |
| chr1 | 233608054 | 233609213 | 1160 | 2.9 | 0.0518 | 0.0829 |
| chr21 | 27297949 | 27298716 | 768 | -1.16 | 0.0518 | 0.083 |
| chr16 | 67782570 | 67783406 | 837 | -0.97 | 0.0519 | 0.083 |
| chr22 | 25760256 | 25761081 | 826 | -1 | 0.0519 | 0.0831 |
| chr16 | 55055012 | 55055612 | 601 | 1.25 | 0.0519 | 0.0831 |
| chr12 | 25801071 | 25802045 | 975 | 3.04 | 0.0519 | 0.0831 |
| chr19 | 15783010 | 15784552 | 1543 | 0.65 | 0.0519 | 0.0831 |
| chr2 | 27433971 | 27451760 | 17790 | -0.18 | 0.0519 | 0.0831 |

| | | | | | | |
|---|---|---|---|---|---|---|
| chr19 | 17447018 | 17469903 | 22886 | 0.27 | 0.0519 | 0.0831 |
| chr1 | 23693933 | 23711399 | 17467 | -0.21 | 0.052 | 0.0831 |
| chr22 | 41346546 | 41349267 | 2722 | 0.38 | 0.052 | 0.0832 |
| chr22 | 30503111 | 30513279 | 10169 | 0.29 | 0.052 | 0.0832 |
| chr14 | 67260832 | 67261414 | 583 | 1.8 | 0.052 | 0.0832 |
| chrX | 16684591 | 16685082 | 492 | -1.21 | 0.052 | 0.0832 |
| chr7 | 158412076 | 158412557 | 482 | -1.35 | 0.052 | 0.0833 |
| chr19 | 52772060 | 52781783 | 9724 | 0.22 | 0.052 | 0.0833 |
| chr17 | 30507539 | 30534129 | 26591 | 0.18 | 0.052 | 0.0833 |
| chr8 | 81118627 | 81119916 | 1290 | 0.7 | 0.052 | 0.0833 |
| chrX | 17303787 | 17304923 | 1137 | -0.77 | 0.052 | 0.0833 |
| chr1 | 20799923 | 20821090 | 21168 | 0.16 | 0.0521 | 0.0833 |
| chr13 | 42293357 | 42295366 | 2010 | -0.56 | 0.0521 | 0.0833 |
| chr18 | 45820930 | 45827421 | 6492 | -1.06 | 0.0521 | 0.0833 |
| chr9 | 126979048 | 126984474 | 5427 | 0.49 | 0.0521 | 0.0833 |
| chr12 | 65188883 | 65191545 | 2663 | 0.31 | 0.0522 | 0.0835 |
| chr16 | 54481682 | 54482679 | 998 | -1.11 | 0.0522 | 0.0835 |
| chr6 | 109653775 | 109671577 | 17803 | 0.5 | 0.0522 | 0.0835 |
| chr14 | 58682338 | 58684663 | 2326 | -0.43 | 0.0523 | 0.0837 |
| chr1 | 35318590 | 35332076 | 13487 | 0.43 | 0.0524 | 0.0837 |
| chr8 | 124428005 | 124430121 | 2117 | 0.38 | 0.0524 | 0.0837 |
| chr9 | 88944580 | 88945234 | 655 | 1.16 | 0.0524 | 0.0838 |
| chr3 | 12688783 | 12721946 | 33164 | 0.2 | 0.0524 | 0.0838 |
| chr1 | 46908148 | 46908445 | 298 | 1.26 | 0.0525 | 0.0839 |
| chr12 | 26392302 | 26392799 | 498 | 1.61 | 0.0525 | 0.084 |
| chr4 | 102755818 | 102757266 | 1449 | 0.51 | 0.0525 | 0.084 |
| chr2 | 152230539 | 152232012 | 1474 | 0.55 | 0.0526 | 0.084 |
| chr22 | 26824571 | 26825006 | 436 | 1.52 | 0.0526 | 0.084 |
| chr5 | 71851556 | 71853455 | 1900 | -0.31 | 0.0526 | 0.084 |
| chr7 | 101278800 | 101279229 | 430 | -2 | 0.0528 | 0.0843 |
| chr14 | 60043023 | 60043530 | 508 | -0.96 | 0.0528 | 0.0844 |
| chr19 | 48610828 | 48674839 | 64012 | -0.15 | 0.0528 | 0.0844 |
| chr2 | 9123465 | 9124449 | 985 | -0.66 | 0.0528 | 0.0844 |
| chr7 | 70036563 | 70037026 | 464 | -1.15 | 0.0529 | 0.0844 |
| chr11 | 118229829 | 118231087 | 1259 | 0.6 | 0.0529 | 0.0845 |
| chr1 | 243687520 | 243689037 | 1518 | -0.51 | 0.0529 | 0.0845 |
| chr6 | 52281617 | 52289762 | 8146 | -0.26 | 0.0529 | 0.0845 |
| chr21 | 40749147 | 40760878 | 11732 | 0.22 | 0.0529 | 0.0846 |
| chr9 | 91258661 | 91273783 | 15123 | -3.42 | 0.0529 | 0.0846 |
| chr4 | 84971196 | 84972495 | 1300 | -0.9 | 0.053 | 0.0846 |
| chr8 | 1949118 | 1950500 | 1383 | -0.71 | 0.053 | 0.0847 |
| chr20 | 30842918 | 30845182 | 2265 | 0.48 | 0.0531 | 0.0847 |
| chr20 | 3075459 | 3122849 | 47391 | -0.17 | 0.0531 | 0.0848 |
| chr5 | 78948946 | 78950017 | 1072 | 1.34 | 0.0531 | 0.0848 |
| chr2 | 70436844 | 70438477 | 1634 | -0.81 | 0.0532 | 0.0849 |
| chr13 | 79199670 | 79200096 | 427 | -1.34 | 0.0532 | 0.0849 |
| chr10 | 30023933 | 30026422 | 2490 | 0.4 | 0.0532 | 0.0849 |
| chr7 | 30347342 | 30362249 | 14908 | 0.3 | 0.0532 | 0.0849 |
| chr3 | 42002184 | 42004672 | 2489 | -0.34 | 0.0532 | 0.0849 |
| chr5 | 32112595 | 32129116 | 16522 | -0.33 | 0.0533 | 0.085 |
| chr13 | 102101254 | 102109442 | 8189 | 0.21 | 0.0533 | 0.085 |
| chr8 | 6586623 | 6606665 | 20043 | 0.24 | 0.0533 | 0.0851 |
| chr10 | 129240093 | 129241031 | 939 | 1.05 | 0.0533 | 0.0851 |
| chr2 | 150468449 | 150469535 | 1087 | 1.2 | 0.0534 | 0.0852 |
| chr2 | 203973362 | 203988220 | 14859 | 0.42 | 0.0534 | 0.0852 |
| chr2 | 86789225 | 86791435 | 2211 | -0.24 | 0.0534 | 0.0852 |
| chr4 | 112432711 | 112434361 | 1651 | -0.42 | 0.0534 | 0.0852 |
| chr3 | 87010330 | 87011085 | 756 | -0.84 | 0.0534 | 0.0852 |
| chr7 | 130903909 | 130939507 | 35599 | 0.1 | 0.0535 | 0.0853 |
| chr5 | 177018904 | 177028038 | 9135 | 0.37 | 0.0535 | 0.0853 |
| chr22 | 37625206 | 37626174 | 969 | -0.82 | 0.0535 | 0.0853 |

| | | | | | | |
|---|---|---|---|---|---|---|
| chr9 | 32425987 | 32428088 | 2102 | -0.88 | 0.0535 | 0.0853 |
| chr14 | 32659075 | 32673743 | 14669 | 0.19 | 0.0535 | 0.0853 |
| chr3 | 126107720 | 126119062 | 11343 | 1.82 | 0.0535 | 0.0853 |
| chr12 | 121340875 | 121342879 | 2005 | 0.6 | 0.0535 | 0.0854 |
| chr17 | 34090959 | 34092190 | 1232 | 0.82 | 0.0536 | 0.0855 |
| chr6 | 151562005 | 151562721 | 717 | -1.16 | 0.0536 | 0.0855 |
| chr16 | 3660959 | 3670016 | 9058 | 0.34 | 0.0536 | 0.0855 |
| chr12 | 107978379 | 107979168 | 790 | -1.19 | 0.0536 | 0.0855 |
| chr12 | 59300410 | 59315049 | 14640 | 0.4 | 0.0537 | 0.0855 |
| chr3 | 155586968 | 155593405 | 6438 | 0.25 | 0.0537 | 0.0855 |
| chr3 | 70898491 | 70900219 | 1729 | 0.54 | 0.0537 | 0.0856 |
| chr1 | 27152528 | 27153014 | 487 | 1.21 | 0.0537 | 0.0856 |
| chr5 | 139118431 | 139119661 | 1231 | -0.95 | 0.0537 | 0.0856 |
| chr20 | 11185595 | 11210982 | 25388 | 0.22 | 0.0538 | 0.0857 |
| chr11 | 31424888 | 31425507 | 620 | 1.1 | 0.0538 | 0.0858 |
| chr12 | 89410651 | 89411051 | 401 | 1.31 | 0.0538 | 0.0858 |
| chr10 | 45454250 | 45455103 | 854 | 0.63 | 0.0539 | 0.0859 |
| chr22 | 47044925 | 47058282 | 13358 | 0.29 | 0.0539 | 0.086 |
| chr9 | 77850238 | 77850912 | 675 | -1.35 | 0.054 | 0.086 |
| chr1 | 22345078 | 22353153 | 8076 | 0.31 | 0.054 | 0.0861 |
| chr22 | 30058635 | 30059068 | 434 | -1.51 | 0.054 | 0.0861 |
| chr1 | 24608793 | 24624288 | 15496 | -0.15 | 0.054 | 0.0861 |
| chr8 | 124991303 | 124995012 | 3710 | 0.28 | 0.0541 | 0.0862 |
| chr16 | 84870889 | 84886092 | 15204 | 0.32 | 0.0541 | 0.0862 |
| chr5 | 52433614 | 52434059 | 446 | 1.9 | 0.0542 | 0.0863 |
| chr1 | 226495946 | 226497909 | 1964 | -0.37 | 0.0542 | 0.0863 |
| chr12 | 119591273 | 119592290 | 1018 | 1.29 | 0.0542 | 0.0863 |
| chr20 | 52293558 | 52308847 | 15290 | 0.21 | 0.0542 | 0.0863 |
| chr3 | 189556660 | 189558017 | 1358 | 0.81 | 0.0542 | 0.0863 |
| chr17 | 35765117 | 35790862 | 25746 | 0.16 | 0.0542 | 0.0863 |
| chr4 | 24643846 | 24662411 | 18566 | 0.22 | 0.0542 | 0.0863 |
| chr12 | 123668070 | 123680549 | 12480 | 0.35 | 0.0542 | 0.0863 |
| chr16 | 70585647 | 70586553 | 907 | -1.03 | 0.0542 | 0.0863 |
| chr22 | 32446356 | 32447241 | 886 | -1.08 | 0.0543 | 0.0865 |
| chr3 | 67546187 | 67547409 | 1223 | 0.9 | 0.0544 | 0.0866 |
| chr1 | 77747654 | 77748513 | 860 | -1.31 | 0.0544 | 0.0866 |
| chr4 | 39589211 | 39642179 | 52969 | -0.12 | 0.0545 | 0.0867 |
| chr19 | 38824926 | 38828183 | 3258 | 0.44 | 0.0545 | 0.0867 |
| chr6 | 75287850 | 75288382 | 533 | -1.26 | 0.0545 | 0.0868 |
| chr10 | 25051044 | 25052744 | 1701 | 0.74 | 0.0545 | 0.0868 |
| chr10 | 126106338 | 126108263 | 1926 | 0.54 | 0.0546 | 0.0869 |
| chr15 | 91444914 | 91447004 | 2091 | 0.52 | 0.0546 | 0.0869 |
| chr21 | 18864183 | 18864916 | 734 | -1.06 | 0.0546 | 0.0869 |
| chr1 | 236755241 | 236768559 | 13319 | 0.36 | 0.0546 | 0.0869 |
| chr16 | 86985361 | 86986768 | 1408 | -0.59 | 0.0546 | 0.0869 |
| chr3 | 49255233 | 49256867 | 1635 | -0.36 | 0.0547 | 0.087 |
| chr14 | 21823861 | 21824532 | 672 | 1.11 | 0.0547 | 0.087 |
| chr13 | 96000918 | 96002048 | 1131 | -1.54 | 0.0547 | 0.0871 |
| chr1 | 9134260 | 9141663 | 7404 | 1.8 | 0.0548 | 0.0872 |
| chr8 | 62892140 | 62893137 | 998 | 3.02 | 0.0548 | 0.0872 |
| chrX | 101914455 | 101915450 | 996 | 0.92 | 0.0548 | 0.0872 |
| chr12 | 122296089 | 122298561 | 2473 | -1.11 | 0.0548 | 0.0872 |
| chr1 | 27425025 | 27449894 | 24870 | -0.27 | 0.0548 | 0.0872 |
| chr16 | 9102372 | 9103188 | 817 | -1.15 | 0.0549 | 0.0872 |
| chr10 | 80166969 | 80167623 | 655 | -1.24 | 0.055 | 0.0875 |
| chr9 | 87862938 | 87864295 | 1358 | 2.99 | 0.055 | 0.0875 |
| chr2 | 9721623 | 9780140 | 58518 | -0.12 | 0.055 | 0.0875 |
| chr2 | 220954766 | 220955272 | 507 | -1.36 | 0.055 | 0.0875 |
| chr4 | 170947076 | 170948925 | 1850 | 0.53 | 0.0551 | 0.0875 |
| chr20 | 33703310 | 33736185 | 32876 | -0.18 | 0.0551 | 0.0875 |
| chr3 | 156337774 | 156338829 | 1056 | -1.13 | 0.0551 | 0.0876 |

| | | | | | | |
|---|---|---|---|---|---|---|
| chr12 | 19835568 | 19837015 | 1448 | 0.49 | 0.0551 | 0.0876 |
| chr8 | 56279849 | 56280380 | 532 | -1.89 | 0.0551 | 0.0876 |
| chr6 | 88211309 | 88212254 | 946 | 1.14 | 0.0551 | 0.0876 |
| chr1 | 29062439 | 29078352 | 15914 | 0.24 | 0.0551 | 0.0876 |
| chr22 | 43069727 | 43126445 | 56719 | -0.19 | 0.0552 | 0.0878 |
| chr19 | 50083127 | 50191486 | 108360 | 0.1 | 0.0552 | 0.0878 |
| chr1 | 41615405 | 41620285 | 4881 | -0.32 | 0.0553 | 0.0879 |
| chr1 | 36744798 | 36750788 | 5991 | -0.49 | 0.0554 | 0.088 |
| chrX | 44749156 | 44777257 | 28102 | -0.3 | 0.0554 | 0.088 |
| chr18 | 60586231 | 60586957 | 727 | 1.25 | 0.0554 | 0.088 |
| chr5 | 157345764 | 157352447 | 6684 | 3.02 | 0.0554 | 0.088 |
| chrX | 73458691 | 73513656 | 54966 | 0.12 | 0.0554 | 0.088 |
| chr5 | 141302186 | 141321799 | 19614 | 0.17 | 0.0554 | 0.088 |
| chr11 | 78157135 | 78157862 | 728 | 1.02 | 0.0554 | 0.0881 |
| chr2 | 242837369 | 242838502 | 1134 | 0.89 | 0.0555 | 0.0881 |
| chr2 | 167267640 | 167268379 | 740 | -1.18 | 0.0555 | 0.0882 |
| chr10 | 93271288 | 93274693 | 3406 | -0.46 | 0.0555 | 0.0882 |
| chr4 | 6942039 | 6950482 | 8444 | 0.25 | 0.0555 | 0.0882 |
| chr2 | 44032833 | 44035940 | 3108 | -0.56 | 0.0556 | 0.0883 |
| chr9 | 125639368 | 125645613 | 6246 | -0.53 | 0.0556 | 0.0883 |
| chr2 | 159824046 | 159827536 | 3491 | 0.33 | 0.0556 | 0.0883 |
| chr21 | 40652926 | 40663525 | 10600 | 0.68 | 0.0556 | 0.0884 |
| chr9 | 86277870 | 86281545 | 3676 | 0.61 | 0.0556 | 0.0884 |
| chr3 | 162481999 | 162482540 | 542 | 1.93 | 0.0557 | 0.0884 |
| chr20 | 55081270 | 55081622 | 353 | 2.08 | 0.0557 | 0.0884 |
| chr1 | 168802920 | 168804065 | 1146 | 0.99 | 0.0557 | 0.0885 |
| chr4 | 83636933 | 83637667 | 735 | 1.17 | 0.0558 | 0.0886 |
| chr7 | 51443581 | 51445009 | 1429 | 0.57 | 0.0558 | 0.0886 |
| chr5 | 65336567 | 65337369 | 803 | 0.86 | 0.0558 | 0.0886 |
| chr4 | 40751654 | 40752733 | 1080 | 0.77 | 0.0559 | 0.0887 |
| chr8 | 62923492 | 62924362 | 871 | 0.84 | 0.0559 | 0.0887 |
| chr15 | 101644963 | 101646192 | 1230 | -0.89 | 0.0559 | 0.0887 |
| chr7 | 82358334 | 82358808 | 475 | -1.4 | 0.0559 | 0.0887 |
| chr3 | 171334352 | 171335391 | 1040 | 1.16 | 0.0559 | 0.0887 |
| chr12 | 6013471 | 6014192 | 722 | -1.41 | 0.0559 | 0.0887 |
| chr20 | 32931292 | 32931861 | 570 | 1.15 | 0.0559 | 0.0888 |
| chr1 | 27830335 | 27832221 | 1887 | 0.54 | 0.056 | 0.0888 |
| chr4 | 69447631 | 69448404 | 774 | -1.62 | 0.056 | 0.0888 |
| chrX | 148621894 | 148623525 | 1632 | 0.36 | 0.056 | 0.0888 |
| chr16 | 11705434 | 11737457 | 32024 | -0.13 | 0.056 | 0.0888 |
| chr1 | 27648072 | 27650052 | 1981 | 0.36 | 0.0561 | 0.0889 |
| chr10 | 120100792 | 120102484 | 1693 | -0.38 | 0.0561 | 0.089 |
| chrX | 119618933 | 119647210 | 28278 | -0.35 | 0.0561 | 0.089 |
| chr2 | 58393559 | 58394068 | 510 | 1.67 | 0.0561 | 0.089 |
| chr14 | 58548273 | 58550039 | 1767 | 0.4 | 0.0561 | 0.089 |
| chr8 | 119351903 | 119361496 | 9594 | 0.4 | 0.0562 | 0.0891 |
| chr6 | 90084852 | 90085148 | 297 | 1.88 | 0.0563 | 0.0892 |
| chr10 | 115503346 | 115507190 | 3845 | -0.45 | 0.0563 | 0.0892 |
| chr8 | 18472186 | 18472557 | 372 | 1.78 | 0.0563 | 0.0892 |
| chr15 | 55781240 | 55791417 | 10178 | -0.27 | 0.0563 | 0.0893 |
| chr8 | 18577227 | 18579866 | 2640 | 0.57 | 0.0563 | 0.0893 |
| chr4 | 24474732 | 24475459 | 728 | 1.32 | 0.0563 | 0.0893 |
| chr8 | 74002301 | 74003131 | 831 | 1.28 | 0.0564 | 0.0894 |
| chr8 | 94849571 | 94859362 | 9792 | 0.65 | 0.0564 | 0.0894 |
| chr11 | 45825275 | 45827915 | 2641 | 0.37 | 0.0564 | 0.0894 |
| chr13 | 50612361 | 50612786 | 426 | -1.35 | 0.0565 | 0.0895 |
| chr1 | 26757929 | 26759875 | 1947 | 0.43 | 0.0565 | 0.0895 |
| chr2 | 233141001 | 233141485 | 485 | -1.72 | 0.0566 | 0.0896 |
| chr8 | 121095352 | 121096944 | 1593 | -0.67 | 0.0566 | 0.0896 |
| chr2 | 129133818 | 129134641 | 824 | -1.46 | 0.0566 | 0.0897 |
| chr1 | 26021134 | 26022897 | 1764 | 0.87 | 0.0566 | 0.0897 |

| | | | | | | |
|---|---|---|---|---|---|---|
| chr10 | 32427728 | 32438928 | 11201 | -0.36 | 0.0566 | 0.0897 |
| chr11 | 74853741 | 74854773 | 1033 | -2.83 | 0.0566 | 0.0897 |
| chrX | 22182795 | 22184494 | 1700 | 2.42 | 0.0567 | 0.0898 |
| chr19 | 21511152 | 21512806 | 1655 | 0.48 | 0.0567 | 0.0898 |
| chr8 | 141671598 | 141672710 | 1113 | -0.94 | 0.0567 | 0.0898 |
| chr6 | 34430271 | 34430747 | 477 | -1.89 | 0.0567 | 0.0899 |
| chr11 | 47995837 | 48192801 | 196965 | 0.07 | 0.0568 | 0.0899 |
| chr21 | 27306730 | 27311688 | 4959 | 0.51 | 0.0568 | 0.0899 |
| chr5 | 41095075 | 41095586 | 512 | -1.86 | 0.0568 | 0.0899 |
| chr17 | 854142 | 872338 | 18197 | -0.27 | 0.0568 | 0.09 |
| chr1 | 87597607 | 87597910 | 304 | -1.89 | 0.0568 | 0.09 |
| chr11 | 33124919 | 33125586 | 668 | -1.42 | 0.0568 | 0.09 |
| chr4 | 108852168 | 108853110 | 943 | 0.95 | 0.0569 | 0.09 |
| chr2 | 65857395 | 65858391 | 997 | -0.84 | 0.0569 | 0.0901 |
| chr12 | 76016985 | 76018362 | 1378 | 0.91 | 0.0569 | 0.0901 |
| chr1 | 64434195 | 64436279 | 2085 | 0.62 | 0.0569 | 0.0901 |
| chr5 | 61671128 | 61675430 | 4303 | -0.33 | 0.057 | 0.0902 |
| chr1 | 113614835 | 113617245 | 2411 | 0.45 | 0.057 | 0.0902 |
| chr1 | 235421738 | 235434731 | 12994 | 0.5 | 0.057 | 0.0902 |
| chr15 | 90293462 | 90408119 | 114658 | 0.12 | 0.057 | 0.0903 |
| chr1 | 182730238 | 182731236 | 999 | -1.41 | 0.0571 | 0.0904 |
| chr12 | 46916510 | 46919126 | 2617 | 0.31 | 0.0571 | 0.0904 |
| chr7 | 17977484 | 17981093 | 3610 | 0.24 | 0.0571 | 0.0904 |
| chr1 | 22298625 | 22299422 | 798 | 1 | 0.0571 | 0.0904 |
| chrX | 135267180 | 135268016 | 837 | 0.78 | 0.0571 | 0.0904 |
| chr16 | 56370032 | 56370494 | 463 | 2 | 0.0571 | 0.0904 |
| chr12 | 448587 | 449588 | 1002 | 0.95 | 0.0571 | 0.0904 |
| chr12 | 111325387 | 111325990 | 604 | 0.97 | 0.0572 | 0.0904 |
| chr6 | 131948316 | 131949326 | 1011 | 0.61 | 0.0572 | 0.0905 |
| chr13 | 45410997 | 45411782 | 786 | 1.9 | 0.0572 | 0.0905 |
| chr5 | 5421460 | 5435240 | 13781 | 0.2 | 0.0572 | 0.0905 |
| chr12 | 47741702 | 47742173 | 472 | -1.92 | 0.0573 | 0.0906 |
| chr11 | 34805110 | 34816578 | 11469 | -3.14 | 0.0573 | 0.0907 |
| chr15 | 67705519 | 67710103 | 4585 | 0.32 | 0.0573 | 0.0907 |
| chr4 | 152329503 | 152330655 | 1153 | 1.1 | 0.0574 | 0.0907 |
| chr7 | 83771612 | 83776894 | 5283 | 0.15 | 0.0574 | 0.0908 |
| chr1 | 25827259 | 25828771 | 1513 | 0.68 | 0.0574 | 0.0908 |
| chr12 | 32812735 | 32813387 | 653 | 1.16 | 0.0574 | 0.0908 |
| chr6 | 135734151 | 135734767 | 617 | -1.48 | 0.0574 | 0.0908 |
| chr10 | 124712777 | 124714285 | 1509 | 0.48 | 0.0575 | 0.0909 |
| chr20 | 5862339 | 5863156 | 818 | 2.84 | 0.0575 | 0.0909 |
| chr10 | 69523146 | 69525742 | 2597 | 0.37 | 0.0575 | 0.0909 |
| chr4 | 139796831 | 139798376 | 1546 | 0.53 | 0.0575 | 0.0909 |
| chr1 | 6784840 | 6785340 | 501 | 1.33 | 0.0575 | 0.0909 |
| chr7 | 151812061 | 151825395 | 13335 | 0.82 | 0.0575 | 0.0909 |
| chr1 | 205512308 | 205513302 | 995 | 1.2 | 0.0575 | 0.0909 |
| chr9 | 136996040 | 137002608 | 6569 | -0.34 | 0.0576 | 0.091 |
| chr7 | 150809777 | 150814455 | 4679 | 0.31 | 0.0576 | 0.091 |
| chr1 | 156280373 | 156309196 | 28824 | 0.21 | 0.0576 | 0.091 |
| chr22 | 39867125 | 39868190 | 1066 | 1.1 | 0.0576 | 0.091 |
| chr7 | 37488046 | 37488459 | 414 | -1.63 | 0.0576 | 0.091 |
| chr15 | 83725675 | 83737974 | 12300 | 0.2 | 0.0576 | 0.0911 |
| chr3 | 119419114 | 119422045 | 2932 | 0.64 | 0.0577 | 0.0912 |
| chr9 | 85930347 | 85931189 | 843 | -0.94 | 0.0577 | 0.0912 |
| chr17 | 9704521 | 9706765 | 2245 | 0.26 | 0.0578 | 0.0914 |
| chr8 | 41840584 | 41840988 | 405 | -1.28 | 0.0579 | 0.0914 |
| chr17 | 79174031 | 79175433 | 1403 | -0.67 | 0.0579 | 0.0914 |
| chr19 | 55554749 | 55557091 | 2343 | -1.25 | 0.058 | 0.0916 |
| chr21 | 33650460 | 33652613 | 2154 | 0.59 | 0.058 | 0.0916 |
| chr22 | 37054657 | 37057351 | 2695 | -0.23 | 0.058 | 0.0917 |
| chr1 | 61946564 | 61963210 | 16647 | 0.53 | 0.058 | 0.0917 |

| | | | | | | |
|---|---|---|---|---|---|---|
| chr9 | 5814207 | 5853674 | 39468 | -0.11 | 0.0581 | 0.0917 |
| chr2 | 44221593 | 44224398 | 2806 | -0.3 | 0.0581 | 0.0917 |
| chr5 | 31470795 | 31476503 | 5709 | -2.38 | 0.0581 | 0.0918 |
| chr14 | 89603367 | 89604361 | 995 | 0.96 | 0.0581 | 0.0918 |
| chrX | 85301243 | 85302995 | 1753 | 0.34 | 0.0581 | 0.0918 |
| chr5 | 31429946 | 31431142 | 1197 | 0.86 | 0.0582 | 0.0918 |
| chr20 | 46113543 | 46294672 | 181130 | 0.07 | 0.0582 | 0.0919 |
| chr17 | 44809090 | 44820767 | 11678 | -0.72 | 0.0582 | 0.0919 |
| chr10 | 70160848 | 70168163 | 7316 | 0.19 | 0.0582 | 0.0919 |
| chr10 | 105785220 | 105795141 | 9922 | 0.38 | 0.0582 | 0.0919 |
| chr20 | 56686192 | 56688360 | 2169 | -0.53 | 0.0583 | 0.092 |
| chr12 | 74668951 | 74669774 | 824 | 1.3 | 0.0583 | 0.092 |
| chr6 | 34191231 | 34227707 | 36477 | -0.14 | 0.0583 | 0.092 |
| chr3 | 32725605 | 32728554 | 2950 | 0.27 | 0.0583 | 0.092 |
| chr2 | 239110787 | 239113129 | 2343 | -0.39 | 0.0583 | 0.0921 |
| chr11 | 113214313 | 113215391 | 1079 | -1.04 | 0.0583 | 0.0921 |
| chr7 | 148562 | 150340 | 1779 | 0.29 | 0.0584 | 0.0921 |
| chr4 | 114468379 | 114483009 | 14631 | -0.31 | 0.0584 | 0.0921 |
| chr6 | 169470444 | 169471644 | 1201 | -0.79 | 0.0584 | 0.0921 |
| chr17 | 30656346 | 30721260 | 64915 | 0.15 | 0.0584 | 0.0922 |
| chr17 | 78426981 | 78429614 | 2634 | 0.23 | 0.0584 | 0.0922 |
| chr8 | 71313746 | 71317481 | 3736 | 0.26 | 0.0584 | 0.0922 |
| chrX | 49686639 | 49687762 | 1124 | 0.75 | 0.0585 | 0.0922 |
| chr1 | 236065943 | 236084260 | 18318 | 0.21 | 0.0585 | 0.0922 |
| chr1 | 28201411 | 28202072 | 662 | 1.2 | 0.0585 | 0.0923 |
| chr2 | 46720117 | 46727736 | 7620 | 0.59 | 0.0585 | 0.0923 |
| chr8 | 96812527 | 96815031 | 2505 | -0.44 | 0.0586 | 0.0923 |
| chr8 | 9412227 | 9414972 | 2746 | 0.24 | 0.0586 | 0.0923 |
| chr5 | 66103885 | 66104279 | 395 | -1.43 | 0.0586 | 0.0923 |
| chr17 | 11981932 | 11982332 | 401 | -1.48 | 0.0586 | 0.0924 |
| chr14 | 53917429 | 53926391 | 8963 | 0.5 | 0.0586 | 0.0924 |
| chr4 | 87959691 | 87960924 | 1234 | -0.96 | 0.0586 | 0.0924 |
| chr10 | 80534752 | 80535339 | 588 | -1.57 | 0.0586 | 0.0924 |
| chr18 | 64177865 | 64191302 | 13438 | 0.27 | 0.0586 | 0.0924 |
| chrX | 41448706 | 41450127 | 1422 | 0.56 | 0.0586 | 0.0924 |
| chr11 | 113730441 | 113747293 | 16853 | 0.26 | 0.0587 | 0.0926 |
| chr12 | 116491157 | 116491866 | 710 | 1 | 0.0587 | 0.0926 |
| chr3 | 23587444 | 23588323 | 880 | 1.88 | 0.0588 | 0.0926 |
| chr10 | 354118 | 355010 | 893 | -1.22 | 0.0588 | 0.0927 |
| chr17 | 2041042 | 2042376 | 1335 | -0.79 | 0.0589 | 0.0928 |
| chr3 | 106926516 | 106936145 | 9630 | 0.89 | 0.0589 | 0.0928 |
| chr1 | 93643451 | 93647127 | 3677 | 0.35 | 0.0589 | 0.0928 |
| chr17 | 9354936 | 9356381 | 1446 | 0.79 | 0.0589 | 0.0928 |
| chr1 | 37979475 | 37981339 | 1865 | 0.43 | 0.059 | 0.0929 |
| chr2 | 181485385 | 181487357 | 1973 | 0.71 | 0.0591 | 0.0931 |
| chr13 | 79219782 | 79234751 | 14970 | 0.36 | 0.0591 | 0.0931 |
| chr5 | 67474200 | 67475076 | 877 | 1.09 | 0.0591 | 0.0931 |
| chr19 | 39997587 | 39998502 | 916 | -0.97 | 0.0591 | 0.0931 |
| chr1 | 12267043 | 12267682 | 640 | -1.18 | 0.0591 | 0.0931 |
| chr11 | 70633946 | 70635555 | 1610 | -0.51 | 0.0591 | 0.0931 |
| chr20 | 50178078 | 50182928 | 4851 | -0.18 | 0.0591 | 0.0931 |
| chr6 | 12403410 | 12424703 | 21294 | -0.19 | 0.0592 | 0.0932 |
| chr4 | 41694188 | 41695413 | 1226 | 1.11 | 0.0592 | 0.0932 |
| chr12 | 26194901 | 26195708 | 808 | 0.85 | 0.0592 | 0.0932 |
| chr11 | 76081901 | 76092935 | 11035 | -0.34 | 0.0592 | 0.0932 |
| chr3 | 153838692 | 153844719 | 6028 | 0.29 | 0.0592 | 0.0932 |
| chr4 | 91278537 | 91279020 | 484 | 1.62 | 0.0592 | 0.0932 |
| chr11 | 110226693 | 110237544 | 10852 | 0.19 | 0.0592 | 0.0932 |
| chr2 | 120122858 | 120126077 | 3220 | -0.34 | 0.0592 | 0.0932 |
| chr19 | 18246891 | 18247544 | 654 | -0.94 | 0.0593 | 0.0933 |
| chr4 | 76837365 | 76837887 | 523 | 0.99 | 0.0593 | 0.0934 |

| | | | | | | |
|---|---|---|---|---|---|---|
| chr5 | 145339957 | 145359271 | 19315 | -0.53 | 0.0594 | 0.0935 |
| chr1 | 235603896 | 235615294 | 11399 | -0.19 | 0.0594 | 0.0935 |
| chr1 | 63987982 | 63990663 | 2682 | -0.24 | 0.0594 | 0.0935 |
| chr3 | 180396995 | 180398020 | 1026 | 0.76 | 0.0594 | 0.0935 |
| chr5 | 145602806 | 145603698 | 893 | 1.07 | 0.0594 | 0.0935 |
| chr1 | 26855802 | 26870604 | 14803 | -0.28 | 0.0594 | 0.0935 |
| chr7 | 101685308 | 101685892 | 585 | -1.85 | 0.0594 | 0.0935 |
| chr5 | 35230143 | 35230636 | 494 | 1.63 | 0.0595 | 0.0936 |
| chr17 | 69544857 | 69545425 | 569 | -1.84 | 0.0595 | 0.0936 |
| chr1 | 100055774 | 100065773 | 10000 | 3.18 | 0.0595 | 0.0936 |
| chr10 | 4849803 | 4908542 | 58740 | 0.12 | 0.0595 | 0.0936 |
| chr11 | 34936516 | 34952143 | 15628 | 0.21 | 0.0595 | 0.0937 |
| chr8 | 28479443 | 28492021 | 12579 | -0.58 | 0.0596 | 0.0937 |
| chr10 | 369519 | 396579 | 27061 | 0.24 | 0.0596 | 0.0938 |
| chr12 | 22677952 | 22679021 | 1070 | 0.93 | 0.0596 | 0.0938 |
| chr15 | 65550540 | 65551475 | 936 | -1.19 | 0.0597 | 0.0938 |
| chr12 | 121788885 | 121791130 | 2246 | 0.29 | 0.0597 | 0.0939 |
| chr4 | 184557166 | 184560335 | 3170 | 0.23 | 0.0597 | 0.0939 |
| chr7 | 102754713 | 102771325 | 16613 | -0.38 | 0.0597 | 0.094 |
| chr17 | 37170419 | 37175530 | 5112 | -0.48 | 0.0598 | 0.094 |
| chr7 | 133500410 | 133500760 | 351 | 1.52 | 0.0598 | 0.094 |
| chr8 | 124642246 | 124643329 | 1084 | -0.66 | 0.0598 | 0.094 |
| chr9 | 117483717 | 117506061 | 22345 | 3.72 | 0.0598 | 0.094 |
| chr14 | 36559345 | 36561663 | 2319 | 0.34 | 0.0598 | 0.0941 |
| chr5 | 74345857 | 74349949 | 4093 | -0.69 | 0.0598 | 0.0941 |
| chr20 | 60876911 | 60886083 | 9173 | 0.32 | 0.0598 | 0.0941 |
| chr2 | 99756700 | 99773026 | 16327 | 0.24 | 0.0599 | 0.0941 |
| chr5 | 180525843 | 180526498 | 656 | 0.95 | 0.0599 | 0.0941 |
| chr10 | 5359190 | 5360278 | 1089 | -0.75 | 0.0599 | 0.0942 |
| chr3 | 102179136 | 102180249 | 1114 | -0.97 | 0.06 | 0.0943 |
| chr2 | 181844054 | 181848944 | 4891 | -0.36 | 0.06 | 0.0944 |
| chr7 | 139256760 | 139257305 | 546 | 1.93 | 0.0601 | 0.0944 |
| chr11 | 86808235 | 86808786 | 552 | -1.14 | 0.0601 | 0.0944 |
| chr13 | 99852044 | 99854628 | 2585 | 0.32 | 0.0601 | 0.0945 |
| chr16 | 27121349 | 27121878 | 530 | 1.26 | 0.0602 | 0.0945 |
| chr11 | 85453657 | 85460238 | 6582 | 1.87 | 0.0602 | 0.0945 |
| chr5 | 91421 | 92364 | 944 | 1.09 | 0.0602 | 0.0946 |
| chr4 | 101269322 | 101270575 | 1254 | -0.62 | 0.0602 | 0.0946 |
| chr12 | 52316149 | 52317582 | 1434 | 0.8 | 0.0602 | 0.0946 |
| chr1 | 150504397 | 150505615 | 1219 | 0.94 | 0.0602 | 0.0946 |
| chr18 | 33194773 | 33195910 | 1138 | 0.77 | 0.0603 | 0.0947 |
| chr5 | 167927645 | 167928273 | 629 | 1.43 | 0.0603 | 0.0947 |
| chrX | 83756353 | 83757797 | 1445 | -0.45 | 0.0603 | 0.0947 |
| chr6 | 27280078 | 27280568 | 491 | 1.25 | 0.0603 | 0.0947 |
| chr1 | 202112137 | 202133168 | 21032 | 0.25 | 0.0603 | 0.0947 |
| chr16 | 25137750 | 25138820 | 1071 | -1.08 | 0.0603 | 0.0947 |
| chr1 | 16674987 | 16679902 | 4916 | 0.32 | 0.0603 | 0.0948 |
| chr19 | 7953024 | 7953976 | 953 | -0.63 | 0.0604 | 0.0948 |
| chr13 | 91570195 | 91586273 | 16079 | 0.46 | 0.0604 | 0.0948 |
| chr3 | 48594651 | 48604597 | 9947 | -0.3 | 0.0604 | 0.0948 |
| chr14 | 100196441 | 100233970 | 37530 | 0.22 | 0.0604 | 0.0948 |
| chr7 | 19938608 | 19940256 | 1649 | -0.44 | 0.0604 | 0.0948 |
| chr12 | 26984897 | 26986154 | 1258 | 0.47 | 0.0604 | 0.0948 |
| chr22 | 31314268 | 31315042 | 775 | -1.01 | 0.0604 | 0.0949 |
| chr1 | 172557509 | 172558548 | 1040 | -1.02 | 0.0605 | 0.0949 |
| chr5 | 78253955 | 78263446 | 9492 | 1.88 | 0.0605 | 0.095 |
| chr6 | 38546879 | 38547210 | 332 | -1.47 | 0.0605 | 0.095 |
| chr7 | 133620293 | 133621375 | 1083 | -1.22 | 0.0606 | 0.095 |
| chr1 | 19793302 | 19813711 | 20410 | 0.21 | 0.0606 | 0.0951 |
| chr2 | 240963735 | 240965743 | 2009 | -0.46 | 0.0607 | 0.0952 |
| chr8 | 95247120 | 95256318 | 9199 | 0.19 | 0.0607 | 0.0952 |

| | | | | | | |
|---|---|---|---|---|---|---|
| chr7 | 155569358 | 155580381 | 11024 | 0.76 | 0.0607 | 0.0952 |
| chr4 | 174290198 | 174294482 | 4285 | 0.34 | 0.0608 | 0.0955 |
| chr5 | 86656631 | 86657185 | 555 | -1.23 | 0.0608 | 0.0955 |
| chr1 | 208063379 | 208064565 | 1187 | -0.81 | 0.0608 | 0.0955 |
| chr7 | 36032576 | 36034062 | 1487 | -0.61 | 0.0609 | 0.0955 |
| chr1 | 80276713 | 80277718 | 1006 | 3.01 | 0.0609 | 0.0955 |
| chr2 | 206947341 | 206953315 | 5975 | -0.2 | 0.0609 | 0.0955 |
| chr14 | 94545818 | 94548311 | 2494 | 0.36 | 0.0609 | 0.0956 |
| chr2 | 228079112 | 228093776 | 14665 | -0.36 | 0.0609 | 0.0956 |
| chr19 | 3141356 | 3156004 | 14649 | -0.4 | 0.0609 | 0.0956 |
| chr3 | 69773949 | 69774564 | 616 | -1.93 | 0.061 | 0.0956 |
| chr5 | 127489037 | 127492147 | 3111 | -0.57 | 0.061 | 0.0956 |
| chr11 | 28323435 | 28326219 | 2785 | 0.36 | 0.061 | 0.0956 |
| chr7 | 45054905 | 45055550 | 646 | -1.09 | 0.061 | 0.0957 |
| chr22 | 18155246 | 18157247 | 2002 | 0.65 | 0.061 | 0.0957 |
| chr9 | 113476307 | 113477183 | 877 | -0.99 | 0.061 | 0.0957 |
| chr11 | 16653963 | 16654658 | 696 | 1.05 | 0.061 | 0.0957 |
| chr11 | 74108225 | 74110517 | 2293 | 0.41 | 0.061 | 0.0957 |
| chr4 | 102057307 | 102062631 | 5325 | 0.51 | 0.0611 | 0.0957 |
| chr16 | 69684792 | 69797673 | 112882 | -0.1 | 0.0611 | 0.0957 |
| chr12 | 116425861 | 116433373 | 7513 | -0.48 | 0.0611 | 0.0957 |
| chr10 | 32201077 | 32251608 | 50532 | 0.08 | 0.0611 | 0.0958 |
| chr16 | 84681546 | 84688927 | 7382 | -0.44 | 0.0611 | 0.0958 |
| chr11 | 116722986 | 116724483 | 1498 | 0.86 | 0.0611 | 0.0958 |
| chr14 | 32312721 | 32313053 | 333 | -1.83 | 0.0612 | 0.0958 |
| chr3 | 29978195 | 29979273 | 1079 | -0.8 | 0.0612 | 0.0959 |
| chr9 | 123581746 | 123641011 | 59266 | 0.16 | 0.0613 | 0.096 |
| chr19 | 3035117 | 3037779 | 2663 | 0.45 | 0.0613 | 0.0961 |
| chr6 | 15638246 | 15638705 | 460 | 1.57 | 0.0613 | 0.0961 |
| chr13 | 67197437 | 67210263 | 12827 | 0.66 | 0.0613 | 0.0961 |
| chr12 | 131272524 | 131272889 | 366 | -1.77 | 0.0614 | 0.0962 |
| chr10 | 95360747 | 95361695 | 949 | -1.02 | 0.0614 | 0.0962 |
| chrX | 133593642 | 133604488 | 10847 | 0.44 | 0.0614 | 0.0962 |
| chr17 | 56591324 | 56597546 | 6223 | 0.27 | 0.0614 | 0.0962 |
| chr6 | 157341216 | 157355054 | 13839 | 0.26 | 0.0614 | 0.0962 |
| chr1 | 175969185 | 175969554 | 370 | -1.91 | 0.0615 | 0.0963 |
| chrX | 11775568 | 11805721 | 30154 | 0.2 | 0.0615 | 0.0963 |
| chr16 | 50361203 | 50372128 | 10926 | -0.24 | 0.0615 | 0.0963 |
| chr18 | 8657042 | 8674164 | 17123 | 0.42 | 0.0615 | 0.0963 |
| chr1 | 33297220 | 33297750 | 531 | -1.5 | 0.0615 | 0.0963 |
| chr1 | 173791863 | 173795510 | 3648 | 0.22 | 0.0616 | 0.0964 |
| chr6 | 34494449 | 34524962 | 30514 | -0.23 | 0.0616 | 0.0964 |
| chr17 | 59521659 | 59522825 | 1167 | 0.87 | 0.0616 | 0.0965 |
| chr2 | 115089481 | 115089877 | 397 | -1.51 | 0.0616 | 0.0965 |
| chr11 | 124823359 | 124824456 | 1098 | 0.66 | 0.0617 | 0.0965 |
| chr6 | 135808729 | 135827521 | 18793 | 0.2 | 0.0617 | 0.0966 |
| chrX | 12156542 | 12169059 | 12518 | 2.86 | 0.0617 | 0.0966 |
| chr2 | 220461246 | 220464304 | 3059 | -0.23 | 0.0617 | 0.0966 |
| chr3 | 44551174 | 44553196 | 2023 | 0.4 | 0.0618 | 0.0967 |
| chr6 | 113001007 | 113013191 | 12185 | -0.44 | 0.0618 | 0.0968 |
| chr8 | 146277048 | 146278970 | 1923 | 0.38 | 0.0618 | 0.0968 |
| chr1 | 24737139 | 24746513 | 9375 | 0.24 | 0.0618 | 0.0968 |
| chr17 | 63543112 | 63544092 | 981 | 0.94 | 0.0619 | 0.0969 |
| chr1 | 166807717 | 166846170 | 38454 | 0.2 | 0.0619 | 0.0969 |
| chr3 | 128929840 | 128950343 | 20504 | 0.17 | 0.0619 | 0.0969 |
| chr7 | 30810283 | 30812229 | 1947 | -0.37 | 0.0619 | 0.0969 |
| chr7 | 133718656 | 133720256 | 1601 | 0.54 | 0.0619 | 0.0969 |
| chr8 | 42310669 | 42311704 | 1036 | -1.25 | 0.0619 | 0.0969 |
| chr5 | 14870997 | 14873256 | 2260 | 0.4 | 0.062 | 0.0969 |
| chr1 | 33546088 | 33546600 | 513 | -1.07 | 0.062 | 0.097 |
| chr20 | 25019827 | 25024419 | 4593 | -0.73 | 0.0621 | 0.0971 |

| | | | | | | |
|---|---|---|---|---|---|---|
| chr11 | 123389415 | 123390577 | 1163 | -0.9 | 0.0621 | 0.0971 |
| chr3 | 118744102 | 118754407 | 10306 | -2.89 | 0.0621 | 0.0972 |
| chr16 | 71840247 | 71851296 | 11050 | 0.17 | 0.0621 | 0.0972 |
| chr6 | 128976761 | 128989193 | 12433 | 0.65 | 0.0621 | 0.0972 |
| chr10 | 17009911 | 17011115 | 1205 | -1.08 | 0.0622 | 0.0972 |
| chr10 | 22937283 | 22949814 | 12532 | 0.39 | 0.0622 | 0.0972 |
| chr19 | 58280422 | 58282348 | 1927 | 0.55 | 0.0623 | 0.0973 |
| chr4 | 114001028 | 114001816 | 789 | 1.34 | 0.0623 | 0.0974 |
| chr10 | 78095761 | 78096263 | 503 | 1.25 | 0.0623 | 0.0975 |
| chr22 | 18293857 | 18294512 | 656 | -1.39 | 0.0624 | 0.0975 |
| chr6 | 24980645 | 24982245 | 1601 | 3 | 0.0624 | 0.0975 |
| chr11 | 64757541 | 64811582 | 54042 | 0.15 | 0.0624 | 0.0976 |
| chr6 | 1678349 | 1678915 | 567 | 1.4 | 0.0625 | 0.0977 |
| chr5 | 137468550 | 137469279 | 730 | 0.89 | 0.0625 | 0.0977 |
| chr5 | 39502144 | 39502819 | 676 | 1.85 | 0.0625 | 0.0977 |
| chrX | 55478147 | 55479234 | 1088 | 0.99 | 0.0626 | 0.0978 |
| chr18 | 7789501 | 7791882 | 2382 | 0.56 | 0.0626 | 0.0978 |
| chr3 | 170715626 | 170721488 | 5863 | 0.26 | 0.0627 | 0.098 |
| chrX | 80063061 | 80066693 | 3633 | 0.3 | 0.0628 | 0.0981 |
| chr5 | 73926106 | 73943253 | 17148 | 0.17 | 0.0628 | 0.0981 |
| chr7 | 17902677 | 17903145 | 469 | -1.24 | 0.0628 | 0.0981 |
| chr10 | 121334538 | 121335866 | 1329 | 0.92 | 0.0628 | 0.0982 |
| chr20 | 20739507 | 20740023 | 517 | 1.06 | 0.0628 | 0.0982 |
| chr2 | 58208294 | 58209071 | 778 | 1 | 0.0629 | 0.0982 |
| chr15 | 89973304 | 89974347 | 1044 | -0.59 | 0.0629 | 0.0982 |
| chr2 | 27472845 | 27473423 | 579 | -1.26 | 0.0629 | 0.0983 |
| chr5 | 76372552 | 76384176 | 11625 | 0.27 | 0.063 | 0.0983 |
| chr6 | 45557684 | 45558837 | 1154 | 0.82 | 0.063 | 0.0984 |
| chr7 | 93758149 | 93760481 | 2333 | 0.48 | 0.063 | 0.0984 |
| chrX | 106216208 | 106218275 | 2068 | -0.79 | 0.063 | 0.0984 |
| chr4 | 100483535 | 100485750 | 2216 | 0.28 | 0.063 | 0.0984 |
| chr18 | 7811095 | 7813589 | 2495 | -0.64 | 0.0631 | 0.0985 |
| chr3 | 151971488 | 151991467 | 19980 | 0.15 | 0.0632 | 0.0987 |
| chr22 | 38900170 | 38903555 | 3386 | 0.21 | 0.0632 | 0.0987 |
| chr7 | 104732661 | 104733734 | 1074 | 1.06 | 0.0632 | 0.0987 |
| chr10 | 121209431 | 121210227 | 797 | -1.38 | 0.0632 | 0.0987 |
| chr11 | 16778944 | 16782012 | 3069 | 0.69 | 0.0633 | 0.0988 |
| chr4 | 79859426 | 79861249 | 1824 | 0.62 | 0.0633 | 0.0988 |
| chr1 | 214296122 | 214296918 | 797 | 1.21 | 0.0633 | 0.0988 |
| chr1 | 92545382 | 92546874 | 1493 | 0.45 | 0.0633 | 0.0988 |
| chr15 | 87135243 | 87136188 | 946 | 2.84 | 0.0633 | 0.0988 |
| chrX | 46259761 | 46260243 | 483 | 1.19 | 0.0633 | 0.0989 |
| chr15 | 85217820 | 85363117 | 145298 | -0.07 | 0.0634 | 0.0989 |
| chr2 | 24895306 | 24895715 | 410 | -1.55 | 0.0634 | 0.0989 |
| chr21 | 43428941 | 43431576 | 2636 | 0.36 | 0.0634 | 0.0989 |
| chr4 | 124939864 | 124940460 | 597 | 1.63 | 0.0634 | 0.0989 |
| chr11 | 128212666 | 128213134 | 469 | 1.29 | 0.0635 | 0.099 |
| chr13 | 44823203 | 44823980 | 778 | 0.99 | 0.0635 | 0.099 |
| chr1 | 227231184 | 227232750 | 1567 | -0.54 | 0.0635 | 0.099 |
| chr14 | 45618544 | 45619170 | 627 | -1.1 | 0.0635 | 0.0991 |
| chr1 | 109693569 | 109695574 | 2006 | 0.71 | 0.0635 | 0.0991 |
| chr8 | 103271382 | 103284472 | 13091 | 0.42 | 0.0635 | 0.0991 |
| chr9 | 103360792 | 103361644 | 853 | 0.88 | 0.0636 | 0.0992 |
| chr1 | 36443691 | 36444751 | 1061 | 0.82 | 0.0636 | 0.0992 |
| chr1 | 3663460 | 3664319 | 860 | 1.04 | 0.0636 | 0.0992 |
| chr17 | 8353776 | 8354472 | 697 | 0.94 | 0.0636 | 0.0993 |
| chr3 | 52624541 | 52625082 | 542 | -1.85 | 0.0637 | 0.0993 |
| chr2 | 109449298 | 109450263 | 966 | 0.82 | 0.0638 | 0.0994 |
| chr6 | 10195711 | 10196334 | 624 | -1.83 | 0.0638 | 0.0995 |
| chr6 | 4401230 | 4403614 | 2385 | 0.35 | 0.0638 | 0.0995 |
| chr2 | 46186214 | 46192502 | 6289 | 0.29 | 0.0638 | 0.0995 |

| | | | | | | |
|---|---|---|---|---|---|---|
| chr5 | 85432946 | 85433505 | 560 | -1.15 | 0.0638 | 0.0995 |
| chr4 | 95524212 | 95525176 | 965 | -1.07 | 0.0639 | 0.0995 |
| chr19 | 11175091 | 11176143 | 1053 | -0.76 | 0.0639 | 0.0996 |
| chr16 | 12594511 | 12611376 | 16866 | 0.49 | 0.0639 | 0.0996 |
| chr14 | 29046513 | 29047722 | 1210 | 0.59 | 0.0639 | 0.0997 |
| chr7 | 93387749 | 93388419 | 671 | 0.98 | 0.064 | 0.0997 |
| chr3 | 193310255 | 193312745 | 2491 | 0.36 | 0.064 | 0.0998 |
| chr18 | 33766658 | 33768332 | 1675 | 0.53 | 0.064 | 0.0998 |
| chrX | 69352454 | 69354635 | 2182 | -0.29 | 0.064 | 0.0998 |
| chr11 | 56948752 | 56949403 | 652 | -1.48 | 0.064 | 0.0998 |
| chr2 | 161850520 | 161851042 | 523 | -1.35 | 0.064 | 0.0998 |
| chr9 | 134374543 | 134378763 | 4221 | 0.64 | 0.0641 | 0.0999 |
| chr2 | 179277677 | 179280323 | 2647 | -0.23 | 0.0641 | 0.0999 |
| chr8 | 22478974 | 22481054 | 2081 | -0.67 | 0.0642 | 0.1 |
| chr3 | 177216234 | 177216913 | 680 | 1.23 | 0.0642 | 0.1 |
| chr19 | 51896863 | 51899108 | 2246 | 0.3 | 0.0642 | 0.1 |
| chr3 | 14629058 | 14630456 | 1399 | -0.75 | 0.0642 | 0.1 |
| chr6 | 16421126 | 16421755 | 630 | 1.08 | 0.0642 | 0.1 |
| chr1 | 111681115 | 111684425 | 3311 | 0.3 | 0.0642 | 0.1 |
| chr5 | 59321985 | 59322404 | 420 | -1.55 | 0.0642 | 0.1 |
| chr21 | 27407383 | 27423666 | 16284 | -0.46 | 0.0642 | 0.1 |
| chr1 | 156051817 | 156117036 | 65220 | 0.08 | 0.0643 | 0.1 |
| chr6 | 28366829 | 28368205 | 1377 | 0.41 | 0.0643 | 0.1 |
| chr1 | 200552020 | 200552483 | 464 | -1.2 | 0.0643 | 0.1 |
| chr22 | 24973977 | 24975333 | 1357 | -0.94 | 0.0643 | 0.1 |
| chr16 | 2800972 | 2806333 | 5362 | 0.19 | 0.0643 | 0.1 |
| chr15 | 90039445 | 90042793 | 3349 | -0.31 | 0.0644 | 0.1 |
| chr2 | 39795162 | 39796263 | 1102 | 1 | 0.0644 | 0.1 |
| chr2 | 216782772 | 216783391 | 620 | -0.95 | 0.0644 | 0.1 |
| chr4 | 106393197 | 106395680 | 2484 | 0.31 | 0.0645 | 0.1 |
| chr1 | 93608578 | 93608990 | 413 | -1.58 | 0.0645 | 0.1 |
| chr21 | 17531130 | 17532452 | 1323 | 0.63 | 0.0646 | 0.101 |
| chr2 | 101177866 | 101180702 | 2837 | -0.28 | 0.0646 | 0.101 |
| chr12 | 117174676 | 117177059 | 2384 | 0.52 | 0.0647 | 0.101 |
| chr10 | 31870817 | 31872656 | 1840 | -0.42 | 0.0647 | 0.101 |
| chr2 | 170789903 | 170790838 | 936 | -1.11 | 0.0648 | 0.101 |
| chr3 | 152975242 | 152976498 | 1257 | 0.8 | 0.0648 | 0.101 |
| chr2 | 159474802 | 159477617 | 2816 | 0.32 | 0.0648 | 0.101 |
| chr3 | 148708704 | 148710394 | 1691 | 0.49 | 0.0648 | 0.101 |
| chr4 | 18021722 | 18024520 | 2799 | -0.36 | 0.0649 | 0.101 |
| chr12 | 48146989 | 48224061 | 77073 | 0.08 | 0.0649 | 0.101 |
| chr3 | 136580163 | 136582776 | 2614 | 0.4 | 0.0649 | 0.101 |
| chr16 | 30499745 | 30504401 | 4657 | -0.39 | 0.0649 | 0.101 |
| chr13 | 115044344 | 115048548 | 4205 | 0.27 | 0.0649 | 0.101 |
| chr11 | 109991978 | 109999566 | 7589 | -0.55 | 0.065 | 0.101 |
| chr20 | 2848570 | 2855288 | 6719 | -0.43 | 0.065 | 0.101 |
| chr6 | 107076594 | 107078888 | 2295 | 0.34 | 0.065 | 0.101 |
| chr4 | 33748467 | 33749163 | 697 | -0.96 | 0.0651 | 0.101 |
| chrX | 47341432 | 47343781 | 2350 | 0.33 | 0.0651 | 0.101 |
| chr16 | 71264110 | 71265068 | 959 | -0.94 | 0.0651 | 0.101 |
| chr13 | 31720479 | 31747586 | 27108 | 0.18 | 0.0652 | 0.101 |
| chr9 | 5875797 | 5894625 | 18829 | -0.37 | 0.0652 | 0.101 |
| chr8 | 144977869 | 144990081 | 12213 | 0.35 | 0.0652 | 0.101 |
| chr21 | 42536793 | 42540801 | 4009 | 0.54 | 0.0652 | 0.101 |
| chr3 | 46919688 | 46921144 | 1457 | 0.66 | 0.0652 | 0.101 |
| chr4 | 78500128 | 78501272 | 1145 | 0.98 | 0.0653 | 0.101 |
| chr6 | 36874170 | 36882874 | 8705 | 0.63 | 0.0653 | 0.102 |
| chr20 | 7960659 | 7970222 | 9564 | 0.92 | 0.0654 | 0.102 |
| chr9 | 139538813 | 139560101 | 21289 | -0.86 | 0.0654 | 0.102 |
| chr14 | 45551571 | 45587641 | 36071 | 0.14 | 0.0654 | 0.102 |
| chr1 | 63832660 | 63834157 | 1498 | 0.51 | 0.0654 | 0.102 |

| | | | | | | |
|---|---|---|---|---|---|---|
| chr17 | 45317036 | 45318045 | 1010 | 0.66 | 0.0655 | 0.102 |
| chr17 | 72375821 | 72376791 | 971 | -1.24 | 0.0655 | 0.102 |
| chr7 | 66766253 | 66769027 | 2775 | 0.33 | 0.0655 | 0.102 |
| chr11 | 11334374 | 11335862 | 1489 | -0.65 | 0.0655 | 0.102 |
| chr2 | 143553385 | 143554281 | 897 | -0.97 | 0.0655 | 0.102 |
| chr8 | 124252426 | 124254027 | 1602 | 0.56 | 0.0656 | 0.102 |
| chr12 | 57984378 | 58005974 | 21597 | 0.26 | 0.0656 | 0.102 |
| chr20 | 18673007 | 18673437 | 431 | -1.46 | 0.0656 | 0.102 |
| chr7 | 93774530 | 93784629 | 10100 | -0.47 | 0.0656 | 0.102 |
| chr2 | 63173803 | 63174937 | 1135 | -1.04 | 0.0656 | 0.102 |
| chr1 | 211209961 | 211211478 | 1518 | -0.43 | 0.0658 | 0.102 |
| chr4 | 159917505 | 159932644 | 15140 | 0.33 | 0.0658 | 0.102 |
| chr10 | 114914354 | 114914924 | 571 | 1.94 | 0.0659 | 0.102 |
| chr3 | 72599049 | 72599666 | 618 | 1.13 | 0.0659 | 0.102 |
| chr11 | 36521792 | 36532554 | 10763 | 0.27 | 0.0659 | 0.102 |
| chr6 | 3156905 | 3164116 | 7212 | 0.3 | 0.066 | 0.102 |
| chr19 | 14800167 | 14801554 | 1388 | 0.55 | 0.066 | 0.102 |
| chr3 | 53230236 | 53231202 | 967 | 0.8 | 0.066 | 0.103 |
| chr4 | 2043070 | 2044079 | 1010 | -0.85 | 0.0661 | 0.103 |
| chr5 | 37738480 | 37739304 | 825 | 0.95 | 0.0661 | 0.103 |
| chr17 | 49512086 | 49517381 | 5296 | 3.27 | 0.0661 | 0.103 |
| chr8 | 105597477 | 105602217 | 4741 | 0.3 | 0.0662 | 0.103 |
| chr11 | 57466492 | 57482718 | 16227 | 0.19 | 0.0662 | 0.103 |
| chr3 | 178068422 | 178071275 | 2854 | 0.45 | 0.0662 | 0.103 |
| chr7 | 105028070 | 105043870 | 15801 | -0.26 | 0.0662 | 0.103 |
| chr19 | 42965997 | 42970641 | 4645 | -2.8 | 0.0662 | 0.103 |
| chr1 | 110105473 | 110106525 | 1053 | -1.12 | 0.0662 | 0.103 |
| chr14 | 67590941 | 67592731 | 1791 | 3.09 | 0.0662 | 0.103 |
| chr18 | 21407145 | 21409556 | 2412 | 0.34 | 0.0662 | 0.103 |
| chr21 | 33260587 | 33264382 | 3796 | 0.23 | 0.0662 | 0.103 |
| chr16 | 717405 | 740674 | 23270 | -0.24 | 0.0662 | 0.103 |
| chr16 | 9030066 | 9031110 | 1045 | -0.93 | 0.0663 | 0.103 |
| chr17 | 44999891 | 45002070 | 2180 | 0.23 | 0.0663 | 0.103 |
| chr4 | 103138148 | 103144590 | 6443 | 0.66 | 0.0663 | 0.103 |
| chr3 | 197193436 | 197196417 | 2982 | -0.68 | 0.0663 | 0.103 |
| chr18 | 30004932 | 30006456 | 1525 | 0.81 | 0.0663 | 0.103 |
| chr13 | 106748400 | 106757361 | 8962 | 0.23 | 0.0663 | 0.103 |
| chr8 | 37935237 | 37935981 | 745 | -1.23 | 0.0663 | 0.103 |
| chr6 | 152587323 | 152593653 | 6331 | -0.47 | 0.0663 | 0.103 |
| chr9 | 105945932 | 105946581 | 650 | -1.32 | 0.0664 | 0.103 |
| chr18 | 24197423 | 24198623 | 1201 | 0.8 | 0.0664 | 0.103 |
| chr1 | 21044651 | 21045214 | 564 | -1.36 | 0.0664 | 0.103 |
| chr19 | 13509481 | 13510696 | 1216 | -0.66 | 0.0665 | 0.103 |
| chr15 | 37473015 | 37474188 | 1174 | 3 | 0.0665 | 0.103 |
| chrX | 23956527 | 23957462 | 936 | -2.75 | 0.0665 | 0.103 |
| chr15 | 34651423 | 34660741 | 9319 | 0.26 | 0.0665 | 0.103 |
| chr2 | 99373890 | 99385956 | 12067 | 0.42 | 0.0665 | 0.103 |
| chr22 | 43200274 | 43203309 | 3036 | 0.35 | 0.0666 | 0.103 |
| chr3 | 61575258 | 61575939 | 682 | -1.48 | 0.0666 | 0.103 |
| chr4 | 129348611 | 129350233 | 1623 | -0.54 | 0.0666 | 0.103 |
| chr1 | 183027952 | 183067231 | 39280 | -0.22 | 0.0666 | 0.103 |
| chr9 | 135752771 | 135761972 | 9202 | -0.37 | 0.0666 | 0.103 |
| chr13 | 30880430 | 30883357 | 2928 | -0.37 | 0.0666 | 0.103 |
| chr14 | 73474843 | 73494608 | 19766 | -0.38 | 0.0666 | 0.103 |
| chrX | 96487623 | 96494748 | 7126 | 0.38 | 0.0667 | 0.103 |
| chr13 | 42802100 | 42803914 | 1815 | 0.52 | 0.0668 | 0.104 |
| chr5 | 150004205 | 150036858 | 32654 | -0.11 | 0.0669 | 0.104 |
| chr7 | 139748860 | 139749636 | 777 | 0.9 | 0.0669 | 0.104 |
| chr1 | 202760334 | 202780754 | 20421 | 0.23 | 0.0669 | 0.104 |
| chr16 | 87379166 | 87379827 | 662 | -1.31 | 0.0669 | 0.104 |
| chr17 | 79255604 | 79284290 | 28687 | -0.2 | 0.067 | 0.104 |

| | | | | | | |
|---|---|---|---|---|---|---|
| chr1 | 31750491 | 31750918 | 428 | -1.59 | 0.067 | 0.104 |
| chr8 | 81446178 | 81447601 | 1424 | 0.8 | 0.067 | 0.104 |
| chr6 | 109380110 | 109381410 | 1301 | 1.04 | 0.067 | 0.104 |
| chr9 | 96792250 | 96793961 | 1712 | -0.36 | 0.0671 | 0.104 |
| chr4 | 48699690 | 48703000 | 3311 | 0.28 | 0.0672 | 0.104 |
| chr19 | 55606769 | 55607708 | 940 | -1.16 | 0.0672 | 0.104 |
| chr10 | 73787487 | 73808371 | 20885 | 0.37 | 0.0672 | 0.104 |
| chr2 | 234155561 | 234169953 | 14393 | -0.2 | 0.0672 | 0.104 |
| chr21 | 30688285 | 30689395 | 1111 | -1.3 | 0.0673 | 0.104 |
| chr2 | 217318167 | 217318819 | 653 | -1.49 | 0.0673 | 0.104 |
| chr8 | 67960634 | 67978175 | 17542 | 0.18 | 0.0673 | 0.104 |
| chr2 | 101222850 | 101223299 | 450 | 1.33 | 0.0673 | 0.104 |
| chr1 | 236029075 | 236031533 | 2459 | 0.47 | 0.0674 | 0.104 |
| chr8 | 28012921 | 28013769 | 849 | -1.01 | 0.0674 | 0.104 |
| chr8 | 37100875 | 37107993 | 7119 | 1.1 | 0.0674 | 0.104 |
| chr9 | 14692121 | 14693795 | 1675 | 0.49 | 0.0674 | 0.104 |
| chr8 | 41674148 | 41686891 | 12744 | -0.29 | 0.0674 | 0.104 |
| chr11 | 74129012 | 74130272 | 1261 | 0.71 | 0.0675 | 0.104 |
| chr1 | 230778204 | 230780767 | 2564 | 0.44 | 0.0675 | 0.104 |
| chr3 | 132227175 | 132227711 | 537 | -1.33 | 0.0675 | 0.104 |
| chr4 | 56304989 | 56311912 | 6924 | 1.11 | 0.0675 | 0.104 |
| chr13 | 65296681 | 65298579 | 1899 | -0.42 | 0.0675 | 0.105 |
| chrX | 84258195 | 84259867 | 1673 | 0.38 | 0.0676 | 0.105 |
| chr12 | 52932420 | 52933412 | 993 | 1.11 | 0.0677 | 0.105 |
| chr9 | 133507766 | 133508964 | 1199 | -1.17 | 0.0677 | 0.105 |
| chr3 | 53074012 | 53080908 | 6897 | 0.54 | 0.0677 | 0.105 |
| chr11 | 19138448 | 19139770 | 1323 | 0.47 | 0.0678 | 0.105 |
| chr2 | 232872969 | 232874795 | 1827 | -0.56 | 0.068 | 0.105 |
| chr6 | 42768215 | 42769195 | 981 | 0.81 | 0.0681 | 0.105 |
| chr14 | 61899790 | 61906773 | 6984 | -0.29 | 0.0681 | 0.105 |
| chr16 | 15149005 | 15156364 | 7360 | -0.48 | 0.0682 | 0.105 |
| chr12 | 127604080 | 127615511 | 11432 | 0.19 | 0.0682 | 0.106 |
| chr5 | 162834137 | 162834862 | 726 | 1.12 | 0.0683 | 0.106 |
| chr2 | 145914324 | 145916663 | 2340 | -0.54 | 0.0683 | 0.106 |
| chr3 | 185890291 | 185891451 | 1161 | -1.12 | 0.0683 | 0.106 |
| chr10 | 95505995 | 95513430 | 7436 | 3.26 | 0.0683 | 0.106 |
| chr2 | 153468063 | 153469487 | 1425 | -0.65 | 0.0683 | 0.106 |
| chr3 | 114102394 | 114103399 | 1006 | -0.81 | 0.0684 | 0.106 |
| chr13 | 94427524 | 94428449 | 926 | 1.1 | 0.0684 | 0.106 |
| chr22 | 21354651 | 21369791 | 15141 | -0.31 | 0.0684 | 0.106 |
| chr5 | 68462002 | 68470454 | 8453 | 0.23 | 0.0684 | 0.106 |
| chr2 | 132250020 | 132251254 | 1235 | -0.61 | 0.0685 | 0.106 |
| chr8 | 143274735 | 143275682 | 948 | 3.02 | 0.0685 | 0.106 |
| chr7 | 13144036 | 13144582 | 547 | -1.32 | 0.0686 | 0.106 |
| chr1 | 201416938 | 201482461 | 65524 | 0.09 | 0.0686 | 0.106 |
| chr10 | 101927419 | 101928273 | 855 | 0.93 | 0.0686 | 0.106 |
| chr1 | 186504464 | 186505883 | 1420 | 0.64 | 0.0686 | 0.106 |
| chr7 | 26437573 | 26439134 | 1562 | 0.92 | 0.0687 | 0.106 |
| chr19 | 3760790 | 3763738 | 2949 | 0.34 | 0.0687 | 0.106 |
| chr7 | 4322173 | 4322981 | 809 | 0.9 | 0.0687 | 0.106 |
| chr2 | 136394694 | 136395251 | 558 | -1.14 | 0.0687 | 0.106 |
| chr18 | 54040139 | 54042803 | 2665 | 0.26 | 0.0688 | 0.106 |
| chr19 | 42578797 | 42593487 | 14691 | 0.13 | 0.0688 | 0.106 |
| chrX | 75391731 | 75394513 | 2783 | 0.28 | 0.069 | 0.107 |
| chr3 | 185462690 | 185464245 | 1556 | 0.85 | 0.069 | 0.107 |
| chr8 | 86088364 | 86091693 | 3330 | -0.22 | 0.069 | 0.107 |
| chr10 | 78915496 | 78926155 | 10660 | 0.2 | 0.0691 | 0.107 |
| chr3 | 185215564 | 185216783 | 1220 | 0.65 | 0.0691 | 0.107 |
| chr7 | 69238473 | 69239427 | 955 | -1.11 | 0.0691 | 0.107 |
| chr4 | 7760195 | 7762825 | 2631 | -0.51 | 0.0692 | 0.107 |
| chr15 | 55512769 | 55514251 | 1483 | 0.63 | 0.0692 | 0.107 |

| | | | | | | |
|---|---|---|---|---|---|---|
| chr10 | 7819310 | 7841877 | 22568 | 0.25 | 0.0692 | 0.107 |
| chr3 | 158188810 | 158190373 | 1564 | 0.64 | 0.0692 | 0.107 |
| chr16 | 66960247 | 66976508 | 16262 | -0.24 | 0.0692 | 0.107 |
| chr7 | 151083428 | 151084023 | 596 | -1.21 | 0.0693 | 0.107 |
| chr9 | 92061094 | 92072956 | 11863 | -3.26 | 0.0693 | 0.107 |
| chr17 | 63556882 | 63557950 | 1069 | -0.45 | 0.0693 | 0.107 |
| chr12 | 21614421 | 21615029 | 609 | -1.39 | 0.0693 | 0.107 |
| chr13 | 25074800 | 25087633 | 12834 | 0.15 | 0.0693 | 0.107 |
| chr6 | 153486846 | 153492894 | 6049 | -0.16 | 0.0693 | 0.107 |
| chr12 | 27615030 | 27615746 | 717 | 0.75 | 0.0693 | 0.107 |
| chr2 | 99796844 | 99799082 | 2239 | -0.39 | 0.0693 | 0.107 |
| chr20 | 37376237 | 37377809 | 1573 | 0.71 | 0.0693 | 0.107 |
| chr9 | 80850000 | 80852026 | 2027 | 0.34 | 0.0694 | 0.107 |
| chr1 | 55322337 | 55350048 | 27712 | 0.32 | 0.0694 | 0.107 |
| chr1 | 38356281 | 38357012 | 732 | 1.2 | 0.0694 | 0.107 |
| chr6 | 75243373 | 75246537 | 3165 | 0.22 | 0.0694 | 0.107 |
| chr13 | 50180811 | 50181329 | 519 | -0.95 | 0.0694 | 0.107 |
| chr12 | 102303003 | 102303525 | 523 | 1.37 | 0.0694 | 0.107 |
| chr6 | 109701789 | 109706239 | 4451 | 0.28 | 0.0694 | 0.107 |
| chr6 | 153298515 | 153306870 | 8356 | 0.26 | 0.0695 | 0.107 |
| chr9 | 4698394 | 4698806 | 413 | -1.84 | 0.0695 | 0.107 |
| chr2 | 179387306 | 179388879 | 1574 | -0.52 | 0.0695 | 0.107 |
| chr12 | 21629066 | 21629594 | 529 | -1.22 | 0.0696 | 0.107 |
| chr19 | 52097083 | 52098399 | 1317 | -0.59 | 0.0696 | 0.107 |
| chr2 | 37310542 | 37312189 | 1648 | 0.47 | 0.0696 | 0.107 |
| chr16 | 55604908 | 55607714 | 2807 | 2.08 | 0.0697 | 0.107 |
| chr2 | 68693474 | 68695460 | 1987 | 0.3 | 0.0697 | 0.107 |
| chr2 | 165697175 | 165698968 | 1794 | 0.56 | 0.0697 | 0.107 |
| chr6 | 10919904 | 10920429 | 526 | -1.09 | 0.0698 | 0.108 |
| chr22 | 46226421 | 46230424 | 4004 | -0.49 | 0.0698 | 0.108 |
| chr9 | 35470301 | 35471743 | 1443 | 0.91 | 0.0699 | 0.108 |
| chr20 | 42375007 | 42375718 | 712 | -1.25 | 0.0699 | 0.108 |
| chr9 | 98979558 | 98980864 | 1307 | -0.69 | 0.0699 | 0.108 |
| chrX | 66763669 | 66785079 | 21411 | -0.22 | 0.0699 | 0.108 |
| chr5 | 155956593 | 155957454 | 862 | 0.92 | 0.0701 | 0.108 |
| chr6 | 135857258 | 135857950 | 693 | -1.06 | 0.0701 | 0.108 |
| chr11 | 27716402 | 27723539 | 7138 | -0.25 | 0.0701 | 0.108 |
| chr11 | 103150131 | 103151361 | 1231 | -0.84 | 0.0702 | 0.108 |
| chr7 | 24612339 | 24620826 | 8488 | 0.51 | 0.0702 | 0.108 |
| chr17 | 67534507 | 67537115 | 2609 | -0.48 | 0.0703 | 0.108 |
| chr22 | 19096691 | 19110659 | 13969 | 0.31 | 0.0703 | 0.108 |
| chr2 | 58860769 | 58862800 | 2032 | 0.4 | 0.0703 | 0.108 |
| chr10 | 5760727 | 5764953 | 4227 | -0.6 | 0.0703 | 0.108 |
| chr1 | 24643308 | 24643740 | 433 | -1.36 | 0.0704 | 0.109 |
| chr7 | 7951466 | 7951837 | 372 | 1.28 | 0.0705 | 0.109 |
| chr3 | 170679741 | 170680075 | 335 | 1.65 | 0.0705 | 0.109 |
| chr10 | 49459458 | 49459987 | 530 | -1.36 | 0.0706 | 0.109 |
| chr20 | 44439690 | 44463312 | 23623 | -0.27 | 0.0706 | 0.109 |
| chr11 | 20427371 | 20429406 | 2036 | -0.55 | 0.0706 | 0.109 |
| chr2 | 160966681 | 160967556 | 876 | -1.18 | 0.0706 | 0.109 |
| chr7 | 68895199 | 68897249 | 2051 | -0.73 | 0.0706 | 0.109 |
| chr14 | 67999331 | 68000621 | 1291 | -0.61 | 0.0706 | 0.109 |
| chr7 | 138389937 | 138395710 | 5774 | 0.67 | 0.0706 | 0.109 |
| chr5 | 112255962 | 112264737 | 8776 | 0.32 | 0.0707 | 0.109 |
| chr17 | 41778088 | 41778386 | 299 | 1.51 | 0.0707 | 0.109 |
| chr3 | 194391899 | 194394132 | 2234 | 0.28 | 0.0707 | 0.109 |
| chr2 | 15773157 | 15773710 | 554 | -1.48 | 0.0708 | 0.109 |
| chr1 | 108114487 | 108115945 | 1459 | 0.8 | 0.0708 | 0.109 |
| chr17 | 21261285 | 21262866 | 1582 | 0.84 | 0.0709 | 0.109 |
| chr3 | 121178503 | 121204006 | 25504 | 0.24 | 0.0709 | 0.109 |
| chr11 | 68775521 | 68777911 | 2391 | 0.54 | 0.0709 | 0.109 |

| | | | | | | |
|---|---|---|---|---|---|---|
| chr3 | 20119475 | 20120349 | 875 | -1.23 | 0.0709 | 0.109 |
| chr8 | 95270874 | 95279279 | 8406 | 0.29 | 0.0709 | 0.109 |
| chr12 | 9564046 | 9564333 | 288 | 1.48 | 0.0709 | 0.109 |
| chr10 | 60679375 | 60681105 | 1731 | 0.54 | 0.0709 | 0.109 |
| chr2 | 224766705 | 224774428 | 7724 | -0.46 | 0.071 | 0.109 |
| chr3 | 181499383 | 181501532 | 2150 | 0.3 | 0.0711 | 0.109 |
| chr17 | 27104560 | 27105154 | 595 | -1.22 | 0.0711 | 0.109 |
| chr17 | 57478068 | 57485549 | 7482 | 0.67 | 0.0711 | 0.109 |
| chr11 | 43851089 | 43853409 | 2321 | 0.45 | 0.0712 | 0.109 |
| chr2 | 190438987 | 190446148 | 7162 | 0.39 | 0.0712 | 0.11 |
| chr2 | 54038644 | 54039252 | 609 | -1.05 | 0.0712 | 0.11 |
| chr8 | 22274614 | 22299958 | 25345 | -0.21 | 0.0712 | 0.11 |
| chr16 | 19696809 | 19730737 | 33929 | 0.16 | 0.0713 | 0.11 |
| chr2 | 26565562 | 26580599 | 15038 | 0.14 | 0.0713 | 0.11 |
| chr4 | 164471147 | 164471560 | 414 | 1.03 | 0.0714 | 0.11 |
| chr13 | 109089781 | 109090349 | 569 | -1.77 | 0.0714 | 0.11 |
| chr4 | 103638219 | 103639355 | 1137 | 1 | 0.0714 | 0.11 |
| chr12 | 112721541 | 112723063 | 1523 | -0.92 | 0.0715 | 0.11 |
| chr8 | 17806250 | 17820924 | 14675 | 0.44 | 0.0715 | 0.11 |
| chr1 | 78143669 | 78154115 | 10447 | 0.25 | 0.0715 | 0.11 |
| chr14 | 50599999 | 50615380 | 15382 | -0.27 | 0.0715 | 0.11 |
| chr11 | 57296959 | 57299255 | 2297 | 0.29 | 0.0716 | 0.11 |
| chr16 | 558650 | 559963 | 1314 | 0.67 | 0.0716 | 0.11 |
| chr19 | 17940560 | 17977055 | 36496 | 0.23 | 0.0717 | 0.11 |
| chr8 | 26101125 | 26101888 | 764 | 1.58 | 0.0717 | 0.11 |
| chr12 | 79406228 | 79407363 | 1136 | 0.7 | 0.0717 | 0.11 |
| chr2 | 56150301 | 56151213 | 913 | 1.14 | 0.0719 | 0.11 |
| chr3 | 186856461 | 186857617 | 1157 | 0.45 | 0.0719 | 0.11 |
| chr8 | 134582085 | 134585465 | 3381 | 0.31 | 0.0719 | 0.111 |
| chr11 | 27255296 | 27256326 | 1031 | -1.06 | 0.0719 | 0.111 |
| chr6 | 41287724 | 41289596 | 1873 | -0.53 | 0.0719 | 0.111 |
| chr8 | 40297838 | 40305848 | 8011 | -0.51 | 0.072 | 0.111 |
| chr2 | 133741682 | 133742704 | 1023 | -1.23 | 0.072 | 0.111 |
| chr2 | 159251384 | 159252491 | 1108 | 0.82 | 0.072 | 0.111 |
| chr1 | 48174572 | 48175770 | 1199 | 1.14 | 0.072 | 0.111 |
| chr17 | 78056246 | 78056882 | 637 | -0.96 | 0.072 | 0.111 |
| chr16 | 89976199 | 89992443 | 16245 | -0.19 | 0.0721 | 0.111 |
| chr9 | 129701699 | 129702176 | 478 | 1.82 | 0.0721 | 0.111 |
| chr13 | 77787175 | 77788528 | 1354 | 0.82 | 0.0721 | 0.111 |
| chr1 | 178340553 | 178341062 | 510 | 1.18 | 0.0721 | 0.111 |
| chr12 | 88421592 | 88436312 | 14721 | 0.34 | 0.0721 | 0.111 |
| chr10 | 121061804 | 121085150 | 23347 | -0.12 | 0.0721 | 0.111 |
| chr7 | 1126155 | 1126699 | 545 | -0.91 | 0.0722 | 0.111 |
| chr11 | 72003116 | 72004481 | 1366 | -1.27 | 0.0723 | 0.111 |
| chr4 | 2262679 | 2265002 | 2324 | -0.52 | 0.0724 | 0.111 |
| chr4 | 189075885 | 189077112 | 1228 | -0.45 | 0.0724 | 0.111 |
| chr1 | 154832263 | 154833142 | 880 | -1.79 | 0.0724 | 0.111 |
| chr17 | 59833998 | 59835474 | 1477 | 0.68 | 0.0724 | 0.111 |
| chr11 | 111554725 | 111555600 | 876 | -1.35 | 0.0724 | 0.111 |
| chr1 | 68697754 | 68698675 | 922 | 1.14 | 0.0724 | 0.111 |
| chr3 | 181465401 | 181465962 | 562 | 1.15 | 0.0725 | 0.111 |
| chr13 | 74656554 | 74657378 | 825 | 1.1 | 0.0725 | 0.111 |
| chr2 | 206809275 | 206810159 | 885 | 1.26 | 0.0726 | 0.111 |
| chr11 | 119564355 | 119567665 | 3311 | -0.65 | 0.0726 | 0.111 |
| chr10 | 111766801 | 111769154 | 2354 | 0.56 | 0.0726 | 0.111 |
| chr11 | 71774115 | 71824920 | 50806 | 0.1 | 0.0727 | 0.112 |
| chr10 | 25303962 | 25307233 | 3272 | -0.24 | 0.0727 | 0.112 |
| chr19 | 10490371 | 10492045 | 1675 | 0.59 | 0.0727 | 0.112 |
| chr4 | 5754572 | 5775588 | 21017 | -0.3 | 0.0727 | 0.112 |
| chr9 | 99344776 | 99345273 | 498 | 1.26 | 0.0727 | 0.112 |
| chr14 | 69468774 | 69481902 | 13129 | -0.42 | 0.0727 | 0.112 |

| | | | | | | |
|---|---|---|---|---|---|---|
| chr11 | 64879433 | 64903316 | 23884 | 0.14 | 0.0728 | 0.112 |
| chr8 | 94737863 | 94768325 | 30463 | -0.14 | 0.0728 | 0.112 |
| chr17 | 33463073 | 33469734 | 6662 | -0.39 | 0.0728 | 0.112 |
| chr12 | 17436708 | 17437534 | 827 | -0.87 | 0.0729 | 0.112 |
| chr17 | 26605690 | 26606074 | 385 | 1.19 | 0.0729 | 0.112 |
| chr16 | 27984049 | 27984470 | 422 | 1.52 | 0.073 | 0.112 |
| chr9 | 107845267 | 107846090 | 824 | 0.95 | 0.073 | 0.112 |
| chr9 | 113112978 | 113126842 | 13865 | -3.07 | 0.073 | 0.112 |
| chr7 | 79763336 | 79789631 | 26296 | 0.16 | 0.073 | 0.112 |
| chr9 | 3920132 | 3921602 | 1471 | 0.53 | 0.073 | 0.112 |
| chr18 | 71901160 | 71902703 | 1544 | 0.76 | 0.073 | 0.112 |
| chr12 | 108055207 | 108056591 | 1385 | -0.8 | 0.073 | 0.112 |
| chr7 | 139575609 | 139576563 | 955 | 1 | 0.0731 | 0.112 |
| chr7 | 91728066 | 91728779 | 714 | 1.65 | 0.0731 | 0.112 |
| chr7 | 99592031 | 99633335 | 41305 | 0.19 | 0.0732 | 0.112 |
| chr4 | 17578101 | 17579899 | 1799 | 0.51 | 0.0732 | 0.112 |
| chr5 | 171366772 | 171408235 | 41464 | -0.18 | 0.0732 | 0.112 |
| chr3 | 99872850 | 99873794 | 945 | -0.8 | 0.0732 | 0.112 |
| chr17 | 78387958 | 78390364 | 2407 | -0.24 | 0.0732 | 0.112 |
| chr4 | 3746840 | 3748177 | 1338 | 0.54 | 0.0732 | 0.112 |
| chr15 | 77196777 | 77198349 | 1573 | 0.47 | 0.0733 | 0.112 |
| chr3 | 178944742 | 178945534 | 793 | 1.25 | 0.0733 | 0.112 |
| chr3 | 136193532 | 136193960 | 429 | 1.48 | 0.0733 | 0.112 |
| chr4 | 95746874 | 95764582 | 17709 | 0.27 | 0.0733 | 0.112 |
| chr7 | 86813659 | 86815190 | 1532 | 0.75 | 0.0733 | 0.112 |
| chr9 | 27018930 | 27020467 | 1538 | -0.52 | 0.0734 | 0.112 |
| chr17 | 48543593 | 48561608 | 18016 | 0.22 | 0.0734 | 0.113 |
| chr22 | 21131054 | 21133417 | 2364 | 0.41 | 0.0735 | 0.113 |
| chr2 | 47685882 | 47686535 | 654 | -0.97 | 0.0735 | 0.113 |
| chr12 | 12223638 | 12236192 | 12555 | 0.31 | 0.0736 | 0.113 |
| chr6 | 149400692 | 149401433 | 742 | -1.05 | 0.0736 | 0.113 |
| chr8 | 29027029 | 29044828 | 17800 | 0.2 | 0.0736 | 0.113 |
| chrX | 47695418 | 47697041 | 1624 | 0.39 | 0.0736 | 0.113 |
| chr2 | 205591403 | 205592080 | 678 | -1.11 | 0.0736 | 0.113 |
| chr1 | 118196997 | 118198353 | 1357 | -0.67 | 0.0736 | 0.113 |
| chr1 | 236957736 | 236959782 | 2047 | 0.39 | 0.0736 | 0.113 |
| chr1 | 185408038 | 185424174 | 16137 | -0.31 | 0.0737 | 0.113 |
| chr6 | 79765443 | 79766284 | 842 | 1.22 | 0.0737 | 0.113 |
| chr1 | 44172169 | 44174612 | 2444 | 0.36 | 0.0737 | 0.113 |
| chr2 | 11968947 | 11971424 | 2478 | 0.35 | 0.0737 | 0.113 |
| chr12 | 41185936 | 41196942 | 11007 | 3.48 | 0.0737 | 0.113 |
| chr1 | 46932065 | 46932505 | 441 | -1.39 | 0.0737 | 0.113 |
| chr14 | 93403674 | 93414312 | 10639 | 0.28 | 0.0737 | 0.113 |
| chr21 | 45225958 | 45226830 | 873 | -1.03 | 0.0738 | 0.113 |
| chr16 | 30064076 | 30108449 | 44374 | 0.11 | 0.0738 | 0.113 |
| chr5 | 18732964 | 18747045 | 14082 | 0.55 | 0.0738 | 0.113 |
| chr17 | 30320279 | 30320747 | 469 | -1.14 | 0.0738 | 0.113 |
| chr10 | 133836173 | 133837120 | 948 | 0.88 | 0.0738 | 0.113 |
| chr2 | 192053992 | 192054649 | 658 | 1.38 | 0.0738 | 0.113 |
| chr12 | 68725128 | 68726771 | 1644 | 0.49 | 0.0738 | 0.113 |
| chr14 | 74398397 | 74401047 | 2651 | -0.66 | 0.0739 | 0.113 |
| chr2 | 70630270 | 70631406 | 1137 | -1.05 | 0.0739 | 0.113 |
| chr6 | 109063199 | 109066333 | 3135 | 0.73 | 0.0739 | 0.113 |
| chr3 | 113665820 | 113666611 | 792 | 0.93 | 0.074 | 0.113 |
| chr13 | 45095824 | 45099515 | 3692 | 0.22 | 0.074 | 0.113 |
| chr2 | 174940345 | 174941263 | 919 | -0.92 | 0.074 | 0.113 |
| chr1 | 156932759 | 156933859 | 1101 | 0.94 | 0.0741 | 0.113 |
| chr1 | 168004196 | 168005415 | 1220 | 0.69 | 0.0741 | 0.113 |
| chr9 | 128651909 | 128652380 | 472 | 1.24 | 0.0741 | 0.113 |
| chr5 | 8456581 | 8458641 | 2061 | 0.46 | 0.0741 | 0.113 |
| chr10 | 127836589 | 127837778 | 1190 | -0.98 | 0.0741 | 0.113 |

| | | | | | | |
|---|---|---|---|---|---|---|
| chr1 | 26146012 | 26151510 | 5499 | -0.48 | 0.0741 | 0.113 |
| chr9 | 128320738 | 128321315 | 578 | 1.23 | 0.0741 | 0.113 |
| chr5 | 61833866 | 61848684 | 14819 | 0.41 | 0.0741 | 0.113 |
| chr14 | 101928494 | 101949594 | 21101 | 0.1 | 0.0742 | 0.113 |
| chr14 | 77474276 | 77474964 | 689 | -1.22 | 0.0742 | 0.113 |
| chr16 | 31062661 | 31087194 | 24534 | -0.15 | 0.0742 | 0.113 |
| chrX | 47020385 | 47021635 | 1251 | -1.11 | 0.0742 | 0.113 |
| chr22 | 47467122 | 47467711 | 590 | -1.27 | 0.0742 | 0.113 |
| chr2 | 66660267 | 66663090 | 2824 | 0.41 | 0.0743 | 0.114 |
| chr11 | 103903290 | 103905100 | 1811 | 0.5 | 0.0743 | 0.114 |
| chr5 | 157344594 | 157345077 | 484 | -1.38 | 0.0743 | 0.114 |
| chr11 | 26641801 | 26645173 | 3373 | 3.43 | 0.0743 | 0.114 |
| chr15 | 101930621 | 101931060 | 440 | -1.6 | 0.0743 | 0.114 |
| chr2 | 54106219 | 54106986 | 768 | -1.08 | 0.0744 | 0.114 |
| chr1 | 54384491 | 54385191 | 701 | 1.28 | 0.0744 | 0.114 |
| chr9 | 14148093 | 14149826 | 1734 | -0.49 | 0.0744 | 0.114 |
| chr16 | 70312761 | 70339351 | 26591 | 0.13 | 0.0744 | 0.114 |
| chr5 | 143094325 | 143094937 | 613 | -1.77 | 0.0745 | 0.114 |
| chr11 | 105892123 | 105893986 | 1864 | 0.46 | 0.0745 | 0.114 |
| chr19 | 28282467 | 28287097 | 4631 | 0.12 | 0.0746 | 0.114 |
| chr2 | 206968821 | 206974161 | 5341 | -0.34 | 0.0746 | 0.114 |
| chr6 | 116691396 | 116693424 | 2029 | 0.43 | 0.0746 | 0.114 |
| chr14 | 105348991 | 105349340 | 350 | 1.46 | 0.0746 | 0.114 |
| chr3 | 171488770 | 171489384 | 615 | 1.23 | 0.0747 | 0.114 |
| chr9 | 130006915 | 130008250 | 1336 | -0.55 | 0.0747 | 0.114 |
| chr17 | 29484949 | 29486461 | 1513 | 0.65 | 0.0747 | 0.114 |
| chr16 | 29167685 | 29169272 | 1588 | 0.61 | 0.0747 | 0.114 |
| chr3 | 134145113 | 134147726 | 2614 | -0.42 | 0.0748 | 0.114 |
| chr22 | 19628979 | 19629615 | 637 | 1.25 | 0.0748 | 0.114 |
| chr9 | 130888732 | 130895260 | 6529 | 0.28 | 0.0748 | 0.114 |
| chr19 | 505980 | 507999 | 2020 | -0.31 | 0.0748 | 0.114 |
| chr10 | 15194147 | 15194822 | 676 | -1.33 | 0.0749 | 0.114 |
| chr2 | 26009049 | 26010630 | 1582 | 0.63 | 0.0749 | 0.114 |
| chr10 | 69539979 | 69615469 | 75491 | 0.08 | 0.0749 | 0.114 |
| chr7 | 2859961 | 2862394 | 2434 | -0.37 | 0.0749 | 0.114 |
| chr2 | 39596107 | 39607461 | 11355 | -0.69 | 0.075 | 0.114 |
| chr3 | 63841067 | 63850943 | 9877 | 0.26 | 0.075 | 0.114 |
| chr3 | 130644191 | 130645214 | 1024 | -0.82 | 0.075 | 0.115 |
| chr7 | 103787329 | 103787705 | 377 | 1.83 | 0.0751 | 0.115 |
| chr3 | 9884414 | 9886378 | 1965 | 0.61 | 0.0751 | 0.115 |
| chr20 | 10073112 | 10075796 | 2685 | -0.4 | 0.0752 | 0.115 |
| chr1 | 161493769 | 161501671 | 7903 | -0.56 | 0.0752 | 0.115 |
| chr5 | 14663737 | 14679668 | 15932 | 0.21 | 0.0753 | 0.115 |
| chr11 | 35817588 | 35820987 | 3400 | -0.64 | 0.0753 | 0.115 |
| chr1 | 103240210 | 103240550 | 341 | 1.26 | 0.0754 | 0.115 |
| chr12 | 77458548 | 77460481 | 1934 | 0.33 | 0.0754 | 0.115 |
| chr13 | 50341777 | 50342407 | 631 | 1.55 | 0.0754 | 0.115 |
| chr8 | 143859040 | 143859699 | 660 | 0.83 | 0.0754 | 0.115 |
| chr9 | 98075131 | 98080824 | 5694 | 0.2 | 0.0754 | 0.115 |
| chr7 | 100886738 | 100896347 | 9610 | 0.27 | 0.0755 | 0.115 |
| chr7 | 135092340 | 135093671 | 1332 | -0.85 | 0.0756 | 0.115 |
| chr8 | 81416340 | 81417280 | 941 | 1.78 | 0.0756 | 0.115 |
| chr11 | 7040507 | 7042110 | 1604 | 0.45 | 0.0756 | 0.115 |
| chr7 | 16459782 | 16462063 | 2282 | 0.24 | 0.0756 | 0.115 |
| chr2 | 19131158 | 19132697 | 1540 | -0.42 | 0.0757 | 0.115 |
| chr10 | 76676546 | 76697515 | 20970 | -0.32 | 0.0757 | 0.115 |
| chr19 | 5902511 | 5915322 | 12812 | 0.27 | 0.0757 | 0.115 |
| chr13 | 113858997 | 113880635 | 21639 | 0.18 | 0.0758 | 0.116 |
| chr6 | 99391619 | 99396539 | 4921 | 0.26 | 0.0758 | 0.116 |
| chr4 | 148119552 | 148120655 | 1104 | 1.03 | 0.0759 | 0.116 |
| chr2 | 39571659 | 39582799 | 11141 | -0.93 | 0.076 | 0.116 |

| | | | | | | |
|---|---|---|---|---|---|---|
| chr4 | 168818044 | 168818637 | 594 | -1.21 | 0.076 | 0.116 |
| chr3 | 69062052 | 69063658 | 1607 | 0.34 | 0.0761 | 0.116 |
| chr4 | 40122608 | 40123956 | 1349 | 0.77 | 0.0761 | 0.116 |
| chr10 | 134404332 | 134405771 | 1440 | 0.67 | 0.0761 | 0.116 |
| chr1 | 63248855 | 63250874 | 2020 | 0.53 | 0.0761 | 0.116 |
| chr8 | 1649100 | 1650004 | 905 | 0.7 | 0.0761 | 0.116 |
| chr3 | 87125932 | 87140688 | 14757 | 0.26 | 0.0762 | 0.116 |
| chrX | 39584616 | 39596175 | 11560 | 0.54 | 0.0762 | 0.116 |
| chr7 | 32789746 | 32790864 | 1119 | -0.64 | 0.0763 | 0.116 |
| chr3 | 157260629 | 157261659 | 1031 | 0.65 | 0.0763 | 0.116 |
| chr6 | 139307912 | 139310166 | 2255 | 0.41 | 0.0763 | 0.116 |
| chr2 | 63925294 | 63927228 | 1935 | -0.35 | 0.0763 | 0.116 |
| chr14 | 102358409 | 102360513 | 2105 | 0.67 | 0.0764 | 0.116 |
| chr8 | 42248154 | 42250737 | 2584 | -0.25 | 0.0764 | 0.116 |
| chr14 | 37866709 | 37867429 | 721 | -1.19 | 0.0764 | 0.116 |
| chr6 | 12061105 | 12072095 | 10991 | 0.54 | 0.0765 | 0.117 |
| chr3 | 177863697 | 177864684 | 988 | 0.82 | 0.0765 | 0.117 |
| chr6 | 167372368 | 167372854 | 487 | 1.67 | 0.0765 | 0.117 |
| chr8 | 133188602 | 133188951 | 350 | -1.74 | 0.0765 | 0.117 |
| chr1 | 199867680 | 199868572 | 893 | 0.8 | 0.0765 | 0.117 |
| chr15 | 43759868 | 43763420 | 3553 | -0.67 | 0.0766 | 0.117 |
| chr3 | 186982128 | 186982600 | 473 | 1.1 | 0.0766 | 0.117 |
| chr8 | 124151491 | 124155469 | 3979 | -0.51 | 0.0766 | 0.117 |
| chr8 | 95625350 | 95625791 | 442 | 1.69 | 0.0766 | 0.117 |
| chr11 | 93583141 | 93583769 | 629 | 0.84 | 0.0767 | 0.117 |
| chr20 | 30823518 | 30824361 | 844 | -0.96 | 0.0767 | 0.117 |
| chr11 | 74344280 | 74349012 | 4733 | -0.42 | 0.0767 | 0.117 |
| chr22 | 29345281 | 29346558 | 1278 | -1.38 | 0.0767 | 0.117 |
| chr4 | 136150792 | 136151855 | 1064 | 0.86 | 0.0767 | 0.117 |
| chr3 | 120460064 | 120469869 | 9806 | 0.18 | 0.0767 | 0.117 |
| chr2 | 30633158 | 30633971 | 814 | -1.03 | 0.0767 | 0.117 |
| chr6 | 170491832 | 170492385 | 554 | 1.2 | 0.0768 | 0.117 |
| chr9 | 19206956 | 19207429 | 474 | 1.84 | 0.0768 | 0.117 |
| chr6 | 43444661 | 43456880 | 12220 | 0.34 | 0.0769 | 0.117 |
| chr6 | 105993834 | 105994898 | 1065 | -0.83 | 0.0769 | 0.117 |
| chr22 | 26907575 | 26909932 | 2358 | 0.28 | 0.0769 | 0.117 |
| chr10 | 129151693 | 129153303 | 1611 | -0.79 | 0.0769 | 0.117 |
| chr17 | 55333019 | 55335353 | 2335 | -0.41 | 0.0769 | 0.117 |
| chr21 | 45759013 | 45760232 | 1220 | 0.39 | 0.077 | 0.117 |
| chr19 | 16553264 | 16553941 | 678 | 1.23 | 0.077 | 0.117 |
| chr3 | 64047456 | 64052138 | 4683 | -0.22 | 0.077 | 0.117 |
| chr1 | 23823763 | 23824334 | 572 | -0.92 | 0.077 | 0.117 |
| chr1 | 24967965 | 24971336 | 3372 | 0.34 | 0.077 | 0.117 |
| chr22 | 24261879 | 24262840 | 962 | -0.9 | 0.0771 | 0.117 |
| chr22 | 29058254 | 29059488 | 1235 | 0.44 | 0.0771 | 0.117 |
| chr3 | 9502368 | 9516489 | 14122 | -0.27 | 0.0771 | 0.117 |
| chr3 | 106143956 | 106144293 | 338 | 1.67 | 0.0771 | 0.117 |
| chr14 | 67397239 | 67401410 | 4172 | 0.2 | 0.0772 | 0.117 |
| chr2 | 169868744 | 169869244 | 501 | -1.66 | 0.0772 | 0.117 |
| chr15 | 101211783 | 101213273 | 1491 | -0.51 | 0.0772 | 0.117 |
| chr4 | 76438311 | 76440896 | 2586 | 0.26 | 0.0772 | 0.117 |
| chr1 | 64023326 | 64023737 | 412 | 1.28 | 0.0772 | 0.117 |
| chr3 | 32533148 | 32544936 | 11789 | 0.4 | 0.0773 | 0.118 |
| chr5 | 52187774 | 52189029 | 1256 | 0.84 | 0.0774 | 0.118 |
| chr7 | 94769155 | 94769596 | 442 | -1.32 | 0.0774 | 0.118 |
| chr12 | 72148048 | 72150051 | 2004 | 0.27 | 0.0775 | 0.118 |
| chr1 | 201923129 | 201926431 | 3303 | 0.26 | 0.0775 | 0.118 |
| chr8 | 30954042 | 30965075 | 11034 | 0.59 | 0.0776 | 0.118 |
| chr7 | 39698779 | 39699515 | 737 | -0.92 | 0.0776 | 0.118 |
| chr13 | 99738177 | 99740647 | 2471 | 0.51 | 0.0776 | 0.118 |
| chr11 | 10387680 | 10388417 | 738 | -0.71 | 0.0776 | 0.118 |

| | | | | | | |
|---|---|---|---|---|---|---|
| chr1 | 149933854 | 149952884 | 19031 | -0.25 | 0.0777 | 0.118 |
| chr15 | 91466000 | 91546977 | 80978 | 0.09 | 0.0778 | 0.118 |
| chrX | 133770847 | 133771335 | 489 | -1.76 | 0.0778 | 0.118 |
| chr7 | 98475703 | 98478221 | 2519 | 0.36 | 0.0778 | 0.118 |
| chr3 | 150344296 | 150345324 | 1029 | -0.95 | 0.0778 | 0.118 |
| chr17 | 77070020 | 77072248 | 2229 | 0.46 | 0.0778 | 0.118 |
| chr15 | 101252356 | 101274494 | 22139 | 0.14 | 0.0778 | 0.118 |
| chr2 | 233893003 | 233894211 | 1209 | -0.95 | 0.0779 | 0.118 |
| chr17 | 79618234 | 79619075 | 842 | -0.92 | 0.0779 | 0.118 |
| chr3 | 181898058 | 181898943 | 886 | 1.79 | 0.0779 | 0.118 |
| chr1 | 42360556 | 42368601 | 8046 | -0.25 | 0.0779 | 0.118 |
| chr10 | 830523 | 835802 | 5280 | 0.49 | 0.078 | 0.118 |
| chr18 | 55148024 | 55148952 | 929 | 0.87 | 0.078 | 0.118 |
| chr1 | 231057409 | 231058500 | 1092 | -0.82 | 0.078 | 0.118 |
| chr14 | 27277488 | 27291341 | 13854 | 3.49 | 0.0781 | 0.119 |
| chr3 | 117591853 | 117595489 | 3637 | 0.32 | 0.0781 | 0.119 |
| chr11 | 93411897 | 93426150 | 14254 | 0.33 | 0.0782 | 0.119 |
| chr2 | 25517411 | 25518452 | 1042 | -0.87 | 0.0782 | 0.119 |
| chr1 | 201138956 | 201140969 | 2014 | 0.31 | 0.0783 | 0.119 |
| chr4 | 68546234 | 68552097 | 5864 | -0.61 | 0.0783 | 0.119 |
| chr22 | 28113362 | 28125895 | 12534 | 0.22 | 0.0784 | 0.119 |
| chr16 | 845903 | 858346 | 12444 | -0.37 | 0.0785 | 0.119 |
| chr11 | 124632060 | 124641181 | 9122 | 0.37 | 0.0785 | 0.119 |
| chr9 | 129773905 | 129774968 | 1064 | 1.17 | 0.0785 | 0.119 |
| chr12 | 46209424 | 46209874 | 451 | -1.42 | 0.0785 | 0.119 |
| chr1 | 91481159 | 91488666 | 7508 | 0.19 | 0.0785 | 0.119 |
| chr1 | 202379495 | 202380427 | 933 | -0.92 | 0.0786 | 0.119 |
| chr19 | 13136624 | 13138249 | 1626 | 0.49 | 0.0786 | 0.119 |
| chr8 | 135652672 | 135654049 | 1378 | 0.67 | 0.0787 | 0.119 |
| chr17 | 65334368 | 65334895 | 528 | -1.66 | 0.0787 | 0.119 |
| chr19 | 5045611 | 5050610 | 5000 | -0.32 | 0.0787 | 0.119 |
| chr14 | 32327698 | 32328831 | 1134 | 0.85 | 0.0787 | 0.119 |
| chr19 | 41168367 | 41285658 | 117292 | -0.07 | 0.0787 | 0.119 |
| chr9 | 95104753 | 95105616 | 864 | -1.07 | 0.0788 | 0.119 |
| chr18 | 72548229 | 72549020 | 792 | 1.02 | 0.0788 | 0.119 |
| chr1 | 28743014 | 28745442 | 2429 | 0.4 | 0.0788 | 0.119 |
| chr2 | 89053411 | 89054181 | 771 | -1.04 | 0.0788 | 0.119 |
| chr1 | 25240916 | 25241965 | 1050 | 0.88 | 0.0788 | 0.12 |
| chr2 | 107131328 | 107132261 | 934 | -0.74 | 0.0788 | 0.12 |
| chr10 | 91173589 | 91175678 | 2090 | 0.35 | 0.0788 | 0.12 |
| chr14 | 24898202 | 24913088 | 14887 | 0.14 | 0.0789 | 0.12 |
| chr17 | 64649015 | 64804245 | 155231 | -0.09 | 0.0789 | 0.12 |
| chr9 | 72938798 | 72939909 | 1112 | 0.89 | 0.0789 | 0.12 |
| chr4 | 121842091 | 121844793 | 2703 | 0.27 | 0.0789 | 0.12 |
| chr1 | 167597053 | 167597803 | 751 | 1.38 | 0.079 | 0.12 |
| chr1 | 81863425 | 81865166 | 1742 | 2.78 | 0.079 | 0.12 |
| chr3 | 159745242 | 159746369 | 1128 | -0.86 | 0.079 | 0.12 |
| chr1 | 119821365 | 119823024 | 1660 | 0.59 | 0.079 | 0.12 |
| chr6 | 116599787 | 116602195 | 2409 | -0.27 | 0.0791 | 0.12 |
| chr16 | 24528565 | 24560041 | 31477 | 0.18 | 0.0791 | 0.12 |
| chr16 | 22199384 | 22257363 | 57980 | 0.11 | 0.0791 | 0.12 |
| chr2 | 239437344 | 239439127 | 1784 | -0.49 | 0.0791 | 0.12 |
| chr3 | 132255131 | 132273405 | 18275 | 0.41 | 0.0792 | 0.12 |
| chr12 | 80481059 | 80482432 | 1374 | -0.7 | 0.0793 | 0.12 |
| chr7 | 5085092 | 5086207 | 1116 | 0.48 | 0.0793 | 0.12 |
| chr1 | 162665757 | 162666790 | 1034 | 0.64 | 0.0793 | 0.12 |
| chr1 | 203856999 | 203885627 | 28629 | -0.15 | 0.0794 | 0.12 |
| chr16 | 2648670 | 2654045 | 5376 | 0.39 | 0.0794 | 0.12 |
| chr7 | 5630744 | 5648062 | 17319 | -0.2 | 0.0794 | 0.12 |
| chr19 | 7069454 | 7070530 | 1077 | 0.59 | 0.0794 | 0.12 |
| chr15 | 75912593 | 75965998 | 53406 | 0.09 | 0.0795 | 0.12 |

| | | | | | | |
|---|---|---|---|---|---|---|
| chr17 | 70115948 | 70131272 | 15325 | 0.33 | 0.0795 | 0.12 |
| chr12 | 48290189 | 48299694 | 9506 | 3.14 | 0.0795 | 0.12 |
| chr7 | 100063626 | 100076909 | 13284 | -0.31 | 0.0795 | 0.12 |
| chr15 | 41185988 | 41199927 | 13940 | 0.27 | 0.0795 | 0.12 |
| chr2 | 175971606 | 175977911 | 6306 | -0.55 | 0.0795 | 0.12 |
| chr20 | 47699817 | 47735907 | 36091 | 0.19 | 0.0796 | 0.121 |
| chr1 | 22378547 | 22381206 | 2660 | 0.34 | 0.0796 | 0.121 |
| chr5 | 41869927 | 41870984 | 1058 | 0.97 | 0.0797 | 0.121 |
| chr21 | 44738989 | 44776219 | 37231 | 0.23 | 0.0797 | 0.121 |
| chr16 | 53195653 | 53196939 | 1287 | 0.88 | 0.0797 | 0.121 |
| chr5 | 61601055 | 61610581 | 9527 | 0.23 | 0.0797 | 0.121 |
| chr20 | 55129328 | 55130943 | 1616 | 0.54 | 0.0798 | 0.121 |
| chr2 | 26464886 | 26468935 | 4050 | -0.24 | 0.0798 | 0.121 |
| chr4 | 54772292 | 54772668 | 377 | -1.77 | 0.0798 | 0.121 |
| chr7 | 46424532 | 46427298 | 2767 | 0.32 | 0.0798 | 0.121 |
| chr3 | 158283606 | 158289471 | 5866 | -0.36 | 0.0798 | 0.121 |
| chr5 | 125984819 | 125985420 | 602 | 0.99 | 0.0798 | 0.121 |
| chr3 | 155507626 | 155509916 | 2291 | -0.51 | 0.0798 | 0.121 |
| chr11 | 28542338 | 28543639 | 1302 | -1.08 | 0.0798 | 0.121 |
| chr2 | 128478781 | 128507947 | 29167 | -0.18 | 0.0799 | 0.121 |
| chr8 | 99076416 | 99082052 | 5637 | 3.03 | 0.0799 | 0.121 |
| chr16 | 325452 | 326336 | 885 | -1.06 | 0.0799 | 0.121 |
| chr8 | 15592352 | 15593052 | 701 | -1.11 | 0.0799 | 0.121 |
| chr3 | 51377240 | 51382588 | 5349 | -0.27 | 0.08 | 0.121 |
| chr5 | 30084184 | 30085438 | 1255 | 0.74 | 0.0801 | 0.121 |
| chr12 | 9122159 | 9123652 | 1494 | 0.42 | 0.0802 | 0.121 |
| chr3 | 162252764 | 162253383 | 620 | 1.77 | 0.0802 | 0.121 |
| chr17 | 8391939 | 8395729 | 3791 | -0.45 | 0.0802 | 0.121 |
| chr14 | 90420023 | 90423697 | 3675 | -0.32 | 0.0803 | 0.121 |
| chr22 | 45944034 | 45950656 | 6623 | -0.23 | 0.0803 | 0.121 |
| chr6 | 36163659 | 36165494 | 1836 | -0.57 | 0.0803 | 0.121 |
| chr13 | 24700875 | 24701704 | 830 | -1.61 | 0.0803 | 0.121 |
| chr18 | 72369093 | 72370142 | 1050 | -0.92 | 0.0803 | 0.121 |
| chr8 | 38175919 | 38177229 | 1311 | 0.89 | 0.0804 | 0.122 |
| chr16 | 2849489 | 2849917 | 429 | 1.76 | 0.0804 | 0.122 |
| chr1 | 241865430 | 241871761 | 6332 | 0.25 | 0.0804 | 0.122 |
| chr15 | 43248122 | 43250759 | 2638 | 0.4 | 0.0804 | 0.122 |
| chr3 | 177622503 | 177624401 | 1899 | -0.73 | 0.0805 | 0.122 |
| chr19 | 34786650 | 34790440 | 3791 | 0.7 | 0.0805 | 0.122 |
| chr16 | 57059853 | 57060473 | 621 | -1.22 | 0.0806 | 0.122 |
| chr12 | 17191450 | 17193074 | 1625 | 0.54 | 0.0806 | 0.122 |
| chr9 | 114611794 | 114612741 | 948 | 0.9 | 0.0806 | 0.122 |
| chr14 | 69526321 | 69569184 | 42864 | -0.15 | 0.0806 | 0.122 |
| chr14 | 99372411 | 99373777 | 1367 | 0.85 | 0.0806 | 0.122 |
| chrX | 4850589 | 4851399 | 811 | 1.1 | 0.0806 | 0.122 |
| chr5 | 15499579 | 15500842 | 1264 | 0.78 | 0.0806 | 0.122 |
| chr6 | 37457082 | 37468030 | 10949 | -0.55 | 0.0808 | 0.122 |
| chr19 | 58144494 | 58145316 | 823 | -0.66 | 0.0808 | 0.122 |
| chr9 | 79940165 | 79941345 | 1181 | 0.84 | 0.0808 | 0.122 |
| chr2 | 47346030 | 47346490 | 461 | 1.27 | 0.0808 | 0.122 |
| chr3 | 5096780 | 5097916 | 1137 | -1.46 | 0.0808 | 0.122 |
| chr2 | 38754963 | 38770682 | 15720 | -0.52 | 0.0808 | 0.122 |
| chr6 | 6781149 | 6782259 | 1111 | 0.91 | 0.0809 | 0.122 |
| chr10 | 29818622 | 29825915 | 7294 | 0.36 | 0.0809 | 0.122 |
| chr1 | 35753411 | 35755096 | 1686 | 0.95 | 0.0809 | 0.122 |
| chr2 | 202444452 | 202445734 | 1283 | -0.8 | 0.081 | 0.122 |
| chr19 | 15666906 | 15667344 | 439 | 1.12 | 0.081 | 0.122 |
| chr8 | 66590295 | 66591375 | 1081 | -0.85 | 0.081 | 0.122 |
| chr6 | 35052471 | 35052918 | 448 | -1.61 | 0.081 | 0.122 |
| chr18 | 52340289 | 52340956 | 668 | 1.24 | 0.081 | 0.122 |
| chr10 | 21906156 | 21907678 | 1523 | 0.73 | 0.0811 | 0.123 |

| | | | | | | |
|---|---|---|---|---|---|---|
| chr2 | 210288063 | 210288550 | 488 | 1.02 | 0.0812 | 0.123 |
| chr6 | 34919810 | 34922626 | 2817 | -0.27 | 0.0812 | 0.123 |
| chr15 | 65300636 | 65350285 | 49650 | -0.16 | 0.0812 | 0.123 |
| chr13 | 80937879 | 80938183 | 305 | 1.11 | 0.0812 | 0.123 |
| chr10 | 70267553 | 70268435 | 883 | -1.11 | 0.0812 | 0.123 |
| chr8 | 133915521 | 133916193 | 673 | -0.89 | 0.0812 | 0.123 |
| chr3 | 133965453 | 133970187 | 4735 | 0.39 | 0.0813 | 0.123 |
| chr10 | 15209484 | 15210626 | 1143 | 0.59 | 0.0813 | 0.123 |
| chr1 | 39623189 | 39680146 | 56958 | 0.09 | 0.0813 | 0.123 |
| chr9 | 134053961 | 134066905 | 12945 | -0.99 | 0.0814 | 0.123 |
| chr22 | 45659435 | 45660248 | 814 | -1.11 | 0.0814 | 0.123 |
| chr1 | 53386523 | 53393939 | 7417 | 0.43 | 0.0814 | 0.123 |
| chr1 | 17865993 | 17919010 | 53018 | 0.15 | 0.0815 | 0.123 |
| chr11 | 85954818 | 85963097 | 8280 | 0.19 | 0.0815 | 0.123 |
| chr2 | 63055613 | 63056114 | 502 | 1.07 | 0.0815 | 0.123 |
| chr2 | 97101316 | 97102105 | 790 | 1.77 | 0.0816 | 0.123 |
| chr10 | 45359839 | 45361423 | 1585 | 0.86 | 0.0816 | 0.123 |
| chr9 | 37119596 | 37121468 | 1873 | 0.35 | 0.0817 | 0.123 |
| chr7 | 138914989 | 138917735 | 2747 | 0.35 | 0.0817 | 0.123 |
| chr6 | 76122512 | 76135447 | 12936 | -0.22 | 0.0817 | 0.123 |
| chr22 | 29661881 | 29672981 | 11101 | -0.19 | 0.0817 | 0.123 |
| chr3 | 52684494 | 52685075 | 582 | 1.03 | 0.0818 | 0.123 |
| chr15 | 43298291 | 43298924 | 634 | 1.41 | 0.0818 | 0.123 |
| chr1 | 228589372 | 228595645 | 6274 | -0.26 | 0.0818 | 0.123 |
| chr17 | 75276731 | 75488547 | 211817 | 0.05 | 0.0818 | 0.123 |
| chr20 | 18859505 | 18860863 | 1359 | 0.84 | 0.0819 | 0.123 |
| chr20 | 2081768 | 2096891 | 15124 | -0.14 | 0.0819 | 0.123 |
| chr1 | 46394448 | 46395701 | 1254 | 0.81 | 0.0819 | 0.123 |
| chrX | 17139219 | 17139977 | 759 | -1.8 | 0.0819 | 0.124 |
| chr12 | 96944577 | 96945154 | 578 | -1.51 | 0.0819 | 0.124 |
| chr10 | 33653956 | 33679548 | 25593 | 0.43 | 0.0819 | 0.124 |
| chr1 | 229642314 | 229645100 | 2787 | 0.25 | 0.082 | 0.124 |
| chr4 | 88295109 | 88295766 | 658 | -1.17 | 0.082 | 0.124 |
| chr5 | 123252160 | 123252940 | 781 | 0.84 | 0.082 | 0.124 |
| chr13 | 46871035 | 46871634 | 600 | -1.07 | 0.082 | 0.124 |
| chr2 | 64789495 | 64790629 | 1135 | 0.87 | 0.0821 | 0.124 |
| chr15 | 80827865 | 80829053 | 1189 | 2.94 | 0.0821 | 0.124 |
| chr7 | 151178819 | 151218020 | 39202 | 0.11 | 0.0821 | 0.124 |
| chr3 | 101421222 | 101426806 | 5585 | -0.68 | 0.0821 | 0.124 |
| chr6 | 132316836 | 132317616 | 781 | 1.13 | 0.0821 | 0.124 |
| chr3 | 10067152 | 10069344 | 2193 | -0.29 | 0.0821 | 0.124 |
| chr1 | 6659890 | 6686009 | 26120 | -0.26 | 0.0822 | 0.124 |
| chr6 | 56406507 | 56409460 | 2954 | 0.29 | 0.0822 | 0.124 |
| chr3 | 12543002 | 12550109 | 7108 | 0.38 | 0.0822 | 0.124 |
| chr9 | 137733326 | 137734533 | 1208 | -0.95 | 0.0822 | 0.124 |
| chr17 | 61483306 | 61484059 | 754 | 0.92 | 0.0823 | 0.124 |
| chr22 | 45873855 | 45874964 | 1110 | 0.72 | 0.0823 | 0.124 |
| chr4 | 84277854 | 84280314 | 2461 | -0.54 | 0.0823 | 0.124 |
| chr12 | 124594479 | 124595028 | 550 | 1.2 | 0.0823 | 0.124 |
| chr14 | 92459916 | 92464686 | 4771 | 1.21 | 0.0823 | 0.124 |
| chr2 | 75503557 | 75504846 | 1290 | -2.92 | 0.0823 | 0.124 |
| chr10 | 104010093 | 104010703 | 611 | -1.1 | 0.0824 | 0.124 |
| chr7 | 55029065 | 55036271 | 7207 | 0.31 | 0.0824 | 0.124 |
| chr1 | 23302449 | 23303000 | 552 | -0.96 | 0.0824 | 0.124 |
| chr2 | 120995702 | 121011467 | 15766 | -0.28 | 0.0824 | 0.124 |
| chr10 | 49716063 | 49716824 | 762 | -1.07 | 0.0824 | 0.124 |
| chr7 | 17672261 | 17673321 | 1061 | 0.68 | 0.0825 | 0.124 |
| chrX | 107757093 | 107761201 | 4109 | -0.35 | 0.0825 | 0.124 |
| chr10 | 101039010 | 101040387 | 1378 | -0.72 | 0.0826 | 0.124 |
| chr11 | 59543981 | 59554955 | 10975 | 3.13 | 0.0826 | 0.124 |
| chr22 | 35694962 | 35698244 | 3283 | 0.39 | 0.0826 | 0.124 |

| | | | | | | |
|---|---|---|---|---|---|---|
| chr5 | 122544195 | 122546003 | 1809 | -0.52 | 0.0827 | 0.124 |
| chr4 | 160947512 | 160960516 | 13005 | -0.77 | 0.0827 | 0.124 |
| chr1 | 39827632 | 39835419 | 7788 | 0.48 | 0.0827 | 0.124 |
| chr14 | 95996770 | 96002892 | 6123 | -0.21 | 0.0827 | 0.125 |
| chr1 | 44748709 | 44750306 | 1598 | 1.99 | 0.0828 | 0.125 |
| chr9 | 128546261 | 128547081 | 821 | 1.74 | 0.0829 | 0.125 |
| chr3 | 196157965 | 196160534 | 2570 | 0.4 | 0.0829 | 0.125 |
| chr2 | 190227764 | 190245634 | 17871 | -0.3 | 0.083 | 0.125 |
| chr12 | 7494242 | 7494829 | 588 | -1.6 | 0.0831 | 0.125 |
| chr5 | 54798028 | 54799193 | 1166 | -1.09 | 0.0831 | 0.125 |
| chr3 | 40546494 | 40548471 | 1978 | 0.54 | 0.0831 | 0.125 |
| chr2 | 191531635 | 191532072 | 438 | -1.21 | 0.0833 | 0.125 |
| chr16 | 46607325 | 46608021 | 697 | 0.92 | 0.0833 | 0.125 |
| chr6 | 89743213 | 89760697 | 17485 | 0.19 | 0.0833 | 0.125 |
| chr6 | 52407696 | 52443217 | 35522 | 0.16 | 0.0833 | 0.125 |
| chr10 | 114922273 | 114923791 | 1519 | -0.73 | 0.0834 | 0.125 |
| chr6 | 43337585 | 43338291 | 707 | 0.78 | 0.0834 | 0.125 |
| chr18 | 13539094 | 13542892 | 3799 | -0.46 | 0.0834 | 0.126 |
| chr7 | 76074714 | 76075045 | 332 | 1.46 | 0.0835 | 0.126 |
| chr4 | 39497320 | 39561110 | 63791 | -0.08 | 0.0835 | 0.126 |
| chr3 | 114166668 | 114177714 | 11047 | 0.6 | 0.0835 | 0.126 |
| chr3 | 182510425 | 182512735 | 2311 | 0.3 | 0.0835 | 0.126 |
| chr5 | 142580311 | 142581044 | 734 | -1.21 | 0.0835 | 0.126 |
| chr18 | 18584312 | 18585303 | 992 | 0.98 | 0.0835 | 0.126 |
| chrX | 16203634 | 16215361 | 11728 | -2.83 | 0.0835 | 0.126 |
| chr18 | 55133221 | 55134508 | 1288 | 0.58 | 0.0836 | 0.126 |
| chr11 | 35965133 | 35966307 | 1175 | -0.57 | 0.0836 | 0.126 |
| chr9 | 99271960 | 99272526 | 567 | -0.98 | 0.0836 | 0.126 |
| chr12 | 114403179 | 114404673 | 1495 | 0.45 | 0.0837 | 0.126 |
| chr15 | 28392921 | 28397454 | 4534 | 0.66 | 0.0837 | 0.126 |
| chr2 | 25474147 | 25475972 | 1826 | -0.41 | 0.0838 | 0.126 |
| chr16 | 4768155 | 4769837 | 1683 | -0.81 | 0.0838 | 0.126 |
| chr6 | 36985782 | 36998122 | 12341 | 0.35 | 0.0838 | 0.126 |
| chr3 | 160224604 | 160227908 | 3305 | -0.44 | 0.0838 | 0.126 |
| chr14 | 89834923 | 89835888 | 966 | -0.97 | 0.0838 | 0.126 |
| chr4 | 123745702 | 123777226 | 31525 | 0.18 | 0.0838 | 0.126 |
| chr6 | 44378733 | 44390359 | 11627 | 0.46 | 0.0838 | 0.126 |
| chr2 | 150884790 | 150885395 | 606 | 1.1 | 0.0838 | 0.126 |
| chr16 | 84209695 | 84221262 | 11568 | -0.31 | 0.0839 | 0.126 |
| chr5 | 43161248 | 43172663 | 11416 | 0.44 | 0.0839 | 0.126 |
| chr2 | 64083900 | 64084414 | 515 | -1.24 | 0.0839 | 0.126 |
| chr22 | 22315580 | 22316526 | 947 | 0.83 | 0.0839 | 0.126 |
| chr19 | 46497811 | 46499530 | 1720 | -0.55 | 0.0839 | 0.126 |
| chr5 | 106198700 | 106199129 | 430 | -1.29 | 0.0841 | 0.126 |
| chr1 | 2186473 | 2188235 | 1763 | 0.76 | 0.0841 | 0.126 |
| chr6 | 111549348 | 111550344 | 997 | 0.99 | 0.0842 | 0.126 |
| chr15 | 56244410 | 56262839 | 18430 | -0.41 | 0.0842 | 0.126 |
| chr16 | 64567746 | 64568637 | 892 | -0.91 | 0.0843 | 0.127 |
| chr20 | 55606354 | 55606803 | 450 | -1.28 | 0.0843 | 0.127 |
| chr13 | 76537305 | 76537911 | 607 | -1.42 | 0.0843 | 0.127 |
| chr15 | 44322141 | 44331320 | 9180 | 2.83 | 0.0844 | 0.127 |
| chr6 | 120326916 | 120327723 | 808 | 0.89 | 0.0844 | 0.127 |
| chr14 | 73429705 | 73430669 | 965 | 1.39 | 0.0844 | 0.127 |
| chr16 | 58425904 | 58434913 | 9010 | -0.26 | 0.0845 | 0.127 |
| chr5 | 92527687 | 92528488 | 802 | 0.88 | 0.0845 | 0.127 |
| chr12 | 110382835 | 110391617 | 8783 | -0.36 | 0.0845 | 0.127 |
| chr7 | 91847416 | 91848342 | 927 | -0.79 | 0.0845 | 0.127 |
| chr10 | 1119617 | 1120123 | 507 | -1.28 | 0.0846 | 0.127 |
| chrX | 118674887 | 118676093 | 1207 | -0.58 | 0.0846 | 0.127 |
| chr10 | 51488757 | 51489117 | 361 | -1.77 | 0.0846 | 0.127 |
| chr1 | 196485689 | 196487328 | 1640 | 1.07 | 0.0847 | 0.127 |

| | | | | | | |
|---|---|---|---|---|---|---|
| chr6 | 17971114 | 17973242 | 2129 | -0.51 | 0.0847 | 0.127 |
| chr20 | 22506844 | 22507561 | 718 | -1.73 | 0.0848 | 0.127 |
| chr7 | 101704905 | 101723177 | 18273 | -0.26 | 0.0848 | 0.127 |
| chr20 | 42331018 | 42337870 | 6853 | -0.55 | 0.0848 | 0.127 |
| chr5 | 142554528 | 142555124 | 597 | -1.58 | 0.0848 | 0.127 |
| chr1 | 203480974 | 203483743 | 2770 | 0.39 | 0.0849 | 0.127 |
| chr3 | 11539943 | 11540791 | 849 | 0.82 | 0.0849 | 0.127 |
| chr16 | 69407481 | 69516805 | 109325 | 0.06 | 0.0849 | 0.127 |
| chr18 | 77770094 | 77771031 | 938 | 1.01 | 0.0849 | 0.127 |
| chr4 | 137426127 | 137426779 | 653 | -0.99 | 0.0849 | 0.127 |
| chr8 | 103595628 | 103598436 | 2809 | -0.49 | 0.0849 | 0.127 |
| chr2 | 109255788 | 109257718 | 1931 | -2.52 | 0.085 | 0.127 |
| chr6 | 139043874 | 139044571 | 698 | -0.97 | 0.085 | 0.127 |
| chr2 | 225502965 | 225503354 | 390 | -1.18 | 0.085 | 0.127 |
| chr2 | 223890296 | 223913067 | 22772 | -0.22 | 0.085 | 0.127 |
| chr17 | 16274751 | 16366266 | 91516 | -0.11 | 0.085 | 0.127 |
| chr17 | 49449558 | 49450058 | 501 | -1.14 | 0.085 | 0.128 |
| chr21 | 16493568 | 16495103 | 1536 | 0.41 | 0.0851 | 0.128 |
| chr10 | 100074051 | 100074887 | 837 | 0.95 | 0.0851 | 0.128 |
| chr1 | 24779558 | 24780087 | 530 | -1.2 | 0.0852 | 0.128 |
| chr7 | 36267221 | 36272855 | 5635 | 0.98 | 0.0852 | 0.128 |
| chr14 | 65501618 | 65522143 | 20526 | -0.41 | 0.0852 | 0.128 |
| chr4 | 1282602 | 1284934 | 2333 | -0.39 | 0.0853 | 0.128 |
| chr10 | 65116243 | 65116642 | 400 | -1.74 | 0.0853 | 0.128 |
| chr8 | 105338036 | 105338346 | 311 | 1.49 | 0.0853 | 0.128 |
| chr16 | 89834278 | 89835292 | 1015 | 1.06 | 0.0853 | 0.128 |
| chr2 | 43764113 | 43768344 | 4232 | -0.49 | 0.0854 | 0.128 |
| chr19 | 6199169 | 6202575 | 3407 | 0.58 | 0.0855 | 0.128 |
| chr14 | 51326274 | 51327336 | 1063 | 0.66 | 0.0855 | 0.128 |
| chr3 | 155523050 | 155525012 | 1963 | 0.45 | 0.0855 | 0.128 |
| chrX | 151138452 | 151155712 | 17261 | 0.19 | 0.0855 | 0.128 |
| chr1 | 231663218 | 231669018 | 5801 | 0.2 | 0.0856 | 0.128 |
| chr17 | 53570502 | 53577745 | 7244 | -0.23 | 0.0856 | 0.128 |
| chr2 | 143439489 | 143451546 | 12058 | -1.88 | 0.0856 | 0.128 |
| chrX | 118886920 | 118888395 | 1476 | 0.77 | 0.0856 | 0.128 |
| chr3 | 149901813 | 149902530 | 718 | -1.28 | 0.0857 | 0.128 |
| chr9 | 83852621 | 83853486 | 866 | -0.62 | 0.0857 | 0.128 |
| chr11 | 69957697 | 69958631 | 935 | -0.98 | 0.0857 | 0.128 |
| chr12 | 26170381 | 26180624 | 10244 | 2.97 | 0.0857 | 0.128 |
| chr4 | 145783091 | 145783694 | 604 | -0.95 | 0.0857 | 0.128 |
| chr12 | 51109396 | 51109869 | 474 | 1.08 | 0.0858 | 0.129 |
| chr2 | 192127754 | 192148338 | 20585 | -3.25 | 0.0859 | 0.129 |
| chr8 | 10137597 | 10138455 | 859 | -0.84 | 0.0859 | 0.129 |
| chr3 | 87904357 | 87905863 | 1507 | -0.59 | 0.0859 | 0.129 |
| chr17 | 47559684 | 47560373 | 690 | -1.57 | 0.086 | 0.129 |
| chr8 | 30039166 | 30039641 | 476 | 1.76 | 0.086 | 0.129 |
| chr1 | 246273391 | 246281224 | 7834 | -0.28 | 0.0861 | 0.129 |
| chr9 | 79432378 | 79434133 | 1756 | -0.39 | 0.0861 | 0.129 |
| chr20 | 11159232 | 11160519 | 1288 | 3.06 | 0.0861 | 0.129 |
| chr6 | 138692104 | 138693773 | 1670 | -0.56 | 0.0861 | 0.129 |
| chrX | 129039233 | 129041497 | 2265 | 0.33 | 0.0861 | 0.129 |
| chr21 | 45527215 | 45527955 | 741 | -0.8 | 0.0862 | 0.129 |
| chr15 | 73877209 | 73878901 | 1693 | -0.66 | 0.0862 | 0.129 |
| chr1 | 65789107 | 65789544 | 438 | -1.76 | 0.0862 | 0.129 |
| chr8 | 80826536 | 80827135 | 600 | 0.96 | 0.0862 | 0.129 |
| chr12 | 123069254 | 123069746 | 493 | 1.33 | 0.0863 | 0.129 |
| chr4 | 87361047 | 87363498 | 2452 | -0.33 | 0.0863 | 0.129 |
| chr20 | 19830019 | 19928804 | 98786 | 0.06 | 0.0864 | 0.129 |
| chr22 | 31002852 | 31006148 | 3297 | 0.39 | 0.0865 | 0.129 |
| chr4 | 129730134 | 129794174 | 64041 | -0.09 | 0.0865 | 0.129 |
| chr18 | 2515617 | 2516973 | 1357 | -0.64 | 0.0865 | 0.129 |

| | | | | | | |
|---|---|---|---|---|---|---|
| chr1 | 183240859 | 183242862 | 2004 | 0.69 | 0.0867 | 0.13 |
| chr10 | 5210382 | 5211568 | 1187 | 1.28 | 0.0868 | 0.13 |
| chr2 | 176994181 | 177005839 | 11659 | -0.33 | 0.0868 | 0.13 |
| chr7 | 5013601 | 5014861 | 1261 | 0.47 | 0.0868 | 0.13 |
| chrX | 118985783 | 118987960 | 2178 | 0.42 | 0.0868 | 0.13 |
| chr4 | 89977543 | 89979124 | 1582 | 0.42 | 0.0869 | 0.13 |
| chr9 | 118111417 | 118116633 | 5217 | 0.85 | 0.0869 | 0.13 |
| chr5 | 137513072 | 137522305 | 9234 | 0.23 | 0.0869 | 0.13 |
| chr3 | 29365268 | 29378341 | 13074 | -0.14 | 0.0869 | 0.13 |
| chr16 | 1231310 | 1234173 | 2864 | 0.36 | 0.087 | 0.13 |
| chr20 | 11241956 | 11248966 | 7011 | 0.39 | 0.087 | 0.13 |
| chr10 | 21920388 | 21968130 | 47743 | -0.19 | 0.087 | 0.13 |
| chr6 | 118845645 | 118846423 | 779 | 1.12 | 0.0871 | 0.13 |
| chr2 | 106438469 | 106439057 | 589 | -1.07 | 0.0872 | 0.13 |
| chr8 | 22624181 | 22628600 | 4420 | 0.17 | 0.0872 | 0.13 |
| chr16 | 11089322 | 11089887 | 566 | -1.41 | 0.0872 | 0.13 |
| chr7 | 68789119 | 68789909 | 791 | -1.23 | 0.0872 | 0.13 |
| chr8 | 71272623 | 71273803 | 1181 | 1.03 | 0.0873 | 0.13 |
| chr5 | 5178043 | 5179136 | 1094 | 0.88 | 0.0873 | 0.131 |
| chr14 | 52924290 | 52925460 | 1171 | 0.68 | 0.0875 | 0.131 |
| chr17 | 77669360 | 77670441 | 1082 | -1.18 | 0.0876 | 0.131 |
| chr2 | 100896037 | 100919582 | 23546 | -0.14 | 0.0876 | 0.131 |
| chr13 | 35730681 | 35731690 | 1010 | -1.08 | 0.0876 | 0.131 |
| chr5 | 91706722 | 91709027 | 2306 | -0.26 | 0.0876 | 0.131 |
| chr5 | 16520923 | 16522362 | 1440 | 1.2 | 0.0876 | 0.131 |
| chr8 | 27183000 | 27199273 | 16274 | 0.14 | 0.0876 | 0.131 |
| chr4 | 158036598 | 158049664 | 13067 | 3.04 | 0.0877 | 0.131 |
| chr2 | 56112344 | 56119685 | 7342 | 0.44 | 0.0878 | 0.131 |
| chr3 | 49129609 | 49143088 | 13480 | 0.2 | 0.0878 | 0.131 |
| chr2 | 138630703 | 138631672 | 970 | -0.8 | 0.0878 | 0.131 |
| chr16 | 68233627 | 68234538 | 912 | -0.85 | 0.0878 | 0.131 |
| chr22 | 31497859 | 31499099 | 1241 | -0.88 | 0.0878 | 0.131 |
| chr8 | 103135836 | 103136378 | 543 | 1.17 | 0.0878 | 0.131 |
| chr5 | 71577277 | 71617120 | 39844 | -0.13 | 0.0878 | 0.131 |
| chrX | 9172612 | 9173482 | 871 | 0.82 | 0.0878 | 0.131 |
| chr10 | 3494676 | 3543718 | 49043 | 0.1 | 0.0878 | 0.131 |
| chr17 | 55638585 | 55639311 | 727 | -0.98 | 0.0879 | 0.131 |
| chr4 | 166222540 | 166223441 | 902 | 0.81 | 0.0879 | 0.131 |
| chr18 | 34909120 | 34909805 | 686 | -1.02 | 0.0879 | 0.131 |
| chr20 | 21105613 | 21107781 | 2169 | 0.36 | 0.088 | 0.131 |
| chr7 | 18125911 | 18126419 | 509 | 1.12 | 0.088 | 0.131 |
| chr12 | 15021661 | 15022198 | 538 | -1.3 | 0.088 | 0.131 |
| chr2 | 122512285 | 122515074 | 2790 | -0.31 | 0.0881 | 0.132 |
| chr12 | 26408917 | 26409621 | 705 | -1.08 | 0.0881 | 0.132 |
| chrX | 110923309 | 110925621 | 2313 | 0.24 | 0.0882 | 0.132 |
| chr4 | 88961736 | 88962210 | 475 | -1.73 | 0.0883 | 0.132 |
| chr1 | 28907273 | 28910033 | 2761 | 0.31 | 0.0883 | 0.132 |
| chr1 | 3712151 | 3713849 | 1699 | 0.58 | 0.0883 | 0.132 |
| chr6 | 35835744 | 35836482 | 739 | -1.42 | 0.0883 | 0.132 |
| chr10 | 106060824 | 106061315 | 492 | 1.24 | 0.0884 | 0.132 |
| chr2 | 238161946 | 238162446 | 501 | -1.21 | 0.0884 | 0.132 |
| chr6 | 25991826 | 25994353 | 2528 | 0.27 | 0.0884 | 0.132 |
| chr5 | 170921407 | 170922541 | 1135 | -0.5 | 0.0884 | 0.132 |
| chr1 | 179555200 | 179561325 | 6126 | 2.68 | 0.0884 | 0.132 |
| chr20 | 56344571 | 56348361 | 3791 | -0.34 | 0.0884 | 0.132 |
| chr1 | 155324699 | 155325156 | 458 | -1.8 | 0.0884 | 0.132 |
| chrX | 44401045 | 44402923 | 1879 | 0.32 | 0.0885 | 0.132 |
| chr19 | 42890652 | 42911086 | 20435 | -0.16 | 0.0885 | 0.132 |
| chr11 | 66277402 | 66288128 | 10727 | -0.42 | 0.0885 | 0.132 |
| chr10 | 17729317 | 17730140 | 824 | -1.15 | 0.0886 | 0.132 |
| chr12 | 105647619 | 105648758 | 1140 | -1.13 | 0.0886 | 0.132 |

| | | | | | | |
|---|---|---|---|---|---|---|
| chr13 | 25483288 | 25498376 | 15089 | -0.18 | 0.0886 | 0.132 |
| chr3 | 180677704 | 180679837 | 2134 | 0.65 | 0.0886 | 0.132 |
| chr3 | 48935560 | 48937106 | 1547 | 0.71 | 0.0887 | 0.132 |
| chrX | 129090833 | 129095969 | 5137 | 0.43 | 0.0888 | 0.132 |
| chr1 | 18092191 | 18095917 | 3727 | 0.61 | 0.0888 | 0.132 |
| chr16 | 3492872 | 3535109 | 42238 | 0.19 | 0.0889 | 0.133 |
| chr22 | 36230066 | 36243997 | 13932 | 0.19 | 0.0889 | 0.133 |
| chr13 | 25245202 | 25246151 | 950 | -1.05 | 0.0889 | 0.133 |
| chr10 | 30610208 | 30638975 | 28768 | 0.18 | 0.0889 | 0.133 |
| chr11 | 47824994 | 47826846 | 1853 | 0.97 | 0.089 | 0.133 |
| chr4 | 38276802 | 38277637 | 836 | 1.3 | 0.089 | 0.133 |
| chr5 | 107996994 | 107998222 | 1229 | 2.55 | 0.089 | 0.133 |
| chr1 | 22220018 | 22266918 | 46901 | 0.1 | 0.0891 | 0.133 |
| chr4 | 3135476 | 3136027 | 552 | -1.1 | 0.0891 | 0.133 |
| chr15 | 77597981 | 77605526 | 7546 | -2.88 | 0.0891 | 0.133 |
| chr15 | 70935632 | 70936976 | 1345 | 0.57 | 0.0891 | 0.133 |
| chr19 | 4399278 | 4417292 | 18015 | 0.24 | 0.0891 | 0.133 |
| chr16 | 1098668 | 1099035 | 368 | -1.42 | 0.0892 | 0.133 |
| chrX | 71681689 | 71682298 | 610 | -1.48 | 0.0893 | 0.133 |
| chr2 | 220024702 | 220044084 | 19383 | -0.2 | 0.0893 | 0.133 |
| chr6 | 111677125 | 111679342 | 2218 | 0.71 | 0.0893 | 0.133 |
| chr8 | 70269678 | 70270066 | 389 | 1.62 | 0.0893 | 0.133 |
| chr1 | 41847394 | 41849091 | 1698 | 0.38 | 0.0893 | 0.133 |
| chr9 | 134405606 | 134422629 | 17024 | 0.41 | 0.0893 | 0.133 |
| chr2 | 44196729 | 44206014 | 9286 | -0.29 | 0.0893 | 0.133 |
| chr14 | 70019740 | 70020571 | 832 | 1.45 | 0.0893 | 0.133 |
| chr3 | 43459437 | 43460944 | 1508 | 2.97 | 0.0893 | 0.133 |
| chr1 | 221785195 | 221795383 | 10189 | 0.25 | 0.0893 | 0.133 |
| chr2 | 232017215 | 232027209 | 9995 | -0.38 | 0.0894 | 0.133 |
| chr13 | 20341496 | 20342528 | 1033 | 1.15 | 0.0894 | 0.133 |
| chr7 | 74123807 | 74124929 | 1123 | -1.05 | 0.0894 | 0.133 |
| chr1 | 230321195 | 230346396 | 25202 | -0.21 | 0.0894 | 0.133 |
| chr15 | 41729675 | 41740537 | 10863 | -0.48 | 0.0894 | 0.133 |
| chr2 | 62533979 | 62535021 | 1043 | -0.67 | 0.0894 | 0.133 |
| chr6 | 8079103 | 8087404 | 8302 | 0.33 | 0.0894 | 0.133 |
| chr18 | 48721156 | 48725440 | 4285 | 0.24 | 0.0895 | 0.133 |
| chr2 | 216161897 | 216163083 | 1187 | -0.77 | 0.0895 | 0.133 |
| chr10 | 11799163 | 11800611 | 1449 | -0.62 | 0.0895 | 0.133 |
| chr13 | 65114745 | 65115540 | 796 | -0.93 | 0.0896 | 0.133 |
| chr14 | 90741421 | 90748089 | 6669 | -0.57 | 0.0896 | 0.133 |
| chr1 | 39563138 | 39583076 | 19939 | -0.22 | 0.0896 | 0.133 |
| chr3 | 41158513 | 41159652 | 1140 | 0.93 | 0.0897 | 0.133 |
| chr15 | 70911321 | 70911949 | 629 | -1.69 | 0.0897 | 0.133 |
| chr2 | 95824400 | 95832385 | 7986 | 0.2 | 0.0897 | 0.133 |
| chr1 | 207509163 | 207511618 | 2456 | 0.54 | 0.0897 | 0.134 |
| chr1 | 51109793 | 51110857 | 1065 | 0.82 | 0.0898 | 0.134 |
| chr22 | 41025874 | 41051125 | 25252 | -0.17 | 0.0898 | 0.134 |
| chr4 | 108910383 | 108912262 | 1880 | 0.48 | 0.0899 | 0.134 |
| chr10 | 104747964 | 104758080 | 10117 | 0.48 | 0.0899 | 0.134 |
| chrX | 46813749 | 46815032 | 1284 | 1.39 | 0.0899 | 0.134 |
| chr6 | 143928322 | 143929105 | 784 | -1.29 | 0.09 | 0.134 |
| chr16 | 77003670 | 77004889 | 1220 | 0.65 | 0.0901 | 0.134 |
| chr1 | 19447199 | 19448325 | 1127 | -0.74 | 0.0901 | 0.134 |
| chr17 | 55483993 | 55484454 | 462 | -1.73 | 0.0903 | 0.134 |
| chr15 | 49315539 | 49339486 | 23948 | 0.29 | 0.0903 | 0.134 |
| chr2 | 242031944 | 242042918 | 10975 | 0.25 | 0.0903 | 0.134 |
| chr9 | 33779280 | 33782991 | 3712 | 0.68 | 0.0903 | 0.134 |
| chr13 | 111657610 | 111659334 | 1725 | 2.86 | 0.0904 | 0.134 |
| chr19 | 42616870 | 42626191 | 9322 | -3.41 | 0.0904 | 0.134 |
| chr1 | 68515575 | 68516635 | 1061 | 1.06 | 0.0904 | 0.134 |
| chr18 | 7440855 | 7441428 | 574 | 1.16 | 0.0904 | 0.134 |

| | | | | | | |
|---|---|---|---|---|---|---|
| chr14 | 50813846 | 50814892 | 1047 | -0.55 | 0.0904 | 0.134 |
| chr17 | 16912156 | 17094805 | 182650 | 0.05 | 0.0905 | 0.135 |
| chr12 | 32398071 | 32398508 | 438 | 1.24 | 0.0905 | 0.135 |
| chr14 | 75016254 | 75017063 | 810 | -1.27 | 0.0905 | 0.135 |
| chr5 | 33147011 | 33154119 | 7109 | 0.21 | 0.0906 | 0.135 |
| chr19 | 11532601 | 11555736 | 23136 | -0.22 | 0.0907 | 0.135 |
| chr3 | 44689783 | 44691132 | 1350 | 0.6 | 0.0907 | 0.135 |
| chr6 | 111334573 | 111335083 | 511 | 1.43 | 0.0907 | 0.135 |
| chr17 | 70627568 | 70637271 | 9704 | 0.35 | 0.0908 | 0.135 |
| chr9 | 85549108 | 85555171 | 6064 | 0.21 | 0.0908 | 0.135 |
| chr9 | 81975391 | 81978337 | 2947 | -0.27 | 0.0909 | 0.135 |
| chr12 | 108908936 | 108910425 | 1490 | 0.42 | 0.0909 | 0.135 |
| chr17 | 25533744 | 25534759 | 1016 | 1.05 | 0.0909 | 0.135 |
| chr15 | 45509371 | 45510463 | 1093 | 0.84 | 0.091 | 0.135 |
| chr13 | 44732224 | 44734872 | 2649 | -0.37 | 0.0911 | 0.135 |
| chr21 | 38070293 | 38073389 | 3097 | -0.36 | 0.0911 | 0.135 |
| chr8 | 39084651 | 39085245 | 595 | -1.02 | 0.0911 | 0.135 |
| chr2 | 85940023 | 85941341 | 1319 | 1.3 | 0.0912 | 0.135 |
| chr10 | 90694797 | 90695241 | 445 | 1.26 | 0.0912 | 0.135 |
| chr10 | 69758454 | 69870584 | 112131 | 0.07 | 0.0912 | 0.136 |
| chr14 | 104376184 | 104388491 | 12308 | 0.35 | 0.0913 | 0.136 |
| chr17 | 58676170 | 58709104 | 32935 | -0.15 | 0.0913 | 0.136 |
| chr20 | 37483134 | 37483812 | 679 | -1.68 | 0.0913 | 0.136 |
| chr6 | 90473045 | 90474009 | 965 | -0.75 | 0.0913 | 0.136 |
| chrX | 16803721 | 16857917 | 54197 | 0.13 | 0.0914 | 0.136 |
| chr1 | 234959569 | 234960163 | 595 | 1.27 | 0.0914 | 0.136 |
| chr11 | 103830161 | 103830554 | 394 | -1.7 | 0.0915 | 0.136 |
| chr7 | 77313646 | 77328961 | 15316 | 0.29 | 0.0915 | 0.136 |
| chr17 | 18247406 | 18267343 | 19938 | 0.27 | 0.0915 | 0.136 |
| chr7 | 101904080 | 101930937 | 26858 | -0.33 | 0.0916 | 0.136 |
| chr2 | 197652546 | 197665032 | 12487 | -0.29 | 0.0916 | 0.136 |
| chr2 | 32921884 | 32930424 | 8541 | -1.22 | 0.0916 | 0.136 |
| chr12 | 95387985 | 95398319 | 10335 | 0.31 | 0.0917 | 0.136 |
| chr1 | 55504635 | 55506040 | 1406 | 0.65 | 0.0917 | 0.136 |
| chr12 | 6945020 | 6945447 | 428 | 0.95 | 0.0918 | 0.136 |
| chr4 | 122632466 | 122633552 | 1087 | 0.38 | 0.0918 | 0.136 |
| chr4 | 79505938 | 79518482 | 12545 | 0.43 | 0.0918 | 0.136 |
| chr19 | 34011780 | 34013346 | 1567 | 0.45 | 0.0919 | 0.136 |
| chr18 | 20957058 | 20966998 | 9941 | -0.25 | 0.0919 | 0.136 |
| chr22 | 18982136 | 18983165 | 1030 | 0.7 | 0.0921 | 0.137 |
| chr6 | 27463010 | 27473345 | 10336 | -0.44 | 0.0921 | 0.137 |
| chr9 | 101864616 | 101868682 | 4067 | 0.34 | 0.0921 | 0.137 |
| chr1 | 59246872 | 59251752 | 4881 | 0.17 | 0.0922 | 0.137 |
| chr1 | 247056120 | 247057125 | 1006 | 0.94 | 0.0922 | 0.137 |
| chr6 | 109442218 | 109443171 | 954 | 0.89 | 0.0923 | 0.137 |
| chr17 | 21217169 | 21219140 | 1972 | 0.73 | 0.0924 | 0.137 |
| chr2 | 55873960 | 55875036 | 1077 | -0.97 | 0.0924 | 0.137 |
| chr5 | 130945864 | 130947195 | 1332 | -0.71 | 0.0925 | 0.137 |
| chr5 | 74531794 | 74533343 | 1550 | -0.53 | 0.0925 | 0.137 |
| chr1 | 41290206 | 41291308 | 1103 | -0.98 | 0.0926 | 0.137 |
| chr6 | 64307680 | 64309051 | 1372 | 0.53 | 0.0926 | 0.137 |
| chr16 | 67122541 | 67123838 | 1298 | -0.67 | 0.0926 | 0.137 |
| chr7 | 3067192 | 3068523 | 1332 | 0.45 | 0.0926 | 0.137 |
| chr12 | 51631868 | 51634932 | 3065 | 0.23 | 0.0926 | 0.137 |
| chr1 | 7357593 | 7364206 | 6614 | -0.21 | 0.0926 | 0.137 |
| chr3 | 33584951 | 33585504 | 554 | 1.02 | 0.0926 | 0.137 |
| chr14 | 23380533 | 23401179 | 20647 | 0.14 | 0.0927 | 0.137 |
| chr6 | 109803512 | 109804446 | 935 | -0.62 | 0.0927 | 0.137 |
| chr19 | 34624780 | 34626302 | 1523 | 0.3 | 0.0927 | 0.137 |
| chr19 | 46635599 | 46638248 | 2650 | 0.28 | 0.0927 | 0.137 |
| chr11 | 9709970 | 9715188 | 5219 | -0.45 | 0.0927 | 0.137 |

| | | | | | | |
|---|---|---|---|---|---|---|
| chr17 | 1613122 | 1640274 | 27153 | -0.11 | 0.0927 | 0.137 |
| chr5 | 64944729 | 64947298 | 2570 | -0.73 | 0.0928 | 0.138 |
| chr5 | 134072262 | 134075959 | 3698 | 0.22 | 0.093 | 0.138 |
| chr4 | 79784491 | 79785020 | 530 | 1.75 | 0.093 | 0.138 |
| chr6 | 19804178 | 19806021 | 1844 | -0.53 | 0.093 | 0.138 |
| chr7 | 130079609 | 130081933 | 2325 | -0.22 | 0.093 | 0.138 |
| chr5 | 1593803 | 1597412 | 3610 | 0.54 | 0.093 | 0.138 |
| chr6 | 25278452 | 25281315 | 2864 | 0.38 | 0.093 | 0.138 |
| chr3 | 124478798 | 124628936 | 150139 | -0.06 | 0.093 | 0.138 |
| chr6 | 22906121 | 22906427 | 307 | 1.78 | 0.093 | 0.138 |
| chr20 | 8995403 | 9007445 | 12043 | -0.34 | 0.093 | 0.138 |
| chr1 | 224644393 | 224645373 | 981 | -0.61 | 0.0931 | 0.138 |
| chrX | 16138205 | 16138927 | 723 | -1.03 | 0.0931 | 0.138 |
| chr3 | 157844876 | 157846876 | 2001 | -0.37 | 0.0932 | 0.138 |
| chr6 | 89070252 | 89070687 | 436 | -1.43 | 0.0932 | 0.138 |
| chr17 | 45971982 | 45983265 | 11284 | 0.21 | 0.0932 | 0.138 |
| chr4 | 74782752 | 74789145 | 6394 | -0.71 | 0.0932 | 0.138 |
| chr1 | 233307806 | 233317582 | 9777 | 0.34 | 0.0933 | 0.138 |
| chr19 | 33661910 | 33669252 | 7343 | -0.17 | 0.0933 | 0.138 |
| chr1 | 53738706 | 53741156 | 2451 | -0.5 | 0.0933 | 0.138 |
| chr2 | 204102638 | 204105560 | 2923 | -0.27 | 0.0933 | 0.138 |
| chr12 | 19726029 | 19726661 | 633 | 0.99 | 0.0934 | 0.138 |
| chr16 | 81410775 | 81412185 | 1411 | -0.77 | 0.0934 | 0.138 |
| chr9 | 139350760 | 139379600 | 28841 | -0.16 | 0.0934 | 0.138 |
| chr6 | 20400875 | 20428304 | 27430 | -0.16 | 0.0934 | 0.138 |
| chr17 | 55355714 | 55373326 | 17613 | -3.42 | 0.0935 | 0.138 |
| chr1 | 40040662 | 40042470 | 1809 | 0.46 | 0.0935 | 0.138 |
| chr9 | 86086372 | 86089907 | 3536 | -0.28 | 0.0935 | 0.138 |
| chr15 | 44954003 | 44956784 | 2782 | 0.25 | 0.0936 | 0.139 |
| chr2 | 164548989 | 164550142 | 1154 | -0.82 | 0.0936 | 0.139 |
| chr10 | 70518710 | 70520233 | 1524 | -0.69 | 0.0936 | 0.139 |
| chr7 | 38766919 | 38767286 | 368 | -1.32 | 0.0936 | 0.139 |
| chr2 | 147631496 | 147632162 | 667 | 1.21 | 0.0937 | 0.139 |
| chr2 | 3370136 | 3384785 | 14650 | 0.21 | 0.0937 | 0.139 |
| chr1 | 51795945 | 51797785 | 1841 | -0.49 | 0.0937 | 0.139 |
| chr1 | 48936984 | 48938794 | 1811 | 0.4 | 0.0937 | 0.139 |
| chr11 | 9633320 | 9636979 | 3660 | 0.18 | 0.0937 | 0.139 |
| chr10 | 118976345 | 118977194 | 850 | -0.75 | 0.0937 | 0.139 |
| chr15 | 64010513 | 64011578 | 1066 | 0.67 | 0.0938 | 0.139 |
| chr19 | 6925287 | 6926646 | 1360 | 0.32 | 0.0939 | 0.139 |
| chr1 | 210800381 | 210801910 | 1530 | -0.58 | 0.0939 | 0.139 |
| chr7 | 128378306 | 128380784 | 2479 | 0.25 | 0.094 | 0.139 |
| chr4 | 5038949 | 5054221 | 15273 | -0.34 | 0.094 | 0.139 |
| chr4 | 41127296 | 41127941 | 646 | 1.13 | 0.094 | 0.139 |
| chr3 | 104105820 | 104106803 | 984 | 0.93 | 0.094 | 0.139 |
| chr6 | 116421107 | 116423233 | 2127 | 0.34 | 0.0941 | 0.139 |
| chr1 | 54586923 | 54587494 | 572 | -0.88 | 0.0941 | 0.139 |
| chr2 | 202506615 | 202508628 | 2014 | -0.26 | 0.0941 | 0.139 |
| chr14 | 56348270 | 56356797 | 8528 | 0.35 | 0.0941 | 0.139 |
| chr3 | 47697814 | 47727334 | 29521 | 0.28 | 0.0941 | 0.139 |
| chr9 | 130123091 | 130124294 | 1204 | -0.84 | 0.0942 | 0.139 |
| chr7 | 30443116 | 30443757 | 642 | -1.09 | 0.0942 | 0.139 |
| chr1 | 150227956 | 150267778 | 39823 | -0.16 | 0.0942 | 0.139 |
| chr18 | 33646192 | 33648028 | 1837 | 0.34 | 0.0942 | 0.139 |
| chrX | 56588890 | 56597022 | 8133 | 0.18 | 0.0944 | 0.14 |
| chr5 | 64433225 | 64434210 | 986 | 0.83 | 0.0944 | 0.14 |
| chr2 | 42841786 | 42847051 | 5266 | 0.43 | 0.0945 | 0.14 |
| chr9 | 116638591 | 116673530 | 34940 | 0.16 | 0.0945 | 0.14 |
| chr4 | 103681424 | 103682742 | 1319 | -0.67 | 0.0945 | 0.14 |
| chr8 | 40958361 | 40959679 | 1319 | 0.85 | 0.0945 | 0.14 |
| chr7 | 129040709 | 129042701 | 1993 | -0.49 | 0.0946 | 0.14 |

| | | | | | | |
|---|---|---|---|---|---|---|
| chr7 | 99715877 | 99725567 | 9691 | -0.2 | 0.0946 | 0.14 |
| chr8 | 42994208 | 42997281 | 3074 | 0.25 | 0.0946 | 0.14 |
| chr6 | 39081998 | 39083452 | 1455 | 0.45 | 0.0946 | 0.14 |
| chr6 | 99962381 | 99970573 | 8193 | 0.19 | 0.0947 | 0.14 |
| chr2 | 119593347 | 119594168 | 822 | -0.6 | 0.0948 | 0.14 |
| chr17 | 57500049 | 57501560 | 1512 | -0.42 | 0.0948 | 0.14 |
| chr4 | 169238971 | 169240595 | 1625 | 0.38 | 0.0948 | 0.14 |
| chr8 | 17545664 | 17560005 | 14342 | -0.28 | 0.0948 | 0.14 |
| chr1 | 176002587 | 176003955 | 1369 | 0.83 | 0.0949 | 0.14 |
| chr15 | 96935273 | 96936145 | 873 | -0.85 | 0.0949 | 0.14 |
| chr1 | 32320950 | 32321717 | 768 | 0.84 | 0.0949 | 0.14 |
| chr11 | 74677737 | 74685094 | 7358 | -0.5 | 0.095 | 0.14 |
| chr7 | 73393102 | 73393516 | 415 | 0.98 | 0.095 | 0.14 |
| chr5 | 17694607 | 17695960 | 1354 | 0.61 | 0.095 | 0.14 |
| chr5 | 17056081 | 17071028 | 14948 | 0.31 | 0.0951 | 0.14 |
| chr16 | 30708883 | 30725141 | 16259 | 0.23 | 0.0951 | 0.14 |
| chrX | 10050376 | 10071484 | 21109 | -0.33 | 0.0951 | 0.14 |
| chr14 | 100020365 | 100021853 | 1489 | 0.7 | 0.0952 | 0.141 |
| chr4 | 667363 | 700019 | 32657 | -0.3 | 0.0953 | 0.141 |
| chr4 | 2585054 | 2585786 | 733 | -0.93 | 0.0953 | 0.141 |
| chr14 | 39517608 | 39520627 | 3020 | 0.49 | 0.0953 | 0.141 |
| chr12 | 66442385 | 66443542 | 1158 | 2.5 | 0.0953 | 0.141 |
| chr16 | 88984665 | 88991774 | 7110 | 0.53 | 0.0954 | 0.141 |
| chr2 | 53994076 | 54015935 | 21860 | -0.16 | 0.0954 | 0.141 |
| chr20 | 60163362 | 60164676 | 1315 | 0.9 | 0.0954 | 0.141 |
| chr17 | 8441698 | 8447447 | 5750 | -0.99 | 0.0954 | 0.141 |
| chr1 | 199963991 | 199966286 | 2296 | 0.66 | 0.0954 | 0.141 |
| chr15 | 40924585 | 40927326 | 2742 | -0.43 | 0.0955 | 0.141 |
| chr2 | 190647276 | 190650820 | 3545 | -0.28 | 0.0955 | 0.141 |
| chr2 | 101033340 | 101034679 | 1340 | -0.4 | 0.0955 | 0.141 |
| chr5 | 170340884 | 170341664 | 781 | 1.15 | 0.0956 | 0.141 |
| chr2 | 174254385 | 174254793 | 409 | -1.62 | 0.0956 | 0.141 |
| chr17 | 73422236 | 73567854 | 145619 | 0.07 | 0.0956 | 0.141 |
| chr6 | 74098200 | 74099884 | 1685 | 0.41 | 0.0956 | 0.141 |
| chr6 | 38122242 | 38124346 | 2105 | -0.52 | 0.0957 | 0.141 |
| chr2 | 236816636 | 236817142 | 507 | -1.12 | 0.0957 | 0.141 |
| chr1 | 232764967 | 232767205 | 2239 | 0.38 | 0.0958 | 0.141 |
| chr5 | 171840904 | 171882207 | 41304 | -0.19 | 0.0958 | 0.141 |
| chr4 | 4290964 | 4292862 | 1899 | 0.6 | 0.0958 | 0.141 |
| chr12 | 68703971 | 68704636 | 666 | 0.9 | 0.0958 | 0.141 |
| chr1 | 75178423 | 75181250 | 2828 | 0.5 | 0.0958 | 0.141 |
| chr20 | 11409841 | 11410259 | 419 | -1.67 | 0.0958 | 0.141 |
| chr11 | 102237758 | 102239419 | 1662 | 0.54 | 0.0959 | 0.141 |
| chr5 | 157502536 | 157503028 | 493 | 0.95 | 0.0959 | 0.141 |
| chr1 | 39451035 | 39469908 | 18874 | 0.19 | 0.0959 | 0.141 |
| chr8 | 6456706 | 6457837 | 1132 | 1.08 | 0.0959 | 0.141 |
| chr3 | 194201614 | 194222693 | 21080 | 0.25 | 0.096 | 0.141 |
| chr1 | 15237491 | 15238019 | 529 | -1.18 | 0.096 | 0.141 |
| chr9 | 86316323 | 86324253 | 7931 | 0.17 | 0.0961 | 0.142 |
| chr17 | 28431557 | 28457606 | 26050 | 0.15 | 0.0961 | 0.142 |
| chr3 | 196225703 | 196231527 | 5825 | 0.21 | 0.0961 | 0.142 |
| chr6 | 151989555 | 151990775 | 1221 | 0.69 | 0.0961 | 0.142 |
| chr1 | 172535145 | 172536250 | 1106 | -1.53 | 0.0962 | 0.142 |
| chr12 | 20799466 | 20799786 | 321 | -1.4 | 0.0962 | 0.142 |
| chr7 | 116659836 | 116660738 | 903 | 0.71 | 0.0963 | 0.142 |
| chr10 | 114344512 | 114344968 | 457 | 0.98 | 0.0963 | 0.142 |
| chr15 | 89708812 | 89717111 | 8300 | -0.21 | 0.0963 | 0.142 |
| chr19 | 51012175 | 51016105 | 3931 | -0.29 | 0.0964 | 0.142 |
| chr11 | 86045822 | 86046915 | 1094 | -0.68 | 0.0964 | 0.142 |
| chr7 | 91808423 | 91808794 | 372 | 1.14 | 0.0964 | 0.142 |
| chr10 | 49196565 | 49197223 | 659 | 0.31 | 0.0964 | 0.142 |

| | | | | | | |
|---|---|---|---|---|---|---|
| chr3 | 128255809 | 128256499 | 691 | 1.11 | 0.0965 | 0.142 |
| chr12 | 96466882 | 96468030 | 1149 | 0.98 | 0.0965 | 0.142 |
| chr6 | 158033596 | 158035794 | 2199 | 0.45 | 0.0965 | 0.142 |
| chr1 | 54354689 | 54356153 | 1465 | 0.53 | 0.0966 | 0.142 |
| chr3 | 4458667 | 4463415 | 4749 | -0.62 | 0.0966 | 0.142 |
| chr2 | 196183086 | 196187772 | 4687 | -0.27 | 0.0967 | 0.142 |
| chr7 | 27940473 | 27941471 | 999 | 0.73 | 0.0967 | 0.142 |
| chr19 | 5218733 | 5230081 | 11349 | -0.41 | 0.0967 | 0.142 |
| chr9 | 80982973 | 80983846 | 874 | -0.82 | 0.0967 | 0.142 |
| chr10 | 116103534 | 116104454 | 921 | 0.94 | 0.0968 | 0.143 |
| chr1 | 196246275 | 196251722 | 5448 | 3.06 | 0.0968 | 0.143 |
| chr11 | 47188589 | 47216461 | 27873 | 0.22 | 0.0968 | 0.143 |
| chr11 | 45376621 | 45377520 | 900 | 0.62 | 0.0968 | 0.143 |
| chr17 | 41131435 | 41208129 | 76695 | 0.08 | 0.0968 | 0.143 |
| chr17 | 33287705 | 33289597 | 1893 | 0.19 | 0.0969 | 0.143 |
| chr2 | 148811295 | 148811909 | 615 | -0.92 | 0.0969 | 0.143 |
| chr7 | 71097967 | 71099282 | 1316 | 2.81 | 0.097 | 0.143 |
| chr22 | 30389634 | 30392075 | 2442 | 0.33 | 0.0971 | 0.143 |
| chr16 | 30609720 | 30622528 | 12809 | 0.23 | 0.0972 | 0.143 |
| chr14 | 57676534 | 57677694 | 1161 | 0.97 | 0.0972 | 0.143 |
| chr3 | 66307055 | 66307695 | 641 | -0.94 | 0.0972 | 0.143 |
| chr8 | 16988542 | 16989267 | 726 | -1.04 | 0.0972 | 0.143 |
| chr6 | 107385790 | 107386383 | 594 | 1.23 | 0.0973 | 0.143 |
| chr4 | 95812771 | 95813652 | 882 | 1.69 | 0.0973 | 0.143 |
| chrX | 40824062 | 40831704 | 7643 | 0.59 | 0.0973 | 0.143 |
| chr20 | 49015086 | 49017603 | 2518 | -0.51 | 0.0973 | 0.143 |
| chr9 | 21415111 | 21416612 | 1502 | 0.51 | 0.0973 | 0.143 |
| chr19 | 6502076 | 6502940 | 865 | -1.04 | 0.0974 | 0.143 |
| chr22 | 31503105 | 31503481 | 377 | 1.03 | 0.0974 | 0.143 |
| chr15 | 26178707 | 26179669 | 963 | -0.67 | 0.0974 | 0.143 |
| chr13 | 44844710 | 44852939 | 8230 | 0.48 | 0.0975 | 0.143 |
| chr3 | 126423106 | 126423459 | 354 | -1.08 | 0.0975 | 0.143 |
| chr5 | 91976099 | 91978085 | 1987 | 0.28 | 0.0975 | 0.143 |
| chr7 | 155249692 | 155250507 | 816 | 0.84 | 0.0976 | 0.144 |
| chr8 | 145114909 | 145116236 | 1328 | -0.49 | 0.0976 | 0.144 |
| chr19 | 50215174 | 50228083 | 12910 | 0.36 | 0.0976 | 0.144 |
| chr20 | 62670071 | 62696157 | 26087 | -0.34 | 0.0977 | 0.144 |
| chr9 | 85497433 | 85498374 | 942 | 1.01 | 0.0977 | 0.144 |
| chr3 | 152176477 | 152190315 | 13839 | 0.3 | 0.0978 | 0.144 |
| chr7 | 104612119 | 104692307 | 80189 | 0.11 | 0.0978 | 0.144 |
| chrX | 135532239 | 135532982 | 744 | 1.26 | 0.0978 | 0.144 |
| chr2 | 224902848 | 224904499 | 1652 | -0.67 | 0.0979 | 0.144 |
| chr10 | 123513518 | 123514655 | 1138 | 1.66 | 0.0979 | 0.144 |
| chr12 | 51142658 | 51143258 | 601 | -1.04 | 0.0979 | 0.144 |
| chr17 | 58041213 | 58042989 | 1777 | 0.31 | 0.098 | 0.144 |
| chr3 | 47138643 | 47146043 | 7401 | -0.48 | 0.098 | 0.144 |
| chr2 | 174982438 | 174985909 | 3472 | 0.54 | 0.098 | 0.144 |
| chr5 | 60893893 | 60922960 | 29068 | 0.12 | 0.0981 | 0.144 |
| chr17 | 35749178 | 35749858 | 681 | 1.1 | 0.0981 | 0.144 |
| chr3 | 196820000 | 196820666 | 667 | -1.04 | 0.0982 | 0.144 |
| chr3 | 185191043 | 185191943 | 901 | 1.28 | 0.0982 | 0.144 |
| chr19 | 18262073 | 18272433 | 10361 | -0.22 | 0.0983 | 0.144 |
| chr4 | 140418682 | 140419606 | 925 | -1.17 | 0.0983 | 0.144 |
| chr22 | 24092697 | 24094315 | 1619 | 0.41 | 0.0983 | 0.144 |
| chr17 | 3860988 | 3869127 | 8140 | -0.36 | 0.0984 | 0.145 |
| chr6 | 33636018 | 33645530 | 9513 | 0.23 | 0.0984 | 0.145 |
| chr10 | 104103699 | 104104455 | 757 | -0.91 | 0.0984 | 0.145 |
| chr11 | 19973317 | 19974436 | 1120 | 0.8 | 0.0985 | 0.145 |
| chr19 | 35600023 | 35601109 | 1087 | 0.69 | 0.0985 | 0.145 |
| chrX | 114876152 | 114877370 | 1219 | -0.75 | 0.0985 | 0.145 |
| chr11 | 13726777 | 13727580 | 804 | 1.29 | 0.0985 | 0.145 |

| | | | | | | |
|---|---|---|---|---|---|---|
| chr1 | 210545156 | 210558586 | 13431 | -0.22 | 0.0986 | 0.145 |
| chr1 | 39943414 | 39945019 | 1606 | -0.71 | 0.0986 | 0.145 |
| chr7 | 131310022 | 131319045 | 9024 | 0.23 | 0.0986 | 0.145 |
| chr17 | 33993453 | 33994115 | 663 | -0.95 | 0.0986 | 0.145 |
| chr7 | 114399170 | 114400441 | 1272 | 0.9 | 0.0986 | 0.145 |
| chr10 | 70358933 | 70361253 | 2321 | 0.25 | 0.0987 | 0.145 |
| chr10 | 105991203 | 105992767 | 1565 | -0.32 | 0.0987 | 0.145 |
| chr10 | 15125813 | 15149170 | 23358 | 0.17 | 0.0988 | 0.145 |
| chr1 | 234401465 | 234402109 | 645 | -0.98 | 0.0988 | 0.145 |
| chr11 | 77592570 | 77593779 | 1210 | -0.83 | 0.0989 | 0.145 |
| chr16 | 82185531 | 82204367 | 18837 | 0.27 | 0.0989 | 0.145 |
| chr14 | 64525985 | 64527496 | 1512 | -0.73 | 0.0989 | 0.145 |
| chr6 | 11343718 | 11345436 | 1719 | 0.45 | 0.099 | 0.145 |
| chr3 | 189110048 | 189111783 | 1736 | -0.55 | 0.099 | 0.145 |
| chr10 | 73155882 | 73156590 | 709 | -0.85 | 0.0991 | 0.145 |
| chr3 | 150295174 | 150303841 | 8668 | 0.44 | 0.0991 | 0.145 |
| chrX | 133970416 | 133971308 | 893 | 0.84 | 0.0991 | 0.145 |
| chr6 | 10885893 | 10888300 | 2408 | 0.25 | 0.0992 | 0.146 |
| chr6 | 97371382 | 97373146 | 1765 | 2.44 | 0.0992 | 0.146 |
| chr12 | 107186462 | 107199418 | 12957 | 0.37 | 0.0992 | 0.146 |
| chr3 | 100369139 | 100370560 | 1422 | 0.48 | 0.0992 | 0.146 |
| chr2 | 173591121 | 173600706 | 9586 | 2.86 | 0.0992 | 0.146 |
| chr19 | 16337257 | 16338062 | 806 | -0.95 | 0.0993 | 0.146 |
| chr14 | 24604638 | 24637207 | 32570 | 0.15 | 0.0993 | 0.146 |
| chr1 | 4327102 | 4328336 | 1235 | 0.56 | 0.0993 | 0.146 |
| chr7 | 65842712 | 65843517 | 806 | -1.03 | 0.0994 | 0.146 |
| chr7 | 5861653 | 5862966 | 1314 | 0.39 | 0.0994 | 0.146 |
| chr17 | 62615059 | 62705974 | 90916 | -0.1 | 0.0994 | 0.146 |
| chr1 | 2573230 | 2575200 | 1971 | -0.47 | 0.0994 | 0.146 |
| chr13 | 20592481 | 20596353 | 3873 | 0.53 | 0.0994 | 0.146 |
| chr11 | 67396365 | 67398781 | 2417 | 0.43 | 0.0994 | 0.146 |
| chr22 | 21976228 | 22021415 | 45188 | -0.16 | 0.0994 | 0.146 |
| chr20 | 44561779 | 44579078 | 17300 | -0.17 | 0.0995 | 0.146 |
| chr10 | 24983243 | 25014412 | 31170 | 0.16 | 0.0996 | 0.146 |
| chr18 | 3581995 | 3723299 | 141305 | -0.07 | 0.0996 | 0.146 |
| chr22 | 42083151 | 42094076 | 10926 | 0.2 | 0.0997 | 0.146 |
| chr20 | 34327107 | 34332897 | 5791 | 0.15 | 0.0998 | 0.146 |
| chr1 | 32670533 | 32707747 | 37215 | -0.15 | 0.0998 | 0.146 |
| chr6 | 97593718 | 97594484 | 767 | 1.05 | 0.0999 | 0.146 |
| chr4 | 173747731 | 173752366 | 4636 | -0.52 | 0.0999 | 0.146 |
| chr9 | 135904979 | 135907142 | 2164 | 0.27 | 0.0999 | 0.146 |
| chrX | 41026924 | 41027639 | 716 | 1.01 | 0.1 | 0.147 |
| chr9 | 131201168 | 131201638 | 471 | -1.09 | 0.1 | 0.147 |
| chr10 | 129704661 | 129705849 | 1189 | 0.92 | 0.1 | 0.147 |
| chr7 | 94432951 | 94433655 | 705 | -1.64 | 0.1 | 0.147 |
| chr7 | 120995638 | 121013532 | 17895 | 1.64 | 0.1 | 0.147 |
| chr5 | 159612555 | 159627304 | 14750 | 0.42 | 0.1 | 0.147 |
| chr7 | 68156188 | 68174689 | 18502 | -0.12 | 0.1 | 0.147 |
| chr3 | 64426206 | 64426730 | 525 | 1.15 | 0.1 | 0.147 |
| chr2 | 223859038 | 223859736 | 699 | 0.94 | 0.1 | 0.147 |
| chr11 | 75945178 | 75947881 | 2704 | -0.38 | 0.1 | 0.147 |
| chr1 | 29471179 | 29474038 | 2860 | 0.31 | 0.1 | 0.147 |
| chr10 | 94632516 | 94633529 | 1014 | 0.84 | 0.1 | 0.147 |
| chr11 | 130391606 | 130393080 | 1475 | 0.57 | 0.1 | 0.147 |
| chr4 | 184319367 | 184329182 | 9816 | 0.52 | 0.1 | 0.147 |
| chr22 | 29412063 | 29441273 | 29211 | -0.16 | 0.101 | 0.147 |
| chr22 | 48533011 | 48534976 | 1966 | 0.51 | 0.101 | 0.147 |
| chr16 | 63421510 | 63422179 | 670 | 0.78 | 0.101 | 0.147 |
| chr2 | 159442596 | 159457550 | 14955 | -0.47 | 0.101 | 0.148 |
| chr7 | 92296583 | 92297378 | 796 | -0.99 | 0.101 | 0.148 |
| chr12 | 21712824 | 21719837 | 7014 | 0.72 | 0.101 | 0.148 |

| | | | | | | |
|---|---|---|---|---|---|---|
| chr11 | 118868002 | 118890663 | 22662 | 0.19 | 0.101 | 0.148 |
| chr20 | 2969528 | 2971147 | 1620 | -0.85 | 0.101 | 0.148 |
| chr2 | 162090210 | 162102274 | 12065 | -0.34 | 0.101 | 0.148 |
| chr22 | 47125101 | 47176675 | 51575 | 0.11 | 0.101 | 0.148 |
| chr7 | 19995119 | 20012297 | 17179 | -0.12 | 0.101 | 0.148 |
| chr15 | 31120715 | 31121386 | 672 | 0.66 | 0.101 | 0.148 |
| chr12 | 12640794 | 12641565 | 772 | -1.09 | 0.101 | 0.148 |
| chr16 | 25244643 | 25245059 | 417 | 1.58 | 0.101 | 0.148 |
| chr16 | 27635516 | 27637480 | 1965 | 0.53 | 0.101 | 0.148 |
| chrX | 107816013 | 107817176 | 1164 | 0.71 | 0.101 | 0.148 |
| chr16 | 4465354 | 4486867 | 21514 | -0.16 | 0.101 | 0.148 |
| chr6 | 154808430 | 154810601 | 2172 | 0.77 | 0.101 | 0.148 |
| chr11 | 117069587 | 117070387 | 801 | 0.95 | 0.101 | 0.148 |
| chrX | 70752028 | 70776942 | 24915 | 0.21 | 0.101 | 0.148 |
| chr2 | 183452141 | 183453250 | 1110 | 0.78 | 0.101 | 0.148 |
| chr11 | 77789517 | 77791846 | 2330 | 0.28 | 0.101 | 0.148 |
| chr1 | 169417920 | 169418808 | 889 | 2.74 | 0.101 | 0.148 |
| chr7 | 131337804 | 131338589 | 786 | -0.98 | 0.101 | 0.148 |
| chr1 | 38454601 | 38496145 | 41545 | -0.16 | 0.101 | 0.149 |
| chr12 | 42800956 | 42801308 | 353 | -0.98 | 0.102 | 0.149 |
| chr16 | 48303442 | 48304441 | 1000 | -1.12 | 0.102 | 0.149 |
| chr5 | 127033741 | 127041540 | 7800 | -0.42 | 0.102 | 0.149 |
| chr7 | 45961999 | 45964536 | 2538 | -0.65 | 0.102 | 0.149 |
| chr15 | 41708287 | 41710604 | 2318 | 0.42 | 0.102 | 0.149 |
| chr6 | 15132780 | 15134070 | 1291 | -0.7 | 0.102 | 0.149 |
| chr5 | 158758457 | 158769684 | 11228 | 0.49 | 0.102 | 0.149 |
| chr17 | 65901029 | 65904583 | 3555 | -0.7 | 0.102 | 0.149 |
| chr22 | 24407074 | 24409058 | 1985 | -0.43 | 0.102 | 0.149 |
| chr11 | 64682698 | 64692744 | 10047 | 0.23 | 0.102 | 0.149 |
| chr8 | 68013176 | 68044703 | 31528 | 0.28 | 0.102 | 0.149 |
| chr9 | 83907076 | 83907608 | 533 | -1.1 | 0.102 | 0.149 |
| chr5 | 172520510 | 172521826 | 1317 | 0.69 | 0.102 | 0.149 |
| chr1 | 46766306 | 46775140 | 8835 | 0.2 | 0.102 | 0.149 |
| chr17 | 28926414 | 28928693 | 2280 | 0.28 | 0.102 | 0.149 |
| chr11 | 85338026 | 85340763 | 2738 | 0.25 | 0.102 | 0.149 |
| chrX | 56755339 | 56757152 | 1814 | 0.34 | 0.102 | 0.15 |
| chr17 | 57591398 | 57592593 | 1196 | 0.79 | 0.102 | 0.15 |
| chr1 | 215301745 | 215302422 | 678 | -0.99 | 0.102 | 0.15 |
| chr3 | 188103012 | 188112927 | 9916 | -0.56 | 0.102 | 0.15 |
| chr15 | 90844207 | 90845042 | 836 | -1.06 | 0.102 | 0.15 |
| chrX | 109511908 | 109530216 | 18309 | 0.31 | 0.102 | 0.15 |
| chr13 | 37632415 | 37634094 | 1680 | 0.31 | 0.103 | 0.15 |
| chr8 | 106788073 | 106789715 | 1643 | 0.99 | 0.103 | 0.15 |
| chr15 | 78568300 | 78569441 | 1142 | -0.61 | 0.103 | 0.15 |
| chr16 | 10579083 | 10579705 | 623 | 2.54 | 0.103 | 0.15 |
| chr8 | 19124617 | 19125240 | 624 | 1.12 | 0.103 | 0.15 |
| chr22 | 29396014 | 29396599 | 586 | 0.98 | 0.103 | 0.15 |
| chr1 | 226588241 | 226596606 | 8366 | -0.29 | 0.103 | 0.15 |
| chr2 | 190014299 | 190015405 | 1107 | -0.75 | 0.103 | 0.15 |
| chr1 | 182118296 | 182118983 | 688 | -1.08 | 0.103 | 0.15 |
| chr14 | 51999285 | 52015360 | 16076 | 0.33 | 0.103 | 0.15 |
| chr7 | 69246074 | 69247061 | 988 | 0.87 | 0.103 | 0.15 |
| chr2 | 187349755 | 187361908 | 12154 | -0.18 | 0.103 | 0.15 |
| chr7 | 16409060 | 16409939 | 880 | 0.86 | 0.103 | 0.15 |
| chr3 | 47767128 | 47767906 | 779 | -0.93 | 0.103 | 0.15 |
| chr1 | 161671340 | 161672861 | 1522 | -0.53 | 0.103 | 0.15 |
| chr6 | 82706728 | 82707706 | 979 | 0.8 | 0.103 | 0.151 |
| chr2 | 238581907 | 238582523 | 617 | -1.11 | 0.103 | 0.151 |
| chr3 | 27352484 | 27353860 | 1377 | -0.4 | 0.103 | 0.151 |
| chr1 | 116544601 | 116545330 | 730 | -1.63 | 0.103 | 0.151 |
| chr22 | 38597338 | 38599549 | 2212 | 0.38 | 0.103 | 0.151 |

| | | | | | | |
|---|---|---|---|---|---|---|
| chr16 | 11918217 | 11919061 | 845 | 1.4 | 0.103 | 0.151 |
| chr1 | 116944136 | 116945026 | 891 | -0.73 | 0.103 | 0.151 |
| chr8 | 17141560 | 17142505 | 946 | -1 | 0.103 | 0.151 |
| chr2 | 136763219 | 136765086 | 1868 | 0.47 | 0.103 | 0.151 |
| chr1 | 61515295 | 61524067 | 8773 | 0.26 | 0.103 | 0.151 |
| chr9 | 113002392 | 113022446 | 20055 | -0.1 | 0.103 | 0.151 |
| chr6 | 11765460 | 11772870 | 7411 | -0.21 | 0.103 | 0.151 |
| chr17 | 21156854 | 21157292 | 439 | 1.67 | 0.103 | 0.151 |
| chr1 | 168591729 | 168593144 | 1416 | 0.51 | 0.104 | 0.151 |
| chr10 | 103909701 | 103930431 | 20731 | -0.16 | 0.104 | 0.151 |
| chr13 | 97680706 | 97681086 | 381 | 0.9 | 0.104 | 0.151 |
| chr5 | 18114283 | 18114677 | 395 | -1.66 | 0.104 | 0.151 |
| chr7 | 51029412 | 51031802 | 2391 | -0.34 | 0.104 | 0.151 |
| chr7 | 50823486 | 50824702 | 1217 | -1.24 | 0.104 | 0.151 |
| chr18 | 519104 | 519620 | 517 | -1.21 | 0.104 | 0.151 |
| chr2 | 43035955 | 43038820 | 2866 | -0.22 | 0.104 | 0.151 |
| chr15 | 39159733 | 39188680 | 28948 | -0.12 | 0.104 | 0.151 |
| chr2 | 20130018 | 20147536 | 17519 | -0.11 | 0.104 | 0.151 |
| chr11 | 120270733 | 120272534 | 1802 | 0.42 | 0.104 | 0.152 |
| chr14 | 68696100 | 68719959 | 23860 | -0.1 | 0.104 | 0.152 |
| chr8 | 101387616 | 101388418 | 803 | 1 | 0.104 | 0.152 |
| chr10 | 119051139 | 119052612 | 1474 | 0.62 | 0.104 | 0.152 |
| chr4 | 141612106 | 141613285 | 1180 | 2.15 | 0.104 | 0.152 |
| chr3 | 52570706 | 52586884 | 16179 | 0.24 | 0.104 | 0.152 |
| chr11 | 66934755 | 66936694 | 1940 | -0.51 | 0.104 | 0.152 |
| chr2 | 65081846 | 65104297 | 22452 | 0.19 | 0.104 | 0.152 |
| chr18 | 188833 | 189650 | 818 | 1.21 | 0.104 | 0.152 |
| chr15 | 32943681 | 32952749 | 9069 | -0.34 | 0.104 | 0.152 |
| chr1 | 235529578 | 235578812 | 49235 | 0.1 | 0.104 | 0.152 |
| chrX | 24071531 | 24104198 | 32668 | 0.11 | 0.104 | 0.152 |
| chr10 | 32308671 | 32355553 | 46883 | 0.15 | 0.104 | 0.152 |
| chr3 | 183265448 | 183266731 | 1284 | 2.74 | 0.104 | 0.152 |
| chr15 | 31195445 | 31197045 | 1601 | 0.43 | 0.104 | 0.152 |
| chr1 | 219444325 | 219455882 | 11558 | -0.4 | 0.104 | 0.152 |
| chr9 | 140082006 | 140173116 | 91111 | -0.11 | 0.105 | 0.152 |
| chr9 | 36566311 | 36625902 | 59592 | 0.12 | 0.105 | 0.152 |
| chr1 | 222987619 | 222999805 | 12187 | 0.33 | 0.105 | 0.153 |
| chr5 | 138513390 | 138515087 | 1698 | -0.61 | 0.105 | 0.153 |
| chr1 | 94607245 | 94609777 | 2533 | 0.59 | 0.105 | 0.153 |
| chr13 | 109421514 | 109432007 | 10494 | 0.25 | 0.105 | 0.153 |
| chr4 | 148890919 | 148892279 | 1361 | 0.71 | 0.105 | 0.153 |
| chr12 | 104849921 | 104857191 | 7271 | 0.3 | 0.105 | 0.153 |
| chr8 | 61135499 | 61139981 | 4483 | 0.32 | 0.105 | 0.153 |
| chr15 | 59396469 | 59681961 | 285493 | 0.04 | 0.105 | 0.153 |
| chr9 | 124675180 | 124675950 | 771 | 1.07 | 0.105 | 0.153 |
| chr17 | 37792601 | 37801607 | 9007 | -0.26 | 0.105 | 0.153 |
| chr8 | 53624999 | 53628953 | 3955 | 0.22 | 0.105 | 0.153 |
| chr17 | 66507083 | 66519672 | 12590 | 0.12 | 0.105 | 0.153 |
| chr11 | 93394191 | 93397110 | 2920 | 0.49 | 0.105 | 0.153 |
| chr5 | 85913060 | 85915652 | 2593 | 0.26 | 0.105 | 0.153 |
| chr11 | 103675026 | 103677968 | 2943 | 0.35 | 0.105 | 0.153 |
| chr16 | 85672521 | 85675779 | 3259 | -0.65 | 0.105 | 0.154 |
| chr15 | 50343468 | 50344086 | 619 | -1.63 | 0.105 | 0.154 |
| chr14 | 34857848 | 34858839 | 992 | -0.92 | 0.105 | 0.154 |
| chr1 | 244004712 | 244015559 | 10848 | 0.15 | 0.105 | 0.154 |
| chr8 | 23574798 | 23579586 | 4789 | -0.33 | 0.106 | 0.154 |
| chr18 | 61088240 | 61090942 | 2703 | 0.21 | 0.106 | 0.154 |
| chr18 | 13037321 | 13037856 | 536 | -1.15 | 0.106 | 0.154 |
| chr1 | 54516980 | 54520184 | 3205 | 0.33 | 0.106 | 0.154 |
| chr16 | 1047023 | 1053638 | 6616 | 0.4 | 0.106 | 0.154 |
| chr3 | 104077671 | 104090977 | 13307 | -0.26 | 0.106 | 0.154 |

| | | | | | | |
|---|---|---|---|---|---|---|
| chr6 | 42837805 | 42859434 | 21630 | 0.17 | 0.106 | 0.154 |
| chr7 | 32728045 | 32728657 | 613 | 0.95 | 0.106 | 0.154 |
| chr9 | 116719238 | 116721243 | 2006 | 0.33 | 0.106 | 0.154 |
| chr19 | 42443984 | 42445070 | 1087 | 0.53 | 0.106 | 0.154 |
| chr1 | 217735315 | 217736481 | 1167 | 0.62 | 0.106 | 0.154 |
| chr15 | 41987968 | 41989113 | 1146 | 0.78 | 0.106 | 0.154 |
| chr2 | 227227478 | 227228318 | 841 | 0.85 | 0.106 | 0.154 |
| chr11 | 75524548 | 75538287 | 13740 | -0.26 | 0.106 | 0.155 |
| chr5 | 64330254 | 64332548 | 2295 | 0.25 | 0.106 | 0.155 |
| chr5 | 52896734 | 52900957 | 4224 | 0.43 | 0.106 | 0.155 |
| chr5 | 176673188 | 176697771 | 24584 | -0.24 | 0.106 | 0.155 |
| chr6 | 8101774 | 8109227 | 7454 | -0.3 | 0.106 | 0.155 |
| chr4 | 106342917 | 106343609 | 693 | 1.27 | 0.106 | 0.155 |
| chr4 | 5709421 | 5726680 | 17260 | -0.19 | 0.107 | 0.155 |
| chrX | 35717813 | 35718779 | 967 | 0.96 | 0.107 | 0.155 |
| chr16 | 22357928 | 22398824 | 40897 | -0.15 | 0.107 | 0.155 |
| chr5 | 140295584 | 140307137 | 11554 | 2.99 | 0.107 | 0.155 |
| chr2 | 139294327 | 139303093 | 8767 | -0.99 | 0.107 | 0.155 |
| chr1 | 54302005 | 54304467 | 2463 | -0.28 | 0.107 | 0.155 |
| chr14 | 75229058 | 75231699 | 2642 | -0.24 | 0.107 | 0.155 |
| chr9 | 110473335 | 110474659 | 1325 | 0.62 | 0.107 | 0.155 |
| chr6 | 41679747 | 41710582 | 30836 | -0.19 | 0.107 | 0.155 |
| chr3 | 123017791 | 123030303 | 12513 | -0.71 | 0.107 | 0.156 |
| chr17 | 66118042 | 66127746 | 9705 | -0.78 | 0.107 | 0.156 |
| chr3 | 156965953 | 156966863 | 911 | 0.8 | 0.107 | 0.156 |
| chr7 | 19155818 | 19157307 | 1490 | 0.59 | 0.107 | 0.156 |
| chr13 | 80585815 | 80586602 | 788 | 0.93 | 0.107 | 0.156 |
| chr11 | 12279394 | 12280933 | 1540 | 0.54 | 0.107 | 0.156 |
| chr18 | 39577130 | 39579261 | 2132 | 0.53 | 0.107 | 0.156 |
| chr17 | 63738573 | 63739440 | 868 | -1.36 | 0.107 | 0.156 |
| chr12 | 87182486 | 87183349 | 864 | 2.28 | 0.107 | 0.156 |
| chr3 | 187568307 | 187571600 | 3294 | 2.78 | 0.107 | 0.156 |
| chr20 | 47376886 | 47377574 | 689 | -0.94 | 0.107 | 0.156 |
| chr7 | 3800379 | 3802185 | 1807 | 0.37 | 0.107 | 0.156 |
| chr2 | 10008087 | 10009581 | 1495 | -0.83 | 0.107 | 0.156 |
| chr18 | 844391 | 848878 | 4488 | 0.37 | 0.107 | 0.156 |
| chr17 | 61655350 | 61656851 | 1502 | -0.93 | 0.107 | 0.156 |
| chr7 | 95932191 | 95933486 | 1296 | 0.8 | 0.108 | 0.156 |
| chr5 | 72900069 | 73006756 | 106688 | -0.09 | 0.108 | 0.156 |
| chr1 | 224075195 | 224076473 | 1279 | 0.54 | 0.108 | 0.156 |
| chr17 | 16380238 | 16390542 | 10305 | -0.52 | 0.108 | 0.156 |
| chr19 | 21203391 | 21206454 | 3064 | 0.48 | 0.108 | 0.156 |
| chr14 | 28750184 | 28751049 | 866 | 0.81 | 0.108 | 0.156 |
| chr4 | 25359033 | 25360872 | 1840 | 0.61 | 0.108 | 0.156 |
| chr6 | 24402754 | 24404369 | 1616 | -0.43 | 0.108 | 0.157 |
| chr9 | 129904996 | 129905958 | 963 | 0.87 | 0.108 | 0.157 |
| chr11 | 3898654 | 3927723 | 29070 | -0.15 | 0.108 | 0.157 |
| chr6 | 147831338 | 147832605 | 1268 | -0.74 | 0.108 | 0.157 |
| chr8 | 125240720 | 125241812 | 1093 | -0.6 | 0.108 | 0.157 |
| chr20 | 45809439 | 45810466 | 1028 | 0.57 | 0.108 | 0.157 |
| chr11 | 31705777 | 31706642 | 866 | 1.36 | 0.108 | 0.157 |
| chr15 | 40212773 | 40235024 | 22252 | 0.27 | 0.108 | 0.157 |
| chr2 | 71785310 | 71786194 | 885 | -0.92 | 0.108 | 0.157 |
| chr1 | 156540099 | 156542393 | 2295 | 0.34 | 0.108 | 0.157 |
| chr19 | 4812099 | 4816401 | 4303 | 0.28 | 0.108 | 0.157 |
| chr10 | 60144081 | 60154188 | 10108 | 0.23 | 0.108 | 0.157 |
| chr5 | 139536172 | 139537508 | 1337 | 0.6 | 0.108 | 0.157 |
| chr2 | 65526650 | 65528853 | 2204 | -0.48 | 0.108 | 0.157 |
| chr12 | 69550413 | 69551329 | 917 | 0.74 | 0.108 | 0.157 |
| chr10 | 114231870 | 114232224 | 355 | -1.09 | 0.108 | 0.157 |
| chr14 | 78157660 | 78158605 | 946 | -0.77 | 0.108 | 0.157 |

| | | | | | | |
|---|---|---|---|---|---|---|
| chr5 | 80540057 | 80542714 | 2658 | -0.61 | 0.108 | 0.157 |
| chr12 | 76083804 | 76085001 | 1198 | -0.56 | 0.109 | 0.157 |
| chr1 | 36183576 | 36185423 | 1848 | -0.3 | 0.109 | 0.158 |
| chr2 | 145209819 | 145211318 | 1500 | -0.4 | 0.109 | 0.158 |
| chr1 | 212320309 | 212321465 | 1157 | 0.49 | 0.109 | 0.158 |
| chr1 | 68084032 | 68085110 | 1079 | 0.67 | 0.109 | 0.158 |
| chr2 | 102842614 | 102857699 | 15086 | -0.17 | 0.109 | 0.158 |
| chr17 | 20902789 | 20956087 | 53299 | 0.12 | 0.109 | 0.158 |
| chr3 | 17473198 | 17475052 | 1855 | -0.36 | 0.109 | 0.158 |
| chr1 | 207911167 | 207926920 | 15754 | 0.35 | 0.109 | 0.158 |
| chr11 | 12694811 | 12945753 | 250943 | -0.04 | 0.109 | 0.158 |
| chr8 | 98684446 | 98703826 | 19381 | -0.26 | 0.109 | 0.158 |
| chr10 | 121301862 | 121303752 | 1891 | -0.46 | 0.109 | 0.158 |
| chr19 | 555556 | 557360 | 1805 | 0.5 | 0.109 | 0.158 |
| chr7 | 50859254 | 50861812 | 2559 | -0.37 | 0.109 | 0.158 |
| chr1 | 59536573 | 59537566 | 994 | 0.92 | 0.109 | 0.158 |
| chr17 | 39894466 | 39900205 | 5740 | -0.32 | 0.109 | 0.158 |
| chr3 | 23531548 | 23532520 | 973 | 0.74 | 0.109 | 0.158 |
| chr20 | 33678820 | 33684975 | 6156 | 0.23 | 0.109 | 0.159 |
| chr15 | 62618701 | 62619293 | 593 | 1.04 | 0.109 | 0.159 |
| chr9 | 133302522 | 133302980 | 459 | -1.02 | 0.109 | 0.159 |
| chr1 | 47815872 | 47817842 | 1971 | -0.4 | 0.109 | 0.159 |
| chrX | 45155050 | 45155668 | 619 | 1.02 | 0.11 | 0.159 |
| chr17 | 71870200 | 71870824 | 625 | -1.59 | 0.11 | 0.159 |
| chr14 | 76506947 | 76507469 | 523 | -1.63 | 0.11 | 0.159 |
| chr1 | 16084798 | 16128784 | 43987 | 0.11 | 0.11 | 0.159 |
| chr3 | 185702750 | 185704414 | 1665 | -0.58 | 0.11 | 0.159 |
| chr9 | 127520513 | 127555707 | 35195 | -0.18 | 0.11 | 0.159 |
| chr1 | 241161140 | 241161609 | 470 | 1.02 | 0.11 | 0.159 |
| chr15 | 41406559 | 41421680 | 15122 | 0.15 | 0.11 | 0.159 |
| chr3 | 109883366 | 109886143 | 2778 | 0.31 | 0.11 | 0.159 |
| chr9 | 138798606 | 138801014 | 2409 | 0.52 | 0.11 | 0.159 |
| chr1 | 201083085 | 201084382 | 1298 | -0.53 | 0.11 | 0.159 |
| chr19 | 8478194 | 8525481 | 47288 | 0.13 | 0.11 | 0.159 |
| chr15 | 65241678 | 65243902 | 2225 | -0.53 | 0.11 | 0.16 |
| chr3 | 72640253 | 72641836 | 1584 | 0.64 | 0.11 | 0.16 |
| chr6 | 44414523 | 44420696 | 6174 | 0.54 | 0.11 | 0.16 |
| chr12 | 49333887 | 49352266 | 18380 | 0.22 | 0.11 | 0.16 |
| chr11 | 8127928 | 8128495 | 568 | -1.62 | 0.11 | 0.16 |
| chr19 | 44644948 | 44646609 | 1662 | 0.36 | 0.11 | 0.16 |
| chr5 | 65438928 | 65443011 | 4084 | 0.17 | 0.11 | 0.16 |
| chr6 | 116954422 | 116955154 | 733 | -1.08 | 0.11 | 0.16 |
| chr15 | 101688865 | 101689423 | 559 | 1.16 | 0.11 | 0.16 |
| chr7 | 4922655 | 4923370 | 716 | -0.88 | 0.11 | 0.16 |
| chr1 | 31374350 | 31376147 | 1798 | 0.56 | 0.11 | 0.16 |
| chr1 | 160093429 | 160098345 | 4917 | 0.24 | 0.11 | 0.16 |
| chr19 | 17716457 | 17717991 | 1535 | 0.44 | 0.11 | 0.16 |
| chr1 | 180531644 | 180532766 | 1123 | -0.64 | 0.11 | 0.16 |
| chr1 | 53703103 | 53716112 | 13010 | 0.24 | 0.11 | 0.16 |
| chr2 | 197790311 | 197792434 | 2124 | -0.32 | 0.11 | 0.16 |
| chr1 | 111667558 | 111668373 | 816 | -0.88 | 0.111 | 0.16 |
| chr5 | 146270155 | 146270825 | 671 | -0.96 | 0.111 | 0.16 |
| chr11 | 111984407 | 111984752 | 346 | 0.94 | 0.111 | 0.16 |
| chr6 | 21281072 | 21282198 | 1127 | 1.63 | 0.111 | 0.16 |
| chr14 | 53018512 | 53019284 | 773 | 0.6 | 0.111 | 0.16 |
| chr22 | 24734901 | 24735269 | 369 | 1.21 | 0.111 | 0.16 |
| chr6 | 75459017 | 75460601 | 1585 | 2.6 | 0.111 | 0.16 |
| chr12 | 107484837 | 107488776 | 3940 | 0.15 | 0.111 | 0.16 |
| chr20 | 46907697 | 46928892 | 21196 | -0.26 | 0.111 | 0.16 |
| chr5 | 55163466 | 55165968 | 2503 | -0.32 | 0.111 | 0.16 |
| chr10 | 133955884 | 133956673 | 790 | -0.97 | 0.111 | 0.161 |

| | | | | | | |
|---|---|---|---|---|---|---|
| chr12 | 77193230 | 77194389 | 1160 | -0.8 | 0.111 | 0.161 |
| chr2 | 188364660 | 188369531 | 4872 | -0.21 | 0.111 | 0.161 |
| chr9 | 100637132 | 100640877 | 3746 | -0.18 | 0.111 | 0.161 |
| chr1 | 184258693 | 184259761 | 1069 | -2.49 | 0.111 | 0.161 |
| chr3 | 25795480 | 25796109 | 630 | 1.15 | 0.111 | 0.161 |
| chr2 | 236735551 | 236737746 | 2196 | -0.45 | 0.111 | 0.161 |
| chr7 | 80240272 | 80241672 | 1401 | 0.47 | 0.111 | 0.161 |
| chr9 | 85895734 | 85897146 | 1413 | -0.53 | 0.111 | 0.161 |
| chr2 | 145088887 | 145091151 | 2265 | -0.3 | 0.111 | 0.161 |
| chr6 | 148089919 | 148104383 | 14465 | 2.96 | 0.111 | 0.161 |
| chr14 | 103216620 | 103217176 | 557 | -1.33 | 0.111 | 0.161 |
| chr17 | 80673736 | 80675737 | 2002 | 0.45 | 0.112 | 0.161 |
| chr8 | 11368505 | 11369179 | 675 | 1.08 | 0.112 | 0.161 |
| chr1 | 151734470 | 151737117 | 2648 | 0.22 | 0.112 | 0.161 |
| chr17 | 71087891 | 71089492 | 1602 | -0.36 | 0.112 | 0.162 |
| chr8 | 66698697 | 66700096 | 1400 | 0.71 | 0.112 | 0.162 |
| chr9 | 140008835 | 140025231 | 16397 | -0.31 | 0.112 | 0.162 |
| chr11 | 130903222 | 130904039 | 818 | 0.99 | 0.112 | 0.162 |
| chr1 | 24165276 | 24165750 | 475 | 1.03 | 0.112 | 0.162 |
| chr17 | 49263112 | 49267768 | 4657 | -0.43 | 0.112 | 0.162 |
| chr5 | 33011531 | 33012425 | 895 | -0.74 | 0.112 | 0.162 |
| chr17 | 29106826 | 29172774 | 65949 | 0.13 | 0.112 | 0.162 |
| chr2 | 108809000 | 108811374 | 2375 | -0.41 | 0.112 | 0.162 |
| chr11 | 13298336 | 13300195 | 1860 | 0.5 | 0.112 | 0.162 |
| chr10 | 115345531 | 115346226 | 696 | -0.99 | 0.112 | 0.162 |
| chr18 | 21082704 | 21129703 | 47000 | -0.13 | 0.112 | 0.162 |
| chr9 | 139516909 | 139520059 | 3151 | -0.53 | 0.112 | 0.162 |
| chr1 | 32341169 | 32341844 | 676 | 0.87 | 0.112 | 0.162 |
| chr3 | 120066463 | 120069920 | 3458 | -0.26 | 0.112 | 0.162 |
| chr4 | 16252426 | 16252759 | 334 | -1.37 | 0.112 | 0.162 |
| chr7 | 140237048 | 140243206 | 6159 | 0.19 | 0.112 | 0.162 |
| chr17 | 11992605 | 11993793 | 1189 | 0.69 | 0.112 | 0.162 |
| chr4 | 174132427 | 174140397 | 7971 | -0.36 | 0.112 | 0.162 |
| chr11 | 8927965 | 8939314 | 11350 | 0.91 | 0.112 | 0.162 |
| chr3 | 64128623 | 64138752 | 10130 | -0.16 | 0.112 | 0.162 |
| chr2 | 106777663 | 106779511 | 1849 | 0.73 | 0.113 | 0.163 |
| chr9 | 111033516 | 111035212 | 1697 | 0.48 | 0.113 | 0.163 |
| chr10 | 126091828 | 126092939 | 1112 | -0.8 | 0.113 | 0.163 |
| chr5 | 92621013 | 92621758 | 746 | -0.77 | 0.113 | 0.163 |
| chr12 | 26424054 | 26429192 | 5139 | 0.16 | 0.113 | 0.163 |
| chr13 | 101015100 | 101016727 | 1628 | 0.69 | 0.113 | 0.163 |
| chr7 | 65540140 | 65542128 | 1989 | 0.35 | 0.113 | 0.163 |
| chr8 | 106796197 | 106796717 | 521 | -1.63 | 0.113 | 0.163 |
| chr17 | 25960401 | 25961278 | 878 | -1.02 | 0.113 | 0.163 |
| chr1 | 118470714 | 118473743 | 3030 | 0.38 | 0.113 | 0.163 |
| chr3 | 20080669 | 20096889 | 16221 | 0.43 | 0.113 | 0.163 |
| chr7 | 117749237 | 117750178 | 942 | 2.49 | 0.113 | 0.163 |
| chr12 | 101852344 | 101854534 | 2191 | 0.24 | 0.113 | 0.163 |
| chr10 | 104087999 | 104089442 | 1444 | -1.06 | 0.113 | 0.164 |
| chr3 | 194875284 | 194876149 | 866 | -0.96 | 0.113 | 0.164 |
| chr16 | 46863101 | 46865880 | 2780 | 0.29 | 0.113 | 0.164 |
| chr10 | 75894090 | 75912261 | 18172 | 0.19 | 0.113 | 0.164 |
| chr1 | 167934976 | 167935688 | 713 | 0.88 | 0.113 | 0.164 |
| chr4 | 101570256 | 101571075 | 820 | 0.83 | 0.113 | 0.164 |
| chr5 | 56433374 | 56434203 | 830 | -1.04 | 0.114 | 0.164 |
| chr12 | 112927525 | 112929192 | 1668 | -0.5 | 0.114 | 0.164 |
| chr12 | 50156397 | 50158279 | 1883 | 0.85 | 0.114 | 0.164 |
| chr2 | 231674151 | 231675021 | 871 | -1.1 | 0.114 | 0.164 |
| chr18 | 18633073 | 18633967 | 895 | -1.06 | 0.114 | 0.164 |
| chr5 | 149144707 | 149145138 | 432 | -1.66 | 0.114 | 0.164 |
| chr12 | 79666794 | 79667271 | 478 | -1.61 | 0.114 | 0.164 |

| | | | | | | |
|---|---|---|---|---|---|---|
| chr2 | 205010202 | 205010895 | 694 | -0.6 | 0.114 | 0.164 |
| chr8 | 125485202 | 125489493 | 4292 | 0.19 | 0.114 | 0.164 |
| chr22 | 31555514 | 31556832 | 1319 | 0.46 | 0.114 | 0.164 |
| chr11 | 41900723 | 41901416 | 694 | -0.94 | 0.114 | 0.164 |
| chr5 | 150787940 | 150788996 | 1057 | 1.04 | 0.114 | 0.164 |
| chr15 | 56440400 | 56441850 | 1451 | 2.52 | 0.114 | 0.164 |
| chr5 | 67563991 | 67564781 | 791 | 0.78 | 0.114 | 0.164 |
| chr17 | 64974 | 66519 | 1546 | -0.5 | 0.114 | 0.164 |
| chr6 | 35063825 | 35064950 | 1126 | 0.81 | 0.114 | 0.164 |
| chr3 | 11375635 | 11376605 | 971 | -0.98 | 0.114 | 0.164 |
| chr5 | 110427555 | 110432488 | 4934 | 0.39 | 0.114 | 0.164 |
| chr18 | 77711287 | 77749572 | 38286 | 0.16 | 0.114 | 0.164 |
| chr19 | 14527591 | 14531225 | 3635 | 0.35 | 0.114 | 0.165 |
| chr20 | 62543834 | 62546972 | 3139 | -0.52 | 0.114 | 0.165 |
| chr1 | 225922748 | 225936546 | 13799 | 0.17 | 0.114 | 0.165 |
| chrX | 149135607 | 149136147 | 541 | -1.38 | 0.114 | 0.165 |
| chr4 | 137290061 | 137290856 | 796 | 0.94 | 0.114 | 0.165 |
| chr19 | 15217401 | 15219473 | 2073 | -0.34 | 0.114 | 0.165 |
| chr2 | 46323683 | 46327628 | 3946 | 0.2 | 0.114 | 0.165 |
| chr1 | 59164256 | 59166381 | 2126 | 0.33 | 0.114 | 0.165 |
| chr2 | 42908632 | 42921713 | 13082 | -1.36 | 0.114 | 0.165 |
| chr5 | 67631147 | 67641195 | 10049 | -0.92 | 0.114 | 0.165 |
| chr2 | 236593689 | 236594207 | 519 | -0.97 | 0.114 | 0.165 |
| chr4 | 131521856 | 131522209 | 354 | -1.16 | 0.114 | 0.165 |
| chr3 | 126372988 | 126374311 | 1324 | 0.59 | 0.114 | 0.165 |
| chr5 | 52404770 | 52406390 | 1621 | 0.47 | 0.114 | 0.165 |
| chr11 | 33394119 | 33394585 | 467 | -1.19 | 0.115 | 0.165 |
| chr11 | 71198934 | 71200531 | 1598 | -0.57 | 0.115 | 0.165 |
| chr8 | 95154457 | 95154988 | 532 | 0.99 | 0.115 | 0.165 |
| chr1 | 222790448 | 222791987 | 1540 | 0.49 | 0.115 | 0.165 |
| chr18 | 55252074 | 55255695 | 3622 | -0.3 | 0.115 | 0.165 |
| chr19 | 52799703 | 52801334 | 1632 | 0.3 | 0.115 | 0.165 |
| chr4 | 77770711 | 77771635 | 925 | 0.81 | 0.115 | 0.165 |
| chr7 | 98569271 | 98569810 | 540 | 1.04 | 0.115 | 0.165 |
| chr13 | 35952735 | 35953965 | 1231 | -0.5 | 0.115 | 0.165 |
| chr6 | 121704584 | 121705177 | 594 | -1.01 | 0.115 | 0.166 |
| chr2 | 228660875 | 228661771 | 897 | -1.05 | 0.115 | 0.166 |
| chrX | 130875281 | 130898397 | 23117 | -0.2 | 0.115 | 0.166 |
| chr4 | 170021083 | 170038957 | 17875 | -0.2 | 0.115 | 0.166 |
| chr10 | 65348264 | 65349341 | 1078 | 0.63 | 0.115 | 0.166 |
| chrX | 122744829 | 122745325 | 497 | -1.23 | 0.115 | 0.166 |
| chr6 | 134608217 | 134618074 | 9858 | 0.2 | 0.115 | 0.166 |
| chr3 | 149625939 | 149626991 | 1053 | 0.62 | 0.115 | 0.166 |
| chr1 | 228802357 | 228803341 | 985 | 0.95 | 0.115 | 0.166 |
| chr1 | 26826534 | 26828694 | 2161 | 0.28 | 0.115 | 0.166 |
| chr11 | 43757396 | 43758568 | 1173 | 0.76 | 0.115 | 0.166 |
| chr10 | 121409962 | 121442317 | 32356 | -0.08 | 0.115 | 0.166 |
| chr17 | 40201885 | 40202919 | 1035 | 0.47 | 0.115 | 0.166 |
| chr8 | 74832743 | 74833415 | 673 | 1.06 | 0.116 | 0.166 |
| chr12 | 92425818 | 92429653 | 3836 | 0.41 | 0.116 | 0.166 |
| chr1 | 154687143 | 154688014 | 872 | 0.83 | 0.116 | 0.166 |
| chr13 | 65502939 | 65503637 | 699 | -1.02 | 0.116 | 0.166 |
| chr6 | 14286074 | 14286971 | 898 | 0.98 | 0.116 | 0.166 |
| chr4 | 47855431 | 47855981 | 551 | 0.96 | 0.116 | 0.166 |
| chr6 | 89632774 | 89633566 | 793 | 0.88 | 0.116 | 0.166 |
| chr4 | 145320815 | 145321159 | 345 | -1.15 | 0.116 | 0.166 |
| chr2 | 112703353 | 112704777 | 1425 | -0.47 | 0.116 | 0.166 |
| chr4 | 184285391 | 184285965 | 575 | 0.96 | 0.116 | 0.166 |
| chr4 | 119512091 | 119512805 | 715 | 0.46 | 0.116 | 0.166 |
| chr5 | 109075040 | 109080274 | 5235 | 0.33 | 0.116 | 0.167 |
| chr17 | 33569049 | 33585586 | 16538 | -0.15 | 0.116 | 0.167 |

| | | | | | | |
|---|---|---|---|---|---|---|
| chr15 | 80397260 | 80398618 | 1359 | 0.56 | 0.116 | 0.167 |
| chr4 | 183837393 | 183839671 | 2279 | 0.27 | 0.116 | 0.167 |
| chr3 | 142572057 | 142572863 | 807 | -0.94 | 0.116 | 0.167 |
| chr5 | 131745684 | 131773836 | 28153 | 0.15 | 0.116 | 0.167 |
| chr3 | 120003713 | 120004535 | 823 | 0.79 | 0.116 | 0.167 |
| chr1 | 78465929 | 78477103 | 11175 | 0.12 | 0.116 | 0.167 |
| chr16 | 59543420 | 59544776 | 1357 | -0.66 | 0.116 | 0.167 |
| chr8 | 125259260 | 125293265 | 34006 | 0.23 | 0.116 | 0.167 |
| chr20 | 61493498 | 61508366 | 14869 | 0.25 | 0.116 | 0.167 |
| chr10 | 14964712 | 14966216 | 1505 | -0.9 | 0.116 | 0.167 |
| chr4 | 131485042 | 131485576 | 535 | -0.99 | 0.116 | 0.167 |
| chr5 | 106551475 | 106551919 | 445 | 1.05 | 0.116 | 0.167 |
| chr2 | 221333541 | 221334256 | 716 | 1.05 | 0.116 | 0.167 |
| chr2 | 150845334 | 150847105 | 1772 | 0.47 | 0.116 | 0.167 |
| chr12 | 31538871 | 31559934 | 21064 | 0.24 | 0.116 | 0.167 |
| chr2 | 225817730 | 225818303 | 574 | -1.08 | 0.116 | 0.167 |
| chr11 | 47502314 | 47601494 | 99181 | -0.09 | 0.116 | 0.167 |
| chr4 | 183896887 | 183897607 | 721 | 0.92 | 0.116 | 0.167 |
| chr4 | 14372309 | 14381586 | 9278 | 0.49 | 0.116 | 0.167 |
| chr5 | 15008104 | 15019252 | 11149 | 0.26 | 0.117 | 0.167 |
| chr1 | 180123034 | 180138199 | 15166 | 0.31 | 0.117 | 0.168 |
| chr8 | 43046146 | 43047199 | 1054 | -0.64 | 0.117 | 0.168 |
| chr2 | 180839348 | 180839801 | 454 | -1.47 | 0.117 | 0.168 |
| chr15 | 62199478 | 62200321 | 844 | 1.41 | 0.117 | 0.168 |
| chr14 | 91857082 | 91862881 | 5800 | -0.88 | 0.117 | 0.168 |
| chr11 | 77810705 | 77813357 | 2653 | 0.25 | 0.117 | 0.168 |
| chr3 | 107808490 | 107810684 | 2195 | 0.36 | 0.117 | 0.168 |
| chr7 | 631639 | 632346 | 708 | -1.2 | 0.117 | 0.168 |
| chr12 | 68856543 | 68857146 | 604 | -0.97 | 0.117 | 0.168 |
| chr10 | 96390028 | 96390546 | 519 | 1.07 | 0.117 | 0.168 |
| chr10 | 128128194 | 128129157 | 964 | 0.7 | 0.117 | 0.168 |
| chr7 | 98619209 | 98651512 | 32304 | 0.16 | 0.117 | 0.168 |
| chr2 | 101417352 | 101418791 | 1440 | 2.39 | 0.117 | 0.168 |
| chr5 | 145233825 | 145235006 | 1182 | 0.46 | 0.117 | 0.168 |
| chr3 | 24517334 | 24517843 | 510 | 1.14 | 0.117 | 0.168 |
| chr16 | 70556561 | 70572020 | 15460 | 0.17 | 0.117 | 0.168 |
| chrX | 118125590 | 118126462 | 873 | 0.72 | 0.117 | 0.168 |
| chr17 | 28492845 | 28505634 | 12790 | -0.31 | 0.117 | 0.168 |
| chr12 | 10284251 | 10285668 | 1418 | 0.46 | 0.117 | 0.168 |
| chr22 | 29278296 | 29280383 | 2088 | -0.57 | 0.117 | 0.169 |
| chr4 | 83954750 | 83957169 | 2420 | 0.28 | 0.117 | 0.169 |
| chr12 | 132513986 | 132515985 | 2000 | 0.66 | 0.117 | 0.169 |
| chr15 | 54104025 | 54104411 | 387 | -1.13 | 0.117 | 0.169 |
| chr4 | 148233409 | 148234450 | 1042 | 0.63 | 0.118 | 0.169 |
| chr2 | 7057547 | 7058771 | 1225 | -0.61 | 0.118 | 0.169 |
| chrX | 23759463 | 23762394 | 2932 | 0.17 | 0.118 | 0.169 |
| chr1 | 92230339 | 92231387 | 1049 | 1.12 | 0.118 | 0.169 |
| chr12 | 107877742 | 107914596 | 36855 | 0.14 | 0.118 | 0.169 |
| chr2 | 128457360 | 128475233 | 17874 | -0.16 | 0.118 | 0.169 |
| chr10 | 120457250 | 120458666 | 1417 | 0.82 | 0.118 | 0.169 |
| chr21 | 38160085 | 38161341 | 1257 | 0.52 | 0.118 | 0.169 |
| chr6 | 5179813 | 5180651 | 839 | -0.79 | 0.118 | 0.169 |
| chr9 | 34636666 | 34667084 | 30419 | 0.19 | 0.118 | 0.169 |
| chr3 | 38288874 | 38289356 | 483 | 1.02 | 0.118 | 0.169 |
| chr15 | 51295977 | 51297004 | 1028 | 0.7 | 0.118 | 0.169 |
| chr18 | 39618023 | 39628992 | 10970 | 0.34 | 0.118 | 0.169 |
| chr2 | 201165244 | 201203913 | 38670 | 0.15 | 0.118 | 0.169 |
| chr4 | 78268357 | 78269172 | 816 | 0.74 | 0.118 | 0.169 |
| chr5 | 37156095 | 37156645 | 551 | -1.04 | 0.118 | 0.169 |
| chr5 | 109094659 | 109106309 | 11651 | 1.07 | 0.118 | 0.169 |
| chr10 | 67330211 | 67331300 | 1090 | -0.39 | 0.118 | 0.169 |

| | | | | | | |
|---|---|---|---|---|---|---|
| chr6 | 16669162 | 16730233 | 61072 | -0.13 | 0.118 | 0.169 |
| chr11 | 34417235 | 34418399 | 1165 | 0.6 | 0.118 | 0.169 |
| chr2 | 17868973 | 17950216 | 81244 | -0.07 | 0.118 | 0.169 |
| chr19 | 51305100 | 51309035 | 3936 | 0.18 | 0.118 | 0.17 |
| chr2 | 72006299 | 72023830 | 17532 | -3.12 | 0.118 | 0.17 |
| chr4 | 181748301 | 181749260 | 960 | 0.75 | 0.118 | 0.17 |
| chr4 | 159794176 | 159795796 | 1621 | -0.49 | 0.118 | 0.17 |
| chr9 | 113995409 | 113996098 | 690 | -1.37 | 0.118 | 0.17 |
| chr6 | 116936919 | 116938406 | 1488 | -0.43 | 0.118 | 0.17 |
| chr3 | 117551396 | 117551876 | 481 | -1 | 0.118 | 0.17 |
| chr17 | 59857364 | 59857815 | 452 | 1.21 | 0.118 | 0.17 |
| chr18 | 21002432 | 21003884 | 1453 | 0.71 | 0.118 | 0.17 |
| chr15 | 44675515 | 44678440 | 2926 | -0.36 | 0.119 | 0.17 |
| chr16 | 77387268 | 77393539 | 6272 | 0.34 | 0.119 | 0.17 |
| chr1 | 116959936 | 116961863 | 1928 | 0.32 | 0.119 | 0.17 |
| chr12 | 57542156 | 57543961 | 1806 | -0.68 | 0.119 | 0.17 |
| chr13 | 49793773 | 49794570 | 798 | -0.81 | 0.119 | 0.17 |
| chr11 | 60665093 | 60693560 | 28468 | -0.15 | 0.119 | 0.17 |
| chr2 | 135116536 | 135118176 | 1641 | 2.73 | 0.119 | 0.17 |
| chr1 | 23157867 | 23159357 | 1491 | -0.35 | 0.119 | 0.17 |
| chr1 | 204542109 | 204542641 | 533 | 1.02 | 0.119 | 0.17 |
| chr11 | 124668948 | 124671268 | 2321 | 0.26 | 0.119 | 0.17 |
| chr1 | 16010282 | 16011788 | 1507 | 0.48 | 0.119 | 0.17 |
| chr11 | 66444524 | 66446730 | 2207 | 0.23 | 0.119 | 0.17 |
| chr6 | 17816299 | 17818896 | 2598 | 0.3 | 0.119 | 0.17 |
| chr2 | 232378234 | 232379877 | 1644 | -0.32 | 0.119 | 0.17 |
| chr10 | 26985715 | 26987567 | 1853 | 0.3 | 0.119 | 0.17 |
| chr3 | 150233135 | 150233718 | 584 | -1.57 | 0.119 | 0.17 |
| chr2 | 171626786 | 171627224 | 439 | 1.03 | 0.119 | 0.17 |
| chr6 | 35382792 | 35383691 | 900 | -0.86 | 0.119 | 0.17 |
| chr20 | 39806429 | 39807040 | 612 | -1.29 | 0.119 | 0.17 |
| chr12 | 122667454 | 122667980 | 527 | -1.12 | 0.119 | 0.171 |
| chr7 | 94536239 | 94538431 | 2193 | 0.2 | 0.119 | 0.171 |
| chr12 | 31515420 | 31522309 | 6890 | 0.53 | 0.119 | 0.171 |
| chr20 | 35871902 | 35872449 | 548 | -1.16 | 0.119 | 0.171 |
| chr11 | 124044866 | 124045394 | 529 | -0.99 | 0.119 | 0.171 |
| chr17 | 32739766 | 32740552 | 787 | -0.91 | 0.119 | 0.171 |
| chr21 | 30775641 | 30776803 | 1163 | -0.87 | 0.119 | 0.171 |
| chr22 | 45697512 | 45706686 | 9175 | 0.29 | 0.119 | 0.171 |
| chr10 | 93410962 | 93420687 | 9726 | 0.34 | 0.12 | 0.171 |
| chr1 | 233210306 | 233223371 | 13066 | 0.28 | 0.12 | 0.171 |
| chr9 | 133760516 | 133762393 | 1878 | -0.52 | 0.12 | 0.171 |
| chr10 | 73922645 | 74125679 | 203035 | 0.05 | 0.12 | 0.171 |
| chr2 | 71969044 | 71975847 | 6804 | -0.25 | 0.12 | 0.171 |
| chr21 | 36598657 | 36599102 | 446 | -1.15 | 0.12 | 0.171 |
| chr21 | 17003867 | 17004683 | 817 | 0.7 | 0.12 | 0.171 |
| chr9 | 125202128 | 125228093 | 25966 | 1.02 | 0.12 | 0.171 |
| chr15 | 75679950 | 75693591 | 13642 | -0.16 | 0.12 | 0.171 |
| chr6 | 111978088 | 111980411 | 2324 | 2.78 | 0.12 | 0.172 |
| chr16 | 47526829 | 47527551 | 723 | -0.92 | 0.12 | 0.172 |
| chr1 | 24562811 | 24563465 | 655 | -1.03 | 0.12 | 0.172 |
| chr3 | 41549289 | 41549821 | 533 | -1 | 0.12 | 0.172 |
| chr4 | 2413390 | 2421280 | 7891 | 0.19 | 0.12 | 0.172 |
| chr11 | 70048628 | 70059132 | 10505 | 0.23 | 0.12 | 0.172 |
| chr16 | 50059097 | 50069533 | 10437 | 0.32 | 0.12 | 0.172 |
| chr5 | 172482718 | 172486146 | 3429 | 0.25 | 0.12 | 0.172 |
| chr9 | 131579922 | 131592113 | 12192 | 0.34 | 0.12 | 0.172 |
| chr3 | 45634261 | 45732826 | 98566 | 0.08 | 0.12 | 0.172 |
| chr3 | 179168423 | 179169879 | 1457 | 0.53 | 0.12 | 0.172 |
| chr5 | 168139209 | 168140618 | 1410 | -0.49 | 0.12 | 0.172 |
| chr9 | 110445892 | 110446304 | 413 | -1.59 | 0.12 | 0.172 |

| | | | | | | |
|---|---|---|---|---|---|---|
| chr2 | 20207927 | 20213402 | 5476 | -0.34 | 0.12 | 0.172 |
| chr8 | 61186067 | 61194491 | 8425 | 0.19 | 0.12 | 0.172 |
| chr2 | 37570803 | 37578438 | 7636 | 0.24 | 0.121 | 0.172 |
| chr5 | 93136670 | 93137087 | 418 | 1.02 | 0.121 | 0.172 |
| chr16 | 19341231 | 19343104 | 1874 | 0.43 | 0.121 | 0.172 |
| chr13 | 65485855 | 65486984 | 1130 | -0.84 | 0.121 | 0.172 |
| chr14 | 62303934 | 62304387 | 454 | -1.12 | 0.121 | 0.172 |
| chr6 | 10306141 | 10306933 | 793 | 0.71 | 0.121 | 0.172 |
| chr20 | 47154585 | 47156056 | 1472 | 0.82 | 0.121 | 0.172 |
| chr8 | 82517408 | 82518676 | 1269 | 0.64 | 0.121 | 0.172 |
| chr1 | 233160986 | 233161644 | 659 | -1.02 | 0.121 | 0.172 |
| chr11 | 85393457 | 85394141 | 685 | 0.68 | 0.121 | 0.173 |
| chr19 | 6108381 | 6112047 | 3667 | 0.3 | 0.121 | 0.173 |
| chr15 | 79011755 | 79021420 | 9666 | -0.45 | 0.121 | 0.173 |
| chr7 | 24516580 | 24518695 | 2116 | -0.22 | 0.121 | 0.173 |
| chr16 | 66636943 | 66645418 | 8476 | 0.2 | 0.121 | 0.173 |
| chr2 | 135848923 | 135852206 | 3284 | 0.42 | 0.121 | 0.173 |
| chr14 | 100304812 | 100305826 | 1015 | 0.69 | 0.121 | 0.173 |
| chr2 | 134325496 | 134326235 | 740 | -0.92 | 0.121 | 0.173 |
| chr9 | 88677720 | 88678958 | 1239 | -0.83 | 0.121 | 0.173 |
| chr6 | 11093044 | 11096627 | 3584 | 0.25 | 0.121 | 0.173 |
| chr5 | 95047957 | 95048845 | 889 | 2.47 | 0.121 | 0.173 |
| chr7 | 156575868 | 156577965 | 2098 | -0.54 | 0.121 | 0.173 |
| chr20 | 62289077 | 62290595 | 1519 | 0.51 | 0.121 | 0.173 |
| chr6 | 4775135 | 4778333 | 3199 | 0.37 | 0.121 | 0.173 |
| chr2 | 28077862 | 28079148 | 1287 | -0.58 | 0.121 | 0.173 |
| chr1 | 79728384 | 79728789 | 406 | 1.17 | 0.122 | 0.174 |
| chr5 | 145552214 | 145563061 | 10848 | 0.2 | 0.122 | 0.174 |
| chr6 | 16651882 | 16652408 | 527 | 1.62 | 0.122 | 0.174 |
| chr9 | 43661476 | 43661793 | 318 | 1.62 | 0.122 | 0.174 |
| chr16 | 11184876 | 11188441 | 3566 | -0.55 | 0.122 | 0.174 |
| chr20 | 32859902 | 32865308 | 5407 | -0.5 | 0.122 | 0.174 |
| chr7 | 64734260 | 64734909 | 650 | -0.68 | 0.122 | 0.174 |
| chr1 | 234081070 | 234082108 | 1039 | 0.71 | 0.122 | 0.174 |
| chr3 | 196943224 | 196948647 | 5424 | 0.53 | 0.122 | 0.174 |
| chr4 | 83565062 | 83566210 | 1149 | 0.84 | 0.122 | 0.174 |
| chr1 | 202425100 | 202425985 | 886 | 0.66 | 0.122 | 0.174 |
| chr11 | 20107219 | 20107623 | 405 | -1.47 | 0.122 | 0.174 |
| chr13 | 73328124 | 73329091 | 968 | -0.72 | 0.122 | 0.174 |
| chr4 | 87845333 | 87860444 | 15112 | 0.16 | 0.122 | 0.174 |
| chr4 | 773959 | 776982 | 3024 | 0.28 | 0.122 | 0.174 |
| chr1 | 169055771 | 169056351 | 581 | 1 | 0.122 | 0.174 |
| chr1 | 204653442 | 204661822 | 8381 | -2.37 | 0.122 | 0.174 |
| chr5 | 175873664 | 175891545 | 17882 | 0.14 | 0.122 | 0.175 |
| chr1 | 167758762 | 167792350 | 33589 | 0.14 | 0.122 | 0.175 |
| chr15 | 73891452 | 73895745 | 4294 | 0.26 | 0.122 | 0.175 |
| chr3 | 53106613 | 53107764 | 1152 | 0.7 | 0.122 | 0.175 |
| chr13 | 80614410 | 80615753 | 1344 | 0.61 | 0.122 | 0.175 |
| chr9 | 35095306 | 35116947 | 21642 | -0.12 | 0.122 | 0.175 |
| chr10 | 95924938 | 95925832 | 895 | -0.84 | 0.122 | 0.175 |
| chr6 | 34351626 | 34360823 | 9198 | -0.32 | 0.122 | 0.175 |
| chr7 | 81696060 | 81698476 | 2417 | 0.26 | 0.123 | 0.175 |
| chr1 | 92763221 | 92768055 | 4835 | 0.22 | 0.123 | 0.175 |
| chr16 | 70284753 | 70294630 | 9878 | -0.3 | 0.123 | 0.175 |
| chr4 | 129241861 | 129245010 | 3150 | -0.32 | 0.123 | 0.175 |
| chr16 | 58298449 | 58300156 | 1708 | 0.56 | 0.123 | 0.175 |
| chr7 | 38947887 | 38948809 | 923 | 0.44 | 0.123 | 0.175 |
| chr3 | 124277637 | 124278757 | 1121 | -0.7 | 0.123 | 0.175 |
| chr22 | 24642933 | 24643449 | 517 | -0.93 | 0.123 | 0.175 |
| chr4 | 110392477 | 110393993 | 1517 | -0.59 | 0.123 | 0.175 |
| chr7 | 69959457 | 69960353 | 897 | 0.93 | 0.123 | 0.175 |

| | | | | | | |
|---|---|---|---|---|---|---|
| chr11 | 15671284 | 15672585 | 1302 | 0.42 | 0.123 | 0.175 |
| chr2 | 38290850 | 38305365 | 14516 | 0.17 | 0.123 | 0.175 |
| chrX | 15508375 | 15512715 | 4341 | -0.2 | 0.123 | 0.175 |
| chr1 | 36272737 | 36274986 | 2250 | 0.47 | 0.123 | 0.175 |
| chr7 | 156516225 | 156516959 | 735 | -1.57 | 0.123 | 0.175 |
| chr5 | 102473536 | 102474365 | 830 | 0.93 | 0.123 | 0.175 |
| chr7 | 18934847 | 18935564 | 718 | 0.89 | 0.123 | 0.175 |
| chr6 | 5752122 | 5753209 | 1088 | -0.88 | 0.123 | 0.175 |
| chr11 | 129011906 | 129018893 | 6988 | -0.35 | 0.123 | 0.175 |
| chr3 | 66052957 | 66054220 | 1264 | 0.51 | 0.123 | 0.176 |
| chr17 | 78793711 | 78808289 | 14579 | -0.19 | 0.123 | 0.176 |
| chr16 | 66838536 | 66839137 | 602 | -1.2 | 0.123 | 0.176 |
| chr8 | 102299769 | 102301428 | 1660 | 0.59 | 0.123 | 0.176 |
| chr6 | 100650242 | 100652155 | 1914 | 0.35 | 0.123 | 0.176 |
| chrX | 107504160 | 107506256 | 2097 | -0.46 | 0.123 | 0.176 |
| chr16 | 30661236 | 30687219 | 25984 | 0.14 | 0.123 | 0.176 |
| chr5 | 121388687 | 121389991 | 1305 | -0.82 | 0.123 | 0.176 |
| chr15 | 102041576 | 102042495 | 920 | -2.43 | 0.123 | 0.176 |
| chr4 | 24974861 | 24975964 | 1104 | -0.74 | 0.124 | 0.176 |
| chr5 | 158721807 | 158722281 | 475 | -1.16 | 0.124 | 0.176 |
| chr12 | 102171218 | 102172381 | 1164 | -0.82 | 0.124 | 0.176 |
| chr7 | 102937197 | 102967063 | 29867 | 0.18 | 0.124 | 0.176 |
| chr13 | 41929792 | 41930434 | 643 | 1.1 | 0.124 | 0.176 |
| chr9 | 6595828 | 6596762 | 935 | 0.93 | 0.124 | 0.176 |
| chr6 | 87863935 | 87867052 | 3118 | 0.3 | 0.124 | 0.176 |
| chr1 | 214587495 | 214587997 | 503 | 1.18 | 0.124 | 0.176 |
| chr9 | 80867067 | 80867401 | 335 | 1.62 | 0.124 | 0.176 |
| chr19 | 7287586 | 7295232 | 7647 | -0.57 | 0.124 | 0.177 |
| chr1 | 17302911 | 17311541 | 8631 | -0.7 | 0.124 | 0.177 |
| chr5 | 123613524 | 123614542 | 1019 | 2.57 | 0.124 | 0.177 |
| chr8 | 24366436 | 24368048 | 1613 | 0.45 | 0.124 | 0.177 |
| chr10 | 17569005 | 17570425 | 1421 | 0.43 | 0.124 | 0.177 |
| chr9 | 98686010 | 98700384 | 14375 | -0.45 | 0.124 | 0.177 |
| chr8 | 11793357 | 11794187 | 831 | 0.88 | 0.124 | 0.177 |
| chr1 | 234906876 | 234908909 | 2034 | 0.2 | 0.124 | 0.177 |
| chr14 | 89679932 | 89688289 | 8358 | -0.41 | 0.124 | 0.177 |
| chr8 | 71227832 | 71228795 | 964 | -0.71 | 0.124 | 0.177 |
| chr22 | 37373677 | 37374184 | 508 | 0.83 | 0.124 | 0.177 |
| chr16 | 69892877 | 69894487 | 1611 | -0.58 | 0.124 | 0.177 |
| chr3 | 29874237 | 29874680 | 444 | -1.06 | 0.124 | 0.177 |
| chr4 | 114879383 | 114883795 | 4413 | 0.22 | 0.124 | 0.177 |
| chr17 | 61505301 | 61524751 | 19451 | -0.16 | 0.124 | 0.177 |
| chr17 | 3538804 | 3543937 | 5134 | 0.28 | 0.124 | 0.177 |
| chr5 | 58710472 | 58726377 | 15906 | -0.22 | 0.124 | 0.177 |
| chr17 | 63583489 | 63585364 | 1876 | -0.61 | 0.124 | 0.177 |
| chr6 | 134684197 | 134695283 | 11087 | -0.45 | 0.124 | 0.177 |
| chr2 | 127808846 | 127874092 | 65247 | 0.09 | 0.125 | 0.177 |
| chr19 | 47000438 | 47001605 | 1168 | -0.74 | 0.125 | 0.177 |
| chr19 | 43335618 | 43337965 | 2348 | 0.33 | 0.125 | 0.177 |
| chr12 | 83409609 | 83410394 | 786 | 0.86 | 0.125 | 0.177 |
| chr1 | 240254873 | 240256989 | 2117 | -0.56 | 0.125 | 0.178 |
| chr6 | 140029650 | 140030108 | 459 | 1.22 | 0.125 | 0.178 |
| chr6 | 49430076 | 49432033 | 1958 | 0.26 | 0.125 | 0.178 |
| chr1 | 153860028 | 153879798 | 19771 | 0.22 | 0.125 | 0.178 |
| chr3 | 117265772 | 117268677 | 2906 | -0.42 | 0.125 | 0.178 |
| chr20 | 40271371 | 40273000 | 1630 | -0.36 | 0.125 | 0.178 |
| chr7 | 28218777 | 28219495 | 719 | 0.67 | 0.125 | 0.178 |
| chr11 | 68444494 | 68445609 | 1116 | 0.8 | 0.125 | 0.178 |
| chr19 | 43354261 | 43356305 | 2045 | 0.37 | 0.125 | 0.178 |
| chr2 | 152190717 | 152191341 | 625 | -0.95 | 0.125 | 0.178 |
| chr3 | 196438034 | 196444590 | 6557 | 0.28 | 0.125 | 0.178 |

| | | | | | | |
|---|---|---|---|---|---|---|
| chr10 | 30440912 | 30441583 | 672 | -1.06 | 0.125 | 0.178 |
| chr16 | 3155763 | 3186186 | 30424 | 0.2 | 0.125 | 0.178 |
| chr5 | 168801522 | 168802354 | 833 | -1.22 | 0.125 | 0.178 |
| chr1 | 39424252 | 39425429 | 1178 | -0.5 | 0.125 | 0.178 |
| chr17 | 41580365 | 41637985 | 57621 | -0.09 | 0.125 | 0.178 |
| chr5 | 175814095 | 175855794 | 41700 | 0.16 | 0.125 | 0.178 |
| chr2 | 55668584 | 55669450 | 867 | 0.79 | 0.125 | 0.178 |
| chr2 | 201979141 | 202023403 | 44263 | -0.07 | 0.125 | 0.178 |
| chr6 | 147608764 | 147609873 | 1110 | 0.6 | 0.126 | 0.178 |
| chr13 | 41768110 | 41769043 | 934 | -0.58 | 0.126 | 0.178 |
| chr15 | 28433135 | 28434458 | 1324 | -0.7 | 0.126 | 0.178 |
| chr8 | 81489529 | 81491362 | 1834 | -0.49 | 0.126 | 0.178 |
| chrX | 55514625 | 55516778 | 2154 | 0.33 | 0.126 | 0.178 |
| chr9 | 6412385 | 6420044 | 7660 | 0.21 | 0.126 | 0.178 |
| chr1 | 28285440 | 28287048 | 1609 | 0.53 | 0.126 | 0.179 |
| chr4 | 57357671 | 57359069 | 1399 | 0.68 | 0.126 | 0.179 |
| chr1 | 112280979 | 112283554 | 2576 | -0.39 | 0.126 | 0.179 |
| chr1 | 145682819 | 145698189 | 15371 | -0.45 | 0.126 | 0.179 |
| chr11 | 85694563 | 85694933 | 371 | -1.33 | 0.126 | 0.179 |
| chr12 | 69909174 | 69922425 | 13252 | 0.69 | 0.126 | 0.179 |
| chr8 | 119633403 | 119634812 | 1410 | 0.67 | 0.126 | 0.179 |
| chr14 | 61412760 | 61413422 | 663 | -0.9 | 0.126 | 0.179 |
| chr3 | 131262173 | 131263575 | 1403 | -0.53 | 0.126 | 0.179 |
| chrX | 27907299 | 27909574 | 2276 | 0.34 | 0.126 | 0.179 |
| chrX | 130036037 | 130037285 | 1249 | 0.36 | 0.126 | 0.179 |
| chr12 | 70609163 | 70610015 | 853 | 0.74 | 0.126 | 0.179 |
| chr11 | 93516920 | 93518955 | 2036 | 0.45 | 0.126 | 0.179 |
| chr15 | 85194509 | 85202580 | 8072 | -0.17 | 0.126 | 0.179 |
| chr4 | 148518453 | 148518951 | 499 | 1.05 | 0.126 | 0.179 |
| chr14 | 55422986 | 55443384 | 20399 | 0.24 | 0.126 | 0.179 |
| chr3 | 129208951 | 129216731 | 7781 | -0.2 | 0.126 | 0.179 |
| chr7 | 73661104 | 73670444 | 9341 | -0.22 | 0.126 | 0.179 |
| chr1 | 145606607 | 145611456 | 4850 | 0.23 | 0.126 | 0.179 |
| chr18 | 10525552 | 10527247 | 1696 | 0.33 | 0.126 | 0.179 |
| chr16 | 88771847 | 88773944 | 2098 | 0.4 | 0.126 | 0.179 |
| chr2 | 164872220 | 164872588 | 369 | -1.2 | 0.126 | 0.179 |
| chr1 | 156181851 | 156214843 | 32993 | -0.16 | 0.126 | 0.179 |
| chr3 | 9424337 | 9484552 | 60216 | 0.09 | 0.126 | 0.179 |
| chr4 | 90460368 | 90471186 | 10819 | 2.71 | 0.126 | 0.179 |
| chr13 | 27705266 | 27705865 | 600 | 0.93 | 0.126 | 0.179 |
| chr20 | 48505465 | 48516774 | 11310 | 0.46 | 0.127 | 0.18 |
| chrX | 45364351 | 45375900 | 11550 | -0.19 | 0.127 | 0.18 |
| chr6 | 76365691 | 76367954 | 2264 | 0.83 | 0.127 | 0.18 |
| chr3 | 176737274 | 176738071 | 798 | -0.84 | 0.127 | 0.18 |
| chr3 | 126753698 | 126755235 | 1538 | 0.66 | 0.127 | 0.18 |
| chr9 | 16191720 | 16222561 | 30842 | 0.21 | 0.127 | 0.18 |
| chr4 | 88156516 | 88162095 | 5580 | 0.26 | 0.127 | 0.18 |
| chr7 | 47292993 | 47731428 | 438436 | -0.04 | 0.127 | 0.18 |
| chr12 | 51009500 | 51010810 | 1311 | 0.56 | 0.127 | 0.18 |
| chr7 | 128200864 | 128201536 | 673 | 0.79 | 0.127 | 0.18 |
| chr4 | 38856711 | 38875903 | 19193 | 0.11 | 0.127 | 0.18 |
| chr2 | 26135838 | 26136853 | 1016 | -0.91 | 0.127 | 0.18 |
| chr7 | 156792335 | 156804091 | 11757 | -0.26 | 0.127 | 0.18 |
| chr6 | 149805854 | 149806855 | 1002 | -0.94 | 0.127 | 0.18 |
| chr2 | 121180171 | 121180593 | 423 | -1.29 | 0.127 | 0.18 |
| chr2 | 220159031 | 220159752 | 722 | -0.7 | 0.127 | 0.18 |
| chr7 | 5794062 | 5821919 | 27858 | -0.18 | 0.127 | 0.18 |
| chr4 | 100642987 | 100645844 | 2858 | 0.3 | 0.127 | 0.18 |
| chr10 | 126449145 | 126464747 | 15603 | 0.41 | 0.127 | 0.181 |
| chrX | 15712846 | 15714034 | 1189 | 0.88 | 0.127 | 0.181 |
| chr9 | 116035965 | 116053197 | 17233 | 0.14 | 0.127 | 0.181 |

| | | | | | | |
|---|---|---|---|---|---|---|
| chr9 | 131037218 | 131039776 | 2559 | 0.34 | 0.127 | 0.181 |
| chr1 | 43813810 | 43839914 | 26105 | 0.13 | 0.128 | 0.181 |
| chr1 | 27136778 | 27137115 | 338 | -1.47 | 0.128 | 0.181 |
| chr2 | 89014751 | 89015557 | 807 | -1.02 | 0.128 | 0.181 |
| chr4 | 119605955 | 119607490 | 1536 | 0.33 | 0.128 | 0.181 |
| chrX | 135960933 | 135963689 | 2757 | 0.21 | 0.128 | 0.181 |
| chr2 | 239143048 | 239149358 | 6311 | -0.64 | 0.128 | 0.181 |
| chr1 | 202555112 | 202569680 | 14569 | -0.17 | 0.128 | 0.181 |
| chr8 | 100510243 | 100511899 | 1657 | -0.39 | 0.128 | 0.181 |
| chr16 | 50401088 | 50409698 | 8611 | -0.34 | 0.128 | 0.181 |
| chr9 | 139607029 | 139607922 | 894 | 0.93 | 0.128 | 0.181 |
| chr2 | 202897225 | 202904621 | 7397 | -0.24 | 0.128 | 0.181 |
| chr3 | 111837715 | 111848943 | 11229 | -0.74 | 0.128 | 0.181 |
| chr17 | 49797 | 50444 | 648 | -1.06 | 0.128 | 0.181 |
| chr20 | 43445894 | 43446574 | 681 | -2.42 | 0.128 | 0.181 |
| chr11 | 36473816 | 36478390 | 4575 | 0.28 | 0.128 | 0.182 |
| chr7 | 34996700 | 34997582 | 883 | -1.13 | 0.128 | 0.182 |
| chr17 | 56735629 | 56775234 | 39606 | 0.17 | 0.128 | 0.182 |
| chr3 | 52803874 | 52805969 | 2096 | -0.28 | 0.128 | 0.182 |
| chr16 | 81128849 | 81148051 | 19203 | -0.28 | 0.128 | 0.182 |
| chr10 | 49598735 | 49600187 | 1453 | 2.69 | 0.128 | 0.182 |
| chr3 | 196864505 | 196865095 | 591 | -1.04 | 0.128 | 0.182 |
| chr6 | 35310377 | 35311002 | 626 | 0.6 | 0.128 | 0.182 |
| chr4 | 11733393 | 11750569 | 17177 | 2.9 | 0.128 | 0.182 |
| chr15 | 99602390 | 99603268 | 879 | -0.44 | 0.129 | 0.182 |
| chr6 | 79576752 | 79578581 | 1830 | -0.28 | 0.129 | 0.182 |
| chr1 | 116914755 | 116921199 | 6445 | 0.22 | 0.129 | 0.182 |
| chr5 | 157002125 | 157003222 | 1098 | -0.73 | 0.129 | 0.182 |
| chr3 | 107369938 | 107370838 | 901 | 0.7 | 0.129 | 0.182 |
| chr1 | 42949902 | 42950654 | 753 | -0.93 | 0.129 | 0.182 |
| chr15 | 91403899 | 91427163 | 23265 | -0.22 | 0.129 | 0.182 |
| chr7 | 128827916 | 128829626 | 1711 | -0.4 | 0.129 | 0.182 |
| chr4 | 126236335 | 126245552 | 9218 | 0.42 | 0.129 | 0.183 |
| chr7 | 129463905 | 129516650 | 52746 | 0.12 | 0.129 | 0.183 |
| chr13 | 35937589 | 35938786 | 1198 | 0.88 | 0.129 | 0.183 |
| chr7 | 127545519 | 127546214 | 696 | 1.09 | 0.129 | 0.183 |
| chr3 | 31740627 | 31747123 | 6497 | -0.27 | 0.129 | 0.183 |
| chr2 | 29419935 | 29420777 | 843 | 0.86 | 0.129 | 0.183 |
| chr10 | 70920387 | 70921167 | 781 | -0.74 | 0.129 | 0.183 |
| chr17 | 35577087 | 35598063 | 20977 | -0.21 | 0.129 | 0.183 |
| chr11 | 122924658 | 122935116 | 10459 | 0.15 | 0.129 | 0.183 |
| chr6 | 659414 | 660165 | 752 | -0.9 | 0.129 | 0.183 |
| chr21 | 40144974 | 40145370 | 397 | -1.51 | 0.13 | 0.183 |
| chr8 | 117560161 | 117560709 | 549 | 0.94 | 0.13 | 0.183 |
| chr15 | 41452175 | 41454660 | 2486 | 0.38 | 0.13 | 0.183 |
| chr4 | 57774093 | 57776433 | 2341 | 0.32 | 0.13 | 0.184 |
| chr2 | 112677581 | 112678071 | 491 | -1.13 | 0.13 | 0.184 |
| chr12 | 49297868 | 49298605 | 738 | -0.53 | 0.13 | 0.184 |
| chr8 | 125641539 | 125642977 | 1439 | 0.57 | 0.13 | 0.184 |
| chr11 | 45938408 | 45953395 | 14988 | -0.28 | 0.13 | 0.184 |
| chr18 | 10578640 | 10579165 | 526 | 1.25 | 0.13 | 0.184 |
| chr11 | 47234756 | 47293023 | 58268 | 0.1 | 0.13 | 0.184 |
| chr7 | 7736077 | 7736796 | 720 | 0.78 | 0.13 | 0.184 |
| chr17 | 79694593 | 79695493 | 901 | 0.87 | 0.13 | 0.184 |
| chrX | 53448121 | 53469462 | 21342 | -0.19 | 0.13 | 0.184 |
| chr6 | 141065921 | 141066569 | 649 | -1.16 | 0.13 | 0.184 |
| chr9 | 35923178 | 35925015 | 1838 | -0.39 | 0.13 | 0.184 |
| chr19 | 14513357 | 14513801 | 445 | 1.45 | 0.13 | 0.184 |
| chr3 | 110132319 | 110140303 | 7985 | 0.2 | 0.13 | 0.184 |
| chr6 | 20932594 | 20939875 | 7282 | 0.4 | 0.13 | 0.184 |
| chr3 | 148673331 | 148674138 | 808 | -0.73 | 0.13 | 0.184 |

| | | | | | | |
|---|---|---|---|---|---|---|
| chr11 | 27860876 | 27861235 | 360 | 0.98 | 0.13 | 0.184 |
| chr8 | 59789018 | 59789713 | 696 | 0.72 | 0.13 | 0.184 |
| chr11 | 68977666 | 68978476 | 811 | -0.9 | 0.13 | 0.184 |
| chr2 | 218517342 | 218518888 | 1547 | -0.47 | 0.131 | 0.184 |
| chr7 | 18466260 | 18466928 | 669 | 0.98 | 0.131 | 0.184 |
| chr6 | 155316465 | 155317253 | 789 | 0.94 | 0.131 | 0.185 |
| chr11 | 47376338 | 47377161 | 824 | -0.68 | 0.131 | 0.185 |
| chr2 | 46427465 | 46428142 | 678 | -0.77 | 0.131 | 0.185 |
| chr2 | 169728410 | 169728791 | 382 | -1.06 | 0.131 | 0.185 |
| chr1 | 229672105 | 229673676 | 1572 | 0.74 | 0.131 | 0.185 |
| chr22 | 39139561 | 39160891 | 21331 | -0.22 | 0.131 | 0.185 |
| chr9 | 125138177 | 125146819 | 8643 | -0.53 | 0.131 | 0.185 |
| chr9 | 129961451 | 129962192 | 742 | -1.57 | 0.131 | 0.185 |
| chr20 | 25676171 | 25678027 | 1857 | 0.26 | 0.131 | 0.185 |
| chr2 | 61371095 | 61373516 | 2422 | 0.22 | 0.131 | 0.185 |
| chr11 | 57173807 | 57174391 | 585 | -1.16 | 0.131 | 0.185 |
| chr3 | 13454951 | 13463086 | 8136 | -0.28 | 0.131 | 0.185 |
| chr5 | 94955894 | 94956298 | 405 | 1.07 | 0.131 | 0.185 |
| chr4 | 114548836 | 114579509 | 30674 | 0.11 | 0.131 | 0.185 |
| chr6 | 16799322 | 16806282 | 6961 | 0.33 | 0.131 | 0.186 |
| chr14 | 101694639 | 101702160 | 7522 | -0.25 | 0.131 | 0.186 |
| chr6 | 15026072 | 15028208 | 2137 | 0.4 | 0.132 | 0.186 |
| chr16 | 22447926 | 22448779 | 854 | 0.43 | 0.132 | 0.186 |
| chr12 | 57212029 | 57213062 | 1034 | -0.6 | 0.132 | 0.186 |
| chr19 | 6218766 | 6219084 | 319 | -1.26 | 0.132 | 0.186 |
| chr9 | 132163279 | 132206292 | 43014 | -0.12 | 0.132 | 0.186 |
| chr12 | 60531441 | 60532629 | 1189 | -0.65 | 0.132 | 0.186 |
| chr19 | 1313316 | 1325648 | 12333 | -0.52 | 0.132 | 0.186 |
| chr4 | 6887337 | 6896547 | 9211 | -0.16 | 0.132 | 0.186 |
| chrX | 39991426 | 39992618 | 1193 | 0.55 | 0.132 | 0.186 |
| chr17 | 78342168 | 78343498 | 1331 | -1.14 | 0.132 | 0.186 |
| chr10 | 76864484 | 76911723 | 47240 | 0.14 | 0.132 | 0.186 |
| chr1 | 10002184 | 10011789 | 9606 | 0.23 | 0.132 | 0.186 |
| chr2 | 196455348 | 196456084 | 737 | 0.94 | 0.132 | 0.186 |
| chr5 | 145581937 | 145597552 | 15616 | -0.18 | 0.132 | 0.186 |
| chr1 | 95582073 | 95583956 | 1884 | 0.43 | 0.132 | 0.186 |
| chr2 | 54242693 | 54244291 | 1599 | -0.4 | 0.132 | 0.186 |
| chr11 | 73896650 | 73897243 | 594 | -1.02 | 0.132 | 0.186 |
| chr3 | 23870850 | 23872345 | 1496 | -0.41 | 0.132 | 0.186 |
| chr9 | 12759785 | 12794896 | 35112 | -0.09 | 0.132 | 0.186 |
| chr17 | 37556595 | 37558770 | 2176 | 0.31 | 0.132 | 0.186 |
| chr3 | 16142356 | 16145905 | 3550 | 0.21 | 0.132 | 0.186 |
| chr20 | 24938216 | 24951775 | 13560 | -0.9 | 0.132 | 0.186 |
| chr22 | 42639185 | 42640385 | 1201 | -0.54 | 0.132 | 0.186 |
| chr1 | 173924949 | 173947747 | 22799 | 0.21 | 0.132 | 0.187 |
| chrX | 118739472 | 118741312 | 1841 | 0.3 | 0.132 | 0.187 |
| chr5 | 60240011 | 60242057 | 2047 | 0.35 | 0.132 | 0.187 |
| chr7 | 33708493 | 33709042 | 550 | 1.29 | 0.132 | 0.187 |
| chr1 | 227456387 | 227456989 | 603 | 1.08 | 0.133 | 0.187 |
| chr20 | 13392756 | 13417720 | 24965 | -0.21 | 0.133 | 0.187 |
| chr15 | 91327522 | 91328333 | 812 | -0.9 | 0.133 | 0.187 |
| chr13 | 105722245 | 105723529 | 1285 | 0.75 | 0.133 | 0.187 |
| chr4 | 48900834 | 48909422 | 8589 | 0.25 | 0.133 | 0.187 |
| chr2 | 25537223 | 25538820 | 1598 | -0.47 | 0.133 | 0.187 |
| chrX | 122781564 | 122783299 | 1736 | 0.56 | 0.133 | 0.187 |
| chr1 | 46418117 | 46423773 | 5657 | 0.14 | 0.133 | 0.187 |
| chr21 | 36528069 | 36532587 | 4519 | 2.7 | 0.133 | 0.187 |
| chr1 | 225997258 | 226014467 | 17210 | -0.16 | 0.133 | 0.188 |
| chr3 | 194353069 | 194355129 | 2061 | 0.36 | 0.133 | 0.188 |
| chr12 | 30878396 | 30891849 | 13454 | 0.3 | 0.133 | 0.188 |
| chr6 | 54200052 | 54200937 | 886 | -0.67 | 0.133 | 0.188 |

| | | | | | | |
|---|---|---|---|---|---|---|
| chr16 | 58065100 | 58065542 | 443 | -1.57 | 0.133 | 0.188 |
| chr11 | 94706054 | 94707536 | 1483 | 0.42 | 0.133 | 0.188 |
| chr22 | 27196602 | 27197125 | 524 | -0.87 | 0.133 | 0.188 |
| chr15 | 78831874 | 78839396 | 7523 | 0.15 | 0.133 | 0.188 |
| chr3 | 15373067 | 15381969 | 8903 | -0.5 | 0.133 | 0.188 |
| chr18 | 42596222 | 42597335 | 1114 | 0.73 | 0.133 | 0.188 |
| chr10 | 29988572 | 29989374 | 803 | 0.93 | 0.133 | 0.188 |
| chr3 | 151739255 | 151745474 | 6220 | -0.26 | 0.134 | 0.188 |
| chr13 | 66976812 | 66977279 | 468 | -1.15 | 0.134 | 0.188 |
| chr16 | 58528173 | 58534805 | 6633 | -0.42 | 0.134 | 0.188 |
| chr16 | 75454857 | 75469179 | 14323 | 0.16 | 0.134 | 0.188 |
| chr13 | 114880601 | 114886889 | 6289 | 0.23 | 0.134 | 0.188 |
| chr11 | 67349003 | 67354617 | 5615 | -0.28 | 0.134 | 0.188 |
| chr12 | 104801805 | 104803886 | 2082 | -0.3 | 0.134 | 0.188 |
| chr7 | 149387409 | 149388533 | 1125 | 0.62 | 0.134 | 0.188 |
| chr1 | 94654515 | 94655136 | 622 | -1.32 | 0.134 | 0.188 |
| chr1 | 92247453 | 92253536 | 6084 | -0.32 | 0.134 | 0.188 |
| chr14 | 63990404 | 63992332 | 1929 | 0.51 | 0.134 | 0.189 |
| chr2 | 113681987 | 113683405 | 1419 | 2.79 | 0.134 | 0.189 |
| chr5 | 108017221 | 108018137 | 917 | -0.58 | 0.134 | 0.189 |
| chr4 | 81105509 | 81105896 | 388 | 1.04 | 0.134 | 0.189 |
| chr15 | 43776123 | 43786209 | 10087 | -0.24 | 0.134 | 0.189 |
| chr16 | 3984542 | 4030990 | 46449 | 0.18 | 0.134 | 0.189 |
| chr19 | 36266444 | 36266820 | 377 | 0.95 | 0.134 | 0.189 |
| chr17 | 78693794 | 78699055 | 5262 | -0.41 | 0.134 | 0.189 |
| chr2 | 198456344 | 198456879 | 536 | 1.28 | 0.134 | 0.189 |
| chr1 | 84542605 | 84544849 | 2245 | 0.26 | 0.134 | 0.189 |
| chr12 | 46233519 | 46234228 | 710 | -0.78 | 0.134 | 0.189 |
| chr18 | 46773718 | 46776251 | 2534 | 0.53 | 0.135 | 0.189 |
| chr19 | 17281733 | 17282080 | 348 | 1.58 | 0.135 | 0.189 |
| chr1 | 243584127 | 243585744 | 1618 | 0.35 | 0.135 | 0.189 |
| chrX | 96374576 | 96375074 | 499 | -0.92 | 0.135 | 0.189 |
| chr3 | 115231545 | 115232043 | 499 | 0.94 | 0.135 | 0.189 |
| chr5 | 57503962 | 57505042 | 1081 | -1.21 | 0.135 | 0.189 |
| chr18 | 77881795 | 77882756 | 962 | 0.93 | 0.135 | 0.19 |
| chr18 | 46986514 | 46987714 | 1201 | -0.58 | 0.135 | 0.19 |
| chr14 | 54655891 | 54656988 | 1098 | 0.55 | 0.135 | 0.19 |
| chr10 | 89768649 | 89769496 | 848 | -0.92 | 0.135 | 0.19 |
| chr14 | 90239692 | 90240973 | 1282 | -0.57 | 0.135 | 0.19 |
| chr11 | 85734286 | 85735158 | 873 | -0.8 | 0.135 | 0.19 |
| chr2 | 3447512 | 3448408 | 897 | -1.4 | 0.135 | 0.19 |
| chr12 | 42580357 | 42588466 | 8110 | -0.45 | 0.135 | 0.19 |
| chr5 | 156692635 | 156693696 | 1062 | 0.84 | 0.135 | 0.19 |
| chr2 | 69649589 | 69650230 | 642 | -0.95 | 0.135 | 0.19 |
| chr19 | 6947547 | 6948444 | 898 | 0.88 | 0.135 | 0.19 |
| chr17 | 76030592 | 76038342 | 7751 | 0.2 | 0.135 | 0.19 |
| chr14 | 100534100 | 100535224 | 1125 | -0.74 | 0.135 | 0.19 |
| chr7 | 151288350 | 151305901 | 17552 | -0.19 | 0.135 | 0.19 |
| chr6 | 37904673 | 37905101 | 429 | -1.01 | 0.135 | 0.19 |
| chr4 | 85791387 | 85792503 | 1117 | -0.62 | 0.135 | 0.19 |
| chrX | 110283378 | 110284255 | 878 | 0.83 | 0.135 | 0.19 |
| chr16 | 187507 | 189589 | 2083 | 0.36 | 0.135 | 0.19 |
| chr22 | 35653027 | 35663054 | 10028 | 0.22 | 0.135 | 0.19 |
| chr11 | 101733103 | 101742085 | 8983 | 0.2 | 0.135 | 0.19 |
| chr3 | 188425399 | 188426200 | 802 | 0.93 | 0.136 | 0.191 |
| chr5 | 13985601 | 13986631 | 1031 | -0.65 | 0.136 | 0.191 |
| chr6 | 4280600 | 4281057 | 458 | 1.57 | 0.136 | 0.191 |
| chr2 | 208392669 | 208400231 | 7563 | 0.19 | 0.136 | 0.191 |
| chr12 | 8057562 | 8060533 | 2972 | 0.41 | 0.136 | 0.191 |
| chr2 | 231967836 | 231968962 | 1127 | 0.87 | 0.136 | 0.191 |
| chr17 | 79116295 | 79117117 | 823 | -1.31 | 0.136 | 0.191 |

| | | | | | | |
|---|---|---|---|---|---|---|
| chr3 | 72460007 | 72461370 | 1364 | -0.71 | 0.136 | 0.191 |
| chr9 | 134000032 | 134002357 | 2326 | 0.22 | 0.136 | 0.191 |
| chr3 | 155432156 | 155433317 | 1162 | 0.74 | 0.136 | 0.191 |
| chr5 | 17819664 | 17839260 | 19597 | 2.58 | 0.136 | 0.191 |
| chr3 | 127770226 | 127772450 | 2225 | 0.28 | 0.136 | 0.191 |
| chrX | 44813734 | 44814538 | 805 | 0.94 | 0.136 | 0.191 |
| chr2 | 131128553 | 131132879 | 4327 | -0.24 | 0.136 | 0.191 |
| chr12 | 121903695 | 121905147 | 1453 | 0.65 | 0.136 | 0.191 |
| chr2 | 216847170 | 216848116 | 947 | -0.58 | 0.136 | 0.191 |
| chr19 | 3434136 | 3435859 | 1724 | 0.5 | 0.136 | 0.191 |
| chr4 | 41472866 | 41478904 | 6039 | -0.27 | 0.136 | 0.191 |
| chr17 | 29323454 | 29333899 | 10446 | -0.5 | 0.136 | 0.191 |
| chr5 | 17365132 | 17379485 | 14354 | -0.15 | 0.136 | 0.192 |
| chr20 | 43921854 | 43922511 | 658 | 0.73 | 0.136 | 0.192 |
| chr14 | 35260637 | 35266890 | 6254 | 0.38 | 0.136 | 0.192 |
| chr3 | 113296395 | 113297576 | 1182 | 0.67 | 0.137 | 0.192 |
| chrX | 10256181 | 10257067 | 887 | 0.81 | 0.137 | 0.192 |
| chr1 | 44898707 | 44899395 | 689 | -0.96 | 0.137 | 0.192 |
| chr13 | 86015296 | 86016221 | 926 | -0.65 | 0.137 | 0.192 |
| chr5 | 1105671 | 1112281 | 6611 | 0.25 | 0.137 | 0.192 |
| chr18 | 53069026 | 53070038 | 1013 | 0.74 | 0.137 | 0.192 |
| chr6 | 10693583 | 10697126 | 3544 | 0.23 | 0.137 | 0.192 |
| chr5 | 53843272 | 53844310 | 1039 | -1.12 | 0.137 | 0.192 |
| chr5 | 42912551 | 42913287 | 737 | 0.92 | 0.137 | 0.192 |
| chr8 | 18504076 | 18505026 | 951 | 0.77 | 0.137 | 0.192 |
| chr12 | 109822934 | 109823454 | 521 | -1.25 | 0.137 | 0.192 |
| chrX | 68062542 | 68063492 | 951 | 0.81 | 0.137 | 0.192 |
| chr7 | 155636126 | 155637601 | 1476 | 0.36 | 0.137 | 0.192 |
| chr2 | 32389326 | 32415387 | 26062 | -0.13 | 0.137 | 0.192 |
| chr7 | 2387070 | 2396659 | 9590 | -0.13 | 0.137 | 0.192 |
| chr9 | 132040440 | 132041144 | 705 | -0.97 | 0.137 | 0.192 |
| chr4 | 152967268 | 152967661 | 394 | 1.48 | 0.137 | 0.192 |
| chr4 | 9693149 | 9693437 | 289 | 1.04 | 0.137 | 0.192 |
| chr17 | 55657308 | 55658827 | 1520 | -2.38 | 0.137 | 0.192 |
| chr1 | 40325293 | 40326335 | 1043 | 2.56 | 0.137 | 0.193 |
| chr2 | 169636266 | 169638563 | 2298 | -0.5 | 0.137 | 0.193 |
| chr6 | 166943919 | 166944843 | 925 | 1.13 | 0.137 | 0.193 |
| chr17 | 40810642 | 40835113 | 24472 | 0.11 | 0.137 | 0.193 |
| chr20 | 23569334 | 23569951 | 618 | 1.54 | 0.138 | 0.193 |
| chr2 | 201390515 | 201391903 | 1389 | -0.45 | 0.138 | 0.193 |
| chr11 | 74659573 | 74661399 | 1827 | -0.25 | 0.138 | 0.193 |
| chr3 | 31355765 | 31356317 | 553 | -1.29 | 0.138 | 0.193 |
| chr1 | 212002413 | 212004913 | 2501 | 0.25 | 0.138 | 0.193 |
| chr1 | 77877205 | 77879068 | 1864 | 0.46 | 0.138 | 0.193 |
| chr11 | 101956601 | 101957227 | 627 | -0.88 | 0.138 | 0.193 |
| chr1 | 192873349 | 192874836 | 1488 | 0.48 | 0.138 | 0.193 |
| chr18 | 11887496 | 11888108 | 613 | -1.04 | 0.138 | 0.193 |
| chr5 | 74700506 | 74708649 | 8144 | 0.35 | 0.138 | 0.193 |
| chr2 | 197688145 | 197689073 | 929 | 0.82 | 0.138 | 0.194 |
| chrX | 41271319 | 41276811 | 5493 | 0.39 | 0.138 | 0.194 |
| chr3 | 116763631 | 116764771 | 1141 | 0.85 | 0.138 | 0.194 |
| chr4 | 103265142 | 103267243 | 2102 | 0.46 | 0.138 | 0.194 |
| chrX | 149737110 | 149738173 | 1064 | 0.56 | 0.138 | 0.194 |
| chrX | 28126401 | 28139122 | 12722 | 2.6 | 0.138 | 0.194 |
| chr6 | 75307997 | 75317231 | 9235 | -0.56 | 0.138 | 0.194 |
| chr16 | 12789138 | 12790403 | 1266 | 0.6 | 0.138 | 0.194 |
| chr3 | 177292235 | 177292990 | 756 | 0.6 | 0.138 | 0.194 |
| chr16 | 12888770 | 12911608 | 22839 | 0.17 | 0.138 | 0.194 |
| chr1 | 54714480 | 54716063 | 1584 | 0.71 | 0.138 | 0.194 |
| chr10 | 103476497 | 103489834 | 13338 | -0.4 | 0.138 | 0.194 |
| chr5 | 176592278 | 176592771 | 494 | 1.55 | 0.138 | 0.194 |

| | | | | | | |
|---|---|---|---|---|---|---|
| chr9 | 21096093 | 21096426 | 334 | 1.27 | 0.139 | 0.194 |
| chr17 | 58114688 | 58115118 | 431 | -1.18 | 0.139 | 0.194 |
| chr1 | 217800428 | 217805462 | 5035 | -0.13 | 0.139 | 0.194 |
| chr17 | 65135376 | 65137363 | 1988 | 0.25 | 0.139 | 0.194 |
| chr5 | 131891363 | 131895540 | 4178 | 0.12 | 0.139 | 0.194 |
| chr16 | 31101605 | 31153591 | 51987 | 0.13 | 0.139 | 0.194 |
| chr9 | 95823114 | 95823693 | 580 | -1.17 | 0.139 | 0.194 |
| chr4 | 164432745 | 164433641 | 897 | -0.78 | 0.139 | 0.194 |
| chr11 | 28636079 | 28654833 | 18755 | 0.26 | 0.139 | 0.194 |
| chr9 | 125861784 | 125874488 | 12705 | 0.35 | 0.139 | 0.195 |
| chr16 | 9940003 | 9941180 | 1178 | -0.61 | 0.139 | 0.195 |
| chr13 | 21139759 | 21142998 | 3240 | 0.21 | 0.139 | 0.195 |
| chr2 | 223620940 | 223622680 | 1741 | -0.43 | 0.139 | 0.195 |
| chr5 | 142413115 | 142413513 | 399 | 1.19 | 0.139 | 0.195 |
| chr1 | 117601010 | 117603890 | 2881 | 0.34 | 0.139 | 0.195 |
| chr5 | 16491862 | 16503810 | 11949 | 2.74 | 0.139 | 0.195 |
| chrX | 149715902 | 149716729 | 828 | 0.68 | 0.139 | 0.195 |
| chr7 | 86798341 | 86799018 | 678 | 0.88 | 0.139 | 0.195 |
| chr3 | 171708885 | 171731322 | 22438 | 0.21 | 0.139 | 0.195 |
| chr12 | 31382120 | 31383184 | 1065 | 0.75 | 0.14 | 0.195 |
| chr22 | 35726604 | 35726944 | 341 | -1.16 | 0.14 | 0.195 |
| chr1 | 228870628 | 228872232 | 1605 | 0.63 | 0.14 | 0.195 |
| chr10 | 30046297 | 30047830 | 1534 | 2.63 | 0.14 | 0.195 |
| chr20 | 23417478 | 23417909 | 432 | 1.09 | 0.14 | 0.195 |
| chr9 | 78830575 | 78831412 | 838 | -1.3 | 0.14 | 0.195 |
| chr4 | 156347709 | 156348094 | 386 | 1.55 | 0.14 | 0.196 |
| chr2 | 47703102 | 47717024 | 13923 | -0.41 | 0.14 | 0.196 |
| chr21 | 18166944 | 18168027 | 1084 | 0.77 | 0.14 | 0.196 |
| chr20 | 37541345 | 37591910 | 50566 | 0.11 | 0.14 | 0.196 |
| chr12 | 49243325 | 49284835 | 41511 | 0.12 | 0.14 | 0.196 |
| chr20 | 62273585 | 62273981 | 397 | 1.11 | 0.14 | 0.196 |
| chr1 | 11216451 | 11217266 | 816 | 0.88 | 0.14 | 0.196 |
| chr12 | 125549540 | 125550735 | 1196 | 0.59 | 0.14 | 0.196 |
| chr20 | 34081034 | 34081674 | 641 | 0.94 | 0.14 | 0.196 |
| chr2 | 19546923 | 19563535 | 16613 | -0.18 | 0.14 | 0.196 |
| chr20 | 34787150 | 34787740 | 591 | 1.12 | 0.14 | 0.196 |
| chr16 | 66508628 | 66522830 | 14203 | 0.21 | 0.14 | 0.196 |
| chr7 | 100181576 | 100184859 | 3284 | 0.35 | 0.14 | 0.196 |
| chr9 | 130186138 | 130187855 | 1718 | 0.34 | 0.14 | 0.196 |
| chr15 | 31342442 | 31342819 | 378 | -1.24 | 0.14 | 0.196 |
| chr1 | 61635556 | 61650843 | 15288 | 0.8 | 0.14 | 0.196 |
| chr20 | 1098416 | 1139611 | 41196 | -0.13 | 0.14 | 0.196 |
| chr1 | 187172254 | 187172705 | 452 | -1.07 | 0.141 | 0.196 |
| chr10 | 35548913 | 35549264 | 352 | -1.01 | 0.141 | 0.197 |
| chr5 | 156799128 | 156818911 | 19784 | 0.14 | 0.141 | 0.197 |
| chr17 | 14098262 | 14099172 | 911 | -1.06 | 0.141 | 0.197 |
| chr5 | 149401306 | 149402383 | 1078 | -0.64 | 0.141 | 0.197 |
| chr1 | 187996045 | 187997490 | 1446 | 0.48 | 0.141 | 0.197 |
| chr17 | 8532725 | 8535465 | 2741 | 0.32 | 0.141 | 0.197 |
| chr1 | 167232844 | 167238276 | 5433 | 0.31 | 0.141 | 0.197 |
| chr5 | 172730701 | 172731260 | 560 | -1.5 | 0.141 | 0.197 |
| chr16 | 88922462 | 88924772 | 2311 | -0.37 | 0.141 | 0.197 |
| chr8 | 39038953 | 39039329 | 377 | -1.09 | 0.141 | 0.197 |
| chr17 | 37817562 | 37860337 | 42776 | 0.14 | 0.141 | 0.197 |
| chr6 | 125378128 | 125379747 | 1620 | -0.48 | 0.141 | 0.197 |
| chr22 | 30901003 | 30902126 | 1124 | -0.45 | 0.141 | 0.197 |
| chr8 | 100023809 | 100031723 | 7915 | 0.2 | 0.141 | 0.197 |
| chr1 | 198465801 | 198467200 | 1400 | -0.7 | 0.141 | 0.197 |
| chr15 | 35102582 | 35103495 | 914 | -0.8 | 0.141 | 0.197 |
| chr2 | 11837232 | 11838015 | 784 | -0.79 | 0.141 | 0.197 |
| chr5 | 57071245 | 57073535 | 2291 | -2.83 | 0.141 | 0.197 |

| | | | | | | |
|---|---|---|---|---|---|---|
| chr5 | 169577408 | 169582188 | 4781 | 0.16 | 0.141 | 0.197 |
| chr6 | 143831517 | 143838480 | 6964 | -0.25 | 0.141 | 0.197 |
| chr16 | 85924166 | 85924789 | 624 | -1.21 | 0.141 | 0.197 |
| chr16 | 19668820 | 19673792 | 4973 | -0.52 | 0.141 | 0.198 |
| chr20 | 4665644 | 4668595 | 2952 | 0.2 | 0.142 | 0.198 |
| chr3 | 48309393 | 48309922 | 530 | 1.15 | 0.142 | 0.198 |
| chr2 | 24297547 | 24308488 | 10942 | 0.14 | 0.142 | 0.198 |
| chr16 | 88099892 | 88100915 | 1024 | -1.03 | 0.142 | 0.198 |
| chr4 | 38805549 | 38806473 | 925 | 0.48 | 0.142 | 0.198 |
| chr19 | 55972497 | 55999577 | 27081 | -0.13 | 0.142 | 0.198 |
| chr2 | 202096372 | 202101701 | 5330 | -0.23 | 0.142 | 0.198 |
| chr2 | 207947011 | 207948019 | 1009 | 0.81 | 0.142 | 0.198 |
| chr12 | 51502990 | 51528451 | 25462 | -0.2 | 0.142 | 0.198 |
| chr3 | 193534521 | 193535062 | 542 | 0.98 | 0.142 | 0.198 |
| chr1 | 116184513 | 116194350 | 9838 | -0.32 | 0.142 | 0.198 |
| chr13 | 110299111 | 110300073 | 963 | 0.68 | 0.142 | 0.198 |
| chr9 | 673703 | 706932 | 33230 | -0.22 | 0.142 | 0.198 |
| chr16 | 57197505 | 57198627 | 1123 | 0.67 | 0.142 | 0.198 |
| chr10 | 77054288 | 77055368 | 1081 | 0.93 | 0.142 | 0.198 |
| chr5 | 55249356 | 55251334 | 1979 | 0.6 | 0.142 | 0.198 |
| chr5 | 140180376 | 140187177 | 6802 | 2.4 | 0.142 | 0.199 |
| chr19 | 49946370 | 50016943 | 70574 | 0.08 | 0.142 | 0.199 |
| chr5 | 65310886 | 65312670 | 1785 | -0.54 | 0.142 | 0.199 |
| chr15 | 77739093 | 77740126 | 1034 | 0.67 | 0.142 | 0.199 |
| chr2 | 42771633 | 42772492 | 860 | 0.79 | 0.142 | 0.199 |
| chr14 | 44175600 | 44177028 | 1429 | -0.43 | 0.143 | 0.199 |
| chr2 | 85803617 | 85845143 | 41527 | 0.12 | 0.143 | 0.199 |
| chr2 | 196602059 | 196602812 | 754 | -0.9 | 0.143 | 0.199 |
| chr20 | 708593 | 709416 | 824 | -0.74 | 0.143 | 0.199 |
| chr6 | 108878543 | 108882320 | 3778 | 0.25 | 0.143 | 0.199 |
| chr12 | 39762003 | 39762817 | 815 | 0.67 | 0.143 | 0.2 |
| chr1 | 189774327 | 189775614 | 1288 | -0.63 | 0.143 | 0.2 |
| chr5 | 131562598 | 131574051 | 11454 | -0.34 | 0.143 | 0.2 |
| chr9 | 107889721 | 107890433 | 713 | 0.96 | 0.143 | 0.2 |
| chr4 | 75096057 | 75098389 | 2333 | -0.34 | 0.143 | 0.2 |
| chr16 | 11874294 | 11904623 | 30330 | 0.15 | 0.143 | 0.2 |
| chr4 | 145276904 | 145278282 | 1379 | 0.56 | 0.144 | 0.2 |
| chr2 | 183579426 | 183584275 | 4850 | -0.16 | 0.144 | 0.2 |
| chr10 | 60271935 | 60292120 | 20186 | -0.1 | 0.144 | 0.2 |
| chr8 | 48980860 | 48989594 | 8735 | -0.27 | 0.144 | 0.2 |
| chr3 | 48741270 | 48755114 | 13845 | -0.2 | 0.144 | 0.2 |
| chr18 | 21544516 | 21544918 | 403 | 1.12 | 0.144 | 0.2 |
| chr6 | 16622892 | 16624060 | 1169 | -0.56 | 0.144 | 0.2 |
| chr4 | 176339584 | 176343854 | 4271 | -0.2 | 0.144 | 0.2 |
| chr1 | 36233583 | 36235956 | 2374 | 0.34 | 0.144 | 0.2 |
| chr8 | 126009584 | 126012322 | 2739 | 0.26 | 0.144 | 0.201 |
| chr1 | 174160031 | 174167988 | 7958 | 0.56 | 0.144 | 0.201 |
| chr11 | 1845993 | 1866652 | 20660 | 0.24 | 0.144 | 0.201 |
| chr1 | 55593356 | 55595886 | 2531 | 0.45 | 0.144 | 0.201 |
| chr3 | 194167619 | 194168482 | 864 | 0.87 | 0.144 | 0.201 |
| chr3 | 128016156 | 128017299 | 1144 | 0.58 | 0.144 | 0.201 |
| chr11 | 94464387 | 94484571 | 20185 | 0.1 | 0.144 | 0.201 |
| chr15 | 89942665 | 89944023 | 1359 | -0.34 | 0.144 | 0.201 |
| chr3 | 63810026 | 63810642 | 617 | 0.74 | 0.144 | 0.201 |
| chr16 | 88600452 | 88601774 | 1323 | 0.64 | 0.144 | 0.201 |
| chr5 | 122180006 | 122186340 | 6335 | 0.18 | 0.144 | 0.201 |
| chr2 | 190626418 | 190628872 | 2455 | -0.23 | 0.144 | 0.201 |
| chr8 | 22999797 | 23004743 | 4947 | -0.32 | 0.144 | 0.201 |
| chr1 | 113008429 | 113009479 | 1051 | 0.73 | 0.144 | 0.201 |
| chr1 | 44619863 | 44621390 | 1528 | 0.37 | 0.145 | 0.201 |
| chr19 | 57945916 | 57947850 | 1935 | 0.34 | 0.145 | 0.201 |

| | | | | | | |
|---|---|---|---|---|---|---|
| chrX | 118170222 | 118171034 | 813 | -0.78 | 0.145 | 0.201 |
| chr15 | 75628423 | 75666095 | 37673 | -0.17 | 0.145 | 0.201 |
| chr17 | 78474783 | 78476712 | 1930 | -0.33 | 0.145 | 0.201 |
| chr6 | 135374871 | 135376898 | 2028 | 0.23 | 0.145 | 0.201 |
| chr3 | 118863974 | 118865021 | 1048 | -0.92 | 0.145 | 0.201 |
| chr12 | 59344322 | 59344936 | 615 | 0.77 | 0.145 | 0.201 |
| chr2 | 191583264 | 191584307 | 1044 | -0.75 | 0.145 | 0.201 |
| chr10 | 78871944 | 78872451 | 508 | -0.87 | 0.145 | 0.201 |
| chr11 | 64490751 | 64491155 | 405 | -0.82 | 0.145 | 0.202 |
| chr11 | 63705026 | 63715056 | 10031 | -0.24 | 0.145 | 0.202 |
| chr14 | 68878621 | 68895200 | 16580 | 0.2 | 0.145 | 0.202 |
| chr6 | 74403895 | 74425687 | 21793 | -0.1 | 0.145 | 0.202 |
| chr7 | 23052707 | 23054544 | 1838 | 0.54 | 0.145 | 0.202 |
| chr8 | 141791600 | 141814640 | 23041 | 0.12 | 0.145 | 0.202 |
| chr1 | 52536115 | 52549718 | 13604 | 0.26 | 0.145 | 0.202 |
| chr1 | 54953368 | 54955292 | 1925 | -0.24 | 0.145 | 0.202 |
| chr7 | 89398 | 89854 | 457 | -0.95 | 0.145 | 0.202 |
| chr8 | 60031789 | 60032297 | 509 | 0.76 | 0.145 | 0.202 |
| chr12 | 30839239 | 30849663 | 10425 | -0.18 | 0.145 | 0.202 |
| chr2 | 236949538 | 236949973 | 436 | -1.37 | 0.146 | 0.202 |
| chr3 | 179109819 | 179110453 | 635 | -0.99 | 0.146 | 0.203 |
| chr1 | 38059885 | 38062575 | 2691 | -0.26 | 0.146 | 0.203 |
| chr20 | 33945144 | 33946882 | 1739 | -0.43 | 0.146 | 0.203 |
| chr12 | 80235343 | 80242365 | 7023 | 0.3 | 0.146 | 0.203 |
| chrX | 102629390 | 102632696 | 3307 | -0.25 | 0.146 | 0.203 |
| chr2 | 171873515 | 171878602 | 5088 | -0.45 | 0.146 | 0.203 |
| chr11 | 43958583 | 43959104 | 522 | 1.07 | 0.146 | 0.203 |
| chr15 | 91388212 | 91389013 | 802 | -0.86 | 0.146 | 0.203 |
| chr4 | 177499945 | 177501256 | 1312 | 0.57 | 0.146 | 0.203 |
| chr1 | 229576268 | 229577302 | 1035 | -0.7 | 0.146 | 0.203 |
| chr20 | 36147235 | 36157437 | 10203 | 0.17 | 0.146 | 0.204 |
| chr2 | 64244326 | 64249516 | 5191 | 0.16 | 0.146 | 0.204 |
| chr22 | 40572661 | 40578060 | 5400 | 0.29 | 0.146 | 0.204 |
| chr2 | 29288032 | 29288779 | 748 | -1.11 | 0.146 | 0.204 |
| chr11 | 76818582 | 76819517 | 936 | 0.82 | 0.147 | 0.204 |
| chr6 | 155184869 | 155185847 | 979 | 0.84 | 0.147 | 0.204 |
| chr4 | 71703103 | 71705610 | 2508 | 0.33 | 0.147 | 0.204 |
| chr9 | 94908774 | 94910454 | 1681 | -0.36 | 0.147 | 0.204 |
| chr14 | 96505706 | 96506152 | 447 | -1 | 0.147 | 0.204 |
| chr4 | 48505538 | 48507429 | 1892 | -0.33 | 0.147 | 0.204 |
| chr9 | 36764866 | 36766721 | 1856 | -0.43 | 0.147 | 0.204 |
| chr10 | 101189597 | 101191897 | 2301 | 0.28 | 0.147 | 0.204 |
| chr4 | 177459507 | 177460414 | 908 | -1.09 | 0.147 | 0.204 |
| chr7 | 151722126 | 151723899 | 1774 | 0.37 | 0.147 | 0.204 |
| chr11 | 82569110 | 82570710 | 1601 | 0.35 | 0.147 | 0.204 |
| chr1 | 203642700 | 203644312 | 1613 | -0.61 | 0.147 | 0.204 |
| chr19 | 9150235 | 9151021 | 787 | -0.86 | 0.147 | 0.204 |
| chr6 | 90993803 | 90995839 | 2037 | 0.45 | 0.147 | 0.204 |
| chr2 | 101358632 | 101359483 | 852 | -0.89 | 0.147 | 0.204 |
| chr7 | 105359046 | 105359813 | 768 | -1.04 | 0.147 | 0.204 |
| chr4 | 34733694 | 34736370 | 2677 | -0.42 | 0.147 | 0.204 |
| chr11 | 11471616 | 11474444 | 2829 | -0.46 | 0.147 | 0.204 |
| chr16 | 56615175 | 56616715 | 1541 | -0.53 | 0.147 | 0.204 |
| chr15 | 79191785 | 79218979 | 27195 | -0.17 | 0.147 | 0.204 |
| chr2 | 232570101 | 232583005 | 12905 | -0.13 | 0.147 | 0.204 |
| chrX | 153307921 | 153323271 | 15351 | -0.18 | 0.147 | 0.204 |
| chr10 | 35675550 | 35680880 | 5331 | 0.48 | 0.147 | 0.204 |
| chr3 | 122328665 | 122329120 | 456 | 2.38 | 0.147 | 0.204 |
| chr2 | 146450599 | 146451508 | 910 | 0.82 | 0.147 | 0.205 |
| chr8 | 66921616 | 66954764 | 33149 | 0.13 | 0.147 | 0.205 |
| chr2 | 61282104 | 61294699 | 12596 | 0.24 | 0.147 | 0.205 |

| | | | | | | |
|---|---|---|---|---|---|---|
| chr6 | 43737405 | 43743261 | 5857 | -0.25 | 0.147 | 0.205 |
| chr8 | 141349308 | 141355810 | 6503 | -0.3 | 0.147 | 0.205 |
| chr9 | 127143811 | 127181948 | 38138 | -0.35 | 0.147 | 0.205 |
| chr12 | 119645669 | 119647514 | 1846 | 2.56 | 0.148 | 0.205 |
| chr1 | 45451610 | 45453338 | 1729 | 0.49 | 0.148 | 0.205 |
| chr4 | 76670482 | 76671831 | 1350 | 0.73 | 0.148 | 0.205 |
| chr3 | 27596375 | 27596743 | 369 | -1.53 | 0.148 | 0.205 |
| chr4 | 84405440 | 84406973 | 1534 | 0.48 | 0.148 | 0.205 |
| chr22 | 21335469 | 21337227 | 1759 | -0.37 | 0.148 | 0.205 |
| chr16 | 15125142 | 15136259 | 11118 | -0.39 | 0.148 | 0.205 |
| chr3 | 29526952 | 29528964 | 2013 | -0.31 | 0.148 | 0.205 |
| chr20 | 31161502 | 31163413 | 1912 | -0.41 | 0.148 | 0.205 |
| chr6 | 42336899 | 42337437 | 539 | 1.25 | 0.148 | 0.205 |
| chr16 | 492439 | 508999 | 16561 | 0.23 | 0.148 | 0.205 |
| chr15 | 38361053 | 38365908 | 4856 | 0.28 | 0.148 | 0.205 |
| chr16 | 8715198 | 8727653 | 12456 | 0.19 | 0.148 | 0.205 |
| chr11 | 36393780 | 36395487 | 1708 | 0.49 | 0.148 | 0.205 |
| chr1 | 28539640 | 28540575 | 936 | -0.89 | 0.148 | 0.206 |
| chr9 | 117919323 | 117933683 | 14361 | 0.95 | 0.148 | 0.206 |
| chr12 | 122014612 | 122028757 | 14146 | 0.19 | 0.148 | 0.206 |
| chr3 | 59003290 | 59003936 | 647 | 1.02 | 0.148 | 0.206 |
| chr6 | 114056005 | 114056993 | 989 | 0.65 | 0.148 | 0.206 |
| chr1 | 230960056 | 230961320 | 1265 | 0.59 | 0.148 | 0.206 |
| chr3 | 133662541 | 133662952 | 412 | 1.24 | 0.148 | 0.206 |
| chr6 | 150389729 | 150393879 | 4151 | 0.44 | 0.148 | 0.206 |
| chrX | 24773638 | 24774308 | 671 | 1.3 | 0.148 | 0.206 |
| chr17 | 10017286 | 10030786 | 13501 | -0.46 | 0.148 | 0.206 |
| chr18 | 30039835 | 30051474 | 11640 | 2.72 | 0.148 | 0.206 |
| chr5 | 121814466 | 121815876 | 1411 | 1.1 | 0.148 | 0.206 |
| chr15 | 45076277 | 45077024 | 748 | 0.66 | 0.149 | 0.206 |
| chr1 | 33004405 | 33005996 | 1592 | 0.45 | 0.149 | 0.206 |
| chr5 | 105733142 | 105735697 | 2556 | 2.55 | 0.149 | 0.206 |
| chr2 | 151259731 | 151260892 | 1162 | 0.72 | 0.149 | 0.206 |
| chr10 | 112562178 | 112564367 | 2190 | 2.6 | 0.149 | 0.206 |
| chr19 | 47392869 | 47462506 | 69638 | -0.09 | 0.149 | 0.206 |
| chr2 | 27903446 | 27908186 | 4741 | 0.22 | 0.149 | 0.206 |
| chr5 | 10333400 | 10334126 | 727 | -0.73 | 0.149 | 0.206 |
| chr6 | 160321097 | 160321553 | 457 | -0.93 | 0.149 | 0.206 |
| chr15 | 43662503 | 43675980 | 13478 | -0.32 | 0.149 | 0.206 |
| chr15 | 50295603 | 50296656 | 1054 | -0.85 | 0.149 | 0.206 |
| chr1 | 219486143 | 219487235 | 1093 | -0.89 | 0.149 | 0.206 |
| chr10 | 115438937 | 115441030 | 2094 | 0.22 | 0.149 | 0.206 |
| chr2 | 205993899 | 205994663 | 765 | -0.82 | 0.149 | 0.206 |
| chr12 | 112883779 | 112885113 | 1335 | -0.7 | 0.149 | 0.207 |
| chr19 | 10443037 | 10445503 | 2467 | -0.41 | 0.149 | 0.207 |
| chr9 | 103064713 | 103068943 | 4231 | 0.68 | 0.149 | 0.207 |
| chr2 | 99723442 | 99723944 | 503 | -0.94 | 0.149 | 0.207 |
| chr9 | 128411238 | 128412481 | 1244 | 0.55 | 0.149 | 0.207 |
| chr11 | 104322498 | 104323842 | 1345 | 0.51 | 0.149 | 0.207 |
| chr13 | 52586506 | 52587422 | 917 | 0.55 | 0.149 | 0.207 |
| chr16 | 4851495 | 4853477 | 1983 | 0.37 | 0.149 | 0.207 |
| chr5 | 96142461 | 96144756 | 2296 | 0.23 | 0.149 | 0.207 |
| chr9 | 116985263 | 116987009 | 1747 | -0.52 | 0.149 | 0.207 |
| chr14 | 36139862 | 36147950 | 8089 | -0.59 | 0.15 | 0.207 |
| chr4 | 175342463 | 175360494 | 18032 | 0.17 | 0.15 | 0.207 |
| chr2 | 20000735 | 20002778 | 2044 | -0.27 | 0.15 | 0.207 |
| chr20 | 43513698 | 43518451 | 4754 | 0.24 | 0.15 | 0.208 |
| chr4 | 185186565 | 185190350 | 3786 | 0.24 | 0.15 | 0.208 |
| chr1 | 24592803 | 24594532 | 1730 | 0.5 | 0.15 | 0.208 |
| chr11 | 15914203 | 15924253 | 10051 | 1.92 | 0.15 | 0.208 |
| chr10 | 30707270 | 30754051 | 46782 | -0.12 | 0.15 | 0.208 |

| | | | | | | |
|---|---|---|---|---|---|---|
| chr11 | 45167534 | 45180244 | 12711 | 0.16 | 0.15 | 0.208 |
| chr2 | 207114841 | 207115528 | 688 | -0.63 | 0.15 | 0.208 |
| chr15 | 43702542 | 43704031 | 1490 | -0.63 | 0.15 | 0.208 |
| chr13 | 98019566 | 98022497 | 2932 | 0.38 | 0.15 | 0.208 |
| chr19 | 48758231 | 48775547 | 17317 | 0.18 | 0.15 | 0.208 |
| chr15 | 56391158 | 56392223 | 1066 | 0.61 | 0.15 | 0.208 |
| chr15 | 77756254 | 77757734 | 1481 | -0.81 | 0.15 | 0.208 |
| chr13 | 50589481 | 50599911 | 10431 | 0.49 | 0.15 | 0.208 |
| chr3 | 145866309 | 145885586 | 19278 | -0.07 | 0.15 | 0.208 |
| chr3 | 172636008 | 172636735 | 728 | 0.99 | 0.15 | 0.208 |
| chr3 | 183852151 | 183854058 | 1908 | 0.34 | 0.151 | 0.208 |
| chr5 | 180454652 | 180455700 | 1049 | 0.55 | 0.151 | 0.208 |
| chrX | 46998036 | 47006101 | 8066 | 0.24 | 0.151 | 0.208 |
| chr4 | 114897034 | 114902139 | 5106 | 0.23 | 0.151 | 0.208 |
| chr4 | 104675673 | 104676976 | 1304 | 0.7 | 0.151 | 0.208 |
| chr13 | 110064503 | 110065331 | 829 | -1.58 | 0.151 | 0.208 |
| chr10 | 52093445 | 52094048 | 604 | 0.85 | 0.151 | 0.208 |
| chr3 | 4763559 | 4765149 | 1591 | 0.55 | 0.151 | 0.208 |
| chr15 | 58903883 | 58905241 | 1359 | 0.48 | 0.151 | 0.208 |
| chr6 | 112364025 | 112364981 | 957 | 0.45 | 0.151 | 0.208 |
| chr3 | 110347518 | 110348081 | 564 | 1.07 | 0.151 | 0.208 |
| chr15 | 70909658 | 70910036 | 379 | 1.2 | 0.151 | 0.208 |
| chrX | 117568093 | 117569001 | 909 | -0.79 | 0.151 | 0.209 |
| chr4 | 22943104 | 22945212 | 2109 | 2.69 | 0.151 | 0.209 |
| chr22 | 46663373 | 46672372 | 9000 | 0.45 | 0.151 | 0.209 |
| chr7 | 100828352 | 100829270 | 919 | -1 | 0.151 | 0.209 |
| chr9 | 16243387 | 16244079 | 693 | 0.8 | 0.151 | 0.209 |
| chr2 | 174313464 | 174314119 | 656 | -0.53 | 0.151 | 0.209 |
| chr19 | 50650818 | 50651543 | 726 | 0.74 | 0.151 | 0.209 |
| chr7 | 149417102 | 149417809 | 708 | -0.82 | 0.151 | 0.209 |
| chr20 | 10385709 | 10429814 | 44106 | -0.09 | 0.151 | 0.209 |
| chr17 | 34274316 | 34274874 | 559 | -0.74 | 0.151 | 0.209 |
| chr6 | 149067322 | 149068809 | 1488 | -0.56 | 0.151 | 0.209 |
| chr10 | 70712946 | 70758008 | 45063 | 0.11 | 0.151 | 0.209 |
| chr2 | 135208735 | 135215909 | 7175 | -2.21 | 0.151 | 0.209 |
| chr2 | 33171533 | 33173650 | 2118 | -0.4 | 0.151 | 0.209 |
| chr4 | 128681352 | 128681824 | 473 | 1.25 | 0.151 | 0.209 |
| chr17 | 29419241 | 29423957 | 4717 | 0.18 | 0.151 | 0.209 |
| chr20 | 32031040 | 32032556 | 1517 | 0.81 | 0.151 | 0.209 |
| chr2 | 86098883 | 86191182 | 92300 | 0.06 | 0.151 | 0.209 |
| chr15 | 69493048 | 69493720 | 673 | 0.78 | 0.152 | 0.209 |
| chr7 | 135167312 | 135168383 | 1072 | -0.85 | 0.152 | 0.21 |
| chr6 | 163974983 | 163975846 | 864 | -0.92 | 0.152 | 0.21 |
| chr18 | 43310304 | 43312579 | 2276 | -0.45 | 0.152 | 0.21 |
| chr21 | 43298288 | 43300282 | 1995 | -0.34 | 0.152 | 0.21 |
| chr1 | 186286120 | 186287109 | 990 | -0.72 | 0.152 | 0.21 |
| chr1 | 220854741 | 220864329 | 9589 | 2.44 | 0.152 | 0.21 |
| chr14 | 75551713 | 75561555 | 9843 | -0.52 | 0.152 | 0.21 |
| chr2 | 54029753 | 54030617 | 865 | -0.71 | 0.152 | 0.21 |
| chr19 | 50706885 | 50707394 | 510 | 0.61 | 0.152 | 0.21 |
| chr4 | 95127960 | 95130159 | 2200 | -0.22 | 0.152 | 0.21 |
| chr8 | 68107099 | 68107946 | 848 | -0.76 | 0.152 | 0.21 |
| chr1 | 116694571 | 116695225 | 655 | -0.65 | 0.152 | 0.21 |
| chr11 | 93932900 | 93934861 | 1962 | -0.48 | 0.152 | 0.211 |
| chr2 | 197268157 | 197269205 | 1049 | 0.66 | 0.153 | 0.211 |
| chr1 | 52284070 | 52285223 | 1154 | 0.8 | 0.153 | 0.211 |
| chr9 | 110217700 | 110228627 | 10928 | 0.67 | 0.153 | 0.211 |
| chr7 | 67778599 | 67795832 | 17234 | 1.19 | 0.153 | 0.211 |
| chr11 | 99196007 | 99197546 | 1540 | 0.37 | 0.153 | 0.211 |
| chr4 | 178150071 | 178150975 | 905 | 0.64 | 0.153 | 0.211 |
| chr9 | 127380215 | 127381262 | 1048 | 0.65 | 0.153 | 0.211 |

| | | | | | | |
|---|---|---|---|---|---|---|
| chr2 | 43954243 | 43954583 | 341 | -1.09 | 0.153 | 0.211 |
| chr12 | 100493481 | 100503691 | 10211 | -0.44 | 0.153 | 0.211 |
| chr15 | 76195153 | 76219626 | 24474 | -0.15 | 0.153 | 0.211 |
| chr17 | 897903 | 1026316 | 128414 | -0.07 | 0.153 | 0.211 |
| chr7 | 50517165 | 50518733 | 1569 | 0.34 | 0.153 | 0.211 |
| chr20 | 16533051 | 16534743 | 1693 | -0.72 | 0.153 | 0.211 |
| chr2 | 69551299 | 69557736 | 6438 | -1.48 | 0.153 | 0.211 |
| chr4 | 106366045 | 106367595 | 1551 | 0.59 | 0.153 | 0.212 |
| chr13 | 76209752 | 76211097 | 1346 | 0.58 | 0.153 | 0.212 |
| chr6 | 156983377 | 156983920 | 544 | 0.88 | 0.153 | 0.212 |
| chr8 | 98288940 | 98290670 | 1731 | 0.38 | 0.153 | 0.212 |
| chr2 | 170715650 | 170716125 | 476 | -1.01 | 0.153 | 0.212 |
| chr4 | 141445294 | 141446424 | 1131 | 0.38 | 0.154 | 0.212 |
| chr1 | 54872197 | 54874982 | 2786 | -0.3 | 0.154 | 0.212 |
| chr6 | 149887365 | 149888599 | 1235 | 0.43 | 0.154 | 0.212 |
| chr5 | 2752025 | 2752580 | 556 | 1 | 0.154 | 0.212 |
| chr5 | 167695990 | 167697731 | 1742 | -0.4 | 0.154 | 0.212 |
| chr7 | 27411781 | 27412724 | 944 | 0.67 | 0.154 | 0.212 |
| chr16 | 66906549 | 66915275 | 8727 | -0.34 | 0.154 | 0.212 |
| chr7 | 128430900 | 128431669 | 770 | 0.83 | 0.154 | 0.212 |
| chr1 | 161067651 | 161068720 | 1070 | 0.67 | 0.154 | 0.212 |
| chr1 | 3049932 | 3050904 | 973 | 0.88 | 0.154 | 0.212 |
| chr4 | 108945084 | 108945800 | 717 | -0.71 | 0.154 | 0.212 |
| chr21 | 35287276 | 35288691 | 1416 | 0.4 | 0.154 | 0.213 |
| chr2 | 25932558 | 25935268 | 2711 | 0.51 | 0.154 | 0.213 |
| chr15 | 40452817 | 40454966 | 2150 | -0.42 | 0.154 | 0.213 |
| chr14 | 56135898 | 56140295 | 4398 | -0.34 | 0.154 | 0.213 |
| chr20 | 13902690 | 13903631 | 942 | -0.69 | 0.154 | 0.213 |
| chr9 | 86519288 | 86519617 | 330 | -1.44 | 0.154 | 0.213 |
| chr2 | 72520732 | 72521713 | 982 | 0.66 | 0.154 | 0.213 |
| chr2 | 219081023 | 219096364 | 15342 | -0.16 | 0.155 | 0.213 |
| chr9 | 139240271 | 139241029 | 759 | -1.34 | 0.155 | 0.213 |
| chr18 | 60986848 | 60987915 | 1068 | 0.7 | 0.155 | 0.213 |
| chr11 | 306801 | 308541 | 1741 | 0.51 | 0.155 | 0.213 |
| chr3 | 14218622 | 14221743 | 3122 | 0.18 | 0.155 | 0.213 |
| chr1 | 94310174 | 94314220 | 4047 | 0.25 | 0.155 | 0.213 |
| chr7 | 112060282 | 112064139 | 3858 | -0.2 | 0.155 | 0.213 |
| chr5 | 65262720 | 65263201 | 482 | 1 | 0.155 | 0.213 |
| chr2 | 220251772 | 220252625 | 854 | -0.45 | 0.155 | 0.214 |
| chr4 | 67355049 | 67356514 | 1466 | 0.83 | 0.155 | 0.214 |
| chr1 | 169818517 | 169819288 | 772 | -0.98 | 0.155 | 0.214 |
| chr3 | 136334790 | 136352427 | 17638 | 0.24 | 0.155 | 0.214 |
| chr1 | 23143283 | 23144553 | 1271 | -2.34 | 0.155 | 0.214 |
| chr20 | 4224248 | 4227255 | 3008 | 0.23 | 0.156 | 0.214 |
| chr2 | 172811168 | 172812174 | 1007 | -0.68 | 0.156 | 0.214 |
| chr11 | 18332078 | 18353138 | 21061 | 0.11 | 0.156 | 0.215 |
| chr2 | 48475204 | 48477032 | 1829 | -0.5 | 0.156 | 0.215 |
| chr2 | 100936826 | 100939381 | 2556 | 0.5 | 0.156 | 0.215 |
| chr4 | 146428646 | 146430141 | 1496 | 0.55 | 0.156 | 0.215 |
| chr1 | 3541624 | 3542073 | 450 | 0.99 | 0.156 | 0.215 |
| chr20 | 44978355 | 44993865 | 15511 | 0.15 | 0.156 | 0.215 |
| chr6 | 170150608 | 170152883 | 2276 | 0.17 | 0.156 | 0.215 |
| chr3 | 57896260 | 57897455 | 1196 | 0.86 | 0.156 | 0.215 |
| chr4 | 1339587 | 1345975 | 6389 | 0.24 | 0.156 | 0.215 |
| chr3 | 106317192 | 106317795 | 604 | 2.29 | 0.156 | 0.215 |
| chr8 | 141779941 | 141781145 | 1205 | -0.67 | 0.156 | 0.216 |
| chr1 | 47999753 | 48000393 | 641 | -0.99 | 0.157 | 0.216 |
| chrX | 24509224 | 24513475 | 4252 | -0.42 | 0.157 | 0.216 |
| chr10 | 76192527 | 76199647 | 7121 | 0.42 | 0.157 | 0.216 |
| chr10 | 79761277 | 79762062 | 786 | -0.94 | 0.157 | 0.216 |
| chr7 | 130125427 | 130127069 | 1643 | 0.42 | 0.157 | 0.216 |

| | | | | | | |
|---|---|---|---|---|---|---|
| chrX | 39753467 | 39769180 | 15714 | 0.19 | 0.157 | 0.216 |
| chr18 | 12862430 | 12863452 | 1023 | -0.83 | 0.157 | 0.216 |
| chr7 | 129141480 | 129143229 | 1750 | -0.32 | 0.157 | 0.216 |
| chr6 | 2952004 | 3025435 | 73432 | -0.09 | 0.157 | 0.216 |
| chrX | 122847032 | 122867433 | 20402 | -0.15 | 0.157 | 0.216 |
| chr15 | 68870149 | 68879300 | 9152 | 0.92 | 0.157 | 0.216 |
| chrX | 21171623 | 21186594 | 14972 | 0.18 | 0.157 | 0.216 |
| chr2 | 70120461 | 70143133 | 22673 | -0.14 | 0.157 | 0.216 |
| chr19 | 18667890 | 18683882 | 15993 | 0.19 | 0.157 | 0.216 |
| chr1 | 48861520 | 48862206 | 687 | -0.71 | 0.157 | 0.217 |
| chr5 | 176920112 | 176924569 | 4458 | 0.27 | 0.157 | 0.217 |
| chr17 | 41754988 | 41755686 | 699 | -1.15 | 0.157 | 0.217 |
| chr9 | 125059736 | 125078827 | 19092 | -0.32 | 0.157 | 0.217 |
| chr12 | 31774865 | 31775602 | 738 | 0.72 | 0.157 | 0.217 |
| chr2 | 20455298 | 20466629 | 11332 | 0.73 | 0.157 | 0.217 |
| chr4 | 106983212 | 106984842 | 1631 | 0.45 | 0.157 | 0.217 |
| chr10 | 28820478 | 28884377 | 63900 | 0.09 | 0.158 | 0.217 |
| chr10 | 114561345 | 114584246 | 22902 | -0.16 | 0.158 | 0.217 |
| chr18 | 700474 | 713416 | 12943 | 0.16 | 0.158 | 0.217 |
| chr6 | 72077067 | 72077920 | 854 | -0.75 | 0.158 | 0.217 |
| chr6 | 107996140 | 107996687 | 548 | 0.56 | 0.158 | 0.217 |
| chr9 | 126803093 | 126803961 | 869 | -0.69 | 0.158 | 0.217 |
| chr14 | 74768227 | 74770521 | 2295 | -0.28 | 0.158 | 0.217 |
| chr17 | 74191052 | 74191590 | 539 | -0.88 | 0.158 | 0.217 |
| chr2 | 134005116 | 134005632 | 517 | -1.02 | 0.158 | 0.217 |
| chr5 | 158406398 | 158408316 | 1919 | 0.25 | 0.158 | 0.217 |
| chr14 | 91777438 | 91784756 | 7319 | -0.59 | 0.158 | 0.217 |
| chr10 | 102487815 | 102499795 | 11981 | 0.16 | 0.158 | 0.217 |
| chr8 | 67831844 | 67839083 | 7240 | 0.17 | 0.158 | 0.217 |
| chr18 | 45687225 | 45688877 | 1653 | 0.5 | 0.158 | 0.217 |
| chr1 | 91965785 | 91968685 | 2901 | 0.25 | 0.158 | 0.217 |
| chrX | 134070268 | 134084647 | 14380 | -0.24 | 0.158 | 0.217 |
| chr2 | 95872549 | 95874103 | 1555 | 0.45 | 0.158 | 0.217 |
| chr11 | 126517376 | 126518387 | 1012 | -0.68 | 0.158 | 0.217 |
| chr9 | 100744471 | 100784837 | 40367 | 0.1 | 0.158 | 0.218 |
| chr14 | 104089851 | 104132860 | 43010 | -0.11 | 0.158 | 0.218 |
| chr22 | 36902570 | 36903732 | 1163 | 0.56 | 0.158 | 0.218 |
| chr1 | 25389627 | 25391049 | 1423 | -0.58 | 0.158 | 0.218 |
| chr3 | 47058352 | 47058993 | 642 | -0.91 | 0.158 | 0.218 |
| chr8 | 134044191 | 134046537 | 2347 | -0.42 | 0.159 | 0.218 |
| chr11 | 10108671 | 10118039 | 9369 | -0.6 | 0.159 | 0.218 |
| chr8 | 10487857 | 10488306 | 450 | -1.01 | 0.159 | 0.218 |
| chrX | 75367970 | 75369013 | 1044 | -0.52 | 0.159 | 0.218 |
| chr12 | 52241818 | 52242343 | 526 | 1.47 | 0.159 | 0.218 |
| chr18 | 56448387 | 56464013 | 15627 | 0.08 | 0.159 | 0.218 |
| chr20 | 59753598 | 59763415 | 9818 | 0.17 | 0.159 | 0.218 |
| chr5 | 131411010 | 131411623 | 614 | -1.07 | 0.159 | 0.218 |
| chr12 | 101750353 | 101750879 | 527 | 0.91 | 0.159 | 0.218 |
| chr3 | 196254562 | 196256393 | 1832 | -0.36 | 0.159 | 0.218 |
| chr15 | 45314705 | 45316302 | 1598 | -0.46 | 0.159 | 0.219 |
| chr8 | 98607625 | 98608776 | 1152 | 0.43 | 0.159 | 0.219 |
| chr13 | 113807715 | 113808228 | 514 | 0.72 | 0.159 | 0.219 |
| chr13 | 45562407 | 45564654 | 2248 | 0.26 | 0.159 | 0.219 |
| chr3 | 139062288 | 139063984 | 1697 | 0.31 | 0.159 | 0.219 |
| chr9 | 138818319 | 138819403 | 1085 | -0.47 | 0.159 | 0.219 |
| chr19 | 5679607 | 5692170 | 12564 | 0.24 | 0.159 | 0.219 |
| chr10 | 54067908 | 54096154 | 28247 | 0.11 | 0.159 | 0.219 |
| chr12 | 48539981 | 48552286 | 12306 | -0.22 | 0.159 | 0.219 |
| chr19 | 8274226 | 8274884 | 659 | 0.75 | 0.159 | 0.219 |
| chr4 | 77342092 | 77342689 | 598 | 0.97 | 0.16 | 0.219 |
| chr15 | 41390519 | 41391537 | 1019 | -0.82 | 0.16 | 0.219 |

| | | | | | | |
|---|---|---|---|---|---|---|
| chr15 | 45693941 | 45712287 | 18347 | 0.24 | 0.16 | 0.219 |
| chr1 | 240548400 | 240550968 | 2569 | 0.21 | 0.16 | 0.219 |
| chr12 | 56827819 | 56829631 | 1813 | -0.58 | 0.16 | 0.219 |
| chr18 | 45633490 | 45633934 | 445 | -1.2 | 0.16 | 0.219 |
| chr12 | 55328182 | 55330006 | 1825 | 0.3 | 0.16 | 0.219 |
| chr1 | 109756059 | 109768940 | 12882 | 0.26 | 0.16 | 0.219 |
| chr1 | 184693393 | 184694050 | 658 | -0.75 | 0.16 | 0.219 |
| chr21 | 44291335 | 44314399 | 23065 | 0.17 | 0.16 | 0.219 |
| chr2 | 175352002 | 175352690 | 689 | -0.54 | 0.16 | 0.22 |
| chr17 | 65774795 | 65776005 | 1211 | 0.42 | 0.16 | 0.22 |
| chr7 | 81833653 | 81835186 | 1534 | -0.45 | 0.16 | 0.22 |
| chr2 | 33275794 | 33299183 | 23390 | -0.14 | 0.16 | 0.22 |
| chr1 | 51068498 | 51070217 | 1720 | 0.44 | 0.16 | 0.22 |
| chr3 | 197005098 | 197006535 | 1438 | 0.73 | 0.16 | 0.22 |
| chr2 | 240306546 | 240307298 | 753 | -1.04 | 0.16 | 0.22 |
| chr2 | 19910686 | 19912124 | 1439 | 0.5 | 0.161 | 0.22 |
| chr9 | 100937394 | 100937743 | 350 | -1.26 | 0.161 | 0.22 |
| chr8 | 133686959 | 133688464 | 1506 | 0.36 | 0.161 | 0.22 |
| chr1 | 229524759 | 229525639 | 881 | 0.77 | 0.161 | 0.22 |
| chrX | 43535636 | 43537183 | 1548 | 0.43 | 0.161 | 0.22 |
| chr10 | 98079588 | 98080615 | 1028 | -2.32 | 0.161 | 0.22 |
| chr1 | 186362289 | 186375386 | 13098 | -0.29 | 0.161 | 0.22 |
| chr20 | 48988461 | 48997748 | 9288 | 0.24 | 0.161 | 0.221 |
| chrX | 51230049 | 51231327 | 1279 | -0.57 | 0.161 | 0.221 |
| chr9 | 36969520 | 36970946 | 1427 | -0.66 | 0.161 | 0.221 |
| chr20 | 61903165 | 61906205 | 3041 | -0.3 | 0.161 | 0.221 |
| chr8 | 58970341 | 58992313 | 21973 | 2.48 | 0.161 | 0.221 |
| chr4 | 56995310 | 56995663 | 354 | -1 | 0.161 | 0.221 |
| chr15 | 29269011 | 29270511 | 1501 | -0.41 | 0.161 | 0.221 |
| chr19 | 18960284 | 18961027 | 744 | -0.81 | 0.161 | 0.221 |
| chr3 | 25276311 | 25276709 | 399 | -1.2 | 0.161 | 0.221 |
| chr2 | 170335537 | 170336747 | 1211 | -0.43 | 0.161 | 0.221 |
| chr11 | 111852089 | 111860438 | 8350 | 0.31 | 0.161 | 0.221 |
| chr3 | 15246918 | 15265566 | 18649 | 0.19 | 0.161 | 0.221 |
| chr11 | 108669835 | 108670801 | 967 | 1.45 | 0.161 | 0.221 |
| chr10 | 56372795 | 56373428 | 634 | -2.25 | 0.161 | 0.221 |
| chr1 | 25009802 | 25011093 | 1292 | 0.57 | 0.162 | 0.221 |
| chr7 | 30386659 | 30388905 | 2247 | 0.29 | 0.162 | 0.222 |
| chr6 | 41783692 | 41794117 | 10426 | 0.21 | 0.162 | 0.222 |
| chr1 | 201717382 | 201718786 | 1405 | 0.5 | 0.162 | 0.222 |
| chr19 | 8146781 | 8147454 | 674 | 0.87 | 0.162 | 0.222 |
| chr3 | 8765947 | 8767159 | 1213 | 2.34 | 0.162 | 0.222 |
| chr19 | 2899944 | 2901104 | 1161 | 0.56 | 0.162 | 0.222 |
| chr3 | 49411017 | 49411616 | 600 | -0.91 | 0.162 | 0.222 |
| chr2 | 187368443 | 187369860 | 1418 | 0.73 | 0.162 | 0.222 |
| chr1 | 155579165 | 155579847 | 683 | 0.48 | 0.162 | 0.222 |
| chr3 | 130935911 | 130936530 | 620 | 0.89 | 0.162 | 0.222 |
| chr21 | 45358408 | 45359439 | 1032 | 0.66 | 0.162 | 0.222 |
| chr5 | 131041722 | 131042831 | 1110 | -0.49 | 0.162 | 0.222 |
| chr2 | 37034207 | 37035336 | 1130 | 0.73 | 0.162 | 0.222 |
| chr3 | 126697750 | 126701770 | 4021 | -0.47 | 0.162 | 0.223 |
| chr14 | 31362572 | 31363939 | 1368 | -0.41 | 0.162 | 0.223 |
| chr5 | 61752666 | 61755009 | 2344 | -0.47 | 0.162 | 0.223 |
| chr2 | 98702450 | 98704372 | 1923 | 0.46 | 0.163 | 0.223 |
| chr13 | 44482475 | 44483973 | 1499 | -0.47 | 0.163 | 0.223 |
| chr17 | 40915059 | 40932321 | 17263 | -0.17 | 0.163 | 0.223 |
| chr2 | 69943466 | 69944156 | 691 | -0.71 | 0.163 | 0.223 |
| chr8 | 132987022 | 132987790 | 769 | 0.6 | 0.163 | 0.223 |
| chr11 | 74318781 | 74324517 | 5737 | -0.2 | 0.163 | 0.223 |
| chr7 | 150671813 | 150672707 | 895 | -0.59 | 0.163 | 0.223 |
| chr9 | 114384410 | 114395250 | 10841 | -0.34 | 0.163 | 0.223 |

| | | | | | | |
|---|---|---|---|---|---|---|
| chr7 | 26191193 | 26248375 | 57183 | 0.11 | 0.163 | 0.223 |
| chr14 | 20914104 | 20941996 | 27893 | 0.12 | 0.163 | 0.223 |
| chr2 | 158146739 | 158149202 | 2464 | -0.31 | 0.163 | 0.223 |
| chr9 | 132627146 | 132627791 | 646 | -0.82 | 0.163 | 0.224 |
| chr9 | 132716601 | 132732704 | 16104 | -0.19 | 0.163 | 0.224 |
| chr9 | 140609477 | 140618899 | 9423 | 0.45 | 0.163 | 0.224 |
| chr8 | 71000584 | 71000960 | 377 | -0.85 | 0.163 | 0.224 |
| chr1 | 149855096 | 149872470 | 17375 | -0.1 | 0.164 | 0.224 |
| chr15 | 96569195 | 96570299 | 1105 | -0.56 | 0.164 | 0.224 |
| chr9 | 72965289 | 72971508 | 6220 | 0.38 | 0.164 | 0.224 |
| chr17 | 60663772 | 60675572 | 11801 | 0.27 | 0.164 | 0.224 |
| chr12 | 64797185 | 64804475 | 7291 | 0.15 | 0.164 | 0.224 |
| chr1 | 214506495 | 214507928 | 1434 | -0.76 | 0.164 | 0.224 |
| chr10 | 124766702 | 124769162 | 2461 | 0.27 | 0.164 | 0.224 |
| chr13 | 21451240 | 21457321 | 6082 | -0.64 | 0.164 | 0.224 |
| chr1 | 208292777 | 208293882 | 1106 | 0.79 | 0.164 | 0.224 |
| chr3 | 184529506 | 184530704 | 1199 | 0.51 | 0.164 | 0.224 |
| chr20 | 31000219 | 31012666 | 12448 | -0.25 | 0.164 | 0.224 |
| chr13 | 91067040 | 91067795 | 756 | -0.79 | 0.164 | 0.224 |
| chr4 | 183817111 | 183817673 | 563 | -0.86 | 0.164 | 0.224 |
| chr1 | 24645829 | 24646507 | 679 | 0.77 | 0.164 | 0.224 |
| chr1 | 197454607 | 197455588 | 982 | 2.09 | 0.164 | 0.225 |
| chr3 | 47653092 | 47653898 | 807 | -0.85 | 0.164 | 0.225 |
| chr22 | 30279191 | 30280017 | 827 | 0.57 | 0.164 | 0.225 |
| chr2 | 20307519 | 20311909 | 4391 | -0.33 | 0.164 | 0.225 |
| chr19 | 44808152 | 44809664 | 1513 | 0.28 | 0.164 | 0.225 |
| chr3 | 66603739 | 66606334 | 2596 | -0.34 | 0.164 | 0.225 |
| chr20 | 43669868 | 43671058 | 1191 | 0.52 | 0.164 | 0.225 |
| chr12 | 17391179 | 17391546 | 368 | 1.13 | 0.165 | 0.225 |
| chr2 | 38812574 | 38831357 | 18784 | -0.16 | 0.165 | 0.225 |
| chr11 | 75459331 | 75480047 | 20717 | 2.62 | 0.165 | 0.225 |
| chr11 | 94527158 | 94528118 | 961 | 1 | 0.165 | 0.225 |
| chr9 | 107730339 | 107731451 | 1113 | -0.47 | 0.165 | 0.225 |
| chr3 | 44924338 | 44936604 | 12267 | 0.19 | 0.165 | 0.225 |
| chr2 | 189063753 | 189064912 | 1160 | 0.57 | 0.165 | 0.225 |
| chr4 | 33091700 | 33092023 | 324 | -1.27 | 0.165 | 0.225 |
| chr1 | 9205191 | 9243275 | 38085 | -0.11 | 0.165 | 0.226 |
| chr1 | 9294253 | 9295970 | 1718 | 0.29 | 0.165 | 0.226 |
| chr5 | 159300570 | 159307245 | 6676 | -0.23 | 0.165 | 0.226 |
| chr11 | 107327649 | 107329227 | 1579 | 0.39 | 0.165 | 0.226 |
| chr13 | 99427013 | 99429634 | 2622 | 0.26 | 0.165 | 0.226 |
| chr12 | 67821825 | 67829316 | 7492 | 2.51 | 0.165 | 0.226 |
| chrX | 134654990 | 134655526 | 537 | 0.85 | 0.165 | 0.226 |
| chr14 | 66584140 | 66584862 | 723 | -0.74 | 0.165 | 0.226 |
| chr20 | 37637432 | 37638715 | 1284 | -0.67 | 0.165 | 0.226 |
| chr8 | 28608999 | 28610194 | 1196 | -1.04 | 0.165 | 0.226 |
| chr8 | 28919072 | 28922125 | 3054 | -0.52 | 0.165 | 0.226 |
| chr11 | 77889972 | 77900691 | 10720 | 0.23 | 0.165 | 0.226 |
| chr4 | 87877025 | 87880738 | 3714 | 0.59 | 0.165 | 0.226 |
| chr6 | 111839304 | 111839912 | 609 | -0.89 | 0.166 | 0.226 |
| chr13 | 44748185 | 44749642 | 1458 | -0.41 | 0.166 | 0.226 |
| chr17 | 29215221 | 29216232 | 1012 | -0.81 | 0.166 | 0.226 |
| chrX | 119867597 | 119871176 | 3580 | -0.22 | 0.166 | 0.226 |
| chr12 | 125534357 | 125534807 | 451 | 0.84 | 0.166 | 0.226 |
| chr6 | 14732675 | 14745087 | 12413 | 0.15 | 0.166 | 0.226 |
| chr2 | 43786598 | 43787671 | 1074 | -0.57 | 0.166 | 0.226 |
| chr5 | 134293879 | 134296726 | 2848 | -0.51 | 0.166 | 0.227 |
| chr16 | 70687877 | 70688342 | 466 | -0.85 | 0.166 | 0.227 |
| chr11 | 72880076 | 72880524 | 449 | -1.02 | 0.166 | 0.227 |
| chr1 | 162039009 | 162039384 | 376 | 1.03 | 0.166 | 0.227 |
| chr5 | 109158898 | 109169985 | 11088 | 0.34 | 0.166 | 0.227 |

| | | | | | | |
|---|---|---|---|---|---|---|
| chr10 | 47655292 | 47656919 | 1628 | -0.36 | 0.167 | 0.227 |
| chr18 | 60381496 | 60385164 | 3669 | 0.29 | 0.167 | 0.227 |
| chr3 | 113414442 | 113418097 | 3656 | 0.24 | 0.167 | 0.227 |
| chr1 | 237068430 | 237068999 | 570 | -1.12 | 0.167 | 0.228 |
| chr2 | 179058379 | 179060289 | 1911 | -0.53 | 0.167 | 0.228 |
| chrX | 110968426 | 110969276 | 851 | -0.88 | 0.167 | 0.228 |
| chr15 | 72562631 | 72565985 | 3355 | 0.19 | 0.167 | 0.228 |
| chr1 | 40915137 | 40916683 | 1547 | 0.51 | 0.167 | 0.228 |
| chrX | 102840064 | 102841333 | 1270 | 0.43 | 0.167 | 0.228 |
| chrX | 77149639 | 77210045 | 60407 | 0.09 | 0.167 | 0.228 |
| chr17 | 64218922 | 64238515 | 19594 | 0.26 | 0.167 | 0.229 |
| chr1 | 16263650 | 16280259 | 16610 | 0.18 | 0.168 | 0.229 |
| chr7 | 110777819 | 110778924 | 1106 | 0.6 | 0.168 | 0.229 |
| chr13 | 91999889 | 92015214 | 15326 | 0.22 | 0.168 | 0.229 |
| chr12 | 51202489 | 51219544 | 17056 | 0.21 | 0.168 | 0.229 |
| chr15 | 67532671 | 67548088 | 15418 | 0.14 | 0.168 | 0.229 |
| chr2 | 202980655 | 202981358 | 704 | -0.81 | 0.168 | 0.229 |
| chr8 | 144716823 | 144719302 | 2480 | 0.23 | 0.168 | 0.229 |
| chr17 | 49368707 | 49370722 | 2016 | 0.54 | 0.168 | 0.229 |
| chr4 | 89921787 | 89922539 | 753 | -0.91 | 0.168 | 0.229 |
| chr1 | 208392475 | 208394813 | 2339 | 0.54 | 0.168 | 0.229 |
| chr2 | 193026064 | 193027255 | 1192 | -0.85 | 0.168 | 0.229 |
| chr17 | 3694683 | 3750420 | 55738 | -0.3 | 0.168 | 0.229 |
| chr2 | 219308105 | 219326516 | 18412 | 0.18 | 0.168 | 0.229 |
| chr13 | 27824355 | 27834489 | 10135 | 0.13 | 0.168 | 0.229 |
| chr14 | 23414276 | 23427164 | 12889 | 0.21 | 0.168 | 0.229 |
| chr6 | 38606717 | 38608715 | 1999 | 0.25 | 0.168 | 0.229 |
| chr4 | 160095538 | 160097981 | 2444 | 0.35 | 0.168 | 0.229 |
| chr4 | 2600245 | 2600818 | 574 | -0.88 | 0.168 | 0.23 |
| chrX | 153218611 | 153219508 | 898 | 0.77 | 0.168 | 0.23 |
| chr6 | 114290363 | 114293017 | 2655 | 0.2 | 0.168 | 0.23 |
| chr2 | 205924561 | 205927226 | 2666 | -0.56 | 0.169 | 0.23 |
| chr1 | 161300058 | 161300787 | 730 | -0.9 | 0.169 | 0.23 |
| chr3 | 193921788 | 193922321 | 534 | 0.88 | 0.169 | 0.23 |
| chr8 | 17692405 | 17723399 | 30995 | 0.09 | 0.169 | 0.23 |
| chr6 | 133118840 | 133121085 | 2246 | 0.31 | 0.169 | 0.23 |
| chr22 | 24988496 | 24990105 | 1610 | -0.43 | 0.169 | 0.23 |
| chr16 | 80574043 | 80575924 | 1882 | -0.21 | 0.169 | 0.23 |
| chr10 | 33084742 | 33085676 | 935 | -0.72 | 0.169 | 0.23 |
| chr16 | 77245696 | 77253978 | 8283 | -0.23 | 0.169 | 0.23 |
| chr3 | 125230270 | 125240056 | 9787 | 0.21 | 0.169 | 0.23 |
| chr11 | 8796025 | 8796400 | 376 | -1.47 | 0.169 | 0.23 |
| chr1 | 20914913 | 20917008 | 2096 | -0.43 | 0.169 | 0.23 |
| chr13 | 99196241 | 99197882 | 1642 | 0.71 | 0.169 | 0.231 |
| chr7 | 12279775 | 12280446 | 672 | 0.7 | 0.169 | 0.231 |
| chr21 | 45425553 | 45433694 | 8142 | 0.26 | 0.169 | 0.231 |
| chr14 | 51794768 | 51795318 | 551 | -0.83 | 0.169 | 0.231 |
| chr1 | 184698633 | 184700791 | 2159 | 0.47 | 0.169 | 0.231 |
| chr2 | 70261828 | 70263135 | 1308 | 0.51 | 0.17 | 0.231 |
| chr2 | 227379272 | 227381306 | 2035 | 0.24 | 0.17 | 0.231 |
| chr8 | 42286836 | 42287609 | 774 | -1.01 | 0.17 | 0.231 |
| chr2 | 24406151 | 24406787 | 637 | 0.8 | 0.17 | 0.231 |
| chr1 | 236305807 | 236306512 | 706 | 0.53 | 0.17 | 0.231 |
| chr11 | 1136148 | 1136660 | 513 | 1.06 | 0.17 | 0.231 |
| chr6 | 135306050 | 135306475 | 426 | -0.96 | 0.17 | 0.231 |
| chr1 | 28878862 | 28881137 | 2276 | 0.33 | 0.17 | 0.231 |
| chr6 | 11315178 | 11320571 | 5394 | -0.14 | 0.17 | 0.231 |
| chr18 | 3498900 | 3499621 | 722 | -0.9 | 0.17 | 0.231 |
| chr8 | 95850163 | 95850691 | 529 | -1.01 | 0.17 | 0.231 |
| chr12 | 118526522 | 118527836 | 1315 | -0.52 | 0.17 | 0.232 |
| chr6 | 4930320 | 4946114 | 15795 | 0.14 | 0.17 | 0.232 |

| | | | | | | |
|---|---|---|---|---|---|---|
| chr10 | 118664833 | 118665746 | 914 | -0.47 | 0.17 | 0.232 |
| chr3 | 150420952 | 150422589 | 1638 | -0.24 | 0.17 | 0.232 |
| chrX | 106398405 | 106399113 | 709 | 0.7 | 0.17 | 0.232 |
| chr4 | 103453004 | 103453819 | 816 | -0.89 | 0.171 | 0.232 |
| chr15 | 66858957 | 66862967 | 4011 | -0.75 | 0.171 | 0.232 |
| chr1 | 57092679 | 57112152 | 19474 | 0.32 | 0.171 | 0.232 |
| chr18 | 24324326 | 24344856 | 20531 | 0.11 | 0.171 | 0.232 |
| chr3 | 151178469 | 151179180 | 712 | 0.72 | 0.171 | 0.232 |
| chr4 | 155470221 | 155472225 | 2005 | 0.35 | 0.171 | 0.232 |
| chr14 | 24656602 | 24667066 | 10465 | -0.14 | 0.171 | 0.232 |
| chr14 | 64658139 | 64662313 | 4175 | -0.28 | 0.171 | 0.232 |
| chr3 | 47801489 | 47802740 | 1252 | -0.68 | 0.171 | 0.232 |
| chr3 | 101497507 | 101512989 | 15483 | -0.15 | 0.171 | 0.233 |
| chr11 | 77849455 | 77851366 | 1912 | 0.2 | 0.171 | 0.233 |
| chr12 | 20029994 | 20037977 | 7984 | 2.54 | 0.171 | 0.233 |
| chr7 | 105331370 | 105332028 | 659 | 0.72 | 0.171 | 0.233 |
| chr5 | 143567012 | 143579442 | 12431 | 0.12 | 0.171 | 0.233 |
| chr14 | 51706308 | 51727471 | 21164 | -0.21 | 0.171 | 0.233 |
| chr18 | 33732571 | 33734218 | 1648 | 0.45 | 0.171 | 0.233 |
| chr9 | 74524222 | 74536646 | 12425 | 0.24 | 0.171 | 0.233 |
| chr8 | 23655580 | 23656820 | 1241 | 0.56 | 0.172 | 0.233 |
| chr2 | 96192248 | 96193076 | 829 | 0.9 | 0.172 | 0.233 |
| chr15 | 70962670 | 70963029 | 360 | -1.42 | 0.172 | 0.234 |
| chr4 | 86699406 | 86718799 | 19394 | 0.33 | 0.172 | 0.234 |
| chr17 | 20811246 | 20812799 | 1554 | 0.27 | 0.172 | 0.234 |
| chr6 | 166442765 | 166447470 | 4706 | -0.22 | 0.172 | 0.234 |
| chr1 | 245254712 | 245255996 | 1285 | 0.38 | 0.172 | 0.234 |
| chr5 | 142533942 | 142535208 | 1267 | 0.67 | 0.172 | 0.234 |
| chr19 | 51338722 | 51339832 | 1111 | 0.54 | 0.172 | 0.234 |
| chr2 | 136661553 | 136662687 | 1135 | -0.54 | 0.172 | 0.234 |
| chr6 | 142795523 | 142799798 | 4276 | -0.63 | 0.172 | 0.234 |
| chrX | 102939666 | 102943406 | 3741 | 0.18 | 0.172 | 0.234 |
| chr2 | 174279094 | 174280103 | 1010 | 0.61 | 0.172 | 0.234 |
| chr2 | 58789327 | 58789839 | 513 | 0.97 | 0.172 | 0.234 |
| chr3 | 47799587 | 47800016 | 430 | 1.04 | 0.172 | 0.234 |
| chr4 | 2443129 | 2443507 | 379 | -1.25 | 0.172 | 0.234 |
| chr11 | 1313465 | 1331699 | 18235 | -0.18 | 0.172 | 0.234 |
| chr13 | 51970931 | 51971802 | 872 | 0.82 | 0.173 | 0.235 |
| chr6 | 105584175 | 105585079 | 905 | -0.73 | 0.173 | 0.235 |
| chr6 | 27705563 | 27721109 | 15547 | 0.28 | 0.173 | 0.235 |
| chr22 | 31199141 | 31202870 | 3730 | 0.56 | 0.173 | 0.235 |
| chr1 | 213223987 | 213225432 | 1446 | 0.28 | 0.173 | 0.235 |
| chr22 | 32266471 | 32267858 | 1388 | 0.63 | 0.173 | 0.235 |
| chr5 | 138235659 | 138243298 | 7640 | 0.33 | 0.173 | 0.235 |
| chr3 | 186819117 | 186820413 | 1297 | 0.57 | 0.173 | 0.235 |
| chr13 | 79183354 | 79183858 | 505 | 1.03 | 0.173 | 0.235 |
| chr11 | 85658670 | 85659403 | 734 | 1.07 | 0.173 | 0.235 |
| chr19 | 40694349 | 40698621 | 4273 | -0.23 | 0.173 | 0.235 |
| chr2 | 38846790 | 38848057 | 1268 | 2.31 | 0.173 | 0.235 |
| chr16 | 85828868 | 85834901 | 6034 | 0.23 | 0.173 | 0.235 |
| chr2 | 61185797 | 61189129 | 3333 | -0.46 | 0.173 | 0.235 |
| chr6 | 64057802 | 64058191 | 390 | -0.91 | 0.173 | 0.236 |
| chr2 | 170122819 | 170123308 | 490 | 1.06 | 0.173 | 0.236 |
| chr5 | 114504867 | 114516296 | 11430 | 2.32 | 0.174 | 0.236 |
| chr2 | 172247443 | 172248318 | 876 | 0.84 | 0.174 | 0.236 |
| chr2 | 239468239 | 239471032 | 2794 | 0.31 | 0.174 | 0.236 |
| chr4 | 90147862 | 90148801 | 940 | -0.88 | 0.174 | 0.236 |
| chr3 | 58222602 | 58224263 | 1662 | 0.45 | 0.174 | 0.236 |
| chr1 | 65639773 | 65640492 | 720 | 1.2 | 0.174 | 0.236 |
| chr17 | 28696926 | 28733131 | 36206 | 0.15 | 0.174 | 0.236 |
| chr6 | 132452243 | 132452931 | 689 | 0.81 | 0.174 | 0.236 |

| | | | | | | |
|---|---|---|---|---|---|---|
| chr8 | 99613606 | 99625740 | 12135 | -2.49 | 0.174 | 0.236 |
| chr22 | 23430444 | 23430957 | 514 | -0.88 | 0.174 | 0.236 |
| chr19 | 41302113 | 41314713 | 12601 | -0.13 | 0.174 | 0.236 |
| chr15 | 71107643 | 71127074 | 19432 | -0.17 | 0.174 | 0.237 |
| chr7 | 85183844 | 85184446 | 603 | 0.68 | 0.174 | 0.237 |
| chr12 | 108930593 | 108961099 | 30507 | 0.11 | 0.174 | 0.237 |
| chr9 | 121080804 | 121094980 | 14177 | 2.44 | 0.174 | 0.237 |
| chr13 | 50570118 | 50573014 | 2897 | 0.23 | 0.174 | 0.237 |
| chr17 | 17438932 | 17443668 | 4737 | -0.2 | 0.174 | 0.237 |
| chr3 | 33318322 | 33320515 | 2194 | 0.25 | 0.174 | 0.237 |
| chr6 | 37849642 | 37859879 | 10238 | 0.29 | 0.175 | 0.237 |
| chr1 | 85154768 | 85157218 | 2451 | 0.32 | 0.175 | 0.237 |
| chr3 | 135860796 | 135869344 | 8549 | 0.33 | 0.175 | 0.237 |
| chr5 | 146940739 | 146941125 | 387 | 0.94 | 0.175 | 0.237 |
| chr19 | 10919123 | 10927813 | 8691 | -0.2 | 0.175 | 0.237 |
| chr4 | 89299843 | 89300774 | 932 | 0.52 | 0.175 | 0.237 |
| chr7 | 4814111 | 4816514 | 2404 | 0.27 | 0.175 | 0.237 |
| chr10 | 16858549 | 16860352 | 1804 | 0.36 | 0.175 | 0.237 |
| chr7 | 128502342 | 128504211 | 1870 | 0.41 | 0.175 | 0.237 |
| chr5 | 372943 | 374658 | 1716 | 0.51 | 0.175 | 0.237 |
| chr6 | 21874437 | 21889113 | 14677 | -0.24 | 0.175 | 0.237 |
| chr5 | 136527605 | 136528875 | 1271 | -0.59 | 0.175 | 0.237 |
| chr12 | 41341715 | 41342044 | 330 | 1.11 | 0.175 | 0.237 |
| chr7 | 77457028 | 77467886 | 10859 | -0.13 | 0.175 | 0.238 |
| chr8 | 23691343 | 23691966 | 624 | -0.9 | 0.175 | 0.238 |
| chr16 | 79673365 | 79674966 | 1602 | 0.57 | 0.175 | 0.238 |
| chr9 | 134851493 | 134852608 | 1116 | 0.54 | 0.175 | 0.238 |
| chr6 | 18042704 | 18054284 | 11581 | 1.19 | 0.175 | 0.238 |
| chr3 | 13438097 | 13439224 | 1128 | -0.68 | 0.175 | 0.238 |
| chr3 | 150515980 | 150516450 | 471 | -1 | 0.175 | 0.238 |
| chr8 | 8105314 | 8106148 | 835 | 0.87 | 0.175 | 0.238 |
| chr5 | 132165426 | 132174172 | 8747 | 0.35 | 0.175 | 0.238 |
| chr10 | 90013338 | 90014385 | 1048 | 0.56 | 0.176 | 0.238 |
| chr6 | 138724295 | 138726718 | 2424 | 0.24 | 0.176 | 0.238 |
| chr8 | 98769002 | 98831239 | 62238 | -0.11 | 0.176 | 0.238 |
| chr15 | 30940389 | 30943883 | 3495 | 0.52 | 0.176 | 0.238 |
| chr5 | 125956455 | 125962171 | 5717 | -0.29 | 0.176 | 0.238 |
| chr2 | 141148339 | 141149078 | 740 | -1.21 | 0.176 | 0.238 |
| chr11 | 87907419 | 87908643 | 1225 | 0.62 | 0.176 | 0.238 |
| chr15 | 76050546 | 76051781 | 1236 | 0.28 | 0.176 | 0.238 |
| chr1 | 155057071 | 155067082 | 10012 | -0.63 | 0.176 | 0.239 |
| chr2 | 54123967 | 54125115 | 1149 | -0.63 | 0.176 | 0.239 |
| chr5 | 102560035 | 102560421 | 387 | 1.04 | 0.176 | 0.239 |
| chr1 | 16532539 | 16534268 | 1730 | 0.38 | 0.176 | 0.239 |
| chr4 | 166423170 | 166424975 | 1806 | -0.43 | 0.176 | 0.239 |
| chr19 | 6392211 | 6394388 | 2178 | -0.21 | 0.176 | 0.239 |
| chr4 | 13428971 | 13430537 | 1567 | 0.71 | 0.176 | 0.239 |
| chr19 | 17790521 | 17791591 | 1071 | 0.71 | 0.176 | 0.239 |
| chr2 | 61023607 | 61024056 | 450 | -0.79 | 0.176 | 0.239 |
| chr13 | 41836967 | 41896565 | 59599 | 0.15 | 0.176 | 0.239 |
| chr11 | 61085931 | 61131422 | 45492 | -0.11 | 0.176 | 0.239 |
| chr19 | 11434319 | 11458177 | 23859 | -0.22 | 0.177 | 0.239 |
| chr8 | 103837260 | 103841914 | 4655 | -0.57 | 0.177 | 0.239 |
| chr2 | 237906635 | 237907916 | 1282 | 0.56 | 0.177 | 0.239 |
| chr12 | 54446609 | 54448360 | 1752 | 0.25 | 0.177 | 0.239 |
| chr2 | 203172713 | 203173125 | 413 | -1.09 | 0.177 | 0.239 |
| chr3 | 20201460 | 20206794 | 5335 | 0.4 | 0.177 | 0.239 |
| chr2 | 153963346 | 153964035 | 690 | 2.2 | 0.177 | 0.239 |
| chr22 | 19793495 | 19794583 | 1089 | -0.66 | 0.177 | 0.239 |
| chr16 | 81347423 | 81349752 | 2330 | 0.35 | 0.177 | 0.24 |
| chr8 | 66860125 | 66861396 | 1272 | 0.65 | 0.177 | 0.24 |

| | | | | | | |
|---|---|---|---|---|---|---|
| chr12 | 58191587 | 58192268 | 682 | 0.78 | 0.177 | 0.24 |
| chr19 | 8635515 | 8682689 | 47175 | -0.16 | 0.177 | 0.24 |
| chr20 | 34129206 | 34143107 | 13902 | -0.28 | 0.177 | 0.24 |
| chr1 | 87378754 | 87381635 | 2882 | 0.25 | 0.177 | 0.24 |
| chr4 | 187527668 | 187528129 | 462 | -0.88 | 0.178 | 0.24 |
| chr20 | 9969675 | 9970197 | 523 | 1.07 | 0.178 | 0.24 |
| chr20 | 5022012 | 5022636 | 625 | -0.76 | 0.178 | 0.24 |
| chr1 | 182088509 | 182090963 | 2455 | 0.22 | 0.178 | 0.241 |
| chr1 | 63941458 | 63942759 | 1302 | 0.6 | 0.178 | 0.241 |
| chr8 | 37378829 | 37379143 | 315 | -1.21 | 0.178 | 0.241 |
| chr20 | 35805917 | 35814710 | 8794 | 0.14 | 0.178 | 0.241 |
| chr8 | 42497407 | 42498432 | 1026 | 2.43 | 0.178 | 0.241 |
| chr3 | 32494621 | 32496240 | 1620 | -2.28 | 0.178 | 0.241 |
| chr16 | 47746744 | 47747601 | 858 | -0.73 | 0.178 | 0.241 |
| chr11 | 35929516 | 35930570 | 1055 | 0.76 | 0.178 | 0.241 |
| chr14 | 34836778 | 34849480 | 12703 | 0.18 | 0.178 | 0.241 |
| chr4 | 1166829 | 1169139 | 2311 | 0.29 | 0.178 | 0.241 |
| chr2 | 163200641 | 163202137 | 1497 | 0.3 | 0.178 | 0.241 |
| chr5 | 158876376 | 158898001 | 21626 | -0.21 | 0.178 | 0.241 |
| chr9 | 115913320 | 115913910 | 591 | 1.18 | 0.179 | 0.241 |
| chr10 | 119804037 | 119807447 | 3411 | -0.15 | 0.179 | 0.241 |
| chr21 | 34977726 | 34978210 | 485 | -0.85 | 0.179 | 0.241 |
| chr6 | 136389797 | 136402331 | 12535 | -2.58 | 0.179 | 0.242 |
| chr13 | 114864160 | 114865010 | 851 | -0.62 | 0.179 | 0.242 |
| chr7 | 152242120 | 152242489 | 370 | 0.99 | 0.179 | 0.242 |
| chr5 | 142654290 | 142676968 | 22679 | 0.15 | 0.179 | 0.242 |
| chr8 | 22772689 | 22774528 | 1840 | -0.39 | 0.179 | 0.242 |
| chr2 | 126156271 | 126164340 | 8070 | 0.39 | 0.179 | 0.242 |
| chr6 | 52476938 | 52478162 | 1225 | -0.49 | 0.179 | 0.242 |
| chrX | 41480250 | 41481830 | 1581 | 0.53 | 0.179 | 0.242 |
| chr9 | 111304821 | 111312078 | 7258 | 0.38 | 0.179 | 0.242 |
| chr8 | 136469041 | 136469512 | 472 | 0.83 | 0.179 | 0.242 |
| chr2 | 172203893 | 172204758 | 866 | -0.81 | 0.179 | 0.242 |
| chr14 | 32805794 | 32815600 | 9807 | -0.38 | 0.179 | 0.242 |
| chr10 | 81107226 | 81109845 | 2620 | -0.28 | 0.179 | 0.242 |
| chr7 | 32132916 | 32133624 | 709 | -1.05 | 0.179 | 0.242 |
| chr4 | 95964045 | 95975951 | 11907 | -0.14 | 0.18 | 0.243 |
| chr11 | 130744281 | 130744817 | 537 | 0.92 | 0.18 | 0.243 |
| chr1 | 45476622 | 45478923 | 2302 | 0.38 | 0.18 | 0.243 |
| chr10 | 99204824 | 99259774 | 54951 | -0.12 | 0.18 | 0.243 |
| chr19 | 3983773 | 3988461 | 4689 | -0.25 | 0.18 | 0.243 |
| chr11 | 17297028 | 17299343 | 2316 | 0.3 | 0.18 | 0.243 |
| chr2 | 225306931 | 225307557 | 627 | -1.03 | 0.18 | 0.243 |
| chr1 | 235242273 | 235303022 | 60750 | -0.1 | 0.18 | 0.243 |
| chr5 | 134093519 | 134099825 | 6307 | -0.17 | 0.18 | 0.243 |
| chr2 | 37498726 | 37499318 | 593 | -1.04 | 0.18 | 0.243 |
| chr17 | 39482076 | 39484554 | 2479 | -0.49 | 0.18 | 0.243 |
| chr14 | 61446351 | 61450263 | 3913 | -0.14 | 0.18 | 0.243 |
| chr17 | 5403987 | 5404605 | 619 | -1 | 0.18 | 0.243 |
| chr9 | 84003601 | 84005555 | 1955 | 0.34 | 0.18 | 0.243 |
| chr5 | 55619518 | 55625173 | 5656 | 0.63 | 0.18 | 0.243 |
| chr14 | 69893667 | 69894393 | 727 | -1.39 | 0.18 | 0.243 |
| chr7 | 103815236 | 103816565 | 1330 | 0.62 | 0.18 | 0.243 |
| chr2 | 8943308 | 8945794 | 2487 | 0.23 | 0.18 | 0.244 |
| chr9 | 14271290 | 14271886 | 597 | 0.89 | 0.18 | 0.244 |
| chr10 | 133877567 | 133881043 | 3477 | -0.18 | 0.181 | 0.244 |
| chr3 | 126629622 | 126631577 | 1956 | -0.29 | 0.181 | 0.244 |
| chr6 | 25489851 | 25490720 | 870 | 0.71 | 0.181 | 0.244 |
| chr1 | 95631224 | 95637127 | 5904 | 0.13 | 0.181 | 0.244 |
| chr3 | 122920000 | 122922081 | 2082 | 0.22 | 0.181 | 0.244 |
| chr2 | 160255942 | 160256930 | 989 | 0.55 | 0.181 | 0.244 |

| | | | | | | |
|---|---|---|---|---|---|---|
| chr2 | 165511383 | 165512413 | 1031 | -0.64 | 0.181 | 0.244 |
| chr9 | 88877116 | 88877725 | 610 | -0.9 | 0.181 | 0.244 |
| chr5 | 38946073 | 38946586 | 514 | 0.84 | 0.181 | 0.244 |
| chr7 | 44495703 | 44531140 | 35438 | 0.12 | 0.181 | 0.244 |
| chr10 | 14878860 | 14884295 | 5436 | 0.22 | 0.181 | 0.244 |
| chr10 | 93623395 | 93624648 | 1254 | 0.61 | 0.181 | 0.244 |
| chr3 | 196514610 | 196520911 | 6302 | -0.24 | 0.181 | 0.244 |
| chr4 | 114594231 | 114630593 | 36363 | -0.14 | 0.181 | 0.244 |
| chr1 | 162680388 | 162680849 | 462 | -0.95 | 0.181 | 0.244 |
| chrX | 130837692 | 130845470 | 7779 | -0.26 | 0.181 | 0.244 |
| chr1 | 24246145 | 24247666 | 1522 | 0.44 | 0.181 | 0.245 |
| chr1 | 45308461 | 45309199 | 739 | -0.66 | 0.181 | 0.245 |
| chr14 | 58636692 | 58638851 | 2160 | 0.26 | 0.181 | 0.245 |
| chr5 | 170946721 | 170962128 | 15408 | -2.61 | 0.181 | 0.245 |
| chr7 | 151122622 | 151124663 | 2042 | -0.27 | 0.181 | 0.245 |
| chr11 | 18720366 | 18720852 | 487 | 1.01 | 0.181 | 0.245 |
| chr8 | 95892254 | 95894317 | 2064 | 0.55 | 0.181 | 0.245 |
| chr7 | 151629908 | 151630470 | 563 | -0.86 | 0.182 | 0.245 |
| chr17 | 7892950 | 7894281 | 1332 | 0.46 | 0.182 | 0.245 |
| chr22 | 19418033 | 19420990 | 2958 | 0.24 | 0.182 | 0.245 |
| chr10 | 134822185 | 134823684 | 1500 | 0.31 | 0.182 | 0.245 |
| chr5 | 138272177 | 138272863 | 687 | -0.86 | 0.182 | 0.245 |
| chr3 | 48469556 | 48489032 | 19477 | 0.17 | 0.182 | 0.245 |
| chr6 | 15365282 | 15366324 | 1043 | 0.67 | 0.182 | 0.245 |
| chr1 | 217448901 | 217449870 | 970 | -0.75 | 0.182 | 0.245 |
| chr19 | 13029726 | 13030728 | 1003 | 0.69 | 0.182 | 0.245 |
| chr12 | 111051274 | 111053589 | 2316 | -0.18 | 0.182 | 0.246 |
| chr6 | 64093771 | 64094198 | 428 | 1.42 | 0.182 | 0.246 |
| chr22 | 47525149 | 47525890 | 742 | -0.73 | 0.182 | 0.246 |
| chr20 | 43035700 | 43036174 | 475 | 0.92 | 0.182 | 0.246 |
| chr3 | 142858328 | 142858826 | 499 | -1.4 | 0.182 | 0.246 |
| chr18 | 52821242 | 52821728 | 487 | 1.03 | 0.182 | 0.246 |
| chr16 | 19565302 | 19568072 | 2771 | 0.2 | 0.182 | 0.246 |
| chr4 | 170637660 | 170638463 | 804 | 1.09 | 0.183 | 0.246 |
| chr3 | 87275289 | 87297996 | 22708 | 0.21 | 0.183 | 0.246 |
| chr14 | 64570407 | 64571002 | 596 | -0.88 | 0.183 | 0.246 |
| chr1 | 23493658 | 23552834 | 59177 | 0.07 | 0.183 | 0.246 |
| chr7 | 94269994 | 94290260 | 20267 | 0.32 | 0.183 | 0.246 |
| chr12 | 8888512 | 8889350 | 839 | 0.88 | 0.183 | 0.246 |
| chr9 | 92137757 | 92138084 | 328 | -1.43 | 0.183 | 0.246 |
| chr6 | 105306902 | 105307750 | 849 | 0.65 | 0.183 | 0.247 |
| chr19 | 8407673 | 8429617 | 21945 | -0.18 | 0.183 | 0.247 |
| chr14 | 104314068 | 104314974 | 907 | 0.55 | 0.183 | 0.247 |
| chr15 | 72664944 | 72669321 | 4378 | 0.22 | 0.183 | 0.247 |
| chr11 | 73450595 | 73452007 | 1413 | 0.62 | 0.183 | 0.247 |
| chr4 | 3076108 | 3078259 | 2152 | 0.46 | 0.183 | 0.247 |
| chrX | 2880623 | 2883776 | 3154 | 0.23 | 0.183 | 0.247 |
| chr13 | 99777398 | 99777804 | 407 | -1 | 0.183 | 0.247 |
| chr5 | 167899624 | 167900097 | 474 | 0.83 | 0.183 | 0.247 |
| chr4 | 122959726 | 122960599 | 874 | 0.77 | 0.184 | 0.247 |
| chr9 | 102732766 | 102733323 | 558 | -0.79 | 0.184 | 0.247 |
| chr12 | 50194129 | 50195238 | 1110 | -0.64 | 0.184 | 0.247 |
| chrX | 35751734 | 35752487 | 754 | -0.88 | 0.184 | 0.248 |
| chr3 | 36688760 | 36691242 | 2483 | 0.36 | 0.184 | 0.248 |
| chr10 | 129214828 | 129215833 | 1006 | -0.72 | 0.184 | 0.248 |
| chr12 | 133148354 | 133148770 | 417 | -1.27 | 0.184 | 0.248 |
| chr22 | 41776495 | 41778247 | 1753 | -0.31 | 0.184 | 0.248 |
| chr14 | 23610654 | 23625043 | 14390 | 0.53 | 0.184 | 0.248 |
| chr12 | 93013298 | 93014781 | 1484 | 0.43 | 0.184 | 0.248 |
| chr7 | 32970814 | 32983100 | 12287 | -0.23 | 0.184 | 0.248 |
| chr4 | 75399608 | 75411569 | 11962 | -0.19 | 0.184 | 0.248 |

| | | | | | | |
|---|---|---|---|---|---|---|
| chr2 | 142942733 | 142948987 | 6255 | 0.23 | 0.184 | 0.248 |
| chr19 | 44905085 | 44906405 | 1321 | 0.5 | 0.184 | 0.248 |
| chr7 | 75820784 | 75821267 | 484 | 1.43 | 0.184 | 0.248 |
| chr12 | 4608288 | 4608715 | 428 | -0.81 | 0.184 | 0.248 |
| chr18 | 34394940 | 34410239 | 15300 | 0.19 | 0.184 | 0.248 |
| chr2 | 55633505 | 55653863 | 20359 | -0.13 | 0.185 | 0.248 |
| chr1 | 220547810 | 220548209 | 400 | -1.1 | 0.185 | 0.249 |
| chr3 | 8524023 | 8621237 | 97215 | -0.07 | 0.185 | 0.249 |
| chr10 | 123659864 | 123735606 | 75743 | 0.06 | 0.185 | 0.249 |
| chr12 | 46271825 | 46286257 | 14433 | 0.27 | 0.185 | 0.249 |
| chr9 | 131285091 | 131286043 | 953 | -0.76 | 0.185 | 0.249 |
| chr15 | 66301014 | 66302298 | 1285 | 0.45 | 0.185 | 0.249 |
| chr10 | 73575729 | 73660462 | 84734 | -0.06 | 0.185 | 0.249 |
| chr4 | 88013137 | 88029779 | 16643 | 0.22 | 0.185 | 0.249 |
| chr9 | 92170534 | 92172078 | 1545 | 0.45 | 0.185 | 0.249 |
| chr15 | 76014470 | 76017234 | 2765 | 0.21 | 0.185 | 0.249 |
| chr8 | 95730993 | 95733878 | 2886 | 0.23 | 0.185 | 0.249 |
| chr15 | 65567528 | 65598390 | 30863 | -0.09 | 0.186 | 0.25 |
| chr20 | 14447914 | 14451769 | 3856 | 0.4 | 0.186 | 0.25 |
| chr3 | 131851850 | 131855940 | 4091 | -0.12 | 0.186 | 0.25 |
| chr14 | 97263677 | 97264574 | 898 | -0.35 | 0.186 | 0.25 |
| chr1 | 150426309 | 150427054 | 746 | -0.73 | 0.186 | 0.25 |
| chr3 | 159706098 | 159710978 | 4881 | -0.46 | 0.186 | 0.25 |
| chr9 | 118757805 | 118795590 | 37786 | 0.13 | 0.186 | 0.25 |
| chr6 | 36088749 | 36089476 | 728 | 0.82 | 0.186 | 0.25 |
| chr9 | 135247846 | 135249488 | 1643 | 0.43 | 0.186 | 0.25 |
| chr16 | 14378838 | 14380352 | 1515 | -0.46 | 0.186 | 0.25 |
| chr17 | 35334733 | 35350378 | 15646 | 0.15 | 0.186 | 0.25 |
| chr20 | 61531220 | 61570655 | 39436 | 0.1 | 0.186 | 0.25 |
| chr5 | 37173161 | 37173693 | 533 | 0.84 | 0.186 | 0.25 |
| chr2 | 85914217 | 85915109 | 893 | -0.68 | 0.186 | 0.25 |
| chr11 | 74494163 | 74495182 | 1020 | 0.56 | 0.186 | 0.25 |
| chr2 | 44455388 | 44474631 | 19244 | -0.08 | 0.186 | 0.25 |
| chr11 | 74699142 | 74700869 | 1728 | 0.31 | 0.186 | 0.25 |
| chr1 | 21443607 | 21444653 | 1047 | 0.7 | 0.186 | 0.25 |
| chr4 | 75149491 | 75149834 | 344 | -1 | 0.186 | 0.25 |
| chr6 | 143770707 | 143772652 | 1946 | 0.32 | 0.186 | 0.25 |
| chr8 | 15397407 | 15399118 | 1712 | 0.34 | 0.186 | 0.251 |
| chr5 | 130838113 | 130851377 | 13265 | -0.39 | 0.187 | 0.251 |
| chr1 | 8498364 | 8499682 | 1319 | 0.75 | 0.187 | 0.251 |
| chr13 | 48668165 | 48669496 | 1332 | 0.33 | 0.187 | 0.251 |
| chr10 | 88470325 | 88472546 | 2222 | 0.19 | 0.187 | 0.251 |
| chr3 | 184016279 | 184018325 | 2047 | 0.4 | 0.187 | 0.251 |
| chr17 | 15382291 | 15383591 | 1301 | -0.56 | 0.187 | 0.251 |
| chr9 | 35488225 | 35564039 | 75815 | -0.05 | 0.187 | 0.251 |
| chr19 | 7717202 | 7717638 | 437 | 0.9 | 0.187 | 0.251 |
| chr2 | 157963566 | 157964519 | 954 | 1.12 | 0.187 | 0.251 |
| chr3 | 170762201 | 170774091 | 11891 | 0.4 | 0.187 | 0.251 |
| chr17 | 33445447 | 33447723 | 2277 | -0.27 | 0.187 | 0.251 |
| chr3 | 183892596 | 183904991 | 12396 | -0.17 | 0.187 | 0.251 |
| chr2 | 164633856 | 164637280 | 3425 | 0.22 | 0.187 | 0.251 |
| chr1 | 42850616 | 42851696 | 1081 | -0.48 | 0.187 | 0.251 |
| chr1 | 212605364 | 212607132 | 1769 | 0.26 | 0.187 | 0.251 |
| chr9 | 97488191 | 97490408 | 2218 | -0.18 | 0.187 | 0.251 |
| chr11 | 4485051 | 4486054 | 1004 | 0.88 | 0.187 | 0.251 |
| chr3 | 176070477 | 176071098 | 622 | 1.38 | 0.187 | 0.251 |
| chr3 | 31945050 | 31946383 | 1334 | -0.86 | 0.187 | 0.251 |
| chr17 | 1156737 | 1173195 | 16459 | -0.17 | 0.187 | 0.251 |
| chr4 | 74024973 | 74025404 | 432 | -1.09 | 0.187 | 0.251 |
| chr7 | 22849958 | 22863106 | 13149 | -0.22 | 0.187 | 0.252 |
| chr12 | 47508261 | 47525499 | 17239 | 0.29 | 0.187 | 0.252 |

| | | | | | | |
|---|---|---|---|---|---|---|
| chr14 | 51451352 | 51452295 | 944 | -0.64 | 0.188 | 0.252 |
| chr17 | 72188864 | 72209537 | 20674 | 0.23 | 0.188 | 0.252 |
| chr6 | 25694417 | 25695684 | 1268 | 0.53 | 0.188 | 0.252 |
| chr3 | 117376189 | 117376713 | 525 | -0.95 | 0.188 | 0.252 |
| chr3 | 67566693 | 67567718 | 1026 | 0.8 | 0.188 | 0.252 |
| chr8 | 19148428 | 19149804 | 1377 | -0.35 | 0.188 | 0.252 |
| chr8 | 18891372 | 18892618 | 1247 | 0.5 | 0.188 | 0.252 |
| chr1 | 171749621 | 171754553 | 4933 | 0.17 | 0.188 | 0.253 |
| chr10 | 69701520 | 69702906 | 1387 | 0.63 | 0.188 | 0.253 |
| chr2 | 169746142 | 169746946 | 805 | -0.58 | 0.188 | 0.253 |
| chr2 | 98278952 | 98281801 | 2850 | -0.18 | 0.188 | 0.253 |
| chr9 | 123659395 | 123659765 | 371 | -1.39 | 0.188 | 0.253 |
| chr11 | 746442 | 778091 | 31650 | -0.11 | 0.189 | 0.253 |
| chr6 | 41471407 | 41472169 | 763 | -0.61 | 0.189 | 0.253 |
| chr3 | 124231668 | 124232647 | 980 | -0.39 | 0.189 | 0.253 |
| chr3 | 53142680 | 53144650 | 1971 | 0.51 | 0.189 | 0.253 |
| chr17 | 41655474 | 41657290 | 1817 | 0.46 | 0.189 | 0.253 |
| chr5 | 56246576 | 56248664 | 2089 | 0.31 | 0.189 | 0.253 |
| chr10 | 51044132 | 51053812 | 9681 | 0.15 | 0.189 | 0.254 |
| chr16 | 57450559 | 57482742 | 32184 | 0.14 | 0.189 | 0.254 |
| chr9 | 13277697 | 13280413 | 2717 | 0.19 | 0.189 | 0.254 |
| chr17 | 55423580 | 55424297 | 718 | -0.57 | 0.189 | 0.254 |
| chr15 | 39695836 | 39697027 | 1192 | 0.58 | 0.189 | 0.254 |
| chr5 | 139868374 | 139869123 | 750 | -0.75 | 0.19 | 0.254 |
| chr2 | 191471651 | 191473623 | 1973 | -0.21 | 0.19 | 0.254 |
| chr13 | 73573720 | 73576862 | 3143 | 0.28 | 0.19 | 0.254 |
| chr1 | 221566913 | 221568074 | 1162 | 2.18 | 0.19 | 0.254 |
| chr9 | 84635695 | 84636938 | 1244 | -0.55 | 0.19 | 0.254 |
| chr6 | 35801264 | 35801810 | 547 | -1.12 | 0.19 | 0.254 |
| chr1 | 236708802 | 236709392 | 591 | -0.81 | 0.19 | 0.254 |
| chr6 | 74508513 | 74514566 | 6054 | -0.3 | 0.19 | 0.254 |
| chr8 | 40227480 | 40235393 | 7914 | 0.1 | 0.19 | 0.255 |
| chr12 | 13394310 | 13395557 | 1248 | 2.44 | 0.19 | 0.255 |
| chrX | 27709738 | 27711207 | 1470 | 2.3 | 0.19 | 0.255 |
| chr14 | 23526409 | 23566259 | 39851 | 0.09 | 0.19 | 0.255 |
| chr20 | 3387263 | 3390244 | 2982 | 0.25 | 0.19 | 0.255 |
| chr2 | 173536232 | 173538978 | 2747 | 0.22 | 0.19 | 0.255 |
| chr7 | 132600087 | 132601038 | 952 | 0.77 | 0.19 | 0.255 |
| chr7 | 75156862 | 75164841 | 7980 | -0.27 | 0.19 | 0.255 |
| chr10 | 62701944 | 62761774 | 59831 | -0.08 | 0.19 | 0.255 |
| chr2 | 234577305 | 234579284 | 1980 | -0.45 | 0.19 | 0.255 |
| chr14 | 42938679 | 42939905 | 1227 | 0.36 | 0.19 | 0.255 |
| chr10 | 28899004 | 28904773 | 5770 | 0.33 | 0.19 | 0.255 |
| chr8 | 144489864 | 144490180 | 317 | 0.97 | 0.19 | 0.255 |
| chr7 | 20230093 | 20239679 | 9587 | 2.43 | 0.191 | 0.255 |
| chr5 | 139925836 | 139956645 | 30810 | -0.1 | 0.191 | 0.255 |
| chr3 | 12837542 | 12838277 | 736 | -0.67 | 0.191 | 0.255 |
| chr1 | 117252981 | 117254404 | 1424 | 2.27 | 0.191 | 0.255 |
| chr12 | 2749855 | 2750604 | 750 | -0.87 | 0.191 | 0.255 |
| chr3 | 101308179 | 101311409 | 3231 | 0.52 | 0.191 | 0.255 |
| chr6 | 12245476 | 12246851 | 1376 | -0.56 | 0.191 | 0.255 |
| chr4 | 106629044 | 106631771 | 2728 | 0.24 | 0.191 | 0.255 |
| chr13 | 53252510 | 53253013 | 504 | -0.82 | 0.191 | 0.255 |
| chr5 | 40550937 | 40553790 | 2854 | -0.2 | 0.191 | 0.255 |
| chr10 | 72259071 | 72259926 | 856 | -0.84 | 0.191 | 0.256 |
| chr1 | 226248117 | 226257880 | 9764 | 0.13 | 0.191 | 0.256 |
| chr13 | 33365396 | 33365988 | 593 | -0.84 | 0.191 | 0.256 |
| chr10 | 916562 | 978280 | 61719 | 0.1 | 0.191 | 0.256 |
| chr19 | 47215980 | 47265257 | 49278 | -0.1 | 0.191 | 0.256 |
| chr2 | 196475066 | 196479423 | 4358 | 0.24 | 0.191 | 0.256 |
| chr2 | 176865978 | 176867672 | 1695 | 0.47 | 0.191 | 0.256 |

| | | | | | | |
|---|---|---|---|---|---|---|
| chr2 | 8877907 | 8878331 | 425 | -0.79 | 0.191 | 0.256 |
| chr12 | 78550731 | 78551659 | 929 | -0.6 | 0.191 | 0.256 |
| chr13 | 91601804 | 91605369 | 3566 | 0.2 | 0.191 | 0.256 |
| chr16 | 30816221 | 30835700 | 19480 | 0.18 | 0.191 | 0.256 |
| chr7 | 134374428 | 134374962 | 535 | -0.78 | 0.191 | 0.256 |
| chr2 | 33201855 | 33202751 | 897 | -0.7 | 0.191 | 0.256 |
| chr14 | 105713917 | 105726123 | 12207 | -0.22 | 0.191 | 0.256 |
| chr15 | 99982908 | 99983205 | 298 | -1.05 | 0.191 | 0.256 |
| chr5 | 114937310 | 114938397 | 1088 | 0.29 | 0.191 | 0.256 |
| chr13 | 96253670 | 96257386 | 3717 | 0.38 | 0.191 | 0.256 |
| chr7 | 131749738 | 131751907 | 2170 | 0.26 | 0.191 | 0.256 |
| chr22 | 50608050 | 50608578 | 529 | -0.65 | 0.192 | 0.256 |
| chr7 | 80914041 | 80914813 | 773 | 0.81 | 0.192 | 0.256 |
| chr3 | 58570691 | 58571168 | 478 | -0.95 | 0.192 | 0.256 |
| chr20 | 16439887 | 16440810 | 924 | -0.61 | 0.192 | 0.257 |
| chr1 | 42998806 | 42999586 | 781 | -0.81 | 0.192 | 0.257 |
| chr4 | 141347720 | 141349454 | 1735 | 0.47 | 0.192 | 0.257 |
| chr7 | 101045130 | 101046778 | 1649 | 0.41 | 0.192 | 0.257 |
| chr6 | 100908852 | 100910309 | 1458 | -0.39 | 0.192 | 0.257 |
| chr10 | 34473720 | 34488569 | 14850 | -0.3 | 0.192 | 0.257 |
| chr16 | 21533641 | 21534215 | 575 | 0.68 | 0.192 | 0.257 |
| chr3 | 140660003 | 140662500 | 2498 | 0.21 | 0.192 | 0.257 |
| chr13 | 20139170 | 20139746 | 577 | 0.77 | 0.192 | 0.257 |
| chr9 | 5783946 | 5785171 | 1226 | 0.74 | 0.193 | 0.257 |
| chr5 | 109201217 | 109201773 | 557 | 0.99 | 0.193 | 0.257 |
| chr16 | 85722053 | 85722636 | 584 | -0.89 | 0.193 | 0.258 |
| chr19 | 50336913 | 50337705 | 793 | 0.8 | 0.193 | 0.258 |
| chr16 | 79793150 | 79804828 | 11679 | 2.33 | 0.193 | 0.258 |
| chr16 | 46947017 | 46947459 | 443 | -0.94 | 0.193 | 0.258 |
| chr6 | 106704109 | 106705614 | 1506 | -0.82 | 0.193 | 0.258 |
| chr1 | 150831724 | 150832149 | 426 | -1.12 | 0.193 | 0.258 |
| chr11 | 122678754 | 122679878 | 1125 | -0.68 | 0.193 | 0.258 |
| chr1 | 205648876 | 205649407 | 532 | -0.86 | 0.193 | 0.258 |
| chr16 | 76979299 | 76989997 | 10699 | -0.19 | 0.193 | 0.258 |
| chr5 | 169094534 | 169096599 | 2066 | 0.28 | 0.193 | 0.258 |
| chr11 | 94225395 | 94228064 | 2670 | 0.27 | 0.193 | 0.258 |
| chr16 | 12069352 | 12072095 | 2744 | 0.23 | 0.193 | 0.258 |
| chr5 | 54710466 | 54719879 | 9414 | -0.23 | 0.194 | 0.259 |
| chr5 | 78429041 | 78440749 | 11709 | -0.4 | 0.194 | 0.259 |
| chr5 | 130578126 | 130579014 | 889 | 0.46 | 0.194 | 0.259 |
| chrX | 65948790 | 65949090 | 301 | 0.64 | 0.194 | 0.259 |
| chr5 | 142811617 | 142812870 | 1254 | -0.45 | 0.194 | 0.259 |
| chr15 | 42499424 | 42501304 | 1881 | -0.21 | 0.194 | 0.259 |
| chr18 | 60517716 | 60518531 | 816 | 0.91 | 0.194 | 0.259 |
| chr2 | 45464581 | 45466031 | 1451 | -0.49 | 0.194 | 0.259 |
| chr2 | 189307245 | 189308742 | 1498 | 0.55 | 0.194 | 0.259 |
| chr11 | 61033592 | 61035377 | 1786 | 0.42 | 0.194 | 0.259 |
| chr8 | 41347222 | 41349413 | 2192 | 0.29 | 0.194 | 0.259 |
| chr10 | 51587756 | 51593689 | 5934 | 0.3 | 0.194 | 0.259 |
| chr12 | 132312385 | 132319768 | 7384 | 0.28 | 0.194 | 0.259 |
| chr7 | 129289545 | 129289916 | 372 | -0.86 | 0.194 | 0.26 |
| chr8 | 98876775 | 98882271 | 5497 | -0.27 | 0.194 | 0.26 |
| chr18 | 77006411 | 77007129 | 719 | -0.53 | 0.194 | 0.26 |
| chr7 | 44083535 | 44085376 | 1842 | 0.32 | 0.194 | 0.26 |
| chr17 | 29468148 | 29469917 | 1770 | 0.62 | 0.195 | 0.26 |
| chr1 | 186461185 | 186488452 | 27268 | -0.11 | 0.195 | 0.26 |
| chr6 | 136603249 | 136612780 | 9532 | 0.12 | 0.195 | 0.26 |
| chr8 | 67332761 | 67351984 | 19224 | 0.17 | 0.195 | 0.26 |
| chr18 | 76828527 | 76830396 | 1870 | 0.41 | 0.195 | 0.26 |
| chr11 | 122752816 | 122754789 | 1974 | 0.21 | 0.195 | 0.26 |
| chr2 | 31456418 | 31457653 | 1236 | -0.57 | 0.195 | 0.26 |

| | | | | | | |
|---|---|---|---|---|---|---|
| chr6 | 82750514 | 82751105 | 592 | 0.81 | 0.195 | 0.26 |
| chr6 | 105164633 | 105165360 | 728 | -0.66 | 0.195 | 0.26 |
| chr5 | 141487709 | 141500516 | 12808 | 0.22 | 0.195 | 0.26 |
| chr15 | 34515913 | 34518443 | 2531 | -0.15 | 0.195 | 0.26 |
| chr4 | 87503125 | 87517104 | 13980 | -0.17 | 0.195 | 0.26 |
| chr18 | 8073720 | 8074380 | 661 | -0.87 | 0.195 | 0.26 |
| chr6 | 53212442 | 53225737 | 13296 | 0.14 | 0.195 | 0.261 |
| chr5 | 38462564 | 38464645 | 2082 | 0.38 | 0.195 | 0.261 |
| chr3 | 161499078 | 161499731 | 654 | 0.7 | 0.195 | 0.261 |
| chr2 | 172010511 | 172018445 | 7935 | -0.24 | 0.195 | 0.261 |
| chr8 | 140866366 | 140868095 | 1730 | 0.39 | 0.195 | 0.261 |
| chr12 | 48357384 | 48358106 | 723 | 0.61 | 0.195 | 0.261 |
| chr3 | 113072591 | 113073913 | 1323 | -0.53 | 0.196 | 0.261 |
| chr22 | 27040700 | 27041374 | 675 | 0.5 | 0.196 | 0.261 |
| chr11 | 129193542 | 129195352 | 1811 | 0.27 | 0.196 | 0.261 |
| chr18 | 25544719 | 25545144 | 426 | 0.95 | 0.196 | 0.261 |
| chr2 | 183515748 | 183516070 | 323 | -1.09 | 0.196 | 0.261 |
| chr17 | 2494980 | 2506648 | 11669 | 0.13 | 0.196 | 0.261 |
| chr14 | 55393898 | 55394236 | 339 | 0.8 | 0.196 | 0.261 |
| chr3 | 93817215 | 93817875 | 661 | -0.67 | 0.196 | 0.261 |
| chr22 | 50246352 | 50248778 | 2427 | 0.27 | 0.196 | 0.261 |
| chr13 | 42428850 | 42429435 | 586 | -0.81 | 0.196 | 0.261 |
| chr7 | 23312421 | 23313924 | 1504 | 0.24 | 0.196 | 0.261 |
| chr10 | 28999179 | 29034112 | 34934 | 0.16 | 0.196 | 0.261 |
| chr3 | 151141849 | 151158055 | 16207 | 0.24 | 0.196 | 0.261 |
| chr11 | 125961612 | 125962364 | 753 | 0.9 | 0.196 | 0.262 |
| chrX | 69509105 | 69528117 | 19013 | 0.17 | 0.196 | 0.262 |
| chr1 | 12079594 | 12080248 | 655 | -0.77 | 0.196 | 0.262 |
| chr8 | 103704778 | 103705750 | 973 | 0.92 | 0.197 | 0.262 |
| chr19 | 18334796 | 18335360 | 565 | 0.62 | 0.197 | 0.262 |
| chr6 | 142732456 | 142736018 | 3563 | -0.21 | 0.197 | 0.262 |
| chr15 | 71922225 | 71924000 | 1776 | -2.27 | 0.197 | 0.262 |
| chr6 | 131422803 | 131424475 | 1673 | 0.39 | 0.197 | 0.262 |
| chr7 | 98864653 | 98872892 | 8240 | -0.22 | 0.197 | 0.262 |
| chr12 | 25486394 | 25487595 | 1202 | 0.5 | 0.197 | 0.262 |
| chr4 | 2844692 | 2846550 | 1859 | 0.38 | 0.197 | 0.262 |
| chr7 | 128654235 | 128671348 | 17114 | -0.22 | 0.197 | 0.263 |
| chr13 | 49637407 | 49638219 | 813 | 0.76 | 0.197 | 0.263 |
| chr10 | 126889615 | 126890221 | 607 | -0.77 | 0.197 | 0.263 |
| chr15 | 101790649 | 101793006 | 2358 | 0.38 | 0.197 | 0.263 |
| chr2 | 47858865 | 47859896 | 1032 | -0.49 | 0.197 | 0.263 |
| chr22 | 24125963 | 24130236 | 4274 | 0.28 | 0.197 | 0.263 |
| chr16 | 23463137 | 23473791 | 10655 | -0.25 | 0.197 | 0.263 |
| chr16 | 85182528 | 85185808 | 3281 | 0.52 | 0.198 | 0.263 |
| chr21 | 45937538 | 45938034 | 497 | 0.86 | 0.198 | 0.263 |
| chr10 | 71008592 | 71013660 | 5069 | -0.47 | 0.198 | 0.263 |
| chr5 | 65181067 | 65182474 | 1408 | 0.48 | 0.198 | 0.263 |
| chr1 | 46804406 | 46814436 | 10031 | 0.16 | 0.198 | 0.264 |
| chr17 | 6425867 | 6427564 | 1698 | 0.53 | 0.198 | 0.264 |
| chr18 | 59894115 | 59894872 | 758 | -0.7 | 0.198 | 0.264 |
| chr8 | 61428441 | 61433854 | 5414 | 0.17 | 0.198 | 0.264 |
| chr16 | 87349792 | 87352350 | 2559 | 0.21 | 0.198 | 0.264 |
| chr3 | 151758173 | 151758842 | 670 | 0.8 | 0.198 | 0.264 |
| chr22 | 39795134 | 39796728 | 1595 | -0.37 | 0.198 | 0.264 |
| chr16 | 9001590 | 9016005 | 14416 | -0.22 | 0.198 | 0.264 |
| chr3 | 38537847 | 38538721 | 875 | 0.44 | 0.198 | 0.264 |
| chr20 | 36651292 | 36663709 | 12418 | -0.16 | 0.198 | 0.264 |
| chr1 | 209419460 | 209420568 | 1109 | -0.69 | 0.198 | 0.264 |
| chr5 | 120587348 | 120588767 | 1420 | 0.34 | 0.198 | 0.264 |
| chr2 | 10975212 | 10985026 | 9815 | -1.04 | 0.198 | 0.264 |
| chr16 | 87179066 | 87179435 | 370 | -0.97 | 0.198 | 0.264 |

| | | | | | | |
|---|---|---|---|---|---|---|
| chr6 | 35778496 | 35779839 | 1344 | 0.45 | 0.199 | 0.264 |
| chr4 | 164058849 | 164059384 | 536 | 1.21 | 0.199 | 0.265 |
| chr3 | 48700321 | 48702357 | 2037 | 0.19 | 0.199 | 0.265 |
| chr12 | 102079379 | 102093244 | 13866 | 0.29 | 0.199 | 0.265 |
| chr7 | 22830677 | 22831317 | 641 | -0.9 | 0.199 | 0.265 |
| chr20 | 4684073 | 4685128 | 1056 | 0.68 | 0.199 | 0.265 |
| chr20 | 62650668 | 62651911 | 1244 | -0.58 | 0.199 | 0.265 |
| chr7 | 100859792 | 100862552 | 2761 | -0.22 | 0.199 | 0.265 |
| chr11 | 75187678 | 75188743 | 1066 | 0.63 | 0.199 | 0.265 |
| chr1 | 146555516 | 146557098 | 1583 | 0.24 | 0.199 | 0.265 |
| chr16 | 11623277 | 11692648 | 69372 | 0.09 | 0.199 | 0.265 |
| chr10 | 101934618 | 101947251 | 12634 | -0.18 | 0.199 | 0.265 |
| chr22 | 18892964 | 18893871 | 908 | 0.3 | 0.199 | 0.265 |
| chr1 | 12290068 | 12291747 | 1680 | 0.36 | 0.199 | 0.265 |
| chr6 | 33739041 | 33757603 | 18563 | 0.13 | 0.199 | 0.265 |
| chr6 | 79675561 | 79678903 | 3343 | 0.4 | 0.2 | 0.265 |
| chr9 | 113505948 | 113516283 | 10336 | 0.14 | 0.2 | 0.265 |
| chr7 | 75193929 | 75205671 | 11743 | 0.24 | 0.2 | 0.266 |
| chr11 | 108027858 | 108032170 | 4313 | 0.5 | 0.2 | 0.266 |
| chr6 | 24717876 | 24722835 | 4960 | 0.12 | 0.2 | 0.266 |
| chr13 | 60737085 | 60739300 | 2216 | 0.22 | 0.2 | 0.266 |
| chr9 | 127596824 | 127597870 | 1047 | 0.79 | 0.2 | 0.266 |
| chr10 | 67178508 | 67180032 | 1525 | -0.38 | 0.2 | 0.266 |
| chr12 | 72258633 | 72259763 | 1131 | -0.75 | 0.2 | 0.266 |
| chr6 | 12119051 | 12152349 | 33299 | -0.11 | 0.2 | 0.266 |
| chr3 | 57260489 | 57263028 | 2540 | 0.31 | 0.2 | 0.266 |
| chr16 | 1520448 | 1526116 | 5669 | 0.25 | 0.2 | 0.266 |
| chr17 | 60769156 | 60794685 | 25530 | -0.25 | 0.2 | 0.266 |
| chr1 | 114084109 | 114085354 | 1246 | -0.57 | 0.2 | 0.266 |
| chr13 | 48403850 | 48404327 | 478 | -1.32 | 0.2 | 0.266 |
| chr12 | 74523517 | 74524038 | 522 | 0.95 | 0.2 | 0.266 |
| chr20 | 45522950 | 45523926 | 977 | -0.68 | 0.2 | 0.266 |
| chr9 | 117159814 | 117161338 | 1525 | -0.37 | 0.2 | 0.266 |
| chr11 | 75201358 | 75248743 | 47386 | 0.13 | 0.2 | 0.266 |
| chr5 | 143203407 | 143207331 | 3925 | -0.34 | 0.2 | 0.266 |
| chr7 | 40126282 | 40126785 | 504 | -0.82 | 0.2 | 0.266 |
| chrX | 18901196 | 18918967 | 17772 | -0.24 | 0.2 | 0.266 |
| chr9 | 109622490 | 109627668 | 5179 | 0.3 | 0.201 | 0.266 |
| chr11 | 95564143 | 95564741 | 599 | 0.67 | 0.201 | 0.267 |
| chr1 | 87792914 | 87801843 | 8930 | 0.16 | 0.201 | 0.267 |
| chr11 | 116642654 | 116644807 | 2154 | 0.24 | 0.201 | 0.267 |
| chr6 | 160930731 | 160942178 | 11448 | 0.32 | 0.201 | 0.267 |
| chr19 | 14247935 | 14249615 | 1681 | 0.43 | 0.201 | 0.267 |
| chr4 | 106123702 | 106124311 | 610 | 0.87 | 0.201 | 0.267 |
| chr10 | 123778544 | 123779852 | 1309 | 0.58 | 0.201 | 0.267 |
| chr10 | 34514250 | 34516006 | 1757 | 0.41 | 0.201 | 0.267 |
| chr10 | 105263971 | 105275798 | 11828 | 2.46 | 0.201 | 0.267 |
| chr7 | 17535193 | 17536058 | 866 | 2.12 | 0.201 | 0.267 |
| chr9 | 92300684 | 92302118 | 1435 | -2.19 | 0.201 | 0.267 |
| chr3 | 32279485 | 32291580 | 12096 | -0.95 | 0.202 | 0.268 |
| chr17 | 80052885 | 80070798 | 17914 | 0.1 | 0.202 | 0.268 |
| chr3 | 73045139 | 73087802 | 42664 | -0.1 | 0.202 | 0.268 |
| chr13 | 21286385 | 21286872 | 488 | -0.89 | 0.202 | 0.268 |
| chr22 | 47013422 | 47030864 | 17443 | 0.25 | 0.202 | 0.268 |
| chr9 | 135544218 | 135549342 | 5125 | 0.19 | 0.202 | 0.268 |
| chr10 | 75159717 | 75174699 | 14983 | 0.16 | 0.202 | 0.268 |
| chr6 | 52793830 | 52795299 | 1470 | 0.38 | 0.202 | 0.268 |
| chrX | 84498790 | 84500048 | 1259 | -0.5 | 0.202 | 0.268 |
| chr10 | 71389377 | 71391197 | 1821 | 0.36 | 0.202 | 0.268 |
| chr17 | 67282970 | 67283878 | 909 | -0.56 | 0.202 | 0.268 |
| chr9 | 35688433 | 35692013 | 3581 | 0.19 | 0.202 | 0.268 |

| | | | | | | |
|---|---|---|---|---|---|---|
| chr1 | 220333933 | 220334572 | 640 | -0.93 | 0.202 | 0.268 |
| chr14 | 71131778 | 71133750 | 1973 | 0.38 | 0.202 | 0.268 |
| chr15 | 35196524 | 35198054 | 1531 | -0.41 | 0.202 | 0.268 |
| chr11 | 1609748 | 1610725 | 978 | 0.65 | 0.202 | 0.268 |
| chr10 | 128147773 | 128149886 | 2114 | -0.51 | 0.202 | 0.269 |
| chr2 | 239298549 | 239299319 | 771 | -0.77 | 0.202 | 0.269 |
| chr17 | 35369539 | 35370035 | 497 | -1.32 | 0.202 | 0.269 |
| chr7 | 106762546 | 106763417 | 872 | -0.62 | 0.203 | 0.269 |
| chr5 | 60315161 | 60316075 | 915 | -0.52 | 0.203 | 0.269 |
| chr1 | 38259493 | 38275099 | 15607 | 0.19 | 0.203 | 0.269 |
| chr1 | 147805771 | 147807688 | 1918 | -0.23 | 0.203 | 0.269 |
| chr22 | 31363911 | 31366670 | 2760 | 0.22 | 0.203 | 0.269 |
| chr3 | 128963988 | 128969394 | 5407 | 0.22 | 0.203 | 0.269 |
| chr11 | 107312253 | 107312680 | 428 | 1.07 | 0.203 | 0.269 |
| chrX | 41042732 | 41053426 | 10695 | -0.28 | 0.203 | 0.269 |
| chr9 | 127985194 | 127986289 | 1096 | -0.67 | 0.203 | 0.269 |
| chr14 | 89506908 | 89507653 | 746 | -0.47 | 0.203 | 0.269 |
| chr12 | 117558198 | 117628670 | 70473 | 0.07 | 0.203 | 0.27 |
| chr8 | 87353206 | 87358095 | 4890 | 0.14 | 0.203 | 0.27 |
| chr4 | 980183 | 988114 | 7932 | 0.18 | 0.204 | 0.27 |
| chrX | 46305744 | 46307453 | 1710 | 0.24 | 0.204 | 0.27 |
| chr8 | 6286646 | 6287852 | 1207 | -0.56 | 0.204 | 0.27 |
| chr5 | 94084750 | 94085572 | 823 | -0.64 | 0.204 | 0.27 |
| chr4 | 2699755 | 2700592 | 838 | -0.85 | 0.204 | 0.27 |
| chr9 | 5626861 | 5633828 | 6968 | 0.18 | 0.204 | 0.27 |
| chr2 | 64836023 | 64837257 | 1235 | -0.26 | 0.204 | 0.271 |
| chr5 | 74615587 | 74618393 | 2807 | -0.23 | 0.204 | 0.271 |
| chr19 | 17829616 | 17830845 | 1230 | -0.43 | 0.204 | 0.271 |
| chr3 | 153942475 | 153942881 | 407 | -1.33 | 0.204 | 0.271 |
| chr15 | 55731367 | 55731979 | 613 | 0.95 | 0.204 | 0.271 |
| chr11 | 65867485 | 65925765 | 58281 | -0.06 | 0.204 | 0.271 |
| chr7 | 81645099 | 81645712 | 614 | -0.67 | 0.204 | 0.271 |
| chr13 | 39580988 | 39581506 | 519 | 0.71 | 0.204 | 0.271 |
| chr1 | 173375823 | 173376381 | 559 | 0.76 | 0.204 | 0.271 |
| chr7 | 44881042 | 44897039 | 15998 | 0.16 | 0.204 | 0.271 |
| chr11 | 68933575 | 68934847 | 1273 | -0.58 | 0.204 | 0.271 |
| chr4 | 154564216 | 154564717 | 502 | 0.75 | 0.204 | 0.271 |
| chr15 | 45906679 | 45907045 | 367 | -0.86 | 0.204 | 0.271 |
| chr8 | 8558651 | 8560110 | 1460 | 0.46 | 0.204 | 0.271 |
| chr2 | 135169103 | 135191894 | 22792 | 0.14 | 0.204 | 0.271 |
| chr21 | 43619292 | 43640403 | 21112 | -0.64 | 0.204 | 0.271 |
| chr18 | 55316997 | 55317773 | 777 | 0.74 | 0.204 | 0.271 |
| chr2 | 204398956 | 204409713 | 10758 | 0.22 | 0.205 | 0.271 |
| chr6 | 13794202 | 13794564 | 363 | -1.24 | 0.205 | 0.271 |
| chr17 | 80762414 | 80790941 | 28528 | -0.21 | 0.205 | 0.271 |
| chr2 | 241189625 | 241196265 | 6641 | 0.21 | 0.205 | 0.271 |
| chr8 | 108497814 | 108519704 | 21891 | 0.19 | 0.205 | 0.271 |
| chr17 | 21102542 | 21132470 | 29929 | -0.15 | 0.205 | 0.271 |
| chr12 | 54828573 | 54829865 | 1293 | -0.45 | 0.205 | 0.271 |
| chr16 | 29801064 | 29843535 | 42472 | 0.07 | 0.205 | 0.271 |
| chr3 | 110417103 | 110417653 | 551 | 0.79 | 0.205 | 0.271 |
| chr10 | 15278632 | 15279310 | 679 | 0.7 | 0.205 | 0.271 |
| chr10 | 104352680 | 104430274 | 77595 | -0.07 | 0.205 | 0.271 |
| chr15 | 45813999 | 45815512 | 1514 | 0.59 | 0.205 | 0.271 |
| chr10 | 77523753 | 77533709 | 9957 | -2.37 | 0.205 | 0.271 |
| chr11 | 12979791 | 12980657 | 867 | 0.63 | 0.205 | 0.271 |
| chrX | 134019937 | 134020903 | 967 | -0.68 | 0.205 | 0.271 |
| chr12 | 17485631 | 17496133 | 10503 | 2.58 | 0.205 | 0.272 |
| chr10 | 70087317 | 70108969 | 21653 | 0.1 | 0.205 | 0.272 |
| chr16 | 56434251 | 56460099 | 25849 | 0.15 | 0.205 | 0.272 |
| chr2 | 127900546 | 127912346 | 11801 | 0.24 | 0.205 | 0.272 |

| | | | | | | |
|---|---|---|---|---|---|---|
| chr1 | 85410633 | 85412583 | 1951 | 0.28 | 0.205 | 0.272 |
| chr13 | 67507074 | 67509878 | 2805 | 0.15 | 0.205 | 0.272 |
| chr1 | 221982889 | 221983398 | 510 | -0.75 | 0.205 | 0.272 |
| chr4 | 140250594 | 140253132 | 2539 | 0.53 | 0.205 | 0.272 |
| chr12 | 46157240 | 46157635 | 396 | -1.35 | 0.205 | 0.272 |
| chr2 | 204443835 | 204447841 | 4007 | -0.4 | 0.205 | 0.272 |
| chr1 | 46740153 | 46740809 | 657 | -0.79 | 0.205 | 0.272 |
| chr4 | 119744565 | 119758359 | 13795 | 0.23 | 0.205 | 0.272 |
| chr5 | 145155055 | 145156922 | 1868 | -0.38 | 0.206 | 0.272 |
| chr22 | 38668270 | 38681303 | 13034 | 0.18 | 0.206 | 0.272 |
| chrX | 69386779 | 69389321 | 2543 | -0.39 | 0.206 | 0.272 |
| chr2 | 175396039 | 175397329 | 1291 | 0.49 | 0.206 | 0.272 |
| chr15 | 74219878 | 74220494 | 617 | 0.74 | 0.206 | 0.272 |
| chr12 | 8891670 | 8891972 | 303 | 0.84 | 0.206 | 0.272 |
| chr5 | 80076798 | 80077620 | 823 | -0.66 | 0.206 | 0.272 |
| chrX | 4912559 | 4913274 | 716 | 0.82 | 0.206 | 0.273 |
| chr17 | 20203872 | 20204358 | 487 | -0.71 | 0.206 | 0.273 |
| chrX | 21901564 | 21902121 | 558 | 1.02 | 0.206 | 0.273 |
| chr7 | 76977582 | 76990287 | 12706 | 1.96 | 0.206 | 0.273 |
| chr4 | 107184918 | 107185730 | 813 | 0.68 | 0.206 | 0.273 |
| chr9 | 33403116 | 33475040 | 71925 | -0.11 | 0.206 | 0.273 |
| chr1 | 86119686 | 86126320 | 6635 | -0.19 | 0.206 | 0.273 |
| chrX | 95939193 | 95941774 | 2582 | 0.27 | 0.206 | 0.273 |
| chr9 | 126031167 | 126031583 | 417 | 0.88 | 0.206 | 0.273 |
| chr20 | 47002809 | 47003127 | 319 | 0.92 | 0.206 | 0.273 |
| chr2 | 172842622 | 172843315 | 694 | -0.96 | 0.207 | 0.273 |
| chr18 | 19282473 | 19285270 | 2798 | 0.24 | 0.207 | 0.273 |
| chr13 | 35321364 | 35321825 | 462 | -0.96 | 0.207 | 0.273 |
| chr16 | 86934890 | 86936205 | 1316 | 0.53 | 0.207 | 0.273 |
| chr15 | 58621783 | 58653472 | 31690 | -0.07 | 0.207 | 0.273 |
| chr4 | 115185428 | 115194067 | 8640 | 2.46 | 0.207 | 0.273 |
| chr18 | 3771021 | 3772509 | 1489 | -0.36 | 0.207 | 0.273 |
| chr13 | 77599710 | 77601882 | 2173 | 0.34 | 0.207 | 0.273 |
| chr9 | 38515723 | 38517105 | 1383 | 0.36 | 0.207 | 0.273 |
| chr12 | 131926354 | 131930113 | 3760 | 0.51 | 0.207 | 0.273 |
| chr5 | 178030433 | 178031296 | 864 | 0.75 | 0.207 | 0.273 |
| chr15 | 45458374 | 45459747 | 1374 | 0.24 | 0.207 | 0.274 |
| chr5 | 122750573 | 122766123 | 15551 | 0.11 | 0.207 | 0.274 |
| chr11 | 118991438 | 118993037 | 1600 | 0.37 | 0.207 | 0.274 |
| chr6 | 105820740 | 105827611 | 6872 | -0.48 | 0.207 | 0.274 |
| chr12 | 105552883 | 105555728 | 2846 | 0.43 | 0.207 | 0.274 |
| chr7 | 19747497 | 19749203 | 1707 | 0.22 | 0.207 | 0.274 |
| chr2 | 148683131 | 148684320 | 1190 | 0.62 | 0.207 | 0.274 |
| chr22 | 30986351 | 30988833 | 2483 | -0.2 | 0.207 | 0.274 |
| chr10 | 71167803 | 71169500 | 1698 | 0.27 | 0.207 | 0.274 |
| chr11 | 35711185 | 35711712 | 528 | 0.81 | 0.207 | 0.274 |
| chr15 | 56467879 | 56480070 | 12192 | -0.48 | 0.208 | 0.274 |
| chr1 | 12221638 | 12222292 | 655 | -0.69 | 0.208 | 0.274 |
| chr14 | 77558900 | 77565659 | 6760 | 0.18 | 0.208 | 0.274 |
| chr14 | 105992018 | 105994850 | 2833 | -0.38 | 0.208 | 0.274 |
| chr5 | 170548815 | 170549865 | 1051 | 0.62 | 0.208 | 0.274 |
| chr17 | 59053345 | 59054660 | 1316 | 1.11 | 0.208 | 0.274 |
| chr11 | 85903867 | 85905061 | 1195 | 0.36 | 0.208 | 0.274 |
| chr1 | 234124197 | 234130422 | 6226 | 0.3 | 0.208 | 0.275 |
| chr2 | 233877026 | 233878055 | 1030 | 0.76 | 0.208 | 0.275 |
| chr14 | 92813989 | 92814491 | 503 | 0.87 | 0.208 | 0.275 |
| chr10 | 14820069 | 14826552 | 6484 | -0.14 | 0.208 | 0.275 |
| chr4 | 48727650 | 48729282 | 1633 | 0.39 | 0.208 | 0.275 |
| chr19 | 50979112 | 50981026 | 1915 | 0.39 | 0.208 | 0.275 |
| chr15 | 64566743 | 64567253 | 511 | 0.9 | 0.208 | 0.275 |
| chr4 | 39881732 | 39882775 | 1044 | -0.66 | 0.208 | 0.275 |

| | | | | | | |
|---|---|---|---|---|---|---|
| chrX | 54466088 | 54467441 | 1354 | 0.44 | 0.208 | 0.275 |
| chr8 | 103849114 | 103852848 | 3735 | 0.51 | 0.209 | 0.275 |
| chr6 | 163732155 | 163754414 | 22260 | 0.14 | 0.209 | 0.275 |
| chr12 | 59120218 | 59121856 | 1639 | -0.46 | 0.209 | 0.275 |
| chr7 | 2995578 | 3003163 | 7586 | 0.21 | 0.209 | 0.275 |
| chr6 | 15662076 | 15663212 | 1137 | 0.55 | 0.209 | 0.275 |
| chr18 | 21374094 | 21374612 | 519 | -1.08 | 0.209 | 0.275 |
| chr22 | 23775875 | 23776288 | 414 | 0.89 | 0.209 | 0.275 |
| chr2 | 216870703 | 216878314 | 7612 | -0.48 | 0.209 | 0.276 |
| chr5 | 90323188 | 90324142 | 955 | 0.63 | 0.209 | 0.276 |
| chr16 | 62259878 | 62260284 | 407 | 0.73 | 0.209 | 0.276 |
| chr11 | 78126867 | 78136070 | 9204 | -0.2 | 0.209 | 0.276 |
| chr22 | 28255502 | 28280279 | 24778 | 0.14 | 0.209 | 0.276 |
| chrX | 83430983 | 83443255 | 12273 | 0.53 | 0.209 | 0.276 |
| chr18 | 58044525 | 58045869 | 1345 | -0.49 | 0.209 | 0.276 |
| chr18 | 13501405 | 13501753 | 349 | 0.88 | 0.209 | 0.276 |
| chr7 | 139184939 | 139185507 | 569 | -0.81 | 0.209 | 0.276 |
| chr15 | 29210031 | 29212439 | 2409 | 0.24 | 0.209 | 0.276 |
| chr10 | 104502818 | 104531238 | 28421 | -0.11 | 0.209 | 0.276 |
| chr22 | 43463032 | 43463543 | 512 | 0.98 | 0.209 | 0.276 |
| chr19 | 16151178 | 16152756 | 1579 | 0.45 | 0.209 | 0.276 |
| chr5 | 139903587 | 139904074 | 488 | 1.02 | 0.209 | 0.276 |
| chr18 | 47900710 | 47902313 | 1604 | 0.37 | 0.209 | 0.276 |
| chr10 | 69910974 | 69915298 | 4325 | -0.21 | 0.21 | 0.276 |
| chr2 | 64925246 | 64926535 | 1290 | -0.54 | 0.21 | 0.276 |
| chr5 | 138532918 | 138534839 | 1922 | 0.21 | 0.21 | 0.277 |
| chr7 | 121783849 | 121784611 | 763 | 0.91 | 0.21 | 0.277 |
| chr10 | 69990229 | 69991613 | 1385 | -0.31 | 0.21 | 0.277 |
| chr1 | 84944534 | 84945974 | 1441 | 0.26 | 0.21 | 0.277 |
| chr3 | 88037115 | 88037663 | 549 | -0.62 | 0.21 | 0.277 |
| chr5 | 17168997 | 17182083 | 13087 | 0.24 | 0.21 | 0.277 |
| chr11 | 122988778 | 122993966 | 5189 | -0.32 | 0.21 | 0.277 |
| chr9 | 114212019 | 114213283 | 1265 | -0.6 | 0.21 | 0.277 |
| chr14 | 69596024 | 69596731 | 708 | -0.67 | 0.21 | 0.277 |
| chr17 | 47723949 | 47724750 | 802 | 0.58 | 0.21 | 0.277 |
| chr11 | 65728218 | 65731727 | 3510 | 0.21 | 0.21 | 0.277 |
| chr1 | 212838557 | 212842754 | 4198 | -0.22 | 0.21 | 0.277 |
| chr17 | 55703149 | 55703481 | 333 | -1.15 | 0.21 | 0.277 |
| chr17 | 56295383 | 56297474 | 2092 | -0.24 | 0.21 | 0.277 |
| chr20 | 21282876 | 21291877 | 9002 | 0.13 | 0.21 | 0.277 |
| chr9 | 134951119 | 134955499 | 4381 | -0.34 | 0.21 | 0.277 |
| chr1 | 16393096 | 16409254 | 16159 | -0.19 | 0.21 | 0.277 |
| chr1 | 93387092 | 93390696 | 3605 | 0.14 | 0.211 | 0.277 |
| chr8 | 52834542 | 52835501 | 960 | 0.82 | 0.211 | 0.278 |
| chr4 | 52708385 | 52720287 | 11903 | -0.16 | 0.211 | 0.278 |
| chr13 | 103451495 | 103453497 | 2003 | 0.44 | 0.211 | 0.278 |
| chr11 | 3787941 | 3789051 | 1111 | 0.68 | 0.211 | 0.278 |
| chr8 | 102137917 | 102172776 | 34860 | 0.11 | 0.211 | 0.278 |
| chr1 | 45768698 | 45770708 | 2011 | 0.29 | 0.211 | 0.278 |
| chr14 | 74550395 | 74552806 | 2412 | 0.22 | 0.211 | 0.278 |
| chr14 | 74269240 | 74270147 | 908 | -0.9 | 0.211 | 0.278 |
| chr7 | 96796417 | 96797800 | 1384 | -0.48 | 0.211 | 0.278 |
| chr6 | 151590236 | 151591307 | 1072 | 0.44 | 0.211 | 0.278 |
| chr7 | 67480115 | 67481374 | 1260 | 0.59 | 0.211 | 0.278 |
| chr1 | 203236585 | 203238313 | 1729 | -0.45 | 0.211 | 0.278 |
| chr17 | 71026313 | 71027252 | 940 | -0.71 | 0.211 | 0.278 |
| chr7 | 117853996 | 117855091 | 1096 | 0.43 | 0.211 | 0.278 |
| chr2 | 74055399 | 74061079 | 5681 | 0.16 | 0.211 | 0.278 |
| chr3 | 13492304 | 13494402 | 2099 | -0.27 | 0.211 | 0.278 |
| chr16 | 67814163 | 67815698 | 1536 | -0.49 | 0.211 | 0.278 |
| chr16 | 67030114 | 67031159 | 1046 | 0.59 | 0.212 | 0.279 |

| | | | | | | |
|---|---|---|---|---|---|---|
| chr19 | 46405169 | 46419390 | 14222 | -0.39 | 0.212 | 0.279 |
| chr6 | 17676839 | 17716044 | 39206 | -0.09 | 0.212 | 0.279 |
| chr2 | 46635099 | 46637955 | 2857 | 0.18 | 0.212 | 0.279 |
| chr6 | 83902214 | 83904168 | 1955 | 0.33 | 0.212 | 0.279 |
| chr3 | 43731697 | 43733538 | 1842 | 0.39 | 0.212 | 0.279 |
| chr3 | 66023372 | 66025809 | 2438 | 0.26 | 0.212 | 0.279 |
| chr19 | 2782111 | 2786454 | 4344 | 0.21 | 0.212 | 0.279 |
| chr11 | 94609278 | 94610938 | 1661 | 0.45 | 0.212 | 0.279 |
| chr10 | 32485774 | 32489430 | 3657 | 2.22 | 0.212 | 0.279 |
| chr4 | 135589022 | 135589997 | 976 | 0.45 | 0.212 | 0.279 |
| chr6 | 42712579 | 42716221 | 3643 | 0.21 | 0.212 | 0.279 |
| chr6 | 138871245 | 138871588 | 344 | 0.95 | 0.212 | 0.279 |
| chr6 | 43174219 | 43175931 | 1713 | 0.42 | 0.212 | 0.279 |
| chr7 | 31014042 | 31015232 | 1191 | 0.72 | 0.212 | 0.279 |
| chr4 | 75693980 | 75695328 | 1349 | 0.54 | 0.213 | 0.28 |
| chr4 | 129609461 | 129609899 | 439 | -0.83 | 0.213 | 0.28 |
| chr10 | 23105575 | 23106527 | 953 | 0.53 | 0.213 | 0.28 |
| chr15 | 64695535 | 64699649 | 4115 | -0.16 | 0.213 | 0.28 |
| chr6 | 111003453 | 111025583 | 22131 | 1.55 | 0.213 | 0.28 |
| chr19 | 18214973 | 18229169 | 14197 | -0.3 | 0.213 | 0.28 |
| chr12 | 57623448 | 57634489 | 11042 | -0.26 | 0.213 | 0.28 |
| chr17 | 42400924 | 42423390 | 22467 | -0.21 | 0.213 | 0.28 |
| chr14 | 54260484 | 54261504 | 1021 | 0.48 | 0.213 | 0.28 |
| chr12 | 70215293 | 70215936 | 644 | 0.73 | 0.213 | 0.281 |
| chr1 | 184020118 | 184022023 | 1906 | -0.19 | 0.213 | 0.281 |
| chr7 | 131164763 | 131168076 | 3314 | 0.26 | 0.213 | 0.281 |
| chr9 | 33676254 | 33688457 | 12204 | 0.45 | 0.213 | 0.281 |
| chr8 | 70906111 | 70907619 | 1509 | -0.44 | 0.213 | 0.281 |
| chr10 | 104180017 | 104192949 | 12933 | 0.21 | 0.213 | 0.281 |
| chr20 | 45854573 | 45855100 | 528 | 1.34 | 0.213 | 0.281 |
| chr5 | 67840561 | 67851361 | 10801 | 0.22 | 0.214 | 0.281 |
| chr9 | 140196777 | 140211487 | 14711 | -0.23 | 0.214 | 0.281 |
| chr2 | 136269495 | 136291126 | 21632 | 0.14 | 0.214 | 0.281 |
| chr18 | 21850851 | 21852796 | 1946 | 0.34 | 0.214 | 0.281 |
| chr10 | 77548028 | 77548741 | 714 | -0.59 | 0.214 | 0.281 |
| chr6 | 17492610 | 17494263 | 1654 | -0.38 | 0.214 | 0.281 |
| chr11 | 118478341 | 118494542 | 16202 | 0.19 | 0.214 | 0.281 |
| chr9 | 20658605 | 20659549 | 945 | 0.4 | 0.214 | 0.281 |
| chr14 | 54976562 | 54977568 | 1007 | 0.37 | 0.214 | 0.281 |
| chr5 | 134271339 | 134272171 | 833 | 0.76 | 0.214 | 0.281 |
| chr4 | 99945467 | 99946610 | 1144 | -0.68 | 0.214 | 0.281 |
| chr9 | 113856460 | 113857578 | 1119 | -0.41 | 0.214 | 0.282 |
| chr2 | 44000801 | 44002544 | 1744 | -0.19 | 0.214 | 0.282 |
| chrX | 152065433 | 152067279 | 1847 | 0.39 | 0.214 | 0.282 |
| chr1 | 152512117 | 152512725 | 609 | -0.9 | 0.214 | 0.282 |
| chr2 | 240677986 | 240681787 | 3802 | -0.19 | 0.214 | 0.282 |
| chr16 | 20412803 | 20413381 | 579 | 0.91 | 0.214 | 0.282 |
| chr16 | 47813274 | 47814179 | 906 | -0.53 | 0.214 | 0.282 |
| chr20 | 49982289 | 49983616 | 1328 | -0.61 | 0.215 | 0.282 |
| chr9 | 4901454 | 4906436 | 4983 | 0.26 | 0.215 | 0.282 |
| chr4 | 22484357 | 22486412 | 2056 | -0.48 | 0.215 | 0.282 |
| chr5 | 150157880 | 150158966 | 1087 | -0.5 | 0.215 | 0.282 |
| chr21 | 37528191 | 37537141 | 8951 | 0.45 | 0.215 | 0.282 |
| chr1 | 154243508 | 154246127 | 2620 | 0.24 | 0.215 | 0.282 |
| chr4 | 153273013 | 153274461 | 1449 | 0.46 | 0.215 | 0.282 |
| chrX | 8922070 | 8923178 | 1109 | 1.9 | 0.215 | 0.282 |
| chr11 | 47735978 | 47755657 | 19680 | 0.16 | 0.215 | 0.282 |
| chr17 | 47326541 | 47338662 | 12122 | -0.4 | 0.215 | 0.282 |
| chr5 | 131109545 | 131111844 | 2300 | 0.27 | 0.215 | 0.282 |
| chr17 | 56612455 | 56613141 | 687 | -1.28 | 0.215 | 0.283 |
| chr10 | 97413184 | 97417316 | 4133 | 0.27 | 0.215 | 0.283 |

| | | | | | | |
|---|---|---|---|---|---|---|
| chr11 | 68197638 | 68255000 | 57363 | 0.11 | 0.215 | 0.283 |
| chr19 | 4208439 | 4208900 | 462 | -0.8 | 0.216 | 0.283 |
| chr10 | 77862551 | 77863780 | 1230 | -0.56 | 0.216 | 0.283 |
| chr10 | 133785546 | 133796185 | 10640 | 0.19 | 0.216 | 0.283 |
| chr4 | 184424127 | 184431426 | 7300 | 0.14 | 0.216 | 0.283 |
| chr4 | 1225869 | 1227597 | 1729 | -0.47 | 0.216 | 0.283 |
| chr15 | 49461673 | 49469137 | 7465 | 0.18 | 0.216 | 0.283 |
| chr5 | 37345890 | 37346318 | 429 | 0.87 | 0.216 | 0.283 |
| chr2 | 208195270 | 208203339 | 8070 | 0.21 | 0.216 | 0.283 |
| chr2 | 192109493 | 192114644 | 5152 | -0.17 | 0.216 | 0.283 |
| chr2 | 65772887 | 65780554 | 7668 | 0.32 | 0.216 | 0.283 |
| chr4 | 1546105 | 1546885 | 781 | 0.52 | 0.216 | 0.283 |
| chr22 | 45554530 | 45566436 | 11907 | 0.15 | 0.216 | 0.283 |
| chr1 | 168889289 | 168891213 | 1925 | 0.21 | 0.216 | 0.283 |
| chr14 | 77278357 | 77279910 | 1554 | -0.32 | 0.216 | 0.283 |
| chr1 | 226520601 | 226521316 | 716 | -0.42 | 0.216 | 0.284 |
| chr16 | 4226789 | 4264284 | 37496 | 0.52 | 0.216 | 0.284 |
| chr10 | 119220868 | 119222106 | 1239 | 0.63 | 0.216 | 0.284 |
| chr1 | 113975850 | 114001393 | 25544 | 0.11 | 0.216 | 0.284 |
| chr7 | 134938384 | 134940306 | 1923 | -0.34 | 0.216 | 0.284 |
| chr10 | 65138774 | 65139345 | 572 | 0.79 | 0.217 | 0.284 |
| chr11 | 12398010 | 12400997 | 2988 | 0.16 | 0.217 | 0.284 |
| chr1 | 151342891 | 151345446 | 2556 | 0.4 | 0.217 | 0.284 |
| chr4 | 13401064 | 13401468 | 405 | 0.77 | 0.217 | 0.284 |
| chr17 | 77881303 | 77910665 | 29363 | 0.11 | 0.217 | 0.284 |
| chr1 | 160202426 | 160233112 | 30687 | 0.12 | 0.217 | 0.284 |
| chr10 | 12139366 | 12142021 | 2656 | 0.31 | 0.217 | 0.284 |
| chr2 | 182521015 | 182522053 | 1039 | -0.49 | 0.217 | 0.284 |
| chr12 | 32519192 | 32520227 | 1036 | 0.57 | 0.217 | 0.284 |
| chr2 | 114633712 | 114662511 | 28800 | -0.1 | 0.217 | 0.284 |
| chr6 | 144583904 | 144608758 | 24855 | 0.14 | 0.217 | 0.285 |
| chr5 | 175906615 | 175930043 | 23429 | 0.13 | 0.217 | 0.285 |
| chr1 | 27288205 | 27289251 | 1047 | -1.06 | 0.217 | 0.285 |
| chr5 | 136553735 | 136554797 | 1063 | 0.57 | 0.217 | 0.285 |
| chr3 | 189381794 | 189383126 | 1333 | -0.47 | 0.217 | 0.285 |
| chr17 | 29684045 | 29686214 | 2170 | -0.29 | 0.217 | 0.285 |
| chr11 | 104033428 | 104035921 | 2494 | 0.26 | 0.217 | 0.285 |
| chr1 | 33077505 | 33078679 | 1175 | 0.38 | 0.218 | 0.285 |
| chr7 | 148435953 | 148436510 | 558 | -0.79 | 0.218 | 0.285 |
| chr18 | 48380371 | 48381156 | 786 | -0.91 | 0.218 | 0.285 |
| chr11 | 46544012 | 46544605 | 594 | 0.96 | 0.218 | 0.286 |
| chr10 | 69894797 | 69897791 | 2995 | -0.16 | 0.218 | 0.286 |
| chr1 | 32965239 | 32966904 | 1666 | 0.31 | 0.218 | 0.286 |
| chr10 | 126067433 | 126078491 | 11059 | 1.32 | 0.218 | 0.286 |
| chr1 | 79085187 | 79085913 | 727 | 0.7 | 0.218 | 0.286 |
| chr20 | 55203559 | 55223574 | 20016 | -0.13 | 0.218 | 0.286 |
| chr10 | 24544772 | 24545465 | 694 | 0.78 | 0.218 | 0.286 |
| chr4 | 175482124 | 175483318 | 1195 | 0.63 | 0.218 | 0.286 |
| chr18 | 72457420 | 72458770 | 1351 | -0.44 | 0.218 | 0.286 |
| chr18 | 56806630 | 56807878 | 1249 | 0.34 | 0.218 | 0.286 |
| chr17 | 35611646 | 35612202 | 557 | -0.68 | 0.218 | 0.286 |
| chr3 | 155352721 | 155353247 | 527 | -0.72 | 0.219 | 0.286 |
| chr8 | 28347001 | 28352716 | 5716 | 0.24 | 0.219 | 0.286 |
| chr2 | 174495276 | 174496652 | 1377 | -0.38 | 0.219 | 0.286 |
| chr2 | 28612208 | 28673659 | 61452 | -0.06 | 0.219 | 0.287 |
| chr1 | 167630409 | 167631102 | 694 | -0.79 | 0.219 | 0.287 |
| chr21 | 17197701 | 17221526 | 23826 | -0.27 | 0.219 | 0.287 |
| chr19 | 15235400 | 15237233 | 1834 | 0.34 | 0.219 | 0.287 |
| chr19 | 4830712 | 4844891 | 14180 | 0.22 | 0.219 | 0.287 |
| chr3 | 12197163 | 12201468 | 4306 | 0.27 | 0.219 | 0.287 |
| chr2 | 136633033 | 136634446 | 1414 | -0.3 | 0.219 | 0.287 |

| | | | | | | |
|---|---|---|---|---|---|---|
| chr17 | 3562016 | 3573550 | 11535 | 0.1 | 0.219 | 0.287 |
| chr4 | 152235823 | 152253689 | 17867 | 0.16 | 0.219 | 0.287 |
| chr10 | 25233420 | 25247759 | 14340 | -0.19 | 0.219 | 0.287 |
| chr1 | 59220864 | 59221716 | 853 | 0.74 | 0.219 | 0.287 |
| chr3 | 93691820 | 93700467 | 8648 | -0.14 | 0.219 | 0.287 |
| chr2 | 70312343 | 70371309 | 58967 | -0.07 | 0.219 | 0.287 |
| chr12 | 1608426 | 1610291 | 1866 | -0.28 | 0.219 | 0.287 |
| chr12 | 132497596 | 132499426 | 1831 | -0.44 | 0.22 | 0.287 |
| chr2 | 127782510 | 127784281 | 1772 | -0.47 | 0.22 | 0.287 |
| chr3 | 66478635 | 66493504 | 14870 | 0.28 | 0.22 | 0.288 |
| chr6 | 121654897 | 121656691 | 1795 | 0.22 | 0.22 | 0.288 |
| chr9 | 119448724 | 119455663 | 6940 | -0.24 | 0.22 | 0.288 |
| chr5 | 124126236 | 124127252 | 1017 | -0.61 | 0.22 | 0.288 |
| chr1 | 110120987 | 110121810 | 824 | -0.63 | 0.22 | 0.288 |
| chr7 | 3475903 | 3477007 | 1105 | 0.47 | 0.22 | 0.288 |
| chr4 | 148928525 | 148929511 | 987 | -0.45 | 0.22 | 0.288 |
| chr4 | 32043838 | 32045403 | 1566 | 0.58 | 0.22 | 0.288 |
| chr3 | 153697823 | 153698467 | 645 | -0.65 | 0.22 | 0.288 |
| chr8 | 42547580 | 42548532 | 953 | -0.58 | 0.22 | 0.288 |
| chr12 | 114252773 | 114253872 | 1100 | -0.55 | 0.22 | 0.288 |
| chr5 | 61785285 | 61785785 | 501 | -0.81 | 0.22 | 0.288 |
| chr10 | 46167186 | 46169256 | 2071 | 0.19 | 0.22 | 0.288 |
| chr3 | 117347484 | 117347848 | 365 | 0.84 | 0.22 | 0.288 |
| chr4 | 114494775 | 114506536 | 11762 | -0.32 | 0.22 | 0.288 |
| chr1 | 225662908 | 225663389 | 482 | -1.01 | 0.221 | 0.288 |
| chr17 | 19351544 | 19352833 | 1290 | 0.52 | 0.221 | 0.289 |
| chr10 | 77228437 | 77229592 | 1156 | -0.49 | 0.221 | 0.289 |
| chr15 | 51894444 | 51896571 | 2128 | 0.26 | 0.221 | 0.289 |
| chrX | 148573712 | 148587320 | 13609 | 0.14 | 0.221 | 0.289 |
| chr3 | 56681480 | 56682213 | 734 | 0.97 | 0.221 | 0.289 |
| chr1 | 1309706 | 1312450 | 2745 | 0.18 | 0.221 | 0.289 |
| chr21 | 36118070 | 36118584 | 515 | -0.71 | 0.221 | 0.289 |
| chr13 | 111805113 | 111807434 | 2322 | 0.18 | 0.221 | 0.289 |
| chr17 | 46770847 | 46771330 | 484 | 0.84 | 0.221 | 0.289 |
| chr2 | 232109558 | 232110713 | 1156 | 0.85 | 0.221 | 0.289 |
| chr4 | 129581211 | 129581907 | 697 | 0.81 | 0.221 | 0.289 |
| chr7 | 117221768 | 117229159 | 7392 | -0.25 | 0.221 | 0.289 |
| chr8 | 110295660 | 110312101 | 16442 | 0.2 | 0.221 | 0.289 |
| chr4 | 78801969 | 78804183 | 2215 | -0.35 | 0.221 | 0.289 |
| chr11 | 47765264 | 47766123 | 860 | -0.63 | 0.221 | 0.289 |
| chr5 | 17995956 | 17996626 | 671 | -0.64 | 0.221 | 0.289 |
| chr20 | 47237262 | 47238835 | 1574 | 0.33 | 0.221 | 0.289 |
| chr6 | 107779750 | 107781600 | 1851 | 0.23 | 0.221 | 0.289 |
| chr5 | 61790410 | 61790931 | 522 | 1.34 | 0.221 | 0.29 |
| chr1 | 23961041 | 23962413 | 1373 | 0.45 | 0.222 | 0.29 |
| chr7 | 141250528 | 141252007 | 1480 | -0.29 | 0.222 | 0.29 |
| chr3 | 32510945 | 32512024 | 1080 | 0.75 | 0.222 | 0.29 |
| chr11 | 22297025 | 22297583 | 559 | 0.66 | 0.222 | 0.29 |
| chr6 | 77743514 | 77744059 | 546 | 0.71 | 0.222 | 0.29 |
| chr5 | 68622276 | 68629650 | 7375 | 0.25 | 0.222 | 0.29 |
| chr21 | 26978900 | 26980686 | 1787 | 0.25 | 0.222 | 0.29 |
| chr3 | 171358321 | 171358999 | 679 | 0.75 | 0.222 | 0.29 |
| chr11 | 119227150 | 119253237 | 26088 | -0.27 | 0.222 | 0.29 |
| chr10 | 73757182 | 73770200 | 13019 | 0.16 | 0.222 | 0.29 |
| chr11 | 82721012 | 82721508 | 497 | 0.75 | 0.222 | 0.29 |
| chr1 | 1195804 | 1217994 | 22191 | 0.31 | 0.222 | 0.29 |
| chr12 | 57379736 | 57380095 | 360 | 0.99 | 0.222 | 0.29 |
| chr4 | 86351381 | 86352598 | 1218 | -0.49 | 0.222 | 0.29 |
| chr8 | 108237978 | 108238702 | 725 | 0.7 | 0.222 | 0.29 |
| chr14 | 50526274 | 50535611 | 9338 | 0.12 | 0.222 | 0.29 |
| chr7 | 95846333 | 95846810 | 478 | -0.77 | 0.222 | 0.29 |

| | | | | | | |
|---|---|---|---|---|---|---|
| chr12 | 31789142 | 31789641 | 500 | -0.75 | 0.222 | 0.29 |
| chr7 | 148532933 | 148537027 | 4095 | 0.36 | 0.222 | 0.29 |
| chr2 | 42721273 | 42722177 | 905 | -0.5 | 0.222 | 0.29 |
| chr15 | 90644979 | 90646370 | 1392 | 0.37 | 0.222 | 0.29 |
| chr7 | 148427202 | 148428028 | 827 | 0.7 | 0.223 | 0.291 |
| chr3 | 125709046 | 125710045 | 1000 | 0.49 | 0.223 | 0.291 |
| chr3 | 101771897 | 101773051 | 1155 | 0.51 | 0.223 | 0.291 |
| chr19 | 55880083 | 55920676 | 40594 | 0.11 | 0.223 | 0.291 |
| chr7 | 152217922 | 152220079 | 2158 | -0.36 | 0.223 | 0.291 |
| chr6 | 5003619 | 5005090 | 1472 | 0.34 | 0.223 | 0.291 |
| chr7 | 109589946 | 109591588 | 1643 | 0.5 | 0.223 | 0.291 |
| chr17 | 40380014 | 40381332 | 1319 | -0.53 | 0.223 | 0.291 |
| chr1 | 38155013 | 38158820 | 3808 | 0.3 | 0.223 | 0.291 |
| chr6 | 144163459 | 144165869 | 2411 | 0.18 | 0.223 | 0.291 |
| chr2 | 99927961 | 99928735 | 775 | -2 | 0.223 | 0.292 |
| chr8 | 124193783 | 124195244 | 1462 | 0.68 | 0.223 | 0.292 |
| chr2 | 98610924 | 98613462 | 2539 | -0.21 | 0.223 | 0.292 |
| chr5 | 135415859 | 135417128 | 1270 | 0.41 | 0.224 | 0.292 |
| chr1 | 204287540 | 204321257 | 33718 | 0.11 | 0.224 | 0.292 |
| chr2 | 65546585 | 65548217 | 1633 | -0.28 | 0.224 | 0.292 |
| chr15 | 81246445 | 81248511 | 2067 | -0.4 | 0.224 | 0.292 |
| chr7 | 135151288 | 135152840 | 1553 | 0.33 | 0.224 | 0.292 |
| chr10 | 35368166 | 35380176 | 12011 | 0.22 | 0.224 | 0.292 |
| chr5 | 159670872 | 159746475 | 75604 | -0.07 | 0.224 | 0.292 |
| chr13 | 45387965 | 45393949 | 5985 | 0.22 | 0.224 | 0.292 |
| chr16 | 11930669 | 11946778 | 16110 | -0.2 | 0.224 | 0.292 |
| chr3 | 111611531 | 111657236 | 45706 | 0.12 | 0.224 | 0.292 |
| chrX | 151998080 | 152000677 | 2598 | -0.18 | 0.224 | 0.292 |
| chr10 | 114286489 | 114287316 | 828 | 0.46 | 0.224 | 0.292 |
| chr8 | 71359237 | 71360055 | 819 | -0.75 | 0.224 | 0.292 |
| chr9 | 84486003 | 84495199 | 9197 | -0.25 | 0.224 | 0.292 |
| chr7 | 129783923 | 129784973 | 1051 | 0.64 | 0.224 | 0.292 |
| chr3 | 57813030 | 57826819 | 13790 | 0.61 | 0.224 | 0.292 |
| chr3 | 44087005 | 44087831 | 827 | 0.8 | 0.224 | 0.293 |
| chr19 | 6799751 | 6803295 | 3545 | -0.16 | 0.224 | 0.293 |
| chr2 | 150780220 | 150781429 | 1210 | 0.41 | 0.224 | 0.293 |
| chr7 | 12536594 | 12537046 | 453 | 0.47 | 0.224 | 0.293 |
| chr1 | 28143167 | 28158874 | 15708 | 0.2 | 0.225 | 0.293 |
| chr19 | 16651728 | 16654502 | 2775 | 0.26 | 0.225 | 0.293 |
| chr3 | 148255610 | 148256011 | 402 | 0.83 | 0.225 | 0.293 |
| chr19 | 45090339 | 45328581 | 238243 | 0.05 | 0.225 | 0.293 |
| chr4 | 161159909 | 161160341 | 433 | -0.86 | 0.225 | 0.293 |
| chr1 | 110745857 | 110753844 | 7988 | 0.48 | 0.225 | 0.293 |
| chr9 | 15463550 | 15465250 | 1701 | 0.56 | 0.225 | 0.293 |
| chrX | 138905576 | 138911222 | 5647 | 0.37 | 0.225 | 0.293 |
| chr19 | 13093881 | 13094459 | 579 | -0.57 | 0.225 | 0.293 |
| chr19 | 1406702 | 1409507 | 2806 | -0.28 | 0.225 | 0.293 |
| chr3 | 179478332 | 179479103 | 772 | -0.61 | 0.225 | 0.294 |
| chr6 | 110529795 | 110531167 | 1373 | -0.48 | 0.225 | 0.294 |
| chr8 | 100416984 | 100417379 | 396 | -1.27 | 0.225 | 0.294 |
| chr12 | 57871464 | 57891909 | 20446 | 0.22 | 0.225 | 0.294 |
| chr5 | 52240712 | 52241670 | 959 | 0.5 | 0.225 | 0.294 |
| chr11 | 111635982 | 111637971 | 1990 | 0.28 | 0.225 | 0.294 |
| chr1 | 34872126 | 34872886 | 761 | -0.87 | 0.226 | 0.294 |
| chr5 | 129630983 | 129631738 | 756 | 2.02 | 0.226 | 0.294 |
| chr6 | 10237940 | 10238275 | 336 | -0.94 | 0.226 | 0.294 |
| chr15 | 64083641 | 64084242 | 602 | 0.62 | 0.226 | 0.295 |
| chr8 | 96279913 | 96283205 | 3293 | 0.17 | 0.226 | 0.295 |
| chr7 | 31914416 | 31941500 | 27085 | -0.08 | 0.227 | 0.295 |
| chr7 | 105415921 | 105418786 | 2866 | -0.16 | 0.227 | 0.295 |
| chr2 | 102615178 | 102616205 | 1028 | 0.67 | 0.227 | 0.295 |

| | | | | | | |
|---|---|---|---|---|---|---|
| chrX | 128674065 | 128675369 | 1305 | 0.47 | 0.227 | 0.296 |
| chr3 | 40841479 | 40841987 | 509 | 0.83 | 0.227 | 0.296 |
| chrX | 70711946 | 70713007 | 1062 | 0.52 | 0.227 | 0.296 |
| chr5 | 177630793 | 177643137 | 12345 | -0.16 | 0.227 | 0.296 |
| chr6 | 41395477 | 41396401 | 925 | 0.86 | 0.227 | 0.296 |
| chr5 | 134367497 | 134370832 | 3336 | 0.2 | 0.227 | 0.296 |
| chr5 | 145876315 | 145877033 | 719 | -1.09 | 0.228 | 0.296 |
| chr10 | 72575019 | 72590456 | 15438 | 0.21 | 0.228 | 0.296 |
| chr9 | 34009279 | 34012542 | 3264 | 0.43 | 0.228 | 0.296 |
| chr10 | 12369859 | 12377191 | 7333 | 0.13 | 0.228 | 0.296 |
| chr17 | 41559919 | 41568361 | 8443 | 0.15 | 0.228 | 0.296 |
| chr18 | 8098826 | 8099393 | 568 | -0.76 | 0.228 | 0.297 |
| chr3 | 64094746 | 64096109 | 1364 | -0.55 | 0.228 | 0.297 |
| chr8 | 106925414 | 106926034 | 621 | -0.72 | 0.228 | 0.297 |
| chr18 | 71814400 | 71816857 | 2458 | -0.26 | 0.228 | 0.297 |
| chr17 | 66532124 | 66540062 | 7939 | 0.28 | 0.228 | 0.297 |
| chr2 | 99524863 | 99534286 | 9424 | -0.22 | 0.228 | 0.297 |
| chr1 | 245748317 | 245748703 | 387 | 1.04 | 0.228 | 0.297 |
| chr6 | 111953113 | 111953695 | 583 | 0.87 | 0.229 | 0.297 |
| chr1 | 182807637 | 182811886 | 4250 | 0.19 | 0.229 | 0.297 |
| chr15 | 36231359 | 36232053 | 695 | 0.49 | 0.229 | 0.298 |
| chr12 | 117029709 | 117039753 | 10045 | -2.26 | 0.229 | 0.298 |
| chr20 | 34041581 | 34057150 | 15570 | -0.13 | 0.229 | 0.298 |
| chr7 | 155497776 | 155498618 | 843 | -0.56 | 0.229 | 0.298 |
| chr9 | 93763764 | 93764169 | 406 | 0.79 | 0.229 | 0.298 |
| chr10 | 433843 | 434654 | 812 | 0.67 | 0.229 | 0.298 |
| chr14 | 69122594 | 69123714 | 1121 | -0.57 | 0.229 | 0.298 |
| chr4 | 78977177 | 78980825 | 3649 | 0.24 | 0.229 | 0.298 |
| chrX | 129064865 | 129066157 | 1293 | 0.3 | 0.229 | 0.298 |
| chr2 | 102241559 | 102242129 | 571 | -0.78 | 0.229 | 0.298 |
| chr21 | 33687647 | 33695755 | 8109 | -0.25 | 0.229 | 0.298 |
| chrX | 46828065 | 46839884 | 11820 | -0.22 | 0.229 | 0.298 |
| chr20 | 9251257 | 9252173 | 917 | -0.46 | 0.23 | 0.299 |
| chr8 | 107057381 | 107073679 | 16299 | 0.12 | 0.23 | 0.299 |
| chr15 | 80986498 | 80989838 | 3341 | 0.22 | 0.23 | 0.299 |
| chr7 | 116715700 | 116716406 | 707 | 0.73 | 0.23 | 0.299 |
| chr11 | 9376639 | 9389397 | 12759 | 0.1 | 0.23 | 0.299 |
| chr15 | 66764627 | 66818498 | 53872 | -0.07 | 0.23 | 0.299 |
| chr1 | 40337583 | 40359451 | 21869 | -0.23 | 0.23 | 0.299 |
| chr8 | 19704146 | 19704685 | 540 | 0.72 | 0.23 | 0.299 |
| chr8 | 8095234 | 8095644 | 411 | -1.02 | 0.23 | 0.299 |
| chr7 | 72394401 | 72396677 | 2277 | 0.21 | 0.23 | 0.299 |
| chr13 | 74760699 | 74762782 | 2084 | 0.34 | 0.23 | 0.299 |
| chr16 | 58584975 | 58586614 | 1640 | -0.73 | 0.23 | 0.299 |
| chr17 | 72390774 | 72391488 | 715 | -0.64 | 0.23 | 0.299 |
| chr6 | 17623751 | 17636924 | 13174 | -0.18 | 0.23 | 0.299 |
| chr4 | 129697472 | 129698760 | 1289 | 0.47 | 0.23 | 0.299 |
| chr5 | 176103007 | 176104070 | 1064 | -0.6 | 0.23 | 0.299 |
| chr20 | 49173787 | 49187200 | 13414 | -0.14 | 0.23 | 0.299 |
| chr4 | 128577399 | 128577794 | 396 | 0.73 | 0.23 | 0.299 |
| chr1 | 147012197 | 147012760 | 564 | -0.9 | 0.23 | 0.299 |
| chr3 | 32610917 | 32613763 | 2847 | -0.17 | 0.23 | 0.299 |
| chr1 | 52493523 | 52504355 | 10833 | -0.14 | 0.23 | 0.299 |
| chr2 | 242156743 | 242158220 | 1478 | -0.44 | 0.23 | 0.299 |
| chr11 | 60897164 | 60898632 | 1469 | -0.44 | 0.231 | 0.3 |
| chr14 | 53111608 | 53112348 | 741 | -0.99 | 0.231 | 0.3 |
| chr22 | 42195766 | 42197276 | 1511 | 0.32 | 0.231 | 0.3 |
| chr12 | 124395035 | 124395425 | 391 | 0.95 | 0.231 | 0.3 |
| chr6 | 4519051 | 4519388 | 338 | 1.02 | 0.231 | 0.3 |
| chrX | 140555882 | 140559101 | 3220 | -0.25 | 0.231 | 0.3 |
| chr12 | 12014149 | 12015566 | 1418 | -0.76 | 0.231 | 0.3 |

| | | | | | | |
|---|---|---|---|---|---|---|
| chr18 | 11322774 | 11326151 | 3378 | -0.36 | 0.231 | 0.3 |
| chr12 | 132530869 | 132531735 | 867 | -0.55 | 0.231 | 0.3 |
| chr2 | 175804035 | 175804445 | 411 | 0.79 | 0.231 | 0.3 |
| chr22 | 30232969 | 30239689 | 6721 | 0.2 | 0.231 | 0.3 |
| chr1 | 171372182 | 171373131 | 950 | 0.74 | 0.231 | 0.3 |
| chr3 | 32744152 | 32745615 | 1464 | -0.52 | 0.231 | 0.3 |
| chr5 | 172086390 | 172087817 | 1428 | 0.42 | 0.231 | 0.3 |
| chr2 | 189652755 | 189653876 | 1122 | 0.55 | 0.231 | 0.3 |
| chr6 | 1267083 | 1268103 | 1021 | 0.57 | 0.231 | 0.3 |
| chr20 | 55434907 | 55435204 | 298 | -1.28 | 0.231 | 0.3 |
| chr3 | 81725250 | 81735372 | 10123 | -0.17 | 0.231 | 0.3 |
| chr11 | 64570827 | 64578733 | 7907 | -0.32 | 0.231 | 0.3 |
| chr6 | 1609545 | 1612598 | 3054 | 0.36 | 0.231 | 0.3 |
| chr3 | 101326541 | 101326992 | 452 | -0.79 | 0.231 | 0.3 |
| chr7 | 7477969 | 7479504 | 1536 | 0.33 | 0.232 | 0.301 |
| chr13 | 35806994 | 35807890 | 897 | -0.58 | 0.232 | 0.301 |
| chr6 | 26520260 | 26578070 | 57811 | 0.08 | 0.232 | 0.301 |
| chr11 | 43665132 | 43666581 | 1450 | 0.27 | 0.232 | 0.301 |
| chr1 | 233102329 | 233123187 | 20859 | -0.22 | 0.232 | 0.301 |
| chr6 | 11920780 | 11922282 | 1503 | -0.36 | 0.232 | 0.301 |
| chr16 | 57728675 | 57739008 | 10334 | -0.27 | 0.232 | 0.301 |
| chr1 | 2029633 | 2030075 | 443 | 1.28 | 0.232 | 0.301 |
| chr20 | 35579131 | 35580849 | 1719 | 0.41 | 0.232 | 0.301 |
| chr1 | 67518274 | 67520402 | 2129 | 0.24 | 0.232 | 0.301 |
| chr20 | 19944154 | 20034036 | 89883 | -0.05 | 0.232 | 0.301 |
| chr8 | 41908334 | 41910838 | 2505 | 0.31 | 0.232 | 0.301 |
| chr22 | 24798226 | 24799327 | 1102 | 0.44 | 0.232 | 0.301 |
| chr7 | 137546660 | 137547487 | 828 | 0.76 | 0.232 | 0.301 |
| chr14 | 64192338 | 64195851 | 3514 | 0.22 | 0.232 | 0.301 |
| chr8 | 435042 | 437628 | 2587 | -0.43 | 0.232 | 0.301 |
| chr1 | 212558271 | 212560476 | 2206 | -0.6 | 0.232 | 0.301 |
| chr8 | 74524715 | 74527349 | 2635 | 0.47 | 0.232 | 0.301 |
| chr10 | 131907103 | 131910722 | 3620 | 0.15 | 0.232 | 0.301 |
| chr15 | 82554609 | 82556346 | 1738 | -0.26 | 0.232 | 0.301 |
| chr17 | 74348904 | 74351420 | 2517 | -0.18 | 0.232 | 0.301 |
| chr10 | 115911174 | 115911967 | 794 | -0.63 | 0.232 | 0.301 |
| chrX | 140000152 | 140001708 | 1557 | -0.33 | 0.232 | 0.301 |
| chr12 | 132459491 | 132473943 | 14453 | -0.17 | 0.232 | 0.301 |
| chr13 | 98252220 | 98253526 | 1307 | 0.91 | 0.232 | 0.301 |
| chr11 | 62193959 | 62208273 | 14315 | -0.35 | 0.232 | 0.301 |
| chr2 | 15731174 | 15754317 | 23144 | -0.12 | 0.232 | 0.301 |
| chr12 | 93893718 | 93894415 | 698 | 0.61 | 0.233 | 0.302 |
| chr15 | 89440899 | 89464455 | 23557 | 0.2 | 0.233 | 0.302 |
| chr9 | 110031736 | 110087796 | 56061 | 0.12 | 0.233 | 0.302 |
| chr2 | 166767546 | 166768111 | 566 | -0.64 | 0.233 | 0.302 |
| chr1 | 55487977 | 55488426 | 450 | 0.65 | 0.233 | 0.302 |
| chr13 | 98627719 | 98631470 | 3752 | 0.19 | 0.233 | 0.302 |
| chr12 | 51034963 | 51036828 | 1866 | 0.32 | 0.233 | 0.302 |
| chr17 | 2689076 | 2699651 | 10576 | 0.5 | 0.233 | 0.302 |
| chr3 | 128061809 | 128063088 | 1280 | -0.41 | 0.233 | 0.302 |
| chr1 | 113408588 | 113409598 | 1011 | 0.68 | 0.233 | 0.302 |
| chr6 | 3196083 | 3210366 | 14284 | -0.28 | 0.233 | 0.302 |
| chrX | 107178684 | 107179326 | 643 | -0.73 | 0.233 | 0.302 |
| chr5 | 163546750 | 163547650 | 901 | -0.56 | 0.233 | 0.302 |
| chr1 | 1981097 | 1981593 | 497 | 0.68 | 0.233 | 0.302 |
| chr1 | 19464628 | 19469010 | 4383 | -0.29 | 0.233 | 0.302 |
| chr3 | 143566942 | 143567565 | 624 | 0.8 | 0.233 | 0.302 |
| chr10 | 33376357 | 33376977 | 621 | 0.74 | 0.233 | 0.302 |
| chr3 | 31397947 | 31412898 | 14952 | 0.22 | 0.233 | 0.302 |
| chr5 | 114701578 | 114708081 | 6504 | 2.03 | 0.233 | 0.302 |
| chr15 | 37301368 | 37302393 | 1026 | 0.61 | 0.233 | 0.302 |

| | | | | | | |
|---|---|---|---|---|---|---|
| chr10 | 44185112 | 44185535 | 424 | 0.75 | 0.233 | 0.302 |
| chr1 | 183687594 | 183693913 | 6320 | 0.23 | 0.234 | 0.302 |
| chr16 | 71371582 | 71373345 | 1764 | 0.43 | 0.234 | 0.303 |
| chr5 | 34866978 | 34867680 | 703 | -0.59 | 0.234 | 0.303 |
| chr6 | 53357955 | 53414685 | 56731 | -0.07 | 0.234 | 0.303 |
| chr10 | 58119762 | 58122033 | 2272 | 0.2 | 0.234 | 0.303 |
| chr8 | 133405352 | 133406728 | 1377 | -2.17 | 0.234 | 0.303 |
| chr4 | 189013708 | 189014605 | 898 | 0.61 | 0.234 | 0.303 |
| chr7 | 67703162 | 67703929 | 768 | 0.74 | 0.234 | 0.303 |
| chr14 | 67634084 | 67635179 | 1096 | -0.64 | 0.234 | 0.303 |
| chr11 | 63888469 | 63888918 | 450 | -0.97 | 0.234 | 0.303 |
| chr17 | 60584231 | 60585100 | 870 | 0.47 | 0.234 | 0.303 |
| chr9 | 35060581 | 35081253 | 20673 | -0.1 | 0.235 | 0.304 |
| chr7 | 105516123 | 105518345 | 2223 | 0.18 | 0.235 | 0.304 |
| chrX | 19685440 | 19701046 | 15607 | 0.1 | 0.235 | 0.304 |
| chr2 | 20488088 | 20499442 | 11355 | -0.18 | 0.235 | 0.304 |
| chr11 | 115248291 | 115249216 | 926 | 0.68 | 0.235 | 0.304 |
| chr12 | 118681818 | 118682236 | 419 | -0.93 | 0.235 | 0.304 |
| chr3 | 196729390 | 196730874 | 1485 | -0.38 | 0.235 | 0.304 |
| chr15 | 63441893 | 63462501 | 20609 | 0.11 | 0.235 | 0.304 |
| chr13 | 63764062 | 63765447 | 1386 | -0.46 | 0.235 | 0.304 |
| chr6 | 20528945 | 20543944 | 15000 | 0.26 | 0.235 | 0.304 |
| chr6 | 150739576 | 150740020 | 445 | 0.9 | 0.235 | 0.304 |
| chr5 | 153963277 | 153967737 | 4461 | 0.38 | 0.235 | 0.304 |
| chr13 | 44119587 | 44120276 | 690 | 0.84 | 0.235 | 0.304 |
| chr17 | 59967669 | 59981792 | 14124 | -0.2 | 0.235 | 0.304 |
| chr13 | 30328401 | 30329313 | 913 | 0.87 | 0.235 | 0.304 |
| chr2 | 54331091 | 54343680 | 12590 | -0.13 | 0.235 | 0.305 |
| chr16 | 75438886 | 75439579 | 694 | 0.67 | 0.235 | 0.305 |
| chr1 | 225878662 | 225889368 | 10707 | -0.1 | 0.236 | 0.305 |
| chr4 | 106310822 | 106311258 | 437 | 0.85 | 0.236 | 0.305 |
| chr2 | 103325359 | 103326479 | 1121 | -0.77 | 0.236 | 0.305 |
| chr1 | 228268932 | 228297748 | 28817 | -0.1 | 0.236 | 0.305 |
| chr7 | 83144217 | 83162451 | 18235 | 0.24 | 0.236 | 0.305 |
| chr1 | 7021926 | 7024134 | 2209 | 0.46 | 0.236 | 0.305 |
| chr22 | 19928238 | 19935350 | 7113 | 0.14 | 0.236 | 0.305 |
| chr16 | 10673771 | 10675025 | 1255 | -0.3 | 0.236 | 0.305 |
| chrX | 135285369 | 135287474 | 2106 | 0.45 | 0.236 | 0.305 |
| chr13 | 30716188 | 30717486 | 1299 | -0.59 | 0.236 | 0.305 |
| chr6 | 53719336 | 53720432 | 1097 | 0.61 | 0.236 | 0.305 |
| chr3 | 171935497 | 171937016 | 1520 | -0.58 | 0.236 | 0.305 |
| chr7 | 40167935 | 40175086 | 7152 | 0.16 | 0.236 | 0.306 |
| chr4 | 39816500 | 39828079 | 11580 | -0.19 | 0.236 | 0.306 |
| chr10 | 94763669 | 94770798 | 7130 | 0.39 | 0.236 | 0.306 |
| chr1 | 77786723 | 77818976 | 32254 | 0.11 | 0.237 | 0.306 |
| chr14 | 81087709 | 81088690 | 982 | -0.63 | 0.237 | 0.306 |
| chrX | 53252680 | 53255269 | 2590 | 0.18 | 0.237 | 0.306 |
| chr13 | 51483346 | 51485438 | 2093 | 0.18 | 0.237 | 0.306 |
| chr9 | 35793405 | 35846316 | 52912 | 0.13 | 0.237 | 0.306 |
| chr1 | 16209247 | 16244020 | 34774 | 0.09 | 0.237 | 0.306 |
| chr9 | 20835553 | 20836164 | 612 | -0.89 | 0.237 | 0.306 |
| chr7 | 32058402 | 32059733 | 1332 | 0.57 | 0.237 | 0.306 |
| chr13 | 36365176 | 36366289 | 1114 | 0.56 | 0.237 | 0.306 |
| chr4 | 124013142 | 124013824 | 683 | 0.96 | 0.237 | 0.306 |
| chr7 | 84027134 | 84043212 | 16079 | -0.17 | 0.237 | 0.306 |
| chr17 | 78282962 | 78309610 | 26649 | -0.16 | 0.237 | 0.307 |
| chr1 | 20478426 | 20480815 | 2390 | 0.44 | 0.237 | 0.307 |
| chr20 | 18567867 | 18569665 | 1799 | 0.24 | 0.237 | 0.307 |
| chr6 | 73945531 | 73946646 | 1116 | -0.47 | 0.237 | 0.307 |
| chr10 | 89675252 | 89675696 | 445 | -1.25 | 0.237 | 0.307 |
| chr7 | 1139972 | 1140637 | 666 | -0.88 | 0.237 | 0.307 |

| | | | | | | |
|---|---|---|---|---|---|---|
| chr5 | 135328992 | 135403253 | 74262 | 0.05 | 0.237 | 0.307 |
| chr3 | 151576386 | 151594395 | 18010 | -0.13 | 0.237 | 0.307 |
| chr4 | 74315903 | 74316705 | 803 | 0.57 | 0.238 | 0.307 |
| chr8 | 43021907 | 43022699 | 793 | 0.64 | 0.238 | 0.307 |
| chr17 | 17203587 | 17207537 | 3951 | -0.23 | 0.238 | 0.307 |
| chr4 | 68410054 | 68412150 | 2097 | 0.23 | 0.238 | 0.307 |
| chr20 | 30696892 | 30698798 | 1907 | -0.19 | 0.238 | 0.307 |
| chr6 | 166308717 | 166309780 | 1064 | 0.62 | 0.238 | 0.307 |
| chr13 | 111903600 | 111905219 | 1620 | 0.43 | 0.238 | 0.307 |
| chr5 | 43193970 | 43194378 | 409 | -0.93 | 0.238 | 0.307 |
| chr12 | 88718344 | 88720000 | 1657 | 0.3 | 0.238 | 0.307 |
| chr22 | 39760385 | 39760759 | 375 | -1.11 | 0.238 | 0.307 |
| chr6 | 110830523 | 110831768 | 1246 | 0.84 | 0.238 | 0.307 |
| chr9 | 84402110 | 84417227 | 15118 | 0.75 | 0.238 | 0.308 |
| chr14 | 101563106 | 101564035 | 930 | -0.58 | 0.238 | 0.308 |
| chrX | 102658815 | 102660550 | 1736 | -0.25 | 0.238 | 0.308 |
| chr6 | 24743100 | 24756127 | 13028 | 0.27 | 0.238 | 0.308 |
| chr14 | 73240693 | 73249373 | 8681 | 0.33 | 0.238 | 0.308 |
| chr2 | 151165618 | 151166811 | 1194 | 0.5 | 0.238 | 0.308 |
| chr10 | 120630253 | 120630629 | 377 | -0.72 | 0.239 | 0.308 |
| chrX | 134048498 | 134049820 | 1323 | 0.38 | 0.239 | 0.308 |
| chr13 | 44977796 | 44978802 | 1007 | -0.78 | 0.239 | 0.308 |
| chr3 | 27508226 | 27526876 | 18651 | -0.12 | 0.239 | 0.308 |
| chr10 | 90495380 | 90496033 | 654 | -0.97 | 0.239 | 0.308 |
| chr7 | 77342182 | 77377091 | 34910 | 0.14 | 0.239 | 0.308 |
| chr7 | 97878222 | 97883054 | 4833 | -0.2 | 0.239 | 0.308 |
| chr15 | 64237048 | 64238087 | 1040 | -0.45 | 0.239 | 0.308 |
| chr1 | 162490369 | 162491340 | 972 | -0.66 | 0.239 | 0.308 |
| chr18 | 54069987 | 54070736 | 750 | -0.8 | 0.239 | 0.308 |
| chr3 | 146130785 | 146140733 | 9949 | 0.54 | 0.239 | 0.308 |
| chr20 | 62151026 | 62153086 | 2061 | 0.24 | 0.239 | 0.308 |
| chr1 | 220211705 | 220220781 | 9077 | 0.15 | 0.239 | 0.309 |
| chr9 | 139221508 | 139223146 | 1639 | -0.44 | 0.239 | 0.309 |
| chr5 | 102628778 | 102630121 | 1344 | 2.13 | 0.239 | 0.309 |
| chr3 | 176753818 | 176759561 | 5744 | -0.23 | 0.239 | 0.309 |
| chr8 | 125371597 | 125385740 | 14144 | -0.19 | 0.239 | 0.309 |
| chr11 | 28984668 | 28998837 | 14170 | 0.29 | 0.239 | 0.309 |
| chr15 | 44718228 | 44734488 | 16261 | 0.11 | 0.239 | 0.309 |
| chr3 | 128878625 | 128881933 | 3309 | 0.21 | 0.239 | 0.309 |
| chr19 | 19549109 | 19601298 | 52190 | -0.12 | 0.239 | 0.309 |
| chr7 | 75795320 | 75808219 | 12900 | 0.32 | 0.239 | 0.309 |
| chr3 | 193988480 | 193989205 | 726 | 0.91 | 0.239 | 0.309 |
| chr6 | 15449427 | 15449847 | 421 | -0.93 | 0.24 | 0.309 |
| chr11 | 73832972 | 73883846 | 50875 | 0.06 | 0.24 | 0.309 |
| chr4 | 88618584 | 88619093 | 510 | 0.92 | 0.24 | 0.309 |
| chr1 | 95390154 | 95409157 | 19004 | 0.12 | 0.24 | 0.309 |
| chr18 | 18689808 | 18692865 | 3058 | 0.23 | 0.24 | 0.309 |
| chr6 | 33610844 | 33611426 | 583 | -0.9 | 0.24 | 0.309 |
| chr17 | 59567865 | 59568315 | 451 | -1.01 | 0.24 | 0.309 |
| chr10 | 27010593 | 27011076 | 484 | 0.85 | 0.24 | 0.309 |
| chr2 | 101850081 | 101889396 | 39316 | 0.12 | 0.24 | 0.309 |
| chr2 | 65453990 | 65495147 | 41158 | -0.08 | 0.24 | 0.309 |
| chr10 | 3796793 | 3798107 | 1315 | 0.48 | 0.24 | 0.309 |
| chr10 | 128593095 | 128595272 | 2178 | 0.22 | 0.24 | 0.309 |
| chr6 | 111563430 | 111563830 | 401 | 0.76 | 0.24 | 0.309 |
| chr1 | 114217622 | 114218143 | 522 | 0.9 | 0.24 | 0.309 |
| chr6 | 2817916 | 2819498 | 1583 | 0.41 | 0.24 | 0.309 |
| chr2 | 58655065 | 58655815 | 751 | -0.64 | 0.24 | 0.309 |
| chr8 | 95856919 | 95858694 | 1776 | 0.57 | 0.24 | 0.309 |
| chr6 | 166327579 | 166337451 | 9873 | 0.27 | 0.24 | 0.31 |
| chr2 | 86348467 | 86349386 | 920 | -0.65 | 0.24 | 0.31 |

| | | | | | | |
|---|---|---|---|---|---|---|
| chr20 | 62249648 | 62260669 | 11022 | 0.19 | 0.24 | 0.31 |
| chr1 | 246545662 | 246546137 | 476 | -0.93 | 0.241 | 0.31 |
| chr12 | 111135660 | 111137103 | 1444 | -0.32 | 0.241 | 0.31 |
| chr17 | 78116328 | 78122290 | 5963 | 0.19 | 0.241 | 0.31 |
| chr3 | 11226005 | 11250316 | 24312 | 0.33 | 0.241 | 0.31 |
| chr6 | 24424810 | 24429383 | 4574 | -0.32 | 0.241 | 0.31 |
| chr4 | 161050075 | 161051397 | 1323 | 0.54 | 0.241 | 0.31 |
| chr6 | 26986846 | 26988576 | 1731 | 0.15 | 0.241 | 0.31 |
| chr3 | 113488098 | 113488775 | 678 | -0.67 | 0.241 | 0.31 |
| chr10 | 100019747 | 100030198 | 10452 | -0.3 | 0.241 | 0.31 |
| chr14 | 71274207 | 71289143 | 14937 | -0.19 | 0.241 | 0.31 |
| chr7 | 64838726 | 64840226 | 1501 | 0.28 | 0.241 | 0.311 |
| chr9 | 111822335 | 111883272 | 60938 | 0.06 | 0.241 | 0.311 |
| chr20 | 10198111 | 10206785 | 8675 | -0.2 | 0.241 | 0.311 |
| chr6 | 6749852 | 6750684 | 833 | -0.55 | 0.241 | 0.311 |
| chr3 | 183601747 | 183603342 | 1596 | 0.37 | 0.241 | 0.311 |
| chr15 | 45985455 | 45998242 | 12788 | 0.44 | 0.242 | 0.311 |
| chr6 | 5084599 | 5085643 | 1045 | -0.54 | 0.242 | 0.311 |
| chr4 | 139903236 | 139903605 | 370 | -0.86 | 0.242 | 0.312 |
| chr2 | 231548648 | 231552981 | 4334 | 0.3 | 0.242 | 0.312 |
| chr11 | 124745786 | 124746962 | 1177 | 0.33 | 0.242 | 0.312 |
| chr20 | 48687424 | 48687968 | 545 | 0.53 | 0.242 | 0.312 |
| chr8 | 67939879 | 67941513 | 1635 | -0.28 | 0.242 | 0.312 |
| chr14 | 55821442 | 55821903 | 462 | -1.24 | 0.242 | 0.312 |
| chr17 | 58326160 | 58326929 | 770 | 1.95 | 0.242 | 0.312 |
| chr8 | 73921122 | 73922016 | 895 | 0.37 | 0.242 | 0.312 |
| chr1 | 186844805 | 186845522 | 718 | -0.66 | 0.242 | 0.312 |
| chr20 | 16387514 | 16404796 | 17283 | 1.42 | 0.242 | 0.312 |
| chr6 | 108580899 | 108583535 | 2637 | 0.2 | 0.242 | 0.312 |
| chr13 | 95672255 | 95697781 | 25527 | 0.17 | 0.242 | 0.312 |
| chr1 | 193155073 | 193156130 | 1058 | 0.51 | 0.242 | 0.312 |
| chr22 | 32979412 | 32980750 | 1339 | 0.49 | 0.243 | 0.312 |
| chr4 | 8421004 | 8443565 | 22562 | -0.18 | 0.243 | 0.312 |
| chr7 | 155949096 | 155949630 | 535 | -1.24 | 0.243 | 0.312 |
| chrX | 48534111 | 48535706 | 1596 | 0.31 | 0.243 | 0.313 |
| chr3 | 131245535 | 131246032 | 498 | -0.81 | 0.243 | 0.313 |
| chr7 | 140211988 | 140218506 | 6519 | 0.55 | 0.243 | 0.313 |
| chr13 | 110521154 | 110522998 | 1845 | 0.39 | 0.243 | 0.313 |
| chr12 | 47411298 | 47412363 | 1066 | 0.44 | 0.243 | 0.313 |
| chr11 | 19940613 | 19941348 | 736 | -0.81 | 0.243 | 0.313 |
| chr10 | 72050029 | 72057231 | 7203 | 0.43 | 0.243 | 0.313 |
| chr12 | 89968034 | 89976631 | 8598 | 0.28 | 0.243 | 0.313 |
| chr18 | 52714708 | 52716901 | 2194 | 0.36 | 0.243 | 0.313 |
| chr3 | 106954208 | 106961018 | 6811 | 0.25 | 0.243 | 0.313 |
| chr20 | 25433818 | 25434402 | 585 | 0.84 | 0.243 | 0.313 |
| chr11 | 8891905 | 8893398 | 1494 | -0.36 | 0.243 | 0.313 |
| chr4 | 187821321 | 187826193 | 4873 | 0.24 | 0.243 | 0.313 |
| chr15 | 91072411 | 91074166 | 1756 | 0.26 | 0.243 | 0.313 |
| chr4 | 94234262 | 94234797 | 536 | -0.85 | 0.243 | 0.313 |
| chr5 | 106275916 | 106277961 | 2046 | -0.32 | 0.243 | 0.313 |
| chr7 | 129690330 | 129691957 | 1628 | 0.23 | 0.243 | 0.313 |
| chr9 | 140636869 | 140656003 | 19135 | -0.14 | 0.243 | 0.313 |
| chr3 | 64008006 | 64029315 | 21310 | -0.17 | 0.243 | 0.313 |
| chrX | 111112160 | 111112740 | 581 | -0.77 | 0.244 | 0.313 |
| chr8 | 142401474 | 142402914 | 1441 | -0.4 | 0.244 | 0.313 |
| chr1 | 27049310 | 27050410 | 1101 | 0.71 | 0.244 | 0.313 |
| chr22 | 35843735 | 35844134 | 400 | 0.79 | 0.244 | 0.313 |
| chr18 | 73129953 | 73131224 | 1272 | -0.6 | 0.244 | 0.314 |
| chr7 | 139899187 | 139900715 | 1529 | -0.42 | 0.244 | 0.314 |
| chr16 | 89938868 | 89949720 | 10853 | 0.16 | 0.244 | 0.314 |
| chr4 | 84883487 | 84884293 | 807 | -0.64 | 0.244 | 0.314 |

| | | | | | | |
|---|---|---|---|---|---|---|
| chr5 | 308787 | 322226 | 13440 | -0.27 | 0.244 | 0.314 |
| chr7 | 148637302 | 148639607 | 2306 | 0.16 | 0.244 | 0.314 |
| chr1 | 168289839 | 168291070 | 1232 | 0.52 | 0.244 | 0.314 |
| chr1 | 9747246 | 9747807 | 562 | -0.59 | 0.244 | 0.314 |
| chrX | 134476793 | 134479520 | 2728 | 0.16 | 0.244 | 0.314 |
| chr6 | 15244655 | 15249997 | 5343 | -0.16 | 0.245 | 0.314 |
| chr13 | 20565430 | 20568435 | 3006 | -0.38 | 0.245 | 0.314 |
| chrX | 53688413 | 53693539 | 5127 | 0.35 | 0.245 | 0.314 |
| chr15 | 91259587 | 91261389 | 1803 | 0.2 | 0.245 | 0.314 |
| chr22 | 21270512 | 21273195 | 2684 | 0.26 | 0.245 | 0.314 |
| chr8 | 110084405 | 110084805 | 401 | -0.99 | 0.245 | 0.315 |
| chr14 | 81302412 | 81311765 | 9354 | -0.35 | 0.245 | 0.315 |
| chr5 | 458889 | 459371 | 483 | 0.87 | 0.245 | 0.315 |
| chr2 | 242822918 | 242823729 | 812 | -0.52 | 0.245 | 0.315 |
| chr11 | 128985405 | 128986712 | 1308 | 0.55 | 0.245 | 0.315 |
| chr2 | 86325278 | 86335379 | 10102 | -0.18 | 0.245 | 0.315 |
| chr15 | 56656940 | 56658578 | 1639 | 0.29 | 0.246 | 0.315 |
| chr1 | 61869761 | 61870444 | 684 | -0.78 | 0.246 | 0.315 |
| chr6 | 127686694 | 127689108 | 2415 | 0.2 | 0.246 | 0.316 |
| chr8 | 144766025 | 144779268 | 13244 | -0.19 | 0.246 | 0.316 |
| chr7 | 84103818 | 84117326 | 13509 | 0.44 | 0.246 | 0.316 |
| chr14 | 24520858 | 24521363 | 506 | -0.71 | 0.246 | 0.316 |
| chr9 | 26928164 | 26957559 | 29396 | 0.1 | 0.246 | 0.316 |
| chr1 | 222638359 | 222640392 | 2034 | 0.18 | 0.246 | 0.316 |
| chr18 | 20839580 | 20840596 | 1017 | -0.57 | 0.246 | 0.316 |
| chr3 | 38387795 | 38389190 | 1396 | -0.36 | 0.246 | 0.316 |
| chr22 | 21212109 | 21214097 | 1989 | 0.32 | 0.246 | 0.316 |
| chr11 | 69707623 | 69716514 | 8892 | -0.34 | 0.246 | 0.316 |
| chr7 | 152028090 | 152028444 | 355 | 1 | 0.246 | 0.316 |
| chr3 | 37083159 | 37098974 | 15816 | 0.19 | 0.246 | 0.316 |
| chr8 | 30084507 | 30086007 | 1501 | -0.49 | 0.246 | 0.316 |
| chr19 | 49163077 | 49164928 | 1852 | 0.29 | 0.246 | 0.316 |
| chr2 | 8551593 | 8552927 | 1335 | 0.52 | 0.246 | 0.316 |
| chr8 | 99502620 | 99503312 | 693 | -0.56 | 0.246 | 0.316 |
| chr6 | 43805943 | 43807132 | 1190 | -0.44 | 0.246 | 0.316 |
| chr14 | 61508496 | 61509317 | 822 | 0.62 | 0.246 | 0.316 |
| chr2 | 99951132 | 99958556 | 7425 | -0.18 | 0.247 | 0.316 |
| chr16 | 11295644 | 11296676 | 1033 | -0.76 | 0.247 | 0.316 |
| chr18 | 77438141 | 77441383 | 3243 | 0.19 | 0.247 | 0.316 |
| chr13 | 33558756 | 33559777 | 1022 | 0.56 | 0.247 | 0.316 |
| chr15 | 80930994 | 80931348 | 355 | 1.27 | 0.247 | 0.317 |
| chr10 | 3355635 | 3356151 | 517 | -0.79 | 0.247 | 0.317 |
| chr1 | 159198877 | 159201023 | 2147 | -0.44 | 0.247 | 0.317 |
| chr1 | 188332498 | 188333440 | 943 | 0.38 | 0.247 | 0.317 |
| chr2 | 17498446 | 17499927 | 1482 | 0.4 | 0.247 | 0.317 |
| chr22 | 36043433 | 36045737 | 2305 | -0.17 | 0.247 | 0.317 |
| chr11 | 82691982 | 82692349 | 368 | -0.78 | 0.247 | 0.317 |
| chr17 | 27168413 | 27182935 | 14523 | 0.19 | 0.247 | 0.317 |
| chr1 | 210000460 | 210023838 | 23379 | -0.13 | 0.247 | 0.317 |
| chr3 | 15899981 | 15902669 | 2689 | -0.23 | 0.247 | 0.317 |
| chrX | 20338545 | 20339309 | 765 | -0.63 | 0.247 | 0.317 |
| chr9 | 135818930 | 135820918 | 1989 | 0.18 | 0.247 | 0.317 |
| chr3 | 124448689 | 124460412 | 11724 | -0.17 | 0.247 | 0.317 |
| chr14 | 35240748 | 35241113 | 366 | 0.82 | 0.248 | 0.317 |
| chr5 | 177579833 | 177581629 | 1797 | 0.32 | 0.248 | 0.317 |
| chr10 | 89621371 | 89628573 | 7203 | 0.13 | 0.248 | 0.318 |
| chr13 | 30688357 | 30689199 | 843 | -0.64 | 0.248 | 0.318 |
| chr4 | 169074474 | 169075102 | 629 | 0.76 | 0.248 | 0.318 |
| chr17 | 28474935 | 28477060 | 2126 | -0.35 | 0.248 | 0.318 |
| chr7 | 128171047 | 128171900 | 854 | -0.56 | 0.248 | 0.318 |
| chr13 | 20531036 | 20534381 | 3346 | -0.18 | 0.248 | 0.318 |

| | | | | | | |
|---|---|---|---|---|---|---|
| chr12 | 42487423 | 42499468 | 12046 | -0.36 | 0.248 | 0.318 |
| chr5 | 40447316 | 40447902 | 587 | 0.8 | 0.248 | 0.318 |
| chr16 | 85636359 | 85656606 | 20248 | 0.13 | 0.248 | 0.318 |
| chr7 | 24946869 | 24961424 | 14556 | 0.26 | 0.248 | 0.318 |
| chr20 | 22817126 | 22818333 | 1208 | -0.56 | 0.248 | 0.318 |
| chr12 | 101691936 | 101693843 | 1908 | -0.4 | 0.248 | 0.318 |
| chr6 | 155447913 | 155449490 | 1578 | 0.47 | 0.249 | 0.319 |
| chr1 | 15648943 | 15652358 | 3416 | 0.31 | 0.249 | 0.319 |
| chr16 | 89625201 | 89653471 | 28271 | 0.11 | 0.249 | 0.319 |
| chr1 | 201562240 | 201575186 | 12947 | -0.18 | 0.249 | 0.319 |
| chr6 | 35885280 | 35889685 | 4406 | 0.17 | 0.249 | 0.319 |
| chr6 | 126300007 | 126308528 | 8522 | 0.21 | 0.249 | 0.319 |
| chrX | 119377890 | 119386062 | 8173 | 0.16 | 0.249 | 0.319 |
| chr14 | 45382384 | 45383282 | 899 | 0.53 | 0.249 | 0.319 |
| chr6 | 157169602 | 157170286 | 685 | -0.84 | 0.249 | 0.319 |
| chr7 | 32697031 | 32711256 | 14226 | 0.12 | 0.249 | 0.319 |
| chr2 | 85959651 | 85981894 | 22244 | -0.1 | 0.249 | 0.319 |
| chr16 | 32265551 | 32265911 | 361 | 0.62 | 0.249 | 0.319 |
| chr2 | 68907046 | 68907720 | 675 | 0.61 | 0.249 | 0.319 |
| chr1 | 40024322 | 40025900 | 1579 | 0.28 | 0.249 | 0.319 |
| chr11 | 46558839 | 46559534 | 696 | -0.69 | 0.25 | 0.32 |
| chr9 | 107707466 | 107707797 | 332 | 0.82 | 0.25 | 0.32 |
| chr12 | 83468027 | 83468708 | 682 | -0.62 | 0.25 | 0.32 |
| chr22 | 39897662 | 39899512 | 1851 | 0.42 | 0.25 | 0.32 |
| chr12 | 1582643 | 1584002 | 1360 | 0.48 | 0.25 | 0.32 |
| chr6 | 38229355 | 38230896 | 1542 | -0.38 | 0.25 | 0.32 |
| chr1 | 46064835 | 46087753 | 22919 | -0.18 | 0.25 | 0.32 |
| chr7 | 6167495 | 6179651 | 12157 | 0.33 | 0.25 | 0.32 |
| chr6 | 35636956 | 35637729 | 774 | -0.69 | 0.25 | 0.32 |
| chr5 | 76534789 | 76535727 | 939 | 0.53 | 0.25 | 0.32 |
| chr11 | 12546552 | 12547505 | 954 | -1.92 | 0.25 | 0.32 |
| chr3 | 56873994 | 56875003 | 1010 | -0.56 | 0.251 | 0.321 |
| chr9 | 26869858 | 26870732 | 875 | 0.56 | 0.251 | 0.321 |
| chr3 | 128838731 | 128840625 | 1895 | 0.37 | 0.251 | 0.321 |
| chr3 | 48282420 | 48283472 | 1053 | 0.43 | 0.251 | 0.321 |
| chr2 | 32936426 | 32937438 | 1013 | 0.57 | 0.251 | 0.321 |
| chr1 | 32131042 | 32136431 | 5390 | -0.22 | 0.251 | 0.321 |
| chr14 | 34904792 | 34935183 | 30392 | 0.09 | 0.251 | 0.321 |
| chr3 | 125926964 | 125932517 | 5554 | 0.24 | 0.251 | 0.322 |
| chr15 | 74243713 | 74244193 | 481 | -0.64 | 0.251 | 0.322 |
| chr14 | 76113724 | 76129000 | 15277 | -0.14 | 0.251 | 0.322 |
| chr8 | 8859721 | 8876079 | 16359 | 0.15 | 0.252 | 0.322 |
| chr4 | 70849430 | 70850162 | 733 | -1.88 | 0.252 | 0.322 |
| chr3 | 159530656 | 159600295 | 69640 | 0.1 | 0.252 | 0.322 |
| chr17 | 7417495 | 7418641 | 1147 | -0.63 | 0.252 | 0.322 |
| chr12 | 105055817 | 105080759 | 24943 | 0.08 | 0.252 | 0.322 |
| chr15 | 44974584 | 44975364 | 781 | 0.86 | 0.252 | 0.322 |
| chr8 | 6404878 | 6405984 | 1107 | 0.54 | 0.252 | 0.322 |
| chr21 | 16357912 | 16379455 | 21544 | 0.18 | 0.252 | 0.322 |
| chr4 | 142723258 | 142724130 | 873 | 0.53 | 0.252 | 0.322 |
| chr2 | 96068013 | 96069713 | 1701 | -0.24 | 0.252 | 0.322 |
| chr1 | 234321608 | 234322502 | 895 | -2.03 | 0.252 | 0.322 |
| chr15 | 90686588 | 90793592 | 107005 | 0.05 | 0.252 | 0.323 |
| chr7 | 85324569 | 85325564 | 996 | 0.52 | 0.252 | 0.323 |
| chr5 | 217825 | 219300 | 1476 | 0.37 | 0.252 | 0.323 |
| chrX | 45181854 | 45182598 | 745 | -0.65 | 0.252 | 0.323 |
| chr6 | 111579746 | 111581595 | 1850 | -0.29 | 0.253 | 0.323 |
| chr3 | 136390536 | 136391715 | 1180 | 0.65 | 0.253 | 0.323 |
| chrX | 154296038 | 154300789 | 4752 | 0.17 | 0.253 | 0.323 |
| chr7 | 73184297 | 73189760 | 5464 | -0.38 | 0.253 | 0.323 |
| chr18 | 34871011 | 34880514 | 9504 | -0.16 | 0.253 | 0.323 |

| | | | | | | |
|---|---|---|---|---|---|---|
| chr3 | 172035115 | 172073744 | 38630 | 0.13 | 0.253 | 0.323 |
| chr12 | 12546939 | 12568936 | 21998 | 0.11 | 0.253 | 0.323 |
| chr18 | 21961047 | 21961529 | 483 | -0.81 | 0.253 | 0.323 |
| chr7 | 35748826 | 35749184 | 359 | 0.69 | 0.253 | 0.324 |
| chr12 | 96160619 | 96161307 | 689 | -0.81 | 0.253 | 0.324 |
| chr12 | 17862097 | 17862848 | 752 | 0.48 | 0.253 | 0.324 |
| chr20 | 44509201 | 44520757 | 11557 | 0.2 | 0.254 | 0.324 |
| chr17 | 30588026 | 30589206 | 1181 | 0.64 | 0.254 | 0.324 |
| chr10 | 94246051 | 94250393 | 4343 | -0.3 | 0.254 | 0.324 |
| chrX | 44287709 | 44291109 | 3401 | -0.31 | 0.254 | 0.324 |
| chr16 | 11961133 | 11975181 | 14049 | -0.19 | 0.254 | 0.324 |
| chr20 | 44959975 | 44961127 | 1153 | 1.98 | 0.254 | 0.324 |
| chr19 | 10974844 | 11022694 | 47851 | -0.12 | 0.254 | 0.325 |
| chr5 | 180227996 | 180239051 | 11056 | 0.12 | 0.254 | 0.325 |
| chr1 | 1333573 | 1344117 | 10545 | -0.18 | 0.254 | 0.325 |
| chr22 | 19047504 | 19048509 | 1006 | -0.55 | 0.254 | 0.325 |
| chr7 | 140267266 | 140267997 | 732 | -0.69 | 0.254 | 0.325 |
| chr10 | 111985376 | 111986147 | 772 | -0.53 | 0.254 | 0.325 |
| chr14 | 74058279 | 74059298 | 1020 | 0.48 | 0.254 | 0.325 |
| chr11 | 74154027 | 74154564 | 538 | 0.85 | 0.254 | 0.325 |
| chr2 | 99988659 | 99989911 | 1253 | -0.52 | 0.254 | 0.325 |
| chrX | 53706053 | 53714070 | 8018 | 0.12 | 0.254 | 0.325 |
| chr1 | 78223726 | 78226094 | 2369 | 0.26 | 0.254 | 0.325 |
| chr9 | 90885372 | 90887109 | 1738 | 0.34 | 0.254 | 0.325 |
| chr19 | 51227609 | 51229343 | 1735 | 0.32 | 0.254 | 0.325 |
| chr1 | 31190316 | 31192218 | 1903 | 0.24 | 0.255 | 0.325 |
| chr4 | 129000141 | 129001010 | 870 | 0.76 | 0.255 | 0.325 |
| chr3 | 49313785 | 49315615 | 1831 | -0.31 | 0.255 | 0.325 |
| chr8 | 74291735 | 74293355 | 1621 | 0.31 | 0.255 | 0.325 |
| chr2 | 175128518 | 175129301 | 784 | 0.79 | 0.255 | 0.325 |
| chr6 | 156102429 | 156105059 | 2631 | -0.18 | 0.255 | 0.325 |
| chr1 | 62587138 | 62588478 | 1341 | 0.47 | 0.255 | 0.325 |
| chr19 | 49215747 | 49250966 | 35220 | 0.11 | 0.255 | 0.326 |
| chr1 | 109233876 | 109237476 | 3601 | 0.16 | 0.255 | 0.326 |
| chr22 | 41272902 | 41273915 | 1014 | 0.62 | 0.255 | 0.326 |
| chr1 | 94928678 | 94930142 | 1465 | -0.58 | 0.255 | 0.326 |
| chr4 | 14836667 | 14837600 | 934 | 0.87 | 0.255 | 0.326 |
| chr10 | 112325850 | 112328754 | 2905 | -0.12 | 0.256 | 0.326 |
| chr6 | 118971151 | 118973521 | 2371 | 0.27 | 0.256 | 0.326 |
| chr4 | 185304559 | 185305846 | 1288 | -0.52 | 0.256 | 0.326 |
| chr7 | 30265332 | 30266092 | 761 | 0.56 | 0.256 | 0.326 |
| chr10 | 32266460 | 32267566 | 1107 | 0.46 | 0.256 | 0.326 |
| chrX | 55025778 | 55035777 | 10000 | 0.14 | 0.256 | 0.326 |
| chr16 | 58209723 | 58231673 | 21951 | -0.17 | 0.256 | 0.326 |
| chr19 | 10341370 | 10342582 | 1213 | 0.48 | 0.256 | 0.326 |
| chr2 | 220385749 | 220386932 | 1184 | -0.61 | 0.256 | 0.327 |
| chr19 | 17579737 | 17587029 | 7293 | -0.24 | 0.256 | 0.327 |
| chr5 | 76345429 | 76346079 | 651 | 0.66 | 0.256 | 0.327 |
| chr7 | 110832386 | 110833338 | 953 | -0.53 | 0.256 | 0.327 |
| chr1 | 53128037 | 53129241 | 1205 | 0.33 | 0.256 | 0.327 |
| chr17 | 63118313 | 63120672 | 2360 | 0.2 | 0.256 | 0.327 |
| chr2 | 215671802 | 215675591 | 3790 | 0.2 | 0.256 | 0.327 |
| chr14 | 60798110 | 60798611 | 502 | 0.79 | 0.257 | 0.327 |
| chr1 | 27094582 | 27099663 | 5082 | 0.21 | 0.257 | 0.327 |
| chr6 | 106757532 | 106758651 | 1120 | 0.94 | 0.257 | 0.327 |
| chr16 | 66666161 | 66666664 | 504 | 0.81 | 0.257 | 0.327 |
| chr11 | 59364093 | 59365140 | 1048 | -0.56 | 0.257 | 0.327 |
| chr7 | 35669695 | 35670716 | 1022 | -0.62 | 0.257 | 0.327 |
| chr8 | 52837403 | 52838191 | 789 | -1.9 | 0.257 | 0.328 |
| chr16 | 56482084 | 56497487 | 15404 | 0.17 | 0.257 | 0.328 |
| chr3 | 135232613 | 135233325 | 713 | -0.78 | 0.257 | 0.328 |

| | | | | | | |
|---|---|---|---|---|---|---|
| chr19 | 19753798 | 19754846 | 1049 | 0.4 | 0.257 | 0.328 |
| chr11 | 63933052 | 63934815 | 1764 | -0.32 | 0.257 | 0.328 |
| chr3 | 171843601 | 171894913 | 51313 | -0.06 | 0.257 | 0.328 |
| chrX | 76911995 | 76927779 | 15785 | 0.14 | 0.257 | 0.328 |
| chr7 | 140304258 | 140305457 | 1200 | -0.6 | 0.257 | 0.328 |
| chr9 | 108418179 | 108419064 | 886 | 0.56 | 0.258 | 0.328 |
| chrX | 41840541 | 41841205 | 665 | -0.7 | 0.258 | 0.328 |
| chr7 | 124537346 | 124538008 | 663 | -0.73 | 0.258 | 0.328 |
| chr10 | 58297613 | 58298563 | 951 | 1.96 | 0.258 | 0.328 |
| chr7 | 143005317 | 143005767 | 451 | -0.86 | 0.258 | 0.328 |
| chr9 | 136241026 | 136243688 | 2663 | 0.2 | 0.258 | 0.329 |
| chr5 | 71682767 | 71699564 | 16798 | -0.63 | 0.258 | 0.329 |
| chr10 | 34164841 | 34166345 | 1505 | -1.62 | 0.258 | 0.329 |
| chr4 | 148521031 | 148521639 | 609 | -0.84 | 0.258 | 0.329 |
| chr12 | 34442605 | 34443826 | 1222 | 0.47 | 0.258 | 0.329 |
| chr15 | 62228532 | 62229026 | 495 | -0.56 | 0.258 | 0.329 |
| chr2 | 227699093 | 227704622 | 5530 | 0.15 | 0.258 | 0.329 |
| chr8 | 57285529 | 57286265 | 737 | -0.56 | 0.258 | 0.329 |
| chr13 | 77900526 | 77904903 | 4378 | 0.23 | 0.259 | 0.329 |
| chr15 | 57884112 | 57884620 | 509 | 0.8 | 0.259 | 0.329 |
| chr2 | 69055863 | 69057075 | 1213 | 0.45 | 0.259 | 0.329 |
| chr20 | 57555661 | 57583033 | 27373 | -0.13 | 0.259 | 0.329 |
| chr7 | 33922936 | 33923492 | 557 | -0.69 | 0.259 | 0.33 |
| chr9 | 34440973 | 34441552 | 580 | -0.69 | 0.259 | 0.33 |
| chr15 | 22832774 | 22835368 | 2595 | 0.18 | 0.259 | 0.33 |
| chr3 | 88080441 | 88081689 | 1249 | 0.3 | 0.259 | 0.33 |
| chr10 | 60094563 | 60108774 | 14212 | -0.27 | 0.259 | 0.33 |
| chr21 | 44580900 | 44599311 | 18412 | 0.11 | 0.26 | 0.33 |
| chr1 | 217833507 | 217834770 | 1264 | 0.47 | 0.26 | 0.331 |
| chr15 | 65742589 | 65753984 | 11396 | 0.24 | 0.26 | 0.331 |
| chr6 | 125444928 | 125445876 | 949 | -0.54 | 0.26 | 0.331 |
| chr3 | 106271218 | 106280603 | 9386 | -0.39 | 0.26 | 0.331 |
| chr7 | 191836 | 192401 | 566 | 0.58 | 0.26 | 0.331 |
| chr7 | 134895737 | 134897139 | 1403 | 0.33 | 0.26 | 0.331 |
| chr3 | 48213814 | 48214400 | 587 | 1.07 | 0.26 | 0.331 |
| chr13 | 40229356 | 40230535 | 1180 | 0.46 | 0.26 | 0.331 |
| chr9 | 88940669 | 88941560 | 892 | -0.63 | 0.26 | 0.331 |
| chr10 | 21869915 | 21871378 | 1464 | -0.47 | 0.26 | 0.331 |
| chr19 | 40501793 | 40504013 | 2221 | 0.13 | 0.26 | 0.331 |
| chr9 | 139257107 | 139267887 | 10781 | -0.25 | 0.26 | 0.331 |
| chr17 | 70818578 | 70819386 | 809 | 1.88 | 0.26 | 0.331 |
| chr9 | 33071729 | 33082214 | 10486 | 0.13 | 0.26 | 0.331 |
| chrX | 118826258 | 118827444 | 1187 | -0.41 | 0.261 | 0.332 |
| chr14 | 102227505 | 102227823 | 319 | 0.78 | 0.261 | 0.332 |
| chr19 | 8570589 | 8591647 | 21059 | 0.2 | 0.261 | 0.332 |
| chr17 | 20189749 | 20191087 | 1339 | -0.56 | 0.261 | 0.332 |
| chr7 | 93716716 | 93718355 | 1640 | 0.33 | 0.261 | 0.332 |
| chr3 | 14410911 | 14415349 | 4439 | 0.26 | 0.261 | 0.332 |
| chr22 | 25844024 | 25845584 | 1561 | 0.37 | 0.261 | 0.332 |
| chr10 | 70205625 | 70244607 | 38983 | -0.13 | 0.261 | 0.332 |
| chr14 | 91090438 | 91091403 | 966 | 0.69 | 0.261 | 0.332 |
| chr14 | 101878495 | 101879986 | 1492 | 0.37 | 0.261 | 0.332 |
| chr1 | 21058162 | 21115575 | 57414 | -0.07 | 0.261 | 0.333 |
| chr10 | 14115963 | 14116490 | 528 | 0.65 | 0.262 | 0.333 |
| chr9 | 86152838 | 86153695 | 858 | -0.61 | 0.262 | 0.333 |
| chr11 | 59577304 | 59578902 | 1599 | 0.25 | 0.262 | 0.333 |
| chr14 | 30395865 | 30397577 | 1713 | -0.5 | 0.262 | 0.333 |
| chr1 | 26560021 | 26566789 | 6769 | 0.19 | 0.262 | 0.333 |
| chr2 | 32053614 | 32066214 | 12601 | -0.43 | 0.262 | 0.333 |
| chr1 | 186291169 | 186292851 | 1683 | 0.69 | 0.262 | 0.333 |
| chr8 | 74378788 | 74379321 | 534 | -0.7 | 0.262 | 0.333 |

| | | | | | | |
|---|---|---|---|---|---|---|
| chr17 | 48276496 | 48288663 | 12168 | 1.29 | 0.262 | 0.334 |
| chr18 | 46693200 | 46693830 | 631 | 0.68 | 0.262 | 0.334 |
| chr20 | 47752543 | 47753311 | 769 | -0.58 | 0.263 | 0.334 |
| chr3 | 121464497 | 121469904 | 5408 | 0.1 | 0.263 | 0.334 |
| chr5 | 112213894 | 112220962 | 7069 | -0.24 | 0.263 | 0.334 |
| chr22 | 27634559 | 27649130 | 14572 | -0.2 | 0.263 | 0.334 |
| chr10 | 22215563 | 22216312 | 750 | 0.65 | 0.263 | 0.334 |
| chr4 | 85200262 | 85201423 | 1162 | 0.43 | 0.263 | 0.334 |
| chrX | 15483299 | 15484275 | 977 | 0.64 | 0.263 | 0.334 |
| chr15 | 44760656 | 44761606 | 951 | -0.63 | 0.263 | 0.334 |
| chr10 | 3290469 | 3291859 | 1391 | 0.23 | 0.263 | 0.334 |
| chrX | 71773419 | 71775164 | 1746 | -0.23 | 0.263 | 0.335 |
| chr9 | 120426518 | 120427274 | 757 | -0.73 | 0.263 | 0.335 |
| chr18 | 21148088 | 21174862 | 26775 | -0.12 | 0.263 | 0.335 |
| chr11 | 110104666 | 110107699 | 3034 | 0.29 | 0.263 | 0.335 |
| chr11 | 94570753 | 94571725 | 973 | 0.54 | 0.264 | 0.335 |
| chr9 | 74559318 | 74560519 | 1202 | 0.66 | 0.264 | 0.335 |
| chr2 | 197179259 | 197181420 | 2162 | 0.26 | 0.264 | 0.335 |
| chr3 | 124750871 | 124752143 | 1273 | -0.39 | 0.264 | 0.335 |
| chr3 | 123602570 | 123603786 | 1217 | -0.65 | 0.264 | 0.335 |
| chr1 | 45792219 | 45807149 | 14931 | -0.15 | 0.264 | 0.335 |
| chr17 | 14664072 | 14664879 | 808 | 0.47 | 0.264 | 0.335 |
| chr2 | 32682016 | 32718648 | 36633 | 0.13 | 0.264 | 0.335 |
| chr20 | 32597024 | 32598378 | 1355 | 0.53 | 0.264 | 0.335 |
| chr6 | 3868832 | 3889779 | 20948 | 0.17 | 0.264 | 0.335 |
| chr6 | 163148074 | 163149887 | 1814 | 0.26 | 0.264 | 0.335 |
| chr3 | 125985593 | 125987000 | 1408 | 0.36 | 0.264 | 0.336 |
| chr4 | 102042894 | 102043771 | 878 | -0.61 | 0.264 | 0.336 |
| chr22 | 18506885 | 18521242 | 14358 | -0.23 | 0.264 | 0.336 |
| chr1 | 100730924 | 100733027 | 2104 | 0.29 | 0.264 | 0.336 |
| chr16 | 420693 | 431706 | 11014 | -0.39 | 0.264 | 0.336 |
| chr11 | 71144658 | 71145222 | 565 | 1.05 | 0.265 | 0.336 |
| chr19 | 14660891 | 14662610 | 1720 | 0.42 | 0.265 | 0.336 |
| chr4 | 136485246 | 136485878 | 633 | -0.76 | 0.265 | 0.336 |
| chr8 | 135697032 | 135698428 | 1397 | 0.53 | 0.265 | 0.336 |
| chr3 | 47130578 | 47131779 | 1202 | 0.56 | 0.265 | 0.336 |
| chr1 | 180959200 | 180960840 | 1641 | -0.46 | 0.265 | 0.336 |
| chr13 | 99473050 | 99478720 | 5671 | 0.29 | 0.265 | 0.337 |
| chr11 | 65478224 | 65488300 | 10077 | 0.15 | 0.265 | 0.337 |
| chr16 | 1592909 | 1593792 | 884 | -0.63 | 0.265 | 0.337 |
| chr12 | 88828120 | 88828611 | 492 | -1.18 | 0.265 | 0.337 |
| chr1 | 44678315 | 44679760 | 1446 | 0.38 | 0.265 | 0.337 |
| chr2 | 234074308 | 234075825 | 1518 | 0.33 | 0.265 | 0.337 |
| chr4 | 102108491 | 102153839 | 45349 | -0.07 | 0.265 | 0.337 |
| chr7 | 66366350 | 66371444 | 5095 | -1.02 | 0.266 | 0.337 |
| chr2 | 106234050 | 106234354 | 305 | 0.95 | 0.266 | 0.337 |
| chr2 | 196440420 | 196441556 | 1137 | -0.4 | 0.266 | 0.337 |
| chr16 | 11752731 | 11753307 | 577 | -0.77 | 0.266 | 0.337 |
| chr1 | 235904385 | 235904684 | 300 | 1.21 | 0.266 | 0.337 |
| chr10 | 94106076 | 94107078 | 1003 | 0.52 | 0.266 | 0.338 |
| chr15 | 57510958 | 57512889 | 1932 | -0.51 | 0.266 | 0.338 |
| chr16 | 11035978 | 11043908 | 7931 | 0.29 | 0.266 | 0.338 |
| chr2 | 240627892 | 240628471 | 580 | 0.73 | 0.266 | 0.338 |
| chr18 | 74205330 | 74205863 | 534 | 0.69 | 0.266 | 0.338 |
| chr19 | 54634457 | 54715756 | 81300 | 0.06 | 0.266 | 0.338 |
| chr19 | 46945680 | 46947496 | 1817 | 0.23 | 0.267 | 0.338 |
| chr11 | 91397768 | 91398439 | 672 | -0.59 | 0.267 | 0.338 |
| chr14 | 59935387 | 59959098 | 23712 | -0.09 | 0.267 | 0.339 |
| chr6 | 10048390 | 10049732 | 1343 | -0.51 | 0.267 | 0.339 |
| chr9 | 128222998 | 128223458 | 461 | -0.73 | 0.267 | 0.339 |
| chrX | 107068992 | 107070538 | 1547 | 0.31 | 0.267 | 0.339 |

| | | | | | | |
|---|---|---|---|---|---|---|
| chr3 | 192536609 | 192537329 | 721 | -0.64 | 0.267 | 0.339 |
| chr7 | 6616223 | 6631033 | 14811 | 0.16 | 0.267 | 0.339 |
| chr15 | 52848626 | 52863928 | 15303 | 0.11 | 0.267 | 0.339 |
| chr10 | 43930335 | 43952341 | 22007 | 0.09 | 0.267 | 0.339 |
| chr17 | 70366807 | 70368637 | 1831 | -0.37 | 0.267 | 0.339 |
| chr5 | 148608350 | 148608694 | 345 | -0.62 | 0.267 | 0.339 |
| chr7 | 148714880 | 148726889 | 12010 | -0.13 | 0.267 | 0.339 |
| chr2 | 39333773 | 39365524 | 31752 | 0.09 | 0.267 | 0.339 |
| chr1 | 11158559 | 11160664 | 2106 | 0.24 | 0.267 | 0.339 |
| chr8 | 128519038 | 128522838 | 3801 | 0.25 | 0.268 | 0.339 |
| chr2 | 2616716 | 2617820 | 1105 | 0.3 | 0.268 | 0.339 |
| chr2 | 84670094 | 84670860 | 767 | -0.71 | 0.268 | 0.339 |
| chr21 | 47742492 | 47746252 | 3761 | 0.23 | 0.268 | 0.339 |
| chr9 | 107592104 | 107593010 | 907 | -0.78 | 0.268 | 0.339 |
| chr19 | 16959065 | 16960815 | 1751 | 0.42 | 0.268 | 0.34 |
| chr22 | 27613903 | 27615030 | 1128 | -0.87 | 0.268 | 0.34 |
| chr1 | 95368636 | 95369432 | 797 | -0.66 | 0.268 | 0.34 |
| chr4 | 185557112 | 185571844 | 14733 | 0.21 | 0.268 | 0.34 |
| chr2 | 103352619 | 103354149 | 1531 | -0.26 | 0.268 | 0.34 |
| chr2 | 44434877 | 44435757 | 881 | 0.72 | 0.268 | 0.34 |
| chr21 | 44393574 | 44407258 | 13685 | 0.16 | 0.268 | 0.34 |
| chr5 | 55111624 | 55118205 | 6582 | -0.51 | 0.268 | 0.34 |
| chr11 | 57169314 | 57170236 | 923 | 1.84 | 0.268 | 0.34 |
| chr7 | 100826023 | 100826376 | 354 | 0.65 | 0.268 | 0.34 |
| chr16 | 50731827 | 50732293 | 467 | 0.74 | 0.268 | 0.34 |
| chr17 | 47392993 | 47394902 | 1910 | -0.31 | 0.268 | 0.34 |
| chr4 | 152125823 | 152128591 | 2769 | 0.35 | 0.268 | 0.34 |
| chr10 | 14918514 | 14922043 | 3530 | 0.19 | 0.268 | 0.34 |
| chr9 | 4831468 | 4835543 | 4076 | -0.26 | 0.269 | 0.34 |
| chr16 | 46823437 | 46824541 | 1105 | 0.51 | 0.269 | 0.34 |
| chr10 | 23216543 | 23217548 | 1006 | -0.25 | 0.269 | 0.34 |
| chr3 | 133524663 | 133525904 | 1242 | 0.43 | 0.269 | 0.34 |
| chr7 | 137276101 | 137277019 | 919 | -0.55 | 0.269 | 0.34 |
| chr1 | 29487074 | 29489520 | 2447 | 0.41 | 0.269 | 0.34 |
| chr16 | 66467276 | 66468720 | 1445 | -0.48 | 0.269 | 0.34 |
| chr21 | 39675087 | 39679374 | 4288 | 0.3 | 0.269 | 0.34 |
| chr4 | 95159508 | 95160759 | 1252 | 0.55 | 0.269 | 0.34 |
| chr10 | 30889339 | 30906723 | 17385 | 0.12 | 0.269 | 0.34 |
| chr2 | 24712771 | 24714544 | 1774 | 0.27 | 0.269 | 0.341 |
| chr1 | 64637163 | 64638721 | 1559 | -0.36 | 0.269 | 0.341 |
| chr20 | 56949440 | 56949998 | 559 | -0.69 | 0.269 | 0.341 |
| chr20 | 50582481 | 50583011 | 531 | 0.69 | 0.269 | 0.341 |
| chr1 | 219380561 | 219383825 | 3265 | -0.23 | 0.269 | 0.341 |
| chr10 | 94076241 | 94089962 | 13722 | -0.36 | 0.269 | 0.341 |
| chr7 | 44864128 | 44865150 | 1023 | 0.61 | 0.269 | 0.341 |
| chr7 | 84412546 | 84413181 | 636 | 0.55 | 0.269 | 0.341 |
| chr15 | 90222871 | 90234524 | 11654 | -0.31 | 0.269 | 0.341 |
| chr11 | 65292548 | 65293434 | 887 | -0.44 | 0.269 | 0.341 |
| chr4 | 164086461 | 164088860 | 2400 | 0.16 | 0.27 | 0.341 |
| chr2 | 207426813 | 207427596 | 784 | -0.76 | 0.27 | 0.341 |
| chr7 | 102714179 | 102717098 | 2920 | 0.15 | 0.27 | 0.341 |
| chr3 | 121241780 | 121242438 | 659 | 0.65 | 0.27 | 0.341 |
| chr11 | 108534935 | 108536593 | 1659 | 0.32 | 0.27 | 0.341 |
| chr8 | 141827659 | 141865782 | 38124 | 0.1 | 0.27 | 0.341 |
| chr12 | 34175430 | 34176029 | 600 | 0.46 | 0.27 | 0.341 |
| chr7 | 31028519 | 31029467 | 949 | 0.94 | 0.27 | 0.341 |
| chr9 | 117313972 | 117315182 | 1211 | -0.55 | 0.27 | 0.341 |
| chr3 | 183103037 | 183103782 | 746 | 0.71 | 0.27 | 0.342 |
| chr2 | 201371576 | 201375280 | 3705 | -0.22 | 0.27 | 0.342 |
| chr4 | 8279934 | 8280686 | 753 | -0.6 | 0.27 | 0.342 |
| chr17 | 4347718 | 4348615 | 898 | 0.59 | 0.27 | 0.342 |

| | | | | | | |
|---|---|---|---|---|---|---|
| chr16 | 25197808 | 25199077 | 1270 | -0.45 | 0.27 | 0.342 |
| chr6 | 128489010 | 128489566 | 557 | 0.6 | 0.27 | 0.342 |
| chr7 | 17224545 | 17225750 | 1206 | 1.49 | 0.27 | 0.342 |
| chr4 | 38932773 | 38933262 | 490 | -0.76 | 0.27 | 0.342 |
| chr11 | 35843224 | 35844285 | 1062 | 0.46 | 0.271 | 0.342 |
| chr4 | 88923393 | 88940060 | 16668 | 0.22 | 0.271 | 0.342 |
| chr2 | 184043615 | 184044379 | 765 | -0.66 | 0.271 | 0.342 |
| chr8 | 141454511 | 141479975 | 25465 | 0.08 | 0.271 | 0.342 |
| chr12 | 40963268 | 40964363 | 1096 | -0.46 | 0.271 | 0.342 |
| chr22 | 30555062 | 30556553 | 1492 | -0.33 | 0.271 | 0.342 |
| chr4 | 108604558 | 108605661 | 1104 | 0.63 | 0.271 | 0.342 |
| chr15 | 85447203 | 85474671 | 27469 | 0.07 | 0.271 | 0.342 |
| chr13 | 33001844 | 33002696 | 853 | 0.9 | 0.271 | 0.342 |
| chr1 | 202308641 | 202319638 | 10998 | -0.14 | 0.271 | 0.343 |
| chr10 | 70553175 | 70554096 | 922 | -0.67 | 0.271 | 0.343 |
| chr15 | 65647805 | 65648384 | 580 | -1.07 | 0.271 | 0.343 |
| chr15 | 71983822 | 71985094 | 1273 | 1.91 | 0.272 | 0.343 |
| chr12 | 105143572 | 105144191 | 620 | 1.02 | 0.272 | 0.343 |
| chr11 | 13282241 | 13284136 | 1896 | -0.24 | 0.272 | 0.343 |
| chr11 | 129764856 | 129765445 | 590 | 0.49 | 0.272 | 0.344 |
| chrX | 67574521 | 67576559 | 2039 | 0.27 | 0.272 | 0.344 |
| chr3 | 138326946 | 138329092 | 2147 | -0.16 | 0.272 | 0.344 |
| chr19 | 34744518 | 34761938 | 17421 | -0.1 | 0.272 | 0.344 |
| chr7 | 107103805 | 107104684 | 880 | -0.57 | 0.272 | 0.344 |
| chr8 | 30914542 | 30915531 | 990 | 0.48 | 0.272 | 0.344 |
| chr5 | 57285210 | 57286159 | 950 | 0.52 | 0.272 | 0.344 |
| chrX | 21023213 | 21023573 | 361 | 0.72 | 0.272 | 0.344 |
| chr12 | 112818788 | 112826052 | 7265 | 0.23 | 0.272 | 0.344 |
| chr1 | 167373158 | 167374064 | 907 | -0.7 | 0.272 | 0.344 |
| chr15 | 99994929 | 99995663 | 735 | 0.64 | 0.273 | 0.344 |
| chr2 | 10913995 | 10914875 | 881 | -0.84 | 0.273 | 0.344 |
| chr11 | 3718352 | 3776115 | 57764 | 0.09 | 0.273 | 0.344 |
| chr4 | 14608992 | 14609962 | 971 | 0.49 | 0.273 | 0.344 |
| chr2 | 178075335 | 178141000 | 65666 | -0.05 | 0.273 | 0.345 |
| chr7 | 76935190 | 76936125 | 936 | -0.45 | 0.273 | 0.345 |
| chr16 | 27297577 | 27298482 | 906 | -0.49 | 0.273 | 0.345 |
| chr1 | 242183134 | 242183788 | 655 | -0.58 | 0.273 | 0.345 |
| chr19 | 12832278 | 12849313 | 17036 | -0.14 | 0.273 | 0.345 |
| chr18 | 56483023 | 56483916 | 894 | -0.67 | 0.273 | 0.345 |
| chr3 | 16014829 | 16015866 | 1038 | -0.56 | 0.273 | 0.345 |
| chr7 | 38861927 | 38862827 | 901 | -0.61 | 0.273 | 0.345 |
| chr7 | 68263233 | 68279584 | 16352 | -0.17 | 0.273 | 0.345 |
| chr16 | 88636035 | 88638001 | 1967 | 0.31 | 0.274 | 0.345 |
| chr7 | 104992256 | 104993667 | 1412 | 0.36 | 0.274 | 0.345 |
| chrX | 118698508 | 118720148 | 21641 | 0.13 | 0.274 | 0.345 |
| chr19 | 46849169 | 46930537 | 81369 | 0.06 | 0.274 | 0.346 |
| chr6 | 128460205 | 128461313 | 1109 | 0.66 | 0.274 | 0.346 |
| chr7 | 116032695 | 116040841 | 8147 | -0.11 | 0.274 | 0.346 |
| chr15 | 58561983 | 58562971 | 989 | 0.49 | 0.274 | 0.346 |
| chr20 | 813429 | 837912 | 24484 | 0.16 | 0.274 | 0.346 |
| chr16 | 53302587 | 53318042 | 15456 | 0.2 | 0.274 | 0.346 |
| chr4 | 169803331 | 169805063 | 1733 | 0.52 | 0.274 | 0.346 |
| chr6 | 116574376 | 116576112 | 1737 | 0.19 | 0.274 | 0.346 |
| chrX | 23970915 | 23972103 | 1189 | -0.32 | 0.274 | 0.346 |
| chr5 | 127397746 | 127398441 | 696 | 0.63 | 0.274 | 0.346 |
| chr5 | 38555966 | 38557926 | 1961 | -0.24 | 0.274 | 0.346 |
| chr21 | 43915632 | 43917420 | 1789 | 0.27 | 0.274 | 0.346 |
| chr5 | 149942043 | 149942525 | 483 | 0.91 | 0.274 | 0.346 |
| chr16 | 27359041 | 27359581 | 541 | -0.84 | 0.274 | 0.346 |
| chr6 | 113340091 | 113340627 | 537 | 0.68 | 0.274 | 0.346 |
| chr14 | 39493136 | 39493908 | 773 | -0.6 | 0.275 | 0.346 |

| | | | | | | |
|---|---|---|---|---|---|---|
| chr11 | 120294374 | 120304027 | 9654 | 0.17 | 0.275 | 0.347 |
| chr11 | 129993677 | 130017315 | 23639 | -0.18 | 0.275 | 0.347 |
| chr1 | 36668871 | 36669885 | 1015 | -0.51 | 0.275 | 0.347 |
| chr2 | 196923853 | 196934266 | 10414 | -0.19 | 0.275 | 0.347 |
| chr22 | 25795703 | 25802090 | 6388 | -0.18 | 0.275 | 0.347 |
| chr11 | 101841730 | 101846117 | 4388 | 0.28 | 0.275 | 0.347 |
| chr3 | 39136752 | 39150178 | 13427 | -0.13 | 0.275 | 0.347 |
| chr19 | 48352173 | 48352982 | 810 | 0.54 | 0.275 | 0.347 |
| chr9 | 130092096 | 130100442 | 8347 | 0.14 | 0.275 | 0.347 |
| chr3 | 101442615 | 101445673 | 3059 | 0.19 | 0.275 | 0.347 |
| chr9 | 86237303 | 86239526 | 2224 | 0.17 | 0.275 | 0.347 |
| chr6 | 139837283 | 139838152 | 870 | 0.43 | 0.275 | 0.347 |
| chr3 | 123371812 | 123372677 | 866 | -0.59 | 0.276 | 0.347 |
| chr6 | 37400221 | 37401824 | 1604 | 0.24 | 0.276 | 0.347 |
| chr9 | 36036600 | 36037878 | 1279 | -0.39 | 0.276 | 0.347 |
| chr5 | 159591217 | 159593969 | 2753 | 0.14 | 0.276 | 0.348 |
| chr19 | 19143191 | 19145658 | 2468 | 0.23 | 0.276 | 0.348 |
| chr15 | 41035971 | 41056875 | 20905 | 0.18 | 0.276 | 0.348 |
| chr15 | 83500042 | 83502362 | 2321 | 0.34 | 0.276 | 0.348 |
| chr5 | 150099726 | 150115128 | 15403 | 0.26 | 0.276 | 0.348 |
| chr19 | 35454127 | 35455405 | 1279 | -0.3 | 0.276 | 0.348 |
| chr18 | 12990307 | 12992844 | 2538 | 0.21 | 0.276 | 0.348 |
| chr8 | 82542738 | 82543434 | 697 | 0.67 | 0.276 | 0.348 |
| chr16 | 27245807 | 27281006 | 35200 | 0.15 | 0.276 | 0.348 |
| chr7 | 135346575 | 135348740 | 2166 | 0.2 | 0.276 | 0.348 |
| chr6 | 52134264 | 52152361 | 18098 | 0.16 | 0.276 | 0.348 |
| chr10 | 103566243 | 103591804 | 25562 | -0.11 | 0.276 | 0.348 |
| chr1 | 180198425 | 180201306 | 2882 | 0.27 | 0.276 | 0.348 |
| chr15 | 70563375 | 70564603 | 1229 | -0.38 | 0.277 | 0.349 |
| chr5 | 16600463 | 16601253 | 791 | -0.65 | 0.277 | 0.349 |
| chr5 | 131088280 | 131089342 | 1063 | -0.49 | 0.277 | 0.349 |
| chr14 | 31070357 | 31071077 | 721 | -0.61 | 0.277 | 0.349 |
| chr12 | 118619627 | 118620173 | 547 | 0.65 | 0.277 | 0.349 |
| chr11 | 19622263 | 19622921 | 659 | -0.94 | 0.277 | 0.349 |
| chr6 | 43111514 | 43111840 | 327 | -0.76 | 0.277 | 0.349 |
| chr11 | 104788694 | 104789271 | 578 | 0.6 | 0.277 | 0.349 |
| chr1 | 234040445 | 234056759 | 16315 | 0.09 | 0.277 | 0.349 |
| chr4 | 77674052 | 77675380 | 1329 | -0.47 | 0.277 | 0.349 |
| chr4 | 94304947 | 94307058 | 2112 | 0.28 | 0.277 | 0.349 |
| chr11 | 118659543 | 118664254 | 4712 | 0.13 | 0.277 | 0.349 |
| chr2 | 135908470 | 135909577 | 1108 | -0.66 | 0.277 | 0.349 |
| chr1 | 26704296 | 26705188 | 893 | 0.67 | 0.277 | 0.349 |
| chr15 | 50963423 | 50980212 | 16790 | 0.12 | 0.277 | 0.349 |
| chr1 | 161195824 | 161196398 | 575 | 0.57 | 0.277 | 0.349 |
| chr1 | 40104921 | 40106316 | 1396 | 0.35 | 0.278 | 0.349 |
| chr3 | 136557309 | 136558897 | 1589 | -0.37 | 0.278 | 0.349 |
| chr1 | 229685889 | 229695686 | 9798 | -0.24 | 0.278 | 0.349 |
| chr1 | 54007038 | 54008650 | 1613 | 0.33 | 0.278 | 0.35 |
| chr7 | 138542862 | 138583868 | 41007 | 0.07 | 0.278 | 0.35 |
| chr3 | 48125970 | 48132108 | 6139 | 0.13 | 0.278 | 0.35 |
| chr11 | 125876815 | 125877511 | 697 | 0.8 | 0.278 | 0.35 |
| chr2 | 121793545 | 121794433 | 889 | 0.58 | 0.278 | 0.35 |
| chr2 | 233213926 | 233214292 | 367 | 1.2 | 0.278 | 0.35 |
| chr14 | 81075330 | 81087264 | 11935 | 1.04 | 0.278 | 0.35 |
| chr17 | 74520946 | 74582301 | 61356 | -0.11 | 0.278 | 0.35 |
| chr13 | 46086035 | 46087118 | 1084 | 0.59 | 0.278 | 0.35 |
| chr6 | 82914866 | 82926416 | 11551 | 0.25 | 0.278 | 0.35 |
| chr6 | 3457532 | 3458024 | 493 | 0.63 | 0.278 | 0.35 |
| chr15 | 63969606 | 63970678 | 1073 | 0.37 | 0.278 | 0.35 |
| chr9 | 3848631 | 3876460 | 27830 | 0.13 | 0.278 | 0.35 |
| chr6 | 133133656 | 133142975 | 9320 | 0.14 | 0.279 | 0.351 |

| | | | | | | |
|---|---|---|---|---|---|---|
| chr8 | 94712271 | 94713589 | 1319 | -0.32 | 0.279 | 0.351 |
| chr12 | 17092713 | 17093755 | 1043 | 0.53 | 0.279 | 0.351 |
| chr1 | 61850827 | 61852905 | 2079 | -0.39 | 0.279 | 0.351 |
| chr3 | 48884257 | 48886102 | 1846 | 0.3 | 0.279 | 0.351 |
| chr6 | 13487154 | 13487779 | 626 | 0.63 | 0.279 | 0.351 |
| chr1 | 234593856 | 234594850 | 995 | 0.47 | 0.279 | 0.351 |
| chr20 | 39860918 | 39861782 | 865 | 0.5 | 0.279 | 0.351 |
| chr5 | 1633564 | 1634627 | 1064 | -0.45 | 0.279 | 0.351 |
| chr22 | 39527202 | 39527737 | 536 | 0.81 | 0.279 | 0.351 |
| chr17 | 42214701 | 42313072 | 98372 | -0.05 | 0.279 | 0.351 |
| chr14 | 62248946 | 62249844 | 899 | -0.56 | 0.279 | 0.352 |
| chr1 | 109463652 | 109464133 | 482 | 0.77 | 0.28 | 0.352 |
| chr12 | 65138287 | 65139346 | 1060 | 0.6 | 0.28 | 0.352 |
| chr13 | 98795479 | 98796378 | 900 | 0.63 | 0.28 | 0.352 |
| chr16 | 68248336 | 68346042 | 97707 | -0.06 | 0.28 | 0.352 |
| chr11 | 47421487 | 47449488 | 28002 | 0.11 | 0.28 | 0.352 |
| chr1 | 115677393 | 115677941 | 549 | -0.75 | 0.28 | 0.352 |
| chr1 | 40741525 | 40743760 | 2236 | -0.35 | 0.28 | 0.352 |
| chr20 | 43567068 | 43567884 | 817 | -0.56 | 0.28 | 0.352 |
| chr2 | 205527947 | 205532479 | 4533 | -0.12 | 0.28 | 0.352 |
| chr22 | 36357316 | 36358272 | 957 | -0.66 | 0.28 | 0.353 |
| chr6 | 82885427 | 82886847 | 1421 | 0.65 | 0.28 | 0.353 |
| chr2 | 203386529 | 203395914 | 9386 | -0.32 | 0.28 | 0.353 |
| chr9 | 113315433 | 113318818 | 3386 | 0.36 | 0.281 | 0.353 |
| chr14 | 43531069 | 43531615 | 547 | 0.76 | 0.281 | 0.353 |
| chr8 | 145100498 | 145100991 | 494 | 1.18 | 0.281 | 0.353 |
| chr3 | 48777199 | 48778380 | 1182 | 0.49 | 0.281 | 0.353 |
| chr2 | 220928553 | 220929976 | 1424 | 0.47 | 0.281 | 0.353 |
| chr3 | 160048648 | 160049825 | 1178 | 0.49 | 0.281 | 0.353 |
| chr3 | 13146719 | 13148786 | 2068 | 0.49 | 0.281 | 0.353 |
| chr1 | 205646683 | 205646978 | 296 | 1.2 | 0.281 | 0.353 |
| chr11 | 65378136 | 65384667 | 6532 | 0.14 | 0.281 | 0.353 |
| chr16 | 84177916 | 84190814 | 12899 | 0.69 | 0.281 | 0.353 |
| chr1 | 28479145 | 28480846 | 1702 | 0.34 | 0.281 | 0.353 |
| chr3 | 99844207 | 99844651 | 445 | 0.76 | 0.281 | 0.354 |
| chr10 | 28591448 | 28591889 | 442 | 1.17 | 0.281 | 0.354 |
| chr8 | 110326424 | 110326925 | 502 | -0.69 | 0.282 | 0.354 |
| chr12 | 131769445 | 131770001 | 557 | 0.71 | 0.282 | 0.354 |
| chr1 | 161014028 | 161017008 | 2981 | 0.14 | 0.282 | 0.354 |
| chr6 | 76310675 | 76324193 | 13519 | 0.11 | 0.282 | 0.354 |
| chr17 | 42572224 | 42581906 | 9683 | 0.12 | 0.282 | 0.354 |
| chr5 | 92914909 | 92921474 | 6566 | -0.33 | 0.282 | 0.354 |
| chr14 | 77227623 | 77228350 | 728 | -0.95 | 0.282 | 0.354 |
| chr12 | 51126143 | 51127148 | 1006 | -0.57 | 0.282 | 0.354 |
| chr22 | 23590201 | 23599910 | 9710 | -0.28 | 0.282 | 0.354 |
| chr2 | 122493025 | 122495414 | 2390 | 0.19 | 0.282 | 0.354 |
| chr16 | 48575696 | 48576223 | 528 | 0.78 | 0.282 | 0.355 |
| chr6 | 34758904 | 34779252 | 20349 | 0.19 | 0.282 | 0.355 |
| chr5 | 78596738 | 78597609 | 872 | 0.5 | 0.283 | 0.355 |
| chr2 | 65387998 | 65398143 | 10146 | -0.25 | 0.283 | 0.355 |
| chr10 | 90961158 | 90961641 | 484 | -0.81 | 0.283 | 0.355 |
| chr2 | 71503764 | 71504883 | 1120 | 0.61 | 0.283 | 0.355 |
| chr12 | 77425735 | 77426714 | 980 | -0.58 | 0.283 | 0.355 |
| chr20 | 10180973 | 10181432 | 460 | 0.57 | 0.283 | 0.355 |
| chr14 | 71066455 | 71068463 | 2009 | 0.19 | 0.283 | 0.355 |
| chr7 | 102104676 | 102106093 | 1418 | -0.29 | 0.283 | 0.356 |
| chr1 | 202731045 | 202731860 | 816 | 0.52 | 0.283 | 0.356 |
| chr19 | 4007790 | 4008241 | 452 | 0.71 | 0.284 | 0.356 |
| chr2 | 161288778 | 161290052 | 1275 | -0.53 | 0.284 | 0.356 |
| chr19 | 19050770 | 19052094 | 1325 | -0.36 | 0.284 | 0.356 |
| chr18 | 1406714 | 1407802 | 1089 | 0.41 | 0.284 | 0.356 |

| | | | | | | |
|---|---|---|---|---|---|---|
| chr4 | 104072116 | 104083423 | 11308 | 0.92 | 0.284 | 0.356 |
| chr7 | 116592521 | 116594974 | 2454 | 0.18 | 0.284 | 0.357 |
| chr7 | 66004703 | 66005600 | 898 | 0.63 | 0.284 | 0.357 |
| chrX | 152863833 | 152864517 | 685 | 0.57 | 0.284 | 0.357 |
| chr5 | 55574616 | 55575797 | 1182 | 0.57 | 0.284 | 0.357 |
| chr10 | 92719752 | 92732251 | 12500 | 0.29 | 0.284 | 0.357 |
| chr13 | 33692893 | 33701685 | 8793 | -0.18 | 0.284 | 0.357 |
| chr4 | 169059047 | 169060117 | 1071 | 0.48 | 0.284 | 0.357 |
| chr10 | 23341753 | 23342935 | 1183 | -0.34 | 0.284 | 0.357 |
| chr5 | 38869885 | 38888423 | 18539 | -0.09 | 0.285 | 0.357 |
| chr1 | 236162268 | 236165348 | 3081 | 0.55 | 0.285 | 0.357 |
| chr20 | 49120931 | 49127672 | 6742 | -0.18 | 0.285 | 0.357 |
| chr17 | 42486075 | 42495594 | 9520 | 0.31 | 0.285 | 0.357 |
| chr5 | 18698239 | 18698779 | 541 | 0.78 | 0.285 | 0.357 |
| chr10 | 31319396 | 31322117 | 2722 | 0.14 | 0.285 | 0.357 |
| chr14 | 64087987 | 64091170 | 3184 | -0.37 | 0.285 | 0.357 |
| chr9 | 136643696 | 136644435 | 740 | -0.61 | 0.285 | 0.358 |
| chr12 | 98745870 | 98747333 | 1464 | 0.35 | 0.285 | 0.358 |
| chr4 | 53885486 | 53886785 | 1300 | -0.51 | 0.285 | 0.358 |
| chr9 | 132671927 | 132673672 | 1746 | 0.44 | 0.285 | 0.358 |
| chr1 | 6473116 | 6475239 | 2124 | 0.42 | 0.285 | 0.358 |
| chr10 | 43781677 | 43784609 | 2933 | 0.2 | 0.285 | 0.358 |
| chr10 | 93714273 | 93715453 | 1181 | 0.48 | 0.285 | 0.358 |
| chr1 | 179256192 | 179283933 | 27742 | -0.15 | 0.285 | 0.358 |
| chr9 | 74289211 | 74383992 | 94782 | 0.05 | 0.286 | 0.358 |
| chr5 | 130923454 | 130923849 | 396 | -1.04 | 0.286 | 0.358 |
| chr22 | 19464732 | 19468130 | 3399 | -0.16 | 0.286 | 0.358 |
| chr1 | 55613057 | 55614003 | 947 | 0.5 | 0.286 | 0.358 |
| chr16 | 1020212 | 1020937 | 726 | 0.42 | 0.286 | 0.358 |
| chr8 | 71243756 | 71258365 | 14610 | 0.22 | 0.286 | 0.358 |
| chr6 | 79456017 | 79456763 | 747 | 0.66 | 0.286 | 0.358 |
| chr7 | 38235384 | 38264571 | 29188 | -0.1 | 0.286 | 0.358 |
| chr16 | 72932169 | 72933236 | 1068 | 0.61 | 0.286 | 0.359 |
| chr8 | 48088538 | 48100690 | 12153 | 2 | 0.286 | 0.359 |
| chr5 | 139553740 | 139562966 | 9227 | 0.1 | 0.286 | 0.359 |
| chr2 | 188312824 | 188313271 | 448 | 0.57 | 0.286 | 0.359 |
| chr7 | 6076521 | 6099809 | 23289 | -0.1 | 0.286 | 0.359 |
| chr5 | 138122251 | 138132749 | 10499 | 0.2 | 0.286 | 0.359 |
| chr10 | 86229143 | 86229821 | 679 | 0.63 | 0.287 | 0.359 |
| chrX | 53603634 | 53604117 | 484 | -0.74 | 0.287 | 0.359 |
| chr4 | 145566899 | 145567624 | 726 | -0.67 | 0.287 | 0.359 |
| chr3 | 172023190 | 172025481 | 2292 | -0.36 | 0.287 | 0.36 |
| chr2 | 238368394 | 238455753 | 87360 | -0.04 | 0.287 | 0.36 |
| chrX | 71351645 | 71352533 | 889 | -0.61 | 0.287 | 0.36 |
| chr18 | 34802556 | 34802915 | 360 | -0.7 | 0.287 | 0.36 |
| chrX | 15272346 | 15289682 | 17337 | 0.17 | 0.287 | 0.36 |
| chr20 | 25335466 | 25345725 | 10260 | -0.17 | 0.287 | 0.36 |
| chr4 | 47464713 | 47466373 | 1661 | 0.37 | 0.287 | 0.36 |
| chr1 | 211937165 | 211959844 | 22680 | -0.18 | 0.287 | 0.36 |
| chr1 | 59125389 | 59127150 | 1762 | 0.5 | 0.287 | 0.36 |
| chr9 | 111235013 | 111241514 | 6502 | -0.19 | 0.287 | 0.36 |
| chr14 | 74982501 | 74984091 | 1591 | -0.41 | 0.288 | 0.36 |
| chr19 | 13943498 | 13992005 | 48508 | 0.06 | 0.288 | 0.36 |
| chr19 | 7660795 | 7661433 | 639 | 0.82 | 0.288 | 0.36 |
| chr11 | 62520640 | 62540128 | 19489 | 0.16 | 0.288 | 0.36 |
| chr15 | 68548876 | 68550057 | 1182 | 0.33 | 0.288 | 0.36 |
| chr19 | 10827562 | 10830545 | 2984 | 0.31 | 0.288 | 0.361 |
| chr16 | 4134768 | 4167306 | 32539 | -0.11 | 0.288 | 0.361 |
| chr1 | 227125705 | 227133541 | 7837 | -0.12 | 0.288 | 0.361 |
| chr6 | 144065123 | 144077457 | 12335 | -0.15 | 0.288 | 0.361 |
| chr5 | 176037284 | 176037754 | 471 | 0.72 | 0.288 | 0.361 |

| | | | | | | |
|---|---|---|---|---|---|---|
| chr7 | 127343990 | 127344496 | 507 | 0.76 | 0.288 | 0.361 |
| chr8 | 30939175 | 30939753 | 579 | -0.64 | 0.288 | 0.361 |
| chr1 | 244815552 | 244823567 | 8016 | -0.19 | 0.288 | 0.361 |
| chr19 | 17529635 | 17531573 | 1939 | -0.25 | 0.288 | 0.361 |
| chr3 | 179116795 | 179117309 | 515 | 0.8 | 0.288 | 0.361 |
| chr4 | 71536440 | 71537599 | 1160 | 0.49 | 0.288 | 0.361 |
| chr1 | 92336841 | 92366957 | 30117 | -0.14 | 0.289 | 0.361 |
| chr7 | 140742142 | 140742929 | 788 | -0.53 | 0.289 | 0.361 |
| chr14 | 51066536 | 51068239 | 1704 | -0.26 | 0.289 | 0.361 |
| chr4 | 7085947 | 7086455 | 509 | -0.67 | 0.289 | 0.361 |
| chr12 | 112607816 | 112630107 | 22292 | -0.23 | 0.289 | 0.361 |
| chr12 | 106327594 | 106328743 | 1150 | 0.37 | 0.289 | 0.361 |
| chr18 | 33708600 | 33711159 | 2560 | 0.15 | 0.289 | 0.362 |
| chr6 | 121951658 | 121952462 | 805 | 1.56 | 0.289 | 0.362 |
| chr9 | 129986898 | 129988052 | 1155 | 0.49 | 0.289 | 0.362 |
| chr20 | 61958080 | 61958520 | 441 | -0.79 | 0.289 | 0.362 |
| chr16 | 69821170 | 69821888 | 719 | -0.74 | 0.289 | 0.362 |
| chr11 | 28678302 | 28680370 | 2069 | 0.31 | 0.289 | 0.362 |
| chr1 | 110260813 | 110262945 | 2133 | -0.24 | 0.289 | 0.362 |
| chr17 | 54962792 | 54995874 | 33083 | 0.1 | 0.289 | 0.362 |
| chr3 | 130445398 | 130446669 | 1272 | 0.38 | 0.289 | 0.362 |
| chr1 | 202926423 | 202936359 | 9937 | 0.23 | 0.29 | 0.362 |
| chr15 | 23060627 | 23062733 | 2107 | 0.4 | 0.29 | 0.362 |
| chr19 | 40595620 | 40597660 | 2041 | -0.12 | 0.29 | 0.362 |
| chr1 | 178367951 | 178369237 | 1287 | -0.45 | 0.29 | 0.362 |
| chr1 | 171809888 | 171810504 | 617 | 0.48 | 0.29 | 0.362 |
| chr18 | 48633822 | 48634695 | 874 | -0.57 | 0.29 | 0.363 |
| chr14 | 95559896 | 95560337 | 442 | -0.8 | 0.29 | 0.363 |
| chr4 | 54568501 | 54569767 | 1267 | -0.37 | 0.29 | 0.363 |
| chrX | 9430543 | 9435058 | 4516 | 0.16 | 0.29 | 0.363 |
| chr3 | 40175079 | 40175678 | 600 | 0.65 | 0.29 | 0.363 |
| chr17 | 49286588 | 49287652 | 1065 | -1.82 | 0.29 | 0.363 |
| chr1 | 240579014 | 240580141 | 1128 | 0.36 | 0.291 | 0.363 |
| chr1 | 211747712 | 211753493 | 5782 | 0.24 | 0.291 | 0.363 |
| chr8 | 82178649 | 82178995 | 347 | -0.78 | 0.291 | 0.363 |
| chr12 | 121438901 | 121454296 | 15396 | 0.16 | 0.291 | 0.363 |
| chr21 | 45552361 | 45554539 | 2179 | 0.27 | 0.291 | 0.363 |
| chr1 | 52606949 | 52609090 | 2142 | 0.21 | 0.291 | 0.363 |
| chr16 | 4109886 | 4111541 | 1656 | 0.55 | 0.291 | 0.363 |
| chr3 | 9973769 | 10029726 | 55958 | -0.07 | 0.291 | 0.363 |
| chr2 | 192435610 | 192459491 | 23882 | 0.35 | 0.291 | 0.363 |
| chr7 | 157225074 | 157225791 | 718 | 0.66 | 0.291 | 0.363 |
| chr8 | 65710879 | 65711649 | 771 | -0.5 | 0.291 | 0.363 |
| chr5 | 121342898 | 121343702 | 805 | -0.61 | 0.291 | 0.364 |
| chr14 | 42599680 | 42600140 | 461 | 0.73 | 0.291 | 0.364 |
| chr7 | 149569362 | 149571640 | 2279 | 0.21 | 0.291 | 0.364 |
| chr2 | 72857290 | 72858444 | 1155 | -0.37 | 0.291 | 0.364 |
| chr2 | 149266443 | 149267544 | 1102 | 0.66 | 0.291 | 0.364 |
| chr4 | 140289586 | 140290721 | 1136 | 0.61 | 0.291 | 0.364 |
| chr11 | 57015912 | 57017250 | 1339 | -0.55 | 0.291 | 0.364 |
| chr22 | 22896539 | 22902208 | 5670 | 0.24 | 0.291 | 0.364 |
| chrX | 3630579 | 3632105 | 1527 | -0.41 | 0.291 | 0.364 |
| chr6 | 135679462 | 135680309 | 848 | -0.55 | 0.291 | 0.364 |
| chr10 | 60227385 | 60235869 | 8485 | 0.28 | 0.291 | 0.364 |
| chr10 | 94788327 | 94791245 | 2919 | 0.2 | 0.292 | 0.364 |
| chr12 | 17741476 | 17742042 | 567 | 0.51 | 0.292 | 0.364 |
| chr2 | 65170488 | 65171762 | 1275 | 0.44 | 0.292 | 0.364 |
| chr12 | 88940025 | 88953832 | 13808 | 0.12 | 0.292 | 0.364 |
| chr14 | 89258463 | 89259870 | 1408 | -0.37 | 0.292 | 0.364 |
| chr19 | 7745756 | 7746365 | 610 | 0.62 | 0.292 | 0.364 |
| chr14 | 105531068 | 105531758 | 691 | -0.76 | 0.292 | 0.364 |

| | | | | | | |
|---|---|---|---|---|---|---|
| chr1 | 7513856 | 7514250 | 395 | 0.75 | 0.292 | 0.364 |
| chr20 | 13618639 | 13620251 | 1613 | 0.19 | 0.292 | 0.365 |
| chr9 | 35945105 | 35945894 | 790 | -0.42 | 0.292 | 0.365 |
| chrX | 122724721 | 122725779 | 1059 | 0.61 | 0.292 | 0.365 |
| chr1 | 8556075 | 8556634 | 560 | -0.74 | 0.292 | 0.365 |
| chr2 | 72899438 | 72905262 | 5825 | 0.16 | 0.292 | 0.365 |
| chr20 | 4741415 | 4742840 | 1426 | 0.33 | 0.292 | 0.365 |
| chr3 | 188300989 | 188303412 | 2424 | -0.19 | 0.292 | 0.365 |
| chr1 | 201678575 | 201686002 | 7428 | -0.12 | 0.292 | 0.365 |
| chr15 | 68713510 | 68714675 | 1166 | -0.42 | 0.293 | 0.365 |
| chr1 | 76251182 | 76257369 | 6188 | 0.16 | 0.293 | 0.365 |
| chr15 | 74092600 | 74094101 | 1502 | 0.45 | 0.293 | 0.365 |
| chr9 | 88420133 | 88420898 | 766 | 0.62 | 0.293 | 0.365 |
| chr6 | 35012183 | 35012753 | 571 | 0.66 | 0.293 | 0.366 |
| chr16 | 72126064 | 72129105 | 3042 | 0.15 | 0.293 | 0.366 |
| chr1 | 188206190 | 188207637 | 1448 | 0.35 | 0.293 | 0.366 |
| chr5 | 53303293 | 53310157 | 6865 | 0.21 | 0.293 | 0.366 |
| chr10 | 116697895 | 116699714 | 1820 | -0.18 | 0.293 | 0.366 |
| chr6 | 71276232 | 71277980 | 1749 | 0.28 | 0.294 | 0.367 |
| chr7 | 39722467 | 39740410 | 17944 | 0.16 | 0.294 | 0.367 |
| chr9 | 131123820 | 131134655 | 10836 | 0.14 | 0.294 | 0.367 |
| chr7 | 94250261 | 94250749 | 489 | 0.85 | 0.294 | 0.367 |
| chr16 | 30032234 | 30033000 | 767 | -0.57 | 0.294 | 0.367 |
| chr12 | 121266360 | 121267539 | 1180 | 0.44 | 0.294 | 0.367 |
| chr14 | 74415798 | 74438786 | 22989 | -0.1 | 0.294 | 0.367 |
| chr16 | 618501 | 620911 | 2411 | -0.34 | 0.294 | 0.367 |
| chr15 | 50930608 | 50948846 | 18239 | -0.12 | 0.294 | 0.367 |
| chr17 | 49270768 | 49278681 | 7914 | 0.25 | 0.294 | 0.367 |
| chr2 | 30549243 | 30549828 | 586 | 0.66 | 0.294 | 0.367 |
| chr2 | 15814359 | 15815100 | 742 | 0.65 | 0.294 | 0.367 |
| chr6 | 55393165 | 55394030 | 866 | 0.56 | 0.295 | 0.367 |
| chr1 | 75986280 | 75986814 | 535 | 0.83 | 0.295 | 0.367 |
| chr11 | 86770864 | 86771630 | 767 | -0.57 | 0.295 | 0.367 |
| chr9 | 15551952 | 15553799 | 1848 | 0.27 | 0.295 | 0.367 |
| chr14 | 92587186 | 92595617 | 8432 | -0.12 | 0.295 | 0.367 |
| chr5 | 146832892 | 146834208 | 1317 | 0.51 | 0.295 | 0.368 |
| chr12 | 12529901 | 12530683 | 783 | -0.69 | 0.295 | 0.368 |
| chr7 | 129249769 | 129260235 | 10467 | -0.14 | 0.295 | 0.368 |
| chr1 | 155211161 | 155243868 | 32708 | 0.1 | 0.295 | 0.368 |
| chr22 | 45808856 | 45811043 | 2188 | -0.22 | 0.295 | 0.368 |
| chrX | 48857927 | 48859677 | 1751 | 0.17 | 0.295 | 0.368 |
| chr5 | 93011099 | 93012456 | 1358 | 1.86 | 0.295 | 0.368 |
| chr14 | 78207634 | 78212913 | 5280 | -0.36 | 0.295 | 0.368 |
| chr8 | 142634534 | 142635405 | 872 | -0.57 | 0.295 | 0.368 |
| chr3 | 196044408 | 196047545 | 3138 | -0.17 | 0.295 | 0.368 |
| chr1 | 53269261 | 53269759 | 499 | -0.84 | 0.295 | 0.368 |
| chr5 | 37472618 | 37473294 | 677 | -1.18 | 0.296 | 0.369 |
| chr11 | 92753144 | 92763032 | 9889 | 0.34 | 0.296 | 0.369 |
| chr17 | 21075683 | 21076113 | 431 | -0.97 | 0.296 | 0.369 |
| chr11 | 43941486 | 43943064 | 1579 | 0.34 | 0.296 | 0.369 |
| chr4 | 155533364 | 155534554 | 1191 | 0.56 | 0.296 | 0.369 |
| chr16 | 23433918 | 23447510 | 13593 | 0.12 | 0.296 | 0.369 |
| chr9 | 34988828 | 34990328 | 1501 | 0.34 | 0.297 | 0.369 |
| chr16 | 15763754 | 15797132 | 33379 | -0.11 | 0.297 | 0.369 |
| chr5 | 80922093 | 80923272 | 1180 | 0.38 | 0.297 | 0.369 |
| chr12 | 95119353 | 95119688 | 336 | 0.71 | 0.297 | 0.369 |
| chr10 | 30115504 | 30116065 | 562 | 0.76 | 0.297 | 0.37 |
| chr6 | 101168039 | 101168568 | 530 | 0.78 | 0.297 | 0.37 |
| chr4 | 95198839 | 95200717 | 1879 | -0.35 | 0.297 | 0.37 |
| chr5 | 133787363 | 133813765 | 26403 | -0.12 | 0.297 | 0.37 |
| chr7 | 139276037 | 139276493 | 457 | -0.75 | 0.297 | 0.37 |

| | | | | | | |
|---|---|---|---|---|---|---|
| chr5 | 79999870 | 80000457 | 588 | 0.52 | 0.297 | 0.37 |
| chr12 | 56074556 | 56076745 | 2190 | 0.27 | 0.297 | 0.37 |
| chr9 | 37784122 | 37786072 | 1951 | 0.21 | 0.297 | 0.37 |
| chr8 | 28860339 | 28860899 | 561 | 0.82 | 0.297 | 0.37 |
| chr7 | 148395048 | 148398007 | 2960 | 0.14 | 0.297 | 0.37 |
| chr2 | 58237500 | 58238757 | 1258 | -0.5 | 0.297 | 0.37 |
| chr5 | 176448357 | 176463785 | 15429 | -0.17 | 0.297 | 0.37 |
| chr17 | 61698696 | 61700988 | 2293 | 0.28 | 0.297 | 0.37 |
| chrX | 44557413 | 44565843 | 8431 | 1.89 | 0.297 | 0.37 |
| chr2 | 69663149 | 69665762 | 2614 | -0.12 | 0.298 | 0.37 |
| chr18 | 23965328 | 23966952 | 1625 | 0.3 | 0.298 | 0.37 |
| chr8 | 104992167 | 104993353 | 1187 | -0.59 | 0.298 | 0.37 |
| chr4 | 151840939 | 151853337 | 12399 | 0.44 | 0.298 | 0.37 |
| chr3 | 13936806 | 13937682 | 877 | 0.56 | 0.298 | 0.37 |
| chr14 | 104544593 | 104545106 | 514 | 0.59 | 0.298 | 0.371 |
| chr20 | 14385980 | 14396454 | 10475 | -0.13 | 0.298 | 0.371 |
| chr14 | 100904649 | 100907082 | 2434 | 0.2 | 0.298 | 0.371 |
| chr2 | 228044693 | 228057330 | 12638 | 0.18 | 0.298 | 0.371 |
| chr16 | 84259278 | 84263708 | 4431 | -0.15 | 0.298 | 0.371 |
| chr4 | 42659255 | 42660011 | 757 | 0.3 | 0.298 | 0.371 |
| chr6 | 20997174 | 21007491 | 10318 | 0.14 | 0.298 | 0.371 |
| chr6 | 163993150 | 164007547 | 14398 | 0.28 | 0.298 | 0.371 |
| chr9 | 17134411 | 17136251 | 1841 | -0.2 | 0.298 | 0.371 |
| chr3 | 100087052 | 100087684 | 633 | 0.84 | 0.299 | 0.372 |
| chr19 | 11707300 | 11709524 | 2225 | 0.2 | 0.299 | 0.372 |
| chr17 | 47467712 | 47493049 | 25338 | -0.13 | 0.299 | 0.372 |
| chr8 | 1770992 | 1791802 | 20811 | 0.16 | 0.299 | 0.372 |
| chr1 | 65532057 | 65534697 | 2641 | -0.18 | 0.299 | 0.372 |
| chr1 | 180988462 | 181014432 | 25971 | 0.12 | 0.299 | 0.372 |
| chr17 | 33431492 | 33432433 | 942 | -0.79 | 0.299 | 0.372 |
| chr17 | 59544374 | 59545750 | 1377 | -0.38 | 0.299 | 0.372 |
| chr10 | 89263893 | 89266144 | 2252 | 0.22 | 0.299 | 0.372 |
| chr6 | 90347076 | 90349458 | 2383 | 0.15 | 0.299 | 0.372 |
| chr7 | 148936092 | 148938369 | 2278 | 0.25 | 0.299 | 0.372 |
| chr19 | 660944 | 680393 | 19450 | 0.11 | 0.299 | 0.372 |
| chrX | 48748012 | 48777058 | 29047 | 0.09 | 0.299 | 0.372 |
| chr17 | 78147997 | 78198849 | 50853 | -0.13 | 0.299 | 0.372 |
| chr13 | 48842235 | 48842815 | 581 | -0.68 | 0.3 | 0.372 |
| chr8 | 68225693 | 68227304 | 1612 | -0.32 | 0.3 | 0.373 |
| chr2 | 108195483 | 108196103 | 621 | -0.73 | 0.3 | 0.373 |
| chr15 | 42999123 | 43000635 | 1513 | -0.41 | 0.3 | 0.373 |
| chr1 | 201951370 | 201954104 | 2735 | 0.23 | 0.3 | 0.373 |
| chr5 | 180212315 | 180213442 | 1128 | 0.52 | 0.3 | 0.373 |
| chr15 | 52673116 | 52673741 | 626 | -0.46 | 0.3 | 0.373 |
| chr15 | 50194139 | 50195770 | 1632 | -2 | 0.3 | 0.373 |
| chr1 | 1840151 | 1841559 | 1409 | 0.36 | 0.3 | 0.373 |
| chr18 | 55758212 | 55758721 | 510 | -0.68 | 0.3 | 0.373 |
| chr1 | 65421088 | 65432901 | 11814 | 0.21 | 0.3 | 0.373 |
| chr11 | 103399781 | 103412359 | 12579 | -2.12 | 0.3 | 0.373 |
| chr9 | 132564753 | 132566453 | 1701 | 0.32 | 0.3 | 0.373 |
| chr20 | 43538304 | 43540640 | 2337 | -0.31 | 0.3 | 0.373 |
| chr16 | 88694263 | 88695441 | 1179 | 0.44 | 0.3 | 0.373 |
| chr2 | 87034787 | 87035547 | 761 | -0.66 | 0.3 | 0.373 |
| chr6 | 169928539 | 169931713 | 3175 | -0.29 | 0.301 | 0.373 |
| chr22 | 38429714 | 38431443 | 1730 | 0.26 | 0.301 | 0.373 |
| chr14 | 68066173 | 68067682 | 1510 | 0.28 | 0.301 | 0.373 |
| chr16 | 23156376 | 23161540 | 5165 | -0.2 | 0.301 | 0.373 |
| chr2 | 139178841 | 139179556 | 716 | 0.67 | 0.301 | 0.373 |
| chrX | 109376991 | 109377745 | 755 | 0.65 | 0.301 | 0.374 |
| chr7 | 97852159 | 97853081 | 923 | -0.56 | 0.301 | 0.374 |
| chr5 | 65082367 | 65082819 | 453 | -1.14 | 0.301 | 0.374 |

| | | | | | | |
|---|---|---|---|---|---|---|
| chr10 | 86119993 | 86121392 | 1400 | 0.51 | 0.301 | 0.374 |
| chr11 | 68267665 | 68271338 | 3674 | -0.35 | 0.301 | 0.374 |
| chr1 | 51762579 | 51780827 | 18249 | -0.16 | 0.301 | 0.374 |
| chr10 | 123572817 | 123573782 | 966 | -0.51 | 0.301 | 0.374 |
| chr5 | 131150327 | 131150786 | 460 | 0.85 | 0.301 | 0.374 |
| chr8 | 42936667 | 42957798 | 21132 | 0.13 | 0.301 | 0.374 |
| chr11 | 18378094 | 18442537 | 64444 | -0.06 | 0.301 | 0.374 |
| chr9 | 17229620 | 17235956 | 6337 | 1.07 | 0.302 | 0.374 |
| chr4 | 1037124 | 1038385 | 1262 | 0.36 | 0.302 | 0.374 |
| chr14 | 35760397 | 35776971 | 16575 | 0.1 | 0.302 | 0.374 |
| chr10 | 96968789 | 97069738 | 100950 | -0.04 | 0.302 | 0.374 |
| chrX | 43513835 | 43515626 | 1792 | 0.25 | 0.302 | 0.375 |
| chr4 | 154407678 | 154408145 | 468 | 0.86 | 0.302 | 0.375 |
| chr6 | 148761649 | 148764013 | 2365 | 0.15 | 0.302 | 0.375 |
| chr19 | 15517430 | 15529816 | 12387 | -0.18 | 0.302 | 0.375 |
| chr5 | 81500392 | 81508219 | 7828 | -1.94 | 0.302 | 0.375 |
| chr2 | 220407768 | 220408566 | 799 | -0.57 | 0.302 | 0.375 |
| chr9 | 94800887 | 94801584 | 698 | 0.68 | 0.302 | 0.375 |
| chr21 | 34886595 | 34898044 | 11450 | -0.21 | 0.302 | 0.375 |
| chr7 | 141675746 | 141676667 | 922 | 0.41 | 0.302 | 0.375 |
| chr22 | 46566195 | 46567780 | 1586 | 0.36 | 0.303 | 0.376 |
| chr7 | 129843434 | 129846280 | 2847 | 0.12 | 0.303 | 0.376 |
| chr1 | 1590071 | 1590791 | 721 | 0.35 | 0.303 | 0.376 |
| chr16 | 2819746 | 2828926 | 9181 | 0.11 | 0.303 | 0.376 |
| chr3 | 195994212 | 196015823 | 21612 | -0.11 | 0.303 | 0.376 |
| chr20 | 44065206 | 44065666 | 461 | 0.51 | 0.303 | 0.376 |
| chr1 | 61827846 | 61828646 | 801 | 0.66 | 0.303 | 0.376 |
| chr14 | 77534243 | 77535185 | 943 | 0.57 | 0.303 | 0.376 |
| chr7 | 38167252 | 38167908 | 657 | -0.53 | 0.303 | 0.376 |
| chr3 | 152812300 | 152812803 | 504 | -0.72 | 0.303 | 0.376 |
| chr2 | 197391899 | 197393120 | 1222 | 0.19 | 0.303 | 0.376 |
| chr5 | 176936634 | 176947000 | 10367 | -0.2 | 0.303 | 0.376 |
| chr13 | 46785557 | 46786275 | 719 | -0.9 | 0.303 | 0.376 |
| chr16 | 23136491 | 23137323 | 833 | -0.54 | 0.303 | 0.376 |
| chr9 | 5509924 | 5510996 | 1073 | -0.54 | 0.303 | 0.376 |
| chr4 | 4542432 | 4544390 | 1959 | -0.18 | 0.303 | 0.376 |
| chr8 | 16606429 | 16607190 | 762 | -0.51 | 0.303 | 0.376 |
| chr16 | 28984874 | 28986834 | 1961 | 0.26 | 0.304 | 0.376 |
| chr20 | 62204633 | 62206219 | 1587 | 0.38 | 0.304 | 0.377 |
| chr3 | 135968655 | 135970406 | 1752 | 0.23 | 0.304 | 0.377 |
| chrX | 10723101 | 10724508 | 1408 | 0.44 | 0.304 | 0.377 |
| chr8 | 17606963 | 17608089 | 1127 | 0.77 | 0.304 | 0.377 |
| chr2 | 128433132 | 128433572 | 441 | -0.77 | 0.304 | 0.377 |
| chr5 | 56594467 | 56595324 | 858 | -0.57 | 0.304 | 0.377 |
| chr1 | 61688531 | 61689070 | 540 | 0.79 | 0.304 | 0.377 |
| chr1 | 161196950 | 161197831 | 882 | -0.63 | 0.304 | 0.377 |
| chr6 | 80380903 | 80381956 | 1054 | 0.59 | 0.304 | 0.377 |
| chr4 | 75572602 | 75609114 | 36513 | -0.06 | 0.304 | 0.377 |
| chr6 | 42514885 | 42515975 | 1091 | 0.34 | 0.305 | 0.377 |
| chr6 | 27355910 | 27357193 | 1284 | 0.31 | 0.305 | 0.378 |
| chr12 | 70836581 | 70837471 | 891 | -0.66 | 0.305 | 0.378 |
| chr1 | 27307071 | 27340679 | 33609 | 0.08 | 0.305 | 0.378 |
| chr17 | 65877856 | 65892377 | 14522 | 0.2 | 0.305 | 0.378 |
| chr18 | 21358974 | 21360635 | 1662 | -0.44 | 0.305 | 0.378 |
| chrX | 129837862 | 129838829 | 968 | -0.47 | 0.305 | 0.378 |
| chr14 | 93566867 | 93583319 | 16453 | -0.12 | 0.305 | 0.378 |
| chrX | 34935442 | 34937192 | 1751 | -0.35 | 0.305 | 0.378 |
| chr22 | 26867068 | 26880933 | 13866 | -0.18 | 0.305 | 0.378 |
| chr3 | 10093409 | 10094001 | 593 | -0.62 | 0.305 | 0.378 |
| chr12 | 21680406 | 21681024 | 619 | -0.76 | 0.305 | 0.378 |
| chr19 | 17241609 | 17247517 | 5909 | 0.16 | 0.306 | 0.379 |

| | | | | | | |
|---|---|---|---|---|---|---|
| chr4 | 108131798 | 108132845 | 1048 | -0.45 | 0.306 | 0.379 |
| chr17 | 73347956 | 73363125 | 15170 | 0.23 | 0.306 | 0.379 |
| chr15 | 59254900 | 59255969 | 1070 | -0.48 | 0.306 | 0.379 |
| chr19 | 11290166 | 11309167 | 19002 | 0.21 | 0.306 | 0.379 |
| chr9 | 72917854 | 72918218 | 365 | -0.83 | 0.306 | 0.379 |
| chr2 | 24082506 | 24083906 | 1401 | -0.36 | 0.306 | 0.379 |
| chr1 | 44215526 | 44223491 | 7966 | 0.33 | 0.306 | 0.379 |
| chr15 | 38543091 | 38559095 | 16005 | 0.14 | 0.306 | 0.379 |
| chr9 | 90588611 | 90590600 | 1990 | -0.19 | 0.306 | 0.379 |
| chr19 | 1566317 | 1567494 | 1178 | 0.43 | 0.306 | 0.38 |
| chr3 | 184078942 | 184082122 | 3181 | 0.15 | 0.306 | 0.38 |
| chr22 | 48588566 | 48589559 | 994 | -0.47 | 0.307 | 0.38 |
| chr4 | 132117446 | 132118322 | 877 | 0.51 | 0.307 | 0.38 |
| chr6 | 161859840 | 161860659 | 820 | 0.54 | 0.307 | 0.38 |
| chr6 | 146284275 | 146286325 | 2051 | 0.23 | 0.307 | 0.38 |
| chr1 | 161734355 | 161737033 | 2679 | 0.12 | 0.307 | 0.38 |
| chr7 | 135242220 | 135273194 | 30975 | 0.12 | 0.307 | 0.38 |
| chr21 | 40127333 | 40129127 | 1795 | -0.39 | 0.307 | 0.38 |
| chr4 | 140186947 | 140188472 | 1526 | -0.22 | 0.307 | 0.38 |
| chr1 | 154598975 | 154601381 | 2407 | 0.14 | 0.307 | 0.38 |
| chr8 | 37619400 | 37621088 | 1689 | 0.19 | 0.307 | 0.38 |
| chr10 | 94290088 | 94298974 | 8887 | 0.28 | 0.307 | 0.38 |
| chr16 | 3199271 | 3241915 | 42645 | -0.13 | 0.307 | 0.381 |
| chr3 | 184971433 | 184972123 | 691 | 0.42 | 0.308 | 0.381 |
| chr9 | 36189585 | 36230618 | 41034 | -0.07 | 0.308 | 0.381 |
| chr18 | 2905622 | 2917556 | 11935 | 0.19 | 0.308 | 0.381 |
| chr3 | 27647854 | 27649603 | 1750 | -0.32 | 0.308 | 0.381 |
| chr11 | 66462411 | 66463238 | 828 | -0.43 | 0.308 | 0.381 |
| chr16 | 48092304 | 48092596 | 293 | 0.98 | 0.308 | 0.381 |
| chr9 | 115512071 | 115514285 | 2215 | 0.26 | 0.308 | 0.381 |
| chr1 | 115322433 | 115323948 | 1516 | 0.2 | 0.308 | 0.381 |
| chr9 | 115163454 | 115164176 | 723 | -0.6 | 0.308 | 0.382 |
| chr20 | 52322303 | 52328590 | 6288 | 0.24 | 0.308 | 0.382 |
| chr20 | 34203196 | 34225900 | 22705 | -0.11 | 0.308 | 0.382 |
| chr15 | 28518717 | 28519150 | 434 | 0.61 | 0.308 | 0.382 |
| chr2 | 224476991 | 224478901 | 1911 | -0.17 | 0.309 | 0.382 |
| chr12 | 42536935 | 42567556 | 30622 | 0.11 | 0.309 | 0.382 |
| chr8 | 87458915 | 87460044 | 1130 | 0.44 | 0.309 | 0.382 |
| chr4 | 162840021 | 162847664 | 7644 | -0.21 | 0.309 | 0.382 |
| chr8 | 60180813 | 60181979 | 1167 | 0.42 | 0.309 | 0.382 |
| chr5 | 134056339 | 134058746 | 2408 | -0.27 | 0.309 | 0.382 |
| chr11 | 74171553 | 74172190 | 638 | -0.55 | 0.309 | 0.382 |
| chr17 | 29839436 | 29861236 | 21801 | -1.64 | 0.309 | 0.382 |
| chr11 | 1714543 | 1715361 | 819 | 0.29 | 0.309 | 0.382 |
| chr3 | 48540910 | 48545679 | 4770 | 0.23 | 0.309 | 0.382 |
| chr7 | 80350003 | 80355393 | 5391 | -0.08 | 0.309 | 0.382 |
| chr20 | 62370439 | 62373287 | 2849 | 0.27 | 0.309 | 0.382 |
| chr3 | 138654311 | 138666590 | 12280 | -0.22 | 0.309 | 0.382 |
| chr6 | 150263103 | 150263609 | 507 | 0.81 | 0.309 | 0.382 |
| chr6 | 45344480 | 45346881 | 2402 | 0.14 | 0.309 | 0.382 |
| chr4 | 41238166 | 41238870 | 705 | -0.57 | 0.309 | 0.383 |
| chrX | 150151040 | 150153808 | 2769 | 0.25 | 0.309 | 0.383 |
| chr8 | 103523761 | 103524314 | 554 | -1.7 | 0.309 | 0.383 |
| chr16 | 56676642 | 56677966 | 1325 | 0.51 | 0.31 | 0.383 |
| chr5 | 173586288 | 173587255 | 968 | 0.59 | 0.31 | 0.383 |
| chr11 | 8815974 | 8836119 | 20146 | 0.22 | 0.31 | 0.383 |
| chr18 | 2772521 | 2772973 | 453 | -0.69 | 0.31 | 0.383 |
| chr11 | 126726810 | 126742721 | 15912 | 0.23 | 0.31 | 0.383 |
| chr15 | 48751891 | 48753136 | 1246 | 0.44 | 0.31 | 0.383 |
| chr6 | 91240651 | 91241067 | 417 | -0.81 | 0.31 | 0.383 |
| chr7 | 76633781 | 76640451 | 6671 | -0.21 | 0.31 | 0.383 |

| | | | | | | |
|---|---|---|---|---|---|---|
| chr6 | 123166963 | 123167747 | 785 | 0.42 | 0.31 | 0.383 |
| chr16 | 11110053 | 11111406 | 1354 | 0.41 | 0.31 | 0.383 |
| chr2 | 32113095 | 32114321 | 1227 | -0.41 | 0.31 | 0.383 |
| chr18 | 33929064 | 33930200 | 1137 | 1.6 | 0.31 | 0.383 |
| chr4 | 189343414 | 189377462 | 34049 | -0.07 | 0.31 | 0.383 |
| chr1 | 15573631 | 15574394 | 764 | -0.45 | 0.31 | 0.383 |
| chr9 | 72434858 | 72441587 | 6730 | 1.93 | 0.31 | 0.383 |
| chr4 | 177419320 | 177421277 | 1958 | 0.36 | 0.31 | 0.383 |
| chr2 | 151128763 | 151129264 | 502 | 0.65 | 0.31 | 0.383 |
| chr6 | 139349049 | 139350816 | 1768 | 0.27 | 0.31 | 0.383 |
| chr4 | 25633629 | 25634361 | 733 | 0.53 | 0.31 | 0.383 |
| chr17 | 67756002 | 67777043 | 21042 | 0.07 | 0.311 | 0.384 |
| chr13 | 33240824 | 33241750 | 927 | 0.47 | 0.311 | 0.384 |
| chr7 | 140712917 | 140715257 | 2341 | -0.17 | 0.311 | 0.384 |
| chr13 | 77873738 | 77881442 | 7705 | -0.36 | 0.311 | 0.384 |
| chr13 | 75904057 | 75904478 | 422 | -0.81 | 0.311 | 0.384 |
| chr8 | 42327717 | 42382002 | 54286 | -0.12 | 0.311 | 0.384 |
| chr9 | 110854853 | 110858215 | 3363 | 0.18 | 0.311 | 0.384 |
| chr16 | 11405659 | 11407234 | 1576 | 0.39 | 0.311 | 0.384 |
| chr16 | 14013258 | 14031601 | 18344 | -0.13 | 0.311 | 0.385 |
| chr12 | 65602447 | 65604376 | 1930 | 0.42 | 0.311 | 0.385 |
| chr10 | 105808871 | 105813186 | 4316 | 0.31 | 0.311 | 0.385 |
| chr2 | 201723401 | 201730411 | 7011 | -0.11 | 0.312 | 0.385 |
| chr4 | 184642069 | 184643554 | 1486 | 0.32 | 0.312 | 0.385 |
| chr14 | 34456298 | 34459924 | 3627 | 0.42 | 0.312 | 0.385 |
| chr6 | 135997854 | 136003033 | 5180 | -0.45 | 0.312 | 0.385 |
| chr8 | 11218172 | 11220648 | 2477 | 0.21 | 0.312 | 0.385 |
| chr17 | 4451744 | 4459569 | 7826 | 0.16 | 0.312 | 0.385 |
| chrX | 138802194 | 138802712 | 519 | -0.8 | 0.312 | 0.385 |
| chr16 | 50214650 | 50245156 | 30507 | 0.07 | 0.312 | 0.385 |
| chr1 | 86895109 | 86896187 | 1079 | 0.52 | 0.312 | 0.385 |
| chr19 | 13884478 | 13909541 | 25064 | -0.13 | 0.312 | 0.385 |
| chr10 | 49863202 | 49864931 | 1730 | -0.26 | 0.312 | 0.385 |
| chr4 | 24528097 | 24561453 | 33357 | 0.1 | 0.312 | 0.385 |
| chr1 | 6408541 | 6428382 | 19842 | -0.14 | 0.312 | 0.385 |
| chr11 | 70950488 | 70965618 | 15131 | -0.11 | 0.312 | 0.386 |
| chr1 | 28558583 | 28587200 | 28618 | -0.13 | 0.312 | 0.386 |
| chr2 | 197560753 | 197563308 | 2556 | 0.35 | 0.312 | 0.386 |
| chr10 | 105085430 | 105085833 | 404 | -0.82 | 0.313 | 0.386 |
| chr15 | 49169835 | 49171167 | 1333 | 0.3 | 0.313 | 0.386 |
| chr18 | 34252265 | 34253306 | 1042 | 0.45 | 0.313 | 0.386 |
| chr14 | 53477119 | 53478374 | 1256 | -0.37 | 0.313 | 0.386 |
| chr2 | 74681332 | 74711887 | 30556 | -0.08 | 0.313 | 0.386 |
| chr16 | 85145732 | 85158160 | 12429 | -0.23 | 0.313 | 0.386 |
| chr1 | 95219100 | 95221852 | 2753 | 0.35 | 0.313 | 0.386 |
| chr18 | 46012463 | 46016733 | 4271 | 0.16 | 0.313 | 0.386 |
| chr2 | 32834806 | 32858169 | 23364 | -0.13 | 0.313 | 0.386 |
| chr17 | 62955837 | 62982167 | 26331 | 0.1 | 0.313 | 0.386 |
| chr1 | 85603125 | 85604244 | 1120 | 1.79 | 0.313 | 0.386 |
| chr4 | 57665852 | 57666551 | 700 | 0.44 | 0.313 | 0.386 |
| chr3 | 93722924 | 93723749 | 826 | 0.4 | 0.313 | 0.386 |
| chr9 | 36136235 | 36163806 | 27572 | 0.18 | 0.313 | 0.386 |
| chr1 | 43312288 | 43313314 | 1027 | -0.53 | 0.313 | 0.386 |
| chr5 | 148960226 | 148962029 | 1804 | 0.25 | 0.313 | 0.386 |
| chr12 | 40225359 | 40226287 | 929 | 0.42 | 0.313 | 0.386 |
| chr19 | 18942779 | 18944275 | 1497 | 0.35 | 0.313 | 0.387 |
| chr5 | 72497456 | 72497784 | 329 | -0.78 | 0.314 | 0.387 |
| chr9 | 86414597 | 86415788 | 1192 | 1.74 | 0.314 | 0.387 |
| chr6 | 54284512 | 54285406 | 895 | 0.82 | 0.314 | 0.387 |
| chr2 | 69518138 | 69518774 | 637 | 0.47 | 0.314 | 0.387 |
| chr9 | 132751340 | 132752434 | 1095 | -0.5 | 0.314 | 0.387 |

| | | | | | | |
|---|---|---|---|---|---|---|
| chr13 | 21745358 | 21755436 | 10079 | 0.15 | 0.314 | 0.387 |
| chr5 | 72857104 | 72875229 | 18126 | 0.13 | 0.314 | 0.387 |
| chr16 | 58748474 | 58769534 | 21061 | -0.12 | 0.314 | 0.388 |
| chr10 | 102746449 | 102773507 | 27059 | 0.08 | 0.315 | 0.388 |
| chr22 | 38289667 | 38303276 | 13610 | 0.14 | 0.315 | 0.388 |
| chr9 | 140706300 | 140711537 | 5238 | -0.32 | 0.315 | 0.388 |
| chrX | 128726711 | 128738637 | 11927 | -0.36 | 0.315 | 0.388 |
| chr3 | 158414013 | 158467016 | 53004 | -0.06 | 0.315 | 0.388 |
| chr17 | 35206935 | 35209547 | 2613 | 0.17 | 0.315 | 0.388 |
| chr2 | 159416508 | 159417414 | 907 | -0.54 | 0.315 | 0.388 |
| chr1 | 54778247 | 54779259 | 1013 | 0.48 | 0.315 | 0.388 |
| chr2 | 197582967 | 197584923 | 1957 | -0.17 | 0.315 | 0.388 |
| chr14 | 95602733 | 95604843 | 2111 | 0.3 | 0.315 | 0.388 |
| chr7 | 157128941 | 157206582 | 77642 | -0.09 | 0.315 | 0.388 |
| chr22 | 20103100 | 20120525 | 17426 | -0.17 | 0.315 | 0.388 |
| chr7 | 73866916 | 73872885 | 5970 | -0.23 | 0.315 | 0.389 |
| chr1 | 224369728 | 224392780 | 23053 | 0.11 | 0.315 | 0.389 |
| chr3 | 140675653 | 140676577 | 925 | 0.61 | 0.315 | 0.389 |
| chr16 | 74944201 | 74945070 | 870 | -0.5 | 0.316 | 0.389 |
| chr4 | 11398553 | 11399728 | 1176 | 1.71 | 0.316 | 0.389 |
| chr10 | 135186910 | 135187710 | 801 | -0.4 | 0.316 | 0.389 |
| chr6 | 13813254 | 13815470 | 2217 | 0.25 | 0.316 | 0.389 |
| chr7 | 128337290 | 128337879 | 590 | -0.57 | 0.316 | 0.389 |
| chr9 | 127926999 | 127963888 | 36890 | 0.11 | 0.316 | 0.389 |
| chr18 | 34974154 | 34974599 | 446 | -0.85 | 0.316 | 0.389 |
| chr1 | 24017495 | 24020032 | 2538 | 0.21 | 0.316 | 0.39 |
| chr9 | 128507870 | 128511209 | 3340 | 0.25 | 0.316 | 0.39 |
| chr7 | 99515659 | 99518147 | 2489 | 0.17 | 0.316 | 0.39 |
| chr9 | 118351517 | 118356646 | 5130 | 0.14 | 0.316 | 0.39 |
| chr21 | 46825146 | 46892263 | 67118 | -0.07 | 0.317 | 0.39 |
| chr1 | 110075133 | 110075630 | 498 | -0.5 | 0.317 | 0.39 |
| chr5 | 139672607 | 139683366 | 10760 | 0.14 | 0.317 | 0.39 |
| chr12 | 27820391 | 27820965 | 575 | -0.59 | 0.317 | 0.39 |
| chr1 | 1549532 | 1551449 | 1918 | -0.26 | 0.317 | 0.39 |
| chr2 | 103220989 | 103221534 | 546 | 0.63 | 0.317 | 0.39 |
| chr17 | 37729546 | 37730781 | 1236 | -0.28 | 0.317 | 0.39 |
| chr6 | 101530818 | 101532392 | 1575 | -0.26 | 0.317 | 0.39 |
| chr15 | 68290064 | 68291892 | 1829 | -0.26 | 0.317 | 0.39 |
| chr14 | 23025463 | 23084608 | 59146 | -0.05 | 0.317 | 0.391 |
| chr12 | 127647777 | 127648357 | 581 | 0.67 | 0.317 | 0.391 |
| chr8 | 144343197 | 144374604 | 31408 | -0.12 | 0.317 | 0.391 |
| chr1 | 196342339 | 196342813 | 475 | -0.53 | 0.318 | 0.391 |
| chr1 | 109504442 | 109506884 | 2443 | 0.15 | 0.318 | 0.391 |
| chr2 | 161521487 | 161524480 | 2994 | -0.12 | 0.318 | 0.391 |
| chr11 | 70880215 | 70880942 | 728 | -0.63 | 0.318 | 0.391 |
| chr11 | 92732894 | 92735481 | 2588 | -0.15 | 0.318 | 0.391 |
| chr2 | 225132579 | 225134155 | 1577 | -0.18 | 0.318 | 0.392 |
| chr18 | 9707751 | 9744830 | 37080 | 0.09 | 0.318 | 0.392 |
| chr17 | 34928917 | 34968769 | 39853 | -0.09 | 0.318 | 0.392 |
| chr6 | 147800835 | 147801475 | 641 | -0.72 | 0.318 | 0.392 |
| chr17 | 65850974 | 65851740 | 767 | -1.06 | 0.318 | 0.392 |
| chr11 | 95521240 | 95525083 | 3844 | 0.11 | 0.319 | 0.392 |
| chr3 | 157052879 | 157054786 | 1908 | 0.25 | 0.319 | 0.392 |
| chr5 | 176073365 | 176075532 | 2168 | -0.23 | 0.319 | 0.392 |
| chr19 | 2426450 | 2429103 | 2654 | -0.21 | 0.319 | 0.392 |
| chr2 | 86205405 | 86226968 | 21564 | -0.17 | 0.319 | 0.392 |
| chr12 | 51734669 | 51735669 | 1001 | -0.49 | 0.319 | 0.392 |
| chr15 | 72754027 | 72788698 | 34672 | 0.06 | 0.319 | 0.392 |
| chr22 | 24585046 | 24606069 | 21024 | -0.08 | 0.319 | 0.392 |
| chr1 | 98403641 | 98404373 | 733 | 0.48 | 0.319 | 0.392 |
| chr6 | 7260830 | 7262366 | 1537 | 0.31 | 0.319 | 0.392 |

| | | | | | | |
|---|---|---|---|---|---|---|
| chr18 | 535385 | 535713 | 329 | 0.82 | 0.319 | 0.392 |
| chr22 | 41204682 | 41216226 | 11545 | -0.21 | 0.319 | 0.392 |
| chr1 | 20091729 | 20093112 | 1384 | 0.4 | 0.319 | 0.392 |
| chr14 | 102675705 | 102702584 | 26880 | -0.14 | 0.319 | 0.392 |
| chr4 | 108664553 | 108665310 | 758 | 0.57 | 0.319 | 0.392 |
| chr2 | 130938647 | 130939751 | 1105 | -0.32 | 0.319 | 0.392 |
| chr14 | 97685039 | 97685505 | 467 | -0.69 | 0.319 | 0.392 |
| chr10 | 118441409 | 118441913 | 505 | 0.55 | 0.319 | 0.392 |
| chr9 | 99636980 | 99647188 | 10209 | 1.96 | 0.319 | 0.392 |
| chr9 | 17344041 | 17344514 | 474 | 0.7 | 0.319 | 0.392 |
| chr7 | 77183915 | 77294943 | 111029 | -0.05 | 0.319 | 0.392 |
| chr3 | 113501743 | 113503247 | 1505 | -0.36 | 0.319 | 0.392 |
| chr6 | 6931660 | 6932072 | 413 | -0.63 | 0.319 | 0.392 |
| chr5 | 137774349 | 137775301 | 953 | -0.5 | 0.319 | 0.392 |
| chr11 | 67374047 | 67375742 | 1696 | 0.26 | 0.319 | 0.392 |
| chr5 | 125689860 | 125690832 | 973 | 0.45 | 0.319 | 0.392 |
| chr2 | 171784295 | 171794625 | 10331 | -0.14 | 0.32 | 0.393 |
| chr15 | 36924001 | 36936361 | 12361 | 0.38 | 0.32 | 0.393 |
| chr15 | 63728626 | 63816139 | 87514 | -0.05 | 0.32 | 0.393 |
| chr1 | 20986998 | 20988964 | 1967 | 0.26 | 0.32 | 0.393 |
| chr3 | 150145370 | 150146152 | 783 | -0.81 | 0.32 | 0.393 |
| chr14 | 54833575 | 54836645 | 3071 | 0.32 | 0.32 | 0.393 |
| chr9 | 124399957 | 124414388 | 14432 | -0.23 | 0.32 | 0.393 |
| chrX | 129535284 | 129536918 | 1635 | 0.28 | 0.32 | 0.394 |
| chr4 | 71858159 | 71860360 | 2202 | 0.19 | 0.321 | 0.394 |
| chr18 | 54282825 | 54283650 | 826 | 0.65 | 0.321 | 0.394 |
| chr2 | 8288174 | 8288950 | 777 | 0.53 | 0.321 | 0.394 |
| chr3 | 186904566 | 186905096 | 531 | 0.63 | 0.321 | 0.394 |
| chr3 | 112248907 | 112250329 | 1423 | 0.36 | 0.321 | 0.394 |
| chr13 | 21099558 | 21100878 | 1321 | 0.37 | 0.321 | 0.394 |
| chr10 | 121276178 | 121282059 | 5882 | -0.41 | 0.321 | 0.394 |
| chr2 | 179360433 | 179361233 | 801 | -0.55 | 0.321 | 0.394 |
| chr12 | 76230146 | 76230880 | 735 | 1.74 | 0.321 | 0.394 |
| chr14 | 91224887 | 91225716 | 830 | -0.24 | 0.321 | 0.394 |
| chr11 | 18771226 | 18771748 | 523 | -0.64 | 0.321 | 0.395 |
| chr4 | 56829561 | 56830072 | 512 | -0.83 | 0.321 | 0.395 |
| chr12 | 123101880 | 123102357 | 478 | 0.68 | 0.322 | 0.395 |
| chr4 | 186713500 | 186714105 | 606 | 0.55 | 0.322 | 0.395 |
| chr3 | 142033833 | 142038840 | 5008 | -0.32 | 0.322 | 0.396 |
| chr12 | 48339794 | 48340777 | 984 | -0.54 | 0.322 | 0.396 |
| chr1 | 211430465 | 211435069 | 4605 | 0.14 | 0.322 | 0.396 |
| chr21 | 38145326 | 38146103 | 778 | 0.59 | 0.323 | 0.396 |
| chr14 | 67157498 | 67157868 | 371 | -0.71 | 0.323 | 0.396 |
| chr12 | 15723726 | 15724649 | 924 | 0.25 | 0.323 | 0.396 |
| chr1 | 197870896 | 197885486 | 14591 | 0.18 | 0.323 | 0.396 |
| chr2 | 30929436 | 30929862 | 427 | 0.74 | 0.323 | 0.397 |
| chr17 | 67263195 | 67264353 | 1159 | -0.43 | 0.323 | 0.397 |
| chr11 | 20132675 | 20135428 | 2754 | 0.17 | 0.323 | 0.397 |
| chrX | 9288749 | 9289939 | 1191 | 0.36 | 0.323 | 0.397 |
| chr16 | 85524837 | 85525884 | 1048 | -0.67 | 0.323 | 0.397 |
| chr10 | 126479199 | 126491451 | 12253 | 0.13 | 0.324 | 0.397 |
| chr10 | 556876 | 558086 | 1211 | -0.32 | 0.324 | 0.397 |
| chr11 | 17096821 | 17100528 | 3708 | -0.12 | 0.324 | 0.397 |
| chr1 | 212201015 | 212221323 | 20309 | 0.1 | 0.324 | 0.397 |
| chr14 | 99945762 | 99949723 | 3962 | 0.12 | 0.324 | 0.398 |
| chrX | 54103245 | 54104095 | 851 | -0.54 | 0.324 | 0.398 |
| chr5 | 59994677 | 59996810 | 2134 | 0.16 | 0.324 | 0.398 |
| chr6 | 155053036 | 155060715 | 7680 | -0.13 | 0.324 | 0.398 |
| chr1 | 145574660 | 145589711 | 15052 | 0.19 | 0.324 | 0.398 |
| chr16 | 67548224 | 67564827 | 16604 | -0.12 | 0.324 | 0.398 |
| chr15 | 100154703 | 100155246 | 544 | 0.82 | 0.324 | 0.398 |

| | | | | | | |
|---|---|---|---|---|---|---|
| chr19 | 49178048 | 49202213 | 24166 | 2.18 | 0.324 | 0.398 |
| chr2 | 216221999 | 216222817 | 819 | -0.57 | 0.324 | 0.398 |
| chr5 | 5270982 | 5279474 | 8493 | 0.3 | 0.325 | 0.398 |
| chr8 | 25863380 | 25863875 | 496 | 0.63 | 0.325 | 0.398 |
| chr7 | 127427112 | 127427856 | 745 | 0.47 | 0.325 | 0.398 |
| chr11 | 58873847 | 58874497 | 651 | 0.54 | 0.325 | 0.399 |
| chr2 | 236809428 | 236810133 | 706 | 0.61 | 0.325 | 0.399 |
| chr6 | 24472157 | 24473107 | 951 | 0.52 | 0.325 | 0.399 |
| chr10 | 80757784 | 80758490 | 707 | -0.66 | 0.325 | 0.399 |
| chr4 | 85612239 | 85612974 | 736 | -0.55 | 0.325 | 0.399 |
| chr11 | 15217479 | 15217896 | 418 | 0.75 | 0.325 | 0.399 |
| chr16 | 248957 | 249514 | 558 | 0.75 | 0.325 | 0.399 |
| chr20 | 23372791 | 23379282 | 6492 | -1.67 | 0.325 | 0.399 |
| chr1 | 110563041 | 110578149 | 15109 | -0.16 | 0.326 | 0.399 |
| chr1 | 59949437 | 59950397 | 961 | 0.49 | 0.326 | 0.399 |
| chr7 | 84314695 | 84332602 | 17908 | 0.12 | 0.326 | 0.399 |
| chr11 | 76771615 | 76785390 | 13776 | 0.19 | 0.326 | 0.399 |
| chrX | 135990747 | 135992918 | 2172 | -0.2 | 0.326 | 0.4 |
| chr7 | 16724412 | 16724889 | 478 | -0.61 | 0.326 | 0.4 |
| chr15 | 62327638 | 62328263 | 626 | 0.61 | 0.326 | 0.4 |
| chr7 | 131830015 | 131830947 | 933 | 0.55 | 0.326 | 0.4 |
| chr7 | 97820004 | 97820607 | 604 | -0.71 | 0.326 | 0.4 |
| chr11 | 85426916 | 85438101 | 11186 | 0.18 | 0.326 | 0.4 |
| chr3 | 28282345 | 28284289 | 1945 | 0.18 | 0.326 | 0.4 |
| chr2 | 172543691 | 172545532 | 1842 | 0.17 | 0.326 | 0.4 |
| chr3 | 117236861 | 117238086 | 1226 | 0.38 | 0.327 | 0.4 |
| chr9 | 118517796 | 118520905 | 3110 | 0.27 | 0.327 | 0.4 |
| chr3 | 151414989 | 151415908 | 920 | -0.35 | 0.327 | 0.4 |
| chr4 | 123794064 | 123850362 | 56299 | -0.07 | 0.327 | 0.4 |
| chr12 | 112179109 | 112179502 | 394 | 0.6 | 0.327 | 0.4 |
| chr17 | 76857160 | 76932989 | 75830 | -0.05 | 0.327 | 0.401 |
| chr1 | 169762959 | 169765113 | 2155 | 0.13 | 0.327 | 0.401 |
| chr7 | 105206188 | 105222596 | 16409 | 0.15 | 0.327 | 0.401 |
| chr3 | 38061029 | 38073579 | 12551 | -0.24 | 0.327 | 0.401 |
| chr1 | 224294046 | 224302984 | 8939 | -0.18 | 0.327 | 0.401 |
| chr2 | 206212247 | 206212932 | 686 | -0.54 | 0.327 | 0.401 |
| chr3 | 50606375 | 50607843 | 1469 | 0.31 | 0.328 | 0.401 |
| chr15 | 74485694 | 74508599 | 22906 | -0.1 | 0.328 | 0.401 |
| chr14 | 103961713 | 104009793 | 48081 | -0.09 | 0.328 | 0.401 |
| chr10 | 77159986 | 77169633 | 9648 | -0.17 | 0.328 | 0.401 |
| chr21 | 17791328 | 17793632 | 2305 | -0.31 | 0.328 | 0.402 |
| chr4 | 154004325 | 154004700 | 376 | 0.67 | 0.328 | 0.402 |
| chr10 | 81741122 | 81742436 | 1315 | 0.29 | 0.328 | 0.402 |
| chr18 | 52547325 | 52548558 | 1234 | 0.51 | 0.328 | 0.402 |
| chr10 | 29841987 | 29843117 | 1131 | -0.47 | 0.328 | 0.402 |
| chr19 | 44630962 | 44631501 | 540 | 1.09 | 0.328 | 0.402 |
| chr10 | 13628717 | 13629947 | 1231 | 0.36 | 0.329 | 0.402 |
| chr1 | 52151916 | 52159937 | 8022 | -0.12 | 0.329 | 0.402 |
| chr3 | 70202623 | 70203185 | 563 | 0.52 | 0.329 | 0.402 |
| chr10 | 114635618 | 114636158 | 541 | -0.54 | 0.329 | 0.402 |
| chr10 | 80059582 | 80063197 | 3616 | 0.22 | 0.329 | 0.402 |
| chr7 | 105007685 | 105009377 | 1693 | -0.35 | 0.329 | 0.403 |
| chr5 | 168586952 | 168592254 | 5303 | 0.14 | 0.329 | 0.403 |
| chr22 | 21234454 | 21242074 | 7621 | -0.21 | 0.329 | 0.403 |
| chr6 | 18122371 | 18123767 | 1397 | 0.28 | 0.329 | 0.403 |
| chr3 | 37313726 | 37315515 | 1790 | -0.43 | 0.329 | 0.403 |
| chr1 | 97432535 | 97432883 | 349 | -0.91 | 0.329 | 0.403 |
| chr11 | 1967686 | 1971001 | 3316 | -0.14 | 0.329 | 0.403 |
| chr4 | 87429260 | 87430377 | 1118 | 0.43 | 0.329 | 0.403 |
| chr8 | 10191617 | 10193042 | 1426 | -0.32 | 0.33 | 0.403 |
| chr6 | 126660701 | 126662198 | 1498 | 0.28 | 0.33 | 0.403 |

| | | | | | | |
|---|---|---|---|---|---|---|
| chr6 | 2902303 | 2903751 | 1449 | 0.33 | 0.33 | 0.403 |
| chr2 | 122204089 | 122204388 | 300 | 0.68 | 0.33 | 0.403 |
| chr13 | 44439742 | 44454667 | 14926 | -0.16 | 0.33 | 0.403 |
| chr2 | 230827773 | 230829094 | 1322 | -0.42 | 0.33 | 0.403 |
| chr11 | 34339333 | 34393919 | 54587 | -0.06 | 0.33 | 0.403 |
| chr10 | 120759612 | 120767648 | 8037 | -0.2 | 0.33 | 0.403 |
| chr15 | 55469389 | 55469746 | 358 | -0.82 | 0.33 | 0.403 |
| chr1 | 229842296 | 229843670 | 1375 | -0.49 | 0.33 | 0.403 |
| chr16 | 27471120 | 27485109 | 13990 | -0.23 | 0.33 | 0.403 |
| chr15 | 86069751 | 86072242 | 2492 | 1.93 | 0.33 | 0.404 |
| chr18 | 64207066 | 64207841 | 776 | 0.52 | 0.33 | 0.404 |
| chr11 | 9025033 | 9025554 | 522 | 0.64 | 0.33 | 0.404 |
| chr16 | 85586563 | 85597865 | 11303 | -0.14 | 0.33 | 0.404 |
| chr6 | 130341306 | 130342123 | 818 | 0.5 | 0.33 | 0.404 |
| chr16 | 637498 | 640800 | 3303 | -0.15 | 0.33 | 0.404 |
| chr1 | 117395544 | 117397481 | 1938 | 0.3 | 0.33 | 0.404 |
| chr16 | 70510205 | 70516703 | 6499 | -0.23 | 0.33 | 0.404 |
| chr1 | 42277878 | 42278158 | 281 | -0.67 | 0.33 | 0.404 |
| chr1 | 244376946 | 244378841 | 1896 | 0.31 | 0.331 | 0.404 |
| chr8 | 42050944 | 42069038 | 18095 | -0.14 | 0.331 | 0.404 |
| chr14 | 94528951 | 94532956 | 4006 | 0.27 | 0.331 | 0.404 |
| chr19 | 13226380 | 13240358 | 13979 | -0.17 | 0.331 | 0.404 |
| chr22 | 29810828 | 29811678 | 851 | 0.59 | 0.331 | 0.404 |
| chrX | 70502108 | 70505478 | 3371 | -0.12 | 0.331 | 0.404 |
| chr8 | 99128696 | 99130890 | 2195 | -0.21 | 0.331 | 0.405 |
| chr7 | 26782640 | 26783515 | 876 | 0.55 | 0.331 | 0.405 |
| chr3 | 129273061 | 129280985 | 7925 | -0.16 | 0.332 | 0.405 |
| chr1 | 41216104 | 41216991 | 888 | 0.54 | 0.332 | 0.405 |
| chr1 | 211785828 | 211788393 | 2566 | 0.23 | 0.332 | 0.405 |
| chr4 | 140894487 | 140910937 | 16451 | 0.21 | 0.332 | 0.405 |
| chr1 | 65837933 | 65844079 | 6147 | -0.33 | 0.332 | 0.405 |
| chr2 | 101924285 | 101925857 | 1573 | -0.26 | 0.332 | 0.405 |
| chr10 | 93751146 | 93755866 | 4721 | 0.22 | 0.332 | 0.406 |
| chr10 | 97089459 | 97095162 | 5704 | -0.24 | 0.332 | 0.406 |
| chr7 | 127775479 | 127775821 | 343 | -0.92 | 0.332 | 0.406 |
| chr1 | 100314946 | 100316538 | 1593 | 0.25 | 0.332 | 0.406 |
| chr6 | 155109482 | 155114293 | 4812 | 0.34 | 0.332 | 0.406 |
| chr20 | 56754354 | 56754707 | 354 | -0.69 | 0.332 | 0.406 |
| chr10 | 45745176 | 45745863 | 688 | 0.53 | 0.332 | 0.406 |
| chr5 | 17852546 | 17861785 | 9240 | -0.19 | 0.332 | 0.406 |
| chr1 | 198596862 | 198597728 | 867 | -0.6 | 0.333 | 0.406 |
| chr12 | 45730663 | 45764249 | 33587 | -0.16 | 0.333 | 0.406 |
| chr2 | 136741167 | 136743865 | 2699 | -0.16 | 0.333 | 0.406 |
| chr19 | 33458425 | 33470159 | 11735 | -0.1 | 0.333 | 0.406 |
| chr4 | 88051673 | 88053292 | 1620 | -0.38 | 0.333 | 0.406 |
| chrX | 53278546 | 53317205 | 38660 | -0.08 | 0.333 | 0.407 |
| chr18 | 59852991 | 59855827 | 2837 | 0.26 | 0.333 | 0.407 |
| chr10 | 98829440 | 98829851 | 412 | -0.68 | 0.333 | 0.407 |
| chr6 | 25006702 | 25007675 | 974 | 0.48 | 0.333 | 0.407 |
| chr22 | 50628320 | 50630124 | 1805 | 0.22 | 0.333 | 0.407 |
| chr8 | 6317918 | 6324282 | 6365 | 0.16 | 0.333 | 0.407 |
| chr19 | 10399153 | 10400266 | 1114 | -0.41 | 0.333 | 0.407 |
| chr5 | 1073524 | 1074028 | 505 | -0.75 | 0.333 | 0.407 |
| chr3 | 48025449 | 48026287 | 839 | 0.72 | 0.333 | 0.407 |
| chr14 | 86053238 | 86054100 | 863 | 0.52 | 0.333 | 0.407 |
| chr19 | 24128632 | 24141154 | 12523 | 1.72 | 0.333 | 0.407 |
| chr2 | 178977059 | 178978136 | 1078 | -0.26 | 0.333 | 0.407 |
| chr3 | 133349054 | 133350026 | 973 | -0.47 | 0.333 | 0.407 |
| chr3 | 149897896 | 149898593 | 698 | 0.63 | 0.334 | 0.407 |
| chr12 | 118453686 | 118473160 | 19475 | 0.11 | 0.334 | 0.407 |
| chr18 | 13374927 | 13383170 | 8244 | 0.33 | 0.334 | 0.407 |

| | | | | | | |
|---|---|---|---|---|---|---|
| chr18 | 51883494 | 51885197 | 1704 | -0.28 | 0.334 | 0.407 |
| chr13 | 35515535 | 35516153 | 619 | -0.66 | 0.334 | 0.407 |
| chr4 | 136195462 | 136196045 | 584 | 0.64 | 0.334 | 0.407 |
| chr1 | 114299506 | 114303104 | 3599 | 0.14 | 0.334 | 0.408 |
| chr14 | 54575071 | 54575839 | 769 | -0.49 | 0.334 | 0.408 |
| chr19 | 19215995 | 19216749 | 755 | -0.4 | 0.334 | 0.408 |
| chr1 | 52867877 | 52879769 | 11893 | -0.14 | 0.335 | 0.408 |
| chr2 | 159979889 | 159982032 | 2144 | 0.19 | 0.335 | 0.408 |
| chr2 | 165476978 | 165478626 | 1649 | -0.3 | 0.335 | 0.408 |
| chr2 | 66722815 | 66724254 | 1440 | -0.4 | 0.335 | 0.408 |
| chr16 | 3930867 | 3931735 | 869 | 0.28 | 0.335 | 0.408 |
| chr3 | 127266867 | 127267379 | 513 | 0.59 | 0.335 | 0.409 |
| chr2 | 101821438 | 101835183 | 13746 | -0.34 | 0.335 | 0.409 |
| chr7 | 2264291 | 2320266 | 55976 | -0.08 | 0.335 | 0.409 |
| chr2 | 44292282 | 44293285 | 1004 | 0.55 | 0.335 | 0.409 |
| chr7 | 73785747 | 73786384 | 638 | 0.7 | 0.335 | 0.409 |
| chr4 | 44679759 | 44683195 | 3437 | 0.16 | 0.335 | 0.409 |
| chr14 | 101131806 | 101132360 | 555 | -0.65 | 0.335 | 0.409 |
| chr9 | 127457216 | 127461311 | 4096 | -1.52 | 0.335 | 0.409 |
| chr13 | 66784512 | 66785377 | 866 | 0.51 | 0.335 | 0.409 |
| chr14 | 42579608 | 42580071 | 464 | -0.66 | 0.335 | 0.409 |
| chr17 | 3485761 | 3487133 | 1373 | -0.56 | 0.335 | 0.409 |
| chr10 | 32292950 | 32293429 | 480 | -0.78 | 0.335 | 0.409 |
| chr17 | 27386615 | 27387113 | 499 | 0.76 | 0.335 | 0.409 |
| chr6 | 2376772 | 2378449 | 1678 | 0.25 | 0.335 | 0.409 |
| chr15 | 91207802 | 91209645 | 1844 | -0.14 | 0.336 | 0.409 |
| chr1 | 27003687 | 27004469 | 783 | 0.44 | 0.336 | 0.409 |
| chr5 | 144966203 | 144968580 | 2378 | 0.17 | 0.336 | 0.409 |
| chr2 | 159846899 | 159867183 | 20285 | -0.18 | 0.336 | 0.409 |
| chr13 | 96365911 | 96367721 | 1811 | 0.28 | 0.336 | 0.409 |
| chr20 | 55286188 | 55293856 | 7669 | 0.2 | 0.336 | 0.409 |
| chr19 | 39520868 | 39545495 | 24628 | 0.16 | 0.336 | 0.409 |
| chr11 | 101780152 | 101787681 | 7530 | 0.11 | 0.336 | 0.41 |
| chr4 | 71832868 | 71833943 | 1076 | -0.41 | 0.336 | 0.41 |
| chr16 | 25267747 | 25270391 | 2645 | 0.16 | 0.336 | 0.41 |
| chr16 | 84081297 | 84090429 | 9133 | -0.23 | 0.336 | 0.41 |
| chrX | 67334214 | 67335715 | 1502 | 0.3 | 0.336 | 0.41 |
| chr1 | 6284082 | 6296677 | 12596 | 0.21 | 0.336 | 0.41 |
| chr9 | 37522656 | 37523498 | 843 | 0.56 | 0.336 | 0.41 |
| chr10 | 91507967 | 91508543 | 577 | -0.7 | 0.336 | 0.41 |
| chr18 | 19319881 | 19322191 | 2311 | 0.21 | 0.336 | 0.41 |
| chr2 | 203735044 | 203737106 | 2063 | -0.16 | 0.337 | 0.41 |
| chr12 | 46796471 | 46797192 | 722 | -0.57 | 0.337 | 0.41 |
| chr18 | 21716524 | 21720348 | 3825 | 0.2 | 0.337 | 0.41 |
| chr1 | 228673248 | 228678781 | 5534 | -0.13 | 0.337 | 0.41 |
| chr9 | 26890074 | 26893767 | 3694 | 0.13 | 0.337 | 0.41 |
| chr7 | 7835394 | 7848465 | 13072 | -0.18 | 0.337 | 0.41 |
| chr10 | 78938817 | 78948068 | 9252 | -0.39 | 0.337 | 0.41 |
| chr16 | 48381304 | 48444507 | 63204 | 0.08 | 0.337 | 0.411 |
| chr1 | 84464354 | 84465520 | 1167 | -0.31 | 0.337 | 0.411 |
| chr15 | 89079767 | 89090636 | 10870 | 0.14 | 0.337 | 0.411 |
| chr10 | 13544742 | 13545127 | 386 | -0.81 | 0.337 | 0.411 |
| chr22 | 18366722 | 18377847 | 11126 | 0.21 | 0.337 | 0.411 |
| chr3 | 56817091 | 56818724 | 1634 | 0.33 | 0.338 | 0.411 |
| chr10 | 124228230 | 124229208 | 979 | -0.49 | 0.338 | 0.411 |
| chr2 | 33780110 | 33780831 | 722 | 0.6 | 0.338 | 0.411 |
| chr14 | 76334209 | 76345849 | 11641 | 0.42 | 0.338 | 0.412 |
| chr1 | 42500804 | 42501555 | 752 | 0.5 | 0.338 | 0.412 |
| chr3 | 136620881 | 136632376 | 11496 | 0.4 | 0.338 | 0.412 |
| chr15 | 75018308 | 75018855 | 548 | -0.69 | 0.338 | 0.412 |
| chr4 | 25971744 | 25972680 | 937 | 0.58 | 0.338 | 0.412 |

| | | | | | | |
|---|---|---|---|---|---|---|
| chr12 | 104981019 | 105000242 | 19224 | 0.09 | 0.338 | 0.412 |
| chr1 | 240042533 | 240057521 | 14989 | -0.24 | 0.338 | 0.412 |
| chr15 | 77645872 | 77647192 | 1321 | 0.32 | 0.339 | 0.412 |
| chr3 | 11641131 | 11645370 | 4240 | 0.14 | 0.339 | 0.412 |
| chr3 | 128335974 | 128337300 | 1327 | 0.35 | 0.339 | 0.412 |
| chr18 | 71955831 | 71960362 | 4532 | -0.24 | 0.339 | 0.412 |
| chr7 | 3961168 | 3962822 | 1655 | -0.23 | 0.339 | 0.412 |
| chr13 | 42400790 | 42412556 | 11767 | 0.16 | 0.339 | 0.412 |
| chr3 | 120135601 | 120137866 | 2266 | -0.19 | 0.339 | 0.412 |
| chr1 | 98314852 | 98316161 | 1310 | 0.32 | 0.339 | 0.412 |
| chr14 | 31961611 | 31962590 | 980 | -0.46 | 0.339 | 0.412 |
| chr5 | 132250585 | 132252242 | 1658 | 0.4 | 0.339 | 0.412 |
| chr14 | 70193506 | 70194033 | 528 | 0.69 | 0.339 | 0.413 |
| chr9 | 98585103 | 98585827 | 725 | -0.62 | 0.339 | 0.413 |
| chr19 | 288236 | 291365 | 3130 | -0.16 | 0.339 | 0.413 |
| chr7 | 33168134 | 33169921 | 1788 | 0.21 | 0.339 | 0.413 |
| chr11 | 100679440 | 100681175 | 1736 | 0.35 | 0.339 | 0.413 |
| chr1 | 155456553 | 155457950 | 1398 | 0.42 | 0.339 | 0.413 |
| chr10 | 131264686 | 131266034 | 1349 | 0.39 | 0.34 | 0.413 |
| chr6 | 15608303 | 15609674 | 1372 | 0.27 | 0.34 | 0.413 |
| chr13 | 24721430 | 24735833 | 14404 | 0.1 | 0.34 | 0.413 |
| chr1 | 36805454 | 36864154 | 58701 | -0.06 | 0.34 | 0.413 |
| chr12 | 102585569 | 102592109 | 6541 | -0.49 | 0.34 | 0.413 |
| chr16 | 69303877 | 69304827 | 951 | -0.53 | 0.34 | 0.413 |
| chr10 | 128945416 | 128948491 | 3076 | -0.15 | 0.34 | 0.413 |
| chr1 | 243376397 | 243377307 | 911 | -0.56 | 0.34 | 0.413 |
| chrX | 153389293 | 153389825 | 533 | -0.58 | 0.34 | 0.413 |
| chr5 | 125929419 | 125938087 | 8669 | 0.14 | 0.34 | 0.413 |
| chr21 | 40816910 | 40818412 | 1503 | 0.2 | 0.34 | 0.413 |
| chr17 | 49300952 | 49308174 | 7223 | -0.22 | 0.34 | 0.413 |
| chr6 | 83007604 | 83008280 | 677 | -0.55 | 0.34 | 0.413 |
| chr1 | 245985927 | 245987027 | 1101 | 0.56 | 0.34 | 0.413 |
| chr6 | 7610802 | 7611623 | 822 | -0.63 | 0.34 | 0.414 |
| chr11 | 100503381 | 100510516 | 7136 | 0.27 | 0.34 | 0.414 |
| chr22 | 50963483 | 50983656 | 20174 | 0.1 | 0.34 | 0.414 |
| chr2 | 170478610 | 170479403 | 794 | -0.57 | 0.341 | 0.414 |
| chr11 | 67182762 | 67197090 | 14329 | 0.18 | 0.341 | 0.414 |
| chr19 | 18698959 | 18701459 | 2501 | -0.2 | 0.341 | 0.414 |
| chr20 | 6020760 | 6021642 | 883 | 0.55 | 0.341 | 0.414 |
| chr2 | 17824254 | 17825313 | 1060 | -0.51 | 0.341 | 0.414 |
| chr3 | 72620104 | 72620486 | 383 | 0.86 | 0.341 | 0.414 |
| chr21 | 34633576 | 34642331 | 8756 | 0.22 | 0.341 | 0.414 |
| chr5 | 41182822 | 41183183 | 362 | -0.73 | 0.341 | 0.414 |
| chr11 | 62135737 | 62136184 | 448 | 0.65 | 0.341 | 0.414 |
| chr4 | 113683624 | 113684053 | 430 | -0.47 | 0.341 | 0.414 |
| chr10 | 105153532 | 105157666 | 4135 | 0.15 | 0.341 | 0.414 |
| chr3 | 150320076 | 150322595 | 2520 | 0.18 | 0.341 | 0.414 |
| chr2 | 118770797 | 118772275 | 1479 | -0.29 | 0.341 | 0.415 |
| chr13 | 114539904 | 114552425 | 12522 | -0.14 | 0.341 | 0.415 |
| chr10 | 105491302 | 105492296 | 995 | 0.46 | 0.341 | 0.415 |
| chr12 | 111284478 | 111285297 | 820 | -0.37 | 0.342 | 0.415 |
| chr22 | 39409672 | 39411425 | 1754 | -0.24 | 0.342 | 0.415 |
| chr4 | 142297024 | 142298041 | 1018 | 0.37 | 0.342 | 0.415 |
| chr6 | 142477138 | 142494802 | 17665 | 0.16 | 0.342 | 0.415 |
| chr12 | 124585255 | 124585821 | 567 | -0.64 | 0.342 | 0.415 |
| chr3 | 184428176 | 184430472 | 2297 | 0.22 | 0.342 | 0.415 |
| chr8 | 144655051 | 144700960 | 45910 | -0.08 | 0.342 | 0.415 |
| chr7 | 158290798 | 158291786 | 989 | -0.52 | 0.342 | 0.415 |
| chr11 | 107728603 | 107730854 | 2252 | 0.22 | 0.342 | 0.416 |
| chr4 | 177191741 | 177192959 | 1219 | -0.35 | 0.342 | 0.416 |
| chr18 | 21050472 | 21050924 | 453 | -0.59 | 0.343 | 0.416 |

| | | | | | | |
|---|---|---|---|---|---|---|
| chr2 | 6321315 | 6327710 | 6396 | -0.17 | 0.343 | 0.416 |
| chr2 | 211340038 | 211342574 | 2537 | -0.23 | 0.343 | 0.416 |
| chr11 | 95245385 | 95246645 | 1261 | 0.32 | 0.343 | 0.416 |
| chr1 | 162792132 | 162792664 | 533 | 0.72 | 0.343 | 0.416 |
| chr6 | 170199925 | 170201957 | 2033 | 0.3 | 0.343 | 0.416 |
| chr10 | 124263573 | 124264523 | 951 | 0.49 | 0.343 | 0.417 |
| chr7 | 123324879 | 123325289 | 411 | -0.64 | 0.343 | 0.417 |
| chr19 | 14016449 | 14017713 | 1265 | 0.37 | 0.343 | 0.417 |
| chr2 | 63635080 | 63636433 | 1354 | -0.31 | 0.344 | 0.417 |
| chr8 | 101678977 | 101689423 | 10447 | 0.12 | 0.344 | 0.417 |
| chr5 | 55007388 | 55009190 | 1803 | 0.23 | 0.344 | 0.417 |
| chr14 | 43004197 | 43004852 | 656 | 0.4 | 0.344 | 0.417 |
| chr7 | 152456213 | 152458207 | 1995 | 0.18 | 0.344 | 0.417 |
| chr2 | 241926325 | 241937989 | 11665 | -0.34 | 0.344 | 0.417 |
| chr8 | 126697983 | 126713076 | 15094 | -0.22 | 0.344 | 0.417 |
| chr1 | 109044898 | 109046313 | 1416 | -0.36 | 0.344 | 0.418 |
| chr19 | 41028333 | 41047676 | 19344 | -0.18 | 0.344 | 0.418 |
| chr14 | 71448248 | 71449239 | 992 | 1.69 | 0.344 | 0.418 |
| chr4 | 25377952 | 25379788 | 1837 | 0.23 | 0.345 | 0.418 |
| chr10 | 92631208 | 92632581 | 1374 | 0.21 | 0.345 | 0.418 |
| chr3 | 56650077 | 56651085 | 1009 | -0.59 | 0.345 | 0.418 |
| chr12 | 116400555 | 116401987 | 1433 | 0.37 | 0.345 | 0.418 |
| chrX | 24482597 | 24484161 | 1565 | 0.34 | 0.345 | 0.419 |
| chr15 | 69452138 | 69466498 | 14361 | 0.2 | 0.345 | 0.419 |
| chr11 | 111943687 | 111958272 | 14586 | -0.11 | 0.345 | 0.419 |
| chr9 | 89020386 | 89021577 | 1192 | -0.39 | 0.345 | 0.419 |
| chr3 | 112612843 | 112613811 | 969 | -0.33 | 0.345 | 0.419 |
| chr9 | 110241933 | 110269844 | 27912 | -0.08 | 0.346 | 0.419 |
| chr6 | 85306767 | 85307502 | 736 | 1.64 | 0.346 | 0.419 |
| chr19 | 1383582 | 1384518 | 937 | 0.41 | 0.346 | 0.42 |
| chr18 | 42753321 | 42753800 | 480 | -1.02 | 0.346 | 0.42 |
| chr8 | 6564033 | 6568276 | 4244 | 0.15 | 0.346 | 0.42 |
| chr9 | 15349539 | 15350990 | 1452 | 0.26 | 0.346 | 0.42 |
| chr19 | 16737800 | 16740421 | 2622 | 0.24 | 0.346 | 0.42 |
| chr5 | 27471197 | 27477293 | 6097 | -0.11 | 0.347 | 0.42 |
| chr3 | 11682001 | 11686383 | 4383 | 0.09 | 0.347 | 0.42 |
| chr12 | 79153041 | 79153975 | 935 | -0.37 | 0.347 | 0.42 |
| chr17 | 4889103 | 4905694 | 16592 | 0.11 | 0.347 | 0.42 |
| chrX | 16167020 | 16179542 | 12523 | -1.98 | 0.347 | 0.42 |
| chr10 | 127500441 | 127541548 | 41108 | -0.08 | 0.347 | 0.421 |
| chr18 | 77866541 | 77868401 | 1861 | 0.19 | 0.347 | 0.421 |
| chr9 | 85833821 | 85835069 | 1249 | 0.45 | 0.347 | 0.421 |
| chr1 | 154377124 | 154452438 | 75315 | -0.06 | 0.347 | 0.421 |
| chr2 | 86667350 | 86670688 | 3339 | 0.12 | 0.348 | 0.421 |
| chr11 | 111382734 | 111384017 | 1284 | -0.3 | 0.348 | 0.421 |
| chr6 | 37888110 | 37899084 | 10975 | 0.34 | 0.348 | 0.422 |
| chr15 | 51199098 | 51205055 | 5958 | -0.13 | 0.348 | 0.422 |
| chr8 | 66513361 | 66520808 | 7448 | 0.29 | 0.348 | 0.422 |
| chr11 | 61879721 | 61905422 | 25702 | -0.18 | 0.348 | 0.422 |
| chr3 | 15160136 | 15160903 | 768 | 0.58 | 0.348 | 0.422 |
| chr10 | 97396004 | 97396542 | 539 | -0.63 | 0.348 | 0.422 |
| chr6 | 37190017 | 37191250 | 1234 | 0.43 | 0.349 | 0.422 |
| chr6 | 130241121 | 130242431 | 1311 | 0.32 | 0.349 | 0.422 |
| chr22 | 17519396 | 17520260 | 865 | 0.47 | 0.349 | 0.423 |
| chr20 | 17485235 | 17485892 | 658 | -0.55 | 0.349 | 0.423 |
| chr1 | 14026130 | 14027290 | 1161 | 0.27 | 0.349 | 0.423 |
| chr5 | 106470691 | 106471898 | 1208 | 0.36 | 0.349 | 0.423 |
| chr12 | 22435030 | 22435541 | 512 | 0.61 | 0.349 | 0.423 |
| chr17 | 74390207 | 74390839 | 633 | -0.64 | 0.349 | 0.423 |
| chr15 | 41877327 | 41878044 | 718 | -0.59 | 0.349 | 0.423 |
| chr18 | 23620408 | 23621090 | 683 | 0.57 | 0.35 | 0.423 |

| | | | | | | |
|---|---|---|---|---|---|---|
| chr10 | 35142645 | 35153256 | 10612 | 0.14 | 0.35 | 0.424 |
| chr5 | 95237356 | 95238183 | 828 | 0.56 | 0.35 | 0.424 |
| chr5 | 141105144 | 141106229 | 1086 | -0.47 | 0.35 | 0.424 |
| chr17 | 27115276 | 27115571 | 296 | 0.91 | 0.35 | 0.424 |
| chr5 | 157375171 | 157376525 | 1355 | 0.41 | 0.35 | 0.424 |
| chr20 | 304117 | 310812 | 6696 | -0.1 | 0.351 | 0.424 |
| chr1 | 217666406 | 217667429 | 1024 | -0.49 | 0.351 | 0.424 |
| chr6 | 3337507 | 3339268 | 1762 | -0.25 | 0.351 | 0.425 |
| chr2 | 204145296 | 204145817 | 522 | -0.5 | 0.351 | 0.425 |
| chr6 | 10032173 | 10034991 | 2819 | -0.24 | 0.351 | 0.425 |
| chrX | 49968425 | 49970033 | 1609 | 0.23 | 0.351 | 0.425 |
| chr1 | 214836040 | 214837141 | 1102 | 0.42 | 0.351 | 0.425 |
| chr8 | 41455340 | 41477146 | 21807 | 0.1 | 0.351 | 0.425 |
| chr6 | 20810815 | 20812413 | 1599 | -0.17 | 0.351 | 0.425 |
| chr20 | 32680937 | 32701136 | 20200 | 0.08 | 0.351 | 0.425 |
| chr5 | 134209344 | 134211393 | 2050 | -0.15 | 0.351 | 0.425 |
| chr8 | 56791898 | 56793794 | 1897 | 0.26 | 0.351 | 0.425 |
| chr7 | 84266185 | 84268246 | 2062 | 0.22 | 0.352 | 0.426 |
| chr1 | 44792681 | 44794741 | 2061 | 0.22 | 0.352 | 0.426 |
| chr9 | 114137395 | 114138222 | 828 | 0.55 | 0.352 | 0.426 |
| chr12 | 120932153 | 120939182 | 7030 | -0.14 | 0.352 | 0.426 |
| chr9 | 79481070 | 79494759 | 13690 | -0.44 | 0.352 | 0.426 |
| chr6 | 35215768 | 35228799 | 13032 | 0.12 | 0.352 | 0.426 |
| chr10 | 127729934 | 127731705 | 1772 | 0.31 | 0.352 | 0.426 |
| chr15 | 63034375 | 63035612 | 1238 | 0.35 | 0.352 | 0.426 |
| chr3 | 27707731 | 27723426 | 15696 | -0.12 | 0.352 | 0.426 |
| chr22 | 38559394 | 38578726 | 19333 | -0.1 | 0.352 | 0.426 |
| chr12 | 102838327 | 102838858 | 532 | -0.59 | 0.352 | 0.426 |
| chr5 | 126852616 | 126854824 | 2209 | 0.18 | 0.352 | 0.426 |
| chr5 | 53605429 | 53607276 | 1848 | 0.21 | 0.352 | 0.426 |
| chr8 | 31829462 | 31830299 | 838 | -0.6 | 0.353 | 0.427 |
| chr1 | 240721066 | 240721619 | 554 | 0.94 | 0.353 | 0.427 |
| chr11 | 129148227 | 129149093 | 867 | 0.51 | 0.353 | 0.427 |
| chr17 | 40108658 | 40178185 | 69528 | 0.05 | 0.353 | 0.427 |
| chr13 | 42679116 | 42680962 | 1847 | 0.26 | 0.353 | 0.427 |
| chr9 | 100326369 | 100327346 | 978 | -0.42 | 0.353 | 0.427 |
| chr3 | 14060907 | 14061512 | 606 | -0.51 | 0.353 | 0.427 |
| chr10 | 94833213 | 94841279 | 8067 | -0.23 | 0.353 | 0.427 |
| chr1 | 244872949 | 244873500 | 552 | -0.86 | 0.354 | 0.428 |
| chr4 | 128338522 | 128339767 | 1246 | 0.33 | 0.354 | 0.428 |
| chr17 | 58635055 | 58636141 | 1087 | -0.46 | 0.354 | 0.428 |
| chrX | 20007891 | 20009968 | 2078 | 0.29 | 0.354 | 0.428 |
| chr19 | 36427065 | 36428058 | 994 | -0.46 | 0.354 | 0.428 |
| chr1 | 155340551 | 155341230 | 680 | 0.93 | 0.354 | 0.428 |
| chr11 | 3691027 | 3698579 | 7553 | -0.21 | 0.354 | 0.428 |
| chr2 | 8441775 | 8448169 | 6395 | -0.19 | 0.354 | 0.428 |
| chr12 | 956544 | 984986 | 28443 | 0.1 | 0.354 | 0.428 |
| chr4 | 1926384 | 1927413 | 1030 | 0.44 | 0.354 | 0.428 |
| chr2 | 216945576 | 216947278 | 1703 | -0.35 | 0.355 | 0.429 |
| chr5 | 14809591 | 14810850 | 1260 | -0.29 | 0.355 | 0.429 |
| chr14 | 23235129 | 23242937 | 7809 | 0.13 | 0.355 | 0.429 |
| chr8 | 74650022 | 74660170 | 10149 | 0.18 | 0.355 | 0.429 |
| chr12 | 26922139 | 26922755 | 617 | 0.57 | 0.355 | 0.429 |
| chr6 | 34565996 | 34579094 | 13099 | -0.19 | 0.355 | 0.429 |
| chr8 | 17459929 | 17473421 | 13493 | 0.1 | 0.355 | 0.429 |
| chrX | 117250201 | 117251409 | 1209 | 0.45 | 0.355 | 0.429 |
| chr1 | 247092806 | 247096091 | 3286 | 0.18 | 0.355 | 0.429 |
| chr3 | 158362097 | 158397878 | 35782 | 0.07 | 0.355 | 0.429 |
| chr3 | 16947217 | 16949099 | 1883 | 0.28 | 0.355 | 0.429 |
| chr4 | 76860406 | 76862114 | 1709 | -0.25 | 0.355 | 0.429 |
| chrX | 67866970 | 67875086 | 8117 | -0.22 | 0.355 | 0.429 |

| | | | | | | |
|---|---|---|---|---|---|---|
| chr9 | 80129903 | 80136509 | 6607 | 0.39 | 0.356 | 0.43 |
| chr2 | 19873088 | 19883029 | 9942 | -1.62 | 0.356 | 0.43 |
| chr3 | 47743954 | 47746324 | 2371 | 0.29 | 0.356 | 0.43 |
| chr9 | 114423474 | 114425213 | 1740 | -0.27 | 0.356 | 0.43 |
| chr3 | 141658461 | 141659638 | 1178 | 0.38 | 0.356 | 0.43 |
| chr5 | 132561209 | 132562351 | 1143 | -0.33 | 0.356 | 0.43 |
| chr3 | 16304925 | 16308165 | 3241 | -0.11 | 0.357 | 0.431 |
| chr7 | 39503837 | 39505448 | 1612 | -0.25 | 0.357 | 0.431 |
| chr3 | 44802393 | 44804737 | 2345 | -0.19 | 0.357 | 0.431 |
| chr9 | 139858658 | 139859332 | 675 | -0.53 | 0.357 | 0.431 |
| chr9 | 118822769 | 118823503 | 735 | 0.58 | 0.357 | 0.431 |
| chr7 | 128629700 | 128631992 | 2293 | -0.45 | 0.357 | 0.431 |
| chr7 | 133983155 | 133985140 | 1986 | -0.26 | 0.357 | 0.431 |
| chr15 | 64477241 | 64508704 | 31464 | -0.12 | 0.357 | 0.431 |
| chr17 | 61351759 | 61354145 | 2387 | 0.16 | 0.357 | 0.431 |
| chr9 | 107912726 | 107917505 | 4780 | 0.27 | 0.357 | 0.431 |
| chr8 | 141319335 | 141333605 | 14271 | -1.43 | 0.357 | 0.432 |
| chr12 | 26348245 | 26349476 | 1232 | -0.3 | 0.357 | 0.432 |
| chr9 | 103187673 | 103207009 | 19337 | 0.06 | 0.358 | 0.432 |
| chr2 | 109234851 | 109238837 | 3987 | -0.17 | 0.358 | 0.432 |
| chr1 | 225789476 | 225815924 | 26449 | -0.11 | 0.358 | 0.432 |
| chr12 | 12672412 | 12673239 | 828 | -0.46 | 0.358 | 0.432 |
| chr17 | 62519788 | 62547705 | 27918 | -0.09 | 0.358 | 0.432 |
| chr5 | 124258065 | 124269052 | 10988 | 1.83 | 0.358 | 0.432 |
| chrX | 70842354 | 70843071 | 718 | 0.53 | 0.358 | 0.432 |
| chr8 | 67782401 | 67783559 | 1159 | -0.25 | 0.358 | 0.432 |
| chr10 | 70660039 | 70662852 | 2814 | 0.12 | 0.358 | 0.433 |
| chr7 | 111832372 | 111852511 | 20140 | -0.14 | 0.359 | 0.433 |
| chr13 | 21402465 | 21403367 | 903 | 0.45 | 0.359 | 0.433 |
| chr3 | 178787309 | 178790324 | 3016 | -0.18 | 0.359 | 0.433 |
| chr20 | 31406985 | 31430619 | 23635 | 0.11 | 0.359 | 0.433 |
| chr2 | 242178253 | 242179014 | 762 | -0.9 | 0.359 | 0.433 |
| chr16 | 75497940 | 75498652 | 713 | 0.29 | 0.359 | 0.433 |
| chr6 | 166510548 | 166511514 | 967 | -0.42 | 0.359 | 0.433 |
| chr4 | 152681548 | 152682190 | 643 | -0.31 | 0.359 | 0.433 |
| chr3 | 182677331 | 182698931 | 21601 | 0.11 | 0.359 | 0.433 |
| chr20 | 1549737 | 1550842 | 1106 | -0.31 | 0.359 | 0.433 |
| chr1 | 44819720 | 44821456 | 1737 | -0.28 | 0.36 | 0.434 |
| chr3 | 145783688 | 145790875 | 7188 | -0.18 | 0.36 | 0.434 |
| chr1 | 167423825 | 167425299 | 1475 | -0.24 | 0.36 | 0.434 |
| chr3 | 174609106 | 174609522 | 417 | 1.02 | 0.36 | 0.434 |
| chr8 | 47187130 | 47187687 | 558 | 1.02 | 0.36 | 0.434 |
| chr9 | 136821375 | 136822299 | 925 | 0.61 | 0.36 | 0.434 |
| chr8 | 101704865 | 101735745 | 30881 | 0.07 | 0.36 | 0.434 |
| chr19 | 45873267 | 45880762 | 7496 | 0.14 | 0.36 | 0.434 |
| chr15 | 65890203 | 65904746 | 14544 | -0.08 | 0.36 | 0.434 |
| chr17 | 56428460 | 56442523 | 14064 | 0.16 | 0.36 | 0.434 |
| chr9 | 131182337 | 131184002 | 1666 | -0.39 | 0.36 | 0.434 |
| chr2 | 81756976 | 81769674 | 12699 | 1.71 | 0.36 | 0.434 |
| chr20 | 45567975 | 45568680 | 706 | -0.56 | 0.361 | 0.435 |
| chr1 | 207405955 | 207415261 | 9307 | 0.18 | 0.361 | 0.435 |
| chr5 | 31529900 | 31534380 | 4481 | 0.13 | 0.361 | 0.435 |
| chr7 | 114529298 | 114530305 | 1008 | -0.35 | 0.361 | 0.435 |
| chr18 | 21692853 | 21693769 | 917 | -0.44 | 0.361 | 0.435 |
| chr2 | 114083134 | 114087611 | 4478 | 0.18 | 0.361 | 0.435 |
| chr6 | 26420965 | 26422227 | 1263 | 0.26 | 0.361 | 0.435 |
| chr3 | 151260961 | 151262509 | 1549 | -0.23 | 0.361 | 0.435 |
| chr7 | 67402047 | 67403753 | 1707 | 0.22 | 0.361 | 0.435 |
| chr2 | 55262226 | 55280977 | 18752 | -0.05 | 0.361 | 0.435 |
| chr5 | 95359676 | 95373534 | 13859 | -0.27 | 0.361 | 0.435 |
| chr12 | 129276170 | 129282811 | 6642 | 0.23 | 0.361 | 0.436 |

| | | | | | | |
|---|---|---|---|---|---|---|
| chr5 | 149979334 | 149986352 | 7019 | -0.28 | 0.361 | 0.436 |
| chr14 | 95898175 | 95898828 | 654 | -0.55 | 0.362 | 0.436 |
| chr9 | 131484632 | 131496066 | 11435 | -0.16 | 0.362 | 0.436 |
| chr12 | 118294680 | 118295148 | 469 | -0.77 | 0.362 | 0.436 |
| chr22 | 45831210 | 45832175 | 966 | 0.42 | 0.362 | 0.436 |
| chr1 | 59346505 | 59370588 | 24084 | -0.12 | 0.362 | 0.436 |
| chr15 | 89786533 | 89788797 | 2265 | -0.18 | 0.362 | 0.436 |
| chr1 | 41898149 | 41899221 | 1073 | -0.5 | 0.362 | 0.436 |
| chr10 | 5275788 | 5277036 | 1249 | 0.37 | 0.363 | 0.437 |
| chr1 | 234886615 | 234887687 | 1073 | 1.64 | 0.363 | 0.437 |
| chr3 | 140949869 | 140951222 | 1354 | 0.33 | 0.363 | 0.437 |
| chr3 | 33137196 | 33140039 | 2844 | 0.12 | 0.363 | 0.437 |
| chr19 | 850084 | 871004 | 20921 | -0.13 | 0.363 | 0.437 |
| chr9 | 5948583 | 5954426 | 5844 | -0.21 | 0.363 | 0.437 |
| chr1 | 206807429 | 206821025 | 13597 | -0.2 | 0.363 | 0.437 |
| chr19 | 15542551 | 15544212 | 1662 | -0.34 | 0.363 | 0.437 |
| chr13 | 24462248 | 24463565 | 1318 | -0.26 | 0.363 | 0.437 |
| chr9 | 107800237 | 107801255 | 1019 | -0.5 | 0.363 | 0.438 |
| chr2 | 102806762 | 102807378 | 617 | 0.7 | 0.364 | 0.438 |
| chr2 | 205888929 | 205890706 | 1778 | -0.27 | 0.364 | 0.438 |
| chr11 | 64936230 | 64950419 | 14190 | -0.18 | 0.364 | 0.438 |
| chr9 | 129088369 | 129098058 | 9690 | -0.19 | 0.364 | 0.438 |
| chr14 | 51861900 | 51866320 | 4421 | -0.1 | 0.364 | 0.438 |
| chr14 | 78107284 | 78108708 | 1425 | 0.54 | 0.364 | 0.438 |
| chr19 | 4086163 | 4087088 | 926 | 0.44 | 0.364 | 0.438 |
| chr1 | 33281147 | 33284740 | 3594 | 0.13 | 0.364 | 0.439 |
| chr1 | 151406986 | 151432827 | 25842 | 0.09 | 0.365 | 0.439 |
| chr11 | 126172991 | 126174761 | 1771 | 0.19 | 0.365 | 0.439 |
| chr8 | 100904112 | 100906124 | 2013 | 0.21 | 0.365 | 0.439 |
| chrX | 44833141 | 44835690 | 2550 | 0.28 | 0.365 | 0.439 |
| chr17 | 45500660 | 45501003 | 344 | -0.66 | 0.365 | 0.439 |
| chr2 | 197195287 | 197200047 | 4761 | 0.32 | 0.365 | 0.439 |
| chr20 | 22492494 | 22493677 | 1184 | 0.32 | 0.365 | 0.439 |
| chr16 | 90112933 | 90114114 | 1182 | 0.34 | 0.365 | 0.44 |
| chr11 | 121972248 | 121972834 | 587 | -0.58 | 0.365 | 0.44 |
| chr10 | 24932389 | 24933112 | 724 | 0.63 | 0.365 | 0.44 |
| chr6 | 108543123 | 108545172 | 2050 | -0.33 | 0.366 | 0.44 |
| chr5 | 167646533 | 167646911 | 379 | 0.49 | 0.366 | 0.44 |
| chr9 | 110803093 | 110822506 | 19414 | -0.08 | 0.366 | 0.44 |
| chr3 | 134235650 | 134236221 | 572 | 0.87 | 0.366 | 0.44 |
| chr1 | 151284238 | 151285063 | 826 | 0.42 | 0.366 | 0.44 |
| chr14 | 54901104 | 54909182 | 8079 | 0.13 | 0.366 | 0.44 |
| chr3 | 196865531 | 196866544 | 1014 | 0.46 | 0.366 | 0.44 |
| chr16 | 71460743 | 71461438 | 696 | -0.38 | 0.366 | 0.441 |
| chr7 | 37122727 | 37123619 | 893 | -0.49 | 0.366 | 0.441 |
| chr8 | 101155719 | 101171670 | 15952 | 0.1 | 0.366 | 0.441 |
| chr1 | 214775501 | 214801468 | 25968 | 0.1 | 0.366 | 0.441 |
| chr14 | 77239662 | 77240818 | 1157 | 0.43 | 0.366 | 0.441 |
| chr18 | 56336776 | 56350342 | 13567 | 0.14 | 0.367 | 0.441 |
| chr4 | 17615274 | 17617339 | 2066 | 0.16 | 0.367 | 0.441 |
| chr13 | 110830186 | 110830711 | 526 | -0.52 | 0.367 | 0.441 |
| chr1 | 109965583 | 109970451 | 4869 | 0.14 | 0.367 | 0.441 |
| chr9 | 88268295 | 88270652 | 2358 | 0.25 | 0.367 | 0.441 |
| chr7 | 24495635 | 24497559 | 1925 | 0.24 | 0.367 | 0.441 |
| chr11 | 78176398 | 78177104 | 707 | 0.53 | 0.367 | 0.441 |
| chr14 | 65146576 | 65149639 | 3064 | 0.23 | 0.367 | 0.441 |
| chr20 | 31026900 | 31073804 | 46905 | 0.05 | 0.367 | 0.441 |
| chr7 | 65401594 | 65417953 | 16360 | -0.15 | 0.367 | 0.441 |
| chr1 | 23727167 | 23731200 | 4034 | 0.22 | 0.367 | 0.441 |
| chr1 | 31734528 | 31735196 | 669 | 0.48 | 0.367 | 0.441 |
| chr5 | 58980010 | 58990200 | 10191 | -0.33 | 0.367 | 0.441 |

| | | | | | | |
|---|---|---|---|---|---|---|
| chr12 | 123108609 | 123109534 | 926 | -0.49 | 0.367 | 0.442 |
| chr1 | 38291110 | 38292204 | 1095 | 0.52 | 0.367 | 0.442 |
| chr10 | 43700366 | 43701405 | 1040 | -0.4 | 0.367 | 0.442 |
| chr2 | 32619567 | 32653958 | 34392 | -0.14 | 0.368 | 0.442 |
| chr6 | 17277835 | 17283448 | 5614 | -0.08 | 0.368 | 0.442 |
| chr3 | 195269134 | 195270166 | 1033 | 0.39 | 0.368 | 0.442 |
| chr4 | 144153091 | 144163584 | 10494 | 0.35 | 0.368 | 0.442 |
| chr19 | 2971606 | 2978226 | 6621 | 0.3 | 0.368 | 0.442 |
| chr20 | 35830197 | 35838699 | 8503 | -0.2 | 0.368 | 0.442 |
| chr6 | 17863682 | 17879576 | 15895 | 0.09 | 0.368 | 0.442 |
| chrX | 25006116 | 25007212 | 1097 | 0.57 | 0.368 | 0.443 |
| chr12 | 465313 | 466491 | 1179 | 0.41 | 0.368 | 0.443 |
| chr2 | 235771647 | 235797220 | 25574 | -0.13 | 0.369 | 0.443 |
| chr2 | 37116027 | 37121666 | 5640 | 0.14 | 0.369 | 0.443 |
| chr4 | 188539711 | 188540737 | 1027 | 0.42 | 0.369 | 0.443 |
| chr20 | 62710407 | 62711272 | 866 | 0.4 | 0.369 | 0.443 |
| chr1 | 8660214 | 8660800 | 587 | 0.51 | 0.369 | 0.443 |
| chr8 | 119208040 | 119222197 | 14158 | -1.6 | 0.369 | 0.443 |
| chr9 | 126122121 | 126138481 | 16361 | 1.83 | 0.369 | 0.443 |
| chr9 | 99095908 | 99108832 | 12925 | 0.08 | 0.369 | 0.444 |
| chr1 | 27215797 | 27227530 | 11734 | 0.15 | 0.369 | 0.444 |
| chr19 | 42202344 | 42207212 | 4869 | 0.14 | 0.369 | 0.444 |
| chr3 | 9746250 | 9746641 | 392 | -0.74 | 0.37 | 0.444 |
| chr6 | 109329670 | 109331665 | 1996 | 0.29 | 0.37 | 0.444 |
| chr17 | 77019880 | 77038530 | 18651 | 0.11 | 0.37 | 0.444 |
| chr21 | 33031237 | 33044066 | 12830 | 0.15 | 0.37 | 0.444 |
| chr1 | 202204417 | 202206870 | 2454 | -0.23 | 0.37 | 0.444 |
| chr10 | 3700924 | 3701610 | 687 | 0.5 | 0.37 | 0.444 |
| chr9 | 114652979 | 114691030 | 38052 | 0.09 | 0.37 | 0.444 |
| chr4 | 6855786 | 6861764 | 5979 | 0.11 | 0.37 | 0.445 |
| chr14 | 99966935 | 99968265 | 1331 | 0.55 | 0.37 | 0.445 |
| chr4 | 5129826 | 5140674 | 10849 | 0.19 | 0.37 | 0.445 |
| chr18 | 19763699 | 19764554 | 856 | -0.56 | 0.371 | 0.445 |
| chr1 | 161336495 | 161338636 | 2142 | 0.24 | 0.371 | 0.445 |
| chr13 | 30495060 | 30500992 | 5933 | 0.08 | 0.371 | 0.445 |
| chr16 | 11142073 | 11144175 | 2103 | 0.24 | 0.371 | 0.445 |
| chr13 | 52733004 | 52734073 | 1070 | 0.26 | 0.371 | 0.445 |
| chr9 | 88635515 | 88645787 | 10273 | -0.2 | 0.371 | 0.445 |
| chr3 | 170586237 | 170589162 | 2926 | -0.11 | 0.371 | 0.446 |
| chr13 | 32888888 | 32891042 | 2155 | -0.13 | 0.371 | 0.446 |
| chr2 | 69869577 | 69879725 | 10149 | -0.13 | 0.371 | 0.446 |
| chr20 | 61334784 | 61335915 | 1132 | -0.53 | 0.371 | 0.446 |
| chr3 | 128996548 | 128998981 | 2434 | 0.15 | 0.371 | 0.446 |
| chr1 | 120255240 | 120255821 | 582 | -0.55 | 0.372 | 0.446 |
| chr16 | 23606370 | 23628497 | 22128 | -0.1 | 0.372 | 0.446 |
| chr2 | 183626966 | 183627951 | 986 | -0.7 | 0.372 | 0.446 |
| chrX | 54555591 | 54557963 | 2373 | 0.15 | 0.372 | 0.446 |
| chr9 | 19315749 | 19317337 | 1589 | -0.35 | 0.372 | 0.446 |
| chr14 | 91643044 | 91643793 | 750 | 0.53 | 0.372 | 0.446 |
| chr3 | 189289834 | 189291127 | 1294 | -0.49 | 0.372 | 0.447 |
| chr18 | 20220420 | 20220845 | 426 | -0.56 | 0.373 | 0.447 |
| chr22 | 32072356 | 32073442 | 1087 | 0.36 | 0.373 | 0.447 |
| chrX | 119805540 | 119807522 | 1983 | 0.16 | 0.373 | 0.448 |
| chr15 | 44887541 | 44891980 | 4440 | -0.24 | 0.373 | 0.448 |
| chr10 | 112356469 | 112356896 | 428 | 0.62 | 0.373 | 0.448 |
| chr7 | 157035121 | 157035651 | 531 | -0.65 | 0.373 | 0.448 |
| chr10 | 80516043 | 80518458 | 2416 | 0.34 | 0.373 | 0.448 |
| chr18 | 39602365 | 39603522 | 1158 | 0.38 | 0.373 | 0.448 |
| chr9 | 108265738 | 108270228 | 4491 | -0.15 | 0.373 | 0.448 |
| chr11 | 26306077 | 26306574 | 498 | 0.51 | 0.374 | 0.448 |
| chr1 | 155478909 | 155481868 | 2960 | -0.29 | 0.374 | 0.448 |

| | | | | | | |
|---|---|---|---|---|---|---|
| chr3 | 11519709 | 11522471 | 2763 | 0.36 | 0.374 | 0.448 |
| chr4 | 87812642 | 87825945 | 13304 | 0.17 | 0.374 | 0.448 |
| chr1 | 183884751 | 183885217 | 467 | 0.85 | 0.374 | 0.448 |
| chr10 | 11726616 | 11727981 | 1366 | -0.23 | 0.374 | 0.448 |
| chr14 | 100031708 | 100072345 | 40638 | -0.07 | 0.374 | 0.448 |
| chr1 | 93789666 | 93791410 | 1745 | 0.28 | 0.374 | 0.448 |
| chr9 | 115164216 | 115174650 | 10435 | 0.29 | 0.374 | 0.449 |
| chr20 | 24965684 | 24986638 | 20955 | -0.14 | 0.374 | 0.449 |
| chr14 | 55136511 | 55144807 | 8297 | 0.2 | 0.374 | 0.449 |
| chr3 | 36999120 | 37000647 | 1528 | 0.31 | 0.374 | 0.449 |
| chr1 | 94187618 | 94189820 | 2203 | 0.37 | 0.375 | 0.449 |
| chr14 | 34567324 | 34568214 | 891 | -0.49 | 0.375 | 0.449 |
| chr11 | 10372933 | 10374020 | 1088 | -0.34 | 0.375 | 0.449 |
| chr2 | 102972189 | 102972987 | 799 | -0.34 | 0.375 | 0.449 |
| chr7 | 156928723 | 156972536 | 43814 | 0.08 | 0.375 | 0.449 |
| chr21 | 34142697 | 34145460 | 2764 | 0.21 | 0.375 | 0.449 |
| chr7 | 75640697 | 75641422 | 726 | -0.67 | 0.375 | 0.449 |
| chr21 | 37753971 | 37758502 | 4532 | 0.21 | 0.375 | 0.449 |
| chr4 | 71767331 | 71768992 | 1662 | 0.24 | 0.375 | 0.449 |
| chr7 | 140372506 | 140373578 | 1073 | 0.39 | 0.375 | 0.449 |
| chr1 | 162467008 | 162468583 | 1576 | 0.18 | 0.375 | 0.45 |
| chr15 | 57209688 | 57221966 | 12279 | 0.11 | 0.375 | 0.45 |
| chr5 | 148723844 | 148738234 | 14391 | -0.13 | 0.375 | 0.45 |
| chr1 | 204215578 | 204274164 | 58587 | -0.07 | 0.375 | 0.45 |
| chrX | 40333224 | 40334356 | 1133 | 0.34 | 0.376 | 0.45 |
| chr17 | 12017761 | 12018858 | 1098 | -0.43 | 0.376 | 0.45 |
| chr1 | 25756158 | 25758286 | 2129 | 0.2 | 0.376 | 0.45 |
| chr16 | 4893276 | 4914251 | 20976 | 0.11 | 0.376 | 0.45 |
| chr7 | 99938359 | 99971201 | 32843 | -0.12 | 0.376 | 0.45 |
| chr10 | 75117704 | 75144533 | 26830 | -0.11 | 0.376 | 0.451 |
| chr12 | 32810746 | 32811150 | 405 | -0.61 | 0.376 | 0.451 |
| chr12 | 52166753 | 52168609 | 1857 | -0.32 | 0.377 | 0.451 |
| chr10 | 72646230 | 72649024 | 2795 | 0.16 | 0.377 | 0.451 |
| chrX | 15436271 | 15437294 | 1024 | 1.49 | 0.377 | 0.451 |
| chr16 | 53946891 | 53947517 | 627 | 0.48 | 0.377 | 0.451 |
| chr9 | 94007960 | 94008823 | 864 | -0.38 | 0.377 | 0.451 |
| chr16 | 51557487 | 51575264 | 17778 | 0.14 | 0.377 | 0.451 |
| chr4 | 78721805 | 78722570 | 766 | -0.52 | 0.377 | 0.452 |
| chr6 | 112947904 | 112948361 | 458 | -0.55 | 0.378 | 0.452 |
| chr1 | 154519977 | 154532541 | 12565 | -0.13 | 0.378 | 0.452 |
| chr11 | 103794028 | 103795351 | 1324 | 0.37 | 0.378 | 0.452 |
| chr6 | 33661169 | 33679795 | 18627 | -0.13 | 0.378 | 0.453 |
| chr2 | 120980018 | 120981764 | 1747 | -0.2 | 0.378 | 0.453 |
| chr22 | 19737663 | 19739760 | 2098 | 0.26 | 0.378 | 0.453 |
| chr6 | 44443704 | 44444310 | 607 | 0.58 | 0.379 | 0.453 |
| chr16 | 67345934 | 67346771 | 838 | 0.46 | 0.379 | 0.453 |
| chr10 | 88077566 | 88078520 | 955 | -0.57 | 0.379 | 0.453 |
| chrX | 10277530 | 10296135 | 18606 | 0.09 | 0.379 | 0.453 |
| chr12 | 93139913 | 93140357 | 445 | -0.51 | 0.379 | 0.453 |
| chr22 | 39988365 | 39989316 | 952 | -0.44 | 0.379 | 0.453 |
| chr2 | 224539337 | 224540345 | 1009 | -0.36 | 0.379 | 0.453 |
| chr16 | 3018985 | 3034257 | 15273 | -0.13 | 0.379 | 0.454 |
| chr11 | 130503386 | 130509257 | 5872 | -0.09 | 0.38 | 0.454 |
| chr3 | 128368108 | 128370566 | 2459 | 0.14 | 0.38 | 0.454 |
| chr1 | 207114785 | 207127710 | 12926 | 0.18 | 0.38 | 0.454 |
| chr8 | 39847037 | 39850521 | 3485 | -0.14 | 0.38 | 0.455 |
| chr10 | 6621692 | 6623984 | 2293 | 0.19 | 0.38 | 0.455 |
| chr6 | 34267536 | 34268513 | 978 | -0.78 | 0.38 | 0.455 |
| chr19 | 47538283 | 47552782 | 14500 | -0.19 | 0.38 | 0.455 |
| chrX | 76816909 | 76817628 | 720 | -0.49 | 0.381 | 0.455 |
| chr16 | 86652324 | 86665804 | 13481 | -1.28 | 0.381 | 0.455 |

| | | | | | | |
|---|---|---|---|---|---|---|
| chr7 | 133455377 | 133456645 | 1269 | -0.4 | 0.381 | 0.456 |
| chr8 | 41819302 | 41819976 | 675 | 0.48 | 0.381 | 0.456 |
| chr10 | 12489344 | 12509347 | 20004 | -0.12 | 0.381 | 0.456 |
| chrX | 38185397 | 38187418 | 2022 | -0.16 | 0.381 | 0.456 |
| chr3 | 34048534 | 34055898 | 7365 | 0.08 | 0.381 | 0.456 |
| chr11 | 29282084 | 29316517 | 34434 | 0.19 | 0.381 | 0.456 |
| chr4 | 58029841 | 58030362 | 522 | -0.46 | 0.381 | 0.456 |
| chr2 | 159925568 | 159956154 | 30587 | 0.09 | 0.382 | 0.456 |
| chr5 | 54572630 | 54573189 | 560 | -0.5 | 0.382 | 0.456 |
| chr1 | 40071595 | 40081543 | 9949 | -0.32 | 0.382 | 0.456 |
| chr17 | 79649554 | 79653062 | 3509 | 0.13 | 0.382 | 0.457 |
| chr1 | 155692223 | 155715428 | 23206 | 0.17 | 0.382 | 0.457 |
| chr2 | 48726737 | 48757954 | 31218 | -0.09 | 0.382 | 0.457 |
| chr1 | 153837731 | 153839105 | 1375 | 0.59 | 0.382 | 0.457 |
| chr6 | 52925624 | 52931498 | 5875 | 0.14 | 0.382 | 0.457 |
| chr14 | 59654784 | 59657354 | 2571 | 0.19 | 0.382 | 0.457 |
| chr9 | 34611264 | 34621170 | 9907 | -0.14 | 0.382 | 0.457 |
| chr8 | 1872413 | 1873270 | 858 | 0.39 | 0.382 | 0.457 |
| chr12 | 12186026 | 12186378 | 353 | 0.51 | 0.382 | 0.457 |
| chr4 | 175590230 | 175591330 | 1101 | 0.41 | 0.383 | 0.457 |
| chrX | 44428813 | 44431212 | 2400 | 0.24 | 0.383 | 0.457 |
| chr6 | 132384498 | 132385536 | 1039 | 0.38 | 0.383 | 0.457 |
| chr6 | 99796958 | 99798314 | 1357 | 0.38 | 0.383 | 0.457 |
| chr2 | 11393405 | 11395057 | 1653 | 0.28 | 0.383 | 0.457 |
| chr4 | 93890530 | 93891997 | 1468 | 0.25 | 0.383 | 0.457 |
| chr10 | 93557276 | 93573669 | 16394 | 0.12 | 0.383 | 0.457 |
| chr17 | 26104050 | 26106080 | 2031 | -0.2 | 0.383 | 0.457 |
| chr13 | 95702259 | 95954571 | 252313 | -0.03 | 0.383 | 0.457 |
| chr20 | 1789572 | 1794436 | 4865 | -0.15 | 0.383 | 0.458 |
| chr17 | 18216783 | 18236638 | 19856 | -0.09 | 0.383 | 0.458 |
| chr1 | 182931880 | 182932490 | 611 | 0.27 | 0.383 | 0.458 |
| chr16 | 57293947 | 57295441 | 1495 | 0.27 | 0.383 | 0.458 |
| chr10 | 74283636 | 74292974 | 9339 | 0.19 | 0.383 | 0.458 |
| chr22 | 46731100 | 46777686 | 46587 | 0.09 | 0.384 | 0.458 |
| chr5 | 142360933 | 142363934 | 3002 | -0.26 | 0.384 | 0.458 |
| chr5 | 179013322 | 179052035 | 38714 | -0.08 | 0.384 | 0.458 |
| chr3 | 17556586 | 17557948 | 1363 | 0.25 | 0.384 | 0.458 |
| chr8 | 145219394 | 145220217 | 824 | -1.5 | 0.384 | 0.459 |
| chr10 | 121030040 | 121032403 | 2364 | 0.18 | 0.384 | 0.459 |
| chr17 | 65988901 | 65992296 | 3396 | 0.16 | 0.384 | 0.459 |
| chr5 | 168306651 | 168307375 | 725 | -0.4 | 0.384 | 0.459 |
| chr10 | 76657139 | 76658796 | 1658 | -0.27 | 0.384 | 0.459 |
| chr14 | 91305556 | 91318132 | 12577 | -0.4 | 0.384 | 0.459 |
| chr17 | 9018280 | 9020120 | 1841 | -0.28 | 0.384 | 0.459 |
| chr6 | 129944247 | 130038874 | 94628 | -0.03 | 0.384 | 0.459 |
| chr3 | 47516068 | 47518420 | 2353 | 0.25 | 0.385 | 0.459 |
| chr17 | 27940122 | 27945617 | 5496 | -0.34 | 0.385 | 0.459 |
| chr2 | 65114542 | 65127611 | 13070 | -0.48 | 0.385 | 0.46 |
| chr22 | 24758107 | 24758790 | 684 | -0.49 | 0.385 | 0.46 |
| chr8 | 63282082 | 63284030 | 1949 | 0.21 | 0.385 | 0.46 |
| chr14 | 102017435 | 102031872 | 14438 | -0.1 | 0.385 | 0.46 |
| chr4 | 148166254 | 148170271 | 4018 | -0.1 | 0.385 | 0.46 |
| chr17 | 47783220 | 47794159 | 10940 | 0.11 | 0.385 | 0.46 |
| chr11 | 35098695 | 35106429 | 7735 | 0.1 | 0.386 | 0.46 |
| chr15 | 30260484 | 30261365 | 882 | -0.45 | 0.386 | 0.46 |
| chr2 | 219133928 | 219158511 | 24584 | -0.07 | 0.386 | 0.46 |
| chr2 | 233497221 | 233497626 | 406 | -0.67 | 0.386 | 0.461 |
| chrX | 56771235 | 56778039 | 6805 | 0.33 | 0.386 | 0.461 |
| chr3 | 32432594 | 32444822 | 12229 | -0.12 | 0.387 | 0.461 |
| chrX | 154280860 | 154281944 | 1085 | 0.4 | 0.387 | 0.461 |
| chr17 | 48045258 | 48047067 | 1810 | 0.16 | 0.387 | 0.461 |

| | | | | | | |
|---|---|---|---|---|---|---|
| chr2 | 71558268 | 71561519 | 3252 | -0.12 | 0.387 | 0.462 |
| chr2 | 173088247 | 173096833 | 8587 | -0.35 | 0.387 | 0.462 |
| chr19 | 45003389 | 45005366 | 1978 | 0.16 | 0.387 | 0.462 |
| chr17 | 9632794 | 9634030 | 1237 | -0.38 | 0.387 | 0.462 |
| chr17 | 65256208 | 65257017 | 810 | 0.48 | 0.387 | 0.462 |
| chr1 | 36762295 | 36772580 | 10286 | 0.14 | 0.387 | 0.462 |
| chr12 | 76552263 | 76552628 | 366 | 0.42 | 0.388 | 0.462 |
| chr2 | 39516855 | 39517224 | 370 | -0.68 | 0.388 | 0.463 |
| chr5 | 54522229 | 54530753 | 8525 | 0.16 | 0.388 | 0.463 |
| chr6 | 125474902 | 125476381 | 1480 | 0.37 | 0.388 | 0.463 |
| chr6 | 26383355 | 26384342 | 988 | 0.31 | 0.388 | 0.463 |
| chrX | 100645453 | 100674062 | 28610 | -0.07 | 0.389 | 0.464 |
| chr9 | 138923574 | 138924656 | 1083 | 0.46 | 0.389 | 0.464 |
| chr6 | 73971864 | 73973482 | 1619 | -0.22 | 0.389 | 0.464 |
| chr5 | 39638857 | 39649483 | 10627 | 1.73 | 0.389 | 0.464 |
| chr17 | 28135003 | 28135801 | 799 | 0.26 | 0.389 | 0.464 |
| chr18 | 18743696 | 18745494 | 1799 | 0.25 | 0.389 | 0.464 |
| chr12 | 112901198 | 112902633 | 1436 | 0.36 | 0.389 | 0.464 |
| chr6 | 7974653 | 7976534 | 1882 | -0.23 | 0.39 | 0.464 |
| chrX | 62549996 | 62550979 | 984 | 0.42 | 0.39 | 0.464 |
| chrX | 24710990 | 24713244 | 2255 | 0.18 | 0.39 | 0.465 |
| chr2 | 204502397 | 204535199 | 32803 | -0.13 | 0.39 | 0.465 |
| chr2 | 236239211 | 236240733 | 1523 | -0.19 | 0.39 | 0.465 |
| chr19 | 2448285 | 2507996 | 59712 | 0.07 | 0.39 | 0.465 |
| chr12 | 62723831 | 62724276 | 446 | -0.57 | 0.39 | 0.465 |
| chr12 | 76705817 | 76706360 | 544 | -0.58 | 0.39 | 0.465 |
| chr4 | 185759727 | 185760117 | 391 | 0.83 | 0.39 | 0.465 |
| chr16 | 53349519 | 53349986 | 468 | -0.59 | 0.39 | 0.465 |
| chr2 | 25263688 | 25266084 | 2397 | -0.22 | 0.39 | 0.465 |
| chr1 | 160067870 | 160079785 | 11916 | 0.22 | 0.391 | 0.466 |
| chr1 | 204332188 | 204348440 | 16253 | 0.1 | 0.391 | 0.466 |
| chr5 | 57447785 | 57449491 | 1707 | -0.29 | 0.391 | 0.466 |
| chr11 | 107432780 | 107436894 | 4115 | -0.22 | 0.391 | 0.466 |
| chr12 | 78392884 | 78393780 | 897 | -0.47 | 0.391 | 0.466 |
| chr6 | 55765908 | 55768057 | 2150 | 0.19 | 0.391 | 0.466 |
| chr16 | 21312149 | 21315068 | 2920 | 0.17 | 0.391 | 0.466 |
| chr7 | 74488398 | 74491659 | 3262 | 0.15 | 0.391 | 0.466 |
| chr10 | 112424231 | 112432766 | 8536 | 0.18 | 0.391 | 0.466 |
| chr2 | 169585222 | 169585623 | 402 | -0.82 | 0.391 | 0.466 |
| chr3 | 165060617 | 165061097 | 481 | 0.53 | 0.392 | 0.467 |
| chr2 | 152116828 | 152118802 | 1975 | -0.11 | 0.392 | 0.467 |
| chr22 | 50764872 | 50766429 | 1558 | 0.2 | 0.392 | 0.467 |
| chr4 | 174450706 | 174451401 | 696 | -0.41 | 0.392 | 0.467 |
| chrX | 14861668 | 14862602 | 935 | -0.43 | 0.392 | 0.467 |
| chr17 | 62914763 | 62915376 | 614 | -0.3 | 0.392 | 0.467 |
| chr10 | 7299092 | 7307877 | 8786 | 0.15 | 0.392 | 0.467 |
| chr5 | 130045777 | 130047551 | 1775 | -0.26 | 0.392 | 0.467 |
| chr21 | 45284068 | 45286426 | 2359 | 0.27 | 0.392 | 0.467 |
| chr16 | 29873372 | 29875516 | 2145 | 0.12 | 0.392 | 0.467 |
| chr2 | 197158594 | 197159662 | 1069 | 0.49 | 0.393 | 0.467 |
| chr17 | 634706 | 656236 | 21531 | -0.12 | 0.393 | 0.467 |
| chr7 | 372234 | 373587 | 1354 | 0.27 | 0.393 | 0.467 |
| chr9 | 87489396 | 87489737 | 342 | 0.58 | 0.393 | 0.468 |
| chr2 | 240782128 | 240783305 | 1178 | -0.41 | 0.393 | 0.468 |
| chr8 | 141717903 | 141758727 | 40825 | 0.1 | 0.393 | 0.468 |
| chr5 | 53379744 | 53387589 | 7846 | -0.18 | 0.393 | 0.468 |
| chr1 | 27675604 | 27676311 | 708 | -0.49 | 0.393 | 0.468 |
| chr10 | 22330744 | 22332847 | 2104 | -0.12 | 0.393 | 0.468 |
| chr1 | 243527881 | 243528589 | 709 | 0.93 | 0.393 | 0.468 |
| chr4 | 48485255 | 48485921 | 667 | 0.51 | 0.393 | 0.468 |
| chr16 | 27688022 | 27689439 | 1418 | 0.36 | 0.393 | 0.468 |

| | | | | | | |
|---|---|---|---|---|---|---|
| chr3 | 107871943 | 107872803 | 861 | -0.49 | 0.393 | 0.468 |
| chr4 | 131343360 | 131344039 | 680 | 1.52 | 0.394 | 0.468 |
| chr2 | 139339505 | 139340319 | 815 | -0.36 | 0.394 | 0.468 |
| chr9 | 111681324 | 111700520 | 19197 | 0.13 | 0.394 | 0.469 |
| chr20 | 49498132 | 49498838 | 707 | -0.58 | 0.394 | 0.469 |
| chr10 | 89917297 | 89918330 | 1034 | 0.35 | 0.394 | 0.469 |
| chr14 | 95579969 | 95581279 | 1311 | 0.35 | 0.394 | 0.469 |
| chr7 | 44951567 | 44953174 | 1608 | -0.35 | 0.394 | 0.469 |
| chr11 | 4137536 | 4138252 | 717 | -0.48 | 0.394 | 0.469 |
| chrX | 102862346 | 102863422 | 1077 | 0.38 | 0.394 | 0.469 |
| chr6 | 89577519 | 89578387 | 869 | -0.46 | 0.394 | 0.469 |
| chr15 | 80726548 | 80730790 | 4243 | 0.11 | 0.395 | 0.469 |
| chr15 | 69803087 | 69803699 | 613 | -0.57 | 0.395 | 0.469 |
| chr17 | 57404917 | 57410255 | 5339 | -0.12 | 0.395 | 0.469 |
| chr1 | 185329556 | 185329973 | 418 | -0.61 | 0.395 | 0.469 |
| chr3 | 108896087 | 108897339 | 1253 | 0.41 | 0.395 | 0.47 |
| chr22 | 32806744 | 32808677 | 1934 | 0.3 | 0.395 | 0.47 |
| chr14 | 64948303 | 64951340 | 3038 | 0.31 | 0.395 | 0.47 |
| chr7 | 149439034 | 149439376 | 343 | 0.66 | 0.395 | 0.47 |
| chr1 | 203456484 | 203457525 | 1042 | -0.33 | 0.395 | 0.47 |
| chr12 | 93349600 | 93368201 | 18602 | 0.12 | 0.395 | 0.47 |
| chr1 | 43636790 | 43638182 | 1393 | -0.21 | 0.395 | 0.47 |
| chr4 | 101014878 | 101015414 | 537 | 0.46 | 0.395 | 0.47 |
| chr6 | 43196416 | 43197194 | 779 | -0.36 | 0.395 | 0.47 |
| chr3 | 107586736 | 107588690 | 1955 | 0.22 | 0.395 | 0.47 |
| chr2 | 74962257 | 74969250 | 6994 | 0.22 | 0.395 | 0.47 |
| chr8 | 98861540 | 98862310 | 771 | 0.52 | 0.396 | 0.47 |
| chr6 | 33950420 | 33956691 | 6272 | 0.36 | 0.396 | 0.47 |
| chr2 | 231276076 | 231281978 | 5903 | -0.15 | 0.396 | 0.47 |
| chr7 | 50773986 | 50783665 | 9680 | 0.15 | 0.396 | 0.47 |
| chr3 | 197280978 | 197282804 | 1827 | 0.27 | 0.396 | 0.47 |
| chr17 | 5317084 | 5343617 | 26534 | 0.07 | 0.396 | 0.471 |
| chr15 | 70818565 | 70819596 | 1032 | 0.33 | 0.396 | 0.471 |
| chr11 | 75186216 | 75187037 | 822 | -0.5 | 0.396 | 0.471 |
| chr5 | 168893590 | 168894636 | 1047 | 0.42 | 0.396 | 0.471 |
| chr14 | 45428595 | 45434370 | 5776 | -0.09 | 0.397 | 0.471 |
| chr3 | 196133103 | 196134237 | 1135 | -0.46 | 0.397 | 0.471 |
| chrX | 44343282 | 44344131 | 850 | -0.45 | 0.397 | 0.471 |
| chr1 | 68315971 | 68317477 | 1507 | -0.29 | 0.397 | 0.471 |
| chr12 | 132194768 | 132206137 | 11370 | 0.12 | 0.397 | 0.471 |
| chr3 | 98311723 | 98313492 | 1770 | 0.28 | 0.397 | 0.472 |
| chr16 | 81962545 | 81975110 | 12566 | -1.01 | 0.397 | 0.472 |
| chr3 | 196377630 | 196378896 | 1267 | 0.27 | 0.397 | 0.472 |
| chr22 | 50311529 | 50312076 | 548 | -0.41 | 0.397 | 0.472 |
| chr3 | 172364070 | 172436050 | 71981 | 0.05 | 0.397 | 0.472 |
| chr6 | 130051314 | 130052461 | 1148 | -0.36 | 0.397 | 0.472 |
| chr14 | 93697422 | 93708132 | 10711 | 0.16 | 0.397 | 0.472 |
| chr5 | 100345831 | 100354658 | 8828 | 1 | 0.398 | 0.472 |
| chr14 | 68764609 | 68765489 | 881 | 0.41 | 0.398 | 0.472 |
| chr1 | 93269980 | 93271272 | 1293 | -0.31 | 0.398 | 0.472 |
| chr7 | 157280951 | 157284467 | 3517 | 0.15 | 0.398 | 0.472 |
| chr10 | 5778030 | 5778808 | 779 | 0.57 | 0.398 | 0.472 |
| chr1 | 84568194 | 84570022 | 1829 | 0.14 | 0.398 | 0.473 |
| chr1 | 110089144 | 110092223 | 3080 | 0.13 | 0.398 | 0.473 |
| chr2 | 19248170 | 19249791 | 1622 | 0.21 | 0.398 | 0.473 |
| chr14 | 21315722 | 21318601 | 2880 | -0.13 | 0.398 | 0.473 |
| chrX | 122798732 | 122810455 | 11724 | 0.22 | 0.398 | 0.473 |
| chr5 | 168435847 | 168436532 | 686 | -0.48 | 0.398 | 0.473 |
| chrX | 18433215 | 18444473 | 11259 | -0.1 | 0.398 | 0.473 |
| chr4 | 33865657 | 33866284 | 628 | -0.44 | 0.398 | 0.473 |
| chr18 | 30081295 | 30081804 | 510 | -0.58 | 0.398 | 0.473 |

| | | | | | | |
|---|---|---|---|---|---|---|
| chr11 | 43501086 | 43501594 | 509 | 0.58 | 0.398 | 0.473 |
| chr19 | 14682164 | 14685137 | 2974 | 0.27 | 0.399 | 0.473 |
| chr15 | 90930479 | 90973880 | 43402 | 0.04 | 0.399 | 0.473 |
| chr17 | 28828471 | 28829842 | 1372 | -0.33 | 0.399 | 0.473 |
| chr14 | 78008693 | 78009700 | 1008 | 0.46 | 0.399 | 0.473 |
| chr4 | 73967643 | 73970921 | 3279 | 0.24 | 0.399 | 0.473 |
| chr8 | 54645250 | 54650568 | 5319 | -1.64 | 0.399 | 0.473 |
| chr15 | 50778362 | 50778800 | 439 | -0.51 | 0.399 | 0.473 |
| chr2 | 145461083 | 145461981 | 899 | -0.35 | 0.399 | 0.473 |
| chr17 | 15982331 | 16010076 | 27746 | 0.11 | 0.399 | 0.474 |
| chr17 | 74684268 | 74685897 | 1630 | -0.22 | 0.399 | 0.474 |
| chr3 | 133460736 | 133462453 | 1718 | 0.3 | 0.399 | 0.474 |
| chr12 | 118727433 | 118745904 | 18472 | -0.21 | 0.399 | 0.474 |
| chr6 | 25875311 | 25884499 | 9189 | 1.65 | 0.399 | 0.474 |
| chr18 | 13725275 | 13727956 | 2682 | 0.12 | 0.399 | 0.474 |
| chr13 | 50782067 | 50784089 | 2023 | 0.16 | 0.4 | 0.474 |
| chr10 | 99496123 | 99517014 | 20892 | -0.11 | 0.4 | 0.474 |
| chr12 | 41332371 | 41332952 | 582 | -0.45 | 0.4 | 0.474 |
| chr7 | 44224081 | 44225303 | 1223 | -0.4 | 0.4 | 0.475 |
| chr2 | 72463204 | 72469077 | 5874 | 0.27 | 0.4 | 0.475 |
| chr16 | 19247254 | 19248840 | 1587 | 0.38 | 0.4 | 0.475 |
| chr8 | 95369109 | 95370064 | 956 | 0.28 | 0.4 | 0.475 |
| chr9 | 111772671 | 111802398 | 29728 | -0.1 | 0.401 | 0.475 |
| chr2 | 235731943 | 235750917 | 18975 | 0.08 | 0.401 | 0.475 |
| chr3 | 52001062 | 52030377 | 29316 | 0.09 | 0.401 | 0.475 |
| chr11 | 61145383 | 61215117 | 69735 | 0.06 | 0.401 | 0.475 |
| chr22 | 22862360 | 22874640 | 12281 | 0.23 | 0.401 | 0.475 |
| chr11 | 67209934 | 67212001 | 2068 | 0.19 | 0.401 | 0.475 |
| chr1 | 9488521 | 9499491 | 10971 | -0.26 | 0.401 | 0.475 |
| chr12 | 51781521 | 51785635 | 4115 | -1.58 | 0.401 | 0.475 |
| chr12 | 88972771 | 88974776 | 2006 | -0.29 | 0.401 | 0.475 |
| chr16 | 85336345 | 85337637 | 1293 | 0.33 | 0.401 | 0.475 |
| chr16 | 56279781 | 56280669 | 889 | 0.56 | 0.401 | 0.476 |
| chr11 | 64068143 | 64127203 | 59061 | -0.07 | 0.401 | 0.476 |
| chr22 | 41601290 | 41602151 | 862 | 0.27 | 0.401 | 0.476 |
| chr1 | 100597659 | 100599689 | 2031 | -0.14 | 0.401 | 0.476 |
| chr5 | 134002200 | 134009823 | 7624 | 0.23 | 0.401 | 0.476 |
| chr17 | 64050242 | 64051039 | 798 | -0.36 | 0.402 | 0.476 |
| chr19 | 747872 | 802285 | 54414 | 0.06 | 0.402 | 0.476 |
| chr2 | 135892217 | 135892607 | 391 | -0.54 | 0.402 | 0.476 |
| chr5 | 130864338 | 130864982 | 645 | -0.49 | 0.402 | 0.476 |
| chr20 | 60011564 | 60012452 | 889 | 0.43 | 0.402 | 0.477 |
| chr1 | 205473265 | 205474408 | 1144 | 0.35 | 0.402 | 0.477 |
| chr4 | 33495432 | 33496483 | 1052 | -0.33 | 0.402 | 0.477 |
| chr17 | 29378869 | 29380023 | 1155 | -0.31 | 0.402 | 0.477 |
| chr9 | 13096647 | 13096949 | 303 | 0.65 | 0.403 | 0.477 |
| chr11 | 33059887 | 33118010 | 58124 | 0.04 | 0.403 | 0.477 |
| chr3 | 100052607 | 100055230 | 2624 | -0.13 | 0.403 | 0.477 |
| chr20 | 60641095 | 60642123 | 1029 | 0.2 | 0.403 | 0.477 |
| chr14 | 99738020 | 99741404 | 3385 | -0.23 | 0.403 | 0.477 |
| chr2 | 203272594 | 203274413 | 1820 | 0.21 | 0.403 | 0.477 |
| chr3 | 121369874 | 121373124 | 3251 | -0.24 | 0.403 | 0.477 |
| chr8 | 41365354 | 41366433 | 1080 | 0.58 | 0.403 | 0.478 |
| chr11 | 70274914 | 70276747 | 1834 | -0.31 | 0.403 | 0.478 |
| chr4 | 185338580 | 185339660 | 1081 | -0.36 | 0.403 | 0.478 |
| chr3 | 136062372 | 136069808 | 7437 | -0.24 | 0.403 | 0.478 |
| chr5 | 106982081 | 107009891 | 27811 | -0.06 | 0.404 | 0.478 |
| chr12 | 15961469 | 15961889 | 421 | 0.56 | 0.404 | 0.478 |
| chr17 | 45725228 | 45772324 | 47097 | 0.06 | 0.404 | 0.478 |
| chr15 | 43212204 | 43213683 | 1480 | -0.38 | 0.404 | 0.478 |
| chr2 | 45539048 | 45539746 | 699 | 0.44 | 0.404 | 0.478 |

| | | | | | | |
|---|---|---|---|---|---|---|
| chr11 | 104404772 | 104405565 | 794 | -0.37 | 0.404 | 0.478 |
| chr6 | 144536284 | 144538454 | 2171 | 0.18 | 0.404 | 0.479 |
| chr5 | 17340941 | 17345471 | 4531 | -0.16 | 0.404 | 0.479 |
| chr11 | 76154376 | 76157257 | 2882 | 0.11 | 0.404 | 0.479 |
| chr19 | 57106366 | 57107498 | 1133 | -0.29 | 0.405 | 0.479 |
| chr16 | 4815683 | 4819429 | 3747 | 0.14 | 0.405 | 0.479 |
| chr4 | 185320797 | 185322622 | 1826 | 0.32 | 0.405 | 0.479 |
| chr2 | 231377689 | 231378511 | 823 | 0.4 | 0.405 | 0.479 |
| chr7 | 79015759 | 79017171 | 1413 | 0.28 | 0.405 | 0.479 |
| chrX | 114924449 | 114924956 | 508 | -0.63 | 0.405 | 0.48 |
| chr1 | 9045497 | 9048245 | 2749 | -0.18 | 0.405 | 0.48 |
| chr6 | 136569899 | 136572337 | 2439 | -0.16 | 0.405 | 0.48 |
| chr1 | 167027975 | 167028570 | 596 | -0.34 | 0.405 | 0.48 |
| chr6 | 139013428 | 139014899 | 1472 | -0.3 | 0.405 | 0.48 |
| chr3 | 155606713 | 155609156 | 2444 | 0.22 | 0.406 | 0.48 |
| chr7 | 73587020 | 73638560 | 51541 | 0.07 | 0.406 | 0.48 |
| chr18 | 74088077 | 74088962 | 886 | -0.38 | 0.406 | 0.48 |
| chr2 | 72921938 | 72922783 | 846 | 0.44 | 0.406 | 0.48 |
| chr20 | 2631748 | 2645760 | 14013 | 0.08 | 0.406 | 0.48 |
| chr15 | 55879598 | 55881423 | 1826 | -0.17 | 0.406 | 0.48 |
| chr14 | 43399074 | 43399797 | 724 | 0.52 | 0.406 | 0.481 |
| chr14 | 75412150 | 75413158 | 1009 | -0.51 | 0.406 | 0.481 |
| chr12 | 112298153 | 112298786 | 634 | -0.52 | 0.406 | 0.481 |
| chr1 | 205343456 | 205344142 | 687 | -0.5 | 0.406 | 0.481 |
| chr17 | 48343921 | 48368152 | 24232 | 0.13 | 0.406 | 0.481 |
| chr13 | 41052215 | 41052786 | 572 | -0.54 | 0.406 | 0.481 |
| chr6 | 21824424 | 21825446 | 1023 | -0.39 | 0.407 | 0.481 |
| chr2 | 32260883 | 32265640 | 4758 | 0.15 | 0.407 | 0.481 |
| chr14 | 105190544 | 105200909 | 10366 | -0.17 | 0.407 | 0.481 |
| chr6 | 42396234 | 42421493 | 25260 | -0.12 | 0.407 | 0.481 |
| chr7 | 26706816 | 26707189 | 374 | 0.72 | 0.407 | 0.481 |
| chr20 | 31252746 | 31253806 | 1061 | -0.48 | 0.407 | 0.481 |
| chr5 | 176587720 | 176588482 | 763 | 0.53 | 0.407 | 0.481 |
| chr1 | 29528076 | 29529439 | 1364 | -0.37 | 0.407 | 0.481 |
| chr19 | 13342527 | 13343003 | 477 | -0.56 | 0.407 | 0.482 |
| chr2 | 48636094 | 48648600 | 12507 | -0.17 | 0.407 | 0.482 |
| chr1 | 78414017 | 78417279 | 3263 | 0.25 | 0.407 | 0.482 |
| chr5 | 126202779 | 126203852 | 1074 | -0.53 | 0.407 | 0.482 |
| chr5 | 149011673 | 149012395 | 723 | -0.5 | 0.407 | 0.482 |
| chr20 | 43587705 | 43608639 | 20935 | 0.1 | 0.408 | 0.482 |
| chr17 | 58264389 | 58265647 | 1259 | 0.28 | 0.408 | 0.482 |
| chr15 | 62795469 | 62799865 | 4397 | 0.09 | 0.408 | 0.482 |
| chr1 | 180067033 | 180068491 | 1459 | -0.26 | 0.408 | 0.482 |
| chr14 | 95941570 | 95950363 | 8794 | 0.27 | 0.408 | 0.482 |
| chr14 | 105218399 | 105222118 | 3720 | -0.16 | 0.408 | 0.482 |
| chr2 | 198419854 | 198422280 | 2427 | -0.23 | 0.408 | 0.482 |
| chr14 | 92925611 | 92940854 | 15244 | 0.11 | 0.408 | 0.482 |
| chr20 | 23400025 | 23402165 | 2141 | 0.16 | 0.408 | 0.483 |
| chr11 | 119036669 | 119040844 | 4176 | -0.14 | 0.409 | 0.483 |
| chr7 | 64125817 | 64127829 | 2013 | 0.2 | 0.409 | 0.483 |
| chr8 | 29934208 | 29961871 | 27664 | -0.07 | 0.409 | 0.483 |
| chr1 | 26126253 | 26126743 | 491 | 0.54 | 0.409 | 0.484 |
| chr20 | 62572264 | 62614088 | 41825 | 0.1 | 0.409 | 0.484 |
| chr13 | 34630688 | 34630952 | 265 | 0.7 | 0.409 | 0.484 |
| chr3 | 17626819 | 17627781 | 963 | -0.3 | 0.409 | 0.484 |
| chr7 | 73026805 | 73062768 | 35964 | -0.13 | 0.409 | 0.484 |
| chr7 | 132937784 | 132938937 | 1154 | 0.18 | 0.409 | 0.484 |
| chr7 | 39772433 | 39774746 | 2314 | 0.12 | 0.409 | 0.484 |
| chrX | 123246890 | 123247990 | 1101 | 0.36 | 0.409 | 0.484 |
| chrX | 3520842 | 3522212 | 1371 | -0.37 | 0.41 | 0.484 |
| chr5 | 93996384 | 94009663 | 13280 | 0.23 | 0.41 | 0.484 |

| | | | | | | |
|---|---|---|---|---|---|---|
| chr22 | 29949040 | 29950951 | 1912 | 0.17 | 0.41 | 0.484 |
| chr19 | 45680288 | 45683306 | 3019 | -0.13 | 0.41 | 0.484 |
| chr3 | 183164802 | 183166095 | 1294 | 0.28 | 0.41 | 0.485 |
| chr2 | 223846801 | 223847559 | 759 | -0.49 | 0.41 | 0.485 |
| chr4 | 147143270 | 147143953 | 684 | 0.59 | 0.41 | 0.485 |
| chr11 | 110300365 | 110302045 | 1681 | 0.22 | 0.41 | 0.485 |
| chr15 | 57998097 | 58000771 | 2675 | 0.1 | 0.411 | 0.485 |
| chr7 | 65515673 | 65517909 | 2237 | -0.25 | 0.411 | 0.485 |
| chr1 | 222059983 | 222061401 | 1419 | -0.29 | 0.411 | 0.485 |
| chr19 | 10304130 | 10305689 | 1560 | -0.29 | 0.411 | 0.485 |
| chr2 | 128990417 | 128991238 | 822 | -0.39 | 0.411 | 0.485 |
| chr4 | 1686328 | 1686802 | 475 | -0.55 | 0.411 | 0.485 |
| chr2 | 203453600 | 203453943 | 344 | -0.68 | 0.411 | 0.486 |
| chr2 | 148585697 | 148587865 | 2169 | -0.15 | 0.411 | 0.486 |
| chr7 | 7091051 | 7091417 | 367 | 0.63 | 0.412 | 0.486 |
| chr6 | 41862614 | 41864363 | 1750 | 0.3 | 0.412 | 0.486 |
| chr10 | 96108952 | 96123399 | 14448 | 0.12 | 0.412 | 0.486 |
| chr17 | 6734642 | 6735481 | 840 | -0.45 | 0.412 | 0.486 |
| chr17 | 3437998 | 3439541 | 1544 | -0.24 | 0.412 | 0.486 |
| chr7 | 107146524 | 107147404 | 881 | 0.49 | 0.412 | 0.486 |
| chr3 | 52231408 | 52232860 | 1453 | 0.29 | 0.412 | 0.486 |
| chr5 | 78091060 | 78092411 | 1352 | 0.32 | 0.412 | 0.486 |
| chr17 | 45302369 | 45303273 | 905 | 0.4 | 0.412 | 0.486 |
| chr16 | 10710440 | 10721332 | 10893 | 0.13 | 0.412 | 0.486 |
| chr5 | 175781227 | 175789242 | 8016 | 0.15 | 0.412 | 0.486 |
| chr6 | 76336808 | 76337529 | 722 | 0.52 | 0.412 | 0.486 |
| chr18 | 77218131 | 77218954 | 824 | 0.48 | 0.412 | 0.486 |
| chr2 | 190816112 | 190817352 | 1241 | -0.26 | 0.412 | 0.486 |
| chr5 | 156526803 | 156528155 | 1353 | -0.43 | 0.412 | 0.487 |
| chr5 | 134112548 | 134128855 | 16308 | 0.15 | 0.413 | 0.487 |
| chr19 | 34992176 | 34992525 | 350 | 0.57 | 0.413 | 0.487 |
| chr7 | 129995723 | 130003994 | 8272 | 0.26 | 0.413 | 0.487 |
| chr1 | 226270403 | 226271949 | 1547 | -0.22 | 0.413 | 0.487 |
| chr10 | 90977016 | 90977566 | 551 | -0.48 | 0.413 | 0.487 |
| chr5 | 150827118 | 150831887 | 4770 | 0.32 | 0.413 | 0.487 |
| chr14 | 55680012 | 55680553 | 542 | -0.42 | 0.413 | 0.487 |
| chr9 | 83408990 | 83409548 | 559 | -0.48 | 0.413 | 0.487 |
| chr9 | 128023275 | 128031542 | 8268 | -0.2 | 0.413 | 0.488 |
| chr4 | 4576767 | 4577530 | 764 | 0.34 | 0.413 | 0.488 |
| chr10 | 93646817 | 93684926 | 38110 | 0.08 | 0.413 | 0.488 |
| chr15 | 40671568 | 40676463 | 4896 | 0.19 | 0.414 | 0.488 |
| chr22 | 43573643 | 43584598 | 10956 | 0.16 | 0.414 | 0.488 |
| chr22 | 41651098 | 41660466 | 9369 | -0.14 | 0.414 | 0.488 |
| chr3 | 176981218 | 176982177 | 960 | -0.32 | 0.414 | 0.488 |
| chr19 | 10216519 | 10230845 | 14327 | -0.12 | 0.414 | 0.488 |
| chr2 | 231729103 | 231730219 | 1117 | -0.4 | 0.414 | 0.488 |
| chr6 | 83585072 | 83586275 | 1204 | -0.33 | 0.414 | 0.488 |
| chr20 | 19715329 | 19718909 | 3581 | -0.11 | 0.414 | 0.488 |
| chr10 | 127747644 | 127752333 | 4690 | -0.2 | 0.414 | 0.488 |
| chr5 | 72733119 | 72745156 | 12038 | -0.18 | 0.414 | 0.488 |
| chr3 | 87549856 | 87553511 | 3656 | -0.22 | 0.414 | 0.488 |
| chr3 | 177161283 | 177161805 | 523 | 0.72 | 0.414 | 0.489 |
| chrX | 28686838 | 28687631 | 794 | 0.37 | 0.414 | 0.489 |
| chr7 | 82177554 | 82190038 | 12485 | -0.16 | 0.415 | 0.489 |
| chr12 | 66500804 | 66501274 | 471 | 0.57 | 0.415 | 0.489 |
| chr20 | 35151260 | 35152191 | 932 | -0.58 | 0.415 | 0.489 |
| chr3 | 141405153 | 141405994 | 842 | 0.5 | 0.415 | 0.489 |
| chr8 | 20054633 | 20055858 | 1226 | 0.23 | 0.415 | 0.489 |
| chr7 | 103863971 | 103865732 | 1762 | 0.17 | 0.415 | 0.489 |
| chr3 | 148803363 | 148804804 | 1442 | 0.25 | 0.415 | 0.489 |
| chr15 | 101864833 | 101865525 | 693 | 0.41 | 0.415 | 0.489 |

| | | | | | | |
|---|---|---|---|---|---|---|
| chr12 | 104438739 | 104439384 | 646 | 1.48 | 0.415 | 0.49 |
| chr1 | 240353928 | 240355060 | 1133 | 0.3 | 0.415 | 0.49 |
| chr12 | 121864534 | 121866143 | 1610 | -0.4 | 0.416 | 0.49 |
| chr2 | 222437860 | 222438249 | 390 | 0.96 | 0.416 | 0.49 |
| chr9 | 97316475 | 97317705 | 1231 | 0.22 | 0.416 | 0.49 |
| chr3 | 136536630 | 136539356 | 2727 | -0.17 | 0.416 | 0.49 |
| chr2 | 173724092 | 173725017 | 926 | 0.42 | 0.416 | 0.49 |
| chr17 | 28776412 | 28779206 | 2795 | -0.12 | 0.416 | 0.49 |
| chr3 | 119298070 | 119299281 | 1212 | 0.22 | 0.416 | 0.49 |
| chr6 | 131455777 | 131457991 | 2215 | 0.15 | 0.416 | 0.49 |
| chr4 | 184039229 | 184052266 | 13038 | 0.33 | 0.416 | 0.49 |
| chr1 | 24578883 | 24579656 | 774 | -0.4 | 0.416 | 0.49 |
| chr8 | 40248176 | 40249791 | 1616 | -0.3 | 0.416 | 0.49 |
| chr17 | 2952390 | 2952896 | 507 | 0.34 | 0.416 | 0.49 |
| chr8 | 33167448 | 33173267 | 5820 | 0.31 | 0.416 | 0.49 |
| chr9 | 73085123 | 73085838 | 716 | 0.47 | 0.416 | 0.49 |
| chr8 | 22550913 | 22561551 | 10639 | 0.22 | 0.416 | 0.49 |
| chr8 | 99819139 | 99820716 | 1578 | 0.21 | 0.416 | 0.491 |
| chr11 | 64322397 | 64334827 | 12431 | -0.17 | 0.417 | 0.491 |
| chr16 | 27760038 | 27760918 | 881 | 0.55 | 0.417 | 0.491 |
| chrX | 62569783 | 62572000 | 2218 | 0.15 | 0.417 | 0.491 |
| chr17 | 62135215 | 62136034 | 820 | 0.5 | 0.417 | 0.491 |
| chrX | 140497235 | 140502945 | 5711 | 0.06 | 0.417 | 0.491 |
| chr22 | 31518445 | 31519396 | 952 | -0.41 | 0.417 | 0.491 |
| chr18 | 53255021 | 53257716 | 2696 | 0.16 | 0.417 | 0.491 |
| chr3 | 170211253 | 170225756 | 14504 | -0.13 | 0.417 | 0.491 |
| chr6 | 26222037 | 26258064 | 36028 | 0.07 | 0.417 | 0.491 |
| chr5 | 40205841 | 40206632 | 792 | -0.47 | 0.417 | 0.491 |
| chr15 | 62280972 | 62283127 | 2156 | -0.22 | 0.417 | 0.491 |
| chr16 | 58194648 | 58195287 | 640 | -0.38 | 0.418 | 0.492 |
| chr8 | 141697487 | 141703709 | 6223 | -0.18 | 0.418 | 0.492 |
| chr3 | 191094296 | 191095376 | 1081 | -0.38 | 0.418 | 0.492 |
| chr2 | 175995460 | 175996263 | 804 | -0.5 | 0.418 | 0.492 |
| chr5 | 122269154 | 122273964 | 4811 | -0.37 | 0.418 | 0.492 |
| chr20 | 33316005 | 33317025 | 1021 | 0.31 | 0.418 | 0.492 |
| chr9 | 111716536 | 111717210 | 675 | -0.5 | 0.418 | 0.492 |
| chr21 | 39159329 | 39160406 | 1078 | 0.46 | 0.418 | 0.492 |
| chr2 | 71203788 | 71204631 | 844 | 0.47 | 0.418 | 0.492 |
| chr16 | 67752130 | 67759425 | 7296 | -0.16 | 0.418 | 0.492 |
| chr15 | 44037799 | 44039757 | 1959 | 0.19 | 0.419 | 0.493 |
| chr22 | 36172574 | 36181009 | 8436 | -0.15 | 0.419 | 0.493 |
| chr7 | 94891218 | 94897819 | 6602 | -0.06 | 0.419 | 0.493 |
| chr1 | 36683437 | 36724901 | 41465 | -0.06 | 0.419 | 0.493 |
| chr21 | 35226872 | 35227928 | 1057 | 0.39 | 0.419 | 0.493 |
| chr1 | 242039464 | 242070482 | 31019 | 0.09 | 0.419 | 0.493 |
| chr6 | 20455571 | 20456292 | 722 | 0.44 | 0.419 | 0.493 |
| chr2 | 118845274 | 118846961 | 1688 | 0.22 | 0.419 | 0.493 |
| chr14 | 99781719 | 99786119 | 4401 | 0.14 | 0.42 | 0.494 |
| chr17 | 77840658 | 77841354 | 697 | 0.68 | 0.42 | 0.494 |
| chr14 | 32720891 | 32729490 | 8600 | 0.15 | 0.42 | 0.494 |
| chr8 | 99713201 | 99713680 | 480 | 0.47 | 0.42 | 0.494 |
| chr19 | 2327109 | 2329327 | 2219 | -0.23 | 0.42 | 0.494 |
| chr15 | 96868254 | 96900900 | 32647 | 0.05 | 0.42 | 0.494 |
| chr1 | 11071625 | 11073684 | 2060 | 0.14 | 0.42 | 0.494 |
| chr19 | 44528894 | 44530203 | 1310 | 0.2 | 0.42 | 0.494 |
| chr18 | 21976861 | 21977812 | 952 | -0.32 | 0.42 | 0.494 |
| chr7 | 100201963 | 100210661 | 8699 | 0.17 | 0.42 | 0.494 |
| chr1 | 8516035 | 8523966 | 7932 | 0.2 | 0.42 | 0.494 |
| chr18 | 13067228 | 13067869 | 642 | -0.47 | 0.42 | 0.494 |
| chr19 | 47851914 | 47892706 | 40793 | -0.1 | 0.421 | 0.495 |
| chr11 | 6439845 | 6441102 | 1258 | 0.32 | 0.421 | 0.495 |

| | | | | | | |
|---|---|---|---|---|---|---|
| chr2 | 221088831 | 221097072 | 8242 | 0.08 | 0.421 | 0.495 |
| chr20 | 13505095 | 13506684 | 1590 | -0.33 | 0.421 | 0.495 |
| chr12 | 51256010 | 51256890 | 881 | -0.39 | 0.421 | 0.495 |
| chr5 | 169906031 | 169907431 | 1401 | 0.3 | 0.421 | 0.495 |
| chr9 | 137561486 | 137562932 | 1447 | -0.35 | 0.421 | 0.495 |
| chr2 | 232645286 | 232659146 | 13861 | -0.13 | 0.421 | 0.495 |
| chr5 | 138597811 | 138651990 | 54180 | -0.05 | 0.421 | 0.495 |
| chr2 | 28339908 | 28340747 | 840 | 0.47 | 0.421 | 0.495 |
| chr7 | 134570282 | 134587534 | 17253 | 0.08 | 0.421 | 0.495 |
| chr3 | 183723471 | 183736424 | 12954 | -0.13 | 0.422 | 0.495 |
| chr11 | 65584940 | 65585825 | 886 | 0.36 | 0.422 | 0.495 |
| chr13 | 22032171 | 22034177 | 2007 | 0.18 | 0.422 | 0.496 |
| chr1 | 234342532 | 234342970 | 439 | -0.57 | 0.422 | 0.496 |
| chr4 | 103996356 | 103998455 | 2100 | -0.22 | 0.422 | 0.496 |
| chr10 | 31891598 | 31893073 | 1476 | -0.28 | 0.422 | 0.496 |
| chr2 | 189231910 | 189232320 | 411 | -0.49 | 0.422 | 0.496 |
| chr19 | 6278912 | 6279881 | 970 | -0.38 | 0.422 | 0.496 |
| chr14 | 88458767 | 88460262 | 1496 | 0.14 | 0.422 | 0.496 |
| chr15 | 101140738 | 101145584 | 4847 | 0.13 | 0.422 | 0.496 |
| chr8 | 103765998 | 103766755 | 758 | -0.39 | 0.423 | 0.496 |
| chr6 | 130628157 | 130630229 | 2073 | 0.22 | 0.423 | 0.497 |
| chr12 | 40340270 | 40341073 | 804 | -0.86 | 0.423 | 0.497 |
| chr16 | 47646470 | 47647317 | 848 | -0.45 | 0.423 | 0.497 |
| chr1 | 85513613 | 85528701 | 15089 | -0.19 | 0.423 | 0.497 |
| chr6 | 79307701 | 79322410 | 14710 | 0.24 | 0.423 | 0.497 |
| chr5 | 134141755 | 134150703 | 8949 | -0.22 | 0.423 | 0.497 |
| chr18 | 9055136 | 9056139 | 1004 | 0.46 | 0.423 | 0.497 |
| chr3 | 152756434 | 152777067 | 20634 | 0.08 | 0.423 | 0.497 |
| chr1 | 153794184 | 153818896 | 24713 | -0.13 | 0.423 | 0.497 |
| chr16 | 76933640 | 76945492 | 11853 | 0.28 | 0.424 | 0.498 |
| chr14 | 67878092 | 67886161 | 8070 | 0.23 | 0.424 | 0.498 |
| chr1 | 29381307 | 29382060 | 754 | -0.41 | 0.424 | 0.498 |
| chr10 | 25367323 | 25368476 | 1154 | 0.32 | 0.424 | 0.498 |
| chr1 | 161718901 | 161721588 | 2688 | -0.15 | 0.424 | 0.498 |
| chr16 | 474017 | 476685 | 2669 | 0.23 | 0.425 | 0.498 |
| chr13 | 78302833 | 78303623 | 791 | -0.54 | 0.425 | 0.498 |
| chr1 | 44394380 | 44413628 | 19249 | 0.1 | 0.425 | 0.498 |
| chr21 | 46265574 | 46266722 | 1149 | 0.4 | 0.425 | 0.499 |
| chr6 | 71166358 | 71166897 | 540 | 0.49 | 0.425 | 0.499 |
| chr1 | 155507002 | 155541584 | 34583 | -0.06 | 0.425 | 0.499 |
| chr20 | 40103972 | 40105690 | 1719 | 0.33 | 0.425 | 0.499 |
| chr15 | 95789673 | 95791311 | 1639 | 0.17 | 0.425 | 0.499 |
| chr3 | 17197115 | 17197820 | 706 | -0.44 | 0.425 | 0.499 |
| chr17 | 7039136 | 7039598 | 463 | -0.86 | 0.425 | 0.499 |
| chr18 | 51914297 | 51915733 | 1437 | 0.27 | 0.425 | 0.499 |
| chr12 | 64955142 | 64955637 | 496 | 0.57 | 0.426 | 0.499 |
| chr12 | 102413303 | 102414245 | 943 | 0.41 | 0.426 | 0.499 |
| chr4 | 86873657 | 86875565 | 1909 | -0.18 | 0.426 | 0.499 |
| chr6 | 16342432 | 16357002 | 14571 | 0.19 | 0.426 | 0.5 |
| chr4 | 6576494 | 6577455 | 962 | 0.35 | 0.426 | 0.5 |
| chr1 | 201503966 | 201521621 | 17656 | -1.64 | 0.426 | 0.5 |
| chr18 | 12307635 | 12328677 | 21043 | 0.11 | 0.426 | 0.5 |
| chr6 | 167411451 | 167413845 | 2395 | -0.22 | 0.426 | 0.5 |
| chr6 | 105238184 | 105238582 | 399 | 0.57 | 0.426 | 0.5 |
| chr4 | 17512920 | 17514360 | 1441 | 0.27 | 0.427 | 0.5 |
| chr3 | 73101493 | 73106251 | 4759 | -0.15 | 0.427 | 0.5 |
| chr7 | 24771975 | 24782867 | 10893 | -0.15 | 0.427 | 0.5 |
| chr1 | 221509073 | 221509887 | 815 | -0.37 | 0.427 | 0.501 |
| chr1 | 66425393 | 66426238 | 846 | 0.43 | 0.427 | 0.501 |
| chr7 | 30543488 | 30545291 | 1804 | 0.17 | 0.427 | 0.501 |
| chr1 | 26249157 | 26249990 | 834 | 0.34 | 0.428 | 0.501 |

| | | | | | | |
|---|---|---|---|---|---|---|
| chr12 | 17169640 | 17170627 | 988 | -0.49 | 0.428 | 0.501 |
| chr1 | 147775214 | 147776486 | 1273 | -0.19 | 0.428 | 0.502 |
| chr3 | 51702669 | 51715271 | 12603 | -0.13 | 0.428 | 0.502 |
| chr1 | 209700938 | 209701868 | 931 | 0.32 | 0.428 | 0.502 |
| chr11 | 115157308 | 115158268 | 961 | 0.37 | 0.428 | 0.502 |
| chr2 | 161341441 | 161351292 | 9852 | -0.13 | 0.428 | 0.502 |
| chr1 | 9903346 | 9904170 | 825 | 0.46 | 0.428 | 0.502 |
| chr14 | 35789556 | 35874847 | 85292 | -0.04 | 0.428 | 0.502 |
| chrX | 153743713 | 153777165 | 33453 | -0.05 | 0.429 | 0.502 |
| chr1 | 19577326 | 19579472 | 2147 | 0.16 | 0.429 | 0.502 |
| chr2 | 3582563 | 3608816 | 26254 | -0.06 | 0.429 | 0.503 |
| chr21 | 34099074 | 34109237 | 10164 | -0.2 | 0.429 | 0.503 |
| chr3 | 12569180 | 12569663 | 484 | 0.57 | 0.429 | 0.503 |
| chr4 | 719952 | 722108 | 2157 | 0.16 | 0.429 | 0.503 |
| chr11 | 123015433 | 123030460 | 15028 | -0.12 | 0.429 | 0.503 |
| chr4 | 89675146 | 89675791 | 646 | 0.52 | 0.429 | 0.503 |
| chr18 | 60478154 | 60483350 | 5197 | -0.15 | 0.429 | 0.503 |
| chr12 | 3382484 | 3382987 | 504 | 0.5 | 0.429 | 0.503 |
| chr10 | 14693175 | 14714226 | 21052 | 0.06 | 0.429 | 0.503 |
| chr6 | 88031833 | 88032932 | 1100 | 0.32 | 0.429 | 0.503 |
| chr9 | 138852083 | 138853079 | 997 | 0.36 | 0.429 | 0.503 |
| chr5 | 93952678 | 93955884 | 3207 | 0.1 | 0.429 | 0.503 |
| chr6 | 110940626 | 110941077 | 452 | -0.53 | 0.429 | 0.503 |
| chr14 | 43634792 | 43635536 | 745 | 0.43 | 0.429 | 0.503 |
| chr2 | 128784332 | 128785691 | 1360 | -0.29 | 0.43 | 0.503 |
| chr10 | 118368909 | 118369771 | 863 | 0.48 | 0.43 | 0.503 |
| chr10 | 135091311 | 135093130 | 1820 | 0.36 | 0.43 | 0.503 |
| chr5 | 124298926 | 124299798 | 873 | 0.42 | 0.43 | 0.504 |
| chr7 | 94867892 | 94876928 | 9037 | -1.56 | 0.43 | 0.504 |
| chr9 | 12810007 | 12820055 | 10049 | 0.05 | 0.43 | 0.504 |
| chr10 | 90675308 | 90675844 | 537 | -0.71 | 0.43 | 0.504 |
| chrX | 52949241 | 52965022 | 15782 | 0.11 | 0.43 | 0.504 |
| chr7 | 33218925 | 33219501 | 577 | -0.54 | 0.431 | 0.504 |
| chr4 | 15957518 | 15957961 | 444 | 0.54 | 0.431 | 0.504 |
| chr2 | 113357378 | 113357716 | 339 | -0.5 | 0.431 | 0.505 |
| chr2 | 242498101 | 242498634 | 534 | -0.44 | 0.431 | 0.505 |
| chr21 | 27089471 | 27090534 | 1064 | -0.34 | 0.431 | 0.505 |
| chr16 | 67080559 | 67103161 | 22603 | -0.13 | 0.431 | 0.505 |
| chr22 | 23281467 | 23282146 | 680 | 0.41 | 0.432 | 0.505 |
| chr7 | 128972076 | 128972764 | 689 | -0.42 | 0.432 | 0.506 |
| chr2 | 217957246 | 217971530 | 14285 | -0.23 | 0.432 | 0.506 |
| chr15 | 57459687 | 57461037 | 1351 | -0.27 | 0.432 | 0.506 |
| chr22 | 50682354 | 50684863 | 2510 | 0.17 | 0.432 | 0.506 |
| chr17 | 47585289 | 47593293 | 8005 | 0.3 | 0.432 | 0.506 |
| chr1 | 200406812 | 200407256 | 445 | 1.35 | 0.433 | 0.506 |
| chr15 | 73050169 | 73050785 | 617 | -0.47 | 0.433 | 0.506 |
| chr7 | 69336084 | 69337858 | 1775 | 0.31 | 0.433 | 0.506 |
| chr10 | 105109837 | 105111647 | 1811 | -0.22 | 0.433 | 0.507 |
| chr1 | 150121651 | 150122770 | 1120 | -0.33 | 0.433 | 0.507 |
| chr17 | 76209942 | 76215599 | 5658 | -0.14 | 0.433 | 0.507 |
| chr3 | 161317302 | 161318022 | 721 | -0.38 | 0.433 | 0.507 |
| chr6 | 138913425 | 138915340 | 1916 | 0.14 | 0.433 | 0.507 |
| chrX | 48554258 | 48563890 | 9633 | 0.13 | 0.433 | 0.507 |
| chr2 | 233264567 | 233265573 | 1007 | 0.43 | 0.433 | 0.507 |
| chr12 | 121714348 | 121754819 | 40472 | 0.09 | 0.433 | 0.507 |
| chr1 | 108669681 | 108669988 | 308 | 0.64 | 0.433 | 0.507 |
| chr3 | 52443013 | 52445656 | 2644 | 0.21 | 0.433 | 0.507 |
| chr9 | 81627419 | 81643414 | 15996 | 0.13 | 0.433 | 0.507 |
| chr16 | 19055380 | 19055761 | 382 | -0.5 | 0.434 | 0.507 |
| chr17 | 7358879 | 7390735 | 31857 | 0.05 | 0.434 | 0.507 |
| chr10 | 96278545 | 96279883 | 1339 | 0.3 | 0.434 | 0.508 |

| | | | | | | |
|---|---|---|---|---|---|---|
| chr8 | 66626754 | 66644026 | 17273 | 0.08 | 0.434 | 0.508 |
| chr1 | 151445471 | 151446310 | 840 | -0.29 | 0.434 | 0.508 |
| chr22 | 44389331 | 44389991 | 661 | 0.54 | 0.435 | 0.508 |
| chr14 | 100258987 | 100269235 | 10249 | -0.22 | 0.435 | 0.509 |
| chr7 | 765434 | 766892 | 1459 | 0.19 | 0.435 | 0.509 |
| chr11 | 102267789 | 102279040 | 11252 | -0.17 | 0.435 | 0.509 |
| chr5 | 70882444 | 70898058 | 15615 | -0.1 | 0.435 | 0.509 |
| chr17 | 30185091 | 30204967 | 19877 | -0.13 | 0.435 | 0.509 |
| chr22 | 45135179 | 45135593 | 415 | 0.6 | 0.436 | 0.509 |
| chr12 | 64539093 | 64541912 | 2820 | 0.33 | 0.436 | 0.509 |
| chr5 | 139973225 | 139973785 | 561 | 0.5 | 0.436 | 0.509 |
| chr5 | 87532887 | 87550768 | 17882 | 0.15 | 0.436 | 0.51 |
| chr11 | 113184747 | 113186427 | 1681 | -0.17 | 0.436 | 0.51 |
| chr20 | 50782205 | 50783280 | 1076 | 0.42 | 0.436 | 0.51 |
| chr2 | 178256637 | 178264582 | 7946 | -0.1 | 0.436 | 0.51 |
| chr8 | 145633582 | 145704920 | 71339 | -0.06 | 0.436 | 0.51 |
| chr3 | 50126323 | 50133949 | 7627 | 0.14 | 0.436 | 0.51 |
| chr13 | 21513405 | 21518376 | 4972 | -0.14 | 0.437 | 0.51 |
| chrX | 70680555 | 70681440 | 886 | 0.4 | 0.437 | 0.51 |
| chr22 | 34249270 | 34250715 | 1446 | -0.2 | 0.437 | 0.51 |
| chr2 | 205316115 | 205319473 | 3359 | 0.12 | 0.437 | 0.51 |
| chrX | 152599632 | 152600474 | 843 | 0.39 | 0.437 | 0.51 |
| chr3 | 194822934 | 194826420 | 3487 | 0.19 | 0.437 | 0.51 |
| chrX | 69653442 | 69655052 | 1611 | -0.21 | 0.437 | 0.511 |
| chr10 | 76584373 | 76587348 | 2976 | -0.14 | 0.437 | 0.511 |
| chr16 | 25117680 | 25123860 | 6181 | -0.1 | 0.437 | 0.511 |
| chr9 | 123786801 | 123818401 | 31601 | -0.06 | 0.437 | 0.511 |
| chr17 | 79632310 | 79635505 | 3196 | -0.12 | 0.437 | 0.511 |
| chr2 | 70297544 | 70298930 | 1387 | -0.25 | 0.437 | 0.511 |
| chr18 | 53457060 | 53458275 | 1216 | 0.4 | 0.437 | 0.511 |
| chr6 | 157148242 | 157149844 | 1603 | -0.28 | 0.437 | 0.511 |
| chr19 | 54341489 | 54398960 | 57472 | 0.04 | 0.438 | 0.511 |
| chr12 | 57270965 | 57274990 | 4026 | 0.21 | 0.438 | 0.511 |
| chr3 | 176867813 | 176879384 | 11572 | -0.21 | 0.438 | 0.511 |
| chr3 | 28992981 | 28994005 | 1025 | -0.32 | 0.438 | 0.511 |
| chr3 | 33154616 | 33175964 | 21349 | -0.09 | 0.438 | 0.511 |
| chr8 | 103299470 | 103300463 | 994 | 0.4 | 0.438 | 0.511 |
| chr20 | 17511022 | 17520073 | 9052 | -0.09 | 0.438 | 0.512 |
| chr15 | 26327593 | 26328872 | 1280 | 0.21 | 0.438 | 0.512 |
| chr1 | 243894618 | 243895085 | 468 | 0.62 | 0.438 | 0.512 |
| chr3 | 10182653 | 10193321 | 10669 | 0.11 | 0.438 | 0.512 |
| chr6 | 107435085 | 107437593 | 2509 | 0.22 | 0.438 | 0.512 |
| chr9 | 129719448 | 129720302 | 855 | 0.4 | 0.439 | 0.512 |
| chrX | 41419896 | 41420986 | 1091 | 0.3 | 0.439 | 0.512 |
| chr14 | 105258393 | 105283274 | 24882 | 0.1 | 0.439 | 0.513 |
| chr7 | 101288012 | 101288661 | 650 | 0.52 | 0.439 | 0.513 |
| chr9 | 134502910 | 134512935 | 10026 | 0.14 | 0.439 | 0.513 |
| chr17 | 2608406 | 2616164 | 7759 | -0.09 | 0.439 | 0.513 |
| chr16 | 15887984 | 15888873 | 890 | -0.44 | 0.439 | 0.513 |
| chr14 | 71514758 | 71516280 | 1523 | -0.31 | 0.439 | 0.513 |
| chr7 | 94717805 | 94735653 | 17849 | -0.18 | 0.439 | 0.513 |
| chr17 | 73256516 | 73286864 | 30349 | 0.06 | 0.439 | 0.513 |
| chr14 | 71688272 | 71689020 | 749 | -0.38 | 0.44 | 0.513 |
| chr5 | 14381081 | 14429221 | 48141 | -0.08 | 0.44 | 0.513 |
| chr4 | 110833787 | 110834478 | 692 | 0.38 | 0.44 | 0.513 |
| chr4 | 169330250 | 169373379 | 43130 | -0.09 | 0.44 | 0.513 |
| chr2 | 58106635 | 58111419 | 4785 | 0.06 | 0.44 | 0.514 |
| chr10 | 135120670 | 135122691 | 2022 | 0.14 | 0.44 | 0.514 |
| chr9 | 88313440 | 88314133 | 694 | 0.35 | 0.44 | 0.514 |
| chrX | 53566666 | 53567892 | 1227 | 0.37 | 0.44 | 0.514 |
| chr1 | 19637880 | 19639472 | 1593 | -0.28 | 0.44 | 0.514 |

| | | | | | | |
|---|---|---|---|---|---|---|
| chr6 | 117163612 | 117164548 | 937 | 1.41 | 0.441 | 0.514 |
| chr6 | 160559500 | 160561114 | 1615 | 0.31 | 0.441 | 0.514 |
| chr1 | 1366842 | 1374134 | 7293 | 0.2 | 0.441 | 0.514 |
| chr4 | 47566648 | 47567716 | 1069 | 0.3 | 0.441 | 0.514 |
| chr1 | 214453574 | 214455873 | 2300 | 0.13 | 0.441 | 0.514 |
| chr9 | 136039070 | 136039712 | 643 | 0.51 | 0.441 | 0.515 |
| chr1 | 184722513 | 184725146 | 2634 | 0.16 | 0.441 | 0.515 |
| chr5 | 39256214 | 39257248 | 1035 | -0.23 | 0.441 | 0.515 |
| chr2 | 40661108 | 40663694 | 2587 | -0.16 | 0.441 | 0.515 |
| chr1 | 223254543 | 223255179 | 637 | 0.38 | 0.441 | 0.515 |
| chr17 | 38789413 | 38805213 | 15801 | -0.07 | 0.441 | 0.515 |
| chr3 | 52252198 | 52252555 | 358 | -0.7 | 0.441 | 0.515 |
| chr11 | 60608883 | 60611889 | 3007 | 0.14 | 0.441 | 0.515 |
| chr1 | 27113523 | 27127161 | 13639 | -0.13 | 0.441 | 0.515 |
| chr3 | 138892415 | 138892892 | 478 | 0.51 | 0.441 | 0.515 |
| chr2 | 64586797 | 64587708 | 912 | -0.28 | 0.442 | 0.515 |
| chr19 | 10283869 | 10284770 | 902 | 0.48 | 0.442 | 0.515 |
| chr2 | 26401565 | 26408560 | 6996 | -0.2 | 0.442 | 0.515 |
| chr14 | 32694157 | 32700487 | 6331 | 0.2 | 0.442 | 0.515 |
| chr15 | 71888447 | 71902801 | 14355 | -0.13 | 0.442 | 0.515 |
| chr19 | 2259191 | 2274662 | 15472 | -0.15 | 0.442 | 0.515 |
| chr12 | 122788725 | 122824203 | 35479 | 0.07 | 0.442 | 0.515 |
| chr1 | 225837630 | 225848900 | 11271 | 0.06 | 0.442 | 0.516 |
| chr17 | 42949298 | 42950461 | 1164 | -1.24 | 0.442 | 0.516 |
| chr18 | 20657236 | 20658334 | 1099 | -0.37 | 0.442 | 0.516 |
| chr6 | 27059727 | 27065568 | 5842 | 0.15 | 0.442 | 0.516 |
| chr19 | 57922042 | 57923182 | 1141 | 0.37 | 0.442 | 0.516 |
| chrX | 24939149 | 24944726 | 5578 | -0.17 | 0.443 | 0.516 |
| chr5 | 81146874 | 81148829 | 1956 | 0.16 | 0.443 | 0.516 |
| chr1 | 182990101 | 183014524 | 24424 | 0.06 | 0.443 | 0.516 |
| chrX | 34892810 | 34903910 | 11101 | -0.16 | 0.443 | 0.517 |
| chr17 | 27949442 | 27949820 | 379 | 0.59 | 0.444 | 0.517 |
| chr13 | 26827964 | 26829673 | 1710 | 0.18 | 0.444 | 0.518 |
| chr1 | 209862296 | 209866388 | 4093 | 0.22 | 0.444 | 0.518 |
| chr9 | 136576152 | 136576901 | 750 | 0.58 | 0.444 | 0.518 |
| chr7 | 129761363 | 129762602 | 1240 | 0.38 | 0.444 | 0.518 |
| chr8 | 8774981 | 8775433 | 453 | -0.48 | 0.444 | 0.518 |
| chr12 | 30945191 | 30950173 | 4983 | 0.09 | 0.444 | 0.518 |
| chr11 | 17426761 | 17428019 | 1259 | -0.17 | 0.444 | 0.518 |
| chr1 | 8377479 | 8378696 | 1218 | 0.36 | 0.445 | 0.518 |
| chr19 | 30364050 | 30364733 | 684 | -0.33 | 0.445 | 0.518 |
| chr6 | 113672433 | 113683415 | 10983 | -0.11 | 0.445 | 0.519 |
| chrX | 129906035 | 129906990 | 956 | 0.39 | 0.445 | 0.519 |
| chr5 | 78530947 | 78533495 | 2549 | 0.18 | 0.445 | 0.519 |
| chr9 | 16824454 | 16824876 | 423 | -0.52 | 0.445 | 0.519 |
| chr8 | 18982556 | 19010306 | 27751 | 0.08 | 0.445 | 0.519 |
| chr18 | 43732026 | 43755198 | 23173 | -0.12 | 0.446 | 0.519 |
| chr2 | 210866498 | 210868878 | 2381 | -0.15 | 0.446 | 0.519 |
| chr1 | 11740510 | 11741720 | 1211 | 0.31 | 0.446 | 0.519 |
| chr8 | 62420254 | 62421889 | 1636 | -0.24 | 0.446 | 0.519 |
| chr7 | 66309017 | 66311561 | 2545 | 0.11 | 0.446 | 0.52 |
| chr6 | 12395917 | 12396704 | 788 | 0.54 | 0.446 | 0.52 |
| chr1 | 161879069 | 161880501 | 1433 | -0.32 | 0.446 | 0.52 |
| chr1 | 172795347 | 172795917 | 571 | -0.42 | 0.446 | 0.52 |
| chr4 | 184826050 | 184827499 | 1450 | 0.36 | 0.446 | 0.52 |
| chr15 | 73073933 | 73089950 | 16018 | -0.11 | 0.447 | 0.52 |
| chr9 | 125761673 | 125766962 | 5290 | 0.19 | 0.447 | 0.52 |
| chrX | 149857159 | 149862838 | 5680 | 0.2 | 0.447 | 0.52 |
| chrX | 106917540 | 106921128 | 3589 | -0.16 | 0.447 | 0.521 |
| chr16 | 30726123 | 30799382 | 73260 | -0.05 | 0.447 | 0.521 |
| chr7 | 83095771 | 83099648 | 3878 | 0.11 | 0.447 | 0.521 |

| | | | | | | |
|---|---|---|---|---|---|---|
| chr1 | 49343346 | 49345330 | 1985 | -0.12 | 0.447 | 0.521 |
| chr20 | 8851161 | 8855848 | 4688 | -0.24 | 0.447 | 0.521 |
| chr1 | 1744030 | 1823502 | 79473 | 0.05 | 0.447 | 0.521 |
| chr6 | 147226128 | 147237470 | 11343 | -0.07 | 0.448 | 0.521 |
| chr15 | 85143135 | 85150659 | 7525 | -0.16 | 0.448 | 0.521 |
| chrX | 20206994 | 20207687 | 694 | -0.82 | 0.448 | 0.521 |
| chr2 | 158185971 | 158186574 | 604 | -0.43 | 0.448 | 0.521 |
| chr22 | 23023890 | 23037828 | 13939 | 0.12 | 0.448 | 0.522 |
| chr6 | 13614364 | 13616739 | 2376 | 0.14 | 0.448 | 0.522 |
| chr6 | 10837837 | 10838864 | 1028 | -0.28 | 0.448 | 0.522 |
| chr17 | 35227075 | 35227994 | 920 | -0.39 | 0.448 | 0.522 |
| chr12 | 1885208 | 1886148 | 941 | -0.48 | 0.448 | 0.522 |
| chr15 | 90894996 | 90914316 | 19321 | -0.11 | 0.448 | 0.522 |
| chr10 | 5425274 | 5426805 | 1532 | -0.27 | 0.448 | 0.522 |
| chr2 | 157255768 | 157258952 | 3185 | -0.14 | 0.448 | 0.522 |
| chr20 | 48702091 | 48814133 | 112043 | -0.03 | 0.449 | 0.522 |
| chr2 | 8711824 | 8713217 | 1394 | -0.36 | 0.449 | 0.522 |
| chr21 | 26946018 | 26947049 | 1032 | 0.41 | 0.449 | 0.522 |
| chr8 | 6502025 | 6506342 | 4318 | 0.28 | 0.449 | 0.522 |
| chr12 | 1905536 | 1914441 | 8906 | -0.57 | 0.449 | 0.522 |
| chr2 | 203958231 | 203958680 | 450 | 0.63 | 0.449 | 0.522 |
| chr2 | 145353401 | 145354576 | 1176 | 0.33 | 0.449 | 0.522 |
| chr1 | 109358405 | 109374621 | 16217 | -0.11 | 0.449 | 0.522 |
| chr2 | 175292313 | 175293663 | 1351 | 1.46 | 0.449 | 0.523 |
| chr12 | 93860460 | 93880299 | 19840 | 0.09 | 0.449 | 0.523 |
| chr15 | 23085274 | 23087955 | 2682 | 0.13 | 0.449 | 0.523 |
| chr9 | 114719982 | 114730018 | 10037 | -0.1 | 0.449 | 0.523 |
| chr1 | 160174501 | 160186454 | 11954 | 0.09 | 0.45 | 0.523 |
| chr7 | 2420924 | 2421376 | 453 | -0.55 | 0.45 | 0.523 |
| chr17 | 2205454 | 2221391 | 15938 | 0.12 | 0.45 | 0.523 |
| chr12 | 51984018 | 51985525 | 1508 | 0.32 | 0.45 | 0.523 |
| chr12 | 45815497 | 45815905 | 409 | -0.57 | 0.45 | 0.523 |
| chr1 | 198826519 | 198826861 | 343 | -0.49 | 0.45 | 0.523 |
| chr17 | 26505457 | 26506878 | 1422 | 0.3 | 0.45 | 0.523 |
| chr12 | 13679737 | 13680860 | 1124 | -0.29 | 0.45 | 0.523 |
| chr11 | 118305835 | 118308337 | 2503 | 0.23 | 0.451 | 0.524 |
| chr11 | 66790141 | 66801429 | 11289 | 0.25 | 0.451 | 0.524 |
| chr7 | 23529399 | 23530580 | 1182 | 0.26 | 0.451 | 0.524 |
| chr7 | 75669351 | 75700976 | 31626 | -0.06 | 0.451 | 0.524 |
| chr8 | 26181176 | 26188693 | 7518 | -0.24 | 0.451 | 0.524 |
| chr12 | 104019918 | 104020328 | 411 | 0.5 | 0.451 | 0.524 |
| chr7 | 30589858 | 30591972 | 2115 | 0.21 | 0.451 | 0.524 |
| chr1 | 242000818 | 242020797 | 19980 | 0.1 | 0.451 | 0.524 |
| chr17 | 80316452 | 80317094 | 643 | -0.38 | 0.451 | 0.525 |
| chr2 | 84742653 | 84744102 | 1450 | -0.26 | 0.451 | 0.525 |
| chr12 | 121287479 | 121295103 | 7625 | 0.23 | 0.452 | 0.525 |
| chr22 | 24456404 | 24457455 | 1052 | 0.34 | 0.452 | 0.525 |
| chr7 | 108128803 | 108130146 | 1344 | -0.33 | 0.452 | 0.525 |
| chr6 | 165722698 | 165723098 | 401 | 0.44 | 0.452 | 0.525 |
| chr15 | 77223440 | 77234145 | 10706 | -0.14 | 0.452 | 0.525 |
| chr14 | 65077758 | 65134246 | 56489 | -0.05 | 0.452 | 0.525 |
| chr4 | 104118162 | 104120218 | 2057 | -0.16 | 0.452 | 0.525 |
| chr20 | 40086488 | 40087147 | 660 | 0.44 | 0.452 | 0.525 |
| chr15 | 44320509 | 44321009 | 501 | -0.47 | 0.452 | 0.525 |
| chr3 | 48074923 | 48111969 | 37047 | 0.08 | 0.452 | 0.526 |
| chr14 | 70882389 | 70884196 | 1808 | 0.16 | 0.452 | 0.526 |
| chr2 | 37416548 | 37460250 | 43703 | -0.06 | 0.453 | 0.526 |
| chr22 | 42709067 | 42717489 | 8423 | 0.17 | 0.453 | 0.526 |
| chr15 | 76134848 | 76137253 | 2406 | 0.12 | 0.453 | 0.526 |
| chr10 | 25103288 | 25115387 | 12100 | -0.21 | 0.453 | 0.526 |
| chr7 | 105469948 | 105491396 | 21449 | -0.13 | 0.453 | 0.526 |

| | | | | | | |
|---|---|---|---|---|---|---|
| chr6 | 35744220 | 35744634 | 415 | -0.53 | 0.453 | 0.526 |
| chr9 | 19284685 | 19287605 | 2921 | -0.25 | 0.453 | 0.526 |
| chr2 | 37616996 | 37618398 | 1403 | 0.26 | 0.453 | 0.526 |
| chr2 | 158316226 | 158318012 | 1787 | 0.27 | 0.453 | 0.526 |
| chr2 | 47997651 | 48039208 | 41558 | 0.07 | 0.453 | 0.526 |
| chr6 | 40862615 | 40880307 | 17693 | 0.1 | 0.453 | 0.526 |
| chr13 | 74507830 | 74513334 | 5505 | -0.18 | 0.453 | 0.527 |
| chr6 | 27486037 | 27487804 | 1768 | 0.22 | 0.453 | 0.527 |
| chr22 | 28774614 | 28775625 | 1012 | -0.36 | 0.454 | 0.527 |
| chr7 | 155743223 | 155745685 | 2463 | -0.1 | 0.454 | 0.527 |
| chr11 | 76902304 | 76903390 | 1087 | -0.14 | 0.454 | 0.527 |
| chr19 | 16682134 | 16697629 | 15496 | -0.1 | 0.454 | 0.527 |
| chr19 | 7479915 | 7483359 | 3445 | -0.15 | 0.454 | 0.527 |
| chr7 | 30174032 | 30183614 | 9583 | 1.6 | 0.454 | 0.527 |
| chr1 | 183512451 | 183537399 | 24949 | 0.12 | 0.454 | 0.527 |
| chr20 | 44596848 | 44604009 | 7162 | -0.08 | 0.455 | 0.528 |
| chr1 | 200579915 | 200591559 | 11645 | -0.08 | 0.455 | 0.528 |
| chr17 | 68164429 | 68165482 | 1054 | -0.35 | 0.455 | 0.528 |
| chr6 | 75184076 | 75184459 | 384 | 0.51 | 0.455 | 0.528 |
| chr11 | 57334824 | 57337375 | 2552 | -0.34 | 0.455 | 0.528 |
| chr8 | 124228956 | 124230519 | 1564 | 0.32 | 0.455 | 0.528 |
| chr18 | 73186730 | 73188059 | 1330 | 0.35 | 0.455 | 0.528 |
| chr1 | 43222427 | 43233693 | 11267 | -0.16 | 0.455 | 0.528 |
| chr13 | 35516479 | 35517577 | 1099 | 0.36 | 0.455 | 0.528 |
| chr16 | 85546474 | 85559166 | 12693 | 0.08 | 0.455 | 0.528 |
| chr6 | 7291483 | 7291857 | 375 | 0.58 | 0.455 | 0.528 |
| chr2 | 183969853 | 183992217 | 22365 | -0.1 | 0.455 | 0.528 |
| chr4 | 153455632 | 153459599 | 3968 | 0.09 | 0.455 | 0.528 |
| chr22 | 38329477 | 38330560 | 1084 | -0.35 | 0.456 | 0.529 |
| chr18 | 68131710 | 68132065 | 356 | 0.66 | 0.456 | 0.529 |
| chr2 | 198174917 | 198176176 | 1260 | 0.33 | 0.456 | 0.529 |
| chr13 | 93922893 | 93924338 | 1446 | -0.29 | 0.457 | 0.53 |
| chr10 | 418329 | 420410 | 2082 | -0.2 | 0.457 | 0.53 |
| chr11 | 10535248 | 10563620 | 28373 | 0.11 | 0.457 | 0.53 |
| chr3 | 120378895 | 120382176 | 3282 | 0.09 | 0.457 | 0.53 |
| chr12 | 76577155 | 76577961 | 807 | 0.44 | 0.457 | 0.53 |
| chr11 | 6947609 | 6948341 | 733 | 0.39 | 0.457 | 0.53 |
| chr14 | 53300884 | 53330433 | 29550 | 0.15 | 0.457 | 0.53 |
| chr4 | 54929203 | 54933531 | 4329 | 0.12 | 0.457 | 0.53 |
| chr2 | 18740703 | 18742609 | 1907 | 0.11 | 0.457 | 0.53 |
| chr1 | 212262869 | 212273175 | 10307 | -0.26 | 0.457 | 0.53 |
| chr6 | 66416169 | 66416966 | 798 | 0.37 | 0.457 | 0.53 |
| chr3 | 4344632 | 4346229 | 1598 | 0.14 | 0.457 | 0.53 |
| chr6 | 56954291 | 56954647 | 357 | 0.54 | 0.458 | 0.53 |
| chr8 | 118826859 | 118828272 | 1414 | 0.28 | 0.458 | 0.531 |
| chr3 | 119862481 | 119867030 | 4550 | -0.2 | 0.458 | 0.531 |
| chr14 | 100484813 | 100504916 | 20104 | -0.09 | 0.458 | 0.531 |
| chr7 | 3312104 | 3312970 | 867 | -0.41 | 0.458 | 0.531 |
| chr2 | 37478096 | 37481913 | 3818 | -0.23 | 0.458 | 0.531 |
| chr16 | 5083217 | 5084295 | 1079 | 0.26 | 0.458 | 0.531 |
| chr1 | 28449500 | 28450816 | 1317 | -0.28 | 0.458 | 0.531 |
| chrX | 69439150 | 69440983 | 1834 | 0.17 | 0.458 | 0.531 |
| chr14 | 24116065 | 24117183 | 1119 | -0.32 | 0.459 | 0.531 |
| chr1 | 38018228 | 38029576 | 11349 | 0.12 | 0.459 | 0.532 |
| chr14 | 34383589 | 34387415 | 3827 | -0.13 | 0.459 | 0.532 |
| chr1 | 161581720 | 161583089 | 1370 | 0.22 | 0.459 | 0.532 |
| chr18 | 21593860 | 21595466 | 1607 | -0.27 | 0.459 | 0.532 |
| chr1 | 44153703 | 44154727 | 1025 | -0.37 | 0.459 | 0.532 |
| chr8 | 97272469 | 97285125 | 12657 | 0.11 | 0.459 | 0.532 |
| chr18 | 54303178 | 54319277 | 16100 | 0.1 | 0.459 | 0.532 |
| chr17 | 5370744 | 5373009 | 2266 | -0.14 | 0.459 | 0.532 |

| | | | | | | |
|---|---|---|---|---|---|---|
| chr20 | 33820482 | 33820855 | 374 | -0.59 | 0.459 | 0.532 |
| chr20 | 30777394 | 30819535 | 42142 | -0.07 | 0.459 | 0.532 |
| chr11 | 124980807 | 124982416 | 1610 | -0.19 | 0.46 | 0.532 |
| chr11 | 46336099 | 46336757 | 659 | 0.44 | 0.46 | 0.533 |
| chr1 | 222000097 | 222019604 | 19508 | -0.24 | 0.46 | 0.533 |
| chr3 | 27260686 | 27261221 | 536 | 0.44 | 0.46 | 0.533 |
| chr13 | 49596313 | 49598339 | 2027 | -0.25 | 0.46 | 0.533 |
| chr10 | 30870025 | 30876822 | 6798 | -0.09 | 0.46 | 0.533 |
| chr10 | 133849635 | 133862628 | 12994 | 0.89 | 0.46 | 0.533 |
| chr12 | 101752694 | 101753685 | 992 | -0.42 | 0.46 | 0.533 |
| chr10 | 34970874 | 35058664 | 87791 | 0.03 | 0.46 | 0.533 |
| chr19 | 50414134 | 50434397 | 20264 | -0.07 | 0.461 | 0.533 |
| chr1 | 22030020 | 22030704 | 685 | -0.32 | 0.461 | 0.534 |
| chr8 | 37637609 | 37644696 | 7088 | -0.17 | 0.461 | 0.534 |
| chr1 | 203259917 | 203260968 | 1052 | 0.4 | 0.461 | 0.534 |
| chr2 | 21021406 | 21023736 | 2331 | 0.08 | 0.461 | 0.534 |
| chr15 | 73976168 | 73977642 | 1475 | 0.27 | 0.461 | 0.534 |
| chr9 | 132769321 | 132836712 | 67392 | -0.05 | 0.461 | 0.534 |
| chr5 | 76325577 | 76327959 | 2383 | 0.16 | 0.461 | 0.534 |
| chr9 | 138391016 | 138397801 | 6786 | -0.14 | 0.461 | 0.534 |
| chr21 | 46521823 | 46522722 | 900 | 0.34 | 0.462 | 0.534 |
| chr14 | 56116830 | 56122658 | 5829 | 0.16 | 0.462 | 0.534 |
| chr11 | 109962613 | 109966014 | 3402 | 0.11 | 0.462 | 0.534 |
| chr16 | 56964631 | 56983853 | 19223 | 0.07 | 0.462 | 0.535 |
| chr10 | 35588930 | 35589420 | 491 | -0.51 | 0.462 | 0.535 |
| chr11 | 36615994 | 36616914 | 921 | 0.2 | 0.462 | 0.535 |
| chr19 | 3552698 | 3573791 | 21094 | -0.07 | 0.462 | 0.535 |
| chr10 | 90692008 | 90692489 | 482 | -0.6 | 0.462 | 0.535 |
| chr16 | 84517569 | 84518384 | 816 | -0.44 | 0.462 | 0.535 |
| chrX | 44702095 | 44702851 | 757 | 0.43 | 0.462 | 0.535 |
| chr2 | 197769561 | 197770439 | 879 | 0.23 | 0.463 | 0.535 |
| chr14 | 37640776 | 37642352 | 1577 | 0.12 | 0.463 | 0.535 |
| chr8 | 134332639 | 134333324 | 686 | 0.82 | 0.463 | 0.535 |
| chr11 | 22813754 | 22814731 | 978 | 0.33 | 0.463 | 0.535 |
| chr7 | 17949635 | 17951308 | 1674 | -0.35 | 0.463 | 0.535 |
| chr2 | 32580303 | 32606137 | 25835 | 0.06 | 0.463 | 0.536 |
| chr6 | 111265429 | 111280915 | 15487 | -0.07 | 0.463 | 0.536 |
| chr6 | 3231135 | 3232599 | 1465 | -0.22 | 0.463 | 0.536 |
| chr17 | 37664756 | 37671504 | 6749 | 0.16 | 0.463 | 0.536 |
| chr19 | 40714890 | 40715485 | 596 | 0.52 | 0.463 | 0.536 |
| chr15 | 40856428 | 40893146 | 36719 | 0.08 | 0.463 | 0.536 |
| chr19 | 11086011 | 11087074 | 1064 | 0.42 | 0.463 | 0.536 |
| chr17 | 49474742 | 49475888 | 1147 | -0.3 | 0.463 | 0.536 |
| chr19 | 51610417 | 51612612 | 2196 | -0.14 | 0.464 | 0.536 |
| chr9 | 131217235 | 131219816 | 2582 | -0.13 | 0.464 | 0.536 |
| chr5 | 107552387 | 107553162 | 776 | -0.46 | 0.464 | 0.536 |
| chr10 | 95632661 | 95633972 | 1312 | -0.23 | 0.464 | 0.536 |
| chr8 | 37961276 | 37964222 | 2947 | 0.09 | 0.464 | 0.536 |
| chr7 | 108290189 | 108291994 | 1806 | 0.18 | 0.464 | 0.537 |
| chr2 | 10115227 | 10116161 | 935 | 1.28 | 0.464 | 0.537 |
| chr9 | 103277681 | 103278560 | 880 | 0.29 | 0.464 | 0.537 |
| chr11 | 67979439 | 67982262 | 2824 | 0.13 | 0.464 | 0.537 |
| chr6 | 153379353 | 153385683 | 6331 | 0.21 | 0.464 | 0.537 |
| chr2 | 25802412 | 25802916 | 505 | -0.41 | 0.465 | 0.537 |
| chr14 | 51466286 | 51474519 | 8234 | 0.06 | 0.465 | 0.537 |
| chr9 | 130460909 | 130505321 | 44413 | -0.06 | 0.465 | 0.537 |
| chr14 | 22383418 | 22385911 | 2494 | 0.21 | 0.465 | 0.537 |
| chr8 | 144947862 | 144961028 | 13167 | -0.16 | 0.465 | 0.537 |
| chr13 | 96272393 | 96273741 | 1349 | -0.3 | 0.465 | 0.538 |
| chrX | 153266949 | 153286967 | 20019 | -0.08 | 0.465 | 0.538 |
| chrX | 11118193 | 11130977 | 12785 | 0.12 | 0.465 | 0.538 |

| | | | | | | |
|---|---|---|---|---|---|---|
| chr20 | 57797137 | 57797680 | 544 | 0.46 | 0.466 | 0.538 |
| chr12 | 20618437 | 20619010 | 574 | -0.44 | 0.466 | 0.538 |
| chr10 | 133992979 | 133993812 | 834 | -0.45 | 0.466 | 0.538 |
| chr10 | 129720188 | 129727197 | 7010 | -0.16 | 0.466 | 0.538 |
| chr14 | 76448034 | 76455902 | 7869 | -0.13 | 0.466 | 0.539 |
| chr14 | 78173210 | 78181993 | 8784 | 0.13 | 0.467 | 0.539 |
| chr10 | 77189770 | 77191892 | 2123 | -0.22 | 0.467 | 0.539 |
| chr4 | 153768117 | 153768584 | 468 | 0.44 | 0.467 | 0.539 |
| chr10 | 113930146 | 113944334 | 14189 | 0.21 | 0.467 | 0.54 |
| chr1 | 39490927 | 39493615 | 2689 | 0.09 | 0.467 | 0.54 |
| chr4 | 110735830 | 110738172 | 2343 | 0.12 | 0.467 | 0.54 |
| chr3 | 78936276 | 78937133 | 858 | 0.38 | 0.467 | 0.54 |
| chr1 | 39900013 | 39917222 | 17210 | 0.15 | 0.467 | 0.54 |
| chr11 | 103196467 | 103197479 | 1013 | -0.31 | 0.468 | 0.54 |
| chr1 | 168355832 | 168362830 | 6999 | 0.32 | 0.468 | 0.54 |
| chr9 | 38608732 | 38623309 | 14578 | -0.09 | 0.468 | 0.54 |
| chr16 | 31213248 | 31227489 | 14242 | -1.47 | 0.468 | 0.54 |
| chr8 | 126372881 | 126393317 | 20437 | 0.07 | 0.468 | 0.54 |
| chr12 | 75723370 | 75729076 | 5707 | -0.14 | 0.468 | 0.54 |
| chr22 | 21921487 | 21927450 | 5964 | 0.12 | 0.468 | 0.54 |
| chr13 | 43578873 | 43581910 | 3038 | 0.2 | 0.468 | 0.541 |
| chr1 | 183482926 | 183494807 | 11882 | -0.13 | 0.468 | 0.541 |
| chr1 | 111770995 | 111771497 | 503 | -0.43 | 0.468 | 0.541 |
| chr5 | 158655199 | 158668102 | 12904 | -0.16 | 0.468 | 0.541 |
| chr18 | 43677044 | 43685419 | 8376 | -0.16 | 0.468 | 0.541 |
| chr4 | 77869908 | 77872657 | 2750 | -0.18 | 0.469 | 0.541 |
| chr6 | 7589996 | 7591308 | 1313 | -0.32 | 0.469 | 0.541 |
| chr3 | 100007448 | 100008053 | 606 | -0.47 | 0.469 | 0.541 |
| chr3 | 197581425 | 197582931 | 1507 | -0.38 | 0.469 | 0.541 |
| chr14 | 35589468 | 35603939 | 14472 | -0.07 | 0.469 | 0.542 |
| chr6 | 158939184 | 158939728 | 545 | 0.44 | 0.469 | 0.542 |
| chr4 | 169951435 | 169952670 | 1236 | 0.18 | 0.469 | 0.542 |
| chr17 | 38391147 | 38392287 | 1141 | -0.33 | 0.469 | 0.542 |
| chr17 | 8775417 | 8776227 | 811 | 0.28 | 0.469 | 0.542 |
| chr5 | 139725610 | 139738618 | 13009 | -0.22 | 0.469 | 0.542 |
| chr10 | 88713534 | 88734956 | 21423 | -0.07 | 0.47 | 0.542 |
| chr6 | 27559756 | 27601896 | 42141 | -0.09 | 0.47 | 0.542 |
| chr12 | 96709285 | 96709785 | 501 | 0.43 | 0.47 | 0.542 |
| chr18 | 55588565 | 55598822 | 10258 | 0.21 | 0.47 | 0.542 |
| chr14 | 93547956 | 93553735 | 5780 | 0.14 | 0.47 | 0.542 |
| chr1 | 178653407 | 178654365 | 959 | 0.4 | 0.47 | 0.542 |
| chr3 | 15033776 | 15034403 | 628 | -0.51 | 0.47 | 0.542 |
| chr2 | 168916734 | 168919096 | 2363 | -0.16 | 0.47 | 0.542 |
| chr3 | 32145859 | 32148639 | 2781 | 0.25 | 0.47 | 0.542 |
| chr3 | 50087111 | 50088814 | 1704 | 0.2 | 0.47 | 0.542 |
| chr12 | 96754255 | 96758760 | 4506 | 0.15 | 0.47 | 0.542 |
| chr1 | 98727539 | 98745390 | 17852 | -0.08 | 0.47 | 0.542 |
| chr14 | 102323896 | 102324537 | 642 | -0.42 | 0.47 | 0.543 |
| chr16 | 21161647 | 21170524 | 8878 | 0.23 | 0.47 | 0.543 |
| chr3 | 169975068 | 169981253 | 6186 | 0.12 | 0.47 | 0.543 |
| chr8 | 23980276 | 23981663 | 1388 | 0.25 | 0.47 | 0.543 |
| chr13 | 44874370 | 44888791 | 14422 | -0.06 | 0.47 | 0.543 |
| chr14 | 52809423 | 52819576 | 10154 | 0.24 | 0.47 | 0.543 |
| chr9 | 2621019 | 2624451 | 3433 | 0.17 | 0.471 | 0.543 |
| chr8 | 90301889 | 90302635 | 747 | -0.39 | 0.471 | 0.543 |
| chr11 | 107267873 | 107268511 | 639 | -0.4 | 0.471 | 0.543 |
| chr6 | 3742856 | 3752946 | 10091 | -0.11 | 0.471 | 0.544 |
| chr1 | 28695561 | 28702295 | 6735 | -0.12 | 0.471 | 0.544 |
| chr11 | 70327995 | 70329196 | 1202 | -0.32 | 0.472 | 0.544 |
| chr9 | 37799947 | 37813839 | 13893 | 0.12 | 0.472 | 0.544 |
| chr21 | 43653227 | 43655485 | 2259 | 0.22 | 0.472 | 0.544 |

| | | | | | | |
|---|---|---|---|---|---|---|
| chr15 | 74945610 | 74946578 | 969 | 1.28 | 0.472 | 0.544 |
| chr3 | 53348750 | 53385446 | 36697 | 0.08 | 0.472 | 0.544 |
| chr4 | 146728118 | 146730704 | 2587 | 0.23 | 0.472 | 0.544 |
| chr1 | 179819075 | 179826772 | 7698 | 0.14 | 0.472 | 0.545 |
| chr8 | 48949801 | 48950601 | 801 | -0.41 | 0.472 | 0.545 |
| chrX | 40125823 | 40126722 | 900 | -0.37 | 0.472 | 0.545 |
| chr14 | 105399352 | 105399875 | 524 | -0.35 | 0.472 | 0.545 |
| chr8 | 40457121 | 40458578 | 1458 | -0.35 | 0.473 | 0.545 |
| chr8 | 41702409 | 41703135 | 727 | 0.43 | 0.473 | 0.545 |
| chr4 | 79695767 | 79698591 | 2825 | 0.11 | 0.473 | 0.545 |
| chr9 | 2096334 | 2096969 | 636 | -0.4 | 0.473 | 0.545 |
| chr6 | 47500992 | 47501424 | 433 | 0.46 | 0.473 | 0.545 |
| chr2 | 180197294 | 180210146 | 12853 | -0.18 | 0.473 | 0.545 |
| chr1 | 235400205 | 235403564 | 3360 | 0.23 | 0.473 | 0.546 |
| chr16 | 50099342 | 50101102 | 1761 | 0.23 | 0.473 | 0.546 |
| chr11 | 108740989 | 108741860 | 872 | -1.29 | 0.474 | 0.546 |
| chr17 | 26918200 | 26927581 | 9382 | -0.1 | 0.474 | 0.546 |
| chr22 | 36519249 | 36520191 | 943 | 0.22 | 0.474 | 0.546 |
| chr1 | 10570796 | 10571768 | 973 | -0.4 | 0.474 | 0.546 |
| chr4 | 185264557 | 185265122 | 566 | 0.45 | 0.474 | 0.546 |
| chr2 | 143503308 | 143512439 | 9132 | 1.3 | 0.474 | 0.546 |
| chr3 | 119180809 | 119188643 | 7835 | 0.08 | 0.474 | 0.546 |
| chr8 | 59323354 | 59324786 | 1433 | 0.15 | 0.474 | 0.546 |
| chr14 | 53618259 | 53620819 | 2561 | -0.14 | 0.474 | 0.546 |
| chr6 | 64345677 | 64347782 | 2106 | -0.15 | 0.474 | 0.546 |
| chr16 | 54319745 | 54321547 | 1803 | -0.23 | 0.474 | 0.546 |
| chr6 | 36390494 | 36413859 | 23366 | 0.06 | 0.475 | 0.547 |
| chr6 | 38180284 | 38181256 | 973 | 0.4 | 0.475 | 0.547 |
| chr20 | 2448847 | 2452596 | 3750 | 0.07 | 0.475 | 0.547 |
| chr22 | 42915409 | 42915778 | 370 | 0.38 | 0.475 | 0.547 |
| chr7 | 150754377 | 150792910 | 38534 | -0.06 | 0.475 | 0.547 |
| chr4 | 140288313 | 140289134 | 822 | -0.44 | 0.475 | 0.547 |
| chr4 | 189942670 | 189943714 | 1045 | 0.32 | 0.475 | 0.547 |
| chr11 | 98940599 | 98941207 | 609 | 0.38 | 0.475 | 0.547 |
| chr15 | 79603457 | 79622217 | 18761 | -0.09 | 0.475 | 0.547 |
| chr22 | 38497928 | 38508400 | 10473 | 0.13 | 0.475 | 0.547 |
| chr20 | 269918 | 282614 | 12697 | -0.07 | 0.475 | 0.547 |
| chrX | 12809504 | 12810424 | 921 | -0.35 | 0.476 | 0.548 |
| chr1 | 16562168 | 16565062 | 2895 | 0.14 | 0.476 | 0.548 |
| chr5 | 55506411 | 55508610 | 2200 | 0.17 | 0.476 | 0.548 |
| chr2 | 150936232 | 150937284 | 1053 | -0.37 | 0.476 | 0.548 |
| chr4 | 162857941 | 162858554 | 614 | 0.4 | 0.476 | 0.548 |
| chr3 | 33952943 | 33958229 | 5287 | 0.22 | 0.476 | 0.548 |
| chr6 | 43939012 | 43939654 | 643 | 0.46 | 0.476 | 0.548 |
| chr7 | 97500913 | 97502446 | 1534 | 0.17 | 0.476 | 0.548 |
| chr22 | 39349640 | 39351122 | 1483 | 0.34 | 0.476 | 0.549 |
| chr13 | 113758974 | 113760528 | 1555 | -0.22 | 0.477 | 0.549 |
| chr7 | 107694583 | 107695731 | 1149 | 1.36 | 0.477 | 0.549 |
| chr1 | 52251374 | 52259523 | 8150 | -0.16 | 0.477 | 0.549 |
| chr18 | 12402489 | 12411011 | 8523 | -0.13 | 0.477 | 0.549 |
| chr16 | 89848965 | 89849734 | 770 | -0.59 | 0.477 | 0.549 |
| chr16 | 46990947 | 47011397 | 20451 | 0.06 | 0.477 | 0.549 |
| chr1 | 47973583 | 47974180 | 598 | 0.46 | 0.477 | 0.549 |
| chr1 | 20929803 | 20945667 | 15865 | 0.07 | 0.478 | 0.55 |
| chr13 | 42031017 | 42032191 | 1175 | 0.33 | 0.478 | 0.55 |
| chr22 | 47326917 | 47331584 | 4668 | -0.24 | 0.478 | 0.55 |
| chr2 | 64652951 | 64654065 | 1115 | 0.37 | 0.478 | 0.55 |
| chr3 | 142217251 | 142229666 | 12416 | 0.13 | 0.478 | 0.55 |
| chr9 | 40632195 | 40632669 | 475 | 0.33 | 0.478 | 0.55 |
| chr1 | 26496371 | 26497320 | 950 | 0.2 | 0.479 | 0.551 |
| chr4 | 79565186 | 79568469 | 3284 | -0.08 | 0.479 | 0.551 |

| | | | | | | |
|---|---|---|---|---|---|---|
| chr14 | 75720905 | 75777920 | 57016 | 0.05 | 0.479 | 0.551 |
| chrX | 7001600 | 7009447 | 7848 | -0.18 | 0.479 | 0.551 |
| chr13 | 44928233 | 44943595 | 15363 | -0.16 | 0.479 | 0.551 |
| chr7 | 148680122 | 148685960 | 5839 | 0.1 | 0.479 | 0.551 |
| chr9 | 115818394 | 115819850 | 1457 | 0.17 | 0.479 | 0.551 |
| chr6 | 86383040 | 86390520 | 7481 | -0.11 | 0.479 | 0.551 |
| chr11 | 32111754 | 32125354 | 13601 | -0.09 | 0.479 | 0.551 |
| chr12 | 132627856 | 132630266 | 2411 | 0.17 | 0.479 | 0.551 |
| chr18 | 61067665 | 61068620 | 956 | 0.49 | 0.48 | 0.552 |
| chr3 | 126190696 | 126206205 | 15510 | 0.16 | 0.48 | 0.552 |
| chr17 | 75083831 | 75100714 | 16884 | 0.06 | 0.48 | 0.552 |
| chr2 | 112831548 | 112847454 | 15907 | -0.13 | 0.48 | 0.552 |
| chr1 | 100434813 | 100436925 | 2113 | 0.13 | 0.48 | 0.552 |
| chr11 | 16856842 | 16860038 | 3197 | -0.22 | 0.48 | 0.552 |
| chr9 | 87465591 | 87468758 | 3168 | -0.09 | 0.48 | 0.552 |
| chr6 | 151644313 | 151648176 | 3864 | -0.19 | 0.48 | 0.552 |
| chrX | 46696336 | 46700989 | 4654 | 0.11 | 0.48 | 0.552 |
| chr2 | 168148998 | 168149409 | 412 | -0.41 | 0.48 | 0.552 |
| chr13 | 99802583 | 99804574 | 1992 | 0.23 | 0.481 | 0.552 |
| chr5 | 155837661 | 155841938 | 4278 | 0.19 | 0.481 | 0.553 |
| chr4 | 184797141 | 184798044 | 904 | 0.41 | 0.481 | 0.553 |
| chr2 | 220832900 | 220834419 | 1520 | -0.22 | 0.481 | 0.553 |
| chr10 | 34010459 | 34011316 | 858 | -0.41 | 0.481 | 0.553 |
| chr14 | 37050332 | 37051898 | 1567 | 0.29 | 0.481 | 0.553 |
| chr17 | 80449966 | 80455804 | 5839 | -0.14 | 0.481 | 0.553 |
| chr10 | 70586551 | 70587421 | 871 | -0.27 | 0.481 | 0.553 |
| chr7 | 19967709 | 19969310 | 1602 | -0.18 | 0.481 | 0.553 |
| chr1 | 78244555 | 78246240 | 1686 | 0.18 | 0.481 | 0.553 |
| chr1 | 33937644 | 33938831 | 1188 | 0.25 | 0.481 | 0.553 |
| chr17 | 78328060 | 78328931 | 872 | 0.47 | 0.481 | 0.553 |
| chr1 | 65883112 | 65896318 | 13207 | -0.07 | 0.481 | 0.553 |
| chr6 | 26086856 | 26129119 | 42264 | -0.04 | 0.481 | 0.553 |
| chr15 | 64644356 | 64681068 | 36713 | 0.05 | 0.482 | 0.554 |
| chr8 | 101880801 | 101894869 | 14069 | 0.24 | 0.482 | 0.554 |
| chr20 | 44650089 | 44650645 | 557 | 0.41 | 0.482 | 0.554 |
| chr3 | 47039313 | 47040022 | 710 | 0.43 | 0.482 | 0.554 |
| chr18 | 57025307 | 57026507 | 1201 | 0.24 | 0.482 | 0.554 |
| chr7 | 43401985 | 43403261 | 1277 | -0.31 | 0.482 | 0.554 |
| chr17 | 78517776 | 78522635 | 4860 | -0.11 | 0.482 | 0.554 |
| chr3 | 100102722 | 100122693 | 19972 | 0.07 | 0.482 | 0.554 |
| chr12 | 19622133 | 19622863 | 731 | 0.36 | 0.482 | 0.554 |
| chr1 | 100714671 | 100715717 | 1047 | 0.2 | 0.483 | 0.554 |
| chr2 | 37938338 | 37939128 | 791 | 0.36 | 0.483 | 0.554 |
| chr20 | 47661890 | 47674846 | 12957 | 0.07 | 0.483 | 0.555 |
| chr10 | 134973558 | 134974375 | 818 | 0.38 | 0.483 | 0.555 |
| chr4 | 40023345 | 40025128 | 1784 | 0.19 | 0.483 | 0.555 |
| chr6 | 113441232 | 113444650 | 3419 | -0.15 | 0.483 | 0.555 |
| chrX | 129397894 | 129402895 | 5002 | -0.1 | 0.483 | 0.555 |
| chr18 | 32956577 | 32957700 | 1124 | 0.25 | 0.483 | 0.555 |
| chr1 | 90158018 | 90158624 | 607 | 0.48 | 0.484 | 0.555 |
| chr4 | 175408261 | 175445842 | 37582 | -0.05 | 0.484 | 0.555 |
| chr11 | 13689164 | 13698082 | 8919 | 0.12 | 0.484 | 0.556 |
| chr16 | 68554518 | 68574269 | 19752 | 0.18 | 0.484 | 0.556 |
| chr20 | 25656344 | 25658088 | 1745 | 0.22 | 0.484 | 0.556 |
| chr2 | 173419854 | 173421805 | 1952 | -0.17 | 0.484 | 0.556 |
| chrX | 128655281 | 128657488 | 2208 | 0.18 | 0.484 | 0.556 |
| chr2 | 172750016 | 172751595 | 1580 | -0.16 | 0.484 | 0.556 |
| chr2 | 61847643 | 61848191 | 549 | -0.42 | 0.484 | 0.556 |
| chr4 | 77172189 | 77173212 | 1024 | 0.39 | 0.485 | 0.556 |
| chr19 | 1652608 | 1653634 | 1027 | 0.21 | 0.485 | 0.556 |
| chr2 | 15499994 | 15501645 | 1652 | 0.29 | 0.485 | 0.556 |

| | | | | | | |
|---|---|---|---|---|---|---|
| chr20 | 4197347 | 4207101 | 9755 | 0.13 | 0.485 | 0.556 |
| chr14 | 23584414 | 23585028 | 615 | -0.54 | 0.485 | 0.557 |
| chr7 | 17691238 | 17692062 | 825 | -0.38 | 0.485 | 0.557 |
| chr14 | 100376574 | 100377579 | 1006 | -0.35 | 0.485 | 0.557 |
| chrX | 109268216 | 109281674 | 13459 | 0.1 | 0.485 | 0.557 |
| chr6 | 47256396 | 47257279 | 884 | 0.36 | 0.485 | 0.557 |
| chr9 | 94181094 | 94196054 | 14961 | 0.12 | 0.485 | 0.557 |
| chr3 | 32355281 | 32355878 | 598 | -0.54 | 0.486 | 0.557 |
| chr19 | 8942076 | 8943870 | 1795 | 0.15 | 0.486 | 0.558 |
| chr1 | 110026503 | 110027020 | 518 | -0.24 | 0.486 | 0.558 |
| chr6 | 14316727 | 14317543 | 817 | -0.35 | 0.486 | 0.558 |
| chr5 | 10392319 | 10393583 | 1265 | -0.31 | 0.487 | 0.559 |
| chr3 | 109251373 | 109252073 | 701 | 0.41 | 0.487 | 0.559 |
| chr4 | 114959588 | 114960904 | 1317 | 0.27 | 0.487 | 0.559 |
| chr6 | 8040913 | 8041622 | 710 | -0.43 | 0.487 | 0.559 |
| chr17 | 47633458 | 47647520 | 14063 | -0.45 | 0.487 | 0.559 |
| chr5 | 141060491 | 141062445 | 1955 | 0.15 | 0.487 | 0.559 |
| chr18 | 48404385 | 48410489 | 6105 | 0.13 | 0.488 | 0.559 |
| chr5 | 153824408 | 153827236 | 2829 | 0.15 | 0.488 | 0.559 |
| chr12 | 133170097 | 133177607 | 7511 | 1.27 | 0.488 | 0.559 |
| chrX | 152126601 | 152127401 | 801 | 0.21 | 0.488 | 0.559 |
| chr1 | 173238883 | 173239809 | 927 | 0.4 | 0.488 | 0.56 |
| chr9 | 124430453 | 124431224 | 772 | -0.3 | 0.488 | 0.56 |
| chr10 | 71454569 | 71455927 | 1359 | 0.24 | 0.488 | 0.56 |
| chr9 | 85914040 | 85915163 | 1124 | -0.39 | 0.488 | 0.56 |
| chrX | 96276829 | 96277366 | 538 | 0.45 | 0.488 | 0.56 |
| chr9 | 102859624 | 102877116 | 17493 | 0.08 | 0.488 | 0.56 |
| chr6 | 53515235 | 53517528 | 2294 | 0.16 | 0.489 | 0.56 |
| chr2 | 177501730 | 177503285 | 1556 | -0.16 | 0.489 | 0.56 |
| chr16 | 59560222 | 59562357 | 2136 | 0.11 | 0.489 | 0.56 |
| chr12 | 51395416 | 51400998 | 5583 | -0.22 | 0.489 | 0.56 |
| chr18 | 55647913 | 55648477 | 565 | -0.41 | 0.489 | 0.56 |
| chr10 | 52343396 | 52346973 | 3578 | -0.25 | 0.489 | 0.56 |
| chr16 | 22307303 | 22342253 | 34951 | -0.09 | 0.489 | 0.561 |
| chr9 | 103173499 | 103174829 | 1331 | 0.17 | 0.489 | 0.561 |
| chr3 | 195807512 | 195809711 | 2200 | -0.16 | 0.489 | 0.561 |
| chr11 | 107408987 | 107409677 | 691 | -0.43 | 0.489 | 0.561 |
| chr15 | 85438243 | 85438601 | 359 | 0.51 | 0.489 | 0.561 |
| chr1 | 12696342 | 12697145 | 804 | -0.37 | 0.49 | 0.561 |
| chr14 | 22024435 | 22026849 | 2415 | -0.13 | 0.49 | 0.561 |
| chr22 | 43031249 | 43058620 | 27372 | -0.11 | 0.49 | 0.561 |
| chr1 | 151118744 | 151119797 | 1054 | 0.35 | 0.49 | 0.561 |
| chr6 | 170122465 | 170125820 | 3356 | -0.16 | 0.49 | 0.562 |
| chr5 | 142691160 | 142694793 | 3634 | 0.19 | 0.49 | 0.562 |
| chr4 | 131214754 | 131215594 | 841 | -0.33 | 0.49 | 0.562 |
| chr8 | 141501464 | 141656647 | 155184 | 0.03 | 0.49 | 0.562 |
| chrX | 71931806 | 71935398 | 3593 | -0.15 | 0.49 | 0.562 |
| chr21 | 30256347 | 30258208 | 1862 | 0.14 | 0.491 | 0.562 |
| chr3 | 179321019 | 179323536 | 2518 | 0.11 | 0.491 | 0.562 |
| chr7 | 128018494 | 128019228 | 735 | 0.42 | 0.491 | 0.562 |
| chrX | 24020520 | 24021495 | 976 | -0.42 | 0.491 | 0.562 |
| chr18 | 51735923 | 51751779 | 15857 | -0.1 | 0.491 | 0.562 |
| chr15 | 52218529 | 52231937 | 13409 | -0.18 | 0.491 | 0.562 |
| chr7 | 107027552 | 107028144 | 593 | -0.45 | 0.491 | 0.562 |
| chr7 | 27737240 | 27738457 | 1218 | 0.25 | 0.491 | 0.563 |
| chr10 | 91312697 | 91313731 | 1035 | 0.32 | 0.491 | 0.563 |
| chr8 | 87568215 | 87569753 | 1539 | -0.28 | 0.492 | 0.563 |
| chr6 | 26457386 | 26458753 | 1368 | 0.12 | 0.492 | 0.563 |
| chr1 | 229431912 | 229432943 | 1032 | -0.37 | 0.492 | 0.563 |
| chr3 | 142496582 | 142499011 | 2430 | 0.22 | 0.492 | 0.563 |
| chr3 | 12386126 | 12387287 | 1162 | 0.39 | 0.492 | 0.563 |

| | | | | | | |
|---|---|---|---|---|---|---|
| chr1 | 15929002 | 15947219 | 18218 | 0.1 | 0.492 | 0.563 |
| chr3 | 17670961 | 17683240 | 12280 | -0.05 | 0.492 | 0.564 |
| chr3 | 49591048 | 49591759 | 712 | -0.34 | 0.492 | 0.564 |
| chr19 | 49371177 | 49382474 | 11298 | -0.05 | 0.492 | 0.564 |
| chr3 | 120153658 | 120170102 | 16445 | 0.1 | 0.492 | 0.564 |
| chr11 | 19798410 | 19799453 | 1044 | -0.32 | 0.492 | 0.564 |
| chr21 | 33983165 | 33986009 | 2845 | -0.1 | 0.492 | 0.564 |
| chr4 | 126310623 | 126311678 | 1056 | 0.38 | 0.493 | 0.564 |
| chr2 | 223228855 | 223229716 | 862 | 0.35 | 0.493 | 0.564 |
| chr7 | 73082266 | 73082729 | 464 | 0.45 | 0.493 | 0.564 |
| chr3 | 136504697 | 136506776 | 2080 | -0.09 | 0.493 | 0.564 |
| chr6 | 149865139 | 149868225 | 3087 | -0.09 | 0.493 | 0.564 |
| chr5 | 141801602 | 141802434 | 833 | 0.34 | 0.493 | 0.564 |
| chr11 | 74022175 | 74022652 | 478 | -0.49 | 0.493 | 0.564 |
| chr3 | 179361677 | 179372630 | 10954 | 0.13 | 0.493 | 0.564 |
| chr1 | 243955030 | 243963501 | 8472 | 0.25 | 0.493 | 0.564 |
| chr22 | 41958934 | 41962324 | 3391 | -0.23 | 0.494 | 0.565 |
| chr1 | 98870926 | 98872199 | 1274 | 0.25 | 0.494 | 0.565 |
| chr14 | 73957093 | 73958893 | 1801 | 0.16 | 0.494 | 0.565 |
| chr7 | 156772157 | 156773172 | 1016 | -0.34 | 0.494 | 0.565 |
| chr2 | 42866062 | 42866997 | 936 | -1.03 | 0.494 | 0.565 |
| chr4 | 147112914 | 147113905 | 992 | 0.38 | 0.494 | 0.566 |
| chr2 | 161592781 | 161611281 | 18501 | 0.11 | 0.495 | 0.566 |
| chr10 | 104209066 | 104221468 | 12403 | -0.2 | 0.495 | 0.566 |
| chr7 | 75471395 | 75473053 | 1659 | 0.23 | 0.495 | 0.566 |
| chr18 | 8328069 | 8345075 | 17007 | 0.07 | 0.495 | 0.566 |
| chr11 | 4114923 | 4117924 | 3002 | 0.09 | 0.495 | 0.566 |
| chr5 | 171314828 | 171315404 | 577 | -0.41 | 0.495 | 0.566 |
| chr3 | 186522610 | 186545348 | 22739 | 0.12 | 0.495 | 0.566 |
| chr2 | 239246662 | 239249043 | 2382 | -0.28 | 0.495 | 0.567 |
| chr17 | 38374216 | 38377094 | 2879 | -0.09 | 0.496 | 0.567 |
| chr11 | 65546785 | 65548710 | 1926 | 0.15 | 0.496 | 0.567 |
| chr11 | 70115844 | 70117785 | 1942 | 0.2 | 0.496 | 0.567 |
| chr6 | 17986767 | 17989269 | 2503 | 0.14 | 0.496 | 0.567 |
| chr9 | 12955796 | 12957104 | 1309 | 0.29 | 0.496 | 0.567 |
| chr14 | 34958261 | 34968418 | 10158 | 0.05 | 0.496 | 0.567 |
| chr8 | 95524758 | 95525539 | 782 | -0.38 | 0.496 | 0.567 |
| chr3 | 119711672 | 119712929 | 1258 | 0.33 | 0.496 | 0.567 |
| chr3 | 14361070 | 14361721 | 652 | -0.35 | 0.496 | 0.567 |
| chr2 | 201513517 | 201513868 | 352 | -0.43 | 0.497 | 0.568 |
| chr16 | 10831182 | 10838780 | 7599 | 0.1 | 0.497 | 0.568 |
| chr6 | 117996041 | 117998042 | 2002 | -0.24 | 0.497 | 0.568 |
| chr5 | 142024297 | 142025107 | 811 | 0.44 | 0.497 | 0.568 |
| chr5 | 141376174 | 141405259 | 29086 | -0.07 | 0.497 | 0.568 |
| chr11 | 27739159 | 27741202 | 2044 | -0.16 | 0.497 | 0.568 |
| chr13 | 34650783 | 34652020 | 1238 | 0.28 | 0.497 | 0.568 |
| chr3 | 14158173 | 14192397 | 34225 | -0.04 | 0.497 | 0.568 |
| chr3 | 195621281 | 195637137 | 15857 | -0.13 | 0.497 | 0.568 |
| chr14 | 73693121 | 73705467 | 12347 | -0.12 | 0.497 | 0.568 |
| chr3 | 134190140 | 134206517 | 16378 | 0.07 | 0.497 | 0.568 |
| chr3 | 13008217 | 13009904 | 1688 | 0.27 | 0.498 | 0.569 |
| chr5 | 149379059 | 149388594 | 9536 | -0.07 | 0.498 | 0.569 |
| chr7 | 23430836 | 23431787 | 952 | 0.34 | 0.498 | 0.569 |
| chr7 | 44834689 | 44859302 | 24614 | 0.06 | 0.498 | 0.569 |
| chr10 | 32097216 | 32097739 | 524 | 0.34 | 0.498 | 0.569 |
| chr5 | 64801957 | 64803337 | 1381 | -0.33 | 0.498 | 0.569 |
| chr6 | 38669913 | 38670952 | 1040 | 0.19 | 0.498 | 0.569 |
| chrX | 22210616 | 22210947 | 332 | 0.54 | 0.498 | 0.569 |
| chr9 | 74261302 | 74263036 | 1735 | -0.2 | 0.498 | 0.569 |
| chr20 | 55965717 | 55966303 | 587 | -0.32 | 0.499 | 0.57 |
| chr14 | 23122478 | 23122979 | 502 | -0.34 | 0.499 | 0.57 |

| | | | | | | |
|---|---|---|---|---|---|---|
| chr7 | 27644936 | 27645471 | 536 | -0.32 | 0.499 | 0.57 |
| chr4 | 39183065 | 39184764 | 1700 | 0.22 | 0.499 | 0.57 |
| chr13 | 31264185 | 31265656 | 1472 | -0.16 | 0.499 | 0.57 |
| chr4 | 140661179 | 140662496 | 1318 | 0.25 | 0.499 | 0.57 |
| chr8 | 24905589 | 24906668 | 1080 | 0.32 | 0.499 | 0.57 |
| chr2 | 55800588 | 55847972 | 47385 | -0.05 | 0.499 | 0.57 |
| chr12 | 17475554 | 17479936 | 4383 | -0.25 | 0.5 | 0.57 |
| chr10 | 64914767 | 64915802 | 1036 | 0.4 | 0.5 | 0.571 |
| chr7 | 2842004 | 2843743 | 1740 | -0.16 | 0.5 | 0.571 |
| chr5 | 65056511 | 65056860 | 350 | 0.48 | 0.5 | 0.571 |
| chr10 | 99277826 | 99278459 | 634 | -0.5 | 0.501 | 0.572 |
| chr15 | 65870800 | 65872778 | 1979 | -0.25 | 0.501 | 0.572 |
| chr14 | 52745981 | 52746848 | 868 | -0.29 | 0.501 | 0.572 |
| chr6 | 105850008 | 105851904 | 1897 | 0.17 | 0.501 | 0.572 |
| chr11 | 2172883 | 2173333 | 451 | 0.53 | 0.501 | 0.572 |
| chr6 | 147305504 | 147322449 | 16946 | 0.19 | 0.501 | 0.572 |
| chr5 | 137622877 | 137624198 | 1322 | -0.24 | 0.501 | 0.572 |
| chr5 | 171643264 | 171644109 | 846 | -0.38 | 0.501 | 0.572 |
| chr19 | 16770266 | 16772074 | 1809 | 0.22 | 0.501 | 0.572 |
| chr11 | 111101333 | 111101970 | 638 | 0.21 | 0.501 | 0.572 |
| chr19 | 6425242 | 6426224 | 983 | 0.28 | 0.502 | 0.573 |
| chr14 | 56035169 | 56091181 | 56013 | 0.04 | 0.502 | 0.573 |
| chr7 | 5408831 | 5409415 | 585 | -0.41 | 0.502 | 0.573 |
| chr7 | 65371372 | 65386493 | 15122 | -0.1 | 0.502 | 0.573 |
| chr7 | 23220705 | 23236592 | 15888 | 0.08 | 0.502 | 0.573 |
| chr11 | 115092446 | 115098304 | 5859 | 0.14 | 0.502 | 0.573 |
| chr13 | 100691732 | 100693265 | 1534 | 0.18 | 0.502 | 0.573 |
| chr14 | 55116751 | 55121853 | 5103 | 0.06 | 0.502 | 0.573 |
| chr16 | 67578370 | 67637103 | 58734 | 0.06 | 0.503 | 0.574 |
| chr8 | 54624713 | 54625326 | 614 | -0.29 | 0.503 | 0.574 |
| chr4 | 3533590 | 3534771 | 1182 | -0.35 | 0.503 | 0.574 |
| chr10 | 94651197 | 94652881 | 1685 | -0.16 | 0.503 | 0.574 |
| chr18 | 57385598 | 57386079 | 482 | -0.42 | 0.503 | 0.574 |
| chr1 | 168720991 | 168722380 | 1390 | 0.22 | 0.503 | 0.574 |
| chr4 | 79749459 | 79750808 | 1350 | 0.28 | 0.503 | 0.574 |
| chr11 | 70226823 | 70262155 | 35333 | 0.05 | 0.504 | 0.575 |
| chr10 | 18831506 | 18832000 | 495 | 0.74 | 0.504 | 0.575 |
| chr11 | 16809814 | 16822378 | 12565 | 0.08 | 0.504 | 0.575 |
| chr4 | 40397473 | 40398745 | 1273 | 0.37 | 0.504 | 0.575 |
| chr15 | 62939108 | 62940479 | 1372 | -0.24 | 0.504 | 0.575 |
| chr22 | 19835855 | 19844244 | 8390 | 0.09 | 0.504 | 0.575 |
| chr1 | 8577136 | 8586677 | 9542 | -0.15 | 0.504 | 0.575 |
| chr15 | 43476801 | 43478902 | 2102 | 0.17 | 0.504 | 0.575 |
| chr5 | 64063204 | 64065577 | 2374 | 0.08 | 0.504 | 0.575 |
| chr16 | 78156478 | 78156811 | 334 | -0.54 | 0.505 | 0.575 |
| chr6 | 159080529 | 159081381 | 853 | 0.3 | 0.505 | 0.575 |
| chr3 | 48054859 | 48062045 | 7187 | 0.1 | 0.505 | 0.575 |
| chr16 | 12927829 | 12928626 | 798 | -0.34 | 0.505 | 0.575 |
| chr13 | 74465474 | 74466593 | 1120 | 0.28 | 0.505 | 0.575 |
| chr13 | 41237970 | 41247698 | 9729 | 0.13 | 0.505 | 0.575 |
| chr10 | 65169384 | 65195463 | 26080 | -0.06 | 0.505 | 0.575 |
| chr19 | 530426 | 534013 | 3588 | 0.16 | 0.505 | 0.575 |
| chr19 | 11638938 | 11646935 | 7998 | -0.17 | 0.505 | 0.576 |
| chr22 | 17555064 | 17566628 | 11565 | -0.14 | 0.505 | 0.576 |
| chr2 | 132284944 | 132285369 | 426 | -0.42 | 0.505 | 0.576 |
| chr6 | 106582432 | 106583876 | 1445 | -0.21 | 0.505 | 0.576 |
| chr4 | 58356864 | 58357194 | 331 | 0.46 | 0.505 | 0.576 |
| chr17 | 56457855 | 56468940 | 11086 | 0.19 | 0.506 | 0.576 |
| chrX | 39955389 | 39969915 | 14527 | 0.07 | 0.506 | 0.576 |
| chr20 | 3586791 | 3587287 | 497 | -0.39 | 0.506 | 0.577 |
| chr5 | 179222607 | 179269140 | 46534 | 0.05 | 0.506 | 0.577 |

| | | | | | | |
|---|---|---|---|---|---|---|
| chr4 | 6827447 | 6828792 | 1346 | 0.33 | 0.506 | 0.577 |
| chr22 | 33040358 | 33041134 | 777 | 0.48 | 0.506 | 0.577 |
| chr5 | 60823117 | 60836792 | 13676 | -0.48 | 0.507 | 0.578 |
| chr9 | 88917998 | 88918891 | 894 | -0.3 | 0.507 | 0.578 |
| chr22 | 19542252 | 19543551 | 1300 | 0.2 | 0.507 | 0.578 |
| chr1 | 52989129 | 52989701 | 573 | 0.42 | 0.507 | 0.578 |
| chr19 | 42539113 | 42539945 | 833 | -0.19 | 0.507 | 0.578 |
| chr10 | 32842670 | 32843306 | 637 | -0.39 | 0.508 | 0.579 |
| chr5 | 60681632 | 60685044 | 3413 | 0.22 | 0.508 | 0.579 |
| chr4 | 76106049 | 76106681 | 633 | 0.33 | 0.508 | 0.579 |
| chr5 | 74657663 | 74658583 | 921 | 0.59 | 0.508 | 0.579 |
| chr6 | 170678312 | 170679731 | 1420 | -0.25 | 0.508 | 0.579 |
| chr10 | 116580572 | 116594078 | 13507 | 0.08 | 0.508 | 0.579 |
| chrX | 20565479 | 20566483 | 1005 | 0.3 | 0.508 | 0.579 |
| chr11 | 126831725 | 126838365 | 6641 | 0.08 | 0.509 | 0.579 |
| chr20 | 9216099 | 9222899 | 6801 | -0.21 | 0.509 | 0.579 |
| chr12 | 27838383 | 27839684 | 1302 | 0.28 | 0.509 | 0.579 |
| chr7 | 44113820 | 44123060 | 9241 | 0.12 | 0.509 | 0.579 |
| chr18 | 43407418 | 43408193 | 776 | -0.35 | 0.509 | 0.58 |
| chr5 | 57412736 | 57431761 | 19026 | -0.06 | 0.509 | 0.58 |
| chr8 | 70854873 | 70855222 | 350 | -0.55 | 0.51 | 0.58 |
| chr5 | 56134503 | 56135680 | 1178 | -0.18 | 0.51 | 0.58 |
| chr1 | 244846424 | 244847333 | 910 | 0.37 | 0.51 | 0.581 |
| chr1 | 114888620 | 114889178 | 559 | 0.41 | 0.51 | 0.581 |
| chr4 | 1911985 | 1913175 | 1191 | -0.31 | 0.51 | 0.581 |
| chr16 | 3687280 | 3712194 | 24915 | -0.09 | 0.51 | 0.581 |
| chrX | 20148835 | 20161001 | 12167 | 0.1 | 0.51 | 0.581 |
| chr7 | 23283151 | 23283666 | 516 | -0.47 | 0.51 | 0.581 |
| chr22 | 22089621 | 22090076 | 456 | 0.4 | 0.51 | 0.581 |
| chr12 | 54613207 | 54635895 | 22689 | 0.11 | 0.51 | 0.581 |
| chr11 | 117048389 | 117050334 | 1946 | 0.23 | 0.51 | 0.581 |
| chr8 | 56710819 | 56712327 | 1509 | -0.38 | 0.511 | 0.581 |
| chr13 | 30814103 | 30815062 | 960 | 0.34 | 0.511 | 0.581 |
| chr16 | 86774390 | 86775720 | 1331 | 0.2 | 0.511 | 0.581 |
| chr14 | 61113819 | 61136252 | 22434 | -0.21 | 0.511 | 0.581 |
| chr7 | 94021082 | 94022394 | 1313 | 0.29 | 0.511 | 0.581 |
| chr1 | 23810218 | 23810669 | 452 | -0.55 | 0.511 | 0.581 |
| chr9 | 112193045 | 112270212 | 77168 | -0.04 | 0.511 | 0.581 |
| chr3 | 72896581 | 72897702 | 1122 | -0.27 | 0.511 | 0.582 |
| chr5 | 170288888 | 170290178 | 1291 | 0.21 | 0.511 | 0.582 |
| chr3 | 49481091 | 49482077 | 987 | -0.31 | 0.511 | 0.582 |
| chr6 | 41734665 | 41756087 | 21423 | -0.09 | 0.511 | 0.582 |
| chr20 | 25731053 | 25732292 | 1240 | 0.22 | 0.511 | 0.582 |
| chr6 | 112651422 | 112652716 | 1295 | 0.41 | 0.511 | 0.582 |
| chr5 | 91887214 | 91899237 | 12024 | -0.05 | 0.511 | 0.582 |
| chr2 | 55449708 | 55467645 | 17938 | -0.14 | 0.512 | 0.582 |
| chr6 | 160845043 | 160846190 | 1148 | 0.32 | 0.512 | 0.582 |
| chr19 | 21264927 | 21265851 | 925 | -0.24 | 0.512 | 0.582 |
| chr6 | 4732819 | 4735471 | 2653 | -0.11 | 0.512 | 0.582 |
| chr22 | 46253420 | 46260763 | 7344 | -0.1 | 0.512 | 0.583 |
| chr7 | 80127769 | 80129573 | 1805 | 0.18 | 0.512 | 0.583 |
| chr1 | 33474593 | 33475394 | 802 | 0.33 | 0.513 | 0.583 |
| chr4 | 8200181 | 8202621 | 2441 | 0.14 | 0.513 | 0.583 |
| chr4 | 76707854 | 76708636 | 783 | -0.36 | 0.513 | 0.583 |
| chr6 | 4913267 | 4914440 | 1174 | 0.33 | 0.513 | 0.583 |
| chrX | 70337906 | 70339348 | 1443 | 0.18 | 0.513 | 0.583 |
| chr2 | 179342517 | 179346352 | 3836 | -0.09 | 0.513 | 0.583 |
| chr4 | 85594858 | 85603833 | 8976 | -0.18 | 0.513 | 0.583 |
| chr4 | 146653657 | 146656479 | 2823 | -0.18 | 0.513 | 0.583 |
| chr7 | 54891302 | 54900686 | 9385 | 0.09 | 0.514 | 0.584 |
| chr5 | 137730773 | 137744187 | 13415 | -0.18 | 0.514 | 0.584 |

| | | | | | | |
|---|---|---|---|---|---|---|
| chr12 | 9825860 | 9827150 | 1291 | -0.22 | 0.514 | 0.584 |
| chr15 | 37350410 | 37352300 | 1891 | 0.25 | 0.514 | 0.584 |
| chr11 | 69356010 | 69356769 | 760 | -0.36 | 0.514 | 0.584 |
| chr12 | 57733014 | 57736298 | 3285 | 0.16 | 0.514 | 0.585 |
| chr15 | 36165983 | 36167658 | 1676 | 0.21 | 0.514 | 0.585 |
| chr19 | 5268319 | 5293884 | 25566 | -0.13 | 0.515 | 0.585 |
| chr17 | 43096077 | 43097171 | 1095 | -0.29 | 0.515 | 0.585 |
| chr14 | 35513845 | 35533297 | 19453 | -0.04 | 0.515 | 0.585 |
| chrX | 133370847 | 133371564 | 718 | 0.36 | 0.515 | 0.585 |
| chr5 | 44809031 | 44813390 | 4360 | -0.1 | 0.515 | 0.585 |
| chr11 | 125438771 | 125440593 | 1823 | 0.14 | 0.515 | 0.585 |
| chr19 | 5717777 | 5721641 | 3865 | -0.1 | 0.515 | 0.585 |
| chr6 | 43020568 | 43028330 | 7763 | 0.1 | 0.515 | 0.585 |
| chr2 | 201217291 | 201289794 | 72504 | 0.04 | 0.515 | 0.585 |
| chr17 | 58806170 | 58808340 | 2171 | -0.14 | 0.515 | 0.585 |
| chr16 | 67180698 | 67242722 | 62025 | -0.04 | 0.515 | 0.585 |
| chr7 | 24692832 | 24693731 | 900 | 0.29 | 0.515 | 0.585 |
| chr2 | 203901578 | 203902655 | 1078 | 0.42 | 0.515 | 0.585 |
| chr5 | 94889865 | 94892105 | 2241 | 0.19 | 0.516 | 0.586 |
| chr13 | 33791127 | 33804754 | 13628 | 0.13 | 0.516 | 0.586 |
| chr6 | 42325361 | 42326004 | 644 | -0.42 | 0.516 | 0.586 |
| chr9 | 104248113 | 104250266 | 2154 | -0.11 | 0.516 | 0.586 |
| chr12 | 54191945 | 54198955 | 7011 | -1.31 | 0.516 | 0.586 |
| chr2 | 150715023 | 150716179 | 1157 | 0.2 | 0.516 | 0.587 |
| chr16 | 68055877 | 68058089 | 2213 | 0.11 | 0.517 | 0.587 |
| chr16 | 80207923 | 80208980 | 1058 | 0.37 | 0.517 | 0.587 |
| chr14 | 66973420 | 66975834 | 2415 | 0.11 | 0.517 | 0.587 |
| chr18 | 71419562 | 71420475 | 914 | 0.34 | 0.517 | 0.587 |
| chr1 | 192757078 | 192757967 | 890 | 0.25 | 0.517 | 0.588 |
| chrX | 106446948 | 106450615 | 3668 | 0.11 | 0.517 | 0.588 |
| chr7 | 73685646 | 73730158 | 44513 | 0.05 | 0.518 | 0.588 |
| chr6 | 18293581 | 18294113 | 533 | 0.42 | 0.518 | 0.588 |
| chr17 | 11965653 | 11966710 | 1058 | 0.37 | 0.518 | 0.588 |
| chr21 | 46728519 | 46767262 | 38744 | 0.04 | 0.518 | 0.588 |
| chr5 | 41391902 | 41419835 | 27934 | 0.06 | 0.518 | 0.588 |
| chr11 | 110158451 | 110168759 | 10309 | -0.09 | 0.518 | 0.588 |
| chr16 | 89808226 | 89809985 | 1760 | 0.26 | 0.518 | 0.588 |
| chr5 | 77001964 | 77011699 | 9736 | 0.15 | 0.518 | 0.588 |
| chr6 | 157502402 | 157503271 | 870 | 0.38 | 0.518 | 0.588 |
| chr6 | 57085979 | 57087739 | 1761 | 0.22 | 0.518 | 0.588 |
| chr16 | 58442603 | 58443929 | 1327 | 0.28 | 0.518 | 0.588 |
| chr6 | 28234288 | 28249859 | 15572 | 0.12 | 0.518 | 0.588 |
| chr1 | 28718075 | 28727709 | 9635 | -0.09 | 0.518 | 0.588 |
| chr2 | 173117441 | 173118741 | 1301 | -0.27 | 0.518 | 0.588 |
| chr14 | 58795927 | 58798547 | 2621 | -0.14 | 0.518 | 0.588 |
| chr5 | 155888533 | 155909300 | 20768 | -0.1 | 0.519 | 0.589 |
| chr8 | 122927222 | 122927820 | 599 | 1.16 | 0.519 | 0.589 |
| chr9 | 650150 | 650850 | 701 | -0.38 | 0.519 | 0.589 |
| chr5 | 141030658 | 141031734 | 1077 | 0.25 | 0.519 | 0.589 |
| chr6 | 11143367 | 11145951 | 2585 | -0.17 | 0.519 | 0.589 |
| chr3 | 107705482 | 107707550 | 2069 | 0.24 | 0.519 | 0.589 |
| chr10 | 7447175 | 7453527 | 6353 | 0.19 | 0.519 | 0.589 |
| chr11 | 13191222 | 13205746 | 14525 | -1.29 | 0.52 | 0.59 |
| chr2 | 25734961 | 25745596 | 10636 | -0.11 | 0.52 | 0.59 |
| chr17 | 10747878 | 10748595 | 718 | 0.33 | 0.52 | 0.59 |
| chr6 | 71890780 | 71892010 | 1231 | 0.26 | 0.52 | 0.59 |
| chr14 | 20686309 | 20688159 | 1851 | 0.2 | 0.521 | 0.591 |
| chr9 | 6645344 | 6716398 | 71055 | 0.04 | 0.521 | 0.591 |
| chr21 | 30848516 | 30848906 | 391 | -0.28 | 0.521 | 0.591 |
| chr14 | 90722326 | 90724540 | 2215 | -0.08 | 0.521 | 0.591 |
| chr1 | 231647802 | 231648964 | 1163 | -0.23 | 0.521 | 0.591 |

| | | | | | | |
|---|---|---|---|---|---|---|
| chr9 | 32500228 | 32503553 | 3326 | 0.14 | 0.521 | 0.591 |
| chr14 | 59911586 | 59921979 | 10394 | 0.16 | 0.521 | 0.591 |
| chr14 | 102966657 | 102997723 | 31067 | -0.06 | 0.521 | 0.591 |
| chr22 | 50945407 | 50947937 | 2531 | 0.13 | 0.522 | 0.592 |
| chr2 | 159758001 | 159758460 | 460 | 0.35 | 0.522 | 0.592 |
| chr17 | 73378592 | 73401759 | 23168 | -0.06 | 0.522 | 0.592 |
| chr1 | 894088 | 911983 | 17896 | -0.17 | 0.522 | 0.592 |
| chr18 | 1292514 | 1292986 | 473 | 0.49 | 0.522 | 0.592 |
| chr16 | 89723642 | 89725826 | 2185 | -0.13 | 0.522 | 0.592 |
| chr1 | 64471117 | 64491030 | 19914 | 0.1 | 0.522 | 0.592 |
| chr17 | 60187930 | 60189826 | 1897 | 0.17 | 0.522 | 0.592 |
| chr11 | 59411992 | 59412799 | 808 | -0.34 | 0.522 | 0.592 |
| chr5 | 145846038 | 145847368 | 1331 | -0.3 | 0.522 | 0.592 |
| chr15 | 41018322 | 41029947 | 11626 | 0.17 | 0.522 | 0.592 |
| chr21 | 47877350 | 47879359 | 2010 | -0.16 | 0.522 | 0.592 |
| chr6 | 4459196 | 4460049 | 854 | 0.55 | 0.522 | 0.592 |
| chr6 | 116329242 | 116359292 | 30051 | -0.05 | 0.522 | 0.592 |
| chr7 | 156431994 | 156434425 | 2432 | 0.09 | 0.523 | 0.592 |
| chr15 | 70442289 | 70442802 | 514 | -0.33 | 0.523 | 0.592 |
| chr2 | 189340258 | 189350965 | 10708 | 0.17 | 0.523 | 0.593 |
| chr17 | 1569641 | 1589611 | 19971 | 0.08 | 0.523 | 0.593 |
| chr3 | 47203302 | 47207255 | 3954 | 0.1 | 0.523 | 0.593 |
| chr2 | 196621175 | 196621747 | 573 | -0.34 | 0.523 | 0.593 |
| chr15 | 49267808 | 49268549 | 742 | -0.35 | 0.523 | 0.593 |
| chr17 | 1519606 | 1553884 | 34279 | 0.1 | 0.524 | 0.593 |
| chr11 | 9284725 | 9287669 | 2945 | -0.1 | 0.524 | 0.593 |
| chr10 | 16500825 | 16508694 | 7870 | 0.16 | 0.524 | 0.593 |
| chrX | 30670265 | 30672166 | 1902 | 0.17 | 0.524 | 0.593 |
| chr6 | 170614890 | 170616976 | 2087 | 0.12 | 0.524 | 0.594 |
| chr15 | 49541512 | 49541923 | 412 | 0.42 | 0.524 | 0.594 |
| chr4 | 3293149 | 3308263 | 15115 | 0.14 | 0.524 | 0.594 |
| chrX | 17393704 | 17396157 | 2454 | -0.21 | 0.524 | 0.594 |
| chr21 | 41879479 | 41880570 | 1092 | 0.26 | 0.525 | 0.594 |
| chr2 | 70070661 | 70071457 | 797 | 0.33 | 0.525 | 0.594 |
| chr5 | 70842186 | 70843062 | 877 | 0.36 | 0.525 | 0.594 |
| chr16 | 19068880 | 19080988 | 12109 | -0.06 | 0.525 | 0.594 |
| chrX | 67707126 | 67730020 | 22895 | -0.06 | 0.525 | 0.594 |
| chr4 | 185646708 | 185656102 | 9395 | 0.17 | 0.525 | 0.594 |
| chr1 | 43123384 | 43125031 | 1648 | 0.16 | 0.525 | 0.594 |
| chr14 | 35249755 | 35250718 | 964 | -0.3 | 0.525 | 0.595 |
| chr1 | 203575344 | 203576705 | 1362 | -0.24 | 0.525 | 0.595 |
| chr3 | 178968421 | 178979669 | 11249 | 0.18 | 0.525 | 0.595 |
| chr17 | 40853061 | 40853447 | 387 | -0.71 | 0.525 | 0.595 |
| chr16 | 25166988 | 25167560 | 573 | -0.4 | 0.526 | 0.595 |
| chr1 | 44705067 | 44707105 | 2039 | -0.25 | 0.526 | 0.595 |
| chr11 | 68713336 | 68714722 | 1387 | -0.33 | 0.526 | 0.595 |
| chr10 | 27026679 | 27045465 | 18787 | 0.11 | 0.526 | 0.595 |
| chr10 | 114205544 | 114208251 | 2708 | -0.07 | 0.526 | 0.595 |
| chr8 | 131770313 | 131783561 | 13249 | 0.04 | 0.526 | 0.595 |
| chr2 | 98261629 | 98263637 | 2009 | -0.11 | 0.526 | 0.596 |
| chr22 | 20849223 | 20891627 | 42405 | 0.04 | 0.526 | 0.596 |
| chr1 | 239976605 | 239981469 | 4865 | -0.06 | 0.527 | 0.596 |
| chr16 | 48326227 | 48326823 | 597 | -0.34 | 0.527 | 0.596 |
| chr7 | 64363649 | 64364943 | 1295 | 0.18 | 0.527 | 0.596 |
| chr20 | 52515603 | 52559214 | 43612 | 0.04 | 0.527 | 0.596 |
| chr5 | 136616173 | 136618615 | 2443 | -0.2 | 0.527 | 0.597 |
| chr2 | 88335828 | 88339040 | 3213 | -0.12 | 0.527 | 0.597 |
| chr5 | 135489328 | 135496811 | 7484 | 0.15 | 0.528 | 0.597 |
| chr10 | 13366406 | 13392199 | 25794 | -0.06 | 0.528 | 0.597 |
| chr6 | 5259774 | 5263295 | 3522 | -0.08 | 0.528 | 0.597 |
| chr5 | 413719 | 425304 | 11586 | 0.19 | 0.528 | 0.597 |

| | | | | | | |
|---|---|---|---|---|---|---|
| chr17 | 29203237 | 29203837 | 601 | 0.38 | 0.528 | 0.598 |
| chr16 | 2254762 | 2319401 | 64640 | 0.04 | 0.528 | 0.598 |
| chr4 | 13656342 | 13659429 | 3088 | -0.25 | 0.528 | 0.598 |
| chr4 | 48270755 | 48272957 | 2203 | 0.13 | 0.529 | 0.598 |
| chr13 | 44361115 | 44361870 | 756 | -0.36 | 0.529 | 0.598 |
| chr17 | 28877558 | 28878766 | 1209 | -0.31 | 0.529 | 0.598 |
| chr12 | 53661400 | 53662772 | 1373 | 0.2 | 0.529 | 0.598 |
| chr3 | 115715209 | 115720018 | 4810 | 0.21 | 0.529 | 0.598 |
| chr17 | 7451662 | 7531553 | 79892 | -0.04 | 0.529 | 0.598 |
| chr1 | 40562367 | 40563214 | 848 | 0.28 | 0.529 | 0.599 |
| chr8 | 61324989 | 61325807 | 819 | -0.35 | 0.529 | 0.599 |
| chr7 | 67889546 | 67903656 | 14111 | 0.84 | 0.529 | 0.599 |
| chr6 | 76203157 | 76210033 | 6877 | 0.14 | 0.53 | 0.599 |
| chr20 | 62311875 | 62340784 | 28910 | 0.05 | 0.53 | 0.599 |
| chr5 | 138897186 | 138898686 | 1501 | -0.14 | 0.53 | 0.599 |
| chr6 | 35941211 | 35942415 | 1205 | 0.24 | 0.53 | 0.599 |
| chr6 | 149637242 | 149659146 | 21905 | 0.07 | 0.53 | 0.599 |
| chr19 | 56028515 | 56076069 | 47555 | 0.04 | 0.53 | 0.599 |
| chr8 | 27348196 | 27349192 | 997 | -0.26 | 0.53 | 0.6 |
| chr3 | 158040104 | 158057840 | 17737 | 0.91 | 0.531 | 0.6 |
| chr22 | 41863560 | 41866192 | 2633 | 0.09 | 0.531 | 0.6 |
| chr5 | 151137312 | 151173046 | 35735 | 0.06 | 0.531 | 0.6 |
| chr3 | 188947048 | 188947899 | 852 | 0.33 | 0.531 | 0.6 |
| chr1 | 226051679 | 226071633 | 19955 | -0.11 | 0.531 | 0.6 |
| chr15 | 22941254 | 22962130 | 20877 | 0.1 | 0.531 | 0.6 |
| chr17 | 9142785 | 9152635 | 9851 | 0.19 | 0.531 | 0.6 |
| chr7 | 74141463 | 74141968 | 506 | -0.44 | 0.531 | 0.6 |
| chr5 | 147822644 | 147824255 | 1612 | -0.22 | 0.531 | 0.6 |
| chr16 | 58717787 | 58719039 | 1253 | -0.2 | 0.531 | 0.6 |
| chr15 | 74907163 | 74909686 | 2524 | 0.11 | 0.531 | 0.6 |
| chr13 | 45371125 | 45372320 | 1196 | 0.32 | 0.531 | 0.6 |
| chr9 | 37575642 | 37593214 | 17573 | -0.08 | 0.532 | 0.601 |
| chr12 | 60591116 | 60591715 | 600 | 0.33 | 0.532 | 0.601 |
| chr2 | 148600825 | 148603167 | 2343 | -0.12 | 0.532 | 0.602 |
| chr16 | 59911566 | 59920563 | 8998 | -0.09 | 0.533 | 0.602 |
| chr1 | 199820236 | 199829664 | 9429 | 0.1 | 0.533 | 0.602 |
| chr7 | 75400886 | 75402517 | 1632 | 0.21 | 0.533 | 0.602 |
| chr2 | 192701543 | 192744471 | 42929 | 0.04 | 0.533 | 0.603 |
| chr5 | 118774507 | 118774956 | 450 | 0.34 | 0.533 | 0.603 |
| chr12 | 53738146 | 53739637 | 1492 | 0.24 | 0.534 | 0.603 |
| chr7 | 45908847 | 45909500 | 654 | -0.47 | 0.534 | 0.603 |
| chr14 | 50971342 | 50972007 | 666 | -0.37 | 0.534 | 0.603 |
| chr15 | 65876717 | 65877012 | 296 | 0.46 | 0.534 | 0.603 |
| chr9 | 38380187 | 38394787 | 14601 | 0.09 | 0.534 | 0.604 |
| chr14 | 30218461 | 30225155 | 6695 | -0.24 | 0.534 | 0.604 |
| chr4 | 184611731 | 184612470 | 740 | 0.37 | 0.535 | 0.604 |
| chr16 | 71495340 | 71496872 | 1533 | 0.13 | 0.535 | 0.604 |
| chr20 | 7999376 | 8001255 | 1880 | -0.12 | 0.535 | 0.604 |
| chr2 | 32772694 | 32774496 | 1803 | 0.26 | 0.535 | 0.604 |
| chr3 | 31677027 | 31678659 | 1633 | -0.28 | 0.535 | 0.604 |
| chr7 | 132467313 | 132469721 | 2409 | -0.12 | 0.535 | 0.605 |
| chr16 | 30533247 | 30551419 | 18173 | -0.09 | 0.535 | 0.605 |
| chr10 | 30146965 | 30147517 | 553 | 0.42 | 0.535 | 0.605 |
| chr8 | 133804485 | 133804997 | 513 | -0.48 | 0.536 | 0.605 |
| chr1 | 101460848 | 101495176 | 34329 | -0.06 | 0.536 | 0.605 |
| chr2 | 150436663 | 150454749 | 18087 | 0.06 | 0.536 | 0.605 |
| chr7 | 81237956 | 81238672 | 717 | 0.35 | 0.536 | 0.605 |
| chr6 | 121987339 | 121993309 | 5971 | 0.18 | 0.536 | 0.605 |
| chr4 | 155489688 | 155490355 | 668 | -0.38 | 0.536 | 0.605 |
| chr22 | 42664657 | 42666643 | 1987 | 0.16 | 0.536 | 0.605 |
| chr6 | 25359374 | 25360083 | 710 | 0.41 | 0.536 | 0.605 |

| | | | | | | |
|---|---|---|---|---|---|---|
| chr3 | 128398907 | 128418105 | 19199 | 0.09 | 0.536 | 0.605 |
| chr10 | 119202064 | 119203007 | 944 | -0.33 | 0.536 | 0.605 |
| chr16 | 53217039 | 53218404 | 1366 | 0.28 | 0.536 | 0.605 |
| chr20 | 33758801 | 33761187 | 2387 | -0.14 | 0.536 | 0.606 |
| chr5 | 173998376 | 173999436 | 1061 | 0.25 | 0.537 | 0.606 |
| chr5 | 41924413 | 41926506 | 2094 | 0.08 | 0.537 | 0.606 |
| chr7 | 138759514 | 138761431 | 1918 | 0.16 | 0.537 | 0.606 |
| chr2 | 10930923 | 10932594 | 1672 | 0.25 | 0.537 | 0.606 |
| chr1 | 117046565 | 117052806 | 6242 | 0.15 | 0.537 | 0.606 |
| chr12 | 89991297 | 89992037 | 741 | 0.3 | 0.537 | 0.606 |
| chr20 | 1757277 | 1757819 | 543 | 0.3 | 0.537 | 0.606 |
| chr11 | 61657926 | 61661962 | 4037 | -0.2 | 0.537 | 0.606 |
| chr3 | 119012611 | 119013944 | 1334 | 0.21 | 0.537 | 0.606 |
| chr17 | 2237637 | 2263875 | 26239 | -0.08 | 0.537 | 0.606 |
| chr17 | 27684016 | 27787574 | 103559 | -0.03 | 0.537 | 0.606 |
| chr12 | 25428618 | 25430257 | 1640 | -0.14 | 0.537 | 0.606 |
| chr7 | 2973482 | 2973931 | 450 | -0.43 | 0.537 | 0.606 |
| chr10 | 71812193 | 71813882 | 1690 | -0.2 | 0.537 | 0.606 |
| chr10 | 101916148 | 101917205 | 1058 | -0.29 | 0.537 | 0.606 |
| chrX | 13272230 | 13273121 | 892 | -0.25 | 0.537 | 0.606 |
| chr19 | 58110810 | 58112141 | 1332 | 0.26 | 0.537 | 0.606 |
| chr17 | 58578719 | 58612629 | 33911 | 0.04 | 0.537 | 0.606 |
| chr6 | 166525723 | 166526909 | 1187 | 0.2 | 0.537 | 0.606 |
| chr3 | 112012439 | 112013047 | 609 | 0.34 | 0.537 | 0.606 |
| chr22 | 33934180 | 33937586 | 3407 | 0.1 | 0.538 | 0.606 |
| chrX | 57021426 | 57022926 | 1501 | -0.13 | 0.538 | 0.606 |
| chr1 | 16784537 | 16786381 | 1845 | 0.42 | 0.538 | 0.606 |
| chr8 | 143557159 | 143557906 | 748 | 0.27 | 0.538 | 0.606 |
| chr18 | 55548812 | 55549159 | 348 | 0.39 | 0.538 | 0.606 |
| chr10 | 3207119 | 3214964 | 7846 | 0.13 | 0.538 | 0.606 |
| chr3 | 162427046 | 162427397 | 352 | 0.43 | 0.538 | 0.607 |
| chr4 | 54325324 | 54326068 | 745 | 0.38 | 0.538 | 0.607 |
| chr1 | 1166042 | 1168272 | 2231 | 0.18 | 0.538 | 0.607 |
| chr2 | 54159314 | 54199234 | 39921 | -0.04 | 0.538 | 0.607 |
| chrX | 47123665 | 47124802 | 1138 | -0.29 | 0.538 | 0.607 |
| chr16 | 69950107 | 69962446 | 12340 | -0.1 | 0.539 | 0.607 |
| chr14 | 90167443 | 90168834 | 1392 | -0.22 | 0.539 | 0.607 |
| chr14 | 35117356 | 35147906 | 30551 | 0.06 | 0.539 | 0.608 |
| chr17 | 79196048 | 79197683 | 1636 | 0.25 | 0.539 | 0.608 |
| chr12 | 64574278 | 64576909 | 2632 | 0.1 | 0.54 | 0.608 |
| chr8 | 125502342 | 125505228 | 2887 | -0.16 | 0.54 | 0.608 |
| chr17 | 36761894 | 36763188 | 1295 | 0.27 | 0.54 | 0.608 |
| chr12 | 102248961 | 102249631 | 671 | -1.07 | 0.54 | 0.608 |
| chr4 | 175525096 | 175525993 | 898 | -0.3 | 0.54 | 0.608 |
| chrX | 46058308 | 46081104 | 22797 | -0.14 | 0.54 | 0.609 |
| chr17 | 55055523 | 55056252 | 730 | 0.29 | 0.54 | 0.609 |
| chr4 | 17495134 | 17496034 | 901 | -0.31 | 0.54 | 0.609 |
| chr1 | 224449321 | 224459357 | 10037 | -0.11 | 0.54 | 0.609 |
| chr6 | 143380510 | 143382724 | 2215 | 0.16 | 0.541 | 0.609 |
| chr15 | 40698036 | 40698514 | 479 | -0.35 | 0.541 | 0.609 |
| chr16 | 2197417 | 2220323 | 22907 | -0.05 | 0.541 | 0.609 |
| chr1 | 116210608 | 116215210 | 4603 | -0.11 | 0.541 | 0.61 |
| chr8 | 67686686 | 67688128 | 1443 | 0.21 | 0.541 | 0.61 |
| chr5 | 178976794 | 178987933 | 11140 | 0.07 | 0.541 | 0.61 |
| chrX | 154443946 | 154446240 | 2295 | 0.11 | 0.542 | 0.61 |
| chr11 | 71933258 | 71949270 | 16013 | 0.08 | 0.542 | 0.61 |
| chr16 | 66984813 | 66985496 | 684 | 0.4 | 0.542 | 0.61 |
| chr1 | 199945552 | 199946636 | 1085 | 0.24 | 0.542 | 0.61 |
| chr19 | 17325396 | 17379187 | 53792 | 0.05 | 0.542 | 0.611 |
| chr15 | 40558660 | 40559709 | 1050 | -0.26 | 0.542 | 0.611 |
| chr16 | 2509429 | 2525023 | 15595 | -0.14 | 0.542 | 0.611 |

| | | | | | | |
|---|---|---|---|---|---|---|
| chr7 | 129709381 | 129711432 | 2052 | -0.14 | 0.542 | 0.611 |
| chr4 | 41936373 | 41938211 | 1839 | 0.14 | 0.542 | 0.611 |
| chr19 | 11750617 | 11751256 | 640 | 0.4 | 0.542 | 0.611 |
| chr13 | 37393386 | 37393996 | 611 | 0.26 | 0.543 | 0.611 |
| chrX | 153361962 | 153363921 | 1960 | 0.16 | 0.543 | 0.611 |
| chr1 | 212540460 | 212541399 | 940 | -0.35 | 0.543 | 0.612 |
| chr2 | 146281738 | 146282714 | 977 | -0.25 | 0.543 | 0.612 |
| chr17 | 74233660 | 74273629 | 39970 | 0.07 | 0.543 | 0.612 |
| chr13 | 98086472 | 98087279 | 808 | 0.25 | 0.544 | 0.612 |
| chr4 | 135286667 | 135287895 | 1229 | -0.33 | 0.544 | 0.612 |
| chr11 | 2620691 | 2621515 | 825 | -0.34 | 0.544 | 0.612 |
| chr15 | 85173565 | 85177509 | 3945 | 0.12 | 0.544 | 0.612 |
| chr17 | 76821698 | 76838352 | 16655 | 0.06 | 0.544 | 0.612 |
| chr13 | 67229658 | 67240925 | 11268 | -0.04 | 0.544 | 0.613 |
| chr2 | 97170763 | 97171833 | 1071 | 0.27 | 0.544 | 0.613 |
| chr2 | 55495083 | 55510525 | 15443 | -0.07 | 0.544 | 0.613 |
| chr2 | 15384299 | 15384828 | 530 | -0.49 | 0.545 | 0.613 |
| chr21 | 43834707 | 43835844 | 1138 | -0.23 | 0.545 | 0.614 |
| chr8 | 40200864 | 40210099 | 9236 | 0.06 | 0.545 | 0.614 |
| chr5 | 145422137 | 145422813 | 677 | 0.32 | 0.545 | 0.614 |
| chr7 | 100290733 | 100305430 | 14698 | -0.11 | 0.545 | 0.614 |
| chr5 | 180257705 | 180259872 | 2168 | 0.09 | 0.545 | 0.614 |
| chr6 | 168228138 | 168228778 | 641 | 0.37 | 0.545 | 0.614 |
| chr3 | 69997138 | 69998382 | 1245 | 0.19 | 0.545 | 0.614 |
| chrX | 49091688 | 49099713 | 8026 | 0.12 | 0.546 | 0.614 |
| chr1 | 116681263 | 116681755 | 493 | 0.36 | 0.546 | 0.614 |
| chr9 | 99212445 | 99213725 | 1281 | -0.24 | 0.546 | 0.614 |
| chr20 | 36497772 | 36499546 | 1775 | 0.26 | 0.546 | 0.614 |
| chr9 | 125298762 | 125299355 | 594 | 0.47 | 0.546 | 0.614 |
| chr1 | 22290061 | 22290524 | 464 | -0.53 | 0.546 | 0.615 |
| chr2 | 121044506 | 121045734 | 1229 | -0.18 | 0.546 | 0.615 |
| chr1 | 17764265 | 17770631 | 6367 | -0.09 | 0.547 | 0.615 |
| chr18 | 13016621 | 13017551 | 931 | -0.28 | 0.547 | 0.615 |
| chr15 | 56124280 | 56124777 | 498 | -0.45 | 0.547 | 0.615 |
| chr1 | 8615061 | 8632250 | 17190 | -0.17 | 0.547 | 0.615 |
| chr3 | 160203182 | 160204267 | 1086 | 0.21 | 0.547 | 0.615 |
| chr1 | 211581492 | 211590637 | 9146 | 1.01 | 0.547 | 0.616 |
| chr19 | 1590931 | 1606588 | 15658 | 0.09 | 0.548 | 0.616 |
| chr3 | 24562691 | 24563776 | 1086 | -0.17 | 0.548 | 0.616 |
| chr5 | 78223313 | 78223827 | 515 | -0.42 | 0.548 | 0.616 |
| chr22 | 18957777 | 18958672 | 896 | 0.23 | 0.548 | 0.616 |
| chr18 | 3401941 | 3469665 | 67725 | -0.03 | 0.548 | 0.616 |
| chr2 | 45870396 | 45879408 | 9013 | -0.16 | 0.548 | 0.616 |
| chr6 | 17413070 | 17433026 | 19957 | -0.05 | 0.548 | 0.616 |
| chr16 | 2552682 | 2587770 | 35089 | -0.08 | 0.548 | 0.616 |
| chr16 | 2478259 | 2480371 | 2113 | 0.19 | 0.548 | 0.616 |
| chr20 | 49511677 | 49576190 | 64514 | -0.05 | 0.548 | 0.616 |
| chr12 | 8261634 | 8262010 | 377 | 0.35 | 0.549 | 0.617 |
| chr1 | 11986663 | 11995464 | 8802 | 0.14 | 0.549 | 0.617 |
| chr7 | 36697715 | 36699162 | 1448 | -0.24 | 0.549 | 0.617 |
| chr7 | 3532927 | 3533786 | 860 | 0.91 | 0.549 | 0.617 |
| chr20 | 56963708 | 57021691 | 57984 | 0.04 | 0.549 | 0.617 |
| chr11 | 129149637 | 129150171 | 535 | -0.35 | 0.549 | 0.617 |
| chr2 | 75184851 | 75186586 | 1736 | -0.11 | 0.549 | 0.617 |
| chr15 | 83620606 | 83621439 | 834 | 0.36 | 0.549 | 0.617 |
| chr14 | 71430864 | 71431663 | 800 | -0.39 | 0.549 | 0.617 |
| chr2 | 218375045 | 218375940 | 896 | 0.31 | 0.549 | 0.618 |
| chr5 | 96501694 | 96502967 | 1274 | -0.25 | 0.55 | 0.618 |
| chr10 | 105358826 | 105359324 | 499 | 0.39 | 0.55 | 0.618 |
| chr6 | 35132851 | 35134744 | 1894 | 0.17 | 0.55 | 0.618 |
| chr17 | 79565147 | 79604908 | 39762 | 0.05 | 0.55 | 0.618 |

| | | | | | | |
|---|---|---|---|---|---|---|
| chr8 | 122841992 | 122842410 | 419 | 0.36 | 0.55 | 0.618 |
| chr8 | 58493232 | 58503712 | 10481 | 0.67 | 0.55 | 0.618 |
| chr12 | 53553051 | 53576234 | 23184 | 0.09 | 0.55 | 0.618 |
| chr2 | 40004955 | 40006473 | 1519 | -0.1 | 0.55 | 0.618 |
| chr4 | 148848752 | 148850246 | 1495 | -0.17 | 0.55 | 0.618 |
| chr11 | 67111155 | 67170083 | 58929 | -0.02 | 0.55 | 0.618 |
| chr5 | 122110220 | 122112326 | 2107 | 0.17 | 0.55 | 0.618 |
| chr1 | 46049241 | 46051698 | 2458 | -0.14 | 0.55 | 0.618 |
| chr1 | 2477813 | 2479368 | 1556 | 0.18 | 0.55 | 0.618 |
| chr8 | 33411238 | 33425360 | 14123 | 0.14 | 0.551 | 0.619 |
| chr5 | 17212860 | 17285736 | 72877 | 0.04 | 0.551 | 0.619 |
| chr2 | 232212363 | 232277798 | 65436 | -0.04 | 0.551 | 0.619 |
| chr9 | 96212017 | 96264507 | 52491 | -0.04 | 0.551 | 0.619 |
| chr4 | 32022047 | 32029727 | 7681 | 0.06 | 0.552 | 0.62 |
| chr11 | 63852116 | 63863063 | 10948 | -0.08 | 0.552 | 0.62 |
| chr7 | 46402713 | 46404242 | 1530 | 0.69 | 0.552 | 0.62 |
| chr6 | 18325927 | 18326608 | 682 | 0.47 | 0.552 | 0.62 |
| chr12 | 9545278 | 9546130 | 853 | -1.04 | 0.552 | 0.62 |
| chr12 | 122387243 | 122389089 | 1847 | -0.15 | 0.552 | 0.62 |
| chr7 | 12724764 | 12730240 | 5477 | -0.07 | 0.553 | 0.621 |
| chr10 | 3566199 | 3581732 | 15534 | 0.24 | 0.553 | 0.621 |
| chr18 | 8020270 | 8020857 | 588 | -0.34 | 0.553 | 0.621 |
| chr19 | 7967671 | 7969539 | 1869 | -0.18 | 0.553 | 0.621 |
| chr19 | 7598042 | 7601130 | 3089 | -0.14 | 0.553 | 0.621 |
| chr16 | 67736847 | 67737671 | 825 | -0.38 | 0.554 | 0.622 |
| chr5 | 77701776 | 77702701 | 926 | -0.24 | 0.554 | 0.622 |
| chr2 | 112812847 | 112815402 | 2556 | -0.1 | 0.554 | 0.622 |
| chr2 | 20098331 | 20112222 | 13892 | 0.05 | 0.554 | 0.622 |
| chr8 | 11626547 | 11627866 | 1320 | -0.22 | 0.554 | 0.622 |
| chr13 | 76444681 | 76445236 | 556 | 0.33 | 0.554 | 0.622 |
| chr19 | 18787261 | 18794952 | 7692 | -0.13 | 0.554 | 0.622 |
| chr9 | 124461646 | 124462012 | 367 | -0.52 | 0.554 | 0.622 |
| chr2 | 73612091 | 73614267 | 2177 | -0.07 | 0.555 | 0.622 |
| chr2 | 27487368 | 27487988 | 621 | -0.37 | 0.555 | 0.622 |
| chr19 | 7405793 | 7406587 | 795 | 0.49 | 0.555 | 0.623 |
| chr11 | 95656115 | 95659196 | 3082 | 0.06 | 0.555 | 0.623 |
| chr14 | 67981509 | 67982156 | 648 | 0.31 | 0.555 | 0.623 |
| chr2 | 61989182 | 61999444 | 10263 | 0.05 | 0.555 | 0.623 |
| chr3 | 44518523 | 44519635 | 1113 | -0.21 | 0.555 | 0.623 |
| chr3 | 185633014 | 185686673 | 53660 | 0.05 | 0.555 | 0.623 |
| chr13 | 30937943 | 30938822 | 880 | -0.36 | 0.556 | 0.623 |
| chr15 | 51051973 | 51060126 | 8154 | 0.06 | 0.556 | 0.623 |
| chr1 | 225951250 | 225969806 | 18557 | -0.06 | 0.556 | 0.623 |
| chr7 | 103765045 | 103765910 | 866 | 0.37 | 0.556 | 0.623 |
| chr5 | 157097647 | 157098389 | 743 | 0.41 | 0.556 | 0.624 |
| chr14 | 34993134 | 34993577 | 444 | -0.46 | 0.556 | 0.624 |
| chr3 | 72957152 | 72957704 | 553 | 0.35 | 0.556 | 0.624 |
| chr18 | 44701375 | 44702759 | 1385 | 0.19 | 0.556 | 0.624 |
| chr14 | 36294830 | 36296628 | 1799 | 0.16 | 0.556 | 0.624 |
| chr10 | 94268166 | 94269486 | 1321 | -0.25 | 0.556 | 0.624 |
| chr1 | 31480065 | 31520829 | 40765 | -0.07 | 0.556 | 0.624 |
| chr17 | 37394085 | 37395493 | 1409 | 0.23 | 0.556 | 0.624 |
| chr12 | 90144037 | 90152039 | 8003 | 0.07 | 0.557 | 0.624 |
| chr17 | 15413325 | 15413958 | 634 | 0.41 | 0.557 | 0.624 |
| chr18 | 46394964 | 46400082 | 5119 | -0.17 | 0.557 | 0.624 |
| chr15 | 68305472 | 68318999 | 13528 | 0.18 | 0.557 | 0.624 |
| chr22 | 32661526 | 32662290 | 765 | -0.31 | 0.557 | 0.625 |
| chr21 | 43373093 | 43378799 | 5707 | -0.11 | 0.557 | 0.625 |
| chr13 | 45693803 | 45696087 | 2285 | 0.14 | 0.557 | 0.625 |
| chr12 | 49204658 | 49209243 | 4586 | 0.16 | 0.557 | 0.625 |
| chr3 | 132377694 | 132380140 | 2447 | -0.09 | 0.557 | 0.625 |

| | | | | | | |
|---|---|---|---|---|---|---|
| chr1 | 7283382 | 7284264 | 883 | 0.34 | 0.557 | 0.625 |
| chr13 | 114047855 | 114048426 | 572 | -0.39 | 0.557 | 0.625 |
| chr5 | 147805003 | 147807637 | 2635 | -0.15 | 0.558 | 0.625 |
| chr6 | 53537619 | 53544283 | 6665 | 0.27 | 0.558 | 0.625 |
| chr12 | 53490876 | 53497169 | 6294 | 0.08 | 0.558 | 0.625 |
| chr4 | 160270114 | 160271222 | 1109 | 0.18 | 0.558 | 0.625 |
| chr4 | 34148174 | 34151472 | 3299 | 0.12 | 0.558 | 0.626 |
| chr9 | 128001804 | 128007007 | 5204 | 0.06 | 0.558 | 0.626 |
| chr11 | 3145429 | 3145907 | 479 | -0.4 | 0.559 | 0.626 |
| chr14 | 68465393 | 68479077 | 13685 | 0.07 | 0.559 | 0.626 |
| chr1 | 213103711 | 213105189 | 1479 | 0.15 | 0.559 | 0.627 |
| chr6 | 43594623 | 43613664 | 19042 | 0.07 | 0.559 | 0.627 |
| chr21 | 38999378 | 39009896 | 10519 | 0.12 | 0.559 | 0.627 |
| chr1 | 212407631 | 212408559 | 929 | -0.35 | 0.559 | 0.627 |
| chr11 | 73662072 | 73667622 | 5551 | -0.24 | 0.559 | 0.627 |
| chr9 | 116137522 | 116174295 | 36774 | 0.06 | 0.559 | 0.627 |
| chr13 | 34874426 | 34875168 | 743 | 0.29 | 0.559 | 0.627 |
| chr6 | 146226204 | 146226565 | 362 | -0.47 | 0.559 | 0.627 |
| chr1 | 12374618 | 12375764 | 1147 | -0.24 | 0.56 | 0.627 |
| chr1 | 196504735 | 196505655 | 921 | 0.33 | 0.56 | 0.627 |
| chr4 | 38905668 | 38906461 | 794 | 0.31 | 0.56 | 0.627 |
| chr6 | 56263551 | 56265726 | 2176 | 0.2 | 0.56 | 0.627 |
| chr19 | 10654127 | 10689227 | 35101 | -0.06 | 0.56 | 0.627 |
| chr7 | 129170136 | 129170793 | 658 | -0.39 | 0.56 | 0.627 |
| chr11 | 46776745 | 46868327 | 91583 | -0.04 | 0.56 | 0.627 |
| chr8 | 120592127 | 120593739 | 1613 | -0.16 | 0.561 | 0.628 |
| chr6 | 151695212 | 151730544 | 35333 | 0.06 | 0.561 | 0.628 |
| chr17 | 18629848 | 18630178 | 331 | 0.45 | 0.561 | 0.628 |
| chr15 | 50644118 | 50651434 | 7317 | 0.04 | 0.561 | 0.628 |
| chr3 | 131747852 | 131753837 | 5986 | 0.21 | 0.561 | 0.628 |
| chr9 | 133320340 | 133320831 | 492 | 0.38 | 0.561 | 0.628 |
| chr22 | 38965686 | 38967407 | 1722 | -0.21 | 0.561 | 0.628 |
| chr1 | 16766322 | 16768151 | 1830 | -0.17 | 0.561 | 0.628 |
| chr5 | 149236787 | 149237750 | 964 | 0.3 | 0.561 | 0.629 |
| chr8 | 41627537 | 41629156 | 1620 | -0.2 | 0.561 | 0.629 |
| chr17 | 4779768 | 4783450 | 3683 | 0.19 | 0.561 | 0.629 |
| chr1 | 94203185 | 94248586 | 45402 | -0.05 | 0.562 | 0.629 |
| chr20 | 50476489 | 50477330 | 842 | 0.32 | 0.562 | 0.629 |
| chrX | 100602151 | 100604451 | 2301 | 0.14 | 0.562 | 0.629 |
| chr2 | 189263565 | 189264489 | 925 | 0.39 | 0.562 | 0.629 |
| chr10 | 31389989 | 31390759 | 771 | 0.31 | 0.562 | 0.629 |
| chr6 | 36952875 | 36954896 | 2022 | 0.12 | 0.562 | 0.629 |
| chr20 | 6102772 | 6104724 | 1953 | 0.07 | 0.562 | 0.629 |
| chr2 | 32502151 | 32529870 | 27720 | 0.05 | 0.562 | 0.629 |
| chr1 | 40254401 | 40265020 | 10620 | -0.21 | 0.562 | 0.629 |
| chr6 | 42595138 | 42596242 | 1105 | -0.27 | 0.563 | 0.63 |
| chr17 | 64637712 | 64638230 | 519 | 0.42 | 0.563 | 0.63 |
| chr2 | 219330247 | 219330910 | 664 | 0.36 | 0.563 | 0.63 |
| chr19 | 31841375 | 31842155 | 781 | -0.27 | 0.563 | 0.63 |
| chr20 | 40245766 | 40247699 | 1934 | -0.16 | 0.563 | 0.63 |
| chr1 | 150158445 | 150159651 | 1207 | -0.26 | 0.563 | 0.63 |
| chr9 | 95709552 | 95710174 | 623 | -0.29 | 0.564 | 0.631 |
| chr22 | 31030666 | 31032645 | 1980 | -0.18 | 0.564 | 0.631 |
| chrX | 152110296 | 152110928 | 633 | 0.28 | 0.564 | 0.631 |
| chr17 | 46276148 | 46277147 | 1000 | -0.2 | 0.564 | 0.631 |
| chr5 | 55464726 | 55466066 | 1341 | -0.28 | 0.565 | 0.632 |
| chr2 | 109335125 | 109336444 | 1320 | -0.12 | 0.565 | 0.632 |
| chr20 | 33057532 | 33057895 | 364 | 0.44 | 0.565 | 0.632 |
| chr7 | 106684492 | 106684989 | 498 | -0.35 | 0.565 | 0.632 |
| chr7 | 1066803 | 1070169 | 3367 | 0.13 | 0.565 | 0.632 |
| chr5 | 57356652 | 57390793 | 34142 | 0.06 | 0.565 | 0.632 |

| | | | | | | |
|---|---|---|---|---|---|---|
| chr1 | 29111836 | 29112272 | 437 | -0.36 | 0.565 | 0.632 |
| chr3 | 137800565 | 137801618 | 1054 | -0.36 | 0.565 | 0.632 |
| chr18 | 77153285 | 77160733 | 7449 | -0.14 | 0.566 | 0.632 |
| chr2 | 202270370 | 202272030 | 1661 | -0.15 | 0.566 | 0.632 |
| chr16 | 20751242 | 20753799 | 2558 | 0.13 | 0.566 | 0.633 |
| chr11 | 9158802 | 9167640 | 8839 | 0.09 | 0.566 | 0.633 |
| chr2 | 73963297 | 73965294 | 1998 | -0.11 | 0.566 | 0.633 |
| chr1 | 108724531 | 108726473 | 1943 | -0.28 | 0.566 | 0.633 |
| chr10 | 131988658 | 131989189 | 532 | -0.37 | 0.566 | 0.633 |
| chr3 | 113549231 | 113558238 | 9008 | 1.22 | 0.566 | 0.633 |
| chr10 | 94448366 | 94460476 | 12111 | 0.06 | 0.566 | 0.633 |
| chr1 | 226104084 | 226112915 | 8832 | -0.08 | 0.566 | 0.633 |
| chr8 | 141760396 | 141761166 | 771 | 1.04 | 0.566 | 0.633 |
| chr19 | 10361895 | 10363785 | 1891 | -0.16 | 0.567 | 0.633 |
| chr10 | 27358226 | 27358983 | 758 | 0.35 | 0.567 | 0.633 |
| chr1 | 86041826 | 86050799 | 8974 | -0.06 | 0.567 | 0.633 |
| chr11 | 18126737 | 18128074 | 1338 | -0.16 | 0.567 | 0.633 |
| chr5 | 139167282 | 139167695 | 414 | -0.37 | 0.567 | 0.633 |
| chr4 | 83754827 | 83755876 | 1050 | 0.37 | 0.567 | 0.634 |
| chr6 | 54146868 | 54147833 | 966 | 0.28 | 0.567 | 0.634 |
| chr1 | 201891238 | 201891801 | 564 | -0.3 | 0.567 | 0.634 |
| chr1 | 21022963 | 21025675 | 2713 | -0.14 | 0.567 | 0.634 |
| chr14 | 62120101 | 62129962 | 9862 | -0.13 | 0.567 | 0.634 |
| chr5 | 179309741 | 179341943 | 32203 | 0.13 | 0.568 | 0.634 |
| chrX | 10422632 | 10488358 | 65727 | -0.05 | 0.568 | 0.634 |
| chr14 | 93672114 | 93683743 | 11630 | -0.08 | 0.568 | 0.634 |
| chr12 | 105022663 | 105023147 | 485 | 0.37 | 0.568 | 0.635 |
| chr9 | 132418791 | 132419271 | 481 | -0.39 | 0.568 | 0.635 |
| chr4 | 151306709 | 151307280 | 572 | 1.01 | 0.568 | 0.635 |
| chr4 | 151635921 | 151644786 | 8866 | 1.14 | 0.568 | 0.635 |
| chr3 | 177569765 | 177570655 | 891 | -0.22 | 0.568 | 0.635 |
| chr20 | 60605850 | 60606804 | 955 | -0.33 | 0.569 | 0.635 |
| chr9 | 32384641 | 32396301 | 11661 | -1.13 | 0.569 | 0.635 |
| chr3 | 33424340 | 33435432 | 11093 | 0.07 | 0.569 | 0.635 |
| chr20 | 21216542 | 21216804 | 263 | 0.41 | 0.569 | 0.636 |
| chr1 | 93296895 | 93300019 | 3125 | 0.09 | 0.57 | 0.636 |
| chr22 | 29137219 | 29139312 | 2094 | -0.14 | 0.57 | 0.636 |
| chr22 | 36418208 | 36425694 | 7487 | 0.15 | 0.57 | 0.636 |
| chr10 | 7859779 | 7870366 | 10588 | 0.13 | 0.57 | 0.636 |
| chr1 | 59897722 | 59899150 | 1429 | 0.21 | 0.57 | 0.636 |
| chr1 | 17674335 | 17675574 | 1240 | -0.23 | 0.57 | 0.636 |
| chr17 | 47267332 | 47276280 | 8949 | -0.06 | 0.57 | 0.637 |
| chr16 | 74872495 | 74873401 | 907 | 0.33 | 0.57 | 0.637 |
| chr14 | 50887021 | 50887661 | 641 | -0.33 | 0.57 | 0.637 |
| chr5 | 32679077 | 32679934 | 858 | -0.46 | 0.57 | 0.637 |
| chr9 | 136113686 | 136115263 | 1578 | -0.22 | 0.571 | 0.637 |
| chr17 | 47409506 | 47412797 | 3292 | -0.11 | 0.571 | 0.637 |
| chr8 | 124338421 | 124339468 | 1048 | 0.33 | 0.571 | 0.637 |
| chr3 | 10305285 | 10306291 | 1007 | -0.27 | 0.571 | 0.637 |
| chr7 | 99677582 | 99700633 | 23052 | -0.05 | 0.571 | 0.637 |
| chr2 | 38151838 | 38153570 | 1733 | -0.08 | 0.571 | 0.637 |
| chr1 | 10554267 | 10556083 | 1817 | 0.21 | 0.571 | 0.638 |
| chr12 | 11898679 | 11899631 | 953 | -0.27 | 0.571 | 0.638 |
| chr1 | 100981008 | 100981745 | 738 | 0.32 | 0.572 | 0.638 |
| chr11 | 66885644 | 66894905 | 9262 | 0.08 | 0.572 | 0.638 |
| chr2 | 28789084 | 28790413 | 1330 | -0.15 | 0.572 | 0.638 |
| chr8 | 103337085 | 103346376 | 9292 | -0.13 | 0.572 | 0.638 |
| chr6 | 34392246 | 34394675 | 2430 | 0.11 | 0.572 | 0.639 |
| chr4 | 47915391 | 47916688 | 1298 | 0.17 | 0.572 | 0.639 |
| chrX | 54521212 | 54522565 | 1354 | -0.23 | 0.572 | 0.639 |
| chr12 | 3101526 | 3127685 | 26160 | -0.09 | 0.572 | 0.639 |

| | | | | | | |
|---|---|---|---|---|---|---|
| chr11 | 126587409 | 126589976 | 2568 | 0.14 | 0.572 | 0.639 |
| chr18 | 46501596 | 46502981 | 1386 | -0.22 | 0.573 | 0.639 |
| chr2 | 58448504 | 58449507 | 1004 | 0.23 | 0.573 | 0.639 |
| chr3 | 110102292 | 110104602 | 2311 | 0.17 | 0.573 | 0.639 |
| chr1 | 231368338 | 231378583 | 10246 | 0.08 | 0.573 | 0.639 |
| chrX | 16607339 | 16607774 | 436 | -0.37 | 0.573 | 0.639 |
| chr9 | 135501175 | 135513142 | 11968 | -0.12 | 0.573 | 0.639 |
| chr7 | 127290825 | 127319711 | 28887 | -0.04 | 0.573 | 0.639 |
| chr6 | 76382998 | 76383799 | 802 | -0.35 | 0.573 | 0.639 |
| chr19 | 2095353 | 2097501 | 2149 | 0.21 | 0.573 | 0.64 |
| chr1 | 234975031 | 235013648 | 38618 | -0.06 | 0.573 | 0.64 |
| chr19 | 45754866 | 45755741 | 876 | -0.32 | 0.574 | 0.64 |
| chr1 | 16338993 | 16340519 | 1527 | -0.19 | 0.574 | 0.64 |
| chr21 | 37801077 | 37803803 | 2727 | -0.08 | 0.574 | 0.64 |
| chr22 | 40741447 | 40744185 | 2739 | 0.11 | 0.574 | 0.64 |
| chr5 | 73980194 | 73981228 | 1035 | 0.18 | 0.574 | 0.64 |
| chr3 | 19926824 | 19943554 | 16731 | -0.06 | 0.574 | 0.64 |
| chr17 | 29516413 | 29517317 | 905 | 0.3 | 0.574 | 0.64 |
| chr3 | 49577357 | 49578019 | 663 | -0.31 | 0.574 | 0.64 |
| chr14 | 74716535 | 74724972 | 8438 | -0.22 | 0.574 | 0.64 |
| chr12 | 93770190 | 93771865 | 1676 | -0.14 | 0.574 | 0.64 |
| chr2 | 191272427 | 191274305 | 1879 | 0.15 | 0.574 | 0.64 |
| chr6 | 155347764 | 155348203 | 440 | -0.34 | 0.575 | 0.641 |
| chr1 | 229099434 | 229100836 | 1403 | -0.17 | 0.575 | 0.641 |
| chr5 | 72110482 | 72118924 | 8443 | -0.05 | 0.575 | 0.642 |
| chr10 | 44690697 | 44691473 | 777 | 0.32 | 0.575 | 0.642 |
| chr19 | 44616815 | 44618292 | 1478 | -0.16 | 0.576 | 0.642 |
| chr6 | 52680262 | 52680862 | 601 | 0.36 | 0.576 | 0.642 |
| chrX | 40855983 | 40908234 | 52252 | 0.03 | 0.576 | 0.642 |
| chr16 | 59357395 | 59359684 | 2290 | 0.1 | 0.576 | 0.642 |
| chr6 | 37811252 | 37811673 | 422 | 0.39 | 0.576 | 0.642 |
| chr9 | 35051035 | 35051546 | 512 | 0.39 | 0.576 | 0.642 |
| chr14 | 42684121 | 42685379 | 1259 | 0.2 | 0.576 | 0.642 |
| chr21 | 47647995 | 47649891 | 1897 | -0.21 | 0.576 | 0.642 |
| chr1 | 63103369 | 63103774 | 406 | -0.31 | 0.576 | 0.642 |
| chr15 | 41392810 | 41394007 | 1198 | 0.4 | 0.576 | 0.642 |
| chr3 | 57632523 | 57633063 | 541 | 0.37 | 0.576 | 0.642 |
| chrX | 77040323 | 77044589 | 4267 | 0.1 | 0.576 | 0.642 |
| chr6 | 170467872 | 170468924 | 1053 | 0.23 | 0.576 | 0.642 |
| chr1 | 55679655 | 55683924 | 4270 | 0.09 | 0.577 | 0.642 |
| chr12 | 49443349 | 49464436 | 21088 | 0.07 | 0.577 | 0.643 |
| chr19 | 44022978 | 44125177 | 102200 | -0.03 | 0.577 | 0.643 |
| chr16 | 48109136 | 48116449 | 7314 | -0.08 | 0.577 | 0.643 |
| chr12 | 3020905 | 3021779 | 875 | -0.29 | 0.577 | 0.643 |
| chr3 | 114403211 | 114404469 | 1259 | 0.27 | 0.577 | 0.643 |
| chr8 | 38938594 | 38962065 | 23472 | 0.09 | 0.577 | 0.643 |
| chr19 | 51142785 | 51143472 | 688 | -0.33 | 0.577 | 0.643 |
| chr13 | 40176395 | 40178177 | 1783 | 0.16 | 0.577 | 0.643 |
| chrX | 62723363 | 62724199 | 837 | 0.31 | 0.577 | 0.643 |
| chr14 | 105646834 | 105647614 | 781 | 0.31 | 0.578 | 0.643 |
| chr22 | 39267015 | 39269568 | 2554 | -0.11 | 0.578 | 0.643 |
| chr2 | 121070304 | 121070822 | 519 | -0.36 | 0.578 | 0.644 |
| chr16 | 19856076 | 19857007 | 932 | 0.23 | 0.578 | 0.644 |
| chr19 | 49339010 | 49340842 | 1833 | 0.15 | 0.578 | 0.644 |
| chr6 | 52462756 | 52463603 | 848 | -0.29 | 0.579 | 0.644 |
| chr1 | 24053616 | 24072110 | 18495 | -0.09 | 0.579 | 0.644 |
| chr9 | 124497464 | 124529202 | 31739 | -0.05 | 0.579 | 0.645 |
| chrX | 79471422 | 79472276 | 855 | -0.27 | 0.579 | 0.645 |
| chr4 | 25401876 | 25403089 | 1214 | -0.26 | 0.579 | 0.645 |
| chr21 | 45719221 | 45721145 | 1925 | -0.18 | 0.579 | 0.645 |
| chr6 | 157694703 | 157696521 | 1819 | 0.16 | 0.579 | 0.645 |

| | | | | | | |
|---|---|---|---|---|---|---|
| chr17 | 13963216 | 13973721 | 10506 | -0.07 | 0.579 | 0.645 |
| chr3 | 152083189 | 152122137 | 38949 | -0.03 | 0.58 | 0.645 |
| chr4 | 159816899 | 159824343 | 7445 | 0.19 | 0.58 | 0.645 |
| chr6 | 166043701 | 166044515 | 815 | -0.32 | 0.58 | 0.645 |
| chr14 | 105552398 | 105561270 | 8873 | -0.05 | 0.58 | 0.646 |
| chr20 | 47562080 | 47562974 | 895 | 0.24 | 0.58 | 0.646 |
| chr11 | 33036816 | 33037896 | 1081 | 0.25 | 0.581 | 0.646 |
| chr18 | 12835713 | 12843787 | 8075 | 0.04 | 0.581 | 0.646 |
| chr20 | 5002909 | 5005385 | 2477 | -0.18 | 0.581 | 0.647 |
| chr11 | 64917061 | 64919596 | 2536 | 0.08 | 0.581 | 0.647 |
| chr15 | 79046977 | 79047516 | 540 | 0.33 | 0.581 | 0.647 |
| chr8 | 22140869 | 22145523 | 4655 | -0.1 | 0.581 | 0.647 |
| chr6 | 121756257 | 121761712 | 5456 | -0.06 | 0.581 | 0.647 |
| chr3 | 100210624 | 100212009 | 1386 | 0.19 | 0.581 | 0.647 |
| chr1 | 154570860 | 154582677 | 11818 | -0.07 | 0.581 | 0.647 |
| chr16 | 11567241 | 11569118 | 1878 | -0.26 | 0.582 | 0.647 |
| chr7 | 6034121 | 6067020 | 32900 | 0.05 | 0.582 | 0.647 |
| chr2 | 192623189 | 192624216 | 1028 | 0.22 | 0.582 | 0.647 |
| chr19 | 30199117 | 30208077 | 8961 | -0.06 | 0.582 | 0.647 |
| chr8 | 58473796 | 58478398 | 4603 | -0.1 | 0.582 | 0.647 |
| chr6 | 152722322 | 152727029 | 4708 | 0.06 | 0.582 | 0.648 |
| chr9 | 131313647 | 131338221 | 24575 | 0.05 | 0.583 | 0.648 |
| chr21 | 38337837 | 38374176 | 36340 | 0.09 | 0.583 | 0.648 |
| chr12 | 240548 | 241819 | 1272 | -0.24 | 0.583 | 0.648 |
| chr2 | 221420864 | 221421836 | 973 | 0.23 | 0.583 | 0.648 |
| chr6 | 105237511 | 105237937 | 427 | -0.34 | 0.583 | 0.648 |
| chr17 | 37308527 | 37312265 | 3739 | 0.11 | 0.583 | 0.648 |
| chr19 | 19302092 | 19314628 | 12537 | -0.1 | 0.583 | 0.648 |
| chr6 | 132129134 | 132129607 | 474 | -0.34 | 0.583 | 0.648 |
| chr5 | 154061515 | 154063984 | 2470 | -0.09 | 0.583 | 0.648 |
| chr20 | 30903877 | 30918676 | 14800 | 0.06 | 0.583 | 0.648 |
| chr11 | 108564552 | 108565658 | 1107 | 0.26 | 0.583 | 0.648 |
| chr11 | 69484752 | 69516103 | 31352 | 0.07 | 0.583 | 0.649 |
| chr2 | 151181770 | 151182921 | 1152 | -0.22 | 0.583 | 0.649 |
| chr4 | 113739079 | 113739834 | 756 | 0.26 | 0.584 | 0.649 |
| chr15 | 40355586 | 40401932 | 46347 | -0.05 | 0.584 | 0.649 |
| chr5 | 123210867 | 123219571 | 8705 | -0.82 | 0.584 | 0.649 |
| chr4 | 38566732 | 38567936 | 1205 | 0.25 | 0.584 | 0.649 |
| chr4 | 79582292 | 79583940 | 1649 | 0.14 | 0.584 | 0.65 |
| chr1 | 89981369 | 89991244 | 9876 | -0.1 | 0.584 | 0.65 |
| chr14 | 92413669 | 92414428 | 760 | 0.27 | 0.585 | 0.65 |
| chr8 | 27256671 | 27258863 | 2193 | -0.15 | 0.585 | 0.65 |
| chr7 | 127944394 | 127944743 | 350 | -0.43 | 0.585 | 0.65 |
| chr10 | 97666631 | 97667888 | 1258 | 0.19 | 0.585 | 0.65 |
| chr17 | 49533818 | 49534554 | 737 | -0.3 | 0.585 | 0.65 |
| chr8 | 122681431 | 122682340 | 910 | 0.28 | 0.585 | 0.65 |
| chr11 | 65321629 | 65356539 | 34911 | 0.04 | 0.585 | 0.65 |
| chr5 | 132111978 | 132113650 | 1673 | -0.17 | 0.585 | 0.65 |
| chr3 | 15139464 | 15141315 | 1852 | 0.14 | 0.585 | 0.65 |
| chr5 | 14775707 | 14776359 | 653 | -0.27 | 0.585 | 0.65 |
| chr19 | 10509217 | 10536300 | 27084 | 0.08 | 0.585 | 0.65 |
| chr2 | 28914524 | 28915422 | 899 | 0.26 | 0.585 | 0.65 |
| chr18 | 20512357 | 20514617 | 2261 | 0.1 | 0.585 | 0.65 |
| chr12 | 62828566 | 62829650 | 1085 | 0.34 | 0.586 | 0.651 |
| chr22 | 38190284 | 38202664 | 12381 | -0.1 | 0.586 | 0.651 |
| chr4 | 79002627 | 79003070 | 444 | 0.38 | 0.586 | 0.651 |
| chr3 | 10129619 | 10135196 | 5578 | 0.15 | 0.586 | 0.651 |
| chr8 | 31706736 | 31707136 | 401 | -0.35 | 0.586 | 0.651 |
| chr6 | 134519358 | 134519818 | 461 | 0.39 | 0.586 | 0.651 |
| chr8 | 63998185 | 63999128 | 944 | -0.27 | 0.586 | 0.651 |
| chr10 | 127887504 | 127893702 | 6199 | 0.17 | 0.586 | 0.651 |

| | | | | | | |
|---|---|---|---|---|---|---|
| chr11 | 36310098 | 36310952 | 855 | 0.24 | 0.586 | 0.651 |
| chr5 | 137483834 | 137484715 | 882 | -0.27 | 0.587 | 0.651 |
| chr9 | 129949124 | 129950321 | 1198 | 0.24 | 0.587 | 0.651 |
| chr3 | 118570818 | 118571388 | 571 | 0.25 | 0.587 | 0.652 |
| chr22 | 24714904 | 24720434 | 5531 | -0.16 | 0.587 | 0.652 |
| chr5 | 92085372 | 92086371 | 1000 | 0.3 | 0.587 | 0.652 |
| chr12 | 123086167 | 123089145 | 2979 | -0.18 | 0.587 | 0.652 |
| chrX | 153233055 | 153239809 | 6755 | -0.05 | 0.587 | 0.652 |
| chr11 | 47313883 | 47314339 | 457 | 0.34 | 0.587 | 0.652 |
| chr12 | 41061572 | 41061928 | 357 | -0.35 | 0.588 | 0.653 |
| chrX | 14046634 | 14049039 | 2406 | 0.09 | 0.588 | 0.653 |
| chr22 | 24666365 | 24668142 | 1778 | 0.15 | 0.588 | 0.653 |
| chr11 | 28030565 | 28031728 | 1164 | -0.17 | 0.588 | 0.653 |
| chr6 | 11366588 | 11369221 | 2634 | -0.09 | 0.588 | 0.653 |
| chr6 | 71997716 | 71999780 | 2065 | -0.16 | 0.588 | 0.653 |
| chr16 | 68876726 | 68882125 | 5400 | -0.09 | 0.589 | 0.654 |
| chr8 | 55046490 | 55078886 | 32397 | -0.06 | 0.589 | 0.654 |
| chr1 | 46597781 | 46599472 | 1692 | -0.12 | 0.589 | 0.654 |
| chr17 | 35912619 | 35931728 | 19110 | -0.1 | 0.589 | 0.654 |
| chr12 | 9486367 | 9487226 | 860 | -1 | 0.589 | 0.654 |
| chr10 | 8762763 | 8769264 | 6502 | -0.1 | 0.589 | 0.654 |
| chr4 | 141107736 | 141108317 | 582 | 0.45 | 0.589 | 0.654 |
| chr3 | 47994397 | 48003980 | 9584 | -0.17 | 0.589 | 0.654 |
| chr14 | 75258995 | 75262403 | 3409 | -0.16 | 0.59 | 0.654 |
| chr10 | 79788015 | 79802359 | 14345 | 0.06 | 0.59 | 0.654 |
| chr16 | 445834 | 453067 | 7234 | -0.09 | 0.59 | 0.654 |
| chr8 | 98673938 | 98674735 | 798 | 0.38 | 0.59 | 0.654 |
| chr4 | 146051946 | 146052864 | 919 | -0.29 | 0.59 | 0.654 |
| chr7 | 99840299 | 99847707 | 7409 | 0.07 | 0.59 | 0.655 |
| chr3 | 114150885 | 114151603 | 719 | -0.3 | 0.59 | 0.655 |
| chr15 | 66319349 | 66320424 | 1076 | -0.23 | 0.59 | 0.655 |
| chr9 | 16335134 | 16345193 | 10060 | 0.07 | 0.59 | 0.655 |
| chr3 | 119121817 | 119122661 | 845 | -0.26 | 0.591 | 0.656 |
| chr16 | 10652897 | 10653334 | 438 | -0.43 | 0.591 | 0.656 |
| chr20 | 5891642 | 5900838 | 9197 | -1.06 | 0.591 | 0.656 |
| chr3 | 132344797 | 132357325 | 12529 | -0.13 | 0.592 | 0.656 |
| chr1 | 25091405 | 25097369 | 5965 | 0.18 | 0.592 | 0.656 |
| chr1 | 32014719 | 32015871 | 1153 | -0.19 | 0.592 | 0.656 |
| chr3 | 122201338 | 122217358 | 16021 | -0.08 | 0.592 | 0.656 |
| chr1 | 55006908 | 55008914 | 2007 | -0.13 | 0.592 | 0.657 |
| chr2 | 189560694 | 189561333 | 640 | -0.36 | 0.592 | 0.657 |
| chrX | 19063700 | 19066042 | 2343 | -0.14 | 0.592 | 0.657 |
| chr16 | 71393873 | 71395570 | 1698 | 0.21 | 0.593 | 0.657 |
| chr20 | 30864779 | 30879102 | 14324 | 0.06 | 0.593 | 0.658 |
| chr14 | 58862107 | 58863326 | 1220 | -0.18 | 0.593 | 0.658 |
| chr1 | 17379334 | 17381354 | 2021 | 0.17 | 0.593 | 0.658 |
| chr1 | 84898887 | 84899656 | 770 | 0.16 | 0.593 | 0.658 |
| chr4 | 78841457 | 78850314 | 8858 | -0.11 | 0.593 | 0.658 |
| chr11 | 534485 | 536434 | 1950 | -0.19 | 0.593 | 0.658 |
| chr1 | 208136224 | 208137497 | 1274 | 0.14 | 0.594 | 0.658 |
| chr1 | 150376224 | 150391555 | 15332 | 0.09 | 0.594 | 0.658 |
| chr3 | 189306167 | 189306924 | 758 | -0.3 | 0.594 | 0.658 |
| chrX | 55185814 | 55188334 | 2521 | 0.1 | 0.594 | 0.658 |
| chr12 | 133262977 | 133264565 | 1589 | 0.14 | 0.594 | 0.658 |
| chr15 | 100272911 | 100274567 | 1657 | 0.12 | 0.594 | 0.658 |
| chr5 | 108373968 | 108374867 | 900 | 0.25 | 0.594 | 0.659 |
| chr20 | 62886418 | 62887617 | 1200 | 0.16 | 0.594 | 0.659 |
| chr1 | 28827318 | 28854704 | 27387 | 0.04 | 0.594 | 0.659 |
| chr12 | 54746654 | 54773538 | 26885 | 0.08 | 0.594 | 0.659 |
| chr4 | 25569462 | 25571450 | 1989 | -0.17 | 0.594 | 0.659 |
| chr11 | 111307393 | 111307865 | 473 | 0.35 | 0.595 | 0.659 |

| | | | | | | |
|---|---|---|---|---|---|---|
| chr21 | 45303406 | 45303937 | 532 | 0.35 | 0.595 | 0.659 |
| chr16 | 23502336 | 23552660 | 50325 | -0.04 | 0.595 | 0.659 |
| chr5 | 141044832 | 141045303 | 472 | 0.38 | 0.595 | 0.659 |
| chr3 | 152144627 | 152162267 | 17641 | -0.13 | 0.595 | 0.66 |
| chr19 | 47196213 | 47196737 | 525 | 0.44 | 0.595 | 0.66 |
| chr1 | 40203913 | 40205465 | 1553 | -0.1 | 0.595 | 0.66 |
| chr7 | 953617 | 954525 | 909 | -0.31 | 0.596 | 0.66 |
| chr9 | 22307824 | 22309160 | 1337 | 0.31 | 0.596 | 0.66 |
| chr7 | 112032164 | 112033170 | 1007 | 0.18 | 0.596 | 0.66 |
| chr9 | 124763891 | 124765453 | 1563 | 0.15 | 0.596 | 0.66 |
| chr19 | 5339573 | 5352575 | 13003 | -0.14 | 0.596 | 0.66 |
| chr11 | 92929078 | 92931535 | 2458 | 0.07 | 0.596 | 0.66 |
| chr17 | 20087232 | 20089174 | 1943 | -0.19 | 0.596 | 0.661 |
| chr3 | 23244514 | 23245653 | 1140 | -0.18 | 0.596 | 0.661 |
| chr16 | 30346311 | 30347191 | 881 | 0.22 | 0.596 | 0.661 |
| chr17 | 21304637 | 21306241 | 1605 | -0.21 | 0.596 | 0.661 |
| chr1 | 40626033 | 40628487 | 2455 | 0.09 | 0.597 | 0.661 |
| chr22 | 33833306 | 33835293 | 1988 | -0.09 | 0.597 | 0.661 |
| chr2 | 220108985 | 220118693 | 9709 | -0.06 | 0.597 | 0.661 |
| chr8 | 118843787 | 118844481 | 695 | -0.32 | 0.597 | 0.661 |
| chr3 | 12851132 | 12852442 | 1311 | -0.19 | 0.597 | 0.661 |
| chr6 | 88374544 | 88443308 | 68765 | 0.03 | 0.597 | 0.661 |
| chr8 | 138128909 | 138131481 | 2573 | -0.16 | 0.597 | 0.661 |
| chr9 | 131843408 | 131844655 | 1248 | 0.19 | 0.597 | 0.662 |
| chr20 | 1206747 | 1207337 | 591 | 0.35 | 0.598 | 0.662 |
| chr8 | 124084865 | 124087755 | 2891 | 0.12 | 0.598 | 0.662 |
| chr4 | 2931764 | 2939889 | 8126 | -0.08 | 0.598 | 0.662 |
| chr2 | 203925425 | 203927216 | 1792 | 0.15 | 0.598 | 0.662 |
| chr15 | 41784996 | 41785948 | 953 | -0.25 | 0.598 | 0.662 |
| chrX | 10320251 | 10321635 | 1385 | 0.18 | 0.598 | 0.662 |
| chr3 | 149434488 | 149436218 | 1731 | -0.28 | 0.599 | 0.663 |
| chr3 | 48922995 | 48923392 | 398 | -0.38 | 0.599 | 0.663 |
| chr7 | 26488822 | 26489710 | 889 | -0.28 | 0.599 | 0.663 |
| chr6 | 108273384 | 108280382 | 6999 | 0.08 | 0.599 | 0.663 |
| chr5 | 132249074 | 132249492 | 419 | -0.41 | 0.599 | 0.663 |
| chr16 | 30932416 | 30970275 | 37860 | -0.05 | 0.599 | 0.663 |
| chr16 | 83960508 | 83994369 | 33862 | -0.05 | 0.599 | 0.663 |
| chr12 | 76630162 | 76631323 | 1162 | 0.29 | 0.599 | 0.663 |
| chr20 | 48962793 | 48963485 | 693 | 0.29 | 0.6 | 0.664 |
| chr8 | 144813622 | 144816689 | 3068 | -0.08 | 0.6 | 0.664 |
| chr6 | 44160550 | 44160922 | 373 | 0.38 | 0.6 | 0.664 |
| chr19 | 50860508 | 50897620 | 37113 | -0.05 | 0.6 | 0.664 |
| chr4 | 41215592 | 41219514 | 3923 | -0.1 | 0.6 | 0.664 |
| chr16 | 3629270 | 3630390 | 1121 | -0.22 | 0.6 | 0.664 |
| chr20 | 13764170 | 13797609 | 33440 | 0.06 | 0.6 | 0.664 |
| chr21 | 35445108 | 35446907 | 1800 | 0.21 | 0.6 | 0.664 |
| chr10 | 25209478 | 25211319 | 1842 | -0.1 | 0.601 | 0.664 |
| chrX | 18369787 | 18373346 | 3560 | -0.14 | 0.601 | 0.664 |
| chr2 | 3652280 | 3654086 | 1807 | 0.14 | 0.601 | 0.664 |
| chr5 | 154235108 | 154250657 | 15550 | -0.04 | 0.601 | 0.665 |
| chr19 | 8543082 | 8543685 | 604 | 0.37 | 0.601 | 0.665 |
| chr16 | 47071316 | 47072473 | 1158 | -0.2 | 0.601 | 0.665 |
| chr1 | 23474737 | 23476403 | 1667 | 0.18 | 0.601 | 0.665 |
| chr2 | 26522409 | 26522960 | 552 | -0.28 | 0.601 | 0.665 |
| chr5 | 98105813 | 98123773 | 17961 | 0.08 | 0.601 | 0.665 |
| chr9 | 16870332 | 16871116 | 785 | -0.31 | 0.601 | 0.665 |
| chr1 | 160369811 | 160370940 | 1130 | -0.22 | 0.601 | 0.665 |
| chr1 | 40399450 | 40400820 | 1371 | 0.23 | 0.601 | 0.665 |
| chr16 | 837736 | 838342 | 607 | 0.4 | 0.601 | 0.665 |
| chr1 | 150321161 | 150362516 | 41356 | 0.04 | 0.601 | 0.665 |
| chr17 | 62339694 | 62341516 | 1823 | 0.19 | 0.602 | 0.665 |

| | | | | | | |
|---|---|---|---|---|---|---|
| chr10 | 129091590 | 129092393 | 804 | -0.31 | 0.602 | 0.665 |
| chr10 | 102044773 | 102047394 | 2622 | 0.09 | 0.602 | 0.665 |
| chr4 | 13598834 | 13599338 | 505 | -0.36 | 0.602 | 0.665 |
| chr14 | 24274648 | 24275484 | 837 | -0.24 | 0.602 | 0.666 |
| chr1 | 243644958 | 243663026 | 18069 | 0.07 | 0.602 | 0.666 |
| chr15 | 72612229 | 72613282 | 1054 | 0.22 | 0.602 | 0.666 |
| chr12 | 109982736 | 109983370 | 635 | 0.32 | 0.602 | 0.666 |
| chr2 | 198017129 | 198026065 | 8937 | -0.08 | 0.602 | 0.666 |
| chr11 | 93917206 | 93917550 | 345 | 0.37 | 0.603 | 0.666 |
| chr10 | 91001590 | 91022820 | 21231 | -0.18 | 0.603 | 0.666 |
| chr9 | 100615486 | 100617327 | 1842 | 0.11 | 0.603 | 0.666 |
| chr9 | 95417115 | 95434558 | 17444 | -0.06 | 0.603 | 0.666 |
| chrX | 44202381 | 44203416 | 1036 | -0.25 | 0.603 | 0.666 |
| chr4 | 184908220 | 184911077 | 2858 | 0.06 | 0.604 | 0.667 |
| chr17 | 26132439 | 26133459 | 1021 | 0.11 | 0.604 | 0.667 |
| chr1 | 218916778 | 218917736 | 959 | 0.27 | 0.604 | 0.667 |
| chr7 | 129303948 | 129305812 | 1865 | 0.14 | 0.604 | 0.668 |
| chr10 | 98479939 | 98480653 | 715 | -0.3 | 0.604 | 0.668 |
| chr2 | 228462506 | 228463337 | 832 | 0.28 | 0.604 | 0.668 |
| chr15 | 50994704 | 50995479 | 776 | -0.27 | 0.604 | 0.668 |
| chr5 | 127873040 | 127875056 | 2017 | -0.2 | 0.605 | 0.668 |
| chr1 | 46015686 | 46023763 | 8078 | -0.08 | 0.605 | 0.668 |
| chr16 | 71960204 | 71961630 | 1427 | 0.2 | 0.605 | 0.668 |
| chr9 | 21031144 | 21032064 | 921 | -0.22 | 0.605 | 0.668 |
| chr20 | 16382342 | 16382974 | 633 | -0.34 | 0.605 | 0.668 |
| chr5 | 58262858 | 58698682 | 435825 | -0.01 | 0.605 | 0.668 |
| chr3 | 62304185 | 62305240 | 1056 | 0.2 | 0.605 | 0.668 |
| chrX | 68758294 | 68758706 | 413 | -0.28 | 0.605 | 0.669 |
| chr16 | 8735639 | 8735968 | 330 | 0.45 | 0.606 | 0.669 |
| chr3 | 69810429 | 69840451 | 30023 | 0.06 | 0.606 | 0.669 |
| chr11 | 114309649 | 114311971 | 2323 | 0.13 | 0.606 | 0.669 |
| chr20 | 60981689 | 60982996 | 1308 | 0.2 | 0.606 | 0.669 |
| chr11 | 76753874 | 76758804 | 4931 | -0.2 | 0.606 | 0.67 |
| chr17 | 62075474 | 62084042 | 8569 | 0.17 | 0.606 | 0.67 |
| chr19 | 17848107 | 17849179 | 1073 | -0.35 | 0.606 | 0.67 |
| chr5 | 118732959 | 118733912 | 954 | -0.25 | 0.607 | 0.67 |
| chr7 | 23444446 | 23488102 | 43657 | -0.07 | 0.607 | 0.67 |
| chr11 | 125932397 | 125933324 | 928 | -0.17 | 0.607 | 0.67 |
| chr18 | 60172922 | 60174612 | 1691 | 0.13 | 0.607 | 0.67 |
| chr20 | 33992760 | 34000604 | 7845 | 0.08 | 0.607 | 0.67 |
| chr14 | 70394527 | 70395347 | 821 | 0.3 | 0.607 | 0.67 |
| chr11 | 76841682 | 76842812 | 1131 | 0.28 | 0.607 | 0.67 |
| chr11 | 85870827 | 85888843 | 18017 | -0.12 | 0.607 | 0.67 |
| chr2 | 208421083 | 208422148 | 1066 | -0.26 | 0.607 | 0.67 |
| chr11 | 57282718 | 57283661 | 944 | 0.32 | 0.607 | 0.67 |
| chr9 | 96056803 | 96057909 | 1107 | 0.24 | 0.607 | 0.67 |
| chr15 | 50914423 | 50915122 | 700 | 0.48 | 0.607 | 0.67 |
| chr8 | 61495051 | 61512874 | 17824 | 0.13 | 0.608 | 0.671 |
| chr11 | 128327889 | 128338722 | 10834 | 0.09 | 0.608 | 0.671 |
| chr1 | 185014041 | 185015803 | 1763 | 0.14 | 0.608 | 0.671 |
| chr3 | 45016353 | 45018829 | 2477 | -0.09 | 0.608 | 0.671 |
| chr10 | 78656860 | 78657248 | 389 | 0.37 | 0.608 | 0.671 |
| chr1 | 229760471 | 229763580 | 3110 | 0.13 | 0.608 | 0.671 |
| chr21 | 33102856 | 33105454 | 2599 | 0.1 | 0.608 | 0.671 |
| chr21 | 40636967 | 40637827 | 861 | -0.28 | 0.609 | 0.672 |
| chr14 | 64215660 | 64216244 | 585 | 0.3 | 0.609 | 0.672 |
| chr13 | 110999295 | 111001171 | 1877 | -0.16 | 0.609 | 0.672 |
| chr20 | 35676555 | 35705590 | 29036 | -0.09 | 0.609 | 0.672 |
| chr12 | 131477998 | 131485028 | 7031 | -0.26 | 0.609 | 0.672 |
| chr1 | 52913926 | 52914354 | 429 | -0.4 | 0.609 | 0.672 |
| chr3 | 44277097 | 44277534 | 438 | -0.3 | 0.609 | 0.672 |

| | | | | | | |
|---|---|---|---|---|---|---|
| chr2 | 106360101 | 106362414 | 2314 | 0.13 | 0.609 | 0.672 |
| chr10 | 3466245 | 3469261 | 3017 | 0.15 | 0.61 | 0.672 |
| chr18 | 21562248 | 21563020 | 773 | -0.3 | 0.61 | 0.673 |
| chr6 | 110499704 | 110502736 | 3033 | 0.09 | 0.61 | 0.673 |
| chr18 | 50023963 | 50027254 | 3292 | 0.08 | 0.61 | 0.673 |
| chr1 | 153755305 | 153756809 | 1505 | 0.1 | 0.61 | 0.673 |
| chr11 | 116657699 | 116671980 | 14282 | 0.09 | 0.61 | 0.673 |
| chr14 | 24422400 | 24423623 | 1224 | -0.16 | 0.611 | 0.674 |
| chr2 | 198396565 | 198402922 | 6358 | -0.12 | 0.611 | 0.674 |
| chr4 | 186795498 | 186802234 | 6737 | 0.19 | 0.611 | 0.674 |
| chr8 | 120867116 | 120887252 | 20137 | 0.1 | 0.611 | 0.674 |
| chr19 | 2028508 | 2070265 | 41758 | 0.03 | 0.611 | 0.674 |
| chr6 | 152667849 | 152668612 | 764 | -0.29 | 0.611 | 0.674 |
| chr10 | 102014832 | 102028393 | 13562 | 0.06 | 0.611 | 0.674 |
| chr6 | 155593839 | 155596185 | 2347 | -0.12 | 0.611 | 0.674 |
| chr3 | 49443055 | 49467738 | 24684 | 0.04 | 0.611 | 0.674 |
| chr13 | 45047879 | 45057964 | 10086 | -0.12 | 0.612 | 0.674 |
| chr15 | 40762017 | 40764131 | 2115 | -0.12 | 0.612 | 0.674 |
| chr4 | 189862326 | 189864144 | 1819 | 0.12 | 0.612 | 0.674 |
| chr8 | 135723893 | 135726971 | 3079 | -0.06 | 0.612 | 0.674 |
| chr2 | 151426891 | 151427856 | 966 | 0.22 | 0.612 | 0.674 |
| chr2 | 180091401 | 180105716 | 14316 | -0.08 | 0.612 | 0.674 |
| chr15 | 65843008 | 65854984 | 11977 | -0.1 | 0.612 | 0.675 |
| chr17 | 74707008 | 74707670 | 663 | 0.28 | 0.612 | 0.675 |
| chr3 | 128897233 | 128915270 | 18038 | 0.05 | 0.612 | 0.675 |
| chr8 | 124357328 | 124358563 | 1236 | -0.25 | 0.612 | 0.675 |
| chr9 | 108058262 | 108062798 | 4537 | 0.06 | 0.612 | 0.675 |
| chr11 | 106239797 | 106240521 | 725 | 0.31 | 0.612 | 0.675 |
| chrX | 100305758 | 100307873 | 2116 | 0.07 | 0.612 | 0.675 |
| chr4 | 110980797 | 110981482 | 686 | 0.3 | 0.613 | 0.675 |
| chr3 | 71581569 | 71586985 | 5417 | -0.12 | 0.613 | 0.675 |
| chr4 | 160023519 | 160025518 | 2000 | -0.1 | 0.613 | 0.675 |
| chr2 | 220492398 | 220492899 | 502 | -0.38 | 0.613 | 0.675 |
| chr8 | 122631727 | 122655949 | 24223 | -0.08 | 0.613 | 0.675 |
| chr6 | 43276268 | 43276689 | 422 | 0.41 | 0.613 | 0.675 |
| chr11 | 86012581 | 86014899 | 2319 | 0.09 | 0.613 | 0.676 |
| chr3 | 127333270 | 127334965 | 1696 | 0.25 | 0.614 | 0.676 |
| chr5 | 32353617 | 32358503 | 4887 | 0.16 | 0.614 | 0.676 |
| chrX | 75648113 | 75648792 | 680 | -0.27 | 0.614 | 0.676 |
| chr19 | 17797219 | 17797853 | 635 | -0.33 | 0.614 | 0.677 |
| chrX | 40261870 | 40263511 | 1642 | -0.13 | 0.614 | 0.677 |
| chr1 | 186648405 | 186652251 | 3847 | -0.06 | 0.614 | 0.677 |
| chr8 | 90334140 | 90335101 | 962 | 0.2 | 0.614 | 0.677 |
| chr14 | 65346187 | 65346998 | 812 | -0.3 | 0.615 | 0.677 |
| chr20 | 57266908 | 57275640 | 8733 | 0.08 | 0.615 | 0.677 |
| chr6 | 17551838 | 17552863 | 1026 | -0.25 | 0.615 | 0.677 |
| chr6 | 155491941 | 155493271 | 1331 | 0.15 | 0.615 | 0.677 |
| chr4 | 39450659 | 39474662 | 24004 | 0.05 | 0.615 | 0.678 |
| chr10 | 127477088 | 127477524 | 437 | 0.37 | 0.616 | 0.678 |
| chr2 | 27877989 | 27887848 | 9860 | 0.06 | 0.616 | 0.678 |
| chr12 | 96910237 | 96910605 | 369 | 0.43 | 0.616 | 0.678 |
| chr16 | 23114864 | 23116523 | 1660 | 0.21 | 0.616 | 0.678 |
| chr3 | 12666659 | 12672684 | 6026 | 0.12 | 0.616 | 0.678 |
| chr1 | 28240117 | 28241775 | 1659 | -0.14 | 0.616 | 0.678 |
| chr13 | 33042365 | 33043104 | 740 | 0.26 | 0.616 | 0.678 |
| chr2 | 174793746 | 174852306 | 58561 | -0.03 | 0.616 | 0.678 |
| chr3 | 148863085 | 148864547 | 1463 | -0.17 | 0.616 | 0.678 |
| chrX | 133540230 | 133540944 | 715 | 0.32 | 0.616 | 0.678 |
| chr8 | 81413398 | 81413883 | 486 | -0.37 | 0.616 | 0.678 |
| chr1 | 11119340 | 11121214 | 1875 | -0.17 | 0.616 | 0.678 |
| chr16 | 3095580 | 3115752 | 20173 | 0.15 | 0.617 | 0.679 |

| | | | | | | |
|---|---|---|---|---|---|---|
| chr17 | 42008686 | 42009262 | 577 | 0.31 | 0.617 | 0.679 |
| chr6 | 111782795 | 111811923 | 29129 | 0.05 | 0.617 | 0.679 |
| chr17 | 1244933 | 1304252 | 59320 | -0.03 | 0.617 | 0.679 |
| chr22 | 30367110 | 30375719 | 8610 | -0.18 | 0.617 | 0.679 |
| chr15 | 79965553 | 79982098 | 16546 | 0.14 | 0.617 | 0.679 |
| chr8 | 144329114 | 144330414 | 1301 | 0.16 | 0.617 | 0.679 |
| chr2 | 204349234 | 204363838 | 14605 | -0.09 | 0.617 | 0.679 |
| chr4 | 53066390 | 53067223 | 834 | 0.19 | 0.617 | 0.679 |
| chr16 | 85779597 | 85785195 | 5599 | -0.13 | 0.617 | 0.679 |
| chr17 | 6616507 | 6617126 | 620 | -0.36 | 0.617 | 0.679 |
| chr22 | 24058718 | 24060182 | 1465 | -0.13 | 0.617 | 0.679 |
| chr8 | 42269838 | 42270583 | 746 | 0.56 | 0.618 | 0.68 |
| chr10 | 93058241 | 93058609 | 369 | 0.37 | 0.618 | 0.68 |
| chr19 | 1437549 | 1439734 | 2186 | -0.14 | 0.618 | 0.68 |
| chr10 | 100205422 | 100207296 | 1875 | 0.11 | 0.618 | 0.68 |
| chr12 | 131238750 | 131239513 | 764 | 0.29 | 0.618 | 0.68 |
| chr1 | 202994941 | 202995802 | 862 | 0.24 | 0.618 | 0.68 |
| chr18 | 73087519 | 73088653 | 1135 | 0.25 | 0.619 | 0.68 |
| chr6 | 100877768 | 100884285 | 6518 | -0.09 | 0.619 | 0.68 |
| chr4 | 57842585 | 57858694 | 16110 | 0.06 | 0.619 | 0.68 |
| chr2 | 10175541 | 10184806 | 9266 | 0.11 | 0.619 | 0.681 |
| chr17 | 58212069 | 58213117 | 1049 | -0.22 | 0.619 | 0.681 |
| chr9 | 26975460 | 26977592 | 2133 | 0.12 | 0.619 | 0.681 |
| chr1 | 235026701 | 235035022 | 8322 | 0.15 | 0.619 | 0.681 |
| chr1 | 87690178 | 87692795 | 2618 | 0.16 | 0.619 | 0.681 |
| chr12 | 12808614 | 12809419 | 806 | -0.48 | 0.619 | 0.681 |
| chr14 | 102621292 | 102621857 | 566 | 0.25 | 0.62 | 0.681 |
| chr3 | 162609129 | 162613047 | 3919 | 0.07 | 0.62 | 0.682 |
| chr2 | 203239072 | 203243837 | 4766 | 0.06 | 0.62 | 0.682 |
| chr6 | 157254290 | 157254724 | 435 | 0.32 | 0.62 | 0.682 |
| chr8 | 101773564 | 101774993 | 1430 | -0.23 | 0.621 | 0.682 |
| chr10 | 126630933 | 126631802 | 870 | 0.28 | 0.621 | 0.682 |
| chr12 | 1628147 | 1629177 | 1031 | -0.27 | 0.621 | 0.683 |
| chr6 | 3259193 | 3260440 | 1248 | -0.21 | 0.621 | 0.683 |
| chr5 | 102486809 | 102487733 | 925 | -0.23 | 0.621 | 0.683 |
| chr15 | 63568653 | 63570500 | 1848 | 0.1 | 0.621 | 0.683 |
| chr1 | 161762724 | 161763339 | 616 | -0.28 | 0.621 | 0.683 |
| chr18 | 67833046 | 67874308 | 41263 | 0.04 | 0.622 | 0.683 |
| chr6 | 47665592 | 47667294 | 1703 | -0.13 | 0.622 | 0.683 |
| chr7 | 104737735 | 104751171 | 13437 | -0.08 | 0.622 | 0.683 |
| chr16 | 20125372 | 20125841 | 470 | -0.29 | 0.622 | 0.684 |
| chr6 | 36830262 | 36854756 | 24495 | 0.07 | 0.622 | 0.684 |
| chr20 | 60663844 | 60664531 | 688 | -0.27 | 0.622 | 0.684 |
| chr5 | 138011929 | 138012838 | 910 | -0.23 | 0.622 | 0.684 |
| chr17 | 683900 | 696168 | 12269 | 0.06 | 0.622 | 0.684 |
| chr10 | 75384450 | 75386979 | 2530 | -0.07 | 0.622 | 0.684 |
| chr8 | 103861638 | 103877906 | 16269 | 0.04 | 0.622 | 0.684 |
| chr2 | 15424026 | 15425754 | 1729 | 0.2 | 0.622 | 0.684 |
| chr20 | 48168578 | 48179275 | 10698 | -0.13 | 0.622 | 0.684 |
| chr16 | 57624247 | 57625422 | 1176 | -0.19 | 0.623 | 0.684 |
| chr18 | 33268125 | 33269230 | 1106 | -0.21 | 0.623 | 0.685 |
| chr1 | 183200495 | 183204676 | 4182 | 0.11 | 0.623 | 0.685 |
| chr16 | 671971 | 700407 | 28437 | 0.05 | 0.623 | 0.685 |
| chr4 | 120446006 | 120446493 | 488 | -0.33 | 0.623 | 0.685 |
| chr20 | 55990048 | 55990618 | 571 | -0.37 | 0.625 | 0.686 |
| chr7 | 56072733 | 56073117 | 385 | -0.46 | 0.625 | 0.686 |
| chr1 | 117098226 | 117114446 | 16221 | 0.07 | 0.625 | 0.686 |
| chr3 | 13063568 | 13064902 | 1335 | -0.22 | 0.625 | 0.686 |
| chr15 | 68393276 | 68407336 | 14061 | 0.19 | 0.625 | 0.687 |
| chr1 | 220959691 | 220960957 | 1267 | 0.18 | 0.625 | 0.687 |
| chr5 | 70750249 | 70806997 | 56749 | 0.03 | 0.626 | 0.687 |

| | | | | | | |
|---|---|---|---|---|---|---|
| chr8 | 39769877 | 39783147 | 13271 | 0.2 | 0.626 | 0.687 |
| chr15 | 50736318 | 50756480 | 20163 | 0.07 | 0.626 | 0.687 |
| chr4 | 76910869 | 76912783 | 1915 | 0.12 | 0.626 | 0.687 |
| chr17 | 70026047 | 70027302 | 1256 | 0.11 | 0.626 | 0.688 |
| chr6 | 152482521 | 152495877 | 13357 | -0.13 | 0.626 | 0.688 |
| chr13 | 26760246 | 26761586 | 1341 | 0.16 | 0.627 | 0.688 |
| chr2 | 46369016 | 46370282 | 1267 | -0.21 | 0.627 | 0.688 |
| chr13 | 41068964 | 41082991 | 14028 | -0.1 | 0.627 | 0.688 |
| chr6 | 146131336 | 146137136 | 5801 | 0.06 | 0.627 | 0.689 |
| chr20 | 46317298 | 46318795 | 1498 | -0.18 | 0.627 | 0.689 |
| chr10 | 118501237 | 118503295 | 2059 | 0.1 | 0.627 | 0.689 |
| chr7 | 45808688 | 45809339 | 652 | -0.22 | 0.628 | 0.689 |
| chr17 | 3767976 | 3769463 | 1488 | -0.25 | 0.628 | 0.689 |
| chr3 | 12871130 | 12884282 | 13153 | 0.05 | 0.628 | 0.689 |
| chr6 | 76421870 | 76422466 | 597 | -0.32 | 0.628 | 0.689 |
| chr6 | 7730870 | 7733118 | 2249 | -0.18 | 0.628 | 0.689 |
| chr2 | 166731092 | 166731619 | 528 | 0.25 | 0.628 | 0.689 |
| chr9 | 139398190 | 139399004 | 815 | -0.26 | 0.628 | 0.689 |
| chr12 | 91330143 | 91331235 | 1093 | 0.26 | 0.628 | 0.689 |
| chr2 | 3854768 | 3856626 | 1859 | 0.21 | 0.628 | 0.69 |
| chr19 | 40022017 | 40030910 | 8894 | 0.07 | 0.628 | 0.69 |
| chr4 | 175373534 | 175383870 | 10337 | -0.08 | 0.629 | 0.69 |
| chr5 | 172158093 | 172237014 | 78922 | 0.02 | 0.629 | 0.69 |
| chr7 | 5375860 | 5381138 | 5279 | -0.16 | 0.629 | 0.69 |
| chr14 | 45459587 | 45465935 | 6349 | 0.14 | 0.629 | 0.69 |
| chr10 | 22886879 | 22888712 | 1834 | -0.17 | 0.629 | 0.69 |
| chr7 | 138469461 | 138471901 | 2441 | 0.09 | 0.629 | 0.69 |
| chr13 | 21346727 | 21348836 | 2110 | 0.09 | 0.63 | 0.691 |
| chr13 | 50264455 | 50266502 | 2048 | -0.13 | 0.63 | 0.691 |
| chr7 | 64699834 | 64713339 | 13506 | 0.18 | 0.63 | 0.691 |
| chr18 | 26727953 | 26728545 | 593 | 0.24 | 0.63 | 0.691 |
| chr22 | 42031602 | 42044991 | 13390 | 0.08 | 0.63 | 0.691 |
| chr12 | 2958460 | 2967057 | 8598 | -0.12 | 0.63 | 0.691 |
| chrX | 23684849 | 23687341 | 2493 | 0.1 | 0.63 | 0.691 |
| chr11 | 77529859 | 77532994 | 3136 | 0.1 | 0.63 | 0.691 |
| chr9 | 107815242 | 107834527 | 19286 | -0.06 | 0.63 | 0.691 |
| chr5 | 68512553 | 68548667 | 36115 | 0.05 | 0.63 | 0.691 |
| chr21 | 15754408 | 15756627 | 2220 | 0.09 | 0.63 | 0.691 |
| chr22 | 22511294 | 22511722 | 429 | -0.26 | 0.63 | 0.691 |
| chrX | 51486058 | 51487239 | 1182 | 0.15 | 0.631 | 0.691 |
| chr1 | 109418999 | 109420839 | 1841 | 0.18 | 0.631 | 0.692 |
| chr7 | 30418241 | 30419308 | 1068 | -0.23 | 0.631 | 0.692 |
| chr7 | 115979482 | 116004288 | 24807 | 0.03 | 0.632 | 0.693 |
| chr1 | 59758055 | 59763647 | 5593 | 0.09 | 0.632 | 0.693 |
| chr5 | 156938042 | 156939552 | 1511 | -0.12 | 0.632 | 0.693 |
| chr11 | 34876118 | 34877147 | 1030 | 0.29 | 0.632 | 0.693 |
| chr11 | 68064802 | 68127014 | 62213 | 0.03 | 0.632 | 0.693 |
| chr1 | 44241295 | 44250987 | 9693 | 0.06 | 0.632 | 0.693 |
| chr2 | 43123768 | 43134584 | 10817 | 0.17 | 0.632 | 0.693 |
| chr7 | 6676150 | 6676895 | 746 | 0.22 | 0.632 | 0.693 |
| chr14 | 91282036 | 91295221 | 13186 | 0.96 | 0.632 | 0.693 |
| chr12 | 28464149 | 28465067 | 919 | -0.25 | 0.633 | 0.694 |
| chr1 | 12078811 | 12079302 | 492 | 0.28 | 0.633 | 0.694 |
| chr17 | 36830066 | 36831827 | 1762 | 0.14 | 0.633 | 0.694 |
| chr2 | 146351206 | 146352034 | 829 | -0.25 | 0.633 | 0.694 |
| chr4 | 156296882 | 156298876 | 1995 | -0.09 | 0.634 | 0.694 |
| chr20 | 42830424 | 42840663 | 10240 | -0.03 | 0.634 | 0.695 |
| chr18 | 45298781 | 45299470 | 690 | -0.26 | 0.634 | 0.695 |
| chr8 | 42867772 | 42868098 | 327 | 0.4 | 0.634 | 0.695 |
| chr2 | 106731230 | 106747551 | 16322 | 0.05 | 0.634 | 0.695 |
| chr3 | 138488329 | 138507146 | 18818 | 0.08 | 0.635 | 0.695 |

| | | | | | | |
|---|---|---|---|---|---|---|
| chr1 | 212261472 | 212262441 | 970 | 0.22 | 0.635 | 0.695 |
| chr11 | 66713584 | 66774740 | 61157 | 0.04 | 0.635 | 0.696 |
| chr4 | 68325248 | 68325622 | 375 | 0.38 | 0.635 | 0.696 |
| chr16 | 69845338 | 69868100 | 22763 | -0.04 | 0.635 | 0.696 |
| chr10 | 60194933 | 60196256 | 1324 | 0.13 | 0.635 | 0.696 |
| chr1 | 114955084 | 114956275 | 1192 | -0.23 | 0.635 | 0.696 |
| chr1 | 93249404 | 93251778 | 2375 | 0.09 | 0.635 | 0.696 |
| chr3 | 187660484 | 187675666 | 15183 | -0.06 | 0.635 | 0.696 |
| chr6 | 106635334 | 106635784 | 451 | 0.33 | 0.636 | 0.696 |
| chr4 | 166128056 | 166129706 | 1651 | -0.13 | 0.636 | 0.696 |
| chr6 | 1554709 | 1555511 | 803 | -0.18 | 0.636 | 0.696 |
| chr17 | 56807935 | 56813097 | 5163 | -0.1 | 0.636 | 0.696 |
| chr8 | 103140736 | 103141265 | 530 | -0.34 | 0.636 | 0.696 |
| chr1 | 4710043 | 4713393 | 3351 | -0.14 | 0.636 | 0.696 |
| chr20 | 55447705 | 55451059 | 3355 | -0.13 | 0.636 | 0.696 |
| chr10 | 93860623 | 93862258 | 1636 | 0.19 | 0.636 | 0.696 |
| chr16 | 88869162 | 88879739 | 10578 | 0.06 | 0.636 | 0.697 |
| chr1 | 3772654 | 3774505 | 1852 | -0.15 | 0.637 | 0.697 |
| chr2 | 8817431 | 8826046 | 8616 | -0.05 | 0.637 | 0.697 |
| chr1 | 27400743 | 27401360 | 618 | 0.31 | 0.637 | 0.697 |
| chr1 | 155140069 | 155147028 | 6960 | -0.1 | 0.637 | 0.697 |
| chr9 | 77473533 | 77474948 | 1416 | 0.21 | 0.637 | 0.698 |
| chr13 | 31618799 | 31620957 | 2159 | -0.09 | 0.637 | 0.698 |
| chr6 | 45623583 | 45624422 | 840 | 0.25 | 0.637 | 0.698 |
| chr6 | 26659031 | 26660511 | 1481 | 0.17 | 0.638 | 0.698 |
| chr5 | 74146820 | 74162948 | 16129 | 0.06 | 0.638 | 0.698 |
| chrX | 134156515 | 134166749 | 10235 | 0.22 | 0.638 | 0.698 |
| chr4 | 122076381 | 122078785 | 2405 | -0.11 | 0.638 | 0.698 |
| chr2 | 135809385 | 135811100 | 1716 | -0.1 | 0.638 | 0.698 |
| chr13 | 50134264 | 50135581 | 1318 | 0.19 | 0.638 | 0.699 |
| chr1 | 94701670 | 94724516 | 22847 | -0.1 | 0.638 | 0.699 |
| chr5 | 175958061 | 175971709 | 13649 | -0.05 | 0.638 | 0.699 |
| chr20 | 61586702 | 61587577 | 876 | 0.25 | 0.639 | 0.699 |
| chr1 | 190400507 | 190401404 | 898 | -0.22 | 0.639 | 0.699 |
| chr9 | 93821636 | 93834649 | 13014 | 0.08 | 0.639 | 0.699 |
| chr16 | 66929387 | 66934552 | 5166 | -0.18 | 0.639 | 0.699 |
| chr8 | 103367173 | 103381102 | 13930 | -0.1 | 0.639 | 0.699 |
| chr11 | 16686460 | 16692109 | 5650 | -0.24 | 0.639 | 0.699 |
| chr10 | 35809858 | 35810620 | 763 | 0.26 | 0.639 | 0.7 |
| chr15 | 93180556 | 93182328 | 1773 | -0.14 | 0.639 | 0.7 |
| chr2 | 164384780 | 164386186 | 1407 | 0.19 | 0.639 | 0.7 |
| chr8 | 62438521 | 62439938 | 1418 | 0.22 | 0.64 | 0.7 |
| chrX | 102022881 | 102025535 | 2655 | -0.05 | 0.64 | 0.7 |
| chr6 | 129017604 | 129018220 | 617 | -0.3 | 0.64 | 0.7 |
| chr5 | 169122661 | 169124198 | 1538 | -0.16 | 0.64 | 0.7 |
| chr18 | 158530 | 160471 | 1942 | 0.1 | 0.64 | 0.7 |
| chr20 | 34286411 | 34303909 | 17499 | 0.05 | 0.64 | 0.7 |
| chr2 | 19337759 | 19342276 | 4518 | -0.05 | 0.64 | 0.7 |
| chr17 | 59868518 | 59868898 | 381 | -0.53 | 0.64 | 0.7 |
| chr15 | 35310570 | 35311331 | 762 | 0.24 | 0.64 | 0.7 |
| chr19 | 11669313 | 11670953 | 1641 | -0.16 | 0.64 | 0.7 |
| chr8 | 145549236 | 145551407 | 2172 | 0.16 | 0.64 | 0.7 |
| chr11 | 70198026 | 70198650 | 625 | -0.27 | 0.64 | 0.7 |
| chr15 | 35240671 | 35262902 | 22232 | 0.05 | 0.641 | 0.7 |
| chr1 | 2071741 | 2072457 | 717 | 0.23 | 0.641 | 0.7 |
| chr11 | 119093179 | 119093623 | 445 | 0.34 | 0.641 | 0.7 |
| chr4 | 88310929 | 88313353 | 2425 | 0.08 | 0.641 | 0.701 |
| chr13 | 50159158 | 50160538 | 1381 | 0.17 | 0.641 | 0.701 |
| chrX | 24041443 | 24045344 | 3902 | -0.06 | 0.641 | 0.701 |
| chr4 | 923781 | 926894 | 3114 | 0.1 | 0.642 | 0.701 |
| chr12 | 123066736 | 123068779 | 2044 | -0.25 | 0.642 | 0.702 |

| | | | | | | |
|---|---|---|---|---|---|---|
| chr17 | 74786874 | 74788144 | 1271 | 0.13 | 0.643 | 0.703 |
| chr17 | 59101426 | 59102095 | 670 | 0.27 | 0.644 | 0.703 |
| chr11 | 19411261 | 19412048 | 788 | 0.27 | 0.644 | 0.703 |
| chr5 | 159435165 | 159451632 | 16468 | 0.06 | 0.644 | 0.703 |
| chr1 | 15587348 | 15588527 | 1180 | 0.15 | 0.644 | 0.704 |
| chr10 | 95653254 | 95654626 | 1373 | -0.13 | 0.644 | 0.704 |
| chr13 | 51796565 | 51796984 | 420 | -0.32 | 0.644 | 0.704 |
| chr11 | 86747082 | 86750552 | 3471 | 0.06 | 0.644 | 0.704 |
| chr12 | 96651558 | 96658210 | 6653 | 0.09 | 0.644 | 0.704 |
| chr7 | 105152051 | 105173943 | 21893 | 0.05 | 0.644 | 0.704 |
| chr4 | 106066995 | 106069146 | 2152 | -0.07 | 0.644 | 0.704 |
| chr3 | 111685850 | 111698634 | 12785 | 0.13 | 0.644 | 0.704 |
| chr5 | 140648631 | 140649398 | 768 | -0.26 | 0.644 | 0.704 |
| chr1 | 235812813 | 235813205 | 393 | 0.36 | 0.645 | 0.704 |
| chr3 | 177526844 | 177548649 | 21806 | 0.04 | 0.645 | 0.705 |
| chr8 | 54672604 | 54681312 | 8709 | -0.17 | 0.645 | 0.705 |
| chr11 | 118628337 | 118642870 | 14534 | 0.09 | 0.645 | 0.705 |
| chr11 | 2720347 | 2722780 | 2434 | -0.11 | 0.645 | 0.705 |
| chr12 | 133067256 | 133067795 | 540 | 0.3 | 0.645 | 0.705 |
| chr20 | 21305295 | 21317980 | 12686 | -0.08 | 0.645 | 0.705 |
| chr2 | 73404225 | 73404600 | 376 | 0.3 | 0.646 | 0.705 |
| chrX | 17517774 | 17519596 | 1823 | -0.15 | 0.646 | 0.705 |
| chr1 | 234444792 | 234446496 | 1705 | 0.12 | 0.646 | 0.706 |
| chr10 | 104144693 | 104162577 | 17885 | -0.06 | 0.646 | 0.706 |
| chr15 | 99948773 | 99976236 | 27464 | 0.17 | 0.646 | 0.706 |
| chr12 | 112315375 | 112325557 | 10183 | -0.16 | 0.646 | 0.706 |
| chr14 | 100806371 | 100808794 | 2424 | 0.12 | 0.647 | 0.706 |
| chr12 | 22642446 | 22643174 | 729 | 0.23 | 0.647 | 0.706 |
| chr18 | 2706384 | 2717438 | 11055 | -0.14 | 0.647 | 0.707 |
| chr14 | 36178566 | 36179875 | 1310 | -0.2 | 0.647 | 0.707 |
| chr12 | 122687784 | 122688373 | 590 | 0.28 | 0.647 | 0.707 |
| chr3 | 131973548 | 131974768 | 1221 | -0.18 | 0.647 | 0.707 |
| chrX | 21391841 | 21393331 | 1491 | -0.19 | 0.647 | 0.707 |
| chr14 | 91883480 | 91885280 | 1801 | 0.13 | 0.647 | 0.707 |
| chr20 | 14824955 | 14827040 | 2086 | 0.11 | 0.648 | 0.707 |
| chr8 | 27584378 | 27585187 | 810 | 0.21 | 0.648 | 0.707 |
| chr3 | 48228041 | 48230730 | 2690 | -0.11 | 0.648 | 0.707 |
| chr4 | 114523300 | 114540000 | 16701 | 0.08 | 0.648 | 0.707 |
| chr7 | 83807070 | 83807541 | 472 | 0.32 | 0.648 | 0.707 |
| chr11 | 73426652 | 73429179 | 2528 | 0.1 | 0.648 | 0.707 |
| chr10 | 27119017 | 27119686 | 670 | -0.33 | 0.648 | 0.707 |
| chr10 | 30471288 | 30475841 | 4554 | -0.17 | 0.648 | 0.708 |
| chr5 | 122241854 | 122243453 | 1600 | 0.19 | 0.648 | 0.708 |
| chr7 | 76256207 | 76256519 | 313 | -0.31 | 0.649 | 0.708 |
| chr5 | 131962530 | 131963210 | 681 | -0.32 | 0.649 | 0.708 |
| chr21 | 30704726 | 30705973 | 1248 | 0.25 | 0.649 | 0.708 |
| chr12 | 75571211 | 75571886 | 676 | -0.25 | 0.649 | 0.708 |
| chr2 | 71164303 | 71164906 | 604 | 0.29 | 0.649 | 0.708 |
| chr2 | 198257448 | 198270473 | 13026 | -0.12 | 0.649 | 0.709 |
| chrX | 15872022 | 15874030 | 2009 | 0.11 | 0.649 | 0.709 |
| chr17 | 11951165 | 11951626 | 462 | -0.31 | 0.65 | 0.709 |
| chr11 | 118926780 | 118939156 | 12377 | -0.07 | 0.65 | 0.709 |
| chr15 | 57440722 | 57441155 | 434 | -0.31 | 0.65 | 0.709 |
| chr2 | 224508011 | 224509250 | 1240 | 0.22 | 0.65 | 0.709 |
| chr18 | 67799957 | 67816052 | 16096 | -0.09 | 0.65 | 0.709 |
| chr13 | 101240017 | 101241359 | 1343 | 0.14 | 0.65 | 0.71 |
| chr12 | 50056761 | 50057143 | 383 | -0.32 | 0.651 | 0.71 |
| chr4 | 25313796 | 25315405 | 1610 | 0.14 | 0.651 | 0.71 |
| chr16 | 70834162 | 70835062 | 901 | 0.13 | 0.651 | 0.71 |
| chr8 | 95542355 | 95544056 | 1702 | 0.2 | 0.651 | 0.71 |
| chr1 | 53307469 | 53310181 | 2713 | -0.1 | 0.651 | 0.71 |

| | | | | | | |
|---|---|---|---|---|---|---|
| chr3 | 48505537 | 48515552 | 10016 | 0.04 | 0.651 | 0.71 |
| chr10 | 15412003 | 15413087 | 1085 | -0.17 | 0.651 | 0.71 |
| chr2 | 171729441 | 171730159 | 719 | 0.23 | 0.652 | 0.711 |
| chr2 | 107198796 | 107199154 | 359 | 0.26 | 0.652 | 0.711 |
| chr3 | 171756792 | 171785214 | 28423 | 0.04 | 0.652 | 0.711 |
| chr7 | 149534770 | 149536546 | 1777 | 0.13 | 0.652 | 0.711 |
| chr17 | 9478101 | 9480899 | 2799 | -0.09 | 0.652 | 0.711 |
| chr16 | 28911373 | 28937644 | 26272 | -0.05 | 0.652 | 0.711 |
| chr1 | 119681648 | 119685224 | 3577 | 0.05 | 0.652 | 0.711 |
| chr12 | 109938266 | 109939099 | 834 | -0.22 | 0.652 | 0.711 |
| chr22 | 22161193 | 22172052 | 10860 | 0.09 | 0.652 | 0.711 |
| chr11 | 66335683 | 66336674 | 992 | -0.22 | 0.652 | 0.711 |
| chr6 | 129918788 | 129934678 | 15891 | 0.07 | 0.653 | 0.711 |
| chr1 | 29240414 | 29241327 | 914 | -0.25 | 0.653 | 0.712 |
| chr9 | 117372738 | 117380124 | 7387 | -0.06 | 0.653 | 0.712 |
| chr17 | 64946894 | 64949703 | 2810 | -0.06 | 0.653 | 0.712 |
| chr1 | 92495275 | 92495944 | 670 | 0.25 | 0.653 | 0.712 |
| chr10 | 179158 | 182641 | 3484 | -0.06 | 0.653 | 0.712 |
| chr13 | 96107956 | 96108574 | 619 | 0.28 | 0.653 | 0.712 |
| chr17 | 33147802 | 33148545 | 744 | -0.24 | 0.653 | 0.712 |
| chr19 | 14473769 | 14475082 | 1314 | -0.14 | 0.653 | 0.712 |
| chr5 | 52083769 | 52106808 | 23040 | 0.05 | 0.654 | 0.712 |
| chr15 | 74832407 | 74833322 | 916 | 0.14 | 0.654 | 0.712 |
| chr2 | 75935875 | 75938769 | 2895 | -0.05 | 0.654 | 0.713 |
| chr17 | 29192410 | 29199954 | 7545 | -0.11 | 0.654 | 0.713 |
| chr2 | 214171139 | 214182295 | 11157 | -0.11 | 0.654 | 0.713 |
| chr10 | 74836507 | 74837635 | 1129 | 0.22 | 0.654 | 0.713 |
| chr5 | 73147730 | 73164254 | 16525 | -0.06 | 0.654 | 0.713 |
| chr5 | 179284679 | 179297986 | 13308 | -0.07 | 0.654 | 0.713 |
| chr17 | 7736901 | 7769990 | 33090 | -0.03 | 0.655 | 0.713 |
| chr22 | 29975900 | 29977945 | 2046 | -0.14 | 0.655 | 0.713 |
| chr12 | 54238423 | 54240011 | 1589 | -0.15 | 0.655 | 0.713 |
| chr9 | 117264062 | 117268671 | 4610 | 0.09 | 0.655 | 0.713 |
| chrX | 23728009 | 23730365 | 2357 | -0.15 | 0.655 | 0.713 |
| chr10 | 34844043 | 34875316 | 31274 | -0.04 | 0.655 | 0.714 |
| chr5 | 118176197 | 118176644 | 448 | 0.51 | 0.655 | 0.714 |
| chr7 | 158189303 | 158196387 | 7085 | -0.17 | 0.656 | 0.714 |
| chr6 | 131614570 | 131614994 | 425 | 0.3 | 0.656 | 0.714 |
| chr1 | 3815734 | 3818294 | 2561 | 0.08 | 0.656 | 0.714 |
| chr17 | 4690358 | 4711183 | 20826 | -0.05 | 0.656 | 0.714 |
| chr17 | 46749206 | 46749769 | 564 | 0.23 | 0.656 | 0.714 |
| chr8 | 8637371 | 8676243 | 38873 | -0.03 | 0.656 | 0.714 |
| chr17 | 39735326 | 39736414 | 1089 | 0.2 | 0.656 | 0.714 |
| chr2 | 214883412 | 214884780 | 1369 | -0.13 | 0.656 | 0.714 |
| chrX | 18270847 | 18271382 | 536 | 0.33 | 0.656 | 0.714 |
| chr7 | 51101594 | 51102945 | 1352 | 0.19 | 0.656 | 0.714 |
| chr2 | 97221949 | 97223898 | 1950 | -0.13 | 0.656 | 0.715 |
| chr1 | 200777752 | 200779165 | 1414 | -0.21 | 0.656 | 0.715 |
| chr17 | 5283136 | 5285723 | 2588 | -0.15 | 0.657 | 0.715 |
| chr7 | 75420524 | 75421441 | 918 | 0.27 | 0.657 | 0.715 |
| chr12 | 48030098 | 48046671 | 16574 | 0.08 | 0.657 | 0.715 |
| chrX | 52004727 | 52005557 | 831 | 0.16 | 0.657 | 0.715 |
| chr2 | 58269416 | 58282131 | 12716 | 0.05 | 0.657 | 0.715 |
| chr2 | 10037676 | 10038131 | 456 | 0.3 | 0.658 | 0.716 |
| chr10 | 76917340 | 76928594 | 11255 | 0.07 | 0.658 | 0.716 |
| chr2 | 196659921 | 196661169 | 1249 | 0.2 | 0.658 | 0.716 |
| chr4 | 87102372 | 87103854 | 1483 | 0.14 | 0.658 | 0.716 |
| chr15 | 83315755 | 83317110 | 1356 | 0.11 | 0.658 | 0.717 |
| chr9 | 14030303 | 14031180 | 878 | -0.21 | 0.659 | 0.717 |
| chr6 | 1636742 | 1637795 | 1054 | -0.2 | 0.659 | 0.717 |
| chr8 | 27507553 | 27567062 | 59510 | 0.05 | 0.659 | 0.717 |

| | | | | | | |
|---|---|---|---|---|---|---|
| chr18 | 33876337 | 33902208 | 25872 | -0.1 | 0.659 | 0.717 |
| chr5 | 140855560 | 140856096 | 537 | 0.24 | 0.659 | 0.717 |
| chr9 | 6501674 | 6503297 | 1624 | -0.16 | 0.659 | 0.717 |
| chr14 | 68400064 | 68412857 | 12794 | -0.04 | 0.659 | 0.717 |
| chr7 | 16683288 | 16711138 | 27851 | -0.03 | 0.66 | 0.718 |
| chr9 | 132585064 | 132598452 | 13389 | -0.07 | 0.66 | 0.718 |
| chr11 | 19681536 | 19702633 | 21098 | -0.16 | 0.66 | 0.718 |
| chr8 | 122240367 | 122240867 | 501 | -0.24 | 0.66 | 0.718 |
| chr14 | 76669178 | 76670786 | 1609 | 0.16 | 0.66 | 0.718 |
| chr3 | 157885060 | 157886947 | 1888 | 0.14 | 0.66 | 0.718 |
| chr8 | 42475907 | 42476966 | 1060 | -0.2 | 0.66 | 0.718 |
| chr17 | 26360970 | 26369826 | 8857 | -0.11 | 0.661 | 0.719 |
| chr13 | 33269502 | 33270896 | 1395 | 0.19 | 0.661 | 0.719 |
| chr7 | 72834676 | 72854170 | 19495 | -0.11 | 0.661 | 0.719 |
| chr9 | 139701698 | 139703661 | 1964 | -0.13 | 0.661 | 0.719 |
| chr22 | 38029436 | 38030009 | 574 | 0.29 | 0.661 | 0.719 |
| chr9 | 84107471 | 84108643 | 1173 | 0.2 | 0.661 | 0.719 |
| chr3 | 185798100 | 185798835 | 736 | 0.29 | 0.661 | 0.719 |
| chr7 | 3185879 | 3187640 | 1762 | -0.1 | 0.661 | 0.719 |
| chr17 | 60046975 | 60048217 | 1243 | -0.18 | 0.661 | 0.719 |
| chr11 | 116759430 | 116759807 | 378 | 0.35 | 0.662 | 0.719 |
| chr22 | 27136235 | 27174690 | 38456 | -0.03 | 0.662 | 0.72 |
| chr2 | 219176894 | 219195165 | 18272 | -0.07 | 0.662 | 0.72 |
| chr2 | 3409538 | 3410721 | 1184 | 0.23 | 0.662 | 0.72 |
| chr11 | 46139985 | 46155381 | 15397 | -0.06 | 0.662 | 0.72 |
| chr17 | 27219715 | 27254156 | 34442 | 0.05 | 0.662 | 0.72 |
| chr22 | 43484530 | 43486933 | 2404 | 0.1 | 0.662 | 0.72 |
| chr3 | 5202298 | 5203139 | 842 | -0.22 | 0.662 | 0.72 |
| chr22 | 36017747 | 36026931 | 9185 | -0.05 | 0.663 | 0.72 |
| chr9 | 98875440 | 98879782 | 4343 | 0.11 | 0.663 | 0.72 |
| chr1 | 29583540 | 29610781 | 27242 | -0.04 | 0.663 | 0.721 |
| chr7 | 94753837 | 94754908 | 1072 | 0.21 | 0.663 | 0.721 |
| chr14 | 71373361 | 71376054 | 2694 | 0.08 | 0.664 | 0.721 |
| chr11 | 123610920 | 123612852 | 1933 | -0.1 | 0.664 | 0.721 |
| chr8 | 38749227 | 38796408 | 47182 | -0.04 | 0.664 | 0.721 |
| chr17 | 30263026 | 30264470 | 1445 | -0.12 | 0.664 | 0.722 |
| chr4 | 99915571 | 99918210 | 2640 | 0.11 | 0.664 | 0.722 |
| chr10 | 97388454 | 97389727 | 1274 | 0.22 | 0.664 | 0.722 |
| chr9 | 15468469 | 15468810 | 342 | 0.28 | 0.664 | 0.722 |
| chrX | 140270121 | 140271911 | 1791 | 0.12 | 0.664 | 0.722 |
| chr7 | 123174057 | 123175080 | 1024 | 0.15 | 0.664 | 0.722 |
| chr1 | 90097730 | 90100342 | 2613 | 0.09 | 0.665 | 0.722 |
| chr3 | 176826337 | 176828707 | 2371 | 0.16 | 0.665 | 0.722 |
| chr1 | 156774184 | 156784277 | 10094 | -0.14 | 0.665 | 0.722 |
| chr1 | 234274172 | 234276851 | 2680 | -0.11 | 0.665 | 0.723 |
| chr17 | 30333791 | 30335501 | 1711 | -0.16 | 0.665 | 0.723 |
| chr1 | 226288278 | 226323804 | 35527 | 0.04 | 0.665 | 0.723 |
| chr15 | 68638727 | 68646568 | 7842 | 0.06 | 0.665 | 0.723 |
| chrX | 22407492 | 22408406 | 915 | 0.25 | 0.665 | 0.723 |
| chr10 | 125813136 | 125824137 | 11002 | -0.1 | 0.665 | 0.723 |
| chr6 | 86069210 | 86072720 | 3511 | 0.09 | 0.666 | 0.723 |
| chr2 | 214859268 | 214861144 | 1877 | 0.09 | 0.666 | 0.723 |
| chrX | 129693910 | 129696141 | 2232 | -0.12 | 0.666 | 0.724 |
| chr7 | 100994987 | 100996469 | 1483 | 0.19 | 0.667 | 0.724 |
| chr2 | 26496396 | 26497535 | 1140 | -0.2 | 0.667 | 0.724 |
| chr7 | 6511798 | 6524557 | 12760 | 0.08 | 0.667 | 0.724 |
| chr11 | 43404216 | 43405225 | 1010 | -0.28 | 0.667 | 0.724 |
| chr19 | 5790468 | 5791855 | 1388 | 0.09 | 0.667 | 0.724 |
| chr19 | 19626309 | 19627971 | 1663 | 0.09 | 0.667 | 0.724 |
| chr4 | 129162279 | 129162978 | 700 | 0.23 | 0.668 | 0.725 |
| chr12 | 71432936 | 71433270 | 335 | -0.34 | 0.668 | 0.725 |

| | | | | | | |
|---|---|---|---|---|---|---|
| chr9 | 133884199 | 133884953 | 755 | 0.28 | 0.668 | 0.725 |
| chr7 | 2899834 | 2916848 | 17015 | 0.06 | 0.668 | 0.725 |
| chr12 | 52890004 | 52890778 | 775 | -0.23 | 0.668 | 0.725 |
| chr19 | 19886323 | 19888141 | 1819 | 0.13 | 0.668 | 0.725 |
| chr14 | 31343061 | 31344459 | 1399 | -0.14 | 0.668 | 0.725 |
| chr21 | 46220697 | 46222794 | 2098 | 0.13 | 0.668 | 0.725 |
| chr3 | 157815423 | 157829029 | 13607 | -0.07 | 0.668 | 0.725 |
| chr9 | 35217756 | 35218236 | 481 | 0.24 | 0.668 | 0.726 |
| chr22 | 31639669 | 31640724 | 1056 | 0.23 | 0.669 | 0.726 |
| chr18 | 21017231 | 21018280 | 1050 | -0.22 | 0.669 | 0.726 |
| chr5 | 131131568 | 131133829 | 2262 | 0.07 | 0.669 | 0.726 |
| chr1 | 52519425 | 52523862 | 4438 | -0.06 | 0.669 | 0.726 |
| chr12 | 117197525 | 117199063 | 1539 | 0.21 | 0.669 | 0.726 |
| chr22 | 31997983 | 31998911 | 929 | -0.19 | 0.669 | 0.726 |
| chr17 | 38404945 | 38415587 | 10643 | 0.08 | 0.669 | 0.726 |
| chr12 | 94017636 | 94019930 | 2295 | 0.06 | 0.669 | 0.726 |
| chr6 | 160866042 | 160877326 | 11285 | 0.08 | 0.669 | 0.726 |
| chr14 | 55238049 | 55242021 | 3973 | -0.05 | 0.67 | 0.726 |
| chr19 | 19248686 | 19249306 | 621 | -0.15 | 0.67 | 0.727 |
| chr7 | 25067612 | 25068956 | 1345 | 0.17 | 0.67 | 0.727 |
| chr9 | 92112164 | 92113689 | 1526 | -0.17 | 0.67 | 0.727 |
| chr11 | 122007084 | 122010299 | 3216 | -0.08 | 0.67 | 0.727 |
| chr2 | 43746501 | 43747808 | 1308 | 0.1 | 0.67 | 0.727 |
| chr11 | 77258598 | 77259445 | 848 | 0.21 | 0.67 | 0.727 |
| chr4 | 111045707 | 111056871 | 11165 | -0.1 | 0.67 | 0.727 |
| chr2 | 65201661 | 65216961 | 15301 | -0.09 | 0.67 | 0.727 |
| chr22 | 29781136 | 29789859 | 8724 | -0.11 | 0.67 | 0.727 |
| chr3 | 39166961 | 39167915 | 955 | -0.19 | 0.671 | 0.727 |
| chr13 | 23893507 | 23896714 | 3208 | -0.08 | 0.671 | 0.727 |
| chr8 | 106860327 | 106862112 | 1786 | -0.16 | 0.671 | 0.727 |
| chr1 | 59478900 | 59486469 | 7570 | -0.28 | 0.671 | 0.727 |
| chr14 | 24727781 | 24741299 | 13519 | -0.08 | 0.671 | 0.728 |
| chr7 | 102787989 | 102790575 | 2587 | -0.07 | 0.671 | 0.728 |
| chr7 | 38866783 | 38867301 | 519 | 0.25 | 0.671 | 0.728 |
| chr16 | 89962377 | 89962782 | 406 | -0.32 | 0.671 | 0.728 |
| chr15 | 43376816 | 43377992 | 1177 | -0.24 | 0.671 | 0.728 |
| chr1 | 27069874 | 27073088 | 3215 | 0.11 | 0.671 | 0.728 |
| chrX | 9637709 | 9639094 | 1386 | -0.18 | 0.671 | 0.728 |
| chr5 | 180559658 | 180562110 | 2453 | 0.07 | 0.672 | 0.728 |
| chr20 | 33411442 | 33414096 | 2655 | 0.09 | 0.672 | 0.728 |
| chr22 | 43689508 | 43690418 | 911 | 0.22 | 0.672 | 0.728 |
| chr19 | 35615119 | 35616019 | 901 | 0.17 | 0.672 | 0.729 |
| chr8 | 11161658 | 11162932 | 1275 | -0.15 | 0.672 | 0.729 |
| chr1 | 40996536 | 40997971 | 1436 | 0.09 | 0.673 | 0.729 |
| chr10 | 121740955 | 121741943 | 989 | -0.17 | 0.673 | 0.729 |
| chr9 | 130590477 | 130591235 | 759 | 0.23 | 0.673 | 0.729 |
| chr3 | 160787835 | 160788362 | 528 | -0.28 | 0.673 | 0.729 |
| chrX | 71400520 | 71402496 | 1977 | 0.09 | 0.673 | 0.729 |
| chr17 | 2786278 | 2796703 | 10426 | 0.08 | 0.673 | 0.73 |
| chr13 | 108868045 | 108868504 | 460 | 0.23 | 0.673 | 0.73 |
| chr8 | 42834116 | 42834866 | 751 | -0.23 | 0.673 | 0.73 |
| chr22 | 38885834 | 38887232 | 1399 | -0.14 | 0.673 | 0.73 |
| chr5 | 70827707 | 70830099 | 2393 | -0.12 | 0.674 | 0.73 |
| chr8 | 143374962 | 143375541 | 580 | -0.36 | 0.674 | 0.73 |
| chr9 | 15485650 | 15487329 | 1680 | 0.13 | 0.674 | 0.73 |
| chr19 | 4791696 | 4792126 | 431 | -0.29 | 0.674 | 0.73 |
| chrX | 36594720 | 36595375 | 656 | 0.25 | 0.674 | 0.731 |
| chr1 | 43854883 | 43856685 | 1803 | 0.11 | 0.674 | 0.731 |
| chr2 | 7117317 | 7117620 | 304 | -0.28 | 0.674 | 0.731 |
| chr3 | 4534057 | 4534910 | 854 | -0.17 | 0.675 | 0.731 |
| chr11 | 73336998 | 73337420 | 423 | 0.35 | 0.675 | 0.731 |

| | | | | | | |
|---|---|---|---|---|---|---|
| chr1 | 246690917 | 246691375 | 459 | -0.31 | 0.675 | 0.731 |
| chrX | 153990104 | 153992384 | 2281 | -0.08 | 0.675 | 0.731 |
| chr3 | 42829623 | 42829992 | 370 | 0.26 | 0.675 | 0.731 |
| chr12 | 109714588 | 109717040 | 2453 | 0.14 | 0.676 | 0.732 |
| chr17 | 72868311 | 72870816 | 2506 | 0.06 | 0.676 | 0.732 |
| chr5 | 131627964 | 131676329 | 48366 | 0.04 | 0.676 | 0.732 |
| chr1 | 64971344 | 64971975 | 632 | -0.24 | 0.676 | 0.732 |
| chr13 | 49323785 | 49324544 | 760 | 0.26 | 0.676 | 0.732 |
| chr8 | 106998280 | 106998848 | 569 | 0.23 | 0.676 | 0.732 |
| chr3 | 10052275 | 10052848 | 574 | 0.28 | 0.676 | 0.732 |
| chr4 | 87606256 | 87606949 | 694 | -0.26 | 0.676 | 0.732 |
| chr10 | 121353283 | 121357480 | 4198 | -0.06 | 0.676 | 0.732 |
| chr19 | 3094705 | 3117485 | 22781 | -0.06 | 0.676 | 0.732 |
| chr19 | 10763049 | 10767335 | 4287 | -0.05 | 0.677 | 0.733 |
| chr11 | 47346728 | 47359627 | 12900 | 0.23 | 0.677 | 0.733 |
| chr2 | 24560552 | 24561093 | 542 | 0.27 | 0.677 | 0.733 |
| chr14 | 105116390 | 105117140 | 751 | 0.26 | 0.677 | 0.733 |
| chr6 | 7347788 | 7348646 | 859 | -0.16 | 0.677 | 0.733 |
| chr5 | 112073598 | 112073941 | 344 | -0.27 | 0.677 | 0.733 |
| chr9 | 37229771 | 37230247 | 477 | 0.24 | 0.677 | 0.733 |
| chr4 | 5501358 | 5502667 | 1310 | -0.17 | 0.677 | 0.733 |
| chr1 | 90437127 | 90474610 | 37484 | -0.03 | 0.677 | 0.733 |
| chr19 | 5961460 | 5990648 | 29189 | 0.06 | 0.677 | 0.733 |
| chr22 | 36276065 | 36319211 | 43147 | -0.05 | 0.678 | 0.733 |
| chr15 | 31749068 | 31757939 | 8872 | -0.15 | 0.678 | 0.733 |
| chr5 | 34499073 | 34499729 | 657 | -0.25 | 0.678 | 0.734 |
| chr10 | 102699499 | 102700418 | 920 | -0.24 | 0.678 | 0.734 |
| chr19 | 15576979 | 15578558 | 1580 | 0.15 | 0.678 | 0.734 |
| chr10 | 30959701 | 30991407 | 31707 | -0.07 | 0.678 | 0.734 |
| chr22 | 31476526 | 31482132 | 5607 | -0.08 | 0.679 | 0.734 |
| chr2 | 70508958 | 70529224 | 20267 | 0.03 | 0.679 | 0.735 |
| chr13 | 43294296 | 43295321 | 1026 | 0.18 | 0.679 | 0.735 |
| chr11 | 44086795 | 44088962 | 2168 | 0.06 | 0.679 | 0.735 |
| chr13 | 114999959 | 115001678 | 1720 | -0.14 | 0.679 | 0.735 |
| chr17 | 42462417 | 42463501 | 1085 | -0.16 | 0.679 | 0.735 |
| chr2 | 37266618 | 37267292 | 675 | -0.2 | 0.679 | 0.735 |
| chr20 | 16348501 | 16349950 | 1450 | 0.13 | 0.679 | 0.735 |
| chr14 | 53212115 | 53212504 | 390 | 0.26 | 0.68 | 0.735 |
| chr11 | 118758203 | 118759239 | 1037 | -0.17 | 0.68 | 0.735 |
| chr16 | 67301914 | 67325689 | 23776 | -0.05 | 0.68 | 0.736 |
| chr12 | 107953896 | 107954461 | 566 | -0.32 | 0.68 | 0.736 |
| chr20 | 13722427 | 13723693 | 1267 | -0.11 | 0.68 | 0.736 |
| chr1 | 150408011 | 150420818 | 12808 | -0.09 | 0.68 | 0.736 |
| chr5 | 127447594 | 127448571 | 978 | 0.23 | 0.68 | 0.736 |
| chr3 | 123751386 | 123752114 | 729 | -0.21 | 0.681 | 0.736 |
| chr1 | 31768475 | 31771057 | 2583 | 0.1 | 0.681 | 0.736 |
| chr1 | 183287238 | 183287733 | 496 | 0.28 | 0.681 | 0.736 |
| chr16 | 69050691 | 69061736 | 11046 | 0.06 | 0.681 | 0.736 |
| chr6 | 158243578 | 158262811 | 19234 | -0.05 | 0.681 | 0.737 |
| chr3 | 49760166 | 49763008 | 2843 | 0.07 | 0.682 | 0.737 |
| chr19 | 39055719 | 39056391 | 673 | 0.22 | 0.682 | 0.737 |
| chr1 | 189552687 | 189606995 | 54309 | 0.05 | 0.682 | 0.737 |
| chr20 | 56915318 | 56918373 | 3056 | 0.1 | 0.682 | 0.738 |
| chr3 | 36985336 | 36986761 | 1426 | -0.17 | 0.682 | 0.738 |
| chr11 | 22279685 | 22280555 | 871 | 0.23 | 0.682 | 0.738 |
| chr10 | 88622044 | 88637061 | 15018 | -0.09 | 0.683 | 0.738 |
| chr13 | 110371743 | 110374015 | 2273 | 0.13 | 0.683 | 0.738 |
| chr11 | 68624215 | 68625778 | 1564 | -0.14 | 0.683 | 0.738 |
| chr8 | 107868999 | 107879195 | 10197 | 0.05 | 0.683 | 0.738 |
| chr7 | 43611220 | 43636847 | 25628 | 0.05 | 0.683 | 0.738 |
| chr17 | 53342328 | 53342680 | 353 | 0.3 | 0.683 | 0.738 |

| | | | | | | |
|---|---|---|---|---|---|---|
| chr17 | 14036177 | 14036846 | 670 | 0.25 | 0.684 | 0.739 |
| chr1 | 88466865 | 88467741 | 877 | 0.2 | 0.684 | 0.739 |
| chr10 | 52152294 | 52191586 | 39293 | -0.04 | 0.684 | 0.739 |
| chr6 | 16851951 | 16852275 | 325 | -0.3 | 0.684 | 0.739 |
| chr8 | 59506983 | 59521812 | 14830 | 0.07 | 0.684 | 0.739 |
| chr14 | 89817595 | 89818378 | 784 | 0.23 | 0.684 | 0.739 |
| chr9 | 71650615 | 71660451 | 9837 | 0.1 | 0.684 | 0.739 |
| chr13 | 24648902 | 24662291 | 13390 | -0.12 | 0.684 | 0.739 |
| chr14 | 51501858 | 51503451 | 1594 | -0.11 | 0.684 | 0.74 |
| chr5 | 173072113 | 173073372 | 1260 | 0.2 | 0.685 | 0.74 |
| chr11 | 34523521 | 34524132 | 612 | 0.29 | 0.685 | 0.74 |
| chr7 | 34562205 | 34570499 | 8295 | 1.09 | 0.685 | 0.74 |
| chr19 | 40736833 | 40793239 | 56407 | -0.03 | 0.685 | 0.74 |
| chr16 | 57768316 | 57772145 | 3830 | 0.06 | 0.685 | 0.74 |
| chr15 | 74428158 | 74437444 | 9287 | -0.15 | 0.685 | 0.74 |
| chr9 | 116101566 | 116103095 | 1530 | -0.09 | 0.685 | 0.74 |
| chrX | 20084894 | 20086710 | 1817 | 0.12 | 0.686 | 0.741 |
| chr12 | 123122803 | 123123284 | 482 | 0.33 | 0.686 | 0.741 |
| chr3 | 179064203 | 179066484 | 2282 | -0.08 | 0.686 | 0.741 |
| chr17 | 35962333 | 35971911 | 9579 | 0.04 | 0.686 | 0.741 |
| chr7 | 151246758 | 151248475 | 1718 | -0.12 | 0.687 | 0.742 |
| chr1 | 27560292 | 27561839 | 1548 | -0.15 | 0.687 | 0.742 |
| chr6 | 35435656 | 35438964 | 3309 | 0.09 | 0.687 | 0.742 |
| chr12 | 29482124 | 29483319 | 1196 | -0.18 | 0.687 | 0.742 |
| chr3 | 197675733 | 197687905 | 12173 | -0.04 | 0.687 | 0.742 |
| chr9 | 132646032 | 132648823 | 2792 | -0.09 | 0.687 | 0.742 |
| chr7 | 97600401 | 97602308 | 1908 | 0.07 | 0.687 | 0.742 |
| chr13 | 31316931 | 31318825 | 1895 | -0.12 | 0.687 | 0.742 |
| chr6 | 141006034 | 141006684 | 651 | -0.21 | 0.688 | 0.742 |
| chr17 | 60635992 | 60645111 | 9120 | 0.05 | 0.688 | 0.742 |
| chr1 | 46712437 | 46714152 | 1716 | 0.14 | 0.688 | 0.742 |
| chr22 | 44019981 | 44021942 | 1962 | 0.09 | 0.688 | 0.743 |
| chr3 | 159429291 | 159430008 | 718 | 0.21 | 0.688 | 0.743 |
| chr1 | 229205378 | 229206818 | 1441 | 0.15 | 0.689 | 0.743 |
| chr22 | 29469080 | 29469342 | 263 | 0.3 | 0.689 | 0.744 |
| chr8 | 8691352 | 8751853 | 60502 | -0.02 | 0.689 | 0.744 |
| chr7 | 74041859 | 74106908 | 65050 | 0.03 | 0.689 | 0.744 |
| chr8 | 144798522 | 144799010 | 489 | -0.24 | 0.689 | 0.744 |
| chr3 | 47553907 | 47555545 | 1639 | 0.09 | 0.689 | 0.744 |
| chr16 | 69535869 | 69536730 | 862 | -0.21 | 0.689 | 0.744 |
| chr4 | 137662944 | 137664552 | 1609 | -0.17 | 0.689 | 0.744 |
| chr6 | 88117079 | 88118833 | 1755 | -0.09 | 0.689 | 0.744 |
| chr10 | 121630681 | 121633434 | 2754 | 0.08 | 0.689 | 0.744 |
| chr16 | 3870116 | 3870611 | 496 | 0.28 | 0.689 | 0.744 |
| chr5 | 176603741 | 176630253 | 26513 | -0.05 | 0.69 | 0.744 |
| chr2 | 118943231 | 118944491 | 1261 | 0.13 | 0.69 | 0.744 |
| chr2 | 71453694 | 71454201 | 508 | -0.21 | 0.69 | 0.745 |
| chr3 | 125069821 | 125077033 | 7213 | -0.09 | 0.69 | 0.745 |
| chr22 | 20747820 | 20748953 | 1134 | -0.14 | 0.69 | 0.745 |
| chr1 | 51899133 | 51900125 | 993 | 0.2 | 0.691 | 0.745 |
| chr1 | 91316383 | 91317031 | 649 | 0.23 | 0.691 | 0.745 |
| chr17 | 75611576 | 75612552 | 977 | -0.2 | 0.691 | 0.746 |
| chr1 | 45096712 | 45098682 | 1971 | -0.07 | 0.691 | 0.746 |
| chrX | 99986550 | 99987160 | 611 | -0.22 | 0.691 | 0.746 |
| chr9 | 134372004 | 134372933 | 930 | -0.24 | 0.691 | 0.746 |
| chr6 | 14210792 | 14212193 | 1402 | 0.14 | 0.691 | 0.746 |
| chr20 | 13661483 | 13662864 | 1382 | -0.13 | 0.692 | 0.746 |
| chr13 | 96295411 | 96297455 | 2045 | -0.12 | 0.692 | 0.746 |
| chr4 | 114758276 | 114759277 | 1002 | -0.17 | 0.692 | 0.746 |
| chr7 | 45019235 | 45041204 | 21970 | 0.04 | 0.692 | 0.746 |
| chr5 | 80262229 | 80264893 | 2665 | -0.07 | 0.692 | 0.747 |

| | | | | | | |
|---|---|---|---|---|---|---|
| chr4 | 7779970 | 7781319 | 1350 | 0.17 | 0.692 | 0.747 |
| chr17 | 60031798 | 60032485 | 688 | -0.28 | 0.692 | 0.747 |
| chr1 | 229477016 | 229479203 | 2188 | 0.1 | 0.692 | 0.747 |
| chr14 | 99844657 | 99850041 | 5385 | -0.1 | 0.692 | 0.747 |
| chr1 | 2322044 | 2345347 | 23304 | 0.05 | 0.692 | 0.747 |
| chr9 | 90094633 | 90095529 | 897 | -0.15 | 0.692 | 0.747 |
| chr18 | 20377534 | 20379019 | 1486 | 0.16 | 0.692 | 0.747 |
| chr20 | 34026860 | 34027469 | 610 | -0.2 | 0.693 | 0.747 |
| chr3 | 171261900 | 171262411 | 512 | -0.26 | 0.693 | 0.747 |
| chr18 | 10797783 | 10800772 | 2990 | 0.07 | 0.693 | 0.747 |
| chr2 | 61547518 | 61585052 | 37535 | -0.04 | 0.693 | 0.747 |
| chr13 | 96435889 | 96437680 | 1792 | 0.11 | 0.693 | 0.747 |
| chr2 | 61470875 | 61475814 | 4940 | -0.09 | 0.693 | 0.747 |
| chr1 | 240655886 | 240663176 | 7291 | -0.08 | 0.693 | 0.747 |
| chr6 | 97730067 | 97731660 | 1594 | 0.12 | 0.694 | 0.748 |
| chr15 | 56282509 | 56287267 | 4759 | -0.05 | 0.694 | 0.748 |
| chr12 | 113536450 | 113536927 | 478 | 0.23 | 0.694 | 0.748 |
| chr1 | 224544157 | 224552003 | 7847 | 0.07 | 0.694 | 0.748 |
| chr6 | 83106263 | 83119713 | 13451 | 0.06 | 0.694 | 0.748 |
| chr12 | 93984121 | 93985872 | 1752 | 0.08 | 0.695 | 0.749 |
| chr1 | 22440531 | 22440949 | 419 | -0.26 | 0.695 | 0.749 |
| chr12 | 103953235 | 103969865 | 16631 | -0.08 | 0.695 | 0.749 |
| chr10 | 81204038 | 81205419 | 1382 | -0.3 | 0.695 | 0.749 |
| chr14 | 74998137 | 74999236 | 1100 | -0.16 | 0.695 | 0.749 |
| chr7 | 127982490 | 127984532 | 2043 | -0.08 | 0.695 | 0.749 |
| chr17 | 9263170 | 9275077 | 11908 | -0.07 | 0.695 | 0.749 |
| chr3 | 133379462 | 133381575 | 2114 | 0.07 | 0.695 | 0.749 |
| chr9 | 6468475 | 6468874 | 400 | -0.29 | 0.695 | 0.749 |
| chr5 | 145614997 | 145615554 | 558 | -0.24 | 0.695 | 0.749 |
| chr22 | 42839900 | 42840570 | 671 | -0.21 | 0.695 | 0.749 |
| chr4 | 88983088 | 88985255 | 2168 | 0.12 | 0.695 | 0.749 |
| chr16 | 56691108 | 56692286 | 1179 | -0.16 | 0.696 | 0.749 |
| chr5 | 89735701 | 89736226 | 526 | 0.2 | 0.696 | 0.749 |
| chr10 | 331307 | 332242 | 936 | -0.2 | 0.696 | 0.75 |
| chr4 | 89376934 | 89378087 | 1154 | 0.16 | 0.696 | 0.75 |
| chr6 | 56443924 | 56444944 | 1021 | -0.19 | 0.697 | 0.75 |
| chr3 | 9790806 | 9797308 | 6503 | 0.07 | 0.697 | 0.75 |
| chr9 | 102774757 | 102777344 | 2588 | -0.11 | 0.697 | 0.75 |
| chr8 | 107182382 | 107183642 | 1261 | -0.14 | 0.697 | 0.75 |
| chr9 | 94115348 | 94124868 | 9521 | -0.1 | 0.697 | 0.75 |
| chr7 | 534670 | 583441 | 48772 | 0.03 | 0.697 | 0.75 |
| chr1 | 20208894 | 20209951 | 1058 | -0.16 | 0.697 | 0.751 |
| chr17 | 76641629 | 76643033 | 1405 | 0.13 | 0.697 | 0.751 |
| chr1 | 12244131 | 12244793 | 663 | 0.21 | 0.697 | 0.751 |
| chr1 | 180639127 | 180650306 | 11180 | 0.11 | 0.698 | 0.751 |
| chr2 | 27988361 | 28003841 | 15481 | -0.05 | 0.698 | 0.751 |
| chr9 | 27053299 | 27055690 | 2392 | 0.12 | 0.698 | 0.751 |
| chr15 | 71145319 | 71159743 | 14425 | -0.08 | 0.698 | 0.751 |
| chr8 | 29855888 | 29859883 | 3996 | 0.08 | 0.698 | 0.752 |
| chr15 | 41860134 | 41860722 | 589 | -0.28 | 0.698 | 0.752 |
| chr13 | 73354694 | 73358388 | 3695 | -0.07 | 0.698 | 0.752 |
| chr17 | 62206431 | 62208624 | 2194 | -0.1 | 0.699 | 0.752 |
| chr12 | 19503959 | 19504516 | 558 | -0.23 | 0.699 | 0.752 |
| chrX | 128799467 | 128812977 | 13511 | 0.06 | 0.699 | 0.752 |
| chr3 | 9811041 | 9812192 | 1152 | -0.12 | 0.699 | 0.752 |
| chr8 | 117586998 | 117588277 | 1280 | -0.12 | 0.699 | 0.752 |
| chr2 | 97078254 | 97079644 | 1391 | 0.09 | 0.699 | 0.752 |
| chr2 | 227291014 | 227294158 | 3145 | -0.11 | 0.699 | 0.752 |
| chr1 | 92269068 | 92271405 | 2338 | -0.14 | 0.7 | 0.753 |
| chr2 | 178337386 | 178340311 | 2926 | 0.12 | 0.7 | 0.753 |
| chr14 | 75178640 | 75181064 | 2425 | 0.05 | 0.7 | 0.753 |

| | | | | | | |
|---|---|---|---|---|---|---|
| chr17 | 19690943 | 19704145 | 13203 | -0.09 | 0.7 | 0.753 |
| chr16 | 12667912 | 12668603 | 692 | -0.23 | 0.7 | 0.754 |
| chr16 | 59938671 | 59940651 | 1981 | -0.11 | 0.701 | 0.754 |
| chr5 | 177556714 | 177558990 | 2277 | 0.07 | 0.701 | 0.754 |
| chr16 | 87096021 | 87096446 | 426 | 0.22 | 0.701 | 0.754 |
| chr4 | 184183273 | 184183686 | 414 | 0.31 | 0.701 | 0.754 |
| chr20 | 43991064 | 44002936 | 11873 | 0.06 | 0.701 | 0.754 |
| chr17 | 30224607 | 30229483 | 4877 | -0.06 | 0.701 | 0.754 |
| chr16 | 68482746 | 68483110 | 365 | -0.21 | 0.701 | 0.754 |
| chr2 | 32803292 | 32803862 | 571 | -0.2 | 0.701 | 0.754 |
| chr12 | 111806062 | 111807509 | 1448 | 0.15 | 0.701 | 0.754 |
| chr4 | 466942 | 468237 | 1296 | 0.1 | 0.701 | 0.754 |
| chr9 | 3745857 | 3760742 | 14886 | 0.06 | 0.701 | 0.754 |
| chr13 | 32726846 | 32738966 | 12121 | -0.12 | 0.701 | 0.754 |
| chr20 | 44621803 | 44622522 | 720 | -0.24 | 0.701 | 0.754 |
| chr22 | 30485174 | 30486319 | 1146 | 0.13 | 0.701 | 0.754 |
| chr3 | 145808291 | 145851747 | 43457 | 0.03 | 0.701 | 0.754 |
| chr16 | 28850886 | 28887492 | 36607 | -0.03 | 0.701 | 0.754 |
| chr4 | 100461362 | 100462155 | 794 | 0.21 | 0.702 | 0.754 |
| chr6 | 105623683 | 105628691 | 5009 | 0.06 | 0.702 | 0.754 |
| chr2 | 27805088 | 27807621 | 2534 | -0.06 | 0.702 | 0.754 |
| chr13 | 20993160 | 20994783 | 1624 | 0.11 | 0.702 | 0.755 |
| chr4 | 71932829 | 71933814 | 986 | -0.18 | 0.702 | 0.755 |
| chr2 | 99347746 | 99348267 | 522 | 0.23 | 0.702 | 0.755 |
| chr18 | 20012265 | 20013299 | 1035 | -0.15 | 0.702 | 0.755 |
| chr4 | 14531052 | 14531948 | 897 | 0.18 | 0.702 | 0.755 |
| chr15 | 83518642 | 83519146 | 505 | -0.25 | 0.703 | 0.755 |
| chr15 | 42120338 | 42120876 | 539 | -0.2 | 0.703 | 0.756 |
| chr18 | 9594770 | 9649552 | 54783 | 0.02 | 0.703 | 0.756 |
| chr13 | 30426300 | 30427580 | 1281 | -0.17 | 0.703 | 0.756 |
| chr1 | 92035202 | 92044653 | 9452 | 0.14 | 0.703 | 0.756 |
| chr3 | 47462772 | 47463155 | 384 | -0.21 | 0.703 | 0.756 |
| chr10 | 126407415 | 126434616 | 27202 | 0.04 | 0.703 | 0.756 |
| chr8 | 95319571 | 95320260 | 690 | -0.21 | 0.703 | 0.756 |
| chr13 | 99892033 | 99892519 | 487 | 0.24 | 0.704 | 0.756 |
| chr20 | 7822064 | 7823161 | 1098 | 0.15 | 0.704 | 0.756 |
| chr22 | 41983430 | 41986882 | 3453 | 0.08 | 0.704 | 0.756 |
| chr2 | 228582269 | 228583570 | 1302 | 0.13 | 0.704 | 0.756 |
| chr17 | 3620541 | 3628476 | 7936 | -0.06 | 0.704 | 0.756 |
| chr3 | 128596853 | 128609209 | 12357 | 0.05 | 0.704 | 0.757 |
| chr17 | 70973017 | 70973828 | 812 | -0.17 | 0.704 | 0.757 |
| chr9 | 86534522 | 86619220 | 84699 | -0.06 | 0.704 | 0.757 |
| chr4 | 8227205 | 8263076 | 35872 | -0.04 | 0.704 | 0.757 |
| chr10 | 12431217 | 12431627 | 411 | 0.3 | 0.704 | 0.757 |
| chr1 | 36915123 | 36931995 | 16873 | 0.04 | 0.705 | 0.757 |
| chr6 | 136132883 | 136134552 | 1670 | 0.12 | 0.705 | 0.757 |
| chr3 | 27762695 | 27764204 | 1510 | 0.12 | 0.705 | 0.757 |
| chr1 | 59509455 | 59524011 | 14557 | -0.13 | 0.705 | 0.757 |
| chr1 | 39324686 | 39340145 | 15460 | 0.07 | 0.705 | 0.757 |
| chr13 | 50284879 | 50286236 | 1358 | -0.14 | 0.706 | 0.758 |
| chr3 | 150755562 | 150756192 | 631 | 0.22 | 0.706 | 0.758 |
| chr11 | 20498663 | 20499064 | 402 | -0.25 | 0.706 | 0.758 |
| chrX | 45329581 | 45341081 | 11501 | 0.04 | 0.706 | 0.758 |
| chr6 | 111301742 | 111304846 | 3105 | 0.05 | 0.706 | 0.758 |
| chr2 | 225905523 | 225918054 | 12532 | 0.06 | 0.706 | 0.758 |
| chr3 | 27609786 | 27610140 | 355 | -0.27 | 0.706 | 0.758 |
| chrX | 73427297 | 73428236 | 940 | -0.22 | 0.706 | 0.759 |
| chr3 | 183406108 | 183417256 | 11149 | -0.08 | 0.706 | 0.759 |
| chr14 | 61189051 | 61202691 | 13641 | 0.08 | 0.707 | 0.759 |
| chr5 | 148650824 | 148652401 | 1578 | 0.14 | 0.707 | 0.759 |
| chr11 | 99110872 | 99112813 | 1942 | -0.09 | 0.707 | 0.759 |

| | | | | | | |
|---|---|---|---|---|---|---|
| chr3 | 186358177 | 186358964 | 788 | -0.2 | 0.707 | 0.759 |
| chr7 | 109957422 | 109958921 | 1500 | -0.12 | 0.707 | 0.759 |
| chr11 | 45183330 | 45183745 | 416 | 0.28 | 0.707 | 0.759 |
| chr6 | 149081918 | 149082638 | 721 | 0.21 | 0.707 | 0.759 |
| chr22 | 38044107 | 38054331 | 10225 | 0.13 | 0.707 | 0.759 |
| chrX | 54204214 | 54210580 | 6367 | 0.06 | 0.708 | 0.76 |
| chr6 | 24494873 | 24495850 | 978 | 0.15 | 0.708 | 0.76 |
| chr10 | 102282142 | 102296735 | 14594 | 0.06 | 0.708 | 0.76 |
| chr6 | 117187231 | 117188220 | 990 | 0.21 | 0.708 | 0.76 |
| chr3 | 33874785 | 33875622 | 838 | 0.17 | 0.708 | 0.76 |
| chr15 | 62681734 | 62684367 | 2634 | 0.06 | 0.708 | 0.76 |
| chr7 | 75115026 | 75115539 | 514 | -0.2 | 0.708 | 0.76 |
| chr15 | 78419555 | 78424908 | 5354 | -0.06 | 0.708 | 0.76 |
| chr3 | 107429680 | 107430277 | 598 | 0.24 | 0.708 | 0.76 |
| chr8 | 126029155 | 126029723 | 569 | -0.19 | 0.709 | 0.76 |
| chr7 | 149318985 | 149322708 | 3724 | 0.07 | 0.709 | 0.761 |
| chr1 | 66671569 | 66674112 | 2544 | 0.09 | 0.709 | 0.761 |
| chr11 | 43373582 | 43389002 | 15421 | 0.05 | 0.709 | 0.761 |
| chr22 | 22336362 | 22337443 | 1082 | -0.1 | 0.709 | 0.761 |
| chr9 | 139158554 | 139159729 | 1176 | 0.17 | 0.709 | 0.761 |
| chr15 | 78729338 | 78761029 | 31692 | -0.03 | 0.709 | 0.761 |
| chr13 | 114909601 | 114909982 | 382 | 0.26 | 0.709 | 0.761 |
| chr6 | 159415967 | 159422169 | 6203 | 0.05 | 0.71 | 0.761 |
| chr1 | 63147786 | 63154700 | 6915 | 0.08 | 0.71 | 0.761 |
| chr1 | 21344138 | 21355178 | 11041 | -0.03 | 0.71 | 0.761 |
| chrX | 13069330 | 13085360 | 16031 | -0.05 | 0.71 | 0.761 |
| chr20 | 44746550 | 44747330 | 781 | -0.21 | 0.71 | 0.762 |
| chr3 | 170625676 | 170627300 | 1625 | -0.11 | 0.71 | 0.762 |
| chr1 | 39592605 | 39599468 | 6864 | 0.08 | 0.71 | 0.762 |
| chr22 | 20904954 | 20924308 | 19355 | -0.05 | 0.711 | 0.762 |
| chr1 | 171544167 | 171559803 | 15637 | 0.08 | 0.711 | 0.762 |
| chr6 | 136042666 | 136045415 | 2750 | -0.07 | 0.711 | 0.762 |
| chr1 | 87239116 | 87240934 | 1819 | 0.13 | 0.711 | 0.762 |
| chr2 | 47915808 | 47916266 | 459 | -0.22 | 0.711 | 0.762 |
| chr2 | 189377112 | 189377774 | 663 | 0.22 | 0.711 | 0.762 |
| chr14 | 94595391 | 94595975 | 585 | 0.2 | 0.711 | 0.763 |
| chr12 | 31158486 | 31159592 | 1107 | 0.13 | 0.712 | 0.763 |
| chr9 | 128170353 | 128171163 | 811 | 0.15 | 0.712 | 0.763 |
| chr17 | 61623119 | 61633556 | 10438 | -0.04 | 0.712 | 0.763 |
| chr15 | 65203613 | 65203994 | 382 | 0.26 | 0.712 | 0.763 |
| chr5 | 157216329 | 157217882 | 1554 | -0.16 | 0.712 | 0.763 |
| chr7 | 95024407 | 95073237 | 48831 | 0.02 | 0.712 | 0.763 |
| chr2 | 30862702 | 30863559 | 858 | 0.16 | 0.712 | 0.763 |
| chr5 | 138051247 | 138052960 | 1714 | 0.11 | 0.712 | 0.763 |
| chr22 | 38701162 | 38724492 | 23331 | 0.04 | 0.712 | 0.764 |
| chr7 | 36191909 | 36202392 | 10484 | -0.09 | 0.713 | 0.764 |
| chr12 | 3277503 | 3292030 | 14528 | -0.12 | 0.713 | 0.764 |
| chr22 | 31575610 | 31575946 | 337 | 0.24 | 0.713 | 0.764 |
| chr1 | 214667828 | 214680973 | 13146 | 0.15 | 0.713 | 0.764 |
| chr16 | 31190324 | 31196074 | 5751 | 0.05 | 0.713 | 0.765 |
| chrX | 153657042 | 153657398 | 357 | -0.24 | 0.713 | 0.765 |
| chr12 | 69445949 | 69463465 | 17517 | 0.05 | 0.714 | 0.765 |
| chr19 | 42927232 | 42938129 | 10898 | -0.15 | 0.714 | 0.765 |
| chr7 | 1193784 | 1216716 | 22933 | -0.04 | 0.715 | 0.766 |
| chr5 | 118787742 | 118789243 | 1502 | 0.07 | 0.715 | 0.766 |
| chr7 | 40071691 | 40074035 | 2345 | 0.16 | 0.715 | 0.766 |
| chr21 | 35075996 | 35089134 | 13139 | 0.1 | 0.715 | 0.766 |
| chr1 | 8456917 | 8458398 | 1482 | -0.16 | 0.715 | 0.767 |
| chr8 | 141885071 | 141885773 | 703 | -0.18 | 0.716 | 0.767 |
| chr3 | 94389763 | 94390579 | 817 | 0.18 | 0.716 | 0.767 |
| chr17 | 8649081 | 8649409 | 329 | 0.27 | 0.716 | 0.767 |

| | | | | | | |
|---|---|---|---|---|---|---|
| chr1 | 117955038 | 117955771 | 734 | 0.22 | 0.716 | 0.767 |
| chr9 | 82185052 | 82189102 | 4051 | 0.05 | 0.717 | 0.768 |
| chr5 | 137827359 | 137845418 | 18060 | -0.11 | 0.717 | 0.768 |
| chr5 | 77583231 | 77591889 | 8659 | 0.05 | 0.717 | 0.768 |
| chr2 | 141381688 | 141382240 | 553 | 0.21 | 0.718 | 0.769 |
| chr1 | 79658608 | 79659881 | 1274 | -0.12 | 0.718 | 0.769 |
| chr1 | 210741375 | 210742378 | 1004 | -0.19 | 0.718 | 0.769 |
| chr1 | 247241336 | 247242220 | 885 | 0.09 | 0.718 | 0.769 |
| chr17 | 13497512 | 13498658 | 1147 | -0.14 | 0.718 | 0.769 |
| chr2 | 220362500 | 220364286 | 1787 | -0.08 | 0.718 | 0.769 |
| chrX | 119076095 | 119078350 | 2256 | -0.07 | 0.718 | 0.769 |
| chr6 | 9515586 | 9533547 | 17962 | 0.07 | 0.718 | 0.769 |
| chr5 | 65459507 | 65471803 | 12297 | -0.06 | 0.719 | 0.77 |
| chr19 | 42828547 | 42830818 | 2272 | -0.08 | 0.719 | 0.77 |
| chr7 | 100398378 | 100436339 | 37962 | 0.03 | 0.719 | 0.77 |
| chr8 | 40263371 | 40264195 | 825 | 0.18 | 0.719 | 0.77 |
| chr12 | 1153322 | 1159955 | 6634 | 0.17 | 0.72 | 0.77 |
| chr8 | 123049962 | 123050961 | 1000 | 0.15 | 0.72 | 0.771 |
| chr7 | 55477177 | 55477881 | 705 | -0.24 | 0.72 | 0.771 |
| chr1 | 229797003 | 229797783 | 781 | -0.19 | 0.72 | 0.771 |
| chr3 | 110280767 | 110303561 | 22795 | -0.03 | 0.72 | 0.771 |
| chr16 | 78132648 | 78140063 | 7416 | -0.06 | 0.72 | 0.771 |
| chr1 | 1508783 | 1511440 | 2658 | 0.05 | 0.721 | 0.771 |
| chr1 | 11724071 | 11724758 | 688 | -0.21 | 0.721 | 0.771 |
| chr10 | 62206400 | 62207129 | 730 | 0.2 | 0.721 | 0.772 |
| chr3 | 48972855 | 48973341 | 487 | -0.24 | 0.721 | 0.772 |
| chr1 | 206730482 | 206731025 | 544 | -0.23 | 0.721 | 0.772 |
| chrX | 76995604 | 77005856 | 10253 | 0.07 | 0.721 | 0.772 |
| chr7 | 87481900 | 87483398 | 1499 | -0.12 | 0.721 | 0.772 |
| chr11 | 35324995 | 35326816 | 1822 | 0.1 | 0.722 | 0.772 |
| chr1 | 85666105 | 85668291 | 2187 | -0.07 | 0.722 | 0.772 |
| chr8 | 38061074 | 38062330 | 1257 | -0.17 | 0.722 | 0.772 |
| chr16 | 31413565 | 31416534 | 2970 | 0.07 | 0.722 | 0.773 |
| chr10 | 70314121 | 70320991 | 6871 | 0.08 | 0.722 | 0.773 |
| chr7 | 155514791 | 155534924 | 20134 | -0.05 | 0.722 | 0.773 |
| chr16 | 85207367 | 85209015 | 1649 | -0.11 | 0.722 | 0.773 |
| chr10 | 54016606 | 54017439 | 834 | -0.17 | 0.723 | 0.773 |
| chr6 | 25660204 | 25661442 | 1239 | -0.19 | 0.723 | 0.773 |
| chr20 | 5166402 | 5167116 | 715 | 0.19 | 0.723 | 0.773 |
| chr9 | 134881132 | 134889030 | 7899 | -0.05 | 0.723 | 0.774 |
| chr2 | 88353084 | 88356342 | 3259 | 0.03 | 0.723 | 0.774 |
| chr14 | 31824125 | 31826217 | 2093 | -0.1 | 0.723 | 0.774 |
| chr3 | 193753146 | 193753789 | 644 | 0.21 | 0.724 | 0.774 |
| chr11 | 32354945 | 32355498 | 554 | 0.2 | 0.724 | 0.774 |
| chr10 | 3268920 | 3269878 | 959 | 0.16 | 0.724 | 0.774 |
| chr6 | 2932683 | 2933509 | 827 | 0.16 | 0.724 | 0.774 |
| chr6 | 139263032 | 139279575 | 16544 | -0.05 | 0.724 | 0.774 |
| chr3 | 47484934 | 47486248 | 1315 | -0.14 | 0.724 | 0.774 |
| chr16 | 2769160 | 2772182 | 3023 | -0.07 | 0.724 | 0.774 |
| chr19 | 17865018 | 17910036 | 45019 | 0.03 | 0.724 | 0.775 |
| chrX | 46722078 | 46725692 | 3615 | 0.07 | 0.724 | 0.775 |
| chr17 | 29394451 | 29405704 | 11254 | 0.08 | 0.724 | 0.775 |
| chr1 | 26731447 | 26732210 | 764 | 0.19 | 0.725 | 0.775 |
| chr20 | 47314165 | 47315211 | 1047 | -0.18 | 0.725 | 0.775 |
| chr18 | 9513530 | 9515001 | 1472 | -0.12 | 0.725 | 0.775 |
| chr2 | 58467157 | 58468995 | 1839 | -0.05 | 0.725 | 0.775 |
| chr16 | 57208528 | 57240653 | 32126 | -0.03 | 0.725 | 0.775 |
| chr8 | 40009246 | 40074809 | 65564 | 0.01 | 0.726 | 0.776 |
| chr15 | 78933069 | 78933817 | 749 | 0.16 | 0.726 | 0.777 |
| chr3 | 120277006 | 120278739 | 1734 | -0.07 | 0.727 | 0.777 |
| chr21 | 40466351 | 40468044 | 1694 | -0.1 | 0.727 | 0.777 |

| | | | | | | |
|---|---|---|---|---|---|---|
| chr1 | 226925709 | 226927630 | 1922 | 0.08 | 0.727 | 0.777 |
| chr1 | 171786408 | 171787751 | 1344 | 0.12 | 0.727 | 0.778 |
| chr8 | 23779598 | 23781168 | 1571 | -0.1 | 0.728 | 0.778 |
| chr4 | 189743588 | 189744360 | 773 | 0.2 | 0.728 | 0.778 |
| chr10 | 104677308 | 104678751 | 1444 | -0.11 | 0.728 | 0.778 |
| chr11 | 32934428 | 32940238 | 5811 | -0.09 | 0.728 | 0.778 |
| chr16 | 4783317 | 4785590 | 2274 | -0.07 | 0.728 | 0.778 |
| chr3 | 17734197 | 17735298 | 1102 | 0.11 | 0.728 | 0.778 |
| chr15 | 63250590 | 63251113 | 524 | -0.2 | 0.728 | 0.778 |
| chr8 | 130648129 | 130648556 | 428 | 0.2 | 0.729 | 0.779 |
| chr15 | 75808317 | 75844114 | 35798 | -0.03 | 0.729 | 0.779 |
| chr13 | 104116234 | 104118272 | 2039 | 0.07 | 0.729 | 0.779 |
| chr4 | 187841591 | 187842185 | 595 | -0.2 | 0.729 | 0.779 |
| chr22 | 30782547 | 30794263 | 11717 | -0.09 | 0.729 | 0.779 |
| chrX | 41955049 | 41955913 | 865 | -0.13 | 0.73 | 0.779 |
| chr6 | 27082561 | 27129989 | 47429 | 0.02 | 0.73 | 0.78 |
| chrX | 102610585 | 102612659 | 2075 | -0.07 | 0.73 | 0.78 |
| chr13 | 45890164 | 45891126 | 963 | 0.13 | 0.73 | 0.78 |
| chr18 | 37378844 | 37380869 | 2026 | 0.12 | 0.731 | 0.781 |
| chr9 | 4714575 | 4715704 | 1130 | -0.21 | 0.731 | 0.781 |
| chr14 | 39733860 | 39751376 | 17517 | 0.03 | 0.731 | 0.781 |
| chr15 | 42338143 | 42350656 | 12514 | -0.08 | 0.732 | 0.781 |
| chr5 | 157157514 | 157171605 | 14092 | 0.06 | 0.732 | 0.782 |
| chr17 | 2828625 | 2830111 | 1487 | -0.13 | 0.733 | 0.783 |
| chr2 | 202296412 | 202298140 | 1729 | -0.12 | 0.733 | 0.783 |
| chr2 | 201935857 | 201937634 | 1778 | 0.09 | 0.733 | 0.783 |
| chr13 | 50985518 | 50986054 | 537 | -0.24 | 0.733 | 0.783 |
| chr3 | 153960631 | 153967104 | 6474 | -0.09 | 0.734 | 0.783 |
| chr15 | 42065740 | 42067775 | 2036 | 0.09 | 0.734 | 0.783 |
| chr12 | 46518355 | 46519015 | 661 | -0.18 | 0.734 | 0.783 |
| chr11 | 33277149 | 33294161 | 17013 | -0.02 | 0.734 | 0.784 |
| chr17 | 41747050 | 41747539 | 490 | 0.2 | 0.734 | 0.784 |
| chr6 | 71959798 | 71960554 | 757 | 0.19 | 0.734 | 0.784 |
| chr10 | 49514068 | 49515944 | 1877 | 0.07 | 0.735 | 0.784 |
| chr16 | 56510460 | 56511981 | 1522 | -0.1 | 0.735 | 0.784 |
| chr6 | 12545415 | 12546782 | 1368 | -0.13 | 0.735 | 0.785 |
| chr4 | 88140529 | 88142794 | 2266 | 0.08 | 0.735 | 0.785 |
| chr1 | 10313415 | 10318302 | 4888 | 0.11 | 0.735 | 0.785 |
| chr12 | 133421686 | 133422560 | 875 | 0.19 | 0.735 | 0.785 |
| chr8 | 132958633 | 132960213 | 1581 | -0.08 | 0.735 | 0.785 |
| chr8 | 28186646 | 28186989 | 344 | 0.31 | 0.735 | 0.785 |
| chr5 | 68389142 | 68403110 | 13969 | 0.05 | 0.735 | 0.785 |
| chr2 | 85131849 | 85294540 | 162692 | -0.01 | 0.735 | 0.785 |
| chr11 | 111230755 | 111231548 | 794 | -0.16 | 0.735 | 0.785 |
| chr11 | 288607 | 289639 | 1033 | 0.18 | 0.735 | 0.785 |
| chr11 | 86472868 | 86473728 | 861 | 0.31 | 0.735 | 0.785 |
| chr10 | 116539081 | 116541146 | 2066 | -0.08 | 0.736 | 0.785 |
| chr19 | 14063333 | 14064629 | 1297 | 0.18 | 0.736 | 0.785 |
| chr19 | 1354164 | 1354813 | 650 | 0.14 | 0.736 | 0.785 |
| chr3 | 118803487 | 118804134 | 648 | -0.2 | 0.736 | 0.785 |
| chr10 | 70683728 | 70685648 | 1921 | -0.12 | 0.736 | 0.785 |
| chr2 | 32742164 | 32751025 | 8862 | -0.07 | 0.736 | 0.785 |
| chr1 | 234613187 | 234615161 | 1975 | 0.1 | 0.736 | 0.786 |
| chr3 | 69697999 | 69699249 | 1251 | -0.14 | 0.736 | 0.786 |
| chr9 | 503656 | 505926 | 2271 | -0.09 | 0.737 | 0.786 |
| chr14 | 77647529 | 77648824 | 1296 | -0.15 | 0.737 | 0.786 |
| chr8 | 95002735 | 95004679 | 1945 | 0.08 | 0.737 | 0.786 |
| chr5 | 60870691 | 60871332 | 642 | 0.19 | 0.737 | 0.786 |
| chr11 | 8349842 | 8351219 | 1378 | -0.18 | 0.737 | 0.786 |
| chr18 | 74187364 | 74190383 | 3020 | 0.05 | 0.737 | 0.786 |
| chr9 | 133802839 | 133822991 | 20153 | -0.08 | 0.737 | 0.786 |

| | | | | | | |
|---|---|---|---|---|---|---|
| chr3 | 64330382 | 64330838 | 457 | -0.22 | 0.737 | 0.786 |
| chr6 | 52216496 | 52227732 | 11237 | 0.06 | 0.738 | 0.787 |
| chr12 | 31719795 | 31745342 | 25548 | 0.03 | 0.738 | 0.787 |
| chr16 | 75627839 | 75629453 | 1615 | 0.1 | 0.738 | 0.787 |
| chr2 | 62132323 | 62133476 | 1154 | 0.1 | 0.738 | 0.787 |
| chr1 | 12039789 | 12042794 | 3006 | 0.08 | 0.738 | 0.787 |
| chr20 | 50211225 | 50211682 | 458 | -0.19 | 0.738 | 0.787 |
| chr4 | 185631769 | 185633763 | 1995 | 0.1 | 0.738 | 0.787 |
| chr10 | 5726135 | 5735322 | 9188 | -0.05 | 0.738 | 0.787 |
| chr2 | 120301800 | 120302527 | 728 | 0.16 | 0.738 | 0.787 |
| chr2 | 197234672 | 197235243 | 572 | 0.2 | 0.738 | 0.787 |
| chr1 | 113476828 | 113480239 | 3412 | -0.06 | 0.738 | 0.787 |
| chr19 | 7098572 | 7099889 | 1318 | 0.1 | 0.739 | 0.787 |
| chr3 | 127730302 | 127730633 | 332 | 0.24 | 0.739 | 0.787 |
| chr1 | 6257001 | 6267395 | 10395 | 0.04 | 0.739 | 0.788 |
| chr13 | 67132013 | 67135004 | 2992 | -0.13 | 0.739 | 0.788 |
| chr8 | 81033033 | 81035316 | 2284 | -0.12 | 0.739 | 0.788 |
| chr19 | 36499926 | 36506270 | 6345 | 0.07 | 0.739 | 0.788 |
| chr3 | 9192479 | 9193856 | 1378 | 0.14 | 0.74 | 0.788 |
| chr3 | 49893301 | 49894663 | 1363 | 0.12 | 0.74 | 0.788 |
| chr9 | 2559156 | 2560496 | 1341 | -0.14 | 0.74 | 0.788 |
| chr6 | 16438325 | 16440555 | 2231 | -0.08 | 0.74 | 0.788 |
| chr4 | 184019049 | 184029042 | 9994 | 0.04 | 0.74 | 0.789 |
| chr9 | 37414529 | 37509066 | 94538 | -0.02 | 0.74 | 0.789 |
| chr1 | 17330207 | 17330905 | 699 | 0.19 | 0.74 | 0.789 |
| chr7 | 6655493 | 6655945 | 453 | 0.21 | 0.741 | 0.789 |
| chr12 | 28947596 | 28949170 | 1575 | 0.1 | 0.741 | 0.789 |
| chr8 | 10696158 | 10698142 | 1985 | 0.07 | 0.741 | 0.79 |
| chr8 | 23103207 | 23107909 | 4703 | 0.06 | 0.741 | 0.79 |
| chr1 | 77060524 | 77061573 | 1050 | 0.17 | 0.741 | 0.79 |
| chr5 | 79949303 | 79952724 | 3422 | 0.05 | 0.742 | 0.79 |
| chr4 | 48803378 | 48804784 | 1407 | 0.15 | 0.742 | 0.79 |
| chr17 | 58179478 | 58180345 | 868 | -0.1 | 0.742 | 0.79 |
| chr2 | 155119579 | 155121293 | 1715 | -0.13 | 0.742 | 0.79 |
| chr10 | 910676 | 911294 | 619 | -0.23 | 0.742 | 0.79 |
| chr3 | 130744759 | 130746895 | 2137 | 0.04 | 0.742 | 0.79 |
| chr9 | 34097873 | 34132528 | 34656 | 0.03 | 0.742 | 0.79 |
| chr19 | 56824576 | 56827717 | 3142 | -0.04 | 0.742 | 0.791 |
| chr3 | 131220584 | 131222288 | 1705 | -0.07 | 0.743 | 0.792 |
| chr1 | 114223645 | 114224584 | 940 | -0.19 | 0.744 | 0.792 |
| chr1 | 246944676 | 246946135 | 1460 | 0.13 | 0.744 | 0.792 |
| chr14 | 35295564 | 35345494 | 49931 | 0.03 | 0.744 | 0.792 |
| chr7 | 100270757 | 100273369 | 2613 | -0.08 | 0.744 | 0.792 |
| chr17 | 65955182 | 65962185 | 7004 | 0.06 | 0.744 | 0.792 |
| chr12 | 127420870 | 127421756 | 887 | -0.19 | 0.744 | 0.793 |
| chr6 | 26272351 | 26366519 | 94169 | -0.02 | 0.744 | 0.793 |
| chr11 | 87784889 | 87785371 | 483 | -0.21 | 0.745 | 0.793 |
| chr1 | 55180970 | 55182668 | 1699 | 0.09 | 0.746 | 0.794 |
| chr20 | 47579621 | 47580438 | 818 | 0.21 | 0.746 | 0.794 |
| chr3 | 49203014 | 49209384 | 6371 | -0.08 | 0.746 | 0.794 |
| chr11 | 86787294 | 86788274 | 981 | -0.16 | 0.746 | 0.794 |
| chr3 | 23927047 | 23970731 | 43685 | -0.03 | 0.746 | 0.795 |
| chr1 | 35853879 | 35854763 | 885 | -0.15 | 0.746 | 0.795 |
| chr1 | 246487608 | 246491522 | 3915 | -0.07 | 0.747 | 0.795 |
| chr15 | 56083384 | 56092692 | 9309 | -0.62 | 0.748 | 0.796 |
| chr8 | 143799262 | 143821867 | 22606 | 0.03 | 0.748 | 0.796 |
| chr14 | 54887167 | 54887974 | 808 | 0.19 | 0.748 | 0.796 |
| chr3 | 190024439 | 190040236 | 15798 | 0.04 | 0.748 | 0.796 |
| chr6 | 125621601 | 125623245 | 1645 | 0.08 | 0.748 | 0.796 |
| chr13 | 43719484 | 43729010 | 9527 | 0.12 | 0.748 | 0.796 |
| chr10 | 121009575 | 121015368 | 5794 | 0.04 | 0.748 | 0.796 |

| | | | | | | |
|---|---|---|---|---|---|---|
| chr5 | 14494717 | 14502676 | 7960 | -0.09 | 0.748 | 0.796 |
| chr8 | 17827038 | 17827570 | 533 | 0.19 | 0.748 | 0.796 |
| chr20 | 11367151 | 11368840 | 1690 | 0.09 | 0.748 | 0.796 |
| chr18 | 45846359 | 45846861 | 503 | -0.27 | 0.749 | 0.797 |
| chr20 | 55926031 | 55927860 | 1830 | 0.05 | 0.749 | 0.797 |
| chr7 | 91747506 | 91748315 | 810 | 0.17 | 0.749 | 0.797 |
| chr10 | 131934068 | 131957315 | 23248 | -0.04 | 0.749 | 0.797 |
| chr20 | 32943113 | 33035797 | 92685 | -0.02 | 0.749 | 0.797 |
| chr8 | 28458027 | 28459560 | 1534 | -0.1 | 0.75 | 0.798 |
| chr5 | 60105017 | 60108536 | 3520 | -0.05 | 0.75 | 0.798 |
| chr10 | 87679436 | 87679923 | 488 | -0.25 | 0.75 | 0.798 |
| chr7 | 17854839 | 17855783 | 945 | -0.18 | 0.75 | 0.798 |
| chr17 | 34216892 | 34217600 | 709 | 0.09 | 0.75 | 0.798 |
| chr1 | 61541789 | 61551009 | 9221 | -0.04 | 0.75 | 0.798 |
| chr9 | 14347428 | 14348042 | 615 | -0.17 | 0.75 | 0.798 |
| chr2 | 55772270 | 55784822 | 12553 | -0.08 | 0.751 | 0.798 |
| chr19 | 58872613 | 58875617 | 3005 | -0.04 | 0.751 | 0.799 |
| chr5 | 118127466 | 118127847 | 382 | 0.19 | 0.751 | 0.799 |
| chr2 | 25193064 | 25198908 | 5845 | -0.02 | 0.751 | 0.799 |
| chr17 | 68132355 | 68135168 | 2814 | -0.05 | 0.751 | 0.799 |
| chr10 | 82167000 | 82296632 | 129633 | -0.01 | 0.751 | 0.799 |
| chr19 | 18653081 | 18654874 | 1794 | 0.12 | 0.752 | 0.799 |
| chr2 | 157110629 | 157111035 | 407 | 0.2 | 0.752 | 0.799 |
| chr10 | 461319 | 462695 | 1377 | 0.12 | 0.752 | 0.799 |
| chr12 | 2061756 | 2063987 | 2232 | 0.1 | 0.752 | 0.799 |
| chr13 | 62903549 | 62904329 | 781 | 0.14 | 0.752 | 0.8 |
| chr16 | 15640813 | 15701163 | 60351 | -0.02 | 0.752 | 0.8 |
| chr20 | 32670282 | 32672094 | 1813 | -0.15 | 0.752 | 0.8 |
| chr16 | 23565463 | 23573272 | 7810 | 0.04 | 0.752 | 0.8 |
| chr17 | 28053322 | 28054347 | 1026 | 0.15 | 0.752 | 0.8 |
| chr18 | 9558380 | 9578331 | 19952 | -0.05 | 0.752 | 0.8 |
| chr11 | 88069946 | 88072152 | 2207 | 0.06 | 0.752 | 0.8 |
| chrX | 106360823 | 106362498 | 1676 | -0.09 | 0.752 | 0.8 |
| chr14 | 67735951 | 67736928 | 978 | 0.17 | 0.753 | 0.8 |
| chr3 | 124338980 | 124339981 | 1002 | -0.12 | 0.753 | 0.8 |
| chr3 | 49021628 | 49029316 | 7689 | -0.04 | 0.753 | 0.8 |
| chr4 | 125098550 | 125099985 | 1436 | -0.11 | 0.753 | 0.8 |
| chr19 | 1939661 | 1942953 | 3293 | -0.06 | 0.753 | 0.801 |
| chr17 | 48585045 | 48586696 | 1652 | 0.08 | 0.753 | 0.801 |
| chr17 | 38254072 | 38337271 | 83200 | 0.02 | 0.753 | 0.801 |
| chr16 | 81069476 | 81071319 | 1844 | 0.09 | 0.754 | 0.801 |
| chr3 | 135889263 | 135917202 | 27940 | 0.03 | 0.754 | 0.801 |
| chr1 | 235838037 | 235839277 | 1241 | 0.15 | 0.754 | 0.801 |
| chr15 | 89614513 | 89614850 | 338 | 0.21 | 0.754 | 0.801 |
| chr5 | 79286367 | 79287624 | 1258 | -0.07 | 0.754 | 0.801 |
| chr3 | 189698911 | 189700024 | 1114 | -0.12 | 0.754 | 0.801 |
| chr19 | 34649830 | 34725377 | 75548 | 0.01 | 0.754 | 0.802 |
| chr2 | 201064148 | 201065124 | 977 | -0.18 | 0.754 | 0.802 |
| chr7 | 83685193 | 83691529 | 6337 | 0.04 | 0.755 | 0.802 |
| chr11 | 68670301 | 68672832 | 2532 | 0.06 | 0.755 | 0.802 |
| chr7 | 77038154 | 77069591 | 31438 | -0.04 | 0.755 | 0.802 |
| chr7 | 130982273 | 130990240 | 7968 | -0.04 | 0.755 | 0.802 |
| chr2 | 231576791 | 231579228 | 2438 | 0.06 | 0.755 | 0.802 |
| chr4 | 126571371 | 126571723 | 353 | 0.28 | 0.755 | 0.802 |
| chr5 | 59491459 | 59493797 | 2339 | 0.05 | 0.755 | 0.802 |
| chr3 | 126614201 | 126615485 | 1285 | -0.14 | 0.756 | 0.803 |
| chr19 | 14182174 | 14203026 | 20853 | -0.04 | 0.756 | 0.803 |
| chr3 | 107940184 | 107942047 | 1864 | -0.05 | 0.756 | 0.803 |
| chr4 | 99799877 | 99809300 | 9424 | -0.07 | 0.756 | 0.803 |
| chr14 | 104152589 | 104182871 | 30283 | -0.04 | 0.756 | 0.803 |
| chr1 | 11538885 | 11539164 | 280 | -0.18 | 0.756 | 0.803 |

| | | | | | | |
|---|---|---|---|---|---|---|
| chr17 | 33996503 | 34040852 | 44350 | -0.03 | 0.757 | 0.803 |
| chr11 | 43901726 | 43903062 | 1337 | 0.08 | 0.757 | 0.803 |
| chr14 | 61091802 | 61092781 | 980 | -0.14 | 0.757 | 0.804 |
| chr20 | 40321198 | 40323315 | 2118 | 0.05 | 0.757 | 0.804 |
| chr3 | 72201644 | 72202557 | 914 | -0.13 | 0.757 | 0.804 |
| chr12 | 121676009 | 121677570 | 1562 | 0.13 | 0.757 | 0.804 |
| chr2 | 45812628 | 45813054 | 427 | -0.15 | 0.757 | 0.804 |
| chr9 | 88355393 | 88359434 | 4042 | 0.04 | 0.758 | 0.804 |
| chr22 | 48863025 | 48863705 | 681 | 0.18 | 0.758 | 0.804 |
| chr2 | 101349902 | 101350658 | 757 | 0.19 | 0.758 | 0.804 |
| chr4 | 140446313 | 140482132 | 35820 | -0.02 | 0.758 | 0.804 |
| chr10 | 8374174 | 8374861 | 688 | 0.16 | 0.758 | 0.805 |
| chr16 | 277486 | 286327 | 8842 | 0.03 | 0.758 | 0.805 |
| chr14 | 100841550 | 100891680 | 50131 | 0.03 | 0.758 | 0.805 |
| chr13 | 98825328 | 98826270 | 943 | 0.16 | 0.758 | 0.805 |
| chr1 | 39406279 | 39408145 | 1867 | 0.06 | 0.758 | 0.805 |
| chr2 | 112374712 | 112375836 | 1125 | 0.13 | 0.759 | 0.805 |
| chr17 | 63779277 | 63782604 | 3328 | -0.04 | 0.759 | 0.805 |
| chr4 | 189028352 | 189033729 | 5378 | -0.03 | 0.759 | 0.805 |
| chr3 | 178156917 | 178158340 | 1424 | 0.12 | 0.759 | 0.805 |
| chr9 | 140471610 | 140486130 | 14521 | -0.07 | 0.759 | 0.805 |
| chr9 | 127904999 | 127905792 | 794 | 0.15 | 0.759 | 0.805 |
| chr11 | 33361010 | 33361887 | 878 | 0.17 | 0.76 | 0.806 |
| chr11 | 43425078 | 43434007 | 8930 | 0.06 | 0.76 | 0.806 |
| chr6 | 551380 | 551828 | 449 | -0.22 | 0.76 | 0.806 |
| chr19 | 12888148 | 12918218 | 30071 | 0.02 | 0.76 | 0.806 |
| chr6 | 35335983 | 35338805 | 2823 | -0.08 | 0.76 | 0.806 |
| chr3 | 42922189 | 42922816 | 628 | 0.17 | 0.76 | 0.807 |
| chr1 | 8916908 | 8973084 | 56177 | -0.02 | 0.76 | 0.807 |
| chr13 | 41753233 | 41754835 | 1603 | -0.12 | 0.76 | 0.807 |
| chr5 | 145662706 | 145663684 | 979 | -0.13 | 0.76 | 0.807 |
| chr13 | 99097749 | 99103527 | 5779 | 0.09 | 0.761 | 0.807 |
| chr5 | 154356579 | 154357821 | 1243 | -0.13 | 0.761 | 0.807 |
| chrX | 69672152 | 69672717 | 566 | 0.19 | 0.761 | 0.807 |
| chr16 | 25077650 | 25079047 | 1398 | 0.08 | 0.761 | 0.807 |
| chr8 | 17855953 | 17867945 | 11993 | -0.06 | 0.761 | 0.807 |
| chrX | 21429488 | 21431362 | 1875 | 0.08 | 0.761 | 0.807 |
| chr12 | 100583542 | 100599019 | 15478 | 0.03 | 0.762 | 0.808 |
| chr7 | 44280369 | 44281221 | 853 | -0.21 | 0.762 | 0.808 |
| chr22 | 37414637 | 37448912 | 34276 | -0.02 | 0.762 | 0.808 |
| chr4 | 170678043 | 170679394 | 1352 | -0.12 | 0.762 | 0.808 |
| chr1 | 150785300 | 150785690 | 391 | 0.25 | 0.762 | 0.808 |
| chr17 | 59433221 | 59435489 | 2269 | -0.05 | 0.762 | 0.808 |
| chr14 | 105759878 | 105781693 | 21816 | -0.05 | 0.763 | 0.809 |
| chr8 | 29813067 | 29816610 | 3544 | -0.06 | 0.763 | 0.809 |
| chr10 | 35619643 | 35627230 | 7588 | 0.04 | 0.763 | 0.809 |
| chr11 | 10144372 | 10153096 | 8725 | 0.13 | 0.763 | 0.809 |
| chr3 | 159449587 | 159451793 | 2207 | -0.08 | 0.763 | 0.809 |
| chr16 | 1822664 | 1834189 | 11526 | 0.05 | 0.764 | 0.809 |
| chr3 | 132035859 | 132036750 | 892 | -0.14 | 0.764 | 0.81 |
| chr1 | 43146890 | 43159611 | 12722 | 0.05 | 0.764 | 0.81 |
| chr22 | 39541822 | 39549366 | 7545 | -0.09 | 0.764 | 0.81 |
| chr22 | 19365863 | 19366267 | 405 | 0.24 | 0.764 | 0.81 |
| chr9 | 132072593 | 132125164 | 52572 | 0.02 | 0.764 | 0.81 |
| chr17 | 75500393 | 75500939 | 547 | 0.25 | 0.765 | 0.81 |
| chr8 | 29227047 | 29244556 | 17510 | 0.04 | 0.765 | 0.81 |
| chr7 | 99035488 | 99037692 | 2205 | 0.07 | 0.765 | 0.81 |
| chrX | 109378481 | 109379619 | 1139 | -0.16 | 0.765 | 0.81 |
| chrX | 140065417 | 140067770 | 2354 | -0.07 | 0.765 | 0.81 |
| chr1 | 72748770 | 72749383 | 614 | 0.15 | 0.765 | 0.81 |
| chr16 | 30904570 | 30915731 | 11162 | -0.04 | 0.765 | 0.81 |

| | | | | | | |
|---|---|---|---|---|---|---|
| chr17 | 42754232 | 42769053 | 14822 | -0.03 | 0.765 | 0.81 |
| chr13 | 86132958 | 86133497 | 540 | -0.21 | 0.765 | 0.811 |
| chr10 | 130008167 | 130009336 | 1170 | 0.11 | 0.765 | 0.811 |
| chr15 | 78614925 | 78615268 | 344 | 0.2 | 0.765 | 0.811 |
| chr9 | 123193442 | 123209834 | 16393 | 0.05 | 0.765 | 0.811 |
| chr1 | 28796884 | 28799438 | 2555 | -0.05 | 0.765 | 0.811 |
| chr10 | 71490766 | 71491737 | 972 | 0.17 | 0.766 | 0.811 |
| chr7 | 92053542 | 92055601 | 2060 | -0.09 | 0.766 | 0.811 |
| chr10 | 105614336 | 105615698 | 1363 | 0.12 | 0.766 | 0.811 |
| chr15 | 83874451 | 83877293 | 2843 | -0.04 | 0.766 | 0.811 |
| chr19 | 35757438 | 35765065 | 7628 | -0.04 | 0.766 | 0.811 |
| chr4 | 54442925 | 54443336 | 412 | -0.22 | 0.766 | 0.811 |
| chr10 | 93717357 | 93717914 | 558 | 0.19 | 0.766 | 0.811 |
| chr17 | 19140911 | 19151710 | 10800 | -0.06 | 0.766 | 0.811 |
| chr10 | 94049842 | 94061042 | 11201 | 0.04 | 0.766 | 0.811 |
| chr7 | 45111761 | 45113727 | 1967 | -0.12 | 0.766 | 0.811 |
| chr17 | 45570188 | 45570695 | 508 | 0.18 | 0.766 | 0.812 |
| chr15 | 50585166 | 50592929 | 7764 | 0.08 | 0.767 | 0.812 |
| chr12 | 79475959 | 79477923 | 1965 | 0.06 | 0.767 | 0.812 |
| chr1 | 8042090 | 8044504 | 2415 | -0.09 | 0.767 | 0.812 |
| chr17 | 29288286 | 29300750 | 12465 | -0.02 | 0.767 | 0.812 |
| chr7 | 56019028 | 56046243 | 27216 | -0.04 | 0.767 | 0.812 |
| chr5 | 127409557 | 127421213 | 11657 | 0.04 | 0.767 | 0.812 |
| chr20 | 2820378 | 2822227 | 1850 | 0.05 | 0.767 | 0.812 |
| chr1 | 237957022 | 237958172 | 1151 | -0.13 | 0.767 | 0.812 |
| chr3 | 138108748 | 138109758 | 1011 | 0.14 | 0.767 | 0.812 |
| chr7 | 95949831 | 95952135 | 2305 | 0.05 | 0.768 | 0.813 |
| chrX | 12964261 | 12976240 | 11980 | 0.04 | 0.768 | 0.813 |
| chr8 | 27672192 | 27696114 | 23923 | 0.03 | 0.768 | 0.813 |
| chr5 | 172067618 | 172068126 | 509 | -0.14 | 0.768 | 0.813 |
| chr7 | 80633764 | 80643284 | 9521 | 0.24 | 0.768 | 0.813 |
| chr9 | 139734425 | 139762260 | 27836 | 0.03 | 0.769 | 0.813 |
| chr2 | 48088500 | 48096112 | 7613 | -0.06 | 0.769 | 0.814 |
| chr7 | 141317017 | 141317427 | 411 | -0.18 | 0.769 | 0.814 |
| chr16 | 47480523 | 47510320 | 29798 | -0.02 | 0.769 | 0.814 |
| chrX | 13406070 | 13418405 | 12336 | 0.03 | 0.77 | 0.815 |
| chr22 | 31450733 | 31452580 | 1848 | 0.06 | 0.77 | 0.815 |
| chr2 | 198345434 | 198382542 | 37109 | 0.02 | 0.77 | 0.815 |
| chr1 | 246794338 | 246796857 | 2520 | -0.08 | 0.771 | 0.815 |
| chrX | 129193943 | 129194992 | 1050 | 0.13 | 0.771 | 0.815 |
| chr2 | 101434687 | 101777553 | 342867 | -0.01 | 0.771 | 0.815 |
| chr13 | 34985580 | 34996841 | 11262 | 0.11 | 0.771 | 0.815 |
| chr5 | 148818270 | 148836693 | 18424 | -0.04 | 0.771 | 0.815 |
| chr10 | 81837474 | 81840099 | 2626 | 0.03 | 0.771 | 0.816 |
| chr8 | 28008340 | 28008913 | 574 | 0.2 | 0.771 | 0.816 |
| chr9 | 34022389 | 34050441 | 28053 | 0.03 | 0.771 | 0.816 |
| chr15 | 63478102 | 63482165 | 4064 | -0.11 | 0.771 | 0.816 |
| chr2 | 176883809 | 176884551 | 743 | 0.18 | 0.771 | 0.816 |
| chr7 | 674882 | 675458 | 577 | -0.22 | 0.771 | 0.816 |
| chr1 | 213403165 | 213403514 | 350 | -0.21 | 0.771 | 0.816 |
| chr11 | 102845601 | 102846116 | 516 | 0.17 | 0.772 | 0.816 |
| chr16 | 74689574 | 74701865 | 12292 | 0.04 | 0.772 | 0.816 |
| chr6 | 14474689 | 14479359 | 4671 | -0.05 | 0.772 | 0.816 |
| chrX | 18692237 | 18693913 | 1677 | 0.07 | 0.772 | 0.816 |
| chr19 | 36484899 | 36487177 | 2279 | 0.05 | 0.772 | 0.817 |
| chr9 | 135089415 | 135090121 | 707 | 0.17 | 0.773 | 0.817 |
| chr5 | 180541953 | 180542810 | 858 | -0.13 | 0.773 | 0.817 |
| chr12 | 80006387 | 80015210 | 8824 | -0.05 | 0.773 | 0.817 |
| chr17 | 68401329 | 68402521 | 1193 | 0.1 | 0.773 | 0.818 |
| chr14 | 55030953 | 55047158 | 16206 | 0.04 | 0.774 | 0.818 |
| chr2 | 30703341 | 30704938 | 1598 | -0.11 | 0.774 | 0.818 |

| | | | | | | |
|---|---|---|---|---|---|---|
| chr10 | 89435536 | 89436444 | 909 | 0.13 | 0.774 | 0.818 |
| chr11 | 77122362 | 77123778 | 1417 | -0.11 | 0.774 | 0.818 |
| chr6 | 2790543 | 2791442 | 900 | 0.13 | 0.774 | 0.818 |
| chr6 | 3518042 | 3519378 | 1337 | -0.1 | 0.774 | 0.819 |
| chr8 | 83462552 | 83482423 | 19872 | -0.03 | 0.774 | 0.819 |
| chr3 | 114324903 | 114328182 | 3280 | 0.06 | 0.774 | 0.819 |
| chr2 | 57987048 | 57987733 | 686 | 0.12 | 0.775 | 0.819 |
| chr1 | 32986247 | 32986780 | 534 | -0.16 | 0.775 | 0.819 |
| chr22 | 41745894 | 41748876 | 2983 | 0.06 | 0.775 | 0.82 |
| chr10 | 889671 | 893934 | 4264 | 0.08 | 0.776 | 0.82 |
| chr6 | 84018130 | 84019894 | 1765 | -0.1 | 0.776 | 0.82 |
| chr20 | 37393080 | 37394816 | 1737 | -0.1 | 0.776 | 0.82 |
| chr19 | 55836189 | 55857674 | 21486 | 0.03 | 0.776 | 0.82 |
| chr1 | 101087160 | 101089488 | 2329 | 0.06 | 0.776 | 0.82 |
| chrX | 149981750 | 149982718 | 969 | 0.11 | 0.776 | 0.82 |
| chr17 | 2381094 | 2416180 | 35087 | 0.02 | 0.776 | 0.82 |
| chr12 | 47945265 | 47946260 | 996 | -0.15 | 0.776 | 0.82 |
| chr3 | 114270386 | 114272086 | 1701 | -0.07 | 0.776 | 0.82 |
| chr6 | 144732756 | 144737151 | 4396 | 0.09 | 0.777 | 0.82 |
| chr2 | 203941548 | 203942644 | 1097 | -0.12 | 0.777 | 0.821 |
| chr21 | 27443150 | 27455989 | 12840 | -0.06 | 0.777 | 0.821 |
| chr2 | 120183431 | 120190022 | 6592 | -0.06 | 0.777 | 0.821 |
| chr15 | 35836948 | 35850396 | 13449 | -0.04 | 0.778 | 0.822 |
| chr17 | 1040025 | 1041538 | 1514 | -0.08 | 0.778 | 0.822 |
| chr21 | 36569938 | 36571315 | 1378 | -0.14 | 0.778 | 0.822 |
| chr2 | 43985642 | 43987201 | 1560 | -0.08 | 0.778 | 0.822 |
| chr2 | 64749953 | 64753433 | 3481 | -0.04 | 0.779 | 0.822 |
| chr1 | 224356601 | 224357184 | 584 | 0.16 | 0.779 | 0.822 |
| chr2 | 122380955 | 122410074 | 29120 | -0.02 | 0.779 | 0.823 |
| chr6 | 86369586 | 86370619 | 1034 | -0.12 | 0.779 | 0.823 |
| chr2 | 48333152 | 48342652 | 9501 | 0.03 | 0.779 | 0.823 |
| chr2 | 109028832 | 109030623 | 1792 | -0.09 | 0.779 | 0.823 |
| chr3 | 51421353 | 51430034 | 8682 | 0.04 | 0.78 | 0.823 |
| chr14 | 78225780 | 78228401 | 2622 | 0.05 | 0.78 | 0.824 |
| chr12 | 32287057 | 32287795 | 739 | -0.17 | 0.781 | 0.824 |
| chr11 | 8289302 | 8295042 | 5741 | -0.13 | 0.781 | 0.824 |
| chr9 | 104295199 | 104298022 | 2824 | -0.04 | 0.781 | 0.825 |
| chr3 | 51975128 | 51977692 | 2565 | 0.06 | 0.781 | 0.825 |
| chr5 | 123145246 | 123156978 | 11733 | 0.04 | 0.781 | 0.825 |
| chr20 | 18446183 | 18449083 | 2901 | -0.04 | 0.781 | 0.825 |
| chr15 | 99619531 | 99620125 | 595 | -0.1 | 0.781 | 0.825 |
| chr2 | 88990115 | 88992430 | 2316 | -0.05 | 0.782 | 0.825 |
| chr7 | 134354237 | 134355216 | 980 | -0.09 | 0.782 | 0.825 |
| chr5 | 175123225 | 175123705 | 481 | 0.18 | 0.782 | 0.825 |
| chr15 | 67295650 | 67296330 | 681 | -0.14 | 0.782 | 0.825 |
| chr21 | 39714513 | 39724595 | 10083 | -0.06 | 0.782 | 0.825 |
| chr16 | 4702657 | 4753067 | 50411 | -0.02 | 0.782 | 0.825 |
| chr6 | 135229316 | 135231183 | 1868 | -0.1 | 0.782 | 0.825 |
| chr3 | 172534012 | 172534628 | 617 | -0.17 | 0.782 | 0.825 |
| chr22 | 23744787 | 23745641 | 855 | -0.11 | 0.782 | 0.825 |
| chr5 | 53813516 | 53814235 | 720 | 0.11 | 0.782 | 0.826 |
| chr15 | 78285774 | 78286821 | 1048 | -0.1 | 0.782 | 0.826 |
| chr8 | 144896017 | 144923232 | 27216 | 0.03 | 0.782 | 0.826 |
| chr9 | 19228448 | 19248512 | 20065 | 0.04 | 0.783 | 0.826 |
| chr13 | 43559554 | 43560352 | 799 | -0.16 | 0.783 | 0.826 |
| chr1 | 88219677 | 88233071 | 13395 | 0.11 | 0.783 | 0.826 |
| chr4 | 148871577 | 148872885 | 1309 | 0.09 | 0.783 | 0.826 |
| chr20 | 50721372 | 50722370 | 999 | 0.11 | 0.783 | 0.826 |
| chr15 | 81321340 | 81322752 | 1413 | 0.11 | 0.783 | 0.827 |
| chr1 | 12585023 | 12680686 | 95664 | -0.01 | 0.784 | 0.827 |
| chr1 | 94848537 | 94861945 | 13409 | 0.03 | 0.784 | 0.827 |

| | | | | | | |
|---|---|---|---|---|---|---|
| chr17 | 74768590 | 74769563 | 974 | -0.11 | 0.784 | 0.827 |
| chr10 | 31417463 | 31424120 | 6658 | -0.08 | 0.785 | 0.828 |
| chr6 | 54357625 | 54358888 | 1264 | 0.09 | 0.785 | 0.828 |
| chr2 | 10394072 | 10395296 | 1225 | 0.12 | 0.785 | 0.828 |
| chr2 | 26099734 | 26102548 | 2815 | -0.03 | 0.785 | 0.828 |
| chr17 | 76250074 | 76250703 | 630 | -0.13 | 0.785 | 0.828 |
| chr19 | 18156640 | 18176370 | 19731 | -0.07 | 0.785 | 0.828 |
| chr3 | 39423996 | 39426529 | 2534 | -0.04 | 0.785 | 0.828 |
| chr2 | 131484561 | 131485190 | 630 | 0.12 | 0.785 | 0.828 |
| chr7 | 99860950 | 99870486 | 9537 | -0.04 | 0.785 | 0.828 |
| chr4 | 25653658 | 25654585 | 928 | 0.11 | 0.785 | 0.828 |
| chr21 | 40159124 | 40193249 | 34126 | -0.02 | 0.785 | 0.828 |
| chr3 | 87881066 | 87883199 | 2134 | 0.04 | 0.786 | 0.828 |
| chr1 | 149782254 | 149782690 | 437 | 0.17 | 0.786 | 0.828 |
| chr5 | 173146646 | 173147177 | 532 | -0.18 | 0.786 | 0.829 |
| chr4 | 186123319 | 186132140 | 8822 | -0.05 | 0.786 | 0.829 |
| chr1 | 1609960 | 1619062 | 9103 | 0.04 | 0.786 | 0.829 |
| chr7 | 809814 | 816478 | 6665 | -0.09 | 0.786 | 0.829 |
| chr8 | 122905955 | 122906525 | 571 | -0.14 | 0.786 | 0.829 |
| chr9 | 4792196 | 4794129 | 1934 | -0.07 | 0.786 | 0.829 |
| chr17 | 77766306 | 77788117 | 21812 | -0.03 | 0.787 | 0.829 |
| chr19 | 1811919 | 1813806 | 1888 | 0.1 | 0.787 | 0.829 |
| chr6 | 169740240 | 169740674 | 435 | 0.17 | 0.787 | 0.829 |
| chrX | 117552649 | 117553570 | 922 | 0.13 | 0.787 | 0.829 |
| chr13 | 50364877 | 50368030 | 3154 | -0.05 | 0.787 | 0.83 |
| chr2 | 203567662 | 203568445 | 784 | 0.18 | 0.787 | 0.83 |
| chr5 | 145825927 | 145831136 | 5210 | -0.04 | 0.787 | 0.83 |
| chr14 | 24020024 | 24039326 | 19303 | -0.03 | 0.787 | 0.83 |
| chr7 | 140486085 | 140486792 | 708 | 0.16 | 0.787 | 0.83 |
| chr1 | 15910874 | 15911927 | 1054 | -0.11 | 0.788 | 0.83 |
| chr1 | 156250738 | 156266012 | 15275 | 0.03 | 0.788 | 0.83 |
| chr2 | 120769849 | 120775210 | 5362 | 0.06 | 0.788 | 0.83 |
| chr15 | 91183550 | 91189793 | 6244 | 0.06 | 0.788 | 0.831 |
| chr2 | 174453070 | 174456341 | 3272 | -0.05 | 0.788 | 0.831 |
| chr13 | 50068706 | 50070937 | 2232 | 0.05 | 0.788 | 0.831 |
| chr3 | 15599036 | 15599748 | 713 | 0.16 | 0.789 | 0.831 |
| chr17 | 78215803 | 78269887 | 54085 | 0.01 | 0.789 | 0.831 |
| chr13 | 110152450 | 110153062 | 613 | -0.15 | 0.789 | 0.831 |
| chr3 | 115376834 | 115378118 | 1285 | 0.06 | 0.789 | 0.831 |
| chr5 | 158998200 | 159013366 | 15167 | 0.02 | 0.789 | 0.831 |
| chr7 | 15546497 | 15548336 | 1840 | 0.08 | 0.789 | 0.831 |
| chr4 | 53524820 | 53526502 | 1683 | 0.07 | 0.789 | 0.832 |
| chr3 | 183352889 | 183355186 | 2298 | 0.05 | 0.79 | 0.832 |
| chr16 | 49788880 | 49789302 | 423 | -0.18 | 0.79 | 0.832 |
| chr11 | 66233564 | 66248734 | 15171 | -0.03 | 0.79 | 0.832 |
| chr1 | 22182011 | 22197393 | 15383 | 0.04 | 0.79 | 0.832 |
| chr9 | 33813216 | 33889942 | 76727 | 0.01 | 0.79 | 0.832 |
| chr18 | 11835683 | 11836145 | 463 | 0.19 | 0.79 | 0.832 |
| chr16 | 66863498 | 66865087 | 1590 | 0.06 | 0.79 | 0.832 |
| chr2 | 45557917 | 45558288 | 372 | -0.17 | 0.791 | 0.833 |
| chr2 | 32794465 | 32795031 | 567 | 0.19 | 0.791 | 0.833 |
| chr17 | 47752896 | 47766251 | 13356 | -0.05 | 0.791 | 0.833 |
| chr7 | 44240272 | 44254683 | 14412 | 0.07 | 0.791 | 0.833 |
| chr8 | 118531874 | 118534400 | 2527 | 0.04 | 0.791 | 0.833 |
| chr1 | 193251969 | 193253475 | 1507 | -0.08 | 0.792 | 0.833 |
| chr12 | 121968046 | 121976794 | 8749 | 0.05 | 0.792 | 0.833 |
| chr3 | 121408532 | 121409206 | 675 | -0.21 | 0.792 | 0.833 |
| chr11 | 29113030 | 29117116 | 4087 | 0.03 | 0.792 | 0.833 |
| chr3 | 196336444 | 196336938 | 495 | -0.15 | 0.792 | 0.833 |
| chr3 | 93560238 | 93560642 | 405 | -0.12 | 0.792 | 0.834 |
| chr3 | 46027171 | 46037613 | 10443 | -0.05 | 0.792 | 0.834 |

| | | | | | | |
|---|---|---|---|---|---|---|
| chr19 | 7552989 | 7554869 | 1881 | -0.06 | 0.792 | 0.834 |
| chr21 | 40365387 | 40365994 | 608 | 0.16 | 0.792 | 0.834 |
| chr11 | 995211 | 1001744 | 6534 | 0.06 | 0.793 | 0.834 |
| chr15 | 59172617 | 59173720 | 1104 | 0.14 | 0.793 | 0.834 |
| chr17 | 40884094 | 40897461 | 13368 | 0.04 | 0.793 | 0.834 |
| chr3 | 27631791 | 27632965 | 1175 | 0.09 | 0.793 | 0.834 |
| chr11 | 77344603 | 77349620 | 5018 | 0.04 | 0.793 | 0.835 |
| chr17 | 7648951 | 7659240 | 10290 | 0.18 | 0.793 | 0.835 |
| chrX | 152735247 | 152736050 | 804 | 0.11 | 0.793 | 0.835 |
| chr3 | 110789158 | 110792064 | 2907 | -0.05 | 0.793 | 0.835 |
| chr1 | 16029647 | 16030584 | 938 | -0.18 | 0.794 | 0.835 |
| chr10 | 14901669 | 14903612 | 1944 | -0.08 | 0.794 | 0.835 |
| chr3 | 138066250 | 138067888 | 1639 | -0.07 | 0.794 | 0.835 |
| chr10 | 70181010 | 70194947 | 13938 | 0.04 | 0.794 | 0.836 |
| chr6 | 4058315 | 4060017 | 1703 | -0.09 | 0.794 | 0.836 |
| chr19 | 983501 | 985209 | 1709 | -0.07 | 0.795 | 0.836 |
| chr3 | 105257109 | 105257797 | 689 | -0.15 | 0.795 | 0.836 |
| chr20 | 32103803 | 32106466 | 2664 | -0.06 | 0.795 | 0.836 |
| chrX | 71477670 | 71497641 | 19972 | -0.03 | 0.795 | 0.836 |
| chr11 | 86167400 | 86175689 | 8290 | 0.05 | 0.795 | 0.836 |
| chr9 | 123128146 | 123129744 | 1599 | 0.06 | 0.795 | 0.836 |
| chr10 | 124738950 | 124741020 | 2071 | 0.05 | 0.795 | 0.836 |
| chr7 | 148841739 | 148870363 | 28625 | -0.03 | 0.795 | 0.836 |
| chr17 | 65820826 | 65837290 | 16465 | -0.04 | 0.796 | 0.837 |
| chr1 | 28966888 | 28976608 | 9721 | 0.03 | 0.796 | 0.837 |
| chr2 | 159779214 | 159783728 | 4515 | 0.09 | 0.796 | 0.837 |
| chr13 | 33768768 | 33769563 | 796 | 0.14 | 0.796 | 0.837 |
| chr7 | 34360795 | 34361391 | 597 | -0.15 | 0.797 | 0.838 |
| chr5 | 10248639 | 10253602 | 4964 | -0.02 | 0.797 | 0.838 |
| chr11 | 12952529 | 12953089 | 561 | 0.15 | 0.797 | 0.838 |
| chr1 | 36500115 | 36500945 | 831 | -0.15 | 0.798 | 0.839 |
| chr19 | 49587762 | 49674315 | 86554 | 0.02 | 0.798 | 0.839 |
| chr15 | 45475312 | 45476120 | 809 | -0.13 | 0.798 | 0.839 |
| chr5 | 76169705 | 76170405 | 701 | -0.16 | 0.798 | 0.839 |
| chr7 | 110753635 | 110754763 | 1129 | -0.13 | 0.798 | 0.839 |
| chr2 | 135207659 | 135208313 | 655 | 0.15 | 0.798 | 0.839 |
| chr11 | 111472258 | 111474547 | 2290 | -0.05 | 0.799 | 0.84 |
| chr12 | 121769015 | 121775885 | 6871 | -0.05 | 0.799 | 0.84 |
| chr19 | 58837843 | 58839929 | 2087 | -0.06 | 0.799 | 0.84 |
| chr15 | 74987279 | 74989108 | 1830 | 0.05 | 0.799 | 0.84 |
| chr19 | 34842144 | 34985543 | 143400 | 0.01 | 0.799 | 0.84 |
| chr4 | 2242783 | 2243779 | 997 | -0.1 | 0.799 | 0.84 |
| chr6 | 150283863 | 150285286 | 1424 | 0.08 | 0.799 | 0.84 |
| chr2 | 190385879 | 190421400 | 35522 | -0.03 | 0.799 | 0.84 |
| chr2 | 85580648 | 85583119 | 2472 | 0.05 | 0.799 | 0.84 |
| chr1 | 180463817 | 180486203 | 22387 | 0.04 | 0.799 | 0.84 |
| chr4 | 128061221 | 128065658 | 4438 | 0.07 | 0.8 | 0.84 |
| chr17 | 16769357 | 16770172 | 816 | 0.1 | 0.8 | 0.84 |
| chr15 | 75511921 | 75512475 | 555 | 0.16 | 0.8 | 0.84 |
| chr1 | 155657742 | 155660355 | 2614 | -0.04 | 0.8 | 0.841 |
| chr1 | 212872021 | 212873918 | 1898 | 0.08 | 0.801 | 0.841 |
| chr6 | 57132570 | 57132973 | 404 | 0.2 | 0.801 | 0.842 |
| chrX | 117494412 | 117495028 | 617 | -0.15 | 0.801 | 0.842 |
| chr18 | 24234719 | 24239010 | 4292 | -0.04 | 0.801 | 0.842 |
| chr2 | 15869803 | 15871158 | 1356 | -0.09 | 0.801 | 0.842 |
| chr2 | 239896011 | 239901271 | 5261 | -0.06 | 0.801 | 0.842 |
| chr2 | 172410168 | 172411328 | 1161 | -0.1 | 0.802 | 0.842 |
| chr1 | 24827571 | 24843082 | 15512 | -0.04 | 0.802 | 0.842 |
| chr19 | 11403865 | 11405244 | 1380 | 0.1 | 0.802 | 0.842 |
| chr8 | 71468786 | 71478442 | 9657 | -0.04 | 0.802 | 0.842 |
| chr8 | 124049761 | 124055704 | 5944 | 0.04 | 0.802 | 0.842 |

| | | | | | | |
|---|---|---|---|---|---|---|
| chr10 | 103542292 | 103544000 | 1709 | -0.06 | 0.802 | 0.843 |
| chr12 | 57399037 | 57403695 | 4659 | -0.05 | 0.803 | 0.843 |
| chr3 | 188664816 | 188670093 | 5278 | -0.03 | 0.803 | 0.843 |
| chr2 | 184013903 | 184014419 | 517 | -0.15 | 0.803 | 0.843 |
| chr6 | 157541958 | 157542870 | 913 | 0.11 | 0.803 | 0.843 |
| chr2 | 9579813 | 9580217 | 405 | 0.17 | 0.803 | 0.843 |
| chr1 | 110282453 | 110283227 | 775 | 0.13 | 0.803 | 0.843 |
| chr7 | 83710600 | 83722299 | 11700 | 0.03 | 0.803 | 0.843 |
| chr17 | 30468308 | 30484459 | 16152 | 0.03 | 0.803 | 0.843 |
| chr11 | 63528580 | 63537112 | 8533 | 0.09 | 0.803 | 0.843 |
| chr17 | 36146672 | 36148299 | 1628 | -0.07 | 0.803 | 0.843 |
| chr3 | 57203758 | 57204547 | 790 | -0.09 | 0.803 | 0.843 |
| chr2 | 158640619 | 158641986 | 1368 | 0.08 | 0.803 | 0.843 |
| chr6 | 1251691 | 1252057 | 367 | 0.23 | 0.803 | 0.843 |
| chr21 | 40684585 | 40694404 | 9820 | -0.04 | 0.803 | 0.843 |
| chr18 | 20159671 | 20161042 | 1372 | -0.08 | 0.804 | 0.844 |
| chr4 | 176451855 | 176453536 | 1682 | -0.09 | 0.804 | 0.844 |
| chr2 | 208632799 | 208694699 | 61901 | -0.02 | 0.804 | 0.844 |
| chr10 | 11779278 | 11785152 | 5875 | 0.07 | 0.804 | 0.844 |
| chr4 | 109605882 | 109606896 | 1015 | 0.1 | 0.804 | 0.844 |
| chrX | 16872029 | 16872538 | 510 | 0.21 | 0.804 | 0.844 |
| chr1 | 237051141 | 237051836 | 696 | -0.15 | 0.804 | 0.844 |
| chr16 | 72697906 | 72699789 | 1884 | -0.04 | 0.804 | 0.844 |
| chr5 | 158437885 | 158445122 | 7238 | -0.07 | 0.804 | 0.844 |
| chr17 | 65486576 | 65497196 | 10621 | -0.12 | 0.804 | 0.844 |
| chr2 | 207022505 | 207026370 | 3866 | 0.03 | 0.805 | 0.844 |
| chr1 | 205179528 | 205181279 | 1752 | 0.06 | 0.805 | 0.845 |
| chr1 | 153735321 | 153742111 | 6791 | -0.07 | 0.805 | 0.845 |
| chr4 | 110933550 | 110935251 | 1702 | -0.1 | 0.805 | 0.845 |
| chr4 | 150907839 | 150929367 | 21529 | -0.03 | 0.806 | 0.845 |
| chr11 | 7694336 | 7695339 | 1004 | -0.1 | 0.806 | 0.845 |
| chr7 | 130352722 | 130354174 | 1453 | -0.09 | 0.806 | 0.845 |
| chr2 | 71355829 | 71358782 | 2954 | 0.03 | 0.806 | 0.845 |
| chr5 | 149336647 | 149341198 | 4552 | 0.03 | 0.806 | 0.845 |
| chr19 | 6361260 | 6373195 | 11936 | 0.05 | 0.806 | 0.845 |
| chr10 | 7986925 | 7999356 | 12432 | 0.05 | 0.806 | 0.845 |
| chr3 | 11296719 | 11297277 | 559 | -0.15 | 0.806 | 0.845 |
| chr6 | 167426481 | 167427187 | 707 | 0.14 | 0.807 | 0.847 |
| chr16 | 56660918 | 56661764 | 847 | -0.1 | 0.807 | 0.847 |
| chr1 | 155803679 | 155808449 | 4771 | 0.09 | 0.807 | 0.847 |
| chr12 | 80276930 | 80282399 | 5470 | -0.09 | 0.807 | 0.847 |
| chr7 | 81876586 | 81877311 | 726 | 0.12 | 0.808 | 0.847 |
| chr5 | 34584404 | 34611073 | 26670 | 0.01 | 0.808 | 0.847 |
| chr5 | 34550570 | 34551179 | 610 | 0.13 | 0.808 | 0.847 |
| chr19 | 41468342 | 41469152 | 811 | -0.11 | 0.808 | 0.848 |
| chr11 | 123407419 | 123408031 | 613 | 0.13 | 0.808 | 0.848 |
| chr16 | 8914707 | 8917329 | 2623 | -0.08 | 0.808 | 0.848 |
| chr8 | 145855399 | 145856362 | 964 | 0.13 | 0.808 | 0.848 |
| chr16 | 11086544 | 11087337 | 794 | 0.16 | 0.809 | 0.848 |
| chrX | 119602043 | 119603237 | 1195 | 0.05 | 0.809 | 0.848 |
| chr3 | 124773184 | 124783257 | 10074 | -0.04 | 0.809 | 0.848 |
| chr12 | 133100215 | 133101839 | 1625 | 0.08 | 0.809 | 0.848 |
| chr21 | 17160938 | 17180074 | 19137 | -0.05 | 0.809 | 0.849 |
| chr1 | 53017223 | 53020092 | 2870 | 0.05 | 0.809 | 0.849 |
| chr18 | 55360649 | 55380684 | 20036 | 0.03 | 0.81 | 0.849 |
| chr3 | 25651582 | 25652279 | 698 | 0.13 | 0.81 | 0.849 |
| chr2 | 153046094 | 153057894 | 11801 | -0.08 | 0.81 | 0.849 |
| chr22 | 33697857 | 33698895 | 1039 | -0.12 | 0.81 | 0.849 |
| chr12 | 77219395 | 77220112 | 718 | 0.14 | 0.81 | 0.849 |
| chr15 | 22891705 | 22927768 | 36064 | 0.02 | 0.81 | 0.849 |
| chr2 | 121310255 | 121310577 | 323 | -0.14 | 0.811 | 0.85 |

| | | | | | | |
|---|---|---|---|---|---|---|
| chr4 | 2536791 | 2546483 | 9693 | 0.04 | 0.811 | 0.85 |
| chr19 | 12806539 | 12808058 | 1520 | -0.06 | 0.811 | 0.85 |
| chr3 | 51571975 | 51584224 | 12250 | -0.04 | 0.811 | 0.85 |
| chr19 | 18302350 | 18316519 | 14170 | 0.04 | 0.811 | 0.85 |
| chr1 | 151537766 | 151613225 | 75460 | 0.02 | 0.811 | 0.85 |
| chr9 | 115950188 | 115950970 | 783 | 0.14 | 0.811 | 0.85 |
| chr10 | 54325817 | 54326787 | 971 | -0.11 | 0.811 | 0.85 |
| chr7 | 34978336 | 34979383 | 1048 | -0.07 | 0.811 | 0.85 |
| chr17 | 1933182 | 1996715 | 63534 | 0.01 | 0.811 | 0.85 |
| chr4 | 160377035 | 160377486 | 452 | 0.16 | 0.812 | 0.85 |
| chr2 | 166213969 | 166215000 | 1032 | 0.12 | 0.812 | 0.85 |
| chr11 | 120195377 | 120210811 | 15435 | -0.03 | 0.812 | 0.85 |
| chr4 | 103634196 | 103634749 | 554 | -0.18 | 0.812 | 0.85 |
| chr2 | 191198597 | 191209555 | 10959 | -0.05 | 0.812 | 0.85 |
| chr14 | 37116861 | 37117593 | 733 | 0.13 | 0.812 | 0.851 |
| chr21 | 27259793 | 27271724 | 11932 | 0.05 | 0.812 | 0.851 |
| chr2 | 200842133 | 200850938 | 8806 | 0.06 | 0.812 | 0.851 |
| chr5 | 270818 | 272324 | 1507 | 0.06 | 0.812 | 0.851 |
| chr8 | 127836896 | 127837539 | 644 | 0.11 | 0.812 | 0.851 |
| chr12 | 105379609 | 105380982 | 1374 | -0.06 | 0.813 | 0.852 |
| chr6 | 88283573 | 88301133 | 17561 | -0.03 | 0.813 | 0.852 |
| chr21 | 35578375 | 35578878 | 504 | 0.16 | 0.813 | 0.852 |
| chr21 | 44526147 | 44528449 | 2303 | 0.04 | 0.813 | 0.852 |
| chr3 | 88197694 | 88214990 | 17297 | 0.03 | 0.813 | 0.852 |
| chr22 | 32018359 | 32059122 | 40764 | -0.02 | 0.813 | 0.852 |
| chr1 | 36614043 | 36653950 | 39908 | -0.02 | 0.813 | 0.852 |
| chrX | 73639835 | 73641158 | 1324 | 0.06 | 0.813 | 0.852 |
| chr22 | 20066446 | 20069603 | 3158 | 0.05 | 0.813 | 0.852 |
| chr18 | 9868023 | 9878053 | 10031 | -0.03 | 0.813 | 0.852 |
| chr17 | 7107411 | 7168432 | 61022 | -0.01 | 0.814 | 0.852 |
| chr3 | 9690524 | 9702734 | 12211 | -0.04 | 0.814 | 0.852 |
| chr13 | 48640164 | 48653062 | 12899 | -0.05 | 0.814 | 0.852 |
| chr16 | 80006147 | 80006596 | 450 | 0.15 | 0.814 | 0.852 |
| chr2 | 37551183 | 37552494 | 1312 | 0.07 | 0.814 | 0.852 |
| chr19 | 32834631 | 32897759 | 63129 | 0.01 | 0.815 | 0.853 |
| chr2 | 64370184 | 64372327 | 2144 | -0.08 | 0.815 | 0.853 |
| chr6 | 132179938 | 132186230 | 6293 | 0.05 | 0.815 | 0.853 |
| chr15 | 42866691 | 42868323 | 1633 | 0.09 | 0.815 | 0.853 |
| chr10 | 125851396 | 125867006 | 15611 | -0.03 | 0.815 | 0.853 |
| chr3 | 114237481 | 114239301 | 1821 | 0.1 | 0.815 | 0.853 |
| chr1 | 65210030 | 65221646 | 11617 | -0.07 | 0.815 | 0.853 |
| chr9 | 124952803 | 124953488 | 686 | -0.12 | 0.815 | 0.853 |
| chr8 | 8799735 | 8800306 | 572 | -0.11 | 0.815 | 0.853 |
| chr19 | 30250611 | 30251570 | 960 | 0.11 | 0.815 | 0.853 |
| chr5 | 61648992 | 61650023 | 1032 | -0.11 | 0.815 | 0.853 |
| chr5 | 171703136 | 171711899 | 8764 | -0.04 | 0.816 | 0.853 |
| chr10 | 71183654 | 71268009 | 84356 | 0.01 | 0.816 | 0.853 |
| chr17 | 15366671 | 15367291 | 621 | -0.15 | 0.816 | 0.853 |
| chr4 | 99824911 | 99825501 | 591 | -0.14 | 0.816 | 0.854 |
| chr14 | 52294878 | 52304945 | 10068 | 0.22 | 0.816 | 0.854 |
| chr6 | 91211673 | 91213317 | 1645 | -0.07 | 0.816 | 0.854 |
| chr11 | 909951 | 911873 | 1923 | -0.07 | 0.816 | 0.854 |
| chr4 | 140513758 | 140531961 | 18204 | 0.02 | 0.816 | 0.854 |
| chr5 | 91170504 | 91171417 | 914 | 0.1 | 0.816 | 0.854 |
| chr2 | 69134472 | 69136719 | 2248 | 0.03 | 0.817 | 0.854 |
| chr17 | 28803754 | 28805932 | 2179 | -0.02 | 0.817 | 0.854 |
| chr10 | 38382616 | 38384331 | 1716 | 0.06 | 0.817 | 0.854 |
| chr14 | 56009662 | 56010205 | 544 | 0.15 | 0.817 | 0.854 |
| chr7 | 92250539 | 92277479 | 26941 | 0.02 | 0.817 | 0.854 |
| chr18 | 12574452 | 12575674 | 1223 | 0.1 | 0.817 | 0.854 |
| chr3 | 39679233 | 39681774 | 2542 | 0.03 | 0.817 | 0.854 |

| | | | | | | |
|---|---|---|---|---|---|---|
| chr9 | 36090917 | 36103780 | 12864 | 0.06 | 0.817 | 0.855 |
| chr1 | 26323281 | 26325304 | 2024 | 0.05 | 0.818 | 0.855 |
| chr4 | 99563967 | 99576440 | 12474 | 0.48 | 0.818 | 0.855 |
| chr17 | 36740081 | 36748250 | 8170 | -0.07 | 0.818 | 0.855 |
| chr2 | 105986211 | 106021326 | 35116 | 0.02 | 0.818 | 0.856 |
| chr9 | 16133162 | 16133948 | 787 | -0.14 | 0.819 | 0.856 |
| chr2 | 70415273 | 70419216 | 3944 | -0.03 | 0.819 | 0.856 |
| chr6 | 134909748 | 134912547 | 2800 | 0.04 | 0.819 | 0.856 |
| chr19 | 19430605 | 19432765 | 2161 | -0.04 | 0.819 | 0.856 |
| chr5 | 155374469 | 155375175 | 707 | 0.16 | 0.819 | 0.856 |
| chr7 | 68853133 | 68854054 | 922 | -0.11 | 0.819 | 0.856 |
| chr1 | 29450540 | 29452521 | 1982 | 0.06 | 0.819 | 0.856 |
| chr8 | 145581484 | 145582891 | 1408 | 0.09 | 0.819 | 0.857 |
| chr11 | 66205399 | 66206829 | 1431 | 0.04 | 0.82 | 0.857 |
| chr9 | 3962029 | 3970756 | 8728 | 0.04 | 0.82 | 0.857 |
| chr1 | 236444113 | 236445911 | 1799 | 0.08 | 0.82 | 0.857 |
| chr15 | 43028421 | 43030149 | 1729 | 0.07 | 0.82 | 0.857 |
| chr5 | 172661096 | 172662737 | 1642 | -0.05 | 0.82 | 0.857 |
| chr16 | 54110874 | 54115927 | 5054 | 0.04 | 0.82 | 0.857 |
| chr6 | 144637562 | 144638063 | 502 | 0.14 | 0.821 | 0.858 |
| chr16 | 19469952 | 19536467 | 66516 | -0.02 | 0.821 | 0.858 |
| chr2 | 191044957 | 191045641 | 685 | -0.12 | 0.821 | 0.858 |
| chr10 | 118595493 | 118610009 | 14517 | -0.05 | 0.821 | 0.858 |
| chr3 | 49822823 | 49840559 | 17737 | 0.05 | 0.821 | 0.858 |
| chr2 | 54140760 | 54143091 | 2332 | -0.09 | 0.821 | 0.858 |
| chr3 | 71150444 | 71151444 | 1001 | 0.12 | 0.821 | 0.858 |
| chr9 | 4021501 | 4024065 | 2565 | 0.06 | 0.822 | 0.859 |
| chr20 | 44713174 | 44719459 | 6286 | -0.03 | 0.822 | 0.859 |
| chr9 | 140301310 | 140336328 | 35019 | -0.03 | 0.822 | 0.859 |
| chr1 | 22090918 | 22092306 | 1389 | 0.09 | 0.822 | 0.859 |
| chr14 | 102828082 | 102830329 | 2248 | 0.04 | 0.822 | 0.859 |
| chr17 | 40598429 | 40742439 | 144011 | 0.01 | 0.822 | 0.859 |
| chr14 | 55347707 | 55348882 | 1176 | -0.09 | 0.823 | 0.859 |
| chr7 | 6387890 | 6389597 | 1708 | 0.06 | 0.823 | 0.859 |
| chr16 | 1876123 | 1879238 | 3116 | -0.06 | 0.823 | 0.859 |
| chr17 | 59805563 | 59806580 | 1018 | 0.08 | 0.823 | 0.86 |
| chr12 | 79737716 | 79738588 | 873 | 0.13 | 0.823 | 0.86 |
| chr13 | 100258239 | 100259688 | 1450 | 0.07 | 0.823 | 0.86 |
| chr5 | 29192033 | 29192754 | 722 | -0.11 | 0.824 | 0.86 |
| chr17 | 60556351 | 60556718 | 368 | 0.15 | 0.824 | 0.86 |
| chr6 | 24381690 | 24390093 | 8404 | -0.04 | 0.824 | 0.86 |
| chr14 | 81635925 | 81667340 | 31416 | 0.02 | 0.824 | 0.86 |
| chr12 | 25585571 | 25589478 | 3908 | -0.05 | 0.824 | 0.86 |
| chr10 | 105766115 | 105766528 | 414 | -0.22 | 0.824 | 0.86 |
| chr20 | 33622363 | 33641659 | 19297 | -0.03 | 0.824 | 0.86 |
| chr10 | 59678906 | 59681098 | 2193 | 0.04 | 0.824 | 0.86 |
| chr7 | 2102838 | 2104089 | 1252 | 0.1 | 0.824 | 0.861 |
| chr14 | 68950487 | 68955301 | 4815 | -0.07 | 0.824 | 0.861 |
| chr11 | 119191722 | 119206113 | 14392 | 0.06 | 0.825 | 0.861 |
| chr10 | 75570573 | 75580102 | 9530 | 0.05 | 0.825 | 0.861 |
| chr6 | 155074950 | 155076303 | 1354 | -0.09 | 0.825 | 0.861 |
| chr22 | 19434252 | 19436228 | 1977 | 0.05 | 0.825 | 0.861 |
| chr9 | 34072277 | 34074215 | 1939 | -0.05 | 0.825 | 0.861 |
| chr1 | 200606800 | 200610655 | 3856 | 0.03 | 0.825 | 0.862 |
| chr14 | 61787728 | 61788612 | 885 | -0.12 | 0.825 | 0.862 |
| chr16 | 70355047 | 70382672 | 27626 | 0.02 | 0.825 | 0.862 |
| chr22 | 41675963 | 41699159 | 23197 | 0.02 | 0.825 | 0.862 |
| chr16 | 47556855 | 47557932 | 1078 | 0.08 | 0.825 | 0.862 |
| chr15 | 50854453 | 50871874 | 17422 | -0.03 | 0.826 | 0.862 |
| chr19 | 48688282 | 48689858 | 1577 | -0.09 | 0.826 | 0.862 |
| chr8 | 143689967 | 143691079 | 1113 | 0.1 | 0.826 | 0.862 |

| | | | | | | |
|---|---|---|---|---|---|---|
| chr10 | 14728504 | 14729858 | 1355 | -0.07 | 0.826 | 0.862 |
| chr2 | 54104072 | 54105122 | 1051 | 0.11 | 0.826 | 0.862 |
| chr17 | 34841899 | 34868000 | 26102 | -0.03 | 0.826 | 0.862 |
| chr5 | 54038067 | 54040867 | 2801 | 0.06 | 0.826 | 0.862 |
| chr1 | 180600300 | 180602497 | 2198 | -0.04 | 0.826 | 0.862 |
| chr5 | 131799752 | 131801226 | 1475 | 0.08 | 0.826 | 0.862 |
| chr12 | 92624336 | 92624774 | 439 | 0.15 | 0.826 | 0.862 |
| chr22 | 19158424 | 19173367 | 14944 | 0.03 | 0.827 | 0.862 |
| chr15 | 39134964 | 39136683 | 1720 | -0.1 | 0.827 | 0.862 |
| chr15 | 30961175 | 30962680 | 1506 | -0.1 | 0.827 | 0.862 |
| chr2 | 43150887 | 43157628 | 6742 | -0.06 | 0.827 | 0.863 |
| chr1 | 244572770 | 244578605 | 5836 | 0.05 | 0.827 | 0.863 |
| chrX | 100234605 | 100250694 | 16090 | -0.03 | 0.827 | 0.863 |
| chr2 | 12148639 | 12149546 | 908 | 0.1 | 0.827 | 0.863 |
| chr9 | 95054341 | 95089063 | 34723 | 0.02 | 0.827 | 0.863 |
| chr7 | 36405735 | 36407306 | 1572 | -0.04 | 0.828 | 0.863 |
| chr5 | 127472591 | 127473447 | 857 | 0.14 | 0.828 | 0.863 |
| chr3 | 11313497 | 11350434 | 36938 | -0.01 | 0.828 | 0.863 |
| chr8 | 125521699 | 125552684 | 30986 | 0.02 | 0.828 | 0.863 |
| chr3 | 97482513 | 97484601 | 2089 | -0.03 | 0.828 | 0.863 |
| chr4 | 73989858 | 73991179 | 1322 | 0.1 | 0.828 | 0.863 |
| chr17 | 62598538 | 62599497 | 960 | 0.1 | 0.828 | 0.863 |
| chr14 | 75347719 | 75356663 | 8945 | -0.03 | 0.828 | 0.863 |
| chr17 | 47865123 | 47867131 | 2009 | 0.04 | 0.828 | 0.863 |
| chr22 | 24945897 | 24953340 | 7444 | 0.03 | 0.828 | 0.864 |
| chr17 | 51220091 | 51220771 | 681 | -0.09 | 0.828 | 0.864 |
| chr11 | 102316337 | 102324462 | 8126 | -0.03 | 0.829 | 0.864 |
| chr4 | 190122944 | 190124028 | 1085 | -0.08 | 0.829 | 0.864 |
| chr3 | 17110582 | 17113099 | 2518 | 0.03 | 0.829 | 0.864 |
| chr16 | 69320184 | 69321575 | 1392 | -0.11 | 0.83 | 0.865 |
| chr8 | 100202972 | 100203437 | 466 | 0.13 | 0.83 | 0.865 |
| chr10 | 101418302 | 101421575 | 3274 | -0.05 | 0.83 | 0.865 |
| chr19 | 5621495 | 5625065 | 3571 | 0.05 | 0.83 | 0.865 |
| chr20 | 43621182 | 43622612 | 1431 | -0.1 | 0.83 | 0.865 |
| chr8 | 81063760 | 81064397 | 638 | 0.15 | 0.83 | 0.865 |
| chr10 | 99599348 | 99600713 | 1366 | -0.08 | 0.83 | 0.865 |
| chr21 | 38628949 | 38641247 | 12299 | 0.02 | 0.83 | 0.865 |
| chr9 | 138882854 | 138922211 | 39358 | -0.02 | 0.83 | 0.865 |
| chr19 | 44007250 | 44009743 | 2494 | -0.07 | 0.83 | 0.865 |
| chr3 | 120343800 | 120344842 | 1043 | 0.13 | 0.831 | 0.865 |
| chrX | 47441086 | 47442235 | 1150 | 0.07 | 0.831 | 0.866 |
| chr1 | 159750082 | 159751570 | 1489 | 0.06 | 0.831 | 0.866 |
| chr5 | 75002342 | 75013272 | 10931 | -0.04 | 0.831 | 0.866 |
| chr7 | 38371507 | 38381385 | 9879 | -0.1 | 0.831 | 0.866 |
| chr17 | 9944331 | 9945122 | 792 | 0.12 | 0.831 | 0.866 |
| chr5 | 180578837 | 180597504 | 18668 | -0.02 | 0.832 | 0.866 |
| chr15 | 50885668 | 50898018 | 12351 | -0.04 | 0.832 | 0.866 |
| chr6 | 52858513 | 52861795 | 3283 | -0.03 | 0.832 | 0.867 |
| chr7 | 48017825 | 48019782 | 1958 | 0.08 | 0.832 | 0.867 |
| chr19 | 4457542 | 4472970 | 15429 | -0.03 | 0.832 | 0.867 |
| chr20 | 1281222 | 1312737 | 31516 | -0.02 | 0.832 | 0.867 |
| chr20 | 35169520 | 35170197 | 678 | 0.14 | 0.832 | 0.867 |
| chr12 | 49854693 | 49856646 | 1954 | -0.07 | 0.832 | 0.867 |
| chr6 | 15284473 | 15285144 | 672 | 0.14 | 0.833 | 0.867 |
| chr1 | 24678561 | 24721039 | 42479 | -0.02 | 0.833 | 0.867 |
| chr21 | 38592653 | 38594245 | 1593 | 0.05 | 0.833 | 0.867 |
| chr3 | 55436432 | 55437832 | 1401 | 0.1 | 0.833 | 0.868 |
| chr21 | 47401056 | 47401438 | 383 | -0.13 | 0.833 | 0.868 |
| chr4 | 52760146 | 52776155 | 16010 | -0.02 | 0.833 | 0.868 |
| chr22 | 41839830 | 41845535 | 5706 | 0.02 | 0.834 | 0.868 |
| chr1 | 29432086 | 29433342 | 1257 | 0.06 | 0.834 | 0.868 |

| | | | | | | |
|---|---|---|---|---|---|---|
| chr14 | 101014080 | 101014731 | 652 | 0.13 | 0.834 | 0.868 |
| chr7 | 114458096 | 114461350 | 3255 | -0.02 | 0.834 | 0.868 |
| chr2 | 32233597 | 32241999 | 8403 | -0.03 | 0.834 | 0.868 |
| chr1 | 205227243 | 205294676 | 67434 | -0.01 | 0.834 | 0.868 |
| chr7 | 105820438 | 105820857 | 420 | -0.14 | 0.834 | 0.868 |
| chr14 | 52455181 | 52458019 | 2839 | 0.03 | 0.834 | 0.868 |
| chr22 | 41486785 | 41503403 | 16619 | -0.03 | 0.834 | 0.868 |
| chr17 | 30453807 | 30455844 | 2038 | -0.05 | 0.834 | 0.868 |
| chr6 | 137113565 | 137114882 | 1318 | 0.06 | 0.834 | 0.868 |
| chr10 | 77290218 | 77290960 | 743 | 0.12 | 0.834 | 0.869 |
| chr17 | 28111254 | 28113956 | 2703 | -0.04 | 0.835 | 0.869 |
| chr1 | 179324395 | 179336059 | 11665 | 0.02 | 0.835 | 0.869 |
| chr5 | 129672487 | 129673935 | 1449 | 0.07 | 0.835 | 0.869 |
| chr6 | 43477426 | 43493525 | 16100 | -0.03 | 0.835 | 0.869 |
| chr2 | 181627506 | 181629243 | 1738 | -0.07 | 0.835 | 0.869 |
| chr19 | 2235023 | 2247724 | 12702 | 0.03 | 0.835 | 0.869 |
| chr17 | 53404436 | 53405961 | 1526 | 0.07 | 0.835 | 0.869 |
| chr11 | 73372229 | 73372622 | 394 | -0.13 | 0.836 | 0.87 |
| chr15 | 72301282 | 72302170 | 889 | 0.1 | 0.836 | 0.87 |
| chr5 | 102090384 | 102092343 | 1960 | 0.06 | 0.836 | 0.87 |
| chr9 | 20786401 | 20787155 | 755 | 0.1 | 0.836 | 0.87 |
| chr14 | 28962190 | 28964498 | 2309 | 0.04 | 0.836 | 0.87 |
| chr4 | 110452891 | 110453374 | 484 | -0.15 | 0.836 | 0.87 |
| chr15 | 73680012 | 73682684 | 2673 | 0.08 | 0.837 | 0.87 |
| chr19 | 58426952 | 58428450 | 1499 | 0.06 | 0.837 | 0.871 |
| chr6 | 42980801 | 42990326 | 9526 | -0.04 | 0.837 | 0.871 |
| chr8 | 37593415 | 37606248 | 12834 | -0.04 | 0.837 | 0.871 |
| chr22 | 45675340 | 45676655 | 1316 | -0.11 | 0.837 | 0.871 |
| chr15 | 79089995 | 79104558 | 14564 | 0.12 | 0.838 | 0.871 |
| chr10 | 126952174 | 126952977 | 804 | -0.1 | 0.838 | 0.871 |
| chr14 | 55877689 | 55879260 | 1572 | -0.05 | 0.838 | 0.871 |
| chr7 | 148888013 | 148907372 | 19360 | 0.03 | 0.838 | 0.872 |
| chr4 | 26584756 | 26586800 | 2045 | -0.04 | 0.838 | 0.872 |
| chr13 | 34618592 | 34619303 | 712 | -0.14 | 0.838 | 0.872 |
| chrX | 117789384 | 117801562 | 12179 | -0.05 | 0.838 | 0.872 |
| chr4 | 169620198 | 169621454 | 1257 | 0.1 | 0.838 | 0.872 |
| chr11 | 9310308 | 9311203 | 896 | -0.12 | 0.838 | 0.872 |
| chr21 | 30517693 | 30518104 | 412 | 0.15 | 0.839 | 0.872 |
| chr15 | 46211132 | 46214005 | 2874 | 0.06 | 0.839 | 0.872 |
| chr2 | 36823757 | 36825719 | 1963 | -0.04 | 0.839 | 0.872 |
| chr2 | 231621729 | 231622173 | 445 | 0.15 | 0.839 | 0.872 |
| chr2 | 25296530 | 25312221 | 15692 | 0.03 | 0.839 | 0.872 |
| chr10 | 69719204 | 69741697 | 22494 | 0.03 | 0.839 | 0.872 |
| chr1 | 20959703 | 20960466 | 764 | -0.11 | 0.839 | 0.872 |
| chr13 | 20653920 | 20655242 | 1323 | 0.09 | 0.839 | 0.872 |
| chr1 | 20833826 | 20835423 | 1598 | 0.05 | 0.839 | 0.872 |
| chr7 | 2490974 | 2499320 | 8347 | -0.05 | 0.84 | 0.873 |
| chr5 | 6631735 | 6634474 | 2740 | 0.03 | 0.84 | 0.873 |
| chr6 | 53297684 | 53302986 | 5303 | -0.06 | 0.84 | 0.873 |
| chr14 | 89881811 | 89884600 | 2790 | -0.04 | 0.84 | 0.873 |
| chr18 | 56302716 | 56307900 | 5185 | -0.05 | 0.84 | 0.873 |
| chr19 | 16293217 | 16309745 | 16529 | -0.03 | 0.84 | 0.874 |
| chr19 | 13043390 | 13076725 | 33336 | -0.02 | 0.841 | 0.874 |
| chr1 | 47644453 | 47658713 | 14261 | -0.03 | 0.841 | 0.874 |
| chr2 | 11333021 | 11355805 | 22785 | -0.03 | 0.841 | 0.874 |
| chr5 | 111605471 | 111606952 | 1482 | 0.07 | 0.841 | 0.874 |
| chr12 | 13197101 | 13197850 | 750 | -0.1 | 0.841 | 0.874 |
| chr4 | 148974931 | 148976295 | 1365 | -0.08 | 0.841 | 0.874 |
| chr20 | 20692524 | 20694091 | 1568 | 0.06 | 0.841 | 0.874 |
| chr3 | 155665920 | 155667678 | 1759 | 0.07 | 0.841 | 0.874 |
| chr2 | 136499199 | 136502404 | 3206 | -0.02 | 0.841 | 0.874 |

| | | | | | | |
|---|---|---|---|---|---|---|
| chr14 | 37465851 | 37466630 | 780 | -0.11 | 0.841 | 0.874 |
| chr13 | 50508316 | 50510474 | 2159 | -0.06 | 0.841 | 0.874 |
| chr20 | 47689434 | 47690491 | 1058 | 0.09 | 0.841 | 0.874 |
| chrX | 16040330 | 16043478 | 3149 | -0.03 | 0.842 | 0.875 |
| chr11 | 65818354 | 65854420 | 36067 | -0.02 | 0.842 | 0.875 |
| chr1 | 1655382 | 1656090 | 709 | -0.08 | 0.842 | 0.875 |
| chr15 | 75436969 | 75452177 | 15209 | 0.03 | 0.842 | 0.875 |
| chr19 | 10425871 | 10426880 | 1010 | -0.09 | 0.842 | 0.875 |
| chrX | 114575543 | 114580992 | 5450 | 0.03 | 0.843 | 0.876 |
| chr2 | 134522890 | 134523356 | 467 | 0.12 | 0.843 | 0.876 |
| chr6 | 28219378 | 28220647 | 1270 | -0.07 | 0.843 | 0.876 |
| chr9 | 129600580 | 129600988 | 409 | -0.2 | 0.843 | 0.876 |
| chr2 | 139258052 | 139274396 | 16345 | 0.02 | 0.843 | 0.876 |
| chrX | 73163302 | 73165187 | 1886 | -0.03 | 0.843 | 0.876 |
| chr5 | 132283273 | 132284311 | 1039 | 0.1 | 0.843 | 0.876 |
| chr22 | 22291040 | 22297617 | 6578 | 0.03 | 0.843 | 0.876 |
| chr1 | 181224457 | 181225472 | 1016 | -0.11 | 0.843 | 0.876 |
| chr22 | 41294682 | 41295378 | 697 | 0.1 | 0.844 | 0.876 |
| chr17 | 58498394 | 58499494 | 1101 | 0.04 | 0.844 | 0.876 |
| chr16 | 4092104 | 4104635 | 12532 | -0.06 | 0.844 | 0.876 |
| chr22 | 30301758 | 30309398 | 7641 | 0.05 | 0.844 | 0.876 |
| chr16 | 30004037 | 30017150 | 13114 | -0.03 | 0.844 | 0.876 |
| chr11 | 69987971 | 69988792 | 822 | 0.11 | 0.844 | 0.876 |
| chr6 | 3848817 | 3850125 | 1309 | 0.08 | 0.844 | 0.876 |
| chr22 | 37517951 | 37518617 | 667 | -0.13 | 0.844 | 0.876 |
| chr16 | 89336279 | 89336824 | 546 | -0.14 | 0.844 | 0.876 |
| chr12 | 120251458 | 120251917 | 460 | -0.13 | 0.845 | 0.877 |
| chr7 | 65431412 | 65450328 | 18917 | -0.02 | 0.845 | 0.877 |
| chr4 | 95331298 | 95334627 | 3330 | 0.04 | 0.845 | 0.877 |
| chr5 | 28189781 | 28190455 | 675 | 0.12 | 0.845 | 0.877 |
| chr12 | 93561808 | 93562264 | 457 | 0.12 | 0.845 | 0.877 |
| chr13 | 50489043 | 50490405 | 1363 | -0.09 | 0.845 | 0.877 |
| chr2 | 23491245 | 23492492 | 1248 | 0.07 | 0.845 | 0.877 |
| chr3 | 9772208 | 9774731 | 2524 | 0.03 | 0.845 | 0.877 |
| chr15 | 89833896 | 89879512 | 45617 | -0.02 | 0.846 | 0.877 |
| chr1 | 25870202 | 25870777 | 576 | -0.12 | 0.846 | 0.878 |
| chr16 | 74590809 | 74593667 | 2859 | 0.06 | 0.846 | 0.878 |
| chr10 | 15367131 | 15368478 | 1348 | 0.06 | 0.846 | 0.878 |
| chr1 | 219103644 | 219119047 | 15404 | 0.05 | 0.846 | 0.878 |
| chr17 | 30289126 | 30289942 | 817 | -0.1 | 0.846 | 0.878 |
| chr5 | 58935996 | 58960894 | 24899 | -0.02 | 0.846 | 0.878 |
| chr1 | 32756256 | 32758735 | 2480 | 0.04 | 0.846 | 0.878 |
| chr3 | 25895395 | 25896639 | 1245 | -0.08 | 0.847 | 0.878 |
| chr18 | 32923020 | 32923992 | 973 | -0.07 | 0.847 | 0.878 |
| chr8 | 121442191 | 121458877 | 16687 | -0.02 | 0.847 | 0.878 |
| chr1 | 41174179 | 41182093 | 7915 | 0.04 | 0.847 | 0.878 |
| chr5 | 72363091 | 72363956 | 866 | 0.1 | 0.847 | 0.878 |
| chr6 | 133451671 | 133453054 | 1384 | -0.07 | 0.847 | 0.879 |
| chr15 | 59336189 | 59374836 | 38648 | 0.02 | 0.847 | 0.879 |
| chr1 | 180922333 | 180926221 | 3889 | 0.05 | 0.847 | 0.879 |
| chr3 | 141867408 | 141869050 | 1643 | -0.04 | 0.847 | 0.879 |
| chr19 | 32809596 | 32810203 | 608 | -0.1 | 0.847 | 0.879 |
| chr1 | 36022060 | 36024079 | 2020 | -0.03 | 0.847 | 0.879 |
| chr20 | 2187748 | 2188890 | 1143 | -0.08 | 0.847 | 0.879 |
| chr9 | 93799054 | 93800943 | 1890 | 0.04 | 0.848 | 0.879 |
| chrX | 149624375 | 149625421 | 1047 | -0.07 | 0.848 | 0.879 |
| chr8 | 90961642 | 90962460 | 819 | -0.11 | 0.848 | 0.879 |
| chr2 | 19737457 | 19744814 | 7358 | -0.02 | 0.848 | 0.88 |
| chr11 | 125949914 | 125950547 | 634 | -0.11 | 0.848 | 0.88 |
| chr10 | 135171014 | 135171509 | 496 | 0.1 | 0.848 | 0.88 |
| chr4 | 57299279 | 57315018 | 15740 | 0.02 | 0.848 | 0.88 |

| | | | | | | |
|---|---|---|---|---|---|---|
| chr2 | 2668809 | 2669549 | 741 | -0.1 | 0.849 | 0.88 |
| chr2 | 114720477 | 114736321 | 15845 | -0.04 | 0.849 | 0.88 |
| chr3 | 122230238 | 122234044 | 3807 | -0.03 | 0.849 | 0.88 |
| chr16 | 29910062 | 29938927 | 28866 | -0.02 | 0.849 | 0.88 |
| chr6 | 143857699 | 143859129 | 1431 | 0.07 | 0.849 | 0.88 |
| chr7 | 134973843 | 134977473 | 3631 | 0.05 | 0.849 | 0.88 |
| chr4 | 44727621 | 44728998 | 1378 | -0.05 | 0.849 | 0.88 |
| chr15 | 83860878 | 83861859 | 982 | -0.07 | 0.849 | 0.88 |
| chr12 | 54150764 | 54151073 | 310 | 0.13 | 0.85 | 0.881 |
| chr3 | 12512550 | 12527151 | 14602 | 0.03 | 0.85 | 0.881 |
| chr3 | 4508124 | 4509482 | 1359 | -0.03 | 0.85 | 0.881 |
| chr2 | 169312070 | 169350117 | 38048 | 0.02 | 0.85 | 0.881 |
| chr5 | 142873390 | 142888614 | 15225 | 0.02 | 0.85 | 0.881 |
| chr14 | 73737323 | 73738271 | 949 | -0.1 | 0.85 | 0.881 |
| chr9 | 72560554 | 72562483 | 1930 | 0.05 | 0.851 | 0.882 |
| chr16 | 68073986 | 68074566 | 581 | -0.13 | 0.851 | 0.882 |
| chr1 | 227915202 | 227924005 | 8804 | -0.03 | 0.851 | 0.882 |
| chr15 | 81616053 | 81617113 | 1061 | -0.07 | 0.851 | 0.882 |
| chr15 | 34330154 | 34331924 | 1771 | -0.03 | 0.852 | 0.883 |
| chr4 | 90114877 | 90119375 | 4499 | -0.04 | 0.852 | 0.883 |
| chr17 | 15209575 | 15210006 | 432 | -0.12 | 0.852 | 0.883 |
| chr3 | 58257559 | 58258558 | 1000 | -0.08 | 0.852 | 0.883 |
| chr22 | 36633804 | 36649642 | 15839 | 0.03 | 0.853 | 0.883 |
| chr17 | 18059750 | 18062209 | 2460 | 0.06 | 0.853 | 0.883 |
| chr14 | 92619584 | 92621085 | 1502 | -0.07 | 0.853 | 0.883 |
| chr14 | 69329380 | 69455577 | 126198 | -0.01 | 0.853 | 0.884 |
| chr9 | 90811481 | 90814430 | 2950 | -0.03 | 0.853 | 0.884 |
| chr6 | 112407629 | 112409557 | 1929 | 0.04 | 0.853 | 0.884 |
| chr4 | 40543023 | 40598915 | 55893 | 0.01 | 0.854 | 0.884 |
| chr13 | 47258239 | 47262947 | 4709 | -0.05 | 0.854 | 0.884 |
| chr11 | 74516715 | 74532157 | 15443 | -0.03 | 0.854 | 0.884 |
| chr14 | 75468527 | 75492826 | 24300 | 0.03 | 0.854 | 0.885 |
| chr17 | 7982723 | 7983928 | 1206 | 0.07 | 0.854 | 0.885 |
| chr2 | 128195431 | 128196522 | 1092 | 0.09 | 0.854 | 0.885 |
| chr2 | 177133764 | 177135406 | 1643 | -0.03 | 0.855 | 0.885 |
| chr20 | 34518201 | 34543615 | 25415 | 0.02 | 0.855 | 0.885 |
| chr1 | 205743859 | 205744634 | 776 | -0.1 | 0.855 | 0.885 |
| chr1 | 41567475 | 41576853 | 9379 | 0.03 | 0.855 | 0.886 |
| chr1 | 151370851 | 151375954 | 5104 | 0.03 | 0.855 | 0.886 |
| chr1 | 160576416 | 160577002 | 587 | 0.1 | 0.855 | 0.886 |
| chr3 | 159842877 | 159852888 | 10012 | 0.06 | 0.856 | 0.886 |
| chr6 | 170653785 | 170654988 | 1204 | 0.08 | 0.856 | 0.886 |
| chr11 | 14395418 | 14403225 | 7808 | -0.06 | 0.856 | 0.886 |
| chr2 | 160353173 | 160354886 | 1714 | -0.04 | 0.856 | 0.886 |
| chr16 | 29972388 | 29988217 | 15830 | -0.03 | 0.856 | 0.886 |
| chr22 | 50181764 | 50183022 | 1259 | 0.07 | 0.856 | 0.886 |
| chrX | 148712202 | 148713648 | 1447 | -0.08 | 0.856 | 0.886 |
| chr7 | 104524485 | 104524878 | 394 | -0.12 | 0.856 | 0.886 |
| chr4 | 33741520 | 33742319 | 800 | 0.09 | 0.857 | 0.887 |
| chr7 | 101386466 | 101398443 | 11978 | -0.06 | 0.857 | 0.887 |
| chr4 | 2469456 | 2477913 | 8458 | 0.03 | 0.857 | 0.887 |
| chr22 | 39436890 | 39437388 | 499 | -0.1 | 0.857 | 0.887 |
| chr17 | 38136195 | 38139215 | 3021 | -0.03 | 0.857 | 0.887 |
| chr19 | 7570409 | 7581733 | 11325 | -0.04 | 0.857 | 0.887 |
| chr2 | 227445483 | 227450166 | 4684 | -0.03 | 0.857 | 0.887 |
| chr6 | 133868486 | 133874650 | 6165 | -0.03 | 0.858 | 0.887 |
| chr4 | 141164023 | 141178248 | 14226 | -0.03 | 0.858 | 0.887 |
| chr2 | 68477047 | 68481652 | 4606 | -0.02 | 0.858 | 0.888 |
| chr8 | 95826340 | 95836875 | 10536 | -0.02 | 0.858 | 0.888 |
| chr12 | 109489606 | 109491420 | 1815 | 0.04 | 0.858 | 0.888 |
| chr17 | 80360677 | 80377108 | 16432 | -0.06 | 0.858 | 0.888 |

| | | | | | | |
|---|---|---|---|---|---|---|
| chr3 | 197517451 | 197519299 | 1849 | 0.04 | 0.859 | 0.888 |
| chr6 | 40961980 | 41009034 | 47055 | 0.01 | 0.859 | 0.888 |
| chr7 | 73355581 | 73357518 | 1938 | -0.06 | 0.859 | 0.888 |
| chr1 | 16301716 | 16303043 | 1328 | 0.07 | 0.859 | 0.889 |
| chr22 | 39377296 | 39381509 | 4214 | -0.03 | 0.859 | 0.889 |
| chr1 | 228326934 | 228338926 | 11993 | 0.02 | 0.86 | 0.889 |
| chr9 | 33972056 | 33985134 | 13079 | -0.04 | 0.86 | 0.889 |
| chr12 | 98960351 | 98972924 | 12574 | -0.16 | 0.86 | 0.889 |
| chr14 | 105885296 | 105898553 | 13258 | 0.02 | 0.86 | 0.889 |
| chr1 | 172686317 | 172687617 | 1301 | 0.06 | 0.86 | 0.889 |
| chr1 | 25031351 | 25032998 | 1648 | 0.04 | 0.86 | 0.889 |
| chr10 | 120512972 | 120515542 | 2571 | 0.03 | 0.86 | 0.889 |
| chr2 | 149339594 | 149341247 | 1654 | 0.05 | 0.86 | 0.889 |
| chr12 | 49318249 | 49325798 | 7550 | 0.05 | 0.86 | 0.89 |
| chr9 | 86297250 | 86300523 | 3274 | -0.05 | 0.861 | 0.89 |
| chr16 | 84494376 | 84495458 | 1083 | 0.08 | 0.861 | 0.89 |
| chr17 | 5729139 | 5741454 | 12316 | 0.02 | 0.861 | 0.89 |
| chr9 | 123687868 | 123705534 | 17667 | -0.03 | 0.861 | 0.89 |
| chr10 | 99142499 | 99143274 | 776 | 0.12 | 0.861 | 0.89 |
| chr22 | 50780525 | 50782001 | 1477 | -0.07 | 0.861 | 0.89 |
| chr10 | 116468511 | 116475532 | 7022 | 0.04 | 0.861 | 0.89 |
| chr17 | 55484812 | 55492116 | 7305 | 0.06 | 0.861 | 0.89 |
| chr14 | 64791368 | 64806628 | 15261 | -0.03 | 0.861 | 0.89 |
| chr7 | 42950490 | 42973230 | 22741 | 0.02 | 0.861 | 0.89 |
| chr17 | 53315420 | 53316619 | 1200 | -0.08 | 0.862 | 0.891 |
| chr15 | 65808258 | 65829552 | 21295 | 0.02 | 0.862 | 0.891 |
| chr1 | 65467886 | 65469179 | 1294 | 0.06 | 0.862 | 0.891 |
| chr1 | 27718323 | 27729789 | 11467 | 0.03 | 0.862 | 0.891 |
| chr1 | 158083001 | 158083725 | 725 | -0.08 | 0.862 | 0.891 |
| chr16 | 2389591 | 2391387 | 1797 | 0.07 | 0.862 | 0.891 |
| chr17 | 79779047 | 79896086 | 117040 | -0.01 | 0.863 | 0.892 |
| chr5 | 145516293 | 145525030 | 8738 | 0.04 | 0.863 | 0.892 |
| chr2 | 175546317 | 175548554 | 2238 | 0.04 | 0.863 | 0.892 |
| chr4 | 95881621 | 95898434 | 16814 | -0.02 | 0.863 | 0.892 |
| chr4 | 13701496 | 13704927 | 3432 | -0.03 | 0.864 | 0.892 |
| chr11 | 17410467 | 17411228 | 762 | -0.09 | 0.864 | 0.892 |
| chr10 | 1155818 | 1157031 | 1214 | 0.06 | 0.864 | 0.892 |
| chr11 | 66312547 | 66314074 | 1528 | -0.05 | 0.864 | 0.893 |
| chr14 | 60573570 | 60578066 | 4497 | 0.06 | 0.864 | 0.893 |
| chr11 | 119599492 | 119601013 | 1522 | -0.06 | 0.864 | 0.893 |
| chr1 | 240395696 | 240396463 | 768 | 0.08 | 0.864 | 0.893 |
| chr16 | 55656833 | 55657342 | 510 | 0.11 | 0.864 | 0.893 |
| chr17 | 66243041 | 66245920 | 2880 | -0.03 | 0.865 | 0.893 |
| chr4 | 154386658 | 154387336 | 679 | -0.09 | 0.865 | 0.893 |
| chr20 | 5510078 | 5516770 | 6693 | 0.03 | 0.865 | 0.893 |
| chr8 | 23018588 | 23040470 | 21883 | -0.02 | 0.865 | 0.893 |
| chr11 | 65756756 | 65789022 | 32267 | -0.02 | 0.865 | 0.893 |
| chr1 | 109583471 | 109584667 | 1197 | -0.07 | 0.865 | 0.894 |
| chr5 | 158369338 | 158370912 | 1575 | -0.06 | 0.865 | 0.894 |
| chr22 | 38452406 | 38454155 | 1750 | -0.04 | 0.866 | 0.894 |
| chr6 | 11684860 | 11687187 | 2328 | -0.04 | 0.866 | 0.894 |
| chr15 | 72426335 | 72428272 | 1938 | -0.04 | 0.866 | 0.894 |
| chr2 | 157188056 | 157200472 | 12417 | -0.01 | 0.866 | 0.894 |
| chr3 | 122153121 | 122153711 | 591 | 0.12 | 0.866 | 0.894 |
| chr10 | 121689843 | 121692474 | 2632 | 0.05 | 0.866 | 0.894 |
| chr3 | 186386932 | 186388452 | 1521 | -0.07 | 0.866 | 0.894 |
| chr3 | 33252982 | 33253559 | 578 | 0.08 | 0.866 | 0.894 |
| chr21 | 27162569 | 27163114 | 546 | 0.09 | 0.866 | 0.894 |
| chr17 | 46793378 | 46807115 | 13738 | -0.04 | 0.866 | 0.895 |
| chr14 | 56799969 | 56807609 | 7641 | 0.03 | 0.866 | 0.895 |
| chr2 | 85515128 | 85516721 | 1594 | 0.04 | 0.867 | 0.895 |

| | | | | | | |
|---|---|---|---|---|---|---|
| chr15 | 82337009 | 82340161 | 3153 | 0.03 | 0.867 | 0.895 |
| chr17 | 25782992 | 25825057 | 42066 | 0.02 | 0.867 | 0.896 |
| chr2 | 226912758 | 226914205 | 1448 | 0.06 | 0.868 | 0.896 |
| chr5 | 137666500 | 137713216 | 46717 | -0.01 | 0.868 | 0.896 |
| chr10 | 126128121 | 126129435 | 1315 | -0.09 | 0.868 | 0.896 |
| chr3 | 50009311 | 50011000 | 1690 | 0.06 | 0.868 | 0.896 |
| chr15 | 70667192 | 70668345 | 1154 | 0.06 | 0.868 | 0.896 |
| chr4 | 11544959 | 11551045 | 6087 | -0.07 | 0.868 | 0.896 |
| chr8 | 141714397 | 141715611 | 1215 | 0.07 | 0.869 | 0.897 |
| chr7 | 27274555 | 27275739 | 1185 | 0.05 | 0.869 | 0.897 |
| chr7 | 150724580 | 150737506 | 12927 | 0.01 | 0.869 | 0.897 |
| chr10 | 121244270 | 121261577 | 17308 | -0.03 | 0.869 | 0.897 |
| chr4 | 75185815 | 75186746 | 932 | -0.08 | 0.87 | 0.897 |
| chr16 | 67513705 | 67516189 | 2485 | 0.04 | 0.87 | 0.897 |
| chr7 | 106299725 | 106302107 | 2383 | -0.03 | 0.87 | 0.898 |
| chr10 | 103732393 | 103767032 | 34640 | 0.02 | 0.87 | 0.898 |
| chr20 | 45825068 | 45840564 | 15497 | 0.02 | 0.871 | 0.898 |
| chr16 | 89159718 | 89166663 | 6946 | -0.04 | 0.871 | 0.898 |
| chr3 | 150460944 | 150482743 | 21800 | -0.02 | 0.871 | 0.898 |
| chr16 | 90085226 | 90090188 | 4963 | 0.02 | 0.871 | 0.898 |
| chr18 | 23668744 | 23671771 | 3028 | 0.02 | 0.871 | 0.898 |
| chr17 | 73223864 | 73237875 | 14012 | 0.02 | 0.871 | 0.898 |
| chr3 | 36532698 | 36533392 | 695 | -0.09 | 0.871 | 0.898 |
| chr1 | 243706395 | 243723617 | 17223 | -0.09 | 0.871 | 0.899 |
| chr1 | 36788696 | 36790162 | 1467 | -0.08 | 0.872 | 0.899 |
| chr14 | 21537854 | 21539888 | 2035 | -0.06 | 0.872 | 0.899 |
| chr3 | 67611444 | 67613303 | 1860 | -0.08 | 0.872 | 0.899 |
| chr7 | 127636861 | 127637987 | 1127 | 0.07 | 0.872 | 0.9 |
| chr2 | 224401410 | 224451554 | 50145 | -0.02 | 0.872 | 0.9 |
| chr19 | 11615544 | 11617264 | 1721 | -0.04 | 0.873 | 0.9 |
| chr2 | 224588256 | 224596424 | 8169 | 0.01 | 0.873 | 0.9 |
| chr19 | 6863005 | 6863650 | 646 | 0.08 | 0.873 | 0.9 |
| chr1 | 212117167 | 212125965 | 8799 | 0.03 | 0.873 | 0.9 |
| chr7 | 55000554 | 55014823 | 14270 | -0.06 | 0.873 | 0.9 |
| chr1 | 212922786 | 212924346 | 1561 | 0.05 | 0.873 | 0.9 |
| chr1 | 27165094 | 27192932 | 27839 | 0.01 | 0.873 | 0.9 |
| chr9 | 129622225 | 129635803 | 13579 | 0.03 | 0.873 | 0.9 |
| chr18 | 9578612 | 9579502 | 891 | 0.1 | 0.874 | 0.901 |
| chr19 | 49858900 | 49867955 | 9056 | 0.02 | 0.874 | 0.901 |
| chr7 | 68877190 | 68878568 | 1379 | 0.07 | 0.874 | 0.901 |
| chr14 | 30193511 | 30194374 | 864 | -0.09 | 0.874 | 0.901 |
| chr2 | 64067356 | 64070454 | 3099 | -0.03 | 0.874 | 0.901 |
| chr7 | 8008430 | 8009354 | 925 | 0.07 | 0.874 | 0.901 |
| chr15 | 59304890 | 59318553 | 13664 | 0.03 | 0.874 | 0.901 |
| chr21 | 43032555 | 43033767 | 1213 | 0.06 | 0.874 | 0.901 |
| chr10 | 134388827 | 134389772 | 946 | -0.08 | 0.874 | 0.901 |
| chr10 | 80216962 | 80219787 | 2826 | -0.05 | 0.874 | 0.901 |
| chr11 | 11958273 | 11960728 | 2456 | 0.03 | 0.874 | 0.901 |
| chr8 | 493757 | 496383 | 2627 | 0.04 | 0.874 | 0.901 |
| chr2 | 130998232 | 131010952 | 12721 | 0.04 | 0.875 | 0.901 |
| chr11 | 85358172 | 85377519 | 19348 | 0.01 | 0.875 | 0.901 |
| chr7 | 65334180 | 65339574 | 5395 | 0.05 | 0.875 | 0.901 |
| chr18 | 44526753 | 44527176 | 424 | -0.08 | 0.875 | 0.901 |
| chr17 | 42966370 | 43038655 | 72286 | 0.01 | 0.875 | 0.901 |
| chr15 | 41835514 | 41836918 | 1405 | 0.04 | 0.875 | 0.901 |
| chrX | 135332897 | 135334078 | 1182 | -0.07 | 0.875 | 0.902 |
| chr20 | 55042569 | 55047389 | 4821 | 0.02 | 0.875 | 0.902 |
| chr1 | 6919378 | 6929320 | 9943 | 0.04 | 0.875 | 0.902 |
| chr2 | 39491232 | 39492367 | 1136 | 0.07 | 0.875 | 0.902 |
| chr7 | 1493934 | 1514298 | 20365 | -0.03 | 0.875 | 0.902 |
| chr2 | 226894964 | 226897608 | 2645 | -0.03 | 0.876 | 0.902 |

| | | | | | | |
|---|---|---|---|---|---|---|
| chr7 | 7531689 | 7532556 | 868 | 0.07 | 0.876 | 0.902 |
| chr6 | 79757958 | 79758506 | 549 | -0.11 | 0.876 | 0.902 |
| chr2 | 8977143 | 8977721 | 579 | 0.07 | 0.876 | 0.903 |
| chr2 | 61607144 | 61767154 | 160011 | 0.01 | 0.877 | 0.903 |
| chr1 | 91995395 | 91996425 | 1031 | -0.07 | 0.877 | 0.903 |
| chr9 | 104351894 | 104352604 | 711 | -0.07 | 0.877 | 0.903 |
| chr16 | 68895383 | 68905790 | 10408 | 0.05 | 0.877 | 0.903 |
| chr9 | 88661219 | 88662272 | 1054 | 0.08 | 0.877 | 0.904 |
| chr7 | 46296575 | 46300917 | 4343 | 0.03 | 0.877 | 0.904 |
| chr12 | 794477 | 794944 | 468 | 0.13 | 0.877 | 0.904 |
| chr1 | 221052380 | 221056532 | 4153 | -0.04 | 0.878 | 0.904 |
| chr11 | 8255628 | 8256716 | 1089 | 0.07 | 0.878 | 0.904 |
| chr5 | 61985491 | 61986666 | 1176 | -0.07 | 0.878 | 0.904 |
| chr6 | 37635429 | 37636128 | 700 | -0.09 | 0.878 | 0.904 |
| chr4 | 159643235 | 159645449 | 2215 | 0.03 | 0.878 | 0.904 |
| chr14 | 50154125 | 50160935 | 6811 | -0.03 | 0.879 | 0.905 |
| chr7 | 55954457 | 55956374 | 1918 | -0.02 | 0.879 | 0.905 |
| chr6 | 99841004 | 99842565 | 1562 | -0.03 | 0.879 | 0.905 |
| chr18 | 32556734 | 32560089 | 3356 | 0.03 | 0.88 | 0.905 |
| chr2 | 160494065 | 160495193 | 1129 | -0.08 | 0.88 | 0.906 |
| chr2 | 19703568 | 19705985 | 2418 | -0.03 | 0.88 | 0.906 |
| chr15 | 32163173 | 32163577 | 405 | 0.11 | 0.88 | 0.906 |
| chr3 | 72531778 | 72537309 | 5532 | 0.04 | 0.88 | 0.906 |
| chr19 | 2739397 | 2740046 | 650 | -0.08 | 0.881 | 0.907 |
| chr12 | 20521477 | 20521923 | 447 | 0.11 | 0.881 | 0.907 |
| chr7 | 131546389 | 131546768 | 380 | -0.1 | 0.881 | 0.907 |
| chr14 | 71786191 | 71788600 | 2410 | 0.04 | 0.881 | 0.907 |
| chr7 | 23569092 | 23572570 | 3479 | -0.02 | 0.881 | 0.907 |
| chr15 | 45740191 | 45751087 | 10897 | 0.02 | 0.881 | 0.907 |
| chr17 | 66817576 | 66819054 | 1479 | -0.06 | 0.881 | 0.907 |
| chr15 | 29126180 | 29142360 | 16181 | 0.04 | 0.881 | 0.907 |
| chr1 | 1441152 | 1447414 | 6263 | -0.04 | 0.882 | 0.907 |
| chr1 | 117452159 | 117453567 | 1409 | -0.06 | 0.882 | 0.907 |
| chr9 | 125580608 | 125591572 | 10965 | -0.02 | 0.882 | 0.907 |
| chr3 | 9957028 | 9959747 | 2720 | -0.03 | 0.882 | 0.907 |
| chr1 | 65347100 | 65407667 | 60568 | 0.01 | 0.882 | 0.907 |
| chr2 | 61862577 | 61863668 | 1092 | -0.07 | 0.882 | 0.907 |
| chr12 | 123802578 | 123803961 | 1384 | -0.07 | 0.882 | 0.907 |
| chr9 | 129484340 | 129485666 | 1327 | 0.04 | 0.882 | 0.907 |
| chr5 | 953125 | 958685 | 5561 | 0.02 | 0.882 | 0.907 |
| chr2 | 191545113 | 191545643 | 531 | -0.11 | 0.883 | 0.908 |
| chr8 | 129323947 | 129339743 | 15797 | -0.02 | 0.883 | 0.908 |
| chr18 | 60448618 | 60449692 | 1075 | -0.06 | 0.883 | 0.908 |
| chr3 | 130464138 | 130467478 | 3341 | 0.02 | 0.884 | 0.909 |
| chr5 | 55554763 | 55555491 | 729 | 0.07 | 0.884 | 0.909 |
| chr6 | 134430367 | 134430782 | 416 | 0.12 | 0.884 | 0.909 |
| chr12 | 109592445 | 109593554 | 1110 | -0.05 | 0.884 | 0.909 |
| chr20 | 23028382 | 23031993 | 3612 | -0.02 | 0.884 | 0.91 |
| chr5 | 140332801 | 140346821 | 14021 | -0.03 | 0.885 | 0.91 |
| chr11 | 130304569 | 130304966 | 398 | 0.08 | 0.885 | 0.91 |
| chr17 | 29654465 | 29656570 | 2106 | 0.05 | 0.885 | 0.91 |
| chr2 | 176019303 | 176047140 | 27838 | -0.02 | 0.885 | 0.91 |
| chr6 | 89826926 | 89827764 | 839 | 0.07 | 0.885 | 0.91 |
| chr17 | 15339845 | 15340694 | 850 | -0.06 | 0.885 | 0.91 |
| chr10 | 29759292 | 29771863 | 12572 | 0.03 | 0.886 | 0.911 |
| chr16 | 95157 | 104414 | 9258 | 0.04 | 0.886 | 0.911 |
| chr2 | 128282809 | 128285418 | 2610 | -0.02 | 0.886 | 0.911 |
| chr21 | 16399452 | 16407688 | 8237 | -0.04 | 0.886 | 0.911 |
| chr14 | 65035164 | 65037253 | 2090 | 0.03 | 0.886 | 0.911 |
| chr16 | 88715924 | 88730482 | 14559 | -0.03 | 0.886 | 0.911 |
| chr8 | 142425002 | 142425644 | 643 | -0.09 | 0.886 | 0.911 |

| | | | | | | |
|---|---|---|---|---|---|---|
| chr8 | 8312816 | 8325201 | 12386 | 0.04 | 0.886 | 0.911 |
| chrX | 53111523 | 53124280 | 12758 | -0.03 | 0.886 | 0.911 |
| chr17 | 46203274 | 46205583 | 2310 | 0.04 | 0.886 | 0.911 |
| chr17 | 29875647 | 29908943 | 33297 | -0.02 | 0.886 | 0.911 |
| chr15 | 101608742 | 101611563 | 2822 | -0.04 | 0.887 | 0.911 |
| chr22 | 30161927 | 30169640 | 7714 | 0.02 | 0.887 | 0.912 |
| chr17 | 5185590 | 5186744 | 1155 | -0.06 | 0.887 | 0.912 |
| chr10 | 94607795 | 94609350 | 1556 | 0.04 | 0.887 | 0.912 |
| chr2 | 60982842 | 60992606 | 9765 | 0.03 | 0.887 | 0.912 |
| chr3 | 63918284 | 63965023 | 46740 | 0.02 | 0.887 | 0.912 |
| chr13 | 42844874 | 42847169 | 2296 | 0.03 | 0.887 | 0.912 |
| chr16 | 30885538 | 30890796 | 5259 | 0.02 | 0.888 | 0.912 |
| chr2 | 23626804 | 23628098 | 1295 | -0.04 | 0.888 | 0.912 |
| chr6 | 160389319 | 160389934 | 616 | 0.06 | 0.888 | 0.912 |
| chr18 | 68574114 | 68576073 | 1960 | 0.02 | 0.888 | 0.913 |
| chr1 | 55229258 | 55231203 | 1946 | 0.03 | 0.888 | 0.913 |
| chr18 | 22006055 | 22007859 | 1805 | -0.03 | 0.888 | 0.913 |
| chr18 | 55332892 | 55333621 | 730 | 0.09 | 0.888 | 0.913 |
| chr8 | 11297345 | 11297898 | 554 | 0.11 | 0.888 | 0.913 |
| chr8 | 95961161 | 95962713 | 1553 | 0.03 | 0.889 | 0.913 |
| chr1 | 114201510 | 114203071 | 1562 | -0.06 | 0.889 | 0.913 |
| chr2 | 23702986 | 23704757 | 1772 | 0.05 | 0.889 | 0.913 |
| chr4 | 85735153 | 85736709 | 1557 | 0.05 | 0.889 | 0.913 |
| chr11 | 69973645 | 69975225 | 1581 | 0.04 | 0.889 | 0.913 |
| chr17 | 48251961 | 48257025 | 5065 | 0.02 | 0.889 | 0.913 |
| chr5 | 172259111 | 172387764 | 128654 | -0.01 | 0.889 | 0.913 |
| chr18 | 12076589 | 12077089 | 501 | 0.07 | 0.89 | 0.914 |
| chr12 | 125437187 | 125450036 | 12850 | -0.04 | 0.89 | 0.914 |
| chr10 | 74531547 | 74533177 | 1631 | 0.07 | 0.89 | 0.914 |
| chr15 | 77327140 | 77329058 | 1919 | 0.06 | 0.89 | 0.914 |
| chr10 | 35784134 | 35785145 | 1012 | -0.06 | 0.891 | 0.915 |
| chr15 | 38746057 | 38747259 | 1203 | 0.04 | 0.891 | 0.915 |
| chr8 | 102192865 | 102193695 | 831 | 0.08 | 0.891 | 0.915 |
| chr3 | 58418645 | 58420420 | 1776 | -0.05 | 0.891 | 0.915 |
| chr7 | 148579865 | 148583348 | 3484 | -0.02 | 0.892 | 0.915 |
| chr17 | 80709714 | 80740379 | 30666 | -0.02 | 0.892 | 0.915 |
| chr6 | 152806872 | 152807640 | 769 | -0.03 | 0.892 | 0.916 |
| chr1 | 110198285 | 110199064 | 780 | 0.06 | 0.892 | 0.916 |
| chr14 | 21438981 | 21440281 | 1301 | 0.05 | 0.892 | 0.916 |
| chr8 | 27167255 | 27169964 | 2710 | 0.03 | 0.892 | 0.916 |
| chr5 | 61770854 | 61771334 | 481 | -0.09 | 0.892 | 0.916 |
| chr7 | 7260128 | 7262036 | 1909 | 0.05 | 0.892 | 0.916 |
| chr22 | 37822954 | 37830032 | 7079 | -0.03 | 0.892 | 0.916 |
| chr9 | 110495168 | 110518660 | 23493 | 0.02 | 0.893 | 0.916 |
| chr10 | 118647280 | 118649494 | 2215 | -0.04 | 0.893 | 0.916 |
| chr10 | 5701893 | 5708701 | 6809 | -0.04 | 0.893 | 0.916 |
| chr19 | 46761845 | 46775843 | 13999 | -0.03 | 0.893 | 0.916 |
| chr8 | 54720462 | 54721397 | 936 | -0.08 | 0.893 | 0.917 |
| chr5 | 33272716 | 33273114 | 399 | -0.11 | 0.893 | 0.917 |
| chr4 | 2009625 | 2012274 | 2650 | -0.03 | 0.893 | 0.917 |
| chr19 | 15433864 | 15443993 | 10130 | -0.02 | 0.893 | 0.917 |
| chr11 | 118122352 | 118123444 | 1093 | 0.04 | 0.894 | 0.917 |
| chr3 | 125092599 | 125095200 | 2602 | 0.02 | 0.894 | 0.918 |
| chr20 | 36321844 | 36323323 | 1480 | -0.05 | 0.894 | 0.918 |
| chrX | 135302560 | 135303386 | 827 | -0.07 | 0.894 | 0.918 |
| chr12 | 116549281 | 116550578 | 1298 | -0.06 | 0.895 | 0.918 |
| chr12 | 105266707 | 105268145 | 1439 | 0.05 | 0.895 | 0.918 |
| chr3 | 179280015 | 179281870 | 1856 | 0.03 | 0.895 | 0.918 |
| chr6 | 144415598 | 144417816 | 2219 | -0.03 | 0.895 | 0.918 |
| chr9 | 107752558 | 107770591 | 18034 | -0.02 | 0.895 | 0.918 |
| chr3 | 53249891 | 53319681 | 69791 | 0.01 | 0.895 | 0.918 |

| | | | | | | |
|---|---|---|---|---|---|---|
| chr6 | 158395456 | 158502122 | 106667 | -0.01 | 0.895 | 0.918 |
| chr3 | 117572076 | 117573732 | 1657 | 0.04 | 0.895 | 0.918 |
| chr3 | 113871471 | 113872043 | 573 | 0.06 | 0.895 | 0.919 |
| chr6 | 161458835 | 161460068 | 1234 | -0.05 | 0.896 | 0.919 |
| chr15 | 99925102 | 99926953 | 1852 | -0.04 | 0.896 | 0.919 |
| chr6 | 24702813 | 24703704 | 892 | -0.07 | 0.896 | 0.919 |
| chr19 | 10811590 | 10812996 | 1407 | -0.05 | 0.896 | 0.919 |
| chr2 | 106646924 | 106647701 | 778 | 0.06 | 0.897 | 0.919 |
| chr9 | 19408429 | 19409729 | 1301 | 0.07 | 0.897 | 0.919 |
| chr7 | 131010020 | 131015745 | 5726 | 0.01 | 0.897 | 0.919 |
| chr1 | 231174804 | 231175873 | 1070 | 0.07 | 0.897 | 0.92 |
| chr15 | 90669309 | 90670398 | 1090 | -0.06 | 0.897 | 0.92 |
| chr1 | 176143212 | 176147906 | 4695 | -0.03 | 0.897 | 0.92 |
| chr8 | 101571251 | 101572042 | 792 | 0.05 | 0.898 | 0.92 |
| chr20 | 55161868 | 55182334 | 20467 | 0.02 | 0.898 | 0.92 |
| chr7 | 2772771 | 2773535 | 765 | 0.07 | 0.898 | 0.921 |
| chr10 | 6000539 | 6001339 | 801 | -0.07 | 0.898 | 0.921 |
| chr5 | 179405650 | 179406346 | 697 | -0.06 | 0.898 | 0.921 |
| chr6 | 11007475 | 11012949 | 5475 | -0.03 | 0.899 | 0.921 |
| chr1 | 189378026 | 189379738 | 1713 | 0.03 | 0.899 | 0.922 |
| chr1 | 91869720 | 91870530 | 811 | -0.06 | 0.899 | 0.922 |
| chr4 | 142689769 | 142693932 | 4164 | 0.03 | 0.899 | 0.922 |
| chr16 | 21986498 | 21987457 | 960 | -0.06 | 0.899 | 0.922 |
| chr22 | 48845942 | 48846333 | 392 | -0.08 | 0.9 | 0.922 |
| chr6 | 1024420 | 1025373 | 954 | -0.06 | 0.9 | 0.922 |
| chrX | 102883211 | 102896128 | 12918 | -0.02 | 0.9 | 0.922 |
| chr14 | 103034231 | 103035050 | 820 | 0.06 | 0.9 | 0.923 |
| chr7 | 128635733 | 128636311 | 579 | 0.07 | 0.9 | 0.923 |
| chr14 | 35041101 | 35099920 | 58820 | 0.01 | 0.9 | 0.923 |
| chr10 | 89515909 | 89536418 | 20510 | -0.02 | 0.901 | 0.923 |
| chr7 | 2795759 | 2801208 | 5450 | 0.01 | 0.901 | 0.924 |
| chr1 | 6843578 | 6861552 | 17975 | -0.02 | 0.901 | 0.924 |
| chr17 | 27263907 | 27280214 | 16308 | 0.02 | 0.902 | 0.924 |
| chr11 | 11835028 | 11836135 | 1108 | 0.06 | 0.902 | 0.924 |
| chr15 | 51149673 | 51150294 | 622 | -0.04 | 0.902 | 0.924 |
| chrX | 57147201 | 57148847 | 1647 | 0.03 | 0.902 | 0.924 |
| chr19 | 7993591 | 7994739 | 1149 | -0.07 | 0.902 | 0.924 |
| chr11 | 35948944 | 35949713 | 770 | 0.07 | 0.902 | 0.924 |
| chr2 | 238769184 | 238775347 | 6164 | -0.03 | 0.902 | 0.924 |
| chr14 | 89055007 | 89084435 | 29429 | 0.02 | 0.902 | 0.924 |
| chr10 | 94515701 | 94521707 | 6007 | -0.02 | 0.903 | 0.925 |
| chr1 | 10433679 | 10491656 | 57978 | 0.01 | 0.903 | 0.925 |
| chr5 | 64103402 | 64104113 | 712 | -0.07 | 0.904 | 0.926 |
| chr14 | 64960402 | 65007982 | 47581 | -0.01 | 0.904 | 0.926 |
| chr5 | 472148 | 473149 | 1002 | 0.04 | 0.904 | 0.926 |
| chr2 | 48126522 | 48135063 | 8542 | 0.01 | 0.904 | 0.926 |
| chr14 | 28849991 | 28874370 | 24380 | -0.02 | 0.904 | 0.926 |
| chr1 | 26798518 | 26800426 | 1909 | -0.03 | 0.904 | 0.926 |
| chr10 | 6185968 | 6225444 | 39477 | 0.02 | 0.904 | 0.926 |
| chr18 | 49935684 | 49936631 | 948 | 0.05 | 0.904 | 0.926 |
| chr10 | 95938026 | 95938702 | 677 | -0.08 | 0.904 | 0.926 |
| chr22 | 24902836 | 24903594 | 759 | -0.05 | 0.905 | 0.926 |
| chr11 | 130033573 | 130034111 | 539 | 0.08 | 0.905 | 0.926 |
| chr17 | 65347781 | 65387583 | 39803 | -0.01 | 0.905 | 0.926 |
| chr19 | 47164820 | 47165702 | 883 | 0.07 | 0.905 | 0.927 |
| chr16 | 18990216 | 19026741 | 36526 | 0.01 | 0.905 | 0.927 |
| chr13 | 46100408 | 46105495 | 5088 | 0.02 | 0.905 | 0.927 |
| chr15 | 50715171 | 50724940 | 9770 | 0.01 | 0.905 | 0.927 |
| chr2 | 9282041 | 9283623 | 1583 | -0.04 | 0.905 | 0.927 |
| chr7 | 148659799 | 148664739 | 4941 | 0.01 | 0.905 | 0.927 |
| chr18 | 19927867 | 19929411 | 1545 | 0.04 | 0.906 | 0.927 |

| | | | | | | |
|---|---|---|---|---|---|---|
| chr14 | 78266020 | 78267536 | 1517 | 0.02 | 0.906 | 0.927 |
| chr15 | 56532154 | 56538704 | 6551 | 0.01 | 0.906 | 0.927 |
| chr14 | 73392571 | 73394717 | 2147 | -0.02 | 0.906 | 0.927 |
| chr20 | 55785062 | 55788305 | 3244 | -0.05 | 0.906 | 0.927 |
| chr5 | 138774676 | 138775795 | 1120 | 0.03 | 0.906 | 0.927 |
| chr19 | 5001965 | 5002532 | 568 | 0.07 | 0.906 | 0.927 |
| chr5 | 53750070 | 53765954 | 15885 | 0.02 | 0.907 | 0.928 |
| chr12 | 7166858 | 7188034 | 21177 | 0.02 | 0.907 | 0.928 |
| chr3 | 172090646 | 172091670 | 1025 | 0.06 | 0.907 | 0.928 |
| chr9 | 98172123 | 98190071 | 17949 | -0.02 | 0.907 | 0.928 |
| chr6 | 76561992 | 76563208 | 1217 | -0.05 | 0.907 | 0.928 |
| chr9 | 20319993 | 20320551 | 559 | -0.09 | 0.907 | 0.928 |
| chr1 | 22046998 | 22048964 | 1967 | 0.04 | 0.907 | 0.928 |
| chr9 | 34155835 | 34164555 | 8721 | 0.01 | 0.907 | 0.928 |
| chr6 | 43824082 | 43824574 | 493 | -0.07 | 0.908 | 0.929 |
| chr9 | 88896156 | 88898452 | 2297 | -0.02 | 0.908 | 0.929 |
| chr1 | 236134726 | 236135832 | 1107 | 0.05 | 0.908 | 0.929 |
| chr20 | 17534812 | 17663970 | 129159 | 0 | 0.908 | 0.929 |
| chr3 | 114340648 | 114345989 | 5342 | 0.01 | 0.908 | 0.929 |
| chr8 | 680224 | 681776 | 1553 | 0.03 | 0.908 | 0.929 |
| chr12 | 6922089 | 6938800 | 16712 | -0.03 | 0.909 | 0.93 |
| chr16 | 28956240 | 28963263 | 7024 | 0.02 | 0.909 | 0.93 |
| chrX | 149650479 | 149651676 | 1198 | -0.05 | 0.909 | 0.93 |
| chr3 | 170695504 | 170697235 | 1732 | 0.03 | 0.909 | 0.93 |
| chr2 | 11412755 | 11413101 | 347 | -0.08 | 0.909 | 0.93 |
| chr22 | 48485532 | 48486643 | 1112 | 0.04 | 0.909 | 0.93 |
| chr9 | 110310075 | 110311846 | 1772 | -0.04 | 0.909 | 0.93 |
| chr8 | 87301894 | 87318708 | 16815 | 0.02 | 0.91 | 0.93 |
| chr1 | 35449615 | 35451744 | 2130 | -0.03 | 0.91 | 0.93 |
| chr6 | 134272513 | 134275289 | 2777 | 0.02 | 0.91 | 0.93 |
| chr17 | 49125253 | 49135125 | 9873 | 0.02 | 0.91 | 0.93 |
| chr11 | 117197917 | 117199640 | 1724 | 0.02 | 0.91 | 0.93 |
| chr12 | 99540192 | 99540612 | 421 | -0.07 | 0.91 | 0.931 |
| chr15 | 72979007 | 72981086 | 2080 | 0.02 | 0.91 | 0.931 |
| chr16 | 89233542 | 89233979 | 438 | -0.12 | 0.911 | 0.931 |
| chr6 | 56389412 | 56391116 | 1705 | -0.06 | 0.911 | 0.931 |
| chr17 | 2715212 | 2715992 | 781 | 0.06 | 0.911 | 0.931 |
| chr20 | 48530834 | 48569865 | 39032 | -0.01 | 0.911 | 0.931 |
| chr3 | 45429604 | 45431275 | 1672 | -0.03 | 0.911 | 0.931 |
| chr15 | 49238090 | 49238891 | 802 | -0.07 | 0.911 | 0.931 |
| chr19 | 52074183 | 52075237 | 1055 | -0.03 | 0.911 | 0.931 |
| chr1 | 183083167 | 183093776 | 10610 | -0.03 | 0.911 | 0.932 |
| chr7 | 29911919 | 29912416 | 498 | -0.07 | 0.912 | 0.932 |
| chr5 | 170763960 | 170764556 | 597 | 0.07 | 0.912 | 0.932 |
| chr5 | 53784822 | 53785150 | 329 | 0.08 | 0.912 | 0.932 |
| chrX | 134134118 | 134139111 | 4994 | 0.03 | 0.912 | 0.932 |
| chr10 | 51561777 | 51573774 | 11998 | 0.01 | 0.912 | 0.932 |
| chr7 | 148611175 | 148612054 | 880 | -0.06 | 0.912 | 0.932 |
| chr8 | 95397694 | 95400143 | 2450 | 0.03 | 0.912 | 0.932 |
| chr1 | 52829870 | 52831849 | 1980 | 0.03 | 0.912 | 0.932 |
| chr2 | 62422447 | 62425288 | 2842 | -0.02 | 0.913 | 0.933 |
| chr12 | 121115433 | 121164483 | 49051 | 0.01 | 0.913 | 0.933 |
| chr11 | 128931133 | 128932302 | 1170 | 0.05 | 0.913 | 0.933 |
| chr14 | 35564537 | 35565930 | 1394 | -0.05 | 0.913 | 0.933 |
| chr10 | 134755675 | 134756353 | 679 | -0.05 | 0.913 | 0.933 |
| chr1 | 3229105 | 3237102 | 7998 | 0.02 | 0.914 | 0.933 |
| chr6 | 152566093 | 152568318 | 2226 | -0.04 | 0.914 | 0.933 |
| chr2 | 38373192 | 38375282 | 2091 | -0.02 | 0.914 | 0.934 |
| chr8 | 135484092 | 135485018 | 927 | -0.05 | 0.914 | 0.934 |
| chr20 | 48358854 | 48365114 | 6261 | -0.03 | 0.914 | 0.934 |
| chr7 | 68189593 | 68199413 | 9821 | -0.02 | 0.915 | 0.934 |

| | | | | | | |
|---|---|---|---|---|---|---|
| chr12 | 45585295 | 45586744 | 1450 | 0.04 | 0.915 | 0.934 |
| chr10 | 116163201 | 116173673 | 10473 | -0.02 | 0.915 | 0.934 |
| chr5 | 94027263 | 94048916 | 21654 | -0.03 | 0.915 | 0.934 |
| chr2 | 47629188 | 47631820 | 2633 | -0.02 | 0.915 | 0.934 |
| chr1 | 111888597 | 111889787 | 1191 | -0.05 | 0.915 | 0.934 |
| chr1 | 213168392 | 213169478 | 1087 | 0.05 | 0.915 | 0.934 |
| chr19 | 892827 | 894068 | 1242 | -0.02 | 0.915 | 0.934 |
| chr4 | 148629089 | 148629586 | 498 | -0.08 | 0.915 | 0.934 |
| chr5 | 76721996 | 76722535 | 540 | -0.08 | 0.915 | 0.935 |
| chr6 | 37207365 | 37226833 | 19469 | -0.01 | 0.916 | 0.935 |
| chr6 | 126341276 | 126386203 | 44928 | -0.01 | 0.916 | 0.935 |
| chr7 | 111513532 | 111514333 | 802 | -0.05 | 0.916 | 0.935 |
| chr2 | 192268964 | 192269954 | 991 | 0.06 | 0.916 | 0.935 |
| chr1 | 205561168 | 205561634 | 467 | 0.05 | 0.916 | 0.936 |
| chr13 | 43858008 | 43858715 | 708 | 0.06 | 0.916 | 0.936 |
| chr6 | 108521561 | 108522751 | 1191 | -0.05 | 0.917 | 0.936 |
| chr15 | 77834738 | 77836399 | 1662 | -0.02 | 0.917 | 0.936 |
| chr14 | 77923026 | 77925612 | 2587 | 0.02 | 0.917 | 0.936 |
| chr12 | 49715596 | 49743552 | 27957 | 0.01 | 0.917 | 0.936 |
| chr7 | 104583469 | 104587105 | 3637 | 0.01 | 0.917 | 0.936 |
| chr1 | 36553862 | 36565301 | 11440 | 0.02 | 0.917 | 0.936 |
| chr2 | 175259282 | 175261030 | 1749 | 0.02 | 0.918 | 0.937 |
| chr15 | 74471405 | 74473166 | 1762 | 0.04 | 0.918 | 0.937 |
| chr20 | 34458283 | 34491064 | 32782 | 0.01 | 0.918 | 0.937 |
| chr17 | 37351977 | 37366099 | 14123 | 0.01 | 0.918 | 0.937 |
| chr12 | 54807073 | 54813661 | 6589 | 0.02 | 0.919 | 0.937 |
| chr3 | 183872461 | 183875481 | 3021 | 0.02 | 0.92 | 0.938 |
| chr4 | 1185509 | 1186858 | 1350 | -0.03 | 0.92 | 0.938 |
| chr10 | 53973254 | 53973569 | 316 | -0.07 | 0.92 | 0.939 |
| chr9 | 96381994 | 96395665 | 13672 | 0.02 | 0.92 | 0.939 |
| chr1 | 54665142 | 54666938 | 1797 | 0.02 | 0.92 | 0.939 |
| chr11 | 110138589 | 110139597 | 1009 | -0.05 | 0.92 | 0.939 |
| chr5 | 86593215 | 86597248 | 4034 | 0.03 | 0.92 | 0.939 |
| chr16 | 4932610 | 4933149 | 540 | 0.07 | 0.92 | 0.939 |
| chr15 | 52820303 | 52822000 | 1698 | 0.02 | 0.921 | 0.939 |
| chr17 | 54831935 | 54832656 | 722 | -0.05 | 0.921 | 0.939 |
| chr6 | 35453216 | 35465670 | 12455 | 0.03 | 0.921 | 0.94 |
| chr6 | 7106538 | 7113431 | 6894 | -0.02 | 0.921 | 0.94 |
| chr4 | 139991087 | 140007177 | 16091 | -0.01 | 0.921 | 0.94 |
| chr2 | 46842838 | 46845553 | 2716 | 0.01 | 0.922 | 0.94 |
| chr19 | 33407851 | 33433156 | 25306 | -0.01 | 0.922 | 0.94 |
| chr17 | 3515763 | 3516789 | 1027 | 0.07 | 0.922 | 0.94 |
| chr9 | 129552829 | 129568705 | 15877 | 0.02 | 0.922 | 0.94 |
| chr11 | 2231195 | 2233190 | 1996 | -0.03 | 0.922 | 0.941 |
| chr10 | 61860179 | 61860812 | 634 | 0.05 | 0.922 | 0.941 |
| chr8 | 143483764 | 143485423 | 1660 | 0.03 | 0.922 | 0.941 |
| chr21 | 45474219 | 45475931 | 1713 | 0.03 | 0.923 | 0.941 |
| chr18 | 50098493 | 50099457 | 965 | 0.03 | 0.923 | 0.941 |
| chr17 | 59588905 | 59589380 | 476 | 0.06 | 0.923 | 0.941 |
| chr5 | 77991785 | 77997538 | 5754 | 0.01 | 0.923 | 0.941 |
| chrX | 117973014 | 117974044 | 1031 | -0.04 | 0.924 | 0.942 |
| chr2 | 7320519 | 7333262 | 12744 | -0.03 | 0.924 | 0.942 |
| chr8 | 145909403 | 145912197 | 2795 | 0.02 | 0.924 | 0.942 |
| chr2 | 58426767 | 58430531 | 3765 | 0.03 | 0.924 | 0.942 |
| chr5 | 108526273 | 108527928 | 1656 | 0.03 | 0.924 | 0.942 |
| chr8 | 30136771 | 30142763 | 5993 | -0.02 | 0.924 | 0.942 |
| chr15 | 89524520 | 89546273 | 21754 | -0.02 | 0.925 | 0.943 |
| chr6 | 52528713 | 52536277 | 7565 | 0.04 | 0.925 | 0.943 |
| chr21 | 46707987 | 46713741 | 5755 | -0.02 | 0.925 | 0.943 |
| chr10 | 126389696 | 126393138 | 3443 | -0.03 | 0.925 | 0.943 |
| chr4 | 133628878 | 133629781 | 904 | 0.05 | 0.925 | 0.943 |

| | | | | | | |
|---|---|---|---|---|---|---|
| chr13 | 21475481 | 21477686 | 2206 | 0.02 | 0.925 | 0.943 |
| chr12 | 74415567 | 74416243 | 677 | 0.05 | 0.925 | 0.943 |
| chr15 | 42525375 | 42526108 | 734 | -0.06 | 0.925 | 0.943 |
| chr2 | 143787420 | 143788058 | 639 | -0.05 | 0.925 | 0.943 |
| chrX | 41124602 | 41136611 | 12010 | 0.02 | 0.926 | 0.943 |
| chr6 | 17467244 | 17467827 | 584 | -0.05 | 0.926 | 0.943 |
| chr22 | 29997173 | 30030216 | 33044 | -0.01 | 0.926 | 0.943 |
| chr14 | 88851019 | 88856429 | 5411 | 0.01 | 0.926 | 0.943 |
| chr12 | 78576843 | 78577920 | 1078 | -0.05 | 0.926 | 0.944 |
| chr21 | 43870313 | 43870939 | 627 | 0.04 | 0.926 | 0.944 |
| chr1 | 246886826 | 246888508 | 1683 | 0.02 | 0.926 | 0.944 |
| chr22 | 22213862 | 22222990 | 9129 | 0.02 | 0.926 | 0.944 |
| chr15 | 65164462 | 65165453 | 992 | -0.06 | 0.926 | 0.944 |
| chr10 | 17552348 | 17555812 | 3465 | -0.01 | 0.927 | 0.944 |
| chr5 | 36150731 | 36157476 | 6746 | -0.02 | 0.927 | 0.944 |
| chr17 | 67170694 | 67172775 | 2082 | 0.01 | 0.927 | 0.944 |
| chr4 | 74591220 | 74607427 | 16208 | -0.01 | 0.927 | 0.945 |
| chr3 | 47865787 | 47867531 | 1745 | -0.02 | 0.927 | 0.945 |
| chr2 | 178937129 | 178938364 | 1236 | -0.03 | 0.927 | 0.945 |
| chr2 | 189493838 | 189495628 | 1791 | -0.04 | 0.928 | 0.945 |
| chr12 | 121221066 | 121221610 | 545 | -0.05 | 0.928 | 0.945 |
| chr7 | 131139890 | 131145034 | 5145 | 0.02 | 0.928 | 0.945 |
| chr7 | 44161477 | 44164218 | 2742 | 0.01 | 0.928 | 0.945 |
| chr6 | 9766075 | 9767212 | 1138 | 0.05 | 0.928 | 0.945 |
| chr16 | 28502560 | 28503939 | 1380 | 0.01 | 0.928 | 0.945 |
| chr1 | 85724294 | 85725776 | 1483 | 0.03 | 0.928 | 0.945 |
| chr15 | 102110238 | 102120844 | 10607 | 0.04 | 0.928 | 0.945 |
| chr17 | 78964846 | 78966562 | 1717 | -0.03 | 0.929 | 0.946 |
| chrX | 39547746 | 39548763 | 1018 | -0.04 | 0.929 | 0.946 |
| chr11 | 107878992 | 107888032 | 9041 | 0.01 | 0.929 | 0.946 |
| chr5 | 137968421 | 137968845 | 425 | -0.05 | 0.929 | 0.946 |
| chr4 | 123718591 | 123720630 | 2040 | -0.02 | 0.929 | 0.946 |
| chr2 | 219867477 | 219867903 | 427 | -0.08 | 0.929 | 0.946 |
| chr6 | 16128678 | 16132451 | 3774 | 0.01 | 0.93 | 0.946 |
| chr2 | 43903144 | 43904090 | 947 | -0.03 | 0.93 | 0.946 |
| chr15 | 67133733 | 67156470 | 22738 | 0.01 | 0.93 | 0.946 |
| chr13 | 67622185 | 67623505 | 1321 | -0.03 | 0.93 | 0.947 |
| chr10 | 118525578 | 118535298 | 9721 | -0.02 | 0.93 | 0.947 |
| chr1 | 25987758 | 25988867 | 1110 | 0.04 | 0.93 | 0.947 |
| chr18 | 656578 | 659331 | 2754 | 0.02 | 0.93 | 0.947 |
| chr10 | 12886233 | 12888182 | 1950 | -0.02 | 0.93 | 0.947 |
| chr14 | 100136410 | 100150287 | 13878 | -0.02 | 0.93 | 0.947 |
| chr6 | 41536127 | 41552497 | 16371 | 0.01 | 0.93 | 0.947 |
| chr6 | 137143061 | 137144536 | 1476 | -0.02 | 0.93 | 0.947 |
| chr15 | 75134683 | 75232018 | 97336 | 0 | 0.931 | 0.947 |
| chr5 | 132620462 | 132621796 | 1335 | -0.04 | 0.931 | 0.947 |
| chr9 | 6442993 | 6450154 | 7162 | -0.02 | 0.931 | 0.947 |
| chr6 | 144999971 | 145000347 | 377 | -0.05 | 0.931 | 0.947 |
| chr5 | 138260089 | 138269595 | 9507 | -0.02 | 0.931 | 0.947 |
| chr11 | 17186368 | 17187105 | 738 | 0.05 | 0.931 | 0.947 |
| chr9 | 5042558 | 5044775 | 2218 | 0.02 | 0.931 | 0.947 |
| chr16 | 14274448 | 14275713 | 1266 | -0.04 | 0.931 | 0.948 |
| chr4 | 76554822 | 76556403 | 1582 | -0.03 | 0.931 | 0.948 |
| chr2 | 62638787 | 62652886 | 14100 | -0.2 | 0.932 | 0.948 |
| chr20 | 30457914 | 30468512 | 10599 | -0.02 | 0.932 | 0.948 |
| chr14 | 75078570 | 75089738 | 11169 | -0.01 | 0.932 | 0.948 |
| chr4 | 123652979 | 123655644 | 2666 | 0.01 | 0.932 | 0.948 |
| chrX | 109245569 | 109247569 | 2001 | -0.02 | 0.932 | 0.948 |
| chr17 | 66201595 | 66202336 | 742 | -0.03 | 0.932 | 0.948 |
| chr11 | 70005134 | 70007225 | 2092 | -0.02 | 0.932 | 0.948 |
| chr2 | 189023026 | 189023600 | 575 | 0.06 | 0.932 | 0.948 |

| | | | | | | |
|---|---|---|---|---|---|---|
| chr3 | 11137970 | 11138932 | 963 | 0.03 | 0.932 | 0.948 |
| chr1 | 212526147 | 212526675 | 529 | -0.05 | 0.932 | 0.948 |
| chr9 | 14301910 | 14327268 | 25359 | 0.01 | 0.933 | 0.948 |
| chr6 | 138187438 | 138195712 | 8275 | -0.02 | 0.933 | 0.948 |
| chr20 | 42264623 | 42265953 | 1331 | 0.04 | 0.933 | 0.948 |
| chr7 | 76134008 | 76144100 | 10093 | 0.01 | 0.933 | 0.949 |
| chr1 | 33257577 | 33258052 | 476 | 0.04 | 0.933 | 0.949 |
| chrX | 70276208 | 70289037 | 12830 | 0.01 | 0.933 | 0.949 |
| chr6 | 170020083 | 170021077 | 995 | 0.04 | 0.934 | 0.95 |
| chr6 | 53166225 | 53194603 | 28379 | -0.01 | 0.934 | 0.95 |
| chr2 | 24269709 | 24272431 | 2723 | 0.03 | 0.934 | 0.95 |
| chr1 | 246160849 | 246161633 | 785 | 0.04 | 0.934 | 0.95 |
| chr14 | 34999624 | 35000835 | 1212 | 0.04 | 0.934 | 0.95 |
| chr2 | 189767626 | 189768157 | 532 | 0.04 | 0.934 | 0.95 |
| chr20 | 1446146 | 1451790 | 5645 | 0.01 | 0.934 | 0.95 |
| chr6 | 143785190 | 143785906 | 717 | -0.05 | 0.935 | 0.95 |
| chr18 | 20084558 | 20085375 | 818 | -0.05 | 0.935 | 0.95 |
| chr13 | 24007175 | 24007912 | 738 | -0.03 | 0.935 | 0.95 |
| chr2 | 216529469 | 216530242 | 774 | -0.04 | 0.935 | 0.95 |
| chr5 | 61572409 | 61573052 | 644 | -0.04 | 0.935 | 0.95 |
| chr4 | 83784429 | 83788476 | 4048 | 0.02 | 0.935 | 0.95 |
| chr1 | 156617654 | 156635876 | 18223 | 0.01 | 0.935 | 0.95 |
| chrX | 106869109 | 106873667 | 4559 | -0.01 | 0.935 | 0.95 |
| chr13 | 31247661 | 31249007 | 1347 | -0.03 | 0.935 | 0.951 |
| chr9 | 91974541 | 91974887 | 347 | -0.06 | 0.935 | 0.951 |
| chr17 | 56707800 | 56710564 | 2765 | 0.01 | 0.935 | 0.951 |
| chr17 | 45068911 | 45079585 | 10675 | -0.09 | 0.935 | 0.951 |
| chr5 | 112910015 | 112919168 | 9154 | 0.03 | 0.936 | 0.951 |
| chr14 | 51297048 | 51298628 | 1581 | 0.04 | 0.936 | 0.951 |
| chr1 | 156663703 | 156675667 | 11965 | -0.05 | 0.936 | 0.951 |
| chr10 | 23002119 | 23004149 | 2031 | 0.02 | 0.936 | 0.951 |
| chr13 | 24152831 | 24153358 | 528 | 0.04 | 0.936 | 0.951 |
| chr8 | 95127367 | 95128527 | 1161 | -0.05 | 0.936 | 0.951 |
| chr3 | 52703832 | 52741158 | 37327 | 0.01 | 0.937 | 0.952 |
| chr9 | 115248033 | 115269888 | 21856 | 0.01 | 0.937 | 0.952 |
| chr2 | 84685459 | 84687351 | 1893 | 0.01 | 0.937 | 0.952 |
| chr5 | 56988288 | 56995375 | 7088 | 0.01 | 0.937 | 0.952 |
| chr3 | 171972248 | 171974865 | 2618 | -0.03 | 0.937 | 0.952 |
| chrX | 53023783 | 53025535 | 1753 | 0.02 | 0.937 | 0.952 |
| chr11 | 8853883 | 8855013 | 1131 | 0.03 | 0.937 | 0.952 |
| chr7 | 38971911 | 38972243 | 333 | -0.06 | 0.938 | 0.952 |
| chr8 | 30504839 | 30516791 | 11953 | 0.01 | 0.938 | 0.952 |
| chr13 | 45150278 | 45164925 | 14648 | 0.01 | 0.938 | 0.952 |
| chr17 | 6962213 | 6965240 | 3028 | -0.01 | 0.938 | 0.952 |
| chr11 | 72245938 | 72246677 | 740 | -0.06 | 0.938 | 0.952 |
| chr3 | 87855748 | 87856569 | 822 | 0.03 | 0.938 | 0.952 |
| chr2 | 205427199 | 205427798 | 600 | -0.05 | 0.938 | 0.952 |
| chr1 | 45731271 | 45733436 | 2166 | 0.01 | 0.938 | 0.952 |
| chrX | 128976845 | 128977891 | 1047 | 0.03 | 0.938 | 0.953 |
| chr7 | 104836863 | 104837648 | 786 | 0.06 | 0.938 | 0.953 |
| chr4 | 95398146 | 95429725 | 31580 | -0.01 | 0.939 | 0.953 |
| chr4 | 95848287 | 95858643 | 10357 | 0.01 | 0.939 | 0.953 |
| chr17 | 55138838 | 55139936 | 1099 | 0.03 | 0.939 | 0.953 |
| chr7 | 95198529 | 95199641 | 1113 | -0.04 | 0.939 | 0.954 |
| chr9 | 114226672 | 114247992 | 21321 | 0.01 | 0.939 | 0.954 |
| chr20 | 9027911 | 9033388 | 5478 | -0.02 | 0.939 | 0.954 |
| chr1 | 212149177 | 212149780 | 604 | 0.04 | 0.94 | 0.954 |
| chr12 | 80579881 | 80581636 | 1756 | -0.02 | 0.94 | 0.954 |
| chr6 | 26401743 | 26402861 | 1119 | -0.03 | 0.94 | 0.954 |
| chr10 | 33353672 | 33357917 | 4246 | -0.02 | 0.94 | 0.954 |
| chr8 | 104383337 | 104383829 | 493 | -0.04 | 0.94 | 0.954 |

| | | | | | | |
|---|---|---|---|---|---|---|
| chr5 | 86417154 | 86417784 | 631 | 0.04 | 0.94 | 0.954 |
| chr12 | 19911964 | 19912878 | 915 | -0.03 | 0.94 | 0.954 |
| chr11 | 10820160 | 10831920 | 11761 | 0.01 | 0.94 | 0.954 |
| chr7 | 135286999 | 135329493 | 42495 | 0.01 | 0.94 | 0.954 |
| chr7 | 99745725 | 99757049 | 11325 | -0.01 | 0.94 | 0.954 |
| chr7 | 6768829 | 6770273 | 1445 | 0.01 | 0.94 | 0.954 |
| chr2 | 30726055 | 30735456 | 9402 | -0.02 | 0.941 | 0.954 |
| chr9 | 136270325 | 136284284 | 13960 | 0.01 | 0.941 | 0.955 |
| chr8 | 21766223 | 21780764 | 14542 | -0.01 | 0.941 | 0.955 |
| chr4 | 1970725 | 1984733 | 14009 | 0.02 | 0.941 | 0.955 |
| chr7 | 134604401 | 134618468 | 14068 | 0.02 | 0.941 | 0.955 |
| chr11 | 58939144 | 58940069 | 926 | 0.03 | 0.941 | 0.955 |
| chr19 | 5948509 | 5948946 | 438 | -0.05 | 0.941 | 0.955 |
| chr14 | 94492033 | 94493237 | 1205 | 0.03 | 0.941 | 0.955 |
| chr4 | 153699852 | 153702613 | 2762 | 0.01 | 0.941 | 0.955 |
| chr6 | 152542441 | 152543777 | 1337 | -0.03 | 0.941 | 0.955 |
| chr8 | 96036241 | 96038658 | 2418 | 0.02 | 0.942 | 0.955 |
| chr17 | 72732265 | 72773230 | 40966 | -0.01 | 0.942 | 0.955 |
| chr7 | 110209686 | 110215443 | 5758 | 0.01 | 0.942 | 0.955 |
| chr4 | 24754471 | 24765815 | 11345 | -0.05 | 0.942 | 0.955 |
| chr1 | 97603766 | 97604558 | 793 | -0.05 | 0.942 | 0.955 |
| chr3 | 87310497 | 87312047 | 1551 | -0.02 | 0.942 | 0.955 |
| chr6 | 136686492 | 136687672 | 1181 | 0.03 | 0.942 | 0.956 |
| chr2 | 227483149 | 227485194 | 2046 | -0.02 | 0.942 | 0.956 |
| chr2 | 43577059 | 43578068 | 1010 | 0.04 | 0.943 | 0.956 |
| chr7 | 44773095 | 44796239 | 23145 | 0.01 | 0.943 | 0.956 |
| chr21 | 37430907 | 37443106 | 12200 | -0.01 | 0.943 | 0.956 |
| chr7 | 96337742 | 96339810 | 2069 | 0.01 | 0.943 | 0.956 |
| chr3 | 129032814 | 129036141 | 3328 | -0.01 | 0.943 | 0.956 |
| chr15 | 99846120 | 99846765 | 646 | -0.03 | 0.943 | 0.956 |
| chr1 | 204719909 | 204721686 | 1778 | 0.02 | 0.943 | 0.956 |
| chr12 | 69044932 | 69045652 | 721 | -0.04 | 0.943 | 0.956 |
| chr2 | 106193061 | 106193720 | 660 | 0.04 | 0.943 | 0.956 |
| chr11 | 93860766 | 93870189 | 9424 | -0.01 | 0.943 | 0.956 |
| chr11 | 107985684 | 108008060 | 22377 | -0.01 | 0.943 | 0.956 |
| chr16 | 19194472 | 19197929 | 3458 | -0.02 | 0.944 | 0.957 |
| chr20 | 14269572 | 14270397 | 826 | 0.03 | 0.944 | 0.957 |
| chr9 | 93924585 | 93927888 | 3304 | -0.02 | 0.944 | 0.957 |
| chr7 | 126340403 | 126341910 | 1508 | 0.02 | 0.944 | 0.957 |
| chr3 | 43147147 | 43147602 | 456 | -0.04 | 0.944 | 0.957 |
| chr14 | 81956909 | 81957225 | 317 | 0.05 | 0.944 | 0.957 |
| chr1 | 32109024 | 32111701 | 2678 | 0.01 | 0.944 | 0.957 |
| chr1 | 154343407 | 154344601 | 1195 | -0.03 | 0.944 | 0.957 |
| chr3 | 193337702 | 193339221 | 1520 | 0.03 | 0.945 | 0.957 |
| chr21 | 33941373 | 33942427 | 1055 | -0.03 | 0.945 | 0.958 |
| chr1 | 65719384 | 65721122 | 1739 | 0.02 | 0.945 | 0.958 |
| chr15 | 76570500 | 76571423 | 924 | 0.03 | 0.945 | 0.958 |
| chr18 | 55287624 | 55302873 | 15250 | -0.01 | 0.945 | 0.958 |
| chr20 | 19737291 | 19740548 | 3258 | 0.01 | 0.945 | 0.958 |
| chr6 | 57036439 | 57041071 | 4633 | 0.01 | 0.945 | 0.958 |
| chr15 | 70958739 | 70959399 | 661 | 0.04 | 0.945 | 0.958 |
| chr20 | 33655689 | 33658531 | 2843 | 0.02 | 0.945 | 0.958 |
| chr1 | 146744876 | 146745528 | 653 | 0.03 | 0.945 | 0.958 |
| chr11 | 130347324 | 130349308 | 1985 | -0.02 | 0.945 | 0.958 |
| chr10 | 114417373 | 114432009 | 14637 | 0.02 | 0.945 | 0.958 |
| chr16 | 58548386 | 58553852 | 5467 | 0.02 | 0.945 | 0.958 |
| chr19 | 51841407 | 51872278 | 30872 | 0 | 0.946 | 0.958 |
| chr2 | 144891336 | 144892182 | 847 | -0.03 | 0.946 | 0.958 |
| chr14 | 94425399 | 94426990 | 1592 | 0.02 | 0.946 | 0.958 |
| chr15 | 72814516 | 72815105 | 590 | 0.06 | 0.946 | 0.958 |
| chr1 | 174758365 | 174759136 | 772 | 0.03 | 0.946 | 0.958 |

| | | | | | | |
|---|---|---|---|---|---|---|
| chr11 | 7592364 | 7601636 | 9273 | 0.01 | 0.946 | 0.959 |
| chr18 | 52923712 | 52924475 | 764 | 0.04 | 0.946 | 0.959 |
| chr3 | 111296852 | 111305216 | 8365 | -0.01 | 0.947 | 0.959 |
| chr5 | 60348431 | 60349306 | 876 | 0.03 | 0.947 | 0.959 |
| chr3 | 52269823 | 52280666 | 10844 | -0.01 | 0.947 | 0.959 |
| chr6 | 39692710 | 39693180 | 471 | 0.03 | 0.947 | 0.959 |
| chr5 | 140010351 | 140052604 | 42254 | -0.01 | 0.947 | 0.959 |
| chr10 | 12390590 | 12392955 | 2366 | 0.02 | 0.947 | 0.959 |
| chr7 | 149249807 | 149250324 | 518 | -0.04 | 0.947 | 0.959 |
| chr13 | 79975431 | 79981499 | 6069 | 0.01 | 0.948 | 0.959 |
| chr13 | 66481914 | 66482406 | 493 | -0.04 | 0.948 | 0.959 |
| chr5 | 132072376 | 132083118 | 10743 | -0.03 | 0.948 | 0.959 |
| chr6 | 126101555 | 126143736 | 42182 | -0.01 | 0.948 | 0.959 |
| chr6 | 157737728 | 157745330 | 7603 | 0.01 | 0.948 | 0.959 |
| chr14 | 63857397 | 63859619 | 2223 | 0.02 | 0.948 | 0.96 |
| chr6 | 27688382 | 27689726 | 1345 | 0.02 | 0.948 | 0.96 |
| chr2 | 114512500 | 114515368 | 2869 | -0.01 | 0.948 | 0.96 |
| chr4 | 2892640 | 2894082 | 1443 | 0.03 | 0.948 | 0.96 |
| chr10 | 114476256 | 114477547 | 1292 | -0.02 | 0.948 | 0.96 |
| chr17 | 21454451 | 21454905 | 455 | -0.03 | 0.948 | 0.96 |
| chr12 | 124195960 | 124197698 | 1739 | -0.02 | 0.95 | 0.961 |
| chr17 | 1849164 | 1849589 | 426 | -0.04 | 0.95 | 0.961 |
| chr3 | 167097609 | 167098102 | 494 | -0.03 | 0.95 | 0.961 |
| chrX | 41159209 | 41162317 | 3109 | -0.01 | 0.95 | 0.961 |
| chr4 | 330805 | 332214 | 1410 | 0.02 | 0.95 | 0.962 |
| chr1 | 228396896 | 228401330 | 4435 | 0.01 | 0.951 | 0.962 |
| chr12 | 40547343 | 40557843 | 10501 | 0.14 | 0.951 | 0.962 |
| chr17 | 61539377 | 61551697 | 12321 | 0.14 | 0.951 | 0.962 |
| chr20 | 12989604 | 13011546 | 21943 | 0.14 | 0.951 | 0.962 |
| chr7 | 14550953 | 14561383 | 10431 | 0.14 | 0.951 | 0.962 |
| chr7 | 55322735 | 55334282 | 11548 | 0.14 | 0.951 | 0.962 |
| chr1 | 11321751 | 11334301 | 12551 | -0.01 | 0.951 | 0.962 |
| chr1 | 6760779 | 6762488 | 1710 | -0.01 | 0.951 | 0.962 |
| chr7 | 127657829 | 127663666 | 5838 | 0.01 | 0.951 | 0.962 |
| chr2 | 28970279 | 28980771 | 10493 | 0.01 | 0.951 | 0.962 |
| chr22 | 39914457 | 39930920 | 16464 | 0.01 | 0.951 | 0.963 |
| chr20 | 50806545 | 50809962 | 3418 | 0.01 | 0.952 | 0.963 |
| chr9 | 131964623 | 131966075 | 1453 | -0.02 | 0.952 | 0.963 |
| chr22 | 24199356 | 24200629 | 1274 | 0.02 | 0.952 | 0.963 |
| chr17 | 34257244 | 34258231 | 988 | 0.03 | 0.952 | 0.963 |
| chr17 | 36713777 | 36716402 | 2626 | -0.03 | 0.952 | 0.963 |
| chr22 | 42998105 | 43012731 | 14627 | 0.01 | 0.952 | 0.963 |
| chr22 | 20007941 | 20010085 | 2145 | 0.02 | 0.953 | 0.964 |
| chr13 | 111527260 | 111527664 | 405 | 0.04 | 0.953 | 0.964 |
| chr2 | 161727119 | 161727964 | 846 | 0.03 | 0.953 | 0.964 |
| chrX | 117735230 | 117735510 | 281 | 0.04 | 0.953 | 0.964 |
| chr3 | 179039458 | 179042762 | 3305 | 0.01 | 0.953 | 0.964 |
| chr8 | 28681352 | 28690809 | 9458 | 0.02 | 0.953 | 0.964 |
| chr1 | 118727464 | 118728559 | 1096 | -0.03 | 0.953 | 0.964 |
| chr3 | 139396180 | 139396781 | 602 | 0.03 | 0.953 | 0.964 |
| chr6 | 152698817 | 152707541 | 8725 | -0.01 | 0.954 | 0.964 |
| chr2 | 74881460 | 74881963 | 504 | -0.04 | 0.954 | 0.964 |
| chr3 | 183511604 | 183512996 | 1393 | 0.03 | 0.954 | 0.964 |
| chr16 | 49654271 | 49655087 | 817 | 0.03 | 0.954 | 0.964 |
| chr16 | 56547016 | 56554638 | 7623 | -0.01 | 0.954 | 0.964 |
| chrX | 129118420 | 129118787 | 368 | -0.04 | 0.954 | 0.964 |
| chr7 | 40097340 | 40108829 | 11490 | 0.01 | 0.954 | 0.964 |
| chr10 | 62233199 | 62242078 | 8880 | 0.01 | 0.954 | 0.964 |
| chr22 | 31607670 | 31608923 | 1254 | -0.03 | 0.954 | 0.965 |
| chr14 | 59001624 | 59002190 | 567 | -0.03 | 0.954 | 0.965 |
| chr12 | 94765889 | 94767001 | 1113 | -0.02 | 0.954 | 0.965 |

| | | | | | | |
|---|---|---|---|---|---|---|
| chr17 | 19409848 | 19412428 | 2581 | 0.01 | 0.955 | 0.965 |
| chr10 | 103811375 | 103829404 | 18030 | -0.01 | 0.955 | 0.965 |
| chr12 | 88524796 | 88537146 | 12351 | -0.01 | 0.955 | 0.965 |
| chr8 | 59554602 | 59573682 | 19081 | 0 | 0.955 | 0.965 |
| chr1 | 150090098 | 150091886 | 1789 | -0.02 | 0.955 | 0.965 |
| chr7 | 65564590 | 65581373 | 16784 | -0.01 | 0.955 | 0.965 |
| chr19 | 50353334 | 50388590 | 35257 | 0 | 0.955 | 0.965 |
| chr14 | 31864856 | 31865489 | 634 | 0.03 | 0.956 | 0.966 |
| chr6 | 2756860 | 2768062 | 11203 | -0.01 | 0.956 | 0.966 |
| chr9 | 80480454 | 80480945 | 492 | -0.03 | 0.956 | 0.966 |
| chr5 | 74058058 | 74076407 | 18350 | -0.01 | 0.956 | 0.966 |
| chr9 | 79982853 | 79983921 | 1069 | 0.02 | 0.956 | 0.966 |
| chr11 | 126686331 | 126687933 | 1603 | -0.02 | 0.956 | 0.966 |
| chr15 | 74890322 | 74891430 | 1109 | 0.02 | 0.956 | 0.966 |
| chr11 | 71158672 | 71165200 | 6529 | 0.01 | 0.956 | 0.966 |
| chr4 | 84109895 | 84111403 | 1509 | 0.03 | 0.957 | 0.966 |
| chr22 | 19131347 | 19132217 | 871 | -0.02 | 0.957 | 0.967 |
| chr17 | 64336902 | 64355718 | 18817 | -0.01 | 0.957 | 0.967 |
| chr3 | 150164479 | 150166493 | 2015 | 0.02 | 0.957 | 0.967 |
| chr3 | 33677822 | 33679325 | 1504 | -0.02 | 0.957 | 0.967 |
| chr7 | 158769387 | 158769858 | 472 | 0.04 | 0.957 | 0.967 |
| chr7 | 35576810 | 35577178 | 369 | 0.04 | 0.958 | 0.967 |
| chr11 | 113643288 | 113644923 | 1636 | 0.01 | 0.958 | 0.967 |
| chr15 | 63549023 | 63549630 | 608 | -0.03 | 0.958 | 0.967 |
| chr1 | 47902248 | 47904059 | 1812 | -0.02 | 0.958 | 0.968 |
| chr20 | 62518336 | 62529277 | 10942 | 0.01 | 0.958 | 0.968 |
| chr12 | 47053285 | 47060056 | 6772 | 0.01 | 0.959 | 0.968 |
| chr6 | 111135047 | 111138187 | 3141 | -0.01 | 0.959 | 0.968 |
| chr5 | 81412899 | 81413739 | 841 | 0.04 | 0.959 | 0.968 |
| chr19 | 12991474 | 12992963 | 1490 | 0.01 | 0.959 | 0.968 |
| chr13 | 41706438 | 41707855 | 1418 | 0.01 | 0.959 | 0.968 |
| chrX | 77320075 | 77327656 | 7582 | 0.01 | 0.959 | 0.969 |
| chr4 | 89718561 | 89719529 | 969 | -0.02 | 0.96 | 0.969 |
| chr1 | 233004674 | 233005669 | 996 | 0.03 | 0.961 | 0.97 |
| chr10 | 44101876 | 44102619 | 744 | 0.02 | 0.961 | 0.97 |
| chr6 | 5032648 | 5033010 | 363 | -0.03 | 0.961 | 0.97 |
| chr12 | 90125190 | 90126263 | 1074 | -0.02 | 0.961 | 0.97 |
| chr10 | 97372078 | 97372747 | 670 | 0.03 | 0.961 | 0.97 |
| chr5 | 39057740 | 39058599 | 860 | -0.02 | 0.961 | 0.97 |
| chr6 | 28072081 | 28073355 | 1275 | 0.01 | 0.961 | 0.97 |
| chr9 | 84448064 | 84459032 | 10969 | 0.01 | 0.961 | 0.97 |
| chr2 | 54949658 | 54951554 | 1897 | -0.01 | 0.962 | 0.971 |
| chr8 | 66751518 | 66755666 | 4149 | 0 | 0.962 | 0.971 |
| chr8 | 23056689 | 23058073 | 1385 | -0.02 | 0.962 | 0.971 |
| chr1 | 187954638 | 187957110 | 2473 | 0.02 | 0.962 | 0.971 |
| chr1 | 193698424 | 193699568 | 1145 | -0.02 | 0.962 | 0.971 |
| chr14 | 24766007 | 24777210 | 11204 | 0.01 | 0.962 | 0.971 |
| chr3 | 152950345 | 152951130 | 786 | 0.02 | 0.962 | 0.971 |
| chr9 | 79791544 | 79795170 | 3627 | -0.01 | 0.962 | 0.971 |
| chr8 | 30478773 | 30488265 | 9493 | 0.02 | 0.962 | 0.971 |
| chr10 | 48325210 | 48355623 | 30414 | 0 | 0.962 | 0.971 |
| chr4 | 126036640 | 126037950 | 1311 | 0.02 | 0.962 | 0.971 |
| chr4 | 83349916 | 83352756 | 2841 | 0.01 | 0.963 | 0.971 |
| chr7 | 158430460 | 158467826 | 37367 | 0.01 | 0.963 | 0.971 |
| chr14 | 68850419 | 68851171 | 753 | 0.03 | 0.963 | 0.972 |
| chr11 | 82610859 | 82613565 | 2707 | -0.01 | 0.963 | 0.972 |
| chr5 | 118747399 | 118747891 | 493 | 0.02 | 0.963 | 0.972 |
| chr5 | 171806300 | 171811158 | 4859 | -0.01 | 0.963 | 0.972 |
| chr3 | 47181665 | 47187901 | 6237 | -0.01 | 0.963 | 0.972 |
| chr6 | 22600010 | 22601023 | 1014 | 0.02 | 0.964 | 0.972 |
| chr3 | 111436647 | 111437064 | 418 | 0.03 | 0.964 | 0.972 |

| | | | | | | |
|---|---|---|---|---|---|---|
| chr9 | 115006450 | 115012503 | 6054 | -0.01 | 0.964 | 0.972 |
| chr7 | 80328615 | 80337111 | 8497 | 0.01 | 0.964 | 0.972 |
| chr12 | 96078398 | 96079264 | 867 | 0.03 | 0.964 | 0.972 |
| chr7 | 4658909 | 4682835 | 23927 | -0.01 | 0.964 | 0.973 |
| chr7 | 48122956 | 48130596 | 7641 | 0.01 | 0.965 | 0.973 |
| chr20 | 30932326 | 30996037 | 63712 | 0 | 0.965 | 0.973 |
| chr6 | 27507153 | 27534400 | 27248 | -0.01 | 0.965 | 0.973 |
| chr5 | 159771827 | 159772487 | 661 | -0.02 | 0.965 | 0.973 |
| chr10 | 115612714 | 115617471 | 4758 | 0 | 0.965 | 0.973 |
| chr17 | 61673962 | 61679831 | 5870 | 0.01 | 0.965 | 0.973 |
| chr10 | 33796796 | 33799648 | 2853 | -0.01 | 0.965 | 0.973 |
| chr5 | 78366067 | 78366528 | 462 | 0.03 | 0.965 | 0.973 |
| chr4 | 120375169 | 120375700 | 532 | -0.01 | 0.965 | 0.973 |
| chr9 | 37753149 | 37754698 | 1550 | -0.01 | 0.965 | 0.973 |
| chr5 | 176657206 | 176658614 | 1409 | -0.02 | 0.965 | 0.973 |
| chr10 | 120798645 | 120807400 | 8756 | 0.01 | 0.965 | 0.973 |
| chr1 | 235054115 | 235070685 | 16571 | 0.01 | 0.965 | 0.973 |
| chr1 | 161535551 | 161536134 | 584 | 0.01 | 0.965 | 0.973 |
| chr19 | 4866170 | 4871234 | 5065 | 0.01 | 0.965 | 0.973 |
| chr2 | 73038804 | 73053255 | 14452 | -0.01 | 0.966 | 0.973 |
| chr5 | 137548186 | 137550282 | 2097 | 0.01 | 0.966 | 0.973 |
| chr17 | 7619750 | 7621626 | 1877 | 0.01 | 0.966 | 0.973 |
| chr8 | 124460206 | 124460564 | 359 | -0.02 | 0.966 | 0.974 |
| chrX | 41780836 | 41783791 | 2956 | -0.01 | 0.966 | 0.974 |
| chr21 | 47608183 | 47609242 | 1060 | 0.02 | 0.967 | 0.974 |
| chr21 | 39648758 | 39650863 | 2106 | 0.01 | 0.967 | 0.974 |
| chr5 | 74905663 | 74920452 | 14790 | -0.01 | 0.967 | 0.974 |
| chr2 | 216362914 | 216363679 | 766 | -0.02 | 0.967 | 0.975 |
| chr9 | 136890601 | 136891348 | 748 | -0.02 | 0.967 | 0.975 |
| chr7 | 55086047 | 55277370 | 191324 | 0 | 0.967 | 0.975 |
| chr8 | 144416040 | 144417628 | 1589 | -0.01 | 0.967 | 0.975 |
| chr20 | 37678118 | 37679133 | 1016 | -0.02 | 0.968 | 0.975 |
| chr11 | 111912761 | 111915995 | 3235 | 0.01 | 0.968 | 0.975 |
| chr15 | 40327346 | 40340379 | 13034 | 0 | 0.968 | 0.975 |
| chr15 | 80695831 | 80697769 | 1939 | -0.01 | 0.968 | 0.975 |
| chr17 | 25951786 | 25952882 | 1097 | -0.02 | 0.968 | 0.975 |
| chr18 | 23805575 | 23807436 | 1862 | -0.01 | 0.968 | 0.975 |
| chr5 | 65156686 | 65159054 | 2369 | 0.01 | 0.969 | 0.976 |
| chr19 | 50934632 | 50935834 | 1203 | 0.02 | 0.969 | 0.976 |
| chr19 | 46683574 | 46730900 | 47327 | 0 | 0.969 | 0.976 |
| chr2 | 74004108 | 74010994 | 6887 | -0.01 | 0.969 | 0.976 |
| chr19 | 3625714 | 3627682 | 1969 | 0.01 | 0.969 | 0.976 |
| chr6 | 90142939 | 90143367 | 429 | 0.02 | 0.969 | 0.976 |
| chr15 | 90574591 | 90605852 | 31262 | 0 | 0.969 | 0.976 |
| chr17 | 41220737 | 41232702 | 11966 | 0.01 | 0.97 | 0.977 |
| chr9 | 97604868 | 97715661 | 110794 | 0 | 0.97 | 0.977 |
| chr4 | 6274391 | 6275682 | 1292 | -0.02 | 0.97 | 0.977 |
| chrX | 17692642 | 17699597 | 6956 | -0.01 | 0.97 | 0.977 |
| chr2 | 264060 | 279250 | 15191 | -0.01 | 0.97 | 0.977 |
| chr16 | 74930557 | 74931737 | 1181 | -0.02 | 0.971 | 0.977 |
| chr6 | 17393029 | 17395281 | 2253 | -0.01 | 0.971 | 0.978 |
| chr18 | 24115769 | 24130465 | 14697 | 0 | 0.971 | 0.978 |
| chr12 | 64061738 | 64062538 | 801 | -0.01 | 0.971 | 0.978 |
| chr16 | 25043032 | 25063935 | 20904 | 0 | 0.971 | 0.978 |
| chr3 | 155625610 | 155639091 | 13482 | -0.01 | 0.972 | 0.978 |
| chr14 | 31125640 | 31126100 | 461 | -0.02 | 0.972 | 0.978 |
| chr6 | 35995575 | 36007648 | 12074 | -0.01 | 0.972 | 0.979 |
| chr1 | 116710596 | 116711420 | 825 | -0.01 | 0.972 | 0.979 |
| chr15 | 31727478 | 31733738 | 6261 | -0.01 | 0.972 | 0.979 |
| chr10 | 3336721 | 3337953 | 1233 | 0.01 | 0.973 | 0.979 |
| chr14 | 99501739 | 99502994 | 1256 | -0.01 | 0.973 | 0.98 |

| | | | | | | |
|---|---|---|---|---|---|---|
| chr2 | 226758462 | 226760397 | 1936 | -0.01 | 0.973 | 0.98 |
| chr6 | 26593786 | 26597925 | 4140 | -0.01 | 0.973 | 0.98 |
| chr4 | 134834002 | 134835078 | 1077 | 0.01 | 0.974 | 0.98 |
| chr12 | 13408557 | 13409644 | 1088 | -0.01 | 0.974 | 0.98 |
| chr5 | 177658124 | 177660545 | 2422 | -0.01 | 0.974 | 0.98 |
| chr3 | 149160607 | 149166899 | 6293 | -0.01 | 0.974 | 0.98 |
| chr13 | 34774328 | 34783142 | 8815 | 0 | 0.974 | 0.98 |
| chr18 | 12375817 | 12378571 | 2755 | 0 | 0.974 | 0.98 |
| chr3 | 11886865 | 11888697 | 1833 | 0.01 | 0.974 | 0.98 |
| chr10 | 33950194 | 33951204 | 1011 | 0.01 | 0.974 | 0.98 |
| chr17 | 16471906 | 16472492 | 587 | -0.02 | 0.974 | 0.98 |
| chr17 | 35560662 | 35561812 | 1151 | -0.01 | 0.974 | 0.98 |
| chr19 | 58038792 | 58039309 | 518 | 0.02 | 0.975 | 0.981 |
| chr3 | 48717520 | 48732847 | 15328 | 0.01 | 0.975 | 0.981 |
| chr4 | 166247701 | 166250012 | 2312 | 0.01 | 0.975 | 0.981 |
| chr19 | 9928606 | 9957862 | 29257 | 0.01 | 0.975 | 0.981 |
| chr2 | 160918444 | 160919908 | 1465 | 0.01 | 0.975 | 0.981 |
| chr13 | 42534237 | 42536257 | 2021 | -0.01 | 0.976 | 0.981 |
| chr9 | 5600191 | 5600617 | 427 | -0.02 | 0.976 | 0.982 |
| chr2 | 88958768 | 88959456 | 689 | -0.02 | 0.976 | 0.982 |
| chr12 | 112903810 | 112912850 | 9041 | -0.01 | 0.976 | 0.982 |
| chr18 | 44496380 | 44498344 | 1965 | 0.01 | 0.976 | 0.982 |
| chr1 | 23337318 | 23347588 | 10271 | 0.01 | 0.976 | 0.982 |
| chr15 | 78857059 | 78864963 | 7905 | -0.01 | 0.976 | 0.982 |
| chr3 | 8483335 | 8500814 | 17480 | 0 | 0.976 | 0.982 |
| chr2 | 178556992 | 178558111 | 1120 | -0.02 | 0.976 | 0.982 |
| chr14 | 56710501 | 56711734 | 1234 | 0.01 | 0.977 | 0.982 |
| chr15 | 28526599 | 28527238 | 640 | -0.01 | 0.977 | 0.982 |
| chr18 | 11797900 | 11808816 | 10917 | 0.01 | 0.977 | 0.983 |
| chr19 | 12777018 | 12793446 | 16429 | 0 | 0.977 | 0.983 |
| chr8 | 41434919 | 41441727 | 6809 | 0 | 0.977 | 0.983 |
| chr4 | 154265131 | 154267154 | 2024 | -0.01 | 0.977 | 0.983 |
| chr6 | 47637485 | 47638560 | 1076 | 0.01 | 0.977 | 0.983 |
| chr17 | 8311211 | 8312215 | 1005 | -0.01 | 0.977 | 0.983 |
| chr11 | 85710689 | 85729815 | 19127 | 0 | 0.978 | 0.983 |
| chr11 | 77299710 | 77324468 | 24759 | 0 | 0.978 | 0.983 |
| chr8 | 107023154 | 107025067 | 1914 | 0.01 | 0.978 | 0.983 |
| chr11 | 111076183 | 111078227 | 2045 | -0.01 | 0.978 | 0.983 |
| chr5 | 16616175 | 16617627 | 1453 | -0.01 | 0.978 | 0.983 |
| chr9 | 35603352 | 35606397 | 3046 | 0 | 0.978 | 0.983 |
| chr5 | 77147797 | 77148673 | 877 | -0.01 | 0.978 | 0.983 |
| chr8 | 103914860 | 103923854 | 8995 | 0 | 0.978 | 0.983 |
| chr1 | 217778043 | 217785730 | 7688 | -0.01 | 0.978 | 0.983 |
| chr21 | 38019737 | 38020589 | 853 | -0.01 | 0.979 | 0.983 |
| chr19 | 15749885 | 15753437 | 3553 | 0 | 0.979 | 0.984 |
| chr11 | 57405058 | 57408176 | 3119 | 0 | 0.979 | 0.984 |
| chr2 | 136702222 | 136719299 | 17078 | 0 | 0.979 | 0.984 |
| chr5 | 32398735 | 32399572 | 838 | 0.02 | 0.979 | 0.984 |
| chr17 | 46078697 | 46189412 | 110716 | 0 | 0.979 | 0.984 |
| chr12 | 113732459 | 113751562 | 19104 | 0 | 0.98 | 0.984 |
| chr10 | 89552595 | 89553942 | 1348 | -0.01 | 0.98 | 0.984 |
| chr1 | 1239756 | 1296138 | 56383 | 0 | 0.98 | 0.984 |
| chr18 | 11499724 | 11502417 | 2694 | 0 | 0.98 | 0.984 |
| chr1 | 6444805 | 6455112 | 10308 | 0 | 0.98 | 0.984 |
| chr17 | 31254360 | 31264605 | 10246 | -0.01 | 0.98 | 0.984 |
| chr19 | 10196598 | 10197335 | 738 | -0.02 | 0.98 | 0.984 |
| chr4 | 129555186 | 129561485 | 6300 | 0 | 0.98 | 0.985 |
| chrX | 70658300 | 70659669 | 1370 | -0.01 | 0.98 | 0.985 |
| chr9 | 116569910 | 116570338 | 429 | -0.02 | 0.98 | 0.985 |
| chr10 | 88534552 | 88555869 | 21318 | 0 | 0.98 | 0.985 |
| chr16 | 12768021 | 12773660 | 5640 | 0 | 0.981 | 0.985 |

| | | | | | | |
|---|---|---|---|---|---|---|
| chr6 | 127838310 | 127841509 | 3200 | 0 | 0.981 | 0.985 |
| chr15 | 51231974 | 51232904 | 931 | 0.01 | 0.981 | 0.985 |
| chr1 | 6639304 | 6643039 | 3736 | 0 | 0.981 | 0.985 |
| chr1 | 231762454 | 231762937 | 484 | -0.02 | 0.981 | 0.985 |
| chr17 | 55036941 | 55038849 | 1909 | 0 | 0.981 | 0.985 |
| chr15 | 34389908 | 34395509 | 5602 | 0 | 0.981 | 0.986 |
| chr3 | 195073378 | 195073995 | 618 | 0.01 | 0.982 | 0.986 |
| chr1 | 53191646 | 53193269 | 1624 | 0.01 | 0.982 | 0.986 |
| chr20 | 33779596 | 33804313 | 24718 | 0.01 | 0.982 | 0.986 |
| chr3 | 125802129 | 125803556 | 1428 | -0.01 | 0.982 | 0.986 |
| chr8 | 20081531 | 20081853 | 323 | -0.01 | 0.983 | 0.987 |
| chr20 | 3450890 | 3465353 | 14464 | 0 | 0.983 | 0.987 |
| chr11 | 121322610 | 121323403 | 794 | 0.01 | 0.983 | 0.987 |
| chr18 | 12701252 | 12704670 | 3419 | 0 | 0.983 | 0.987 |
| chr7 | 139105516 | 139119886 | 14371 | 0 | 0.983 | 0.987 |
| chrX | 69548709 | 69549702 | 994 | 0.01 | 0.983 | 0.987 |
| chr3 | 23769906 | 23803309 | 33404 | 0 | 0.983 | 0.987 |
| chr2 | 153914243 | 153917427 | 3185 | 0 | 0.983 | 0.987 |
| chr3 | 97731163 | 97733810 | 2648 | 0 | 0.984 | 0.988 |
| chr19 | 55622870 | 55630572 | 7703 | 0 | 0.984 | 0.988 |
| chr4 | 15304939 | 15305516 | 578 | -0.01 | 0.984 | 0.988 |
| chr7 | 55980078 | 55981196 | 1119 | 0.01 | 0.984 | 0.988 |
| chr5 | 77049032 | 77072821 | 23790 | 0 | 0.984 | 0.988 |
| chr22 | 28471277 | 28471871 | 595 | 0.01 | 0.984 | 0.988 |
| chr1 | 201645182 | 201656465 | 11284 | 0.01 | 0.984 | 0.988 |
| chr7 | 135607508 | 135628956 | 21449 | 0 | 0.985 | 0.988 |
| chr6 | 161678415 | 161679743 | 1329 | -0.01 | 0.985 | 0.989 |
| chr5 | 138671844 | 138679303 | 7460 | 0 | 0.985 | 0.989 |
| chr17 | 6914357 | 6940293 | 25937 | 0 | 0.985 | 0.989 |
| chr11 | 101917586 | 101919144 | 1559 | 0 | 0.986 | 0.989 |
| chr10 | 93771361 | 93771740 | 380 | 0.01 | 0.986 | 0.989 |
| chr10 | 69670010 | 69696195 | 26186 | 0 | 0.986 | 0.989 |
| chr20 | 42085468 | 42095925 | 10458 | 0 | 0.986 | 0.989 |
| chr11 | 14464988 | 14476660 | 11673 | 0 | 0.986 | 0.989 |
| chr2 | 201908011 | 201909267 | 1257 | -0.01 | 0.986 | 0.989 |
| chr7 | 42926859 | 42929768 | 2910 | 0 | 0.986 | 0.989 |
| chr7 | 101331415 | 101335988 | 4574 | 0.01 | 0.986 | 0.989 |
| chr18 | 19444752 | 19445912 | 1161 | -0.01 | 0.986 | 0.989 |
| chr9 | 137536766 | 137537553 | 788 | -0.01 | 0.986 | 0.99 |
| chr4 | 76633087 | 76653841 | 20755 | 0 | 0.987 | 0.99 |
| chr1 | 165566713 | 165568239 | 1527 | 0 | 0.987 | 0.99 |
| chr7 | 67926009 | 67926963 | 955 | -0.01 | 0.987 | 0.99 |
| chr6 | 4351208 | 4384263 | 33056 | 0 | 0.987 | 0.99 |
| chr14 | 69847678 | 69866807 | 19130 | 0 | 0.987 | 0.99 |
| chr22 | 45898057 | 45899365 | 1309 | -0.01 | 0.987 | 0.99 |
| chr22 | 40765647 | 40771000 | 5354 | 0 | 0.987 | 0.99 |
| chr9 | 131258389 | 131268051 | 9663 | 0 | 0.987 | 0.99 |
| chr3 | 109641270 | 109649816 | 8547 | 0 | 0.987 | 0.99 |
| chr9 | 115975889 | 115996025 | 20137 | 0 | 0.987 | 0.99 |
| chr14 | 21847986 | 21853664 | 5679 | 0 | 0.987 | 0.99 |
| chr11 | 16839475 | 16840263 | 789 | -0.01 | 0.988 | 0.99 |
| chr22 | 41416929 | 41419158 | 2230 | 0 | 0.988 | 0.99 |
| chr6 | 108169699 | 108170140 | 442 | 0.01 | 0.988 | 0.991 |
| chr19 | 8372028 | 8387356 | 15329 | 0 | 0.988 | 0.991 |
| chr2 | 68288941 | 68291099 | 2159 | 0 | 0.989 | 0.991 |
| chr11 | 74459174 | 74466950 | 7777 | 0 | 0.989 | 0.991 |
| chr17 | 16049644 | 16095967 | 46324 | 0 | 0.989 | 0.991 |
| chr10 | 86299518 | 86300832 | 1315 | -0.01 | 0.989 | 0.991 |
| chr1 | 160782481 | 160797942 | 15462 | 0 | 0.989 | 0.991 |
| chr15 | 57307754 | 57308660 | 907 | 0 | 0.989 | 0.991 |
| chr3 | 52785729 | 52786803 | 1075 | 0 | 0.989 | 0.991 |

| | | | | | | |
|---|---|---|---|---|---|---|
| chr20 | 2731323 | 2733493 | 2171 | 0 | 0.989 | 0.991 |
| chr4 | 176390704 | 176391568 | 865 | -0.01 | 0.989 | 0.992 |
| chr6 | 139693544 | 139697907 | 4364 | 0 | 0.99 | 0.992 |
| chr17 | 69971446 | 69972778 | 1333 | -0.01 | 0.99 | 0.992 |
| chr20 | 56193057 | 56193516 | 460 | -0.01 | 0.99 | 0.992 |
| chr20 | 61733458 | 61734132 | 675 | -0.01 | 0.99 | 0.992 |
| chr9 | 136201980 | 136224288 | 22309 | 0 | 0.99 | 0.992 |
| chr6 | 37321198 | 37322787 | 1590 | 0 | 0.99 | 0.992 |
| chr1 | 204120934 | 204121905 | 972 | -0.01 | 0.991 | 0.993 |
| chr13 | 76248817 | 76249854 | 1038 | 0 | 0.991 | 0.993 |
| chr9 | 96024874 | 96025617 | 744 | -0.01 | 0.991 | 0.993 |
| chr22 | 27598061 | 27609303 | 11243 | 0 | 0.991 | 0.993 |
| chr21 | 37581755 | 37582986 | 1232 | 0 | 0.991 | 0.993 |
| chr11 | 8615248 | 8626425 | 11178 | 0.01 | 0.991 | 0.993 |
| chr3 | 160032007 | 160032557 | 551 | 0 | 0.991 | 0.993 |
| chr4 | 145587600 | 145589357 | 1758 | 0 | 0.991 | 0.993 |
| chr11 | 44117227 | 44118377 | 1151 | -0.01 | 0.992 | 0.993 |
| chr5 | 31855032 | 31855537 | 506 | 0.01 | 0.992 | 0.993 |
| chr6 | 158980440 | 158982407 | 1968 | 0 | 0.992 | 0.993 |
| chr1 | 115282639 | 115301618 | 18980 | 0 | 0.992 | 0.993 |
| chr4 | 5159314 | 5160812 | 1499 | 0 | 0.992 | 0.994 |
| chr22 | 46645746 | 46647073 | 1328 | 0 | 0.992 | 0.994 |
| chr3 | 195573727 | 195574846 | 1120 | 0 | 0.992 | 0.994 |
| chr11 | 13961787 | 13965056 | 3270 | 0 | 0.993 | 0.994 |
| chr9 | 135978297 | 136025508 | 47212 | 0 | 0.993 | 0.994 |
| chr17 | 7283959 | 7309100 | 25142 | 0 | 0.993 | 0.994 |
| chr10 | 100296107 | 100296982 | 876 | 0 | 0.993 | 0.994 |
| chr12 | 22428448 | 22429375 | 928 | 0 | 0.993 | 0.994 |
| chr17 | 30771093 | 30790659 | 19567 | 0 | 0.993 | 0.994 |
| chr15 | 66192152 | 66209064 | 16913 | 0 | 0.993 | 0.995 |
| chr13 | 67404887 | 67409789 | 4903 | 0 | 0.993 | 0.995 |
| chr1 | 120189860 | 120191708 | 1849 | 0 | 0.993 | 0.995 |
| chr14 | 104024182 | 104055608 | 31427 | 0 | 0.993 | 0.995 |
| chr3 | 118706142 | 118706497 | 356 | 0 | 0.993 | 0.995 |
| chrX | 71447295 | 71459758 | 12464 | 0 | 0.994 | 0.995 |
| chr8 | 81165274 | 81169910 | 4637 | 0 | 0.994 | 0.995 |
| chr6 | 25962269 | 25972413 | 10145 | 0 | 0.994 | 0.995 |
| chr3 | 56590930 | 56592327 | 1398 | 0 | 0.994 | 0.995 |
| chr15 | 26123104 | 26149114 | 26011 | 0 | 0.995 | 0.996 |
| chr9 | 103113889 | 103115908 | 2020 | 0 | 0.995 | 0.996 |
| chr9 | 99615818 | 99617301 | 1484 | 0 | 0.995 | 0.996 |
| chr11 | 10306365 | 10349263 | 42899 | 0 | 0.995 | 0.996 |
| chr5 | 171613510 | 171615844 | 2335 | 0 | 0.996 | 0.996 |
| chr7 | 517100 | 517875 | 776 | 0 | 0.996 | 0.996 |
| chr15 | 57714073 | 57720857 | 6785 | 0 | 0.996 | 0.996 |
| chr2 | 187558343 | 187559188 | 846 | 0 | 0.996 | 0.997 |
| chrX | 71420765 | 71422971 | 2207 | 0 | 0.996 | 0.997 |
| chr6 | 134095213 | 134096344 | 1132 | 0 | 0.996 | 0.997 |
| chr10 | 88344409 | 88345769 | 1361 | 0 | 0.997 | 0.997 |
| chr9 | 88771704 | 88772118 | 415 | 0 | 0.997 | 0.998 |
| chr6 | 27739974 | 27745584 | 5611 | 0 | 0.997 | 0.998 |
| chr4 | 74844753 | 74845250 | 498 | 0 | 0.997 | 0.998 |
| chr11 | 111894736 | 111897008 | 2273 | 0 | 0.997 | 0.998 |
| chr2 | 181934296 | 181935871 | 1576 | 0 | 0.997 | 0.998 |
| chr8 | 103936288 | 103942905 | 6618 | 0 | 0.998 | 0.998 |
| chr7 | 27238557 | 27250742 | 12186 | 0 | 0.998 | 0.998 |
| chr8 | 26370859 | 26372278 | 1420 | 0 | 0.998 | 0.998 |
| chr10 | 103371515 | 103372155 | 641 | 0 | 0.998 | 0.998 |
| chr17 | 29057728 | 29059105 | 1378 | 0 | 0.998 | 0.998 |
| chr17 | 73304622 | 73323064 | 18443 | 0 | 0.998 | 0.999 |
| chr21 | 38419024 | 38460308 | 41285 | 0 | 0.998 | 0.999 |

| | | | | | | |
|---|---|---|---|---|---|---|
| chr16 | 88518674 | 88522601 | 3928 | 0 | 0.999 | 0.999 |
| chr11 | 28898805 | 28900199 | 1395 | 0 | 0.999 | 0.999 |
| chr2 | 122092167 | 122093992 | 1826 | 0 | 0.999 | 0.999 |
| chr4 | 39320853 | 39327324 | 6472 | 0 | 0.999 | 1 |
| chr2 | 44314091 | 44328359 | 14269 | 0 | 0.999 | 1 |
| chr17 | 65524761 | 65525752 | 992 | 0 | 1 | 1 |
| chr1 | 109796578 | 109797922 | 1345 | 0 | 1 | 1 |
| chr5 | 150399640 | 150400873 | 1234 | 0 | 1 | 1 |
| chr10 | 121161016 | 121162244 | 1229 | 0 | 1 | 1 |

**Supplementary Table 4C, H3K9me3 differential binding analysis, Related to Figure 4D, Supplementary Figure 4G, H, I**

| chrom | start | end | width | fold | pvalue | FDR |
|---|---|---|---|---|---|---|
| chr10 | 65641215 | 67330169 | 1688955 | 3.75 | 0 | 0 |
| chr10 | 86410049 | 87227464 | 817416 | 4.11 | 0 | 0 |
| chr13 | 37658722 | 38686314 | 1027593 | 4.38 | 0 | 0 |
| chr16 | 5334817 | 8589427 | 3254611 | 1.96 | 0 | 0 |
| chr18 | 30529777 | 32068684 | 1538908 | 3.48 | 0 | 0 |
| chr18 | 37581043 | 39481749 | 1900707 | 2.86 | 0 | 0 |
| chr18 | 39673654 | 41189629 | 1515976 | 2.73 | 0 | 0 |
| chr18 | 50228573 | 51638568 | 1409996 | 3.33 | 0 | 0 |
| chr2 | 137223841 | 138567273 | 1343433 | 3.5 | 0 | 0 |
| chr2 | 147646467 | 148524377 | 877911 | 4.45 | 0 | 0 |
| chr21 | 30885238 | 32434053 | 1548816 | 2.43 | 0 | 0 |
| chr5 | 96583268 | 98084371 | 1501104 | 3.05 | 0 | 0 |
| chr8 | 131942039 | 132776535 | 834497 | 2.94 | 0 | 0 |
| chr9 | 22549240 | 23362671 | 813432 | 5.45 | 0 | 0 |
| chr9 | 120312646 | 120924708 | 612063 | 3.6 | 0 | 0 |
| chrX | 14144972 | 14817664 | 672693 | 4.07 | 0 | 0 |
| chr6 | 49825232 | 50773867 | 948636 | 3.62 | 9.41E-304 | 2.10E-300 |
| chr5 | 7906721 | 10124227 | 2217507 | 2.59 | 2.28E-303 | 4.79E-300 |
| chr8 | 134688253 | 135480504 | 792252 | 3.24 | 3.74E-302 | 7.46E-299 |
| chr20 | 37789938 | 39301742 | 1511805 | 2.79 | 7.16E-289 | 1.36E-285 |
| chr16 | 25706595 | 27056748 | 1350154 | 2.29 | 9.52E-289 | 1.72E-285 |
| chr4 | 181757894 | 183057040 | 1299147 | 2.87 | 5.76E-288 | 9.91E-285 |
| chr4 | 36426607 | 37290347 | 863741 | 2.95 | 1.31E-284 | 2.15E-281 |
| chr8 | 126905949 | 127297216 | 391268 | 3.66 | 5.11E-276 | 8.05E-273 |
| chr12 | 44401337 | 45321133 | 919797 | 3.92 | 4.42E-272 | 6.70E-269 |
| chr11 | 131568741 | 132938331 | 1369591 | 2.08 | 1.83E-262 | 2.66E-259 |
| chr20 | 40331688 | 41949588 | 1617901 | 2.78 | 2.06E-261 | 2.88E-258 |
| chr18 | 65197306 | 66316027 | 1118722 | 2.8 | 8.33E-259 | 1.13E-255 |
| chr12 | 22877532 | 25138810 | 2261279 | 3.24 | 1.55E-255 | 2.02E-252 |
| chr20 | 6120065 | 6742422 | 622358 | 2.38 | 1.88E-253 | 2.37E-250 |
| chr5 | 88215170 | 89301434 | 1086265 | 3.39 | 6.25E-253 | 7.63E-250 |
| chr1 | 95912397 | 96668248 | 755852 | 4.18 | 4.52E-249 | 5.35E-246 |
| chrX | 141312511 | 142589506 | 1276996 | 2.41 | 6.40E-249 | 7.35E-246 |
| chr20 | 11918012 | 12888851 | 970840 | 2.64 | 9.99E-244 | 1.11E-240 |
| chr20 | 53301550 | 54889346 | 1587797 | 2.11 | 2.88E-239 | 3.11E-236 |
| chr4 | 42712500 | 44006412 | 1293913 | 2.46 | 1.46E-236 | 1.54E-233 |
| chr1 | 72812329 | 74646625 | 1834297 | 2.94 | 4.61E-234 | 4.72E-231 |
| chr5 | 109218833 | 110048984 | 830152 | 5.17 | 1.35E-228 | 1.35E-225 |
| chrX | 50763216 | 51055749 | 292534 | 4.06 | 6.55E-228 | 6.36E-225 |
| chr12 | 81170033 | 82715597 | 1545565 | 3.19 | 4.85E-227 | 4.59E-224 |
| chr3 | 25915586 | 26426096 | 510511 | 3.66 | 2.31E-221 | 2.14E-218 |
| chr3 | 73181214 | 75286888 | 2105675 | 2.14 | 9.58E-216 | 8.64E-213 |
| chr11 | 55661342 | 56385536 | 724195 | 3.66 | 4.70E-215 | 4.14E-212 |
| chr3 | 21478435 | 22789590 | 1311156 | 4.99 | 3.38E-213 | 2.91E-210 |

| | | | | | | |
|---|---|---|---|---|---|---|
| chr8 | 64220161 | 65474772 | 1254612 | 3.46 | 8.95E-213 | 7.41E-210 |
| chr6 | 120490653 | 121326897 | 836245 | 3.6 | 9.00E-213 | 7.41E-210 |
| chrX | 147263901 | 148517383 | 1253483 | 4.32 | 1.57E-207 | 1.26E-204 |
| chr20 | 50857094 | 52020249 | 1163156 | 2.4 | 3.41E-206 | 2.69E-203 |
| chr11 | 40743101 | 43295772 | 2552672 | 2.02 | 3.50E-203 | 2.71E-200 |
| chr16 | 82274608 | 83841275 | 1566668 | 2.28 | 2.38E-200 | 1.80E-197 |
| chr11 | 132956022 | 133684661 | 728640 | 2.71 | 3.13E-199 | 2.32E-196 |
| chr13 | 72129497 | 72427610 | 298114 | 4.49 | 1.13E-198 | 8.19E-196 |
| chr17 | 10687141 | 11834733 | 1147593 | 1.96 | 3.02E-198 | 2.16E-195 |
| chr8 | 15622427 | 16461094 | 838668 | 2.1 | 4.70E-197 | 3.29E-194 |
| chr12 | 33209976 | 34083991 | 874016 | 3.35 | 6.99E-194 | 4.81E-191 |
| chr1 | 181489308 | 181973987 | 484680 | 3.3 | 7.75E-193 | 5.24E-190 |
| chr8 | 57361539 | 57832649 | 471111 | 2.55 | 2.84E-192 | 1.89E-189 |
| chr6 | 96069252 | 96953130 | 883879 | 2.99 | 2.89E-185 | 1.88E-182 |
| chr14 | 29246929 | 30014897 | 767969 | 4.36 | 1.02E-182 | 6.51E-180 |
| chr8 | 2343733 | 6248114 | 3904382 | 1.94 | 3.25E-182 | 2.05E-179 |
| chr4 | 11813514 | 12714888 | 901375 | 2.63 | 1.66E-181 | 1.03E-178 |
| chr12 | 127656685 | 129260799 | 1604115 | 2.07 | 2.85E-181 | 1.74E-178 |
| chrX | 97183034 | 99614143 | 2431110 | 3.27 | 1.07E-179 | 6.40E-177 |
| chr18 | 4700779 | 5225399 | 524621 | 2.78 | 2.00E-175 | 1.18E-172 |
| chr7 | 48342117 | 48881629 | 539513 | 2.31 | 6.71E-175 | 3.91E-172 |
| chr2 | 58999883 | 60747278 | 1747396 | 2.85 | 1.36E-174 | 7.82E-172 |
| chr7 | 135690754 | 137264911 | 1574158 | 4.08 | 1.03E-171 | 5.81E-169 |
| chr7 | 48977825 | 50087740 | 1109916 | 2.49 | 1.97E-171 | 1.10E-168 |
| chr18 | 22946548 | 23496336 | 549789 | 2.41 | 2.82E-167 | 1.55E-164 |
| chr13 | 47558057 | 48286295 | 728239 | 4.22 | 8.26E-167 | 4.46E-164 |
| chr11 | 15102001 | 15659413 | 557413 | 3.34 | 3.21E-164 | 1.71E-161 |
| chr3 | 97007093 | 97368164 | 361072 | 4.51 | 1.92E-163 | 1.01E-160 |
| chr3 | 6525635 | 7331088 | 805454 | 2.73 | 2.05E-163 | 1.06E-160 |
| chr16 | 64494229 | 65147858 | 653630 | 1.98 | 1.20E-160 | 6.12E-158 |
| chr3 | 60187476 | 61158182 | 970707 | 2.47 | 1.79E-159 | 9.02E-157 |
| chr3 | 3246635 | 4307066 | 1060432 | 3.72 | 1.53E-157 | 7.64E-155 |
| chr2 | 34342046 | 34692426 | 350381 | 2.38 | 2.05E-157 | 1.01E-154 |
| chr18 | 35412634 | 36483241 | 1070608 | 2.56 | 5.87E-156 | 2.85E-153 |
| chr7 | 51503832 | 53189439 | 1685608 | 2.01 | 3.28E-155 | 1.57E-152 |
| chr7 | 96831530 | 97348887 | 517358 | 1.97 | 5.10E-154 | 2.41E-151 |
| chr10 | 1782121 | 2845363 | 1063243 | 2.06 | 2.96E-149 | 1.39E-146 |
| chr13 | 103650086 | 103827392 | 177307 | 5.88 | 5.20E-147 | 2.40E-144 |
| chr18 | 24804450 | 25156824 | 352375 | 5.09 | 2.11E-145 | 9.63E-143 |
| chr7 | 53238479 | 54286067 | 1047589 | 2.08 | 5.40E-145 | 2.43E-142 |
| chr11 | 20705889 | 21313107 | 607219 | 3.94 | 6.18E-145 | 2.75E-142 |
| chr2 | 129595659 | 130514559 | 918901 | 1.84 | 1.08E-144 | 4.77E-142 |
| chr6 | 101854441 | 104967631 | 3113191 | 4.4 | 1.49E-144 | 6.47E-142 |
| chr5 | 50261739 | 52022066 | 1760328 | 2.41 | 2.25E-144 | 9.69E-142 |
| chr8 | 97360008 | 97643343 | 283336 | 3.8 | 8.13E-143 | 3.46E-140 |
| chr12 | 29550735 | 30638049 | 1087315 | 1.96 | 1.97E-138 | 8.29E-136 |
| chr5 | 2764111 | 4859303 | 2095193 | 1.45 | 1.13E-137 | 4.70E-135 |
| chr8 | 87725702 | 88127831 | 402130 | 4.65 | 1.44E-137 | 5.94E-135 |
| chr12 | 43932677 | 44057130 | 124454 | 3.76 | 3.39E-133 | 1.38E-130 |
| chr18 | 41203423 | 41929319 | 725897 | 2.43 | 2.31E-132 | 9.30E-130 |
| chr13 | 107587963 | 107853959 | 265997 | 3.23 | 2.38E-132 | 9.50E-130 |
| chr11 | 127652669 | 127889203 | 236535 | 4.52 | 7.60E-132 | 3.00E-129 |
| chr16 | 65163667 | 66250305 | 1086639 | 2.5 | 1.42E-131 | 5.54E-129 |
| chr10 | 106705626 | 111593356 | 4887731 | 2.27 | 5.21E-131 | 2.01E-128 |
| chr3 | 64990685 | 65146267 | 155583 | 4.4 | 4.42E-130 | 1.69E-127 |
| chrX | 30928528 | 34481018 | 3552491 | 2.14 | 7.28E-128 | 2.76E-125 |
| chr3 | 88258961 | 90203431 | 1944471 | 2.12 | 6.62E-127 | 2.48E-124 |
| chr10 | 91719714 | 92073366 | 353653 | 3.27 | 6.87E-125 | 2.55E-122 |
| chrX | 97059005 | 97170215 | 111211 | 6.32 | 1.07E-124 | 3.92E-122 |

| | | | | | | |
|---|---|---|---|---|---|---|
| chr14 | 41656151 | 42065746 | 409596 | 2.27 | 3.07E-124 | 1.12E-121 |
| chr12 | 60942880 | 62510184 | 1567305 | 2.83 | 2.16E-123 | 7.80E-121 |
| chrX | 95319594 | 95563310 | 243717 | 3.91 | 2.28E-123 | 8.15E-121 |
| chr14 | 97652171 | 99364148 | 1711978 | 1.69 | 7.30E-123 | 2.58E-120 |
| chr14 | 45795632 | 47043097 | 1247466 | 2.21 | 9.18E-123 | 3.22E-120 |
| chr9 | 23378002 | 23812522 | 434521 | 3.84 | 2.19E-122 | 7.59E-120 |
| chr14 | 62401216 | 62588626 | 187411 | 3.1 | 2.89E-122 | 9.95E-120 |
| chr3 | 20318230 | 21367593 | 1049364 | 3.39 | 3.69E-122 | 1.26E-119 |
| chr11 | 59969406 | 60495237 | 525832 | 2.16 | 5.31E-122 | 1.79E-119 |
| chr4 | 16307030 | 17469284 | 1162255 | 1.82 | 1.37E-121 | 4.60E-119 |
| chrX | 11565925 | 11724787 | 158863 | 4.09 | 8.25E-121 | 2.74E-118 |
| chr14 | 25480606 | 26778417 | 1297812 | 1.78 | 3.59E-120 | 1.18E-117 |
| chr12 | 114447827 | 114908240 | 460414 | 2.25 | 2.37E-118 | 7.72E-116 |
| chr17 | 49628685 | 51630960 | 2002276 | 2.23 | 3.27E-118 | 1.06E-115 |
| chr11 | 79159497 | 82387493 | 3227997 | 2.31 | 5.72E-118 | 1.84E-115 |
| chr8 | 33904934 | 36990801 | 3085868 | 1.43 | 1.24E-117 | 3.96E-115 |
| chr12 | 4970132 | 6018923 | 1048792 | 3.55 | 2.19E-117 | 6.92E-115 |
| chr5 | 7311948 | 7738251 | 426304 | 3.45 | 5.31E-116 | 1.66E-113 |
| chr17 | 31284273 | 32558924 | 1274652 | 3.31 | 1.08E-115 | 3.35E-113 |
| chrX | 136524023 | 138286215 | 1762193 | 4.72 | 2.73E-113 | 8.40E-111 |
| chr13 | 58210847 | 60085519 | 1874673 | 2.53 | 4.42E-113 | 1.35E-110 |
| chr18 | 58239304 | 59209805 | 970502 | 3.63 | 1.12E-112 | 3.40E-110 |
| chr1 | 118637156 | 119394122 | 756967 | 3.3 | 1.13E-112 | 3.40E-110 |
| chr13 | 26084455 | 26614175 | 529721 | 2.35 | 1.86E-112 | 5.55E-110 |
| chr6 | 23454950 | 24037410 | 582461 | 2.26 | 1.26E-111 | 3.74E-109 |
| chr7 | 85493137 | 86561557 | 1068421 | 3.16 | 2.47E-111 | 7.25E-109 |
| chr12 | 125891104 | 127227089 | 1335986 | 2.07 | 2.93E-111 | 8.54E-109 |
| chr1 | 101829476 | 103338127 | 1508652 | 2.43 | 1.45E-110 | 4.18E-108 |
| chr5 | 62086464 | 63454297 | 1367834 | 2.63 | 1.89E-110 | 5.43E-108 |
| chr5 | 15017141 | 16413620 | 1396480 | 2.48 | 2.18E-109 | 6.19E-107 |
| chr8 | 136472898 | 140019889 | 3546992 | 2.21 | 4.00E-109 | 1.13E-106 |
| chr8 | 65477717 | 66032986 | 555270 | 2.21 | 2.69E-108 | 7.54E-106 |
| chr1 | 71565448 | 72741679 | 1176232 | 2.78 | 9.07E-108 | 2.52E-105 |
| chr18 | 25809032 | 26572893 | 763862 | 2.29 | 9.88E-108 | 2.73E-105 |
| chr11 | 130965176 | 131537235 | 572060 | 2.26 | 8.46E-107 | 2.32E-104 |
| chr19 | 31846091 | 32706366 | 860276 | 1.46 | 1.15E-106 | 3.14E-104 |
| chr11 | 112844914 | 113145521 | 300608 | 2.85 | 1.12E-105 | 3.04E-103 |
| chr3 | 662988 | 3157905 | 2494918 | 2.36 | 2.84E-105 | 7.63E-103 |
| chr2 | 241200484 | 241223655 | 23172 | -4.02 | 2.50E-104 | 6.68E-102 |
| chr8 | 12708209 | 15393391 | 2685183 | 2.16 | 2.75E-104 | 7.27E-102 |
| chr2 | 12853546 | 13514437 | 660892 | 2.21 | 6.04E-104 | 1.59E-101 |
| chr15 | 61552871 | 62120544 | 567674 | 2.08 | 9.14E-104 | 2.39E-101 |
| chr8 | 50348300 | 52435432 | 2087133 | 1.89 | 1.25E-103 | 3.24E-101 |
| chr6 | 91345484 | 94113553 | 2768070 | 2.52 | 1.68E-103 | 4.32E-101 |
| chr18 | 49226598 | 49801901 | 575304 | 2.03 | 6.15E-103 | 1.57E-100 |
| chr5 | 163005901 | 165788983 | 2783083 | 2.76 | 1.58E-102 | 4.01E-100 |
| chr5 | 10945171 | 13326677 | 2381507 | 2.51 | 5.59E-102 | 1.41E-99 |
| chr5 | 165805258 | 166109885 | 304628 | 2.96 | 7.87E-102 | 1.97E-99 |
| chr9 | 77823471 | 78493638 | 670168 | 3.9 | 2.07E-101 | 5.15E-99 |
| chr16 | 59956011 | 63163468 | 3207458 | 1.85 | 2.23E-101 | 5.52E-99 |
| chr4 | 44771547 | 47382512 | 2610966 | 2.61 | 2.71E-101 | 6.66E-99 |
| chr15 | 47404130 | 48382488 | 978359 | 2.11 | 6.78E-101 | 1.66E-98 |
| chr18 | 5888485 | 6687310 | 798826 | 1.72 | 2.30E-100 | 5.57E-98 |
| chr12 | 129311656 | 130493745 | 1182090 | 2 | 5.78E-99 | 1.39E-96 |
| chr21 | 29434574 | 30206548 | 771975 | 2.45 | 1.23E-98 | 2.94E-96 |
| chr14 | 86577626 | 87057771 | 480146 | 1.98 | 7.30E-98 | 1.74E-95 |
| chr10 | 83582234 | 84100819 | 518586 | 1.92 | 1.45E-97 | 3.43E-95 |
| chr13 | 37014955 | 37361103 | 346149 | 2.18 | 4.48E-97 | 1.05E-94 |
| chr18 | 75772376 | 76210287 | 437912 | 1.7 | 1.66E-96 | 3.87E-94 |
| chrX | 8149370 | 8430292 | 280923 | 2.62 | 1.71E-96 | 3.98E-94 |
| chr4 | 27122587 | 27668560 | 545974 | 3.7 | 2.75E-96 | 6.35E-94 |

| | | | | | | |
|---|---|---|---|---|---|---|
| chr3 | 143725570 | 144813764 | 1088195 | 2.97 | 1.33E-94 | 3.05E-92 |
| chr2 | 48840110 | 53083553 | 4243444 | 2.09 | 2.02E-94 | 4.61E-92 |
| chr22 | 34325868 | 35397032 | 1071165 | 2.05 | 1.44E-93 | 3.26E-91 |
| chr4 | 81369198 | 81880525 | 511328 | 5.76 | 5.29E-92 | 1.19E-89 |
| chr13 | 74865382 | 75784100 | 918719 | 2.61 | 7.67E-92 | 1.72E-89 |
| chr3 | 98660 | 612068 | 513409 | 2.14 | 1.30E-91 | 2.89E-89 |
| chr20 | 21506063 | 22227379 | 721317 | 2.94 | 1.95E-91 | 4.33E-89 |
| chrX | 7119150 | 7623449 | 504300 | 2.03 | 3.23E-91 | 7.11E-89 |
| chr10 | 23746199 | 24475633 | 729435 | 2.15 | 3.26E-91 | 7.13E-89 |
| chrX | 111929767 | 112078596 | 148830 | 3.29 | 5.97E-91 | 1.30E-88 |
| chr6 | 86737113 | 87367128 | 630016 | 2.38 | 8.31E-91 | 1.80E-88 |
| chr13 | 29483983 | 29907792 | 423810 | 1.62 | 6.87E-90 | 1.48E-87 |
| chr15 | 93620570 | 95795094 | 2174525 | 1.8 | 3.34E-88 | 7.15E-86 |
| chr1 | 83255265 | 83598195 | 342931 | 2.44 | 3.03E-87 | 6.45E-85 |
| chr16 | 9405561 | 10271338 | 865778 | 2.59 | 1.86E-86 | 3.93E-84 |
| chr4 | 58437519 | 58760430 | 322912 | 2.54 | 1.07E-85 | 2.25E-83 |
| chr10 | 67446472 | 68820579 | 1374108 | 2.33 | 1.42E-85 | 2.97E-83 |
| chrX | 5221897 | 6130241 | 908345 | 1.85 | 2.05E-85 | 4.27E-83 |
| chr14 | 39996140 | 41628591 | 1632452 | 2.63 | 2.59E-85 | 5.36E-83 |
| chr5 | 115968431 | 118119155 | 2150725 | 2.81 | 3.06E-85 | 6.29E-83 |
| chr14 | 87078400 | 87879569 | 801170 | 2.21 | 8.68E-85 | 1.78E-82 |
| chr6 | 70231827 | 70370113 | 138287 | 3.1 | 8.86E-85 | 1.80E-82 |
| chr14 | 87895659 | 88395144 | 499486 | 2.73 | 1.06E-84 | 2.15E-82 |
| chr18 | 4139606 | 4678633 | 539028 | 4.39 | 1.23E-84 | 2.48E-82 |
| chr6 | 76821372 | 77192945 | 371574 | 3.18 | 4.28E-84 | 8.57E-82 |
| chr2 | 18075981 | 18553797 | 477817 | 3.33 | 8.44E-84 | 1.68E-81 |
| chr8 | 68429120 | 70734261 | 2305142 | 3.51 | 1.08E-83 | 2.14E-81 |
| chr7 | 54301640 | 54373729 | 72090 | 2.92 | 2.13E-83 | 4.19E-81 |
| chr17 | 10234537 | 10515094 | 280558 | 1.59 | 9.67E-83 | 1.90E-80 |
| chr15 | 24823609 | 25775452 | 951844 | -1.99 | 1.71E-82 | 3.34E-80 |
| chrX | 26675552 | 27301913 | 626362 | 2.01 | 1.04E-81 | 2.02E-79 |
| chr2 | 122776218 | 127429712 | 4653495 | 1.71 | 3.99E-81 | 7.70E-79 |
| chr10 | 100277312 | 100806996 | 529685 | 2.81 | 9.42E-81 | 1.81E-78 |
| chr4 | 173886995 | 173931882 | 44888 | -4.41 | 1.81E-80 | 3.46E-78 |
| chr8 | 49918735 | 50332576 | 413842 | 1.95 | 2.61E-80 | 4.97E-78 |
| chr8 | 89346784 | 90018314 | 671531 | 2.72 | 2.67E-80 | 5.05E-78 |
| chr14 | 82033602 | 85553525 | 3519924 | 2.24 | 5.74E-80 | 1.08E-77 |
| chr10 | 54544771 | 57375719 | 2830949 | 2.19 | 7.39E-80 | 1.39E-77 |
| chr21 | 47990631 | 48015686 | 25056 | -5.9 | 1.63E-79 | 3.03E-77 |
| chr20 | 6758265 | 7904758 | 1146494 | 1.52 | 3.27E-79 | 6.07E-77 |
| chr5 | 44139394 | 44723255 | 583862 | 2.77 | 4.79E-79 | 8.84E-77 |
| chr3 | 76000243 | 78628298 | 2628056 | 1.77 | 2.34E-78 | 4.30E-76 |
| chr14 | 33417648 | 34255096 | 837449 | 5.12 | 6.53E-78 | 1.19E-75 |
| chr9 | 26343258 | 26653736 | 310479 | 2.19 | 2.14E-77 | 3.89E-75 |
| chr7 | 130450292 | 130466077 | 15786 | -4.91 | 1.05E-76 | 1.90E-74 |
| chr10 | 125732340 | 125751033 | 18694 | -6.06 | 1.69E-76 | 3.05E-74 |
| chrX | 75813520 | 76255342 | 441823 | 2.12 | 1.79E-76 | 3.21E-74 |
| chr10 | 84557801 | 85851777 | 1293977 | 2.21 | 1.34E-75 | 2.39E-73 |
| chr7 | 153521518 | 153655086 | 133569 | 2.34 | 3.74E-75 | 6.64E-73 |
| chr5 | 66547873 | 67435724 | 887852 | 3.1 | 6.07E-75 | 1.07E-72 |
| chr15 | 27139399 | 28347910 | 1208512 | 1.21 | 6.93E-75 | 1.22E-72 |
| chr8 | 135891096 | 136461178 | 570083 | 2.56 | 2.46E-74 | 4.31E-72 |
| chr3 | 146536175 | 147657884 | 1121710 | 3.02 | 8.84E-74 | 1.54E-71 |
| chr3 | 34460642 | 35974052 | 1513411 | 3.73 | 1.47E-73 | 2.56E-71 |
| chr13 | 61552754 | 62407272 | 854519 | 2.82 | 2.90E-73 | 5.01E-71 |
| chr2 | 168157075 | 168700902 | 543828 | 3.59 | 6.14E-73 | 1.06E-70 |
| chr15 | 81635089 | 82321837 | 686749 | 1.66 | 7.55E-73 | 1.29E-70 |
| chr18 | 6708018 | 6872336 | 164319 | 3.12 | 9.75E-73 | 1.66E-70 |
| chr18 | 64875638 | 65177073 | 301436 | 4.33 | 2.31E-72 | 3.92E-70 |
| chr1 | 34135549 | 34622672 | 487124 | 2.83 | 3.99E-72 | 6.75E-70 |
| chrX | 25338915 | 26659712 | 1320798 | 2.44 | 7.61E-72 | 1.28E-69 |

| | | | | | | |
|---|---|---|---|---|---|---|
| chr2 | 114835946 | 118520527 | 3684582 | 1.73 | 9.59E-72 | 1.61E-69 |
| chr11 | 39399271 | 40301796 | 902526 | 2.1 | 9.73E-72 | 1.62E-69 |
| chr1 | 49423142 | 50785232 | 1362091 | 2.29 | 1.80E-71 | 2.99E-69 |
| chrX | 8513945 | 8692970 | 179026 | 2.73 | 2.98E-71 | 4.92E-69 |
| chr18 | 61691718 | 64157625 | 2465908 | 2.29 | 5.86E-71 | 9.64E-69 |
| chr19 | 56676609 | 58869217 | 2192609 | -18.26 | 6.86E-71 | 1.12E-68 |
| chr15 | 96926335 | 98446221 | 1519887 | 2.15 | 1.61E-70 | 2.62E-68 |
| chr14 | 62601808 | 63137561 | 535754 | 1.8 | 1.89E-70 | 3.06E-68 |
| chr1 | 208479778 | 209150261 | 670484 | 1.9 | 2.01E-70 | 3.24E-68 |
| chr8 | 86411230 | 86546967 | 135738 | 4.23 | 1.09E-69 | 1.76E-67 |
| chr4 | 73199227 | 73426163 | 226937 | 5.47 | 1.47E-69 | 2.35E-67 |
| chr14 | 30476645 | 30986195 | 509551 | 1.72 | 4.81E-69 | 7.68E-67 |
| chrX | 132115717 | 132293833 | 178117 | 2.82 | 5.93E-69 | 9.44E-67 |
| chrX | 112914591 | 113162269 | 247679 | 2.74 | 1.36E-68 | 2.16E-66 |
| chrX | 124570374 | 125260863 | 690490 | 3.28 | 2.17E-68 | 3.42E-66 |
| chrX | 116920218 | 117029976 | 109759 | 3.66 | 5.16E-68 | 8.11E-66 |
| chr19 | 51925599 | 54300627 | 2375029 | -17.85 | 6.21E-68 | 9.71E-66 |
| chr18 | 980662 | 1194816 | 214155 | 3.34 | 1.36E-67 | 2.12E-65 |
| chr19 | 36807874 | 37752663 | 944790 | -17.72 | 6.25E-67 | 9.70E-65 |
| chr21 | 41165852 | 41836704 | 670853 | 2.22 | 1.30E-66 | 2.00E-64 |
| chr2 | 21189211 | 23417331 | 2228121 | 2.67 | 2.04E-65 | 3.14E-63 |
| chr15 | 54131728 | 55452819 | 1321092 | 2.78 | 2.65E-65 | 4.05E-63 |
| chrX | 111597417 | 111916962 | 319546 | 3.36 | 3.50E-65 | 5.34E-63 |
| chr7 | 42289576 | 42852892 | 563317 | 2.49 | 4.43E-63 | 6.73E-61 |
| chr5 | 119810860 | 120123579 | 312720 | 3.51 | 4.50E-63 | 6.81E-61 |
| chrX | 78346734 | 78568033 | 221300 | 2.86 | 5.20E-63 | 7.84E-61 |
| chr15 | 46263069 | 47339016 | 1075948 | 1.76 | 6.31E-63 | 9.48E-61 |
| chr19 | 44292942 | 45065483 | 772542 | -17.12 | 1.10E-62 | 1.64E-60 |
| chr11 | 48555429 | 48626361 | 70933 | 3.07 | 1.44E-62 | 2.14E-60 |
| chr19 | 31184070 | 31827722 | 643653 | 1.36 | 1.76E-62 | 2.62E-60 |
| chr20 | 20098167 | 20171088 | 72922 | -3.23 | 3.65E-62 | 5.40E-60 |
| chr4 | 10234708 | 10398405 | 163698 | 2.76 | 4.89E-62 | 7.21E-60 |
| chr19 | 37795150 | 38387254 | 592105 | -17.02 | 6.26E-62 | 9.19E-60 |
| chr6 | 15695435 | 15723288 | 27854 | -3.53 | 8.36E-62 | 1.22E-59 |
| chrX | 6150054 | 6921990 | 771937 | 1.63 | 4.62E-61 | 6.72E-59 |
| chr5 | 7080729 | 7281993 | 201265 | 2.98 | 1.19E-60 | 1.73E-58 |
| chr14 | 23136660 | 23193726 | 57067 | -3.74 | 1.78E-60 | 2.57E-58 |
| chr7 | 124581919 | 126513796 | 1931878 | 3.37 | 1.81E-60 | 2.60E-58 |
| chr13 | 107867008 | 108517817 | 650810 | 3.37 | 2.61E-60 | 3.74E-58 |
| chr12 | 103354793 | 103877286 | 522494 | 1.99 | 3.82E-60 | 5.46E-58 |
| chr3 | 117909988 | 118329768 | 419781 | 2.73 | 4.29E-60 | 6.11E-58 |
| chr13 | 54195891 | 57713074 | 3517184 | 2.1 | 4.43E-60 | 6.28E-58 |
| chr7 | 28794051 | 28943607 | 149557 | 2.5 | 5.52E-60 | 7.80E-58 |
| chr10 | 9390290 | 11476309 | 2086020 | 1.72 | 5.62E-60 | 7.90E-58 |
| chr7 | 38446868 | 38757262 | 310395 | 1.82 | 8.76E-60 | 1.23E-57 |
| chr10 | 19660050 | 20844598 | 1184549 | 1.97 | 2.49E-59 | 3.48E-57 |
| chr6 | 69352819 | 70190326 | 837508 | 3.21 | 2.94E-59 | 4.09E-57 |
| chr7 | 119581170 | 120416465 | 835296 | 5.22 | 8.03E-59 | 1.11E-56 |
| chr8 | 20243531 | 20893227 | 649697 | 2.03 | 1.08E-58 | 1.49E-56 |
| chrX | 125818210 | 127866790 | 2048581 | 2.66 | 1.62E-58 | 2.23E-56 |
| chr21 | 19529792 | 26928065 | 7398274 | 1.81 | 1.72E-58 | 2.36E-56 |
| chr8 | 23452016 | 23476455 | 24440 | -3.17 | 1.83E-58 | 2.50E-56 |
| chrX | 143559184 | 145883385 | 2324202 | 2.05 | 2.40E-58 | 3.27E-56 |
| chr9 | 7817986 | 12679223 | 4861238 | 2.3 | 2.18E-57 | 2.96E-55 |
| chr9 | 30791101 | 32336859 | 1545759 | 2.13 | 2.47E-57 | 3.34E-55 |
| chr1 | 210451622 | 210460222 | 8601 | -3.69 | 4.85E-57 | 6.54E-55 |
| chr18 | 36539645 | 36949100 | 409456 | 3.11 | 6.43E-57 | 8.64E-55 |
| chr4 | 10592750 | 11236703 | 643954 | 2.44 | 7.66E-57 | 1.02E-54 |
| chr10 | 28042525 | 28463200 | 420676 | 2.41 | 2.58E-56 | 3.43E-54 |
| chr11 | 4432355 | 4511141 | 78787 | 3.23 | 4.19E-56 | 5.56E-54 |
| chr4 | 44025844 | 44443126 | 417283 | 2.89 | 4.93E-56 | 6.53E-54 |

| | | | | | | |
|---|---|---|---|---|---|---|
| chr1 | 237252710 | 237537514 | 284805 | 2.53 | 9.49E-56 | 1.25E-53 |
| chrX | 77992974 | 78327668 | 334695 | 3.31 | 1.10E-55 | 1.45E-53 |
| chr19 | 22861748 | 24182605 | 1320858 | -16.16 | 2.00E-55 | 2.62E-53 |
| chr17 | 53863762 | 54667123 | 803362 | 16.02 | 2.79E-55 | 3.65E-53 |
| chr8 | 71661391 | 72712788 | 1051398 | 16.02 | 2.83E-55 | 3.69E-53 |
| chr5 | 177689638 | 178831448 | 1141811 | -16.05 | 2.96E-55 | 3.84E-53 |
| chr7 | 63294186 | 64467901 | 1173716 | -16.02 | 4.37E-55 | 5.65E-53 |
| chr7 | 152644736 | 153402731 | 757996 | 2.34 | 8.50E-55 | 1.09E-52 |
| chr11 | 35496803 | 35542007 | 45205 | -4.37 | 9.81E-55 | 1.26E-52 |
| chr6 | 51087423 | 51818065 | 730643 | 2.96 | 1.23E-54 | 1.58E-52 |
| chrX | 104173877 | 105644787 | 1470911 | 15.89 | 2.03E-54 | 2.59E-52 |
| chr9 | 7275571 | 7780392 | 504822 | 2.37 | 2.83E-54 | 3.60E-52 |
| chr16 | 78714023 | 79612374 | 898352 | 1.62 | 3.31E-54 | 4.19E-52 |
| chr6 | 97860759 | 99265824 | 1405066 | 15.87 | 3.56E-54 | 4.49E-52 |
| chr12 | 55470709 | 55586163 | 115455 | 2.39 | 4.29E-54 | 5.40E-52 |
| chr7 | 77774884 | 78809614 | 1034731 | 15.84 | 4.46E-54 | 5.59E-52 |
| chr19 | 11709372 | 12760281 | 1050910 | -15.92 | 4.52E-54 | 5.65E-52 |
| chr11 | 36775911 | 39387390 | 2611480 | 1.88 | 1.12E-53 | 1.40E-51 |
| chr19 | 9195476 | 9927963 | 732488 | -15.83 | 1.13E-53 | 1.40E-51 |
| chr16 | 25346301 | 25692941 | 346641 | 1.74 | 2.31E-53 | 2.85E-51 |
| chr6 | 78316416 | 78395947 | 79532 | 3.14 | 2.36E-53 | 2.91E-51 |
| chr2 | 158801117 | 158844778 | 43662 | -3.63 | 4.25E-53 | 5.22E-51 |
| chr5 | 143620090 | 144715510 | 1095421 | 15.68 | 6.78E-53 | 8.31E-51 |
| chr1 | 215815989 | 216767627 | 951639 | 15.7 | 7.72E-53 | 9.43E-51 |
| chr11 | 29466830 | 30321095 | 854266 | 15.62 | 1.68E-52 | 2.04E-50 |
| chr8 | 56023582 | 56615157 | 591576 | 3.59 | 2.17E-52 | 2.63E-50 |
| chrX | 42871032 | 43152224 | 281193 | 2.18 | 4.42E-52 | 5.34E-50 |
| chrX | 51081912 | 51159445 | 77534 | 4.48 | 4.67E-52 | 5.63E-50 |
| chr1 | 176203977 | 177117992 | 914016 | 15.54 | 4.70E-52 | 5.64E-50 |
| chr1 | 103373947 | 103784953 | 411007 | 2.99 | 5.13E-52 | 6.14E-50 |
| chr6 | 114768838 | 116157864 | 1389027 | 15.56 | 6.68E-52 | 7.97E-50 |
| chr19 | 40434921 | 40643818 | 208898 | -15.53 | 7.41E-52 | 8.82E-50 |
| chr1 | 158180994 | 159097001 | 916008 | 15.51 | 7.92E-52 | 9.40E-50 |
| chr8 | 55472736 | 56010424 | 537689 | 4.9 | 9.46E-52 | 1.12E-49 |
| chr7 | 18679289 | 18892383 | 213095 | 3.93 | 9.54E-52 | 1.13E-49 |
| chrX | 92375476 | 92916665 | 541190 | 2.44 | 9.83E-52 | 1.16E-49 |
| chr11 | 83550005 | 84260231 | 710227 | 15.49 | 1.40E-51 | 1.64E-49 |
| chr20 | 8221301 | 8367243 | 145943 | 3.65 | 1.44E-51 | 1.68E-49 |
| chr21 | 47068174 | 47117233 | 49060 | -3 | 1.75E-51 | 2.04E-49 |
| chr10 | 52754685 | 53445003 | 690319 | 15.45 | 1.95E-51 | 2.27E-49 |
| chr20 | 11626204 | 11870580 | 244377 | 1.84 | 2.20E-51 | 2.55E-49 |
| chrX | 145905406 | 146962707 | 1057302 | 2.32 | 2.27E-51 | 2.62E-49 |
| chr10 | 58306048 | 59091779 | 785732 | 2.04 | 2.52E-51 | 2.89E-49 |
| chr4 | 69687 | 666382 | 596696 | -15.46 | 2.52E-51 | 2.89E-49 |
| chr21 | 28234138 | 29216635 | 982498 | 1.95 | 3.01E-51 | 3.44E-49 |
| chr7 | 112769545 | 113646756 | 877212 | 15.43 | 4.32E-51 | 4.93E-49 |
| chrX | 65406479 | 65773379 | 366901 | 2.32 | 4.34E-51 | 4.93E-49 |
| chr15 | 79979397 | 80045462 | 66066 | -9.33 | 8.10E-51 | 9.18E-49 |
| chr2 | 40819426 | 41708717 | 889292 | 15.39 | 8.22E-51 | 9.29E-49 |
| chr10 | 37487185 | 38452095 | 964911 | -15.41 | 1.00E-50 | 1.13E-48 |
| chr9 | 27603941 | 28344428 | 740488 | 15.31 | 1.62E-50 | 1.82E-48 |
| chr17 | 52351910 | 52958141 | 606232 | 1.9 | 1.78E-50 | 1.99E-48 |
| chr5 | 146840058 | 147742183 | 902126 | 15.33 | 1.85E-50 | 2.06E-48 |
| chr11 | 96443550 | 98188978 | 1745429 | 2.5 | 2.79E-50 | 3.11E-48 |
| chr17 | 8710068 | 8729749 | 19682 | -3.22 | 2.84E-50 | 3.15E-48 |
| chr12 | 28998956 | 29288958 | 290003 | 1.46 | 3.22E-50 | 3.56E-48 |
| chr16 | 3255305 | 3502767 | 247463 | -15.28 | 3.28E-50 | 3.62E-48 |
| chr15 | 69126352 | 69164597 | 38246 | -9.29 | 5.18E-50 | 5.70E-48 |
| chr3 | 144926138 | 145737430 | 811293 | 15.31 | 5.72E-50 | 6.27E-48 |
| chr1 | 247059684 | 247470403 | 410720 | -15.24 | 6.15E-50 | 6.73E-48 |
| chr3 | 137512989 | 137621423 | 108435 | 4.53 | 6.33E-50 | 6.90E-48 |

| | | | | | | |
|---|---|---|---|---|---|---|
| chr19 | 20957497 | 21812350 | 854854 | -15.24 | 7.28E-50 | 7.92E-48 |
| chr5 | 135646039 | 136524567 | 878529 | 15.21 | 7.49E-50 | 8.12E-48 |
| chr14 | 38688904 | 39289881 | 600978 | 15.19 | 8.95E-50 | 9.67E-48 |
| chr6 | 64636317 | 66011043 | 1374727 | 2.55 | 1.00E-49 | 1.08E-47 |
| chr13 | 38721229 | 38906271 | 185043 | 3.04 | 1.17E-49 | 1.26E-47 |
| chr21 | 41893553 | 42478033 | 584481 | 1.64 | 1.20E-49 | 1.29E-47 |
| chr1 | 5027567 | 5325124 | 297558 | 1.59 | 1.28E-49 | 1.37E-47 |
| chr11 | 4654962 | 5282632 | 627671 | 15.19 | 1.53E-49 | 1.64E-47 |
| chr6 | 124131223 | 125027793 | 896571 | 15.2 | 1.55E-49 | 1.64E-47 |
| chr8 | 79656013 | 80505862 | 849850 | 15.19 | 1.56E-49 | 1.65E-47 |
| chr3 | 19194118 | 19894656 | 700539 | 15.17 | 1.81E-49 | 1.91E-47 |
| chr4 | 171074136 | 171173238 | 99103 | 4.32 | 1.81E-49 | 1.91E-47 |
| chr9 | 23819584 | 25661108 | 1841525 | 2.02 | 1.85E-49 | 1.95E-47 |
| chr14 | 33084421 | 33398421 | 314001 | 4.71 | 2.04E-49 | 2.13E-47 |
| chrX | 142607306 | 143161020 | 553715 | 1.84 | 2.94E-49 | 3.07E-47 |
| chr8 | 73111765 | 73831892 | 720128 | 15.14 | 2.97E-49 | 3.10E-47 |
| chr9 | 1059485 | 2002146 | 942662 | 2.43 | 3.03E-49 | 3.15E-47 |
| chr3 | 154056763 | 154777259 | 720497 | 15.15 | 4.02E-49 | 4.17E-47 |
| chrX | 80224295 | 80282460 | 58166 | 3.87 | 4.33E-49 | 4.48E-47 |
| chr9 | 76411159 | 77101013 | 689855 | 15.11 | 4.58E-49 | 4.72E-47 |
| chr5 | 20293120 | 20695358 | 402239 | 2.69 | 4.65E-49 | 4.78E-47 |
| chr20 | 29856145 | 29892103 | 35959 | 3.3 | 1.15E-48 | 1.18E-46 |
| chr5 | 41948123 | 42577386 | 629264 | 15.02 | 1.31E-48 | 1.34E-46 |
| chr8 | 146038240 | 146277301 | 239062 | -15.08 | 1.36E-48 | 1.38E-46 |
| chr5 | 140423647 | 140872131 | 448485 | -15.04 | 1.71E-48 | 1.74E-46 |
| chr3 | 139661958 | 140511049 | 849092 | 15 | 2.00E-48 | 2.03E-46 |
| chr4 | 79968231 | 80805346 | 837116 | 15.02 | 2.15E-48 | 2.17E-46 |
| chr17 | 16427902 | 16582476 | 154575 | -14.98 | 2.29E-48 | 2.31E-46 |
| chr5 | 128475323 | 129097172 | 621850 | 15.01 | 2.69E-48 | 2.71E-46 |
| chr13 | 72454780 | 73184265 | 729486 | 14.96 | 2.80E-48 | 2.81E-46 |
| chr3 | 18532854 | 19179452 | 646599 | 14.97 | 3.64E-48 | 3.65E-46 |
| chr18 | 1372777 | 1506973 | 134197 | 3.34 | 3.70E-48 | 3.69E-46 |
| chr1 | 58037012 | 58860303 | 823292 | 14.98 | 3.87E-48 | 3.86E-46 |
| chr9 | 73353138 | 73862156 | 509019 | 14.93 | 4.57E-48 | 4.54E-46 |
| chr13 | 53484421 | 54170040 | 685620 | 3.4 | 6.48E-48 | 6.42E-46 |
| chr3 | 62613066 | 63328336 | 715271 | 14.9 | 8.10E-48 | 8.00E-46 |
| chr9 | 99420685 | 99657093 | 236409 | -14.89 | 9.13E-48 | 9.00E-46 |
| chr3 | 44462812 | 44836478 | 373667 | -14.9 | 9.27E-48 | 9.12E-46 |
| chr1 | 60449884 | 61295049 | 845166 | 14.87 | 1.16E-47 | 1.13E-45 |
| chr19 | 36665172 | 36759488 | 94317 | -14.87 | 1.18E-47 | 1.15E-45 |
| chr1 | 57309678 | 58023034 | 713357 | 14.87 | 1.39E-47 | 1.36E-45 |
| chr7 | 121793105 | 122515397 | 722293 | 14.94 | 1.46E-47 | 1.42E-45 |
| chr1 | 175171856 | 175821865 | 650010 | 14.9 | 1.86E-47 | 1.80E-45 |
| chr1 | 14185301 | 14923911 | 738611 | 1.17 | 2.23E-47 | 2.16E-45 |
| chr4 | 82403468 | 83094889 | 691422 | 14.83 | 2.28E-47 | 2.20E-45 |
| chr4 | 70563076 | 71378928 | 815853 | 14.84 | 2.73E-47 | 2.63E-45 |
| chrX | 112098789 | 112897591 | 798803 | 2.49 | 3.31E-47 | 3.18E-45 |
| chr1 | 177229405 | 177834009 | 604605 | 14.8 | 3.54E-47 | 3.39E-45 |
| chr11 | 28534346 | 28998699 | 464354 | -14.8 | 3.89E-47 | 3.71E-45 |
| chrX | 50026059 | 50749635 | 723577 | 14.79 | 3.90E-47 | 3.71E-45 |
| chr2 | 228915088 | 230113704 | 1198617 | 2.53 | 4.93E-47 | 4.69E-45 |
| chr7 | 122538066 | 123116938 | 578873 | 14.78 | 5.72E-47 | 5.43E-45 |
| chr2 | 199188790 | 199986659 | 797870 | 14.83 | 5.95E-47 | 5.63E-45 |
| chr13 | 69775647 | 71803939 | 2028293 | 2.21 | 6.24E-47 | 5.89E-45 |
| chr11 | 83035802 | 83525510 | 489709 | 14.76 | 6.81E-47 | 6.41E-45 |
| chr2 | 18743476 | 18768099 | 24624 | -6.25 | 1.05E-46 | 9.86E-45 |
| chr6 | 123321748 | 124115379 | 793632 | 14.75 | 1.27E-46 | 1.19E-44 |
| chr10 | 5042509 | 5379121 | 336613 | -14.73 | 1.29E-46 | 1.20E-44 |
| chr13 | 38946910 | 39477612 | 530703 | 14.7 | 1.52E-46 | 1.42E-44 |
| chrX | 123607755 | 124307034 | 699280 | 14.69 | 1.69E-46 | 1.57E-44 |
| chrX | 125279484 | 125524365 | 244882 | 2.68 | 1.78E-46 | 1.65E-44 |

| | | | | | | |
|---|---|---|---|---|---|---|
| chr4 | 1524370 | 1531586 | 7217 | -4.39 | 2.08E-46 | 1.93E-44 |
| chr11 | 89153215 | 89284353 | 131139 | 4.22 | 3.13E-46 | 2.89E-44 |
| chr1 | 106492592 | 107966703 | 1474112 | 2.17 | 3.87E-46 | 3.57E-44 |
| chr10 | 53470077 | 53914033 | 443957 | 14.61 | 5.08E-46 | 4.66E-44 |
| chr12 | 133682323 | 133821588 | 139266 | -14.64 | 5.87E-46 | 5.38E-44 |
| chr12 | 71222304 | 71807955 | 585652 | 4.3 | 5.93E-46 | 5.43E-44 |
| chr13 | 96799352 | 97453078 | 653727 | 14.61 | 6.14E-46 | 5.60E-44 |
| chr19 | 19775760 | 20402350 | 626591 | -14.66 | 6.90E-46 | 6.28E-44 |
| chr19 | 24202069 | 24501274 | 299206 | -14.63 | 8.19E-46 | 7.43E-44 |
| chr20 | 21384608 | 21408758 | 24151 | -8.89 | 8.39E-46 | 7.60E-44 |
| chr10 | 112881014 | 113881694 | 1000681 | 5.04 | 8.60E-46 | 7.77E-44 |
| chr4 | 16276194 | 16302286 | 26093 | -6.85 | 9.36E-46 | 8.43E-44 |
| chr7 | 121267370 | 121771169 | 503800 | 14.58 | 9.55E-46 | 8.59E-44 |
| chr12 | 72664386 | 74012651 | 1348266 | 2.26 | 1.05E-45 | 9.43E-44 |
| chr16 | 16856653 | 17557899 | 701247 | 2.09 | 1.36E-45 | 1.22E-43 |
| chr11 | 55076330 | 55627855 | 551526 | 1.65 | 2.57E-45 | 2.29E-43 |
| chr19 | 21827248 | 22560632 | 733385 | -14.65 | 2.76E-45 | 2.46E-43 |
| chr14 | 47066248 | 48588317 | 1522070 | 1.67 | 3.39E-45 | 3.01E-43 |
| chr8 | 46959224 | 47476324 | 517101 | -14.54 | 3.93E-45 | 3.49E-43 |
| chr8 | 96363382 | 96797105 | 433724 | 14.47 | 4.05E-45 | 3.58E-43 |
| chr6 | 119704549 | 120324194 | 619646 | 14.44 | 6.75E-45 | 5.95E-43 |
| chr10 | 92086721 | 92495002 | 408282 | 14.44 | 7.49E-45 | 6.59E-43 |
| chr6 | 164455649 | 165190987 | 735339 | 2.7 | 7.59E-45 | 6.67E-43 |
| chr5 | 159947918 | 162729453 | 2781536 | 4.72 | 9.26E-45 | 8.12E-43 |
| chr2 | 99846069 | 99852521 | 6453 | -5.18 | 9.60E-45 | 8.39E-43 |
| chr10 | 100813505 | 100912611 | 99107 | 4.96 | 1.05E-44 | 9.19E-43 |
| chr18 | 30185930 | 30272684 | 86755 | 2.65 | 1.15E-44 | 9.99E-43 |
| chr2 | 95779443 | 95966065 | 186623 | -14.41 | 1.27E-44 | 1.10E-42 |
| chrX | 147044584 | 147248726 | 204143 | 4.43 | 1.64E-44 | 1.42E-42 |
| chr1 | 227636200 | 227915214 | 279015 | -14.4 | 1.89E-44 | 1.64E-42 |
| chr12 | 77728010 | 78073347 | 345338 | 14.39 | 1.94E-44 | 1.67E-42 |
| chr1 | 216782942 | 217243970 | 461029 | 14.38 | 2.02E-44 | 1.74E-42 |
| chrX | 92935346 | 93634925 | 699580 | 2.12 | 2.07E-44 | 1.78E-42 |
| chr5 | 98878091 | 99381056 | 502966 | 3.12 | 2.12E-44 | 1.81E-42 |
| chr4 | 109711239 | 110279425 | 568187 | 14.41 | 2.64E-44 | 2.25E-42 |
| chr3 | 147671225 | 148065306 | 394082 | 14.4 | 2.69E-44 | 2.30E-42 |
| chr4 | 18606255 | 19194684 | 588430 | 1.95 | 2.70E-44 | 2.30E-42 |
| chrX | 75198042 | 75288686 | 90645 | 2.56 | 2.92E-44 | 2.48E-42 |
| chr5 | 100787957 | 101979046 | 1191090 | 2.75 | 3.00E-44 | 2.54E-42 |
| chr18 | 59295478 | 59555874 | 260397 | 2.61 | 3.43E-44 | 2.90E-42 |
| chrX | 65996543 | 66141786 | 145244 | 2.32 | 3.63E-44 | 3.06E-42 |
| chr2 | 234386693 | 234695641 | 308949 | -14.3 | 4.39E-44 | 3.69E-42 |
| chr11 | 7128029 | 7524001 | 395973 | 14.29 | 6.22E-44 | 5.22E-42 |
| chr14 | 63516079 | 63739938 | 223860 | 1.99 | 7.21E-44 | 6.04E-42 |
| chr3 | 179659510 | 180273709 | 614200 | 14.3 | 8.97E-44 | 7.50E-42 |
| chr1 | 218564329 | 218890881 | 326553 | -14.38 | 1.00E-43 | 8.34E-42 |
| chr3 | 98905046 | 99348098 | 443053 | 14.25 | 1.08E-43 | 8.99E-42 |
| chr1 | 76649853 | 77214247 | 564395 | 14.28 | 1.09E-43 | 9.03E-42 |
| chr3 | 26446744 | 27122684 | 675941 | 4.91 | 1.15E-43 | 9.56E-42 |
| chr11 | 78361259 | 78749062 | 387804 | 14.33 | 1.19E-43 | 9.87E-42 |
| chr9 | 81396344 | 81757930 | 361587 | -14.26 | 1.30E-43 | 1.07E-41 |
| chr6 | 77206374 | 77384524 | 178151 | 2.59 | 1.41E-43 | 1.16E-41 |
| chr2 | 144020537 | 144545411 | 524875 | 14.23 | 1.56E-43 | 1.28E-41 |
| chr1 | 111601735 | 111652916 | 51182 | -3.17 | 1.97E-43 | 1.61E-41 |
| chr15 | 98518944 | 98931973 | 413030 | 1.8 | 2.24E-43 | 1.83E-41 |
| chr8 | 43289946 | 43756784 | 466839 | -14.29 | 2.34E-43 | 1.91E-41 |
| chr8 | 129122053 | 129464420 | 342368 | -14.32 | 2.50E-43 | 2.04E-41 |
| chr10 | 43968739 | 44191659 | 222921 | -14.19 | 2.64E-43 | 2.14E-41 |
| chr1 | 162833344 | 163216870 | 383527 | 14.18 | 2.78E-43 | 2.25E-41 |
| chr13 | 79260836 | 79791094 | 530259 | 14.18 | 3.15E-43 | 2.55E-41 |
| chr10 | 130078118 | 131676453 | 1598336 | 1.5 | 3.30E-43 | 2.66E-41 |

| | | | | | | |
|---|---|---|---|---|---|---|
| chr11 | 87142352 | 87578900 | 436549 | 14.16 | 3.64E-43 | 2.93E-41 |
| chr1 | 69472257 | 70024167 | 551911 | 14.24 | 3.76E-43 | 3.02E-41 |
| chr3 | 6374718 | 6521567 | 146850 | 2.85 | 3.83E-43 | 3.07E-41 |
| chr5 | 98566741 | 98726583 | 159843 | 3.18 | 4.30E-43 | 3.44E-41 |
| chr6 | 72877009 | 73319221 | 442213 | 14.18 | 4.48E-43 | 3.58E-41 |
| chr12 | 91395870 | 91715849 | 319980 | 14.16 | 5.07E-43 | 4.04E-41 |
| chr8 | 88595914 | 89326336 | 730423 | 2.22 | 8.40E-43 | 6.68E-41 |
| chr3 | 30226905 | 30645837 | 418933 | -14.33 | 8.45E-43 | 6.71E-41 |
| chr5 | 113023074 | 113378610 | 355537 | 14.1 | 1.04E-42 | 8.22E-41 |
| chr9 | 78519690 | 78876378 | 356689 | 14.09 | 1.12E-42 | 8.87E-41 |
| chr7 | 126563629 | 126883437 | 319809 | 14.1 | 1.21E-42 | 9.52E-41 |
| chr10 | 22457666 | 22472875 | 15210 | -4.9 | 1.27E-42 | 1.00E-40 |
| chrX | 87162662 | 88456257 | 1293596 | 1.94 | 1.33E-42 | 1.04E-40 |
| chr5 | 22217330 | 23004902 | 787573 | 1.79 | 1.42E-42 | 1.11E-40 |
| chr5 | 87085915 | 87415696 | 329782 | 14.09 | 1.63E-42 | 1.27E-40 |
| chr5 | 86729405 | 87068855 | 339451 | 14.1 | 1.64E-42 | 1.28E-40 |
| chr1 | 71066851 | 71504659 | 437809 | 14.04 | 2.14E-42 | 1.66E-40 |
| chr18 | 66409717 | 67475756 | 1066040 | 4.06 | 2.18E-42 | 1.69E-40 |
| chr1 | 152663772 | 153148743 | 484972 | 14.07 | 2.22E-42 | 1.72E-40 |
| chr6 | 6072986 | 6544979 | 471994 | 2.26 | 2.39E-42 | 1.85E-40 |
| chr7 | 28969453 | 29069232 | 99780 | -2.26 | 2.62E-42 | 2.02E-40 |
| chr11 | 32302975 | 32315604 | 12630 | -3.86 | 2.73E-42 | 2.10E-40 |
| chrX | 94610347 | 95214004 | 603658 | 2.68 | 2.75E-42 | 2.12E-40 |
| chr4 | 111662094 | 112315672 | 653579 | 14.13 | 2.82E-42 | 2.16E-40 |
| chr8 | 129482718 | 129881996 | 399279 | -14.19 | 2.90E-42 | 2.23E-40 |
| chr2 | 1909116 | 2699817 | 790702 | 1.58 | 2.97E-42 | 2.27E-40 |
| chr11 | 5839702 | 6184126 | 344425 | 14.04 | 3.09E-42 | 2.36E-40 |
| chr8 | 108798821 | 109174028 | 375208 | 14.03 | 3.60E-42 | 2.74E-40 |
| chr8 | 85103700 | 85985941 | 882242 | 2.59 | 4.00E-42 | 3.04E-40 |
| chr10 | 122221526 | 122565914 | 344389 | 14.02 | 4.47E-42 | 3.39E-40 |
| chr13 | 78717223 | 79086729 | 369507 | 13.98 | 5.13E-42 | 3.88E-40 |
| chr7 | 33507747 | 33933581 | 425835 | 13.98 | 5.58E-42 | 4.21E-40 |
| chr7 | 6688967 | 6775464 | 86498 | -13.96 | 6.63E-42 | 5.00E-40 |
| chr7 | 18268872 | 18306463 | 37592 | -2.42 | 8.20E-42 | 6.17E-40 |
| chr6 | 133123510 | 133128262 | 4753 | -5 | 9.77E-42 | 7.33E-40 |
| chr10 | 106347602 | 106684538 | 336937 | 13.95 | 9.78E-42 | 7.33E-40 |
| chr5 | 87596868 | 87925107 | 328240 | 13.94 | 1.02E-41 | 7.60E-40 |
| chr7 | 90322828 | 90703097 | 380270 | 13.99 | 1.14E-41 | 8.54E-40 |
| chr17 | 12936384 | 13921341 | 984958 | 0.77 | 1.15E-41 | 8.58E-40 |
| chr12 | 98373581 | 98689787 | 316207 | 13.93 | 1.17E-41 | 8.69E-40 |
| chr17 | 4967607 | 5110779 | 143173 | -13.97 | 1.21E-41 | 8.95E-40 |
| chr11 | 22359534 | 22655467 | 295934 | 13.91 | 1.55E-41 | 1.15E-39 |
| chr6 | 73341772 | 73680076 | 338305 | 13.9 | 1.65E-41 | 1.22E-39 |
| chrX | 47717636 | 47963470 | 245835 | -13.88 | 1.74E-41 | 1.28E-39 |
| chr18 | 11430995 | 11462008 | 31014 | -4.03 | 1.81E-41 | 1.33E-39 |
| chr5 | 179641705 | 179716625 | 74921 | -13.88 | 2.12E-41 | 1.56E-39 |
| chr18 | 68624813 | 69086912 | 462100 | 13.95 | 2.12E-41 | 1.56E-39 |
| chr11 | 3358407 | 3408933 | 50527 | -13.88 | 2.14E-41 | 1.56E-39 |
| chr10 | 57392036 | 58285587 | 893552 | 1.97 | 2.35E-41 | 1.72E-39 |
| chr12 | 55588228 | 55863550 | 275323 | 13.93 | 2.63E-41 | 1.92E-39 |
| chr14 | 24993461 | 25344802 | 351342 | 13.88 | 2.64E-41 | 1.92E-39 |
| chr1 | 215337981 | 215688745 | 350765 | 13.89 | 2.78E-41 | 2.02E-39 |
| chr14 | 104949233 | 105017082 | 67850 | -9.06 | 2.99E-41 | 2.17E-39 |
| chrX | 12472241 | 12548889 | 76649 | 3.19 | 3.15E-41 | 2.28E-39 |
| chr3 | 178223084 | 178647425 | 424342 | 13.85 | 3.37E-41 | 2.44E-39 |
| chr18 | 32818721 | 32923381 | 104661 | -13.8 | 5.18E-41 | 3.73E-39 |
| chr6 | 27272094 | 27439418 | 167325 | -13.79 | 6.46E-41 | 4.65E-39 |
| chr3 | 63346200 | 63746512 | 400313 | 13.81 | 6.77E-41 | 4.86E-39 |
| chr15 | 93022519 | 93052989 | 30471 | -3.07 | 7.75E-41 | 5.56E-39 |
| chr7 | 142595640 | 142700478 | 104839 | 2.34 | 8.83E-41 | 6.32E-39 |
| chr7 | 145740697 | 147950681 | 2209985 | 1.8 | 1.71E-40 | 1.22E-38 |

| | | | | | | |
|---|---|---|---|---|---|---|
| chr1 | 40893213 | 41088022 | 194810 | -13.77 | 1.75E-40 | 1.25E-38 |
| chr5 | 151528992 | 151865739 | 336748 | 13.72 | 2.24E-40 | 1.59E-38 |
| chr4 | 158161695 | 158612891 | 451197 | 13.8 | 2.77E-40 | 1.97E-38 |
| chr9 | 118636021 | 118835354 | 199334 | -13.91 | 2.80E-40 | 1.98E-38 |
| chr14 | 20304065 | 20434264 | 130200 | 13.7 | 2.83E-40 | 2.01E-38 |
| chr2 | 198732982 | 199139390 | 406409 | 13.68 | 3.51E-40 | 2.48E-38 |
| chr19 | 30142670 | 30146926 | 4257 | -5.96 | 4.70E-40 | 3.31E-38 |
| chr3 | 172768673 | 173099786 | 331114 | 13.69 | 4.74E-40 | 3.34E-38 |
| chr5 | 110109440 | 110396577 | 287138 | 13.69 | 5.10E-40 | 3.58E-38 |
| chr13 | 102615152 | 103030184 | 415033 | 13.67 | 5.43E-40 | 3.81E-38 |
| chrX | 7957225 | 8124938 | 167714 | 2.68 | 5.47E-40 | 3.83E-38 |
| chr1 | 74688442 | 75043849 | 355408 | 13.64 | 5.78E-40 | 4.04E-38 |
| chr6 | 118229799 | 118598005 | 368207 | 13.64 | 6.66E-40 | 4.64E-38 |
| chr5 | 140106930 | 140303957 | 197028 | -13.67 | 7.18E-40 | 5.00E-38 |
| chr19 | 35651787 | 35741885 | 90099 | -13.63 | 7.54E-40 | 5.24E-38 |
| chr4 | 156920582 | 157355700 | 435119 | 13.68 | 8.02E-40 | 5.56E-38 |
| chr1 | 170905390 | 171187261 | 281872 | 13.61 | 9.16E-40 | 6.34E-38 |
| chr18 | 10215169 | 10231674 | 16506 | -4.37 | 1.16E-39 | 8.02E-38 |
| chr1 | 70213602 | 70581187 | 367586 | 13.61 | 1.28E-39 | 8.81E-38 |
| chr9 | 113543236 | 113728587 | 185352 | -13.61 | 1.28E-39 | 8.81E-38 |
| chr20 | 23985166 | 24565097 | 579932 | 2.72 | 1.32E-39 | 9.06E-38 |
| chrX | 11205032 | 11536399 | 331368 | 13.59 | 1.38E-39 | 9.49E-38 |
| chr7 | 96359768 | 96612302 | 252535 | 13.59 | 1.43E-39 | 9.82E-38 |
| chr9 | 101656692 | 101661506 | 4815 | -6.05 | 1.46E-39 | 1.00E-37 |
| chr6 | 18605362 | 19786221 | 1180860 | 2.17 | 1.54E-39 | 1.05E-37 |
| chr2 | 227204112 | 227562973 | 358862 | -13.63 | 1.60E-39 | 1.09E-37 |
| chr18 | 75621213 | 75721482 | 100270 | 2.25 | 1.70E-39 | 1.15E-37 |
| chr1 | 237549568 | 237949522 | 399955 | 3.15 | 1.76E-39 | 1.20E-37 |
| chrX | 78716025 | 79295192 | 579168 | 2.39 | 1.93E-39 | 1.31E-37 |
| chr19 | 54736320 | 54778110 | 41791 | -7.54 | 1.98E-39 | 1.34E-37 |
| chr8 | 78016038 | 79564175 | 1548138 | 2.47 | 1.98E-39 | 1.34E-37 |
| chr11 | 26788708 | 27058101 | 269394 | 13.56 | 2.08E-39 | 1.40E-37 |
| chr8 | 145911725 | 145979358 | 67634 | -13.55 | 2.38E-39 | 1.60E-37 |
| chr14 | 49653594 | 49902478 | 248885 | 13.54 | 2.49E-39 | 1.67E-37 |
| chr8 | 130699023 | 130832257 | 133235 | -13.51 | 2.79E-39 | 1.87E-37 |
| chr11 | 56591702 | 56769928 | 178227 | -13.56 | 2.87E-39 | 1.92E-37 |
| chr16 | 68555279 | 68673999 | 118721 | -13.56 | 4.04E-39 | 2.70E-37 |
| chr19 | 2824229 | 2941167 | 116939 | -13.52 | 4.42E-39 | 2.94E-37 |
| chr9 | 107178028 | 107427694 | 249667 | 13.49 | 5.07E-39 | 3.37E-37 |
| chrX | 90287382 | 90398534 | 111153 | 1.92 | 5.09E-39 | 3.38E-37 |
| chr13 | 92076313 | 93868500 | 1792188 | 3.23 | 5.89E-39 | 3.90E-37 |
| chr1 | 238072493 | 239305144 | 1232652 | 1.88 | 6.13E-39 | 4.06E-37 |
| chr10 | 47070161 | 47148782 | 78622 | -13.47 | 7.15E-39 | 4.72E-37 |
| chr6 | 70582328 | 70860047 | 277720 | 13.46 | 7.33E-39 | 4.83E-37 |
| chr2 | 133567023 | 133923886 | 356864 | 13.46 | 8.02E-39 | 5.28E-37 |
| chr9 | 22123451 | 22273799 | 150349 | -13.46 | 8.08E-39 | 5.31E-37 |
| chr8 | 96829568 | 97147845 | 318278 | 4.26 | 9.45E-39 | 6.20E-37 |
| chr5 | 111185474 | 111472727 | 287254 | 13.46 | 9.56E-39 | 6.26E-37 |
| chr2 | 226932942 | 227177288 | 244347 | -13.43 | 9.91E-39 | 6.48E-37 |
| chr2 | 182528756 | 182547877 | 19122 | -4.14 | 1.15E-38 | 7.51E-37 |
| chr14 | 68530199 | 68723428 | 193230 | -13.53 | 1.30E-38 | 8.46E-37 |
| chr1 | 166221236 | 166441784 | 220549 | 13.44 | 1.33E-38 | 8.64E-37 |
| chrX | 127942161 | 128316504 | 374344 | 13.45 | 1.33E-38 | 8.65E-37 |
| chr8 | 109542187 | 109791806 | 249620 | 13.42 | 1.37E-38 | 8.91E-37 |
| chr11 | 29012655 | 29205981 | 193327 | -13.4 | 1.38E-38 | 8.92E-37 |
| chr2 | 113701768 | 113826034 | 124267 | -2.16 | 1.39E-38 | 8.99E-37 |
| chr10 | 42947938 | 43189496 | 241559 | -13.43 | 1.47E-38 | 9.45E-37 |
| chr11 | 123952081 | 124260905 | 308825 | 13.42 | 1.50E-38 | 9.63E-37 |
| chr11 | 5321547 | 5568617 | 247071 | 13.41 | 1.94E-38 | 1.24E-36 |
| chr12 | 41761212 | 41907676 | 146465 | 2.43 | 2.54E-38 | 1.63E-36 |
| chr14 | 50902431 | 50978367 | 75937 | -13.35 | 2.56E-38 | 1.64E-36 |

| | | | | | | |
|---|---|---|---|---|---|---|
| chr8 | 106339970 | 106659639 | 319670 | 13.53 | 2.60E-38 | 1.66E-36 |
| chrX | 139311556 | 139578933 | 267378 | 13.37 | 2.60E-38 | 1.66E-36 |
| chr9 | 104375517 | 106813481 | 2437965 | 2.38 | 2.76E-38 | 1.76E-36 |
| chr3 | 158576959 | 158846444 | 269486 | 13.36 | 3.15E-38 | 2.00E-36 |
| chr10 | 68914365 | 69217214 | 302850 | 13.37 | 3.86E-38 | 2.45E-36 |
| chr20 | 53025995 | 53255829 | 229835 | 13.34 | 4.25E-38 | 2.69E-36 |
| chr3 | 97794580 | 98043111 | 248532 | 13.35 | 5.60E-38 | 3.54E-36 |
| chr8 | 104812833 | 105121801 | 308969 | 13.48 | 6.37E-38 | 4.03E-36 |
| chr1 | 219640582 | 219712378 | 71797 | 9.13 | 7.53E-38 | 4.75E-36 |
| chr8 | 91136144 | 91401248 | 265105 | 13.31 | 8.24E-38 | 5.19E-36 |
| chr2 | 73864547 | 73898125 | 33579 | -2.65 | 8.34E-38 | 5.24E-36 |
| chr6 | 72604732 | 72863698 | 258967 | 13.28 | 9.45E-38 | 5.93E-36 |
| chrX | 103834171 | 104126807 | 292637 | 13.27 | 9.76E-38 | 6.12E-36 |
| chr17 | 26228671 | 26236766 | 8096 | -3.35 | 1.01E-37 | 6.29E-36 |
| chr9 | 103511782 | 103786598 | 274817 | 13.32 | 1.08E-37 | 6.72E-36 |
| chr19 | 50519372 | 50571343 | 51972 | -13.23 | 1.30E-37 | 8.11E-36 |
| chr9 | 104107095 | 104216318 | 109224 | -13.23 | 1.33E-37 | 8.29E-36 |
| chrX | 136218991 | 136504455 | 285465 | 13.27 | 1.36E-37 | 8.43E-36 |
| chr5 | 146394578 | 146596272 | 201695 | 13.24 | 1.61E-37 | 9.98E-36 |
| chr1 | 182065527 | 182105786 | 40260 | -3.7 | 1.90E-37 | 1.18E-35 |
| chr16 | 75174981 | 75231904 | 56924 | -13.21 | 1.95E-37 | 1.21E-35 |
| chr9 | 71411962 | 71611349 | 199388 | 4.23 | 2.01E-37 | 1.24E-35 |
| chr7 | 8412747 | 10918949 | 2506203 | 2.22 | 2.86E-37 | 1.76E-35 |
| chr16 | 63843545 | 64087373 | 243829 | 2.09 | 3.25E-37 | 2.00E-35 |
| chr10 | 72954786 | 72961603 | 6818 | -4.77 | 3.33E-37 | 2.05E-35 |
| chrX | 116757391 | 116830853 | 73463 | 3.1 | 3.35E-37 | 2.05E-35 |
| chrX | 77494081 | 77804614 | 310534 | 13.19 | 3.35E-37 | 2.05E-35 |
| chr7 | 5089326 | 5155360 | 66035 | -13.17 | 3.70E-37 | 2.26E-35 |
| chrX | 132313190 | 132538712 | 225523 | 2.14 | 3.85E-37 | 2.35E-35 |
| chr1 | 83955646 | 84225189 | 269544 | 13.16 | 4.75E-37 | 2.90E-35 |
| chrX | 22982029 | 23320690 | 338662 | 2.19 | 5.53E-37 | 3.37E-35 |
| chr10 | 20861831 | 21088436 | 226606 | 2.37 | 6.19E-37 | 3.76E-35 |
| chr10 | 26018289 | 26671057 | 652769 | 3.3 | 7.05E-37 | 4.28E-35 |
| chr6 | 122366587 | 122656279 | 289693 | 13.2 | 8.37E-37 | 5.07E-35 |
| chr18 | 27879860 | 28398869 | 519010 | 2.14 | 9.15E-37 | 5.53E-35 |
| chr5 | 110579538 | 110807367 | 227830 | 13.14 | 1.05E-36 | 6.33E-35 |
| chr4 | 165200861 | 165465773 | 264913 | 2.32 | 1.07E-36 | 6.46E-35 |
| chr3 | 33005374 | 33009392 | 4019 | -5.55 | 1.16E-36 | 7.00E-35 |
| chr5 | 147870157 | 148080118 | 209962 | 13.09 | 1.38E-36 | 8.28E-35 |
| chr3 | 132770701 | 133035271 | 264571 | 13.07 | 1.46E-36 | 8.79E-35 |
| chr3 | 55558287 | 56043119 | 484833 | 1.98 | 1.47E-36 | 8.79E-35 |
| chr5 | 99442738 | 99666656 | 223919 | 3.16 | 1.57E-36 | 9.38E-35 |
| chr2 | 154045597 | 154310266 | 264670 | 13.09 | 1.58E-36 | 9.43E-35 |
| chr5 | 72625276 | 72659336 | 34061 | -2.49 | 1.67E-36 | 9.94E-35 |
| chr11 | 25099248 | 26289834 | 1190587 | 1.82 | 1.68E-36 | 9.99E-35 |
| chr3 | 86424431 | 86696972 | 272542 | 2.18 | 1.78E-36 | 1.06E-34 |
| chr21 | 33575514 | 33624219 | 48706 | -2.2 | 1.86E-36 | 1.10E-34 |
| chr22 | 44810865 | 44818174 | 7310 | -8.69 | 1.90E-36 | 1.12E-34 |
| chr1 | 165922038 | 166124845 | 202808 | 13.06 | 2.29E-36 | 1.35E-34 |
| chr20 | 42729192 | 42738428 | 9237 | -3.28 | 2.29E-36 | 1.35E-34 |
| chr14 | 92201060 | 92231253 | 30194 | -8.78 | 2.36E-36 | 1.39E-34 |
| chr8 | 53045571 | 53240595 | 195025 | 13.02 | 2.99E-36 | 1.76E-34 |
| chr14 | 59989455 | 59997821 | 8367 | -4.38 | 3.40E-36 | 2.00E-34 |
| chr3 | 167155275 | 167379500 | 224226 | 13.03 | 3.43E-36 | 2.01E-34 |
| chr14 | 20455684 | 20626379 | 170696 | 13 | 3.54E-36 | 2.08E-34 |
| chr18 | 77647524 | 77665061 | 17538 | -8.93 | 3.62E-36 | 2.12E-34 |
| chr8 | 29455092 | 29630122 | 175031 | -13.09 | 4.01E-36 | 2.34E-34 |
| chr18 | 67971386 | 68068466 | 97081 | -12.97 | 4.08E-36 | 2.38E-34 |
| chr9 | 114271288 | 114333125 | 61838 | -12.97 | 4.41E-36 | 2.57E-34 |
| chr12 | 34555477 | 34761228 | 205752 | -13.12 | 4.61E-36 | 2.68E-34 |
| chr7 | 155976301 | 155986450 | 10150 | -2.88 | 4.62E-36 | 2.68E-34 |

| | | | | | | |
|---|---|---|---|---|---|---|
| chr9 | 116454906 | 116627566 | 172661 | -13.16 | 4.67E-36 | 2.71E-34 |
| chr7 | 30225337 | 30238750 | 13414 | -3.78 | 5.38E-36 | 3.11E-34 |
| chr10 | 77553625 | 77780919 | 227295 | 13.01 | 5.51E-36 | 3.18E-34 |
| chr10 | 62032140 | 62046478 | 14339 | -3.01 | 5.61E-36 | 3.24E-34 |
| chr1 | 78679431 | 78944195 | 264765 | 12.97 | 5.63E-36 | 3.25E-34 |
| chr5 | 33381328 | 33399553 | 18226 | -3.13 | 5.87E-36 | 3.38E-34 |
| chr2 | 209770084 | 210064628 | 294545 | 13.05 | 6.19E-36 | 3.55E-34 |
| chr8 | 39995083 | 40149066 | 153984 | -13 | 7.61E-36 | 4.37E-34 |
| chr5 | 57315300 | 57582118 | 266819 | -12.98 | 8.46E-36 | 4.85E-34 |
| chr4 | 13816489 | 13941887 | 125399 | -12.92 | 1.05E-35 | 5.97E-34 |
| chr4 | 44458978 | 44657190 | 198213 | 12.93 | 1.07E-35 | 6.10E-34 |
| chr9 | 95575152 | 95634415 | 59264 | -12.88 | 1.34E-35 | 7.62E-34 |
| chr2 | 111745870 | 111786383 | 40514 | -2.86 | 1.54E-35 | 8.77E-34 |
| chr2 | 239613512 | 239749415 | 135904 | -12.91 | 1.68E-35 | 9.56E-34 |
| chr3 | 67174292 | 67385097 | 210806 | 12.89 | 1.69E-35 | 9.59E-34 |
| chr2 | 134027910 | 134303126 | 275217 | 12.92 | 1.86E-35 | 1.05E-33 |
| chr8 | 104555043 | 104787005 | 231963 | 12.94 | 1.86E-35 | 1.05E-33 |
| chr8 | 11189749 | 11196445 | 6697 | -8.08 | 2.13E-35 | 1.21E-33 |
| chr5 | 146125251 | 146338010 | 212760 | 3.64 | 2.44E-35 | 1.38E-33 |
| chr22 | 49987835 | 50008485 | 20651 | -8.1 | 2.56E-35 | 1.44E-33 |
| chrX | 110516691 | 110768827 | 252137 | 12.83 | 3.42E-35 | 1.92E-33 |
| chr6 | 97016236 | 97240572 | 224337 | 12.83 | 3.44E-35 | 1.93E-33 |
| chr12 | 133438605 | 133502618 | 64014 | -12.85 | 5.15E-35 | 2.89E-33 |
| chrX | 46315865 | 46383715 | 67851 | -12.78 | 5.39E-35 | 3.02E-33 |
| chr4 | 126932501 | 127468411 | 535911 | 2.53 | 6.27E-35 | 3.51E-33 |
| chr2 | 173801620 | 173809337 | 7718 | -4.05 | 7.00E-35 | 3.91E-33 |
| chr10 | 102599017 | 102618100 | 19084 | -3.17 | 7.44E-35 | 4.14E-33 |
| chr10 | 60249670 | 60366746 | 117077 | -12.94 | 7.63E-35 | 4.25E-33 |
| chr3 | 154843490 | 154999817 | 156328 | 12.77 | 8.18E-35 | 4.54E-33 |
| chr19 | 58922605 | 58962411 | 39807 | -12.75 | 1.10E-34 | 6.12E-33 |
| chr2 | 33909406 | 34316353 | 406948 | 2.21 | 1.22E-34 | 6.76E-33 |
| chr3 | 167479985 | 167704438 | 224454 | 12.88 | 1.25E-34 | 6.93E-33 |
| chr10 | 13045522 | 13086176 | 40655 | -2.22 | 1.27E-34 | 7.03E-33 |
| chr3 | 146376630 | 146515587 | 138958 | 12.87 | 1.31E-34 | 7.25E-33 |
| chrX | 113179423 | 113389548 | 210126 | 1.8 | 1.43E-34 | 7.87E-33 |
| chr13 | 34709566 | 34880215 | 170650 | -12.83 | 1.67E-34 | 9.19E-33 |
| chr8 | 131682925 | 131792778 | 109854 | -12.83 | 1.68E-34 | 9.22E-33 |
| chr8 | 109283158 | 109427887 | 144730 | 12.77 | 1.83E-34 | 1.00E-32 |
| chr11 | 22674402 | 22806845 | 132444 | 12.71 | 1.93E-34 | 1.06E-32 |
| chr6 | 26611716 | 26667368 | 55653 | -12.68 | 1.94E-34 | 1.06E-32 |
| chr2 | 133122440 | 133268439 | 146000 | -12.7 | 1.95E-34 | 1.06E-32 |
| chr5 | 6797450 | 6923992 | 126543 | -12.67 | 1.97E-34 | 1.08E-32 |
| chr8 | 123972233 | 124012444 | 40212 | -8.56 | 2.02E-34 | 1.10E-32 |
| chr5 | 151314681 | 151507578 | 192898 | 12.71 | 2.05E-34 | 1.11E-32 |
| chr15 | 29657661 | 29665651 | 7991 | -7.9 | 2.17E-34 | 1.18E-32 |
| chr20 | 39601781 | 39608104 | 6324 | -4.95 | 2.60E-34 | 1.41E-32 |
| chrX | 90504090 | 90734139 | 230050 | 2.31 | 2.66E-34 | 1.44E-32 |
| chr6 | 12292452 | 12438864 | 146413 | -12.74 | 3.29E-34 | 1.78E-32 |
| chr16 | 87629051 | 87634314 | 5264 | -4.12 | 3.53E-34 | 1.91E-32 |
| chr5 | 121654898 | 121808749 | 153852 | 12.65 | 3.64E-34 | 1.96E-32 |
| chr10 | 44892500 | 45345560 | 453061 | 1.31 | 3.88E-34 | 2.09E-32 |
| chr10 | 14499433 | 14507880 | 8448 | -3.6 | 3.91E-34 | 2.10E-32 |
| chr13 | 77192873 | 77386979 | 194107 | 12.65 | 3.98E-34 | 2.13E-32 |
| chr12 | 81034743 | 81152020 | 117278 | 12.76 | 4.10E-34 | 2.20E-32 |
| chr2 | 56429271 | 56629065 | 199795 | 12.67 | 4.19E-34 | 2.25E-32 |
| chr8 | 122683312 | 122799961 | 116650 | -12.73 | 4.21E-34 | 2.25E-32 |
| chr8 | 22890389 | 22921939 | 31551 | -12.63 | 4.24E-34 | 2.26E-32 |
| chrX | 41991774 | 42000004 | 8231 | -4.24 | 4.43E-34 | 2.36E-32 |
| chr11 | 89835939 | 89879294 | 43356 | 3.13 | 5.27E-34 | 2.81E-32 |
| chr7 | 90722298 | 90888211 | 165914 | 12.61 | 6.01E-34 | 3.19E-32 |
| chr8 | 55095706 | 55122210 | 26505 | -7.96 | 6.74E-34 | 3.58E-32 |

| | | | | | | |
|---|---|---|---|---|---|---|
| chr21 | 46023804 | 46115018 | 91215 | -12.57 | 7.42E-34 | 3.93E-32 |
| chr7 | 144560304 | 145693286 | 1132983 | 1.87 | 7.72E-34 | 4.08E-32 |
| chr4 | 21202585 | 21502644 | 300060 | 1.51 | 7.75E-34 | 4.10E-32 |
| chr1 | 83055749 | 83238621 | 182873 | 2.02 | 8.23E-34 | 4.34E-32 |
| chr2 | 35313690 | 35968987 | 655298 | 3.85 | 8.26E-34 | 4.35E-32 |
| chr6 | 133877437 | 133984534 | 107098 | -12.8 | 8.27E-34 | 4.36E-32 |
| chr7 | 103468151 | 103622852 | 154702 | 12.59 | 8.33E-34 | 4.38E-32 |
| chr4 | 13957349 | 14111034 | 153686 | -12.59 | 8.63E-34 | 4.53E-32 |
| chr12 | 74705529 | 74806436 | 100908 | -12.67 | 9.05E-34 | 4.74E-32 |
| chr2 | 211379737 | 212896820 | 1517084 | 2.5 | 9.23E-34 | 4.83E-32 |
| chr4 | 175259897 | 175438824 | 178928 | -12.54 | 1.14E-33 | 5.94E-32 |
| chr18 | 70849486 | 70929543 | 80058 | -12.56 | 1.16E-33 | 6.06E-32 |
| chr2 | 65894773 | 66608534 | 713762 | 2.75 | 1.23E-33 | 6.43E-32 |
| chr2 | 36691181 | 36711842 | 20662 | -12.53 | 1.24E-33 | 6.48E-32 |
| chr11 | 22872082 | 23924769 | 1052688 | 1.81 | 1.25E-33 | 6.50E-32 |
| chr6 | 73704702 | 73867382 | 162681 | 12.56 | 1.27E-33 | 6.60E-32 |
| chr1 | 70049739 | 70197456 | 147718 | 12.6 | 1.34E-33 | 6.94E-32 |
| chr5 | 89397435 | 89556551 | 159117 | 12.58 | 1.34E-33 | 6.94E-32 |
| chr19 | 7063374 | 7098671 | 35298 | -12.52 | 1.35E-33 | 6.98E-32 |
| chr10 | 116975869 | 117110906 | 135038 | 12.62 | 1.35E-33 | 6.98E-32 |
| chr3 | 6176440 | 6252611 | 76172 | 4.35 | 1.60E-33 | 8.25E-32 |
| chr5 | 143454229 | 143500321 | 46093 | -7.99 | 1.74E-33 | 8.94E-32 |
| chr9 | 27343512 | 27517515 | 174004 | 12.55 | 1.84E-33 | 9.46E-32 |
| chr4 | 75493425 | 75610277 | 116853 | -12.51 | 1.84E-33 | 9.46E-32 |
| chr20 | 60642739 | 60696064 | 53326 | -12.55 | 1.90E-33 | 9.73E-32 |
| chr2 | 176360160 | 176546353 | 186194 | 12.57 | 2.00E-33 | 1.02E-31 |
| chr3 | 127211248 | 127247160 | 35913 | -5.2 | 2.07E-33 | 1.06E-31 |
| chr20 | 57050689 | 57140968 | 90280 | -12.5 | 2.13E-33 | 1.09E-31 |
| chr7 | 84848139 | 85041684 | 193546 | 12.68 | 2.44E-33 | 1.24E-31 |
| chrX | 4192249 | 4406702 | 214454 | 2.09 | 2.55E-33 | 1.30E-31 |
| chr16 | 59599194 | 59625233 | 26040 | -7.89 | 2.67E-33 | 1.36E-31 |
| chr13 | 103850759 | 104043754 | 192996 | 12.5 | 2.75E-33 | 1.40E-31 |
| chr3 | 189011716 | 189131837 | 120122 | -12.5 | 3.04E-33 | 1.54E-31 |
| chrX | 47249625 | 47330895 | 81271 | -12.47 | 3.19E-33 | 1.62E-31 |
| chr2 | 5797384 | 7030354 | 1232971 | 1.05 | 3.33E-33 | 1.68E-31 |
| chr12 | 14479562 | 14595671 | 116110 | -12.47 | 4.09E-33 | 2.07E-31 |
| chrX | 11849408 | 12033098 | 183691 | -12.44 | 4.31E-33 | 2.17E-31 |
| chr1 | 18532545 | 18561687 | 29143 | -3.62 | 4.44E-33 | 2.24E-31 |
| chr7 | 111063617 | 111181762 | 118146 | -12.43 | 4.88E-33 | 2.46E-31 |
| chr13 | 32042361 | 32297512 | 255152 | 1.51 | 5.34E-33 | 2.69E-31 |
| chr8 | 42952277 | 42987702 | 35426 | -12.41 | 5.96E-33 | 2.99E-31 |
| chr6 | 133998176 | 134158389 | 160214 | -12.58 | 6.07E-33 | 3.05E-31 |
| chr8 | 40194723 | 40328750 | 134028 | -12.44 | 7.06E-33 | 3.53E-31 |
| chr7 | 153864192 | 154270518 | 406327 | 3.22 | 7.44E-33 | 3.72E-31 |
| chr7 | 99117873 | 99148686 | 30814 | -12.4 | 7.99E-33 | 3.99E-31 |
| chr16 | 50442428 | 50596073 | 153646 | -12.37 | 1.05E-32 | 5.23E-31 |
| chr10 | 77849802 | 77861507 | 11706 | -4.1 | 1.11E-32 | 5.54E-31 |
| chr8 | 86220008 | 86354443 | 134436 | 12.38 | 1.11E-32 | 5.54E-31 |
| chr11 | 34664754 | 34760346 | 95593 | -12.37 | 1.15E-32 | 5.70E-31 |
| chr7 | 45524870 | 45602524 | 77655 | -2.33 | 1.23E-32 | 6.08E-31 |
| chr5 | 135010222 | 135018212 | 7991 | -8.22 | 1.26E-32 | 6.25E-31 |
| chr7 | 68505996 | 68667161 | 161166 | -12.5 | 1.35E-32 | 6.68E-31 |
| chr6 | 72344735 | 72495062 | 150328 | 12.41 | 1.47E-32 | 7.26E-31 |
| chr3 | 72270671 | 72281703 | 11033 | -3.12 | 1.49E-32 | 7.34E-31 |
| chrX | 139595070 | 139754303 | 159234 | 12.39 | 1.52E-32 | 7.49E-31 |
| chr22 | 26394724 | 26554589 | 159866 | 3.12 | 1.52E-32 | 7.50E-31 |
| chr7 | 99657399 | 99678111 | 20713 | -12.33 | 1.53E-32 | 7.53E-31 |
| chr4 | 13639028 | 13800679 | 161652 | -12.36 | 1.58E-32 | 7.76E-31 |
| chr2 | 195423844 | 195673103 | 249260 | 12.6 | 1.71E-32 | 8.36E-31 |
| chr2 | 587135 | 705577 | 118443 | -12.32 | 1.79E-32 | 8.75E-31 |
| chr17 | 5379685 | 5403260 | 23576 | -12.32 | 1.79E-32 | 8.75E-31 |

| | | | | | | |
|---|---|---|---|---|---|---|
| chr3 | 173314179 | 173861559 | 547381 | 2.92 | 1.94E-32 | 9.46E-31 |
| chr8 | 144721644 | 144757760 | 36117 | -12.34 | 2.03E-32 | 9.91E-31 |
| chr3 | 87335697 | 87417013 | 81317 | -12.3 | 2.18E-32 | 1.06E-30 |
| chr2 | 221553886 | 221756452 | 202567 | -12.31 | 2.27E-32 | 1.10E-30 |
| chr2 | 82904555 | 83218338 | 313784 | -12.48 | 2.29E-32 | 1.12E-30 |
| chr14 | 61768269 | 61793290 | 25022 | -8.57 | 2.57E-32 | 1.25E-30 |
| chr10 | 54177125 | 54274649 | 97525 | -12.29 | 2.58E-32 | 1.25E-30 |
| chr8 | 108618530 | 108774795 | 156266 | 12.37 | 2.68E-32 | 1.30E-30 |
| chr1 | 227988806 | 228068581 | 79776 | -12.28 | 2.79E-32 | 1.35E-30 |
| chr13 | 86910568 | 88221318 | 1310751 | 2.06 | 3.50E-32 | 1.69E-30 |
| chr2 | 242852818 | 242862845 | 10028 | -7.83 | 3.61E-32 | 1.74E-30 |
| chr3 | 169067963 | 169203093 | 135131 | 12.29 | 3.74E-32 | 1.80E-30 |
| chr6 | 28116391 | 28175317 | 58927 | -12.27 | 3.91E-32 | 1.88E-30 |
| chr4 | 110709269 | 110719519 | 10251 | -4.11 | 4.46E-32 | 2.14E-30 |
| chr16 | 75846152 | 76678576 | 832425 | 2.54 | 4.56E-32 | 2.19E-30 |
| chr1 | 231380321 | 231404792 | 24472 | -12.26 | 4.73E-32 | 2.27E-30 |
| chr3 | 137385936 | 137469371 | 83436 | 3.98 | 5.17E-32 | 2.48E-30 |
| chr8 | 66667683 | 66741654 | 73972 | -12.27 | 5.21E-32 | 2.49E-30 |
| chr7 | 40379207 | 40789653 | 410447 | 1.47 | 5.23E-32 | 2.49E-30 |
| chr22 | 25110305 | 25121618 | 11314 | -8.48 | 5.37E-32 | 2.56E-30 |
| chr11 | 11210125 | 11233429 | 23305 | -2.6 | 5.63E-32 | 2.68E-30 |
| chr2 | 144561489 | 144618611 | 57123 | 2.56 | 5.80E-32 | 2.76E-30 |
| chr12 | 13757065 | 13887887 | 130823 | 12.26 | 6.73E-32 | 3.20E-30 |
| chr9 | 97547793 | 97655441 | 107649 | -12.26 | 7.87E-32 | 3.73E-30 |
| chrX | 66157245 | 66465281 | 308037 | 2.14 | 8.10E-32 | 3.84E-30 |
| chr16 | 71431597 | 71512857 | 81261 | -12.29 | 8.19E-32 | 3.88E-30 |
| chr8 | 99328122 | 99335498 | 7377 | -3.86 | 8.58E-32 | 4.05E-30 |
| chr4 | 121179645 | 121328971 | 149327 | 12.22 | 9.32E-32 | 4.40E-30 |
| chr17 | 5658087 | 5733145 | 75059 | -12.19 | 9.34E-32 | 4.40E-30 |
| chr5 | 155500082 | 155659228 | 159147 | -12.28 | 9.39E-32 | 4.42E-30 |
| chr8 | 8208659 | 8225490 | 16832 | -12.22 | 1.23E-31 | 5.77E-30 |
| chr15 | 34443307 | 34469354 | 26048 | -12.17 | 1.24E-31 | 5.83E-30 |
| chr3 | 173138461 | 173296266 | 157806 | 12.32 | 1.25E-31 | 5.86E-30 |
| chr12 | 80712771 | 80795078 | 82308 | 12.24 | 1.28E-31 | 6.01E-30 |
| chr7 | 112598736 | 112720409 | 121674 | 12.31 | 1.29E-31 | 6.06E-30 |
| chr8 | 186000 | 240654 | 54655 | -12.16 | 1.49E-31 | 6.96E-30 |
| chr7 | 19911072 | 20034612 | 123541 | -12.2 | 1.52E-31 | 7.09E-30 |
| chr7 | 22647933 | 22729234 | 81302 | -12.18 | 1.52E-31 | 7.09E-30 |
| chr7 | 134234676 | 134313497 | 78822 | -12.27 | 1.58E-31 | 7.36E-30 |
| chr4 | 18116061 | 18589778 | 473718 | 3.92 | 1.66E-31 | 7.72E-30 |
| chr4 | 172742590 | 172940930 | 198341 | 12.21 | 1.70E-31 | 7.90E-30 |
| chr2 | 163233061 | 163435304 | 202244 | 12.25 | 1.73E-31 | 8.04E-30 |
| chr2 | 65673878 | 65752728 | 78851 | -12.25 | 1.78E-31 | 8.23E-30 |
| chr3 | 180724133 | 180823979 | 99847 | 2.69 | 1.83E-31 | 8.45E-30 |
| chr5 | 58745567 | 58877072 | 131506 | -12.15 | 1.83E-31 | 8.46E-30 |
| chr10 | 27873339 | 28024261 | 150923 | 1.83 | 1.85E-31 | 8.55E-30 |
| chr2 | 239421693 | 239536520 | 114828 | -12.23 | 1.99E-31 | 9.15E-30 |
| chr9 | 90821906 | 90829741 | 7836 | -7.75 | 2.01E-31 | 9.24E-30 |
| chr10 | 80182744 | 80190066 | 7323 | -7.71 | 2.24E-31 | 1.03E-29 |
| chr10 | 14947054 | 14968354 | 21301 | -12.14 | 2.28E-31 | 1.05E-29 |
| chr12 | 101225809 | 101348274 | 122466 | 12.17 | 2.28E-31 | 1.05E-29 |
| chr14 | 66899725 | 66908329 | 8605 | -3.52 | 2.38E-31 | 1.09E-29 |
| chr8 | 102327766 | 102420402 | 92637 | -12.32 | 2.54E-31 | 1.16E-29 |
| chr7 | 17361778 | 17495966 | 134189 | -12.18 | 2.60E-31 | 1.19E-29 |
| chr7 | 64841052 | 64913390 | 72339 | -12.11 | 2.66E-31 | 1.21E-29 |
| chr2 | 62934892 | 62950544 | 15653 | -12.16 | 2.78E-31 | 1.27E-29 |
| chr6 | 166761687 | 166773464 | 11778 | -3.62 | 2.86E-31 | 1.30E-29 |
| chr13 | 94764571 | 94975212 | 210642 | 12.21 | 3.06E-31 | 1.39E-29 |
| chrX | 51522598 | 51630095 | 107498 | -12.12 | 3.24E-31 | 1.47E-29 |
| chr12 | 46857292 | 46923454 | 66163 | -12.12 | 3.34E-31 | 1.52E-29 |
| chr15 | 93486337 | 93507355 | 21019 | -12.09 | 3.58E-31 | 1.62E-29 |

| | | | | | | |
|---|---|---|---|---|---|---|
| chr7 | 68367813 | 68487858 | 120046 | -12.23 | 3.72E-31 | 1.68E-29 |
| chr5 | 52609691 | 52726425 | 116735 | -12.19 | 3.93E-31 | 1.78E-29 |
| chr5 | 180256606 | 180286043 | 29438 | -12.1 | 4.12E-31 | 1.86E-29 |
| chr7 | 115872540 | 116011327 | 138788 | -12.1 | 4.15E-31 | 1.87E-29 |
| chr19 | 13422708 | 13564268 | 141561 | -12.07 | 4.30E-31 | 1.94E-29 |
| chr5 | 154543021 | 154588468 | 45448 | -2.34 | 4.32E-31 | 1.94E-29 |
| chr14 | 105583851 | 105592047 | 8197 | -2.96 | 4.46E-31 | 2.01E-29 |
| chr4 | 107881578 | 107952061 | 70484 | 4.69 | 4.86E-31 | 2.18E-29 |
| chr9 | 13456744 | 13906991 | 450248 | 1.16 | 5.02E-31 | 2.25E-29 |
| chr10 | 60412817 | 60505983 | 93167 | -12.19 | 5.32E-31 | 2.38E-29 |
| chr3 | 123540361 | 123614595 | 74235 | -12.06 | 5.40E-31 | 2.42E-29 |
| chr17 | 68157680 | 68615459 | 457780 | 2.65 | 5.79E-31 | 2.59E-29 |
| chr1 | 90875174 | 91033038 | 157865 | 12.15 | 6.14E-31 | 2.74E-29 |
| chr10 | 6277128 | 6440210 | 163083 | -12.43 | 6.23E-31 | 2.78E-29 |
| chr2 | 70800359 | 70804421 | 4063 | -4.92 | 6.42E-31 | 2.86E-29 |
| chr8 | 105224731 | 105451286 | 226556 | 1.99 | 6.74E-31 | 3.00E-29 |
| chr4 | 165718387 | 165847949 | 129563 | 12.05 | 7.53E-31 | 3.35E-29 |
| chr2 | 75962962 | 77894276 | 1931315 | 1.43 | 7.77E-31 | 3.45E-29 |
| chr7 | 138183355 | 138263221 | 79867 | -12.03 | 8.10E-31 | 3.59E-29 |
| chr14 | 104795247 | 104884758 | 89512 | -12.06 | 8.40E-31 | 3.72E-29 |
| chr9 | 108645184 | 109571656 | 926473 | 2.41 | 8.72E-31 | 3.85E-29 |
| chr5 | 1424555 | 1466853 | 42299 | -12 | 9.42E-31 | 4.16E-29 |
| chr19 | 46540507 | 46616852 | 76346 | -12.06 | 9.97E-31 | 4.40E-29 |
| chr9 | 28359344 | 28457550 | 98207 | 12.09 | 1.01E-30 | 4.44E-29 |
| chr9 | 96946948 | 97008495 | 61548 | -12.04 | 1.08E-30 | 4.74E-29 |
| chr12 | 113940481 | 114071249 | 130769 | -11.98 | 1.23E-30 | 5.42E-29 |
| chrX | 45316971 | 45408386 | 91416 | -12.03 | 1.31E-30 | 5.76E-29 |
| chr8 | 58273347 | 58286845 | 13499 | -8.32 | 1.38E-30 | 6.05E-29 |
| chr2 | 102689633 | 102790527 | 100895 | -12.22 | 1.43E-30 | 6.26E-29 |
| chr2 | 149939548 | 150122475 | 182928 | 3.91 | 1.57E-30 | 6.86E-29 |
| chr7 | 155418128 | 155452919 | 34792 | -12.03 | 1.58E-30 | 6.92E-29 |
| chr17 | 59774808 | 59850682 | 75875 | -11.99 | 1.63E-30 | 7.11E-29 |
| chr7 | 86575711 | 86669189 | 93479 | 11.97 | 1.85E-30 | 8.05E-29 |
| chr5 | 102647296 | 105437785 | 2790490 | 2.67 | 1.89E-30 | 8.23E-29 |
| chr9 | 2772540 | 2803633 | 31094 | -11.94 | 1.95E-30 | 8.48E-29 |
| chr13 | 91909764 | 91932135 | 22372 | 3.49 | 2.05E-30 | 8.92E-29 |
| chr5 | 123455896 | 123587099 | 131204 | -12.28 | 2.12E-30 | 9.21E-29 |
| chr7 | 157693275 | 157764930 | 71656 | -11.93 | 2.20E-30 | 9.53E-29 |
| chrX | 46099230 | 46261477 | 162248 | -12.02 | 2.23E-30 | 9.68E-29 |
| chr15 | 99354997 | 99384863 | 29867 | -12.07 | 2.24E-30 | 9.68E-29 |
| chr2 | 37422942 | 37445525 | 22584 | -11.92 | 2.49E-30 | 1.07E-28 |
| chr6 | 146384594 | 146540347 | 155754 | 12.01 | 2.57E-30 | 1.11E-28 |
| chr8 | 17654282 | 17705485 | 51204 | -12.32 | 2.57E-30 | 1.11E-28 |
| chr12 | 14373218 | 14464652 | 91435 | -12.02 | 2.60E-30 | 1.12E-28 |
| chr9 | 21519861 | 21545782 | 25922 | -11.92 | 2.64E-30 | 1.14E-28 |
| chr10 | 22274511 | 22409932 | 135422 | -11.99 | 2.69E-30 | 1.15E-28 |
| chr2 | 67745100 | 67824903 | 79804 | -11.92 | 2.82E-30 | 1.21E-28 |
| chr1 | 233288139 | 233335007 | 46869 | -11.93 | 2.85E-30 | 1.22E-28 |
| chr17 | 29946207 | 30003390 | 57184 | -11.98 | 2.85E-30 | 1.22E-28 |
| chr2 | 200383086 | 200522843 | 139758 | 11.96 | 2.95E-30 | 1.26E-28 |
| chr3 | 110046039 | 110143927 | 97889 | -11.99 | 3.14E-30 | 1.34E-28 |
| chr1 | 4532713 | 4553005 | 20293 | -3.35 | 3.78E-30 | 1.61E-28 |
| chr2 | 162576804 | 162720612 | 143809 | 11.97 | 4.08E-30 | 1.74E-28 |
| chrX | 139196801 | 139292593 | 95793 | 11.91 | 4.34E-30 | 1.85E-28 |
| chrX | 55574334 | 55654079 | 79746 | 2.95 | 4.35E-30 | 1.85E-28 |
| chr14 | 71705635 | 71782228 | 76594 | -11.89 | 4.41E-30 | 1.87E-28 |
| chr5 | 34984013 | 35615461 | 631449 | 2.68 | 4.63E-30 | 1.97E-28 |
| chr8 | 118149031 | 118298414 | 149384 | 12.07 | 4.67E-30 | 1.98E-28 |
| chr17 | 48301229 | 48329624 | 28396 | -11.93 | 4.87E-30 | 2.06E-28 |
| chr12 | 48594087 | 48604806 | 10720 | -3.99 | 5.53E-30 | 2.34E-28 |
| chr15 | 72041218 | 72065189 | 23972 | -11.88 | 6.56E-30 | 2.77E-28 |

| | | | | | | |
|---|---|---|---|---|---|---|
| chr15 | 29835078 | 29878968 | 43891 | -11.83 | 7.03E-30 | 2.97E-28 |
| chr2 | 31881322 | 31987643 | 106322 | -11.84 | 7.23E-30 | 3.05E-28 |
| chr1 | 228411066 | 228450619 | 39554 | -11.91 | 7.55E-30 | 3.18E-28 |
| chr11 | 28322050 | 28408401 | 86352 | -11.82 | 7.82E-30 | 3.29E-28 |
| chr7 | 123557286 | 123667585 | 110300 | 11.87 | 8.05E-30 | 3.38E-28 |
| chr2 | 150980637 | 151053412 | 72776 | -11.82 | 8.84E-30 | 3.71E-28 |
| chr10 | 5413104 | 5474655 | 61552 | -11.84 | 8.91E-30 | 3.74E-28 |
| chr5 | 178848798 | 178921986 | 73189 | -11.82 | 9.18E-30 | 3.84E-28 |
| chr8 | 98481766 | 98542398 | 60633 | -3.41 | 9.25E-30 | 3.87E-28 |
| chr11 | 104984853 | 105725859 | 741007 | 3.73 | 9.57E-30 | 4.00E-28 |
| chr16 | 27370071 | 27465094 | 95024 | -11.83 | 9.63E-30 | 4.02E-28 |
| chr1 | 233887463 | 233985977 | 98515 | -11.84 | 1.03E-29 | 4.29E-28 |
| chr1 | 91047167 | 91143423 | 96257 | 1.93 | 1.05E-29 | 4.39E-28 |
| chr10 | 117895995 | 118036746 | 140752 | 11.83 | 1.06E-29 | 4.39E-28 |
| chr11 | 133997281 | 134044226 | 46946 | -11.91 | 1.08E-29 | 4.48E-28 |
| chr1 | 163340850 | 164442311 | 1101462 | 1.63 | 1.10E-29 | 4.56E-28 |
| chrX | 3896317 | 4167196 | 270880 | 2.45 | 1.16E-29 | 4.82E-28 |
| chr12 | 84405592 | 84653789 | 248198 | -11.79 | 1.20E-29 | 4.97E-28 |
| chr10 | 122665207 | 122782151 | 116945 | -11.85 | 1.21E-29 | 5.00E-28 |
| chr11 | 78288101 | 78346990 | 58890 | -2.61 | 1.22E-29 | 5.03E-28 |
| chr19 | 54791248 | 54926196 | 134949 | -11.94 | 1.28E-29 | 5.30E-28 |
| chr2 | 235040485 | 235221719 | 181235 | -11.79 | 1.32E-29 | 5.43E-28 |
| chr10 | 119598341 | 119718956 | 120616 | 1.83 | 1.36E-29 | 5.58E-28 |
| chr2 | 13879211 | 15298586 | 1419376 | 1.76 | 1.37E-29 | 5.65E-28 |
| chr17 | 9734596 | 9790978 | 56383 | -11.82 | 1.38E-29 | 5.68E-28 |
| chr11 | 112474469 | 112831714 | 357246 | 3.46 | 1.41E-29 | 5.80E-28 |
| chr19 | 14812556 | 14854447 | 41892 | -11.79 | 1.50E-29 | 6.14E-28 |
| chrX | 115006287 | 115092033 | 85747 | -11.86 | 1.56E-29 | 6.37E-28 |
| chr2 | 221050928 | 221198129 | 147202 | -12 | 1.56E-29 | 6.39E-28 |
| chr6 | 135791972 | 135815593 | 23622 | -11.76 | 1.81E-29 | 7.41E-28 |
| chr7 | 55963219 | 56019395 | 56177 | -11.78 | 1.85E-29 | 7.56E-28 |
| chr1 | 113985953 | 114112367 | 126415 | -11.75 | 1.90E-29 | 7.76E-28 |
| chr9 | 72619224 | 72719400 | 100177 | 11.79 | 1.95E-29 | 7.96E-28 |
| chr11 | 103465203 | 103599681 | 134479 | -11.86 | 2.02E-29 | 8.22E-28 |
| chr11 | 74570053 | 74613945 | 43893 | -11.75 | 2.03E-29 | 8.27E-28 |
| chr5 | 18764931 | 20279450 | 1514520 | 2.68 | 2.05E-29 | 8.33E-28 |
| chr8 | 32876013 | 32924919 | 48907 | -11.81 | 2.12E-29 | 8.62E-28 |
| chr19 | 30475366 | 30492265 | 16900 | -11.74 | 2.20E-29 | 8.90E-28 |
| chr5 | 43827082 | 43938268 | 111187 | 11.79 | 2.22E-29 | 8.98E-28 |
| chr15 | 71452778 | 71476825 | 24048 | -11.8 | 2.32E-29 | 9.37E-28 |
| chr8 | 108116041 | 108170845 | 54805 | -11.74 | 2.38E-29 | 9.60E-28 |
| chr13 | 78376379 | 78487980 | 111602 | 11.76 | 2.42E-29 | 9.75E-28 |
| chr9 | 138786601 | 138812819 | 26219 | -11.73 | 2.42E-29 | 9.77E-28 |
| chr2 | 46319341 | 46340208 | 20868 | -11.83 | 2.50E-29 | 1.01E-27 |
| chr12 | 38375969 | 38442953 | 66985 | -11.92 | 2.51E-29 | 1.01E-27 |
| chr12 | 111382264 | 111458495 | 76232 | -11.75 | 2.67E-29 | 1.07E-27 |
| chr7 | 3767028 | 3824772 | 57745 | -11.72 | 2.76E-29 | 1.11E-27 |
| chr4 | 188617703 | 188631006 | 13304 | -5.79 | 2.87E-29 | 1.15E-27 |
| chrX | 45173023 | 45285195 | 112173 | -11.72 | 2.88E-29 | 1.15E-27 |
| chr10 | 18636510 | 18755671 | 119162 | 11.79 | 2.89E-29 | 1.15E-27 |
| chr2 | 58291430 | 58333083 | 41654 | -11.77 | 3.27E-29 | 1.31E-27 |
| chr13 | 34900106 | 34983914 | 83809 | -11.81 | 3.46E-29 | 1.38E-27 |
| chr16 | 27819943 | 27909312 | 89370 | -11.76 | 3.49E-29 | 1.39E-27 |
| chr9 | 73126998 | 73207955 | 80958 | 11.78 | 3.54E-29 | 1.41E-27 |
| chr2 | 63431357 | 63438056 | 6700 | -7.61 | 3.60E-29 | 1.43E-27 |
| chr12 | 101552775 | 101636600 | 83826 | 11.78 | 3.62E-29 | 1.44E-27 |
| chr9 | 77198990 | 77295022 | 96033 | 11.79 | 3.78E-29 | 1.50E-27 |
| chr15 | 80199816 | 80217941 | 18126 | -11.71 | 3.79E-29 | 1.50E-27 |
| chr12 | 38262994 | 38330265 | 67272 | -11.96 | 3.80E-29 | 1.51E-27 |
| chr19 | 33803214 | 33851043 | 47830 | -11.79 | 3.81E-29 | 1.51E-27 |
| chrX | 67804828 | 67830685 | 25858 | -8.85 | 3.81E-29 | 1.51E-27 |

| | | | | | | |
|---|---|---|---|---|---|---|
| chr14 | 86026516 | 86539915 | 513400 | 4.11 | 3.81E-29 | 1.51E-27 |
| chr2 | 240324270 | 240403703 | 79434 | -11.77 | 3.82E-29 | 1.51E-27 |
| chr2 | 171738519 | 171784771 | 46253 | -11.71 | 3.83E-29 | 1.51E-27 |
| chr14 | 58376645 | 58418755 | 42111 | 2.34 | 3.87E-29 | 1.52E-27 |
| chr7 | 30966468 | 31002731 | 36264 | -11.71 | 3.90E-29 | 1.53E-27 |
| chr3 | 172525030 | 172545759 | 20730 | -11.76 | 3.90E-29 | 1.53E-27 |
| chr7 | 92468042 | 92532731 | 64690 | -11.72 | 3.98E-29 | 1.56E-27 |
| chr12 | 124677220 | 124733273 | 56054 | -11.88 | 4.14E-29 | 1.63E-27 |
| chr16 | 52692835 | 52773793 | 80959 | -11.73 | 4.26E-29 | 1.67E-27 |
| chrX | 73189913 | 73274037 | 84125 | -11.77 | 4.33E-29 | 1.69E-27 |
| chr17 | 56104261 | 56122266 | 18006 | -7.49 | 4.45E-29 | 1.74E-27 |
| chrX | 134397486 | 134467661 | 70176 | -11.73 | 4.58E-29 | 1.79E-27 |
| chr2 | 11172398 | 11181854 | 9457 | -3.45 | 4.71E-29 | 1.84E-27 |
| chr7 | 55848234 | 55949892 | 101659 | -11.73 | 4.96E-29 | 1.93E-27 |
| chr8 | 71530793 | 71589113 | 58321 | -11.67 | 5.12E-29 | 1.99E-27 |
| chr14 | 22784605 | 22837034 | 52430 | -11.67 | 5.14E-29 | 2.00E-27 |
| chr6 | 40324822 | 40428833 | 104012 | 11.73 | 5.32E-29 | 2.07E-27 |
| chr9 | 32622983 | 32702624 | 79642 | -11.71 | 5.32E-29 | 2.07E-27 |
| chr2 | 72269687 | 72326485 | 56799 | -11.67 | 5.36E-29 | 2.08E-27 |
| chr17 | 15607784 | 15616939 | 9156 | -11.69 | 5.50E-29 | 2.13E-27 |
| chr7 | 92406368 | 92452649 | 46282 | -11.66 | 5.59E-29 | 2.16E-27 |
| chr12 | 130950370 | 130954515 | 4146 | -6.17 | 5.60E-29 | 2.16E-27 |
| chr2 | 30156989 | 30239707 | 82719 | 1.91 | 5.65E-29 | 2.18E-27 |
| chr22 | 31361854 | 31426674 | 64821 | -11.67 | 5.65E-29 | 2.18E-27 |
| chr15 | 38427228 | 38450075 | 22848 | -8.13 | 5.71E-29 | 2.20E-27 |
| chr15 | 72108277 | 72125288 | 17012 | -11.75 | 5.74E-29 | 2.21E-27 |
| chr1 | 30074990 | 30089743 | 14754 | -3.91 | 5.82E-29 | 2.24E-27 |
| chr19 | 39770623 | 39782179 | 11557 | -11.68 | 6.06E-29 | 2.33E-27 |
| chrX | 41055483 | 41076441 | 20959 | -11.69 | 6.13E-29 | 2.35E-27 |
| chr1 | 68320278 | 68391046 | 70769 | -11.65 | 6.16E-29 | 2.36E-27 |
| chr8 | 80748142 | 80761664 | 13523 | -7.81 | 6.71E-29 | 2.57E-27 |
| chr9 | 118851332 | 118927955 | 76624 | -11.86 | 6.90E-29 | 2.64E-27 |
| chr4 | 15023787 | 15092805 | 69019 | -11.66 | 7.13E-29 | 2.72E-27 |
| chrX | 85425250 | 86544285 | 1119036 | 2.27 | 7.15E-29 | 2.73E-27 |
| chr13 | 43070347 | 43083861 | 13515 | -3.44 | 7.21E-29 | 2.75E-27 |
| chr3 | 184212235 | 184231091 | 18857 | -9.79 | 7.21E-29 | 2.75E-27 |
| chr12 | 20626254 | 20703198 | 76945 | -11.8 | 7.36E-29 | 2.80E-27 |
| chr4 | 183078143 | 183338461 | 260319 | 2.43 | 7.36E-29 | 2.80E-27 |
| chr16 | 57244391 | 57270910 | 26520 | -11.64 | 7.42E-29 | 2.82E-27 |
| chr17 | 3925809 | 3935494 | 9686 | -11.64 | 7.54E-29 | 2.86E-27 |
| chr12 | 47622007 | 47732689 | 110683 | -11.81 | 7.56E-29 | 2.87E-27 |
| chr1 | 90475650 | 90496925 | 21276 | -11.7 | 7.61E-29 | 2.88E-27 |
| chr10 | 83201401 | 83451668 | 250268 | -11.65 | 7.65E-29 | 2.90E-27 |
| chr21 | 19204319 | 19260678 | 56360 | 2.97 | 7.77E-29 | 2.94E-27 |
| chr17 | 56954090 | 57010569 | 56480 | -11.74 | 8.07E-29 | 3.05E-27 |
| chr3 | 94922677 | 95232308 | 309632 | -11.98 | 8.09E-29 | 3.05E-27 |
| chr3 | 119142091 | 119164006 | 21916 | -11.65 | 8.20E-29 | 3.09E-27 |
| chrX | 106290501 | 106356783 | 66283 | -11.62 | 8.78E-29 | 3.31E-27 |
| chr1 | 7135689 | 7142167 | 6479 | -3.6 | 8.78E-29 | 3.31E-27 |
| chr19 | 38420845 | 38430606 | 9762 | -11.7 | 8.89E-29 | 3.34E-27 |
| chr5 | 33102037 | 33195530 | 93494 | -11.62 | 8.96E-29 | 3.37E-27 |
| chr5 | 123248324 | 123358434 | 110111 | -11.87 | 9.03E-29 | 3.39E-27 |
| chr6 | 44483827 | 44492873 | 9047 | -3.35 | 9.08E-29 | 3.40E-27 |
| chr1 | 244669387 | 244755485 | 86099 | -11.66 | 9.47E-29 | 3.54E-27 |
| chr5 | 92273096 | 92345152 | 72057 | -11.67 | 9.64E-29 | 3.61E-27 |
| chr11 | 7054663 | 7114226 | 59564 | 11.67 | 9.73E-29 | 3.63E-27 |
| chr2 | 63031697 | 63081495 | 49799 | -11.74 | 9.82E-29 | 3.67E-27 |
| chr8 | 32771596 | 32862493 | 90898 | -11.83 | 1.02E-28 | 3.79E-27 |
| chr7 | 105976796 | 106058125 | 81330 | -11.86 | 1.05E-28 | 3.90E-27 |
| chr7 | 149200449 | 149240444 | 39996 | -11.61 | 1.07E-28 | 3.98E-27 |
| chr15 | 43648215 | 43662134 | 13920 | -11.62 | 1.10E-28 | 4.09E-27 |

| | | | | | | |
|---|---|---|---|---|---|---|
| chr6 | 117441539 | 117565973 | 124435 | 11.66 | 1.11E-28 | 4.12E-27 |
| chr7 | 120791460 | 120889822 | 98363 | 11.68 | 1.13E-28 | 4.20E-27 |
| chr15 | 49792861 | 49919167 | 126307 | -11.72 | 1.16E-28 | 4.30E-27 |
| chr19 | 49752415 | 49773968 | 21554 | -11.6 | 1.24E-28 | 4.59E-27 |
| chr2 | 162450117 | 162564180 | 114064 | 11.66 | 1.24E-28 | 4.60E-27 |
| chr3 | 98623813 | 98678358 | 54546 | -11.6 | 1.26E-28 | 4.64E-27 |
| chr16 | 10431159 | 10510693 | 79535 | -11.68 | 1.29E-28 | 4.77E-27 |
| chr12 | 17433334 | 17498001 | 64668 | -11.69 | 1.31E-28 | 4.82E-27 |
| chr2 | 96484396 | 96500100 | 15705 | -3.34 | 1.31E-28 | 4.83E-27 |
| chr20 | 8383134 | 8478860 | 95727 | 11.64 | 1.32E-28 | 4.85E-27 |
| chr14 | 72315605 | 72900834 | 585230 | 0.79 | 1.34E-28 | 4.94E-27 |
| chr2 | 63094895 | 63117373 | 22479 | -11.63 | 1.39E-28 | 5.12E-27 |
| chr7 | 16926689 | 17037631 | 110943 | -11.91 | 1.40E-28 | 5.15E-27 |
| chr2 | 89188121 | 89276840 | 88720 | 11.61 | 1.43E-28 | 5.23E-27 |
| chrX | 9920550 | 10005509 | 84960 | -11.66 | 1.44E-28 | 5.28E-27 |
| chrX | 38428859 | 38511928 | 83070 | 11.61 | 1.45E-28 | 5.31E-27 |
| chr4 | 164499204 | 165180764 | 681561 | 2.81 | 1.47E-28 | 5.36E-27 |
| chr1 | 98680524 | 98806061 | 125538 | -11.6 | 1.51E-28 | 5.53E-27 |
| chr16 | 80152136 | 80176535 | 24400 | -3.67 | 1.53E-28 | 5.58E-27 |
| chr3 | 30738898 | 30779293 | 40396 | -11.57 | 1.57E-28 | 5.73E-27 |
| chr10 | 82373160 | 82420439 | 47280 | -2.63 | 1.57E-28 | 5.73E-27 |
| chr2 | 239559246 | 239599772 | 40527 | -11.58 | 1.60E-28 | 5.84E-27 |
| chr7 | 127161658 | 127188618 | 26961 | -4.05 | 1.63E-28 | 5.93E-27 |
| chrX | 120146184 | 120174107 | 27924 | 3.44 | 1.68E-28 | 6.09E-27 |
| chr17 | 57835533 | 57890682 | 55150 | -11.58 | 1.69E-28 | 6.13E-27 |
| chr14 | 28322712 | 28617319 | 294608 | 1.65 | 1.70E-28 | 6.16E-27 |
| chr10 | 52643091 | 52730754 | 87664 | 11.59 | 1.77E-28 | 6.40E-27 |
| chr4 | 73448459 | 73545741 | 97283 | 11.61 | 1.78E-28 | 6.43E-27 |
| chr3 | 176391292 | 176520621 | 129330 | 11.67 | 1.89E-28 | 6.82E-27 |
| chr2 | 167283159 | 168142597 | 859439 | 2.29 | 1.91E-28 | 6.88E-27 |
| chr6 | 169324662 | 169349827 | 25166 | -7.9 | 1.95E-28 | 7.03E-27 |
| chr12 | 94381035 | 94458552 | 77518 | -11.58 | 1.97E-28 | 7.11E-27 |
| chr11 | 114511205 | 114942074 | 430870 | 1.93 | 2.00E-28 | 7.22E-27 |
| chr2 | 216536338 | 216698472 | 162135 | -11.94 | 2.17E-28 | 7.82E-27 |
| chr17 | 67738344 | 67794561 | 56218 | -11.73 | 2.25E-28 | 8.10E-27 |
| chr15 | 39397474 | 39512924 | 115451 | -11.71 | 2.26E-28 | 8.11E-27 |
| chr12 | 55876916 | 55992126 | 115211 | 4.01 | 2.26E-28 | 8.11E-27 |
| chr6 | 119262039 | 119272063 | 10025 | -4.78 | 2.31E-28 | 8.28E-27 |
| chr1 | 23679651 | 23693886 | 14236 | -11.55 | 2.31E-28 | 8.29E-27 |
| chr8 | 31357462 | 31484120 | 126659 | -1.63 | 2.54E-28 | 9.10E-27 |
| chr15 | 36655359 | 36736112 | 80754 | -11.6 | 2.58E-28 | 9.22E-27 |
| chr2 | 184004787 | 184075138 | 70352 | -11.55 | 2.59E-28 | 9.25E-27 |
| chr20 | 16732613 | 17442236 | 709624 | 3.92 | 2.62E-28 | 9.35E-27 |
| chr7 | 283928 | 333185 | 49258 | -11.66 | 2.65E-28 | 9.44E-27 |
| chr17 | 58533138 | 58574431 | 41294 | -11.57 | 2.72E-28 | 9.67E-27 |
| chr5 | 172985595 | 173032994 | 47400 | -11.59 | 2.75E-28 | 9.78E-27 |
| chr5 | 21581682 | 21797493 | 215812 | 2.21 | 2.77E-28 | 9.85E-27 |
| chr6 | 19843368 | 19935476 | 92109 | 11.62 | 2.84E-28 | 1.01E-26 |
| chr8 | 6501229 | 6569851 | 68623 | -11.56 | 2.86E-28 | 1.02E-26 |
| chr13 | 46427599 | 46441994 | 14396 | -3.42 | 2.89E-28 | 1.02E-26 |
| chr5 | 136689039 | 136795699 | 106661 | 11.76 | 2.90E-28 | 1.03E-26 |
| chr7 | 98067727 | 98166445 | 98719 | -11.73 | 3.04E-28 | 1.08E-26 |
| chr2 | 144625543 | 144633161 | 7619 | -3.06 | 3.09E-28 | 1.09E-26 |
| chr4 | 173610189 | 173704853 | 94665 | -11.51 | 3.10E-28 | 1.10E-26 |
| chr15 | 56411443 | 56448108 | 36666 | -11.51 | 3.15E-28 | 1.11E-26 |
| chr2 | 226101916 | 226194064 | 92149 | -11.51 | 3.21E-28 | 1.13E-26 |
| chr11 | 58530475 | 58550642 | 20168 | 4.48 | 3.21E-28 | 1.13E-26 |
| chr9 | 74471903 | 74500430 | 28528 | -11.51 | 3.22E-28 | 1.13E-26 |
| chr9 | 81060084 | 81161109 | 101026 | -11.74 | 3.22E-28 | 1.13E-26 |
| chr1 | 68409238 | 68475054 | 65817 | -11.52 | 3.27E-28 | 1.15E-26 |
| chr11 | 91094022 | 91177893 | 83872 | -11.74 | 3.30E-28 | 1.16E-26 |

| | | | | | | |
|---|---|---|---|---|---|---|
| chr8 | 446674 | 482470 | 35797 | -11.51 | 3.35E-28 | 1.17E-26 |
| chr3 | 141035569 | 141138460 | 102892 | -11.51 | 3.35E-28 | 1.17E-26 |
| chr20 | 13954793 | 14002382 | 47590 | -11.51 | 3.40E-28 | 1.19E-26 |
| chr11 | 5739740 | 5826301 | 86562 | 11.59 | 3.45E-28 | 1.20E-26 |
| chr4 | 55409637 | 55823957 | 414321 | 3.18 | 3.48E-28 | 1.22E-26 |
| chr17 | 32774565 | 32842593 | 68029 | 11.64 | 3.57E-28 | 1.25E-26 |
| chr22 | 26573337 | 26611885 | 38549 | 2.95 | 3.62E-28 | 1.26E-26 |
| chr4 | 55201824 | 55402048 | 200225 | 1.61 | 3.76E-28 | 1.31E-26 |
| chr7 | 120648353 | 120736438 | 88086 | 11.54 | 3.86E-28 | 1.34E-26 |
| chr2 | 102518161 | 102590553 | 72393 | -11.51 | 4.05E-28 | 1.41E-26 |
| chr20 | 49832072 | 49947213 | 115142 | -11.59 | 4.09E-28 | 1.42E-26 |
| chr17 | 6811168 | 6839733 | 28566 | -11.54 | 4.13E-28 | 1.43E-26 |
| chr12 | 7316032 | 7335178 | 19147 | -2.44 | 4.15E-28 | 1.44E-26 |
| chr7 | 7476044 | 7541888 | 65845 | -11.49 | 4.29E-28 | 1.49E-26 |
| chr2 | 83559652 | 83762303 | 202652 | -11.5 | 4.31E-28 | 1.49E-26 |
| chr3 | 110234342 | 110315340 | 80999 | -11.49 | 4.32E-28 | 1.49E-26 |
| chr19 | 54546432 | 54598404 | 51973 | -11.61 | 4.35E-28 | 1.50E-26 |
| chr18 | 76641569 | 76661556 | 19988 | -2.44 | 4.39E-28 | 1.51E-26 |
| chr11 | 74762302 | 74841105 | 78804 | -11.51 | 4.45E-28 | 1.53E-26 |
| chr7 | 422321 | 459227 | 36907 | -11.52 | 4.46E-28 | 1.54E-26 |
| chr3 | 173875084 | 173915283 | 40200 | 2.42 | 4.47E-28 | 1.54E-26 |
| chr6 | 72513810 | 72591983 | 78174 | 11.51 | 4.59E-28 | 1.58E-26 |
| chr16 | 16055327 | 16079103 | 23777 | -11.49 | 4.79E-28 | 1.64E-26 |
| chr2 | 101665294 | 101721310 | 56017 | -11.53 | 4.95E-28 | 1.70E-26 |
| chr5 | 170978220 | 171037343 | 59124 | -11.47 | 4.99E-28 | 1.71E-26 |
| chr1 | 75431805 | 75452275 | 20471 | 4.82 | 5.04E-28 | 1.73E-26 |
| chr16 | 72415383 | 72499474 | 84092 | -11.56 | 5.12E-28 | 1.75E-26 |
| chr9 | 115657566 | 115822130 | 164565 | -3.62 | 5.20E-28 | 1.78E-26 |
| chr5 | 58283309 | 58384856 | 101548 | -11.55 | 5.27E-28 | 1.80E-26 |
| chr7 | 99080064 | 99099639 | 19576 | -11.47 | 5.68E-28 | 1.94E-26 |
| chr9 | 73929356 | 74022556 | 93201 | 11.49 | 5.81E-28 | 1.98E-26 |
| chr3 | 192331096 | 192435387 | 104292 | 11.55 | 5.91E-28 | 2.01E-26 |
| chr10 | 777809 | 817535 | 39727 | -11.47 | 5.94E-28 | 2.02E-26 |
| chr2 | 57915789 | 57919702 | 3914 | -5.86 | 6.09E-28 | 2.07E-26 |
| chr4 | 74911509 | 75013724 | 102216 | -11.54 | 6.13E-28 | 2.08E-26 |
| chr3 | 10930804 | 10967609 | 36806 | -11.46 | 6.15E-28 | 2.09E-26 |
| chr2 | 205076158 | 205082393 | 6236 | -2.68 | 6.20E-28 | 2.10E-26 |
| chr2 | 98924170 | 98987246 | 63077 | -11.45 | 6.26E-28 | 2.12E-26 |
| chr3 | 31684019 | 31702104 | 18086 | -11.47 | 6.28E-28 | 2.13E-26 |
| chr9 | 118427510 | 118499207 | 71698 | -11.46 | 6.41E-28 | 2.17E-26 |
| chr8 | 41286130 | 41311033 | 24904 | -2.77 | 6.45E-28 | 2.18E-26 |
| chr20 | 14898561 | 14983198 | 84638 | -11.87 | 6.53E-28 | 2.21E-26 |
| chr6 | 77974509 | 78021586 | 47078 | -11.49 | 6.74E-28 | 2.27E-26 |
| chr9 | 129000487 | 129049897 | 49411 | -11.45 | 6.75E-28 | 2.28E-26 |
| chr2 | 83234194 | 83409821 | 175628 | -11.62 | 7.01E-28 | 2.36E-26 |
| chrX | 10080741 | 10115377 | 34637 | -11.5 | 7.33E-28 | 2.47E-26 |
| chr7 | 46462195 | 46515571 | 53377 | -11.48 | 7.42E-28 | 2.49E-26 |
| chr9 | 72734928 | 72830479 | 95552 | 11.49 | 8.08E-28 | 2.71E-26 |
| chr19 | 20769003 | 20929644 | 160642 | -11.67 | 8.29E-28 | 2.78E-26 |
| chr11 | 69857716 | 69903377 | 45662 | -11.47 | 8.32E-28 | 2.79E-26 |
| chr12 | 32116840 | 32151233 | 34394 | -11.48 | 8.44E-28 | 2.83E-26 |
| chr2 | 28171239 | 28226480 | 55242 | -11.44 | 9.07E-28 | 3.04E-26 |
| chr7 | 158719544 | 158795572 | 76029 | -11.69 | 9.21E-28 | 3.08E-26 |
| chr15 | 56503950 | 56527842 | 23893 | -11.43 | 9.24E-28 | 3.09E-26 |
| chr3 | 70122005 | 70129197 | 7193 | -3.25 | 9.37E-28 | 3.13E-26 |
| chr11 | 35888246 | 35928615 | 40370 | -11.42 | 9.38E-28 | 3.13E-26 |
| chr20 | 45692908 | 45743981 | 51074 | -11.48 | 9.39E-28 | 3.13E-26 |
| chr19 | 38447442 | 38466527 | 19086 | -11.42 | 9.61E-28 | 3.20E-26 |
| chr6 | 39542781 | 39680217 | 137437 | 1.73 | 9.76E-28 | 3.25E-26 |
| chrX | 22422180 | 22964647 | 542468 | 2.94 | 9.77E-28 | 3.25E-26 |
| chr12 | 79669084 | 79738209 | 69126 | -11.62 | 9.80E-28 | 3.25E-26 |

| | | | | | | |
|---|---|---|---|---|---|---|
| chr17 | 55272575 | 55307322 | 34748 | -11.43 | 9.99E-28 | 3.31E-26 |
| chr3 | 140524098 | 140598978 | 74881 | 2.3 | 1.02E-27 | 3.37E-26 |
| chr20 | 4507789 | 4578443 | 70655 | -11.42 | 1.03E-27 | 3.40E-26 |
| chr7 | 34245761 | 34339647 | 93887 | 11.44 | 1.05E-27 | 3.46E-26 |
| chr7 | 95582204 | 95658466 | 76263 | 11.48 | 1.05E-27 | 3.46E-26 |
| chr12 | 79885669 | 79969467 | 83799 | -11.51 | 1.05E-27 | 3.46E-26 |
| chr2 | 121378648 | 121449873 | 71226 | -11.65 | 1.09E-27 | 3.60E-26 |
| chr13 | 42997258 | 43046969 | 49712 | -2.33 | 1.16E-27 | 3.84E-26 |
| chr1 | 55836344 | 55947997 | 111654 | -11.51 | 1.19E-27 | 3.91E-26 |
| chr9 | 73258496 | 73316803 | 58308 | 11.46 | 1.19E-27 | 3.92E-26 |
| chr11 | 65497648 | 65537329 | 39682 | -11.41 | 1.20E-27 | 3.94E-26 |
| chr12 | 43327635 | 43439916 | 112282 | -11.41 | 1.20E-27 | 3.94E-26 |
| chr2 | 129086793 | 129114794 | 28002 | -11.4 | 1.21E-27 | 3.97E-26 |
| chr18 | 22318386 | 22917728 | 599343 | 4.84 | 1.21E-27 | 3.98E-26 |
| chr4 | 129400232 | 129457762 | 57531 | -11.44 | 1.22E-27 | 3.98E-26 |
| chr5 | 174977500 | 175078794 | 101295 | -11.65 | 1.25E-27 | 4.09E-26 |
| chr7 | 46531468 | 46615604 | 84137 | -11.44 | 1.30E-27 | 4.26E-26 |
| chr17 | 80898295 | 80915433 | 17139 | -11.41 | 1.32E-27 | 4.32E-26 |
| chr10 | 123057100 | 123187888 | 130789 | -11.39 | 1.34E-27 | 4.37E-26 |
| chr12 | 13905004 | 13973219 | 68216 | 11.41 | 1.42E-27 | 4.64E-26 |
| chr6 | 125053071 | 125153338 | 100268 | 11.42 | 1.43E-27 | 4.67E-26 |
| chr10 | 49872360 | 50594948 | 722589 | 2.36 | 1.51E-27 | 4.92E-26 |
| chrX | 62614889 | 62720628 | 105740 | -11.4 | 1.51E-27 | 4.92E-26 |
| chr5 | 93965044 | 93995645 | 30602 | -11.5 | 1.53E-27 | 4.99E-26 |
| chr8 | 129043691 | 129094774 | 51084 | -11.38 | 1.53E-27 | 4.99E-26 |
| chr2 | 143702592 | 143725035 | 22444 | -11.38 | 1.54E-27 | 4.99E-26 |
| chr11 | 92226592 | 92331098 | 104507 | 11.43 | 1.56E-27 | 5.07E-26 |
| chr3 | 179553459 | 179642023 | 88565 | 11.41 | 1.59E-27 | 5.16E-26 |
| chr19 | 47050775 | 47085371 | 34597 | -11.38 | 1.61E-27 | 5.20E-26 |
| chr11 | 2227762 | 2276965 | 49204 | -11.44 | 1.64E-27 | 5.30E-26 |
| chr5 | 35708991 | 36093149 | 384159 | 4.01 | 1.65E-27 | 5.32E-26 |
| chr2 | 62782591 | 62885953 | 103363 | -11.6 | 1.66E-27 | 5.36E-26 |
| chr3 | 180824671 | 180920990 | 96320 | 11.4 | 1.67E-27 | 5.38E-26 |
| chr9 | 132268614 | 132305108 | 36495 | -11.74 | 1.68E-27 | 5.40E-26 |
| chr2 | 71948805 | 72000723 | 51919 | -11.46 | 1.68E-27 | 5.40E-26 |
| chr21 | 45960548 | 46007360 | 46813 | -11.37 | 1.68E-27 | 5.41E-26 |
| chr12 | 72144766 | 72165402 | 20637 | -11.37 | 1.68E-27 | 5.41E-26 |
| chr1 | 12442401 | 12474028 | 31628 | -11.38 | 1.72E-27 | 5.51E-26 |
| chr2 | 171173615 | 171195416 | 21802 | -3.67 | 1.76E-27 | 5.64E-26 |
| chr20 | 52714925 | 52789365 | 74441 | -11.4 | 1.78E-27 | 5.70E-26 |
| chr7 | 110092397 | 110213680 | 121284 | -11.5 | 1.82E-27 | 5.82E-26 |
| chr11 | 114140909 | 114218421 | 77513 | -11.45 | 1.82E-27 | 5.83E-26 |
| chr18 | 42692092 | 43122311 | 430220 | 1.92 | 1.82E-27 | 5.83E-26 |
| chr17 | 16850957 | 16881318 | 30362 | -11.39 | 1.83E-27 | 5.85E-26 |
| chr18 | 14091795 | 14124970 | 33176 | -11.38 | 1.86E-27 | 5.95E-26 |
| chr2 | 224071146 | 224077715 | 6570 | -7.57 | 1.88E-27 | 6.00E-26 |
| chr3 | 151418230 | 151527470 | 109241 | -11.5 | 1.89E-27 | 6.02E-26 |
| chr4 | 156455229 | 156563349 | 108121 | 11.41 | 1.89E-27 | 6.02E-26 |
| chr2 | 155322343 | 155427563 | 105221 | 11.69 | 1.95E-27 | 6.20E-26 |
| chr12 | 88166826 | 88249111 | 82286 | -11.63 | 2.00E-27 | 6.37E-26 |
| chr5 | 171936635 | 171965388 | 28754 | -11.36 | 2.02E-27 | 6.42E-26 |
| chr2 | 202365643 | 202404439 | 38797 | -11.36 | 2.05E-27 | 6.51E-26 |
| chr13 | 106921148 | 107019607 | 98460 | -11.35 | 2.07E-27 | 6.58E-26 |
| chr20 | 58589553 | 58683721 | 94169 | -11.36 | 2.10E-27 | 6.67E-26 |
| chr8 | 87238831 | 87330816 | 91986 | -11.71 | 2.16E-27 | 6.84E-26 |
| chr6 | 41227779 | 41233751 | 5973 | -2.96 | 2.18E-27 | 6.88E-26 |
| chr8 | 19424072 | 19538700 | 114629 | -11.36 | 2.19E-27 | 6.92E-26 |
| chr17 | 66833539 | 67134693 | 301155 | 2.58 | 2.19E-27 | 6.93E-26 |
| chr19 | 16048127 | 16108660 | 60534 | -11.35 | 2.23E-27 | 7.02E-26 |
| chr7 | 1801621 | 1841824 | 40204 | -11.34 | 2.29E-27 | 7.21E-26 |
| chr14 | 60073389 | 60344103 | 270715 | 1.89 | 2.39E-27 | 7.53E-26 |

| | | | | | | |
|---|---|---|---|---|---|---|
| chr1 | 166141916 | 166207201 | 65286 | 11.43 | 2.40E-27 | 7.54E-26 |
| chr7 | 3893544 | 3932775 | 39232 | -11.34 | 2.40E-27 | 7.54E-26 |
| chr5 | 17890885 | 17981335 | 90451 | -11.53 | 2.42E-27 | 7.59E-26 |
| chr3 | 125910656 | 125963296 | 52641 | -11.35 | 2.44E-27 | 7.67E-26 |
| chr1 | 111454790 | 111523917 | 69128 | -11.34 | 2.48E-27 | 7.78E-26 |
| chr13 | 109875195 | 109932429 | 57235 | -11.35 | 2.48E-27 | 7.78E-26 |
| chr11 | 9950189 | 10039513 | 89325 | -11.62 | 2.61E-27 | 8.16E-26 |
| chr1 | 178698378 | 178748203 | 49826 | -11.54 | 2.64E-27 | 8.27E-26 |
| chr5 | 89573137 | 89659206 | 86070 | 11.49 | 2.68E-27 | 8.37E-26 |
| chr2 | 131529606 | 131542415 | 12810 | -7.65 | 2.68E-27 | 8.38E-26 |
| chr1 | 7321330 | 7399928 | 78599 | -11.41 | 2.87E-27 | 8.95E-26 |
| chr20 | 2696980 | 2717446 | 20467 | -11.35 | 2.90E-27 | 9.05E-26 |
| chr11 | 31271052 | 31366267 | 95216 | 11.44 | 3.02E-27 | 9.42E-26 |
| chr6 | 101588359 | 101618525 | 30167 | 7.39 | 3.08E-27 | 9.58E-26 |
| chr12 | 120045624 | 120058056 | 12433 | -7.37 | 3.37E-27 | 1.05E-25 |
| chr12 | 21098843 | 21164334 | 65492 | -11.31 | 3.41E-27 | 1.06E-25 |
| chr7 | 24004757 | 24319992 | 315236 | 2 | 3.51E-27 | 1.09E-25 |
| chr5 | 166419614 | 166472115 | 52502 | 3.84 | 3.52E-27 | 1.09E-25 |
| chr8 | 59249337 | 59282729 | 33393 | -4.31 | 3.58E-27 | 1.11E-25 |
| chr14 | 50277857 | 50312336 | 34480 | -11.35 | 3.65E-27 | 1.13E-25 |
| chr16 | 89289750 | 89307024 | 17275 | -11.34 | 3.73E-27 | 1.16E-25 |
| chr11 | 31085433 | 31246600 | 161168 | 5.95 | 3.76E-27 | 1.16E-25 |
| chr3 | 181067835 | 181157843 | 90009 | 11.34 | 3.80E-27 | 1.18E-25 |
| chr9 | 84025791 | 84075248 | 49458 | -11.48 | 3.92E-27 | 1.21E-25 |
| chr11 | 3255385 | 3268216 | 12832 | -8.52 | 3.95E-27 | 1.22E-25 |
| chrX | 113590854 | 113692450 | 101597 | 3.78 | 3.97E-27 | 1.23E-25 |
| chr2 | 211125436 | 211206744 | 81309 | 11.35 | 4.05E-27 | 1.25E-25 |
| chr7 | 131017387 | 131036508 | 19122 | -11.33 | 4.08E-27 | 1.26E-25 |
| chr2 | 227626357 | 227653321 | 26965 | -11.31 | 4.10E-27 | 1.26E-25 |
| chr18 | 32138377 | 32206044 | 67668 | 11.34 | 4.32E-27 | 1.33E-25 |
| chr3 | 56438647 | 56464845 | 26199 | 3.26 | 4.34E-27 | 1.33E-25 |
| chr3 | 143402205 | 143479133 | 76929 | 11.37 | 4.44E-27 | 1.36E-25 |
| chr19 | 41364765 | 41383394 | 18630 | -4.35 | 4.65E-27 | 1.43E-25 |
| chr8 | 122149446 | 122241946 | 92501 | -11.36 | 4.66E-27 | 1.43E-25 |
| chr20 | 751075 | 791654 | 40580 | -11.33 | 4.77E-27 | 1.46E-25 |
| chr8 | 2195731 | 2324266 | 128536 | 2.29 | 4.83E-27 | 1.48E-25 |
| chr14 | 39518118 | 39543471 | 25354 | -11.4 | 4.84E-27 | 1.48E-25 |
| chr5 | 58696448 | 58717575 | 21128 | -11.3 | 5.03E-27 | 1.54E-25 |
| chr11 | 34598960 | 34625171 | 26212 | -11.28 | 5.10E-27 | 1.56E-25 |
| chr2 | 17748566 | 17813701 | 65136 | 11.34 | 5.11E-27 | 1.56E-25 |
| chr10 | 77041632 | 77148413 | 106782 | -11.51 | 5.18E-27 | 1.58E-25 |
| chr13 | 21785306 | 21798189 | 12884 | -3.48 | 5.22E-27 | 1.59E-25 |
| chr6 | 66036996 | 66313234 | 276239 | 3.38 | 5.27E-27 | 1.60E-25 |
| chr11 | 43478513 | 43506834 | 28322 | -11.33 | 5.33E-27 | 1.62E-25 |
| chr7 | 21659122 | 21722778 | 63657 | -11.31 | 5.52E-27 | 1.68E-25 |
| chr6 | 49659537 | 49752612 | 93076 | 11.3 | 5.64E-27 | 1.71E-25 |
| chr8 | 28061645 | 28098875 | 37231 | -11.28 | 5.66E-27 | 1.72E-25 |
| chr16 | 80605157 | 80775838 | 170682 | 1.6 | 5.78E-27 | 1.75E-25 |
| chr17 | 15228273 | 15241507 | 13235 | -7.91 | 5.79E-27 | 1.75E-25 |
| chr2 | 202149850 | 202204597 | 54748 | -11.3 | 5.86E-27 | 1.77E-25 |
| chr5 | 123398552 | 123432712 | 34161 | -7.97 | 5.96E-27 | 1.80E-25 |
| chr14 | 102199085 | 102221353 | 22269 | -9.63 | 6.10E-27 | 1.84E-25 |
| chr8 | 120115341 | 120137357 | 22017 | 2.86 | 6.27E-27 | 1.89E-25 |
| chr16 | 87555980 | 87580381 | 24402 | -11.32 | 6.28E-27 | 1.89E-25 |
| chr10 | 102908019 | 102941468 | 33450 | -11.25 | 6.42E-27 | 1.93E-25 |
| chr12 | 131392239 | 131435627 | 43389 | -11.32 | 6.46E-27 | 1.95E-25 |
| chr2 | 231636313 | 231661142 | 24830 | -11.25 | 6.51E-27 | 1.96E-25 |
| chr8 | 32939476 | 33024109 | 84634 | -11.46 | 6.66E-27 | 2.00E-25 |
| chr2 | 40540184 | 40543367 | 3184 | -7.88 | 6.77E-27 | 2.03E-25 |
| chr6 | 136279892 | 136357345 | 77454 | -11.3 | 7.29E-27 | 2.19E-25 |
| chr16 | 86035116 | 86065285 | 30170 | -11.24 | 7.34E-27 | 2.20E-25 |

| | | | | | | |
|---|---|---|---|---|---|---|
| chr3 | 5676850 | 5734691 | 57842 | -11.24 | 7.53E-27 | 2.26E-25 |
| chr7 | 99508844 | 99561236 | 52393 | -11.31 | 7.54E-27 | 2.26E-25 |
| chr3 | 160975061 | 161046705 | 71645 | -11.24 | 7.64E-27 | 2.29E-25 |
| chr11 | 11071487 | 11081904 | 10418 | -3.33 | 7.67E-27 | 2.29E-25 |
| chr20 | 499676 | 531695 | 32020 | -11.27 | 7.98E-27 | 2.38E-25 |
| chrX | 107712905 | 107785461 | 72557 | -11.24 | 8.00E-27 | 2.39E-25 |
| chr5 | 71113066 | 71123716 | 10651 | -7.33 | 8.07E-27 | 2.41E-25 |
| chr2 | 197487991 | 197523436 | 35446 | -11.26 | 8.24E-27 | 2.46E-25 |
| chr4 | 7549942 | 7592268 | 42327 | -11.26 | 8.43E-27 | 2.51E-25 |
| chr1 | 108710223 | 108740538 | 30316 | -11.23 | 8.61E-27 | 2.56E-25 |
| chr17 | 59692977 | 59731290 | 38314 | -11.27 | 8.66E-27 | 2.57E-25 |
| chr4 | 189332963 | 189387357 | 54395 | -11.27 | 9.02E-27 | 2.68E-25 |
| chr9 | 114026462 | 114033123 | 6662 | -5.93 | 9.02E-27 | 2.68E-25 |
| chr12 | 80195939 | 80209742 | 13804 | -11.22 | 9.21E-27 | 2.73E-25 |
| chr19 | 33183921 | 33202891 | 18971 | -11.22 | 9.56E-27 | 2.83E-25 |
| chrX | 128553137 | 128564969 | 11833 | -4.04 | 9.60E-27 | 2.84E-25 |
| chr15 | 56868238 | 56911717 | 43480 | -11.27 | 9.72E-27 | 2.88E-25 |
| chr9 | 135564696 | 135583948 | 19253 | -11.22 | 1.01E-26 | 2.98E-25 |
| chr15 | 91756264 | 92387932 | 631669 | 1.01 | 1.02E-26 | 3.01E-25 |
| chr10 | 37317935 | 37446487 | 128553 | -11.32 | 1.02E-26 | 3.01E-25 |
| chr7 | 95003041 | 95019719 | 16679 | -8.47 | 1.03E-26 | 3.05E-25 |
| chr4 | 6483941 | 6515630 | 31690 | -11.21 | 1.04E-26 | 3.06E-25 |
| chr4 | 90175582 | 90215296 | 39715 | -11.21 | 1.06E-26 | 3.11E-25 |
| chr2 | 31631005 | 31700833 | 69829 | -11.23 | 1.08E-26 | 3.18E-25 |
| chrX | 140487054 | 140560910 | 73857 | -11.36 | 1.10E-26 | 3.22E-25 |
| chr1 | 97194521 | 97226611 | 32091 | -11.24 | 1.13E-26 | 3.31E-25 |
| chr4 | 56148141 | 56163604 | 15464 | -4.34 | 1.13E-26 | 3.32E-25 |
| chr12 | 46672881 | 46745345 | 72465 | -11.41 | 1.17E-26 | 3.43E-25 |
| chr19 | 34054813 | 34105503 | 50691 | -4 | 1.18E-26 | 3.45E-25 |
| chr12 | 47502652 | 47567692 | 65041 | -11.36 | 1.19E-26 | 3.50E-25 |
| chr3 | 100726708 | 100812646 | 85939 | -11.45 | 1.20E-26 | 3.51E-25 |
| chr4 | 74865560 | 74896805 | 31246 | -11.21 | 1.23E-26 | 3.60E-25 |
| chr9 | 126914424 | 126960309 | 45886 | -11.2 | 1.24E-26 | 3.63E-25 |
| chr5 | 117544 | 151242 | 33699 | -11.21 | 1.27E-26 | 3.72E-25 |
| chr1 | 172873847 | 172933163 | 59317 | -11.19 | 1.29E-26 | 3.75E-25 |
| chr15 | 59465280 | 59482220 | 16941 | -11.21 | 1.34E-26 | 3.92E-25 |
| chr18 | 48842757 | 49086327 | 243571 | 3.15 | 1.38E-26 | 4.02E-25 |
| chr8 | 130050147 | 130172100 | 121954 | -11.42 | 1.41E-26 | 4.11E-25 |
| chr14 | 57027649 | 57035834 | 8186 | -3.13 | 1.47E-26 | 4.28E-25 |
| chr7 | 67700088 | 67767646 | 67559 | -11.35 | 1.51E-26 | 4.40E-25 |
| chr3 | 9605611 | 9668022 | 62412 | -11.3 | 1.52E-26 | 4.42E-25 |
| chr15 | 56651073 | 56655585 | 4513 | -7.49 | 1.53E-26 | 4.43E-25 |
| chr2 | 139632569 | 141028301 | 1395733 | 1.97 | 1.69E-26 | 4.89E-25 |
| chr12 | 41723920 | 41733499 | 9580 | 7.73 | 1.70E-26 | 4.93E-25 |
| chr15 | 32332353 | 32445362 | 113010 | 1.5 | 1.71E-26 | 4.94E-25 |
| chr22 | 47622689 | 47662554 | 39866 | -2.6 | 1.71E-26 | 4.95E-25 |
| chr9 | 130200568 | 130213091 | 12524 | -11.25 | 1.75E-26 | 5.07E-25 |
| chr8 | 127443273 | 127509836 | 66564 | -11.19 | 1.77E-26 | 5.13E-25 |
| chr12 | 58699111 | 58750836 | 51726 | -11.21 | 1.79E-26 | 5.17E-25 |
| chr11 | 58883134 | 58903482 | 20349 | -11.26 | 1.83E-26 | 5.29E-25 |
| chr9 | 35128803 | 35160339 | 31537 | -11.2 | 1.89E-26 | 5.44E-25 |
| chr4 | 23899866 | 24280256 | 380391 | 3.53 | 1.89E-26 | 5.44E-25 |
| chr5 | 127429318 | 127455537 | 26220 | -11.16 | 1.92E-26 | 5.52E-25 |
| chr18 | 70461530 | 70543286 | 81757 | 2.68 | 1.92E-26 | 5.53E-25 |
| chr3 | 151544289 | 151575952 | 31664 | -11.36 | 1.94E-26 | 5.59E-25 |
| chr11 | 34779410 | 34888559 | 109150 | -11.52 | 1.96E-26 | 5.63E-25 |
| chr12 | 57242344 | 57282250 | 39907 | -11.17 | 1.97E-26 | 5.67E-25 |
| chr22 | 19299069 | 19333837 | 34769 | -11.16 | 1.99E-26 | 5.70E-25 |
| chr5 | 13339624 | 13424363 | 84740 | 11.35 | 2.05E-26 | 5.87E-25 |
| chr1 | 86084953 | 86131912 | 46960 | -11.19 | 2.07E-26 | 5.93E-25 |
| chr14 | 69062138 | 69114664 | 52527 | -11.33 | 2.09E-26 | 5.97E-25 |

| | | | | | | |
|---|---|---|---|---|---|---|
| chr11 | 58565395 | 58674120 | 108726 | 2.1 | 2.10E-26 | 5.99E-25 |
| chr8 | 131812613 | 131893435 | 80823 | -11.62 | 2.14E-26 | 6.12E-25 |
| chr11 | 110476555 | 110563407 | 86853 | 11.26 | 2.25E-26 | 6.41E-25 |
| chr7 | 35325553 | 35334048 | 8496 | -2.67 | 2.25E-26 | 6.41E-25 |
| chr3 | 102642170 | 102834624 | 192455 | -11.38 | 2.25E-26 | 6.42E-25 |
| chr3 | 16098143 | 16212058 | 113916 | -11.83 | 2.29E-26 | 6.53E-25 |
| chr5 | 98454168 | 98515066 | 60899 | 11.18 | 2.30E-26 | 6.55E-25 |
| chr11 | 10063557 | 10139168 | 75612 | -11.41 | 2.36E-26 | 6.70E-25 |
| chrX | 73410128 | 73450765 | 40638 | -11.19 | 2.38E-26 | 6.76E-25 |
| chr8 | 144831841 | 144880036 | 48196 | -11.28 | 2.46E-26 | 6.99E-25 |
| chr5 | 38988029 | 39048753 | 60725 | -11.19 | 2.52E-26 | 7.16E-25 |
| chr2 | 136509152 | 136560377 | 51226 | -11.32 | 2.53E-26 | 7.17E-25 |
| chr20 | 36693017 | 36710698 | 17682 | -11.19 | 2.57E-26 | 7.28E-25 |
| chr8 | 54828476 | 54857168 | 28693 | -11.14 | 2.57E-26 | 7.28E-25 |
| chr20 | 62342897 | 62358230 | 15334 | -11.18 | 2.60E-26 | 7.36E-25 |
| chr7 | 103810988 | 103863992 | 53005 | -11.17 | 2.61E-26 | 7.38E-25 |
| chr5 | 38890450 | 38936137 | 45688 | -11.22 | 2.63E-26 | 7.43E-25 |
| chr3 | 188723742 | 188787553 | 63812 | -11.15 | 2.68E-26 | 7.56E-25 |
| chr19 | 30028787 | 30082607 | 53821 | -11.29 | 2.70E-26 | 7.63E-25 |
| chr7 | 67117528 | 67186973 | 69446 | -11.41 | 2.71E-26 | 7.65E-25 |
| chr10 | 123376250 | 123380310 | 4061 | -7.81 | 2.77E-26 | 7.79E-25 |
| chr21 | 46982647 | 47024710 | 42064 | -11.12 | 2.78E-26 | 7.81E-25 |
| chr7 | 80517847 | 80583830 | 65984 | -11.24 | 2.78E-26 | 7.83E-25 |
| chr13 | 77097039 | 77178990 | 81952 | 11.19 | 2.80E-26 | 7.86E-25 |
| chrX | 39896281 | 39914959 | 18679 | -11.13 | 2.87E-26 | 8.05E-25 |
| chr5 | 140918553 | 140944863 | 26311 | -11.15 | 2.87E-26 | 8.06E-25 |
| chr2 | 238008048 | 238058615 | 50568 | -11.14 | 2.90E-26 | 8.13E-25 |
| chr10 | 77411067 | 77502681 | 91615 | 11.17 | 2.91E-26 | 8.15E-25 |
| chr8 | 25062070 | 25121803 | 59734 | -11.19 | 2.93E-26 | 8.20E-25 |
| chr9 | 81934609 | 82052970 | 118362 | -11.29 | 2.93E-26 | 8.20E-25 |
| chr17 | 35621029 | 35641092 | 20064 | -11.14 | 2.98E-26 | 8.34E-25 |
| chr4 | 10440949 | 10469408 | 28460 | -11.11 | 3.05E-26 | 8.51E-25 |
| chr20 | 18021443 | 18109440 | 87998 | -11.31 | 3.16E-26 | 8.81E-25 |
| chr15 | 51242740 | 51267972 | 25233 | -11.24 | 3.17E-26 | 8.84E-25 |
| chr15 | 80909476 | 80947676 | 38201 | -11.13 | 3.27E-26 | 9.11E-25 |
| chr21 | 39544653 | 39628089 | 83437 | -11.31 | 3.29E-26 | 9.16E-25 |
| chr7 | 1720532 | 1754828 | 34297 | -11.11 | 3.33E-26 | 9.27E-25 |
| chr3 | 25524520 | 25574769 | 50250 | -11.12 | 3.34E-26 | 9.28E-25 |
| chr8 | 19223416 | 19252662 | 29247 | -11.12 | 3.43E-26 | 9.54E-25 |
| chr10 | 95783764 | 95804573 | 20810 | -11.24 | 3.46E-26 | 9.62E-25 |
| chr12 | 25152019 | 25271866 | 119848 | 2.21 | 3.54E-26 | 9.83E-25 |
| chr6 | 166402071 | 166484415 | 82345 | -11.4 | 3.57E-26 | 9.89E-25 |
| chr20 | 36082216 | 36089688 | 7473 | -6.88 | 3.67E-26 | 1.02E-24 |
| chr14 | 73088751 | 73109707 | 20957 | -3.34 | 3.68E-26 | 1.02E-24 |
| chr17 | 48084711 | 48102523 | 17813 | -11.1 | 3.70E-26 | 1.02E-24 |
| chr4 | 160536595 | 160626111 | 89517 | 3.24 | 3.78E-26 | 1.04E-24 |
| chr15 | 24112441 | 24351972 | 239532 | 1.35 | 3.82E-26 | 1.06E-24 |
| chr7 | 45643148 | 45674339 | 31192 | -11.14 | 3.90E-26 | 1.08E-24 |
| chr1 | 203920526 | 203945822 | 25297 | -11.1 | 3.94E-26 | 1.09E-24 |
| chr5 | 5246963 | 5315398 | 68436 | -11.21 | 3.96E-26 | 1.09E-24 |
| chr4 | 14654682 | 14756832 | 102151 | -11.23 | 4.03E-26 | 1.11E-24 |
| chr7 | 93787109 | 93828447 | 41339 | -11.09 | 4.06E-26 | 1.12E-24 |
| chr12 | 84669953 | 84859265 | 189313 | -11.29 | 4.10E-26 | 1.13E-24 |
| chr14 | 89447117 | 89510400 | 63284 | -11.13 | 4.16E-26 | 1.14E-24 |
| chr16 | 27107061 | 27206751 | 99691 | -11.11 | 4.20E-26 | 1.15E-24 |
| chrX | 93930318 | 94595670 | 665353 | 1.88 | 4.27E-26 | 1.17E-24 |
| chr6 | 106003519 | 106082733 | 79215 | -11.08 | 4.36E-26 | 1.20E-24 |
| chr1 | 234176698 | 234226836 | 50139 | -11.12 | 4.54E-26 | 1.24E-24 |
| chr3 | 121533913 | 121552408 | 18496 | -11.11 | 4.57E-26 | 1.25E-24 |
| chr5 | 124728670 | 124801218 | 72549 | -11.4 | 4.59E-26 | 1.26E-24 |
| chr13 | 95741460 | 95763408 | 21949 | -11.08 | 4.64E-26 | 1.27E-24 |

| | | | | | | |
|---|---|---|---|---|---|---|
| chrX | 115185325 | 115265864 | 80540 | 2.45 | 4.80E-26 | 1.31E-24 |
| chr8 | 145783268 | 145842390 | 59123 | -11.34 | 4.81E-26 | 1.31E-24 |
| chr20 | 33767365 | 33794873 | 27509 | -11.11 | 4.86E-26 | 1.33E-24 |
| chr3 | 118854979 | 118922853 | 67875 | -11.22 | 4.87E-26 | 1.33E-24 |
| chr20 | 58161678 | 58171537 | 9860 | -3.01 | 4.91E-26 | 1.34E-24 |
| chr14 | 67395758 | 67428428 | 32671 | -11.07 | 5.06E-26 | 1.38E-24 |
| chr11 | 1331444 | 1357037 | 25594 | -11.15 | 5.12E-26 | 1.39E-24 |
| chr4 | 149522263 | 150151089 | 628827 | 3.4 | 5.57E-26 | 1.51E-24 |
| chr9 | 126586630 | 126610848 | 24219 | -11.08 | 5.66E-26 | 1.54E-24 |
| chr7 | 79786150 | 79814305 | 28156 | -11.19 | 5.67E-26 | 1.54E-24 |
| chr3 | 194110007 | 194113889 | 3883 | -7.26 | 5.73E-26 | 1.56E-24 |
| chrX | 115134092 | 115164854 | 30763 | 3.57 | 5.75E-26 | 1.56E-24 |
| chr2 | 75462362 | 75487516 | 25155 | -11.12 | 5.79E-26 | 1.57E-24 |
| chr7 | 32040339 | 32059211 | 18873 | -11.07 | 5.87E-26 | 1.59E-24 |
| chr12 | 94736791 | 94759928 | 23138 | -11.06 | 5.98E-26 | 1.62E-24 |
| chr5 | 65382034 | 65411152 | 29119 | -11.11 | 6.11E-26 | 1.65E-24 |
| chr9 | 96732138 | 96798458 | 66321 | -11.15 | 6.14E-26 | 1.66E-24 |
| chr6 | 104982818 | 105049642 | 66825 | 11.09 | 6.15E-26 | 1.66E-24 |
| chr10 | 33254313 | 33306395 | 52083 | -11.07 | 6.42E-26 | 1.73E-24 |
| chr3 | 127048341 | 127103494 | 55154 | -11.05 | 6.43E-26 | 1.73E-24 |
| chr9 | 75613705 | 75648950 | 35246 | -11.16 | 6.77E-26 | 1.82E-24 |
| chr2 | 225920427 | 226000102 | 79676 | -11.09 | 6.95E-26 | 1.87E-24 |
| chr7 | 97720533 | 97732840 | 12308 | -11.04 | 7.01E-26 | 1.89E-24 |
| chr20 | 18309820 | 18313889 | 4070 | -8.48 | 7.25E-26 | 1.95E-24 |
| chr5 | 54149381 | 54158734 | 9354 | -3.48 | 7.35E-26 | 1.98E-24 |
| chr8 | 19591638 | 19611223 | 19586 | -11.05 | 7.51E-26 | 2.02E-24 |
| chr2 | 36426079 | 36439041 | 12963 | -4.77 | 7.52E-26 | 2.02E-24 |
| chr20 | 36575049 | 36598541 | 23493 | -11.08 | 7.55E-26 | 2.02E-24 |
| chr17 | 21228958 | 21241292 | 12335 | -11.04 | 7.55E-26 | 2.02E-24 |
| chr2 | 39880040 | 39931173 | 51134 | -11.06 | 7.57E-26 | 2.03E-24 |
| chr9 | 98157924 | 98194711 | 36788 | -11.09 | 7.63E-26 | 2.04E-24 |
| chr4 | 121093403 | 121163514 | 70112 | 11.07 | 7.69E-26 | 2.05E-24 |
| chr2 | 86165255 | 86200486 | 35232 | -11.03 | 7.93E-26 | 2.12E-24 |
| chr16 | 58304003 | 58349807 | 45805 | -11.1 | 7.94E-26 | 2.12E-24 |
| chr8 | 554913 | 608711 | 53799 | -11.12 | 8.00E-26 | 2.13E-24 |
| chr5 | 17250812 | 17339193 | 88382 | -11.06 | 8.14E-26 | 2.17E-24 |
| chr17 | 81087349 | 81095441 | 8093 | -4.37 | 8.19E-26 | 2.18E-24 |
| chr5 | 81660670 | 81702404 | 41735 | -11.03 | 8.34E-26 | 2.22E-24 |
| chr2 | 103161668 | 103203742 | 42075 | -11.04 | 8.34E-26 | 2.22E-24 |
| chr5 | 32746285 | 32778681 | 32397 | -11.04 | 8.43E-26 | 2.24E-24 |
| chr5 | 142742287 | 142768778 | 26492 | -11.03 | 8.48E-26 | 2.25E-24 |
| chr10 | 80637135 | 80658375 | 21241 | -11.05 | 8.73E-26 | 2.32E-24 |
| chr20 | 18259994 | 18281311 | 21318 | -11.02 | 8.87E-26 | 2.35E-24 |
| chr11 | 2803435 | 2852723 | 49289 | -11.12 | 8.98E-26 | 2.38E-24 |
| chr2 | 220516106 | 220585228 | 69123 | -2.13 | 9.11E-26 | 2.41E-24 |
| chr6 | 12452718 | 12525361 | 72644 | -11.19 | 9.20E-26 | 2.44E-24 |
| chr9 | 91844718 | 91891887 | 47170 | -11.02 | 9.21E-26 | 2.44E-24 |
| chr3 | 98566756 | 98602278 | 35523 | -11.07 | 9.57E-26 | 2.53E-24 |
| chr6 | 78584616 | 78834756 | 250141 | 1.98 | 9.67E-26 | 2.55E-24 |
| chr11 | 56827336 | 56885570 | 58235 | -11.09 | 9.77E-26 | 2.58E-24 |
| chr22 | 46771160 | 46773174 | 2015 | -7.47 | 9.86E-26 | 2.60E-24 |
| chr2 | 242518359 | 242529181 | 10823 | -11.01 | 9.96E-26 | 2.62E-24 |
| chr9 | 18262370 | 18293473 | 31104 | -11.01 | 1.01E-25 | 2.67E-24 |
| chr7 | 130087156 | 130133211 | 46056 | -11 | 1.06E-25 | 2.78E-24 |
| chr2 | 26438921 | 26453363 | 14443 | -11.07 | 1.09E-25 | 2.86E-24 |
| chr12 | 47061347 | 47158306 | 96960 | -11.26 | 1.11E-25 | 2.92E-24 |
| chr12 | 41311504 | 41315760 | 4257 | -7.79 | 1.12E-25 | 2.94E-24 |
| chr10 | 118784697 | 118819090 | 34394 | -11.03 | 1.16E-25 | 3.04E-24 |
| chr5 | 100383872 | 100479491 | 95620 | -11.24 | 1.16E-25 | 3.04E-24 |
| chr21 | 38684025 | 38693521 | 9497 | -8.41 | 1.17E-25 | 3.07E-24 |
| chr2 | 188353058 | 188403570 | 50513 | -11.53 | 1.17E-25 | 3.07E-24 |

| | | | | | | |
|---|---|---|---|---|---|---|
| chr15 | 80165725 | 80174182 | 8458 | -11.06 | 1.18E-25 | 3.08E-24 |
| chr7 | 158076135 | 158122514 | 46380 | -11.01 | 1.18E-25 | 3.09E-24 |
| chr2 | 101513091 | 101544067 | 30977 | -11.02 | 1.21E-25 | 3.17E-24 |
| chr10 | 133492720 | 133506799 | 14080 | -3.85 | 1.22E-25 | 3.19E-24 |
| chr15 | 53139829 | 53471109 | 331281 | 2.01 | 1.23E-25 | 3.20E-24 |
| chr19 | 43791830 | 43834549 | 42720 | -11.01 | 1.27E-25 | 3.32E-24 |
| chr3 | 40560255 | 40593990 | 33736 | -11 | 1.28E-25 | 3.32E-24 |
| chr2 | 109423905 | 109468939 | 45035 | -11.07 | 1.28E-25 | 3.32E-24 |
| chr5 | 146743973 | 146823587 | 79615 | 11.14 | 1.29E-25 | 3.35E-24 |
| chr2 | 241461394 | 241484142 | 22749 | -11.01 | 1.31E-25 | 3.40E-24 |
| chr15 | 101333568 | 101378500 | 44933 | -11.08 | 1.31E-25 | 3.42E-24 |
| chr18 | 7269436 | 7315558 | 46123 | -10.99 | 1.37E-25 | 3.55E-24 |
| chr2 | 29555856 | 30134535 | 578680 | 1.25 | 1.43E-25 | 3.72E-24 |
| chr14 | 51002486 | 51045239 | 42754 | -11.04 | 1.46E-25 | 3.78E-24 |
| chr19 | 6147458 | 6196649 | 49192 | -11.13 | 1.51E-25 | 3.92E-24 |
| chr3 | 70294127 | 70334164 | 40038 | -11.01 | 1.53E-25 | 3.97E-24 |
| chr20 | 22618877 | 22696195 | 77319 | -11.2 | 1.53E-25 | 3.97E-24 |
| chr6 | 106100966 | 106164205 | 63240 | -11.22 | 1.53E-25 | 3.97E-24 |
| chr7 | 149374095 | 149411104 | 37010 | -11.03 | 1.54E-25 | 3.99E-24 |
| chr9 | 6134488 | 6186257 | 51770 | -10.97 | 1.55E-25 | 4.01E-24 |
| chr2 | 240621712 | 240638631 | 16920 | -10.99 | 1.56E-25 | 4.02E-24 |
| chr7 | 130539386 | 130551302 | 11917 | -11.03 | 1.56E-25 | 4.03E-24 |
| chr7 | 37962513 | 37978648 | 16136 | -11.01 | 1.69E-25 | 4.37E-24 |
| chr7 | 116036659 | 116119675 | 83017 | -11.02 | 1.70E-25 | 4.39E-24 |
| chr2 | 240821501 | 240875382 | 53882 | -11.03 | 1.71E-25 | 4.39E-24 |
| chr9 | 19545424 | 19905975 | 360552 | 3.91 | 1.72E-25 | 4.44E-24 |
| chr4 | 129975731 | 130007116 | 31386 | -10.96 | 1.74E-25 | 4.46E-24 |
| chr8 | 25376452 | 25386002 | 9551 | -7.38 | 1.75E-25 | 4.50E-24 |
| chr5 | 179571573 | 179593503 | 21931 | -10.96 | 1.77E-25 | 4.55E-24 |
| chr5 | 129827976 | 129865199 | 37224 | -10.97 | 1.80E-25 | 4.63E-24 |
| chr10 | 112780390 | 112828959 | 48570 | -10.96 | 1.87E-25 | 4.79E-24 |
| chr7 | 90008310 | 90031175 | 22866 | -10.96 | 1.94E-25 | 4.97E-24 |
| chr10 | 75064470 | 75094388 | 29919 | -11.05 | 1.96E-25 | 5.02E-24 |
| chr12 | 14746570 | 14791683 | 45114 | -10.96 | 2.01E-25 | 5.14E-24 |
| chr17 | 55450264 | 55472436 | 22173 | -10.97 | 2.04E-25 | 5.22E-24 |
| chr4 | 190717811 | 190747106 | 29296 | -3.02 | 2.10E-25 | 5.37E-24 |
| chr2 | 48563774 | 48583330 | 19557 | -10.94 | 2.11E-25 | 5.38E-24 |
| chr2 | 46427547 | 46454621 | 27075 | -10.96 | 2.11E-25 | 5.38E-24 |
| chr2 | 7879037 | 7905488 | 26452 | -11.08 | 2.14E-25 | 5.46E-24 |
| chr2 | 31034748 | 31101811 | 67064 | -2.32 | 2.21E-25 | 5.64E-24 |
| chr20 | 13028948 | 13076869 | 47922 | -2.25 | 2.22E-25 | 5.66E-24 |
| chr17 | 78272824 | 78280754 | 7931 | -10.94 | 2.23E-25 | 5.67E-24 |
| chr2 | 168713812 | 168771079 | 57268 | 10.99 | 2.25E-25 | 5.73E-24 |
| chr9 | 98698822 | 98723040 | 24219 | -10.98 | 2.28E-25 | 5.81E-24 |
| chr9 | 94677055 | 94758930 | 81876 | -11.21 | 2.34E-25 | 5.95E-24 |
| chr2 | 157561132 | 157654977 | 93846 | 10.96 | 2.50E-25 | 6.34E-24 |
| chr2 | 241289 | 264243 | 22955 | -10.93 | 2.51E-25 | 6.36E-24 |
| chr2 | 174180082 | 174212640 | 32559 | -10.95 | 2.53E-25 | 6.40E-24 |
| chr7 | 81243686 | 81303866 | 60181 | -11.2 | 2.55E-25 | 6.47E-24 |
| chr22 | 43967784 | 44018954 | 51171 | -10.95 | 2.60E-25 | 6.58E-24 |
| chr4 | 155502634 | 155522866 | 20233 | -8.35 | 2.61E-25 | 6.60E-24 |
| chr1 | 183929375 | 183982620 | 53246 | 10.96 | 2.62E-25 | 6.64E-24 |
| chr7 | 347031 | 358934 | 11904 | -11 | 2.67E-25 | 6.74E-24 |
| chr3 | 46500659 | 46518604 | 17946 | -3.5 | 2.67E-25 | 6.75E-24 |
| chr9 | 118072902 | 118115904 | 43003 | -10.92 | 2.68E-25 | 6.76E-24 |
| chr2 | 190829885 | 190886941 | 57057 | -11.16 | 2.68E-25 | 6.76E-24 |
| chr14 | 44763818 | 45292478 | 528661 | 1.27 | 2.73E-25 | 6.88E-24 |
| chr4 | 82331661 | 82380999 | 49339 | 10.97 | 2.76E-25 | 6.94E-24 |
| chr7 | 4092269 | 4134494 | 42226 | -10.97 | 2.80E-25 | 7.04E-24 |
| chr2 | 69226511 | 69231538 | 5028 | -3.85 | 2.80E-25 | 7.05E-24 |
| chr9 | 140230033 | 140271000 | 40968 | -11.06 | 2.84E-25 | 7.13E-24 |

| | | | | | | |
|---|---|---|---|---|---|---|
| chr11 | 88892723 | 88972933 | 80211 | 3.51 | 2.85E-25 | 7.15E-24 |
| chr2 | 44533712 | 44542062 | 8351 | -10.95 | 2.88E-25 | 7.23E-24 |
| chr9 | 35493107 | 35510700 | 17594 | -10.91 | 2.90E-25 | 7.27E-24 |
| chr9 | 116773873 | 116788141 | 14269 | -10.95 | 2.97E-25 | 7.46E-24 |
| chr9 | 83874414 | 83959586 | 85173 | -11.02 | 2.98E-25 | 7.46E-24 |
| chr11 | 9885335 | 9934644 | 49310 | -11.04 | 2.98E-25 | 7.47E-24 |
| chr7 | 89978789 | 89995373 | 16585 | -10.94 | 3.00E-25 | 7.51E-24 |
| chr5 | 148474314 | 148482596 | 8283 | -2.68 | 3.02E-25 | 7.54E-24 |
| chr2 | 30919407 | 30953518 | 34112 | -10.98 | 3.02E-25 | 7.55E-24 |
| chr12 | 43716431 | 43784107 | 67677 | -10.91 | 3.12E-25 | 7.79E-24 |
| chr2 | 137018334 | 137032668 | 14335 | -7.51 | 3.13E-25 | 7.82E-24 |
| chr15 | 58315304 | 58319770 | 4467 | -3.15 | 3.19E-25 | 7.95E-24 |
| chr17 | 75642271 | 75671125 | 28855 | -10.91 | 3.23E-25 | 8.05E-24 |
| chr5 | 126579219 | 126625509 | 46291 | -10.97 | 3.26E-25 | 8.11E-24 |
| chr7 | 67546069 | 67625221 | 79153 | -11.12 | 3.27E-25 | 8.13E-24 |
| chr4 | 95545943 | 95616490 | 70548 | -10.9 | 3.31E-25 | 8.24E-24 |
| chr12 | 78179202 | 78184008 | 4807 | -7.33 | 3.39E-25 | 8.41E-24 |
| chr2 | 163467162 | 163571366 | 104205 | 11.25 | 3.39E-25 | 8.41E-24 |
| chrX | 35442762 | 35503364 | 60603 | -10.93 | 3.41E-25 | 8.46E-24 |
| chr3 | 169284633 | 169371870 | 87238 | 11.09 | 3.44E-25 | 8.53E-24 |
| chr11 | 98254115 | 98831261 | 577147 | 1.92 | 3.45E-25 | 8.55E-24 |
| chr12 | 76456820 | 76475495 | 18676 | -10.93 | 3.50E-25 | 8.68E-24 |
| chr7 | 46010884 | 46058295 | 47412 | -10.91 | 3.57E-25 | 8.85E-24 |
| chr2 | 174367019 | 174422887 | 55869 | -10.89 | 3.61E-25 | 8.93E-24 |
| chr15 | 85652379 | 85676184 | 23806 | -10.89 | 3.63E-25 | 8.98E-24 |
| chr2 | 5119308 | 5764740 | 645433 | 0.82 | 3.71E-25 | 9.16E-24 |
| chr6 | 114521019 | 114593760 | 72742 | 11 | 3.77E-25 | 9.32E-24 |
| chr14 | 104586858 | 104597649 | 10792 | -10.89 | 3.83E-25 | 9.45E-24 |
| chr14 | 73214207 | 73246215 | 32009 | -10.9 | 3.83E-25 | 9.45E-24 |
| chr8 | 2052018 | 2103112 | 51095 | -3.02 | 3.85E-25 | 9.49E-24 |
| chr9 | 21814272 | 21832340 | 18069 | -10.94 | 3.88E-25 | 9.56E-24 |
| chr6 | 79624567 | 79664869 | 40303 | -10.89 | 3.94E-25 | 9.70E-24 |
| chr9 | 4218110 | 4258477 | 40368 | -10.97 | 3.98E-25 | 9.79E-24 |
| chr7 | 68157347 | 68240837 | 83491 | -10.91 | 4.04E-25 | 9.92E-24 |
| chr12 | 87014024 | 87064618 | 50595 | -10.89 | 4.09E-25 | 1.01E-23 |
| chr3 | 149538454 | 149557680 | 19227 | -10.9 | 4.10E-25 | 1.01E-23 |
| chr9 | 113477571 | 113523516 | 45946 | -11.01 | 4.19E-25 | 1.03E-23 |
| chr8 | 122812543 | 122859409 | 46867 | -11.13 | 4.19E-25 | 1.03E-23 |
| chr14 | 99510613 | 99581368 | 70756 | -10.88 | 4.20E-25 | 1.03E-23 |
| chr9 | 118129593 | 118198659 | 69067 | -11.03 | 4.20E-25 | 1.03E-23 |
| chr12 | 10605642 | 10641323 | 35682 | -10.88 | 4.26E-25 | 1.04E-23 |
| chr6 | 17130121 | 17165839 | 35719 | -10.91 | 4.30E-25 | 1.05E-23 |
| chr10 | 29537911 | 29604091 | 66181 | -11.08 | 4.32E-25 | 1.06E-23 |
| chr7 | 141445767 | 141454182 | 8416 | -10.98 | 4.38E-25 | 1.07E-23 |
| chr19 | 33717612 | 33721862 | 4251 | -10.88 | 4.41E-25 | 1.08E-23 |
| chr20 | 36643859 | 36649377 | 5519 | -10.88 | 4.50E-25 | 1.10E-23 |
| chrX | 9672314 | 9725197 | 52884 | -10.87 | 4.50E-25 | 1.10E-23 |
| chr3 | 30958323 | 31008414 | 50092 | -11.03 | 4.57E-25 | 1.11E-23 |
| chr7 | 91797521 | 91837579 | 40059 | -10.87 | 4.59E-25 | 1.12E-23 |
| chr8 | 143709662 | 143754865 | 45204 | -10.98 | 4.69E-25 | 1.14E-23 |
| chr15 | 102054449 | 102061399 | 6951 | -10.87 | 4.79E-25 | 1.16E-23 |
| chr7 | 158295223 | 158328582 | 33360 | -10.87 | 4.79E-25 | 1.16E-23 |
| chr13 | 23903750 | 23926325 | 22576 | -10.92 | 4.86E-25 | 1.18E-23 |
| chr11 | 14021437 | 14029293 | 7857 | -4.94 | 4.88E-25 | 1.18E-23 |
| chr20 | 2584850 | 2624909 | 40060 | -10.99 | 4.88E-25 | 1.18E-23 |
| chr4 | 22965963 | 23040784 | 74822 | -10.87 | 4.91E-25 | 1.19E-23 |
| chr3 | 10735011 | 10780009 | 44999 | -10.89 | 4.91E-25 | 1.19E-23 |
| chr10 | 31738154 | 31799509 | 61356 | -10.87 | 5.02E-25 | 1.22E-23 |
| chrX | 10226657 | 10293379 | 66723 | -10.9 | 5.14E-25 | 1.24E-23 |
| chr1 | 234000111 | 234066667 | 66557 | -10.86 | 5.17E-25 | 1.25E-23 |
| chr3 | 189197388 | 189276781 | 79394 | -10.86 | 5.17E-25 | 1.25E-23 |

| | | | | | | |
|---|---|---|---|---|---|---|
| chr14 | 93049653 | 93095988 | 46336 | -10.88 | 5.19E-25 | 1.25E-23 |
| chr2 | 218949102 | 219023090 | 73989 | -3.05 | 5.22E-25 | 1.26E-23 |
| chr12 | 450790 | 476282 | 25493 | -10.86 | 5.26E-25 | 1.27E-23 |
| chr6 | 101283410 | 101313608 | 30199 | -10.89 | 5.26E-25 | 1.27E-23 |
| chr16 | 14555584 | 14576025 | 20442 | -10.88 | 5.35E-25 | 1.29E-23 |
| chr3 | 149855558 | 149936338 | 80781 | -10.92 | 5.39E-25 | 1.30E-23 |
| chr8 | 17844853 | 17855417 | 10565 | -10.91 | 5.40E-25 | 1.30E-23 |
| chr3 | 31065835 | 31114851 | 49017 | -10.86 | 5.50E-25 | 1.32E-23 |
| chr13 | 95068936 | 95074518 | 5583 | -5.03 | 5.54E-25 | 1.33E-23 |
| chr12 | 46939049 | 47011504 | 72456 | -11.1 | 5.65E-25 | 1.36E-23 |
| chr3 | 133796709 | 133858972 | 62264 | -10.85 | 5.66E-25 | 1.36E-23 |
| chr19 | 30382596 | 30429533 | 46938 | -10.98 | 5.72E-25 | 1.37E-23 |
| chr4 | 25200669 | 25235452 | 34784 | -10.91 | 5.83E-25 | 1.40E-23 |
| chr7 | 103369415 | 103438998 | 69584 | 10.93 | 5.86E-25 | 1.40E-23 |
| chr19 | 42074134 | 42101415 | 27282 | -10.85 | 5.89E-25 | 1.41E-23 |
| chr12 | 87079499 | 87126773 | 47275 | -10.9 | 5.94E-25 | 1.42E-23 |
| chr20 | 8866628 | 8906366 | 39739 | -10.85 | 5.96E-25 | 1.42E-23 |
| chr17 | 9490276 | 9547784 | 57509 | -10.86 | 5.99E-25 | 1.43E-23 |
| chr2 | 122424823 | 122441794 | 16972 | -11.05 | 6.04E-25 | 1.44E-23 |
| chr19 | 32841930 | 32867748 | 25819 | -10.84 | 6.06E-25 | 1.45E-23 |
| chr8 | 102858657 | 102994994 | 136338 | -11.37 | 6.12E-25 | 1.46E-23 |
| chr10 | 8967823 | 9040669 | 72847 | -10.85 | 6.22E-25 | 1.48E-23 |
| chr7 | 107832205 | 107883997 | 51793 | -10.92 | 6.23E-25 | 1.48E-23 |
| chr2 | 129211645 | 129251840 | 40196 | -10.87 | 6.26E-25 | 1.49E-23 |
| chr2 | 107206914 | 107249032 | 42119 | -1.92 | 6.30E-25 | 1.50E-23 |
| chr9 | 95741366 | 95763897 | 22532 | -10.88 | 6.33E-25 | 1.50E-23 |
| chr17 | 80689632 | 80708374 | 18743 | -10.87 | 6.33E-25 | 1.50E-23 |
| chr15 | 31660724 | 31720433 | 59710 | -10.98 | 6.39E-25 | 1.52E-23 |
| chr20 | 23193893 | 23222309 | 28417 | -10.84 | 6.40E-25 | 1.52E-23 |
| chr14 | 94533516 | 94552696 | 19181 | -10.84 | 6.40E-25 | 1.52E-23 |
| chr16 | 87111823 | 87121423 | 9601 | -3.28 | 6.44E-25 | 1.52E-23 |
| chr5 | 127356619 | 127392112 | 35494 | -10.86 | 6.55E-25 | 1.55E-23 |
| chr8 | 60062572 | 60781470 | 718899 | 1.59 | 6.60E-25 | 1.56E-23 |
| chr5 | 13488913 | 13572771 | 83859 | -10.84 | 6.63E-25 | 1.57E-23 |
| chr3 | 70398709 | 70440047 | 41339 | -10.85 | 6.67E-25 | 1.58E-23 |
| chr11 | 68875214 | 68956440 | 81227 | -11.04 | 6.71E-25 | 1.58E-23 |
| chr3 | 14260561 | 14311350 | 50790 | -10.9 | 6.72E-25 | 1.59E-23 |
| chr15 | 99568155 | 99587006 | 18852 | -10.83 | 6.76E-25 | 1.60E-23 |
| chr10 | 129695010 | 129702083 | 7074 | -8.21 | 6.84E-25 | 1.61E-23 |
| chr12 | 51943948 | 51953426 | 9479 | -2.8 | 6.86E-25 | 1.62E-23 |
| chr8 | 120160064 | 120245377 | 85314 | 10.93 | 7.06E-25 | 1.66E-23 |
| chr2 | 205237924 | 205312647 | 74724 | -10.93 | 7.19E-25 | 1.69E-23 |
| chr14 | 32590840 | 32611205 | 20366 | -10.83 | 7.20E-25 | 1.69E-23 |
| chr7 | 3135887 | 3178687 | 42801 | -10.84 | 7.23E-25 | 1.70E-23 |
| chr3 | 32024752 | 32052259 | 27508 | -11.02 | 7.23E-25 | 1.70E-23 |
| chr2 | 224193059 | 224276371 | 83313 | -10.83 | 7.38E-25 | 1.73E-23 |
| chr2 | 238978913 | 238990587 | 11675 | -10.83 | 7.39E-25 | 1.73E-23 |
| chr22 | 49774870 | 49817368 | 42499 | -10.82 | 7.61E-25 | 1.78E-23 |
| chr9 | 137875392 | 137897063 | 21672 | -10.84 | 8.21E-25 | 1.92E-23 |
| chr1 | 233090841 | 233112010 | 21170 | -10.9 | 8.37E-25 | 1.96E-23 |
| chr5 | 60828135 | 60852557 | 24423 | -10.82 | 8.39E-25 | 1.96E-23 |
| chr11 | 130598293 | 130621514 | 23222 | -10.83 | 8.43E-25 | 1.97E-23 |
| chr6 | 117323690 | 117408160 | 84471 | 10.93 | 8.57E-25 | 2.00E-23 |
| chr10 | 101496226 | 101503285 | 7060 | -10.97 | 8.73E-25 | 2.04E-23 |
| chr17 | 55224568 | 55258168 | 33601 | -10.9 | 8.74E-25 | 2.04E-23 |
| chr14 | 68883946 | 68956032 | 72087 | -10.82 | 8.95E-25 | 2.09E-23 |
| chr12 | 58902192 | 58933048 | 30857 | -10.83 | 8.95E-25 | 2.09E-23 |
| chr15 | 85592524 | 85605588 | 13065 | -10.83 | 8.98E-25 | 2.09E-23 |
| chr22 | 50076603 | 50106121 | 29519 | -10.84 | 9.18E-25 | 2.14E-23 |
| chr22 | 46089069 | 46139379 | 50311 | -10.87 | 9.20E-25 | 2.14E-23 |
| chr5 | 156426197 | 156447158 | 20962 | -10.82 | 9.23E-25 | 2.15E-23 |

| | | | | | | |
|---|---|---|---|---|---|---|
| chr9 | 91026769 | 91040526 | 13758 | -10.85 | 9.32E-25 | 2.17E-23 |
| chr1 | 40566119 | 40597321 | 31203 | -10.81 | 9.56E-25 | 2.22E-23 |
| chr12 | 131201777 | 131250866 | 49090 | -10.89 | 9.62E-25 | 2.23E-23 |
| chr5 | 154242874 | 154269224 | 26351 | -10.82 | 9.65E-25 | 2.24E-23 |
| chr2 | 9994409 | 10002440 | 8032 | -10.83 | 9.78E-25 | 2.27E-23 |
| chr2 | 19315425 | 19385698 | 70274 | -10.99 | 9.80E-25 | 2.27E-23 |
| chr2 | 64126696 | 64135186 | 8491 | -10.81 | 9.81E-25 | 2.27E-23 |
| chr8 | 145992419 | 146006848 | 14430 | -10.92 | 9.91E-25 | 2.29E-23 |
| chr2 | 119761819 | 119823964 | 62146 | -10.82 | 9.91E-25 | 2.29E-23 |
| chr12 | 17182945 | 17279151 | 96207 | -11.39 | 9.96E-25 | 2.30E-23 |
| chr2 | 114681557 | 114703679 | 22123 | -10.8 | 1.01E-24 | 2.32E-23 |
| chr3 | 140703229 | 140726482 | 23254 | -7.72 | 1.01E-24 | 2.32E-23 |
| chr9 | 103066543 | 103081087 | 14545 | -10.8 | 1.01E-24 | 2.34E-23 |
| chr10 | 77865218 | 78616227 | 751010 | 1.71 | 1.03E-24 | 2.38E-23 |
| chr2 | 159183126 | 159192600 | 9475 | -4.34 | 1.05E-24 | 2.42E-23 |
| chr14 | 81310294 | 81360354 | 50061 | -10.83 | 1.05E-24 | 2.42E-23 |
| chr18 | 77837365 | 77859774 | 22410 | -10.79 | 1.06E-24 | 2.43E-23 |
| chr13 | 77469265 | 77475241 | 5977 | -4.27 | 1.08E-24 | 2.48E-23 |
| chr1 | 233691496 | 233722389 | 30894 | -10.79 | 1.08E-24 | 2.48E-23 |
| chr10 | 13576858 | 13624914 | 48057 | -10.95 | 1.09E-24 | 2.51E-23 |
| chr1 | 166461357 | 166775839 | 314483 | 3.33 | 1.10E-24 | 2.52E-23 |
| chr4 | 143522955 | 143591454 | 68500 | -10.79 | 1.10E-24 | 2.52E-23 |
| chr3 | 103022753 | 103188523 | 165771 | -11.08 | 1.11E-24 | 2.54E-23 |
| chr2 | 18607499 | 18675963 | 68465 | 10.91 | 1.11E-24 | 2.54E-23 |
| chr4 | 171612311 | 172729176 | 1116866 | 2.57 | 1.12E-24 | 2.56E-23 |
| chr12 | 87804702 | 87849395 | 44694 | -2.06 | 1.12E-24 | 2.56E-23 |
| chr13 | 57748216 | 58055132 | 306917 | 2.14 | 1.17E-24 | 2.67E-23 |
| chr4 | 3573154 | 3653142 | 79989 | -10.79 | 1.18E-24 | 2.70E-23 |
| chr5 | 5372291 | 5433386 | 61096 | -10.92 | 1.22E-24 | 2.79E-23 |
| chr12 | 130509437 | 130614681 | 105245 | -10.97 | 1.22E-24 | 2.79E-23 |
| chr11 | 76485014 | 76492371 | 7358 | -10.87 | 1.22E-24 | 2.79E-23 |
| chr7 | 31969106 | 32025922 | 56817 | -10.82 | 1.23E-24 | 2.81E-23 |
| chr5 | 129742076 | 129814076 | 72001 | -10.87 | 1.27E-24 | 2.89E-23 |
| chr1 | 33950817 | 33974202 | 23386 | -10.77 | 1.28E-24 | 2.93E-23 |
| chr14 | 104692783 | 104728592 | 35810 | -10.79 | 1.32E-24 | 3.01E-23 |
| chr4 | 129332777 | 129384793 | 52017 | -11.06 | 1.33E-24 | 3.03E-23 |
| chr20 | 45042447 | 45176327 | 133881 | -2.19 | 1.38E-24 | 3.13E-23 |
| chr4 | 1243779 | 1265262 | 21484 | -10.81 | 1.40E-24 | 3.18E-23 |
| chr5 | 76794997 | 76807799 | 12803 | -3.37 | 1.41E-24 | 3.21E-23 |
| chr5 | 143031726 | 143076628 | 44903 | -10.84 | 1.44E-24 | 3.28E-23 |
| chr16 | 17578513 | 18167512 | 589000 | 1.52 | 1.45E-24 | 3.30E-23 |
| chr2 | 131683913 | 131714474 | 30562 | -10.81 | 1.48E-24 | 3.35E-23 |
| chr20 | 13477109 | 13494854 | 17746 | -10.8 | 1.49E-24 | 3.38E-23 |
| chr7 | 16141086 | 16168285 | 27200 | -10.8 | 1.51E-24 | 3.41E-23 |
| chr18 | 64233803 | 64857990 | 624188 | 2.75 | 1.55E-24 | 3.52E-23 |
| chr3 | 129615334 | 129652043 | 36710 | -10.94 | 1.57E-24 | 3.54E-23 |
| chr13 | 62420212 | 62869861 | 449650 | 2.71 | 1.57E-24 | 3.54E-23 |
| chr12 | 58608364 | 58677888 | 69525 | -10.76 | 1.58E-24 | 3.57E-23 |
| chr7 | 90924489 | 91177589 | 253101 | 2.57 | 1.59E-24 | 3.58E-23 |
| chr19 | 30513055 | 30532322 | 19268 | -10.76 | 1.59E-24 | 3.59E-23 |
| chr10 | 94863174 | 94945559 | 82386 | -11.13 | 1.60E-24 | 3.60E-23 |
| chr11 | 68462186 | 68489681 | 27496 | -10.77 | 1.61E-24 | 3.63E-23 |
| chr8 | 59420708 | 59438981 | 18274 | -9.22 | 1.62E-24 | 3.66E-23 |
| chr22 | 46243733 | 46270787 | 27055 | -10.75 | 1.63E-24 | 3.68E-23 |
| chr8 | 144083515 | 144092278 | 8764 | -8 | 1.65E-24 | 3.71E-23 |
| chr18 | 20229287 | 20280007 | 50721 | -10.76 | 1.67E-24 | 3.76E-23 |
| chr9 | 75459308 | 75505646 | 46339 | -10.82 | 1.71E-24 | 3.83E-23 |
| chr6 | 108233910 | 108254603 | 20694 | -10.75 | 1.76E-24 | 3.94E-23 |
| chr11 | 92910426 | 92925127 | 14702 | -10.82 | 1.79E-24 | 4.01E-23 |
| chr2 | 37981794 | 38049031 | 67238 | -10.96 | 1.80E-24 | 4.04E-23 |
| chr1 | 228173661 | 228187157 | 13497 | -10.76 | 1.82E-24 | 4.07E-23 |

| | | | | | | |
|---|---|---|---|---|---|---|
| chr8 | 19268266 | 19309470 | 41205 | -10.9 | 1.82E-24 | 4.07E-23 |
| chr2 | 29341984 | 29375511 | 33528 | -10.85 | 1.83E-24 | 4.09E-23 |
| chr4 | 102779097 | 102813129 | 34033 | -10.76 | 1.83E-24 | 4.10E-23 |
| chr2 | 143798276 | 143841015 | 42740 | -10.83 | 1.84E-24 | 4.12E-23 |
| chr10 | 34637466 | 34662160 | 24695 | -10.83 | 1.85E-24 | 4.13E-23 |
| chr10 | 87555170 | 87617255 | 62086 | -10.76 | 1.85E-24 | 4.14E-23 |
| chr11 | 48276708 | 48398262 | 121555 | 9.28 | 1.86E-24 | 4.15E-23 |
| chr11 | 34391319 | 34438339 | 47021 | -10.84 | 1.86E-24 | 4.15E-23 |
| chr6 | 12542477 | 12601430 | 58954 | -10.75 | 1.87E-24 | 4.18E-23 |
| chr2 | 189247408 | 189286556 | 39149 | -10.83 | 1.89E-24 | 4.20E-23 |
| chr6 | 40584405 | 40594834 | 10430 | -4.86 | 1.90E-24 | 4.24E-23 |
| chr19 | 17021184 | 17048281 | 27098 | -10.74 | 1.91E-24 | 4.24E-23 |
| chr2 | 240907666 | 240927212 | 19547 | -10.77 | 1.91E-24 | 4.24E-23 |
| chrX | 82239298 | 82303885 | 64588 | -10.74 | 1.97E-24 | 4.37E-23 |
| chr21 | 44952744 | 44988072 | 35329 | -10.85 | 1.97E-24 | 4.37E-23 |
| chr4 | 21556656 | 21616711 | 60056 | 1.89 | 1.99E-24 | 4.42E-23 |
| chr11 | 86419018 | 86474573 | 55556 | -10.8 | 1.99E-24 | 4.42E-23 |
| chr14 | 32425820 | 32496153 | 70334 | -10.93 | 2.03E-24 | 4.51E-23 |
| chr9 | 84130289 | 84180745 | 50457 | -10.75 | 2.04E-24 | 4.51E-23 |
| chr19 | 46792357 | 46834086 | 41730 | -10.83 | 2.04E-24 | 4.51E-23 |
| chr9 | 83735743 | 83808142 | 72400 | -10.87 | 2.04E-24 | 4.52E-23 |
| chr17 | 27603429 | 27613058 | 9630 | -10.95 | 2.04E-24 | 4.52E-23 |
| chr16 | 11638672 | 11653894 | 15223 | -10.73 | 2.05E-24 | 4.53E-23 |
| chr17 | 56369826 | 56377343 | 7518 | -7.31 | 2.06E-24 | 4.56E-23 |
| chr5 | 43443251 | 43473306 | 30056 | -10.75 | 2.13E-24 | 4.71E-23 |
| chr2 | 223046148 | 223106180 | 60033 | 4.12 | 2.14E-24 | 4.73E-23 |
| chr3 | 5787514 | 5837897 | 50384 | -10.76 | 2.17E-24 | 4.78E-23 |
| chr6 | 161402644 | 161406771 | 4128 | -4.44 | 2.20E-24 | 4.85E-23 |
| chr8 | 120493685 | 120556102 | 62418 | 10.78 | 2.20E-24 | 4.86E-23 |
| chr8 | 62228575 | 62239759 | 11185 | -8.21 | 2.21E-24 | 4.87E-23 |
| chr6 | 39916236 | 40303827 | 387592 | 1.44 | 2.23E-24 | 4.90E-23 |
| chr14 | 101948115 | 101983314 | 35200 | -10.79 | 2.28E-24 | 5.01E-23 |
| chr22 | 44894963 | 44923360 | 28398 | -10.72 | 2.28E-24 | 5.01E-23 |
| chr10 | 93359914 | 93418978 | 59065 | -10.78 | 2.32E-24 | 5.09E-23 |
| chr2 | 234318705 | 234340470 | 21766 | -10.84 | 2.37E-24 | 5.21E-23 |
| chr3 | 165579463 | 167138270 | 1558808 | 2.49 | 2.45E-24 | 5.38E-23 |
| chr17 | 70696533 | 70710415 | 13883 | -10.74 | 2.46E-24 | 5.39E-23 |
| chr13 | 94693430 | 94751507 | 58078 | 3.31 | 2.48E-24 | 5.43E-23 |
| chr11 | 6938781 | 7023948 | 85168 | 10.87 | 2.48E-24 | 5.43E-23 |
| chr1 | 71002522 | 71054090 | 51569 | 10.85 | 2.49E-24 | 5.45E-23 |
| chr7 | 54998017 | 55060916 | 62900 | -10.73 | 2.53E-24 | 5.53E-23 |
| chr2 | 151130703 | 151174369 | 43667 | -10.77 | 2.57E-24 | 5.61E-23 |
| chr12 | 51181165 | 51197153 | 15989 | -10.73 | 2.57E-24 | 5.61E-23 |
| chr4 | 38775862 | 38855365 | 79504 | -10.71 | 2.58E-24 | 5.64E-23 |
| chr8 | 93982240 | 94630855 | 648616 | 3.43 | 2.61E-24 | 5.71E-23 |
| chr15 | 71554318 | 71585601 | 31284 | -10.77 | 2.66E-24 | 5.80E-23 |
| chr13 | 49317011 | 49357578 | 40568 | -10.73 | 2.67E-24 | 5.81E-23 |
| chr1 | 31915887 | 31964506 | 48620 | -10.88 | 2.67E-24 | 5.81E-23 |
| chr9 | 81199490 | 81228279 | 28790 | -10.78 | 2.70E-24 | 5.88E-23 |
| chr20 | 6092320 | 6099130 | 6811 | -10.76 | 2.71E-24 | 5.89E-23 |
| chr3 | 43364654 | 43400121 | 35468 | -10.71 | 2.74E-24 | 5.96E-23 |
| chr11 | 92986357 | 93003036 | 16680 | -2.93 | 2.76E-24 | 6.01E-23 |
| chr13 | 109238765 | 109309342 | 70578 | -10.88 | 2.77E-24 | 6.02E-23 |
| chr4 | 15265509 | 15279083 | 13575 | -3.8 | 2.78E-24 | 6.04E-23 |
| chr12 | 14616369 | 14634891 | 18523 | -10.71 | 2.83E-24 | 6.14E-23 |
| chr15 | 91589487 | 91642618 | 53132 | -10.82 | 2.83E-24 | 6.14E-23 |
| chr16 | 27927204 | 28005768 | 78565 | -11 | 2.87E-24 | 6.22E-23 |
| chr5 | 23360744 | 23504089 | 143346 | -10.75 | 2.89E-24 | 6.26E-23 |
| chr2 | 100905175 | 100936679 | 31505 | -10.75 | 2.89E-24 | 6.26E-23 |
| chr20 | 19839802 | 19881270 | 41469 | -10.75 | 2.91E-24 | 6.29E-23 |
| chr2 | 58437038 | 58442133 | 5096 | -7.36 | 2.94E-24 | 6.36E-23 |

| | | | | | | |
|---|---|---|---|---|---|---|
| chr1 | 243976253 | 244003079 | 26827 | -10.7 | 2.97E-24 | 6.42E-23 |
| chr6 | 28331630 | 28361605 | 29976 | -10.77 | 2.97E-24 | 6.43E-23 |
| chr5 | 127261186 | 127335142 | 73957 | -10.89 | 2.98E-24 | 6.44E-23 |
| chr8 | 120332606 | 120388455 | 55850 | 10.76 | 3.00E-24 | 6.47E-23 |
| chr3 | 39338447 | 39362399 | 23953 | -10.7 | 3.11E-24 | 6.71E-23 |
| chr2 | 136079931 | 136111523 | 31593 | -10.7 | 3.12E-24 | 6.73E-23 |
| chr3 | 156903126 | 156914760 | 11635 | -2.84 | 3.12E-24 | 6.73E-23 |
| chr8 | 11107177 | 11170922 | 63746 | -10.75 | 3.13E-24 | 6.74E-23 |
| chr5 | 6620665 | 6631238 | 10574 | -10.79 | 3.14E-24 | 6.75E-23 |
| chr12 | 41092619 | 41153203 | 60585 | -10.74 | 3.22E-24 | 6.92E-23 |
| chr2 | 161980518 | 162015615 | 35098 | -10.71 | 3.23E-24 | 6.95E-23 |
| chr5 | 167937930 | 167943972 | 6043 | -10.73 | 3.24E-24 | 6.95E-23 |
| chrX | 45090871 | 45147175 | 56305 | -9.07 | 3.24E-24 | 6.95E-23 |
| chr10 | 29381103 | 29437225 | 56123 | -10.91 | 3.25E-24 | 6.97E-23 |
| chr2 | 178313154 | 178345037 | 31884 | -10.79 | 3.30E-24 | 7.08E-23 |
| chr20 | 14364045 | 14392470 | 28426 | -10.79 | 3.31E-24 | 7.10E-23 |
| chr12 | 46129467 | 46138061 | 8595 | -10.69 | 3.33E-24 | 7.13E-23 |
| chr8 | 53692594 | 53739877 | 47284 | -10.73 | 3.34E-24 | 7.16E-23 |
| chr15 | 67195013 | 67242583 | 47571 | -11.02 | 3.37E-24 | 7.21E-23 |
| chr15 | 90170823 | 90186394 | 15572 | -10.69 | 3.38E-24 | 7.23E-23 |
| chr17 | 61348734 | 61389717 | 40984 | -10.69 | 3.39E-24 | 7.23E-23 |
| chr2 | 63493289 | 63531478 | 38190 | -10.83 | 3.40E-24 | 7.27E-23 |
| chr15 | 85160149 | 85168324 | 8176 | -10.7 | 3.42E-24 | 7.30E-23 |
| chr17 | 10094570 | 10185757 | 91188 | -2.23 | 3.53E-24 | 7.52E-23 |
| chr3 | 96185163 | 96304748 | 119586 | -10.7 | 3.60E-24 | 7.67E-23 |
| chr7 | 134141465 | 134173060 | 31596 | -10.72 | 3.62E-24 | 7.72E-23 |
| chr2 | 26300654 | 26308166 | 7513 | -10.76 | 3.63E-24 | 7.74E-23 |
| chr3 | 161087866 | 161139819 | 51954 | -10.78 | 3.65E-24 | 7.78E-23 |
| chr7 | 126900135 | 126952982 | 52848 | 10.74 | 3.76E-24 | 7.99E-23 |
| chr9 | 5704908 | 5723028 | 18121 | -10.68 | 3.83E-24 | 8.14E-23 |
| chr9 | 2907162 | 3137154 | 229993 | 1.72 | 3.83E-24 | 8.14E-23 |
| chr3 | 67986009 | 68905155 | 919147 | 5.95 | 3.88E-24 | 8.24E-23 |
| chr1 | 15131226 | 15181922 | 50697 | -10.67 | 3.89E-24 | 8.26E-23 |
| chr2 | 239360013 | 239390683 | 30671 | -10.69 | 3.93E-24 | 8.34E-23 |
| chr2 | 46937189 | 46958726 | 21538 | -10.7 | 3.94E-24 | 8.35E-23 |
| chr20 | 11392194 | 11439398 | 47205 | -10.86 | 3.97E-24 | 8.41E-23 |
| chr2 | 230256190 | 230301929 | 45740 | -10.77 | 3.99E-24 | 8.45E-23 |
| chr20 | 721784 | 737108 | 15325 | -10.77 | 3.99E-24 | 8.45E-23 |
| chr2 | 217998339 | 218044122 | 45784 | -10.67 | 4.05E-24 | 8.57E-23 |
| chr9 | 102222607 | 102532305 | 309699 | 4.15 | 4.09E-24 | 8.64E-23 |
| chr2 | 221905675 | 221994778 | 89104 | -10.89 | 4.11E-24 | 8.68E-23 |
| chr2 | 101939900 | 101943726 | 3827 | -7.08 | 4.11E-24 | 8.69E-23 |
| chr11 | 72207003 | 72227927 | 20925 | -3.08 | 4.18E-24 | 8.83E-23 |
| chr2 | 151954087 | 151989552 | 35466 | -10.7 | 4.25E-24 | 8.96E-23 |
| chr2 | 83815072 | 83923727 | 108656 | -10.74 | 4.28E-24 | 9.02E-23 |
| chr3 | 113130665 | 113147816 | 17152 | -10.67 | 4.28E-24 | 9.03E-23 |
| chr2 | 178691625 | 178759071 | 67447 | 10.7 | 4.43E-24 | 9.34E-23 |
| chr3 | 95247970 | 95375582 | 127613 | -10.72 | 4.49E-24 | 9.46E-23 |
| chr9 | 81316824 | 81368646 | 51823 | -10.84 | 4.53E-24 | 9.53E-23 |
| chr1 | 230946380 | 230954003 | 7624 | -5.41 | 4.68E-24 | 9.84E-23 |
| chr14 | 101725745 | 101794575 | 68831 | -10.77 | 4.71E-24 | 9.89E-23 |
| chr2 | 39584889 | 39614369 | 29481 | -10.76 | 4.72E-24 | 9.91E-23 |
| chr2 | 241942383 | 241968828 | 26446 | -10.65 | 4.80E-24 | 1.01E-22 |
| chr9 | 80772004 | 80789653 | 17650 | -10.68 | 4.92E-24 | 1.03E-22 |
| chr2 | 112864476 | 112886992 | 22517 | -10.66 | 5.00E-24 | 1.05E-22 |
| chr20 | 22747487 | 22789375 | 41889 | -10.66 | 5.01E-24 | 1.05E-22 |
| chr2 | 240512680 | 240578824 | 66145 | -10.82 | 5.05E-24 | 1.06E-22 |
| chr15 | 71311502 | 71433733 | 122232 | -11.23 | 5.08E-24 | 1.06E-22 |
| chr11 | 13206991 | 13223884 | 16894 | -10.81 | 5.16E-24 | 1.08E-22 |
| chr17 | 19578410 | 19594901 | 16492 | -10.67 | 5.21E-24 | 1.09E-22 |
| chr2 | 182069488 | 182077478 | 7991 | -8.16 | 5.33E-24 | 1.11E-22 |

| | | | | | | |
|---|---|---|---|---|---|---|
| chr9 | 85774403 | 85793988 | 19586 | -10.65 | 5.37E-24 | 1.12E-22 |
| chr21 | 17252231 | 17278914 | 26684 | -10.7 | 5.47E-24 | 1.14E-22 |
| chr8 | 23994708 | 24032320 | 37613 | -10.8 | 5.51E-24 | 1.15E-22 |
| chr1 | 170514920 | 170551571 | 36652 | -10.7 | 5.56E-24 | 1.16E-22 |
| chr8 | 17491845 | 17513776 | 21932 | -10.79 | 5.65E-24 | 1.18E-22 |
| chr20 | 22368928 | 22443315 | 74388 | -10.87 | 5.74E-24 | 1.20E-22 |
| chr12 | 125683992 | 125749058 | 65067 | -10.81 | 5.79E-24 | 1.21E-22 |
| chr12 | 69689452 | 69719875 | 30424 | -10.64 | 5.81E-24 | 1.21E-22 |
| chr8 | 127525900 | 127549410 | 23511 | -10.66 | 5.82E-24 | 1.21E-22 |
| chr12 | 103937397 | 103965478 | 28082 | -10.88 | 5.87E-24 | 1.22E-22 |
| chr7 | 73337718 | 73380484 | 42767 | -10.78 | 5.92E-24 | 1.23E-22 |
| chr2 | 241411983 | 241436324 | 24342 | -10.69 | 5.99E-24 | 1.24E-22 |
| chr5 | 56580530 | 56690426 | 109897 | -10.72 | 6.12E-24 | 1.27E-22 |
| chr6 | 67949153 | 69338752 | 1389600 | 1.71 | 6.13E-24 | 1.27E-22 |
| chr15 | 50026756 | 50162272 | 135517 | -1.69 | 6.18E-24 | 1.28E-22 |
| chr18 | 53281099 | 53287953 | 6855 | -7 | 6.20E-24 | 1.28E-22 |
| chr15 | 62601580 | 62616379 | 14800 | -10.82 | 6.23E-24 | 1.29E-22 |
| chr22 | 49156078 | 49189658 | 33581 | -10.63 | 6.30E-24 | 1.30E-22 |
| chr4 | 14207082 | 14307363 | 100282 | -10.69 | 6.37E-24 | 1.32E-22 |
| chr12 | 105913502 | 105962200 | 48699 | -10.67 | 6.44E-24 | 1.33E-22 |
| chr17 | 20206941 | 20213521 | 6581 | -10.62 | 6.65E-24 | 1.37E-22 |
| chr7 | 132425033 | 132455340 | 30308 | -10.63 | 6.71E-24 | 1.39E-22 |
| chr17 | 54964187 | 55011726 | 47540 | -10.69 | 6.71E-24 | 1.39E-22 |
| chr3 | 11142129 | 11167766 | 25638 | -10.63 | 6.77E-24 | 1.40E-22 |
| chr9 | 120112173 | 120160803 | 48631 | 10.69 | 6.78E-24 | 1.40E-22 |
| chr4 | 86106278 | 86210512 | 104235 | -10.82 | 6.92E-24 | 1.43E-22 |
| chr2 | 72985412 | 73013180 | 27769 | -10.65 | 7.01E-24 | 1.44E-22 |
| chr15 | 86274148 | 86279829 | 5682 | -10.74 | 7.05E-24 | 1.45E-22 |
| chr9 | 114491567 | 114531207 | 39641 | -10.76 | 7.09E-24 | 1.46E-22 |
| chr3 | 31624231 | 31649517 | 25287 | -10.64 | 7.10E-24 | 1.46E-22 |
| chr9 | 136935896 | 136972319 | 36424 | -10.74 | 7.10E-24 | 1.46E-22 |
| chr6 | 62782940 | 62991020 | 208081 | 1.87 | 7.11E-24 | 1.46E-22 |
| chr16 | 15889329 | 15915055 | 25727 | -10.7 | 7.21E-24 | 1.48E-22 |
| chr3 | 46231630 | 46240638 | 9009 | -2.96 | 7.22E-24 | 1.48E-22 |
| chr2 | 99501503 | 99505380 | 3878 | -7.01 | 7.32E-24 | 1.50E-22 |
| chr12 | 75315634 | 75365932 | 50299 | -11.16 | 7.42E-24 | 1.52E-22 |
| chr10 | 133431644 | 133478887 | 47244 | -10.65 | 7.45E-24 | 1.53E-22 |
| chr4 | 185025867 | 185063546 | 37680 | -10.73 | 7.75E-24 | 1.59E-22 |
| chr5 | 40114678 | 40179369 | 64692 | -10.69 | 7.75E-24 | 1.59E-22 |
| chr17 | 5212470 | 5227563 | 15094 | -10.68 | 7.79E-24 | 1.59E-22 |
| chr21 | 34530122 | 34541410 | 11289 | -2.78 | 7.79E-24 | 1.59E-22 |
| chr14 | 70319894 | 70366416 | 46523 | -10.8 | 7.80E-24 | 1.59E-22 |
| chr3 | 112565655 | 112636577 | 70923 | -10.94 | 7.94E-24 | 1.62E-22 |
| chr11 | 10892542 | 10941558 | 49017 | -10.67 | 7.95E-24 | 1.62E-22 |
| chr20 | 61768555 | 61781760 | 13206 | -10.67 | 7.99E-24 | 1.63E-22 |
| chr9 | 38165485 | 38188634 | 23150 | -10.61 | 8.00E-24 | 1.63E-22 |
| chr12 | 94317458 | 94366296 | 48839 | -10.61 | 8.00E-24 | 1.63E-22 |
| chr11 | 84368695 | 84424539 | 55845 | 10.65 | 8.00E-24 | 1.63E-22 |
| chr2 | 238398270 | 238413082 | 14813 | -10.64 | 8.03E-24 | 1.63E-22 |
| chr2 | 191602510 | 191611306 | 8797 | -7.07 | 8.08E-24 | 1.64E-22 |
| chr10 | 131827928 | 131896265 | 68338 | -10.72 | 8.23E-24 | 1.67E-22 |
| chr5 | 82858053 | 82877017 | 18965 | -10.6 | 8.29E-24 | 1.68E-22 |
| chr8 | 39497477 | 39510445 | 12969 | -7.05 | 8.29E-24 | 1.68E-22 |
| chr6 | 123244986 | 123307396 | 62411 | 10.64 | 8.31E-24 | 1.69E-22 |
| chr12 | 10786431 | 10823930 | 37500 | -10.62 | 8.44E-24 | 1.71E-22 |
| chr20 | 905842 | 930984 | 25143 | -10.61 | 8.45E-24 | 1.71E-22 |
| chr12 | 58073598 | 58106765 | 33168 | -10.63 | 8.49E-24 | 1.72E-22 |
| chr16 | 89086897 | 89099472 | 12576 | -10.61 | 8.54E-24 | 1.73E-22 |
| chr1 | 243390135 | 243404309 | 14175 | -10.64 | 8.72E-24 | 1.76E-22 |
| chr12 | 113419160 | 113451206 | 32047 | -10.6 | 8.76E-24 | 1.77E-22 |
| chr5 | 32849726 | 32921749 | 72024 | -10.66 | 8.84E-24 | 1.79E-22 |

| | | | | | | |
|---|---|---|---|---|---|---|
| chr2 | 26880375 | 26901582 | 21208 | -10.78 | 8.92E-24 | 1.80E-22 |
| chr2 | 10388296 | 10393204 | 4909 | -7.25 | 8.98E-24 | 1.81E-22 |
| chr2 | 83430320 | 83540355 | 110036 | -10.81 | 9.03E-24 | 1.82E-22 |
| chr2 | 204753644 | 204769165 | 15522 | -3.08 | 9.21E-24 | 1.86E-22 |
| chr19 | 34173345 | 34225454 | 52110 | -10.61 | 9.22E-24 | 1.86E-22 |
| chr5 | 67630428 | 67710745 | 80318 | -10.88 | 9.23E-24 | 1.86E-22 |
| chr2 | 95716413 | 95766166 | 49754 | -10.81 | 9.25E-24 | 1.86E-22 |
| chrX | 7675515 | 7803284 | 127770 | 2.61 | 9.39E-24 | 1.89E-22 |
| chr8 | 144030260 | 144063134 | 32875 | -10.72 | 9.40E-24 | 1.89E-22 |
| chr16 | 76956376 | 77003649 | 47274 | -10.61 | 9.54E-24 | 1.92E-22 |
| chr17 | 38678677 | 38692297 | 13621 | -10.59 | 9.58E-24 | 1.92E-22 |
| chr10 | 22800018 | 22825579 | 25562 | -10.59 | 9.58E-24 | 1.92E-22 |
| chr3 | 149737655 | 149763256 | 25602 | -10.59 | 9.69E-24 | 1.95E-22 |
| chr14 | 101606599 | 101649244 | 42646 | -10.6 | 9.72E-24 | 1.95E-22 |
| chr22 | 22447045 | 22455633 | 8589 | -2.81 | 9.83E-24 | 1.97E-22 |
| chr11 | 29330898 | 29362803 | 31906 | -10.62 | 9.84E-24 | 1.97E-22 |
| chr18 | 60337382 | 60367524 | 30143 | -10.62 | 9.85E-24 | 1.97E-22 |
| chr8 | 128476966 | 128588922 | 111957 | -10.82 | 9.94E-24 | 1.99E-22 |
| chr2 | 112934465 | 112980617 | 46153 | -10.6 | 9.94E-24 | 1.99E-22 |
| chr6 | 17891528 | 17903888 | 12361 | -10.6 | 9.95E-24 | 1.99E-22 |
| chr8 | 46909061 | 46946452 | 37392 | -10.59 | 9.98E-24 | 2.00E-22 |
| chr3 | 170626624 | 170661366 | 34743 | -10.58 | 1.01E-23 | 2.01E-22 |
| chr17 | 6484649 | 6490962 | 6314 | -10.64 | 1.01E-23 | 2.02E-22 |
| chr17 | 15904110 | 15908185 | 4076 | -10.58 | 1.03E-23 | 2.05E-22 |
| chr12 | 66529918 | 66538995 | 9078 | -10.58 | 1.04E-23 | 2.08E-22 |
| chr2 | 11198468 | 11204990 | 6523 | -10.65 | 1.05E-23 | 2.09E-22 |
| chr2 | 128355304 | 128372778 | 17475 | -10.6 | 1.07E-23 | 2.13E-22 |
| chr7 | 106598172 | 106660186 | 62015 | 10.62 | 1.08E-23 | 2.15E-22 |
| chr19 | 14726988 | 14755835 | 28848 | -10.58 | 1.10E-23 | 2.20E-22 |
| chr20 | 45238904 | 45260797 | 21894 | -10.57 | 1.12E-23 | 2.23E-22 |
| chr13 | 97707153 | 97754643 | 47491 | -10.6 | 1.13E-23 | 2.25E-22 |
| chr10 | 43222841 | 43276764 | 53924 | -10.64 | 1.13E-23 | 2.25E-22 |
| chr7 | 758959 | 776603 | 17645 | -10.58 | 1.14E-23 | 2.26E-22 |
| chrX | 108204167 | 108250279 | 46113 | 3.84 | 1.14E-23 | 2.27E-22 |
| chr12 | 94810975 | 94850998 | 40024 | -10.6 | 1.15E-23 | 2.29E-22 |
| chr13 | 47514907 | 47540694 | 25788 | 2.56 | 1.16E-23 | 2.30E-22 |
| chr15 | 31471624 | 31486918 | 15295 | -10.58 | 1.16E-23 | 2.30E-22 |
| chr3 | 87632790 | 87681550 | 48761 | -10.61 | 1.18E-23 | 2.34E-22 |
| chr2 | 143524939 | 143589817 | 64879 | -10.74 | 1.18E-23 | 2.34E-22 |
| chr19 | 38650027 | 38654499 | 4473 | -10.57 | 1.20E-23 | 2.37E-22 |
| chr1 | 154003917 | 154060626 | 56710 | -10.6 | 1.21E-23 | 2.38E-22 |
| chr7 | 67794095 | 67813211 | 19117 | -10.63 | 1.22E-23 | 2.41E-22 |
| chr17 | 52990648 | 52996390 | 5743 | -10.79 | 1.22E-23 | 2.41E-22 |
| chrX | 154806859 | 154861090 | 54232 | -10.56 | 1.23E-23 | 2.43E-22 |
| chr18 | 45381506 | 45388504 | 6999 | -10.68 | 1.24E-23 | 2.45E-22 |
| chr8 | 143381285 | 143412288 | 31004 | -10.79 | 1.24E-23 | 2.45E-22 |
| chr5 | 123654762 | 123727848 | 73087 | -11.01 | 1.26E-23 | 2.48E-22 |
| chr20 | 14837946 | 14884906 | 46961 | -10.97 | 1.29E-23 | 2.54E-22 |
| chr20 | 18797003 | 19699134 | 902132 | 2.18 | 1.30E-23 | 2.56E-22 |
| chr9 | 5634024 | 5671221 | 37198 | -10.73 | 1.30E-23 | 2.56E-22 |
| chr13 | 114291724 | 114318010 | 26287 | -10.56 | 1.31E-23 | 2.58E-22 |
| chr2 | 192721165 | 192768612 | 47448 | -10.62 | 1.33E-23 | 2.62E-22 |
| chr15 | 29492993 | 29548163 | 55171 | -10.6 | 1.34E-23 | 2.63E-22 |
| chr2 | 211225262 | 211281234 | 55973 | 10.62 | 1.35E-23 | 2.65E-22 |
| chr5 | 31336122 | 31371798 | 35677 | 8.06 | 1.36E-23 | 2.68E-22 |
| chr2 | 29167768 | 29173474 | 5707 | -10.6 | 1.37E-23 | 2.68E-22 |
| chr5 | 115611349 | 115674560 | 63212 | -10.56 | 1.37E-23 | 2.68E-22 |
| chr8 | 143775619 | 143800745 | 25127 | -10.57 | 1.40E-23 | 2.74E-22 |
| chr12 | 84035394 | 84131678 | 96285 | -10.56 | 1.40E-23 | 2.74E-22 |
| chr5 | 566020 | 596776 | 30757 | -10.63 | 1.41E-23 | 2.76E-22 |
| chr3 | 119000283 | 119008219 | 7937 | -3.1 | 1.41E-23 | 2.76E-22 |

| | | | | | | |
|---|---|---|---|---|---|---|
| chr4 | 38293165 | 38356798 | 63634 | -10.67 | 1.44E-23 | 2.82E-22 |
| chr15 | 68386939 | 68397104 | 10166 | -10.55 | 1.48E-23 | 2.90E-22 |
| chr16 | 74825906 | 74863892 | 37987 | -10.66 | 1.51E-23 | 2.95E-22 |
| chr19 | 15985083 | 16029828 | 44746 | -10.62 | 1.51E-23 | 2.96E-22 |
| chr9 | 95451323 | 95463532 | 12210 | -10.56 | 1.52E-23 | 2.96E-22 |
| chr10 | 31448718 | 31469900 | 21183 | -10.54 | 1.53E-23 | 2.98E-22 |
| chr3 | 196961477 | 196978583 | 17107 | -10.56 | 1.53E-23 | 2.98E-22 |
| chr8 | 30785995 | 30832243 | 46249 | -10.56 | 1.56E-23 | 3.04E-22 |
| chr5 | 38806654 | 38869217 | 62564 | -10.54 | 1.57E-23 | 3.06E-22 |
| chr10 | 19233027 | 19328939 | 95913 | -10.66 | 1.60E-23 | 3.13E-22 |
| chr3 | 135337634 | 135374735 | 37102 | 10.58 | 1.61E-23 | 3.14E-22 |
| chr1 | 233238817 | 233259383 | 20567 | -10.54 | 1.63E-23 | 3.17E-22 |
| chr20 | 10317975 | 10371344 | 53370 | -10.8 | 1.65E-23 | 3.21E-22 |
| chr11 | 31982032 | 32004097 | 22066 | 7.61 | 1.66E-23 | 3.23E-22 |
| chr9 | 6308639 | 6350324 | 41686 | -10.54 | 1.67E-23 | 3.24E-22 |
| chr1 | 245299139 | 245336968 | 37830 | -10.68 | 1.69E-23 | 3.27E-22 |
| chr1 | 179360194 | 179406384 | 46191 | -10.66 | 1.69E-23 | 3.28E-22 |
| chr9 | 35174235 | 35226893 | 52659 | -10.83 | 1.69E-23 | 3.28E-22 |
| chr20 | 18201217 | 18227473 | 26257 | -10.58 | 1.71E-23 | 3.31E-22 |
| chr6 | 166278634 | 166320181 | 41548 | -10.53 | 1.71E-23 | 3.32E-22 |
| chr6 | 20994976 | 21029581 | 34606 | -10.57 | 1.72E-23 | 3.34E-22 |
| chr1 | 75073197 | 75132001 | 58805 | 10.6 | 1.75E-23 | 3.38E-22 |
| chr12 | 120071577 | 120100242 | 28666 | -10.53 | 1.76E-23 | 3.40E-22 |
| chr16 | 31284637 | 31336970 | 52334 | -10.55 | 1.77E-23 | 3.42E-22 |
| chr2 | 175031406 | 175042144 | 10739 | -10.58 | 1.77E-23 | 3.42E-22 |
| chr3 | 34003691 | 34060708 | 57018 | -10.56 | 1.78E-23 | 3.44E-22 |
| chr14 | 56263704 | 56304709 | 41006 | -10.6 | 1.80E-23 | 3.47E-22 |
| chr1 | 243739879 | 243768238 | 28360 | -10.53 | 1.80E-23 | 3.48E-22 |
| chr13 | 110210002 | 110257965 | 47964 | -10.56 | 1.81E-23 | 3.49E-22 |
| chr2 | 210672555 | 210735697 | 63143 | 10.57 | 1.82E-23 | 3.50E-22 |
| chr2 | 191401310 | 191434944 | 33635 | -10.67 | 1.82E-23 | 3.50E-22 |
| chr19 | 13782672 | 13827455 | 44784 | -10.65 | 1.85E-23 | 3.57E-22 |
| chr8 | 29334319 | 29368608 | 34290 | -10.69 | 1.88E-23 | 3.62E-22 |
| chr9 | 102933441 | 102963060 | 29620 | -10.57 | 1.91E-23 | 3.68E-22 |
| chr7 | 130921758 | 130973454 | 51697 | -10.64 | 1.95E-23 | 3.74E-22 |
| chr2 | 238460227 | 238465235 | 5009 | -10.57 | 1.95E-23 | 3.74E-22 |
| chr18 | 75367492 | 75602614 | 235123 | 1.11 | 1.95E-23 | 3.74E-22 |
| chr9 | 14922272 | 14978657 | 56386 | 4.72 | 1.97E-23 | 3.79E-22 |
| chr8 | 122008754 | 122044095 | 35342 | -10.53 | 2.00E-23 | 3.83E-22 |
| chr14 | 88864052 | 88887004 | 22953 | -10.52 | 2.00E-23 | 3.84E-22 |
| chrX | 149977596 | 150004144 | 26549 | -10.53 | 2.03E-23 | 3.90E-22 |
| chr8 | 408578 | 428656 | 20079 | -10.52 | 2.06E-23 | 3.95E-22 |
| chr2 | 210556146 | 210602433 | 46288 | 10.57 | 2.07E-23 | 3.97E-22 |
| chr15 | 58613016 | 58646007 | 32992 | -10.77 | 2.10E-23 | 4.03E-22 |
| chr4 | 148162174 | 148204187 | 42014 | -10.62 | 2.11E-23 | 4.03E-22 |
| chr5 | 67847155 | 67875234 | 28080 | -10.52 | 2.14E-23 | 4.09E-22 |
| chr17 | 70392026 | 70416275 | 24250 | -10.54 | 2.15E-23 | 4.10E-22 |
| chr5 | 966677 | 989291 | 22615 | -10.52 | 2.17E-23 | 4.14E-22 |
| chr7 | 43701410 | 43767531 | 66122 | -10.79 | 2.17E-23 | 4.15E-22 |
| chr1 | 220417959 | 220442538 | 24580 | -10.52 | 2.21E-23 | 4.22E-22 |
| chr17 | 75162205 | 75172636 | 10432 | -10.54 | 2.21E-23 | 4.22E-22 |
| chr2 | 11973939 | 12229909 | 255971 | -2.27 | 2.23E-23 | 4.24E-22 |
| chr16 | 47791580 | 47831423 | 39844 | -10.51 | 2.23E-23 | 4.24E-22 |
| chrX | 110356371 | 110413481 | 57111 | 10.57 | 2.23E-23 | 4.24E-22 |
| chr20 | 55650516 | 55740428 | 89913 | -10.89 | 2.23E-23 | 4.24E-22 |
| chr5 | 52980449 | 53017205 | 36757 | -10.66 | 2.25E-23 | 4.28E-22 |
| chr7 | 50998398 | 51045417 | 47020 | -10.51 | 2.30E-23 | 4.38E-22 |
| chr2 | 118892689 | 118945134 | 52446 | -10.53 | 2.37E-23 | 4.50E-22 |
| chr8 | 87657512 | 87696028 | 38517 | 10.54 | 2.38E-23 | 4.51E-22 |
| chr15 | 96322161 | 96352212 | 30052 | -10.5 | 2.39E-23 | 4.53E-22 |
| chr4 | 128072351 | 128085093 | 12743 | -4.27 | 2.39E-23 | 4.53E-22 |

| | | | | | | |
|---|---|---|---|---|---|---|
| chr12 | 30680395 | 30763234 | 82840 | -10.87 | 2.40E-23 | 4.56E-22 |
| chr2 | 108808986 | 108866857 | 57872 | -10.5 | 2.41E-23 | 4.57E-22 |
| chr10 | 74720924 | 74741327 | 20404 | -10.54 | 2.46E-23 | 4.66E-22 |
| chr2 | 207147346 | 207184826 | 37481 | -10.5 | 2.46E-23 | 4.67E-22 |
| chr14 | 68974140 | 69048524 | 74385 | -10.78 | 2.47E-23 | 4.68E-22 |
| chr9 | 114162384 | 114169266 | 6883 | -10.57 | 2.47E-23 | 4.68E-22 |
| chr10 | 34339871 | 34389631 | 49761 | -10.82 | 2.48E-23 | 4.70E-22 |
| chr1 | 244562592 | 244569055 | 6464 | -10.54 | 2.53E-23 | 4.78E-22 |
| chr4 | 161229078 | 161263135 | 34058 | -10.5 | 2.60E-23 | 4.90E-22 |
| chr2 | 25114548 | 25134994 | 20447 | -10.49 | 2.60E-23 | 4.91E-22 |
| chr2 | 120254699 | 120274224 | 19526 | -10.55 | 2.60E-23 | 4.91E-22 |
| chr7 | 67462922 | 67508739 | 45818 | -10.63 | 2.61E-23 | 4.93E-22 |
| chr2 | 11786496 | 11816303 | 29808 | -10.67 | 2.63E-23 | 4.95E-22 |
| chr19 | 315480 | 356199 | 40720 | -10.5 | 2.63E-23 | 4.95E-22 |
| chr12 | 31038547 | 31054620 | 16074 | -4.43 | 2.63E-23 | 4.96E-22 |
| chr12 | 70827924 | 70916998 | 89075 | -10.7 | 2.64E-23 | 4.96E-22 |
| chr18 | 3355655 | 3421201 | 65547 | -10.59 | 2.67E-23 | 5.02E-22 |
| chr2 | 35997501 | 36102301 | 104801 | 4.35 | 2.68E-23 | 5.04E-22 |
| chr7 | 86979323 | 87025138 | 45816 | -10.55 | 2.70E-23 | 5.08E-22 |
| chr19 | 39024101 | 39034495 | 10395 | -10.51 | 2.70E-23 | 5.08E-22 |
| chr20 | 11488835 | 11523070 | 34236 | -10.5 | 2.71E-23 | 5.09E-22 |
| chr20 | 57528748 | 57550976 | 22229 | -10.59 | 2.74E-23 | 5.14E-22 |
| chr3 | 136592455 | 136640790 | 48336 | -10.66 | 2.80E-23 | 5.26E-22 |
| chr9 | 83469820 | 83511462 | 41643 | -10.5 | 2.82E-23 | 5.28E-22 |
| chr10 | 134652751 | 134670695 | 17945 | -10.55 | 2.84E-23 | 5.32E-22 |
| chr8 | 52448408 | 52708837 | 260430 | 1.66 | 2.87E-23 | 5.38E-22 |
| chr5 | 118589039 | 118604002 | 14964 | -10.56 | 2.89E-23 | 5.42E-22 |
| chr9 | 96634432 | 96663093 | 28662 | -10.48 | 2.90E-23 | 5.42E-22 |
| chr4 | 56641903 | 56644420 | 2518 | -5.92 | 2.92E-23 | 5.47E-22 |
| chr1 | 225049596 | 225100853 | 51258 | -10.78 | 2.95E-23 | 5.52E-22 |
| chr8 | 17082642 | 17090128 | 7487 | -10.54 | 2.96E-23 | 5.53E-22 |
| chr11 | 92351557 | 92407124 | 55568 | 10.52 | 2.99E-23 | 5.58E-22 |
| chr8 | 115801968 | 115930537 | 128570 | -10.77 | 2.99E-23 | 5.59E-22 |
| chr13 | 65168133 | 65236123 | 67991 | 10.56 | 3.00E-23 | 5.60E-22 |
| chr1 | 246856160 | 246910944 | 54785 | -10.85 | 3.01E-23 | 5.61E-22 |
| chr2 | 237129874 | 237145275 | 15402 | -10.52 | 3.04E-23 | 5.67E-22 |
| chr10 | 80246892 | 80265928 | 19037 | -10.53 | 3.06E-23 | 5.71E-22 |
| chr2 | 103068713 | 103087823 | 19111 | -10.48 | 3.09E-23 | 5.76E-22 |
| chr2 | 173049792 | 173067596 | 17805 | -10.48 | 3.10E-23 | 5.77E-22 |
| chr8 | 23892837 | 23951993 | 59157 | -10.59 | 3.11E-23 | 5.79E-22 |
| chrX | 108492403 | 108551581 | 59179 | 10.52 | 3.13E-23 | 5.82E-22 |
| chr17 | 39266817 | 39302648 | 35832 | -10.48 | 3.16E-23 | 5.88E-22 |
| chr7 | 132790180 | 132799661 | 9482 | -3.86 | 3.17E-23 | 5.89E-22 |
| chr2 | 69165878 | 69212773 | 46896 | -10.62 | 3.27E-23 | 6.08E-22 |
| chr12 | 62683832 | 62701950 | 18119 | -10.56 | 3.27E-23 | 6.08E-22 |
| chr11 | 34903132 | 34936033 | 32902 | -10.47 | 3.28E-23 | 6.08E-22 |
| chr8 | 130494520 | 130547533 | 53014 | -10.5 | 3.29E-23 | 6.10E-22 |
| chr17 | 63793955 | 63807376 | 13422 | -10.5 | 3.29E-23 | 6.10E-22 |
| chr11 | 13137477 | 13180383 | 42907 | -10.86 | 3.29E-23 | 6.10E-22 |
| chr7 | 149260479 | 149282165 | 21687 | -10.48 | 3.31E-23 | 6.13E-22 |
| chr2 | 241091233 | 241147811 | 56579 | -10.64 | 3.35E-23 | 6.21E-22 |
| chr12 | 108878685 | 108904237 | 25553 | -10.52 | 3.42E-23 | 6.33E-22 |
| chr16 | 81983521 | 82011036 | 27516 | -10.47 | 3.54E-23 | 6.55E-22 |
| chr3 | 95690337 | 95807731 | 117395 | -10.64 | 3.60E-23 | 6.66E-22 |
| chr8 | 115300083 | 115455104 | 155022 | -10.76 | 3.69E-23 | 6.82E-22 |
| chr7 | 17593549 | 17650425 | 56877 | -10.82 | 3.73E-23 | 6.88E-22 |
| chr19 | 13628732 | 13672891 | 44160 | -10.47 | 3.76E-23 | 6.94E-22 |
| chr11 | 71024869 | 71073018 | 48150 | -10.59 | 3.78E-23 | 6.97E-22 |
| chr10 | 7486791 | 7507313 | 20523 | -10.47 | 3.79E-23 | 6.98E-22 |
| chr2 | 110055671 | 110063031 | 7361 | -10.51 | 3.84E-23 | 7.08E-22 |
| chr9 | 29122235 | 29152681 | 30447 | -10.54 | 3.86E-23 | 7.12E-22 |

| | | | | | | |
|---|---|---|---|---|---|---|
| chr2 | 214187464 | 214214180 | 26717 | -10.46 | 3.87E-23 | 7.12E-22 |
| chr13 | 25355199 | 25360902 | 5704 | -4.18 | 3.89E-23 | 7.15E-22 |
| chr1 | 228483044 | 228514215 | 31172 | -10.46 | 3.90E-23 | 7.17E-22 |
| chr3 | 33738391 | 33775594 | 37204 | -10.51 | 3.92E-23 | 7.21E-22 |
| chr20 | 55558277 | 55575606 | 17330 | -10.48 | 3.92E-23 | 7.21E-22 |
| chr20 | 36862391 | 36875050 | 12660 | -10.58 | 3.94E-23 | 7.23E-22 |
| chr12 | 110620120 | 110633050 | 12931 | -10.46 | 3.98E-23 | 7.31E-22 |
| chr7 | 54391679 | 54605438 | 213760 | 7.07 | 4.02E-23 | 7.37E-22 |
| chr17 | 80992267 | 81006227 | 13961 | -10.46 | 4.06E-23 | 7.44E-22 |
| chr11 | 26730225 | 26759653 | 29429 | 10.51 | 4.07E-23 | 7.45E-22 |
| chr19 | 48784993 | 48792003 | 7011 | -10.47 | 4.09E-23 | 7.49E-22 |
| chr15 | 36430558 | 36486571 | 56014 | -10.88 | 4.09E-23 | 7.49E-22 |
| chr14 | 89343972 | 89364817 | 20846 | -10.46 | 4.16E-23 | 7.61E-22 |
| chr4 | 142448312 | 142514653 | 66342 | -10.5 | 4.16E-23 | 7.61E-22 |
| chr3 | 112652146 | 112685372 | 33227 | -10.79 | 4.27E-23 | 7.80E-22 |
| chr14 | 59848248 | 59883116 | 34869 | -10.45 | 4.38E-23 | 8.00E-22 |
| chr20 | 14138505 | 14168332 | 29828 | -10.45 | 4.43E-23 | 8.09E-22 |
| chr20 | 23353013 | 23368660 | 15648 | -10.5 | 4.45E-23 | 8.13E-22 |
| chr16 | 51511144 | 51572322 | 61179 | -10.53 | 4.46E-23 | 8.13E-22 |
| chr3 | 38692966 | 38715684 | 22719 | -10.44 | 4.47E-23 | 8.16E-22 |
| chr20 | 36260748 | 36296717 | 35970 | -10.61 | 4.49E-23 | 8.19E-22 |
| chr4 | 189884282 | 189935010 | 50729 | -10.44 | 4.50E-23 | 8.20E-22 |
| chr10 | 31822761 | 31883227 | 60467 | -2.97 | 4.54E-23 | 8.28E-22 |
| chr5 | 54718435 | 54733185 | 14751 | -10.45 | 4.55E-23 | 8.28E-22 |
| chr8 | 67860765 | 67891799 | 31035 | -10.58 | 4.58E-23 | 8.33E-22 |
| chr10 | 128030212 | 128082474 | 52263 | -10.45 | 4.58E-23 | 8.33E-22 |
| chr4 | 143005762 | 143050270 | 44509 | -10.63 | 4.66E-23 | 8.47E-22 |
| chr8 | 22726116 | 22728713 | 2598 | -6.58 | 4.70E-23 | 8.54E-22 |
| chr4 | 88251194 | 88275070 | 23877 | -10.44 | 4.70E-23 | 8.54E-22 |
| chr8 | 41910766 | 41931614 | 20849 | -10.5 | 4.76E-23 | 8.65E-22 |
| chr1 | 211973915 | 211994888 | 20974 | -10.44 | 4.77E-23 | 8.66E-22 |
| chr3 | 5608261 | 5662405 | 54145 | -10.44 | 4.80E-23 | 8.70E-22 |
| chr2 | 88331758 | 88339755 | 7998 | -10.44 | 4.83E-23 | 8.75E-22 |
| chr3 | 13841563 | 13873476 | 31914 | -10.44 | 4.85E-23 | 8.78E-22 |
| chr2 | 48020837 | 48043316 | 22480 | -10.55 | 4.89E-23 | 8.86E-22 |
| chr20 | 2366503 | 2407602 | 41100 | -10.57 | 4.91E-23 | 8.89E-22 |
| chr8 | 102220742 | 102233252 | 12511 | -10.5 | 4.92E-23 | 8.90E-22 |
| chr7 | 44375690 | 44409711 | 34022 | -10.51 | 4.98E-23 | 9.01E-22 |
| chr8 | 142471327 | 142513173 | 41847 | -10.46 | 5.06E-23 | 9.14E-22 |
| chr2 | 47769546 | 47783943 | 14398 | -10.43 | 5.09E-23 | 9.19E-22 |
| chr20 | 62015061 | 62024249 | 9189 | -10.44 | 5.10E-23 | 9.22E-22 |
| chr6 | 169859015 | 169962347 | 103333 | -10.66 | 5.16E-23 | 9.31E-22 |
| chr5 | 120550632 | 120613989 | 63358 | -10.49 | 5.16E-23 | 9.31E-22 |
| chr6 | 155785642 | 155825390 | 39749 | -10.43 | 5.17E-23 | 9.32E-22 |
| chrX | 49558103 | 49587871 | 29769 | -4.01 | 5.20E-23 | 9.36E-22 |
| chr6 | 136219109 | 136267214 | 48106 | -10.43 | 5.21E-23 | 9.39E-22 |
| chr13 | 44641044 | 44680282 | 39239 | -10.43 | 5.22E-23 | 9.40E-22 |
| chr4 | 24910826 | 24968392 | 57567 | -3.06 | 5.23E-23 | 9.41E-22 |
| chr11 | 2496844 | 2525979 | 29136 | -4.08 | 5.28E-23 | 9.51E-22 |
| chr10 | 8735432 | 8771997 | 36566 | -10.47 | 5.32E-23 | 9.56E-22 |
| chr7 | 127766704 | 127785117 | 18414 | -10.44 | 5.39E-23 | 9.69E-22 |
| chrX | 68173594 | 68196345 | 22752 | -10.57 | 5.51E-23 | 9.89E-22 |
| chr10 | 61671398 | 61691207 | 19810 | -10.44 | 5.51E-23 | 9.90E-22 |
| chr5 | 55627949 | 55659268 | 31320 | -10.52 | 5.61E-23 | 1.01E-21 |
| chr15 | 99528041 | 99544540 | 16500 | -10.5 | 5.66E-23 | 1.01E-21 |
| chr6 | 39807806 | 39826002 | 18197 | 3.1 | 5.81E-23 | 1.04E-21 |
| chr15 | 83884113 | 83915261 | 31149 | -10.48 | 5.81E-23 | 1.04E-21 |
| chr3 | 5483701 | 5518259 | 34559 | -10.43 | 5.83E-23 | 1.04E-21 |
| chr8 | 76469515 | 77272382 | 802868 | 2.01 | 5.85E-23 | 1.05E-21 |
| chr15 | 83737715 | 83750889 | 13175 | -10.42 | 5.92E-23 | 1.06E-21 |
| chr1 | 219356337 | 219374070 | 17734 | -10.53 | 5.99E-23 | 1.07E-21 |

| | | | | | | |
|---|---|---|---|---|---|---|
| chr1 | 147148724 | 147217917 | 69194 | -10.41 | 6.08E-23 | 1.09E-21 |
| chr17 | 69486446 | 69836876 | 350431 | 2.32 | 6.12E-23 | 1.09E-21 |
| chr3 | 149007611 | 149071765 | 64155 | -10.56 | 6.29E-23 | 1.12E-21 |
| chr1 | 154600885 | 154614396 | 13512 | -10.41 | 6.39E-23 | 1.14E-21 |
| chr3 | 87242196 | 87286166 | 43971 | -10.41 | 6.46E-23 | 1.15E-21 |
| chr2 | 111967852 | 111991439 | 23588 | -10.44 | 6.56E-23 | 1.17E-21 |
| chr11 | 31025391 | 31072031 | 46641 | 10.49 | 6.66E-23 | 1.19E-21 |
| chr6 | 142892846 | 142935447 | 42602 | -1.82 | 6.74E-23 | 1.20E-21 |
| chr5 | 179308590 | 179314642 | 6053 | -10.41 | 6.74E-23 | 1.20E-21 |
| chr6 | 2254118 | 2300248 | 46131 | -10.55 | 6.77E-23 | 1.20E-21 |
| chr11 | 57372877 | 57400193 | 27317 | -7.68 | 6.81E-23 | 1.21E-21 |
| chr5 | 134310719 | 134328096 | 17378 | -10.4 | 6.82E-23 | 1.21E-21 |
| chr7 | 99628760 | 99636167 | 7408 | -10.41 | 6.83E-23 | 1.21E-21 |
| chr12 | 63792637 | 63828212 | 35576 | -10.43 | 6.84E-23 | 1.22E-21 |
| chr10 | 25628741 | 25683247 | 54507 | -10.42 | 6.85E-23 | 1.22E-21 |
| chr2 | 99609582 | 99671772 | 62191 | -10.55 | 6.89E-23 | 1.22E-21 |
| chr10 | 133535880 | 133573853 | 37974 | -10.42 | 6.97E-23 | 1.24E-21 |
| chr7 | 32537582 | 32563107 | 25526 | -10.54 | 6.97E-23 | 1.24E-21 |
| chr4 | 189813798 | 189870321 | 56524 | -10.49 | 7.03E-23 | 1.25E-21 |
| chr9 | 79252322 | 79260180 | 7859 | 5.3 | 7.17E-23 | 1.27E-21 |
| chr1 | 205513064 | 205532308 | 19245 | -10.48 | 7.19E-23 | 1.27E-21 |
| chr21 | 39504794 | 39530784 | 25991 | -10.49 | 7.22E-23 | 1.28E-21 |
| chr15 | 79910247 | 79963853 | 53607 | -10.54 | 7.27E-23 | 1.29E-21 |
| chr3 | 105099616 | 105137607 | 37992 | -10.49 | 7.31E-23 | 1.29E-21 |
| chr13 | 52634322 | 52652519 | 18198 | -10.4 | 7.33E-23 | 1.30E-21 |
| chr7 | 127349111 | 127355146 | 6036 | -10.47 | 7.37E-23 | 1.30E-21 |
| chr17 | 63230612 | 63233098 | 2487 | -7.91 | 7.39E-23 | 1.30E-21 |
| chr2 | 120367352 | 120408227 | 40876 | -10.41 | 7.39E-23 | 1.30E-21 |
| chr8 | 145872866 | 145894587 | 21722 | -10.53 | 7.39E-23 | 1.30E-21 |
| chr3 | 149644872 | 149673496 | 28625 | -10.49 | 7.43E-23 | 1.31E-21 |
| chr9 | 76362656 | 76398536 | 35881 | 10.53 | 7.45E-23 | 1.31E-21 |
| chr7 | 23375179 | 23385593 | 10415 | -10.52 | 7.47E-23 | 1.32E-21 |
| chr11 | 125660837 | 125688841 | 28005 | -3.84 | 7.48E-23 | 1.32E-21 |
| chr10 | 25735602 | 25816264 | 80663 | -10.61 | 7.48E-23 | 1.32E-21 |
| chr12 | 28355501 | 28379386 | 23886 | -10.4 | 7.52E-23 | 1.32E-21 |
| chr8 | 140680169 | 140706291 | 26123 | -10.4 | 7.59E-23 | 1.34E-21 |
| chr3 | 149106609 | 149123807 | 17199 | -10.39 | 7.63E-23 | 1.34E-21 |
| chr8 | 110281254 | 110299957 | 18704 | -10.44 | 7.78E-23 | 1.37E-21 |
| chr9 | 126423227 | 126445248 | 22022 | -10.45 | 7.80E-23 | 1.37E-21 |
| chr9 | 94802963 | 94827371 | 24409 | -10.47 | 7.82E-23 | 1.37E-21 |
| chr9 | 110086189 | 110095102 | 8914 | -10.4 | 7.90E-23 | 1.39E-21 |
| chr2 | 241739357 | 241770357 | 31001 | -10.45 | 7.90E-23 | 1.39E-21 |
| chr5 | 133647793 | 133677791 | 29999 | -10.46 | 8.04E-23 | 1.41E-21 |
| chrX | 10389740 | 10427136 | 37397 | -10.4 | 8.25E-23 | 1.45E-21 |
| chr3 | 142191603 | 142203935 | 12333 | -10.41 | 8.34E-23 | 1.46E-21 |
| chr7 | 31880024 | 31887908 | 7885 | -10.45 | 8.39E-23 | 1.47E-21 |
| chr16 | 50677109 | 50742916 | 65808 | -10.75 | 8.42E-23 | 1.47E-21 |
| chr8 | 29812950 | 29913779 | 100830 | -10.91 | 8.58E-23 | 1.50E-21 |
| chr10 | 92666631 | 92692747 | 26117 | -10.46 | 8.77E-23 | 1.53E-21 |
| chr16 | 74159881 | 74217254 | 57374 | -10.5 | 8.80E-23 | 1.54E-21 |
| chr5 | 32996558 | 33017950 | 21393 | -10.38 | 8.81E-23 | 1.54E-21 |
| chr3 | 190636151 | 190691303 | 55153 | 10.43 | 8.82E-23 | 1.54E-21 |
| chr14 | 101669648 | 101680519 | 10872 | -10.38 | 8.84E-23 | 1.54E-21 |
| chr11 | 61699472 | 61710674 | 11203 | -10.4 | 8.85E-23 | 1.54E-21 |
| chr16 | 80024719 | 80086345 | 61627 | -10.49 | 8.92E-23 | 1.55E-21 |
| chr14 | 91933630 | 91942480 | 8851 | -10.6 | 8.92E-23 | 1.55E-21 |
| chr12 | 32034279 | 32079083 | 44805 | -10.43 | 9.05E-23 | 1.58E-21 |
| chr2 | 118774604 | 118801637 | 27034 | -2.93 | 9.24E-23 | 1.61E-21 |
| chr1 | 2728448 | 2764058 | 35611 | -10.37 | 9.31E-23 | 1.62E-21 |
| chrX | 150021300 | 150049204 | 27905 | -10.41 | 9.39E-23 | 1.63E-21 |
| chr2 | 192044429 | 192109024 | 64596 | -10.39 | 9.40E-23 | 1.63E-21 |

| | | | | | | |
|---|---|---|---|---|---|---|
| chr7 | 101213111 | 101240590 | 27480 | -10.38 | 9.48E-23 | 1.65E-21 |
| chr15 | 33008350 | 33096775 | 88426 | -10.8 | 9.51E-23 | 1.65E-21 |
| chr12 | 96137597 | 96155750 | 18154 | -10.39 | 9.61E-23 | 1.67E-21 |
| chr10 | 82488155 | 82662311 | 174157 | 1.37 | 9.68E-23 | 1.68E-21 |
| chr8 | 130606718 | 130638774 | 32057 | -10.37 | 9.72E-23 | 1.69E-21 |
| chr11 | 69663496 | 69674676 | 11181 | -10.38 | 9.81E-23 | 1.70E-21 |
| chr18 | 32082614 | 32114240 | 31627 | 10.43 | 9.85E-23 | 1.71E-21 |
| chr2 | 84305668 | 84389359 | 83692 | -10.42 | 9.89E-23 | 1.71E-21 |
| chr5 | 151897625 | 153327286 | 1429662 | 4.89 | 9.92E-23 | 1.72E-21 |
| chr9 | 114787096 | 114827446 | 40351 | -10.37 | 1.00E-22 | 1.73E-21 |
| chr2 | 111802610 | 111847427 | 44818 | -2.49 | 1.01E-22 | 1.74E-21 |
| chr10 | 45698327 | 45740660 | 42334 | -10.41 | 1.02E-22 | 1.76E-21 |
| chr12 | 79334163 | 79352539 | 18377 | -10.38 | 1.02E-22 | 1.76E-21 |
| chr12 | 28504135 | 28527260 | 23126 | -10.37 | 1.02E-22 | 1.76E-21 |
| chr8 | 39523178 | 39561135 | 37958 | -3.21 | 1.03E-22 | 1.78E-21 |
| chr10 | 96862544 | 96902530 | 39987 | -10.37 | 1.05E-22 | 1.80E-21 |
| chr5 | 30071645 | 30151794 | 80150 | -10.73 | 1.05E-22 | 1.81E-21 |
| chr7 | 81674964 | 81700748 | 25785 | -10.42 | 1.07E-22 | 1.84E-21 |
| chr6 | 72062868 | 72085206 | 22339 | -10.39 | 1.08E-22 | 1.87E-21 |
| chr16 | 58163744 | 58179894 | 16151 | -10.37 | 1.08E-22 | 1.87E-21 |
| chr1 | 229469388 | 229502246 | 32859 | -10.38 | 1.10E-22 | 1.89E-21 |
| chr22 | 27077181 | 27105349 | 28169 | -10.36 | 1.10E-22 | 1.89E-21 |
| chr8 | 66798118 | 66851802 | 53685 | -10.71 | 1.12E-22 | 1.92E-21 |
| chr8 | 115654377 | 115785422 | 131046 | -10.51 | 1.13E-22 | 1.94E-21 |
| chr13 | 48529339 | 48551822 | 22484 | -10.37 | 1.13E-22 | 1.94E-21 |
| chr9 | 138278191 | 138326547 | 48357 | -10.37 | 1.13E-22 | 1.95E-21 |
| chr1 | 11553139 | 11574252 | 21114 | -10.42 | 1.15E-22 | 1.97E-21 |
| chr5 | 176179308 | 176195343 | 16036 | -10.48 | 1.15E-22 | 1.97E-21 |
| chr4 | 4954245 | 4980592 | 26348 | -10.35 | 1.15E-22 | 1.98E-21 |
| chr1 | 242920869 | 242924652 | 3784 | -5.14 | 1.15E-22 | 1.98E-21 |
| chr21 | 46238603 | 46250411 | 11809 | -10.35 | 1.16E-22 | 2.00E-21 |
| chr13 | 47384871 | 47388463 | 3593 | -7.89 | 1.17E-22 | 2.00E-21 |
| chr15 | 58890657 | 58923895 | 33239 | -10.35 | 1.17E-22 | 2.01E-21 |
| chr9 | 22347752 | 22374330 | 26579 | -10.38 | 1.18E-22 | 2.02E-21 |
| chr9 | 4170114 | 4204284 | 34171 | -10.62 | 1.20E-22 | 2.05E-21 |
| chr10 | 126922654 | 126942796 | 20143 | -10.43 | 1.20E-22 | 2.05E-21 |
| chr2 | 153791497 | 153844537 | 53041 | 10.46 | 1.21E-22 | 2.08E-21 |
| chr11 | 76764740 | 76773955 | 9216 | -10.41 | 1.22E-22 | 2.09E-21 |
| chr5 | 98770393 | 98803750 | 33358 | 3 | 1.22E-22 | 2.09E-21 |
| chr8 | 110417600 | 110482055 | 64456 | 10.42 | 1.22E-22 | 2.09E-21 |
| chr7 | 21739739 | 21782233 | 42495 | -10.45 | 1.24E-22 | 2.11E-21 |
| chr1 | 242377222 | 242408077 | 30856 | -10.46 | 1.24E-22 | 2.12E-21 |
| chr10 | 19349618 | 19413094 | 63477 | -10.47 | 1.25E-22 | 2.12E-21 |
| chr14 | 92260734 | 92294429 | 33696 | -10.43 | 1.26E-22 | 2.14E-21 |
| chr9 | 100163319 | 100173210 | 9892 | -10.41 | 1.26E-22 | 2.14E-21 |
| chr3 | 169226902 | 169270145 | 43244 | 10.6 | 1.26E-22 | 2.15E-21 |
| chr19 | 13739730 | 13767005 | 27276 | -10.35 | 1.27E-22 | 2.16E-21 |
| chr8 | 48242017 | 48281364 | 39348 | -10.38 | 1.29E-22 | 2.20E-21 |
| chr12 | 43553610 | 43596300 | 42691 | -10.34 | 1.31E-22 | 2.23E-21 |
| chr5 | 173664553 | 173721379 | 56827 | -2.11 | 1.31E-22 | 2.23E-21 |
| chr20 | 61669689 | 61713487 | 43799 | -10.41 | 1.32E-22 | 2.24E-21 |
| chr14 | 31224736 | 31270911 | 46176 | -1.81 | 1.33E-22 | 2.26E-21 |
| chr7 | 55524522 | 55536286 | 11765 | -10.34 | 1.34E-22 | 2.27E-21 |
| chr3 | 118792465 | 118841522 | 49058 | -10.37 | 1.34E-22 | 2.27E-21 |
| chr7 | 142893544 | 142959688 | 66145 | -10.4 | 1.34E-22 | 2.27E-21 |
| chr8 | 118069574 | 118100065 | 30492 | 10.4 | 1.34E-22 | 2.28E-21 |
| chr8 | 143445664 | 143471406 | 25743 | -10.45 | 1.34E-22 | 2.28E-21 |
| chr5 | 31446931 | 31467320 | 20390 | -10.56 | 1.35E-22 | 2.28E-21 |
| chrX | 154740702 | 154784183 | 43482 | -10.37 | 1.40E-22 | 2.38E-21 |
| chr5 | 168968090 | 168994651 | 26562 | -10.33 | 1.43E-22 | 2.41E-21 |
| chr11 | 70764327 | 70791812 | 27486 | -10.39 | 1.43E-22 | 2.41E-21 |

| | | | | | | |
|---|---|---|---|---|---|---|
| chr9 | 100042488 | 100068492 | 26005 | -10.44 | 1.44E-22 | 2.44E-21 |
| chr20 | 2327439 | 2352394 | 24956 | -10.4 | 1.46E-22 | 2.46E-21 |
| chr5 | 153668618 | 153687703 | 19086 | -10.44 | 1.48E-22 | 2.50E-21 |
| chr8 | 28027045 | 28034523 | 7479 | -10.33 | 1.48E-22 | 2.50E-21 |
| chr11 | 44040339 | 44063870 | 23532 | -10.34 | 1.49E-22 | 2.51E-21 |
| chr5 | 155204312 | 155230284 | 25973 | -10.45 | 1.49E-22 | 2.51E-21 |
| chr3 | 161991626 | 162021250 | 29625 | -10.62 | 1.49E-22 | 2.51E-21 |
| chr10 | 74524237 | 74540026 | 15790 | -10.46 | 1.49E-22 | 2.51E-21 |
| chr10 | 25554840 | 25578937 | 24098 | -10.33 | 1.49E-22 | 2.51E-21 |
| chr5 | 151205915 | 151225482 | 19568 | -10.35 | 1.49E-22 | 2.52E-21 |
| chr4 | 14773236 | 14821748 | 48513 | -10.56 | 1.50E-22 | 2.52E-21 |
| chr5 | 60229799 | 60290368 | 60570 | -10.51 | 1.50E-22 | 2.53E-21 |
| chr10 | 4891277 | 4909522 | 18246 | -10.32 | 1.57E-22 | 2.64E-21 |
| chr5 | 93440288 | 93482837 | 42550 | -10.33 | 1.57E-22 | 2.64E-21 |
| chr7 | 7192905 | 7246847 | 53943 | -10.42 | 1.59E-22 | 2.67E-21 |
| chr8 | 122069059 | 122103833 | 34775 | -10.34 | 1.60E-22 | 2.68E-21 |
| chr12 | 120178297 | 120187251 | 8955 | -10.34 | 1.60E-22 | 2.69E-21 |
| chr7 | 16846745 | 16893831 | 47087 | -10.33 | 1.62E-22 | 2.71E-21 |
| chr20 | 42516154 | 42542710 | 26557 | -10.32 | 1.62E-22 | 2.72E-21 |
| chr3 | 43866938 | 43903214 | 36277 | -10.33 | 1.62E-22 | 2.72E-21 |
| chr16 | 90055294 | 90065039 | 9746 | -10.38 | 1.64E-22 | 2.74E-21 |
| chr6 | 107206514 | 107230943 | 24430 | -10.32 | 1.64E-22 | 2.75E-21 |
| chr1 | 167490252 | 167494195 | 3944 | -3.07 | 1.64E-22 | 2.75E-21 |
| chr11 | 40347297 | 40385395 | 38099 | -10.36 | 1.68E-22 | 2.81E-21 |
| chr5 | 56991725 | 57035499 | 43775 | -10.59 | 1.69E-22 | 2.83E-21 |
| chr8 | 57846068 | 57887585 | 41518 | -10.33 | 1.69E-22 | 2.83E-21 |
| chr2 | 77916741 | 82885919 | 4969179 | 0.97 | 1.71E-22 | 2.85E-21 |
| chr5 | 146351630 | 146380622 | 28993 | 10.38 | 1.72E-22 | 2.87E-21 |
| chrX | 65808045 | 65830244 | 22200 | -10.36 | 1.72E-22 | 2.88E-21 |
| chr2 | 207639222 | 207646177 | 6956 | -10.43 | 1.73E-22 | 2.88E-21 |
| chr17 | 78204279 | 78209465 | 5187 | -10.31 | 1.73E-22 | 2.88E-21 |
| chr8 | 52773777 | 52784113 | 10337 | -10.31 | 1.73E-22 | 2.88E-21 |
| chr6 | 86646143 | 86694448 | 48306 | 10.47 | 1.73E-22 | 2.89E-21 |
| chr9 | 93249021 | 93268500 | 19480 | -10.32 | 1.74E-22 | 2.90E-21 |
| chr1 | 219105171 | 219148159 | 42989 | -10.49 | 1.75E-22 | 2.91E-21 |
| chr2 | 240090918 | 240111647 | 20730 | -10.33 | 1.76E-22 | 2.93E-21 |
| chr12 | 65613021 | 65625335 | 12315 | -10.31 | 1.77E-22 | 2.94E-21 |
| chr8 | 123660453 | 123726643 | 66191 | -10.42 | 1.77E-22 | 2.94E-21 |
| chrX | 62749369 | 62808826 | 59458 | -10.32 | 1.78E-22 | 2.96E-21 |
| chr3 | 30664691 | 30698415 | 33725 | -10.34 | 1.78E-22 | 2.96E-21 |
| chr8 | 27706726 | 27728892 | 22167 | -10.44 | 1.81E-22 | 3.00E-21 |
| chr5 | 124574964 | 124615650 | 40687 | -10.58 | 1.81E-22 | 3.00E-21 |
| chr7 | 116136068 | 116159339 | 23272 | -10.33 | 1.81E-22 | 3.00E-21 |
| chr1 | 22620333 | 22649402 | 29070 | -10.4 | 1.81E-22 | 3.01E-21 |
| chr5 | 180478509 | 180515518 | 37010 | -10.35 | 1.83E-22 | 3.03E-21 |
| chr5 | 74267351 | 74310615 | 43265 | -10.38 | 1.84E-22 | 3.05E-21 |
| chr17 | 47497959 | 47508214 | 10256 | -10.32 | 1.87E-22 | 3.09E-21 |
| chr2 | 223848170 | 223864030 | 15861 | -7.01 | 1.88E-22 | 3.11E-21 |
| chrX | 9117743 | 9150206 | 32464 | -10.35 | 1.92E-22 | 3.17E-21 |
| chr7 | 137846761 | 137875981 | 29221 | -10.54 | 1.92E-22 | 3.17E-21 |
| chr10 | 123735016 | 123773055 | 38040 | -10.61 | 1.92E-22 | 3.18E-21 |
| chr12 | 43638256 | 43698640 | 60385 | -10.3 | 1.93E-22 | 3.18E-21 |
| chr12 | 119684733 | 119963072 | 278340 | 1.72 | 1.93E-22 | 3.19E-21 |
| chr2 | 133516470 | 133556539 | 40070 | 10.34 | 1.93E-22 | 3.19E-21 |
| chr7 | 47790850 | 47834218 | 43369 | -10.34 | 1.93E-22 | 3.19E-21 |
| chr12 | 118182459 | 118235889 | 53431 | -10.37 | 1.93E-22 | 3.19E-21 |
| chr11 | 123584293 | 123934954 | 350662 | 1.62 | 1.94E-22 | 3.20E-21 |
| chr1 | 29694426 | 29707801 | 13376 | -10.31 | 1.94E-22 | 3.20E-21 |
| chr11 | 27070646 | 27126256 | 55611 | 10.47 | 1.95E-22 | 3.22E-21 |
| chr14 | 104445249 | 104458017 | 12769 | -10.32 | 1.96E-22 | 3.23E-21 |
| chr8 | 24321636 | 24352478 | 30843 | 3.7 | 1.97E-22 | 3.25E-21 |

| | | | | | | |
|---|---|---|---|---|---|---|
| chr7 | 17291645 | 17315013 | 23369 | -10.37 | 1.98E-22 | 3.26E-21 |
| chr1 | 109325435 | 109336915 | 11481 | -10.32 | 1.99E-22 | 3.28E-21 |
| chr19 | 48681973 | 48694181 | 12209 | -10.3 | 2.01E-22 | 3.31E-21 |
| chr3 | 156587803 | 156614321 | 26519 | -10.31 | 2.01E-22 | 3.31E-21 |
| chr7 | 77433981 | 77466039 | 32059 | -10.44 | 2.09E-22 | 3.44E-21 |
| chr14 | 48746541 | 48858201 | 111661 | -10.49 | 2.11E-22 | 3.47E-21 |
| chr2 | 175364729 | 175393907 | 29179 | -10.46 | 2.11E-22 | 3.47E-21 |
| chr2 | 158557185 | 158586700 | 29516 | -10.4 | 2.13E-22 | 3.49E-21 |
| chr9 | 118349394 | 118386402 | 37009 | -10.31 | 2.13E-22 | 3.49E-21 |
| chr9 | 17240835 | 17263029 | 22195 | -10.29 | 2.14E-22 | 3.50E-21 |
| chr12 | 103013537 | 103073348 | 59812 | -10.45 | 2.14E-22 | 3.51E-21 |
| chr21 | 35531039 | 35551455 | 20417 | -10.29 | 2.16E-22 | 3.54E-21 |
| chr8 | 48125215 | 48162934 | 37720 | -10.48 | 2.19E-22 | 3.59E-21 |
| chr6 | 95831122 | 96007021 | 175900 | 3.94 | 2.21E-22 | 3.61E-21 |
| chr1 | 42752895 | 42771738 | 18844 | -10.37 | 2.22E-22 | 3.63E-21 |
| chr2 | 72688444 | 72708058 | 19615 | -10.44 | 2.26E-22 | 3.69E-21 |
| chr1 | 180345350 | 180391949 | 46600 | -10.33 | 2.26E-22 | 3.69E-21 |
| chr15 | 52151112 | 52164999 | 13888 | -10.29 | 2.26E-22 | 3.69E-21 |
| chr7 | 100528493 | 100543411 | 14919 | -10.34 | 2.26E-22 | 3.70E-21 |
| chr4 | 160973727 | 161036317 | 62591 | -10.36 | 2.28E-22 | 3.72E-21 |
| chr2 | 207204059 | 207260405 | 56347 | -10.41 | 2.31E-22 | 3.77E-21 |
| chr7 | 46807325 | 46879043 | 71719 | -10.45 | 2.33E-22 | 3.80E-21 |
| chr20 | 1326013 | 1342683 | 16671 | -10.28 | 2.39E-22 | 3.89E-21 |
| chr14 | 107107571 | 107124180 | 16610 | -3.82 | 2.39E-22 | 3.90E-21 |
| chr1 | 45689878 | 45729445 | 39568 | -10.48 | 2.39E-22 | 3.90E-21 |
| chr1 | 172296028 | 172339898 | 43871 | -10.28 | 2.41E-22 | 3.92E-21 |
| chr5 | 125138248 | 125200109 | 61862 | -10.44 | 2.43E-22 | 3.95E-21 |
| chr14 | 106398993 | 106410743 | 11751 | -10.28 | 2.44E-22 | 3.96E-21 |
| chr20 | 61630646 | 61653585 | 22940 | -10.3 | 2.44E-22 | 3.96E-21 |
| chr11 | 61785722 | 61817708 | 31987 | -10.49 | 2.44E-22 | 3.97E-21 |
| chr3 | 125026531 | 125071348 | 44818 | -10.35 | 2.46E-22 | 4.00E-21 |
| chr20 | 688425 | 696726 | 8302 | -10.28 | 2.46E-22 | 4.00E-21 |
| chr14 | 102107967 | 102118783 | 10817 | -10.3 | 2.52E-22 | 4.10E-21 |
| chr9 | 96170792 | 96189210 | 18419 | -3.92 | 2.55E-22 | 4.14E-21 |
| chr4 | 129845626 | 129871348 | 25723 | -10.33 | 2.57E-22 | 4.16E-21 |
| chr10 | 59775387 | 59803019 | 27633 | -10.34 | 2.58E-22 | 4.19E-21 |
| chr12 | 20347999 | 20392471 | 44473 | -10.75 | 2.60E-22 | 4.22E-21 |
| chr8 | 104259225 | 104304028 | 44804 | -10.29 | 2.60E-22 | 4.22E-21 |
| chr19 | 36095421 | 36100787 | 5367 | -10.27 | 2.62E-22 | 4.25E-21 |
| chr15 | 70880969 | 70917029 | 36061 | -10.35 | 2.63E-22 | 4.26E-21 |
| chr9 | 30252833 | 30351726 | 98894 | -10.31 | 2.64E-22 | 4.27E-21 |
| chr2 | 218150055 | 218174927 | 24873 | -10.29 | 2.66E-22 | 4.31E-21 |
| chr2 | 86230906 | 86244243 | 13338 | -10.39 | 2.68E-22 | 4.33E-21 |
| chr2 | 221265491 | 221288777 | 23287 | -10.29 | 2.71E-22 | 4.37E-21 |
| chr4 | 6703369 | 6717498 | 14130 | -10.3 | 2.72E-22 | 4.39E-21 |
| chr13 | 85517104 | 86484586 | 967483 | 4.49 | 2.73E-22 | 4.40E-21 |
| chr5 | 32716460 | 32733556 | 17097 | -10.29 | 2.74E-22 | 4.43E-21 |
| chr8 | 29400412 | 29440826 | 40415 | -10.3 | 2.76E-22 | 4.45E-21 |
| chr9 | 93286012 | 93326043 | 40032 | -10.27 | 2.78E-22 | 4.48E-21 |
| chr8 | 140460814 | 140593619 | 132806 | 1.85 | 2.80E-22 | 4.51E-21 |
| chr7 | 16410974 | 16444528 | 33555 | -10.3 | 2.85E-22 | 4.58E-21 |
| chr8 | 143500295 | 143518410 | 18116 | -10.32 | 2.87E-22 | 4.63E-21 |
| chr5 | 95563860 | 95607816 | 43957 | -10.36 | 2.88E-22 | 4.64E-21 |
| chr9 | 99934510 | 99983497 | 48988 | -10.28 | 2.91E-22 | 4.69E-21 |
| chr5 | 175128681 | 175161128 | 32448 | -10.4 | 2.92E-22 | 4.70E-21 |
| chr13 | 74003085 | 74035498 | 32414 | -10.41 | 2.96E-22 | 4.76E-21 |
| chr3 | 10578952 | 10587725 | 8774 | -3.62 | 2.99E-22 | 4.81E-21 |
| chr20 | 56679567 | 56703264 | 23698 | -10.29 | 3.00E-22 | 4.82E-21 |
| chr1 | 212562141 | 212581897 | 19757 | -10.28 | 3.03E-22 | 4.86E-21 |
| chr3 | 8969690 | 8988431 | 18742 | -10.43 | 3.04E-22 | 4.88E-21 |
| chr2 | 234919164 | 234978558 | 59395 | -10.37 | 3.05E-22 | 4.90E-21 |

| | | | | | | |
|---|---|---|---|---|---|---|
| chr16 | 47847745 | 47872707 | 24963 | -10.27 | 3.07E-22 | 4.92E-21 |
| chr17 | 11846062 | 11882834 | 36773 | -10.26 | 3.09E-22 | 4.96E-21 |
| chr1 | 151857929 | 151885979 | 28051 | -10.26 | 3.10E-22 | 4.97E-21 |
| chr12 | 41394211 | 41406904 | 12694 | -10.26 | 3.11E-22 | 4.98E-21 |
| chr8 | 39886781 | 39940436 | 53656 | -10.72 | 3.11E-22 | 4.98E-21 |
| chr12 | 43110905 | 43150252 | 39348 | -10.26 | 3.12E-22 | 5.00E-21 |
| chr8 | 53773843 | 53796351 | 22509 | -10.26 | 3.13E-22 | 5.00E-21 |
| chr2 | 102980279 | 103003261 | 22983 | -10.26 | 3.15E-22 | 5.04E-21 |
| chrX | 40164672 | 40225593 | 60922 | -10.45 | 3.15E-22 | 5.04E-21 |
| chr4 | 169335603 | 169416197 | 80595 | -10.53 | 3.17E-22 | 5.06E-21 |
| chr9 | 28899593 | 28934259 | 34667 | -10.31 | 3.18E-22 | 5.08E-21 |
| chr10 | 72374432 | 72393666 | 19235 | -10.32 | 3.22E-22 | 5.14E-21 |
| chr1 | 2854678 | 2876310 | 21633 | -10.28 | 3.25E-22 | 5.18E-21 |
| chr2 | 143756442 | 143776227 | 19786 | -10.36 | 3.29E-22 | 5.25E-21 |
| chr14 | 95256888 | 95323640 | 66753 | -10.31 | 3.41E-22 | 5.44E-21 |
| chr9 | 38201224 | 38269207 | 67984 | -10.29 | 3.44E-22 | 5.48E-21 |
| chr11 | 12318274 | 12340323 | 22050 | -10.25 | 3.47E-22 | 5.52E-21 |
| chr9 | 125035080 | 125041108 | 6029 | -10.32 | 3.49E-22 | 5.55E-21 |
| chr4 | 5068450 | 5099935 | 31486 | -10.3 | 3.49E-22 | 5.56E-21 |
| chr2 | 215638895 | 215660705 | 21811 | -10.26 | 3.51E-22 | 5.58E-21 |
| chr1 | 55286646 | 55298810 | 12165 | -10.26 | 3.52E-22 | 5.60E-21 |
| chr4 | 58176884 | 58227123 | 50240 | 10.28 | 3.55E-22 | 5.65E-21 |
| chr4 | 162702715 | 162730676 | 27962 | -10.25 | 3.62E-22 | 5.75E-21 |
| chr19 | 20595689 | 20673307 | 77619 | -10.39 | 3.62E-22 | 5.75E-21 |
| chr22 | 42395004 | 42411116 | 16113 | -10.32 | 3.65E-22 | 5.79E-21 |
| chr17 | 37150263 | 37165592 | 15330 | -10.37 | 3.68E-22 | 5.85E-21 |
| chr10 | 29450811 | 29465417 | 14607 | -10.24 | 3.69E-22 | 5.85E-21 |
| chr14 | 101846057 | 101860319 | 14263 | -10.24 | 3.70E-22 | 5.87E-21 |
| chr12 | 111738531 | 111746873 | 8343 | -10.25 | 3.72E-22 | 5.89E-21 |
| chr1 | 42958957 | 42989930 | 30974 | -10.43 | 3.74E-22 | 5.92E-21 |
| chr9 | 120258303 | 120299522 | 41220 | 10.32 | 3.74E-22 | 5.92E-21 |
| chr5 | 31282131 | 31320339 | 38209 | 10.29 | 3.78E-22 | 5.98E-21 |
| chr18 | 33296597 | 33338947 | 42351 | -10.25 | 3.83E-22 | 6.05E-21 |
| chrX | 57263193 | 57321987 | 58795 | -10.39 | 3.83E-22 | 6.06E-21 |
| chr15 | 93431744 | 93442904 | 11161 | -7.2 | 3.84E-22 | 6.07E-21 |
| chrX | 10313757 | 10370373 | 56617 | -10.4 | 3.86E-22 | 6.09E-21 |
| chr7 | 721257 | 744518 | 23262 | -10.29 | 3.87E-22 | 6.11E-21 |
| chr12 | 88596725 | 88619774 | 23050 | -10.33 | 3.88E-22 | 6.13E-21 |
| chr1 | 75318400 | 75373702 | 55303 | 2.33 | 3.89E-22 | 6.14E-21 |
| chrX | 151165592 | 151236889 | 71298 | -10.24 | 3.90E-22 | 6.15E-21 |
| chr10 | 114604533 | 114672450 | 67918 | -10.48 | 3.93E-22 | 6.20E-21 |
| chrX | 128336023 | 128374880 | 38858 | 10.29 | 3.95E-22 | 6.23E-21 |
| chr12 | 124299016 | 124320745 | 21730 | -10.47 | 3.97E-22 | 6.26E-21 |
| chr9 | 102684408 | 102700389 | 15982 | -10.31 | 3.98E-22 | 6.27E-21 |
| chr14 | 92666497 | 92716746 | 50250 | -2.18 | 4.01E-22 | 6.31E-21 |
| chr10 | 47015270 | 47055298 | 40029 | -10.6 | 4.05E-22 | 6.37E-21 |
| chr16 | 80214029 | 80275976 | 61948 | -10.25 | 4.11E-22 | 6.45E-21 |
| chr8 | 143634776 | 143644390 | 9615 | -10.25 | 4.11E-22 | 6.45E-21 |
| chr12 | 124757786 | 124774042 | 16257 | -10.37 | 4.12E-22 | 6.48E-21 |
| chr15 | 68674630 | 68697215 | 22586 | -10.23 | 4.13E-22 | 6.49E-21 |
| chr13 | 40869285 | 40882305 | 13021 | -8.68 | 4.20E-22 | 6.60E-21 |
| chr14 | 68396416 | 68437221 | 40806 | -10.42 | 4.22E-22 | 6.62E-21 |
| chr19 | 38587930 | 38592690 | 4761 | -10.24 | 4.22E-22 | 6.62E-21 |
| chr10 | 43503718 | 43531274 | 27557 | -10.24 | 4.24E-22 | 6.65E-21 |
| chr14 | 72211757 | 72300067 | 88311 | -4.57 | 4.26E-22 | 6.67E-21 |
| chr1 | 169530988 | 169534924 | 3937 | -7.14 | 4.26E-22 | 6.67E-21 |
| chr2 | 86974867 | 86980246 | 5380 | -10.23 | 4.27E-22 | 6.68E-21 |
| chr12 | 47990677 | 48009253 | 18577 | -10.24 | 4.28E-22 | 6.71E-21 |
| chr7 | 80747643 | 80811686 | 64044 | -10.23 | 4.29E-22 | 6.72E-21 |
| chr10 | 124714018 | 124731079 | 17062 | -10.22 | 4.33E-22 | 6.77E-21 |
| chr2 | 136267775 | 136295727 | 27953 | -10.23 | 4.34E-22 | 6.78E-21 |

| | | | | | | |
|---|---|---|---|---|---|---|
| chr12 | 40677864 | 40692450 | 14587 | -10.23 | 4.34E-22 | 6.78E-21 |
| chr12 | 59536387 | 59555439 | 19053 | -10.24 | 4.37E-22 | 6.83E-21 |
| chr2 | 112383922 | 112406715 | 22794 | -10.31 | 4.41E-22 | 6.88E-21 |
| chr1 | 41601367 | 41616051 | 14685 | -10.33 | 4.41E-22 | 6.88E-21 |
| chr2 | 220977462 | 221011284 | 33823 | -10.36 | 4.55E-22 | 7.10E-21 |
| chr10 | 69728695 | 69735347 | 6653 | -10.27 | 4.55E-22 | 7.10E-21 |
| chr10 | 49460990 | 49494794 | 33805 | -10.3 | 4.58E-22 | 7.14E-21 |
| chr14 | 105865299 | 105873676 | 8378 | -10.24 | 4.63E-22 | 7.21E-21 |
| chr5 | 37254167 | 37288705 | 34539 | -10.38 | 4.66E-22 | 7.27E-21 |
| chr15 | 30012547 | 30026201 | 13655 | -10.29 | 4.67E-22 | 7.27E-21 |
| chr7 | 157824585 | 157855012 | 30428 | -10.22 | 4.72E-22 | 7.34E-21 |
| chr12 | 16543987 | 16595213 | 51227 | -10.39 | 4.81E-22 | 7.48E-21 |
| chr3 | 189613040 | 189666063 | 53024 | -10.28 | 4.83E-22 | 7.51E-21 |
| chr10 | 1043035 | 1049613 | 6579 | -10.29 | 4.84E-22 | 7.53E-21 |
| chr2 | 241271055 | 241301638 | 30584 | -10.31 | 4.84E-22 | 7.53E-21 |
| chr1 | 228211766 | 228247765 | 36000 | -10.25 | 4.87E-22 | 7.56E-21 |
| chr7 | 108000406 | 108013956 | 13551 | -10.49 | 4.89E-22 | 7.59E-21 |
| chr1 | 31028937 | 31040727 | 11791 | -7.09 | 4.93E-22 | 7.65E-21 |
| chr7 | 45396668 | 45453496 | 56829 | -8.16 | 4.94E-22 | 7.67E-21 |
| chr3 | 37062645 | 37078877 | 16233 | -10.27 | 4.96E-22 | 7.69E-21 |
| chr2 | 7930185 | 7995697 | 65513 | -10.35 | 5.01E-22 | 7.77E-21 |
| chr2 | 231694041 | 231724943 | 30903 | -10.3 | 5.04E-22 | 7.80E-21 |
| chr2 | 3396624 | 3402333 | 5710 | -10.22 | 5.04E-22 | 7.81E-21 |
| chr6 | 144521721 | 144569511 | 47791 | -10.23 | 5.05E-22 | 7.81E-21 |
| chr10 | 135222716 | 135227222 | 4507 | -10.25 | 5.09E-22 | 7.88E-21 |
| chr12 | 70142537 | 70166455 | 23919 | -10.21 | 5.10E-22 | 7.89E-21 |
| chr14 | 76467911 | 76492073 | 24163 | -10.22 | 5.12E-22 | 7.91E-21 |
| chr2 | 72576311 | 72648390 | 72080 | -10.42 | 5.24E-22 | 8.10E-21 |
| chr2 | 235540569 | 235587842 | 47274 | -10.47 | 5.26E-22 | 8.13E-21 |
| chr9 | 82089228 | 82116257 | 27030 | -10.26 | 5.31E-22 | 8.19E-21 |
| chr7 | 31677452 | 31680744 | 3293 | -3.41 | 5.37E-22 | 8.29E-21 |
| chr7 | 48037648 | 48064913 | 27266 | -10.32 | 5.38E-22 | 8.30E-21 |
| chr1 | 185538057 | 185558310 | 20254 | -10.22 | 5.43E-22 | 8.37E-21 |
| chr2 | 86721017 | 86730919 | 9903 | -10.25 | 5.47E-22 | 8.43E-21 |
| chr11 | 59785964 | 59860076 | 74113 | 2.01 | 5.49E-22 | 8.45E-21 |
| chr1 | 10451936 | 10457556 | 5621 | -10.2 | 5.55E-22 | 8.55E-21 |
| chr13 | 94444374 | 94513116 | 68743 | 10.51 | 5.57E-22 | 8.58E-21 |
| chr4 | 171190518 | 171598740 | 408223 | 1.98 | 5.58E-22 | 8.59E-21 |
| chr2 | 40025682 | 40043278 | 17597 | -2.32 | 5.58E-22 | 8.59E-21 |
| chr2 | 191106537 | 191143042 | 36506 | -10.24 | 5.60E-22 | 8.61E-21 |
| chr1 | 104008200 | 104045972 | 37773 | 10.33 | 5.69E-22 | 8.76E-21 |
| chr9 | 80128936 | 80161910 | 32975 | -10.29 | 5.70E-22 | 8.76E-21 |
| chr9 | 98622911 | 98636377 | 13467 | -10.2 | 5.75E-22 | 8.83E-21 |
| chr12 | 64427826 | 64449896 | 22071 | -10.21 | 5.78E-22 | 8.87E-21 |
| chr9 | 94461957 | 94479430 | 17474 | -10.2 | 5.82E-22 | 8.94E-21 |
| chr7 | 81106677 | 81144284 | 37608 | -10.4 | 5.84E-22 | 8.95E-21 |
| chr10 | 8643271 | 8685455 | 42185 | -10.26 | 5.87E-22 | 9.00E-21 |
| chr15 | 86412879 | 87032182 | 619304 | 3.71 | 5.87E-22 | 9.00E-21 |
| chr14 | 70795032 | 70824838 | 29807 | -10.21 | 5.89E-22 | 9.03E-21 |
| chr2 | 149353963 | 149362241 | 8279 | -10.2 | 5.93E-22 | 9.09E-21 |
| chr3 | 88114142 | 88154517 | 40376 | -10.22 | 6.02E-22 | 9.22E-21 |
| chr10 | 4967068 | 5012841 | 45774 | -10.19 | 6.03E-22 | 9.23E-21 |
| chr5 | 37135162 | 37146271 | 11110 | -10.2 | 6.04E-22 | 9.24E-21 |
| chr8 | 144912025 | 144940730 | 28706 | -10.27 | 6.05E-22 | 9.25E-21 |
| chr8 | 11795326 | 11806558 | 11233 | -10.22 | 6.05E-22 | 9.25E-21 |
| chr13 | 21138091 | 21159899 | 21809 | -10.22 | 6.08E-22 | 9.29E-21 |
| chr22 | 50258161 | 50291971 | 33811 | -10.2 | 6.09E-22 | 9.30E-21 |
| chr3 | 126886123 | 126923143 | 37021 | -10.22 | 6.13E-22 | 9.36E-21 |
| chr1 | 225119078 | 225135130 | 16053 | -10.25 | 6.14E-22 | 9.38E-21 |
| chr2 | 30881132 | 30896495 | 15364 | -10.28 | 6.16E-22 | 9.39E-21 |
| chr15 | 99502349 | 99508633 | 6285 | -10.2 | 6.20E-22 | 9.46E-21 |

| | | | | | | |
|---|---|---|---|---|---|---|
| chrX | 83025052 | 83163094 | 138043 | 2.62 | 6.23E-22 | 9.49E-21 |
| chrX | 20251807 | 20273887 | 22081 | -10.19 | 6.30E-22 | 9.59E-21 |
| chr7 | 134090091 | 134127865 | 37775 | -10.24 | 6.39E-22 | 9.74E-21 |
| chr17 | 70785196 | 70836122 | 50927 | -10.54 | 6.43E-22 | 9.78E-21 |
| chr8 | 110224493 | 110241177 | 16685 | -10.19 | 6.49E-22 | 9.87E-21 |
| chr4 | 37307438 | 37339314 | 31877 | 2.83 | 6.50E-22 | 9.89E-21 |
| chr14 | 53946861 | 53974284 | 27424 | -10.19 | 6.51E-22 | 9.90E-21 |
| chr14 | 92135267 | 92177739 | 42473 | -10.4 | 6.52E-22 | 9.91E-21 |
| chr8 | 9217908 | 9259242 | 41335 | -10.2 | 6.56E-22 | 9.96E-21 |
| chr5 | 14252345 | 14260569 | 8225 | -10.26 | 6.56E-22 | 9.96E-21 |
| chr2 | 8340059 | 8354876 | 14818 | -10.18 | 6.57E-22 | 9.97E-21 |
| chr8 | 49628186 | 49666106 | 37921 | -10.27 | 6.65E-22 | 1.01E-20 |
| chr17 | 35352571 | 35360322 | 7752 | -10.39 | 6.66E-22 | 1.01E-20 |
| chr9 | 137492856 | 137508692 | 15837 | -10.22 | 6.68E-22 | 1.01E-20 |
| chr6 | 153276497 | 153296336 | 19840 | -10.26 | 6.71E-22 | 1.02E-20 |
| chrX | 105665764 | 105712632 | 46869 | 10.39 | 6.76E-22 | 1.02E-20 |
| chr2 | 36850399 | 36888742 | 38344 | -10.24 | 6.78E-22 | 1.03E-20 |
| chr12 | 20988835 | 21008425 | 19591 | -10.47 | 6.78E-22 | 1.03E-20 |
| chr2 | 239881053 | 239925784 | 44732 | -10.33 | 6.87E-22 | 1.04E-20 |
| chr11 | 68990687 | 69028150 | 37464 | -10.22 | 6.89E-22 | 1.04E-20 |
| chr17 | 21039274 | 21060324 | 21051 | -10.33 | 6.91E-22 | 1.05E-20 |
| chr22 | 33205114 | 33374376 | 169263 | 1.55 | 6.92E-22 | 1.05E-20 |
| chr17 | 77634219 | 77650219 | 16001 | -10.32 | 6.93E-22 | 1.05E-20 |
| chr12 | 90500926 | 90537383 | 36458 | -10.22 | 7.00E-22 | 1.06E-20 |
| chr8 | 77843610 | 77858544 | 14935 | 4.38 | 7.04E-22 | 1.06E-20 |
| chr4 | 19685830 | 19743008 | 57179 | -10.18 | 7.08E-22 | 1.07E-20 |
| chr11 | 29269703 | 29317703 | 48001 | -10.2 | 7.10E-22 | 1.07E-20 |
| chr12 | 58349376 | 58373123 | 23748 | -10.18 | 7.13E-22 | 1.07E-20 |
| chr5 | 38617314 | 38648374 | 31061 | -8.81 | 7.13E-22 | 1.07E-20 |
| chrX | 140389101 | 140458004 | 68904 | -10.57 | 7.14E-22 | 1.08E-20 |
| chr19 | 5500486 | 5536394 | 35909 | -10.28 | 7.14E-22 | 1.08E-20 |
| chr16 | 72338521 | 72376049 | 37529 | -10.18 | 7.15E-22 | 1.08E-20 |
| chr1 | 90386956 | 90411320 | 24365 | -10.29 | 7.21E-22 | 1.08E-20 |
| chr12 | 85321966 | 85374700 | 52735 | 10.44 | 7.25E-22 | 1.09E-20 |
| chr7 | 3090322 | 3122770 | 32449 | -10.24 | 7.26E-22 | 1.09E-20 |
| chr4 | 22519567 | 22572493 | 52927 | -10.18 | 7.34E-22 | 1.10E-20 |
| chr6 | 9845203 | 9889761 | 44559 | -10.4 | 7.34E-22 | 1.10E-20 |
| chrX | 13068841 | 13095045 | 26205 | -10.19 | 7.42E-22 | 1.11E-20 |
| chr2 | 62495048 | 62514090 | 19043 | -10.32 | 7.47E-22 | 1.12E-20 |
| chr18 | 77807172 | 77819256 | 12085 | -10.22 | 7.47E-22 | 1.12E-20 |
| chr3 | 95908224 | 96013704 | 105481 | -10.49 | 7.49E-22 | 1.12E-20 |
| chr4 | 69927374 | 69970050 | 42677 | -10.18 | 7.53E-22 | 1.13E-20 |
| chr4 | 41713401 | 41742128 | 28728 | -10.18 | 7.62E-22 | 1.14E-20 |
| chr3 | 96072929 | 96166367 | 93439 | -10.18 | 7.64E-22 | 1.14E-20 |
| chr18 | 56413152 | 56418377 | 5226 | -10.29 | 7.67E-22 | 1.15E-20 |
| chr20 | 19767303 | 19801299 | 33997 | -10.4 | 7.79E-22 | 1.17E-20 |
| chr14 | 97513451 | 97565717 | 52267 | 10.38 | 7.80E-22 | 1.17E-20 |
| chr7 | 4149973 | 4167399 | 17427 | -10.18 | 7.99E-22 | 1.20E-20 |
| chr12 | 63482340 | 63525393 | 43054 | -10.22 | 8.00E-22 | 1.20E-20 |
| chr7 | 137955597 | 138039114 | 83518 | -10.65 | 8.05E-22 | 1.20E-20 |
| chr17 | 15288431 | 15299322 | 10892 | -2.8 | 8.11E-22 | 1.21E-20 |
| chr1 | 35980601 | 35999618 | 19018 | -10.17 | 8.14E-22 | 1.22E-20 |
| chr3 | 15678907 | 15709765 | 30859 | -10.19 | 8.23E-22 | 1.23E-20 |
| chr7 | 140715629 | 140732481 | 16853 | -10.18 | 8.24E-22 | 1.23E-20 |
| chr11 | 121156701 | 121162296 | 5596 | -3.6 | 8.27E-22 | 1.23E-20 |
| chr11 | 57767523 | 58271985 | 504463 | 6.47 | 8.28E-22 | 1.23E-20 |
| chr4 | 123961165 | 123974529 | 13365 | -10.18 | 8.28E-22 | 1.23E-20 |
| chr2 | 241866675 | 241882157 | 15483 | -10.17 | 8.36E-22 | 1.25E-20 |
| chr12 | 91911570 | 91943930 | 32361 | -10.16 | 8.38E-22 | 1.25E-20 |
| chr2 | 161662150 | 161701920 | 39771 | -10.16 | 8.50E-22 | 1.27E-20 |
| chrX | 73287664 | 73341433 | 53770 | -10.32 | 8.51E-22 | 1.27E-20 |

| | | | | | | |
|---|---|---|---|---|---|---|
| chr13 | 20580586 | 20591923 | 11338 | -10.16 | 8.52E-22 | 1.27E-20 |
| chr9 | 87735365 | 87762197 | 26833 | -10.24 | 8.52E-22 | 1.27E-20 |
| chr6 | 155640052 | 155688645 | 48594 | -10.17 | 8.59E-22 | 1.28E-20 |
| chr2 | 47078656 | 47114662 | 36007 | -10.17 | 8.60E-22 | 1.28E-20 |
| chr11 | 86295158 | 86361962 | 66805 | -10.17 | 8.76E-22 | 1.30E-20 |
| chr2 | 222969069 | 223009265 | 40197 | 10.25 | 8.90E-22 | 1.32E-20 |
| chr5 | 68211999 | 68238648 | 26650 | -10.15 | 8.98E-22 | 1.33E-20 |
| chr13 | 34593147 | 34629724 | 36578 | -10.31 | 8.99E-22 | 1.33E-20 |
| chr7 | 3523938 | 3552217 | 28280 | -10.18 | 9.07E-22 | 1.35E-20 |
| chr8 | 109937122 | 110000572 | 63451 | -10.3 | 9.25E-22 | 1.37E-20 |
| chr6 | 126677349 | 126734240 | 56892 | -10.15 | 9.39E-22 | 1.39E-20 |
| chr20 | 2781087 | 2794256 | 13170 | -10.21 | 9.41E-22 | 1.39E-20 |
| chr10 | 94312222 | 94321169 | 8948 | -10.15 | 9.45E-22 | 1.40E-20 |
| chr17 | 15676607 | 15686742 | 10136 | -10.16 | 9.55E-22 | 1.41E-20 |
| chr21 | 16394398 | 16420936 | 26539 | -10.15 | 9.62E-22 | 1.42E-20 |
| chr20 | 33067277 | 33089464 | 22188 | -10.16 | 9.67E-22 | 1.43E-20 |
| chr4 | 103004742 | 103056995 | 52254 | -10.18 | 9.70E-22 | 1.43E-20 |
| chr5 | 23705442 | 23801745 | 96304 | -10.16 | 9.72E-22 | 1.44E-20 |
| chr15 | 59417017 | 59427317 | 10301 | -10.15 | 9.73E-22 | 1.44E-20 |
| chr22 | 32402375 | 32417874 | 15500 | -10.15 | 9.74E-22 | 1.44E-20 |
| chr9 | 103046041 | 103052506 | 6466 | -10.15 | 9.74E-22 | 1.44E-20 |
| chr8 | 33038758 | 33072311 | 33554 | -10.15 | 9.82E-22 | 1.45E-20 |
| chr3 | 5547017 | 5593415 | 46399 | -10.26 | 9.83E-22 | 1.45E-20 |
| chrX | 74603904 | 74619462 | 15559 | 2.7 | 9.87E-22 | 1.45E-20 |
| chr6 | 170080060 | 170097891 | 17832 | -10.16 | 9.91E-22 | 1.46E-20 |
| chr7 | 68070498 | 68084286 | 13789 | -10.2 | 9.95E-22 | 1.47E-20 |
| chr3 | 190707403 | 190759721 | 52319 | 10.18 | 9.95E-22 | 1.47E-20 |
| chr1 | 2695715 | 2712980 | 17266 | -10.17 | 1.00E-21 | 1.48E-20 |
| chr5 | 139465186 | 139472141 | 6956 | -7.7 | 1.00E-21 | 1.48E-20 |
| chr3 | 194603174 | 194626213 | 23040 | -10.15 | 1.01E-21 | 1.48E-20 |
| chr10 | 127562705 | 127567416 | 4712 | -10.17 | 1.01E-21 | 1.48E-20 |
| chr1 | 4247587 | 4322168 | 74582 | -1.77 | 1.01E-21 | 1.48E-20 |
| chr15 | 55842080 | 55865807 | 23728 | -10.14 | 1.01E-21 | 1.49E-20 |
| chr8 | 143107707 | 143137534 | 29828 | -10.23 | 1.03E-21 | 1.51E-20 |
| chr17 | 17535080 | 17563948 | 28869 | -10.26 | 1.03E-21 | 1.51E-20 |
| chr11 | 102414368 | 102468451 | 54084 | -10.23 | 1.04E-21 | 1.52E-20 |
| chr11 | 34485826 | 34491259 | 5434 | -10.14 | 1.04E-21 | 1.53E-20 |
| chr12 | 94255279 | 94271308 | 16030 | -10.14 | 1.04E-21 | 1.53E-20 |
| chr7 | 103283278 | 103289925 | 6648 | -2.78 | 1.05E-21 | 1.53E-20 |
| chr17 | 22173835 | 22203425 | 29591 | -10.3 | 1.05E-21 | 1.53E-20 |
| chr2 | 86915995 | 86936154 | 20160 | -10.29 | 1.06E-21 | 1.55E-20 |
| chr2 | 15895431 | 15902987 | 7557 | -3.63 | 1.06E-21 | 1.55E-20 |
| chr13 | 45534842 | 45540392 | 5551 | -10.18 | 1.07E-21 | 1.56E-20 |
| chr10 | 93203143 | 93232887 | 29745 | -10.26 | 1.09E-21 | 1.59E-20 |
| chr15 | 35183692 | 35187814 | 4123 | -10.23 | 1.09E-21 | 1.59E-20 |
| chr20 | 24955703 | 24963802 | 8100 | -10.13 | 1.10E-21 | 1.61E-20 |
| chr4 | 154591455 | 154618935 | 27481 | -10.14 | 1.10E-21 | 1.61E-20 |
| chr7 | 32106777 | 32202364 | 95588 | -10.44 | 1.11E-21 | 1.62E-20 |
| chr9 | 98928724 | 98993664 | 64941 | -10.94 | 1.11E-21 | 1.62E-20 |
| chr10 | 22102277 | 22123486 | 21210 | -10.13 | 1.11E-21 | 1.62E-20 |
| chr9 | 120191752 | 120226986 | 35235 | 10.31 | 1.11E-21 | 1.62E-20 |
| chr18 | 44448362 | 44471212 | 22851 | -10.14 | 1.12E-21 | 1.63E-20 |
| chr9 | 103890131 | 103940182 | 50052 | 10.17 | 1.13E-21 | 1.65E-20 |
| chr7 | 31904077 | 31946307 | 42231 | -10.38 | 1.13E-21 | 1.65E-20 |
| chr15 | 73058056 | 73062132 | 4077 | -10.22 | 1.14E-21 | 1.66E-20 |
| chr9 | 72307485 | 72319668 | 12184 | -10.25 | 1.14E-21 | 1.66E-20 |
| chr9 | 765646 | 795680 | 30035 | -10.28 | 1.15E-21 | 1.67E-20 |
| chr10 | 125943518 | 125968184 | 24667 | -10.15 | 1.15E-21 | 1.67E-20 |
| chr7 | 127823543 | 127845406 | 21864 | -10.19 | 1.16E-21 | 1.69E-20 |
| chr11 | 2300006 | 2311470 | 11465 | -10.13 | 1.16E-21 | 1.69E-20 |
| chr7 | 4014600 | 4074860 | 60261 | -10.26 | 1.17E-21 | 1.69E-20 |

| | | | | | | |
|---|---|---|---|---|---|---|
| chr3 | 162599236 | 162635535 | 36300 | -10.27 | 1.17E-21 | 1.70E-20 |
| chr5 | 175558595 | 175607182 | 48588 | -10.14 | 1.17E-21 | 1.70E-20 |
| chr10 | 119397792 | 119418249 | 20458 | -10.17 | 1.17E-21 | 1.70E-20 |
| chr4 | 8824643 | 8855412 | 30770 | -10.13 | 1.18E-21 | 1.71E-20 |
| chr1 | 206202061 | 206222298 | 20238 | -10.13 | 1.19E-21 | 1.73E-20 |
| chr3 | 142814056 | 142833026 | 18971 | -3.7 | 1.20E-21 | 1.74E-20 |
| chr7 | 41162266 | 41214770 | 52505 | -10.15 | 1.20E-21 | 1.74E-20 |
| chr15 | 71197081 | 71202685 | 5605 | -10.14 | 1.22E-21 | 1.77E-20 |
| chr5 | 174072277 | 174112316 | 40040 | -10.38 | 1.22E-21 | 1.77E-20 |
| chr2 | 19500391 | 19521754 | 21364 | -10.25 | 1.22E-21 | 1.77E-20 |
| chr12 | 45530855 | 45585347 | 54493 | -10.12 | 1.23E-21 | 1.78E-20 |
| chr9 | 117719860 | 117723864 | 4005 | -3.73 | 1.23E-21 | 1.78E-20 |
| chr7 | 134837750 | 134849076 | 11327 | -10.12 | 1.24E-21 | 1.79E-20 |
| chr8 | 8464848 | 8520634 | 55787 | -10.12 | 1.24E-21 | 1.80E-20 |
| chr15 | 85619781 | 85632074 | 12294 | -10.16 | 1.25E-21 | 1.81E-20 |
| chr6 | 28031074 | 28065420 | 34347 | -10.14 | 1.25E-21 | 1.81E-20 |
| chr18 | 69822172 | 69880520 | 58349 | -10.42 | 1.27E-21 | 1.84E-20 |
| chr1 | 240538554 | 240557662 | 19109 | -10.14 | 1.28E-21 | 1.85E-20 |
| chr8 | 101479012 | 101497428 | 18417 | -10.12 | 1.28E-21 | 1.85E-20 |
| chr5 | 67888026 | 67918730 | 30705 | -10.17 | 1.28E-21 | 1.85E-20 |
| chr8 | 128160774 | 128195563 | 34790 | -10.34 | 1.30E-21 | 1.87E-20 |
| chr7 | 105869293 | 105893666 | 24374 | -10.12 | 1.30E-21 | 1.88E-20 |
| chr4 | 20716141 | 20740597 | 24457 | -10.12 | 1.31E-21 | 1.88E-20 |
| chr11 | 117951576 | 117987431 | 35856 | -10.32 | 1.31E-21 | 1.88E-20 |
| chr8 | 98290728 | 98331021 | 40294 | 10.16 | 1.31E-21 | 1.89E-20 |
| chr1 | 2908742 | 2927689 | 18948 | -10.12 | 1.32E-21 | 1.90E-20 |
| chr5 | 17856308 | 17873066 | 16759 | -10.21 | 1.32E-21 | 1.90E-20 |
| chr6 | 64049151 | 64090627 | 41477 | -10.18 | 1.33E-21 | 1.91E-20 |
| chr16 | 14634495 | 14639415 | 4921 | -10.15 | 1.33E-21 | 1.91E-20 |
| chr4 | 84512399 | 84545837 | 33439 | -10.13 | 1.33E-21 | 1.91E-20 |
| chr1 | 185577319 | 185626652 | 49334 | -10.39 | 1.33E-21 | 1.91E-20 |
| chr6 | 28287667 | 28309361 | 21695 | -10.11 | 1.33E-21 | 1.91E-20 |
| chr17 | 41420495 | 41431158 | 10664 | -10.13 | 1.34E-21 | 1.92E-20 |
| chr20 | 10262114 | 10278153 | 16040 | -10.14 | 1.35E-21 | 1.93E-20 |
| chr14 | 43445244 | 43502896 | 57653 | -10.42 | 1.36E-21 | 1.95E-20 |
| chr2 | 190672163 | 190677774 | 5612 | -10.11 | 1.37E-21 | 1.96E-20 |
| chr16 | 72672175 | 72712724 | 40550 | -10.16 | 1.37E-21 | 1.97E-20 |
| chr21 | 46656422 | 46672017 | 15596 | -10.14 | 1.39E-21 | 2.00E-20 |
| chr5 | 173579976 | 173604731 | 24756 | -10.11 | 1.40E-21 | 2.00E-20 |
| chr1 | 25952036 | 25967373 | 15338 | -6.92 | 1.41E-21 | 2.02E-20 |
| chr10 | 127698597 | 127723907 | 25311 | -10.31 | 1.44E-21 | 2.05E-20 |
| chr11 | 45135232 | 45153750 | 18519 | -10.17 | 1.46E-21 | 2.08E-20 |
| chr11 | 74531900 | 74540926 | 9027 | -10.1 | 1.47E-21 | 2.11E-20 |
| chr18 | 8925598 | 8975240 | 49643 | -10.31 | 1.49E-21 | 2.12E-20 |
| chr9 | 77154070 | 77182955 | 28886 | 10.17 | 1.49E-21 | 2.13E-20 |
| chr3 | 150384343 | 150411125 | 26783 | -10.21 | 1.50E-21 | 2.15E-20 |
| chr8 | 125798562 | 125834659 | 36098 | -10.23 | 1.52E-21 | 2.17E-20 |
| chr18 | 53491900 | 53522880 | 30981 | -10.1 | 1.54E-21 | 2.20E-20 |
| chr9 | 86143935 | 86177514 | 33580 | -10.1 | 1.54E-21 | 2.20E-20 |
| chr18 | 55321473 | 55342525 | 21053 | -10.12 | 1.54E-21 | 2.20E-20 |
| chr5 | 58173972 | 58208183 | 34212 | -10.28 | 1.54E-21 | 2.20E-20 |
| chr2 | 97605273 | 97612208 | 6936 | -10.11 | 1.56E-21 | 2.22E-20 |
| chr5 | 169030938 | 169068489 | 37552 | -10.15 | 1.56E-21 | 2.22E-20 |
| chr7 | 95044464 | 95113186 | 68723 | -10.4 | 1.56E-21 | 2.22E-20 |
| chr18 | 68091438 | 68113820 | 22383 | -10.1 | 1.57E-21 | 2.24E-20 |
| chr4 | 133878573 | 133945338 | 66766 | 10.39 | 1.58E-21 | 2.24E-20 |
| chr10 | 93491557 | 93524292 | 32736 | -10.24 | 1.58E-21 | 2.25E-20 |
| chr3 | 5318607 | 5338837 | 20231 | -10.11 | 1.59E-21 | 2.26E-20 |
| chr10 | 118050111 | 118084641 | 34531 | 10.3 | 1.59E-21 | 2.27E-20 |
| chr8 | 101591299 | 101613487 | 22189 | -10.2 | 1.60E-21 | 2.28E-20 |
| chr22 | 44604148 | 44647804 | 43657 | -10.14 | 1.61E-21 | 2.28E-20 |

| | | | | | | |
|---|---|---|---|---|---|---|
| chr5 | 39081746 | 39106360 | 24615 | -10.42 | 1.62E-21 | 2.30E-20 |
| chr1 | 239381346 | 239421294 | 39949 | -10.1 | 1.63E-21 | 2.31E-20 |
| chr2 | 62982042 | 63011681 | 29640 | -10.12 | 1.64E-21 | 2.32E-20 |
| chr7 | 157556236 | 157568547 | 12312 | -6.92 | 1.66E-21 | 2.36E-20 |
| chr3 | 122944301 | 122960128 | 15828 | -10.1 | 1.66E-21 | 2.36E-20 |
| chr5 | 130114622 | 130145594 | 30973 | -10.24 | 1.69E-21 | 2.40E-20 |
| chr8 | 24827361 | 24847892 | 20532 | -8.57 | 1.69E-21 | 2.40E-20 |
| chr3 | 172284600 | 172290254 | 5655 | -7.46 | 1.69E-21 | 2.40E-20 |
| chr9 | 130285136 | 130289476 | 4341 | -10.09 | 1.71E-21 | 2.42E-20 |
| chr20 | 1231221 | 1244277 | 13057 | -10.09 | 1.71E-21 | 2.42E-20 |
| chr6 | 113573456 | 113611818 | 38363 | -10.18 | 1.72E-21 | 2.43E-20 |
| chr12 | 92959295 | 93001626 | 42332 | -10.14 | 1.73E-21 | 2.45E-20 |
| chr5 | 68078093 | 68081988 | 3896 | -3.5 | 1.75E-21 | 2.47E-20 |
| chr6 | 149023259 | 149053466 | 30208 | -4.17 | 1.75E-21 | 2.47E-20 |
| chr5 | 81603987 | 81628234 | 24248 | -2.49 | 1.75E-21 | 2.47E-20 |
| chr1 | 234312070 | 234348424 | 36355 | -10.09 | 1.75E-21 | 2.47E-20 |
| chrX | 116131217 | 116183508 | 52292 | 2.33 | 1.76E-21 | 2.48E-20 |
| chr3 | 14201701 | 14206340 | 4640 | -10.24 | 1.76E-21 | 2.49E-20 |
| chr8 | 58301808 | 58350231 | 48424 | -10.09 | 1.77E-21 | 2.49E-20 |
| chr5 | 171186120 | 171209204 | 23085 | -10.1 | 1.77E-21 | 2.49E-20 |
| chr8 | 90891784 | 90925860 | 34077 | -10.1 | 1.79E-21 | 2.52E-20 |
| chr19 | 44193660 | 44214596 | 20937 | -10.17 | 1.79E-21 | 2.52E-20 |
| chr2 | 64099124 | 64103211 | 4088 | -10.12 | 1.81E-21 | 2.55E-20 |
| chr4 | 114873859 | 114908780 | 34922 | -10.2 | 1.82E-21 | 2.56E-20 |
| chr3 | 109403128 | 109458646 | 55519 | -10.09 | 1.83E-21 | 2.57E-20 |
| chr9 | 27168943 | 27180036 | 11094 | -2.74 | 1.83E-21 | 2.58E-20 |
| chr6 | 94770094 | 95516331 | 746238 | 1.88 | 1.86E-21 | 2.61E-20 |
| chr19 | 46630198 | 46668352 | 38155 | -10.15 | 1.86E-21 | 2.61E-20 |
| chr13 | 79106272 | 79139041 | 32770 | 10.14 | 1.86E-21 | 2.62E-20 |
| chr3 | 59086430 | 59736829 | 650400 | 1.88 | 1.88E-21 | 2.63E-20 |
| chr5 | 130034831 | 130084949 | 50119 | -10.11 | 1.89E-21 | 2.65E-20 |
| chr9 | 115254106 | 115259153 | 5048 | -10.11 | 1.90E-21 | 2.67E-20 |
| chr10 | 120616373 | 120653204 | 36832 | -10.13 | 1.91E-21 | 2.68E-20 |
| chr3 | 27478150 | 27490549 | 12400 | -10.09 | 1.93E-21 | 2.71E-20 |
| chr12 | 67698449 | 67735230 | 36782 | -10.16 | 1.94E-21 | 2.72E-20 |
| chr8 | 41560167 | 41575354 | 15188 | -10.09 | 1.95E-21 | 2.74E-20 |
| chr7 | 46397886 | 46442282 | 44397 | -10.2 | 1.97E-21 | 2.76E-20 |
| chr2 | 173436874 | 173445992 | 9119 | -10.11 | 1.98E-21 | 2.78E-20 |
| chr11 | 87956190 | 87974338 | 18149 | -10.08 | 2.00E-21 | 2.80E-20 |
| chr8 | 9948940 | 9977955 | 29016 | -10.07 | 2.00E-21 | 2.80E-20 |
| chr2 | 184191521 | 186273687 | 2082167 | 2 | 2.01E-21 | 2.81E-20 |
| chr1 | 219193323 | 219247097 | 53775 | -10.44 | 2.04E-21 | 2.85E-20 |
| chr18 | 56653630 | 56688552 | 34923 | -10.16 | 2.05E-21 | 2.86E-20 |
| chr1 | 57145065 | 57170684 | 25620 | -10.07 | 2.05E-21 | 2.87E-20 |
| chr9 | 117989235 | 118027612 | 38378 | -10.14 | 2.08E-21 | 2.90E-20 |
| chr1 | 19075774 | 19085928 | 10155 | -10.16 | 2.08E-21 | 2.90E-20 |
| chr1 | 210749007 | 210787496 | 38490 | -10.12 | 2.10E-21 | 2.93E-20 |
| chr16 | 14361338 | 14384991 | 23654 | -10.09 | 2.11E-21 | 2.94E-20 |
| chr19 | 30356139 | 30360369 | 4231 | -10.07 | 2.11E-21 | 2.94E-20 |
| chr18 | 30362894 | 30464299 | 101406 | 3 | 2.12E-21 | 2.96E-20 |
| chr12 | 7361629 | 7402167 | 40539 | -10.2 | 2.13E-21 | 2.97E-20 |
| chr5 | 76186507 | 76212073 | 25567 | -10.16 | 2.14E-21 | 2.98E-20 |
| chr10 | 7346112 | 7386268 | 40157 | -10.11 | 2.14E-21 | 2.98E-20 |
| chr9 | 97041223 | 97054268 | 13046 | -10.08 | 2.15E-21 | 2.99E-20 |
| chr8 | 55026130 | 55034482 | 8353 | -10.09 | 2.16E-21 | 3.01E-20 |
| chr2 | 181179285 | 181214002 | 34718 | -10.09 | 2.18E-21 | 3.03E-20 |
| chr9 | 93014582 | 93044924 | 30343 | -10.1 | 2.19E-21 | 3.04E-20 |
| chr7 | 151601399 | 151613532 | 12134 | -7.66 | 2.20E-21 | 3.06E-20 |
| chr1 | 93685095 | 93731324 | 46230 | -10.11 | 2.23E-21 | 3.10E-20 |
| chr3 | 48354126 | 48367624 | 13499 | -10.08 | 2.24E-21 | 3.11E-20 |
| chr14 | 70951203 | 70999966 | 48764 | -10.22 | 2.25E-21 | 3.12E-20 |

| | | | | | | |
|---|---|---|---|---|---|---|
| chr4 | 173738612 | 173785789 | 47178 | -10.23 | 2.26E-21 | 3.14E-20 |
| chr3 | 17755103 | 17773722 | 18620 | -10.08 | 2.26E-21 | 3.14E-20 |
| chr1 | 90631692 | 90654904 | 23213 | 6.9 | 2.27E-21 | 3.14E-20 |
| chr12 | 100058766 | 100092329 | 33564 | -10.06 | 2.28E-21 | 3.16E-20 |
| chr18 | 73157887 | 73197729 | 39843 | -10.12 | 2.28E-21 | 3.16E-20 |
| chr11 | 26412911 | 26460719 | 47809 | 10.34 | 2.29E-21 | 3.17E-20 |
| chr11 | 120022238 | 120048223 | 25986 | -10.25 | 2.34E-21 | 3.24E-20 |
| chr2 | 32788617 | 32793212 | 4596 | -10.07 | 2.35E-21 | 3.26E-20 |
| chr6 | 55938866 | 55982129 | 43264 | -10.24 | 2.36E-21 | 3.26E-20 |
| chr10 | 24550677 | 24599964 | 49288 | -10.21 | 2.36E-21 | 3.27E-20 |
| chr17 | 47245039 | 47264222 | 19184 | -10.28 | 2.37E-21 | 3.27E-20 |
| chr17 | 19519577 | 19527327 | 7751 | -10.13 | 2.38E-21 | 3.29E-20 |
| chr3 | 141253373 | 141267969 | 14597 | -10.12 | 2.39E-21 | 3.31E-20 |
| chr5 | 105859175 | 105906837 | 47663 | -10.06 | 2.40E-21 | 3.31E-20 |
| chr3 | 64933919 | 64972423 | 38505 | 10.12 | 2.40E-21 | 3.31E-20 |
| chr3 | 16332810 | 16339908 | 7099 | -10.06 | 2.42E-21 | 3.35E-20 |
| chr4 | 145113739 | 145205776 | 92038 | 2.39 | 2.44E-21 | 3.37E-20 |
| chr1 | 178931074 | 178966428 | 35355 | -10.05 | 2.45E-21 | 3.38E-20 |
| chr6 | 166337522 | 166372888 | 35367 | -10.15 | 2.46E-21 | 3.39E-20 |
| chr11 | 17639247 | 17649208 | 9962 | -7.45 | 2.49E-21 | 3.43E-20 |
| chr7 | 139764077 | 139771922 | 7846 | -10.05 | 2.50E-21 | 3.44E-20 |
| chr2 | 86566241 | 86578498 | 12258 | -10.12 | 2.52E-21 | 3.47E-20 |
| chr20 | 13745903 | 13748840 | 2938 | -10.14 | 2.55E-21 | 3.51E-20 |
| chr14 | 100162681 | 100180044 | 17364 | -10.1 | 2.55E-21 | 3.51E-20 |
| chr21 | 16115727 | 16186325 | 70599 | -10.44 | 2.55E-21 | 3.51E-20 |
| chr9 | 129111761 | 129126849 | 15089 | -10.06 | 2.56E-21 | 3.51E-20 |
| chr11 | 120594564 | 120604795 | 10232 | -4.37 | 2.58E-21 | 3.54E-20 |
| chr4 | 1481339 | 1490073 | 8735 | -10.05 | 2.62E-21 | 3.60E-20 |
| chr9 | 128790264 | 128809256 | 18993 | -10.06 | 2.62E-21 | 3.60E-20 |
| chr2 | 220813137 | 220864184 | 51048 | -10.39 | 2.62E-21 | 3.60E-20 |
| chr16 | 55748119 | 55765923 | 17805 | -7.69 | 2.63E-21 | 3.61E-20 |
| chrX | 53873138 | 53883412 | 10275 | -2.9 | 2.63E-21 | 3.62E-20 |
| chr12 | 99990004 | 100038468 | 48465 | -2.59 | 2.66E-21 | 3.65E-20 |
| chr19 | 49871089 | 49887519 | 16431 | -10.15 | 2.71E-21 | 3.72E-20 |
| chr5 | 55712292 | 55749374 | 37083 | -10.25 | 2.72E-21 | 3.73E-20 |
| chr12 | 110588159 | 110596366 | 8208 | -10.07 | 2.73E-21 | 3.75E-20 |
| chr3 | 126020536 | 126050987 | 30452 | -10.07 | 2.74E-21 | 3.75E-20 |
| chr7 | 84145636 | 84164027 | 18392 | -10.17 | 2.76E-21 | 3.78E-20 |
| chr2 | 73924397 | 73944262 | 19866 | -10.05 | 2.77E-21 | 3.79E-20 |
| chr12 | 62809191 | 62856881 | 47691 | -10.19 | 2.77E-21 | 3.79E-20 |
| chr7 | 17729095 | 17743765 | 14671 | -10.24 | 2.77E-21 | 3.79E-20 |
| chr9 | 75561681 | 75591421 | 29741 | -10.04 | 2.79E-21 | 3.82E-20 |
| chr22 | 31292268 | 31313487 | 21220 | -10.05 | 2.80E-21 | 3.82E-20 |
| chrX | 130631457 | 130658763 | 27307 | -10.06 | 2.81E-21 | 3.84E-20 |
| chr12 | 111236359 | 111244713 | 8355 | -10.07 | 2.85E-21 | 3.89E-20 |
| chr6 | 138496653 | 138501363 | 4711 | -5.43 | 2.88E-21 | 3.93E-20 |
| chr11 | 59445334 | 59463949 | 18616 | -10.04 | 2.91E-21 | 3.97E-20 |
| chr12 | 124949236 | 124959398 | 10163 | -10.04 | 2.92E-21 | 3.98E-20 |
| chr18 | 4021198 | 4070147 | 48950 | -2.02 | 2.92E-21 | 3.98E-20 |
| chr7 | 33155812 | 33183170 | 27359 | -10.25 | 2.92E-21 | 3.98E-20 |
| chr9 | 118314798 | 118331434 | 16637 | -10.04 | 2.93E-21 | 3.99E-20 |
| chr7 | 20519960 | 20541622 | 21663 | -10.04 | 2.94E-21 | 4.00E-20 |
| chr5 | 125022931 | 125072828 | 49898 | -10.35 | 2.94E-21 | 4.00E-20 |
| chr3 | 151732205 | 151769909 | 37705 | -10.1 | 2.94E-21 | 4.01E-20 |
| chr15 | 67965337 | 67992342 | 27006 | -10.14 | 2.94E-21 | 4.01E-20 |
| chr9 | 113876128 | 113902493 | 26366 | -10.16 | 2.94E-21 | 4.01E-20 |
| chr17 | 37190716 | 37212846 | 22131 | -10.18 | 2.96E-21 | 4.02E-20 |
| chr5 | 142910634 | 142928890 | 18257 | -10.17 | 2.96E-21 | 4.03E-20 |
| chr5 | 42597686 | 42632084 | 34399 | 10.16 | 2.97E-21 | 4.04E-20 |
| chr14 | 107238011 | 107289530 | 51520 | -10.2 | 2.99E-21 | 4.06E-20 |
| chr7 | 138106456 | 138121087 | 14632 | -10.24 | 2.99E-21 | 4.06E-20 |

| | | | | | | |
|---|---|---|---|---|---|---|
| chr14 | 105369522 | 105391356 | 21835 | -10.2 | 3.01E-21 | 4.09E-20 |
| chr4 | 142886114 | 142944112 | 57999 | -10.18 | 3.03E-21 | 4.12E-20 |
| chr7 | 33984251 | 34202880 | 218630 | 1.43 | 3.04E-21 | 4.12E-20 |
| chr7 | 97786 | 128412 | 30627 | -10.18 | 3.04E-21 | 4.12E-20 |
| chr12 | 94016558 | 94036162 | 19605 | -10.07 | 3.06E-21 | 4.15E-20 |
| chr19 | 33517843 | 33538599 | 20757 | -10.04 | 3.08E-21 | 4.17E-20 |
| chr20 | 24667506 | 24671272 | 3767 | -5.61 | 3.08E-21 | 4.17E-20 |
| chrX | 19331809 | 19356770 | 24962 | -10.05 | 3.12E-21 | 4.22E-20 |
| chr11 | 124551023 | 124581087 | 30065 | -10.09 | 3.12E-21 | 4.23E-20 |
| chr21 | 46937541 | 46942968 | 5428 | -10.12 | 3.16E-21 | 4.27E-20 |
| chr7 | 156646295 | 156657596 | 11302 | -10.03 | 3.20E-21 | 4.33E-20 |
| chr7 | 94448447 | 94508565 | 60119 | -10.34 | 3.23E-21 | 4.37E-20 |
| chr10 | 134683229 | 134689856 | 6628 | -8.52 | 3.24E-21 | 4.39E-20 |
| chr10 | 29196483 | 29253816 | 57334 | -10.45 | 3.27E-21 | 4.42E-20 |
| chr2 | 240452438 | 240499436 | 46999 | -10.13 | 3.27E-21 | 4.42E-20 |
| chr3 | 87887312 | 87930770 | 43459 | -10.05 | 3.28E-21 | 4.43E-20 |
| chr16 | 48612201 | 48630017 | 17817 | -10.14 | 3.30E-21 | 4.45E-20 |
| chr3 | 15879931 | 15913870 | 33940 | -10.54 | 3.30E-21 | 4.46E-20 |
| chr5 | 33511244 | 33551128 | 39885 | 10.07 | 3.31E-21 | 4.47E-20 |
| chr2 | 226812133 | 226847817 | 35685 | -10.02 | 3.33E-21 | 4.49E-20 |
| chr12 | 72045729 | 72051287 | 5559 | -10.08 | 3.35E-21 | 4.51E-20 |
| chr19 | 51135482 | 51180793 | 45312 | -10.17 | 3.38E-21 | 4.56E-20 |
| chr19 | 43387222 | 43429921 | 42700 | -10.12 | 3.39E-21 | 4.56E-20 |
| chr12 | 47024792 | 47045356 | 20565 | -10.11 | 3.40E-21 | 4.57E-20 |
| chr8 | 49207972 | 49226856 | 18885 | -10.02 | 3.40E-21 | 4.57E-20 |
| chr6 | 46214967 | 46313203 | 98237 | 3.11 | 3.41E-21 | 4.58E-20 |
| chr5 | 82747861 | 82775656 | 27796 | -10.07 | 3.42E-21 | 4.60E-20 |
| chr15 | 51775414 | 51780487 | 5074 | -10.19 | 3.43E-21 | 4.62E-20 |
| chr3 | 126986452 | 127016250 | 29799 | -2.66 | 3.46E-21 | 4.65E-20 |
| chr8 | 32086714 | 32131242 | 44529 | -10.02 | 3.48E-21 | 4.67E-20 |
| chr21 | 16802544 | 16823992 | 21449 | -10.13 | 3.49E-21 | 4.68E-20 |
| chr11 | 86221334 | 86258412 | 37079 | -10.15 | 3.49E-21 | 4.68E-20 |
| chr15 | 67042141 | 67050636 | 8496 | -10.11 | 3.49E-21 | 4.68E-20 |
| chr4 | 178246685 | 178254155 | 7471 | -10.02 | 3.56E-21 | 4.78E-20 |
| chr5 | 60487576 | 60528965 | 41390 | -10.12 | 3.57E-21 | 4.79E-20 |
| chr7 | 4685992 | 4715601 | 29610 | -10.19 | 3.59E-21 | 4.81E-20 |
| chr1 | 150900950 | 150907412 | 6463 | -10.13 | 3.59E-21 | 4.82E-20 |
| chr3 | 95437787 | 95518823 | 81037 | -10.08 | 3.60E-21 | 4.82E-20 |
| chr4 | 14565693 | 14637896 | 72204 | -10.16 | 3.60E-21 | 4.83E-20 |
| chr7 | 19859304 | 19890634 | 31331 | -10.06 | 3.61E-21 | 4.83E-20 |
| chr6 | 149698751 | 149718939 | 20189 | -10.13 | 3.62E-21 | 4.84E-20 |
| chr1 | 192843123 | 192848281 | 5159 | -7 | 3.64E-21 | 4.88E-20 |
| chr9 | 94274050 | 94318135 | 44086 | -10.03 | 3.65E-21 | 4.89E-20 |
| chr4 | 177746274 | 177809556 | 63283 | -10.12 | 3.66E-21 | 4.89E-20 |
| chr4 | 85963881 | 86021741 | 57861 | -10.07 | 3.66E-21 | 4.89E-20 |
| chr5 | 59919010 | 59955077 | 36068 | -10.53 | 3.67E-21 | 4.90E-20 |
| chr8 | 59296901 | 59321320 | 24420 | -10.02 | 3.68E-21 | 4.92E-20 |
| chr15 | 69840034 | 69846087 | 6054 | -10.03 | 3.71E-21 | 4.95E-20 |
| chr4 | 20623064 | 20667553 | 44490 | 10.08 | 3.73E-21 | 4.98E-20 |
| chr14 | 95066526 | 95098743 | 32218 | -10.17 | 3.74E-21 | 4.99E-20 |
| chr3 | 119752836 | 119774265 | 21430 | -10.01 | 3.74E-21 | 4.99E-20 |
| chr7 | 14767730 | 14841464 | 73735 | -10.01 | 3.76E-21 | 5.01E-20 |
| chr7 | 121143287 | 121155176 | 11890 | -10.09 | 3.76E-21 | 5.02E-20 |
| chr2 | 67883288 | 67890442 | 7155 | -3.12 | 3.86E-21 | 5.15E-20 |
| chr18 | 33163990 | 33184760 | 20771 | -10.05 | 3.87E-21 | 5.15E-20 |
| chr8 | 48357900 | 48383159 | 25260 | -10.21 | 3.87E-21 | 5.15E-20 |
| chr9 | 14651642 | 14658940 | 7299 | -10.03 | 3.91E-21 | 5.21E-20 |
| chr3 | 184950008 | 184978598 | 28591 | -10.09 | 3.91E-21 | 5.21E-20 |
| chr7 | 150312185 | 150369045 | 56861 | 2.49 | 3.92E-21 | 5.22E-20 |
| chr16 | 22347138 | 22353478 | 6341 | -10.03 | 3.95E-21 | 5.25E-20 |
| chr6 | 144300823 | 144335467 | 34645 | -10.1 | 3.96E-21 | 5.26E-20 |

| | | | | | | |
|---|---|---|---|---|---|---|
| chr3 | 139469101 | 139646846 | 177746 | 9.23 | 3.96E-21 | 5.26E-20 |
| chr2 | 121897144 | 121929679 | 32536 | -10.15 | 3.98E-21 | 5.29E-20 |
| chr17 | 25696493 | 25705436 | 8944 | -10.06 | 4.01E-21 | 5.33E-20 |
| chr6 | 12877684 | 12893583 | 15900 | -7.47 | 4.01E-21 | 5.33E-20 |
| chr4 | 123015959 | 123061888 | 45930 | -10.07 | 4.02E-21 | 5.34E-20 |
| chr16 | 54407963 | 54429625 | 21663 | -2.55 | 4.05E-21 | 5.37E-20 |
| chr13 | 28654203 | 28679219 | 25017 | -10.02 | 4.06E-21 | 5.39E-20 |
| chr12 | 90018231 | 90047957 | 29727 | -10.13 | 4.08E-21 | 5.40E-20 |
| chr6 | 41356987 | 41370944 | 13958 | -10 | 4.08E-21 | 5.40E-20 |
| chr11 | 108577339 | 108596753 | 19415 | -10.14 | 4.08E-21 | 5.41E-20 |
| chrX | 103012145 | 103014806 | 2662 | -4.17 | 4.15E-21 | 5.50E-20 |
| chr20 | 1382890 | 1415336 | 32447 | -10 | 4.18E-21 | 5.53E-20 |
| chr4 | 74550566 | 74620583 | 70018 | -10.05 | 4.20E-21 | 5.56E-20 |
| chr4 | 153386652 | 153405945 | 19294 | -10.02 | 4.21E-21 | 5.57E-20 |
| chr3 | 128637847 | 128649913 | 12067 | -10 | 4.26E-21 | 5.64E-20 |
| chr17 | 269855 | 288157 | 18303 | -10.19 | 4.27E-21 | 5.65E-20 |
| chr10 | 21737108 | 21751919 | 14812 | -10.05 | 4.28E-21 | 5.65E-20 |
| chr16 | 53453094 | 53465068 | 11975 | -10.04 | 4.30E-21 | 5.67E-20 |
| chrX | 81732048 | 81777483 | 45436 | -10.02 | 4.30E-21 | 5.67E-20 |
| chr3 | 110661168 | 110709005 | 47838 | -10.31 | 4.32E-21 | 5.71E-20 |
| chr2 | 33003442 | 33017749 | 14308 | -10.04 | 4.32E-21 | 5.71E-20 |
| chr1 | 234469640 | 234505405 | 35766 | -10.01 | 4.34E-21 | 5.72E-20 |
| chr5 | 158717392 | 158732110 | 14719 | -10.01 | 4.35E-21 | 5.74E-20 |
| chr9 | 108321209 | 108338423 | 17215 | -10 | 4.37E-21 | 5.76E-20 |
| chr20 | 10820583 | 10872266 | 51684 | -10.02 | 4.38E-21 | 5.77E-20 |
| chr1 | 162794889 | 162811957 | 17069 | -2.18 | 4.40E-21 | 5.80E-20 |
| chr2 | 47725452 | 47737176 | 11725 | -10 | 4.42E-21 | 5.82E-20 |
| chr12 | 17536768 | 17571423 | 34656 | -10.21 | 4.43E-21 | 5.83E-20 |
| chr15 | 68181745 | 68243962 | 62218 | -10.68 | 4.43E-21 | 5.83E-20 |
| chr7 | 12095037 | 12153270 | 58234 | -10.08 | 4.47E-21 | 5.88E-20 |
| chr7 | 41752005 | 41976865 | 224861 | 1.45 | 4.52E-21 | 5.94E-20 |
| chr3 | 79138427 | 80870711 | 1732285 | 1.35 | 4.52E-21 | 5.94E-20 |
| chr12 | 31865576 | 31922022 | 56447 | -10.06 | 4.53E-21 | 5.95E-20 |
| chr2 | 120946541 | 120958004 | 11464 | -9.99 | 4.54E-21 | 5.96E-20 |
| chr7 | 41080457 | 41113646 | 33190 | -10.01 | 4.55E-21 | 5.97E-20 |
| chr5 | 79756807 | 79780933 | 24127 | -9.99 | 4.55E-21 | 5.98E-20 |
| chr9 | 117055062 | 117059463 | 4402 | -10.09 | 4.58E-21 | 6.00E-20 |
| chr10 | 23369531 | 23386691 | 17161 | -10.06 | 4.59E-21 | 6.02E-20 |
| chr1 | 39250225 | 39281048 | 30824 | -10.04 | 4.60E-21 | 6.03E-20 |
| chr1 | 197048784 | 197055933 | 7150 | -10.02 | 4.63E-21 | 6.07E-20 |
| chr19 | 47958394 | 47960797 | 2404 | -9.99 | 4.64E-21 | 6.08E-20 |
| chr12 | 74417866 | 74487796 | 69931 | -10.28 | 4.65E-21 | 6.09E-20 |
| chr1 | 3995384 | 4012913 | 17530 | -10.08 | 4.67E-21 | 6.11E-20 |
| chr10 | 127630469 | 127652955 | 22487 | -10.1 | 4.69E-21 | 6.13E-20 |
| chr11 | 121921490 | 121937929 | 16440 | -10.03 | 4.69E-21 | 6.14E-20 |
| chr7 | 102613392 | 102645696 | 32305 | -10.01 | 4.69E-21 | 6.14E-20 |
| chr4 | 88988384 | 89006504 | 18121 | -9.99 | 4.74E-21 | 6.20E-20 |
| chr19 | 49026732 | 49033047 | 6316 | -10.11 | 4.75E-21 | 6.21E-20 |
| chr1 | 213070913 | 213092144 | 21232 | -9.99 | 4.77E-21 | 6.23E-20 |
| chr12 | 69579992 | 69602444 | 22453 | -10.09 | 4.77E-21 | 6.23E-20 |
| chr4 | 89026816 | 89050224 | 23409 | -10.04 | 4.77E-21 | 6.23E-20 |
| chr3 | 32259896 | 32304443 | 44548 | -10.14 | 4.77E-21 | 6.23E-20 |
| chr5 | 57114431 | 57167592 | 53162 | -10.03 | 4.85E-21 | 6.33E-20 |
| chr14 | 64178035 | 64188029 | 9995 | -10.03 | 4.85E-21 | 6.33E-20 |
| chr16 | 24972424 | 24977158 | 4735 | -10.02 | 4.91E-21 | 6.40E-20 |
| chr15 | 99113278 | 99146574 | 33297 | -10.21 | 4.96E-21 | 6.46E-20 |
| chr21 | 32598496 | 32618930 | 20435 | -10.06 | 5.04E-21 | 6.57E-20 |
| chr16 | 12750875 | 12774886 | 24012 | -10.02 | 5.12E-21 | 6.67E-20 |
| chr7 | 101696771 | 101733421 | 36651 | -10.04 | 5.14E-21 | 6.70E-20 |
| chr6 | 77575577 | 77605972 | 30396 | -10 | 5.18E-21 | 6.74E-20 |
| chr6 | 165420009 | 165442316 | 22308 | 3.7 | 5.20E-21 | 6.76E-20 |

| | | | | | | |
|---|---|---|---|---|---|---|
| chr20 | 56623756 | 56639286 | 15531 | -10.11 | 5.21E-21 | 6.78E-20 |
| chr10 | 33350692 | 33407203 | 56512 | -10.17 | 5.24E-21 | 6.81E-20 |
| chr12 | 20427691 | 20471310 | 43620 | -10.24 | 5.27E-21 | 6.85E-20 |
| chr15 | 57074811 | 57103139 | 28329 | -10.24 | 5.27E-21 | 6.85E-20 |
| chr12 | 12346349 | 12364567 | 18219 | -9.98 | 5.27E-21 | 6.85E-20 |
| chr4 | 142615735 | 142639725 | 23991 | -10.02 | 5.30E-21 | 6.88E-20 |
| chr6 | 77702268 | 77749130 | 46863 | -10.04 | 5.37E-21 | 6.97E-20 |
| chr1 | 171978722 | 171987976 | 9255 | -10 | 5.38E-21 | 6.99E-20 |
| chr2 | 55246413 | 55267228 | 20816 | -9.99 | 5.40E-21 | 7.01E-20 |
| chr1 | 215703517 | 215735231 | 31715 | -9.97 | 5.57E-21 | 7.23E-20 |
| chr15 | 96474710 | 96519007 | 44298 | -2.24 | 5.62E-21 | 7.28E-20 |
| chr7 | 1656178 | 1678266 | 22089 | -10.04 | 5.63E-21 | 7.29E-20 |
| chr7 | 39752381 | 39756731 | 4351 | -7 | 5.65E-21 | 7.32E-20 |
| chr3 | 5907425 | 5947315 | 39891 | -10.33 | 5.67E-21 | 7.34E-20 |
| chr3 | 33564252 | 33574232 | 9981 | -10.01 | 5.67E-21 | 7.34E-20 |
| chr16 | 56411542 | 56425411 | 13870 | -9.97 | 5.74E-21 | 7.43E-20 |
| chr1 | 183760692 | 183773857 | 13166 | -10.12 | 5.75E-21 | 7.44E-20 |
| chr2 | 28368546 | 28381515 | 12970 | -9.99 | 5.79E-21 | 7.49E-20 |
| chr12 | 117077379 | 117092426 | 15048 | -10.01 | 5.80E-21 | 7.50E-20 |
| chr1 | 98541112 | 98575792 | 34681 | -10.17 | 5.85E-21 | 7.55E-20 |
| chr12 | 14872634 | 14884890 | 12257 | -10 | 5.86E-21 | 7.57E-20 |
| chr2 | 83940994 | 83995712 | 54719 | -10.22 | 5.88E-21 | 7.59E-20 |
| chrX | 154324225 | 154364310 | 40086 | -9.97 | 5.88E-21 | 7.60E-20 |
| chr15 | 73442997 | 73460279 | 17283 | -10.11 | 5.91E-21 | 7.63E-20 |
| chr7 | 3592029 | 3620554 | 28526 | -9.98 | 5.92E-21 | 7.64E-20 |
| chr3 | 64042591 | 64081515 | 38925 | -10.07 | 5.93E-21 | 7.65E-20 |
| chr3 | 105688065 | 105719674 | 31610 | -10.04 | 5.95E-21 | 7.68E-20 |
| chr12 | 107731486 | 107748756 | 17271 | -9.97 | 5.96E-21 | 7.69E-20 |
| chrX | 25199097 | 25325240 | 126144 | 2.93 | 5.98E-21 | 7.71E-20 |
| chr3 | 109586323 | 109645506 | 59184 | -10.02 | 6.02E-21 | 7.76E-20 |
| chr6 | 56132199 | 56173814 | 41616 | -10.04 | 6.11E-21 | 7.87E-20 |
| chr12 | 105887301 | 105892669 | 5369 | -6.78 | 6.14E-21 | 7.90E-20 |
| chr5 | 115148303 | 115165774 | 17472 | -9.99 | 6.21E-21 | 7.99E-20 |
| chr12 | 43258753 | 43306874 | 48122 | -10.03 | 6.25E-21 | 8.04E-20 |
| chr15 | 59752588 | 59782922 | 30335 | -9.96 | 6.27E-21 | 8.06E-20 |
| chr16 | 10323431 | 10328881 | 5451 | -4.22 | 6.34E-21 | 8.14E-20 |
| chr4 | 32007229 | 32057781 | 50553 | -9.96 | 6.34E-21 | 8.15E-20 |
| chr1 | 85252620 | 85275291 | 22672 | -10.01 | 6.38E-21 | 8.19E-20 |
| chr11 | 109560365 | 109613026 | 52662 | -9.97 | 6.42E-21 | 8.24E-20 |
| chr4 | 109142697 | 109160929 | 18233 | 7.75 | 6.42E-21 | 8.24E-20 |
| chr7 | 32698231 | 32747903 | 49673 | -10.06 | 6.43E-21 | 8.25E-20 |
| chr17 | 57024024 | 57031721 | 7698 | -10 | 6.44E-21 | 8.26E-20 |
| chr5 | 73817944 | 73833621 | 15678 | -10.03 | 6.46E-21 | 8.29E-20 |
| chr7 | 86827902 | 86838302 | 10401 | -9.96 | 6.51E-21 | 8.34E-20 |
| chrX | 154499322 | 154509624 | 10303 | -4.35 | 6.51E-21 | 8.34E-20 |
| chr7 | 6159238 | 6173686 | 14449 | -9.98 | 6.55E-21 | 8.39E-20 |
| chr9 | 5777114 | 5784174 | 7061 | -10 | 6.64E-21 | 8.51E-20 |
| chr1 | 204705410 | 204710010 | 4601 | -9.97 | 6.66E-21 | 8.53E-20 |
| chr5 | 32791622 | 32836794 | 45173 | -10.01 | 6.68E-21 | 8.55E-20 |
| chr2 | 69126743 | 69136495 | 9753 | -10.01 | 6.70E-21 | 8.57E-20 |
| chr10 | 35896758 | 35920968 | 24211 | -10.06 | 6.70E-21 | 8.57E-20 |
| chr22 | 27736680 | 27753556 | 16877 | -9.95 | 6.74E-21 | 8.62E-20 |
| chr2 | 99739353 | 99756292 | 16940 | -9.97 | 6.76E-21 | 8.63E-20 |
| chr5 | 71200619 | 71246387 | 45769 | -10.16 | 6.76E-21 | 8.64E-20 |
| chr4 | 12764291 | 13135623 | 371333 | 1.59 | 6.79E-21 | 8.67E-20 |
| chr8 | 115141027 | 115253657 | 112631 | -10.41 | 6.80E-21 | 8.68E-20 |
| chr15 | 99888215 | 99934739 | 46525 | -10.26 | 6.81E-21 | 8.69E-20 |
| chr13 | 27770712 | 27807477 | 36766 | -10 | 6.93E-21 | 8.84E-20 |
| chr7 | 3265013 | 3311617 | 46605 | -9.96 | 6.94E-21 | 8.86E-20 |
| chr5 | 58409064 | 58451467 | 42404 | -9.96 | 6.96E-21 | 8.88E-20 |
| chr2 | 196208031 | 196234200 | 26170 | -10.12 | 6.96E-21 | 8.88E-20 |

| | | | | | | |
|---|---|---|---|---|---|---|
| chr12 | 6749392 | 6757822 | 8431 | -10.08 | 6.97E-21 | 8.88E-20 |
| chr20 | 45319509 | 45354016 | 34508 | -10.34 | 6.99E-21 | 8.90E-20 |
| chr1 | 94246737 | 94277902 | 31166 | -9.96 | 7.02E-21 | 8.94E-20 |
| chr12 | 90061288 | 90081055 | 19768 | -9.99 | 7.06E-21 | 8.98E-20 |
| chr20 | 10290903 | 10302508 | 11606 | -9.97 | 7.15E-21 | 9.09E-20 |
| chr7 | 46921795 | 46966160 | 44366 | -9.96 | 7.15E-21 | 9.10E-20 |
| chr1 | 92849865 | 92871092 | 21228 | -10.16 | 7.18E-21 | 9.13E-20 |
| chr3 | 95822980 | 95895666 | 72687 | -10.09 | 7.20E-21 | 9.15E-20 |
| chr17 | 17261715 | 17270050 | 8336 | -9.96 | 7.22E-21 | 9.17E-20 |
| chr10 | 32933012 | 32949106 | 16095 | -9.96 | 7.27E-21 | 9.23E-20 |
| chr21 | 16216095 | 16256444 | 40350 | -9.97 | 7.27E-21 | 9.24E-20 |
| chr2 | 3122781 | 3145845 | 23065 | -9.94 | 7.29E-21 | 9.26E-20 |
| chr12 | 102921769 | 102965956 | 44188 | -9.94 | 7.35E-21 | 9.32E-20 |
| chr9 | 75676830 | 75687765 | 10936 | -8.54 | 7.46E-21 | 9.46E-20 |
| chr18 | 77484342 | 77499475 | 15134 | -9.99 | 7.47E-21 | 9.48E-20 |
| chr19 | 6285662 | 6324045 | 38384 | -10.04 | 7.53E-21 | 9.54E-20 |
| chr13 | 107362521 | 107405991 | 43471 | 2.1 | 7.56E-21 | 9.58E-20 |
| chr11 | 4156426 | 4163210 | 6785 | -10.02 | 7.58E-21 | 9.60E-20 |
| chr17 | 8941948 | 8953164 | 11217 | -10.06 | 7.59E-21 | 9.61E-20 |
| chr2 | 242631459 | 242663460 | 32002 | -9.95 | 7.59E-21 | 9.61E-20 |
| chr21 | 42942820 | 42952030 | 9211 | -9.94 | 7.61E-21 | 9.63E-20 |
| chr2 | 219387878 | 219393920 | 6043 | -10.03 | 7.65E-21 | 9.67E-20 |
| chr10 | 94606434 | 94625460 | 19027 | -9.94 | 7.68E-21 | 9.71E-20 |
| chr2 | 161402815 | 161449363 | 46549 | -10.05 | 7.69E-21 | 9.72E-20 |
| chr2 | 69529418 | 69546867 | 17450 | -9.97 | 7.72E-21 | 9.75E-20 |
| chr17 | 56905771 | 56933534 | 27764 | -9.98 | 7.77E-21 | 9.81E-20 |
| chr2 | 109482657 | 109507752 | 25096 | -9.96 | 7.80E-21 | 9.85E-20 |
| chr4 | 15135313 | 15175743 | 40431 | -10.24 | 7.81E-21 | 9.85E-20 |
| chrX | 139849894 | 139897264 | 47371 | -10.2 | 7.81E-21 | 9.85E-20 |
| chr15 | 71713806 | 71732003 | 18198 | -10.2 | 7.81E-21 | 9.86E-20 |
| chr9 | 21460061 | 21470593 | 10533 | -10 | 7.83E-21 | 9.87E-20 |
| chr2 | 224310308 | 224333417 | 23110 | -9.98 | 7.88E-21 | 9.94E-20 |
| chr4 | 52776701 | 52795391 | 18691 | -9.99 | 7.91E-21 | 9.97E-20 |
| chr16 | 55544469 | 55557192 | 12724 | -9.97 | 7.99E-21 | 1.01E-19 |
| chr12 | 96118913 | 96124495 | 5583 | -9.94 | 8.04E-21 | 1.01E-19 |
| chr7 | 11565652 | 11608189 | 42538 | -9.95 | 8.14E-21 | 1.03E-19 |
| chr14 | 42082091 | 42622093 | 540003 | 2.13 | 8.17E-21 | 1.03E-19 |
| chr4 | 152963497 | 152981228 | 17732 | -10.01 | 8.19E-21 | 1.03E-19 |
| chr12 | 13663141 | 13681569 | 18429 | -9.96 | 8.21E-21 | 1.03E-19 |
| chr7 | 39034577 | 39408284 | 373708 | 3.02 | 8.23E-21 | 1.03E-19 |
| chrX | 72847892 | 72891493 | 43602 | -10.02 | 8.26E-21 | 1.04E-19 |
| chr17 | 15177000 | 15204457 | 27458 | -10.24 | 8.28E-21 | 1.04E-19 |
| chr16 | 88607665 | 88623785 | 16121 | -10 | 8.37E-21 | 1.05E-19 |
| chr2 | 30457560 | 30490266 | 32707 | -10 | 8.37E-21 | 1.05E-19 |
| chr17 | 63823736 | 63841063 | 17328 | -10.09 | 8.37E-21 | 1.05E-19 |
| chr2 | 40058362 | 40072315 | 13954 | -9.94 | 8.41E-21 | 1.06E-19 |
| chrX | 45884343 | 45963618 | 79276 | -10.14 | 8.48E-21 | 1.06E-19 |
| chrX | 58433665 | 58460674 | 27010 | -10.16 | 8.52E-21 | 1.07E-19 |
| chr5 | 1738148 | 1744370 | 6223 | -9.97 | 8.55E-21 | 1.07E-19 |
| chr8 | 100031784 | 100046509 | 14726 | -9.94 | 8.57E-21 | 1.07E-19 |
| chr11 | 35684837 | 35690545 | 5709 | -9.93 | 8.64E-21 | 1.08E-19 |
| chr11 | 92721393 | 92752245 | 30853 | -9.93 | 8.66E-21 | 1.09E-19 |
| chr12 | 83950448 | 84021964 | 71517 | -9.95 | 8.67E-21 | 1.09E-19 |
| chr2 | 200561432 | 200618216 | 56785 | -10.28 | 8.67E-21 | 1.09E-19 |
| chr2 | 208225323 | 208270557 | 45235 | -9.93 | 8.68E-21 | 1.09E-19 |
| chr1 | 229215825 | 229233174 | 17350 | -10.03 | 8.69E-21 | 1.09E-19 |
| chr20 | 25489643 | 25497679 | 8037 | -9.93 | 8.75E-21 | 1.09E-19 |
| chr3 | 134388813 | 134396360 | 7548 | -2.66 | 8.75E-21 | 1.09E-19 |
| chr12 | 104751779 | 104768406 | 16628 | -9.93 | 8.76E-21 | 1.09E-19 |
| chr15 | 59888923 | 59914455 | 25533 | -10.07 | 8.79E-21 | 1.10E-19 |
| chr17 | 37914994 | 37925692 | 10699 | -7.74 | 8.79E-21 | 1.10E-19 |

| | | | | | | |
|---|---|---|---|---|---|---|
| chr5 | 82192617 | 82213943 | 21327 | -9.94 | 8.81E-21 | 1.10E-19 |
| chr2 | 162214544 | 162221543 | 7000 | -9.95 | 8.81E-21 | 1.10E-19 |
| chr2 | 109071775 | 109082713 | 10939 | -9.98 | 9.03E-21 | 1.13E-19 |
| chr3 | 168891872 | 168941632 | 49761 | 10.11 | 9.08E-21 | 1.13E-19 |
| chr1 | 43942339 | 43958979 | 16641 | -9.94 | 9.10E-21 | 1.13E-19 |
| chr8 | 105566579 | 105603936 | 37358 | -9.93 | 9.19E-21 | 1.15E-19 |
| chr5 | 158538925 | 158555338 | 16414 | -9.99 | 9.22E-21 | 1.15E-19 |
| chr2 | 46642477 | 46649250 | 6774 | -9.94 | 9.22E-21 | 1.15E-19 |
| chr1 | 92203331 | 92215906 | 12576 | -9.94 | 9.29E-21 | 1.16E-19 |
| chr10 | 124019290 | 124023175 | 3886 | -9.93 | 9.29E-21 | 1.16E-19 |
| chr22 | 46219127 | 46227949 | 8823 | -9.92 | 9.29E-21 | 1.16E-19 |
| chr2 | 178288421 | 178300373 | 11953 | -9.93 | 9.30E-21 | 1.16E-19 |
| chr6 | 3551536 | 3556298 | 4763 | -8.53 | 9.38E-21 | 1.17E-19 |
| chr9 | 35874004 | 35878738 | 4735 | -9.93 | 9.54E-21 | 1.19E-19 |
| chr12 | 46635302 | 46646897 | 11596 | -9.93 | 9.57E-21 | 1.19E-19 |
| chr16 | 50030646 | 50044136 | 13491 | -9.92 | 9.59E-21 | 1.19E-19 |
| chr5 | 67777649 | 67797814 | 20166 | -9.92 | 9.68E-21 | 1.20E-19 |
| chr21 | 38507441 | 38511037 | 3597 | -7.8 | 9.70E-21 | 1.20E-19 |
| chr7 | 3428639 | 3441698 | 13060 | -9.94 | 9.83E-21 | 1.22E-19 |
| chr2 | 214167078 | 214171933 | 4856 | -9.96 | 9.87E-21 | 1.22E-19 |
| chr5 | 22015487 | 22124963 | 109477 | 2.16 | 9.90E-21 | 1.23E-19 |
| chr2 | 9087814 | 9097124 | 9311 | -9.91 | 9.99E-21 | 1.24E-19 |
| chr12 | 96254365 | 96257560 | 3196 | -9.91 | 1.00E-20 | 1.24E-19 |
| chr20 | 17665177 | 17677615 | 12439 | -9.91 | 1.02E-20 | 1.26E-19 |
| chr9 | 33319062 | 33341748 | 22687 | -10.07 | 1.03E-20 | 1.28E-19 |
| chr11 | 12630738 | 12652095 | 21358 | -9.97 | 1.04E-20 | 1.28E-19 |
| chr16 | 48341228 | 48348098 | 6871 | -9.92 | 1.04E-20 | 1.28E-19 |
| chr7 | 109608929 | 109623867 | 14939 | -9.94 | 1.04E-20 | 1.28E-19 |
| chr1 | 180624483 | 180641282 | 16800 | -9.92 | 1.04E-20 | 1.29E-19 |
| chr7 | 150979436 | 151037998 | 58563 | -10.25 | 1.05E-20 | 1.30E-19 |
| chr4 | 83115105 | 83156678 | 41574 | 9.96 | 1.05E-20 | 1.30E-19 |
| chr20 | 20354094 | 20365653 | 11560 | -2.17 | 1.06E-20 | 1.31E-19 |
| chr12 | 40098949 | 40139980 | 41032 | -9.94 | 1.06E-20 | 1.31E-19 |
| chr2 | 225577772 | 225603830 | 26059 | -10.11 | 1.06E-20 | 1.31E-19 |
| chr7 | 84021685 | 84077562 | 55878 | -10.31 | 1.07E-20 | 1.32E-19 |
| chr19 | 50786984 | 50792969 | 5986 | -9.91 | 1.07E-20 | 1.32E-19 |
| chr5 | 78893309 | 78921267 | 27959 | -9.95 | 1.08E-20 | 1.33E-19 |
| chr20 | 14737300 | 14801754 | 64455 | -10.73 | 1.08E-20 | 1.33E-19 |
| chr2 | 4009763 | 5018149 | 1008387 | 1.32 | 1.08E-20 | 1.34E-19 |
| chr1 | 116107865 | 116132454 | 24590 | -9.91 | 1.10E-20 | 1.35E-19 |
| chr7 | 89796759 | 89824047 | 27289 | -10.1 | 1.10E-20 | 1.35E-19 |
| chr3 | 164936936 | 165353980 | 417045 | 2.23 | 1.11E-20 | 1.37E-19 |
| chr20 | 24825213 | 24848323 | 23111 | -2.29 | 1.12E-20 | 1.37E-19 |
| chr8 | 24992790 | 25017381 | 24592 | -10.07 | 1.12E-20 | 1.38E-19 |
| chr1 | 163298511 | 163306746 | 8236 | -10.07 | 1.13E-20 | 1.39E-19 |
| chr2 | 25176700 | 25180107 | 3408 | -10.02 | 1.13E-20 | 1.39E-19 |
| chr5 | 157430820 | 157434325 | 3506 | -7.58 | 1.13E-20 | 1.39E-19 |
| chr11 | 74962443 | 74967358 | 4916 | -9.91 | 1.14E-20 | 1.40E-19 |
| chr12 | 11250882 | 11279601 | 28720 | -9.92 | 1.14E-20 | 1.40E-19 |
| chr10 | 79403996 | 79420861 | 16866 | -9.98 | 1.15E-20 | 1.41E-19 |
| chr10 | 29926268 | 29947516 | 21249 | -9.9 | 1.17E-20 | 1.44E-19 |
| chr1 | 189563190 | 189587870 | 24681 | -9.91 | 1.20E-20 | 1.48E-19 |
| chr3 | 53685242 | 53700769 | 15528 | -10.3 | 1.22E-20 | 1.50E-19 |
| chr5 | 87421295 | 87484655 | 63361 | -10.02 | 1.22E-20 | 1.50E-19 |
| chr17 | 64303145 | 64319871 | 16727 | -9.9 | 1.23E-20 | 1.51E-19 |
| chr17 | 33696049 | 33714183 | 18135 | -9.9 | 1.23E-20 | 1.51E-19 |
| chr7 | 46996925 | 47014460 | 17536 | -9.95 | 1.24E-20 | 1.52E-19 |
| chr3 | 126299798 | 126323384 | 23587 | -9.97 | 1.24E-20 | 1.52E-19 |
| chr20 | 23435359 | 23448889 | 13531 | -10 | 1.24E-20 | 1.52E-19 |
| chr21 | 32845451 | 32865996 | 20546 | -10.01 | 1.24E-20 | 1.52E-19 |
| chr1 | 7452919 | 7461576 | 8658 | -9.99 | 1.25E-20 | 1.53E-19 |

| | | | | | | |
|---|---|---|---|---|---|---|
| chr11 | 119944615 | 119973022 | 28408 | -9.89 | 1.26E-20 | 1.55E-19 |
| chr13 | 50425700 | 50455171 | 29472 | -9.98 | 1.27E-20 | 1.55E-19 |
| chr4 | 177492116 | 177520160 | 28045 | -9.99 | 1.27E-20 | 1.55E-19 |
| chr12 | 94710175 | 94720386 | 10212 | -9.91 | 1.27E-20 | 1.55E-19 |
| chr2 | 141639510 | 142851203 | 1211694 | 5.01 | 1.27E-20 | 1.56E-19 |
| chr21 | 44603255 | 44625830 | 22576 | -10.04 | 1.28E-20 | 1.56E-19 |
| chr2 | 39765173 | 39793974 | 28802 | -10.2 | 1.30E-20 | 1.58E-19 |
| chr18 | 45520585 | 45537265 | 16681 | -9.93 | 1.30E-20 | 1.59E-19 |
| chr4 | 181633634 | 181738740 | 105107 | 2.63 | 1.32E-20 | 1.61E-19 |
| chr9 | 116684448 | 116703452 | 19005 | -9.99 | 1.32E-20 | 1.61E-19 |
| chr16 | 19398500 | 19433517 | 35018 | -9.89 | 1.32E-20 | 1.62E-19 |
| chr9 | 93750966 | 93791641 | 40676 | -9.95 | 1.32E-20 | 1.62E-19 |
| chr1 | 182881522 | 182905792 | 24271 | -9.88 | 1.33E-20 | 1.62E-19 |
| chr21 | 39291941 | 39319943 | 28003 | -10.03 | 1.35E-20 | 1.65E-19 |
| chr9 | 121485446 | 121533958 | 48513 | -9.92 | 1.36E-20 | 1.66E-19 |
| chr12 | 40722158 | 40726540 | 4383 | -9.89 | 1.37E-20 | 1.67E-19 |
| chr19 | 16321837 | 16333909 | 12073 | -9.9 | 1.37E-20 | 1.67E-19 |
| chr18 | 76562851 | 76625219 | 62369 | -2.48 | 1.38E-20 | 1.67E-19 |
| chr2 | 63327684 | 63333832 | 6149 | -9.88 | 1.39E-20 | 1.69E-19 |
| chr14 | 104411136 | 104421102 | 9967 | -9.88 | 1.40E-20 | 1.71E-19 |
| chr2 | 11150558 | 11154514 | 3957 | -7.58 | 1.40E-20 | 1.71E-19 |
| chr6 | 105856149 | 105883950 | 27802 | -9.99 | 1.41E-20 | 1.71E-19 |
| chr1 | 12525821 | 12552044 | 26224 | -9.88 | 1.41E-20 | 1.71E-19 |
| chr1 | 234084222 | 234101715 | 17494 | -9.95 | 1.42E-20 | 1.72E-19 |
| chr3 | 52761565 | 52775353 | 13789 | -9.99 | 1.42E-20 | 1.72E-19 |
| chr12 | 102496753 | 102505221 | 8469 | -10 | 1.42E-20 | 1.72E-19 |
| chr12 | 58810849 | 58831015 | 20167 | -9.88 | 1.42E-20 | 1.73E-19 |
| chr6 | 50815017 | 50856665 | 41649 | 2.87 | 1.42E-20 | 1.73E-19 |
| chr14 | 32726836 | 32759576 | 32741 | -10.01 | 1.42E-20 | 1.73E-19 |
| chr6 | 136034724 | 136052091 | 17368 | -9.88 | 1.43E-20 | 1.73E-19 |
| chr6 | 77631203 | 77668656 | 37454 | -9.93 | 1.44E-20 | 1.75E-19 |
| chr7 | 67202443 | 67234721 | 32279 | -10.14 | 1.44E-20 | 1.75E-19 |
| chr20 | 11289543 | 11304772 | 15230 | -9.88 | 1.45E-20 | 1.75E-19 |
| chr6 | 170495935 | 170520688 | 24754 | -9.88 | 1.45E-20 | 1.76E-19 |
| chr3 | 38469465 | 38481322 | 11858 | -9.9 | 1.46E-20 | 1.77E-19 |
| chr14 | 101273635 | 101279958 | 6324 | -9.95 | 1.46E-20 | 1.77E-19 |
| chr16 | 84492129 | 84506559 | 14431 | -9.88 | 1.46E-20 | 1.77E-19 |
| chr16 | 54372974 | 54391709 | 18736 | -9.98 | 1.47E-20 | 1.77E-19 |
| chr14 | 101875025 | 101897893 | 22869 | -9.88 | 1.48E-20 | 1.79E-19 |
| chr3 | 14720929 | 14756433 | 35505 | -9.88 | 1.48E-20 | 1.79E-19 |
| chr15 | 65304079 | 65307814 | 3736 | -9.91 | 1.50E-20 | 1.81E-19 |
| chr2 | 86746349 | 86775467 | 29119 | -10.09 | 1.50E-20 | 1.81E-19 |
| chr22 | 42429480 | 42457336 | 27857 | -9.95 | 1.50E-20 | 1.81E-19 |
| chr8 | 8288216 | 8306370 | 18155 | -10 | 1.51E-20 | 1.83E-19 |
| chr1 | 176112742 | 176140391 | 27650 | -9.89 | 1.52E-20 | 1.83E-19 |
| chr15 | 102108667 | 102145664 | 36998 | -9.87 | 1.52E-20 | 1.83E-19 |
| chr18 | 67762862 | 67798478 | 35617 | -9.99 | 1.52E-20 | 1.84E-19 |
| chr5 | 114487419 | 114521129 | 33711 | -10.05 | 1.53E-20 | 1.84E-19 |
| chr11 | 68749284 | 68756106 | 6823 | -9.87 | 1.56E-20 | 1.88E-19 |
| chr1 | 225249135 | 225284044 | 34910 | -10 | 1.56E-20 | 1.88E-19 |
| chr5 | 141718139 | 141728634 | 10496 | -9.91 | 1.58E-20 | 1.90E-19 |
| chr15 | 101702974 | 101709098 | 6125 | -9.88 | 1.59E-20 | 1.91E-19 |
| chr14 | 102863176 | 102871132 | 7957 | -9.87 | 1.59E-20 | 1.91E-19 |
| chr1 | 218904466 | 218946101 | 41636 | -10.3 | 1.59E-20 | 1.91E-19 |
| chr16 | 59907807 | 59933907 | 26101 | -9.87 | 1.59E-20 | 1.92E-19 |
| chr7 | 8291875 | 8320598 | 28724 | -9.89 | 1.60E-20 | 1.92E-19 |
| chr1 | 67531458 | 67540649 | 9192 | -2.72 | 1.60E-20 | 1.92E-19 |
| chr4 | 23070988 | 23105214 | 34227 | -9.88 | 1.61E-20 | 1.94E-19 |
| chrX | 41699177 | 41709422 | 10246 | -9.87 | 1.62E-20 | 1.95E-19 |
| chr11 | 12964699 | 12977995 | 13297 | -9.87 | 1.62E-20 | 1.95E-19 |
| chr3 | 189362689 | 189397391 | 34703 | -9.87 | 1.63E-20 | 1.96E-19 |

| | | | | | | |
|---|---|---|---|---|---|---|
| chr2 | 169895568 | 169914407 | 18840 | -9.89 | 1.63E-20 | 1.96E-19 |
| chr10 | 23413348 | 23430855 | 17508 | -2.4 | 1.64E-20 | 1.96E-19 |
| chr7 | 30528721 | 30548068 | 19348 | -9.86 | 1.64E-20 | 1.97E-19 |
| chr12 | 17058560 | 17107772 | 49213 | -10.12 | 1.64E-20 | 1.97E-19 |
| chr7 | 83606039 | 83627556 | 21518 | -10.08 | 1.65E-20 | 1.97E-19 |
| chr12 | 89580259 | 89588993 | 8735 | -9.87 | 1.65E-20 | 1.98E-19 |
| chr4 | 14128175 | 14185158 | 56984 | -9.96 | 1.66E-20 | 1.99E-19 |
| chr2 | 58154378 | 58165218 | 10841 | -9.86 | 1.67E-20 | 2.00E-19 |
| chr4 | 21744329 | 21752889 | 8561 | 5.07 | 1.67E-20 | 2.00E-19 |
| chr14 | 101807776 | 101824228 | 16453 | -9.86 | 1.67E-20 | 2.00E-19 |
| chr2 | 109561790 | 109571411 | 9622 | -9.87 | 1.69E-20 | 2.02E-19 |
| chr12 | 124529801 | 124552796 | 22996 | -10.09 | 1.70E-20 | 2.03E-19 |
| chr12 | 64731563 | 64775944 | 44382 | -10.1 | 1.70E-20 | 2.04E-19 |
| chr10 | 48565290 | 48589720 | 24431 | -2 | 1.73E-20 | 2.06E-19 |
| chr3 | 157961512 | 157996456 | 34945 | -9.91 | 1.76E-20 | 2.10E-19 |
| chr1 | 199501683 | 199511784 | 10102 | -9.86 | 1.76E-20 | 2.11E-19 |
| chrX | 71636860 | 71657105 | 20246 | -9.86 | 1.77E-20 | 2.11E-19 |
| chr4 | 22134708 | 22154905 | 20198 | 7.61 | 1.77E-20 | 2.11E-19 |
| chr8 | 90663041 | 90706381 | 43341 | -9.9 | 1.77E-20 | 2.11E-19 |
| chr10 | 22427244 | 22445240 | 17997 | -2 | 1.77E-20 | 2.11E-19 |
| chr13 | 21322099 | 21345669 | 23571 | -9.86 | 1.77E-20 | 2.11E-19 |
| chr2 | 150814342 | 150855726 | 41385 | -9.88 | 1.77E-20 | 2.11E-19 |
| chr6 | 169811533 | 169837483 | 25951 | -9.87 | 1.77E-20 | 2.11E-19 |
| chr10 | 123514278 | 123529637 | 15360 | -9.9 | 1.78E-20 | 2.12E-19 |
| chr9 | 100537872 | 100556920 | 19049 | -9.86 | 1.78E-20 | 2.12E-19 |
| chr4 | 184992946 | 185010492 | 17547 | -8.63 | 1.79E-20 | 2.13E-19 |
| chrX | 63634941 | 64193519 | 558579 | 1.5 | 1.79E-20 | 2.13E-19 |
| chr6 | 97443663 | 97477791 | 34129 | 9.93 | 1.79E-20 | 2.13E-19 |
| chr12 | 107003533 | 107015304 | 11772 | -9.87 | 1.80E-20 | 2.14E-19 |
| chr5 | 124403115 | 124432974 | 29860 | -9.98 | 1.80E-20 | 2.14E-19 |
| chr5 | 126482705 | 126518397 | 35693 | -9.99 | 1.81E-20 | 2.16E-19 |
| chr5 | 91883899 | 91914619 | 30721 | -9.88 | 1.82E-20 | 2.16E-19 |
| chr3 | 31149996 | 31193407 | 43412 | -10.05 | 1.82E-20 | 2.16E-19 |
| chr9 | 96885374 | 96894684 | 9311 | -9.85 | 1.82E-20 | 2.16E-19 |
| chr10 | 67373425 | 67376554 | 3130 | -4.64 | 1.82E-20 | 2.17E-19 |
| chr12 | 13699365 | 13722784 | 23420 | 10 | 1.83E-20 | 2.17E-19 |
| chrX | 14993068 | 15035721 | 42654 | 2.47 | 1.84E-20 | 2.18E-19 |
| chr1 | 243785767 | 243823202 | 37436 | -9.87 | 1.86E-20 | 2.21E-19 |
| chr4 | 4708708 | 4752225 | 43518 | -4.58 | 1.88E-20 | 2.23E-19 |
| chr14 | 94867829 | 94924050 | 56222 | -1.78 | 1.88E-20 | 2.23E-19 |
| chr4 | 19760302 | 19799044 | 38743 | -9.87 | 1.88E-20 | 2.23E-19 |
| chr15 | 29436591 | 29456792 | 20202 | -9.87 | 1.88E-20 | 2.23E-19 |
| chr3 | 72218227 | 72253204 | 34978 | -2.57 | 1.89E-20 | 2.24E-19 |
| chr2 | 38555871 | 38573020 | 17150 | -9.87 | 1.89E-20 | 2.24E-19 |
| chr5 | 123788678 | 123802597 | 13920 | -10.15 | 1.89E-20 | 2.24E-19 |
| chr18 | 45427248 | 45434865 | 7618 | -9.88 | 1.90E-20 | 2.25E-19 |
| chr3 | 181792020 | 181828868 | 36849 | -9.87 | 1.92E-20 | 2.27E-19 |
| chr10 | 115429308 | 115455268 | 25961 | -9.87 | 1.93E-20 | 2.28E-19 |
| chr10 | 29118895 | 29151162 | 32268 | -10.14 | 1.94E-20 | 2.29E-19 |
| chr5 | 30238681 | 30318585 | 79905 | -10.1 | 1.94E-20 | 2.29E-19 |
| chr9 | 129576361 | 129583543 | 7183 | -9.86 | 1.94E-20 | 2.29E-19 |
| chr17 | 61013461 | 61027011 | 13551 | -9.88 | 1.95E-20 | 2.31E-19 |
| chr10 | 3645253 | 3666231 | 20979 | -9.97 | 1.95E-20 | 2.31E-19 |
| chr14 | 71613608 | 71637692 | 24085 | -9.91 | 1.96E-20 | 2.31E-19 |
| chr12 | 65962347 | 65975428 | 13082 | -4.95 | 1.97E-20 | 2.32E-19 |
| chr17 | 49371181 | 49399126 | 27946 | -9.85 | 1.98E-20 | 2.33E-19 |
| chr5 | 41386052 | 41417663 | 31612 | -9.84 | 2.02E-20 | 2.38E-19 |
| chr12 | 45824858 | 45902288 | 77431 | -10.04 | 2.02E-20 | 2.38E-19 |
| chrX | 62898508 | 62919950 | 21443 | -9.95 | 2.02E-20 | 2.38E-19 |
| chr2 | 242767296 | 242779789 | 12494 | -9.85 | 2.02E-20 | 2.38E-19 |
| chr2 | 174004214 | 174028497 | 24284 | -9.85 | 2.03E-20 | 2.39E-19 |

| | | | | | | |
|---|---|---|---|---|---|---|
| chr14 | 103452669 | 103458645 | 5977 | -9.84 | 2.04E-20 | 2.40E-19 |
| chr10 | 8562724 | 8604633 | 41910 | -9.94 | 2.04E-20 | 2.40E-19 |
| chr5 | 102560559 | 102585419 | 24861 | -9.87 | 2.04E-20 | 2.40E-19 |
| chr5 | 54590148 | 54612664 | 22517 | -9.85 | 2.05E-20 | 2.41E-19 |
| chr12 | 40658278 | 40659812 | 1535 | -7.08 | 2.06E-20 | 2.42E-19 |
| chr2 | 207084561 | 207118465 | 33905 | -9.96 | 2.06E-20 | 2.42E-19 |
| chr5 | 172930525 | 172949858 | 19334 | -9.89 | 2.06E-20 | 2.42E-19 |
| chr18 | 3459561 | 3475750 | 16190 | -9.85 | 2.07E-20 | 2.43E-19 |
| chr15 | 29612131 | 29620538 | 8408 | -9.85 | 2.07E-20 | 2.43E-19 |
| chr9 | 113390573 | 113421403 | 30831 | -10.02 | 2.08E-20 | 2.44E-19 |
| chr10 | 118936662 | 118973859 | 37198 | -9.94 | 2.09E-20 | 2.46E-19 |
| chr3 | 29717732 | 29745363 | 27632 | -9.84 | 2.10E-20 | 2.46E-19 |
| chr15 | 68323150 | 68333629 | 10480 | -10.16 | 2.10E-20 | 2.47E-19 |
| chr19 | 33245519 | 33273828 | 28310 | -10.2 | 2.11E-20 | 2.47E-19 |
| chr4 | 112331363 | 112379228 | 47866 | 9.99 | 2.12E-20 | 2.48E-19 |
| chr8 | 54999017 | 55004302 | 5286 | -9.85 | 2.15E-20 | 2.52E-19 |
| chr8 | 141895876 | 141916727 | 20852 | -9.98 | 2.16E-20 | 2.54E-19 |
| chr3 | 126766812 | 126781000 | 14189 | -9.86 | 2.17E-20 | 2.54E-19 |
| chr3 | 33350643 | 33391827 | 41185 | -9.92 | 2.19E-20 | 2.56E-19 |
| chr13 | 80494465 | 80522568 | 28104 | -9.92 | 2.19E-20 | 2.56E-19 |
| chr5 | 131842863 | 131868523 | 25661 | -10.22 | 2.20E-20 | 2.57E-19 |
| chr5 | 73057631 | 73078074 | 20444 | -9.86 | 2.20E-20 | 2.57E-19 |
| chr13 | 44863294 | 44902382 | 39089 | -9.94 | 2.21E-20 | 2.59E-19 |
| chr2 | 208454080 | 208459395 | 5316 | -9.83 | 2.22E-20 | 2.60E-19 |
| chr10 | 15442471 | 15465540 | 23070 | -9.86 | 2.24E-20 | 2.61E-19 |
| chr15 | 64869919 | 64874904 | 4986 | -9.84 | 2.27E-20 | 2.65E-19 |
| chr8 | 58711589 | 58762323 | 50735 | -9.84 | 2.27E-20 | 2.65E-19 |
| chr7 | 51157571 | 51167605 | 10035 | -9.84 | 2.28E-20 | 2.67E-19 |
| chr12 | 31977123 | 32006706 | 29584 | -10.06 | 2.28E-20 | 2.67E-19 |
| chr3 | 123270715 | 123276956 | 6242 | -9.91 | 2.29E-20 | 2.67E-19 |
| chr1 | 88251209 | 88277056 | 25848 | -9.91 | 2.35E-20 | 2.74E-19 |
| chr10 | 80000805 | 80033392 | 32588 | -9.92 | 2.36E-20 | 2.75E-19 |
| chr13 | 58068716 | 58196276 | 127561 | 2.43 | 2.36E-20 | 2.75E-19 |
| chr7 | 66808494 | 66846116 | 37623 | -10.09 | 2.38E-20 | 2.78E-19 |
| chr12 | 75397790 | 75426606 | 28817 | -10.22 | 2.38E-20 | 2.78E-19 |
| chr10 | 88017798 | 88040606 | 22809 | -9.83 | 2.39E-20 | 2.78E-19 |
| chr12 | 63647323 | 63669517 | 22195 | -9.84 | 2.39E-20 | 2.79E-19 |
| chr5 | 136967172 | 136982471 | 15300 | -9.83 | 2.41E-20 | 2.80E-19 |
| chr1 | 247676940 | 248584800 | 907861 | 1.47 | 2.42E-20 | 2.81E-19 |
| chr16 | 84350968 | 84372363 | 21396 | -10.18 | 2.42E-20 | 2.82E-19 |
| chr10 | 12050887 | 12055607 | 4721 | -9.87 | 2.44E-20 | 2.84E-19 |
| chr10 | 96435746 | 96479428 | 43683 | -9.83 | 2.45E-20 | 2.85E-19 |
| chrX | 101869854 | 101899998 | 30145 | -9.83 | 2.46E-20 | 2.86E-19 |
| chr3 | 23169143 | 23220852 | 51710 | -9.82 | 2.47E-20 | 2.87E-19 |
| chr15 | 39157221 | 39220609 | 63389 | -10.28 | 2.49E-20 | 2.89E-19 |
| chr8 | 22751639 | 22754336 | 2698 | -3.15 | 2.49E-20 | 2.89E-19 |
| chr7 | 130729816 | 130776692 | 46877 | -9.86 | 2.50E-20 | 2.90E-19 |
| chr19 | 33381941 | 33398241 | 16301 | -9.84 | 2.50E-20 | 2.90E-19 |
| chr4 | 13151113 | 13191351 | 40239 | 2.17 | 2.50E-20 | 2.90E-19 |
| chrX | 75398118 | 75414765 | 16648 | -9.85 | 2.50E-20 | 2.90E-19 |
| chr2 | 10653434 | 10658167 | 4734 | -9.83 | 2.52E-20 | 2.92E-19 |
| chr2 | 120891108 | 120927773 | 36666 | -9.99 | 2.53E-20 | 2.93E-19 |
| chr5 | 125095339 | 125124507 | 29169 | -9.84 | 2.54E-20 | 2.94E-19 |
| chr10 | 90076393 | 90084297 | 7905 | -9.82 | 2.55E-20 | 2.95E-19 |
| chr22 | 44033807 | 44051184 | 17378 | -9.84 | 2.56E-20 | 2.96E-19 |
| chr18 | 57712374 | 57762485 | 50112 | -10.01 | 2.56E-20 | 2.97E-19 |
| chr12 | 21038881 | 21059472 | 20592 | -10.3 | 2.60E-20 | 3.01E-19 |
| chr5 | 18350335 | 18728000 | 377666 | 2.74 | 2.62E-20 | 3.03E-19 |
| chr12 | 107316653 | 107344512 | 27860 | -9.85 | 2.63E-20 | 3.04E-19 |
| chr1 | 199721870 | 199739686 | 17817 | -9.85 | 2.66E-20 | 3.07E-19 |
| chr15 | 70656357 | 70723078 | 66722 | -10.53 | 2.66E-20 | 3.08E-19 |

| | | | | | | |
|---|---|---|---|---|---|---|
| chr1 | 221025172 | 221041243 | 16072 | -9.82 | 2.71E-20 | 3.13E-19 |
| chr4 | 183964995 | 183993292 | 28298 | -9.87 | 2.72E-20 | 3.14E-19 |
| chr4 | 129929431 | 129944966 | 15536 | -9.85 | 2.73E-20 | 3.15E-19 |
| chr4 | 8712641 | 8737393 | 24753 | -9.84 | 2.76E-20 | 3.18E-19 |
| chr5 | 114994367 | 115026543 | 32177 | -9.83 | 2.76E-20 | 3.19E-19 |
| chr5 | 142661363 | 142674845 | 13483 | -9.81 | 2.79E-20 | 3.22E-19 |
| chr9 | 101500944 | 101534399 | 33456 | -10 | 2.80E-20 | 3.23E-19 |
| chr13 | 114223470 | 114232415 | 8946 | -9.81 | 2.81E-20 | 3.24E-19 |
| chr2 | 88980046 | 88990310 | 10265 | -9.88 | 2.83E-20 | 3.27E-19 |
| chr2 | 47010767 | 47058075 | 47309 | -9.99 | 2.85E-20 | 3.29E-19 |
| chr21 | 45823456 | 45833230 | 9775 | -7.04 | 2.87E-20 | 3.30E-19 |
| chr8 | 49440692 | 49451088 | 10397 | -9.81 | 2.91E-20 | 3.35E-19 |
| chr7 | 80871716 | 80895342 | 23627 | -9.82 | 2.91E-20 | 3.35E-19 |
| chr11 | 35747689 | 35759127 | 11439 | -9.8 | 2.94E-20 | 3.38E-19 |
| chr11 | 1230617 | 1239055 | 8439 | -10.04 | 2.94E-20 | 3.38E-19 |
| chr11 | 128045312 | 128094122 | 48811 | -9.94 | 2.96E-20 | 3.40E-19 |
| chr3 | 112894821 | 112918127 | 23307 | -9.92 | 2.96E-20 | 3.40E-19 |
| chr13 | 80962155 | 80999536 | 37382 | -9.8 | 2.99E-20 | 3.44E-19 |
| chr5 | 131576564 | 131584436 | 7873 | -9.81 | 3.00E-20 | 3.44E-19 |
| chr1 | 156987379 | 156994774 | 7396 | -9.88 | 3.00E-20 | 3.45E-19 |
| chr10 | 8197805 | 8213825 | 16021 | -9.8 | 3.04E-20 | 3.49E-19 |
| chr13 | 67678371 | 67691336 | 12966 | -9.8 | 3.04E-20 | 3.49E-19 |
| chr21 | 46443061 | 46469116 | 26056 | -9.84 | 3.04E-20 | 3.49E-19 |
| chr10 | 73170047 | 73179274 | 9228 | -9.86 | 3.05E-20 | 3.50E-19 |
| chr7 | 109708344 | 109743074 | 34731 | -9.89 | 3.05E-20 | 3.50E-19 |
| chr16 | 88315349 | 88377694 | 62346 | -10.18 | 3.05E-20 | 3.50E-19 |
| chr1 | 191223035 | 191314441 | 91407 | -10.03 | 3.06E-20 | 3.51E-19 |
| chr12 | 2092078 | 2098917 | 6840 | -9.85 | 3.06E-20 | 3.51E-19 |
| chr1 | 228142025 | 228146614 | 4590 | -9.81 | 3.07E-20 | 3.51E-19 |
| chr18 | 3285464 | 3302464 | 17001 | -9.9 | 3.07E-20 | 3.52E-19 |
| chr1 | 226168192 | 226172255 | 4064 | -9.94 | 3.09E-20 | 3.54E-19 |
| chr21 | 47506535 | 47535645 | 29111 | -9.8 | 3.09E-20 | 3.54E-19 |
| chr16 | 16208998 | 16219837 | 10840 | -9.84 | 3.09E-20 | 3.54E-19 |
| chr7 | 65721863 | 65738966 | 17104 | -9.95 | 3.12E-20 | 3.57E-19 |
| chr5 | 17690158 | 17715549 | 25392 | -10.01 | 3.15E-20 | 3.61E-19 |
| chr18 | 69918259 | 69961869 | 43611 | -9.81 | 3.16E-20 | 3.61E-19 |
| chr12 | 133186628 | 133194764 | 8137 | -9.8 | 3.16E-20 | 3.61E-19 |
| chr20 | 40212465 | 40232040 | 19576 | -9.99 | 3.18E-20 | 3.63E-19 |
| chr10 | 71936860 | 71942045 | 5186 | -9.82 | 3.20E-20 | 3.66E-19 |
| chr7 | 30031158 | 30039182 | 8025 | -9.93 | 3.23E-20 | 3.69E-19 |
| chr6 | 53460465 | 53488099 | 27635 | -9.93 | 3.24E-20 | 3.70E-19 |
| chrX | 74721722 | 74769605 | 47884 | 2.05 | 3.26E-20 | 3.72E-19 |
| chr11 | 14356684 | 14370742 | 14059 | -9.8 | 3.26E-20 | 3.72E-19 |
| chr7 | 130291521 | 130334706 | 43186 | -9.79 | 3.29E-20 | 3.75E-19 |
| chr13 | 23296517 | 23350821 | 54305 | -9.85 | 3.31E-20 | 3.78E-19 |
| chr12 | 20812233 | 20832145 | 19913 | -9.89 | 3.32E-20 | 3.78E-19 |
| chr18 | 77684276 | 77697919 | 13644 | -9.8 | 3.32E-20 | 3.78E-19 |
| chr11 | 1432140 | 1444222 | 12083 | -9.81 | 3.34E-20 | 3.80E-19 |
| chr22 | 48778040 | 48807493 | 29454 | -10.07 | 3.34E-20 | 3.80E-19 |
| chr12 | 9105705 | 9127437 | 21733 | -9.92 | 3.34E-20 | 3.80E-19 |
| chr12 | 51225042 | 51260568 | 35527 | -10.08 | 3.36E-20 | 3.82E-19 |
| chr3 | 40524643 | 40545665 | 21023 | -9.81 | 3.37E-20 | 3.83E-19 |
| chr20 | 17703170 | 17746567 | 43398 | -10.22 | 3.37E-20 | 3.84E-19 |
| chr1 | 77616829 | 77645268 | 28440 | -9.79 | 3.38E-20 | 3.85E-19 |
| chr19 | 45217513 | 45220389 | 2877 | -9.79 | 3.44E-20 | 3.91E-19 |
| chr7 | 45467716 | 45486976 | 19261 | -9.85 | 3.45E-20 | 3.92E-19 |
| chr12 | 80229985 | 80247944 | 17960 | -9.79 | 3.45E-20 | 3.92E-19 |
| chr12 | 41044597 | 41073396 | 28800 | -9.96 | 3.47E-20 | 3.94E-19 |
| chr10 | 129752848 | 129757705 | 4858 | -6.76 | 3.48E-20 | 3.96E-19 |
| chr20 | 46146523 | 46156576 | 10054 | -9.8 | 3.49E-20 | 3.96E-19 |
| chr3 | 102847608 | 102919769 | 72162 | -9.87 | 3.50E-20 | 3.97E-19 |

| | | | | | | |
|---|---|---|---|---|---|---|
| chr7 | 3640370 | 3672413 | 32044 | -9.79 | 3.52E-20 | 3.99E-19 |
| chr2 | 72889412 | 72896154 | 6743 | -9.81 | 3.52E-20 | 3.99E-19 |
| chr3 | 25346476 | 25382824 | 36349 | 9.97 | 3.52E-20 | 3.99E-19 |
| chr8 | 24198673 | 24230092 | 31420 | -9.87 | 3.56E-20 | 4.03E-19 |
| chr7 | 30743091 | 30776813 | 33723 | -9.81 | 3.57E-20 | 4.04E-19 |
| chr17 | 4157803 | 4164178 | 6376 | -9.8 | 3.59E-20 | 4.06E-19 |
| chrX | 82063751 | 82095216 | 31466 | -9.79 | 3.60E-20 | 4.08E-19 |
| chr8 | 135679343 | 135687953 | 8611 | -6.71 | 3.62E-20 | 4.09E-19 |
| chr9 | 93950255 | 93968951 | 18697 | -9.97 | 3.62E-20 | 4.09E-19 |
| chr14 | 53882809 | 53901627 | 18819 | -9.82 | 3.63E-20 | 4.10E-19 |
| chr2 | 183221939 | 183250213 | 28275 | -9.85 | 3.65E-20 | 4.13E-19 |
| chr22 | 46002908 | 46022031 | 19124 | -9.8 | 3.65E-20 | 4.13E-19 |
| chr18 | 74338610 | 74344511 | 5902 | -7.03 | 3.67E-20 | 4.14E-19 |
| chr5 | 124813788 | 124877295 | 63508 | -10.14 | 3.67E-20 | 4.15E-19 |
| chr6 | 131371326 | 131380148 | 8823 | -9.78 | 3.72E-20 | 4.21E-19 |
| chr5 | 27462811 | 27490960 | 28150 | -9.79 | 3.74E-20 | 4.22E-19 |
| chr12 | 117431494 | 117455017 | 23524 | -10.21 | 3.76E-20 | 4.24E-19 |
| chr12 | 103170833 | 103201150 | 30318 | 5 | 3.77E-20 | 4.25E-19 |
| chr10 | 116740228 | 116747788 | 7561 | -9.79 | 3.83E-20 | 4.32E-19 |
| chr13 | 80331995 | 80365206 | 33212 | -9.79 | 3.84E-20 | 4.33E-19 |
| chr15 | 37777167 | 38042350 | 265184 | 1.39 | 3.85E-20 | 4.34E-19 |
| chr2 | 172212432 | 172239324 | 26893 | -9.84 | 3.87E-20 | 4.36E-19 |
| chr2 | 84179287 | 84241565 | 62279 | -9.78 | 3.90E-20 | 4.39E-19 |
| chr2 | 179568615 | 179642241 | 73627 | 1.72 | 3.91E-20 | 4.40E-19 |
| chr3 | 183365695 | 183376526 | 10832 | -9.93 | 3.91E-20 | 4.41E-19 |
| chr8 | 58981969 | 59039338 | 57370 | -9.78 | 3.92E-20 | 4.42E-19 |
| chrX | 41747415 | 41768299 | 20885 | -9.8 | 3.93E-20 | 4.42E-19 |
| chr5 | 149392575 | 149396929 | 4355 | -9.9 | 3.98E-20 | 4.48E-19 |
| chr2 | 71874294 | 71888206 | 13913 | -9.79 | 4.01E-20 | 4.51E-19 |
| chr8 | 94654171 | 94680013 | 25843 | -3.36 | 4.04E-20 | 4.55E-19 |
| chr20 | 22914127 | 22981564 | 67438 | -9.86 | 4.05E-20 | 4.55E-19 |
| chr7 | 85357634 | 85381352 | 23719 | -9.83 | 4.07E-20 | 4.58E-19 |
| chr9 | 38435727 | 38477949 | 42223 | -9.87 | 4.08E-20 | 4.58E-19 |
| chr10 | 115629201 | 115647359 | 18159 | -10.15 | 4.14E-20 | 4.65E-19 |
| chr8 | 140603771 | 140633406 | 29636 | 9.85 | 4.14E-20 | 4.65E-19 |
| chr3 | 104810028 | 104852151 | 42124 | -9.8 | 4.15E-20 | 4.65E-19 |
| chr17 | 75328644 | 75340882 | 12239 | -9.88 | 4.16E-20 | 4.67E-19 |
| chr8 | 144242250 | 144246304 | 4055 | -9.78 | 4.18E-20 | 4.68E-19 |
| chr15 | 101273977 | 101298278 | 24302 | -9.8 | 4.18E-20 | 4.68E-19 |
| chr12 | 31599887 | 31623933 | 24047 | -9.97 | 4.18E-20 | 4.68E-19 |
| chr15 | 89504918 | 89520393 | 15476 | -9.96 | 4.18E-20 | 4.69E-19 |
| chr16 | 58388420 | 58435738 | 47319 | -9.88 | 4.24E-20 | 4.75E-19 |
| chr18 | 33531735 | 33543184 | 11450 | -9.77 | 4.24E-20 | 4.75E-19 |
| chr4 | 153838906 | 153843056 | 4151 | -6.61 | 4.25E-20 | 4.76E-19 |
| chr1 | 231135339 | 231145659 | 10321 | -9.9 | 4.33E-20 | 4.85E-19 |
| chr2 | 103106536 | 103129768 | 23233 | -9.76 | 4.40E-20 | 4.93E-19 |
| chr15 | 33334475 | 33345281 | 10807 | -9.87 | 4.42E-20 | 4.95E-19 |
| chr7 | 148489744 | 148496489 | 6746 | -9.77 | 4.43E-20 | 4.95E-19 |
| chr20 | 50701749 | 50706623 | 4875 | -9.89 | 4.43E-20 | 4.96E-19 |
| chr6 | 4976483 | 4981051 | 4569 | -6.84 | 4.44E-20 | 4.97E-19 |
| chr11 | 17689963 | 17726045 | 36083 | -1.95 | 4.44E-20 | 4.97E-19 |
| chr11 | 36728703 | 36763339 | 34637 | 10.03 | 4.45E-20 | 4.97E-19 |
| chr13 | 67476351 | 67517585 | 41235 | -9.86 | 4.47E-20 | 4.99E-19 |
| chr5 | 72043324 | 72077663 | 34340 | -9.82 | 4.48E-20 | 5.00E-19 |
| chr9 | 126829388 | 126850060 | 20673 | -9.77 | 4.51E-20 | 5.04E-19 |
| chr16 | 2404758 | 2410056 | 5299 | -9.79 | 4.56E-20 | 5.09E-19 |
| chr5 | 40920857 | 40928312 | 7456 | -2.51 | 4.57E-20 | 5.10E-19 |
| chr12 | 124620874 | 124654476 | 33603 | -10.35 | 4.58E-20 | 5.10E-19 |
| chr3 | 133620712 | 133641494 | 20783 | -9.76 | 4.58E-20 | 5.11E-19 |
| chr10 | 81122924 | 81132548 | 9625 | -9.76 | 4.61E-20 | 5.14E-19 |
| chr12 | 114090803 | 114097891 | 7089 | -9.76 | 4.67E-20 | 5.20E-19 |

| | | | | | | |
|---|---|---|---|---|---|---|
| chr5 | 125715750 | 125748592 | 32843 | -9.8 | 4.69E-20 | 5.22E-19 |
| chr2 | 196991324 | 197001134 | 9811 | -9.77 | 4.70E-20 | 5.24E-19 |
| chr4 | 153727822 | 153775965 | 48144 | -9.78 | 4.70E-20 | 5.24E-19 |
| chr14 | 104904910 | 104920555 | 15646 | -4.45 | 4.71E-20 | 5.24E-19 |
| chr10 | 23057481 | 23091703 | 34223 | -9.77 | 4.71E-20 | 5.24E-19 |
| chr7 | 149161914 | 149183337 | 21424 | -9.77 | 4.74E-20 | 5.27E-19 |
| chr1 | 181113918 | 181143643 | 29726 | -9.85 | 4.76E-20 | 5.29E-19 |
| chr14 | 51147226 | 51170194 | 22969 | -9.96 | 4.77E-20 | 5.30E-19 |
| chr22 | 50417655 | 50430858 | 13204 | -9.77 | 4.79E-20 | 5.33E-19 |
| chr7 | 80313052 | 80376579 | 63528 | -9.86 | 4.80E-20 | 5.34E-19 |
| chr2 | 36926 | 83250 | 46325 | -9.92 | 4.81E-20 | 5.34E-19 |
| chr18 | 9445305 | 9470648 | 25344 | -9.87 | 4.81E-20 | 5.34E-19 |
| chr14 | 71531125 | 71563959 | 32835 | -9.75 | 4.96E-20 | 5.51E-19 |
| chr12 | 47470548 | 47486914 | 16367 | -9.85 | 4.97E-20 | 5.51E-19 |
| chr7 | 158371025 | 158386867 | 15843 | -9.78 | 4.99E-20 | 5.54E-19 |
| chr10 | 114945305 | 114983765 | 38461 | -10.02 | 5.00E-20 | 5.54E-19 |
| chr14 | 95842743 | 95861720 | 18978 | -9.83 | 5.02E-20 | 5.56E-19 |
| chr2 | 47138771 | 47159548 | 20778 | -9.87 | 5.05E-20 | 5.60E-19 |
| chr9 | 100574968 | 100603458 | 28491 | -9.77 | 5.06E-20 | 5.61E-19 |
| chr16 | 87663374 | 87666652 | 3279 | -9.75 | 5.08E-20 | 5.63E-19 |
| chr1 | 175949598 | 175978714 | 29117 | -9.84 | 5.09E-20 | 5.64E-19 |
| chr2 | 54541952 | 54553982 | 12031 | -9.78 | 5.10E-20 | 5.65E-19 |
| chr2 | 88402292 | 88504297 | 102006 | -1.79 | 5.15E-20 | 5.70E-19 |
| chr1 | 35345650 | 35361220 | 15571 | -9.8 | 5.19E-20 | 5.74E-19 |
| chr8 | 24926795 | 24959350 | 32556 | -9.9 | 5.20E-20 | 5.75E-19 |
| chr21 | 45015645 | 45050124 | 34480 | -9.87 | 5.22E-20 | 5.77E-19 |
| chr5 | 172792091 | 172821973 | 29883 | -9.75 | 5.22E-20 | 5.77E-19 |
| chr14 | 77632688 | 77639091 | 6404 | -9.75 | 5.25E-20 | 5.80E-19 |
| chr6 | 145793900 | 145848880 | 54981 | -9.78 | 5.25E-20 | 5.81E-19 |
| chr7 | 158835988 | 158855619 | 19632 | -9.91 | 5.26E-20 | 5.81E-19 |
| chr15 | 83764743 | 83772680 | 7938 | -4.17 | 5.26E-20 | 5.81E-19 |
| chr5 | 64892940 | 64903011 | 10072 | -9.78 | 5.30E-20 | 5.85E-19 |
| chr5 | 79534388 | 79540929 | 6542 | -9.8 | 5.30E-20 | 5.85E-19 |
| chr6 | 2321821 | 2343610 | 21790 | -9.75 | 5.31E-20 | 5.86E-19 |
| chr1 | 33871444 | 33891414 | 19971 | -9.76 | 5.42E-20 | 5.97E-19 |
| chr19 | 20710995 | 20750440 | 39446 | -9.94 | 5.42E-20 | 5.98E-19 |
| chr7 | 93503451 | 93528180 | 24730 | -9.92 | 5.45E-20 | 6.00E-19 |
| chr13 | 73494727 | 73523428 | 28702 | -10.03 | 5.49E-20 | 6.05E-19 |
| chr4 | 5307160 | 5354777 | 47618 | -9.95 | 5.50E-20 | 6.05E-19 |
| chr12 | 121536438 | 121539417 | 2980 | -8.27 | 5.50E-20 | 6.06E-19 |
| chr9 | 113748919 | 113770141 | 21223 | -9.75 | 5.51E-20 | 6.06E-19 |
| chr3 | 188982052 | 188999021 | 16970 | -9.78 | 5.51E-20 | 6.06E-19 |
| chr6 | 4449463 | 4502131 | 52669 | -9.88 | 5.52E-20 | 6.08E-19 |
| chr9 | 132056144 | 132067168 | 11025 | -9.88 | 5.54E-20 | 6.10E-19 |
| chr15 | 58583222 | 58594081 | 10860 | -9.88 | 5.57E-20 | 6.12E-19 |
| chr13 | 110732326 | 110755833 | 23508 | -9.86 | 5.59E-20 | 6.15E-19 |
| chr7 | 72794200 | 72806604 | 12405 | -9.84 | 5.60E-20 | 6.15E-19 |
| chr6 | 84132981 | 84137680 | 4700 | -9.81 | 5.67E-20 | 6.23E-19 |
| chr17 | 22068426 | 22117438 | 49013 | 1.72 | 5.72E-20 | 6.29E-19 |
| chr16 | 10534103 | 10542598 | 8496 | -9.74 | 5.73E-20 | 6.29E-19 |
| chr12 | 58032975 | 58056619 | 23645 | -10.01 | 5.73E-20 | 6.29E-19 |
| chr12 | 107979438 | 107991798 | 12361 | -9.85 | 5.76E-20 | 6.32E-19 |
| chr12 | 70526403 | 70556124 | 29722 | -9.9 | 5.76E-20 | 6.32E-19 |
| chr18 | 67929538 | 67950975 | 21438 | -9.85 | 5.82E-20 | 6.38E-19 |
| chr1 | 225465916 | 225484998 | 19083 | -9.76 | 5.89E-20 | 6.46E-19 |
| chr12 | 118249769 | 118305920 | 56152 | -9.74 | 5.94E-20 | 6.51E-19 |
| chr6 | 163875193 | 163895122 | 19930 | -9.73 | 5.94E-20 | 6.51E-19 |
| chr4 | 144503416 | 144605385 | 101970 | 2.39 | 5.99E-20 | 6.56E-19 |
| chr17 | 59490534 | 59518068 | 27535 | -9.9 | 6.00E-20 | 6.57E-19 |
| chr1 | 173136674 | 173148565 | 11892 | -9.78 | 6.03E-20 | 6.60E-19 |
| chr5 | 159099980 | 159121323 | 21344 | -9.73 | 6.06E-20 | 6.63E-19 |

| | | | | | | |
|---|---|---|---|---|---|---|
| chr12 | 99407579 | 99447553 | 39975 | -9.83 | 6.06E-20 | 6.63E-19 |
| chr13 | 51494745 | 51512603 | 17859 | -9.73 | 6.07E-20 | 6.64E-19 |
| chr8 | 131527182 | 131565432 | 38251 | -9.87 | 6.11E-20 | 6.68E-19 |
| chr1 | 155355563 | 155362378 | 6816 | -9.73 | 6.11E-20 | 6.68E-19 |
| chr8 | 62511100 | 62526382 | 15283 | -9.73 | 6.15E-20 | 6.73E-19 |
| chr22 | 42470489 | 42513258 | 42770 | -9.99 | 6.18E-20 | 6.76E-19 |
| chr5 | 156819365 | 156842022 | 22658 | -9.74 | 6.20E-20 | 6.77E-19 |
| chr1 | 186678727 | 186701215 | 22489 | -9.76 | 6.22E-20 | 6.79E-19 |
| chr14 | 71089829 | 71104316 | 14488 | -9.76 | 6.24E-20 | 6.81E-19 |
| chr2 | 207888177 | 207899925 | 11749 | -9.73 | 6.26E-20 | 6.83E-19 |
| chr9 | 77575270 | 77592878 | 17609 | -9.73 | 6.27E-20 | 6.85E-19 |
| chr5 | 130369230 | 130435577 | 66348 | -9.79 | 6.29E-20 | 6.87E-19 |
| chr14 | 85782350 | 85787022 | 4673 | -7 | 6.30E-20 | 6.87E-19 |
| chr15 | 63392173 | 63413708 | 21536 | -9.84 | 6.31E-20 | 6.88E-19 |
| chrX | 113414879 | 113516621 | 101743 | 1.76 | 6.32E-20 | 6.88E-19 |
| chr8 | 140732503 | 140749368 | 16866 | -9.78 | 6.36E-20 | 6.93E-19 |
| chr2 | 48383573 | 48389395 | 5823 | -3.8 | 6.42E-20 | 7.00E-19 |
| chr19 | 743826 | 761955 | 18130 | -9.73 | 6.45E-20 | 7.02E-19 |
| chr11 | 102224322 | 102231414 | 7093 | -9.86 | 6.45E-20 | 7.02E-19 |
| chr19 | 50648257 | 50662933 | 14677 | -9.83 | 6.49E-20 | 7.07E-19 |
| chr4 | 121995540 | 122008577 | 13038 | -9.74 | 6.51E-20 | 7.08E-19 |
| chr20 | 60165453 | 60193338 | 27886 | -9.77 | 6.52E-20 | 7.10E-19 |
| chr20 | 17868605 | 17880285 | 11681 | -9.76 | 6.57E-20 | 7.15E-19 |
| chr15 | 35564418 | 35577770 | 13353 | -9.73 | 6.60E-20 | 7.17E-19 |
| chr15 | 101573165 | 101582136 | 8972 | -9.88 | 6.61E-20 | 7.19E-19 |
| chr6 | 4760287 | 4785117 | 24831 | -9.75 | 6.63E-20 | 7.20E-19 |
| chr7 | 3950158 | 3997580 | 47423 | -9.93 | 6.69E-20 | 7.26E-19 |
| chr12 | 18202545 | 18259888 | 57344 | -9.8 | 6.76E-20 | 7.34E-19 |
| chr17 | 8579949 | 8585669 | 5721 | -2.84 | 6.78E-20 | 7.36E-19 |
| chr13 | 108931472 | 108971021 | 39550 | -9.72 | 6.79E-20 | 7.37E-19 |
| chr1 | 168124004 | 168130884 | 6881 | -9.74 | 6.79E-20 | 7.37E-19 |
| chr6 | 151840864 | 151862659 | 21796 | -9.81 | 6.87E-20 | 7.45E-19 |
| chr3 | 167390570 | 167398349 | 7780 | -9.84 | 6.90E-20 | 7.48E-19 |
| chr1 | 228375260 | 228390151 | 14892 | -9.91 | 6.98E-20 | 7.56E-19 |
| chr16 | 883568 | 907332 | 23765 | -9.76 | 7.01E-20 | 7.60E-19 |
| chr5 | 179529886 | 179536080 | 6195 | -9.73 | 7.01E-20 | 7.60E-19 |
| chr1 | 55751401 | 55812239 | 60839 | -9.94 | 7.01E-20 | 7.60E-19 |
| chr2 | 190128936 | 190161704 | 32769 | -9.73 | 7.04E-20 | 7.62E-19 |
| chr1 | 232810758 | 232857090 | 46333 | -10.08 | 7.05E-20 | 7.63E-19 |
| chr10 | 88662074 | 88668173 | 6100 | -9.83 | 7.07E-20 | 7.65E-19 |
| chr6 | 163919468 | 163930653 | 11186 | -9.72 | 7.08E-20 | 7.66E-19 |
| chr5 | 134612621 | 134638261 | 25641 | -9.74 | 7.08E-20 | 7.66E-19 |
| chr17 | 13959286 | 13971123 | 11838 | -9.72 | 7.09E-20 | 7.66E-19 |
| chr11 | 70635593 | 70643575 | 7983 | -9.76 | 7.12E-20 | 7.70E-19 |
| chr13 | 99715839 | 99730307 | 14469 | -9.83 | 7.14E-20 | 7.72E-19 |
| chr4 | 152863820 | 152900530 | 36711 | -2.23 | 7.20E-20 | 7.78E-19 |
| chr2 | 163033938 | 163074833 | 40896 | 9.76 | 7.20E-20 | 7.78E-19 |
| chr18 | 21255726 | 21301310 | 45585 | -9.73 | 7.21E-20 | 7.79E-19 |
| chr7 | 98794421 | 98806716 | 12296 | -9.75 | 7.25E-20 | 7.82E-19 |
| chr3 | 122298933 | 122304512 | 5580 | -9.72 | 7.27E-20 | 7.85E-19 |
| chrX | 119830701 | 119879672 | 48972 | -9.86 | 7.34E-20 | 7.92E-19 |
| chr12 | 59856021 | 59956176 | 100156 | -9.72 | 7.37E-20 | 7.94E-19 |
| chr12 | 53529084 | 53534900 | 5817 | -9.71 | 7.37E-20 | 7.94E-19 |
| chr6 | 160789704 | 160805311 | 15608 | -9.76 | 7.37E-20 | 7.94E-19 |
| chr2 | 231384422 | 231436675 | 52254 | -9.95 | 7.43E-20 | 8.01E-19 |
| chr9 | 81832025 | 81849475 | 17451 | -9.74 | 7.50E-20 | 8.07E-19 |
| chrX | 9751282 | 9769604 | 18323 | -9.71 | 7.51E-20 | 8.09E-19 |
| chr2 | 119053031 | 119121179 | 68149 | -2.55 | 7.52E-20 | 8.10E-19 |
| chr9 | 38098897 | 38116650 | 17754 | -9.91 | 7.53E-20 | 8.11E-19 |
| chr16 | 81369591 | 81383998 | 14408 | -9.71 | 7.56E-20 | 8.13E-19 |
| chr17 | 28678656 | 28691666 | 13011 | -9.73 | 7.60E-20 | 8.18E-19 |

| | | | | | | |
|---|---|---|---|---|---|---|
| chr17 | 65042923 | 65046827 | 3905 | -9.71 | 7.61E-20 | 8.19E-19 |
| chr5 | 169541641 | 169559674 | 18034 | -9.72 | 7.65E-20 | 8.23E-19 |
| chr11 | 121269262 | 121274290 | 5029 | -4.33 | 7.68E-20 | 8.25E-19 |
| chr21 | 17115174 | 17135944 | 20771 | -9.8 | 7.69E-20 | 8.26E-19 |
| chr7 | 141504444 | 141668706 | 164263 | 1.37 | 7.74E-20 | 8.31E-19 |
| chr14 | 91213755 | 91236195 | 22441 | -9.73 | 7.76E-20 | 8.33E-19 |
| chr10 | 29960383 | 29968356 | 7974 | -9.86 | 7.80E-20 | 8.37E-19 |
| chr19 | 38669077 | 38672080 | 3004 | -9.82 | 7.80E-20 | 8.37E-19 |
| chr7 | 31033305 | 31053010 | 19706 | -9.76 | 7.82E-20 | 8.39E-19 |
| chr9 | 82140983 | 82184187 | 43205 | -9.94 | 7.82E-20 | 8.39E-19 |
| chr2 | 37380856 | 37394655 | 13800 | -9.77 | 7.86E-20 | 8.42E-19 |
| chr2 | 153631541 | 153672601 | 41061 | -9.7 | 7.96E-20 | 8.54E-19 |
| chr2 | 84619012 | 84639349 | 20338 | 5.02 | 7.97E-20 | 8.54E-19 |
| chr16 | 54448652 | 54479507 | 30856 | -9.81 | 7.97E-20 | 8.54E-19 |
| chr15 | 90553373 | 90557908 | 4536 | -9.7 | 7.99E-20 | 8.56E-19 |
| chr9 | 136453942 | 136465265 | 11324 | -9.76 | 7.99E-20 | 8.56E-19 |
| chr4 | 5402955 | 5433151 | 30197 | -9.73 | 8.11E-20 | 8.68E-19 |
| chr8 | 102760874 | 102771410 | 10537 | -9.8 | 8.12E-20 | 8.69E-19 |
| chr20 | 9110417 | 9133695 | 23279 | -9.71 | 8.14E-20 | 8.71E-19 |
| chr11 | 69173287 | 69184999 | 11713 | -9.7 | 8.15E-20 | 8.71E-19 |
| chr2 | 202584057 | 202603525 | 19469 | -9.88 | 8.15E-20 | 8.71E-19 |
| chr1 | 193663973 | 193682434 | 18462 | -9.79 | 8.19E-20 | 8.75E-19 |
| chr12 | 28744609 | 28776922 | 32314 | -9.73 | 8.20E-20 | 8.76E-19 |
| chr2 | 86864831 | 86878496 | 13666 | -9.7 | 8.24E-20 | 8.81E-19 |
| chr6 | 136771765 | 136780976 | 9212 | -9.73 | 8.25E-20 | 8.81E-19 |
| chr14 | 96569389 | 96587360 | 17972 | -9.9 | 8.25E-20 | 8.81E-19 |
| chr11 | 120636879 | 120645487 | 8609 | -3.21 | 8.25E-20 | 8.81E-19 |
| chr9 | 88702307 | 88708612 | 6306 | -9.73 | 8.31E-20 | 8.86E-19 |
| chr11 | 44647741 | 44672540 | 24800 | -9.71 | 8.32E-20 | 8.88E-19 |
| chr2 | 148978473 | 149018203 | 39731 | -9.72 | 8.32E-20 | 8.88E-19 |
| chr20 | 15437020 | 16222562 | 785543 | 0.75 | 8.35E-20 | 8.90E-19 |
| chr2 | 127936296 | 128000352 | 64057 | -9.81 | 8.40E-20 | 8.95E-19 |
| chr1 | 190979625 | 191096553 | 116929 | -10.24 | 8.40E-20 | 8.95E-19 |
| chr10 | 124589826 | 124605299 | 15474 | -9.82 | 8.42E-20 | 8.97E-19 |
| chr5 | 71694334 | 71703087 | 8754 | -9.7 | 8.45E-20 | 9.00E-19 |
| chr2 | 9195568 | 9223527 | 27960 | -9.71 | 8.52E-20 | 9.08E-19 |
| chrX | 13407101 | 13421800 | 14700 | -9.7 | 8.56E-20 | 9.11E-19 |
| chr7 | 78898833 | 78932237 | 33405 | -9.75 | 8.56E-20 | 9.11E-19 |
| chr1 | 64754276 | 64765562 | 11287 | -9.69 | 8.61E-20 | 9.16E-19 |
| chr12 | 74148201 | 74197404 | 49204 | -9.69 | 8.68E-20 | 9.23E-19 |
| chr12 | 1852526 | 1870095 | 17570 | -9.74 | 8.69E-20 | 9.23E-19 |
| chr2 | 197544920 | 197553444 | 8525 | -9.69 | 8.77E-20 | 9.32E-19 |
| chr7 | 150135245 | 150155139 | 19895 | -9.7 | 8.80E-20 | 9.35E-19 |
| chrX | 73098716 | 73128077 | 29362 | -9.69 | 8.81E-20 | 9.36E-19 |
| chr3 | 31465633 | 31480472 | 14840 | -9.74 | 8.82E-20 | 9.36E-19 |
| chr20 | 61738032 | 61755622 | 17591 | -9.71 | 8.85E-20 | 9.40E-19 |
| chr21 | 32520293 | 32532035 | 11743 | -9.73 | 8.86E-20 | 9.40E-19 |
| chr10 | 102363140 | 102375045 | 11906 | -3.05 | 8.86E-20 | 9.40E-19 |
| chr3 | 129554344 | 129580973 | 26630 | -9.7 | 8.88E-20 | 9.42E-19 |
| chr1 | 54385802 | 54402499 | 16698 | -9.73 | 8.89E-20 | 9.43E-19 |
| chr1 | 48064555 | 48076030 | 11476 | -9.71 | 8.93E-20 | 9.46E-19 |
| chr2 | 241000972 | 241059875 | 58904 | -3.77 | 8.95E-20 | 9.49E-19 |
| chrX | 17273249 | 17283284 | 10036 | -2.8 | 8.96E-20 | 9.49E-19 |
| chr5 | 57672192 | 57702689 | 30498 | -9.75 | 9.02E-20 | 9.55E-19 |
| chr10 | 125630245 | 125637813 | 7569 | -2.48 | 9.05E-20 | 9.58E-19 |
| chr11 | 76642905 | 76657186 | 14282 | -9.74 | 9.06E-20 | 9.59E-19 |
| chr15 | 67403116 | 67405401 | 2286 | -9.74 | 9.06E-20 | 9.59E-19 |
| chr14 | 42654658 | 42674133 | 19476 | -10.02 | 9.06E-20 | 9.59E-19 |
| chr1 | 233515643 | 233558650 | 43008 | -10.14 | 9.08E-20 | 9.61E-19 |
| chr15 | 31575608 | 31586886 | 11279 | -9.82 | 9.18E-20 | 9.71E-19 |
| chr5 | 170271326 | 170302310 | 30985 | -9.85 | 9.18E-20 | 9.71E-19 |

| | | | | | | |
|---|---|---|---|---|---|---|
| chr1 | 154131625 | 154135609 | 3985 | -9.72 | 9.20E-20 | 9.72E-19 |
| chr7 | 135460509 | 135464816 | 4308 | -9.7 | 9.21E-20 | 9.73E-19 |
| chr11 | 68690145 | 68695073 | 4929 | -9.69 | 9.21E-20 | 9.73E-19 |
| chr4 | 156403703 | 156439940 | 36238 | 9.95 | 9.21E-20 | 9.73E-19 |
| chr3 | 153975806 | 153988274 | 12469 | -9.76 | 9.29E-20 | 9.80E-19 |
| chr3 | 123305904 | 123311559 | 5656 | -9.7 | 9.30E-20 | 9.81E-19 |
| chr18 | 9292408 | 9329270 | 36863 | -9.88 | 9.30E-20 | 9.82E-19 |
| chr3 | 96472166 | 96497276 | 25111 | -6.65 | 9.30E-20 | 9.82E-19 |
| chr1 | 14001492 | 14026260 | 24769 | -9.71 | 9.31E-20 | 9.82E-19 |
| chr9 | 81781885 | 81812939 | 31055 | -9.69 | 9.34E-20 | 9.84E-19 |
| chr6 | 66714490 | 67777450 | 1062961 | 1.94 | 9.34E-20 | 9.84E-19 |
| chr14 | 106007846 | 106017755 | 9910 | -9.69 | 9.34E-20 | 9.85E-19 |
| chr10 | 104921304 | 104926562 | 5259 | -9.7 | 9.35E-20 | 9.85E-19 |
| chr9 | 4053993 | 4062891 | 8899 | -9.78 | 9.36E-20 | 9.86E-19 |
| chr12 | 43018794 | 43092885 | 74092 | -9.97 | 9.39E-20 | 9.88E-19 |
| chr18 | 72788629 | 72814701 | 26073 | -9.7 | 9.47E-20 | 9.97E-19 |
| chr7 | 48095311 | 48112538 | 17228 | -9.73 | 9.53E-20 | 1.00E-18 |
| chr1 | 167284360 | 167324324 | 39965 | -9.85 | 9.55E-20 | 1.00E-18 |
| chr13 | 32907367 | 32915867 | 8501 | -9.76 | 9.56E-20 | 1.01E-18 |
| chr12 | 117055766 | 117064096 | 8331 | -9.84 | 9.61E-20 | 1.01E-18 |
| chr2 | 103412828 | 103422535 | 9708 | -9.74 | 9.70E-20 | 1.02E-18 |
| chr9 | 16085910 | 16095825 | 9916 | -3.84 | 9.72E-20 | 1.02E-18 |
| chr16 | 12804411 | 12840865 | 36455 | -10.03 | 9.74E-20 | 1.02E-18 |
| chr10 | 21986246 | 21991158 | 4913 | -9.68 | 9.76E-20 | 1.03E-18 |
| chr6 | 166960038 | 166974660 | 14623 | -9.72 | 9.81E-20 | 1.03E-18 |
| chr8 | 54967947 | 54985779 | 17833 | -9.94 | 9.85E-20 | 1.03E-18 |
| chr2 | 121754914 | 121787598 | 32685 | -9.85 | 9.86E-20 | 1.04E-18 |
| chr5 | 92516236 | 92536218 | 19983 | -9.72 | 9.91E-20 | 1.04E-18 |
| chr8 | 63014315 | 63031398 | 17084 | -9.8 | 1.01E-19 | 1.06E-18 |
| chrX | 69318725 | 69341964 | 23240 | -9.69 | 1.02E-19 | 1.06E-18 |
| chr8 | 82099305 | 82138350 | 39046 | -4.71 | 1.02E-19 | 1.07E-18 |
| chr4 | 13272032 | 13301294 | 29263 | -6.9 | 1.03E-19 | 1.08E-18 |
| chr2 | 148579715 | 148600993 | 21279 | -9.71 | 1.03E-19 | 1.08E-18 |
| chr20 | 60565124 | 60578167 | 13044 | -9.68 | 1.04E-19 | 1.08E-18 |
| chr8 | 64098819 | 64106530 | 7712 | -9.9 | 1.04E-19 | 1.09E-18 |
| chr9 | 115400072 | 115412844 | 12773 | -9.68 | 1.05E-19 | 1.10E-18 |
| chr10 | 210744 | 254848 | 44105 | -9.67 | 1.05E-19 | 1.10E-18 |
| chrX | 143295716 | 143504041 | 208326 | 1.97 | 1.05E-19 | 1.10E-18 |
| chr19 | 49820693 | 49827854 | 7162 | -9.67 | 1.06E-19 | 1.11E-18 |
| chr20 | 13555912 | 13571943 | 16032 | -9.74 | 1.06E-19 | 1.11E-18 |
| chrX | 64209727 | 64669630 | 459904 | 1.8 | 1.06E-19 | 1.11E-18 |
| chr2 | 20606814 | 20611169 | 4356 | -9.76 | 1.07E-19 | 1.11E-18 |
| chr1 | 77874667 | 77912535 | 37869 | -9.73 | 1.07E-19 | 1.12E-18 |
| chr12 | 12798022 | 12807002 | 8981 | -9.75 | 1.07E-19 | 1.12E-18 |
| chr6 | 24334562 | 24347871 | 13310 | -9.74 | 1.08E-19 | 1.13E-18 |
| chr3 | 149703643 | 149717576 | 13934 | -9.69 | 1.08E-19 | 1.13E-18 |
| chr2 | 36902731 | 36911168 | 8438 | -9.71 | 1.08E-19 | 1.13E-18 |
| chr11 | 85540124 | 85570858 | 30735 | -9.89 | 1.10E-19 | 1.14E-18 |
| chr11 | 48134402 | 48138097 | 3696 | -9.85 | 1.10E-19 | 1.15E-18 |
| chr9 | 100626665 | 100644017 | 17353 | -9.67 | 1.11E-19 | 1.16E-18 |
| chr12 | 127241650 | 127272294 | 30645 | -9.68 | 1.11E-19 | 1.16E-18 |
| chr3 | 113359594 | 113368210 | 8617 | -9.7 | 1.12E-19 | 1.16E-18 |
| chr14 | 103666284 | 103685447 | 19164 | -9.83 | 1.12E-19 | 1.16E-18 |
| chr7 | 38158598 | 38197157 | 38560 | -10.16 | 1.13E-19 | 1.17E-18 |
| chr2 | 110889521 | 110914684 | 25164 | -9.71 | 1.13E-19 | 1.17E-18 |
| chr9 | 88617962 | 88621592 | 3631 | -9.87 | 1.13E-19 | 1.18E-18 |
| chr5 | 55905903 | 55926730 | 20828 | -9.82 | 1.13E-19 | 1.18E-18 |
| chrX | 153819413 | 153836235 | 16823 | -9.74 | 1.14E-19 | 1.18E-18 |
| chr3 | 99517154 | 99536305 | 19152 | -9.75 | 1.14E-19 | 1.18E-18 |
| chr9 | 7182141 | 7234850 | 52710 | -9.7 | 1.14E-19 | 1.18E-18 |
| chr9 | 110801975 | 110855598 | 53624 | -9.93 | 1.14E-19 | 1.19E-18 |

| | | | | | | |
|---|---|---|---|---|---|---|
| chr21 | 19364558 | 19406174 | 41617 | 1.86 | 1.15E-19 | 1.19E-18 |
| chr2 | 98589203 | 98606560 | 17358 | -9.73 | 1.16E-19 | 1.20E-18 |
| chrX | 69894904 | 69948266 | 53363 | -9.7 | 1.16E-19 | 1.21E-18 |
| chr2 | 225620894 | 225632894 | 12001 | -9.66 | 1.16E-19 | 1.21E-18 |
| chrX | 4576571 | 4590183 | 13613 | -9.68 | 1.17E-19 | 1.21E-18 |
| chr1 | 43705710 | 43730249 | 24540 | -9.8 | 1.17E-19 | 1.22E-18 |
| chr3 | 62130957 | 62164677 | 33721 | 9.76 | 1.18E-19 | 1.22E-18 |
| chr2 | 181242198 | 181261841 | 19644 | -9.66 | 1.18E-19 | 1.23E-18 |
| chr11 | 9056316 | 9064950 | 8635 | -9.67 | 1.19E-19 | 1.23E-18 |
| chr10 | 60183229 | 60223533 | 40305 | -10.16 | 1.19E-19 | 1.24E-18 |
| chr4 | 143375492 | 143417630 | 42139 | -9.67 | 1.20E-19 | 1.24E-18 |
| chr6 | 106730761 | 106740983 | 10223 | -9.78 | 1.20E-19 | 1.25E-18 |
| chr7 | 101120173 | 101134761 | 14589 | -9.76 | 1.21E-19 | 1.25E-18 |
| chr14 | 99386984 | 99389654 | 2671 | -6.8 | 1.21E-19 | 1.25E-18 |
| chr2 | 201404313 | 201423048 | 18736 | -9.67 | 1.21E-19 | 1.25E-18 |
| chr2 | 200998897 | 201032172 | 33276 | -9.83 | 1.21E-19 | 1.26E-18 |
| chr4 | 142268542 | 142299034 | 30493 | -9.68 | 1.22E-19 | 1.26E-18 |
| chr2 | 86052632 | 86083599 | 30968 | -9.66 | 1.22E-19 | 1.27E-18 |
| chr14 | 103817479 | 103832515 | 15037 | -9.89 | 1.23E-19 | 1.27E-18 |
| chr2 | 172311759 | 172316978 | 5220 | -9.7 | 1.23E-19 | 1.27E-18 |
| chr8 | 102571795 | 102585656 | 13862 | -9.72 | 1.23E-19 | 1.27E-18 |
| chr2 | 100958566 | 100989074 | 30509 | -9.74 | 1.24E-19 | 1.28E-18 |
| chr17 | 59640338 | 59651004 | 10667 | -9.77 | 1.24E-19 | 1.28E-18 |
| chr8 | 10594399 | 10611259 | 16861 | -3.31 | 1.25E-19 | 1.29E-18 |
| chr10 | 29483685 | 29516381 | 32697 | -9.69 | 1.25E-19 | 1.29E-18 |
| chr4 | 8301187 | 8340512 | 39326 | -9.78 | 1.25E-19 | 1.29E-18 |
| chr3 | 37901271 | 37919110 | 17840 | -9.68 | 1.25E-19 | 1.29E-18 |
| chr15 | 41351972 | 41358160 | 6189 | -9.67 | 1.25E-19 | 1.29E-18 |
| chr2 | 102174541 | 102206404 | 31864 | -9.94 | 1.27E-19 | 1.31E-18 |
| chr3 | 190053818 | 190074325 | 20508 | -9.65 | 1.27E-19 | 1.31E-18 |
| chr9 | 90649786 | 90661621 | 11836 | -9.7 | 1.28E-19 | 1.31E-18 |
| chr16 | 68495250 | 68537640 | 42391 | -10.04 | 1.28E-19 | 1.32E-18 |
| chr7 | 111260961 | 111303759 | 42799 | -9.73 | 1.29E-19 | 1.32E-18 |
| chr2 | 173608432 | 173627930 | 19499 | -9.67 | 1.29E-19 | 1.33E-18 |
| chr11 | 44207773 | 44218661 | 10889 | -9.65 | 1.30E-19 | 1.34E-18 |
| chr7 | 156341381 | 156386089 | 44709 | -9.78 | 1.30E-19 | 1.34E-18 |
| chr10 | 35978474 | 37017721 | 1039248 | 1.47 | 1.30E-19 | 1.34E-18 |
| chr12 | 96948683 | 96968890 | 20208 | -9.65 | 1.31E-19 | 1.34E-18 |
| chr3 | 126377124 | 126394782 | 17659 | -9.66 | 1.31E-19 | 1.34E-18 |
| chr10 | 8824970 | 8833500 | 8531 | -9.65 | 1.31E-19 | 1.35E-18 |
| chr4 | 58774719 | 58935222 | 160504 | 1.62 | 1.32E-19 | 1.36E-18 |
| chr9 | 77491454 | 77501112 | 9659 | -9.66 | 1.32E-19 | 1.36E-18 |
| chr9 | 3639139 | 3641401 | 2263 | -4.06 | 1.33E-19 | 1.37E-18 |
| chr12 | 104611054 | 104631935 | 20882 | -9.85 | 1.33E-19 | 1.37E-18 |
| chr14 | 78400442 | 78416617 | 16176 | -9.7 | 1.33E-19 | 1.37E-18 |
| chr7 | 27799086 | 27802596 | 3511 | -9.69 | 1.33E-19 | 1.37E-18 |
| chr7 | 26520659 | 26526862 | 6204 | -9.66 | 1.34E-19 | 1.37E-18 |
| chr8 | 75078750 | 75094391 | 15642 | -2.06 | 1.34E-19 | 1.38E-18 |
| chr8 | 29754756 | 29783377 | 28622 | -9.89 | 1.35E-19 | 1.38E-18 |
| chr9 | 136631239 | 136641385 | 10147 | -9.66 | 1.35E-19 | 1.38E-18 |
| chr1 | 41798918 | 41806897 | 7980 | -9.65 | 1.36E-19 | 1.39E-18 |
| chr11 | 67060523 | 67064972 | 4450 | -9.75 | 1.36E-19 | 1.39E-18 |
| chr8 | 117731150 | 117736473 | 5324 | -9.71 | 1.36E-19 | 1.39E-18 |
| chr7 | 131122545 | 131126933 | 4389 | -9.81 | 1.36E-19 | 1.39E-18 |
| chr15 | 69771338 | 69785196 | 13859 | -9.82 | 1.36E-19 | 1.39E-18 |
| chr6 | 12268848 | 12279273 | 10426 | -9.65 | 1.37E-19 | 1.40E-18 |
| chr7 | 106072672 | 106096167 | 23496 | -9.79 | 1.38E-19 | 1.41E-18 |
| chr9 | 93190013 | 93223874 | 33862 | -9.72 | 1.38E-19 | 1.41E-18 |
| chr8 | 87449730 | 87457058 | 7329 | -9.78 | 1.38E-19 | 1.41E-18 |
| chr12 | 59069918 | 59095698 | 25781 | -9.91 | 1.38E-19 | 1.42E-18 |
| chr5 | 155718944 | 155746078 | 27135 | -9.96 | 1.39E-19 | 1.42E-18 |

| | | | | | | |
|---|---|---|---|---|---|---|
| chr13 | 41718335 | 41727954 | 9620 | -9.66 | 1.39E-19 | 1.42E-18 |
| chr8 | 23045157 | 23069048 | 23892 | -9.7 | 1.40E-19 | 1.43E-18 |
| chr3 | 117569238 | 117597334 | 28097 | -9.71 | 1.40E-19 | 1.43E-18 |
| chr3 | 85146083 | 85220028 | 73946 | -9.84 | 1.40E-19 | 1.43E-18 |
| chr17 | 62145295 | 62149587 | 4293 | -9.82 | 1.41E-19 | 1.44E-18 |
| chr9 | 96332084 | 96349186 | 17103 | -9.73 | 1.41E-19 | 1.44E-18 |
| chr19 | 5381056 | 5409013 | 27958 | -9.69 | 1.41E-19 | 1.44E-18 |
| chr8 | 59741086 | 59770121 | 29036 | -10.05 | 1.41E-19 | 1.44E-18 |
| chr12 | 88978406 | 88982389 | 3984 | -6.59 | 1.42E-19 | 1.45E-18 |
| chr12 | 58387765 | 58397611 | 9847 | -9.64 | 1.42E-19 | 1.45E-18 |
| chr5 | 100572451 | 100600473 | 28023 | -10.05 | 1.43E-19 | 1.45E-18 |
| chr12 | 114923392 | 115090118 | 166727 | 1.17 | 1.43E-19 | 1.46E-18 |
| chr9 | 90968519 | 90993295 | 24777 | -9.79 | 1.44E-19 | 1.46E-18 |
| chr7 | 43297263 | 43312733 | 15471 | -9.65 | 1.44E-19 | 1.46E-18 |
| chr1 | 91334124 | 91349370 | 15247 | -9.77 | 1.45E-19 | 1.48E-18 |
| chr3 | 182844867 | 182859608 | 14742 | -9.66 | 1.45E-19 | 1.48E-18 |
| chr1 | 66720060 | 66736595 | 16536 | -9.64 | 1.45E-19 | 1.48E-18 |
| chr4 | 11602678 | 11646813 | 44136 | -9.85 | 1.46E-19 | 1.48E-18 |
| chr5 | 179609762 | 179626154 | 16393 | -9.71 | 1.46E-19 | 1.49E-18 |
| chr14 | 55777036 | 55783538 | 6503 | -9.66 | 1.47E-19 | 1.49E-18 |
| chr10 | 37187851 | 37240506 | 52656 | -9.79 | 1.47E-19 | 1.50E-18 |
| chr10 | 60384339 | 60395202 | 10864 | -9.79 | 1.48E-19 | 1.50E-18 |
| chr14 | 63153251 | 63502605 | 349355 | 1.27 | 1.49E-19 | 1.51E-18 |
| chr2 | 221387965 | 221409033 | 21069 | -9.65 | 1.49E-19 | 1.51E-18 |
| chr5 | 32682345 | 32701664 | 19320 | -9.69 | 1.49E-19 | 1.52E-18 |
| chr7 | 134949574 | 134952700 | 3127 | -6.71 | 1.49E-19 | 1.52E-18 |
| chr1 | 197070750 | 197078874 | 8125 | -9.7 | 1.50E-19 | 1.52E-18 |
| chr19 | 1769674 | 1791308 | 21635 | -9.65 | 1.51E-19 | 1.53E-18 |
| chr20 | 37730365 | 37754774 | 24410 | -9.65 | 1.51E-19 | 1.54E-18 |
| chr4 | 141491120 | 141494003 | 2884 | -6.77 | 1.52E-19 | 1.54E-18 |
| chr21 | 38234115 | 38253247 | 19133 | -9.73 | 1.52E-19 | 1.54E-18 |
| chr8 | 40459527 | 40483955 | 24429 | -9.81 | 1.52E-19 | 1.54E-18 |
| chr2 | 190800637 | 190809316 | 8680 | -9.74 | 1.53E-19 | 1.55E-18 |
| chr17 | 70602889 | 70626290 | 23402 | -9.75 | 1.53E-19 | 1.55E-18 |
| chr5 | 168660802 | 168668893 | 8092 | -9.82 | 1.54E-19 | 1.55E-18 |
| chr10 | 33488038 | 33522406 | 34369 | -9.79 | 1.54E-19 | 1.55E-18 |
| chr3 | 113639528 | 113648245 | 8718 | -2.15 | 1.54E-19 | 1.56E-18 |
| chr3 | 82222670 | 82377566 | 154897 | 2.1 | 1.54E-19 | 1.56E-18 |
| chr3 | 111617224 | 111654849 | 37626 | -10.14 | 1.56E-19 | 1.58E-18 |
| chr2 | 135825412 | 135847394 | 21983 | -9.71 | 1.56E-19 | 1.58E-18 |
| chr6 | 62384054 | 62770069 | 386016 | 1.82 | 1.56E-19 | 1.58E-18 |
| chr13 | 111433586 | 111444766 | 11181 | -2.48 | 1.57E-19 | 1.58E-18 |
| chr8 | 130657862 | 130684524 | 26663 | -9.73 | 1.58E-19 | 1.60E-18 |
| chr10 | 16751863 | 16782017 | 30155 | -9.63 | 1.59E-19 | 1.60E-18 |
| chr2 | 159551988 | 159572165 | 20178 | -9.63 | 1.59E-19 | 1.61E-18 |
| chr6 | 107101887 | 107108810 | 6924 | -9.65 | 1.59E-19 | 1.61E-18 |
| chr2 | 107635669 | 107671604 | 35936 | -2.51 | 1.60E-19 | 1.61E-18 |
| chr11 | 32627034 | 32657741 | 30708 | -9.64 | 1.61E-19 | 1.62E-18 |
| chr19 | 16877592 | 16907501 | 29910 | -9.67 | 1.61E-19 | 1.62E-18 |
| chr10 | 80056920 | 80099234 | 42315 | -9.66 | 1.61E-19 | 1.63E-18 |
| chr5 | 38735235 | 38747497 | 12263 | -9.63 | 1.63E-19 | 1.64E-18 |
| chr8 | 8113028 | 8175922 | 62895 | -9.83 | 1.63E-19 | 1.64E-18 |
| chr2 | 86289937 | 86298569 | 8633 | -9.92 | 1.63E-19 | 1.64E-18 |
| chr11 | 111343655 | 111360649 | 16995 | -9.64 | 1.64E-19 | 1.65E-18 |
| chr17 | 75614932 | 75624974 | 10043 | -9.72 | 1.65E-19 | 1.66E-18 |
| chr7 | 82978185 | 82983277 | 5093 | -2.88 | 1.65E-19 | 1.66E-18 |
| chr2 | 99471198 | 99480058 | 8861 | -9.69 | 1.66E-19 | 1.67E-18 |
| chr8 | 48215496 | 48223953 | 8458 | -9.7 | 1.66E-19 | 1.68E-18 |
| chr6 | 51837216 | 51893151 | 55936 | 9.68 | 1.67E-19 | 1.68E-18 |
| chr2 | 112792100 | 112807953 | 15854 | -9.63 | 1.67E-19 | 1.68E-18 |
| chr2 | 224977064 | 225003712 | 26649 | -9.64 | 1.68E-19 | 1.69E-18 |

| | | | | | | |
|---|---|---|---|---|---|---|
| chr2 | 189133553 | 189186327 | 52775 | -10.09 | 1.69E-19 | 1.70E-18 |
| chr2 | 99036733 | 99051871 | 15139 | -9.65 | 1.69E-19 | 1.70E-18 |
| chr10 | 133361348 | 133381250 | 19903 | -9.65 | 1.70E-19 | 1.70E-18 |
| chr3 | 30106930 | 30178586 | 71657 | -10.45 | 1.70E-19 | 1.71E-18 |
| chr1 | 5819726 | 5838789 | 19064 | -9.63 | 1.70E-19 | 1.71E-18 |
| chr5 | 84007259 | 84934375 | 927117 | 1.58 | 1.70E-19 | 1.71E-18 |
| chr16 | 90119633 | 90146586 | 26954 | -9.68 | 1.71E-19 | 1.72E-18 |
| chr5 | 10738788 | 10755928 | 17141 | -9.66 | 1.71E-19 | 1.72E-18 |
| chr21 | 32789117 | 32817821 | 28705 | -9.67 | 1.72E-19 | 1.72E-18 |
| chr15 | 39251227 | 39277255 | 26029 | -9.91 | 1.72E-19 | 1.73E-18 |
| chr6 | 169773007 | 169786273 | 13267 | -9.73 | 1.73E-19 | 1.74E-18 |
| chr10 | 63931015 | 63945950 | 14936 | -9.65 | 1.75E-19 | 1.75E-18 |
| chr10 | 32709211 | 32734051 | 24841 | -9.67 | 1.75E-19 | 1.76E-18 |
| chr18 | 3643623 | 3688348 | 44726 | -9.94 | 1.76E-19 | 1.76E-18 |
| chr12 | 47429149 | 47450656 | 21508 | -9.72 | 1.79E-19 | 1.79E-18 |
| chr2 | 129180588 | 129190218 | 9631 | -9.62 | 1.79E-19 | 1.79E-18 |
| chr17 | 20150635 | 20162225 | 11591 | -9.7 | 1.80E-19 | 1.81E-18 |
| chr15 | 101542811 | 101547024 | 4214 | -9.62 | 1.81E-19 | 1.81E-18 |
| chr1 | 221115139 | 221144298 | 29160 | -9.66 | 1.82E-19 | 1.82E-18 |
| chr6 | 147616059 | 147626525 | 10467 | -9.62 | 1.82E-19 | 1.82E-18 |
| chr2 | 1552968 | 1741023 | 188056 | -1.83 | 1.82E-19 | 1.82E-18 |
| chr17 | 9117049 | 9121488 | 4440 | -9.62 | 1.83E-19 | 1.83E-18 |
| chr7 | 55387003 | 55427653 | 40651 | -9.89 | 1.83E-19 | 1.83E-18 |
| chr1 | 219263417 | 219288275 | 24859 | -9.7 | 1.83E-19 | 1.83E-18 |
| chr15 | 52377228 | 52384094 | 6867 | -9.62 | 1.84E-19 | 1.84E-18 |
| chr2 | 164117692 | 164141849 | 24158 | -9.62 | 1.85E-19 | 1.84E-18 |
| chr1 | 177981904 | 177988211 | 6308 | -5.04 | 1.85E-19 | 1.85E-18 |
| chr12 | 118397803 | 118415586 | 17784 | -9.62 | 1.85E-19 | 1.85E-18 |
| chr9 | 95847285 | 95867904 | 20620 | -9.66 | 1.86E-19 | 1.85E-18 |
| chr22 | 49940109 | 49957698 | 17590 | -2.12 | 1.87E-19 | 1.86E-18 |
| chr6 | 22233150 | 22258539 | 25390 | -9.72 | 1.87E-19 | 1.86E-18 |
| chr10 | 33633035 | 33667417 | 34383 | -9.74 | 1.88E-19 | 1.87E-18 |
| chr3 | 56574492 | 56587693 | 13202 | -6.91 | 1.88E-19 | 1.88E-18 |
| chr4 | 1588554 | 1594230 | 5677 | -9.62 | 1.91E-19 | 1.90E-18 |
| chr4 | 47992752 | 48012637 | 19886 | -9.61 | 1.91E-19 | 1.90E-18 |
| chr3 | 114483690 | 114496410 | 12721 | -9.85 | 1.93E-19 | 1.93E-18 |
| chr7 | 34728223 | 34738186 | 9964 | -9.62 | 1.93E-19 | 1.93E-18 |
| chr6 | 137498344 | 137512481 | 14138 | -9.61 | 1.94E-19 | 1.93E-18 |
| chr2 | 133977873 | 134014324 | 36452 | 9.85 | 1.94E-19 | 1.94E-18 |
| chr2 | 171952968 | 171958012 | 5045 | -9.61 | 1.95E-19 | 1.94E-18 |
| chr9 | 4992030 | 5042302 | 50273 | -9.91 | 1.96E-19 | 1.95E-18 |
| chr17 | 61048877 | 61063680 | 14804 | -9.71 | 1.96E-19 | 1.95E-18 |
| chr8 | 52876577 | 52890156 | 13580 | -9.62 | 1.97E-19 | 1.96E-18 |
| chr1 | 185473235 | 185493782 | 20548 | -9.76 | 1.97E-19 | 1.96E-18 |
| chr3 | 65177501 | 65209612 | 32112 | 9.66 | 1.97E-19 | 1.96E-18 |
| chr19 | 30569269 | 30573045 | 3777 | -9.63 | 1.97E-19 | 1.96E-18 |
| chr2 | 198346954 | 198366843 | 19890 | -9.61 | 1.98E-19 | 1.97E-18 |
| chr8 | 39144579 | 39170202 | 25624 | -9.61 | 1.98E-19 | 1.97E-18 |
| chr10 | 60919280 | 60931014 | 11735 | -9.61 | 1.98E-19 | 1.97E-18 |
| chr14 | 55037835 | 55041325 | 3491 | -9.61 | 2.00E-19 | 1.99E-18 |
| chr5 | 173771714 | 173844715 | 73002 | -2.16 | 2.01E-19 | 1.99E-18 |
| chr17 | 8966756 | 9006821 | 40066 | -10.02 | 2.01E-19 | 2.00E-18 |
| chr2 | 9738345 | 9742386 | 4042 | -9.86 | 2.02E-19 | 2.00E-18 |
| chr15 | 57442359 | 57446303 | 3945 | -9.61 | 2.02E-19 | 2.01E-18 |
| chr1 | 147779692 | 147804871 | 25180 | -9.68 | 2.03E-19 | 2.01E-18 |
| chr3 | 34161219 | 34184312 | 23094 | -9.66 | 2.03E-19 | 2.01E-18 |
| chr11 | 19265567 | 19276661 | 11095 | -9.85 | 2.05E-19 | 2.03E-18 |
| chr5 | 34758401 | 34787936 | 29536 | -9.61 | 2.06E-19 | 2.04E-18 |
| chr20 | 32790569 | 32802307 | 11739 | -9.64 | 2.06E-19 | 2.04E-18 |
| chr10 | 33207014 | 33217600 | 10587 | -9.63 | 2.06E-19 | 2.04E-18 |
| chr9 | 107982849 | 107989323 | 6475 | -9.62 | 2.07E-19 | 2.04E-18 |

| | | | | | | |
|---|---|---|---|---|---|---|
| chr21 | 43151746 | 43160860 | 9115 | -9.62 | 2.10E-19 | 2.08E-18 |
| chr5 | 68099921 | 68110510 | 10590 | -9.6 | 2.10E-19 | 2.08E-18 |
| chr2 | 67649311 | 67661826 | 12516 | -9.61 | 2.12E-19 | 2.09E-18 |
| chr9 | 16007648 | 16038442 | 30795 | -9.61 | 2.13E-19 | 2.11E-18 |
| chr4 | 38441431 | 38478314 | 36884 | -9.83 | 2.13E-19 | 2.11E-18 |
| chr3 | 99937390 | 99958466 | 21077 | -9.6 | 2.13E-19 | 2.11E-18 |
| chr2 | 196158753 | 196195166 | 36414 | -9.81 | 2.14E-19 | 2.12E-18 |
| chr14 | 81087042 | 81118555 | 31514 | -9.78 | 2.14E-19 | 2.12E-18 |
| chr9 | 135399843 | 135412359 | 12517 | -9.7 | 2.16E-19 | 2.13E-18 |
| chr9 | 128110782 | 128115304 | 4523 | -9.6 | 2.16E-19 | 2.14E-18 |
| chr3 | 135387934 | 135611090 | 223157 | 3.61 | 2.16E-19 | 2.14E-18 |
| chr16 | 58798615 | 58803211 | 4597 | -4.43 | 2.17E-19 | 2.14E-18 |
| chr7 | 128754086 | 128762688 | 8603 | -9.62 | 2.17E-19 | 2.14E-18 |
| chr21 | 18427450 | 18453562 | 26113 | 9.67 | 2.17E-19 | 2.14E-18 |
| chr7 | 74125646 | 74128963 | 3318 | -9.75 | 2.18E-19 | 2.15E-18 |
| chr9 | 116650874 | 116658894 | 8021 | -9.6 | 2.18E-19 | 2.15E-18 |
| chr13 | 34254989 | 34295615 | 40627 | -9.96 | 2.19E-19 | 2.15E-18 |
| chr1 | 232878721 | 232901685 | 22965 | -9.96 | 2.19E-19 | 2.16E-18 |
| chr11 | 20248614 | 20265123 | 16510 | -9.7 | 2.20E-19 | 2.17E-18 |
| chr13 | 34644237 | 34684445 | 40209 | -9.91 | 2.20E-19 | 2.17E-18 |
| chr22 | 47702584 | 47732137 | 29554 | -9.6 | 2.21E-19 | 2.17E-18 |
| chr15 | 96552128 | 96570261 | 18134 | -4.86 | 2.21E-19 | 2.18E-18 |
| chr16 | 19815912 | 19827101 | 11190 | -9.6 | 2.22E-19 | 2.18E-18 |
| chr7 | 3707491 | 3752372 | 44882 | -9.85 | 2.22E-19 | 2.19E-18 |
| chr4 | 66867067 | 66897808 | 30742 | -9.66 | 2.23E-19 | 2.19E-18 |
| chr6 | 10028199 | 10055001 | 26803 | -9.66 | 2.23E-19 | 2.19E-18 |
| chr1 | 179013069 | 179029903 | 16835 | -9.6 | 2.23E-19 | 2.19E-18 |
| chr7 | 83794823 | 83816038 | 21216 | -9.86 | 2.24E-19 | 2.20E-18 |
| chr22 | 33743109 | 33793449 | 50341 | -10 | 2.24E-19 | 2.20E-18 |
| chr13 | 49536769 | 49549597 | 12829 | -9.71 | 2.25E-19 | 2.21E-18 |
| chr2 | 241357055 | 241366435 | 9381 | -9.69 | 2.25E-19 | 2.21E-18 |
| chr11 | 77543576 | 77568411 | 24836 | -9.7 | 2.26E-19 | 2.22E-18 |
| chr5 | 57217687 | 57272155 | 54469 | -9.97 | 2.29E-19 | 2.25E-18 |
| chr18 | 7988471 | 8004573 | 16103 | -9.6 | 2.30E-19 | 2.26E-18 |
| chr17 | 39354174 | 39396909 | 42736 | -9.88 | 2.31E-19 | 2.27E-18 |
| chr8 | 49321488 | 49329514 | 8027 | -9.72 | 2.31E-19 | 2.27E-18 |
| chr7 | 137549247 | 137592487 | 43241 | -9.69 | 2.32E-19 | 2.28E-18 |
| chr5 | 158804742 | 158818625 | 13884 | -9.64 | 2.34E-19 | 2.29E-18 |
| chr17 | 8385806 | 8389806 | 4001 | -9.7 | 2.35E-19 | 2.31E-18 |
| chr6 | 159989203 | 160008945 | 19743 | -2.64 | 2.36E-19 | 2.31E-18 |
| chr1 | 52999400 | 53026169 | 26770 | -9.66 | 2.38E-19 | 2.33E-18 |
| chr16 | 85863797 | 85903582 | 39786 | -9.7 | 2.38E-19 | 2.33E-18 |
| chr3 | 141375539 | 141399068 | 23530 | -9.62 | 2.39E-19 | 2.34E-18 |
| chr12 | 69788673 | 69801048 | 12376 | -9.63 | 2.39E-19 | 2.34E-18 |
| chr19 | 33215648 | 33219871 | 4224 | -9.59 | 2.40E-19 | 2.34E-18 |
| chr16 | 19208844 | 19260023 | 51180 | -9.87 | 2.40E-19 | 2.35E-18 |
| chr2 | 95978627 | 96008640 | 30014 | -9.69 | 2.40E-19 | 2.35E-18 |
| chr7 | 92365583 | 92393754 | 28172 | -9.59 | 2.41E-19 | 2.35E-18 |
| chr11 | 28211100 | 28219349 | 8250 | -9.59 | 2.41E-19 | 2.35E-18 |
| chr3 | 148741722 | 148744488 | 2767 | -9.65 | 2.42E-19 | 2.36E-18 |
| chr12 | 80142384 | 80153352 | 10969 | -9.77 | 2.42E-19 | 2.37E-18 |
| chr5 | 142977000 | 142996404 | 19405 | -9.79 | 2.44E-19 | 2.38E-18 |
| chr7 | 94881354 | 94904862 | 23509 | -9.63 | 2.45E-19 | 2.39E-18 |
| chr5 | 95539896 | 95549922 | 10027 | -9.81 | 2.45E-19 | 2.39E-18 |
| chr10 | 115173503 | 115273864 | 100362 | -1.71 | 2.45E-19 | 2.39E-18 |
| chr2 | 73282436 | 73285847 | 3412 | -9.64 | 2.45E-19 | 2.39E-18 |
| chr2 | 97038305 | 97044963 | 6659 | -9.59 | 2.45E-19 | 2.39E-18 |
| chr7 | 50959229 | 50980519 | 21291 | -9.59 | 2.46E-19 | 2.40E-18 |
| chr15 | 70798902 | 70805899 | 6998 | -9.65 | 2.47E-19 | 2.41E-18 |
| chr3 | 65265652 | 65293870 | 28219 | 9.65 | 2.48E-19 | 2.41E-18 |
| chr3 | 124533230 | 124560342 | 27113 | -9.6 | 2.48E-19 | 2.42E-18 |

| | | | | | | |
|---|---|---|---|---|---|---|
| chr8 | 144515083 | 144539495 | 24413 | -9.59 | 2.49E-19 | 2.43E-18 |
| chr1 | 222767642 | 222787617 | 19976 | -9.61 | 2.51E-19 | 2.44E-18 |
| chr2 | 217572926 | 217575190 | 2265 | -5.58 | 2.51E-19 | 2.45E-18 |
| chr1 | 168223316 | 168229432 | 6117 | -9.62 | 2.53E-19 | 2.46E-18 |
| chr10 | 74299042 | 74320672 | 21631 | -9.63 | 2.53E-19 | 2.47E-18 |
| chr10 | 116053496 | 116058269 | 4774 | -9.76 | 2.54E-19 | 2.47E-18 |
| chr15 | 58990011 | 58994181 | 4171 | -9.6 | 2.54E-19 | 2.47E-18 |
| chr14 | 65176755 | 65182049 | 5295 | -9.59 | 2.54E-19 | 2.47E-18 |
| chr2 | 42138116 | 42153161 | 15046 | -9.76 | 2.56E-19 | 2.49E-18 |
| chr12 | 12553539 | 12591060 | 37522 | -9.59 | 2.57E-19 | 2.50E-18 |
| chr20 | 14305613 | 14324627 | 19015 | -9.81 | 2.57E-19 | 2.50E-18 |
| chr9 | 123789344 | 123833314 | 43971 | -9.58 | 2.58E-19 | 2.50E-18 |
| chr1 | 183711608 | 183746049 | 34442 | -9.88 | 2.58E-19 | 2.51E-18 |
| chr2 | 73625160 | 73632634 | 7475 | -9.6 | 2.61E-19 | 2.53E-18 |
| chr21 | 38715988 | 38726115 | 10128 | -9.58 | 2.62E-19 | 2.54E-18 |
| chr4 | 106648131 | 106661353 | 13223 | -9.64 | 2.64E-19 | 2.56E-18 |
| chr3 | 4985716 | 5024131 | 38416 | -9.83 | 2.64E-19 | 2.56E-18 |
| chr9 | 136165280 | 136175036 | 9757 | -9.62 | 2.70E-19 | 2.62E-18 |
| chr2 | 178780080 | 178803833 | 23754 | 9.62 | 2.70E-19 | 2.62E-18 |
| chr19 | 46031823 | 46042948 | 11126 | -9.61 | 2.70E-19 | 2.62E-18 |
| chr11 | 33010247 | 33021647 | 11401 | -9.58 | 2.72E-19 | 2.63E-18 |
| chr3 | 87187789 | 87199635 | 11847 | -9.58 | 2.72E-19 | 2.64E-18 |
| chr3 | 53439372 | 53451902 | 12531 | -9.71 | 2.74E-19 | 2.65E-18 |
| chr16 | 63675858 | 63725299 | 49442 | -9.65 | 2.75E-19 | 2.66E-18 |
| chr13 | 81078300 | 81118686 | 40387 | -9.6 | 2.80E-19 | 2.71E-18 |
| chrX | 35169396 | 35205638 | 36243 | -9.63 | 2.82E-19 | 2.72E-18 |
| chr6 | 36047897 | 36055339 | 7443 | -9.64 | 2.82E-19 | 2.73E-18 |
| chr7 | 151154092 | 151163212 | 9121 | -9.64 | 2.83E-19 | 2.74E-18 |
| chr21 | 43839372 | 43857789 | 18418 | -9.59 | 2.83E-19 | 2.74E-18 |
| chr19 | 34026521 | 34038053 | 11533 | -9.71 | 2.84E-19 | 2.75E-18 |
| chr7 | 33950425 | 33969224 | 18800 | 9.62 | 2.84E-19 | 2.75E-18 |
| chr8 | 121483151 | 121493570 | 10420 | -9.6 | 2.85E-19 | 2.75E-18 |
| chr3 | 180978880 | 181018275 | 39396 | 9.91 | 2.86E-19 | 2.77E-18 |
| chr2 | 8695106 | 8703569 | 8464 | -9.65 | 2.93E-19 | 2.83E-18 |
| chr4 | 90345662 | 90379078 | 33417 | -9.83 | 2.94E-19 | 2.84E-18 |
| chr7 | 25281319 | 25600111 | 318793 | 3.81 | 2.95E-19 | 2.85E-18 |
| chrX | 22354215 | 22386103 | 31889 | 9.67 | 2.96E-19 | 2.86E-18 |
| chr15 | 57226841 | 57249875 | 23035 | -9.68 | 2.96E-19 | 2.86E-18 |
| chr1 | 5341113 | 5460554 | 119442 | 1.56 | 2.98E-19 | 2.87E-18 |
| chr9 | 18233120 | 18245978 | 12859 | -9.66 | 2.99E-19 | 2.88E-18 |
| chr18 | 13193248 | 13202300 | 9053 | -9.57 | 3.01E-19 | 2.90E-18 |
| chr7 | 157263580 | 157268916 | 5337 | -9.56 | 3.01E-19 | 2.90E-18 |
| chr2 | 232406042 | 232419243 | 13202 | -9.62 | 3.02E-19 | 2.91E-18 |
| chr8 | 124881396 | 124926443 | 45048 | -10 | 3.02E-19 | 2.91E-18 |
| chr12 | 101401474 | 101424593 | 23120 | -9.59 | 3.04E-19 | 2.93E-18 |
| chr22 | 19242634 | 19257777 | 15144 | -9.57 | 3.04E-19 | 2.93E-18 |
| chr5 | 76022627 | 76027209 | 4583 | -9.58 | 3.04E-19 | 2.93E-18 |
| chr5 | 81210264 | 81241532 | 31269 | -9.68 | 3.07E-19 | 2.95E-18 |
| chr12 | 90671633 | 90684884 | 13252 | -9.57 | 3.07E-19 | 2.95E-18 |
| chr8 | 87419897 | 87434329 | 14433 | -9.57 | 3.07E-19 | 2.96E-18 |
| chr20 | 4073907 | 4079939 | 6033 | -9.57 | 3.08E-19 | 2.96E-18 |
| chr3 | 128571247 | 128581893 | 10647 | -9.58 | 3.08E-19 | 2.96E-18 |
| chr11 | 70593840 | 70621304 | 27465 | -9.66 | 3.09E-19 | 2.97E-18 |
| chr8 | 100949957 | 100980608 | 30652 | -9.56 | 3.12E-19 | 3.00E-18 |
| chr1 | 25475532 | 25489482 | 13951 | -9.73 | 3.13E-19 | 3.01E-18 |
| chr10 | 16862354 | 16867167 | 4814 | -9.56 | 3.14E-19 | 3.01E-18 |
| chr6 | 137556950 | 137591800 | 34851 | -9.69 | 3.14E-19 | 3.01E-18 |
| chr8 | 33214563 | 33239280 | 24718 | -9.59 | 3.15E-19 | 3.03E-18 |
| chr8 | 80662940 | 80667271 | 4332 | -2.66 | 3.15E-19 | 3.03E-18 |
| chr18 | 60677519 | 60711220 | 33702 | -9.57 | 3.16E-19 | 3.04E-18 |
| chr14 | 54555902 | 54565823 | 9922 | -9.65 | 3.17E-19 | 3.04E-18 |

| | | | | | | |
|---|---|---|---|---|---|---|
| chr7 | 44336159 | 44356367 | 20209 | -9.6 | 3.20E-19 | 3.07E-18 |
| chr1 | 69396349 | 69423716 | 27368 | 9.62 | 3.21E-19 | 3.07E-18 |
| chr5 | 166343559 | 166400893 | 57335 | -9.95 | 3.21E-19 | 3.07E-18 |
| chr18 | 50004226 | 50031967 | 27742 | -9.59 | 3.22E-19 | 3.08E-18 |
| chr9 | 72905150 | 72925276 | 20127 | -9.67 | 3.22E-19 | 3.08E-18 |
| chr6 | 121444474 | 121447712 | 3239 | -5.12 | 3.23E-19 | 3.09E-18 |
| chr12 | 113349181 | 113368299 | 19119 | -9.56 | 3.23E-19 | 3.09E-18 |
| chr5 | 134532075 | 134539987 | 7913 | -9.56 | 3.24E-19 | 3.11E-18 |
| chr7 | 90156741 | 90185201 | 28461 | 9.69 | 3.25E-19 | 3.11E-18 |
| chr11 | 8203786 | 8209097 | 5312 | -9.55 | 3.33E-19 | 3.18E-18 |
| chr3 | 151627538 | 151648765 | 21228 | -9.82 | 3.33E-19 | 3.18E-18 |
| chr10 | 84408917 | 84456912 | 47996 | -9.56 | 3.33E-19 | 3.18E-18 |
| chr6 | 47475533 | 47495492 | 19960 | -9.58 | 3.33E-19 | 3.18E-18 |
| chr6 | 151347404 | 151359146 | 11743 | -9.76 | 3.34E-19 | 3.19E-18 |
| chrX | 45464104 | 45490355 | 26252 | -9.59 | 3.34E-19 | 3.20E-18 |
| chr12 | 76817193 | 76828894 | 11702 | -9.63 | 3.35E-19 | 3.20E-18 |
| chr2 | 113603402 | 113630882 | 27481 | -9.64 | 3.35E-19 | 3.20E-18 |
| chr12 | 117617984 | 117640325 | 22342 | -9.56 | 3.37E-19 | 3.22E-18 |
| chr2 | 149456513 | 149463196 | 6684 | -9.56 | 3.40E-19 | 3.25E-18 |
| chr7 | 11702858 | 11726499 | 23642 | -9.57 | 3.42E-19 | 3.27E-18 |
| chr13 | 90029616 | 90056947 | 27332 | -9.91 | 3.43E-19 | 3.27E-18 |
| chr11 | 65956776 | 65972447 | 15672 | -9.56 | 3.44E-19 | 3.28E-18 |
| chr3 | 148888275 | 148896938 | 8664 | -9.55 | 3.45E-19 | 3.29E-18 |
| chr2 | 8110795 | 8133028 | 22234 | -9.69 | 3.45E-19 | 3.29E-18 |
| chr1 | 111395500 | 111412094 | 16595 | -2.79 | 3.45E-19 | 3.29E-18 |
| chr9 | 115224069 | 115241402 | 17334 | -9.66 | 3.46E-19 | 3.29E-18 |
| chr3 | 172715837 | 172738684 | 22848 | 9.7 | 3.48E-19 | 3.32E-18 |
| chr9 | 139761417 | 139768679 | 7263 | -9.66 | 3.48E-19 | 3.32E-18 |
| chrX | 81853437 | 81887788 | 34352 | -9.58 | 3.49E-19 | 3.32E-18 |
| chr3 | 145834051 | 145849310 | 15260 | -9.73 | 3.49E-19 | 3.33E-18 |
| chr21 | 16293186 | 16322336 | 29151 | -9.55 | 3.50E-19 | 3.33E-18 |
| chr10 | 35725788 | 35735253 | 9466 | -9.56 | 3.52E-19 | 3.35E-18 |
| chr1 | 245367576 | 245372844 | 5269 | -9.55 | 3.53E-19 | 3.35E-18 |
| chr3 | 112410634 | 112450964 | 40331 | -9.61 | 3.53E-19 | 3.35E-18 |
| chr6 | 151382951 | 151400307 | 17357 | -9.55 | 3.54E-19 | 3.37E-18 |
| chr14 | 64056506 | 64067233 | 10728 | -9.55 | 3.55E-19 | 3.37E-18 |
| chr2 | 61579536 | 61590970 | 11435 | -9.58 | 3.56E-19 | 3.38E-18 |
| chr10 | 79709418 | 79726192 | 16775 | -9.59 | 3.57E-19 | 3.39E-18 |
| chr12 | 31817289 | 31847160 | 29872 | -9.57 | 3.58E-19 | 3.40E-18 |
| chr12 | 79603929 | 79648678 | 44750 | -9.95 | 3.59E-19 | 3.41E-18 |
| chrX | 123560967 | 123593414 | 32448 | 9.84 | 3.59E-19 | 3.41E-18 |
| chr6 | 168408669 | 168421573 | 12905 | -9.63 | 3.59E-19 | 3.41E-18 |
| chr2 | 73034368 | 73049156 | 14789 | -9.56 | 3.62E-19 | 3.44E-18 |
| chr2 | 26325265 | 26327947 | 2683 | -9.57 | 3.64E-19 | 3.45E-18 |
| chr7 | 155799393 | 155827451 | 28059 | -9.66 | 3.66E-19 | 3.47E-18 |
| chr1 | 241734920 | 241758154 | 23235 | -9.56 | 3.68E-19 | 3.49E-18 |
| chr1 | 152437458 | 152462363 | 24906 | -9.54 | 3.73E-19 | 3.53E-18 |
| chr4 | 13601343 | 13609489 | 8147 | -9.64 | 3.75E-19 | 3.55E-18 |
| chr2 | 120712760 | 120729029 | 16270 | -9.65 | 3.75E-19 | 3.56E-18 |
| chr12 | 120392643 | 120399158 | 6516 | -9.64 | 3.76E-19 | 3.56E-18 |
| chr12 | 58570418 | 58594084 | 23667 | -9.64 | 3.76E-19 | 3.56E-18 |
| chr20 | 34951953 | 34956191 | 4239 | -9.55 | 3.78E-19 | 3.58E-18 |
| chr1 | 42373897 | 42417764 | 43868 | -9.55 | 3.82E-19 | 3.62E-18 |
| chr7 | 94198668 | 94219498 | 20831 | -9.68 | 3.82E-19 | 3.62E-18 |
| chr5 | 176139322 | 176144246 | 4925 | -9.54 | 3.83E-19 | 3.62E-18 |
| chr6 | 169689683 | 169708318 | 18636 | -9.55 | 3.85E-19 | 3.64E-18 |
| chr11 | 67298941 | 67304729 | 5789 | -9.54 | 3.85E-19 | 3.64E-18 |
| chr14 | 48992353 | 49037706 | 45354 | -9.63 | 3.87E-19 | 3.66E-18 |
| chr4 | 41023994 | 41061942 | 37949 | -9.7 | 3.89E-19 | 3.67E-18 |
| chr7 | 87850894 | 87861542 | 10649 | -9.58 | 3.93E-19 | 3.71E-18 |
| chr3 | 127513211 | 127558414 | 45204 | -9.7 | 3.93E-19 | 3.71E-18 |

| | | | | | | |
|---|---|---|---|---|---|---|
| chr15 | 89779737 | 89798421 | 18685 | -9.68 | 3.94E-19 | 3.72E-18 |
| chr15 | 31399770 | 31416857 | 17088 | -9.6 | 3.94E-19 | 3.72E-18 |
| chr4 | 142653944 | 142664921 | 10978 | -9.59 | 3.95E-19 | 3.73E-18 |
| chr22 | 48504131 | 48539266 | 35136 | -9.7 | 3.95E-19 | 3.73E-18 |
| chr11 | 125412602 | 125426571 | 13970 | -9.54 | 3.96E-19 | 3.73E-18 |
| chr9 | 15891904 | 15918531 | 26628 | -9.55 | 3.96E-19 | 3.74E-18 |
| chr14 | 68468313 | 68499072 | 30760 | -9.83 | 3.99E-19 | 3.76E-18 |
| chr11 | 43881941 | 43902209 | 20269 | -9.54 | 4.00E-19 | 3.77E-18 |
| chr17 | 41013876 | 41025598 | 11723 | -9.54 | 4.01E-19 | 3.78E-18 |
| chr12 | 84933241 | 85230173 | 296933 | 1.92 | 4.05E-19 | 3.81E-18 |
| chr10 | 14551449 | 14559528 | 8080 | -9.55 | 4.05E-19 | 3.82E-18 |
| chr2 | 143304771 | 143319231 | 14461 | -9.57 | 4.06E-19 | 3.83E-18 |
| chr8 | 119558569 | 119585927 | 27359 | -9.58 | 4.08E-19 | 3.84E-18 |
| chr22 | 46056041 | 46072886 | 16846 | -9.53 | 4.10E-19 | 3.86E-18 |
| chr6 | 170415655 | 170439258 | 23604 | -9.57 | 4.15E-19 | 3.90E-18 |
| chr18 | 13142583 | 13171595 | 29013 | -9.54 | 4.16E-19 | 3.91E-18 |
| chr17 | 75000629 | 75008775 | 8147 | -9.66 | 4.17E-19 | 3.92E-18 |
| chr15 | 56178222 | 56183137 | 4916 | -9.53 | 4.17E-19 | 3.92E-18 |
| chr1 | 246675487 | 246695675 | 20189 | -9.65 | 4.19E-19 | 3.94E-18 |
| chr20 | 39815995 | 39829062 | 13068 | -9.53 | 4.19E-19 | 3.94E-18 |
| chr10 | 29688927 | 29697693 | 8767 | -9.53 | 4.20E-19 | 3.94E-18 |
| chr2 | 210105323 | 210146726 | 41404 | 9.58 | 4.20E-19 | 3.94E-18 |
| chr10 | 89321999 | 89341563 | 19565 | -9.78 | 4.21E-19 | 3.96E-18 |
| chr2 | 216480328 | 216509257 | 28930 | -10.18 | 4.22E-19 | 3.97E-18 |
| chr3 | 111861509 | 111885706 | 24198 | -9.54 | 4.23E-19 | 3.97E-18 |
| chr4 | 8055275 | 8068470 | 13196 | -9.58 | 4.23E-19 | 3.97E-18 |
| chr17 | 57819456 | 57822554 | 3099 | -9.65 | 4.23E-19 | 3.97E-18 |
| chr12 | 90164880 | 90185417 | 20538 | -9.6 | 4.25E-19 | 3.99E-18 |
| chr4 | 15893405 | 15919485 | 26081 | -9.59 | 4.26E-19 | 3.99E-18 |
| chr10 | 87281331 | 87314284 | 32954 | -9.53 | 4.26E-19 | 3.99E-18 |
| chr12 | 90809534 | 90862832 | 53299 | -9.63 | 4.27E-19 | 4.00E-18 |
| chr8 | 41532327 | 41537758 | 5432 | -9.56 | 4.31E-19 | 4.04E-18 |
| chr4 | 39856027 | 39860069 | 4043 | -9.55 | 4.31E-19 | 4.04E-18 |
| chr10 | 70131035 | 70135617 | 4583 | -9.53 | 4.33E-19 | 4.06E-18 |
| chr4 | 74031020 | 74042140 | 11121 | -9.56 | 4.39E-19 | 4.11E-18 |
| chr17 | 75870379 | 75875043 | 4665 | -9.53 | 4.39E-19 | 4.11E-18 |
| chr7 | 101478340 | 101494370 | 16031 | -9.61 | 4.40E-19 | 4.12E-18 |
| chr1 | 110468025 | 110521783 | 53759 | -9.75 | 4.40E-19 | 4.12E-18 |
| chr5 | 168889640 | 168918106 | 28467 | -9.53 | 4.40E-19 | 4.12E-18 |
| chr14 | 32647056 | 32655873 | 8818 | -9.55 | 4.42E-19 | 4.13E-18 |
| chr8 | 100893445 | 100907731 | 14287 | -9.52 | 4.44E-19 | 4.15E-18 |
| chr6 | 12115856 | 12140621 | 24766 | -9.53 | 4.48E-19 | 4.19E-18 |
| chr11 | 14960477 | 14983398 | 22922 | 9.58 | 4.48E-19 | 4.19E-18 |
| chr12 | 11293659 | 11310831 | 17173 | -9.55 | 4.49E-19 | 4.20E-18 |
| chr10 | 12007316 | 12010551 | 3236 | -9.67 | 4.52E-19 | 4.22E-18 |
| chrX | 84823197 | 85002146 | 178950 | 2.77 | 4.52E-19 | 4.22E-18 |
| chr3 | 67791150 | 67815763 | 24614 | -9.6 | 4.54E-19 | 4.24E-18 |
| chr10 | 5581792 | 5587147 | 5356 | -9.52 | 4.55E-19 | 4.25E-18 |
| chr9 | 86191495 | 86205501 | 14007 | -9.52 | 4.56E-19 | 4.25E-18 |
| chr7 | 34816023 | 34853901 | 37879 | -9.57 | 4.56E-19 | 4.26E-18 |
| chr3 | 150425140 | 150435794 | 10655 | -9.53 | 4.57E-19 | 4.26E-18 |
| chr7 | 134011858 | 134072472 | 60615 | -9.78 | 4.58E-19 | 4.27E-18 |
| chr8 | 102473317 | 102493200 | 19884 | -9.55 | 4.59E-19 | 4.28E-18 |
| chr12 | 62536558 | 62556753 | 20196 | 9.58 | 4.59E-19 | 4.28E-18 |
| chr17 | 72470662 | 72486353 | 15692 | -9.55 | 4.60E-19 | 4.29E-18 |
| chr12 | 22492821 | 22502252 | 9432 | -2.88 | 4.63E-19 | 4.31E-18 |
| chr13 | 79968171 | 79972503 | 4333 | -9.54 | 4.65E-19 | 4.33E-18 |
| chr18 | 48449075 | 48454458 | 5384 | -9.55 | 4.65E-19 | 4.33E-18 |
| chr1 | 44273352 | 44288519 | 15168 | -9.52 | 4.66E-19 | 4.34E-18 |
| chr4 | 8888450 | 8906367 | 17918 | -9.62 | 4.68E-19 | 4.35E-18 |
| chr8 | 33493188 | 33517974 | 24787 | -9.62 | 4.70E-19 | 4.37E-18 |

| | | | | | | |
|---|---|---|---|---|---|---|
| chr5 | 77514463 | 77521202 | 6740 | -9.6 | 4.74E-19 | 4.41E-18 |
| chr8 | 73950585 | 73974855 | 24271 | -9.65 | 4.74E-19 | 4.41E-18 |
| chr10 | 105850366 | 105869417 | 19052 | -9.76 | 4.75E-19 | 4.42E-18 |
| chr4 | 169848786 | 169863097 | 14312 | -9.6 | 4.75E-19 | 4.42E-18 |
| chr12 | 104413093 | 104453290 | 40198 | -9.85 | 4.80E-19 | 4.46E-18 |
| chr15 | 70520491 | 70526364 | 5874 | -9.57 | 4.83E-19 | 4.48E-18 |
| chr6 | 14166447 | 14173608 | 7162 | -9.53 | 4.83E-19 | 4.49E-18 |
| chr20 | 23941255 | 23947657 | 6403 | -9.52 | 4.84E-19 | 4.50E-18 |
| chr13 | 44718488 | 44765671 | 47184 | -9.52 | 4.84E-19 | 4.50E-18 |
| chr15 | 99851219 | 99868880 | 17662 | -9.65 | 4.85E-19 | 4.50E-18 |
| chr9 | 97510414 | 97514299 | 3886 | -9.53 | 4.85E-19 | 4.50E-18 |
| chr10 | 86142168 | 86181676 | 39509 | -9.53 | 4.91E-19 | 4.55E-18 |
| chr7 | 111195697 | 111219353 | 23657 | -9.52 | 4.94E-19 | 4.58E-18 |
| chr4 | 28284023 | 28312490 | 28468 | -9.61 | 4.95E-19 | 4.59E-18 |
| chr17 | 30490914 | 30496171 | 5258 | -9.56 | 4.96E-19 | 4.60E-18 |
| chr12 | 69018430 | 69022855 | 4426 | -9.54 | 4.97E-19 | 4.60E-18 |
| chr14 | 96724892 | 96745901 | 21010 | -9.61 | 4.97E-19 | 4.60E-18 |
| chr7 | 75337218 | 75340755 | 3538 | -9.53 | 4.97E-19 | 4.60E-18 |
| chrX | 10523892 | 10553108 | 29217 | -9.52 | 4.97E-19 | 4.60E-18 |
| chr8 | 110055183 | 110063771 | 8589 | -9.53 | 4.99E-19 | 4.62E-18 |
| chr1 | 88395150 | 88429206 | 34057 | -9.55 | 4.99E-19 | 4.62E-18 |
| chr1 | 168418780 | 168444770 | 25991 | -9.58 | 5.01E-19 | 4.64E-18 |
| chr2 | 74947549 | 74965308 | 17760 | -9.52 | 5.01E-19 | 4.64E-18 |
| chr9 | 14045165 | 14055649 | 10485 | -9.7 | 5.04E-19 | 4.66E-18 |
| chr8 | 23658159 | 23680770 | 22612 | -9.52 | 5.07E-19 | 4.69E-18 |
| chr17 | 80638462 | 80654313 | 15852 | -9.8 | 5.08E-19 | 4.70E-18 |
| chr2 | 20213259 | 20231189 | 17931 | -9.81 | 5.09E-19 | 4.70E-18 |
| chr10 | 7263565 | 7284570 | 21006 | -9.57 | 5.12E-19 | 4.73E-18 |
| chr8 | 42845208 | 42849538 | 4331 | -9.52 | 5.13E-19 | 4.74E-18 |
| chr2 | 98487514 | 98504903 | 17390 | -9.61 | 5.15E-19 | 4.75E-18 |
| chr1 | 231773071 | 231808128 | 35058 | -9.63 | 5.16E-19 | 4.77E-18 |
| chr12 | 1166735 | 1173932 | 7198 | -9.51 | 5.18E-19 | 4.79E-18 |
| chr4 | 187770543 | 187796224 | 25682 | -9.72 | 5.18E-19 | 4.79E-18 |
| chr5 | 91842809 | 91857235 | 14427 | -9.63 | 5.19E-19 | 4.79E-18 |
| chr17 | 43014349 | 43023262 | 8914 | -9.66 | 5.19E-19 | 4.79E-18 |
| chr14 | 50871600 | 50885312 | 13713 | -9.7 | 5.24E-19 | 4.83E-18 |
| chr18 | 2684220 | 2690249 | 6030 | -9.53 | 5.25E-19 | 4.84E-18 |
| chr5 | 143313736 | 143332675 | 18940 | -9.68 | 5.27E-19 | 4.86E-18 |
| chr3 | 37458001 | 37462304 | 4304 | -9.57 | 5.28E-19 | 4.86E-18 |
| chr9 | 87516489 | 87542792 | 26304 | -9.53 | 5.32E-19 | 4.91E-18 |
| chr10 | 7900489 | 7906709 | 6221 | -9.53 | 5.34E-19 | 4.92E-18 |
| chr9 | 134411982 | 134420481 | 8500 | -9.69 | 5.36E-19 | 4.94E-18 |
| chr6 | 158368336 | 158382212 | 13877 | -9.6 | 5.37E-19 | 4.94E-18 |
| chr2 | 198545006 | 198559934 | 14929 | -9.62 | 5.39E-19 | 4.97E-18 |
| chr12 | 109283578 | 109321355 | 37778 | -9.6 | 5.41E-19 | 4.98E-18 |
| chr6 | 111592042 | 111600383 | 8342 | -9.54 | 5.51E-19 | 5.07E-18 |
| chr9 | 103838956 | 103872702 | 33747 | 9.72 | 5.52E-19 | 5.08E-18 |
| chr4 | 153622957 | 153639430 | 16474 | -9.67 | 5.54E-19 | 5.10E-18 |
| chr4 | 99109024 | 99122953 | 13930 | -9.51 | 5.54E-19 | 5.10E-18 |
| chr11 | 3195953 | 3201702 | 5750 | -4.32 | 5.56E-19 | 5.11E-18 |
| chr7 | 66175965 | 66202009 | 26045 | -9.64 | 5.56E-19 | 5.11E-18 |
| chr6 | 5047543 | 5068400 | 20858 | -9.63 | 5.58E-19 | 5.13E-18 |
| chr4 | 120610069 | 120650648 | 40580 | -9.66 | 5.59E-19 | 5.13E-18 |
| chr1 | 235831817 | 235839818 | 8002 | -9.54 | 5.61E-19 | 5.15E-18 |
| chr3 | 43936085 | 43969115 | 33031 | -9.83 | 5.62E-19 | 5.16E-18 |
| chr6 | 127669377 | 127693801 | 24425 | -9.54 | 5.65E-19 | 5.19E-18 |
| chr10 | 33144354 | 33165848 | 21495 | -9.53 | 5.68E-19 | 5.21E-18 |
| chr1 | 77696996 | 77722082 | 25087 | -9.5 | 5.70E-19 | 5.23E-18 |
| chrX | 17398608 | 17428012 | 29405 | -9.69 | 5.71E-19 | 5.24E-18 |
| chr10 | 33423636 | 33454864 | 31229 | -9.58 | 5.75E-19 | 5.27E-18 |
| chr1 | 85044258 | 85080953 | 36696 | -9.61 | 5.75E-19 | 5.27E-18 |

| | | | | | | |
|---|---|---|---|---|---|---|
| chr9 | 81250304 | 81286137 | 35834 | -9.77 | 5.75E-19 | 5.27E-18 |
| chr10 | 96646446 | 96696120 | 49675 | -9.55 | 5.80E-19 | 5.32E-18 |
| chr10 | 54069792 | 54091422 | 21631 | -9.6 | 5.85E-19 | 5.36E-18 |
| chr5 | 154042222 | 154073672 | 31451 | -9.92 | 5.87E-19 | 5.37E-18 |
| chr4 | 87553217 | 87579659 | 26443 | -9.58 | 5.87E-19 | 5.37E-18 |
| chr11 | 32750775 | 32770676 | 19902 | -2.82 | 5.91E-19 | 5.41E-18 |
| chr3 | 120643908 | 121063673 | 419766 | 4.3 | 5.92E-19 | 5.42E-18 |
| chr10 | 125985357 | 126004285 | 18929 | -9.71 | 5.93E-19 | 5.43E-18 |
| chr7 | 2987356 | 3006622 | 19267 | -9.49 | 5.95E-19 | 5.44E-18 |
| chr13 | 21018606 | 21025623 | 7018 | -9.5 | 5.95E-19 | 5.44E-18 |
| chr3 | 8378210 | 8393009 | 14800 | -9.52 | 5.95E-19 | 5.44E-18 |
| chr10 | 96717529 | 96753076 | 35548 | -9.54 | 5.95E-19 | 5.44E-18 |
| chr10 | 123395244 | 123402881 | 7638 | -9.65 | 5.95E-19 | 5.44E-18 |
| chr16 | 87155488 | 87206460 | 50973 | -9.69 | 5.96E-19 | 5.44E-18 |
| chr18 | 57786304 | 57808424 | 22121 | -9.55 | 5.98E-19 | 5.46E-18 |
| chr3 | 87513043 | 87554282 | 41240 | -9.61 | 6.00E-19 | 5.48E-18 |
| chr12 | 96729321 | 96740347 | 11027 | -9.49 | 6.01E-19 | 5.49E-18 |
| chr12 | 17684668 | 17703594 | 18927 | -9.55 | 6.03E-19 | 5.50E-18 |
| chr7 | 116410151 | 116415421 | 5271 | -9.55 | 6.06E-19 | 5.53E-18 |
| chr10 | 54499938 | 54527151 | 27214 | 10 | 6.06E-19 | 5.53E-18 |
| chr3 | 156513620 | 156529190 | 15571 | -9.53 | 6.15E-19 | 5.61E-18 |
| chr2 | 241983219 | 241986721 | 3503 | -9.51 | 6.26E-19 | 5.70E-18 |
| chr1 | 233183908 | 233211247 | 27340 | -9.52 | 6.29E-19 | 5.74E-18 |
| chr2 | 73738490 | 73765793 | 27304 | -9.65 | 6.31E-19 | 5.75E-18 |
| chr6 | 83375220 | 83398734 | 23515 | -9.55 | 6.32E-19 | 5.76E-18 |
| chr10 | 45433360 | 45442234 | 8875 | -9.54 | 6.33E-19 | 5.77E-18 |
| chr5 | 74736909 | 74745658 | 8750 | -9.53 | 6.36E-19 | 5.80E-18 |
| chr3 | 64160011 | 64183690 | 23680 | -9.64 | 6.37E-19 | 5.80E-18 |
| chr6 | 117174331 | 117195597 | 21267 | -9.52 | 6.37E-19 | 5.80E-18 |
| chr17 | 30755521 | 30770383 | 14863 | -9.59 | 6.40E-19 | 5.83E-18 |
| chr6 | 161540129 | 161550706 | 10578 | -9.49 | 6.41E-19 | 5.83E-18 |
| chr12 | 26459113 | 26474804 | 15692 | -9.5 | 6.41E-19 | 5.83E-18 |
| chr22 | 46189009 | 46196593 | 7585 | -9.5 | 6.45E-19 | 5.87E-18 |
| chr12 | 59360474 | 59392538 | 32065 | -9.48 | 6.48E-19 | 5.89E-18 |
| chr18 | 18586055 | 18596758 | 10704 | -9.56 | 6.49E-19 | 5.89E-18 |
| chr3 | 69956958 | 69966930 | 9973 | -9.62 | 6.49E-19 | 5.89E-18 |
| chr13 | 96309457 | 96324728 | 15272 | -9.55 | 6.51E-19 | 5.91E-18 |
| chrX | 35319938 | 35343438 | 23501 | -9.6 | 6.53E-19 | 5.93E-18 |
| chr6 | 144171452 | 144210831 | 39380 | -9.51 | 6.55E-19 | 5.95E-18 |
| chr10 | 123277777 | 123324965 | 47189 | -9.86 | 6.59E-19 | 5.98E-18 |
| chr4 | 128661562 | 128680535 | 18974 | -9.53 | 6.59E-19 | 5.98E-18 |
| chr9 | 16990271 | 16999800 | 9530 | -9.59 | 6.60E-19 | 5.99E-18 |
| chr2 | 46740426 | 46745414 | 4989 | -9.49 | 6.60E-19 | 5.99E-18 |
| chr1 | 18815499 | 18850002 | 34504 | -3.42 | 6.62E-19 | 6.00E-18 |
| chr4 | 101672808 | 101700639 | 27832 | -9.56 | 6.63E-19 | 6.01E-18 |
| chr8 | 141665139 | 141668333 | 3195 | -9.55 | 6.65E-19 | 6.03E-18 |
| chr4 | 41290287 | 41326370 | 36084 | -9.78 | 6.67E-19 | 6.05E-18 |
| chr16 | 23713514 | 23723090 | 9577 | -9.49 | 6.70E-19 | 6.07E-18 |
| chr15 | 44640325 | 44651959 | 11635 | -9.48 | 6.75E-19 | 6.11E-18 |
| chr17 | 9637490 | 9658665 | 21176 | -9.58 | 6.75E-19 | 6.12E-18 |
| chr13 | 73880054 | 73921684 | 41631 | -9.61 | 6.85E-19 | 6.20E-18 |
| chr1 | 53820646 | 53825154 | 4509 | -9.49 | 6.86E-19 | 6.21E-18 |
| chr11 | 9668727 | 9702454 | 33728 | -9.58 | 6.89E-19 | 6.24E-18 |
| chr11 | 126758108 | 126772558 | 14451 | -9.58 | 6.97E-19 | 6.31E-18 |
| chr11 | 93440267 | 93445054 | 4788 | -9.48 | 6.99E-19 | 6.33E-18 |
| chr3 | 102565789 | 102625962 | 60174 | -9.52 | 7.07E-19 | 6.40E-18 |
| chr10 | 34784485 | 34822703 | 38219 | -9.7 | 7.09E-19 | 6.41E-18 |
| chr20 | 56954708 | 56958437 | 3730 | -8.09 | 7.11E-19 | 6.43E-18 |
| chr1 | 31094955 | 31102297 | 7343 | -9.48 | 7.12E-19 | 6.43E-18 |
| chr8 | 54883450 | 54918836 | 35387 | -9.6 | 7.24E-19 | 6.54E-18 |
| chr10 | 118101763 | 118123204 | 21442 | 9.62 | 7.25E-19 | 6.55E-18 |

| | | | | | | |
|---|---|---|---|---|---|---|
| chr18 | 73063062 | 73095148 | 32087 | -9.59 | 7.27E-19 | 6.56E-18 |
| chr8 | 32709423 | 32745647 | 36225 | -9.65 | 7.28E-19 | 6.57E-18 |
| chr5 | 143377032 | 143396898 | 19867 | -9.54 | 7.29E-19 | 6.58E-18 |
| chr11 | 94342360 | 94363517 | 21158 | -9.56 | 7.31E-19 | 6.60E-18 |
| chr19 | 16155291 | 16173995 | 18705 | -9.54 | 7.35E-19 | 6.63E-18 |
| chr5 | 60807076 | 60813588 | 6513 | -9.49 | 7.36E-19 | 6.64E-18 |
| chr3 | 180453172 | 180483166 | 29995 | 9.52 | 7.40E-19 | 6.68E-18 |
| chr7 | 68312145 | 68350672 | 38528 | -9.61 | 7.41E-19 | 6.68E-18 |
| chr13 | 27263496 | 27289820 | 26325 | -9.5 | 7.41E-19 | 6.68E-18 |
| chr20 | 56326135 | 56351206 | 25072 | -9.51 | 7.41E-19 | 6.68E-18 |
| chr15 | 52323501 | 52327762 | 4262 | -9.48 | 7.44E-19 | 6.70E-18 |
| chr10 | 4480496 | 4489801 | 9306 | -9.49 | 7.45E-19 | 6.71E-18 |
| chr3 | 148344378 | 148387457 | 43080 | -9.72 | 7.46E-19 | 6.72E-18 |
| chr16 | 16174488 | 16177967 | 3480 | -9.49 | 7.46E-19 | 6.72E-18 |
| chr19 | 20466546 | 20503846 | 37301 | -9.67 | 7.48E-19 | 6.73E-18 |
| chr18 | 47352787 | 47358760 | 5974 | -9.56 | 7.49E-19 | 6.74E-18 |
| chr10 | 98350619 | 98360707 | 10089 | -3.82 | 7.56E-19 | 6.80E-18 |
| chr2 | 39668762 | 39687963 | 19202 | -9.51 | 7.57E-19 | 6.80E-18 |
| chr11 | 47318843 | 47325591 | 6749 | -9.5 | 7.57E-19 | 6.80E-18 |
| chr12 | 75818370 | 75832973 | 14604 | -9.48 | 7.58E-19 | 6.81E-18 |
| chr9 | 87014370 | 87050613 | 36244 | -9.56 | 7.58E-19 | 6.81E-18 |
| chr5 | 49594530 | 49634342 | 39813 | -9.47 | 7.64E-19 | 6.87E-18 |
| chr9 | 128767952 | 128773900 | 5949 | -9.51 | 7.66E-19 | 6.88E-18 |
| chr4 | 100851022 | 100865676 | 14655 | -9.58 | 7.68E-19 | 6.90E-18 |
| chr1 | 192962465 | 192975753 | 13289 | -9.49 | 7.70E-19 | 6.91E-18 |
| chr11 | 103160868 | 103168831 | 7964 | -9.47 | 7.71E-19 | 6.92E-18 |
| chr20 | 4106213 | 4123089 | 16877 | -9.47 | 7.73E-19 | 6.94E-18 |
| chr12 | 76588806 | 76604255 | 15450 | -9.47 | 7.74E-19 | 6.94E-18 |
| chr7 | 23868140 | 23887367 | 19228 | -2.27 | 7.75E-19 | 6.95E-18 |
| chr10 | 77201598 | 77207112 | 5515 | -9.46 | 7.77E-19 | 6.97E-18 |
| chr9 | 115033137 | 115054555 | 21419 | -9.46 | 7.77E-19 | 6.97E-18 |
| chr4 | 74820242 | 74841720 | 21479 | -9.69 | 7.80E-19 | 6.99E-18 |
| chr13 | 78515846 | 78544588 | 28743 | 9.57 | 7.80E-19 | 6.99E-18 |
| chr10 | 105909595 | 105916453 | 6859 | -9.47 | 7.82E-19 | 7.01E-18 |
| chr1 | 227269678 | 227279173 | 9496 | -9.52 | 7.83E-19 | 7.01E-18 |
| chr3 | 141310165 | 141323009 | 12845 | -9.59 | 7.86E-19 | 7.04E-18 |
| chr2 | 99394456 | 99401763 | 7308 | -9.47 | 7.87E-19 | 7.04E-18 |
| chr4 | 190038914 | 190074004 | 35091 | -9.58 | 7.90E-19 | 7.07E-18 |
| chr14 | 94259324 | 94354782 | 95459 | -3.17 | 8.02E-19 | 7.18E-18 |
| chr12 | 102875350 | 102902500 | 27151 | -9.46 | 8.04E-19 | 7.19E-18 |
| chr6 | 77868102 | 77913411 | 45310 | -9.72 | 8.04E-19 | 7.19E-18 |
| chrX | 29663901 | 29686376 | 22476 | -9.72 | 8.22E-19 | 7.35E-18 |
| chr1 | 212022358 | 212055930 | 33573 | -9.76 | 8.23E-19 | 7.36E-18 |
| chr10 | 1017899 | 1028832 | 10934 | -9.54 | 8.23E-19 | 7.36E-18 |
| chr2 | 43316302 | 43347986 | 31685 | -9.67 | 8.27E-19 | 7.39E-18 |
| chr3 | 119969894 | 119979565 | 9672 | -9.46 | 8.27E-19 | 7.39E-18 |
| chr3 | 14350354 | 14363818 | 13465 | -3.03 | 8.35E-19 | 7.46E-18 |
| chr11 | 125713254 | 125727449 | 14196 | -9.49 | 8.45E-19 | 7.55E-18 |
| chr2 | 105767831 | 105782386 | 14556 | -2.2 | 8.46E-19 | 7.55E-18 |
| chr3 | 94837434 | 94896432 | 58999 | -9.61 | 8.46E-19 | 7.55E-18 |
| chr19 | 33657048 | 33660769 | 3722 | -9.48 | 8.47E-19 | 7.56E-18 |
| chr2 | 97579516 | 97586618 | 7103 | -9.49 | 8.48E-19 | 7.57E-18 |
| chr17 | 8555273 | 8561696 | 6424 | -9.46 | 8.53E-19 | 7.61E-18 |
| chr7 | 151112762 | 151120851 | 8090 | -9.52 | 8.54E-19 | 7.62E-18 |
| chr1 | 75190282 | 75206025 | 15744 | -9.63 | 8.54E-19 | 7.62E-18 |
| chr7 | 27655250 | 27661764 | 6515 | -9.46 | 8.55E-19 | 7.62E-18 |
| chr1 | 191333059 | 191404266 | 71208 | -9.58 | 8.56E-19 | 7.62E-18 |
| chr15 | 68821981 | 68839764 | 17784 | -9.58 | 8.56E-19 | 7.62E-18 |
| chr2 | 8158066 | 8183560 | 25495 | -9.51 | 8.56E-19 | 7.62E-18 |
| chr3 | 14945153 | 14949761 | 4609 | -9.53 | 8.59E-19 | 7.65E-18 |
| chr10 | 43734410 | 43744431 | 10022 | -9.57 | 8.60E-19 | 7.66E-18 |

| | | | | | | |
|---|---|---|---|---|---|---|
| chr17 | 64902388 | 64947886 | 45499 | -9.68 | 8.63E-19 | 7.68E-18 |
| chrX | 80539523 | 80555815 | 16293 | -9.47 | 8.64E-19 | 7.69E-18 |
| chr14 | 58446818 | 58477603 | 30786 | 9.67 | 8.66E-19 | 7.70E-18 |
| chr4 | 108692520 | 108710676 | 18157 | -9.45 | 8.66E-19 | 7.71E-18 |
| chr12 | 44137204 | 44149199 | 11996 | -9.5 | 8.76E-19 | 7.79E-18 |
| chr1 | 30899976 | 30911534 | 11559 | -2.4 | 8.77E-19 | 7.80E-18 |
| chr17 | 77008850 | 77012260 | 3411 | -9.56 | 8.81E-19 | 7.83E-18 |
| chr12 | 60025429 | 60090511 | 65083 | -9.66 | 8.82E-19 | 7.84E-18 |
| chr2 | 47345213 | 47376612 | 31400 | -9.47 | 8.85E-19 | 7.87E-18 |
| chr14 | 78094423 | 78113892 | 19470 | -9.54 | 8.87E-19 | 7.88E-18 |
| chr6 | 160729117 | 160744473 | 15357 | -9.51 | 8.87E-19 | 7.88E-18 |
| chr6 | 163625557 | 163647711 | 22155 | -9.5 | 8.99E-19 | 7.98E-18 |
| chr7 | 68925700 | 68945702 | 20003 | -9.59 | 9.03E-19 | 8.02E-18 |
| chr12 | 74887923 | 74919155 | 31233 | -9.56 | 9.04E-19 | 8.02E-18 |
| chr5 | 74763158 | 74786311 | 23154 | -9.58 | 9.07E-19 | 8.04E-18 |
| chr3 | 138392656 | 138396976 | 4321 | -9.51 | 9.11E-19 | 8.08E-18 |
| chr13 | 49370818 | 49414760 | 43943 | -9.74 | 9.18E-19 | 8.14E-18 |
| chr10 | 30585833 | 30592334 | 6502 | -9.47 | 9.22E-19 | 8.18E-18 |
| chr1 | 57113784 | 57124477 | 10694 | -9.45 | 9.23E-19 | 8.18E-18 |
| chr6 | 22102798 | 22136275 | 33478 | -9.52 | 9.26E-19 | 8.21E-18 |
| chr2 | 218240868 | 218259851 | 18984 | -9.49 | 9.29E-19 | 8.23E-18 |
| chr16 | 84051707 | 84070460 | 18754 | -9.54 | 9.29E-19 | 8.23E-18 |
| chr1 | 224317053 | 224320073 | 3021 | -9.52 | 9.32E-19 | 8.26E-18 |
| chr17 | 45395359 | 45399666 | 4308 | -9.48 | 9.34E-19 | 8.27E-18 |
| chr7 | 14406582 | 14452121 | 45540 | -9.45 | 9.39E-19 | 8.31E-18 |
| chr11 | 43721148 | 43747204 | 26057 | -9.49 | 9.40E-19 | 8.32E-18 |
| chr22 | 25664010 | 25665754 | 1745 | -7.41 | 9.41E-19 | 8.33E-18 |
| chr3 | 81795337 | 81808952 | 13616 | -9.51 | 9.45E-19 | 8.36E-18 |
| chr11 | 69266911 | 69289514 | 22604 | -9.48 | 9.46E-19 | 8.37E-18 |
| chr1 | 19843043 | 19879488 | 36446 | -9.48 | 9.48E-19 | 8.38E-18 |
| chr11 | 19135948 | 19156892 | 20945 | -9.54 | 9.49E-19 | 8.39E-18 |
| chr3 | 138359926 | 138371666 | 11741 | -9.46 | 9.68E-19 | 8.55E-18 |
| chr9 | 137092177 | 137105304 | 13128 | -9.46 | 9.71E-19 | 8.58E-18 |
| chr6 | 70529891 | 70557625 | 27735 | 9.59 | 9.71E-19 | 8.58E-18 |
| chr10 | 139009 | 144716 | 5708 | -9.45 | 9.75E-19 | 8.61E-18 |
| chr1 | 62015250 | 62034015 | 18766 | -9.46 | 9.76E-19 | 8.61E-18 |
| chr4 | 20556378 | 20589050 | 32673 | 9.61 | 9.79E-19 | 8.64E-18 |
| chr16 | 23144844 | 23152930 | 8087 | -9.55 | 9.80E-19 | 8.65E-18 |
| chr4 | 87129930 | 87144259 | 14330 | -9.45 | 9.83E-19 | 8.68E-18 |
| chr4 | 173800294 | 173821405 | 21112 | -9.55 | 9.86E-19 | 8.70E-18 |
| chr2 | 202226153 | 202234223 | 8071 | -9.44 | 9.90E-19 | 8.73E-18 |
| chr8 | 74910771 | 74951627 | 40857 | -9.52 | 9.91E-19 | 8.74E-18 |
| chr2 | 238076469 | 238109867 | 33399 | -9.49 | 9.96E-19 | 8.78E-18 |
| chr8 | 25259238 | 25273643 | 14406 | -9.56 | 9.99E-19 | 8.80E-18 |
| chr10 | 61734343 | 61749221 | 14879 | -9.53 | 1.00E-18 | 8.81E-18 |
| chr3 | 124573036 | 124601254 | 28219 | -9.5 | 1.00E-18 | 8.84E-18 |
| chr17 | 57728230 | 57730528 | 2299 | -9.5 | 1.00E-18 | 8.84E-18 |
| chr8 | 99794238 | 99805169 | 10932 | -9.54 | 1.01E-18 | 8.86E-18 |
| chr4 | 6534148 | 6567488 | 33341 | -9.45 | 1.01E-18 | 8.86E-18 |
| chr2 | 120781901 | 120835818 | 53918 | -9.94 | 1.01E-18 | 8.89E-18 |
| chr1 | 242310691 | 242332746 | 22056 | -9.78 | 1.02E-18 | 8.95E-18 |
| chr12 | 131468526 | 131486448 | 17923 | -9.59 | 1.02E-18 | 8.95E-18 |
| chr12 | 98824035 | 98830321 | 6287 | -9.45 | 1.02E-18 | 8.96E-18 |
| chr12 | 14087225 | 14108980 | 21756 | -9.55 | 1.02E-18 | 8.96E-18 |
| chr5 | 158994974 | 159030368 | 35395 | -9.46 | 1.02E-18 | 8.99E-18 |
| chr2 | 97180673 | 97187536 | 6864 | -9.48 | 1.03E-18 | 9.02E-18 |
| chr20 | 13352218 | 13368462 | 16245 | -9.44 | 1.03E-18 | 9.03E-18 |
| chr7 | 92317130 | 92336690 | 19561 | -9.46 | 1.03E-18 | 9.05E-18 |
| chr8 | 38543296 | 38556115 | 12820 | -9.44 | 1.03E-18 | 9.07E-18 |
| chr12 | 66593173 | 66615420 | 22248 | -9.45 | 1.04E-18 | 9.09E-18 |
| chr12 | 95292279 | 95309626 | 17348 | -9.47 | 1.04E-18 | 9.11E-18 |

| | | | | | | |
|---|---|---|---|---|---|---|
| chr6 | 71212067 | 71215465 | 3399 | -9.58 | 1.04E-18 | 9.14E-18 |
| chr4 | 84042428 | 84061848 | 19421 | -9.43 | 1.05E-18 | 9.18E-18 |
| chr21 | 40661187 | 40674979 | 13793 | -9.46 | 1.05E-18 | 9.21E-18 |
| chr2 | 101483073 | 101491752 | 8680 | -9.46 | 1.05E-18 | 9.22E-18 |
| chr13 | 80130955 | 80141245 | 10291 | -9.43 | 1.05E-18 | 9.24E-18 |
| chr2 | 38626708 | 38641252 | 14545 | -9.49 | 1.06E-18 | 9.29E-18 |
| chr15 | 31290170 | 31306647 | 16478 | -9.5 | 1.06E-18 | 9.29E-18 |
| chr18 | 33109145 | 33120286 | 11142 | -9.46 | 1.06E-18 | 9.32E-18 |
| chr1 | 232265670 | 232302538 | 36869 | -9.47 | 1.06E-18 | 9.32E-18 |
| chr6 | 148453016 | 148466318 | 13303 | -9.46 | 1.07E-18 | 9.35E-18 |
| chr1 | 193045792 | 193049664 | 3873 | -9.59 | 1.07E-18 | 9.35E-18 |
| chr1 | 69059814 | 69088518 | 28705 | -9.45 | 1.07E-18 | 9.35E-18 |
| chr4 | 69889015 | 69907958 | 18944 | -9.59 | 1.07E-18 | 9.36E-18 |
| chr1 | 114587609 | 114617305 | 29697 | -9.45 | 1.08E-18 | 9.41E-18 |
| chr8 | 32348963 | 32386096 | 37134 | -10.09 | 1.08E-18 | 9.44E-18 |
| chr13 | 114545188 | 114552967 | 7780 | -9.43 | 1.08E-18 | 9.45E-18 |
| chr6 | 2506714 | 2533738 | 27025 | -9.97 | 1.08E-18 | 9.48E-18 |
| chr12 | 107083913 | 107098308 | 14396 | -9.44 | 1.09E-18 | 9.49E-18 |
| chr11 | 35250930 | 35266704 | 15775 | -9.52 | 1.10E-18 | 9.58E-18 |
| chr1 | 221264718 | 221305199 | 40482 | -9.52 | 1.10E-18 | 9.62E-18 |
| chr9 | 136068359 | 136072409 | 4051 | -9.43 | 1.10E-18 | 9.63E-18 |
| chr9 | 95538687 | 95550020 | 11334 | -9.53 | 1.11E-18 | 9.68E-18 |
| chr9 | 111068325 | 111094117 | 25793 | -9.66 | 1.12E-18 | 9.75E-18 |
| chr9 | 555348 | 568680 | 13333 | -9.43 | 1.12E-18 | 9.78E-18 |
| chr11 | 12667333 | 12689681 | 22349 | -9.47 | 1.13E-18 | 9.83E-18 |
| chr22 | 48615634 | 48631620 | 15987 | -9.59 | 1.13E-18 | 9.86E-18 |
| chr18 | 47944843 | 47975622 | 30780 | -9.48 | 1.13E-18 | 9.86E-18 |
| chr1 | 114232048 | 114240062 | 8015 | -9.54 | 1.13E-18 | 9.89E-18 |
| chr8 | 38891827 | 38923387 | 31561 | -9.45 | 1.13E-18 | 9.89E-18 |
| chr11 | 17389952 | 17400846 | 10895 | -9.47 | 1.13E-18 | 9.89E-18 |
| chr16 | 69030921 | 69039642 | 8722 | -9.51 | 1.14E-18 | 9.97E-18 |
| chr8 | 81854675 | 81869239 | 14565 | -9.43 | 1.15E-18 | 1.00E-17 |
| chr11 | 34980279 | 35006822 | 26544 | -9.52 | 1.15E-18 | 1.01E-17 |
| chr7 | 44521777 | 44524728 | 2952 | -9.44 | 1.16E-18 | 1.01E-17 |
| chr20 | 13283758 | 13323068 | 39311 | -9.49 | 1.16E-18 | 1.01E-17 |
| chr15 | 82566121 | 82581553 | 15433 | -9.46 | 1.16E-18 | 1.01E-17 |
| chr17 | 16016837 | 16021079 | 4243 | -9.42 | 1.16E-18 | 1.01E-17 |
| chr7 | 95727227 | 95744335 | 17109 | -9.47 | 1.16E-18 | 1.01E-17 |
| chr10 | 88364620 | 88382559 | 17940 | -9.42 | 1.17E-18 | 1.02E-17 |
| chr7 | 22776451 | 22805840 | 29390 | -9.5 | 1.17E-18 | 1.02E-17 |
| chr9 | 32585529 | 32608627 | 23099 | -9.46 | 1.18E-18 | 1.03E-17 |
| chr2 | 158207336 | 158245376 | 38041 | -9.43 | 1.19E-18 | 1.03E-17 |
| chrX | 47454802 | 47473455 | 18654 | -9.42 | 1.19E-18 | 1.03E-17 |
| chr10 | 19072299 | 19146666 | 74368 | 1.86 | 1.19E-18 | 1.03E-17 |
| chr4 | 184710558 | 184717150 | 6593 | -9.44 | 1.19E-18 | 1.04E-17 |
| chr5 | 139318129 | 139324273 | 6145 | -9.42 | 1.19E-18 | 1.04E-17 |
| chr16 | 80128473 | 80137050 | 8578 | -9.42 | 1.19E-18 | 1.04E-17 |
| chr20 | 25381290 | 25387577 | 6288 | -9.52 | 1.20E-18 | 1.04E-17 |
| chr15 | 28485878 | 28495257 | 9380 | -9.5 | 1.21E-18 | 1.05E-17 |
| chr12 | 116357659 | 116378894 | 21236 | -9.61 | 1.23E-18 | 1.07E-17 |
| chr13 | 81565255 | 81598712 | 33458 | -9.43 | 1.24E-18 | 1.08E-17 |
| chr5 | 158955089 | 158979169 | 24081 | -9.42 | 1.24E-18 | 1.08E-17 |
| chr4 | 189291853 | 189314562 | 22710 | -9.49 | 1.25E-18 | 1.08E-17 |
| chr5 | 21044198 | 21105793 | 61596 | -9.65 | 1.26E-18 | 1.09E-17 |
| chr3 | 160731923 | 160751452 | 19530 | -9.43 | 1.26E-18 | 1.09E-17 |
| chr7 | 137900649 | 137941722 | 41074 | -10.04 | 1.27E-18 | 1.10E-17 |
| chr12 | 45743787 | 45764807 | 21021 | -9.54 | 1.27E-18 | 1.10E-17 |
| chr14 | 23664178 | 23700688 | 36511 | -9.62 | 1.28E-18 | 1.11E-17 |
| chr17 | 10668405 | 10684234 | 15830 | -9.48 | 1.29E-18 | 1.11E-17 |
| chr10 | 96958329 | 97000504 | 42176 | -9.43 | 1.29E-18 | 1.12E-17 |
| chr11 | 77273118 | 77294804 | 21687 | -9.55 | 1.30E-18 | 1.13E-17 |

| | | | | | | |
|---|---|---|---|---|---|---|
| chr16 | 49643010 | 49684175 | 41166 | -9.58 | 1.31E-18 | 1.13E-17 |
| chr2 | 28097676 | 28105911 | 8236 | -9.51 | 1.31E-18 | 1.13E-17 |
| chr10 | 5713443 | 5726344 | 12902 | -9.41 | 1.32E-18 | 1.14E-17 |
| chr3 | 185022476 | 185030215 | 7740 | -9.42 | 1.32E-18 | 1.14E-17 |
| chr17 | 21439647 | 21461659 | 22013 | -9.43 | 1.32E-18 | 1.14E-17 |
| chr2 | 165015695 | 165057499 | 41805 | -9.63 | 1.33E-18 | 1.15E-17 |
| chr6 | 159385578 | 159391855 | 6278 | -9.5 | 1.34E-18 | 1.16E-17 |
| chr2 | 238559347 | 238567423 | 8077 | -9.41 | 1.34E-18 | 1.16E-17 |
| chr2 | 118639211 | 118670819 | 31609 | -2.78 | 1.34E-18 | 1.16E-17 |
| chr2 | 159861125 | 159874065 | 12941 | -9.54 | 1.35E-18 | 1.17E-17 |
| chr7 | 83647997 | 83663940 | 15944 | -9.66 | 1.36E-18 | 1.17E-17 |
| chr5 | 138223823 | 138234715 | 10893 | -9.41 | 1.36E-18 | 1.17E-17 |
| chr21 | 33715580 | 33719058 | 3479 | -9.43 | 1.37E-18 | 1.18E-17 |
| chr1 | 203851359 | 203884671 | 33313 | -9.57 | 1.38E-18 | 1.19E-17 |
| chr5 | 23216719 | 23278139 | 61421 | -9.56 | 1.38E-18 | 1.19E-17 |
| chr1 | 227468466 | 227485563 | 17098 | -9.46 | 1.39E-18 | 1.20E-17 |
| chr4 | 102870775 | 102906891 | 36117 | -9.53 | 1.40E-18 | 1.20E-17 |
| chr7 | 109223505 | 109227954 | 4450 | -6.98 | 1.40E-18 | 1.21E-17 |
| chr4 | 74718872 | 74743617 | 24746 | -9.49 | 1.40E-18 | 1.21E-17 |
| chr2 | 19573099 | 19580900 | 7802 | -3.29 | 1.40E-18 | 1.21E-17 |
| chr8 | 131020307 | 131024469 | 4163 | -9.4 | 1.41E-18 | 1.22E-17 |
| chr1 | 235141871 | 235176424 | 34554 | -9.71 | 1.44E-18 | 1.24E-17 |
| chr22 | 47747151 | 47761344 | 14194 | -9.41 | 1.44E-18 | 1.24E-17 |
| chr16 | 89062102 | 89068331 | 6230 | -9.42 | 1.44E-18 | 1.24E-17 |
| chr15 | 59943584 | 59946032 | 2449 | -9.4 | 1.44E-18 | 1.24E-17 |
| chr2 | 215983736 | 215998828 | 15093 | -9.41 | 1.45E-18 | 1.25E-17 |
| chr15 | 55896877 | 55903067 | 6191 | -9.45 | 1.45E-18 | 1.25E-17 |
| chr10 | 97015202 | 97018958 | 3757 | -9.51 | 1.45E-18 | 1.25E-17 |
| chr14 | 92629498 | 92633339 | 3842 | -9.56 | 1.45E-18 | 1.25E-17 |
| chr7 | 30610166 | 30628160 | 17995 | -9.44 | 1.45E-18 | 1.25E-17 |
| chr19 | 43313005 | 43340278 | 27274 | -9.4 | 1.46E-18 | 1.26E-17 |
| chr7 | 27037033 | 27060039 | 23007 | -9.44 | 1.46E-18 | 1.26E-17 |
| chr18 | 45284154 | 45290453 | 6300 | -9.46 | 1.46E-18 | 1.26E-17 |
| chr5 | 56717596 | 56740701 | 23106 | -9.41 | 1.47E-18 | 1.26E-17 |
| chr16 | 86680276 | 86714541 | 34266 | -9.4 | 1.47E-18 | 1.26E-17 |
| chr14 | 70410411 | 70418186 | 7776 | -9.49 | 1.48E-18 | 1.27E-17 |
| chr11 | 35021704 | 35093816 | 72113 | -10.06 | 1.48E-18 | 1.27E-17 |
| chr1 | 66847864 | 66856610 | 8747 | -9.44 | 1.49E-18 | 1.28E-17 |
| chr1 | 231674324 | 231678372 | 4049 | -9.41 | 1.49E-18 | 1.28E-17 |
| chr6 | 38837178 | 38850388 | 13211 | -3.13 | 1.50E-18 | 1.28E-17 |
| chr2 | 48502212 | 48506766 | 4555 | -9.4 | 1.51E-18 | 1.29E-17 |
| chr3 | 183802276 | 183814888 | 12613 | -9.48 | 1.51E-18 | 1.29E-17 |
| chr2 | 86790845 | 86818705 | 27861 | -9.39 | 1.54E-18 | 1.32E-17 |
| chr6 | 89542871 | 89554133 | 11263 | -9.4 | 1.54E-18 | 1.32E-17 |
| chr1 | 212225397 | 212232600 | 7204 | -9.41 | 1.54E-18 | 1.32E-17 |
| chr13 | 33186330 | 33199177 | 12848 | -9.44 | 1.55E-18 | 1.32E-17 |
| chr7 | 129297909 | 129316191 | 18283 | -9.5 | 1.56E-18 | 1.34E-17 |
| chr9 | 83577283 | 83599340 | 22058 | -9.46 | 1.57E-18 | 1.34E-17 |
| chr14 | 35913020 | 35937850 | 24831 | -9.43 | 1.57E-18 | 1.35E-17 |
| chr20 | 50826957 | 50839831 | 12875 | -9.6 | 1.58E-18 | 1.35E-17 |
| chr4 | 78557922 | 78569237 | 11316 | -9.41 | 1.58E-18 | 1.35E-17 |
| chrX | 35923182 | 35944059 | 20878 | -9.39 | 1.58E-18 | 1.35E-17 |
| chr18 | 48610748 | 48616556 | 5809 | -9.45 | 1.58E-18 | 1.35E-17 |
| chr3 | 31980617 | 31999060 | 18444 | -9.45 | 1.58E-18 | 1.35E-17 |
| chr2 | 102127743 | 102141050 | 13308 | -9.39 | 1.59E-18 | 1.36E-17 |
| chr7 | 111947983 | 111967628 | 19646 | -9.44 | 1.59E-18 | 1.36E-17 |
| chr14 | 97426952 | 97444461 | 17510 | -9.39 | 1.60E-18 | 1.36E-17 |
| chr18 | 53600991 | 53614724 | 13734 | -9.39 | 1.60E-18 | 1.37E-17 |
| chr5 | 93335947 | 93359449 | 23503 | -9.39 | 1.61E-18 | 1.37E-17 |
| chr10 | 60599568 | 60643160 | 43593 | -9.58 | 1.61E-18 | 1.38E-17 |
| chr4 | 15618914 | 15636972 | 18059 | -9.39 | 1.62E-18 | 1.38E-17 |

| | | | | | | |
|---|---|---|---|---|---|---|
| chr5 | 139821175 | 139831915 | 10741 | -9.4 | 1.62E-18 | 1.38E-17 |
| chr6 | 41314973 | 41332030 | 17058 | -9.41 | 1.62E-18 | 1.38E-17 |
| chrX | 108320762 | 108354222 | 33461 | 9.5 | 1.63E-18 | 1.39E-17 |
| chr11 | 68194108 | 68198174 | 4067 | -9.39 | 1.63E-18 | 1.39E-17 |
| chr12 | 113321836 | 113325389 | 3554 | -9.42 | 1.63E-18 | 1.39E-17 |
| chr7 | 37056660 | 37100839 | 44180 | -9.69 | 1.63E-18 | 1.39E-17 |
| chr10 | 86380119 | 86397237 | 17119 | 9.44 | 1.64E-18 | 1.40E-17 |
| chr11 | 15951641 | 15976506 | 24866 | -9.59 | 1.65E-18 | 1.41E-17 |
| chr3 | 29097215 | 29173384 | 76170 | -9.66 | 1.66E-18 | 1.41E-17 |
| chr2 | 198611122 | 198634194 | 23073 | -9.5 | 1.66E-18 | 1.41E-17 |
| chr9 | 113286403 | 113298910 | 12508 | -9.44 | 1.67E-18 | 1.42E-17 |
| chr20 | 32322965 | 32326766 | 3802 | -9.38 | 1.67E-18 | 1.43E-17 |
| chr17 | 47452826 | 47461689 | 8864 | -9.7 | 1.68E-18 | 1.43E-17 |
| chr15 | 61350901 | 61386486 | 35586 | 9.43 | 1.68E-18 | 1.43E-17 |
| chr4 | 11677882 | 11729996 | 52115 | -9.72 | 1.69E-18 | 1.44E-17 |
| chr20 | 16662826 | 16671160 | 8335 | 3.35 | 1.70E-18 | 1.45E-17 |
| chr22 | 44516480 | 44534830 | 18351 | -9.38 | 1.70E-18 | 1.45E-17 |
| chr8 | 129011269 | 129024418 | 13150 | -9.38 | 1.71E-18 | 1.45E-17 |
| chr1 | 215186684 | 215222895 | 36212 | 7.56 | 1.71E-18 | 1.46E-17 |
| chr7 | 70168567 | 70198247 | 29681 | -9.41 | 1.72E-18 | 1.46E-17 |
| chr8 | 121186640 | 121226038 | 39399 | 9.44 | 1.72E-18 | 1.47E-17 |
| chr17 | 30906368 | 30911344 | 4977 | -9.42 | 1.73E-18 | 1.47E-17 |
| chrX | 17521634 | 17536629 | 14996 | -9.39 | 1.73E-18 | 1.47E-17 |
| chr9 | 12779039 | 12783974 | 4936 | -9.38 | 1.74E-18 | 1.48E-17 |
| chr5 | 133496079 | 133499117 | 3039 | -9.44 | 1.74E-18 | 1.48E-17 |
| chr3 | 141159270 | 141165700 | 6431 | -9.41 | 1.76E-18 | 1.50E-17 |
| chr9 | 121388847 | 121412062 | 23216 | -9.43 | 1.77E-18 | 1.50E-17 |
| chr18 | 60131517 | 60150920 | 19404 | -9.48 | 1.77E-18 | 1.50E-17 |
| chr10 | 31049386 | 31066922 | 17537 | -9.4 | 1.77E-18 | 1.50E-17 |
| chr3 | 5389944 | 5402262 | 12319 | -9.39 | 1.78E-18 | 1.51E-17 |
| chr14 | 38080293 | 38104308 | 24016 | -9.41 | 1.78E-18 | 1.51E-17 |
| chr10 | 103619498 | 103627637 | 8140 | -9.38 | 1.80E-18 | 1.53E-17 |
| chr9 | 35009403 | 35031956 | 22554 | -9.41 | 1.80E-18 | 1.53E-17 |
| chr4 | 103779054 | 103799633 | 20580 | -9.46 | 1.81E-18 | 1.53E-17 |
| chr1 | 99747702 | 99783853 | 36152 | 9.69 | 1.81E-18 | 1.54E-17 |
| chr11 | 71155940 | 71158530 | 2591 | -9.41 | 1.82E-18 | 1.54E-17 |
| chr9 | 130654254 | 130658694 | 4441 | -9.38 | 1.82E-18 | 1.54E-17 |
| chr3 | 17473236 | 17481630 | 8395 | -9.47 | 1.82E-18 | 1.54E-17 |
| chr3 | 169377615 | 169395097 | 17483 | -9.42 | 1.82E-18 | 1.54E-17 |
| chr9 | 20133153 | 20156965 | 23813 | 9.49 | 1.83E-18 | 1.55E-17 |
| chr7 | 151210454 | 151247486 | 37033 | -9.54 | 1.84E-18 | 1.55E-17 |
| chr2 | 150456039 | 150466315 | 10277 | -9.38 | 1.84E-18 | 1.55E-17 |
| chr7 | 46334919 | 46354736 | 19818 | -9.44 | 1.84E-18 | 1.55E-17 |
| chr6 | 134230636 | 134237430 | 6795 | -9.54 | 1.85E-18 | 1.57E-17 |
| chr7 | 23743090 | 23748753 | 5664 | -9.37 | 1.85E-18 | 1.57E-17 |
| chr2 | 169805672 | 169813233 | 7562 | -9.4 | 1.85E-18 | 1.57E-17 |
| chr6 | 53414088 | 53428481 | 14394 | -9.5 | 1.85E-18 | 1.57E-17 |
| chr7 | 100943440 | 100955760 | 12321 | -9.42 | 1.86E-18 | 1.57E-17 |
| chr2 | 24669718 | 24674848 | 5131 | -9.37 | 1.87E-18 | 1.58E-17 |
| chr5 | 123757048 | 123773410 | 16363 | -9.46 | 1.88E-18 | 1.59E-17 |
| chr5 | 39748724 | 39759299 | 10576 | -9.37 | 1.88E-18 | 1.59E-17 |
| chr2 | 97144192 | 97161964 | 17773 | -9.41 | 1.88E-18 | 1.59E-17 |
| chr6 | 44651992 | 44657853 | 5862 | -3.77 | 1.89E-18 | 1.59E-17 |
| chr11 | 28019564 | 28032829 | 13266 | -9.47 | 1.89E-18 | 1.59E-17 |
| chr19 | 19576785 | 19581427 | 4643 | -9.43 | 1.90E-18 | 1.60E-17 |
| chr5 | 71879300 | 71901140 | 21841 | -9.38 | 1.92E-18 | 1.62E-17 |
| chr16 | 53602068 | 53608915 | 6848 | -9.37 | 1.93E-18 | 1.63E-17 |
| chr2 | 237979223 | 237991981 | 12759 | -9.55 | 1.93E-18 | 1.63E-17 |
| chr1 | 207324688 | 207338886 | 14199 | -9.38 | 1.95E-18 | 1.65E-17 |
| chrX | 149194335 | 149215443 | 21109 | -9.43 | 1.96E-18 | 1.65E-17 |
| chr13 | 73440267 | 73473760 | 33494 | -9.43 | 1.96E-18 | 1.65E-17 |

| | | | | | | |
|---|---|---|---|---|---|---|
| chr2 | 240965513 | 240981230 | 15718 | -6.69 | 1.96E-18 | 1.65E-17 |
| chr19 | 33778318 | 33781544 | 3227 | -9.38 | 1.97E-18 | 1.65E-17 |
| chr5 | 156026967 | 156040598 | 13632 | -9.38 | 1.97E-18 | 1.66E-17 |
| chr1 | 38742089 | 38746438 | 4350 | -2.82 | 1.98E-18 | 1.66E-17 |
| chr15 | 58945714 | 58953820 | 8107 | -9.37 | 1.98E-18 | 1.66E-17 |
| chr19 | 33099440 | 33105215 | 5776 | -9.38 | 1.99E-18 | 1.67E-17 |
| chr16 | 64413986 | 64457130 | 43145 | -9.39 | 1.99E-18 | 1.67E-17 |
| chr5 | 107707456 | 107713965 | 6510 | -9.36 | 2.00E-18 | 1.68E-17 |
| chr9 | 6200454 | 6213525 | 13072 | -9.38 | 2.00E-18 | 1.68E-17 |
| chr11 | 130546449 | 130557274 | 10826 | -9.44 | 2.00E-18 | 1.68E-17 |
| chr18 | 1297453 | 1324804 | 27352 | 9.42 | 2.03E-18 | 1.70E-17 |
| chr3 | 105187411 | 105211714 | 24304 | -9.63 | 2.03E-18 | 1.70E-17 |
| chr21 | 48064095 | 48077125 | 13031 | -9.54 | 2.04E-18 | 1.71E-17 |
| chr3 | 169020048 | 169052827 | 32780 | 9.45 | 2.05E-18 | 1.72E-17 |
| chr12 | 16353465 | 16355746 | 2282 | -5.26 | 2.05E-18 | 1.72E-17 |
| chr15 | 93002457 | 93016526 | 14070 | 9.51 | 2.05E-18 | 1.72E-17 |
| chrX | 107686906 | 107698410 | 11505 | -9.36 | 2.05E-18 | 1.72E-17 |
| chr2 | 97081743 | 97126403 | 44661 | -9.6 | 2.05E-18 | 1.72E-17 |
| chr7 | 79983549 | 79996120 | 12572 | -9.52 | 2.06E-18 | 1.72E-17 |
| chr16 | 54061597 | 54076264 | 14668 | -9.38 | 2.06E-18 | 1.73E-17 |
| chr1 | 207504918 | 207523968 | 19051 | -9.52 | 2.06E-18 | 1.73E-17 |
| chrX | 16101675 | 16138726 | 37052 | -9.37 | 2.06E-18 | 1.73E-17 |
| chr2 | 63355790 | 63370297 | 14508 | -9.37 | 2.06E-18 | 1.73E-17 |
| chr21 | 46605484 | 46611592 | 6109 | -9.39 | 2.06E-18 | 1.73E-17 |
| chr1 | 224529209 | 224535693 | 6485 | -9.49 | 2.07E-18 | 1.73E-17 |
| chr5 | 118469265 | 118477392 | 8128 | -9.37 | 2.09E-18 | 1.75E-17 |
| chr12 | 1497922 | 1503843 | 5922 | -9.38 | 2.11E-18 | 1.76E-17 |
| chrX | 76768852 | 76800365 | 31514 | -9.36 | 2.11E-18 | 1.77E-17 |
| chr10 | 82697877 | 82768144 | 70268 | 2.68 | 2.12E-18 | 1.78E-17 |
| chr15 | 73153969 | 73194096 | 40128 | -1.69 | 2.13E-18 | 1.78E-17 |
| chr3 | 102355286 | 102386993 | 31708 | 9.42 | 2.13E-18 | 1.78E-17 |
| chr2 | 70859380 | 70862617 | 3238 | -9.41 | 2.13E-18 | 1.78E-17 |
| chr18 | 2808981 | 2824953 | 15973 | -9.48 | 2.14E-18 | 1.79E-17 |
| chr2 | 40465665 | 40469472 | 3808 | -8 | 2.15E-18 | 1.80E-17 |
| chr13 | 80541211 | 80574054 | 32844 | -9.4 | 2.15E-18 | 1.80E-17 |
| chr4 | 169891021 | 169904212 | 13192 | -9.39 | 2.15E-18 | 1.80E-17 |
| chr1 | 64190431 | 64211462 | 21032 | -9.37 | 2.16E-18 | 1.80E-17 |
| chr10 | 747576 | 762209 | 14634 | -3.79 | 2.18E-18 | 1.82E-17 |
| chr2 | 161562895 | 161592685 | 29791 | -9.55 | 2.18E-18 | 1.82E-17 |
| chr1 | 19458657 | 19463573 | 4917 | -9.43 | 2.19E-18 | 1.82E-17 |
| chr2 | 174437993 | 174482547 | 44555 | -9.4 | 2.19E-18 | 1.83E-17 |
| chr8 | 37503391 | 37527858 | 24468 | -9.71 | 2.20E-18 | 1.83E-17 |
| chr18 | 9597417 | 9602670 | 5254 | -9.36 | 2.20E-18 | 1.84E-17 |
| chr11 | 45456613 | 45462984 | 6372 | -9.35 | 2.22E-18 | 1.85E-17 |
| chr4 | 184961970 | 184970122 | 8153 | -9.36 | 2.23E-18 | 1.86E-17 |
| chr11 | 1126824 | 1140184 | 13361 | -9.56 | 2.23E-18 | 1.86E-17 |
| chr3 | 136037630 | 136044954 | 7325 | -9.35 | 2.24E-18 | 1.86E-17 |
| chr7 | 14316700 | 14376443 | 59744 | -9.35 | 2.24E-18 | 1.86E-17 |
| chr5 | 21125204 | 21189715 | 64512 | -9.61 | 2.26E-18 | 1.88E-17 |
| chr3 | 141950624 | 141983642 | 33019 | -9.56 | 2.26E-18 | 1.88E-17 |
| chr6 | 170658345 | 170664082 | 5738 | -7.97 | 2.26E-18 | 1.89E-17 |
| chr8 | 115947309 | 115999864 | 52556 | -9.49 | 2.27E-18 | 1.89E-17 |
| chr8 | 40497526 | 40537521 | 39996 | -9.48 | 2.27E-18 | 1.89E-17 |
| chr9 | 118518772 | 118590639 | 71868 | -10.08 | 2.28E-18 | 1.90E-17 |
| chr20 | 9572197 | 9588282 | 16086 | -9.46 | 2.28E-18 | 1.90E-17 |
| chr9 | 94920793 | 94961614 | 40822 | -9.48 | 2.28E-18 | 1.90E-17 |
| chr7 | 130685934 | 130690700 | 4767 | -9.4 | 2.29E-18 | 1.91E-17 |
| chr17 | 63999759 | 64021941 | 22183 | -9.39 | 2.30E-18 | 1.91E-17 |
| chr10 | 124941211 | 124946420 | 5210 | -9.37 | 2.30E-18 | 1.91E-17 |
| chr7 | 17880465 | 17906995 | 26531 | -9.43 | 2.31E-18 | 1.92E-17 |
| chr8 | 126024551 | 126031893 | 7343 | -9.36 | 2.31E-18 | 1.92E-17 |

| | | | | | | |
|---|---|---|---|---|---|---|
| chr3 | 70864481 | 70868191 | 3711 | -6.42 | 2.31E-18 | 1.92E-17 |
| chr5 | 111818268 | 111849612 | 31345 | -2.55 | 2.32E-18 | 1.92E-17 |
| chr18 | 74501108 | 74511306 | 10199 | -9.37 | 2.32E-18 | 1.92E-17 |
| chr5 | 114338372 | 114356227 | 17856 | 9.43 | 2.32E-18 | 1.93E-17 |
| chr15 | 61305356 | 61322304 | 16949 | 9.41 | 2.33E-18 | 1.94E-17 |
| chr2 | 44477982 | 44482218 | 4237 | -9.44 | 2.35E-18 | 1.95E-17 |
| chr13 | 101027839 | 101065005 | 37167 | -9.57 | 2.37E-18 | 1.96E-17 |
| chr22 | 45293878 | 45308280 | 14403 | -9.38 | 2.37E-18 | 1.96E-17 |
| chr5 | 60735392 | 60768170 | 32779 | -9.79 | 2.38E-18 | 1.97E-17 |
| chr15 | 79325486 | 79336996 | 11511 | -9.37 | 2.38E-18 | 1.97E-17 |
| chr20 | 11233353 | 11240527 | 7175 | -9.35 | 2.39E-18 | 1.98E-17 |
| chr22 | 44769262 | 44775525 | 6264 | -9.34 | 2.40E-18 | 1.99E-17 |
| chr11 | 71251620 | 71275256 | 23637 | -6.98 | 2.41E-18 | 2.00E-17 |
| chr6 | 83553431 | 83573830 | 20400 | -6.81 | 2.41E-18 | 2.00E-17 |
| chr10 | 74602999 | 74613557 | 10559 | -9.42 | 2.41E-18 | 2.00E-17 |
| chr8 | 128333112 | 128386148 | 53037 | -9.62 | 2.42E-18 | 2.01E-17 |
| chr7 | 18639185 | 18647163 | 7979 | -9.41 | 2.43E-18 | 2.01E-17 |
| chr9 | 5344988 | 5351393 | 6406 | -9.35 | 2.43E-18 | 2.01E-17 |
| chr6 | 146099488 | 146128186 | 28699 | -9.43 | 2.43E-18 | 2.01E-17 |
| chr16 | 66438613 | 66451896 | 13284 | -9.35 | 2.46E-18 | 2.04E-17 |
| chr6 | 85677144 | 85703407 | 26264 | -9.41 | 2.46E-18 | 2.04E-17 |
| chr4 | 174539855 | 174562503 | 22649 | -9.36 | 2.48E-18 | 2.05E-17 |
| chr3 | 100906245 | 100940071 | 33827 | -9.4 | 2.49E-18 | 2.06E-17 |
| chr3 | 182793232 | 182800252 | 7021 | -9.35 | 2.49E-18 | 2.06E-17 |
| chr20 | 49661786 | 49702923 | 41138 | -9.68 | 2.49E-18 | 2.06E-17 |
| chr18 | 51862420 | 51879439 | 17020 | -6.69 | 2.49E-18 | 2.06E-17 |
| chr7 | 45202175 | 45210407 | 8233 | -9.34 | 2.50E-18 | 2.07E-17 |
| chr2 | 55146669 | 55166073 | 19405 | -9.47 | 2.50E-18 | 2.07E-17 |
| chr14 | 53603894 | 53610786 | 6893 | -9.36 | 2.51E-18 | 2.07E-17 |
| chr16 | 89728393 | 89752154 | 23762 | -9.49 | 2.51E-18 | 2.07E-17 |
| chr10 | 73941044 | 73945770 | 4727 | -9.36 | 2.51E-18 | 2.07E-17 |
| chr8 | 108071362 | 108093920 | 22559 | -9.5 | 2.52E-18 | 2.08E-17 |
| chr2 | 61248080 | 61263329 | 15250 | -9.51 | 2.52E-18 | 2.08E-17 |
| chr15 | 58542092 | 58566744 | 24653 | -9.88 | 2.52E-18 | 2.08E-17 |
| chr1 | 90685560 | 90743858 | 58299 | 2.65 | 2.55E-18 | 2.10E-17 |
| chr12 | 69527123 | 69537323 | 10201 | -9.35 | 2.55E-18 | 2.10E-17 |
| chr12 | 105817576 | 105837952 | 20377 | -9.47 | 2.56E-18 | 2.11E-17 |
| chr1 | 50961046 | 50969981 | 8936 | -9.43 | 2.57E-18 | 2.12E-17 |
| chr9 | 99789720 | 99830287 | 40568 | -9.78 | 2.57E-18 | 2.12E-17 |
| chr7 | 16371098 | 16390857 | 19760 | -9.44 | 2.58E-18 | 2.13E-17 |
| chr7 | 35853930 | 35862326 | 8397 | -9.38 | 2.58E-18 | 2.13E-17 |
| chr19 | 7865113 | 7885097 | 19985 | -9.44 | 2.60E-18 | 2.14E-17 |
| chr10 | 7551501 | 7586386 | 34886 | -9.52 | 2.60E-18 | 2.14E-17 |
| chr7 | 149348313 | 149358886 | 10574 | -9.34 | 2.60E-18 | 2.14E-17 |
| chr4 | 88591466 | 88620865 | 29400 | -7.46 | 2.60E-18 | 2.14E-17 |
| chr12 | 111005189 | 111011918 | 6730 | -9.34 | 2.63E-18 | 2.16E-17 |
| chr1 | 222935462 | 222964002 | 28541 | -9.34 | 2.64E-18 | 2.17E-17 |
| chr12 | 46356122 | 46369269 | 13148 | -9.48 | 2.65E-18 | 2.18E-17 |
| chr5 | 40444064 | 40461915 | 17852 | -9.39 | 2.65E-18 | 2.18E-17 |
| chr1 | 22761769 | 22771179 | 9411 | -9.33 | 2.67E-18 | 2.19E-17 |
| chr12 | 102597217 | 102603833 | 6617 | -9.34 | 2.67E-18 | 2.20E-17 |
| chr2 | 172686596 | 172690481 | 3886 | -9.35 | 2.68E-18 | 2.20E-17 |
| chr7 | 7321040 | 7328010 | 6971 | -9.33 | 2.69E-18 | 2.21E-17 |
| chr8 | 27537167 | 27567046 | 29880 | -9.81 | 2.69E-18 | 2.21E-17 |
| chr17 | 79929439 | 79933939 | 4501 | -9.4 | 2.70E-18 | 2.22E-17 |
| chr2 | 187535936 | 187538974 | 3039 | -9.47 | 2.71E-18 | 2.23E-17 |
| chr2 | 190196904 | 190227957 | 31054 | -9.48 | 2.71E-18 | 2.23E-17 |
| chr2 | 148721096 | 148737465 | 16370 | -9.5 | 2.72E-18 | 2.23E-17 |
| chr1 | 243455838 | 243465916 | 10079 | -9.33 | 2.72E-18 | 2.23E-17 |
| chr14 | 88823576 | 88843770 | 20195 | -2.82 | 2.73E-18 | 2.24E-17 |
| chr10 | 62416966 | 62471838 | 54873 | -9.61 | 2.73E-18 | 2.24E-17 |

| | | | | | | |
|---|---|---|---|---|---|---|
| chr2 | 192340527 | 192349253 | 8727 | -9.44 | 2.73E-18 | 2.24E-17 |
| chr17 | 72432234 | 72436696 | 4463 | -9.4 | 2.74E-18 | 2.25E-17 |
| chr22 | 51083959 | 51090187 | 6229 | -9.38 | 2.75E-18 | 2.25E-17 |
| chr15 | 69360064 | 69365907 | 5844 | -9.49 | 2.75E-18 | 2.25E-17 |
| chr5 | 33040661 | 33063020 | 22360 | -9.53 | 2.75E-18 | 2.25E-17 |
| chr2 | 102818515 | 102835761 | 17247 | -9.54 | 2.76E-18 | 2.26E-17 |
| chr1 | 172164021 | 172177995 | 13975 | -9.51 | 2.80E-18 | 2.29E-17 |
| chr5 | 173407844 | 173419935 | 12092 | -9.58 | 2.81E-18 | 2.30E-17 |
| chr16 | 78655829 | 78699478 | 43650 | -9.4 | 2.82E-18 | 2.31E-17 |
| chr10 | 65249318 | 65280807 | 31490 | -9.37 | 2.82E-18 | 2.31E-17 |
| chr18 | 53699337 | 53719686 | 20350 | -9.33 | 2.83E-18 | 2.31E-17 |
| chr5 | 95624744 | 95662140 | 37397 | -9.5 | 2.84E-18 | 2.32E-17 |
| chr12 | 106772324 | 106776826 | 4503 | -9.33 | 2.84E-18 | 2.32E-17 |
| chr2 | 28079168 | 28082026 | 2859 | -9.33 | 2.85E-18 | 2.33E-17 |
| chr17 | 75504900 | 75513629 | 8730 | -9.4 | 2.86E-18 | 2.34E-17 |
| chr20 | 13713790 | 13724416 | 10627 | -9.35 | 2.89E-18 | 2.36E-17 |
| chr15 | 71267759 | 71293957 | 26199 | -9.62 | 2.92E-18 | 2.39E-17 |
| chr3 | 189458279 | 189480366 | 22088 | -9.69 | 2.93E-18 | 2.39E-17 |
| chr6 | 74610423 | 74657717 | 47295 | -9.32 | 2.93E-18 | 2.39E-17 |
| chr10 | 29995825 | 30011878 | 16054 | -9.38 | 2.93E-18 | 2.39E-17 |
| chr22 | 44251541 | 44271805 | 20265 | -9.36 | 2.95E-18 | 2.40E-17 |
| chrX | 2926030 | 2950148 | 24119 | -9.45 | 2.97E-18 | 2.43E-17 |
| chr7 | 148293404 | 148315973 | 22570 | -9.39 | 2.98E-18 | 2.43E-17 |
| chr12 | 22683251 | 22686210 | 2960 | -9.32 | 2.98E-18 | 2.43E-17 |
| chr3 | 157916243 | 157921119 | 4877 | -9.32 | 2.98E-18 | 2.43E-17 |
| chr16 | 54328475 | 54346729 | 18255 | -9.62 | 2.98E-18 | 2.43E-17 |
| chr1 | 19323252 | 19332975 | 9724 | -9.42 | 2.99E-18 | 2.43E-17 |
| chrX | 107878562 | 107885886 | 7325 | -9.33 | 3.00E-18 | 2.44E-17 |
| chr7 | 44696896 | 44702576 | 5681 | -9.37 | 3.01E-18 | 2.45E-17 |
| chr2 | 223883259 | 223905773 | 22515 | -9.41 | 3.01E-18 | 2.45E-17 |
| chr1 | 29854412 | 29862070 | 7659 | -9.33 | 3.01E-18 | 2.45E-17 |
| chr10 | 13120922 | 13128757 | 7836 | -9.37 | 3.02E-18 | 2.46E-17 |
| chr3 | 99378222 | 99395857 | 17636 | 9.51 | 3.03E-18 | 2.47E-17 |
| chr12 | 65919644 | 65938545 | 18902 | -9.68 | 3.04E-18 | 2.47E-17 |
| chr9 | 99034812 | 99039574 | 4763 | -9.33 | 3.06E-18 | 2.49E-17 |
| chrX | 9216109 | 9233086 | 16978 | -9.47 | 3.06E-18 | 2.49E-17 |
| chr12 | 125650027 | 125666017 | 15991 | -9.39 | 3.07E-18 | 2.50E-17 |
| chr5 | 135540662 | 135561200 | 20539 | -9.34 | 3.09E-18 | 2.51E-17 |
| chr10 | 84350265 | 84393603 | 43339 | -9.34 | 3.09E-18 | 2.51E-17 |
| chr4 | 38250704 | 38259405 | 8702 | -9.36 | 3.10E-18 | 2.52E-17 |
| chr15 | 23937293 | 24108738 | 171446 | 0.85 | 3.10E-18 | 2.52E-17 |
| chr2 | 198388650 | 198393656 | 5007 | -9.37 | 3.11E-18 | 2.52E-17 |
| chr9 | 131948664 | 131958743 | 10080 | -9.42 | 3.11E-18 | 2.53E-17 |
| chr3 | 13325187 | 13330302 | 5116 | -9.35 | 3.14E-18 | 2.55E-17 |
| chr2 | 128444056 | 128449489 | 5434 | -9.32 | 3.17E-18 | 2.57E-17 |
| chr5 | 130734357 | 130747213 | 12857 | -9.38 | 3.20E-18 | 2.60E-17 |
| chrX | 46037046 | 46057375 | 20330 | -9.42 | 3.21E-18 | 2.60E-17 |
| chr4 | 188016879 | 188020722 | 3844 | -3.51 | 3.21E-18 | 2.61E-17 |
| chrX | 13111130 | 13140938 | 29809 | -9.36 | 3.22E-18 | 2.61E-17 |
| chr3 | 127485747 | 127498330 | 12584 | -9.33 | 3.22E-18 | 2.61E-17 |
| chr15 | 44855506 | 44858751 | 3246 | -9.32 | 3.23E-18 | 2.62E-17 |
| chr12 | 75613823 | 75660806 | 46984 | -9.33 | 3.27E-18 | 2.65E-17 |
| chr3 | 169616062 | 169634979 | 18918 | -9.44 | 3.27E-18 | 2.65E-17 |
| chr1 | 200750798 | 200762924 | 12127 | -9.32 | 3.28E-18 | 2.66E-17 |
| chr17 | 53493263 | 53497503 | 4241 | -9.34 | 3.29E-18 | 2.67E-17 |
| chr6 | 83628293 | 83659289 | 30997 | -9.31 | 3.34E-18 | 2.71E-17 |
| chr4 | 14373438 | 14406152 | 32715 | -9.37 | 3.36E-18 | 2.72E-17 |
| chr3 | 189290001 | 189330281 | 40281 | -9.51 | 3.37E-18 | 2.73E-17 |
| chr3 | 120056553 | 120060597 | 4045 | -9.33 | 3.37E-18 | 2.73E-17 |
| chr14 | 53746106 | 53757462 | 11357 | -9.32 | 3.38E-18 | 2.73E-17 |
| chr2 | 228294905 | 228301065 | 6161 | -9.48 | 3.38E-18 | 2.73E-17 |

| | | | | | | |
|---|---|---|---|---|---|---|
| chr10 | 33970888 | 33986872 | 15985 | -9.39 | 3.38E-18 | 2.73E-17 |
| chr1 | 240321840 | 240329909 | 8070 | -9.33 | 3.38E-18 | 2.73E-17 |
| chr7 | 150039310 | 150046104 | 6795 | -9.32 | 3.38E-18 | 2.74E-17 |
| chr17 | 39622711 | 39629095 | 6385 | -9.35 | 3.39E-18 | 2.74E-17 |
| chr13 | 106831159 | 106858767 | 27609 | -9.31 | 3.40E-18 | 2.75E-17 |
| chr8 | 135778191 | 135787462 | 9272 | -9.37 | 3.40E-18 | 2.75E-17 |
| chr6 | 87448778 | 87512238 | 63461 | 1.7 | 3.41E-18 | 2.75E-17 |
| chr10 | 120732169 | 120762705 | 30537 | -9.7 | 3.41E-18 | 2.75E-17 |
| chr20 | 31577997 | 31593946 | 15950 | -9.46 | 3.42E-18 | 2.76E-17 |
| chr14 | 48931331 | 48979689 | 48359 | -9.32 | 3.43E-18 | 2.77E-17 |
| chr10 | 23139559 | 23243145 | 103587 | -3.1 | 3.43E-18 | 2.77E-17 |
| chr7 | 3484338 | 3497732 | 13395 | -9.37 | 3.43E-18 | 2.77E-17 |
| chr2 | 165363021 | 165386694 | 23674 | -9.32 | 3.44E-18 | 2.78E-17 |
| chr10 | 119092569 | 119096674 | 4106 | -9.38 | 3.47E-18 | 2.80E-17 |
| chr15 | 100124083 | 100134443 | 10361 | -9.31 | 3.50E-18 | 2.83E-17 |
| chr5 | 31826801 | 31840501 | 13701 | -9.3 | 3.51E-18 | 2.83E-17 |
| chr17 | 57915963 | 57941139 | 25177 | -9.43 | 3.51E-18 | 2.83E-17 |
| chr11 | 61752756 | 61768366 | 15611 | -9.48 | 3.52E-18 | 2.84E-17 |
| chr6 | 8965793 | 8979688 | 13896 | -9.47 | 3.52E-18 | 2.84E-17 |
| chr1 | 115779430 | 115803033 | 23604 | -9.35 | 3.53E-18 | 2.84E-17 |
| chr6 | 162854558 | 163113815 | 259258 | 2.23 | 3.53E-18 | 2.84E-17 |
| chr1 | 119468239 | 119498628 | 30390 | 7.42 | 3.55E-18 | 2.86E-17 |
| chr12 | 64868201 | 64871300 | 3100 | -9.42 | 3.55E-18 | 2.86E-17 |
| chr4 | 102278966 | 102301349 | 22384 | -9.35 | 3.55E-18 | 2.86E-17 |
| chr9 | 140747001 | 140764280 | 17280 | -9.38 | 3.58E-18 | 2.88E-17 |
| chr12 | 57758899 | 57762344 | 3446 | -9.33 | 3.61E-18 | 2.90E-17 |
| chr1 | 63098776 | 63111514 | 12739 | -9.39 | 3.61E-18 | 2.91E-17 |
| chr4 | 139442044 | 139457826 | 15783 | -9.5 | 3.64E-18 | 2.93E-17 |
| chr15 | 72869149 | 72872404 | 3256 | -9.31 | 3.67E-18 | 2.95E-17 |
| chr3 | 27237931 | 27277951 | 40021 | -9.74 | 3.68E-18 | 2.95E-17 |
| chr16 | 15677167 | 15683980 | 6814 | -9.34 | 3.68E-18 | 2.96E-17 |
| chr3 | 45825144 | 45832435 | 7292 | -9.3 | 3.70E-18 | 2.97E-17 |
| chr7 | 6504655 | 6508587 | 3933 | -9.3 | 3.72E-18 | 2.99E-17 |
| chr9 | 83671497 | 83682354 | 10858 | -9.44 | 3.72E-18 | 2.99E-17 |
| chr13 | 27697075 | 27717009 | 19935 | -9.34 | 3.72E-18 | 2.99E-17 |
| chr9 | 82785380 | 82801659 | 16280 | -9.34 | 3.72E-18 | 2.99E-17 |
| chr7 | 47307226 | 47326657 | 19432 | -9.32 | 3.74E-18 | 3.00E-17 |
| chr5 | 58890203 | 58899713 | 9511 | -9.41 | 3.75E-18 | 3.01E-17 |
| chr10 | 80367024 | 80372734 | 5711 | -9.37 | 3.76E-18 | 3.01E-17 |
| chr2 | 120643825 | 120662392 | 18568 | -9.3 | 3.77E-18 | 3.03E-17 |
| chr4 | 123882527 | 123910734 | 28208 | -9.3 | 3.80E-18 | 3.05E-17 |
| chr20 | 18331729 | 18350761 | 19033 | -9.36 | 3.80E-18 | 3.05E-17 |
| chr5 | 118839351 | 118848899 | 9549 | -9.36 | 3.80E-18 | 3.05E-17 |
| chrX | 99721365 | 99728069 | 6705 | 4 | 3.81E-18 | 3.05E-17 |
| chr8 | 116018259 | 116071823 | 53565 | -9.44 | 3.81E-18 | 3.05E-17 |
| chr3 | 193670818 | 193675159 | 4342 | -7.39 | 3.82E-18 | 3.06E-17 |
| chr5 | 106468077 | 106479429 | 11353 | -9.32 | 3.83E-18 | 3.06E-17 |
| chr8 | 63069868 | 63087170 | 17303 | -9.32 | 3.84E-18 | 3.07E-17 |
| chr3 | 25809929 | 25819139 | 9211 | -9.31 | 3.85E-18 | 3.08E-17 |
| chr14 | 92004923 | 92035319 | 30397 | -9.65 | 3.85E-18 | 3.08E-17 |
| chr12 | 48451423 | 48455337 | 3915 | -9.48 | 3.88E-18 | 3.11E-17 |
| chr8 | 116365458 | 116434993 | 69536 | -9.3 | 3.88E-18 | 3.11E-17 |
| chr20 | 30535205 | 30537723 | 2519 | -9.29 | 3.89E-18 | 3.11E-17 |
| chrX | 79693544 | 79712632 | 19089 | 9.4 | 3.89E-18 | 3.11E-17 |
| chr5 | 82407999 | 82423622 | 15624 | -9.33 | 3.90E-18 | 3.12E-17 |
| chr1 | 154262207 | 154281834 | 19628 | -9.5 | 3.94E-18 | 3.15E-17 |
| chr2 | 150591335 | 150613601 | 22267 | -9.34 | 3.96E-18 | 3.16E-17 |
| chr18 | 9816222 | 9823138 | 6917 | -9.33 | 3.96E-18 | 3.17E-17 |
| chr20 | 25678859 | 25683009 | 4151 | -9.29 | 3.97E-18 | 3.17E-17 |
| chr6 | 39126588 | 39147344 | 20757 | -9.51 | 3.97E-18 | 3.17E-17 |
| chr3 | 44949074 | 44957916 | 8843 | -9.35 | 3.97E-18 | 3.17E-17 |

| | | | | | | |
|---|---|---|---|---|---|---|
| chr18 | 12297889 | 12313510 | 15622 | -9.42 | 3.98E-18 | 3.18E-17 |
| chr22 | 35577711 | 35587282 | 9572 | -9.29 | 3.99E-18 | 3.19E-17 |
| chr6 | 109533072 | 109547646 | 14575 | -9.29 | 3.99E-18 | 3.19E-17 |
| chr1 | 42700925 | 42717786 | 16862 | -9.29 | 4.02E-18 | 3.21E-17 |
| chr12 | 90127203 | 90144140 | 16938 | -9.53 | 4.02E-18 | 3.21E-17 |
| chr9 | 75799581 | 75805437 | 5857 | -9.35 | 4.02E-18 | 3.21E-17 |
| chr15 | 85388848 | 85392599 | 3752 | -9.49 | 4.06E-18 | 3.23E-17 |
| chr22 | 19632916 | 19640882 | 7967 | -3.93 | 4.07E-18 | 3.24E-17 |
| chr7 | 8347271 | 8360237 | 12967 | -9.29 | 4.08E-18 | 3.25E-17 |
| chr1 | 99025404 | 99030887 | 5484 | -6.49 | 4.09E-18 | 3.26E-17 |
| chr11 | 34963212 | 34967378 | 4167 | -9.3 | 4.13E-18 | 3.29E-17 |
| chr7 | 2063763 | 2083310 | 19548 | -9.32 | 4.16E-18 | 3.31E-17 |
| chr15 | 59052994 | 59058876 | 5883 | -9.3 | 4.20E-18 | 3.34E-17 |
| chr22 | 31233272 | 31251474 | 18203 | -9.29 | 4.20E-18 | 3.35E-17 |
| chr3 | 176837345 | 176854863 | 17519 | -9.32 | 4.21E-18 | 3.35E-17 |
| chr3 | 107994278 | 108011415 | 17138 | -9.29 | 4.21E-18 | 3.36E-17 |
| chr8 | 86151836 | 86163608 | 11773 | -9.3 | 4.23E-18 | 3.36E-17 |
| chr7 | 110970473 | 110986220 | 15748 | -9.29 | 4.23E-18 | 3.37E-17 |
| chr2 | 138919937 | 138931338 | 11402 | -9.37 | 4.23E-18 | 3.37E-17 |
| chr8 | 71269452 | 71272659 | 3208 | -9.31 | 4.24E-18 | 3.38E-17 |
| chr15 | 77686955 | 77693309 | 6355 | -9.28 | 4.25E-18 | 3.38E-17 |
| chr1 | 222236703 | 222270176 | 33474 | -9.39 | 4.27E-18 | 3.40E-17 |
| chr15 | 42544266 | 42551344 | 7079 | -9.29 | 4.29E-18 | 3.41E-17 |
| chr5 | 85951635 | 85970892 | 19258 | 9.34 | 4.29E-18 | 3.41E-17 |
| chr3 | 134346109 | 134353494 | 7386 | -9.28 | 4.30E-18 | 3.41E-17 |
| chr11 | 106012258 | 106039430 | 27173 | -9.48 | 4.31E-18 | 3.42E-17 |
| chr17 | 26512054 | 26514674 | 2621 | -9.47 | 4.31E-18 | 3.43E-17 |
| chr13 | 28221750 | 28226521 | 4772 | -9.49 | 4.34E-18 | 3.45E-17 |
| chr18 | 69732561 | 69754982 | 22422 | -9.32 | 4.35E-18 | 3.46E-17 |
| chr8 | 11377653 | 11389148 | 11496 | -9.41 | 4.36E-18 | 3.46E-17 |
| chr15 | 83832644 | 83839315 | 6672 | -9.29 | 4.37E-18 | 3.47E-17 |
| chr17 | 3877692 | 3902094 | 24403 | -9.49 | 4.38E-18 | 3.48E-17 |
| chr11 | 109880045 | 109914245 | 34201 | -9.63 | 4.39E-18 | 3.48E-17 |
| chr5 | 91686306 | 91717833 | 31528 | -9.3 | 4.41E-18 | 3.50E-17 |
| chr16 | 48176637 | 48187656 | 11020 | -9.32 | 4.42E-18 | 3.50E-17 |
| chr1 | 221893279 | 221899156 | 5878 | -9.33 | 4.44E-18 | 3.52E-17 |
| chr19 | 4241297 | 4257082 | 15786 | -9.3 | 4.44E-18 | 3.52E-17 |
| chr5 | 153601626 | 153608298 | 6673 | -9.37 | 4.44E-18 | 3.52E-17 |
| chr4 | 744171 | 772160 | 27990 | -9.35 | 4.44E-18 | 3.52E-17 |
| chr11 | 117759754 | 117768046 | 8293 | -9.29 | 4.45E-18 | 3.52E-17 |
| chr3 | 95592419 | 95645015 | 52597 | -9.37 | 4.47E-18 | 3.54E-17 |
| chr3 | 105747639 | 105784648 | 37010 | -9.4 | 4.48E-18 | 3.55E-17 |
| chr19 | 15229445 | 15278218 | 48774 | -9.53 | 4.50E-18 | 3.56E-17 |
| chr8 | 53418195 | 53440674 | 22480 | -9.4 | 4.51E-18 | 3.57E-17 |
| chr15 | 84353707 | 84750942 | 397236 | 1.91 | 4.51E-18 | 3.57E-17 |
| chr5 | 32424270 | 32429401 | 5132 | -9.28 | 4.52E-18 | 3.58E-17 |
| chr2 | 133283498 | 133311158 | 27661 | -9.28 | 4.53E-18 | 3.58E-17 |
| chr21 | 44233534 | 44250590 | 17057 | -9.51 | 4.54E-18 | 3.59E-17 |
| chr8 | 61506263 | 61518738 | 12476 | -9.3 | 4.55E-18 | 3.60E-17 |
| chr7 | 29936295 | 29948571 | 12277 | -9.32 | 4.55E-18 | 3.60E-17 |
| chr17 | 18796127 | 18802831 | 6705 | -9.32 | 4.56E-18 | 3.61E-17 |
| chrX | 150623446 | 150647880 | 24435 | 2.7 | 4.58E-18 | 3.61E-17 |
| chr2 | 241166230 | 241181468 | 15239 | -9.28 | 4.59E-18 | 3.62E-17 |
| chr6 | 22412818 | 22425260 | 12443 | -9.28 | 4.59E-18 | 3.63E-17 |
| chr3 | 16749765 | 16785662 | 35898 | -9.65 | 4.61E-18 | 3.64E-17 |
| chr1 | 165840843 | 165848709 | 7867 | -9.33 | 4.61E-18 | 3.64E-17 |
| chr6 | 106247936 | 106273322 | 25387 | -9.38 | 4.62E-18 | 3.64E-17 |
| chr2 | 159925253 | 159954426 | 29174 | -9.53 | 4.65E-18 | 3.67E-17 |
| chr5 | 129878348 | 129901256 | 22909 | -9.28 | 4.66E-18 | 3.67E-17 |
| chr12 | 116686023 | 116691147 | 5125 | -9.28 | 4.66E-18 | 3.68E-17 |
| chr11 | 2774865 | 2789535 | 14671 | -9.29 | 4.70E-18 | 3.70E-17 |

| | | | | | | |
|---|---|---|---|---|---|---|
| chr12 | 29298753 | 29457949 | 159197 | 2.59 | 4.70E-18 | 3.71E-17 |
| chr2 | 158020922 | 158043984 | 23063 | -9.32 | 4.70E-18 | 3.71E-17 |
| chr6 | 4297684 | 4340018 | 42335 | -9.29 | 4.71E-18 | 3.71E-17 |
| chr4 | 186404779 | 186423782 | 19004 | -9.28 | 4.76E-18 | 3.75E-17 |
| chr9 | 93163566 | 93174817 | 11252 | -9.33 | 4.78E-18 | 3.76E-17 |
| chr5 | 76487294 | 76495303 | 8010 | -9.28 | 4.79E-18 | 3.78E-17 |
| chr19 | 50755375 | 50761443 | 6069 | -9.47 | 4.80E-18 | 3.78E-17 |
| chr10 | 76101843 | 76122484 | 20642 | -9.28 | 4.81E-18 | 3.79E-17 |
| chr3 | 196233054 | 196250208 | 17155 | -9.37 | 4.82E-18 | 3.79E-17 |
| chr16 | 52833192 | 52850398 | 17207 | -9.35 | 4.82E-18 | 3.79E-17 |
| chr6 | 20879404 | 20888417 | 9014 | -9.33 | 4.82E-18 | 3.79E-17 |
| chr3 | 160297951 | 160312981 | 15031 | -6.65 | 4.83E-18 | 3.80E-17 |
| chr12 | 131657928 | 131684656 | 26729 | -9.43 | 4.87E-18 | 3.83E-17 |
| chr17 | 6536080 | 6538330 | 2251 | -9.32 | 4.89E-18 | 3.84E-17 |
| chr12 | 11095044 | 11109985 | 14942 | -9.46 | 4.89E-18 | 3.85E-17 |
| chr9 | 127222684 | 127226903 | 4220 | -9.42 | 4.91E-18 | 3.86E-17 |
| chr10 | 75673972 | 75689421 | 15450 | -9.73 | 4.92E-18 | 3.87E-17 |
| chr3 | 44016872 | 44035315 | 18444 | -9.63 | 4.95E-18 | 3.89E-17 |
| chr17 | 18898043 | 18910686 | 12644 | -9.3 | 4.98E-18 | 3.91E-17 |
| chr2 | 44961813 | 44966300 | 4488 | -9.34 | 4.98E-18 | 3.91E-17 |
| chr2 | 221452730 | 221487922 | 35193 | -9.35 | 4.99E-18 | 3.92E-17 |
| chr2 | 178863457 | 178892678 | 29222 | 9.5 | 5.00E-18 | 3.92E-17 |
| chrX | 124532177 | 124553183 | 21007 | 9.35 | 5.00E-18 | 3.92E-17 |
| chr6 | 1329134 | 1354790 | 25657 | -9.27 | 5.00E-18 | 3.93E-17 |
| chr16 | 19004099 | 19033771 | 29673 | -9.35 | 5.01E-18 | 3.93E-17 |
| chrX | 21068040 | 21092400 | 24361 | -9.4 | 5.03E-18 | 3.95E-17 |
| chr13 | 76752561 | 76785024 | 32464 | 2.25 | 5.03E-18 | 3.95E-17 |
| chr2 | 33111532 | 33123327 | 11796 | -9.27 | 5.05E-18 | 3.96E-17 |
| chr4 | 147076263 | 147084476 | 8214 | -9.27 | 5.05E-18 | 3.96E-17 |
| chr8 | 19556658 | 19573914 | 17257 | -9.56 | 5.05E-18 | 3.96E-17 |
| chr2 | 55688746 | 55704347 | 15602 | -9.27 | 5.05E-18 | 3.96E-17 |
| chr1 | 152104656 | 152123476 | 18821 | -9.28 | 5.07E-18 | 3.97E-17 |
| chr7 | 157282944 | 157302692 | 19749 | -9.45 | 5.09E-18 | 3.99E-17 |
| chr20 | 36936980 | 36968642 | 31663 | -9.27 | 5.11E-18 | 4.00E-17 |
| chr3 | 186859806 | 186879179 | 19374 | -9.27 | 5.13E-18 | 4.01E-17 |
| chr14 | 93252416 | 93258205 | 5790 | -9.27 | 5.13E-18 | 4.01E-17 |
| chr5 | 135302624 | 135354769 | 52146 | -9.29 | 5.15E-18 | 4.03E-17 |
| chr14 | 31797766 | 31803786 | 6021 | -9.33 | 5.18E-18 | 4.05E-17 |
| chr7 | 54882908 | 54898234 | 15327 | -9.3 | 5.18E-18 | 4.06E-17 |
| chr10 | 54290309 | 54325377 | 35069 | -9.41 | 5.19E-18 | 4.06E-17 |
| chr10 | 134709848 | 134719669 | 9822 | -9.29 | 5.20E-18 | 4.07E-17 |
| chr8 | 91528703 | 91549907 | 21205 | 9.37 | 5.23E-18 | 4.09E-17 |
| chr22 | 43852209 | 43861114 | 8906 | -9.26 | 5.23E-18 | 4.09E-17 |
| chr11 | 126830013 | 126854927 | 24915 | -9.39 | 5.23E-18 | 4.09E-17 |
| chr15 | 72082261 | 72095691 | 13431 | -9.32 | 5.26E-18 | 4.11E-17 |
| chr1 | 169298040 | 169335516 | 37477 | -9.37 | 5.29E-18 | 4.13E-17 |
| chr22 | 45945982 | 45968967 | 22986 | -9.33 | 5.32E-18 | 4.15E-17 |
| chr2 | 160196066 | 160200790 | 4725 | -9.27 | 5.32E-18 | 4.15E-17 |
| chr12 | 47382394 | 47411044 | 28651 | -9.27 | 5.35E-18 | 4.17E-17 |
| chr3 | 134107888 | 134134486 | 26599 | -9.28 | 5.42E-18 | 4.23E-17 |
| chr2 | 43509537 | 43514202 | 4666 | -9.33 | 5.49E-18 | 4.28E-17 |
| chr2 | 74977810 | 74986071 | 8262 | -9.26 | 5.49E-18 | 4.28E-17 |
| chr2 | 98899551 | 98909932 | 10382 | -3.46 | 5.52E-18 | 4.31E-17 |
| chr10 | 21895887 | 21899043 | 3157 | -9.32 | 5.55E-18 | 4.33E-17 |
| chr10 | 119138330 | 119150358 | 12029 | -7.89 | 5.56E-18 | 4.34E-17 |
| chr4 | 151031613 | 151044255 | 12643 | -9.29 | 5.57E-18 | 4.34E-17 |
| chr12 | 132946444 | 132952010 | 5567 | -9.26 | 5.57E-18 | 4.34E-17 |
| chr9 | 124923674 | 124932054 | 8381 | -9.28 | 5.60E-18 | 4.36E-17 |
| chr1 | 241084561 | 241116601 | 32041 | -9.49 | 5.61E-18 | 4.37E-17 |
| chr11 | 56786709 | 56813519 | 26811 | -9.42 | 5.62E-18 | 4.38E-17 |
| chr7 | 156291419 | 156306229 | 14811 | -9.28 | 5.62E-18 | 4.38E-17 |

| | | | | | | |
|---|---|---|---|---|---|---|
| chr18 | 54827538 | 54855837 | 28300 | -2.38 | 5.63E-18 | 4.38E-17 |
| chr15 | 33559896 | 33570267 | 10372 | -9.25 | 5.64E-18 | 4.39E-17 |
| chr3 | 131813948 | 131852658 | 38711 | -9.27 | 5.77E-18 | 4.49E-17 |
| chr4 | 79186282 | 79188463 | 2182 | -6.71 | 5.82E-18 | 4.53E-17 |
| chr4 | 184193868 | 184198114 | 4247 | -9.37 | 5.83E-18 | 4.53E-17 |
| chr13 | 44566939 | 44581933 | 14995 | -9.26 | 5.86E-18 | 4.56E-17 |
| chr8 | 143343836 | 143349490 | 5655 | -8.09 | 5.87E-18 | 4.56E-17 |
| chr20 | 50773172 | 50783557 | 10386 | -9.33 | 5.87E-18 | 4.57E-17 |
| chr4 | 5769589 | 5779757 | 10169 | -9.25 | 5.88E-18 | 4.57E-17 |
| chr7 | 11453359 | 11481487 | 28129 | -9.32 | 5.89E-18 | 4.58E-17 |
| chr10 | 64346955 | 64374609 | 27655 | -9.28 | 5.89E-18 | 4.58E-17 |
| chr3 | 113258932 | 113268365 | 9434 | -9.25 | 5.89E-18 | 4.58E-17 |
| chr8 | 93949700 | 93965959 | 16260 | -9.28 | 5.92E-18 | 4.60E-17 |
| chr12 | 3171544 | 3176892 | 5349 | -9.35 | 5.93E-18 | 4.60E-17 |
| chr16 | 87318184 | 87326704 | 8521 | -9.25 | 5.94E-18 | 4.61E-17 |
| chr13 | 35594574 | 35608973 | 14400 | -9.27 | 5.95E-18 | 4.62E-17 |
| chr9 | 119276961 | 119301595 | 24635 | -9.31 | 5.96E-18 | 4.62E-17 |
| chr5 | 82907119 | 82925692 | 18574 | -9.29 | 5.96E-18 | 4.62E-17 |
| chr9 | 126336894 | 126341406 | 4513 | -9.27 | 5.97E-18 | 4.63E-17 |
| chr7 | 70262542 | 71989970 | 1727429 | 0.59 | 5.99E-18 | 4.65E-17 |
| chr7 | 21530077 | 21550172 | 20096 | -9.66 | 6.01E-18 | 4.66E-17 |
| chr19 | 15626199 | 15650512 | 24314 | -9.41 | 6.03E-18 | 4.67E-17 |
| chr10 | 13438504 | 13470594 | 32091 | -9.69 | 6.03E-18 | 4.67E-17 |
| chr6 | 167375265 | 167385632 | 10368 | -9.28 | 6.04E-18 | 4.68E-17 |
| chr3 | 146070585 | 146098505 | 27921 | -9.25 | 6.06E-18 | 4.69E-17 |
| chr1 | 25299503 | 25306378 | 6876 | -9.29 | 6.10E-18 | 4.72E-17 |
| chr1 | 198179267 | 198187490 | 8224 | -9.24 | 6.12E-18 | 4.74E-17 |
| chr4 | 133814098 | 133843411 | 29314 | 9.3 | 6.18E-18 | 4.78E-17 |
| chr6 | 132480414 | 132502677 | 22264 | -9.24 | 6.19E-18 | 4.79E-17 |
| chrX | 83448711 | 83455561 | 6851 | -6.53 | 6.19E-18 | 4.79E-17 |
| chr2 | 209291402 | 209299624 | 8223 | -3.69 | 6.21E-18 | 4.81E-17 |
| chr18 | 7832703 | 7852518 | 19816 | -9.24 | 6.21E-18 | 4.81E-17 |
| chr9 | 125620172 | 125625378 | 5207 | -9.3 | 6.24E-18 | 4.83E-17 |
| chr10 | 61909888 | 61923100 | 13213 | -9.32 | 6.24E-18 | 4.83E-17 |
| chr9 | 75831293 | 75846419 | 15127 | -9.3 | 6.26E-18 | 4.84E-17 |
| chr8 | 87505628 | 87510798 | 5171 | -9.26 | 6.28E-18 | 4.85E-17 |
| chr18 | 70114368 | 70132968 | 18601 | -9.27 | 6.28E-18 | 4.85E-17 |
| chr6 | 122291001 | 122327164 | 36164 | 9.42 | 6.31E-18 | 4.88E-17 |
| chr2 | 228348855 | 228352522 | 3668 | -9.3 | 6.32E-18 | 4.88E-17 |
| chr14 | 67122324 | 67148976 | 26653 | -9.26 | 6.37E-18 | 4.92E-17 |
| chr2 | 150712916 | 150736046 | 23131 | -9.28 | 6.39E-18 | 4.94E-17 |
| chr2 | 151229541 | 151257392 | 27852 | -9.5 | 6.42E-18 | 4.96E-17 |
| chr2 | 232715567 | 232736101 | 20535 | -9.4 | 6.43E-18 | 4.96E-17 |
| chr13 | 33433644 | 33445304 | 11661 | -4.35 | 6.43E-18 | 4.96E-17 |
| chr1 | 40792176 | 40808980 | 16805 | -9.28 | 6.44E-18 | 4.97E-17 |
| chr5 | 74048970 | 74053548 | 4579 | -9.24 | 6.49E-18 | 5.00E-17 |
| chr19 | 27995271 | 29728403 | 1733133 | 0.65 | 6.52E-18 | 5.03E-17 |
| chr14 | 80973492 | 80988980 | 15489 | -9.32 | 6.55E-18 | 5.05E-17 |
| chr7 | 93927438 | 93952836 | 25399 | -9.45 | 6.56E-18 | 5.06E-17 |
| chr2 | 55786794 | 55790510 | 3717 | -9.27 | 6.58E-18 | 5.07E-17 |
| chr15 | 31240833 | 31244172 | 3340 | -9.3 | 6.59E-18 | 5.08E-17 |
| chr12 | 60644851 | 60695057 | 50207 | -9.57 | 6.59E-18 | 5.08E-17 |
| chr9 | 137136860 | 137141903 | 5044 | -9.28 | 6.59E-18 | 5.08E-17 |
| chr12 | 89183117 | 89194979 | 11863 | -9.24 | 6.63E-18 | 5.11E-17 |
| chr10 | 841427 | 848099 | 6673 | -9.24 | 6.66E-18 | 5.13E-17 |
| chr1 | 112691948 | 112764068 | 72121 | 2.29 | 6.67E-18 | 5.13E-17 |
| chr8 | 90348910 | 90365136 | 16227 | -9.27 | 6.68E-18 | 5.14E-17 |
| chr6 | 22814914 | 22843423 | 28510 | -9.29 | 6.68E-18 | 5.14E-17 |
| chr4 | 129261556 | 129299872 | 38317 | -9.63 | 6.70E-18 | 5.15E-17 |
| chr12 | 110650251 | 110653162 | 2912 | -9.3 | 6.70E-18 | 5.16E-17 |
| chr16 | 51158328 | 51188927 | 30600 | -9.63 | 6.71E-18 | 5.16E-17 |

| | | | | | | |
|---|---|---|---|---|---|---|
| chr14 | 106799604 | 106805247 | 5644 | 9.29 | 6.72E-18 | 5.17E-17 |
| chr4 | 3877488 | 3881490 | 4003 | -9.25 | 6.79E-18 | 5.22E-17 |
| chr2 | 180053233 | 180077122 | 23890 | -9.36 | 6.80E-18 | 5.23E-17 |
| chr8 | 58625054 | 58650642 | 25589 | -9.33 | 6.81E-18 | 5.23E-17 |
| chr7 | 12437190 | 12467541 | 30352 | -9.28 | 6.85E-18 | 5.26E-17 |
| chr18 | 67879146 | 67909688 | 30543 | -9.47 | 6.86E-18 | 5.27E-17 |
| chr20 | 8982756 | 9007622 | 24867 | -9.24 | 6.90E-18 | 5.30E-17 |
| chr16 | 17571698 | 17576699 | 5002 | -3.13 | 6.90E-18 | 5.30E-17 |
| chr8 | 58813305 | 58835406 | 22102 | -9.27 | 6.93E-18 | 5.32E-17 |
| chr2 | 96022342 | 96037440 | 15099 | -9.24 | 6.93E-18 | 5.32E-17 |
| chr11 | 119635756 | 119640359 | 4604 | -9.34 | 7.01E-18 | 5.38E-17 |
| chr9 | 90275455 | 90284110 | 8656 | -9.24 | 7.05E-18 | 5.41E-17 |
| chr18 | 52631194 | 52656875 | 25682 | -9.25 | 7.07E-18 | 5.42E-17 |
| chr5 | 124492879 | 124514978 | 22100 | -9.55 | 7.07E-18 | 5.42E-17 |
| chr12 | 46548946 | 46563080 | 14135 | -9.24 | 7.08E-18 | 5.43E-17 |
| chr18 | 32961058 | 33000738 | 39681 | -9.58 | 7.09E-18 | 5.43E-17 |
| chr5 | 126638149 | 126653667 | 15519 | -9.3 | 7.10E-18 | 5.44E-17 |
| chr2 | 224392264 | 224431502 | 39239 | -9.25 | 7.11E-18 | 5.45E-17 |
| chr10 | 28805527 | 28815381 | 9855 | -9.34 | 7.12E-18 | 5.45E-17 |
| chrX | 48573398 | 48589780 | 16383 | -9.71 | 7.16E-18 | 5.49E-17 |
| chr20 | 18181632 | 18186853 | 5222 | -9.24 | 7.22E-18 | 5.53E-17 |
| chr7 | 120449257 | 120472483 | 23227 | 9.32 | 7.22E-18 | 5.53E-17 |
| chr12 | 48350751 | 48354319 | 3569 | -3.45 | 7.22E-18 | 5.53E-17 |
| chr7 | 68708710 | 68752602 | 43893 | -9.56 | 7.24E-18 | 5.54E-17 |
| chr11 | 16073312 | 16097603 | 24292 | -9.48 | 7.26E-18 | 5.55E-17 |
| chr1 | 152493670 | 152502337 | 8668 | -9.26 | 7.28E-18 | 5.57E-17 |
| chr11 | 64187360 | 64208034 | 20675 | -9.5 | 7.29E-18 | 5.58E-17 |
| chr2 | 113546868 | 113570177 | 23310 | -9.23 | 7.30E-18 | 5.58E-17 |
| chr13 | 23615330 | 23662105 | 46776 | -9.25 | 7.33E-18 | 5.61E-17 |
| chr3 | 152192641 | 152202767 | 10127 | -9.32 | 7.33E-18 | 5.61E-17 |
| chr13 | 109018681 | 109033135 | 14455 | -9.3 | 7.34E-18 | 5.61E-17 |
| chr7 | 151274980 | 151279412 | 4433 | -9.24 | 7.34E-18 | 5.61E-17 |
| chr19 | 17619459 | 17640343 | 20885 | -9.23 | 7.40E-18 | 5.65E-17 |
| chr9 | 90862036 | 90871150 | 9115 | -9.27 | 7.42E-18 | 5.66E-17 |
| chr16 | 28246144 | 28265153 | 19010 | -9.39 | 7.42E-18 | 5.67E-17 |
| chr9 | 98085313 | 98095505 | 10193 | -9.35 | 7.44E-18 | 5.68E-17 |
| chr18 | 76702909 | 76724530 | 21622 | -9.24 | 7.45E-18 | 5.68E-17 |
| chr4 | 101138261 | 101162432 | 24172 | -9.29 | 7.49E-18 | 5.72E-17 |
| chr8 | 144785547 | 144793033 | 7487 | -9.29 | 7.51E-18 | 5.73E-17 |
| chr5 | 146681862 | 146695268 | 13407 | -9.26 | 7.52E-18 | 5.73E-17 |
| chr1 | 172946824 | 172972117 | 25294 | -9.5 | 7.53E-18 | 5.74E-17 |
| chr20 | 32534268 | 32540251 | 5984 | -9.25 | 7.54E-18 | 5.75E-17 |
| chr6 | 160612707 | 160648260 | 35554 | -9.53 | 7.56E-18 | 5.76E-17 |
| chr2 | 28525372 | 28532338 | 6967 | -9.25 | 7.67E-18 | 5.85E-17 |
| chr12 | 109690250 | 109708983 | 18734 | -9.22 | 7.68E-18 | 5.85E-17 |
| chr2 | 157506627 | 157538696 | 32070 | 9.31 | 7.69E-18 | 5.86E-17 |
| chr6 | 140328592 | 140359787 | 31196 | -9.22 | 7.73E-18 | 5.89E-17 |
| chr4 | 8514591 | 8546129 | 31539 | -9.22 | 7.74E-18 | 5.89E-17 |
| chr10 | 33683074 | 33697736 | 14663 | -9.61 | 7.79E-18 | 5.93E-17 |
| chr6 | 25132687 | 25158682 | 25996 | -9.23 | 7.83E-18 | 5.96E-17 |
| chr13 | 33556674 | 33618176 | 61503 | -1.73 | 7.84E-18 | 5.97E-17 |
| chr2 | 109790671 | 109799179 | 8509 | -9.22 | 7.88E-18 | 6.00E-17 |
| chr22 | 40965766 | 40980683 | 14918 | -9.22 | 7.89E-18 | 6.00E-17 |
| chr7 | 46685894 | 46699585 | 13692 | -6.57 | 7.89E-18 | 6.00E-17 |
| chr3 | 158968466 | 158972840 | 4375 | -9.22 | 7.89E-18 | 6.00E-17 |
| chr8 | 118119156 | 118132693 | 13538 | 9.27 | 7.90E-18 | 6.01E-17 |
| chr6 | 86129431 | 86153003 | 23573 | -9.3 | 7.91E-18 | 6.01E-17 |
| chr7 | 23816693 | 23851758 | 35066 | -9.22 | 7.91E-18 | 6.01E-17 |
| chr7 | 118731629 | 118794153 | 62525 | -9.48 | 7.93E-18 | 6.02E-17 |
| chr10 | 33583989 | 33618494 | 34506 | -9.51 | 7.94E-18 | 6.03E-17 |
| chr13 | 95909459 | 95919511 | 10053 | -9.22 | 7.95E-18 | 6.04E-17 |

| | | | | | | |
|---|---|---|---|---|---|---|
| chr1 | 219480438 | 219501568 | 21131 | -9.26 | 7.98E-18 | 6.06E-17 |
| chr13 | 49563402 | 49575317 | 11916 | -9.32 | 7.98E-18 | 6.06E-17 |
| chr12 | 67949588 | 67964138 | 14551 | -7.02 | 8.07E-18 | 6.12E-17 |
| chr5 | 115306740 | 115327566 | 20827 | -9.34 | 8.09E-18 | 6.14E-17 |
| chr17 | 66390637 | 66413823 | 23187 | -9.39 | 8.13E-18 | 6.17E-17 |
| chr3 | 119234673 | 119240415 | 5743 | -9.3 | 8.17E-18 | 6.20E-17 |
| chr1 | 118408503 | 118419449 | 10947 | -9.21 | 8.17E-18 | 6.20E-17 |
| chr6 | 145478406 | 145506876 | 28471 | -9.36 | 8.22E-18 | 6.23E-17 |
| chr1 | 234277085 | 234284581 | 7497 | -9.56 | 8.23E-18 | 6.24E-17 |
| chr17 | 9581693 | 9599858 | 18166 | -9.25 | 8.29E-18 | 6.29E-17 |
| chr12 | 21470169 | 21487495 | 17327 | -9.31 | 8.30E-18 | 6.29E-17 |
| chr2 | 224106555 | 224127325 | 20771 | -9.21 | 8.32E-18 | 6.31E-17 |
| chr10 | 123895200 | 123905952 | 10753 | -9.21 | 8.37E-18 | 6.34E-17 |
| chr11 | 4044277 | 4059438 | 15162 | -9.28 | 8.44E-18 | 6.39E-17 |
| chr14 | 71027264 | 71047566 | 20303 | -9.33 | 8.50E-18 | 6.44E-17 |
| chr14 | 66813469 | 66828767 | 15299 | -9.34 | 8.50E-18 | 6.44E-17 |
| chr15 | 34340653 | 34346809 | 6157 | -9.23 | 8.55E-18 | 6.47E-17 |
| chr5 | 150164143 | 150381387 | 217245 | -3.1 | 8.55E-18 | 6.47E-17 |
| chr11 | 130210275 | 130230100 | 19826 | -9.21 | 8.56E-18 | 6.48E-17 |
| chr14 | 76249970 | 76267261 | 17292 | -9.23 | 8.61E-18 | 6.52E-17 |
| chr4 | 88162187 | 88184737 | 22551 | -9.29 | 8.65E-18 | 6.54E-17 |
| chr1 | 35134034 | 35169512 | 35479 | -9.23 | 8.67E-18 | 6.56E-17 |
| chr10 | 103942995 | 103953784 | 10790 | -9.42 | 8.70E-18 | 6.58E-17 |
| chr4 | 146013669 | 146017250 | 3582 | -9.34 | 8.73E-18 | 6.60E-17 |
| chr17 | 66463049 | 66482209 | 19161 | -9.49 | 8.74E-18 | 6.60E-17 |
| chrX | 56763963 | 56774068 | 10106 | -9.23 | 8.75E-18 | 6.61E-17 |
| chr10 | 122791435 | 122842142 | 50708 | -9.21 | 8.75E-18 | 6.61E-17 |
| chr10 | 90141688 | 90160531 | 18844 | -9.32 | 8.79E-18 | 6.64E-17 |
| chr12 | 19853217 | 19879909 | 26693 | -9.2 | 8.83E-18 | 6.67E-17 |
| chr12 | 64152509 | 64168177 | 15669 | -9.26 | 8.83E-18 | 6.67E-17 |
| chr6 | 125601738 | 125610814 | 9077 | -9.33 | 8.83E-18 | 6.67E-17 |
| chr4 | 169267931 | 169291752 | 23822 | -9.21 | 8.84E-18 | 6.67E-17 |
| chr16 | 72742788 | 72750635 | 7848 | -9.25 | 8.88E-18 | 6.70E-17 |
| chr17 | 64079134 | 64118758 | 39625 | -9.5 | 8.91E-18 | 6.72E-17 |
| chr14 | 27072186 | 27096622 | 24437 | -9.46 | 8.97E-18 | 6.77E-17 |
| chr19 | 17078724 | 17110459 | 31736 | -9.48 | 8.98E-18 | 6.77E-17 |
| chr17 | 58991634 | 58995270 | 3637 | -9.21 | 8.99E-18 | 6.78E-17 |
| chr17 | 78760654 | 78782728 | 22075 | -9.54 | 9.04E-18 | 6.82E-17 |
| chr2 | 231292487 | 231302987 | 10501 | -9.21 | 9.05E-18 | 6.82E-17 |
| chr2 | 188072131 | 188110283 | 38153 | 9.26 | 9.06E-18 | 6.83E-17 |
| chr18 | 43669078 | 43672548 | 3471 | -9.21 | 9.08E-18 | 6.84E-17 |
| chr6 | 48389650 | 48420865 | 31216 | -9.35 | 9.10E-18 | 6.85E-17 |
| chr16 | 70835638 | 70839905 | 4268 | -9.21 | 9.13E-18 | 6.88E-17 |
| chr5 | 169142478 | 169161930 | 19453 | -9.21 | 9.13E-18 | 6.88E-17 |
| chr15 | 85701516 | 85717548 | 16033 | -9.47 | 9.14E-18 | 6.88E-17 |
| chr7 | 141257760 | 141260936 | 3177 | -9.25 | 9.17E-18 | 6.90E-17 |
| chr8 | 131381999 | 131384617 | 2619 | -9.28 | 9.18E-18 | 6.91E-17 |
| chr15 | 67636536 | 67646585 | 10050 | -9.21 | 9.22E-18 | 6.93E-17 |
| chr4 | 81026227 | 81040252 | 14026 | -9.3 | 9.23E-18 | 6.94E-17 |
| chr14 | 34876656 | 34902144 | 25489 | -9.51 | 9.34E-18 | 7.03E-17 |
| chr18 | 55538875 | 55568615 | 29741 | -9.54 | 9.37E-18 | 7.04E-17 |
| chr16 | 23751012 | 23778452 | 27441 | -9.28 | 9.38E-18 | 7.05E-17 |
| chr10 | 21329254 | 21349911 | 20658 | -9.25 | 9.48E-18 | 7.12E-17 |
| chr4 | 6058182 | 6071243 | 13062 | -9.22 | 9.58E-18 | 7.20E-17 |
| chr6 | 142517070 | 142536031 | 18962 | -9.2 | 9.59E-18 | 7.20E-17 |
| chr4 | 119992252 | 120035270 | 43019 | -9.31 | 9.69E-18 | 7.28E-17 |
| chr1 | 221391762 | 221410797 | 19036 | -9.22 | 9.70E-18 | 7.29E-17 |
| chr15 | 71242552 | 71254618 | 12067 | -9.38 | 9.71E-18 | 7.29E-17 |
| chr5 | 126909424 | 126918119 | 8696 | -9.19 | 9.75E-18 | 7.32E-17 |
| chr1 | 240173309 | 240183568 | 10260 | -9.26 | 9.76E-18 | 7.33E-17 |
| chr18 | 7710756 | 7747730 | 36975 | -9.26 | 9.77E-18 | 7.33E-17 |

| | | | | | | |
|---|---|---|---|---|---|---|
| chrX | 58312241 | 58329381 | 17141 | -9.2 | 9.80E-18 | 7.36E-17 |
| chr14 | 54220250 | 54254508 | 34259 | -9.61 | 9.81E-18 | 7.36E-17 |
| chr10 | 77309530 | 77343894 | 34365 | -2.46 | 9.82E-18 | 7.36E-17 |
| chr11 | 13446773 | 13452168 | 5396 | -9.28 | 9.82E-18 | 7.36E-17 |
| chr1 | 209245584 | 209266959 | 21376 | -9.33 | 9.86E-18 | 7.39E-17 |
| chr6 | 11340818 | 11396992 | 56175 | -9.33 | 1.00E-17 | 7.50E-17 |
| chr17 | 413631 | 432265 | 18635 | -9.44 | 1.01E-17 | 7.54E-17 |
| chr16 | 88003339 | 88007738 | 4400 | -9.19 | 1.01E-17 | 7.58E-17 |
| chr3 | 38145468 | 38148464 | 2997 | -9.2 | 1.02E-17 | 7.67E-17 |
| chr19 | 36414733 | 36417710 | 2978 | -9.19 | 1.02E-17 | 7.67E-17 |
| chr14 | 37623541 | 37631619 | 8079 | -9.22 | 1.03E-17 | 7.69E-17 |
| chr4 | 67417301 | 67439716 | 22416 | -9.19 | 1.03E-17 | 7.73E-17 |
| chrX | 57201665 | 57222317 | 20653 | -9.22 | 1.04E-17 | 7.75E-17 |
| chr7 | 21617016 | 21645388 | 28373 | -9.46 | 1.04E-17 | 7.75E-17 |
| chr9 | 123565131 | 123573210 | 8080 | -9.19 | 1.04E-17 | 7.77E-17 |
| chr5 | 52506857 | 52527022 | 20166 | -9.35 | 1.04E-17 | 7.78E-17 |
| chr4 | 147714823 | 147757098 | 42276 | 9.45 | 1.05E-17 | 7.86E-17 |
| chr18 | 11601275 | 11603996 | 2722 | -3.77 | 1.05E-17 | 7.88E-17 |
| chr8 | 25984158 | 26021063 | 36906 | -9.25 | 1.06E-17 | 7.95E-17 |
| chr8 | 58392066 | 58411429 | 19364 | -9.28 | 1.08E-17 | 8.06E-17 |
| chr5 | 162813560 | 162822086 | 8527 | -9.22 | 1.08E-17 | 8.07E-17 |
| chr4 | 76884462 | 76899359 | 14898 | -9.26 | 1.08E-17 | 8.09E-17 |
| chr3 | 110196819 | 110219775 | 22957 | -9.44 | 1.09E-17 | 8.11E-17 |
| chr2 | 30720008 | 30774931 | 54924 | -9.37 | 1.09E-17 | 8.11E-17 |
| chr21 | 27665962 | 27677811 | 11850 | -9.26 | 1.09E-17 | 8.12E-17 |
| chr12 | 74629033 | 74640492 | 11460 | -9.18 | 1.09E-17 | 8.15E-17 |
| chr4 | 177904666 | 177935592 | 30927 | -9.22 | 1.10E-17 | 8.19E-17 |
| chr5 | 134482551 | 134497657 | 15107 | -9.21 | 1.10E-17 | 8.22E-17 |
| chr4 | 140935073 | 140938365 | 3293 | -9.18 | 1.11E-17 | 8.25E-17 |
| chr5 | 32336039 | 32342340 | 6302 | -9.28 | 1.11E-17 | 8.27E-17 |
| chr9 | 140596401 | 140612716 | 16316 | -9.18 | 1.11E-17 | 8.31E-17 |
| chr4 | 145795230 | 145822744 | 27515 | 9.24 | 1.12E-17 | 8.36E-17 |
| chr10 | 77515792 | 77545967 | 30176 | 9.3 | 1.13E-17 | 8.40E-17 |
| chr12 | 62902691 | 62909750 | 7060 | -9.18 | 1.14E-17 | 8.51E-17 |
| chr2 | 240680734 | 240723634 | 42901 | -3.59 | 1.15E-17 | 8.55E-17 |
| chr11 | 73850671 | 73865654 | 14984 | -9.19 | 1.15E-17 | 8.56E-17 |
| chr20 | 10436443 | 10452873 | 16431 | -9.36 | 1.15E-17 | 8.56E-17 |
| chr20 | 24887063 | 24890760 | 3698 | -9.17 | 1.16E-17 | 8.63E-17 |
| chr7 | 12730320 | 12744853 | 14534 | -9.38 | 1.16E-17 | 8.66E-17 |
| chr6 | 95540240 | 95679750 | 139511 | 1.96 | 1.16E-17 | 8.67E-17 |
| chr20 | 4417512 | 4440716 | 23205 | -9.4 | 1.17E-17 | 8.68E-17 |
| chr1 | 236520437 | 236552148 | 31712 | -9.65 | 1.17E-17 | 8.68E-17 |
| chr2 | 119014375 | 119018710 | 4336 | -7.93 | 1.17E-17 | 8.69E-17 |
| chr15 | 99831366 | 99836186 | 4821 | -9.18 | 1.17E-17 | 8.73E-17 |
| chr3 | 160879831 | 160895096 | 15266 | -9.19 | 1.18E-17 | 8.74E-17 |
| chr9 | 106895341 | 106898569 | 3229 | -9.45 | 1.18E-17 | 8.77E-17 |
| chr7 | 111579657 | 111585869 | 6213 | -9.19 | 1.18E-17 | 8.77E-17 |
| chr1 | 42482958 | 42493748 | 10791 | -9.27 | 1.18E-17 | 8.80E-17 |
| chr10 | 74636888 | 74639747 | 2860 | -9.17 | 1.19E-17 | 8.82E-17 |
| chr10 | 8884104 | 8894128 | 10025 | -9.21 | 1.19E-17 | 8.82E-17 |
| chr9 | 38623485 | 38626778 | 3294 | -9.25 | 1.19E-17 | 8.86E-17 |
| chr16 | 49738160 | 49766956 | 28797 | -9.39 | 1.20E-17 | 8.88E-17 |
| chr11 | 60921216 | 60925285 | 4070 | -9.19 | 1.20E-17 | 8.89E-17 |
| chr14 | 27349684 | 27371149 | 21466 | -9.24 | 1.20E-17 | 8.92E-17 |
| chr4 | 1380644 | 1393035 | 12392 | -9.21 | 1.20E-17 | 8.93E-17 |
| chr12 | 101526566 | 101539861 | 13296 | 2.4 | 1.20E-17 | 8.93E-17 |
| chr3 | 171420534 | 171436811 | 16278 | -9.18 | 1.21E-17 | 8.96E-17 |
| chr16 | 79876180 | 79896498 | 20319 | -9.2 | 1.21E-17 | 8.96E-17 |
| chr2 | 218343106 | 218359413 | 16308 | -9.34 | 1.21E-17 | 8.98E-17 |
| chr6 | 141851759 | 141876518 | 24760 | -9.28 | 1.21E-17 | 9.00E-17 |
| chr14 | 55289173 | 55299514 | 10342 | -9.2 | 1.22E-17 | 9.02E-17 |

| | | | | | | |
|---|---|---|---|---|---|---|
| chr6 | 137991717 | 138013355 | 21639 | -9.51 | 1.22E-17 | 9.05E-17 |
| chr16 | 69668352 | 69672304 | 3953 | -9.28 | 1.22E-17 | 9.05E-17 |
| chr10 | 87348406 | 87371399 | 22994 | -9.43 | 1.23E-17 | 9.13E-17 |
| chr15 | 72222630 | 72226886 | 4257 | -9.24 | 1.23E-17 | 9.14E-17 |
| chr4 | 109439796 | 109482172 | 42377 | 1.96 | 1.24E-17 | 9.20E-17 |
| chr3 | 145952581 | 145957631 | 5051 | -9.17 | 1.24E-17 | 9.21E-17 |
| chr11 | 116310734 | 116323006 | 12273 | -9.17 | 1.25E-17 | 9.23E-17 |
| chr10 | 13681834 | 13687148 | 5315 | -9.18 | 1.25E-17 | 9.27E-17 |
| chr3 | 150194040 | 150236560 | 42521 | -9.17 | 1.25E-17 | 9.27E-17 |
| chr5 | 172235718 | 172254577 | 18860 | -9.35 | 1.25E-17 | 9.28E-17 |
| chrX | 4740144 | 4758243 | 18100 | -9.24 | 1.26E-17 | 9.30E-17 |
| chr3 | 127655780 | 127686201 | 30422 | -9.17 | 1.26E-17 | 9.30E-17 |
| chr8 | 67149249 | 67178778 | 29530 | -9.27 | 1.26E-17 | 9.32E-17 |
| chrX | 130592072 | 130617987 | 25916 | -9.2 | 1.26E-17 | 9.33E-17 |
| chrX | 80297344 | 80319405 | 22062 | 9.32 | 1.26E-17 | 9.33E-17 |
| chr7 | 17240226 | 17278084 | 37859 | -9.25 | 1.26E-17 | 9.34E-17 |
| chr8 | 92026235 | 92030141 | 3907 | -9.32 | 1.27E-17 | 9.37E-17 |
| chr16 | 4109476 | 4114761 | 5286 | -9.16 | 1.27E-17 | 9.39E-17 |
| chr15 | 80889590 | 80894587 | 4998 | -9.29 | 1.27E-17 | 9.40E-17 |
| chr1 | 91444313 | 91470916 | 26604 | -9.2 | 1.28E-17 | 9.43E-17 |
| chr1 | 18730214 | 18762848 | 32635 | -9.5 | 1.28E-17 | 9.47E-17 |
| chr8 | 59352011 | 59356168 | 4158 | -9.33 | 1.29E-17 | 9.52E-17 |
| chr2 | 11077132 | 11094863 | 17732 | -9.16 | 1.29E-17 | 9.53E-17 |
| chr1 | 193286774 | 193304291 | 17518 | -9.24 | 1.29E-17 | 9.56E-17 |
| chr11 | 57064092 | 57069527 | 5436 | -9.17 | 1.30E-17 | 9.62E-17 |
| chr14 | 90911545 | 90915692 | 4148 | -9.17 | 1.31E-17 | 9.63E-17 |
| chr16 | 31427662 | 31437696 | 10035 | -9.39 | 1.31E-17 | 9.63E-17 |
| chr7 | 152400210 | 152404160 | 3951 | -9.18 | 1.32E-17 | 9.72E-17 |
| chr16 | 12682818 | 12711544 | 28727 | -9.26 | 1.32E-17 | 9.74E-17 |
| chr2 | 65270215 | 65279888 | 9674 | -9.21 | 1.32E-17 | 9.74E-17 |
| chr15 | 101451991 | 101457173 | 5183 | -9.19 | 1.32E-17 | 9.74E-17 |
| chr5 | 43358612 | 43377279 | 18668 | -9.32 | 1.32E-17 | 9.76E-17 |
| chr10 | 50633260 | 50672097 | 38838 | -9.37 | 1.33E-17 | 9.77E-17 |
| chr6 | 14199986 | 14280811 | 80826 | -9.57 | 1.33E-17 | 9.77E-17 |
| chr17 | 64175152 | 64181496 | 6345 | -9.23 | 1.33E-17 | 9.78E-17 |
| chr13 | 73264776 | 73290906 | 26131 | -9.16 | 1.33E-17 | 9.79E-17 |
| chr16 | 53887557 | 53906449 | 18893 | -9.17 | 1.34E-17 | 9.87E-17 |
| chr15 | 80072913 | 80131399 | 58487 | -9.4 | 1.35E-17 | 9.91E-17 |
| chr7 | 12263331 | 12279055 | 15725 | -9.32 | 1.35E-17 | 9.92E-17 |
| chr10 | 105622990 | 105626806 | 3817 | -9.16 | 1.35E-17 | 9.93E-17 |
| chr20 | 2198455 | 2201782 | 3328 | -9.18 | 1.35E-17 | 9.93E-17 |
| chr4 | 187635891 | 187643336 | 7446 | -9.18 | 1.36E-17 | 9.97E-17 |
| chr8 | 17250611 | 17351852 | 101242 | -2.24 | 1.36E-17 | 9.97E-17 |
| chrX | 56061153 | 56086604 | 25452 | -9.17 | 1.36E-17 | 9.97E-17 |
| chr11 | 70936790 | 70941580 | 4791 | -9.18 | 1.36E-17 | 9.98E-17 |
| chr15 | 76382254 | 76418868 | 36615 | -9.7 | 1.36E-17 | 1.00E-16 |
| chr7 | 123448216 | 123452558 | 4343 | -9.17 | 1.37E-17 | 1.00E-16 |
| chr8 | 100382545 | 100395004 | 12460 | -9.17 | 1.37E-17 | 1.01E-16 |
| chr5 | 42949146 | 42955085 | 5940 | -9.16 | 1.38E-17 | 1.01E-16 |
| chr15 | 36919169 | 36929851 | 10683 | -9.16 | 1.38E-17 | 1.01E-16 |
| chr8 | 141980428 | 141990685 | 10258 | -9.27 | 1.38E-17 | 1.01E-16 |
| chr2 | 173375003 | 173380447 | 5445 | -9.17 | 1.39E-17 | 1.02E-16 |
| chr6 | 75211916 | 75226994 | 15079 | -9.37 | 1.39E-17 | 1.02E-16 |
| chr7 | 129883925 | 129900166 | 16242 | -9.2 | 1.39E-17 | 1.02E-16 |
| chr17 | 44917251 | 44921814 | 4564 | -9.16 | 1.39E-17 | 1.02E-16 |
| chr3 | 138716115 | 138732097 | 15983 | -9.19 | 1.40E-17 | 1.03E-16 |
| chr17 | 7876783 | 7879852 | 3070 | -9.17 | 1.40E-17 | 1.03E-16 |
| chr3 | 45689457 | 45702887 | 13431 | -9.15 | 1.41E-17 | 1.03E-16 |
| chr17 | 41866250 | 41877750 | 11501 | -9.16 | 1.41E-17 | 1.04E-16 |
| chr6 | 22760321 | 22796688 | 36368 | -9.33 | 1.41E-17 | 1.04E-16 |
| chr17 | 61581288 | 61584733 | 3446 | -2.53 | 1.42E-17 | 1.04E-16 |

| | | | | | | |
|---|---|---|---|---|---|---|
| chr6 | 11436433 | 11452493 | 16061 | -9.18 | 1.42E-17 | 1.04E-16 |
| chr4 | 5941718 | 5962826 | 21109 | -9.37 | 1.43E-17 | 1.05E-16 |
| chr1 | 213197663 | 213210323 | 12661 | -9.15 | 1.43E-17 | 1.05E-16 |
| chr3 | 6265848 | 6365901 | 100054 | 2.32 | 1.44E-17 | 1.05E-16 |
| chr10 | 30262594 | 30276441 | 13848 | -9.17 | 1.44E-17 | 1.06E-16 |
| chr13 | 47056163 | 47062231 | 6069 | -9.15 | 1.45E-17 | 1.06E-16 |
| chr10 | 22155454 | 22193238 | 37785 | -9.15 | 1.45E-17 | 1.06E-16 |
| chr15 | 30071989 | 30110690 | 38702 | -9.17 | 1.45E-17 | 1.06E-16 |
| chr2 | 84927672 | 84931329 | 3658 | -3.45 | 1.45E-17 | 1.06E-16 |
| chr3 | 177523540 | 177539726 | 16187 | -9.19 | 1.46E-17 | 1.07E-16 |
| chr7 | 75432297 | 75438020 | 5724 | -9.21 | 1.48E-17 | 1.08E-16 |
| chr6 | 28250441 | 28261354 | 10914 | -9.16 | 1.48E-17 | 1.08E-16 |
| chr4 | 47927361 | 47972980 | 45620 | -9.22 | 1.48E-17 | 1.08E-16 |
| chr4 | 159216046 | 159237655 | 21610 | -9.15 | 1.48E-17 | 1.08E-16 |
| chr5 | 169275868 | 169285838 | 9971 | -9.2 | 1.49E-17 | 1.09E-16 |
| chr18 | 53641156 | 53656588 | 15433 | -9.15 | 1.49E-17 | 1.09E-16 |
| chr12 | 17325011 | 17358582 | 33572 | -9.48 | 1.49E-17 | 1.09E-16 |
| chr10 | 14698759 | 14709367 | 10609 | -9.28 | 1.49E-17 | 1.09E-16 |
| chr3 | 110812671 | 110838578 | 25908 | -9.35 | 1.50E-17 | 1.10E-16 |
| chr8 | 144491911 | 144499215 | 7305 | -9.31 | 1.50E-17 | 1.10E-16 |
| chr6 | 99386195 | 99398794 | 12600 | -9.15 | 1.51E-17 | 1.10E-16 |
| chr2 | 133475953 | 133495983 | 20031 | 9.21 | 1.52E-17 | 1.11E-16 |
| chr12 | 18710741 | 18760289 | 49549 | -9.24 | 1.52E-17 | 1.11E-16 |
| chr19 | 40296512 | 40312288 | 15777 | -9.2 | 1.55E-17 | 1.13E-16 |
| chr14 | 58159584 | 58188665 | 29082 | -9.28 | 1.55E-17 | 1.13E-16 |
| chr8 | 126200146 | 126203562 | 3417 | -9.15 | 1.57E-17 | 1.14E-16 |
| chr11 | 60547220 | 60585992 | 38773 | -9.61 | 1.57E-17 | 1.14E-16 |
| chr18 | 12824946 | 12830079 | 5134 | -9.2 | 1.57E-17 | 1.15E-16 |
| chr3 | 126828559 | 126837344 | 8786 | -9.16 | 1.58E-17 | 1.15E-16 |
| chr6 | 9351936 | 9366829 | 14894 | -9.14 | 1.59E-17 | 1.16E-16 |
| chr11 | 28185740 | 28197169 | 11430 | -9.14 | 1.60E-17 | 1.17E-16 |
| chr4 | 123789193 | 123808461 | 19269 | -9.24 | 1.61E-17 | 1.17E-16 |
| chr14 | 76211483 | 76215385 | 3903 | -9.14 | 1.61E-17 | 1.17E-16 |
| chr16 | 86903594 | 86910466 | 6873 | -9.15 | 1.61E-17 | 1.17E-16 |
| chr1 | 180698121 | 180728035 | 29915 | -9.25 | 1.62E-17 | 1.18E-16 |
| chr2 | 128196432 | 128205459 | 9028 | -9.32 | 1.62E-17 | 1.18E-16 |
| chr15 | 75365961 | 75380112 | 14152 | -9.28 | 1.62E-17 | 1.18E-16 |
| chr15 | 69497138 | 69544888 | 47751 | -9.72 | 1.62E-17 | 1.18E-16 |
| chr13 | 91866410 | 91886530 | 20121 | 9.2 | 1.63E-17 | 1.18E-16 |
| chr11 | 13920996 | 13961323 | 40328 | -9.14 | 1.63E-17 | 1.18E-16 |
| chr9 | 130891456 | 130897940 | 6485 | -9.15 | 1.64E-17 | 1.19E-16 |
| chr7 | 34920706 | 34936200 | 15495 | -9.28 | 1.64E-17 | 1.19E-16 |
| chr18 | 77331670 | 77355262 | 23593 | -9.25 | 1.66E-17 | 1.20E-16 |
| chr8 | 27054465 | 27094107 | 39643 | -2.5 | 1.66E-17 | 1.20E-16 |
| chr8 | 39094472 | 39103509 | 9038 | -9.2 | 1.67E-17 | 1.21E-16 |
| chr2 | 28041743 | 28054778 | 13036 | -9.29 | 1.69E-17 | 1.22E-16 |
| chr9 | 117864723 | 117889039 | 24317 | -9.13 | 1.69E-17 | 1.23E-16 |
| chr19 | 36321070 | 36334665 | 13596 | -9.37 | 1.69E-17 | 1.23E-16 |
| chr12 | 94086581 | 94117547 | 30967 | -9.31 | 1.69E-17 | 1.23E-16 |
| chr10 | 8924667 | 8938620 | 13954 | -9.14 | 1.70E-17 | 1.23E-16 |
| chr3 | 183159500 | 183163443 | 3944 | -9.14 | 1.70E-17 | 1.23E-16 |
| chr7 | 138888749 | 138895825 | 7077 | -9.15 | 1.70E-17 | 1.24E-16 |
| chr15 | 79610966 | 79623609 | 12644 | -9.13 | 1.70E-17 | 1.24E-16 |
| chr2 | 28684090 | 28687262 | 3173 | -9.13 | 1.71E-17 | 1.24E-16 |
| chr1 | 89972511 | 89989610 | 17100 | -9.13 | 1.71E-17 | 1.24E-16 |
| chr1 | 43569361 | 43601552 | 32192 | -9.16 | 1.71E-17 | 1.24E-16 |
| chr1 | 101112178 | 101116556 | 4379 | -5.17 | 1.72E-17 | 1.24E-16 |
| chr10 | 4080902 | 4087024 | 6123 | -9.14 | 1.72E-17 | 1.24E-16 |
| chr8 | 100471211 | 100476015 | 4805 | -9.18 | 1.73E-17 | 1.26E-16 |
| chr2 | 220275685 | 220279226 | 3542 | -9.13 | 1.74E-17 | 1.26E-16 |
| chr11 | 45436834 | 45442548 | 5715 | -3.05 | 1.74E-17 | 1.26E-16 |

| | | | | | | |
|---|---|---|---|---|---|---|
| chr17 | 28278917 | 28343201 | 64285 | -9.6 | 1.74E-17 | 1.26E-16 |
| chr2 | 167103709 | 167137220 | 33512 | 9.23 | 1.74E-17 | 1.26E-16 |
| chr4 | 5983318 | 5989015 | 5698 | -9.13 | 1.74E-17 | 1.26E-16 |
| chr3 | 197288529 | 197306790 | 18262 | -9.31 | 1.74E-17 | 1.26E-16 |
| chr1 | 173430877 | 173442072 | 11196 | -9.24 | 1.74E-17 | 1.26E-16 |
| chr22 | 26202280 | 26227360 | 25081 | -9.18 | 1.75E-17 | 1.26E-16 |
| chr14 | 93439831 | 93457042 | 17212 | -9.16 | 1.75E-17 | 1.26E-16 |
| chr8 | 126538045 | 126551067 | 13023 | -9.36 | 1.75E-17 | 1.27E-16 |
| chr15 | 80766839 | 80769989 | 3151 | -9.13 | 1.76E-17 | 1.27E-16 |
| chr6 | 153466602 | 153491227 | 24626 | -9.44 | 1.77E-17 | 1.28E-16 |
| chr2 | 120546211 | 120590095 | 43885 | -9.3 | 1.77E-17 | 1.28E-16 |
| chr11 | 110598341 | 111096781 | 498441 | 3.86 | 1.77E-17 | 1.28E-16 |
| chr14 | 91903060 | 91913598 | 10539 | -9.41 | 1.79E-17 | 1.29E-16 |
| chr1 | 245887220 | 245897535 | 10316 | -9.13 | 1.79E-17 | 1.30E-16 |
| chr2 | 210234463 | 210247328 | 12866 | -9.16 | 1.80E-17 | 1.30E-16 |
| chr1 | 12617003 | 12637666 | 20664 | -9.27 | 1.80E-17 | 1.30E-16 |
| chr2 | 10026241 | 10029029 | 2789 | -9.3 | 1.80E-17 | 1.30E-16 |
| chrX | 46594254 | 46602105 | 7852 | -9.13 | 1.81E-17 | 1.30E-16 |
| chr2 | 121481801 | 121486471 | 4671 | -9.13 | 1.81E-17 | 1.31E-16 |
| chr1 | 233354633 | 233358228 | 3596 | -9.16 | 1.81E-17 | 1.31E-16 |
| chr21 | 47258685 | 47264072 | 5388 | -9.13 | 1.81E-17 | 1.31E-16 |
| chr13 | 48639927 | 48647449 | 7523 | -9.15 | 1.82E-17 | 1.31E-16 |
| chr16 | 75630650 | 75637953 | 7304 | -9.17 | 1.83E-17 | 1.32E-16 |
| chr1 | 33524765 | 33536703 | 11939 | -9.16 | 1.83E-17 | 1.32E-16 |
| chr11 | 56431920 | 56453326 | 21407 | 9.45 | 1.83E-17 | 1.32E-16 |
| chr13 | 66868275 | 66894105 | 25831 | -9.28 | 1.83E-17 | 1.32E-16 |
| chr9 | 22440344 | 22446047 | 5704 | -9.12 | 1.83E-17 | 1.32E-16 |
| chr8 | 88260611 | 88291761 | 31151 | -9.13 | 1.84E-17 | 1.32E-16 |
| chr20 | 48237747 | 48246767 | 9021 | -9.16 | 1.84E-17 | 1.32E-16 |
| chr5 | 39832334 | 39849371 | 17038 | -9.23 | 1.84E-17 | 1.32E-16 |
| chr2 | 161609710 | 161619991 | 10282 | -9.13 | 1.84E-17 | 1.33E-16 |
| chr1 | 207555475 | 207567018 | 11544 | -9.27 | 1.84E-17 | 1.33E-16 |
| chr4 | 157649723 | 157665405 | 15683 | -9.14 | 1.86E-17 | 1.34E-16 |
| chr14 | 37673886 | 37695650 | 21765 | -9.4 | 1.86E-17 | 1.34E-16 |
| chr12 | 89302237 | 89319024 | 16788 | -9.2 | 1.86E-17 | 1.34E-16 |
| chr11 | 87044413 | 87051054 | 6642 | -9.15 | 1.87E-17 | 1.34E-16 |
| chr3 | 123447570 | 123479353 | 31784 | -9.36 | 1.87E-17 | 1.34E-16 |
| chr12 | 622402 | 642756 | 20355 | -9.45 | 1.87E-17 | 1.34E-16 |
| chr6 | 153384316 | 153410059 | 25744 | -9.13 | 1.87E-17 | 1.35E-16 |
| chr2 | 190091558 | 190110449 | 18892 | -9.12 | 1.87E-17 | 1.35E-16 |
| chr1 | 246417609 | 246425878 | 8270 | -9.21 | 1.88E-17 | 1.35E-16 |
| chr12 | 14672001 | 14679937 | 7937 | -9.12 | 1.88E-17 | 1.35E-16 |
| chr14 | 22757813 | 22770927 | 13115 | -9.18 | 1.88E-17 | 1.35E-16 |
| chr20 | 56198569 | 56222555 | 23987 | -9.14 | 1.88E-17 | 1.35E-16 |
| chr9 | 86016060 | 86050546 | 34487 | -9.17 | 1.90E-17 | 1.36E-16 |
| chr4 | 106988224 | 107012687 | 24464 | -9.13 | 1.90E-17 | 1.36E-16 |
| chr17 | 39497807 | 39515138 | 17332 | -9.28 | 1.90E-17 | 1.37E-16 |
| chr7 | 36469315 | 36486861 | 17547 | -9.58 | 1.91E-17 | 1.37E-16 |
| chr17 | 55088720 | 55107580 | 18861 | -9.34 | 1.91E-17 | 1.37E-16 |
| chr6 | 97607051 | 97610860 | 3810 | -9.31 | 1.91E-17 | 1.37E-16 |
| chr13 | 80292544 | 80311531 | 18988 | -9.15 | 1.91E-17 | 1.37E-16 |
| chr21 | 40944263 | 40968413 | 24151 | -9.13 | 1.92E-17 | 1.38E-16 |
| chr1 | 234574069 | 234577436 | 3368 | -9.12 | 1.93E-17 | 1.38E-16 |
| chr1 | 212938786 | 212945441 | 6656 | -9.13 | 1.93E-17 | 1.38E-16 |
| chr6 | 140705016 | 140738728 | 33713 | -9.44 | 1.94E-17 | 1.39E-16 |
| chr11 | 31485393 | 31490223 | 4831 | -9.12 | 1.94E-17 | 1.39E-16 |
| chr1 | 48299051 | 48302603 | 3553 | -7.8 | 1.95E-17 | 1.39E-16 |
| chr8 | 21868438 | 21872503 | 4066 | -9.16 | 1.95E-17 | 1.39E-16 |
| chr14 | 61303340 | 61324336 | 20997 | -9.33 | 1.95E-17 | 1.39E-16 |
| chr3 | 160266229 | 160270012 | 3784 | -9.17 | 1.95E-17 | 1.40E-16 |
| chr2 | 225424568 | 225433241 | 8674 | -9.31 | 1.95E-17 | 1.40E-16 |

| | | | | | | |
|---|---|---|---|---|---|---|
| chr16 | 53283905 | 53295240 | 11336 | -9.28 | 1.95E-17 | 1.40E-16 |
| chr3 | 45846662 | 45863651 | 16990 | -6.46 | 1.96E-17 | 1.40E-16 |
| chr12 | 19479667 | 19503442 | 23776 | -9.16 | 1.96E-17 | 1.40E-16 |
| chr8 | 134174258 | 134182132 | 7875 | -9.2 | 1.96E-17 | 1.41E-16 |
| chr1 | 165766513 | 165770870 | 4358 | -9.12 | 1.97E-17 | 1.41E-16 |
| chr14 | 96603886 | 96622080 | 18195 | -9.44 | 1.97E-17 | 1.41E-16 |
| chr6 | 148366180 | 148394531 | 28352 | -9.45 | 1.97E-17 | 1.41E-16 |
| chr12 | 112342151 | 112348939 | 6789 | -9.18 | 1.97E-17 | 1.41E-16 |
| chr2 | 110970858 | 110982025 | 11168 | -2.89 | 1.97E-17 | 1.41E-16 |
| chr5 | 56360773 | 56367672 | 6900 | -9.23 | 1.98E-17 | 1.41E-16 |
| chr22 | 49225038 | 49229826 | 4789 | -9.18 | 1.98E-17 | 1.41E-16 |
| chr1 | 187906859 | 187923776 | 16918 | -9.14 | 1.98E-17 | 1.42E-16 |
| chr10 | 15101466 | 15110940 | 9475 | -9.12 | 1.99E-17 | 1.42E-16 |
| chr3 | 189144357 | 189181761 | 37405 | -9.36 | 1.99E-17 | 1.42E-16 |
| chr12 | 6516780 | 6527514 | 10735 | -9.16 | 2.00E-17 | 1.43E-16 |
| chr18 | 29551747 | 29559879 | 8133 | -9.21 | 2.01E-17 | 1.43E-16 |
| chr4 | 64192069 | 64225879 | 33811 | -9.12 | 2.01E-17 | 1.43E-16 |
| chr3 | 38505625 | 38513035 | 7411 | -9.16 | 2.02E-17 | 1.44E-16 |
| chr13 | 109422982 | 109438153 | 15172 | -9.2 | 2.02E-17 | 1.44E-16 |
| chr7 | 106180462 | 106205926 | 25465 | 9.26 | 2.02E-17 | 1.44E-16 |
| chr5 | 157079251 | 157089914 | 10664 | -9.11 | 2.03E-17 | 1.44E-16 |
| chr12 | 83874964 | 83918156 | 43193 | -9.15 | 2.03E-17 | 1.44E-16 |
| chr3 | 186978233 | 186991514 | 13282 | -9.13 | 2.03E-17 | 1.44E-16 |
| chr6 | 109676606 | 109685381 | 8776 | -9.16 | 2.03E-17 | 1.45E-16 |
| chr3 | 95389320 | 95424823 | 35504 | -9.14 | 2.03E-17 | 1.45E-16 |
| chr3 | 70185227 | 70197880 | 12654 | -9.12 | 2.03E-17 | 1.45E-16 |
| chr7 | 34651568 | 34691125 | 39558 | -9.29 | 2.03E-17 | 1.45E-16 |
| chr14 | 57095163 | 57106503 | 11341 | -9.13 | 2.03E-17 | 1.45E-16 |
| chr4 | 102673995 | 102721569 | 47575 | -9.23 | 2.04E-17 | 1.46E-16 |
| chr7 | 67882603 | 67933151 | 50549 | -9.51 | 2.05E-17 | 1.46E-16 |
| chr17 | 2924171 | 2928310 | 4140 | -9.28 | 2.06E-17 | 1.46E-16 |
| chr11 | 49915158 | 50036455 | 121298 | 1.36 | 2.07E-17 | 1.48E-16 |
| chr4 | 68463733 | 68480518 | 16786 | -9.11 | 2.10E-17 | 1.49E-16 |
| chr5 | 74802972 | 74831014 | 28043 | -9.16 | 2.11E-17 | 1.50E-16 |
| chr3 | 188897342 | 188915699 | 18358 | -9.19 | 2.11E-17 | 1.50E-16 |
| chr9 | 73222574 | 73240225 | 17652 | 9.17 | 2.11E-17 | 1.50E-16 |
| chr18 | 44771434 | 44776306 | 4873 | -4.88 | 2.11E-17 | 1.50E-16 |
| chr5 | 73004119 | 73007217 | 3099 | -9.11 | 2.13E-17 | 1.51E-16 |
| chr3 | 4780429 | 4809983 | 29555 | -9.26 | 2.13E-17 | 1.51E-16 |
| chr18 | 13594374 | 13601861 | 7488 | -9.11 | 2.14E-17 | 1.52E-16 |
| chr12 | 54270298 | 54277340 | 7043 | -3.23 | 2.14E-17 | 1.52E-16 |
| chr4 | 176320209 | 176345562 | 25354 | -9.16 | 2.14E-17 | 1.52E-16 |
| chr12 | 117942937 | 117993605 | 50669 | -9.48 | 2.14E-17 | 1.52E-16 |
| chr2 | 106818228 | 106854203 | 35976 | -5.13 | 2.14E-17 | 1.52E-16 |
| chr17 | 18568929 | 18582183 | 13255 | -9.19 | 2.14E-17 | 1.52E-16 |
| chr18 | 13700405 | 13718859 | 18455 | -9.23 | 2.16E-17 | 1.53E-16 |
| chr5 | 124638222 | 124656492 | 18271 | -9.32 | 2.16E-17 | 1.53E-16 |
| chr2 | 72180729 | 72227067 | 46339 | -1.87 | 2.16E-17 | 1.53E-16 |
| chr8 | 83251814 | 83290596 | 38783 | -9.12 | 2.17E-17 | 1.54E-16 |
| chr14 | 35579379 | 35582732 | 3354 | -9.11 | 2.18E-17 | 1.54E-16 |
| chr7 | 68258209 | 68297916 | 39708 | -9.65 | 2.18E-17 | 1.54E-16 |
| chr11 | 103655288 | 103707995 | 52708 | -9.49 | 2.19E-17 | 1.55E-16 |
| chr20 | 56792152 | 56796510 | 4359 | -9.1 | 2.20E-17 | 1.56E-16 |
| chr4 | 66058738 | 66082759 | 24022 | -7.08 | 2.22E-17 | 1.57E-16 |
| chr5 | 91748036 | 91772027 | 23992 | -9.43 | 2.23E-17 | 1.58E-16 |
| chr3 | 49251188 | 49276569 | 25382 | -9.12 | 2.23E-17 | 1.58E-16 |
| chr2 | 152324221 | 152330799 | 6579 | -9.13 | 2.23E-17 | 1.58E-16 |
| chr14 | 76592492 | 76617937 | 25446 | -9.3 | 2.23E-17 | 1.58E-16 |
| chr13 | 25379595 | 25398387 | 18793 | -9.13 | 2.23E-17 | 1.58E-16 |
| chrX | 4608400 | 4645265 | 36866 | -9.35 | 2.25E-17 | 1.59E-16 |
| chr12 | 82926928 | 82946950 | 20023 | -9.2 | 2.26E-17 | 1.60E-16 |

| | | | | | | |
|---|---|---|---|---|---|---|
| chr12 | 31759251 | 31765421 | 6171 | -9.17 | 2.27E-17 | 1.61E-16 |
| chr3 | 98353160 | 98400088 | 46929 | -9.53 | 2.27E-17 | 1.61E-16 |
| chr11 | 45299664 | 45310997 | 11334 | -9.11 | 2.27E-17 | 1.61E-16 |
| chr6 | 117752248 | 117767336 | 15089 | -9.1 | 2.29E-17 | 1.62E-16 |
| chr15 | 89555428 | 89569853 | 14426 | -9.1 | 2.29E-17 | 1.62E-16 |
| chr3 | 180946916 | 180962276 | 15361 | 9.26 | 2.30E-17 | 1.62E-16 |
| chr9 | 91355678 | 91394334 | 38657 | -9.32 | 2.31E-17 | 1.63E-16 |
| chrX | 4941989 | 4969935 | 27947 | -9.11 | 2.31E-17 | 1.63E-16 |
| chr7 | 21969854 | 21978878 | 9025 | -9.13 | 2.33E-17 | 1.65E-16 |
| chr9 | 139645050 | 139651514 | 6465 | -9.23 | 2.34E-17 | 1.65E-16 |
| chr10 | 33872694 | 33909577 | 36884 | -9.46 | 2.34E-17 | 1.66E-16 |
| chr1 | 189944177 | 189971929 | 27753 | -9.44 | 2.35E-17 | 1.66E-16 |
| chr4 | 129508475 | 129520709 | 12235 | -9.17 | 2.35E-17 | 1.66E-16 |
| chr7 | 46174775 | 46196389 | 21615 | -9.1 | 2.37E-17 | 1.67E-16 |
| chr7 | 48924312 | 48953477 | 29166 | -9.16 | 2.38E-17 | 1.68E-16 |
| chr7 | 130989698 | 131002776 | 13079 | -9.42 | 2.38E-17 | 1.68E-16 |
| chr12 | 52896486 | 52906176 | 9691 | -9.1 | 2.38E-17 | 1.68E-16 |
| chr4 | 170132982 | 170137983 | 5002 | -9.1 | 2.38E-17 | 1.68E-16 |
| chr3 | 73031348 | 73036853 | 5506 | -9.1 | 2.38E-17 | 1.68E-16 |
| chr5 | 6958609 | 7047186 | 88578 | 1.93 | 2.38E-17 | 1.68E-16 |
| chr5 | 140322456 | 140335549 | 13094 | -9.1 | 2.39E-17 | 1.69E-16 |
| chr6 | 11500782 | 11520229 | 19448 | -9.11 | 2.40E-17 | 1.69E-16 |
| chr1 | 30312465 | 30317322 | 4858 | -9.09 | 2.41E-17 | 1.70E-16 |
| chr1 | 63965376 | 63981820 | 16445 | -9.13 | 2.41E-17 | 1.70E-16 |
| chr17 | 4362488 | 4365296 | 2809 | -9.12 | 2.41E-17 | 1.70E-16 |
| chr14 | 60729051 | 60733951 | 4901 | -9.16 | 2.41E-17 | 1.70E-16 |
| chr20 | 11158353 | 11168529 | 10177 | -9.1 | 2.41E-17 | 1.70E-16 |
| chr18 | 36517469 | 36523571 | 6103 | -3.43 | 2.42E-17 | 1.70E-16 |
| chr2 | 128077267 | 128080117 | 2851 | -9.22 | 2.42E-17 | 1.70E-16 |
| chr3 | 102950993 | 103001943 | 50951 | -9.19 | 2.46E-17 | 1.73E-16 |
| chr15 | 79737653 | 79741980 | 4328 | -9.1 | 2.46E-17 | 1.73E-16 |
| chr2 | 237779307 | 237794562 | 15256 | -6.58 | 2.46E-17 | 1.73E-16 |
| chr12 | 46781558 | 46843122 | 61565 | -9.93 | 2.47E-17 | 1.73E-16 |
| chr12 | 114223035 | 114229869 | 6835 | -9.09 | 2.47E-17 | 1.74E-16 |
| chr4 | 56423943 | 56456033 | 32091 | -9.1 | 2.48E-17 | 1.74E-16 |
| chr2 | 143333312 | 143356708 | 23397 | -9.43 | 2.48E-17 | 1.74E-16 |
| chr16 | 14113412 | 14133855 | 20444 | -9.09 | 2.50E-17 | 1.76E-16 |
| chr4 | 52684059 | 52699904 | 15846 | -9.09 | 2.50E-17 | 1.76E-16 |
| chr5 | 150567216 | 150573191 | 5976 | -9.09 | 2.50E-17 | 1.76E-16 |
| chr18 | 70010006 | 70047708 | 37703 | -9.1 | 2.50E-17 | 1.76E-16 |
| chr10 | 101067114 | 101073017 | 5904 | -9.09 | 2.51E-17 | 1.76E-16 |
| chr15 | 81320546 | 81324298 | 3753 | -9.09 | 2.51E-17 | 1.76E-16 |
| chr6 | 151093688 | 151097305 | 3618 | -5.31 | 2.51E-17 | 1.76E-16 |
| chr14 | 76678651 | 76691741 | 13091 | -9.12 | 2.51E-17 | 1.76E-16 |
| chr12 | 30955560 | 30963639 | 8080 | -9.11 | 2.51E-17 | 1.76E-16 |
| chr6 | 1293020 | 1306911 | 13892 | -9.18 | 2.52E-17 | 1.77E-16 |
| chr7 | 82579398 | 82585809 | 6412 | -6.61 | 2.52E-17 | 1.77E-16 |
| chr5 | 124669611 | 124706220 | 36610 | -9.55 | 2.53E-17 | 1.77E-16 |
| chr5 | 179270788 | 179274670 | 3883 | -9.14 | 2.55E-17 | 1.78E-16 |
| chr14 | 92809741 | 92811653 | 1913 | -7.03 | 2.57E-17 | 1.80E-16 |
| chr20 | 45578407 | 45587650 | 9244 | -9.09 | 2.57E-17 | 1.80E-16 |
| chr5 | 124975824 | 124998000 | 22177 | -9.16 | 2.58E-17 | 1.81E-16 |
| chr2 | 69284221 | 69292787 | 8567 | -9.11 | 2.58E-17 | 1.81E-16 |
| chr12 | 57301687 | 57315700 | 14014 | -9.09 | 2.59E-17 | 1.81E-16 |
| chrX | 151026253 | 151058260 | 32008 | -9.09 | 2.60E-17 | 1.82E-16 |
| chr12 | 26670479 | 26680592 | 10114 | -9.14 | 2.61E-17 | 1.83E-16 |
| chr6 | 83004998 | 83024865 | 19868 | -9.13 | 2.61E-17 | 1.83E-16 |
| chr9 | 21626617 | 21661884 | 35268 | -9.42 | 2.62E-17 | 1.83E-16 |
| chrX | 154263147 | 154268509 | 5363 | -9.09 | 2.62E-17 | 1.84E-16 |
| chr12 | 106397477 | 106420071 | 22595 | -9.17 | 2.62E-17 | 1.84E-16 |
| chr10 | 101029967 | 101046244 | 16278 | -9.4 | 2.63E-17 | 1.84E-16 |

| | | | | | | |
|---|---|---|---|---|---|---|
| chrX | 61920351 | 62055352 | 135002 | -8.34 | 2.64E-17 | 1.84E-16 |
| chr1 | 169030156 | 169038104 | 7949 | -9.12 | 2.65E-17 | 1.86E-16 |
| chr2 | 139156883 | 139174360 | 17478 | -9.09 | 2.66E-17 | 1.86E-16 |
| chr6 | 22442918 | 22486269 | 43352 | -9.16 | 2.66E-17 | 1.86E-16 |
| chr9 | 135202409 | 135211247 | 8839 | -9.11 | 2.66E-17 | 1.86E-16 |
| chr2 | 191902340 | 191911917 | 9578 | -9.16 | 2.66E-17 | 1.86E-16 |
| chr1 | 2567637 | 2581124 | 13488 | -9.14 | 2.67E-17 | 1.86E-16 |
| chr1 | 95705348 | 95720798 | 15451 | -9.13 | 2.67E-17 | 1.86E-16 |
| chr3 | 159695374 | 159709929 | 14556 | -9.1 | 2.67E-17 | 1.87E-16 |
| chr5 | 59658034 | 59686064 | 28031 | 9.33 | 2.69E-17 | 1.88E-16 |
| chr13 | 48923212 | 48929720 | 6509 | -9.09 | 2.71E-17 | 1.89E-16 |
| chr1 | 94902010 | 94925551 | 23542 | -9.1 | 2.72E-17 | 1.90E-16 |
| chr10 | 27546024 | 27566131 | 20108 | -9.11 | 2.72E-17 | 1.90E-16 |
| chr15 | 26159209 | 26176107 | 16899 | -9.14 | 2.73E-17 | 1.91E-16 |
| chr1 | 64418305 | 64424862 | 6558 | -9.08 | 2.74E-17 | 1.91E-16 |
| chr3 | 94388694 | 94407405 | 18712 | -9.12 | 2.74E-17 | 1.91E-16 |
| chr1 | 97493842 | 97532603 | 38762 | -9.14 | 2.74E-17 | 1.91E-16 |
| chr2 | 219589384 | 219594764 | 5381 | -9.09 | 2.74E-17 | 1.91E-16 |
| chr15 | 76922317 | 76930361 | 8045 | -9.11 | 2.75E-17 | 1.91E-16 |
| chr17 | 65725438 | 65728327 | 2890 | -9.16 | 2.76E-17 | 1.93E-16 |
| chr1 | 11406552 | 11414908 | 8357 | -9.1 | 2.77E-17 | 1.93E-16 |
| chr3 | 94421003 | 94432481 | 11479 | -9.08 | 2.78E-17 | 1.94E-16 |
| chr1 | 56211215 | 56241463 | 30249 | -9.18 | 2.79E-17 | 1.95E-16 |
| chr11 | 69569239 | 69584444 | 15206 | -9.13 | 2.80E-17 | 1.95E-16 |
| chr11 | 65911586 | 65916701 | 5116 | -9.08 | 2.82E-17 | 1.96E-16 |
| chr13 | 111389697 | 111418671 | 28975 | -9.23 | 2.82E-17 | 1.97E-16 |
| chr7 | 109782677 | 109805277 | 22601 | -9.1 | 2.82E-17 | 1.97E-16 |
| chr1 | 211574754 | 211592136 | 17383 | -9.15 | 2.83E-17 | 1.97E-16 |
| chr21 | 34186125 | 34211625 | 25501 | -9.09 | 2.85E-17 | 1.99E-16 |
| chr19 | 33735739 | 33750135 | 14397 | -9.08 | 2.85E-17 | 1.99E-16 |
| chr4 | 20461882 | 20495557 | 33676 | 9.57 | 2.86E-17 | 1.99E-16 |
| chr9 | 96387732 | 96402829 | 15098 | -9.3 | 2.87E-17 | 1.99E-16 |
| chr6 | 85655619 | 85661290 | 5672 | -6.64 | 2.90E-17 | 2.02E-16 |
| chr6 | 89268022 | 89302664 | 34643 | -9.08 | 2.91E-17 | 2.02E-16 |
| chr11 | 70885196 | 70890208 | 5013 | -9.07 | 2.91E-17 | 2.02E-16 |
| chr10 | 6811198 | 6820487 | 9290 | -9.09 | 2.92E-17 | 2.03E-16 |
| chr20 | 4156155 | 4159586 | 3432 | -9.09 | 2.92E-17 | 2.03E-16 |
| chr2 | 146700518 | 146719788 | 19271 | -9.09 | 2.94E-17 | 2.04E-16 |
| chr1 | 47113153 | 47116652 | 3500 | -9.11 | 2.94E-17 | 2.04E-16 |
| chr16 | 80986048 | 80995890 | 9843 | -9.3 | 2.95E-17 | 2.05E-16 |
| chr19 | 13686851 | 13691711 | 4861 | -9.08 | 2.95E-17 | 2.05E-16 |
| chr3 | 27339291 | 27352075 | 12785 | -9.07 | 2.96E-17 | 2.05E-16 |
| chr10 | 18221729 | 18414254 | 192526 | 1.08 | 2.96E-17 | 2.05E-16 |
| chr16 | 20757356 | 20784501 | 27146 | -9.42 | 2.97E-17 | 2.06E-16 |
| chr14 | 81275063 | 81292081 | 17019 | -9.24 | 2.98E-17 | 2.06E-16 |
| chr6 | 152542194 | 152545985 | 3792 | -9.28 | 2.98E-17 | 2.06E-16 |
| chr2 | 175148695 | 175159100 | 10406 | -9.15 | 2.98E-17 | 2.07E-16 |
| chr18 | 45998414 | 46035266 | 36853 | -9.56 | 3.00E-17 | 2.08E-16 |
| chr2 | 177230517 | 177245423 | 14907 | -9.1 | 3.00E-17 | 2.08E-16 |
| chr7 | 111338321 | 111374590 | 36270 | -9.46 | 3.00E-17 | 2.08E-16 |
| chr1 | 89379860 | 89386545 | 6686 | -9.07 | 3.01E-17 | 2.08E-16 |
| chr13 | 109477293 | 109495459 | 18167 | -9.24 | 3.01E-17 | 2.09E-16 |
| chr21 | 16497957 | 16508091 | 10135 | -9.07 | 3.02E-17 | 2.09E-16 |
| chr3 | 169532271 | 169536340 | 4070 | -9.08 | 3.02E-17 | 2.09E-16 |
| chr11 | 44481754 | 44488545 | 6792 | -9.1 | 3.02E-17 | 2.09E-16 |
| chr5 | 115803841 | 115818125 | 14285 | -9.07 | 3.02E-17 | 2.09E-16 |
| chr7 | 46116642 | 46159282 | 42641 | -9.47 | 3.04E-17 | 2.10E-16 |
| chr6 | 4627594 | 4653447 | 25854 | -9.21 | 3.05E-17 | 2.11E-16 |
| chr20 | 60916197 | 60924650 | 8454 | -9.27 | 3.05E-17 | 2.11E-16 |
| chr18 | 46561612 | 46567464 | 5853 | -9.07 | 3.06E-17 | 2.12E-16 |
| chr14 | 38298230 | 38305377 | 7148 | -9.17 | 3.06E-17 | 2.12E-16 |

| | | | | | | |
|---|---|---|---|---|---|---|
| chr8 | 30721648 | 30751008 | 29361 | -1.98 | 3.06E-17 | 2.12E-16 |
| chr12 | 124808188 | 124811392 | 3205 | -9.07 | 3.07E-17 | 2.12E-16 |
| chr5 | 83615982 | 83632146 | 16165 | -9.19 | 3.07E-17 | 2.12E-16 |
| chr5 | 41196581 | 41222164 | 25584 | 9.44 | 3.07E-17 | 2.12E-16 |
| chr4 | 184464625 | 184467452 | 2828 | -6.43 | 3.07E-17 | 2.12E-16 |
| chr3 | 109184990 | 109217593 | 32604 | -9.21 | 3.07E-17 | 2.12E-16 |
| chr12 | 65023010 | 65027855 | 4846 | -9.11 | 3.08E-17 | 2.13E-16 |
| chr20 | 14562374 | 14573354 | 10981 | -9.22 | 3.09E-17 | 2.13E-16 |
| chrX | 66833132 | 66845714 | 12583 | -9.07 | 3.09E-17 | 2.13E-16 |
| chr6 | 41579622 | 41591804 | 12183 | -9.08 | 3.09E-17 | 2.14E-16 |
| chr10 | 95188587 | 95203794 | 15208 | -9.19 | 3.10E-17 | 2.14E-16 |
| chr10 | 4762741 | 4773549 | 10809 | -9.1 | 3.12E-17 | 2.15E-16 |
| chr17 | 59617009 | 59627410 | 10402 | -9.58 | 3.13E-17 | 2.16E-16 |
| chr10 | 134026272 | 134032654 | 6383 | -9.19 | 3.13E-17 | 2.16E-16 |
| chr1 | 177153825 | 177188300 | 34476 | -9.17 | 3.14E-17 | 2.16E-16 |
| chr7 | 55293603 | 55299749 | 6147 | -2.89 | 3.14E-17 | 2.17E-16 |
| chr20 | 13448740 | 13463856 | 15117 | -9.11 | 3.15E-17 | 2.17E-16 |
| chr10 | 120529135 | 120537930 | 8796 | -9.12 | 3.15E-17 | 2.18E-16 |
| chr5 | 94816535 | 94826079 | 9545 | -9.38 | 3.16E-17 | 2.18E-16 |
| chr17 | 39529745 | 39544555 | 14811 | -9.09 | 3.16E-17 | 2.18E-16 |
| chr18 | 34255519 | 34265520 | 10002 | -9.07 | 3.19E-17 | 2.20E-16 |
| chr9 | 74682821 | 74718444 | 35624 | -9.14 | 3.21E-17 | 2.21E-16 |
| chr4 | 128767475 | 128779160 | 11686 | -9.17 | 3.22E-17 | 2.22E-16 |
| chr9 | 135828686 | 135836069 | 7384 | -9.11 | 3.22E-17 | 2.22E-16 |
| chr18 | 9851535 | 9869304 | 17770 | -9.06 | 3.23E-17 | 2.23E-16 |
| chr7 | 133020042 | 133038659 | 18618 | -9.08 | 3.25E-17 | 2.24E-16 |
| chr14 | 81134553 | 81154028 | 19476 | -9.26 | 3.28E-17 | 2.26E-16 |
| chr7 | 67426603 | 67447668 | 21066 | -9.3 | 3.30E-17 | 2.27E-16 |
| chr5 | 142843057 | 142894578 | 51522 | -9.44 | 3.31E-17 | 2.28E-16 |
| chr6 | 127482372 | 127504712 | 22341 | -9.11 | 3.31E-17 | 2.28E-16 |
| chr3 | 180603674 | 180624799 | 21126 | -9.19 | 3.32E-17 | 2.28E-16 |
| chrX | 110238938 | 110338351 | 99414 | 1.69 | 3.33E-17 | 2.29E-16 |
| chr7 | 75299624 | 75305300 | 5677 | -9.09 | 3.33E-17 | 2.29E-16 |
| chr20 | 22828256 | 22850219 | 21964 | -9.08 | 3.34E-17 | 2.30E-16 |
| chr2 | 165175611 | 165192625 | 17015 | -9.06 | 3.35E-17 | 2.30E-16 |
| chr1 | 18079514 | 18086872 | 7359 | -9.09 | 3.35E-17 | 2.30E-16 |
| chr12 | 116825355 | 116840305 | 14951 | -9.19 | 3.35E-17 | 2.30E-16 |
| chr4 | 72472312 | 72494778 | 22467 | -9.27 | 3.36E-17 | 2.31E-16 |
| chr4 | 71903355 | 71914657 | 11303 | -9.17 | 3.38E-17 | 2.32E-16 |
| chr1 | 180911388 | 180931157 | 19770 | -9.56 | 3.38E-17 | 2.32E-16 |
| chr1 | 31682250 | 31687927 | 5678 | -9.07 | 3.38E-17 | 2.32E-16 |
| chr7 | 39583541 | 39590907 | 7367 | -9.07 | 3.38E-17 | 2.32E-16 |
| chr5 | 17438465 | 17463161 | 24697 | -9.21 | 3.38E-17 | 2.32E-16 |
| chr12 | 16460881 | 16464889 | 4009 | -6.26 | 3.38E-17 | 2.32E-16 |
| chr8 | 87200767 | 87207812 | 7046 | -9.06 | 3.39E-17 | 2.32E-16 |
| chr18 | 72256805 | 72264259 | 7455 | -9.31 | 3.39E-17 | 2.33E-16 |
| chr14 | 53093390 | 53109259 | 15870 | -9.06 | 3.39E-17 | 2.33E-16 |
| chr14 | 75970439 | 76001166 | 30728 | -9.28 | 3.41E-17 | 2.34E-16 |
| chr1 | 230196132 | 230200923 | 4792 | -9.07 | 3.41E-17 | 2.34E-16 |
| chr17 | 70529278 | 70543283 | 14006 | -9.06 | 3.42E-17 | 2.34E-16 |
| chr22 | 39273899 | 39305789 | 31891 | -9.28 | 3.42E-17 | 2.34E-16 |
| chr15 | 68160991 | 68166601 | 5611 | -9.4 | 3.42E-17 | 2.34E-16 |
| chr1 | 160629362 | 160641629 | 12268 | -2.07 | 3.44E-17 | 2.36E-16 |
| chr14 | 75513128 | 75528318 | 15191 | -9.08 | 3.44E-17 | 2.36E-16 |
| chrX | 40697239 | 40704226 | 6988 | -9.1 | 3.44E-17 | 2.36E-16 |
| chr12 | 100461492 | 100477779 | 16288 | -9.06 | 3.44E-17 | 2.36E-16 |
| chr5 | 170002118 | 170011240 | 9123 | -9.06 | 3.44E-17 | 2.36E-16 |
| chr2 | 18940213 | 18950796 | 10584 | -9.17 | 3.45E-17 | 2.36E-16 |
| chr12 | 103986542 | 104008928 | 22387 | -9.06 | 3.45E-17 | 2.36E-16 |
| chr20 | 321937 | 325684 | 3748 | -9.06 | 3.45E-17 | 2.36E-16 |
| chr3 | 43046142 | 43065242 | 19101 | -2.34 | 3.45E-17 | 2.36E-16 |

| | | | | | | |
|---|---|---|---|---|---|---|
| chr9 | 2481321 | 2509630 | 28310 | 9.31 | 3.46E-17 | 2.37E-16 |
| chr3 | 109161489 | 109171362 | 9874 | -7.75 | 3.47E-17 | 2.37E-16 |
| chr3 | 8856739 | 8865578 | 8840 | -9.07 | 3.48E-17 | 2.38E-16 |
| chr3 | 152387241 | 152517853 | 130613 | 3.66 | 3.49E-17 | 2.39E-16 |
| chr7 | 20879281 | 21285854 | 406574 | 1.85 | 3.53E-17 | 2.41E-16 |
| chr21 | 45871402 | 45874868 | 3467 | -6.64 | 3.54E-17 | 2.42E-16 |
| chr2 | 61217405 | 61220814 | 3410 | -9.1 | 3.55E-17 | 2.43E-16 |
| chr9 | 101769113 | 101775897 | 6785 | -9.08 | 3.56E-17 | 2.43E-16 |
| chr21 | 16837284 | 16853955 | 16672 | -9.19 | 3.58E-17 | 2.44E-16 |
| chr15 | 36567699 | 36578242 | 10544 | -9.07 | 3.58E-17 | 2.45E-16 |
| chr15 | 69872637 | 69900117 | 27481 | -9.48 | 3.58E-17 | 2.45E-16 |
| chr1 | 93836381 | 93853181 | 16801 | -9.13 | 3.59E-17 | 2.45E-16 |
| chr3 | 11880223 | 11884320 | 4098 | -9.06 | 3.60E-17 | 2.46E-16 |
| chr2 | 148841526 | 148850942 | 9417 | -9.2 | 3.61E-17 | 2.46E-16 |
| chr8 | 143154394 | 143168004 | 13611 | -9.06 | 3.63E-17 | 2.48E-16 |
| chr2 | 145009345 | 145017211 | 7867 | -9.2 | 3.64E-17 | 2.48E-16 |
| chr6 | 163176887 | 163486659 | 309773 | 1.64 | 3.64E-17 | 2.48E-16 |
| chr6 | 72171716 | 72180578 | 8863 | -9.13 | 3.67E-17 | 2.50E-16 |
| chr5 | 114584405 | 114599767 | 15363 | -9.05 | 3.67E-17 | 2.50E-16 |
| chr2 | 72344192 | 72352402 | 8211 | -9.25 | 3.67E-17 | 2.50E-16 |
| chr4 | 1172943 | 1185553 | 12611 | -9.16 | 3.68E-17 | 2.51E-16 |
| chr11 | 1383297 | 1402093 | 18797 | -9.15 | 3.69E-17 | 2.51E-16 |
| chr14 | 76192346 | 76194346 | 2001 | -9.14 | 3.69E-17 | 2.51E-16 |
| chr11 | 102287880 | 102316000 | 28121 | -9.06 | 3.73E-17 | 2.54E-16 |
| chr1 | 246989953 | 246996168 | 6216 | -9.05 | 3.73E-17 | 2.54E-16 |
| chr7 | 155719370 | 155740045 | 20676 | -9.09 | 3.74E-17 | 2.54E-16 |
| chr14 | 56084494 | 56094818 | 10325 | -9.1 | 3.76E-17 | 2.56E-16 |
| chr15 | 52111605 | 52115516 | 3912 | -9.14 | 3.76E-17 | 2.56E-16 |
| chr11 | 947365 | 963858 | 16494 | -9.05 | 3.76E-17 | 2.56E-16 |
| chr2 | 221221983 | 221246966 | 24984 | -9.14 | 3.77E-17 | 2.57E-16 |
| chr2 | 100075669 | 100088026 | 12358 | -9.04 | 3.78E-17 | 2.57E-16 |
| chr17 | 59901620 | 59906440 | 4821 | -9.08 | 3.78E-17 | 2.57E-16 |
| chr7 | 101050524 | 101054431 | 3908 | -9.17 | 3.79E-17 | 2.57E-16 |
| chr2 | 235643589 | 235671058 | 27470 | -9.46 | 3.79E-17 | 2.58E-16 |
| chr19 | 34340961 | 34357288 | 16328 | -9.13 | 3.80E-17 | 2.58E-16 |
| chr13 | 44415135 | 44439965 | 24831 | -9.11 | 3.81E-17 | 2.59E-16 |
| chr22 | 23386813 | 23399941 | 13129 | -9.05 | 3.82E-17 | 2.60E-16 |
| chr12 | 18148186 | 18155882 | 7697 | -9.1 | 3.83E-17 | 2.61E-16 |
| chr12 | 28167165 | 28200028 | 32864 | -9.09 | 3.85E-17 | 2.61E-16 |
| chr20 | 596856 | 607029 | 10174 | -9.29 | 3.86E-17 | 2.62E-16 |
| chr3 | 8743401 | 8750351 | 6951 | -9.11 | 3.88E-17 | 2.63E-16 |
| chr6 | 126236951 | 126252953 | 16003 | -3.77 | 3.89E-17 | 2.64E-16 |
| chr13 | 35821874 | 35844950 | 23077 | -9.23 | 3.89E-17 | 2.64E-16 |
| chrX | 108604466 | 108630369 | 25904 | 9.22 | 3.93E-17 | 2.67E-16 |
| chr2 | 161493304 | 161498490 | 5187 | -9.07 | 3.96E-17 | 2.69E-16 |
| chr13 | 96455500 | 96463588 | 8089 | -9.26 | 3.96E-17 | 2.69E-16 |
| chr1 | 183625088 | 183643892 | 18805 | -9.36 | 3.97E-17 | 2.69E-16 |
| chr9 | 134653662 | 134671910 | 18249 | -9.4 | 3.97E-17 | 2.69E-16 |
| chr5 | 106504336 | 106517446 | 13111 | -9.06 | 3.98E-17 | 2.70E-16 |
| chr11 | 32380473 | 32399996 | 19524 | -9.2 | 3.98E-17 | 2.70E-16 |
| chr8 | 49394432 | 49420778 | 26347 | -9.34 | 3.98E-17 | 2.70E-16 |
| chr7 | 128413242 | 128419571 | 6330 | -9.08 | 4.00E-17 | 2.71E-16 |
| chr3 | 99860662 | 99877269 | 16608 | -9.05 | 4.00E-17 | 2.71E-16 |
| chr8 | 31664734 | 31690246 | 25513 | -9.44 | 4.01E-17 | 2.72E-16 |
| chr6 | 71580344 | 71588318 | 7975 | -9.04 | 4.02E-17 | 2.72E-16 |
| chr2 | 68300071 | 68305044 | 4974 | -4.23 | 4.05E-17 | 2.74E-16 |
| chrX | 69048987 | 69072630 | 23644 | 9.1 | 4.05E-17 | 2.74E-16 |
| chr15 | 90655970 | 90660362 | 4393 | -9.21 | 4.07E-17 | 2.75E-16 |
| chr6 | 100766777 | 100802396 | 35620 | -9.19 | 4.09E-17 | 2.77E-16 |
| chr8 | 86967164 | 86975310 | 8147 | -3.76 | 4.09E-17 | 2.77E-16 |
| chr12 | 63882985 | 63904892 | 21908 | -9.08 | 4.09E-17 | 2.77E-16 |

| | | | | | | |
|---|---|---|---|---|---|---|
| chr5 | 39906871 | 39940295 | 33425 | -9.09 | 4.10E-17 | 2.77E-16 |
| chr2 | 61900967 | 61906607 | 5641 | -9.15 | 4.10E-17 | 2.77E-16 |
| chr14 | 45726479 | 45733699 | 7221 | -9.05 | 4.11E-17 | 2.78E-16 |
| chr15 | 50333028 | 50346955 | 13928 | -9.04 | 4.12E-17 | 2.79E-16 |
| chr11 | 2460932 | 2477522 | 16591 | -9.12 | 4.13E-17 | 2.79E-16 |
| chr11 | 71671793 | 71690720 | 18928 | -9.18 | 4.13E-17 | 2.79E-16 |
| chr5 | 143101166 | 143109522 | 8357 | -9.08 | 4.14E-17 | 2.80E-16 |
| chr2 | 165504070 | 165511626 | 7557 | -9.04 | 4.15E-17 | 2.80E-16 |
| chr22 | 46599844 | 46609818 | 9975 | -9.05 | 4.16E-17 | 2.81E-16 |
| chr3 | 100675550 | 100684088 | 8539 | -9.03 | 4.17E-17 | 2.82E-16 |
| chr16 | 69074639 | 69076827 | 2189 | -9.04 | 4.18E-17 | 2.82E-16 |
| chr8 | 99386393 | 99407561 | 21169 | -9.04 | 4.18E-17 | 2.83E-16 |
| chr10 | 98765130 | 98770091 | 4962 | -9.05 | 4.19E-17 | 2.83E-16 |
| chr2 | 146808837 | 146830517 | 21681 | -9.04 | 4.19E-17 | 2.83E-16 |
| chr5 | 108847585 | 108979427 | 131843 | 3.36 | 4.20E-17 | 2.83E-16 |
| chr1 | 225201653 | 225215540 | 13888 | -9.19 | 4.21E-17 | 2.84E-16 |
| chr17 | 76945234 | 76948719 | 3486 | -9.16 | 4.22E-17 | 2.85E-16 |
| chr6 | 9504795 | 9534070 | 29276 | -9.03 | 4.22E-17 | 2.85E-16 |
| chr15 | 85500318 | 85511944 | 11627 | -9.35 | 4.23E-17 | 2.85E-16 |
| chr5 | 71813633 | 71844516 | 30884 | -9.2 | 4.24E-17 | 2.86E-16 |
| chr5 | 52787229 | 52823014 | 35786 | -9.45 | 4.24E-17 | 2.86E-16 |
| chr5 | 179499243 | 179509743 | 10501 | -9.12 | 4.24E-17 | 2.86E-16 |
| chr16 | 89190383 | 89198077 | 7695 | -9.03 | 4.25E-17 | 2.86E-16 |
| chr11 | 75393471 | 75420671 | 27201 | -9.38 | 4.25E-17 | 2.87E-16 |
| chr19 | 14778065 | 14789852 | 11788 | -2.54 | 4.27E-17 | 2.88E-16 |
| chr1 | 161016111 | 161021351 | 5241 | -9.17 | 4.28E-17 | 2.88E-16 |
| chr10 | 3395223 | 3406682 | 11460 | -9.27 | 4.28E-17 | 2.88E-16 |
| chr10 | 31560385 | 31573220 | 12836 | -9.22 | 4.29E-17 | 2.89E-16 |
| chr12 | 105448253 | 105451887 | 3635 | -9.05 | 4.29E-17 | 2.89E-16 |
| chr7 | 131086301 | 131089564 | 3264 | -9.04 | 4.29E-17 | 2.89E-16 |
| chr2 | 164351355 | 164384508 | 33154 | -9.46 | 4.30E-17 | 2.89E-16 |
| chr14 | 92932497 | 92947791 | 15295 | -9.04 | 4.30E-17 | 2.90E-16 |
| chr2 | 196329114 | 196354720 | 25607 | -9.12 | 4.31E-17 | 2.90E-16 |
| chr16 | 71581967 | 71597346 | 15380 | -9.26 | 4.33E-17 | 2.91E-16 |
| chr5 | 717556 | 720734 | 3179 | -6.66 | 4.33E-17 | 2.91E-16 |
| chr6 | 88738160 | 88748527 | 10368 | -3.41 | 4.33E-17 | 2.91E-16 |
| chr5 | 122688151 | 122694204 | 6054 | -9.03 | 4.33E-17 | 2.91E-16 |
| chr5 | 98140001 | 98162881 | 22881 | -9.03 | 4.34E-17 | 2.92E-16 |
| chr11 | 57603713 | 57629564 | 25852 | -9.08 | 4.36E-17 | 2.93E-16 |
| chr3 | 111531909 | 111540080 | 8172 | -9.03 | 4.37E-17 | 2.94E-16 |
| chr2 | 135548347 | 135571349 | 23003 | -9.09 | 4.41E-17 | 2.96E-16 |
| chr12 | 17382358 | 17401865 | 19508 | -9.36 | 4.42E-17 | 2.97E-16 |
| chr1 | 236255782 | 236262713 | 6932 | -9.04 | 4.44E-17 | 2.98E-16 |
| chr12 | 43469803 | 43487906 | 18104 | -9.19 | 4.44E-17 | 2.98E-16 |
| chr6 | 155844368 | 155858144 | 13777 | -9.12 | 4.44E-17 | 2.98E-16 |
| chr6 | 13668471 | 13681865 | 13395 | -9.04 | 4.45E-17 | 2.99E-16 |
| chr7 | 143585830 | 143596296 | 10467 | -9.03 | 4.47E-17 | 3.00E-16 |
| chr4 | 112641044 | 112677280 | 36237 | 9.19 | 4.48E-17 | 3.01E-16 |
| chr2 | 46056334 | 46059882 | 3549 | -9.04 | 4.49E-17 | 3.01E-16 |
| chr12 | 109338189 | 109343923 | 5735 | -9.1 | 4.50E-17 | 3.02E-16 |
| chr6 | 110850690 | 110881110 | 30421 | -9.13 | 4.51E-17 | 3.02E-16 |
| chr2 | 107691947 | 107718994 | 27048 | 9.09 | 4.52E-17 | 3.03E-16 |
| chr15 | 68645074 | 68651447 | 6374 | -9.15 | 4.53E-17 | 3.04E-16 |
| chr3 | 50757329 | 50793955 | 36627 | -9.19 | 4.56E-17 | 3.05E-16 |
| chr8 | 56995864 | 57004589 | 8726 | -9.08 | 4.57E-17 | 3.06E-16 |
| chr2 | 37708361 | 37727263 | 18903 | -9.04 | 4.57E-17 | 3.06E-16 |
| chr22 | 29630006 | 29640259 | 10254 | -9.16 | 4.57E-17 | 3.06E-16 |
| chr15 | 63875735 | 63878263 | 2529 | -9.05 | 4.57E-17 | 3.06E-16 |
| chr3 | 31386217 | 31446518 | 60302 | -9.65 | 4.58E-17 | 3.06E-16 |
| chr1 | 191143063 | 191195464 | 52402 | -9.1 | 4.58E-17 | 3.06E-16 |
| chr1 | 31145663 | 31153548 | 7886 | -9.03 | 4.58E-17 | 3.07E-16 |

| | | | | | | |
|---|---|---|---|---|---|---|
| chr2 | 235247632 | 235269749 | 22118 | -9.12 | 4.59E-17 | 3.07E-16 |
| chr8 | 119887405 | 119902820 | 15416 | -9.07 | 4.59E-17 | 3.07E-16 |
| chr2 | 98526818 | 98538431 | 11614 | -9.21 | 4.60E-17 | 3.08E-16 |
| chr7 | 143667892 | 143689303 | 21412 | -9.05 | 4.60E-17 | 3.08E-16 |
| chr10 | 87398258 | 87433415 | 35158 | -9.26 | 4.60E-17 | 3.08E-16 |
| chr15 | 50598912 | 50601298 | 2387 | -9.02 | 4.60E-17 | 3.08E-16 |
| chr4 | 183804399 | 183808581 | 4183 | -9.03 | 4.62E-17 | 3.09E-16 |
| chr13 | 110645459 | 110675005 | 29547 | -9.14 | 4.63E-17 | 3.09E-16 |
| chr11 | 69201723 | 69218538 | 16816 | -9.14 | 4.64E-17 | 3.10E-16 |
| chr5 | 72392330 | 72409843 | 17514 | -9.1 | 4.64E-17 | 3.10E-16 |
| chr17 | 16771350 | 16811381 | 40032 | -9.35 | 4.65E-17 | 3.11E-16 |
| chr7 | 158135374 | 158175429 | 40056 | -9.46 | 4.66E-17 | 3.12E-16 |
| chr5 | 6654626 | 6658379 | 3754 | -9.18 | 4.68E-17 | 3.12E-16 |
| chr2 | 69252718 | 69256361 | 3644 | -9.07 | 4.68E-17 | 3.13E-16 |
| chr3 | 123638729 | 123642517 | 3789 | -9.04 | 4.70E-17 | 3.14E-16 |
| chrX | 56605712 | 56636596 | 30885 | -9.02 | 4.72E-17 | 3.15E-16 |
| chr4 | 168871348 | 168901138 | 29791 | -9.03 | 4.73E-17 | 3.16E-16 |
| chr11 | 4010914 | 4018168 | 7255 | -9.12 | 4.74E-17 | 3.16E-16 |
| chr4 | 62250478 | 62294340 | 43863 | -9.06 | 4.75E-17 | 3.17E-16 |
| chr2 | 172278540 | 172294099 | 15560 | -9.02 | 4.75E-17 | 3.17E-16 |
| chr2 | 225044003 | 225061040 | 17038 | -9.1 | 4.76E-17 | 3.17E-16 |
| chr2 | 48521690 | 48541143 | 19454 | -9.14 | 4.77E-17 | 3.18E-16 |
| chr8 | 29642752 | 29653625 | 10874 | -9.11 | 4.77E-17 | 3.18E-16 |
| chr1 | 229400468 | 229418977 | 18510 | -9.02 | 4.78E-17 | 3.19E-16 |
| chr8 | 116150908 | 116185394 | 34487 | -9.06 | 4.79E-17 | 3.19E-16 |
| chr8 | 30532256 | 30536504 | 4249 | -9.03 | 4.81E-17 | 3.20E-16 |
| chr7 | 23133600 | 23142990 | 9391 | -9.02 | 4.82E-17 | 3.21E-16 |
| chr1 | 17477406 | 17486330 | 8925 | -9.08 | 4.86E-17 | 3.24E-16 |
| chr3 | 14893482 | 14899369 | 5888 | -9.02 | 4.86E-17 | 3.24E-16 |
| chr10 | 123433319 | 123438962 | 5644 | -9.02 | 4.91E-17 | 3.27E-16 |
| chr10 | 21467718 | 21522232 | 54515 | -9.68 | 4.91E-17 | 3.27E-16 |
| chr3 | 146299095 | 146302630 | 3536 | -9.09 | 4.95E-17 | 3.30E-16 |
| chr1 | 201737971 | 201743245 | 5275 | -9.07 | 4.95E-17 | 3.30E-16 |
| chr1 | 24201943 | 24207049 | 5107 | -9.05 | 4.96E-17 | 3.30E-16 |
| chr13 | 42498182 | 42509534 | 11353 | -9.02 | 4.97E-17 | 3.31E-16 |
| chr7 | 79360069 | 79387207 | 27139 | -9.23 | 4.99E-17 | 3.32E-16 |
| chr5 | 60165312 | 60179263 | 13952 | -9.02 | 5.00E-17 | 3.33E-16 |
| chr12 | 39904470 | 39920964 | 16495 | -9.04 | 5.01E-17 | 3.33E-16 |
| chr10 | 95277475 | 95281086 | 3612 | -9.18 | 5.01E-17 | 3.33E-16 |
| chr4 | 131828260 | 131860890 | 32631 | -9.17 | 5.04E-17 | 3.35E-16 |
| chr3 | 100106954 | 100110074 | 3121 | -9.02 | 5.06E-17 | 3.36E-16 |
| chr4 | 54993741 | 54999852 | 6112 | -4.75 | 5.07E-17 | 3.37E-16 |
| chr18 | 8889330 | 8910891 | 21562 | -9.04 | 5.13E-17 | 3.41E-16 |
| chr5 | 130229471 | 130259890 | 30420 | -9.18 | 5.14E-17 | 3.41E-16 |
| chr8 | 62785430 | 62817025 | 31596 | -9.42 | 5.19E-17 | 3.45E-16 |
| chr1 | 30490665 | 30506158 | 15494 | -9.1 | 5.19E-17 | 3.45E-16 |
| chr14 | 36310029 | 36322187 | 12159 | -9.01 | 5.19E-17 | 3.45E-16 |
| chr18 | 75066633 | 75084506 | 17874 | -2.05 | 5.21E-17 | 3.46E-16 |
| chr10 | 32887343 | 32917372 | 30030 | -9.45 | 5.21E-17 | 3.46E-16 |
| chr8 | 53837289 | 53864287 | 26999 | -9.09 | 5.23E-17 | 3.47E-16 |
| chrX | 123278873 | 123304065 | 25193 | -9.01 | 5.23E-17 | 3.47E-16 |
| chr3 | 28253963 | 28262567 | 8605 | -9.01 | 5.23E-17 | 3.47E-16 |
| chr7 | 1373417 | 1383557 | 10141 | -9.28 | 5.24E-17 | 3.47E-16 |
| chr7 | 96729543 | 96771873 | 42331 | -9.03 | 5.25E-17 | 3.48E-16 |
| chr2 | 171040828 | 171045959 | 5132 | -9.02 | 5.25E-17 | 3.48E-16 |
| chr10 | 97409663 | 97412687 | 3025 | -9.09 | 5.26E-17 | 3.49E-16 |
| chr12 | 104575815 | 104595109 | 19295 | -9.27 | 5.26E-17 | 3.49E-16 |
| chr6 | 105964348 | 105976733 | 12386 | -9.01 | 5.26E-17 | 3.49E-16 |
| chr7 | 138052228 | 138057299 | 5072 | -9.08 | 5.28E-17 | 3.50E-16 |
| chr3 | 5755747 | 5772461 | 16715 | -9.01 | 5.29E-17 | 3.50E-16 |
| chr7 | 77404769 | 77416456 | 11688 | -9.02 | 5.29E-17 | 3.50E-16 |

| | | | | | | |
|---|---|---|---|---|---|---|
| chr9 | 97924356 | 97930344 | 5989 | -9.01 | 5.29E-17 | 3.51E-16 |
| chr5 | 108135972 | 108158221 | 22250 | -9.31 | 5.32E-17 | 3.52E-16 |
| chr15 | 22804175 | 22819801 | 15627 | -9.03 | 5.32E-17 | 3.52E-16 |
| chr10 | 4166532 | 4185562 | 19031 | -9.02 | 5.34E-17 | 3.53E-16 |
| chr6 | 149556515 | 149565809 | 9295 | -9.01 | 5.35E-17 | 3.54E-16 |
| chr4 | 142400023 | 142411226 | 11204 | -9.07 | 5.36E-17 | 3.55E-16 |
| chr18 | 52329246 | 52374139 | 44894 | -9.01 | 5.37E-17 | 3.55E-16 |
| chr12 | 92121900 | 92155610 | 33711 | -9.32 | 5.38E-17 | 3.56E-16 |
| chr4 | 77727486 | 77741100 | 13615 | -9.16 | 5.39E-17 | 3.56E-16 |
| chr11 | 23939637 | 25080382 | 1140746 | 1.38 | 5.40E-17 | 3.57E-16 |
| chr3 | 36711363 | 36724116 | 12754 | -9.01 | 5.41E-17 | 3.58E-16 |
| chr7 | 1767456 | 1771360 | 3905 | -9.01 | 5.43E-17 | 3.59E-16 |
| chr18 | 18714355 | 18737616 | 23262 | -9.03 | 5.44E-17 | 3.60E-16 |
| chr18 | 61525178 | 61553522 | 28345 | 4.22 | 5.46E-17 | 3.61E-16 |
| chr20 | 15232581 | 15268381 | 35801 | -9.63 | 5.47E-17 | 3.61E-16 |
| chr5 | 79822297 | 79832681 | 10385 | -9.03 | 5.49E-17 | 3.62E-16 |
| chr9 | 103213476 | 103216355 | 2880 | -9.03 | 5.50E-17 | 3.63E-16 |
| chr19 | 51902286 | 51910187 | 7902 | -9.22 | 5.50E-17 | 3.63E-16 |
| chr11 | 43784966 | 43797135 | 12170 | -9 | 5.52E-17 | 3.65E-16 |
| chr17 | 78979315 | 79001701 | 22387 | -9.24 | 5.52E-17 | 3.65E-16 |
| chr20 | 21300394 | 21317720 | 17327 | -9.01 | 5.54E-17 | 3.65E-16 |
| chr3 | 107902914 | 107914587 | 11674 | -9.01 | 5.58E-17 | 3.68E-16 |
| chr12 | 28223271 | 28236720 | 13450 | -9.11 | 5.59E-17 | 3.69E-16 |
| chr9 | 80065334 | 80106589 | 41256 | -9.22 | 5.61E-17 | 3.70E-16 |
| chr5 | 450423 | 453420 | 2998 | -9.12 | 5.64E-17 | 3.72E-16 |
| chr1 | 88205608 | 88238101 | 32494 | -9 | 5.64E-17 | 3.72E-16 |
| chr11 | 119420791 | 119464634 | 43844 | -9.15 | 5.64E-17 | 3.72E-16 |
| chr2 | 198426491 | 198440398 | 13908 | -9.01 | 5.64E-17 | 3.72E-16 |
| chr5 | 13995904 | 14004767 | 8864 | -9 | 5.65E-17 | 3.72E-16 |
| chr12 | 48027375 | 48039463 | 12089 | -9.08 | 5.66E-17 | 3.73E-16 |
| chr2 | 196371116 | 196408799 | 37684 | -9.32 | 5.67E-17 | 3.73E-16 |
| chr10 | 97523039 | 97543241 | 20203 | -2.28 | 5.67E-17 | 3.74E-16 |
| chr5 | 167882597 | 167885534 | 2938 | -9.02 | 5.70E-17 | 3.75E-16 |
| chr21 | 18383523 | 18403085 | 19563 | 9.16 | 5.71E-17 | 3.76E-16 |
| chr22 | 47470072 | 47492336 | 22265 | -9.04 | 5.71E-17 | 3.76E-16 |
| chr17 | 60672359 | 60675228 | 2870 | -9.05 | 5.72E-17 | 3.77E-16 |
| chr8 | 83076931 | 83153391 | 76461 | -9.31 | 5.74E-17 | 3.77E-16 |
| chr7 | 40965610 | 40988123 | 22514 | -1.95 | 5.74E-17 | 3.78E-16 |
| chr8 | 100928707 | 100936724 | 8018 | -9.05 | 5.77E-17 | 3.79E-16 |
| chr5 | 96489320 | 96496056 | 6737 | -9 | 5.77E-17 | 3.79E-16 |
| chr10 | 106005474 | 106008874 | 3401 | -9.01 | 5.80E-17 | 3.81E-16 |
| chr4 | 23379612 | 23422032 | 42421 | -9.15 | 5.81E-17 | 3.82E-16 |
| chr19 | 50252549 | 50260708 | 8160 | -9.24 | 5.82E-17 | 3.83E-16 |
| chr5 | 74845130 | 74861618 | 16489 | -9.05 | 5.83E-17 | 3.83E-16 |
| chr18 | 33867658 | 33873814 | 6157 | -9.04 | 5.83E-17 | 3.83E-16 |
| chr1 | 212541599 | 212547659 | 6061 | -9 | 5.84E-17 | 3.84E-16 |
| chr10 | 35271622 | 35276591 | 4970 | -9.18 | 5.85E-17 | 3.84E-16 |
| chr3 | 101073714 | 101080357 | 6644 | -9 | 5.88E-17 | 3.86E-16 |
| chr19 | 35557325 | 35574041 | 16717 | -9.51 | 5.88E-17 | 3.86E-16 |
| chr2 | 111487846 | 111589598 | 101753 | -2.56 | 5.89E-17 | 3.86E-16 |
| chrX | 76719623 | 76744135 | 24513 | -9 | 5.89E-17 | 3.87E-16 |
| chr4 | 106346002 | 106358753 | 12752 | -9.19 | 5.94E-17 | 3.90E-16 |
| chr3 | 96383812 | 96424026 | 40215 | -8.99 | 5.95E-17 | 3.91E-16 |
| chr2 | 42640170 | 42644537 | 4368 | -8.99 | 5.98E-17 | 3.92E-16 |
| chr12 | 111671297 | 111685120 | 13824 | -9.13 | 6.00E-17 | 3.93E-16 |
| chr2 | 128024532 | 128043887 | 19356 | -9.14 | 6.00E-17 | 3.93E-16 |
| chr8 | 37845092 | 37852430 | 7339 | -9.13 | 6.01E-17 | 3.94E-16 |
| chr4 | 6202816 | 6224679 | 21864 | -9 | 6.02E-17 | 3.95E-16 |
| chr21 | 44015395 | 44032807 | 17413 | -9.13 | 6.03E-17 | 3.95E-16 |
| chr1 | 240362937 | 240367010 | 4074 | -9 | 6.05E-17 | 3.97E-16 |
| chr4 | 25396985 | 25399796 | 2812 | -9.06 | 6.05E-17 | 3.97E-16 |

| | | | | | | |
|---|---|---|---|---|---|---|
| chr7 | 101163698 | 101169233 | 5536 | -9.01 | 6.05E-17 | 3.97E-16 |
| chr4 | 145932016 | 145948894 | 16879 | -8.99 | 6.07E-17 | 3.97E-16 |
| chr1 | 64129962 | 64149213 | 19252 | -9.04 | 6.08E-17 | 3.98E-16 |
| chr5 | 13594415 | 13620445 | 26031 | -9.01 | 6.08E-17 | 3.98E-16 |
| chr21 | 46864801 | 46882601 | 17801 | -8.99 | 6.11E-17 | 4.00E-16 |
| chr16 | 14420279 | 14440838 | 20560 | -9.15 | 6.13E-17 | 4.02E-16 |
| chr7 | 33104259 | 33117651 | 13393 | -2.71 | 6.14E-17 | 4.02E-16 |
| chr16 | 57359563 | 57382895 | 23333 | -9.14 | 6.17E-17 | 4.04E-16 |
| chr1 | 7870439 | 7876482 | 6044 | -9.09 | 6.17E-17 | 4.04E-16 |
| chr11 | 124710518 | 124737136 | 26619 | -9.18 | 6.18E-17 | 4.04E-16 |
| chr17 | 2945457 | 2949755 | 4299 | -9.25 | 6.21E-17 | 4.06E-16 |
| chr3 | 139119776 | 139135698 | 15923 | -9.15 | 6.22E-17 | 4.07E-16 |
| chr5 | 92204770 | 92230292 | 25523 | -9.25 | 6.23E-17 | 4.08E-16 |
| chr20 | 8516953 | 8544228 | 27276 | 2.24 | 6.24E-17 | 4.08E-16 |
| chr17 | 69406228 | 69426035 | 19808 | -9.21 | 6.26E-17 | 4.09E-16 |
| chr1 | 240466001 | 240481315 | 15315 | -8.99 | 6.27E-17 | 4.10E-16 |
| chr5 | 18119153 | 18287209 | 168057 | -1.7 | 6.29E-17 | 4.11E-16 |
| chrX | 51176365 | 51191755 | 15391 | 9.09 | 6.31E-17 | 4.12E-16 |
| chr14 | 30155710 | 30192556 | 36847 | 9.57 | 6.32E-17 | 4.13E-16 |
| chr2 | 73085421 | 73101180 | 15760 | -9.05 | 6.32E-17 | 4.13E-16 |
| chr18 | 48486353 | 48505191 | 18839 | -9.04 | 6.37E-17 | 4.16E-16 |
| chr1 | 228279577 | 228284052 | 4476 | -9.03 | 6.40E-17 | 4.18E-16 |
| chr21 | 43208724 | 43214477 | 5754 | -9.05 | 6.43E-17 | 4.20E-16 |
| chr7 | 97691391 | 97705793 | 14403 | -9.05 | 6.44E-17 | 4.20E-16 |
| chr2 | 224579783 | 224598469 | 18687 | -9.44 | 6.44E-17 | 4.20E-16 |
| chr12 | 9300973 | 9324714 | 23742 | 3.13 | 6.47E-17 | 4.22E-16 |
| chr15 | 100164000 | 100188297 | 24298 | -9.02 | 6.47E-17 | 4.22E-16 |
| chr18 | 9516158 | 9530433 | 14276 | -9.03 | 6.49E-17 | 4.23E-16 |
| chr7 | 7564822 | 7579577 | 14756 | -9.03 | 6.49E-17 | 4.23E-16 |
| chr11 | 129299172 | 129302880 | 3709 | -3.01 | 6.54E-17 | 4.27E-16 |
| chr9 | 118253354 | 118299958 | 46605 | -9 | 6.57E-17 | 4.28E-16 |
| chr18 | 51814130 | 51828121 | 13992 | -8.98 | 6.57E-17 | 4.28E-16 |
| chr2 | 150766377 | 150779122 | 12746 | -8.98 | 6.58E-17 | 4.29E-16 |
| chr9 | 135969483 | 135974011 | 4529 | -8.99 | 6.63E-17 | 4.32E-16 |
| chr1 | 39048997 | 39063002 | 14006 | -9.03 | 6.66E-17 | 4.34E-16 |
| chr4 | 41584610 | 41587869 | 3260 | -9.03 | 6.66E-17 | 4.34E-16 |
| chr7 | 109824444 | 109840206 | 15763 | -8.99 | 6.67E-17 | 4.34E-16 |
| chr6 | 97810347 | 97831934 | 21588 | 6.87 | 6.69E-17 | 4.35E-16 |
| chr1 | 68260857 | 68283367 | 22511 | -9.07 | 6.71E-17 | 4.36E-16 |
| chr2 | 98293498 | 98311601 | 18104 | -8.98 | 6.72E-17 | 4.37E-16 |
| chr3 | 108793892 | 108809688 | 15797 | -8.99 | 6.73E-17 | 4.38E-16 |
| chr10 | 76465164 | 76494044 | 28881 | -9.23 | 6.74E-17 | 4.39E-16 |
| chr18 | 23650252 | 23658079 | 7828 | -9.01 | 6.74E-17 | 4.39E-16 |
| chr14 | 69265119 | 69272862 | 7744 | -9.01 | 6.78E-17 | 4.41E-16 |
| chr12 | 12922256 | 12928021 | 5766 | -9.04 | 6.78E-17 | 4.41E-16 |
| chr7 | 84747996 | 84790342 | 42347 | 9.69 | 6.80E-17 | 4.42E-16 |
| chr12 | 88395501 | 88417164 | 21664 | -9.03 | 6.82E-17 | 4.43E-16 |
| chr8 | 22375346 | 22379262 | 3917 | -8.99 | 6.83E-17 | 4.44E-16 |
| chrX | 73505516 | 73509343 | 3828 | -9.04 | 6.86E-17 | 4.46E-16 |
| chr7 | 120902963 | 120918518 | 15556 | 9.13 | 6.89E-17 | 4.48E-16 |
| chr21 | 37465100 | 37468099 | 3000 | -6.99 | 7.01E-17 | 4.55E-16 |
| chr3 | 122883445 | 122887051 | 3607 | -9.04 | 7.02E-17 | 4.56E-16 |
| chr12 | 117309661 | 117313784 | 4124 | -9 | 7.03E-17 | 4.57E-16 |
| chr2 | 164627648 | 164645920 | 18273 | -9.01 | 7.04E-17 | 4.57E-16 |
| chr4 | 101805405 | 101814610 | 9206 | -8.98 | 7.07E-17 | 4.59E-16 |
| chr1 | 43316007 | 43346965 | 30959 | -8.99 | 7.09E-17 | 4.60E-16 |
| chrX | 84522843 | 84528952 | 6110 | -9.01 | 7.09E-17 | 4.60E-16 |
| chr20 | 10102622 | 10109271 | 6650 | -9.05 | 7.12E-17 | 4.62E-16 |
| chr3 | 106315298 | 106778514 | 463217 | 2.6 | 7.15E-17 | 4.64E-16 |
| chr10 | 116938372 | 116959446 | 21075 | 9.08 | 7.17E-17 | 4.65E-16 |
| chr14 | 54370654 | 54383585 | 12932 | -9.33 | 7.17E-17 | 4.65E-16 |

| | | | | | | |
|---|---|---|---|---|---|---|
| chr7 | 14253010 | 14303374 | 50365 | -8.98 | 7.19E-17 | 4.66E-16 |
| chr14 | 67167360 | 67208018 | 40659 | -9.05 | 7.24E-17 | 4.69E-16 |
| chr17 | 4226875 | 4230763 | 3889 | -9.05 | 7.24E-17 | 4.69E-16 |
| chr12 | 87881969 | 87902774 | 20806 | -9.16 | 7.25E-17 | 4.70E-16 |
| chr15 | 63020553 | 63024436 | 3884 | -9.07 | 7.25E-17 | 4.70E-16 |
| chr9 | 37108093 | 37120135 | 12043 | -8.99 | 7.29E-17 | 4.72E-16 |
| chr2 | 164738183 | 164762641 | 24459 | -9.14 | 7.31E-17 | 4.74E-16 |
| chr16 | 70334108 | 70337408 | 3301 | -8.98 | 7.34E-17 | 4.75E-16 |
| chr17 | 19227114 | 19230129 | 3016 | -9 | 7.36E-17 | 4.76E-16 |
| chr6 | 6563570 | 6578636 | 15067 | -2.25 | 7.41E-17 | 4.80E-16 |
| chr10 | 72407059 | 72420116 | 13058 | -8.97 | 7.41E-17 | 4.80E-16 |
| chr15 | 52343283 | 52345704 | 2422 | -9.19 | 7.47E-17 | 4.83E-16 |
| chr3 | 4857106 | 4866211 | 9106 | -9.03 | 7.49E-17 | 4.84E-16 |
| chrX | 81792818 | 81819229 | 26412 | -9.07 | 7.49E-17 | 4.84E-16 |
| chr7 | 44563754 | 44569081 | 5328 | -9.04 | 7.49E-17 | 4.84E-16 |
| chr4 | 125615682 | 125622623 | 6942 | -9.14 | 7.54E-17 | 4.87E-16 |
| chr2 | 113271341 | 113274083 | 2743 | -9.06 | 7.57E-17 | 4.90E-16 |
| chr4 | 15486447 | 15499608 | 13162 | -8.97 | 7.62E-17 | 4.92E-16 |
| chr10 | 134919896 | 134934059 | 14164 | -9.05 | 7.63E-17 | 4.93E-16 |
| chr15 | 65192415 | 65199589 | 7175 | -9.05 | 7.63E-17 | 4.93E-16 |
| chr11 | 74023594 | 74029776 | 6183 | -9.03 | 7.63E-17 | 4.93E-16 |
| chr9 | 12799637 | 12812814 | 13178 | -9.05 | 7.66E-17 | 4.95E-16 |
| chr15 | 90153008 | 90156828 | 3821 | -9.26 | 7.66E-17 | 4.95E-16 |
| chr21 | 36176075 | 36179788 | 3714 | -9.02 | 7.67E-17 | 4.95E-16 |
| chr11 | 28052891 | 28069071 | 16181 | -9.12 | 7.68E-17 | 4.96E-16 |
| chr10 | 126796111 | 126799293 | 3183 | -8.98 | 7.73E-17 | 4.99E-16 |
| chr8 | 83795960 | 83850346 | 54387 | -9.31 | 7.77E-17 | 5.02E-16 |
| chr9 | 111865099 | 111868254 | 3156 | -8.98 | 7.79E-17 | 5.03E-16 |
| chr20 | 25406662 | 25411585 | 4924 | -9.11 | 7.80E-17 | 5.03E-16 |
| chr3 | 15830444 | 15834153 | 3710 | -9 | 7.81E-17 | 5.04E-16 |
| chr14 | 72108280 | 72122163 | 13884 | -9.12 | 7.84E-17 | 5.06E-16 |
| chr18 | 77526345 | 77530262 | 3918 | -9.01 | 7.85E-17 | 5.06E-16 |
| chr6 | 118893880 | 118902059 | 8180 | -9.06 | 7.85E-17 | 5.06E-16 |
| chrX | 82119022 | 82142037 | 23016 | -9.23 | 7.85E-17 | 5.06E-16 |
| chr3 | 98691694 | 98695817 | 4124 | -8.98 | 7.86E-17 | 5.07E-16 |
| chr16 | 4588923 | 4591619 | 2697 | -9.03 | 7.87E-17 | 5.07E-16 |
| chr1 | 239590106 | 239776089 | 185984 | 1.42 | 7.87E-17 | 5.07E-16 |
| chr7 | 139562715 | 139567432 | 4718 | -8.99 | 7.88E-17 | 5.08E-16 |
| chrX | 81901374 | 81926687 | 25314 | -8.96 | 7.88E-17 | 5.08E-16 |
| chr9 | 80939823 | 80969078 | 29256 | -9.11 | 7.89E-17 | 5.08E-16 |
| chr4 | 146429504 | 146436756 | 7253 | -8.97 | 7.90E-17 | 5.08E-16 |
| chr1 | 246965774 | 246975655 | 9882 | -9.2 | 7.91E-17 | 5.09E-16 |
| chr9 | 91307193 | 91316660 | 9468 | -8.99 | 7.91E-17 | 5.09E-16 |
| chr3 | 177992259 | 178006632 | 14374 | -9.31 | 7.96E-17 | 5.13E-16 |
| chr13 | 46899432 | 46938853 | 39422 | -8.96 | 7.96E-17 | 5.13E-16 |
| chr10 | 17489906 | 17524719 | 34814 | -9.04 | 8.02E-17 | 5.16E-16 |
| chr5 | 56934010 | 56955496 | 21487 | -9.33 | 8.04E-17 | 5.17E-16 |
| chr11 | 86842348 | 86862488 | 20141 | -9.19 | 8.08E-17 | 5.20E-16 |
| chr2 | 200737140 | 200750565 | 13426 | -9.07 | 8.10E-17 | 5.21E-16 |
| chr5 | 17195835 | 17211854 | 16020 | -8.96 | 8.11E-17 | 5.21E-16 |
| chr8 | 48446504 | 48468011 | 21508 | -9.14 | 8.11E-17 | 5.22E-16 |
| chr6 | 142662822 | 142673564 | 10743 | -9.04 | 8.15E-17 | 5.24E-16 |
| chr12 | 52337106 | 52360641 | 23536 | -8.98 | 8.16E-17 | 5.24E-16 |
| chr2 | 200864390 | 200885602 | 21213 | -8.96 | 8.20E-17 | 5.27E-16 |
| chr3 | 171570175 | 171587807 | 17633 | -9.18 | 8.20E-17 | 5.27E-16 |
| chr12 | 59133575 | 59186233 | 52659 | -9.29 | 8.29E-17 | 5.33E-16 |
| chr7 | 36433395 | 36436614 | 3220 | -9.03 | 8.31E-17 | 5.34E-16 |
| chr15 | 39297702 | 39307797 | 10096 | -9.04 | 8.33E-17 | 5.35E-16 |
| chr2 | 3274527 | 3281437 | 6911 | -8.96 | 8.35E-17 | 5.36E-16 |
| chr10 | 62665682 | 62689170 | 23489 | -8.96 | 8.38E-17 | 5.38E-16 |
| chr9 | 140625267 | 140630162 | 4896 | -8.96 | 8.41E-17 | 5.39E-16 |

| | | | | | | |
|---|---|---|---|---|---|---|
| chr21 | 38049074 | 38062041 | 12968 | -9.09 | 8.41E-17 | 5.39E-16 |
| chr2 | 131477400 | 131493345 | 15946 | -8.96 | 8.43E-17 | 5.41E-16 |
| chrX | 66760093 | 66777424 | 17332 | -9.01 | 8.44E-17 | 5.42E-16 |
| chr1 | 165644606 | 165657702 | 13097 | -9.06 | 8.47E-17 | 5.43E-16 |
| chr9 | 111632590 | 111635711 | 3122 | -9.07 | 8.47E-17 | 5.43E-16 |
| chr4 | 15195566 | 15221235 | 25670 | -9.32 | 8.48E-17 | 5.43E-16 |
| chr20 | 45285308 | 45302726 | 17419 | -3.52 | 8.49E-17 | 5.44E-16 |
| chr8 | 99357287 | 99366920 | 9634 | -3.32 | 8.53E-17 | 5.47E-16 |
| chr3 | 13218485 | 13226547 | 8063 | -8.96 | 8.54E-17 | 5.47E-16 |
| chrX | 130843025 | 130875712 | 32688 | -8.98 | 8.54E-17 | 5.47E-16 |
| chr3 | 101097138 | 101127609 | 30472 | -8.98 | 8.54E-17 | 5.47E-16 |
| chr12 | 46224009 | 46229283 | 5275 | -9.07 | 8.57E-17 | 5.49E-16 |
| chr20 | 627996 | 630834 | 2839 | -8.97 | 8.59E-17 | 5.50E-16 |
| chr7 | 65367092 | 65370916 | 3825 | -9.09 | 8.63E-17 | 5.53E-16 |
| chr16 | 22076919 | 22079997 | 3079 | -8.97 | 8.65E-17 | 5.53E-16 |
| chr16 | 25258306 | 25261228 | 2923 | -9.02 | 8.66E-17 | 5.54E-16 |
| chr7 | 18117403 | 18123191 | 5789 | -8.95 | 8.66E-17 | 5.54E-16 |
| chr4 | 169123086 | 169165110 | 42025 | -8.96 | 8.76E-17 | 5.61E-16 |
| chr15 | 81564220 | 81571655 | 7436 | -8.98 | 8.79E-17 | 5.63E-16 |
| chr8 | 123558126 | 123589970 | 31845 | -9.08 | 8.80E-17 | 5.63E-16 |
| chr7 | 11875848 | 11919532 | 43685 | -9.06 | 8.80E-17 | 5.63E-16 |
| chr16 | 23255117 | 23276799 | 21683 | -9.14 | 8.81E-17 | 5.63E-16 |
| chr12 | 90777096 | 90792742 | 15647 | -9.42 | 8.81E-17 | 5.63E-16 |
| chr6 | 137192240 | 137197577 | 5338 | -8.99 | 8.84E-17 | 5.65E-16 |
| chr21 | 16652852 | 16690068 | 37217 | -9.32 | 8.87E-17 | 5.67E-16 |
| chr21 | 47173231 | 47178137 | 4907 | -8.96 | 8.87E-17 | 5.67E-16 |
| chr20 | 14417947 | 14421130 | 3184 | -8.97 | 8.88E-17 | 5.67E-16 |
| chr11 | 8322953 | 8327582 | 4630 | -8.97 | 8.90E-17 | 5.68E-16 |
| chr1 | 100455595 | 100471629 | 16035 | -9 | 8.90E-17 | 5.69E-16 |
| chr3 | 135997161 | 136001043 | 3883 | -8.95 | 8.91E-17 | 5.69E-16 |
| chr10 | 48300662 | 48311032 | 10371 | -9.17 | 8.93E-17 | 5.70E-16 |
| chr9 | 84351210 | 84362518 | 11309 | -8.98 | 8.93E-17 | 5.70E-16 |
| chr3 | 33248701 | 33252662 | 3962 | -8.97 | 8.93E-17 | 5.70E-16 |
| chr13 | 37546766 | 37551388 | 4623 | -9 | 8.94E-17 | 5.71E-16 |
| chr18 | 70068811 | 70099070 | 30260 | -8.96 | 8.98E-17 | 5.73E-16 |
| chr8 | 62453913 | 62458186 | 4274 | -8.98 | 8.99E-17 | 5.73E-16 |
| chr14 | 54925390 | 54935903 | 10514 | -8.97 | 8.99E-17 | 5.73E-16 |
| chr1 | 66408086 | 66410026 | 1941 | -3.15 | 9.00E-17 | 5.74E-16 |
| chr20 | 59718318 | 59721204 | 2887 | -3.78 | 9.01E-17 | 5.74E-16 |
| chr5 | 153930262 | 153942110 | 11849 | -9.08 | 9.01E-17 | 5.74E-16 |
| chr5 | 81382881 | 81390114 | 7234 | -8.95 | 9.01E-17 | 5.74E-16 |
| chr19 | 6563863 | 6572646 | 8784 | -8.99 | 9.03E-17 | 5.76E-16 |
| chr4 | 146474187 | 146493714 | 19528 | -8.96 | 9.09E-17 | 5.79E-16 |
| chr22 | 47229227 | 47235736 | 6510 | -8.95 | 9.10E-17 | 5.80E-16 |
| chr13 | 50287433 | 50291787 | 4355 | -9.21 | 9.11E-17 | 5.80E-16 |
| chr12 | 68982654 | 68989259 | 6606 | -8.96 | 9.11E-17 | 5.80E-16 |
| chrX | 71891772 | 71904402 | 12631 | -9.09 | 9.12E-17 | 5.80E-16 |
| chr7 | 45941624 | 45949195 | 7572 | -8.95 | 9.12E-17 | 5.81E-16 |
| chr13 | 34462182 | 34475583 | 13402 | -8.96 | 9.12E-17 | 5.81E-16 |
| chr2 | 23944431 | 23949377 | 4947 | -9.07 | 9.18E-17 | 5.84E-16 |
| chr11 | 69362461 | 69377420 | 14960 | -9.12 | 9.21E-17 | 5.86E-16 |
| chr6 | 131629533 | 131637011 | 7479 | -8.95 | 9.27E-17 | 5.90E-16 |
| chr5 | 14162873 | 14165345 | 2473 | -8.96 | 9.30E-17 | 5.92E-16 |
| chr7 | 152427932 | 152432940 | 5009 | -9.07 | 9.31E-17 | 5.92E-16 |
| chr14 | 101064379 | 101068738 | 4360 | -8.96 | 9.31E-17 | 5.92E-16 |
| chr12 | 90257248 | 90266920 | 9673 | -9.11 | 9.35E-17 | 5.94E-16 |
| chr8 | 77882047 | 77889829 | 7783 | -8.99 | 9.35E-17 | 5.94E-16 |
| chr5 | 139994596 | 140009403 | 14808 | -9.39 | 9.40E-17 | 5.97E-16 |
| chr14 | 20668462 | 20698875 | 30414 | -9.22 | 9.40E-17 | 5.97E-16 |
| chr10 | 76265205 | 76275901 | 10697 | -8.98 | 9.40E-17 | 5.97E-16 |
| chr3 | 138258060 | 138293085 | 35026 | -9.18 | 9.41E-17 | 5.98E-16 |

| | | | | | | |
|---|---|---|---|---|---|---|
| chr4 | 130166736 | 130177503 | 10768 | -8.95 | 9.42E-17 | 5.98E-16 |
| chr2 | 220916676 | 220964511 | 47836 | -9.55 | 9.45E-17 | 6.00E-16 |
| chr6 | 11647495 | 11698280 | 50786 | -9.09 | 9.45E-17 | 6.00E-16 |
| chr7 | 135504653 | 135556467 | 51815 | 1.74 | 9.47E-17 | 6.01E-16 |
| chr1 | 42518839 | 42568924 | 50086 | -2.76 | 9.47E-17 | 6.01E-16 |
| chr15 | 96840041 | 96843361 | 3321 | -8.96 | 9.48E-17 | 6.02E-16 |
| chr19 | 8158680 | 8164434 | 5755 | -8.94 | 9.60E-17 | 6.09E-16 |
| chr9 | 72462561 | 72477064 | 14504 | -8.95 | 9.62E-17 | 6.10E-16 |
| chr18 | 33616626 | 33634586 | 17961 | -8.94 | 9.66E-17 | 6.12E-16 |
| chr11 | 27827737 | 27873401 | 45665 | -9.27 | 9.66E-17 | 6.12E-16 |
| chr14 | 93330753 | 93333042 | 2290 | -3.03 | 9.66E-17 | 6.13E-16 |
| chr1 | 8236468 | 8241606 | 5139 | -9 | 9.67E-17 | 6.13E-16 |
| chr17 | 78331264 | 78335611 | 4348 | -8.95 | 9.67E-17 | 6.13E-16 |
| chrX | 74787378 | 74804038 | 16661 | 9 | 9.68E-17 | 6.14E-16 |
| chr1 | 117965691 | 117971069 | 5379 | -8.96 | 9.69E-17 | 6.14E-16 |
| chr1 | 43172008 | 43181244 | 9237 | -8.98 | 9.69E-17 | 6.14E-16 |
| chr16 | 22265719 | 22272484 | 6766 | -8.99 | 9.74E-17 | 6.17E-16 |
| chr2 | 27364904 | 27367118 | 2215 | -8.98 | 9.79E-17 | 6.20E-16 |
| chr16 | 77439252 | 77443561 | 4310 | -8.96 | 9.85E-17 | 6.24E-16 |
| chr12 | 13368344 | 13373550 | 5207 | -9.01 | 9.85E-17 | 6.24E-16 |
| chr17 | 45439903 | 45465323 | 25421 | -9.07 | 9.89E-17 | 6.26E-16 |
| chr1 | 97123015 | 97137805 | 14791 | -8.94 | 9.90E-17 | 6.26E-16 |
| chr3 | 66284592 | 66294475 | 9884 | -8.94 | 9.90E-17 | 6.26E-16 |
| chr8 | 24431717 | 24707331 | 275615 | 3.18 | 9.92E-17 | 6.28E-16 |
| chr16 | 86357496 | 86360924 | 3429 | -8.94 | 9.93E-17 | 6.28E-16 |
| chr7 | 55450173 | 55458515 | 8343 | -8.95 | 9.95E-17 | 6.29E-16 |
| chr13 | 109094685 | 109125049 | 30365 | -9.23 | 9.95E-17 | 6.29E-16 |
| chr8 | 26351072 | 26368504 | 17433 | -9 | 9.97E-17 | 6.30E-16 |
| chr7 | 156420408 | 156446300 | 25893 | -9.38 | 9.99E-17 | 6.31E-16 |
| chr16 | 48318665 | 48322020 | 3356 | -8.94 | 1.00E-16 | 6.33E-16 |
| chr14 | 65552150 | 65556441 | 4292 | -9 | 1.01E-16 | 6.36E-16 |
| chr12 | 25649717 | 25667233 | 17517 | -9.01 | 1.01E-16 | 6.37E-16 |
| chr19 | 8658756 | 8671060 | 12305 | -8.94 | 1.01E-16 | 6.38E-16 |
| chr17 | 63761474 | 63779614 | 18141 | -9.25 | 1.01E-16 | 6.38E-16 |
| chr13 | 105802105 | 105824866 | 22762 | -8.98 | 1.02E-16 | 6.41E-16 |
| chr12 | 11051408 | 11058496 | 7089 | -9.01 | 1.02E-16 | 6.42E-16 |
| chr1 | 180409312 | 180448724 | 39413 | -9.33 | 1.02E-16 | 6.45E-16 |
| chr8 | 131433221 | 131452283 | 19063 | -9.24 | 1.03E-16 | 6.48E-16 |
| chr8 | 122649609 | 122669652 | 20044 | -9.26 | 1.03E-16 | 6.51E-16 |
| chr1 | 174157217 | 174202466 | 45250 | -9.5 | 1.03E-16 | 6.51E-16 |
| chr2 | 120606371 | 120612121 | 5751 | -8.96 | 1.03E-16 | 6.52E-16 |
| chr2 | 85061822 | 85065438 | 3617 | -8.93 | 1.03E-16 | 6.52E-16 |
| chr13 | 42654834 | 42679591 | 24758 | -8.95 | 1.04E-16 | 6.53E-16 |
| chr2 | 178232226 | 178243474 | 11249 | -8.93 | 1.04E-16 | 6.53E-16 |
| chr20 | 46440880 | 46444770 | 3891 | -8.94 | 1.04E-16 | 6.56E-16 |
| chr5 | 134754272 | 134766768 | 12497 | -8.96 | 1.04E-16 | 6.57E-16 |
| chr6 | 131748516 | 131760441 | 11926 | -8.96 | 1.05E-16 | 6.62E-16 |
| chr2 | 169228443 | 169237375 | 8933 | -6.67 | 1.06E-16 | 6.67E-16 |
| chr6 | 75090793 | 75094407 | 3615 | -3.49 | 1.06E-16 | 6.67E-16 |
| chr2 | 69759431 | 69775254 | 15824 | -8.93 | 1.06E-16 | 6.68E-16 |
| chr16 | 57582285 | 57595996 | 13712 | -9.4 | 1.06E-16 | 6.70E-16 |
| chr11 | 28431784 | 28457273 | 25490 | -8.96 | 1.06E-16 | 6.70E-16 |
| chr11 | 26336370 | 26352190 | 15821 | 9.08 | 1.07E-16 | 6.71E-16 |
| chr4 | 153172379 | 153185195 | 12817 | -8.94 | 1.07E-16 | 6.72E-16 |
| chr2 | 233665622 | 233668904 | 3283 | -9.05 | 1.08E-16 | 6.77E-16 |
| chr5 | 72678706 | 72683181 | 4476 | -8.94 | 1.08E-16 | 6.78E-16 |
| chr1 | 235644662 | 235647713 | 3052 | -8.98 | 1.08E-16 | 6.78E-16 |
| chr1 | 151322871 | 151336635 | 13765 | -8.96 | 1.09E-16 | 6.85E-16 |
| chr12 | 7257321 | 7260050 | 2730 | -8.95 | 1.10E-16 | 6.89E-16 |
| chr1 | 115651172 | 115668208 | 17037 | -8.99 | 1.10E-16 | 6.89E-16 |
| chr3 | 16594877 | 16608706 | 13830 | -9.13 | 1.10E-16 | 6.89E-16 |

| | | | | | | |
|---|---|---|---|---|---|---|
| chr15 | 76284499 | 76296493 | 11995 | -8.93 | 1.10E-16 | 6.90E-16 |
| chr2 | 176937064 | 176949969 | 12906 | -9.06 | 1.10E-16 | 6.90E-16 |
| chr5 | 102407780 | 102424886 | 17107 | -3.56 | 1.10E-16 | 6.90E-16 |
| chr7 | 2227508 | 2236426 | 8919 | -8.93 | 1.10E-16 | 6.92E-16 |
| chr3 | 113032700 | 113043906 | 11207 | -8.97 | 1.11E-16 | 6.96E-16 |
| chr4 | 95337771 | 95349810 | 12040 | -9.01 | 1.11E-16 | 6.96E-16 |
| chr3 | 146019493 | 146041593 | 22101 | -8.98 | 1.11E-16 | 6.96E-16 |
| chrX | 15435399 | 15443077 | 7679 | -8.94 | 1.11E-16 | 6.97E-16 |
| chr6 | 42399996 | 42404512 | 4517 | -8.93 | 1.11E-16 | 6.98E-16 |
| chr1 | 118289211 | 118292488 | 3278 | -2.89 | 1.11E-16 | 6.98E-16 |
| chr9 | 114623034 | 114644801 | 21768 | -9.06 | 1.11E-16 | 6.99E-16 |
| chr2 | 28790748 | 28796997 | 6250 | -9.13 | 1.11E-16 | 7.00E-16 |
| chr8 | 108586467 | 108605066 | 18600 | -9.3 | 1.12E-16 | 7.00E-16 |
| chr3 | 56193732 | 56214974 | 21243 | -8.93 | 1.12E-16 | 7.00E-16 |
| chr12 | 132224376 | 132239056 | 14681 | -8.97 | 1.12E-16 | 7.01E-16 |
| chr7 | 5045312 | 5052253 | 6942 | -8.94 | 1.12E-16 | 7.01E-16 |
| chr4 | 55096981 | 55109882 | 12902 | -8.92 | 1.12E-16 | 7.05E-16 |
| chr2 | 39745925 | 39750222 | 4298 | -8.96 | 1.13E-16 | 7.07E-16 |
| chr5 | 58466267 | 58499260 | 32994 | -8.96 | 1.13E-16 | 7.10E-16 |
| chr14 | 49055816 | 49106180 | 50365 | -9.1 | 1.14E-16 | 7.15E-16 |
| chr14 | 103206044 | 103210822 | 4779 | -8.93 | 1.14E-16 | 7.16E-16 |
| chr9 | 83703350 | 83716147 | 12798 | -8.98 | 1.14E-16 | 7.17E-16 |
| chr4 | 85439853 | 85477790 | 37938 | -9.04 | 1.14E-16 | 7.17E-16 |
| chr9 | 74790758 | 74807883 | 17126 | -9 | 1.15E-16 | 7.19E-16 |
| chr4 | 90028772 | 90036783 | 8012 | -8.92 | 1.15E-16 | 7.19E-16 |
| chr10 | 99951614 | 99954900 | 3287 | -8.92 | 1.15E-16 | 7.19E-16 |
| chr14 | 71408767 | 71418594 | 9828 | -8.96 | 1.15E-16 | 7.22E-16 |
| chr8 | 141822442 | 141826919 | 4478 | -8.97 | 1.16E-16 | 7.26E-16 |
| chr5 | 171396748 | 171409079 | 12332 | -8.94 | 1.16E-16 | 7.26E-16 |
| chr19 | 5199538 | 5211628 | 12091 | -8.92 | 1.17E-16 | 7.29E-16 |
| chr17 | 39198302 | 39214270 | 15969 | -8.96 | 1.17E-16 | 7.29E-16 |
| chr5 | 34540755 | 34548309 | 7555 | -9 | 1.17E-16 | 7.30E-16 |
| chr11 | 77440482 | 77444545 | 4064 | -8.92 | 1.17E-16 | 7.32E-16 |
| chr8 | 59206960 | 59234009 | 27050 | -8.94 | 1.17E-16 | 7.33E-16 |
| chr9 | 93931288 | 93936189 | 4902 | -8.95 | 1.17E-16 | 7.34E-16 |
| chr12 | 70601371 | 70609013 | 7643 | -9.03 | 1.18E-16 | 7.36E-16 |
| chr2 | 53581912 | 53608020 | 26109 | -8.92 | 1.18E-16 | 7.37E-16 |
| chr12 | 116990464 | 117009070 | 18607 | -9.31 | 1.18E-16 | 7.38E-16 |
| chr10 | 63881716 | 63903970 | 22255 | -9.16 | 1.18E-16 | 7.39E-16 |
| chr1 | 167583155 | 167595199 | 12045 | -9.1 | 1.18E-16 | 7.40E-16 |
| chr14 | 35992486 | 35996431 | 3946 | -8.95 | 1.19E-16 | 7.41E-16 |
| chr2 | 68447069 | 68450298 | 3230 | -8.92 | 1.19E-16 | 7.42E-16 |
| chr12 | 105165358 | 105174825 | 9468 | -9.02 | 1.19E-16 | 7.46E-16 |
| chr6 | 116618304 | 116620823 | 2520 | -6.52 | 1.20E-16 | 7.48E-16 |
| chr10 | 6945613 | 6952168 | 6556 | -9 | 1.20E-16 | 7.49E-16 |
| chr2 | 8196140 | 8217901 | 21762 | -8.92 | 1.20E-16 | 7.50E-16 |
| chr6 | 20420899 | 20441679 | 20781 | -9.04 | 1.20E-16 | 7.50E-16 |
| chr16 | 24368291 | 24505819 | 137529 | -2.06 | 1.20E-16 | 7.51E-16 |
| chr15 | 73472963 | 73478302 | 5340 | -8.93 | 1.20E-16 | 7.51E-16 |
| chr12 | 28422163 | 28444189 | 22027 | -9.24 | 1.21E-16 | 7.54E-16 |
| chr2 | 3251471 | 3255666 | 4196 | -8.92 | 1.22E-16 | 7.62E-16 |
| chr9 | 139824975 | 139829271 | 4297 | -8.98 | 1.22E-16 | 7.62E-16 |
| chr2 | 155206044 | 155224757 | 18714 | 8.98 | 1.23E-16 | 7.64E-16 |
| chr7 | 81534124 | 81545896 | 11773 | -8.93 | 1.23E-16 | 7.66E-16 |
| chr20 | 9417866 | 9420897 | 3032 | -9.18 | 1.23E-16 | 7.66E-16 |
| chr5 | 10918843 | 10933437 | 14595 | -1.91 | 1.24E-16 | 7.73E-16 |
| chr2 | 235602063 | 235608410 | 6348 | -9.16 | 1.24E-16 | 7.73E-16 |
| chr9 | 135640168 | 135666835 | 26668 | -9.09 | 1.24E-16 | 7.74E-16 |
| chr20 | 35620972 | 35625410 | 4439 | -8.91 | 1.24E-16 | 7.74E-16 |
| chr4 | 17983105 | 17991848 | 8744 | -8.93 | 1.25E-16 | 7.76E-16 |
| chr10 | 127764392 | 127782342 | 17951 | -8.98 | 1.25E-16 | 7.77E-16 |

| | | | | | | |
|---|---|---|---|---|---|---|
| chr12 | 111780443 | 111788736 | 8294 | -8.92 | 1.25E-16 | 7.78E-16 |
| chr7 | 94984218 | 94990003 | 5786 | -8.91 | 1.25E-16 | 7.78E-16 |
| chr15 | 56223791 | 56232428 | 8638 | -8.92 | 1.26E-16 | 7.82E-16 |
| chr22 | 45005867 | 45011584 | 5718 | -8.94 | 1.26E-16 | 7.83E-16 |
| chr8 | 63401257 | 63425239 | 23983 | 2.84 | 1.26E-16 | 7.84E-16 |
| chr2 | 23562976 | 23569876 | 6901 | -9.03 | 1.26E-16 | 7.85E-16 |
| chr13 | 50783899 | 50794189 | 10291 | -8.95 | 1.26E-16 | 7.86E-16 |
| chr7 | 35600510 | 35611588 | 11079 | -2.66 | 1.27E-16 | 7.86E-16 |
| chr4 | 120400482 | 120406547 | 6066 | -8.92 | 1.27E-16 | 7.86E-16 |
| chr22 | 24265452 | 24274581 | 9130 | -8.97 | 1.27E-16 | 7.90E-16 |
| chr10 | 104128354 | 104131710 | 3357 | -9.03 | 1.27E-16 | 7.91E-16 |
| chr20 | 25624603 | 25627501 | 2899 | -8.94 | 1.28E-16 | 7.92E-16 |
| chr16 | 8773215 | 8786807 | 13593 | -9.08 | 1.28E-16 | 7.93E-16 |
| chr6 | 145760989 | 145769384 | 8396 | -8.96 | 1.28E-16 | 7.94E-16 |
| chr2 | 165256865 | 165272100 | 15236 | -8.91 | 1.28E-16 | 7.94E-16 |
| chr3 | 46958803 | 46971140 | 12338 | -8.92 | 1.28E-16 | 7.96E-16 |
| chr8 | 17603001 | 17607653 | 4653 | -8.91 | 1.28E-16 | 7.96E-16 |
| chr8 | 19395169 | 19407408 | 12240 | -8.96 | 1.28E-16 | 7.96E-16 |
| chr1 | 243724654 | 243726974 | 2321 | -8.94 | 1.28E-16 | 7.97E-16 |
| chr8 | 64190354 | 64205240 | 14887 | 9.03 | 1.29E-16 | 8.00E-16 |
| chrX | 109925950 | 109944620 | 18671 | 9.03 | 1.29E-16 | 8.00E-16 |
| chr4 | 3667568 | 3675667 | 8100 | -8.95 | 1.29E-16 | 8.02E-16 |
| chr1 | 27505285 | 27512382 | 7098 | -9.11 | 1.30E-16 | 8.04E-16 |
| chr8 | 66562617 | 66572909 | 10293 | -8.92 | 1.30E-16 | 8.05E-16 |
| chr18 | 33213727 | 33229632 | 15906 | -8.91 | 1.30E-16 | 8.08E-16 |
| chr10 | 134960201 | 134964382 | 4182 | -9.08 | 1.30E-16 | 8.08E-16 |
| chr5 | 159164963 | 159181213 | 16251 | -9.12 | 1.30E-16 | 8.08E-16 |
| chr8 | 11438197 | 11468377 | 30181 | -9.2 | 1.31E-16 | 8.10E-16 |
| chr3 | 185791059 | 185804649 | 13591 | -8.99 | 1.31E-16 | 8.10E-16 |
| chr5 | 41466689 | 41478102 | 11414 | -8.91 | 1.31E-16 | 8.11E-16 |
| chr2 | 226737631 | 226762751 | 25121 | -9.13 | 1.31E-16 | 8.11E-16 |
| chr11 | 61853609 | 61885521 | 31913 | -9.13 | 1.31E-16 | 8.13E-16 |
| chr17 | 47539770 | 47554080 | 14311 | -9.04 | 1.31E-16 | 8.13E-16 |
| chr17 | 20718622 | 20727059 | 8438 | -8.93 | 1.32E-16 | 8.16E-16 |
| chr9 | 115127270 | 115139567 | 12298 | -9.19 | 1.33E-16 | 8.20E-16 |
| chr4 | 122274021 | 122277400 | 3380 | -3.19 | 1.33E-16 | 8.21E-16 |
| chr5 | 90336350 | 90350876 | 14527 | -8.95 | 1.33E-16 | 8.23E-16 |
| chr1 | 22679778 | 22709485 | 29708 | -8.91 | 1.33E-16 | 8.25E-16 |
| chr19 | 968994 | 972397 | 3404 | -8.9 | 1.34E-16 | 8.27E-16 |
| chr11 | 72617814 | 72625405 | 7592 | -8.93 | 1.34E-16 | 8.27E-16 |
| chr12 | 89079307 | 89102526 | 23220 | -8.9 | 1.34E-16 | 8.31E-16 |
| chr8 | 90147816 | 90175550 | 27735 | -8.91 | 1.35E-16 | 8.32E-16 |
| chr10 | 23452490 | 23454664 | 2175 | -4.12 | 1.35E-16 | 8.37E-16 |
| chr4 | 28341993 | 28355461 | 13469 | -9.12 | 1.36E-16 | 8.40E-16 |
| chr10 | 14757630 | 14790177 | 32548 | -9.72 | 1.36E-16 | 8.40E-16 |
| chr14 | 27290960 | 27321221 | 30262 | -9.16 | 1.36E-16 | 8.41E-16 |
| chr20 | 62135406 | 62155859 | 20454 | -9.02 | 1.37E-16 | 8.47E-16 |
| chr9 | 111913321 | 111917822 | 4502 | -8.93 | 1.37E-16 | 8.49E-16 |
| chr19 | 43020065 | 43021769 | 1705 | -9.04 | 1.38E-16 | 8.50E-16 |
| chr19 | 46340959 | 46345154 | 4196 | -8.92 | 1.38E-16 | 8.50E-16 |
| chr3 | 101865672 | 101906580 | 40909 | -9.81 | 1.38E-16 | 8.52E-16 |
| chrX | 3162764 | 3220845 | 58082 | -1.76 | 1.38E-16 | 8.53E-16 |
| chr3 | 6131731 | 6154831 | 23101 | 2.21 | 1.38E-16 | 8.53E-16 |
| chr1 | 205443170 | 205445925 | 2756 | -9.06 | 1.39E-16 | 8.57E-16 |
| chr7 | 62374502 | 62678407 | 303906 | -2.57 | 1.39E-16 | 8.57E-16 |
| chr15 | 59609175 | 59614173 | 4999 | -8.94 | 1.39E-16 | 8.59E-16 |
| chr1 | 32768988 | 32771773 | 2786 | -8.97 | 1.39E-16 | 8.60E-16 |
| chr10 | 121724403 | 121729092 | 4690 | -8.9 | 1.39E-16 | 8.60E-16 |
| chr15 | 45532700 | 45539239 | 6540 | -8.91 | 1.40E-16 | 8.64E-16 |
| chr6 | 4112280 | 4133632 | 21353 | -8.93 | 1.40E-16 | 8.65E-16 |
| chr5 | 83243582 | 83249334 | 5753 | 9.08 | 1.41E-16 | 8.69E-16 |

| | | | | | | |
|---|---|---|---|---|---|---|
| chr14 | 20739846 | 20748006 | 8161 | -8.96 | 1.41E-16 | 8.69E-16 |
| chr10 | 33042504 | 33061918 | 19415 | -8.98 | 1.41E-16 | 8.71E-16 |
| chr15 | 50500377 | 50508042 | 7666 | -9.03 | 1.42E-16 | 8.76E-16 |
| chr12 | 127619064 | 127633389 | 14326 | -9 | 1.43E-16 | 8.82E-16 |
| chr2 | 45117982 | 45139500 | 21519 | -8.96 | 1.43E-16 | 8.83E-16 |
| chr10 | 76039643 | 76048221 | 8579 | -9.04 | 1.44E-16 | 8.84E-16 |
| chr11 | 78087368 | 78099992 | 12625 | -9.08 | 1.44E-16 | 8.84E-16 |
| chr21 | 17475686 | 17496838 | 21153 | -8.98 | 1.44E-16 | 8.87E-16 |
| chr20 | 61351114 | 61354123 | 3010 | -8.9 | 1.45E-16 | 8.89E-16 |
| chr1 | 1995122 | 2011937 | 16816 | -8.92 | 1.45E-16 | 8.89E-16 |
| chr12 | 62584792 | 62599015 | 14224 | -8.91 | 1.45E-16 | 8.92E-16 |
| chr7 | 118014396 | 118667128 | 652733 | 1.99 | 1.45E-16 | 8.94E-16 |
| chr8 | 91844939 | 91863287 | 18349 | 9.12 | 1.46E-16 | 8.95E-16 |
| chr21 | 40137541 | 40165534 | 27994 | -8.97 | 1.46E-16 | 8.96E-16 |
| chr11 | 65830159 | 65833442 | 3284 | -8.97 | 1.46E-16 | 8.97E-16 |
| chr10 | 95612598 | 95646694 | 34097 | -9.18 | 1.46E-16 | 8.98E-16 |
| chr1 | 63926050 | 63929336 | 3287 | -8.95 | 1.46E-16 | 9.00E-16 |
| chr3 | 102302911 | 102307592 | 4682 | 7.08 | 1.46E-16 | 9.00E-16 |
| chr17 | 17318319 | 17334296 | 15978 | -9.25 | 1.47E-16 | 9.06E-16 |
| chr7 | 30138367 | 30155702 | 17336 | -9.03 | 1.48E-16 | 9.08E-16 |
| chr2 | 238362487 | 238370354 | 7868 | -8.89 | 1.48E-16 | 9.09E-16 |
| chr9 | 98653303 | 98656139 | 2837 | -8.9 | 1.48E-16 | 9.11E-16 |
| chr8 | 21729968 | 21736574 | 6607 | -8.96 | 1.49E-16 | 9.12E-16 |
| chrX | 116852390 | 116884279 | 31890 | 2.02 | 1.49E-16 | 9.13E-16 |
| chr1 | 6768818 | 6773812 | 4995 | -8.96 | 1.49E-16 | 9.13E-16 |
| chr11 | 122053386 | 122058409 | 5024 | -8.96 | 1.49E-16 | 9.14E-16 |
| chr3 | 169927462 | 169937471 | 10010 | -8.89 | 1.49E-16 | 9.14E-16 |
| chr5 | 134288526 | 134291582 | 3057 | -8.89 | 1.49E-16 | 9.14E-16 |
| chr17 | 7431128 | 7442523 | 11396 | -9.12 | 1.49E-16 | 9.14E-16 |
| chr5 | 171357958 | 171363032 | 5075 | -8.9 | 1.51E-16 | 9.23E-16 |
| chr10 | 96497247 | 96519706 | 22460 | -9.01 | 1.51E-16 | 9.27E-16 |
| chrX | 56569580 | 56574005 | 4426 | -6.71 | 1.52E-16 | 9.30E-16 |
| chr9 | 18983303 | 19006658 | 23356 | -7.04 | 1.52E-16 | 9.30E-16 |
| chr18 | 3755351 | 3769683 | 14333 | -9.18 | 1.52E-16 | 9.30E-16 |
| chr7 | 130800624 | 130805037 | 4414 | -8.92 | 1.52E-16 | 9.31E-16 |
| chr2 | 58460822 | 58470631 | 9810 | -9.02 | 1.52E-16 | 9.32E-16 |
| chr17 | 70872312 | 70877011 | 4700 | -8.9 | 1.52E-16 | 9.33E-16 |
| chr11 | 117322746 | 117326453 | 3708 | -8.89 | 1.52E-16 | 9.33E-16 |
| chr1 | 193399222 | 193406835 | 7614 | -8.96 | 1.52E-16 | 9.33E-16 |
| chr6 | 168951057 | 168968118 | 17062 | -8.94 | 1.53E-16 | 9.38E-16 |
| chr11 | 86942038 | 86945406 | 3369 | -8.9 | 1.53E-16 | 9.38E-16 |
| chr1 | 67343281 | 67346207 | 2927 | -6.97 | 1.53E-16 | 9.39E-16 |
| chr9 | 80302505 | 80327465 | 24961 | -9.12 | 1.54E-16 | 9.40E-16 |
| chr7 | 130509285 | 130513676 | 4392 | -8.9 | 1.54E-16 | 9.41E-16 |
| chr1 | 233599883 | 233654686 | 54804 | -9.5 | 1.54E-16 | 9.41E-16 |
| chr9 | 75145087 | 75153625 | 8539 | -9.03 | 1.54E-16 | 9.43E-16 |
| chr10 | 45392272 | 45419399 | 27128 | -9.18 | 1.54E-16 | 9.44E-16 |
| chr10 | 8044337 | 8048797 | 4461 | -8.93 | 1.54E-16 | 9.44E-16 |
| chr17 | 4519153 | 4526766 | 7614 | -8.97 | 1.55E-16 | 9.45E-16 |
| chr5 | 64257744 | 64262094 | 4351 | -9.08 | 1.55E-16 | 9.47E-16 |
| chr17 | 55323190 | 55329706 | 6517 | -9.13 | 1.55E-16 | 9.48E-16 |
| chr10 | 93013676 | 93019230 | 5555 | -8.89 | 1.55E-16 | 9.48E-16 |
| chr4 | 185891069 | 185931013 | 39945 | -9.23 | 1.56E-16 | 9.52E-16 |
| chr2 | 218708212 | 218712255 | 4044 | -8.95 | 1.56E-16 | 9.53E-16 |
| chr14 | 89270297 | 89277333 | 7037 | -8.91 | 1.56E-16 | 9.54E-16 |
| chr13 | 52597361 | 52605422 | 8062 | -8.94 | 1.56E-16 | 9.54E-16 |
| chrX | 64840497 | 64848998 | 8502 | -8.89 | 1.56E-16 | 9.54E-16 |
| chr12 | 88666933 | 88698676 | 31744 | -8.93 | 1.57E-16 | 9.57E-16 |
| chr5 | 40239797 | 40250985 | 11189 | -8.9 | 1.57E-16 | 9.57E-16 |
| chr1 | 185417223 | 185428960 | 11738 | -8.96 | 1.57E-16 | 9.57E-16 |
| chr1 | 54306568 | 54315350 | 8783 | -8.97 | 1.57E-16 | 9.61E-16 |

| | | | | | | |
|---|---|---|---|---|---|---|
| chr14 | 35950483 | 35971931 | 21449 | -9.11 | 1.59E-16 | 9.67E-16 |
| chr4 | 95699617 | 95705262 | 5646 | -8.89 | 1.59E-16 | 9.70E-16 |
| chr10 | 127825471 | 127841686 | 16216 | -8.92 | 1.59E-16 | 9.72E-16 |
| chr9 | 87701685 | 87721853 | 20169 | -8.89 | 1.59E-16 | 9.73E-16 |
| chr6 | 35875876 | 35880228 | 4353 | -8.99 | 1.60E-16 | 9.75E-16 |
| chr5 | 126855873 | 126861208 | 5336 | -8.89 | 1.60E-16 | 9.77E-16 |
| chr2 | 228012717 | 228033163 | 20447 | -9.14 | 1.61E-16 | 9.81E-16 |
| chr3 | 45522254 | 45526989 | 4736 | -8.93 | 1.61E-16 | 9.84E-16 |
| chr18 | 39494547 | 39504380 | 9834 | 7.73 | 1.62E-16 | 9.87E-16 |
| chr6 | 81329108 | 81356286 | 27179 | -8.9 | 1.62E-16 | 9.89E-16 |
| chr18 | 8408105 | 8415653 | 7549 | -8.88 | 1.63E-16 | 9.92E-16 |
| chr1 | 211616445 | 211636520 | 20076 | -9.09 | 1.63E-16 | 9.95E-16 |
| chr6 | 127829523 | 127834177 | 4655 | -8.92 | 1.64E-16 | 9.97E-16 |
| chr22 | 33543866 | 33649451 | 105586 | -1.1 | 1.64E-16 | 9.99E-16 |
| chr3 | 54242424 | 54282085 | 39662 | -8.98 | 1.65E-16 | 1.00E-15 |
| chr12 | 64966105 | 64970561 | 4457 | -8.99 | 1.65E-16 | 1.00E-15 |
| chr7 | 24771806 | 24779778 | 7973 | -8.91 | 1.66E-16 | 1.01E-15 |
| chr17 | 75132565 | 75135974 | 3410 | -9.02 | 1.66E-16 | 1.01E-15 |
| chr1 | 109872290 | 109876440 | 4151 | -8.89 | 1.66E-16 | 1.01E-15 |
| chr1 | 200900443 | 200917934 | 17492 | -8.88 | 1.67E-16 | 1.01E-15 |
| chr13 | 49721218 | 49725877 | 4660 | -8.89 | 1.67E-16 | 1.02E-15 |
| chr12 | 19175725 | 19189774 | 14050 | -8.88 | 1.67E-16 | 1.02E-15 |
| chrX | 68224991 | 68232383 | 7393 | -9.05 | 1.67E-16 | 1.02E-15 |
| chr5 | 10195480 | 10200553 | 5074 | -8.88 | 1.68E-16 | 1.02E-15 |
| chr2 | 62727828 | 62768458 | 40631 | -9.2 | 1.68E-16 | 1.02E-15 |
| chr17 | 34832891 | 34846820 | 13930 | -8.89 | 1.69E-16 | 1.03E-15 |
| chr1 | 22010733 | 22014032 | 3300 | -8.88 | 1.69E-16 | 1.03E-15 |
| chr4 | 139152180 | 139172067 | 19888 | -8.95 | 1.70E-16 | 1.03E-15 |
| chr14 | 43372342 | 43422566 | 50225 | -9.31 | 1.71E-16 | 1.04E-15 |
| chr10 | 106192489 | 106212549 | 20061 | 9.07 | 1.71E-16 | 1.04E-15 |
| chr3 | 5856710 | 5877372 | 20663 | -9.39 | 1.72E-16 | 1.04E-15 |
| chr11 | 74205855 | 74228992 | 23138 | -9.01 | 1.73E-16 | 1.05E-15 |
| chr12 | 64525412 | 64534338 | 8927 | -9.02 | 1.73E-16 | 1.05E-15 |
| chr15 | 101127068 | 101132589 | 5522 | -8.9 | 1.74E-16 | 1.06E-15 |
| chr8 | 41176251 | 41178028 | 1778 | -6.51 | 1.75E-16 | 1.06E-15 |
| chr5 | 20950652 | 20999546 | 48895 | -9.4 | 1.75E-16 | 1.06E-15 |
| chr8 | 74851291 | 74866564 | 15274 | -8.87 | 1.75E-16 | 1.06E-15 |
| chr6 | 130617849 | 130628516 | 10668 | -8.98 | 1.75E-16 | 1.06E-15 |
| chr5 | 172843229 | 172852198 | 8970 | -8.88 | 1.76E-16 | 1.07E-15 |
| chr4 | 151186633 | 151193624 | 6992 | -8.87 | 1.76E-16 | 1.07E-15 |
| chr14 | 59501533 | 59533294 | 31762 | -3.46 | 1.76E-16 | 1.07E-15 |
| chr10 | 895295 | 907582 | 12288 | -9.02 | 1.78E-16 | 1.08E-15 |
| chr1 | 212806056 | 212814759 | 8704 | -8.91 | 1.78E-16 | 1.08E-15 |
| chr7 | 23690844 | 23699897 | 9054 | -8.88 | 1.78E-16 | 1.08E-15 |
| chr1 | 61851006 | 61856257 | 5252 | -8.94 | 1.78E-16 | 1.08E-15 |
| chr22 | 35827795 | 35830922 | 3128 | -8.91 | 1.78E-16 | 1.08E-15 |
| chr14 | 99434204 | 99449040 | 14837 | -8.93 | 1.78E-16 | 1.08E-15 |
| chr17 | 70664359 | 70677777 | 13419 | -9.05 | 1.79E-16 | 1.08E-15 |
| chr1 | 49366721 | 49396111 | 29391 | -8.88 | 1.79E-16 | 1.09E-15 |
| chr1 | 110663520 | 110668909 | 5390 | -8.87 | 1.81E-16 | 1.09E-15 |
| chrX | 130054411 | 130059761 | 5351 | -8.89 | 1.81E-16 | 1.09E-15 |
| chr4 | 131113348 | 131126909 | 13562 | -8.9 | 1.81E-16 | 1.09E-15 |
| chr2 | 64613694 | 64619007 | 5314 | -8.87 | 1.82E-16 | 1.10E-15 |
| chr4 | 14839731 | 14859498 | 19768 | -8.87 | 1.83E-16 | 1.10E-15 |
| chr4 | 165480706 | 165648086 | 167381 | 2.98 | 1.84E-16 | 1.11E-15 |
| chr3 | 149793065 | 149820344 | 27280 | -8.92 | 1.85E-16 | 1.12E-15 |
| chr9 | 113947769 | 113990881 | 43113 | -9.37 | 1.85E-16 | 1.12E-15 |
| chr7 | 139501250 | 139549143 | 47894 | -9 | 1.85E-16 | 1.12E-15 |
| chr8 | 33551365 | 33839447 | 288083 | 1.91 | 1.86E-16 | 1.12E-15 |
| chr1 | 101502913 | 101507657 | 4745 | -8.93 | 1.86E-16 | 1.12E-15 |
| chr3 | 124974331 | 124978720 | 4390 | -8.87 | 1.86E-16 | 1.13E-15 |

| | | | | | | |
|---|---|---|---|---|---|---|
| chr9 | 119348713 | 119352369 | 3657 | -8.94 | 1.87E-16 | 1.13E-15 |
| chr13 | 35028896 | 35048549 | 19654 | -9.12 | 1.87E-16 | 1.13E-15 |
| chr20 | 4776912 | 4780580 | 3669 | -8.88 | 1.87E-16 | 1.13E-15 |
| chr13 | 90677221 | 90701075 | 23855 | -8.89 | 1.88E-16 | 1.14E-15 |
| chr3 | 10285315 | 10288516 | 3202 | -8.96 | 1.89E-16 | 1.14E-15 |
| chr9 | 29215285 | 29230885 | 15601 | -8.87 | 1.89E-16 | 1.14E-15 |
| chr16 | 11574509 | 11585896 | 11388 | -9.18 | 1.89E-16 | 1.14E-15 |
| chr20 | 10077700 | 10089139 | 11440 | -9.12 | 1.90E-16 | 1.15E-15 |
| chr6 | 56080650 | 56097466 | 16817 | -8.87 | 1.90E-16 | 1.15E-15 |
| chr11 | 113518401 | 113538869 | 20469 | -2.09 | 1.90E-16 | 1.15E-15 |
| chr5 | 14086616 | 14112063 | 25448 | -8.89 | 1.90E-16 | 1.15E-15 |
| chr7 | 86951474 | 86964845 | 13372 | -9.17 | 1.90E-16 | 1.15E-15 |
| chr10 | 128854408 | 128859113 | 4706 | -8.97 | 1.90E-16 | 1.15E-15 |
| chr14 | 61420043 | 61426375 | 6333 | -8.93 | 1.90E-16 | 1.15E-15 |
| chr3 | 164282389 | 164337717 | 55329 | -9.35 | 1.91E-16 | 1.15E-15 |
| chr2 | 158611784 | 158615725 | 3942 | -9 | 1.91E-16 | 1.15E-15 |
| chr4 | 67462677 | 67486478 | 23802 | -8.91 | 1.91E-16 | 1.15E-15 |
| chr1 | 4727108 | 4741651 | 14544 | -8.98 | 1.93E-16 | 1.16E-15 |
| chr1 | 218504367 | 218509065 | 4699 | -8.86 | 1.93E-16 | 1.17E-15 |
| chr7 | 3052773 | 3066431 | 13659 | -9.01 | 1.95E-16 | 1.17E-15 |
| chr17 | 45377966 | 45381547 | 3582 | -8.88 | 1.95E-16 | 1.17E-15 |
| chr15 | 34779911 | 34787871 | 7961 | -9.2 | 1.95E-16 | 1.17E-15 |
| chr8 | 90258583 | 90264096 | 5514 | -8.95 | 1.95E-16 | 1.18E-15 |
| chr5 | 174830665 | 174869690 | 39026 | -3.04 | 1.95E-16 | 1.18E-15 |
| chrX | 36611477 | 36620420 | 8944 | -6.19 | 1.96E-16 | 1.18E-15 |
| chr8 | 32211405 | 32238641 | 27237 | -9.01 | 1.96E-16 | 1.18E-15 |
| chr20 | 3918157 | 3922594 | 4438 | -9.08 | 1.96E-16 | 1.18E-15 |
| chrX | 74527097 | 74553738 | 26642 | -8.88 | 1.96E-16 | 1.18E-15 |
| chr10 | 119452662 | 119488926 | 36265 | -9 | 1.97E-16 | 1.18E-15 |
| chr8 | 100161504 | 100164320 | 2817 | -8.86 | 1.98E-16 | 1.19E-15 |
| chr12 | 34427548 | 34440181 | 12634 | -8.86 | 1.98E-16 | 1.19E-15 |
| chr4 | 126636981 | 126665004 | 28024 | -8.91 | 2.01E-16 | 1.21E-15 |
| chr19 | 7124666 | 7147523 | 22858 | -9.14 | 2.01E-16 | 1.21E-15 |
| chr19 | 3780159 | 3797862 | 17704 | -9.1 | 2.01E-16 | 1.21E-15 |
| chr1 | 37206365 | 37229288 | 22924 | -8.87 | 2.02E-16 | 1.21E-15 |
| chr6 | 403716 | 422331 | 18616 | -9.01 | 2.03E-16 | 1.22E-15 |
| chr6 | 953213 | 993410 | 40198 | -1.76 | 2.03E-16 | 1.22E-15 |
| chr15 | 29573154 | 29589526 | 16373 | -9.1 | 2.03E-16 | 1.22E-15 |
| chr9 | 92158840 | 92163173 | 4334 | -8.86 | 2.03E-16 | 1.22E-15 |
| chr9 | 90067795 | 90073712 | 5918 | -9.02 | 2.04E-16 | 1.22E-15 |
| chr16 | 11301032 | 11313747 | 12716 | -8.87 | 2.04E-16 | 1.22E-15 |
| chr17 | 60736474 | 60740385 | 3912 | -8.86 | 2.04E-16 | 1.22E-15 |
| chr2 | 197357951 | 197366274 | 8324 | -8.88 | 2.04E-16 | 1.23E-15 |
| chr3 | 178691352 | 178713034 | 21683 | -8.9 | 2.04E-16 | 1.23E-15 |
| chrX | 54367107 | 54405879 | 38773 | -9.05 | 2.05E-16 | 1.23E-15 |
| chr17 | 60784692 | 60788189 | 3498 | -8.93 | 2.05E-16 | 1.23E-15 |
| chr14 | 34605825 | 34628109 | 22285 | -8.91 | 2.05E-16 | 1.23E-15 |
| chrX | 45435219 | 45449048 | 13830 | -8.89 | 2.06E-16 | 1.23E-15 |
| chr8 | 74072684 | 74082908 | 10225 | -8.95 | 2.06E-16 | 1.24E-15 |
| chr4 | 57487930 | 57494038 | 6109 | -7.6 | 2.07E-16 | 1.24E-15 |
| chr12 | 121231669 | 121235015 | 3347 | -8.95 | 2.07E-16 | 1.24E-15 |
| chr17 | 66272552 | 66278091 | 5540 | -9.04 | 2.08E-16 | 1.24E-15 |
| chr2 | 151433372 | 151442973 | 9602 | -8.86 | 2.08E-16 | 1.24E-15 |
| chr7 | 31170778 | 31196321 | 25544 | -8.88 | 2.08E-16 | 1.25E-15 |
| chr1 | 87980643 | 88012217 | 31575 | -8.92 | 2.08E-16 | 1.25E-15 |
| chr2 | 45376277 | 45379235 | 2959 | -8.92 | 2.08E-16 | 1.25E-15 |
| chr15 | 66554266 | 66568799 | 14534 | -8.93 | 2.09E-16 | 1.25E-15 |
| chr19 | 33417608 | 33420336 | 2729 | -8.9 | 2.09E-16 | 1.25E-15 |
| chr13 | 112843763 | 112848540 | 4778 | -2.85 | 2.10E-16 | 1.26E-15 |
| chr12 | 109474640 | 109481064 | 6425 | -8.9 | 2.11E-16 | 1.26E-15 |
| chr2 | 72833680 | 72845558 | 11879 | -8.89 | 2.11E-16 | 1.26E-15 |

| | | | | | | |
|---|---|---|---|---|---|---|
| chr20 | 42107304 | 42131668 | 24365 | -9.03 | 2.12E-16 | 1.27E-15 |
| chr4 | 176031051 | 176046316 | 15266 | -9.04 | 2.12E-16 | 1.27E-15 |
| chr2 | 45798110 | 45800501 | 2392 | -8.93 | 2.13E-16 | 1.28E-15 |
| chr15 | 75405016 | 75414779 | 9764 | -9.16 | 2.14E-16 | 1.28E-15 |
| chr1 | 94674428 | 94677267 | 2840 | -8.87 | 2.14E-16 | 1.28E-15 |
| chr7 | 87482217 | 87502330 | 20114 | -8.86 | 2.14E-16 | 1.28E-15 |
| chr13 | 66189216 | 66204582 | 15367 | 8.98 | 2.15E-16 | 1.28E-15 |
| chrX | 25069199 | 25090343 | 21145 | -9.06 | 2.15E-16 | 1.29E-15 |
| chr11 | 101712048 | 101750193 | 38146 | -8.85 | 2.16E-16 | 1.29E-15 |
| chr8 | 32143760 | 32179678 | 35919 | -8.98 | 2.16E-16 | 1.29E-15 |
| chr1 | 43655429 | 43670664 | 15236 | -8.85 | 2.16E-16 | 1.29E-15 |
| chr14 | 70769421 | 70780764 | 11344 | -8.86 | 2.16E-16 | 1.29E-15 |
| chr18 | 3201374 | 3206584 | 5211 | -8.85 | 2.17E-16 | 1.29E-15 |
| chr8 | 82625775 | 82637027 | 11253 | -8.97 | 2.17E-16 | 1.30E-15 |
| chr3 | 38426366 | 38429787 | 3422 | -8.88 | 2.18E-16 | 1.30E-15 |
| chr1 | 191475476 | 191550180 | 74705 | -8.96 | 2.18E-16 | 1.30E-15 |
| chr14 | 105110961 | 105129520 | 18560 | -8.96 | 2.18E-16 | 1.30E-15 |
| chr13 | 109353506 | 109373544 | 20039 | -8.97 | 2.18E-16 | 1.30E-15 |
| chr6 | 56034473 | 56042761 | 8289 | -8.87 | 2.19E-16 | 1.31E-15 |
| chr2 | 235361628 | 235397144 | 35517 | -9.24 | 2.20E-16 | 1.31E-15 |
| chr1 | 169248768 | 169256746 | 7979 | -8.93 | 2.21E-16 | 1.32E-15 |
| chrX | 24422263 | 24452230 | 29968 | -6.56 | 2.21E-16 | 1.32E-15 |
| chrX | 8444435 | 8454562 | 10128 | 2.9 | 2.22E-16 | 1.32E-15 |
| chr9 | 36222132 | 36243145 | 21014 | -8.96 | 2.22E-16 | 1.32E-15 |
| chr1 | 120075668 | 120102055 | 26388 | 8.92 | 2.22E-16 | 1.33E-15 |
| chr14 | 94617784 | 94639888 | 22105 | -8.98 | 2.23E-16 | 1.33E-15 |
| chr14 | 50485828 | 50493898 | 8071 | -8.97 | 2.23E-16 | 1.33E-15 |
| chr6 | 160268313 | 160289543 | 21231 | -8.85 | 2.23E-16 | 1.33E-15 |
| chr12 | 77283997 | 77299718 | 15722 | -8.98 | 2.24E-16 | 1.33E-15 |
| chr10 | 99976016 | 99987449 | 11434 | -8.85 | 2.24E-16 | 1.33E-15 |
| chr4 | 8751412 | 8759531 | 8120 | -8.91 | 2.24E-16 | 1.34E-15 |
| chr11 | 92774212 | 92798315 | 24104 | -8.89 | 2.25E-16 | 1.34E-15 |
| chrX | 109962947 | 109978397 | 15451 | 8.94 | 2.27E-16 | 1.35E-15 |
| chr11 | 85435111 | 85440152 | 5042 | -8.88 | 2.27E-16 | 1.35E-15 |
| chr8 | 124383929 | 124386176 | 2248 | -9 | 2.28E-16 | 1.36E-15 |
| chr11 | 1598190 | 1601211 | 3022 | -8.92 | 2.28E-16 | 1.36E-15 |
| chr10 | 61949293 | 61952010 | 2718 | -8.91 | 2.28E-16 | 1.36E-15 |
| chr17 | 80776250 | 80779711 | 3462 | -8.85 | 2.28E-16 | 1.36E-15 |
| chr2 | 147393806 | 147415199 | 21394 | -8.85 | 2.29E-16 | 1.36E-15 |
| chr3 | 58437764 | 58443488 | 5725 | -9.09 | 2.29E-16 | 1.36E-15 |
| chr2 | 44689915 | 44695302 | 5388 | -8.89 | 2.31E-16 | 1.38E-15 |
| chr6 | 169533857 | 169552705 | 18849 | -8.86 | 2.32E-16 | 1.38E-15 |
| chr2 | 101149215 | 101153633 | 4419 | -6.51 | 2.32E-16 | 1.38E-15 |
| chr13 | 106778761 | 106807785 | 29025 | -8.86 | 2.32E-16 | 1.38E-15 |
| chr4 | 109526603 | 109538792 | 12190 | -8.84 | 2.33E-16 | 1.38E-15 |
| chr7 | 80161621 | 80192576 | 30956 | -9.27 | 2.34E-16 | 1.39E-15 |
| chr5 | 171797083 | 171800198 | 3116 | -8.84 | 2.34E-16 | 1.39E-15 |
| chr2 | 128575484 | 128586407 | 10924 | -8.85 | 2.34E-16 | 1.39E-15 |
| chr2 | 182874870 | 182889617 | 14748 | -8.89 | 2.34E-16 | 1.39E-15 |
| chr9 | 140715289 | 140721525 | 6237 | -8.85 | 2.35E-16 | 1.40E-15 |
| chr10 | 71345544 | 71388950 | 43407 | -2.22 | 2.36E-16 | 1.40E-15 |
| chr3 | 107604145 | 107615339 | 11195 | -2.49 | 2.37E-16 | 1.40E-15 |
| chrX | 44802454 | 44806695 | 4242 | -8.85 | 2.38E-16 | 1.41E-15 |
| chr7 | 115754508 | 115770883 | 16376 | -9.01 | 2.39E-16 | 1.42E-15 |
| chr3 | 193466288 | 193477981 | 11694 | -8.98 | 2.40E-16 | 1.42E-15 |
| chr4 | 39016818 | 39026506 | 9689 | -8.84 | 2.40E-16 | 1.43E-15 |
| chr2 | 110938573 | 110946865 | 8293 | -8.88 | 2.41E-16 | 1.43E-15 |
| chr12 | 72104921 | 72109811 | 4891 | -8.84 | 2.41E-16 | 1.43E-15 |
| chr7 | 32628626 | 32655099 | 26474 | -8.86 | 2.43E-16 | 1.44E-15 |
| chr2 | 8646099 | 8655006 | 8908 | -8.89 | 2.43E-16 | 1.44E-15 |
| chr9 | 101848697 | 101864432 | 15736 | -9 | 2.44E-16 | 1.45E-15 |

| | | | | | | |
|---|---|---|---|---|---|---|
| chr2 | 102335927 | 102342125 | 6199 | -9.05 | 2.45E-16 | 1.45E-15 |
| chr3 | 11445408 | 11456768 | 11361 | -8.91 | 2.45E-16 | 1.45E-15 |
| chr1 | 185100427 | 185103743 | 3317 | -8.88 | 2.45E-16 | 1.45E-15 |
| chr5 | 40076298 | 40092292 | 15995 | -9.17 | 2.46E-16 | 1.45E-15 |
| chr6 | 49317878 | 49336153 | 18276 | -8.85 | 2.47E-16 | 1.46E-15 |
| chr12 | 94777764 | 94792229 | 14466 | -8.85 | 2.48E-16 | 1.47E-15 |
| chr12 | 13994493 | 14003647 | 9155 | 8.92 | 2.49E-16 | 1.47E-15 |
| chr12 | 107771444 | 107784523 | 13080 | -8.84 | 2.50E-16 | 1.48E-15 |
| chr13 | 76327254 | 76368706 | 41453 | -8.89 | 2.50E-16 | 1.48E-15 |
| chr7 | 109175663 | 109199604 | 23942 | -8.94 | 2.51E-16 | 1.48E-15 |
| chr18 | 43273459 | 43291226 | 17768 | -8.85 | 2.51E-16 | 1.49E-15 |
| chrX | 80867524 | 80939293 | 71770 | 1.91 | 2.51E-16 | 1.49E-15 |
| chrX | 89260927 | 89350840 | 89914 | 1.83 | 2.52E-16 | 1.49E-15 |
| chr13 | 60687209 | 60702680 | 15472 | -9 | 2.52E-16 | 1.49E-15 |
| chr6 | 118196614 | 118212626 | 16013 | 8.93 | 2.52E-16 | 1.49E-15 |
| chr9 | 116862738 | 116866876 | 4139 | -9.01 | 2.52E-16 | 1.49E-15 |
| chr4 | 3053573 | 3067264 | 13692 | -9.1 | 2.53E-16 | 1.50E-15 |
| chr4 | 100286037 | 100320200 | 34164 | -9.12 | 2.55E-16 | 1.51E-15 |
| chr2 | 231800207 | 231803548 | 3342 | -9.02 | 2.55E-16 | 1.51E-15 |
| chr14 | 51527764 | 51540310 | 12547 | -8.83 | 2.55E-16 | 1.51E-15 |
| chr1 | 45535981 | 45542898 | 6918 | -8.88 | 2.58E-16 | 1.52E-15 |
| chr1 | 47233080 | 47251925 | 18846 | -8.86 | 2.59E-16 | 1.53E-15 |
| chr6 | 135681631 | 135693280 | 11650 | -8.96 | 2.59E-16 | 1.53E-15 |
| chr4 | 136686022 | 136716467 | 30446 | -8.91 | 2.60E-16 | 1.53E-15 |
| chr2 | 71543399 | 71550225 | 6827 | -8.97 | 2.61E-16 | 1.54E-15 |
| chr7 | 134578753 | 134582661 | 3909 | -8.83 | 2.63E-16 | 1.55E-15 |
| chrX | 152919045 | 152932334 | 13290 | -8.86 | 2.63E-16 | 1.55E-15 |
| chr3 | 113942180 | 113948059 | 5880 | -8.87 | 2.63E-16 | 1.55E-15 |
| chr17 | 65503294 | 65515457 | 12164 | -8.89 | 2.65E-16 | 1.56E-15 |
| chr5 | 80142808 | 80153046 | 10239 | -8.83 | 2.65E-16 | 1.57E-15 |
| chr12 | 94570623 | 94588726 | 18104 | -8.84 | 2.65E-16 | 1.57E-15 |
| chr2 | 119227356 | 119534537 | 307182 | -2.1 | 2.66E-16 | 1.57E-15 |
| chr2 | 233826176 | 233831535 | 5360 | -8.87 | 2.67E-16 | 1.57E-15 |
| chr6 | 12967583 | 12996108 | 28526 | -3.56 | 2.67E-16 | 1.57E-15 |
| chr13 | 96270569 | 96281388 | 10820 | -8.93 | 2.67E-16 | 1.57E-15 |
| chr1 | 1698701 | 1704563 | 5863 | -8.9 | 2.68E-16 | 1.58E-15 |
| chr2 | 223215106 | 223235434 | 20329 | -8.85 | 2.68E-16 | 1.58E-15 |
| chr6 | 12620563 | 12651892 | 31330 | -9.28 | 2.68E-16 | 1.58E-15 |
| chr3 | 54468362 | 54836659 | 368298 | 1.35 | 2.70E-16 | 1.59E-15 |
| chr13 | 110594718 | 110624561 | 29844 | -8.85 | 2.70E-16 | 1.59E-15 |
| chr1 | 164848505 | 165315925 | 467421 | 3.09 | 2.72E-16 | 1.60E-15 |
| chr13 | 95376244 | 95381228 | 4985 | -8.82 | 2.73E-16 | 1.61E-15 |
| chr5 | 145256827 | 145280651 | 23825 | -8.89 | 2.74E-16 | 1.61E-15 |
| chr3 | 105008931 | 105026666 | 17736 | -8.89 | 2.74E-16 | 1.61E-15 |
| chr2 | 147042827 | 147065937 | 23111 | -8.9 | 2.74E-16 | 1.61E-15 |
| chr7 | 117207720 | 117227436 | 19717 | -8.9 | 2.74E-16 | 1.61E-15 |
| chr14 | 67075002 | 67090379 | 15378 | -8.91 | 2.75E-16 | 1.62E-15 |
| chr3 | 150024437 | 150031139 | 6703 | -8.88 | 2.76E-16 | 1.62E-15 |
| chrX | 85199135 | 85210017 | 10883 | -8.88 | 2.76E-16 | 1.62E-15 |
| chr6 | 48350169 | 48373633 | 23465 | -8.92 | 2.77E-16 | 1.63E-15 |
| chr10 | 73875579 | 73889416 | 13838 | -8.85 | 2.77E-16 | 1.63E-15 |
| chr3 | 42710355 | 42719489 | 9135 | -8.92 | 2.78E-16 | 1.63E-15 |
| chr13 | 20879316 | 20902393 | 23078 | -2.77 | 2.78E-16 | 1.63E-15 |
| chr12 | 12178473 | 12181833 | 3361 | -8.87 | 2.78E-16 | 1.63E-15 |
| chr21 | 37971159 | 38010426 | 39268 | -9.23 | 2.78E-16 | 1.63E-15 |
| chr11 | 36133288 | 36137309 | 4022 | -8.89 | 2.79E-16 | 1.64E-15 |
| chr9 | 128217565 | 128220696 | 3132 | -8.82 | 2.79E-16 | 1.64E-15 |
| chr6 | 82061797 | 82132333 | 70537 | 2.34 | 2.79E-16 | 1.64E-15 |
| chr14 | 56325134 | 56331976 | 6843 | -8.9 | 2.79E-16 | 1.64E-15 |
| chr2 | 187633895 | 187656133 | 22239 | -9.21 | 2.81E-16 | 1.65E-15 |
| chr14 | 20718220 | 20722025 | 3806 | -8.82 | 2.81E-16 | 1.65E-15 |

| | | | | | | |
|---|---|---|---|---|---|---|
| chr1 | 231017769 | 231022713 | 4945 | -8.96 | 2.82E-16 | 1.65E-15 |
| chr10 | 59526123 | 59539741 | 13619 | -8.82 | 2.82E-16 | 1.65E-15 |
| chr2 | 27961714 | 27972863 | 11150 | -9.15 | 2.83E-16 | 1.66E-15 |
| chr5 | 71141523 | 71167043 | 25521 | -7.18 | 2.84E-16 | 1.66E-15 |
| chr13 | 22203269 | 22236939 | 33671 | -9.17 | 2.84E-16 | 1.67E-15 |
| chr19 | 38520034 | 38533625 | 13592 | -8.97 | 2.85E-16 | 1.67E-15 |
| chr3 | 133507819 | 133515416 | 7598 | -8.96 | 2.85E-16 | 1.67E-15 |
| chr21 | 16702785 | 16740024 | 37240 | -9.43 | 2.86E-16 | 1.67E-15 |
| chr9 | 101952907 | 101975077 | 22171 | -9.19 | 2.86E-16 | 1.68E-15 |
| chr10 | 49618359 | 49621768 | 3410 | -8.82 | 2.87E-16 | 1.68E-15 |
| chr14 | 25408475 | 25439231 | 30757 | 8.88 | 2.87E-16 | 1.68E-15 |
| chr7 | 20642869 | 20680444 | 37576 | -8.9 | 2.90E-16 | 1.70E-15 |
| chr17 | 39177202 | 39181370 | 4169 | -8.9 | 2.91E-16 | 1.70E-15 |
| chr12 | 39173660 | 39187425 | 13766 | -8.94 | 2.91E-16 | 1.70E-15 |
| chr8 | 38510384 | 38522565 | 12182 | -3.67 | 2.91E-16 | 1.71E-15 |
| chr3 | 71538859 | 71542652 | 3794 | -8.82 | 2.92E-16 | 1.71E-15 |
| chr9 | 85700890 | 85714295 | 13406 | -8.92 | 2.92E-16 | 1.71E-15 |
| chr2 | 39085928 | 39100666 | 14739 | -9.02 | 2.93E-16 | 1.71E-15 |
| chr6 | 159324438 | 159329899 | 5462 | -8.83 | 2.93E-16 | 1.71E-15 |
| chr8 | 111071143 | 111804607 | 733465 | 1.94 | 2.93E-16 | 1.71E-15 |
| chr11 | 96258217 | 96278415 | 20199 | 8.88 | 2.93E-16 | 1.72E-15 |
| chr2 | 218126686 | 218134235 | 7550 | -8.83 | 2.94E-16 | 1.72E-15 |
| chr5 | 55596743 | 55606245 | 9503 | -8.82 | 2.94E-16 | 1.72E-15 |
| chr11 | 70293066 | 70298317 | 5252 | -8.88 | 2.95E-16 | 1.72E-15 |
| chr5 | 93304273 | 93314417 | 10145 | -8.89 | 2.95E-16 | 1.72E-15 |
| chr2 | 205601419 | 205631598 | 30180 | -9.47 | 2.95E-16 | 1.73E-15 |
| chr12 | 66557965 | 66578958 | 20994 | -8.82 | 2.95E-16 | 1.73E-15 |
| chr13 | 40672747 | 40706660 | 33914 | -9.1 | 2.95E-16 | 1.73E-15 |
| chr6 | 43205485 | 43210457 | 4973 | -8.82 | 2.96E-16 | 1.73E-15 |
| chr9 | 137551550 | 137568092 | 16543 | -8.82 | 2.96E-16 | 1.73E-15 |
| chr10 | 3834219 | 3841536 | 7318 | -8.84 | 2.97E-16 | 1.73E-15 |
| chr5 | 40876428 | 40888027 | 11600 | -8.91 | 2.97E-16 | 1.74E-15 |
| chr6 | 88148116 | 88168335 | 20220 | -8.94 | 2.98E-16 | 1.74E-15 |
| chr11 | 86136817 | 86143635 | 6819 | -8.81 | 2.98E-16 | 1.74E-15 |
| chr11 | 36680465 | 36695389 | 14925 | -8.91 | 2.99E-16 | 1.74E-15 |
| chr5 | 14688795 | 14691325 | 2531 | -8.95 | 2.99E-16 | 1.74E-15 |
| chr2 | 106265096 | 106347768 | 82673 | -3.36 | 2.99E-16 | 1.75E-15 |
| chr10 | 76569395 | 76578732 | 9338 | -9.02 | 2.99E-16 | 1.75E-15 |
| chr5 | 55685015 | 55693206 | 8192 | -9.03 | 3.00E-16 | 1.75E-15 |
| chr16 | 21100278 | 21107309 | 7032 | -8.92 | 3.01E-16 | 1.75E-15 |
| chr5 | 118802803 | 118806145 | 3343 | -8.83 | 3.01E-16 | 1.75E-15 |
| chr17 | 12037681 | 12886392 | 848712 | 2.86 | 3.01E-16 | 1.75E-15 |
| chr10 | 26800485 | 26810521 | 10037 | -8.99 | 3.01E-16 | 1.76E-15 |
| chr2 | 139331579 | 139338885 | 7307 | -8.93 | 3.04E-16 | 1.77E-15 |
| chr13 | 36860392 | 36865525 | 5134 | -8.81 | 3.05E-16 | 1.78E-15 |
| chr5 | 114870999 | 114882785 | 11787 | -8.81 | 3.05E-16 | 1.78E-15 |
| chr2 | 153557256 | 153568345 | 11090 | -8.98 | 3.06E-16 | 1.78E-15 |
| chr3 | 34093833 | 34105385 | 11553 | -8.81 | 3.06E-16 | 1.78E-15 |
| chr7 | 139602359 | 139625208 | 22850 | -8.92 | 3.07E-16 | 1.79E-15 |
| chr1 | 114260036 | 114265158 | 5123 | -8.81 | 3.07E-16 | 1.79E-15 |
| chr8 | 41682484 | 41690847 | 8364 | -8.84 | 3.08E-16 | 1.79E-15 |
| chr8 | 683128 | 691545 | 8418 | -8.83 | 3.09E-16 | 1.80E-15 |
| chr3 | 149406443 | 149417440 | 10998 | -8.96 | 3.10E-16 | 1.80E-15 |
| chr4 | 153948885 | 153978921 | 30037 | -8.93 | 3.10E-16 | 1.81E-15 |
| chr14 | 38020389 | 38026976 | 6588 | -8.95 | 3.11E-16 | 1.81E-15 |
| chr19 | 38561627 | 38571912 | 10286 | -8.87 | 3.11E-16 | 1.81E-15 |
| chr2 | 54343280 | 54348641 | 5362 | -8.82 | 3.11E-16 | 1.81E-15 |
| chr1 | 108400705 | 108428229 | 27525 | 8.93 | 3.11E-16 | 1.81E-15 |
| chr9 | 37473315 | 37477042 | 3728 | -8.81 | 3.12E-16 | 1.81E-15 |
| chr20 | 4481902 | 4492375 | 10474 | -8.82 | 3.13E-16 | 1.82E-15 |
| chr20 | 11357918 | 11371896 | 13979 | -8.84 | 3.14E-16 | 1.82E-15 |

| | | | | | | |
|---|---|---|---|---|---|---|
| chr10 | 89640447 | 89657857 | 17411 | -8.81 | 3.14E-16 | 1.83E-15 |
| chr3 | 47698270 | 47701225 | 2956 | -8.83 | 3.14E-16 | 1.83E-15 |
| chr7 | 50581767 | 50604097 | 22331 | -8.82 | 3.14E-16 | 1.83E-15 |
| chr15 | 67161964 | 67165094 | 3131 | -8.81 | 3.15E-16 | 1.83E-15 |
| chr7 | 99242462 | 99262566 | 20105 | -8.95 | 3.15E-16 | 1.83E-15 |
| chr9 | 36771378 | 36813168 | 41791 | -9.23 | 3.17E-16 | 1.84E-15 |
| chr17 | 30133390 | 30141299 | 7910 | -9.22 | 3.17E-16 | 1.84E-15 |
| chr5 | 58233607 | 58259393 | 25787 | -9.07 | 3.18E-16 | 1.84E-15 |
| chr11 | 8135512 | 8162482 | 26971 | -8.82 | 3.20E-16 | 1.86E-15 |
| chr2 | 11471701 | 11478474 | 6774 | -8.85 | 3.20E-16 | 1.86E-15 |
| chr8 | 126837253 | 126892761 | 55509 | 2.35 | 3.20E-16 | 1.86E-15 |
| chr2 | 176091219 | 176106434 | 15216 | -6.59 | 3.20E-16 | 1.86E-15 |
| chr7 | 74104788 | 74111919 | 7132 | -8.88 | 3.22E-16 | 1.87E-15 |
| chr13 | 21958703 | 21962051 | 3349 | -8.85 | 3.22E-16 | 1.87E-15 |
| chr9 | 134434429 | 134440625 | 6197 | -8.88 | 3.23E-16 | 1.88E-15 |
| chr16 | 89933202 | 89935608 | 2407 | -8.81 | 3.23E-16 | 1.88E-15 |
| chr12 | 13124878 | 13128069 | 3192 | -8.81 | 3.25E-16 | 1.88E-15 |
| chr4 | 176499192 | 176532039 | 32848 | -8.86 | 3.25E-16 | 1.89E-15 |
| chr4 | 56395780 | 56405073 | 9294 | -8.82 | 3.26E-16 | 1.89E-15 |
| chr5 | 175265722 | 175268341 | 2620 | -8.8 | 3.27E-16 | 1.89E-15 |
| chr2 | 43114228 | 43118673 | 4446 | -8.93 | 3.27E-16 | 1.90E-15 |
| chr1 | 64045867 | 64050072 | 4206 | -8.81 | 3.27E-16 | 1.90E-15 |
| chr17 | 43075184 | 43078275 | 3092 | -8.9 | 3.28E-16 | 1.90E-15 |
| chr2 | 72081830 | 72084336 | 2507 | -8.8 | 3.28E-16 | 1.90E-15 |
| chr4 | 83239087 | 83253287 | 14201 | -8.8 | 3.28E-16 | 1.90E-15 |
| chr1 | 164519168 | 164541403 | 22236 | -8.81 | 3.29E-16 | 1.91E-15 |
| chr12 | 15009222 | 15022037 | 12816 | -8.91 | 3.29E-16 | 1.91E-15 |
| chr10 | 119203050 | 119236978 | 33929 | -9.01 | 3.30E-16 | 1.91E-15 |
| chr5 | 52196285 | 52200809 | 4525 | -8.81 | 3.30E-16 | 1.91E-15 |
| chr10 | 32797325 | 32827299 | 29975 | -9.02 | 3.31E-16 | 1.92E-15 |
| chr15 | 63435989 | 63459283 | 23295 | -9.02 | 3.31E-16 | 1.92E-15 |
| chr6 | 22861985 | 22887826 | 25842 | -9.02 | 3.31E-16 | 1.92E-15 |
| chr9 | 114883929 | 114897124 | 13196 | -8.91 | 3.32E-16 | 1.92E-15 |
| chr17 | 80949268 | 80957482 | 8215 | -8.81 | 3.32E-16 | 1.92E-15 |
| chr10 | 98529835 | 98545386 | 15552 | -8.88 | 3.35E-16 | 1.94E-15 |
| chr2 | 58235873 | 58244604 | 8732 | -8.89 | 3.36E-16 | 1.94E-15 |
| chr17 | 64692436 | 64696983 | 4548 | -8.84 | 3.37E-16 | 1.95E-15 |
| chr12 | 97955842 | 97988869 | 33028 | -8.87 | 3.37E-16 | 1.95E-15 |
| chr11 | 27942038 | 27980197 | 38160 | -8.82 | 3.38E-16 | 1.95E-15 |
| chr20 | 46045100 | 46058859 | 13760 | -8.93 | 3.38E-16 | 1.95E-15 |
| chr3 | 134237686 | 134244792 | 7107 | -8.82 | 3.38E-16 | 1.95E-15 |
| chr21 | 40865801 | 40868204 | 2404 | -3.64 | 3.39E-16 | 1.96E-15 |
| chr3 | 28042042 | 28046934 | 4893 | -3.52 | 3.40E-16 | 1.96E-15 |
| chr1 | 3258848 | 3262476 | 3629 | -8.82 | 3.41E-16 | 1.97E-15 |
| chr3 | 193746217 | 193771309 | 25093 | -9.11 | 3.41E-16 | 1.97E-15 |
| chr8 | 53457847 | 53484054 | 26208 | -8.86 | 3.41E-16 | 1.97E-15 |
| chr8 | 41594110 | 41599212 | 5103 | -8.87 | 3.42E-16 | 1.97E-15 |
| chr2 | 102286540 | 102294425 | 7886 | -8.82 | 3.42E-16 | 1.97E-15 |
| chr5 | 75358723 | 75372398 | 13676 | 8.87 | 3.43E-16 | 1.98E-15 |
| chr1 | 219423797 | 219436841 | 13045 | -8.89 | 3.44E-16 | 1.98E-15 |
| chr16 | 77284989 | 77306754 | 21766 | -8.94 | 3.45E-16 | 1.99E-15 |
| chr8 | 22183952 | 22187635 | 3684 | -8.8 | 3.45E-16 | 1.99E-15 |
| chr11 | 130781507 | 130788201 | 6695 | -8.84 | 3.46E-16 | 2.00E-15 |
| chr4 | 75399236 | 75419309 | 20074 | -9.1 | 3.47E-16 | 2.00E-15 |
| chr15 | 35203815 | 35206339 | 2525 | -9.03 | 3.48E-16 | 2.00E-15 |
| chr1 | 234423300 | 234436214 | 12915 | -8.8 | 3.48E-16 | 2.01E-15 |
| chrX | 56979431 | 56989560 | 10130 | -8.8 | 3.50E-16 | 2.02E-15 |
| chr12 | 133091796 | 133113769 | 21974 | -9 | 3.51E-16 | 2.02E-15 |
| chr7 | 4216210 | 4240955 | 24746 | -8.92 | 3.51E-16 | 2.02E-15 |
| chr5 | 172173282 | 172197160 | 23879 | -8.81 | 3.52E-16 | 2.03E-15 |
| chr5 | 37470235 | 37489196 | 18962 | -8.82 | 3.55E-16 | 2.05E-15 |

| | | | | | | |
|---|---|---|---|---|---|---|
| chr8 | 91876820 | 91899466 | 22647 | 9.16 | 3.56E-16 | 2.05E-15 |
| chr4 | 112599800 | 112622745 | 22946 | 8.95 | 3.56E-16 | 2.05E-15 |
| chr4 | 85861004 | 85865562 | 4559 | -8.8 | 3.58E-16 | 2.06E-15 |
| chr2 | 118969151 | 118974041 | 4891 | -9.04 | 3.61E-16 | 2.08E-15 |
| chr1 | 48129748 | 48139423 | 9676 | -8.9 | 3.61E-16 | 2.08E-15 |
| chr8 | 102817428 | 102839545 | 22118 | -9.19 | 3.62E-16 | 2.08E-15 |
| chr7 | 7420122 | 7447972 | 27851 | -8.79 | 3.62E-16 | 2.08E-15 |
| chr8 | 58856982 | 58861677 | 4696 | -6.3 | 3.62E-16 | 2.08E-15 |
| chr12 | 117807036 | 117818627 | 11592 | -8.85 | 3.63E-16 | 2.09E-15 |
| chr6 | 168438251 | 168442835 | 4585 | -8.81 | 3.63E-16 | 2.09E-15 |
| chr4 | 75115969 | 75121524 | 5556 | -8.8 | 3.63E-16 | 2.09E-15 |
| chr1 | 35854793 | 35875160 | 20368 | -8.91 | 3.63E-16 | 2.09E-15 |
| chr8 | 121265878 | 121280533 | 14656 | 8.89 | 3.64E-16 | 2.09E-15 |
| chr3 | 17529502 | 17536651 | 7150 | -9.01 | 3.64E-16 | 2.09E-15 |
| chr2 | 225347926 | 225355951 | 8026 | -8.82 | 3.66E-16 | 2.10E-15 |
| chr1 | 43790060 | 43794632 | 4573 | -8.83 | 3.67E-16 | 2.11E-15 |
| chr2 | 64172362 | 64179982 | 7621 | -9 | 3.67E-16 | 2.11E-15 |
| chr17 | 6711622 | 6793247 | 81626 | -8.31 | 3.67E-16 | 2.11E-15 |
| chr9 | 132702953 | 132705800 | 2848 | -8.86 | 3.68E-16 | 2.11E-15 |
| chr2 | 164518077 | 164527832 | 9756 | -8.86 | 3.68E-16 | 2.11E-15 |
| chr6 | 122757962 | 122762614 | 4653 | -8.93 | 3.68E-16 | 2.11E-15 |
| chr19 | 17124481 | 17135265 | 10785 | -8.79 | 3.69E-16 | 2.12E-15 |
| chr21 | 34578031 | 34595266 | 17236 | -8.79 | 3.71E-16 | 2.13E-15 |
| chr5 | 81101414 | 81104716 | 3303 | -8.83 | 3.71E-16 | 2.13E-15 |
| chr4 | 90638170 | 90649224 | 11055 | -8.79 | 3.73E-16 | 2.14E-15 |
| chr2 | 44856478 | 44863184 | 6707 | -8.83 | 3.74E-16 | 2.15E-15 |
| chr7 | 96340908 | 96347696 | 6789 | -8.82 | 3.75E-16 | 2.15E-15 |
| chr2 | 236240352 | 236257750 | 17399 | -8.85 | 3.75E-16 | 2.15E-15 |
| chr14 | 59746027 | 59753127 | 7101 | -8.79 | 3.76E-16 | 2.16E-15 |
| chr7 | 137407674 | 137453785 | 46112 | 1.8 | 3.78E-16 | 2.17E-15 |
| chr2 | 86596547 | 86605097 | 8551 | -8.81 | 3.79E-16 | 2.17E-15 |
| chrX | 76900098 | 76902413 | 2316 | -8.92 | 3.82E-16 | 2.19E-15 |
| chr1 | 240386827 | 240417006 | 30180 | -8.79 | 3.82E-16 | 2.19E-15 |
| chr9 | 119637563 | 119641140 | 3578 | -8.79 | 3.83E-16 | 2.19E-15 |
| chr13 | 111309889 | 111314377 | 4489 | -8.79 | 3.83E-16 | 2.20E-15 |
| chr7 | 38031642 | 38089711 | 58070 | -8.79 | 3.83E-16 | 2.20E-15 |
| chr2 | 46900736 | 46905415 | 4680 | -8.82 | 3.83E-16 | 2.20E-15 |
| chr2 | 190939167 | 190943905 | 4739 | -8.9 | 3.84E-16 | 2.20E-15 |
| chr11 | 19456911 | 19458853 | 1943 | -2.94 | 3.84E-16 | 2.20E-15 |
| chr14 | 56784104 | 56802846 | 18743 | -8.88 | 3.87E-16 | 2.21E-15 |
| chrX | 110993638 | 110996821 | 3184 | -6.61 | 3.87E-16 | 2.21E-15 |
| chr8 | 91073173 | 91074977 | 1805 | -6.16 | 3.88E-16 | 2.22E-15 |
| chr21 | 27343206 | 27346956 | 3751 | -8.81 | 3.89E-16 | 2.23E-15 |
| chr6 | 48795916 | 48811878 | 15963 | -8.8 | 3.89E-16 | 2.23E-15 |
| chr11 | 128952715 | 128977885 | 25171 | -8.78 | 3.89E-16 | 2.23E-15 |
| chr3 | 124425896 | 124429613 | 3718 | -2.77 | 3.91E-16 | 2.24E-15 |
| chr18 | 33808328 | 33829201 | 20874 | -8.78 | 3.91E-16 | 2.24E-15 |
| chr11 | 19630894 | 19658165 | 27272 | -8.79 | 3.92E-16 | 2.24E-15 |
| chr5 | 173231009 | 173256304 | 25296 | -9.42 | 3.92E-16 | 2.24E-15 |
| chr2 | 164051531 | 164074223 | 22693 | -8.91 | 3.92E-16 | 2.24E-15 |
| chr3 | 188521855 | 188528577 | 6723 | -8.81 | 3.93E-16 | 2.24E-15 |
| chr3 | 108291324 | 108295157 | 3834 | -9.15 | 3.93E-16 | 2.25E-15 |
| chr3 | 112522328 | 112544596 | 22269 | -8.95 | 3.93E-16 | 2.25E-15 |
| chr4 | 121360114 | 121381896 | 21783 | 8.89 | 3.93E-16 | 2.25E-15 |
| chr10 | 101569987 | 101575322 | 5336 | -8.81 | 3.94E-16 | 2.25E-15 |
| chr5 | 138197173 | 138200676 | 3504 | -8.8 | 3.94E-16 | 2.25E-15 |
| chr12 | 121354001 | 121357563 | 3563 | -8.83 | 3.95E-16 | 2.26E-15 |
| chr14 | 96453535 | 96493208 | 39674 | -2.39 | 3.96E-16 | 2.26E-15 |
| chr3 | 110367114 | 110396099 | 28986 | -8.99 | 3.99E-16 | 2.28E-15 |
| chr5 | 33290342 | 33317750 | 27409 | -8.87 | 3.99E-16 | 2.28E-15 |
| chr5 | 82998549 | 83005209 | 6661 | -8.79 | 4.00E-16 | 2.28E-15 |

| | | | | | | |
|---|---|---|---|---|---|---|
| chr5 | 55414543 | 55432841 | 18299 | -9.01 | 4.04E-16 | 2.31E-15 |
| chr8 | 498752 | 518064 | 19313 | -4.07 | 4.04E-16 | 2.31E-15 |
| chr4 | 2325740 | 2331573 | 5834 | -8.79 | 4.05E-16 | 2.31E-15 |
| chr2 | 171471241 | 171486881 | 15641 | -2.38 | 4.06E-16 | 2.31E-15 |
| chr5 | 58669325 | 58674621 | 5297 | -8.9 | 4.06E-16 | 2.32E-15 |
| chr3 | 152136417 | 152153799 | 17383 | -8.83 | 4.07E-16 | 2.32E-15 |
| chr1 | 197692035 | 197696775 | 4741 | -8.89 | 4.07E-16 | 2.32E-15 |
| chr3 | 14541298 | 14545134 | 3837 | -8.87 | 4.08E-16 | 2.32E-15 |
| chr3 | 151881264 | 151896761 | 15498 | -8.98 | 4.08E-16 | 2.33E-15 |
| chr8 | 106902571 | 106936448 | 33878 | -9.05 | 4.08E-16 | 2.33E-15 |
| chr9 | 90303243 | 90308744 | 5502 | -8.87 | 4.09E-16 | 2.33E-15 |
| chr2 | 159473317 | 159477585 | 4269 | -8.79 | 4.10E-16 | 2.34E-15 |
| chr22 | 17435387 | 17448337 | 12951 | 8.86 | 4.10E-16 | 2.34E-15 |
| chr18 | 44983935 | 45001690 | 17756 | 8.89 | 4.11E-16 | 2.34E-15 |
| chr3 | 183312961 | 183318452 | 5492 | -8.78 | 4.11E-16 | 2.34E-15 |
| chr4 | 57838591 | 57840941 | 2351 | -8.8 | 4.15E-16 | 2.36E-15 |
| chr3 | 30921491 | 30934739 | 13249 | -2.08 | 4.15E-16 | 2.36E-15 |
| chr11 | 31562538 | 31565867 | 3330 | -8.85 | 4.17E-16 | 2.37E-15 |
| chr10 | 80570454 | 80604345 | 33892 | -9.21 | 4.17E-16 | 2.37E-15 |
| chr11 | 75728129 | 75763774 | 35646 | -8.81 | 4.17E-16 | 2.37E-15 |
| chr3 | 11828271 | 11845581 | 17311 | -8.78 | 4.17E-16 | 2.38E-15 |
| chr4 | 88850849 | 88854578 | 3730 | -8.8 | 4.17E-16 | 2.38E-15 |
| chr2 | 121164939 | 121169776 | 4838 | -8.8 | 4.18E-16 | 2.38E-15 |
| chr11 | 58919073 | 58934999 | 15927 | -8.95 | 4.18E-16 | 2.38E-15 |
| chrX | 9046208 | 9064504 | 18297 | -8.92 | 4.19E-16 | 2.38E-15 |
| chr20 | 25702774 | 25718184 | 15411 | -8.79 | 4.21E-16 | 2.40E-15 |
| chr16 | 47370138 | 47388626 | 18489 | -8.92 | 4.26E-16 | 2.42E-15 |
| chr12 | 38646401 | 38657162 | 10762 | -8.77 | 4.27E-16 | 2.43E-15 |
| chr8 | 128273249 | 128278009 | 4761 | -8.78 | 4.27E-16 | 2.43E-15 |
| chr6 | 170898000 | 170902337 | 4338 | -8.77 | 4.27E-16 | 2.43E-15 |
| chr10 | 128161793 | 128177875 | 16083 | -8.78 | 4.27E-16 | 2.43E-15 |
| chr2 | 97374341 | 97376866 | 2526 | -8.8 | 4.27E-16 | 2.43E-15 |
| chr5 | 110520927 | 110535185 | 14259 | 8.91 | 4.28E-16 | 2.43E-15 |
| chr1 | 244035437 | 244042188 | 6752 | -8.82 | 4.29E-16 | 2.44E-15 |
| chr5 | 167102474 | 167123413 | 20940 | -8.83 | 4.30E-16 | 2.44E-15 |
| chr5 | 64404194 | 64414291 | 10098 | -8.78 | 4.30E-16 | 2.44E-15 |
| chr9 | 114109228 | 114118367 | 9140 | -8.86 | 4.30E-16 | 2.44E-15 |
| chr14 | 36072806 | 36083368 | 10563 | -8.88 | 4.30E-16 | 2.44E-15 |
| chr14 | 101516439 | 101526081 | 9643 | -8.8 | 4.31E-16 | 2.44E-15 |
| chr2 | 63147691 | 63153412 | 5722 | -8.78 | 4.32E-16 | 2.45E-15 |
| chr2 | 31258350 | 31262284 | 3935 | -8.81 | 4.34E-16 | 2.46E-15 |
| chr11 | 72720096 | 72740166 | 20071 | -8.96 | 4.35E-16 | 2.47E-15 |
| chr21 | 35721677 | 35732087 | 10411 | -8.78 | 4.36E-16 | 2.48E-15 |
| chr16 | 69460760 | 69464559 | 3800 | -8.78 | 4.38E-16 | 2.48E-15 |
| chr10 | 114212475 | 114225776 | 13302 | -8.98 | 4.41E-16 | 2.50E-15 |
| chr15 | 80796297 | 80810176 | 13880 | -8.94 | 4.45E-16 | 2.52E-15 |
| chr3 | 113228721 | 113243446 | 14726 | -8.85 | 4.45E-16 | 2.52E-15 |
| chr14 | 62152676 | 62157806 | 5131 | -8.85 | 4.46E-16 | 2.53E-15 |
| chr12 | 116910336 | 116933304 | 22969 | -9.11 | 4.46E-16 | 2.53E-15 |
| chr14 | 68804949 | 68840210 | 35262 | -9.15 | 4.46E-16 | 2.53E-15 |
| chr5 | 1398120 | 1400135 | 2016 | -8.77 | 4.46E-16 | 2.53E-15 |
| chr8 | 32498786 | 32511314 | 12529 | -8.84 | 4.47E-16 | 2.53E-15 |
| chr18 | 52488239 | 52501791 | 13553 | -8.84 | 4.48E-16 | 2.54E-15 |
| chr2 | 29016590 | 29029090 | 12501 | -8.79 | 4.49E-16 | 2.54E-15 |
| chr5 | 179446743 | 179464984 | 18242 | -8.95 | 4.50E-16 | 2.55E-15 |
| chr1 | 241796632 | 241839000 | 42369 | -9.17 | 4.51E-16 | 2.55E-15 |
| chr6 | 135969597 | 135999382 | 29786 | -9.29 | 4.53E-16 | 2.56E-15 |
| chr8 | 95101750 | 95120641 | 18892 | -8.84 | 4.53E-16 | 2.56E-15 |
| chr7 | 38211316 | 38242768 | 31453 | -8.91 | 4.54E-16 | 2.57E-15 |
| chr8 | 142697704 | 142706284 | 8581 | -8.94 | 4.55E-16 | 2.57E-15 |
| chr8 | 94723194 | 94726501 | 3308 | -8.78 | 4.58E-16 | 2.59E-15 |

| | | | | | | |
|---|---|---|---|---|---|---|
| chr4 | 160840261 | 160864306 | 24046 | -8.95 | 4.59E-16 | 2.60E-15 |
| chr3 | 45492968 | 45503389 | 10422 | -8.78 | 4.59E-16 | 2.60E-15 |
| chr12 | 63622109 | 63634478 | 12370 | -8.78 | 4.60E-16 | 2.60E-15 |
| chr1 | 212709039 | 212712619 | 3581 | -8.8 | 4.60E-16 | 2.60E-15 |
| chr1 | 211132966 | 211148852 | 15887 | -8.77 | 4.63E-16 | 2.62E-15 |
| chr3 | 172661493 | 172680860 | 19368 | 8.86 | 4.65E-16 | 2.63E-15 |
| chr12 | 47944539 | 47951208 | 6670 | -8.8 | 4.66E-16 | 2.63E-15 |
| chr11 | 117721042 | 117725021 | 3980 | -8.78 | 4.67E-16 | 2.64E-15 |
| chr2 | 181079816 | 181109917 | 30102 | -8.8 | 4.69E-16 | 2.65E-15 |
| chr1 | 234820413 | 234823517 | 3105 | -8.8 | 4.71E-16 | 2.66E-15 |
| chr3 | 125225739 | 125229898 | 4160 | -8.85 | 4.72E-16 | 2.67E-15 |
| chr14 | 25357676 | 25394892 | 37217 | 8.97 | 4.75E-16 | 2.68E-15 |
| chr12 | 87556673 | 87565402 | 8730 | -8.76 | 4.75E-16 | 2.68E-15 |
| chr9 | 71883433 | 71909942 | 26510 | -8.77 | 4.75E-16 | 2.68E-15 |
| chr9 | 123427446 | 123430328 | 2883 | -8.8 | 4.75E-16 | 2.68E-15 |
| chr1 | 167110953 | 167137236 | 26284 | -8.81 | 4.76E-16 | 2.69E-15 |
| chr5 | 82129866 | 82144066 | 14201 | -6.45 | 4.76E-16 | 2.69E-15 |
| chr12 | 59511639 | 59523544 | 11906 | -8.76 | 4.77E-16 | 2.69E-15 |
| chr10 | 53939161 | 53951715 | 12555 | 8.83 | 4.77E-16 | 2.69E-15 |
| chr1 | 160554371 | 160575379 | 21009 | -2.98 | 4.77E-16 | 2.69E-15 |
| chr7 | 31441987 | 31451513 | 9527 | -8.94 | 4.80E-16 | 2.71E-15 |
| chr9 | 74225167 | 74233194 | 8028 | -8.94 | 4.81E-16 | 2.71E-15 |
| chr5 | 59850784 | 59886855 | 36072 | -8.77 | 4.82E-16 | 2.72E-15 |
| chr2 | 174817081 | 174826800 | 9720 | -8.76 | 4.82E-16 | 2.72E-15 |
| chr11 | 9131217 | 9136166 | 4950 | -8.76 | 4.82E-16 | 2.72E-15 |
| chr8 | 38071835 | 38074373 | 2539 | -8.76 | 4.83E-16 | 2.72E-15 |
| chr13 | 112827524 | 112829420 | 1897 | -3.05 | 4.83E-16 | 2.72E-15 |
| chr5 | 155940926 | 155953261 | 12336 | -8.76 | 4.83E-16 | 2.72E-15 |
| chr19 | 47148860 | 47151708 | 2849 | -8.81 | 4.86E-16 | 2.73E-15 |
| chr5 | 72466260 | 72482389 | 16130 | -8.77 | 4.87E-16 | 2.74E-15 |
| chr18 | 59875439 | 59904560 | 29122 | -9.08 | 4.88E-16 | 2.75E-15 |
| chr4 | 184293469 | 184306504 | 13036 | -8.79 | 4.88E-16 | 2.75E-15 |
| chr2 | 226882014 | 226904585 | 22572 | -8.88 | 4.89E-16 | 2.75E-15 |
| chr14 | 71501694 | 71510883 | 9190 | -8.78 | 4.90E-16 | 2.76E-15 |
| chr7 | 156485318 | 156528564 | 43247 | -8.77 | 4.94E-16 | 2.78E-15 |
| chr16 | 46847205 | 46850226 | 3022 | -8.76 | 4.95E-16 | 2.79E-15 |
| chr13 | 50035136 | 50039898 | 4763 | -8.78 | 4.95E-16 | 2.79E-15 |
| chr3 | 40983505 | 41006899 | 23395 | -8.85 | 4.96E-16 | 2.79E-15 |
| chr1 | 244248088 | 244263486 | 15399 | -8.76 | 4.97E-16 | 2.79E-15 |
| chr1 | 110321711 | 110325152 | 3442 | -8.76 | 4.97E-16 | 2.80E-15 |
| chr4 | 154220379 | 154227728 | 7350 | -8.83 | 4.98E-16 | 2.80E-15 |
| chr11 | 8569067 | 8584605 | 15539 | -8.76 | 5.00E-16 | 2.81E-15 |
| chr5 | 72895740 | 72899554 | 3815 | -8.77 | 5.04E-16 | 2.83E-15 |
| chr16 | 81543673 | 81546272 | 2600 | -8.83 | 5.04E-16 | 2.83E-15 |
| chr11 | 119267231 | 119272969 | 5739 | -8.83 | 5.04E-16 | 2.83E-15 |
| chr13 | 98160870 | 98173738 | 12869 | -2.13 | 5.07E-16 | 2.85E-15 |
| chr4 | 82283349 | 82314353 | 31005 | 7.33 | 5.08E-16 | 2.85E-15 |
| chr8 | 61402069 | 61421576 | 19508 | -8.9 | 5.08E-16 | 2.85E-15 |
| chr2 | 165401163 | 165408781 | 7619 | -8.76 | 5.08E-16 | 2.85E-15 |
| chr5 | 53863618 | 53934173 | 70556 | 1.41 | 5.13E-16 | 2.88E-15 |
| chr1 | 242643965 | 242661884 | 17920 | -8.96 | 5.13E-16 | 2.88E-15 |
| chr10 | 30770454 | 30774574 | 4121 | -8.8 | 5.15E-16 | 2.89E-15 |
| chr6 | 155558516 | 155569503 | 10988 | -8.83 | 5.17E-16 | 2.90E-15 |
| chr9 | 109944590 | 109971258 | 26669 | -8.81 | 5.18E-16 | 2.91E-15 |
| chr6 | 169488493 | 169504630 | 16138 | -8.77 | 5.19E-16 | 2.91E-15 |
| chrX | 20032099 | 20036376 | 4278 | -8.81 | 5.19E-16 | 2.91E-15 |
| chr19 | 39391913 | 39399518 | 7606 | -8.88 | 5.21E-16 | 2.92E-15 |
| chr4 | 99957080 | 99960776 | 3697 | -8.8 | 5.21E-16 | 2.92E-15 |
| chr8 | 124313041 | 124321896 | 8856 | -8.76 | 5.24E-16 | 2.94E-15 |
| chr2 | 173207438 | 173213082 | 5645 | -8.75 | 5.25E-16 | 2.94E-15 |
| chr17 | 77047718 | 77052506 | 4789 | -8.85 | 5.25E-16 | 2.94E-15 |

| | | | | | | |
|---|---|---|---|---|---|---|
| chr11 | 126373959 | 126377806 | 3848 | -8.75 | 5.25E-16 | 2.94E-15 |
| chr3 | 9562507 | 9591341 | 28835 | -8.86 | 5.26E-16 | 2.95E-15 |
| chr12 | 95397910 | 95403289 | 5380 | -8.81 | 5.27E-16 | 2.96E-15 |
| chr12 | 115145471 | 116341934 | 1196464 | 3.51 | 5.29E-16 | 2.96E-15 |
| chr6 | 75241064 | 75255824 | 14761 | -8.84 | 5.29E-16 | 2.96E-15 |
| chr3 | 37019524 | 37026856 | 7333 | -8.94 | 5.30E-16 | 2.97E-15 |
| chr5 | 53578631 | 53585827 | 7197 | -9.02 | 5.31E-16 | 2.98E-15 |
| chr2 | 135897866 | 135902005 | 4140 | -8.8 | 5.36E-16 | 3.00E-15 |
| chr1 | 41632135 | 41637415 | 5281 | -8.76 | 5.37E-16 | 3.00E-15 |
| chr4 | 140351705 | 140362579 | 10875 | -8.85 | 5.38E-16 | 3.01E-15 |
| chr7 | 116379958 | 116388733 | 8776 | -8.75 | 5.38E-16 | 3.01E-15 |
| chr11 | 35228882 | 35233424 | 4543 | -8.87 | 5.38E-16 | 3.01E-15 |
| chr17 | 63619508 | 63652804 | 33297 | -9.26 | 5.38E-16 | 3.01E-15 |
| chr2 | 154852587 | 154867187 | 14601 | -8.78 | 5.38E-16 | 3.01E-15 |
| chr1 | 173586648 | 173625843 | 39196 | -9.14 | 5.40E-16 | 3.02E-15 |
| chr10 | 79126821 | 79130850 | 4030 | -8.75 | 5.41E-16 | 3.02E-15 |
| chr2 | 178819304 | 178837266 | 17963 | 9.02 | 5.41E-16 | 3.03E-15 |
| chr8 | 19132992 | 19142701 | 9710 | -8.75 | 5.44E-16 | 3.04E-15 |
| chr14 | 27170498 | 27220380 | 49883 | -9.45 | 5.44E-16 | 3.04E-15 |
| chr5 | 14935172 | 14951114 | 15943 | -9.67 | 5.45E-16 | 3.05E-15 |
| chr12 | 63751429 | 63772660 | 21232 | -8.77 | 5.45E-16 | 3.05E-15 |
| chr2 | 20566794 | 20572392 | 5599 | -8.76 | 5.46E-16 | 3.05E-15 |
| chr5 | 179828076 | 179842114 | 14039 | -8.85 | 5.46E-16 | 3.05E-15 |
| chr5 | 90524091 | 90540953 | 16863 | -8.82 | 5.48E-16 | 3.06E-15 |
| chr4 | 186948135 | 186959379 | 11245 | -8.78 | 5.50E-16 | 3.07E-15 |
| chr15 | 39564132 | 39613571 | 49440 | -9.14 | 5.50E-16 | 3.07E-15 |
| chr12 | 132676684 | 132680063 | 3380 | -8.76 | 5.50E-16 | 3.07E-15 |
| chr4 | 189402223 | 189418142 | 15920 | -8.76 | 5.50E-16 | 3.07E-15 |
| chr7 | 80675000 | 80683850 | 8851 | -8.83 | 5.51E-16 | 3.07E-15 |
| chr18 | 13305111 | 13318202 | 13092 | -8.75 | 5.53E-16 | 3.09E-15 |
| chr4 | 58313676 | 58417288 | 103613 | 2.98 | 5.53E-16 | 3.09E-15 |
| chr19 | 15702550 | 15707948 | 5399 | -2.72 | 5.54E-16 | 3.09E-15 |
| chr11 | 120990271 | 120999604 | 9334 | -2.42 | 5.56E-16 | 3.10E-15 |
| chr2 | 170923850 | 170928615 | 4766 | -8.94 | 5.56E-16 | 3.10E-15 |
| chr9 | 87777351 | 87786187 | 8837 | -8.81 | 5.56E-16 | 3.10E-15 |
| chr8 | 126330428 | 126334069 | 3642 | -8.76 | 5.56E-16 | 3.10E-15 |
| chr12 | 11777667 | 11799035 | 21369 | -8.9 | 5.57E-16 | 3.10E-15 |
| chr9 | 111764241 | 111767654 | 3414 | -8.8 | 5.57E-16 | 3.10E-15 |
| chr12 | 96201716 | 96207250 | 5535 | -8.75 | 5.62E-16 | 3.13E-15 |
| chr9 | 98796899 | 98823065 | 26167 | -9.25 | 5.62E-16 | 3.13E-15 |
| chr19 | 6617781 | 6628536 | 10756 | -8.78 | 5.64E-16 | 3.14E-15 |
| chr10 | 4858376 | 4868151 | 9776 | -8.98 | 5.67E-16 | 3.16E-15 |
| chr9 | 123259888 | 123275180 | 15293 | -8.86 | 5.67E-16 | 3.16E-15 |
| chr4 | 152294241 | 152321842 | 27602 | -8.75 | 5.72E-16 | 3.19E-15 |
| chr1 | 95038188 | 95048740 | 10553 | -8.83 | 5.72E-16 | 3.19E-15 |
| chr12 | 48706851 | 48714181 | 7331 | -8.79 | 5.74E-16 | 3.19E-15 |
| chr2 | 26375880 | 26380504 | 4625 | -8.77 | 5.74E-16 | 3.19E-15 |
| chr1 | 225533481 | 225537908 | 4428 | -8.74 | 5.76E-16 | 3.21E-15 |
| chr18 | 29357011 | 29360971 | 3961 | -8.74 | 5.76E-16 | 3.21E-15 |
| chr3 | 146202445 | 146227845 | 25401 | -8.86 | 5.77E-16 | 3.21E-15 |
| chr12 | 90570783 | 90580538 | 9756 | -8.74 | 5.77E-16 | 3.21E-15 |
| chr3 | 87443012 | 87465671 | 22660 | -8.75 | 5.80E-16 | 3.22E-15 |
| chr4 | 64632243 | 64663622 | 31380 | -8.85 | 5.80E-16 | 3.23E-15 |
| chr6 | 158945149 | 158954879 | 9731 | -8.81 | 5.81E-16 | 3.23E-15 |
| chr12 | 21188344 | 21197608 | 9265 | -8.82 | 5.83E-16 | 3.24E-15 |
| chr3 | 103300501 | 104222490 | 921990 | 1.15 | 5.84E-16 | 3.25E-15 |
| chr18 | 45116842 | 45139292 | 22451 | -2.32 | 5.85E-16 | 3.25E-15 |
| chr4 | 175200083 | 175215152 | 15070 | -8.88 | 5.86E-16 | 3.26E-15 |
| chr16 | 57889169 | 57892547 | 3379 | -8.78 | 5.86E-16 | 3.26E-15 |
| chr1 | 9088910 | 9096551 | 7642 | -8.84 | 5.87E-16 | 3.26E-15 |
| chr1 | 204490581 | 204494061 | 3481 | -8.74 | 5.92E-16 | 3.29E-15 |

| | | | | | | |
|---|---|---|---|---|---|---|
| chr8 | 54679683 | 54687691 | 8009 | -8.74 | 5.92E-16 | 3.29E-15 |
| chr1 | 77758847 | 77812410 | 53564 | -8.8 | 5.93E-16 | 3.29E-15 |
| chr1 | 193774712 | 193801628 | 26917 | -8.82 | 5.94E-16 | 3.30E-15 |
| chr6 | 134172403 | 134191439 | 19037 | -9.14 | 5.94E-16 | 3.30E-15 |
| chr14 | 34438096 | 34441080 | 2985 | -8.76 | 5.96E-16 | 3.31E-15 |
| chr4 | 99930290 | 99943562 | 13273 | -8.74 | 5.97E-16 | 3.31E-15 |
| chr3 | 102438829 | 102479792 | 40964 | -8.81 | 5.97E-16 | 3.31E-15 |
| chr17 | 8800849 | 8904422 | 103574 | -3.36 | 5.97E-16 | 3.31E-15 |
| chr10 | 118766461 | 118769234 | 2774 | -8.75 | 5.98E-16 | 3.32E-15 |
| chr3 | 160167830 | 160192440 | 24611 | -9.08 | 5.99E-16 | 3.32E-15 |
| chr3 | 196497497 | 196523669 | 26173 | -8.88 | 5.99E-16 | 3.32E-15 |
| chr15 | 52074252 | 52080376 | 6125 | -8.87 | 6.00E-16 | 3.33E-15 |
| chr11 | 71919710 | 71926953 | 7244 | -8.74 | 6.00E-16 | 3.33E-15 |
| chr10 | 121743135 | 122204629 | 461495 | 3.66 | 6.00E-16 | 3.33E-15 |
| chr2 | 106769150 | 106774266 | 5117 | -8.91 | 6.03E-16 | 3.34E-15 |
| chr12 | 47747993 | 47795941 | 47949 | -9.58 | 6.03E-16 | 3.34E-15 |
| chrX | 35661116 | 35689995 | 28880 | -8.74 | 6.06E-16 | 3.36E-15 |
| chr10 | 101519092 | 101534306 | 15215 | -8.83 | 6.08E-16 | 3.37E-15 |
| chr7 | 138671899 | 138674816 | 2918 | -8.79 | 6.08E-16 | 3.37E-15 |
| chr15 | 40304399 | 40308589 | 4191 | -8.76 | 6.09E-16 | 3.37E-15 |
| chr12 | 75217047 | 75232114 | 15068 | -9.33 | 6.10E-16 | 3.38E-15 |
| chr2 | 214904478 | 214912167 | 7690 | -8.76 | 6.16E-16 | 3.41E-15 |
| chr5 | 129961960 | 129981739 | 19780 | -8.81 | 6.16E-16 | 3.41E-15 |
| chrX | 106076696 | 106103236 | 26541 | -8.9 | 6.17E-16 | 3.41E-15 |
| chr2 | 143484269 | 143511335 | 27067 | -9.08 | 6.18E-16 | 3.42E-15 |
| chr2 | 24742789 | 24790005 | 47217 | -9.17 | 6.20E-16 | 3.43E-15 |
| chr12 | 131907179 | 131915149 | 7971 | -8.87 | 6.21E-16 | 3.43E-15 |
| chr2 | 217927996 | 217933834 | 5839 | -8.77 | 6.22E-16 | 3.44E-15 |
| chr20 | 4004270 | 4007060 | 2791 | -8.78 | 6.23E-16 | 3.45E-15 |
| chr1 | 99812533 | 99831048 | 18516 | 8.87 | 6.26E-16 | 3.46E-15 |
| chr3 | 174165773 | 174187877 | 22105 | -8.73 | 6.26E-16 | 3.46E-15 |
| chr3 | 162033819 | 162055492 | 21674 | -8.73 | 6.27E-16 | 3.47E-15 |
| chr8 | 124572620 | 124577122 | 4503 | -3 | 6.28E-16 | 3.47E-15 |
| chr6 | 140785522 | 140810407 | 24886 | -8.88 | 6.29E-16 | 3.48E-15 |
| chr3 | 181692729 | 181703397 | 10669 | -8.89 | 6.29E-16 | 3.48E-15 |
| chr2 | 145847700 | 145877289 | 29590 | -8.9 | 6.30E-16 | 3.48E-15 |
| chr13 | 95224754 | 95237650 | 12897 | -8.76 | 6.30E-16 | 3.48E-15 |
| chr4 | 113365830 | 113385143 | 19314 | -8.75 | 6.31E-16 | 3.49E-15 |
| chr15 | 59856174 | 59859367 | 3194 | -8.88 | 6.32E-16 | 3.49E-15 |
| chr2 | 174747231 | 174756285 | 9055 | -8.78 | 6.33E-16 | 3.49E-15 |
| chr12 | 114137274 | 114148796 | 11523 | -8.73 | 6.38E-16 | 3.52E-15 |
| chr3 | 61610011 | 61633275 | 23265 | -8.74 | 6.40E-16 | 3.53E-15 |
| chr7 | 31785860 | 31806259 | 20400 | -8.8 | 6.40E-16 | 3.53E-15 |
| chr2 | 32004394 | 32020634 | 16241 | -8.88 | 6.41E-16 | 3.54E-15 |
| chr11 | 16155732 | 16190344 | 34613 | -8.99 | 6.41E-16 | 3.54E-15 |
| chr8 | 74636984 | 74644552 | 7569 | -8.82 | 6.45E-16 | 3.56E-15 |
| chr13 | 61068941 | 61087795 | 18855 | -8.73 | 6.46E-16 | 3.57E-15 |
| chr3 | 4901510 | 4905891 | 4382 | -8.76 | 6.49E-16 | 3.58E-15 |
| chr2 | 175505741 | 175522218 | 16478 | -8.74 | 6.51E-16 | 3.59E-15 |
| chr7 | 21491412 | 21496943 | 5532 | -8.95 | 6.51E-16 | 3.59E-15 |
| chr4 | 34114100 | 34219480 | 105381 | -1.76 | 6.51E-16 | 3.59E-15 |
| chr8 | 8808912 | 8836778 | 27867 | -9.05 | 6.51E-16 | 3.59E-15 |
| chr2 | 26824194 | 26828416 | 4223 | -8.99 | 6.52E-16 | 3.59E-15 |
| chr1 | 218528113 | 218544009 | 15897 | -8.81 | 6.54E-16 | 3.60E-15 |
| chr13 | 52320453 | 52323759 | 3307 | -8.73 | 6.58E-16 | 3.63E-15 |
| chr12 | 84356917 | 84388756 | 31840 | -8.73 | 6.60E-16 | 3.64E-15 |
| chr11 | 44724887 | 44732549 | 7663 | -8.96 | 6.61E-16 | 3.64E-15 |
| chrX | 103645968 | 103664223 | 18256 | -8.75 | 6.63E-16 | 3.65E-15 |
| chr3 | 5358436 | 5370035 | 11600 | -8.81 | 6.68E-16 | 3.68E-15 |
| chr4 | 24756337 | 24767522 | 11186 | -8.74 | 6.73E-16 | 3.71E-15 |
| chr2 | 39808014 | 39817315 | 9302 | -8.86 | 6.75E-16 | 3.71E-15 |

| | | | | | | |
|---|---|---|---|---|---|---|
| chr14 | 34540048 | 34558104 | 18057 | -9.05 | 6.75E-16 | 3.71E-15 |
| chr11 | 130366375 | 130379368 | 12994 | -8.95 | 6.76E-16 | 3.72E-15 |
| chr7 | 139362109 | 139372496 | 10388 | -8.75 | 6.78E-16 | 3.73E-15 |
| chr11 | 28155427 | 28163046 | 7620 | -8.81 | 6.81E-16 | 3.75E-15 |
| chr1 | 19617671 | 19623070 | 5400 | -8.73 | 6.81E-16 | 3.75E-15 |
| chr9 | 71925833 | 71932674 | 6842 | -8.72 | 6.85E-16 | 3.77E-15 |
| chr2 | 28989222 | 28992963 | 3742 | -8.74 | 6.87E-16 | 3.78E-15 |
| chr3 | 100519334 | 100551803 | 32470 | -3.24 | 6.89E-16 | 3.79E-15 |
| chr12 | 68828344 | 68844673 | 16330 | -8.93 | 6.90E-16 | 3.79E-15 |
| chrX | 30298950 | 30314257 | 15308 | -8.74 | 6.90E-16 | 3.79E-15 |
| chrX | 128800941 | 128808813 | 7873 | -8.76 | 6.90E-16 | 3.79E-15 |
| chr5 | 55533832 | 55545738 | 11907 | -8.88 | 6.94E-16 | 3.82E-15 |
| chr2 | 84049144 | 84080197 | 31054 | -8.73 | 6.95E-16 | 3.82E-15 |
| chr3 | 115704259 | 115730057 | 25799 | -8.78 | 6.95E-16 | 3.82E-15 |
| chr4 | 177546880 | 177563637 | 16758 | -8.77 | 6.96E-16 | 3.82E-15 |
| chr6 | 57028555 | 57031278 | 2724 | -8.82 | 7.03E-16 | 3.86E-15 |
| chr14 | 90521614 | 90526575 | 4962 | -8.73 | 7.05E-16 | 3.87E-15 |
| chr2 | 163993396 | 164003479 | 10084 | -8.98 | 7.05E-16 | 3.87E-15 |
| chr1 | 14990334 | 15052157 | 61824 | -3.33 | 7.07E-16 | 3.88E-15 |
| chr7 | 27602258 | 27607834 | 5577 | -8.86 | 7.08E-16 | 3.89E-15 |
| chr8 | 48662402 | 48674500 | 12099 | -8.86 | 7.08E-16 | 3.89E-15 |
| chr20 | 10677315 | 10694868 | 17554 | -8.84 | 7.08E-16 | 3.89E-15 |
| chr1 | 97382042 | 97394231 | 12190 | -8.81 | 7.11E-16 | 3.90E-15 |
| chr1 | 215149204 | 215167575 | 18372 | 8.82 | 7.11E-16 | 3.90E-15 |
| chr19 | 10154614 | 10157965 | 3352 | -5.1 | 7.11E-16 | 3.90E-15 |
| chr3 | 94633450 | 94667234 | 33785 | -8.76 | 7.12E-16 | 3.91E-15 |
| chr2 | 233774158 | 233778464 | 4307 | -8.75 | 7.13E-16 | 3.91E-15 |
| chr15 | 23799143 | 23812827 | 13685 | -8.82 | 7.13E-16 | 3.91E-15 |
| chr6 | 170609029 | 170634471 | 25443 | -8.74 | 7.15E-16 | 3.92E-15 |
| chr14 | 35410951 | 35414893 | 3943 | -8.75 | 7.16E-16 | 3.93E-15 |
| chrX | 77310340 | 77332086 | 21747 | -8.79 | 7.19E-16 | 3.94E-15 |
| chr22 | 50462576 | 50492594 | 30019 | -8.78 | 7.19E-16 | 3.94E-15 |
| chr7 | 36517961 | 36534550 | 16590 | -8.88 | 7.21E-16 | 3.95E-15 |
| chr16 | 20846679 | 20849172 | 2494 | -8.76 | 7.23E-16 | 3.96E-15 |
| chr2 | 58016608 | 58027451 | 10844 | -8.95 | 7.24E-16 | 3.97E-15 |
| chr21 | 46690516 | 46695614 | 5099 | -8.71 | 7.28E-16 | 3.99E-15 |
| chr2 | 139000991 | 139044834 | 43844 | -9.05 | 7.28E-16 | 3.99E-15 |
| chr2 | 98269142 | 98273000 | 3859 | -8.73 | 7.31E-16 | 4.00E-15 |
| chr3 | 164077889 | 164145839 | 67951 | -9.1 | 7.32E-16 | 4.01E-15 |
| chr8 | 47894501 | 47930409 | 35909 | -8.74 | 7.34E-16 | 4.02E-15 |
| chr10 | 135004398 | 135008256 | 3859 | -8.71 | 7.35E-16 | 4.02E-15 |
| chr6 | 14090492 | 14108615 | 18124 | -8.8 | 7.35E-16 | 4.02E-15 |
| chr20 | 61931938 | 61934220 | 2283 | -8.81 | 7.35E-16 | 4.02E-15 |
| chr20 | 34069956 | 34073273 | 3318 | -8.97 | 7.35E-16 | 4.02E-15 |
| chr3 | 187727882 | 187753666 | 25785 | -9.27 | 7.36E-16 | 4.03E-15 |
| chr6 | 162396765 | 162422881 | 26117 | -8.82 | 7.36E-16 | 4.03E-15 |
| chr3 | 9449401 | 9453008 | 3608 | -8.72 | 7.37E-16 | 4.03E-15 |
| chr18 | 29277112 | 29322364 | 45253 | -8.98 | 7.37E-16 | 4.03E-15 |
| chr5 | 170446617 | 170452538 | 5922 | -8.73 | 7.37E-16 | 4.03E-15 |
| chr2 | 235735941 | 235805186 | 69246 | -9.36 | 7.39E-16 | 4.04E-15 |
| chr17 | 6333245 | 6346391 | 13147 | -8.75 | 7.43E-16 | 4.06E-15 |
| chr13 | 40352560 | 40370207 | 17648 | -8.78 | 7.45E-16 | 4.07E-15 |
| chr20 | 57941606 | 57952758 | 11153 | 8.78 | 7.48E-16 | 4.09E-15 |
| chr2 | 181721050 | 181750430 | 29381 | -9 | 7.50E-16 | 4.10E-15 |
| chr2 | 101784373 | 101806567 | 22195 | -9.11 | 7.52E-16 | 4.11E-15 |
| chr2 | 201699759 | 201703348 | 3590 | -8.83 | 7.56E-16 | 4.13E-15 |
| chr13 | 40896313 | 40904916 | 8604 | -8.72 | 7.56E-16 | 4.13E-15 |
| chr20 | 6004177 | 6006695 | 2519 | -8.78 | 7.62E-16 | 4.16E-15 |
| chr7 | 34777918 | 34784846 | 6929 | -8.73 | 7.62E-16 | 4.16E-15 |
| chr7 | 39919861 | 39924045 | 4185 | -8.78 | 7.64E-16 | 4.17E-15 |
| chr1 | 4664600 | 4672273 | 7674 | -8.71 | 7.65E-16 | 4.18E-15 |

| | | | | | | |
|---|---|---|---|---|---|---|
| chrX | 36525023 | 36542162 | 17140 | -8.82 | 7.66E-16 | 4.18E-15 |
| chr6 | 116484577 | 116497693 | 13117 | -8.75 | 7.69E-16 | 4.19E-15 |
| chr2 | 230600323 | 230609650 | 9328 | -8.74 | 7.70E-16 | 4.20E-15 |
| chr6 | 28197267 | 28207145 | 9879 | -8.71 | 7.71E-16 | 4.20E-15 |
| chr2 | 170536494 | 170548170 | 11677 | -8.71 | 7.71E-16 | 4.20E-15 |
| chr12 | 31653643 | 31667879 | 14237 | -8.73 | 7.72E-16 | 4.21E-15 |
| chr4 | 830362 | 836740 | 6379 | -8.72 | 7.73E-16 | 4.21E-15 |
| chr2 | 223193714 | 223200505 | 6792 | -8.76 | 7.73E-16 | 4.22E-15 |
| chr3 | 177474888 | 177488251 | 13364 | -8.71 | 7.75E-16 | 4.22E-15 |
| chr18 | 57821860 | 57837963 | 16104 | -8.71 | 7.75E-16 | 4.22E-15 |
| chr5 | 73237128 | 73241114 | 3987 | -8.71 | 7.75E-16 | 4.22E-15 |
| chrX | 14884354 | 14887124 | 2771 | -8.71 | 7.75E-16 | 4.22E-15 |
| chr6 | 109861733 | 109880175 | 18443 | -2.7 | 7.76E-16 | 4.22E-15 |
| chr11 | 16446950 | 16454991 | 8042 | -8.75 | 7.76E-16 | 4.23E-15 |
| chr6 | 97404804 | 97424314 | 19511 | 8.92 | 7.79E-16 | 4.24E-15 |
| chr7 | 109454304 | 109462164 | 7861 | -8.73 | 7.79E-16 | 4.24E-15 |
| chr8 | 143220809 | 143244257 | 23449 | -8.79 | 7.80E-16 | 4.25E-15 |
| chr9 | 93849236 | 93917949 | 68714 | -9.62 | 7.80E-16 | 4.25E-15 |
| chr18 | 77258146 | 77262156 | 4011 | -8.8 | 7.82E-16 | 4.26E-15 |
| chr13 | 103309823 | 103323163 | 13341 | -8.72 | 7.83E-16 | 4.26E-15 |
| chr5 | 146717440 | 146732729 | 15290 | 8.82 | 7.88E-16 | 4.29E-15 |
| chr7 | 18378643 | 18383590 | 4948 | -8.7 | 7.88E-16 | 4.29E-15 |
| chr10 | 99557591 | 99559991 | 2401 | -8.71 | 7.88E-16 | 4.29E-15 |
| chr8 | 53391611 | 53403416 | 11806 | -8.72 | 7.90E-16 | 4.30E-15 |
| chr5 | 166850709 | 166903971 | 53263 | 2.53 | 7.91E-16 | 4.30E-15 |
| chr8 | 107093951 | 107114273 | 20323 | -8.7 | 7.92E-16 | 4.31E-15 |
| chr5 | 120924746 | 120938457 | 13712 | -8.75 | 7.92E-16 | 4.31E-15 |
| chr7 | 128299906 | 128303848 | 3943 | -8.76 | 8.00E-16 | 4.35E-15 |
| chr8 | 100442897 | 100447479 | 4583 | -8.99 | 8.04E-16 | 4.37E-15 |
| chr12 | 19898623 | 19907129 | 8507 | -8.82 | 8.05E-16 | 4.37E-15 |
| chr10 | 48498977 | 48538478 | 39502 | -4.04 | 8.05E-16 | 4.37E-15 |
| chr2 | 65181056 | 65183580 | 2525 | -8.74 | 8.05E-16 | 4.38E-15 |
| chr7 | 23288590 | 23293519 | 4930 | -2.92 | 8.07E-16 | 4.38E-15 |
| chr1 | 205361808 | 205376576 | 14769 | -8.78 | 8.09E-16 | 4.39E-15 |
| chr16 | 52114837 | 52182977 | 68141 | 1.4 | 8.13E-16 | 4.41E-15 |
| chr3 | 149437143 | 149440960 | 3818 | -8.72 | 8.14E-16 | 4.42E-15 |
| chr8 | 123901586 | 123912144 | 10559 | -8.73 | 8.15E-16 | 4.42E-15 |
| chr1 | 117435589 | 117442441 | 6853 | -8.86 | 8.19E-16 | 4.45E-15 |
| chr12 | 83398600 | 83405324 | 6725 | -8.73 | 8.20E-16 | 4.45E-15 |
| chr6 | 5013359 | 5021308 | 7950 | -8.75 | 8.21E-16 | 4.46E-15 |
| chr3 | 101825287 | 101843277 | 17991 | -9.31 | 8.24E-16 | 4.47E-15 |
| chr11 | 9755641 | 9765704 | 10064 | -8.71 | 8.34E-16 | 4.52E-15 |
| chr10 | 115072956 | 115109899 | 36944 | -8.89 | 8.34E-16 | 4.52E-15 |
| chr20 | 24755107 | 24758536 | 3430 | -8.72 | 8.36E-16 | 4.54E-15 |
| chr1 | 150825035 | 150828726 | 3692 | -8.71 | 8.36E-16 | 4.54E-15 |
| chr7 | 98397303 | 98408437 | 11135 | -8.94 | 8.39E-16 | 4.55E-15 |
| chr5 | 93394465 | 93403966 | 9502 | -8.71 | 8.41E-16 | 4.56E-15 |
| chr4 | 152468168 | 152474645 | 6478 | -8.7 | 8.41E-16 | 4.56E-15 |
| chr2 | 40605832 | 40610219 | 4388 | -8.76 | 8.42E-16 | 4.56E-15 |
| chr21 | 33792440 | 33795506 | 3067 | -8.76 | 8.45E-16 | 4.58E-15 |
| chr21 | 38276090 | 38288732 | 12643 | -8.71 | 8.53E-16 | 4.62E-15 |
| chr8 | 134616278 | 134629713 | 13436 | -8.85 | 8.54E-16 | 4.63E-15 |
| chr4 | 108623412 | 108629819 | 6408 | -8.7 | 8.57E-16 | 4.64E-15 |
| chr1 | 226803682 | 226811146 | 7465 | -8.69 | 8.59E-16 | 4.65E-15 |
| chr10 | 94679511 | 94690040 | 10530 | -8.77 | 8.63E-16 | 4.67E-15 |
| chr1 | 242994401 | 243014947 | 20547 | -8.74 | 8.63E-16 | 4.67E-15 |
| chr12 | 84884228 | 84918836 | 34609 | -8.7 | 8.64E-16 | 4.68E-15 |
| chr1 | 22160120 | 22163353 | 3234 | -8.75 | 8.67E-16 | 4.69E-15 |
| chr9 | 13921904 | 13927726 | 5823 | -1.96 | 8.68E-16 | 4.70E-15 |
| chr9 | 77544430 | 77549060 | 4631 | -8.7 | 8.71E-16 | 4.71E-15 |
| chr16 | 72555304 | 72566427 | 11124 | -8.71 | 8.72E-16 | 4.72E-15 |

| | | | | | | |
|---|---|---|---|---|---|---|
| chr9 | 538587 | 541353 | 2767 | -8.69 | 8.72E-16 | 4.72E-15 |
| chr7 | 109933059 | 109957543 | 24485 | -8.71 | 8.76E-16 | 4.74E-15 |
| chr4 | 187250426 | 187263339 | 12914 | -8.75 | 8.77E-16 | 4.74E-15 |
| chr11 | 12043369 | 12047893 | 4525 | -8.69 | 8.80E-16 | 4.76E-15 |
| chr5 | 151309136 | 151312267 | 3132 | -6.24 | 8.84E-16 | 4.78E-15 |
| chr4 | 130797950 | 130831334 | 33385 | -8.72 | 8.85E-16 | 4.78E-15 |
| chrX | 119468832 | 119486979 | 18148 | -2.2 | 8.85E-16 | 4.79E-15 |
| chr3 | 121402477 | 121405470 | 2994 | -8.76 | 8.86E-16 | 4.79E-15 |
| chr8 | 106682149 | 106697201 | 15053 | 8.76 | 8.87E-16 | 4.79E-15 |
| chr9 | 108226132 | 108232493 | 6362 | -8.81 | 8.89E-16 | 4.80E-15 |
| chr1 | 204136761 | 204140858 | 4098 | -8.7 | 8.94E-16 | 4.83E-15 |
| chr21 | 39468135 | 39480663 | 12529 | -8.77 | 8.94E-16 | 4.83E-15 |
| chr7 | 129207033 | 129228553 | 21521 | -8.76 | 8.94E-16 | 4.83E-15 |
| chr5 | 58920486 | 58928302 | 7817 | -8.75 | 8.94E-16 | 4.83E-15 |
| chr10 | 60803706 | 60830947 | 27242 | -9.18 | 8.99E-16 | 4.86E-15 |
| chrX | 102223795 | 102230182 | 6388 | -2.69 | 9.00E-16 | 4.86E-15 |
| chr20 | 11003671 | 11014687 | 11017 | -8.69 | 9.01E-16 | 4.86E-15 |
| chr2 | 164088143 | 164100991 | 12849 | -8.71 | 9.01E-16 | 4.86E-15 |
| chr10 | 113944696 | 113958031 | 13336 | -8.69 | 9.03E-16 | 4.87E-15 |
| chr10 | 22578026 | 22581593 | 3568 | -2.87 | 9.03E-16 | 4.87E-15 |
| chr3 | 148610594 | 148612206 | 1613 | -3.83 | 9.06E-16 | 4.89E-15 |
| chr8 | 63289536 | 63294751 | 5216 | -8.71 | 9.07E-16 | 4.89E-15 |
| chr22 | 49674857 | 49691030 | 16174 | -8.72 | 9.08E-16 | 4.89E-15 |
| chr13 | 107229686 | 107240334 | 10649 | -8.71 | 9.09E-16 | 4.90E-15 |
| chr9 | 110959765 | 110967326 | 7562 | -8.75 | 9.13E-16 | 4.92E-15 |
| chr2 | 69629997 | 69632531 | 2535 | -8.69 | 9.15E-16 | 4.93E-15 |
| chr8 | 102617409 | 102643970 | 26562 | -8.92 | 9.16E-16 | 4.94E-15 |
| chr12 | 18843684 | 18870999 | 27316 | -8.79 | 9.21E-16 | 4.96E-15 |
| chr10 | 62729645 | 62752244 | 22600 | -8.73 | 9.25E-16 | 4.99E-15 |
| chr12 | 344011 | 365656 | 21646 | -8.73 | 9.27E-16 | 4.99E-15 |
| chr1 | 218483063 | 218486233 | 3171 | -8.72 | 9.28E-16 | 5.00E-15 |
| chr14 | 94584363 | 94602958 | 18596 | -8.85 | 9.29E-16 | 5.00E-15 |
| chr11 | 3230816 | 3233902 | 3087 | -8.75 | 9.30E-16 | 5.01E-15 |
| chr11 | 35590329 | 35602747 | 12419 | -8.73 | 9.34E-16 | 5.03E-15 |
| chr14 | 105564250 | 105571259 | 7010 | -8.92 | 9.38E-16 | 5.05E-15 |
| chr12 | 12017168 | 12020673 | 3506 | -8.69 | 9.40E-16 | 5.06E-15 |
| chr10 | 101099066 | 101101264 | 2199 | -8.7 | 9.40E-16 | 5.06E-15 |
| chrX | 84264482 | 84294940 | 30459 | -8.95 | 9.41E-16 | 5.06E-15 |
| chr7 | 152340517 | 152346613 | 6097 | -8.85 | 9.43E-16 | 5.07E-15 |
| chr6 | 37433094 | 37437281 | 4188 | -8.69 | 9.43E-16 | 5.07E-15 |
| chr4 | 139294292 | 139299735 | 5444 | -7.05 | 9.43E-16 | 5.07E-15 |
| chr18 | 33973010 | 33977926 | 4917 | -8.69 | 9.44E-16 | 5.08E-15 |
| chr4 | 95878596 | 95886129 | 7534 | -8.79 | 9.49E-16 | 5.10E-15 |
| chr4 | 7875576 | 7892122 | 16547 | -8.68 | 9.51E-16 | 5.11E-15 |
| chr10 | 3421664 | 3437278 | 15615 | -8.77 | 9.52E-16 | 5.12E-15 |
| chr4 | 188550808 | 188567995 | 17188 | -8.71 | 9.52E-16 | 5.12E-15 |
| chr11 | 95663824 | 95682196 | 18373 | -8.7 | 9.52E-16 | 5.12E-15 |
| chr1 | 236360203 | 236368636 | 8434 | -8.99 | 9.54E-16 | 5.13E-15 |
| chr2 | 101044657 | 101126902 | 82246 | -2.92 | 9.60E-16 | 5.16E-15 |
| chr14 | 48875822 | 48910683 | 34862 | -8.78 | 9.64E-16 | 5.18E-15 |
| chr1 | 229811714 | 229822548 | 10835 | -8.68 | 9.64E-16 | 5.18E-15 |
| chr9 | 6264200 | 6295618 | 31419 | -8.69 | 9.65E-16 | 5.18E-15 |
| chr10 | 25287923 | 25294337 | 6415 | -8.69 | 9.66E-16 | 5.19E-15 |
| chr3 | 109309497 | 109329892 | 20396 | -8.75 | 9.68E-16 | 5.20E-15 |
| chr2 | 70243553 | 70246941 | 3389 | -8.68 | 9.71E-16 | 5.21E-15 |
| chr12 | 97829611 | 97847950 | 18340 | 8.77 | 9.71E-16 | 5.21E-15 |
| chr1 | 233738346 | 233741930 | 3585 | -8.7 | 9.74E-16 | 5.23E-15 |
| chr11 | 72787375 | 72790215 | 2841 | -8.73 | 9.75E-16 | 5.23E-15 |
| chr2 | 84666829 | 84669085 | 2257 | -8.81 | 9.75E-16 | 5.23E-15 |
| chr1 | 66234159 | 66257488 | 23330 | -1.66 | 9.76E-16 | 5.24E-15 |
| chr5 | 96183674 | 96186346 | 2673 | -8.68 | 9.77E-16 | 5.24E-15 |

| | | | | | | |
|---|---|---|---|---|---|---|
| chr2 | 172833499 | 172839047 | 5549 | -8.74 | 9.81E-16 | 5.26E-15 |
| chr7 | 105591271 | 105595626 | 4356 | -8.82 | 9.82E-16 | 5.26E-15 |
| chr14 | 31466704 | 31469387 | 2684 | -8.7 | 9.82E-16 | 5.27E-15 |
| chr5 | 137046784 | 137075245 | 28462 | -9.15 | 9.82E-16 | 5.27E-15 |
| chr11 | 100482275 | 100495162 | 12888 | -8.86 | 9.83E-16 | 5.27E-15 |
| chr1 | 113017752 | 113024646 | 6895 | -8.69 | 9.87E-16 | 5.29E-15 |
| chr15 | 33396455 | 33401864 | 5410 | -5.57 | 9.89E-16 | 5.30E-15 |
| chr4 | 87434325 | 87452783 | 18459 | -8.94 | 9.90E-16 | 5.31E-15 |
| chr1 | 223601195 | 223663568 | 62374 | -9.04 | 9.92E-16 | 5.31E-15 |
| chr1 | 59623035 | 59634490 | 11456 | -8.68 | 9.96E-16 | 5.34E-15 |
| chr6 | 161355823 | 161359402 | 3580 | -6.14 | 9.97E-16 | 5.34E-15 |
| chr7 | 3403224 | 3407434 | 4211 | -8.69 | 9.97E-16 | 5.34E-15 |
| chr9 | 17023543 | 17045178 | 21636 | -8.72 | 9.98E-16 | 5.34E-15 |
| chr7 | 78865029 | 78872810 | 7782 | -5.25 | 1.00E-15 | 5.36E-15 |
| chr4 | 15739201 | 15748767 | 9567 | -8.87 | 1.00E-15 | 5.37E-15 |
| chr16 | 71950956 | 71954407 | 3452 | -8.73 | 1.01E-15 | 5.39E-15 |
| chr4 | 13489055 | 13503781 | 14727 | -8.68 | 1.01E-15 | 5.39E-15 |
| chr4 | 58262126 | 58286998 | 24873 | 9.07 | 1.01E-15 | 5.40E-15 |
| chr4 | 161343646 | 161362388 | 18743 | -8.73 | 1.01E-15 | 5.42E-15 |
| chr4 | 163590026 | 163602766 | 12741 | -8.77 | 1.02E-15 | 5.44E-15 |
| chr10 | 29328325 | 29333164 | 4840 | -8.8 | 1.02E-15 | 5.45E-15 |
| chr20 | 50670731 | 50675351 | 4621 | -8.69 | 1.02E-15 | 5.46E-15 |
| chr9 | 134881853 | 134889017 | 7165 | -8.7 | 1.02E-15 | 5.46E-15 |
| chr7 | 28733227 | 28759715 | 26489 | 9.17 | 1.02E-15 | 5.48E-15 |
| chr1 | 8790820 | 8801729 | 10910 | -8.81 | 1.03E-15 | 5.50E-15 |
| chr17 | 63904416 | 63908900 | 4485 | -8.69 | 1.03E-15 | 5.50E-15 |
| chr21 | 35580920 | 35588055 | 7136 | -8.7 | 1.03E-15 | 5.51E-15 |
| chr17 | 61279931 | 61283709 | 3779 | -8.72 | 1.03E-15 | 5.51E-15 |
| chr2 | 21062885 | 21096802 | 33918 | -3.18 | 1.03E-15 | 5.51E-15 |
| chr13 | 39859640 | 39871883 | 12244 | -8.77 | 1.04E-15 | 5.54E-15 |
| chrX | 153021103 | 153024168 | 3066 | -8.67 | 1.04E-15 | 5.54E-15 |
| chrX | 16010880 | 16020598 | 9719 | -8.87 | 1.04E-15 | 5.54E-15 |
| chrX | 40266870 | 40288680 | 21811 | -8.84 | 1.04E-15 | 5.55E-15 |
| chr22 | 45529625 | 45542652 | 13028 | -8.87 | 1.04E-15 | 5.56E-15 |
| chr4 | 129149293 | 129174567 | 25275 | -8.9 | 1.04E-15 | 5.56E-15 |
| chr6 | 33954105 | 33992311 | 38207 | -9.16 | 1.05E-15 | 5.59E-15 |
| chr7 | 18037714 | 18061472 | 23759 | -8.67 | 1.05E-15 | 5.60E-15 |
| chr3 | 36265002 | 36279396 | 14395 | -8.67 | 1.05E-15 | 5.60E-15 |
| chr1 | 225825276 | 225834240 | 8965 | -8.92 | 1.05E-15 | 5.60E-15 |
| chr3 | 16370523 | 16384111 | 13589 | -8.72 | 1.05E-15 | 5.60E-15 |
| chr7 | 105350136 | 105355100 | 4965 | -8.69 | 1.05E-15 | 5.60E-15 |
| chr20 | 52616627 | 52620015 | 3389 | -8.67 | 1.05E-15 | 5.60E-15 |
| chr12 | 45918854 | 45937356 | 18503 | -8.73 | 1.05E-15 | 5.60E-15 |
| chr22 | 46154792 | 46167915 | 13124 | -8.67 | 1.05E-15 | 5.62E-15 |
| chr2 | 205201604 | 205215206 | 13603 | -8.75 | 1.05E-15 | 5.62E-15 |
| chr19 | 34142080 | 34145063 | 2984 | -8.76 | 1.06E-15 | 5.62E-15 |
| chr9 | 92739678 | 92759084 | 19407 | -9.07 | 1.06E-15 | 5.65E-15 |
| chr5 | 40738731 | 40742326 | 3596 | -8.79 | 1.06E-15 | 5.65E-15 |
| chr17 | 71999154 | 72012352 | 13199 | -8.83 | 1.07E-15 | 5.68E-15 |
| chr14 | 93803062 | 93830404 | 27343 | -8.75 | 1.07E-15 | 5.69E-15 |
| chr11 | 109124410 | 109143339 | 18930 | -8.67 | 1.07E-15 | 5.70E-15 |
| chr3 | 180303951 | 180319793 | 15843 | -8.75 | 1.07E-15 | 5.70E-15 |
| chr19 | 3898050 | 3923679 | 25630 | -8.68 | 1.07E-15 | 5.71E-15 |
| chr6 | 3586973 | 3603057 | 16085 | -8.82 | 1.07E-15 | 5.72E-15 |
| chr7 | 82156429 | 82186075 | 29647 | -8.91 | 1.08E-15 | 5.73E-15 |
| chr18 | 74771739 | 74777320 | 5582 | -8.67 | 1.08E-15 | 5.73E-15 |
| chr1 | 116680719 | 116688115 | 7397 | -8.76 | 1.08E-15 | 5.73E-15 |
| chr12 | 66040169 | 66060573 | 20405 | -8.74 | 1.08E-15 | 5.76E-15 |
| chr3 | 36881929 | 36886466 | 4538 | -8.69 | 1.09E-15 | 5.80E-15 |
| chr4 | 124990404 | 125002959 | 12556 | -8.78 | 1.09E-15 | 5.81E-15 |
| chr12 | 86428428 | 86437063 | 8636 | -7.02 | 1.09E-15 | 5.81E-15 |

| | | | | | | |
|---|---|---|---|---|---|---|
| chr4 | 56563057 | 56582967 | 19911 | -6.36 | 1.10E-15 | 5.84E-15 |
| chr15 | 49024560 | 49029293 | 4734 | -8.91 | 1.10E-15 | 5.85E-15 |
| chr16 | 52652926 | 52670867 | 17942 | -8.77 | 1.10E-15 | 5.86E-15 |
| chr12 | 74213979 | 74221970 | 7992 | -6.28 | 1.10E-15 | 5.86E-15 |
| chr19 | 19233669 | 19244133 | 10465 | -8.7 | 1.11E-15 | 5.87E-15 |
| chr1 | 193607854 | 193625373 | 17520 | -8.67 | 1.11E-15 | 5.88E-15 |
| chr2 | 72556898 | 72561601 | 4704 | -8.67 | 1.11E-15 | 5.89E-15 |
| chr18 | 19588946 | 19596617 | 7672 | -8.67 | 1.11E-15 | 5.90E-15 |
| chr6 | 24876189 | 24878779 | 2591 | -6.85 | 1.12E-15 | 5.93E-15 |
| chr12 | 125216257 | 125251038 | 34782 | -9 | 1.12E-15 | 5.93E-15 |
| chr13 | 74783094 | 74800012 | 16919 | 8.96 | 1.12E-15 | 5.93E-15 |
| chr1 | 10622596 | 10645951 | 23356 | -8.76 | 1.12E-15 | 5.94E-15 |
| chr2 | 106031469 | 106040279 | 8811 | -8.76 | 1.12E-15 | 5.94E-15 |
| chr1 | 113130053 | 113134122 | 4070 | -8.68 | 1.12E-15 | 5.96E-15 |
| chr11 | 11887307 | 11890820 | 3514 | -8.68 | 1.12E-15 | 5.96E-15 |
| chr7 | 8384244 | 8399558 | 15315 | -8.69 | 1.13E-15 | 5.97E-15 |
| chr10 | 23498393 | 23575879 | 77487 | 1.46 | 1.13E-15 | 5.98E-15 |
| chr7 | 117087442 | 117104313 | 16872 | -8.89 | 1.14E-15 | 6.03E-15 |
| chr19 | 43052208 | 43056197 | 3990 | -3.13 | 1.14E-15 | 6.04E-15 |
| chr2 | 11556562 | 11585315 | 28754 | -9.01 | 1.14E-15 | 6.05E-15 |
| chr12 | 93947031 | 93953427 | 6397 | -8.67 | 1.14E-15 | 6.05E-15 |
| chrX | 72448292 | 72463116 | 14825 | 6.5 | 1.15E-15 | 6.07E-15 |
| chr18 | 69980363 | 69996151 | 15789 | -8.72 | 1.15E-15 | 6.07E-15 |
| chr7 | 33476831 | 33494615 | 17785 | 8.77 | 1.15E-15 | 6.11E-15 |
| chr6 | 109138282 | 109149992 | 11711 | -8.78 | 1.15E-15 | 6.11E-15 |
| chr2 | 224908394 | 224924120 | 15727 | -8.77 | 1.16E-15 | 6.15E-15 |
| chr2 | 131652604 | 131666736 | 14133 | -8.72 | 1.16E-15 | 6.16E-15 |
| chr8 | 11343664 | 11349117 | 5454 | -8.88 | 1.16E-15 | 6.17E-15 |
| chr12 | 120773820 | 120777130 | 3311 | -8.67 | 1.17E-15 | 6.18E-15 |
| chr7 | 110301012 | 110315211 | 14200 | -8.67 | 1.17E-15 | 6.18E-15 |
| chr16 | 75053470 | 75056930 | 3461 | -8.76 | 1.17E-15 | 6.18E-15 |
| chr5 | 166485508 | 166669945 | 184438 | 1.32 | 1.17E-15 | 6.19E-15 |
| chrX | 84545737 | 84557841 | 12105 | -8.67 | 1.17E-15 | 6.19E-15 |
| chr7 | 35461389 | 35501350 | 39962 | -2.28 | 1.18E-15 | 6.23E-15 |
| chr15 | 67945018 | 67948659 | 3642 | -8.69 | 1.18E-15 | 6.25E-15 |
| chr6 | 56244768 | 56268533 | 23766 | -8.75 | 1.19E-15 | 6.28E-15 |
| chr3 | 126413258 | 126421122 | 7865 | -8.66 | 1.19E-15 | 6.28E-15 |
| chr4 | 4400235 | 4402925 | 2691 | -8.68 | 1.19E-15 | 6.29E-15 |
| chr4 | 177263580 | 177295745 | 32166 | -8.68 | 1.19E-15 | 6.31E-15 |
| chr12 | 86044571 | 86054285 | 9715 | -8.76 | 1.20E-15 | 6.34E-15 |
| chr8 | 101085641 | 101088451 | 2811 | -8.71 | 1.20E-15 | 6.35E-15 |
| chr4 | 28033467 | 28047508 | 14042 | -8.66 | 1.20E-15 | 6.35E-15 |
| chr3 | 160835246 | 160844348 | 9103 | -8.66 | 1.20E-15 | 6.35E-15 |
| chr1 | 1126004 | 1131242 | 5239 | -8.72 | 1.21E-15 | 6.39E-15 |
| chr11 | 129019723 | 129037475 | 17753 | -8.67 | 1.21E-15 | 6.39E-15 |
| chr6 | 56297979 | 56305907 | 7929 | -8.66 | 1.21E-15 | 6.39E-15 |
| chr9 | 32358652 | 32371992 | 13341 | -8.75 | 1.21E-15 | 6.41E-15 |
| chr6 | 40938827 | 40948538 | 9712 | -8.75 | 1.21E-15 | 6.41E-15 |
| chr3 | 196881691 | 196885383 | 3693 | -8.66 | 1.21E-15 | 6.41E-15 |
| chr6 | 15885862 | 15903735 | 17874 | -8.71 | 1.21E-15 | 6.41E-15 |
| chr3 | 113687547 | 113694487 | 6941 | -8.67 | 1.22E-15 | 6.43E-15 |
| chr4 | 81918388 | 81932302 | 13915 | 8.94 | 1.22E-15 | 6.43E-15 |
| chr20 | 32464906 | 32480475 | 15570 | -8.78 | 1.22E-15 | 6.45E-15 |
| chr17 | 20032697 | 20035427 | 2731 | -8.65 | 1.22E-15 | 6.45E-15 |
| chr1 | 169424881 | 169428541 | 3661 | -8.65 | 1.23E-15 | 6.47E-15 |
| chr6 | 144760952 | 144764608 | 3657 | -8.66 | 1.23E-15 | 6.47E-15 |
| chr17 | 57174789 | 57177835 | 3047 | -8.65 | 1.23E-15 | 6.48E-15 |
| chrX | 40134736 | 40150092 | 15357 | -8.73 | 1.23E-15 | 6.50E-15 |
| chr4 | 83684696 | 83689629 | 4934 | -8.72 | 1.23E-15 | 6.51E-15 |
| chr11 | 69547343 | 69554302 | 6960 | -8.93 | 1.24E-15 | 6.52E-15 |
| chr11 | 8909736 | 8924571 | 14836 | -8.83 | 1.24E-15 | 6.53E-15 |

| | | | | | | |
|---|---|---|---|---|---|---|
| chr3 | 184772969 | 184779461 | 6493 | -8.65 | 1.24E-15 | 6.53E-15 |
| chr19 | 38484116 | 38486678 | 2563 | -8.69 | 1.25E-15 | 6.57E-15 |
| chr12 | 50855678 | 50859741 | 4064 | -8.67 | 1.25E-15 | 6.57E-15 |
| chr18 | 53368958 | 53382442 | 13485 | -8.68 | 1.26E-15 | 6.62E-15 |
| chr5 | 53241627 | 53254546 | 12920 | -8.67 | 1.26E-15 | 6.62E-15 |
| chr6 | 90570651 | 90577657 | 7007 | -8.69 | 1.26E-15 | 6.63E-15 |
| chr11 | 3134538 | 3138276 | 3739 | -8.66 | 1.26E-15 | 6.64E-15 |
| chr6 | 147793710 | 147821452 | 27743 | -8.74 | 1.26E-15 | 6.65E-15 |
| chr1 | 182005654 | 182017426 | 11773 | -8.74 | 1.26E-15 | 6.65E-15 |
| chr14 | 77357239 | 77367002 | 9764 | -8.75 | 1.27E-15 | 6.67E-15 |
| chr5 | 92857157 | 92873962 | 16806 | -8.75 | 1.27E-15 | 6.67E-15 |
| chr8 | 18486070 | 18508042 | 21973 | -8.65 | 1.27E-15 | 6.70E-15 |
| chr20 | 62774322 | 62780935 | 6614 | -8.7 | 1.28E-15 | 6.73E-15 |
| chr14 | 31293238 | 31304979 | 11742 | -8.69 | 1.28E-15 | 6.74E-15 |
| chr5 | 126699892 | 126725104 | 25213 | -8.67 | 1.28E-15 | 6.75E-15 |
| chr3 | 10865446 | 10885963 | 20518 | -9.02 | 1.28E-15 | 6.75E-15 |
| chr1 | 178155432 | 178182338 | 26907 | -8.76 | 1.29E-15 | 6.76E-15 |
| chr8 | 53348635 | 53370011 | 21377 | -8.86 | 1.29E-15 | 6.77E-15 |
| chr4 | 28071445 | 28097646 | 26202 | -8.65 | 1.29E-15 | 6.79E-15 |
| chr3 | 87790046 | 87826653 | 36608 | -9.24 | 1.30E-15 | 6.83E-15 |
| chr2 | 230473007 | 230488527 | 15521 | 8.74 | 1.30E-15 | 6.84E-15 |
| chr4 | 168828212 | 168844028 | 15817 | -8.68 | 1.30E-15 | 6.84E-15 |
| chr8 | 109906004 | 109912063 | 6060 | -8.65 | 1.31E-15 | 6.86E-15 |
| chr14 | 69296882 | 69304005 | 7124 | -8.69 | 1.31E-15 | 6.86E-15 |
| chr20 | 4750816 | 4754539 | 3724 | -8.72 | 1.31E-15 | 6.88E-15 |
| chr9 | 101480467 | 101487567 | 7101 | -8.72 | 1.31E-15 | 6.88E-15 |
| chr12 | 122678790 | 122681811 | 3022 | -8.68 | 1.31E-15 | 6.88E-15 |
| chr7 | 81840760 | 81858734 | 17975 | -8.81 | 1.31E-15 | 6.89E-15 |
| chr9 | 26854728 | 26895792 | 41065 | -8.79 | 1.31E-15 | 6.90E-15 |
| chr2 | 237058557 | 237063419 | 4863 | -8.76 | 1.31E-15 | 6.90E-15 |
| chr14 | 37792479 | 37801612 | 9134 | -8.65 | 1.32E-15 | 6.91E-15 |
| chrX | 71028518 | 71036567 | 8050 | -3.65 | 1.32E-15 | 6.92E-15 |
| chr10 | 129084337 | 129088913 | 4577 | -8.67 | 1.32E-15 | 6.92E-15 |
| chr5 | 143288649 | 143300264 | 11616 | -9.02 | 1.33E-15 | 6.98E-15 |
| chr2 | 54589654 | 54597131 | 7478 | -3.43 | 1.33E-15 | 7.00E-15 |
| chr12 | 65787108 | 65799090 | 11983 | -8.72 | 1.34E-15 | 7.01E-15 |
| chr18 | 33921124 | 33944553 | 23430 | -8.73 | 1.34E-15 | 7.03E-15 |
| chr11 | 10219623 | 10266958 | 47336 | -9.61 | 1.34E-15 | 7.04E-15 |
| chr11 | 78117416 | 78122675 | 5260 | -8.65 | 1.34E-15 | 7.05E-15 |
| chr2 | 159143047 | 159157357 | 14311 | -2.61 | 1.35E-15 | 7.06E-15 |
| chr20 | 22714503 | 22729817 | 15315 | -8.84 | 1.35E-15 | 7.07E-15 |
| chr14 | 55692254 | 55703483 | 11230 | -3.14 | 1.35E-15 | 7.08E-15 |
| chr12 | 50884617 | 50895959 | 11343 | -8.76 | 1.35E-15 | 7.08E-15 |
| chr3 | 155013553 | 155016246 | 2694 | -4.15 | 1.36E-15 | 7.11E-15 |
| chr19 | 33329789 | 33332119 | 2331 | -8.86 | 1.36E-15 | 7.12E-15 |
| chr20 | 18158925 | 18162446 | 3522 | -8.7 | 1.37E-15 | 7.16E-15 |
| chr6 | 17240592 | 17246790 | 6199 | -8.73 | 1.37E-15 | 7.16E-15 |
| chr7 | 81032969 | 81045815 | 12847 | -8.7 | 1.37E-15 | 7.16E-15 |
| chr14 | 61070070 | 61080954 | 10885 | -8.64 | 1.37E-15 | 7.16E-15 |
| chr7 | 132690080 | 132694525 | 4446 | -8.7 | 1.37E-15 | 7.17E-15 |
| chr15 | 60114477 | 60117828 | 3352 | -8.66 | 1.37E-15 | 7.18E-15 |
| chr2 | 202103955 | 202115683 | 11729 | -8.66 | 1.37E-15 | 7.19E-15 |
| chr12 | 86951099 | 86963877 | 12779 | -8.8 | 1.38E-15 | 7.23E-15 |
| chr9 | 85519299 | 85533501 | 14203 | -8.99 | 1.38E-15 | 7.23E-15 |
| chr5 | 155053948 | 155058137 | 4190 | -3.9 | 1.38E-15 | 7.23E-15 |
| chr12 | 80403801 | 80430998 | 27198 | -8.64 | 1.38E-15 | 7.24E-15 |
| chr12 | 96596020 | 96600798 | 4779 | -8.78 | 1.39E-15 | 7.25E-15 |
| chr1 | 235076787 | 235087628 | 10842 | -8.98 | 1.39E-15 | 7.27E-15 |
| chr6 | 144616174 | 144620503 | 4330 | -8.65 | 1.40E-15 | 7.30E-15 |
| chr9 | 129076502 | 129087930 | 11429 | -8.67 | 1.40E-15 | 7.30E-15 |
| chr8 | 59619437 | 59623829 | 4393 | -8.73 | 1.40E-15 | 7.31E-15 |

| | | | | | | |
|---|---|---|---|---|---|---|
| chr4 | 75323921 | 75329632 | 5712 | -8.66 | 1.40E-15 | 7.31E-15 |
| chr19 | 42867361 | 42872003 | 4643 | -8.68 | 1.40E-15 | 7.33E-15 |
| chr9 | 98040256 | 98045394 | 5139 | -8.69 | 1.41E-15 | 7.35E-15 |
| chr14 | 99473392 | 99494244 | 20853 | -8.64 | 1.41E-15 | 7.36E-15 |
| chr12 | 117379619 | 117382222 | 2604 | -8.64 | 1.41E-15 | 7.39E-15 |
| chr15 | 96143035 | 96175539 | 32505 | -8.71 | 1.42E-15 | 7.42E-15 |
| chr6 | 105335023 | 105353149 | 18127 | -8.71 | 1.42E-15 | 7.42E-15 |
| chr16 | 15560370 | 15591071 | 30702 | -8.75 | 1.42E-15 | 7.42E-15 |
| chr8 | 120812565 | 120814822 | 2258 | -8.66 | 1.42E-15 | 7.43E-15 |
| chr12 | 78968303 | 78981358 | 13056 | -8.72 | 1.42E-15 | 7.43E-15 |
| chr3 | 141925436 | 141928614 | 3179 | -8.64 | 1.43E-15 | 7.45E-15 |
| chr3 | 99414248 | 99431520 | 17273 | 8.86 | 1.43E-15 | 7.45E-15 |
| chr3 | 86804267 | 86819428 | 15162 | -8.64 | 1.43E-15 | 7.45E-15 |
| chr2 | 8895302 | 8899214 | 3913 | -8.68 | 1.43E-15 | 7.45E-15 |
| chr1 | 145635697 | 145639562 | 3866 | -8.64 | 1.43E-15 | 7.47E-15 |
| chr13 | 50532646 | 50540283 | 7638 | -8.64 | 1.43E-15 | 7.47E-15 |
| chr7 | 27308132 | 27314077 | 5946 | -8.64 | 1.43E-15 | 7.47E-15 |
| chr2 | 129030924 | 129033678 | 2755 | -8.64 | 1.44E-15 | 7.49E-15 |
| chr9 | 29809595 | 29830079 | 20485 | -8.64 | 1.44E-15 | 7.49E-15 |
| chr6 | 82714281 | 82722586 | 8306 | -9 | 1.44E-15 | 7.50E-15 |
| chr5 | 127049708 | 127069512 | 19805 | -8.76 | 1.44E-15 | 7.52E-15 |
| chr15 | 36771812 | 36789460 | 17649 | -8.64 | 1.44E-15 | 7.52E-15 |
| chr11 | 11168431 | 11189447 | 21017 | -2.95 | 1.44E-15 | 7.52E-15 |
| chr10 | 30828417 | 30862123 | 33707 | -9.42 | 1.45E-15 | 7.53E-15 |
| chr2 | 225518362 | 225530571 | 12210 | -8.68 | 1.45E-15 | 7.54E-15 |
| chr7 | 65761900 | 65765595 | 3696 | -8.66 | 1.45E-15 | 7.55E-15 |
| chr2 | 10963762 | 10983533 | 19772 | -8.85 | 1.45E-15 | 7.56E-15 |
| chr2 | 17871115 | 17882785 | 11671 | -8.78 | 1.45E-15 | 7.56E-15 |
| chr12 | 39950020 | 39954311 | 4292 | -6.72 | 1.46E-15 | 7.60E-15 |
| chr1 | 205846917 | 205868789 | 21873 | -8.63 | 1.46E-15 | 7.62E-15 |
| chr2 | 172257656 | 172260720 | 3065 | -8.67 | 1.46E-15 | 7.62E-15 |
| chr11 | 58409349 | 58414729 | 5381 | -2.35 | 1.47E-15 | 7.65E-15 |
| chr22 | 48585245 | 48599081 | 13837 | -8.8 | 1.48E-15 | 7.68E-15 |
| chr6 | 130092296 | 130103206 | 10911 | -8.65 | 1.48E-15 | 7.70E-15 |
| chr3 | 116806609 | 116822791 | 16183 | -8.8 | 1.48E-15 | 7.70E-15 |
| chr9 | 28725013 | 28747729 | 22717 | -8.7 | 1.48E-15 | 7.71E-15 |
| chr15 | 101667860 | 101675681 | 7822 | -8.72 | 1.49E-15 | 7.73E-15 |
| chr6 | 20806119 | 20815324 | 9206 | -8.69 | 1.49E-15 | 7.76E-15 |
| chr16 | 1133596 | 1137299 | 3704 | -8.63 | 1.49E-15 | 7.76E-15 |
| chr9 | 98879727 | 98884789 | 5063 | -8.79 | 1.49E-15 | 7.76E-15 |
| chr4 | 156190354 | 156196565 | 6212 | -8.78 | 1.49E-15 | 7.76E-15 |
| chr2 | 200766060 | 200784086 | 18027 | -8.74 | 1.49E-15 | 7.77E-15 |
| chr1 | 222743545 | 222746669 | 3125 | -8.71 | 1.51E-15 | 7.83E-15 |
| chr3 | 178188666 | 178204631 | 15966 | 8.77 | 1.51E-15 | 7.84E-15 |
| chr2 | 70762747 | 70766213 | 3467 | -8.66 | 1.51E-15 | 7.85E-15 |
| chr22 | 45862114 | 45886803 | 24690 | -8.75 | 1.52E-15 | 7.90E-15 |
| chr17 | 59162359 | 59172626 | 10268 | -8.78 | 1.52E-15 | 7.90E-15 |
| chr5 | 34420879 | 34469767 | 48889 | -8.92 | 1.52E-15 | 7.91E-15 |
| chr1 | 120474540 | 120487599 | 13060 | -8.63 | 1.53E-15 | 7.94E-15 |
| chr5 | 58520271 | 58522438 | 2168 | -8.68 | 1.53E-15 | 7.94E-15 |
| chr20 | 16280655 | 16285190 | 4536 | -8.63 | 1.53E-15 | 7.95E-15 |
| chr5 | 141525322 | 141528031 | 2710 | -8.84 | 1.53E-15 | 7.95E-15 |
| chr3 | 141181950 | 141191830 | 9881 | -8.86 | 1.54E-15 | 7.99E-15 |
| chr3 | 114664475 | 114675047 | 10573 | -8.88 | 1.54E-15 | 7.99E-15 |
| chr5 | 146103159 | 146111143 | 7985 | -8.67 | 1.54E-15 | 7.99E-15 |
| chr1 | 165581092 | 165584624 | 3533 | -8.63 | 1.55E-15 | 8.03E-15 |
| chr14 | 28928161 | 28937311 | 9151 | -4.26 | 1.55E-15 | 8.03E-15 |
| chr7 | 5218334 | 5227667 | 9334 | -8.89 | 1.55E-15 | 8.05E-15 |
| chr15 | 73735751 | 73748485 | 12735 | -8.73 | 1.56E-15 | 8.08E-15 |
| chr14 | 54981401 | 54984766 | 3366 | -8.63 | 1.56E-15 | 8.08E-15 |
| chr17 | 57098 | 62247 | 5150 | -8.65 | 1.56E-15 | 8.08E-15 |

| | | | | | | |
|---|---|---|---|---|---|---|
| chr3 | 17032790 | 17045424 | 12635 | -8.92 | 1.56E-15 | 8.09E-15 |
| chr16 | 30158901 | 30162808 | 3908 | -8.79 | 1.56E-15 | 8.11E-15 |
| chr2 | 106697620 | 106709502 | 11883 | -8.64 | 1.57E-15 | 8.13E-15 |
| chr2 | 120324975 | 120343160 | 18186 | -8.64 | 1.57E-15 | 8.13E-15 |
| chr6 | 87637863 | 87706960 | 69098 | 3.13 | 1.57E-15 | 8.13E-15 |
| chr6 | 146557938 | 146573747 | 15810 | 7.61 | 1.57E-15 | 8.15E-15 |
| chr5 | 67500502 | 67517757 | 17256 | -8.63 | 1.58E-15 | 8.16E-15 |
| chr1 | 86023042 | 86028212 | 5171 | -8.62 | 1.58E-15 | 8.17E-15 |
| chr8 | 122495371 | 122511179 | 15809 | -8.69 | 1.58E-15 | 8.18E-15 |
| chr10 | 127399734 | 127407474 | 7741 | -8.8 | 1.58E-15 | 8.19E-15 |
| chr17 | 55768440 | 55774061 | 5622 | -8.62 | 1.58E-15 | 8.19E-15 |
| chr12 | 16431092 | 16446418 | 15327 | 2.78 | 1.59E-15 | 8.21E-15 |
| chr4 | 11486687 | 11511411 | 24725 | -8.82 | 1.59E-15 | 8.22E-15 |
| chr2 | 45211073 | 45225739 | 14667 | -2.36 | 1.59E-15 | 8.23E-15 |
| chrX | 118150349 | 118152709 | 2361 | -8.67 | 1.59E-15 | 8.24E-15 |
| chr6 | 26982289 | 26996916 | 14628 | -8.67 | 1.60E-15 | 8.27E-15 |
| chr7 | 131568338 | 131572951 | 4614 | -8.65 | 1.60E-15 | 8.27E-15 |
| chr12 | 74666359 | 74672898 | 6540 | -8.65 | 1.60E-15 | 8.27E-15 |
| chr7 | 75255202 | 75282635 | 27434 | -9.06 | 1.60E-15 | 8.27E-15 |
| chrX | 140991501 | 140997084 | 5584 | -4.84 | 1.61E-15 | 8.30E-15 |
| chr6 | 154703998 | 154736788 | 32791 | -8.7 | 1.61E-15 | 8.31E-15 |
| chr8 | 124938608 | 124942424 | 3817 | -8.84 | 1.61E-15 | 8.33E-15 |
| chr7 | 131870453 | 131873497 | 3045 | -8.67 | 1.61E-15 | 8.34E-15 |
| chr8 | 119321715 | 119329305 | 7591 | -8.62 | 1.62E-15 | 8.35E-15 |
| chr9 | 117951358 | 117954426 | 3069 | -8.62 | 1.62E-15 | 8.35E-15 |
| chr7 | 114460596 | 114469092 | 8497 | -8.62 | 1.62E-15 | 8.38E-15 |
| chr15 | 78482293 | 78484847 | 2555 | -8.67 | 1.62E-15 | 8.38E-15 |
| chr3 | 23939393 | 23946877 | 7485 | -8.67 | 1.63E-15 | 8.40E-15 |
| chr8 | 101387691 | 101414322 | 26632 | -8.91 | 1.63E-15 | 8.42E-15 |
| chr2 | 164667307 | 164676146 | 8840 | -8.92 | 1.63E-15 | 8.43E-15 |
| chr4 | 156758528 | 156813736 | 55209 | -9.03 | 1.64E-15 | 8.44E-15 |
| chr6 | 10638384 | 10667835 | 29452 | -8.69 | 1.64E-15 | 8.48E-15 |
| chr18 | 534158 | 553939 | 19782 | -2.56 | 1.65E-15 | 8.49E-15 |
| chr9 | 137392277 | 137425361 | 33085 | -8.81 | 1.65E-15 | 8.50E-15 |
| chr10 | 30431713 | 30457664 | 25952 | -8.82 | 1.65E-15 | 8.51E-15 |
| chr1 | 111868772 | 111878018 | 9247 | -8.62 | 1.65E-15 | 8.51E-15 |
| chr17 | 7576285 | 7578796 | 2512 | -8.65 | 1.65E-15 | 8.52E-15 |
| chr2 | 67593746 | 67606819 | 13074 | -8.81 | 1.66E-15 | 8.56E-15 |
| chr5 | 67821545 | 67827316 | 5772 | -8.62 | 1.66E-15 | 8.56E-15 |
| chr15 | 51818711 | 51824500 | 5790 | -8.66 | 1.66E-15 | 8.57E-15 |
| chr5 | 59008097 | 59012429 | 4333 | -8.64 | 1.66E-15 | 8.58E-15 |
| chr12 | 65875388 | 65879867 | 4480 | -8.62 | 1.67E-15 | 8.59E-15 |
| chr2 | 161284158 | 161288407 | 4250 | -8.62 | 1.67E-15 | 8.59E-15 |
| chr14 | 63830079 | 63838095 | 8017 | -8.62 | 1.67E-15 | 8.61E-15 |
| chr8 | 118387646 | 118404593 | 16948 | -8.62 | 1.67E-15 | 8.63E-15 |
| chr2 | 46221908 | 46226140 | 4233 | -8.63 | 1.67E-15 | 8.63E-15 |
| chr7 | 117287082 | 117321631 | 34550 | -8.84 | 1.68E-15 | 8.63E-15 |
| chr3 | 100059952 | 100082093 | 22142 | -8.63 | 1.68E-15 | 8.63E-15 |
| chr1 | 234773573 | 234805188 | 31616 | -9.16 | 1.68E-15 | 8.66E-15 |
| chr9 | 97224834 | 97229960 | 5127 | -8.71 | 1.69E-15 | 8.69E-15 |
| chr14 | 23229369 | 23234741 | 5373 | -8.65 | 1.69E-15 | 8.69E-15 |
| chr1 | 30235700 | 30253757 | 18058 | -6.86 | 1.69E-15 | 8.70E-15 |
| chr2 | 43085827 | 43097847 | 12021 | -8.94 | 1.69E-15 | 8.70E-15 |
| chr7 | 12200781 | 12213151 | 12371 | -8.62 | 1.71E-15 | 8.78E-15 |
| chr3 | 121163634 | 121178019 | 14386 | -8.62 | 1.71E-15 | 8.79E-15 |
| chr12 | 94939752 | 94952342 | 12591 | -8.62 | 1.71E-15 | 8.79E-15 |
| chrX | 102479291 | 102481348 | 2058 | -4.45 | 1.71E-15 | 8.82E-15 |
| chr5 | 74184193 | 74214439 | 30247 | -8.82 | 1.72E-15 | 8.82E-15 |
| chr8 | 86062022 | 86072227 | 10206 | -8.62 | 1.72E-15 | 8.83E-15 |
| chr20 | 43709703 | 43712169 | 2467 | -8.63 | 1.72E-15 | 8.85E-15 |
| chr15 | 93259851 | 93266643 | 6793 | -8.88 | 1.73E-15 | 8.87E-15 |

| | | | | | | |
|---|---|---|---|---|---|---|
| chr18 | 9552364 | 9554850 | 2487 | -8.72 | 1.73E-15 | 8.90E-15 |
| chr2 | 28398646 | 28413432 | 14787 | -8.89 | 1.74E-15 | 8.92E-15 |
| chr13 | 41833076 | 41846795 | 13720 | -8.74 | 1.74E-15 | 8.94E-15 |
| chr7 | 148368490 | 148380392 | 11903 | -8.76 | 1.75E-15 | 8.97E-15 |
| chr3 | 171860577 | 171864953 | 4377 | -8.66 | 1.75E-15 | 8.97E-15 |
| chr18 | 24166083 | 24186128 | 20046 | -8.73 | 1.76E-15 | 9.02E-15 |
| chrX | 53376106 | 53380214 | 4109 | -8.62 | 1.76E-15 | 9.05E-15 |
| chr20 | 47816955 | 47829108 | 12154 | -8.79 | 1.76E-15 | 9.05E-15 |
| chr11 | 93093543 | 93116730 | 23188 | -8.64 | 1.77E-15 | 9.08E-15 |
| chr17 | 30975442 | 30979000 | 3559 | -8.63 | 1.77E-15 | 9.08E-15 |
| chr2 | 178122295 | 178124687 | 2393 | -8.62 | 1.78E-15 | 9.12E-15 |
| chr11 | 130705570 | 130719142 | 13573 | -8.65 | 1.78E-15 | 9.12E-15 |
| chr22 | 26774719 | 26785080 | 10362 | -2.46 | 1.78E-15 | 9.12E-15 |
| chr21 | 17322207 | 17328569 | 6363 | -2.75 | 1.79E-15 | 9.17E-15 |
| chr17 | 14069738 | 14074135 | 4398 | -8.77 | 1.79E-15 | 9.18E-15 |
| chr6 | 9202721 | 9217926 | 15206 | -8.61 | 1.79E-15 | 9.18E-15 |
| chr4 | 124210279 | 124216587 | 6309 | -8.61 | 1.79E-15 | 9.18E-15 |
| chr9 | 112028384 | 112031511 | 3128 | -8.61 | 1.79E-15 | 9.18E-15 |
| chr3 | 9329026 | 9336014 | 6989 | -8.67 | 1.79E-15 | 9.20E-15 |
| chr10 | 61422021 | 61428552 | 6532 | -8.72 | 1.80E-15 | 9.21E-15 |
| chr1 | 97354717 | 97369231 | 14515 | -8.62 | 1.80E-15 | 9.21E-15 |
| chr1 | 199593778 | 199616733 | 22956 | -8.83 | 1.80E-15 | 9.21E-15 |
| chr6 | 105802227 | 105817870 | 15644 | -8.61 | 1.80E-15 | 9.21E-15 |
| chr22 | 45761329 | 45764391 | 3063 | -8.62 | 1.80E-15 | 9.23E-15 |
| chr2 | 173517799 | 173554283 | 36485 | -8.83 | 1.80E-15 | 9.24E-15 |
| chrX | 47674284 | 47687327 | 13044 | -6.9 | 1.81E-15 | 9.27E-15 |
| chr7 | 68767904 | 68809999 | 42096 | -9.29 | 1.81E-15 | 9.28E-15 |
| chr7 | 29549497 | 29564081 | 14585 | -8.65 | 1.81E-15 | 9.29E-15 |
| chr3 | 70358900 | 70378234 | 19335 | -8.88 | 1.82E-15 | 9.31E-15 |
| chr2 | 169208662 | 169211238 | 2577 | -8.62 | 1.82E-15 | 9.34E-15 |
| chr5 | 74446785 | 74459757 | 12973 | -8.61 | 1.83E-15 | 9.39E-15 |
| chr13 | 74689577 | 74696295 | 6719 | -8.66 | 1.84E-15 | 9.43E-15 |
| chr2 | 68983206 | 68986000 | 2795 | -3.87 | 1.84E-15 | 9.43E-15 |
| chr4 | 187010359 | 187017289 | 6931 | -8.62 | 1.85E-15 | 9.45E-15 |
| chr19 | 48395544 | 48403072 | 7529 | -8.78 | 1.85E-15 | 9.47E-15 |
| chr12 | 31709054 | 31715160 | 6107 | -8.73 | 1.85E-15 | 9.47E-15 |
| chr20 | 22335034 | 22345147 | 10114 | -8.65 | 1.86E-15 | 9.51E-15 |
| chr12 | 31690429 | 31696101 | 5673 | -8.69 | 1.86E-15 | 9.52E-15 |
| chr1 | 98969464 | 98988577 | 19114 | -8.61 | 1.86E-15 | 9.52E-15 |
| chr10 | 77003284 | 77021214 | 17931 | -8.84 | 1.86E-15 | 9.52E-15 |
| chr6 | 150085728 | 150088936 | 3209 | -8.64 | 1.86E-15 | 9.53E-15 |
| chrX | 154617335 | 154637455 | 20121 | -8.61 | 1.87E-15 | 9.54E-15 |
| chr2 | 222375120 | 222394318 | 19199 | -8.61 | 1.88E-15 | 9.59E-15 |
| chr8 | 144393622 | 144396909 | 3288 | -8.62 | 1.88E-15 | 9.59E-15 |
| chrX | 154034803 | 154045953 | 11151 | -2.07 | 1.88E-15 | 9.63E-15 |
| chr7 | 148826452 | 148829272 | 2821 | -8.6 | 1.89E-15 | 9.63E-15 |
| chr2 | 31371349 | 31374300 | 2952 | -8.77 | 1.89E-15 | 9.63E-15 |
| chr1 | 66882032 | 66898827 | 16796 | -8.8 | 1.90E-15 | 9.68E-15 |
| chr3 | 102325081 | 102341018 | 15938 | 8.77 | 1.90E-15 | 9.69E-15 |
| chr4 | 100366832 | 100375683 | 8852 | -8.89 | 1.90E-15 | 9.70E-15 |
| chr12 | 9849056 | 9860363 | 11308 | -8.63 | 1.90E-15 | 9.70E-15 |
| chr15 | 100891640 | 100901900 | 10261 | -2.68 | 1.90E-15 | 9.71E-15 |
| chr5 | 44879206 | 44901376 | 22171 | -8.67 | 1.90E-15 | 9.71E-15 |
| chr4 | 134225575 | 134259673 | 34099 | 8.71 | 1.90E-15 | 9.71E-15 |
| chr4 | 95120545 | 95125093 | 4549 | -8.61 | 1.91E-15 | 9.75E-15 |
| chr12 | 125881555 | 125888905 | 7351 | -8.63 | 1.91E-15 | 9.75E-15 |
| chr22 | 49058778 | 49069713 | 10936 | -8.72 | 1.91E-15 | 9.76E-15 |
| chr15 | 90329667 | 90332717 | 3051 | -8.76 | 1.91E-15 | 9.76E-15 |
| chr18 | 56241265 | 56265865 | 24601 | -8.91 | 1.91E-15 | 9.76E-15 |
| chr12 | 131932665 | 131951881 | 19217 | -8.96 | 1.92E-15 | 9.77E-15 |
| chr13 | 27196824 | 27227241 | 30418 | -1.83 | 1.92E-15 | 9.79E-15 |

| | | | | | | |
|---|---|---|---|---|---|---|
| chr12 | 17301898 | 17308301 | 6404 | -8.71 | 1.92E-15 | 9.80E-15 |
| chr9 | 87464951 | 87476529 | 11579 | -8.63 | 1.92E-15 | 9.80E-15 |
| chr1 | 66433705 | 66455083 | 21379 | -8.91 | 1.93E-15 | 9.82E-15 |
| chr8 | 24868864 | 24889367 | 20504 | -2.22 | 1.93E-15 | 9.82E-15 |
| chr1 | 93119357 | 93123738 | 4382 | -8.61 | 1.93E-15 | 9.84E-15 |
| chrX | 46477318 | 46480002 | 2685 | -8.62 | 1.94E-15 | 9.87E-15 |
| chrX | 101138864 | 101154060 | 15197 | 8.68 | 1.94E-15 | 9.87E-15 |
| chr2 | 56128532 | 56132064 | 3533 | -8.64 | 1.96E-15 | 9.97E-15 |
| chr15 | 31604751 | 31614103 | 9353 | -8.9 | 1.96E-15 | 9.97E-15 |
| chr5 | 130168457 | 130182956 | 14500 | -8.65 | 1.96E-15 | 9.97E-15 |
| chr20 | 49026411 | 49039146 | 12736 | -8.7 | 1.96E-15 | 9.99E-15 |
| chr1 | 68516751 | 68541453 | 24703 | -9.06 | 1.98E-15 | 1.01E-14 |
| chr5 | 130456706 | 130475337 | 18632 | -8.79 | 1.99E-15 | 1.01E-14 |
| chr11 | 59549434 | 59552760 | 3327 | -3.85 | 1.99E-15 | 1.01E-14 |
| chr4 | 184894223 | 184900323 | 6101 | -8.61 | 1.99E-15 | 1.01E-14 |
| chr19 | 42304797 | 42307029 | 2233 | -8.61 | 2.00E-15 | 1.02E-14 |
| chr2 | 98785547 | 98885914 | 100368 | -1.94 | 2.00E-15 | 1.02E-14 |
| chr17 | 7281065 | 7284002 | 2938 | -8.61 | 2.01E-15 | 1.02E-14 |
| chr22 | 29569811 | 29576666 | 6856 | -8.61 | 2.01E-15 | 1.02E-14 |
| chr3 | 179208154 | 179217313 | 9160 | -7.21 | 2.01E-15 | 1.02E-14 |
| chr11 | 69320401 | 69334572 | 14172 | -8.61 | 2.01E-15 | 1.02E-14 |
| chr20 | 5814652 | 5824035 | 9384 | -8.63 | 2.01E-15 | 1.02E-14 |
| chr11 | 44272678 | 44276572 | 3895 | -8.65 | 2.02E-15 | 1.03E-14 |
| chr5 | 43328681 | 43344629 | 15949 | -8.6 | 2.02E-15 | 1.03E-14 |
| chr2 | 202452888 | 202476617 | 23730 | -8.68 | 2.02E-15 | 1.03E-14 |
| chr3 | 64099540 | 64111605 | 12066 | -8.68 | 2.02E-15 | 1.03E-14 |
| chr1 | 203126563 | 203130366 | 3804 | -8.68 | 2.02E-15 | 1.03E-14 |
| chr17 | 27779382 | 27781765 | 2384 | -8.64 | 2.03E-15 | 1.03E-14 |
| chr11 | 101940759 | 101945995 | 5237 | -8.6 | 2.03E-15 | 1.03E-14 |
| chr9 | 127135844 | 127137956 | 2113 | -8.6 | 2.03E-15 | 1.03E-14 |
| chr21 | 16883330 | 16905789 | 22460 | -8.8 | 2.03E-15 | 1.03E-14 |
| chr1 | 220842603 | 220852178 | 9576 | -8.64 | 2.03E-15 | 1.03E-14 |
| chr12 | 52664852 | 52668440 | 3589 | -8.64 | 2.04E-15 | 1.04E-14 |
| chr12 | 111551785 | 111586265 | 34481 | -8.64 | 2.04E-15 | 1.04E-14 |
| chr21 | 43773839 | 43780583 | 6745 | -8.65 | 2.05E-15 | 1.04E-14 |
| chr14 | 101564087 | 101574509 | 10423 | -8.59 | 2.05E-15 | 1.04E-14 |
| chr9 | 92458224 | 92486833 | 28610 | 8.98 | 2.05E-15 | 1.04E-14 |
| chr6 | 17185912 | 17192153 | 6242 | -8.62 | 2.06E-15 | 1.04E-14 |
| chr18 | 73036181 | 73041556 | 5376 | -8.6 | 2.07E-15 | 1.05E-14 |
| chr4 | 6164804 | 6189557 | 24754 | -8.8 | 2.07E-15 | 1.05E-14 |
| chr5 | 43048891 | 43059468 | 10578 | -8.59 | 2.07E-15 | 1.05E-14 |
| chr3 | 97386717 | 97397815 | 11099 | 8.7 | 2.07E-15 | 1.05E-14 |
| chr9 | 126291564 | 126301869 | 10306 | -8.61 | 2.08E-15 | 1.05E-14 |
| chr2 | 98634630 | 98640278 | 5649 | -8.6 | 2.09E-15 | 1.06E-14 |
| chr7 | 29581806 | 29584852 | 3047 | -8.62 | 2.10E-15 | 1.06E-14 |
| chr11 | 104751045 | 104775015 | 23971 | 8.68 | 2.10E-15 | 1.06E-14 |
| chr2 | 135590084 | 135605727 | 15644 | -8.73 | 2.10E-15 | 1.07E-14 |
| chr6 | 47109090 | 47134097 | 25008 | -8.74 | 2.11E-15 | 1.07E-14 |
| chr4 | 68903639 | 68933118 | 29480 | 9.04 | 2.11E-15 | 1.07E-14 |
| chr11 | 70357378 | 70361317 | 3940 | -8.6 | 2.11E-15 | 1.07E-14 |
| chr21 | 47928910 | 47936955 | 8046 | -8.71 | 2.12E-15 | 1.07E-14 |
| chr5 | 171079764 | 171102583 | 22820 | -9.03 | 2.13E-15 | 1.08E-14 |
| chr2 | 18039532 | 18053050 | 13519 | 8.99 | 2.13E-15 | 1.08E-14 |
| chr2 | 19037806 | 19067964 | 30159 | -9.06 | 2.13E-15 | 1.08E-14 |
| chr9 | 124005276 | 124015481 | 10206 | -8.61 | 2.14E-15 | 1.08E-14 |
| chr7 | 131158019 | 131160713 | 2695 | -8.64 | 2.14E-15 | 1.08E-14 |
| chr3 | 36986279 | 36991863 | 5585 | -8.61 | 2.14E-15 | 1.08E-14 |
| chr5 | 131685539 | 131698654 | 13116 | -8.65 | 2.16E-15 | 1.09E-14 |
| chr10 | 8424246 | 8439518 | 15273 | -8.67 | 2.17E-15 | 1.10E-14 |
| chr9 | 88771447 | 88811014 | 39568 | -9.24 | 2.18E-15 | 1.10E-14 |
| chr18 | 7341113 | 7357430 | 16318 | -4.63 | 2.18E-15 | 1.10E-14 |

| | | | | | | |
|---|---|---|---|---|---|---|
| chr10 | 134631480 | 134633785 | 2306 | -8.65 | 2.18E-15 | 1.10E-14 |
| chr12 | 113689773 | 113692873 | 3101 | -8.61 | 2.18E-15 | 1.10E-14 |
| chr20 | 47056828 | 47076677 | 19850 | -2.6 | 2.19E-15 | 1.11E-14 |
| chr13 | 106532025 | 106551832 | 19808 | -8.71 | 2.19E-15 | 1.11E-14 |
| chr7 | 23777873 | 23794486 | 16614 | -8.75 | 2.19E-15 | 1.11E-14 |
| chr18 | 13332399 | 13350290 | 17892 | -8.62 | 2.19E-15 | 1.11E-14 |
| chr11 | 108284946 | 108309293 | 24348 | -9.05 | 2.20E-15 | 1.11E-14 |
| chr6 | 8902852 | 8914365 | 11514 | -8.76 | 2.21E-15 | 1.11E-14 |
| chr3 | 53241895 | 53246295 | 4401 | -8.64 | 2.21E-15 | 1.12E-14 |
| chr1 | 193953053 | 193970903 | 17851 | -8.78 | 2.21E-15 | 1.12E-14 |
| chr2 | 225081491 | 225098972 | 17482 | -8.59 | 2.22E-15 | 1.12E-14 |
| chr8 | 119984561 | 119999850 | 15290 | -8.6 | 2.22E-15 | 1.12E-14 |
| chr13 | 42607460 | 42612036 | 4577 | -8.59 | 2.22E-15 | 1.12E-14 |
| chr12 | 108596392 | 108598548 | 2157 | -7.34 | 2.22E-15 | 1.12E-14 |
| chr1 | 188026543 | 188040710 | 14168 | -8.63 | 2.23E-15 | 1.12E-14 |
| chr6 | 149416328 | 149422087 | 5760 | -8.6 | 2.23E-15 | 1.13E-14 |
| chr10 | 93760238 | 93765024 | 4787 | -8.59 | 2.24E-15 | 1.13E-14 |
| chr8 | 30099883 | 30131929 | 32047 | -8.74 | 2.25E-15 | 1.13E-14 |
| chr7 | 22012871 | 22015442 | 2572 | -8.59 | 2.25E-15 | 1.14E-14 |
| chr6 | 107714129 | 107727544 | 13416 | -8.59 | 2.25E-15 | 1.14E-14 |
| chr2 | 197723232 | 197726459 | 3228 | -8.6 | 2.26E-15 | 1.14E-14 |
| chrX | 44212633 | 44225066 | 12434 | -8.6 | 2.26E-15 | 1.14E-14 |
| chr2 | 136208916 | 136215819 | 6904 | -8.6 | 2.26E-15 | 1.14E-14 |
| chr6 | 169984531 | 170002308 | 17778 | -8.68 | 2.26E-15 | 1.14E-14 |
| chr3 | 196854602 | 196863768 | 9167 | -8.64 | 2.26E-15 | 1.14E-14 |
| chr11 | 76615530 | 76619847 | 4318 | -8.59 | 2.27E-15 | 1.14E-14 |
| chr11 | 18241954 | 18266302 | 24349 | -8.81 | 2.27E-15 | 1.14E-14 |
| chr11 | 40671745 | 40737152 | 65408 | 1.46 | 2.27E-15 | 1.15E-14 |
| chr2 | 61662097 | 61678777 | 16681 | -8.83 | 2.28E-15 | 1.15E-14 |
| chr14 | 35501682 | 35505850 | 4169 | -8.6 | 2.28E-15 | 1.15E-14 |
| chrX | 13438527 | 13452624 | 14098 | -8.6 | 2.28E-15 | 1.15E-14 |
| chr5 | 173077454 | 173107430 | 29977 | -9.26 | 2.28E-15 | 1.15E-14 |
| chr2 | 73662338 | 73664603 | 2266 | -8.58 | 2.28E-15 | 1.15E-14 |
| chr5 | 23302527 | 23342123 | 39597 | -8.85 | 2.29E-15 | 1.15E-14 |
| chr9 | 32404872 | 32407248 | 2377 | -8.66 | 2.29E-15 | 1.15E-14 |
| chr5 | 130580247 | 130584509 | 4263 | -8.63 | 2.29E-15 | 1.16E-14 |
| chr3 | 98547732 | 98553078 | 5347 | -8.77 | 2.30E-15 | 1.16E-14 |
| chr1 | 168821059 | 168828897 | 7839 | -8.65 | 2.30E-15 | 1.16E-14 |
| chr5 | 17794548 | 17843701 | 49154 | -9.4 | 2.30E-15 | 1.16E-14 |
| chr15 | 58779966 | 58823360 | 43395 | -9.47 | 2.31E-15 | 1.16E-14 |
| chr11 | 1061172 | 1068772 | 7601 | -8.89 | 2.31E-15 | 1.16E-14 |
| chr13 | 33372679 | 33393365 | 20687 | -8.68 | 2.31E-15 | 1.16E-14 |
| chr12 | 43217370 | 43240761 | 23392 | -8.62 | 2.32E-15 | 1.17E-14 |
| chr11 | 67897489 | 67902834 | 5346 | -8.69 | 2.32E-15 | 1.17E-14 |
| chr4 | 15686355 | 15700837 | 14483 | -8.63 | 2.33E-15 | 1.17E-14 |
| chr9 | 74271421 | 74292631 | 21211 | -8.81 | 2.33E-15 | 1.17E-14 |
| chrX | 80437343 | 80447553 | 10211 | -8.6 | 2.33E-15 | 1.17E-14 |
| chr7 | 31742205 | 31758356 | 16152 | 8.77 | 2.33E-15 | 1.17E-14 |
| chr6 | 826669 | 841499 | 14831 | -8.75 | 2.33E-15 | 1.17E-14 |
| chr1 | 231633288 | 231639517 | 6230 | -8.81 | 2.34E-15 | 1.17E-14 |
| chr9 | 91671074 | 91687539 | 16466 | 8.65 | 2.34E-15 | 1.18E-14 |
| chr4 | 90463808 | 90470330 | 6523 | -8.61 | 2.34E-15 | 1.18E-14 |
| chr1 | 55107217 | 55125900 | 18684 | -8.86 | 2.35E-15 | 1.18E-14 |
| chr17 | 2871305 | 2879376 | 8072 | -8.7 | 2.35E-15 | 1.18E-14 |
| chr15 | 53107958 | 53139756 | 31799 | -9.05 | 2.36E-15 | 1.18E-14 |
| chr15 | 62184416 | 62195100 | 10685 | -8.61 | 2.36E-15 | 1.19E-14 |
| chr22 | 46652466 | 46671836 | 19371 | -8.75 | 2.36E-15 | 1.19E-14 |
| chr22 | 39193883 | 39207628 | 13746 | -8.58 | 2.37E-15 | 1.19E-14 |
| chr19 | 8737420 | 8753668 | 16249 | -9.03 | 2.38E-15 | 1.20E-14 |
| chr2 | 171371526 | 171415786 | 44261 | -2.23 | 2.38E-15 | 1.20E-14 |
| chr3 | 97589019 | 97595380 | 6362 | -6.49 | 2.39E-15 | 1.20E-14 |

| | | | | | | |
|---|---|---|---|---|---|---|
| chr1 | 229048138 | 229067020 | 18883 | -8.75 | 2.39E-15 | 1.20E-14 |
| chr2 | 33266792 | 33291297 | 24506 | -8.82 | 2.39E-15 | 1.20E-14 |
| chr2 | 234851888 | 234870666 | 18779 | -7.75 | 2.39E-15 | 1.20E-14 |
| chr14 | 60475376 | 60483484 | 8109 | -8.6 | 2.40E-15 | 1.20E-14 |
| chr11 | 12507662 | 12512141 | 4480 | -8.65 | 2.40E-15 | 1.20E-14 |
| chr6 | 74025654 | 74030621 | 4968 | -8.58 | 2.40E-15 | 1.20E-14 |
| chr11 | 69711522 | 69714449 | 2928 | -8.67 | 2.40E-15 | 1.20E-14 |
| chr7 | 46761233 | 46774857 | 13625 | -8.87 | 2.40E-15 | 1.20E-14 |
| chr18 | 9974143 | 9983594 | 9452 | -8.67 | 2.41E-15 | 1.21E-14 |
| chr1 | 116209902 | 116212548 | 2647 | -8.6 | 2.42E-15 | 1.21E-14 |
| chr6 | 43912265 | 43926942 | 14678 | -8.8 | 2.42E-15 | 1.21E-14 |
| chr12 | 13257842 | 13265783 | 7942 | -8.63 | 2.42E-15 | 1.21E-14 |
| chr6 | 24671171 | 24675084 | 3914 | -8.63 | 2.42E-15 | 1.21E-14 |
| chr1 | 166969082 | 166979411 | 10330 | -8.63 | 2.42E-15 | 1.21E-14 |
| chr5 | 133599256 | 133631193 | 31938 | -9.29 | 2.44E-15 | 1.22E-14 |
| chr1 | 172776877 | 172799817 | 22941 | -8.62 | 2.44E-15 | 1.22E-14 |
| chr1 | 205123725 | 205127072 | 3348 | -8.66 | 2.45E-15 | 1.22E-14 |
| chr1 | 233435962 | 233447238 | 11277 | -9.1 | 2.45E-15 | 1.22E-14 |
| chr2 | 106446838 | 106452960 | 6123 | -8.58 | 2.45E-15 | 1.23E-14 |
| chrX | 154668132 | 154680759 | 12628 | -8.63 | 2.46E-15 | 1.23E-14 |
| chr5 | 55156064 | 55175262 | 19199 | -8.69 | 2.46E-15 | 1.23E-14 |
| chr2 | 178472113 | 178481737 | 9625 | -8.98 | 2.46E-15 | 1.23E-14 |
| chr2 | 86107738 | 86126265 | 18528 | -8.59 | 2.46E-15 | 1.23E-14 |
| chr5 | 155906925 | 155918448 | 11524 | -8.57 | 2.47E-15 | 1.23E-14 |
| chr2 | 183042057 | 183050060 | 8004 | -8.7 | 2.48E-15 | 1.24E-14 |
| chr11 | 73823263 | 73828395 | 5133 | -8.65 | 2.48E-15 | 1.24E-14 |
| chr16 | 47770550 | 47776989 | 6440 | -8.57 | 2.48E-15 | 1.24E-14 |
| chr5 | 93874114 | 93887214 | 13101 | -8.73 | 2.49E-15 | 1.24E-14 |
| chr21 | 32944520 | 32947764 | 3245 | -8.66 | 2.49E-15 | 1.24E-14 |
| chr4 | 169695550 | 169713982 | 18433 | -8.64 | 2.49E-15 | 1.24E-14 |
| chr4 | 131975366 | 131994009 | 18644 | -8.65 | 2.50E-15 | 1.25E-14 |
| chr1 | 182705866 | 182744682 | 38817 | -8.61 | 2.50E-15 | 1.25E-14 |
| chr9 | 96367063 | 96373514 | 6452 | -8.6 | 2.51E-15 | 1.25E-14 |
| chr12 | 20749315 | 20760350 | 11036 | -8.76 | 2.51E-15 | 1.26E-14 |
| chr8 | 126686275 | 126690603 | 4329 | -8.59 | 2.52E-15 | 1.26E-14 |
| chr3 | 33794295 | 33798791 | 4497 | -8.59 | 2.52E-15 | 1.26E-14 |
| chr16 | 23796096 | 23797816 | 1721 | -4.67 | 2.52E-15 | 1.26E-14 |
| chr8 | 115614053 | 115640426 | 26374 | -8.61 | 2.53E-15 | 1.26E-14 |
| chr1 | 93744521 | 93759936 | 15416 | -8.81 | 2.54E-15 | 1.27E-14 |
| chr2 | 33619651 | 33636431 | 16781 | -9.7 | 2.56E-15 | 1.28E-14 |
| chr5 | 37096996 | 37105234 | 8239 | -8.58 | 2.56E-15 | 1.28E-14 |
| chr4 | 160383644 | 160396330 | 12687 | -8.58 | 2.56E-15 | 1.28E-14 |
| chr4 | 146729905 | 146733605 | 3701 | -8.58 | 2.56E-15 | 1.28E-14 |
| chr7 | 32231497 | 32236494 | 4998 | -8.6 | 2.57E-15 | 1.28E-14 |
| chr20 | 46801308 | 46822559 | 21252 | -8.83 | 2.57E-15 | 1.28E-14 |
| chr1 | 11440421 | 11442964 | 2544 | -5.84 | 2.58E-15 | 1.28E-14 |
| chr13 | 40204677 | 40212554 | 7878 | -8.59 | 2.58E-15 | 1.28E-14 |
| chrX | 45650329 | 45653144 | 2816 | -8.58 | 2.58E-15 | 1.29E-14 |
| chr19 | 51034734 | 51037337 | 2604 | -8.58 | 2.58E-15 | 1.29E-14 |
| chr1 | 109534831 | 109538402 | 3572 | -8.76 | 2.58E-15 | 1.29E-14 |
| chr7 | 134358025 | 134360090 | 2066 | -8.57 | 2.58E-15 | 1.29E-14 |
| chr10 | 118616229 | 118620374 | 4146 | -8.59 | 2.58E-15 | 1.29E-14 |
| chr8 | 108212840 | 108226246 | 13407 | -8.58 | 2.59E-15 | 1.29E-14 |
| chr4 | 5588880 | 5604929 | 16050 | -8.64 | 2.61E-15 | 1.30E-14 |
| chr10 | 80542633 | 80554390 | 11758 | -8.57 | 2.61E-15 | 1.30E-14 |
| chr3 | 48400523 | 48407976 | 7454 | -6.66 | 2.61E-15 | 1.30E-14 |
| chr3 | 9293481 | 9300533 | 7053 | -8.65 | 2.61E-15 | 1.30E-14 |
| chr5 | 52918390 | 52922065 | 3676 | -8.66 | 2.61E-15 | 1.30E-14 |
| chr7 | 44545709 | 44550559 | 4851 | -8.58 | 2.61E-15 | 1.30E-14 |
| chr2 | 187974901 | 187996503 | 21603 | 8.76 | 2.62E-15 | 1.30E-14 |
| chr2 | 101443245 | 101445992 | 2748 | -8.57 | 2.62E-15 | 1.31E-14 |

| | | | | | | |
|---|---|---|---|---|---|---|
| chr1 | 219515362 | 219520862 | 5501 | -8.57 | 2.63E-15 | 1.31E-14 |
| chr11 | 112012516 | 112036184 | 23669 | -9.12 | 2.63E-15 | 1.31E-14 |
| chr10 | 35370381 | 35382744 | 12364 | -8.94 | 2.63E-15 | 1.31E-14 |
| chr18 | 19982277 | 20001556 | 19280 | -8.62 | 2.64E-15 | 1.31E-14 |
| chr3 | 164354367 | 164385124 | 30758 | -8.57 | 2.66E-15 | 1.32E-14 |
| chr9 | 94583025 | 94586238 | 3214 | -8.76 | 2.67E-15 | 1.33E-14 |
| chr1 | 228118720 | 228126232 | 7513 | -8.57 | 2.67E-15 | 1.33E-14 |
| chr7 | 95973015 | 96267376 | 294362 | 3.73 | 2.67E-15 | 1.33E-14 |
| chr4 | 77236880 | 77282915 | 46036 | -7.74 | 2.68E-15 | 1.33E-14 |
| chr15 | 62689590 | 62693621 | 4032 | -8.58 | 2.68E-15 | 1.33E-14 |
| chr8 | 134535616 | 134560653 | 25038 | -8.79 | 2.68E-15 | 1.33E-14 |
| chr22 | 44096052 | 44110986 | 14935 | -8.59 | 2.68E-15 | 1.33E-14 |
| chr3 | 158502736 | 158513278 | 10543 | -8.75 | 2.69E-15 | 1.33E-14 |
| chr18 | 77297133 | 77300418 | 3286 | -6.32 | 2.69E-15 | 1.34E-14 |
| chr17 | 65343818 | 65346768 | 2951 | -8.68 | 2.69E-15 | 1.34E-14 |
| chr6 | 117943186 | 117961835 | 18650 | -8.87 | 2.69E-15 | 1.34E-14 |
| chr8 | 56698534 | 56700963 | 2430 | -8.63 | 2.70E-15 | 1.34E-14 |
| chr18 | 57007321 | 57014096 | 6776 | -8.57 | 2.71E-15 | 1.34E-14 |
| chr3 | 39635828 | 39663094 | 27267 | 3.63 | 2.71E-15 | 1.34E-14 |
| chr4 | 6851885 | 6855391 | 3507 | -8.58 | 2.71E-15 | 1.35E-14 |
| chr3 | 12212116 | 12216240 | 4125 | -8.56 | 2.71E-15 | 1.35E-14 |
| chr6 | 15642591 | 15648971 | 6381 | -8.56 | 2.72E-15 | 1.35E-14 |
| chr1 | 152181342 | 152196222 | 14881 | -8.57 | 2.72E-15 | 1.35E-14 |
| chr12 | 8148822 | 8161249 | 12428 | -8.63 | 2.72E-15 | 1.35E-14 |
| chr7 | 127476070 | 127479003 | 2934 | -8.69 | 2.72E-15 | 1.35E-14 |
| chr20 | 23152659 | 23178370 | 25712 | -2.49 | 2.73E-15 | 1.35E-14 |
| chr7 | 39693698 | 39717030 | 23333 | -8.79 | 2.73E-15 | 1.35E-14 |
| chr7 | 43388531 | 43395997 | 7467 | -8.58 | 2.74E-15 | 1.36E-14 |
| chr4 | 173841692 | 173866291 | 24600 | -8.99 | 2.76E-15 | 1.37E-14 |
| chr12 | 93547464 | 93556572 | 9109 | -8.57 | 2.76E-15 | 1.37E-14 |
| chr19 | 19343371 | 19347526 | 4156 | -8.56 | 2.76E-15 | 1.37E-14 |
| chr9 | 113207429 | 113221794 | 14366 | -8.7 | 2.77E-15 | 1.37E-14 |
| chr13 | 21977713 | 21986472 | 8760 | -8.57 | 2.77E-15 | 1.37E-14 |
| chr2 | 145893929 | 145911703 | 17775 | -8.69 | 2.77E-15 | 1.37E-14 |
| chr8 | 25222344 | 25236252 | 13909 | -8.7 | 2.78E-15 | 1.38E-14 |
| chr5 | 40786753 | 40789432 | 2680 | -8.57 | 2.78E-15 | 1.38E-14 |
| chr1 | 78073059 | 78077341 | 4283 | -8.57 | 2.80E-15 | 1.39E-14 |
| chr12 | 40468754 | 40477002 | 8249 | -8.79 | 2.82E-15 | 1.40E-14 |
| chr5 | 68342186 | 68356633 | 14448 | -8.57 | 2.82E-15 | 1.40E-14 |
| chr5 | 39349217 | 39352557 | 3341 | -3.03 | 2.82E-15 | 1.40E-14 |
| chr4 | 113439311 | 113442729 | 3419 | -8.67 | 2.82E-15 | 1.40E-14 |
| chr8 | 62492449 | 62496191 | 3743 | -8.74 | 2.83E-15 | 1.40E-14 |
| chr7 | 109333031 | 109340754 | 7724 | -8.58 | 2.83E-15 | 1.40E-14 |
| chr10 | 8846804 | 8857686 | 10883 | -8.77 | 2.83E-15 | 1.40E-14 |
| chr1 | 54074746 | 54078213 | 3468 | -8.57 | 2.84E-15 | 1.40E-14 |
| chr22 | 20082855 | 20085278 | 2424 | -8.74 | 2.84E-15 | 1.41E-14 |
| chr18 | 61035959 | 61051850 | 15892 | -8.56 | 2.84E-15 | 1.41E-14 |
| chr20 | 58706837 | 59563698 | 856862 | 0.56 | 2.85E-15 | 1.41E-14 |
| chr4 | 37655108 | 37660328 | 5221 | -8.56 | 2.86E-15 | 1.42E-14 |
| chr2 | 181382097 | 181411934 | 29838 | -8.7 | 2.87E-15 | 1.42E-14 |
| chr10 | 69705765 | 69710462 | 4698 | -8.61 | 2.87E-15 | 1.42E-14 |
| chr14 | 49393061 | 49638244 | 245184 | 2.66 | 2.87E-15 | 1.42E-14 |
| chr3 | 120010931 | 120014607 | 3677 | -8.56 | 2.88E-15 | 1.43E-14 |
| chr1 | 231246936 | 231256463 | 9528 | -8.67 | 2.90E-15 | 1.43E-14 |
| chr2 | 201354384 | 201357379 | 2996 | -8.58 | 2.90E-15 | 1.43E-14 |
| chr12 | 90881371 | 90910842 | 29472 | -8.64 | 2.90E-15 | 1.43E-14 |
| chr21 | 40190728 | 40199880 | 9153 | -8.64 | 2.92E-15 | 1.44E-14 |
| chr12 | 107672189 | 107678288 | 6100 | -8.59 | 2.92E-15 | 1.44E-14 |
| chr4 | 124238689 | 124245910 | 7222 | -8.56 | 2.92E-15 | 1.44E-14 |
| chr7 | 143543667 | 143547265 | 3599 | -8.6 | 2.92E-15 | 1.44E-14 |
| chr10 | 3947813 | 3966863 | 19051 | -8.93 | 2.93E-15 | 1.44E-14 |

| | | | | | | |
|---|---|---|---|---|---|---|
| chr12 | 92511726 | 92517732 | 6007 | -8.57 | 2.94E-15 | 1.45E-14 |
| chrX | 34878730 | 34890783 | 12054 | -8.59 | 2.94E-15 | 1.45E-14 |
| chrX | 18934065 | 18937143 | 3079 | -8.56 | 2.95E-15 | 1.45E-14 |
| chr1 | 16295227 | 16310488 | 15262 | -8.59 | 2.95E-15 | 1.46E-14 |
| chr15 | 70286513 | 70309899 | 23387 | -9.02 | 2.96E-15 | 1.46E-14 |
| chr22 | 48144629 | 48197360 | 52732 | -2.76 | 2.96E-15 | 1.46E-14 |
| chr3 | 148992173 | 148994770 | 2598 | -8.59 | 2.96E-15 | 1.46E-14 |
| chr16 | 55784217 | 55788441 | 4225 | -8.59 | 2.96E-15 | 1.46E-14 |
| chr3 | 64017127 | 64023593 | 6467 | -8.63 | 2.96E-15 | 1.46E-14 |
| chr4 | 4350772 | 4377038 | 26267 | -8.63 | 2.96E-15 | 1.46E-14 |
| chr1 | 30764764 | 30771254 | 6491 | -2.7 | 2.96E-15 | 1.46E-14 |
| chr1 | 115705203 | 115717817 | 12615 | -8.6 | 2.96E-15 | 1.46E-14 |
| chr13 | 44795352 | 44805081 | 9730 | -8.55 | 2.97E-15 | 1.46E-14 |
| chr9 | 3781659 | 3790434 | 8776 | -8.61 | 2.98E-15 | 1.47E-14 |
| chr2 | 189751500 | 189755865 | 4366 | -8.7 | 2.98E-15 | 1.47E-14 |
| chr20 | 37696313 | 37715532 | 19220 | -8.61 | 2.98E-15 | 1.47E-14 |
| chr2 | 137123779 | 137128069 | 4291 | -8.55 | 3.00E-15 | 1.48E-14 |
| chr3 | 131797498 | 131800808 | 3311 | -6.28 | 3.00E-15 | 1.48E-14 |
| chr7 | 84259873 | 84266759 | 6887 | -8.69 | 3.00E-15 | 1.48E-14 |
| chr5 | 92408325 | 92437540 | 29216 | -8.62 | 3.00E-15 | 1.48E-14 |
| chr5 | 82063584 | 82089093 | 25510 | -8.64 | 3.01E-15 | 1.48E-14 |
| chr3 | 111598129 | 111600947 | 2819 | -8.58 | 3.01E-15 | 1.48E-14 |
| chr12 | 1528748 | 1532017 | 3270 | -8.57 | 3.02E-15 | 1.49E-14 |
| chr5 | 134415910 | 134419370 | 3461 | -8.58 | 3.02E-15 | 1.49E-14 |
| chr12 | 85979799 | 86007176 | 27378 | -8.74 | 3.04E-15 | 1.50E-14 |
| chr18 | 8740837 | 8762297 | 21461 | -8.6 | 3.04E-15 | 1.50E-14 |
| chr2 | 10459929 | 10475952 | 16024 | -8.91 | 3.04E-15 | 1.50E-14 |
| chrX | 21178609 | 21191967 | 13359 | -8.57 | 3.06E-15 | 1.50E-14 |
| chr1 | 212997202 | 213001750 | 4549 | -8.7 | 3.06E-15 | 1.51E-14 |
| chr10 | 14522758 | 14534966 | 12209 | -8.95 | 3.06E-15 | 1.51E-14 |
| chr6 | 133016391 | 133028490 | 12100 | -8.58 | 3.07E-15 | 1.51E-14 |
| chr22 | 49111183 | 49124668 | 13486 | -8.57 | 3.07E-15 | 1.51E-14 |
| chr1 | 183219008 | 183225518 | 6511 | -8.6 | 3.08E-15 | 1.51E-14 |
| chr12 | 17628899 | 17637733 | 8835 | -8.6 | 3.08E-15 | 1.51E-14 |
| chr10 | 35212385 | 35215930 | 3546 | -8.68 | 3.08E-15 | 1.52E-14 |
| chr10 | 61581595 | 61585926 | 4332 | -8.66 | 3.09E-15 | 1.52E-14 |
| chr8 | 134377091 | 134381196 | 4106 | -8.62 | 3.09E-15 | 1.52E-14 |
| chr1 | 192708525 | 192728831 | 20307 | -8.62 | 3.10E-15 | 1.52E-14 |
| chr6 | 2481523 | 2489318 | 7796 | -8.62 | 3.10E-15 | 1.52E-14 |
| chr7 | 106441673 | 106459591 | 17919 | 8.73 | 3.10E-15 | 1.52E-14 |
| chr3 | 119638678 | 119644659 | 5982 | -8.69 | 3.11E-15 | 1.53E-14 |
| chr20 | 57493918 | 57513036 | 19119 | -8.89 | 3.11E-15 | 1.53E-14 |
| chr7 | 133604185 | 133608168 | 3984 | -8.55 | 3.12E-15 | 1.53E-14 |
| chr14 | 61164456 | 61169400 | 4945 | -8.56 | 3.12E-15 | 1.53E-14 |
| chr1 | 164778260 | 164798761 | 20502 | 8.75 | 3.13E-15 | 1.54E-14 |
| chr5 | 1234332 | 1237245 | 2914 | -8.55 | 3.13E-15 | 1.54E-14 |
| chr2 | 239020022 | 239022725 | 2704 | -8.56 | 3.14E-15 | 1.54E-14 |
| chr10 | 25596369 | 25612833 | 16465 | -8.55 | 3.14E-15 | 1.54E-14 |
| chr8 | 8358491 | 8378391 | 19901 | -9.12 | 3.15E-15 | 1.55E-14 |
| chr8 | 140863117 | 140875311 | 12195 | -8.63 | 3.16E-15 | 1.55E-14 |
| chr5 | 380132 | 386701 | 6570 | -8.57 | 3.18E-15 | 1.56E-14 |
| chr11 | 94241321 | 94246652 | 5332 | -8.56 | 3.20E-15 | 1.57E-14 |
| chr5 | 68000779 | 68019629 | 18851 | -8.56 | 3.20E-15 | 1.57E-14 |
| chr12 | 107570578 | 107588410 | 17833 | -8.6 | 3.20E-15 | 1.57E-14 |
| chr14 | 35051422 | 35062163 | 10742 | -8.65 | 3.21E-15 | 1.57E-14 |
| chr5 | 17646409 | 17674893 | 28485 | -8.65 | 3.21E-15 | 1.57E-14 |
| chr1 | 155440320 | 155443189 | 2870 | -8.56 | 3.22E-15 | 1.58E-14 |
| chr1 | 53669014 | 53673355 | 4342 | -8.56 | 3.22E-15 | 1.58E-14 |
| chr17 | 26396780 | 26399648 | 2869 | -8.55 | 3.22E-15 | 1.58E-14 |
| chr4 | 148812096 | 148825426 | 13331 | -8.62 | 3.23E-15 | 1.58E-14 |
| chr14 | 61348561 | 61354414 | 5854 | -8.55 | 3.25E-15 | 1.59E-14 |

| | | | | | | |
|---|---|---|---|---|---|---|
| chr20 | 36892383 | 36919250 | 26868 | -8.68 | 3.26E-15 | 1.60E-14 |
| chr1 | 223532823 | 223550827 | 18005 | -4.28 | 3.26E-15 | 1.60E-14 |
| chr17 | 55374188 | 55376257 | 2070 | -8.61 | 3.26E-15 | 1.60E-14 |
| chr2 | 182656975 | 182660801 | 3827 | -8.64 | 3.27E-15 | 1.60E-14 |
| chr13 | 96363220 | 96366468 | 3249 | -8.64 | 3.27E-15 | 1.60E-14 |
| chr16 | 69909963 | 69920344 | 10382 | -8.57 | 3.28E-15 | 1.61E-14 |
| chr13 | 51586463 | 51599031 | 12569 | -8.55 | 3.28E-15 | 1.61E-14 |
| chr17 | 71784640 | 71816443 | 31804 | -9.19 | 3.28E-15 | 1.61E-14 |
| chr16 | 52788966 | 52809613 | 20648 | -6.16 | 3.29E-15 | 1.61E-14 |
| chr9 | 114953352 | 114973400 | 20049 | -8.93 | 3.30E-15 | 1.61E-14 |
| chr4 | 89100005 | 89107015 | 7011 | -8.62 | 3.30E-15 | 1.62E-14 |
| chr5 | 153624649 | 153643082 | 18434 | -8.54 | 3.31E-15 | 1.62E-14 |
| chr6 | 140646494 | 140664873 | 18380 | -8.55 | 3.31E-15 | 1.62E-14 |
| chr17 | 56878854 | 56880753 | 1900 | -8.54 | 3.31E-15 | 1.62E-14 |
| chr15 | 53537122 | 53552504 | 15383 | 8.67 | 3.33E-15 | 1.63E-14 |
| chr1 | 167912793 | 167947705 | 34913 | -8.82 | 3.33E-15 | 1.63E-14 |
| chr20 | 14343964 | 14350212 | 6249 | -8.74 | 3.34E-15 | 1.63E-14 |
| chr10 | 4701209 | 4732230 | 31022 | -8.59 | 3.34E-15 | 1.63E-14 |
| chr16 | 3881645 | 3884224 | 2580 | -8.54 | 3.36E-15 | 1.64E-14 |
| chr12 | 78111435 | 78132420 | 20986 | -8.55 | 3.37E-15 | 1.65E-14 |
| chr1 | 230785825 | 230790374 | 4550 | -8.54 | 3.38E-15 | 1.65E-14 |
| chr2 | 8511152 | 8528770 | 17619 | -8.61 | 3.38E-15 | 1.65E-14 |
| chrX | 19535624 | 19543598 | 7975 | -8.56 | 3.38E-15 | 1.65E-14 |
| chr12 | 97315473 | 97318551 | 3079 | -8.55 | 3.39E-15 | 1.66E-14 |
| chr6 | 132410394 | 132425417 | 15024 | -8.54 | 3.39E-15 | 1.66E-14 |
| chr17 | 55693457 | 55695939 | 2483 | -8.54 | 3.40E-15 | 1.66E-14 |
| chr2 | 197438365 | 197445585 | 7221 | -8.56 | 3.40E-15 | 1.66E-14 |
| chr2 | 106202006 | 106207581 | 5576 | -8.69 | 3.42E-15 | 1.67E-14 |
| chr2 | 58099111 | 58107507 | 8397 | -8.77 | 3.42E-15 | 1.67E-14 |
| chr3 | 131602139 | 131611598 | 9460 | -8.54 | 3.42E-15 | 1.67E-14 |
| chr8 | 61723826 | 61727166 | 3341 | -8.74 | 3.43E-15 | 1.68E-14 |
| chr12 | 66097357 | 66101222 | 3866 | -8.6 | 3.44E-15 | 1.68E-14 |
| chr10 | 112448905 | 112464261 | 15357 | -8.89 | 3.44E-15 | 1.68E-14 |
| chr1 | 207591229 | 207603676 | 12448 | -8.53 | 3.44E-15 | 1.68E-14 |
| chr7 | 11527067 | 11544044 | 16978 | -8.57 | 3.44E-15 | 1.68E-14 |
| chr9 | 140367320 | 140371689 | 4370 | -8.61 | 3.45E-15 | 1.68E-14 |
| chrX | 149702124 | 149708025 | 5902 | -8.61 | 3.45E-15 | 1.68E-14 |
| chr7 | 142252243 | 142268186 | 15944 | 8.61 | 3.45E-15 | 1.68E-14 |
| chr12 | 18058604 | 18086658 | 28055 | -8.58 | 3.47E-15 | 1.69E-14 |
| chr21 | 34648357 | 34653613 | 5257 | -8.56 | 3.49E-15 | 1.70E-14 |
| chr5 | 73130546 | 73133852 | 3307 | -8.54 | 3.49E-15 | 1.70E-14 |
| chr9 | 77777755 | 77783135 | 5381 | -8.55 | 3.50E-15 | 1.70E-14 |
| chr7 | 108196246 | 108200486 | 4241 | -8.54 | 3.50E-15 | 1.71E-14 |
| chr2 | 150136199 | 150150896 | 14698 | 8.61 | 3.51E-15 | 1.71E-14 |
| chr2 | 53301032 | 53329786 | 28755 | -8.6 | 3.53E-15 | 1.72E-14 |
| chr1 | 65254946 | 65259053 | 4108 | -8.66 | 3.54E-15 | 1.72E-14 |
| chr12 | 112285117 | 112310535 | 25419 | -8.67 | 3.55E-15 | 1.73E-14 |
| chr20 | 383456 | 393367 | 9912 | -8.77 | 3.55E-15 | 1.73E-14 |
| chr7 | 43007594 | 43052354 | 44761 | -9.19 | 3.56E-15 | 1.73E-14 |
| chr12 | 89231341 | 89249676 | 18336 | -8.57 | 3.56E-15 | 1.73E-14 |
| chr8 | 144259827 | 144263300 | 3474 | -8.64 | 3.57E-15 | 1.74E-14 |
| chr6 | 1562827 | 1584648 | 21822 | -8.67 | 3.57E-15 | 1.74E-14 |
| chr14 | 34583943 | 34587526 | 3584 | -8.56 | 3.58E-15 | 1.74E-14 |
| chr6 | 116311064 | 116314400 | 3337 | -8.55 | 3.59E-15 | 1.74E-14 |
| chr7 | 14623675 | 14667592 | 43918 | -8.7 | 3.59E-15 | 1.74E-14 |
| chr4 | 6819817 | 6823586 | 3770 | -8.54 | 3.60E-15 | 1.75E-14 |
| chr4 | 83944461 | 83948367 | 3907 | -8.68 | 3.60E-15 | 1.75E-14 |
| chr20 | 44015356 | 44031447 | 16092 | -8.55 | 3.61E-15 | 1.75E-14 |
| chr4 | 108475041 | 108491267 | 16227 | -8.55 | 3.61E-15 | 1.75E-14 |
| chr2 | 36131794 | 36148446 | 16653 | 8.84 | 3.61E-15 | 1.75E-14 |
| chr12 | 64394977 | 64401606 | 6630 | -8.53 | 3.62E-15 | 1.76E-14 |

| | | | | | | |
|---|---|---|---|---|---|---|
| chr3 | 37816998 | 37820615 | 3618 | -8.56 | 3.63E-15 | 1.76E-14 |
| chr2 | 181952075 | 181966611 | 14537 | -8.56 | 3.63E-15 | 1.76E-14 |
| chr4 | 24590665 | 24602026 | 11362 | -8.53 | 3.64E-15 | 1.77E-14 |
| chr14 | 95669846 | 95673438 | 3593 | -8.53 | 3.64E-15 | 1.77E-14 |
| chr5 | 179417862 | 179425822 | 7961 | -8.78 | 3.64E-15 | 1.77E-14 |
| chr1 | 67948537 | 67954517 | 5981 | -8.77 | 3.65E-15 | 1.77E-14 |
| chr5 | 158900488 | 158904008 | 3521 | -8.59 | 3.65E-15 | 1.77E-14 |
| chr3 | 197038992 | 197049852 | 10861 | -8.54 | 3.66E-15 | 1.78E-14 |
| chr12 | 100815641 | 100823713 | 8073 | -8.55 | 3.66E-15 | 1.78E-14 |
| chr11 | 94546501 | 94560391 | 13891 | -8.53 | 3.67E-15 | 1.78E-14 |
| chr4 | 48280960 | 48288488 | 7529 | -4.11 | 3.68E-15 | 1.79E-14 |
| chr1 | 77442555 | 77446918 | 4364 | -8.56 | 3.69E-15 | 1.79E-14 |
| chr22 | 49327398 | 49333370 | 5973 | -3.23 | 3.69E-15 | 1.79E-14 |
| chr12 | 42337133 | 42348704 | 11572 | -8.59 | 3.69E-15 | 1.79E-14 |
| chr5 | 133056163 | 133082953 | 26791 | -8.78 | 3.72E-15 | 1.80E-14 |
| chr1 | 64522375 | 64525745 | 3371 | -8.61 | 3.73E-15 | 1.81E-14 |
| chr2 | 235690226 | 235693983 | 3758 | -8.53 | 3.73E-15 | 1.81E-14 |
| chr14 | 95112866 | 95116842 | 3977 | -8.55 | 3.73E-15 | 1.81E-14 |
| chr11 | 95373818 | 95388094 | 14277 | 8.6 | 3.74E-15 | 1.81E-14 |
| chr13 | 96608022 | 96616143 | 8122 | -8.62 | 3.75E-15 | 1.82E-14 |
| chr8 | 126469878 | 126472730 | 2853 | -8.6 | 3.77E-15 | 1.82E-14 |
| chr2 | 169770951 | 169773623 | 2673 | -8.54 | 3.77E-15 | 1.83E-14 |
| chr18 | 8338523 | 8353002 | 14480 | -8.57 | 3.77E-15 | 1.83E-14 |
| chr10 | 133736373 | 133740161 | 3789 | -8.55 | 3.78E-15 | 1.83E-14 |
| chr12 | 76316625 | 76334270 | 17646 | -8.58 | 3.78E-15 | 1.83E-14 |
| chr10 | 60871227 | 60881909 | 10683 | -8.62 | 3.79E-15 | 1.83E-14 |
| chr14 | 104075646 | 104082108 | 6463 | -8.55 | 3.79E-15 | 1.84E-14 |
| chr14 | 104774316 | 104779605 | 5290 | -8.53 | 3.82E-15 | 1.85E-14 |
| chr20 | 2755717 | 2768366 | 12650 | -8.73 | 3.83E-15 | 1.86E-14 |
| chr6 | 66605863 | 66622875 | 17013 | -4.09 | 3.84E-15 | 1.86E-14 |
| chr4 | 27707286 | 27806989 | 99704 | 1.94 | 3.84E-15 | 1.86E-14 |
| chr1 | 88372355 | 88381871 | 9517 | -8.6 | 3.85E-15 | 1.86E-14 |
| chr1 | 29724903 | 29732905 | 8003 | -8.61 | 3.86E-15 | 1.87E-14 |
| chr16 | 23175615 | 23179202 | 3588 | -8.65 | 3.86E-15 | 1.87E-14 |
| chr3 | 149165651 | 149188280 | 22630 | -8.73 | 3.89E-15 | 1.88E-14 |
| chr10 | 124105400 | 124128208 | 22809 | -8.85 | 3.90E-15 | 1.88E-14 |
| chr9 | 136610948 | 136617690 | 6743 | -8.53 | 3.90E-15 | 1.89E-14 |
| chrX | 117866667 | 117872519 | 5853 | -8.53 | 3.93E-15 | 1.90E-14 |
| chr3 | 187941361 | 187945676 | 4316 | -8.54 | 3.93E-15 | 1.90E-14 |
| chr2 | 183826482 | 183837869 | 11388 | -8.52 | 3.94E-15 | 1.90E-14 |
| chr4 | 104076068 | 104091697 | 15630 | -8.52 | 3.94E-15 | 1.90E-14 |
| chr8 | 90515861 | 90520335 | 4475 | -8.56 | 3.95E-15 | 1.91E-14 |
| chr8 | 105791769 | 106323133 | 531365 | 5.29 | 3.95E-15 | 1.91E-14 |
| chr6 | 80053006 | 80080991 | 27986 | -8.58 | 3.95E-15 | 1.91E-14 |
| chr4 | 141659303 | 141673533 | 14231 | -8.53 | 3.95E-15 | 1.91E-14 |
| chr2 | 238685196 | 238718630 | 33435 | -8.52 | 3.96E-15 | 1.92E-14 |
| chr19 | 51263124 | 51282915 | 19792 | -8.53 | 3.97E-15 | 1.92E-14 |
| chr4 | 185524565 | 185526358 | 1794 | -4.38 | 3.98E-15 | 1.92E-14 |
| chr2 | 160397525 | 160400357 | 2833 | -8.52 | 3.99E-15 | 1.93E-14 |
| chr7 | 66860016 | 67107369 | 247354 | -2.7 | 4.00E-15 | 1.93E-14 |
| chr12 | 92292201 | 92301810 | 9610 | -8.66 | 4.00E-15 | 1.93E-14 |
| chr10 | 61517827 | 61556960 | 39134 | -8.53 | 4.02E-15 | 1.94E-14 |
| chr22 | 27618143 | 27626792 | 8650 | -8.53 | 4.02E-15 | 1.94E-14 |
| chrX | 45841020 | 45857596 | 16577 | -8.53 | 4.02E-15 | 1.94E-14 |
| chr9 | 116403361 | 116406098 | 2738 | -8.6 | 4.02E-15 | 1.94E-14 |
| chrX | 8858763 | 8864816 | 6054 | -8.53 | 4.03E-15 | 1.95E-14 |
| chr16 | 3966543 | 3971900 | 5358 | -8.56 | 4.03E-15 | 1.95E-14 |
| chr16 | 75717373 | 75727806 | 10434 | -8.52 | 4.03E-15 | 1.95E-14 |
| chr5 | 129648170 | 129666817 | 18648 | -8.58 | 4.04E-15 | 1.95E-14 |
| chr9 | 116730083 | 116734779 | 4697 | -8.55 | 4.04E-15 | 1.95E-14 |
| chr3 | 114050187 | 114053331 | 3145 | -8.52 | 4.05E-15 | 1.95E-14 |

| | | | | | | |
|---|---|---|---|---|---|---|
| chr4 | 133604144 | 133619014 | 14871 | -8.57 | 4.07E-15 | 1.96E-14 |
| chr12 | 84193848 | 84221764 | 27917 | -8.66 | 4.08E-15 | 1.97E-14 |
| chr3 | 134308765 | 134316216 | 7452 | -8.52 | 4.09E-15 | 1.97E-14 |
| chr5 | 173273342 | 173277287 | 3946 | -8.57 | 4.09E-15 | 1.97E-14 |
| chr20 | 15327282 | 15333544 | 6263 | -8.65 | 4.10E-15 | 1.98E-14 |
| chr2 | 223800566 | 223803948 | 3383 | -8.98 | 4.10E-15 | 1.98E-14 |
| chr21 | 27754059 | 27771603 | 17545 | -8.52 | 4.10E-15 | 1.98E-14 |
| chr6 | 105901814 | 105935709 | 33896 | -9.09 | 4.11E-15 | 1.98E-14 |
| chr8 | 120259854 | 120287719 | 27866 | 9.05 | 4.11E-15 | 1.98E-14 |
| chr8 | 49251903 | 49278488 | 26586 | -8.67 | 4.14E-15 | 1.99E-14 |
| chr1 | 7752539 | 7754443 | 1905 | -6.13 | 4.15E-15 | 2.00E-14 |
| chr7 | 130657708 | 130661185 | 3478 | -8.51 | 4.15E-15 | 2.00E-14 |
| chr8 | 128224374 | 128240552 | 16179 | -8.75 | 4.17E-15 | 2.01E-14 |
| chr12 | 133282034 | 133286778 | 4745 | -8.54 | 4.18E-15 | 2.01E-14 |
| chr11 | 101558074 | 101565660 | 7587 | -7.3 | 4.19E-15 | 2.02E-14 |
| chr15 | 81188037 | 81202661 | 14625 | -8.78 | 4.20E-15 | 2.02E-14 |
| chr16 | 53066837 | 53081856 | 15020 | -8.68 | 4.21E-15 | 2.03E-14 |
| chr1 | 226929371 | 226932869 | 3499 | -8.51 | 4.23E-15 | 2.03E-14 |
| chr7 | 109428084 | 109435366 | 7283 | -8.52 | 4.23E-15 | 2.04E-14 |
| chr14 | 38123877 | 38137811 | 13935 | -8.52 | 4.24E-15 | 2.04E-14 |
| chr3 | 104415428 | 104432263 | 16836 | 8.64 | 4.24E-15 | 2.04E-14 |
| chr2 | 170947614 | 170954890 | 7277 | -8.51 | 4.24E-15 | 2.04E-14 |
| chr20 | 48815027 | 48821277 | 6251 | -8.69 | 4.24E-15 | 2.04E-14 |
| chr12 | 93366905 | 93387710 | 20806 | -8.81 | 4.24E-15 | 2.04E-14 |
| chr16 | 63638878 | 63647707 | 8830 | -8.52 | 4.25E-15 | 2.04E-14 |
| chr1 | 87842980 | 87857272 | 14293 | -8.64 | 4.26E-15 | 2.04E-14 |
| chr6 | 4380648 | 4386763 | 6116 | -8.62 | 4.26E-15 | 2.05E-14 |
| chr3 | 186803601 | 186817919 | 14319 | -8.85 | 4.27E-15 | 2.05E-14 |
| chr13 | 111724957 | 111728241 | 3285 | -8.59 | 4.28E-15 | 2.05E-14 |
| chr13 | 33025676 | 33035774 | 10099 | -8.59 | 4.28E-15 | 2.05E-14 |
| chr3 | 39666078 | 39682691 | 16614 | -8.57 | 4.29E-15 | 2.06E-14 |
| chr9 | 30167178 | 30200755 | 33578 | -8.57 | 4.30E-15 | 2.06E-14 |
| chr15 | 38714605 | 38732384 | 17780 | -8.51 | 4.31E-15 | 2.07E-14 |
| chr10 | 69569187 | 69575309 | 6123 | -8.63 | 4.31E-15 | 2.07E-14 |
| chr1 | 18135935 | 18142611 | 6677 | -8.75 | 4.32E-15 | 2.07E-14 |
| chr3 | 8443007 | 8449448 | 6442 | -8.55 | 4.34E-15 | 2.08E-14 |
| chr18 | 72129231 | 72142889 | 13659 | -2.4 | 4.35E-15 | 2.09E-14 |
| chr13 | 73660076 | 73664361 | 4286 | -8.51 | 4.38E-15 | 2.10E-14 |
| chr21 | 32702105 | 32733724 | 31620 | -8.67 | 4.39E-15 | 2.10E-14 |
| chr12 | 13511756 | 13531501 | 19746 | -8.53 | 4.39E-15 | 2.11E-14 |
| chr4 | 141798455 | 141805679 | 7225 | -8.59 | 4.40E-15 | 2.11E-14 |
| chr3 | 25404089 | 25420108 | 16020 | 8.65 | 4.41E-15 | 2.11E-14 |
| chr4 | 124487010 | 124504237 | 17228 | -8.81 | 4.41E-15 | 2.12E-14 |
| chr14 | 52839315 | 52864577 | 25263 | -8.64 | 4.41E-15 | 2.12E-14 |
| chr11 | 13017444 | 13034839 | 17396 | -9.05 | 4.42E-15 | 2.12E-14 |
| chr3 | 167894719 | 167913093 | 18375 | 8.67 | 4.42E-15 | 2.12E-14 |
| chr16 | 27774789 | 27776924 | 2136 | -8.51 | 4.44E-15 | 2.13E-14 |
| chr20 | 3465836 | 3503400 | 37565 | -9.13 | 4.45E-15 | 2.14E-14 |
| chr15 | 57026091 | 57034451 | 8361 | -8.84 | 4.46E-15 | 2.14E-14 |
| chr5 | 118918725 | 119787517 | 868793 | 4.15 | 4.47E-15 | 2.14E-14 |
| chr1 | 214747645 | 214752080 | 4436 | -8.53 | 4.47E-15 | 2.14E-14 |
| chr3 | 43261366 | 43288374 | 27009 | -1.59 | 4.47E-15 | 2.14E-14 |
| chr13 | 44599749 | 44623228 | 23480 | -8.71 | 4.50E-15 | 2.15E-14 |
| chr1 | 98613649 | 98622889 | 9241 | -8.51 | 4.50E-15 | 2.15E-14 |
| chr7 | 29870284 | 29872584 | 2301 | -8.51 | 4.50E-15 | 2.15E-14 |
| chr6 | 143365246 | 143371590 | 6345 | -8.55 | 4.50E-15 | 2.15E-14 |
| chr15 | 90063919 | 90112799 | 48881 | -9.43 | 4.51E-15 | 2.16E-14 |
| chr2 | 45442118 | 45455840 | 13723 | -8.5 | 4.52E-15 | 2.16E-14 |
| chr10 | 120124275 | 120127635 | 3361 | -8.58 | 4.52E-15 | 2.16E-14 |
| chr10 | 31897283 | 32027677 | 130395 | -3.89 | 4.53E-15 | 2.17E-14 |
| chr22 | 48028796 | 48097452 | 68657 | -2.59 | 4.54E-15 | 2.17E-14 |

| | | | | | | |
|---|---|---|---|---|---|---|
| chr7 | 7039763 | 7135127 | 95365 | -2.82 | 4.54E-15 | 2.17E-14 |
| chr3 | 168964197 | 168978790 | 14594 | 8.59 | 4.56E-15 | 2.18E-14 |
| chr6 | 47393100 | 47403595 | 10496 | -8.74 | 4.56E-15 | 2.18E-14 |
| chr13 | 100120829 | 100122829 | 2001 | -4.02 | 4.57E-15 | 2.18E-14 |
| chr2 | 230958363 | 230973214 | 14852 | -8.88 | 4.58E-15 | 2.19E-14 |
| chr8 | 142950638 | 142952832 | 2195 | -8.5 | 4.59E-15 | 2.19E-14 |
| chr14 | 57608593 | 57622424 | 13832 | 8.62 | 4.59E-15 | 2.19E-14 |
| chr19 | 48933140 | 48935665 | 2526 | -8.5 | 4.61E-15 | 2.20E-14 |
| chr6 | 66525076 | 66534353 | 9278 | -8.5 | 4.65E-15 | 2.22E-14 |
| chr11 | 86802177 | 86804876 | 2700 | -8.56 | 4.68E-15 | 2.23E-14 |
| chr1 | 167066257 | 167084269 | 18013 | -8.57 | 4.70E-15 | 2.24E-14 |
| chr6 | 160034554 | 160040464 | 5911 | -8.5 | 4.71E-15 | 2.25E-14 |
| chr4 | 103397053 | 103420333 | 23281 | -8.57 | 4.71E-15 | 2.25E-14 |
| chr16 | 10994562 | 11004512 | 9951 | -8.56 | 4.71E-15 | 2.25E-14 |
| chr4 | 2098063 | 2106892 | 8830 | -8.6 | 4.72E-15 | 2.25E-14 |
| chrX | 37245984 | 37257133 | 11150 | -8.52 | 4.74E-15 | 2.26E-14 |
| chr1 | 170045906 | 170049414 | 3509 | -8.52 | 4.74E-15 | 2.26E-14 |
| chr13 | 112028662 | 112033859 | 5198 | -3.39 | 4.74E-15 | 2.26E-14 |
| chr18 | 19268942 | 19272253 | 3312 | -8.5 | 4.76E-15 | 2.27E-14 |
| chr8 | 24049598 | 24085976 | 36379 | -9.15 | 4.76E-15 | 2.27E-14 |
| chr10 | 93276291 | 93279334 | 3044 | -8.5 | 4.77E-15 | 2.28E-14 |
| chr7 | 47715209 | 47719227 | 4019 | -8.61 | 4.78E-15 | 2.28E-14 |
| chr1 | 35744263 | 35751298 | 7036 | -8.5 | 4.79E-15 | 2.29E-14 |
| chr20 | 15137059 | 15155735 | 18677 | -9.05 | 4.80E-15 | 2.29E-14 |
| chr2 | 112821593 | 112832316 | 10724 | -8.54 | 4.81E-15 | 2.29E-14 |
| chr14 | 71878425 | 71894539 | 16115 | -8.59 | 4.81E-15 | 2.29E-14 |
| chr4 | 124575944 | 124580079 | 4136 | -4.88 | 4.81E-15 | 2.29E-14 |
| chr10 | 5621882 | 5628432 | 6551 | -8.54 | 4.82E-15 | 2.30E-14 |
| chr9 | 125254036 | 125262764 | 8729 | -8.65 | 4.82E-15 | 2.30E-14 |
| chr20 | 40131854 | 40135964 | 4111 | -8.6 | 4.82E-15 | 2.30E-14 |
| chr7 | 67946102 | 67975229 | 29128 | -8.97 | 4.85E-15 | 2.31E-14 |
| chr4 | 108839087 | 108843856 | 4770 | -8.51 | 4.86E-15 | 2.32E-14 |
| chr6 | 130594058 | 130600183 | 6126 | -8.52 | 4.86E-15 | 2.32E-14 |
| chr2 | 163858432 | 163877615 | 19184 | 8.59 | 4.87E-15 | 2.32E-14 |
| chr12 | 112725594 | 112728263 | 2670 | -8.5 | 4.89E-15 | 2.33E-14 |
| chr3 | 111378782 | 111383696 | 4915 | -8.53 | 4.89E-15 | 2.33E-14 |
| chr16 | 81431898 | 81434307 | 2410 | -8.49 | 4.90E-15 | 2.33E-14 |
| chr2 | 207839236 | 207864161 | 24926 | -8.68 | 4.90E-15 | 2.33E-14 |
| chr1 | 59291604 | 59295376 | 3773 | -8.5 | 4.91E-15 | 2.34E-14 |
| chr3 | 151389023 | 151402155 | 13133 | -8.66 | 4.91E-15 | 2.34E-14 |
| chr3 | 17885741 | 17904364 | 18624 | -8.74 | 4.91E-15 | 2.34E-14 |
| chr7 | 94343259 | 94360415 | 17157 | -8.5 | 4.91E-15 | 2.34E-14 |
| chr17 | 35388271 | 35390471 | 2201 | -8.51 | 4.91E-15 | 2.34E-14 |
| chr12 | 70770996 | 70786060 | 15065 | -8.56 | 4.93E-15 | 2.34E-14 |
| chr22 | 45507499 | 45515797 | 8299 | -8.5 | 4.93E-15 | 2.35E-14 |
| chrX | 12805944 | 12814314 | 8371 | -8.56 | 4.94E-15 | 2.35E-14 |
| chr6 | 7848566 | 7854512 | 5947 | -8.53 | 4.94E-15 | 2.35E-14 |
| chr4 | 88945118 | 88948176 | 3059 | -8.51 | 4.96E-15 | 2.36E-14 |
| chr18 | 45569052 | 45573084 | 4033 | -8.51 | 4.98E-15 | 2.37E-14 |
| chr10 | 22136725 | 22140639 | 3915 | -8.49 | 4.99E-15 | 2.37E-14 |
| chr1 | 244794224 | 244824828 | 30605 | -8.82 | 4.99E-15 | 2.37E-14 |
| chr2 | 85307902 | 85315337 | 7436 | -4.29 | 5.00E-15 | 2.38E-14 |
| chr6 | 6950201 | 6953532 | 3332 | -8.52 | 5.00E-15 | 2.38E-14 |
| chr7 | 45965183 | 45971193 | 6011 | -8.54 | 5.02E-15 | 2.38E-14 |
| chr2 | 203543162 | 203548878 | 5717 | -8.57 | 5.02E-15 | 2.38E-14 |
| chr12 | 111101555 | 111123433 | 21879 | -8.79 | 5.02E-15 | 2.38E-14 |
| chr8 | 91974994 | 91979549 | 4556 | -8.49 | 5.05E-15 | 2.40E-14 |
| chr10 | 121110698 | 121117096 | 6399 | -8.5 | 5.05E-15 | 2.40E-14 |
| chr12 | 93117174 | 93120680 | 3507 | -8.57 | 5.06E-15 | 2.40E-14 |
| chr3 | 128675089 | 128678770 | 3682 | -8.49 | 5.06E-15 | 2.41E-14 |
| chr11 | 108492806 | 108501705 | 8900 | -8.55 | 5.08E-15 | 2.41E-14 |

| | | | | | | |
|---|---|---|---|---|---|---|
| chr5 | 71057885 | 71070919 | 13035 | -8.83 | 5.10E-15 | 2.42E-14 |
| chr7 | 138337331 | 138353859 | 16529 | -8.53 | 5.10E-15 | 2.42E-14 |
| chr7 | 132470500 | 132474608 | 4109 | -8.53 | 5.11E-15 | 2.43E-14 |
| chr6 | 165731669 | 165736934 | 5266 | -8.54 | 5.12E-15 | 2.43E-14 |
| chr11 | 29393958 | 29410966 | 17009 | -8.62 | 5.13E-15 | 2.43E-14 |
| chr12 | 14812970 | 14828716 | 15747 | -8.56 | 5.14E-15 | 2.44E-14 |
| chr4 | 52931728 | 52949358 | 17631 | -2.58 | 5.17E-15 | 2.45E-14 |
| chrX | 19427654 | 19449465 | 21812 | -8.49 | 5.17E-15 | 2.45E-14 |
| chr17 | 58888518 | 58892390 | 3873 | -8.61 | 5.20E-15 | 2.46E-14 |
| chr1 | 244096736 | 244104642 | 7907 | -8.57 | 5.20E-15 | 2.47E-14 |
| chr4 | 126900037 | 126918007 | 17971 | 8.56 | 5.21E-15 | 2.47E-14 |
| chr2 | 217312255 | 217319679 | 7425 | -8.51 | 5.22E-15 | 2.47E-14 |
| chr11 | 2103522 | 2115665 | 12144 | -8.5 | 5.22E-15 | 2.47E-14 |
| chr2 | 44430592 | 44438556 | 7965 | -8.61 | 5.22E-15 | 2.47E-14 |
| chr4 | 105526330 | 105535796 | 9467 | -8.55 | 5.23E-15 | 2.48E-14 |
| chr3 | 136371094 | 136375051 | 3958 | -8.72 | 5.23E-15 | 2.48E-14 |
| chr1 | 185075401 | 185079544 | 4144 | -8.5 | 5.26E-15 | 2.49E-14 |
| chr2 | 58257418 | 58271810 | 14393 | -8.49 | 5.27E-15 | 2.50E-14 |
| chr5 | 132311739 | 132316235 | 4497 | -8.51 | 5.29E-15 | 2.50E-14 |
| chr14 | 69841724 | 69844811 | 3088 | -8.54 | 5.29E-15 | 2.51E-14 |
| chr13 | 113012730 | 113036195 | 23466 | -2.34 | 5.29E-15 | 2.51E-14 |
| chr12 | 90107163 | 90113076 | 5914 | -8.54 | 5.30E-15 | 2.51E-14 |
| chr7 | 20196705 | 20211039 | 14335 | -8.49 | 5.32E-15 | 2.52E-14 |
| chrX | 64970448 | 64974601 | 4154 | -5.07 | 5.33E-15 | 2.52E-14 |
| chr3 | 156462901 | 156479731 | 16831 | -8.5 | 5.34E-15 | 2.53E-14 |
| chr1 | 50873584 | 50879611 | 6028 | -8.51 | 5.35E-15 | 2.53E-14 |
| chr2 | 205805886 | 205814835 | 8950 | -8.6 | 5.36E-15 | 2.54E-14 |
| chr12 | 26401725 | 26420965 | 19241 | -8.8 | 5.37E-15 | 2.54E-14 |
| chr5 | 56046225 | 56049471 | 3247 | -8.48 | 5.40E-15 | 2.55E-14 |
| chr10 | 8709165 | 8721429 | 12265 | -8.5 | 5.40E-15 | 2.56E-14 |
| chr12 | 88077367 | 88103358 | 25992 | -8.5 | 5.41E-15 | 2.56E-14 |
| chr1 | 44607114 | 44613150 | 6037 | -8.48 | 5.41E-15 | 2.56E-14 |
| chr12 | 79237148 | 79249648 | 12501 | -8.73 | 5.43E-15 | 2.57E-14 |
| chr1 | 55648124 | 55652425 | 4302 | -8.5 | 5.43E-15 | 2.57E-14 |
| chr4 | 1150630 | 1153573 | 2944 | -8.48 | 5.45E-15 | 2.58E-14 |
| chr6 | 122139552 | 122148939 | 9388 | -8.53 | 5.45E-15 | 2.58E-14 |
| chr5 | 168001161 | 168026381 | 25221 | -8.66 | 5.46E-15 | 2.58E-14 |
| chr3 | 67832157 | 67854509 | 22353 | -8.5 | 5.47E-15 | 2.58E-14 |
| chr9 | 140649019 | 140665784 | 16766 | -8.49 | 5.48E-15 | 2.59E-14 |
| chr12 | 117908264 | 117922761 | 14498 | -8.49 | 5.50E-15 | 2.60E-14 |
| chr21 | 46633956 | 46638657 | 4702 | -8.55 | 5.51E-15 | 2.60E-14 |
| chrX | 135191006 | 135215060 | 24055 | -8.54 | 5.51E-15 | 2.60E-14 |
| chr1 | 199679052 | 199684329 | 5278 | -8.48 | 5.52E-15 | 2.61E-14 |
| chr5 | 142449696 | 142454662 | 4967 | -8.48 | 5.52E-15 | 2.61E-14 |
| chr2 | 36367132 | 36385700 | 18569 | -8.51 | 5.55E-15 | 2.62E-14 |
| chr6 | 34466981 | 34478047 | 11067 | -8.48 | 5.55E-15 | 2.62E-14 |
| chr16 | 1847223 | 1851112 | 3890 | -8.49 | 5.56E-15 | 2.63E-14 |
| chr14 | 51959474 | 51971197 | 11724 | -8.59 | 5.56E-15 | 2.63E-14 |
| chr1 | 115948380 | 115970792 | 22413 | -8.53 | 5.59E-15 | 2.64E-14 |
| chr4 | 175663935 | 175669609 | 5675 | -8.48 | 5.59E-15 | 2.64E-14 |
| chr20 | 15285094 | 15300969 | 15876 | -9.09 | 5.62E-15 | 2.65E-14 |
| chr3 | 150854274 | 150858169 | 3896 | -8.5 | 5.63E-15 | 2.66E-14 |
| chr1 | 30287329 | 30293538 | 6210 | -2.7 | 5.63E-15 | 2.66E-14 |
| chr10 | 123344710 | 123348359 | 3650 | -3.51 | 5.64E-15 | 2.66E-14 |
| chr3 | 42471335 | 42497048 | 25714 | -8.78 | 5.67E-15 | 2.68E-14 |
| chr8 | 91818665 | 91829610 | 10946 | 8.6 | 5.69E-15 | 2.68E-14 |
| chr11 | 13490949 | 13535847 | 44899 | -9.09 | 5.70E-15 | 2.69E-14 |
| chr13 | 65363659 | 65387416 | 23758 | 8.86 | 5.71E-15 | 2.69E-14 |
| chr21 | 32465572 | 32479435 | 13864 | -2.48 | 5.75E-15 | 2.71E-14 |
| chr20 | 21157451 | 21159444 | 1994 | -8.52 | 5.75E-15 | 2.71E-14 |
| chr17 | 56465921 | 56472703 | 6783 | -2.01 | 5.75E-15 | 2.71E-14 |

| | | | | | | |
|---|---|---|---|---|---|---|
| chr1 | 30563320 | 30567144 | 3825 | -8.49 | 5.76E-15 | 2.71E-14 |
| chr21 | 34703322 | 34708227 | 4906 | -8.52 | 5.78E-15 | 2.72E-14 |
| chr16 | 51585653 | 51610844 | 25192 | -8.81 | 5.80E-15 | 2.73E-14 |
| chr10 | 82210238 | 82212356 | 2119 | -8.48 | 5.80E-15 | 2.73E-14 |
| chr4 | 28176219 | 28188989 | 12771 | -8.49 | 5.81E-15 | 2.73E-14 |
| chr3 | 114636700 | 114642144 | 5445 | -8.62 | 5.81E-15 | 2.73E-14 |
| chr16 | 19135706 | 19138769 | 3064 | -8.65 | 5.82E-15 | 2.74E-14 |
| chr1 | 108675098 | 108691421 | 16324 | -8.56 | 5.82E-15 | 2.74E-14 |
| chr11 | 86753694 | 86758630 | 4937 | -8.48 | 5.84E-15 | 2.75E-14 |
| chr7 | 13628830 | 13661123 | 32294 | -8.52 | 5.85E-15 | 2.75E-14 |
| chr12 | 69936407 | 69939902 | 3496 | -8.7 | 5.87E-15 | 2.76E-14 |
| chr12 | 80682595 | 80691966 | 9372 | 7.13 | 5.87E-15 | 2.76E-14 |
| chr5 | 150987558 | 150999345 | 11788 | -8.56 | 5.87E-15 | 2.76E-14 |
| chr3 | 37379710 | 37381687 | 1978 | -8.48 | 5.88E-15 | 2.77E-14 |
| chr3 | 110890084 | 110894706 | 4623 | -8.51 | 5.89E-15 | 2.77E-14 |
| chr9 | 85994319 | 85999685 | 5367 | -8.59 | 5.89E-15 | 2.77E-14 |
| chr2 | 172481548 | 172516777 | 35230 | -3.81 | 5.90E-15 | 2.78E-14 |
| chr4 | 189436532 | 189445543 | 9012 | -8.48 | 5.93E-15 | 2.79E-14 |
| chr1 | 88161698 | 88170464 | 8767 | -8.47 | 5.94E-15 | 2.79E-14 |
| chr18 | 55681796 | 55684897 | 3102 | -8.47 | 5.96E-15 | 2.80E-14 |
| chr1 | 172696829 | 172726377 | 29549 | -8.86 | 5.98E-15 | 2.81E-14 |
| chr5 | 167804811 | 167820768 | 15958 | -8.55 | 5.99E-15 | 2.81E-14 |
| chr6 | 140853081 | 140867683 | 14603 | -8.55 | 6.01E-15 | 2.82E-14 |
| chr21 | 38166963 | 38172153 | 5191 | -8.48 | 6.01E-15 | 2.82E-14 |
| chr20 | 17638317 | 17641240 | 2924 | -8.57 | 6.02E-15 | 2.83E-14 |
| chr18 | 57667644 | 57680748 | 13105 | -8.49 | 6.05E-15 | 2.84E-14 |
| chr9 | 117496734 | 117523791 | 27058 | -8.71 | 6.05E-15 | 2.84E-14 |
| chr13 | 105715741 | 105749783 | 34043 | -8.5 | 6.07E-15 | 2.85E-14 |
| chr19 | 50981419 | 50994316 | 12898 | -8.58 | 6.11E-15 | 2.87E-14 |
| chr8 | 67031734 | 67044708 | 12975 | -8.58 | 6.11E-15 | 2.87E-14 |
| chr2 | 32121318 | 32137032 | 15715 | -8.62 | 6.15E-15 | 2.89E-14 |
| chr1 | 92790451 | 92795732 | 5282 | -8.49 | 6.20E-15 | 2.91E-14 |
| chr5 | 78702569 | 78706819 | 4251 | -8.57 | 6.20E-15 | 2.91E-14 |
| chr8 | 32654586 | 32678596 | 24011 | -8.64 | 6.20E-15 | 2.91E-14 |
| chr20 | 5677414 | 5709030 | 31617 | -9.1 | 6.21E-15 | 2.91E-14 |
| chr19 | 2569118 | 2585958 | 16841 | -8.69 | 6.23E-15 | 2.92E-14 |
| chr10 | 95745499 | 95770630 | 25132 | -8.48 | 6.30E-15 | 2.95E-14 |
| chr15 | 71680846 | 71682807 | 1962 | -8.47 | 6.33E-15 | 2.97E-14 |
| chr1 | 29782965 | 29823173 | 40209 | -8.85 | 6.33E-15 | 2.97E-14 |
| chr3 | 56781471 | 56786013 | 4543 | -8.61 | 6.33E-15 | 2.97E-14 |
| chr1 | 233128180 | 233138151 | 9972 | -8.57 | 6.34E-15 | 2.97E-14 |
| chr8 | 59637500 | 59648545 | 11046 | -8.48 | 6.37E-15 | 2.99E-14 |
| chr14 | 103166776 | 103173970 | 7195 | -8.61 | 6.38E-15 | 2.99E-14 |
| chr9 | 131609632 | 131614361 | 4730 | -8.47 | 6.39E-15 | 2.99E-14 |
| chr8 | 95991992 | 95997690 | 5699 | -8.46 | 6.40E-15 | 3.00E-14 |
| chr9 | 117361423 | 117365232 | 3810 | -8.47 | 6.41E-15 | 3.00E-14 |
| chr11 | 118155599 | 118165603 | 10005 | -8.57 | 6.43E-15 | 3.01E-14 |
| chr10 | 112079769 | 112086312 | 6544 | -8.47 | 6.46E-15 | 3.02E-14 |
| chr6 | 53652346 | 53656211 | 3866 | -8.47 | 6.46E-15 | 3.02E-14 |
| chr7 | 109385698 | 109415483 | 29786 | -8.47 | 6.51E-15 | 3.05E-14 |
| chr2 | 55578401 | 55580094 | 1694 | -8.49 | 6.52E-15 | 3.05E-14 |
| chrX | 13161834 | 13173305 | 11472 | -8.61 | 6.55E-15 | 3.06E-14 |
| chr20 | 50592357 | 50595712 | 3356 | -8.58 | 6.55E-15 | 3.06E-14 |
| chrX | 102835115 | 102857016 | 21902 | -8.47 | 6.56E-15 | 3.07E-14 |
| chr14 | 54449957 | 54455610 | 5654 | -8.78 | 6.57E-15 | 3.07E-14 |
| chr2 | 54247071 | 54281516 | 34446 | -8.87 | 6.58E-15 | 3.08E-14 |
| chr14 | 66353971 | 66414370 | 60400 | 2.21 | 6.60E-15 | 3.09E-14 |
| chr4 | 85179989 | 85185268 | 5280 | -8.49 | 6.61E-15 | 3.09E-14 |
| chr5 | 129534754 | 129541685 | 6932 | -8.49 | 6.61E-15 | 3.09E-14 |
| chr12 | 12256553 | 12262715 | 6163 | -8.47 | 6.62E-15 | 3.10E-14 |
| chr3 | 170693647 | 170730637 | 36991 | -8.78 | 6.63E-15 | 3.10E-14 |

| | | | | | | |
|---|---|---|---|---|---|---|
| chr2 | 128232251 | 128235303 | 3053 | -8.47 | 6.68E-15 | 3.12E-14 |
| chr10 | 95839614 | 95842319 | 2706 | -8.54 | 6.71E-15 | 3.14E-14 |
| chr18 | 12727364 | 12740773 | 13410 | -8.48 | 6.73E-15 | 3.14E-14 |
| chr5 | 21241513 | 21268093 | 26581 | -8.53 | 6.75E-15 | 3.15E-14 |
| chr12 | 131717584 | 131742016 | 24433 | -8.74 | 6.76E-15 | 3.16E-14 |
| chr15 | 22900945 | 22903827 | 2883 | -8.51 | 6.79E-15 | 3.17E-14 |
| chr6 | 147721257 | 147733967 | 12711 | -8.49 | 6.79E-15 | 3.17E-14 |
| chr18 | 19388349 | 19390635 | 2287 | -8.46 | 6.79E-15 | 3.17E-14 |
| chr5 | 17388608 | 17398405 | 9798 | -8.72 | 6.79E-15 | 3.17E-14 |
| chr15 | 64037625 | 64039851 | 2227 | -8.49 | 6.81E-15 | 3.18E-14 |
| chr4 | 175462771 | 175488010 | 25240 | -9.12 | 6.81E-15 | 3.18E-14 |
| chrX | 41659999 | 41675992 | 15994 | -8.59 | 6.81E-15 | 3.18E-14 |
| chr16 | 53182392 | 53186119 | 3728 | -8.48 | 6.83E-15 | 3.19E-14 |
| chr13 | 100403189 | 100421081 | 17893 | -8.57 | 6.86E-15 | 3.20E-14 |
| chr9 | 137201091 | 137205251 | 4161 | -8.46 | 6.88E-15 | 3.21E-14 |
| chr21 | 16919731 | 16956827 | 37097 | -8.79 | 6.88E-15 | 3.21E-14 |
| chr3 | 109758070 | 109765247 | 7178 | -8.57 | 6.89E-15 | 3.21E-14 |
| chr4 | 170286299 | 170303245 | 16947 | -8.77 | 6.90E-15 | 3.22E-14 |
| chr4 | 178581565 | 178586670 | 5106 | -8.45 | 6.91E-15 | 3.22E-14 |
| chrX | 20532376 | 20563990 | 31615 | -8.85 | 6.92E-15 | 3.23E-14 |
| chr2 | 148752226 | 148766994 | 14769 | -8.46 | 6.95E-15 | 3.24E-14 |
| chr7 | 54916017 | 54926474 | 10458 | -8.58 | 6.97E-15 | 3.25E-14 |
| chr4 | 95235803 | 95250272 | 14470 | -8.45 | 6.97E-15 | 3.25E-14 |
| chr2 | 231201058 | 231222993 | 21936 | -8.64 | 6.98E-15 | 3.26E-14 |
| chr3 | 41509764 | 41516703 | 6940 | -8.46 | 6.99E-15 | 3.26E-14 |
| chr5 | 32185954 | 32189995 | 4042 | -8.47 | 7.00E-15 | 3.26E-14 |
| chr6 | 54413260 | 54429224 | 15965 | -8.59 | 7.00E-15 | 3.26E-14 |
| chr1 | 3361996 | 3367495 | 5500 | -8.46 | 7.02E-15 | 3.27E-14 |
| chr5 | 36272796 | 36675756 | 402961 | 3.13 | 7.02E-15 | 3.27E-14 |
| chr16 | 53106706 | 53112507 | 5802 | -8.46 | 7.02E-15 | 3.27E-14 |
| chr9 | 138090960 | 138104557 | 13598 | -8.45 | 7.03E-15 | 3.27E-14 |
| chr7 | 91715600 | 91720411 | 4812 | -8.49 | 7.04E-15 | 3.28E-14 |
| chr1 | 223564572 | 223580339 | 15768 | -8.53 | 7.04E-15 | 3.28E-14 |
| chr1 | 84531439 | 84537708 | 6270 | -8.45 | 7.07E-15 | 3.29E-14 |
| chr16 | 28281351 | 28283877 | 2527 | -8.49 | 7.08E-15 | 3.29E-14 |
| chr10 | 48462374 | 48477165 | 14792 | -2.7 | 7.08E-15 | 3.29E-14 |
| chr7 | 151819844 | 151822721 | 2878 | -8.47 | 7.08E-15 | 3.30E-14 |
| chr4 | 84445500 | 84450243 | 4744 | -8.52 | 7.08E-15 | 3.30E-14 |
| chr2 | 28959477 | 28962377 | 2901 | -8.49 | 7.08E-15 | 3.30E-14 |
| chr5 | 177517141 | 177521725 | 4585 | -8.46 | 7.10E-15 | 3.31E-14 |
| chr4 | 22915640 | 22938829 | 23190 | -8.54 | 7.12E-15 | 3.31E-14 |
| chr6 | 110366357 | 110379547 | 13191 | -8.49 | 7.13E-15 | 3.32E-14 |
| chr11 | 115315896 | 115320139 | 4244 | -8.46 | 7.15E-15 | 3.33E-14 |
| chr19 | 38606820 | 38611754 | 4935 | -8.56 | 7.19E-15 | 3.35E-14 |
| chr19 | 16990563 | 16994871 | 4309 | -8.47 | 7.21E-15 | 3.35E-14 |
| chr18 | 8223526 | 8244075 | 20550 | -8.57 | 7.22E-15 | 3.36E-14 |
| chr1 | 92619640 | 92627010 | 7371 | -8.47 | 7.26E-15 | 3.37E-14 |
| chr2 | 156243999 | 156731618 | 487620 | 2.1 | 7.26E-15 | 3.37E-14 |
| chr6 | 4531253 | 4543108 | 11856 | -8.48 | 7.26E-15 | 3.38E-14 |
| chr10 | 126019408 | 126025521 | 6114 | -8.49 | 7.28E-15 | 3.38E-14 |
| chr12 | 90385693 | 90391951 | 6259 | -8.5 | 7.28E-15 | 3.38E-14 |
| chr20 | 24916459 | 24930142 | 13684 | -8.5 | 7.31E-15 | 3.39E-14 |
| chr5 | 60095927 | 60105253 | 9327 | -8.46 | 7.31E-15 | 3.39E-14 |
| chr11 | 117267576 | 117277607 | 10032 | -8.45 | 7.31E-15 | 3.40E-14 |
| chr11 | 125533494 | 125537452 | 3959 | -8.45 | 7.32E-15 | 3.40E-14 |
| chr15 | 101509255 | 101521158 | 11904 | -8.53 | 7.37E-15 | 3.42E-14 |
| chr3 | 164222062 | 164265639 | 43578 | -8.83 | 7.38E-15 | 3.43E-14 |
| chr4 | 153575507 | 153580586 | 5080 | -8.54 | 7.39E-15 | 3.43E-14 |
| chr20 | 55366857 | 55393518 | 26662 | -8.85 | 7.41E-15 | 3.44E-14 |
| chr11 | 119159827 | 119163406 | 3580 | -8.56 | 7.46E-15 | 3.46E-14 |
| chr13 | 49124481 | 49130984 | 6504 | -8.46 | 7.46E-15 | 3.46E-14 |

| | | | | | | |
|---|---|---|---|---|---|---|
| chr15 | 29678753 | 29685937 | 7185 | -8.44 | 7.47E-15 | 3.47E-14 |
| chr12 | 46268466 | 46280204 | 11739 | -8.46 | 7.47E-15 | 3.47E-14 |
| chr17 | 48415535 | 48419805 | 4271 | -8.46 | 7.47E-15 | 3.47E-14 |
| chr7 | 138842398 | 138845193 | 2796 | -8.5 | 7.49E-15 | 3.48E-14 |
| chr3 | 108027742 | 108029594 | 1853 | -6.21 | 7.49E-15 | 3.48E-14 |
| chr10 | 100104388 | 100137728 | 33341 | -8.77 | 7.51E-15 | 3.48E-14 |
| chrX | 68388319 | 68396214 | 7896 | -8.45 | 7.51E-15 | 3.48E-14 |
| chr2 | 38965982 | 38967938 | 1957 | -8.48 | 7.54E-15 | 3.50E-14 |
| chr15 | 67618826 | 67621518 | 2693 | -8.57 | 7.58E-15 | 3.52E-14 |
| chr6 | 165459485 | 165464934 | 5450 | -7.36 | 7.59E-15 | 3.52E-14 |
| chr1 | 46848235 | 46860594 | 12360 | -8.44 | 7.61E-15 | 3.53E-14 |
| chr6 | 10964882 | 10968353 | 3472 | -8.44 | 7.61E-15 | 3.53E-14 |
| chr6 | 71714039 | 71732497 | 18459 | -8.44 | 7.61E-15 | 3.53E-14 |
| chr4 | 7630666 | 7647694 | 17029 | -8.44 | 7.66E-15 | 3.55E-14 |
| chr8 | 9271173 | 9284264 | 13092 | -8.45 | 7.66E-15 | 3.55E-14 |
| chr2 | 169940722 | 169943517 | 2796 | -8.44 | 7.71E-15 | 3.57E-14 |
| chr5 | 85817727 | 85924094 | 106368 | 2.92 | 7.71E-15 | 3.57E-14 |
| chr7 | 1404198 | 1443510 | 39313 | -9.04 | 7.72E-15 | 3.57E-14 |
| chr5 | 90883859 | 90890977 | 7119 | -8.44 | 7.73E-15 | 3.58E-14 |
| chr12 | 16194578 | 16208065 | 13488 | -8.51 | 7.75E-15 | 3.59E-14 |
| chr3 | 69748612 | 69753254 | 4643 | -8.44 | 7.75E-15 | 3.59E-14 |
| chr2 | 162933949 | 162939971 | 6023 | -8.44 | 7.75E-15 | 3.59E-14 |
| chr15 | 68086852 | 68089542 | 2691 | -8.51 | 7.76E-15 | 3.59E-14 |
| chr20 | 19821403 | 19824291 | 2889 | -8.54 | 7.77E-15 | 3.60E-14 |
| chr5 | 142641462 | 142648370 | 6909 | -8.62 | 7.77E-15 | 3.60E-14 |
| chr18 | 56583471 | 56593880 | 10410 | -8.45 | 7.77E-15 | 3.60E-14 |
| chr17 | 64811725 | 64823645 | 11921 | -8.54 | 7.79E-15 | 3.60E-14 |
| chr5 | 72827638 | 72832602 | 4965 | -8.44 | 7.79E-15 | 3.60E-14 |
| chr5 | 137245321 | 137253849 | 8529 | -8.67 | 7.81E-15 | 3.61E-14 |
| chr17 | 63928651 | 63968121 | 39471 | -8.76 | 7.81E-15 | 3.61E-14 |
| chr1 | 217331582 | 217405059 | 73478 | 2.09 | 7.84E-15 | 3.62E-14 |
| chr6 | 167660295 | 167671616 | 11322 | -8.44 | 7.86E-15 | 3.64E-14 |
| chr18 | 10081531 | 10084848 | 3318 | -6.08 | 7.88E-15 | 3.64E-14 |
| chr1 | 221562288 | 221576952 | 14665 | -8.46 | 7.89E-15 | 3.65E-14 |
| chr12 | 121873226 | 121876160 | 2935 | -8.48 | 7.94E-15 | 3.67E-14 |
| chr14 | 31112619 | 31115677 | 3059 | -8.51 | 7.99E-15 | 3.69E-14 |
| chr12 | 50101722 | 50122367 | 20646 | -8.59 | 8.02E-15 | 3.71E-14 |
| chr3 | 177560489 | 177582019 | 21531 | -8.94 | 8.03E-15 | 3.71E-14 |
| chr3 | 17629960 | 17633952 | 3993 | -8.44 | 8.03E-15 | 3.71E-14 |
| chr14 | 37249132 | 37251909 | 2778 | -8.49 | 8.06E-15 | 3.72E-14 |
| chr3 | 159378855 | 159381968 | 3114 | -8.51 | 8.08E-15 | 3.73E-14 |
| chr10 | 35677468 | 35680952 | 3485 | -8.51 | 8.09E-15 | 3.74E-14 |
| chr9 | 139054628 | 139059195 | 4568 | -8.53 | 8.10E-15 | 3.74E-14 |
| chr3 | 164161668 | 164200853 | 39186 | -8.94 | 8.11E-15 | 3.74E-14 |
| chr14 | 31052237 | 31054913 | 2677 | -8.49 | 8.12E-15 | 3.75E-14 |
| chr4 | 114397079 | 114399921 | 2843 | -8.47 | 8.12E-15 | 3.75E-14 |
| chr7 | 111435144 | 111444513 | 9370 | -8.65 | 8.13E-15 | 3.75E-14 |
| chr16 | 46749121 | 46760411 | 11291 | -8.6 | 8.16E-15 | 3.77E-14 |
| chr6 | 109108538 | 109113569 | 5032 | -8.54 | 8.17E-15 | 3.77E-14 |
| chr12 | 56785544 | 56794277 | 8734 | -8.62 | 8.19E-15 | 3.78E-14 |
| chr11 | 7587346 | 7591393 | 4048 | -8.45 | 8.22E-15 | 3.79E-14 |
| chr12 | 104214356 | 104226539 | 12184 | -8.7 | 8.24E-15 | 3.80E-14 |
| chr11 | 67992162 | 68017669 | 25508 | -8.6 | 8.26E-15 | 3.81E-14 |
| chr7 | 111713315 | 111716587 | 3273 | -8.45 | 8.26E-15 | 3.81E-14 |
| chr4 | 93159232 | 93162785 | 3554 | -4.41 | 8.26E-15 | 3.81E-14 |
| chr5 | 10618824 | 10622515 | 3692 | -8.5 | 8.27E-15 | 3.81E-14 |
| chr2 | 101752509 | 101763963 | 11455 | -8.54 | 8.27E-15 | 3.81E-14 |
| chr3 | 122367131 | 122395224 | 28094 | -8.51 | 8.28E-15 | 3.81E-14 |
| chr3 | 152682144 | 152698707 | 16564 | -8.48 | 8.28E-15 | 3.81E-14 |
| chr1 | 33574131 | 33577166 | 3036 | -8.44 | 8.28E-15 | 3.81E-14 |
| chr8 | 49712421 | 49722812 | 10392 | -8.46 | 8.28E-15 | 3.82E-14 |

| | | | | | | |
|---|---|---|---|---|---|---|
| chr21 | 38907663 | 38916073 | 8411 | -8.45 | 8.32E-15 | 3.83E-14 |
| chr1 | 40821824 | 40831400 | 9577 | -8.59 | 8.32E-15 | 3.83E-14 |
| chr8 | 99756910 | 99759815 | 2906 | -8.45 | 8.37E-15 | 3.85E-14 |
| chr19 | 45786655 | 45800428 | 13774 | -8.48 | 8.45E-15 | 3.89E-14 |
| chr11 | 45058961 | 45062298 | 3338 | -8.45 | 8.47E-15 | 3.90E-14 |
| chr9 | 123118210 | 123143134 | 24925 | -8.61 | 8.48E-15 | 3.90E-14 |
| chr16 | 10559775 | 10591336 | 31562 | -8.55 | 8.48E-15 | 3.91E-14 |
| chr18 | 47830018 | 47832647 | 2630 | -8.49 | 8.52E-15 | 3.92E-14 |
| chr2 | 224354204 | 224379698 | 25495 | -8.49 | 8.54E-15 | 3.93E-14 |
| chrX | 85118515 | 85122882 | 4368 | -8.43 | 8.55E-15 | 3.93E-14 |
| chr1 | 35497772 | 35500668 | 2897 | -8.44 | 8.55E-15 | 3.93E-14 |
| chr6 | 76118645 | 76140616 | 21972 | -8.43 | 8.56E-15 | 3.94E-14 |
| chr4 | 155783748 | 155805240 | 21493 | -8.46 | 8.57E-15 | 3.94E-14 |
| chr3 | 125791773 | 125811098 | 19326 | -8.43 | 8.61E-15 | 3.96E-14 |
| chr12 | 34358152 | 34366403 | 8252 | -8.61 | 8.62E-15 | 3.96E-14 |
| chr17 | 53422842 | 53452090 | 29249 | -9.6 | 8.63E-15 | 3.97E-14 |
| chr13 | 90161633 | 90172247 | 10615 | -8.55 | 8.65E-15 | 3.98E-14 |
| chr17 | 77872388 | 77874538 | 2151 | -8.46 | 8.66E-15 | 3.98E-14 |
| chrX | 41824102 | 41837862 | 13761 | -7.23 | 8.71E-15 | 4.00E-14 |
| chr7 | 134691097 | 134710847 | 19751 | -8.46 | 8.72E-15 | 4.01E-14 |
| chr18 | 32504346 | 32521370 | 17025 | -8.91 | 8.73E-15 | 4.01E-14 |
| chr18 | 46425125 | 46432502 | 7378 | -8.47 | 8.75E-15 | 4.02E-14 |
| chr9 | 123396439 | 123404013 | 7575 | -8.44 | 8.77E-15 | 4.03E-14 |
| chr15 | 90264290 | 90268624 | 4335 | -8.43 | 8.78E-15 | 4.03E-14 |
| chr3 | 54906295 | 54924356 | 18062 | -8.45 | 8.83E-15 | 4.06E-14 |
| chr14 | 72922714 | 73018744 | 96031 | -1.3 | 8.84E-15 | 4.06E-14 |
| chr1 | 198549206 | 198561050 | 11845 | -8.45 | 8.85E-15 | 4.06E-14 |
| chr8 | 72746157 | 72749969 | 3813 | -8.43 | 8.85E-15 | 4.06E-14 |
| chr17 | 59030217 | 59047881 | 17665 | -9.01 | 8.87E-15 | 4.07E-14 |
| chr7 | 142863356 | 142878207 | 14852 | -8.94 | 8.89E-15 | 4.08E-14 |
| chr3 | 192462942 | 192476394 | 13453 | 8.52 | 8.90E-15 | 4.08E-14 |
| chr1 | 179289499 | 179299783 | 10285 | -8.43 | 8.90E-15 | 4.08E-14 |
| chr3 | 110005335 | 110019022 | 13688 | -8.72 | 8.93E-15 | 4.10E-14 |
| chr5 | 141913334 | 141916872 | 3539 | -8.48 | 8.95E-15 | 4.10E-14 |
| chr6 | 161973797 | 161987044 | 13248 | -4.33 | 8.98E-15 | 4.12E-14 |
| chr2 | 172994316 | 173026913 | 32598 | -8.75 | 8.99E-15 | 4.12E-14 |
| chr20 | 14619435 | 14623572 | 4138 | -8.56 | 9.00E-15 | 4.13E-14 |
| chr9 | 36989677 | 36992642 | 2966 | -8.48 | 9.01E-15 | 4.13E-14 |
| chr4 | 74183697 | 74192197 | 8501 | -6.09 | 9.02E-15 | 4.14E-14 |
| chrX | 103683896 | 103697487 | 13592 | -8.49 | 9.02E-15 | 4.14E-14 |
| chr6 | 15417164 | 15421435 | 4272 | -8.43 | 9.05E-15 | 4.15E-14 |
| chr2 | 151841835 | 151849624 | 7790 | -8.48 | 9.05E-15 | 4.15E-14 |
| chr5 | 82251464 | 82275791 | 24328 | -8.42 | 9.06E-15 | 4.15E-14 |
| chr1 | 37359996 | 37381498 | 21503 | -8.53 | 9.06E-15 | 4.15E-14 |
| chr11 | 16125258 | 16128850 | 3593 | -8.44 | 9.07E-15 | 4.16E-14 |
| chr3 | 190002007 | 190011749 | 9743 | -8.49 | 9.08E-15 | 4.16E-14 |
| chr5 | 65905935 | 65970812 | 64878 | 1.82 | 9.10E-15 | 4.17E-14 |
| chr6 | 137150597 | 137154046 | 3450 | -8.56 | 9.13E-15 | 4.18E-14 |
| chr8 | 9509981 | 9522449 | 12469 | -8.55 | 9.15E-15 | 4.19E-14 |
| chrX | 72807469 | 72811347 | 3879 | -8.44 | 9.16E-15 | 4.19E-14 |
| chr12 | 113583810 | 113586843 | 3034 | -8.45 | 9.16E-15 | 4.19E-14 |
| chr2 | 63564321 | 63572788 | 8468 | -8.62 | 9.17E-15 | 4.20E-14 |
| chr6 | 63347614 | 63365058 | 17445 | -8.46 | 9.20E-15 | 4.21E-14 |
| chrX | 62583861 | 62595544 | 11684 | -8.43 | 9.20E-15 | 4.21E-14 |
| chr11 | 72075114 | 72079383 | 4270 | -8.42 | 9.21E-15 | 4.22E-14 |
| chr11 | 1830466 | 1841539 | 11074 | -8.94 | 9.22E-15 | 4.22E-14 |
| chr10 | 45989374 | 45994245 | 4872 | -8.42 | 9.30E-15 | 4.25E-14 |
| chr17 | 56183919 | 56185925 | 2007 | -8.42 | 9.31E-15 | 4.26E-14 |
| chr17 | 45036463 | 45039812 | 3350 | -8.42 | 9.33E-15 | 4.27E-14 |
| chr12 | 64505361 | 64509319 | 3959 | -8.47 | 9.34E-15 | 4.27E-14 |
| chr1 | 86185747 | 86191128 | 5382 | -8.65 | 9.36E-15 | 4.28E-14 |

| | | | | | | |
|---|---|---|---|---|---|---|
| chr16 | 53622189 | 53631252 | 9064 | -8.69 | 9.39E-15 | 4.29E-14 |
| chr1 | 205819276 | 205826775 | 7500 | -8.47 | 9.41E-15 | 4.30E-14 |
| chr14 | 71300697 | 71304663 | 3967 | -8.47 | 9.43E-15 | 4.31E-14 |
| chr16 | 47924155 | 47944215 | 20061 | -8.46 | 9.45E-15 | 4.32E-14 |
| chr3 | 140850260 | 140863973 | 13714 | -3.09 | 9.47E-15 | 4.33E-14 |
| chr5 | 80014713 | 80017399 | 2687 | -8.42 | 9.50E-15 | 4.34E-14 |
| chr18 | 8131889 | 8159210 | 27322 | -8.5 | 9.51E-15 | 4.34E-14 |
| chr16 | 68138392 | 68145375 | 6984 | -8.6 | 9.51E-15 | 4.34E-14 |
| chr1 | 93571803 | 93574781 | 2979 | -8.42 | 9.53E-15 | 4.35E-14 |
| chr6 | 24287897 | 24316771 | 28875 | -8.84 | 9.54E-15 | 4.36E-14 |
| chr9 | 93109935 | 93124204 | 14270 | -8.44 | 9.56E-15 | 4.37E-14 |
| chr7 | 79903119 | 79924468 | 21350 | -9.12 | 9.57E-15 | 4.37E-14 |
| chr6 | 79761486 | 79763970 | 2485 | -8.43 | 9.58E-15 | 4.38E-14 |
| chr1 | 191720515 | 191745000 | 24486 | -8.42 | 9.59E-15 | 4.38E-14 |
| chr8 | 81929010 | 81945566 | 16557 | -8.43 | 9.60E-15 | 4.38E-14 |
| chr7 | 101572294 | 101582999 | 10706 | -8.47 | 9.60E-15 | 4.38E-14 |
| chr9 | 96092899 | 96103204 | 10306 | -8.78 | 9.62E-15 | 4.39E-14 |
| chr17 | 39474808 | 39477960 | 3153 | -8.58 | 9.64E-15 | 4.40E-14 |
| chr10 | 89559598 | 89570121 | 10524 | -8.42 | 9.64E-15 | 4.40E-14 |
| chr12 | 38773031 | 38800284 | 27254 | -8.41 | 9.64E-15 | 4.40E-14 |
| chrX | 9170378 | 9199099 | 28722 | -8.66 | 9.64E-15 | 4.40E-14 |
| chr2 | 240178309 | 240181217 | 2909 | -8.41 | 9.65E-15 | 4.40E-14 |
| chr3 | 149581023 | 149587375 | 6353 | -8.41 | 9.66E-15 | 4.40E-14 |
| chr3 | 41310523 | 41316397 | 5875 | -8.42 | 9.66E-15 | 4.40E-14 |
| chr3 | 120398377 | 120423359 | 24983 | -8.78 | 9.66E-15 | 4.41E-14 |
| chr7 | 102682986 | 102693986 | 11001 | -8.58 | 9.67E-15 | 4.41E-14 |
| chr3 | 27305248 | 27310411 | 5164 | -8.43 | 9.69E-15 | 4.42E-14 |
| chr20 | 56421875 | 56447927 | 26053 | -8.55 | 9.69E-15 | 4.42E-14 |
| chr14 | 54303785 | 54320542 | 16758 | -8.6 | 9.70E-15 | 4.42E-14 |
| chr1 | 176066627 | 176069880 | 3254 | -8.41 | 9.70E-15 | 4.42E-14 |
| chr12 | 106448126 | 106463897 | 15772 | -8.41 | 9.71E-15 | 4.42E-14 |
| chr3 | 69876131 | 69886082 | 9952 | -8.41 | 9.71E-15 | 4.42E-14 |
| chr18 | 45469692 | 45475578 | 5887 | -7.34 | 9.73E-15 | 4.43E-14 |
| chr8 | 83730595 | 83740250 | 9656 | -8.44 | 9.74E-15 | 4.43E-14 |
| chr2 | 70823966 | 70826612 | 2647 | -8.49 | 9.74E-15 | 4.43E-14 |
| chr12 | 102628757 | 102642543 | 13787 | -8.42 | 9.74E-15 | 4.43E-14 |
| chr16 | 53217984 | 53221994 | 4011 | -8.49 | 9.74E-15 | 4.43E-14 |
| chr8 | 108040891 | 108054692 | 13802 | -8.48 | 9.77E-15 | 4.45E-14 |
| chr3 | 157258750 | 157280146 | 21397 | -8.51 | 9.78E-15 | 4.45E-14 |
| chrX | 106380447 | 106391497 | 11051 | -8.61 | 9.79E-15 | 4.45E-14 |
| chr6 | 43697539 | 43700519 | 2981 | -8.44 | 9.83E-15 | 4.47E-14 |
| chr5 | 81332059 | 81343767 | 11709 | -8.47 | 9.89E-15 | 4.50E-14 |
| chr1 | 42585837 | 42592998 | 7162 | -8.42 | 9.90E-15 | 4.51E-14 |
| chr10 | 64284504 | 64301661 | 17158 | -8.41 | 9.94E-15 | 4.52E-14 |
| chr18 | 74603557 | 74613602 | 10046 | -8.45 | 9.95E-15 | 4.53E-14 |
| chr12 | 84246875 | 84257453 | 10579 | -8.42 | 9.96E-15 | 4.53E-14 |
| chr13 | 35526300 | 35530122 | 3823 | -8.42 | 9.99E-15 | 4.54E-14 |
| chr13 | 80837514 | 80861062 | 23549 | -8.42 | 1.00E-14 | 4.56E-14 |
| chr3 | 23720500 | 23748163 | 27664 | -8.56 | 1.00E-14 | 4.56E-14 |
| chr15 | 37460689 | 37488352 | 27664 | -4.28 | 1.01E-14 | 4.59E-14 |
| chr5 | 159813600 | 159816607 | 3008 | -8.54 | 1.01E-14 | 4.59E-14 |
| chr3 | 115632857 | 115670640 | 37784 | -8.9 | 1.01E-14 | 4.60E-14 |
| chr19 | 1693522 | 1705316 | 11795 | -8.41 | 1.01E-14 | 4.60E-14 |
| chr12 | 30913021 | 30938295 | 25275 | -9.01 | 1.01E-14 | 4.61E-14 |
| chr8 | 124595742 | 124598649 | 2908 | -8.41 | 1.02E-14 | 4.61E-14 |
| chr9 | 86090281 | 86097082 | 6802 | -8.6 | 1.02E-14 | 4.62E-14 |
| chr2 | 210279357 | 210285207 | 5851 | -8.41 | 1.02E-14 | 4.62E-14 |
| chr3 | 104906023 | 104921963 | 15941 | -8.41 | 1.02E-14 | 4.62E-14 |
| chr18 | 10698957 | 10715010 | 16054 | 8.51 | 1.02E-14 | 4.62E-14 |
| chr4 | 8093966 | 8097580 | 3615 | -8.42 | 1.02E-14 | 4.62E-14 |
| chr15 | 30321579 | 30330713 | 9135 | -8.49 | 1.02E-14 | 4.63E-14 |

| | | | | | | |
|---|---|---|---|---|---|---|
| chr3 | 131122276 | 131135231 | 12956 | -8.49 | 1.02E-14 | 4.63E-14 |
| chr3 | 193431258 | 193438944 | 7687 | -8.44 | 1.03E-14 | 4.66E-14 |
| chr6 | 157809889 | 157813370 | 3482 | -8.43 | 1.03E-14 | 4.66E-14 |
| chr7 | 155682465 | 155706294 | 23830 | -8.72 | 1.03E-14 | 4.69E-14 |
| chr9 | 116950778 | 116953525 | 2748 | -8.48 | 1.04E-14 | 4.71E-14 |
| chr14 | 94754092 | 94805122 | 51031 | -8.95 | 1.04E-14 | 4.71E-14 |
| chr4 | 24711385 | 24720861 | 9477 | -8.43 | 1.04E-14 | 4.71E-14 |
| chr18 | 10663263 | 10678045 | 14783 | 8.48 | 1.04E-14 | 4.73E-14 |
| chr2 | 23829281 | 23847911 | 18631 | -8.41 | 1.04E-14 | 4.73E-14 |
| chr4 | 63944446 | 63976390 | 31945 | -8.68 | 1.04E-14 | 4.73E-14 |
| chr2 | 135727172 | 135745235 | 18064 | -8.53 | 1.05E-14 | 4.74E-14 |
| chr5 | 134594209 | 134598582 | 4374 | -8.42 | 1.05E-14 | 4.75E-14 |
| chr6 | 149314153 | 149324276 | 10124 | -8.43 | 1.05E-14 | 4.75E-14 |
| chr1 | 203404063 | 203416433 | 12371 | -8.43 | 1.05E-14 | 4.77E-14 |
| chr9 | 102773063 | 102809736 | 36674 | -8.51 | 1.05E-14 | 4.78E-14 |
| chr12 | 78651784 | 78685442 | 33659 | -9.02 | 1.05E-14 | 4.78E-14 |
| chr6 | 129112431 | 129125320 | 12890 | -2.56 | 1.05E-14 | 4.78E-14 |
| chr1 | 186508050 | 186522362 | 14313 | -8.4 | 1.06E-14 | 4.79E-14 |
| chr3 | 119871671 | 119875656 | 3986 | -8.4 | 1.06E-14 | 4.79E-14 |
| chr7 | 55202551 | 55207054 | 4504 | -8.42 | 1.06E-14 | 4.79E-14 |
| chr9 | 132235770 | 132238569 | 2800 | -8.44 | 1.06E-14 | 4.80E-14 |
| chr12 | 14911795 | 14920509 | 8715 | -8.57 | 1.07E-14 | 4.83E-14 |
| chr8 | 63485370 | 63489067 | 3698 | -2.82 | 1.07E-14 | 4.83E-14 |
| chr18 | 67638986 | 67656986 | 18001 | -8.4 | 1.07E-14 | 4.84E-14 |
| chr6 | 144105203 | 144115026 | 9824 | -8.44 | 1.07E-14 | 4.84E-14 |
| chr6 | 113430092 | 113440821 | 10730 | -8.47 | 1.07E-14 | 4.85E-14 |
| chr22 | 40063792 | 40069659 | 5868 | -8.45 | 1.07E-14 | 4.86E-14 |
| chr13 | 111672185 | 111675785 | 3601 | -8.46 | 1.08E-14 | 4.86E-14 |
| chr10 | 101892920 | 101896999 | 4080 | -8.4 | 1.08E-14 | 4.87E-14 |
| chr7 | 11760816 | 11788632 | 27817 | -8.56 | 1.08E-14 | 4.88E-14 |
| chr7 | 148780855 | 148784659 | 3805 | -8.41 | 1.09E-14 | 4.91E-14 |
| chr1 | 12584261 | 12599927 | 15667 | -8.59 | 1.09E-14 | 4.91E-14 |
| chr22 | 48965440 | 48969184 | 3745 | -8.4 | 1.09E-14 | 4.92E-14 |
| chr20 | 39381970 | 39393587 | 11618 | 8.65 | 1.09E-14 | 4.92E-14 |
| chr20 | 31136428 | 31138514 | 2087 | -8.41 | 1.09E-14 | 4.95E-14 |
| chr5 | 60642600 | 60656403 | 13804 | -8.44 | 1.10E-14 | 4.97E-14 |
| chr9 | 90844464 | 90847995 | 3532 | -8.4 | 1.10E-14 | 4.98E-14 |
| chr22 | 48676488 | 48680910 | 4423 | -8.41 | 1.11E-14 | 5.00E-14 |
| chr18 | 77619217 | 77623297 | 4081 | -3.64 | 1.11E-14 | 5.00E-14 |
| chr3 | 105045610 | 105053066 | 7457 | -8.4 | 1.11E-14 | 5.01E-14 |
| chr4 | 26444720 | 26465857 | 21138 | -8.55 | 1.11E-14 | 5.03E-14 |
| chr1 | 179736669 | 179743531 | 6863 | -8.41 | 1.11E-14 | 5.03E-14 |
| chr5 | 82171196 | 82177266 | 6071 | -8.66 | 1.11E-14 | 5.03E-14 |
| chr12 | 10527671 | 10538880 | 11210 | -8.59 | 1.12E-14 | 5.04E-14 |
| chr6 | 149505414 | 149521940 | 16527 | -8.63 | 1.12E-14 | 5.04E-14 |
| chr22 | 46948348 | 46953962 | 5615 | -8.44 | 1.12E-14 | 5.04E-14 |
| chr6 | 116261868 | 116274678 | 12811 | -8.41 | 1.12E-14 | 5.05E-14 |
| chr1 | 160757923 | 160779690 | 21768 | -8.62 | 1.12E-14 | 5.05E-14 |
| chr5 | 53307457 | 53311309 | 3853 | -8.41 | 1.12E-14 | 5.05E-14 |
| chr21 | 34738000 | 34743010 | 5011 | -8.48 | 1.12E-14 | 5.06E-14 |
| chr8 | 100065487 | 100075612 | 10126 | -8.43 | 1.12E-14 | 5.06E-14 |
| chr2 | 24062790 | 24068775 | 5986 | -8.42 | 1.12E-14 | 5.07E-14 |
| chr6 | 57609403 | 57686385 | 76983 | 1.48 | 1.12E-14 | 5.07E-14 |
| chr14 | 96676541 | 96679534 | 2994 | -8.46 | 1.13E-14 | 5.08E-14 |
| chr3 | 131982420 | 132007801 | 25382 | -8.72 | 1.13E-14 | 5.09E-14 |
| chr5 | 108703365 | 108705389 | 2025 | -8.41 | 1.13E-14 | 5.10E-14 |
| chr4 | 75707113 | 75714836 | 7724 | -8.58 | 1.14E-14 | 5.12E-14 |
| chr12 | 118875672 | 119414359 | 538688 | 4.52 | 1.14E-14 | 5.15E-14 |
| chr12 | 10552144 | 10558561 | 6418 | -8.55 | 1.14E-14 | 5.15E-14 |
| chr7 | 44449343 | 44452150 | 2808 | -8.41 | 1.14E-14 | 5.16E-14 |
| chr13 | 104130744 | 104146946 | 16203 | -8.53 | 1.15E-14 | 5.16E-14 |

| | | | | | | |
|---|---|---|---|---|---|---|
| chrX | 28314824 | 28337181 | 22358 | -8.54 | 1.15E-14 | 5.16E-14 |
| chr2 | 207991879 | 207995779 | 3901 | -8.5 | 1.15E-14 | 5.17E-14 |
| chrX | 122281068 | 122308251 | 27184 | -8.69 | 1.15E-14 | 5.18E-14 |
| chr20 | 3123400 | 3126158 | 2759 | -8.39 | 1.15E-14 | 5.19E-14 |
| chr6 | 13970046 | 13975070 | 5025 | -8.4 | 1.15E-14 | 5.19E-14 |
| chr1 | 146644978 | 146651588 | 6611 | -8.4 | 1.15E-14 | 5.19E-14 |
| chr7 | 659160 | 662351 | 3192 | -8.42 | 1.15E-14 | 5.20E-14 |
| chr2 | 200051796 | 200066565 | 14770 | 8.49 | 1.16E-14 | 5.20E-14 |
| chr12 | 70976571 | 70981090 | 4520 | -8.43 | 1.16E-14 | 5.21E-14 |
| chr1 | 231578390 | 231595050 | 16661 | -8.56 | 1.16E-14 | 5.22E-14 |
| chr1 | 168491326 | 168494983 | 3658 | -8.4 | 1.16E-14 | 5.22E-14 |
| chr12 | 10478126 | 10481770 | 3645 | -8.43 | 1.16E-14 | 5.24E-14 |
| chr22 | 19990190 | 19993231 | 3042 | -8.39 | 1.17E-14 | 5.25E-14 |
| chr7 | 67521778 | 67526064 | 4287 | -8.43 | 1.17E-14 | 5.26E-14 |
| chr1 | 186489689 | 186495336 | 5648 | -8.39 | 1.17E-14 | 5.26E-14 |
| chr6 | 8456625 | 8462977 | 6353 | -8.46 | 1.17E-14 | 5.26E-14 |
| chr2 | 189789430 | 189795798 | 6369 | -8.41 | 1.17E-14 | 5.26E-14 |
| chr10 | 97337766 | 97357468 | 19703 | -8.66 | 1.17E-14 | 5.27E-14 |
| chr12 | 87597330 | 87603919 | 6590 | -8.41 | 1.17E-14 | 5.28E-14 |
| chr2 | 11417958 | 11425531 | 7574 | -8.49 | 1.18E-14 | 5.28E-14 |
| chrX | 3656657 | 3670058 | 13402 | -8.48 | 1.18E-14 | 5.28E-14 |
| chr4 | 90143389 | 90158511 | 15123 | -8.77 | 1.18E-14 | 5.29E-14 |
| chrX | 105770633 | 105779526 | 8894 | 8.49 | 1.18E-14 | 5.31E-14 |
| chr6 | 78070969 | 78077360 | 6392 | -8.54 | 1.18E-14 | 5.31E-14 |
| chr10 | 76377222 | 76382133 | 4912 | -8.39 | 1.18E-14 | 5.31E-14 |
| chr14 | 22871036 | 22889899 | 18864 | -8.86 | 1.19E-14 | 5.33E-14 |
| chr4 | 127755032 | 127773128 | 18097 | -8.5 | 1.19E-14 | 5.33E-14 |
| chr10 | 62311518 | 62337867 | 26350 | -8.41 | 1.19E-14 | 5.34E-14 |
| chr4 | 120142202 | 120146930 | 4729 | -8.51 | 1.19E-14 | 5.34E-14 |
| chr17 | 34436741 | 34458251 | 21511 | -1.98 | 1.19E-14 | 5.35E-14 |
| chr19 | 44150090 | 44165204 | 15115 | -8.43 | 1.19E-14 | 5.36E-14 |
| chr8 | 10558981 | 10567603 | 8623 | -8.53 | 1.19E-14 | 5.36E-14 |
| chr20 | 47422492 | 47426348 | 3857 | -8.39 | 1.20E-14 | 5.39E-14 |
| chr12 | 10654692 | 10661602 | 6911 | -8.4 | 1.20E-14 | 5.40E-14 |
| chrX | 107134226 | 107137461 | 3236 | -8.39 | 1.21E-14 | 5.42E-14 |
| chr5 | 16858918 | 16876435 | 17518 | -8.42 | 1.21E-14 | 5.43E-14 |
| chr12 | 32344712 | 32359127 | 14416 | -8.41 | 1.21E-14 | 5.43E-14 |
| chr11 | 91998647 | 92011308 | 12662 | -8.39 | 1.21E-14 | 5.44E-14 |
| chr1 | 85194773 | 85201408 | 6636 | -3.9 | 1.21E-14 | 5.44E-14 |
| chr5 | 60442063 | 60460389 | 18327 | -8.4 | 1.22E-14 | 5.46E-14 |
| chr5 | 122116468 | 122124407 | 7940 | -8.41 | 1.22E-14 | 5.46E-14 |
| chr9 | 37867257 | 37878584 | 11328 | -8.5 | 1.22E-14 | 5.46E-14 |
| chr2 | 60950821 | 60966434 | 15614 | -8.54 | 1.22E-14 | 5.47E-14 |
| chr12 | 95890945 | 95893885 | 2941 | -8.39 | 1.22E-14 | 5.47E-14 |
| chr14 | 58882117 | 58884143 | 2027 | -8.44 | 1.22E-14 | 5.49E-14 |
| chr18 | 72214457 | 72218522 | 4066 | -8.42 | 1.22E-14 | 5.49E-14 |
| chr12 | 88997683 | 89006097 | 8415 | -8.57 | 1.23E-14 | 5.50E-14 |
| chr14 | 69240090 | 69243623 | 3534 | -8.46 | 1.23E-14 | 5.52E-14 |
| chr1 | 245789420 | 245815723 | 26304 | -1.9 | 1.23E-14 | 5.53E-14 |
| chr1 | 20188224 | 20192506 | 4283 | -8.38 | 1.24E-14 | 5.54E-14 |
| chr2 | 181030354 | 181036724 | 6371 | -8.42 | 1.24E-14 | 5.54E-14 |
| chr1 | 88323816 | 88335077 | 11262 | -8.4 | 1.24E-14 | 5.54E-14 |
| chrX | 73362222 | 73394701 | 32480 | -8.81 | 1.24E-14 | 5.55E-14 |
| chr8 | 49471329 | 49476018 | 4690 | -8.57 | 1.24E-14 | 5.55E-14 |
| chr7 | 82810436 | 82815986 | 5551 | -8.38 | 1.24E-14 | 5.55E-14 |
| chr13 | 63843191 | 63862073 | 18883 | -3.9 | 1.24E-14 | 5.56E-14 |
| chr12 | 18913386 | 18945357 | 31972 | -8.52 | 1.24E-14 | 5.56E-14 |
| chr6 | 20315872 | 20335907 | 20036 | -8.52 | 1.24E-14 | 5.56E-14 |
| chrX | 83279691 | 83285309 | 5619 | 3.55 | 1.24E-14 | 5.57E-14 |
| chr7 | 127262613 | 127265616 | 3004 | -8.38 | 1.25E-14 | 5.58E-14 |
| chr6 | 138233928 | 138239504 | 5577 | -3.61 | 1.25E-14 | 5.59E-14 |

| | | | | | | |
|---|---|---|---|---|---|---|
| chr5 | 135403223 | 135405535 | 2313 | -8.39 | 1.25E-14 | 5.60E-14 |
| chr6 | 2709286 | 2723638 | 14353 | -8.44 | 1.25E-14 | 5.60E-14 |
| chr9 | 84094003 | 84102726 | 8724 | -8.55 | 1.25E-14 | 5.60E-14 |
| chr8 | 8396008 | 8400270 | 4263 | -8.39 | 1.26E-14 | 5.62E-14 |
| chr8 | 10214954 | 10228561 | 13608 | -8.57 | 1.26E-14 | 5.62E-14 |
| chr5 | 106679959 | 106699606 | 19648 | -8.38 | 1.26E-14 | 5.64E-14 |
| chr4 | 183873263 | 183889261 | 15999 | -8.46 | 1.26E-14 | 5.65E-14 |
| chr8 | 17404847 | 17417522 | 12676 | -8.48 | 1.27E-14 | 5.66E-14 |
| chr11 | 64355117 | 64357696 | 2580 | -8.38 | 1.27E-14 | 5.66E-14 |
| chr8 | 124848271 | 124860724 | 12454 | -8.51 | 1.27E-14 | 5.68E-14 |
| chr4 | 3719106 | 3721526 | 2421 | -8.41 | 1.27E-14 | 5.69E-14 |
| chr7 | 3361988 | 3382091 | 20104 | -8.65 | 1.27E-14 | 5.69E-14 |
| chr6 | 39340598 | 39352210 | 11613 | -8.44 | 1.28E-14 | 5.70E-14 |
| chr6 | 106202334 | 106226561 | 24228 | -8.83 | 1.28E-14 | 5.70E-14 |
| chr6 | 17064137 | 17091270 | 27134 | -2.06 | 1.28E-14 | 5.71E-14 |
| chr1 | 229558699 | 229561230 | 2532 | -8.38 | 1.28E-14 | 5.71E-14 |
| chr14 | 81054467 | 81062931 | 8465 | -8.4 | 1.28E-14 | 5.71E-14 |
| chr19 | 10051171 | 10054606 | 3436 | -8.45 | 1.28E-14 | 5.71E-14 |
| chr3 | 87856707 | 87863498 | 6792 | -8.38 | 1.28E-14 | 5.71E-14 |
| chr10 | 33536644 | 33565038 | 28395 | -8.61 | 1.28E-14 | 5.72E-14 |
| chr5 | 172625163 | 172649998 | 24836 | -8.4 | 1.28E-14 | 5.73E-14 |
| chr9 | 90037346 | 90041540 | 4195 | -8.43 | 1.29E-14 | 5.76E-14 |
| chr12 | 104025475 | 104034799 | 9325 | -8.38 | 1.29E-14 | 5.77E-14 |
| chr15 | 49632445 | 49646638 | 14194 | -8.76 | 1.29E-14 | 5.78E-14 |
| chr21 | 35330726 | 35334452 | 3727 | -8.38 | 1.30E-14 | 5.78E-14 |
| chr16 | 47521553 | 47526054 | 4502 | -8.4 | 1.30E-14 | 5.79E-14 |
| chr9 | 71642763 | 71646250 | 3488 | -8.44 | 1.30E-14 | 5.80E-14 |
| chrX | 42757076 | 42850452 | 93377 | 1.3 | 1.30E-14 | 5.80E-14 |
| chr2 | 219214292 | 219218708 | 4417 | -8.68 | 1.30E-14 | 5.80E-14 |
| chr15 | 76856970 | 76875810 | 18841 | -8.51 | 1.30E-14 | 5.81E-14 |
| chr2 | 153117506 | 153157242 | 39737 | -9.02 | 1.30E-14 | 5.81E-14 |
| chr6 | 158329915 | 158340392 | 10478 | -8.39 | 1.31E-14 | 5.82E-14 |
| chr8 | 39033992 | 39057116 | 23125 | -8.39 | 1.31E-14 | 5.82E-14 |
| chr14 | 67490855 | 67504175 | 13321 | -8.38 | 1.31E-14 | 5.82E-14 |
| chrX | 83338058 | 83347303 | 9246 | -8.38 | 1.31E-14 | 5.82E-14 |
| chr2 | 204927303 | 204959907 | 32605 | -1.49 | 1.31E-14 | 5.85E-14 |
| chr11 | 106910429 | 107143458 | 233030 | 3.44 | 1.31E-14 | 5.85E-14 |
| chr12 | 10947699 | 10957271 | 9573 | 1.85 | 1.32E-14 | 5.86E-14 |
| chr5 | 39489548 | 39493115 | 3568 | -8.42 | 1.32E-14 | 5.87E-14 |
| chrX | 150803752 | 150817908 | 14157 | 8.46 | 1.32E-14 | 5.88E-14 |
| chr3 | 8829268 | 8844140 | 14873 | -8.44 | 1.33E-14 | 5.90E-14 |
| chr16 | 56607605 | 56610530 | 2926 | -8.44 | 1.33E-14 | 5.91E-14 |
| chr3 | 122623634 | 122626029 | 2396 | -8.38 | 1.33E-14 | 5.92E-14 |
| chr2 | 99865735 | 99899107 | 33373 | -8.51 | 1.33E-14 | 5.93E-14 |
| chr11 | 125377622 | 125399791 | 22170 | -8.66 | 1.33E-14 | 5.93E-14 |
| chr6 | 3914710 | 3919506 | 4797 | -8.44 | 1.33E-14 | 5.93E-14 |
| chr9 | 116802242 | 116808714 | 6473 | -8.42 | 1.33E-14 | 5.93E-14 |
| chrX | 69864952 | 69880851 | 15900 | -8.42 | 1.34E-14 | 5.95E-14 |
| chr7 | 50855154 | 50867347 | 12194 | -8.49 | 1.34E-14 | 5.96E-14 |
| chr3 | 64871752 | 64897914 | 26163 | 3.41 | 1.34E-14 | 5.98E-14 |
| chr22 | 36192510 | 36211699 | 19190 | -8.58 | 1.35E-14 | 6.00E-14 |
| chr10 | 31267736 | 31285445 | 17710 | -8.69 | 1.35E-14 | 6.00E-14 |
| chr1 | 108153429 | 108159055 | 5627 | -8.38 | 1.35E-14 | 6.01E-14 |
| chr3 | 44875856 | 44880656 | 4801 | -8.39 | 1.35E-14 | 6.02E-14 |
| chr16 | 87221855 | 87225566 | 3712 | -8.41 | 1.36E-14 | 6.03E-14 |
| chr9 | 123587308 | 123589812 | 2505 | -8.53 | 1.36E-14 | 6.03E-14 |
| chr8 | 104367932 | 104382503 | 14572 | -8.56 | 1.36E-14 | 6.03E-14 |
| chr8 | 126136349 | 126138743 | 2395 | -8.39 | 1.36E-14 | 6.03E-14 |
| chr18 | 37432118 | 37501352 | 69235 | 2.1 | 1.36E-14 | 6.03E-14 |
| chr20 | 20493482 | 20496827 | 3346 | -8.39 | 1.36E-14 | 6.04E-14 |
| chr12 | 21236264 | 21275802 | 39539 | -8.44 | 1.36E-14 | 6.05E-14 |

| | | | | | | |
|---|---|---|---|---|---|---|
| chr1 | 114855959 | 114872531 | 16573 | -8.44 | 1.36E-14 | 6.06E-14 |
| chr5 | 77977399 | 78005770 | 28372 | -8.57 | 1.37E-14 | 6.07E-14 |
| chr4 | 68523986 | 68542200 | 18215 | -8.4 | 1.37E-14 | 6.09E-14 |
| chr3 | 187641411 | 187645974 | 4564 | -8.41 | 1.37E-14 | 6.10E-14 |
| chr7 | 21297171 | 21307939 | 10769 | -5.95 | 1.38E-14 | 6.11E-14 |
| chr10 | 79300410 | 79304978 | 4569 | -8.42 | 1.38E-14 | 6.13E-14 |
| chr2 | 120162868 | 120180816 | 17949 | -8.94 | 1.38E-14 | 6.14E-14 |
| chr11 | 11837832 | 11858799 | 20968 | -8.4 | 1.38E-14 | 6.14E-14 |
| chr21 | 42523867 | 42532001 | 8135 | -8.39 | 1.39E-14 | 6.15E-14 |
| chr5 | 82642106 | 82657102 | 14997 | -8.45 | 1.39E-14 | 6.16E-14 |
| chr9 | 16932298 | 16940995 | 8698 | -8.46 | 1.39E-14 | 6.16E-14 |
| chr11 | 63286615 | 63292463 | 5849 | -8.4 | 1.39E-14 | 6.17E-14 |
| chr15 | 71779041 | 71799386 | 20346 | -8.94 | 1.39E-14 | 6.17E-14 |
| chr7 | 76915041 | 76932563 | 17523 | -8.38 | 1.39E-14 | 6.17E-14 |
| chr8 | 17145584 | 17168239 | 22656 | -8.37 | 1.39E-14 | 6.18E-14 |
| chr6 | 17464878 | 17468562 | 3685 | -8.45 | 1.40E-14 | 6.19E-14 |
| chr5 | 78367798 | 78370884 | 3087 | -8.37 | 1.40E-14 | 6.22E-14 |
| chr20 | 45655828 | 45658571 | 2744 | -8.37 | 1.41E-14 | 6.27E-14 |
| chr9 | 80390725 | 80394269 | 3545 | -8.37 | 1.41E-14 | 6.27E-14 |
| chr2 | 202325082 | 202327430 | 2349 | -8.38 | 1.42E-14 | 6.27E-14 |
| chr18 | 9252891 | 9265240 | 12350 | -8.53 | 1.42E-14 | 6.28E-14 |
| chr17 | 58324967 | 58327037 | 2071 | -8.38 | 1.43E-14 | 6.33E-14 |
| chr4 | 7179951 | 7183291 | 3341 | -8.39 | 1.43E-14 | 6.33E-14 |
| chr2 | 238869884 | 238872910 | 3027 | -8.37 | 1.43E-14 | 6.35E-14 |
| chr12 | 55163951 | 55182553 | 18603 | 2.59 | 1.44E-14 | 6.37E-14 |
| chr12 | 32846497 | 32849284 | 2788 | -8.38 | 1.44E-14 | 6.37E-14 |
| chr1 | 72758394 | 72765624 | 7231 | 8.44 | 1.44E-14 | 6.37E-14 |
| chr4 | 125758098 | 125765034 | 6937 | -8.37 | 1.44E-14 | 6.38E-14 |
| chr8 | 22006474 | 22009072 | 2599 | -8.41 | 1.44E-14 | 6.39E-14 |
| chr12 | 88637565 | 88648739 | 11175 | -8.4 | 1.45E-14 | 6.43E-14 |
| chr2 | 130564778 | 130614922 | 50145 | -1.94 | 1.45E-14 | 6.44E-14 |
| chr4 | 42272519 | 42277086 | 4568 | -8.37 | 1.46E-14 | 6.44E-14 |
| chr2 | 127766058 | 127782031 | 15974 | -8.64 | 1.46E-14 | 6.45E-14 |
| chr1 | 236420943 | 236426519 | 5577 | -8.44 | 1.46E-14 | 6.45E-14 |
| chr21 | 40094276 | 40101343 | 7068 | -3.07 | 1.46E-14 | 6.46E-14 |
| chr4 | 95765735 | 95768727 | 2993 | -8.4 | 1.46E-14 | 6.46E-14 |
| chr4 | 115161675 | 115179048 | 17374 | -8.51 | 1.46E-14 | 6.46E-14 |
| chr1 | 173095973 | 173122985 | 27013 | -8.43 | 1.47E-14 | 6.48E-14 |
| chr5 | 56526268 | 56529125 | 2858 | -8.38 | 1.47E-14 | 6.48E-14 |
| chr12 | 63069115 | 63086468 | 17354 | -8.61 | 1.47E-14 | 6.51E-14 |
| chr8 | 61570484 | 61574386 | 3903 | -8.4 | 1.48E-14 | 6.54E-14 |
| chrX | 69962487 | 69975660 | 13174 | -8.39 | 1.48E-14 | 6.54E-14 |
| chr1 | 236602018 | 236609355 | 7338 | -8.37 | 1.48E-14 | 6.54E-14 |
| chr15 | 68804998 | 68806929 | 1932 | -8.38 | 1.48E-14 | 6.54E-14 |
| chr1 | 204879335 | 204888351 | 9017 | -8.42 | 1.48E-14 | 6.55E-14 |
| chr9 | 21593757 | 21596775 | 3019 | -8.4 | 1.48E-14 | 6.55E-14 |
| chr18 | 2838865 | 2841099 | 2235 | -5.96 | 1.49E-14 | 6.57E-14 |
| chr18 | 18660361 | 18665400 | 5040 | -8.37 | 1.49E-14 | 6.57E-14 |
| chr1 | 24456114 | 24458604 | 2491 | -8.39 | 1.49E-14 | 6.58E-14 |
| chr11 | 14714852 | 14729890 | 15039 | -8.36 | 1.49E-14 | 6.59E-14 |
| chr16 | 14316579 | 14321698 | 5120 | -8.52 | 1.50E-14 | 6.60E-14 |
| chr6 | 1189652 | 1197070 | 7419 | -2.43 | 1.50E-14 | 6.60E-14 |
| chr20 | 3841829 | 3843574 | 1746 | -8.43 | 1.50E-14 | 6.60E-14 |
| chr2 | 15779754 | 15782571 | 2818 | -8.37 | 1.50E-14 | 6.61E-14 |
| chr8 | 54800246 | 54804238 | 3993 | -8.37 | 1.50E-14 | 6.62E-14 |
| chr14 | 33053170 | 33058162 | 4993 | -8.4 | 1.50E-14 | 6.62E-14 |
| chr20 | 31755605 | 31772119 | 16515 | -8.73 | 1.51E-14 | 6.65E-14 |
| chr1 | 63894633 | 63897546 | 2914 | -8.38 | 1.51E-14 | 6.67E-14 |
| chr15 | 48997737 | 49010089 | 12353 | -8.36 | 1.51E-14 | 6.68E-14 |
| chr3 | 18055977 | 18063580 | 7604 | -8.4 | 1.52E-14 | 6.68E-14 |
| chr2 | 85630082 | 85632194 | 2113 | -8.37 | 1.52E-14 | 6.69E-14 |

| | | | | | | |
|---|---|---|---|---|---|---|
| chr7 | 32840655 | 32843472 | 2818 | -8.4 | 1.52E-14 | 6.71E-14 |
| chr3 | 31593567 | 31603217 | 9651 | -8.44 | 1.53E-14 | 6.73E-14 |
| chr3 | 158438902 | 158463311 | 24410 | -8.44 | 1.53E-14 | 6.73E-14 |
| chr6 | 111779068 | 111782497 | 3430 | -8.45 | 1.53E-14 | 6.73E-14 |
| chr9 | 19951631 | 20129846 | 178216 | 2.14 | 1.53E-14 | 6.75E-14 |
| chr9 | 84566744 | 84572936 | 6193 | -2.86 | 1.53E-14 | 6.75E-14 |
| chr3 | 170293903 | 170302811 | 8909 | 8.48 | 1.53E-14 | 6.76E-14 |
| chr3 | 24071764 | 24075946 | 4183 | -8.36 | 1.53E-14 | 6.76E-14 |
| chrX | 54594202 | 54597297 | 3096 | -8.46 | 1.53E-14 | 6.76E-14 |
| chr6 | 55509309 | 55529025 | 19717 | -8.52 | 1.53E-14 | 6.76E-14 |
| chr1 | 213112532 | 213116355 | 3824 | -8.46 | 1.54E-14 | 6.76E-14 |
| chr7 | 20560861 | 20577306 | 16446 | -8.42 | 1.54E-14 | 6.77E-14 |
| chr2 | 38699044 | 38709170 | 10127 | -8.8 | 1.54E-14 | 6.78E-14 |
| chr14 | 72066295 | 72071923 | 5629 | -8.38 | 1.55E-14 | 6.82E-14 |
| chr2 | 181290946 | 181297537 | 6592 | -8.39 | 1.56E-14 | 6.87E-14 |
| chr4 | 57987814 | 57995484 | 7671 | -8.41 | 1.56E-14 | 6.88E-14 |
| chr22 | 25300575 | 25305062 | 4488 | -8.36 | 1.56E-14 | 6.88E-14 |
| chr4 | 1131683 | 1136540 | 4858 | -8.44 | 1.56E-14 | 6.88E-14 |
| chr12 | 805499 | 809076 | 3578 | -8.44 | 1.56E-14 | 6.88E-14 |
| chr16 | 51863098 | 51887263 | 24166 | -8.44 | 1.56E-14 | 6.88E-14 |
| chr8 | 83975595 | 85097805 | 1122211 | 1.33 | 1.57E-14 | 6.92E-14 |
| chr2 | 26053940 | 26056853 | 2914 | -8.49 | 1.58E-14 | 6.96E-14 |
| chr15 | 58850245 | 58876772 | 26528 | -8.76 | 1.58E-14 | 6.96E-14 |
| chr7 | 68021402 | 68055410 | 34009 | -8.56 | 1.58E-14 | 6.97E-14 |
| chr5 | 108725896 | 108773558 | 47663 | -8.49 | 1.59E-14 | 6.98E-14 |
| chr8 | 22339992 | 22342225 | 2234 | -8.37 | 1.59E-14 | 6.99E-14 |
| chr9 | 90006719 | 90010144 | 3426 | -8.47 | 1.59E-14 | 6.99E-14 |
| chr2 | 214979405 | 214984238 | 4834 | -8.35 | 1.59E-14 | 6.99E-14 |
| chr5 | 121535847 | 121651548 | 115702 | 1.83 | 1.60E-14 | 7.04E-14 |
| chr6 | 36121742 | 36128550 | 6809 | -8.38 | 1.60E-14 | 7.05E-14 |
| chr4 | 38164660 | 38174564 | 9905 | -8.48 | 1.61E-14 | 7.06E-14 |
| chr12 | 19062027 | 19066893 | 4867 | -8.35 | 1.62E-14 | 7.10E-14 |
| chr22 | 39370926 | 39381349 | 10424 | -8.49 | 1.62E-14 | 7.11E-14 |
| chr1 | 246506639 | 246509819 | 3181 | -8.37 | 1.62E-14 | 7.11E-14 |
| chr10 | 27083759 | 27088686 | 4928 | -8.38 | 1.62E-14 | 7.11E-14 |
| chr17 | 27064040 | 27069113 | 5074 | -8.52 | 1.62E-14 | 7.13E-14 |
| chrX | 40557597 | 40560115 | 2519 | -8.36 | 1.63E-14 | 7.15E-14 |
| chr4 | 5662547 | 5671445 | 8899 | -8.45 | 1.63E-14 | 7.16E-14 |
| chr5 | 73298326 | 73302863 | 4538 | -8.36 | 1.64E-14 | 7.19E-14 |
| chr1 | 236484863 | 236489311 | 4449 | -8.51 | 1.64E-14 | 7.19E-14 |
| chr13 | 24396506 | 24400318 | 3813 | -8.35 | 1.64E-14 | 7.19E-14 |
| chr12 | 105732645 | 105736075 | 3431 | -8.56 | 1.64E-14 | 7.20E-14 |
| chr1 | 207300673 | 207304023 | 3351 | -8.41 | 1.64E-14 | 7.21E-14 |
| chr1 | 211930615 | 211933352 | 2738 | -8.49 | 1.64E-14 | 7.22E-14 |
| chr2 | 190962258 | 190969047 | 6790 | -8.68 | 1.65E-14 | 7.25E-14 |
| chrX | 20981461 | 20988592 | 7132 | -6.61 | 1.65E-14 | 7.25E-14 |
| chr7 | 131691996 | 131701046 | 9051 | -8.55 | 1.66E-14 | 7.27E-14 |
| chr8 | 96114263 | 96118125 | 3863 | -8.46 | 1.66E-14 | 7.27E-14 |
| chr14 | 100823785 | 100826093 | 2309 | -8.52 | 1.67E-14 | 7.33E-14 |
| chr6 | 22181393 | 22187520 | 6128 | -8.36 | 1.67E-14 | 7.33E-14 |
| chr6 | 8996101 | 9005168 | 9068 | -8.36 | 1.67E-14 | 7.34E-14 |
| chr15 | 49499847 | 49520782 | 20936 | -8.52 | 1.67E-14 | 7.34E-14 |
| chr9 | 119731602 | 119736830 | 5229 | -8.39 | 1.68E-14 | 7.36E-14 |
| chr5 | 74414956 | 74427196 | 12241 | -8.6 | 1.68E-14 | 7.36E-14 |
| chr4 | 1737843 | 1740911 | 3069 | -8.35 | 1.68E-14 | 7.37E-14 |
| chr14 | 23920291 | 23922191 | 1901 | -8.43 | 1.68E-14 | 7.37E-14 |
| chr4 | 6245142 | 6249030 | 3889 | -8.42 | 1.68E-14 | 7.38E-14 |
| chr2 | 203768561 | 203771526 | 2966 | -8.45 | 1.69E-14 | 7.38E-14 |
| chrX | 57105975 | 57118768 | 12794 | -8.35 | 1.69E-14 | 7.39E-14 |
| chr20 | 4932375 | 4940576 | 8202 | -8.38 | 1.69E-14 | 7.40E-14 |
| chr18 | 59714764 | 59728398 | 13635 | -8.37 | 1.69E-14 | 7.41E-14 |

| | | | | | | |
|---|---|---|---|---|---|---|
| chr16 | 78586273 | 78612665 | 26393 | -8.35 | 1.70E-14 | 7.45E-14 |
| chr8 | 140997197 | 141010277 | 13081 | -8.37 | 1.70E-14 | 7.45E-14 |
| chr12 | 28466900 | 28483935 | 17036 | -8.35 | 1.70E-14 | 7.45E-14 |
| chr16 | 4975761 | 4978800 | 3040 | -8.35 | 1.71E-14 | 7.47E-14 |
| chr14 | 43527716 | 43563426 | 35711 | -8.81 | 1.71E-14 | 7.47E-14 |
| chr1 | 47979083 | 48008432 | 29350 | -8.7 | 1.71E-14 | 7.48E-14 |
| chr5 | 72100542 | 72109075 | 8534 | -8.5 | 1.71E-14 | 7.48E-14 |
| chr10 | 82781160 | 82865089 | 83930 | 1.79 | 1.71E-14 | 7.49E-14 |
| chrX | 130520668 | 130524181 | 3514 | -8.36 | 1.71E-14 | 7.49E-14 |
| chr3 | 194689522 | 194703780 | 14259 | -8.38 | 1.71E-14 | 7.49E-14 |
| chr17 | 15954493 | 15958940 | 4448 | -8.37 | 1.72E-14 | 7.52E-14 |
| chr16 | 3649153 | 3651296 | 2144 | -8.36 | 1.72E-14 | 7.53E-14 |
| chr6 | 27077964 | 27083331 | 5368 | -8.36 | 1.73E-14 | 7.54E-14 |
| chr6 | 165763091 | 165777846 | 14756 | -8.44 | 1.73E-14 | 7.57E-14 |
| chr15 | 53923327 | 53925850 | 2524 | -8.35 | 1.73E-14 | 7.57E-14 |
| chr2 | 37911594 | 37950547 | 38954 | -8.35 | 1.73E-14 | 7.57E-14 |
| chr11 | 12078316 | 12106192 | 27877 | -8.67 | 1.73E-14 | 7.57E-14 |
| chr1 | 61309774 | 61318759 | 8986 | -8.34 | 1.73E-14 | 7.57E-14 |
| chr2 | 69832479 | 69838461 | 5983 | -8.35 | 1.74E-14 | 7.59E-14 |
| chr3 | 148240885 | 148246387 | 5503 | -8.59 | 1.74E-14 | 7.60E-14 |
| chr18 | 11648679 | 11662849 | 14171 | -8.38 | 1.74E-14 | 7.60E-14 |
| chr9 | 19072225 | 19074744 | 2520 | -8.59 | 1.74E-14 | 7.62E-14 |
| chr12 | 53045389 | 53074831 | 29443 | 8.81 | 1.75E-14 | 7.63E-14 |
| chr2 | 42956134 | 42963200 | 7067 | -8.34 | 1.75E-14 | 7.64E-14 |
| chr13 | 78223761 | 78230901 | 7141 | -3.32 | 1.75E-14 | 7.65E-14 |
| chr1 | 202962413 | 202981013 | 18601 | -8.35 | 1.75E-14 | 7.66E-14 |
| chr4 | 168572379 | 168585205 | 12827 | -8.36 | 1.76E-14 | 7.69E-14 |
| chr15 | 31499913 | 31504755 | 4843 | -8.52 | 1.76E-14 | 7.69E-14 |
| chr2 | 236081226 | 236096436 | 15211 | -8.43 | 1.77E-14 | 7.71E-14 |
| chr1 | 91420239 | 91425811 | 5573 | -8.36 | 1.77E-14 | 7.71E-14 |
| chr19 | 35812475 | 35814178 | 1704 | -8.39 | 1.77E-14 | 7.73E-14 |
| chr17 | 47615120 | 47619698 | 4579 | -8.34 | 1.78E-14 | 7.76E-14 |
| chr5 | 38518558 | 38532705 | 14148 | -8.47 | 1.78E-14 | 7.77E-14 |
| chr11 | 22071805 | 22104342 | 32538 | -8.43 | 1.78E-14 | 7.77E-14 |
| chr2 | 238134374 | 238158779 | 24406 | -8.37 | 1.79E-14 | 7.80E-14 |
| chr4 | 180813719 | 180825645 | 11927 | -8.36 | 1.79E-14 | 7.82E-14 |
| chr16 | 5022428 | 5039005 | 16578 | -8.54 | 1.79E-14 | 7.82E-14 |
| chr21 | 37439779 | 37442095 | 2317 | -8.38 | 1.79E-14 | 7.82E-14 |
| chr22 | 21093869 | 21096299 | 2431 | -8.35 | 1.80E-14 | 7.83E-14 |
| chr7 | 129455494 | 129458714 | 3221 | -8.34 | 1.80E-14 | 7.83E-14 |
| chr3 | 100186250 | 100209128 | 22879 | -8.53 | 1.80E-14 | 7.84E-14 |
| chr1 | 93354137 | 93386294 | 32158 | -8.53 | 1.80E-14 | 7.86E-14 |
| chr12 | 94476773 | 94481817 | 5045 | -8.35 | 1.81E-14 | 7.87E-14 |
| chr2 | 23659385 | 23665826 | 6442 | -8.39 | 1.81E-14 | 7.87E-14 |
| chr9 | 86792722 | 86818295 | 25574 | -8.44 | 1.82E-14 | 7.91E-14 |
| chr14 | 105464461 | 105467574 | 3114 | -8.37 | 1.82E-14 | 7.91E-14 |
| chr6 | 107183052 | 107192312 | 9261 | -8.36 | 1.82E-14 | 7.92E-14 |
| chr8 | 1741553 | 1744472 | 2920 | -8.39 | 1.82E-14 | 7.92E-14 |
| chrX | 73555475 | 73560626 | 5152 | -8.35 | 1.82E-14 | 7.92E-14 |
| chr2 | 53177105 | 53202290 | 25186 | -8.34 | 1.84E-14 | 8.01E-14 |
| chr2 | 64982282 | 64984405 | 2124 | -8.34 | 1.85E-14 | 8.04E-14 |
| chr2 | 164019977 | 164023635 | 3659 | -8.34 | 1.85E-14 | 8.04E-14 |
| chr4 | 25093892 | 25104359 | 10468 | -8.53 | 1.85E-14 | 8.04E-14 |
| chr11 | 19088303 | 19091815 | 3513 | -8.34 | 1.85E-14 | 8.05E-14 |
| chr16 | 50086098 | 50092379 | 6282 | -8.34 | 1.85E-14 | 8.05E-14 |
| chr4 | 103343754 | 103357889 | 14136 | -8.37 | 1.85E-14 | 8.05E-14 |
| chr1 | 213456064 | 213473573 | 17510 | -8.42 | 1.85E-14 | 8.06E-14 |
| chr12 | 19686498 | 19690631 | 4134 | -8.34 | 1.85E-14 | 8.07E-14 |
| chr16 | 87263732 | 87267461 | 3730 | -8.34 | 1.87E-14 | 8.13E-14 |
| chr14 | 66173682 | 66187244 | 13563 | -8.37 | 1.87E-14 | 8.13E-14 |
| chr2 | 137089887 | 137109119 | 19233 | -8.34 | 1.87E-14 | 8.14E-14 |

| | | | | | | |
|---|---|---|---|---|---|---|
| chr3 | 50835782 | 50840419 | 4638 | -8.5 | 1.87E-14 | 8.14E-14 |
| chr14 | 32248438 | 32253301 | 4864 | -8.45 | 1.88E-14 | 8.17E-14 |
| chr2 | 113019757 | 113026657 | 6901 | -8.36 | 1.88E-14 | 8.18E-14 |
| chr3 | 188219737 | 188223003 | 3267 | -8.33 | 1.88E-14 | 8.18E-14 |
| chr15 | 59719692 | 59722524 | 2833 | -8.34 | 1.89E-14 | 8.22E-14 |
| chrX | 107534235 | 107537144 | 2910 | -8.41 | 1.89E-14 | 8.23E-14 |
| chr18 | 23923686 | 23931439 | 7754 | -8.43 | 1.90E-14 | 8.26E-14 |
| chr9 | 86266380 | 86270512 | 4133 | -8.61 | 1.90E-14 | 8.27E-14 |
| chr3 | 142486551 | 142490323 | 3773 | -8.5 | 1.91E-14 | 8.31E-14 |
| chr2 | 180789617 | 180796071 | 6455 | -8.33 | 1.92E-14 | 8.34E-14 |
| chr14 | 76631481 | 76634316 | 2836 | -8.45 | 1.92E-14 | 8.34E-14 |
| chr4 | 9959439 | 9988123 | 28685 | -8.82 | 1.92E-14 | 8.35E-14 |
| chr8 | 67072368 | 67078575 | 6208 | -8.37 | 1.92E-14 | 8.35E-14 |
| chr3 | 125346498 | 125357323 | 10826 | -8.35 | 1.92E-14 | 8.35E-14 |
| chr4 | 151831834 | 151835187 | 3354 | -8.63 | 1.93E-14 | 8.36E-14 |
| chr4 | 40151113 | 40154462 | 3350 | -8.47 | 1.93E-14 | 8.37E-14 |
| chr10 | 48338886 | 48353916 | 15031 | -8.34 | 1.94E-14 | 8.40E-14 |
| chr12 | 16165962 | 16175319 | 9358 | -8.46 | 1.95E-14 | 8.46E-14 |
| chr7 | 44210555 | 44223383 | 12829 | -8.33 | 1.95E-14 | 8.48E-14 |
| chr7 | 7636400 | 7640106 | 3707 | -8.4 | 1.96E-14 | 8.48E-14 |
| chr2 | 46599304 | 46600998 | 1695 | -8.49 | 1.96E-14 | 8.49E-14 |
| chr7 | 27563948 | 27582414 | 18467 | -8.43 | 1.96E-14 | 8.49E-14 |
| chr3 | 99478346 | 99490312 | 11967 | 8.53 | 1.96E-14 | 8.49E-14 |
| chr2 | 174161871 | 174167205 | 5335 | -8.4 | 1.96E-14 | 8.50E-14 |
| chr4 | 14956957 | 14976498 | 19542 | -8.44 | 1.96E-14 | 8.52E-14 |
| chr8 | 29710848 | 29714937 | 4090 | -8.37 | 1.97E-14 | 8.54E-14 |
| chr16 | 48097365 | 48118359 | 20995 | -8.61 | 1.97E-14 | 8.56E-14 |
| chr20 | 35920361 | 35930849 | 10489 | -8.34 | 1.98E-14 | 8.60E-14 |
| chr12 | 131801235 | 131808512 | 7278 | -8.42 | 1.99E-14 | 8.65E-14 |
| chr5 | 166308186 | 166327864 | 19679 | -8.47 | 2.00E-14 | 8.65E-14 |
| chr6 | 55839854 | 55860008 | 20155 | -8.53 | 2.00E-14 | 8.65E-14 |
| chr1 | 172582810 | 172587449 | 4640 | -8.34 | 2.00E-14 | 8.67E-14 |
| chr7 | 133713225 | 133722541 | 9317 | -8.37 | 2.01E-14 | 8.72E-14 |
| chr1 | 7237222 | 7242688 | 5467 | -8.36 | 2.01E-14 | 8.72E-14 |
| chr6 | 79452405 | 79459439 | 7035 | -8.39 | 2.01E-14 | 8.72E-14 |
| chr2 | 171096079 | 171153699 | 57621 | -1.99 | 2.02E-14 | 8.75E-14 |
| chr7 | 55076225 | 55083284 | 7060 | -8.47 | 2.02E-14 | 8.77E-14 |
| chr13 | 109586065 | 109591421 | 5357 | -8.34 | 2.03E-14 | 8.80E-14 |
| chr6 | 97712401 | 97715371 | 2971 | -8.38 | 2.04E-14 | 8.83E-14 |
| chr2 | 74524691 | 74527276 | 2586 | -8.32 | 2.05E-14 | 8.87E-14 |
| chr3 | 35992547 | 36001095 | 8549 | -8.37 | 2.05E-14 | 8.87E-14 |
| chr14 | 95610547 | 95613602 | 3056 | -8.48 | 2.06E-14 | 8.92E-14 |
| chr12 | 9504603 | 9508453 | 3851 | -8.33 | 2.06E-14 | 8.93E-14 |
| chr9 | 138345145 | 138347876 | 2732 | -8.38 | 2.07E-14 | 8.95E-14 |
| chrX | 38527759 | 38567786 | 40028 | 7.37 | 2.07E-14 | 8.96E-14 |
| chr18 | 8087363 | 8104194 | 16832 | -8.32 | 2.08E-14 | 8.99E-14 |
| chr1 | 240018064 | 240028391 | 10328 | -8.58 | 2.08E-14 | 8.99E-14 |
| chr12 | 93585341 | 93588427 | 3087 | -8.34 | 2.08E-14 | 9.00E-14 |
| chr2 | 109098979 | 109101420 | 2442 | -8.39 | 2.09E-14 | 9.04E-14 |
| chrX | 45819798 | 45828024 | 8227 | -8.45 | 2.09E-14 | 9.05E-14 |
| chr1 | 113714444 | 113728676 | 14233 | -8.41 | 2.10E-14 | 9.07E-14 |
| chr22 | 27640010 | 27652916 | 12907 | -8.36 | 2.12E-14 | 9.14E-14 |
| chr7 | 20246202 | 20250355 | 4154 | -8.35 | 2.12E-14 | 9.17E-14 |
| chr10 | 115774067 | 115776867 | 2801 | -8.33 | 2.13E-14 | 9.19E-14 |
| chr3 | 146128988 | 146147445 | 18458 | -8.46 | 2.14E-14 | 9.23E-14 |
| chr10 | 54152539 | 54163845 | 11307 | -8.34 | 2.14E-14 | 9.24E-14 |
| chr4 | 99286342 | 99291058 | 4717 | -8.33 | 2.14E-14 | 9.25E-14 |
| chr7 | 4282861 | 4478961 | 196101 | -3.26 | 2.14E-14 | 9.26E-14 |
| chr16 | 20075762 | 20080935 | 5174 | -6.68 | 2.14E-14 | 9.26E-14 |
| chr7 | 43328313 | 43331646 | 3334 | -8.4 | 2.15E-14 | 9.28E-14 |
| chr4 | 124718957 | 124721282 | 2326 | -4.13 | 2.15E-14 | 9.28E-14 |

| | | | | | | |
|---|---|---|---|---|---|---|
| chr5 | 91504212 | 91521487 | 17276 | -6.49 | 2.16E-14 | 9.31E-14 |
| chr3 | 71895323 | 71921001 | 25679 | -2.18 | 2.16E-14 | 9.32E-14 |
| chr1 | 150076853 | 150088610 | 11758 | -8.32 | 2.17E-14 | 9.35E-14 |
| chr1 | 64631699 | 64635300 | 3602 | -8.33 | 2.17E-14 | 9.37E-14 |
| chr5 | 155689936 | 155698051 | 8116 | -8.5 | 2.17E-14 | 9.38E-14 |
| chr15 | 43247994 | 43266709 | 18716 | -8.32 | 2.18E-14 | 9.39E-14 |
| chr12 | 10494578 | 10511211 | 16634 | -8.33 | 2.18E-14 | 9.41E-14 |
| chr19 | 11327036 | 11333056 | 6021 | -8.32 | 2.19E-14 | 9.45E-14 |
| chrX | 106204616 | 106216212 | 11597 | -8.32 | 2.19E-14 | 9.46E-14 |
| chrX | 109736651 | 109745811 | 9161 | 8.39 | 2.20E-14 | 9.47E-14 |
| chr11 | 127242861 | 127638826 | 395966 | 4.79 | 2.20E-14 | 9.48E-14 |
| chr17 | 15145196 | 15146926 | 1731 | -8.32 | 2.20E-14 | 9.48E-14 |
| chr4 | 5203975 | 5225639 | 21665 | -8.32 | 2.20E-14 | 9.48E-14 |
| chr4 | 67028821 | 67041483 | 12663 | -8.37 | 2.20E-14 | 9.50E-14 |
| chr17 | 70191993 | 70272823 | 80831 | -9.83 | 2.20E-14 | 9.50E-14 |
| chr9 | 137336733 | 137354076 | 17344 | -8.39 | 2.21E-14 | 9.51E-14 |
| chr10 | 24614822 | 24619166 | 4345 | -8.39 | 2.21E-14 | 9.53E-14 |
| chr15 | 57542499 | 57544226 | 1728 | -8.39 | 2.22E-14 | 9.55E-14 |
| chr14 | 57291518 | 57522262 | 230745 | 1.27 | 2.22E-14 | 9.57E-14 |
| chr4 | 187130049 | 187147238 | 17190 | -8.35 | 2.23E-14 | 9.59E-14 |
| chr3 | 136541178 | 136547052 | 5875 | -8.33 | 2.23E-14 | 9.60E-14 |
| chr4 | 164104860 | 164120610 | 15751 | -6.9 | 2.24E-14 | 9.64E-14 |
| chr4 | 96265361 | 96280591 | 15231 | -8.35 | 2.25E-14 | 9.68E-14 |
| chr12 | 131333073 | 131347021 | 13949 | -8.48 | 2.26E-14 | 9.72E-14 |
| chr15 | 79794409 | 79803591 | 9183 | 8.44 | 2.26E-14 | 9.72E-14 |
| chr13 | 63578438 | 63592379 | 13942 | -8.41 | 2.26E-14 | 9.72E-14 |
| chr17 | 35933637 | 35935569 | 1933 | -8.41 | 2.26E-14 | 9.73E-14 |
| chr10 | 77797201 | 77800872 | 3672 | -8.36 | 2.26E-14 | 9.75E-14 |
| chr2 | 57429769 | 57685086 | 255318 | 1.76 | 2.27E-14 | 9.78E-14 |
| chr11 | 94158142 | 94162950 | 4809 | -8.31 | 2.28E-14 | 9.83E-14 |
| chr2 | 69890033 | 69905848 | 15816 | -8.58 | 2.28E-14 | 9.83E-14 |
| chr2 | 169300212 | 169311924 | 11713 | -8.59 | 2.29E-14 | 9.86E-14 |
| chr5 | 37772276 | 37776450 | 4175 | -8.33 | 2.29E-14 | 9.86E-14 |
| chr9 | 83972751 | 83983178 | 10428 | -8.51 | 2.29E-14 | 9.86E-14 |
| chr10 | 79951843 | 79954249 | 2407 | -8.35 | 2.30E-14 | 9.91E-14 |
| chrX | 66899783 | 66903367 | 3585 | -8.37 | 2.30E-14 | 9.91E-14 |
| chr6 | 17358261 | 17371516 | 13256 | -8.36 | 2.30E-14 | 9.91E-14 |
| chr14 | 39371259 | 39382335 | 11077 | 8.59 | 2.31E-14 | 9.91E-14 |
| chr20 | 7973292 | 7977307 | 4016 | -8.43 | 2.31E-14 | 9.92E-14 |
| chr2 | 89297293 | 89303816 | 6524 | 8.38 | 2.31E-14 | 9.93E-14 |
| chr11 | 34462152 | 34464705 | 2554 | -8.39 | 2.31E-14 | 9.94E-14 |
| chr9 | 90491697 | 90495372 | 3676 | -8.32 | 2.32E-14 | 9.95E-14 |
| chr13 | 46365371 | 46393042 | 27672 | -8.41 | 2.32E-14 | 9.96E-14 |
| chrX | 140924773 | 140950621 | 25849 | -8.33 | 2.32E-14 | 9.98E-14 |
| chr14 | 102134516 | 102140544 | 6029 | -6.51 | 2.33E-14 | 1.00E-13 |
| chr19 | 30448550 | 30452189 | 3640 | -8.31 | 2.33E-14 | 1.00E-13 |
| chr9 | 115438631 | 115441685 | 3055 | -8.33 | 2.33E-14 | 1.00E-13 |
| chr5 | 115446782 | 115456675 | 9894 | -8.34 | 2.34E-14 | 1.00E-13 |
| chr4 | 185832950 | 185870274 | 37325 | -3.61 | 2.34E-14 | 1.00E-13 |
| chr18 | 29459825 | 29462551 | 2727 | -8.32 | 2.35E-14 | 1.01E-13 |
| chr15 | 99608317 | 99627462 | 19146 | -8.5 | 2.35E-14 | 1.01E-13 |
| chr3 | 120456591 | 120459631 | 3041 | -8.32 | 2.36E-14 | 1.01E-13 |
| chr4 | 5715682 | 5717847 | 2166 | -8.31 | 2.36E-14 | 1.01E-13 |
| chr2 | 207274957 | 207284082 | 9126 | -6.27 | 2.36E-14 | 1.01E-13 |
| chr14 | 54472643 | 54487458 | 14816 | -8.8 | 2.36E-14 | 1.01E-13 |
| chr15 | 91276013 | 91278303 | 2291 | -8.37 | 2.37E-14 | 1.02E-13 |
| chr16 | 47572999 | 47586681 | 13683 | -8.37 | 2.37E-14 | 1.02E-13 |
| chr3 | 192304934 | 192315010 | 10077 | 8.38 | 2.37E-14 | 1.02E-13 |
| chr4 | 24619205 | 24623745 | 4541 | -8.32 | 2.37E-14 | 1.02E-13 |
| chr8 | 131600007 | 131629200 | 29194 | -8.33 | 2.38E-14 | 1.02E-13 |
| chr3 | 138767398 | 138774686 | 7289 | -8.45 | 2.38E-14 | 1.02E-13 |

| | | | | | | |
|---|---|---|---|---|---|---|
| chr8 | 18082618 | 18369723 | 287106 | -1.13 | 2.38E-14 | 1.02E-13 |
| chr3 | 112468061 | 112492347 | 24287 | -8.64 | 2.38E-14 | 1.02E-13 |
| chr2 | 173136895 | 173142909 | 6015 | -8.46 | 2.39E-14 | 1.03E-13 |
| chr3 | 126501782 | 126512917 | 11136 | -8.31 | 2.39E-14 | 1.03E-13 |
| chr7 | 26922651 | 26926189 | 3539 | -8.37 | 2.39E-14 | 1.03E-13 |
| chr9 | 37067183 | 37079569 | 12387 | -8.31 | 2.40E-14 | 1.03E-13 |
| chr2 | 162159060 | 162163367 | 4308 | -8.31 | 2.40E-14 | 1.03E-13 |
| chr4 | 39372506 | 39383104 | 10599 | -8.31 | 2.40E-14 | 1.03E-13 |
| chr11 | 2020338 | 2033532 | 13195 | -8.51 | 2.41E-14 | 1.03E-13 |
| chr1 | 221235668 | 221248387 | 12720 | -8.32 | 2.41E-14 | 1.03E-13 |
| chr11 | 45084405 | 45088800 | 4396 | -8.32 | 2.42E-14 | 1.03E-13 |
| chr13 | 83645869 | 83670066 | 24198 | -8.63 | 2.42E-14 | 1.04E-13 |
| chr5 | 129924739 | 129931889 | 7151 | -8.36 | 2.42E-14 | 1.04E-13 |
| chr1 | 28004690 | 28010750 | 6061 | -8.33 | 2.43E-14 | 1.04E-13 |
| chr12 | 107475016 | 107491726 | 16711 | -8.38 | 2.43E-14 | 1.04E-13 |
| chr17 | 62368079 | 62370431 | 2353 | -8.38 | 2.43E-14 | 1.04E-13 |
| chr1 | 162445708 | 162451145 | 5438 | -8.35 | 2.44E-14 | 1.04E-13 |
| chrX | 43925892 | 43940723 | 14832 | -8.33 | 2.44E-14 | 1.04E-13 |
| chr1 | 79521532 | 79536241 | 14710 | -8.61 | 2.45E-14 | 1.05E-13 |
| chr9 | 14575851 | 14609851 | 34001 | -8.51 | 2.45E-14 | 1.05E-13 |
| chr5 | 68045433 | 68059497 | 14065 | -3.08 | 2.45E-14 | 1.05E-13 |
| chr7 | 42902880 | 42909909 | 7030 | -8.42 | 2.46E-14 | 1.05E-13 |
| chr14 | 105605164 | 105608016 | 2853 | -8.32 | 2.46E-14 | 1.05E-13 |
| chr18 | 56534155 | 56536413 | 2259 | -8.32 | 2.46E-14 | 1.05E-13 |
| chr12 | 64371595 | 64374454 | 2860 | -8.31 | 2.47E-14 | 1.06E-13 |
| chr3 | 52134889 | 52138343 | 3455 | -8.32 | 2.47E-14 | 1.06E-13 |
| chr2 | 131632450 | 131637810 | 5361 | -8.31 | 2.48E-14 | 1.06E-13 |
| chr3 | 126144115 | 126148635 | 4521 | -8.31 | 2.48E-14 | 1.06E-13 |
| chr6 | 77821821 | 77839480 | 17660 | -8.31 | 2.49E-14 | 1.06E-13 |
| chr12 | 60808889 | 60817728 | 8840 | -8.46 | 2.49E-14 | 1.06E-13 |
| chr2 | 128833864 | 128842048 | 8185 | -8.35 | 2.50E-14 | 1.07E-13 |
| chr2 | 89042298 | 89064288 | 21991 | -8.67 | 2.50E-14 | 1.07E-13 |
| chr13 | 19502380 | 19728731 | 226352 | 0.98 | 2.51E-14 | 1.07E-13 |
| chr14 | 36362968 | 36366245 | 3278 | -8.31 | 2.51E-14 | 1.07E-13 |
| chr7 | 17771831 | 17803190 | 31360 | -8.56 | 2.52E-14 | 1.08E-13 |
| chr2 | 36535979 | 36556199 | 20221 | -8.32 | 2.52E-14 | 1.08E-13 |
| chr16 | 10362916 | 10373590 | 10675 | -8.42 | 2.53E-14 | 1.08E-13 |
| chr5 | 52256411 | 52274803 | 18393 | -8.31 | 2.54E-14 | 1.08E-13 |
| chr2 | 204667541 | 204671847 | 4307 | -8.3 | 2.54E-14 | 1.08E-13 |
| chr6 | 138257170 | 138261189 | 4020 | -8.42 | 2.55E-14 | 1.09E-13 |
| chr22 | 27152752 | 27176422 | 23671 | -8.43 | 2.55E-14 | 1.09E-13 |
| chr12 | 19028759 | 19045100 | 16342 | -8.39 | 2.55E-14 | 1.09E-13 |
| chr2 | 53218237 | 53235764 | 17528 | -8.39 | 2.56E-14 | 1.09E-13 |
| chr12 | 57151557 | 57153694 | 2138 | -8.33 | 2.57E-14 | 1.09E-13 |
| chr12 | 17586730 | 17597247 | 10518 | -8.37 | 2.57E-14 | 1.10E-13 |
| chr4 | 2027431 | 2033567 | 6137 | -8.37 | 2.57E-14 | 1.10E-13 |
| chr5 | 79717843 | 79724101 | 6259 | -8.3 | 2.57E-14 | 1.10E-13 |
| chr2 | 173716194 | 173720486 | 4293 | -8.39 | 2.58E-14 | 1.10E-13 |
| chr2 | 235850862 | 235853630 | 2769 | -8.41 | 2.59E-14 | 1.10E-13 |
| chr17 | 63727027 | 63736624 | 9598 | -8.72 | 2.59E-14 | 1.10E-13 |
| chr20 | 61493628 | 61502076 | 8449 | -8.35 | 2.59E-14 | 1.11E-13 |
| chr3 | 157830141 | 157846715 | 16575 | -8.4 | 2.60E-14 | 1.11E-13 |
| chr10 | 25195076 | 25212020 | 16945 | -8.46 | 2.60E-14 | 1.11E-13 |
| chr12 | 92017391 | 92027506 | 10116 | -8.3 | 2.61E-14 | 1.11E-13 |
| chr12 | 117526978 | 117528840 | 1863 | -8.33 | 2.62E-14 | 1.11E-13 |
| chr2 | 74834234 | 74841108 | 6875 | -2.22 | 2.62E-14 | 1.12E-13 |
| chr8 | 53510998 | 53517522 | 6525 | -8.38 | 2.62E-14 | 1.12E-13 |
| chr2 | 55302462 | 55308887 | 6426 | -8.38 | 2.62E-14 | 1.12E-13 |
| chr9 | 84195799 | 84222192 | 26394 | -8.33 | 2.62E-14 | 1.12E-13 |
| chr3 | 124949829 | 124955529 | 5701 | -8.46 | 2.63E-14 | 1.12E-13 |
| chr5 | 179869394 | 179890600 | 21207 | -8.68 | 2.64E-14 | 1.12E-13 |

| | | | | | | |
|---|---|---|---|---|---|---|
| chr5 | 88045644 | 88050627 | 4984 | -8.31 | 2.64E-14 | 1.12E-13 |
| chr13 | 66806482 | 66820925 | 14444 | -8.53 | 2.64E-14 | 1.12E-13 |
| chr7 | 23445947 | 23449284 | 3338 | -8.3 | 2.64E-14 | 1.13E-13 |
| chr4 | 62333752 | 62365000 | 31249 | -8.31 | 2.65E-14 | 1.13E-13 |
| chr8 | 119836886 | 119854057 | 17172 | -8.29 | 2.65E-14 | 1.13E-13 |
| chr7 | 112467345 | 112471587 | 4243 | -8.3 | 2.65E-14 | 1.13E-13 |
| chr5 | 92566360 | 92570555 | 4196 | -8.45 | 2.66E-14 | 1.13E-13 |
| chr4 | 186762847 | 186779966 | 17120 | -8.31 | 2.66E-14 | 1.13E-13 |
| chr8 | 30895139 | 30899128 | 3990 | -8.3 | 2.66E-14 | 1.13E-13 |
| chr15 | 73768132 | 73775657 | 7526 | -8.39 | 2.67E-14 | 1.13E-13 |
| chr11 | 94619562 | 94639896 | 20335 | -8.33 | 2.67E-14 | 1.14E-13 |
| chr6 | 82800924 | 82820578 | 19655 | -8.47 | 2.67E-14 | 1.14E-13 |
| chr12 | 4826025 | 4953754 | 127730 | 2.48 | 2.68E-14 | 1.14E-13 |
| chr13 | 65286265 | 65316594 | 30330 | -8.72 | 2.68E-14 | 1.14E-13 |
| chr21 | 33826174 | 33835026 | 8853 | -8.3 | 2.68E-14 | 1.14E-13 |
| chr2 | 181134226 | 181138574 | 4349 | -8.33 | 2.68E-14 | 1.14E-13 |
| chr4 | 102627089 | 102641441 | 14353 | -8.38 | 2.68E-14 | 1.14E-13 |
| chr2 | 218061266 | 218076595 | 15330 | -8.5 | 2.69E-14 | 1.14E-13 |
| chr2 | 62269523 | 62273399 | 3877 | -8.29 | 2.69E-14 | 1.14E-13 |
| chr7 | 29615478 | 29630065 | 14588 | -8.37 | 2.69E-14 | 1.14E-13 |
| chr12 | 93233941 | 93236146 | 2206 | -8.48 | 2.69E-14 | 1.14E-13 |
| chr14 | 102079807 | 102085394 | 5588 | -8.29 | 2.71E-14 | 1.15E-13 |
| chr20 | 19724951 | 19727993 | 3043 | -8.31 | 2.71E-14 | 1.15E-13 |
| chr15 | 99731450 | 99747415 | 15966 | -8.34 | 2.71E-14 | 1.15E-13 |
| chr3 | 13449177 | 13452712 | 3536 | -8.29 | 2.73E-14 | 1.16E-13 |
| chr20 | 43922763 | 43924994 | 2232 | -8.29 | 2.73E-14 | 1.16E-13 |
| chr1 | 40488355 | 40492639 | 4285 | -8.41 | 2.73E-14 | 1.16E-13 |
| chr10 | 105673560 | 105675876 | 2317 | -8.32 | 2.74E-14 | 1.16E-13 |
| chr12 | 20928120 | 20943704 | 15585 | -8.63 | 2.75E-14 | 1.17E-13 |
| chr15 | 64548255 | 64550063 | 1809 | -8.3 | 2.75E-14 | 1.17E-13 |
| chr16 | 84603111 | 84605839 | 2729 | -8.3 | 2.75E-14 | 1.17E-13 |
| chr5 | 158919998 | 158937980 | 17983 | -8.3 | 2.76E-14 | 1.17E-13 |
| chr3 | 116923689 | 116932333 | 8645 | -8.47 | 2.76E-14 | 1.17E-13 |
| chr17 | 18553341 | 18556261 | 2921 | -8.39 | 2.76E-14 | 1.17E-13 |
| chr9 | 21483913 | 21487313 | 3401 | -8.39 | 2.76E-14 | 1.17E-13 |
| chr2 | 119866818 | 119871209 | 4392 | -8.35 | 2.77E-14 | 1.18E-13 |
| chr5 | 92796360 | 92816921 | 20562 | -8.7 | 2.77E-14 | 1.18E-13 |
| chr8 | 29674538 | 29694716 | 20179 | -8.96 | 2.79E-14 | 1.18E-13 |
| chr8 | 123024548 | 123029607 | 5060 | -8.55 | 2.79E-14 | 1.19E-13 |
| chr9 | 135074385 | 135081592 | 7208 | -8.41 | 2.80E-14 | 1.19E-13 |
| chr6 | 49279839 | 49305044 | 25206 | -8.34 | 2.81E-14 | 1.19E-13 |
| chr17 | 35703297 | 35705517 | 2221 | -8.29 | 2.81E-14 | 1.19E-13 |
| chr9 | 119218061 | 119220916 | 2856 | -8.31 | 2.81E-14 | 1.19E-13 |
| chr6 | 139439200 | 139442255 | 3056 | -5.95 | 2.82E-14 | 1.20E-13 |
| chr13 | 101469712 | 101481987 | 12276 | -8.36 | 2.83E-14 | 1.20E-13 |
| chr14 | 56720755 | 56726588 | 5834 | -8.3 | 2.84E-14 | 1.20E-13 |
| chr10 | 123453215 | 123480540 | 27326 | -8.51 | 2.84E-14 | 1.20E-13 |
| chr3 | 31493408 | 31502759 | 9352 | -7.11 | 2.84E-14 | 1.20E-13 |
| chr1 | 221597539 | 221604768 | 7230 | -8.4 | 2.85E-14 | 1.21E-13 |
| chr8 | 41346425 | 41361786 | 15362 | -8.44 | 2.85E-14 | 1.21E-13 |
| chr11 | 356704 | 361813 | 5110 | -8.3 | 2.86E-14 | 1.21E-13 |
| chr14 | 93032087 | 93034509 | 2423 | -8.34 | 2.86E-14 | 1.21E-13 |
| chr2 | 70877782 | 70880508 | 2727 | -8.29 | 2.86E-14 | 1.21E-13 |
| chr1 | 185698866 | 185720753 | 21888 | -8.44 | 2.87E-14 | 1.21E-13 |
| chr8 | 37777367 | 37789947 | 12581 | -8.54 | 2.87E-14 | 1.22E-13 |
| chr10 | 116721712 | 116724021 | 2310 | -8.29 | 2.87E-14 | 1.22E-13 |
| chr4 | 103708834 | 103714665 | 5832 | -8.29 | 2.87E-14 | 1.22E-13 |
| chr6 | 1442197 | 1452909 | 10713 | -8.31 | 2.88E-14 | 1.22E-13 |
| chr5 | 657569 | 661175 | 3607 | -8.32 | 2.89E-14 | 1.22E-13 |
| chr16 | 14148954 | 14152170 | 3217 | -8.29 | 2.89E-14 | 1.22E-13 |
| chrX | 21588826 | 21610726 | 21901 | -8.46 | 2.89E-14 | 1.22E-13 |

| | | | | | | |
|---|---|---|---|---|---|---|
| chr2 | 242983745 | 243003603 | 19859 | -8.48 | 2.90E-14 | 1.23E-13 |
| chr22 | 25690961 | 25695608 | 4648 | -6.19 | 2.91E-14 | 1.23E-13 |
| chr15 | 35775442 | 35781374 | 5933 | -8.32 | 2.91E-14 | 1.23E-13 |
| chr1 | 218453766 | 218457306 | 3541 | -8.29 | 2.91E-14 | 1.23E-13 |
| chr5 | 180009086 | 180013224 | 4139 | -8.47 | 2.91E-14 | 1.23E-13 |
| chr11 | 120103380 | 120104893 | 1514 | -6.12 | 2.91E-14 | 1.23E-13 |
| chr14 | 106681284 | 106775280 | 93997 | 5.25 | 2.92E-14 | 1.23E-13 |
| chr3 | 14819299 | 14828755 | 9457 | -8.58 | 2.92E-14 | 1.23E-13 |
| chr5 | 66234203 | 66237567 | 3365 | -8.33 | 2.93E-14 | 1.24E-13 |
| chr3 | 120376083 | 120382615 | 6533 | -8.5 | 2.93E-14 | 1.24E-13 |
| chr3 | 33540291 | 33548788 | 8498 | -8.3 | 2.93E-14 | 1.24E-13 |
| chr2 | 129265014 | 129294357 | 29344 | -8.55 | 2.94E-14 | 1.24E-13 |
| chr11 | 8295175 | 8303268 | 8094 | -8.35 | 2.94E-14 | 1.24E-13 |
| chr13 | 41780569 | 41786564 | 5996 | -8.32 | 2.95E-14 | 1.25E-13 |
| chr8 | 30042290 | 30053904 | 11615 | -8.28 | 2.95E-14 | 1.25E-13 |
| chr5 | 73504310 | 73521248 | 16939 | -8.29 | 2.96E-14 | 1.25E-13 |
| chr3 | 101006909 | 101029504 | 22596 | -8.3 | 2.97E-14 | 1.25E-13 |
| chr18 | 43382838 | 43386464 | 3627 | -8.3 | 2.99E-14 | 1.26E-13 |
| chrX | 13261929 | 13273561 | 11633 | -8.31 | 3.00E-14 | 1.26E-13 |
| chr11 | 68371085 | 68374088 | 3004 | -8.55 | 3.00E-14 | 1.27E-13 |
| chr12 | 1623687 | 1626650 | 2964 | -8.48 | 3.00E-14 | 1.27E-13 |
| chr19 | 41647804 | 41664070 | 16267 | -9 | 3.01E-14 | 1.27E-13 |
| chr6 | 155038184 | 155043188 | 5005 | -8.28 | 3.01E-14 | 1.27E-13 |
| chr16 | 9107084 | 9110168 | 3085 | -8.29 | 3.01E-14 | 1.27E-13 |
| chr3 | 67620495 | 67625015 | 4521 | -8.35 | 3.02E-14 | 1.28E-13 |
| chr13 | 36988803 | 37004375 | 15573 | -8.3 | 3.03E-14 | 1.28E-13 |
| chr4 | 188120180 | 188129826 | 9647 | -8.28 | 3.03E-14 | 1.28E-13 |
| chr6 | 83740806 | 83745059 | 4254 | -8.28 | 3.03E-14 | 1.28E-13 |
| chr7 | 3325882 | 3331292 | 5411 | -8.32 | 3.04E-14 | 1.28E-13 |
| chr4 | 42041526 | 42044064 | 2539 | -6.49 | 3.04E-14 | 1.28E-13 |
| chr20 | 32344368 | 32346730 | 2363 | -8.36 | 3.05E-14 | 1.28E-13 |
| chr8 | 27900338 | 27903857 | 3520 | -7.53 | 3.05E-14 | 1.28E-13 |
| chr5 | 156280639 | 156283960 | 3322 | -8.41 | 3.05E-14 | 1.29E-13 |
| chr2 | 196108277 | 196119730 | 11454 | -8.3 | 3.06E-14 | 1.29E-13 |
| chr6 | 170144440 | 170147730 | 3291 | -8.28 | 3.06E-14 | 1.29E-13 |
| chr1 | 38238086 | 38241064 | 2979 | -8.31 | 3.07E-14 | 1.29E-13 |
| chr12 | 124257437 | 124278490 | 21054 | -8.64 | 3.07E-14 | 1.29E-13 |
| chr12 | 71207091 | 71214852 | 7762 | -8.33 | 3.08E-14 | 1.30E-13 |
| chr2 | 71109037 | 71111262 | 2226 | -4.19 | 3.08E-14 | 1.30E-13 |
| chr3 | 192729820 | 192732008 | 2189 | -5.93 | 3.08E-14 | 1.30E-13 |
| chr17 | 9236394 | 9241068 | 4675 | -8.31 | 3.09E-14 | 1.30E-13 |
| chr2 | 179661830 | 179665197 | 3368 | -8.28 | 3.09E-14 | 1.30E-13 |
| chr7 | 144533574 | 144549677 | 16104 | -8.29 | 3.09E-14 | 1.30E-13 |
| chr1 | 160860901 | 160907029 | 46129 | -8.91 | 3.10E-14 | 1.30E-13 |
| chr2 | 75763281 | 75765861 | 2581 | -5.89 | 3.10E-14 | 1.30E-13 |
| chr20 | 33992506 | 33996810 | 4305 | -8.47 | 3.11E-14 | 1.31E-13 |
| chr2 | 112450839 | 112455141 | 4303 | -8.33 | 3.11E-14 | 1.31E-13 |
| chrX | 149401770 | 149416465 | 14696 | -8.28 | 3.12E-14 | 1.31E-13 |
| chr1 | 115000177 | 115003712 | 3536 | -8.36 | 3.12E-14 | 1.31E-13 |
| chr3 | 64506933 | 64510481 | 3549 | -8.28 | 3.12E-14 | 1.31E-13 |
| chr8 | 146292209 | 146300816 | 8608 | -8.36 | 3.12E-14 | 1.31E-13 |
| chr7 | 82749964 | 82753122 | 3159 | -8.36 | 3.12E-14 | 1.31E-13 |
| chr3 | 108710564 | 108733112 | 22549 | 2.6 | 3.14E-14 | 1.32E-13 |
| chr1 | 17500636 | 17515908 | 15273 | -8.34 | 3.14E-14 | 1.32E-13 |
| chr11 | 72154474 | 72157003 | 2530 | -8.3 | 3.15E-14 | 1.33E-13 |
| chr11 | 130174853 | 130193494 | 18642 | -8.59 | 3.15E-14 | 1.33E-13 |
| chr2 | 48108851 | 48111547 | 2697 | -8.31 | 3.17E-14 | 1.33E-13 |
| chr9 | 74448316 | 74451219 | 2904 | -8.31 | 3.17E-14 | 1.33E-13 |
| chr5 | 13933139 | 13935916 | 2778 | -8.33 | 3.18E-14 | 1.33E-13 |
| chr2 | 56003619 | 56006386 | 2768 | -8.29 | 3.19E-14 | 1.34E-13 |
| chr2 | 73412348 | 73429528 | 17181 | -8.87 | 3.19E-14 | 1.34E-13 |

| | | | | | | |
|---|---|---|---|---|---|---|
| chr5 | 139961905 | 139965893 | 3989 | -8.36 | 3.20E-14 | 1.35E-13 |
| chr11 | 95701268 | 95719669 | 18402 | -8.36 | 3.20E-14 | 1.35E-13 |
| chr4 | 23206584 | 23211395 | 4812 | -7.1 | 3.20E-14 | 1.35E-13 |
| chr1 | 221622442 | 221642501 | 20060 | -8.27 | 3.22E-14 | 1.35E-13 |
| chr20 | 14819549 | 14825234 | 5686 | -8.43 | 3.22E-14 | 1.35E-13 |
| chr4 | 170948815 | 170951030 | 2216 | -8.28 | 3.23E-14 | 1.35E-13 |
| chr22 | 43947745 | 43954962 | 7218 | -8.37 | 3.23E-14 | 1.36E-13 |
| chr9 | 20538872 | 20542180 | 3309 | -8.41 | 3.24E-14 | 1.36E-13 |
| chr21 | 19179009 | 19183367 | 4359 | -8.33 | 3.25E-14 | 1.36E-13 |
| chr16 | 46820350 | 46830499 | 10150 | -8.28 | 3.25E-14 | 1.36E-13 |
| chr5 | 159231618 | 159236096 | 4479 | -8.27 | 3.27E-14 | 1.37E-13 |
| chr5 | 130640813 | 130658104 | 17292 | -8.42 | 3.28E-14 | 1.38E-13 |
| chr18 | 277181 | 280814 | 3634 | -8.31 | 3.28E-14 | 1.38E-13 |
| chr5 | 53117010 | 53138562 | 21553 | -8.99 | 3.28E-14 | 1.38E-13 |
| chr12 | 109572753 | 109575053 | 2301 | -8.28 | 3.30E-14 | 1.38E-13 |
| chr21 | 30621528 | 30628314 | 6787 | -8.28 | 3.30E-14 | 1.38E-13 |
| chr2 | 121287275 | 121290276 | 3002 | -8.27 | 3.30E-14 | 1.39E-13 |
| chr17 | 40523361 | 40525210 | 1850 | -8.58 | 3.31E-14 | 1.39E-13 |
| chr14 | 99834215 | 99838933 | 4719 | -8.33 | 3.31E-14 | 1.39E-13 |
| chr17 | 54728021 | 54751046 | 23026 | -8.54 | 3.31E-14 | 1.39E-13 |
| chr15 | 101219370 | 101243620 | 24251 | -8.33 | 3.32E-14 | 1.39E-13 |
| chr2 | 159614683 | 159625545 | 10863 | -8.47 | 3.33E-14 | 1.40E-13 |
| chr6 | 97736213 | 97744655 | 8443 | -8.27 | 3.33E-14 | 1.40E-13 |
| chr18 | 77414404 | 77419806 | 5403 | -8.27 | 3.34E-14 | 1.40E-13 |
| chr4 | 111453431 | 111459969 | 6539 | -8.27 | 3.34E-14 | 1.40E-13 |
| chr2 | 43287832 | 43291750 | 3919 | -8.28 | 3.34E-14 | 1.40E-13 |
| chr17 | 19856030 | 19858287 | 2258 | -8.31 | 3.34E-14 | 1.40E-13 |
| chrX | 107628611 | 107637386 | 8776 | -8.29 | 3.35E-14 | 1.40E-13 |
| chr4 | 106324759 | 106328922 | 4164 | -8.27 | 3.36E-14 | 1.41E-13 |
| chr12 | 106554061 | 106558566 | 4506 | -8.3 | 3.37E-14 | 1.41E-13 |
| chr4 | 103435401 | 103438759 | 3359 | -8.32 | 3.37E-14 | 1.41E-13 |
| chr2 | 65151259 | 65164206 | 12948 | -3.18 | 3.37E-14 | 1.41E-13 |
| chr14 | 100468825 | 100472837 | 4013 | -8.27 | 3.38E-14 | 1.42E-13 |
| chr12 | 98154728 | 98172380 | 17653 | -8.27 | 3.38E-14 | 1.42E-13 |
| chrX | 39823163 | 39832372 | 9210 | -8.37 | 3.38E-14 | 1.42E-13 |
| chr15 | 60845129 | 60846769 | 1641 | -8.3 | 3.42E-14 | 1.43E-13 |
| chr14 | 89379583 | 89425992 | 46410 | -2.34 | 3.43E-14 | 1.43E-13 |
| chr3 | 182050526 | 182063166 | 12641 | -8.26 | 3.43E-14 | 1.43E-13 |
| chr16 | 21299789 | 21302892 | 3104 | -8.34 | 3.44E-14 | 1.44E-13 |
| chr5 | 2258014 | 2756294 | 498281 | 0.76 | 3.45E-14 | 1.44E-13 |
| chr5 | 156546604 | 156555512 | 8909 | -8.36 | 3.46E-14 | 1.45E-13 |
| chr4 | 162858705 | 162879578 | 20874 | -8.29 | 3.47E-14 | 1.45E-13 |
| chr6 | 132736728 | 132744380 | 7653 | -8.26 | 3.47E-14 | 1.45E-13 |
| chr8 | 115545932 | 115582716 | 36785 | -8.45 | 3.48E-14 | 1.45E-13 |
| chr2 | 71026054 | 71032452 | 6399 | -8.26 | 3.48E-14 | 1.45E-13 |
| chr16 | 78191695 | 78209779 | 18085 | -8.51 | 3.49E-14 | 1.46E-13 |
| chr10 | 70075018 | 70077908 | 2891 | -8.37 | 3.49E-14 | 1.46E-13 |
| chr14 | 76530341 | 76533076 | 2736 | -8.32 | 3.49E-14 | 1.46E-13 |
| chrX | 74697342 | 74706314 | 8973 | 8.33 | 3.49E-14 | 1.46E-13 |
| chr12 | 98792955 | 98804938 | 11984 | -8.4 | 3.50E-14 | 1.46E-13 |
| chr4 | 185629540 | 185633875 | 4336 | -8.6 | 3.50E-14 | 1.46E-13 |
| chr2 | 8670003 | 8673409 | 3407 | -8.27 | 3.51E-14 | 1.47E-13 |
| chr8 | 128091637 | 128104483 | 12847 | -8.41 | 3.51E-14 | 1.47E-13 |
| chr10 | 101699237 | 101713232 | 13996 | -8.28 | 3.52E-14 | 1.47E-13 |
| chr2 | 136241622 | 136245999 | 4378 | -8.46 | 3.52E-14 | 1.47E-13 |
| chr4 | 38999478 | 39003296 | 3819 | -8.27 | 3.53E-14 | 1.47E-13 |
| chr17 | 33734602 | 33743366 | 8765 | -3 | 3.53E-14 | 1.47E-13 |
| chr11 | 74247796 | 74273571 | 25776 | -8.52 | 3.53E-14 | 1.47E-13 |
| chr2 | 161632903 | 161638574 | 5672 | -8.26 | 3.53E-14 | 1.47E-13 |
| chr10 | 92526304 | 92541984 | 15681 | 8.34 | 3.54E-14 | 1.48E-13 |
| chr7 | 14504357 | 14526868 | 22512 | -8.62 | 3.54E-14 | 1.48E-13 |

| | | | | | | |
|---|---|---|---|---|---|---|
| chr7 | 123748063 | 123757695 | 9633 | 2.72 | 3.55E-14 | 1.48E-13 |
| chr1 | 43919664 | 43927997 | 8334 | -8.36 | 3.56E-14 | 1.48E-13 |
| chr2 | 175786321 | 175800300 | 13980 | -8.26 | 3.57E-14 | 1.49E-13 |
| chr6 | 14487292 | 14498207 | 10916 | -8.37 | 3.58E-14 | 1.49E-13 |
| chr5 | 149941240 | 149944115 | 2876 | -8.26 | 3.59E-14 | 1.50E-13 |
| chr3 | 71638720 | 71661358 | 22639 | -8.3 | 3.60E-14 | 1.50E-13 |
| chr1 | 101353986 | 101371924 | 17939 | -8.38 | 3.60E-14 | 1.50E-13 |
| chr12 | 60313387 | 60340454 | 27068 | -8.27 | 3.60E-14 | 1.50E-13 |
| chr2 | 96742075 | 96747564 | 5490 | -8.44 | 3.60E-14 | 1.50E-13 |
| chr11 | 10998538 | 11028010 | 29473 | -8.38 | 3.61E-14 | 1.51E-13 |
| chr7 | 32430956 | 32438821 | 7866 | -8.28 | 3.61E-14 | 1.51E-13 |
| chr7 | 139022263 | 139024949 | 2687 | -8.28 | 3.61E-14 | 1.51E-13 |
| chr4 | 95952108 | 95958164 | 6057 | -8.4 | 3.62E-14 | 1.51E-13 |
| chr17 | 75021467 | 75029095 | 7629 | -8.69 | 3.63E-14 | 1.51E-13 |
| chr15 | 36373094 | 36379856 | 6763 | -8.28 | 3.63E-14 | 1.51E-13 |
| chr3 | 18209285 | 18239597 | 30313 | -8.39 | 3.63E-14 | 1.51E-13 |
| chr6 | 86420154 | 86438516 | 18363 | -8.34 | 3.64E-14 | 1.51E-13 |
| chr4 | 140616267 | 140631083 | 14817 | -8.31 | 3.64E-14 | 1.52E-13 |
| chr20 | 39914032 | 39917049 | 3018 | -8.26 | 3.65E-14 | 1.52E-13 |
| chr6 | 20262180 | 20265476 | 3297 | -8.34 | 3.66E-14 | 1.52E-13 |
| chr8 | 58914913 | 58923600 | 8688 | -8.28 | 3.67E-14 | 1.53E-13 |
| chr14 | 53776021 | 53784992 | 8972 | -8.32 | 3.67E-14 | 1.53E-13 |
| chr18 | 53336331 | 53345658 | 9328 | -8.25 | 3.67E-14 | 1.53E-13 |
| chr6 | 38686898 | 38703997 | 17100 | -8.3 | 3.68E-14 | 1.53E-13 |
| chr1 | 79629839 | 79646289 | 16451 | -8.31 | 3.68E-14 | 1.53E-13 |
| chr1 | 41335471 | 41343758 | 8288 | -8.3 | 3.68E-14 | 1.53E-13 |
| chr10 | 96391210 | 96417118 | 25909 | -8.65 | 3.68E-14 | 1.53E-13 |
| chrX | 114374090 | 114379559 | 5470 | -2.67 | 3.69E-14 | 1.54E-13 |
| chr12 | 40926729 | 40946816 | 20088 | -8.37 | 3.69E-14 | 1.54E-13 |
| chr3 | 101509668 | 101513166 | 3499 | -8.25 | 3.73E-14 | 1.55E-13 |
| chr22 | 30538770 | 30546260 | 7491 | -8.34 | 3.73E-14 | 1.55E-13 |
| chr5 | 30499515 | 30517841 | 18327 | -8.26 | 3.74E-14 | 1.55E-13 |
| chr2 | 27007542 | 27014562 | 7021 | -8.33 | 3.74E-14 | 1.55E-13 |
| chr4 | 131185579 | 131190428 | 4850 | -8.27 | 3.74E-14 | 1.56E-13 |
| chr11 | 109972161 | 109979531 | 7371 | -8.41 | 3.76E-14 | 1.56E-13 |
| chr5 | 167162603 | 167176763 | 14161 | -2.39 | 3.76E-14 | 1.56E-13 |
| chr2 | 62551355 | 62553549 | 2195 | -5.92 | 3.76E-14 | 1.56E-13 |
| chr15 | 78792608 | 78794888 | 2281 | -8.29 | 3.77E-14 | 1.57E-13 |
| chr12 | 38538839 | 38542647 | 3809 | -8.31 | 3.77E-14 | 1.57E-13 |
| chr3 | 17077801 | 17091757 | 13957 | -8.35 | 3.78E-14 | 1.57E-13 |
| chr6 | 113077630 | 113090460 | 12831 | -8.31 | 3.79E-14 | 1.57E-13 |
| chr1 | 202645672 | 202648273 | 2602 | -8.36 | 3.81E-14 | 1.58E-13 |
| chr3 | 194910132 | 194913996 | 3865 | -8.26 | 3.82E-14 | 1.59E-13 |
| chr1 | 114536506 | 114548969 | 12464 | -8.25 | 3.83E-14 | 1.59E-13 |
| chr14 | 93109323 | 93112449 | 3127 | -8.25 | 3.83E-14 | 1.59E-13 |
| chr10 | 8513631 | 8520224 | 6594 | -8.31 | 3.83E-14 | 1.59E-13 |
| chr9 | 93645944 | 93650320 | 4377 | -8.25 | 3.84E-14 | 1.59E-13 |
| chr3 | 133436277 | 133451665 | 15389 | -3.22 | 3.86E-14 | 1.60E-13 |
| chr4 | 77760724 | 77770631 | 9908 | -8.36 | 3.86E-14 | 1.60E-13 |
| chr5 | 54668063 | 54689533 | 21471 | -8.39 | 3.86E-14 | 1.60E-13 |
| chr6 | 94138826 | 94518897 | 380072 | 1.32 | 3.87E-14 | 1.61E-13 |
| chr4 | 65039268 | 65076229 | 36962 | -8.77 | 3.87E-14 | 1.61E-13 |
| chrX | 133794064 | 133809055 | 14992 | -8.25 | 3.89E-14 | 1.61E-13 |
| chr10 | 90865898 | 90873416 | 7519 | -8.37 | 3.90E-14 | 1.62E-13 |
| chr7 | 16910517 | 16913158 | 2642 | -8.3 | 3.90E-14 | 1.62E-13 |
| chr9 | 112405806 | 112422475 | 16670 | -8.47 | 3.90E-14 | 1.62E-13 |
| chr7 | 95290314 | 95293485 | 3172 | -8.28 | 3.90E-14 | 1.62E-13 |
| chr9 | 99058163 | 99093038 | 34876 | -9 | 3.97E-14 | 1.65E-13 |
| chr1 | 227511363 | 227531590 | 20228 | -8.96 | 3.98E-14 | 1.65E-13 |
| chr8 | 133512288 | 133521658 | 9371 | -2.5 | 3.99E-14 | 1.65E-13 |
| chr2 | 107441234 | 107445763 | 4530 | -3 | 4.00E-14 | 1.66E-13 |

| | | | | | | |
|---|---|---|---|---|---|---|
| chr16 | 74311742 | 74327494 | 15753 | -8.61 | 4.03E-14 | 1.67E-13 |
| chr19 | 54512883 | 54529125 | 16243 | -8.27 | 4.03E-14 | 1.67E-13 |
| chr4 | 170010752 | 170020222 | 9471 | -8.27 | 4.03E-14 | 1.67E-13 |
| chr17 | 71130446 | 71149083 | 18638 | -8.87 | 4.05E-14 | 1.68E-13 |
| chr7 | 148437494 | 148441317 | 3824 | -8.35 | 4.05E-14 | 1.68E-13 |
| chr14 | 102350860 | 102353528 | 2669 | -8.25 | 4.07E-14 | 1.69E-13 |
| chr21 | 38772883 | 38776461 | 3579 | -8.24 | 4.07E-14 | 1.69E-13 |
| chr8 | 40345093 | 40379918 | 34826 | -8.56 | 4.09E-14 | 1.69E-13 |
| chr8 | 59117044 | 59126852 | 9809 | -8.39 | 4.09E-14 | 1.69E-13 |
| chr5 | 37804091 | 37814273 | 10183 | 8.31 | 4.09E-14 | 1.69E-13 |
| chr12 | 32194873 | 32203118 | 8246 | -8.6 | 4.09E-14 | 1.69E-13 |
| chr22 | 34182948 | 34188287 | 5340 | -8.44 | 4.10E-14 | 1.70E-13 |
| chr3 | 33499194 | 33523865 | 24672 | -8.49 | 4.10E-14 | 1.70E-13 |
| chr22 | 30479760 | 30492085 | 12326 | -8.24 | 4.10E-14 | 1.70E-13 |
| chr5 | 124321754 | 124340009 | 18256 | -9.11 | 4.12E-14 | 1.70E-13 |
| chr8 | 102186387 | 102195523 | 9137 | -8.24 | 4.12E-14 | 1.70E-13 |
| chr20 | 22456707 | 22468999 | 12293 | -8.76 | 4.13E-14 | 1.71E-13 |
| chr18 | 55381932 | 55394542 | 12611 | -8.41 | 4.13E-14 | 1.71E-13 |
| chr4 | 105255779 | 105277226 | 21448 | -8.37 | 4.14E-14 | 1.71E-13 |
| chr6 | 147284438 | 147289031 | 4594 | -8.27 | 4.14E-14 | 1.71E-13 |
| chr16 | 19158763 | 19177152 | 18390 | -8.53 | 4.14E-14 | 1.71E-13 |
| chr11 | 70999348 | 71008920 | 9573 | -8.44 | 4.15E-14 | 1.72E-13 |
| chr9 | 86620638 | 86623334 | 2697 | -8.29 | 4.15E-14 | 1.72E-13 |
| chr1 | 87386850 | 87391848 | 4999 | -8.24 | 4.16E-14 | 1.72E-13 |
| chr7 | 23713687 | 23715962 | 2276 | -8.24 | 4.17E-14 | 1.72E-13 |
| chr4 | 111515532 | 111520872 | 5341 | -8.26 | 4.18E-14 | 1.73E-13 |
| chr18 | 20021052 | 20031388 | 10337 | -8.25 | 4.19E-14 | 1.73E-13 |
| chr19 | 7813106 | 7850516 | 37411 | -7.3 | 4.19E-14 | 1.73E-13 |
| chr2 | 238193477 | 238214264 | 20788 | -8.25 | 4.20E-14 | 1.73E-13 |
| chr8 | 38683485 | 38686093 | 2609 | -8.24 | 4.20E-14 | 1.73E-13 |
| chr20 | 42685641 | 42690839 | 5199 | -8.27 | 4.20E-14 | 1.74E-13 |
| chr5 | 99740380 | 99762388 | 22009 | 8.84 | 4.24E-14 | 1.75E-13 |
| chr7 | 4932360 | 4963303 | 30944 | -8.44 | 4.24E-14 | 1.75E-13 |
| chr11 | 78969826 | 78977308 | 7483 | 8.53 | 4.24E-14 | 1.75E-13 |
| chr20 | 1111937 | 1114173 | 2237 | -8.24 | 4.24E-14 | 1.75E-13 |
| chr8 | 90118315 | 90125334 | 7020 | -8.32 | 4.25E-14 | 1.75E-13 |
| chr17 | 19345670 | 19349251 | 3582 | -8.31 | 4.25E-14 | 1.76E-13 |
| chr11 | 101626354 | 101655307 | 28954 | -8.35 | 4.28E-14 | 1.77E-13 |
| chr5 | 13965179 | 13974104 | 8926 | -3.26 | 4.28E-14 | 1.77E-13 |
| chr1 | 85715613 | 85730245 | 14633 | -8.4 | 4.28E-14 | 1.77E-13 |
| chr7 | 93754574 | 93761397 | 6824 | -8.38 | 4.29E-14 | 1.77E-13 |
| chr10 | 81951340 | 81953988 | 2649 | -8.26 | 4.29E-14 | 1.77E-13 |
| chr7 | 112544804 | 112552694 | 7891 | -8.49 | 4.29E-14 | 1.77E-13 |
| chr2 | 241554999 | 241558854 | 3856 | -8.24 | 4.29E-14 | 1.77E-13 |
| chr6 | 41635654 | 41638734 | 3081 | -8.24 | 4.29E-14 | 1.77E-13 |
| chr8 | 28212463 | 28214002 | 1540 | -8.28 | 4.30E-14 | 1.77E-13 |
| chr12 | 49112315 | 49116070 | 3756 | -8.24 | 4.30E-14 | 1.77E-13 |
| chr2 | 46102376 | 46104849 | 2474 | -8.31 | 4.31E-14 | 1.78E-13 |
| chr17 | 64266334 | 64268582 | 2249 | -8.29 | 4.31E-14 | 1.78E-13 |
| chr20 | 56827515 | 56830526 | 3012 | -8.25 | 4.33E-14 | 1.78E-13 |
| chr1 | 225441210 | 225449445 | 8236 | -8.24 | 4.33E-14 | 1.78E-13 |
| chr7 | 141273520 | 141288609 | 15090 | -8.44 | 4.33E-14 | 1.79E-13 |
| chr4 | 156256496 | 156260374 | 3879 | -8.28 | 4.33E-14 | 1.79E-13 |
| chr1 | 219176868 | 219180269 | 3402 | -8.29 | 4.34E-14 | 1.79E-13 |
| chr2 | 73684671 | 73687915 | 3245 | -8.3 | 4.34E-14 | 1.79E-13 |
| chr16 | 72081778 | 72084688 | 2911 | -8.25 | 4.35E-14 | 1.79E-13 |
| chr5 | 35631037 | 35635998 | 4962 | -8.26 | 4.35E-14 | 1.79E-13 |
| chr6 | 34126044 | 34136027 | 9984 | -8.27 | 4.36E-14 | 1.80E-13 |
| chr1 | 67152289 | 67158406 | 6118 | -8.34 | 4.37E-14 | 1.80E-13 |
| chr13 | 104210464 | 104226868 | 16405 | -8.23 | 4.37E-14 | 1.80E-13 |
| chr17 | 28483975 | 28487517 | 3543 | -8.45 | 4.38E-14 | 1.80E-13 |

| | | | | | | |
|---|---|---|---|---|---|---|
| chr7 | 978788 | 981513 | 2726 | -8.25 | 4.38E-14 | 1.80E-13 |
| chr20 | 35702842 | 35704647 | 1806 | -8.23 | 4.38E-14 | 1.80E-13 |
| chr12 | 56765179 | 56768441 | 3263 | -8.35 | 4.39E-14 | 1.81E-13 |
| chr22 | 45227419 | 45241192 | 13774 | -8.24 | 4.40E-14 | 1.81E-13 |
| chr5 | 175186098 | 175221007 | 34910 | -9.37 | 4.40E-14 | 1.81E-13 |
| chr9 | 139105326 | 139107526 | 2201 | -8.23 | 4.40E-14 | 1.81E-13 |
| chr5 | 93177917 | 93181622 | 3706 | -8.24 | 4.40E-14 | 1.81E-13 |
| chr2 | 105389095 | 105441481 | 52387 | 2.04 | 4.40E-14 | 1.81E-13 |
| chr16 | 24874497 | 24878431 | 3935 | -8.23 | 4.41E-14 | 1.81E-13 |
| chr16 | 56719696 | 56732448 | 12753 | -8.48 | 4.41E-14 | 1.81E-13 |
| chr14 | 21665106 | 21668986 | 3881 | -8.32 | 4.44E-14 | 1.82E-13 |
| chr7 | 34953990 | 34974266 | 20277 | -8.36 | 4.44E-14 | 1.83E-13 |
| chr4 | 5549422 | 5560459 | 11038 | -8.26 | 4.45E-14 | 1.83E-13 |
| chr2 | 87046375 | 87050855 | 4481 | -8.24 | 4.50E-14 | 1.85E-13 |
| chr3 | 160055038 | 160057263 | 2226 | -8.24 | 4.50E-14 | 1.85E-13 |
| chr4 | 180091807 | 180110100 | 18294 | -8.3 | 4.50E-14 | 1.85E-13 |
| chr1 | 228303583 | 228307962 | 4380 | -8.46 | 4.50E-14 | 1.85E-13 |
| chr2 | 218210473 | 218215816 | 5344 | -8.29 | 4.51E-14 | 1.85E-13 |
| chr1 | 219030116 | 219059433 | 29318 | -8.91 | 4.52E-14 | 1.86E-13 |
| chr3 | 30195899 | 30208914 | 13016 | -8.63 | 4.54E-14 | 1.86E-13 |
| chr5 | 123817684 | 123829841 | 12158 | -8.41 | 4.55E-14 | 1.87E-13 |
| chr8 | 95325088 | 95331656 | 6569 | -8.25 | 4.56E-14 | 1.87E-13 |
| chr5 | 133208856 | 133211719 | 2864 | -8.35 | 4.56E-14 | 1.87E-13 |
| chr1 | 174809650 | 174820286 | 10637 | -8.26 | 4.56E-14 | 1.87E-13 |
| chr11 | 29238721 | 29248232 | 9512 | -8.27 | 4.57E-14 | 1.88E-13 |
| chr15 | 59999114 | 60001512 | 2399 | -8.36 | 4.58E-14 | 1.88E-13 |
| chrX | 76817514 | 76823762 | 6249 | -8.28 | 4.58E-14 | 1.88E-13 |
| chr22 | 24506277 | 24509242 | 2966 | -8.32 | 4.59E-14 | 1.88E-13 |
| chr1 | 50804134 | 50821504 | 17371 | -8.23 | 4.59E-14 | 1.88E-13 |
| chr8 | 67437030 | 67440444 | 3415 | -8.3 | 4.61E-14 | 1.89E-13 |
| chr9 | 97750438 | 97752852 | 2415 | -8.37 | 4.63E-14 | 1.90E-13 |
| chr2 | 43775872 | 43777558 | 1687 | -8.29 | 4.64E-14 | 1.90E-13 |
| chr13 | 106507093 | 106512728 | 5636 | -8.24 | 4.65E-14 | 1.91E-13 |
| chr6 | 64262691 | 64278618 | 15928 | -8.28 | 4.67E-14 | 1.91E-13 |
| chr13 | 27729845 | 27734991 | 5147 | -8.23 | 4.67E-14 | 1.92E-13 |
| chr4 | 26407088 | 26409159 | 2072 | -8.23 | 4.67E-14 | 1.92E-13 |
| chr2 | 191642918 | 191649831 | 6914 | -8.25 | 4.67E-14 | 1.92E-13 |
| chr14 | 31013432 | 31017493 | 4062 | -8.23 | 4.67E-14 | 1.92E-13 |
| chr20 | 47212657 | 47219995 | 7339 | -8.23 | 4.67E-14 | 1.92E-13 |
| chr3 | 169474480 | 169484311 | 9832 | -8.26 | 4.67E-14 | 1.92E-13 |
| chr8 | 29304258 | 29315417 | 11160 | -8.26 | 4.67E-14 | 1.92E-13 |
| chr9 | 22310724 | 22331380 | 20657 | -8.41 | 4.68E-14 | 1.92E-13 |
| chr16 | 52279879 | 52294765 | 14887 | -8.4 | 4.68E-14 | 1.92E-13 |
| chr1 | 173532902 | 173537953 | 5052 | -8.25 | 4.68E-14 | 1.92E-13 |
| chr17 | 75897240 | 75918680 | 21441 | -8.57 | 4.69E-14 | 1.92E-13 |
| chr6 | 22039106 | 22057158 | 18053 | -8.32 | 4.71E-14 | 1.93E-13 |
| chr12 | 60606061 | 60621646 | 15586 | -8.22 | 4.72E-14 | 1.93E-13 |
| chr9 | 72495114 | 72498720 | 3607 | -8.23 | 4.73E-14 | 1.94E-13 |
| chr20 | 30464973 | 30482260 | 17288 | -8.46 | 4.74E-14 | 1.94E-13 |
| chr20 | 55777150 | 55784246 | 7097 | -8.25 | 4.75E-14 | 1.94E-13 |
| chr2 | 40003006 | 40004568 | 1563 | -8.37 | 4.76E-14 | 1.95E-13 |
| chr4 | 169436767 | 169455052 | 18286 | -8.48 | 4.77E-14 | 1.95E-13 |
| chr20 | 43250842 | 43253929 | 3088 | -8.37 | 4.77E-14 | 1.95E-13 |
| chr5 | 171889545 | 171896034 | 6490 | -8.25 | 4.77E-14 | 1.95E-13 |
| chr7 | 106959250 | 106982539 | 23290 | -8.26 | 4.77E-14 | 1.95E-13 |
| chr6 | 21878947 | 21886004 | 7058 | -8.42 | 4.78E-14 | 1.96E-13 |
| chr7 | 28288104 | 28313870 | 25767 | -3.12 | 4.79E-14 | 1.96E-13 |
| chr16 | 81878907 | 81881471 | 2565 | -6.48 | 4.80E-14 | 1.96E-13 |
| chr10 | 69780103 | 69783380 | 3278 | -8.22 | 4.80E-14 | 1.96E-13 |
| chr2 | 151073267 | 151105921 | 32655 | -8.69 | 4.80E-14 | 1.97E-13 |
| chr5 | 177043656 | 177047113 | 3458 | -8.22 | 4.81E-14 | 1.97E-13 |

| | | | | | | |
|---|---|---|---|---|---|---|
| chr2 | 70655867 | 70661254 | 5388 | -8.22 | 4.82E-14 | 1.97E-13 |
| chr8 | 134315452 | 134319956 | 4505 | -8.26 | 4.84E-14 | 1.98E-13 |
| chr7 | 83848850 | 83851624 | 2775 | -8.29 | 4.85E-14 | 1.98E-13 |
| chr7 | 51213736 | 51217326 | 3591 | -8.22 | 4.86E-14 | 1.99E-13 |
| chr3 | 196445722 | 196449482 | 3761 | -8.31 | 4.89E-14 | 2.00E-13 |
| chr2 | 181151359 | 181164270 | 12912 | -8.31 | 4.90E-14 | 2.00E-13 |
| chr10 | 4017245 | 4027263 | 10019 | -8.3 | 4.90E-14 | 2.00E-13 |
| chr7 | 115806403 | 115828752 | 22350 | -8.48 | 4.91E-14 | 2.01E-13 |
| chr14 | 73285256 | 73289084 | 3829 | -8.25 | 4.92E-14 | 2.01E-13 |
| chr1 | 217990006 | 218002192 | 12187 | -8.23 | 4.92E-14 | 2.01E-13 |
| chr3 | 28311329 | 28315940 | 4612 | -8.23 | 4.93E-14 | 2.02E-13 |
| chr15 | 100068453 | 100074893 | 6441 | -8.37 | 4.94E-14 | 2.02E-13 |
| chr11 | 59350971 | 59354270 | 3300 | -8.24 | 4.94E-14 | 2.02E-13 |
| chr2 | 197599089 | 197605806 | 6718 | -8.28 | 4.95E-14 | 2.02E-13 |
| chr18 | 9782745 | 9795641 | 12897 | -8.37 | 4.95E-14 | 2.02E-13 |
| chr8 | 62768180 | 62771392 | 3213 | -8.22 | 4.96E-14 | 2.03E-13 |
| chr17 | 29477565 | 29480249 | 2685 | -8.28 | 4.97E-14 | 2.03E-13 |
| chr4 | 84258986 | 84276264 | 17279 | -8.32 | 4.97E-14 | 2.03E-13 |
| chr9 | 15931175 | 15933676 | 2502 | -8.23 | 4.98E-14 | 2.03E-13 |
| chr3 | 136654418 | 136656776 | 2359 | -8.25 | 4.98E-14 | 2.03E-13 |
| chr11 | 87796437 | 87811182 | 14746 | -8.22 | 5.01E-14 | 2.04E-13 |
| chr1 | 223271407 | 223275694 | 4288 | -8.22 | 5.01E-14 | 2.05E-13 |
| chr2 | 7223012 | 7229209 | 6198 | -2.91 | 5.01E-14 | 2.05E-13 |
| chr1 | 243536032 | 243538977 | 2946 | -8.3 | 5.05E-14 | 2.06E-13 |
| chr21 | 39000005 | 39007224 | 7220 | -8.24 | 5.08E-14 | 2.07E-13 |
| chr12 | 72221953 | 72258147 | 36195 | -8.74 | 5.08E-14 | 2.07E-13 |
| chr8 | 127969584 | 127972367 | 2784 | -8.58 | 5.08E-14 | 2.07E-13 |
| chr2 | 135663975 | 135673010 | 9036 | -8.31 | 5.09E-14 | 2.08E-13 |
| chr1 | 59655491 | 59684409 | 28919 | -8.3 | 5.09E-14 | 2.08E-13 |
| chr3 | 23051490 | 23069557 | 18068 | -8.32 | 5.10E-14 | 2.08E-13 |
| chr8 | 118780581 | 118793476 | 12896 | -8.26 | 5.12E-14 | 2.09E-13 |
| chr3 | 124752243 | 124755839 | 3597 | -8.24 | 5.13E-14 | 2.09E-13 |
| chr10 | 98646322 | 98653079 | 6758 | -8.21 | 5.14E-14 | 2.10E-13 |
| chrX | 14053023 | 14067644 | 14622 | -8.23 | 5.15E-14 | 2.10E-13 |
| chr14 | 77546239 | 77559359 | 13121 | -8.22 | 5.15E-14 | 2.10E-13 |
| chr15 | 36298938 | 36307609 | 8672 | -8.27 | 5.15E-14 | 2.10E-13 |
| chr7 | 150686103 | 150689365 | 3263 | -8.38 | 5.15E-14 | 2.10E-13 |
| chr17 | 26136087 | 26142011 | 5925 | -8.21 | 5.17E-14 | 2.11E-13 |
| chr4 | 8694098 | 8699996 | 5899 | -8.23 | 5.17E-14 | 2.11E-13 |
| chr12 | 83828557 | 83857104 | 28548 | -8.22 | 5.19E-14 | 2.11E-13 |
| chr2 | 174043477 | 174046785 | 3309 | -8.45 | 5.20E-14 | 2.12E-13 |
| chr18 | 24201679 | 24206928 | 5250 | -8.29 | 5.23E-14 | 2.13E-13 |
| chr9 | 111802953 | 111812981 | 10029 | -8.31 | 5.24E-14 | 2.13E-13 |
| chr2 | 64998539 | 65004021 | 5483 | -8.21 | 5.26E-14 | 2.14E-13 |
| chr7 | 152212399 | 152218105 | 5707 | -8.24 | 5.27E-14 | 2.14E-13 |
| chr4 | 87221109 | 87227001 | 5893 | -8.22 | 5.28E-14 | 2.15E-13 |
| chr6 | 166930121 | 166939932 | 9812 | -8.22 | 5.28E-14 | 2.15E-13 |
| chr2 | 105330810 | 105349473 | 18664 | 2.02 | 5.28E-14 | 2.15E-13 |
| chr7 | 44821717 | 44825403 | 3687 | -8.23 | 5.29E-14 | 2.15E-13 |
| chr2 | 172099500 | 172112107 | 12608 | -8.33 | 5.30E-14 | 2.16E-13 |
| chr10 | 101787059 | 101789339 | 2281 | -8.22 | 5.30E-14 | 2.16E-13 |
| chr18 | 9230612 | 9233882 | 3271 | -8.22 | 5.30E-14 | 2.16E-13 |
| chr8 | 105617084 | 105644510 | 27427 | -1.6 | 5.31E-14 | 2.16E-13 |
| chr2 | 189972703 | 189999377 | 26675 | -8.67 | 5.31E-14 | 2.16E-13 |
| chr9 | 100244379 | 100251905 | 7527 | -8.41 | 5.31E-14 | 2.16E-13 |
| chr18 | 2386387 | 2402050 | 15664 | -8.21 | 5.34E-14 | 2.17E-13 |
| chr10 | 123695807 | 123701825 | 6019 | -8.24 | 5.34E-14 | 2.17E-13 |
| chr6 | 36224586 | 36234543 | 9958 | -8.23 | 5.34E-14 | 2.17E-13 |
| chr9 | 132988871 | 132991363 | 2493 | -8.22 | 5.35E-14 | 2.17E-13 |
| chr2 | 109008730 | 109023660 | 14931 | -8.51 | 5.36E-14 | 2.18E-13 |
| chr4 | 154657875 | 154685008 | 27134 | -8.61 | 5.36E-14 | 2.18E-13 |

| | | | | | | |
|---|---|---|---|---|---|---|
| chr5 | 57632149 | 57647325 | 15177 | -8.23 | 5.36E-14 | 2.18E-13 |
| chr12 | 69996488 | 70012742 | 16255 | -8.25 | 5.36E-14 | 2.18E-13 |
| chr17 | 57486585 | 57489035 | 2451 | -8.26 | 5.38E-14 | 2.19E-13 |
| chr10 | 72524490 | 72527984 | 3495 | -8.25 | 5.38E-14 | 2.19E-13 |
| chr6 | 79262047 | 79279022 | 16976 | -8.23 | 5.39E-14 | 2.19E-13 |
| chr10 | 133796204 | 133802477 | 6274 | -8.35 | 5.40E-14 | 2.19E-13 |
| chr12 | 108017446 | 108026507 | 9062 | -8.3 | 5.40E-14 | 2.19E-13 |
| chr12 | 16019840 | 16024637 | 4798 | -8.23 | 5.40E-14 | 2.19E-13 |
| chr15 | 57477720 | 57479614 | 1895 | -8.3 | 5.40E-14 | 2.19E-13 |
| chr5 | 91807134 | 91810396 | 3263 | -8.27 | 5.41E-14 | 2.19E-13 |
| chr3 | 72330711 | 72333896 | 3186 | -8.21 | 5.43E-14 | 2.20E-13 |
| chr17 | 63983558 | 63986980 | 3423 | -8.33 | 5.43E-14 | 2.20E-13 |
| chr1 | 54207390 | 54210363 | 2974 | -8.24 | 5.43E-14 | 2.20E-13 |
| chr14 | 39877732 | 39886681 | 8950 | -8.23 | 5.44E-14 | 2.21E-13 |
| chr8 | 73873022 | 73887790 | 14769 | 8.39 | 5.44E-14 | 2.21E-13 |
| chr3 | 169850011 | 169853307 | 3297 | -8.22 | 5.45E-14 | 2.21E-13 |
| chr10 | 76678154 | 76680678 | 2525 | -8.21 | 5.45E-14 | 2.21E-13 |
| chr1 | 75287933 | 75301714 | 13782 | 8.45 | 5.46E-14 | 2.21E-13 |
| chr9 | 96121261 | 96124726 | 3466 | -8.36 | 5.48E-14 | 2.22E-13 |
| chr3 | 187136357 | 187147755 | 11399 | -8.23 | 5.49E-14 | 2.23E-13 |
| chr3 | 42615684 | 42618185 | 2502 | -8.38 | 5.50E-14 | 2.23E-13 |
| chrX | 35595034 | 35616078 | 21045 | -8.29 | 5.50E-14 | 2.23E-13 |
| chr2 | 10551817 | 10553801 | 1985 | -8.21 | 5.51E-14 | 2.24E-13 |
| chr16 | 5270394 | 5283190 | 12797 | -8.21 | 5.52E-14 | 2.24E-13 |
| chr7 | 80603646 | 80611375 | 7730 | -8.55 | 5.52E-14 | 2.24E-13 |
| chr10 | 64212981 | 64237758 | 24778 | -8.21 | 5.54E-14 | 2.25E-13 |
| chr1 | 162584769 | 162588713 | 3945 | -8.47 | 5.55E-14 | 2.25E-13 |
| chr2 | 166705534 | 166710938 | 5405 | -8.27 | 5.57E-14 | 2.26E-13 |
| chr1 | 56357154 | 56367184 | 10031 | -8.24 | 5.58E-14 | 2.26E-13 |
| chr6 | 12014519 | 12031536 | 17018 | -8.23 | 5.60E-14 | 2.27E-13 |
| chr5 | 54542850 | 54545379 | 2530 | -8.2 | 5.61E-14 | 2.27E-13 |
| chr2 | 159266192 | 159268799 | 2608 | -8.31 | 5.62E-14 | 2.27E-13 |
| chr6 | 3010067 | 3013328 | 3262 | -8.31 | 5.62E-14 | 2.28E-13 |
| chr14 | 99766413 | 99777811 | 11399 | -8.47 | 5.63E-14 | 2.28E-13 |
| chr1 | 207180966 | 207187224 | 6259 | -8.37 | 5.64E-14 | 2.28E-13 |
| chr10 | 72009631 | 72045501 | 35871 | -8.66 | 5.65E-14 | 2.29E-13 |
| chr9 | 90781983 | 90784747 | 2765 | -8.22 | 5.65E-14 | 2.29E-13 |
| chr1 | 244468228 | 244496438 | 28211 | -8.8 | 5.67E-14 | 2.30E-13 |
| chr12 | 27627815 | 27633183 | 5369 | -8.25 | 5.69E-14 | 2.30E-13 |
| chr1 | 244433184 | 244453194 | 20011 | -8.46 | 5.70E-14 | 2.31E-13 |
| chr2 | 204609432 | 204633020 | 23589 | -8.22 | 5.70E-14 | 2.31E-13 |
| chr7 | 111685800 | 111689068 | 3269 | -8.36 | 5.70E-14 | 2.31E-13 |
| chr19 | 33968715 | 33974619 | 5905 | -8.24 | 5.71E-14 | 2.31E-13 |
| chr11 | 121203135 | 121204910 | 1776 | -5.92 | 5.71E-14 | 2.31E-13 |
| chr10 | 30896811 | 30900167 | 3357 | -8.22 | 5.71E-14 | 2.31E-13 |
| chr1 | 169397293 | 169410048 | 12756 | -1.77 | 5.71E-14 | 2.31E-13 |
| chr1 | 179245055 | 179249796 | 4742 | -8.4 | 5.73E-14 | 2.32E-13 |
| chr2 | 235519351 | 235524693 | 5343 | -8.2 | 5.77E-14 | 2.33E-13 |
| chr6 | 9259220 | 9270979 | 11760 | -8.29 | 5.80E-14 | 2.35E-13 |
| chr9 | 119926545 | 119930297 | 3753 | 4.76 | 5.81E-14 | 2.35E-13 |
| chr1 | 2503546 | 2506096 | 2551 | -8.2 | 5.83E-14 | 2.36E-13 |
| chr7 | 157631885 | 157643862 | 11978 | -8.36 | 5.83E-14 | 2.36E-13 |
| chr1 | 178804053 | 178806964 | 2912 | -8.32 | 5.85E-14 | 2.36E-13 |
| chr1 | 112217925 | 112224044 | 6120 | -8.2 | 5.86E-14 | 2.37E-13 |
| chr15 | 73672498 | 73676158 | 3661 | -8.23 | 5.86E-14 | 2.37E-13 |
| chr8 | 131583831 | 131587363 | 3533 | -8.23 | 5.88E-14 | 2.38E-13 |
| chr6 | 109021794 | 109025437 | 3644 | -8.2 | 5.88E-14 | 2.38E-13 |
| chr9 | 4891125 | 4907749 | 16625 | -8.24 | 5.90E-14 | 2.38E-13 |
| chr10 | 191298 | 196760 | 5463 | -8.22 | 5.91E-14 | 2.39E-13 |
| chr5 | 54629579 | 54643577 | 13999 | -8.25 | 5.92E-14 | 2.39E-13 |
| chr14 | 56628849 | 56634132 | 5284 | -8.41 | 5.93E-14 | 2.40E-13 |

| | | | | | | |
|---|---|---|---|---|---|---|
| chr1 | 182588886 | 182591786 | 2901 | -8.2 | 5.94E-14 | 2.40E-13 |
| chr4 | 121394400 | 121407425 | 13026 | 8.27 | 5.95E-14 | 2.40E-13 |
| chr3 | 149998886 | 150010221 | 11336 | -8.2 | 5.95E-14 | 2.40E-13 |
| chr1 | 64492870 | 64495648 | 2779 | -8.2 | 5.95E-14 | 2.40E-13 |
| chr6 | 9624767 | 9632511 | 7745 | -8.27 | 5.96E-14 | 2.41E-13 |
| chr18 | 60178927 | 60183593 | 4667 | -8.48 | 5.97E-14 | 2.41E-13 |
| chr1 | 56088174 | 56103532 | 15359 | -8.2 | 5.97E-14 | 2.41E-13 |
| chr2 | 139389760 | 139391906 | 2147 | -8.3 | 5.99E-14 | 2.42E-13 |
| chr6 | 161630566 | 161641640 | 11075 | -8.27 | 5.99E-14 | 2.42E-13 |
| chr8 | 42093981 | 42112196 | 18216 | -8.81 | 6.00E-14 | 2.42E-13 |
| chr3 | 9368291 | 9374158 | 5868 | -8.37 | 6.00E-14 | 2.42E-13 |
| chr17 | 35142327 | 35157245 | 14919 | -8.58 | 6.03E-14 | 2.43E-13 |
| chr3 | 51938880 | 51945491 | 6612 | -8.24 | 6.04E-14 | 2.43E-13 |
| chr10 | 95027487 | 95031565 | 4079 | -8.31 | 6.04E-14 | 2.43E-13 |
| chr1 | 147351740 | 147354436 | 2697 | -8.25 | 6.04E-14 | 2.43E-13 |
| chr4 | 104048863 | 104051466 | 2604 | -8.21 | 6.05E-14 | 2.44E-13 |
| chr2 | 242494565 | 242497329 | 2765 | -8.24 | 6.06E-14 | 2.44E-13 |
| chr3 | 53351839 | 53372170 | 20332 | -8.81 | 6.07E-14 | 2.45E-13 |
| chr14 | 103257850 | 103260865 | 3016 | -8.2 | 6.10E-14 | 2.46E-13 |
| chr6 | 35072398 | 35080490 | 8093 | -8.21 | 6.13E-14 | 2.47E-13 |
| chr5 | 137554551 | 137576198 | 21648 | -8.63 | 6.14E-14 | 2.47E-13 |
| chr9 | 113927171 | 113935257 | 8087 | -8.26 | 6.18E-14 | 2.49E-13 |
| chr7 | 141310473 | 141312418 | 1946 | -8.54 | 6.18E-14 | 2.49E-13 |
| chr13 | 111041691 | 111046488 | 4798 | -8.19 | 6.20E-14 | 2.50E-13 |
| chr18 | 41947458 | 41992508 | 45051 | 1.54 | 6.21E-14 | 2.50E-13 |
| chr11 | 73991268 | 74010049 | 18782 | -8.19 | 6.21E-14 | 2.50E-13 |
| chr15 | 76228410 | 76233801 | 5392 | -8.19 | 6.21E-14 | 2.50E-13 |
| chr3 | 64254790 | 64268838 | 14049 | -8.23 | 6.22E-14 | 2.51E-13 |
| chr1 | 220636978 | 220648426 | 11449 | -8.2 | 6.26E-14 | 2.52E-13 |
| chr8 | 119522195 | 119525180 | 2986 | -8.2 | 6.27E-14 | 2.52E-13 |
| chr1 | 92816946 | 92819225 | 2280 | -8.23 | 6.27E-14 | 2.52E-13 |
| chr22 | 27672515 | 27679973 | 7459 | -8.28 | 6.28E-14 | 2.53E-13 |
| chr15 | 58752428 | 58761040 | 8613 | -8.29 | 6.30E-14 | 2.54E-13 |
| chr4 | 148386061 | 148401379 | 15319 | -8.24 | 6.34E-14 | 2.55E-13 |
| chr8 | 119937888 | 119962817 | 24930 | -8.4 | 6.37E-14 | 2.56E-13 |
| chr4 | 57033706 | 57043682 | 9977 | -8.21 | 6.38E-14 | 2.56E-13 |
| chr8 | 142256372 | 142261200 | 4829 | -8.25 | 6.39E-14 | 2.57E-13 |
| chr17 | 79197657 | 79201576 | 3920 | -8.2 | 6.39E-14 | 2.57E-13 |
| chr7 | 137718938 | 137737691 | 18754 | -8.63 | 6.39E-14 | 2.57E-13 |
| chr3 | 113622095 | 113624983 | 2889 | -8.21 | 6.40E-14 | 2.57E-13 |
| chr9 | 114222842 | 114225671 | 2830 | -8.19 | 6.40E-14 | 2.57E-13 |
| chr14 | 37859030 | 37865390 | 6361 | -8.19 | 6.41E-14 | 2.58E-13 |
| chr1 | 244293593 | 244296426 | 2834 | -4.87 | 6.42E-14 | 2.58E-13 |
| chr10 | 118988821 | 118991566 | 2746 | -2.78 | 6.45E-14 | 2.59E-13 |
| chr6 | 136482534 | 136488891 | 6358 | -8.21 | 6.46E-14 | 2.60E-13 |
| chr14 | 105653745 | 105656130 | 2386 | -8.26 | 6.48E-14 | 2.60E-13 |
| chr9 | 15234909 | 15239036 | 4128 | -8.38 | 6.48E-14 | 2.60E-13 |
| chr13 | 95584268 | 95596083 | 11816 | -8.19 | 6.49E-14 | 2.61E-13 |
| chr3 | 31338137 | 31341124 | 2988 | -8.23 | 6.49E-14 | 2.61E-13 |
| chrX | 129861351 | 129867446 | 6096 | -8.2 | 6.49E-14 | 2.61E-13 |
| chr16 | 52604048 | 52609525 | 5478 | -8.22 | 6.50E-14 | 2.61E-13 |
| chr6 | 22307727 | 22313777 | 6051 | -8.23 | 6.51E-14 | 2.61E-13 |
| chr12 | 90958893 | 90973101 | 14209 | -8.76 | 6.54E-14 | 2.63E-13 |
| chr5 | 92608343 | 92625664 | 17322 | -8.31 | 6.56E-14 | 2.63E-13 |
| chr7 | 83684736 | 83688272 | 3537 | -8.2 | 6.60E-14 | 2.65E-13 |
| chr12 | 22021595 | 22027942 | 6348 | -8.26 | 6.60E-14 | 2.65E-13 |
| chr17 | 74869772 | 74872244 | 2473 | -8.2 | 6.62E-14 | 2.66E-13 |
| chr2 | 54943565 | 54946437 | 2873 | -8.28 | 6.62E-14 | 2.66E-13 |
| chr6 | 140449625 | 140457508 | 7884 | -8.18 | 6.66E-14 | 2.67E-13 |
| chr18 | 45761107 | 45785145 | 24039 | -4.08 | 6.68E-14 | 2.68E-13 |
| chr20 | 57630073 | 57633460 | 3388 | -8.24 | 6.69E-14 | 2.68E-13 |

| | | | | | | |
|---|---|---|---|---|---|---|
| chr8 | 23214550 | 23218417 | 3868 | -8.45 | 6.70E-14 | 2.69E-13 |
| chr17 | 33835195 | 33845721 | 10527 | -8.23 | 6.71E-14 | 2.69E-13 |
| chr2 | 219701537 | 219707331 | 5795 | -8.19 | 6.72E-14 | 2.69E-13 |
| chr15 | 64246771 | 64258286 | 11516 | -8.24 | 6.72E-14 | 2.70E-13 |
| chr21 | 46791655 | 46797927 | 6273 | -8.4 | 6.73E-14 | 2.70E-13 |
| chr10 | 119862117 | 119994568 | 132452 | 2.71 | 6.73E-14 | 2.70E-13 |
| chr17 | 74641680 | 74649730 | 8051 | -8.35 | 6.77E-14 | 2.71E-13 |
| chr1 | 97335054 | 97339542 | 4489 | -8.18 | 6.78E-14 | 2.72E-13 |
| chr6 | 152208673 | 152228028 | 19356 | -8.51 | 6.80E-14 | 2.73E-13 |
| chr17 | 53611952 | 53627094 | 15143 | -8.19 | 6.81E-14 | 2.73E-13 |
| chr15 | 64928735 | 64938433 | 9699 | -8.47 | 6.82E-14 | 2.73E-13 |
| chr10 | 30967815 | 30970634 | 2820 | -8.23 | 6.83E-14 | 2.74E-13 |
| chr3 | 11085073 | 11087320 | 2248 | -8.32 | 6.85E-14 | 2.74E-13 |
| chr14 | 65647502 | 65650418 | 2917 | -8.18 | 6.87E-14 | 2.75E-13 |
| chr2 | 190540447 | 190544373 | 3927 | -8.3 | 6.88E-14 | 2.76E-13 |
| chr4 | 161051487 | 161055886 | 4400 | -8.21 | 6.89E-14 | 2.76E-13 |
| chrX | 19092964 | 19117764 | 24801 | -8.24 | 6.89E-14 | 2.76E-13 |
| chr8 | 10090616 | 10103956 | 13341 | -8.18 | 6.91E-14 | 2.77E-13 |
| chr6 | 40977503 | 40989037 | 11535 | -8.3 | 6.93E-14 | 2.77E-13 |
| chr13 | 96500885 | 96504081 | 3197 | -8.33 | 6.94E-14 | 2.78E-13 |
| chr1 | 7261211 | 7280891 | 19681 | -8.2 | 6.95E-14 | 2.78E-13 |
| chr20 | 5610977 | 5621439 | 10463 | -8.56 | 6.97E-14 | 2.79E-13 |
| chr11 | 2061939 | 2092370 | 30432 | 1.5 | 6.97E-14 | 2.79E-13 |
| chr5 | 127830568 | 127844817 | 14250 | -8.63 | 7.01E-14 | 2.81E-13 |
| chr4 | 140731120 | 140742975 | 11856 | -8.31 | 7.02E-14 | 2.81E-13 |
| chr19 | 48088777 | 48092778 | 4002 | -8.41 | 7.02E-14 | 2.81E-13 |
| chr11 | 16266067 | 16270693 | 4627 | -8.25 | 7.03E-14 | 2.81E-13 |
| chr2 | 159376109 | 159379416 | 3308 | -8.48 | 7.03E-14 | 2.81E-13 |
| chr2 | 197323504 | 197329900 | 6397 | -8.36 | 7.05E-14 | 2.82E-13 |
| chr1 | 60389800 | 60392194 | 2395 | -8.25 | 7.05E-14 | 2.82E-13 |
| chr6 | 27936850 | 27940203 | 3354 | -8.25 | 7.06E-14 | 2.82E-13 |
| chr7 | 33326078 | 33343106 | 17029 | -8.32 | 7.06E-14 | 2.82E-13 |
| chr22 | 48706634 | 48726325 | 19692 | -8.49 | 7.07E-14 | 2.83E-13 |
| chr2 | 38433299 | 38462403 | 29105 | -8.57 | 7.07E-14 | 2.83E-13 |
| chr3 | 36505952 | 36521811 | 15860 | -8.42 | 7.08E-14 | 2.83E-13 |
| chr9 | 138622684 | 138624807 | 2124 | -8.21 | 7.09E-14 | 2.83E-13 |
| chr13 | 33285817 | 33288235 | 2419 | -8.29 | 7.10E-14 | 2.84E-13 |
| chr4 | 101863156 | 101867312 | 4157 | -8.33 | 7.12E-14 | 2.84E-13 |
| chr1 | 212196480 | 212198909 | 2430 | -8.18 | 7.12E-14 | 2.84E-13 |
| chr6 | 83499842 | 83512211 | 12370 | -2.39 | 7.14E-14 | 2.85E-13 |
| chr12 | 18307282 | 18315472 | 8191 | -8.2 | 7.14E-14 | 2.85E-13 |
| chr11 | 116401090 | 116422063 | 20974 | -3.43 | 7.15E-14 | 2.86E-13 |
| chrX | 40780318 | 40789079 | 8762 | -8.46 | 7.16E-14 | 2.86E-13 |
| chr22 | 44955568 | 44966980 | 11413 | -8.18 | 7.17E-14 | 2.86E-13 |
| chr7 | 109565336 | 109593665 | 28330 | -8.26 | 7.17E-14 | 2.86E-13 |
| chr17 | 56520642 | 56533400 | 12759 | -8.71 | 7.18E-14 | 2.86E-13 |
| chr6 | 79833530 | 79846620 | 13091 | -6.02 | 7.18E-14 | 2.87E-13 |
| chr17 | 48145853 | 48149475 | 3623 | -8.18 | 7.18E-14 | 2.87E-13 |
| chrX | 96209688 | 96224855 | 15168 | -8.27 | 7.18E-14 | 2.87E-13 |
| chr2 | 150930041 | 150943128 | 13088 | -8.22 | 7.19E-14 | 2.87E-13 |
| chr3 | 48653488 | 48656645 | 3158 | -8.28 | 7.19E-14 | 2.87E-13 |
| chr9 | 101559547 | 101587051 | 27505 | -8.78 | 7.19E-14 | 2.87E-13 |
| chr5 | 108540195 | 108584496 | 44302 | -2.97 | 7.24E-14 | 2.89E-13 |
| chr6 | 74671179 | 74692095 | 20917 | -8.39 | 7.26E-14 | 2.89E-13 |
| chr22 | 39553683 | 39563124 | 9442 | -8.18 | 7.27E-14 | 2.90E-13 |
| chr2 | 19645149 | 19665792 | 20644 | -8.42 | 7.33E-14 | 2.92E-13 |
| chr20 | 15182718 | 15193775 | 11058 | -8.64 | 7.36E-14 | 2.93E-13 |
| chr5 | 52577272 | 52585712 | 8441 | -8.22 | 7.36E-14 | 2.94E-13 |
| chr1 | 234143558 | 234151486 | 7929 | -8.26 | 7.37E-14 | 2.94E-13 |
| chr2 | 233931860 | 233938237 | 6378 | -8.17 | 7.37E-14 | 2.94E-13 |
| chr5 | 124952704 | 124960789 | 8086 | -8.27 | 7.38E-14 | 2.94E-13 |

| | | | | | | |
|---|---|---|---|---|---|---|
| chr7 | 76837336 | 76843432 | 6097 | -8.17 | 7.41E-14 | 2.95E-13 |
| chr18 | 7813314 | 7816566 | 3253 | -8.28 | 7.43E-14 | 2.96E-13 |
| chr20 | 48192692 | 48199947 | 7256 | -8.23 | 7.45E-14 | 2.97E-13 |
| chr2 | 30345618 | 30347800 | 2183 | -8.2 | 7.45E-14 | 2.97E-13 |
| chr5 | 80370999 | 80378419 | 7421 | -3.07 | 7.45E-14 | 2.97E-13 |
| chr17 | 6442017 | 6444053 | 2037 | -8.21 | 7.46E-14 | 2.97E-13 |
| chr13 | 77504552 | 77507169 | 2618 | -6.6 | 7.46E-14 | 2.97E-13 |
| chr6 | 86101211 | 86108535 | 7325 | -8.18 | 7.48E-14 | 2.98E-13 |
| chr16 | 75802474 | 75805660 | 3187 | -8.18 | 7.49E-14 | 2.98E-13 |
| chr1 | 8145180 | 8187142 | 41963 | -8.8 | 7.50E-14 | 2.98E-13 |
| chr5 | 118533675 | 118539437 | 5763 | -8.29 | 7.50E-14 | 2.99E-13 |
| chr2 | 186957805 | 186984676 | 26872 | 8.58 | 7.54E-14 | 3.00E-13 |
| chr1 | 116586390 | 116602530 | 16141 | -6.08 | 7.55E-14 | 3.01E-13 |
| chr5 | 17354651 | 17373675 | 19025 | -8.21 | 7.55E-14 | 3.01E-13 |
| chr20 | 56107867 | 56115258 | 7392 | -8.36 | 7.56E-14 | 3.01E-13 |
| chr16 | 72117621 | 72120173 | 2553 | -8.24 | 7.56E-14 | 3.01E-13 |
| chr5 | 113507029 | 113521564 | 14536 | -8.25 | 7.57E-14 | 3.01E-13 |
| chr18 | 7905532 | 7919113 | 13582 | -8.45 | 7.58E-14 | 3.01E-13 |
| chr7 | 19776943 | 19794809 | 17867 | -8.45 | 7.58E-14 | 3.01E-13 |
| chr9 | 127098837 | 127101046 | 2210 | -8.25 | 7.58E-14 | 3.01E-13 |
| chrX | 24910516 | 24912821 | 2306 | -8.36 | 7.61E-14 | 3.03E-13 |
| chr13 | 99524752 | 99527512 | 2761 | -8.18 | 7.61E-14 | 3.03E-13 |
| chr3 | 190275981 | 190285107 | 9127 | -8.18 | 7.62E-14 | 3.03E-13 |
| chr1 | 198204734 | 198206650 | 1917 | -8.2 | 7.66E-14 | 3.04E-13 |
| chr3 | 188325298 | 188327687 | 2390 | -8.2 | 7.69E-14 | 3.06E-13 |
| chr20 | 42457276 | 42460822 | 3547 | -8.17 | 7.70E-14 | 3.06E-13 |
| chr3 | 4413946 | 4417225 | 3280 | -8.21 | 7.70E-14 | 3.06E-13 |
| chr6 | 138315693 | 138321042 | 5350 | -8.22 | 7.71E-14 | 3.06E-13 |
| chr14 | 39820869 | 39834678 | 13810 | -8.45 | 7.73E-14 | 3.07E-13 |
| chr10 | 111744290 | 111749684 | 5395 | -8.37 | 7.73E-14 | 3.07E-13 |
| chr20 | 30632355 | 30651442 | 19088 | -8.19 | 7.74E-14 | 3.07E-13 |
| chr11 | 68428974 | 68442295 | 13322 | -8.47 | 7.74E-14 | 3.08E-13 |
| chr11 | 129605991 | 129624055 | 18065 | -8.25 | 7.77E-14 | 3.09E-13 |
| chr3 | 159460371 | 159475925 | 15555 | -8.45 | 7.78E-14 | 3.09E-13 |
| chr7 | 157235314 | 157249382 | 14069 | -8.43 | 7.79E-14 | 3.10E-13 |
| chr6 | 11141316 | 11160977 | 19662 | -8.26 | 7.80E-14 | 3.10E-13 |
| chr2 | 196545401 | 196566529 | 21129 | -8.24 | 7.81E-14 | 3.10E-13 |
| chr3 | 58463909 | 58467590 | 3682 | -8.35 | 7.86E-14 | 3.12E-13 |
| chr6 | 75300463 | 75315714 | 15252 | -8.22 | 7.87E-14 | 3.12E-13 |
| chr2 | 3910310 | 3993269 | 82960 | 1.32 | 7.90E-14 | 3.14E-13 |
| chr20 | 45518065 | 45520641 | 2577 | -8.17 | 7.91E-14 | 3.14E-13 |
| chr9 | 108496629 | 108515105 | 18477 | -8.33 | 7.91E-14 | 3.14E-13 |
| chr3 | 72780907 | 72784644 | 3738 | -8.29 | 7.93E-14 | 3.14E-13 |
| chr20 | 43457385 | 43484294 | 26910 | -7.16 | 7.97E-14 | 3.16E-13 |
| chr6 | 125429850 | 125434287 | 4438 | -8.2 | 7.98E-14 | 3.17E-13 |
| chr1 | 117353999 | 117357015 | 3017 | -8.17 | 8.01E-14 | 3.17E-13 |
| chr1 | 240430622 | 240433732 | 3111 | -8.17 | 8.01E-14 | 3.17E-13 |
| chr4 | 96486757 | 96495826 | 9070 | -8.49 | 8.01E-14 | 3.18E-13 |
| chr6 | 48656773 | 48666072 | 9300 | -8.17 | 8.03E-14 | 3.18E-13 |
| chr1 | 114167927 | 114172143 | 4217 | -8.19 | 8.06E-14 | 3.19E-13 |
| chr6 | 78411799 | 78426262 | 14464 | 2.11 | 8.07E-14 | 3.20E-13 |
| chr19 | 50682692 | 50693435 | 10744 | -8.46 | 8.08E-14 | 3.20E-13 |
| chr14 | 102739796 | 102754978 | 15183 | -8.55 | 8.09E-14 | 3.21E-13 |
| chr1 | 187336121 | 187352810 | 16690 | 8.32 | 8.11E-14 | 3.21E-13 |
| chr3 | 19947259 | 19955960 | 8702 | -8.54 | 8.11E-14 | 3.21E-13 |
| chr14 | 70297023 | 70300697 | 3675 | -8.2 | 8.12E-14 | 3.22E-13 |
| chr7 | 148933947 | 148941200 | 7254 | -8.16 | 8.15E-14 | 3.23E-13 |
| chr10 | 4039974 | 4044139 | 4166 | -8.16 | 8.16E-14 | 3.23E-13 |
| chrX | 47498968 | 47500996 | 2029 | -8.2 | 8.18E-14 | 3.24E-13 |
| chr11 | 10572805 | 10574649 | 1845 | -6.15 | 8.19E-14 | 3.24E-13 |
| chr13 | 75970449 | 75974891 | 4443 | -8.21 | 8.23E-14 | 3.26E-13 |

| | | | | | | |
|---|---|---|---|---|---|---|
| chr5 | 127879066 | 127890372 | 11307 | -8.2 | 8.26E-14 | 3.27E-13 |
| chr7 | 33248944 | 33254942 | 5999 | -8.2 | 8.32E-14 | 3.29E-13 |
| chr4 | 78050532 | 78053737 | 3206 | -8.16 | 8.33E-14 | 3.30E-13 |
| chr8 | 90077718 | 90086397 | 8680 | -8.28 | 8.35E-14 | 3.30E-13 |
| chr12 | 125060575 | 125084496 | 23922 | -8.65 | 8.35E-14 | 3.30E-13 |
| chr7 | 4890284 | 4908878 | 18595 | -8.54 | 8.36E-14 | 3.31E-13 |
| chr7 | 68118200 | 68123223 | 5024 | -8.31 | 8.37E-14 | 3.31E-13 |
| chr4 | 3795488 | 3799216 | 3729 | -8.27 | 8.39E-14 | 3.32E-13 |
| chr2 | 167157854 | 167174878 | 17025 | 8.23 | 8.40E-14 | 3.32E-13 |
| chr6 | 133789551 | 133791304 | 1754 | -8.2 | 8.44E-14 | 3.34E-13 |
| chr17 | 9294358 | 9297605 | 3248 | -8.25 | 8.44E-14 | 3.34E-13 |
| chr5 | 60979064 | 61083203 | 104140 | -1.03 | 8.49E-14 | 3.36E-13 |
| chr8 | 66755959 | 66758843 | 2885 | -8.16 | 8.52E-14 | 3.37E-13 |
| chr14 | 22645190 | 22651681 | 6492 | 2.46 | 8.53E-14 | 3.37E-13 |
| chr6 | 153656675 | 153667903 | 11229 | -8.27 | 8.53E-14 | 3.37E-13 |
| chr8 | 60802879 | 60958862 | 155984 | 3.61 | 8.53E-14 | 3.37E-13 |
| chr11 | 6602931 | 6606665 | 3735 | -8.16 | 8.53E-14 | 3.37E-13 |
| chr7 | 157322989 | 157326723 | 3735 | -8.18 | 8.54E-14 | 3.38E-13 |
| chr6 | 159151243 | 159161733 | 10491 | -8.32 | 8.55E-14 | 3.38E-13 |
| chr11 | 4386579 | 4396348 | 9770 | 8.27 | 8.55E-14 | 3.38E-13 |
| chr1 | 82483642 | 82487834 | 4193 | -6.02 | 8.57E-14 | 3.39E-13 |
| chr9 | 6370022 | 6374510 | 4489 | -8.16 | 8.62E-14 | 3.41E-13 |
| chr16 | 1015088 | 1018352 | 3265 | -8.2 | 8.62E-14 | 3.41E-13 |
| chr1 | 184994946 | 184999318 | 4373 | -2.85 | 8.63E-14 | 3.41E-13 |
| chr3 | 182919596 | 182923238 | 3643 | -8.19 | 8.63E-14 | 3.41E-13 |
| chr11 | 16910989 | 16913609 | 2621 | -8.23 | 8.66E-14 | 3.42E-13 |
| chr20 | 15208821 | 15211766 | 2946 | -8.23 | 8.67E-14 | 3.42E-13 |
| chr4 | 169078877 | 169083297 | 4421 | -8.16 | 8.67E-14 | 3.42E-13 |
| chr2 | 106648858 | 106652007 | 3150 | -8.26 | 8.69E-14 | 3.43E-13 |
| chr7 | 107720799 | 107724054 | 3256 | -8.22 | 8.70E-14 | 3.43E-13 |
| chr21 | 47435302 | 47451276 | 15975 | -8.15 | 8.72E-14 | 3.44E-13 |
| chr5 | 159047697 | 159064993 | 17297 | -8.77 | 8.72E-14 | 3.44E-13 |
| chr6 | 55187269 | 55205494 | 18226 | -8.32 | 8.75E-14 | 3.45E-13 |
| chr2 | 742339 | 842172 | 99834 | -1.42 | 8.78E-14 | 3.46E-13 |
| chr14 | 70106727 | 70109277 | 2551 | -8.16 | 8.78E-14 | 3.46E-13 |
| chr1 | 41684010 | 41687012 | 3003 | -8.2 | 8.81E-14 | 3.48E-13 |
| chr6 | 11099411 | 11102959 | 3549 | -8.19 | 8.84E-14 | 3.49E-13 |
| chr13 | 50941372 | 50956674 | 15303 | -8.37 | 8.84E-14 | 3.49E-13 |
| chr2 | 170971771 | 170993357 | 21587 | -8.3 | 8.86E-14 | 3.49E-13 |
| chr1 | 178892445 | 178894966 | 2522 | -8.16 | 8.87E-14 | 3.50E-13 |
| chr6 | 13940174 | 13944027 | 3854 | -8.18 | 8.88E-14 | 3.50E-13 |
| chr1 | 67849378 | 67853509 | 4132 | -8.18 | 8.91E-14 | 3.51E-13 |
| chr2 | 3660347 | 3671635 | 11289 | -8.46 | 8.92E-14 | 3.52E-13 |
| chr3 | 119950216 | 119954712 | 4497 | -8.18 | 8.93E-14 | 3.52E-13 |
| chr14 | 99699261 | 99704503 | 5243 | -8.29 | 8.94E-14 | 3.52E-13 |
| chr20 | 14714288 | 14717615 | 3328 | -8.26 | 8.95E-14 | 3.53E-13 |
| chr7 | 80638333 | 80642193 | 3861 | -8.17 | 8.95E-14 | 3.53E-13 |
| chr12 | 95942579 | 95944287 | 1709 | -8.15 | 8.98E-14 | 3.54E-13 |
| chr10 | 34514537 | 34517658 | 3122 | -8.26 | 8.98E-14 | 3.54E-13 |
| chr16 | 27318764 | 27321038 | 2275 | -8.2 | 9.04E-14 | 3.56E-13 |
| chr4 | 34855127 | 34865659 | 10533 | 2.84 | 9.10E-14 | 3.59E-13 |
| chr4 | 71860118 | 71878100 | 17983 | -8.34 | 9.11E-14 | 3.59E-13 |
| chr9 | 17136362 | 17171473 | 35112 | -8.37 | 9.12E-14 | 3.59E-13 |
| chr9 | 29292590 | 29300701 | 8112 | -8.16 | 9.16E-14 | 3.61E-13 |
| chr9 | 114358109 | 114366144 | 8036 | -8.15 | 9.17E-14 | 3.61E-13 |
| chr6 | 164011638 | 164018621 | 6984 | -8.16 | 9.18E-14 | 3.61E-13 |
| chr11 | 76044576 | 76048170 | 3595 | -8.19 | 9.19E-14 | 3.62E-13 |
| chr4 | 105647055 | 105655768 | 8714 | -8.19 | 9.19E-14 | 3.62E-13 |
| chr5 | 55281710 | 55300210 | 18501 | -8.61 | 9.20E-14 | 3.62E-13 |
| chr11 | 117812910 | 117828793 | 15884 | -8.14 | 9.20E-14 | 3.62E-13 |
| chr2 | 62578357 | 62614675 | 36319 | -8.86 | 9.21E-14 | 3.62E-13 |

| | | | | | | |
|---|---|---|---|---|---|---|
| chr5 | 39868183 | 39882311 | 14129 | -8.45 | 9.23E-14 | 3.63E-13 |
| chr18 | 54246945 | 54253324 | 6380 | -8.17 | 9.23E-14 | 3.63E-13 |
| chr1 | 197844104 | 197848370 | 4267 | -8.24 | 9.24E-14 | 3.64E-13 |
| chr3 | 111463732 | 111467637 | 3906 | -8.25 | 9.25E-14 | 3.64E-13 |
| chr12 | 60125268 | 60145008 | 19741 | -8.33 | 9.26E-14 | 3.64E-13 |
| chr3 | 67692705 | 67727864 | 35160 | -8.41 | 9.26E-14 | 3.64E-13 |
| chr10 | 101931557 | 101933594 | 2038 | -8.25 | 9.27E-14 | 3.64E-13 |
| chr2 | 157276964 | 157279365 | 2402 | -8.31 | 9.29E-14 | 3.65E-13 |
| chr12 | 96566039 | 96571627 | 5589 | -8.16 | 9.32E-14 | 3.66E-13 |
| chr5 | 176111311 | 176113619 | 2309 | -8.16 | 9.33E-14 | 3.67E-13 |
| chr6 | 162483536 | 162502915 | 19380 | -8.15 | 9.37E-14 | 3.68E-13 |
| chrX | 74116884 | 74132465 | 15582 | 2.33 | 9.38E-14 | 3.69E-13 |
| chr16 | 50366694 | 50390223 | 23530 | -8.15 | 9.38E-14 | 3.69E-13 |
| chrX | 107495273 | 107511021 | 15749 | -8.24 | 9.41E-14 | 3.70E-13 |
| chr1 | 41241212 | 41243585 | 2374 | -8.14 | 9.43E-14 | 3.70E-13 |
| chr2 | 65854478 | 65860817 | 6340 | -8.16 | 9.50E-14 | 3.73E-13 |
| chr9 | 79293780 | 79296020 | 2241 | -8.17 | 9.50E-14 | 3.73E-13 |
| chr1 | 92485136 | 92490571 | 5436 | -8.2 | 9.55E-14 | 3.75E-13 |
| chr10 | 96266966 | 96269596 | 2631 | -8.24 | 9.56E-14 | 3.76E-13 |
| chr13 | 91817494 | 91823632 | 6139 | 8.27 | 9.59E-14 | 3.77E-13 |
| chr12 | 14702421 | 14707017 | 4597 | -8.17 | 9.60E-14 | 3.77E-13 |
| chr10 | 5809279 | 5828462 | 19184 | -8.16 | 9.61E-14 | 3.77E-13 |
| chrX | 44832354 | 44838557 | 6204 | -8.14 | 9.61E-14 | 3.77E-13 |
| chr10 | 13711098 | 13715923 | 4826 | -8.53 | 9.64E-14 | 3.78E-13 |
| chr12 | 69623404 | 69627920 | 4517 | -8.44 | 9.64E-14 | 3.78E-13 |
| chr12 | 18438836 | 18456682 | 17847 | -8.15 | 9.64E-14 | 3.78E-13 |
| chr6 | 36144520 | 36148279 | 3760 | -8.17 | 9.65E-14 | 3.79E-13 |
| chr19 | 41229129 | 41234041 | 4913 | -8.19 | 9.68E-14 | 3.80E-13 |
| chr1 | 189229649 | 189239959 | 10311 | -8.32 | 9.70E-14 | 3.80E-13 |
| chr1 | 162059409 | 162079120 | 19712 | -8.43 | 9.71E-14 | 3.81E-13 |
| chr1 | 42625358 | 42627550 | 2193 | -8.19 | 9.72E-14 | 3.81E-13 |
| chr7 | 80428762 | 80439739 | 10978 | -8.19 | 9.72E-14 | 3.81E-13 |
| chr2 | 163826027 | 163841304 | 15278 | 8.25 | 9.72E-14 | 3.81E-13 |
| chr21 | 38949248 | 38953268 | 4021 | -8.25 | 9.73E-14 | 3.81E-13 |
| chr17 | 44975753 | 44977513 | 1761 | -8.14 | 9.73E-14 | 3.81E-13 |
| chr1 | 221970238 | 221974370 | 4133 | -8.19 | 9.76E-14 | 3.82E-13 |
| chr1 | 19702944 | 19704809 | 1866 | -8.14 | 9.76E-14 | 3.83E-13 |
| chr9 | 72547876 | 72558172 | 10297 | -8.14 | 9.76E-14 | 3.83E-13 |
| chr5 | 169087483 | 169091744 | 4262 | -8.14 | 9.79E-14 | 3.84E-13 |
| chr15 | 29975129 | 29978970 | 3842 | -8.14 | 9.81E-14 | 3.84E-13 |
| chr4 | 161190868 | 161203940 | 13073 | -8.26 | 9.81E-14 | 3.85E-13 |
| chr3 | 123073128 | 123088829 | 15702 | -8.15 | 9.82E-14 | 3.85E-13 |
| chr5 | 141857769 | 141860808 | 3040 | -8.15 | 9.83E-14 | 3.85E-13 |
| chr17 | 46310618 | 46313544 | 2927 | -8.22 | 9.89E-14 | 3.88E-13 |
| chr13 | 51377930 | 51381755 | 3826 | -8.23 | 9.90E-14 | 3.88E-13 |
| chr7 | 98906532 | 98909048 | 2517 | -8.14 | 9.90E-14 | 3.88E-13 |
| chr3 | 152042470 | 152044990 | 2521 | -8.24 | 9.91E-14 | 3.88E-13 |
| chr12 | 99373383 | 99380815 | 7433 | -8.23 | 9.93E-14 | 3.89E-13 |
| chr15 | 76511758 | 76514272 | 2515 | -8.13 | 9.99E-14 | 3.91E-13 |
| chr2 | 99708911 | 99712192 | 3282 | -8.14 | 1.00E-13 | 3.92E-13 |
| chr2 | 153716973 | 153721208 | 4236 | -8.18 | 1.01E-13 | 3.93E-13 |
| chr9 | 5402204 | 5409910 | 7707 | -8.28 | 1.01E-13 | 3.94E-13 |
| chr10 | 132009800 | 132892615 | 882816 | 0.71 | 1.01E-13 | 3.95E-13 |
| chr3 | 28133942 | 28157459 | 23518 | -2.39 | 1.01E-13 | 3.95E-13 |
| chr17 | 542974 | 545450 | 2477 | -8.14 | 1.01E-13 | 3.97E-13 |
| chr3 | 184726562 | 184738383 | 11822 | -8.13 | 1.02E-13 | 3.98E-13 |
| chr1 | 224763473 | 224771964 | 8492 | -2.06 | 1.02E-13 | 3.98E-13 |
| chr6 | 134732257 | 134743011 | 10755 | -8.13 | 1.02E-13 | 3.99E-13 |
| chr9 | 91410641 | 91419301 | 8661 | -8.14 | 1.02E-13 | 3.99E-13 |
| chr3 | 125161232 | 125165699 | 4468 | -8.16 | 1.02E-13 | 4.00E-13 |
| chrX | 53036509 | 53039383 | 2875 | -8.16 | 1.02E-13 | 4.00E-13 |

| | | | | | | |
|---|---|---|---|---|---|---|
| chr13 | 76404555 | 76417628 | 13074 | -8.23 | 1.02E-13 | 4.00E-13 |
| chr16 | 68886855 | 68892237 | 5383 | -8.24 | 1.02E-13 | 4.00E-13 |
| chr12 | 45718097 | 45723056 | 4960 | -8.4 | 1.03E-13 | 4.01E-13 |
| chr2 | 54379323 | 54383113 | 3791 | -8.24 | 1.03E-13 | 4.01E-13 |
| chr5 | 95878096 | 95900066 | 21971 | -6.73 | 1.03E-13 | 4.03E-13 |
| chr17 | 26536529 | 26539154 | 2626 | -8.14 | 1.03E-13 | 4.03E-13 |
| chr4 | 42314730 | 42326478 | 11749 | -8.16 | 1.03E-13 | 4.03E-13 |
| chr5 | 363640 | 366974 | 3335 | -8.31 | 1.03E-13 | 4.03E-13 |
| chr4 | 3168787 | 3172093 | 3307 | -8.13 | 1.03E-13 | 4.04E-13 |
| chr7 | 139644895 | 139648610 | 3716 | -8.19 | 1.04E-13 | 4.04E-13 |
| chrX | 62511789 | 62529517 | 17729 | -8.44 | 1.04E-13 | 4.05E-13 |
| chr5 | 100341803 | 100370658 | 28856 | -8.39 | 1.04E-13 | 4.06E-13 |
| chr15 | 65428198 | 65431517 | 3320 | -8.13 | 1.04E-13 | 4.07E-13 |
| chr7 | 38120337 | 38140438 | 20102 | -8.15 | 1.04E-13 | 4.07E-13 |
| chr12 | 92443651 | 92471590 | 27940 | -8.32 | 1.04E-13 | 4.07E-13 |
| chr4 | 170402549 | 170424527 | 21979 | -8.13 | 1.04E-13 | 4.07E-13 |
| chr3 | 142005250 | 142009162 | 3913 | -8.23 | 1.05E-13 | 4.10E-13 |
| chr3 | 105968684 | 105973486 | 4803 | -8.21 | 1.05E-13 | 4.10E-13 |
| chr3 | 24665923 | 24678370 | 12448 | -8.16 | 1.05E-13 | 4.11E-13 |
| chr2 | 162328229 | 162339639 | 11411 | -3.44 | 1.06E-13 | 4.12E-13 |
| chr16 | 8810450 | 8813067 | 2618 | -8.24 | 1.06E-13 | 4.13E-13 |
| chr3 | 3226321 | 3230307 | 3987 | -8.2 | 1.06E-13 | 4.14E-13 |
| chr10 | 77362103 | 77377932 | 15830 | 8.35 | 1.06E-13 | 4.14E-13 |
| chr5 | 64524221 | 64527778 | 3558 | -3.41 | 1.06E-13 | 4.14E-13 |
| chr7 | 51267024 | 51273057 | 6034 | -8.25 | 1.06E-13 | 4.15E-13 |
| chr5 | 148988619 | 148991247 | 2629 | -8.13 | 1.06E-13 | 4.15E-13 |
| chrX | 82549309 | 82566955 | 17647 | -2.8 | 1.06E-13 | 4.15E-13 |
| chr4 | 19436202 | 19454554 | 18353 | -8.13 | 1.06E-13 | 4.15E-13 |
| chr9 | 119793967 | 119798446 | 4480 | 8.25 | 1.06E-13 | 4.15E-13 |
| chr12 | 11141534 | 11154645 | 13112 | -8.16 | 1.07E-13 | 4.16E-13 |
| chr5 | 92647403 | 92651622 | 4220 | -8.13 | 1.07E-13 | 4.17E-13 |
| chr4 | 76423469 | 76425585 | 2117 | -8.19 | 1.07E-13 | 4.18E-13 |
| chr14 | 43586532 | 43606659 | 20128 | -8.4 | 1.07E-13 | 4.19E-13 |
| chr2 | 242109667 | 242112223 | 2557 | -8.18 | 1.07E-13 | 4.19E-13 |
| chr15 | 71599149 | 71612328 | 13180 | -8.34 | 1.08E-13 | 4.19E-13 |
| chr2 | 225677651 | 225681433 | 3783 | -8.25 | 1.08E-13 | 4.19E-13 |
| chr11 | 134520236 | 134545222 | 24987 | -3.88 | 1.08E-13 | 4.21E-13 |
| chr1 | 211039015 | 211046249 | 7235 | -8.16 | 1.08E-13 | 4.22E-13 |
| chr13 | 67251523 | 67261678 | 10156 | -8.23 | 1.08E-13 | 4.22E-13 |
| chr1 | 220476821 | 220484608 | 7788 | -8.13 | 1.08E-13 | 4.22E-13 |
| chr2 | 154756691 | 154766074 | 9384 | -8.31 | 1.09E-13 | 4.24E-13 |
| chr17 | 17347160 | 17373283 | 26124 | -8.81 | 1.09E-13 | 4.26E-13 |
| chr10 | 62942975 | 62960376 | 17402 | -8.41 | 1.09E-13 | 4.26E-13 |
| chr4 | 93886684 | 93894544 | 7861 | -8.22 | 1.10E-13 | 4.27E-13 |
| chr7 | 22963255 | 22974529 | 11275 | -8.27 | 1.10E-13 | 4.27E-13 |
| chr6 | 539333 | 543386 | 4054 | -8.12 | 1.10E-13 | 4.28E-13 |
| chr17 | 59268143 | 59271060 | 2918 | -8.12 | 1.10E-13 | 4.29E-13 |
| chr13 | 110354025 | 110379801 | 25777 | -9.02 | 1.10E-13 | 4.29E-13 |
| chr4 | 124634310 | 124656112 | 21803 | -8.48 | 1.11E-13 | 4.31E-13 |
| chr3 | 189957172 | 189965409 | 8238 | -8.33 | 1.11E-13 | 4.31E-13 |
| chr11 | 63557359 | 63560700 | 3342 | -8.22 | 1.11E-13 | 4.32E-13 |
| chr20 | 57026656 | 57030936 | 4281 | -8.3 | 1.11E-13 | 4.32E-13 |
| chr9 | 16287253 | 16299294 | 12042 | -8.38 | 1.11E-13 | 4.33E-13 |
| chr2 | 44390546 | 44393589 | 3044 | -8.26 | 1.11E-13 | 4.33E-13 |
| chr1 | 223408961 | 223411598 | 2638 | -8.18 | 1.12E-13 | 4.34E-13 |
| chr4 | 102477954 | 102505210 | 27257 | -8.62 | 1.12E-13 | 4.35E-13 |
| chr10 | 118692070 | 118700090 | 8021 | -8.12 | 1.12E-13 | 4.35E-13 |
| chr15 | 90921499 | 90923733 | 2235 | -6.8 | 1.12E-13 | 4.36E-13 |
| chr4 | 161120790 | 161146117 | 25328 | -8.29 | 1.12E-13 | 4.36E-13 |
| chr13 | 36504618 | 36518189 | 13572 | -8.12 | 1.13E-13 | 4.39E-13 |
| chr22 | 25732588 | 25741085 | 8498 | -8.12 | 1.13E-13 | 4.39E-13 |

| | | | | | | |
|---|---|---|---|---|---|---|
| chr12 | 21740029 | 21745799 | 5771 | -8.15 | 1.13E-13 | 4.39E-13 |
| chr6 | 170457346 | 170459823 | 2478 | -5.92 | 1.13E-13 | 4.39E-13 |
| chr6 | 161781644 | 161786094 | 4451 | -8.12 | 1.13E-13 | 4.40E-13 |
| chr14 | 68763217 | 68770940 | 7724 | -8.38 | 1.13E-13 | 4.40E-13 |
| chr4 | 56194878 | 56198665 | 3788 | -8.28 | 1.13E-13 | 4.40E-13 |
| chr19 | 50841935 | 50849031 | 7097 | -8.4 | 1.14E-13 | 4.42E-13 |
| chrX | 29574470 | 29580976 | 6507 | -8.22 | 1.14E-13 | 4.43E-13 |
| chr14 | 61519658 | 61528564 | 8907 | -8.17 | 1.14E-13 | 4.43E-13 |
| chr2 | 136348167 | 136351156 | 2990 | -8.13 | 1.14E-13 | 4.44E-13 |
| chr1 | 221791748 | 221800889 | 9142 | -8.15 | 1.15E-13 | 4.45E-13 |
| chr5 | 74150996 | 74153887 | 2892 | -8.23 | 1.15E-13 | 4.45E-13 |
| chr2 | 11011531 | 11014452 | 2922 | -8.16 | 1.15E-13 | 4.45E-13 |
| chr14 | 52370127 | 52374226 | 4100 | -8.12 | 1.15E-13 | 4.45E-13 |
| chr3 | 118758312 | 118773840 | 15529 | -8.26 | 1.15E-13 | 4.46E-13 |
| chr2 | 181353382 | 181364953 | 11572 | -8.18 | 1.15E-13 | 4.46E-13 |
| chr16 | 19457534 | 19459178 | 1645 | -8.13 | 1.15E-13 | 4.47E-13 |
| chr1 | 230987842 | 230990736 | 2895 | -8.24 | 1.15E-13 | 4.47E-13 |
| chr7 | 127616933 | 127619780 | 2848 | -8.18 | 1.15E-13 | 4.47E-13 |
| chr10 | 63194399 | 63203241 | 8843 | 8.28 | 1.15E-13 | 4.47E-13 |
| chr2 | 204261811 | 204283937 | 22127 | -8.58 | 1.15E-13 | 4.47E-13 |
| chr3 | 61946631 | 61960779 | 14149 | 8.27 | 1.15E-13 | 4.48E-13 |
| chr8 | 48309176 | 48314766 | 5591 | -8.12 | 1.16E-13 | 4.48E-13 |
| chr9 | 114843918 | 114851481 | 7564 | -8.35 | 1.16E-13 | 4.48E-13 |
| chrX | 131852164 | 132076881 | 224718 | 2.79 | 1.16E-13 | 4.48E-13 |
| chr4 | 81325562 | 81335986 | 10425 | 8.3 | 1.16E-13 | 4.49E-13 |
| chr11 | 45707036 | 45716327 | 9292 | 8.27 | 1.16E-13 | 4.49E-13 |
| chr4 | 3850796 | 3853266 | 2471 | -7.04 | 1.16E-13 | 4.50E-13 |
| chrX | 109316193 | 109320647 | 4455 | -8.12 | 1.16E-13 | 4.51E-13 |
| chr20 | 52906743 | 52924817 | 18075 | 8.71 | 1.17E-13 | 4.52E-13 |
| chr2 | 233176473 | 233179072 | 2600 | -8.13 | 1.17E-13 | 4.52E-13 |
| chr4 | 101362057 | 101374095 | 12039 | -8.15 | 1.17E-13 | 4.53E-13 |
| chr3 | 36292151 | 36306387 | 14237 | -8.22 | 1.18E-13 | 4.55E-13 |
| chr16 | 75154368 | 75158056 | 3689 | -8.4 | 1.18E-13 | 4.55E-13 |
| chr2 | 61953153 | 61956085 | 2933 | -8.19 | 1.18E-13 | 4.56E-13 |
| chr6 | 131327815 | 131344037 | 16223 | -8.13 | 1.18E-13 | 4.56E-13 |
| chr20 | 10659361 | 10662534 | 3174 | -8.12 | 1.18E-13 | 4.56E-13 |
| chr3 | 138469063 | 138488830 | 19768 | -8.14 | 1.18E-13 | 4.58E-13 |
| chr6 | 128695682 | 128699848 | 4167 | -8.11 | 1.18E-13 | 4.58E-13 |
| chr1 | 117944605 | 117947141 | 2537 | -8.16 | 1.18E-13 | 4.58E-13 |
| chr3 | 98734296 | 98753916 | 19621 | -8.18 | 1.19E-13 | 4.59E-13 |
| chr5 | 177605720 | 177622032 | 16313 | -8.52 | 1.19E-13 | 4.61E-13 |
| chr22 | 29341166 | 29344741 | 3576 | -8.2 | 1.20E-13 | 4.65E-13 |
| chr3 | 87744847 | 87757229 | 12383 | -8.16 | 1.20E-13 | 4.66E-13 |
| chr2 | 202704589 | 202706960 | 2372 | -8.15 | 1.21E-13 | 4.67E-13 |
| chr16 | 24723018 | 24733561 | 10544 | -8.18 | 1.21E-13 | 4.69E-13 |
| chr12 | 16795699 | 16993853 | 198155 | 2.45 | 1.21E-13 | 4.69E-13 |
| chr2 | 19895482 | 19902851 | 7370 | -8.11 | 1.22E-13 | 4.71E-13 |
| chr14 | 68010406 | 68012852 | 2447 | -8.21 | 1.22E-13 | 4.71E-13 |
| chr7 | 133834688 | 133837370 | 2683 | -8.11 | 1.22E-13 | 4.72E-13 |
| chr3 | 185573717 | 185576242 | 2526 | -8.11 | 1.22E-13 | 4.72E-13 |
| chr7 | 129795511 | 129801677 | 6167 | -8.23 | 1.22E-13 | 4.72E-13 |
| chrX | 81650692 | 81662588 | 11897 | -8.22 | 1.23E-13 | 4.74E-13 |
| chr3 | 138682620 | 138685811 | 3192 | -8.11 | 1.23E-13 | 4.74E-13 |
| chr2 | 190996860 | 191011253 | 14394 | -8.5 | 1.23E-13 | 4.74E-13 |
| chr10 | 43343853 | 43347194 | 3342 | -8.12 | 1.23E-13 | 4.75E-13 |
| chr10 | 94175292 | 94183913 | 8622 | -8.15 | 1.23E-13 | 4.75E-13 |
| chr7 | 107177290 | 107186874 | 9585 | -8.12 | 1.23E-13 | 4.75E-13 |
| chr8 | 87383784 | 87386505 | 2722 | -8.23 | 1.23E-13 | 4.76E-13 |
| chr20 | 56719448 | 56724387 | 4940 | -8.32 | 1.23E-13 | 4.76E-13 |
| chr5 | 13445546 | 13450987 | 5442 | 8.19 | 1.23E-13 | 4.76E-13 |
| chr2 | 145720978 | 145732115 | 11138 | -6.26 | 1.23E-13 | 4.77E-13 |

| | | | | | | |
|---|---|---|---|---|---|---|
| chr4 | 189640628 | 189651534 | 10907 | -8.11 | 1.24E-13 | 4.77E-13 |
| chr5 | 126531438 | 126550615 | 19178 | -8.43 | 1.24E-13 | 4.77E-13 |
| chr1 | 94023764 | 94027364 | 3601 | -8.4 | 1.24E-13 | 4.77E-13 |
| chr4 | 13547959 | 13554886 | 6928 | -8.11 | 1.24E-13 | 4.80E-13 |
| chr20 | 46019453 | 46030308 | 10856 | -8.22 | 1.25E-13 | 4.81E-13 |
| chrX | 35883184 | 35893134 | 9951 | -8.18 | 1.25E-13 | 4.81E-13 |
| chr14 | 25452876 | 25464741 | 11866 | 8.42 | 1.25E-13 | 4.81E-13 |
| chr3 | 98523679 | 98527642 | 3964 | -8.12 | 1.25E-13 | 4.82E-13 |
| chr11 | 49155597 | 49166317 | 10721 | 2.8 | 1.25E-13 | 4.83E-13 |
| chr11 | 36087029 | 36098464 | 11436 | -8.14 | 1.25E-13 | 4.84E-13 |
| chr8 | 74500436 | 74503575 | 3140 | -8.1 | 1.26E-13 | 4.85E-13 |
| chr4 | 1055886 | 1066307 | 10422 | -8.13 | 1.26E-13 | 4.85E-13 |
| chr5 | 75944049 | 75946410 | 2362 | -8.15 | 1.26E-13 | 4.86E-13 |
| chr17 | 26774221 | 26780273 | 6053 | -8.14 | 1.27E-13 | 4.88E-13 |
| chr7 | 13207520 | 13228632 | 21113 | -8.28 | 1.27E-13 | 4.88E-13 |
| chrX | 152692502 | 152697372 | 4871 | -8.12 | 1.27E-13 | 4.88E-13 |
| chr3 | 11750262 | 11753028 | 2767 | -8.11 | 1.27E-13 | 4.91E-13 |
| chr8 | 67092644 | 67096892 | 4249 | -8.11 | 1.27E-13 | 4.91E-13 |
| chr12 | 121295534 | 121297824 | 2291 | -8.11 | 1.27E-13 | 4.91E-13 |
| chr5 | 169512123 | 169526844 | 14722 | -5.99 | 1.27E-13 | 4.91E-13 |
| chr1 | 3752926 | 3755161 | 2236 | -8.13 | 1.27E-13 | 4.91E-13 |
| chr1 | 207933264 | 207951774 | 18511 | -8.34 | 1.28E-13 | 4.92E-13 |
| chr7 | 3460198 | 3462979 | 2782 | -8.11 | 1.28E-13 | 4.93E-13 |
| chr3 | 188828052 | 188838118 | 10067 | -8.22 | 1.28E-13 | 4.93E-13 |
| chr3 | 161635713 | 161646464 | 10752 | -3.08 | 1.28E-13 | 4.94E-13 |
| chr16 | 84433615 | 84436645 | 3031 | -8.43 | 1.28E-13 | 4.94E-13 |
| chr6 | 16809011 | 16814974 | 5964 | -8.17 | 1.28E-13 | 4.94E-13 |
| chrX | 122640445 | 122673899 | 33455 | -7.01 | 1.29E-13 | 4.97E-13 |
| chr6 | 37800942 | 37803240 | 2299 | -8.11 | 1.30E-13 | 5.00E-13 |
| chr2 | 86027559 | 86030042 | 2484 | -8.15 | 1.30E-13 | 5.02E-13 |
| chr14 | 67958504 | 67961642 | 3139 | -8.1 | 1.30E-13 | 5.02E-13 |
| chr2 | 209257228 | 209260675 | 3448 | -8.1 | 1.30E-13 | 5.02E-13 |
| chr12 | 109813380 | 109868489 | 55110 | -9.07 | 1.30E-13 | 5.02E-13 |
| chr9 | 124609151 | 124612306 | 3156 | -8.13 | 1.31E-13 | 5.02E-13 |
| chr15 | 100224063 | 100228081 | 4019 | -8.42 | 1.31E-13 | 5.02E-13 |
| chr1 | 40372750 | 40376019 | 3270 | -8.15 | 1.31E-13 | 5.03E-13 |
| chr9 | 79005919 | 79007816 | 1898 | -8.15 | 1.31E-13 | 5.03E-13 |
| chr2 | 105972525 | 105974698 | 2174 | -8.13 | 1.31E-13 | 5.04E-13 |
| chr4 | 164043759 | 164046520 | 2762 | -8.1 | 1.31E-13 | 5.04E-13 |
| chr7 | 47211197 | 47218635 | 7439 | -8.21 | 1.31E-13 | 5.05E-13 |
| chr16 | 85957992 | 85975505 | 17514 | -8.5 | 1.31E-13 | 5.06E-13 |
| chr20 | 25344568 | 25355553 | 10986 | -8.27 | 1.32E-13 | 5.07E-13 |
| chr6 | 76348573 | 76351802 | 3230 | -8.27 | 1.32E-13 | 5.07E-13 |
| chr12 | 121595039 | 121597416 | 2378 | -8.11 | 1.32E-13 | 5.08E-13 |
| chr12 | 77431004 | 77433470 | 2467 | -8.28 | 1.32E-13 | 5.09E-13 |
| chr19 | 5097506 | 5101193 | 3688 | -8.27 | 1.32E-13 | 5.09E-13 |
| chr6 | 151531259 | 151536551 | 5293 | -8.42 | 1.33E-13 | 5.11E-13 |
| chr4 | 157908177 | 157919563 | 11387 | -8.23 | 1.33E-13 | 5.13E-13 |
| chr5 | 172700780 | 172703598 | 2819 | -8.11 | 1.34E-13 | 5.14E-13 |
| chr1 | 196634921 | 196640125 | 5205 | -8.13 | 1.34E-13 | 5.15E-13 |
| chr7 | 75558892 | 75587958 | 29067 | -8.24 | 1.34E-13 | 5.15E-13 |
| chr2 | 64072849 | 64085882 | 13034 | -8.57 | 1.34E-13 | 5.15E-13 |
| chr11 | 109660722 | 109678687 | 17966 | -8.11 | 1.34E-13 | 5.15E-13 |
| chr1 | 225363051 | 225368377 | 5327 | -8.15 | 1.34E-13 | 5.16E-13 |
| chr13 | 40777212 | 40792638 | 15427 | -8.31 | 1.34E-13 | 5.16E-13 |
| chr8 | 38447367 | 38450482 | 3116 | -8.19 | 1.34E-13 | 5.16E-13 |
| chr2 | 3165855 | 3170010 | 4156 | -8.15 | 1.35E-13 | 5.16E-13 |
| chrX | 45035953 | 45040630 | 4678 | -8.13 | 1.35E-13 | 5.17E-13 |
| chr3 | 94266455 | 94278137 | 11683 | -8.2 | 1.35E-13 | 5.17E-13 |
| chr1 | 58880805 | 58892723 | 11919 | 8.48 | 1.35E-13 | 5.18E-13 |
| chr11 | 77072840 | 77078169 | 5330 | -8.1 | 1.35E-13 | 5.18E-13 |

| | | | | | | |
|---|---|---|---|---|---|---|
| chr12 | 52729614 | 52738411 | 8798 | -8.21 | 1.35E-13 | 5.20E-13 |
| chr9 | 74582100 | 74586263 | 4164 | -8.16 | 1.36E-13 | 5.21E-13 |
| chr9 | 100482534 | 100520300 | 37767 | -8.36 | 1.36E-13 | 5.21E-13 |
| chr5 | 39594844 | 39610258 | 15415 | -8.28 | 1.36E-13 | 5.22E-13 |
| chr2 | 86140521 | 86148284 | 7764 | -8.4 | 1.36E-13 | 5.22E-13 |
| chr16 | 87033822 | 87043143 | 9322 | -8.14 | 1.36E-13 | 5.22E-13 |
| chr5 | 79244924 | 79263711 | 18788 | 8.17 | 1.36E-13 | 5.23E-13 |
| chr2 | 209142691 | 209156269 | 13579 | -8.45 | 1.37E-13 | 5.24E-13 |
| chr16 | 86970122 | 86975281 | 5160 | -8.33 | 1.37E-13 | 5.24E-13 |
| chr17 | 2459241 | 2465539 | 6299 | -8.4 | 1.37E-13 | 5.25E-13 |
| chr8 | 124117142 | 124119195 | 2054 | -8.16 | 1.37E-13 | 5.26E-13 |
| chr3 | 88013852 | 88030374 | 16523 | -8.32 | 1.38E-13 | 5.27E-13 |
| chr4 | 90688725 | 90700798 | 12074 | -8.1 | 1.38E-13 | 5.28E-13 |
| chr2 | 224497323 | 224505118 | 7796 | -8.18 | 1.38E-13 | 5.30E-13 |
| chr17 | 71066886 | 71080989 | 14104 | -8.13 | 1.38E-13 | 5.30E-13 |
| chr5 | 83352507 | 83358296 | 5790 | 8.28 | 1.39E-13 | 5.32E-13 |
| chr11 | 103242738 | 103247718 | 4981 | -8.11 | 1.40E-13 | 5.36E-13 |
| chr11 | 92675762 | 92684571 | 8810 | -8.1 | 1.41E-13 | 5.41E-13 |
| chr8 | 81768476 | 81784496 | 16021 | -8.09 | 1.42E-13 | 5.44E-13 |
| chr1 | 28026906 | 28044360 | 17455 | -8.18 | 1.42E-13 | 5.44E-13 |
| chr16 | 19101663 | 19111906 | 10244 | -8.18 | 1.42E-13 | 5.44E-13 |
| chr4 | 169801676 | 169804859 | 3184 | -8.22 | 1.43E-13 | 5.47E-13 |
| chr19 | 48829934 | 48832530 | 2597 | -8.12 | 1.43E-13 | 5.47E-13 |
| chr11 | 122436834 | 122460414 | 23581 | -8.31 | 1.43E-13 | 5.48E-13 |
| chr7 | 140683029 | 140684958 | 1930 | -8.21 | 1.43E-13 | 5.49E-13 |
| chr2 | 33194451 | 33207018 | 12568 | -8.44 | 1.44E-13 | 5.50E-13 |
| chr8 | 31565599 | 31570272 | 4674 | -8.13 | 1.44E-13 | 5.52E-13 |
| chr10 | 45493712 | 45521764 | 28053 | -7.39 | 1.45E-13 | 5.54E-13 |
| chr3 | 72469071 | 72472440 | 3370 | -8.13 | 1.45E-13 | 5.55E-13 |
| chr3 | 134063749 | 134067267 | 3519 | -8.14 | 1.45E-13 | 5.55E-13 |
| chr3 | 29675783 | 29679030 | 3248 | -8.28 | 1.45E-13 | 5.56E-13 |
| chr7 | 103761181 | 103787503 | 26323 | -8.11 | 1.46E-13 | 5.57E-13 |
| chr12 | 42429079 | 42433214 | 4136 | -8.22 | 1.46E-13 | 5.57E-13 |
| chr19 | 5254460 | 5256980 | 2521 | -8.09 | 1.46E-13 | 5.57E-13 |
| chr13 | 100160139 | 100162643 | 2505 | -8.09 | 1.46E-13 | 5.58E-13 |
| chr9 | 86318937 | 86349443 | 30507 | -8.39 | 1.46E-13 | 5.58E-13 |
| chr1 | 15638055 | 15648520 | 10466 | -8.09 | 1.46E-13 | 5.58E-13 |
| chr1 | 66698777 | 66706381 | 7605 | -8.09 | 1.46E-13 | 5.59E-13 |
| chr18 | 5311705 | 5862493 | 550789 | 5.54 | 1.46E-13 | 5.59E-13 |
| chr2 | 235887579 | 235888982 | 1404 | -8.14 | 1.46E-13 | 5.60E-13 |
| chr3 | 151785858 | 151812372 | 26515 | -8.66 | 1.46E-13 | 5.60E-13 |
| chr11 | 65802776 | 65805646 | 2871 | -8.24 | 1.48E-13 | 5.66E-13 |
| chr10 | 7796908 | 7799005 | 2098 | -8.14 | 1.48E-13 | 5.67E-13 |
| chr9 | 136190266 | 136193703 | 3438 | -8.09 | 1.48E-13 | 5.67E-13 |
| chr16 | 51466110 | 51471577 | 5468 | -8.13 | 1.49E-13 | 5.67E-13 |
| chr12 | 39247901 | 39255149 | 7249 | -8.25 | 1.49E-13 | 5.69E-13 |
| chr2 | 228446002 | 228455145 | 9144 | -8.26 | 1.50E-13 | 5.72E-13 |
| chr3 | 64646867 | 64649378 | 2512 | -8.09 | 1.50E-13 | 5.72E-13 |
| chr1 | 7090879 | 7094865 | 3987 | -8.09 | 1.50E-13 | 5.73E-13 |
| chr20 | 47149423 | 47161137 | 11715 | -8.22 | 1.50E-13 | 5.73E-13 |
| chr12 | 101741596 | 101744344 | 2749 | -8.13 | 1.50E-13 | 5.74E-13 |
| chr20 | 62639083 | 62640967 | 1885 | -8.12 | 1.51E-13 | 5.75E-13 |
| chr3 | 13195840 | 13198602 | 2763 | -8.16 | 1.51E-13 | 5.76E-13 |
| chr20 | 47926669 | 47935267 | 8599 | -8.23 | 1.51E-13 | 5.77E-13 |
| chr20 | 37449594 | 37451703 | 2110 | -8.08 | 1.51E-13 | 5.78E-13 |
| chr4 | 131472990 | 131479773 | 6784 | -8.08 | 1.52E-13 | 5.78E-13 |
| chr1 | 197765318 | 197782659 | 17342 | -8.09 | 1.52E-13 | 5.79E-13 |
| chr1 | 93336904 | 93340092 | 3189 | -8.15 | 1.52E-13 | 5.80E-13 |
| chr10 | 51013124 | 51016269 | 3146 | -8.15 | 1.52E-13 | 5.80E-13 |
| chr5 | 171684180 | 171686687 | 2508 | -8.11 | 1.52E-13 | 5.80E-13 |
| chr4 | 166392315 | 166396743 | 4429 | -8.08 | 1.52E-13 | 5.80E-13 |

| | | | | | | |
|---|---|---|---|---|---|---|
| chr9 | 132116785 | 132134144 | 17360 | -8.47 | 1.52E-13 | 5.81E-13 |
| chr5 | 159419263 | 159421623 | 2361 | -8.08 | 1.53E-13 | 5.82E-13 |
| chr13 | 46603391 | 46606274 | 2884 | -8.09 | 1.53E-13 | 5.82E-13 |
| chr11 | 69083978 | 69086699 | 2722 | -8.24 | 1.53E-13 | 5.82E-13 |
| chr9 | 77669731 | 77674267 | 4537 | -8.08 | 1.53E-13 | 5.83E-13 |
| chr21 | 43260254 | 43263309 | 3056 | -8.14 | 1.53E-13 | 5.84E-13 |
| chr5 | 613123 | 616133 | 3011 | -8.1 | 1.53E-13 | 5.84E-13 |
| chr5 | 13634240 | 13655330 | 21091 | -8.38 | 1.53E-13 | 5.84E-13 |
| chr11 | 92825634 | 92836955 | 11322 | -8.09 | 1.53E-13 | 5.84E-13 |
| chr2 | 33052836 | 33056614 | 3779 | -8.26 | 1.54E-13 | 5.85E-13 |
| chr3 | 87212308 | 87216415 | 4108 | -8.1 | 1.54E-13 | 5.87E-13 |
| chr20 | 47391744 | 47396948 | 5205 | -8.11 | 1.54E-13 | 5.87E-13 |
| chr11 | 27162110 | 27164819 | 2710 | -8.1 | 1.54E-13 | 5.88E-13 |
| chr7 | 120556272 | 120564747 | 8476 | 8.43 | 1.54E-13 | 5.88E-13 |
| chr7 | 69630724 | 70045285 | 414562 | 0.85 | 1.54E-13 | 5.88E-13 |
| chr11 | 120453077 | 120455948 | 2872 | -8.08 | 1.55E-13 | 5.89E-13 |
| chr12 | 104310349 | 104313277 | 2929 | -8.16 | 1.55E-13 | 5.89E-13 |
| chr6 | 48988341 | 48994386 | 6046 | -8.08 | 1.55E-13 | 5.90E-13 |
| chr14 | 70208568 | 70212605 | 4038 | -8.12 | 1.55E-13 | 5.90E-13 |
| chr18 | 67827589 | 67831210 | 3622 | -8.08 | 1.55E-13 | 5.90E-13 |
| chrX | 62838825 | 62853782 | 14958 | -8.11 | 1.55E-13 | 5.91E-13 |
| chr1 | 88703599 | 88711378 | 7780 | -8.1 | 1.55E-13 | 5.92E-13 |
| chr17 | 31227461 | 31231330 | 3870 | -8.28 | 1.56E-13 | 5.93E-13 |
| chr18 | 20149056 | 20170342 | 21287 | -8.33 | 1.56E-13 | 5.94E-13 |
| chr9 | 6063392 | 6071573 | 8182 | -8.08 | 1.56E-13 | 5.94E-13 |
| chr9 | 119056253 | 119058725 | 2473 | -8.12 | 1.57E-13 | 5.96E-13 |
| chr5 | 56199417 | 56201341 | 1925 | -8.08 | 1.57E-13 | 5.96E-13 |
| chr6 | 170777521 | 170803754 | 26234 | -2.21 | 1.58E-13 | 6.01E-13 |
| chr1 | 233672291 | 233676991 | 4701 | -8.1 | 1.58E-13 | 6.01E-13 |
| chr18 | 19496279 | 19500231 | 3953 | -8.11 | 1.58E-13 | 6.02E-13 |
| chr8 | 28856865 | 28859133 | 2269 | -8.08 | 1.59E-13 | 6.04E-13 |
| chr9 | 133012512 | 133022134 | 9623 | -8.09 | 1.59E-13 | 6.06E-13 |
| chr16 | 9169298 | 9173993 | 4696 | -8.1 | 1.60E-13 | 6.07E-13 |
| chr10 | 87944915 | 87947441 | 2527 | -8.09 | 1.60E-13 | 6.10E-13 |
| chr1 | 63699463 | 63702219 | 2757 | -4.77 | 1.61E-13 | 6.12E-13 |
| chr7 | 159102766 | 159124639 | 21874 | -1.34 | 1.61E-13 | 6.14E-13 |
| chr21 | 41860885 | 41866521 | 5637 | -3.76 | 1.62E-13 | 6.14E-13 |
| chr9 | 133649279 | 133705103 | 55825 | -8.98 | 1.62E-13 | 6.15E-13 |
| chr4 | 23287930 | 23296372 | 8443 | -8.1 | 1.62E-13 | 6.15E-13 |
| chr2 | 146461949 | 146469711 | 7763 | -8.08 | 1.62E-13 | 6.15E-13 |
| chr15 | 69651157 | 69665529 | 14373 | -8.15 | 1.62E-13 | 6.16E-13 |
| chr10 | 17092567 | 17096372 | 3806 | -8.14 | 1.62E-13 | 6.16E-13 |
| chr12 | 21312545 | 21322791 | 10247 | -8.25 | 1.62E-13 | 6.16E-13 |
| chr7 | 134978309 | 134984350 | 6042 | -8.19 | 1.62E-13 | 6.17E-13 |
| chr1 | 15430055 | 15432760 | 2706 | -8.08 | 1.63E-13 | 6.17E-13 |
| chr2 | 88798551 | 88814419 | 15869 | -8.33 | 1.63E-13 | 6.17E-13 |
| chr14 | 68311879 | 68317038 | 5160 | -8.11 | 1.63E-13 | 6.19E-13 |
| chr12 | 111627228 | 111628963 | 1736 | -8.08 | 1.63E-13 | 6.20E-13 |
| chr13 | 91093319 | 91101529 | 8211 | -8.23 | 1.64E-13 | 6.21E-13 |
| chr19 | 18936418 | 18939010 | 2593 | -8.11 | 1.64E-13 | 6.21E-13 |
| chr5 | 55500200 | 55517420 | 17221 | -8.46 | 1.64E-13 | 6.22E-13 |
| chr17 | 62651149 | 62653086 | 1938 | -8.08 | 1.65E-13 | 6.24E-13 |
| chrX | 152621521 | 152626496 | 4976 | -8.08 | 1.65E-13 | 6.26E-13 |
| chr5 | 60008066 | 60014030 | 5965 | -8.09 | 1.65E-13 | 6.27E-13 |
| chr2 | 226716830 | 226722250 | 5421 | -8.2 | 1.66E-13 | 6.29E-13 |
| chr12 | 76761553 | 76763555 | 2003 | -8.09 | 1.66E-13 | 6.30E-13 |
| chr19 | 7518148 | 7521301 | 3154 | -3.14 | 1.66E-13 | 6.31E-13 |
| chr3 | 14669184 | 14680648 | 11465 | -8.07 | 1.67E-13 | 6.34E-13 |
| chr1 | 202900984 | 202902889 | 1906 | -8.07 | 1.67E-13 | 6.35E-13 |
| chr5 | 170674460 | 170677559 | 3100 | -8.07 | 1.67E-13 | 6.35E-13 |
| chr4 | 2132683 | 2136010 | 3328 | -8.08 | 1.67E-13 | 6.35E-13 |

| | | | | | | |
|---|---|---|---|---|---|---|
| chr7 | 20058367 | 20068061 | 9695 | -8.1 | 1.67E-13 | 6.35E-13 |
| chr9 | 127464459 | 127469931 | 5473 | -8.21 | 1.67E-13 | 6.35E-13 |
| chr12 | 96072200 | 96087628 | 15429 | -8.45 | 1.68E-13 | 6.36E-13 |
| chr10 | 118729015 | 118732315 | 3301 | -8.09 | 1.68E-13 | 6.37E-13 |
| chr1 | 222872386 | 222879912 | 7527 | -8.07 | 1.68E-13 | 6.38E-13 |
| chr3 | 112337662 | 112341389 | 3728 | -8.13 | 1.69E-13 | 6.40E-13 |
| chr17 | 60847742 | 60860843 | 13102 | -8.42 | 1.70E-13 | 6.42E-13 |
| chr1 | 208161271 | 208180191 | 18921 | -8.45 | 1.70E-13 | 6.42E-13 |
| chr14 | 74378384 | 74380940 | 2557 | -8.28 | 1.70E-13 | 6.42E-13 |
| chr20 | 46199036 | 46201076 | 2041 | -8.13 | 1.70E-13 | 6.43E-13 |
| chr14 | 101375899 | 101400221 | 24323 | -8.38 | 1.70E-13 | 6.45E-13 |
| chr1 | 4371854 | 4375660 | 3807 | -2.87 | 1.72E-13 | 6.50E-13 |
| chr18 | 33244976 | 33248493 | 3518 | -8.11 | 1.73E-13 | 6.55E-13 |
| chrX | 44429904 | 44433877 | 3974 | -8.15 | 1.73E-13 | 6.55E-13 |
| chrX | 86975703 | 87149836 | 174134 | 2.14 | 1.74E-13 | 6.57E-13 |
| chr9 | 128707533 | 128709751 | 2219 | -8.12 | 1.74E-13 | 6.58E-13 |
| chr3 | 172595665 | 172617830 | 22166 | -8.09 | 1.74E-13 | 6.58E-13 |
| chr6 | 145221898 | 145225575 | 3678 | -8.18 | 1.74E-13 | 6.59E-13 |
| chr16 | 14668246 | 14671329 | 3084 | -8.1 | 1.74E-13 | 6.59E-13 |
| chr10 | 49847274 | 49849715 | 2442 | -8.09 | 1.74E-13 | 6.59E-13 |
| chr16 | 50003014 | 50012523 | 9510 | -8.08 | 1.74E-13 | 6.59E-13 |
| chr13 | 35282465 | 35296046 | 13582 | -8.29 | 1.74E-13 | 6.60E-13 |
| chr12 | 69879686 | 69881998 | 2313 | -8.11 | 1.74E-13 | 6.60E-13 |
| chr18 | 7597826 | 7600328 | 2503 | -8.07 | 1.75E-13 | 6.60E-13 |
| chr2 | 189686929 | 189706633 | 19705 | -8.21 | 1.75E-13 | 6.61E-13 |
| chr3 | 131872206 | 131904305 | 32100 | -8.08 | 1.75E-13 | 6.62E-13 |
| chr18 | 22173387 | 22303907 | 130521 | 1.26 | 1.75E-13 | 6.62E-13 |
| chr9 | 114460704 | 114464272 | 3569 | -8.09 | 1.76E-13 | 6.63E-13 |
| chr2 | 101236428 | 101271614 | 35187 | -8.81 | 1.76E-13 | 6.64E-13 |
| chr1 | 12665123 | 12668882 | 3760 | -8.07 | 1.76E-13 | 6.64E-13 |
| chr11 | 84941062 | 84951392 | 10331 | -8.36 | 1.76E-13 | 6.66E-13 |
| chr5 | 78676180 | 78679598 | 3419 | -8.09 | 1.77E-13 | 6.70E-13 |
| chr19 | 41174195 | 41176462 | 2268 | -8.49 | 1.78E-13 | 6.71E-13 |
| chr7 | 8031157 | 8035311 | 4155 | -8.31 | 1.79E-13 | 6.75E-13 |
| chr9 | 80836249 | 80839709 | 3461 | -6.18 | 1.79E-13 | 6.77E-13 |
| chr7 | 128701025 | 128703929 | 2905 | -8.07 | 1.80E-13 | 6.78E-13 |
| chr1 | 162270767 | 162279106 | 8340 | -8.14 | 1.80E-13 | 6.79E-13 |
| chr10 | 30117347 | 30120969 | 3623 | -6.44 | 1.80E-13 | 6.79E-13 |
| chr11 | 19590775 | 19593302 | 2528 | -8.07 | 1.80E-13 | 6.80E-13 |
| chr3 | 105633520 | 105640521 | 7002 | -8.33 | 1.80E-13 | 6.80E-13 |
| chr7 | 7669295 | 7672231 | 2937 | -8.1 | 1.80E-13 | 6.80E-13 |
| chr7 | 40175199 | 40188743 | 13545 | -8.07 | 1.81E-13 | 6.83E-13 |
| chr7 | 105266054 | 105267473 | 1420 | -8.08 | 1.81E-13 | 6.83E-13 |
| chr4 | 173512618 | 173532754 | 20137 | -8.25 | 1.81E-13 | 6.83E-13 |
| chr2 | 11626694 | 11645000 | 18307 | -8.71 | 1.82E-13 | 6.85E-13 |
| chr7 | 108033775 | 108036232 | 2458 | -3.69 | 1.83E-13 | 6.89E-13 |
| chr6 | 89515825 | 89519630 | 3806 | -8.07 | 1.83E-13 | 6.91E-13 |
| chr14 | 102575068 | 102579142 | 4075 | -8.1 | 1.83E-13 | 6.91E-13 |
| chr20 | 1801260 | 1804980 | 3721 | -8.22 | 1.83E-13 | 6.91E-13 |
| chr5 | 82040645 | 82049010 | 8366 | -8.06 | 1.83E-13 | 6.92E-13 |
| chr13 | 113110423 | 113135234 | 24812 | -8.32 | 1.83E-13 | 6.92E-13 |
| chr19 | 39705489 | 39721384 | 15896 | -8.38 | 1.84E-13 | 6.94E-13 |
| chrX | 134545754 | 134552520 | 6767 | -8.09 | 1.84E-13 | 6.94E-13 |
| chrX | 117435962 | 117446680 | 10719 | -6.36 | 1.84E-13 | 6.94E-13 |
| chr4 | 173988924 | 174011341 | 22418 | -8.26 | 1.84E-13 | 6.95E-13 |
| chr7 | 116902144 | 116909079 | 6936 | -3.58 | 1.85E-13 | 6.96E-13 |
| chr2 | 187426188 | 187434161 | 7974 | -8.21 | 1.85E-13 | 6.97E-13 |
| chr20 | 10709887 | 10714403 | 4517 | -8.18 | 1.85E-13 | 6.98E-13 |
| chr8 | 101849299 | 101851539 | 2241 | -8.06 | 1.85E-13 | 6.98E-13 |
| chr22 | 19539653 | 19552464 | 12812 | -8.06 | 1.85E-13 | 6.99E-13 |
| chr2 | 192187884 | 192189913 | 2030 | -8.14 | 1.86E-13 | 7.00E-13 |

| | | | | | | |
|---|---|---|---|---|---|---|
| chr12 | 116858813 | 116870797 | 11985 | -8.06 | 1.86E-13 | 7.00E-13 |
| chr18 | 44176620 | 44179975 | 3356 | -5.86 | 1.86E-13 | 7.01E-13 |
| chr7 | 137758194 | 137762873 | 4680 | -8.06 | 1.86E-13 | 7.02E-13 |
| chr1 | 114137990 | 114141758 | 3769 | -8.07 | 1.87E-13 | 7.04E-13 |
| chr2 | 138746330 | 138753634 | 7305 | -2.71 | 1.87E-13 | 7.04E-13 |
| chr12 | 27555553 | 27557878 | 2326 | -8.1 | 1.87E-13 | 7.04E-13 |
| chr5 | 176891789 | 176893527 | 1739 | -8.06 | 1.87E-13 | 7.06E-13 |
| chr2 | 214139916 | 214144054 | 4139 | -8.22 | 1.88E-13 | 7.08E-13 |
| chr20 | 44900178 | 44903891 | 3714 | -8.07 | 1.88E-13 | 7.09E-13 |
| chr14 | 65718210 | 65728989 | 10780 | -8.07 | 1.89E-13 | 7.10E-13 |
| chr3 | 148637069 | 148642320 | 5252 | -8.11 | 1.89E-13 | 7.11E-13 |
| chr10 | 96917970 | 96937344 | 19375 | -8.26 | 1.89E-13 | 7.11E-13 |
| chr6 | 4569491 | 4573027 | 3537 | -8.07 | 1.89E-13 | 7.12E-13 |
| chr11 | 86272247 | 86276377 | 4131 | -8.24 | 1.89E-13 | 7.13E-13 |
| chr15 | 35161028 | 35162830 | 1803 | -8.13 | 1.89E-13 | 7.13E-13 |
| chr19 | 40682613 | 40684638 | 2026 | -8.13 | 1.89E-13 | 7.13E-13 |
| chr14 | 99804537 | 99807430 | 2894 | -8.17 | 1.90E-13 | 7.14E-13 |
| chr9 | 75246960 | 75251065 | 4106 | -8.12 | 1.90E-13 | 7.16E-13 |
| chr19 | 30178682 | 30182545 | 3864 | -8.43 | 1.91E-13 | 7.18E-13 |
| chr14 | 54028651 | 54086045 | 57395 | -9.01 | 1.91E-13 | 7.19E-13 |
| chr3 | 119328925 | 119331213 | 2289 | -8.11 | 1.92E-13 | 7.21E-13 |
| chrX | 81682568 | 81696257 | 13690 | -8.05 | 1.92E-13 | 7.22E-13 |
| chr20 | 34159644 | 34162974 | 3331 | -8.07 | 1.92E-13 | 7.22E-13 |
| chr2 | 139132915 | 139137895 | 4981 | -8.05 | 1.92E-13 | 7.23E-13 |
| chr9 | 117477563 | 117483563 | 6001 | -8.11 | 1.93E-13 | 7.24E-13 |
| chr1 | 234909314 | 234912525 | 3212 | -8.3 | 1.93E-13 | 7.27E-13 |
| chr4 | 154292025 | 154294945 | 2921 | -8.16 | 1.94E-13 | 7.27E-13 |
| chr11 | 9735926 | 9742666 | 6741 | -8.12 | 1.94E-13 | 7.30E-13 |
| chr12 | 84271914 | 84285761 | 13848 | -8.06 | 1.95E-13 | 7.32E-13 |
| chr10 | 23588572 | 23595415 | 6844 | 3.77 | 1.95E-13 | 7.34E-13 |
| chr5 | 155820225 | 155826173 | 5949 | -8.12 | 1.97E-13 | 7.38E-13 |
| chr4 | 78930909 | 78934667 | 3759 | -8.06 | 1.97E-13 | 7.38E-13 |
| chr18 | 7422443 | 7427806 | 5364 | -8.06 | 1.97E-13 | 7.39E-13 |
| chr9 | 110262058 | 110266211 | 4154 | -8.21 | 1.97E-13 | 7.41E-13 |
| chr11 | 68052018 | 68072300 | 20283 | -8.26 | 1.98E-13 | 7.42E-13 |
| chrX | 138780044 | 138786468 | 6425 | 8.13 | 1.98E-13 | 7.42E-13 |
| chr7 | 143705005 | 143708351 | 3347 | -2.87 | 1.98E-13 | 7.42E-13 |
| chrX | 8929531 | 8941019 | 11489 | -8.1 | 1.99E-13 | 7.47E-13 |
| chr1 | 182208939 | 182216584 | 7646 | 8.12 | 1.99E-13 | 7.48E-13 |
| chr2 | 201636500 | 201639511 | 3012 | -8.05 | 2.00E-13 | 7.49E-13 |
| chr6 | 131713783 | 131732358 | 18576 | -8.05 | 2.00E-13 | 7.49E-13 |
| chr8 | 135760839 | 135763928 | 3090 | -8.06 | 2.00E-13 | 7.51E-13 |
| chr5 | 80450504 | 80453535 | 3032 | -8.08 | 2.01E-13 | 7.55E-13 |
| chr22 | 45266520 | 45274128 | 7609 | -8.05 | 2.01E-13 | 7.55E-13 |
| chr12 | 87959302 | 87963470 | 4169 | -8.27 | 2.02E-13 | 7.56E-13 |
| chr18 | 54027997 | 54031421 | 3425 | -8.05 | 2.02E-13 | 7.56E-13 |
| chr2 | 38352933 | 38356128 | 3196 | -8.05 | 2.02E-13 | 7.57E-13 |
| chr15 | 50451735 | 50462628 | 10894 | -8.2 | 2.02E-13 | 7.59E-13 |
| chr9 | 82257927 | 82260782 | 2856 | -8.11 | 2.03E-13 | 7.62E-13 |
| chr2 | 146968271 | 146975413 | 7143 | -8.05 | 2.04E-13 | 7.65E-13 |
| chr19 | 55613983 | 55615843 | 1861 | -8.05 | 2.04E-13 | 7.65E-13 |
| chr5 | 115124323 | 115127219 | 2897 | -8.05 | 2.04E-13 | 7.65E-13 |
| chrX | 39359040 | 39370320 | 11281 | -4.68 | 2.04E-13 | 7.66E-13 |
| chr22 | 33916707 | 33918956 | 2250 | -8.07 | 2.04E-13 | 7.66E-13 |
| chr1 | 241851517 | 241868516 | 17000 | -8.35 | 2.05E-13 | 7.66E-13 |
| chr8 | 141029230 | 141040875 | 11646 | -8.21 | 2.05E-13 | 7.67E-13 |
| chr10 | 78964164 | 78971617 | 7454 | -8.04 | 2.07E-13 | 7.75E-13 |
| chr8 | 59943115 | 59947384 | 4270 | -8.15 | 2.07E-13 | 7.76E-13 |
| chrX | 18103177 | 18114000 | 10824 | -2.35 | 2.07E-13 | 7.76E-13 |
| chr2 | 71329328 | 71334124 | 4797 | -8.27 | 2.07E-13 | 7.77E-13 |
| chr6 | 100387013 | 100399202 | 12190 | 8.28 | 2.07E-13 | 7.77E-13 |

| | | | | | | |
|---|---|---|---|---|---|---|
| chr1 | 198471404 | 198475401 | 3998 | -8.16 | 2.08E-13 | 7.79E-13 |
| chr15 | 37105510 | 37118845 | 13336 | -8.51 | 2.10E-13 | 7.87E-13 |
| chr10 | 74172942 | 74174938 | 1997 | -8.18 | 2.11E-13 | 7.89E-13 |
| chr3 | 192973251 | 193173190 | 199940 | 6.43 | 2.11E-13 | 7.89E-13 |
| chr16 | 58213392 | 58216868 | 3477 | -8.04 | 2.11E-13 | 7.91E-13 |
| chrX | 119159220 | 119163045 | 3826 | -8.1 | 2.12E-13 | 7.92E-13 |
| chr6 | 146217683 | 146220345 | 2663 | -8.1 | 2.12E-13 | 7.93E-13 |
| chr4 | 56495286 | 56511916 | 16631 | -8.15 | 2.12E-13 | 7.94E-13 |
| chr5 | 71276667 | 71279931 | 3265 | -8.06 | 2.13E-13 | 7.95E-13 |
| chrX | 35298753 | 35306714 | 7962 | -8.11 | 2.13E-13 | 7.95E-13 |
| chr9 | 137742837 | 137767218 | 24382 | -8.28 | 2.13E-13 | 7.96E-13 |
| chr19 | 55722661 | 55732932 | 10272 | -8.11 | 2.13E-13 | 7.97E-13 |
| chr12 | 76712069 | 76725308 | 13240 | -8.06 | 2.14E-13 | 8.01E-13 |
| chrX | 73947736 | 74100246 | 152511 | 4.5 | 2.14E-13 | 8.01E-13 |
| chr7 | 23655392 | 23659149 | 3758 | -8.13 | 2.14E-13 | 8.02E-13 |
| chr1 | 64272523 | 64292236 | 19714 | -8.17 | 2.15E-13 | 8.02E-13 |
| chr8 | 99536023 | 99545582 | 9560 | -8.04 | 2.15E-13 | 8.05E-13 |
| chr3 | 38575770 | 38578810 | 3041 | -8.11 | 2.15E-13 | 8.05E-13 |
| chr10 | 63217954 | 63333220 | 115267 | 2.22 | 2.15E-13 | 8.05E-13 |
| chr15 | 78924778 | 78926843 | 2066 | -8.13 | 2.18E-13 | 8.13E-13 |
| chr2 | 170418835 | 170421541 | 2707 | -8.1 | 2.18E-13 | 8.14E-13 |
| chr18 | 72387949 | 72390753 | 2805 | -8.04 | 2.18E-13 | 8.15E-13 |
| chr6 | 56805364 | 56817230 | 11867 | -8.1 | 2.18E-13 | 8.16E-13 |
| chr1 | 198764772 | 198767436 | 2665 | -6.91 | 2.18E-13 | 8.16E-13 |
| chr3 | 94761476 | 94787254 | 25779 | -8.11 | 2.19E-13 | 8.16E-13 |
| chr2 | 75328405 | 75330511 | 2107 | -8.06 | 2.19E-13 | 8.18E-13 |
| chr18 | 24271071 | 24275278 | 4208 | -8.05 | 2.19E-13 | 8.19E-13 |
| chr6 | 90265969 | 90268810 | 2842 | -8.07 | 2.20E-13 | 8.21E-13 |
| chr22 | 37380801 | 37387684 | 6884 | -8.36 | 2.20E-13 | 8.21E-13 |
| chr1 | 234391339 | 234394804 | 3466 | -8.06 | 2.20E-13 | 8.23E-13 |
| chr11 | 34148920 | 34150831 | 1912 | -8.07 | 2.21E-13 | 8.26E-13 |
| chr17 | 5466534 | 5483184 | 16651 | -1.68 | 2.23E-13 | 8.32E-13 |
| chr10 | 24740655 | 24745189 | 4535 | -8.39 | 2.23E-13 | 8.32E-13 |
| chr15 | 35517693 | 35525425 | 7733 | -2.47 | 2.23E-13 | 8.34E-13 |
| chr10 | 74911237 | 74927181 | 15945 | -8.04 | 2.24E-13 | 8.36E-13 |
| chr13 | 73936622 | 73950574 | 13953 | -8.12 | 2.24E-13 | 8.37E-13 |
| chr3 | 53640272 | 53644711 | 4440 | -8.2 | 2.24E-13 | 8.37E-13 |
| chr12 | 42568653 | 42571232 | 2580 | -8.03 | 2.24E-13 | 8.37E-13 |
| chr2 | 74391753 | 74404316 | 12564 | -8.08 | 2.25E-13 | 8.38E-13 |
| chr7 | 110864469 | 110876675 | 12207 | -8.22 | 2.25E-13 | 8.39E-13 |
| chr11 | 112178696 | 112192101 | 13406 | -8.06 | 2.26E-13 | 8.44E-13 |
| chr2 | 146847936 | 146853495 | 5560 | -8.18 | 2.26E-13 | 8.44E-13 |
| chr10 | 116892884 | 116908951 | 16068 | -8.09 | 2.27E-13 | 8.47E-13 |
| chr22 | 40839854 | 40842394 | 2541 | -8.04 | 2.27E-13 | 8.47E-13 |
| chr12 | 21076098 | 21081680 | 5583 | -8.27 | 2.28E-13 | 8.49E-13 |
| chr7 | 8063989 | 8070403 | 6415 | -8.09 | 2.28E-13 | 8.49E-13 |
| chr1 | 53562239 | 53578712 | 16474 | -8.07 | 2.28E-13 | 8.50E-13 |
| chr5 | 171457154 | 171460487 | 3334 | -8.04 | 2.28E-13 | 8.52E-13 |
| chr1 | 7897560 | 7900394 | 2835 | -8.2 | 2.29E-13 | 8.54E-13 |
| chr8 | 39800653 | 39805805 | 5153 | -8.17 | 2.30E-13 | 8.57E-13 |
| chr4 | 140225064 | 140237870 | 12807 | -8.25 | 2.30E-13 | 8.58E-13 |
| chr8 | 143860369 | 143862519 | 2151 | -8.12 | 2.30E-13 | 8.59E-13 |
| chrX | 102162132 | 102177673 | 15542 | -8.04 | 2.31E-13 | 8.59E-13 |
| chr12 | 66732867 | 66739360 | 6494 | -8.21 | 2.31E-13 | 8.60E-13 |
| chr8 | 110955518 | 110969052 | 13535 | -8.04 | 2.31E-13 | 8.61E-13 |
| chr13 | 100772468 | 100775267 | 2800 | -8.03 | 2.31E-13 | 8.61E-13 |
| chr3 | 115173256 | 115179010 | 5755 | -8.21 | 2.31E-13 | 8.62E-13 |
| chr10 | 101815166 | 101819343 | 4178 | -8.11 | 2.32E-13 | 8.62E-13 |
| chr4 | 10042586 | 10046054 | 3469 | -8.1 | 2.32E-13 | 8.65E-13 |
| chr8 | 68397393 | 68399281 | 1889 | -8.03 | 2.33E-13 | 8.66E-13 |
| chr2 | 122394884 | 122397478 | 2595 | -8.03 | 2.33E-13 | 8.67E-13 |

| | | | | | | |
|---|---|---|---|---|---|---|
| chr1 | 44918575 | 44938161 | 19587 | -8.18 | 2.33E-13 | 8.67E-13 |
| chr20 | 9437993 | 9442374 | 4382 | -8.1 | 2.33E-13 | 8.68E-13 |
| chr22 | 32984962 | 32987929 | 2968 | -2.42 | 2.33E-13 | 8.68E-13 |
| chr7 | 156551192 | 156554177 | 2986 | -8.15 | 2.33E-13 | 8.69E-13 |
| chr10 | 11758645 | 11781390 | 22746 | -8.4 | 2.34E-13 | 8.71E-13 |
| chr4 | 17933559 | 17938599 | 5041 | -8.05 | 2.35E-13 | 8.73E-13 |
| chrX | 68262001 | 68284410 | 22410 | -8.09 | 2.35E-13 | 8.75E-13 |
| chr7 | 7892722 | 7897841 | 5120 | -8.21 | 2.35E-13 | 8.75E-13 |
| chr1 | 169841095 | 169843164 | 2070 | -5.93 | 2.35E-13 | 8.76E-13 |
| chr12 | 32957486 | 32962469 | 4984 | -8.08 | 2.36E-13 | 8.79E-13 |
| chr12 | 15881012 | 15897776 | 16765 | -8.04 | 2.37E-13 | 8.80E-13 |
| chr12 | 78739324 | 78742273 | 2950 | -8.06 | 2.38E-13 | 8.85E-13 |
| chr13 | 42376444 | 42390313 | 13870 | -8.2 | 2.38E-13 | 8.86E-13 |
| chr12 | 117235811 | 117238271 | 2461 | -8.03 | 2.39E-13 | 8.88E-13 |
| chr15 | 44762495 | 44764710 | 2216 | -8.13 | 2.39E-13 | 8.90E-13 |
| chr10 | 30692564 | 30695020 | 2457 | -8.03 | 2.39E-13 | 8.90E-13 |
| chr10 | 114567336 | 114571384 | 4049 | -8.14 | 2.40E-13 | 8.90E-13 |
| chr2 | 25403958 | 25405733 | 1776 | -8.09 | 2.40E-13 | 8.91E-13 |
| chr6 | 136178783 | 136186711 | 7929 | -8.31 | 2.40E-13 | 8.91E-13 |
| chr7 | 20095953 | 20103721 | 7769 | -8.24 | 2.40E-13 | 8.92E-13 |
| chr9 | 85586357 | 85593859 | 7503 | -8.2 | 2.41E-13 | 8.94E-13 |
| chr10 | 101404201 | 101410852 | 6652 | -2.84 | 2.41E-13 | 8.96E-13 |
| chr6 | 127902728 | 127913990 | 11263 | -8.04 | 2.42E-13 | 8.98E-13 |
| chr18 | 61503723 | 61507048 | 3326 | -8.09 | 2.43E-13 | 9.02E-13 |
| chr18 | 55478322 | 55481209 | 2888 | -8.03 | 2.43E-13 | 9.04E-13 |
| chr1 | 226446936 | 226450568 | 3633 | -8.08 | 2.44E-13 | 9.05E-13 |
| chr5 | 94311002 | 94314852 | 3851 | -8.08 | 2.44E-13 | 9.06E-13 |
| chr9 | 20675162 | 20677506 | 2345 | -8.02 | 2.44E-13 | 9.06E-13 |
| chr1 | 171333578 | 171351983 | 18406 | -8.12 | 2.44E-13 | 9.08E-13 |
| chr8 | 48988604 | 49012857 | 24254 | -8.67 | 2.45E-13 | 9.09E-13 |
| chr6 | 44376614 | 44378690 | 2077 | -8.22 | 2.45E-13 | 9.10E-13 |
| chr10 | 59404438 | 59412676 | 8239 | -8.04 | 2.45E-13 | 9.11E-13 |
| chr3 | 65631695 | 65636991 | 5297 | -8.03 | 2.45E-13 | 9.11E-13 |
| chr17 | 39761100 | 39765142 | 4043 | -8.17 | 2.46E-13 | 9.13E-13 |
| chr2 | 196302252 | 196308313 | 6062 | -8.25 | 2.46E-13 | 9.13E-13 |
| chr3 | 100502753 | 100505747 | 2995 | -8.17 | 2.46E-13 | 9.13E-13 |
| chr2 | 102154018 | 102160109 | 6092 | -8.02 | 2.47E-13 | 9.15E-13 |
| chr13 | 48887792 | 48897306 | 9515 | -8.03 | 2.47E-13 | 9.15E-13 |
| chr3 | 195873990 | 195878848 | 4859 | -8.03 | 2.47E-13 | 9.15E-13 |
| chr3 | 183213125 | 183224528 | 11404 | -8.06 | 2.47E-13 | 9.18E-13 |
| chr10 | 43672426 | 43683321 | 10896 | -8.02 | 2.49E-13 | 9.22E-13 |
| chr1 | 231277586 | 231288918 | 11333 | -8.42 | 2.49E-13 | 9.22E-13 |
| chr2 | 192218328 | 192221999 | 3672 | -8.02 | 2.49E-13 | 9.23E-13 |
| chr2 | 160550639 | 160552529 | 1891 | -8.03 | 2.50E-13 | 9.27E-13 |
| chrX | 4835711 | 4839720 | 4010 | -8.04 | 2.51E-13 | 9.29E-13 |
| chr2 | 211005136 | 211012357 | 7222 | -8.05 | 2.51E-13 | 9.29E-13 |
| chr2 | 236965720 | 236968416 | 2697 | -8.02 | 2.51E-13 | 9.31E-13 |
| chr12 | 100325143 | 100346268 | 21126 | -8.67 | 2.52E-13 | 9.33E-13 |
| chr2 | 143669404 | 143672146 | 2743 | -8.07 | 2.53E-13 | 9.37E-13 |
| chr18 | 33509184 | 33513293 | 4110 | -8.05 | 2.53E-13 | 9.38E-13 |
| chr2 | 209215073 | 209231651 | 16579 | -8.03 | 2.53E-13 | 9.38E-13 |
| chr6 | 76228898 | 76251087 | 22190 | -8.78 | 2.54E-13 | 9.40E-13 |
| chr3 | 7472823 | 7479119 | 6297 | -8.02 | 2.54E-13 | 9.41E-13 |
| chr1 | 191645892 | 191673810 | 27919 | -8.17 | 2.55E-13 | 9.42E-13 |
| chr14 | 93193258 | 93199194 | 5937 | -8.02 | 2.55E-13 | 9.45E-13 |
| chr22 | 44064921 | 44076782 | 11862 | -8.03 | 2.56E-13 | 9.48E-13 |
| chr2 | 43752204 | 43753870 | 1667 | -8.07 | 2.56E-13 | 9.48E-13 |
| chrX | 154067656 | 154078284 | 10629 | -8.22 | 2.56E-13 | 9.49E-13 |
| chr8 | 109465277 | 109471871 | 6595 | -8.1 | 2.57E-13 | 9.52E-13 |
| chr4 | 78681573 | 78684684 | 3112 | -8.02 | 2.58E-13 | 9.55E-13 |
| chr2 | 173233169 | 173245392 | 12224 | -8.31 | 2.59E-13 | 9.58E-13 |

| | | | | | | |
|---|---|---|---|---|---|---|
| chr14 | 50028356 | 50040855 | 12500 | -8.1 | 2.59E-13 | 9.59E-13 |
| chr9 | 26937384 | 26940290 | 2907 | -8.17 | 2.59E-13 | 9.59E-13 |
| chr1 | 214354945 | 214357970 | 3026 | -8.09 | 2.59E-13 | 9.59E-13 |
| chr4 | 178117046 | 178130031 | 12986 | -8.12 | 2.60E-13 | 9.61E-13 |
| chr4 | 175824060 | 175830026 | 5967 | -8.03 | 2.60E-13 | 9.63E-13 |
| chr5 | 58974470 | 58977203 | 2734 | -8.03 | 2.60E-13 | 9.63E-13 |
| chr3 | 176760833 | 176762990 | 2158 | -8.08 | 2.62E-13 | 9.68E-13 |
| chr6 | 141639189 | 141662882 | 23694 | -8.04 | 2.62E-13 | 9.69E-13 |
| chr3 | 4730698 | 4733178 | 2481 | -8.02 | 2.62E-13 | 9.69E-13 |
| chr8 | 81061316 | 81063923 | 2608 | -8.05 | 2.63E-13 | 9.71E-13 |
| chr3 | 133074443 | 133076960 | 2518 | -3.51 | 2.63E-13 | 9.72E-13 |
| chr12 | 28143245 | 28145697 | 2453 | -8.02 | 2.63E-13 | 9.72E-13 |
| chr7 | 93244703 | 93257954 | 13252 | -8.19 | 2.63E-13 | 9.72E-13 |
| chr2 | 217875738 | 217892661 | 16924 | -8.08 | 2.64E-13 | 9.74E-13 |
| chr17 | 61097336 | 61100546 | 3211 | -8.07 | 2.64E-13 | 9.74E-13 |
| chr6 | 150308460 | 150310505 | 2046 | -8.05 | 2.64E-13 | 9.76E-13 |
| chr2 | 128788317 | 128812795 | 24479 | -4.66 | 2.64E-13 | 9.76E-13 |
| chr5 | 34493622 | 34509343 | 15722 | -8.17 | 2.64E-13 | 9.76E-13 |
| chr10 | 118508052 | 118525645 | 17594 | -8.02 | 2.66E-13 | 9.81E-13 |
| chr16 | 28577432 | 28579809 | 2378 | -8.02 | 2.66E-13 | 9.81E-13 |
| chr14 | 100312042 | 100314429 | 2388 | -8.01 | 2.66E-13 | 9.84E-13 |
| chr9 | 138959524 | 138962385 | 2862 | -8.1 | 2.67E-13 | 9.85E-13 |
| chr20 | 48929866 | 48942293 | 12428 | -8.24 | 2.67E-13 | 9.86E-13 |
| chr2 | 99109244 | 99114257 | 5014 | -8.09 | 2.67E-13 | 9.87E-13 |
| chr4 | 151064004 | 151066003 | 2000 | -8.02 | 2.69E-13 | 9.94E-13 |
| chr2 | 58053164 | 58056751 | 3588 | -8.08 | 2.70E-13 | 9.97E-13 |
| chr11 | 5661077 | 5676329 | 15253 | -8.23 | 2.70E-13 | 9.98E-13 |
| chr2 | 235922392 | 235925535 | 3144 | -8.01 | 2.70E-13 | 9.98E-13 |
| chr2 | 3354161 | 3357557 | 3397 | -8.03 | 2.71E-13 | 9.99E-13 |
| chr11 | 44778248 | 44790431 | 12184 | -8.07 | 2.71E-13 | 1.00E-12 |
| chr17 | 167851 | 169611 | 1761 | -8.03 | 2.72E-13 | 1.00E-12 |
| chr2 | 39483290 | 39485056 | 1767 | -8.01 | 2.72E-13 | 1.00E-12 |
| chrX | 63443692 | 63558727 | 115036 | 1.5 | 2.73E-13 | 1.01E-12 |
| chr9 | 22393939 | 22412752 | 18814 | -8.01 | 2.74E-13 | 1.01E-12 |
| chr1 | 193215955 | 193218177 | 2223 | -8.06 | 2.74E-13 | 1.01E-12 |
| chr5 | 100679516 | 100685421 | 5906 | -8.23 | 2.75E-13 | 1.01E-12 |
| chr6 | 107298866 | 107309391 | 10526 | -8.18 | 2.75E-13 | 1.01E-12 |
| chr9 | 135367898 | 135371570 | 3673 | -8.18 | 2.75E-13 | 1.01E-12 |
| chr8 | 106750578 | 106758600 | 8023 | 8.21 | 2.75E-13 | 1.01E-12 |
| chr17 | 8162729 | 8164861 | 2133 | -8.04 | 2.75E-13 | 1.01E-12 |
| chr11 | 27311685 | 27328953 | 17269 | -8.09 | 2.75E-13 | 1.01E-12 |
| chr18 | 9047110 | 9053798 | 6689 | -8.01 | 2.76E-13 | 1.02E-12 |
| chr22 | 27332509 | 27335187 | 2679 | -8.03 | 2.76E-13 | 1.02E-12 |
| chr12 | 86915111 | 86920629 | 5519 | -8.25 | 2.76E-13 | 1.02E-12 |
| chr12 | 48470843 | 48472674 | 1832 | -8.09 | 2.77E-13 | 1.02E-12 |
| chr8 | 61159846 | 61162673 | 2828 | -8.11 | 2.77E-13 | 1.02E-12 |
| chr6 | 4865726 | 4869650 | 3925 | -8.07 | 2.78E-13 | 1.02E-12 |
| chr5 | 77914485 | 77929892 | 15408 | -8.01 | 2.78E-13 | 1.02E-12 |
| chr3 | 99787008 | 99791308 | 4301 | -8.13 | 2.78E-13 | 1.02E-12 |
| chr6 | 78153506 | 78169207 | 15702 | -8.01 | 2.79E-13 | 1.03E-12 |
| chr7 | 80850686 | 80854429 | 3744 | -8.01 | 2.80E-13 | 1.03E-12 |
| chr2 | 108977111 | 108982862 | 5752 | -8.14 | 2.80E-13 | 1.03E-12 |
| chr1 | 169367365 | 169377543 | 10179 | -8.07 | 2.80E-13 | 1.03E-12 |
| chr9 | 101177181 | 101197176 | 19996 | -8.03 | 2.80E-13 | 1.03E-12 |
| chr11 | 78802853 | 78805478 | 2626 | -8.03 | 2.81E-13 | 1.03E-12 |
| chr6 | 168721407 | 168726089 | 4683 | -8.19 | 2.83E-13 | 1.04E-12 |
| chr4 | 5898249 | 5917874 | 19626 | -4.36 | 2.83E-13 | 1.04E-12 |
| chr8 | 23092696 | 23096430 | 3735 | -8.15 | 2.83E-13 | 1.04E-12 |
| chr1 | 21153873 | 21172031 | 18159 | -8.17 | 2.83E-13 | 1.04E-12 |
| chr12 | 74247758 | 74253291 | 5534 | -8.03 | 2.84E-13 | 1.04E-12 |
| chr4 | 116935523 | 116944199 | 8677 | -8.11 | 2.84E-13 | 1.04E-12 |

| | | | | | | |
|---|---|---|---|---|---|---|
| chr3 | 39517168 | 39530093 | 12926 | -2.61 | 2.84E-13 | 1.04E-12 |
| chr13 | 78280234 | 78289265 | 9032 | -8.01 | 2.85E-13 | 1.05E-12 |
| chr3 | 148696652 | 148701835 | 5184 | -8.22 | 2.85E-13 | 1.05E-12 |
| chr18 | 53041102 | 53051232 | 10131 | -8.12 | 2.86E-13 | 1.05E-12 |
| chr16 | 53779966 | 53782541 | 2576 | -8.02 | 2.87E-13 | 1.05E-12 |
| chr4 | 184621883 | 184624462 | 2580 | -8.1 | 2.87E-13 | 1.06E-12 |
| chr6 | 53811215 | 53816584 | 5370 | -8.04 | 2.87E-13 | 1.06E-12 |
| chr5 | 23674306 | 23687380 | 13075 | -8.11 | 2.87E-13 | 1.06E-12 |
| chr8 | 86092253 | 86096505 | 4253 | -8.08 | 2.88E-13 | 1.06E-12 |
| chr7 | 69549261 | 69567279 | 18019 | -8.18 | 2.88E-13 | 1.06E-12 |
| chr6 | 165968305 | 165984929 | 16625 | -8.04 | 2.88E-13 | 1.06E-12 |
| chr18 | 660130 | 666726 | 6597 | -8.19 | 2.89E-13 | 1.06E-12 |
| chr20 | 62755807 | 62761383 | 5577 | -6.33 | 2.89E-13 | 1.06E-12 |
| chr4 | 39608046 | 39610008 | 1963 | -8 | 2.89E-13 | 1.06E-12 |
| chr10 | 5525115 | 5535700 | 10586 | -8.03 | 2.89E-13 | 1.06E-12 |
| chr2 | 158281343 | 158284186 | 2844 | -8.03 | 2.89E-13 | 1.06E-12 |
| chr22 | 39998352 | 40009032 | 10681 | -8.01 | 2.89E-13 | 1.06E-12 |
| chr5 | 67748068 | 67759342 | 11275 | -8.17 | 2.90E-13 | 1.06E-12 |
| chr2 | 47439198 | 47441807 | 2610 | -8.21 | 2.90E-13 | 1.07E-12 |
| chr9 | 128925725 | 128985610 | 59886 | -3.34 | 2.91E-13 | 1.07E-12 |
| chr1 | 182126640 | 182130893 | 4254 | -2.83 | 2.91E-13 | 1.07E-12 |
| chr8 | 31797158 | 31814183 | 17026 | -8.2 | 2.94E-13 | 1.08E-12 |
| chr7 | 41037788 | 41054270 | 16483 | -8.01 | 2.95E-13 | 1.08E-12 |
| chr1 | 53086477 | 53100962 | 14486 | -8.26 | 2.95E-13 | 1.08E-12 |
| chr7 | 73309768 | 73312304 | 2537 | -8 | 2.95E-13 | 1.08E-12 |
| chr5 | 77483309 | 77500582 | 17274 | -8.06 | 2.96E-13 | 1.09E-12 |
| chr7 | 2713057 | 2715650 | 2594 | -8.05 | 2.96E-13 | 1.09E-12 |
| chr15 | 67849236 | 67851696 | 2461 | -8.02 | 3.00E-13 | 1.10E-12 |
| chr7 | 32983036 | 32988610 | 5575 | -8.26 | 3.01E-13 | 1.10E-12 |
| chr2 | 8383309 | 8392516 | 9208 | -6.64 | 3.01E-13 | 1.10E-12 |
| chr17 | 32856015 | 32866117 | 10103 | 8.1 | 3.01E-13 | 1.11E-12 |
| chr11 | 47672584 | 47675054 | 2471 | -8.07 | 3.02E-13 | 1.11E-12 |
| chr14 | 65587488 | 65589728 | 2241 | -8.02 | 3.02E-13 | 1.11E-12 |
| chr5 | 170491176 | 170505910 | 14735 | -8.02 | 3.03E-13 | 1.11E-12 |
| chr6 | 89185818 | 89195752 | 9935 | -8.06 | 3.04E-13 | 1.11E-12 |
| chr8 | 116480182 | 116515297 | 35116 | -8.26 | 3.04E-13 | 1.11E-12 |
| chr14 | 32198737 | 32201550 | 2814 | -8.01 | 3.05E-13 | 1.12E-12 |
| chr5 | 57605506 | 57617346 | 11841 | -8.03 | 3.06E-13 | 1.12E-12 |
| chr14 | 103708604 | 103724800 | 16197 | -8.39 | 3.06E-13 | 1.12E-12 |
| chr4 | 130504824 | 130516500 | 11677 | -8.16 | 3.06E-13 | 1.12E-12 |
| chr6 | 130645037 | 130658738 | 13702 | -2.84 | 3.06E-13 | 1.12E-12 |
| chr20 | 50069854 | 50071877 | 2024 | -7.99 | 3.08E-13 | 1.13E-12 |
| chr8 | 30676406 | 30680429 | 4024 | -8.29 | 3.08E-13 | 1.13E-12 |
| chr5 | 159134390 | 159136846 | 2457 | -8.09 | 3.09E-13 | 1.13E-12 |
| chr2 | 201441084 | 201444574 | 3491 | -8.04 | 3.09E-13 | 1.13E-12 |
| chr19 | 45099161 | 45103461 | 4301 | -8.39 | 3.09E-13 | 1.13E-12 |
| chr1 | 2352806 | 2361768 | 8963 | -8 | 3.10E-13 | 1.13E-12 |
| chr3 | 52633139 | 52636327 | 3189 | -8.08 | 3.10E-13 | 1.13E-12 |
| chr17 | 25719749 | 25726895 | 7147 | -3.21 | 3.11E-13 | 1.14E-12 |
| chr1 | 109116400 | 109124882 | 8483 | -8.11 | 3.11E-13 | 1.14E-12 |
| chr17 | 70349549 | 70354572 | 5024 | -8.11 | 3.11E-13 | 1.14E-12 |
| chr4 | 157834196 | 157858274 | 24079 | -8.12 | 3.15E-13 | 1.15E-12 |
| chr8 | 115105513 | 115127351 | 21839 | -8.36 | 3.15E-13 | 1.15E-12 |
| chr9 | 90232485 | 90245359 | 12875 | -7.99 | 3.16E-13 | 1.16E-12 |
| chr9 | 96449346 | 96617252 | 167907 | -2.91 | 3.16E-13 | 1.16E-12 |
| chr1 | 52681066 | 52684069 | 3004 | -8 | 3.17E-13 | 1.16E-12 |
| chr3 | 27174017 | 27183917 | 9901 | -8.04 | 3.19E-13 | 1.17E-12 |
| chr8 | 42422956 | 42425182 | 2227 | -6.87 | 3.19E-13 | 1.17E-12 |
| chrX | 82035195 | 82043389 | 8195 | -8.02 | 3.20E-13 | 1.17E-12 |
| chr1 | 10902791 | 10915860 | 13070 | -8.16 | 3.20E-13 | 1.17E-12 |
| chr3 | 33328429 | 33337863 | 9435 | -7.99 | 3.21E-13 | 1.17E-12 |

| | | | | | | |
|---|---|---|---|---|---|---|
| chr14 | 95630330 | 95634784 | 4455 | -8.04 | 3.21E-13 | 1.17E-12 |
| chr10 | 51028957 | 51031651 | 2695 | -8 | 3.22E-13 | 1.18E-12 |
| chr12 | 111502433 | 111510727 | 8295 | -8.15 | 3.23E-13 | 1.18E-12 |
| chr4 | 161159884 | 161167168 | 7285 | -7.99 | 3.23E-13 | 1.18E-12 |
| chr3 | 95548220 | 95558124 | 9905 | -8.02 | 3.24E-13 | 1.18E-12 |
| chr5 | 10164523 | 10173655 | 9133 | -7.2 | 3.26E-13 | 1.19E-12 |
| chr20 | 2662047 | 2663575 | 1529 | -3.95 | 3.27E-13 | 1.19E-12 |
| chr1 | 43975755 | 43985588 | 9834 | -7.99 | 3.28E-13 | 1.20E-12 |
| chr1 | 99106199 | 99116451 | 10253 | -8.04 | 3.28E-13 | 1.20E-12 |
| chr3 | 5418029 | 5426463 | 8435 | -8.21 | 3.29E-13 | 1.20E-12 |
| chr3 | 33843881 | 33846313 | 2433 | -8.02 | 3.31E-13 | 1.21E-12 |
| chr10 | 59733742 | 59738085 | 4344 | -8.11 | 3.31E-13 | 1.21E-12 |
| chr5 | 16945369 | 16948467 | 3099 | -8.11 | 3.31E-13 | 1.21E-12 |
| chr1 | 113698168 | 113701196 | 3029 | -8.04 | 3.33E-13 | 1.22E-12 |
| chr12 | 113273479 | 113276523 | 3045 | -8.02 | 3.34E-13 | 1.22E-12 |
| chrX | 34576500 | 34587499 | 11000 | -6.24 | 3.35E-13 | 1.22E-12 |
| chr3 | 38186490 | 38189354 | 2865 | -8.22 | 3.36E-13 | 1.23E-12 |
| chr2 | 172441343 | 172448065 | 6723 | -8 | 3.36E-13 | 1.23E-12 |
| chr9 | 34354841 | 34358508 | 3668 | -8.02 | 3.37E-13 | 1.23E-12 |
| chr9 | 101459830 | 101467795 | 7966 | -7.99 | 3.37E-13 | 1.23E-12 |
| chr6 | 153028033 | 153035218 | 7186 | -7.98 | 3.38E-13 | 1.23E-12 |
| chr7 | 13039138 | 13051210 | 12073 | -8.17 | 3.38E-13 | 1.23E-12 |
| chr12 | 10450896 | 10457319 | 6424 | -8.02 | 3.39E-13 | 1.24E-12 |
| chr16 | 19274815 | 19290169 | 15355 | -8.43 | 3.40E-13 | 1.24E-12 |
| chr21 | 43995851 | 43998515 | 2665 | -7.98 | 3.41E-13 | 1.24E-12 |
| chr4 | 148352064 | 148355601 | 3538 | -7.98 | 3.41E-13 | 1.24E-12 |
| chr1 | 3675952 | 3677947 | 1996 | -8 | 3.42E-13 | 1.25E-12 |
| chr9 | 2170788 | 2176076 | 5289 | -8.05 | 3.42E-13 | 1.25E-12 |
| chr2 | 128150611 | 128152750 | 2140 | -7.98 | 3.43E-13 | 1.25E-12 |
| chr10 | 134412082 | 134415449 | 3368 | -8.04 | 3.43E-13 | 1.25E-12 |
| chr7 | 87875033 | 87892575 | 17543 | -4.46 | 3.43E-13 | 1.25E-12 |
| chr9 | 82215907 | 82223761 | 7855 | -8.11 | 3.43E-13 | 1.25E-12 |
| chr12 | 11007576 | 11011025 | 3450 | -8.04 | 3.43E-13 | 1.25E-12 |
| chr12 | 69913249 | 69917913 | 4665 | -7.98 | 3.44E-13 | 1.25E-12 |
| chr7 | 20483182 | 20485754 | 2573 | -7.98 | 3.44E-13 | 1.25E-12 |
| chr12 | 65813581 | 65826519 | 12939 | -8.03 | 3.44E-13 | 1.25E-12 |
| chr5 | 76059342 | 76062142 | 2801 | -8.04 | 3.44E-13 | 1.25E-12 |
| chr9 | 137843748 | 137850785 | 7038 | -8.28 | 3.46E-13 | 1.26E-12 |
| chr6 | 136718505 | 136723715 | 5211 | -8.07 | 3.46E-13 | 1.26E-12 |
| chr11 | 9210473 | 9214468 | 3996 | -8 | 3.46E-13 | 1.26E-12 |
| chr1 | 6176748 | 6193584 | 16837 | -7.99 | 3.47E-13 | 1.26E-12 |
| chr5 | 37627382 | 37630376 | 2995 | -8 | 3.47E-13 | 1.26E-12 |
| chr1 | 69268775 | 69276908 | 8134 | 7.02 | 3.48E-13 | 1.27E-12 |
| chr11 | 45211623 | 45225351 | 13729 | -7.99 | 3.48E-13 | 1.27E-12 |
| chr3 | 61470131 | 61476701 | 6571 | 6.64 | 3.48E-13 | 1.27E-12 |
| chr17 | 9892404 | 9899500 | 7097 | -8 | 3.50E-13 | 1.27E-12 |
| chr9 | 114048678 | 114060753 | 12076 | -8.38 | 3.50E-13 | 1.28E-12 |
| chrX | 53547630 | 53555848 | 8219 | -8.01 | 3.51E-13 | 1.28E-12 |
| chr6 | 108445517 | 108452545 | 7029 | -7.98 | 3.51E-13 | 1.28E-12 |
| chr8 | 141143256 | 141145429 | 2174 | -8.13 | 3.51E-13 | 1.28E-12 |
| chr5 | 76970876 | 76974171 | 3296 | -8 | 3.51E-13 | 1.28E-12 |
| chrX | 77885595 | 77904213 | 18619 | -2.16 | 3.52E-13 | 1.28E-12 |
| chrX | 12143176 | 12150864 | 7689 | -2.1 | 3.53E-13 | 1.28E-12 |
| chr5 | 30332679 | 30368588 | 35910 | -8.18 | 3.53E-13 | 1.29E-12 |
| chr16 | 54163389 | 54191345 | 27957 | -8.41 | 3.56E-13 | 1.29E-12 |
| chr6 | 8425528 | 8437002 | 11475 | -7.99 | 3.56E-13 | 1.30E-12 |
| chr8 | 37019347 | 37034001 | 14655 | -1.93 | 3.56E-13 | 1.30E-12 |
| chr2 | 142911070 | 142945934 | 34865 | -8.2 | 3.57E-13 | 1.30E-12 |
| chr2 | 28482325 | 28484409 | 2085 | -7.99 | 3.58E-13 | 1.30E-12 |
| chr15 | 67447582 | 67449413 | 1832 | -8 | 3.58E-13 | 1.30E-12 |
| chr8 | 73016824 | 73027923 | 11100 | -8.07 | 3.58E-13 | 1.30E-12 |

| | | | | | | |
|---|---|---|---|---|---|---|
| chr2 | 55001234 | 55004286 | 3053 | -7.99 | 3.59E-13 | 1.30E-12 |
| chr8 | 74476250 | 74481249 | 5000 | -7.98 | 3.59E-13 | 1.30E-12 |
| chr12 | 19238106 | 19250243 | 12138 | -8.17 | 3.60E-13 | 1.31E-12 |
| chr6 | 35815927 | 35817818 | 1892 | -8 | 3.60E-13 | 1.31E-12 |
| chr16 | 1562558 | 1565666 | 3109 | -8 | 3.61E-13 | 1.31E-12 |
| chr10 | 81908605 | 81910852 | 2248 | -8.02 | 3.61E-13 | 1.31E-12 |
| chr5 | 93523715 | 93525367 | 1653 | -3.6 | 3.62E-13 | 1.31E-12 |
| chr4 | 174225503 | 174227907 | 2405 | -7.99 | 3.62E-13 | 1.32E-12 |
| chr3 | 134219841 | 134224131 | 4291 | -8.02 | 3.62E-13 | 1.32E-12 |
| chr9 | 108569691 | 108586594 | 16904 | -8.04 | 3.62E-13 | 1.32E-12 |
| chr21 | 34440615 | 34442273 | 1659 | -8.1 | 3.63E-13 | 1.32E-12 |
| chrX | 100973119 | 100990775 | 17657 | 8.47 | 3.63E-13 | 1.32E-12 |
| chr1 | 181306566 | 181309137 | 2572 | -8.04 | 3.64E-13 | 1.32E-12 |
| chr4 | 106776517 | 106781576 | 5060 | -7.98 | 3.66E-13 | 1.33E-12 |
| chr7 | 158190478 | 158193154 | 2677 | -7.97 | 3.67E-13 | 1.33E-12 |
| chr3 | 176809256 | 176813651 | 4396 | -8.03 | 3.67E-13 | 1.33E-12 |
| chr1 | 222105277 | 222120335 | 15059 | -8.38 | 3.68E-13 | 1.34E-12 |
| chr14 | 65860131 | 65864108 | 3978 | -8.38 | 3.68E-13 | 1.34E-12 |
| chr8 | 33096029 | 33121269 | 25241 | -8.35 | 3.70E-13 | 1.34E-12 |
| chr10 | 105713579 | 105716096 | 2518 | -8 | 3.71E-13 | 1.34E-12 |
| chr2 | 45973855 | 45975290 | 1436 | -8.01 | 3.71E-13 | 1.35E-12 |
| chr16 | 12930513 | 12935376 | 4864 | -8.17 | 3.72E-13 | 1.35E-12 |
| chr1 | 37745020 | 37751047 | 6028 | -8.02 | 3.72E-13 | 1.35E-12 |
| chr21 | 27642425 | 27650684 | 8260 | -2.3 | 3.72E-13 | 1.35E-12 |
| chr8 | 127954402 | 127957030 | 2629 | -8.2 | 3.72E-13 | 1.35E-12 |
| chr3 | 68918478 | 68933982 | 15505 | -6.78 | 3.73E-13 | 1.35E-12 |
| chr6 | 86024460 | 86028907 | 4448 | -8.02 | 3.73E-13 | 1.35E-12 |
| chr2 | 165217360 | 165232161 | 14802 | -8.27 | 3.74E-13 | 1.35E-12 |
| chr7 | 7834673 | 7837665 | 2993 | -8.17 | 3.75E-13 | 1.36E-12 |
| chr4 | 169496271 | 169504676 | 8406 | -8.13 | 3.75E-13 | 1.36E-12 |
| chr7 | 13288410 | 13292338 | 3929 | -8.04 | 3.77E-13 | 1.37E-12 |
| chr2 | 165561889 | 165572343 | 10455 | -8.09 | 3.77E-13 | 1.37E-12 |
| chr3 | 64360240 | 64362803 | 2564 | -7.99 | 3.78E-13 | 1.37E-12 |
| chr20 | 56490642 | 56505017 | 14376 | -8.11 | 3.79E-13 | 1.37E-12 |
| chrX | 11105618 | 11110330 | 4713 | -8.11 | 3.79E-13 | 1.37E-12 |
| chr2 | 232749704 | 232751908 | 2205 | -7.97 | 3.80E-13 | 1.38E-12 |
| chr7 | 84184138 | 84187769 | 3632 | -7.99 | 3.81E-13 | 1.38E-12 |
| chr8 | 52829905 | 52842378 | 12474 | -8.07 | 3.81E-13 | 1.38E-12 |
| chr1 | 186567423 | 186574830 | 7408 | -7.98 | 3.81E-13 | 1.38E-12 |
| chr20 | 33510113 | 33512859 | 2747 | -7.97 | 3.81E-13 | 1.38E-12 |
| chr20 | 10985737 | 10988760 | 3024 | -7.97 | 3.83E-13 | 1.39E-12 |
| chr7 | 17949035 | 17957908 | 8874 | -8.09 | 3.84E-13 | 1.39E-12 |
| chr14 | 101477867 | 101492179 | 14313 | -7.97 | 3.85E-13 | 1.39E-12 |
| chr14 | 89558380 | 89603772 | 45393 | -4.11 | 3.85E-13 | 1.39E-12 |
| chrX | 15745527 | 15750153 | 4627 | -7.97 | 3.85E-13 | 1.39E-12 |
| chr4 | 47834323 | 47836697 | 2375 | -7.98 | 3.86E-13 | 1.40E-12 |
| chr16 | 4804766 | 4806601 | 1836 | -8.01 | 3.88E-13 | 1.40E-12 |
| chr20 | 11051307 | 11053822 | 2516 | -7.97 | 3.88E-13 | 1.40E-12 |
| chr1 | 186853744 | 186856260 | 2517 | -7.98 | 3.88E-13 | 1.41E-12 |
| chr19 | 6453144 | 6455734 | 2591 | -7.97 | 3.89E-13 | 1.41E-12 |
| chr2 | 160101192 | 160104325 | 3134 | -7.97 | 3.89E-13 | 1.41E-12 |
| chr8 | 87146012 | 87148101 | 2090 | -8.35 | 3.89E-13 | 1.41E-12 |
| chr9 | 124257607 | 124259657 | 2051 | -8.08 | 3.89E-13 | 1.41E-12 |
| chr5 | 168935845 | 168954884 | 19040 | -8.01 | 3.90E-13 | 1.41E-12 |
| chr15 | 33114257 | 33163112 | 48856 | -9.25 | 3.91E-13 | 1.41E-12 |
| chr1 | 162228214 | 162232632 | 4419 | -8 | 3.92E-13 | 1.42E-12 |
| chr19 | 45121726 | 45139301 | 17576 | -8.05 | 3.92E-13 | 1.42E-12 |
| chr6 | 41386201 | 41388596 | 2396 | -7.97 | 3.92E-13 | 1.42E-12 |
| chr19 | 51833611 | 51837341 | 3731 | -8.11 | 3.93E-13 | 1.42E-12 |
| chr8 | 135713258 | 135715441 | 2184 | -7.98 | 3.94E-13 | 1.42E-12 |
| chr9 | 77127273 | 77135351 | 8079 | 8.18 | 3.95E-13 | 1.43E-12 |

| | | | | | | |
|---|---|---|---|---|---|---|
| chr5 | 30456948 | 30476202 | 19255 | -8.01 | 3.95E-13 | 1.43E-12 |
| chr21 | 34682178 | 34687415 | 5238 | -8.12 | 3.95E-13 | 1.43E-12 |
| chr21 | 36979123 | 37139637 | 160515 | 1.15 | 3.97E-13 | 1.43E-12 |
| chrX | 17297672 | 17341335 | 43664 | -1.6 | 3.97E-13 | 1.43E-12 |
| chr3 | 94362982 | 94372025 | 9044 | -7.99 | 3.98E-13 | 1.44E-12 |
| chr10 | 47601085 | 47614867 | 13783 | -8.33 | 3.98E-13 | 1.44E-12 |
| chr9 | 128078223 | 128081156 | 2934 | -8.14 | 3.99E-13 | 1.44E-12 |
| chr4 | 155851132 | 155855844 | 4713 | -8.01 | 4.00E-13 | 1.45E-12 |
| chr6 | 14734105 | 14746641 | 12537 | -8.05 | 4.02E-13 | 1.45E-12 |
| chr10 | 30806975 | 30809453 | 2479 | -8.1 | 4.02E-13 | 1.45E-12 |
| chr3 | 38597715 | 38616060 | 18346 | -8.38 | 4.04E-13 | 1.46E-12 |
| chr6 | 100650205 | 100653799 | 3595 | -7.99 | 4.04E-13 | 1.46E-12 |
| chr11 | 86622445 | 86625527 | 3083 | -7.96 | 4.05E-13 | 1.46E-12 |
| chr14 | 67040634 | 67055010 | 14377 | -7.97 | 4.05E-13 | 1.46E-12 |
| chrX | 133853110 | 133856989 | 3880 | -7.97 | 4.06E-13 | 1.47E-12 |
| chr7 | 43643218 | 43645267 | 2050 | -8 | 4.07E-13 | 1.47E-12 |
| chr4 | 24863803 | 24867394 | 3592 | -8.08 | 4.07E-13 | 1.47E-12 |
| chrX | 7020889 | 7022530 | 1642 | -6.22 | 4.07E-13 | 1.47E-12 |
| chr1 | 184645813 | 184652360 | 6548 | -7.98 | 4.08E-13 | 1.47E-12 |
| chr6 | 82680929 | 82690915 | 9987 | -8.27 | 4.08E-13 | 1.47E-12 |
| chr9 | 119324257 | 119332500 | 8244 | -7.97 | 4.09E-13 | 1.48E-12 |
| chr1 | 62674718 | 62678991 | 4274 | -8.23 | 4.11E-13 | 1.48E-12 |
| chr10 | 29282187 | 29308189 | 26003 | -8.25 | 4.12E-13 | 1.49E-12 |
| chr6 | 46124285 | 46157975 | 33691 | 2.08 | 4.12E-13 | 1.49E-12 |
| chr4 | 158949078 | 158962467 | 13390 | -8.09 | 4.13E-13 | 1.49E-12 |
| chr11 | 128244226 | 128252432 | 8207 | -8.01 | 4.14E-13 | 1.49E-12 |
| chr7 | 47663998 | 47685939 | 21942 | -8.33 | 4.14E-13 | 1.49E-12 |
| chr7 | 30869818 | 30874568 | 4751 | -7.96 | 4.16E-13 | 1.50E-12 |
| chr3 | 151662226 | 151668271 | 6046 | -8.04 | 4.16E-13 | 1.50E-12 |
| chrX | 110783250 | 110799117 | 15868 | 8.07 | 4.18E-13 | 1.51E-12 |
| chr2 | 99579564 | 99594666 | 15103 | -2.94 | 4.18E-13 | 1.51E-12 |
| chr2 | 37868346 | 37876191 | 7846 | -8.11 | 4.18E-13 | 1.51E-12 |
| chr16 | 68855925 | 68864677 | 8753 | -8.36 | 4.19E-13 | 1.51E-12 |
| chr4 | 9841114 | 9858679 | 17566 | -2.43 | 4.20E-13 | 1.51E-12 |
| chr1 | 20718984 | 20723321 | 4338 | -3.54 | 4.20E-13 | 1.51E-12 |
| chr14 | 27138357 | 27157408 | 19052 | -7.96 | 4.20E-13 | 1.51E-12 |
| chr4 | 106004907 | 106025121 | 20215 | -1.85 | 4.21E-13 | 1.52E-12 |
| chr11 | 78220275 | 78222431 | 2157 | -7.97 | 4.22E-13 | 1.52E-12 |
| chr12 | 90601554 | 90605330 | 3777 | -8.02 | 4.23E-13 | 1.52E-12 |
| chr10 | 119059300 | 119065867 | 6568 | -7.95 | 4.24E-13 | 1.53E-12 |
| chr11 | 76210694 | 76213287 | 2594 | -7.98 | 4.24E-13 | 1.53E-12 |
| chr12 | 8977392 | 8979983 | 2592 | -7.98 | 4.25E-13 | 1.53E-12 |
| chr8 | 47866013 | 47877891 | 11879 | -8.01 | 4.26E-13 | 1.53E-12 |
| chr6 | 89746363 | 89749845 | 3483 | -7.96 | 4.26E-13 | 1.53E-12 |
| chr10 | 115522270 | 115524213 | 1944 | -7.96 | 4.27E-13 | 1.54E-12 |
| chr9 | 124565768 | 124568292 | 2525 | -8.09 | 4.27E-13 | 1.54E-12 |
| chr10 | 129872168 | 129874724 | 2557 | -7.95 | 4.27E-13 | 1.54E-12 |
| chr1 | 153964037 | 153968867 | 4831 | -5.92 | 4.27E-13 | 1.54E-12 |
| chr9 | 129356465 | 129364064 | 7600 | -7.98 | 4.28E-13 | 1.54E-12 |
| chr2 | 46531465 | 46533066 | 1602 | -8.13 | 4.28E-13 | 1.54E-12 |
| chr12 | 68207176 | 68211863 | 4688 | -8.1 | 4.29E-13 | 1.54E-12 |
| chr11 | 112113171 | 112117357 | 4187 | -8.1 | 4.29E-13 | 1.54E-12 |
| chr6 | 86291194 | 86301294 | 10101 | -8.1 | 4.30E-13 | 1.54E-12 |
| chr4 | 180372972 | 180387428 | 14457 | -8 | 4.31E-13 | 1.55E-12 |
| chr5 | 73412930 | 73416893 | 3964 | -7.95 | 4.31E-13 | 1.55E-12 |
| chrX | 9345019 | 9360074 | 15056 | -8.2 | 4.33E-13 | 1.56E-12 |
| chr12 | 119657378 | 119669442 | 12065 | -8.11 | 4.33E-13 | 1.56E-12 |
| chrX | 125539994 | 125591484 | 51491 | 1.74 | 4.33E-13 | 1.56E-12 |
| chr20 | 35657585 | 35668800 | 11216 | -8.06 | 4.34E-13 | 1.56E-12 |
| chr12 | 18981164 | 18985307 | 4144 | -8.03 | 4.35E-13 | 1.56E-12 |
| chr16 | 14028236 | 14049524 | 21289 | -8.06 | 4.35E-13 | 1.56E-12 |

| | | | | | | |
|---|---|---|---|---|---|---|
| chr5 | 45978800 | 46072235 | 93436 | -1.77 | 4.37E-13 | 1.57E-12 |
| chr4 | 187946435 | 187955347 | 8913 | -7.96 | 4.37E-13 | 1.57E-12 |
| chr21 | 35117977 | 35121454 | 3478 | -7.99 | 4.37E-13 | 1.57E-12 |
| chr1 | 12281406 | 12284985 | 3580 | -7.96 | 4.38E-13 | 1.57E-12 |
| chr1 | 227293902 | 227297323 | 3422 | -8.08 | 4.40E-13 | 1.58E-12 |
| chr21 | 42806023 | 42808503 | 2481 | -7.96 | 4.41E-13 | 1.58E-12 |
| chr4 | 73902057 | 73904899 | 2843 | -7.95 | 4.42E-13 | 1.59E-12 |
| chr19 | 18012629 | 18028432 | 15804 | -8.03 | 4.43E-13 | 1.59E-12 |
| chr2 | 46472962 | 46475985 | 3024 | -8.12 | 4.44E-13 | 1.59E-12 |
| chr6 | 138363083 | 138379570 | 16488 | -8.23 | 4.44E-13 | 1.59E-12 |
| chr17 | 58630294 | 58636290 | 5997 | -7.97 | 4.44E-13 | 1.60E-12 |
| chr15 | 37275101 | 37278895 | 3795 | -7.95 | 4.45E-13 | 1.60E-12 |
| chr7 | 81732486 | 81739953 | 7468 | -7.95 | 4.46E-13 | 1.60E-12 |
| chr3 | 148268045 | 148287684 | 19640 | -8.19 | 4.47E-13 | 1.60E-12 |
| chr17 | 74890118 | 74891848 | 1731 | -7.96 | 4.48E-13 | 1.61E-12 |
| chr1 | 118312034 | 118322977 | 10944 | -1.86 | 4.49E-13 | 1.61E-12 |
| chr6 | 21300079 | 21304996 | 4918 | -8.11 | 4.50E-13 | 1.62E-12 |
| chr6 | 132807478 | 132816180 | 8703 | -8.11 | 4.50E-13 | 1.62E-12 |
| chr3 | 15007657 | 15042437 | 34781 | -7.98 | 4.50E-13 | 1.62E-12 |
| chr22 | 44870051 | 44874375 | 4325 | -2.21 | 4.51E-13 | 1.62E-12 |
| chrX | 35371568 | 35376294 | 4727 | -7.97 | 4.52E-13 | 1.62E-12 |
| chr3 | 142890222 | 142900988 | 10767 | -3.47 | 4.52E-13 | 1.62E-12 |
| chr4 | 120483648 | 120487629 | 3982 | -7.95 | 4.54E-13 | 1.63E-12 |
| chr10 | 28936701 | 28938848 | 2148 | -8.12 | 4.54E-13 | 1.63E-12 |
| chr2 | 182592396 | 182612966 | 20571 | -7.95 | 4.56E-13 | 1.63E-12 |
| chr21 | 43413852 | 43417937 | 4086 | -7.95 | 4.57E-13 | 1.64E-12 |
| chr3 | 8555397 | 8561039 | 5643 | -8.11 | 4.57E-13 | 1.64E-12 |
| chr2 | 136771499 | 136775753 | 4255 | -8.21 | 4.58E-13 | 1.64E-12 |
| chr2 | 177965563 | 177968941 | 3379 | -7.94 | 4.60E-13 | 1.65E-12 |
| chr11 | 18598969 | 18600778 | 1810 | -7.94 | 4.60E-13 | 1.65E-12 |
| chr8 | 31607744 | 31616381 | 8638 | -8.18 | 4.61E-13 | 1.65E-12 |
| chr2 | 237543006 | 237592245 | 49240 | -1.55 | 4.61E-13 | 1.65E-12 |
| chr1 | 197656936 | 197658967 | 2032 | -7.94 | 4.61E-13 | 1.65E-12 |
| chr3 | 23663683 | 23667916 | 4234 | -8.07 | 4.61E-13 | 1.65E-12 |
| chr4 | 39114716 | 39125992 | 11277 | -8.01 | 4.62E-13 | 1.65E-12 |
| chr13 | 110112379 | 110117221 | 4843 | -8.07 | 4.62E-13 | 1.66E-12 |
| chr12 | 107941623 | 107943878 | 2256 | -8.09 | 4.63E-13 | 1.66E-12 |
| chr1 | 172228405 | 172241272 | 12868 | -8.09 | 4.64E-13 | 1.66E-12 |
| chr6 | 39394162 | 39405624 | 11463 | -7.95 | 4.65E-13 | 1.67E-12 |
| chr20 | 60324263 | 60327131 | 2869 | -8.1 | 4.67E-13 | 1.67E-12 |
| chr20 | 16313871 | 16316662 | 2792 | -7.94 | 4.67E-13 | 1.67E-12 |
| chr6 | 141087788 | 141094006 | 6219 | -7.94 | 4.67E-13 | 1.67E-12 |
| chr1 | 223026429 | 223032846 | 6418 | -8.11 | 4.68E-13 | 1.68E-12 |
| chr16 | 46619558 | 46625562 | 6005 | -8.05 | 4.69E-13 | 1.68E-12 |
| chr19 | 43294423 | 43300347 | 5925 | -7.94 | 4.69E-13 | 1.68E-12 |
| chr8 | 121798247 | 121802952 | 4706 | -3.99 | 4.70E-13 | 1.68E-12 |
| chr6 | 42434970 | 42438273 | 3304 | -7.94 | 4.70E-13 | 1.68E-12 |
| chr4 | 64819221 | 64840575 | 21355 | -7.95 | 4.70E-13 | 1.68E-12 |
| chr5 | 90229877 | 90251833 | 21957 | -8.06 | 4.71E-13 | 1.68E-12 |
| chr9 | 27218807 | 27260100 | 41294 | -1.36 | 4.71E-13 | 1.68E-12 |
| chr6 | 99438726 | 99559869 | 121144 | 4.39 | 4.71E-13 | 1.68E-12 |
| chr4 | 181495790 | 181502588 | 6799 | 8.15 | 4.71E-13 | 1.68E-12 |
| chr13 | 40310576 | 40315472 | 4897 | -7.94 | 4.72E-13 | 1.69E-12 |
| chr15 | 64086945 | 64090639 | 3695 | -8.02 | 4.72E-13 | 1.69E-12 |
| chr17 | 71879102 | 71930239 | 51138 | -2.82 | 4.73E-13 | 1.69E-12 |
| chr9 | 83996763 | 84008406 | 11644 | -7.94 | 4.73E-13 | 1.69E-12 |
| chr14 | 61608511 | 61612460 | 3950 | -8.26 | 4.73E-13 | 1.69E-12 |
| chr1 | 176045017 | 176048612 | 3596 | -8.01 | 4.75E-13 | 1.70E-12 |
| chr7 | 87242775 | 87266801 | 24027 | -7.96 | 4.75E-13 | 1.70E-12 |
| chr6 | 25204052 | 25207015 | 2964 | -8 | 4.75E-13 | 1.70E-12 |
| chr5 | 93924776 | 93943449 | 18674 | -7.96 | 4.76E-13 | 1.70E-12 |

| | | | | | | |
|---|---|---|---|---|---|---|
| chrX | 37776253 | 37779686 | 3434 | -8 | 4.76E-13 | 1.70E-12 |
| chr20 | 43439974 | 43441736 | 1763 | -7.94 | 4.77E-13 | 1.70E-12 |
| chr5 | 39065894 | 39068659 | 2766 | -7.95 | 4.80E-13 | 1.71E-12 |
| chr12 | 131268748 | 131271156 | 2409 | -7.99 | 4.85E-13 | 1.73E-12 |
| chr10 | 100208294 | 100210284 | 1991 | -7.99 | 4.86E-13 | 1.73E-12 |
| chr12 | 91838596 | 91860590 | 21995 | -7.99 | 4.86E-13 | 1.73E-12 |
| chr9 | 82066133 | 82069530 | 3398 | -7.99 | 4.86E-13 | 1.73E-12 |
| chr15 | 66665468 | 66667620 | 2153 | -7.97 | 4.87E-13 | 1.74E-12 |
| chr12 | 110411816 | 110417078 | 5263 | -7.96 | 4.88E-13 | 1.74E-12 |
| chr10 | 72730913 | 72733326 | 2414 | -8.06 | 4.90E-13 | 1.75E-12 |
| chr10 | 51536921 | 51539491 | 2571 | -8.11 | 4.97E-13 | 1.77E-12 |
| chr12 | 70626270 | 70632076 | 5807 | -8.03 | 4.99E-13 | 1.78E-12 |
| chr18 | 13021390 | 13023681 | 2292 | -7.99 | 4.99E-13 | 1.78E-12 |
| chr16 | 66365244 | 66386598 | 21355 | -8.4 | 5.01E-13 | 1.79E-12 |
| chr17 | 66103688 | 66105742 | 2055 | -8.06 | 5.04E-13 | 1.80E-12 |
| chr9 | 85547173 | 85552943 | 5771 | -8.02 | 5.04E-13 | 1.80E-12 |
| chr13 | 24052250 | 24062657 | 10408 | -8.02 | 5.05E-13 | 1.80E-12 |
| chr5 | 77208591 | 77252300 | 43710 | -7.98 | 5.05E-13 | 1.80E-12 |
| chr1 | 214876696 | 214880588 | 3893 | -2.8 | 5.06E-13 | 1.80E-12 |
| chr1 | 243680283 | 243682018 | 1736 | -7.94 | 5.06E-13 | 1.81E-12 |
| chr1 | 199071069 | 199076536 | 5468 | -8.01 | 5.07E-13 | 1.81E-12 |
| chr4 | 160942583 | 160959389 | 16807 | -8.01 | 5.07E-13 | 1.81E-12 |
| chr2 | 114663016 | 114664834 | 1819 | -7.96 | 5.07E-13 | 1.81E-12 |
| chr10 | 114547326 | 114550989 | 3664 | -7.93 | 5.08E-13 | 1.81E-12 |
| chr4 | 88204062 | 88208930 | 4869 | -7.97 | 5.09E-13 | 1.81E-12 |
| chr4 | 78888640 | 78891681 | 3042 | -7.93 | 5.11E-13 | 1.82E-12 |
| chr8 | 117442933 | 117450107 | 7175 | -7.97 | 5.12E-13 | 1.82E-12 |
| chr3 | 117038830 | 117052659 | 13830 | -7.96 | 5.12E-13 | 1.82E-12 |
| chr1 | 212169857 | 212171913 | 2057 | -8.13 | 5.13E-13 | 1.83E-12 |
| chr14 | 32890628 | 32893602 | 2975 | -7.95 | 5.13E-13 | 1.83E-12 |
| chr7 | 45334938 | 45383730 | 48793 | -2.04 | 5.13E-13 | 1.83E-12 |
| chr10 | 96242483 | 96245438 | 2956 | -8.03 | 5.14E-13 | 1.83E-12 |
| chr6 | 9428377 | 9444192 | 15816 | -7.98 | 5.15E-13 | 1.83E-12 |
| chr12 | 76200177 | 76203518 | 3342 | -7.93 | 5.16E-13 | 1.84E-12 |
| chr16 | 19590307 | 19592898 | 2592 | -7.93 | 5.16E-13 | 1.84E-12 |
| chr13 | 104193006 | 104197843 | 4838 | -7.93 | 5.17E-13 | 1.84E-12 |
| chr12 | 78337499 | 78344490 | 6992 | -8.38 | 5.18E-13 | 1.84E-12 |
| chr16 | 16027247 | 16030209 | 2963 | -7.97 | 5.18E-13 | 1.84E-12 |
| chr1 | 194676174 | 195720533 | 1044360 | 1.69 | 5.18E-13 | 1.84E-12 |
| chr9 | 16189033 | 16191216 | 2184 | -6.83 | 5.22E-13 | 1.86E-12 |
| chr3 | 33719304 | 33721322 | 2019 | -7.98 | 5.22E-13 | 1.86E-12 |
| chr3 | 193783861 | 193787727 | 3867 | -7.94 | 5.23E-13 | 1.86E-12 |
| chr15 | 50291741 | 50293110 | 1370 | -7.93 | 5.24E-13 | 1.86E-12 |
| chr4 | 5375099 | 5380382 | 5284 | -7.93 | 5.25E-13 | 1.87E-12 |
| chr5 | 81721580 | 81737911 | 16332 | -7.96 | 5.26E-13 | 1.87E-12 |
| chr2 | 210815962 | 210818246 | 2285 | -3.67 | 5.27E-13 | 1.87E-12 |
| chr6 | 113222230 | 113226664 | 4435 | -8 | 5.27E-13 | 1.88E-12 |
| chr1 | 169005120 | 169007966 | 2847 | -4.46 | 5.28E-13 | 1.88E-12 |
| chr1 | 173560934 | 173566281 | 5348 | -7.93 | 5.28E-13 | 1.88E-12 |
| chr3 | 82390542 | 82764012 | 373471 | 1.21 | 5.28E-13 | 1.88E-12 |
| chr10 | 75049315 | 75051617 | 2303 | -8.05 | 5.29E-13 | 1.88E-12 |
| chr20 | 43315283 | 43321166 | 5884 | -7.93 | 5.29E-13 | 1.88E-12 |
| chr2 | 234734384 | 234746408 | 12025 | -7.93 | 5.30E-13 | 1.89E-12 |
| chr1 | 219569033 | 219573344 | 4312 | -8.03 | 5.31E-13 | 1.89E-12 |
| chr12 | 15937366 | 15946490 | 9125 | -7.98 | 5.31E-13 | 1.89E-12 |
| chr17 | 67537017 | 67550654 | 13638 | -8.14 | 5.31E-13 | 1.89E-12 |
| chr1 | 94399762 | 94423433 | 23672 | -3.68 | 5.31E-13 | 1.89E-12 |
| chr3 | 107634306 | 107656541 | 22236 | -3.71 | 5.32E-13 | 1.89E-12 |
| chr3 | 105477354 | 105483078 | 5725 | -8.08 | 5.32E-13 | 1.89E-12 |
| chr3 | 57297083 | 57300518 | 3436 | -7.97 | 5.33E-13 | 1.89E-12 |
| chr8 | 63150105 | 63157612 | 7508 | -8.1 | 5.33E-13 | 1.89E-12 |

| | | | | | | |
|---|---|---|---|---|---|---|
| chr22 | 36882477 | 36885088 | 2612 | -7.93 | 5.34E-13 | 1.90E-12 |
| chr4 | 25488297 | 25496147 | 7851 | -8.11 | 5.35E-13 | 1.90E-12 |
| chr18 | 9161984 | 9183895 | 21912 | -8.15 | 5.36E-13 | 1.90E-12 |
| chr5 | 94948072 | 94951400 | 3329 | -8.05 | 5.36E-13 | 1.90E-12 |
| chr10 | 31400651 | 31418919 | 18269 | -8.33 | 5.39E-13 | 1.91E-12 |
| chr19 | 51634124 | 51658544 | 24421 | -3.23 | 5.39E-13 | 1.91E-12 |
| chr15 | 66102846 | 66106284 | 3439 | -7.99 | 5.41E-13 | 1.92E-12 |
| chr20 | 50093008 | 50095676 | 2669 | -7.92 | 5.43E-13 | 1.93E-12 |
| chr8 | 27229874 | 27232528 | 2655 | -8.13 | 5.43E-13 | 1.93E-12 |
| chr16 | 2800947 | 2815755 | 14809 | -8.13 | 5.44E-13 | 1.93E-12 |
| chr19 | 43605453 | 43620728 | 15276 | -8.08 | 5.44E-13 | 1.93E-12 |
| chr8 | 82379417 | 82392394 | 12978 | 8.26 | 5.45E-13 | 1.93E-12 |
| chr16 | 11525440 | 11549651 | 24212 | -8.54 | 5.46E-13 | 1.94E-12 |
| chr3 | 7544593 | 7546657 | 2065 | -8 | 5.48E-13 | 1.94E-12 |
| chr5 | 16494370 | 16497536 | 3167 | -8 | 5.49E-13 | 1.95E-12 |
| chr7 | 133979866 | 133982060 | 2195 | -8.04 | 5.50E-13 | 1.95E-12 |
| chr2 | 190057400 | 190062829 | 5430 | -7.92 | 5.52E-13 | 1.96E-12 |
| chr1 | 230084848 | 230119592 | 34745 | -2.82 | 5.53E-13 | 1.96E-12 |
| chr8 | 103799940 | 103814664 | 14725 | -8.08 | 5.53E-13 | 1.96E-12 |
| chr3 | 23695821 | 23698696 | 2876 | -7.92 | 5.54E-13 | 1.96E-12 |
| chr12 | 83769346 | 83783759 | 14414 | -7.94 | 5.54E-13 | 1.96E-12 |
| chr5 | 107042658 | 107059976 | 17319 | -7.92 | 5.58E-13 | 1.98E-12 |
| chr14 | 52001767 | 52006054 | 4288 | -7.95 | 5.59E-13 | 1.98E-12 |
| chr2 | 19477498 | 19483478 | 5981 | -8.18 | 5.60E-13 | 1.99E-12 |
| chr13 | 35562204 | 35565981 | 3778 | -7.97 | 5.61E-13 | 1.99E-12 |
| chr5 | 132752262 | 132755730 | 3469 | -5.82 | 5.62E-13 | 1.99E-12 |
| chr6 | 164101764 | 164107258 | 5495 | -3.17 | 5.62E-13 | 1.99E-12 |
| chr2 | 73981653 | 73984040 | 2388 | -7.92 | 5.64E-13 | 2.00E-12 |
| chr8 | 17921720 | 17923718 | 1999 | -8.07 | 5.65E-13 | 2.00E-12 |
| chr3 | 9528150 | 9531146 | 2997 | -7.94 | 5.66E-13 | 2.00E-12 |
| chr14 | 36929428 | 36932925 | 3498 | 6.37 | 5.66E-13 | 2.00E-12 |
| chr18 | 52391935 | 52408635 | 16701 | -7.94 | 5.67E-13 | 2.01E-12 |
| chr9 | 107795561 | 107831192 | 35632 | -8.17 | 5.67E-13 | 2.01E-12 |
| chr6 | 77928615 | 77933692 | 5078 | -7.99 | 5.67E-13 | 2.01E-12 |
| chr10 | 49721508 | 49737666 | 16159 | -7.92 | 5.69E-13 | 2.01E-12 |
| chr17 | 41303750 | 41306330 | 2581 | -7.92 | 5.70E-13 | 2.02E-12 |
| chr6 | 162558533 | 162563521 | 4989 | -7.92 | 5.72E-13 | 2.03E-12 |
| chr6 | 154752558 | 154754678 | 2121 | -7.91 | 5.74E-13 | 2.03E-12 |
| chr3 | 163988393 | 163997667 | 9275 | -7.97 | 5.79E-13 | 2.05E-12 |
| chr13 | 114971614 | 114974025 | 2412 | -7.93 | 5.80E-13 | 2.05E-12 |
| chr3 | 58530583 | 58538665 | 8083 | -7.92 | 5.80E-13 | 2.05E-12 |
| chr2 | 11844680 | 11847196 | 2517 | -8.02 | 5.82E-13 | 2.06E-12 |
| chr10 | 50974176 | 50986774 | 12599 | -7.99 | 5.84E-13 | 2.06E-12 |
| chr10 | 94118625 | 94121333 | 2709 | -7.99 | 5.84E-13 | 2.06E-12 |
| chr1 | 174951409 | 174953532 | 2124 | -8.03 | 5.84E-13 | 2.07E-12 |
| chr10 | 133885715 | 133888873 | 3159 | -7.93 | 5.87E-13 | 2.07E-12 |
| chr11 | 63336498 | 63348561 | 12064 | -8.01 | 5.88E-13 | 2.08E-12 |
| chr4 | 135608238 | 135632213 | 23976 | 8.58 | 5.89E-13 | 2.08E-12 |
| chr3 | 33268396 | 33285273 | 16878 | -8.12 | 5.90E-13 | 2.09E-12 |
| chr3 | 72542381 | 72553261 | 10881 | -7.91 | 5.90E-13 | 2.09E-12 |
| chr5 | 125979719 | 125982994 | 3276 | -7.98 | 5.90E-13 | 2.09E-12 |
| chr4 | 129054997 | 129089836 | 34840 | -8.04 | 5.91E-13 | 2.09E-12 |
| chr5 | 173745681 | 173749235 | 3555 | -7.95 | 5.91E-13 | 2.09E-12 |
| chr7 | 14587928 | 14605472 | 17545 | -7.98 | 5.92E-13 | 2.09E-12 |
| chr7 | 24641546 | 24666400 | 24855 | -7.96 | 5.94E-13 | 2.10E-12 |
| chr3 | 112504927 | 112508459 | 3533 | -7.94 | 5.94E-13 | 2.10E-12 |
| chr4 | 77042524 | 77044821 | 2298 | -8.08 | 5.94E-13 | 2.10E-12 |
| chr7 | 18588441 | 18604467 | 16027 | -8.84 | 5.94E-13 | 2.10E-12 |
| chr14 | 75167933 | 75170883 | 2951 | -7.93 | 5.96E-13 | 2.11E-12 |
| chr11 | 134058521 | 134062436 | 3916 | -8.25 | 5.96E-13 | 2.11E-12 |
| chr2 | 159323068 | 159330086 | 7019 | -8.08 | 5.97E-13 | 2.11E-12 |

| | | | | | | |
|---|---|---|---|---|---|---|
| chr2 | 232815344 | 232816801 | 1458 | -7.92 | 5.97E-13 | 2.11E-12 |
| chr4 | 100686085 | 100701388 | 15304 | -3.02 | 5.98E-13 | 2.11E-12 |
| chr11 | 44172631 | 44175148 | 2518 | -7.97 | 5.99E-13 | 2.12E-12 |
| chr20 | 60444494 | 60448254 | 3761 | -8.01 | 6.00E-13 | 2.12E-12 |
| chr13 | 22042179 | 22046054 | 3876 | -8.01 | 6.00E-13 | 2.12E-12 |
| chr1 | 173035325 | 173055922 | 20598 | -7.99 | 6.01E-13 | 2.12E-12 |
| chr6 | 82484059 | 82507196 | 23138 | -8.77 | 6.03E-13 | 2.13E-12 |
| chr15 | 52666370 | 52667751 | 1382 | -7.95 | 6.03E-13 | 2.13E-12 |
| chr14 | 91535812 | 91538117 | 2306 | -7.92 | 6.03E-13 | 2.13E-12 |
| chr3 | 56147873 | 56161846 | 13974 | -7.92 | 6.03E-13 | 2.13E-12 |
| chr16 | 84277923 | 84293352 | 15430 | -8.34 | 6.05E-13 | 2.14E-12 |
| chr6 | 58764237 | 58770947 | 6711 | -8.04 | 6.07E-13 | 2.14E-12 |
| chr22 | 47151245 | 47153497 | 2253 | -7.92 | 6.07E-13 | 2.14E-12 |
| chr4 | 64876040 | 64896543 | 20504 | -8.04 | 6.08E-13 | 2.14E-12 |
| chr3 | 126692483 | 126695096 | 2614 | -7.91 | 6.08E-13 | 2.15E-12 |
| chr19 | 42697350 | 42698701 | 1352 | -7.91 | 6.10E-13 | 2.15E-12 |
| chr6 | 112259047 | 112264436 | 5390 | -5.7 | 6.11E-13 | 2.15E-12 |
| chr15 | 62945458 | 62947392 | 1935 | -8 | 6.11E-13 | 2.16E-12 |
| chr3 | 71521873 | 71525979 | 4107 | -8.07 | 6.12E-13 | 2.16E-12 |
| chr13 | 21195994 | 21198084 | 2091 | -7.99 | 6.13E-13 | 2.16E-12 |
| chr10 | 74430738 | 74437618 | 6881 | -8.39 | 6.13E-13 | 2.16E-12 |
| chr1 | 66749318 | 66755177 | 5860 | -7.94 | 6.15E-13 | 2.17E-12 |
| chr13 | 100985238 | 100988376 | 3139 | -7.93 | 6.15E-13 | 2.17E-12 |
| chrX | 35630638 | 35639434 | 8797 | -8.05 | 6.16E-13 | 2.17E-12 |
| chr12 | 18560451 | 18585403 | 24953 | -8.38 | 6.16E-13 | 2.17E-12 |
| chr18 | 7529789 | 7532700 | 2912 | -7.95 | 6.16E-13 | 2.17E-12 |
| chr11 | 89961876 | 91083417 | 1121542 | 2.73 | 6.16E-13 | 2.17E-12 |
| chr6 | 167010300 | 167019208 | 8909 | -1.79 | 6.17E-13 | 2.17E-12 |
| chr13 | 33735310 | 33739067 | 3758 | -7.94 | 6.18E-13 | 2.18E-12 |
| chr8 | 91676832 | 91679689 | 2858 | -7.91 | 6.21E-13 | 2.19E-12 |
| chr2 | 57799921 | 57814350 | 14430 | 8.03 | 6.25E-13 | 2.20E-12 |
| chr19 | 3846694 | 3853162 | 6469 | -7.9 | 6.26E-13 | 2.20E-12 |
| chr9 | 94776567 | 94785175 | 8609 | -7.9 | 6.26E-13 | 2.20E-12 |
| chr15 | 77055029 | 77083936 | 28908 | -8.13 | 6.26E-13 | 2.20E-12 |
| chr18 | 70811113 | 70818798 | 7686 | -7.92 | 6.31E-13 | 2.22E-12 |
| chr1 | 214999734 | 215002868 | 3135 | -7.91 | 6.32E-13 | 2.22E-12 |
| chr2 | 241318748 | 241321104 | 2357 | -7.91 | 6.32E-13 | 2.22E-12 |
| chr9 | 124864652 | 124868147 | 3496 | -8.1 | 6.32E-13 | 2.23E-12 |
| chr4 | 110424966 | 110427410 | 2445 | -8.12 | 6.34E-13 | 2.23E-12 |
| chr15 | 58511209 | 58513027 | 1819 | -7.91 | 6.34E-13 | 2.23E-12 |
| chr12 | 64295674 | 64302450 | 6777 | -7.92 | 6.35E-13 | 2.24E-12 |
| chr1 | 19897292 | 19900045 | 2754 | -7.95 | 6.36E-13 | 2.24E-12 |
| chr22 | 24766559 | 24773181 | 6623 | -7.92 | 6.36E-13 | 2.24E-12 |
| chr4 | 170724512 | 170727859 | 3348 | -7.9 | 6.39E-13 | 2.25E-12 |
| chr6 | 144002793 | 144005830 | 3038 | -7.98 | 6.40E-13 | 2.25E-12 |
| chr4 | 131221253 | 131233307 | 12055 | -8.21 | 6.40E-13 | 2.25E-12 |
| chr12 | 25955912 | 25971054 | 15143 | -7.9 | 6.41E-13 | 2.25E-12 |
| chrX | 35545787 | 35573772 | 27986 | -8.18 | 6.41E-13 | 2.26E-12 |
| chr4 | 125497329 | 125503798 | 6470 | -7.93 | 6.42E-13 | 2.26E-12 |
| chr7 | 110701735 | 110706662 | 4928 | -7.9 | 6.43E-13 | 2.26E-12 |
| chr7 | 121178020 | 121186731 | 8712 | -7.92 | 6.44E-13 | 2.26E-12 |
| chr7 | 129998049 | 130001548 | 3500 | -7.9 | 6.45E-13 | 2.27E-12 |
| chr21 | 45910849 | 45914319 | 3471 | -7.91 | 6.45E-13 | 2.27E-12 |
| chr2 | 153351766 | 153354554 | 2789 | -8.14 | 6.45E-13 | 2.27E-12 |
| chr5 | 153532126 | 153536393 | 4268 | -6.02 | 6.45E-13 | 2.27E-12 |
| chrX | 4537438 | 4558270 | 20833 | -8.13 | 6.46E-13 | 2.27E-12 |
| chr2 | 99428066 | 99430960 | 2895 | -5.93 | 6.47E-13 | 2.27E-12 |
| chr14 | 20648067 | 20657451 | 9385 | 8 | 6.47E-13 | 2.27E-12 |
| chr3 | 122037032 | 122039865 | 2834 | -7.91 | 6.47E-13 | 2.27E-12 |
| chrX | 65351459 | 65393869 | 42411 | 3.05 | 6.51E-13 | 2.29E-12 |
| chr9 | 18217507 | 18219681 | 2175 | -7.96 | 6.53E-13 | 2.29E-12 |

| | | | | | | |
|---|---|---|---|---|---|---|
| chr7 | 95191334 | 95203891 | 12558 | -8.04 | 6.53E-13 | 2.29E-12 |
| chr1 | 158077772 | 158080656 | 2885 | -7.92 | 6.55E-13 | 2.30E-12 |
| chr11 | 515095 | 517145 | 2051 | -7.91 | 6.55E-13 | 2.30E-12 |
| chr2 | 139538416 | 139553471 | 15056 | -7.9 | 6.55E-13 | 2.30E-12 |
| chr14 | 32984805 | 32986980 | 2176 | -3.81 | 6.58E-13 | 2.31E-12 |
| chr8 | 103142205 | 103208504 | 66300 | -2.52 | 6.58E-13 | 2.31E-12 |
| chr9 | 110710450 | 110716134 | 5685 | -7.95 | 6.58E-13 | 2.31E-12 |
| chr3 | 142058489 | 142061964 | 3476 | -7.92 | 6.58E-13 | 2.31E-12 |
| chr9 | 79902243 | 79904227 | 1985 | -7.91 | 6.58E-13 | 2.31E-12 |
| chr10 | 60749478 | 60753124 | 3647 | -8.08 | 6.58E-13 | 2.31E-12 |
| chr17 | 2575305 | 2579389 | 4085 | -7.93 | 6.59E-13 | 2.31E-12 |
| chr5 | 56392830 | 56397878 | 5049 | -8.1 | 6.59E-13 | 2.31E-12 |
| chr22 | 24701132 | 24703128 | 1997 | -7.99 | 6.61E-13 | 2.32E-12 |
| chr2 | 242265020 | 242267127 | 2108 | -7.98 | 6.62E-13 | 2.32E-12 |
| chr1 | 178766537 | 178768681 | 2145 | -7.92 | 6.62E-13 | 2.32E-12 |
| chr22 | 24748825 | 24750602 | 1778 | -7.9 | 6.64E-13 | 2.33E-12 |
| chr7 | 19486080 | 19701514 | 215435 | 1.18 | 6.64E-13 | 2.33E-12 |
| chr2 | 191528815 | 191531113 | 2299 | -7.92 | 6.65E-13 | 2.33E-12 |
| chr16 | 50797063 | 50801210 | 4148 | -7.93 | 6.68E-13 | 2.34E-12 |
| chr14 | 52962289 | 52966048 | 3760 | -7.94 | 6.69E-13 | 2.34E-12 |
| chr7 | 30418939 | 30424307 | 5369 | -7.94 | 6.69E-13 | 2.34E-12 |
| chr9 | 126996203 | 126998662 | 2460 | -7.92 | 6.71E-13 | 2.35E-12 |
| chr7 | 93115833 | 93128204 | 12372 | 8.07 | 6.71E-13 | 2.35E-12 |
| chr10 | 12694867 | 12712378 | 17512 | -8.04 | 6.75E-13 | 2.37E-12 |
| chr14 | 102534362 | 102541612 | 7251 | -7.9 | 6.76E-13 | 2.37E-12 |
| chr11 | 111491837 | 111494249 | 2413 | -7.93 | 6.78E-13 | 2.38E-12 |
| chr14 | 94086663 | 94171617 | 84955 | -1.7 | 6.78E-13 | 2.38E-12 |
| chr13 | 33247086 | 33249660 | 2575 | -8.15 | 6.78E-13 | 2.38E-12 |
| chr7 | 38363796 | 38366699 | 2904 | -7.99 | 6.79E-13 | 2.38E-12 |
| chr8 | 17560037 | 17567732 | 7696 | -7.98 | 6.80E-13 | 2.38E-12 |
| chr2 | 20779738 | 20782278 | 2541 | -8 | 6.81E-13 | 2.38E-12 |
| chr21 | 36607568 | 36633720 | 26153 | -8.01 | 6.81E-13 | 2.39E-12 |
| chr21 | 16774742 | 16785215 | 10474 | -8.01 | 6.82E-13 | 2.39E-12 |
| chr3 | 157695448 | 157772307 | 76860 | -1.61 | 6.82E-13 | 2.39E-12 |
| chrX | 135926970 | 135930048 | 3079 | -7.97 | 6.82E-13 | 2.39E-12 |
| chr2 | 38369137 | 38375218 | 6082 | -7.9 | 6.84E-13 | 2.39E-12 |
| chr6 | 111222370 | 111226037 | 3668 | -8.05 | 6.85E-13 | 2.40E-12 |
| chr18 | 51669154 | 51671740 | 2587 | -7.89 | 6.85E-13 | 2.40E-12 |
| chr17 | 78615911 | 78633093 | 17183 | -8.46 | 6.86E-13 | 2.40E-12 |
| chr20 | 10542361 | 10544861 | 2501 | -7.99 | 6.87E-13 | 2.40E-12 |
| chr11 | 69238709 | 69240731 | 2023 | -7.89 | 6.88E-13 | 2.41E-12 |
| chr13 | 98142571 | 98144877 | 2307 | -7.93 | 6.88E-13 | 2.41E-12 |
| chr7 | 105539652 | 105568460 | 28809 | -7.9 | 6.88E-13 | 2.41E-12 |
| chr5 | 172733773 | 172736184 | 2412 | -7.9 | 6.88E-13 | 2.41E-12 |
| chr1 | 924875 | 929057 | 4183 | -7.89 | 6.88E-13 | 2.41E-12 |
| chr3 | 141518875 | 141521183 | 2309 | -7.94 | 6.90E-13 | 2.41E-12 |
| chr4 | 77857749 | 77864479 | 6731 | -8.01 | 6.90E-13 | 2.41E-12 |
| chr6 | 147569560 | 147573014 | 3455 | -7.9 | 6.92E-13 | 2.42E-12 |
| chr14 | 57900418 | 57902423 | 2006 | -6.24 | 6.92E-13 | 2.42E-12 |
| chr6 | 151879187 | 151885064 | 5878 | -3.54 | 6.95E-13 | 2.43E-12 |
| chr17 | 194348 | 224037 | 29690 | -8.43 | 6.96E-13 | 2.43E-12 |
| chrX | 111055076 | 111063282 | 8207 | 7.97 | 6.96E-13 | 2.43E-12 |
| chr2 | 48322265 | 48327892 | 5628 | -3.9 | 6.98E-13 | 2.44E-12 |
| chr3 | 121422336 | 121424816 | 2481 | -7.9 | 6.99E-13 | 2.44E-12 |
| chr6 | 139257157 | 139259164 | 2008 | -7.9 | 6.99E-13 | 2.44E-12 |
| chr7 | 1216930 | 1218508 | 1579 | -7.89 | 7.00E-13 | 2.45E-12 |
| chr11 | 72377821 | 72380071 | 2251 | -7.89 | 7.01E-13 | 2.45E-12 |
| chr1 | 227112032 | 227160580 | 48549 | -8.34 | 7.01E-13 | 2.45E-12 |
| chr10 | 72092243 | 72094616 | 2374 | -7.98 | 7.01E-13 | 2.45E-12 |
| chr16 | 3618472 | 3622591 | 4120 | -7.94 | 7.01E-13 | 2.45E-12 |
| chrX | 96783925 | 96787511 | 3587 | -8 | 7.03E-13 | 2.45E-12 |

| | | | | | | |
|---|---|---|---|---|---|---|
| chr6 | 143670789 | 143677701 | 6913 | -8.01 | 7.04E-13 | 2.46E-12 |
| chr11 | 67317494 | 67320650 | 3157 | -7.92 | 7.05E-13 | 2.46E-12 |
| chr4 | 152411188 | 152416632 | 5445 | -7.96 | 7.06E-13 | 2.47E-12 |
| chr9 | 95692630 | 95701999 | 9370 | -7.94 | 7.06E-13 | 2.47E-12 |
| chr5 | 101992801 | 102004494 | 11694 | 7.99 | 7.08E-13 | 2.47E-12 |
| chr6 | 109176070 | 109179638 | 3569 | -8.04 | 7.08E-13 | 2.47E-12 |
| chr21 | 43048800 | 43059727 | 10928 | -7.89 | 7.11E-13 | 2.48E-12 |
| chr7 | 116299239 | 116310537 | 11299 | -8.1 | 7.12E-13 | 2.48E-12 |
| chr1 | 47191630 | 47193376 | 1747 | -7.9 | 7.13E-13 | 2.49E-12 |
| chr11 | 10380556 | 10414588 | 34033 | -8.3 | 7.13E-13 | 2.49E-12 |
| chr6 | 168751091 | 168762709 | 11619 | -8.37 | 7.16E-13 | 2.50E-12 |
| chr1 | 114388011 | 114390962 | 2952 | -7.89 | 7.17E-13 | 2.50E-12 |
| chr2 | 127896348 | 127910646 | 14299 | -8.12 | 7.17E-13 | 2.50E-12 |
| chr2 | 242880758 | 242883472 | 2715 | -3.07 | 7.17E-13 | 2.50E-12 |
| chr11 | 123148292 | 123159773 | 11482 | -7.9 | 7.18E-13 | 2.50E-12 |
| chr12 | 131634101 | 131637875 | 3775 | -8.16 | 7.18E-13 | 2.50E-12 |
| chr3 | 189929268 | 189938137 | 8870 | -7.94 | 7.21E-13 | 2.51E-12 |
| chr15 | 101160389 | 101163095 | 2707 | -8.01 | 7.22E-13 | 2.52E-12 |
| chr9 | 113789673 | 113793495 | 3823 | -7.98 | 7.22E-13 | 2.52E-12 |
| chrX | 66480400 | 66521278 | 40879 | 1.62 | 7.23E-13 | 2.52E-12 |
| chr3 | 41704490 | 41713952 | 9463 | -7.9 | 7.23E-13 | 2.52E-12 |
| chr4 | 102402168 | 102412567 | 10400 | -7.92 | 7.23E-13 | 2.52E-12 |
| chr4 | 84620813 | 84624234 | 3422 | -7.89 | 7.25E-13 | 2.53E-12 |
| chr6 | 11224111 | 11227222 | 3112 | -7.91 | 7.28E-13 | 2.54E-12 |
| chr16 | 9135979 | 9141347 | 5369 | -7.93 | 7.29E-13 | 2.54E-12 |
| chr7 | 28082102 | 28099680 | 17579 | -8.38 | 7.30E-13 | 2.54E-12 |
| chr21 | 46819446 | 46822834 | 3389 | -7.89 | 7.30E-13 | 2.54E-12 |
| chr3 | 23086408 | 23102417 | 16010 | -8.3 | 7.31E-13 | 2.55E-12 |
| chr3 | 142642352 | 142644380 | 2029 | -7.9 | 7.33E-13 | 2.55E-12 |
| chr9 | 126482080 | 126491235 | 9156 | -8.11 | 7.33E-13 | 2.55E-12 |
| chr18 | 72046059 | 72107728 | 61670 | -1.82 | 7.35E-13 | 2.56E-12 |
| chr4 | 150945936 | 150958655 | 12720 | -8.02 | 7.36E-13 | 2.56E-12 |
| chr16 | 14231317 | 14234115 | 2799 | -7.94 | 7.37E-13 | 2.57E-12 |
| chr4 | 55024221 | 55028121 | 3901 | -7.94 | 7.39E-13 | 2.57E-12 |
| chr15 | 43835272 | 43840472 | 5201 | -7.92 | 7.40E-13 | 2.58E-12 |
| chr11 | 10872959 | 10877131 | 4173 | -7.94 | 7.42E-13 | 2.58E-12 |
| chr17 | 36157942 | 36172809 | 14868 | -7.89 | 7.46E-13 | 2.60E-12 |
| chr2 | 157807120 | 157816665 | 9546 | -7.91 | 7.48E-13 | 2.60E-12 |
| chr4 | 132553035 | 132562603 | 9569 | -7.93 | 7.50E-13 | 2.61E-12 |
| chr12 | 76082977 | 76093722 | 10746 | -8 | 7.50E-13 | 2.61E-12 |
| chr2 | 54440276 | 54443990 | 3715 | -7.88 | 7.51E-13 | 2.61E-12 |
| chr20 | 16685009 | 16710570 | 25562 | -1.2 | 7.53E-13 | 2.62E-12 |
| chrX | 95228538 | 95304815 | 76278 | 2.79 | 7.55E-13 | 2.63E-12 |
| chr3 | 109567601 | 109571247 | 3647 | -7.9 | 7.55E-13 | 2.63E-12 |
| chr2 | 214862736 | 214886861 | 24126 | -7.94 | 7.59E-13 | 2.64E-12 |
| chr7 | 17119699 | 17124759 | 5061 | -8.07 | 7.61E-13 | 2.65E-12 |
| chr3 | 110590858 | 110594831 | 3974 | -7.88 | 7.62E-13 | 2.65E-12 |
| chr1 | 94202313 | 94214580 | 12268 | -7.94 | 7.63E-13 | 2.65E-12 |
| chr3 | 196462650 | 196484491 | 21842 | -7.89 | 7.65E-13 | 2.66E-12 |
| chr16 | 81062529 | 81084833 | 22305 | -8.1 | 7.67E-13 | 2.67E-12 |
| chr4 | 161071060 | 161093538 | 22479 | -8.23 | 7.68E-13 | 2.67E-12 |
| chr7 | 79672565 | 79675618 | 3054 | -7.9 | 7.70E-13 | 2.67E-12 |
| chr3 | 81969771 | 81972304 | 2534 | -6.18 | 7.70E-13 | 2.68E-12 |
| chr3 | 8464593 | 8468128 | 3536 | -7.96 | 7.70E-13 | 2.68E-12 |
| chr4 | 184766750 | 184769716 | 2967 | -7.91 | 7.71E-13 | 2.68E-12 |
| chr2 | 25697828 | 25699564 | 1737 | -7.88 | 7.72E-13 | 2.68E-12 |
| chr2 | 174608071 | 174611994 | 3924 | -8 | 7.73E-13 | 2.69E-12 |
| chr1 | 225744288 | 225747003 | 2716 | -7.98 | 7.73E-13 | 2.69E-12 |
| chr3 | 32659404 | 32676035 | 16632 | -7.91 | 7.80E-13 | 2.71E-12 |
| chr14 | 32214613 | 32220248 | 5636 | -8.42 | 7.80E-13 | 2.71E-12 |
| chr12 | 17040556 | 17043987 | 3432 | -7.94 | 7.83E-13 | 2.72E-12 |

| | | | | | | |
|---|---|---|---|---|---|---|
| chr4 | 184246619 | 184257564 | 10946 | -7.94 | 7.84E-13 | 2.72E-12 |
| chr2 | 165116891 | 165136479 | 19589 | -8.21 | 7.84E-13 | 2.72E-12 |
| chr9 | 6830709 | 6832403 | 1695 | -7.87 | 7.86E-13 | 2.73E-12 |
| chr3 | 157345577 | 157359464 | 13888 | -1.7 | 7.88E-13 | 2.73E-12 |
| chr1 | 193999839 | 194004886 | 5048 | -7.89 | 7.88E-13 | 2.74E-12 |
| chr2 | 172166999 | 172170519 | 3521 | -7.89 | 7.89E-13 | 2.74E-12 |
| chr7 | 1627471 | 1629864 | 2394 | -7.93 | 7.90E-13 | 2.74E-12 |
| chr2 | 105863304 | 105866182 | 2879 | -8.05 | 7.90E-13 | 2.74E-12 |
| chr3 | 96042255 | 96060239 | 17985 | -8.07 | 7.91E-13 | 2.74E-12 |
| chrX | 24928524 | 24931534 | 3011 | -7.9 | 7.92E-13 | 2.75E-12 |
| chr6 | 79392065 | 79404352 | 12288 | -7.91 | 7.93E-13 | 2.75E-12 |
| chr2 | 159117006 | 159119121 | 2116 | -5.75 | 7.94E-13 | 2.75E-12 |
| chr11 | 120247985 | 120266977 | 18993 | -7.98 | 7.94E-13 | 2.76E-12 |
| chr10 | 59333287 | 59344573 | 11287 | -7.99 | 7.96E-13 | 2.76E-12 |
| chr2 | 220254980 | 220259194 | 4215 | -7.87 | 7.97E-13 | 2.76E-12 |
| chr12 | 11190955 | 11217737 | 26783 | -8.06 | 7.97E-13 | 2.76E-12 |
| chr13 | 41762597 | 41767276 | 4680 | -7.89 | 7.98E-13 | 2.77E-12 |
| chr3 | 153275418 | 153283700 | 8283 | -3.06 | 7.98E-13 | 2.77E-12 |
| chr12 | 15324877 | 15329210 | 4334 | -7.95 | 7.99E-13 | 2.77E-12 |
| chr4 | 67543255 | 67565156 | 21902 | -7.95 | 7.99E-13 | 2.77E-12 |
| chr2 | 75885180 | 75887686 | 2507 | -8.09 | 7.99E-13 | 2.77E-12 |
| chrX | 35999695 | 36008523 | 8829 | -8.03 | 7.99E-13 | 2.77E-12 |
| chr12 | 41929673 | 41981349 | 51677 | 7.55 | 7.99E-13 | 2.77E-12 |
| chr4 | 64958673 | 64988962 | 30290 | -8.34 | 8.00E-13 | 2.77E-12 |
| chr1 | 244897422 | 244900232 | 2811 | -7.88 | 8.01E-13 | 2.78E-12 |
| chr7 | 1557316 | 1559879 | 2564 | -7.92 | 8.02E-13 | 2.78E-12 |
| chr3 | 136173892 | 136176880 | 2989 | -7.89 | 8.03E-13 | 2.78E-12 |
| chr2 | 228244898 | 228257025 | 12128 | -7.92 | 8.03E-13 | 2.78E-12 |
| chr1 | 45742097 | 45750465 | 8369 | -8.08 | 8.06E-13 | 2.79E-12 |
| chrX | 110469151 | 110478044 | 8894 | 8.16 | 8.06E-13 | 2.79E-12 |
| chr5 | 173626203 | 173646989 | 20787 | -2.34 | 8.07E-13 | 2.79E-12 |
| chr3 | 153483767 | 153489161 | 5395 | -7.91 | 8.07E-13 | 2.79E-12 |
| chr3 | 113710455 | 113725599 | 15145 | -7.87 | 8.13E-13 | 2.81E-12 |
| chr5 | 118210495 | 118215297 | 4803 | -7.93 | 8.13E-13 | 2.82E-12 |
| chr6 | 3735031 | 3736432 | 1402 | -7.87 | 8.15E-13 | 2.82E-12 |
| chr17 | 9220081 | 9223511 | 3431 | -7.88 | 8.15E-13 | 2.82E-12 |
| chr3 | 145746969 | 145762518 | 15550 | -7.9 | 8.17E-13 | 2.83E-12 |
| chr2 | 107474438 | 107477732 | 3295 | -3.64 | 8.18E-13 | 2.83E-12 |
| chr1 | 26593365 | 26596313 | 2949 | -7.88 | 8.23E-13 | 2.85E-12 |
| chr1 | 84760876 | 84804211 | 43336 | -7.88 | 8.24E-13 | 2.85E-12 |
| chr17 | 78713367 | 78716796 | 3430 | -7.88 | 8.24E-13 | 2.85E-12 |
| chr1 | 169988740 | 169998201 | 9462 | -7.87 | 8.24E-13 | 2.85E-12 |
| chr2 | 165929917 | 165940796 | 10880 | -7.93 | 8.25E-13 | 2.85E-12 |
| chr8 | 657816 | 663837 | 6022 | -7.87 | 8.26E-13 | 2.86E-12 |
| chr16 | 10640717 | 10657940 | 17224 | -8 | 8.28E-13 | 2.86E-12 |
| chr10 | 46976973 | 46982143 | 5171 | -8.34 | 8.28E-13 | 2.86E-12 |
| chr10 | 59677326 | 59697771 | 20446 | -8.26 | 8.30E-13 | 2.87E-12 |
| chr6 | 37934211 | 37943780 | 9570 | -8.02 | 8.30E-13 | 2.87E-12 |
| chr2 | 192318623 | 192325020 | 6398 | -7.87 | 8.32E-13 | 2.87E-12 |
| chr3 | 64281893 | 64284470 | 2578 | -7.93 | 8.32E-13 | 2.88E-12 |
| chrX | 150201179 | 150228956 | 27778 | -8.72 | 8.33E-13 | 2.88E-12 |
| chr7 | 91469648 | 91486168 | 16521 | -8.04 | 8.34E-13 | 2.88E-12 |
| chr7 | 139428015 | 139434743 | 6729 | -7.87 | 8.34E-13 | 2.88E-12 |
| chr17 | 80379759 | 80381766 | 2008 | -7.89 | 8.37E-13 | 2.89E-12 |
| chr13 | 107419203 | 107554331 | 135129 | 1.81 | 8.37E-13 | 2.89E-12 |
| chr7 | 157660507 | 157678753 | 18247 | -8.35 | 8.37E-13 | 2.89E-12 |
| chr7 | 131537592 | 131540684 | 3093 | -7.9 | 8.39E-13 | 2.90E-12 |
| chr12 | 90303857 | 90308018 | 4162 | -8.01 | 8.40E-13 | 2.90E-12 |
| chr6 | 18167562 | 18169414 | 1853 | -7.88 | 8.42E-13 | 2.91E-12 |
| chr2 | 200908570 | 200918589 | 10020 | -8.19 | 8.43E-13 | 2.91E-12 |
| chrX | 81085073 | 81088114 | 3042 | 4.13 | 8.44E-13 | 2.91E-12 |

| | | | | | | |
|---|---|---|---|---|---|---|
| chr9 | 108618567 | 108622078 | 3512 | 6 | 8.44E-13 | 2.91E-12 |
| chr2 | 18691414 | 18693364 | 1951 | -7.9 | 8.44E-13 | 2.91E-12 |
| chr10 | 124282219 | 124298592 | 16374 | -2.12 | 8.47E-13 | 2.92E-12 |
| chr5 | 156297212 | 156299971 | 2760 | -7.87 | 8.48E-13 | 2.93E-12 |
| chr18 | 68260443 | 68281300 | 20858 | 8.15 | 8.52E-13 | 2.94E-12 |
| chr2 | 241912331 | 241914896 | 2566 | -7.87 | 8.53E-13 | 2.94E-12 |
| chr1 | 185753422 | 185770687 | 17266 | -8.07 | 8.53E-13 | 2.94E-12 |
| chr9 | 141016729 | 141026214 | 9486 | -7.86 | 8.53E-13 | 2.94E-12 |
| chr19 | 15212517 | 15216115 | 3599 | -7.89 | 8.54E-13 | 2.95E-12 |
| chrX | 100681708 | 100684149 | 2442 | -7.93 | 8.55E-13 | 2.95E-12 |
| chr1 | 80852681 | 80857228 | 4548 | -7.91 | 8.55E-13 | 2.95E-12 |
| chr12 | 102522195 | 102539997 | 17803 | -8.23 | 8.56E-13 | 2.95E-12 |
| chr7 | 65785440 | 65812880 | 27441 | -8.28 | 8.56E-13 | 2.95E-12 |
| chr1 | 41358665 | 41362623 | 3959 | -8 | 8.57E-13 | 2.96E-12 |
| chr5 | 180562834 | 180565838 | 3005 | -8.03 | 8.57E-13 | 2.96E-12 |
| chr17 | 30460482 | 30463071 | 2590 | -7.93 | 8.59E-13 | 2.96E-12 |
| chr1 | 84684406 | 84690055 | 5650 | -7.87 | 8.63E-13 | 2.98E-12 |
| chr2 | 218833686 | 218835677 | 1992 | -7.89 | 8.63E-13 | 2.98E-12 |
| chr22 | 43217131 | 43218801 | 1671 | -7.92 | 8.64E-13 | 2.98E-12 |
| chr10 | 52556446 | 52563466 | 7021 | 7.94 | 8.65E-13 | 2.98E-12 |
| chr1 | 59767739 | 59772557 | 4819 | -7.86 | 8.65E-13 | 2.98E-12 |
| chr14 | 75218019 | 75220964 | 2946 | -7.92 | 8.67E-13 | 2.99E-12 |
| chr4 | 26246795 | 26261866 | 15072 | -8.06 | 8.68E-13 | 2.99E-12 |
| chr12 | 59014082 | 59030706 | 16625 | -7.92 | 8.70E-13 | 3.00E-12 |
| chr7 | 94099629 | 94103055 | 3427 | -7.86 | 8.70E-13 | 3.00E-12 |
| chr1 | 44690729 | 44694212 | 3484 | -7.91 | 8.72E-13 | 3.01E-12 |
| chr3 | 159586491 | 159588240 | 1750 | -7.95 | 8.72E-13 | 3.01E-12 |
| chr4 | 15801115 | 15804603 | 3489 | -7.95 | 8.73E-13 | 3.01E-12 |
| chr11 | 69839669 | 69854450 | 14782 | -7.94 | 8.75E-13 | 3.01E-12 |
| chr1 | 173725746 | 173746910 | 21165 | -7.91 | 8.75E-13 | 3.01E-12 |
| chr19 | 16500112 | 16502198 | 2087 | -8.32 | 8.75E-13 | 3.01E-12 |
| chr6 | 54932856 | 54954848 | 21993 | -7.87 | 8.80E-13 | 3.03E-12 |
| chr20 | 35008414 | 35010582 | 2169 | -7.86 | 8.83E-13 | 3.04E-12 |
| chr2 | 146247954 | 146251893 | 3940 | -7.9 | 8.84E-13 | 3.04E-12 |
| chr11 | 101449263 | 101452765 | 3503 | -7.87 | 8.84E-13 | 3.04E-12 |
| chr11 | 68630323 | 68632366 | 2044 | -7.86 | 8.85E-13 | 3.05E-12 |
| chr2 | 173861841 | 173866333 | 4493 | -7.86 | 8.85E-13 | 3.05E-12 |
| chr12 | 53476695 | 53479822 | 3128 | -8.07 | 8.86E-13 | 3.05E-12 |
| chr1 | 68705004 | 68718249 | 13246 | -7.86 | 8.86E-13 | 3.05E-12 |
| chr6 | 126910308 | 126912545 | 2238 | -5.81 | 8.87E-13 | 3.05E-12 |
| chr10 | 116316036 | 116318988 | 2953 | -7.93 | 8.87E-13 | 3.05E-12 |
| chr17 | 16397644 | 16403515 | 5872 | -8.07 | 8.89E-13 | 3.06E-12 |
| chr12 | 43163640 | 43168026 | 4387 | -7.87 | 8.89E-13 | 3.06E-12 |
| chr7 | 7597428 | 7611007 | 13580 | -8.03 | 8.90E-13 | 3.06E-12 |
| chr5 | 174027924 | 174042733 | 14810 | -8.15 | 8.90E-13 | 3.06E-12 |
| chr13 | 33798083 | 33815521 | 17439 | -7.92 | 8.92E-13 | 3.07E-12 |
| chr3 | 99897181 | 99900086 | 2906 | -7.88 | 8.92E-13 | 3.07E-12 |
| chr1 | 84560291 | 84573951 | 13661 | -8.12 | 8.94E-13 | 3.07E-12 |
| chr3 | 139206168 | 139216001 | 9834 | -7.9 | 8.95E-13 | 3.08E-12 |
| chr8 | 120048569 | 120052827 | 4259 | 7.93 | 8.96E-13 | 3.08E-12 |
| chr9 | 16383322 | 16385449 | 2128 | -7.9 | 9.00E-13 | 3.09E-12 |
| chr2 | 28704624 | 28707688 | 3065 | -7.87 | 9.00E-13 | 3.09E-12 |
| chr9 | 132010395 | 132013534 | 3140 | -8.01 | 9.01E-13 | 3.10E-12 |
| chr1 | 186333566 | 186339853 | 6288 | -7.95 | 9.06E-13 | 3.11E-12 |
| chr11 | 113694431 | 113696748 | 2318 | -7.86 | 9.06E-13 | 3.11E-12 |
| chr18 | 33659231 | 33665094 | 5864 | -7.86 | 9.11E-13 | 3.13E-12 |
| chr6 | 23365214 | 23441105 | 75892 | 1.43 | 9.15E-13 | 3.14E-12 |
| chr14 | 94236694 | 94243003 | 6310 | -2.36 | 9.15E-13 | 3.14E-12 |
| chr12 | 15764572 | 15769546 | 4975 | -8.13 | 9.16E-13 | 3.15E-12 |
| chr8 | 99280331 | 99288935 | 8605 | -7.87 | 9.17E-13 | 3.15E-12 |
| chr19 | 41787974 | 41791421 | 3448 | -8.25 | 9.18E-13 | 3.15E-12 |

| | | | | | | |
|---|---|---|---|---|---|---|
| chr3 | 96776362 | 96991421 | 215060 | 1.44 | 9.18E-13 | 3.15E-12 |
| chr3 | 162709881 | 162713394 | 3514 | -7.86 | 9.19E-13 | 3.16E-12 |
| chr2 | 205761235 | 205764689 | 3455 | -7.88 | 9.20E-13 | 3.16E-12 |
| chr4 | 149043578 | 149047500 | 3923 | -7.9 | 9.20E-13 | 3.16E-12 |
| chr18 | 2427433 | 2442243 | 14811 | -7.92 | 9.20E-13 | 3.16E-12 |
| chr18 | 27020953 | 27439227 | 418275 | 1.46 | 9.23E-13 | 3.17E-12 |
| chr15 | 44369003 | 44372501 | 3499 | -7.86 | 9.26E-13 | 3.18E-12 |
| chr10 | 35544579 | 35548040 | 3462 | -7.91 | 9.27E-13 | 3.18E-12 |
| chr8 | 107871762 | 107875092 | 3331 | -7.86 | 9.28E-13 | 3.19E-12 |
| chr18 | 2212552 | 2217857 | 5306 | -7.87 | 9.29E-13 | 3.19E-12 |
| chr7 | 6031038 | 6034442 | 3405 | -7.87 | 9.29E-13 | 3.19E-12 |
| chr16 | 21025079 | 21027439 | 2361 | -7.87 | 9.30E-13 | 3.19E-12 |
| chr4 | 185404054 | 185407159 | 3106 | -7.95 | 9.30E-13 | 3.19E-12 |
| chr6 | 83163931 | 83169602 | 5672 | -8.03 | 9.31E-13 | 3.19E-12 |
| chr20 | 9382430 | 9384416 | 1987 | -7.88 | 9.31E-13 | 3.19E-12 |
| chr8 | 107183358 | 107191302 | 7945 | -7.85 | 9.31E-13 | 3.19E-12 |
| chr5 | 113392950 | 113410376 | 17427 | 8.74 | 9.34E-13 | 3.20E-12 |
| chr9 | 91587083 | 91598492 | 11410 | -7.93 | 9.35E-13 | 3.20E-12 |
| chrX | 102800980 | 102802904 | 1925 | -7.87 | 9.37E-13 | 3.21E-12 |
| chr8 | 41411169 | 41414023 | 2855 | -7.86 | 9.40E-13 | 3.22E-12 |
| chr17 | 28054754 | 28056951 | 2198 | -7.92 | 9.41E-13 | 3.23E-12 |
| chr3 | 11896890 | 11975908 | 79019 | -3.12 | 9.41E-13 | 3.23E-12 |
| chr5 | 93732913 | 93744970 | 12058 | 7.93 | 9.43E-13 | 3.23E-12 |
| chr21 | 27317192 | 27319142 | 1951 | -7.93 | 9.46E-13 | 3.24E-12 |
| chrX | 24884505 | 24890321 | 5817 | -7.97 | 9.48E-13 | 3.25E-12 |
| chr10 | 129777381 | 129780029 | 2649 | -7.97 | 9.49E-13 | 3.25E-12 |
| chrX | 16037554 | 16055861 | 18308 | -7.93 | 9.50E-13 | 3.25E-12 |
| chr3 | 129353967 | 129356586 | 2620 | -7.86 | 9.51E-13 | 3.26E-12 |
| chr4 | 84904085 | 84911518 | 7434 | -7.85 | 9.55E-13 | 3.27E-12 |
| chr5 | 76457006 | 76462411 | 5406 | -7.94 | 9.60E-13 | 3.29E-12 |
| chr3 | 129470910 | 129473516 | 2607 | -7.95 | 9.60E-13 | 3.29E-12 |
| chr7 | 139158125 | 139160021 | 1897 | -7.9 | 9.61E-13 | 3.29E-12 |
| chr6 | 154575375 | 154578121 | 2747 | -7.85 | 9.61E-13 | 3.29E-12 |
| chr4 | 121799805 | 121803628 | 3824 | -7.85 | 9.64E-13 | 3.30E-12 |
| chr6 | 150192333 | 150194985 | 2653 | -7.85 | 9.64E-13 | 3.30E-12 |
| chr5 | 21822545 | 21935785 | 113241 | 2.68 | 9.66E-13 | 3.31E-12 |
| chr2 | 160579977 | 160592618 | 12642 | -7.99 | 9.66E-13 | 3.31E-12 |
| chr9 | 132092300 | 132094503 | 2204 | -7.92 | 9.66E-13 | 3.31E-12 |
| chr12 | 26802925 | 26804880 | 1956 | -7.97 | 9.67E-13 | 3.31E-12 |
| chr12 | 30984373 | 30998499 | 14127 | -8.04 | 9.67E-13 | 3.31E-12 |
| chr3 | 182356541 | 182359223 | 2683 | -6.77 | 9.71E-13 | 3.32E-12 |
| chr15 | 68576291 | 68577762 | 1472 | -7.86 | 9.72E-13 | 3.33E-12 |
| chr4 | 5147505 | 5157820 | 10316 | -7.85 | 9.75E-13 | 3.33E-12 |
| chr10 | 5500630 | 5510965 | 10336 | -7.91 | 9.76E-13 | 3.34E-12 |
| chr5 | 61862266 | 61864837 | 2572 | -7.87 | 9.78E-13 | 3.34E-12 |
| chr4 | 24786205 | 24791145 | 4941 | -7.88 | 9.78E-13 | 3.35E-12 |
| chr21 | 35800298 | 35877033 | 76736 | -3.11 | 9.79E-13 | 3.35E-12 |
| chr11 | 95328338 | 95342606 | 14269 | 7.93 | 9.81E-13 | 3.35E-12 |
| chrX | 19835035 | 19840979 | 5945 | -7.93 | 9.83E-13 | 3.36E-12 |
| chr19 | 2366144 | 2381273 | 15130 | -8.16 | 9.86E-13 | 3.37E-12 |
| chr3 | 193707737 | 193715266 | 7530 | -8.11 | 9.88E-13 | 3.38E-12 |
| chr5 | 71656915 | 71671372 | 14458 | -7.9 | 9.89E-13 | 3.38E-12 |
| chr2 | 63737788 | 63764211 | 26424 | -8.25 | 9.89E-13 | 3.38E-12 |
| chr11 | 120411770 | 120414542 | 2773 | -7.9 | 9.92E-13 | 3.39E-12 |
| chr17 | 25505715 | 25517431 | 11717 | -7.96 | 9.95E-13 | 3.40E-12 |
| chrX | 45064233 | 45072640 | 8408 | -7.84 | 9.95E-13 | 3.40E-12 |
| chr18 | 73017843 | 73023514 | 5672 | -7.89 | 9.97E-13 | 3.41E-12 |
| chrX | 134493627 | 134503500 | 9874 | -7.87 | 9.98E-13 | 3.41E-12 |
| chr6 | 4842644 | 4846440 | 3797 | -8.01 | 9.98E-13 | 3.41E-12 |
| chr10 | 79098149 | 79100956 | 2808 | -7.91 | 9.99E-13 | 3.41E-12 |
| chr16 | 24991923 | 24994056 | 2134 | -7.87 | 1.00E-12 | 3.43E-12 |

| | | | | | | |
|---|---|---|---|---|---|---|
| chr7 | 51350890 | 51353510 | 2621 | -7.85 | 1.01E-12 | 3.44E-12 |
| chr2 | 43003909 | 43006843 | 2935 | -7.98 | 1.01E-12 | 3.44E-12 |
| chrX | 82384390 | 82393758 | 9369 | -8 | 1.01E-12 | 3.45E-12 |
| chr4 | 156037859 | 156054065 | 16207 | -7.86 | 1.02E-12 | 3.47E-12 |
| chr2 | 63783572 | 63796678 | 13107 | -8.04 | 1.02E-12 | 3.48E-12 |
| chr14 | 69131751 | 69168347 | 36597 | -8.07 | 1.02E-12 | 3.49E-12 |
| chr1 | 190024022 | 190031424 | 7403 | -7.84 | 1.02E-12 | 3.49E-12 |
| chr4 | 131033228 | 131042034 | 8807 | -7.94 | 1.02E-12 | 3.49E-12 |
| chr13 | 102098832 | 102115203 | 16372 | -8.21 | 1.02E-12 | 3.49E-12 |
| chr6 | 3479279 | 3481561 | 2283 | -7.84 | 1.02E-12 | 3.50E-12 |
| chr18 | 24022681 | 24026403 | 3723 | -7.89 | 1.03E-12 | 3.50E-12 |
| chr1 | 41908342 | 41912446 | 4105 | -8.07 | 1.03E-12 | 3.50E-12 |
| chr2 | 203968103 | 203978307 | 10205 | -7.96 | 1.03E-12 | 3.51E-12 |
| chrX | 90136890 | 90145226 | 8337 | 1.91 | 1.03E-12 | 3.51E-12 |
| chr20 | 45609839 | 45613515 | 3677 | -8.03 | 1.03E-12 | 3.51E-12 |
| chr6 | 21072030 | 21078045 | 6016 | -7.84 | 1.03E-12 | 3.52E-12 |
| chr17 | 15798292 | 15826501 | 28210 | -8.59 | 1.03E-12 | 3.52E-12 |
| chrX | 101187202 | 101197843 | 10642 | -7.89 | 1.03E-12 | 3.52E-12 |
| chr22 | 40915834 | 40918165 | 2332 | -7.98 | 1.04E-12 | 3.55E-12 |
| chr6 | 145183412 | 145195515 | 12104 | -8.03 | 1.04E-12 | 3.55E-12 |
| chrX | 101398886 | 101408990 | 10105 | 2.42 | 1.04E-12 | 3.56E-12 |
| chr5 | 67438118 | 67440873 | 2756 | -3.93 | 1.04E-12 | 3.56E-12 |
| chr7 | 85233437 | 85245802 | 12366 | -7.9 | 1.05E-12 | 3.57E-12 |
| chr22 | 37401059 | 37403604 | 2546 | -7.84 | 1.05E-12 | 3.57E-12 |
| chr20 | 50626385 | 50629317 | 2933 | -7.86 | 1.05E-12 | 3.58E-12 |
| chr10 | 92848897 | 92867953 | 19057 | -6.78 | 1.05E-12 | 3.58E-12 |
| chr3 | 187568260 | 187572113 | 3854 | -7.99 | 1.05E-12 | 3.58E-12 |
| chr22 | 18964185 | 18969524 | 5340 | -7.87 | 1.06E-12 | 3.60E-12 |
| chr11 | 75079119 | 75082450 | 3332 | -8.23 | 1.06E-12 | 3.60E-12 |
| chr12 | 96771401 | 96777461 | 6061 | -7.88 | 1.06E-12 | 3.60E-12 |
| chr12 | 75275334 | 75287557 | 12224 | -7.95 | 1.06E-12 | 3.61E-12 |
| chr16 | 87476751 | 87479539 | 2789 | -7.87 | 1.06E-12 | 3.61E-12 |
| chrX | 153370325 | 153383723 | 13399 | -7.87 | 1.06E-12 | 3.61E-12 |
| chr2 | 242361891 | 242370574 | 8684 | -7.84 | 1.06E-12 | 3.61E-12 |
| chr13 | 77864090 | 77866312 | 2223 | -7.84 | 1.06E-12 | 3.62E-12 |
| chr16 | 4680397 | 4682707 | 2311 | -7.84 | 1.06E-12 | 3.62E-12 |
| chr6 | 56008366 | 56020279 | 11914 | -7.84 | 1.06E-12 | 3.62E-12 |
| chr2 | 10433736 | 10440159 | 6424 | -8.05 | 1.07E-12 | 3.63E-12 |
| chr12 | 120244508 | 120246130 | 1623 | -7.87 | 1.07E-12 | 3.63E-12 |
| chr2 | 8266884 | 8291505 | 24622 | -8.37 | 1.07E-12 | 3.63E-12 |
| chr1 | 171518679 | 171529230 | 10552 | -7.92 | 1.07E-12 | 3.63E-12 |
| chr2 | 218580097 | 218587177 | 7081 | -7.84 | 1.07E-12 | 3.65E-12 |
| chr8 | 24358141 | 24360832 | 2692 | -5.7 | 1.07E-12 | 3.66E-12 |
| chr16 | 22042064 | 22044135 | 2072 | -7.84 | 1.08E-12 | 3.66E-12 |
| chr14 | 38160205 | 38164382 | 4178 | -7.86 | 1.08E-12 | 3.66E-12 |
| chr3 | 171958130 | 171961117 | 2988 | -7.86 | 1.08E-12 | 3.66E-12 |
| chr6 | 16912343 | 16915526 | 3184 | -7.84 | 1.08E-12 | 3.66E-12 |
| chr2 | 36209185 | 36222244 | 13060 | -5.82 | 1.08E-12 | 3.66E-12 |
| chr2 | 101825218 | 101828551 | 3334 | -8.22 | 1.08E-12 | 3.67E-12 |
| chr5 | 74583013 | 74589907 | 6895 | -8.07 | 1.08E-12 | 3.68E-12 |
| chr20 | 24729704 | 24742243 | 12540 | -7.9 | 1.08E-12 | 3.68E-12 |
| chr2 | 37689700 | 37693632 | 3933 | -8.04 | 1.08E-12 | 3.68E-12 |
| chr4 | 159411519 | 159429254 | 17736 | -7.83 | 1.09E-12 | 3.69E-12 |
| chr6 | 4691236 | 4694266 | 3031 | -7.85 | 1.09E-12 | 3.70E-12 |
| chr2 | 54299412 | 54302129 | 2718 | -7.86 | 1.09E-12 | 3.70E-12 |
| chr10 | 59101212 | 59328063 | 226852 | 5.21 | 1.09E-12 | 3.70E-12 |
| chr2 | 208826190 | 208828341 | 2152 | -7.92 | 1.09E-12 | 3.71E-12 |
| chr20 | 36195716 | 36199025 | 3310 | -8.33 | 1.09E-12 | 3.71E-12 |
| chr13 | 41280000 | 41288271 | 8272 | -7.92 | 1.09E-12 | 3.71E-12 |
| chr1 | 8603670 | 8605652 | 1983 | -7.84 | 1.09E-12 | 3.72E-12 |
| chr9 | 100868441 | 100870064 | 1624 | -7.85 | 1.09E-12 | 3.72E-12 |

| | | | | | | |
|---|---|---|---|---|---|---|
| chr3 | 28392955 | 28412218 | 19264 | -6.52 | 1.10E-12 | 3.73E-12 |
| chr2 | 37792928 | 37834219 | 41292 | -8.3 | 1.10E-12 | 3.73E-12 |
| chr8 | 71011418 | 71024439 | 13022 | -7.95 | 1.10E-12 | 3.73E-12 |
| chr2 | 197895109 | 197899783 | 4675 | -7.91 | 1.10E-12 | 3.73E-12 |
| chr3 | 25455035 | 25460254 | 5220 | -7.96 | 1.10E-12 | 3.73E-12 |
| chr18 | 45488102 | 45502880 | 14779 | -8.14 | 1.10E-12 | 3.74E-12 |
| chr18 | 55971089 | 55974137 | 3049 | -7.85 | 1.10E-12 | 3.74E-12 |
| chr19 | 7169735 | 7171919 | 2185 | -7.85 | 1.10E-12 | 3.74E-12 |
| chr11 | 77525430 | 77527589 | 2160 | -7.85 | 1.10E-12 | 3.75E-12 |
| chr1 | 6905439 | 6915587 | 10149 | -7.85 | 1.10E-12 | 3.75E-12 |
| chr4 | 129470988 | 129486958 | 15971 | -8.07 | 1.11E-12 | 3.75E-12 |
| chr10 | 21610468 | 21657894 | 47427 | -7.88 | 1.11E-12 | 3.76E-12 |
| chr5 | 175282759 | 175292954 | 10196 | -7.92 | 1.11E-12 | 3.76E-12 |
| chr13 | 28076254 | 28103121 | 26868 | -8.12 | 1.11E-12 | 3.76E-12 |
| chr17 | 42623079 | 42624793 | 1715 | -7.83 | 1.11E-12 | 3.77E-12 |
| chr19 | 8133673 | 8135964 | 2292 | -7.85 | 1.11E-12 | 3.77E-12 |
| chr3 | 14487586 | 14493719 | 6134 | -8.03 | 1.11E-12 | 3.77E-12 |
| chr8 | 42678607 | 42687709 | 9103 | -7.97 | 1.11E-12 | 3.77E-12 |
| chr7 | 116540587 | 116553417 | 12831 | -8.19 | 1.11E-12 | 3.77E-12 |
| chr6 | 71039695 | 71050504 | 10810 | -8.05 | 1.11E-12 | 3.78E-12 |
| chr14 | 36110063 | 36114447 | 4385 | -7.91 | 1.11E-12 | 3.78E-12 |
| chr4 | 176886694 | 176890850 | 4157 | -7.83 | 1.12E-12 | 3.78E-12 |
| chr9 | 125181733 | 125197309 | 15577 | -8.01 | 1.12E-12 | 3.79E-12 |
| chr20 | 46483743 | 46495039 | 11297 | -8.46 | 1.12E-12 | 3.79E-12 |
| chr1 | 8099329 | 8104977 | 5649 | -8.14 | 1.12E-12 | 3.79E-12 |
| chr2 | 10206871 | 10209017 | 2147 | -7.98 | 1.12E-12 | 3.80E-12 |
| chr17 | 72658480 | 72668208 | 9729 | -3.89 | 1.12E-12 | 3.80E-12 |
| chr10 | 62892879 | 62904789 | 11911 | -7.94 | 1.13E-12 | 3.82E-12 |
| chr4 | 77383490 | 77399727 | 16238 | -7.99 | 1.13E-12 | 3.83E-12 |
| chr9 | 133315469 | 133318337 | 2869 | -7.89 | 1.13E-12 | 3.83E-12 |
| chr7 | 47176548 | 47182824 | 6277 | -7.83 | 1.13E-12 | 3.83E-12 |
| chr15 | 77431711 | 77434505 | 2795 | -7.89 | 1.13E-12 | 3.84E-12 |
| chr12 | 560705 | 568597 | 7893 | -7.89 | 1.14E-12 | 3.84E-12 |
| chr1 | 201492459 | 201494570 | 2112 | -7.9 | 1.14E-12 | 3.84E-12 |
| chr2 | 236383484 | 236400867 | 17384 | -8.06 | 1.14E-12 | 3.85E-12 |
| chr8 | 32430702 | 32432781 | 2080 | -7.85 | 1.14E-12 | 3.85E-12 |
| chr4 | 130886583 | 130891869 | 5287 | -7.88 | 1.14E-12 | 3.86E-12 |
| chr15 | 28527446 | 28537095 | 9650 | -8.02 | 1.14E-12 | 3.87E-12 |
| chrX | 51741530 | 51753787 | 12258 | 7.9 | 1.14E-12 | 3.87E-12 |
| chr4 | 13246573 | 13250568 | 3996 | -2.76 | 1.14E-12 | 3.87E-12 |
| chr8 | 32281686 | 32296285 | 14600 | -8.21 | 1.14E-12 | 3.87E-12 |
| chr1 | 39169223 | 39189236 | 20014 | -2.02 | 1.14E-12 | 3.87E-12 |
| chr12 | 121618468 | 121620760 | 2293 | -7.83 | 1.15E-12 | 3.88E-12 |
| chr1 | 88125532 | 88145966 | 20435 | -7.85 | 1.15E-12 | 3.89E-12 |
| chr2 | 175057073 | 175059776 | 2704 | -7.85 | 1.15E-12 | 3.89E-12 |
| chr1 | 30165282 | 30167438 | 2157 | -7.83 | 1.15E-12 | 3.89E-12 |
| chr11 | 72686151 | 72688858 | 2708 | -7.83 | 1.15E-12 | 3.89E-12 |
| chr3 | 8521000 | 8528052 | 7053 | -8.09 | 1.15E-12 | 3.90E-12 |
| chr4 | 54619010 | 54650811 | 31802 | -8.14 | 1.15E-12 | 3.90E-12 |
| chr15 | 74755467 | 74759600 | 4134 | -7.84 | 1.16E-12 | 3.91E-12 |
| chr1 | 246553365 | 246555876 | 2512 | -7.83 | 1.16E-12 | 3.94E-12 |
| chr22 | 30453010 | 30461678 | 8669 | -7.83 | 1.16E-12 | 3.94E-12 |
| chr12 | 105676807 | 105693329 | 16523 | -7.91 | 1.17E-12 | 3.94E-12 |
| chr18 | 24338750 | 24348052 | 9303 | -7.94 | 1.17E-12 | 3.95E-12 |
| chr11 | 109375894 | 109388987 | 13094 | 2.54 | 1.17E-12 | 3.95E-12 |
| chr11 | 57039717 | 57042411 | 2695 | -7.84 | 1.17E-12 | 3.95E-12 |
| chr9 | 119371467 | 119374465 | 2999 | -7.82 | 1.17E-12 | 3.96E-12 |
| chr12 | 87749431 | 87755013 | 5583 | -7.83 | 1.18E-12 | 3.98E-12 |
| chr2 | 20806628 | 20808390 | 1763 | -7.84 | 1.18E-12 | 3.98E-12 |
| chr8 | 108491808 | 108495809 | 4002 | -8.18 | 1.18E-12 | 3.99E-12 |
| chr18 | 53869596 | 53874342 | 4747 | -5.82 | 1.18E-12 | 4.00E-12 |

| | | | | | | |
|---|---|---|---|---|---|---|
| chr6 | 37977533 | 38004250 | 26718 | -8.1 | 1.19E-12 | 4.00E-12 |
| chr4 | 8168972 | 8171859 | 2888 | -7.86 | 1.19E-12 | 4.01E-12 |
| chr2 | 207697863 | 207699519 | 1657 | -5.01 | 1.19E-12 | 4.01E-12 |
| chr15 | 89828941 | 89830150 | 1210 | -7.83 | 1.19E-12 | 4.01E-12 |
| chr18 | 24054088 | 24056705 | 2618 | -8.23 | 1.19E-12 | 4.01E-12 |
| chr19 | 43549275 | 43560936 | 11662 | -7.83 | 1.19E-12 | 4.01E-12 |
| chr22 | 37835561 | 37850744 | 15184 | -7.85 | 1.19E-12 | 4.03E-12 |
| chr1 | 161956498 | 161962847 | 6350 | -7.83 | 1.20E-12 | 4.04E-12 |
| chrX | 84617879 | 84629935 | 12057 | -3.55 | 1.20E-12 | 4.05E-12 |
| chr15 | 44990566 | 44996480 | 5915 | -4.03 | 1.20E-12 | 4.05E-12 |
| chr4 | 1998130 | 2002232 | 4103 | -7.83 | 1.20E-12 | 4.05E-12 |
| chr22 | 27822774 | 27828892 | 6119 | -8.01 | 1.20E-12 | 4.06E-12 |
| chr7 | 73769527 | 73771578 | 2052 | -7.86 | 1.20E-12 | 4.06E-12 |
| chr2 | 242684293 | 242686739 | 2447 | -7.9 | 1.20E-12 | 4.06E-12 |
| chr22 | 28132720 | 28134644 | 1925 | -7.82 | 1.21E-12 | 4.07E-12 |
| chr6 | 113491362 | 113517092 | 25731 | -8.12 | 1.21E-12 | 4.07E-12 |
| chr15 | 30962971 | 30967995 | 5025 | -7.82 | 1.21E-12 | 4.08E-12 |
| chr14 | 56666886 | 56669089 | 2204 | -7.89 | 1.22E-12 | 4.10E-12 |
| chr5 | 52469016 | 52472460 | 3445 | -7.89 | 1.22E-12 | 4.10E-12 |
| chr3 | 196563298 | 196576642 | 13345 | -8.03 | 1.22E-12 | 4.10E-12 |
| chr12 | 74966058 | 74969645 | 3588 | -7.82 | 1.22E-12 | 4.10E-12 |
| chr11 | 27709068 | 27712529 | 3462 | -7.83 | 1.22E-12 | 4.11E-12 |
| chr10 | 32971187 | 33003120 | 31934 | -7.99 | 1.22E-12 | 4.12E-12 |
| chr3 | 153063645 | 153082473 | 18829 | -8.09 | 1.22E-12 | 4.12E-12 |
| chr17 | 65111931 | 65113738 | 1808 | -7.82 | 1.23E-12 | 4.13E-12 |
| chr12 | 87391300 | 87412165 | 20866 | -8.21 | 1.23E-12 | 4.13E-12 |
| chr18 | 23991633 | 23996236 | 4604 | -7.84 | 1.23E-12 | 4.14E-12 |
| chr14 | 39449778 | 39454489 | 4712 | -2.64 | 1.23E-12 | 4.15E-12 |
| chr12 | 78827607 | 78833080 | 5474 | -7.97 | 1.23E-12 | 4.15E-12 |
| chr1 | 21705017 | 21708094 | 3078 | -8 | 1.24E-12 | 4.19E-12 |
| chr4 | 187276477 | 187287446 | 10970 | -7.83 | 1.24E-12 | 4.19E-12 |
| chr2 | 134985217 | 134988432 | 3216 | -7.82 | 1.25E-12 | 4.19E-12 |
| chr18 | 35168260 | 35171342 | 3083 | -2.97 | 1.25E-12 | 4.20E-12 |
| chr1 | 232329087 | 232339474 | 10388 | -7.9 | 1.25E-12 | 4.20E-12 |
| chr1 | 44559024 | 44563944 | 4921 | -7.82 | 1.26E-12 | 4.23E-12 |
| chr16 | 47105095 | 47107023 | 1929 | -7.82 | 1.26E-12 | 4.24E-12 |
| chr8 | 9641158 | 9650866 | 9709 | -7.83 | 1.26E-12 | 4.24E-12 |
| chr14 | 97294462 | 97297426 | 2965 | -7.89 | 1.26E-12 | 4.25E-12 |
| chr4 | 180486603 | 180495931 | 9329 | -7.88 | 1.26E-12 | 4.25E-12 |
| chr10 | 123017245 | 123039132 | 21888 | -7.88 | 1.27E-12 | 4.27E-12 |
| chr7 | 95962523 | 95965213 | 2691 | -7.88 | 1.27E-12 | 4.27E-12 |
| chr12 | 96892498 | 96895957 | 3460 | -7.81 | 1.27E-12 | 4.27E-12 |
| chr4 | 141120234 | 141124551 | 4318 | -7.81 | 1.27E-12 | 4.27E-12 |
| chr1 | 34640454 | 35085892 | 445439 | 2.26 | 1.27E-12 | 4.27E-12 |
| chr11 | 76189253 | 76193690 | 4438 | -7.81 | 1.27E-12 | 4.28E-12 |
| chr13 | 24704792 | 24707144 | 2353 | -7.83 | 1.27E-12 | 4.28E-12 |
| chr3 | 182372171 | 182392965 | 20795 | -7.96 | 1.27E-12 | 4.28E-12 |
| chr15 | 85990119 | 85995109 | 4991 | -8.08 | 1.27E-12 | 4.28E-12 |
| chr2 | 110867163 | 110870892 | 3730 | -7.87 | 1.27E-12 | 4.29E-12 |
| chr2 | 58712466 | 58716042 | 3577 | -7.81 | 1.28E-12 | 4.29E-12 |
| chr13 | 76157398 | 76160342 | 2945 | -7.96 | 1.28E-12 | 4.29E-12 |
| chr1 | 24583063 | 24608745 | 25683 | -8.22 | 1.28E-12 | 4.30E-12 |
| chr1 | 89056423 | 89063738 | 7316 | -7.82 | 1.28E-12 | 4.30E-12 |
| chr2 | 217095827 | 217098522 | 2696 | -7.82 | 1.28E-12 | 4.30E-12 |
| chr7 | 16267237 | 16272881 | 5645 | -7.97 | 1.28E-12 | 4.30E-12 |
| chrX | 69370331 | 69372907 | 2577 | -7.82 | 1.28E-12 | 4.31E-12 |
| chr2 | 84133372 | 84150105 | 16734 | -7.87 | 1.28E-12 | 4.31E-12 |
| chr12 | 93456049 | 93480848 | 24800 | -7.82 | 1.29E-12 | 4.32E-12 |
| chr2 | 8411100 | 8417298 | 6199 | -2.21 | 1.29E-12 | 4.33E-12 |
| chr7 | 87453243 | 87458460 | 5218 | -8 | 1.29E-12 | 4.33E-12 |
| chr3 | 154791455 | 154796199 | 4745 | -2.3 | 1.29E-12 | 4.33E-12 |

| | | | | | | |
|---|---|---|---|---|---|---|
| chr4 | 96208788 | 96222145 | 13358 | -7.94 | 1.29E-12 | 4.35E-12 |
| chr5 | 40540337 | 40568385 | 28049 | -7.82 | 1.30E-12 | 4.37E-12 |
| chr6 | 142371574 | 142375508 | 3935 | -4.25 | 1.31E-12 | 4.39E-12 |
| chr3 | 112151431 | 112170568 | 19138 | -2.26 | 1.31E-12 | 4.39E-12 |
| chr5 | 162983676 | 162987509 | 3834 | -6.9 | 1.31E-12 | 4.40E-12 |
| chr5 | 77860439 | 77862454 | 2016 | -7.84 | 1.31E-12 | 4.41E-12 |
| chr15 | 83445157 | 83463902 | 18746 | -8.17 | 1.31E-12 | 4.41E-12 |
| chr13 | 110397166 | 110406942 | 9777 | -8.07 | 1.32E-12 | 4.42E-12 |
| chr3 | 56358217 | 56366721 | 8505 | 7.92 | 1.32E-12 | 4.44E-12 |
| chr1 | 210183327 | 210184793 | 1467 | -7.81 | 1.32E-12 | 4.44E-12 |
| chr19 | 51441726 | 51444688 | 2963 | -7.83 | 1.32E-12 | 4.44E-12 |
| chr20 | 60839614 | 60841762 | 2149 | -7.95 | 1.32E-12 | 4.45E-12 |
| chr14 | 91591597 | 91595236 | 3640 | -7.85 | 1.33E-12 | 4.45E-12 |
| chr2 | 30820654 | 30823725 | 3072 | -7.89 | 1.33E-12 | 4.45E-12 |
| chr8 | 103071119 | 103097240 | 26122 | -8.58 | 1.33E-12 | 4.46E-12 |
| chr11 | 9032356 | 9034537 | 2182 | -6.3 | 1.33E-12 | 4.47E-12 |
| chr8 | 48576428 | 48583388 | 6961 | -8 | 1.34E-12 | 4.48E-12 |
| chr2 | 198528874 | 198532201 | 3328 | -8 | 1.34E-12 | 4.50E-12 |
| chr16 | 21318715 | 21320854 | 2140 | -7.81 | 1.34E-12 | 4.50E-12 |
| chr21 | 39645992 | 39653962 | 7971 | -7.84 | 1.34E-12 | 4.51E-12 |
| chr19 | 47007882 | 47030072 | 22191 | -8.02 | 1.34E-12 | 4.51E-12 |
| chr20 | 34590463 | 34592834 | 2372 | -8 | 1.34E-12 | 4.51E-12 |
| chrX | 152588556 | 152592801 | 4246 | -7.82 | 1.35E-12 | 4.51E-12 |
| chr17 | 41687550 | 41690292 | 2743 | -7.81 | 1.35E-12 | 4.51E-12 |
| chr20 | 37401888 | 37403456 | 1569 | -7.81 | 1.35E-12 | 4.51E-12 |
| chr7 | 134201890 | 134216962 | 15073 | -7.83 | 1.35E-12 | 4.52E-12 |
| chr5 | 41570875 | 41639936 | 69062 | 1.8 | 1.35E-12 | 4.52E-12 |
| chr2 | 131765580 | 131786611 | 21032 | -7.81 | 1.35E-12 | 4.54E-12 |
| chrX | 150171920 | 150183024 | 11105 | -8.18 | 1.36E-12 | 4.54E-12 |
| chr4 | 103081677 | 103083796 | 2120 | -5.79 | 1.36E-12 | 4.56E-12 |
| chrX | 49657757 | 49670671 | 12915 | -6.51 | 1.36E-12 | 4.56E-12 |
| chr13 | 19857807 | 19903044 | 45238 | 1.59 | 1.36E-12 | 4.57E-12 |
| chr21 | 45847702 | 45851534 | 3833 | -7.88 | 1.37E-12 | 4.57E-12 |
| chr6 | 83049894 | 83056279 | 6386 | -7.8 | 1.37E-12 | 4.60E-12 |
| chr9 | 135287277 | 135288753 | 1477 | -7.93 | 1.38E-12 | 4.61E-12 |
| chr1 | 210368668 | 210372413 | 3746 | -7.81 | 1.38E-12 | 4.61E-12 |
| chr2 | 157232502 | 157235767 | 3266 | -7.82 | 1.38E-12 | 4.61E-12 |
| chr12 | 121508559 | 121511551 | 2993 | -7.88 | 1.38E-12 | 4.61E-12 |
| chr14 | 42689971 | 42708060 | 18090 | -7.97 | 1.38E-12 | 4.62E-12 |
| chr11 | 8599320 | 8601845 | 2526 | -7.81 | 1.38E-12 | 4.62E-12 |
| chr14 | 45524669 | 45540005 | 15337 | -7.85 | 1.38E-12 | 4.63E-12 |
| chr10 | 22223395 | 22225855 | 2461 | -7.81 | 1.38E-12 | 4.63E-12 |
| chr1 | 232543752 | 232546901 | 3150 | -7.99 | 1.39E-12 | 4.65E-12 |
| chr9 | 16205016 | 16212768 | 7753 | -7.81 | 1.39E-12 | 4.65E-12 |
| chr18 | 3977743 | 3981695 | 3953 | -7.81 | 1.39E-12 | 4.66E-12 |
| chr9 | 30066651 | 30081556 | 14906 | -7.81 | 1.39E-12 | 4.66E-12 |
| chr12 | 76530995 | 76533779 | 2785 | -7.82 | 1.40E-12 | 4.67E-12 |
| chr2 | 208504656 | 208509179 | 4524 | -7.82 | 1.40E-12 | 4.67E-12 |
| chr19 | 47100960 | 47103237 | 2278 | -7.8 | 1.40E-12 | 4.67E-12 |
| chr9 | 135692127 | 135695336 | 3210 | -7.98 | 1.40E-12 | 4.67E-12 |
| chr9 | 71963713 | 71985064 | 21352 | -7.8 | 1.40E-12 | 4.68E-12 |
| chr5 | 31951074 | 31959118 | 8045 | -7.95 | 1.40E-12 | 4.69E-12 |
| chr5 | 52073699 | 52082270 | 8572 | -7.81 | 1.41E-12 | 4.71E-12 |
| chr16 | 87690674 | 87692793 | 2120 | -7.84 | 1.41E-12 | 4.71E-12 |
| chr12 | 52275073 | 52279642 | 4570 | -7.86 | 1.41E-12 | 4.72E-12 |
| chr6 | 145637506 | 145640134 | 2629 | -5.62 | 1.41E-12 | 4.73E-12 |
| chr2 | 237455839 | 237472152 | 16314 | -3.26 | 1.42E-12 | 4.73E-12 |
| chr5 | 31901280 | 31905899 | 4620 | -7.81 | 1.42E-12 | 4.73E-12 |
| chr12 | 56152251 | 56154588 | 2338 | -7.91 | 1.42E-12 | 4.73E-12 |
| chr2 | 161058326 | 161075217 | 16892 | -8.1 | 1.42E-12 | 4.74E-12 |
| chr4 | 88513435 | 88521594 | 8160 | -3.22 | 1.42E-12 | 4.76E-12 |

| | | | | | | |
|---|---|---|---|---|---|---|
| chr15 | 40591391 | 40593504 | 2114 | -7.81 | 1.42E-12 | 4.76E-12 |
| chr9 | 15856695 | 15863300 | 6606 | -8.03 | 1.43E-12 | 4.77E-12 |
| chr8 | 119440141 | 119455153 | 15013 | -7.93 | 1.43E-12 | 4.77E-12 |
| chr8 | 134456855 | 134459228 | 2374 | -7.93 | 1.43E-12 | 4.78E-12 |
| chr7 | 114517084 | 114520823 | 3740 | -7.82 | 1.43E-12 | 4.78E-12 |
| chr1 | 151642442 | 151644338 | 1897 | -7.82 | 1.44E-12 | 4.81E-12 |
| chr7 | 84322911 | 84340135 | 17225 | -8.34 | 1.44E-12 | 4.81E-12 |
| chr1 | 66630991 | 66639770 | 8780 | -8.04 | 1.44E-12 | 4.82E-12 |
| chr9 | 99162463 | 99165359 | 2897 | -7.8 | 1.44E-12 | 4.82E-12 |
| chr1 | 12365829 | 12376708 | 10880 | -7.8 | 1.44E-12 | 4.82E-12 |
| chr14 | 53572764 | 53575330 | 2567 | -7.8 | 1.45E-12 | 4.83E-12 |
| chr3 | 49595635 | 49597872 | 2238 | -7.8 | 1.45E-12 | 4.83E-12 |
| chr3 | 119470872 | 119495975 | 25104 | -4.98 | 1.45E-12 | 4.84E-12 |
| chr3 | 113476276 | 113478152 | 1877 | -7.8 | 1.45E-12 | 4.84E-12 |
| chr17 | 33240073 | 33242799 | 2727 | -7.8 | 1.45E-12 | 4.85E-12 |
| chr6 | 126365284 | 126366940 | 1657 | -6.73 | 1.46E-12 | 4.86E-12 |
| chr2 | 11930071 | 11931827 | 1757 | -7.83 | 1.46E-12 | 4.86E-12 |
| chr7 | 81921162 | 81932643 | 11482 | -7.91 | 1.46E-12 | 4.88E-12 |
| chrX | 44984168 | 44988287 | 4120 | -7.85 | 1.47E-12 | 4.90E-12 |
| chr1 | 230375630 | 230377529 | 1900 | -7.81 | 1.47E-12 | 4.91E-12 |
| chr11 | 12800509 | 12803093 | 2585 | -7.92 | 1.47E-12 | 4.91E-12 |
| chr20 | 16403105 | 16420950 | 17846 | -7.88 | 1.48E-12 | 4.92E-12 |
| chr2 | 170826018 | 170833641 | 7624 | -7.8 | 1.48E-12 | 4.93E-12 |
| chrX | 21146011 | 21165810 | 19800 | -6.48 | 1.48E-12 | 4.95E-12 |
| chr1 | 208430373 | 208437888 | 7516 | -1.88 | 1.48E-12 | 4.95E-12 |
| chr4 | 190111390 | 190117135 | 5746 | -7.84 | 1.48E-12 | 4.95E-12 |
| chr2 | 63993801 | 63995503 | 1703 | -7.8 | 1.49E-12 | 4.96E-12 |
| chr17 | 25574986 | 25577732 | 2747 | -7.86 | 1.49E-12 | 4.96E-12 |
| chr4 | 75733664 | 75754770 | 21107 | -7.84 | 1.49E-12 | 4.97E-12 |
| chr12 | 86386049 | 86395828 | 9780 | -7.81 | 1.49E-12 | 4.97E-12 |
| chr9 | 99113123 | 99115499 | 2377 | -7.81 | 1.49E-12 | 4.98E-12 |
| chr13 | 111581743 | 111586312 | 4570 | -7.84 | 1.50E-12 | 4.98E-12 |
| chr9 | 28686428 | 28699280 | 12853 | -8 | 1.50E-12 | 4.99E-12 |
| chr6 | 148246012 | 148251251 | 5240 | -8.13 | 1.50E-12 | 5.01E-12 |
| chr3 | 8349631 | 8352152 | 2522 | -7.81 | 1.51E-12 | 5.03E-12 |
| chr8 | 27447215 | 27449727 | 2513 | -7.88 | 1.51E-12 | 5.04E-12 |
| chr20 | 14690912 | 14699972 | 9061 | -7.91 | 1.52E-12 | 5.05E-12 |
| chr21 | 17562305 | 17573700 | 11396 | -7.81 | 1.52E-12 | 5.06E-12 |
| chr6 | 138394906 | 138402483 | 7578 | -7.91 | 1.52E-12 | 5.06E-12 |
| chr4 | 134154049 | 134171943 | 17895 | 7.86 | 1.52E-12 | 5.06E-12 |
| chr9 | 79444215 | 79462221 | 18007 | -7.79 | 1.52E-12 | 5.07E-12 |
| chrX | 70544391 | 70547953 | 3563 | -7.79 | 1.52E-12 | 5.07E-12 |
| chr6 | 107835563 | 107838695 | 3133 | -7.96 | 1.52E-12 | 5.08E-12 |
| chr5 | 129117082 | 129124560 | 7479 | 7.89 | 1.52E-12 | 5.08E-12 |
| chr12 | 80481320 | 80484167 | 2848 | -5.64 | 1.53E-12 | 5.08E-12 |
| chr2 | 128424141 | 128428002 | 3862 | -7.8 | 1.53E-12 | 5.09E-12 |
| chr6 | 132762168 | 132765629 | 3462 | -7.82 | 1.54E-12 | 5.12E-12 |
| chr3 | 121075190 | 121093020 | 17831 | 2.76 | 1.54E-12 | 5.12E-12 |
| chr4 | 77691611 | 77698131 | 6521 | -7.81 | 1.54E-12 | 5.13E-12 |
| chr12 | 7895684 | 7899591 | 3908 | -7.85 | 1.54E-12 | 5.13E-12 |
| chr4 | 38196169 | 38207171 | 11003 | -7.85 | 1.54E-12 | 5.13E-12 |
| chr13 | 30657868 | 30660276 | 2409 | -7.79 | 1.54E-12 | 5.13E-12 |
| chr15 | 23082343 | 23084617 | 2275 | -7.93 | 1.54E-12 | 5.13E-12 |
| chr5 | 89739706 | 89743673 | 3968 | -7.8 | 1.54E-12 | 5.14E-12 |
| chr7 | 140657210 | 140663077 | 5868 | -7.84 | 1.54E-12 | 5.14E-12 |
| chrX | 43996280 | 43999158 | 2879 | -8.11 | 1.55E-12 | 5.15E-12 |
| chr18 | 11955273 | 11958543 | 3271 | -7.82 | 1.55E-12 | 5.15E-12 |
| chr1 | 20321878 | 20377868 | 55991 | -1.89 | 1.55E-12 | 5.15E-12 |
| chr13 | 30925843 | 30928284 | 2442 | -7.79 | 1.55E-12 | 5.16E-12 |
| chr3 | 155129326 | 155147820 | 18495 | -7.94 | 1.56E-12 | 5.18E-12 |
| chr4 | 6436114 | 6438911 | 2798 | -7.85 | 1.56E-12 | 5.20E-12 |

| | | | | | | |
|---|---|---|---|---|---|---|
| chr11 | 87858346 | 87865915 | 7570 | -7.83 | 1.57E-12 | 5.20E-12 |
| chr20 | 60285647 | 60292498 | 6852 | -7.91 | 1.57E-12 | 5.21E-12 |
| chr17 | 19970431 | 19987484 | 17054 | -7.91 | 1.57E-12 | 5.22E-12 |
| chr2 | 7315896 | 7335869 | 19974 | -7.98 | 1.58E-12 | 5.24E-12 |
| chr11 | 126803374 | 126806659 | 3286 | -7.83 | 1.58E-12 | 5.24E-12 |
| chr3 | 12905761 | 12908270 | 2510 | -7.83 | 1.58E-12 | 5.24E-12 |
| chr13 | 74158382 | 74162487 | 4106 | -7.79 | 1.58E-12 | 5.25E-12 |
| chr1 | 26914156 | 26916424 | 2269 | -7.8 | 1.58E-12 | 5.26E-12 |
| chr16 | 1743238 | 1747229 | 3992 | -7.94 | 1.58E-12 | 5.26E-12 |
| chr8 | 100596520 | 100599246 | 2727 | -7.94 | 1.58E-12 | 5.26E-12 |
| chr11 | 130524774 | 130528600 | 3827 | -7.96 | 1.59E-12 | 5.26E-12 |
| chr14 | 32379615 | 32390658 | 11044 | -7.92 | 1.59E-12 | 5.27E-12 |
| chr8 | 74420986 | 74426417 | 5432 | -7.78 | 1.60E-12 | 5.30E-12 |
| chr10 | 8400305 | 8411503 | 11199 | -7.88 | 1.60E-12 | 5.31E-12 |
| chr15 | 56462831 | 56477893 | 15063 | -7.87 | 1.60E-12 | 5.31E-12 |
| chr12 | 21410062 | 21413704 | 3643 | -7.8 | 1.60E-12 | 5.32E-12 |
| chr12 | 59297661 | 59299780 | 2120 | -7.83 | 1.60E-12 | 5.32E-12 |
| chr12 | 32092996 | 32101214 | 8219 | -8.09 | 1.60E-12 | 5.32E-12 |
| chr3 | 194417187 | 194429367 | 12181 | -7.97 | 1.61E-12 | 5.33E-12 |
| chr18 | 19532267 | 19535878 | 3612 | -7.84 | 1.61E-12 | 5.33E-12 |
| chr8 | 16892498 | 16918487 | 25990 | -8.01 | 1.61E-12 | 5.33E-12 |
| chr12 | 11390603 | 11394300 | 3698 | 7.11 | 1.61E-12 | 5.33E-12 |
| chr10 | 124694447 | 124697199 | 2753 | -7.86 | 1.61E-12 | 5.35E-12 |
| chr1 | 108348008 | 108360729 | 12722 | 3.96 | 1.62E-12 | 5.36E-12 |
| chr17 | 66185444 | 66187885 | 2442 | -7.83 | 1.62E-12 | 5.37E-12 |
| chr6 | 137758792 | 137765521 | 6730 | -7.78 | 1.63E-12 | 5.39E-12 |
| chr2 | 206356066 | 206385100 | 29035 | -8.14 | 1.63E-12 | 5.40E-12 |
| chr3 | 153921229 | 153925112 | 3884 | -7.92 | 1.63E-12 | 5.40E-12 |
| chr11 | 70984025 | 70986079 | 2055 | -7.78 | 1.63E-12 | 5.42E-12 |
| chr11 | 16557424 | 16571635 | 14212 | -7.8 | 1.64E-12 | 5.43E-12 |
| chr13 | 66575315 | 66579849 | 4535 | -8.11 | 1.64E-12 | 5.44E-12 |
| chr9 | 99880754 | 99883815 | 3062 | -7.8 | 1.64E-12 | 5.44E-12 |
| chr5 | 159538330 | 159540214 | 1885 | -7.84 | 1.64E-12 | 5.44E-12 |
| chr8 | 99681995 | 99683989 | 1995 | -7.84 | 1.64E-12 | 5.44E-12 |
| chr6 | 132242487 | 132245571 | 3085 | -7.84 | 1.65E-12 | 5.46E-12 |
| chr5 | 168628007 | 168636722 | 8716 | -8.12 | 1.65E-12 | 5.46E-12 |
| chrX | 40043120 | 40046246 | 3127 | -7.81 | 1.65E-12 | 5.46E-12 |
| chr1 | 187086975 | 187102248 | 15274 | 8.18 | 1.65E-12 | 5.47E-12 |
| chr13 | 80177554 | 80196506 | 18953 | -7.96 | 1.65E-12 | 5.48E-12 |
| chr9 | 35530265 | 35544294 | 14030 | -7.79 | 1.65E-12 | 5.48E-12 |
| chr2 | 181999213 | 182033130 | 33918 | -8.12 | 1.66E-12 | 5.49E-12 |
| chr11 | 66217143 | 66231298 | 14156 | -8.05 | 1.66E-12 | 5.49E-12 |
| chr5 | 13669820 | 13676075 | 6256 | -7.78 | 1.66E-12 | 5.50E-12 |
| chr6 | 4798354 | 4809564 | 11211 | -7.87 | 1.67E-12 | 5.52E-12 |
| chr15 | 51410449 | 51418478 | 8030 | -7.89 | 1.67E-12 | 5.53E-12 |
| chr17 | 1872318 | 1874812 | 2495 | -7.86 | 1.67E-12 | 5.53E-12 |
| chr7 | 128912971 | 128915018 | 2048 | -7.82 | 1.68E-12 | 5.56E-12 |
| chr3 | 72604541 | 72611989 | 7449 | -7.9 | 1.68E-12 | 5.56E-12 |
| chr10 | 63489224 | 63504714 | 15491 | -8.38 | 1.68E-12 | 5.56E-12 |
| chr22 | 37658740 | 37661634 | 2895 | -7.82 | 1.69E-12 | 5.58E-12 |
| chr2 | 101344543 | 101347958 | 3416 | -7.85 | 1.69E-12 | 5.59E-12 |
| chr10 | 47538631 | 47543258 | 4628 | -7.81 | 1.70E-12 | 5.61E-12 |
| chr8 | 90796461 | 90835643 | 39183 | -8.04 | 1.70E-12 | 5.61E-12 |
| chr2 | 203606461 | 203610380 | 3920 | -7.78 | 1.70E-12 | 5.62E-12 |
| chrX | 47349638 | 47382370 | 32733 | -8.31 | 1.70E-12 | 5.62E-12 |
| chr7 | 97959677 | 97981300 | 21624 | -7.92 | 1.70E-12 | 5.63E-12 |
| chr16 | 48528559 | 48542733 | 14175 | -8.02 | 1.70E-12 | 5.64E-12 |
| chr12 | 125777207 | 125788567 | 11361 | -7.83 | 1.71E-12 | 5.66E-12 |
| chr3 | 40122124 | 40125278 | 3155 | -7.78 | 1.71E-12 | 5.67E-12 |
| chr12 | 75131780 | 75153124 | 21345 | -8.06 | 1.71E-12 | 5.67E-12 |
| chr5 | 80431355 | 80433765 | 2411 | -7.77 | 1.72E-12 | 5.68E-12 |

| | | | | | | |
|---|---|---|---|---|---|---|
| chr1 | 84973592 | 84976340 | 2749 | -7.96 | 1.72E-12 | 5.69E-12 |
| chr3 | 101745571 | 101768823 | 23253 | -8.27 | 1.72E-12 | 5.70E-12 |
| chr5 | 168584324 | 168599864 | 15541 | -7.83 | 1.72E-12 | 5.70E-12 |
| chr15 | 91231749 | 91233665 | 1917 | -7.86 | 1.73E-12 | 5.71E-12 |
| chr3 | 114353826 | 114356546 | 2721 | -7.83 | 1.73E-12 | 5.72E-12 |
| chr2 | 218187828 | 218195689 | 7862 | -7.78 | 1.73E-12 | 5.72E-12 |
| chr20 | 55122174 | 55131163 | 8990 | -7.85 | 1.74E-12 | 5.74E-12 |
| chr1 | 169808014 | 169810867 | 2854 | -7.78 | 1.74E-12 | 5.76E-12 |
| chr17 | 65137659 | 65139148 | 1490 | -7.89 | 1.74E-12 | 5.76E-12 |
| chr5 | 168795565 | 168803148 | 7584 | -7.82 | 1.75E-12 | 5.76E-12 |
| chr11 | 44103202 | 44107919 | 4718 | -7.78 | 1.75E-12 | 5.79E-12 |
| chr4 | 99222398 | 99226060 | 3663 | -7.85 | 1.75E-12 | 5.79E-12 |
| chr4 | 4587995 | 4597801 | 9807 | -8.04 | 1.75E-12 | 5.79E-12 |
| chr5 | 168823526 | 168826887 | 3362 | -7.8 | 1.75E-12 | 5.79E-12 |
| chr9 | 83619617 | 83647468 | 27852 | -7.97 | 1.76E-12 | 5.81E-12 |
| chr5 | 10473636 | 10478535 | 4900 | -7.82 | 1.76E-12 | 5.82E-12 |
| chr16 | 56786120 | 56789889 | 3770 | -8.15 | 1.78E-12 | 5.86E-12 |
| chr1 | 227231463 | 227237467 | 6005 | -8.13 | 1.78E-12 | 5.87E-12 |
| chr1 | 43113570 | 43115842 | 2273 | -7.87 | 1.78E-12 | 5.87E-12 |
| chrX | 82734383 | 82739921 | 5539 | 3.82 | 1.78E-12 | 5.88E-12 |
| chr2 | 68647871 | 68650145 | 2275 | 4.89 | 1.78E-12 | 5.88E-12 |
| chrX | 2820101 | 2823243 | 3143 | -7.93 | 1.79E-12 | 5.90E-12 |
| chr7 | 14541153 | 14558052 | 16900 | -8.15 | 1.79E-12 | 5.91E-12 |
| chr3 | 131958386 | 131960866 | 2481 | -2.77 | 1.79E-12 | 5.91E-12 |
| chr6 | 113896676 | 113902700 | 6025 | -7.78 | 1.80E-12 | 5.93E-12 |
| chr9 | 35412629 | 35429177 | 16549 | -7.78 | 1.80E-12 | 5.93E-12 |
| chr14 | 92868348 | 92912602 | 44255 | -8.92 | 1.80E-12 | 5.93E-12 |
| chr2 | 202945127 | 202947566 | 2440 | -8.03 | 1.81E-12 | 5.97E-12 |
| chr2 | 224712768 | 224716224 | 3457 | -7.86 | 1.81E-12 | 5.97E-12 |
| chr6 | 169281497 | 169308094 | 26598 | -8.45 | 1.81E-12 | 5.98E-12 |
| chr5 | 81541862 | 81544312 | 2451 | -7.78 | 1.81E-12 | 5.98E-12 |
| chr4 | 84650055 | 84666508 | 16454 | -7.97 | 1.82E-12 | 5.99E-12 |
| chr13 | 80157744 | 80163515 | 5772 | -7.89 | 1.82E-12 | 5.99E-12 |
| chr2 | 55552710 | 55554612 | 1903 | -7.89 | 1.82E-12 | 6.00E-12 |
| chr7 | 35032136 | 35042577 | 10442 | -7.83 | 1.82E-12 | 6.00E-12 |
| chr16 | 85217091 | 85219215 | 2125 | -7.94 | 1.82E-12 | 6.01E-12 |
| chr1 | 196524373 | 196534650 | 10278 | -7.92 | 1.83E-12 | 6.02E-12 |
| chr9 | 135182863 | 135184892 | 2030 | -7.94 | 1.83E-12 | 6.02E-12 |
| chr21 | 30557737 | 30575549 | 17813 | -8.02 | 1.83E-12 | 6.02E-12 |
| chr4 | 101060281 | 101063300 | 3020 | -7.9 | 1.83E-12 | 6.03E-12 |
| chr15 | 39634277 | 39637586 | 3310 | -7.78 | 1.83E-12 | 6.04E-12 |
| chr20 | 60889965 | 60892872 | 2908 | -7.91 | 1.84E-12 | 6.07E-12 |
| chr15 | 25993615 | 25999236 | 5622 | -7.76 | 1.84E-12 | 6.07E-12 |
| chr16 | 52588533 | 52590948 | 2416 | -7.76 | 1.84E-12 | 6.07E-12 |
| chr17 | 38633528 | 38636330 | 2803 | -7.8 | 1.84E-12 | 6.07E-12 |
| chr21 | 33936440 | 33939371 | 2932 | -7.78 | 1.85E-12 | 6.08E-12 |
| chr13 | 30540016 | 30542920 | 2905 | -7.77 | 1.85E-12 | 6.09E-12 |
| chr1 | 218962954 | 218983890 | 20937 | -8.75 | 1.85E-12 | 6.09E-12 |
| chr3 | 182957290 | 182961879 | 4590 | -7.8 | 1.85E-12 | 6.09E-12 |
| chr6 | 53314133 | 53321260 | 7128 | -7.84 | 1.85E-12 | 6.09E-12 |
| chr21 | 40396045 | 40400956 | 4912 | -7.89 | 1.86E-12 | 6.12E-12 |
| chr1 | 235459098 | 235461706 | 2609 | -7.81 | 1.86E-12 | 6.12E-12 |
| chr16 | 70663880 | 70666443 | 2564 | -7.81 | 1.86E-12 | 6.12E-12 |
| chr22 | 27203273 | 27216134 | 12862 | -7.98 | 1.86E-12 | 6.13E-12 |
| chr5 | 23601946 | 23617922 | 15977 | -7.93 | 1.86E-12 | 6.13E-12 |
| chr15 | 80826340 | 80828674 | 2335 | -7.83 | 1.86E-12 | 6.13E-12 |
| chr16 | 12869252 | 12905765 | 36514 | -7.99 | 1.87E-12 | 6.15E-12 |
| chr5 | 86291204 | 86298047 | 6844 | 7.9 | 1.87E-12 | 6.16E-12 |
| chr1 | 184003380 | 184029584 | 26205 | -8.32 | 1.87E-12 | 6.17E-12 |
| chr5 | 108455603 | 108458978 | 3376 | -7.76 | 1.88E-12 | 6.17E-12 |
| chr1 | 20895810 | 20899381 | 3572 | -7.76 | 1.88E-12 | 6.18E-12 |

| | | | | | | |
|---|---|---|---|---|---|---|
| chr1 | 183398162 | 183408490 | 10329 | -7.79 | 1.89E-12 | 6.22E-12 |
| chrX | 119720746 | 119733767 | 13022 | -7.82 | 1.89E-12 | 6.22E-12 |
| chr1 | 193183704 | 193198148 | 14445 | -8.09 | 1.90E-12 | 6.24E-12 |
| chr12 | 17908219 | 17910691 | 2473 | -7.79 | 1.90E-12 | 6.24E-12 |
| chr10 | 61613507 | 61621603 | 8097 | -8.17 | 1.90E-12 | 6.24E-12 |
| chr4 | 129107552 | 129135161 | 27610 | -7.92 | 1.90E-12 | 6.24E-12 |
| chr22 | 38937581 | 38941755 | 4175 | -7.82 | 1.90E-12 | 6.24E-12 |
| chr2 | 191029512 | 191039626 | 10115 | -7.77 | 1.90E-12 | 6.24E-12 |
| chrX | 68710354 | 68720684 | 10331 | -7.97 | 1.90E-12 | 6.24E-12 |
| chr2 | 236823593 | 236825484 | 1892 | -7.97 | 1.90E-12 | 6.25E-12 |
| chrX | 100571968 | 100575238 | 3271 | -7.76 | 1.90E-12 | 6.25E-12 |
| chr22 | 23769911 | 23773390 | 3480 | -7.85 | 1.90E-12 | 6.25E-12 |
| chr5 | 33263547 | 33266228 | 2682 | -7.83 | 1.91E-12 | 6.27E-12 |
| chr12 | 27750521 | 27758432 | 7912 | -7.83 | 1.91E-12 | 6.28E-12 |
| chr1 | 182860133 | 182865095 | 4963 | -7.8 | 1.91E-12 | 6.28E-12 |
| chr18 | 24726198 | 24729989 | 3792 | -7.77 | 1.92E-12 | 6.30E-12 |
| chr8 | 142210289 | 142215208 | 4920 | -7.93 | 1.92E-12 | 6.30E-12 |
| chr1 | 59379652 | 59382470 | 2819 | -7.77 | 1.92E-12 | 6.30E-12 |
| chr4 | 101080938 | 101084383 | 3446 | -2.2 | 1.92E-12 | 6.31E-12 |
| chr17 | 5506115 | 5643268 | 137154 | -2.08 | 1.92E-12 | 6.31E-12 |
| chr11 | 66736494 | 66748824 | 12331 | -7.89 | 1.92E-12 | 6.32E-12 |
| chr2 | 191781625 | 191788374 | 6750 | -8.05 | 1.93E-12 | 6.32E-12 |
| chr6 | 137328940 | 137364155 | 35216 | -3.32 | 1.94E-12 | 6.35E-12 |
| chrX | 13186082 | 13194918 | 8837 | -7.78 | 1.94E-12 | 6.37E-12 |
| chr15 | 38935683 | 38942492 | 6810 | -7.82 | 1.95E-12 | 6.38E-12 |
| chr6 | 127370451 | 127385380 | 14930 | -8.12 | 1.95E-12 | 6.39E-12 |
| chr18 | 74459533 | 74462483 | 2951 | -7.76 | 1.95E-12 | 6.40E-12 |
| chr5 | 137333231 | 137342222 | 8992 | -7.78 | 1.95E-12 | 6.41E-12 |
| chr11 | 27692367 | 27695406 | 3040 | -7.76 | 1.95E-12 | 6.41E-12 |
| chr7 | 33206420 | 33221073 | 14654 | -7.82 | 1.95E-12 | 6.41E-12 |
| chr9 | 28989556 | 29010935 | 21380 | -7.81 | 1.96E-12 | 6.42E-12 |
| chrX | 13991568 | 14000841 | 9274 | -2.05 | 1.96E-12 | 6.43E-12 |
| chr1 | 88503409 | 88507616 | 4208 | -7.77 | 1.97E-12 | 6.45E-12 |
| chr8 | 42767858 | 42769525 | 1668 | -7.85 | 1.97E-12 | 6.46E-12 |
| chr5 | 141827591 | 141837044 | 9454 | -8.1 | 1.97E-12 | 6.46E-12 |
| chr11 | 64310545 | 64320392 | 9848 | -7.92 | 1.97E-12 | 6.47E-12 |
| chr11 | 35852698 | 35865239 | 12542 | -7.77 | 1.97E-12 | 6.47E-12 |
| chr3 | 34127133 | 34143359 | 16227 | -7.97 | 1.97E-12 | 6.47E-12 |
| chr6 | 33866194 | 33870346 | 4153 | -5.86 | 1.98E-12 | 6.47E-12 |
| chr3 | 69422015 | 69428123 | 6109 | -7.82 | 2.00E-12 | 6.55E-12 |
| chr13 | 83165287 | 83181662 | 16376 | -8.07 | 2.00E-12 | 6.55E-12 |
| chr12 | 65215548 | 65227589 | 12042 | -8.02 | 2.00E-12 | 6.55E-12 |
| chr2 | 45727518 | 45730104 | 2587 | -7.82 | 2.00E-12 | 6.56E-12 |
| chr8 | 133980098 | 133986211 | 6114 | 7.98 | 2.01E-12 | 6.58E-12 |
| chr2 | 10705067 | 10717891 | 12825 | -8 | 2.01E-12 | 6.58E-12 |
| chr1 | 65552397 | 65561976 | 9580 | -7.8 | 2.01E-12 | 6.59E-12 |
| chr12 | 32389346 | 32391553 | 2208 | -7.77 | 2.01E-12 | 6.60E-12 |
| chr6 | 21577554 | 21584129 | 6576 | -8 | 2.02E-12 | 6.61E-12 |
| chr5 | 33427835 | 33431031 | 3197 | -7.81 | 2.02E-12 | 6.62E-12 |
| chr12 | 69654643 | 69657622 | 2980 | -7.78 | 2.02E-12 | 6.63E-12 |
| chr15 | 70155864 | 70231726 | 75863 | -2.99 | 2.03E-12 | 6.63E-12 |
| chr2 | 53934275 | 53958044 | 23770 | -7.94 | 2.03E-12 | 6.64E-12 |
| chr3 | 124655074 | 124656882 | 1809 | -7.81 | 2.03E-12 | 6.66E-12 |
| chr10 | 96776340 | 96790807 | 14468 | -7.75 | 2.03E-12 | 6.66E-12 |
| chr3 | 100153013 | 100156125 | 3113 | -7.75 | 2.03E-12 | 6.66E-12 |
| chr4 | 141438401 | 141449475 | 11075 | -7.85 | 2.03E-12 | 6.66E-12 |
| chr5 | 40348372 | 40381302 | 32931 | -8.32 | 2.04E-12 | 6.67E-12 |
| chr3 | 70991374 | 70994093 | 2720 | 3.25 | 2.04E-12 | 6.69E-12 |
| chrX | 35128461 | 35134173 | 5713 | -7.77 | 2.05E-12 | 6.70E-12 |
| chr9 | 33724424 | 33728992 | 4569 | -7.8 | 2.05E-12 | 6.70E-12 |
| chr12 | 100361885 | 100369803 | 7919 | -7.88 | 2.05E-12 | 6.71E-12 |

| | | | | | | |
|---|---|---|---|---|---|---|
| chrX | 108580792 | 108589388 | 8597 | 8.17 | 2.05E-12 | 6.71E-12 |
| chr7 | 51182765 | 51185523 | 2759 | -7.75 | 2.06E-12 | 6.72E-12 |
| chr17 | 29748765 | 29750562 | 1798 | -7.84 | 2.06E-12 | 6.72E-12 |
| chr5 | 105988360 | 105991629 | 3270 | -7.76 | 2.06E-12 | 6.74E-12 |
| chr1 | 233580204 | 233587051 | 6848 | -7.82 | 2.06E-12 | 6.74E-12 |
| chr13 | 51989424 | 51991508 | 2085 | -7.87 | 2.06E-12 | 6.75E-12 |
| chr4 | 39484509 | 39488328 | 3820 | -7.75 | 2.07E-12 | 6.76E-12 |
| chr3 | 16655147 | 16671319 | 16173 | -8.22 | 2.08E-12 | 6.82E-12 |
| chr4 | 7809813 | 7811890 | 2078 | -7.79 | 2.09E-12 | 6.82E-12 |
| chr2 | 188936417 | 188952765 | 16349 | -7.78 | 2.09E-12 | 6.82E-12 |
| chr9 | 129833446 | 129835182 | 1737 | -7.75 | 2.10E-12 | 6.85E-12 |
| chrX | 74951658 | 74968838 | 17181 | -2.01 | 2.10E-12 | 6.85E-12 |
| chr16 | 86953458 | 86956423 | 2966 | -7.77 | 2.10E-12 | 6.86E-12 |
| chr2 | 43893436 | 43895490 | 2055 | -7.75 | 2.10E-12 | 6.87E-12 |
| chr2 | 159287907 | 159297281 | 9375 | -7.85 | 2.10E-12 | 6.87E-12 |
| chr4 | 72443306 | 72449441 | 6136 | -7.84 | 2.10E-12 | 6.87E-12 |
| chr17 | 75581173 | 75586929 | 5757 | -3.28 | 2.11E-12 | 6.89E-12 |
| chr13 | 51303353 | 51325138 | 21786 | -7.83 | 2.11E-12 | 6.89E-12 |
| chr1 | 75829535 | 75841349 | 11815 | -7.79 | 2.11E-12 | 6.89E-12 |
| chr2 | 152913949 | 152917422 | 3474 | -7.78 | 2.12E-12 | 6.92E-12 |
| chr10 | 91521270 | 91536086 | 14817 | -8.37 | 2.12E-12 | 6.93E-12 |
| chr6 | 2815339 | 2817584 | 2246 | -7.8 | 2.13E-12 | 6.94E-12 |
| chr2 | 15877317 | 15882857 | 5541 | -2.31 | 2.13E-12 | 6.94E-12 |
| chr6 | 14061574 | 14063929 | 2356 | -7.8 | 2.13E-12 | 6.96E-12 |
| chr13 | 24197082 | 24201578 | 4497 | -7.82 | 2.13E-12 | 6.96E-12 |
| chr8 | 110177313 | 110180242 | 2930 | -7.76 | 2.13E-12 | 6.96E-12 |
| chr7 | 109261810 | 109266795 | 4986 | -7.75 | 2.14E-12 | 6.97E-12 |
| chr5 | 131055345 | 131066115 | 10771 | -7.75 | 2.14E-12 | 6.97E-12 |
| chr6 | 20904064 | 20917167 | 13104 | -7.81 | 2.14E-12 | 6.98E-12 |
| chr16 | 14190409 | 14192642 | 2234 | -7.85 | 2.14E-12 | 6.99E-12 |
| chr6 | 144717400 | 144720098 | 2699 | -7.77 | 2.15E-12 | 7.02E-12 |
| chr21 | 33888948 | 33892443 | 3496 | -7.78 | 2.16E-12 | 7.05E-12 |
| chr1 | 55395871 | 55400638 | 4768 | -8.07 | 2.17E-12 | 7.07E-12 |
| chr8 | 67105761 | 67125648 | 19888 | -1.44 | 2.17E-12 | 7.09E-12 |
| chr14 | 103505803 | 103509410 | 3608 | -7.9 | 2.17E-12 | 7.09E-12 |
| chr9 | 125912589 | 125914869 | 2281 | -7.81 | 2.17E-12 | 7.09E-12 |
| chr18 | 48138492 | 48142200 | 3709 | -2.99 | 2.18E-12 | 7.11E-12 |
| chr8 | 23251893 | 23253615 | 1723 | -7.81 | 2.18E-12 | 7.12E-12 |
| chr6 | 141824514 | 141828985 | 4472 | -7.76 | 2.18E-12 | 7.12E-12 |
| chr11 | 86682068 | 86685196 | 3129 | -7.77 | 2.19E-12 | 7.13E-12 |
| chr11 | 93840936 | 93844495 | 3560 | -7.75 | 2.19E-12 | 7.14E-12 |
| chr20 | 56158804 | 56163606 | 4803 | -8.04 | 2.19E-12 | 7.14E-12 |
| chr4 | 187460428 | 187466585 | 6158 | -7.87 | 2.19E-12 | 7.14E-12 |
| chr1 | 35616825 | 35631621 | 14797 | -7.79 | 2.19E-12 | 7.14E-12 |
| chr5 | 148435620 | 148439105 | 3486 | -7.78 | 2.19E-12 | 7.15E-12 |
| chr12 | 127285209 | 127500998 | 215790 | 1.38 | 2.20E-12 | 7.16E-12 |
| chr1 | 159665635 | 159673342 | 7708 | 7.82 | 2.20E-12 | 7.16E-12 |
| chrX | 110444238 | 110451187 | 6950 | 7.84 | 2.21E-12 | 7.19E-12 |
| chr1 | 226501537 | 226509591 | 8055 | -7.76 | 2.21E-12 | 7.19E-12 |
| chr12 | 78997022 | 79002470 | 5449 | -7.76 | 2.21E-12 | 7.19E-12 |
| chr9 | 90564378 | 90566411 | 2034 | -7.75 | 2.21E-12 | 7.19E-12 |
| chr3 | 159859966 | 159862761 | 2796 | -7.78 | 2.21E-12 | 7.20E-12 |
| chr17 | 48723013 | 48730505 | 7493 | -7.88 | 2.21E-12 | 7.20E-12 |
| chr20 | 20044095 | 20045825 | 1731 | -6.23 | 2.22E-12 | 7.21E-12 |
| chr3 | 42276022 | 42280849 | 4828 | -2.83 | 2.22E-12 | 7.21E-12 |
| chrX | 46778544 | 46780416 | 1873 | -7.77 | 2.23E-12 | 7.25E-12 |
| chr12 | 92321271 | 92334385 | 13115 | -8.28 | 2.23E-12 | 7.27E-12 |
| chr22 | 26648780 | 26655559 | 6780 | 7.81 | 2.24E-12 | 7.28E-12 |
| chr1 | 154750570 | 154752664 | 2095 | -7.8 | 2.24E-12 | 7.28E-12 |
| chr9 | 100130029 | 100133228 | 3200 | -7.8 | 2.24E-12 | 7.28E-12 |
| chr12 | 54022894 | 54026521 | 3628 | -7.99 | 2.24E-12 | 7.29E-12 |

| | | | | | | |
|---|---|---|---|---|---|---|
| chr9 | 139453252 | 139455638 | 2387 | -7.8 | 2.24E-12 | 7.29E-12 |
| chr18 | 52435605 | 52446889 | 11285 | -7.79 | 2.24E-12 | 7.30E-12 |
| chr6 | 52979268 | 52981228 | 1961 | -5.84 | 2.24E-12 | 7.30E-12 |
| chr1 | 4774790 | 4783058 | 8269 | -5.88 | 2.25E-12 | 7.31E-12 |
| chr16 | 57622712 | 57632821 | 10110 | -8.09 | 2.25E-12 | 7.31E-12 |
| chr3 | 152959369 | 152970107 | 10739 | -7.74 | 2.25E-12 | 7.31E-12 |
| chr2 | 223276798 | 223281691 | 4894 | -7.76 | 2.25E-12 | 7.33E-12 |
| chr11 | 85412562 | 85415540 | 2979 | -7.95 | 2.26E-12 | 7.34E-12 |
| chr2 | 196265563 | 196269172 | 3610 | -7.74 | 2.26E-12 | 7.35E-12 |
| chr6 | 83195125 | 83200616 | 5492 | -7.74 | 2.26E-12 | 7.36E-12 |
| chr11 | 103827964 | 103833565 | 5602 | -7.74 | 2.27E-12 | 7.37E-12 |
| chr5 | 55127047 | 55143426 | 16380 | -8.13 | 2.28E-12 | 7.40E-12 |
| chr2 | 63681674 | 63689216 | 7543 | -7.8 | 2.29E-12 | 7.44E-12 |
| chr16 | 19694700 | 19717245 | 22546 | -7.92 | 2.30E-12 | 7.49E-12 |
| chr20 | 56758496 | 56763046 | 4551 | -7.89 | 2.31E-12 | 7.52E-12 |
| chr7 | 151783895 | 151787361 | 3467 | -7.89 | 2.32E-12 | 7.54E-12 |
| chr2 | 203936473 | 203938936 | 2464 | -7.76 | 2.33E-12 | 7.56E-12 |
| chr16 | 87836512 | 87853161 | 16650 | -8.44 | 2.33E-12 | 7.57E-12 |
| chr9 | 121336541 | 121341181 | 4641 | -7.73 | 2.33E-12 | 7.57E-12 |
| chr1 | 219307754 | 219312520 | 4767 | -7.85 | 2.33E-12 | 7.57E-12 |
| chrX | 109428218 | 109431319 | 3102 | -7.84 | 2.33E-12 | 7.57E-12 |
| chr4 | 188163397 | 188166642 | 3246 | -7.73 | 2.33E-12 | 7.57E-12 |
| chr1 | 51504190 | 51526721 | 22532 | -8.26 | 2.33E-12 | 7.58E-12 |
| chr1 | 178377644 | 178387072 | 9429 | -7.74 | 2.33E-12 | 7.58E-12 |
| chr2 | 151458747 | 151479982 | 21236 | -8.35 | 2.34E-12 | 7.58E-12 |
| chr2 | 189620752 | 189624671 | 3920 | -7.93 | 2.34E-12 | 7.58E-12 |
| chr14 | 90288731 | 90291244 | 2514 | -7.81 | 2.35E-12 | 7.62E-12 |
| chr6 | 106799530 | 106802040 | 2511 | -7.77 | 2.35E-12 | 7.63E-12 |
| chr15 | 70621742 | 70630460 | 8719 | -7.48 | 2.36E-12 | 7.66E-12 |
| chr4 | 67376457 | 67393197 | 16741 | -7.75 | 2.37E-12 | 7.69E-12 |
| chr14 | 37580997 | 37589846 | 8850 | -7.75 | 2.37E-12 | 7.71E-12 |
| chr1 | 210312915 | 210315771 | 2857 | -7.8 | 2.38E-12 | 7.72E-12 |
| chr10 | 74581344 | 74583053 | 1710 | -7.73 | 2.38E-12 | 7.72E-12 |
| chr12 | 105572835 | 105580649 | 7815 | -7.95 | 2.38E-12 | 7.73E-12 |
| chr11 | 46979922 | 46983692 | 3771 | -7.75 | 2.38E-12 | 7.73E-12 |
| chr16 | 49923308 | 49925471 | 2164 | -7.73 | 2.38E-12 | 7.73E-12 |
| chr7 | 43616874 | 43619894 | 3021 | -7.73 | 2.39E-12 | 7.75E-12 |
| chr10 | 61449352 | 61481753 | 32402 | -8.11 | 2.39E-12 | 7.77E-12 |
| chr4 | 142792173 | 142798110 | 5938 | -7.79 | 2.40E-12 | 7.78E-12 |
| chr1 | 232939330 | 232944674 | 5345 | -7.94 | 2.40E-12 | 7.79E-12 |
| chr20 | 61310564 | 61312645 | 2082 | -7.83 | 2.41E-12 | 7.81E-12 |
| chr21 | 30770073 | 30773742 | 3670 | -5.56 | 2.41E-12 | 7.81E-12 |
| chr6 | 107257322 | 107262401 | 5080 | -8.03 | 2.41E-12 | 7.81E-12 |
| chr4 | 39566313 | 39568624 | 2312 | -5.68 | 2.41E-12 | 7.81E-12 |
| chr1 | 233831638 | 233854343 | 22706 | -7.84 | 2.41E-12 | 7.82E-12 |
| chr14 | 89636683 | 89639853 | 3171 | -7.74 | 2.42E-12 | 7.84E-12 |
| chr18 | 46681845 | 46692589 | 10745 | -7.73 | 2.42E-12 | 7.84E-12 |
| chr11 | 21699643 | 21707096 | 7454 | -7.78 | 2.42E-12 | 7.84E-12 |
| chr12 | 85626484 | 85629496 | 3013 | -7.73 | 2.42E-12 | 7.84E-12 |
| chr3 | 130638250 | 130643608 | 5359 | -7.74 | 2.43E-12 | 7.87E-12 |
| chr6 | 141729601 | 141748218 | 18618 | -7.8 | 2.43E-12 | 7.88E-12 |
| chr6 | 75979878 | 75983816 | 3939 | -7.99 | 2.43E-12 | 7.89E-12 |
| chr7 | 134770015 | 134774668 | 4654 | -5.05 | 2.44E-12 | 7.89E-12 |
| chr2 | 153095602 | 153098282 | 2681 | -7.74 | 2.44E-12 | 7.89E-12 |
| chr2 | 114004918 | 114013883 | 8966 | -7.75 | 2.44E-12 | 7.90E-12 |
| chr14 | 85895674 | 85909285 | 13612 | -8.11 | 2.45E-12 | 7.93E-12 |
| chr5 | 112866087 | 112868937 | 2851 | -7.9 | 2.45E-12 | 7.94E-12 |
| chr8 | 77326246 | 77345775 | 19530 | -8.17 | 2.46E-12 | 7.97E-12 |
| chr3 | 182649380 | 182651750 | 2371 | -7.73 | 2.47E-12 | 8.00E-12 |
| chr4 | 168679363 | 168691172 | 11810 | -7.75 | 2.48E-12 | 8.02E-12 |
| chr3 | 66715933 | 66734752 | 18820 | -2.05 | 2.48E-12 | 8.02E-12 |

| | | | | | | |
|---|---|---|---|---|---|---|
| chr6 | 56840619 | 56844232 | 3614 | -7.79 | 2.48E-12 | 8.03E-12 |
| chr1 | 157940388 | 157946657 | 6270 | -7.77 | 2.49E-12 | 8.07E-12 |
| chr18 | 53894338 | 53900866 | 6529 | -7.73 | 2.49E-12 | 8.07E-12 |
| chr13 | 60127221 | 60129808 | 2588 | -6.64 | 2.50E-12 | 8.08E-12 |
| chr1 | 145466415 | 145468863 | 2449 | -7.76 | 2.50E-12 | 8.09E-12 |
| chr20 | 24614156 | 24626744 | 12589 | -8.03 | 2.50E-12 | 8.10E-12 |
| chr12 | 19279187 | 19281743 | 2557 | -7.73 | 2.51E-12 | 8.12E-12 |
| chr8 | 142006200 | 142008064 | 1865 | -7.73 | 2.51E-12 | 8.12E-12 |
| chr9 | 115527950 | 115529375 | 1426 | -7.8 | 2.52E-12 | 8.14E-12 |
| chr6 | 15929016 | 15939244 | 10229 | -7.72 | 2.52E-12 | 8.14E-12 |
| chr6 | 4242288 | 4248093 | 5806 | -7.74 | 2.52E-12 | 8.14E-12 |
| chr3 | 133202504 | 133206280 | 3777 | -7.8 | 2.52E-12 | 8.16E-12 |
| chr8 | 116788811 | 116804534 | 15724 | -7.72 | 2.52E-12 | 8.16E-12 |
| chr20 | 15031145 | 15034863 | 3719 | -7.99 | 2.52E-12 | 8.16E-12 |
| chr9 | 15312614 | 15326454 | 13841 | -7.86 | 2.53E-12 | 8.18E-12 |
| chr2 | 165756414 | 165758139 | 1726 | -7.73 | 2.54E-12 | 8.21E-12 |
| chr1 | 168694254 | 168699942 | 5689 | -7.88 | 2.56E-12 | 8.27E-12 |
| chr7 | 38855257 | 38864141 | 8885 | -7.88 | 2.56E-12 | 8.27E-12 |
| chr11 | 33406414 | 33408759 | 2346 | -7.85 | 2.56E-12 | 8.28E-12 |
| chr3 | 40049697 | 40063076 | 13380 | 7.83 | 2.56E-12 | 8.28E-12 |
| chr1 | 49218945 | 49226181 | 7237 | 7.8 | 2.57E-12 | 8.31E-12 |
| chr3 | 127940695 | 127943334 | 2640 | -7.72 | 2.57E-12 | 8.31E-12 |
| chr2 | 220876828 | 220901335 | 24508 | -8.09 | 2.58E-12 | 8.32E-12 |
| chr3 | 186844484 | 186846573 | 2090 | -7.77 | 2.58E-12 | 8.33E-12 |
| chr8 | 100178104 | 100179643 | 1540 | -7.74 | 2.58E-12 | 8.33E-12 |
| chr12 | 66476659 | 66495510 | 18852 | -7.78 | 2.58E-12 | 8.34E-12 |
| chr5 | 138134907 | 138136662 | 1756 | -7.79 | 2.59E-12 | 8.37E-12 |
| chr10 | 8791277 | 8797957 | 6681 | -7.8 | 2.59E-12 | 8.37E-12 |
| chr5 | 124356671 | 124360201 | 3531 | -7.8 | 2.59E-12 | 8.37E-12 |
| chr5 | 82385783 | 82389956 | 4174 | -7.77 | 2.59E-12 | 8.37E-12 |
| chr11 | 76788604 | 76790970 | 2367 | -7.84 | 2.59E-12 | 8.37E-12 |
| chr2 | 177592973 | 177605822 | 12850 | 8.54 | 2.60E-12 | 8.38E-12 |
| chr4 | 148556299 | 148558187 | 1889 | -7.78 | 2.60E-12 | 8.38E-12 |
| chr14 | 68367444 | 68372955 | 5512 | -7.72 | 2.60E-12 | 8.39E-12 |
| chr5 | 14133174 | 14136570 | 3397 | -7.73 | 2.61E-12 | 8.41E-12 |
| chr4 | 41378850 | 41381662 | 2813 | -7.72 | 2.61E-12 | 8.42E-12 |
| chr3 | 190534784 | 190537616 | 2833 | -7.72 | 2.61E-12 | 8.42E-12 |
| chr7 | 73934255 | 73937113 | 2859 | -7.76 | 2.62E-12 | 8.47E-12 |
| chr15 | 33222502 | 33254236 | 31735 | -8.87 | 2.63E-12 | 8.48E-12 |
| chr7 | 131659071 | 131674326 | 15256 | -2.23 | 2.63E-12 | 8.49E-12 |
| chr18 | 19244220 | 19247521 | 3302 | -7.72 | 2.64E-12 | 8.50E-12 |
| chr11 | 21601425 | 21618021 | 16597 | -7.81 | 2.65E-12 | 8.55E-12 |
| chr12 | 108093401 | 108099742 | 6342 | -8 | 2.65E-12 | 8.56E-12 |
| chr12 | 18023112 | 18044574 | 21463 | -7.72 | 2.65E-12 | 8.56E-12 |
| chr12 | 113559991 | 113561727 | 1737 | -7.77 | 2.66E-12 | 8.58E-12 |
| chr8 | 48396182 | 48398359 | 2178 | -7.77 | 2.66E-12 | 8.58E-12 |
| chr5 | 171731644 | 171733814 | 2171 | -7.73 | 2.67E-12 | 8.60E-12 |
| chr17 | 9319587 | 9323028 | 3442 | -7.72 | 2.67E-12 | 8.60E-12 |
| chr3 | 177614443 | 177619082 | 4640 | -8.04 | 2.67E-12 | 8.60E-12 |
| chr6 | 127627309 | 127631248 | 3940 | -7.88 | 2.67E-12 | 8.60E-12 |
| chr3 | 7499019 | 7509334 | 10316 | -7.79 | 2.67E-12 | 8.61E-12 |
| chr4 | 125364061 | 125369777 | 5717 | -7.76 | 2.67E-12 | 8.62E-12 |
| chr8 | 124765998 | 124770002 | 4005 | -7.76 | 2.67E-12 | 8.62E-12 |
| chr6 | 90931222 | 90935054 | 3833 | -7.72 | 2.68E-12 | 8.62E-12 |
| chrX | 16874791 | 16876530 | 1740 | -8.12 | 2.68E-12 | 8.62E-12 |
| chr12 | 40960686 | 40977096 | 16411 | -7.88 | 2.68E-12 | 8.62E-12 |
| chr10 | 73747064 | 73754663 | 7600 | -7.75 | 2.68E-12 | 8.64E-12 |
| chr12 | 78505845 | 78508744 | 2900 | -7.72 | 2.68E-12 | 8.64E-12 |
| chr7 | 128962561 | 128965236 | 2676 | -7.91 | 2.68E-12 | 8.64E-12 |
| chr10 | 92928569 | 92941635 | 13067 | -8.28 | 2.69E-12 | 8.65E-12 |
| chr3 | 67469544 | 67472088 | 2545 | -7.96 | 2.69E-12 | 8.67E-12 |

| | | | | | | |
|---|---|---|---|---|---|---|
| chr11 | 108830167 | 108843330 | 13164 | -7.79 | 2.70E-12 | 8.68E-12 |
| chr17 | 29299092 | 29301450 | 2359 | -7.94 | 2.70E-12 | 8.69E-12 |
| chr12 | 110542135 | 110550823 | 8689 | -7.74 | 2.70E-12 | 8.70E-12 |
| chr7 | 12572792 | 12577255 | 4464 | -7.74 | 2.70E-12 | 8.70E-12 |
| chr20 | 46609294 | 46788307 | 179014 | -3.33 | 2.71E-12 | 8.72E-12 |
| chr6 | 88546825 | 88548830 | 2006 | -7.73 | 2.71E-12 | 8.73E-12 |
| chr9 | 82442414 | 82466489 | 24076 | -2.17 | 2.72E-12 | 8.74E-12 |
| chr19 | 17063634 | 17065394 | 1761 | -6.03 | 2.73E-12 | 8.79E-12 |
| chr13 | 45475405 | 45487344 | 11940 | -7.87 | 2.73E-12 | 8.79E-12 |
| chr14 | 69392426 | 69394779 | 2354 | -7.77 | 2.75E-12 | 8.84E-12 |
| chr1 | 55419723 | 55424448 | 4726 | -7.71 | 2.75E-12 | 8.84E-12 |
| chr4 | 78831526 | 78847737 | 16212 | -8.02 | 2.75E-12 | 8.85E-12 |
| chr18 | 34594739 | 34608856 | 14118 | -7.72 | 2.75E-12 | 8.85E-12 |
| chr7 | 73408669 | 73411202 | 2534 | -7.72 | 2.75E-12 | 8.85E-12 |
| chr14 | 64101550 | 64104760 | 3211 | -7.83 | 2.76E-12 | 8.89E-12 |
| chr7 | 25894347 | 25898330 | 3984 | -7.8 | 2.77E-12 | 8.90E-12 |
| chr14 | 60644077 | 60648478 | 4402 | -7.83 | 2.77E-12 | 8.90E-12 |
| chr1 | 189885811 | 189894911 | 9101 | -7.83 | 2.77E-12 | 8.91E-12 |
| chr8 | 81114561 | 81118006 | 3446 | -7.83 | 2.77E-12 | 8.91E-12 |
| chrX | 24836249 | 24838343 | 2095 | -7.95 | 2.77E-12 | 8.92E-12 |
| chr4 | 141762050 | 141766531 | 4482 | -7.79 | 2.77E-12 | 8.92E-12 |
| chr7 | 83869801 | 83876041 | 6241 | -7.88 | 2.78E-12 | 8.93E-12 |
| chr3 | 156640856 | 156664657 | 23802 | -7.98 | 2.78E-12 | 8.93E-12 |
| chr5 | 134912539 | 134916858 | 4320 | -8.33 | 2.78E-12 | 8.94E-12 |
| chr20 | 24983080 | 24984708 | 1629 | -7.73 | 2.79E-12 | 8.97E-12 |
| chr13 | 91172765 | 91176681 | 3917 | -7.73 | 2.80E-12 | 8.99E-12 |
| chr14 | 95732736 | 95734902 | 2167 | -7.71 | 2.80E-12 | 8.99E-12 |
| chr21 | 47480362 | 47483567 | 3206 | -7.71 | 2.80E-12 | 8.99E-12 |
| chr9 | 138028548 | 138032627 | 4080 | -7.71 | 2.80E-12 | 8.99E-12 |
| chr11 | 16524011 | 16542112 | 18102 | -7.71 | 2.82E-12 | 9.05E-12 |
| chr5 | 142393389 | 142398129 | 4741 | -7.74 | 2.82E-12 | 9.06E-12 |
| chr4 | 121932347 | 121942366 | 10020 | 7.85 | 2.83E-12 | 9.07E-12 |
| chr2 | 149483659 | 149485909 | 2251 | -7.74 | 2.83E-12 | 9.10E-12 |
| chr6 | 21218118 | 21254784 | 36667 | -8.76 | 2.84E-12 | 9.13E-12 |
| chr7 | 42250935 | 42266270 | 15336 | -7.92 | 2.84E-12 | 9.13E-12 |
| chr16 | 59715954 | 59730291 | 14338 | -2.48 | 2.85E-12 | 9.14E-12 |
| chr10 | 64187269 | 64199819 | 12551 | -7.75 | 2.85E-12 | 9.14E-12 |
| chr10 | 79854168 | 79916686 | 62519 | -1.21 | 2.85E-12 | 9.15E-12 |
| chr6 | 130943578 | 130946081 | 2504 | -3.49 | 2.85E-12 | 9.15E-12 |
| chr3 | 51006373 | 51010446 | 4074 | -7.71 | 2.85E-12 | 9.15E-12 |
| chr12 | 52766802 | 52768978 | 2177 | -7.71 | 2.85E-12 | 9.16E-12 |
| chr14 | 50757954 | 50760154 | 2201 | -7.99 | 2.86E-12 | 9.17E-12 |
| chr1 | 85363997 | 85372174 | 8178 | -7.82 | 2.86E-12 | 9.17E-12 |
| chr19 | 49443289 | 49445170 | 1882 | -7.8 | 2.86E-12 | 9.17E-12 |
| chr11 | 69795895 | 69808421 | 12527 | -7.79 | 2.87E-12 | 9.21E-12 |
| chr5 | 96519759 | 96528291 | 8533 | -7.79 | 2.87E-12 | 9.21E-12 |
| chrX | 84140859 | 84149120 | 8262 | 7.78 | 2.88E-12 | 9.24E-12 |
| chr4 | 99519497 | 99522803 | 3307 | -7.78 | 2.88E-12 | 9.24E-12 |
| chr8 | 17631077 | 17638979 | 7903 | -7.78 | 2.89E-12 | 9.26E-12 |
| chrX | 34951697 | 34955606 | 3910 | -7.71 | 2.89E-12 | 9.27E-12 |
| chr21 | 33161028 | 33163637 | 2610 | -7.8 | 2.89E-12 | 9.27E-12 |
| chr8 | 54425633 | 54438652 | 13020 | -7.8 | 2.89E-12 | 9.27E-12 |
| chr1 | 15724933 | 15727667 | 2735 | -7.78 | 2.89E-12 | 9.28E-12 |
| chr1 | 61419762 | 61433567 | 13806 | -4.02 | 2.90E-12 | 9.30E-12 |
| chr2 | 19733673 | 19745881 | 12209 | -7.75 | 2.90E-12 | 9.31E-12 |
| chr7 | 11801579 | 11813240 | 11662 | -7.97 | 2.91E-12 | 9.32E-12 |
| chr6 | 89593882 | 89596503 | 2622 | -7.84 | 2.92E-12 | 9.35E-12 |
| chr12 | 70656033 | 70669635 | 13603 | -7.73 | 2.92E-12 | 9.36E-12 |
| chr4 | 84077136 | 84094296 | 17161 | -7.86 | 2.93E-12 | 9.37E-12 |
| chr15 | 32019531 | 32155922 | 136392 | 2.52 | 2.93E-12 | 9.38E-12 |
| chr8 | 128914361 | 128916409 | 2049 | -7.73 | 2.95E-12 | 9.44E-12 |

| | | | | | | |
|---|---|---|---|---|---|---|
| chr14 | 32628610 | 32632049 | 3440 | -7.86 | 2.95E-12 | 9.44E-12 |
| chr12 | 10384690 | 10388483 | 3794 | -7.74 | 2.95E-12 | 9.44E-12 |
| chr20 | 18579841 | 18582369 | 2529 | -7.89 | 2.96E-12 | 9.46E-12 |
| chr16 | 52256617 | 52265526 | 8910 | -7.7 | 2.96E-12 | 9.49E-12 |
| chr2 | 161530830 | 161538428 | 7599 | -7.76 | 2.96E-12 | 9.49E-12 |
| chr3 | 57381088 | 57398718 | 17631 | -7.82 | 2.97E-12 | 9.50E-12 |
| chr2 | 240275800 | 240279762 | 3963 | -8.03 | 2.97E-12 | 9.50E-12 |
| chr2 | 196737611 | 196742631 | 5021 | -7.72 | 2.98E-12 | 9.53E-12 |
| chr1 | 66107806 | 66108986 | 1181 | -3.65 | 2.98E-12 | 9.53E-12 |
| chr1 | 46284472 | 46288011 | 3540 | -7.71 | 2.98E-12 | 9.55E-12 |
| chr6 | 3950851 | 3954614 | 3764 | -7.77 | 3.00E-12 | 9.59E-12 |
| chr6 | 117918215 | 117919933 | 1719 | -7.74 | 3.00E-12 | 9.59E-12 |
| chr6 | 56343964 | 56345132 | 1169 | -6.7 | 3.00E-12 | 9.60E-12 |
| chr13 | 101232507 | 101235413 | 2907 | -7.7 | 3.01E-12 | 9.65E-12 |
| chr11 | 43826437 | 43828254 | 1818 | -7.81 | 3.02E-12 | 9.65E-12 |
| chr13 | 89871595 | 89875753 | 4159 | -7.91 | 3.02E-12 | 9.65E-12 |
| chr10 | 86201145 | 86202768 | 1624 | -7.75 | 3.02E-12 | 9.65E-12 |
| chr6 | 136402900 | 136417101 | 14202 | -7.96 | 3.02E-12 | 9.66E-12 |
| chr2 | 33676114 | 33679643 | 3530 | -8.15 | 3.02E-12 | 9.66E-12 |
| chr1 | 214583612 | 214585368 | 1757 | -8.04 | 3.02E-12 | 9.66E-12 |
| chr14 | 27434102 | 27439494 | 5393 | -7.91 | 3.04E-12 | 9.73E-12 |
| chrX | 63150754 | 63394454 | 243701 | 1.16 | 3.05E-12 | 9.74E-12 |
| chr7 | 15179010 | 15194196 | 15187 | -7.7 | 3.05E-12 | 9.74E-12 |
| chr4 | 166057900 | 166062369 | 4470 | -7.71 | 3.05E-12 | 9.75E-12 |
| chr10 | 124064427 | 124081194 | 16768 | -7.74 | 3.05E-12 | 9.76E-12 |
| chr6 | 167055822 | 167066343 | 10522 | -7.74 | 3.05E-12 | 9.76E-12 |
| chr4 | 169647038 | 169650409 | 3372 | -7.92 | 3.05E-12 | 9.76E-12 |
| chr13 | 27624313 | 27638486 | 14174 | -8.04 | 3.06E-12 | 9.78E-12 |
| chr2 | 202031505 | 202053585 | 22081 | -7.78 | 3.07E-12 | 9.81E-12 |
| chr1 | 161779451 | 161781058 | 1608 | -7.71 | 3.08E-12 | 9.84E-12 |
| chr7 | 45992793 | 45996415 | 3623 | -7.77 | 3.09E-12 | 9.87E-12 |
| chr4 | 152656945 | 152660028 | 3084 | -7.71 | 3.09E-12 | 9.88E-12 |
| chr18 | 71389311 | 71399001 | 9691 | -7.73 | 3.09E-12 | 9.89E-12 |
| chr11 | 1756407 | 1763349 | 6943 | -3.48 | 3.10E-12 | 9.90E-12 |
| chr10 | 94579893 | 94583875 | 3983 | -7.7 | 3.10E-12 | 9.90E-12 |
| chr6 | 20741759 | 20755230 | 13472 | -7.87 | 3.10E-12 | 9.92E-12 |
| chr12 | 79806844 | 79822912 | 16069 | -7.93 | 3.11E-12 | 9.92E-12 |
| chr5 | 151086182 | 151092765 | 6584 | -7.71 | 3.12E-12 | 9.95E-12 |
| chr15 | 69802100 | 69819042 | 16943 | -8.89 | 3.12E-12 | 9.96E-12 |
| chr10 | 86218075 | 86219771 | 1697 | -7.71 | 3.12E-12 | 9.96E-12 |
| chr10 | 134284640 | 134286431 | 1792 | -7.7 | 3.12E-12 | 9.97E-12 |
| chr21 | 44300148 | 44305159 | 5012 | -7.7 | 3.14E-12 | 1.00E-11 |
| chr4 | 106224378 | 106226926 | 2549 | -7.69 | 3.14E-12 | 1.00E-11 |
| chr1 | 228531143 | 228533354 | 2212 | -7.86 | 3.14E-12 | 1.00E-11 |
| chr9 | 72941956 | 72944209 | 2254 | -7.7 | 3.14E-12 | 1.00E-11 |
| chr2 | 155157539 | 155174016 | 16478 | 7.85 | 3.15E-12 | 1.01E-11 |
| chr1 | 7047830 | 7054698 | 6869 | -7.71 | 3.15E-12 | 1.01E-11 |
| chr3 | 155767451 | 155780371 | 12921 | -7.7 | 3.16E-12 | 1.01E-11 |
| chr2 | 153169999 | 153174038 | 4040 | -7.81 | 3.17E-12 | 1.01E-11 |
| chr9 | 5834277 | 5854832 | 20556 | -8.07 | 3.19E-12 | 1.02E-11 |
| chrX | 46848147 | 46858797 | 10651 | -7.88 | 3.19E-12 | 1.02E-11 |
| chr1 | 169501852 | 169511959 | 10108 | -6.68 | 3.20E-12 | 1.02E-11 |
| chr10 | 17136009 | 17138191 | 2183 | -7.83 | 3.20E-12 | 1.02E-11 |
| chr3 | 142735984 | 142737555 | 1572 | -7.71 | 3.20E-12 | 1.02E-11 |
| chr1 | 18257793 | 18259721 | 1929 | -7.7 | 3.21E-12 | 1.02E-11 |
| chr14 | 73555012 | 73558715 | 3704 | -7.74 | 3.21E-12 | 1.02E-11 |
| chr2 | 61532735 | 61537036 | 4302 | -7.74 | 3.21E-12 | 1.02E-11 |
| chr18 | 68460709 | 68548403 | 87695 | 1.64 | 3.21E-12 | 1.02E-11 |
| chr22 | 41016346 | 41019648 | 3303 | -7.73 | 3.22E-12 | 1.03E-11 |
| chr8 | 28005295 | 28007971 | 2677 | -7.7 | 3.22E-12 | 1.03E-11 |
| chr4 | 87798989 | 87803030 | 4042 | -7.7 | 3.23E-12 | 1.03E-11 |

| | | | | | | |
|---|---|---|---|---|---|---|
| chr10 | 62926380 | 62929083 | 2704 | -7.72 | 3.24E-12 | 1.03E-11 |
| chr16 | 31410449 | 31413035 | 2587 | -7.72 | 3.24E-12 | 1.03E-11 |
| chr5 | 122292318 | 122294655 | 2338 | -7.69 | 3.24E-12 | 1.03E-11 |
| chr9 | 95134786 | 95138541 | 3756 | -7.69 | 3.25E-12 | 1.04E-11 |
| chr6 | 140104019 | 140109141 | 5123 | -7.85 | 3.25E-12 | 1.04E-11 |
| chr18 | 37328493 | 37330846 | 2354 | -7.75 | 3.26E-12 | 1.04E-11 |
| chr13 | 66531387 | 66552107 | 20721 | -7.95 | 3.26E-12 | 1.04E-11 |
| chr2 | 56021407 | 56026206 | 4800 | -7.85 | 3.28E-12 | 1.04E-11 |
| chr4 | 41087129 | 41090161 | 3033 | -7.71 | 3.28E-12 | 1.04E-11 |
| chr6 | 107767336 | 107772054 | 4719 | -7.7 | 3.28E-12 | 1.04E-11 |
| chr2 | 160942184 | 160951345 | 9162 | -8.15 | 3.28E-12 | 1.04E-11 |
| chr5 | 549515 | 551805 | 2291 | -6 | 3.29E-12 | 1.05E-11 |
| chr2 | 46665205 | 46673936 | 8732 | -7.77 | 3.29E-12 | 1.05E-11 |
| chr5 | 106774282 | 106788088 | 13807 | -7.72 | 3.30E-12 | 1.05E-11 |
| chrX | 130703199 | 130705088 | 1890 | -7.77 | 3.30E-12 | 1.05E-11 |
| chr18 | 47726566 | 47728335 | 1770 | -6.73 | 3.30E-12 | 1.05E-11 |
| chr13 | 44696300 | 44700418 | 4119 | -7.77 | 3.30E-12 | 1.05E-11 |
| chr7 | 79412172 | 79414443 | 2272 | -7.71 | 3.30E-12 | 1.05E-11 |
| chr11 | 107371786 | 107375005 | 3220 | -7.87 | 3.31E-12 | 1.05E-11 |
| chr21 | 40995330 | 40999081 | 3752 | -7.71 | 3.31E-12 | 1.05E-11 |
| chr20 | 45769147 | 45770753 | 1607 | -7.77 | 3.31E-12 | 1.05E-11 |
| chr2 | 43844849 | 43867182 | 22334 | -8.39 | 3.31E-12 | 1.05E-11 |
| chr11 | 35387431 | 35389217 | 1787 | -7.69 | 3.32E-12 | 1.06E-11 |
| chr10 | 69890040 | 69912257 | 22218 | -8.3 | 3.32E-12 | 1.06E-11 |
| chr11 | 50106024 | 50283297 | 177274 | 0.92 | 3.32E-12 | 1.06E-11 |
| chr9 | 4804513 | 4807920 | 3408 | -7.69 | 3.33E-12 | 1.06E-11 |
| chr8 | 133664627 | 133666896 | 2270 | -7.73 | 3.33E-12 | 1.06E-11 |
| chr20 | 49262854 | 49265568 | 2715 | -7.69 | 3.33E-12 | 1.06E-11 |
| chr5 | 113719227 | 114324814 | 605588 | 5.14 | 3.33E-12 | 1.06E-11 |
| chr3 | 136477036 | 136483771 | 6736 | -7.83 | 3.33E-12 | 1.06E-11 |
| chr1 | 198923505 | 198925371 | 1867 | -7.69 | 3.34E-12 | 1.06E-11 |
| chr10 | 24660402 | 24668471 | 8070 | -7.83 | 3.34E-12 | 1.06E-11 |
| chr16 | 88112263 | 88115058 | 2796 | -7.69 | 3.37E-12 | 1.07E-11 |
| chr1 | 231948111 | 231962723 | 14613 | -7.69 | 3.37E-12 | 1.07E-11 |
| chr4 | 64065710 | 64093205 | 27496 | -8.06 | 3.37E-12 | 1.07E-11 |
| chr3 | 101922994 | 101934376 | 11383 | -8.13 | 3.38E-12 | 1.07E-11 |
| chr6 | 17736884 | 17739482 | 2599 | -7.69 | 3.39E-12 | 1.08E-11 |
| chr6 | 40468133 | 40474173 | 6041 | 7.76 | 3.39E-12 | 1.08E-11 |
| chr13 | 22003172 | 22013219 | 10048 | -7.79 | 3.39E-12 | 1.08E-11 |
| chr13 | 49503854 | 49509789 | 5936 | -7.71 | 3.40E-12 | 1.08E-11 |
| chr1 | 196503942 | 196507871 | 3930 | -7.7 | 3.41E-12 | 1.08E-11 |
| chr15 | 67565964 | 67581439 | 15476 | -8.1 | 3.41E-12 | 1.08E-11 |
| chr8 | 83478751 | 83500535 | 21785 | -7.7 | 3.41E-12 | 1.08E-11 |
| chr5 | 40711203 | 40724875 | 13673 | -7.74 | 3.42E-12 | 1.09E-11 |
| chr6 | 139984347 | 139988736 | 4390 | -7.69 | 3.43E-12 | 1.09E-11 |
| chr14 | 32859044 | 32863100 | 4057 | -2.62 | 3.43E-12 | 1.09E-11 |
| chr17 | 53250795 | 53252632 | 1838 | -7.68 | 3.43E-12 | 1.09E-11 |
| chr20 | 3365900 | 3377405 | 11506 | -8.14 | 3.44E-12 | 1.09E-11 |
| chr8 | 59059519 | 59066948 | 7430 | -7.68 | 3.44E-12 | 1.09E-11 |
| chr7 | 68988631 | 69006675 | 18045 | -8.28 | 3.44E-12 | 1.09E-11 |
| chr1 | 229089868 | 229091769 | 1902 | -8 | 3.45E-12 | 1.09E-11 |
| chr1 | 154865008 | 154867027 | 2020 | -7.89 | 3.45E-12 | 1.09E-11 |
| chr16 | 30107851 | 30110804 | 2954 | -7.69 | 3.45E-12 | 1.09E-11 |
| chr10 | 134326385 | 134346295 | 19911 | -7.69 | 3.45E-12 | 1.09E-11 |
| chr4 | 19657743 | 19665683 | 7941 | -7.77 | 3.45E-12 | 1.09E-11 |
| chr17 | 31049321 | 31053398 | 4078 | -7.75 | 3.45E-12 | 1.09E-11 |
| chr3 | 45883789 | 45916577 | 32789 | -2.54 | 3.45E-12 | 1.10E-11 |
| chr11 | 126020990 | 126023026 | 2037 | -7.69 | 3.46E-12 | 1.10E-11 |
| chr2 | 75944367 | 75947551 | 3185 | -7.73 | 3.46E-12 | 1.10E-11 |
| chr6 | 158751992 | 158754825 | 2834 | -7.68 | 3.47E-12 | 1.10E-11 |
| chr2 | 43222768 | 43250949 | 28182 | -8.64 | 3.47E-12 | 1.10E-11 |

| | | | | | | |
|---|---|---|---|---|---|---|
| chr4 | 101277125 | 101281021 | 3897 | -7.69 | 3.47E-12 | 1.10E-11 |
| chr5 | 127407050 | 127409799 | 2750 | -7.69 | 3.47E-12 | 1.10E-11 |
| chr5 | 34826713 | 34835167 | 8455 | -7.7 | 3.48E-12 | 1.10E-11 |
| chr10 | 21769505 | 21773572 | 4068 | -7.74 | 3.48E-12 | 1.10E-11 |
| chr20 | 49576663 | 49610134 | 33472 | -8.04 | 3.50E-12 | 1.11E-11 |
| chr3 | 67659048 | 67661345 | 2298 | -7.69 | 3.50E-12 | 1.11E-11 |
| chr21 | 33115700 | 33119444 | 3745 | -7.71 | 3.50E-12 | 1.11E-11 |
| chr8 | 103688456 | 103690486 | 2031 | -7.71 | 3.50E-12 | 1.11E-11 |
| chr19 | 20419635 | 20428198 | 8564 | -7.68 | 3.51E-12 | 1.11E-11 |
| chrX | 4721429 | 4726673 | 5245 | -7.79 | 3.51E-12 | 1.11E-11 |
| chr14 | 66920857 | 66937411 | 16555 | -7.77 | 3.52E-12 | 1.11E-11 |
| chr2 | 99988878 | 99991510 | 2633 | -7.77 | 3.52E-12 | 1.11E-11 |
| chr13 | 45000663 | 45003098 | 2436 | -7.77 | 3.52E-12 | 1.12E-11 |
| chr12 | 112942378 | 112944698 | 2321 | -7.74 | 3.52E-12 | 1.12E-11 |
| chr6 | 39783779 | 39788829 | 5051 | 7.81 | 3.52E-12 | 1.12E-11 |
| chr14 | 23329017 | 23333078 | 4062 | -8.11 | 3.52E-12 | 1.12E-11 |
| chr17 | 7931157 | 7932799 | 1643 | -7.88 | 3.53E-12 | 1.12E-11 |
| chr12 | 6136118 | 6138757 | 2640 | -5.14 | 3.54E-12 | 1.12E-11 |
| chr7 | 68879600 | 68910291 | 30692 | -8.26 | 3.54E-12 | 1.12E-11 |
| chr19 | 55533112 | 55538091 | 4980 | -7.68 | 3.55E-12 | 1.12E-11 |
| chr21 | 19454612 | 19511108 | 56497 | 1.5 | 3.56E-12 | 1.13E-11 |
| chr14 | 24274050 | 24288640 | 14591 | -7.69 | 3.56E-12 | 1.13E-11 |
| chr4 | 154182273 | 154190919 | 8647 | -7.69 | 3.56E-12 | 1.13E-11 |
| chr5 | 68435669 | 68437670 | 2002 | -7.68 | 3.57E-12 | 1.13E-11 |
| chr5 | 133806110 | 133810928 | 4819 | -8.01 | 3.57E-12 | 1.13E-11 |
| chr7 | 11120567 | 11124101 | 3535 | -7.73 | 3.58E-12 | 1.13E-11 |
| chr15 | 43052007 | 43055118 | 3112 | -7.7 | 3.59E-12 | 1.13E-11 |
| chr3 | 178075752 | 178087935 | 12184 | -8.01 | 3.59E-12 | 1.14E-11 |
| chr4 | 142976138 | 142979148 | 3011 | -7.68 | 3.59E-12 | 1.14E-11 |
| chr14 | 74882590 | 74888600 | 6011 | -7.96 | 3.60E-12 | 1.14E-11 |
| chr10 | 119179119 | 119181763 | 2645 | -7.98 | 3.60E-12 | 1.14E-11 |
| chr8 | 83685859 | 83713773 | 27915 | -8.03 | 3.60E-12 | 1.14E-11 |
| chr7 | 17673698 | 17700530 | 26833 | -8.71 | 3.60E-12 | 1.14E-11 |
| chr11 | 113578415 | 113582565 | 4151 | -7.71 | 3.60E-12 | 1.14E-11 |
| chr3 | 195039322 | 195042054 | 2733 | -7.86 | 3.61E-12 | 1.14E-11 |
| chr19 | 49851940 | 49854248 | 2309 | -7.68 | 3.61E-12 | 1.14E-11 |
| chrX | 69527422 | 69529489 | 2068 | -7.68 | 3.61E-12 | 1.14E-11 |
| chr11 | 2957668 | 2961525 | 3858 | -7.89 | 3.61E-12 | 1.14E-11 |
| chr10 | 116838261 | 116873631 | 35371 | -4.36 | 3.62E-12 | 1.14E-11 |
| chr22 | 25608610 | 25612856 | 4247 | -7.72 | 3.62E-12 | 1.14E-11 |
| chr10 | 83464248 | 83488592 | 24345 | -7.87 | 3.62E-12 | 1.15E-11 |
| chr19 | 14109628 | 14111251 | 1624 | -7.68 | 3.63E-12 | 1.15E-11 |
| chr22 | 27512956 | 27517464 | 4509 | -7.68 | 3.63E-12 | 1.15E-11 |
| chr22 | 33041257 | 33043377 | 2121 | -3.73 | 3.63E-12 | 1.15E-11 |
| chr6 | 170348081 | 170358699 | 10619 | -7.77 | 3.64E-12 | 1.15E-11 |
| chr10 | 114421136 | 114422971 | 1836 | -7.69 | 3.64E-12 | 1.15E-11 |
| chr1 | 213556577 | 214109936 | 553360 | 3.31 | 3.64E-12 | 1.15E-11 |
| chr11 | 8488590 | 8491935 | 3346 | -7.68 | 3.65E-12 | 1.15E-11 |
| chr17 | 48956133 | 48957756 | 1624 | -7.87 | 3.65E-12 | 1.15E-11 |
| chr5 | 168844879 | 168866186 | 21308 | -7.72 | 3.65E-12 | 1.15E-11 |
| chr1 | 35534817 | 35537483 | 2667 | -7.74 | 3.66E-12 | 1.15E-11 |
| chr21 | 39716652 | 39722869 | 6218 | -7.75 | 3.66E-12 | 1.16E-11 |
| chr17 | 48899251 | 48903152 | 3902 | -2.27 | 3.66E-12 | 1.16E-11 |
| chr13 | 106300513 | 106316807 | 16295 | -7.75 | 3.68E-12 | 1.16E-11 |
| chr11 | 129095187 | 129097848 | 2662 | -7.68 | 3.68E-12 | 1.16E-11 |
| chr9 | 33230154 | 33233398 | 3245 | -7.68 | 3.69E-12 | 1.16E-11 |
| chr4 | 184669810 | 184674085 | 4276 | -7.81 | 3.69E-12 | 1.16E-11 |
| chr12 | 57834459 | 57836650 | 2192 | -7.67 | 3.70E-12 | 1.17E-11 |
| chr1 | 85848049 | 85854057 | 6009 | -7.72 | 3.71E-12 | 1.17E-11 |
| chr7 | 17140113 | 17156555 | 16443 | -7.82 | 3.72E-12 | 1.17E-11 |
| chr3 | 51560276 | 51563596 | 3321 | -7.74 | 3.72E-12 | 1.18E-11 |

| | | | | | | |
|---|---|---|---|---|---|---|
| chr7 | 45501622 | 45512154 | 10533 | -7.67 | 3.73E-12 | 1.18E-11 |
| chr12 | 90209686 | 90224288 | 14603 | -7.72 | 3.73E-12 | 1.18E-11 |
| chr5 | 92019552 | 92026729 | 7178 | -7.72 | 3.74E-12 | 1.18E-11 |
| chr8 | 95153289 | 95156675 | 3387 | -7.73 | 3.74E-12 | 1.18E-11 |
| chr6 | 76176126 | 76180257 | 4132 | -7.68 | 3.74E-12 | 1.18E-11 |
| chr15 | 36954113 | 36956593 | 2481 | -6.15 | 3.74E-12 | 1.18E-11 |
| chr7 | 123129165 | 123154410 | 25246 | -8.03 | 3.74E-12 | 1.18E-11 |
| chr11 | 15835426 | 15843617 | 8192 | -7.78 | 3.75E-12 | 1.18E-11 |
| chr10 | 126188911 | 126190277 | 1367 | -7.67 | 3.75E-12 | 1.18E-11 |
| chr17 | 38650237 | 38653555 | 3319 | -7.71 | 3.75E-12 | 1.18E-11 |
| chr7 | 42932120 | 42936948 | 4829 | -7.73 | 3.77E-12 | 1.19E-11 |
| chr9 | 135144395 | 135148002 | 3608 | -7.97 | 3.78E-12 | 1.19E-11 |
| chr14 | 88482144 | 88542319 | 60176 | -1.39 | 3.78E-12 | 1.19E-11 |
| chr20 | 61010835 | 61012545 | 1711 | -7.76 | 3.78E-12 | 1.19E-11 |
| chr15 | 33542330 | 33545565 | 3236 | -5.82 | 3.78E-12 | 1.19E-11 |
| chr1 | 44307229 | 44309569 | 2341 | -7.84 | 3.78E-12 | 1.19E-11 |
| chr15 | 49403697 | 49405547 | 1851 | -7.74 | 3.79E-12 | 1.20E-11 |
| chr12 | 112389198 | 112410209 | 21012 | -7.75 | 3.80E-12 | 1.20E-11 |
| chr17 | 62058413 | 62060846 | 2434 | -7.81 | 3.81E-12 | 1.20E-11 |
| chr12 | 22754423 | 22770032 | 15610 | -8.14 | 3.81E-12 | 1.20E-11 |
| chr10 | 50760552 | 50779272 | 18721 | -2.09 | 3.81E-12 | 1.20E-11 |
| chr4 | 99188482 | 99197451 | 8970 | -7.86 | 3.84E-12 | 1.21E-11 |
| chr16 | 84883923 | 84895882 | 11960 | -7.95 | 3.85E-12 | 1.21E-11 |
| chr9 | 85632458 | 85634890 | 2433 | -7.73 | 3.86E-12 | 1.21E-11 |
| chr2 | 88830141 | 88839003 | 8863 | -7.71 | 3.87E-12 | 1.22E-11 |
| chr4 | 113281737 | 113289949 | 8213 | -7.89 | 3.89E-12 | 1.22E-11 |
| chr15 | 30138981 | 30140973 | 1993 | -7.7 | 3.90E-12 | 1.23E-11 |
| chr14 | 64284952 | 64290051 | 5100 | -7.77 | 3.90E-12 | 1.23E-11 |
| chr3 | 106998622 | 107015049 | 16428 | -7.77 | 3.90E-12 | 1.23E-11 |
| chr6 | 159599768 | 159609511 | 9744 | -2.18 | 3.90E-12 | 1.23E-11 |
| chr14 | 107000195 | 107076124 | 75930 | 1.52 | 3.90E-12 | 1.23E-11 |
| chr19 | 50385460 | 50386968 | 1509 | -7.76 | 3.92E-12 | 1.23E-11 |
| chr3 | 81866655 | 81869271 | 2617 | -7.67 | 3.92E-12 | 1.23E-11 |
| chr6 | 165998770 | 166008164 | 9395 | -7.67 | 3.92E-12 | 1.23E-11 |
| chr2 | 183111376 | 183123020 | 11645 | 8.04 | 3.92E-12 | 1.23E-11 |
| chr1 | 56809488 | 56814645 | 5158 | -7.72 | 3.93E-12 | 1.23E-11 |
| chr6 | 137973697 | 137977704 | 4008 | -7.87 | 3.93E-12 | 1.23E-11 |
| chr6 | 132218283 | 132221284 | 3002 | -7.67 | 3.93E-12 | 1.24E-11 |
| chr2 | 228403643 | 228405279 | 1637 | -7.79 | 3.93E-12 | 1.24E-11 |
| chr7 | 92252349 | 92273040 | 20692 | -8 | 3.94E-12 | 1.24E-11 |
| chr1 | 119729362 | 119743472 | 14111 | -7.69 | 3.96E-12 | 1.25E-11 |
| chr2 | 47996896 | 48000212 | 3317 | -7.68 | 3.96E-12 | 1.25E-11 |
| chr2 | 160355937 | 160358783 | 2847 | -7.67 | 3.97E-12 | 1.25E-11 |
| chr12 | 129288352 | 129290397 | 2046 | -7.68 | 3.98E-12 | 1.25E-11 |
| chr7 | 15314135 | 15323900 | 9766 | -5.94 | 3.99E-12 | 1.25E-11 |
| chr3 | 192257596 | 192263502 | 5907 | 7.8 | 3.99E-12 | 1.25E-11 |
| chr4 | 141228450 | 141258923 | 30474 | -8.32 | 3.99E-12 | 1.25E-11 |
| chr20 | 46399008 | 46406566 | 7559 | -7.82 | 4.00E-12 | 1.26E-11 |
| chrX | 79549586 | 79561739 | 12154 | -8 | 4.00E-12 | 1.26E-11 |
| chr6 | 127555618 | 127558269 | 2652 | -7.66 | 4.01E-12 | 1.26E-11 |
| chr9 | 95389054 | 95392828 | 3775 | -7.7 | 4.03E-12 | 1.27E-11 |
| chr2 | 237674720 | 237755659 | 80940 | -1.36 | 4.03E-12 | 1.27E-11 |
| chr17 | 46852584 | 46857325 | 4742 | -7.86 | 4.04E-12 | 1.27E-11 |
| chr4 | 123387099 | 123391753 | 4655 | -7.75 | 4.05E-12 | 1.27E-11 |
| chr6 | 142598704 | 142606660 | 7957 | -7.83 | 4.06E-12 | 1.27E-11 |
| chr12 | 93301598 | 93303807 | 2210 | -7.79 | 4.06E-12 | 1.27E-11 |
| chr1 | 61572652 | 61574828 | 2177 | -7.66 | 4.06E-12 | 1.28E-11 |
| chr22 | 27703355 | 27718203 | 14849 | -8.01 | 4.06E-12 | 1.28E-11 |
| chr6 | 26105353 | 26108742 | 3390 | -7.68 | 4.07E-12 | 1.28E-11 |
| chr12 | 93249989 | 93252036 | 2048 | -7.68 | 4.08E-12 | 1.28E-11 |
| chr17 | 76542095 | 76544048 | 1954 | -7.84 | 4.08E-12 | 1.28E-11 |

| | | | | | | |
|---|---|---|---|---|---|---|
| chr2 | 57707250 | 57715209 | 7960 | 7.74 | 4.09E-12 | 1.28E-11 |
| chr6 | 100739370 | 100751575 | 12206 | -7.83 | 4.09E-12 | 1.28E-11 |
| chr3 | 102536276 | 102551045 | 14770 | -7.66 | 4.09E-12 | 1.28E-11 |
| chr8 | 107051715 | 107054087 | 2373 | -7.77 | 4.09E-12 | 1.28E-11 |
| chr9 | 15492412 | 15494957 | 2546 | -7.71 | 4.10E-12 | 1.29E-11 |
| chr15 | 67106163 | 67107897 | 1735 | -7.68 | 4.11E-12 | 1.29E-11 |
| chr16 | 48561618 | 48564796 | 3179 | -8.02 | 4.11E-12 | 1.29E-11 |
| chr1 | 190126319 | 190129350 | 3032 | -7.67 | 4.12E-12 | 1.29E-11 |
| chr1 | 231881749 | 231889873 | 8125 | -7.66 | 4.12E-12 | 1.29E-11 |
| chr7 | 47347498 | 47352828 | 5331 | -7.84 | 4.12E-12 | 1.29E-11 |
| chr9 | 35579053 | 35594962 | 15910 | -8.42 | 4.13E-12 | 1.30E-11 |
| chr6 | 39034519 | 39040643 | 6125 | -2.64 | 4.14E-12 | 1.30E-11 |
| chr1 | 79493481 | 79504619 | 11139 | -7.94 | 4.15E-12 | 1.30E-11 |
| chr4 | 99619973 | 99627880 | 7908 | -5.9 | 4.15E-12 | 1.30E-11 |
| chr12 | 76654851 | 76658334 | 3484 | -7.68 | 4.15E-12 | 1.30E-11 |
| chr6 | 126455349 | 126462221 | 6873 | -7.73 | 4.15E-12 | 1.30E-11 |
| chr6 | 26196726 | 26200355 | 3630 | -7.72 | 4.16E-12 | 1.30E-11 |
| chr2 | 28655752 | 28666409 | 10658 | -7.88 | 4.17E-12 | 1.31E-11 |
| chr8 | 41128806 | 41131551 | 2746 | -7.66 | 4.18E-12 | 1.31E-11 |
| chr20 | 25240587 | 25246550 | 5964 | -7.68 | 4.18E-12 | 1.31E-11 |
| chr21 | 38873122 | 38875885 | 2764 | -7.71 | 4.18E-12 | 1.31E-11 |
| chr10 | 16467665 | 16471898 | 4234 | -7.66 | 4.18E-12 | 1.31E-11 |
| chr4 | 131438349 | 131446725 | 8377 | -7.66 | 4.18E-12 | 1.31E-11 |
| chr4 | 87099989 | 87102212 | 2224 | -7.66 | 4.18E-12 | 1.31E-11 |
| chr12 | 47187278 | 47308819 | 121542 | -1.53 | 4.18E-12 | 1.31E-11 |
| chr19 | 41338104 | 41348095 | 9992 | -8.01 | 4.19E-12 | 1.31E-11 |
| chr9 | 32716690 | 32729202 | 12513 | -7.77 | 4.19E-12 | 1.31E-11 |
| chr15 | 71501431 | 71506629 | 5199 | -7.72 | 4.19E-12 | 1.31E-11 |
| chr6 | 6015436 | 6018693 | 3258 | -5.34 | 4.19E-12 | 1.31E-11 |
| chr11 | 46174260 | 46181033 | 6774 | -7.79 | 4.21E-12 | 1.32E-11 |
| chr2 | 15530899 | 15537569 | 6671 | -7.67 | 4.22E-12 | 1.32E-11 |
| chr2 | 234768612 | 234795673 | 27062 | -7.53 | 4.22E-12 | 1.32E-11 |
| chr16 | 21684218 | 21685954 | 1737 | -7.73 | 4.22E-12 | 1.32E-11 |
| chr3 | 56717437 | 56719208 | 1772 | -7.66 | 4.22E-12 | 1.32E-11 |
| chr10 | 71031136 | 71033926 | 2791 | -7.69 | 4.23E-12 | 1.32E-11 |
| chr5 | 90308608 | 90312507 | 3900 | -7.79 | 4.24E-12 | 1.33E-11 |
| chr15 | 40235869 | 40238928 | 3060 | -7.71 | 4.24E-12 | 1.33E-11 |
| chr20 | 8946927 | 8949988 | 3062 | -7.65 | 4.25E-12 | 1.33E-11 |
| chr13 | 25149198 | 25152550 | 3353 | -7.66 | 4.25E-12 | 1.33E-11 |
| chr5 | 127214180 | 127219774 | 5595 | -7.97 | 4.26E-12 | 1.33E-11 |
| chr20 | 18597146 | 18599160 | 2015 | -7.66 | 4.26E-12 | 1.33E-11 |
| chrX | 35147458 | 35156204 | 8747 | -2.53 | 4.26E-12 | 1.33E-11 |
| chr7 | 151059047 | 151061001 | 1955 | -7.89 | 4.27E-12 | 1.34E-11 |
| chr20 | 36314148 | 36316364 | 2217 | -7.75 | 4.28E-12 | 1.34E-11 |
| chr19 | 12993860 | 12995996 | 2137 | -7.67 | 4.29E-12 | 1.34E-11 |
| chr2 | 210616978 | 210621603 | 4626 | 7.74 | 4.29E-12 | 1.34E-11 |
| chr1 | 7849595 | 7851935 | 2341 | -7.68 | 4.29E-12 | 1.34E-11 |
| chr1 | 37789432 | 37807626 | 18195 | -8.02 | 4.30E-12 | 1.34E-11 |
| chr20 | 34667283 | 34672822 | 5540 | -7.71 | 4.31E-12 | 1.35E-11 |
| chr11 | 85480173 | 85497551 | 17379 | -8.11 | 4.31E-12 | 1.35E-11 |
| chr15 | 53592003 | 53602058 | 10056 | -7.66 | 4.32E-12 | 1.35E-11 |
| chr13 | 80616676 | 80635840 | 19165 | -7.73 | 4.32E-12 | 1.35E-11 |
| chrX | 107032755 | 107046644 | 13890 | -1.64 | 4.32E-12 | 1.35E-11 |
| chr15 | 73634870 | 73636689 | 1820 | -7.67 | 4.32E-12 | 1.35E-11 |
| chr2 | 161098992 | 161117062 | 18071 | -7.79 | 4.33E-12 | 1.35E-11 |
| chr20 | 49979236 | 49981544 | 2309 | -7.71 | 4.33E-12 | 1.35E-11 |
| chr7 | 15578879 | 15587006 | 8128 | -7.68 | 4.33E-12 | 1.35E-11 |
| chr22 | 49904027 | 49907533 | 3507 | -7.75 | 4.33E-12 | 1.35E-11 |
| chr7 | 12035476 | 12037622 | 2147 | -7.74 | 4.34E-12 | 1.36E-11 |
| chr12 | 3097729 | 3099586 | 1858 | -7.66 | 4.34E-12 | 1.36E-11 |
| chr5 | 168711734 | 168714245 | 2512 | -7.67 | 4.36E-12 | 1.36E-11 |

| | | | | | | |
|---|---|---|---|---|---|---|
| chr6 | 91077081 | 91090178 | 13098 | -7.67 | 4.37E-12 | 1.37E-11 |
| chr9 | 90474377 | 90478982 | 4606 | -7.66 | 4.40E-12 | 1.37E-11 |
| chr3 | 182033524 | 182037815 | 4292 | -7.71 | 4.41E-12 | 1.38E-11 |
| chr5 | 158614464 | 158617992 | 3529 | -7.65 | 4.41E-12 | 1.38E-11 |
| chr10 | 8498765 | 8500993 | 2229 | -7.83 | 4.42E-12 | 1.38E-11 |
| chr11 | 36240302 | 36242444 | 2143 | -7.65 | 4.43E-12 | 1.38E-11 |
| chr1 | 167771079 | 167786663 | 15585 | -8.22 | 4.43E-12 | 1.38E-11 |
| chr6 | 574291 | 576571 | 2281 | -7.69 | 4.43E-12 | 1.38E-11 |
| chr10 | 24956586 | 24968811 | 12226 | -8.25 | 4.46E-12 | 1.39E-11 |
| chr6 | 78034718 | 78045728 | 11011 | -7.81 | 4.48E-12 | 1.40E-11 |
| chr3 | 24358826 | 24361921 | 3096 | -7.74 | 4.48E-12 | 1.40E-11 |
| chr1 | 38574185 | 38576619 | 2435 | -5.89 | 4.48E-12 | 1.40E-11 |
| chr10 | 37131001 | 37152752 | 21752 | -8.17 | 4.48E-12 | 1.40E-11 |
| chr15 | 72374636 | 72378228 | 3593 | -7.87 | 4.49E-12 | 1.40E-11 |
| chr1 | 111577991 | 111582377 | 4387 | -7.65 | 4.49E-12 | 1.40E-11 |
| chr8 | 58775488 | 58795916 | 20429 | -7.76 | 4.49E-12 | 1.40E-11 |
| chr1 | 229855169 | 229973653 | 118485 | -1.95 | 4.49E-12 | 1.40E-11 |
| chr6 | 54848299 | 54859214 | 10916 | -7.74 | 4.50E-12 | 1.40E-11 |
| chr19 | 8295260 | 8309188 | 13929 | -7.74 | 4.50E-12 | 1.40E-11 |
| chr4 | 2188371 | 2202328 | 13958 | -7.68 | 4.50E-12 | 1.40E-11 |
| chr4 | 6100512 | 6105792 | 5281 | -5.81 | 4.50E-12 | 1.40E-11 |
| chr2 | 155039914 | 155043637 | 3724 | -7.83 | 4.51E-12 | 1.41E-11 |
| chr1 | 55732859 | 55735846 | 2988 | -7.78 | 4.53E-12 | 1.41E-11 |
| chr4 | 135160263 | 135176305 | 16043 | 7.83 | 4.53E-12 | 1.41E-11 |
| chr18 | 25266134 | 25271518 | 5385 | -7.66 | 4.55E-12 | 1.42E-11 |
| chr13 | 112046960 | 112054088 | 7129 | -7.65 | 4.56E-12 | 1.42E-11 |
| chr12 | 114243939 | 114250153 | 6215 | -7.95 | 4.56E-12 | 1.42E-11 |
| chr2 | 138856753 | 138863886 | 7134 | -7.84 | 4.57E-12 | 1.42E-11 |
| chr20 | 36727582 | 36731358 | 3777 | -7.86 | 4.59E-12 | 1.43E-11 |
| chr21 | 43398151 | 43400904 | 2754 | -7.71 | 4.59E-12 | 1.43E-11 |
| chr3 | 160438958 | 160443904 | 4947 | -2.88 | 4.60E-12 | 1.43E-11 |
| chr9 | 96832529 | 96834979 | 2451 | -7.65 | 4.60E-12 | 1.43E-11 |
| chr20 | 10738036 | 10740140 | 2105 | -7.7 | 4.61E-12 | 1.43E-11 |
| chr5 | 90947156 | 90950730 | 3575 | -7.8 | 4.63E-12 | 1.44E-11 |
| chr14 | 57774741 | 57780467 | 5727 | -7.64 | 4.64E-12 | 1.45E-11 |
| chr16 | 72399328 | 72402089 | 2762 | -7.64 | 4.65E-12 | 1.45E-11 |
| chr7 | 109130386 | 109142720 | 12335 | -7.75 | 4.65E-12 | 1.45E-11 |
| chr9 | 85740762 | 85744495 | 3734 | -7.64 | 4.65E-12 | 1.45E-11 |
| chr3 | 107127556 | 107129568 | 2013 | -7.74 | 4.65E-12 | 1.45E-11 |
| chr9 | 134346324 | 134347796 | 1473 | -7.66 | 4.65E-12 | 1.45E-11 |
| chr17 | 26340596 | 26364880 | 24285 | -8.05 | 4.65E-12 | 1.45E-11 |
| chr6 | 159408910 | 159414451 | 5542 | -7.74 | 4.65E-12 | 1.45E-11 |
| chr6 | 84115548 | 84119846 | 4299 | -7.65 | 4.65E-12 | 1.45E-11 |
| chr4 | 75194832 | 75211158 | 16327 | -8 | 4.67E-12 | 1.45E-11 |
| chr7 | 82629418 | 82634407 | 4990 | -7.96 | 4.68E-12 | 1.46E-11 |
| chr10 | 98562688 | 98564371 | 1684 | -7.67 | 4.71E-12 | 1.46E-11 |
| chr1 | 8213345 | 8219969 | 6625 | -7.72 | 4.71E-12 | 1.46E-11 |
| chr9 | 3960329 | 4002515 | 42187 | -8.24 | 4.72E-12 | 1.47E-11 |
| chr5 | 80207292 | 80209846 | 2555 | -7.64 | 4.73E-12 | 1.47E-11 |
| chr6 | 142747777 | 142750253 | 2477 | -7.64 | 4.73E-12 | 1.47E-11 |
| chr11 | 82483641 | 82493746 | 10106 | -7.66 | 4.73E-12 | 1.47E-11 |
| chr14 | 100244161 | 100246880 | 2720 | -7.86 | 4.73E-12 | 1.47E-11 |
| chr22 | 46564901 | 46577304 | 12404 | -7.67 | 4.75E-12 | 1.48E-11 |
| chr5 | 132346865 | 132353961 | 7097 | -7.79 | 4.75E-12 | 1.48E-11 |
| chr4 | 183773567 | 183775929 | 2363 | -7.68 | 4.75E-12 | 1.48E-11 |
| chr11 | 33593059 | 33595389 | 2331 | -7.71 | 4.77E-12 | 1.48E-11 |
| chr1 | 66494814 | 66497848 | 3035 | -7.64 | 4.78E-12 | 1.49E-11 |
| chr12 | 10732380 | 10735262 | 2883 | -7.7 | 4.78E-12 | 1.49E-11 |
| chr1 | 240833806 | 240837990 | 4185 | -7.8 | 4.79E-12 | 1.49E-11 |
| chr18 | 43443178 | 43456059 | 12882 | -7.74 | 4.82E-12 | 1.50E-11 |
| chr12 | 107047924 | 107062674 | 14751 | -1.86 | 4.82E-12 | 1.50E-11 |

| | | | | | | |
|---|---|---|---|---|---|---|
| chr5 | 73393966 | 73396615 | 2650 | -7.84 | 4.83E-12 | 1.50E-11 |
| chr9 | 110418015 | 110421137 | 3123 | -7.76 | 4.83E-12 | 1.50E-11 |
| chr19 | 43109804 | 43224884 | 115081 | -2.12 | 4.84E-12 | 1.50E-11 |
| chr6 | 87965374 | 87970511 | 5138 | -7.64 | 4.84E-12 | 1.50E-11 |
| chr9 | 93814037 | 93831980 | 17944 | -8.05 | 4.84E-12 | 1.50E-11 |
| chr12 | 18274066 | 18280400 | 6335 | -7.77 | 4.85E-12 | 1.51E-11 |
| chr22 | 41102024 | 41104709 | 2686 | -7.65 | 4.85E-12 | 1.51E-11 |
| chr4 | 89563628 | 89565964 | 2337 | -7.65 | 4.86E-12 | 1.51E-11 |
| chr12 | 80347469 | 80349897 | 2429 | -7.64 | 4.86E-12 | 1.51E-11 |
| chr3 | 8489983 | 8492223 | 2241 | -7.64 | 4.86E-12 | 1.51E-11 |
| chrX | 36907993 | 36924401 | 16409 | -7.66 | 4.87E-12 | 1.51E-11 |
| chr6 | 40846494 | 40847866 | 1373 | -6.58 | 4.88E-12 | 1.51E-11 |
| chr9 | 12835461 | 12848751 | 13291 | -8.03 | 4.88E-12 | 1.52E-11 |
| chr4 | 166811690 | 166818089 | 6400 | -7.74 | 4.88E-12 | 1.52E-11 |
| chr12 | 76397560 | 76399472 | 1913 | -7.64 | 4.89E-12 | 1.52E-11 |
| chr2 | 17986223 | 17995083 | 8861 | 7.81 | 4.89E-12 | 1.52E-11 |
| chr12 | 10145408 | 10148614 | 3207 | 7.73 | 4.90E-12 | 1.52E-11 |
| chr11 | 4190541 | 4198641 | 8101 | -7.67 | 4.91E-12 | 1.52E-11 |
| chr12 | 50003457 | 50010415 | 6959 | -7.67 | 4.91E-12 | 1.52E-11 |
| chr4 | 93350836 | 93352761 | 1926 | -6.05 | 4.91E-12 | 1.52E-11 |
| chrX | 117268486 | 117272835 | 4350 | -7.64 | 4.91E-12 | 1.52E-11 |
| chr1 | 217637313 | 217639711 | 2399 | -7.66 | 4.93E-12 | 1.53E-11 |
| chr13 | 112221064 | 112234156 | 13093 | -2.56 | 4.93E-12 | 1.53E-11 |
| chr3 | 72723674 | 72732162 | 8489 | -7.64 | 4.94E-12 | 1.53E-11 |
| chr9 | 81865055 | 81879117 | 14063 | -8.16 | 4.94E-12 | 1.53E-11 |
| chr7 | 29348509 | 29351326 | 2818 | -2.59 | 4.94E-12 | 1.53E-11 |
| chr15 | 52436580 | 52438888 | 2309 | -7.71 | 4.94E-12 | 1.53E-11 |
| chr3 | 17258956 | 17264219 | 5264 | -7.82 | 4.95E-12 | 1.54E-11 |
| chr9 | 112462307 | 112463740 | 1434 | -3.65 | 4.96E-12 | 1.54E-11 |
| chr5 | 129319769 | 129323363 | 3595 | -2.56 | 4.96E-12 | 1.54E-11 |
| chr15 | 41547187 | 41555774 | 8588 | -7.64 | 4.96E-12 | 1.54E-11 |
| chr17 | 67332471 | 67334594 | 2124 | -7.64 | 4.96E-12 | 1.54E-11 |
| chr2 | 201296677 | 201302515 | 5839 | -7.89 | 4.97E-12 | 1.54E-11 |
| chr10 | 82235763 | 82238818 | 3056 | -7.65 | 4.98E-12 | 1.54E-11 |
| chr8 | 48632413 | 48635729 | 3317 | -7.68 | 4.98E-12 | 1.54E-11 |
| chr3 | 13711438 | 13720591 | 9154 | -7.74 | 4.99E-12 | 1.55E-11 |
| chr3 | 159403779 | 159411779 | 8001 | -7.64 | 5.00E-12 | 1.55E-11 |
| chr16 | 53034177 | 53049892 | 15716 | -7.72 | 5.01E-12 | 1.55E-11 |
| chrX | 16543070 | 16555301 | 12232 | -7.99 | 5.01E-12 | 1.55E-11 |
| chr7 | 102492500 | 102526851 | 34352 | -8.2 | 5.01E-12 | 1.55E-11 |
| chr1 | 115817694 | 115821518 | 3825 | -7.65 | 5.01E-12 | 1.55E-11 |
| chr14 | 64778730 | 64782012 | 3283 | -7.63 | 5.01E-12 | 1.55E-11 |
| chr6 | 109497434 | 109515221 | 17788 | -7.84 | 5.02E-12 | 1.55E-11 |
| chr5 | 168154806 | 168158147 | 3342 | -7.64 | 5.02E-12 | 1.55E-11 |
| chr7 | 139291852 | 139297455 | 5604 | -8.11 | 5.02E-12 | 1.55E-11 |
| chr7 | 99277874 | 99280775 | 2902 | -7.71 | 5.03E-12 | 1.56E-11 |
| chr9 | 126878393 | 126885011 | 6619 | -7.65 | 5.03E-12 | 1.56E-11 |
| chr9 | 85670258 | 85685369 | 15112 | -7.8 | 5.06E-12 | 1.57E-11 |
| chr14 | 55570562 | 55573662 | 3101 | -7.7 | 5.08E-12 | 1.57E-11 |
| chr18 | 9744697 | 9747723 | 3027 | -7.82 | 5.09E-12 | 1.57E-11 |
| chr3 | 70947745 | 70951524 | 3780 | -3.31 | 5.10E-12 | 1.58E-11 |
| chr10 | 33930237 | 33953965 | 23729 | -8.53 | 5.11E-12 | 1.58E-11 |
| chr3 | 33588594 | 33591213 | 2620 | -7.74 | 5.11E-12 | 1.58E-11 |
| chr2 | 216165589 | 216167994 | 2406 | -7.63 | 5.13E-12 | 1.59E-11 |
| chr2 | 189218022 | 189221957 | 3936 | -7.82 | 5.14E-12 | 1.59E-11 |
| chr8 | 40853991 | 40856843 | 2853 | -7.94 | 5.14E-12 | 1.59E-11 |
| chr4 | 150977457 | 150982122 | 4666 | -7.69 | 5.15E-12 | 1.59E-11 |
| chr13 | 43951867 | 43955493 | 3627 | -7.77 | 5.16E-12 | 1.60E-11 |
| chr5 | 129559913 | 129561978 | 2066 | -7.64 | 5.17E-12 | 1.60E-11 |
| chr7 | 94652717 | 94657089 | 4373 | -7.72 | 5.18E-12 | 1.60E-11 |
| chr4 | 169319513 | 169322067 | 2555 | -7.66 | 5.19E-12 | 1.60E-11 |

| | | | | | | |
|---|---|---|---|---|---|---|
| chr8 | 10700036 | 10790596 | 90561 | -1.64 | 5.20E-12 | 1.61E-11 |
| chr17 | 16949157 | 16950905 | 1749 | -7.67 | 5.20E-12 | 1.61E-11 |
| chr10 | 126619420 | 126621509 | 2090 | -7.63 | 5.21E-12 | 1.61E-11 |
| chr1 | 45348794 | 45356065 | 7272 | -7.76 | 5.21E-12 | 1.61E-11 |
| chr4 | 40168443 | 40170944 | 2502 | -7.64 | 5.21E-12 | 1.61E-11 |
| chr2 | 67565064 | 67578120 | 13057 | -7.65 | 5.23E-12 | 1.62E-11 |
| chr15 | 71696138 | 71697907 | 1770 | -7.86 | 5.23E-12 | 1.62E-11 |
| chr19 | 3299118 | 3308766 | 9649 | -7.77 | 5.24E-12 | 1.62E-11 |
| chrX | 4502197 | 4521471 | 19275 | -7.91 | 5.25E-12 | 1.62E-11 |
| chr12 | 10405852 | 10409722 | 3871 | -7.79 | 5.25E-12 | 1.62E-11 |
| chr10 | 14825489 | 14827977 | 2489 | -7.77 | 5.27E-12 | 1.63E-11 |
| chr15 | 75033923 | 75036127 | 2205 | -7.65 | 5.27E-12 | 1.63E-11 |
| chr4 | 180727553 | 180741511 | 13959 | -7.65 | 5.27E-12 | 1.63E-11 |
| chr10 | 103016896 | 103019087 | 2192 | -7.63 | 5.28E-12 | 1.63E-11 |
| chr12 | 56271185 | 56273138 | 1954 | -7.64 | 5.28E-12 | 1.63E-11 |
| chr17 | 72572457 | 72593988 | 21532 | -8.07 | 5.29E-12 | 1.63E-11 |
| chr10 | 89671498 | 89673939 | 2442 | -7.84 | 5.31E-12 | 1.64E-11 |
| chr2 | 39423251 | 39428049 | 4799 | -7.87 | 5.32E-12 | 1.64E-11 |
| chr2 | 62178305 | 62188725 | 10421 | -7.89 | 5.33E-12 | 1.65E-11 |
| chr8 | 76306948 | 76319409 | 12462 | -7.96 | 5.33E-12 | 1.65E-11 |
| chr11 | 71953945 | 71966416 | 12472 | -8.01 | 5.34E-12 | 1.65E-11 |
| chr9 | 94126552 | 94140605 | 14054 | -7.68 | 5.34E-12 | 1.65E-11 |
| chr22 | 50640990 | 50642534 | 1545 | -7.66 | 5.35E-12 | 1.65E-11 |
| chr17 | 36767673 | 36769089 | 1417 | -7.66 | 5.38E-12 | 1.66E-11 |
| chr22 | 23011619 | 23014333 | 2715 | -7.71 | 5.38E-12 | 1.66E-11 |
| chr12 | 32575361 | 32583541 | 8181 | -7.75 | 5.39E-12 | 1.66E-11 |
| chr12 | 9832088 | 9835603 | 3516 | -7.7 | 5.40E-12 | 1.67E-11 |
| chr4 | 5010720 | 5031487 | 20768 | -7.98 | 5.40E-12 | 1.67E-11 |
| chr19 | 6794258 | 6821039 | 26782 | -7.83 | 5.41E-12 | 1.67E-11 |
| chrX | 107204555 | 107212576 | 8022 | -7.76 | 5.41E-12 | 1.67E-11 |
| chr2 | 40703024 | 40719248 | 16225 | -7.7 | 5.42E-12 | 1.67E-11 |
| chr2 | 181332284 | 181335003 | 2720 | -7.7 | 5.42E-12 | 1.67E-11 |
| chr1 | 90110402 | 90113146 | 2745 | -7.62 | 5.42E-12 | 1.67E-11 |
| chr14 | 66120428 | 66124289 | 3862 | -7.63 | 5.44E-12 | 1.68E-11 |
| chr3 | 37969488 | 37971509 | 2022 | -7.69 | 5.45E-12 | 1.68E-11 |
| chr11 | 77500969 | 77504625 | 3657 | -7.65 | 5.45E-12 | 1.68E-11 |
| chr11 | 30331664 | 30344347 | 12684 | -7.77 | 5.46E-12 | 1.68E-11 |
| chr2 | 75241475 | 75243141 | 1667 | -7.63 | 5.47E-12 | 1.69E-11 |
| chr14 | 36668291 | 36673820 | 5530 | -2.32 | 5.47E-12 | 1.69E-11 |
| chr5 | 30383283 | 30406030 | 22748 | -8.4 | 5.48E-12 | 1.69E-11 |
| chr5 | 52487337 | 52489409 | 2073 | -7.63 | 5.48E-12 | 1.69E-11 |
| chr3 | 112818062 | 112821568 | 3507 | -7.68 | 5.48E-12 | 1.69E-11 |
| chr14 | 76022822 | 76025213 | 2392 | -5.52 | 5.48E-12 | 1.69E-11 |
| chr13 | 33228682 | 33230989 | 2308 | -7.63 | 5.49E-12 | 1.69E-11 |
| chr5 | 90400728 | 90402953 | 2226 | -7.65 | 5.49E-12 | 1.69E-11 |
| chr18 | 19313140 | 19318870 | 5731 | -7.64 | 5.50E-12 | 1.70E-11 |
| chr12 | 17837941 | 17842639 | 4699 | -7.78 | 5.54E-12 | 1.71E-11 |
| chr2 | 152088144 | 152090707 | 2564 | -7.73 | 5.56E-12 | 1.71E-11 |
| chr13 | 73843568 | 73852628 | 9061 | -7.65 | 5.56E-12 | 1.71E-11 |
| chr2 | 152750389 | 152752563 | 2175 | -7.62 | 5.56E-12 | 1.71E-11 |
| chr11 | 123295199 | 123300317 | 5119 | -7.71 | 5.56E-12 | 1.71E-11 |
| chr2 | 98343176 | 98360300 | 17125 | -7.73 | 5.56E-12 | 1.71E-11 |
| chr10 | 96184064 | 96198303 | 14240 | -7.74 | 5.58E-12 | 1.72E-11 |
| chr3 | 116583402 | 116594934 | 11533 | -7.96 | 5.58E-12 | 1.72E-11 |
| chr1 | 65027252 | 65029843 | 2592 | -7.67 | 5.60E-12 | 1.72E-11 |
| chr4 | 95092102 | 95094358 | 2257 | -6.55 | 5.60E-12 | 1.72E-11 |
| chr16 | 67732357 | 67734137 | 1781 | -7.63 | 5.60E-12 | 1.72E-11 |
| chr10 | 54125860 | 54136189 | 10330 | -7.82 | 5.60E-12 | 1.72E-11 |
| chr11 | 33614876 | 33619082 | 4207 | -7.69 | 5.62E-12 | 1.73E-11 |
| chr21 | 47038226 | 47040829 | 2604 | -7.62 | 5.62E-12 | 1.73E-11 |
| chr8 | 107682564 | 107687462 | 4899 | -7.76 | 5.64E-12 | 1.74E-11 |

| | | | | | | |
|---|---|---|---|---|---|---|
| chrX | 140247358 | 140260446 | 13089 | -8.02 | 5.68E-12 | 1.75E-11 |
| chr18 | 1348640 | 1358068 | 9429 | 7.8 | 5.68E-12 | 1.75E-11 |
| chr13 | 35384534 | 35388815 | 4282 | -5.94 | 5.71E-12 | 1.76E-11 |
| chr6 | 47257790 | 47260384 | 2595 | -7.62 | 5.71E-12 | 1.76E-11 |
| chr11 | 126066723 | 126068860 | 2138 | -7.62 | 5.72E-12 | 1.76E-11 |
| chr11 | 108556119 | 108558274 | 2156 | -7.62 | 5.72E-12 | 1.76E-11 |
| chr2 | 121214430 | 121230251 | 15822 | -7.84 | 5.74E-12 | 1.77E-11 |
| chr7 | 114844388 | 114847756 | 3369 | -7.62 | 5.75E-12 | 1.77E-11 |
| chr8 | 74736548 | 74738545 | 1998 | -7.64 | 5.75E-12 | 1.77E-11 |
| chr2 | 178421835 | 178423436 | 1602 | -6 | 5.76E-12 | 1.77E-11 |
| chr5 | 95337844 | 95374441 | 36598 | -8.15 | 5.76E-12 | 1.77E-11 |
| chr2 | 54464158 | 54465629 | 1472 | -7.68 | 5.76E-12 | 1.77E-11 |
| chr3 | 108044571 | 108047121 | 2551 | -5.66 | 5.76E-12 | 1.77E-11 |
| chr1 | 219847165 | 219963905 | 116741 | 1.21 | 5.76E-12 | 1.77E-11 |
| chr3 | 23263777 | 23266724 | 2948 | -7.73 | 5.77E-12 | 1.77E-11 |
| chr12 | 100694279 | 100698566 | 4288 | -7.62 | 5.77E-12 | 1.77E-11 |
| chr2 | 206880089 | 206882304 | 2216 | -7.62 | 5.77E-12 | 1.77E-11 |
| chr12 | 104536782 | 104540124 | 3343 | -7.74 | 5.80E-12 | 1.78E-11 |
| chr10 | 37276005 | 37285336 | 9332 | -7.82 | 5.81E-12 | 1.78E-11 |
| chr22 | 26160953 | 26163894 | 2942 | -7.69 | 5.81E-12 | 1.79E-11 |
| chr7 | 130859094 | 130873370 | 14277 | -7.96 | 5.81E-12 | 1.79E-11 |
| chr6 | 64118567 | 64133833 | 15267 | -7.65 | 5.82E-12 | 1.79E-11 |
| chr7 | 140878367 | 140893577 | 15211 | 7.91 | 5.83E-12 | 1.79E-11 |
| chr4 | 142686339 | 142705133 | 18795 | -7.62 | 5.85E-12 | 1.80E-11 |
| chr20 | 34417731 | 34431674 | 13944 | -8.07 | 5.86E-12 | 1.80E-11 |
| chr4 | 108768521 | 108771490 | 2970 | -7.62 | 5.86E-12 | 1.80E-11 |
| chr20 | 53007949 | 53009598 | 1650 | -5.88 | 5.86E-12 | 1.80E-11 |
| chr6 | 132715097 | 132717633 | 2537 | -7.73 | 5.86E-12 | 1.80E-11 |
| chr5 | 129590592 | 129604597 | 14006 | -7.64 | 5.86E-12 | 1.80E-11 |
| chr18 | 77883774 | 77885609 | 1836 | -7.74 | 5.87E-12 | 1.80E-11 |
| chrX | 104142026 | 104147260 | 5235 | 7.74 | 5.89E-12 | 1.81E-11 |
| chr2 | 72664080 | 72673543 | 9464 | -7.76 | 5.89E-12 | 1.81E-11 |
| chr5 | 171551071 | 171565024 | 13954 | -7.8 | 5.90E-12 | 1.81E-11 |
| chr3 | 155873192 | 155874412 | 1221 | -4.08 | 5.90E-12 | 1.81E-11 |
| chr13 | 100898544 | 100901619 | 3076 | -7.61 | 5.91E-12 | 1.81E-11 |
| chr2 | 61312213 | 61313393 | 1181 | -7.68 | 5.93E-12 | 1.82E-11 |
| chrX | 3561234 | 3563120 | 1887 | -7.64 | 5.94E-12 | 1.82E-11 |
| chr12 | 26155141 | 26157335 | 2195 | -7.65 | 5.94E-12 | 1.82E-11 |
| chr1 | 223450973 | 223459430 | 8458 | -6.63 | 5.95E-12 | 1.82E-11 |
| chr7 | 14222464 | 14239241 | 16778 | -7.77 | 5.95E-12 | 1.82E-11 |
| chr7 | 20333404 | 20336001 | 2598 | -7.62 | 5.96E-12 | 1.83E-11 |
| chr14 | 56447269 | 56478183 | 30915 | -3.21 | 5.97E-12 | 1.83E-11 |
| chr19 | 3250538 | 3255278 | 4741 | -7.81 | 5.97E-12 | 1.83E-11 |
| chr18 | 32783799 | 32794846 | 11048 | -7.72 | 5.97E-12 | 1.83E-11 |
| chr12 | 51308432 | 51311984 | 3553 | -7.63 | 5.99E-12 | 1.84E-11 |
| chrX | 77211332 | 77215248 | 3917 | -7.65 | 6.00E-12 | 1.84E-11 |
| chr4 | 115102279 | 115117529 | 15251 | -7.74 | 6.00E-12 | 1.84E-11 |
| chr12 | 104788198 | 104801349 | 13152 | -7.61 | 6.00E-12 | 1.84E-11 |
| chrX | 12099934 | 12108531 | 8598 | -7.74 | 6.01E-12 | 1.84E-11 |
| chr1 | 56061907 | 56075087 | 13181 | -7.79 | 6.02E-12 | 1.84E-11 |
| chr1 | 89265752 | 89267927 | 2176 | -7.61 | 6.02E-12 | 1.85E-11 |
| chr6 | 133836970 | 133839999 | 3030 | -7.74 | 6.03E-12 | 1.85E-11 |
| chr15 | 49942931 | 49948006 | 5076 | -5.47 | 6.03E-12 | 1.85E-11 |
| chr1 | 199358987 | 199363438 | 4452 | -2.9 | 6.03E-12 | 1.85E-11 |
| chr10 | 4580507 | 4585160 | 4654 | -7.85 | 6.06E-12 | 1.85E-11 |
| chr1 | 214782577 | 214784857 | 2281 | -7.61 | 6.08E-12 | 1.86E-11 |
| chr3 | 87147253 | 87171819 | 24567 | -7.62 | 6.12E-12 | 1.88E-11 |
| chr1 | 237064756 | 237094795 | 30040 | -4.11 | 6.13E-12 | 1.88E-11 |
| chr4 | 134373078 | 134383246 | 10169 | 7.69 | 6.13E-12 | 1.88E-11 |
| chr2 | 25738521 | 25740529 | 2009 | -7.7 | 6.13E-12 | 1.88E-11 |
| chr15 | 49999668 | 50003185 | 3518 | -7.61 | 6.14E-12 | 1.88E-11 |

| | | | | | | |
|---|---|---|---|---|---|---|
| chr6 | 118808657 | 118811742 | 3086 | -7.63 | 6.14E-12 | 1.88E-11 |
| chr8 | 83170179 | 83192915 | 22737 | -8 | 6.14E-12 | 1.88E-11 |
| chr9 | 136876001 | 136881561 | 5561 | -7.61 | 6.15E-12 | 1.88E-11 |
| chr13 | 44520949 | 44523326 | 2378 | -7.69 | 6.16E-12 | 1.88E-11 |
| chr2 | 38329455 | 38332855 | 3401 | -7.77 | 6.18E-12 | 1.89E-11 |
| chr14 | 51724766 | 51727276 | 2511 | -7.68 | 6.19E-12 | 1.89E-11 |
| chr3 | 114876633 | 114878982 | 2350 | -6.11 | 6.20E-12 | 1.90E-11 |
| chr9 | 114751649 | 114754475 | 2827 | -7.72 | 6.20E-12 | 1.90E-11 |
| chr7 | 19729011 | 19743708 | 14698 | -7.62 | 6.21E-12 | 1.90E-11 |
| chr13 | 76472602 | 76475625 | 3024 | -7.64 | 6.22E-12 | 1.90E-11 |
| chr19 | 44042445 | 44043924 | 1480 | -7.63 | 6.23E-12 | 1.91E-11 |
| chr9 | 85934241 | 85955911 | 21671 | -7.99 | 6.23E-12 | 1.91E-11 |
| chr18 | 21345443 | 21348916 | 3474 | -7.81 | 6.24E-12 | 1.91E-11 |
| chr19 | 15797256 | 15836333 | 39078 | -7.09 | 6.24E-12 | 1.91E-11 |
| chr19 | 42628394 | 42629452 | 1059 | -7.66 | 6.24E-12 | 1.91E-11 |
| chr8 | 115266744 | 115286689 | 19946 | -7.78 | 6.26E-12 | 1.91E-11 |
| chr6 | 108468902 | 108470690 | 1789 | -7.61 | 6.27E-12 | 1.92E-11 |
| chr13 | 67224189 | 67229158 | 4970 | -7.64 | 6.30E-12 | 1.92E-11 |
| chr15 | 24570253 | 24624375 | 54123 | 1.7 | 6.30E-12 | 1.92E-11 |
| chr11 | 111425513 | 111428052 | 2540 | -7.6 | 6.33E-12 | 1.93E-11 |
| chr4 | 99570669 | 99573661 | 2993 | -7.64 | 6.33E-12 | 1.93E-11 |
| chr12 | 86821514 | 86826153 | 4640 | -7.85 | 6.33E-12 | 1.93E-11 |
| chr11 | 94393129 | 94396990 | 3862 | -7.61 | 6.35E-12 | 1.94E-11 |
| chr11 | 109819258 | 109822499 | 3242 | -7.6 | 6.36E-12 | 1.94E-11 |
| chr6 | 105640849 | 105645781 | 4933 | -7.62 | 6.39E-12 | 1.95E-11 |
| chr10 | 12252441 | 12254529 | 2089 | -7.71 | 6.40E-12 | 1.96E-11 |
| chr3 | 40609153 | 40612107 | 2955 | -7.65 | 6.40E-12 | 1.96E-11 |
| chr15 | 68954453 | 68964053 | 9601 | -7.66 | 6.40E-12 | 1.96E-11 |
| chr6 | 90996307 | 90998404 | 2098 | -7.61 | 6.40E-12 | 1.96E-11 |
| chr3 | 104968290 | 104981923 | 13634 | -7.64 | 6.41E-12 | 1.96E-11 |
| chr6 | 142316927 | 142331408 | 14482 | -7.69 | 6.43E-12 | 1.96E-11 |
| chr12 | 63053580 | 63055359 | 1780 | -7.61 | 6.43E-12 | 1.96E-11 |
| chr6 | 17671488 | 17674599 | 3112 | -7.68 | 6.43E-12 | 1.96E-11 |
| chr10 | 111920342 | 111921770 | 1429 | -7.73 | 6.44E-12 | 1.96E-11 |
| chrX | 44027715 | 44036021 | 8307 | -7.8 | 6.44E-12 | 1.97E-11 |
| chr8 | 145177772 | 145187649 | 9878 | -7.96 | 6.44E-12 | 1.97E-11 |
| chr7 | 150399666 | 150403402 | 3737 | -2.44 | 6.46E-12 | 1.97E-11 |
| chr7 | 45709506 | 45718063 | 8558 | -2.09 | 6.46E-12 | 1.97E-11 |
| chr6 | 21929872 | 21936051 | 6180 | -7.78 | 6.47E-12 | 1.97E-11 |
| chr1 | 215313778 | 215317047 | 3270 | -7.68 | 6.47E-12 | 1.97E-11 |
| chr10 | 79038642 | 79040795 | 2154 | -7.61 | 6.48E-12 | 1.98E-11 |
| chr21 | 32635861 | 32646792 | 10932 | -7.74 | 6.49E-12 | 1.98E-11 |
| chr14 | 61930799 | 61932684 | 1886 | -7.74 | 6.49E-12 | 1.98E-11 |
| chr20 | 11457606 | 11460290 | 2685 | -7.63 | 6.50E-12 | 1.98E-11 |
| chr9 | 107013830 | 107021082 | 7253 | -8.04 | 6.50E-12 | 1.98E-11 |
| chr7 | 36359605 | 36386411 | 26807 | -7.84 | 6.51E-12 | 1.98E-11 |
| chr2 | 38728652 | 38730179 | 1528 | -7.65 | 6.51E-12 | 1.98E-11 |
| chr8 | 133317254 | 133321630 | 4377 | -4.15 | 6.51E-12 | 1.98E-11 |
| chr4 | 79724873 | 79729059 | 4187 | -7.64 | 6.51E-12 | 1.99E-11 |
| chr20 | 36834160 | 36836281 | 2122 | -7.61 | 6.53E-12 | 1.99E-11 |
| chr17 | 38041584 | 38048509 | 6926 | -7.84 | 6.53E-12 | 1.99E-11 |
| chr1 | 12497584 | 12504967 | 7384 | -7.62 | 6.55E-12 | 2.00E-11 |
| chr16 | 19572060 | 19576007 | 3948 | -7.6 | 6.57E-12 | 2.00E-11 |
| chr15 | 36513797 | 36519213 | 5417 | -7.63 | 6.58E-12 | 2.01E-11 |
| chr20 | 13624881 | 13641311 | 16431 | -8.08 | 6.59E-12 | 2.01E-11 |
| chr14 | 71653409 | 71688204 | 34796 | -8.67 | 6.61E-12 | 2.01E-11 |
| chr14 | 95653433 | 95656028 | 2596 | -7.64 | 6.61E-12 | 2.01E-11 |
| chr8 | 18568560 | 18572588 | 4029 | -7.62 | 6.61E-12 | 2.02E-11 |
| chr21 | 36268623 | 36270843 | 2221 | -7.6 | 6.62E-12 | 2.02E-11 |
| chr16 | 14065652 | 14075442 | 9791 | -7.6 | 6.64E-12 | 2.02E-11 |
| chr3 | 32710811 | 32716003 | 5193 | -7.61 | 6.66E-12 | 2.03E-11 |

| | | | | | | |
|---|---|---|---|---|---|---|
| chr6 | 131836091 | 131840128 | 4038 | -7.69 | 6.67E-12 | 2.03E-11 |
| chr1 | 66522366 | 66525945 | 3580 | -3.29 | 6.68E-12 | 2.03E-11 |
| chr1 | 54242605 | 54244677 | 2073 | -7.62 | 6.68E-12 | 2.03E-11 |
| chr4 | 39202955 | 39205106 | 2152 | -7.62 | 6.68E-12 | 2.03E-11 |
| chr2 | 83784388 | 83795256 | 10869 | -7.72 | 6.69E-12 | 2.04E-11 |
| chr9 | 130610464 | 130613068 | 2605 | -7.77 | 6.71E-12 | 2.04E-11 |
| chr2 | 112733077 | 112735925 | 2849 | -7.6 | 6.72E-12 | 2.04E-11 |
| chr12 | 107186036 | 107188598 | 2563 | -7.72 | 6.72E-12 | 2.05E-11 |
| chr15 | 36082479 | 36086826 | 4348 | -7.61 | 6.74E-12 | 2.05E-11 |
| chr4 | 38959847 | 38967322 | 7476 | -7.74 | 6.76E-12 | 2.06E-11 |
| chr12 | 110188593 | 110201773 | 13181 | -7.73 | 6.78E-12 | 2.06E-11 |
| chr13 | 99584693 | 99589002 | 4310 | -7.63 | 6.78E-12 | 2.06E-11 |
| chr15 | 39902240 | 39912594 | 10355 | -8 | 6.81E-12 | 2.07E-11 |
| chr3 | 42116895 | 42120637 | 3743 | -7.64 | 6.82E-12 | 2.08E-11 |
| chr7 | 131929545 | 131933488 | 3944 | -7.6 | 6.85E-12 | 2.08E-11 |
| chr21 | 28099191 | 28133617 | 34427 | 1.83 | 6.85E-12 | 2.08E-11 |
| chr3 | 100982868 | 100984658 | 1791 | -7.63 | 6.87E-12 | 2.09E-11 |
| chr12 | 64467247 | 64469759 | 2513 | -7.59 | 6.87E-12 | 2.09E-11 |
| chr9 | 90898005 | 90901026 | 3022 | -7.62 | 6.88E-12 | 2.09E-11 |
| chr14 | 24923054 | 24926641 | 3588 | -7.6 | 6.88E-12 | 2.09E-11 |
| chr10 | 96616780 | 96630141 | 13362 | -7.76 | 6.89E-12 | 2.10E-11 |
| chr5 | 169174642 | 169177834 | 3193 | -7.71 | 6.92E-12 | 2.10E-11 |
| chr1 | 146862674 | 146955788 | 93115 | -1.66 | 6.93E-12 | 2.11E-11 |
| chr22 | 27121086 | 27124065 | 2980 | -7.65 | 6.93E-12 | 2.11E-11 |
| chr9 | 91643975 | 91664872 | 20898 | -2.11 | 6.95E-12 | 2.11E-11 |
| chr2 | 192000081 | 192005273 | 5193 | -7.72 | 6.95E-12 | 2.11E-11 |
| chr4 | 79145890 | 79149141 | 3252 | -7.64 | 6.97E-12 | 2.12E-11 |
| chr10 | 97457231 | 97466553 | 9323 | -7.67 | 6.97E-12 | 2.12E-11 |
| chr5 | 74512838 | 74517220 | 4383 | -7.59 | 6.97E-12 | 2.12E-11 |
| chr6 | 56888591 | 56890992 | 2402 | -7.63 | 6.98E-12 | 2.12E-11 |
| chr15 | 71127291 | 71128604 | 1314 | -7.59 | 6.99E-12 | 2.12E-11 |
| chr10 | 24762848 | 24769481 | 6634 | -7.62 | 7.00E-12 | 2.13E-11 |
| chr7 | 121216460 | 121224631 | 8172 | 8.05 | 7.00E-12 | 2.13E-11 |
| chr8 | 123743961 | 123746025 | 2065 | -7.61 | 7.00E-12 | 2.13E-11 |
| chr13 | 98251984 | 98255339 | 3356 | -7.62 | 7.00E-12 | 2.13E-11 |
| chr5 | 52894667 | 52897141 | 2475 | -7.59 | 7.00E-12 | 2.13E-11 |
| chr10 | 609532 | 616979 | 7448 | -7.66 | 7.01E-12 | 2.13E-11 |
| chr1 | 65701397 | 65704056 | 2660 | -7.61 | 7.01E-12 | 2.13E-11 |
| chr5 | 1302336 | 1311304 | 8969 | -8.02 | 7.03E-12 | 2.14E-11 |
| chr3 | 177174306 | 177176791 | 2486 | -7.59 | 7.04E-12 | 2.14E-11 |
| chr10 | 6650427 | 6657496 | 7070 | -1.88 | 7.07E-12 | 2.15E-11 |
| chr12 | 75076950 | 75079897 | 2948 | -7.63 | 7.08E-12 | 2.15E-11 |
| chr7 | 47289299 | 47292379 | 3081 | -7.59 | 7.08E-12 | 2.15E-11 |
| chr18 | 59765206 | 59767935 | 2730 | -7.65 | 7.09E-12 | 2.15E-11 |
| chr2 | 226573826 | 226576746 | 2921 | -5.71 | 7.12E-12 | 2.16E-11 |
| chr18 | 34621424 | 34624702 | 3279 | -7.59 | 7.13E-12 | 2.16E-11 |
| chr11 | 125741279 | 125756464 | 15186 | -7.6 | 7.14E-12 | 2.17E-11 |
| chr4 | 32290392 | 32292839 | 2448 | -6.01 | 7.14E-12 | 2.17E-11 |
| chr2 | 191726014 | 191734031 | 8018 | -7.82 | 7.15E-12 | 2.17E-11 |
| chr10 | 3219822 | 3225087 | 5266 | -7.59 | 7.16E-12 | 2.17E-11 |
| chr5 | 175846892 | 175849358 | 2467 | -7.64 | 7.20E-12 | 2.18E-11 |
| chr8 | 66995750 | 67012339 | 16590 | -8.12 | 7.20E-12 | 2.18E-11 |
| chr1 | 174054493 | 174057490 | 2998 | -7.75 | 7.21E-12 | 2.19E-11 |
| chr20 | 2298683 | 2306193 | 7511 | -8.08 | 7.22E-12 | 2.19E-11 |
| chr1 | 92670599 | 92673057 | 2459 | -7.6 | 7.23E-12 | 2.19E-11 |
| chr21 | 39741346 | 39745068 | 3723 | -7.75 | 7.26E-12 | 2.20E-11 |
| chr10 | 89391872 | 89407334 | 15463 | -7.89 | 7.27E-12 | 2.20E-11 |
| chr4 | 38390183 | 38402242 | 12060 | -8.18 | 7.27E-12 | 2.21E-11 |
| chr10 | 116682573 | 116692974 | 10402 | -7.59 | 7.28E-12 | 2.21E-11 |
| chr20 | 18436211 | 18438167 | 1957 | -7.58 | 7.28E-12 | 2.21E-11 |
| chr6 | 49095833 | 49100103 | 4271 | -7.64 | 7.28E-12 | 2.21E-11 |

| | | | | | | |
|---|---|---|---|---|---|---|
| chrX | 139999181 | 140007831 | 8651 | -7.77 | 7.31E-12 | 2.21E-11 |
| chr3 | 24334777 | 24337352 | 2576 | -7.59 | 7.32E-12 | 2.22E-11 |
| chr1 | 87140336 | 87144761 | 4426 | -7.59 | 7.32E-12 | 2.22E-11 |
| chr7 | 29259138 | 29285076 | 25939 | -7.77 | 7.33E-12 | 2.22E-11 |
| chr6 | 13085733 | 13087960 | 2228 | -7.59 | 7.33E-12 | 2.22E-11 |
| chr17 | 60638653 | 60644759 | 6107 | -7.7 | 7.35E-12 | 2.23E-11 |
| chr20 | 1201929 | 1204866 | 2938 | -5.7 | 7.36E-12 | 2.23E-11 |
| chr2 | 174920414 | 174924456 | 4043 | -7.58 | 7.36E-12 | 2.23E-11 |
| chr15 | 66323597 | 66337912 | 14316 | -7.9 | 7.38E-12 | 2.24E-11 |
| chr4 | 159278363 | 159293743 | 15381 | -7.58 | 7.39E-12 | 2.24E-11 |
| chr2 | 164780084 | 164783215 | 3132 | -7.62 | 7.39E-12 | 2.24E-11 |
| chr9 | 632681 | 635092 | 2412 | -7.62 | 7.39E-12 | 2.24E-11 |
| chr1 | 207425739 | 207430620 | 4882 | -7.6 | 7.40E-12 | 2.24E-11 |
| chr1 | 94707351 | 94709493 | 2143 | -7.66 | 7.40E-12 | 2.24E-11 |
| chr22 | 44306467 | 44309704 | 3238 | -4.19 | 7.42E-12 | 2.25E-11 |
| chr3 | 32212188 | 32214480 | 2293 | -7.64 | 7.42E-12 | 2.25E-11 |
| chr12 | 88147953 | 88151911 | 3959 | -7.64 | 7.43E-12 | 2.25E-11 |
| chr1 | 2786920 | 2789363 | 2444 | -7.58 | 7.44E-12 | 2.25E-11 |
| chr12 | 63921400 | 63922779 | 1380 | -7.65 | 7.44E-12 | 2.25E-11 |
| chr19 | 36471087 | 36481021 | 9935 | -7.71 | 7.46E-12 | 2.26E-11 |
| chr3 | 149347673 | 149352249 | 4577 | -7.62 | 7.46E-12 | 2.26E-11 |
| chr7 | 140010993 | 140020392 | 9400 | -7.76 | 7.50E-12 | 2.27E-11 |
| chr3 | 13582089 | 13584735 | 2647 | -7.6 | 7.50E-12 | 2.27E-11 |
| chr10 | 43442455 | 43452936 | 10482 | -7.72 | 7.51E-12 | 2.27E-11 |
| chr4 | 90807668 | 90811355 | 3688 | -7.85 | 7.56E-12 | 2.29E-11 |
| chr5 | 30034656 | 30057874 | 23219 | -8.41 | 7.56E-12 | 2.29E-11 |
| chr5 | 142576276 | 142582270 | 5995 | -7.89 | 7.57E-12 | 2.29E-11 |
| chr6 | 111806961 | 111809082 | 2122 | -7.77 | 7.58E-12 | 2.29E-11 |
| chr2 | 101874394 | 101876015 | 1622 | -7.59 | 7.59E-12 | 2.29E-11 |
| chr6 | 35396091 | 35398496 | 2406 | -7.59 | 7.60E-12 | 2.30E-11 |
| chr8 | 11258875 | 11266095 | 7221 | -5.95 | 7.63E-12 | 2.31E-11 |
| chr1 | 188670076 | 188675653 | 5578 | -7.79 | 7.64E-12 | 2.31E-11 |
| chr7 | 130825047 | 130839279 | 14233 | -7.58 | 7.68E-12 | 2.32E-11 |
| chr10 | 73242327 | 73250666 | 8340 | -7.68 | 7.68E-12 | 2.32E-11 |
| chr11 | 64961251 | 64962977 | 1727 | -7.58 | 7.69E-12 | 2.32E-11 |
| chr14 | 105305948 | 105314373 | 8426 | -7.81 | 7.69E-12 | 2.33E-11 |
| chr4 | 14476081 | 14482488 | 6408 | -7.58 | 7.73E-12 | 2.34E-11 |
| chr17 | 5791623 | 5945575 | 153953 | -1.95 | 7.77E-12 | 2.35E-11 |
| chr20 | 4884633 | 4890007 | 5375 | -7.58 | 7.79E-12 | 2.35E-11 |
| chr6 | 117807433 | 117809448 | 2016 | -7.57 | 7.79E-12 | 2.35E-11 |
| chr10 | 114688821 | 114691840 | 3020 | -7.59 | 7.80E-12 | 2.36E-11 |
| chr19 | 17407903 | 17409864 | 1962 | -7.58 | 7.81E-12 | 2.36E-11 |
| chr1 | 242570806 | 242574836 | 4031 | -7.79 | 7.81E-12 | 2.36E-11 |
| chr6 | 114401423 | 114408344 | 6922 | -7.58 | 7.83E-12 | 2.37E-11 |
| chr4 | 189458867 | 189462549 | 3683 | -7.64 | 7.85E-12 | 2.37E-11 |
| chr8 | 49547599 | 49551971 | 4373 | -7.64 | 7.87E-12 | 2.38E-11 |
| chr8 | 6339469 | 6341197 | 1729 | -7.64 | 7.88E-12 | 2.38E-11 |
| chr1 | 35431513 | 35435998 | 4486 | -7.57 | 7.88E-12 | 2.38E-11 |
| chr4 | 38571414 | 38604701 | 33288 | -8.14 | 7.89E-12 | 2.38E-11 |
| chr9 | 112743952 | 112758156 | 14205 | -7.58 | 7.89E-12 | 2.38E-11 |
| chr8 | 67820829 | 67824841 | 4013 | -7.59 | 7.92E-12 | 2.39E-11 |
| chr11 | 49106563 | 49114390 | 7828 | 7.87 | 7.94E-12 | 2.40E-11 |
| chr2 | 236204454 | 236220686 | 16233 | -7.8 | 7.94E-12 | 2.40E-11 |
| chr6 | 23200804 | 23352596 | 151793 | 1.12 | 7.94E-12 | 2.40E-11 |
| chr5 | 137147007 | 137163355 | 16349 | -7.93 | 7.95E-12 | 2.40E-11 |
| chr7 | 68138082 | 68141520 | 3439 | -7.58 | 7.96E-12 | 2.40E-11 |
| chrX | 96985933 | 96994148 | 8216 | -7.58 | 7.96E-12 | 2.40E-11 |
| chr12 | 93014675 | 93017278 | 2604 | -7.59 | 7.96E-12 | 2.40E-11 |
| chr14 | 27452530 | 27471590 | 19061 | -8.41 | 7.96E-12 | 2.40E-11 |
| chr13 | 44778778 | 44782753 | 3976 | -7.67 | 7.99E-12 | 2.41E-11 |
| chr1 | 108217149 | 108228498 | 11350 | -7.59 | 8.01E-12 | 2.42E-11 |

| | | | | | | |
|---|---|---|---|---|---|---|
| chr1 | 151562663 | 151564801 | 2139 | -7.57 | 8.02E-12 | 2.42E-11 |
| chr19 | 7780552 | 7782628 | 2077 | -7.59 | 8.06E-12 | 2.43E-11 |
| chr5 | 115501112 | 115510546 | 9435 | -7.78 | 8.07E-12 | 2.43E-11 |
| chr11 | 120487589 | 120490528 | 2940 | -7.61 | 8.07E-12 | 2.43E-11 |
| chr11 | 76524757 | 76526518 | 1762 | -7.62 | 8.08E-12 | 2.44E-11 |
| chr12 | 75956526 | 75960968 | 4443 | -7.62 | 8.08E-12 | 2.44E-11 |
| chr1 | 3794559 | 3796900 | 2342 | -7.57 | 8.09E-12 | 2.44E-11 |
| chr3 | 94807013 | 94822008 | 14996 | -7.75 | 8.09E-12 | 2.44E-11 |
| chr2 | 189856477 | 189866529 | 10053 | -7.58 | 8.09E-12 | 2.44E-11 |
| chrX | 68455590 | 68468262 | 12673 | -5.61 | 8.10E-12 | 2.44E-11 |
| chr2 | 31574033 | 31577148 | 3116 | -7.7 | 8.11E-12 | 2.44E-11 |
| chr5 | 142199937 | 142201673 | 1737 | -7.62 | 8.12E-12 | 2.45E-11 |
| chr18 | 19293482 | 19298864 | 5383 | -7.72 | 8.14E-12 | 2.45E-11 |
| chr15 | 74438247 | 74441455 | 3209 | -7.58 | 8.14E-12 | 2.45E-11 |
| chr10 | 7527377 | 7529783 | 2407 | -7.57 | 8.15E-12 | 2.46E-11 |
| chr17 | 21925147 | 22034201 | 109055 | 1 | 8.16E-12 | 2.46E-11 |
| chr19 | 7413381 | 7418196 | 4816 | -7.79 | 8.20E-12 | 2.47E-11 |
| chr10 | 3752744 | 3755420 | 2677 | -7.76 | 8.21E-12 | 2.47E-11 |
| chr2 | 109689268 | 109695659 | 6392 | -7.63 | 8.22E-12 | 2.48E-11 |
| chr8 | 103947025 | 103949409 | 2385 | -7.61 | 8.23E-12 | 2.48E-11 |
| chr2 | 179942088 | 179949170 | 7083 | -7.66 | 8.30E-12 | 2.50E-11 |
| chr3 | 9103723 | 9105419 | 1697 | -7.56 | 8.31E-12 | 2.50E-11 |
| chr2 | 151378518 | 151396089 | 17572 | -7.6 | 8.31E-12 | 2.50E-11 |
| chr6 | 122120024 | 122124684 | 4661 | -7.69 | 8.32E-12 | 2.50E-11 |
| chr4 | 95983826 | 96001580 | 17755 | -8.09 | 8.33E-12 | 2.51E-11 |
| chr11 | 130127518 | 130130317 | 2800 | -7.62 | 8.34E-12 | 2.51E-11 |
| chr16 | 47662104 | 47669426 | 7323 | -7.65 | 8.35E-12 | 2.51E-11 |
| chr10 | 116534726 | 116538136 | 3411 | -7.62 | 8.37E-12 | 2.52E-11 |
| chr7 | 131625344 | 131642071 | 16728 | -8.09 | 8.38E-12 | 2.52E-11 |
| chr12 | 9941141 | 9943851 | 2711 | -2.53 | 8.41E-12 | 2.53E-11 |
| chr6 | 7491916 | 7493471 | 1556 | -7.57 | 8.42E-12 | 2.53E-11 |
| chr9 | 116228829 | 116235002 | 6174 | -7.65 | 8.47E-12 | 2.55E-11 |
| chr6 | 125748379 | 125763243 | 14865 | 7.81 | 8.49E-12 | 2.56E-11 |
| chrX | 57641182 | 57656049 | 14868 | -2.59 | 8.50E-12 | 2.56E-11 |
| chrX | 16961365 | 16963257 | 1893 | -7.56 | 8.51E-12 | 2.56E-11 |
| chr4 | 3695858 | 3699580 | 3723 | -7.85 | 8.51E-12 | 2.56E-11 |
| chr12 | 82972139 | 82980715 | 8577 | -7.7 | 8.52E-12 | 2.56E-11 |
| chr21 | 17185058 | 17188937 | 3880 | -7.58 | 8.53E-12 | 2.57E-11 |
| chr1 | 52406165 | 52408515 | 2351 | -7.61 | 8.56E-12 | 2.57E-11 |
| chr12 | 64605838 | 64607023 | 1186 | -7.58 | 8.60E-12 | 2.59E-11 |
| chr5 | 124380557 | 124387889 | 7333 | -7.58 | 8.60E-12 | 2.59E-11 |
| chr12 | 4778220 | 4781722 | 3503 | -2.65 | 8.62E-12 | 2.59E-11 |
| chr4 | 25073960 | 25080970 | 7011 | -7.57 | 8.62E-12 | 2.59E-11 |
| chr6 | 76327143 | 76329033 | 1891 | -7.57 | 8.62E-12 | 2.59E-11 |
| chr13 | 64423617 | 65153291 | 729675 | 3.54 | 8.64E-12 | 2.60E-11 |
| chr6 | 163523858 | 163560842 | 36985 | -2.16 | 8.65E-12 | 2.60E-11 |
| chr8 | 91628377 | 91631042 | 2666 | -7.59 | 8.65E-12 | 2.60E-11 |
| chr1 | 174281550 | 174297010 | 15461 | -8.19 | 8.66E-12 | 2.60E-11 |
| chr1 | 8056181 | 8059555 | 3375 | -7.56 | 8.67E-12 | 2.60E-11 |
| chr1 | 152395120 | 152409659 | 14540 | -7.64 | 8.68E-12 | 2.61E-11 |
| chr7 | 14166627 | 14185220 | 18594 | -7.58 | 8.70E-12 | 2.61E-11 |
| chr10 | 115547958 | 115551067 | 3110 | -7.56 | 8.70E-12 | 2.61E-11 |
| chr1 | 71547207 | 71549847 | 2641 | -5.69 | 8.75E-12 | 2.63E-11 |
| chr5 | 52554074 | 52563979 | 9906 | -7.58 | 8.75E-12 | 2.63E-11 |
| chr2 | 146749851 | 146754263 | 4413 | -7.73 | 8.75E-12 | 2.63E-11 |
| chr6 | 109471505 | 109473447 | 1943 | -7.61 | 8.76E-12 | 2.63E-11 |
| chr22 | 39013846 | 39016104 | 2259 | -7.58 | 8.77E-12 | 2.64E-11 |
| chr2 | 171270826 | 171273136 | 2311 | -3.26 | 8.78E-12 | 2.64E-11 |
| chr6 | 157345909 | 157347935 | 2027 | -7.58 | 8.79E-12 | 2.64E-11 |
| chr15 | 56071625 | 56088144 | 16520 | -7.98 | 8.80E-12 | 2.64E-11 |
| chr3 | 36961035 | 36964580 | 3546 | -7.81 | 8.81E-12 | 2.65E-11 |

| | | | | | | |
|---|---|---|---|---|---|---|
| chr1 | 100775325 | 100783554 | 8230 | -7.56 | 8.83E-12 | 2.65E-11 |
| chr2 | 166635163 | 166637628 | 2466 | -7.67 | 8.83E-12 | 2.65E-11 |
| chr6 | 17395477 | 17397358 | 1882 | -7.56 | 8.83E-12 | 2.65E-11 |
| chr14 | 31571165 | 31572929 | 1765 | -7.56 | 8.83E-12 | 2.65E-11 |
| chr17 | 40476737 | 40480686 | 3950 | -7.61 | 8.83E-12 | 2.65E-11 |
| chr2 | 216314554 | 216316730 | 2177 | -7.57 | 8.84E-12 | 2.65E-11 |
| chr2 | 27117364 | 27120799 | 3436 | -3.97 | 8.85E-12 | 2.66E-11 |
| chr4 | 180218527 | 180233868 | 15342 | -7.81 | 8.85E-12 | 2.66E-11 |
| chr5 | 149144901 | 149146669 | 1769 | -7.56 | 8.85E-12 | 2.66E-11 |
| chr1 | 245351155 | 245354325 | 3171 | -7.57 | 8.86E-12 | 2.66E-11 |
| chr6 | 24253032 | 24256250 | 3219 | -7.57 | 8.86E-12 | 2.66E-11 |
| chr1 | 176090985 | 176093266 | 2282 | -7.57 | 8.86E-12 | 2.66E-11 |
| chr8 | 31025202 | 31027486 | 2285 | -7.68 | 8.86E-12 | 2.66E-11 |
| chr17 | 19955900 | 19957425 | 1526 | -7.56 | 8.87E-12 | 2.66E-11 |
| chr8 | 142318836 | 142320508 | 1673 | -7.81 | 8.89E-12 | 2.67E-11 |
| chrX | 141013207 | 141015492 | 2286 | -3.44 | 8.93E-12 | 2.68E-11 |
| chrX | 100156676 | 100159891 | 3216 | -7.68 | 8.94E-12 | 2.68E-11 |
| chr16 | 89441435 | 89443732 | 2298 | -7.68 | 8.95E-12 | 2.68E-11 |
| chr3 | 126443242 | 126459219 | 15978 | -7.61 | 8.95E-12 | 2.68E-11 |
| chr2 | 122257107 | 122259267 | 2161 | -7.7 | 8.96E-12 | 2.69E-11 |
| chr2 | 219765104 | 219766702 | 1599 | -7.56 | 8.96E-12 | 2.69E-11 |
| chr7 | 91964165 | 91965851 | 1687 | -7.58 | 8.98E-12 | 2.69E-11 |
| chr14 | 36249075 | 36251804 | 2730 | -7.57 | 8.99E-12 | 2.69E-11 |
| chr20 | 9999562 | 10008718 | 9157 | -8.01 | 9.01E-12 | 2.70E-11 |
| chrX | 35107295 | 35114034 | 6740 | -7.56 | 9.02E-12 | 2.70E-11 |
| chr6 | 51896418 | 51899753 | 3336 | -2.84 | 9.02E-12 | 2.70E-11 |
| chr3 | 102506324 | 102521830 | 15507 | -7.58 | 9.03E-12 | 2.71E-11 |
| chr12 | 26569801 | 26571871 | 2071 | -7.56 | 9.05E-12 | 2.71E-11 |
| chrX | 14921526 | 14923658 | 2133 | -7.62 | 9.07E-12 | 2.72E-11 |
| chr6 | 16893127 | 16898138 | 5012 | -6.09 | 9.09E-12 | 2.72E-11 |
| chr22 | 28352043 | 28368446 | 16404 | -7.56 | 9.09E-12 | 2.72E-11 |
| chr5 | 43133732 | 43137046 | 3315 | -7.55 | 9.10E-12 | 2.72E-11 |
| chr5 | 126875036 | 126876784 | 1749 | -7.61 | 9.12E-12 | 2.73E-11 |
| chr15 | 52746662 | 52748740 | 2079 | -7.75 | 9.16E-12 | 2.74E-11 |
| chr2 | 7245756 | 7301784 | 56029 | -2.92 | 9.18E-12 | 2.75E-11 |
| chr6 | 84674484 | 84677121 | 2638 | -7.55 | 9.19E-12 | 2.75E-11 |
| chr4 | 130597244 | 130613583 | 16340 | -7.59 | 9.20E-12 | 2.75E-11 |
| chr2 | 170468877 | 170473662 | 4786 | -7.56 | 9.21E-12 | 2.76E-11 |
| chr5 | 141958167 | 141960575 | 2409 | -7.72 | 9.23E-12 | 2.76E-11 |
| chr4 | 151646853 | 151663192 | 16340 | -7.58 | 9.26E-12 | 2.77E-11 |
| chr1 | 150367950 | 150373199 | 5250 | -7.64 | 9.27E-12 | 2.77E-11 |
| chr2 | 164162257 | 164173448 | 11192 | -7.64 | 9.28E-12 | 2.78E-11 |
| chr6 | 39419933 | 39422580 | 2648 | -7.63 | 9.31E-12 | 2.79E-11 |
| chr5 | 158133545 | 158136325 | 2781 | -7.55 | 9.33E-12 | 2.79E-11 |
| chr15 | 46141316 | 46166191 | 24876 | -1.91 | 9.34E-12 | 2.80E-11 |
| chr10 | 13228177 | 13229641 | 1465 | -7.63 | 9.36E-12 | 2.80E-11 |
| chr3 | 187969331 | 187971024 | 1694 | -7.56 | 9.37E-12 | 2.80E-11 |
| chr10 | 4213453 | 4216092 | 2640 | -7.76 | 9.39E-12 | 2.81E-11 |
| chr4 | 63722968 | 63739742 | 16775 | -7.57 | 9.39E-12 | 2.81E-11 |
| chr16 | 54936039 | 54939669 | 3631 | -7.61 | 9.46E-12 | 2.83E-11 |
| chr16 | 955677 | 958240 | 2564 | -7.55 | 9.54E-12 | 2.85E-11 |
| chr6 | 135205781 | 135214750 | 8970 | -7.55 | 9.58E-12 | 2.86E-11 |
| chr12 | 113614456 | 113616681 | 2226 | -7.55 | 9.59E-12 | 2.87E-11 |
| chr4 | 101445341 | 101458553 | 13213 | -7.55 | 9.60E-12 | 2.87E-11 |
| chr2 | 58895320 | 58899256 | 3937 | -7.66 | 9.62E-12 | 2.88E-11 |
| chr2 | 170439209 | 170440788 | 1580 | -7.63 | 9.64E-12 | 2.88E-11 |
| chr9 | 836262 | 843827 | 7566 | -7.63 | 9.64E-12 | 2.88E-11 |
| chr12 | 101650878 | 101662294 | 11417 | -3.37 | 9.66E-12 | 2.89E-11 |
| chr5 | 58601689 | 58605990 | 4302 | -7.56 | 9.67E-12 | 2.89E-11 |
| chr2 | 210911079 | 210921208 | 10130 | -7.63 | 9.67E-12 | 2.89E-11 |
| chr2 | 239953334 | 239955348 | 2015 | -7.55 | 9.67E-12 | 2.89E-11 |

| | | | | | | |
|---|---|---|---|---|---|---|
| chr4 | 169008335 | 169026702 | 18368 | -7.83 | 9.68E-12 | 2.89E-11 |
| chr4 | 95390916 | 95392965 | 2050 | -7.64 | 9.70E-12 | 2.90E-11 |
| chr18 | 46252131 | 46254753 | 2623 | -7.56 | 9.71E-12 | 2.90E-11 |
| chr12 | 94175351 | 94182912 | 7562 | -7.66 | 9.71E-12 | 2.90E-11 |
| chr2 | 136437265 | 136439059 | 1795 | -7.55 | 9.71E-12 | 2.90E-11 |
| chr3 | 133669480 | 133671806 | 2327 | -3.41 | 9.71E-12 | 2.90E-11 |
| chr7 | 82091007 | 82094909 | 3903 | -7.73 | 9.73E-12 | 2.91E-11 |
| chr3 | 69607834 | 69691391 | 83558 | -1.02 | 9.75E-12 | 2.91E-11 |
| chr7 | 84522856 | 84526289 | 3434 | -7.54 | 9.75E-12 | 2.91E-11 |
| chr1 | 46563970 | 46566455 | 2486 | -7.56 | 9.75E-12 | 2.91E-11 |
| chr16 | 21053149 | 21077937 | 24789 | -8.06 | 9.81E-12 | 2.93E-11 |
| chr12 | 43807006 | 43811817 | 4812 | -7.81 | 9.86E-12 | 2.94E-11 |
| chr13 | 111196004 | 111199249 | 3246 | -7.56 | 9.86E-12 | 2.94E-11 |
| chr8 | 33859110 | 33864045 | 4936 | -4.37 | 9.86E-12 | 2.94E-11 |
| chr2 | 225734246 | 225743317 | 9072 | -7.74 | 9.86E-12 | 2.94E-11 |
| chr1 | 51376627 | 51390833 | 14207 | -7.56 | 9.89E-12 | 2.95E-11 |
| chr11 | 84323918 | 84338439 | 14522 | 8.34 | 9.92E-12 | 2.96E-11 |
| chr18 | 45331566 | 45335292 | 3727 | -7.6 | 9.95E-12 | 2.97E-11 |
| chr11 | 46069464 | 46070801 | 1338 | -7.56 | 9.97E-12 | 2.97E-11 |
| chr12 | 77248682 | 77251189 | 2508 | -7.55 | 9.98E-12 | 2.98E-11 |
| chr2 | 180322212 | 180329950 | 7739 | -7.55 | 9.98E-12 | 2.98E-11 |
| chr17 | 60095927 | 60097624 | 1698 | -7.71 | 9.98E-12 | 2.98E-11 |
| chr8 | 28831813 | 28833695 | 1883 | -7.71 | 1.00E-11 | 2.98E-11 |
| chr6 | 161745910 | 161750135 | 4226 | -7.67 | 1.00E-11 | 2.99E-11 |
| chr3 | 49412860 | 49418559 | 5700 | -7.97 | 1.00E-11 | 2.99E-11 |
| chr3 | 114752359 | 114754943 | 2585 | -7.56 | 1.00E-11 | 2.99E-11 |
| chr15 | 78516975 | 78518944 | 1970 | -7.55 | 1.01E-11 | 3.00E-11 |
| chr6 | 1138368 | 1141116 | 2749 | -7.56 | 1.01E-11 | 3.02E-11 |
| chr3 | 183135039 | 183139381 | 4343 | -7.56 | 1.01E-11 | 3.02E-11 |
| chr2 | 64776530 | 64777698 | 1169 | -7.54 | 1.01E-11 | 3.02E-11 |
| chr5 | 80113501 | 80125662 | 12162 | -7.54 | 1.01E-11 | 3.02E-11 |
| chr3 | 56739200 | 56751700 | 12501 | -7.76 | 1.01E-11 | 3.02E-11 |
| chr16 | 653702 | 655972 | 2271 | -7.83 | 1.02E-11 | 3.02E-11 |
| chr5 | 52853561 | 52866515 | 12955 | -7.79 | 1.02E-11 | 3.03E-11 |
| chrX | 41575979 | 41579735 | 3757 | -7.81 | 1.02E-11 | 3.03E-11 |
| chr6 | 17564992 | 17578192 | 13201 | -7.69 | 1.02E-11 | 3.03E-11 |
| chr6 | 147699764 | 147702557 | 2794 | -7.76 | 1.02E-11 | 3.03E-11 |
| chr1 | 116288962 | 116303526 | 14565 | -7.55 | 1.02E-11 | 3.03E-11 |
| chr15 | 76902660 | 76907662 | 5003 | -7.55 | 1.02E-11 | 3.04E-11 |
| chr2 | 151307482 | 151309878 | 2397 | -7.55 | 1.02E-11 | 3.04E-11 |
| chr5 | 44838646 | 44857284 | 18639 | -7.6 | 1.02E-11 | 3.04E-11 |
| chr1 | 46436368 | 46458369 | 22002 | -8.1 | 1.02E-11 | 3.04E-11 |
| chr1 | 51618526 | 51638450 | 19925 | -7.81 | 1.02E-11 | 3.04E-11 |
| chr9 | 85829625 | 85871247 | 41623 | -8.45 | 1.02E-11 | 3.04E-11 |
| chr6 | 136799682 | 136804601 | 4920 | -7.58 | 1.02E-11 | 3.05E-11 |
| chr17 | 61453653 | 61455589 | 1937 | -7.56 | 1.02E-11 | 3.05E-11 |
| chr6 | 24635160 | 24639280 | 4121 | -7.6 | 1.03E-11 | 3.05E-11 |
| chr8 | 140938195 | 140942608 | 4414 | -7.74 | 1.03E-11 | 3.06E-11 |
| chr13 | 68859239 | 68869139 | 9901 | 7.8 | 1.03E-11 | 3.06E-11 |
| chr16 | 86624846 | 86639948 | 15103 | -7.61 | 1.03E-11 | 3.06E-11 |
| chr13 | 106479768 | 106483365 | 3598 | -7.55 | 1.03E-11 | 3.06E-11 |
| chr2 | 194889120 | 195399493 | 510374 | 2.94 | 1.03E-11 | 3.07E-11 |
| chr7 | 4987605 | 4994787 | 7183 | -7.58 | 1.03E-11 | 3.07E-11 |
| chr5 | 90118050 | 90133059 | 15010 | -7.86 | 1.03E-11 | 3.07E-11 |
| chr5 | 79412996 | 79423438 | 10443 | -7.84 | 1.04E-11 | 3.08E-11 |
| chr2 | 171557848 | 171566977 | 9130 | -6.32 | 1.04E-11 | 3.09E-11 |
| chr1 | 69310711 | 69319692 | 8982 | 7.8 | 1.04E-11 | 3.09E-11 |
| chrX | 44307268 | 44325759 | 18492 | -7.54 | 1.04E-11 | 3.09E-11 |
| chr3 | 39592015 | 39612399 | 20385 | 2.23 | 1.04E-11 | 3.09E-11 |
| chr19 | 4775591 | 4781898 | 6308 | -7.62 | 1.04E-11 | 3.09E-11 |
| chr22 | 25920926 | 25931190 | 10265 | -7.62 | 1.04E-11 | 3.09E-11 |

| | | | | | | |
|---|---|---|---|---|---|---|
| chr1 | 172848389 | 172859066 | 10678 | -7.66 | 1.04E-11 | 3.09E-11 |
| chr2 | 38946967 | 38948581 | 1615 | -7.54 | 1.04E-11 | 3.10E-11 |
| chr1 | 23318603 | 23321990 | 3388 | -7.82 | 1.04E-11 | 3.10E-11 |
| chr5 | 68536063 | 68538685 | 2623 | -7.58 | 1.04E-11 | 3.10E-11 |
| chr17 | 45511512 | 45516883 | 5372 | -7.67 | 1.04E-11 | 3.10E-11 |
| chr6 | 133514250 | 133516829 | 2580 | -7.58 | 1.05E-11 | 3.11E-11 |
| chrX | 39021714 | 39030682 | 8969 | 7.98 | 1.05E-11 | 3.11E-11 |
| chr3 | 39024836 | 39052546 | 27711 | -7.67 | 1.05E-11 | 3.11E-11 |
| chr7 | 49883 | 53902 | 4020 | -7.83 | 1.05E-11 | 3.11E-11 |
| chr8 | 95699688 | 95701064 | 1377 | -5.78 | 1.05E-11 | 3.12E-11 |
| chr2 | 105665568 | 105668904 | 3337 | -8.33 | 1.05E-11 | 3.12E-11 |
| chr7 | 151709680 | 151711389 | 1710 | -7.61 | 1.05E-11 | 3.12E-11 |
| chr4 | 37403696 | 37421680 | 17985 | -7.57 | 1.05E-11 | 3.12E-11 |
| chr10 | 103709590 | 103711267 | 1678 | -7.53 | 1.05E-11 | 3.13E-11 |
| chr5 | 53790053 | 53805812 | 15760 | -3.57 | 1.05E-11 | 3.13E-11 |
| chr9 | 411041 | 414261 | 3221 | -7.78 | 1.06E-11 | 3.15E-11 |
| chr1 | 186715592 | 186718549 | 2958 | -7.76 | 1.06E-11 | 3.15E-11 |
| chr1 | 209180314 | 209207468 | 27155 | -2.68 | 1.06E-11 | 3.16E-11 |
| chr1 | 9347225 | 9349732 | 2508 | -7.54 | 1.06E-11 | 3.16E-11 |
| chr12 | 13579950 | 13582294 | 2345 | -7.64 | 1.06E-11 | 3.16E-11 |
| chrX | 29444403 | 29455766 | 11364 | -7.55 | 1.07E-11 | 3.16E-11 |
| chr6 | 71185381 | 71187768 | 2388 | -7.63 | 1.07E-11 | 3.16E-11 |
| chr1 | 67499435 | 67503113 | 3679 | -7.56 | 1.07E-11 | 3.16E-11 |
| chr4 | 151737144 | 151739292 | 2149 | -7.66 | 1.07E-11 | 3.17E-11 |
| chr11 | 57117737 | 57131174 | 13438 | -8.23 | 1.07E-11 | 3.17E-11 |
| chr8 | 134144056 | 134148487 | 4432 | -7.88 | 1.07E-11 | 3.17E-11 |
| chr13 | 103457557 | 103464481 | 6925 | -7.66 | 1.07E-11 | 3.17E-11 |
| chr2 | 221533224 | 221536849 | 3626 | -7.62 | 1.07E-11 | 3.17E-11 |
| chr20 | 25223534 | 25225390 | 1857 | -7.58 | 1.08E-11 | 3.19E-11 |
| chr10 | 15842091 | 15844545 | 2455 | -7.59 | 1.08E-11 | 3.19E-11 |
| chr2 | 8493506 | 8497225 | 3720 | -7.59 | 1.08E-11 | 3.19E-11 |
| chr13 | 94225955 | 94233816 | 7862 | 7.68 | 1.08E-11 | 3.19E-11 |
| chr5 | 106384495 | 106386913 | 2419 | -7.62 | 1.08E-11 | 3.20E-11 |
| chr3 | 51483569 | 51485262 | 1694 | -7.56 | 1.08E-11 | 3.20E-11 |
| chr2 | 9927572 | 9929726 | 2155 | -7.62 | 1.08E-11 | 3.20E-11 |
| chr7 | 51066697 | 51069288 | 2592 | -7.53 | 1.08E-11 | 3.21E-11 |
| chr10 | 71958132 | 71959988 | 1857 | -7.68 | 1.09E-11 | 3.22E-11 |
| chr3 | 130968327 | 130982499 | 14173 | -7.58 | 1.09E-11 | 3.22E-11 |
| chrX | 106167695 | 106169715 | 2021 | -7.53 | 1.09E-11 | 3.23E-11 |
| chr1 | 221503730 | 221506125 | 2396 | -7.56 | 1.09E-11 | 3.24E-11 |
| chr22 | 33856586 | 33859434 | 2849 | -7.56 | 1.09E-11 | 3.24E-11 |
| chr18 | 44356597 | 44367401 | 10805 | -7.75 | 1.09E-11 | 3.24E-11 |
| chr21 | 40917494 | 40931335 | 13842 | -7.55 | 1.10E-11 | 3.25E-11 |
| chr12 | 78217370 | 78219701 | 2332 | -7.53 | 1.10E-11 | 3.25E-11 |
| chr7 | 5472417 | 5477960 | 5544 | -7.53 | 1.10E-11 | 3.25E-11 |
| chr4 | 159871733 | 159875687 | 3955 | -7.75 | 1.10E-11 | 3.25E-11 |
| chr22 | 39577440 | 39597310 | 19871 | -7.93 | 1.10E-11 | 3.26E-11 |
| chr1 | 242701484 | 242702961 | 1478 | -7.53 | 1.10E-11 | 3.26E-11 |
| chr13 | 80385983 | 80390572 | 4590 | -7.55 | 1.10E-11 | 3.26E-11 |
| chr8 | 6275941 | 6278231 | 2291 | -7.53 | 1.10E-11 | 3.26E-11 |
| chr3 | 118574346 | 118591614 | 17269 | -3.28 | 1.10E-11 | 3.27E-11 |
| chr10 | 98804508 | 98814650 | 10143 | -7.55 | 1.10E-11 | 3.27E-11 |
| chr9 | 124661697 | 124663716 | 2020 | -7.57 | 1.10E-11 | 3.27E-11 |
| chr11 | 4575806 | 4581461 | 5656 | 2.05 | 1.11E-11 | 3.27E-11 |
| chr19 | 33077850 | 33085684 | 7835 | -7.53 | 1.11E-11 | 3.27E-11 |
| chr8 | 9543039 | 9555673 | 12635 | -7.59 | 1.11E-11 | 3.27E-11 |
| chr12 | 25598359 | 25619249 | 20891 | -7.69 | 1.11E-11 | 3.28E-11 |
| chr5 | 75402341 | 75405605 | 3265 | 5.65 | 1.11E-11 | 3.28E-11 |
| chr2 | 55031892 | 55033729 | 1838 | -6.26 | 1.11E-11 | 3.29E-11 |
| chr4 | 173426873 | 173435014 | 8142 | -7.7 | 1.11E-11 | 3.30E-11 |
| chr5 | 61114487 | 61115640 | 1154 | -7.55 | 1.11E-11 | 3.30E-11 |

| | | | | | | |
|---|---|---|---|---|---|---|
| chr6 | 80706898 | 80708877 | 1980 | -7.63 | 1.12E-11 | 3.30E-11 |
| chr6 | 157925910 | 157928242 | 2333 | -7.63 | 1.12E-11 | 3.30E-11 |
| chr2 | 31114510 | 31116687 | 2178 | -7.53 | 1.12E-11 | 3.30E-11 |
| chr3 | 12401967 | 12408420 | 6454 | -7.53 | 1.12E-11 | 3.30E-11 |
| chr3 | 117247102 | 117274304 | 27203 | -8.4 | 1.12E-11 | 3.30E-11 |
| chr13 | 27449010 | 27462447 | 13438 | -1.91 | 1.12E-11 | 3.31E-11 |
| chr16 | 1108642 | 1110198 | 1557 | -7.52 | 1.12E-11 | 3.33E-11 |
| chr6 | 139178687 | 139181815 | 3129 | -7.52 | 1.12E-11 | 3.33E-11 |
| chr8 | 81682130 | 81684324 | 2195 | -7.6 | 1.13E-11 | 3.33E-11 |
| chr19 | 46447278 | 46457275 | 9998 | -7.55 | 1.13E-11 | 3.33E-11 |
| chr12 | 47958607 | 47967794 | 9188 | 7.6 | 1.13E-11 | 3.33E-11 |
| chr19 | 10478654 | 10480661 | 2008 | -7.53 | 1.13E-11 | 3.33E-11 |
| chr12 | 87767532 | 87770812 | 3281 | -7.6 | 1.13E-11 | 3.35E-11 |
| chr9 | 126685838 | 126699450 | 13613 | -7.53 | 1.13E-11 | 3.35E-11 |
| chr3 | 197436676 | 197451006 | 14331 | -7.55 | 1.13E-11 | 3.35E-11 |
| chr20 | 56848623 | 56850788 | 2166 | -7.59 | 1.14E-11 | 3.37E-11 |
| chrX | 110838805 | 110845410 | 6606 | 7.61 | 1.14E-11 | 3.38E-11 |
| chr3 | 55028812 | 55039058 | 10247 | -7.59 | 1.14E-11 | 3.38E-11 |
| chr9 | 127843035 | 127845926 | 2892 | -7.59 | 1.14E-11 | 3.38E-11 |
| chr1 | 220513018 | 220519426 | 6409 | 7.65 | 1.15E-11 | 3.39E-11 |
| chr8 | 48702939 | 48705443 | 2505 | -7.53 | 1.15E-11 | 3.39E-11 |
| chr8 | 122473417 | 122475308 | 1892 | -7.53 | 1.15E-11 | 3.39E-11 |
| chr4 | 102101940 | 102104776 | 2837 | -7.53 | 1.15E-11 | 3.39E-11 |
| chr4 | 40614569 | 40616817 | 2249 | -7.72 | 1.15E-11 | 3.39E-11 |
| chr10 | 32507025 | 32513669 | 6645 | -7.69 | 1.15E-11 | 3.40E-11 |
| chr16 | 990566 | 992354 | 1789 | -7.54 | 1.15E-11 | 3.40E-11 |
| chr2 | 196903901 | 196907212 | 3312 | -7.69 | 1.15E-11 | 3.41E-11 |
| chr1 | 245200843 | 245202322 | 1480 | -7.69 | 1.16E-11 | 3.42E-11 |
| chr13 | 45705372 | 45708889 | 3518 | -7.52 | 1.16E-11 | 3.42E-11 |
| chr6 | 108329402 | 108331782 | 2381 | -7.64 | 1.16E-11 | 3.43E-11 |
| chr6 | 125560277 | 125565125 | 4849 | -7.56 | 1.16E-11 | 3.43E-11 |
| chr7 | 7299141 | 7303424 | 4284 | -7.54 | 1.16E-11 | 3.44E-11 |
| chr7 | 12542816 | 12557788 | 14973 | -7.65 | 1.16E-11 | 3.44E-11 |
| chr6 | 49185126 | 49197626 | 12501 | -7.57 | 1.17E-11 | 3.45E-11 |
| chr2 | 109709893 | 109713813 | 3921 | -7.7 | 1.17E-11 | 3.45E-11 |
| chr16 | 53519363 | 53521145 | 1783 | -7.52 | 1.17E-11 | 3.45E-11 |
| chr14 | 37929379 | 37933890 | 4512 | -7.6 | 1.17E-11 | 3.45E-11 |
| chr6 | 4149007 | 4152309 | 3303 | -5.97 | 1.17E-11 | 3.46E-11 |
| chr5 | 94367761 | 94376558 | 8798 | -7.76 | 1.17E-11 | 3.46E-11 |
| chr2 | 31611455 | 31616488 | 5034 | -7.65 | 1.18E-11 | 3.47E-11 |
| chr5 | 133752877 | 133757296 | 4420 | -7.9 | 1.18E-11 | 3.47E-11 |
| chr3 | 171707947 | 171723065 | 15119 | -7.92 | 1.18E-11 | 3.47E-11 |
| chr6 | 88937717 | 88944384 | 6668 | -2.4 | 1.18E-11 | 3.47E-11 |
| chr11 | 13634716 | 13688198 | 53483 | -3.23 | 1.18E-11 | 3.47E-11 |
| chr4 | 7952564 | 7956968 | 4405 | -7.55 | 1.18E-11 | 3.48E-11 |
| chr12 | 87457412 | 87471100 | 13689 | -7.57 | 1.18E-11 | 3.48E-11 |
| chr5 | 91954236 | 91966832 | 12597 | -8.08 | 1.18E-11 | 3.48E-11 |
| chr13 | 44064193 | 44080394 | 16202 | -7.52 | 1.18E-11 | 3.48E-11 |
| chr12 | 16635418 | 16638359 | 2942 | -7.54 | 1.18E-11 | 3.48E-11 |
| chr5 | 100288790 | 100299362 | 10573 | -5.98 | 1.19E-11 | 3.49E-11 |
| chr6 | 11285452 | 11299302 | 13851 | -7.8 | 1.19E-11 | 3.50E-11 |
| chr11 | 116620701 | 116622627 | 1927 | -7.53 | 1.19E-11 | 3.50E-11 |
| chr7 | 151473299 | 151476367 | 3069 | -7.52 | 1.19E-11 | 3.52E-11 |
| chr1 | 109559047 | 109561053 | 2007 | -7.56 | 1.20E-11 | 3.54E-11 |
| chr12 | 125191201 | 125193346 | 2146 | -7.56 | 1.20E-11 | 3.54E-11 |
| chr12 | 92425404 | 92427841 | 2438 | -7.62 | 1.20E-11 | 3.55E-11 |
| chr13 | 24112464 | 24116284 | 3821 | -7.63 | 1.20E-11 | 3.55E-11 |
| chr11 | 27206864 | 27209017 | 2154 | -7.8 | 1.20E-11 | 3.55E-11 |
| chr3 | 177322401 | 177347704 | 25304 | -8.4 | 1.21E-11 | 3.55E-11 |
| chr6 | 48510652 | 48534391 | 23740 | -7.57 | 1.21E-11 | 3.55E-11 |
| chr2 | 139220004 | 139223996 | 3993 | -7.53 | 1.21E-11 | 3.55E-11 |

| | | | | | | |
|---|---|---|---|---|---|---|
| chr10 | 117518105 | 117525932 | 7828 | 7.78 | 1.21E-11 | 3.56E-11 |
| chr1 | 41541950 | 41545516 | 3567 | -7.69 | 1.21E-11 | 3.56E-11 |
| chr2 | 44747369 | 44764239 | 16871 | -7.87 | 1.21E-11 | 3.57E-11 |
| chr11 | 102171161 | 102174958 | 3798 | -7.55 | 1.22E-11 | 3.59E-11 |
| chr12 | 88357043 | 88370968 | 13926 | -7.73 | 1.22E-11 | 3.60E-11 |
| chr10 | 45971357 | 45976198 | 4842 | -7.55 | 1.23E-11 | 3.61E-11 |
| chr14 | 50533849 | 50536991 | 3143 | -7.55 | 1.23E-11 | 3.61E-11 |
| chr3 | 130666114 | 130668082 | 1969 | -7.63 | 1.23E-11 | 3.61E-11 |
| chr12 | 9037170 | 9041180 | 4011 | -7.6 | 1.23E-11 | 3.62E-11 |
| chr4 | 101254772 | 101258445 | 3674 | -6.55 | 1.23E-11 | 3.62E-11 |
| chr14 | 75462763 | 75468532 | 5770 | -7.51 | 1.23E-11 | 3.63E-11 |
| chr6 | 20941647 | 20945212 | 3566 | -7.59 | 1.24E-11 | 3.64E-11 |
| chr8 | 55314466 | 55322283 | 7818 | -7.74 | 1.24E-11 | 3.64E-11 |
| chr5 | 78435651 | 78437900 | 2250 | -7.52 | 1.24E-11 | 3.65E-11 |
| chr3 | 194652387 | 194659213 | 6827 | -7.77 | 1.24E-11 | 3.65E-11 |
| chr1 | 10983940 | 10989664 | 5725 | -7.54 | 1.24E-11 | 3.66E-11 |
| chr20 | 1176712 | 1177922 | 1211 | -7.66 | 1.24E-11 | 3.66E-11 |
| chr16 | 22220978 | 22222879 | 1902 | -7.68 | 1.25E-11 | 3.67E-11 |
| chr3 | 31960716 | 31963566 | 2851 | -7.55 | 1.25E-11 | 3.67E-11 |
| chr15 | 87221801 | 88982623 | 1760823 | 4.49 | 1.25E-11 | 3.67E-11 |
| chr13 | 99680010 | 99688740 | 8731 | -7.62 | 1.25E-11 | 3.68E-11 |
| chr8 | 53550400 | 53552146 | 1747 | -7.53 | 1.25E-11 | 3.68E-11 |
| chr4 | 20858102 | 20878859 | 20758 | -7.63 | 1.25E-11 | 3.68E-11 |
| chrX | 13299352 | 13304855 | 5504 | -7.64 | 1.25E-11 | 3.69E-11 |
| chr18 | 50130189 | 50132812 | 2624 | -5.61 | 1.26E-11 | 3.70E-11 |
| chr2 | 54068549 | 54070303 | 1755 | -7.53 | 1.26E-11 | 3.71E-11 |
| chr5 | 92165283 | 92172504 | 7222 | -7.65 | 1.27E-11 | 3.72E-11 |
| chr9 | 90356603 | 90369394 | 12792 | -7.7 | 1.27E-11 | 3.73E-11 |
| chr9 | 119660429 | 119663829 | 3401 | -7.57 | 1.27E-11 | 3.73E-11 |
| chr4 | 190771150 | 190776804 | 5655 | -7.51 | 1.27E-11 | 3.73E-11 |
| chr4 | 139352166 | 139365634 | 13469 | -7.51 | 1.27E-11 | 3.73E-11 |
| chr5 | 77632791 | 77638556 | 5766 | -7.7 | 1.27E-11 | 3.74E-11 |
| chr12 | 88578145 | 88579633 | 1489 | -7.85 | 1.27E-11 | 3.74E-11 |
| chr15 | 36535076 | 36539994 | 4919 | -6.27 | 1.28E-11 | 3.76E-11 |
| chrX | 101935710 | 101938950 | 3241 | -7.6 | 1.29E-11 | 3.78E-11 |
| chr6 | 157060574 | 157063717 | 3144 | -7.53 | 1.29E-11 | 3.79E-11 |
| chr1 | 236150975 | 236153982 | 3008 | -7.52 | 1.29E-11 | 3.80E-11 |
| chr11 | 120542137 | 120547875 | 5739 | -2.46 | 1.30E-11 | 3.81E-11 |
| chr9 | 101718434 | 101720773 | 2340 | -7.51 | 1.30E-11 | 3.82E-11 |
| chr20 | 50394410 | 50397042 | 2633 | -7.59 | 1.30E-11 | 3.82E-11 |
| chr14 | 35839735 | 35842320 | 2586 | -7.53 | 1.30E-11 | 3.82E-11 |
| chr5 | 118628394 | 118636742 | 8349 | -7.58 | 1.30E-11 | 3.83E-11 |
| chr16 | 87003328 | 87009086 | 5759 | -7.51 | 1.31E-11 | 3.83E-11 |
| chr12 | 4467504 | 4488633 | 21130 | -2.5 | 1.31E-11 | 3.84E-11 |
| chr11 | 31880228 | 31884404 | 4177 | 7.73 | 1.31E-11 | 3.85E-11 |
| chr14 | 30124123 | 30135119 | 10997 | 7.77 | 1.31E-11 | 3.85E-11 |
| chr10 | 81826756 | 81829308 | 2553 | -7.65 | 1.32E-11 | 3.86E-11 |
| chr7 | 6870435 | 6877794 | 7360 | -7.8 | 1.32E-11 | 3.87E-11 |
| chr4 | 76456246 | 76462385 | 6140 | -7.65 | 1.32E-11 | 3.88E-11 |
| chr12 | 49570593 | 49572421 | 1829 | -7.5 | 1.32E-11 | 3.88E-11 |
| chr18 | 60095472 | 60098235 | 2764 | -7.55 | 1.32E-11 | 3.88E-11 |
| chr4 | 105569666 | 105573423 | 3758 | -7.51 | 1.33E-11 | 3.89E-11 |
| chr17 | 74973147 | 74975381 | 2235 | -7.78 | 1.33E-11 | 3.89E-11 |
| chr14 | 37211856 | 37223882 | 12027 | -4.94 | 1.33E-11 | 3.90E-11 |
| chr3 | 194007633 | 194010521 | 2889 | -7.5 | 1.33E-11 | 3.90E-11 |
| chr10 | 134506414 | 134520662 | 14249 | -7.76 | 1.33E-11 | 3.91E-11 |
| chr12 | 91283424 | 91287543 | 4120 | 6.64 | 1.33E-11 | 3.91E-11 |
| chr15 | 24768354 | 24809790 | 41437 | 1.9 | 1.33E-11 | 3.91E-11 |
| chr1 | 18311525 | 18314311 | 2787 | -5.57 | 1.33E-11 | 3.91E-11 |
| chr18 | 29239603 | 29241871 | 2269 | -7.51 | 1.34E-11 | 3.92E-11 |
| chrX | 116517001 | 116546515 | 29515 | 2.13 | 1.34E-11 | 3.92E-11 |

| | | | | | | |
|---|---|---|---|---|---|---|
| chr6 | 71147787 | 71150026 | 2240 | -7.5 | 1.34E-11 | 3.92E-11 |
| chr2 | 215279833 | 215290637 | 10805 | 7.57 | 1.34E-11 | 3.92E-11 |
| chr3 | 7564588 | 7569422 | 4835 | -7.55 | 1.34E-11 | 3.93E-11 |
| chr6 | 111748669 | 111751078 | 2410 | -7.56 | 1.34E-11 | 3.94E-11 |
| chr7 | 33442377 | 33449528 | 7152 | -7.56 | 1.34E-11 | 3.94E-11 |
| chr2 | 128962515 | 128964687 | 2173 | -7.6 | 1.35E-11 | 3.95E-11 |
| chr1 | 234969384 | 235002067 | 32684 | -8.47 | 1.35E-11 | 3.95E-11 |
| chr11 | 72819891 | 72836348 | 16458 | -7.51 | 1.35E-11 | 3.96E-11 |
| chr7 | 109758372 | 109762870 | 4499 | -7.6 | 1.35E-11 | 3.96E-11 |
| chr14 | 21717814 | 21720332 | 2519 | -7.62 | 1.35E-11 | 3.96E-11 |
| chr11 | 101994066 | 102009507 | 15442 | -7.62 | 1.35E-11 | 3.96E-11 |
| chr9 | 28948566 | 28952878 | 4313 | -7.5 | 1.35E-11 | 3.97E-11 |
| chr17 | 969453 | 971852 | 2400 | -7.76 | 1.36E-11 | 3.98E-11 |
| chr20 | 60485440 | 60494798 | 9359 | -7.82 | 1.36E-11 | 3.98E-11 |
| chr6 | 11612757 | 11625870 | 13114 | -7.63 | 1.36E-11 | 3.98E-11 |
| chr20 | 47018908 | 47022698 | 3791 | -7.64 | 1.36E-11 | 3.99E-11 |
| chr11 | 106479661 | 106550129 | 70469 | 3.1 | 1.36E-11 | 3.99E-11 |
| chr3 | 195757306 | 195759489 | 2184 | -7.53 | 1.37E-11 | 4.00E-11 |
| chr3 | 152320957 | 152323590 | 2634 | -2.92 | 1.37E-11 | 4.01E-11 |
| chr12 | 42794720 | 42796734 | 2015 | -7.74 | 1.38E-11 | 4.03E-11 |
| chr2 | 25549110 | 25550956 | 1847 | -7.5 | 1.38E-11 | 4.03E-11 |
| chr9 | 34668343 | 34670556 | 2214 | -7.51 | 1.38E-11 | 4.04E-11 |
| chr10 | 61772603 | 61776027 | 3425 | -7.51 | 1.38E-11 | 4.04E-11 |
| chr1 | 218173847 | 218178027 | 4181 | 7.58 | 1.38E-11 | 4.05E-11 |
| chr11 | 31587386 | 31595929 | 8544 | -7.75 | 1.38E-11 | 4.05E-11 |
| chr3 | 186548211 | 186553107 | 4897 | -6.54 | 1.39E-11 | 4.06E-11 |
| chr4 | 91963636 | 91978002 | 14367 | -7.56 | 1.39E-11 | 4.07E-11 |
| chr1 | 116539159 | 116542030 | 2872 | -7.51 | 1.40E-11 | 4.08E-11 |
| chr4 | 91423069 | 91427876 | 4808 | -7.52 | 1.40E-11 | 4.09E-11 |
| chr4 | 79315884 | 79318390 | 2507 | -7.51 | 1.40E-11 | 4.09E-11 |
| chr9 | 112131661 | 112141918 | 10258 | -7.61 | 1.40E-11 | 4.10E-11 |
| chr2 | 2754693 | 2849330 | 94638 | -1.83 | 1.40E-11 | 4.10E-11 |
| chr12 | 106084595 | 106095876 | 11282 | -2.91 | 1.41E-11 | 4.11E-11 |
| chr21 | 30254775 | 30275757 | 20983 | -7.52 | 1.41E-11 | 4.12E-11 |
| chr2 | 218372165 | 218374991 | 2827 | -7.5 | 1.41E-11 | 4.12E-11 |
| chr7 | 55798008 | 55802861 | 4854 | -7.49 | 1.41E-11 | 4.13E-11 |
| chr15 | 36193878 | 36211023 | 17146 | -7.79 | 1.41E-11 | 4.13E-11 |
| chr19 | 2313680 | 2316008 | 2329 | -7.51 | 1.42E-11 | 4.14E-11 |
| chr11 | 45260940 | 45262522 | 1583 | -7.51 | 1.42E-11 | 4.14E-11 |
| chr8 | 55228546 | 55242969 | 14424 | 7.79 | 1.42E-11 | 4.15E-11 |
| chr5 | 72340643 | 72342057 | 1415 | -7.49 | 1.42E-11 | 4.16E-11 |
| chr1 | 204859767 | 204873521 | 13755 | 3.06 | 1.43E-11 | 4.18E-11 |
| chr14 | 51479011 | 51503086 | 24076 | -7.86 | 1.43E-11 | 4.19E-11 |
| chr10 | 103170224 | 103172279 | 2056 | -7.5 | 1.43E-11 | 4.19E-11 |
| chr4 | 14327586 | 14337000 | 9415 | -7.69 | 1.44E-11 | 4.19E-11 |
| chr2 | 46863374 | 46869124 | 5751 | -7.53 | 1.44E-11 | 4.20E-11 |
| chr3 | 67770331 | 67773207 | 2877 | -7.53 | 1.44E-11 | 4.20E-11 |
| chr17 | 80244016 | 80245881 | 1866 | -7.5 | 1.44E-11 | 4.20E-11 |
| chrX | 4994051 | 4999927 | 5877 | -7.49 | 1.44E-11 | 4.20E-11 |
| chr1 | 222306436 | 222374689 | 68254 | -1.95 | 1.44E-11 | 4.20E-11 |
| chr19 | 1099284 | 1101086 | 1803 | -7.49 | 1.44E-11 | 4.20E-11 |
| chr4 | 114338509 | 114343889 | 5381 | -7.49 | 1.44E-11 | 4.20E-11 |
| chr2 | 210358964 | 210364996 | 6033 | -7.51 | 1.45E-11 | 4.23E-11 |
| chr3 | 176722678 | 176725371 | 2694 | -7.61 | 1.46E-11 | 4.26E-11 |
| chr3 | 14570437 | 14572029 | 1593 | -7.5 | 1.46E-11 | 4.26E-11 |
| chr6 | 24987202 | 24994720 | 7519 | -7.67 | 1.46E-11 | 4.26E-11 |
| chrX | 24242950 | 24244699 | 1750 | -7.49 | 1.46E-11 | 4.27E-11 |
| chr4 | 90908724 | 90912880 | 4157 | -5.71 | 1.46E-11 | 4.27E-11 |
| chr9 | 14374988 | 14376797 | 1810 | -7.52 | 1.47E-11 | 4.28E-11 |
| chr4 | 795258 | 814174 | 18917 | -8.02 | 1.47E-11 | 4.28E-11 |
| chr19 | 17491155 | 17500917 | 9763 | -7.5 | 1.48E-11 | 4.30E-11 |

| | | | | | | |
|---|---|---|---|---|---|---|
| chrX | 17795880 | 17799395 | 3516 | -7.56 | 1.48E-11 | 4.32E-11 |
| chr2 | 143386295 | 143412981 | 26687 | -8.1 | 1.48E-11 | 4.32E-11 |
| chr2 | 88372461 | 88374189 | 1729 | -7.51 | 1.48E-11 | 4.32E-11 |
| chr7 | 91632613 | 91641459 | 8847 | -7.49 | 1.48E-11 | 4.33E-11 |
| chr2 | 64411948 | 64421665 | 9718 | 7.56 | 1.48E-11 | 4.33E-11 |
| chr3 | 18135241 | 18139819 | 4579 | -7.6 | 1.48E-11 | 4.33E-11 |
| chr12 | 74932582 | 74936085 | 3504 | -7.52 | 1.49E-11 | 4.34E-11 |
| chr6 | 2650377 | 2663886 | 13510 | -7.65 | 1.49E-11 | 4.35E-11 |
| chr7 | 151341392 | 151343492 | 2101 | -7.62 | 1.49E-11 | 4.35E-11 |
| chr18 | 54955707 | 54958127 | 2421 | -5.7 | 1.50E-11 | 4.36E-11 |
| chr14 | 91403301 | 91407045 | 3745 | -7.69 | 1.50E-11 | 4.36E-11 |
| chr5 | 129480413 | 129489748 | 9336 | -7.8 | 1.50E-11 | 4.37E-11 |
| chr5 | 139656440 | 139658471 | 2032 | -7.49 | 1.50E-11 | 4.38E-11 |
| chr9 | 37563532 | 37565522 | 1991 | -7.49 | 1.50E-11 | 4.38E-11 |
| chr4 | 2155040 | 2157282 | 2243 | -7.49 | 1.51E-11 | 4.40E-11 |
| chr22 | 27586095 | 27601329 | 15235 | -7.66 | 1.51E-11 | 4.41E-11 |
| chr22 | 47974683 | 47979809 | 5127 | -7.66 | 1.51E-11 | 4.41E-11 |
| chr4 | 106931766 | 106937360 | 5595 | -7.69 | 1.51E-11 | 4.41E-11 |
| chr19 | 59039735 | 59043551 | 3817 | -7.52 | 1.51E-11 | 4.41E-11 |
| chr15 | 39707432 | 39710357 | 2926 | -7.54 | 1.52E-11 | 4.42E-11 |
| chr3 | 15789466 | 15793095 | 3630 | -7.61 | 1.52E-11 | 4.43E-11 |
| chr2 | 96665575 | 96667662 | 2088 | -3.16 | 1.52E-11 | 4.44E-11 |
| chr2 | 190726356 | 190736147 | 9792 | -7.51 | 1.52E-11 | 4.44E-11 |
| chr17 | 25814141 | 25829875 | 15735 | -7.78 | 1.52E-11 | 4.44E-11 |
| chr13 | 106023672 | 106033973 | 10302 | -7.65 | 1.53E-11 | 4.44E-11 |
| chr5 | 176420637 | 176422659 | 2023 | -7.52 | 1.53E-11 | 4.44E-11 |
| chr6 | 143978301 | 143981986 | 3686 | -7.53 | 1.53E-11 | 4.45E-11 |
| chr10 | 96581600 | 96602373 | 20774 | -7.62 | 1.53E-11 | 4.45E-11 |
| chr12 | 132996849 | 133003176 | 6328 | -7.56 | 1.53E-11 | 4.46E-11 |
| chr7 | 140516886 | 140518519 | 1634 | -7.53 | 1.54E-11 | 4.47E-11 |
| chr1 | 9782405 | 9784679 | 2275 | -7.48 | 1.54E-11 | 4.47E-11 |
| chrX | 16240638 | 16243788 | 3151 | -7.48 | 1.54E-11 | 4.47E-11 |
| chr3 | 97488915 | 97491044 | 2130 | -7.54 | 1.54E-11 | 4.47E-11 |
| chr18 | 57937400 | 57954786 | 17387 | -7.49 | 1.54E-11 | 4.48E-11 |
| chr20 | 23280387 | 23324836 | 44450 | -8.64 | 1.54E-11 | 4.48E-11 |
| chr1 | 91902974 | 91907465 | 4492 | -7.55 | 1.54E-11 | 4.48E-11 |
| chr6 | 125325403 | 125328556 | 3154 | -7.53 | 1.54E-11 | 4.49E-11 |
| chr6 | 74551462 | 74554177 | 2716 | -7.51 | 1.55E-11 | 4.50E-11 |
| chrX | 44063125 | 44080261 | 17137 | -7.52 | 1.55E-11 | 4.51E-11 |
| chr3 | 151155107 | 151167907 | 12801 | -7.74 | 1.55E-11 | 4.51E-11 |
| chr9 | 91284268 | 91286960 | 2693 | -6.1 | 1.55E-11 | 4.52E-11 |
| chr7 | 93033472 | 93035929 | 2458 | -7.64 | 1.55E-11 | 4.52E-11 |
| chr8 | 43081319 | 43083159 | 1841 | -7.57 | 1.56E-11 | 4.54E-11 |
| chr9 | 80227801 | 80252682 | 24882 | -8.07 | 1.56E-11 | 4.55E-11 |
| chr5 | 127943126 | 127946431 | 3306 | -3.56 | 1.57E-11 | 4.56E-11 |
| chr12 | 98007010 | 98014047 | 7038 | -7.58 | 1.57E-11 | 4.57E-11 |
| chr14 | 102010066 | 102013258 | 3193 | -7.59 | 1.57E-11 | 4.57E-11 |
| chr1 | 16428648 | 16436321 | 7674 | -7.58 | 1.57E-11 | 4.57E-11 |
| chr19 | 22587107 | 22835185 | 248079 | -2.13 | 1.57E-11 | 4.58E-11 |
| chr13 | 96338167 | 96350350 | 12184 | -7.57 | 1.58E-11 | 4.58E-11 |
| chr13 | 28519779 | 28522184 | 2406 | -7.49 | 1.58E-11 | 4.59E-11 |
| chr6 | 136541996 | 136545225 | 3230 | -7.61 | 1.58E-11 | 4.59E-11 |
| chr11 | 93686550 | 93689112 | 2563 | -3.8 | 1.58E-11 | 4.60E-11 |
| chr1 | 51848457 | 51851557 | 3101 | -7.63 | 1.58E-11 | 4.60E-11 |
| chr18 | 46911584 | 46923185 | 11602 | -7.7 | 1.59E-11 | 4.61E-11 |
| chr4 | 160306907 | 160309656 | 2750 | -7.48 | 1.59E-11 | 4.61E-11 |
| chr12 | 125150498 | 125152323 | 1826 | -7.49 | 1.59E-11 | 4.63E-11 |
| chr16 | 54279612 | 54306697 | 27086 | -8.03 | 1.59E-11 | 4.63E-11 |
| chr20 | 60041992 | 60044462 | 2471 | -7.48 | 1.60E-11 | 4.64E-11 |
| chr9 | 139667109 | 139669222 | 2114 | -7.49 | 1.60E-11 | 4.66E-11 |
| chr17 | 65453186 | 65468423 | 15238 | -7.87 | 1.60E-11 | 4.66E-11 |

| | | | | | | |
|---|---|---|---|---|---|---|
| chr21 | 36052265 | 36067996 | 15732 | -7.54 | 1.61E-11 | 4.66E-11 |
| chr6 | 81527512 | 81557480 | 29969 | 1.94 | 1.61E-11 | 4.68E-11 |
| chr12 | 87677970 | 87690309 | 12340 | -8.02 | 1.61E-11 | 4.68E-11 |
| chr1 | 194460098 | 194480914 | 20817 | -7.76 | 1.61E-11 | 4.68E-11 |
| chr1 | 165376530 | 165549297 | 172768 | 4.01 | 1.62E-11 | 4.69E-11 |
| chr6 | 131070294 | 131083451 | 13158 | -7.54 | 1.62E-11 | 4.69E-11 |
| chr12 | 79563505 | 79575134 | 11630 | -8.04 | 1.62E-11 | 4.69E-11 |
| chr13 | 66849596 | 66853119 | 3524 | -7.49 | 1.62E-11 | 4.71E-11 |
| chr9 | 128536782 | 128545801 | 9020 | -7.48 | 1.62E-11 | 4.71E-11 |
| chrX | 55436191 | 55442038 | 5848 | 7.56 | 1.63E-11 | 4.73E-11 |
| chr13 | 28055242 | 28057235 | 1994 | -7.58 | 1.63E-11 | 4.74E-11 |
| chr1 | 214682123 | 214685215 | 3093 | -7.59 | 1.63E-11 | 4.74E-11 |
| chr16 | 973245 | 977213 | 3969 | -7.57 | 1.63E-11 | 4.74E-11 |
| chr4 | 177836484 | 177840776 | 4293 | -7.57 | 1.63E-11 | 4.74E-11 |
| chr14 | 62128834 | 62130600 | 1767 | -7.57 | 1.63E-11 | 4.74E-11 |
| chr8 | 58885501 | 58893316 | 7816 | -4.6 | 1.63E-11 | 4.74E-11 |
| chr2 | 97329030 | 97330717 | 1688 | -7.48 | 1.64E-11 | 4.75E-11 |
| chr4 | 168968138 | 168972841 | 4704 | -7.48 | 1.64E-11 | 4.75E-11 |
| chr2 | 128169680 | 128171397 | 1718 | -7.6 | 1.64E-11 | 4.76E-11 |
| chr2 | 73379450 | 73380815 | 1366 | -7.47 | 1.64E-11 | 4.76E-11 |
| chr6 | 94127302 | 94131257 | 3956 | -7.56 | 1.64E-11 | 4.77E-11 |
| chr6 | 88248956 | 88251200 | 2245 | -7.49 | 1.65E-11 | 4.78E-11 |
| chr4 | 28124785 | 28132532 | 7748 | -7.49 | 1.65E-11 | 4.78E-11 |
| chr3 | 180644628 | 180646640 | 2013 | -7.52 | 1.65E-11 | 4.79E-11 |
| chr2 | 204789158 | 204791548 | 2391 | -6.41 | 1.66E-11 | 4.80E-11 |
| chr12 | 93609252 | 93634031 | 24780 | -7.47 | 1.66E-11 | 4.80E-11 |
| chr16 | 56837456 | 56839937 | 2482 | -7.48 | 1.66E-11 | 4.81E-11 |
| chr8 | 103715788 | 103749875 | 34088 | -8.07 | 1.66E-11 | 4.82E-11 |
| chr4 | 101653775 | 101658254 | 4480 | -7.47 | 1.66E-11 | 4.82E-11 |
| chr13 | 40003085 | 40014900 | 11816 | -7.54 | 1.66E-11 | 4.82E-11 |
| chr20 | 47524910 | 47526809 | 1900 | -7.54 | 1.66E-11 | 4.82E-11 |
| chr3 | 131707924 | 131712820 | 4897 | -7.63 | 1.67E-11 | 4.84E-11 |
| chr13 | 95009358 | 95023958 | 14601 | 7.61 | 1.67E-11 | 4.84E-11 |
| chr11 | 105864078 | 105880463 | 16386 | -8.22 | 1.67E-11 | 4.85E-11 |
| chr10 | 25125360 | 25127640 | 2281 | -7.49 | 1.68E-11 | 4.86E-11 |
| chr9 | 86942131 | 86945263 | 3133 | -7.8 | 1.68E-11 | 4.86E-11 |
| chr11 | 70968854 | 70971265 | 2412 | -7.63 | 1.68E-11 | 4.86E-11 |
| chr6 | 127009640 | 127014180 | 4541 | -7.51 | 1.68E-11 | 4.86E-11 |
| chr3 | 17194240 | 17196747 | 2508 | -7.48 | 1.68E-11 | 4.87E-11 |
| chr18 | 23858790 | 23860392 | 1603 | -7.48 | 1.68E-11 | 4.88E-11 |
| chr6 | 9737699 | 9740968 | 3270 | -7.71 | 1.68E-11 | 4.88E-11 |
| chr12 | 39507254 | 39516155 | 8902 | -7.72 | 1.68E-11 | 4.88E-11 |
| chr6 | 127967415 | 127970193 | 2779 | -7.47 | 1.69E-11 | 4.88E-11 |
| chr12 | 111934736 | 111936284 | 1549 | -7.47 | 1.69E-11 | 4.89E-11 |
| chr10 | 95958038 | 95967697 | 9660 | -7.51 | 1.69E-11 | 4.89E-11 |
| chr12 | 66000333 | 66002766 | 2434 | -7.49 | 1.69E-11 | 4.89E-11 |
| chr4 | 68557256 | 68573973 | 16718 | -7.69 | 1.69E-11 | 4.90E-11 |
| chr10 | 134101479 | 134103417 | 1939 | -7.49 | 1.70E-11 | 4.91E-11 |
| chr21 | 44714234 | 44716563 | 2330 | -7.58 | 1.70E-11 | 4.91E-11 |
| chr1 | 53602665 | 53604737 | 2073 | -7.5 | 1.70E-11 | 4.92E-11 |
| chrX | 45792868 | 45798211 | 5344 | -7.5 | 1.70E-11 | 4.92E-11 |
| chr14 | 24054435 | 24063885 | 9451 | -7.74 | 1.71E-11 | 4.93E-11 |
| chr4 | 53558685 | 53561067 | 2383 | -7.54 | 1.72E-11 | 4.96E-11 |
| chr21 | 45598024 | 45611773 | 13750 | -7.84 | 1.72E-11 | 4.98E-11 |
| chr4 | 178217238 | 178221089 | 3852 | -7.62 | 1.73E-11 | 5.00E-11 |
| chr13 | 67203110 | 67205665 | 2556 | -7.47 | 1.73E-11 | 5.00E-11 |
| chr19 | 50714136 | 50728871 | 14736 | -7.47 | 1.73E-11 | 5.00E-11 |
| chr4 | 74334764 | 74342001 | 7238 | -1.69 | 1.73E-11 | 5.01E-11 |
| chr14 | 71818478 | 71821849 | 3372 | -7.47 | 1.73E-11 | 5.01E-11 |
| chr11 | 5696750 | 5699750 | 3001 | -7.47 | 1.74E-11 | 5.02E-11 |
| chr13 | 50851309 | 50864334 | 13026 | -7.47 | 1.74E-11 | 5.02E-11 |

| | | | | | | |
|---|---|---|---|---|---|---|
| chr7 | 108917759 | 108929054 | 11296 | 8.1 | 1.74E-11 | 5.04E-11 |
| chr2 | 69307736 | 69314583 | 6848 | -5.94 | 1.74E-11 | 5.04E-11 |
| chr1 | 197535440 | 197538378 | 2939 | -7.54 | 1.75E-11 | 5.04E-11 |
| chrX | 109526170 | 109545698 | 19529 | -7.51 | 1.75E-11 | 5.05E-11 |
| chr8 | 135830430 | 135831510 | 1081 | -7.53 | 1.75E-11 | 5.06E-11 |
| chr21 | 44908562 | 44917510 | 8949 | -7.49 | 1.75E-11 | 5.06E-11 |
| chr7 | 36588948 | 36600138 | 11191 | -7.49 | 1.75E-11 | 5.07E-11 |
| chr14 | 57801169 | 57804199 | 3031 | -7.48 | 1.76E-11 | 5.08E-11 |
| chr10 | 32375896 | 32388803 | 12908 | -7.67 | 1.76E-11 | 5.08E-11 |
| chr18 | 7868004 | 7876724 | 8721 | -7.67 | 1.76E-11 | 5.08E-11 |
| chr7 | 87382825 | 87384858 | 2034 | 5.51 | 1.76E-11 | 5.09E-11 |
| chr15 | 77017158 | 77018447 | 1290 | -7.46 | 1.76E-11 | 5.09E-11 |
| chr7 | 33357136 | 33360914 | 3779 | -7.47 | 1.76E-11 | 5.09E-11 |
| chr16 | 53677545 | 53679070 | 1526 | -7.49 | 1.76E-11 | 5.09E-11 |
| chr18 | 60259226 | 60267716 | 8491 | -7.46 | 1.76E-11 | 5.09E-11 |
| chr12 | 16084422 | 16092420 | 7999 | -7.57 | 1.76E-11 | 5.09E-11 |
| chr6 | 144701633 | 144704000 | 2368 | -7.58 | 1.77E-11 | 5.10E-11 |
| chr12 | 21885081 | 21898546 | 13466 | -7.46 | 1.77E-11 | 5.10E-11 |
| chr12 | 132610698 | 132613840 | 3143 | -7.48 | 1.77E-11 | 5.10E-11 |
| chr5 | 55485373 | 55487507 | 2135 | -7.46 | 1.77E-11 | 5.11E-11 |
| chr12 | 105292027 | 105293887 | 1861 | -7.55 | 1.78E-11 | 5.13E-11 |
| chr17 | 62274193 | 62275710 | 1518 | -7.46 | 1.78E-11 | 5.13E-11 |
| chr20 | 46911627 | 46947940 | 36314 | -7.54 | 1.78E-11 | 5.14E-11 |
| chr3 | 174656728 | 174662187 | 5460 | -7.54 | 1.78E-11 | 5.14E-11 |
| chr12 | 12534245 | 12536062 | 1818 | -7.6 | 1.78E-11 | 5.14E-11 |
| chr5 | 175753999 | 175761837 | 7839 | -7.46 | 1.79E-11 | 5.15E-11 |
| chr1 | 97739706 | 97751441 | 11736 | -7.9 | 1.79E-11 | 5.16E-11 |
| chr8 | 144323236 | 144325456 | 2221 | -7.69 | 1.80E-11 | 5.18E-11 |
| chr13 | 91479618 | 91490232 | 10615 | -7.65 | 1.80E-11 | 5.19E-11 |
| chr15 | 77527328 | 77529402 | 2075 | -7.46 | 1.80E-11 | 5.20E-11 |
| chr6 | 126878051 | 126881436 | 3386 | -7.48 | 1.80E-11 | 5.20E-11 |
| chr8 | 91728319 | 91730679 | 2361 | -7.48 | 1.80E-11 | 5.20E-11 |
| chr15 | 89105302 | 89106556 | 1255 | -7.48 | 1.81E-11 | 5.21E-11 |
| chr16 | 69808894 | 69810939 | 2046 | -7.46 | 1.81E-11 | 5.21E-11 |
| chr3 | 41047808 | 41051238 | 3431 | -7.46 | 1.81E-11 | 5.22E-11 |
| chr2 | 173916204 | 173924379 | 8176 | -7.66 | 1.81E-11 | 5.23E-11 |
| chr11 | 108802986 | 108807830 | 4845 | -7.65 | 1.82E-11 | 5.23E-11 |
| chr1 | 233805900 | 233816044 | 10145 | -7.63 | 1.82E-11 | 5.24E-11 |
| chr10 | 43302218 | 43307732 | 5515 | -7.46 | 1.82E-11 | 5.24E-11 |
| chr22 | 50222824 | 50225951 | 3128 | -7.5 | 1.82E-11 | 5.24E-11 |
| chr12 | 74991413 | 74995330 | 3918 | -7.49 | 1.82E-11 | 5.24E-11 |
| chr7 | 21565155 | 21597905 | 32751 | -6.75 | 1.82E-11 | 5.25E-11 |
| chr6 | 140588198 | 140597417 | 9220 | -7.63 | 1.82E-11 | 5.25E-11 |
| chr2 | 170664885 | 170674610 | 9726 | -7.61 | 1.83E-11 | 5.27E-11 |
| chr12 | 46092576 | 46096059 | 3484 | -5.57 | 1.83E-11 | 5.28E-11 |
| chr11 | 88167239 | 88171262 | 4024 | -7.73 | 1.83E-11 | 5.28E-11 |
| chr8 | 68412782 | 68415200 | 2419 | -7.48 | 1.83E-11 | 5.28E-11 |
| chr12 | 133036298 | 133039782 | 3485 | -7.67 | 1.84E-11 | 5.29E-11 |
| chr3 | 70025620 | 70028838 | 3219 | -4.11 | 1.84E-11 | 5.29E-11 |
| chr5 | 124166659 | 124170127 | 3469 | -7.83 | 1.84E-11 | 5.29E-11 |
| chr15 | 36752579 | 36756666 | 4088 | -7.55 | 1.84E-11 | 5.30E-11 |
| chr13 | 67559608 | 67564052 | 4445 | -7.46 | 1.84E-11 | 5.31E-11 |
| chr7 | 99299764 | 99317114 | 17351 | -7.46 | 1.84E-11 | 5.31E-11 |
| chr5 | 148524799 | 148527659 | 2861 | -7.46 | 1.85E-11 | 5.34E-11 |
| chr7 | 1917346 | 1919261 | 1916 | -7.46 | 1.86E-11 | 5.34E-11 |
| chr4 | 96018668 | 96022035 | 3368 | -7.49 | 1.86E-11 | 5.36E-11 |
| chr4 | 130665309 | 130670177 | 4869 | -7.46 | 1.86E-11 | 5.37E-11 |
| chr17 | 19916305 | 19917891 | 1587 | -7.61 | 1.87E-11 | 5.37E-11 |
| chr3 | 94563545 | 94594424 | 30880 | -7.96 | 1.87E-11 | 5.37E-11 |
| chr7 | 15003087 | 15029282 | 26196 | -7.79 | 1.87E-11 | 5.38E-11 |
| chr12 | 32441652 | 32443199 | 1548 | -7.46 | 1.88E-11 | 5.40E-11 |

| | | | | | | |
|---|---|---|---|---|---|---|
| chr7 | 108218770 | 108221183 | 2414 | -7.46 | 1.88E-11 | 5.41E-11 |
| chr9 | 37272538 | 37274277 | 1740 | -7.45 | 1.89E-11 | 5.43E-11 |
| chr2 | 55428660 | 55432106 | 3447 | -7.45 | 1.89E-11 | 5.43E-11 |
| chr3 | 11017287 | 11033153 | 15867 | -7.66 | 1.89E-11 | 5.44E-11 |
| chr16 | 84010487 | 84012874 | 2388 | -7.46 | 1.89E-11 | 5.45E-11 |
| chr3 | 38444821 | 38447086 | 2266 | -7.5 | 1.89E-11 | 5.45E-11 |
| chr7 | 110573459 | 110577443 | 3985 | -7.47 | 1.90E-11 | 5.48E-11 |
| chr17 | 5136356 | 5143198 | 6843 | -8.03 | 1.91E-11 | 5.48E-11 |
| chr2 | 24384892 | 24386350 | 1459 | -7.67 | 1.91E-11 | 5.49E-11 |
| chr4 | 55841411 | 55973329 | 131919 | 1.7 | 1.91E-11 | 5.49E-11 |
| chr13 | 96148308 | 96155883 | 7576 | -7.55 | 1.91E-11 | 5.49E-11 |
| chr10 | 86036367 | 86047203 | 10837 | -7.97 | 1.92E-11 | 5.51E-11 |
| chr9 | 114082001 | 114089127 | 7127 | -7.58 | 1.92E-11 | 5.51E-11 |
| chr11 | 125020428 | 125026132 | 5705 | -7.48 | 1.92E-11 | 5.51E-11 |
| chr15 | 45038044 | 45050321 | 12278 | -2.41 | 1.92E-11 | 5.52E-11 |
| chr2 | 37469426 | 37470900 | 1475 | -7.45 | 1.92E-11 | 5.52E-11 |
| chr4 | 81896367 | 81903257 | 6891 | 7.87 | 1.93E-11 | 5.53E-11 |
| chr5 | 64215436 | 64218266 | 2831 | -7.45 | 1.93E-11 | 5.55E-11 |
| chr15 | 67537296 | 67540235 | 2940 | -7.7 | 1.93E-11 | 5.55E-11 |
| chr2 | 30691000 | 30693736 | 2737 | -7.89 | 1.94E-11 | 5.57E-11 |
| chr5 | 42984384 | 42986380 | 1997 | -7.6 | 1.94E-11 | 5.58E-11 |
| chr6 | 35507739 | 35520664 | 12926 | -7.45 | 1.95E-11 | 5.60E-11 |
| chr11 | 56553221 | 56555314 | 2094 | -7.45 | 1.96E-11 | 5.62E-11 |
| chrX | 102069950 | 102074941 | 4992 | -7.48 | 1.96E-11 | 5.63E-11 |
| chr15 | 71105876 | 71113140 | 7265 | -8.07 | 1.96E-11 | 5.63E-11 |
| chr4 | 171052430 | 171057338 | 4909 | -3.2 | 1.97E-11 | 5.65E-11 |
| chr11 | 86888078 | 86890067 | 1990 | -7.45 | 1.97E-11 | 5.65E-11 |
| chr2 | 143154253 | 143161087 | 6835 | -2.61 | 1.97E-11 | 5.66E-11 |
| chr18 | 44023251 | 44025711 | 2461 | -2.8 | 1.98E-11 | 5.68E-11 |
| chr4 | 21111298 | 21127253 | 15956 | -3.04 | 1.98E-11 | 5.69E-11 |
| chrX | 12943860 | 12956721 | 12862 | -7.55 | 1.98E-11 | 5.69E-11 |
| chr2 | 219639324 | 219653056 | 13733 | -7.5 | 1.98E-11 | 5.69E-11 |
| chr7 | 22593024 | 22634966 | 41943 | -7.82 | 1.99E-11 | 5.70E-11 |
| chr8 | 122129093 | 122134091 | 4999 | -7.57 | 1.99E-11 | 5.70E-11 |
| chr8 | 41005776 | 41018490 | 12715 | -7.46 | 1.99E-11 | 5.70E-11 |
| chr14 | 100143450 | 100145850 | 2401 | -7.57 | 1.99E-11 | 5.70E-11 |
| chr4 | 108806987 | 108811607 | 4621 | -7.54 | 1.99E-11 | 5.71E-11 |
| chr9 | 4129165 | 4133569 | 4405 | -7.57 | 1.99E-11 | 5.71E-11 |
| chr11 | 2338442 | 2344679 | 6238 | -7.95 | 1.99E-11 | 5.71E-11 |
| chr16 | 70752590 | 70754350 | 1761 | -7.47 | 1.99E-11 | 5.72E-11 |
| chr11 | 35800542 | 35802087 | 1546 | -7.44 | 2.00E-11 | 5.73E-11 |
| chr10 | 123812249 | 123827242 | 14994 | -8.09 | 2.00E-11 | 5.74E-11 |
| chr20 | 42148325 | 42153677 | 5353 | -7.75 | 2.00E-11 | 5.75E-11 |
| chr14 | 94499288 | 94501966 | 2679 | -7.59 | 2.00E-11 | 5.75E-11 |
| chr20 | 23243906 | 23245527 | 1622 | -7.45 | 2.01E-11 | 5.76E-11 |
| chr1 | 86469742 | 86472553 | 2812 | -7.48 | 2.01E-11 | 5.76E-11 |
| chr1 | 221426812 | 221437555 | 10744 | -7.45 | 2.01E-11 | 5.76E-11 |
| chr8 | 134226430 | 134228046 | 1617 | -7.49 | 2.01E-11 | 5.76E-11 |
| chr10 | 53460150 | 53463293 | 3144 | -7.46 | 2.01E-11 | 5.77E-11 |
| chr16 | 70700766 | 70705081 | 4316 | -7.46 | 2.01E-11 | 5.77E-11 |
| chr7 | 94703973 | 94705859 | 1887 | -7.46 | 2.02E-11 | 5.78E-11 |
| chr22 | 49265578 | 49273107 | 7530 | -7.52 | 2.02E-11 | 5.78E-11 |
| chr14 | 61045040 | 61049824 | 4785 | -7.49 | 2.02E-11 | 5.79E-11 |
| chr12 | 131557469 | 131566269 | 8801 | -7.6 | 2.03E-11 | 5.82E-11 |
| chrX | 62470596 | 62474396 | 3801 | -7.6 | 2.03E-11 | 5.83E-11 |
| chr18 | 21621940 | 21628895 | 6956 | -7.49 | 2.03E-11 | 5.83E-11 |
| chr2 | 69379460 | 69380563 | 1104 | -3.4 | 2.04E-11 | 5.83E-11 |
| chr1 | 232011064 | 232013134 | 2071 | -7.44 | 2.04E-11 | 5.83E-11 |
| chr5 | 80570061 | 80590846 | 20786 | -7.92 | 2.04E-11 | 5.84E-11 |
| chr8 | 42808613 | 42809894 | 1282 | -7.5 | 2.05E-11 | 5.86E-11 |
| chr9 | 102714170 | 102716306 | 2137 | -7.5 | 2.05E-11 | 5.88E-11 |

| | | | | | | |
|---|---|---|---|---|---|---|
| chr13 | 52421160 | 52430810 | 9651 | -7.45 | 2.05E-11 | 5.88E-11 |
| chr20 | 52589396 | 52598565 | 9170 | -8.15 | 2.06E-11 | 5.89E-11 |
| chr5 | 153473674 | 153475121 | 1448 | -7.47 | 2.06E-11 | 5.89E-11 |
| chrX | 53995368 | 53996864 | 1497 | -7.48 | 2.06E-11 | 5.90E-11 |
| chr6 | 82611991 | 82614052 | 2062 | -7.55 | 2.06E-11 | 5.90E-11 |
| chr6 | 113921278 | 113935486 | 14209 | -7.54 | 2.06E-11 | 5.91E-11 |
| chr7 | 7263446 | 7272171 | 8726 | -7.79 | 2.06E-11 | 5.91E-11 |
| chr14 | 64307462 | 64309607 | 2146 | -7.54 | 2.07E-11 | 5.91E-11 |
| chr6 | 165576667 | 165596053 | 19387 | -8.04 | 2.07E-11 | 5.92E-11 |
| chr9 | 90601201 | 90624945 | 23745 | -7.72 | 2.07E-11 | 5.92E-11 |
| chr1 | 1465463 | 1467643 | 2181 | -7.44 | 2.07E-11 | 5.92E-11 |
| chr11 | 261158 | 277488 | 16331 | -8.11 | 2.07E-11 | 5.92E-11 |
| chr9 | 110649667 | 110687650 | 37984 | -8.43 | 2.08E-11 | 5.94E-11 |
| chr3 | 129665287 | 129666895 | 1609 | -7.46 | 2.08E-11 | 5.94E-11 |
| chr2 | 46830463 | 46838084 | 7622 | -7.49 | 2.08E-11 | 5.95E-11 |
| chr10 | 82047298 | 82092724 | 45427 | -3.86 | 2.08E-11 | 5.95E-11 |
| chr7 | 20226248 | 20230244 | 3997 | -7.52 | 2.09E-11 | 5.97E-11 |
| chr11 | 60957576 | 60967059 | 9484 | -7.69 | 2.09E-11 | 5.98E-11 |
| chr17 | 78240257 | 78246297 | 6041 | -7.49 | 2.09E-11 | 5.98E-11 |
| chr14 | 93145520 | 93149472 | 3953 | -7.51 | 2.10E-11 | 6.00E-11 |
| chr4 | 103377869 | 103383838 | 5970 | -1.98 | 2.10E-11 | 6.00E-11 |
| chr4 | 3737683 | 3745432 | 7750 | -7.46 | 2.10E-11 | 6.00E-11 |
| chr14 | 38549674 | 38571601 | 21928 | -7.72 | 2.10E-11 | 6.00E-11 |
| chr2 | 33580415 | 33582092 | 1678 | -7.45 | 2.11E-11 | 6.02E-11 |
| chr3 | 142467601 | 142469731 | 2131 | -7.64 | 2.11E-11 | 6.02E-11 |
| chr1 | 185999102 | 186001519 | 2418 | -7.58 | 2.11E-11 | 6.03E-11 |
| chr8 | 27867961 | 27870275 | 2315 | -6.38 | 2.11E-11 | 6.03E-11 |
| chr9 | 135306475 | 135318545 | 12071 | -7.44 | 2.11E-11 | 6.03E-11 |
| chr7 | 54940843 | 54947687 | 6845 | -7.6 | 2.11E-11 | 6.03E-11 |
| chr17 | 19155675 | 19157035 | 1361 | -7.51 | 2.12E-11 | 6.04E-11 |
| chr3 | 18082248 | 18098430 | 16183 | -7.68 | 2.12E-11 | 6.04E-11 |
| chr3 | 133050829 | 133057946 | 7118 | 7.8 | 2.12E-11 | 6.05E-11 |
| chr4 | 185343084 | 185344664 | 1581 | -7.5 | 2.12E-11 | 6.06E-11 |
| chr10 | 6552328 | 6553928 | 1601 | -5.71 | 2.12E-11 | 6.06E-11 |
| chr6 | 42678768 | 42681050 | 2283 | -7.46 | 2.14E-11 | 6.12E-11 |
| chr22 | 43105170 | 43107221 | 2052 | -7.48 | 2.15E-11 | 6.13E-11 |
| chr4 | 98327603 | 98338704 | 11102 | -7.45 | 2.15E-11 | 6.14E-11 |
| chr2 | 210977796 | 210991468 | 13673 | -7.53 | 2.15E-11 | 6.14E-11 |
| chr20 | 9066610 | 9068356 | 1747 | -7.43 | 2.16E-11 | 6.16E-11 |
| chr3 | 159102812 | 159162657 | 59846 | -1.79 | 2.16E-11 | 6.16E-11 |
| chr6 | 4898334 | 4900696 | 2363 | -7.49 | 2.16E-11 | 6.16E-11 |
| chr7 | 28949379 | 28953653 | 4275 | -7.44 | 2.16E-11 | 6.17E-11 |
| chr9 | 12731483 | 12742758 | 11276 | -7.49 | 2.16E-11 | 6.18E-11 |
| chr6 | 169001129 | 169006220 | 5092 | -7.51 | 2.17E-11 | 6.18E-11 |
| chr12 | 132187774 | 132190857 | 3084 | -7.49 | 2.17E-11 | 6.19E-11 |
| chr2 | 101631611 | 101634103 | 2493 | -7.44 | 2.17E-11 | 6.20E-11 |
| chr1 | 56259353 | 56266713 | 7361 | -7.51 | 2.18E-11 | 6.21E-11 |
| chr7 | 2521961 | 2523686 | 1726 | -7.43 | 2.18E-11 | 6.22E-11 |
| chr2 | 220605374 | 220678122 | 72749 | -1.56 | 2.18E-11 | 6.22E-11 |
| chr8 | 90587327 | 90599535 | 12209 | -7.46 | 2.18E-11 | 6.23E-11 |
| chr22 | 36061156 | 36078234 | 17079 | -7.43 | 2.18E-11 | 6.23E-11 |
| chr8 | 18461116 | 18463592 | 2477 | -7.45 | 2.19E-11 | 6.24E-11 |
| chr2 | 183738170 | 183746533 | 8364 | -7.5 | 2.20E-11 | 6.27E-11 |
| chr11 | 13422012 | 13424479 | 2468 | -7.56 | 2.20E-11 | 6.27E-11 |
| chr2 | 134422518 | 134428918 | 6401 | -2.65 | 2.20E-11 | 6.28E-11 |
| chr1 | 23265943 | 23278379 | 12437 | -8.01 | 2.20E-11 | 6.28E-11 |
| chr6 | 149114524 | 149136668 | 22145 | -7.8 | 2.21E-11 | 6.29E-11 |
| chr2 | 228048118 | 228057024 | 8907 | -7.77 | 2.21E-11 | 6.29E-11 |
| chr5 | 114774546 | 114777392 | 2847 | -7.46 | 2.21E-11 | 6.30E-11 |
| chr9 | 4104084 | 4109075 | 4992 | -7.43 | 2.22E-11 | 6.33E-11 |
| chr12 | 80057176 | 80069480 | 12305 | -7.49 | 2.22E-11 | 6.34E-11 |

| | | | | | | |
|---|---|---|---|---|---|---|
| chr7 | 43585593 | 43587599 | 2007 | -7.53 | 2.23E-11 | 6.37E-11 |
| chr4 | 133336819 | 133341089 | 4271 | -7.45 | 2.24E-11 | 6.39E-11 |
| chr18 | 56321599 | 56324191 | 2593 | -7.56 | 2.25E-11 | 6.40E-11 |
| chr15 | 37043529 | 37052461 | 8933 | -7.44 | 2.25E-11 | 6.41E-11 |
| chr10 | 70024226 | 70039003 | 14778 | -7.9 | 2.25E-11 | 6.41E-11 |
| chr1 | 224684092 | 224685378 | 1287 | -7.47 | 2.25E-11 | 6.41E-11 |
| chr7 | 56503299 | 56797946 | 294648 | -1.6 | 2.25E-11 | 6.41E-11 |
| chr8 | 118353071 | 118360027 | 6957 | -2.05 | 2.25E-11 | 6.41E-11 |
| chr6 | 132313784 | 132326761 | 12978 | -7.46 | 2.25E-11 | 6.41E-11 |
| chr10 | 59649021 | 59662191 | 13171 | -8.07 | 2.26E-11 | 6.45E-11 |
| chr17 | 2524403 | 2531023 | 6621 | -7.43 | 2.27E-11 | 6.46E-11 |
| chr2 | 161471693 | 161473770 | 2078 | -7.45 | 2.27E-11 | 6.47E-11 |
| chr20 | 60727375 | 60729356 | 1982 | -7.44 | 2.27E-11 | 6.47E-11 |
| chr9 | 124159137 | 124168728 | 9592 | -8.07 | 2.28E-11 | 6.49E-11 |
| chr1 | 24733827 | 24735810 | 1984 | -7.43 | 2.28E-11 | 6.49E-11 |
| chr14 | 75880554 | 75893172 | 12619 | -7.43 | 2.28E-11 | 6.49E-11 |
| chr13 | 99411847 | 99423342 | 11496 | -7.43 | 2.29E-11 | 6.51E-11 |
| chr6 | 132656701 | 132676929 | 20229 | -7.43 | 2.29E-11 | 6.51E-11 |
| chr7 | 94955356 | 94968136 | 12781 | -7.75 | 2.30E-11 | 6.54E-11 |
| chrX | 36185624 | 36188391 | 2768 | -7.47 | 2.31E-11 | 6.57E-11 |
| chr2 | 177443225 | 177446971 | 3747 | -7.55 | 2.31E-11 | 6.58E-11 |
| chr15 | 73083611 | 73084626 | 1016 | -7.52 | 2.31E-11 | 6.58E-11 |
| chr17 | 3662138 | 3664515 | 2378 | -7.48 | 2.31E-11 | 6.59E-11 |
| chr7 | 40074876 | 40077167 | 2292 | -7.56 | 2.32E-11 | 6.59E-11 |
| chr18 | 60598322 | 60601878 | 3557 | -7.42 | 2.32E-11 | 6.59E-11 |
| chr9 | 137636191 | 137640418 | 4228 | -7.42 | 2.32E-11 | 6.61E-11 |
| chr19 | 19147707 | 19149539 | 1833 | -7.52 | 2.32E-11 | 6.61E-11 |
| chr5 | 125861414 | 125864543 | 3130 | -7.43 | 2.32E-11 | 6.61E-11 |
| chr3 | 151600670 | 151612836 | 12167 | -7.48 | 2.33E-11 | 6.61E-11 |
| chr19 | 50351076 | 50353138 | 2063 | -7.47 | 2.33E-11 | 6.63E-11 |
| chr6 | 11855668 | 11857660 | 1993 | -7.43 | 2.33E-11 | 6.63E-11 |
| chr7 | 94325170 | 94330506 | 5337 | -7.44 | 2.33E-11 | 6.64E-11 |
| chr9 | 131537442 | 131540407 | 2966 | -7.56 | 2.34E-11 | 6.65E-11 |
| chr4 | 122807579 | 122829809 | 22231 | -7.68 | 2.34E-11 | 6.65E-11 |
| chr10 | 135242540 | 135372816 | 130277 | -2.85 | 2.34E-11 | 6.65E-11 |
| chr2 | 232274604 | 232276221 | 1618 | -7.45 | 2.34E-11 | 6.66E-11 |
| chr14 | 27273729 | 27276013 | 2285 | -7.57 | 2.34E-11 | 6.67E-11 |
| chr12 | 13002957 | 13023611 | 20655 | -7.71 | 2.35E-11 | 6.67E-11 |
| chr14 | 97373482 | 97389677 | 16196 | -7.71 | 2.36E-11 | 6.71E-11 |
| chr11 | 103983412 | 103989406 | 5995 | -7.68 | 2.36E-11 | 6.71E-11 |
| chr12 | 118431978 | 118434135 | 2158 | -7.43 | 2.36E-11 | 6.71E-11 |
| chr7 | 25610633 | 25865746 | 255114 | 4.64 | 2.36E-11 | 6.71E-11 |
| chr11 | 101296417 | 101299731 | 3315 | -7.44 | 2.36E-11 | 6.72E-11 |
| chr6 | 24396192 | 24398402 | 2211 | -7.62 | 2.36E-11 | 6.72E-11 |
| chr14 | 76705543 | 76719210 | 13668 | -7.85 | 2.37E-11 | 6.72E-11 |
| chr1 | 70732628 | 70734522 | 1895 | -7.42 | 2.37E-11 | 6.73E-11 |
| chr8 | 107946446 | 107953943 | 7498 | -7.79 | 2.37E-11 | 6.73E-11 |
| chr9 | 96138606 | 96147646 | 9041 | -7.79 | 2.37E-11 | 6.73E-11 |
| chr2 | 203803576 | 203813673 | 10098 | -7.69 | 2.37E-11 | 6.74E-11 |
| chr4 | 129606387 | 129608634 | 2248 | -6.61 | 2.38E-11 | 6.75E-11 |
| chr2 | 188436152 | 188439081 | 2930 | -7.86 | 2.38E-11 | 6.75E-11 |
| chr16 | 51401269 | 51436497 | 35229 | -8.39 | 2.38E-11 | 6.77E-11 |
| chrX | 67192306 | 67195658 | 3353 | -5.68 | 2.39E-11 | 6.79E-11 |
| chr8 | 140271283 | 140415314 | 144032 | 1.99 | 2.40E-11 | 6.80E-11 |
| chr13 | 112096955 | 112119393 | 22439 | -3.15 | 2.40E-11 | 6.80E-11 |
| chr8 | 63453146 | 63456536 | 3391 | 7.6 | 2.40E-11 | 6.80E-11 |
| chr7 | 4659177 | 4662879 | 3703 | -7.63 | 2.40E-11 | 6.81E-11 |
| chr11 | 13112900 | 13121210 | 8311 | -7.6 | 2.40E-11 | 6.82E-11 |
| chrX | 133832913 | 133836222 | 3310 | -7.46 | 2.40E-11 | 6.82E-11 |
| chr4 | 133407372 | 133411307 | 3936 | -7.43 | 2.41E-11 | 6.83E-11 |
| chr6 | 44913585 | 44918367 | 4783 | -7.43 | 2.41E-11 | 6.85E-11 |

| | | | | | | |
|---|---|---|---|---|---|---|
| chr6 | 82533203 | 82545892 | 12690 | -8.12 | 2.41E-11 | 6.85E-11 |
| chr6 | 47432779 | 47439237 | 6459 | -7.45 | 2.42E-11 | 6.86E-11 |
| chr8 | 6627288 | 6641976 | 14689 | -7.6 | 2.42E-11 | 6.86E-11 |
| chr1 | 53954363 | 53956606 | 2244 | -7.46 | 2.42E-11 | 6.87E-11 |
| chr12 | 20969520 | 20970958 | 1439 | -7.53 | 2.42E-11 | 6.87E-11 |
| chr8 | 144433137 | 144435900 | 2764 | -7.86 | 2.43E-11 | 6.89E-11 |
| chr1 | 223821313 | 223837457 | 16145 | -7.84 | 2.43E-11 | 6.89E-11 |
| chr11 | 86006423 | 86009968 | 3546 | -7.55 | 2.43E-11 | 6.89E-11 |
| chr18 | 28940945 | 28948305 | 7361 | -7.55 | 2.43E-11 | 6.89E-11 |
| chr7 | 6607868 | 6610395 | 2528 | -7.43 | 2.43E-11 | 6.89E-11 |
| chr2 | 198496643 | 198499346 | 2704 | -7.42 | 2.43E-11 | 6.90E-11 |
| chr3 | 38226629 | 38228571 | 1943 | -7.62 | 2.44E-11 | 6.91E-11 |
| chr3 | 170555520 | 170563304 | 7785 | -7.56 | 2.44E-11 | 6.92E-11 |
| chr5 | 107549368 | 107552052 | 2685 | -7.5 | 2.44E-11 | 6.93E-11 |
| chr11 | 11953291 | 11956032 | 2742 | -7.46 | 2.44E-11 | 6.93E-11 |
| chr7 | 85301260 | 85326796 | 25537 | -7.42 | 2.46E-11 | 6.96E-11 |
| chr3 | 99733197 | 99735202 | 2006 | -7.43 | 2.46E-11 | 6.97E-11 |
| chr12 | 64550978 | 64560793 | 9816 | -7.73 | 2.46E-11 | 6.97E-11 |
| chr4 | 77825177 | 77831744 | 6568 | -2.55 | 2.46E-11 | 6.97E-11 |
| chr9 | 4436400 | 4453763 | 17364 | -1.38 | 2.47E-11 | 6.99E-11 |
| chr11 | 92656673 | 92661244 | 4572 | -7.42 | 2.47E-11 | 6.99E-11 |
| chr2 | 26231938 | 26233266 | 1329 | -7.44 | 2.47E-11 | 7.00E-11 |
| chr3 | 134014115 | 134015851 | 1737 | -7.41 | 2.47E-11 | 7.01E-11 |
| chrX | 9839519 | 9855117 | 15599 | -8.02 | 2.48E-11 | 7.03E-11 |
| chr9 | 84628342 | 84633103 | 4762 | -7.58 | 2.48E-11 | 7.04E-11 |
| chr8 | 123000493 | 123005852 | 5360 | -6.05 | 2.49E-11 | 7.05E-11 |
| chr14 | 101456053 | 101463498 | 7446 | -8.07 | 2.49E-11 | 7.06E-11 |
| chr18 | 20075960 | 20082014 | 6055 | -7.48 | 2.50E-11 | 7.07E-11 |
| chr7 | 34485227 | 34489146 | 3920 | -7.47 | 2.51E-11 | 7.10E-11 |
| chr7 | 35691430 | 35696723 | 5294 | -7.47 | 2.51E-11 | 7.10E-11 |
| chr2 | 187879999 | 187887838 | 7840 | 7.48 | 2.51E-11 | 7.11E-11 |
| chr7 | 68847410 | 68850252 | 2843 | -7.52 | 2.51E-11 | 7.11E-11 |
| chr10 | 74263188 | 74265473 | 2286 | -7.53 | 2.51E-11 | 7.12E-11 |
| chr21 | 33860381 | 33862146 | 1766 | -7.42 | 2.52E-11 | 7.12E-11 |
| chr2 | 42347617 | 42349394 | 1778 | -7.43 | 2.52E-11 | 7.12E-11 |
| chr20 | 10232880 | 10236632 | 3753 | -7.47 | 2.52E-11 | 7.13E-11 |
| chr2 | 75353630 | 75377727 | 24098 | -8.16 | 2.52E-11 | 7.14E-11 |
| chr2 | 120421129 | 120424032 | 2904 | -3.02 | 2.52E-11 | 7.14E-11 |
| chr1 | 193440898 | 193460444 | 19547 | -7.6 | 2.52E-11 | 7.14E-11 |
| chr3 | 195188472 | 195198408 | 9937 | -7.5 | 2.52E-11 | 7.14E-11 |
| chr4 | 85507696 | 85510708 | 3013 | -7.49 | 2.54E-11 | 7.18E-11 |
| chr1 | 151709378 | 151711637 | 2260 | -7.43 | 2.55E-11 | 7.20E-11 |
| chr9 | 461002 | 463458 | 2457 | -7.43 | 2.55E-11 | 7.20E-11 |
| chr7 | 156828277 | 156830006 | 1730 | -7.49 | 2.55E-11 | 7.20E-11 |
| chr20 | 34867344 | 34880587 | 13244 | -2.78 | 2.55E-11 | 7.20E-11 |
| chr13 | 24998006 | 24999694 | 1689 | -7.46 | 2.55E-11 | 7.22E-11 |
| chr8 | 82840810 | 82844523 | 3714 | 7.75 | 2.55E-11 | 7.22E-11 |
| chr5 | 5171208 | 5176398 | 5191 | -7.46 | 2.56E-11 | 7.22E-11 |
| chr5 | 53337346 | 53356489 | 19144 | -7.45 | 2.56E-11 | 7.24E-11 |
| chr6 | 141367104 | 141370352 | 3249 | -7.45 | 2.56E-11 | 7.24E-11 |
| chr1 | 5648753 | 5652805 | 4053 | -2.73 | 2.56E-11 | 7.25E-11 |
| chr16 | 72247687 | 72250119 | 2433 | -7.41 | 2.56E-11 | 7.25E-11 |
| chr14 | 69895167 | 69899672 | 4506 | -7.42 | 2.57E-11 | 7.26E-11 |
| chr9 | 133369580 | 133371785 | 2206 | -7.43 | 2.58E-11 | 7.28E-11 |
| chr3 | 66470522 | 66474290 | 3769 | -7.48 | 2.58E-11 | 7.28E-11 |
| chr1 | 230393223 | 230395607 | 2385 | -7.56 | 2.58E-11 | 7.30E-11 |
| chr3 | 169972302 | 169974341 | 2040 | -7.45 | 2.59E-11 | 7.31E-11 |
| chr4 | 31848204 | 31856327 | 8124 | -7.45 | 2.59E-11 | 7.31E-11 |
| chr19 | 1634732 | 1638567 | 3836 | -7.42 | 2.60E-11 | 7.35E-11 |
| chr11 | 120934914 | 120936857 | 1944 | -7.43 | 2.62E-11 | 7.40E-11 |
| chr8 | 125319082 | 125320821 | 1740 | -7.41 | 2.62E-11 | 7.40E-11 |

| | | | | | | |
|---|---|---|---|---|---|---|
| chr16 | 74557650 | 74559359 | 1710 | -7.44 | 2.62E-11 | 7.41E-11 |
| chr4 | 59796714 | 59804888 | 8175 | -7.46 | 2.62E-11 | 7.41E-11 |
| chr2 | 110475273 | 110486555 | 11283 | -7.5 | 2.62E-11 | 7.41E-11 |
| chr8 | 107916706 | 107918769 | 2064 | -7.59 | 2.62E-11 | 7.41E-11 |
| chr7 | 108900550 | 108904484 | 3935 | 7.47 | 2.63E-11 | 7.42E-11 |
| chr4 | 17643622 | 17645946 | 2325 | -7.41 | 2.63E-11 | 7.43E-11 |
| chr9 | 33355534 | 33357119 | 1586 | -7.63 | 2.63E-11 | 7.43E-11 |
| chr5 | 127111267 | 127129037 | 17771 | -6.36 | 2.65E-11 | 7.47E-11 |
| chr17 | 58509389 | 58511773 | 2385 | -7.48 | 2.65E-11 | 7.48E-11 |
| chr12 | 21792911 | 21795428 | 2518 | -5.65 | 2.65E-11 | 7.49E-11 |
| chr2 | 162291349 | 162293125 | 1777 | -7.4 | 2.66E-11 | 7.51E-11 |
| chr2 | 24833883 | 24835861 | 1979 | -7.43 | 2.67E-11 | 7.55E-11 |
| chr5 | 141285014 | 141286604 | 1591 | -7.43 | 2.67E-11 | 7.55E-11 |
| chr19 | 19394961 | 19404711 | 9751 | -7.45 | 2.68E-11 | 7.55E-11 |
| chr17 | 76614142 | 76617099 | 2958 | -7.89 | 2.68E-11 | 7.57E-11 |
| chrX | 30634095 | 30641528 | 7434 | -7.42 | 2.68E-11 | 7.57E-11 |
| chr17 | 60888664 | 60892227 | 3564 | -7.4 | 2.69E-11 | 7.59E-11 |
| chr15 | 67177647 | 67180209 | 2563 | -7.47 | 2.69E-11 | 7.60E-11 |
| chr4 | 162567682 | 162571344 | 3663 | -7.41 | 2.69E-11 | 7.60E-11 |
| chr12 | 101780941 | 101786513 | 5573 | -7.68 | 2.70E-11 | 7.61E-11 |
| chr13 | 103515757 | 103517643 | 1887 | -7.61 | 2.70E-11 | 7.62E-11 |
| chr4 | 154204137 | 154207653 | 3517 | -7.62 | 2.72E-11 | 7.66E-11 |
| chr11 | 67790927 | 67792964 | 2038 | -7.89 | 2.72E-11 | 7.67E-11 |
| chr8 | 8612661 | 8623391 | 10731 | -7.58 | 2.72E-11 | 7.67E-11 |
| chr9 | 86836775 | 86855336 | 18562 | -7.89 | 2.73E-11 | 7.68E-11 |
| chr17 | 32638206 | 32743729 | 105524 | 4.03 | 2.73E-11 | 7.68E-11 |
| chr10 | 127953243 | 127955920 | 2678 | -7.4 | 2.73E-11 | 7.69E-11 |
| chr15 | 79418550 | 79443469 | 24920 | -2.01 | 2.73E-11 | 7.69E-11 |
| chr3 | 40705460 | 40709142 | 3683 | 6.49 | 2.73E-11 | 7.69E-11 |
| chr3 | 193871765 | 193873516 | 1752 | -3.69 | 2.73E-11 | 7.69E-11 |
| chr12 | 28551077 | 28554661 | 3585 | -7.41 | 2.73E-11 | 7.70E-11 |
| chr15 | 101399998 | 101410428 | 10431 | -8.24 | 2.73E-11 | 7.71E-11 |
| chr12 | 40271574 | 40274067 | 2494 | -7.41 | 2.74E-11 | 7.71E-11 |
| chr5 | 137686372 | 137688139 | 1768 | -7.41 | 2.76E-11 | 7.77E-11 |
| chr5 | 133548405 | 133550356 | 1952 | -7.57 | 2.76E-11 | 7.78E-11 |
| chr4 | 189587967 | 189593084 | 5118 | -7.4 | 2.76E-11 | 7.79E-11 |
| chr7 | 132933070 | 132934961 | 1892 | -7.45 | 2.77E-11 | 7.80E-11 |
| chr19 | 46882747 | 46893456 | 10710 | -7.42 | 2.78E-11 | 7.82E-11 |
| chr13 | 40428333 | 40461275 | 32943 | -2.52 | 2.78E-11 | 7.83E-11 |
| chr1 | 167541377 | 167544802 | 3426 | -7.47 | 2.78E-11 | 7.84E-11 |
| chr7 | 83770286 | 83773055 | 2770 | -7.47 | 2.78E-11 | 7.84E-11 |
| chr2 | 165533244 | 165535673 | 2430 | -7.44 | 2.79E-11 | 7.85E-11 |
| chr12 | 46193042 | 46195892 | 2851 | -7.67 | 2.79E-11 | 7.85E-11 |
| chr6 | 64361475 | 64363939 | 2465 | -7.41 | 2.79E-11 | 7.85E-11 |
| chr8 | 126380949 | 126383054 | 2106 | -7.41 | 2.79E-11 | 7.85E-11 |
| chr1 | 180665529 | 180672647 | 7119 | -7.67 | 2.79E-11 | 7.86E-11 |
| chr12 | 17768772 | 17782000 | 13229 | -7.75 | 2.79E-11 | 7.86E-11 |
| chr6 | 56060754 | 56063296 | 2543 | -7.43 | 2.79E-11 | 7.86E-11 |
| chr4 | 166647170 | 166659936 | 12767 | -7.63 | 2.80E-11 | 7.88E-11 |
| chr11 | 12060679 | 12065469 | 4791 | -7.4 | 2.81E-11 | 7.89E-11 |
| chr2 | 144675123 | 144678710 | 3588 | -4.18 | 2.81E-11 | 7.92E-11 |
| chr12 | 68404833 | 68409564 | 4732 | 7.74 | 2.81E-11 | 7.92E-11 |
| chr12 | 108115154 | 108116723 | 1570 | -7.4 | 2.82E-11 | 7.92E-11 |
| chr11 | 98916422 | 98926007 | 9586 | 7.51 | 2.82E-11 | 7.92E-11 |
| chr11 | 11738607 | 11752268 | 13662 | -1.72 | 2.82E-11 | 7.93E-11 |
| chr13 | 46072066 | 46073790 | 1725 | -7.48 | 2.83E-11 | 7.96E-11 |
| chr11 | 17953735 | 17966290 | 12556 | -7.4 | 2.83E-11 | 7.96E-11 |
| chr10 | 33739665 | 33741679 | 2015 | -7.43 | 2.83E-11 | 7.96E-11 |
| chr11 | 91952274 | 91954885 | 2612 | -7.52 | 2.84E-11 | 7.98E-11 |
| chr2 | 152667539 | 152669841 | 2303 | -7.97 | 2.85E-11 | 8.02E-11 |
| chr11 | 66349095 | 66351188 | 2094 | -7.45 | 2.85E-11 | 8.02E-11 |

| | | | | | | |
|---|---|---|---|---|---|---|
| chr4 | 158723442 | 158733513 | 10072 | 7.49 | 2.85E-11 | 8.02E-11 |
| chrX | 51479136 | 51508868 | 29733 | -2 | 2.86E-11 | 8.03E-11 |
| chr1 | 222629806 | 222632729 | 2924 | -7.43 | 2.87E-11 | 8.06E-11 |
| chr10 | 73806018 | 73810393 | 4376 | -7.43 | 2.87E-11 | 8.06E-11 |
| chr10 | 100191354 | 100193385 | 2032 | -7.47 | 2.88E-11 | 8.09E-11 |
| chr1 | 77531887 | 77540969 | 9083 | -7.41 | 2.89E-11 | 8.11E-11 |
| chr5 | 131946556 | 131948013 | 1458 | -7.42 | 2.89E-11 | 8.12E-11 |
| chr7 | 92651542 | 92662427 | 10886 | -7.74 | 2.89E-11 | 8.12E-11 |
| chr2 | 172627296 | 172635595 | 8300 | -7.56 | 2.90E-11 | 8.14E-11 |
| chr18 | 8438588 | 8443198 | 4611 | -7.42 | 2.90E-11 | 8.16E-11 |
| chr2 | 157155722 | 157157509 | 1788 | -7.4 | 2.91E-11 | 8.16E-11 |
| chr11 | 118117794 | 118130462 | 12669 | -2.39 | 2.92E-11 | 8.20E-11 |
| chr3 | 56631151 | 56640881 | 9731 | -7.4 | 2.92E-11 | 8.20E-11 |
| chr15 | 37086466 | 37092755 | 6290 | -7.57 | 2.92E-11 | 8.21E-11 |
| chr13 | 27569699 | 27578229 | 8531 | -7.42 | 2.93E-11 | 8.24E-11 |
| chr7 | 128077072 | 128078755 | 1684 | -7.57 | 2.94E-11 | 8.25E-11 |
| chr12 | 100121747 | 100124296 | 2550 | -5.58 | 2.94E-11 | 8.27E-11 |
| chr3 | 56601202 | 56603469 | 2268 | -7.4 | 2.95E-11 | 8.29E-11 |
| chr21 | 18242508 | 18251697 | 9190 | 5.75 | 2.95E-11 | 8.29E-11 |
| chr6 | 148319827 | 148333488 | 13662 | -8.15 | 2.96E-11 | 8.31E-11 |
| chr22 | 30006022 | 30007838 | 1817 | -7.51 | 2.96E-11 | 8.31E-11 |
| chr4 | 125881046 | 125883969 | 2924 | -7.41 | 2.96E-11 | 8.31E-11 |
| chr20 | 53001487 | 53004929 | 3443 | 7.47 | 2.96E-11 | 8.31E-11 |
| chr3 | 32485795 | 32488595 | 2801 | -7.42 | 2.96E-11 | 8.32E-11 |
| chrX | 10163224 | 10165004 | 1781 | -7.48 | 2.97E-11 | 8.33E-11 |
| chr5 | 67979510 | 67987670 | 8161 | -7.48 | 2.97E-11 | 8.34E-11 |
| chr5 | 139089781 | 139100663 | 10883 | -7.39 | 2.99E-11 | 8.40E-11 |
| chr1 | 4066170 | 4068534 | 2365 | -7.46 | 3.00E-11 | 8.41E-11 |
| chr16 | 66744333 | 66746871 | 2539 | -7.46 | 3.00E-11 | 8.41E-11 |
| chr1 | 191110510 | 191122518 | 12009 | -7.39 | 3.00E-11 | 8.41E-11 |
| chr10 | 95227417 | 95237071 | 9655 | -7.46 | 3.00E-11 | 8.43E-11 |
| chr6 | 111427491 | 111429523 | 2033 | -7.44 | 3.00E-11 | 8.43E-11 |
| chr9 | 135126111 | 135127847 | 1737 | -7.46 | 3.01E-11 | 8.43E-11 |
| chr2 | 197782147 | 197783608 | 1462 | -7.46 | 3.01E-11 | 8.44E-11 |
| chr12 | 13597745 | 13599313 | 1569 | -7.4 | 3.02E-11 | 8.46E-11 |
| chr15 | 50953616 | 50955269 | 1654 | -7.55 | 3.02E-11 | 8.47E-11 |
| chr10 | 9242124 | 9244775 | 2652 | -7.56 | 3.03E-11 | 8.50E-11 |
| chr5 | 56452818 | 56454855 | 2038 | -7.42 | 3.03E-11 | 8.50E-11 |
| chr9 | 125804251 | 125806318 | 2068 | -7.42 | 3.03E-11 | 8.50E-11 |
| chr8 | 115018386 | 115049502 | 31117 | -2.58 | 3.04E-11 | 8.53E-11 |
| chr3 | 157198260 | 157200752 | 2493 | -3.4 | 3.04E-11 | 8.53E-11 |
| chr4 | 86719171 | 86721254 | 2084 | -7.45 | 3.06E-11 | 8.56E-11 |
| chrX | 35056719 | 35060369 | 3651 | -7.41 | 3.06E-11 | 8.57E-11 |
| chr2 | 225269684 | 225271824 | 2141 | -7.44 | 3.07E-11 | 8.60E-11 |
| chr3 | 170592096 | 170603114 | 11019 | -7.49 | 3.07E-11 | 8.61E-11 |
| chr2 | 162912151 | 162915209 | 3059 | -7.48 | 3.08E-11 | 8.62E-11 |
| chr17 | 77934628 | 77936346 | 1719 | -7.4 | 3.08E-11 | 8.62E-11 |
| chr15 | 78463929 | 78464905 | 977 | -7.39 | 3.08E-11 | 8.62E-11 |
| chr2 | 47864842 | 47866635 | 1794 | -7.51 | 3.08E-11 | 8.63E-11 |
| chr2 | 9873013 | 9882326 | 9314 | -7.88 | 3.08E-11 | 8.63E-11 |
| chr6 | 8088232 | 8091803 | 3572 | -7.81 | 3.08E-11 | 8.63E-11 |
| chr3 | 112374772 | 112385125 | 10354 | -7.52 | 3.09E-11 | 8.66E-11 |
| chr3 | 64416231 | 64422737 | 6507 | -7.44 | 3.09E-11 | 8.66E-11 |
| chr1 | 189823888 | 189829227 | 5340 | -7.53 | 3.11E-11 | 8.71E-11 |
| chr14 | 45563577 | 45564977 | 1401 | -7.38 | 3.11E-11 | 8.71E-11 |
| chr5 | 60339591 | 60340826 | 1236 | -7.51 | 3.12E-11 | 8.73E-11 |
| chr10 | 21450525 | 21452213 | 1689 | -7.4 | 3.12E-11 | 8.73E-11 |
| chr3 | 170942929 | 170944866 | 1938 | -7.38 | 3.13E-11 | 8.75E-11 |
| chr11 | 102065306 | 102067198 | 1893 | -7.38 | 3.13E-11 | 8.76E-11 |
| chr6 | 166110878 | 166113801 | 2924 | -7.47 | 3.13E-11 | 8.77E-11 |
| chr6 | 86588460 | 86595075 | 6616 | 7.61 | 3.14E-11 | 8.79E-11 |

| | | | | | | |
|---|---|---|---|---|---|---|
| chr2 | 164931819 | 164953153 | 21335 | -7.91 | 3.14E-11 | 8.79E-11 |
| chr13 | 49471993 | 49477271 | 5279 | -7.46 | 3.14E-11 | 8.80E-11 |
| chr3 | 127608741 | 127611059 | 2319 | -7.38 | 3.15E-11 | 8.81E-11 |
| chr1 | 112581988 | 112668569 | 86582 | 2.66 | 3.16E-11 | 8.85E-11 |
| chr12 | 10691639 | 10693870 | 2232 | -7.41 | 3.18E-11 | 8.88E-11 |
| chrX | 149134147 | 149136481 | 2335 | -7.59 | 3.18E-11 | 8.89E-11 |
| chr1 | 187993994 | 188008801 | 14808 | -7.45 | 3.18E-11 | 8.90E-11 |
| chr13 | 45182890 | 45186330 | 3441 | -7.44 | 3.18E-11 | 8.90E-11 |
| chr7 | 131586701 | 131589295 | 2595 | -7.64 | 3.19E-11 | 8.92E-11 |
| chr4 | 185216332 | 185226254 | 9923 | -7.45 | 3.19E-11 | 8.92E-11 |
| chr12 | 64210265 | 64224803 | 14539 | -8.05 | 3.19E-11 | 8.93E-11 |
| chr8 | 94747539 | 94751196 | 3658 | -7.38 | 3.19E-11 | 8.93E-11 |
| chr12 | 57336458 | 57338369 | 1912 | -7.38 | 3.20E-11 | 8.94E-11 |
| chr1 | 56146426 | 56149391 | 2966 | -7.44 | 3.21E-11 | 8.97E-11 |
| chr21 | 27710953 | 27723321 | 12369 | -6.18 | 3.21E-11 | 8.98E-11 |
| chr1 | 161723541 | 161726210 | 2670 | -7.4 | 3.22E-11 | 8.99E-11 |
| chr1 | 169116487 | 169120154 | 3668 | -7.56 | 3.22E-11 | 8.99E-11 |
| chr1 | 242101395 | 242102974 | 1580 | -7.52 | 3.23E-11 | 9.02E-11 |
| chr10 | 7236591 | 7238834 | 2244 | -7.42 | 3.23E-11 | 9.02E-11 |
| chr1 | 151365894 | 151367789 | 1896 | -7.57 | 3.23E-11 | 9.02E-11 |
| chr1 | 172560152 | 172567117 | 6966 | -7.38 | 3.23E-11 | 9.02E-11 |
| chr4 | 95805903 | 95838293 | 32391 | -7.91 | 3.23E-11 | 9.03E-11 |
| chr9 | 15020289 | 15062733 | 42445 | 3.26 | 3.24E-11 | 9.05E-11 |
| chr13 | 91599696 | 91606991 | 7296 | -7.56 | 3.24E-11 | 9.06E-11 |
| chr7 | 803935 | 806286 | 2352 | -7.39 | 3.25E-11 | 9.09E-11 |
| chr16 | 22659383 | 22808553 | 149171 | -2.31 | 3.26E-11 | 9.12E-11 |
| chr2 | 19701143 | 19717808 | 16666 | -6.62 | 3.27E-11 | 9.14E-11 |
| chr21 | 18906766 | 18912551 | 5786 | -7.58 | 3.28E-11 | 9.15E-11 |
| chr11 | 75984598 | 75986651 | 2054 | -7.38 | 3.28E-11 | 9.16E-11 |
| chr17 | 16082992 | 16084305 | 1314 | -7.42 | 3.28E-11 | 9.17E-11 |
| chr3 | 67418378 | 67421680 | 3303 | -7.47 | 3.29E-11 | 9.18E-11 |
| chr1 | 53852146 | 53854166 | 2021 | -7.62 | 3.29E-11 | 9.18E-11 |
| chr8 | 132859949 | 132861725 | 1777 | -7.41 | 3.29E-11 | 9.18E-11 |
| chr2 | 28927548 | 28930177 | 2630 | -7.38 | 3.30E-11 | 9.20E-11 |
| chr6 | 100715685 | 100723984 | 8300 | -7.38 | 3.30E-11 | 9.21E-11 |
| chr3 | 179014716 | 179017797 | 3082 | -5.96 | 3.31E-11 | 9.23E-11 |
| chr11 | 109944868 | 109951103 | 6236 | -7.45 | 3.31E-11 | 9.24E-11 |
| chr19 | 6117020 | 6128557 | 11538 | -7.63 | 3.31E-11 | 9.25E-11 |
| chr16 | 58855223 | 58858321 | 3099 | -7.43 | 3.31E-11 | 9.25E-11 |
| chr12 | 109610186 | 109612476 | 2291 | -7.38 | 3.31E-11 | 9.25E-11 |
| chr10 | 61878961 | 61880960 | 2000 | -7.4 | 3.32E-11 | 9.28E-11 |
| chr10 | 60659753 | 60663845 | 4093 | -7.96 | 3.33E-11 | 9.28E-11 |
| chr11 | 17235481 | 17236823 | 1343 | -7.49 | 3.33E-11 | 9.29E-11 |
| chr3 | 31819418 | 31824366 | 4949 | -7.4 | 3.33E-11 | 9.29E-11 |
| chr6 | 107745931 | 107749689 | 3759 | -7.37 | 3.34E-11 | 9.31E-11 |
| chr1 | 231704750 | 231715702 | 10953 | -7.37 | 3.34E-11 | 9.33E-11 |
| chr15 | 74295184 | 74305986 | 10803 | -7.54 | 3.34E-11 | 9.33E-11 |
| chr22 | 28052197 | 28099206 | 47010 | -8.24 | 3.35E-11 | 9.33E-11 |
| chr5 | 172606848 | 172609460 | 2613 | -7.46 | 3.35E-11 | 9.33E-11 |
| chr21 | 38480013 | 38481663 | 1651 | -7.42 | 3.35E-11 | 9.33E-11 |
| chr2 | 231052964 | 231058902 | 5939 | -7.51 | 3.35E-11 | 9.33E-11 |
| chr9 | 113848308 | 113856217 | 7910 | -7.51 | 3.35E-11 | 9.33E-11 |
| chr8 | 119736354 | 119741340 | 4987 | 2.92 | 3.36E-11 | 9.36E-11 |
| chr10 | 115914222 | 115930259 | 16038 | -7.69 | 3.38E-11 | 9.42E-11 |
| chr1 | 94099618 | 94101562 | 1945 | -7.38 | 3.39E-11 | 9.46E-11 |
| chr8 | 82608772 | 82610921 | 2150 | -7.38 | 3.39E-11 | 9.46E-11 |
| chr6 | 90653880 | 90655746 | 1867 | -7.39 | 3.41E-11 | 9.50E-11 |
| chr15 | 71824859 | 71862242 | 37384 | -7.37 | 3.41E-11 | 9.50E-11 |
| chr2 | 63596982 | 63599414 | 2433 | -7.37 | 3.41E-11 | 9.50E-11 |
| chr1 | 162043711 | 162045483 | 1773 | -7.42 | 3.44E-11 | 9.57E-11 |
| chr10 | 96210874 | 96223433 | 12560 | -7.43 | 3.44E-11 | 9.57E-11 |

| | | | | | | |
|---|---|---|---|---|---|---|
| chr12 | 60004560 | 60008529 | 3970 | -7.41 | 3.44E-11 | 9.59E-11 |
| chr14 | 85730618 | 85734674 | 4057 | -7.48 | 3.44E-11 | 9.59E-11 |
| chr6 | 82849782 | 82853113 | 3332 | -4.2 | 3.45E-11 | 9.60E-11 |
| chr3 | 16499091 | 16541368 | 42278 | -8.7 | 3.45E-11 | 9.61E-11 |
| chr1 | 161226418 | 161233202 | 6785 | -7.43 | 3.45E-11 | 9.61E-11 |
| chr16 | 19046295 | 19048896 | 2602 | -7.37 | 3.46E-11 | 9.64E-11 |
| chr1 | 24173393 | 24175068 | 1676 | -7.37 | 3.46E-11 | 9.65E-11 |
| chr10 | 32081053 | 32083017 | 1965 | -7.4 | 3.46E-11 | 9.65E-11 |
| chr11 | 27535993 | 27537994 | 2002 | -7.47 | 3.46E-11 | 9.65E-11 |
| chr4 | 130272603 | 130283682 | 11080 | -7.42 | 3.47E-11 | 9.66E-11 |
| chr10 | 87995915 | 87999376 | 3462 | -7.41 | 3.47E-11 | 9.66E-11 |
| chr5 | 34725912 | 34728165 | 2254 | -7.37 | 3.47E-11 | 9.66E-11 |
| chr6 | 161806614 | 161811274 | 4661 | -7.37 | 3.47E-11 | 9.66E-11 |
| chr22 | 36546676 | 36549231 | 2556 | -7.37 | 3.47E-11 | 9.67E-11 |
| chr4 | 63671181 | 63684134 | 12954 | -7.63 | 3.48E-11 | 9.69E-11 |
| chr11 | 122539649 | 122540811 | 1163 | -7.37 | 3.48E-11 | 9.69E-11 |
| chr2 | 17583813 | 17675900 | 92088 | 2.54 | 3.48E-11 | 9.69E-11 |
| chrX | 83578988 | 83591370 | 12383 | 7.62 | 3.48E-11 | 9.69E-11 |
| chr10 | 4533234 | 4534988 | 1755 | -7.37 | 3.49E-11 | 9.71E-11 |
| chr11 | 47849605 | 47851242 | 1638 | -7.37 | 3.49E-11 | 9.71E-11 |
| chr1 | 215785911 | 215787538 | 1628 | -7.52 | 3.50E-11 | 9.72E-11 |
| chr12 | 69270854 | 69276350 | 5497 | -7.42 | 3.50E-11 | 9.73E-11 |
| chr11 | 123128251 | 123130767 | 2517 | -7.39 | 3.50E-11 | 9.73E-11 |
| chrX | 116042053 | 116050797 | 8745 | -7.36 | 3.50E-11 | 9.75E-11 |
| chr4 | 47874509 | 47877180 | 2672 | -7.58 | 3.51E-11 | 9.76E-11 |
| chr4 | 131724295 | 131738066 | 13772 | -7.6 | 3.51E-11 | 9.77E-11 |
| chr20 | 44587007 | 44588546 | 1540 | -7.38 | 3.51E-11 | 9.77E-11 |
| chr5 | 137092583 | 137096817 | 4235 | -7.36 | 3.53E-11 | 9.80E-11 |
| chr10 | 112508320 | 112509852 | 1533 | -7.41 | 3.54E-11 | 9.84E-11 |
| chrX | 84353884 | 84356517 | 2634 | -7.38 | 3.55E-11 | 9.86E-11 |
| chr6 | 13108661 | 13113687 | 5027 | -4.98 | 3.55E-11 | 9.88E-11 |
| chr5 | 57068794 | 57080516 | 11723 | -7.36 | 3.55E-11 | 9.88E-11 |
| chr6 | 10118602 | 10147093 | 28492 | -8.18 | 3.56E-11 | 9.89E-11 |
| chr8 | 81645541 | 81647392 | 1852 | -7.36 | 3.56E-11 | 9.90E-11 |
| chr7 | 98713779 | 98731658 | 17880 | -7.39 | 3.57E-11 | 9.93E-11 |
| chr7 | 69375222 | 69391414 | 16193 | -7.36 | 3.58E-11 | 9.94E-11 |
| chr5 | 86585817 | 86588170 | 2354 | -7.39 | 3.58E-11 | 9.94E-11 |
| chr2 | 230231435 | 230234984 | 3550 | -7.54 | 3.58E-11 | 9.95E-11 |
| chr11 | 89094543 | 89097449 | 2907 | 7.48 | 3.58E-11 | 9.95E-11 |
| chr2 | 87021383 | 87025463 | 4081 | -6.37 | 3.59E-11 | 9.97E-11 |
| chr3 | 23892535 | 23894480 | 1946 | -7.36 | 3.59E-11 | 9.98E-11 |
| chr4 | 131298068 | 131300424 | 2357 | -7.36 | 3.59E-11 | 9.98E-11 |
| chr9 | 77621893 | 77624661 | 2769 | -7.58 | 3.60E-11 | 9.99E-11 |
| chr4 | 127976583 | 127991185 | 14603 | -7.55 | 3.61E-11 | 1.00E-10 |
| chr3 | 164678451 | 164893693 | 215243 | 2.22 | 3.61E-11 | 1.00E-10 |
| chr12 | 69442022 | 69453234 | 11213 | -7.48 | 3.61E-11 | 1.00E-10 |
| chr8 | 99624818 | 99626953 | 2136 | -7.46 | 3.62E-11 | 1.00E-10 |
| chr4 | 104911029 | 104914536 | 3508 | -7.49 | 3.62E-11 | 1.00E-10 |
| chr3 | 126637119 | 126639861 | 2743 | -7.47 | 3.62E-11 | 1.01E-10 |
| chr2 | 197918336 | 197920944 | 2609 | -7.48 | 3.63E-11 | 1.01E-10 |
| chr5 | 100259245 | 100272804 | 13560 | -7.78 | 3.63E-11 | 1.01E-10 |
| chr17 | 66498555 | 66514951 | 16397 | -7.45 | 3.63E-11 | 1.01E-10 |
| chr12 | 86321898 | 86341022 | 19125 | -7.7 | 3.63E-11 | 1.01E-10 |
| chr6 | 138418996 | 138421443 | 2448 | -7.36 | 3.64E-11 | 1.01E-10 |
| chr14 | 89659597 | 89663848 | 4252 | -7.37 | 3.64E-11 | 1.01E-10 |
| chr8 | 141787210 | 141789873 | 2664 | -7.41 | 3.64E-11 | 1.01E-10 |
| chr1 | 173693127 | 173706035 | 12909 | -7.37 | 3.64E-11 | 1.01E-10 |
| chr21 | 36789903 | 36825344 | 35442 | -1.2 | 3.65E-11 | 1.01E-10 |
| chr13 | 111623596 | 111630102 | 6507 | -7.45 | 3.65E-11 | 1.01E-10 |
| chr6 | 145254589 | 145266515 | 11927 | -7.38 | 3.65E-11 | 1.01E-10 |
| chr6 | 140946135 | 140950636 | 4502 | -7.51 | 3.66E-11 | 1.01E-10 |

| | | | | | | |
|---|---|---|---|---|---|---|
| chr2 | 216392986 | 216399875 | 6890 | -7.41 | 3.67E-11 | 1.02E-10 |
| chr7 | 15978547 | 15994226 | 15680 | 7.55 | 3.67E-11 | 1.02E-10 |
| chr16 | 49713041 | 49715025 | 1985 | -7.39 | 3.67E-11 | 1.02E-10 |
| chr12 | 38515810 | 38524264 | 8455 | -7.43 | 3.67E-11 | 1.02E-10 |
| chr4 | 184156518 | 184159033 | 2516 | -7.37 | 3.68E-11 | 1.02E-10 |
| chr1 | 86365531 | 86367429 | 1899 | -7.36 | 3.68E-11 | 1.02E-10 |
| chr2 | 241783609 | 241794741 | 11133 | -7.38 | 3.69E-11 | 1.02E-10 |
| chr12 | 83797150 | 83809935 | 12786 | -7.37 | 3.69E-11 | 1.02E-10 |
| chr7 | 109493821 | 109496738 | 2918 | -7.39 | 3.69E-11 | 1.02E-10 |
| chr9 | 19266324 | 19271476 | 5153 | -7.39 | 3.69E-11 | 1.02E-10 |
| chr8 | 143289794 | 143291369 | 1576 | -7.38 | 3.70E-11 | 1.03E-10 |
| chr8 | 76419443 | 76455421 | 35979 | -7.88 | 3.70E-11 | 1.03E-10 |
| chrX | 100185382 | 100187787 | 2406 | -7.36 | 3.72E-11 | 1.03E-10 |
| chr1 | 186469157 | 186475604 | 6448 | -7.4 | 3.73E-11 | 1.03E-10 |
| chr12 | 110216250 | 110220384 | 4135 | -7.36 | 3.73E-11 | 1.03E-10 |
| chr5 | 56417843 | 56420503 | 2661 | -7.36 | 3.75E-11 | 1.04E-10 |
| chrX | 21118040 | 21120520 | 2481 | -7.37 | 3.75E-11 | 1.04E-10 |
| chr6 | 15796426 | 15799959 | 3534 | -7.36 | 3.75E-11 | 1.04E-10 |
| chr3 | 95666661 | 95672702 | 6042 | -6.08 | 3.75E-11 | 1.04E-10 |
| chr8 | 9593198 | 9594224 | 1027 | -7.35 | 3.76E-11 | 1.04E-10 |
| chr5 | 53275221 | 53278006 | 2786 | -7.46 | 3.76E-11 | 1.04E-10 |
| chr8 | 127859580 | 127861484 | 1905 | -7.37 | 3.76E-11 | 1.04E-10 |
| chr1 | 22524830 | 22527056 | 2227 | -7.38 | 3.77E-11 | 1.04E-10 |
| chr4 | 39351279 | 39353162 | 1884 | -7.41 | 3.78E-11 | 1.05E-10 |
| chr7 | 64738862 | 64747544 | 8683 | -7.45 | 3.78E-11 | 1.05E-10 |
| chr2 | 75133796 | 75135626 | 1831 | -3.98 | 3.78E-11 | 1.05E-10 |
| chr22 | 28904321 | 28908864 | 4544 | -7.37 | 3.78E-11 | 1.05E-10 |
| chr4 | 174204643 | 174208781 | 4139 | -7.44 | 3.78E-11 | 1.05E-10 |
| chr10 | 103272951 | 103274212 | 1262 | -7.38 | 3.78E-11 | 1.05E-10 |
| chr9 | 119433491 | 119435147 | 1657 | -7.38 | 3.78E-11 | 1.05E-10 |
| chr22 | 49971579 | 49975124 | 3546 | -4.21 | 3.79E-11 | 1.05E-10 |
| chr22 | 24846751 | 24848658 | 1908 | -7.36 | 3.80E-11 | 1.05E-10 |
| chr12 | 25886529 | 25892866 | 6338 | -2.86 | 3.82E-11 | 1.06E-10 |
| chr4 | 6145545 | 6147360 | 1816 | -7.4 | 3.82E-11 | 1.06E-10 |
| chr12 | 21541370 | 21543993 | 2624 | 7.44 | 3.82E-11 | 1.06E-10 |
| chrX | 150665042 | 150669080 | 4039 | 7.44 | 3.82E-11 | 1.06E-10 |
| chr4 | 122356485 | 122361189 | 4705 | -7.39 | 3.83E-11 | 1.06E-10 |
| chr14 | 58580099 | 58594384 | 14286 | -5.32 | 3.84E-11 | 1.06E-10 |
| chr7 | 102543256 | 102548638 | 5383 | -7.36 | 3.84E-11 | 1.06E-10 |
| chr1 | 199385199 | 199389347 | 4149 | -7.37 | 3.85E-11 | 1.06E-10 |
| chr8 | 25308446 | 25310133 | 1688 | -7.5 | 3.85E-11 | 1.07E-10 |
| chr17 | 61925185 | 61927062 | 1878 | -7.43 | 3.86E-11 | 1.07E-10 |
| chr6 | 121883139 | 121886657 | 3519 | -7.57 | 3.87E-11 | 1.07E-10 |
| chrX | 128841556 | 128844778 | 3223 | -5.8 | 3.87E-11 | 1.07E-10 |
| chr8 | 95011111 | 95014326 | 3216 | -7.36 | 3.87E-11 | 1.07E-10 |
| chr5 | 135079825 | 135082581 | 2757 | -7.42 | 3.87E-11 | 1.07E-10 |
| chr8 | 39628255 | 39667280 | 39026 | -1.37 | 3.89E-11 | 1.07E-10 |
| chr1 | 84449259 | 84453607 | 4349 | -7.38 | 3.89E-11 | 1.08E-10 |
| chr1 | 113854572 | 113860554 | 5983 | -7.53 | 3.90E-11 | 1.08E-10 |
| chr1 | 168179588 | 168194157 | 14570 | -7.57 | 3.92E-11 | 1.08E-10 |
| chr3 | 69705789 | 69730696 | 24908 | -7.81 | 3.92E-11 | 1.08E-10 |
| chr11 | 129425821 | 129445248 | 19428 | -2.56 | 3.92E-11 | 1.08E-10 |
| chr15 | 83488238 | 83489475 | 1238 | -7.37 | 3.92E-11 | 1.08E-10 |
| chr3 | 189575666 | 189591224 | 15559 | -7.48 | 3.92E-11 | 1.08E-10 |
| chr6 | 76292979 | 76295106 | 2128 | -7.54 | 3.92E-11 | 1.08E-10 |
| chr3 | 190820241 | 190827109 | 6869 | 7.44 | 3.93E-11 | 1.09E-10 |
| chr7 | 79307372 | 79313473 | 6102 | -7.44 | 3.93E-11 | 1.09E-10 |
| chr1 | 114485828 | 114488612 | 2785 | -7.38 | 3.93E-11 | 1.09E-10 |
| chr14 | 62201310 | 62202997 | 1688 | -7.38 | 3.93E-11 | 1.09E-10 |
| chr5 | 80690569 | 80691589 | 1021 | -7.38 | 3.93E-11 | 1.09E-10 |
| chr9 | 98748661 | 98750859 | 2199 | -7.44 | 3.94E-11 | 1.09E-10 |

| | | | | | | |
|---|---|---|---|---|---|---|
| chr17 | 10062448 | 10077050 | 14603 | -7.86 | 3.94E-11 | 1.09E-10 |
| chr2 | 180231573 | 180243502 | 11930 | -7.4 | 3.94E-11 | 1.09E-10 |
| chr5 | 74477725 | 74481364 | 3640 | -5.92 | 3.96E-11 | 1.09E-10 |
| chr12 | 112915269 | 112917541 | 2273 | -7.37 | 3.97E-11 | 1.10E-10 |
| chr17 | 8564757 | 8567012 | 2256 | 7.55 | 3.97E-11 | 1.10E-10 |
| chr12 | 96478836 | 96485999 | 7164 | -7.5 | 3.97E-11 | 1.10E-10 |
| chr5 | 56478871 | 56480522 | 1652 | -7.36 | 3.99E-11 | 1.10E-10 |
| chr8 | 66051331 | 66466951 | 415621 | 3.51 | 3.99E-11 | 1.10E-10 |
| chr13 | 98856153 | 98859010 | 2858 | -7.8 | 3.99E-11 | 1.10E-10 |
| chr9 | 20574877 | 20576981 | 2105 | -7.35 | 4.00E-11 | 1.10E-10 |
| chr2 | 145669820 | 145699474 | 29655 | -7.59 | 4.00E-11 | 1.10E-10 |
| chr10 | 35603083 | 35623911 | 20829 | -7.37 | 4.00E-11 | 1.10E-10 |
| chr11 | 33784466 | 33786008 | 1543 | -7.46 | 4.02E-11 | 1.11E-10 |
| chr15 | 31741915 | 31748203 | 6289 | -7.59 | 4.03E-11 | 1.11E-10 |
| chr6 | 152578244 | 152580591 | 2348 | -7.48 | 4.03E-11 | 1.11E-10 |
| chr6 | 166222894 | 166226162 | 3269 | -7.36 | 4.04E-11 | 1.11E-10 |
| chr15 | 34492268 | 34494859 | 2592 | -7.48 | 4.04E-11 | 1.11E-10 |
| chr11 | 63122225 | 63269731 | 147507 | -2.36 | 4.05E-11 | 1.12E-10 |
| chr3 | 141607124 | 141632204 | 25081 | -7.41 | 4.05E-11 | 1.12E-10 |
| chr12 | 28302829 | 28305469 | 2641 | -7.35 | 4.05E-11 | 1.12E-10 |
| chr15 | 92539100 | 92541751 | 2652 | -7.34 | 4.06E-11 | 1.12E-10 |
| chr2 | 40088577 | 40433426 | 344850 | 1.12 | 4.07E-11 | 1.12E-10 |
| chr4 | 124010862 | 124013178 | 2317 | -7.36 | 4.07E-11 | 1.12E-10 |
| chr8 | 27970124 | 27978735 | 8612 | -7.36 | 4.07E-11 | 1.12E-10 |
| chr8 | 134425817 | 134434038 | 8222 | -7.92 | 4.08E-11 | 1.13E-10 |
| chr2 | 109908905 | 109912293 | 3389 | -7.47 | 4.08E-11 | 1.13E-10 |
| chr16 | 89268907 | 89270208 | 1302 | -7.36 | 4.10E-11 | 1.13E-10 |
| chr1 | 44998052 | 45001983 | 3932 | -7.72 | 4.11E-11 | 1.13E-10 |
| chr14 | 100970309 | 100981923 | 11615 | -7.34 | 4.11E-11 | 1.13E-10 |
| chr16 | 78425211 | 78427439 | 2229 | -7.39 | 4.11E-11 | 1.13E-10 |
| chr12 | 80177489 | 80180404 | 2916 | -7.41 | 4.13E-11 | 1.14E-10 |
| chr8 | 141065567 | 141066712 | 1146 | -7.39 | 4.13E-11 | 1.14E-10 |
| chr4 | 86470946 | 86475644 | 4699 | -7.5 | 4.14E-11 | 1.14E-10 |
| chr12 | 90429119 | 90431025 | 1907 | -7.34 | 4.15E-11 | 1.14E-10 |
| chr4 | 31915438 | 31919876 | 4439 | -7.34 | 4.15E-11 | 1.14E-10 |
| chr6 | 9695483 | 9697430 | 1948 | -7.34 | 4.16E-11 | 1.15E-10 |
| chr13 | 86502478 | 86760071 | 257594 | 1.98 | 4.17E-11 | 1.15E-10 |
| chr11 | 35952907 | 35954579 | 1673 | -7.34 | 4.17E-11 | 1.15E-10 |
| chr18 | 42459623 | 42472156 | 12534 | -7.62 | 4.17E-11 | 1.15E-10 |
| chr14 | 95196465 | 95199611 | 3147 | -7.39 | 4.18E-11 | 1.15E-10 |
| chr8 | 9925896 | 9931374 | 5479 | -7.41 | 4.19E-11 | 1.15E-10 |
| chr11 | 10720073 | 10722360 | 2288 | -5.65 | 4.20E-11 | 1.16E-10 |
| chr3 | 141346571 | 141349220 | 2650 | -7.59 | 4.20E-11 | 1.16E-10 |
| chr12 | 1422666 | 1424507 | 1842 | -7.34 | 4.22E-11 | 1.16E-10 |
| chr5 | 167745913 | 167750429 | 4517 | -7.34 | 4.22E-11 | 1.16E-10 |
| chr18 | 34129842 | 34137253 | 7412 | -7.66 | 4.23E-11 | 1.17E-10 |
| chr18 | 68252598 | 68257584 | 4987 | -7.34 | 4.23E-11 | 1.17E-10 |
| chr3 | 145887212 | 145902055 | 14844 | -7.34 | 4.24E-11 | 1.17E-10 |
| chr6 | 56862283 | 56865878 | 3596 | -7.6 | 4.24E-11 | 1.17E-10 |
| chr16 | 21127253 | 21131576 | 4324 | -7.8 | 4.25E-11 | 1.17E-10 |
| chr20 | 47179781 | 47182780 | 3000 | -7.47 | 4.26E-11 | 1.17E-10 |
| chr10 | 134781886 | 134783744 | 1859 | -7.43 | 4.26E-11 | 1.17E-10 |
| chr5 | 167711306 | 167713269 | 1964 | -7.47 | 4.28E-11 | 1.18E-10 |
| chr4 | 54723720 | 54727309 | 3590 | -7.43 | 4.28E-11 | 1.18E-10 |
| chr7 | 111476027 | 111478115 | 2089 | -7.42 | 4.28E-11 | 1.18E-10 |
| chr11 | 108075869 | 108082891 | 7023 | -7.57 | 4.28E-11 | 1.18E-10 |
| chr4 | 53640386 | 53649448 | 9063 | -2.65 | 4.29E-11 | 1.18E-10 |
| chr5 | 75617418 | 75620908 | 3491 | -7.48 | 4.29E-11 | 1.18E-10 |
| chr3 | 31292969 | 31302699 | 9731 | -7.86 | 4.30E-11 | 1.18E-10 |
| chr10 | 133862426 | 133869810 | 7385 | -6.72 | 4.30E-11 | 1.18E-10 |
| chr7 | 16184561 | 16186948 | 2388 | -7.39 | 4.30E-11 | 1.18E-10 |

| | | | | | | |
|---|---|---|---|---|---|---|
| chr2 | 201459801 | 201461557 | 1757 | -7.34 | 4.30E-11 | 1.18E-10 |
| chr4 | 124282406 | 124284175 | 1770 | -7.34 | 4.30E-11 | 1.18E-10 |
| chr3 | 142512919 | 142514897 | 1979 | -7.42 | 4.31E-11 | 1.18E-10 |
| chr3 | 10983777 | 10985611 | 1835 | -7.35 | 4.32E-11 | 1.19E-10 |
| chr20 | 14594757 | 14597326 | 2570 | -7.33 | 4.33E-11 | 1.19E-10 |
| chr9 | 20758245 | 20771360 | 13116 | -7.8 | 4.34E-11 | 1.19E-10 |
| chr2 | 234111002 | 234113422 | 2421 | -7.34 | 4.34E-11 | 1.19E-10 |
| chr6 | 151732927 | 151736139 | 3213 | -7.34 | 4.34E-11 | 1.19E-10 |
| chr1 | 85798119 | 85800970 | 2852 | -7.33 | 4.34E-11 | 1.19E-10 |
| chr10 | 127888539 | 127895832 | 7294 | -7.41 | 4.35E-11 | 1.19E-10 |
| chr3 | 161174201 | 161182910 | 8710 | -7.64 | 4.36E-11 | 1.20E-10 |
| chr16 | 15538257 | 15539907 | 1651 | -7.34 | 4.36E-11 | 1.20E-10 |
| chr13 | 33353043 | 33354894 | 1852 | -7.36 | 4.37E-11 | 1.20E-10 |
| chr19 | 34123897 | 34128717 | 4821 | -7.37 | 4.37E-11 | 1.20E-10 |
| chr19 | 48329572 | 48331040 | 1469 | -7.34 | 4.38E-11 | 1.20E-10 |
| chr5 | 141399212 | 141400806 | 1595 | -7.34 | 4.38E-11 | 1.20E-10 |
| chr8 | 66978667 | 66981770 | 3104 | -7.34 | 4.39E-11 | 1.20E-10 |
| chr1 | 174498672 | 174502375 | 3704 | -7.62 | 4.39E-11 | 1.21E-10 |
| chr4 | 64240104 | 64256293 | 16190 | -7.35 | 4.40E-11 | 1.21E-10 |
| chr12 | 15140443 | 15145193 | 4751 | -7.33 | 4.43E-11 | 1.22E-10 |
| chr7 | 66539681 | 66541791 | 2111 | -7.56 | 4.44E-11 | 1.22E-10 |
| chr3 | 127734409 | 127736855 | 2447 | -7.39 | 4.44E-11 | 1.22E-10 |
| chrX | 35085665 | 35091065 | 5401 | -7.33 | 4.46E-11 | 1.22E-10 |
| chr15 | 69481506 | 69483600 | 2095 | -7.47 | 4.46E-11 | 1.22E-10 |
| chr1 | 38202739 | 38213275 | 10537 | -7.38 | 4.47E-11 | 1.23E-10 |
| chr1 | 168011456 | 168012970 | 1515 | -7.81 | 4.47E-11 | 1.23E-10 |
| chr17 | 67572854 | 67577799 | 4946 | -5.15 | 4.47E-11 | 1.23E-10 |
| chr3 | 79039799 | 79048888 | 9090 | -7.47 | 4.47E-11 | 1.23E-10 |
| chr12 | 2168367 | 2206632 | 38266 | -1.97 | 4.48E-11 | 1.23E-10 |
| chr10 | 65149741 | 65151750 | 2010 | -7.38 | 4.49E-11 | 1.23E-10 |
| chr10 | 33075195 | 33096640 | 21446 | -8.11 | 4.51E-11 | 1.24E-10 |
| chr1 | 151453403 | 151456043 | 2641 | -7.66 | 4.51E-11 | 1.24E-10 |
| chr7 | 139212623 | 139214393 | 1771 | -5.75 | 4.51E-11 | 1.24E-10 |
| chr15 | 88982857 | 88990252 | 7396 | -6.63 | 4.52E-11 | 1.24E-10 |
| chr18 | 28518239 | 28522536 | 4298 | -7.33 | 4.53E-11 | 1.24E-10 |
| chr8 | 70880366 | 70882741 | 2376 | -7.36 | 4.55E-11 | 1.25E-10 |
| chr9 | 5736470 | 5738977 | 2508 | -7.36 | 4.55E-11 | 1.25E-10 |
| chr2 | 236777739 | 236780172 | 2434 | -7.37 | 4.56E-11 | 1.25E-10 |
| chr20 | 55144957 | 55147126 | 2170 | -7.34 | 4.57E-11 | 1.25E-10 |
| chr18 | 24396828 | 24399976 | 3149 | -7.35 | 4.57E-11 | 1.25E-10 |
| chr15 | 71982484 | 71988793 | 6310 | -7.34 | 4.57E-11 | 1.25E-10 |
| chr1 | 197862777 | 197867909 | 5133 | -7.37 | 4.57E-11 | 1.25E-10 |
| chr2 | 20256776 | 20279323 | 22548 | -8.12 | 4.58E-11 | 1.25E-10 |
| chr10 | 95901767 | 95906274 | 4508 | -7.33 | 4.58E-11 | 1.26E-10 |
| chr7 | 110275419 | 110279208 | 3790 | -7.35 | 4.58E-11 | 1.26E-10 |
| chr18 | 13499805 | 13519799 | 19995 | -7.39 | 4.59E-11 | 1.26E-10 |
| chr18 | 69784287 | 69787690 | 3404 | -7.34 | 4.59E-11 | 1.26E-10 |
| chr3 | 171108244 | 171122922 | 14679 | -7.77 | 4.61E-11 | 1.26E-10 |
| chr11 | 61138847 | 61141114 | 2268 | -7.45 | 4.62E-11 | 1.27E-10 |
| chr4 | 186553789 | 186556064 | 2276 | -7.45 | 4.62E-11 | 1.27E-10 |
| chr20 | 5844707 | 5847844 | 3138 | -7.62 | 4.62E-11 | 1.27E-10 |
| chr12 | 113882777 | 113884909 | 2133 | -7.46 | 4.63E-11 | 1.27E-10 |
| chr6 | 128225708 | 128233128 | 7421 | -7.33 | 4.64E-11 | 1.27E-10 |
| chr8 | 11580867 | 11589597 | 8731 | -7.33 | 4.64E-11 | 1.27E-10 |
| chr5 | 14620343 | 14630779 | 10437 | -7.62 | 4.64E-11 | 1.27E-10 |
| chr6 | 164402719 | 164442216 | 39498 | 1.53 | 4.65E-11 | 1.27E-10 |
| chr4 | 155932477 | 155943019 | 10543 | -7.61 | 4.66E-11 | 1.28E-10 |
| chr18 | 24378523 | 24381202 | 2680 | -7.38 | 4.66E-11 | 1.28E-10 |
| chr20 | 48423913 | 48437021 | 13109 | -7.56 | 4.67E-11 | 1.28E-10 |
| chr22 | 20912660 | 20914344 | 1685 | -7.36 | 4.68E-11 | 1.28E-10 |
| chr15 | 66264298 | 66265521 | 1224 | -7.38 | 4.69E-11 | 1.28E-10 |

| | | | | | | |
|---|---|---|---|---|---|---|
| chr1 | 30465433 | 30468423 | 2991 | -7.33 | 4.69E-11 | 1.28E-10 |
| chr2 | 36807538 | 36809634 | 2097 | -7.43 | 4.69E-11 | 1.28E-10 |
| chr16 | 23110203 | 23122099 | 11897 | -7.63 | 4.69E-11 | 1.28E-10 |
| chr5 | 23647694 | 23658244 | 10551 | -7.6 | 4.70E-11 | 1.29E-10 |
| chr6 | 3699026 | 3716254 | 17229 | -7.85 | 4.70E-11 | 1.29E-10 |
| chr3 | 118726134 | 118729009 | 2876 | -7.33 | 4.71E-11 | 1.29E-10 |
| chr20 | 43668475 | 43669815 | 1341 | -7.33 | 4.72E-11 | 1.29E-10 |
| chr5 | 37599839 | 37601965 | 2127 | -7.39 | 4.72E-11 | 1.29E-10 |
| chr20 | 60609459 | 60614114 | 4656 | -7.37 | 4.74E-11 | 1.30E-10 |
| chr1 | 11666233 | 11674133 | 7901 | -7.32 | 4.75E-11 | 1.30E-10 |
| chr5 | 76992047 | 76993315 | 1269 | -7.32 | 4.75E-11 | 1.30E-10 |
| chr5 | 133299084 | 133300557 | 1474 | -7.32 | 4.75E-11 | 1.30E-10 |
| chr9 | 79582016 | 79597281 | 15266 | -1.71 | 4.75E-11 | 1.30E-10 |
| chr4 | 82050440 | 82054157 | 3718 | -7.48 | 4.75E-11 | 1.30E-10 |
| chr2 | 30565016 | 30567496 | 2481 | -7.57 | 4.76E-11 | 1.30E-10 |
| chr7 | 11431058 | 11438781 | 7724 | -7.33 | 4.77E-11 | 1.30E-10 |
| chr9 | 137290241 | 137291491 | 1251 | -7.33 | 4.77E-11 | 1.30E-10 |
| chr19 | 5728444 | 5730437 | 1994 | -7.37 | 4.77E-11 | 1.30E-10 |
| chr8 | 125782394 | 125784529 | 2136 | -7.37 | 4.77E-11 | 1.30E-10 |
| chr11 | 75449299 | 75467219 | 17921 | -7.5 | 4.77E-11 | 1.30E-10 |
| chr22 | 28758966 | 28766060 | 7095 | -7.34 | 4.77E-11 | 1.30E-10 |
| chr6 | 99346298 | 99347789 | 1492 | -7.39 | 4.79E-11 | 1.31E-10 |
| chr6 | 42463127 | 42469784 | 6658 | -7.34 | 4.79E-11 | 1.31E-10 |
| chr9 | 599373 | 602247 | 2875 | -7.34 | 4.79E-11 | 1.31E-10 |
| chr9 | 74615257 | 74617438 | 2182 | -7.35 | 4.80E-11 | 1.31E-10 |
| chr17 | 60986351 | 60998031 | 11681 | -7.34 | 4.80E-11 | 1.31E-10 |
| chr18 | 21069927 | 21073398 | 3472 | -7.36 | 4.81E-11 | 1.31E-10 |
| chr1 | 51357891 | 51362709 | 4819 | -7.42 | 4.82E-11 | 1.32E-10 |
| chr6 | 11248594 | 11250877 | 2284 | -7.42 | 4.82E-11 | 1.32E-10 |
| chr21 | 45084906 | 45087100 | 2195 | -7.39 | 4.82E-11 | 1.32E-10 |
| chr3 | 143524719 | 143538317 | 13599 | 2.59 | 4.82E-11 | 1.32E-10 |
| chrX | 121437905 | 121441349 | 3445 | 7.38 | 4.83E-11 | 1.32E-10 |
| chr15 | 92476591 | 92478489 | 1899 | -7.33 | 4.84E-11 | 1.32E-10 |
| chr18 | 46348549 | 46350758 | 2210 | -7.33 | 4.84E-11 | 1.32E-10 |
| chr17 | 15843240 | 15846933 | 3694 | -7.84 | 4.84E-11 | 1.32E-10 |
| chr4 | 53674515 | 53689912 | 15398 | -7.46 | 4.85E-11 | 1.32E-10 |
| chr5 | 14042349 | 14072959 | 30611 | -7.8 | 4.85E-11 | 1.32E-10 |
| chr10 | 114375469 | 114377115 | 1647 | -7.35 | 4.85E-11 | 1.32E-10 |
| chr20 | 48155329 | 48163441 | 8113 | -7.72 | 4.85E-11 | 1.32E-10 |
| chr15 | 63128780 | 63151646 | 22867 | -7.82 | 4.86E-11 | 1.33E-10 |
| chr12 | 26835713 | 26837769 | 2057 | -7.38 | 4.87E-11 | 1.33E-10 |
| chr9 | 110516318 | 110536813 | 20496 | -7.7 | 4.87E-11 | 1.33E-10 |
| chr6 | 12816471 | 12840621 | 24151 | -2.07 | 4.87E-11 | 1.33E-10 |
| chr2 | 139294667 | 139296549 | 1883 | -7.33 | 4.87E-11 | 1.33E-10 |
| chr10 | 3589453 | 3591743 | 2291 | -7.36 | 4.88E-11 | 1.33E-10 |
| chr2 | 44319525 | 44321688 | 2164 | -7.34 | 4.89E-11 | 1.34E-10 |
| chr11 | 103615294 | 103618653 | 3360 | -7.48 | 4.90E-11 | 1.34E-10 |
| chr7 | 81783862 | 81791527 | 7666 | -7.62 | 4.92E-11 | 1.34E-10 |
| chr11 | 70735530 | 70746370 | 10841 | -7.8 | 4.93E-11 | 1.34E-10 |
| chr2 | 57356010 | 57414678 | 58669 | 1.89 | 4.94E-11 | 1.35E-10 |
| chr9 | 102884384 | 102886143 | 1760 | -7.56 | 4.94E-11 | 1.35E-10 |
| chr1 | 169684761 | 169687669 | 2909 | -7.43 | 4.94E-11 | 1.35E-10 |
| chr8 | 9335688 | 9339334 | 3647 | -7.43 | 4.94E-11 | 1.35E-10 |
| chr5 | 65584212 | 65587333 | 3122 | -7.32 | 4.95E-11 | 1.35E-10 |
| chr17 | 78460135 | 78461633 | 1499 | -7.35 | 4.96E-11 | 1.35E-10 |
| chr20 | 14655098 | 14657919 | 2822 | -7.35 | 4.96E-11 | 1.35E-10 |
| chr21 | 16982936 | 16985617 | 2682 | -7.35 | 4.96E-11 | 1.35E-10 |
| chr10 | 17682343 | 17696883 | 14541 | -7.71 | 4.99E-11 | 1.36E-10 |
| chr7 | 397548 | 399832 | 2285 | -7.32 | 5.00E-11 | 1.36E-10 |
| chr13 | 34030455 | 34035395 | 4941 | -7.5 | 5.00E-11 | 1.36E-10 |
| chr15 | 92580678 | 92583509 | 2832 | -7.34 | 5.01E-11 | 1.37E-10 |

| | | | | | | |
|---|---|---|---|---|---|---|
| chr15 | 72205460 | 72207272 | 1813 | -7.34 | 5.01E-11 | 1.37E-10 |
| chr15 | 93280227 | 93290744 | 10518 | -7.74 | 5.01E-11 | 1.37E-10 |
| chr16 | 74760560 | 74762310 | 1751 | -7.34 | 5.02E-11 | 1.37E-10 |
| chr8 | 30003216 | 30009675 | 6460 | -5.52 | 5.02E-11 | 1.37E-10 |
| chr12 | 47593378 | 47599328 | 5951 | -7.95 | 5.02E-11 | 1.37E-10 |
| chr7 | 18003236 | 18005243 | 2008 | -7.32 | 5.02E-11 | 1.37E-10 |
| chr6 | 146677216 | 146694541 | 17326 | -7.31 | 5.02E-11 | 1.37E-10 |
| chr5 | 37233859 | 37235839 | 1981 | -7.33 | 5.03E-11 | 1.37E-10 |
| chr5 | 162783920 | 162796318 | 12399 | -7.61 | 5.04E-11 | 1.37E-10 |
| chr21 | 40252636 | 40266114 | 13479 | -7.79 | 5.05E-11 | 1.37E-10 |
| chr3 | 120034780 | 120038736 | 3957 | -7.55 | 5.05E-11 | 1.38E-10 |
| chr12 | 67425720 | 67428578 | 2859 | -7.45 | 5.05E-11 | 1.38E-10 |
| chr3 | 117615357 | 117618062 | 2706 | -7.45 | 5.05E-11 | 1.38E-10 |
| chr10 | 31002152 | 31003951 | 1800 | -7.4 | 5.05E-11 | 1.38E-10 |
| chr2 | 170007095 | 170026260 | 19166 | -2.62 | 5.06E-11 | 1.38E-10 |
| chr16 | 2903436 | 2912064 | 8629 | -7.31 | 5.09E-11 | 1.39E-10 |
| chr8 | 90059490 | 90062922 | 3433 | -7.31 | 5.09E-11 | 1.39E-10 |
| chr16 | 16269901 | 16274084 | 4184 | -7.39 | 5.10E-11 | 1.39E-10 |
| chr11 | 86787494 | 86788967 | 1474 | -7.31 | 5.11E-11 | 1.39E-10 |
| chr14 | 32821674 | 32824097 | 2424 | -7.31 | 5.11E-11 | 1.39E-10 |
| chr7 | 110346023 | 110356324 | 10302 | -7.67 | 5.11E-11 | 1.39E-10 |
| chr9 | 29928138 | 29940595 | 12458 | -7.48 | 5.12E-11 | 1.39E-10 |
| chr22 | 46405053 | 46406777 | 1725 | -7.32 | 5.13E-11 | 1.40E-10 |
| chr22 | 21013498 | 21018000 | 4503 | -7.31 | 5.17E-11 | 1.41E-10 |
| chr9 | 110885113 | 110888329 | 3217 | -7.49 | 5.17E-11 | 1.41E-10 |
| chr11 | 47079023 | 47081244 | 2222 | -7.31 | 5.17E-11 | 1.41E-10 |
| chr8 | 96170797 | 96172672 | 1876 | -7.33 | 5.17E-11 | 1.41E-10 |
| chr10 | 101856226 | 101865146 | 8921 | -7.78 | 5.17E-11 | 1.41E-10 |
| chr13 | 63755376 | 63770460 | 15085 | -7.82 | 5.19E-11 | 1.41E-10 |
| chr16 | 50822777 | 50824565 | 1789 | -7.36 | 5.20E-11 | 1.41E-10 |
| chr18 | 3522894 | 3525013 | 2120 | -7.33 | 5.20E-11 | 1.41E-10 |
| chr1 | 187257012 | 187261985 | 4974 | 7.39 | 5.20E-11 | 1.41E-10 |
| chr20 | 46852581 | 46865741 | 13161 | -6.94 | 5.22E-11 | 1.42E-10 |
| chr11 | 125831557 | 125833931 | 2375 | -7.41 | 5.23E-11 | 1.42E-10 |
| chr2 | 32642180 | 32643990 | 1811 | -7.4 | 5.25E-11 | 1.43E-10 |
| chr4 | 186258738 | 186260760 | 2023 | -7.36 | 5.26E-11 | 1.43E-10 |
| chr6 | 10091483 | 10094910 | 3428 | -7.4 | 5.26E-11 | 1.43E-10 |
| chr3 | 117102401 | 117104457 | 2057 | -7.31 | 5.30E-11 | 1.44E-10 |
| chr15 | 30263738 | 30265149 | 1412 | -7.31 | 5.30E-11 | 1.44E-10 |
| chr3 | 153789696 | 153800748 | 11053 | -7.31 | 5.30E-11 | 1.44E-10 |
| chr3 | 114734073 | 114736906 | 2834 | -7.46 | 5.30E-11 | 1.44E-10 |
| chr7 | 24452238 | 24478702 | 26465 | -7.46 | 5.30E-11 | 1.44E-10 |
| chr21 | 39080644 | 39100684 | 20041 | -8.01 | 5.32E-11 | 1.44E-10 |
| chr1 | 67431253 | 67433457 | 2205 | -7.38 | 5.32E-11 | 1.44E-10 |
| chr15 | 99253516 | 99255643 | 2128 | -7.47 | 5.33E-11 | 1.45E-10 |
| chrX | 53694464 | 53696177 | 1714 | -7.38 | 5.33E-11 | 1.45E-10 |
| chr1 | 93220175 | 93237369 | 17195 | -8.11 | 5.34E-11 | 1.45E-10 |
| chr1 | 221988481 | 221993789 | 5309 | -7.37 | 5.35E-11 | 1.45E-10 |
| chr1 | 212306736 | 212309551 | 2816 | -7.32 | 5.36E-11 | 1.46E-10 |
| chr1 | 25421880 | 25444511 | 22632 | -7.31 | 5.36E-11 | 1.46E-10 |
| chr5 | 91601805 | 91619423 | 17619 | -7.62 | 5.37E-11 | 1.46E-10 |
| chr15 | 81464049 | 81469668 | 5620 | 5.78 | 5.39E-11 | 1.46E-10 |
| chr4 | 125278525 | 125290571 | 12047 | -7.52 | 5.40E-11 | 1.47E-10 |
| chr2 | 48402494 | 48406349 | 3856 | -7.65 | 5.40E-11 | 1.47E-10 |
| chr5 | 65173224 | 65178745 | 5522 | -7.31 | 5.41E-11 | 1.47E-10 |
| chr7 | 95369872 | 95371654 | 1783 | 3.87 | 5.42E-11 | 1.47E-10 |
| chr8 | 127909787 | 127913906 | 4120 | -7.3 | 5.43E-11 | 1.47E-10 |
| chr13 | 33406662 | 33417405 | 10744 | -7.49 | 5.44E-11 | 1.48E-10 |
| chr2 | 46973107 | 46974251 | 1145 | -7.53 | 5.44E-11 | 1.48E-10 |
| chr1 | 91930435 | 91933919 | 3485 | -3.13 | 5.45E-11 | 1.48E-10 |
| chr6 | 130997773 | 131000217 | 2445 | -5.52 | 5.45E-11 | 1.48E-10 |

| | | | | | | |
|---|---|---|---|---|---|---|
| chr7 | 130583183 | 130593741 | 10559 | -7.75 | 5.45E-11 | 1.48E-10 |
| chr7 | 2469467 | 2471634 | 2168 | -7.42 | 5.45E-11 | 1.48E-10 |
| chr9 | 124539527 | 124541287 | 1761 | -7.32 | 5.46E-11 | 1.48E-10 |
| chr2 | 191290941 | 191294210 | 3270 | -7.39 | 5.48E-11 | 1.49E-10 |
| chr1 | 64842683 | 64850860 | 8178 | -7.36 | 5.48E-11 | 1.49E-10 |
| chr12 | 112218122 | 112219995 | 1874 | -7.42 | 5.49E-11 | 1.49E-10 |
| chr1 | 70955041 | 70957049 | 2009 | -7.39 | 5.51E-11 | 1.49E-10 |
| chr1 | 87921996 | 87926574 | 4579 | -5.77 | 5.51E-11 | 1.49E-10 |
| chr14 | 23206835 | 23212573 | 5739 | -7.45 | 5.52E-11 | 1.50E-10 |
| chr12 | 58794396 | 58796988 | 2593 | -7.32 | 5.53E-11 | 1.50E-10 |
| chr4 | 28646008 | 28710402 | 64395 | 1.28 | 5.53E-11 | 1.50E-10 |
| chr8 | 111014947 | 111024563 | 9617 | -7.35 | 5.54E-11 | 1.50E-10 |
| chr4 | 96787540 | 96791613 | 4074 | -7.46 | 5.54E-11 | 1.50E-10 |
| chr5 | 34578002 | 34585523 | 7522 | -7.51 | 5.56E-11 | 1.51E-10 |
| chr1 | 32379254 | 32386500 | 7247 | -7.37 | 5.57E-11 | 1.51E-10 |
| chr5 | 39239831 | 39242126 | 2296 | -7.37 | 5.57E-11 | 1.51E-10 |
| chr20 | 35994020 | 36004439 | 10420 | -7.63 | 5.57E-11 | 1.51E-10 |
| chr10 | 82255860 | 82257393 | 1534 | -7.38 | 5.58E-11 | 1.51E-10 |
| chr6 | 141507970 | 141514964 | 6995 | -7.31 | 5.59E-11 | 1.51E-10 |
| chr9 | 3477514 | 3479917 | 2404 | -7.3 | 5.59E-11 | 1.52E-10 |
| chr3 | 45315627 | 45358196 | 42570 | -1.76 | 5.60E-11 | 1.52E-10 |
| chrX | 12239675 | 12244933 | 5259 | -7.37 | 5.60E-11 | 1.52E-10 |
| chr4 | 89141362 | 89157219 | 15858 | -7.64 | 5.60E-11 | 1.52E-10 |
| chr1 | 243440496 | 243442194 | 1699 | -7.34 | 5.60E-11 | 1.52E-10 |
| chr10 | 70404903 | 70408481 | 3579 | -7.48 | 5.62E-11 | 1.52E-10 |
| chr7 | 16659351 | 16662001 | 2651 | -7.71 | 5.64E-11 | 1.53E-10 |
| chr18 | 12948870 | 12959135 | 10266 | -7.3 | 5.64E-11 | 1.53E-10 |
| chr4 | 132020129 | 132028675 | 8547 | -7.36 | 5.66E-11 | 1.53E-10 |
| chr2 | 20985893 | 20994966 | 9074 | -7.31 | 5.67E-11 | 1.54E-10 |
| chr7 | 93987308 | 93989148 | 1841 | -7.3 | 5.68E-11 | 1.54E-10 |
| chr12 | 132369322 | 132371089 | 1768 | -7.41 | 5.69E-11 | 1.54E-10 |
| chr7 | 105673102 | 105675023 | 1922 | -7.41 | 5.69E-11 | 1.54E-10 |
| chr14 | 53542290 | 53545515 | 3226 | -7.35 | 5.69E-11 | 1.54E-10 |
| chr16 | 63360500 | 63374242 | 13743 | -7.31 | 5.69E-11 | 1.54E-10 |
| chr12 | 74597345 | 74613946 | 16602 | -7.65 | 5.69E-11 | 1.54E-10 |
| chr10 | 126560769 | 126571837 | 11069 | -7.39 | 5.70E-11 | 1.54E-10 |
| chr10 | 43643820 | 43645932 | 2113 | -7.32 | 5.70E-11 | 1.54E-10 |
| chr6 | 26416738 | 26419535 | 2798 | -7.36 | 5.71E-11 | 1.54E-10 |
| chr1 | 91497008 | 91508663 | 11656 | -7.49 | 5.71E-11 | 1.54E-10 |
| chr19 | 39172318 | 39180236 | 7919 | -7.65 | 5.71E-11 | 1.55E-10 |
| chr11 | 130646839 | 130670553 | 23715 | -7.73 | 5.74E-11 | 1.55E-10 |
| chr3 | 107085130 | 107087889 | 2760 | -7.33 | 5.74E-11 | 1.55E-10 |
| chr20 | 14282359 | 14284005 | 1647 | -7.45 | 5.76E-11 | 1.56E-10 |
| chr18 | 9409298 | 9411894 | 2597 | -7.35 | 5.76E-11 | 1.56E-10 |
| chr9 | 97701082 | 97706909 | 5828 | -7.3 | 5.77E-11 | 1.56E-10 |
| chr9 | 3352581 | 3354789 | 2209 | -7.5 | 5.77E-11 | 1.56E-10 |
| chr1 | 210495741 | 210500224 | 4484 | -7.37 | 5.77E-11 | 1.56E-10 |
| chr11 | 67104645 | 67110457 | 5813 | -7.31 | 5.77E-11 | 1.56E-10 |
| chr11 | 68728998 | 68732751 | 3754 | -7.57 | 5.78E-11 | 1.56E-10 |
| chr2 | 65055370 | 65057460 | 2091 | -7.3 | 5.78E-11 | 1.56E-10 |
| chr2 | 209126450 | 209127951 | 1502 | -7.37 | 5.79E-11 | 1.57E-10 |
| chr12 | 45340818 | 45344559 | 3742 | -7.29 | 5.80E-11 | 1.57E-10 |
| chr14 | 67548728 | 67550143 | 1416 | -7.29 | 5.80E-11 | 1.57E-10 |
| chr15 | 44313664 | 44316703 | 3040 | -7.29 | 5.80E-11 | 1.57E-10 |
| chr2 | 239337838 | 239340269 | 2432 | -7.29 | 5.80E-11 | 1.57E-10 |
| chr16 | 58039589 | 58041099 | 1511 | -7.31 | 5.82E-11 | 1.57E-10 |
| chrX | 133735815 | 133737295 | 1481 | -7.3 | 5.83E-11 | 1.58E-10 |
| chrX | 122794674 | 122798096 | 3423 | -7.31 | 5.84E-11 | 1.58E-10 |
| chr1 | 19347834 | 19350391 | 2558 | -7.3 | 5.84E-11 | 1.58E-10 |
| chr3 | 47399068 | 47402323 | 3256 | -7.3 | 5.84E-11 | 1.58E-10 |
| chr1 | 101159174 | 101164794 | 5621 | -7.51 | 5.87E-11 | 1.59E-10 |

| | | | | | | |
|---|---|---|---|---|---|---|
| chr7 | 113850500 | 113882634 | 32135 | 8.75 | 5.88E-11 | 1.59E-10 |
| chr4 | 85021896 | 85036839 | 14944 | -7.45 | 5.88E-11 | 1.59E-10 |
| chr17 | 71597377 | 71618398 | 21022 | -8.53 | 5.89E-11 | 1.59E-10 |
| chr2 | 86328933 | 86330124 | 1192 | -7.46 | 5.92E-11 | 1.60E-10 |
| chr11 | 21993518 | 21996044 | 2527 | -7.29 | 5.93E-11 | 1.60E-10 |
| chr6 | 26260036 | 26272855 | 12820 | -7.4 | 5.93E-11 | 1.60E-10 |
| chr18 | 32755658 | 32757685 | 2028 | -7.31 | 5.93E-11 | 1.60E-10 |
| chr11 | 28271266 | 28303151 | 31886 | -7.52 | 5.94E-11 | 1.60E-10 |
| chr16 | 30605901 | 30612721 | 6821 | -7.32 | 5.95E-11 | 1.61E-10 |
| chr1 | 86272731 | 86276105 | 3375 | -7.59 | 5.96E-11 | 1.61E-10 |
| chr7 | 138071819 | 138073798 | 1980 | -7.31 | 5.96E-11 | 1.61E-10 |
| chr2 | 24185662 | 24187755 | 2094 | -7.62 | 5.96E-11 | 1.61E-10 |
| chr6 | 78090314 | 78099988 | 9675 | -7.43 | 5.97E-11 | 1.61E-10 |
| chr2 | 175227831 | 175230797 | 2967 | -7.49 | 5.97E-11 | 1.61E-10 |
| chr12 | 89211804 | 89214461 | 2658 | -7.52 | 5.99E-11 | 1.62E-10 |
| chr3 | 151682248 | 151696887 | 14640 | -7.53 | 5.99E-11 | 1.62E-10 |
| chr8 | 33198844 | 33200939 | 2096 | -7.53 | 5.99E-11 | 1.62E-10 |
| chr12 | 58484619 | 58503535 | 18917 | -7.83 | 6.00E-11 | 1.62E-10 |
| chr6 | 45565614 | 45568949 | 3336 | -7.35 | 6.01E-11 | 1.62E-10 |
| chr9 | 100736363 | 100738286 | 1924 | -7.37 | 6.02E-11 | 1.62E-10 |
| chr15 | 71956901 | 71958958 | 2058 | -7.57 | 6.02E-11 | 1.62E-10 |
| chr8 | 30201859 | 30203473 | 1615 | -7.34 | 6.03E-11 | 1.63E-10 |
| chr5 | 31523023 | 31524891 | 1869 | -7.57 | 6.04E-11 | 1.63E-10 |
| chr9 | 98853115 | 98859203 | 6089 | -7.73 | 6.05E-11 | 1.63E-10 |
| chr4 | 161437544 | 161450824 | 13281 | -7.48 | 6.05E-11 | 1.63E-10 |
| chr9 | 95662200 | 95673626 | 11427 | -7.63 | 6.05E-11 | 1.63E-10 |
| chr10 | 15768785 | 15776990 | 8206 | -2.85 | 6.06E-11 | 1.63E-10 |
| chr18 | 1522185 | 1885078 | 362894 | 5.28 | 6.07E-11 | 1.64E-10 |
| chr7 | 135477856 | 135479454 | 1599 | -7.46 | 6.07E-11 | 1.64E-10 |
| chrX | 149026314 | 149029258 | 2945 | -2.53 | 6.08E-11 | 1.64E-10 |
| chr4 | 106285503 | 106288473 | 2971 | -7.32 | 6.08E-11 | 1.64E-10 |
| chr6 | 148207787 | 148210637 | 2851 | -7.46 | 6.09E-11 | 1.64E-10 |
| chr6 | 14525593 | 14538572 | 12980 | -7.86 | 6.10E-11 | 1.64E-10 |
| chr13 | 47094427 | 47097300 | 2874 | -7.3 | 6.11E-11 | 1.65E-10 |
| chr1 | 117925296 | 117927566 | 2271 | -7.29 | 6.11E-11 | 1.65E-10 |
| chr6 | 145442803 | 145451428 | 8626 | -7.32 | 6.12E-11 | 1.65E-10 |
| chr3 | 197224585 | 197229080 | 4496 | -7.3 | 6.12E-11 | 1.65E-10 |
| chr12 | 931444 | 933145 | 1702 | -7.35 | 6.13E-11 | 1.65E-10 |
| chr12 | 54231727 | 54235044 | 3318 | -7.35 | 6.13E-11 | 1.65E-10 |
| chr2 | 99456320 | 99458553 | 2234 | -7.47 | 6.15E-11 | 1.66E-10 |
| chr17 | 61252252 | 61253809 | 1558 | -7.32 | 6.15E-11 | 1.66E-10 |
| chr11 | 108767777 | 108769884 | 2108 | -7.43 | 6.15E-11 | 1.66E-10 |
| chr12 | 68230084 | 68234433 | 4350 | -3.59 | 6.15E-11 | 1.66E-10 |
| chr2 | 145771111 | 145793461 | 22351 | -7.35 | 6.16E-11 | 1.66E-10 |
| chr12 | 13553275 | 13565337 | 12063 | -7.32 | 6.17E-11 | 1.66E-10 |
| chr3 | 108412183 | 108434944 | 22762 | -4.38 | 6.18E-11 | 1.67E-10 |
| chr3 | 33129992 | 33131327 | 1336 | -7.29 | 6.19E-11 | 1.67E-10 |
| chr3 | 125184431 | 125187028 | 2598 | -7.29 | 6.21E-11 | 1.67E-10 |
| chr19 | 2660713 | 2663780 | 3068 | -7.28 | 6.21E-11 | 1.67E-10 |
| chr6 | 7969582 | 7971910 | 2329 | -7.28 | 6.21E-11 | 1.67E-10 |
| chr2 | 187918051 | 187923181 | 5131 | 7.42 | 6.21E-11 | 1.67E-10 |
| chr1 | 42667176 | 42669720 | 2545 | -7.36 | 6.22E-11 | 1.67E-10 |
| chr21 | 47130420 | 47137682 | 7263 | -4.44 | 6.22E-11 | 1.67E-10 |
| chr3 | 137841594 | 137843632 | 2039 | -7.34 | 6.23E-11 | 1.68E-10 |
| chr12 | 58859089 | 58872248 | 13160 | -7.29 | 6.23E-11 | 1.68E-10 |
| chr13 | 50895473 | 50898479 | 3007 | -7.55 | 6.23E-11 | 1.68E-10 |
| chr11 | 124873419 | 124884667 | 11249 | -7.43 | 6.24E-11 | 1.68E-10 |
| chr2 | 88938460 | 88943972 | 5513 | -7.43 | 6.25E-11 | 1.68E-10 |
| chr15 | 53515519 | 53523672 | 8154 | 7.77 | 6.26E-11 | 1.68E-10 |
| chr3 | 170255815 | 170261105 | 5291 | 3.09 | 6.27E-11 | 1.69E-10 |
| chr20 | 9283717 | 9295748 | 12032 | -7.31 | 6.28E-11 | 1.69E-10 |

| | | | | | | |
|---|---|---|---|---|---|---|
| chr1 | 78114393 | 78117299 | 2907 | -7.3 | 6.28E-11 | 1.69E-10 |
| chr2 | 102925067 | 102965065 | 39999 | -2.13 | 6.29E-11 | 1.69E-10 |
| chr9 | 95976782 | 95978950 | 2169 | -7.33 | 6.29E-11 | 1.69E-10 |
| chr8 | 49297978 | 49302579 | 4602 | -7.36 | 6.30E-11 | 1.69E-10 |
| chr2 | 181575748 | 181578870 | 3123 | -7.28 | 6.30E-11 | 1.69E-10 |
| chr4 | 120703570 | 120713978 | 10409 | -7.59 | 6.31E-11 | 1.70E-10 |
| chr3 | 109233017 | 109241228 | 8212 | -5.81 | 6.32E-11 | 1.70E-10 |
| chr9 | 75732124 | 75736097 | 3974 | -7.34 | 6.32E-11 | 1.70E-10 |
| chr12 | 76173488 | 76176855 | 3368 | -7.35 | 6.32E-11 | 1.70E-10 |
| chr4 | 64584425 | 64596954 | 12530 | -7.33 | 6.33E-11 | 1.70E-10 |
| chr5 | 170405826 | 170408946 | 3121 | -7.29 | 6.34E-11 | 1.70E-10 |
| chr2 | 151646292 | 151650104 | 3813 | -7.6 | 6.35E-11 | 1.71E-10 |
| chrX | 84115264 | 84122822 | 7559 | 3.84 | 6.35E-11 | 1.71E-10 |
| chr9 | 114441467 | 114443575 | 2109 | -7.28 | 6.36E-11 | 1.71E-10 |
| chr18 | 71099144 | 71100784 | 1641 | -6.27 | 6.38E-11 | 1.71E-10 |
| chr10 | 101624438 | 101628079 | 3642 | -7.3 | 6.38E-11 | 1.71E-10 |
| chr3 | 115027864 | 115031117 | 3254 | -7.32 | 6.39E-11 | 1.72E-10 |
| chr1 | 43052649 | 43067171 | 14523 | -7.6 | 6.41E-11 | 1.72E-10 |
| chr8 | 101782874 | 101799729 | 16856 | -7.54 | 6.42E-11 | 1.73E-10 |
| chr4 | 75049096 | 75059795 | 10700 | -7.39 | 6.43E-11 | 1.73E-10 |
| chr2 | 55617613 | 55619269 | 1657 | -7.4 | 6.43E-11 | 1.73E-10 |
| chr1 | 181027908 | 181040266 | 12359 | -7.6 | 6.44E-11 | 1.73E-10 |
| chr8 | 116199763 | 116217065 | 17303 | -7.69 | 6.44E-11 | 1.73E-10 |
| chr2 | 8031395 | 8041926 | 10532 | -7.33 | 6.45E-11 | 1.73E-10 |
| chr10 | 13185833 | 13192459 | 6627 | -7.4 | 6.46E-11 | 1.73E-10 |
| chr7 | 50555286 | 50558398 | 3113 | -7.4 | 6.46E-11 | 1.73E-10 |
| chr5 | 138689225 | 138690785 | 1561 | -7.42 | 6.48E-11 | 1.74E-10 |
| chr14 | 36184006 | 36185979 | 1974 | -7.33 | 6.48E-11 | 1.74E-10 |
| chr7 | 151741284 | 151744198 | 2915 | -7.36 | 6.51E-11 | 1.75E-10 |
| chr2 | 114565759 | 114567724 | 1966 | -6.42 | 6.52E-11 | 1.75E-10 |
| chr5 | 124220375 | 124222382 | 2008 | -7.43 | 6.52E-11 | 1.75E-10 |
| chr1 | 223089108 | 223091252 | 2145 | -7.33 | 6.53E-11 | 1.75E-10 |
| chr1 | 116628826 | 116639911 | 11086 | -4.39 | 6.54E-11 | 1.75E-10 |
| chr6 | 42288564 | 42293489 | 4926 | -7.27 | 6.55E-11 | 1.76E-10 |
| chr8 | 102707843 | 102709258 | 1416 | -7.32 | 6.55E-11 | 1.76E-10 |
| chr13 | 98392410 | 98445558 | 53149 | -1.94 | 6.57E-11 | 1.76E-10 |
| chr11 | 10601529 | 10605381 | 3853 | 7.4 | 6.57E-11 | 1.76E-10 |
| chr5 | 49949494 | 49957660 | 8167 | -7.29 | 6.58E-11 | 1.76E-10 |
| chr20 | 4214091 | 4221279 | 7189 | -7.44 | 6.59E-11 | 1.77E-10 |
| chr3 | 23327748 | 23329426 | 1679 | -7.44 | 6.59E-11 | 1.77E-10 |
| chr2 | 108157834 | 108183896 | 26063 | -1.33 | 6.60E-11 | 1.77E-10 |
| chr2 | 188192211 | 188194799 | 2589 | -5.75 | 6.61E-11 | 1.77E-10 |
| chr13 | 23437253 | 23448029 | 10777 | -7.3 | 6.63E-11 | 1.78E-10 |
| chr12 | 107697680 | 107702671 | 4992 | -3.94 | 6.63E-11 | 1.78E-10 |
| chr5 | 155407163 | 155414334 | 7172 | -7.7 | 6.66E-11 | 1.79E-10 |
| chr6 | 135837599 | 135841029 | 3431 | -7.46 | 6.67E-11 | 1.79E-10 |
| chr2 | 234887970 | 234894651 | 6682 | -7.29 | 6.67E-11 | 1.79E-10 |
| chr13 | 40288460 | 40290922 | 2463 | -7.31 | 6.68E-11 | 1.79E-10 |
| chrX | 15562745 | 15564194 | 1450 | -6.29 | 6.68E-11 | 1.79E-10 |
| chr17 | 76040310 | 76042722 | 2413 | -7.28 | 6.68E-11 | 1.79E-10 |
| chr3 | 110171742 | 110178050 | 6309 | -7.28 | 6.68E-11 | 1.79E-10 |
| chr20 | 45673134 | 45676621 | 3488 | -7.33 | 6.71E-11 | 1.80E-10 |
| chr11 | 64157429 | 64172717 | 15289 | -7.73 | 6.71E-11 | 1.80E-10 |
| chr9 | 32517467 | 32532489 | 15023 | -7.71 | 6.74E-11 | 1.80E-10 |
| chr2 | 204402206 | 204404965 | 2760 | -7.38 | 6.74E-11 | 1.81E-10 |
| chr2 | 145354954 | 145356853 | 1900 | -7.28 | 6.76E-11 | 1.81E-10 |
| chr15 | 57872748 | 57876797 | 4050 | -6.37 | 6.77E-11 | 1.81E-10 |
| chr4 | 88786531 | 88789707 | 3177 | -7.33 | 6.77E-11 | 1.81E-10 |
| chr6 | 48269796 | 48272430 | 2635 | -7.33 | 6.77E-11 | 1.81E-10 |
| chr6 | 40630273 | 40636044 | 5772 | -5.89 | 6.78E-11 | 1.81E-10 |
| chr12 | 94145900 | 94149410 | 3511 | -7.41 | 6.78E-11 | 1.82E-10 |

| | | | | | | |
|---|---|---|---|---|---|---|
| chr4 | 163466423 | 163470710 | 4288 | -7.39 | 6.78E-11 | 1.82E-10 |
| chr1 | 239906713 | 239916202 | 9490 | 7.52 | 6.79E-11 | 1.82E-10 |
| chr2 | 204567459 | 204568489 | 1031 | -7.27 | 6.80E-11 | 1.82E-10 |
| chr5 | 148400133 | 148402814 | 2682 | -7.37 | 6.80E-11 | 1.82E-10 |
| chr2 | 223647583 | 223652090 | 4508 | -7.51 | 6.80E-11 | 1.82E-10 |
| chr13 | 36644631 | 36686327 | 41697 | 1.51 | 6.80E-11 | 1.82E-10 |
| chr4 | 13454921 | 13468376 | 13456 | -7.32 | 6.81E-11 | 1.82E-10 |
| chr4 | 143290936 | 143293594 | 2659 | -7.41 | 6.82E-11 | 1.83E-10 |
| chr16 | 19779867 | 19782118 | 2252 | -7.28 | 6.83E-11 | 1.83E-10 |
| chr9 | 5973553 | 5976417 | 2865 | -7.27 | 6.85E-11 | 1.83E-10 |
| chr1 | 65075177 | 65078317 | 3141 | -7.32 | 6.86E-11 | 1.84E-10 |
| chr11 | 104534998 | 104546115 | 11118 | -2.08 | 6.86E-11 | 1.84E-10 |
| chr14 | 100436808 | 100454484 | 17677 | -7.58 | 6.86E-11 | 1.84E-10 |
| chr18 | 5234918 | 5237545 | 2628 | -5.47 | 6.86E-11 | 1.84E-10 |
| chr9 | 94549390 | 94551362 | 1973 | -7.27 | 6.87E-11 | 1.84E-10 |
| chr12 | 118665579 | 118667932 | 2354 | -7.35 | 6.89E-11 | 1.84E-10 |
| chrX | 35767424 | 35778131 | 10708 | -7.44 | 6.91E-11 | 1.85E-10 |
| chr15 | 84237833 | 84240856 | 3024 | -7.49 | 6.94E-11 | 1.86E-10 |
| chr4 | 38713227 | 38715703 | 2477 | -7.27 | 6.95E-11 | 1.86E-10 |
| chr4 | 3274140 | 3284385 | 10246 | -7.64 | 6.95E-11 | 1.86E-10 |
| chr16 | 20269075 | 20272373 | 3299 | -5.43 | 6.95E-11 | 1.86E-10 |
| chr3 | 172135706 | 172151737 | 16032 | -7.27 | 6.99E-11 | 1.87E-10 |
| chr21 | 47912275 | 47914426 | 2152 | -7.55 | 7.01E-11 | 1.88E-10 |
| chr7 | 80268010 | 80270515 | 2506 | -7.31 | 7.02E-11 | 1.88E-10 |
| chr1 | 47668408 | 47669981 | 1574 | -7.39 | 7.03E-11 | 1.88E-10 |
| chr7 | 44738118 | 44740130 | 2013 | -7.4 | 7.04E-11 | 1.88E-10 |
| chr6 | 146180778 | 146183250 | 2473 | -7.26 | 7.05E-11 | 1.88E-10 |
| chr4 | 135573105 | 135586949 | 13845 | 7.78 | 7.06E-11 | 1.89E-10 |
| chr14 | 56577617 | 56580770 | 3154 | -7.4 | 7.06E-11 | 1.89E-10 |
| chr5 | 132825513 | 132827421 | 1909 | -7.3 | 7.06E-11 | 1.89E-10 |
| chr11 | 13257976 | 13259686 | 1711 | -7.27 | 7.07E-11 | 1.89E-10 |
| chr6 | 44323535 | 44327046 | 3512 | -7.32 | 7.09E-11 | 1.89E-10 |
| chr9 | 116889209 | 116899869 | 10661 | -7.32 | 7.09E-11 | 1.89E-10 |
| chr18 | 7648678 | 7651613 | 2936 | -7.27 | 7.09E-11 | 1.90E-10 |
| chr20 | 4718752 | 4731760 | 13009 | -7.91 | 7.12E-11 | 1.90E-10 |
| chr11 | 74306172 | 74308139 | 1968 | -7.29 | 7.15E-11 | 1.91E-10 |
| chr7 | 46294880 | 46297108 | 2229 | -7.27 | 7.16E-11 | 1.91E-10 |
| chr22 | 26732083 | 26747076 | 14994 | -1.73 | 7.17E-11 | 1.91E-10 |
| chr4 | 155899269 | 155902041 | 2773 | -7.39 | 7.17E-11 | 1.91E-10 |
| chr13 | 74063794 | 74069793 | 6000 | -7.27 | 7.17E-11 | 1.91E-10 |
| chrX | 109096430 | 109098362 | 1933 | -7.27 | 7.17E-11 | 1.91E-10 |
| chr1 | 221749680 | 221760638 | 10959 | -7.35 | 7.19E-11 | 1.92E-10 |
| chr2 | 138689511 | 138693624 | 4114 | -7.26 | 7.20E-11 | 1.92E-10 |
| chrX | 107554554 | 107562286 | 7733 | -7.28 | 7.22E-11 | 1.93E-10 |
| chr13 | 106340622 | 106348067 | 7446 | -7.48 | 7.24E-11 | 1.93E-10 |
| chr15 | 66005755 | 66007253 | 1499 | -7.48 | 7.24E-11 | 1.93E-10 |
| chr5 | 142234207 | 142236739 | 2533 | -7.33 | 7.25E-11 | 1.93E-10 |
| chr15 | 64598338 | 64600050 | 1713 | -7.27 | 7.25E-11 | 1.94E-10 |
| chr1 | 221170115 | 221183200 | 13086 | -7.31 | 7.26E-11 | 1.94E-10 |
| chr6 | 84029065 | 84030457 | 1393 | -7.31 | 7.26E-11 | 1.94E-10 |
| chr4 | 99238679 | 99241833 | 3155 | -7.26 | 7.27E-11 | 1.94E-10 |
| chr1 | 90424021 | 90426292 | 2272 | -7.26 | 7.28E-11 | 1.94E-10 |
| chr17 | 66778131 | 66786716 | 8586 | -7.28 | 7.28E-11 | 1.94E-10 |
| chr10 | 13882430 | 13884649 | 2220 | -7.26 | 7.30E-11 | 1.95E-10 |
| chr4 | 143662227 | 143667530 | 5304 | -7.3 | 7.30E-11 | 1.95E-10 |
| chr3 | 159759054 | 159761877 | 2824 | -7.28 | 7.32E-11 | 1.95E-10 |
| chr1 | 194127139 | 194128507 | 1369 | -7.29 | 7.32E-11 | 1.95E-10 |
| chr17 | 60964750 | 60967184 | 2435 | -7.29 | 7.32E-11 | 1.95E-10 |
| chr1 | 21425161 | 21426779 | 1619 | -7.26 | 7.33E-11 | 1.96E-10 |
| chr10 | 91012631 | 91015822 | 3192 | -7.37 | 7.33E-11 | 1.96E-10 |
| chr19 | 56014508 | 56017444 | 2937 | -7.38 | 7.33E-11 | 1.96E-10 |

| | | | | | | |
|---|---|---|---|---|---|---|
| chr10 | 115683099 | 115696971 | 13873 | -7.72 | 7.34E-11 | 1.96E-10 |
| chr6 | 12154902 | 12159572 | 4671 | -7.59 | 7.35E-11 | 1.96E-10 |
| chr19 | 17274056 | 17274943 | 888 | -7.33 | 7.36E-11 | 1.96E-10 |
| chr8 | 9460359 | 9462843 | 2485 | -7.33 | 7.36E-11 | 1.96E-10 |
| chr17 | 71036164 | 71038383 | 2220 | -7.39 | 7.36E-11 | 1.96E-10 |
| chr1 | 182616831 | 182619945 | 3115 | 7.32 | 7.38E-11 | 1.97E-10 |
| chr18 | 46800725 | 46802946 | 2222 | -7.27 | 7.38E-11 | 1.97E-10 |
| chr2 | 221024251 | 221026771 | 2521 | -7.32 | 7.38E-11 | 1.97E-10 |
| chr1 | 65997594 | 66000360 | 2767 | -7.28 | 7.41E-11 | 1.97E-10 |
| chr3 | 101591491 | 101596258 | 4768 | -7.54 | 7.44E-11 | 1.98E-10 |
| chr15 | 69915171 | 69950476 | 35306 | -3.3 | 7.45E-11 | 1.98E-10 |
| chr6 | 5752979 | 5769082 | 16104 | -7.6 | 7.47E-11 | 1.99E-10 |
| chrX | 103422520 | 103424037 | 1518 | -7.26 | 7.48E-11 | 1.99E-10 |
| chr3 | 137711777 | 137716409 | 4633 | -5.68 | 7.49E-11 | 1.99E-10 |
| chr7 | 110065626 | 110079503 | 13878 | -7.47 | 7.50E-11 | 2.00E-10 |
| chr7 | 81073167 | 81081447 | 8281 | -7.33 | 7.50E-11 | 2.00E-10 |
| chr3 | 127718311 | 127719806 | 1496 | -7.29 | 7.51E-11 | 2.00E-10 |
| chr12 | 14991471 | 14993445 | 1975 | -7.28 | 7.52E-11 | 2.00E-10 |
| chr1 | 108035212 | 108044317 | 9106 | 7.32 | 7.52E-11 | 2.00E-10 |
| chrX | 130006341 | 130010932 | 4592 | -7.3 | 7.53E-11 | 2.00E-10 |
| chr2 | 32682051 | 32684017 | 1967 | -7.76 | 7.54E-11 | 2.01E-10 |
| chr2 | 20358663 | 20360246 | 1584 | -7.29 | 7.54E-11 | 2.01E-10 |
| chr15 | 70602563 | 70604141 | 1579 | -7.26 | 7.55E-11 | 2.01E-10 |
| chr6 | 158998225 | 159008131 | 9907 | -7.38 | 7.56E-11 | 2.01E-10 |
| chr4 | 107799940 | 107856255 | 56316 | 2.01 | 7.56E-11 | 2.01E-10 |
| chr12 | 41177419 | 41179743 | 2325 | -7.35 | 7.57E-11 | 2.01E-10 |
| chr5 | 151188339 | 151191013 | 2675 | -7.4 | 7.59E-11 | 2.02E-10 |
| chr1 | 193723595 | 193725681 | 2087 | -7.26 | 7.63E-11 | 2.03E-10 |
| chr14 | 69973979 | 69983085 | 9107 | -7.77 | 7.63E-11 | 2.03E-10 |
| chr5 | 155469042 | 155472399 | 3358 | -7.25 | 7.64E-11 | 2.03E-10 |
| chr1 | 207269564 | 207274040 | 4477 | -7.37 | 7.65E-11 | 2.04E-10 |
| chr21 | 42703550 | 42716561 | 13012 | -2.64 | 7.65E-11 | 2.04E-10 |
| chrX | 95972930 | 95975239 | 2310 | -7.36 | 7.65E-11 | 2.04E-10 |
| chr1 | 36997488 | 36999751 | 2264 | -7.28 | 7.66E-11 | 2.04E-10 |
| chr16 | 84418023 | 84420324 | 2302 | -7.36 | 7.66E-11 | 2.04E-10 |
| chr4 | 53527163 | 53531584 | 4422 | -7.31 | 7.68E-11 | 2.04E-10 |
| chr3 | 191126139 | 191128812 | 2674 | -7.35 | 7.69E-11 | 2.05E-10 |
| chr6 | 54389700 | 54394879 | 5180 | -7.35 | 7.69E-11 | 2.05E-10 |
| chr13 | 84758802 | 84771531 | 12730 | -7.39 | 7.69E-11 | 2.05E-10 |
| chr9 | 12891683 | 12899859 | 8177 | -7.39 | 7.70E-11 | 2.05E-10 |
| chr2 | 9035865 | 9052096 | 16232 | -7.67 | 7.70E-11 | 2.05E-10 |
| chr3 | 126280232 | 126282422 | 2191 | -7.29 | 7.71E-11 | 2.05E-10 |
| chr6 | 107151987 | 107154354 | 2368 | -7.29 | 7.71E-11 | 2.05E-10 |
| chr5 | 131088367 | 131104286 | 15920 | -7.56 | 7.71E-11 | 2.05E-10 |
| chr10 | 62615710 | 62618809 | 3100 | -7.28 | 7.72E-11 | 2.05E-10 |
| chr6 | 114249761 | 114252491 | 2731 | -7.27 | 7.74E-11 | 2.06E-10 |
| chr7 | 3021237 | 3035937 | 14701 | -7.27 | 7.74E-11 | 2.06E-10 |
| chr4 | 142863598 | 142871467 | 7870 | -7.43 | 7.75E-11 | 2.06E-10 |
| chr1 | 55020340 | 55022387 | 2048 | -7.33 | 7.78E-11 | 2.07E-10 |
| chr1 | 173414631 | 173416617 | 1987 | -7.25 | 7.79E-11 | 2.07E-10 |
| chr6 | 55612062 | 55616542 | 4481 | -7.25 | 7.81E-11 | 2.07E-10 |
| chr10 | 93677988 | 93680605 | 2618 | -7.32 | 7.81E-11 | 2.07E-10 |
| chr1 | 154712432 | 154713937 | 1506 | -7.29 | 7.82E-11 | 2.08E-10 |
| chr2 | 237859741 | 237865534 | 5794 | -2.1 | 7.83E-11 | 2.08E-10 |
| chr8 | 100686671 | 100698578 | 11908 | -7.26 | 7.83E-11 | 2.08E-10 |
| chr12 | 100147060 | 100186592 | 39533 | -7.63 | 7.85E-11 | 2.08E-10 |
| chr6 | 9479217 | 9482205 | 2989 | -7.29 | 7.86E-11 | 2.09E-10 |
| chr5 | 95511087 | 95523432 | 12346 | -8.01 | 7.88E-11 | 2.09E-10 |
| chr11 | 95537576 | 95540429 | 2854 | -7.29 | 7.88E-11 | 2.09E-10 |
| chr18 | 77978261 | 77980844 | 2584 | -7.26 | 7.90E-11 | 2.10E-10 |
| chr14 | 64480922 | 64482209 | 1288 | -7.31 | 7.90E-11 | 2.10E-10 |

| | | | | | | |
|---|---|---|---|---|---|---|
| chr20 | 47371100 | 47373353 | 2254 | -7.29 | 7.92E-11 | 2.10E-10 |
| chr14 | 97113454 | 97166958 | 53505 | -3.47 | 7.96E-11 | 2.11E-10 |
| chr10 | 90987607 | 90989838 | 2232 | -7.25 | 7.96E-11 | 2.11E-10 |
| chr12 | 111709475 | 111719923 | 10449 | -7.61 | 7.96E-11 | 2.11E-10 |
| chr19 | 49686001 | 49694723 | 8723 | -7.79 | 7.97E-11 | 2.11E-10 |
| chr5 | 80545659 | 80547968 | 2310 | -7.37 | 7.97E-11 | 2.12E-10 |
| chr18 | 34517868 | 34522328 | 4461 | -7.25 | 7.98E-11 | 2.12E-10 |
| chr1 | 22736013 | 22738211 | 2199 | -7.36 | 7.99E-11 | 2.12E-10 |
| chr2 | 194449077 | 194460671 | 11595 | -7.36 | 7.99E-11 | 2.12E-10 |
| chr7 | 105411805 | 105413546 | 1742 | -7.35 | 7.99E-11 | 2.12E-10 |
| chr8 | 49828541 | 49829960 | 1420 | -7.27 | 7.99E-11 | 2.12E-10 |
| chr3 | 87573319 | 87577121 | 3803 | -7.34 | 8.03E-11 | 2.13E-10 |
| chr15 | 92413049 | 92414516 | 1468 | -7.28 | 8.05E-11 | 2.14E-10 |
| chr1 | 46406984 | 46414811 | 7828 | -7.33 | 8.06E-11 | 2.14E-10 |
| chr6 | 131413964 | 131422762 | 8799 | -7.33 | 8.08E-11 | 2.14E-10 |
| chrX | 81628343 | 81634446 | 6104 | -7.33 | 8.08E-11 | 2.14E-10 |
| chr14 | 24668969 | 24671308 | 2340 | -7.25 | 8.08E-11 | 2.14E-10 |
| chr6 | 49350369 | 49368612 | 18244 | -7.41 | 8.09E-11 | 2.15E-10 |
| chr21 | 39679259 | 39693750 | 14492 | -7.41 | 8.12E-11 | 2.15E-10 |
| chr2 | 158392718 | 158422987 | 30270 | -7.28 | 8.12E-11 | 2.15E-10 |
| chr2 | 45511328 | 45514025 | 2698 | -7.29 | 8.13E-11 | 2.16E-10 |
| chr7 | 35731226 | 35737585 | 6360 | -7.26 | 8.14E-11 | 2.16E-10 |
| chrX | 74502796 | 74509527 | 6732 | -7.65 | 8.16E-11 | 2.16E-10 |
| chr21 | 45367851 | 45369768 | 1918 | -7.27 | 8.17E-11 | 2.17E-10 |
| chr4 | 139418012 | 139425494 | 7483 | -7.26 | 8.19E-11 | 2.17E-10 |
| chr11 | 95609175 | 95611944 | 2770 | -7.24 | 8.19E-11 | 2.17E-10 |
| chrX | 102703025 | 102707262 | 4238 | -7.24 | 8.19E-11 | 2.17E-10 |
| chr2 | 161212770 | 161214333 | 1564 | -7.24 | 8.21E-11 | 2.18E-10 |
| chr22 | 27967851 | 27969206 | 1356 | -7.25 | 8.21E-11 | 2.18E-10 |
| chr4 | 179842223 | 179860233 | 18011 | -7.63 | 8.28E-11 | 2.19E-10 |
| chr5 | 59633798 | 59642252 | 8455 | 7.76 | 8.28E-11 | 2.19E-10 |
| chr2 | 58397397 | 58415192 | 17796 | -7.3 | 8.29E-11 | 2.20E-10 |
| chr4 | 103901843 | 103920388 | 18546 | -7.66 | 8.29E-11 | 2.20E-10 |
| chr7 | 151763407 | 151765618 | 2212 | -7.25 | 8.29E-11 | 2.20E-10 |
| chr20 | 35589686 | 35591865 | 2180 | -7.25 | 8.32E-11 | 2.20E-10 |
| chr16 | 21639100 | 21641125 | 2026 | -7.27 | 8.34E-11 | 2.21E-10 |
| chr22 | 28320948 | 28324226 | 3279 | -7.34 | 8.35E-11 | 2.21E-10 |
| chr16 | 80380552 | 80386865 | 6314 | -7.24 | 8.35E-11 | 2.21E-10 |
| chr6 | 113242152 | 113245173 | 3022 | -7.38 | 8.37E-11 | 2.22E-10 |
| chr12 | 27323894 | 27327370 | 3477 | -7.48 | 8.38E-11 | 2.22E-10 |
| chr3 | 17940720 | 17944427 | 3708 | -7.26 | 8.39E-11 | 2.22E-10 |
| chr12 | 70329525 | 70332389 | 2865 | -7.33 | 8.40E-11 | 2.22E-10 |
| chr9 | 29343205 | 29344947 | 1743 | -7.35 | 8.41E-11 | 2.23E-10 |
| chr8 | 121337506 | 121340424 | 2919 | 2.88 | 8.42E-11 | 2.23E-10 |
| chrX | 82001251 | 82010448 | 9198 | -7.27 | 8.43E-11 | 2.23E-10 |
| chr4 | 168481062 | 168484975 | 3914 | -7.25 | 8.44E-11 | 2.23E-10 |
| chr22 | 50815143 | 50817308 | 2166 | -7.34 | 8.44E-11 | 2.23E-10 |
| chr13 | 63207241 | 63219718 | 12478 | -7.38 | 8.44E-11 | 2.23E-10 |
| chr1 | 201409611 | 201410887 | 1277 | -7.26 | 8.46E-11 | 2.24E-10 |
| chr13 | 68297410 | 68302116 | 4707 | -6.26 | 8.46E-11 | 2.24E-10 |
| chr6 | 108802324 | 108805041 | 2718 | -7.26 | 8.47E-11 | 2.24E-10 |
| chr4 | 155759618 | 155763027 | 3410 | -7.27 | 8.47E-11 | 2.24E-10 |
| chr6 | 138838196 | 138839972 | 1777 | -7.27 | 8.47E-11 | 2.24E-10 |
| chr2 | 101850039 | 101853672 | 3634 | -7.32 | 8.48E-11 | 2.24E-10 |
| chr10 | 34433544 | 34435267 | 1724 | -7.25 | 8.50E-11 | 2.25E-10 |
| chr8 | 95475711 | 95479426 | 3716 | -7.46 | 8.53E-11 | 2.25E-10 |
| chr1 | 172605525 | 172612134 | 6610 | -7.46 | 8.55E-11 | 2.26E-10 |
| chr6 | 78554529 | 78572069 | 17541 | 2 | 8.55E-11 | 2.26E-10 |
| chr2 | 179795474 | 179801388 | 5915 | -7.41 | 8.56E-11 | 2.26E-10 |
| chr15 | 57773358 | 57776700 | 3343 | -7.28 | 8.56E-11 | 2.26E-10 |
| chr9 | 134539156 | 134541097 | 1942 | -7.27 | 8.56E-11 | 2.26E-10 |

| | | | | | | |
|---|---|---|---|---|---|---|
| chr1 | 211524687 | 211526790 | 2104 | -7.24 | 8.56E-11 | 2.26E-10 |
| chr12 | 87645997 | 87648348 | 2352 | -7.28 | 8.58E-11 | 2.27E-10 |
| chr5 | 90580428 | 90583579 | 3152 | -7.41 | 8.60E-11 | 2.27E-10 |
| chr1 | 120031917 | 120056458 | 24542 | 1.59 | 8.60E-11 | 2.27E-10 |
| chr10 | 60011461 | 60019593 | 8133 | -7.56 | 8.62E-11 | 2.28E-10 |
| chr18 | 35089031 | 35091753 | 2723 | -2.92 | 8.63E-11 | 2.28E-10 |
| chrX | 57431870 | 57435314 | 3445 | -7.27 | 8.63E-11 | 2.28E-10 |
| chr9 | 3701338 | 3724654 | 23317 | -2.05 | 8.66E-11 | 2.29E-10 |
| chr4 | 145219022 | 145224963 | 5942 | 7.31 | 8.69E-11 | 2.30E-10 |
| chr5 | 127749457 | 127752531 | 3075 | -7.29 | 8.70E-11 | 2.30E-10 |
| chr12 | 8772792 | 8774301 | 1510 | -7.36 | 8.71E-11 | 2.30E-10 |
| chr6 | 122359460 | 122361868 | 2409 | -7.36 | 8.72E-11 | 2.30E-10 |
| chr15 | 55812882 | 55819846 | 6965 | -6.56 | 8.72E-11 | 2.30E-10 |
| chr7 | 34877818 | 34894553 | 16736 | -7.92 | 8.73E-11 | 2.31E-10 |
| chr4 | 64918390 | 64926095 | 7706 | -7.26 | 8.74E-11 | 2.31E-10 |
| chr9 | 133824606 | 133840399 | 15794 | -5.89 | 8.75E-11 | 2.31E-10 |
| chr19 | 18467206 | 18477540 | 10335 | -7.79 | 8.75E-11 | 2.31E-10 |
| chr18 | 48111445 | 48124613 | 13169 | -2.2 | 8.78E-11 | 2.32E-10 |
| chr10 | 1320154 | 1338538 | 18385 | -2.98 | 8.79E-11 | 2.32E-10 |
| chr14 | 61839092 | 61843712 | 4621 | -6.09 | 8.80E-11 | 2.32E-10 |
| chr1 | 26284197 | 26285807 | 1611 | -7.25 | 8.82E-11 | 2.33E-10 |
| chr17 | 1915524 | 1917958 | 2435 | -7.27 | 8.87E-11 | 2.34E-10 |
| chr2 | 238542885 | 238545968 | 3084 | -7.35 | 8.88E-11 | 2.34E-10 |
| chr21 | 17153557 | 17156852 | 3296 | -7.34 | 8.88E-11 | 2.34E-10 |
| chr2 | 233999826 | 234001692 | 1867 | -7.58 | 8.90E-11 | 2.35E-10 |
| chr12 | 8254325 | 8259409 | 5085 | -7.4 | 8.91E-11 | 2.35E-10 |
| chr2 | 106173318 | 106188313 | 14996 | -7.4 | 8.91E-11 | 2.35E-10 |
| chr15 | 36974518 | 36980540 | 6023 | -7.33 | 8.91E-11 | 2.35E-10 |
| chr2 | 121687198 | 121689608 | 2411 | -7.37 | 8.91E-11 | 2.35E-10 |
| chr1 | 187431381 | 187435398 | 4018 | -3.05 | 8.92E-11 | 2.35E-10 |
| chr2 | 201051749 | 201053183 | 1435 | -7.28 | 8.92E-11 | 2.35E-10 |
| chr2 | 201376786 | 201383099 | 6314 | -2.07 | 8.93E-11 | 2.36E-10 |
| chr6 | 46612070 | 46614121 | 2052 | -6.42 | 8.95E-11 | 2.36E-10 |
| chr16 | 68706891 | 68709844 | 2954 | -7.27 | 8.96E-11 | 2.36E-10 |
| chr11 | 95081502 | 95096731 | 15230 | -7.61 | 8.96E-11 | 2.36E-10 |
| chr2 | 20437466 | 20438614 | 1149 | -7.23 | 8.99E-11 | 2.37E-10 |
| chr11 | 119481262 | 119484336 | 3075 | -7.25 | 8.99E-11 | 2.37E-10 |
| chr11 | 85730927 | 85733016 | 2090 | -7.23 | 9.00E-11 | 2.37E-10 |
| chr12 | 84166157 | 84172443 | 6287 | -7.31 | 9.01E-11 | 2.38E-10 |
| chr5 | 133005621 | 133008322 | 2702 | -7.26 | 9.04E-11 | 2.38E-10 |
| chr6 | 74914664 | 74916512 | 1849 | -5.02 | 9.05E-11 | 2.39E-10 |
| chr16 | 72790499 | 72793507 | 3009 | -7.3 | 9.05E-11 | 2.39E-10 |
| chr5 | 77702731 | 77704765 | 2035 | -7.4 | 9.07E-11 | 2.39E-10 |
| chr8 | 101979980 | 101982241 | 2262 | -7.42 | 9.08E-11 | 2.39E-10 |
| chr3 | 159636798 | 159653856 | 17059 | -7.88 | 9.09E-11 | 2.39E-10 |
| chr4 | 65319079 | 65330305 | 11227 | -7.35 | 9.09E-11 | 2.40E-10 |
| chr11 | 60939755 | 60942993 | 3239 | -7.34 | 9.09E-11 | 2.40E-10 |
| chr8 | 39818756 | 39829465 | 10710 | -7.24 | 9.10E-11 | 2.40E-10 |
| chr9 | 79276189 | 79277577 | 1389 | -7.27 | 9.11E-11 | 2.40E-10 |
| chr12 | 57740708 | 57742809 | 2102 | -7.23 | 9.13E-11 | 2.41E-10 |
| chr20 | 45930082 | 45946633 | 16552 | -7.3 | 9.14E-11 | 2.41E-10 |
| chr12 | 51274823 | 51277465 | 2643 | -7.43 | 9.14E-11 | 2.41E-10 |
| chr20 | 60400692 | 60404284 | 3593 | -7.23 | 9.17E-11 | 2.42E-10 |
| chr6 | 161884440 | 161888481 | 4042 | -7.23 | 9.17E-11 | 2.42E-10 |
| chr11 | 77950726 | 77954115 | 3390 | -7.24 | 9.18E-11 | 2.42E-10 |
| chr4 | 38421946 | 38428855 | 6910 | -7.26 | 9.18E-11 | 2.42E-10 |
| chr15 | 69008234 | 69013901 | 5668 | -7.31 | 9.20E-11 | 2.42E-10 |
| chr6 | 33932009 | 33940268 | 8260 | -7.48 | 9.21E-11 | 2.43E-10 |
| chr6 | 145117170 | 145131867 | 14698 | -7.59 | 9.21E-11 | 2.43E-10 |
| chr4 | 135069637 | 135081530 | 11894 | 7.48 | 9.23E-11 | 2.43E-10 |
| chr4 | 19519910 | 19533176 | 13267 | -7.39 | 9.24E-11 | 2.43E-10 |

| | | | | | | |
|---|---|---|---|---|---|---|
| chrX | 37958191 | 37960387 | 2197 | -7.28 | 9.24E-11 | 2.43E-10 |
| chr7 | 123684817 | 123728179 | 43363 | 1.83 | 9.24E-11 | 2.43E-10 |
| chr2 | 100411204 | 100421048 | 9845 | -2.39 | 9.24E-11 | 2.43E-10 |
| chr18 | 67850826 | 67854197 | 3372 | -7.29 | 9.24E-11 | 2.43E-10 |
| chr8 | 38169416 | 38170907 | 1492 | -7.42 | 9.25E-11 | 2.43E-10 |
| chr2 | 209673365 | 209676319 | 2955 | -7.24 | 9.27E-11 | 2.44E-10 |
| chr6 | 136891655 | 136894121 | 2467 | -5.71 | 9.32E-11 | 2.45E-10 |
| chr14 | 89779576 | 89787332 | 7757 | -7.29 | 9.32E-11 | 2.45E-10 |
| chr4 | 79782593 | 79784278 | 1686 | -7.28 | 9.32E-11 | 2.45E-10 |
| chr4 | 2981981 | 2989450 | 7470 | -7.45 | 9.33E-11 | 2.45E-10 |
| chr7 | 23061285 | 23073815 | 12531 | -7.4 | 9.36E-11 | 2.46E-10 |
| chr14 | 102161797 | 102165628 | 3832 | -1.99 | 9.36E-11 | 2.46E-10 |
| chr19 | 14430396 | 14432224 | 1829 | -7.24 | 9.38E-11 | 2.47E-10 |
| chr3 | 197777489 | 197780147 | 2659 | -7.24 | 9.38E-11 | 2.47E-10 |
| chr9 | 16358233 | 16360452 | 2220 | -7.23 | 9.39E-11 | 2.47E-10 |
| chr4 | 17792957 | 17795033 | 2077 | -7.24 | 9.39E-11 | 2.47E-10 |
| chr11 | 95423140 | 95426028 | 2889 | 6.37 | 9.41E-11 | 2.47E-10 |
| chr5 | 172527307 | 172529490 | 2184 | -7.72 | 9.41E-11 | 2.47E-10 |
| chr7 | 11983617 | 11987010 | 3394 | -7.27 | 9.42E-11 | 2.48E-10 |
| chr7 | 76995175 | 76997481 | 2307 | -7.41 | 9.42E-11 | 2.48E-10 |
| chr6 | 82338515 | 82342402 | 3888 | -7.36 | 9.42E-11 | 2.48E-10 |
| chr11 | 18522576 | 18524084 | 1509 | -7.23 | 9.42E-11 | 2.48E-10 |
| chr1 | 199294063 | 199297754 | 3692 | -7.37 | 9.43E-11 | 2.48E-10 |
| chr1 | 160703275 | 160707441 | 4167 | -7.24 | 9.43E-11 | 2.48E-10 |
| chr11 | 108323771 | 108326024 | 2254 | -7.44 | 9.44E-11 | 2.48E-10 |
| chr15 | 79774678 | 79784802 | 10125 | -7.22 | 9.46E-11 | 2.48E-10 |
| chr20 | 58364112 | 58379830 | 15719 | -2.88 | 9.46E-11 | 2.49E-10 |
| chr4 | 186802695 | 186815214 | 12520 | -7.33 | 9.50E-11 | 2.50E-10 |
| chr7 | 54976586 | 54978638 | 2053 | -7.32 | 9.51E-11 | 2.50E-10 |
| chrX | 48079305 | 48087164 | 7860 | -7.32 | 9.51E-11 | 2.50E-10 |
| chr3 | 113203614 | 113205613 | 2000 | -7.25 | 9.53E-11 | 2.50E-10 |
| chr7 | 37937497 | 37945727 | 8231 | -7.38 | 9.53E-11 | 2.50E-10 |
| chr5 | 39281695 | 39299914 | 18220 | -8.02 | 9.55E-11 | 2.51E-10 |
| chr2 | 235432300 | 235448723 | 16424 | -7.5 | 9.57E-11 | 2.51E-10 |
| chr1 | 95096772 | 95106404 | 9633 | -7.47 | 9.57E-11 | 2.51E-10 |
| chr20 | 223489 | 235252 | 11764 | -5.3 | 9.57E-11 | 2.51E-10 |
| chr16 | 18846224 | 18847730 | 1507 | -7.22 | 9.59E-11 | 2.52E-10 |
| chr3 | 186215895 | 186219394 | 3500 | -7.27 | 9.61E-11 | 2.52E-10 |
| chr6 | 35661798 | 35671918 | 10121 | -7.23 | 9.62E-11 | 2.52E-10 |
| chrX | 151801962 | 151803534 | 1573 | -7.23 | 9.62E-11 | 2.52E-10 |
| chr17 | 64996556 | 65006042 | 9487 | -7.86 | 9.62E-11 | 2.53E-10 |
| chr2 | 165785442 | 165789195 | 3754 | -7.38 | 9.62E-11 | 2.53E-10 |
| chr11 | 34033934 | 34045232 | 11299 | -7.3 | 9.64E-11 | 2.53E-10 |
| chr4 | 158626364 | 158708467 | 82104 | 3.84 | 9.65E-11 | 2.53E-10 |
| chr10 | 116341566 | 116354553 | 12988 | -7.47 | 9.66E-11 | 2.54E-10 |
| chr5 | 14515219 | 14516855 | 1637 | -7.26 | 9.67E-11 | 2.54E-10 |
| chr5 | 127666093 | 127667835 | 1743 | -7.26 | 9.67E-11 | 2.54E-10 |
| chr20 | 3954886 | 3956274 | 1389 | -7.25 | 9.67E-11 | 2.54E-10 |
| chrX | 35739837 | 35748547 | 8711 | -7.24 | 9.68E-11 | 2.54E-10 |
| chr11 | 19343680 | 19347351 | 3672 | -6 | 9.68E-11 | 2.54E-10 |
| chr11 | 78764294 | 78770149 | 5856 | 7.29 | 9.69E-11 | 2.54E-10 |
| chr9 | 117432132 | 117451668 | 19537 | -7.88 | 9.71E-11 | 2.55E-10 |
| chr12 | 118329076 | 118350351 | 21276 | -7.76 | 9.72E-11 | 2.55E-10 |
| chr10 | 73691562 | 73698563 | 7002 | -7.44 | 9.74E-11 | 2.56E-10 |
| chr21 | 44206718 | 44208712 | 1995 | -7.23 | 9.76E-11 | 2.56E-10 |
| chr5 | 172545570 | 172548004 | 2435 | -7.6 | 9.77E-11 | 2.56E-10 |
| chr19 | 32944267 | 32946018 | 1752 | -7.41 | 9.78E-11 | 2.56E-10 |
| chr20 | 9397625 | 9398907 | 1283 | -7.42 | 9.81E-11 | 2.57E-10 |
| chr6 | 88898602 | 88914806 | 16205 | -2.16 | 9.81E-11 | 2.57E-10 |
| chr13 | 35543234 | 35545557 | 2324 | -7.49 | 9.85E-11 | 2.58E-10 |
| chr6 | 154900367 | 154903486 | 3120 | -7.25 | 9.86E-11 | 2.58E-10 |

| | | | | | | |
|---|---|---|---|---|---|---|
| chr1 | 168244645 | 168247289 | 2645 | -7.45 | 9.91E-11 | 2.60E-10 |
| chr6 | 141908792 | 141914269 | 5478 | -7.35 | 9.91E-11 | 2.60E-10 |
| chr7 | 96310587 | 96315052 | 4466 | -7.21 | 9.91E-11 | 2.60E-10 |
| chr3 | 5455863 | 5457568 | 1706 | -5.33 | 9.92E-11 | 2.60E-10 |
| chr16 | 56567602 | 56576427 | 8826 | -7.33 | 9.92E-11 | 2.60E-10 |
| chr14 | 32331797 | 32335630 | 3834 | -7.27 | 9.92E-11 | 2.60E-10 |
| chr13 | 31886898 | 31891034 | 4137 | -7.41 | 9.94E-11 | 2.60E-10 |
| chr11 | 126705234 | 126707666 | 2433 | -7.23 | 9.95E-11 | 2.61E-10 |
| chr3 | 188346939 | 188353947 | 7009 | -7.23 | 9.95E-11 | 2.61E-10 |
| chr18 | 9390726 | 9392920 | 2195 | -7.31 | 9.96E-11 | 2.61E-10 |
| chr13 | 67785051 | 67801964 | 16914 | -7.43 | 9.96E-11 | 2.61E-10 |
| chr11 | 82875910 | 82879627 | 3718 | -7.31 | 9.97E-11 | 2.61E-10 |
| chr20 | 31706954 | 31709368 | 2415 | -7.36 | 9.98E-11 | 2.61E-10 |
| chr12 | 101722300 | 101723916 | 1617 | -7.3 | 9.99E-11 | 2.62E-10 |
| chr2 | 65407067 | 65408781 | 1715 | -7.3 | 9.99E-11 | 2.62E-10 |
| chr4 | 189550035 | 189555004 | 4970 | -7.36 | 9.99E-11 | 2.62E-10 |
| chr18 | 2989101 | 3002260 | 13160 | -7.25 | 1.00E-10 | 2.63E-10 |
| chr17 | 40897189 | 40898307 | 1119 | -7.27 | 1.00E-10 | 2.63E-10 |
| chr3 | 12473068 | 12475223 | 2156 | -7.27 | 1.00E-10 | 2.63E-10 |
| chr2 | 164809290 | 164811197 | 1908 | -7.37 | 1.01E-10 | 2.63E-10 |
| chr2 | 151552519 | 151556680 | 4162 | -7.21 | 1.01E-10 | 2.63E-10 |
| chr15 | 53737150 | 53762229 | 25080 | -8.02 | 1.01E-10 | 2.63E-10 |
| chr9 | 29983636 | 30015302 | 31667 | -8.05 | 1.01E-10 | 2.64E-10 |
| chr20 | 32422376 | 32423906 | 1531 | -7.23 | 1.01E-10 | 2.64E-10 |
| chr3 | 36391914 | 36401995 | 10082 | -7.23 | 1.01E-10 | 2.64E-10 |
| chr3 | 122567739 | 122570396 | 2658 | -7.59 | 1.01E-10 | 2.65E-10 |
| chr7 | 86919186 | 86930641 | 11456 | -1.71 | 1.02E-10 | 2.66E-10 |
| chr7 | 80244640 | 80249676 | 5037 | -7.39 | 1.02E-10 | 2.66E-10 |
| chr3 | 163270640 | 163883065 | 612426 | 1.85 | 1.02E-10 | 2.67E-10 |
| chr5 | 789921 | 794036 | 4116 | -7.24 | 1.02E-10 | 2.68E-10 |
| chr2 | 29390532 | 29391669 | 1138 | -7.25 | 1.03E-10 | 2.68E-10 |
| chr6 | 1420375 | 1421839 | 1465 | -7.27 | 1.03E-10 | 2.69E-10 |
| chr9 | 74855265 | 74866514 | 11250 | -7.21 | 1.03E-10 | 2.69E-10 |
| chr5 | 174012507 | 174014882 | 2376 | -7.41 | 1.03E-10 | 2.70E-10 |
| chr12 | 80040487 | 80042437 | 1951 | -7.22 | 1.04E-10 | 2.71E-10 |
| chr22 | 47316851 | 47318634 | 1784 | -7.22 | 1.04E-10 | 2.71E-10 |
| chr1 | 3332952 | 3334664 | 1713 | -7.23 | 1.04E-10 | 2.71E-10 |
| chr22 | 20034397 | 20036738 | 2342 | -7.23 | 1.04E-10 | 2.71E-10 |
| chr5 | 64933539 | 64934993 | 1455 | -7.33 | 1.04E-10 | 2.71E-10 |
| chrX | 131596940 | 131598837 | 1898 | -7.41 | 1.04E-10 | 2.71E-10 |
| chr4 | 106093814 | 106098441 | 4628 | -7.45 | 1.04E-10 | 2.72E-10 |
| chr7 | 106883940 | 106887585 | 3646 | -7.3 | 1.04E-10 | 2.72E-10 |
| chr11 | 108638891 | 108656431 | 17541 | -7.41 | 1.04E-10 | 2.72E-10 |
| chr11 | 104224652 | 104227011 | 2360 | -4.81 | 1.04E-10 | 2.73E-10 |
| chr4 | 74099212 | 74102622 | 3411 | -7.35 | 1.04E-10 | 2.73E-10 |
| chr2 | 72111267 | 72150422 | 39156 | -3.88 | 1.05E-10 | 2.73E-10 |
| chr9 | 87589372 | 87596378 | 7007 | -7.23 | 1.05E-10 | 2.73E-10 |
| chrX | 100721154 | 100726587 | 5434 | -7.23 | 1.05E-10 | 2.73E-10 |
| chr11 | 74061396 | 74062724 | 1329 | -7.31 | 1.05E-10 | 2.74E-10 |
| chr3 | 138995160 | 138997610 | 2451 | -7.24 | 1.05E-10 | 2.74E-10 |
| chr4 | 95287250 | 95289546 | 2297 | -7.24 | 1.05E-10 | 2.74E-10 |
| chr17 | 53363414 | 53369551 | 6138 | -7.42 | 1.05E-10 | 2.74E-10 |
| chr9 | 111158796 | 111160677 | 1882 | -7.3 | 1.05E-10 | 2.74E-10 |
| chr7 | 15350441 | 15382041 | 31601 | -7.8 | 1.06E-10 | 2.76E-10 |
| chr1 | 23348062 | 23350177 | 2116 | -7.21 | 1.06E-10 | 2.76E-10 |
| chr5 | 169873628 | 169875655 | 2028 | -7.21 | 1.06E-10 | 2.76E-10 |
| chr2 | 222892730 | 222895327 | 2598 | -7.29 | 1.06E-10 | 2.76E-10 |
| chr1 | 27410373 | 27412834 | 2462 | -7.28 | 1.06E-10 | 2.77E-10 |
| chr13 | 95954472 | 95969154 | 14683 | -7.38 | 1.06E-10 | 2.78E-10 |
| chr14 | 58983891 | 58993759 | 9869 | -7.53 | 1.06E-10 | 2.78E-10 |
| chr1 | 35590859 | 35593182 | 2324 | -7.28 | 1.07E-10 | 2.78E-10 |

| | | | | | | |
|---|---|---|---|---|---|---|
| chr2 | 189334620 | 189336417 | 1798 | -7.25 | 1.07E-10 | 2.79E-10 |
| chr7 | 6901694 | 6916469 | 14776 | -7.27 | 1.07E-10 | 2.80E-10 |
| chr4 | 114601891 | 114603409 | 1519 | -7.21 | 1.07E-10 | 2.80E-10 |
| chr13 | 24620167 | 24622426 | 2260 | -7.32 | 1.08E-10 | 2.83E-10 |
| chr5 | 122802721 | 122806181 | 3461 | -7.32 | 1.08E-10 | 2.83E-10 |
| chr2 | 37101259 | 37102566 | 1308 | -7.45 | 1.09E-10 | 2.83E-10 |
| chr3 | 105154227 | 105165386 | 11160 | -7.61 | 1.09E-10 | 2.84E-10 |
| chrX | 138835609 | 138838307 | 2699 | -7.2 | 1.09E-10 | 2.85E-10 |
| chr9 | 2900210 | 2903553 | 3344 | -3.87 | 1.09E-10 | 2.85E-10 |
| chr6 | 109293119 | 109296243 | 3125 | -7.35 | 1.09E-10 | 2.85E-10 |
| chr1 | 37606750 | 37614533 | 7784 | 7.95 | 1.09E-10 | 2.85E-10 |
| chr13 | 109946608 | 109949113 | 2506 | -7.29 | 1.09E-10 | 2.85E-10 |
| chr4 | 118777707 | 118779612 | 1906 | 5.35 | 1.09E-10 | 2.85E-10 |
| chr4 | 99685543 | 99691535 | 5993 | -6.09 | 1.09E-10 | 2.85E-10 |
| chr18 | 54129052 | 54135447 | 6396 | -3.06 | 1.10E-10 | 2.86E-10 |
| chr17 | 36960040 | 36961508 | 1469 | -7.24 | 1.10E-10 | 2.86E-10 |
| chr17 | 28827793 | 28829124 | 1332 | -7.25 | 1.10E-10 | 2.86E-10 |
| chr6 | 113668453 | 113672383 | 3931 | -7.4 | 1.10E-10 | 2.86E-10 |
| chr5 | 131982436 | 131984988 | 2553 | -7.22 | 1.10E-10 | 2.87E-10 |
| chr7 | 31830155 | 31844903 | 14749 | -7.21 | 1.10E-10 | 2.88E-10 |
| chr9 | 90714615 | 90724360 | 9746 | -7.31 | 1.11E-10 | 2.88E-10 |
| chr10 | 11626534 | 11628166 | 1633 | -7.22 | 1.11E-10 | 2.89E-10 |
| chr5 | 175900205 | 175909208 | 9004 | -7.22 | 1.11E-10 | 2.89E-10 |
| chr7 | 23581965 | 23583441 | 1477 | -7.29 | 1.11E-10 | 2.91E-10 |
| chr6 | 5436870 | 5438355 | 1486 | -7.35 | 1.12E-10 | 2.91E-10 |
| chr13 | 113842622 | 113844464 | 1843 | -7.35 | 1.12E-10 | 2.91E-10 |
| chr10 | 90504793 | 90510258 | 5466 | -5.27 | 1.12E-10 | 2.91E-10 |
| chr17 | 53474442 | 53476636 | 2195 | -7.21 | 1.12E-10 | 2.91E-10 |
| chr15 | 66306769 | 66308757 | 1989 | -7.2 | 1.13E-10 | 2.94E-10 |
| chr16 | 50855953 | 50869090 | 13138 | -1.5 | 1.13E-10 | 2.95E-10 |
| chr1 | 241942360 | 241945372 | 3013 | -7.32 | 1.13E-10 | 2.95E-10 |
| chr1 | 100165957 | 100167791 | 1835 | -7.3 | 1.13E-10 | 2.95E-10 |
| chr7 | 682990 | 693681 | 10692 | -7.32 | 1.13E-10 | 2.95E-10 |
| chr3 | 160504541 | 160506501 | 1961 | -7.3 | 1.13E-10 | 2.95E-10 |
| chr7 | 17846949 | 17853059 | 6111 | -7.29 | 1.13E-10 | 2.95E-10 |
| chr17 | 47357531 | 47359804 | 2274 | -7.48 | 1.14E-10 | 2.97E-10 |
| chr20 | 36419315 | 36421365 | 2051 | -7.38 | 1.14E-10 | 2.97E-10 |
| chr10 | 13857120 | 13859847 | 2728 | -7.2 | 1.14E-10 | 2.98E-10 |
| chr20 | 57460089 | 57461381 | 1293 | -7.19 | 1.14E-10 | 2.98E-10 |
| chr4 | 125107040 | 125116574 | 9535 | -7.23 | 1.15E-10 | 2.98E-10 |
| chr3 | 125713362 | 125716186 | 2825 | -7.26 | 1.15E-10 | 2.99E-10 |
| chr11 | 133863958 | 133867074 | 3117 | -5.58 | 1.15E-10 | 2.99E-10 |
| chr3 | 124673934 | 124676100 | 2167 | -7.21 | 1.15E-10 | 2.99E-10 |
| chr1 | 218236782 | 218239374 | 2593 | -7.39 | 1.15E-10 | 2.99E-10 |
| chr19 | 56604697 | 56607096 | 2400 | -7.2 | 1.15E-10 | 2.99E-10 |
| chr9 | 81043419 | 81046789 | 3371 | -7.27 | 1.15E-10 | 2.99E-10 |
| chr15 | 50809408 | 50810797 | 1390 | -7.25 | 1.15E-10 | 3.00E-10 |
| chr13 | 44504246 | 44507374 | 3129 | -7.27 | 1.15E-10 | 3.00E-10 |
| chrX | 120178170 | 120182997 | 4828 | -7.19 | 1.16E-10 | 3.01E-10 |
| chr9 | 6092475 | 6097808 | 5334 | -7.19 | 1.16E-10 | 3.01E-10 |
| chrX | 41630166 | 41633920 | 3755 | -7.2 | 1.16E-10 | 3.01E-10 |
| chr6 | 132623487 | 132639032 | 15546 | -7.22 | 1.16E-10 | 3.02E-10 |
| chr6 | 106181383 | 106185719 | 4337 | -7.37 | 1.16E-10 | 3.03E-10 |
| chrX | 86861982 | 86865994 | 4013 | -7.2 | 1.17E-10 | 3.03E-10 |
| chrX | 29726416 | 29728498 | 2083 | -7.21 | 1.17E-10 | 3.03E-10 |
| chr6 | 74197900 | 74200755 | 2856 | -7.3 | 1.17E-10 | 3.04E-10 |
| chr4 | 76696115 | 76699013 | 2899 | -7.32 | 1.17E-10 | 3.04E-10 |
| chr3 | 180435367 | 180440170 | 4804 | 7.28 | 1.17E-10 | 3.04E-10 |
| chr4 | 124323986 | 124327082 | 3097 | -7.46 | 1.17E-10 | 3.05E-10 |
| chr8 | 111849055 | 111851768 | 2714 | -7.25 | 1.17E-10 | 3.05E-10 |
| chr18 | 73111169 | 73117060 | 5892 | -3.06 | 1.17E-10 | 3.05E-10 |

| | | | | | | |
|---|---|---|---|---|---|---|
| chr3 | 152840040 | 152843066 | 3027 | -7.38 | 1.18E-10 | 3.06E-10 |
| chr5 | 168773209 | 168781096 | 7888 | -7.34 | 1.18E-10 | 3.06E-10 |
| chr4 | 88638899 | 88649568 | 10670 | -2.15 | 1.18E-10 | 3.06E-10 |
| chr12 | 106674773 | 106676886 | 2114 | -7.3 | 1.18E-10 | 3.07E-10 |
| chr3 | 55419880 | 55459760 | 39881 | -2.9 | 1.19E-10 | 3.08E-10 |
| chr6 | 26578559 | 26580487 | 1929 | -7.34 | 1.19E-10 | 3.09E-10 |
| chr20 | 42585710 | 42587360 | 1651 | -7.24 | 1.19E-10 | 3.10E-10 |
| chr12 | 91213907 | 91217961 | 4055 | 7.28 | 1.19E-10 | 3.10E-10 |
| chr16 | 2437804 | 2440429 | 2626 | -7.47 | 1.19E-10 | 3.10E-10 |
| chr2 | 203447771 | 203450001 | 2231 | -7.19 | 1.19E-10 | 3.10E-10 |
| chrX | 129396945 | 129417898 | 20954 | -7.68 | 1.20E-10 | 3.11E-10 |
| chr1 | 68573461 | 68576399 | 2939 | -7.21 | 1.20E-10 | 3.11E-10 |
| chr1 | 63033123 | 63038440 | 5318 | -7.22 | 1.20E-10 | 3.12E-10 |
| chr8 | 81379601 | 81384484 | 4884 | -7.22 | 1.20E-10 | 3.12E-10 |
| chr19 | 55919873 | 55922193 | 2321 | -7.35 | 1.20E-10 | 3.12E-10 |
| chr19 | 39638994 | 39640191 | 1198 | -7.35 | 1.20E-10 | 3.12E-10 |
| chr6 | 153053446 | 153055646 | 2201 | -7.19 | 1.20E-10 | 3.12E-10 |
| chr5 | 139407275 | 139415029 | 7755 | -7.2 | 1.20E-10 | 3.13E-10 |
| chr12 | 2224021 | 2628634 | 404614 | 1.61 | 1.20E-10 | 3.13E-10 |
| chr6 | 108563149 | 108565309 | 2161 | -7.25 | 1.21E-10 | 3.13E-10 |
| chr1 | 46314038 | 46316430 | 2393 | -7.21 | 1.21E-10 | 3.13E-10 |
| chr12 | 98185154 | 98192529 | 7376 | 2.05 | 1.21E-10 | 3.14E-10 |
| chr20 | 35193464 | 35194418 | 955 | -7.19 | 1.21E-10 | 3.14E-10 |
| chr2 | 175284194 | 175295110 | 10917 | -7.19 | 1.21E-10 | 3.14E-10 |
| chr1 | 87477442 | 87480521 | 3080 | -7.22 | 1.21E-10 | 3.14E-10 |
| chr2 | 103258117 | 103265483 | 7367 | -7.41 | 1.21E-10 | 3.15E-10 |
| chr10 | 12563504 | 12567645 | 4142 | -7.19 | 1.21E-10 | 3.15E-10 |
| chr9 | 91122918 | 91124784 | 1867 | -7.19 | 1.21E-10 | 3.15E-10 |
| chr12 | 100978176 | 100983164 | 4989 | -7.25 | 1.21E-10 | 3.15E-10 |
| chr2 | 38168116 | 38183454 | 15339 | -7.83 | 1.22E-10 | 3.16E-10 |
| chr11 | 69442945 | 69445179 | 2235 | -7.19 | 1.22E-10 | 3.17E-10 |
| chr6 | 22615602 | 22619568 | 3967 | -7.19 | 1.22E-10 | 3.17E-10 |
| chr13 | 102245193 | 102247240 | 2048 | -7.19 | 1.23E-10 | 3.18E-10 |
| chr8 | 95085475 | 95088263 | 2789 | -7.38 | 1.23E-10 | 3.19E-10 |
| chr4 | 139497809 | 139504144 | 6336 | -7.72 | 1.23E-10 | 3.19E-10 |
| chr8 | 122890054 | 122903984 | 13931 | -7.72 | 1.23E-10 | 3.19E-10 |
| chr3 | 40172939 | 40184092 | 11154 | -7.4 | 1.23E-10 | 3.20E-10 |
| chr2 | 216073669 | 216075679 | 2011 | -7.18 | 1.24E-10 | 3.20E-10 |
| chr4 | 168113311 | 168116749 | 3439 | -7.18 | 1.24E-10 | 3.20E-10 |
| chr4 | 57203273 | 57205107 | 1835 | -7.29 | 1.24E-10 | 3.20E-10 |
| chr4 | 148293434 | 148315257 | 21824 | -7.72 | 1.24E-10 | 3.20E-10 |
| chr20 | 58433893 | 58436302 | 2410 | -7.29 | 1.24E-10 | 3.20E-10 |
| chr3 | 72176149 | 72180096 | 3948 | -7.32 | 1.24E-10 | 3.21E-10 |
| chr2 | 172463882 | 172466332 | 2451 | -7.23 | 1.24E-10 | 3.21E-10 |
| chr12 | 58277350 | 58279799 | 2450 | -7.53 | 1.24E-10 | 3.21E-10 |
| chr9 | 134117870 | 134120042 | 2173 | -7.28 | 1.24E-10 | 3.22E-10 |
| chr11 | 33453826 | 33464458 | 10633 | -7.27 | 1.24E-10 | 3.22E-10 |
| chr2 | 209441197 | 209443628 | 2432 | -7.49 | 1.24E-10 | 3.22E-10 |
| chr12 | 96614498 | 96616987 | 2490 | -7.19 | 1.24E-10 | 3.22E-10 |
| chr11 | 111255279 | 111257588 | 2310 | -7.24 | 1.25E-10 | 3.23E-10 |
| chr1 | 196596568 | 196599132 | 2565 | -7.27 | 1.25E-10 | 3.23E-10 |
| chr6 | 160565245 | 160575940 | 10696 | -7.34 | 1.25E-10 | 3.24E-10 |
| chr5 | 40974040 | 41175508 | 201469 | 3.95 | 1.25E-10 | 3.25E-10 |
| chr6 | 127418833 | 127432027 | 13195 | -7.26 | 1.26E-10 | 3.26E-10 |
| chr3 | 121368259 | 121370358 | 2100 | -7.18 | 1.26E-10 | 3.26E-10 |
| chr1 | 169274288 | 169278346 | 4059 | -7.26 | 1.26E-10 | 3.26E-10 |
| chr18 | 46103694 | 46105221 | 1528 | -7.2 | 1.26E-10 | 3.27E-10 |
| chr7 | 80107547 | 80109828 | 2282 | -7.18 | 1.26E-10 | 3.27E-10 |
| chr6 | 86355377 | 86377569 | 22193 | -7.25 | 1.27E-10 | 3.28E-10 |
| chr20 | 42604075 | 42605740 | 1666 | -7.25 | 1.27E-10 | 3.29E-10 |
| chr13 | 98271562 | 98285199 | 13638 | -7.41 | 1.27E-10 | 3.29E-10 |

| | | | | | | |
|---|---|---|---|---|---|---|
| chrX | 15394405 | 15396603 | 2199 | -7.2 | 1.27E-10 | 3.30E-10 |
| chr18 | 45310505 | 45314387 | 3883 | -6.08 | 1.27E-10 | 3.30E-10 |
| chr6 | 24114711 | 24119135 | 4425 | 7.45 | 1.28E-10 | 3.31E-10 |
| chr21 | 36565912 | 36583516 | 17605 | -7.66 | 1.28E-10 | 3.31E-10 |
| chr5 | 170365862 | 170367668 | 1807 | -7.22 | 1.28E-10 | 3.31E-10 |
| chr2 | 118994157 | 118997513 | 3357 | -7.37 | 1.28E-10 | 3.31E-10 |
| chr15 | 63529026 | 63536616 | 7591 | -7.52 | 1.28E-10 | 3.31E-10 |
| chr3 | 177304290 | 177306790 | 2501 | -7.24 | 1.28E-10 | 3.32E-10 |
| chr6 | 108725065 | 108727455 | 2391 | -7.18 | 1.28E-10 | 3.32E-10 |
| chr1 | 193743122 | 193746443 | 3322 | -7.18 | 1.29E-10 | 3.33E-10 |
| chr14 | 102617929 | 102619837 | 1909 | -7.18 | 1.29E-10 | 3.33E-10 |
| chr15 | 46060160 | 46074139 | 13980 | -1.54 | 1.29E-10 | 3.33E-10 |
| chr6 | 9549359 | 9554356 | 4998 | -3.1 | 1.29E-10 | 3.33E-10 |
| chr5 | 53459503 | 53462068 | 2566 | -7.24 | 1.29E-10 | 3.33E-10 |
| chr21 | 16337972 | 16357047 | 19076 | -7.61 | 1.29E-10 | 3.33E-10 |
| chr7 | 148970214 | 148972326 | 2113 | -7.19 | 1.29E-10 | 3.33E-10 |
| chr2 | 18965581 | 18967233 | 1653 | -7.22 | 1.29E-10 | 3.34E-10 |
| chr2 | 103216742 | 103228610 | 11869 | -7.18 | 1.29E-10 | 3.34E-10 |
| chr21 | 36446351 | 36458468 | 12118 | -7.29 | 1.29E-10 | 3.34E-10 |
| chr16 | 85661901 | 85663998 | 2098 | -7.29 | 1.29E-10 | 3.34E-10 |
| chr18 | 77760475 | 77762618 | 2144 | -7.31 | 1.30E-10 | 3.35E-10 |
| chr15 | 99442712 | 99447277 | 4566 | -7.32 | 1.30E-10 | 3.35E-10 |
| chr6 | 13341430 | 13343466 | 2037 | -7.18 | 1.30E-10 | 3.36E-10 |
| chr14 | 91420739 | 91435009 | 14271 | -8.06 | 1.30E-10 | 3.36E-10 |
| chr22 | 43925362 | 43934138 | 8777 | -7.32 | 1.30E-10 | 3.36E-10 |
| chr13 | 33933971 | 33937840 | 3870 | -7.27 | 1.30E-10 | 3.36E-10 |
| chr11 | 61209231 | 61210981 | 1751 | -7.18 | 1.30E-10 | 3.36E-10 |
| chr7 | 157600310 | 157602867 | 2558 | -7.18 | 1.30E-10 | 3.37E-10 |
| chr7 | 15623371 | 15626166 | 2796 | -7.26 | 1.30E-10 | 3.37E-10 |
| chr3 | 107863988 | 107866153 | 2166 | -7.2 | 1.31E-10 | 3.37E-10 |
| chr6 | 5639798 | 5642122 | 2325 | -7.2 | 1.31E-10 | 3.37E-10 |
| chr14 | 50780341 | 50782360 | 2020 | -7.23 | 1.31E-10 | 3.37E-10 |
| chr16 | 68470140 | 68472961 | 2822 | -5.96 | 1.31E-10 | 3.38E-10 |
| chr5 | 71398868 | 71400504 | 1637 | -7.19 | 1.31E-10 | 3.38E-10 |
| chr9 | 36320823 | 36322838 | 2016 | -7.19 | 1.31E-10 | 3.38E-10 |
| chr4 | 126459660 | 126461950 | 2291 | -7.17 | 1.31E-10 | 3.38E-10 |
| chr3 | 194253412 | 194268359 | 14948 | -7.81 | 1.31E-10 | 3.39E-10 |
| chr13 | 44178336 | 44191998 | 13663 | -7.23 | 1.31E-10 | 3.39E-10 |
| chr16 | 85572653 | 85574210 | 1558 | -7.18 | 1.31E-10 | 3.39E-10 |
| chr4 | 134532001 | 134538059 | 6059 | 7.28 | 1.32E-10 | 3.40E-10 |
| chr17 | 30088847 | 30104007 | 15161 | -7.43 | 1.32E-10 | 3.40E-10 |
| chr3 | 94221422 | 94224323 | 2902 | -7.18 | 1.32E-10 | 3.41E-10 |
| chr2 | 128720506 | 128722286 | 1781 | -7.2 | 1.32E-10 | 3.41E-10 |
| chr20 | 42175146 | 42177088 | 1943 | -7.25 | 1.32E-10 | 3.42E-10 |
| chr2 | 176805597 | 176807862 | 2266 | -7.34 | 1.32E-10 | 3.42E-10 |
| chr8 | 61349074 | 61360326 | 11253 | -7.35 | 1.33E-10 | 3.42E-10 |
| chr4 | 103954147 | 103957306 | 3160 | -7.21 | 1.33E-10 | 3.42E-10 |
| chr17 | 64606410 | 64608045 | 1636 | -7.22 | 1.33E-10 | 3.43E-10 |
| chrX | 151832011 | 151835725 | 3715 | -7.17 | 1.33E-10 | 3.43E-10 |
| chr6 | 43942763 | 43944833 | 2071 | -7.4 | 1.33E-10 | 3.43E-10 |
| chr2 | 89003234 | 89005614 | 2381 | -7.18 | 1.33E-10 | 3.43E-10 |
| chr18 | 69668583 | 69695592 | 27010 | -7.41 | 1.34E-10 | 3.45E-10 |
| chr8 | 41989485 | 41990768 | 1284 | -7.19 | 1.34E-10 | 3.45E-10 |
| chr3 | 136247871 | 136252351 | 4481 | -7.44 | 1.34E-10 | 3.45E-10 |
| chr20 | 56461775 | 56463815 | 2041 | -7.17 | 1.34E-10 | 3.46E-10 |
| chr18 | 28870706 | 28872822 | 2117 | -7.23 | 1.34E-10 | 3.46E-10 |
| chr20 | 39508898 | 39510820 | 1923 | -7.36 | 1.34E-10 | 3.46E-10 |
| chr7 | 114874760 | 114886329 | 11570 | -7.48 | 1.35E-10 | 3.47E-10 |
| chr1 | 44242703 | 44246219 | 3517 | -7.17 | 1.35E-10 | 3.48E-10 |
| chr6 | 159297560 | 159300191 | 2632 | -7.29 | 1.35E-10 | 3.48E-10 |
| chr11 | 122814001 | 122816459 | 2459 | -7.21 | 1.35E-10 | 3.48E-10 |

| | | | | | | |
|---|---|---|---|---|---|---|
| chr6 | 56943004 | 56945660 | 2657 | -7.17 | 1.35E-10 | 3.49E-10 |
| chr16 | 76927950 | 76929835 | 1886 | -7.27 | 1.35E-10 | 3.49E-10 |
| chr11 | 32688967 | 32691563 | 2597 | -7.46 | 1.36E-10 | 3.50E-10 |
| chr7 | 128818586 | 128821616 | 3031 | -7.38 | 1.36E-10 | 3.50E-10 |
| chr4 | 132355519 | 132360653 | 5135 | -7.26 | 1.36E-10 | 3.50E-10 |
| chr1 | 223994661 | 223996291 | 1631 | -7.62 | 1.36E-10 | 3.51E-10 |
| chr20 | 4182436 | 4186008 | 3573 | -7.38 | 1.36E-10 | 3.51E-10 |
| chr3 | 114822281 | 114824029 | 1749 | -7.32 | 1.36E-10 | 3.51E-10 |
| chr20 | 14544139 | 14548599 | 4461 | -7.69 | 1.36E-10 | 3.51E-10 |
| chr8 | 75062256 | 75065295 | 3040 | -5.56 | 1.36E-10 | 3.52E-10 |
| chr3 | 128298369 | 128309922 | 11554 | -7.25 | 1.37E-10 | 3.53E-10 |
| chr10 | 95686755 | 95694350 | 7596 | -7.17 | 1.37E-10 | 3.53E-10 |
| chr12 | 85667193 | 85669099 | 1907 | -7.18 | 1.37E-10 | 3.54E-10 |
| chr12 | 105219008 | 105223107 | 4100 | -7.2 | 1.37E-10 | 3.54E-10 |
| chr2 | 160278440 | 160279912 | 1473 | -7.17 | 1.37E-10 | 3.54E-10 |
| chr22 | 50158646 | 50162082 | 3437 | -7.17 | 1.37E-10 | 3.54E-10 |
| chr9 | 36883784 | 36885025 | 1242 | -7.38 | 1.37E-10 | 3.54E-10 |
| chr4 | 96388025 | 96391110 | 3086 | -6.5 | 1.38E-10 | 3.56E-10 |
| chr2 | 143441764 | 143445265 | 3502 | -7.39 | 1.38E-10 | 3.56E-10 |
| chr9 | 138720426 | 138721967 | 1542 | -7.27 | 1.38E-10 | 3.57E-10 |
| chr3 | 194454342 | 194456728 | 2387 | -7.18 | 1.39E-10 | 3.59E-10 |
| chr9 | 96033216 | 96047300 | 14085 | -7.18 | 1.39E-10 | 3.59E-10 |
| chr17 | 28231949 | 28234308 | 2360 | -7.17 | 1.40E-10 | 3.59E-10 |
| chr14 | 107147116 | 107150572 | 3457 | -7.16 | 1.40E-10 | 3.60E-10 |
| chr4 | 74691764 | 74694796 | 3033 | -7.21 | 1.40E-10 | 3.60E-10 |
| chr13 | 22059957 | 22064009 | 4053 | -7.16 | 1.40E-10 | 3.61E-10 |
| chr10 | 116384152 | 116385991 | 1840 | -7.17 | 1.40E-10 | 3.61E-10 |
| chr13 | 50878178 | 50882563 | 4386 | -7.29 | 1.41E-10 | 3.64E-10 |
| chr1 | 179117585 | 179119906 | 2322 | -7.27 | 1.41E-10 | 3.64E-10 |
| chr2 | 109323354 | 109327691 | 4338 | -7.27 | 1.41E-10 | 3.64E-10 |
| chr5 | 124446623 | 124455451 | 8829 | -7.54 | 1.41E-10 | 3.64E-10 |
| chr7 | 29643510 | 29653245 | 9736 | -7.51 | 1.41E-10 | 3.64E-10 |
| chr16 | 4637633 | 4639601 | 1969 | -7.3 | 1.41E-10 | 3.64E-10 |
| chr8 | 82678830 | 82690145 | 11316 | -7.3 | 1.41E-10 | 3.64E-10 |
| chr18 | 20471432 | 20484555 | 13124 | -7.48 | 1.42E-10 | 3.64E-10 |
| chr15 | 54058871 | 54089930 | 31060 | 1.94 | 1.42E-10 | 3.64E-10 |
| chr2 | 190039648 | 190041013 | 1366 | -7.17 | 1.42E-10 | 3.65E-10 |
| chr17 | 59193662 | 59195543 | 1882 | -7.41 | 1.42E-10 | 3.65E-10 |
| chr15 | 75459719 | 75462723 | 3005 | -7.31 | 1.42E-10 | 3.66E-10 |
| chr4 | 168733597 | 168736460 | 2864 | -7.22 | 1.43E-10 | 3.67E-10 |
| chr4 | 54469303 | 54479932 | 10630 | -7.38 | 1.43E-10 | 3.67E-10 |
| chr9 | 29357699 | 29360980 | 3282 | -5.32 | 1.43E-10 | 3.67E-10 |
| chr18 | 20848850 | 20858223 | 9374 | -7.32 | 1.43E-10 | 3.68E-10 |
| chr14 | 56143711 | 56179304 | 35594 | -7.35 | 1.43E-10 | 3.68E-10 |
| chr6 | 74814811 | 74819129 | 4319 | 2.88 | 1.43E-10 | 3.69E-10 |
| chr13 | 66480285 | 66485597 | 5313 | -5.29 | 1.44E-10 | 3.69E-10 |
| chr6 | 163151122 | 163152457 | 1336 | -5.43 | 1.44E-10 | 3.70E-10 |
| chr10 | 102203940 | 102206550 | 2611 | -7.18 | 1.44E-10 | 3.70E-10 |
| chr1 | 66656402 | 66665658 | 9257 | -7.33 | 1.44E-10 | 3.70E-10 |
| chr13 | 110139557 | 110142859 | 3303 | -7.17 | 1.44E-10 | 3.70E-10 |
| chr1 | 58976233 | 58978380 | 2148 | -7.21 | 1.44E-10 | 3.71E-10 |
| chr12 | 108325778 | 108333762 | 7985 | 2.92 | 1.44E-10 | 3.71E-10 |
| chr4 | 167875255 | 167892230 | 16976 | -7.33 | 1.44E-10 | 3.71E-10 |
| chr11 | 121395780 | 121401071 | 5292 | -7.2 | 1.44E-10 | 3.71E-10 |
| chr1 | 98109863 | 98112813 | 2951 | -7.23 | 1.45E-10 | 3.72E-10 |
| chr5 | 36977618 | 36979533 | 1916 | -7.37 | 1.45E-10 | 3.72E-10 |
| chr5 | 138975147 | 138978138 | 2992 | -7.16 | 1.45E-10 | 3.72E-10 |
| chr5 | 115687545 | 115691583 | 4039 | -7.34 | 1.45E-10 | 3.73E-10 |
| chrX | 46001268 | 46012269 | 11002 | -7.39 | 1.45E-10 | 3.73E-10 |
| chr11 | 44919597 | 44921320 | 1724 | -7.2 | 1.47E-10 | 3.76E-10 |
| chr6 | 160818824 | 160820955 | 2132 | -7.18 | 1.47E-10 | 3.76E-10 |

| | | | | | | |
|---|---|---|---|---|---|---|
| chr9 | 121369474 | 121375186 | 5713 | -7.16 | 1.47E-10 | 3.77E-10 |
| chr15 | 93589021 | 93590233 | 1213 | -7.19 | 1.47E-10 | 3.77E-10 |
| chr1 | 155677973 | 155679936 | 1964 | -7.16 | 1.47E-10 | 3.77E-10 |
| chr5 | 134949898 | 134976580 | 26683 | -1.36 | 1.47E-10 | 3.79E-10 |
| chr1 | 45885173 | 45888839 | 3667 | -7.28 | 1.48E-10 | 3.79E-10 |
| chr6 | 163828580 | 163830587 | 2008 | -7.16 | 1.48E-10 | 3.79E-10 |
| chr8 | 8415333 | 8429305 | 13973 | -7.44 | 1.48E-10 | 3.79E-10 |
| chr14 | 35559357 | 35560594 | 1238 | -7.27 | 1.48E-10 | 3.79E-10 |
| chr15 | 41578438 | 41579720 | 1283 | -5.33 | 1.48E-10 | 3.79E-10 |
| chr7 | 47094491 | 47117053 | 22563 | -7.68 | 1.48E-10 | 3.80E-10 |
| chr3 | 185547548 | 185549909 | 2362 | -7.19 | 1.48E-10 | 3.80E-10 |
| chr5 | 123375307 | 123380815 | 5509 | -7.65 | 1.48E-10 | 3.80E-10 |
| chr15 | 78815214 | 78817458 | 2245 | -7.2 | 1.48E-10 | 3.81E-10 |
| chr6 | 64615849 | 64620643 | 4795 | 7.22 | 1.48E-10 | 3.81E-10 |
| chr3 | 4954601 | 4958043 | 3443 | -7.44 | 1.48E-10 | 3.81E-10 |
| chr7 | 140796493 | 140807261 | 10769 | 7.31 | 1.49E-10 | 3.83E-10 |
| chr15 | 62235429 | 62236900 | 1472 | -7.17 | 1.49E-10 | 3.83E-10 |
| chr11 | 103263087 | 103272537 | 9451 | -7.33 | 1.49E-10 | 3.83E-10 |
| chr17 | 37727063 | 37728243 | 1181 | -7.33 | 1.49E-10 | 3.83E-10 |
| chr4 | 64750276 | 64762669 | 12394 | -7.37 | 1.50E-10 | 3.83E-10 |
| chr4 | 70032395 | 70035492 | 3098 | -7.31 | 1.50E-10 | 3.83E-10 |
| chr5 | 39709762 | 39720964 | 11203 | -7.6 | 1.50E-10 | 3.84E-10 |
| chr10 | 94527438 | 94529130 | 1693 | -7.41 | 1.50E-10 | 3.84E-10 |
| chr2 | 192813105 | 192816094 | 2990 | -7.17 | 1.50E-10 | 3.84E-10 |
| chr4 | 7406309 | 7410772 | 4464 | -7.2 | 1.50E-10 | 3.84E-10 |
| chr1 | 245234741 | 245241134 | 6394 | -7.37 | 1.50E-10 | 3.85E-10 |
| chr11 | 94716446 | 94721430 | 4985 | -7.19 | 1.51E-10 | 3.86E-10 |
| chr5 | 123989385 | 124002842 | 13458 | -7.49 | 1.51E-10 | 3.88E-10 |
| chr10 | 4274721 | 4280225 | 5505 | -7.18 | 1.52E-10 | 3.89E-10 |
| chr2 | 74753370 | 74754705 | 1336 | -7.18 | 1.52E-10 | 3.89E-10 |
| chr13 | 26766755 | 26770419 | 3665 | -7.15 | 1.52E-10 | 3.89E-10 |
| chr10 | 21682433 | 21685645 | 3213 | -7.38 | 1.52E-10 | 3.89E-10 |
| chr16 | 75070742 | 75083560 | 12819 | -7.42 | 1.52E-10 | 3.90E-10 |
| chr7 | 12851796 | 12887482 | 35687 | -8.51 | 1.52E-10 | 3.90E-10 |
| chr6 | 17409895 | 17411626 | 1732 | -7.22 | 1.53E-10 | 3.91E-10 |
| chr1 | 46702109 | 46703617 | 1509 | -7.17 | 1.53E-10 | 3.91E-10 |
| chr8 | 116318676 | 116323414 | 4739 | -7.15 | 1.53E-10 | 3.92E-10 |
| chr10 | 6762225 | 6770317 | 8093 | -7.17 | 1.53E-10 | 3.92E-10 |
| chr5 | 100075388 | 100077406 | 2019 | -4.03 | 1.53E-10 | 3.93E-10 |
| chr2 | 152490222 | 152492529 | 2308 | -7.19 | 1.53E-10 | 3.93E-10 |
| chr14 | 35288552 | 35289896 | 1345 | -7.16 | 1.53E-10 | 3.93E-10 |
| chr12 | 67658176 | 67660737 | 2562 | -7.21 | 1.53E-10 | 3.93E-10 |
| chr14 | 77521579 | 77523815 | 2237 | -7.15 | 1.54E-10 | 3.94E-10 |
| chr12 | 55137764 | 55153197 | 15434 | -7.26 | 1.55E-10 | 3.96E-10 |
| chr2 | 149171727 | 149175483 | 3757 | -7.16 | 1.55E-10 | 3.97E-10 |
| chr3 | 17494294 | 17501275 | 6982 | -7.44 | 1.55E-10 | 3.97E-10 |
| chr17 | 18527390 | 18531948 | 4559 | -7.53 | 1.55E-10 | 3.98E-10 |
| chr4 | 26075818 | 26135559 | 59742 | -2.57 | 1.56E-10 | 4.00E-10 |
| chr15 | 74976692 | 74978006 | 1315 | -7.23 | 1.57E-10 | 4.01E-10 |
| chr18 | 33681151 | 33683577 | 2427 | -7.31 | 1.57E-10 | 4.01E-10 |
| chr3 | 116971410 | 116974594 | 3185 | -7.47 | 1.57E-10 | 4.01E-10 |
| chr14 | 37353291 | 37355628 | 2338 | -7.16 | 1.57E-10 | 4.02E-10 |
| chr3 | 57797269 | 57799128 | 1860 | -7.18 | 1.57E-10 | 4.02E-10 |
| chr5 | 148646871 | 148649095 | 2225 | -7.18 | 1.57E-10 | 4.02E-10 |
| chr7 | 105523023 | 105524719 | 1697 | -7.22 | 1.57E-10 | 4.02E-10 |
| chr4 | 187320575 | 187325482 | 4908 | -7.22 | 1.58E-10 | 4.03E-10 |
| chr9 | 88323364 | 88324954 | 1591 | -7.15 | 1.58E-10 | 4.04E-10 |
| chr7 | 139949339 | 139958064 | 8726 | -7.69 | 1.58E-10 | 4.04E-10 |
| chr12 | 97941108 | 97943121 | 2014 | -7.33 | 1.58E-10 | 4.05E-10 |
| chr6 | 81998449 | 82043374 | 44926 | 1.85 | 1.58E-10 | 4.05E-10 |
| chr7 | 73511202 | 73512745 | 1544 | -7.14 | 1.58E-10 | 4.05E-10 |

| | | | | | | |
|---|---|---|---|---|---|---|
| chr4 | 160035384 | 160037061 | 1678 | -7.15 | 1.58E-10 | 4.05E-10 |
| chrX | 19695574 | 19697638 | 2065 | -7.22 | 1.59E-10 | 4.06E-10 |
| chr15 | 96711700 | 96713459 | 1760 | -5.63 | 1.59E-10 | 4.06E-10 |
| chr14 | 50402739 | 50410703 | 7965 | -7.52 | 1.59E-10 | 4.06E-10 |
| chr4 | 128614938 | 128617067 | 2130 | -7.21 | 1.59E-10 | 4.07E-10 |
| chr3 | 27631919 | 27664130 | 32212 | -8.27 | 1.59E-10 | 4.07E-10 |
| chr21 | 30501253 | 30502818 | 1566 | -7.18 | 1.59E-10 | 4.07E-10 |
| chr3 | 57590456 | 57592834 | 2379 | -7.15 | 1.60E-10 | 4.09E-10 |
| chr12 | 51291790 | 51293537 | 1748 | -7.15 | 1.60E-10 | 4.09E-10 |
| chr1 | 185329920 | 185332661 | 2742 | -7.16 | 1.61E-10 | 4.11E-10 |
| chr5 | 82795419 | 82797546 | 2128 | -7.34 | 1.61E-10 | 4.12E-10 |
| chr8 | 9318645 | 9320563 | 1919 | -7.15 | 1.61E-10 | 4.12E-10 |
| chr4 | 170442577 | 170445656 | 3080 | -7.25 | 1.62E-10 | 4.14E-10 |
| chr1 | 154477875 | 154479600 | 1726 | -7.24 | 1.62E-10 | 4.14E-10 |
| chr13 | 96658543 | 96661164 | 2622 | -7.45 | 1.62E-10 | 4.14E-10 |
| chr17 | 18070029 | 18071831 | 1803 | -7.24 | 1.62E-10 | 4.15E-10 |
| chr2 | 191868026 | 191880811 | 12786 | -7.65 | 1.62E-10 | 4.15E-10 |
| chr3 | 155670207 | 155674090 | 3884 | -7.16 | 1.63E-10 | 4.16E-10 |
| chr6 | 122228118 | 122230909 | 2792 | -7.23 | 1.63E-10 | 4.17E-10 |
| chr9 | 115299421 | 115301703 | 2283 | -7.23 | 1.63E-10 | 4.17E-10 |
| chr11 | 108610746 | 108612434 | 1689 | -7.19 | 1.64E-10 | 4.17E-10 |
| chr18 | 19633348 | 19636131 | 2784 | -7.19 | 1.64E-10 | 4.17E-10 |
| chr6 | 83724574 | 83727938 | 3365 | -7.25 | 1.64E-10 | 4.17E-10 |
| chr12 | 68782718 | 68784578 | 1861 | -7.44 | 1.64E-10 | 4.19E-10 |
| chr18 | 19699257 | 19702047 | 2791 | -7.44 | 1.64E-10 | 4.19E-10 |
| chr8 | 102165555 | 102171194 | 5640 | -7.14 | 1.64E-10 | 4.19E-10 |
| chr13 | 90749370 | 90753509 | 4140 | -7.29 | 1.65E-10 | 4.21E-10 |
| chr11 | 87886741 | 87894685 | 7945 | -6.05 | 1.65E-10 | 4.21E-10 |
| chr2 | 206306841 | 206308796 | 1956 | -7.15 | 1.65E-10 | 4.21E-10 |
| chr8 | 62675416 | 62677138 | 1723 | -7.42 | 1.65E-10 | 4.21E-10 |
| chr6 | 55752573 | 55765573 | 13001 | -7.19 | 1.65E-10 | 4.22E-10 |
| chr1 | 86382095 | 86387201 | 5107 | -7.14 | 1.66E-10 | 4.23E-10 |
| chr7 | 90093201 | 90096244 | 3044 | -5.21 | 1.66E-10 | 4.23E-10 |
| chr1 | 53347429 | 53348697 | 1269 | -7.15 | 1.66E-10 | 4.24E-10 |
| chr4 | 84127153 | 84137813 | 10661 | -7.31 | 1.66E-10 | 4.24E-10 |
| chr15 | 56669666 | 56692597 | 22932 | -7.71 | 1.66E-10 | 4.24E-10 |
| chr2 | 33513961 | 33518714 | 4754 | -7.17 | 1.67E-10 | 4.25E-10 |
| chr15 | 50228909 | 50259348 | 30440 | -2.39 | 1.67E-10 | 4.26E-10 |
| chr12 | 42878339 | 42879559 | 1221 | -7.14 | 1.67E-10 | 4.26E-10 |
| chr8 | 87345139 | 87349215 | 4077 | -7.2 | 1.67E-10 | 4.26E-10 |
| chr2 | 119849235 | 119853393 | 4159 | -7.17 | 1.67E-10 | 4.27E-10 |
| chr12 | 69466502 | 69475122 | 8621 | -7.14 | 1.68E-10 | 4.27E-10 |
| chr13 | 25176808 | 25179825 | 3018 | -6.17 | 1.68E-10 | 4.27E-10 |
| chr5 | 131349587 | 131351496 | 1910 | -7.21 | 1.68E-10 | 4.28E-10 |
| chr11 | 32783340 | 32790920 | 7581 | -7.14 | 1.68E-10 | 4.28E-10 |
| chr13 | 80072710 | 80074226 | 1517 | -7.14 | 1.68E-10 | 4.28E-10 |
| chr13 | 22156394 | 22158198 | 1805 | -7.15 | 1.68E-10 | 4.28E-10 |
| chr16 | 73243015 | 73251175 | 8161 | -7.2 | 1.68E-10 | 4.28E-10 |
| chr16 | 58250196 | 58255010 | 4815 | -3.38 | 1.68E-10 | 4.29E-10 |
| chr11 | 70919427 | 70921257 | 1831 | -7.18 | 1.68E-10 | 4.29E-10 |
| chr6 | 67863090 | 67878033 | 14944 | -2.55 | 1.68E-10 | 4.29E-10 |
| chr16 | 56545861 | 56546878 | 1018 | -7.16 | 1.68E-10 | 4.29E-10 |
| chr17 | 3600579 | 3602794 | 2216 | -7.16 | 1.68E-10 | 4.29E-10 |
| chr20 | 35950423 | 35951549 | 1127 | -7.14 | 1.69E-10 | 4.29E-10 |
| chr10 | 111807448 | 111809692 | 2245 | -7.14 | 1.69E-10 | 4.30E-10 |
| chr6 | 48605302 | 48611967 | 6666 | -7.25 | 1.69E-10 | 4.31E-10 |
| chr9 | 93089046 | 93092925 | 3880 | -7.39 | 1.69E-10 | 4.31E-10 |
| chr20 | 43505957 | 43507302 | 1346 | -7.14 | 1.70E-10 | 4.33E-10 |
| chr6 | 77407991 | 77412975 | 4985 | -7.15 | 1.70E-10 | 4.33E-10 |
| chr1 | 86343138 | 86346399 | 3262 | -7.13 | 1.70E-10 | 4.33E-10 |
| chr15 | 62892573 | 62894177 | 1605 | -7.15 | 1.70E-10 | 4.33E-10 |

| | | | | | | |
|---|---|---|---|---|---|---|
| chr7 | 151501615 | 151503320 | 1706 | -5.27 | 1.71E-10 | 4.35E-10 |
| chr1 | 80417787 | 80427630 | 9844 | -7.17 | 1.71E-10 | 4.36E-10 |
| chr2 | 134867455 | 134869687 | 2233 | -7.17 | 1.71E-10 | 4.36E-10 |
| chr3 | 16855871 | 16862545 | 6675 | -7.19 | 1.71E-10 | 4.36E-10 |
| chr3 | 33023117 | 33025663 | 2547 | -7.35 | 1.72E-10 | 4.37E-10 |
| chrX | 78679109 | 78691852 | 12744 | 1.71 | 1.72E-10 | 4.38E-10 |
| chr4 | 16063074 | 16065434 | 2361 | -7.17 | 1.72E-10 | 4.38E-10 |
| chr11 | 3633125 | 3634246 | 1122 | -7.15 | 1.72E-10 | 4.38E-10 |
| chr13 | 46872866 | 46886018 | 13153 | -3.53 | 1.72E-10 | 4.38E-10 |
| chr1 | 62528525 | 62533101 | 4577 | -7.13 | 1.72E-10 | 4.39E-10 |
| chr4 | 25777731 | 25779610 | 1880 | -7.17 | 1.73E-10 | 4.40E-10 |
| chr13 | 43618172 | 43633966 | 15795 | -7.85 | 1.73E-10 | 4.40E-10 |
| chr6 | 26480202 | 26488464 | 8263 | -7.14 | 1.73E-10 | 4.41E-10 |
| chr8 | 27394139 | 27396347 | 2209 | -7.14 | 1.73E-10 | 4.41E-10 |
| chr5 | 82610836 | 82612298 | 1463 | -6.14 | 1.74E-10 | 4.41E-10 |
| chr15 | 99413080 | 99415163 | 2084 | -7.13 | 1.74E-10 | 4.42E-10 |
| chr9 | 75397237 | 75419177 | 21941 | -8.28 | 1.74E-10 | 4.43E-10 |
| chr3 | 125999709 | 126002377 | 2669 | -7.13 | 1.74E-10 | 4.43E-10 |
| chr9 | 93663134 | 93681514 | 18381 | -7.83 | 1.74E-10 | 4.43E-10 |
| chr4 | 69133355 | 69137591 | 4237 | -7.32 | 1.74E-10 | 4.44E-10 |
| chr11 | 69144999 | 69147180 | 2182 | -7.22 | 1.75E-10 | 4.44E-10 |
| chrX | 103602083 | 103612288 | 10206 | -7.22 | 1.75E-10 | 4.44E-10 |
| chr5 | 138367802 | 138370261 | 2460 | -7.13 | 1.75E-10 | 4.44E-10 |
| chr22 | 48412768 | 48418941 | 6174 | -3.29 | 1.75E-10 | 4.44E-10 |
| chr10 | 13025357 | 13027489 | 2133 | -7.41 | 1.75E-10 | 4.45E-10 |
| chrX | 120218532 | 120220908 | 2377 | -5.84 | 1.75E-10 | 4.45E-10 |
| chr16 | 15056964 | 15058282 | 1319 | -7.13 | 1.75E-10 | 4.46E-10 |
| chr1 | 240199062 | 240203126 | 4065 | -7.18 | 1.75E-10 | 4.46E-10 |
| chr9 | 126664544 | 126665773 | 1230 | -7.14 | 1.76E-10 | 4.47E-10 |
| chr11 | 117491657 | 117493543 | 1887 | -7.16 | 1.76E-10 | 4.48E-10 |
| chr17 | 29178940 | 29180930 | 1991 | -7.16 | 1.76E-10 | 4.48E-10 |
| chr6 | 28083015 | 28084856 | 1842 | -7.16 | 1.76E-10 | 4.48E-10 |
| chr18 | 57185573 | 57193714 | 8142 | -7.38 | 1.76E-10 | 4.48E-10 |
| chr5 | 166918240 | 166986256 | 68017 | 1.78 | 1.76E-10 | 4.48E-10 |
| chr2 | 19457618 | 19459236 | 1619 | -7.42 | 1.77E-10 | 4.49E-10 |
| chrX | 37482337 | 37485863 | 3527 | 7.22 | 1.77E-10 | 4.49E-10 |
| chr10 | 124197857 | 124199992 | 2136 | -7.25 | 1.77E-10 | 4.49E-10 |
| chr6 | 9231753 | 9233910 | 2158 | -7.14 | 1.77E-10 | 4.50E-10 |
| chr21 | 35079980 | 35082694 | 2715 | -7.17 | 1.78E-10 | 4.51E-10 |
| chr5 | 61660361 | 61661835 | 1475 | -7.16 | 1.78E-10 | 4.51E-10 |
| chr1 | 196378942 | 196381280 | 2339 | -7.27 | 1.78E-10 | 4.52E-10 |
| chr1 | 195806699 | 195949729 | 143031 | 2.2 | 1.78E-10 | 4.53E-10 |
| chr1 | 185375642 | 185377751 | 2110 | -7.22 | 1.79E-10 | 4.53E-10 |
| chr10 | 94786846 | 94788221 | 1376 | -7.13 | 1.79E-10 | 4.53E-10 |
| chr2 | 241571425 | 241575140 | 3716 | -5.69 | 1.79E-10 | 4.54E-10 |
| chr15 | 29714772 | 29720937 | 6166 | -5.28 | 1.79E-10 | 4.54E-10 |
| chr7 | 123372050 | 123374468 | 2419 | -7.15 | 1.79E-10 | 4.54E-10 |
| chr10 | 34131833 | 34136129 | 4297 | -3.02 | 1.79E-10 | 4.54E-10 |
| chr15 | 91659437 | 91665805 | 6369 | -1.85 | 1.79E-10 | 4.54E-10 |
| chr15 | 56999767 | 57002528 | 2762 | -7.13 | 1.79E-10 | 4.54E-10 |
| chr18 | 48425066 | 48433888 | 8823 | -7.14 | 1.79E-10 | 4.55E-10 |
| chr5 | 150139101 | 150145605 | 6505 | -7.39 | 1.79E-10 | 4.55E-10 |
| chr8 | 124678504 | 124680703 | 2200 | -7.32 | 1.80E-10 | 4.56E-10 |
| chr5 | 1928875 | 1953766 | 24892 | -7.58 | 1.80E-10 | 4.56E-10 |
| chr21 | 44417873 | 44420869 | 2997 | -7.13 | 1.80E-10 | 4.56E-10 |
| chr19 | 46222908 | 46227082 | 4175 | -7.3 | 1.80E-10 | 4.56E-10 |
| chr13 | 109508477 | 109516182 | 7706 | -7.73 | 1.80E-10 | 4.57E-10 |
| chr6 | 18105609 | 18108514 | 2906 | -7.2 | 1.80E-10 | 4.57E-10 |
| chr6 | 79900193 | 79902539 | 2347 | -7.13 | 1.80E-10 | 4.57E-10 |
| chr5 | 23528090 | 23545407 | 17318 | -7.39 | 1.80E-10 | 4.57E-10 |
| chr1 | 186550756 | 186554037 | 3282 | -7.35 | 1.80E-10 | 4.58E-10 |

| | | | | | | |
|---|---|---|---|---|---|---|
| chr4 | 14524250 | 14529701 | 5452 | -7.17 | 1.81E-10 | 4.60E-10 |
| chr1 | 6439828 | 6444893 | 5066 | -7.18 | 1.82E-10 | 4.61E-10 |
| chr4 | 63907997 | 63915720 | 7724 | -7.36 | 1.82E-10 | 4.61E-10 |
| chr4 | 132206369 | 132210236 | 3868 | -7.12 | 1.82E-10 | 4.62E-10 |
| chr9 | 26964383 | 26966957 | 2575 | -7.13 | 1.82E-10 | 4.62E-10 |
| chr19 | 35971755 | 35973226 | 1472 | -7.36 | 1.83E-10 | 4.63E-10 |
| chr9 | 108403652 | 108405471 | 1820 | -7.26 | 1.84E-10 | 4.66E-10 |
| chr18 | 48001410 | 48003346 | 1937 | -2.8 | 1.84E-10 | 4.66E-10 |
| chr4 | 176441871 | 176445048 | 3178 | -7.26 | 1.84E-10 | 4.66E-10 |
| chr12 | 60582247 | 60585337 | 3091 | -7.14 | 1.84E-10 | 4.67E-10 |
| chr12 | 100762287 | 100763846 | 1560 | -7.14 | 1.84E-10 | 4.67E-10 |
| chr13 | 90553354 | 90558195 | 4842 | -7.37 | 1.84E-10 | 4.67E-10 |
| chr7 | 101769673 | 101771038 | 1366 | -7.32 | 1.84E-10 | 4.67E-10 |
| chr20 | 58554757 | 58557126 | 2370 | -7.17 | 1.85E-10 | 4.68E-10 |
| chr6 | 156960591 | 156964252 | 3662 | -7.13 | 1.85E-10 | 4.68E-10 |
| chr5 | 155078069 | 155081403 | 3335 | -7.14 | 1.85E-10 | 4.69E-10 |
| chr10 | 29021648 | 29023787 | 2140 | -7.24 | 1.85E-10 | 4.69E-10 |
| chrX | 53395831 | 53399005 | 3175 | -7.24 | 1.85E-10 | 4.69E-10 |
| chr1 | 110406600 | 110409506 | 2907 | -7.24 | 1.85E-10 | 4.69E-10 |
| chr1 | 153170179 | 153188087 | 17909 | 1.91 | 1.85E-10 | 4.69E-10 |
| chr8 | 145763939 | 145766156 | 2218 | -7.23 | 1.85E-10 | 4.69E-10 |
| chr3 | 124892428 | 124894009 | 1582 | -7.22 | 1.86E-10 | 4.70E-10 |
| chr18 | 73132637 | 73135291 | 2655 | -7.26 | 1.86E-10 | 4.70E-10 |
| chr13 | 78088451 | 78112564 | 24114 | -7.33 | 1.86E-10 | 4.70E-10 |
| chr3 | 177026796 | 177035507 | 8712 | -7.54 | 1.86E-10 | 4.70E-10 |
| chr1 | 186214829 | 186218280 | 3452 | -7.2 | 1.86E-10 | 4.71E-10 |
| chr9 | 91907390 | 91909582 | 2193 | -7.16 | 1.86E-10 | 4.71E-10 |
| chrX | 54788781 | 54811589 | 22809 | -4.83 | 1.86E-10 | 4.72E-10 |
| chr3 | 66694570 | 66696791 | 2222 | -7.13 | 1.87E-10 | 4.72E-10 |
| chr3 | 131165610 | 131167764 | 2155 | -7.13 | 1.87E-10 | 4.72E-10 |
| chr11 | 43700580 | 43701849 | 1270 | -7.29 | 1.87E-10 | 4.72E-10 |
| chr13 | 96172997 | 96181036 | 8040 | -7.29 | 1.87E-10 | 4.72E-10 |
| chr3 | 138787300 | 138818002 | 30703 | -3.12 | 1.87E-10 | 4.73E-10 |
| chr13 | 43775716 | 43778411 | 2696 | -7.21 | 1.87E-10 | 4.73E-10 |
| chr4 | 65190746 | 65207233 | 16488 | -7.48 | 1.87E-10 | 4.73E-10 |
| chr6 | 54796516 | 54799647 | 3132 | -7.15 | 1.88E-10 | 4.75E-10 |
| chr2 | 216848403 | 216854189 | 5787 | -7.38 | 1.88E-10 | 4.75E-10 |
| chr8 | 1961107 | 1963947 | 2841 | -7.12 | 1.88E-10 | 4.75E-10 |
| chr5 | 133097691 | 133100281 | 2591 | 4.28 | 1.88E-10 | 4.75E-10 |
| chr16 | 48499612 | 48510061 | 10450 | -7.29 | 1.88E-10 | 4.76E-10 |
| chr11 | 20014289 | 20015625 | 1337 | -7.13 | 1.88E-10 | 4.77E-10 |
| chr4 | 41344078 | 41351748 | 7671 | -7.17 | 1.89E-10 | 4.77E-10 |
| chr14 | 97210868 | 97213106 | 2239 | -7.14 | 1.89E-10 | 4.78E-10 |
| chr10 | 93925535 | 93935797 | 10263 | -7.49 | 1.89E-10 | 4.78E-10 |
| chr16 | 47136170 | 47138775 | 2606 | -7.12 | 1.89E-10 | 4.78E-10 |
| chr3 | 170880326 | 170882704 | 2379 | -7.3 | 1.89E-10 | 4.79E-10 |
| chr17 | 71013562 | 71015245 | 1684 | -7.15 | 1.90E-10 | 4.79E-10 |
| chr2 | 194276156 | 194284659 | 8504 | -7.15 | 1.90E-10 | 4.79E-10 |
| chr19 | 34900982 | 34902055 | 1074 | -7.16 | 1.90E-10 | 4.81E-10 |
| chr5 | 17773535 | 17775160 | 1626 | -7.46 | 1.90E-10 | 4.81E-10 |
| chr12 | 132154650 | 132160094 | 5445 | -7.12 | 1.90E-10 | 4.81E-10 |
| chr16 | 72765154 | 72769251 | 4098 | -7.19 | 1.91E-10 | 4.82E-10 |
| chr13 | 110578472 | 110580378 | 1907 | -7.13 | 1.91E-10 | 4.83E-10 |
| chr5 | 167050924 | 167053325 | 2402 | -7.12 | 1.91E-10 | 4.84E-10 |
| chr3 | 24696353 | 24698207 | 1855 | -7.16 | 1.91E-10 | 4.84E-10 |
| chr2 | 176559330 | 176755131 | 195802 | 1.9 | 1.92E-10 | 4.85E-10 |
| chr3 | 54348165 | 54449252 | 101088 | 1.27 | 1.92E-10 | 4.85E-10 |
| chr8 | 145142437 | 145144407 | 1971 | -7.14 | 1.93E-10 | 4.87E-10 |
| chr17 | 74753784 | 74755296 | 1513 | -7.43 | 1.93E-10 | 4.87E-10 |
| chr19 | 43363269 | 43368077 | 4809 | -7.12 | 1.93E-10 | 4.87E-10 |
| chr13 | 113248690 | 113264975 | 16286 | -7.59 | 1.93E-10 | 4.88E-10 |

| | | | | | | |
|---|---|---|---|---|---|---|
| chr1 | 115446427 | 115448692 | 2266 | -2.94 | 1.94E-10 | 4.89E-10 |
| chr4 | 64535066 | 64548934 | 13869 | -7.24 | 1.94E-10 | 4.89E-10 |
| chr5 | 76563767 | 76566616 | 2850 | -7.14 | 1.94E-10 | 4.90E-10 |
| chrX | 130826071 | 130828510 | 2440 | -7.14 | 1.94E-10 | 4.90E-10 |
| chr3 | 155192876 | 155196073 | 3198 | -7.24 | 1.94E-10 | 4.90E-10 |
| chr6 | 37621393 | 37628987 | 7595 | -7.25 | 1.94E-10 | 4.90E-10 |
| chr1 | 231063815 | 231064919 | 1105 | -7.39 | 1.94E-10 | 4.90E-10 |
| chr12 | 7877875 | 7881431 | 3557 | -7.54 | 1.94E-10 | 4.90E-10 |
| chr9 | 103134842 | 103147761 | 12920 | -7.51 | 1.94E-10 | 4.91E-10 |
| chr3 | 134505081 | 134507046 | 1966 | -3.04 | 1.94E-10 | 4.91E-10 |
| chr22 | 47998394 | 48006782 | 8389 | -7.26 | 1.94E-10 | 4.91E-10 |
| chr15 | 71533238 | 71535942 | 2705 | -7.21 | 1.95E-10 | 4.91E-10 |
| chr16 | 73118476 | 73120381 | 1906 | -7.37 | 1.95E-10 | 4.91E-10 |
| chr21 | 36650784 | 36660988 | 10205 | -7.26 | 1.95E-10 | 4.92E-10 |
| chr5 | 100504722 | 100513624 | 8903 | -7.57 | 1.95E-10 | 4.93E-10 |
| chr10 | 125855467 | 125860960 | 5494 | -7.41 | 1.95E-10 | 4.93E-10 |
| chr7 | 106930280 | 106931962 | 1683 | -7.17 | 1.96E-10 | 4.96E-10 |
| chr11 | 33698670 | 33709550 | 10881 | -7.38 | 1.97E-10 | 4.96E-10 |
| chr5 | 17736465 | 17742695 | 6231 | -7.72 | 1.98E-10 | 4.98E-10 |
| chr2 | 182567288 | 182574520 | 7233 | -7.29 | 1.98E-10 | 4.98E-10 |
| chrX | 128529081 | 128533050 | 3970 | -7.29 | 1.98E-10 | 5.00E-10 |
| chr13 | 82786997 | 82790598 | 3602 | -7.29 | 1.99E-10 | 5.01E-10 |
| chr3 | 112061504 | 112065138 | 3635 | -2.23 | 1.99E-10 | 5.02E-10 |
| chr3 | 45131909 | 45133026 | 1118 | -6.23 | 1.99E-10 | 5.03E-10 |
| chr8 | 68091552 | 68092950 | 1399 | -7.45 | 1.99E-10 | 5.03E-10 |
| chr5 | 950070 | 952420 | 2351 | -7.15 | 2.01E-10 | 5.06E-10 |
| chr4 | 76043343 | 76047358 | 4016 | -7.15 | 2.01E-10 | 5.06E-10 |
| chr4 | 99344223 | 99346787 | 2565 | -7.17 | 2.01E-10 | 5.06E-10 |
| chr19 | 30341031 | 30342679 | 1649 | -7.11 | 2.02E-10 | 5.08E-10 |
| chr1 | 68956093 | 68978451 | 22359 | -7.17 | 2.02E-10 | 5.09E-10 |
| chr9 | 100349180 | 100351839 | 2660 | -7.15 | 2.02E-10 | 5.10E-10 |
| chr3 | 66435066 | 66437201 | 2136 | -7.15 | 2.02E-10 | 5.10E-10 |
| chr2 | 10631509 | 10632800 | 1292 | -7.22 | 2.03E-10 | 5.11E-10 |
| chr11 | 130576486 | 130579655 | 3170 | -7.21 | 2.03E-10 | 5.12E-10 |
| chr12 | 54944264 | 54955234 | 10971 | -2.35 | 2.04E-10 | 5.13E-10 |
| chrX | 7916425 | 7942417 | 25993 | 3.33 | 2.04E-10 | 5.13E-10 |
| chr4 | 177727759 | 177730542 | 2784 | -7.14 | 2.04E-10 | 5.13E-10 |
| chr9 | 100418262 | 100420778 | 2517 | -3.94 | 2.04E-10 | 5.13E-10 |
| chr2 | 226693748 | 226696281 | 2534 | -7.16 | 2.04E-10 | 5.15E-10 |
| chr5 | 153771153 | 153772772 | 1620 | -7.16 | 2.04E-10 | 5.15E-10 |
| chr3 | 45191564 | 45194042 | 2479 | -7.4 | 2.05E-10 | 5.15E-10 |
| chr2 | 146066488 | 146071231 | 4744 | -7.5 | 2.05E-10 | 5.15E-10 |
| chr20 | 40165229 | 40178987 | 13759 | -7.37 | 2.05E-10 | 5.16E-10 |
| chr2 | 69482100 | 69491998 | 9899 | -3.21 | 2.05E-10 | 5.16E-10 |
| chrX | 18397519 | 18399308 | 1790 | -7.22 | 2.05E-10 | 5.17E-10 |
| chr10 | 115034006 | 115036635 | 2630 | -7.22 | 2.05E-10 | 5.17E-10 |
| chr7 | 101366332 | 101368066 | 1735 | -7.16 | 2.06E-10 | 5.18E-10 |
| chr16 | 77023400 | 77025824 | 2425 | -7.19 | 2.06E-10 | 5.19E-10 |
| chr7 | 158227753 | 158229811 | 2059 | -7.25 | 2.06E-10 | 5.19E-10 |
| chr13 | 23415858 | 23422979 | 7122 | -7.11 | 2.07E-10 | 5.20E-10 |
| chr2 | 219073573 | 219075397 | 1825 | -7.11 | 2.07E-10 | 5.20E-10 |
| chr5 | 129428430 | 129434756 | 6327 | -7.11 | 2.07E-10 | 5.20E-10 |
| chr1 | 28241418 | 28242487 | 1070 | -7.15 | 2.07E-10 | 5.21E-10 |
| chr10 | 79245784 | 79247742 | 1959 | -7.11 | 2.07E-10 | 5.21E-10 |
| chr12 | 81015637 | 81019524 | 3888 | 7.17 | 2.07E-10 | 5.21E-10 |
| chr19 | 48295322 | 48305092 | 9771 | -7.34 | 2.08E-10 | 5.23E-10 |
| chr12 | 10165511 | 10179046 | 13536 | -7.34 | 2.08E-10 | 5.23E-10 |
| chr12 | 102659675 | 102662896 | 3222 | 7.17 | 2.08E-10 | 5.23E-10 |
| chr14 | 77733605 | 77735207 | 1603 | -7.18 | 2.08E-10 | 5.24E-10 |
| chr22 | 27138196 | 27140230 | 2035 | -7.19 | 2.09E-10 | 5.25E-10 |
| chr17 | 27254829 | 27256240 | 1412 | -7.11 | 2.09E-10 | 5.26E-10 |

| | | | | | | |
|---|---|---|---|---|---|---|
| chr9 | 87555676 | 87575552 | 19877 | -7.52 | 2.09E-10 | 5.26E-10 |
| chr5 | 141424062 | 141425237 | 1176 | -7.17 | 2.09E-10 | 5.27E-10 |
| chr5 | 76317655 | 76325917 | 8263 | -3.46 | 2.10E-10 | 5.27E-10 |
| chr2 | 239267534 | 239268718 | 1185 | -7.15 | 2.10E-10 | 5.28E-10 |
| chr4 | 93821321 | 93824764 | 3444 | -7.35 | 2.10E-10 | 5.29E-10 |
| chr6 | 101169305 | 101170769 | 1465 | -7.1 | 2.10E-10 | 5.29E-10 |
| chr6 | 166175655 | 166177946 | 2292 | -7.15 | 2.11E-10 | 5.29E-10 |
| chr15 | 53618750 | 53621651 | 2902 | 7.17 | 2.11E-10 | 5.30E-10 |
| chr6 | 111108758 | 111131848 | 23091 | -7.79 | 2.11E-10 | 5.30E-10 |
| chr6 | 48830430 | 48832998 | 2569 | -7.1 | 2.11E-10 | 5.31E-10 |
| chr12 | 87303795 | 87308306 | 4512 | -7.35 | 2.11E-10 | 5.31E-10 |
| chr7 | 131741619 | 131745305 | 3687 | -7.1 | 2.12E-10 | 5.34E-10 |
| chr10 | 94440575 | 94447701 | 7127 | -7.22 | 2.12E-10 | 5.34E-10 |
| chr4 | 84298132 | 84300487 | 2356 | -7.23 | 2.12E-10 | 5.34E-10 |
| chr9 | 134933463 | 134935704 | 2242 | -7.1 | 2.13E-10 | 5.35E-10 |
| chr13 | 89835246 | 89838461 | 3216 | -5.38 | 2.13E-10 | 5.35E-10 |
| chr8 | 98979793 | 98983512 | 3720 | -7.2 | 2.13E-10 | 5.35E-10 |
| chr16 | 53833680 | 53836337 | 2658 | -7.2 | 2.14E-10 | 5.37E-10 |
| chr6 | 47149017 | 47151851 | 2835 | -3.38 | 2.14E-10 | 5.38E-10 |
| chr1 | 191417966 | 191441822 | 23857 | -7.68 | 2.15E-10 | 5.39E-10 |
| chr12 | 107516662 | 107518195 | 1534 | -7.1 | 2.15E-10 | 5.39E-10 |
| chr2 | 150685946 | 150696536 | 10591 | -7.26 | 2.15E-10 | 5.41E-10 |
| chr4 | 54110469 | 54114162 | 3694 | -7.27 | 2.16E-10 | 5.42E-10 |
| chr18 | 10932146 | 11001850 | 69705 | -1.66 | 2.17E-10 | 5.44E-10 |
| chrX | 39542141 | 39556909 | 14769 | -7.47 | 2.17E-10 | 5.44E-10 |
| chr7 | 148896219 | 148897631 | 1413 | -7.32 | 2.17E-10 | 5.45E-10 |
| chr12 | 120265965 | 120289952 | 23988 | -7.57 | 2.17E-10 | 5.46E-10 |
| chr6 | 25732223 | 25739870 | 7648 | -3.51 | 2.18E-10 | 5.47E-10 |
| chr4 | 89544588 | 89546244 | 1657 | -7.13 | 2.18E-10 | 5.48E-10 |
| chr1 | 175920604 | 175922100 | 1497 | -7.15 | 2.19E-10 | 5.49E-10 |
| chr19 | 30591180 | 30593934 | 2755 | -2.36 | 2.19E-10 | 5.49E-10 |
| chr4 | 102236615 | 102241644 | 5030 | -7.29 | 2.19E-10 | 5.50E-10 |
| chr9 | 3424887 | 3427116 | 2230 | -7.29 | 2.19E-10 | 5.50E-10 |
| chr5 | 173151569 | 173154424 | 2856 | -7.45 | 2.19E-10 | 5.50E-10 |
| chr6 | 44939499 | 44942951 | 3453 | -7.16 | 2.20E-10 | 5.53E-10 |
| chr4 | 98217545 | 98223569 | 6025 | -7.13 | 2.21E-10 | 5.53E-10 |
| chr9 | 36145272 | 36158555 | 13284 | -7.61 | 2.21E-10 | 5.54E-10 |
| chr13 | 19793568 | 19843510 | 49943 | 1.79 | 2.21E-10 | 5.54E-10 |
| chr15 | 38677487 | 38686037 | 8551 | -7.27 | 2.21E-10 | 5.55E-10 |
| chr13 | 25564912 | 25567285 | 2374 | -7.1 | 2.21E-10 | 5.55E-10 |
| chr20 | 14070714 | 14072415 | 1702 | -7.1 | 2.21E-10 | 5.55E-10 |
| chrX | 135061332 | 135064185 | 2854 | -7.55 | 2.21E-10 | 5.55E-10 |
| chr12 | 89723807 | 89726034 | 2228 | -7.3 | 2.21E-10 | 5.55E-10 |
| chr4 | 140389102 | 140394289 | 5188 | -7.14 | 2.22E-10 | 5.57E-10 |
| chr11 | 16286880 | 16291963 | 5084 | -7.21 | 2.22E-10 | 5.58E-10 |
| chr18 | 486499 | 488245 | 1747 | -7.22 | 2.22E-10 | 5.58E-10 |
| chr4 | 180863099 | 180869252 | 6154 | -7.09 | 2.23E-10 | 5.58E-10 |
| chr1 | 55344530 | 55347504 | 2975 | -7.43 | 2.23E-10 | 5.59E-10 |
| chr2 | 163798380 | 163801526 | 3147 | -6.14 | 2.23E-10 | 5.59E-10 |
| chr6 | 48203245 | 48205277 | 2033 | -7.2 | 2.23E-10 | 5.59E-10 |
| chr9 | 124742827 | 124744464 | 1638 | -7.31 | 2.23E-10 | 5.59E-10 |
| chr6 | 66640299 | 66648563 | 8265 | -7.35 | 2.23E-10 | 5.60E-10 |
| chr2 | 131734687 | 131736188 | 1502 | -7.11 | 2.23E-10 | 5.60E-10 |
| chr3 | 18337546 | 18339626 | 2081 | -7.11 | 2.23E-10 | 5.60E-10 |
| chr7 | 26840932 | 26843599 | 2668 | -7.1 | 2.23E-10 | 5.60E-10 |
| chr4 | 32171731 | 32176531 | 4801 | -7.39 | 2.24E-10 | 5.61E-10 |
| chr1 | 184780935 | 184783429 | 2495 | -7.13 | 2.24E-10 | 5.62E-10 |
| chr7 | 123278596 | 123287176 | 8581 | -7.19 | 2.24E-10 | 5.62E-10 |
| chr16 | 11122243 | 11123495 | 1253 | -7.25 | 2.25E-10 | 5.64E-10 |
| chr8 | 8884804 | 8893965 | 9162 | -7.18 | 2.25E-10 | 5.64E-10 |
| chr3 | 15954652 | 15956423 | 1772 | -7.13 | 2.25E-10 | 5.65E-10 |

| | | | | | | |
|---|---|---|---|---|---|---|
| chr3 | 184139396 | 184141556 | 2161 | -7.32 | 2.26E-10 | 5.66E-10 |
| chr2 | 64830795 | 64832213 | 1419 | -7.13 | 2.26E-10 | 5.67E-10 |
| chr7 | 91883800 | 91894139 | 10340 | -7.1 | 2.27E-10 | 5.67E-10 |
| chr7 | 157893311 | 157923814 | 30504 | -3.36 | 2.27E-10 | 5.69E-10 |
| chr1 | 207122677 | 207124365 | 1689 | -7.09 | 2.28E-10 | 5.70E-10 |
| chr5 | 135372818 | 135387162 | 14345 | -7.36 | 2.28E-10 | 5.70E-10 |
| chr3 | 110948384 | 110962397 | 14014 | -7.5 | 2.28E-10 | 5.70E-10 |
| chr12 | 28946300 | 28963850 | 17551 | -6.15 | 2.28E-10 | 5.71E-10 |
| chr8 | 95415780 | 95417409 | 1630 | -7.1 | 2.28E-10 | 5.71E-10 |
| chr6 | 88994615 | 89001775 | 7161 | -5.68 | 2.28E-10 | 5.71E-10 |
| chr4 | 32081745 | 32083930 | 2186 | -7.1 | 2.30E-10 | 5.75E-10 |
| chr8 | 107145926 | 107154023 | 8098 | -7.37 | 2.30E-10 | 5.76E-10 |
| chr4 | 188649607 | 188679231 | 29625 | -6.6 | 2.31E-10 | 5.77E-10 |
| chr2 | 48156458 | 48158211 | 1754 | -7.15 | 2.31E-10 | 5.78E-10 |
| chr3 | 149600301 | 149602145 | 1845 | -7.22 | 2.31E-10 | 5.78E-10 |
| chr2 | 221504681 | 221509577 | 4897 | -7.11 | 2.31E-10 | 5.78E-10 |
| chr8 | 142555656 | 142567035 | 11380 | -7.12 | 2.31E-10 | 5.78E-10 |
| chr2 | 233331422 | 233338100 | 6679 | -3.44 | 2.32E-10 | 5.80E-10 |
| chr1 | 38106312 | 38109159 | 2848 | -7.15 | 2.32E-10 | 5.81E-10 |
| chrX | 103091320 | 103106768 | 15449 | -5.86 | 2.33E-10 | 5.82E-10 |
| chr3 | 138317017 | 138322915 | 5899 | -7.13 | 2.33E-10 | 5.82E-10 |
| chr14 | 70042483 | 70045130 | 2648 | -7.2 | 2.33E-10 | 5.83E-10 |
| chr1 | 95490012 | 95495221 | 5210 | -7.1 | 2.33E-10 | 5.83E-10 |
| chr17 | 28626594 | 28628166 | 1573 | -7.1 | 2.33E-10 | 5.83E-10 |
| chr1 | 235958414 | 235960150 | 1737 | -7.13 | 2.34E-10 | 5.84E-10 |
| chr5 | 61579996 | 61587437 | 7442 | -7.19 | 2.34E-10 | 5.84E-10 |
| chr16 | 66540324 | 66541909 | 1586 | -7.19 | 2.34E-10 | 5.85E-10 |
| chr20 | 45217883 | 45222123 | 4241 | -7.11 | 2.34E-10 | 5.86E-10 |
| chrX | 16999727 | 17006748 | 7022 | -7.09 | 2.35E-10 | 5.86E-10 |
| chr4 | 106499424 | 106502646 | 3223 | -7.17 | 2.35E-10 | 5.88E-10 |
| chr11 | 45848383 | 45849760 | 1378 | -7.31 | 2.35E-10 | 5.89E-10 |
| chr6 | 141067500 | 141071469 | 3970 | -7.25 | 2.36E-10 | 5.89E-10 |
| chr14 | 43206274 | 43208862 | 2589 | 6.38 | 2.36E-10 | 5.90E-10 |
| chr20 | 9027742 | 9030765 | 3024 | -7.11 | 2.36E-10 | 5.90E-10 |
| chr12 | 10009206 | 10010858 | 1653 | 7.26 | 2.37E-10 | 5.92E-10 |
| chr14 | 67638987 | 67646237 | 7251 | -7.26 | 2.37E-10 | 5.92E-10 |
| chr8 | 90543343 | 90555109 | 11767 | -7.38 | 2.37E-10 | 5.92E-10 |
| chr10 | 119310220 | 119312113 | 1894 | -7.13 | 2.38E-10 | 5.95E-10 |
| chr2 | 223821158 | 223832522 | 11365 | -7.26 | 2.38E-10 | 5.95E-10 |
| chr12 | 107810551 | 107821334 | 10784 | -7.11 | 2.39E-10 | 5.97E-10 |
| chr1 | 18869675 | 18871896 | 2222 | -7.1 | 2.39E-10 | 5.97E-10 |
| chr6 | 136130354 | 136134935 | 4582 | -7.1 | 2.39E-10 | 5.97E-10 |
| chr16 | 22624981 | 22637370 | 12390 | -7.15 | 2.40E-10 | 5.99E-10 |
| chr4 | 11752740 | 11755664 | 2925 | -7.15 | 2.40E-10 | 5.99E-10 |
| chr3 | 192620983 | 192622566 | 1584 | -7.09 | 2.40E-10 | 5.99E-10 |
| chr12 | 68918691 | 68921628 | 2938 | -7.09 | 2.40E-10 | 6.00E-10 |
| chr7 | 32344681 | 32347516 | 2836 | -2.87 | 2.41E-10 | 6.01E-10 |
| chrX | 78631333 | 78663006 | 31674 | 1.8 | 2.41E-10 | 6.01E-10 |
| chr1 | 162384246 | 162394961 | 10716 | -7.4 | 2.41E-10 | 6.02E-10 |
| chr22 | 35629640 | 35632033 | 2394 | -7.09 | 2.41E-10 | 6.02E-10 |
| chr13 | 104574189 | 104589671 | 15483 | -7.11 | 2.41E-10 | 6.02E-10 |
| chr13 | 29931278 | 29955602 | 24325 | -3.79 | 2.42E-10 | 6.05E-10 |
| chr4 | 11531588 | 11533351 | 1764 | -7.1 | 2.42E-10 | 6.05E-10 |
| chr4 | 75636521 | 75641964 | 5444 | -7.33 | 2.43E-10 | 6.06E-10 |
| chr10 | 22488476 | 22498070 | 9595 | -3.42 | 2.43E-10 | 6.07E-10 |
| chr5 | 59355752 | 59384990 | 29239 | -1.57 | 2.44E-10 | 6.08E-10 |
| chr8 | 104538352 | 104546609 | 8258 | -7.08 | 2.44E-10 | 6.08E-10 |
| chr11 | 22147000 | 22155660 | 8661 | -7.2 | 2.44E-10 | 6.09E-10 |
| chr6 | 144993600 | 144996724 | 3125 | -7.21 | 2.44E-10 | 6.09E-10 |
| chr7 | 138472736 | 138477064 | 4329 | -7.21 | 2.44E-10 | 6.09E-10 |
| chr3 | 47479314 | 47491036 | 11723 | -7.22 | 2.45E-10 | 6.11E-10 |

| | | | | | | |
|---|---|---|---|---|---|---|
| chr5 | 77059488 | 77067771 | 8284 | -7.4 | 2.45E-10 | 6.11E-10 |
| chr15 | 58668625 | 58670071 | 1447 | -7.08 | 2.45E-10 | 6.11E-10 |
| chr6 | 10718298 | 10720573 | 2276 | -7.08 | 2.45E-10 | 6.12E-10 |
| chr17 | 6866783 | 6894052 | 27270 | -8.28 | 2.46E-10 | 6.13E-10 |
| chr1 | 223391372 | 223393936 | 2565 | -7.34 | 2.46E-10 | 6.14E-10 |
| chr4 | 74142918 | 74145492 | 2575 | -7.24 | 2.46E-10 | 6.14E-10 |
| chr10 | 128585895 | 128588684 | 2790 | -7.1 | 2.46E-10 | 6.14E-10 |
| chr4 | 188297151 | 188312518 | 15368 | -7.3 | 2.47E-10 | 6.15E-10 |
| chr4 | 123486624 | 123489229 | 2606 | -7.08 | 2.48E-10 | 6.18E-10 |
| chr4 | 178048912 | 178052005 | 3094 | -7.16 | 2.48E-10 | 6.18E-10 |
| chr6 | 152393949 | 152399885 | 5937 | -2.65 | 2.48E-10 | 6.18E-10 |
| chr9 | 13213238 | 13214714 | 1477 | -7.11 | 2.49E-10 | 6.20E-10 |
| chr15 | 45588259 | 45597177 | 8919 | -5.89 | 2.49E-10 | 6.21E-10 |
| chr2 | 24642542 | 24643737 | 1196 | -7.1 | 2.49E-10 | 6.21E-10 |
| chr4 | 41430850 | 41437456 | 6607 | -7.09 | 2.50E-10 | 6.22E-10 |
| chr7 | 98583275 | 98584405 | 1131 | -7.15 | 2.50E-10 | 6.23E-10 |
| chr5 | 125225351 | 125229812 | 4462 | -4.73 | 2.50E-10 | 6.24E-10 |
| chr2 | 161043673 | 161045631 | 1959 | -7.15 | 2.51E-10 | 6.25E-10 |
| chr4 | 26679903 | 26682140 | 2238 | -7.15 | 2.51E-10 | 6.25E-10 |
| chr6 | 109946220 | 109961890 | 15671 | -7.11 | 2.51E-10 | 6.26E-10 |
| chr15 | 85865438 | 85867069 | 1632 | -7.09 | 2.52E-10 | 6.28E-10 |
| chr6 | 81283770 | 81297680 | 13911 | -7.08 | 2.53E-10 | 6.31E-10 |
| chr12 | 90339266 | 90355433 | 16168 | -7.14 | 2.53E-10 | 6.32E-10 |
| chr9 | 129673287 | 129674471 | 1185 | -7.1 | 2.54E-10 | 6.33E-10 |
| chr12 | 93346888 | 93350621 | 3734 | -7.28 | 2.54E-10 | 6.33E-10 |
| chr16 | 9266369 | 9270645 | 4277 | -5.35 | 2.55E-10 | 6.35E-10 |
| chr9 | 3462350 | 3464680 | 2331 | -7.09 | 2.55E-10 | 6.36E-10 |
| chr6 | 45045510 | 45048588 | 3079 | -7.11 | 2.55E-10 | 6.36E-10 |
| chr6 | 75331950 | 75336146 | 4197 | -4.63 | 2.56E-10 | 6.36E-10 |
| chr6 | 81243978 | 81247122 | 3145 | -7.47 | 2.56E-10 | 6.36E-10 |
| chr2 | 175417577 | 175419440 | 1864 | -7.11 | 2.56E-10 | 6.37E-10 |
| chr1 | 93632503 | 93634674 | 2172 | -7.21 | 2.56E-10 | 6.38E-10 |
| chr1 | 35955594 | 35959267 | 3674 | -7.13 | 2.56E-10 | 6.39E-10 |
| chr2 | 19815950 | 19828180 | 12231 | -7.35 | 2.58E-10 | 6.42E-10 |
| chr15 | 77143975 | 77145833 | 1859 | -7.07 | 2.58E-10 | 6.42E-10 |
| chr4 | 189734125 | 189736955 | 2831 | -7.08 | 2.58E-10 | 6.43E-10 |
| chr14 | 56194732 | 56203996 | 9265 | -7.45 | 2.59E-10 | 6.44E-10 |
| chr13 | 21813479 | 21815844 | 2366 | -7.07 | 2.59E-10 | 6.44E-10 |
| chr7 | 105719700 | 105729094 | 9395 | -7.51 | 2.60E-10 | 6.46E-10 |
| chr2 | 237169045 | 237206551 | 37507 | -2.41 | 2.61E-10 | 6.49E-10 |
| chr15 | 100052849 | 100054255 | 1407 | -7.1 | 2.61E-10 | 6.49E-10 |
| chr12 | 133405979 | 133408992 | 3014 | -7.08 | 2.61E-10 | 6.50E-10 |
| chr4 | 183418363 | 183688903 | 270541 | 3.52 | 2.61E-10 | 6.50E-10 |
| chr1 | 45488423 | 45490031 | 1609 | -7.16 | 2.61E-10 | 6.50E-10 |
| chr18 | 56225632 | 56228587 | 2956 | -7.16 | 2.61E-10 | 6.50E-10 |
| chr19 | 43658593 | 43665307 | 6715 | -7.16 | 2.61E-10 | 6.50E-10 |
| chr2 | 177791027 | 177791993 | 967 | -7.07 | 2.62E-10 | 6.51E-10 |
| chr12 | 93200129 | 93201380 | 1252 | -7.16 | 2.62E-10 | 6.51E-10 |
| chr1 | 101061222 | 101071129 | 9908 | -7.14 | 2.62E-10 | 6.51E-10 |
| chrX | 69236256 | 69239536 | 3281 | -7.25 | 2.62E-10 | 6.51E-10 |
| chr13 | 52704801 | 52706323 | 1523 | -7.08 | 2.63E-10 | 6.55E-10 |
| chr10 | 94421541 | 94424267 | 2727 | -7.42 | 2.64E-10 | 6.55E-10 |
| chr5 | 10510337 | 10523852 | 13516 | -7.42 | 2.64E-10 | 6.55E-10 |
| chr13 | 105765602 | 105767403 | 1802 | -7.15 | 2.64E-10 | 6.56E-10 |
| chr4 | 5811323 | 5813236 | 1914 | -7.07 | 2.64E-10 | 6.56E-10 |
| chr11 | 72919854 | 72921465 | 1612 | -7.1 | 2.64E-10 | 6.57E-10 |
| chr6 | 18072994 | 18092202 | 19209 | -8.03 | 2.64E-10 | 6.57E-10 |
| chr6 | 88494395 | 88497311 | 2917 | -7.49 | 2.66E-10 | 6.60E-10 |
| chr17 | 35426127 | 35427446 | 1320 | -7.07 | 2.66E-10 | 6.61E-10 |
| chr6 | 40651229 | 40653515 | 2287 | -7.07 | 2.66E-10 | 6.61E-10 |
| chr10 | 45931607 | 45933936 | 2330 | -7.59 | 2.66E-10 | 6.61E-10 |

| | | | | | | |
|---|---|---|---|---|---|---|
| chr4 | 121671169 | 121673377 | 2209 | -7.14 | 2.66E-10 | 6.62E-10 |
| chr12 | 76218818 | 76241344 | 22527 | -7.97 | 2.66E-10 | 6.62E-10 |
| chr6 | 34525380 | 34526897 | 1518 | -7.07 | 2.66E-10 | 6.62E-10 |
| chr1 | 93929400 | 93933207 | 3808 | -7.08 | 2.67E-10 | 6.62E-10 |
| chr18 | 59654179 | 59657188 | 3010 | 7.14 | 2.67E-10 | 6.62E-10 |
| chrX | 152878990 | 152880826 | 1837 | -7.07 | 2.67E-10 | 6.63E-10 |
| chr2 | 119710189 | 119718582 | 8394 | -2.75 | 2.67E-10 | 6.63E-10 |
| chr9 | 74897900 | 74906046 | 8147 | -7.53 | 2.67E-10 | 6.63E-10 |
| chr1 | 173276012 | 173278006 | 1995 | -5.84 | 2.67E-10 | 6.64E-10 |
| chr4 | 57966257 | 57974618 | 8362 | -7.5 | 2.68E-10 | 6.65E-10 |
| chr8 | 30145463 | 30147352 | 1890 | -7.28 | 2.68E-10 | 6.66E-10 |
| chr7 | 44968796 | 44970480 | 1685 | -7.2 | 2.68E-10 | 6.66E-10 |
| chr9 | 14724828 | 14909690 | 184863 | 2.46 | 2.69E-10 | 6.67E-10 |
| chr6 | 106482534 | 106484750 | 2217 | -7.16 | 2.69E-10 | 6.68E-10 |
| chr4 | 90266673 | 90269706 | 3034 | -7.28 | 2.70E-10 | 6.69E-10 |
| chr18 | 25419964 | 25426594 | 6631 | -3.19 | 2.70E-10 | 6.71E-10 |
| chrX | 55734686 | 55737754 | 3069 | -7.08 | 2.71E-10 | 6.72E-10 |
| chr5 | 21015634 | 21024537 | 8904 | -7.44 | 2.71E-10 | 6.73E-10 |
| chr6 | 126577696 | 126582793 | 5098 | -7.06 | 2.71E-10 | 6.73E-10 |
| chr1 | 172483228 | 172485215 | 1988 | -7.06 | 2.72E-10 | 6.76E-10 |
| chr14 | 66983699 | 66984697 | 999 | -7.06 | 2.72E-10 | 6.76E-10 |
| chr3 | 190178025 | 190190317 | 12293 | -7.14 | 2.72E-10 | 6.76E-10 |
| chr22 | 44983265 | 44985994 | 2730 | -7.07 | 2.73E-10 | 6.77E-10 |
| chr15 | 51299469 | 51313102 | 13634 | -7.06 | 2.74E-10 | 6.81E-10 |
| chr12 | 40063957 | 40074234 | 10278 | -7.1 | 2.74E-10 | 6.81E-10 |
| chr2 | 198019873 | 198021705 | 1833 | -7.08 | 2.75E-10 | 6.81E-10 |
| chr2 | 149315069 | 149321875 | 6807 | -6.22 | 2.75E-10 | 6.81E-10 |
| chr3 | 167949386 | 167955164 | 5779 | 5.35 | 2.75E-10 | 6.82E-10 |
| chr11 | 77684465 | 77685587 | 1123 | -7.12 | 2.75E-10 | 6.82E-10 |
| chr3 | 119573980 | 119578098 | 4119 | -7.59 | 2.76E-10 | 6.84E-10 |
| chr5 | 73201673 | 73202977 | 1305 | -7.07 | 2.76E-10 | 6.84E-10 |
| chr1 | 187562962 | 187565187 | 2226 | -3.37 | 2.76E-10 | 6.84E-10 |
| chr17 | 34311086 | 34426218 | 115133 | -1.43 | 2.76E-10 | 6.85E-10 |
| chr7 | 35894831 | 35896731 | 1901 | -7.21 | 2.76E-10 | 6.85E-10 |
| chr10 | 46079902 | 46082908 | 3007 | -7.61 | 2.76E-10 | 6.85E-10 |
| chr14 | 90804542 | 90806544 | 2003 | -6.2 | 2.76E-10 | 6.85E-10 |
| chr10 | 74374288 | 74376746 | 2459 | -7.11 | 2.76E-10 | 6.85E-10 |
| chr12 | 7512651 | 7524020 | 11370 | -7.11 | 2.76E-10 | 6.85E-10 |
| chr1 | 96992893 | 96995418 | 2526 | -7.16 | 2.76E-10 | 6.85E-10 |
| chr3 | 64137613 | 64140304 | 2692 | -7.09 | 2.77E-10 | 6.86E-10 |
| chr12 | 79475852 | 79478416 | 2565 | -7.15 | 2.77E-10 | 6.88E-10 |
| chr1 | 168375601 | 168377033 | 1433 | -7.14 | 2.79E-10 | 6.91E-10 |
| chr12 | 132692662 | 132706716 | 14055 | -7.26 | 2.79E-10 | 6.92E-10 |
| chr6 | 40872733 | 40877280 | 4548 | -7.09 | 2.79E-10 | 6.92E-10 |
| chr7 | 156957806 | 156960204 | 2399 | -7.3 | 2.79E-10 | 6.92E-10 |
| chr15 | 81288307 | 81289916 | 1610 | -7.06 | 2.80E-10 | 6.94E-10 |
| chr10 | 114499934 | 114501150 | 1217 | -7.06 | 2.80E-10 | 6.94E-10 |
| chr7 | 132190512 | 132193729 | 3218 | 2.81 | 2.81E-10 | 6.96E-10 |
| chr1 | 47567174 | 47570032 | 2859 | -5.22 | 2.82E-10 | 6.98E-10 |
| chr5 | 55339413 | 55350702 | 11290 | -8.16 | 2.82E-10 | 6.99E-10 |
| chr7 | 13245102 | 13246412 | 1311 | -5.33 | 2.83E-10 | 7.01E-10 |
| chr21 | 36522696 | 36538060 | 15365 | -7.15 | 2.83E-10 | 7.01E-10 |
| chr9 | 123969013 | 123970752 | 1740 | -7.15 | 2.83E-10 | 7.01E-10 |
| chr8 | 61252828 | 61284897 | 32070 | -3.01 | 2.83E-10 | 7.02E-10 |
| chr15 | 22972237 | 22973791 | 1555 | -7.06 | 2.85E-10 | 7.05E-10 |
| chr4 | 155738703 | 155740706 | 2004 | -7.41 | 2.86E-10 | 7.09E-10 |
| chr11 | 32725832 | 32731303 | 5472 | -7.66 | 2.87E-10 | 7.10E-10 |
| chr1 | 222029839 | 222041573 | 11735 | -7.34 | 2.90E-10 | 7.18E-10 |
| chr3 | 16902973 | 16905501 | 2529 | -7.09 | 2.90E-10 | 7.18E-10 |
| chr1 | 15590763 | 15592800 | 2038 | -7.09 | 2.90E-10 | 7.18E-10 |
| chr1 | 3214623 | 3219473 | 4851 | -7.34 | 2.91E-10 | 7.21E-10 |

| | | | | | | |
|---|---|---|---|---|---|---|
| chr9 | 30134357 | 30147521 | 13165 | -7.34 | 2.91E-10 | 7.21E-10 |
| chr10 | 105509326 | 105513054 | 3729 | -7.12 | 2.92E-10 | 7.23E-10 |
| chr12 | 59219940 | 59230453 | 10514 | -7.05 | 2.92E-10 | 7.23E-10 |
| chr2 | 161912205 | 161914258 | 2054 | -7.31 | 2.92E-10 | 7.23E-10 |
| chr1 | 232112468 | 232114795 | 2328 | -7.09 | 2.92E-10 | 7.24E-10 |
| chr2 | 166940367 | 166955452 | 15086 | 7.62 | 2.93E-10 | 7.24E-10 |
| chr13 | 73705560 | 73707720 | 2161 | -7.15 | 2.93E-10 | 7.25E-10 |
| chr2 | 111634698 | 111689547 | 54850 | -3.38 | 2.93E-10 | 7.26E-10 |
| chr2 | 225704026 | 225706257 | 2232 | -7.63 | 2.93E-10 | 7.26E-10 |
| chr15 | 42626329 | 42628110 | 1782 | -7.08 | 2.94E-10 | 7.26E-10 |
| chr7 | 51110767 | 51113349 | 2583 | -7.06 | 2.94E-10 | 7.27E-10 |
| chr7 | 24422358 | 24423694 | 1337 | -7.07 | 2.94E-10 | 7.27E-10 |
| chr12 | 20330154 | 20333698 | 3545 | -7.42 | 2.94E-10 | 7.27E-10 |
| chr15 | 96810287 | 96811846 | 1560 | -7.1 | 2.94E-10 | 7.28E-10 |
| chr6 | 71891627 | 71894848 | 3222 | -7.22 | 2.95E-10 | 7.31E-10 |
| chr6 | 134875798 | 134879828 | 4031 | -7.13 | 2.96E-10 | 7.32E-10 |
| chr20 | 37165564 | 37167811 | 2248 | -7.22 | 2.96E-10 | 7.32E-10 |
| chr5 | 32962505 | 32966539 | 4035 | -7.22 | 2.96E-10 | 7.32E-10 |
| chr14 | 62287793 | 62292312 | 4520 | -7.32 | 2.96E-10 | 7.33E-10 |
| chr10 | 104894019 | 104896378 | 2360 | -7.39 | 2.96E-10 | 7.33E-10 |
| chr13 | 105926707 | 105929864 | 3158 | -7.07 | 2.96E-10 | 7.33E-10 |
| chr9 | 79350696 | 79364117 | 13422 | -1.45 | 2.97E-10 | 7.34E-10 |
| chr17 | 54763551 | 54765396 | 1846 | -2.93 | 2.97E-10 | 7.34E-10 |
| chr11 | 106265326 | 106268180 | 2855 | -5.35 | 2.98E-10 | 7.37E-10 |
| chr11 | 88537890 | 88550169 | 12280 | -7.22 | 2.98E-10 | 7.37E-10 |
| chrX | 36376510 | 36380884 | 4375 | -7.24 | 2.99E-10 | 7.40E-10 |
| chr1 | 65463478 | 65465372 | 1895 | -7.06 | 2.99E-10 | 7.40E-10 |
| chr10 | 45358798 | 45359945 | 1148 | -7.06 | 3.00E-10 | 7.40E-10 |
| chr10 | 51511483 | 51522930 | 11448 | -7.28 | 3.00E-10 | 7.40E-10 |
| chr15 | 100037153 | 100040054 | 2902 | -7.28 | 3.00E-10 | 7.40E-10 |
| chr9 | 18374837 | 18381805 | 6969 | -7.28 | 3.00E-10 | 7.40E-10 |
| chr20 | 56578637 | 56610559 | 31923 | -7.05 | 3.00E-10 | 7.40E-10 |
| chr18 | 2454861 | 2459094 | 4234 | -7.05 | 3.00E-10 | 7.42E-10 |
| chr6 | 13519396 | 13520767 | 1372 | -5.65 | 3.01E-10 | 7.44E-10 |
| chr8 | 128143309 | 128145997 | 2689 | -7.09 | 3.02E-10 | 7.45E-10 |
| chr10 | 90389387 | 90399167 | 9781 | -3.05 | 3.03E-10 | 7.47E-10 |
| chr5 | 121379943 | 121381851 | 1909 | -7.05 | 3.03E-10 | 7.48E-10 |
| chr11 | 75328990 | 75331274 | 2285 | -7.24 | 3.03E-10 | 7.49E-10 |
| chr10 | 93744489 | 93746378 | 1890 | -7.29 | 3.03E-10 | 7.49E-10 |
| chr1 | 242679686 | 242681209 | 1524 | -7.07 | 3.04E-10 | 7.51E-10 |
| chr5 | 127493752 | 127508269 | 14518 | -7.08 | 3.05E-10 | 7.52E-10 |
| chr7 | 139902315 | 139904232 | 1918 | -7.05 | 3.05E-10 | 7.54E-10 |
| chr12 | 39055156 | 39056390 | 1235 | -7.05 | 3.06E-10 | 7.54E-10 |
| chrX | 39638221 | 39640219 | 1999 | -7.05 | 3.06E-10 | 7.54E-10 |
| chr4 | 28962759 | 29018358 | 55600 | 1.57 | 3.06E-10 | 7.56E-10 |
| chr9 | 20856263 | 20858466 | 2204 | -7.05 | 3.07E-10 | 7.58E-10 |
| chr7 | 137387345 | 137389922 | 2578 | -5.29 | 3.07E-10 | 7.58E-10 |
| chr4 | 54799393 | 54807767 | 8375 | -7.26 | 3.08E-10 | 7.59E-10 |
| chr2 | 113901788 | 113903595 | 1808 | -7.07 | 3.08E-10 | 7.60E-10 |
| chr6 | 81384997 | 81387404 | 2408 | -7.16 | 3.08E-10 | 7.60E-10 |
| chr6 | 146034786 | 146037194 | 2409 | -7.05 | 3.08E-10 | 7.60E-10 |
| chr3 | 40682542 | 40692157 | 9616 | 4.11 | 3.08E-10 | 7.60E-10 |
| chr20 | 25143445 | 25147422 | 3978 | -7.16 | 3.08E-10 | 7.60E-10 |
| chrX | 24401048 | 24405490 | 4443 | -7.16 | 3.08E-10 | 7.60E-10 |
| chr10 | 61970373 | 61976133 | 5761 | -7.42 | 3.08E-10 | 7.61E-10 |
| chr19 | 18986644 | 18988642 | 1999 | -7.04 | 3.09E-10 | 7.61E-10 |
| chr9 | 82287156 | 82293534 | 6379 | -7.19 | 3.10E-10 | 7.64E-10 |
| chr20 | 61579856 | 61581308 | 1453 | -7.19 | 3.10E-10 | 7.65E-10 |
| chr7 | 80697937 | 80716473 | 18537 | -7.64 | 3.11E-10 | 7.66E-10 |
| chr7 | 109853247 | 109859878 | 6632 | -7.07 | 3.11E-10 | 7.67E-10 |
| chr4 | 79475685 | 79479547 | 3863 | -7.14 | 3.12E-10 | 7.68E-10 |

| | | | | | | |
|---|---|---|---|---|---|---|
| chr16 | 88248821 | 88253590 | 4770 | -5.37 | 3.12E-10 | 7.68E-10 |
| chrX | 71426649 | 71430666 | 4018 | -7.05 | 3.12E-10 | 7.68E-10 |
| chr3 | 10109395 | 10110587 | 1193 | -7.07 | 3.12E-10 | 7.69E-10 |
| chr19 | 43850961 | 43857001 | 6041 | -7.1 | 3.12E-10 | 7.69E-10 |
| chr16 | 47901165 | 47903053 | 1889 | -7.27 | 3.13E-10 | 7.71E-10 |
| chr22 | 46380600 | 46382409 | 1810 | -7.06 | 3.13E-10 | 7.72E-10 |
| chr12 | 76967213 | 76974001 | 6789 | -7.22 | 3.15E-10 | 7.76E-10 |
| chr5 | 142690431 | 142692787 | 2357 | -7.22 | 3.15E-10 | 7.76E-10 |
| chr2 | 232248441 | 232250644 | 2204 | -7.04 | 3.15E-10 | 7.76E-10 |
| chr4 | 90121473 | 90129866 | 8394 | -7.53 | 3.15E-10 | 7.77E-10 |
| chr12 | 105791198 | 105793139 | 1942 | -7.12 | 3.15E-10 | 7.77E-10 |
| chr1 | 212730079 | 212736864 | 6786 | -7.09 | 3.16E-10 | 7.79E-10 |
| chr2 | 25288781 | 25290199 | 1419 | -7.11 | 3.17E-10 | 7.82E-10 |
| chr6 | 38797795 | 38807058 | 9264 | 5.64 | 3.18E-10 | 7.83E-10 |
| chr17 | 42642059 | 42643878 | 1820 | -7.11 | 3.18E-10 | 7.83E-10 |
| chr15 | 93069967 | 93129378 | 59412 | -1.56 | 3.19E-10 | 7.85E-10 |
| chr3 | 111273470 | 111275103 | 1634 | -7.24 | 3.19E-10 | 7.86E-10 |
| chr6 | 158229157 | 158231542 | 2386 | -7.08 | 3.19E-10 | 7.86E-10 |
| chr6 | 56217402 | 56219748 | 2347 | -7.4 | 3.19E-10 | 7.86E-10 |
| chr20 | 61962885 | 61966163 | 3279 | -7.24 | 3.20E-10 | 7.88E-10 |
| chr21 | 16524277 | 16534563 | 10287 | -7.44 | 3.20E-10 | 7.89E-10 |
| chrX | 6962664 | 6964595 | 1932 | -4.14 | 3.21E-10 | 7.90E-10 |
| chr3 | 182276721 | 182292332 | 15612 | -7.08 | 3.22E-10 | 7.92E-10 |
| chr5 | 169485833 | 169487685 | 1853 | -7.04 | 3.22E-10 | 7.93E-10 |
| chr21 | 27911488 | 27965638 | 54151 | -1.42 | 3.22E-10 | 7.93E-10 |
| chr11 | 44574035 | 44575465 | 1431 | -7.1 | 3.23E-10 | 7.94E-10 |
| chr7 | 91768143 | 91771559 | 3417 | -7.41 | 3.23E-10 | 7.95E-10 |
| chr1 | 4049380 | 4051717 | 2338 | -7.16 | 3.23E-10 | 7.95E-10 |
| chr4 | 138770691 | 138772542 | 1852 | -7.04 | 3.23E-10 | 7.95E-10 |
| chr14 | 35729323 | 35730988 | 1666 | -7.04 | 3.23E-10 | 7.95E-10 |
| chr1 | 162512789 | 162514207 | 1419 | -7.1 | 3.23E-10 | 7.95E-10 |
| chr22 | 46821250 | 46823705 | 2456 | -7.1 | 3.23E-10 | 7.95E-10 |
| chr6 | 70463204 | 70467235 | 4032 | -7.1 | 3.23E-10 | 7.95E-10 |
| chr12 | 123042023 | 123044069 | 2047 | -7.25 | 3.23E-10 | 7.95E-10 |
| chr5 | 111787374 | 111791692 | 4319 | 7.3 | 3.23E-10 | 7.95E-10 |
| chr3 | 114526093 | 114528217 | 2125 | -7.25 | 3.24E-10 | 7.96E-10 |
| chr8 | 124191000 | 124192802 | 1803 | -7.04 | 3.24E-10 | 7.96E-10 |
| chrX | 129181630 | 129183310 | 1681 | -7.45 | 3.24E-10 | 7.96E-10 |
| chr3 | 138138473 | 138153238 | 14766 | -7.14 | 3.24E-10 | 7.97E-10 |
| chr10 | 52242796 | 52245739 | 2944 | -7.08 | 3.25E-10 | 7.99E-10 |
| chr4 | 28395863 | 28401483 | 5621 | -7.3 | 3.26E-10 | 8.00E-10 |
| chr5 | 53033096 | 53064563 | 31468 | -8.01 | 3.26E-10 | 8.01E-10 |
| chr6 | 75586198 | 75588851 | 2654 | 5.24 | 3.26E-10 | 8.02E-10 |
| chr3 | 187797569 | 187799135 | 1567 | -7.2 | 3.26E-10 | 8.02E-10 |
| chr5 | 44933214 | 44944441 | 11228 | -7.52 | 3.27E-10 | 8.03E-10 |
| chr10 | 46130566 | 46133844 | 3279 | -7.2 | 3.27E-10 | 8.04E-10 |
| chr22 | 29242265 | 29244762 | 2498 | -7.2 | 3.27E-10 | 8.04E-10 |
| chr3 | 137862203 | 137874092 | 11890 | -7.12 | 3.27E-10 | 8.04E-10 |
| chr1 | 188274811 | 188277228 | 2418 | -7.04 | 3.27E-10 | 8.04E-10 |
| chr9 | 13419456 | 13428592 | 9137 | -7.38 | 3.27E-10 | 8.04E-10 |
| chr5 | 917871 | 919669 | 1799 | -7.15 | 3.28E-10 | 8.06E-10 |
| chr12 | 88742055 | 88756241 | 14187 | -7.04 | 3.28E-10 | 8.06E-10 |
| chr11 | 12846070 | 12858902 | 12833 | -7.07 | 3.28E-10 | 8.06E-10 |
| chr19 | 55778000 | 55779501 | 1502 | -7.14 | 3.29E-10 | 8.07E-10 |
| chr9 | 113026242 | 113028133 | 1892 | -7.04 | 3.29E-10 | 8.07E-10 |
| chr3 | 182587161 | 182588338 | 1178 | -7.14 | 3.29E-10 | 8.08E-10 |
| chr1 | 219013531 | 219016723 | 3193 | -7.21 | 3.30E-10 | 8.11E-10 |
| chr9 | 98138996 | 98142404 | 3409 | -7.42 | 3.30E-10 | 8.11E-10 |
| chr4 | 6460302 | 6462187 | 1886 | -7.03 | 3.30E-10 | 8.11E-10 |
| chr11 | 94418972 | 94431800 | 12829 | -7.43 | 3.32E-10 | 8.16E-10 |
| chr10 | 62250603 | 62252267 | 1665 | -7.06 | 3.33E-10 | 8.17E-10 |

| | | | | | | |
|---|---|---|---|---|---|---|
| chr18 | 9664461 | 9678091 | 13631 | -7.39 | 3.33E-10 | 8.17E-10 |
| chrX | 135386155 | 135387251 | 1097 | -4.02 | 3.33E-10 | 8.18E-10 |
| chr18 | 34381677 | 34383682 | 2006 | -7.21 | 3.33E-10 | 8.18E-10 |
| chr4 | 2736672 | 2738994 | 2323 | -7.21 | 3.33E-10 | 8.18E-10 |
| chr7 | 84218328 | 84234887 | 16560 | -8.19 | 3.34E-10 | 8.19E-10 |
| chr15 | 91679911 | 91712866 | 32956 | -3.28 | 3.34E-10 | 8.20E-10 |
| chr5 | 106211340 | 106213648 | 2309 | -7.08 | 3.35E-10 | 8.22E-10 |
| chr2 | 162312026 | 162313850 | 1825 | -7.04 | 3.35E-10 | 8.22E-10 |
| chr6 | 150567799 | 150570031 | 2233 | -7.04 | 3.35E-10 | 8.22E-10 |
| chr11 | 27912375 | 27929338 | 16964 | -7.47 | 3.35E-10 | 8.22E-10 |
| chr12 | 22175520 | 22177841 | 2322 | -7.1 | 3.35E-10 | 8.22E-10 |
| chr5 | 31938929 | 31941988 | 3060 | 3.84 | 3.35E-10 | 8.23E-10 |
| chr9 | 76236570 | 76238093 | 1524 | -7.04 | 3.35E-10 | 8.23E-10 |
| chrX | 18728385 | 18731249 | 2865 | -7.36 | 3.36E-10 | 8.24E-10 |
| chr2 | 70599133 | 70601058 | 1926 | -3.11 | 3.36E-10 | 8.25E-10 |
| chr8 | 53608211 | 53611021 | 2811 | -7.11 | 3.36E-10 | 8.25E-10 |
| chrX | 70821096 | 70823477 | 2382 | -7.22 | 3.37E-10 | 8.27E-10 |
| chr7 | 115280856 | 115283057 | 2202 | -7.34 | 3.37E-10 | 8.27E-10 |
| chr8 | 141485978 | 141513607 | 27630 | -8.16 | 3.37E-10 | 8.27E-10 |
| chr7 | 141685825 | 141704016 | 18192 | -7.86 | 3.39E-10 | 8.31E-10 |
| chr12 | 76004364 | 76010840 | 6477 | -7.14 | 3.39E-10 | 8.32E-10 |
| chr5 | 108113506 | 108121119 | 7614 | -7.14 | 3.39E-10 | 8.32E-10 |
| chr12 | 102352186 | 102366664 | 14479 | -7.54 | 3.40E-10 | 8.32E-10 |
| chr6 | 74949342 | 74958151 | 8810 | -3.33 | 3.40E-10 | 8.33E-10 |
| chr3 | 140777344 | 140779870 | 2527 | -7.03 | 3.40E-10 | 8.34E-10 |
| chr6 | 158839851 | 158841687 | 1837 | -7.09 | 3.41E-10 | 8.37E-10 |
| chr14 | 69330875 | 69332817 | 1943 | -7.05 | 3.43E-10 | 8.41E-10 |
| chr2 | 147214935 | 147218705 | 3771 | -7.08 | 3.44E-10 | 8.42E-10 |
| chr3 | 20130815 | 20132175 | 1361 | -7.08 | 3.44E-10 | 8.42E-10 |
| chr5 | 172575152 | 172577069 | 1918 | -7.08 | 3.44E-10 | 8.42E-10 |
| chr4 | 163827082 | 163839077 | 11996 | -7.06 | 3.45E-10 | 8.44E-10 |
| chr6 | 149463163 | 149464893 | 1731 | -7.06 | 3.45E-10 | 8.44E-10 |
| chr12 | 86148514 | 86150822 | 2309 | -7.03 | 3.45E-10 | 8.44E-10 |
| chr7 | 102592078 | 102594770 | 2693 | -7.03 | 3.45E-10 | 8.44E-10 |
| chr11 | 69497741 | 69507915 | 10175 | -7.3 | 3.45E-10 | 8.44E-10 |
| chr1 | 62930828 | 62934092 | 3265 | -7.11 | 3.45E-10 | 8.46E-10 |
| chr11 | 86074145 | 86088400 | 14256 | -7.04 | 3.46E-10 | 8.47E-10 |
| chr7 | 5760843 | 5762325 | 1483 | -7.04 | 3.46E-10 | 8.47E-10 |
| chr1 | 232178015 | 232182915 | 4901 | 7.36 | 3.46E-10 | 8.47E-10 |
| chr8 | 10521627 | 10526642 | 5016 | -4.08 | 3.46E-10 | 8.49E-10 |
| chr14 | 21295662 | 21297951 | 2290 | -7.1 | 3.47E-10 | 8.49E-10 |
| chr15 | 59364695 | 59366596 | 1902 | -7.03 | 3.47E-10 | 8.49E-10 |
| chrX | 109596254 | 109599258 | 3005 | -5.14 | 3.48E-10 | 8.52E-10 |
| chr1 | 168292942 | 168294925 | 1984 | -7.1 | 3.48E-10 | 8.52E-10 |
| chr6 | 86205834 | 86208591 | 2758 | -7.27 | 3.50E-10 | 8.56E-10 |
| chr4 | 130562674 | 130565207 | 2534 | -7.09 | 3.50E-10 | 8.56E-10 |
| chr8 | 42288892 | 42291361 | 2470 | -7.12 | 3.50E-10 | 8.56E-10 |
| chr8 | 126622433 | 126624124 | 1692 | -7.02 | 3.50E-10 | 8.57E-10 |
| chr15 | 45611583 | 45613671 | 2089 | -7.22 | 3.51E-10 | 8.58E-10 |
| chr13 | 83308680 | 83313102 | 4423 | -7.09 | 3.52E-10 | 8.61E-10 |
| chr1 | 185451441 | 185455099 | 3659 | -7.04 | 3.52E-10 | 8.62E-10 |
| chrX | 13004011 | 13005872 | 1862 | -7.05 | 3.53E-10 | 8.63E-10 |
| chrX | 36751766 | 36802544 | 50779 | 4.34 | 3.53E-10 | 8.63E-10 |
| chr18 | 74853924 | 74856910 | 2987 | -7.03 | 3.53E-10 | 8.63E-10 |
| chr20 | 37060146 | 37062151 | 2006 | -7.07 | 3.53E-10 | 8.63E-10 |
| chrX | 76419960 | 76474820 | 54861 | 1.65 | 3.53E-10 | 8.64E-10 |
| chr7 | 98446113 | 98447868 | 1756 | -7.03 | 3.54E-10 | 8.66E-10 |
| chr1 | 146761435 | 146846928 | 85494 | -2.01 | 3.55E-10 | 8.68E-10 |
| chr21 | 45782103 | 45784003 | 1901 | -7.14 | 3.55E-10 | 8.69E-10 |
| chr10 | 133606279 | 133634582 | 28304 | -3.85 | 3.56E-10 | 8.71E-10 |
| chr4 | 159313136 | 159326809 | 13674 | -7.29 | 3.56E-10 | 8.71E-10 |

| | | | | | | |
|---|---|---|---|---|---|---|
| chr9 | 38383790 | 38386136 | 2347 | -7.17 | 3.56E-10 | 8.72E-10 |
| chr8 | 62706533 | 62710149 | 3617 | -7.33 | 3.57E-10 | 8.74E-10 |
| chr12 | 88920954 | 88922583 | 1630 | -7.37 | 3.58E-10 | 8.75E-10 |
| chr13 | 32563675 | 32569642 | 5968 | -7.06 | 3.58E-10 | 8.76E-10 |
| chr3 | 70160243 | 70170717 | 10475 | -7.4 | 3.59E-10 | 8.77E-10 |
| chr4 | 105017298 | 105021696 | 4399 | -7.07 | 3.61E-10 | 8.82E-10 |
| chr10 | 93793946 | 93795807 | 1862 | -7.1 | 3.61E-10 | 8.82E-10 |
| chr7 | 31486621 | 31488678 | 2058 | -7.05 | 3.62E-10 | 8.84E-10 |
| chr8 | 38653664 | 38660913 | 7250 | -7.2 | 3.62E-10 | 8.85E-10 |
| chr12 | 133386413 | 133389880 | 3468 | -7.06 | 3.63E-10 | 8.87E-10 |
| chr2 | 8855489 | 8857233 | 1745 | -7.23 | 3.63E-10 | 8.88E-10 |
| chr8 | 142595191 | 142597374 | 2184 | -7.26 | 3.65E-10 | 8.92E-10 |
| chr2 | 135772642 | 135774433 | 1792 | -7.09 | 3.65E-10 | 8.93E-10 |
| chr13 | 101439020 | 101449346 | 10327 | -2.03 | 3.68E-10 | 8.98E-10 |
| chr9 | 109620868 | 109624329 | 3462 | -7.22 | 3.68E-10 | 8.99E-10 |
| chr5 | 95829371 | 95837545 | 8175 | -7.06 | 3.69E-10 | 9.01E-10 |
| chr5 | 39639472 | 39643022 | 3551 | -7.08 | 3.69E-10 | 9.01E-10 |
| chr16 | 82187719 | 82189262 | 1544 | -7.13 | 3.70E-10 | 9.03E-10 |
| chr1 | 156282543 | 156284998 | 2456 | -7.15 | 3.70E-10 | 9.04E-10 |
| chr11 | 31514561 | 31517482 | 2922 | -7.02 | 3.70E-10 | 9.05E-10 |
| chr15 | 62279964 | 62282281 | 2318 | -7.12 | 3.71E-10 | 9.05E-10 |
| chr3 | 181395766 | 181397318 | 1553 | -7.12 | 3.71E-10 | 9.05E-10 |
| chr20 | 49100976 | 49103060 | 2085 | -7.32 | 3.71E-10 | 9.05E-10 |
| chrX | 140302924 | 140314300 | 11377 | -7.42 | 3.71E-10 | 9.06E-10 |
| chr7 | 47614530 | 47625978 | 11449 | -7.05 | 3.72E-10 | 9.09E-10 |
| chr4 | 151553509 | 151555544 | 2036 | -7.06 | 3.72E-10 | 9.09E-10 |
| chr2 | 162033270 | 162034806 | 1537 | -7.11 | 3.73E-10 | 9.11E-10 |
| chr11 | 108894857 | 108897456 | 2600 | -7.02 | 3.73E-10 | 9.11E-10 |
| chr6 | 158272993 | 158275142 | 2150 | -7.07 | 3.73E-10 | 9.12E-10 |
| chr1 | 204361110 | 204366595 | 5486 | -7.4 | 3.74E-10 | 9.14E-10 |
| chr3 | 72196564 | 72203945 | 7382 | -7.16 | 3.74E-10 | 9.14E-10 |
| chr10 | 30133523 | 30135192 | 1670 | -7.17 | 3.75E-10 | 9.15E-10 |
| chr1 | 33310260 | 33312134 | 1875 | -7.31 | 3.75E-10 | 9.16E-10 |
| chr3 | 162734143 | 162737893 | 3751 | -7.03 | 3.75E-10 | 9.16E-10 |
| chr9 | 114548294 | 114550110 | 1817 | -7.1 | 3.76E-10 | 9.19E-10 |
| chr15 | 29376532 | 29377906 | 1375 | -7.09 | 3.78E-10 | 9.22E-10 |
| chr5 | 118313489 | 118316839 | 3351 | -7.03 | 3.78E-10 | 9.23E-10 |
| chr8 | 49518933 | 49528149 | 9217 | -7.2 | 3.78E-10 | 9.23E-10 |
| chr2 | 179498208 | 179501860 | 3653 | 5.28 | 3.78E-10 | 9.23E-10 |
| chr1 | 84990634 | 84994016 | 3383 | -7.02 | 3.79E-10 | 9.24E-10 |
| chr15 | 34426686 | 34428384 | 1699 | -5.42 | 3.79E-10 | 9.25E-10 |
| chr4 | 180634738 | 180640188 | 5451 | -7.01 | 3.80E-10 | 9.26E-10 |
| chr1 | 182256184 | 182266581 | 10398 | 7.13 | 3.80E-10 | 9.27E-10 |
| chr2 | 101426923 | 101428047 | 1125 | -5.24 | 3.80E-10 | 9.28E-10 |
| chr1 | 60342279 | 60354434 | 12156 | -7.02 | 3.81E-10 | 9.30E-10 |
| chr13 | 111751055 | 111765596 | 14542 | -1.87 | 3.82E-10 | 9.31E-10 |
| chr4 | 90846833 | 90848891 | 2059 | -7.02 | 3.82E-10 | 9.31E-10 |
| chrX | 67588883 | 67590872 | 1990 | -7.02 | 3.82E-10 | 9.31E-10 |
| chr11 | 48172880 | 48175551 | 2672 | -7.08 | 3.82E-10 | 9.32E-10 |
| chr3 | 23823505 | 23826440 | 2936 | -7.41 | 3.83E-10 | 9.33E-10 |
| chr2 | 160924651 | 160927170 | 2520 | -7.25 | 3.83E-10 | 9.33E-10 |
| chr5 | 39172113 | 39174499 | 2387 | -7.25 | 3.83E-10 | 9.33E-10 |
| chr1 | 193639863 | 193643725 | 3863 | -7.03 | 3.83E-10 | 9.33E-10 |
| chrX | 71702949 | 71704607 | 1659 | -7.21 | 3.83E-10 | 9.34E-10 |
| chr1 | 236323928 | 236326067 | 2140 | -7.04 | 3.84E-10 | 9.36E-10 |
| chr1 | 116329493 | 116331365 | 1873 | -7.22 | 3.85E-10 | 9.39E-10 |
| chr7 | 53213463 | 53217844 | 4382 | 1.97 | 3.85E-10 | 9.39E-10 |
| chr3 | 15426997 | 15440093 | 13097 | -7.38 | 3.86E-10 | 9.40E-10 |
| chr7 | 95923690 | 95934378 | 10689 | -7.33 | 3.86E-10 | 9.42E-10 |
| chr1 | 39646815 | 39648850 | 2036 | -7.01 | 3.87E-10 | 9.42E-10 |
| chr18 | 29575998 | 29577632 | 1635 | -7.01 | 3.87E-10 | 9.42E-10 |

| | | | | | | |
|---|---|---|---|---|---|---|
| chr5 | 33473468 | 33475482 | 2015 | -7.01 | 3.87E-10 | 9.43E-10 |
| chr6 | 145684013 | 145699150 | 15138 | -7.07 | 3.87E-10 | 9.44E-10 |
| chr7 | 16708891 | 16712737 | 3847 | -7.29 | 3.88E-10 | 9.46E-10 |
| chr17 | 15643965 | 15659739 | 15775 | -7.45 | 3.88E-10 | 9.46E-10 |
| chrX | 56955188 | 56961993 | 6806 | -4.08 | 3.89E-10 | 9.46E-10 |
| chr17 | 73349936 | 73351344 | 1409 | -7.04 | 3.89E-10 | 9.47E-10 |
| chr14 | 104494576 | 104541989 | 47414 | -7.01 | 3.89E-10 | 9.48E-10 |
| chr5 | 52308723 | 52310230 | 1508 | -7.13 | 3.92E-10 | 9.55E-10 |
| chr12 | 132427410 | 132429667 | 2258 | -7.11 | 3.92E-10 | 9.56E-10 |
| chr6 | 158812535 | 158815646 | 3112 | -7.15 | 3.93E-10 | 9.56E-10 |
| chr8 | 40432855 | 40434777 | 1923 | -7.15 | 3.93E-10 | 9.56E-10 |
| chr2 | 44358291 | 44361406 | 3116 | -7.62 | 3.93E-10 | 9.57E-10 |
| chr6 | 24506890 | 24508819 | 1930 | -7.34 | 3.93E-10 | 9.57E-10 |
| chr4 | 64359686 | 64365779 | 6094 | -7.16 | 3.93E-10 | 9.57E-10 |
| chr2 | 61154459 | 61155557 | 1099 | -7.16 | 3.95E-10 | 9.61E-10 |
| chr17 | 77987867 | 77989962 | 2096 | -7.03 | 3.95E-10 | 9.61E-10 |
| chr12 | 10300262 | 10311487 | 11226 | -7.17 | 3.96E-10 | 9.63E-10 |
| chr22 | 17863020 | 17865886 | 2867 | -7.09 | 3.96E-10 | 9.64E-10 |
| chr11 | 129642216 | 129644131 | 1916 | -7.01 | 3.97E-10 | 9.66E-10 |
| chr1 | 76410519 | 76416085 | 5567 | -7.09 | 3.97E-10 | 9.67E-10 |
| chr1 | 204298139 | 204299787 | 1649 | -7.17 | 3.98E-10 | 9.68E-10 |
| chr10 | 79016612 | 79018394 | 1783 | -7.17 | 3.98E-10 | 9.68E-10 |
| chr8 | 58451909 | 58453772 | 1864 | -7.02 | 3.98E-10 | 9.69E-10 |
| chr9 | 16128817 | 16131108 | 2292 | -7.02 | 3.98E-10 | 9.69E-10 |
| chr2 | 218861910 | 218864340 | 2431 | -7.18 | 3.98E-10 | 9.69E-10 |
| chr6 | 57084103 | 57085578 | 1476 | -7.04 | 3.99E-10 | 9.71E-10 |
| chrX | 13233019 | 13245907 | 12889 | -7.09 | 4.00E-10 | 9.72E-10 |
| chr8 | 91932548 | 91936499 | 3952 | 7.53 | 4.00E-10 | 9.74E-10 |
| chr11 | 92637012 | 92638480 | 1469 | -4.16 | 4.01E-10 | 9.76E-10 |
| chr8 | 61098523 | 61100619 | 2097 | -7.02 | 4.02E-10 | 9.77E-10 |
| chr11 | 9581066 | 9584203 | 3138 | -7.2 | 4.03E-10 | 9.80E-10 |
| chr5 | 60716934 | 60718418 | 1485 | -7.2 | 4.03E-10 | 9.80E-10 |
| chr1 | 221708904 | 221710848 | 1945 | -7 | 4.03E-10 | 9.81E-10 |
| chr4 | 184820339 | 184821827 | 1489 | -4.59 | 4.04E-10 | 9.83E-10 |
| chr4 | 6606517 | 6607579 | 1063 | -7.07 | 4.04E-10 | 9.83E-10 |
| chr12 | 46014839 | 46016413 | 1575 | -7.02 | 4.05E-10 | 9.83E-10 |
| chr4 | 188326359 | 188336683 | 10325 | -7.02 | 4.05E-10 | 9.83E-10 |
| chrX | 23437658 | 23441470 | 3813 | -3.76 | 4.05E-10 | 9.84E-10 |
| chr1 | 88069885 | 88086317 | 16433 | -7.34 | 4.06E-10 | 9.86E-10 |
| chr6 | 141800991 | 141808567 | 7577 | -7.34 | 4.06E-10 | 9.86E-10 |
| chr6 | 140683513 | 140687030 | 3518 | -7.07 | 4.07E-10 | 9.88E-10 |
| chr6 | 56582514 | 56584289 | 1776 | -7 | 4.07E-10 | 9.89E-10 |
| chr1 | 145672209 | 145674283 | 2075 | -7.01 | 4.07E-10 | 9.90E-10 |
| chr17 | 6970764 | 6972465 | 1702 | -7 | 4.08E-10 | 9.90E-10 |
| chr14 | 88927830 | 88929508 | 1679 | -7.21 | 4.09E-10 | 9.92E-10 |
| chr7 | 30478037 | 30485431 | 7395 | -7.03 | 4.09E-10 | 9.93E-10 |
| chr3 | 121442308 | 121443271 | 964 | -7.07 | 4.09E-10 | 9.93E-10 |
| chr19 | 34836062 | 34841390 | 5329 | -7.26 | 4.09E-10 | 9.93E-10 |
| chr2 | 54649512 | 54651233 | 1722 | -7 | 4.10E-10 | 9.96E-10 |
| chr12 | 18002792 | 18008763 | 5972 | -7.13 | 4.10E-10 | 9.96E-10 |
| chr4 | 179240250 | 179243876 | 3627 | -7.13 | 4.10E-10 | 9.96E-10 |
| chrX | 13333726 | 13340354 | 6629 | -7.06 | 4.11E-10 | 9.97E-10 |
| chr11 | 14672485 | 14674651 | 2167 | -7.14 | 4.11E-10 | 9.98E-10 |
| chr8 | 124506937 | 124508639 | 1703 | -7.06 | 4.12E-10 | 9.99E-10 |
| chr11 | 68975011 | 68977230 | 2220 | -7.16 | 4.13E-10 | 1.00E-09 |
| chr20 | 17911860 | 17913371 | 1512 | -7 | 4.14E-10 | 1.00E-09 |
| chrX | 18708454 | 18714196 | 5743 | -7.23 | 4.14E-10 | 1.01E-09 |
| chr6 | 54598897 | 54611230 | 12334 | -7.23 | 4.16E-10 | 1.01E-09 |
| chr6 | 125298797 | 125309660 | 10864 | -7.12 | 4.17E-10 | 1.01E-09 |
| chrX | 21224382 | 21263907 | 39526 | -6.38 | 4.17E-10 | 1.01E-09 |
| chr12 | 39275727 | 39285935 | 10209 | -7.28 | 4.17E-10 | 1.01E-09 |

| | | | | | | |
|---|---|---|---|---|---|---|
| chr10 | 115501884 | 115505421 | 3538 | -7 | 4.18E-10 | 1.01E-09 |
| chrX | 69610506 | 69614536 | 4031 | -7 | 4.18E-10 | 1.01E-09 |
| chr12 | 59625125 | 59672139 | 47015 | 1.95 | 4.18E-10 | 1.01E-09 |
| chr1 | 118349587 | 118351225 | 1639 | -3.35 | 4.19E-10 | 1.02E-09 |
| chr3 | 169788801 | 169791134 | 2334 | -7 | 4.19E-10 | 1.02E-09 |
| chr11 | 71988160 | 71993811 | 5652 | -7.24 | 4.19E-10 | 1.02E-09 |
| chr13 | 50624970 | 50626928 | 1959 | -7.05 | 4.20E-10 | 1.02E-09 |
| chr14 | 36387718 | 36389674 | 1957 | -7.05 | 4.20E-10 | 1.02E-09 |
| chr4 | 140912496 | 140917267 | 4772 | -7.53 | 4.21E-10 | 1.02E-09 |
| chr12 | 59048624 | 59055832 | 7209 | -7.16 | 4.21E-10 | 1.02E-09 |
| chr18 | 9762055 | 9764093 | 2039 | -7.01 | 4.21E-10 | 1.02E-09 |
| chr16 | 46701506 | 46702994 | 1489 | -7.16 | 4.21E-10 | 1.02E-09 |
| chr6 | 154791307 | 154792454 | 1148 | -7 | 4.22E-10 | 1.02E-09 |
| chrX | 20355010 | 20364304 | 9295 | -7.29 | 4.23E-10 | 1.03E-09 |
| chr7 | 123783949 | 123785893 | 1945 | 5.44 | 4.24E-10 | 1.03E-09 |
| chr1 | 207895668 | 207896803 | 1136 | -7.03 | 4.24E-10 | 1.03E-09 |
| chr19 | 10176317 | 10185502 | 9186 | -7.03 | 4.24E-10 | 1.03E-09 |
| chr17 | 19736541 | 19738241 | 1701 | -7.08 | 4.24E-10 | 1.03E-09 |
| chr5 | 82594591 | 82596353 | 1763 | -7.17 | 4.25E-10 | 1.03E-09 |
| chr2 | 62438830 | 62448367 | 9538 | -7.08 | 4.25E-10 | 1.03E-09 |
| chrX | 149489369 | 149492965 | 3597 | -7.08 | 4.25E-10 | 1.03E-09 |
| chr4 | 123296175 | 123310131 | 13957 | -7.04 | 4.28E-10 | 1.04E-09 |
| chr16 | 19523784 | 19525852 | 2069 | -7.44 | 4.28E-10 | 1.04E-09 |
| chr13 | 83785669 | 83789312 | 3644 | -7.01 | 4.28E-10 | 1.04E-09 |
| chr10 | 74352041 | 74353357 | 1317 | -6.99 | 4.29E-10 | 1.04E-09 |
| chr21 | 36198278 | 36201526 | 3249 | -7.19 | 4.29E-10 | 1.04E-09 |
| chr5 | 114554546 | 114555582 | 1037 | -5.61 | 4.29E-10 | 1.04E-09 |
| chr12 | 93662272 | 93666338 | 4067 | -7.01 | 4.29E-10 | 1.04E-09 |
| chr11 | 121095073 | 121099524 | 4452 | -2.35 | 4.29E-10 | 1.04E-09 |
| chr2 | 147351354 | 147353981 | 2628 | -7.21 | 4.29E-10 | 1.04E-09 |
| chr3 | 32460130 | 32464573 | 4444 | -7.01 | 4.30E-10 | 1.04E-09 |
| chr7 | 116218365 | 116227648 | 9284 | -7.3 | 4.30E-10 | 1.04E-09 |
| chr3 | 125746382 | 125748128 | 1747 | -6.99 | 4.31E-10 | 1.04E-09 |
| chrX | 13504609 | 13511181 | 6573 | -7.03 | 4.31E-10 | 1.04E-09 |
| chr2 | 196629022 | 196631119 | 2098 | -7.1 | 4.32E-10 | 1.05E-09 |
| chr5 | 110905878 | 110953123 | 47246 | 1.52 | 4.32E-10 | 1.05E-09 |
| chrX | 131325892 | 131328026 | 2135 | -7.09 | 4.32E-10 | 1.05E-09 |
| chr1 | 89304851 | 89306787 | 1937 | -7 | 4.33E-10 | 1.05E-09 |
| chr5 | 13827700 | 13829343 | 1644 | -6.03 | 4.33E-10 | 1.05E-09 |
| chr17 | 35229658 | 35240573 | 10916 | -7 | 4.35E-10 | 1.05E-09 |
| chr7 | 54663145 | 54671587 | 8443 | 2.07 | 4.35E-10 | 1.05E-09 |
| chr10 | 64083474 | 64115742 | 32269 | -4.22 | 4.36E-10 | 1.06E-09 |
| chr1 | 97025544 | 97028558 | 3015 | -7 | 4.36E-10 | 1.06E-09 |
| chr2 | 187177329 | 187184911 | 7583 | -7.01 | 4.37E-10 | 1.06E-09 |
| chr12 | 68292996 | 68301832 | 8837 | -7.05 | 4.38E-10 | 1.06E-09 |
| chr7 | 50803543 | 50806439 | 2897 | -7.49 | 4.38E-10 | 1.06E-09 |
| chrX | 40965644 | 40967434 | 1791 | -6.99 | 4.39E-10 | 1.06E-09 |
| chr12 | 113758899 | 113760983 | 2085 | -7 | 4.40E-10 | 1.06E-09 |
| chr13 | 29216778 | 29219600 | 2823 | -6.99 | 4.40E-10 | 1.06E-09 |
| chr1 | 30193614 | 30195982 | 2369 | -7.02 | 4.41E-10 | 1.07E-09 |
| chr4 | 135489941 | 135499578 | 9638 | 7.25 | 4.41E-10 | 1.07E-09 |
| chr1 | 246766415 | 246768280 | 1866 | -7.04 | 4.41E-10 | 1.07E-09 |
| chr10 | 87742204 | 87745807 | 3604 | 7.09 | 4.42E-10 | 1.07E-09 |
| chr14 | 92050185 | 92052503 | 2319 | -7.01 | 4.42E-10 | 1.07E-09 |
| chr17 | 48217480 | 48219819 | 2340 | -7.01 | 4.42E-10 | 1.07E-09 |
| chr13 | 95784145 | 95798723 | 14579 | -7.18 | 4.43E-10 | 1.07E-09 |
| chr8 | 81815454 | 81818497 | 3044 | -7.22 | 4.43E-10 | 1.07E-09 |
| chr15 | 49113448 | 49126483 | 13036 | -7.24 | 4.44E-10 | 1.07E-09 |
| chr3 | 99915191 | 99924582 | 9392 | -6.99 | 4.44E-10 | 1.07E-09 |
| chr5 | 61949031 | 61951712 | 2682 | -7.14 | 4.45E-10 | 1.08E-09 |
| chr9 | 136829193 | 136831008 | 1816 | -6.99 | 4.45E-10 | 1.08E-09 |

| | | | | | | |
|---|---|---|---|---|---|---|
| chr7 | 73075937 | 73078409 | 2473 | -7.02 | 4.45E-10 | 1.08E-09 |
| chr12 | 1657813 | 1666834 | 9022 | -7.02 | 4.46E-10 | 1.08E-09 |
| chr2 | 224958590 | 224963070 | 4481 | -7.23 | 4.46E-10 | 1.08E-09 |
| chr1 | 84894631 | 84897436 | 2806 | -6.99 | 4.48E-10 | 1.08E-09 |
| chr3 | 45045209 | 45046967 | 1759 | -6.99 | 4.48E-10 | 1.08E-09 |
| chr2 | 169107543 | 169116230 | 8688 | -7.25 | 4.48E-10 | 1.08E-09 |
| chr15 | 57314209 | 57316332 | 2124 | -6.99 | 4.49E-10 | 1.08E-09 |
| chr4 | 114712452 | 114726063 | 13612 | -6.99 | 4.49E-10 | 1.08E-09 |
| chr12 | 12039630 | 12041663 | 2034 | -7.02 | 4.49E-10 | 1.09E-09 |
| chr9 | 120058767 | 120062650 | 3884 | 7.07 | 4.50E-10 | 1.09E-09 |
| chr7 | 13414330 | 13424718 | 10389 | -7.37 | 4.50E-10 | 1.09E-09 |
| chr7 | 8219084 | 8221901 | 2818 | -2.82 | 4.52E-10 | 1.09E-09 |
| chrX | 138920444 | 138921921 | 1478 | -7.11 | 4.52E-10 | 1.09E-09 |
| chr11 | 130410704 | 130413869 | 3166 | -7.1 | 4.53E-10 | 1.09E-09 |
| chr14 | 103029063 | 103030906 | 1844 | -7.13 | 4.53E-10 | 1.09E-09 |
| chr4 | 122398079 | 122408469 | 10391 | -6.99 | 4.53E-10 | 1.09E-09 |
| chr9 | 124139853 | 124141235 | 1383 | -7.03 | 4.53E-10 | 1.09E-09 |
| chr18 | 33786979 | 33788623 | 1645 | -7.09 | 4.54E-10 | 1.10E-09 |
| chr3 | 98204432 | 98207121 | 2690 | 7.05 | 4.54E-10 | 1.10E-09 |
| chr19 | 18151358 | 18152736 | 1379 | -6.99 | 4.54E-10 | 1.10E-09 |
| chr3 | 29612585 | 29614334 | 1750 | -6.99 | 4.55E-10 | 1.10E-09 |
| chr19 | 17841346 | 17842611 | 1266 | -6.98 | 4.55E-10 | 1.10E-09 |
| chr1 | 171856841 | 171862055 | 5215 | -7.44 | 4.55E-10 | 1.10E-09 |
| chr1 | 51272395 | 51274953 | 2559 | -7.01 | 4.56E-10 | 1.10E-09 |
| chr11 | 2568307 | 2575355 | 7049 | -7.26 | 4.57E-10 | 1.10E-09 |
| chrX | 38575932 | 38579362 | 3431 | -3 | 4.58E-10 | 1.10E-09 |
| chr2 | 190575782 | 190579835 | 4054 | -7.07 | 4.58E-10 | 1.11E-09 |
| chr20 | 40107197 | 40108492 | 1296 | -7 | 4.59E-10 | 1.11E-09 |
| chr5 | 39133976 | 39142030 | 8055 | -7.02 | 4.60E-10 | 1.11E-09 |
| chr4 | 97291289 | 97306686 | 15398 | -6.99 | 4.62E-10 | 1.11E-09 |
| chr17 | 734134 | 736220 | 2087 | -6.99 | 4.62E-10 | 1.11E-09 |
| chr6 | 10009180 | 10011404 | 2225 | -6.99 | 4.62E-10 | 1.11E-09 |
| chr8 | 126641142 | 126643353 | 2212 | -7.01 | 4.62E-10 | 1.12E-09 |
| chr19 | 3272421 | 3273903 | 1483 | -7.02 | 4.63E-10 | 1.12E-09 |
| chr12 | 87517609 | 87532772 | 15164 | -7.17 | 4.63E-10 | 1.12E-09 |
| chr8 | 142666311 | 142669188 | 2878 | 5.81 | 4.63E-10 | 1.12E-09 |
| chrX | 16922833 | 16925731 | 2899 | -7.03 | 4.63E-10 | 1.12E-09 |
| chr13 | 109068553 | 109071204 | 2652 | -7.02 | 4.64E-10 | 1.12E-09 |
| chr1 | 84623229 | 84628644 | 5416 | -6.98 | 4.64E-10 | 1.12E-09 |
| chr6 | 165856074 | 165858082 | 2009 | -7 | 4.64E-10 | 1.12E-09 |
| chr7 | 107458161 | 107481179 | 23019 | -7.66 | 4.65E-10 | 1.12E-09 |
| chr12 | 93699456 | 93701175 | 1720 | -6.98 | 4.66E-10 | 1.12E-09 |
| chr1 | 153591775 | 153593461 | 1687 | -7.05 | 4.66E-10 | 1.12E-09 |
| chr10 | 103332007 | 103333444 | 1438 | -7 | 4.67E-10 | 1.12E-09 |
| chr3 | 136411785 | 136415759 | 3975 | -7.03 | 4.67E-10 | 1.13E-09 |
| chr6 | 14561586 | 14566533 | 4948 | -7 | 4.68E-10 | 1.13E-09 |
| chr22 | 39078923 | 39080195 | 1273 | -6.99 | 4.68E-10 | 1.13E-09 |
| chr8 | 110319672 | 110323591 | 3920 | -7 | 4.69E-10 | 1.13E-09 |
| chr9 | 136584912 | 136589626 | 4715 | -7.43 | 4.70E-10 | 1.13E-09 |
| chr6 | 135099945 | 135113926 | 13982 | -7.28 | 4.71E-10 | 1.14E-09 |
| chr16 | 87455043 | 87456729 | 1687 | -6.99 | 4.72E-10 | 1.14E-09 |
| chr17 | 68641856 | 68644112 | 2257 | -7.02 | 4.72E-10 | 1.14E-09 |
| chr8 | 101041399 | 101044096 | 2698 | -7.02 | 4.72E-10 | 1.14E-09 |
| chr12 | 42317947 | 42320112 | 2166 | -7.09 | 4.72E-10 | 1.14E-09 |
| chr6 | 63460549 | 63467315 | 6767 | -7.24 | 4.73E-10 | 1.14E-09 |
| chr8 | 124962476 | 124965082 | 2607 | -7.1 | 4.74E-10 | 1.14E-09 |
| chr4 | 117837419 | 117842874 | 5456 | -7.11 | 4.74E-10 | 1.14E-09 |
| chr22 | 29065170 | 29067364 | 2195 | -7.03 | 4.75E-10 | 1.14E-09 |
| chr22 | 27532036 | 27533454 | 1419 | -7.01 | 4.76E-10 | 1.15E-09 |
| chr5 | 114912199 | 114914273 | 2075 | -7.14 | 4.76E-10 | 1.15E-09 |
| chr4 | 108509991 | 108512984 | 2994 | -7.02 | 4.76E-10 | 1.15E-09 |

| | | | | | | |
|---|---|---|---|---|---|---|
| chr2 | 53624754 | 53638325 | 13572 | -7.08 | 4.77E-10 | 1.15E-09 |
| chr6 | 160703687 | 160707026 | 3340 | -7.2 | 4.77E-10 | 1.15E-09 |
| chr11 | 21457621 | 21479225 | 21605 | -7.89 | 4.77E-10 | 1.15E-09 |
| chr11 | 77982503 | 77985291 | 2789 | -7.08 | 4.77E-10 | 1.15E-09 |
| chr15 | 45861949 | 45868047 | 6099 | -7.07 | 4.79E-10 | 1.15E-09 |
| chr6 | 75126035 | 75132198 | 6164 | -7.07 | 4.79E-10 | 1.15E-09 |
| chr19 | 3467285 | 3468632 | 1348 | -6.98 | 4.79E-10 | 1.15E-09 |
| chr9 | 107947903 | 107951811 | 3909 | -6.98 | 4.80E-10 | 1.16E-09 |
| chr10 | 64504566 | 64509953 | 5388 | 7.07 | 4.81E-10 | 1.16E-09 |
| chr13 | 100283751 | 100286035 | 2285 | -7.01 | 4.81E-10 | 1.16E-09 |
| chr13 | 109791444 | 109805734 | 14291 | -7.01 | 4.81E-10 | 1.16E-09 |
| chr20 | 48271829 | 48273241 | 1413 | -7.05 | 4.82E-10 | 1.16E-09 |
| chr9 | 29098401 | 29099930 | 1530 | -7.05 | 4.82E-10 | 1.16E-09 |
| chr2 | 193876065 | 193909526 | 33462 | -7.71 | 4.84E-10 | 1.16E-09 |
| chr10 | 21559622 | 21580416 | 20795 | -7.68 | 4.84E-10 | 1.16E-09 |
| chr9 | 90119783 | 90133011 | 13229 | -7.41 | 4.84E-10 | 1.17E-09 |
| chr9 | 113807031 | 113827523 | 20493 | -7.72 | 4.87E-10 | 1.17E-09 |
| chr4 | 130106828 | 130118006 | 11179 | -7.22 | 4.87E-10 | 1.17E-09 |
| chr1 | 79789696 | 79792805 | 3110 | -7.02 | 4.87E-10 | 1.17E-09 |
| chr12 | 1570214 | 1571731 | 1518 | -7.02 | 4.87E-10 | 1.17E-09 |
| chr18 | 11679687 | 11684816 | 5130 | -2.57 | 4.87E-10 | 1.17E-09 |
| chr4 | 173561830 | 173569902 | 8073 | -6.97 | 4.89E-10 | 1.17E-09 |
| chr4 | 111139508 | 111141360 | 1853 | -6.97 | 4.89E-10 | 1.18E-09 |
| chr2 | 238815317 | 238816907 | 1591 | -6.98 | 4.89E-10 | 1.18E-09 |
| chr2 | 71855791 | 71857183 | 1393 | -7 | 4.90E-10 | 1.18E-09 |
| chr2 | 61338767 | 61340769 | 2003 | -7.3 | 4.90E-10 | 1.18E-09 |
| chr13 | 36709025 | 36764580 | 55556 | 1.28 | 4.91E-10 | 1.18E-09 |
| chr11 | 116290737 | 116292111 | 1375 | -6.97 | 4.91E-10 | 1.18E-09 |
| chr18 | 20828018 | 20829946 | 1929 | -7.01 | 4.91E-10 | 1.18E-09 |
| chr3 | 182342157 | 182343889 | 1733 | -7.01 | 4.91E-10 | 1.18E-09 |
| chr8 | 40396724 | 40411074 | 14351 | -7.27 | 4.92E-10 | 1.18E-09 |
| chr1 | 41669070 | 41670839 | 1770 | -7.04 | 4.92E-10 | 1.18E-09 |
| chr18 | 60398635 | 60400632 | 1998 | -6.99 | 4.93E-10 | 1.18E-09 |
| chr5 | 90290103 | 90293308 | 3206 | -6.99 | 4.93E-10 | 1.18E-09 |
| chr18 | 71187318 | 71196111 | 8794 | -7.18 | 4.94E-10 | 1.19E-09 |
| chr9 | 6862873 | 6865486 | 2614 | -7.03 | 4.95E-10 | 1.19E-09 |
| chr3 | 177810389 | 177813529 | 3141 | -7.19 | 4.96E-10 | 1.19E-09 |
| chr5 | 155852999 | 155857236 | 4238 | -7.46 | 4.97E-10 | 1.19E-09 |
| chr3 | 110503916 | 110506293 | 2378 | -7.09 | 4.98E-10 | 1.20E-09 |
| chr14 | 65974121 | 65976949 | 2829 | -7.09 | 4.98E-10 | 1.20E-09 |
| chr7 | 131350722 | 131352208 | 1487 | -5.1 | 4.98E-10 | 1.20E-09 |
| chr3 | 138205973 | 138208427 | 2455 | -7.08 | 4.99E-10 | 1.20E-09 |
| chr1 | 86699228 | 86702941 | 3714 | -2.58 | 4.99E-10 | 1.20E-09 |
| chr3 | 57633856 | 57636095 | 2240 | -6.97 | 4.99E-10 | 1.20E-09 |
| chr17 | 17517243 | 17519573 | 2331 | -6.98 | 4.99E-10 | 1.20E-09 |
| chr17 | 7768322 | 7769394 | 1073 | -7.03 | 5.01E-10 | 1.20E-09 |
| chr6 | 27913405 | 27917132 | 3728 | -7.11 | 5.01E-10 | 1.20E-09 |
| chr1 | 111240801 | 111242245 | 1445 | -6.98 | 5.02E-10 | 1.20E-09 |
| chr10 | 81767192 | 81769555 | 2364 | -6.97 | 5.02E-10 | 1.20E-09 |
| chr20 | 60126267 | 60128297 | 2031 | -7.2 | 5.03E-10 | 1.21E-09 |
| chr15 | 89621685 | 89623461 | 1777 | -7.06 | 5.03E-10 | 1.21E-09 |
| chr4 | 3024542 | 3026017 | 1476 | -7.06 | 5.03E-10 | 1.21E-09 |
| chr10 | 71194391 | 71204856 | 10466 | -6.99 | 5.04E-10 | 1.21E-09 |
| chr6 | 161701109 | 161703192 | 2084 | -7 | 5.04E-10 | 1.21E-09 |
| chr3 | 27517658 | 27519654 | 1997 | -6.98 | 5.05E-10 | 1.21E-09 |
| chr15 | 31067072 | 31074682 | 7611 | -7.14 | 5.05E-10 | 1.21E-09 |
| chr6 | 35848547 | 35849758 | 1212 | -7.36 | 5.05E-10 | 1.21E-09 |
| chr5 | 129395301 | 129396521 | 1221 | -6.97 | 5.06E-10 | 1.21E-09 |
| chr5 | 57187203 | 57190348 | 3146 | -6.97 | 5.06E-10 | 1.21E-09 |
| chr9 | 97483435 | 97484745 | 1311 | -6.97 | 5.06E-10 | 1.21E-09 |
| chr3 | 82019330 | 82041537 | 22208 | 6.13 | 5.07E-10 | 1.22E-09 |

| | | | | | | |
|---|---|---|---|---|---|---|
| chr9 | 119901963 | 119905940 | 3978 | 7.45 | 5.08E-10 | 1.22E-09 |
| chr20 | 18381472 | 18383331 | 1860 | -7.02 | 5.09E-10 | 1.22E-09 |
| chr4 | 93716621 | 93718517 | 1897 | -6.02 | 5.09E-10 | 1.22E-09 |
| chr5 | 6779138 | 6781574 | 2437 | -6.97 | 5.10E-10 | 1.22E-09 |
| chr13 | 84967022 | 84974077 | 7056 | -7.21 | 5.11E-10 | 1.23E-09 |
| chr13 | 84915280 | 84928173 | 12894 | -7.04 | 5.12E-10 | 1.23E-09 |
| chr7 | 95903473 | 95905592 | 2120 | -6.98 | 5.12E-10 | 1.23E-09 |
| chr13 | 94609718 | 94614484 | 4767 | -7.01 | 5.12E-10 | 1.23E-09 |
| chr7 | 27357921 | 27373887 | 15967 | -7.26 | 5.14E-10 | 1.23E-09 |
| chr20 | 16245509 | 16247582 | 2074 | -7.04 | 5.14E-10 | 1.23E-09 |
| chrX | 8903974 | 8909285 | 5312 | -7.34 | 5.17E-10 | 1.24E-09 |
| chr10 | 3607518 | 3609070 | 1553 | -6.97 | 5.18E-10 | 1.24E-09 |
| chr9 | 124838804 | 124841028 | 2225 | -7.17 | 5.18E-10 | 1.24E-09 |
| chr4 | 74668804 | 74671657 | 2854 | -6.99 | 5.18E-10 | 1.24E-09 |
| chr2 | 203112404 | 203114165 | 1762 | -6.97 | 5.19E-10 | 1.24E-09 |
| chr13 | 33879959 | 33893708 | 13750 | -6.98 | 5.19E-10 | 1.24E-09 |
| chr18 | 48018412 | 48097098 | 78687 | -2.13 | 5.20E-10 | 1.25E-09 |
| chr7 | 79690345 | 79716976 | 26632 | -7.66 | 5.21E-10 | 1.25E-09 |
| chr2 | 191572434 | 191581837 | 9404 | -7.45 | 5.21E-10 | 1.25E-09 |
| chr9 | 37319694 | 37320959 | 1266 | -7.17 | 5.21E-10 | 1.25E-09 |
| chr2 | 75529105 | 75536161 | 7057 | -7.27 | 5.22E-10 | 1.25E-09 |
| chr4 | 8498162 | 8500589 | 2428 | -7.09 | 5.23E-10 | 1.25E-09 |
| chr17 | 53164438 | 53166164 | 1727 | -7.08 | 5.23E-10 | 1.25E-09 |
| chr3 | 100125796 | 100130894 | 5099 | -7.11 | 5.25E-10 | 1.26E-09 |
| chr16 | 77460830 | 77463574 | 2745 | -5.96 | 5.26E-10 | 1.26E-09 |
| chr12 | 40556928 | 40559462 | 2535 | -7.24 | 5.26E-10 | 1.26E-09 |
| chr7 | 13148942 | 13151503 | 2562 | -7.02 | 5.27E-10 | 1.26E-09 |
| chr9 | 20839584 | 20842371 | 2788 | -7.06 | 5.28E-10 | 1.26E-09 |
| chr14 | 96010443 | 96018242 | 7800 | -7.3 | 5.29E-10 | 1.27E-09 |
| chr10 | 100928397 | 100932676 | 4280 | 7.34 | 5.30E-10 | 1.27E-09 |
| chr5 | 72193734 | 72200349 | 6616 | -7.44 | 5.30E-10 | 1.27E-09 |
| chr1 | 12120135 | 12121481 | 1347 | -6.96 | 5.31E-10 | 1.27E-09 |
| chr14 | 85568560 | 85612685 | 44126 | 1.94 | 5.32E-10 | 1.27E-09 |
| chr12 | 18115319 | 18119085 | 3767 | -7.2 | 5.32E-10 | 1.27E-09 |
| chr7 | 116668809 | 116671423 | 2615 | -7.14 | 5.33E-10 | 1.28E-09 |
| chr11 | 69421957 | 69424136 | 2180 | -7.04 | 5.33E-10 | 1.28E-09 |
| chr22 | 42945580 | 42947938 | 2359 | -7.04 | 5.33E-10 | 1.28E-09 |
| chr9 | 88299430 | 88301094 | 1665 | -7.1 | 5.33E-10 | 1.28E-09 |
| chr18 | 46831976 | 46833310 | 1335 | -6.96 | 5.34E-10 | 1.28E-09 |
| chr6 | 126094496 | 126095879 | 1384 | -7.11 | 5.34E-10 | 1.28E-09 |
| chr7 | 67677373 | 67679913 | 2541 | -6.96 | 5.35E-10 | 1.28E-09 |
| chr19 | 49790156 | 49793812 | 3657 | -7.04 | 5.35E-10 | 1.28E-09 |
| chr14 | 76137608 | 76139428 | 1821 | -6.98 | 5.36E-10 | 1.28E-09 |
| chr11 | 103411301 | 103413919 | 2619 | -5.16 | 5.37E-10 | 1.28E-09 |
| chr5 | 127912625 | 127919752 | 7128 | -7.31 | 5.39E-10 | 1.29E-09 |
| chr2 | 12249237 | 12847687 | 598451 | 0.79 | 5.41E-10 | 1.29E-09 |
| chr2 | 178637942 | 178640874 | 2933 | -5.31 | 5.42E-10 | 1.30E-09 |
| chr2 | 152105162 | 152106839 | 1678 | -7 | 5.44E-10 | 1.30E-09 |
| chr1 | 173076073 | 173078778 | 2706 | -6.99 | 5.45E-10 | 1.30E-09 |
| chr5 | 136818035 | 136827338 | 9304 | 7.7 | 5.45E-10 | 1.30E-09 |
| chr10 | 33113747 | 33115342 | 1596 | -7.02 | 5.46E-10 | 1.31E-09 |
| chr1 | 86543830 | 86545588 | 1759 | -3.61 | 5.47E-10 | 1.31E-09 |
| chr1 | 224234665 | 224236098 | 1434 | -7.22 | 5.47E-10 | 1.31E-09 |
| chr18 | 29728342 | 29731113 | 2772 | -6.96 | 5.47E-10 | 1.31E-09 |
| chr4 | 159004072 | 159013424 | 9353 | -7 | 5.48E-10 | 1.31E-09 |
| chr6 | 143441433 | 143459523 | 18091 | -7.77 | 5.49E-10 | 1.31E-09 |
| chr12 | 63264157 | 63265520 | 1364 | -6.97 | 5.49E-10 | 1.31E-09 |
| chr2 | 63651680 | 63653810 | 2131 | -7.23 | 5.50E-10 | 1.31E-09 |
| chr4 | 81297780 | 81301913 | 4134 | 7.23 | 5.50E-10 | 1.31E-09 |
| chr16 | 47641307 | 47648512 | 7206 | -7.1 | 5.51E-10 | 1.32E-09 |
| chr6 | 19954139 | 20032146 | 78008 | 2.82 | 5.51E-10 | 1.32E-09 |

| | | | | | | |
|---|---|---|---|---|---|---|
| chr4 | 185196781 | 185199622 | 2842 | -7.06 | 5.51E-10 | 1.32E-09 |
| chr6 | 44832279 | 44838298 | 6020 | -6.98 | 5.52E-10 | 1.32E-09 |
| chr6 | 75170590 | 75173440 | 2851 | -7.42 | 5.52E-10 | 1.32E-09 |
| chr16 | 86170104 | 86222730 | 52627 | -1.94 | 5.52E-10 | 1.32E-09 |
| chr4 | 69072461 | 69078990 | 6530 | -2.56 | 5.52E-10 | 1.32E-09 |
| chr2 | 150562861 | 150564210 | 1350 | -7.11 | 5.52E-10 | 1.32E-09 |
| chr2 | 237408996 | 237432806 | 23811 | -3.05 | 5.54E-10 | 1.32E-09 |
| chr10 | 104981115 | 104985376 | 4262 | -7.6 | 5.56E-10 | 1.33E-09 |
| chr2 | 160495352 | 160497818 | 2467 | -6.95 | 5.57E-10 | 1.33E-09 |
| chr1 | 241607732 | 241609449 | 1718 | -7.23 | 5.58E-10 | 1.33E-09 |
| chr6 | 126172936 | 126174791 | 1856 | -6.98 | 5.58E-10 | 1.33E-09 |
| chr6 | 140753910 | 140756425 | 2516 | -7.04 | 5.58E-10 | 1.33E-09 |
| chr3 | 104786598 | 104794473 | 7876 | -7.36 | 5.59E-10 | 1.33E-09 |
| chr1 | 180971607 | 180973044 | 1438 | -6.95 | 5.60E-10 | 1.34E-09 |
| chr2 | 72471380 | 72472586 | 1207 | -6.95 | 5.60E-10 | 1.34E-09 |
| chr2 | 179853303 | 179861234 | 7932 | -6.99 | 5.60E-10 | 1.34E-09 |
| chr15 | 42407212 | 42409752 | 2541 | -6.96 | 5.60E-10 | 1.34E-09 |
| chr4 | 133072838 | 133079008 | 6171 | -6.96 | 5.60E-10 | 1.34E-09 |
| chr1 | 65223743 | 65237747 | 14005 | -7.29 | 5.62E-10 | 1.34E-09 |
| chr13 | 102511611 | 102513216 | 1606 | 5.41 | 5.62E-10 | 1.34E-09 |
| chr5 | 142707221 | 142709581 | 2361 | -6.97 | 5.63E-10 | 1.34E-09 |
| chr8 | 122428445 | 122430333 | 1889 | -6.95 | 5.63E-10 | 1.34E-09 |
| chr5 | 40218048 | 40220416 | 2369 | -3.05 | 5.64E-10 | 1.35E-09 |
| chr14 | 52600255 | 52675237 | 74983 | -2.36 | 5.65E-10 | 1.35E-09 |
| chr3 | 128766683 | 128768487 | 1805 | -7.03 | 5.65E-10 | 1.35E-09 |
| chr9 | 127873659 | 127877189 | 3531 | -6.99 | 5.66E-10 | 1.35E-09 |
| chr17 | 8495635 | 8498350 | 2716 | -7.08 | 5.66E-10 | 1.35E-09 |
| chr5 | 40423434 | 40427013 | 3580 | -7.15 | 5.66E-10 | 1.35E-09 |
| chr1 | 168855203 | 168859310 | 4108 | -7.4 | 5.66E-10 | 1.35E-09 |
| chr12 | 69391229 | 69395270 | 4042 | -7.09 | 5.67E-10 | 1.35E-09 |
| chr6 | 135267686 | 135282529 | 14844 | -7.02 | 5.68E-10 | 1.35E-09 |
| chr5 | 136656724 | 136658709 | 1986 | -7.06 | 5.68E-10 | 1.35E-09 |
| chr14 | 45438768 | 45440203 | 1436 | -7.1 | 5.69E-10 | 1.36E-09 |
| chr12 | 45674327 | 45675802 | 1476 | -7 | 5.70E-10 | 1.36E-09 |
| chr11 | 94957315 | 94958863 | 1549 | -6.95 | 5.73E-10 | 1.37E-09 |
| chr17 | 4590856 | 4593374 | 2519 | -6.95 | 5.73E-10 | 1.37E-09 |
| chr1 | 98910742 | 98913788 | 3047 | -6.95 | 5.74E-10 | 1.37E-09 |
| chrX | 129790961 | 129793067 | 2107 | -6.95 | 5.74E-10 | 1.37E-09 |
| chr12 | 98956441 | 98958317 | 1877 | -7 | 5.74E-10 | 1.37E-09 |
| chr12 | 72302381 | 72304501 | 2121 | -7.12 | 5.74E-10 | 1.37E-09 |
| chr1 | 194108760 | 194110482 | 1723 | -6.97 | 5.74E-10 | 1.37E-09 |
| chr1 | 98877560 | 98885667 | 8108 | -7.01 | 5.74E-10 | 1.37E-09 |
| chr8 | 119171547 | 119173784 | 2238 | -6.96 | 5.75E-10 | 1.37E-09 |
| chr12 | 14147609 | 14223572 | 75964 | 0.97 | 5.75E-10 | 1.37E-09 |
| chr6 | 128865059 | 128867449 | 2391 | -7.13 | 5.76E-10 | 1.37E-09 |
| chr4 | 183730241 | 183734147 | 3907 | -6.95 | 5.76E-10 | 1.37E-09 |
| chr9 | 76297688 | 76298888 | 1201 | -6.95 | 5.77E-10 | 1.37E-09 |
| chr2 | 96762310 | 96772605 | 10296 | -7.22 | 5.77E-10 | 1.37E-09 |
| chr1 | 194523591 | 194529485 | 5895 | -7.07 | 5.77E-10 | 1.38E-09 |
| chr8 | 104399168 | 104401131 | 1964 | -7.26 | 5.79E-10 | 1.38E-09 |
| chr7 | 135119597 | 135120972 | 1376 | -7.13 | 5.79E-10 | 1.38E-09 |
| chr7 | 94150432 | 94151911 | 1480 | -7.03 | 5.80E-10 | 1.38E-09 |
| chr9 | 80472895 | 80479435 | 6541 | -7.05 | 5.80E-10 | 1.38E-09 |
| chr12 | 50916178 | 50918129 | 1952 | -6.95 | 5.80E-10 | 1.38E-09 |
| chr8 | 53669678 | 53671268 | 1591 | -5.47 | 5.81E-10 | 1.38E-09 |
| chr7 | 26075420 | 26100312 | 24893 | -2.85 | 5.81E-10 | 1.38E-09 |
| chr5 | 78140012 | 78151418 | 11407 | -6.95 | 5.81E-10 | 1.38E-09 |
| chr12 | 50684348 | 50686648 | 2301 | -7.04 | 5.82E-10 | 1.38E-09 |
| chr1 | 171420713 | 171423141 | 2429 | -6.96 | 5.82E-10 | 1.39E-09 |
| chr7 | 35438627 | 35441286 | 2660 | -6.97 | 5.83E-10 | 1.39E-09 |
| chr3 | 187694442 | 187696268 | 1827 | -7.04 | 5.85E-10 | 1.39E-09 |

| | | | | | | |
|---|---|---|---|---|---|---|
| chr13 | 23882938 | 23886012 | 3075 | -6.95 | 5.86E-10 | 1.39E-09 |
| chr5 | 169859317 | 169860730 | 1414 | -6.95 | 5.86E-10 | 1.39E-09 |
| chrX | 40822840 | 40838168 | 15329 | -7.46 | 5.87E-10 | 1.40E-09 |
| chr2 | 72727858 | 72729565 | 1708 | -6.97 | 5.87E-10 | 1.40E-09 |
| chr19 | 16285677 | 16288548 | 2872 | -7.27 | 5.88E-10 | 1.40E-09 |
| chr18 | 28888614 | 28891167 | 2554 | -5.2 | 5.89E-10 | 1.40E-09 |
| chr20 | 8836920 | 8838268 | 1349 | -6.98 | 5.89E-10 | 1.40E-09 |
| chr21 | 46202668 | 46204415 | 1748 | -6.97 | 5.89E-10 | 1.40E-09 |
| chr2 | 177113780 | 177118693 | 4914 | -7 | 5.91E-10 | 1.41E-09 |
| chr3 | 114280177 | 114283567 | 3391 | -7.02 | 5.91E-10 | 1.41E-09 |
| chr2 | 190398495 | 190400848 | 2354 | -6.96 | 5.93E-10 | 1.41E-09 |
| chr3 | 193540432 | 193542729 | 2298 | -6.95 | 5.94E-10 | 1.41E-09 |
| chr1 | 110861638 | 110873148 | 11511 | -7.2 | 5.95E-10 | 1.41E-09 |
| chr12 | 117564447 | 117567046 | 2600 | -6.99 | 5.95E-10 | 1.42E-09 |
| chr17 | 19660683 | 19664578 | 3896 | -7.01 | 5.97E-10 | 1.42E-09 |
| chr3 | 69800741 | 69803807 | 3067 | -7.15 | 5.97E-10 | 1.42E-09 |
| chr2 | 191485142 | 191490151 | 5010 | -6.95 | 5.97E-10 | 1.42E-09 |
| chr2 | 48778524 | 48780330 | 1807 | -6.96 | 5.98E-10 | 1.42E-09 |
| chr12 | 86617931 | 86620719 | 2789 | -7 | 5.98E-10 | 1.42E-09 |
| chr4 | 148483912 | 148488821 | 4910 | -6.96 | 6.03E-10 | 1.43E-09 |
| chr2 | 204117008 | 204118999 | 1992 | -7.06 | 6.03E-10 | 1.43E-09 |
| chr13 | 35227182 | 35235173 | 7992 | -7 | 6.04E-10 | 1.44E-09 |
| chr6 | 138280940 | 138282769 | 1830 | -7.16 | 6.04E-10 | 1.44E-09 |
| chr9 | 125775843 | 125777304 | 1462 | -7.09 | 6.04E-10 | 1.44E-09 |
| chr12 | 16480152 | 16497795 | 17644 | -5.98 | 6.05E-10 | 1.44E-09 |
| chr18 | 42567795 | 42570375 | 2581 | -7.05 | 6.05E-10 | 1.44E-09 |
| chr4 | 14442579 | 14449109 | 6531 | -7.48 | 6.06E-10 | 1.44E-09 |
| chr17 | 80471158 | 80473535 | 2378 | -7.21 | 6.06E-10 | 1.44E-09 |
| chr5 | 56791637 | 56797075 | 5439 | -7.1 | 6.06E-10 | 1.44E-09 |
| chr19 | 5481112 | 5482528 | 1417 | -7.16 | 6.06E-10 | 1.44E-09 |
| chr1 | 111120626 | 111121945 | 1320 | -7.07 | 6.07E-10 | 1.44E-09 |
| chr21 | 44121053 | 44122893 | 1841 | -7.07 | 6.07E-10 | 1.44E-09 |
| chr1 | 232975776 | 232978276 | 2501 | -7.08 | 6.07E-10 | 1.44E-09 |
| chr3 | 177241865 | 177243754 | 1890 | -7.05 | 6.07E-10 | 1.44E-09 |
| chr4 | 103490931 | 103492028 | 1098 | -6.94 | 6.08E-10 | 1.44E-09 |
| chr10 | 32353591 | 32354918 | 1328 | -7.05 | 6.08E-10 | 1.44E-09 |
| chr4 | 122787614 | 122789217 | 1604 | -7.1 | 6.08E-10 | 1.44E-09 |
| chr22 | 48115292 | 48130113 | 14822 | -2.23 | 6.09E-10 | 1.45E-09 |
| chr12 | 101979506 | 101981846 | 2341 | -2.98 | 6.09E-10 | 1.45E-09 |
| chr6 | 97657148 | 97659878 | 2731 | -6.95 | 6.10E-10 | 1.45E-09 |
| chr4 | 158823232 | 158832417 | 9186 | 7.07 | 6.10E-10 | 1.45E-09 |
| chr20 | 60546933 | 60550102 | 3170 | -6.97 | 6.11E-10 | 1.45E-09 |
| chr6 | 157630262 | 157631617 | 1356 | -6.98 | 6.13E-10 | 1.45E-09 |
| chr20 | 20708991 | 21099445 | 390455 | 1.28 | 6.14E-10 | 1.46E-09 |
| chr15 | 41970779 | 41972153 | 1375 | -6.94 | 6.14E-10 | 1.46E-09 |
| chr14 | 31490001 | 31491616 | 1616 | -7.02 | 6.14E-10 | 1.46E-09 |
| chr19 | 2627954 | 2631280 | 3327 | -6.95 | 6.15E-10 | 1.46E-09 |
| chr3 | 12326402 | 12327345 | 944 | -6.95 | 6.15E-10 | 1.46E-09 |
| chr6 | 25945290 | 25961338 | 16049 | -7.48 | 6.16E-10 | 1.46E-09 |
| chr4 | 107082636 | 107085174 | 2539 | -6.99 | 6.17E-10 | 1.46E-09 |
| chr1 | 210584847 | 210598580 | 13734 | -7.63 | 6.18E-10 | 1.47E-09 |
| chr5 | 78957243 | 78958838 | 1596 | -7.01 | 6.18E-10 | 1.47E-09 |
| chrX | 46621374 | 46623754 | 2381 | -6.94 | 6.19E-10 | 1.47E-09 |
| chr20 | 32554795 | 32566553 | 11759 | -7.63 | 6.20E-10 | 1.47E-09 |
| chr16 | 87087166 | 87088912 | 1747 | -6.94 | 6.21E-10 | 1.47E-09 |
| chr11 | 85933808 | 85936729 | 2922 | -3.14 | 6.21E-10 | 1.47E-09 |
| chr10 | 63690580 | 63693731 | 3152 | -7.23 | 6.22E-10 | 1.47E-09 |
| chr8 | 19631314 | 19656720 | 25407 | -8.04 | 6.22E-10 | 1.48E-09 |
| chr3 | 66149688 | 66153232 | 3545 | -2.8 | 6.23E-10 | 1.48E-09 |
| chr6 | 144046943 | 144050224 | 3282 | -6.94 | 6.23E-10 | 1.48E-09 |
| chr18 | 34478871 | 34480710 | 1840 | -6.98 | 6.26E-10 | 1.48E-09 |

| | | | | | | |
|---|---|---|---|---|---|---|
| chr11 | 36009612 | 36027604 | 17993 | -7.46 | 6.26E-10 | 1.48E-09 |
| chrX | 153999088 | 154000996 | 1909 | -7 | 6.31E-10 | 1.50E-09 |
| chr3 | 43569317 | 43571070 | 1754 | -6.94 | 6.31E-10 | 1.50E-09 |
| chr3 | 57349889 | 57368333 | 18445 | -2.5 | 6.35E-10 | 1.50E-09 |
| chr11 | 118545136 | 118548466 | 3331 | -7.25 | 6.35E-10 | 1.51E-09 |
| chr2 | 172062613 | 172081505 | 18893 | -7.65 | 6.36E-10 | 1.51E-09 |
| chr6 | 42486392 | 42488480 | 2089 | -6.95 | 6.36E-10 | 1.51E-09 |
| chr2 | 114602469 | 114604979 | 2511 | -7.06 | 6.38E-10 | 1.51E-09 |
| chr3 | 13643712 | 13647712 | 4001 | -6.94 | 6.38E-10 | 1.51E-09 |
| chr9 | 101406119 | 101417910 | 11792 | -7.07 | 6.39E-10 | 1.51E-09 |
| chr6 | 158118596 | 158120802 | 2207 | -6.93 | 6.39E-10 | 1.51E-09 |
| chr1 | 206797148 | 206808471 | 11324 | -7.08 | 6.41E-10 | 1.52E-09 |
| chr6 | 148007152 | 148010120 | 2969 | -7.09 | 6.42E-10 | 1.52E-09 |
| chr19 | 36251059 | 36252102 | 1044 | -6.93 | 6.44E-10 | 1.53E-09 |
| chr2 | 121339948 | 121341297 | 1350 | -7.17 | 6.45E-10 | 1.53E-09 |
| chr14 | 90785770 | 90787446 | 1677 | -7.03 | 6.47E-10 | 1.53E-09 |
| chr19 | 46706889 | 46708849 | 1961 | -7.03 | 6.47E-10 | 1.53E-09 |
| chr6 | 21425609 | 21427465 | 1857 | -7.11 | 6.47E-10 | 1.53E-09 |
| chr18 | 53552608 | 53554650 | 2043 | -7.1 | 6.48E-10 | 1.53E-09 |
| chr4 | 24738245 | 24740821 | 2577 | -7.02 | 6.48E-10 | 1.54E-09 |
| chr4 | 102379016 | 102382589 | 3574 | -5.33 | 6.49E-10 | 1.54E-09 |
| chr4 | 174123595 | 174125823 | 2229 | -6.95 | 6.51E-10 | 1.54E-09 |
| chr2 | 162256409 | 162259692 | 3284 | -7.01 | 6.51E-10 | 1.54E-09 |
| chr5 | 167484490 | 167486056 | 1567 | -6.98 | 6.51E-10 | 1.54E-09 |
| chr6 | 147898444 | 147900740 | 2297 | -6.94 | 6.51E-10 | 1.54E-09 |
| chr2 | 190781248 | 190783412 | 2165 | -6.94 | 6.51E-10 | 1.54E-09 |
| chr4 | 143358813 | 143360996 | 2184 | -7.01 | 6.54E-10 | 1.55E-09 |
| chr4 | 78959232 | 78961702 | 2471 | -6.96 | 6.55E-10 | 1.55E-09 |
| chr9 | 28837069 | 28840592 | 3524 | -6.93 | 6.57E-10 | 1.55E-09 |
| chr13 | 95694706 | 95696433 | 1728 | -3.37 | 6.57E-10 | 1.55E-09 |
| chr10 | 32212079 | 32213033 | 955 | -7.19 | 6.58E-10 | 1.56E-09 |
| chrX | 99836519 | 99848211 | 11693 | 7.46 | 6.60E-10 | 1.56E-09 |
| chr20 | 52647817 | 52666872 | 19056 | -7.83 | 6.61E-10 | 1.56E-09 |
| chr12 | 93037030 | 93039503 | 2474 | -7.25 | 6.61E-10 | 1.56E-09 |
| chr10 | 126268489 | 126281120 | 12632 | -7.39 | 6.63E-10 | 1.57E-09 |
| chr4 | 186483231 | 186487073 | 3843 | -6.97 | 6.65E-10 | 1.57E-09 |
| chr5 | 74551390 | 74564402 | 13013 | -6.97 | 6.65E-10 | 1.57E-09 |
| chr7 | 117016328 | 117034871 | 18544 | -7.24 | 6.67E-10 | 1.58E-09 |
| chr2 | 44648567 | 44649787 | 1221 | -6.99 | 6.68E-10 | 1.58E-09 |
| chr21 | 38448310 | 38452334 | 4025 | -6.99 | 6.68E-10 | 1.58E-09 |
| chr9 | 126464570 | 126465969 | 1400 | -6.94 | 6.69E-10 | 1.58E-09 |
| chrX | 149958474 | 149960505 | 2032 | -6.94 | 6.69E-10 | 1.58E-09 |
| chr2 | 219668746 | 219670134 | 1389 | -6.94 | 6.70E-10 | 1.58E-09 |
| chr2 | 135933523 | 135952176 | 18654 | -6.94 | 6.70E-10 | 1.59E-09 |
| chr11 | 86400990 | 86402906 | 1917 | -6.93 | 6.72E-10 | 1.59E-09 |
| chr12 | 118130369 | 118132864 | 2496 | -7.03 | 6.72E-10 | 1.59E-09 |
| chr4 | 101388995 | 101392351 | 3357 | -6.92 | 6.72E-10 | 1.59E-09 |
| chr2 | 30840215 | 30841850 | 1636 | -7.08 | 6.75E-10 | 1.59E-09 |
| chr19 | 42329843 | 42333054 | 3212 | -7.29 | 6.75E-10 | 1.60E-09 |
| chr6 | 108120438 | 108121798 | 1361 | -5.29 | 6.75E-10 | 1.60E-09 |
| chr12 | 27192493 | 27194368 | 1876 | -7.02 | 6.78E-10 | 1.60E-09 |
| chr19 | 10947383 | 10950800 | 3418 | -7.21 | 6.79E-10 | 1.61E-09 |
| chr4 | 178340829 | 178343379 | 2551 | -6.93 | 6.80E-10 | 1.61E-09 |
| chr13 | 49928503 | 49930506 | 2004 | -6.93 | 6.80E-10 | 1.61E-09 |
| chr20 | 30862382 | 30863878 | 1497 | -6.94 | 6.81E-10 | 1.61E-09 |
| chr4 | 17862945 | 17865745 | 2801 | -6.94 | 6.81E-10 | 1.61E-09 |
| chr18 | 26809833 | 26814264 | 4432 | -6.96 | 6.82E-10 | 1.61E-09 |
| chrX | 9480503 | 9482249 | 1747 | -7.01 | 6.82E-10 | 1.61E-09 |
| chr2 | 134539515 | 134572836 | 33322 | -1.66 | 6.85E-10 | 1.62E-09 |
| chr18 | 60057533 | 60059242 | 1710 | -6.93 | 6.86E-10 | 1.62E-09 |
| chr2 | 181595930 | 181598922 | 2993 | -7.02 | 6.87E-10 | 1.62E-09 |

| | | | | | | |
|---|---|---|---|---|---|---|
| chr8 | 26462481 | 26464009 | 1529 | -7.23 | 6.87E-10 | 1.62E-09 |
| chr1 | 199639662 | 199641824 | 2163 | -7 | 6.88E-10 | 1.62E-09 |
| chr2 | 20528447 | 20531784 | 3338 | -7 | 6.88E-10 | 1.62E-09 |
| chr5 | 89764172 | 89766427 | 2256 | -6.95 | 6.88E-10 | 1.62E-09 |
| chr13 | 75856889 | 75858525 | 1637 | -7.01 | 6.91E-10 | 1.63E-09 |
| chr13 | 79827547 | 79836880 | 9334 | 4.78 | 6.91E-10 | 1.63E-09 |
| chr1 | 110358015 | 110359538 | 1524 | -6.97 | 6.91E-10 | 1.63E-09 |
| chr1 | 11013172 | 11014779 | 1608 | -6.94 | 6.92E-10 | 1.63E-09 |
| chr12 | 112832433 | 112834250 | 1818 | -6.94 | 6.92E-10 | 1.63E-09 |
| chr3 | 32134289 | 32136491 | 2203 | -6.94 | 6.92E-10 | 1.63E-09 |
| chr5 | 96130840 | 96132416 | 1577 | -6.92 | 6.92E-10 | 1.63E-09 |
| chr5 | 16658113 | 16659758 | 1646 | -6.94 | 6.93E-10 | 1.63E-09 |
| chr13 | 46344317 | 46346291 | 1975 | -6.99 | 6.94E-10 | 1.64E-09 |
| chr8 | 28450021 | 28451445 | 1425 | -6.92 | 6.96E-10 | 1.64E-09 |
| chr2 | 151579337 | 151583782 | 4446 | -7.18 | 6.96E-10 | 1.64E-09 |
| chr8 | 40945010 | 40946581 | 1572 | -7.18 | 6.97E-10 | 1.64E-09 |
| chr18 | 70392667 | 70398782 | 6116 | -7.23 | 6.98E-10 | 1.65E-09 |
| chr4 | 112400377 | 112403658 | 3282 | 3.88 | 6.99E-10 | 1.65E-09 |
| chr2 | 15763868 | 15765261 | 1394 | -7.11 | 7.01E-10 | 1.65E-09 |
| chr1 | 233028172 | 233030973 | 2802 | -6.94 | 7.01E-10 | 1.65E-09 |
| chr14 | 69186006 | 69187696 | 1691 | -6.94 | 7.01E-10 | 1.65E-09 |
| chr21 | 46732075 | 46736461 | 4387 | -7.35 | 7.03E-10 | 1.66E-09 |
| chr5 | 1757671 | 1774211 | 16541 | -6.93 | 7.05E-10 | 1.66E-09 |
| chr5 | 144917014 | 144918432 | 1419 | -6.93 | 7.05E-10 | 1.66E-09 |
| chr12 | 42381511 | 42383901 | 2391 | -6.92 | 7.06E-10 | 1.66E-09 |
| chr11 | 14588999 | 14596165 | 7167 | -7.03 | 7.07E-10 | 1.67E-09 |
| chr14 | 78237266 | 78240799 | 3534 | -7.03 | 7.07E-10 | 1.67E-09 |
| chr4 | 163960269 | 163962745 | 2477 | -6.99 | 7.07E-10 | 1.67E-09 |
| chr1 | 87230208 | 87233952 | 3745 | -6.98 | 7.08E-10 | 1.67E-09 |
| chr11 | 5585370 | 5595126 | 9757 | 7.95 | 7.08E-10 | 1.67E-09 |
| chr18 | 9367422 | 9376451 | 9030 | -7.13 | 7.08E-10 | 1.67E-09 |
| chr15 | 71876871 | 71879380 | 2510 | -7.15 | 7.09E-10 | 1.67E-09 |
| chr1 | 201765004 | 201766836 | 1833 | -6.92 | 7.10E-10 | 1.67E-09 |
| chr1 | 161982701 | 161997405 | 14705 | -6.92 | 7.10E-10 | 1.67E-09 |
| chr1 | 236205594 | 236219089 | 13496 | -6.92 | 7.10E-10 | 1.67E-09 |
| chr6 | 81406397 | 81412303 | 5907 | -6.92 | 7.10E-10 | 1.67E-09 |
| chr6 | 118151677 | 118153501 | 1825 | -6.93 | 7.11E-10 | 1.68E-09 |
| chr4 | 168497917 | 168502677 | 4761 | -7.08 | 7.12E-10 | 1.68E-09 |
| chr12 | 75250796 | 75253576 | 2781 | -7.01 | 7.13E-10 | 1.68E-09 |
| chr4 | 132414982 | 132416956 | 1975 | -6.94 | 7.13E-10 | 1.68E-09 |
| chr2 | 171700865 | 171720425 | 19561 | -7.73 | 7.14E-10 | 1.68E-09 |
| chr14 | 58008690 | 58023371 | 14682 | -1.9 | 7.15E-10 | 1.68E-09 |
| chr3 | 120348902 | 120363278 | 14377 | -7.58 | 7.15E-10 | 1.69E-09 |
| chr1 | 212630924 | 212633097 | 2174 | -7.1 | 7.17E-10 | 1.69E-09 |
| chr17 | 38812675 | 38814873 | 2199 | -6.92 | 7.17E-10 | 1.69E-09 |
| chr12 | 85597582 | 85601405 | 3824 | -6.93 | 7.18E-10 | 1.69E-09 |
| chrX | 20501288 | 20508022 | 6735 | -7.21 | 7.18E-10 | 1.69E-09 |
| chr13 | 68629589 | 68631532 | 1944 | -6.93 | 7.18E-10 | 1.69E-09 |
| chr4 | 186625809 | 186628608 | 2800 | -6.93 | 7.18E-10 | 1.69E-09 |
| chr16 | 14477285 | 14479419 | 2135 | -6.97 | 7.18E-10 | 1.69E-09 |
| chr16 | 58725300 | 58727195 | 1896 | -6.97 | 7.18E-10 | 1.69E-09 |
| chr8 | 82971824 | 82978286 | 6463 | -7.16 | 7.19E-10 | 1.69E-09 |
| chr8 | 42459986 | 42461944 | 1959 | -6.92 | 7.19E-10 | 1.69E-09 |
| chr15 | 93219365 | 93243129 | 23765 | -8.3 | 7.20E-10 | 1.70E-09 |
| chr3 | 197697986 | 197699818 | 1833 | -6.97 | 7.21E-10 | 1.70E-09 |
| chr11 | 17880146 | 17891934 | 11789 | -7.1 | 7.21E-10 | 1.70E-09 |
| chr7 | 24339653 | 24346449 | 6797 | -2.04 | 7.22E-10 | 1.70E-09 |
| chr15 | 82370758 | 82386192 | 15435 | -7.33 | 7.22E-10 | 1.70E-09 |
| chr13 | 79841578 | 79846626 | 5049 | -5.56 | 7.25E-10 | 1.71E-09 |
| chr17 | 37297281 | 37298484 | 1204 | -6.99 | 7.27E-10 | 1.71E-09 |
| chr16 | 52872148 | 53011079 | 138932 | -7.39 | 7.27E-10 | 1.71E-09 |

| | | | | | | |
|---|---|---|---|---|---|---|
| chr2 | 42114452 | 42116839 | 2388 | -7.03 | 7.29E-10 | 1.71E-09 |
| chr11 | 22120206 | 22121689 | 1484 | -5.99 | 7.29E-10 | 1.71E-09 |
| chr15 | 71936846 | 71938084 | 1239 | -6.91 | 7.29E-10 | 1.72E-09 |
| chr2 | 179222816 | 179224853 | 2038 | -5.28 | 7.30E-10 | 1.72E-09 |
| chr6 | 149346883 | 149348360 | 1478 | -7.06 | 7.30E-10 | 1.72E-09 |
| chr12 | 13483668 | 13495607 | 11940 | -6.92 | 7.30E-10 | 1.72E-09 |
| chr5 | 27391141 | 27400381 | 9241 | -6.91 | 7.30E-10 | 1.72E-09 |
| chr3 | 170055389 | 170058201 | 2813 | -7.18 | 7.32E-10 | 1.72E-09 |
| chr5 | 52123369 | 52124615 | 1247 | -7.07 | 7.32E-10 | 1.72E-09 |
| chr6 | 21272357 | 21275033 | 2677 | -7.22 | 7.33E-10 | 1.72E-09 |
| chrX | 35031028 | 35033564 | 2537 | -7.13 | 7.37E-10 | 1.73E-09 |
| chrX | 68941259 | 69035787 | 94529 | 3.98 | 7.37E-10 | 1.73E-09 |
| chr11 | 82622170 | 82623609 | 1440 | -6.92 | 7.40E-10 | 1.74E-09 |
| chr6 | 1153842 | 1160177 | 6336 | -2.34 | 7.40E-10 | 1.74E-09 |
| chr11 | 66663214 | 66664949 | 1736 | -6.91 | 7.40E-10 | 1.74E-09 |
| chr17 | 5169463 | 5171752 | 2290 | -7.23 | 7.41E-10 | 1.74E-09 |
| chr11 | 49483485 | 49522339 | 38855 | 1.83 | 7.41E-10 | 1.74E-09 |
| chr14 | 62219843 | 62223393 | 3551 | -7.04 | 7.41E-10 | 1.74E-09 |
| chr2 | 223602105 | 223605050 | 2946 | -7.04 | 7.41E-10 | 1.74E-09 |
| chr7 | 47568999 | 47573662 | 4664 | -7.42 | 7.42E-10 | 1.74E-09 |
| chr14 | 65901641 | 65903041 | 1401 | -7.02 | 7.42E-10 | 1.74E-09 |
| chr11 | 11388988 | 11391112 | 2125 | -7.23 | 7.43E-10 | 1.75E-09 |
| chr11 | 35720612 | 35724552 | 3941 | -6.95 | 7.44E-10 | 1.75E-09 |
| chr6 | 53704156 | 53705568 | 1413 | -7.01 | 7.45E-10 | 1.75E-09 |
| chr8 | 81584825 | 81587748 | 2924 | -6.94 | 7.45E-10 | 1.75E-09 |
| chr2 | 23993676 | 23995098 | 1423 | -7.33 | 7.45E-10 | 1.75E-09 |
| chr13 | 61342984 | 61367109 | 24126 | 1.82 | 7.47E-10 | 1.75E-09 |
| chr14 | 27573625 | 28221238 | 647614 | 1.21 | 7.47E-10 | 1.76E-09 |
| chr19 | 1995926 | 1996970 | 1045 | -6.98 | 7.49E-10 | 1.76E-09 |
| chr20 | 48341666 | 48343284 | 1619 | -6.91 | 7.49E-10 | 1.76E-09 |
| chr10 | 420402 | 421968 | 1567 | -7 | 7.50E-10 | 1.76E-09 |
| chr9 | 100384391 | 100386520 | 2130 | -7 | 7.50E-10 | 1.76E-09 |
| chr4 | 101781766 | 101784138 | 2373 | -5.22 | 7.50E-10 | 1.76E-09 |
| chr1 | 75854062 | 75857971 | 3910 | -6.93 | 7.51E-10 | 1.76E-09 |
| chr4 | 128899235 | 128900767 | 1533 | -7.33 | 7.51E-10 | 1.76E-09 |
| chr17 | 78015359 | 78016855 | 1497 | -6.91 | 7.52E-10 | 1.77E-09 |
| chr14 | 97329975 | 97331207 | 1233 | -6.91 | 7.52E-10 | 1.77E-09 |
| chr9 | 108276006 | 108302508 | 26503 | -7.85 | 7.53E-10 | 1.77E-09 |
| chr4 | 128472347 | 128474555 | 2209 | -5.09 | 7.54E-10 | 1.77E-09 |
| chr6 | 2009756 | 2016101 | 6346 | -7.2 | 7.55E-10 | 1.77E-09 |
| chr12 | 124596913 | 124598981 | 2069 | -7.15 | 7.56E-10 | 1.77E-09 |
| chr10 | 70611191 | 70613194 | 2004 | -6.96 | 7.56E-10 | 1.77E-09 |
| chr11 | 101503614 | 101507150 | 3537 | -6.96 | 7.56E-10 | 1.77E-09 |
| chr1 | 241058141 | 241067699 | 9559 | -6.92 | 7.56E-10 | 1.77E-09 |
| chr3 | 116905110 | 116906722 | 1613 | -6.97 | 7.58E-10 | 1.78E-09 |
| chr17 | 70446066 | 70502356 | 56291 | -7.9 | 7.61E-10 | 1.79E-09 |
| chr5 | 82679319 | 82681771 | 2453 | -6.94 | 7.61E-10 | 1.79E-09 |
| chr5 | 111897950 | 111901555 | 3606 | -4.02 | 7.61E-10 | 1.79E-09 |
| chr16 | 31521646 | 31711453 | 189808 | -1.54 | 7.62E-10 | 1.79E-09 |
| chr9 | 128263454 | 128265232 | 1779 | -7.16 | 7.63E-10 | 1.79E-09 |
| chr12 | 111814585 | 111816795 | 2211 | -6.95 | 7.63E-10 | 1.79E-09 |
| chr11 | 93035574 | 93063160 | 27587 | -1.99 | 7.65E-10 | 1.79E-09 |
| chr12 | 88821250 | 88823686 | 2437 | -7.16 | 7.66E-10 | 1.80E-09 |
| chrX | 109567528 | 109570082 | 2555 | -7.16 | 7.66E-10 | 1.80E-09 |
| chr7 | 93430392 | 93433097 | 2706 | -6.95 | 7.68E-10 | 1.80E-09 |
| chr2 | 105509214 | 105527496 | 18283 | -1.68 | 7.69E-10 | 1.80E-09 |
| chr3 | 136692118 | 136694123 | 2006 | -6.97 | 7.69E-10 | 1.80E-09 |
| chr16 | 69614543 | 69617383 | 2841 | -7.25 | 7.69E-10 | 1.80E-09 |
| chr4 | 123700257 | 123702349 | 2093 | -6.93 | 7.70E-10 | 1.81E-09 |
| chr11 | 75801368 | 75803001 | 1634 | -6.96 | 7.71E-10 | 1.81E-09 |
| chr4 | 75885331 | 75887567 | 2237 | -6.96 | 7.71E-10 | 1.81E-09 |

| | | | | | | |
|---|---|---|---|---|---|---|
| chr1 | 160814425 | 160823152 | 8728 | -5.13 | 7.72E-10 | 1.81E-09 |
| chr1 | 198399433 | 198402089 | 2657 | -6.91 | 7.75E-10 | 1.82E-09 |
| chr2 | 196795495 | 196821435 | 25941 | -7.87 | 7.75E-10 | 1.82E-09 |
| chr17 | 78545358 | 78546668 | 1311 | -6.9 | 7.76E-10 | 1.82E-09 |
| chr1 | 188737297 | 188740928 | 3632 | -6.96 | 7.78E-10 | 1.82E-09 |
| chr16 | 54044511 | 54047511 | 3001 | -6.96 | 7.78E-10 | 1.82E-09 |
| chr9 | 127825521 | 127827218 | 1698 | -7.03 | 7.80E-10 | 1.83E-09 |
| chr1 | 64354744 | 64356552 | 1809 | -6.91 | 7.80E-10 | 1.83E-09 |
| chr5 | 172466818 | 172468263 | 1446 | -7.01 | 7.81E-10 | 1.83E-09 |
| chr13 | 21104901 | 21112571 | 7671 | -7.01 | 7.82E-10 | 1.83E-09 |
| chrX | 19994770 | 20003977 | 9208 | -7.01 | 7.82E-10 | 1.83E-09 |
| chr7 | 46628727 | 46634671 | 5945 | -7.07 | 7.82E-10 | 1.83E-09 |
| chr6 | 128739037 | 128741160 | 2124 | -7.01 | 7.83E-10 | 1.83E-09 |
| chr1 | 11059519 | 11061715 | 2197 | -7.23 | 7.83E-10 | 1.83E-09 |
| chr2 | 151722760 | 151767474 | 44715 | -7.03 | 7.84E-10 | 1.84E-09 |
| chr4 | 40779249 | 40786706 | 7458 | -7.13 | 7.84E-10 | 1.84E-09 |
| chr17 | 34891540 | 34894619 | 3080 | -7.45 | 7.85E-10 | 1.84E-09 |
| chr16 | 8633718 | 8636093 | 2376 | -6.99 | 7.86E-10 | 1.84E-09 |
| chr2 | 75924723 | 75926255 | 1533 | -7.36 | 7.86E-10 | 1.84E-09 |
| chr5 | 130197886 | 130201351 | 3466 | -7.27 | 7.88E-10 | 1.84E-09 |
| chr1 | 91819007 | 91832388 | 13382 | -1.86 | 7.88E-10 | 1.84E-09 |
| chr7 | 107073448 | 107076538 | 3091 | -6.99 | 7.89E-10 | 1.85E-09 |
| chr12 | 118515337 | 118516556 | 1220 | -6.93 | 7.90E-10 | 1.85E-09 |
| chr4 | 122704544 | 122705921 | 1378 | -6.95 | 7.91E-10 | 1.85E-09 |
| chr7 | 13385051 | 13392588 | 7538 | -6.99 | 7.91E-10 | 1.85E-09 |
| chr1 | 51296642 | 51327073 | 30432 | -7.53 | 7.93E-10 | 1.86E-09 |
| chr14 | 51613797 | 51617150 | 3354 | -6.98 | 7.96E-10 | 1.86E-09 |
| chr2 | 227688268 | 227707615 | 19348 | -7.28 | 7.97E-10 | 1.86E-09 |
| chr3 | 108150876 | 108168046 | 17171 | -6.98 | 7.97E-10 | 1.87E-09 |
| chr11 | 74871452 | 74872975 | 1524 | -6.93 | 7.99E-10 | 1.87E-09 |
| chr3 | 29645934 | 29655835 | 9902 | -6.93 | 7.99E-10 | 1.87E-09 |
| chr6 | 100041330 | 100045170 | 3841 | -2.16 | 8.02E-10 | 1.88E-09 |
| chr7 | 131725696 | 131727626 | 1931 | -6.93 | 8.02E-10 | 1.88E-09 |
| chr17 | 64663842 | 64675598 | 11757 | -7.47 | 8.03E-10 | 1.88E-09 |
| chr12 | 78758928 | 78774277 | 15350 | -7.1 | 8.05E-10 | 1.88E-09 |
| chr5 | 87972923 | 87991482 | 18560 | -7.1 | 8.05E-10 | 1.88E-09 |
| chr4 | 138860438 | 138862865 | 2428 | -6.9 | 8.05E-10 | 1.88E-09 |
| chr12 | 34098027 | 34133685 | 35659 | 2.84 | 8.07E-10 | 1.89E-09 |
| chr12 | 3955394 | 3989945 | 34552 | -4.32 | 8.07E-10 | 1.89E-09 |
| chr10 | 49555010 | 49557237 | 2228 | -6.9 | 8.09E-10 | 1.89E-09 |
| chrX | 102436838 | 102442405 | 5568 | -6.9 | 8.10E-10 | 1.89E-09 |
| chr6 | 56452692 | 56454449 | 1758 | -7.12 | 8.12E-10 | 1.90E-09 |
| chr13 | 63970106 | 64049266 | 79161 | 1.88 | 8.15E-10 | 1.91E-09 |
| chr2 | 157176146 | 157179838 | 3693 | -6.94 | 8.17E-10 | 1.91E-09 |
| chr1 | 209958087 | 209968502 | 10416 | -2.38 | 8.18E-10 | 1.91E-09 |
| chr18 | 12805245 | 12807332 | 2088 | -6.9 | 8.20E-10 | 1.92E-09 |
| chr3 | 127422197 | 127424348 | 2152 | -6.9 | 8.20E-10 | 1.92E-09 |
| chr3 | 16263064 | 16264667 | 1604 | -7.01 | 8.21E-10 | 1.92E-09 |
| chr15 | 35220778 | 35222575 | 1798 | -6.9 | 8.22E-10 | 1.92E-09 |
| chrX | 75017052 | 75031817 | 14766 | 2.03 | 8.24E-10 | 1.93E-09 |
| chrX | 108646702 | 108687212 | 40511 | 1.51 | 8.27E-10 | 1.93E-09 |
| chr7 | 5783099 | 5784686 | 1588 | -7.06 | 8.28E-10 | 1.94E-09 |
| chr6 | 81587241 | 81608610 | 21370 | 2.4 | 8.29E-10 | 1.94E-09 |
| chr3 | 113182086 | 113183516 | 1431 | -7 | 8.29E-10 | 1.94E-09 |
| chr9 | 88067708 | 88072818 | 5111 | 7.14 | 8.31E-10 | 1.94E-09 |
| chr7 | 155070709 | 155073987 | 3279 | -6.9 | 8.31E-10 | 1.94E-09 |
| chr2 | 88528106 | 88619157 | 91052 | -2.4 | 8.32E-10 | 1.94E-09 |
| chr7 | 141827295 | 141839291 | 11997 | -6.9 | 8.32E-10 | 1.94E-09 |
| chr1 | 199792772 | 199794820 | 2049 | -7.31 | 8.34E-10 | 1.95E-09 |
| chr10 | 133841019 | 133842751 | 1733 | -7.07 | 8.34E-10 | 1.95E-09 |
| chr3 | 46142827 | 46144488 | 1662 | -6.89 | 8.37E-10 | 1.95E-09 |

| | | | | | | |
|---|---|---|---|---|---|---|
| chr21 | 38983648 | 38986477 | 2830 | -6.89 | 8.37E-10 | 1.95E-09 |
| chr6 | 170015802 | 170017893 | 2092 | -6.89 | 8.37E-10 | 1.95E-09 |
| chr4 | 141199032 | 141201902 | 2871 | -6.94 | 8.38E-10 | 1.96E-09 |
| chr12 | 76914743 | 76917628 | 2886 | -7.19 | 8.39E-10 | 1.96E-09 |
| chr2 | 232945596 | 232957235 | 11640 | -7.08 | 8.39E-10 | 1.96E-09 |
| chr5 | 122351020 | 122353124 | 2105 | -6.89 | 8.39E-10 | 1.96E-09 |
| chr14 | 67328354 | 67340485 | 12132 | -7.35 | 8.40E-10 | 1.96E-09 |
| chr17 | 75529801 | 75531595 | 1795 | -6.91 | 8.40E-10 | 1.96E-09 |
| chr2 | 152123584 | 152125183 | 1600 | -6.97 | 8.42E-10 | 1.97E-09 |
| chr2 | 238894913 | 238909642 | 14730 | -6.91 | 8.44E-10 | 1.97E-09 |
| chr7 | 128352373 | 128366331 | 13959 | -6.91 | 8.44E-10 | 1.97E-09 |
| chr1 | 12137000 | 12143539 | 6540 | -6.93 | 8.47E-10 | 1.98E-09 |
| chr7 | 76049425 | 76050462 | 1038 | -6.97 | 8.48E-10 | 1.98E-09 |
| chr2 | 53118284 | 53130654 | 12371 | -3.02 | 8.49E-10 | 1.98E-09 |
| chr2 | 206461751 | 206480272 | 18522 | -2.68 | 8.50E-10 | 1.98E-09 |
| chrX | 57757909 | 57770703 | 12795 | 2.1 | 8.53E-10 | 1.99E-09 |
| chr3 | 114458289 | 114462120 | 3832 | -7.09 | 8.53E-10 | 1.99E-09 |
| chr8 | 80934077 | 80935227 | 1151 | -6.92 | 8.55E-10 | 2.00E-09 |
| chr1 | 224358349 | 224360324 | 1976 | -6.9 | 8.55E-10 | 2.00E-09 |
| chr4 | 23735790 | 23885208 | 149419 | 1.04 | 8.55E-10 | 2.00E-09 |
| chr11 | 88413489 | 88418969 | 5481 | -6.91 | 8.57E-10 | 2.00E-09 |
| chr12 | 28778730 | 28790772 | 12043 | 7.3 | 8.58E-10 | 2.00E-09 |
| chrX | 45018856 | 45021581 | 2726 | -7.16 | 8.60E-10 | 2.01E-09 |
| chr10 | 126963560 | 127030006 | 66447 | -2.84 | 8.60E-10 | 2.01E-09 |
| chr17 | 6666263 | 6704794 | 38532 | -1.63 | 8.61E-10 | 2.01E-09 |
| chr3 | 177945303 | 177959640 | 14338 | -6.89 | 8.62E-10 | 2.01E-09 |
| chr3 | 157891896 | 157894126 | 2231 | -7.01 | 8.63E-10 | 2.01E-09 |
| chr13 | 32992091 | 32994532 | 2442 | -6.89 | 8.64E-10 | 2.01E-09 |
| chr6 | 14372977 | 14379335 | 6359 | -6.89 | 8.64E-10 | 2.01E-09 |
| chr6 | 35164486 | 35166490 | 2005 | -7.02 | 8.64E-10 | 2.01E-09 |
| chr2 | 45857307 | 45858447 | 1141 | -7.02 | 8.64E-10 | 2.01E-09 |
| chr5 | 169828400 | 169831692 | 3293 | -7.02 | 8.64E-10 | 2.01E-09 |
| chr12 | 105085768 | 105087217 | 1450 | -7 | 8.64E-10 | 2.01E-09 |
| chrX | 12065693 | 12067813 | 2121 | -6.99 | 8.66E-10 | 2.02E-09 |
| chr17 | 38961170 | 38962319 | 1150 | -6.91 | 8.67E-10 | 2.02E-09 |
| chr10 | 8288158 | 8300319 | 12162 | -6.99 | 8.68E-10 | 2.02E-09 |
| chr4 | 139234592 | 139237079 | 2488 | -6.06 | 8.68E-10 | 2.02E-09 |
| chr4 | 189487643 | 189495451 | 7809 | -7.05 | 8.70E-10 | 2.03E-09 |
| chr8 | 95588561 | 95590664 | 2104 | -7.05 | 8.70E-10 | 2.03E-09 |
| chr2 | 42302652 | 42304593 | 1942 | -6.89 | 8.70E-10 | 2.03E-09 |
| chr6 | 45198932 | 45211343 | 12412 | -6.89 | 8.70E-10 | 2.03E-09 |
| chr4 | 14937828 | 14942434 | 4607 | -6.9 | 8.71E-10 | 2.03E-09 |
| chr13 | 32947089 | 32949434 | 2346 | -6.98 | 8.71E-10 | 2.03E-09 |
| chr7 | 109544600 | 109546230 | 1631 | -5.95 | 8.73E-10 | 2.03E-09 |
| chr3 | 9431985 | 9433317 | 1333 | -5.15 | 8.74E-10 | 2.04E-09 |
| chr6 | 21543792 | 21550520 | 6729 | -7.13 | 8.75E-10 | 2.04E-09 |
| chr6 | 42562982 | 42564475 | 1494 | -6.91 | 8.75E-10 | 2.04E-09 |
| chr10 | 4937903 | 4941287 | 3385 | -6.88 | 8.75E-10 | 2.04E-09 |
| chr3 | 113108527 | 113110396 | 1870 | -6.88 | 8.75E-10 | 2.04E-09 |
| chr3 | 128008500 | 128010564 | 2065 | -6.89 | 8.77E-10 | 2.04E-09 |
| chr12 | 77125677 | 77131591 | 5915 | -7.18 | 8.77E-10 | 2.04E-09 |
| chr2 | 30507845 | 30509921 | 2077 | -7.06 | 8.78E-10 | 2.04E-09 |
| chr17 | 56130790 | 56133144 | 2355 | 5.41 | 8.79E-10 | 2.05E-09 |
| chr1 | 214428404 | 214429927 | 1524 | -6.91 | 8.79E-10 | 2.05E-09 |
| chr18 | 24317838 | 24320577 | 2740 | -7.07 | 8.80E-10 | 2.05E-09 |
| chr15 | 91178000 | 91187403 | 9404 | -7.13 | 8.81E-10 | 2.05E-09 |
| chr6 | 46519432 | 46526801 | 7370 | 6.95 | 8.81E-10 | 2.05E-09 |
| chr9 | 125567992 | 125569717 | 1726 | -6.92 | 8.82E-10 | 2.05E-09 |
| chr5 | 95480407 | 95482850 | 2444 | -7.04 | 8.85E-10 | 2.06E-09 |
| chr12 | 15969902 | 15971556 | 1655 | -7.08 | 8.85E-10 | 2.06E-09 |
| chr4 | 132185516 | 132188592 | 3077 | -6.88 | 8.86E-10 | 2.06E-09 |

| | | | | | | |
|---|---|---|---|---|---|---|
| chr17 | 72262826 | 72264303 | 1478 | -6.9 | 8.86E-10 | 2.06E-09 |
| chr4 | 91733587 | 91736882 | 3296 | -6.9 | 8.86E-10 | 2.06E-09 |
| chr6 | 43980063 | 43986316 | 6254 | -6.93 | 8.87E-10 | 2.06E-09 |
| chrX | 55221750 | 55224903 | 3154 | -6.88 | 8.88E-10 | 2.07E-09 |
| chr4 | 189769023 | 189772068 | 3046 | -6.88 | 8.88E-10 | 2.07E-09 |
| chr16 | 70814875 | 70816465 | 1591 | -6.91 | 8.91E-10 | 2.07E-09 |
| chr8 | 132845888 | 132847226 | 1339 | -6.97 | 8.92E-10 | 2.07E-09 |
| chr4 | 149093957 | 149095956 | 2000 | -5.43 | 8.94E-10 | 2.08E-09 |
| chr1 | 193253089 | 193255436 | 2348 | -6.88 | 8.96E-10 | 2.08E-09 |
| chr5 | 139255135 | 139256243 | 1109 | -6.88 | 8.96E-10 | 2.08E-09 |
| chr9 | 2749281 | 2751179 | 1899 | -6.89 | 8.96E-10 | 2.08E-09 |
| chr17 | 19696521 | 19697760 | 1240 | -7.07 | 8.97E-10 | 2.09E-09 |
| chr4 | 168650062 | 168653573 | 3512 | -6.93 | 9.00E-10 | 2.09E-09 |
| chr7 | 12812462 | 12814502 | 2041 | -6.96 | 9.02E-10 | 2.10E-09 |
| chr13 | 29260952 | 29262591 | 1640 | -6.88 | 9.03E-10 | 2.10E-09 |
| chr9 | 132334206 | 132343451 | 9246 | -7.54 | 9.03E-10 | 2.10E-09 |
| chrX | 15272705 | 15275192 | 2488 | -7.15 | 9.04E-10 | 2.10E-09 |
| chr2 | 31766712 | 31854370 | 87659 | -3.11 | 9.05E-10 | 2.10E-09 |
| chr20 | 3390856 | 3392268 | 1413 | -6.89 | 9.07E-10 | 2.11E-09 |
| chr1 | 68499055 | 68502697 | 3643 | -6.92 | 9.07E-10 | 2.11E-09 |
| chr20 | 5038957 | 5040246 | 1290 | -6.88 | 9.08E-10 | 2.11E-09 |
| chr6 | 117057043 | 117067280 | 10238 | -7.01 | 9.09E-10 | 2.11E-09 |
| chr1 | 207386256 | 207388877 | 2622 | -6.95 | 9.09E-10 | 2.11E-09 |
| chr12 | 42909079 | 42910892 | 1814 | -7 | 9.10E-10 | 2.11E-09 |
| chr18 | 56178530 | 56205296 | 26767 | -6.9 | 9.10E-10 | 2.11E-09 |
| chr1 | 179590630 | 179592247 | 1618 | -6.89 | 9.10E-10 | 2.11E-09 |
| chr15 | 52830824 | 52832968 | 2145 | -7.44 | 9.12E-10 | 2.12E-09 |
| chr18 | 9343077 | 9344351 | 1275 | -6.92 | 9.13E-10 | 2.12E-09 |
| chr7 | 6528468 | 6530469 | 2002 | -7.16 | 9.14E-10 | 2.12E-09 |
| chr15 | 56769269 | 56854926 | 85658 | -3.07 | 9.16E-10 | 2.13E-09 |
| chr1 | 241883264 | 241885726 | 2463 | -7.25 | 9.16E-10 | 2.13E-09 |
| chr4 | 146632028 | 146634728 | 2701 | -6.88 | 9.18E-10 | 2.13E-09 |
| chr2 | 106494002 | 106630257 | 136256 | -7.52 | 9.22E-10 | 2.14E-09 |
| chr13 | 83990802 | 84001901 | 11100 | -6.89 | 9.22E-10 | 2.14E-09 |
| chr1 | 168639958 | 168642750 | 2793 | -6.93 | 9.23E-10 | 2.14E-09 |
| chr19 | 9107369 | 9108617 | 1249 | -6.96 | 9.23E-10 | 2.14E-09 |
| chr2 | 45538762 | 45555352 | 16591 | -7.6 | 9.24E-10 | 2.14E-09 |
| chr3 | 17659070 | 17661149 | 2080 | -5.34 | 9.24E-10 | 2.15E-09 |
| chr4 | 18005342 | 18007695 | 2354 | -6.88 | 9.26E-10 | 2.15E-09 |
| chr9 | 111132080 | 111136504 | 4425 | -6.88 | 9.26E-10 | 2.15E-09 |
| chr7 | 13683575 | 13696297 | 12723 | -7.22 | 9.26E-10 | 2.15E-09 |
| chrX | 74142651 | 74144268 | 1618 | -6.96 | 9.27E-10 | 2.15E-09 |
| chr1 | 7300517 | 7303967 | 3451 | -6.91 | 9.27E-10 | 2.15E-09 |
| chr5 | 31857301 | 31863433 | 6133 | -6.37 | 9.30E-10 | 2.16E-09 |
| chr8 | 133019117 | 133020054 | 938 | -6.95 | 9.31E-10 | 2.16E-09 |
| chr2 | 120002654 | 120004136 | 1483 | -6.87 | 9.35E-10 | 2.17E-09 |
| chr1 | 246245145 | 246247621 | 2477 | -6.89 | 9.36E-10 | 2.17E-09 |
| chr2 | 162198609 | 162200439 | 1831 | -6.89 | 9.36E-10 | 2.17E-09 |
| chr10 | 97886942 | 97889413 | 2472 | -2.7 | 9.38E-10 | 2.18E-09 |
| chr11 | 92206797 | 92210106 | 3310 | 6.97 | 9.39E-10 | 2.18E-09 |
| chr11 | 73924680 | 73934021 | 9342 | -6.88 | 9.40E-10 | 2.18E-09 |
| chr10 | 101729468 | 101731588 | 2121 | -7.01 | 9.40E-10 | 2.18E-09 |
| chrX | 10914467 | 10925746 | 11280 | -6.88 | 9.41E-10 | 2.18E-09 |
| chr18 | 24563265 | 24570699 | 7435 | -3.37 | 9.42E-10 | 2.18E-09 |
| chr4 | 101411893 | 101414777 | 2885 | -7.02 | 9.42E-10 | 2.18E-09 |
| chr8 | 29929053 | 29930054 | 1002 | -6.9 | 9.46E-10 | 2.19E-09 |
| chr4 | 160911067 | 160913597 | 2531 | -6.92 | 9.50E-10 | 2.20E-09 |
| chr19 | 49213971 | 49215167 | 1197 | -6.9 | 9.50E-10 | 2.20E-09 |
| chr9 | 26793129 | 26795565 | 2437 | -6.9 | 9.50E-10 | 2.20E-09 |
| chr4 | 170363926 | 170375175 | 11250 | -6.94 | 9.51E-10 | 2.20E-09 |
| chr11 | 68245990 | 68252553 | 6564 | -6.96 | 9.53E-10 | 2.21E-09 |

| | | | | | | |
|---|---|---|---|---|---|---|
| chr2 | 174246116 | 174249485 | 3370 | -6.96 | 9.53E-10 | 2.21E-09 |
| chr12 | 14057356 | 14059441 | 2086 | -7.1 | 9.53E-10 | 2.21E-09 |
| chr7 | 84485730 | 84488022 | 2293 | -7.15 | 9.56E-10 | 2.22E-09 |
| chr1 | 188224751 | 188226578 | 1828 | -5.58 | 9.57E-10 | 2.22E-09 |
| chr3 | 51318420 | 51357712 | 39293 | -6.58 | 9.58E-10 | 2.22E-09 |
| chr6 | 134523503 | 134525816 | 2314 | -6.99 | 9.58E-10 | 2.22E-09 |
| chr4 | 176805705 | 176809552 | 3848 | -6.95 | 9.58E-10 | 2.22E-09 |
| chr11 | 94448541 | 94451694 | 3154 | -6.95 | 9.61E-10 | 2.23E-09 |
| chr6 | 143337003 | 143349038 | 12036 | -7.16 | 9.61E-10 | 2.23E-09 |
| chr2 | 169687086 | 169688446 | 1361 | -5.44 | 9.63E-10 | 2.23E-09 |
| chr4 | 101604281 | 101607519 | 3239 | -6.87 | 9.63E-10 | 2.23E-09 |
| chr13 | 35944172 | 35947168 | 2997 | -7.03 | 9.63E-10 | 2.23E-09 |
| chr12 | 105479322 | 105481049 | 1728 | -7.11 | 9.69E-10 | 2.24E-09 |
| chr13 | 100853656 | 100866909 | 13254 | -7.11 | 9.69E-10 | 2.24E-09 |
| chr15 | 29113054 | 29117678 | 4625 | -7.11 | 9.69E-10 | 2.24E-09 |
| chr1 | 117476131 | 117477944 | 1814 | -6.91 | 9.70E-10 | 2.25E-09 |
| chr9 | 29078765 | 29080189 | 1425 | -6.88 | 9.70E-10 | 2.25E-09 |
| chr6 | 18459243 | 18460738 | 1496 | -5.9 | 9.71E-10 | 2.25E-09 |
| chr11 | 128145031 | 128147070 | 2040 | -7.21 | 9.74E-10 | 2.26E-09 |
| chr2 | 200934486 | 200944854 | 10369 | -7.05 | 9.75E-10 | 2.26E-09 |
| chr7 | 134905377 | 134908551 | 3175 | -6.87 | 9.75E-10 | 2.26E-09 |
| chr2 | 135634974 | 135637635 | 2662 | -6.89 | 9.75E-10 | 2.26E-09 |
| chr7 | 140745465 | 140761658 | 16194 | -7.37 | 9.76E-10 | 2.26E-09 |
| chr2 | 15367339 | 15368612 | 1274 | -6.94 | 9.77E-10 | 2.26E-09 |
| chr6 | 53196220 | 53197570 | 1351 | -6.94 | 9.77E-10 | 2.26E-09 |
| chr7 | 34531209 | 34534622 | 3414 | 7.19 | 9.77E-10 | 2.26E-09 |
| chr20 | 19958526 | 19959767 | 1242 | -6.91 | 9.78E-10 | 2.26E-09 |
| chr15 | 39329819 | 39343144 | 13326 | -7.45 | 9.78E-10 | 2.26E-09 |
| chr1 | 39370993 | 39373041 | 2049 | -6.88 | 9.79E-10 | 2.26E-09 |
| chr4 | 56697977 | 56708010 | 10034 | -2.72 | 9.81E-10 | 2.27E-09 |
| chr4 | 96575859 | 96578487 | 2629 | -6.93 | 9.82E-10 | 2.27E-09 |
| chr14 | 68175532 | 68194349 | 18818 | -7.82 | 9.87E-10 | 2.28E-09 |
| chr11 | 86918512 | 86920370 | 1859 | -5.52 | 9.88E-10 | 2.29E-09 |
| chr11 | 66466827 | 66468468 | 1642 | -6.86 | 9.88E-10 | 2.29E-09 |
| chr4 | 148505214 | 148507059 | 1846 | -6.9 | 9.89E-10 | 2.29E-09 |
| chr19 | 6246093 | 6248065 | 1973 | -6.94 | 9.90E-10 | 2.29E-09 |
| chr11 | 96080746 | 96082783 | 2038 | -6.87 | 9.96E-10 | 2.30E-09 |
| chr13 | 32879695 | 32881054 | 1360 | -6.87 | 9.96E-10 | 2.30E-09 |
| chr10 | 25151411 | 25165288 | 13878 | -7.46 | 9.99E-10 | 2.31E-09 |
| chr4 | 147860845 | 147862488 | 1644 | -5.55 | 1.00E-09 | 2.31E-09 |
| chr3 | 10653101 | 10658311 | 5211 | -6.42 | 1.00E-09 | 2.32E-09 |
| chr3 | 67743339 | 67745804 | 2466 | -6.93 | 1.00E-09 | 2.32E-09 |
| chr6 | 54087133 | 54088799 | 1667 | -6.89 | 1.00E-09 | 2.32E-09 |
| chr4 | 24677078 | 24696052 | 18975 | -7.62 | 1.00E-09 | 2.32E-09 |
| chr3 | 165360572 | 165516173 | 155602 | 2.12 | 1.00E-09 | 2.32E-09 |
| chr3 | 114371321 | 114397593 | 26273 | -7.46 | 1.00E-09 | 2.32E-09 |
| chr8 | 10256812 | 10259112 | 2301 | -6.86 | 1.00E-09 | 2.32E-09 |
| chr2 | 71000895 | 71003858 | 2964 | -6.88 | 1.01E-09 | 2.33E-09 |
| chr5 | 77549147 | 77554889 | 5743 | -6.88 | 1.01E-09 | 2.33E-09 |
| chr4 | 124093665 | 124095772 | 2108 | -7.02 | 1.01E-09 | 2.33E-09 |
| chr1 | 246596786 | 246598623 | 1838 | -7 | 1.01E-09 | 2.33E-09 |
| chr4 | 15967164 | 15969582 | 2419 | -6.98 | 1.01E-09 | 2.33E-09 |
| chr9 | 13294109 | 13321189 | 27081 | -3.58 | 1.01E-09 | 2.33E-09 |
| chr3 | 101615366 | 101616507 | 1142 | -6.97 | 1.01E-09 | 2.33E-09 |
| chr9 | 120010134 | 120042643 | 32510 | 6.71 | 1.01E-09 | 2.34E-09 |
| chr1 | 200681940 | 200684663 | 2724 | -6.97 | 1.01E-09 | 2.34E-09 |
| chr1 | 9913362 | 9915055 | 1694 | -7.03 | 1.01E-09 | 2.34E-09 |
| chr14 | 54128712 | 54129844 | 1133 | -6.96 | 1.02E-09 | 2.34E-09 |
| chr1 | 192398696 | 192400923 | 2228 | -6.89 | 1.02E-09 | 2.35E-09 |
| chr4 | 119305708 | 119309360 | 3653 | -6.86 | 1.02E-09 | 2.35E-09 |
| chr2 | 164722641 | 164724763 | 2123 | -6.9 | 1.02E-09 | 2.35E-09 |

| | | | | | | |
|---|---|---|---|---|---|---|
| chr9 | 29269234 | 29277200 | 7967 | -7.24 | 1.02E-09 | 2.35E-09 |
| chr13 | 100063429 | 100065244 | 1816 | -6.86 | 1.02E-09 | 2.36E-09 |
| chr8 | 76353378 | 76365081 | 11704 | -6.95 | 1.02E-09 | 2.36E-09 |
| chr6 | 82444230 | 82446210 | 1981 | -6.91 | 1.02E-09 | 2.37E-09 |
| chr6 | 9983590 | 9985284 | 1695 | -7.05 | 1.03E-09 | 2.38E-09 |
| chr16 | 59701583 | 59703186 | 1604 | -5.22 | 1.03E-09 | 2.38E-09 |
| chr1 | 86317787 | 86322491 | 4705 | -3.4 | 1.03E-09 | 2.38E-09 |
| chr1 | 25924681 | 25926582 | 1902 | -6.86 | 1.03E-09 | 2.38E-09 |
| chr1 | 62493900 | 62495216 | 1317 | -6.86 | 1.03E-09 | 2.38E-09 |
| chr10 | 75821552 | 75823625 | 2074 | -7.06 | 1.03E-09 | 2.38E-09 |
| chr8 | 27313398 | 27315276 | 1879 | 7.09 | 1.03E-09 | 2.38E-09 |
| chr12 | 15728232 | 15735410 | 7179 | -7.12 | 1.03E-09 | 2.39E-09 |
| chr16 | 71531912 | 71556779 | 24868 | -7.48 | 1.03E-09 | 2.39E-09 |
| chr11 | 10348960 | 10361251 | 12292 | -7.56 | 1.03E-09 | 2.39E-09 |
| chr22 | 27255581 | 27272626 | 17046 | -7.17 | 1.04E-09 | 2.40E-09 |
| chr2 | 193337344 | 193341445 | 4102 | -7.06 | 1.04E-09 | 2.40E-09 |
| chr21 | 18877145 | 18878456 | 1312 | -6.9 | 1.04E-09 | 2.41E-09 |
| chr21 | 46716363 | 46718178 | 1816 | -6.87 | 1.04E-09 | 2.41E-09 |
| chr1 | 3404187 | 3405910 | 1724 | -6.86 | 1.04E-09 | 2.41E-09 |
| chr3 | 122916653 | 122918252 | 1600 | -6.9 | 1.05E-09 | 2.42E-09 |
| chr2 | 210781756 | 210785878 | 4123 | 6.94 | 1.05E-09 | 2.43E-09 |
| chr13 | 99241982 | 99243676 | 1695 | -7.08 | 1.06E-09 | 2.43E-09 |
| chr5 | 10419164 | 10420838 | 1675 | -7.09 | 1.06E-09 | 2.43E-09 |
| chr6 | 36366007 | 36382401 | 16395 | -6.87 | 1.06E-09 | 2.45E-09 |
| chrX | 20605644 | 20607992 | 2349 | -6.89 | 1.06E-09 | 2.45E-09 |
| chr11 | 73769973 | 73779308 | 9336 | -7 | 1.06E-09 | 2.45E-09 |
| chr8 | 8543735 | 8545456 | 1722 | -7.09 | 1.06E-09 | 2.45E-09 |
| chr1 | 67578286 | 67587607 | 9322 | -7.1 | 1.07E-09 | 2.46E-09 |
| chr10 | 105591799 | 105593139 | 1341 | -6.96 | 1.07E-09 | 2.46E-09 |
| chr18 | 32575326 | 32577245 | 1920 | -6.96 | 1.07E-09 | 2.46E-09 |
| chr7 | 82470884 | 82472471 | 1588 | -7.19 | 1.07E-09 | 2.46E-09 |
| chrX | 72792110 | 72794447 | 2338 | -6.89 | 1.07E-09 | 2.46E-09 |
| chr4 | 92751931 | 92756928 | 4998 | -6.95 | 1.07E-09 | 2.46E-09 |
| chr10 | 89284959 | 89287311 | 2353 | -7.02 | 1.07E-09 | 2.47E-09 |
| chr2 | 73129563 | 73134988 | 5426 | -6.68 | 1.07E-09 | 2.47E-09 |
| chr5 | 107116379 | 107118065 | 1687 | -6.97 | 1.07E-09 | 2.47E-09 |
| chr12 | 52050317 | 52052289 | 1973 | -6.86 | 1.07E-09 | 2.47E-09 |
| chr16 | 28195804 | 28197329 | 1526 | -7.01 | 1.08E-09 | 2.48E-09 |
| chr7 | 91858032 | 91858947 | 916 | -6.96 | 1.08E-09 | 2.48E-09 |
| chr8 | 41048715 | 41050502 | 1788 | -6.85 | 1.08E-09 | 2.48E-09 |
| chr12 | 69376241 | 69378409 | 2169 | -6.85 | 1.08E-09 | 2.48E-09 |
| chr14 | 52298098 | 52300090 | 1993 | -6.85 | 1.08E-09 | 2.48E-09 |
| chr9 | 92851798 | 92853596 | 1799 | -2.57 | 1.08E-09 | 2.48E-09 |
| chr3 | 54220147 | 54222894 | 2748 | 5.26 | 1.08E-09 | 2.48E-09 |
| chr1 | 57052397 | 57076115 | 23719 | -7.36 | 1.08E-09 | 2.49E-09 |
| chr4 | 130973246 | 130975624 | 2379 | -7.04 | 1.08E-09 | 2.49E-09 |
| chr2 | 134850725 | 134853032 | 2308 | -6.85 | 1.08E-09 | 2.49E-09 |
| chr22 | 45026596 | 45050088 | 23493 | -7.67 | 1.08E-09 | 2.49E-09 |
| chr2 | 189116884 | 189118813 | 1930 | -6.93 | 1.08E-09 | 2.49E-09 |
| chr9 | 3805849 | 3809460 | 3612 | -6.93 | 1.08E-09 | 2.49E-09 |
| chr5 | 114680778 | 114682337 | 1560 | -6.9 | 1.08E-09 | 2.49E-09 |
| chr3 | 11567922 | 11569689 | 1768 | -7.04 | 1.08E-09 | 2.49E-09 |
| chr2 | 178909535 | 178914073 | 4539 | 7.04 | 1.08E-09 | 2.49E-09 |
| chr21 | 15447215 | 15451897 | 4683 | -6.88 | 1.09E-09 | 2.50E-09 |
| chr2 | 171008552 | 171010926 | 2375 | -7.03 | 1.09E-09 | 2.50E-09 |
| chr17 | 76301920 | 76303783 | 1864 | -6.9 | 1.09E-09 | 2.51E-09 |
| chr2 | 233850424 | 233858662 | 8239 | -7.56 | 1.09E-09 | 2.51E-09 |
| chr4 | 146159185 | 146160855 | 1671 | -6.01 | 1.09E-09 | 2.51E-09 |
| chr3 | 171605107 | 171607854 | 2748 | -7.11 | 1.09E-09 | 2.52E-09 |
| chr1 | 200805293 | 200806731 | 1439 | -7.12 | 1.10E-09 | 2.53E-09 |
| chr2 | 215521553 | 215545184 | 23632 | -7.82 | 1.10E-09 | 2.53E-09 |

| | | | | | | |
|---|---|---|---|---|---|---|
| chr6 | 109840567 | 109846429 | 5863 | -5.99 | 1.10E-09 | 2.53E-09 |
| chr12 | 94062387 | 94064287 | 1901 | -6.88 | 1.10E-09 | 2.53E-09 |
| chr3 | 149207834 | 149208958 | 1125 | -6.88 | 1.10E-09 | 2.53E-09 |
| chr16 | 31385728 | 31387527 | 1800 | -6.87 | 1.10E-09 | 2.54E-09 |
| chr4 | 189981607 | 189984656 | 3050 | -6.87 | 1.10E-09 | 2.54E-09 |
| chr3 | 142103819 | 142109447 | 5629 | -6.86 | 1.10E-09 | 2.54E-09 |
| chr7 | 51421226 | 51422770 | 1545 | -6.86 | 1.10E-09 | 2.54E-09 |
| chr8 | 108396191 | 108398386 | 2196 | -7.12 | 1.10E-09 | 2.54E-09 |
| chr12 | 45784640 | 45786074 | 1435 | -6.85 | 1.10E-09 | 2.54E-09 |
| chr21 | 43743573 | 43753401 | 9829 | -4.1 | 1.11E-09 | 2.55E-09 |
| chr16 | 81455119 | 81457085 | 1967 | -6.89 | 1.11E-09 | 2.55E-09 |
| chr4 | 124885831 | 124888728 | 2898 | -6.85 | 1.11E-09 | 2.55E-09 |
| chr7 | 57204408 | 57282153 | 77746 | -1.99 | 1.11E-09 | 2.56E-09 |
| chr12 | 117680069 | 117682556 | 2488 | 6.97 | 1.11E-09 | 2.56E-09 |
| chr1 | 175061023 | 175065134 | 4112 | 6.92 | 1.12E-09 | 2.57E-09 |
| chr6 | 18234532 | 18235975 | 1444 | -6.85 | 1.12E-09 | 2.57E-09 |
| chr8 | 120415211 | 120418958 | 3748 | 7.2 | 1.12E-09 | 2.57E-09 |
| chr9 | 129531015 | 129532550 | 1536 | -6.98 | 1.12E-09 | 2.57E-09 |
| chr16 | 19492379 | 19493707 | 1329 | -6.86 | 1.12E-09 | 2.57E-09 |
| chr6 | 169020269 | 169036511 | 16243 | -3.95 | 1.12E-09 | 2.57E-09 |
| chr4 | 110780110 | 110783375 | 3266 | -6.99 | 1.12E-09 | 2.57E-09 |
| chr10 | 29005045 | 29008367 | 3323 | -6.88 | 1.12E-09 | 2.57E-09 |
| chr12 | 111488589 | 111489815 | 1227 | -7.08 | 1.12E-09 | 2.57E-09 |
| chr2 | 208051171 | 208077039 | 25869 | -7.34 | 1.12E-09 | 2.58E-09 |
| chr1 | 100058478 | 100074687 | 16210 | -7.14 | 1.13E-09 | 2.59E-09 |
| chr10 | 79536381 | 79538318 | 1938 | -6.88 | 1.13E-09 | 2.59E-09 |
| chr7 | 24800146 | 24801524 | 1379 | -6.84 | 1.13E-09 | 2.59E-09 |
| chr7 | 102748414 | 102751606 | 3193 | -6.87 | 1.13E-09 | 2.59E-09 |
| chr15 | 51169967 | 51187386 | 17420 | -7.75 | 1.13E-09 | 2.59E-09 |
| chrX | 45981109 | 45984081 | 2973 | -6.9 | 1.13E-09 | 2.59E-09 |
| chr2 | 187948211 | 187952154 | 3944 | 6.95 | 1.14E-09 | 2.61E-09 |
| chr5 | 108811157 | 108822744 | 11588 | -7.39 | 1.14E-09 | 2.62E-09 |
| chr16 | 82204660 | 82207009 | 2350 | -6.85 | 1.14E-09 | 2.62E-09 |
| chrX | 37924887 | 37929077 | 4191 | -6.93 | 1.14E-09 | 2.62E-09 |
| chr6 | 1933161 | 1935786 | 2626 | -6.97 | 1.14E-09 | 2.62E-09 |
| chr6 | 150535945 | 150546077 | 10133 | -7.24 | 1.14E-09 | 2.62E-09 |
| chr2 | 89175352 | 89176760 | 1409 | -6.98 | 1.14E-09 | 2.62E-09 |
| chr3 | 175695507 | 175708055 | 12549 | -6.85 | 1.14E-09 | 2.62E-09 |
| chr8 | 132825632 | 132830577 | 4946 | -7.09 | 1.14E-09 | 2.63E-09 |
| chrX | 17814482 | 17816391 | 1910 | -6.9 | 1.15E-09 | 2.63E-09 |
| chrX | 34665602 | 34668169 | 2568 | -3.6 | 1.15E-09 | 2.63E-09 |
| chr2 | 9577192 | 9578546 | 1355 | -6.95 | 1.15E-09 | 2.63E-09 |
| chr2 | 99276552 | 99277897 | 1346 | -6.88 | 1.15E-09 | 2.63E-09 |
| chr4 | 84808976 | 84810859 | 1884 | -7 | 1.15E-09 | 2.64E-09 |
| chr9 | 37930614 | 37931912 | 1299 | -7 | 1.15E-09 | 2.64E-09 |
| chr12 | 122383806 | 122391584 | 7779 | -7.28 | 1.15E-09 | 2.64E-09 |
| chr12 | 55326328 | 55330635 | 4308 | -7.01 | 1.15E-09 | 2.64E-09 |
| chr6 | 129912688 | 129914301 | 1614 | -6.85 | 1.15E-09 | 2.64E-09 |
| chr6 | 13576545 | 13578575 | 2031 | -6.92 | 1.15E-09 | 2.64E-09 |
| chr4 | 134769526 | 134781597 | 12072 | 7.56 | 1.15E-09 | 2.65E-09 |
| chr17 | 77388474 | 77389492 | 1019 | -6.86 | 1.16E-09 | 2.65E-09 |
| chr14 | 61639441 | 61641196 | 1756 | -7.02 | 1.16E-09 | 2.65E-09 |
| chr14 | 105232417 | 105238863 | 6447 | -7.05 | 1.16E-09 | 2.65E-09 |
| chr15 | 53037589 | 53039950 | 2362 | -7.11 | 1.16E-09 | 2.65E-09 |
| chr1 | 101139412 | 101141097 | 1686 | -6.89 | 1.16E-09 | 2.66E-09 |
| chr11 | 121898336 | 121901168 | 2833 | -6.87 | 1.16E-09 | 2.66E-09 |
| chr19 | 5061638 | 5063498 | 1861 | -6.85 | 1.16E-09 | 2.66E-09 |
| chr20 | 60529316 | 60533213 | 3898 | -7.11 | 1.16E-09 | 2.67E-09 |
| chr4 | 32223987 | 32227024 | 3038 | -7.03 | 1.16E-09 | 2.67E-09 |
| chr17 | 28729590 | 28731211 | 1622 | -7.03 | 1.17E-09 | 2.67E-09 |
| chr3 | 151186539 | 151201583 | 15045 | -7.73 | 1.17E-09 | 2.67E-09 |

| | | | | | | |
|---|---|---|---|---|---|---|
| chrX | 53205170 | 53209519 | 4350 | -6.91 | 1.17E-09 | 2.67E-09 |
| chr10 | 60961347 | 60970539 | 9193 | -6.84 | 1.17E-09 | 2.67E-09 |
| chr14 | 91348732 | 91350744 | 2013 | -6.84 | 1.17E-09 | 2.67E-09 |
| chr12 | 75747966 | 75749996 | 2031 | -7.12 | 1.17E-09 | 2.68E-09 |
| chr5 | 55009318 | 55015476 | 6159 | -7.33 | 1.17E-09 | 2.68E-09 |
| chr14 | 51215854 | 51216966 | 1113 | -6.85 | 1.17E-09 | 2.68E-09 |
| chr4 | 25910048 | 25912034 | 1987 | -6.86 | 1.17E-09 | 2.68E-09 |
| chr6 | 122936700 | 122938633 | 1934 | -6.84 | 1.17E-09 | 2.69E-09 |
| chr13 | 45295588 | 45297767 | 2180 | -6.84 | 1.17E-09 | 2.69E-09 |
| chr3 | 57425746 | 57427682 | 1937 | -7.04 | 1.18E-09 | 2.70E-09 |
| chr21 | 34610303 | 34612923 | 2621 | -6.97 | 1.18E-09 | 2.70E-09 |
| chr1 | 42055341 | 42067977 | 12637 | -6.98 | 1.18E-09 | 2.70E-09 |
| chr3 | 153337572 | 153339581 | 2010 | -6.98 | 1.18E-09 | 2.70E-09 |
| chr12 | 117581928 | 117584721 | 2794 | -6.95 | 1.18E-09 | 2.70E-09 |
| chr15 | 52770469 | 52771567 | 1099 | -6.85 | 1.18E-09 | 2.71E-09 |
| chr1 | 230811771 | 230813071 | 1301 | -6.84 | 1.18E-09 | 2.71E-09 |
| chr8 | 121366616 | 121415804 | 49189 | -3.57 | 1.19E-09 | 2.72E-09 |
| chr7 | 25074220 | 25076664 | 2445 | -6.94 | 1.19E-09 | 2.72E-09 |
| chr6 | 160901044 | 160902921 | 1878 | -6.85 | 1.19E-09 | 2.72E-09 |
| chr9 | 136128769 | 136140110 | 11342 | -7.26 | 1.19E-09 | 2.72E-09 |
| chr2 | 75843948 | 75845055 | 1108 | -5.89 | 1.19E-09 | 2.72E-09 |
| chr2 | 196947605 | 196950109 | 2505 | -6.89 | 1.19E-09 | 2.73E-09 |
| chr7 | 168724 | 171306 | 2583 | -7.34 | 1.20E-09 | 2.74E-09 |
| chr9 | 91330057 | 91332969 | 2913 | -7.09 | 1.20E-09 | 2.74E-09 |
| chr3 | 95572481 | 95579323 | 6843 | -6.89 | 1.20E-09 | 2.75E-09 |
| chr12 | 26233925 | 26235862 | 1938 | -6.83 | 1.20E-09 | 2.75E-09 |
| chr20 | 48104229 | 48115642 | 11414 | -7.03 | 1.20E-09 | 2.75E-09 |
| chr15 | 57962419 | 57964129 | 1711 | -6.85 | 1.20E-09 | 2.75E-09 |
| chr6 | 34692934 | 34694616 | 1683 | -6.85 | 1.21E-09 | 2.76E-09 |
| chr1 | 210127954 | 210129353 | 1400 | -5.16 | 1.21E-09 | 2.76E-09 |
| chr19 | 30547174 | 30548290 | 1117 | -6.87 | 1.21E-09 | 2.77E-09 |
| chr8 | 75280617 | 75283993 | 3377 | -6.83 | 1.21E-09 | 2.77E-09 |
| chr5 | 170897923 | 170901171 | 3249 | -7.04 | 1.21E-09 | 2.77E-09 |
| chr7 | 129933537 | 129935632 | 2096 | -6.89 | 1.21E-09 | 2.77E-09 |
| chr3 | 126943186 | 126971217 | 28032 | -6.91 | 1.21E-09 | 2.78E-09 |
| chr2 | 24407666 | 24422441 | 14776 | -7.1 | 1.22E-09 | 2.79E-09 |
| chr10 | 32190226 | 32191430 | 1205 | -7.08 | 1.22E-09 | 2.79E-09 |
| chr11 | 11621648 | 11623522 | 1875 | -6.89 | 1.22E-09 | 2.79E-09 |
| chr17 | 34005100 | 34005998 | 899 | -6.89 | 1.22E-09 | 2.79E-09 |
| chr9 | 110590739 | 110593161 | 2423 | -6.96 | 1.22E-09 | 2.79E-09 |
| chr2 | 70529503 | 70530448 | 946 | -6.83 | 1.22E-09 | 2.79E-09 |
| chr6 | 83916856 | 83918693 | 1838 | -6.98 | 1.22E-09 | 2.79E-09 |
| chr7 | 140946259 | 141219048 | 272790 | 2.77 | 1.22E-09 | 2.80E-09 |
| chr13 | 51678057 | 51705384 | 27328 | -7.52 | 1.23E-09 | 2.80E-09 |
| chr4 | 68076478 | 68090238 | 13761 | 2.01 | 1.23E-09 | 2.80E-09 |
| chr10 | 32462776 | 32464607 | 1832 | -7.05 | 1.23E-09 | 2.81E-09 |
| chr22 | 30587660 | 30598932 | 11273 | -7.26 | 1.23E-09 | 2.81E-09 |
| chr17 | 54839331 | 54840149 | 819 | -4.93 | 1.23E-09 | 2.81E-09 |
| chr4 | 13437373 | 13440032 | 2660 | -6.83 | 1.23E-09 | 2.81E-09 |
| chr18 | 12540886 | 12555537 | 14652 | -7.06 | 1.23E-09 | 2.82E-09 |
| chr3 | 127968770 | 127980115 | 11346 | -7.25 | 1.24E-09 | 2.83E-09 |
| chr7 | 157416199 | 157438597 | 22399 | -2.78 | 1.24E-09 | 2.83E-09 |
| chr3 | 183455688 | 183459000 | 3313 | -6.83 | 1.24E-09 | 2.84E-09 |
| chr4 | 64397493 | 64408421 | 10929 | -6.95 | 1.24E-09 | 2.84E-09 |
| chr4 | 132899586 | 132903496 | 3911 | -6.97 | 1.24E-09 | 2.84E-09 |
| chr1 | 17390636 | 17392417 | 1782 | -6.95 | 1.24E-09 | 2.84E-09 |
| chr8 | 90451314 | 90458281 | 6968 | -7.02 | 1.24E-09 | 2.84E-09 |
| chr12 | 76267495 | 76269940 | 2446 | -6.94 | 1.24E-09 | 2.84E-09 |
| chr11 | 75669346 | 75685805 | 16460 | -7.29 | 1.24E-09 | 2.84E-09 |
| chr12 | 21395761 | 21397370 | 1610 | -6.88 | 1.25E-09 | 2.85E-09 |
| chr9 | 112997946 | 112999273 | 1328 | -5.93 | 1.25E-09 | 2.85E-09 |

| | | | | | | |
|---|---|---|---|---|---|---|
| chr5 | 154022166 | 154024841 | 2676 | -6.85 | 1.25E-09 | 2.85E-09 |
| chr4 | 87030831 | 87034207 | 3377 | -7.12 | 1.25E-09 | 2.85E-09 |
| chr11 | 44232560 | 44234492 | 1933 | -6.83 | 1.25E-09 | 2.86E-09 |
| chr2 | 11766402 | 11768268 | 1867 | -7.29 | 1.25E-09 | 2.86E-09 |
| chr2 | 226339471 | 226343386 | 3916 | -6.93 | 1.25E-09 | 2.86E-09 |
| chr9 | 130344904 | 130346454 | 1551 | -7.03 | 1.25E-09 | 2.86E-09 |
| chr3 | 40011723 | 40014536 | 2814 | 6.9 | 1.25E-09 | 2.86E-09 |
| chr5 | 53622787 | 53654168 | 31382 | -2.65 | 1.26E-09 | 2.87E-09 |
| chr12 | 64314999 | 64323014 | 8016 | -7.38 | 1.26E-09 | 2.87E-09 |
| chr5 | 105697540 | 105708445 | 10906 | -6.87 | 1.26E-09 | 2.87E-09 |
| chr1 | 199222791 | 199226282 | 3492 | -2.96 | 1.26E-09 | 2.87E-09 |
| chr19 | 14203876 | 14205161 | 1286 | -7.01 | 1.26E-09 | 2.87E-09 |
| chr5 | 31564259 | 31567401 | 3143 | -7.01 | 1.26E-09 | 2.87E-09 |
| chr11 | 102921858 | 102923790 | 1933 | -7.04 | 1.26E-09 | 2.87E-09 |
| chr3 | 24125978 | 24145074 | 19097 | -7.58 | 1.26E-09 | 2.87E-09 |
| chrX | 22026000 | 22028206 | 2207 | -6.9 | 1.26E-09 | 2.87E-09 |
| chr3 | 138026491 | 138032383 | 5893 | -2.46 | 1.26E-09 | 2.87E-09 |
| chr20 | 52568829 | 52570272 | 1444 | -6.86 | 1.26E-09 | 2.88E-09 |
| chrX | 153154670 | 153161990 | 7321 | -6.84 | 1.26E-09 | 2.88E-09 |
| chr1 | 214517129 | 214520429 | 3301 | -6.87 | 1.26E-09 | 2.88E-09 |
| chr6 | 61967021 | 62060707 | 93687 | 1.14 | 1.27E-09 | 2.89E-09 |
| chr4 | 7440321 | 7442122 | 1802 | -6.83 | 1.27E-09 | 2.90E-09 |
| chr7 | 102467392 | 102469091 | 1700 | -7.05 | 1.27E-09 | 2.91E-09 |
| chr4 | 147535000 | 147539977 | 4978 | -6.83 | 1.27E-09 | 2.91E-09 |
| chr11 | 74728856 | 74731931 | 3076 | -7.03 | 1.28E-09 | 2.91E-09 |
| chr4 | 128945796 | 128946958 | 1163 | -6.83 | 1.28E-09 | 2.91E-09 |
| chr5 | 132537264 | 132538729 | 1466 | -6.83 | 1.28E-09 | 2.92E-09 |
| chr9 | 130941007 | 130942726 | 1720 | -7.06 | 1.28E-09 | 2.92E-09 |
| chr5 | 76441067 | 76444500 | 3434 | -6.83 | 1.29E-09 | 2.93E-09 |
| chr6 | 53074936 | 53078797 | 3862 | -6.83 | 1.29E-09 | 2.94E-09 |
| chr2 | 155443730 | 156220695 | 776966 | 3.38 | 1.29E-09 | 2.94E-09 |
| chr17 | 62087001 | 62088079 | 1079 | -6.89 | 1.29E-09 | 2.94E-09 |
| chr1 | 49246648 | 49279134 | 32487 | 1.97 | 1.29E-09 | 2.94E-09 |
| chr14 | 28758585 | 28913443 | 154859 | -2.44 | 1.29E-09 | 2.94E-09 |
| chr21 | 16602071 | 16606853 | 4783 | -6.85 | 1.29E-09 | 2.95E-09 |
| chr18 | 54803201 | 54814094 | 10894 | -1.96 | 1.30E-09 | 2.95E-09 |
| chr4 | 60342318 | 60349768 | 7451 | -6.84 | 1.30E-09 | 2.96E-09 |
| chrX | 35390632 | 35403138 | 12507 | -6.96 | 1.30E-09 | 2.97E-09 |
| chr12 | 27395708 | 27397019 | 1312 | -5.45 | 1.30E-09 | 2.97E-09 |
| chr13 | 80471850 | 80476271 | 4422 | -6.82 | 1.30E-09 | 2.97E-09 |
| chr8 | 133625234 | 133629967 | 4734 | -7.16 | 1.30E-09 | 2.97E-09 |
| chr1 | 193700318 | 193701794 | 1477 | -6.88 | 1.31E-09 | 2.97E-09 |
| chr1 | 225169934 | 225173955 | 4022 | -6.88 | 1.31E-09 | 2.97E-09 |
| chr9 | 86701052 | 86707174 | 6123 | -1.74 | 1.31E-09 | 2.97E-09 |
| chr6 | 144583521 | 144593203 | 9683 | -7.06 | 1.31E-09 | 2.98E-09 |
| chr21 | 40777366 | 40778781 | 1416 | -6.93 | 1.31E-09 | 2.98E-09 |
| chr7 | 20713243 | 20716068 | 2826 | -4.37 | 1.31E-09 | 2.98E-09 |
| chr6 | 24181860 | 24184351 | 2492 | -7.2 | 1.31E-09 | 2.99E-09 |
| chr5 | 25384755 | 27019666 | 1634912 | 1.16 | 1.31E-09 | 2.99E-09 |
| chr6 | 100002539 | 100004168 | 1630 | -6.92 | 1.31E-09 | 2.99E-09 |
| chr13 | 83513011 | 83520517 | 7507 | -6.88 | 1.32E-09 | 2.99E-09 |
| chr9 | 4856695 | 4857889 | 1195 | -6.88 | 1.32E-09 | 2.99E-09 |
| chrX | 149346025 | 149348120 | 2096 | -6.99 | 1.32E-09 | 3.00E-09 |
| chr2 | 113084169 | 113086409 | 2241 | -6.83 | 1.32E-09 | 3.01E-09 |
| chr3 | 109126660 | 109129146 | 2487 | -6.83 | 1.32E-09 | 3.01E-09 |
| chr2 | 190380211 | 190383007 | 2797 | -7.37 | 1.32E-09 | 3.01E-09 |
| chr3 | 94194535 | 94205007 | 10473 | -7.36 | 1.33E-09 | 3.03E-09 |
| chr1 | 32996129 | 32998696 | 2568 | -7.01 | 1.33E-09 | 3.03E-09 |
| chr3 | 193975514 | 193976947 | 1434 | -6.9 | 1.33E-09 | 3.03E-09 |
| chr5 | 60034857 | 60037426 | 2570 | -6.87 | 1.34E-09 | 3.04E-09 |
| chr18 | 25307272 | 25310657 | 3386 | -6.9 | 1.34E-09 | 3.04E-09 |

| | | | | | | |
|---|---|---|---|---|---|---|
| chr5 | 157056717 | 157058344 | 1628 | -7.31 | 1.34E-09 | 3.05E-09 |
| chr10 | 25376197 | 25395829 | 19633 | -6.48 | 1.34E-09 | 3.05E-09 |
| chrX | 28079249 | 28090741 | 11493 | -7.22 | 1.34E-09 | 3.05E-09 |
| chr11 | 134270207 | 134271565 | 1359 | -5.37 | 1.35E-09 | 3.06E-09 |
| chr9 | 126720543 | 126731050 | 10508 | -7.3 | 1.35E-09 | 3.07E-09 |
| chr19 | 14031452 | 14032420 | 969 | -6.89 | 1.36E-09 | 3.08E-09 |
| chr6 | 150228671 | 150231177 | 2507 | -6.84 | 1.36E-09 | 3.09E-09 |
| chr2 | 230799154 | 230800574 | 1421 | -6.82 | 1.36E-09 | 3.09E-09 |
| chr7 | 80482019 | 80504960 | 22942 | -7.63 | 1.36E-09 | 3.09E-09 |
| chrX | 29225624 | 29230017 | 4394 | -7.09 | 1.36E-09 | 3.09E-09 |
| chr2 | 196843802 | 196846210 | 2409 | -7.04 | 1.36E-09 | 3.10E-09 |
| chr8 | 59550556 | 59563476 | 12921 | -7.17 | 1.37E-09 | 3.11E-09 |
| chr20 | 1772335 | 1774563 | 2229 | -7.05 | 1.37E-09 | 3.12E-09 |
| chr2 | 24331416 | 24372214 | 40799 | -8.33 | 1.37E-09 | 3.12E-09 |
| chr19 | 5346881 | 5348083 | 1203 | -6.84 | 1.37E-09 | 3.12E-09 |
| chr2 | 61819877 | 61821886 | 2010 | -6.85 | 1.37E-09 | 3.12E-09 |
| chr9 | 110907645 | 110909797 | 2153 | -7.05 | 1.37E-09 | 3.13E-09 |
| chr20 | 20082466 | 20084147 | 1682 | -5.07 | 1.38E-09 | 3.13E-09 |
| chr5 | 95307355 | 95308883 | 1529 | -6.93 | 1.38E-09 | 3.14E-09 |
| chr6 | 149084193 | 149086662 | 2470 | -6.93 | 1.38E-09 | 3.14E-09 |
| chr4 | 168163502 | 168167027 | 3526 | -6.92 | 1.39E-09 | 3.15E-09 |
| chr10 | 11597529 | 11598927 | 1399 | -6.85 | 1.39E-09 | 3.16E-09 |
| chr4 | 146393196 | 146395781 | 2586 | -6.98 | 1.39E-09 | 3.16E-09 |
| chr12 | 132873113 | 132889558 | 16446 | -6.98 | 1.39E-09 | 3.17E-09 |
| chr11 | 122986158 | 122987874 | 1717 | -6.81 | 1.40E-09 | 3.17E-09 |
| chr1 | 221473801 | 221475680 | 1880 | -6.97 | 1.40E-09 | 3.18E-09 |
| chr4 | 176959761 | 176963716 | 3956 | -4.27 | 1.40E-09 | 3.19E-09 |
| chr5 | 142347671 | 142348982 | 1312 | -6.91 | 1.40E-09 | 3.19E-09 |
| chr17 | 61222467 | 61223563 | 1097 | -6.81 | 1.40E-09 | 3.19E-09 |
| chr1 | 174115083 | 174118836 | 3754 | -7.21 | 1.40E-09 | 3.19E-09 |
| chr4 | 131147636 | 131150373 | 2738 | -6.82 | 1.41E-09 | 3.20E-09 |
| chr1 | 59253237 | 59255050 | 1814 | -6.9 | 1.41E-09 | 3.20E-09 |
| chr4 | 146525770 | 146528138 | 2369 | -6.81 | 1.41E-09 | 3.21E-09 |
| chr1 | 182316798 | 182319985 | 3188 | 6.93 | 1.41E-09 | 3.21E-09 |
| chrX | 135643171 | 135655423 | 12253 | -7.13 | 1.42E-09 | 3.22E-09 |
| chr1 | 117221655 | 117224520 | 2866 | -7.08 | 1.42E-09 | 3.22E-09 |
| chr4 | 161282090 | 161284579 | 2490 | -6.89 | 1.42E-09 | 3.22E-09 |
| chr4 | 186880804 | 186882793 | 1990 | -6.82 | 1.42E-09 | 3.22E-09 |
| chr12 | 15800244 | 15801403 | 1160 | -6.81 | 1.42E-09 | 3.22E-09 |
| chr2 | 58968032 | 58982190 | 14159 | -1.7 | 1.42E-09 | 3.23E-09 |
| chr7 | 94235334 | 94239412 | 4079 | -6.81 | 1.42E-09 | 3.23E-09 |
| chr8 | 83775187 | 83780399 | 5213 | -6.83 | 1.42E-09 | 3.23E-09 |
| chr14 | 65007927 | 65009580 | 1654 | -6.85 | 1.43E-09 | 3.23E-09 |
| chr2 | 192979338 | 192980394 | 1057 | -6.81 | 1.43E-09 | 3.23E-09 |
| chr2 | 24120378 | 24122565 | 2188 | -6.84 | 1.43E-09 | 3.24E-09 |
| chrX | 16191420 | 16203410 | 11991 | -6.84 | 1.43E-09 | 3.24E-09 |
| chr5 | 107617264 | 107620807 | 3544 | -7.02 | 1.43E-09 | 3.24E-09 |
| chr7 | 99481075 | 99488564 | 7490 | -7.02 | 1.43E-09 | 3.24E-09 |
| chr2 | 63456820 | 63460140 | 3321 | -6.81 | 1.43E-09 | 3.24E-09 |
| chr1 | 60188345 | 60203645 | 15301 | -6.81 | 1.43E-09 | 3.25E-09 |
| chr1 | 201308833 | 201313312 | 4480 | -6.45 | 1.43E-09 | 3.25E-09 |
| chrX | 102870845 | 102873544 | 2700 | -6.85 | 1.44E-09 | 3.26E-09 |
| chr22 | 31710769 | 31712431 | 1663 | -7.03 | 1.44E-09 | 3.26E-09 |
| chr7 | 141237795 | 141240758 | 2964 | -6.85 | 1.44E-09 | 3.26E-09 |
| chr6 | 10225049 | 10228922 | 3874 | -1.9 | 1.44E-09 | 3.26E-09 |
| chr4 | 129554723 | 129560093 | 5371 | -6.81 | 1.44E-09 | 3.27E-09 |
| chr4 | 93084306 | 93089283 | 4978 | -6.81 | 1.44E-09 | 3.27E-09 |
| chr12 | 74525629 | 74527442 | 1814 | -6.83 | 1.44E-09 | 3.27E-09 |
| chrX | 110150029 | 110223645 | 73617 | 4.08 | 1.44E-09 | 3.27E-09 |
| chr3 | 173931832 | 173968002 | 36171 | 2.07 | 1.44E-09 | 3.27E-09 |
| chrX | 68011065 | 68034679 | 23615 | -6.03 | 1.45E-09 | 3.28E-09 |

| | | | | | | |
|---|---|---|---|---|---|---|
| chr1 | 198371560 | 198373792 | 2233 | -6.88 | 1.45E-09 | 3.28E-09 |
| chr8 | 23498206 | 23533002 | 34797 | -2.04 | 1.45E-09 | 3.28E-09 |
| chr7 | 66716278 | 66718363 | 2086 | -6.82 | 1.45E-09 | 3.28E-09 |
| chr16 | 47316753 | 47332836 | 16084 | -7.64 | 1.45E-09 | 3.28E-09 |
| chr3 | 11786410 | 11787941 | 1532 | -6.85 | 1.45E-09 | 3.29E-09 |
| chr2 | 35044378 | 35237747 | 193370 | 1.64 | 1.45E-09 | 3.29E-09 |
| chr11 | 61987233 | 62030774 | 43542 | -2.8 | 1.45E-09 | 3.30E-09 |
| chr3 | 135802886 | 135811666 | 8781 | -6.95 | 1.46E-09 | 3.30E-09 |
| chr6 | 143394776 | 143396241 | 1466 | -6.95 | 1.46E-09 | 3.30E-09 |
| chr5 | 10779417 | 10781657 | 2241 | -5.05 | 1.46E-09 | 3.30E-09 |
| chr13 | 77958727 | 77961313 | 2587 | -6.92 | 1.46E-09 | 3.31E-09 |
| chr2 | 136184530 | 136186425 | 1896 | -7.03 | 1.46E-09 | 3.31E-09 |
| chr6 | 4204672 | 4215790 | 11119 | -7.19 | 1.46E-09 | 3.31E-09 |
| chr12 | 75045334 | 75046902 | 1569 | -6.83 | 1.46E-09 | 3.31E-09 |
| chr2 | 220068747 | 220070042 | 1296 | -6.83 | 1.46E-09 | 3.31E-09 |
| chr2 | 227810620 | 227826033 | 15414 | -7.44 | 1.46E-09 | 3.31E-09 |
| chr4 | 25472911 | 25474539 | 1629 | -6.97 | 1.46E-09 | 3.31E-09 |
| chr5 | 59079650 | 59081593 | 1944 | -6.8 | 1.47E-09 | 3.33E-09 |
| chr5 | 21216381 | 21226012 | 9632 | -6.81 | 1.47E-09 | 3.33E-09 |
| chr6 | 51983910 | 51985687 | 1778 | -6.81 | 1.47E-09 | 3.33E-09 |
| chr17 | 1650746 | 1652950 | 2205 | -6.98 | 1.47E-09 | 3.33E-09 |
| chr8 | 63983991 | 63990267 | 6277 | -6.9 | 1.47E-09 | 3.33E-09 |
| chr15 | 45888478 | 45890244 | 1767 | -6.81 | 1.48E-09 | 3.34E-09 |
| chr10 | 6722976 | 6725195 | 2220 | -6.81 | 1.48E-09 | 3.34E-09 |
| chr3 | 160091069 | 160092065 | 997 | -6.81 | 1.48E-09 | 3.34E-09 |
| chr6 | 90882530 | 90884397 | 1868 | -6.86 | 1.48E-09 | 3.34E-09 |
| chr2 | 25658165 | 25659092 | 928 | -6.89 | 1.48E-09 | 3.35E-09 |
| chr6 | 7255467 | 7257432 | 1966 | -6.89 | 1.48E-09 | 3.36E-09 |
| chr15 | 77490390 | 77499923 | 9534 | -7.28 | 1.49E-09 | 3.36E-09 |
| chr4 | 175610136 | 175612369 | 2234 | -7.06 | 1.49E-09 | 3.37E-09 |
| chr22 | 25322398 | 25326745 | 4348 | -7 | 1.49E-09 | 3.37E-09 |
| chr14 | 61707022 | 61709043 | 2022 | -6.85 | 1.49E-09 | 3.37E-09 |
| chr14 | 27417633 | 27420803 | 3171 | -6.91 | 1.49E-09 | 3.38E-09 |
| chr1 | 94853827 | 94856561 | 2735 | -6.82 | 1.50E-09 | 3.38E-09 |
| chr5 | 42845879 | 42863452 | 17574 | -7.21 | 1.50E-09 | 3.39E-09 |
| chr12 | 70250936 | 70254277 | 3342 | -6.88 | 1.50E-09 | 3.39E-09 |
| chr8 | 27691142 | 27692455 | 1314 | -6.97 | 1.50E-09 | 3.39E-09 |
| chr1 | 231179797 | 231192457 | 12661 | -7.41 | 1.50E-09 | 3.40E-09 |
| chr8 | 9304227 | 9305639 | 1413 | -6.83 | 1.50E-09 | 3.40E-09 |
| chr8 | 122966806 | 122974771 | 7966 | -7.01 | 1.51E-09 | 3.40E-09 |
| chr7 | 13595811 | 13615164 | 19354 | -7.17 | 1.51E-09 | 3.41E-09 |
| chr7 | 102049646 | 102051062 | 1417 | -6.81 | 1.51E-09 | 3.41E-09 |
| chr6 | 48177320 | 48178888 | 1569 | -6.85 | 1.51E-09 | 3.41E-09 |
| chr2 | 205478908 | 205480485 | 1578 | -6.8 | 1.51E-09 | 3.41E-09 |
| chr3 | 11393240 | 11395593 | 2354 | -6.8 | 1.51E-09 | 3.41E-09 |
| chr1 | 180259731 | 180302932 | 43202 | -7.95 | 1.51E-09 | 3.42E-09 |
| chr4 | 26499776 | 26502228 | 2453 | -6.98 | 1.51E-09 | 3.42E-09 |
| chr18 | 70751231 | 70758649 | 7419 | -6.87 | 1.51E-09 | 3.42E-09 |
| chr6 | 113376560 | 113392810 | 16251 | -7.17 | 1.51E-09 | 3.42E-09 |
| chr4 | 104135865 | 104138515 | 2651 | -7.07 | 1.52E-09 | 3.43E-09 |
| chr1 | 172365370 | 172367969 | 2600 | -6.85 | 1.52E-09 | 3.43E-09 |
| chr2 | 44838077 | 44839238 | 1162 | -6.83 | 1.52E-09 | 3.44E-09 |
| chr4 | 120585757 | 120587767 | 2011 | -6.83 | 1.52E-09 | 3.44E-09 |
| chr2 | 138638304 | 138640327 | 2024 | -6.8 | 1.52E-09 | 3.45E-09 |
| chr9 | 125221593 | 125224641 | 3049 | -6.88 | 1.53E-09 | 3.46E-09 |
| chr6 | 35715011 | 35716661 | 1651 | -6.8 | 1.53E-09 | 3.46E-09 |
| chrX | 18790433 | 18793063 | 2631 | -6.84 | 1.53E-09 | 3.46E-09 |
| chr4 | 125832927 | 125835344 | 2418 | -7.03 | 1.53E-09 | 3.47E-09 |
| chr11 | 87766551 | 87769383 | 2833 | -6.92 | 1.54E-09 | 3.47E-09 |
| chr13 | 89800516 | 89803318 | 2803 | -6.92 | 1.54E-09 | 3.47E-09 |
| chr2 | 206815880 | 206817691 | 1812 | -6.82 | 1.54E-09 | 3.47E-09 |

| | | | | | | |
|---|---|---|---|---|---|---|
| chr9 | 36179952 | 36181547 | 1596 | -6.95 | 1.54E-09 | 3.48E-09 |
| chr4 | 180682355 | 180689652 | 7298 | -6.91 | 1.54E-09 | 3.48E-09 |
| chr14 | 53689297 | 53711149 | 21853 | -7.39 | 1.54E-09 | 3.48E-09 |
| chr7 | 133149511 | 133151768 | 2258 | -6.85 | 1.55E-09 | 3.49E-09 |
| chr7 | 28464562 | 28705853 | 241292 | 2.03 | 1.55E-09 | 3.49E-09 |
| chr7 | 17211700 | 17213765 | 2066 | -6.83 | 1.55E-09 | 3.50E-09 |
| chr20 | 11329533 | 11344132 | 14600 | -6.89 | 1.55E-09 | 3.50E-09 |
| chr5 | 36901110 | 36902917 | 1808 | -6.79 | 1.55E-09 | 3.51E-09 |
| chr6 | 90809799 | 90826969 | 17171 | -1.74 | 1.56E-09 | 3.51E-09 |
| chr2 | 159775978 | 159794739 | 18762 | -6.79 | 1.56E-09 | 3.51E-09 |
| chr15 | 99281787 | 99283243 | 1457 | -6.81 | 1.56E-09 | 3.52E-09 |
| chr13 | 89681176 | 89687318 | 6143 | -6.88 | 1.56E-09 | 3.52E-09 |
| chr11 | 58695922 | 58792122 | 96201 | 2.4 | 1.56E-09 | 3.52E-09 |
| chr22 | 42633488 | 42634771 | 1284 | -6.79 | 1.56E-09 | 3.53E-09 |
| chr4 | 91822257 | 91824889 | 2633 | -6.79 | 1.56E-09 | 3.53E-09 |
| chr6 | 109044217 | 109046425 | 2209 | -6.79 | 1.56E-09 | 3.53E-09 |
| chr12 | 102258416 | 102263559 | 5144 | -6.99 | 1.57E-09 | 3.54E-09 |
| chr1 | 51489649 | 51491561 | 1913 | -7 | 1.58E-09 | 3.55E-09 |
| chr3 | 15807440 | 15813817 | 6378 | -6.87 | 1.58E-09 | 3.57E-09 |
| chr3 | 113402370 | 113404849 | 2480 | -7 | 1.58E-09 | 3.57E-09 |
| chr2 | 233142981 | 233144547 | 1567 | -7.06 | 1.58E-09 | 3.57E-09 |
| chr6 | 4013488 | 4014567 | 1080 | -6.8 | 1.58E-09 | 3.57E-09 |
| chr3 | 17163834 | 17179609 | 15776 | -7.21 | 1.59E-09 | 3.58E-09 |
| chr4 | 163184521 | 163186813 | 2293 | -6.84 | 1.59E-09 | 3.58E-09 |
| chr6 | 85608640 | 85610562 | 1923 | -6.82 | 1.59E-09 | 3.58E-09 |
| chr9 | 124188924 | 124191022 | 2099 | -6.86 | 1.59E-09 | 3.58E-09 |
| chr1 | 59957377 | 59962152 | 4776 | -6.8 | 1.59E-09 | 3.59E-09 |
| chr2 | 224522862 | 224527632 | 4771 | -6.8 | 1.59E-09 | 3.59E-09 |
| chr3 | 10795648 | 10815667 | 20020 | -7.5 | 1.60E-09 | 3.60E-09 |
| chr7 | 82849549 | 82853277 | 3729 | -6.79 | 1.60E-09 | 3.60E-09 |
| chrX | 107924693 | 107927243 | 2551 | -6.79 | 1.60E-09 | 3.60E-09 |
| chr5 | 87560580 | 87572928 | 12349 | -6.95 | 1.60E-09 | 3.61E-09 |
| chr7 | 65844205 | 65846038 | 1834 | -6.79 | 1.60E-09 | 3.61E-09 |
| chr4 | 107261327 | 107262978 | 1652 | -6.84 | 1.61E-09 | 3.62E-09 |
| chr15 | 59545566 | 59547140 | 1575 | -7.12 | 1.61E-09 | 3.62E-09 |
| chr1 | 150732820 | 150735162 | 2343 | -7.02 | 1.61E-09 | 3.63E-09 |
| chr3 | 99712166 | 99713536 | 1371 | -5.18 | 1.62E-09 | 3.65E-09 |
| chr3 | 177097169 | 177099136 | 1968 | -6.83 | 1.62E-09 | 3.65E-09 |
| chr1 | 81196749 | 81204842 | 8094 | -6.93 | 1.62E-09 | 3.65E-09 |
| chr10 | 88307317 | 88310299 | 2983 | -6.85 | 1.62E-09 | 3.65E-09 |
| chr16 | 24658333 | 24660102 | 1770 | -6.94 | 1.62E-09 | 3.66E-09 |
| chr11 | 12002376 | 12014725 | 12350 | -6.9 | 1.62E-09 | 3.66E-09 |
| chr1 | 167707083 | 167708497 | 1415 | -6.83 | 1.63E-09 | 3.68E-09 |
| chr10 | 35047616 | 35049320 | 1705 | -6.79 | 1.64E-09 | 3.68E-09 |
| chr7 | 79329286 | 79331574 | 2289 | -6.82 | 1.64E-09 | 3.69E-09 |
| chr6 | 117303782 | 117305103 | 1322 | -6.1 | 1.65E-09 | 3.71E-09 |
| chr10 | 102858820 | 102861148 | 2329 | -6.84 | 1.65E-09 | 3.71E-09 |
| chr15 | 68268526 | 68272100 | 3575 | -6.79 | 1.65E-09 | 3.72E-09 |
| chr4 | 177875240 | 177877064 | 1825 | -6.79 | 1.65E-09 | 3.72E-09 |
| chr3 | 100342874 | 100345660 | 2787 | -6.79 | 1.65E-09 | 3.73E-09 |
| chr8 | 32318918 | 32320948 | 2031 | -6.85 | 1.65E-09 | 3.73E-09 |
| chr12 | 50657402 | 50658535 | 1134 | -6.79 | 1.66E-09 | 3.73E-09 |
| chr3 | 110464066 | 110466108 | 2043 | -7.1 | 1.66E-09 | 3.74E-09 |
| chr10 | 71060475 | 71062539 | 2065 | -6.83 | 1.66E-09 | 3.74E-09 |
| chr5 | 153443842 | 153444950 | 1109 | -6.79 | 1.67E-09 | 3.75E-09 |
| chr2 | 57748477 | 57750026 | 1550 | -6.85 | 1.67E-09 | 3.76E-09 |
| chr8 | 42635221 | 42637425 | 2205 | -5.13 | 1.67E-09 | 3.77E-09 |
| chr1 | 187176318 | 187181783 | 5466 | 7.08 | 1.67E-09 | 3.77E-09 |
| chr9 | 27204875 | 27207266 | 2392 | 6.99 | 1.68E-09 | 3.77E-09 |
| chr3 | 37858660 | 37872103 | 13444 | -7.63 | 1.68E-09 | 3.78E-09 |
| chr14 | 67258492 | 67260737 | 2246 | -6.84 | 1.68E-09 | 3.78E-09 |

| | | | | | | |
|---|---|---|---|---|---|---|
| chr7 | 138317756 | 138318918 | 1163 | -6.79 | 1.68E-09 | 3.79E-09 |
| chr13 | 47548684 | 47550133 | 1450 | -6.78 | 1.69E-09 | 3.81E-09 |
| chr12 | 79523770 | 79526800 | 3031 | -6.78 | 1.70E-09 | 3.82E-09 |
| chr11 | 61157058 | 61163394 | 6337 | -6.8 | 1.70E-09 | 3.82E-09 |
| chr4 | 86413917 | 86421826 | 7910 | -6.78 | 1.70E-09 | 3.82E-09 |
| chr7 | 140626759 | 140627931 | 1173 | -6.78 | 1.70E-09 | 3.82E-09 |
| chr1 | 213362959 | 213365148 | 2190 | -6.81 | 1.70E-09 | 3.83E-09 |
| chr5 | 156392484 | 156394340 | 1857 | -6.81 | 1.70E-09 | 3.83E-09 |
| chr7 | 158480519 | 158482067 | 1549 | -6.81 | 1.70E-09 | 3.83E-09 |
| chr3 | 115449478 | 115473749 | 24272 | -1.57 | 1.71E-09 | 3.84E-09 |
| chr9 | 37846500 | 37848465 | 1966 | -6.93 | 1.71E-09 | 3.84E-09 |
| chr20 | 55936264 | 55937504 | 1241 | -6.84 | 1.71E-09 | 3.85E-09 |
| chr4 | 139928346 | 139929848 | 1503 | -6.79 | 1.71E-09 | 3.85E-09 |
| chr5 | 139893449 | 139895324 | 1876 | -6.89 | 1.71E-09 | 3.86E-09 |
| chr12 | 111962037 | 111963795 | 1759 | -6.95 | 1.72E-09 | 3.87E-09 |
| chr12 | 52030785 | 52032583 | 1799 | -6.78 | 1.72E-09 | 3.88E-09 |
| chr20 | 36810126 | 36811680 | 1555 | -6.78 | 1.72E-09 | 3.88E-09 |
| chr8 | 24104759 | 24106831 | 2073 | -6.78 | 1.72E-09 | 3.88E-09 |
| chr2 | 184152020 | 184155202 | 3183 | 6.91 | 1.72E-09 | 3.88E-09 |
| chr20 | 61141515 | 61142907 | 1393 | -6.88 | 1.73E-09 | 3.88E-09 |
| chr9 | 126191271 | 126192757 | 1487 | -6.8 | 1.73E-09 | 3.88E-09 |
| chr3 | 32844080 | 32845463 | 1384 | -6.87 | 1.73E-09 | 3.89E-09 |
| chr3 | 169878874 | 169880748 | 1875 | -6.87 | 1.73E-09 | 3.89E-09 |
| chr12 | 60209697 | 60215851 | 6155 | -6.81 | 1.73E-09 | 3.89E-09 |
| chr8 | 49738551 | 49752696 | 14146 | -7.12 | 1.73E-09 | 3.89E-09 |
| chrX | 28830574 | 28832988 | 2415 | -6.96 | 1.73E-09 | 3.89E-09 |
| chr18 | 37279530 | 37282139 | 2610 | -6.86 | 1.74E-09 | 3.91E-09 |
| chr7 | 13349780 | 13355207 | 5428 | -7.48 | 1.74E-09 | 3.91E-09 |
| chr16 | 25289888 | 25332976 | 43089 | -3.7 | 1.74E-09 | 3.91E-09 |
| chr13 | 110794555 | 110796274 | 1720 | -6.79 | 1.74E-09 | 3.91E-09 |
| chr7 | 47592812 | 47595339 | 2528 | -6.79 | 1.74E-09 | 3.91E-09 |
| chr2 | 99142790 | 99145036 | 2247 | -6.97 | 1.74E-09 | 3.91E-09 |
| chrX | 148900410 | 148902882 | 2473 | -4.97 | 1.74E-09 | 3.92E-09 |
| chr2 | 42695125 | 42696386 | 1262 | -6.78 | 1.75E-09 | 3.92E-09 |
| chr12 | 3614334 | 3898883 | 284550 | 4.25 | 1.75E-09 | 3.92E-09 |
| chr12 | 66020557 | 66024047 | 3491 | -6.85 | 1.75E-09 | 3.94E-09 |
| chr7 | 82649894 | 82651255 | 1362 | -6.8 | 1.75E-09 | 3.94E-09 |
| chr10 | 71010414 | 71011969 | 1556 | -6.77 | 1.76E-09 | 3.95E-09 |
| chr4 | 121458109 | 121468667 | 10559 | -6.77 | 1.76E-09 | 3.95E-09 |
| chrX | 49901604 | 49911165 | 9562 | 5.51 | 1.76E-09 | 3.95E-09 |
| chr18 | 72281372 | 72282862 | 1491 | -6.78 | 1.76E-09 | 3.95E-09 |
| chr3 | 138898092 | 138900189 | 2098 | -6.85 | 1.76E-09 | 3.95E-09 |
| chr14 | 56815480 | 56817108 | 1629 | -6.8 | 1.76E-09 | 3.96E-09 |
| chr10 | 22980770 | 22982280 | 1511 | -7.15 | 1.76E-09 | 3.96E-09 |
| chr13 | 105975028 | 105988440 | 13413 | -7.62 | 1.77E-09 | 3.96E-09 |
| chr5 | 54478904 | 54500217 | 21314 | -6.78 | 1.77E-09 | 3.96E-09 |
| chr15 | 92664438 | 92666251 | 1814 | -6.81 | 1.77E-09 | 3.96E-09 |
| chr1 | 115588030 | 115589771 | 1742 | -6.78 | 1.77E-09 | 3.97E-09 |
| chr4 | 124420658 | 124425253 | 4596 | -6.8 | 1.77E-09 | 3.98E-09 |
| chr19 | 39992282 | 40002644 | 10363 | -7.11 | 1.78E-09 | 3.99E-09 |
| chr8 | 23008335 | 23010424 | 2090 | -7 | 1.78E-09 | 3.99E-09 |
| chr13 | 30287125 | 30307162 | 20038 | -1.72 | 1.78E-09 | 4.00E-09 |
| chr15 | 50384134 | 50385322 | 1189 | -6.81 | 1.78E-09 | 4.00E-09 |
| chr8 | 17114589 | 17117095 | 2507 | -6.84 | 1.78E-09 | 4.00E-09 |
| chr5 | 95282311 | 95288038 | 5728 | -6.8 | 1.78E-09 | 4.00E-09 |
| chr1 | 225395478 | 225422261 | 26784 | -7.68 | 1.78E-09 | 4.00E-09 |
| chr10 | 103436058 | 103437416 | 1359 | -6.78 | 1.79E-09 | 4.00E-09 |
| chr19 | 13367177 | 13368255 | 1079 | -6.78 | 1.79E-09 | 4.00E-09 |
| chr14 | 81843431 | 81847271 | 3841 | -7.06 | 1.79E-09 | 4.01E-09 |
| chr8 | 130926735 | 130930068 | 3334 | -7.5 | 1.79E-09 | 4.01E-09 |
| chr6 | 75023764 | 75027223 | 3460 | -6.83 | 1.80E-09 | 4.03E-09 |

| | | | | | | |
|---|---|---|---|---|---|---|
| chr2 | 231756666 | 231757752 | 1087 | -6.8 | 1.80E-09 | 4.04E-09 |
| chr2 | 75184621 | 75197222 | 12602 | -6.78 | 1.80E-09 | 4.04E-09 |
| chr7 | 143952510 | 143953766 | 1257 | 6.84 | 1.80E-09 | 4.05E-09 |
| chr1 | 45510560 | 45512912 | 2353 | -6.92 | 1.81E-09 | 4.05E-09 |
| chr7 | 138578448 | 138586501 | 8054 | -6.88 | 1.81E-09 | 4.06E-09 |
| chr13 | 20552363 | 20554049 | 1687 | -6.83 | 1.82E-09 | 4.08E-09 |
| chr4 | 156218530 | 156220933 | 2404 | -6.83 | 1.82E-09 | 4.08E-09 |
| chr13 | 31908968 | 31912644 | 3677 | -6.89 | 1.83E-09 | 4.09E-09 |
| chr4 | 66375375 | 66382419 | 7045 | -6.79 | 1.83E-09 | 4.09E-09 |
| chr7 | 156769429 | 156772791 | 3363 | -6.82 | 1.83E-09 | 4.10E-09 |
| chr3 | 192138718 | 192147498 | 8781 | -6.88 | 1.83E-09 | 4.10E-09 |
| chr2 | 226780487 | 226784035 | 3549 | -6.86 | 1.84E-09 | 4.11E-09 |
| chr14 | 23803528 | 23813027 | 9500 | -2.75 | 1.84E-09 | 4.13E-09 |
| chr14 | 102934072 | 102935808 | 1737 | -6.78 | 1.85E-09 | 4.14E-09 |
| chr3 | 51026823 | 51030345 | 3523 | -2.66 | 1.85E-09 | 4.15E-09 |
| chr18 | 23765698 | 23781880 | 16183 | -6.8 | 1.85E-09 | 4.15E-09 |
| chr16 | 48021053 | 48041236 | 20184 | -1.51 | 1.85E-09 | 4.15E-09 |
| chr10 | 96805687 | 96807833 | 2147 | -6.78 | 1.85E-09 | 4.15E-09 |
| chr13 | 40600940 | 40616217 | 15278 | -7.27 | 1.85E-09 | 4.15E-09 |
| chr2 | 164195630 | 164219246 | 23617 | -7.39 | 1.86E-09 | 4.16E-09 |
| chr12 | 91745044 | 91746946 | 1903 | -6.96 | 1.86E-09 | 4.16E-09 |
| chr3 | 107068680 | 107071596 | 2917 | -6.85 | 1.86E-09 | 4.16E-09 |
| chr4 | 66766614 | 66771980 | 5367 | -6.84 | 1.86E-09 | 4.17E-09 |
| chr11 | 74135461 | 74137885 | 2425 | -7.27 | 1.86E-09 | 4.17E-09 |
| chr8 | 39585033 | 39586224 | 1192 | -2.76 | 1.86E-09 | 4.17E-09 |
| chr3 | 42933017 | 43021085 | 88069 | -3.47 | 1.87E-09 | 4.19E-09 |
| chr11 | 129922495 | 129923962 | 1468 | -6.77 | 1.87E-09 | 4.19E-09 |
| chr3 | 31246960 | 31251082 | 4123 | -6.98 | 1.87E-09 | 4.19E-09 |
| chr7 | 158211627 | 158213910 | 2284 | -6.97 | 1.87E-09 | 4.19E-09 |
| chr8 | 130979395 | 130982177 | 2783 | -6.97 | 1.87E-09 | 4.19E-09 |
| chr3 | 28228479 | 28240032 | 11554 | 2.01 | 1.88E-09 | 4.20E-09 |
| chr4 | 176264286 | 176266896 | 2611 | -6.79 | 1.88E-09 | 4.20E-09 |
| chr12 | 82767764 | 82770186 | 2423 | -6.78 | 1.88E-09 | 4.21E-09 |
| chr3 | 15213697 | 15228103 | 14407 | -3.41 | 1.88E-09 | 4.21E-09 |
| chr11 | 1044705 | 1045759 | 1055 | -6.76 | 1.89E-09 | 4.22E-09 |
| chr6 | 10256681 | 10288650 | 31970 | -8 | 1.89E-09 | 4.22E-09 |
| chr7 | 150168443 | 150294467 | 126025 | 2.15 | 1.89E-09 | 4.23E-09 |
| chrX | 49610168 | 49614865 | 4698 | -2.33 | 1.90E-09 | 4.25E-09 |
| chr4 | 119798733 | 119800307 | 1575 | -5.13 | 1.90E-09 | 4.25E-09 |
| chr4 | 22443015 | 22444303 | 1289 | -5.6 | 1.90E-09 | 4.25E-09 |
| chr9 | 98601291 | 98607015 | 5725 | -6.9 | 1.91E-09 | 4.26E-09 |
| chr20 | 25427999 | 25429232 | 1234 | -6.76 | 1.91E-09 | 4.26E-09 |
| chr8 | 112366522 | 112791292 | 424771 | 1.48 | 1.91E-09 | 4.26E-09 |
| chr13 | 21575116 | 21577226 | 2111 | -6.76 | 1.91E-09 | 4.27E-09 |
| chr8 | 101430954 | 101432248 | 1295 | -6.76 | 1.91E-09 | 4.27E-09 |
| chr3 | 196274738 | 196277857 | 3120 | -6.92 | 1.91E-09 | 4.27E-09 |
| chr14 | 90992491 | 90994085 | 1595 | -6.76 | 1.91E-09 | 4.28E-09 |
| chr6 | 41171717 | 41189424 | 17708 | -3.61 | 1.91E-09 | 4.28E-09 |
| chr17 | 47716079 | 47717681 | 1603 | -6.93 | 1.91E-09 | 4.28E-09 |
| chr4 | 184412407 | 184414321 | 1915 | -6.93 | 1.91E-09 | 4.28E-09 |
| chr2 | 121849189 | 121850768 | 1580 | -6.87 | 1.92E-09 | 4.28E-09 |
| chr16 | 81753739 | 81755699 | 1961 | -6.77 | 1.92E-09 | 4.29E-09 |
| chr1 | 161466282 | 161468489 | 2208 | 5.11 | 1.92E-09 | 4.29E-09 |
| chr9 | 133520900 | 133523501 | 2602 | -7.07 | 1.92E-09 | 4.30E-09 |
| chr20 | 17773272 | 17780761 | 7490 | -6.93 | 1.92E-09 | 4.30E-09 |
| chr3 | 153450557 | 153454579 | 4023 | -7.07 | 1.93E-09 | 4.31E-09 |
| chr1 | 210675897 | 210686156 | 10260 | 7.21 | 1.93E-09 | 4.31E-09 |
| chr2 | 194328864 | 194341376 | 12513 | -6.78 | 1.93E-09 | 4.32E-09 |
| chr2 | 109850100 | 109851647 | 1548 | -6.76 | 1.93E-09 | 4.32E-09 |
| chr8 | 135569475 | 135570918 | 1444 | -6.76 | 1.93E-09 | 4.32E-09 |
| chr22 | 25575206 | 25576327 | 1122 | -6.76 | 1.93E-09 | 4.32E-09 |

| | | | | | | |
|---|---|---|---|---|---|---|
| chr6 | 149446046 | 149449043 | 2998 | -6.85 | 1.94E-09 | 4.33E-09 |
| chr21 | 15572562 | 15573863 | 1302 | -3.19 | 1.94E-09 | 4.33E-09 |
| chr2 | 139071189 | 139074775 | 3587 | -6.82 | 1.94E-09 | 4.34E-09 |
| chr2 | 236131920 | 236133172 | 1253 | -6.81 | 1.94E-09 | 4.34E-09 |
| chr8 | 55158176 | 55159965 | 1790 | -3.75 | 1.95E-09 | 4.35E-09 |
| chr2 | 61354818 | 61356539 | 1722 | -6.76 | 1.95E-09 | 4.35E-09 |
| chr12 | 1306737 | 1307920 | 1184 | -6.77 | 1.95E-09 | 4.36E-09 |
| chr12 | 32821823 | 32823917 | 2095 | -7.03 | 1.95E-09 | 4.36E-09 |
| chr14 | 100263318 | 100264537 | 1220 | -6.96 | 1.95E-09 | 4.36E-09 |
| chrX | 16068525 | 16077642 | 9118 | -6.88 | 1.95E-09 | 4.36E-09 |
| chr3 | 122547324 | 122548635 | 1312 | -6.84 | 1.96E-09 | 4.37E-09 |
| chr2 | 191847489 | 191848773 | 1285 | -6.88 | 1.96E-09 | 4.37E-09 |
| chr13 | 95875618 | 95877164 | 1547 | -6.76 | 1.96E-09 | 4.37E-09 |
| chr22 | 19769237 | 19779907 | 10671 | -6.76 | 1.96E-09 | 4.38E-09 |
| chr13 | 80048793 | 80050284 | 1492 | -6.81 | 1.96E-09 | 4.38E-09 |
| chr18 | 32942526 | 32945967 | 3442 | -6.81 | 1.96E-09 | 4.38E-09 |
| chrX | 85228747 | 85230759 | 2013 | -6.87 | 1.96E-09 | 4.38E-09 |
| chr13 | 73325239 | 73326847 | 1609 | -6.86 | 1.96E-09 | 4.38E-09 |
| chr11 | 44693782 | 44711479 | 17698 | -7.78 | 1.97E-09 | 4.39E-09 |
| chr1 | 101091962 | 101095822 | 3861 | -6.77 | 1.97E-09 | 4.39E-09 |
| chr13 | 35764208 | 35765764 | 1557 | -7.04 | 1.97E-09 | 4.40E-09 |
| chr11 | 35108144 | 35111134 | 2991 | -6.79 | 1.98E-09 | 4.42E-09 |
| chrX | 37216640 | 37219960 | 3321 | -6.79 | 1.98E-09 | 4.42E-09 |
| chrX | 65114557 | 65121435 | 6879 | -6.79 | 1.98E-09 | 4.42E-09 |
| chr9 | 18319094 | 18326356 | 7263 | -6.76 | 1.98E-09 | 4.42E-09 |
| chr1 | 15816573 | 15825428 | 8856 | -6.75 | 1.98E-09 | 4.42E-09 |
| chr20 | 14183372 | 14200742 | 17371 | -7.48 | 1.98E-09 | 4.42E-09 |
| chr14 | 67353447 | 67356191 | 2745 | -6.76 | 1.99E-09 | 4.43E-09 |
| chr6 | 28392265 | 28393716 | 1452 | -6.76 | 1.99E-09 | 4.43E-09 |
| chr17 | 9411206 | 9412918 | 1713 | -6.77 | 1.99E-09 | 4.44E-09 |
| chr13 | 63650664 | 63657644 | 6981 | -6.99 | 1.99E-09 | 4.44E-09 |
| chr2 | 43637061 | 43638400 | 1340 | -6.99 | 1.99E-09 | 4.44E-09 |
| chr3 | 11103927 | 11107048 | 3122 | -6.95 | 1.99E-09 | 4.44E-09 |
| chr4 | 114934576 | 114940273 | 5698 | -7.21 | 1.99E-09 | 4.44E-09 |
| chr14 | 45406759 | 45417083 | 10325 | -6.96 | 2.00E-09 | 4.45E-09 |
| chr3 | 114794376 | 114795959 | 1584 | -6.8 | 2.00E-09 | 4.45E-09 |
| chr11 | 18168913 | 18189401 | 20489 | -1.83 | 2.00E-09 | 4.45E-09 |
| chr1 | 221853096 | 221867895 | 14800 | -7.45 | 2.00E-09 | 4.46E-09 |
| chr12 | 70311548 | 70313036 | 1489 | -6.77 | 2.00E-09 | 4.46E-09 |
| chr1 | 41422891 | 41424896 | 2006 | -6.78 | 2.01E-09 | 4.49E-09 |
| chr5 | 106190649 | 106192391 | 1743 | -6.78 | 2.01E-09 | 4.49E-09 |
| chr3 | 30820411 | 30822411 | 2001 | -7.11 | 2.01E-09 | 4.49E-09 |
| chr7 | 21801776 | 21815680 | 13905 | -7.37 | 2.01E-09 | 4.49E-09 |
| chr20 | 49237258 | 49238715 | 1458 | -6.9 | 2.01E-09 | 4.49E-09 |
| chr6 | 51053402 | 51059517 | 6116 | -6.87 | 2.02E-09 | 4.49E-09 |
| chr4 | 151313424 | 151315154 | 1731 | -6.75 | 2.02E-09 | 4.50E-09 |
| chr12 | 64576830 | 64579432 | 2603 | -6.83 | 2.02E-09 | 4.50E-09 |
| chrX | 79471893 | 79474815 | 2923 | -6.83 | 2.02E-09 | 4.50E-09 |
| chr9 | 120243510 | 120246880 | 3371 | 6.82 | 2.02E-09 | 4.50E-09 |
| chr5 | 76356842 | 76365447 | 8606 | -6.91 | 2.02E-09 | 4.50E-09 |
| chr10 | 32643222 | 32644901 | 1680 | -6.92 | 2.02E-09 | 4.50E-09 |
| chr12 | 117781316 | 117783147 | 1832 | -6.92 | 2.02E-09 | 4.50E-09 |
| chrX | 135845658 | 135854206 | 8549 | -6.92 | 2.03E-09 | 4.51E-09 |
| chr9 | 28971248 | 28974484 | 3237 | -6.77 | 2.03E-09 | 4.52E-09 |
| chr5 | 30172970 | 30176920 | 3951 | -6.76 | 2.03E-09 | 4.52E-09 |
| chr1 | 174259614 | 174260995 | 1382 | -6.82 | 2.04E-09 | 4.54E-09 |
| chr5 | 94660528 | 94685605 | 25078 | 1.64 | 2.04E-09 | 4.54E-09 |
| chr12 | 89692077 | 89693970 | 1894 | -6.75 | 2.05E-09 | 4.56E-09 |
| chr12 | 80824790 | 80830650 | 5861 | 7.44 | 2.05E-09 | 4.57E-09 |
| chr4 | 28015214 | 28018176 | 2963 | -6.75 | 2.05E-09 | 4.57E-09 |
| chr1 | 30815953 | 30823392 | 7440 | -6.25 | 2.05E-09 | 4.57E-09 |

| | | | | | | |
|---|---|---|---|---|---|---|
| chr4 | 78230411 | 78267741 | 37331 | -1.09 | 2.06E-09 | 4.58E-09 |
| chr12 | 38624587 | 38626630 | 2044 | -6.81 | 2.06E-09 | 4.59E-09 |
| chr5 | 131908663 | 131911686 | 3024 | -7 | 2.06E-09 | 4.59E-09 |
| chr6 | 16474528 | 16484051 | 9524 | -6.76 | 2.06E-09 | 4.59E-09 |
| chr4 | 94740768 | 94749337 | 8570 | -6.8 | 2.07E-09 | 4.61E-09 |
| chr11 | 60841402 | 60851930 | 10529 | -7.35 | 2.08E-09 | 4.63E-09 |
| chr5 | 131229144 | 131269329 | 40186 | -1.92 | 2.08E-09 | 4.63E-09 |
| chr1 | 12407554 | 12410291 | 2738 | -6.75 | 2.08E-09 | 4.64E-09 |
| chr12 | 20237007 | 20238986 | 1980 | -6.78 | 2.09E-09 | 4.65E-09 |
| chr3 | 46610846 | 46612368 | 1523 | -5.05 | 2.09E-09 | 4.65E-09 |
| chr4 | 85905193 | 85908555 | 3363 | -6.87 | 2.09E-09 | 4.66E-09 |
| chr15 | 66740528 | 66742303 | 1776 | -6.78 | 2.10E-09 | 4.67E-09 |
| chr7 | 111039271 | 111047451 | 8181 | -6.98 | 2.10E-09 | 4.67E-09 |
| chr9 | 115474228 | 115476093 | 1866 | -6.76 | 2.10E-09 | 4.68E-09 |
| chr11 | 125564875 | 125566164 | 1290 | -6.76 | 2.11E-09 | 4.69E-09 |
| chr3 | 102223608 | 102225762 | 2155 | -6.85 | 2.11E-09 | 4.69E-09 |
| chr12 | 27033638 | 27037304 | 3667 | -6.75 | 2.11E-09 | 4.70E-09 |
| chr9 | 111968826 | 111970524 | 1699 | -6.75 | 2.11E-09 | 4.70E-09 |
| chr6 | 121955462 | 121958565 | 3104 | -5.07 | 2.12E-09 | 4.71E-09 |
| chr5 | 127171034 | 127174611 | 3578 | -7.05 | 2.12E-09 | 4.72E-09 |
| chr11 | 7729739 | 7988018 | 258280 | 3.31 | 2.12E-09 | 4.72E-09 |
| chr4 | 40408077 | 40422176 | 14100 | -6.87 | 2.13E-09 | 4.73E-09 |
| chr4 | 93458568 | 93461242 | 2675 | -6.83 | 2.13E-09 | 4.73E-09 |
| chr4 | 173160555 | 173162654 | 2100 | -6.89 | 2.13E-09 | 4.73E-09 |
| chr3 | 23357519 | 23360183 | 2665 | -6.9 | 2.13E-09 | 4.73E-09 |
| chr2 | 143639245 | 143640695 | 1451 | -6.94 | 2.13E-09 | 4.73E-09 |
| chr10 | 134126966 | 134139193 | 12228 | -6.86 | 2.13E-09 | 4.74E-09 |
| chr13 | 91198575 | 91201316 | 2742 | -6.86 | 2.13E-09 | 4.74E-09 |
| chr9 | 16331797 | 16335843 | 4047 | -6.83 | 2.13E-09 | 4.74E-09 |
| chr14 | 60419178 | 60427142 | 7965 | -3.24 | 2.14E-09 | 4.75E-09 |
| chr8 | 101634427 | 101635912 | 1486 | -6.74 | 2.14E-09 | 4.76E-09 |
| chrX | 40072421 | 40080004 | 7584 | -6.76 | 2.15E-09 | 4.77E-09 |
| chr6 | 137698554 | 137702608 | 4055 | -3.79 | 2.15E-09 | 4.77E-09 |
| chr11 | 88196970 | 88200569 | 3600 | -6.84 | 2.15E-09 | 4.78E-09 |
| chrX | 77826604 | 77828753 | 2150 | 6.89 | 2.15E-09 | 4.78E-09 |
| chr4 | 127792096 | 127794846 | 2751 | -6.75 | 2.16E-09 | 4.79E-09 |
| chr8 | 59147901 | 59152836 | 4936 | -7.04 | 2.16E-09 | 4.79E-09 |
| chr1 | 61501262 | 61502808 | 1547 | -6.77 | 2.16E-09 | 4.80E-09 |
| chr14 | 77696568 | 77698158 | 1591 | -6.77 | 2.16E-09 | 4.80E-09 |
| chrX | 77230169 | 77232156 | 1988 | -6.83 | 2.16E-09 | 4.81E-09 |
| chr2 | 163108399 | 163116384 | 7986 | 6.88 | 2.17E-09 | 4.83E-09 |
| chr16 | 11204138 | 11205636 | 1499 | -6.82 | 2.17E-09 | 4.83E-09 |
| chr5 | 79457533 | 79459225 | 1693 | -6.95 | 2.18E-09 | 4.84E-09 |
| chr4 | 77672275 | 77673391 | 1117 | -6.74 | 2.18E-09 | 4.84E-09 |
| chrX | 15373590 | 15374636 | 1047 | -6.74 | 2.18E-09 | 4.84E-09 |
| chr7 | 92170475 | 92173247 | 2773 | -7.02 | 2.19E-09 | 4.86E-09 |
| chrX | 29549172 | 29559853 | 10682 | -7.52 | 2.19E-09 | 4.86E-09 |
| chr4 | 87749717 | 87752014 | 2298 | -6.77 | 2.19E-09 | 4.86E-09 |
| chr20 | 33544520 | 33547034 | 2515 | -6.81 | 2.20E-09 | 4.88E-09 |
| chr8 | 57238570 | 57306091 | 67522 | -2.35 | 2.20E-09 | 4.89E-09 |
| chr2 | 75500306 | 75502468 | 2163 | -6.96 | 2.20E-09 | 4.89E-09 |
| chr1 | 84657903 | 84660213 | 2311 | -6.77 | 2.21E-09 | 4.90E-09 |
| chr5 | 141999068 | 142000720 | 1653 | -6.74 | 2.21E-09 | 4.90E-09 |
| chr12 | 32727801 | 32731920 | 4120 | -6.96 | 2.21E-09 | 4.91E-09 |
| chr14 | 35273909 | 35275031 | 1123 | -7.12 | 2.21E-09 | 4.91E-09 |
| chr8 | 40608082 | 40609421 | 1340 | -5.82 | 2.21E-09 | 4.91E-09 |
| chr9 | 119029222 | 119034186 | 4965 | -5.25 | 2.22E-09 | 4.92E-09 |
| chr3 | 46679450 | 46697118 | 17669 | -3.74 | 2.22E-09 | 4.92E-09 |
| chr7 | 14123268 | 14126371 | 3104 | -7.13 | 2.22E-09 | 4.93E-09 |
| chr6 | 108860857 | 108862182 | 1326 | -6.74 | 2.22E-09 | 4.94E-09 |
| chr1 | 186815616 | 186827879 | 12264 | -7.13 | 2.23E-09 | 4.95E-09 |

| | | | | | | |
|---|---|---|---|---|---|---|
| chrX | 30117801 | 30120109 | 2309 | -6.75 | 2.23E-09 | 4.95E-09 |
| chr9 | 3287197 | 3289013 | 1817 | -6.75 | 2.23E-09 | 4.95E-09 |
| chr10 | 93319947 | 93341713 | 21767 | -7.18 | 2.23E-09 | 4.95E-09 |
| chrX | 106856826 | 106866680 | 9855 | -7.18 | 2.23E-09 | 4.95E-09 |
| chr21 | 33768744 | 33770305 | 1562 | -6.78 | 2.23E-09 | 4.95E-09 |
| chr16 | 14291285 | 14292572 | 1288 | -6.8 | 2.24E-09 | 4.97E-09 |
| chr3 | 161716507 | 161727348 | 10842 | -2.39 | 2.24E-09 | 4.97E-09 |
| chr5 | 67612715 | 67614890 | 2176 | -6.75 | 2.24E-09 | 4.97E-09 |
| chr12 | 105267199 | 105278558 | 11360 | -6.87 | 2.24E-09 | 4.97E-09 |
| chr4 | 48888193 | 48893145 | 4953 | -6.88 | 2.24E-09 | 4.97E-09 |
| chr20 | 5189974 | 5191165 | 1192 | -6.86 | 2.24E-09 | 4.97E-09 |
| chr1 | 62742917 | 62758459 | 15543 | -6.87 | 2.25E-09 | 4.98E-09 |
| chr13 | 83339217 | 83342354 | 3138 | -6.87 | 2.25E-09 | 4.98E-09 |
| chr3 | 186752255 | 186754498 | 2244 | -6.86 | 2.25E-09 | 4.98E-09 |
| chr5 | 77362806 | 77364386 | 1581 | -6.89 | 2.25E-09 | 4.99E-09 |
| chr16 | 66984743 | 66986052 | 1310 | -6.79 | 2.26E-09 | 5.00E-09 |
| chr3 | 121833703 | 121902552 | 68850 | -2.25 | 2.26E-09 | 5.01E-09 |
| chr5 | 37791561 | 37798922 | 7362 | -3.03 | 2.26E-09 | 5.01E-09 |
| chr1 | 89432830 | 89434340 | 1511 | -6.73 | 2.26E-09 | 5.01E-09 |
| chr14 | 59902836 | 59904083 | 1248 | -6.91 | 2.26E-09 | 5.01E-09 |
| chr7 | 82326229 | 82329092 | 2864 | -7.14 | 2.26E-09 | 5.01E-09 |
| chr7 | 84424681 | 84427070 | 2390 | -6.99 | 2.27E-09 | 5.02E-09 |
| chr14 | 51281109 | 51282756 | 1648 | -6.73 | 2.27E-09 | 5.03E-09 |
| chr3 | 153735848 | 153739097 | 3250 | -6.83 | 2.27E-09 | 5.03E-09 |
| chr4 | 101044806 | 101047137 | 2332 | -6.83 | 2.28E-09 | 5.04E-09 |
| chr1 | 2027959 | 2030028 | 2070 | -6.83 | 2.28E-09 | 5.05E-09 |
| chr14 | 55261349 | 55268539 | 7191 | -6.83 | 2.28E-09 | 5.05E-09 |
| chr7 | 155194540 | 155195740 | 1201 | -6.73 | 2.28E-09 | 5.06E-09 |
| chrX | 43700183 | 43702070 | 1888 | -6.73 | 2.28E-09 | 5.06E-09 |
| chr10 | 105017495 | 105030397 | 12903 | -7.77 | 2.29E-09 | 5.07E-09 |
| chr2 | 233796431 | 233807054 | 10624 | -6.76 | 2.29E-09 | 5.07E-09 |
| chr9 | 28821840 | 28824297 | 2458 | -6.73 | 2.29E-09 | 5.07E-09 |
| chr11 | 12485782 | 12487525 | 1744 | -6.74 | 2.29E-09 | 5.08E-09 |
| chr10 | 16489625 | 16491949 | 2325 | -6.73 | 2.30E-09 | 5.08E-09 |
| chr18 | 57405716 | 57415089 | 9374 | -7.02 | 2.30E-09 | 5.08E-09 |
| chr1 | 232196666 | 232214351 | 17686 | -7.37 | 2.30E-09 | 5.08E-09 |
| chr6 | 155480706 | 155482799 | 2094 | -6.82 | 2.30E-09 | 5.09E-09 |
| chr2 | 205526229 | 205533948 | 7720 | -6.81 | 2.30E-09 | 5.09E-09 |
| chr9 | 130719126 | 130720518 | 1393 | -6.81 | 2.30E-09 | 5.09E-09 |
| chr16 | 82040069 | 82060947 | 20879 | -2.21 | 2.30E-09 | 5.09E-09 |
| chr8 | 126361519 | 126363737 | 2219 | -6.74 | 2.30E-09 | 5.09E-09 |
| chr15 | 93182586 | 93200429 | 17844 | -7.85 | 2.30E-09 | 5.09E-09 |
| chr5 | 138009404 | 138021162 | 11759 | -7.06 | 2.30E-09 | 5.10E-09 |
| chr13 | 35170367 | 35173330 | 2964 | -7.16 | 2.31E-09 | 5.12E-09 |
| chr12 | 78618520 | 78624543 | 6024 | -7.43 | 2.32E-09 | 5.13E-09 |
| chr13 | 89322714 | 89334778 | 12065 | -6.78 | 2.32E-09 | 5.13E-09 |
| chr15 | 31172453 | 31173474 | 1022 | -6.76 | 2.32E-09 | 5.13E-09 |
| chr14 | 106828563 | 106852955 | 24393 | 2.97 | 2.32E-09 | 5.13E-09 |
| chr14 | 32090415 | 32092911 | 2497 | -6.81 | 2.32E-09 | 5.14E-09 |
| chr15 | 24439702 | 24475952 | 36251 | 2 | 2.33E-09 | 5.14E-09 |
| chr1 | 94631452 | 94633602 | 2151 | -6.95 | 2.33E-09 | 5.15E-09 |
| chr9 | 26811890 | 26815511 | 3622 | -6.75 | 2.33E-09 | 5.15E-09 |
| chr3 | 8660307 | 8662204 | 1898 | -6.73 | 2.34E-09 | 5.17E-09 |
| chr17 | 61634151 | 61635222 | 1072 | -6.8 | 2.34E-09 | 5.18E-09 |
| chr1 | 43379448 | 43381882 | 2435 | -7.02 | 2.34E-09 | 5.18E-09 |
| chr7 | 90299343 | 90305046 | 5704 | 6.93 | 2.36E-09 | 5.21E-09 |
| chr4 | 158754384 | 158757135 | 2752 | 6.82 | 2.36E-09 | 5.23E-09 |
| chr3 | 159674998 | 159676484 | 1487 | -6.85 | 2.37E-09 | 5.25E-09 |
| chr3 | 12607331 | 12608972 | 1642 | -6.76 | 2.38E-09 | 5.26E-09 |
| chr3 | 27369621 | 27379495 | 9875 | -7.09 | 2.38E-09 | 5.26E-09 |
| chr12 | 17131245 | 17135002 | 3758 | -6.73 | 2.38E-09 | 5.27E-09 |

| | | | | | | |
|---|---|---|---|---|---|---|
| chr16 | 22171185 | 22179033 | 7849 | -6.98 | 2.38E-09 | 5.27E-09 |
| chr2 | 42424469 | 42426110 | 1642 | -6.84 | 2.38E-09 | 5.27E-09 |
| chr10 | 17182158 | 17184248 | 2091 | -6.89 | 2.38E-09 | 5.27E-09 |
| chr4 | 67071672 | 67075661 | 3990 | -6.89 | 2.38E-09 | 5.27E-09 |
| chr1 | 189094013 | 189095452 | 1440 | -6.76 | 2.39E-09 | 5.29E-09 |
| chr5 | 59595664 | 59597204 | 1541 | -6.76 | 2.39E-09 | 5.29E-09 |
| chr14 | 70267612 | 70269052 | 1441 | -6.75 | 2.39E-09 | 5.29E-09 |
| chrX | 40750143 | 40764304 | 14162 | -7.18 | 2.40E-09 | 5.30E-09 |
| chr10 | 42490265 | 42524068 | 33804 | -3.69 | 2.40E-09 | 5.30E-09 |
| chr6 | 167238979 | 167261946 | 22968 | -2.23 | 2.40E-09 | 5.31E-09 |
| chr3 | 48330832 | 48334166 | 3335 | -6.82 | 2.40E-09 | 5.31E-09 |
| chr11 | 21736504 | 21752377 | 15874 | -6.99 | 2.40E-09 | 5.31E-09 |
| chr15 | 89068737 | 89079893 | 11157 | -6.88 | 2.41E-09 | 5.31E-09 |
| chr13 | 34151820 | 34153370 | 1551 | -5.36 | 2.41E-09 | 5.32E-09 |
| chr10 | 34049855 | 34052070 | 2216 | -6.82 | 2.41E-09 | 5.32E-09 |
| chr8 | 109529972 | 109532948 | 2977 | -3.06 | 2.41E-09 | 5.33E-09 |
| chr8 | 134600643 | 134601523 | 881 | -6.83 | 2.42E-09 | 5.33E-09 |
| chr4 | 139253023 | 139255525 | 2503 | -6.81 | 2.42E-09 | 5.34E-09 |
| chr11 | 117598059 | 117600182 | 2124 | -6.72 | 2.42E-09 | 5.35E-09 |
| chr3 | 82045128 | 82099830 | 54703 | 6.77 | 2.42E-09 | 5.35E-09 |
| chr1 | 171208648 | 171211476 | 2829 | -6.75 | 2.42E-09 | 5.35E-09 |
| chr4 | 118870441 | 118871908 | 1468 | 3.17 | 2.43E-09 | 5.35E-09 |
| chr6 | 101147255 | 101150617 | 3363 | -7 | 2.43E-09 | 5.36E-09 |
| chr1 | 49317605 | 49329469 | 11865 | -2.12 | 2.43E-09 | 5.36E-09 |
| chr7 | 96795592 | 96797378 | 1787 | -6.77 | 2.43E-09 | 5.36E-09 |
| chr6 | 105713488 | 105714942 | 1455 | -6.12 | 2.44E-09 | 5.38E-09 |
| chr3 | 47596797 | 47598790 | 1994 | -7 | 2.44E-09 | 5.38E-09 |
| chr11 | 8416759 | 8426931 | 10173 | -5.47 | 2.44E-09 | 5.39E-09 |
| chr19 | 7239053 | 7247384 | 8332 | -6.8 | 2.44E-09 | 5.39E-09 |
| chr7 | 149108360 | 149110454 | 2095 | -6.8 | 2.44E-09 | 5.39E-09 |
| chr5 | 136569197 | 136574787 | 5591 | 7.16 | 2.45E-09 | 5.42E-09 |
| chr7 | 137814269 | 137818410 | 4142 | -6.75 | 2.45E-09 | 5.42E-09 |
| chr18 | 8068964 | 8070837 | 1874 | -6.73 | 2.45E-09 | 5.42E-09 |
| chr10 | 99594933 | 99596407 | 1475 | -6.8 | 2.46E-09 | 5.42E-09 |
| chr20 | 450596 | 452155 | 1560 | -6.95 | 2.48E-09 | 5.46E-09 |
| chr21 | 35032636 | 35034112 | 1477 | -6.76 | 2.48E-09 | 5.47E-09 |
| chr5 | 148626226 | 148627132 | 907 | -6.76 | 2.48E-09 | 5.47E-09 |
| chr10 | 54036066 | 54039027 | 2962 | -7.25 | 2.48E-09 | 5.48E-09 |
| chr7 | 148677776 | 148678904 | 1129 | -6.79 | 2.49E-09 | 5.49E-09 |
| chr13 | 30898983 | 30901240 | 2258 | -6.72 | 2.49E-09 | 5.49E-09 |
| chr13 | 49238154 | 49241811 | 3658 | -2.33 | 2.49E-09 | 5.49E-09 |
| chr11 | 82392116 | 82393370 | 1255 | -6.72 | 2.49E-09 | 5.49E-09 |
| chr5 | 155798721 | 155800605 | 1885 | -6.72 | 2.50E-09 | 5.51E-09 |
| chr3 | 122018836 | 122019980 | 1145 | -6.96 | 2.50E-09 | 5.51E-09 |
| chr21 | 42625405 | 42626961 | 1557 | -6.72 | 2.50E-09 | 5.52E-09 |
| chr14 | 39610527 | 39612149 | 1623 | -6.73 | 2.50E-09 | 5.52E-09 |
| chr5 | 123918897 | 123944937 | 26041 | -7.9 | 2.51E-09 | 5.53E-09 |
| chr4 | 106413527 | 106415482 | 1956 | -6.84 | 2.51E-09 | 5.53E-09 |
| chr14 | 97348748 | 97350371 | 1624 | -6.72 | 2.52E-09 | 5.56E-09 |
| chr15 | 51877534 | 51892745 | 15212 | -7.4 | 2.53E-09 | 5.57E-09 |
| chr6 | 81131611 | 81136528 | 4918 | -6.82 | 2.53E-09 | 5.57E-09 |
| chr15 | 44269258 | 44271058 | 1801 | -6.75 | 2.53E-09 | 5.58E-09 |
| chr3 | 184992154 | 184996819 | 4666 | -6.9 | 2.53E-09 | 5.59E-09 |
| chr16 | 52366079 | 52367606 | 1528 | -4.98 | 2.54E-09 | 5.60E-09 |
| chr16 | 87137234 | 87140365 | 3132 | -6.73 | 2.55E-09 | 5.62E-09 |
| chr4 | 180595481 | 180598812 | 3332 | -6.98 | 2.55E-09 | 5.63E-09 |
| chr6 | 22721907 | 22724744 | 2838 | -6.91 | 2.55E-09 | 5.63E-09 |
| chr4 | 102967262 | 102971581 | 4320 | -6.71 | 2.55E-09 | 5.63E-09 |
| chr2 | 147371350 | 147374204 | 2855 | -5.09 | 2.56E-09 | 5.63E-09 |
| chr10 | 690355 | 691546 | 1192 | -6.71 | 2.56E-09 | 5.64E-09 |
| chr8 | 75303687 | 76062670 | 758984 | 5.82 | 2.56E-09 | 5.65E-09 |

| | | | | | | |
|---|---|---|---|---|---|---|
| chr7 | 79137814 | 79140809 | 2996 | -6.99 | 2.57E-09 | 5.65E-09 |
| chr8 | 95393976 | 95396619 | 2644 | -6.71 | 2.57E-09 | 5.66E-09 |
| chr11 | 105896463 | 105898401 | 1939 | -6.74 | 2.57E-09 | 5.66E-09 |
| chr4 | 187030918 | 187039544 | 8627 | -6.74 | 2.57E-09 | 5.66E-09 |
| chr4 | 79623188 | 79625270 | 2083 | -6.72 | 2.57E-09 | 5.66E-09 |
| chr1 | 168565110 | 168567214 | 2105 | -6.76 | 2.57E-09 | 5.66E-09 |
| chr2 | 129535829 | 129538143 | 2315 | -6.8 | 2.57E-09 | 5.66E-09 |
| chr9 | 113092903 | 113094073 | 1171 | -5.25 | 2.57E-09 | 5.67E-09 |
| chr3 | 118469150 | 118471518 | 2369 | -2.53 | 2.58E-09 | 5.67E-09 |
| chr5 | 64712119 | 64718108 | 5990 | -2.39 | 2.58E-09 | 5.68E-09 |
| chr1 | 81395380 | 81399149 | 3770 | -6.73 | 2.58E-09 | 5.69E-09 |
| chr6 | 126404375 | 126406248 | 1874 | -6.83 | 2.58E-09 | 5.69E-09 |
| chr7 | 5544584 | 5545888 | 1305 | -6.83 | 2.58E-09 | 5.69E-09 |
| chr2 | 146028870 | 146031868 | 2999 | -5.07 | 2.59E-09 | 5.71E-09 |
| chr12 | 92786253 | 92788828 | 2576 | -6.72 | 2.60E-09 | 5.72E-09 |
| chr7 | 154867185 | 154993828 | 126644 | -2.1 | 2.60E-09 | 5.72E-09 |
| chr2 | 191681043 | 191694956 | 13914 | -7.18 | 2.61E-09 | 5.75E-09 |
| chr12 | 114163888 | 114191220 | 27333 | -7.06 | 2.61E-09 | 5.75E-09 |
| chr17 | 30159822 | 30163719 | 3898 | -7.01 | 2.62E-09 | 5.76E-09 |
| chrX | 28952330 | 28957952 | 5623 | -7.01 | 2.62E-09 | 5.76E-09 |
| chr4 | 97630616 | 97643504 | 12889 | -6.72 | 2.62E-09 | 5.76E-09 |
| chr2 | 439150 | 448159 | 9010 | -6.94 | 2.62E-09 | 5.76E-09 |
| chr20 | 35479026 | 35480158 | 1133 | -6.91 | 2.62E-09 | 5.76E-09 |
| chr12 | 51512250 | 51513368 | 1119 | -6.74 | 2.62E-09 | 5.77E-09 |
| chr5 | 1351590 | 1380380 | 28791 | -7.69 | 2.62E-09 | 5.77E-09 |
| chr5 | 170127402 | 170166529 | 39128 | -1.95 | 2.63E-09 | 5.78E-09 |
| chr10 | 28685862 | 28687007 | 1146 | -4.95 | 2.63E-09 | 5.79E-09 |
| chr20 | 37532813 | 37534775 | 1963 | -6.78 | 2.63E-09 | 5.79E-09 |
| chr11 | 76674321 | 76683396 | 9076 | -7.2 | 2.63E-09 | 5.79E-09 |
| chr1 | 162480704 | 162482075 | 1372 | -6.71 | 2.64E-09 | 5.81E-09 |
| chr10 | 21870962 | 21872834 | 1873 | -6.75 | 2.65E-09 | 5.82E-09 |
| chr19 | 3684185 | 3685418 | 1234 | -6.73 | 2.65E-09 | 5.84E-09 |
| chr8 | 131233495 | 131234490 | 996 | -6.73 | 2.65E-09 | 5.84E-09 |
| chr17 | 60913360 | 60915667 | 2308 | -6.86 | 2.66E-09 | 5.84E-09 |
| chr5 | 73865466 | 73868040 | 2575 | -6.72 | 2.66E-09 | 5.84E-09 |
| chr6 | 15434566 | 15435898 | 1333 | -6.88 | 2.67E-09 | 5.87E-09 |
| chr18 | 20939665 | 20941002 | 1338 | -6.71 | 2.68E-09 | 5.89E-09 |
| chr18 | 48337605 | 48338982 | 1378 | -6.71 | 2.68E-09 | 5.89E-09 |
| chr4 | 1000097 | 1001322 | 1226 | -6.71 | 2.68E-09 | 5.89E-09 |
| chr8 | 82175549 | 82177042 | 1494 | -6.71 | 2.68E-09 | 5.89E-09 |
| chr4 | 132338508 | 132341266 | 2759 | -5.81 | 2.68E-09 | 5.89E-09 |
| chr17 | 64046157 | 64057106 | 10950 | -6.78 | 2.68E-09 | 5.89E-09 |
| chr11 | 101329051 | 101330878 | 1828 | -6.71 | 2.68E-09 | 5.89E-09 |
| chr5 | 100018057 | 100019795 | 1739 | -4.93 | 2.69E-09 | 5.90E-09 |
| chr7 | 11057085 | 11064457 | 7373 | -6.9 | 2.70E-09 | 5.93E-09 |
| chr3 | 158891301 | 158893579 | 2279 | -6.72 | 2.70E-09 | 5.93E-09 |
| chr2 | 235832986 | 235834586 | 1601 | -6.72 | 2.70E-09 | 5.94E-09 |
| chr11 | 62792018 | 62904227 | 112210 | -2 | 2.70E-09 | 5.94E-09 |
| chr10 | 5397356 | 5400594 | 3239 | -7.4 | 2.71E-09 | 5.94E-09 |
| chr15 | 69041204 | 69043419 | 2216 | -7.04 | 2.71E-09 | 5.96E-09 |
| chr3 | 43465863 | 43468311 | 2449 | -6.7 | 2.72E-09 | 5.97E-09 |
| chr1 | 61711328 | 61712969 | 1642 | -6.7 | 2.72E-09 | 5.97E-09 |
| chr2 | 154803624 | 154819507 | 15884 | -6.71 | 2.73E-09 | 5.99E-09 |
| chr19 | 47935584 | 47937018 | 1435 | -6.92 | 2.73E-09 | 5.99E-09 |
| chr6 | 9820192 | 9823851 | 3660 | -7.04 | 2.73E-09 | 5.99E-09 |
| chr3 | 109498037 | 109513952 | 15916 | -7.16 | 2.74E-09 | 6.01E-09 |
| chr12 | 52125778 | 52128155 | 2378 | -6.71 | 2.75E-09 | 6.03E-09 |
| chr2 | 62207909 | 62210058 | 2150 | -6.93 | 2.75E-09 | 6.03E-09 |
| chr1 | 79107711 | 79113473 | 5763 | -2.61 | 2.75E-09 | 6.04E-09 |
| chr7 | 13073567 | 13092312 | 18746 | -7.1 | 2.76E-09 | 6.06E-09 |
| chr3 | 69109935 | 69111468 | 1534 | -6.84 | 2.76E-09 | 6.07E-09 |

| | | | | | | |
|---|---|---|---|---|---|---|
| chr11 | 74150792 | 74152726 | 1935 | -6.7 | 2.76E-09 | 6.07E-09 |
| chr5 | 121289994 | 121294178 | 4185 | -5.3 | 2.77E-09 | 6.07E-09 |
| chr9 | 18576098 | 18974903 | 398806 | 1.79 | 2.77E-09 | 6.08E-09 |
| chr2 | 88631537 | 88640748 | 9212 | -1.78 | 2.77E-09 | 6.08E-09 |
| chr4 | 119170421 | 119171989 | 1569 | -6.87 | 2.78E-09 | 6.10E-09 |
| chrX | 19308935 | 19320981 | 12047 | 3.15 | 2.78E-09 | 6.10E-09 |
| chr15 | 95809486 | 95819548 | 10063 | -2.03 | 2.78E-09 | 6.11E-09 |
| chr13 | 91557900 | 91571841 | 13942 | -6.81 | 2.78E-09 | 6.11E-09 |
| chr8 | 11509854 | 11545195 | 35342 | -6.71 | 2.79E-09 | 6.13E-09 |
| chr6 | 8479909 | 8482083 | 2175 | -6.73 | 2.80E-09 | 6.15E-09 |
| chr13 | 74674173 | 74676456 | 2284 | -6.83 | 2.81E-09 | 6.16E-09 |
| chr5 | 155987583 | 155989622 | 2040 | -6.83 | 2.81E-09 | 6.16E-09 |
| chr21 | 40435596 | 40437870 | 2275 | -6.83 | 2.81E-09 | 6.16E-09 |
| chr1 | 178217458 | 178220294 | 2837 | -6.82 | 2.81E-09 | 6.16E-09 |
| chr9 | 97432038 | 97439226 | 7189 | -6.82 | 2.81E-09 | 6.16E-09 |
| chr8 | 58671481 | 58697049 | 25569 | -2.66 | 2.81E-09 | 6.16E-09 |
| chr2 | 56172056 | 56174116 | 2061 | -6.95 | 2.81E-09 | 6.17E-09 |
| chr10 | 12426454 | 12427910 | 1457 | -6.79 | 2.81E-09 | 6.17E-09 |
| chr12 | 109633336 | 109634730 | 1395 | -6.79 | 2.81E-09 | 6.17E-09 |
| chr20 | 55474930 | 55478725 | 3796 | -6.79 | 2.81E-09 | 6.17E-09 |
| chr4 | 39283530 | 39285142 | 1613 | -6.7 | 2.82E-09 | 6.18E-09 |
| chr19 | 41396194 | 41623389 | 227196 | -2.36 | 2.82E-09 | 6.18E-09 |
| chr12 | 21215067 | 21216991 | 1925 | -6.81 | 2.82E-09 | 6.18E-09 |
| chr1 | 228970550 | 228992470 | 21921 | -6.7 | 2.82E-09 | 6.19E-09 |
| chr2 | 242159051 | 242160263 | 1213 | -6.7 | 2.83E-09 | 6.20E-09 |
| chr2 | 189894829 | 189895979 | 1151 | -6.91 | 2.83E-09 | 6.20E-09 |
| chr14 | 37832965 | 37834864 | 1900 | -6.76 | 2.83E-09 | 6.20E-09 |
| chr2 | 73706184 | 73707162 | 979 | -6.78 | 2.83E-09 | 6.21E-09 |
| chr8 | 38764070 | 38765243 | 1174 | -6.78 | 2.83E-09 | 6.21E-09 |
| chr15 | 55749891 | 55753625 | 3735 | -6.88 | 2.83E-09 | 6.21E-09 |
| chr11 | 92516879 | 92519337 | 2459 | -6.72 | 2.83E-09 | 6.21E-09 |
| chr16 | 84624317 | 84625524 | 1208 | -6.72 | 2.83E-09 | 6.21E-09 |
| chr5 | 153911557 | 153913036 | 1480 | -5.89 | 2.83E-09 | 6.21E-09 |
| chr22 | 51129504 | 51131157 | 1654 | -6.72 | 2.84E-09 | 6.21E-09 |
| chr8 | 10177163 | 10179830 | 2668 | -6.72 | 2.84E-09 | 6.21E-09 |
| chr20 | 52448753 | 52461027 | 12275 | -7.35 | 2.84E-09 | 6.22E-09 |
| chr7 | 79212574 | 79214347 | 1774 | -6.88 | 2.84E-09 | 6.22E-09 |
| chr2 | 131092342 | 131093799 | 1458 | -6.7 | 2.84E-09 | 6.23E-09 |
| chr4 | 38747795 | 38750206 | 2412 | -6.7 | 2.84E-09 | 6.23E-09 |
| chr7 | 140112214 | 140113500 | 1287 | -6.7 | 2.84E-09 | 6.23E-09 |
| chr7 | 43674220 | 43675719 | 1500 | -7 | 2.84E-09 | 6.23E-09 |
| chr1 | 220744629 | 220749891 | 5263 | -3.97 | 2.85E-09 | 6.24E-09 |
| chr4 | 84879170 | 84887229 | 8060 | -7.17 | 2.85E-09 | 6.24E-09 |
| chr7 | 18214572 | 18227882 | 13311 | -5.87 | 2.86E-09 | 6.25E-09 |
| chr5 | 44050622 | 44126543 | 75922 | 2.74 | 2.86E-09 | 6.26E-09 |
| chr14 | 60366713 | 60371010 | 4298 | -2.3 | 2.86E-09 | 6.26E-09 |
| chr3 | 194515879 | 194517848 | 1970 | -6.7 | 2.88E-09 | 6.30E-09 |
| chr1 | 231325029 | 231326974 | 1946 | -6.98 | 2.88E-09 | 6.30E-09 |
| chr22 | 46966996 | 46968893 | 1898 | -6.71 | 2.88E-09 | 6.31E-09 |
| chr6 | 35973977 | 35975518 | 1542 | -6.7 | 2.88E-09 | 6.31E-09 |
| chr11 | 116341589 | 116343620 | 2032 | -4.93 | 2.89E-09 | 6.31E-09 |
| chr9 | 116011317 | 116013587 | 2271 | -6.69 | 2.89E-09 | 6.32E-09 |
| chr2 | 58877215 | 58881490 | 4276 | -2.57 | 2.89E-09 | 6.32E-09 |
| chr12 | 96168414 | 96169285 | 872 | -6.7 | 2.89E-09 | 6.33E-09 |
| chr16 | 47186040 | 47187726 | 1687 | -6.7 | 2.89E-09 | 6.33E-09 |
| chr6 | 84403020 | 84404370 | 1351 | -6.72 | 2.91E-09 | 6.36E-09 |
| chr13 | 41100969 | 41102958 | 1990 | -6.74 | 2.91E-09 | 6.37E-09 |
| chr11 | 21972408 | 21976814 | 4407 | -5.79 | 2.91E-09 | 6.37E-09 |
| chr5 | 17999139 | 18017524 | 18386 | -7.51 | 2.92E-09 | 6.38E-09 |
| chr2 | 63929194 | 63936043 | 6850 | -6.72 | 2.92E-09 | 6.38E-09 |
| chr20 | 8777352 | 8778704 | 1353 | -6.72 | 2.92E-09 | 6.38E-09 |

| | | | | | | |
|---|---|---|---|---|---|---|
| chrX | 44452428 | 44464274 | 11847 | -7.09 | 2.92E-09 | 6.38E-09 |
| chr10 | 1424873 | 1429012 | 4140 | -2.48 | 2.92E-09 | 6.39E-09 |
| chr1 | 108647567 | 108649413 | 1847 | -6.7 | 2.93E-09 | 6.40E-09 |
| chr8 | 141408524 | 141410660 | 2137 | -6.7 | 2.93E-09 | 6.40E-09 |
| chr19 | 3425761 | 3427431 | 1671 | -6.7 | 2.93E-09 | 6.40E-09 |
| chr11 | 113666043 | 113667478 | 1436 | -6.76 | 2.93E-09 | 6.40E-09 |
| chr3 | 17116908 | 17118231 | 1324 | -6.71 | 2.93E-09 | 6.40E-09 |
| chr5 | 93810112 | 93812278 | 2167 | 6.77 | 2.93E-09 | 6.41E-09 |
| chr3 | 129447494 | 129448624 | 1131 | -6.75 | 2.94E-09 | 6.42E-09 |
| chr7 | 126981808 | 126983952 | 2145 | -6.73 | 2.94E-09 | 6.43E-09 |
| chr9 | 126040731 | 126042105 | 1375 | -6.7 | 2.95E-09 | 6.44E-09 |
| chrX | 29887877 | 29891193 | 3317 | -6.76 | 2.95E-09 | 6.44E-09 |
| chr5 | 171171190 | 171173746 | 2557 | 6.9 | 2.95E-09 | 6.45E-09 |
| chr12 | 47851425 | 47934533 | 83109 | -1.27 | 2.95E-09 | 6.45E-09 |
| chr7 | 13454842 | 13462258 | 7417 | -7.29 | 2.96E-09 | 6.46E-09 |
| chr2 | 233576152 | 233582347 | 6196 | -6.69 | 2.96E-09 | 6.47E-09 |
| chr5 | 127150675 | 127154328 | 3654 | -6.83 | 2.97E-09 | 6.49E-09 |
| chr18 | 21676222 | 21677751 | 1530 | -6.82 | 2.97E-09 | 6.49E-09 |
| chr3 | 187817720 | 187819553 | 1834 | -6.69 | 2.97E-09 | 6.50E-09 |
| chr17 | 42604059 | 42604994 | 936 | -6.95 | 2.98E-09 | 6.50E-09 |
| chr12 | 82995955 | 82998354 | 2400 | -6.7 | 2.98E-09 | 6.50E-09 |
| chr2 | 58073440 | 58074946 | 1507 | -6.74 | 2.98E-09 | 6.50E-09 |
| chr2 | 226464077 | 226467219 | 3143 | -6.81 | 2.98E-09 | 6.51E-09 |
| chr5 | 153703954 | 153705747 | 1794 | -6.8 | 2.98E-09 | 6.51E-09 |
| chr14 | 38530706 | 38532669 | 1964 | -6.69 | 2.98E-09 | 6.51E-09 |
| chr5 | 92390462 | 92392749 | 2288 | -7.01 | 2.98E-09 | 6.52E-09 |
| chr12 | 15074641 | 15075841 | 1201 | -6.86 | 2.99E-09 | 6.52E-09 |
| chr4 | 128251566 | 128255860 | 4295 | 3.55 | 2.99E-09 | 6.52E-09 |
| chr7 | 158914155 | 158916322 | 2168 | -6.8 | 2.99E-09 | 6.53E-09 |
| chr21 | 44736854 | 44738744 | 1891 | -6.87 | 3.00E-09 | 6.55E-09 |
| chr12 | 112596063 | 112598017 | 1955 | -6.96 | 3.00E-09 | 6.55E-09 |
| chr13 | 106567914 | 106571245 | 3332 | -6.69 | 3.00E-09 | 6.56E-09 |
| chr14 | 67280439 | 67283192 | 2754 | -6.69 | 3.01E-09 | 6.57E-09 |
| chr11 | 43621924 | 43640406 | 18483 | -4.86 | 3.02E-09 | 6.58E-09 |
| chr14 | 57215421 | 57277236 | 61816 | 2.47 | 3.02E-09 | 6.59E-09 |
| chr12 | 113219278 | 113220487 | 1210 | -6.78 | 3.02E-09 | 6.59E-09 |
| chr1 | 221344320 | 221345856 | 1537 | -6.89 | 3.02E-09 | 6.59E-09 |
| chr9 | 33613401 | 33615310 | 1910 | -6.74 | 3.02E-09 | 6.60E-09 |
| chr4 | 7081092 | 7083128 | 2037 | -6.7 | 3.03E-09 | 6.61E-09 |
| chr7 | 116943565 | 116945349 | 1785 | -6.89 | 3.03E-09 | 6.61E-09 |
| chr7 | 13730502 | 13738602 | 8101 | -6.96 | 3.03E-09 | 6.61E-09 |
| chr21 | 39138260 | 39155039 | 16780 | -7.07 | 3.03E-09 | 6.62E-09 |
| chr15 | 70483235 | 70485086 | 1852 | -6.97 | 3.04E-09 | 6.64E-09 |
| chr6 | 111301199 | 111302555 | 1357 | -6.69 | 3.05E-09 | 6.66E-09 |
| chr12 | 21840628 | 21843344 | 2717 | -6.9 | 3.06E-09 | 6.68E-09 |
| chr7 | 114369061 | 114371064 | 2004 | -6.69 | 3.07E-09 | 6.70E-09 |
| chr17 | 67149396 | 67153156 | 3761 | 6.84 | 3.07E-09 | 6.70E-09 |
| chr1 | 30035135 | 30037777 | 2643 | -6.69 | 3.07E-09 | 6.71E-09 |
| chr12 | 112812639 | 112813557 | 919 | -6.69 | 3.07E-09 | 6.71E-09 |
| chr3 | 184754063 | 184757125 | 3063 | -6.73 | 3.08E-09 | 6.71E-09 |
| chr5 | 143439136 | 143441173 | 2038 | -6.68 | 3.08E-09 | 6.71E-09 |
| chr12 | 21338996 | 21349029 | 10034 | -6.76 | 3.08E-09 | 6.72E-09 |
| chr7 | 94777044 | 94778959 | 1916 | -6.76 | 3.08E-09 | 6.72E-09 |
| chr15 | 33282424 | 33303402 | 20979 | -7.91 | 3.08E-09 | 6.73E-09 |
| chr1 | 2419420 | 2421040 | 1621 | -6.69 | 3.08E-09 | 6.73E-09 |
| chr8 | 68159069 | 68160989 | 1921 | -6.69 | 3.09E-09 | 6.75E-09 |
| chr7 | 155221210 | 155222705 | 1496 | -6.69 | 3.10E-09 | 6.76E-09 |
| chr1 | 189859817 | 189861842 | 2026 | -6.71 | 3.10E-09 | 6.77E-09 |
| chr14 | 96803649 | 96804561 | 913 | -6.7 | 3.11E-09 | 6.77E-09 |
| chr2 | 139445190 | 139446604 | 1415 | -4.41 | 3.11E-09 | 6.77E-09 |
| chr7 | 141371303 | 141375531 | 4229 | -7.32 | 3.11E-09 | 6.79E-09 |

| | | | | | | |
|---|---|---|---|---|---|---|
| chr3 | 152591512 | 152593167 | 1656 | -4.89 | 3.12E-09 | 6.80E-09 |
| chr8 | 121439789 | 121442043 | 2255 | -6.69 | 3.12E-09 | 6.81E-09 |
| chr5 | 61130850 | 61218204 | 87355 | -1.45 | 3.12E-09 | 6.81E-09 |
| chr7 | 30007210 | 30008464 | 1255 | -6.93 | 3.13E-09 | 6.82E-09 |
| chr2 | 173488263 | 173490017 | 1755 | -6.82 | 3.14E-09 | 6.85E-09 |
| chr10 | 88169237 | 88171153 | 1917 | -6.69 | 3.14E-09 | 6.85E-09 |
| chr4 | 138797234 | 138799513 | 2280 | -6.74 | 3.14E-09 | 6.85E-09 |
| chr2 | 10181574 | 10183156 | 1583 | -6.74 | 3.15E-09 | 6.85E-09 |
| chr10 | 35447303 | 35457162 | 9860 | -7 | 3.16E-09 | 6.87E-09 |
| chr3 | 193624525 | 193627235 | 2711 | -4.9 | 3.16E-09 | 6.89E-09 |
| chr4 | 124347048 | 124350391 | 3344 | -6.69 | 3.17E-09 | 6.90E-09 |
| chr17 | 57641946 | 57642836 | 891 | -6.68 | 3.17E-09 | 6.91E-09 |
| chr4 | 104279386 | 104281873 | 2488 | -6.68 | 3.17E-09 | 6.91E-09 |
| chr3 | 13530367 | 13532221 | 1855 | -6.86 | 3.17E-09 | 6.91E-09 |
| chr17 | 2348724 | 2349751 | 1028 | -6.78 | 3.17E-09 | 6.91E-09 |
| chr11 | 45285374 | 45286813 | 1440 | -6.69 | 3.18E-09 | 6.91E-09 |
| chr11 | 128309514 | 128311421 | 1908 | -6.69 | 3.18E-09 | 6.91E-09 |
| chr12 | 60518779 | 60521637 | 2859 | -6.69 | 3.18E-09 | 6.91E-09 |
| chr3 | 101464318 | 101467898 | 3581 | -6.68 | 3.18E-09 | 6.92E-09 |
| chrX | 2848236 | 2849860 | 1625 | -6.68 | 3.18E-09 | 6.92E-09 |
| chr17 | 40809907 | 40810845 | 939 | -6.8 | 3.19E-09 | 6.95E-09 |
| chr2 | 62918767 | 62920569 | 1803 | -6.68 | 3.19E-09 | 6.95E-09 |
| chr8 | 23978496 | 23981691 | 3196 | -6.79 | 3.20E-09 | 6.97E-09 |
| chr12 | 84320267 | 84329110 | 8844 | -7.21 | 3.22E-09 | 7.01E-09 |
| chr1 | 54961984 | 54963883 | 1900 | -6.69 | 3.23E-09 | 7.03E-09 |
| chr11 | 43681098 | 43683471 | 2374 | -6.69 | 3.23E-09 | 7.03E-09 |
| chr3 | 97572389 | 97573829 | 1441 | -6.69 | 3.23E-09 | 7.03E-09 |
| chr7 | 148733176 | 148735019 | 1844 | -6.73 | 3.23E-09 | 7.03E-09 |
| chr4 | 180912279 | 180919355 | 7077 | -6.75 | 3.24E-09 | 7.06E-09 |
| chr2 | 190012369 | 190013860 | 1492 | -5.75 | 3.25E-09 | 7.06E-09 |
| chr1 | 244862951 | 244865001 | 2051 | -6.75 | 3.26E-09 | 7.10E-09 |
| chr19 | 6491142 | 6492751 | 1610 | -6.67 | 3.26E-09 | 7.10E-09 |
| chr9 | 5196862 | 5246396 | 49535 | -2.08 | 3.27E-09 | 7.11E-09 |
| chr9 | 117737252 | 117743111 | 5860 | -2.57 | 3.28E-09 | 7.13E-09 |
| chr5 | 115580912 | 115583266 | 2355 | -7.03 | 3.28E-09 | 7.13E-09 |
| chr1 | 15279590 | 15281117 | 1528 | -6.67 | 3.28E-09 | 7.13E-09 |
| chr6 | 27608905 | 27609932 | 1028 | -6.72 | 3.29E-09 | 7.16E-09 |
| chr6 | 133553290 | 133557258 | 3969 | -7.14 | 3.30E-09 | 7.19E-09 |
| chr9 | 108552904 | 108555792 | 2889 | -6.74 | 3.31E-09 | 7.19E-09 |
| chr7 | 37992227 | 37994027 | 1801 | -6.7 | 3.32E-09 | 7.21E-09 |
| chr8 | 42006047 | 42007923 | 1877 | -6.68 | 3.32E-09 | 7.22E-09 |
| chr16 | 21154686 | 21155992 | 1307 | -6.71 | 3.32E-09 | 7.22E-09 |
| chr3 | 110479051 | 110487673 | 8623 | -6.81 | 3.33E-09 | 7.23E-09 |
| chr1 | 160437609 | 160508530 | 70922 | -1.5 | 3.33E-09 | 7.23E-09 |
| chr4 | 121481802 | 121484547 | 2746 | -6.7 | 3.33E-09 | 7.23E-09 |
| chr6 | 16660708 | 16662910 | 2203 | -6.8 | 3.33E-09 | 7.24E-09 |
| chr6 | 52251965 | 52253640 | 1676 | -6.83 | 3.33E-09 | 7.25E-09 |
| chr15 | 77633069 | 77634487 | 1419 | -6.92 | 3.34E-09 | 7.27E-09 |
| chr1 | 198705648 | 198708475 | 2828 | -2.22 | 3.35E-09 | 7.27E-09 |
| chr4 | 105321743 | 105323509 | 1767 | -6.74 | 3.35E-09 | 7.28E-09 |
| chr7 | 117270675 | 117272481 | 1807 | -5.2 | 3.35E-09 | 7.28E-09 |
| chr3 | 4598727 | 4600790 | 2064 | -6.71 | 3.35E-09 | 7.28E-09 |
| chr17 | 58486761 | 58488350 | 1590 | -6.94 | 3.36E-09 | 7.30E-09 |
| chr15 | 53821775 | 53825425 | 3651 | -2.5 | 3.36E-09 | 7.30E-09 |
| chr7 | 144423367 | 144428368 | 5002 | -6.77 | 3.36E-09 | 7.30E-09 |
| chr11 | 45027517 | 45033591 | 6075 | -7.05 | 3.37E-09 | 7.32E-09 |
| chr2 | 220699015 | 220728907 | 29893 | -1.81 | 3.37E-09 | 7.32E-09 |
| chr4 | 137518597 | 137521581 | 2985 | -6.77 | 3.37E-09 | 7.33E-09 |
| chr2 | 228147094 | 228148309 | 1216 | -6.7 | 3.38E-09 | 7.35E-09 |
| chr4 | 85945835 | 85948453 | 2619 | -6.68 | 3.39E-09 | 7.36E-09 |
| chr1 | 180478893 | 180481018 | 2126 | -6.87 | 3.39E-09 | 7.37E-09 |

| | | | | | | |
|---|---|---|---|---|---|---|
| chr6 | 161612854 | 161615177 | 2324 | -6.87 | 3.39E-09 | 7.37E-09 |
| chr16 | 49945622 | 49984268 | 38647 | -7.82 | 3.40E-09 | 7.38E-09 |
| chr14 | 56381894 | 56385083 | 3190 | -6.76 | 3.40E-09 | 7.38E-09 |
| chr3 | 141830349 | 141831956 | 1608 | -6.69 | 3.41E-09 | 7.40E-09 |
| chr10 | 95118284 | 95119251 | 968 | -6.94 | 3.41E-09 | 7.41E-09 |
| chr1 | 16634106 | 16635228 | 1123 | -6.82 | 3.42E-09 | 7.43E-09 |
| chr12 | 53977361 | 53978872 | 1512 | -6.82 | 3.42E-09 | 7.43E-09 |
| chr8 | 38971922 | 38987146 | 15225 | -7.61 | 3.42E-09 | 7.43E-09 |
| chr22 | 47181010 | 47182782 | 1773 | -6.83 | 3.43E-09 | 7.45E-09 |
| chr5 | 139568059 | 139569355 | 1297 | -6.72 | 3.44E-09 | 7.46E-09 |
| chr6 | 140164566 | 140173065 | 8500 | -6.78 | 3.44E-09 | 7.46E-09 |
| chrX | 45602217 | 45605324 | 3108 | -6.78 | 3.44E-09 | 7.46E-09 |
| chr7 | 82250086 | 82251552 | 1467 | -6.67 | 3.44E-09 | 7.47E-09 |
| chr15 | 53877671 | 53879490 | 1820 | -7.02 | 3.44E-09 | 7.48E-09 |
| chr4 | 68051159 | 68062649 | 11491 | 2.43 | 3.45E-09 | 7.48E-09 |
| chr7 | 87737384 | 87738828 | 1445 | -6.68 | 3.45E-09 | 7.49E-09 |
| chr3 | 134163167 | 134165150 | 1984 | -6.7 | 3.46E-09 | 7.51E-09 |
| chr7 | 75219761 | 75221035 | 1275 | -6.7 | 3.46E-09 | 7.51E-09 |
| chr3 | 170768134 | 170772320 | 4187 | -6.77 | 3.46E-09 | 7.51E-09 |
| chr13 | 106226636 | 106232860 | 6225 | -7.26 | 3.46E-09 | 7.51E-09 |
| chr9 | 109668520 | 109669746 | 1227 | -6.66 | 3.46E-09 | 7.52E-09 |
| chr5 | 81311530 | 81312953 | 1424 | -6.9 | 3.47E-09 | 7.52E-09 |
| chr21 | 40494518 | 40517403 | 22886 | -7.52 | 3.48E-09 | 7.55E-09 |
| chr8 | 68221184 | 68225924 | 4741 | -6.96 | 3.48E-09 | 7.55E-09 |
| chr20 | 30676094 | 30677267 | 1174 | -6.67 | 3.49E-09 | 7.56E-09 |
| chr9 | 513517 | 515014 | 1498 | -6.67 | 3.49E-09 | 7.56E-09 |
| chr9 | 108025558 | 108027897 | 2340 | -6.67 | 3.49E-09 | 7.57E-09 |
| chr18 | 33054223 | 33056138 | 1916 | -6.75 | 3.49E-09 | 7.57E-09 |
| chr2 | 202799229 | 202801150 | 1922 | -5.55 | 3.49E-09 | 7.58E-09 |
| chr1 | 181974757 | 181991369 | 16613 | -3.9 | 3.50E-09 | 7.58E-09 |
| chr12 | 63686146 | 63690205 | 4060 | -6.68 | 3.50E-09 | 7.60E-09 |
| chr12 | 44324072 | 44325870 | 1799 | -6.73 | 3.50E-09 | 7.60E-09 |
| chrX | 100269892 | 100270728 | 837 | -6.88 | 3.50E-09 | 7.60E-09 |
| chr5 | 41493957 | 41496471 | 2515 | -6.75 | 3.50E-09 | 7.60E-09 |
| chr1 | 25206749 | 25219634 | 12886 | -7.03 | 3.51E-09 | 7.60E-09 |
| chr3 | 120213639 | 120215780 | 2142 | -6.74 | 3.52E-09 | 7.63E-09 |
| chr22 | 17278210 | 17332208 | 53999 | 1.43 | 3.53E-09 | 7.64E-09 |
| chr1 | 229378821 | 229380324 | 1504 | -6.82 | 3.53E-09 | 7.64E-09 |
| chr1 | 246580200 | 246582756 | 2557 | -6.79 | 3.53E-09 | 7.64E-09 |
| chr4 | 142597796 | 142600117 | 2322 | -6.66 | 3.53E-09 | 7.64E-09 |
| chrX | 100825680 | 100827508 | 1829 | -6.72 | 3.53E-09 | 7.65E-09 |
| chr10 | 112229027 | 112231454 | 2428 | -6.84 | 3.55E-09 | 7.68E-09 |
| chr16 | 15656280 | 15658029 | 1750 | -6.85 | 3.56E-09 | 7.72E-09 |
| chr8 | 1308026 | 1316547 | 8522 | -6.85 | 3.56E-09 | 7.72E-09 |
| chr1 | 29602711 | 29603938 | 1228 | -6.66 | 3.57E-09 | 7.74E-09 |
| chrX | 123213890 | 123227567 | 13678 | -6.9 | 3.57E-09 | 7.74E-09 |
| chr20 | 10893004 | 10898394 | 5391 | -6.69 | 3.58E-09 | 7.76E-09 |
| chr9 | 2220786 | 2223623 | 2838 | -6.7 | 3.58E-09 | 7.76E-09 |
| chr10 | 52108569 | 52123414 | 14846 | -6.66 | 3.59E-09 | 7.77E-09 |
| chrX | 68842033 | 68844835 | 2803 | -5.26 | 3.60E-09 | 7.79E-09 |
| chr18 | 8723925 | 8727493 | 3569 | -7.18 | 3.61E-09 | 7.83E-09 |
| chr8 | 37716229 | 37717520 | 1292 | -6.67 | 3.62E-09 | 7.83E-09 |
| chr7 | 155549460 | 155550849 | 1390 | -6.66 | 3.62E-09 | 7.84E-09 |
| chr6 | 167524045 | 167534711 | 10667 | -7.39 | 3.62E-09 | 7.84E-09 |
| chr3 | 145793792 | 145794684 | 893 | -6.72 | 3.63E-09 | 7.86E-09 |
| chr21 | 30820164 | 30822588 | 2425 | -6.66 | 3.64E-09 | 7.88E-09 |
| chr1 | 63754247 | 63756185 | 1939 | -6.68 | 3.64E-09 | 7.88E-09 |
| chr13 | 61243031 | 61328095 | 85065 | 7.09 | 3.65E-09 | 7.90E-09 |
| chr3 | 38968993 | 38998112 | 29120 | -1.58 | 3.66E-09 | 7.92E-09 |
| chr4 | 121730504 | 121732089 | 1586 | -6.78 | 3.68E-09 | 7.97E-09 |
| chr9 | 118215634 | 118220336 | 4703 | -7.02 | 3.69E-09 | 7.98E-09 |

| | | | | | | |
|---|---|---|---|---|---|---|
| chr20 | 33933477 | 33934697 | 1221 | -6.82 | 3.69E-09 | 7.98E-09 |
| chr1 | 239785142 | 239786395 | 1254 | -5.22 | 3.69E-09 | 7.99E-09 |
| chr14 | 100620400 | 100622055 | 1656 | -6.9 | 3.70E-09 | 8.00E-09 |
| chr6 | 85356173 | 85360359 | 4187 | -6.83 | 3.70E-09 | 8.02E-09 |
| chr11 | 116769913 | 116771115 | 1203 | -6.67 | 3.71E-09 | 8.03E-09 |
| chr8 | 81280581 | 81281908 | 1328 | -6.67 | 3.71E-09 | 8.03E-09 |
| chr16 | 29983866 | 29984782 | 917 | -6.66 | 3.72E-09 | 8.05E-09 |
| chr16 | 47606868 | 47609123 | 2256 | -6.66 | 3.72E-09 | 8.05E-09 |
| chr10 | 88400848 | 88402076 | 1229 | -6.7 | 3.74E-09 | 8.09E-09 |
| chr21 | 35899863 | 35901615 | 1753 | -6.7 | 3.74E-09 | 8.09E-09 |
| chr6 | 63595686 | 63599170 | 3485 | -6.82 | 3.74E-09 | 8.10E-09 |
| chr10 | 114531223 | 114532349 | 1127 | -6.77 | 3.74E-09 | 8.10E-09 |
| chr4 | 64270931 | 64276761 | 5831 | -6.77 | 3.74E-09 | 8.10E-09 |
| chr8 | 74894249 | 74896202 | 1954 | -6.77 | 3.74E-09 | 8.10E-09 |
| chr3 | 109816310 | 109817683 | 1374 | -6.66 | 3.75E-09 | 8.10E-09 |
| chr6 | 85741712 | 85745021 | 3310 | -5.09 | 3.75E-09 | 8.10E-09 |
| chr15 | 78997291 | 79008050 | 10760 | -6.92 | 3.76E-09 | 8.14E-09 |
| chr8 | 108379920 | 108382179 | 2260 | -6.68 | 3.79E-09 | 8.20E-09 |
| chr20 | 43365168 | 43366959 | 1792 | -6.85 | 3.79E-09 | 8.20E-09 |
| chr13 | 76838695 | 76860134 | 21440 | -2.33 | 3.79E-09 | 8.20E-09 |
| chr8 | 135731847 | 135738290 | 6444 | -7.09 | 3.79E-09 | 8.20E-09 |
| chr9 | 82746375 | 82771285 | 24911 | -1.35 | 3.80E-09 | 8.21E-09 |
| chr7 | 105484700 | 105487526 | 2827 | -6.65 | 3.80E-09 | 8.21E-09 |
| chr2 | 203706610 | 203708647 | 2038 | -6.93 | 3.80E-09 | 8.21E-09 |
| chr15 | 36599548 | 36602191 | 2644 | -6.74 | 3.80E-09 | 8.21E-09 |
| chr2 | 112469160 | 112470565 | 1406 | -6.66 | 3.80E-09 | 8.22E-09 |
| chr11 | 126902695 | 126913940 | 11246 | -6.85 | 3.80E-09 | 8.22E-09 |
| chr11 | 43861859 | 43862743 | 885 | -6.65 | 3.80E-09 | 8.22E-09 |
| chrX | 42421058 | 42458674 | 37617 | 2.28 | 3.80E-09 | 8.22E-09 |
| chr14 | 58505421 | 58508163 | 2743 | -5.54 | 3.81E-09 | 8.22E-09 |
| chr4 | 133266583 | 133277683 | 11101 | -6.65 | 3.81E-09 | 8.23E-09 |
| chr5 | 34877367 | 34879259 | 1893 | -6.66 | 3.82E-09 | 8.24E-09 |
| chr6 | 148110785 | 148112538 | 1754 | -6.66 | 3.82E-09 | 8.24E-09 |
| chr3 | 51205467 | 51224044 | 18578 | -2.95 | 3.84E-09 | 8.29E-09 |
| chr22 | 49286757 | 49288055 | 1299 | -6.65 | 3.84E-09 | 8.30E-09 |
| chr8 | 59335065 | 59336478 | 1414 | -6.65 | 3.84E-09 | 8.30E-09 |
| chr8 | 41491796 | 41493016 | 1221 | -6.87 | 3.84E-09 | 8.30E-09 |
| chr3 | 13383887 | 13385332 | 1446 | -6.72 | 3.85E-09 | 8.31E-09 |
| chrX | 53901296 | 53928401 | 27106 | -4.34 | 3.85E-09 | 8.32E-09 |
| chrX | 118064442 | 118066021 | 1580 | -6.68 | 3.85E-09 | 8.32E-09 |
| chr22 | 49889360 | 49891288 | 1929 | -6.65 | 3.85E-09 | 8.32E-09 |
| chr5 | 78241183 | 78243521 | 2339 | -6.65 | 3.85E-09 | 8.32E-09 |
| chrX | 4474886 | 4486685 | 11800 | -7 | 3.86E-09 | 8.33E-09 |
| chr3 | 112355101 | 112356558 | 1458 | -6.71 | 3.87E-09 | 8.35E-09 |
| chr6 | 85497838 | 85506623 | 8786 | -6.71 | 3.87E-09 | 8.35E-09 |
| chr14 | 77341024 | 77342468 | 1445 | -6.65 | 3.89E-09 | 8.40E-09 |
| chr7 | 115736543 | 115738504 | 1962 | -6.65 | 3.89E-09 | 8.40E-09 |
| chr1 | 231821064 | 231823786 | 2723 | -3.22 | 3.89E-09 | 8.40E-09 |
| chr2 | 228747246 | 228749612 | 2367 | -6.71 | 3.89E-09 | 8.40E-09 |
| chr4 | 105456425 | 105458232 | 1808 | -6.67 | 3.91E-09 | 8.44E-09 |
| chr17 | 53280839 | 53282091 | 1253 | -5.09 | 3.92E-09 | 8.46E-09 |
| chr14 | 101091543 | 101093529 | 1987 | -6.64 | 3.93E-09 | 8.48E-09 |
| chr3 | 7364842 | 7396365 | 31524 | 6.44 | 3.93E-09 | 8.49E-09 |
| chrX | 52076484 | 52079206 | 2723 | -2.69 | 3.93E-09 | 8.49E-09 |
| chr11 | 94762044 | 94764183 | 2140 | -5.74 | 3.94E-09 | 8.50E-09 |
| chr7 | 104820253 | 104821955 | 1703 | -6.65 | 3.94E-09 | 8.51E-09 |
| chr8 | 122576108 | 122579590 | 3483 | -6.65 | 3.94E-09 | 8.51E-09 |
| chr2 | 153940674 | 153949738 | 9065 | -2.48 | 3.95E-09 | 8.51E-09 |
| chr8 | 61382503 | 61384027 | 1525 | -6.8 | 3.95E-09 | 8.53E-09 |
| chr8 | 23844410 | 23846461 | 2052 | -6.77 | 3.95E-09 | 8.53E-09 |
| chr2 | 183885924 | 183887131 | 1208 | -6.81 | 3.96E-09 | 8.54E-09 |

| | | | | | | |
|---|---|---|---|---|---|---|
| chr9 | 29509183 | 29513435 | 4253 | -6.68 | 3.96E-09 | 8.54E-09 |
| chr5 | 31986807 | 31988532 | 1726 | -6.81 | 3.96E-09 | 8.55E-09 |
| chr14 | 65922444 | 65923715 | 1272 | -6.81 | 3.97E-09 | 8.56E-09 |
| chr4 | 129576155 | 129591706 | 15552 | -7.17 | 3.97E-09 | 8.57E-09 |
| chr8 | 70734731 | 70741455 | 6725 | -6.76 | 3.97E-09 | 8.57E-09 |
| chr12 | 17865453 | 17866952 | 1500 | -6.7 | 3.98E-09 | 8.57E-09 |
| chr11 | 6534700 | 6536442 | 1743 | -6.82 | 3.98E-09 | 8.58E-09 |
| chr16 | 24846657 | 24856830 | 10174 | -6.75 | 3.98E-09 | 8.58E-09 |
| chr21 | 15711370 | 15713601 | 2232 | 3.53 | 3.99E-09 | 8.59E-09 |
| chr11 | 20555858 | 20557330 | 1473 | -6.65 | 4.00E-09 | 8.62E-09 |
| chr1 | 165681503 | 165683358 | 1856 | -6.92 | 4.00E-09 | 8.63E-09 |
| chr15 | 101479570 | 101480941 | 1372 | -6.7 | 4.00E-09 | 8.63E-09 |
| chr2 | 224157942 | 224160642 | 2701 | -6.7 | 4.00E-09 | 8.63E-09 |
| chr4 | 126618432 | 126620174 | 1743 | -6.84 | 4.00E-09 | 8.63E-09 |
| chr7 | 22942249 | 22943583 | 1335 | -6.67 | 4.01E-09 | 8.63E-09 |
| chr4 | 101558628 | 101560642 | 2015 | -6.84 | 4.01E-09 | 8.64E-09 |
| chr2 | 189317999 | 189320368 | 2370 | -6.92 | 4.02E-09 | 8.66E-09 |
| chr4 | 7683077 | 7685434 | 2358 | -6.73 | 4.03E-09 | 8.68E-09 |
| chr9 | 17434045 | 17435943 | 1899 | -6.65 | 4.03E-09 | 8.69E-09 |
| chr10 | 12522443 | 12523981 | 1539 | -6.67 | 4.03E-09 | 8.69E-09 |
| chr10 | 62004694 | 62007995 | 3302 | -6.73 | 4.04E-09 | 8.70E-09 |
| chr7 | 138771397 | 138773898 | 2502 | -6.73 | 4.04E-09 | 8.70E-09 |
| chr5 | 13880186 | 13882604 | 2419 | -6.64 | 4.04E-09 | 8.71E-09 |
| chr6 | 96039361 | 96040714 | 1354 | -6.64 | 4.05E-09 | 8.72E-09 |
| chr2 | 226434527 | 226441919 | 7393 | -2.32 | 4.05E-09 | 8.72E-09 |
| chrX | 56020954 | 56024105 | 3152 | 2.33 | 4.06E-09 | 8.74E-09 |
| chr20 | 1651825 | 1707943 | 56119 | -1.32 | 4.07E-09 | 8.76E-09 |
| chr2 | 15585801 | 15586729 | 929 | -6.72 | 4.07E-09 | 8.76E-09 |
| chr3 | 184401557 | 184403026 | 1470 | -6.64 | 4.07E-09 | 8.76E-09 |
| chr17 | 70316171 | 70322052 | 5882 | -1.94 | 4.07E-09 | 8.76E-09 |
| chr13 | 102559108 | 102560272 | 1165 | 5.41 | 4.08E-09 | 8.78E-09 |
| chr4 | 24495606 | 24497539 | 1934 | -6.68 | 4.08E-09 | 8.78E-09 |
| chr1 | 119407144 | 119455093 | 47950 | 3.15 | 4.08E-09 | 8.78E-09 |
| chr4 | 107315658 | 107319382 | 3725 | -6.95 | 4.08E-09 | 8.79E-09 |
| chr5 | 172890557 | 172894504 | 3948 | -6.69 | 4.08E-09 | 8.79E-09 |
| chr2 | 68828297 | 68934658 | 106362 | -2.74 | 4.09E-09 | 8.80E-09 |
| chr1 | 45662500 | 45664733 | 2234 | -6.67 | 4.09E-09 | 8.80E-09 |
| chr2 | 114521440 | 114523338 | 1899 | -6.87 | 4.10E-09 | 8.82E-09 |
| chr3 | 9122177 | 9124921 | 2745 | -6.65 | 4.10E-09 | 8.82E-09 |
| chr3 | 97729366 | 97730668 | 1303 | -6.65 | 4.10E-09 | 8.82E-09 |
| chr9 | 100093427 | 100096320 | 2894 | -6.71 | 4.12E-09 | 8.86E-09 |
| chr2 | 235988581 | 236005990 | 17410 | -7.35 | 4.12E-09 | 8.87E-09 |
| chr11 | 56935628 | 56938501 | 2874 | -7.11 | 4.13E-09 | 8.89E-09 |
| chr6 | 35130720 | 35132479 | 1760 | -6.64 | 4.14E-09 | 8.91E-09 |
| chr6 | 125654225 | 125657360 | 3136 | -2.19 | 4.14E-09 | 8.91E-09 |
| chr12 | 57894150 | 57895431 | 1282 | -6.66 | 4.15E-09 | 8.93E-09 |
| chr7 | 5686406 | 5689093 | 2688 | -6.66 | 4.16E-09 | 8.96E-09 |
| chr11 | 102524619 | 102527162 | 2544 | -6.96 | 4.16E-09 | 8.96E-09 |
| chr11 | 92056120 | 92057315 | 1196 | -6.63 | 4.18E-09 | 8.98E-09 |
| chr1 | 188193445 | 188195279 | 1835 | -6.68 | 4.18E-09 | 8.99E-09 |
| chr8 | 116244334 | 116252005 | 7672 | -6.68 | 4.18E-09 | 8.99E-09 |
| chr12 | 79409128 | 79412047 | 2920 | -6.65 | 4.18E-09 | 8.99E-09 |
| chr17 | 5196085 | 5197731 | 1647 | -6.78 | 4.19E-09 | 9.00E-09 |
| chr12 | 114321441 | 114322954 | 1514 | -6.78 | 4.19E-09 | 9.00E-09 |
| chr3 | 109377650 | 109384060 | 6411 | -7.07 | 4.20E-09 | 9.03E-09 |
| chr1 | 2294255 | 2296038 | 1784 | -6.69 | 4.21E-09 | 9.05E-09 |
| chr7 | 19823308 | 19825681 | 2374 | -6.69 | 4.21E-09 | 9.05E-09 |
| chr2 | 165289167 | 165290569 | 1403 | -6.63 | 4.22E-09 | 9.07E-09 |
| chr18 | 20350754 | 20352525 | 1772 | -6.66 | 4.22E-09 | 9.07E-09 |
| chr2 | 176837841 | 176839242 | 1402 | -6.65 | 4.22E-09 | 9.08E-09 |
| chr5 | 697550 | 699579 | 2030 | -6.65 | 4.22E-09 | 9.08E-09 |

| | | | | | | |
|---|---|---|---|---|---|---|
| chr13 | 73983199 | 73986724 | 3526 | -6.67 | 4.22E-09 | 9.08E-09 |
| chr3 | 189431705 | 189434453 | 2749 | -6.67 | 4.22E-09 | 9.08E-09 |
| chr14 | 65109783 | 65119308 | 9526 | -7.22 | 4.23E-09 | 9.08E-09 |
| chr6 | 135529979 | 135531368 | 1390 | -6.64 | 4.23E-09 | 9.09E-09 |
| chr9 | 108352734 | 108354526 | 1793 | -6.69 | 4.25E-09 | 9.13E-09 |
| chr3 | 24017102 | 24018299 | 1198 | -6.73 | 4.25E-09 | 9.14E-09 |
| chr4 | 86232375 | 86235036 | 2662 | -3.22 | 4.26E-09 | 9.14E-09 |
| chr9 | 137915786 | 137920155 | 4370 | -1.86 | 4.26E-09 | 9.16E-09 |
| chr3 | 30015096 | 30017320 | 2225 | -6.69 | 4.26E-09 | 9.16E-09 |
| chr16 | 86426973 | 86525045 | 98073 | -1.77 | 4.26E-09 | 9.16E-09 |
| chr4 | 90938980 | 90941357 | 2378 | -6.8 | 4.26E-09 | 9.16E-09 |
| chr9 | 117215529 | 117217258 | 1730 | -6.74 | 4.28E-09 | 9.19E-09 |
| chr2 | 149837908 | 149842203 | 4296 | -6.64 | 4.29E-09 | 9.20E-09 |
| chr4 | 15878701 | 15880267 | 1567 | -6.82 | 4.30E-09 | 9.23E-09 |
| chr8 | 8326285 | 8328479 | 2195 | -6.73 | 4.31E-09 | 9.25E-09 |
| chr12 | 122436058 | 122457744 | 21687 | -6.71 | 4.31E-09 | 9.26E-09 |
| chr4 | 40362428 | 40364199 | 1772 | -6.71 | 4.31E-09 | 9.26E-09 |
| chr5 | 5328236 | 5342703 | 14468 | -7.24 | 4.31E-09 | 9.26E-09 |
| chr12 | 69321458 | 69323048 | 1591 | -6.68 | 4.33E-09 | 9.30E-09 |
| chr7 | 155646483 | 155662522 | 16040 | -7.24 | 4.33E-09 | 9.31E-09 |
| chr18 | 57891587 | 57893610 | 2024 | -6.72 | 4.34E-09 | 9.31E-09 |
| chr12 | 87200893 | 87203860 | 2968 | -6.86 | 4.34E-09 | 9.32E-09 |
| chr17 | 49312515 | 49329460 | 16946 | -6.66 | 4.34E-09 | 9.32E-09 |
| chr5 | 92730727 | 92741470 | 10744 | -7.09 | 4.34E-09 | 9.32E-09 |
| chr1 | 232480399 | 232482097 | 1699 | -2.49 | 4.35E-09 | 9.35E-09 |
| chr1 | 192352446 | 192354525 | 2080 | -6.87 | 4.36E-09 | 9.35E-09 |
| chr11 | 28483274 | 28485601 | 2328 | -6.67 | 4.37E-09 | 9.38E-09 |
| chr1 | 77396259 | 77397980 | 1722 | -3.29 | 4.37E-09 | 9.38E-09 |
| chr4 | 79866180 | 79869260 | 3081 | -3.04 | 4.37E-09 | 9.38E-09 |
| chr15 | 43135166 | 43149526 | 14361 | -6.88 | 4.39E-09 | 9.42E-09 |
| chr4 | 137133967 | 137140409 | 6443 | -6.69 | 4.40E-09 | 9.43E-09 |
| chr6 | 47581094 | 47583484 | 2391 | -6.7 | 4.41E-09 | 9.46E-09 |
| chr8 | 119283130 | 119284848 | 1719 | -6.7 | 4.41E-09 | 9.46E-09 |
| chr21 | 32659936 | 32661760 | 1825 | -6.67 | 4.41E-09 | 9.46E-09 |
| chr16 | 28024885 | 28061524 | 36640 | -6.78 | 4.42E-09 | 9.48E-09 |
| chr1 | 111695702 | 111697462 | 1761 | -6.86 | 4.42E-09 | 9.48E-09 |
| chrX | 18807246 | 18810590 | 3345 | -6.69 | 4.43E-09 | 9.50E-09 |
| chr20 | 32815151 | 32828467 | 13317 | -3.21 | 4.43E-09 | 9.50E-09 |
| chrX | 57359053 | 57377237 | 18185 | -5.89 | 4.43E-09 | 9.51E-09 |
| chr1 | 245153440 | 245155093 | 1654 | -6.87 | 4.43E-09 | 9.51E-09 |
| chr3 | 149151123 | 149152739 | 1617 | -6.67 | 4.43E-09 | 9.51E-09 |
| chr7 | 101625930 | 101627069 | 1140 | -6.67 | 4.43E-09 | 9.51E-09 |
| chr10 | 116487362 | 116488686 | 1325 | -6.63 | 4.43E-09 | 9.51E-09 |
| chr15 | 53079330 | 53081201 | 1872 | -6.62 | 4.44E-09 | 9.51E-09 |
| chr6 | 131512824 | 131515106 | 2283 | -6.62 | 4.44E-09 | 9.51E-09 |
| chr4 | 170837284 | 170839568 | 2285 | -6.78 | 4.44E-09 | 9.52E-09 |
| chrX | 81472079 | 81474671 | 2593 | -6.64 | 4.44E-09 | 9.53E-09 |
| chr3 | 33309123 | 33314576 | 5454 | -6.75 | 4.45E-09 | 9.53E-09 |
| chr6 | 137118265 | 137120513 | 2249 | -6.75 | 4.45E-09 | 9.53E-09 |
| chrX | 64881665 | 64883256 | 1592 | -6.74 | 4.45E-09 | 9.54E-09 |
| chr7 | 87230296 | 87232376 | 2081 | -6.63 | 4.45E-09 | 9.54E-09 |
| chr14 | 63865234 | 63866209 | 976 | -6.69 | 4.45E-09 | 9.55E-09 |
| chr12 | 28327501 | 28342753 | 15253 | -6.74 | 4.46E-09 | 9.55E-09 |
| chr10 | 62062211 | 62063216 | 1006 | -6.73 | 4.48E-09 | 9.60E-09 |
| chrX | 74355925 | 74367534 | 11610 | -6.73 | 4.48E-09 | 9.60E-09 |
| chr3 | 163948342 | 163955048 | 6707 | -6.96 | 4.48E-09 | 9.60E-09 |
| chr4 | 13525500 | 13526991 | 1492 | -6.68 | 4.51E-09 | 9.65E-09 |
| chr2 | 38838517 | 38839694 | 1178 | -6.72 | 4.51E-09 | 9.66E-09 |
| chr2 | 197392443 | 197394852 | 2410 | -6.62 | 4.52E-09 | 9.68E-09 |
| chr3 | 106790574 | 106827689 | 37116 | 1.9 | 4.52E-09 | 9.68E-09 |
| chr22 | 40885683 | 40890780 | 5098 | -6.89 | 4.52E-09 | 9.69E-09 |

| | | | | | | |
|---|---|---|---|---|---|---|
| chr7 | 83258840 | 83262385 | 3546 | -6.71 | 4.53E-09 | 9.70E-09 |
| chr4 | 94524389 | 94526628 | 2240 | -6.63 | 4.53E-09 | 9.71E-09 |
| chr12 | 97917613 | 97920262 | 2650 | -6.68 | 4.54E-09 | 9.72E-09 |
| chr10 | 49823278 | 49824905 | 1628 | -6.65 | 4.54E-09 | 9.72E-09 |
| chr6 | 84465081 | 84467513 | 2433 | -6.65 | 4.54E-09 | 9.72E-09 |
| chr16 | 59216891 | 59263941 | 47051 | -2.69 | 4.55E-09 | 9.74E-09 |
| chr16 | 55410599 | 55449563 | 38965 | 2.49 | 4.55E-09 | 9.74E-09 |
| chr13 | 66993239 | 66996173 | 2935 | -6.71 | 4.55E-09 | 9.74E-09 |
| chr3 | 121234797 | 121236188 | 1392 | -6.64 | 4.56E-09 | 9.76E-09 |
| chr14 | 21034147 | 21038048 | 3902 | -3.69 | 4.56E-09 | 9.77E-09 |
| chrX | 57623461 | 57625541 | 2081 | -6.77 | 4.57E-09 | 9.79E-09 |
| chr4 | 142210682 | 142213395 | 2714 | -6.77 | 4.57E-09 | 9.79E-09 |
| chr20 | 45368403 | 45375055 | 6653 | -6.92 | 4.58E-09 | 9.80E-09 |
| chr5 | 176563078 | 176565001 | 1924 | -6.62 | 4.58E-09 | 9.81E-09 |
| chr10 | 112320587 | 112321662 | 1076 | -6.74 | 4.58E-09 | 9.81E-09 |
| chr17 | 65410038 | 65411160 | 1123 | -6.62 | 4.59E-09 | 9.82E-09 |
| chr6 | 3865036 | 3866718 | 1683 | -6.62 | 4.59E-09 | 9.82E-09 |
| chr7 | 68005877 | 68007303 | 1427 | -6.65 | 4.59E-09 | 9.83E-09 |
| chr7 | 109092466 | 109094544 | 2079 | -6.65 | 4.59E-09 | 9.83E-09 |
| chr2 | 75430358 | 75448135 | 17778 | -6.92 | 4.60E-09 | 9.84E-09 |
| chr14 | 43861895 | 44744629 | 882735 | 1.1 | 4.62E-09 | 9.88E-09 |
| chr22 | 22863589 | 22865544 | 1956 | -6.62 | 4.62E-09 | 9.88E-09 |
| chr16 | 14525641 | 14526954 | 1314 | -6.63 | 4.63E-09 | 9.90E-09 |
| chr16 | 71804609 | 71805894 | 1286 | -6.82 | 4.66E-09 | 9.97E-09 |
| chr4 | 108543543 | 108545496 | 1954 | -6.82 | 4.66E-09 | 9.97E-09 |
| chr3 | 150357927 | 150364043 | 6117 | -7.35 | 4.67E-09 | 9.98E-09 |
| chr10 | 76078995 | 76080755 | 1761 | -6.83 | 4.67E-09 | 9.99E-09 |
| chr1 | 207462678 | 207466382 | 3705 | -5.5 | 4.67E-09 | 9.99E-09 |
| chr10 | 70864287 | 70865442 | 1156 | -6.64 | 4.68E-09 | 1.00E-08 |
| chr2 | 19135458 | 19151574 | 16117 | -7.66 | 4.68E-09 | 1.00E-08 |
| chr3 | 141235345 | 141236575 | 1231 | -7.13 | 4.68E-09 | 1.00E-08 |
| chrX | 27519605 | 27522882 | 3278 | -6.66 | 4.71E-09 | 1.01E-08 |
| chr19 | 45306868 | 45308031 | 1164 | -6.78 | 4.71E-09 | 1.01E-08 |
| chr20 | 47665601 | 47667911 | 2311 | -7.07 | 4.72E-09 | 1.01E-08 |
| chr7 | 50412180 | 50441973 | 29794 | -6.58 | 4.72E-09 | 1.01E-08 |
| chr5 | 145372004 | 145374347 | 2344 | -6.78 | 4.72E-09 | 1.01E-08 |
| chrX | 95768530 | 95770518 | 1989 | -4.97 | 4.72E-09 | 1.01E-08 |
| chr3 | 155290774 | 155291878 | 1105 | -6.61 | 4.73E-09 | 1.01E-08 |
| chr7 | 36100492 | 36111925 | 11434 | -6.72 | 4.74E-09 | 1.01E-08 |
| chr14 | 77446762 | 77458338 | 11577 | -7.21 | 4.75E-09 | 1.01E-08 |
| chr1 | 154366541 | 154368961 | 2421 | -6.72 | 4.75E-09 | 1.02E-08 |
| chr3 | 122441995 | 122443486 | 1492 | -6.71 | 4.77E-09 | 1.02E-08 |
| chr3 | 7526657 | 7529361 | 2705 | -6.62 | 4.79E-09 | 1.02E-08 |
| chr4 | 14998307 | 15000308 | 2002 | -6.96 | 4.80E-09 | 1.03E-08 |
| chr1 | 185032746 | 185034279 | 1534 | -6.64 | 4.83E-09 | 1.03E-08 |
| chr10 | 77274387 | 77275516 | 1130 | -6.64 | 4.83E-09 | 1.03E-08 |
| chr10 | 134474318 | 134475584 | 1267 | -6.64 | 4.83E-09 | 1.03E-08 |
| chr11 | 28250598 | 28251914 | 1317 | -6.62 | 4.83E-09 | 1.03E-08 |
| chr15 | 50746541 | 50747725 | 1185 | -6.62 | 4.83E-09 | 1.03E-08 |
| chr2 | 225016494 | 225018441 | 1948 | -6.63 | 4.83E-09 | 1.03E-08 |
| chr12 | 92493413 | 92498521 | 5109 | -7.08 | 4.84E-09 | 1.03E-08 |
| chr8 | 117807687 | 117812398 | 4712 | -2.21 | 4.84E-09 | 1.03E-08 |
| chr3 | 156364453 | 156365565 | 1113 | -6.62 | 4.85E-09 | 1.03E-08 |
| chr11 | 78941666 | 78945222 | 3557 | 6.79 | 4.85E-09 | 1.04E-08 |
| chr1 | 212914360 | 212915831 | 1472 | -6.84 | 4.86E-09 | 1.04E-08 |
| chr1 | 110816594 | 110817803 | 1210 | -4.42 | 4.87E-09 | 1.04E-08 |
| chr6 | 52472980 | 52474806 | 1827 | -5.94 | 4.87E-09 | 1.04E-08 |
| chr1 | 204688081 | 204689604 | 1524 | -6.67 | 4.87E-09 | 1.04E-08 |
| chrX | 96643804 | 96645355 | 1552 | -6.67 | 4.87E-09 | 1.04E-08 |
| chr2 | 167204131 | 167207104 | 2974 | 6.91 | 4.87E-09 | 1.04E-08 |
| chr10 | 90359881 | 90361466 | 1586 | -6.61 | 4.89E-09 | 1.04E-08 |

| | | | | | | |
|---|---|---|---|---|---|---|
| chr14 | 39721413 | 39743195 | 21783 | -7.23 | 4.89E-09 | 1.04E-08 |
| chr1 | 47054510 | 47056399 | 1890 | -6.64 | 4.90E-09 | 1.05E-08 |
| chr15 | 78071550 | 78073765 | 2216 | -6.92 | 4.90E-09 | 1.05E-08 |
| chr4 | 47529267 | 47537618 | 8352 | -6.63 | 4.90E-09 | 1.05E-08 |
| chr9 | 29658838 | 29673969 | 15132 | -6.68 | 4.91E-09 | 1.05E-08 |
| chr16 | 73191863 | 73193544 | 1682 | -6.85 | 4.91E-09 | 1.05E-08 |
| chr2 | 62479923 | 62481701 | 1779 | -6.62 | 4.92E-09 | 1.05E-08 |
| chr1 | 190166537 | 190171812 | 5276 | -6.61 | 4.93E-09 | 1.05E-08 |
| chr15 | 64379733 | 64380629 | 897 | -6.73 | 4.94E-09 | 1.05E-08 |
| chr6 | 34397499 | 34399777 | 2279 | -6.65 | 4.94E-09 | 1.05E-08 |
| chr4 | 66935637 | 66944355 | 8719 | -6.66 | 4.94E-09 | 1.05E-08 |
| chr1 | 197977882 | 197980323 | 2442 | -7.04 | 4.95E-09 | 1.06E-08 |
| chr18 | 22059246 | 22074910 | 15665 | -7.3 | 4.95E-09 | 1.06E-08 |
| chr9 | 95366359 | 95368455 | 2097 | -6.61 | 4.97E-09 | 1.06E-08 |
| chr10 | 90531393 | 90533644 | 2252 | -5.46 | 4.97E-09 | 1.06E-08 |
| chr10 | 35879224 | 35883142 | 3919 | -6.07 | 4.97E-09 | 1.06E-08 |
| chr1 | 87679611 | 87683004 | 3394 | -3.44 | 4.98E-09 | 1.06E-08 |
| chr20 | 37032721 | 37041384 | 8664 | -6.91 | 4.99E-09 | 1.06E-08 |
| chr14 | 30439275 | 30460484 | 21210 | 1.92 | 4.99E-09 | 1.06E-08 |
| chr14 | 97190526 | 97191927 | 1402 | -6.67 | 4.99E-09 | 1.06E-08 |
| chr20 | 50153993 | 50156070 | 2078 | -6.75 | 4.99E-09 | 1.06E-08 |
| chr3 | 54945611 | 54947381 | 1771 | -6.75 | 4.99E-09 | 1.06E-08 |
| chrX | 70494683 | 70497505 | 2823 | -6.74 | 4.99E-09 | 1.06E-08 |
| chr11 | 120212918 | 120214498 | 1581 | -6.77 | 5.00E-09 | 1.07E-08 |
| chr10 | 34980309 | 34981599 | 1291 | -6.7 | 5.02E-09 | 1.07E-08 |
| chr7 | 110536849 | 110538131 | 1283 | -6.79 | 5.03E-09 | 1.07E-08 |
| chr11 | 17026931 | 17028399 | 1469 | -6.82 | 5.03E-09 | 1.07E-08 |
| chr7 | 38928401 | 38930575 | 2175 | -6.82 | 5.03E-09 | 1.07E-08 |
| chr1 | 10514490 | 10515722 | 1233 | -6.61 | 5.03E-09 | 1.07E-08 |
| chr13 | 98730272 | 98743595 | 13324 | -6.61 | 5.03E-09 | 1.07E-08 |
| chr10 | 31212808 | 31214014 | 1207 | -6.6 | 5.04E-09 | 1.07E-08 |
| chr4 | 98367473 | 98371837 | 4365 | -6.62 | 5.05E-09 | 1.08E-08 |
| chr8 | 19363794 | 19370254 | 6461 | -7.16 | 5.05E-09 | 1.08E-08 |
| chr11 | 100309261 | 100395831 | 86571 | 2.75 | 5.05E-09 | 1.08E-08 |
| chr7 | 31525012 | 31564779 | 39768 | -1.98 | 5.06E-09 | 1.08E-08 |
| chr6 | 51907948 | 51909428 | 1481 | 5.75 | 5.06E-09 | 1.08E-08 |
| chr2 | 26691453 | 26692974 | 1522 | -6.8 | 5.07E-09 | 1.08E-08 |
| chr13 | 95679770 | 95681048 | 1279 | -6.83 | 5.07E-09 | 1.08E-08 |
| chr4 | 140519810 | 140521175 | 1366 | -6.7 | 5.08E-09 | 1.08E-08 |
| chr3 | 184692069 | 184693632 | 1564 | -6.66 | 5.08E-09 | 1.08E-08 |
| chr8 | 64213537 | 64216130 | 2594 | -6.66 | 5.08E-09 | 1.08E-08 |
| chr5 | 71414013 | 71415509 | 1497 | -7.06 | 5.08E-09 | 1.08E-08 |
| chr12 | 127564224 | 127566767 | 2544 | -6.89 | 5.09E-09 | 1.08E-08 |
| chr2 | 42481070 | 42482213 | 1144 | -6.61 | 5.10E-09 | 1.08E-08 |
| chr15 | 67372398 | 67374469 | 2072 | -6.84 | 5.10E-09 | 1.09E-08 |
| chr11 | 73971694 | 73973304 | 1611 | -6.63 | 5.11E-09 | 1.09E-08 |
| chr2 | 64350670 | 64351498 | 829 | -6.69 | 5.12E-09 | 1.09E-08 |
| chr3 | 67640744 | 67642970 | 2227 | -6.61 | 5.12E-09 | 1.09E-08 |
| chr8 | 29098561 | 29106184 | 7624 | -7.02 | 5.13E-09 | 1.09E-08 |
| chr1 | 198783507 | 198786235 | 2729 | -4.94 | 5.15E-09 | 1.10E-08 |
| chr7 | 1316259 | 1318333 | 2075 | -6.62 | 5.15E-09 | 1.10E-08 |
| chr6 | 37741157 | 37742500 | 1344 | -6.61 | 5.16E-09 | 1.10E-08 |
| chr4 | 133097175 | 133103165 | 5991 | -6.85 | 5.17E-09 | 1.10E-08 |
| chr13 | 50706420 | 50711057 | 4638 | -6.91 | 5.17E-09 | 1.10E-08 |
| chr8 | 26888093 | 26890745 | 2653 | -6.64 | 5.17E-09 | 1.10E-08 |
| chr2 | 128287030 | 128288958 | 1929 | -6.65 | 5.18E-09 | 1.10E-08 |
| chr4 | 48059908 | 48062184 | 2277 | -6.63 | 5.22E-09 | 1.11E-08 |
| chr7 | 132370666 | 132371629 | 964 | -5.24 | 5.22E-09 | 1.11E-08 |
| chr13 | 43435588 | 43437147 | 1560 | -6.85 | 5.23E-09 | 1.11E-08 |
| chr3 | 53480624 | 53482472 | 1849 | -6.85 | 5.23E-09 | 1.11E-08 |
| chr5 | 16480117 | 16481207 | 1091 | -6.61 | 5.24E-09 | 1.11E-08 |

| | | | | | | |
|---|---|---|---|---|---|---|
| chrX | 12641729 | 12643629 | 1901 | -6.61 | 5.24E-09 | 1.11E-08 |
| chr13 | 101859723 | 101865392 | 5670 | -6.87 | 5.24E-09 | 1.11E-08 |
| chr7 | 135587732 | 135592845 | 5114 | 6.73 | 5.25E-09 | 1.12E-08 |
| chrX | 108367858 | 108384000 | 16143 | 3.29 | 5.25E-09 | 1.12E-08 |
| chr2 | 174515783 | 174517070 | 1288 | -6.6 | 5.27E-09 | 1.12E-08 |
| chr4 | 150724169 | 150725869 | 1701 | -6.6 | 5.27E-09 | 1.12E-08 |
| chr1 | 56016379 | 56034913 | 18535 | -6.93 | 5.27E-09 | 1.12E-08 |
| chr19 | 1739927 | 1740967 | 1041 | -6.74 | 5.29E-09 | 1.13E-08 |
| chr13 | 82745453 | 82747926 | 2474 | -6.75 | 5.30E-09 | 1.13E-08 |
| chr5 | 130817487 | 130824424 | 6938 | -6.73 | 5.31E-09 | 1.13E-08 |
| chr21 | 42491542 | 42501827 | 10286 | -6.94 | 5.33E-09 | 1.13E-08 |
| chr11 | 76973049 | 76974580 | 1532 | -6.61 | 5.33E-09 | 1.13E-08 |
| chr12 | 13320563 | 13322872 | 2310 | -6.61 | 5.33E-09 | 1.13E-08 |
| chr2 | 189508268 | 189509628 | 1361 | -6.61 | 5.34E-09 | 1.13E-08 |
| chr1 | 167444173 | 167445492 | 1320 | -6.59 | 5.37E-09 | 1.14E-08 |
| chr7 | 81995587 | 81996849 | 1263 | -6.62 | 5.38E-09 | 1.14E-08 |
| chr4 | 188581997 | 188583419 | 1423 | -5.77 | 5.38E-09 | 1.14E-08 |
| chr19 | 48376960 | 48379657 | 2698 | -6.59 | 5.39E-09 | 1.14E-08 |
| chr2 | 226068759 | 226070231 | 1473 | -6.59 | 5.39E-09 | 1.14E-08 |
| chr4 | 124805212 | 124807007 | 1796 | -6.59 | 5.39E-09 | 1.14E-08 |
| chr3 | 13311042 | 13312213 | 1172 | -6.69 | 5.39E-09 | 1.14E-08 |
| chr2 | 39704745 | 39706125 | 1381 | -6.59 | 5.39E-09 | 1.15E-08 |
| chrX | 122849725 | 122851019 | 1295 | -6.59 | 5.39E-09 | 1.15E-08 |
| chr18 | 28612867 | 28614148 | 1282 | -6.59 | 5.40E-09 | 1.15E-08 |
| chr10 | 124965004 | 125062770 | 97767 | -1.83 | 5.40E-09 | 1.15E-08 |
| chr8 | 103776706 | 103779103 | 2398 | -6.89 | 5.42E-09 | 1.15E-08 |
| chr1 | 92309076 | 92310438 | 1363 | -6.81 | 5.43E-09 | 1.15E-08 |
| chr12 | 123732190 | 123734347 | 2158 | -6.64 | 5.45E-09 | 1.16E-08 |
| chr12 | 106013623 | 106015414 | 1792 | -4.93 | 5.48E-09 | 1.16E-08 |
| chr7 | 138534372 | 138543211 | 8840 | -6.63 | 5.49E-09 | 1.17E-08 |
| chr15 | 34319683 | 34321165 | 1483 | -6.59 | 5.50E-09 | 1.17E-08 |
| chr2 | 161752182 | 161753574 | 1393 | -6.83 | 5.50E-09 | 1.17E-08 |
| chr5 | 120666548 | 120671568 | 5021 | -7.28 | 5.51E-09 | 1.17E-08 |
| chr6 | 142691652 | 142692503 | 852 | -6.59 | 5.52E-09 | 1.17E-08 |
| chr10 | 5560680 | 5562511 | 1832 | -6.62 | 5.53E-09 | 1.17E-08 |
| chr7 | 114806341 | 114807867 | 1527 | -6.62 | 5.53E-09 | 1.17E-08 |
| chr20 | 10941110 | 10955005 | 13896 | -7.51 | 5.53E-09 | 1.17E-08 |
| chr2 | 155307326 | 155310007 | 2682 | -2.63 | 5.54E-09 | 1.18E-08 |
| chr8 | 17985719 | 18065275 | 79557 | -2.99 | 5.54E-09 | 1.18E-08 |
| chr6 | 74133556 | 74135017 | 1462 | -6.61 | 5.55E-09 | 1.18E-08 |
| chr6 | 158455733 | 158457715 | 1983 | -6.61 | 5.55E-09 | 1.18E-08 |
| chr7 | 69285282 | 69294245 | 8964 | -6.84 | 5.55E-09 | 1.18E-08 |
| chr5 | 60387138 | 60388490 | 1353 | -6.85 | 5.57E-09 | 1.18E-08 |
| chr2 | 228195792 | 228196740 | 949 | -6.6 | 5.57E-09 | 1.18E-08 |
| chr6 | 100977353 | 100979028 | 1676 | -6.6 | 5.57E-09 | 1.18E-08 |
| chr8 | 82352734 | 82354949 | 2216 | 6.7 | 5.58E-09 | 1.18E-08 |
| chr3 | 121103284 | 121114091 | 10808 | -2.12 | 5.60E-09 | 1.19E-08 |
| chr9 | 2245470 | 2247504 | 2035 | -6.58 | 5.60E-09 | 1.19E-08 |
| chr16 | 31245431 | 31247069 | 1639 | -6.74 | 5.62E-09 | 1.19E-08 |
| chr2 | 121867973 | 121871303 | 3331 | -5.25 | 5.62E-09 | 1.19E-08 |
| chr3 | 106932753 | 106945913 | 13161 | -6.59 | 5.63E-09 | 1.19E-08 |
| chr1 | 210398177 | 210404793 | 6617 | -5.62 | 5.63E-09 | 1.19E-08 |
| chr3 | 73146779 | 73147898 | 1120 | -6.59 | 5.63E-09 | 1.19E-08 |
| chr3 | 33075125 | 33078535 | 3411 | -6.86 | 5.63E-09 | 1.19E-08 |
| chr17 | 45829748 | 45853915 | 24168 | -4.46 | 5.63E-09 | 1.19E-08 |
| chr6 | 146645079 | 146647518 | 2440 | -6.59 | 5.63E-09 | 1.19E-08 |
| chr6 | 5918542 | 5963437 | 44896 | -1.43 | 5.64E-09 | 1.20E-08 |
| chr1 | 63858170 | 63860673 | 2504 | -6.86 | 5.64E-09 | 1.20E-08 |
| chr3 | 124061077 | 124062174 | 1098 | -5.22 | 5.65E-09 | 1.20E-08 |
| chr1 | 192501969 | 192507844 | 5876 | -6.93 | 5.65E-09 | 1.20E-08 |
| chr2 | 31171072 | 31172127 | 1056 | -5.77 | 5.66E-09 | 1.20E-08 |

| | | | | | | |
|---|---|---|---|---|---|---|
| chr12 | 26535454 | 26536881 | 1428 | -6.58 | 5.67E-09 | 1.20E-08 |
| chr19 | 6872159 | 7015903 | 143745 | -4.39 | 5.69E-09 | 1.21E-08 |
| chr22 | 43769338 | 43778300 | 8963 | -6.78 | 5.69E-09 | 1.21E-08 |
| chr17 | 2552234 | 2554257 | 2024 | -6.59 | 5.69E-09 | 1.21E-08 |
| chr5 | 60669122 | 60671055 | 1934 | -6.59 | 5.69E-09 | 1.21E-08 |
| chr5 | 91258651 | 91261247 | 2597 | -6.64 | 5.70E-09 | 1.21E-08 |
| chr8 | 100349523 | 100351777 | 2255 | -6.79 | 5.70E-09 | 1.21E-08 |
| chr8 | 68345222 | 68347090 | 1869 | -6.62 | 5.71E-09 | 1.21E-08 |
| chr1 | 17690652 | 17691601 | 950 | -6.6 | 5.71E-09 | 1.21E-08 |
| chr18 | 68342345 | 68364180 | 21836 | 2.86 | 5.72E-09 | 1.21E-08 |
| chr2 | 193807954 | 193821129 | 13176 | -6.61 | 5.72E-09 | 1.21E-08 |
| chr5 | 118768818 | 118771076 | 2259 | -6.68 | 5.72E-09 | 1.21E-08 |
| chr16 | 54632376 | 54920934 | 288559 | -1.85 | 5.72E-09 | 1.21E-08 |
| chr7 | 50703712 | 50705024 | 1313 | -6.58 | 5.73E-09 | 1.21E-08 |
| chr22 | 23569960 | 23571872 | 1913 | -6.76 | 5.74E-09 | 1.22E-08 |
| chr14 | 74592203 | 74593345 | 1143 | -6.58 | 5.74E-09 | 1.22E-08 |
| chr18 | 13632058 | 13634459 | 2402 | -6.58 | 5.74E-09 | 1.22E-08 |
| chr1 | 207829294 | 207841915 | 12622 | -2.6 | 5.74E-09 | 1.22E-08 |
| chr12 | 78383967 | 78385184 | 1218 | -6.7 | 5.74E-09 | 1.22E-08 |
| chr5 | 56157443 | 56158544 | 1102 | -6.67 | 5.74E-09 | 1.22E-08 |
| chr7 | 118808134 | 118817041 | 8908 | -6.88 | 5.76E-09 | 1.22E-08 |
| chr3 | 10008829 | 10010049 | 1221 | -6.77 | 5.76E-09 | 1.22E-08 |
| chr15 | 70456070 | 70457155 | 1086 | -6.58 | 5.76E-09 | 1.22E-08 |
| chr3 | 98086028 | 98191721 | 105694 | 7.17 | 5.77E-09 | 1.22E-08 |
| chr7 | 14040858 | 14044481 | 3624 | -6.67 | 5.77E-09 | 1.22E-08 |
| chr4 | 10178204 | 10211177 | 32974 | 2.42 | 5.77E-09 | 1.22E-08 |
| chr7 | 91253348 | 91395697 | 142350 | 1.85 | 5.78E-09 | 1.22E-08 |
| chr5 | 81777661 | 81933774 | 156114 | -2.59 | 5.79E-09 | 1.22E-08 |
| chr2 | 219095795 | 219097431 | 1637 | -6.61 | 5.79E-09 | 1.23E-08 |
| chr1 | 201670541 | 201671479 | 939 | -6.59 | 5.79E-09 | 1.23E-08 |
| chr7 | 30929657 | 30931369 | 1713 | -6.59 | 5.79E-09 | 1.23E-08 |
| chr1 | 193031438 | 193032474 | 1037 | -6.78 | 5.80E-09 | 1.23E-08 |
| chr1 | 92105064 | 92107735 | 2672 | -6.89 | 5.81E-09 | 1.23E-08 |
| chr12 | 22827824 | 22829122 | 1299 | -7.04 | 5.82E-09 | 1.23E-08 |
| chr19 | 7738143 | 7739518 | 1376 | -6.58 | 5.82E-09 | 1.23E-08 |
| chr5 | 80039109 | 80040550 | 1442 | -6.58 | 5.82E-09 | 1.23E-08 |
| chr9 | 97993078 | 97994698 | 1621 | -6.79 | 5.82E-09 | 1.23E-08 |
| chr16 | 63594193 | 63596458 | 2266 | -6.66 | 5.84E-09 | 1.24E-08 |
| chr4 | 81078814 | 81085509 | 6696 | -6.9 | 5.87E-09 | 1.24E-08 |
| chr1 | 207785992 | 207787295 | 1304 | -5.76 | 5.87E-09 | 1.24E-08 |
| chr11 | 77999170 | 78000483 | 1314 | -6.61 | 5.88E-09 | 1.24E-08 |
| chr1 | 117800855 | 117802939 | 2085 | -6.58 | 5.88E-09 | 1.24E-08 |
| chr11 | 32868868 | 32869838 | 971 | -6.65 | 5.88E-09 | 1.24E-08 |
| chr15 | 101904878 | 101906068 | 1191 | -6.58 | 5.90E-09 | 1.25E-08 |
| chr15 | 39950717 | 39951795 | 1079 | -6.62 | 5.92E-09 | 1.25E-08 |
| chr3 | 192806392 | 192809955 | 3564 | 5.49 | 5.93E-09 | 1.25E-08 |
| chr6 | 14448101 | 14473877 | 25777 | -7.53 | 5.93E-09 | 1.25E-08 |
| chr4 | 5132768 | 5134100 | 1333 | -6.59 | 5.93E-09 | 1.25E-08 |
| chr7 | 158638091 | 158639867 | 1777 | -6.59 | 5.93E-09 | 1.25E-08 |
| chr3 | 107327792 | 107331252 | 3461 | -6.84 | 5.94E-09 | 1.25E-08 |
| chr13 | 74451370 | 74453129 | 1760 | -6.62 | 5.95E-09 | 1.26E-08 |
| chr3 | 53496829 | 53508367 | 11539 | -6.73 | 5.97E-09 | 1.26E-08 |
| chr5 | 168500279 | 168501784 | 1506 | -6.6 | 5.98E-09 | 1.26E-08 |
| chr6 | 80752459 | 80753912 | 1454 | -6.6 | 5.98E-09 | 1.26E-08 |
| chr9 | 125114826 | 125116102 | 1277 | -6.6 | 5.98E-09 | 1.26E-08 |
| chrX | 54264312 | 54265930 | 1619 | -6.6 | 5.98E-09 | 1.26E-08 |
| chr13 | 48421896 | 48425273 | 3378 | -3.09 | 5.99E-09 | 1.27E-08 |
| chr12 | 56485810 | 56487521 | 1712 | -6.98 | 6.00E-09 | 1.27E-08 |
| chr4 | 151797147 | 151802369 | 5223 | -6.64 | 6.00E-09 | 1.27E-08 |
| chr4 | 130710733 | 130712410 | 1678 | -6.61 | 6.01E-09 | 1.27E-08 |
| chr2 | 236476564 | 236477771 | 1208 | -6.58 | 6.01E-09 | 1.27E-08 |

| | | | | | | |
|---|---|---|---|---|---|---|
| chr2 | 100876015 | 100888600 | 12586 | -6.86 | 6.02E-09 | 1.27E-08 |
| chr6 | 83424328 | 83435459 | 11132 | -6.86 | 6.02E-09 | 1.27E-08 |
| chr3 | 187300526 | 187323083 | 22558 | 1.4 | 6.03E-09 | 1.27E-08 |
| chr4 | 177367169 | 177378552 | 11384 | -6.59 | 6.04E-09 | 1.28E-08 |
| chrX | 40338374 | 40340591 | 2218 | -6.67 | 6.05E-09 | 1.28E-08 |
| chr10 | 76595159 | 76596205 | 1047 | -6.57 | 6.06E-09 | 1.28E-08 |
| chr16 | 81790957 | 81792859 | 1903 | -6.99 | 6.07E-09 | 1.28E-08 |
| chr3 | 44894636 | 44895462 | 827 | -6.93 | 6.07E-09 | 1.28E-08 |
| chr13 | 112602515 | 112607436 | 4922 | -2.17 | 6.07E-09 | 1.28E-08 |
| chr10 | 98828701 | 98830679 | 1979 | -7.09 | 6.07E-09 | 1.28E-08 |
| chr20 | 3786275 | 3791230 | 4956 | -6.64 | 6.08E-09 | 1.28E-08 |
| chr2 | 177298887 | 177302329 | 3443 | -6.58 | 6.09E-09 | 1.29E-08 |
| chr1 | 115364368 | 115369176 | 4809 | -2.7 | 6.09E-09 | 1.29E-08 |
| chr9 | 127341690 | 127342769 | 1080 | -6.66 | 6.13E-09 | 1.29E-08 |
| chr7 | 137513162 | 137520295 | 7134 | -1.51 | 6.14E-09 | 1.30E-08 |
| chr4 | 137996574 | 137998394 | 1821 | -6.6 | 6.14E-09 | 1.30E-08 |
| chr1 | 97057910 | 97059445 | 1536 | -6.57 | 6.16E-09 | 1.30E-08 |
| chr6 | 9945834 | 9948511 | 2678 | -6.6 | 6.16E-09 | 1.30E-08 |
| chr11 | 110310048 | 110311886 | 1839 | -6.7 | 6.17E-09 | 1.30E-08 |
| chr16 | 11614150 | 11615978 | 1829 | -6.7 | 6.17E-09 | 1.30E-08 |
| chr7 | 75059725 | 75061059 | 1335 | -6.59 | 6.19E-09 | 1.31E-08 |
| chr9 | 110162540 | 110164778 | 2239 | -6.59 | 6.19E-09 | 1.31E-08 |
| chr1 | 56706838 | 56708650 | 1813 | -6.58 | 6.20E-09 | 1.31E-08 |
| chr16 | 10676815 | 10678089 | 1275 | -6.65 | 6.21E-09 | 1.31E-08 |
| chr18 | 71143443 | 71145498 | 2056 | -5.23 | 6.21E-09 | 1.31E-08 |
| chr1 | 51222427 | 51224164 | 1738 | -6.59 | 6.21E-09 | 1.31E-08 |
| chr9 | 3918122 | 3919929 | 1808 | -6.76 | 6.22E-09 | 1.31E-08 |
| chr13 | 90352231 | 90354862 | 2632 | -6.57 | 6.22E-09 | 1.31E-08 |
| chr2 | 65430570 | 65431638 | 1069 | -6.57 | 6.22E-09 | 1.31E-08 |
| chr5 | 127785511 | 127787028 | 1518 | -6.57 | 6.22E-09 | 1.31E-08 |
| chr10 | 118578786 | 118581558 | 2773 | -6.67 | 6.22E-09 | 1.31E-08 |
| chr21 | 18065514 | 18069023 | 3510 | -5.07 | 6.24E-09 | 1.32E-08 |
| chr11 | 73234101 | 73236006 | 1906 | -6.82 | 6.25E-09 | 1.32E-08 |
| chr4 | 103142960 | 103150013 | 7054 | -6.9 | 6.26E-09 | 1.32E-08 |
| chr9 | 90114289 | 90115492 | 1204 | 6.63 | 6.28E-09 | 1.33E-08 |
| chr3 | 113295550 | 113296842 | 1293 | -6.57 | 6.29E-09 | 1.33E-08 |
| chr18 | 8996696 | 9004895 | 8200 | -6.83 | 6.30E-09 | 1.33E-08 |
| chr3 | 107504039 | 107506131 | 2093 | -6.79 | 6.30E-09 | 1.33E-08 |
| chr1 | 54537084 | 54538066 | 983 | -6.58 | 6.30E-09 | 1.33E-08 |
| chr3 | 98325637 | 98329173 | 3537 | -6.56 | 6.30E-09 | 1.33E-08 |
| chr18 | 23573586 | 23579866 | 6281 | -6.91 | 6.32E-09 | 1.33E-08 |
| chr11 | 26575958 | 26576911 | 954 | 5.32 | 6.33E-09 | 1.33E-08 |
| chr13 | 83759402 | 83762356 | 2955 | -6.56 | 6.33E-09 | 1.33E-08 |
| chr8 | 31704692 | 31716420 | 11729 | -6.72 | 6.33E-09 | 1.33E-08 |
| chr12 | 21676649 | 21678016 | 1368 | -6.7 | 6.33E-09 | 1.34E-08 |
| chr10 | 112422942 | 112424282 | 1341 | -6.63 | 6.35E-09 | 1.34E-08 |
| chr9 | 29871618 | 29877698 | 6081 | -6.63 | 6.35E-09 | 1.34E-08 |
| chr3 | 66080775 | 66082354 | 1580 | -6.69 | 6.36E-09 | 1.34E-08 |
| chr7 | 108157923 | 108159065 | 1143 | -6.6 | 6.38E-09 | 1.34E-08 |
| chrX | 75502816 | 75504973 | 2158 | -6.6 | 6.38E-09 | 1.34E-08 |
| chr20 | 52475725 | 52478746 | 3022 | -6.75 | 6.38E-09 | 1.34E-08 |
| chr3 | 148176561 | 148179294 | 2734 | -5.2 | 6.38E-09 | 1.35E-08 |
| chr1 | 84715446 | 84725857 | 10412 | -6.56 | 6.39E-09 | 1.35E-08 |
| chr18 | 9722922 | 9725125 | 2204 | -6.9 | 6.39E-09 | 1.35E-08 |
| chr17 | 77849361 | 77850677 | 1317 | -6.76 | 6.39E-09 | 1.35E-08 |
| chr20 | 44221499 | 44270255 | 48757 | -1.27 | 6.39E-09 | 1.35E-08 |
| chr6 | 8657290 | 8662355 | 5066 | 2.42 | 6.40E-09 | 1.35E-08 |
| chr5 | 120995174 | 121003747 | 8574 | -6.67 | 6.40E-09 | 1.35E-08 |
| chr12 | 66367415 | 66383320 | 15906 | -6.62 | 6.41E-09 | 1.35E-08 |
| chr15 | 86096568 | 86097435 | 868 | -6.64 | 6.42E-09 | 1.35E-08 |
| chr12 | 107910408 | 107915866 | 5459 | -6.56 | 6.43E-09 | 1.35E-08 |

| | | | | | | |
|---|---|---|---|---|---|---|
| chr2 | 44817046 | 44821338 | 4293 | -6.56 | 6.43E-09 | 1.35E-08 |
| chr7 | 16478941 | 16480684 | 1744 | -6.58 | 6.43E-09 | 1.35E-08 |
| chr11 | 2614184 | 2690922 | 76739 | -9.63 | 6.43E-09 | 1.35E-08 |
| chr3 | 176230905 | 176234175 | 3271 | -6.57 | 6.45E-09 | 1.36E-08 |
| chrX | 133488753 | 133490205 | 1453 | -6.63 | 6.46E-09 | 1.36E-08 |
| chr3 | 101150373 | 101162833 | 12461 | -6.58 | 6.47E-09 | 1.36E-08 |
| chr8 | 37811989 | 37813789 | 1801 | -6.86 | 6.48E-09 | 1.36E-08 |
| chr4 | 151760781 | 151762127 | 1347 | -5.19 | 6.48E-09 | 1.36E-08 |
| chr1 | 8901410 | 8903926 | 2517 | -6.99 | 6.51E-09 | 1.37E-08 |
| chr13 | 114856569 | 114858992 | 2424 | -6.79 | 6.51E-09 | 1.37E-08 |
| chr6 | 108933142 | 108934593 | 1452 | -6.65 | 6.52E-09 | 1.37E-08 |
| chr2 | 231319967 | 231321513 | 1547 | -6.56 | 6.52E-09 | 1.37E-08 |
| chr8 | 31782942 | 31784523 | 1582 | -6.57 | 6.52E-09 | 1.37E-08 |
| chrX | 85243476 | 85245518 | 2043 | -6.57 | 6.52E-09 | 1.37E-08 |
| chr14 | 37522513 | 37524403 | 1891 | -6.56 | 6.52E-09 | 1.37E-08 |
| chr14 | 36444527 | 36445302 | 776 | -6.64 | 6.55E-09 | 1.38E-08 |
| chr15 | 78898444 | 78900086 | 1643 | -6.64 | 6.55E-09 | 1.38E-08 |
| chrX | 84571971 | 84575162 | 3192 | -6.64 | 6.55E-09 | 1.38E-08 |
| chr1 | 176007144 | 176008080 | 937 | -6.56 | 6.56E-09 | 1.38E-08 |
| chr5 | 168753317 | 168757176 | 3860 | -6.8 | 6.58E-09 | 1.38E-08 |
| chr5 | 83924518 | 83993190 | 68673 | 1.18 | 6.58E-09 | 1.38E-08 |
| chr8 | 18922917 | 18924254 | 1338 | -6.59 | 6.59E-09 | 1.39E-08 |
| chr12 | 89483502 | 89488365 | 4864 | -2.05 | 6.60E-09 | 1.39E-08 |
| chr6 | 46715147 | 46717041 | 1895 | -6.7 | 6.64E-09 | 1.40E-08 |
| chr14 | 76809563 | 76811383 | 1821 | -6.57 | 6.64E-09 | 1.40E-08 |
| chr8 | 61751511 | 61752869 | 1359 | -6.63 | 6.64E-09 | 1.40E-08 |
| chr3 | 44909043 | 44910377 | 1335 | -6.72 | 6.64E-09 | 1.40E-08 |
| chr5 | 72582588 | 72583753 | 1166 | -6.73 | 6.65E-09 | 1.40E-08 |
| chr1 | 35268881 | 35270258 | 1378 | -6.59 | 6.65E-09 | 1.40E-08 |
| chr1 | 80607888 | 80611260 | 3373 | -6.67 | 6.68E-09 | 1.40E-08 |
| chr16 | 4188257 | 4199237 | 10981 | -7.17 | 6.70E-09 | 1.41E-08 |
| chr1 | 200948765 | 200949843 | 1079 | -6.71 | 6.72E-09 | 1.41E-08 |
| chr9 | 30103831 | 30117557 | 13727 | -6.71 | 6.72E-09 | 1.41E-08 |
| chr1 | 92124601 | 92127199 | 2599 | -7.02 | 6.73E-09 | 1.41E-08 |
| chr19 | 15729708 | 15745825 | 16118 | -7.02 | 6.73E-09 | 1.41E-08 |
| chr3 | 139418245 | 139421045 | 2801 | 6.63 | 6.73E-09 | 1.41E-08 |
| chr13 | 109743807 | 109745318 | 1512 | -6.61 | 6.74E-09 | 1.42E-08 |
| chr8 | 80968812 | 80983228 | 14417 | -6.96 | 6.75E-09 | 1.42E-08 |
| chr10 | 80777115 | 80778478 | 1364 | -6.62 | 6.75E-09 | 1.42E-08 |
| chr4 | 152265482 | 152267125 | 1644 | -6.73 | 6.75E-09 | 1.42E-08 |
| chr21 | 47231775 | 47233824 | 2050 | -6.55 | 6.76E-09 | 1.42E-08 |
| chr2 | 65089537 | 65090412 | 876 | -6.65 | 6.76E-09 | 1.42E-08 |
| chr7 | 13752000 | 13762355 | 10356 | -7.02 | 6.77E-09 | 1.42E-08 |
| chr10 | 72813842 | 72815385 | 1544 | -6.56 | 6.77E-09 | 1.42E-08 |
| chr6 | 140537820 | 140539829 | 2010 | -6.66 | 6.78E-09 | 1.42E-08 |
| chr14 | 90611468 | 90727973 | 116506 | -2.94 | 6.79E-09 | 1.43E-08 |
| chr7 | 15145713 | 15154303 | 8591 | -5.22 | 6.80E-09 | 1.43E-08 |
| chr3 | 110980649 | 110987891 | 7243 | -2.39 | 6.80E-09 | 1.43E-08 |
| chr15 | 50409747 | 50417491 | 7745 | -6.75 | 6.81E-09 | 1.43E-08 |
| chr11 | 92889085 | 92891351 | 2267 | -6.65 | 6.83E-09 | 1.43E-08 |
| chr8 | 81603634 | 81606263 | 2630 | -5.06 | 6.84E-09 | 1.44E-08 |
| chr5 | 146003234 | 146067185 | 63952 | 3.91 | 6.85E-09 | 1.44E-08 |
| chr2 | 143937067 | 143939637 | 2571 | -2.96 | 6.86E-09 | 1.44E-08 |
| chr1 | 213386589 | 213387597 | 1009 | -4.81 | 6.86E-09 | 1.44E-08 |
| chr3 | 187926591 | 187928066 | 1476 | -6.59 | 6.87E-09 | 1.44E-08 |
| chr4 | 150796167 | 150797602 | 1436 | -6.6 | 6.87E-09 | 1.44E-08 |
| chr3 | 24508681 | 24510623 | 1943 | -6.63 | 6.88E-09 | 1.44E-08 |
| chr1 | 50988514 | 50989718 | 1205 | -6.55 | 6.88E-09 | 1.44E-08 |
| chr11 | 82761098 | 82762711 | 1614 | -6.55 | 6.88E-09 | 1.44E-08 |
| chr12 | 91992338 | 91994166 | 1829 | -6.55 | 6.88E-09 | 1.44E-08 |
| chr3 | 153251895 | 153253041 | 1147 | -4.51 | 6.88E-09 | 1.44E-08 |

| | | | | | | |
|---|---|---|---|---|---|---|
| chr9 | 86375923 | 86377702 | 1780 | -6.61 | 6.88E-09 | 1.44E-08 |
| chr8 | 22969770 | 22971493 | 1724 | -6.57 | 6.89E-09 | 1.45E-08 |
| chr10 | 83171907 | 83172876 | 970 | 5.04 | 6.89E-09 | 1.45E-08 |
| chr2 | 7639501 | 7649945 | 10445 | 2.16 | 6.90E-09 | 1.45E-08 |
| chr5 | 115096188 | 115100891 | 4704 | -6.98 | 6.90E-09 | 1.45E-08 |
| chr6 | 76391606 | 76393084 | 1479 | -6.63 | 6.91E-09 | 1.45E-08 |
| chr13 | 76586689 | 76598992 | 12304 | -6.89 | 6.93E-09 | 1.45E-08 |
| chr6 | 134646243 | 134647604 | 1362 | -6.57 | 6.93E-09 | 1.45E-08 |
| chr8 | 26166070 | 26177680 | 11611 | -7.32 | 6.94E-09 | 1.46E-08 |
| chr9 | 88138268 | 88142555 | 4288 | -2.44 | 6.94E-09 | 1.46E-08 |
| chr11 | 96101125 | 96102323 | 1199 | -6.55 | 6.94E-09 | 1.46E-08 |
| chr17 | 78294509 | 78297129 | 2621 | -5.37 | 6.97E-09 | 1.46E-08 |
| chr11 | 120571674 | 120579596 | 7923 | -4.07 | 6.98E-09 | 1.46E-08 |
| chr7 | 20627714 | 20630049 | 2336 | -6.55 | 7.00E-09 | 1.47E-08 |
| chr14 | 30347686 | 30349444 | 1759 | -5.97 | 7.01E-09 | 1.47E-08 |
| chr13 | 114594365 | 114601358 | 6994 | -1.71 | 7.02E-09 | 1.47E-08 |
| chr2 | 119632615 | 119706852 | 74238 | -1.99 | 7.03E-09 | 1.47E-08 |
| chr3 | 150046755 | 150056575 | 9821 | -6.57 | 7.03E-09 | 1.47E-08 |
| chr19 | 17732293 | 17733385 | 1093 | -6.59 | 7.03E-09 | 1.47E-08 |
| chr6 | 36818297 | 36819462 | 1166 | -6.59 | 7.03E-09 | 1.47E-08 |
| chr6 | 147307094 | 147309163 | 2070 | -6.55 | 7.04E-09 | 1.48E-08 |
| chr4 | 2473024 | 2474143 | 1120 | -6.55 | 7.05E-09 | 1.48E-08 |
| chr8 | 92163698 | 93872307 | 1708610 | 11.57 | 7.08E-09 | 1.48E-08 |
| chr4 | 167080286 | 167106997 | 26712 | -6.99 | 7.08E-09 | 1.48E-08 |
| chr16 | 88131644 | 88132914 | 1271 | -6.55 | 7.10E-09 | 1.49E-08 |
| chr13 | 106104840 | 106108662 | 3823 | -6.83 | 7.11E-09 | 1.49E-08 |
| chr20 | 46124270 | 46126598 | 2329 | -6.56 | 7.11E-09 | 1.49E-08 |
| chr2 | 206678343 | 206704446 | 26104 | -7.88 | 7.14E-09 | 1.50E-08 |
| chrX | 122219033 | 122235322 | 16290 | -5.06 | 7.14E-09 | 1.50E-08 |
| chr6 | 54453934 | 54456125 | 2192 | -6.56 | 7.16E-09 | 1.50E-08 |
| chr17 | 6369864 | 6371314 | 1451 | -6.72 | 7.16E-09 | 1.50E-08 |
| chr3 | 10258637 | 10259726 | 1090 | -6.92 | 7.17E-09 | 1.50E-08 |
| chr2 | 148539819 | 148542334 | 2516 | 6.61 | 7.17E-09 | 1.50E-08 |
| chr5 | 154462282 | 154464157 | 1876 | -6.7 | 7.17E-09 | 1.50E-08 |
| chr12 | 88786999 | 88796409 | 9411 | -7.34 | 7.18E-09 | 1.50E-08 |
| chr9 | 100295731 | 100297092 | 1362 | -6.68 | 7.18E-09 | 1.50E-08 |
| chr7 | 51199856 | 51201138 | 1283 | -6.54 | 7.18E-09 | 1.50E-08 |
| chr7 | 97945111 | 97946672 | 1562 | -6.54 | 7.18E-09 | 1.50E-08 |
| chr3 | 150781478 | 150782940 | 1463 | -6.61 | 7.18E-09 | 1.50E-08 |
| chr2 | 122502946 | 122506665 | 3720 | -6.6 | 7.19E-09 | 1.51E-08 |
| chr13 | 111964244 | 111965939 | 1696 | -6.66 | 7.20E-09 | 1.51E-08 |
| chr20 | 948606 | 949894 | 1289 | -6.57 | 7.20E-09 | 1.51E-08 |
| chrX | 16608615 | 16609982 | 1368 | -6.54 | 7.21E-09 | 1.51E-08 |
| chr4 | 177473046 | 177474959 | 1914 | -6.96 | 7.24E-09 | 1.51E-08 |
| chr4 | 48247247 | 48249667 | 2421 | -6.65 | 7.25E-09 | 1.52E-08 |
| chr4 | 162351552 | 162355440 | 3889 | -6.57 | 7.25E-09 | 1.52E-08 |
| chr2 | 205420303 | 205423180 | 2878 | -6.75 | 7.25E-09 | 1.52E-08 |
| chr11 | 129176330 | 129178030 | 1701 | -5.69 | 7.25E-09 | 1.52E-08 |
| chr2 | 10763695 | 10764931 | 1237 | -6.64 | 7.26E-09 | 1.52E-08 |
| chrX | 16274415 | 16276184 | 1770 | -6.75 | 7.26E-09 | 1.52E-08 |
| chr1 | 225693801 | 225695206 | 1406 | -6.65 | 7.27E-09 | 1.52E-08 |
| chr20 | 24690450 | 24696577 | 6128 | -1.4 | 7.29E-09 | 1.53E-08 |
| chr18 | 20641743 | 20644526 | 2784 | -6.54 | 7.29E-09 | 1.53E-08 |
| chr18 | 70230581 | 70232472 | 1892 | -5.03 | 7.30E-09 | 1.53E-08 |
| chr5 | 59532990 | 59555759 | 22770 | -2.49 | 7.30E-09 | 1.53E-08 |
| chr10 | 3551344 | 3555361 | 4018 | -6.58 | 7.31E-09 | 1.53E-08 |
| chr6 | 139554626 | 139556500 | 1875 | -6.58 | 7.31E-09 | 1.53E-08 |
| chr5 | 112883179 | 112885179 | 2001 | -6.76 | 7.31E-09 | 1.53E-08 |
| chr4 | 189273400 | 189275630 | 2231 | -6.64 | 7.31E-09 | 1.53E-08 |
| chr11 | 117142412 | 117144658 | 2247 | -6.94 | 7.32E-09 | 1.53E-08 |
| chr12 | 27987173 | 28011343 | 24171 | -7.67 | 7.32E-09 | 1.53E-08 |

| | | | | | | |
|---|---|---|---|---|---|---|
| chr1 | 192673052 | 192674976 | 1925 | -6.76 | 7.33E-09 | 1.53E-08 |
| chr10 | 72780549 | 72782019 | 1471 | -6.56 | 7.33E-09 | 1.53E-08 |
| chr1 | 109066971 | 109070800 | 3830 | -6.77 | 7.34E-09 | 1.53E-08 |
| chr13 | 65245700 | 65267747 | 22048 | -3.48 | 7.34E-09 | 1.54E-08 |
| chr12 | 105040361 | 105041853 | 1493 | -6.54 | 7.36E-09 | 1.54E-08 |
| chr2 | 55351433 | 55352715 | 1283 | -6.54 | 7.36E-09 | 1.54E-08 |
| chr7 | 26549615 | 26562982 | 13368 | -7.14 | 7.36E-09 | 1.54E-08 |
| chr5 | 112014915 | 112035727 | 20813 | -7.51 | 7.37E-09 | 1.54E-08 |
| chr10 | 22553850 | 22555428 | 1579 | -3.24 | 7.38E-09 | 1.54E-08 |
| chr5 | 96284290 | 96285586 | 1297 | -6.78 | 7.39E-09 | 1.54E-08 |
| chr2 | 9411732 | 9413314 | 1583 | -6.59 | 7.40E-09 | 1.55E-08 |
| chr12 | 41624996 | 41627148 | 2153 | -6.62 | 7.41E-09 | 1.55E-08 |
| chr6 | 112052876 | 112055522 | 2647 | -3.31 | 7.41E-09 | 1.55E-08 |
| chr6 | 137041911 | 137043051 | 1141 | -6.54 | 7.41E-09 | 1.55E-08 |
| chr18 | 9633879 | 9636433 | 2555 | -6.56 | 7.44E-09 | 1.56E-08 |
| chr11 | 122693459 | 122695125 | 1667 | -6.62 | 7.44E-09 | 1.56E-08 |
| chr10 | 91113077 | 91114623 | 1547 | -6.55 | 7.46E-09 | 1.56E-08 |
| chr15 | 52259454 | 52260445 | 992 | -6.55 | 7.46E-09 | 1.56E-08 |
| chr6 | 2547271 | 2549119 | 1849 | -6.56 | 7.47E-09 | 1.56E-08 |
| chrX | 113960860 | 113970553 | 9694 | 2.91 | 7.49E-09 | 1.57E-08 |
| chr21 | 17074188 | 17078143 | 3956 | -7.05 | 7.50E-09 | 1.57E-08 |
| chr3 | 170104485 | 170116693 | 12209 | -7.1 | 7.51E-09 | 1.57E-08 |
| chr20 | 17546801 | 17547736 | 936 | -6.61 | 7.51E-09 | 1.57E-08 |
| chr4 | 113401588 | 113414588 | 13001 | -5.84 | 7.51E-09 | 1.57E-08 |
| chr5 | 87589866 | 87592685 | 2820 | -6.94 | 7.51E-09 | 1.57E-08 |
| chr18 | 12490559 | 12492771 | 2213 | -6.8 | 7.52E-09 | 1.57E-08 |
| chr4 | 133146606 | 133148543 | 1938 | -6.54 | 7.53E-09 | 1.57E-08 |
| chr1 | 63540136 | 63544191 | 4056 | -1.88 | 7.54E-09 | 1.58E-08 |
| chr3 | 53591663 | 53593827 | 2165 | -6.54 | 7.55E-09 | 1.58E-08 |
| chr8 | 18813665 | 18814791 | 1127 | -6.54 | 7.55E-09 | 1.58E-08 |
| chr5 | 60357214 | 60359458 | 2245 | -6.88 | 7.56E-09 | 1.58E-08 |
| chr10 | 112855527 | 112858043 | 2517 | 6.67 | 7.59E-09 | 1.58E-08 |
| chr1 | 225320424 | 225322825 | 2402 | -6.69 | 7.59E-09 | 1.59E-08 |
| chr16 | 48808481 | 49304626 | 496146 | 10.55 | 7.61E-09 | 1.59E-08 |
| chr10 | 106170175 | 106171998 | 1824 | -7.05 | 7.63E-09 | 1.59E-08 |
| chr4 | 48029746 | 48030925 | 1180 | -6.54 | 7.64E-09 | 1.59E-08 |
| chrX | 75729235 | 75762981 | 33747 | 3.25 | 7.66E-09 | 1.60E-08 |
| chr22 | 30241743 | 30243357 | 1615 | -6.73 | 7.66E-09 | 1.60E-08 |
| chr1 | 168674811 | 168677730 | 2920 | -6.64 | 7.67E-09 | 1.60E-08 |
| chr1 | 172831170 | 172832771 | 1602 | -6.56 | 7.67E-09 | 1.60E-08 |
| chr10 | 18427451 | 18488663 | 61213 | 1.19 | 7.68E-09 | 1.60E-08 |
| chr1 | 77983880 | 77985623 | 1744 | -6.57 | 7.68E-09 | 1.60E-08 |
| chr10 | 82465567 | 82466820 | 1254 | -6.54 | 7.71E-09 | 1.61E-08 |
| chr15 | 66846878 | 66848807 | 1930 | -6.83 | 7.71E-09 | 1.61E-08 |
| chr6 | 82949283 | 82950206 | 924 | -6.53 | 7.72E-09 | 1.61E-08 |
| chr6 | 148411880 | 148413520 | 1641 | -6.53 | 7.72E-09 | 1.61E-08 |
| chr4 | 156898905 | 156901959 | 3055 | -6.63 | 7.72E-09 | 1.61E-08 |
| chr10 | 69496441 | 69521787 | 25347 | -1.5 | 7.72E-09 | 1.61E-08 |
| chr1 | 93986075 | 93987514 | 1440 | -6.59 | 7.74E-09 | 1.61E-08 |
| chr6 | 38093747 | 38095206 | 1460 | -6.67 | 7.74E-09 | 1.61E-08 |
| chr4 | 115272645 | 115274449 | 1805 | -6.67 | 7.74E-09 | 1.61E-08 |
| chr2 | 61375427 | 61379252 | 3826 | -6.66 | 7.76E-09 | 1.62E-08 |
| chrX | 114435665 | 114440308 | 4644 | -2.26 | 7.78E-09 | 1.62E-08 |
| chr8 | 117284506 | 117288694 | 4189 | -6.56 | 7.78E-09 | 1.62E-08 |
| chr10 | 95989654 | 95990487 | 834 | -6.56 | 7.78E-09 | 1.62E-08 |
| chr2 | 213439772 | 213740633 | 300862 | 2.23 | 7.79E-09 | 1.62E-08 |
| chr6 | 151294293 | 151295541 | 1249 | -6.53 | 7.80E-09 | 1.63E-08 |
| chr4 | 89956255 | 89958585 | 2331 | -6.55 | 7.81E-09 | 1.63E-08 |
| chr20 | 48452101 | 48459680 | 7580 | -6.64 | 7.81E-09 | 1.63E-08 |
| chr2 | 108232569 | 108264506 | 31938 | -3.58 | 7.85E-09 | 1.64E-08 |
| chr9 | 86224673 | 86226016 | 1344 | -6.74 | 7.85E-09 | 1.64E-08 |

| | | | | | | |
|---|---|---|---|---|---|---|
| chr3 | 5285184 | 5285798 | 615 | -6.58 | 7.87E-09 | 1.64E-08 |
| chr6 | 44994703 | 44996013 | 1311 | -6.58 | 7.87E-09 | 1.64E-08 |
| chr9 | 89027028 | 89028152 | 1125 | -6.58 | 7.87E-09 | 1.64E-08 |
| chr7 | 151363074 | 151365766 | 2693 | -6.61 | 7.88E-09 | 1.64E-08 |
| chr2 | 203576581 | 203579101 | 2521 | -6.53 | 7.88E-09 | 1.64E-08 |
| chr18 | 61221747 | 61223741 | 1995 | 6.65 | 7.89E-09 | 1.64E-08 |
| chr4 | 136631546 | 136633645 | 2100 | -6.53 | 7.91E-09 | 1.65E-08 |
| chr1 | 100560734 | 100562987 | 2254 | -6.86 | 7.93E-09 | 1.65E-08 |
| chr6 | 78124147 | 78125348 | 1202 | -6.6 | 7.93E-09 | 1.65E-08 |
| chr4 | 120518771 | 120534392 | 15622 | -6.98 | 7.94E-09 | 1.65E-08 |
| chr11 | 13795204 | 13796580 | 1377 | -6.62 | 7.95E-09 | 1.66E-08 |
| chr8 | 99728222 | 99729702 | 1481 | -6.53 | 7.96E-09 | 1.66E-08 |
| chr10 | 90907725 | 90909037 | 1313 | -5.02 | 7.97E-09 | 1.66E-08 |
| chr6 | 45548813 | 45551080 | 2268 | -6.77 | 8.01E-09 | 1.67E-08 |
| chr14 | 91666164 | 91677643 | 11480 | -6.87 | 8.02E-09 | 1.67E-08 |
| chr4 | 177978493 | 177980541 | 2049 | -6.59 | 8.05E-09 | 1.68E-08 |
| chr12 | 79023869 | 79025418 | 1550 | -6.67 | 8.07E-09 | 1.68E-08 |
| chr20 | 31075671 | 31077373 | 1703 | -6.67 | 8.07E-09 | 1.68E-08 |
| chr20 | 23405781 | 23413757 | 7977 | -6.68 | 8.08E-09 | 1.68E-08 |
| chr22 | 40947935 | 40951047 | 3113 | -6.81 | 8.08E-09 | 1.68E-08 |
| chr10 | 60526702 | 60528182 | 1481 | -6.78 | 8.08E-09 | 1.68E-08 |
| chr12 | 89334605 | 89338046 | 3442 | -6.78 | 8.08E-09 | 1.68E-08 |
| chr4 | 42556392 | 42557860 | 1469 | -6.53 | 8.09E-09 | 1.68E-08 |
| chr2 | 131612768 | 131614239 | 1472 | -6.66 | 8.09E-09 | 1.68E-08 |
| chr4 | 96158913 | 96167521 | 8609 | -6.88 | 8.11E-09 | 1.69E-08 |
| chr5 | 156249292 | 156256163 | 6872 | -6.59 | 8.12E-09 | 1.69E-08 |
| chr14 | 56248989 | 56250698 | 1710 | -6.71 | 8.13E-09 | 1.69E-08 |
| chr4 | 3404974 | 3406695 | 1722 | -6.71 | 8.13E-09 | 1.69E-08 |
| chr2 | 95583746 | 95588737 | 4992 | -2.32 | 8.13E-09 | 1.69E-08 |
| chr3 | 99695191 | 99697529 | 2339 | -6.59 | 8.15E-09 | 1.70E-08 |
| chr6 | 79703073 | 79704423 | 1351 | -6.59 | 8.15E-09 | 1.70E-08 |
| chr4 | 162390495 | 162398088 | 7594 | -6.55 | 8.17E-09 | 1.70E-08 |
| chr4 | 178198753 | 178201171 | 2419 | -6.63 | 8.18E-09 | 1.70E-08 |
| chr4 | 178318121 | 178320074 | 1954 | -6.52 | 8.18E-09 | 1.70E-08 |
| chr15 | 75001815 | 75003001 | 1187 | -4.9 | 8.19E-09 | 1.70E-08 |
| chr13 | 100434421 | 100436225 | 1805 | -6.62 | 8.21E-09 | 1.71E-08 |
| chr1 | 14943804 | 14950248 | 6445 | -2.6 | 8.22E-09 | 1.71E-08 |
| chrX | 53666120 | 53667394 | 1275 | -6.54 | 8.22E-09 | 1.71E-08 |
| chr1 | 37085980 | 37088302 | 2323 | -5.25 | 8.23E-09 | 1.71E-08 |
| chr6 | 24068992 | 24095743 | 26752 | 1.9 | 8.25E-09 | 1.72E-08 |
| chr7 | 40822653 | 40951277 | 128625 | -1.46 | 8.28E-09 | 1.72E-08 |
| chr13 | 27658670 | 27659988 | 1319 | -6.55 | 8.29E-09 | 1.72E-08 |
| chr5 | 39520558 | 39522165 | 1608 | -6.55 | 8.29E-09 | 1.72E-08 |
| chr1 | 220116632 | 220121228 | 4597 | -6.57 | 8.29E-09 | 1.72E-08 |
| chr8 | 31876017 | 31892108 | 16092 | -7.24 | 8.29E-09 | 1.72E-08 |
| chr22 | 32741477 | 32742545 | 1069 | -6.54 | 8.32E-09 | 1.73E-08 |
| chr2 | 28895497 | 28896780 | 1284 | -4.9 | 8.34E-09 | 1.73E-08 |
| chr12 | 117753420 | 117754985 | 1566 | -6.66 | 8.36E-09 | 1.74E-08 |
| chr7 | 34203008 | 34222916 | 19909 | -7.31 | 8.37E-09 | 1.74E-08 |
| chr7 | 117932616 | 117981790 | 49175 | 2.52 | 8.39E-09 | 1.74E-08 |
| chr12 | 116468390 | 116470305 | 1916 | -6.64 | 8.39E-09 | 1.74E-08 |
| chr3 | 133951723 | 133953067 | 1345 | -6.63 | 8.41E-09 | 1.75E-08 |
| chr19 | 15533987 | 15535032 | 1046 | -6.71 | 8.41E-09 | 1.75E-08 |
| chr3 | 119681279 | 119682873 | 1595 | -6.55 | 8.42E-09 | 1.75E-08 |
| chr3 | 115524151 | 115526418 | 2268 | -6.63 | 8.43E-09 | 1.75E-08 |
| chr6 | 44409492 | 44411249 | 1758 | -6.62 | 8.45E-09 | 1.76E-08 |
| chr13 | 24371990 | 24373459 | 1470 | -6.52 | 8.46E-09 | 1.76E-08 |
| chr7 | 104471031 | 104471902 | 872 | -6.57 | 8.46E-09 | 1.76E-08 |
| chr18 | 55424205 | 55426479 | 2275 | -6.55 | 8.47E-09 | 1.76E-08 |
| chr4 | 130222882 | 130225134 | 2253 | -6.53 | 8.49E-09 | 1.76E-08 |
| chrX | 131018805 | 131020573 | 1769 | -6.52 | 8.49E-09 | 1.76E-08 |

| | | | | | | |
|---|---|---|---|---|---|---|
| chr1 | 218032789 | 218034132 | 1344 | -6.51 | 8.50E-09 | 1.76E-08 |
| chr22 | 25809106 | 25810541 | 1436 | -6.51 | 8.50E-09 | 1.76E-08 |
| chr1 | 196048107 | 196051173 | 3067 | -6.52 | 8.50E-09 | 1.77E-08 |
| chrX | 43820394 | 43822353 | 1960 | -6.52 | 8.50E-09 | 1.77E-08 |
| chr2 | 100731364 | 100750926 | 19563 | -2.37 | 8.52E-09 | 1.77E-08 |
| chr10 | 93076562 | 93092024 | 15463 | -2.28 | 8.56E-09 | 1.78E-08 |
| chr8 | 117079791 | 117082427 | 2637 | -6.99 | 8.57E-09 | 1.78E-08 |
| chr12 | 120868485 | 120869458 | 974 | -6.66 | 8.58E-09 | 1.78E-08 |
| chr12 | 59340730 | 59344713 | 3984 | -6.67 | 8.58E-09 | 1.78E-08 |
| chr12 | 20001222 | 20003367 | 2146 | -6.67 | 8.59E-09 | 1.78E-08 |
| chr14 | 55151890 | 55153025 | 1136 | -6.68 | 8.59E-09 | 1.78E-08 |
| chr19 | 30867355 | 31167873 | 300519 | 1.01 | 8.61E-09 | 1.79E-08 |
| chr1 | 212074226 | 212090680 | 16455 | -6.94 | 8.66E-09 | 1.80E-08 |
| chr14 | 91189366 | 91192175 | 2810 | -6.88 | 8.66E-09 | 1.80E-08 |
| chr4 | 106208135 | 106210080 | 1946 | -6.53 | 8.67E-09 | 1.80E-08 |
| chr3 | 176277759 | 176302191 | 24433 | 2.26 | 8.67E-09 | 1.80E-08 |
| chr20 | 46882304 | 46893526 | 11223 | -2.69 | 8.67E-09 | 1.80E-08 |
| chr10 | 14989148 | 14990113 | 966 | -6.59 | 8.69E-09 | 1.80E-08 |
| chr5 | 76764672 | 76777012 | 12341 | -6.59 | 8.69E-09 | 1.80E-08 |
| chr1 | 4813570 | 4826461 | 12892 | -2.67 | 8.69E-09 | 1.80E-08 |
| chr7 | 110889375 | 110891986 | 2612 | -6.52 | 8.69E-09 | 1.80E-08 |
| chr1 | 227612762 | 227614349 | 1588 | -6.55 | 8.70E-09 | 1.80E-08 |
| chrX | 102635721 | 102637669 | 1949 | -4.88 | 8.71E-09 | 1.81E-08 |
| chr14 | 51126990 | 51129227 | 2238 | -6.54 | 8.71E-09 | 1.81E-08 |
| chr17 | 45170026 | 45171089 | 1064 | -6.81 | 8.71E-09 | 1.81E-08 |
| chr15 | 90839995 | 90841327 | 1333 | -6.57 | 8.73E-09 | 1.81E-08 |
| chr6 | 10842936 | 10844542 | 1607 | -6.57 | 8.73E-09 | 1.81E-08 |
| chr2 | 108557935 | 108605033 | 47099 | -3.21 | 8.73E-09 | 1.81E-08 |
| chr12 | 77224911 | 77226783 | 1873 | -6.95 | 8.74E-09 | 1.81E-08 |
| chr12 | 55201615 | 55274207 | 72593 | 1.3 | 8.75E-09 | 1.81E-08 |
| chr10 | 22736925 | 22739871 | 2947 | -6.51 | 8.76E-09 | 1.82E-08 |
| chr2 | 209095563 | 209097802 | 2240 | -6.51 | 8.76E-09 | 1.82E-08 |
| chr1 | 100296883 | 100306510 | 9628 | -6.74 | 8.78E-09 | 1.82E-08 |
| chr11 | 35353173 | 35359977 | 6805 | -6.95 | 8.80E-09 | 1.83E-08 |
| chr1 | 78250096 | 78254610 | 4515 | -6.51 | 8.82E-09 | 1.83E-08 |
| chr6 | 45175981 | 45177788 | 1808 | -6.51 | 8.82E-09 | 1.83E-08 |
| chr7 | 65532979 | 65533951 | 973 | -6.51 | 8.82E-09 | 1.83E-08 |
| chr7 | 117553393 | 117584342 | 30950 | -3.85 | 8.83E-09 | 1.83E-08 |
| chr9 | 102750418 | 102757032 | 6615 | -6.52 | 8.84E-09 | 1.83E-08 |
| chrX | 66796911 | 66800534 | 3624 | -6.58 | 8.84E-09 | 1.83E-08 |
| chr2 | 224788955 | 224805333 | 16379 | -6.9 | 8.85E-09 | 1.83E-08 |
| chr19 | 48217655 | 48219097 | 1443 | -6.51 | 8.85E-09 | 1.83E-08 |
| chr6 | 27136969 | 27138656 | 1688 | -6.51 | 8.85E-09 | 1.83E-08 |
| chr10 | 14607997 | 14613274 | 5278 | -6.96 | 8.85E-09 | 1.83E-08 |
| chr4 | 175548484 | 175552721 | 4238 | -7.01 | 8.86E-09 | 1.84E-08 |
| chr18 | 74274252 | 74288049 | 13798 | -6.59 | 8.87E-09 | 1.84E-08 |
| chr13 | 64062886 | 64290798 | 227913 | 3.26 | 8.87E-09 | 1.84E-08 |
| chr21 | 41024686 | 41026422 | 1737 | -6.53 | 8.87E-09 | 1.84E-08 |
| chr10 | 97287078 | 97290044 | 2967 | -7.31 | 8.89E-09 | 1.84E-08 |
| chrX | 17355108 | 17371612 | 16505 | -5.98 | 8.89E-09 | 1.84E-08 |
| chr7 | 50737808 | 50740813 | 3006 | -7.13 | 8.89E-09 | 1.84E-08 |
| chr4 | 41790938 | 41845930 | 54993 | -1.41 | 8.90E-09 | 1.84E-08 |
| chr1 | 90082789 | 90084601 | 1813 | -6.76 | 8.90E-09 | 1.84E-08 |
| chr22 | 43074433 | 43076250 | 1818 | -6.59 | 8.92E-09 | 1.85E-08 |
| chr19 | 2714913 | 2716121 | 1209 | -6.52 | 8.94E-09 | 1.85E-08 |
| chr17 | 64540552 | 64578791 | 38240 | -7.63 | 8.94E-09 | 1.85E-08 |
| chr1 | 173463992 | 173466899 | 2908 | -6.54 | 8.97E-09 | 1.86E-08 |
| chr1 | 183304208 | 183305946 | 1739 | -6.54 | 8.97E-09 | 1.86E-08 |
| chr6 | 517595 | 522050 | 4456 | -6.81 | 8.99E-09 | 1.86E-08 |
| chr2 | 36786445 | 36793965 | 7521 | -6.9 | 9.01E-09 | 1.87E-08 |
| chr3 | 142239837 | 142252217 | 12381 | -6.9 | 9.01E-09 | 1.87E-08 |

| | | | | | | |
|---|---|---|---|---|---|---|
| chr6 | 10889901 | 10891599 | 1699 | -6.67 | 9.03E-09 | 1.87E-08 |
| chr10 | 35084915 | 35086243 | 1329 | -6.5 | 9.04E-09 | 1.87E-08 |
| chr12 | 38846432 | 38848402 | 1971 | 6.6 | 9.04E-09 | 1.87E-08 |
| chr16 | 12337077 | 12338510 | 1434 | -6.52 | 9.05E-09 | 1.87E-08 |
| chr2 | 28120233 | 28121792 | 1560 | -6.52 | 9.05E-09 | 1.87E-08 |
| chr5 | 79305702 | 79307736 | 2035 | -2.93 | 9.05E-09 | 1.87E-08 |
| chr2 | 175001720 | 175002869 | 1150 | -6.58 | 9.05E-09 | 1.87E-08 |
| chr2 | 208898667 | 208900217 | 1551 | -6.58 | 9.05E-09 | 1.87E-08 |
| chr9 | 138657162 | 138659248 | 2087 | -6.58 | 9.05E-09 | 1.87E-08 |
| chr10 | 89757280 | 89771655 | 14376 | -6.85 | 9.06E-09 | 1.88E-08 |
| chrX | 77816063 | 77817288 | 1226 | -5.87 | 9.08E-09 | 1.88E-08 |
| chr11 | 121450207 | 121451898 | 1692 | -6.5 | 9.09E-09 | 1.88E-08 |
| chrX | 9099913 | 9101513 | 1601 | -6.5 | 9.09E-09 | 1.88E-08 |
| chr3 | 134287261 | 134290501 | 3241 | -6.51 | 9.11E-09 | 1.88E-08 |
| chr8 | 40171882 | 40174564 | 2683 | -6.51 | 9.11E-09 | 1.88E-08 |
| chr20 | 25543654 | 25544889 | 1236 | -6.52 | 9.11E-09 | 1.89E-08 |
| chr3 | 184515393 | 184516922 | 1530 | -6.53 | 9.12E-09 | 1.89E-08 |
| chr1 | 169220298 | 169228113 | 7816 | -6.66 | 9.13E-09 | 1.89E-08 |
| chr12 | 19373174 | 19374598 | 1425 | -6.66 | 9.13E-09 | 1.89E-08 |
| chr7 | 22885288 | 22888489 | 3202 | -6.66 | 9.13E-09 | 1.89E-08 |
| chr16 | 80342462 | 80348989 | 6528 | -6.63 | 9.17E-09 | 1.90E-08 |
| chr2 | 225483572 | 225486567 | 2996 | -6.6 | 9.18E-09 | 1.90E-08 |
| chr20 | 23607373 | 23609540 | 2168 | -6.62 | 9.18E-09 | 1.90E-08 |
| chr17 | 73462386 | 73463323 | 938 | -6.55 | 9.20E-09 | 1.90E-08 |
| chr10 | 116295805 | 116296885 | 1081 | -6.72 | 9.20E-09 | 1.90E-08 |
| chr6 | 24753542 | 24755516 | 1975 | -6.7 | 9.21E-09 | 1.90E-08 |
| chr16 | 22969620 | 23070165 | 100546 | -1.35 | 9.22E-09 | 1.91E-08 |
| chr1 | 85347288 | 85348603 | 1316 | -6.53 | 9.23E-09 | 1.91E-08 |
| chr5 | 166718091 | 166720695 | 2605 | -6.93 | 9.25E-09 | 1.91E-08 |
| chr1 | 78091746 | 78093268 | 1523 | -6.5 | 9.26E-09 | 1.91E-08 |
| chr3 | 127856748 | 127858066 | 1319 | -6.5 | 9.26E-09 | 1.91E-08 |
| chr1 | 217702393 | 217703759 | 1367 | -6.51 | 9.27E-09 | 1.92E-08 |
| chr21 | 36495840 | 36498181 | 2342 | -6.51 | 9.27E-09 | 1.92E-08 |
| chr14 | 51371528 | 51372546 | 1019 | -6.51 | 9.30E-09 | 1.92E-08 |
| chr18 | 3240611 | 3242966 | 2356 | -6.6 | 9.30E-09 | 1.92E-08 |
| chr19 | 11177683 | 11178760 | 1078 | -6.6 | 9.30E-09 | 1.92E-08 |
| chr5 | 90000021 | 90001928 | 1908 | 4.84 | 9.30E-09 | 1.92E-08 |
| chr4 | 179081274 | 179083398 | 2125 | -5.37 | 9.30E-09 | 1.92E-08 |
| chr3 | 5141650 | 5144195 | 2546 | -7 | 9.30E-09 | 1.92E-08 |
| chr2 | 47648752 | 47650235 | 1484 | -6.74 | 9.31E-09 | 1.92E-08 |
| chr14 | 57937394 | 57938411 | 1018 | -5.18 | 9.33E-09 | 1.93E-08 |
| chr1 | 152229615 | 152230934 | 1320 | -6.75 | 9.34E-09 | 1.93E-08 |
| chrX | 65138268 | 65139714 | 1447 | -6.53 | 9.35E-09 | 1.93E-08 |
| chr10 | 22529903 | 22531526 | 1624 | -6.5 | 9.36E-09 | 1.93E-08 |
| chr13 | 22317067 | 22318600 | 1534 | 5.02 | 9.37E-09 | 1.93E-08 |
| chr2 | 33708795 | 33714389 | 5595 | -6.43 | 9.39E-09 | 1.94E-08 |
| chr2 | 232679138 | 232687364 | 8227 | -6.76 | 9.40E-09 | 1.94E-08 |
| chr7 | 109313256 | 109314921 | 1666 | -6.5 | 9.41E-09 | 1.94E-08 |
| chr13 | 33488807 | 33538390 | 49584 | -2.08 | 9.43E-09 | 1.95E-08 |
| chr3 | 162572778 | 162585607 | 12830 | -7.31 | 9.46E-09 | 1.95E-08 |
| chr14 | 75600230 | 75601025 | 796 | -6.52 | 9.46E-09 | 1.95E-08 |
| chr1 | 177222075 | 177224381 | 2307 | -6.52 | 9.49E-09 | 1.96E-08 |
| chr19 | 16791500 | 16793074 | 1575 | -6.5 | 9.49E-09 | 1.96E-08 |
| chr1 | 29263479 | 29265755 | 2277 | -6.75 | 9.51E-09 | 1.96E-08 |
| chr11 | 62939208 | 62942167 | 2960 | -6.57 | 9.53E-09 | 1.97E-08 |
| chr11 | 8773234 | 8775438 | 2205 | -6.78 | 9.55E-09 | 1.97E-08 |
| chr7 | 88003710 | 89723481 | 1719772 | 5.51 | 9.56E-09 | 1.97E-08 |
| chr6 | 133811145 | 133814195 | 3051 | -6.51 | 9.59E-09 | 1.98E-08 |
| chr8 | 104439453 | 104440678 | 1226 | -6.52 | 9.64E-09 | 1.99E-08 |
| chr10 | 17567822 | 17586846 | 19025 | -6.49 | 9.66E-09 | 1.99E-08 |
| chr1 | 186938785 | 186940554 | 1770 | -6.56 | 9.66E-09 | 1.99E-08 |

| | | | | | | |
|---|---|---|---|---|---|---|
| chr10 | 5939441 | 5940690 | 1250 | -6.65 | 9.72E-09 | 2.01E-08 |
| chr7 | 138372705 | 138375281 | 2577 | -6.65 | 9.72E-09 | 2.01E-08 |
| chr2 | 25022670 | 25023275 | 606 | -6.49 | 9.72E-09 | 2.01E-08 |
| chr3 | 108117861 | 108119937 | 2077 | -6.64 | 9.72E-09 | 2.01E-08 |
| chr4 | 100071409 | 100073284 | 1876 | -6.52 | 9.72E-09 | 2.01E-08 |
| chr4 | 176668989 | 176671479 | 2491 | -6.55 | 9.74E-09 | 2.01E-08 |
| chr1 | 76448915 | 76450398 | 1484 | -6.63 | 9.74E-09 | 2.01E-08 |
| chr4 | 188821226 | 188824489 | 3264 | -6.65 | 9.75E-09 | 2.01E-08 |
| chr11 | 62973814 | 62984176 | 10363 | -6.64 | 9.76E-09 | 2.01E-08 |
| chr5 | 173964859 | 173966559 | 1701 | -6.68 | 9.76E-09 | 2.01E-08 |
| chr1 | 44642064 | 44644313 | 2250 | -6.63 | 9.77E-09 | 2.01E-08 |
| chr13 | 36389178 | 36390696 | 1519 | -6.66 | 9.77E-09 | 2.01E-08 |
| chr6 | 114625945 | 114627264 | 1320 | -6.63 | 9.78E-09 | 2.02E-08 |
| chr18 | 21836487 | 21837987 | 1501 | -6.5 | 9.78E-09 | 2.02E-08 |
| chr6 | 137529487 | 137531621 | 2135 | -6.5 | 9.78E-09 | 2.02E-08 |
| chr10 | 67339659 | 67342343 | 2685 | -2.73 | 9.79E-09 | 2.02E-08 |
| chr2 | 38290739 | 38291371 | 633 | -6.61 | 9.79E-09 | 2.02E-08 |
| chr4 | 151006073 | 151007971 | 1899 | -6.52 | 9.80E-09 | 2.02E-08 |
| chr15 | 77035751 | 77037188 | 1438 | -6.79 | 9.80E-09 | 2.02E-08 |
| chr18 | 13411072 | 13413152 | 2081 | -6.62 | 9.80E-09 | 2.02E-08 |
| chr1 | 87659186 | 87662595 | 3410 | -4.31 | 9.80E-09 | 2.02E-08 |
| chr6 | 79885323 | 79886609 | 1287 | -6.49 | 9.82E-09 | 2.02E-08 |
| chr6 | 147263440 | 147265712 | 2273 | -6.69 | 9.82E-09 | 2.02E-08 |
| chr21 | 16277762 | 16280241 | 2480 | -6.6 | 9.82E-09 | 2.02E-08 |
| chr7 | 36040920 | 36043592 | 2673 | -6.86 | 9.84E-09 | 2.03E-08 |
| chr6 | 127341022 | 127346646 | 5625 | -7.05 | 9.85E-09 | 2.03E-08 |
| chr6 | 39710588 | 39767793 | 57206 | 1.2 | 9.86E-09 | 2.03E-08 |
| chr4 | 161305652 | 161307347 | 1696 | -5.22 | 9.86E-09 | 2.03E-08 |
| chr8 | 102034489 | 102049313 | 14825 | -6.99 | 9.88E-09 | 2.03E-08 |
| chr10 | 73376724 | 73377851 | 1128 | -6.53 | 9.88E-09 | 2.04E-08 |
| chr2 | 225288540 | 225291194 | 2655 | -6.7 | 9.89E-09 | 2.04E-08 |
| chr3 | 72589931 | 72591252 | 1322 | -6.5 | 9.90E-09 | 2.04E-08 |
| chr13 | 40952063 | 40953534 | 1472 | -6.55 | 9.91E-09 | 2.04E-08 |
| chr6 | 147599231 | 147600302 | 1072 | -6.59 | 9.91E-09 | 2.04E-08 |
| chr8 | 48930764 | 48933304 | 2541 | -6.58 | 9.94E-09 | 2.05E-08 |
| chr4 | 147655906 | 147673240 | 17335 | 1.87 | 9.94E-09 | 2.05E-08 |
| chr7 | 133950060 | 133957002 | 6943 | -3.84 | 9.96E-09 | 2.05E-08 |
| chr8 | 10498623 | 10499835 | 1213 | -6.5 | 9.97E-09 | 2.05E-08 |
| chr7 | 116960138 | 116971797 | 11660 | -3.16 | 9.98E-09 | 2.05E-08 |
| chr8 | 63119768 | 63121435 | 1668 | -6.58 | 1.00E-08 | 2.06E-08 |
| chr6 | 44736982 | 44754450 | 17469 | -5.88 | 1.00E-08 | 2.06E-08 |
| chr5 | 52035323 | 52037834 | 2512 | 6.57 | 1.00E-08 | 2.06E-08 |
| chr3 | 4581462 | 4583492 | 2031 | -6.49 | 1.00E-08 | 2.06E-08 |
| chr1 | 63769950 | 63774947 | 4998 | -4.18 | 1.00E-08 | 2.06E-08 |
| chr6 | 140024756 | 140027450 | 2695 | -6.81 | 1.00E-08 | 2.07E-08 |
| chrX | 133892001 | 133893887 | 1887 | -6.5 | 1.01E-08 | 2.07E-08 |
| chr6 | 126806562 | 126814704 | 8143 | -7.01 | 1.01E-08 | 2.08E-08 |
| chr10 | 100065297 | 100066957 | 1661 | -6.74 | 1.01E-08 | 2.08E-08 |
| chr3 | 65902785 | 65904256 | 1472 | -6.5 | 1.01E-08 | 2.09E-08 |
| chr2 | 54706478 | 54708299 | 1822 | -6.53 | 1.01E-08 | 2.09E-08 |
| chrX | 135422820 | 135440853 | 18034 | 1.78 | 1.01E-08 | 2.09E-08 |
| chr12 | 92348369 | 92350874 | 2506 | -6.95 | 1.02E-08 | 2.09E-08 |
| chr13 | 33134894 | 33136826 | 1933 | -6.49 | 1.02E-08 | 2.09E-08 |
| chr3 | 43820482 | 43826595 | 6114 | -6.49 | 1.02E-08 | 2.09E-08 |
| chr3 | 151130409 | 151131814 | 1406 | -6.49 | 1.02E-08 | 2.10E-08 |
| chr2 | 33249233 | 33250377 | 1145 | -6.51 | 1.02E-08 | 2.10E-08 |
| chr22 | 50524989 | 50536439 | 11451 | -7.25 | 1.02E-08 | 2.10E-08 |
| chr17 | 79148915 | 79150132 | 1218 | -6.75 | 1.02E-08 | 2.10E-08 |
| chr15 | 59580045 | 59581533 | 1489 | -6.53 | 1.02E-08 | 2.10E-08 |
| chr20 | 46362971 | 46364312 | 1342 | -6.53 | 1.02E-08 | 2.10E-08 |
| chr4 | 17957653 | 17958712 | 1060 | -6.53 | 1.02E-08 | 2.10E-08 |

| | | | | | | |
|---|---|---|---|---|---|---|
| chr1 | 173343894 | 173345743 | 1850 | -6.51 | 1.02E-08 | 2.10E-08 |
| chr14 | 73849279 | 73851225 | 1947 | -6.84 | 1.02E-08 | 2.11E-08 |
| chrX | 38845658 | 38947974 | 102317 | 1.4 | 1.03E-08 | 2.11E-08 |
| chr12 | 110109049 | 110111230 | 2182 | -6.76 | 1.03E-08 | 2.11E-08 |
| chrX | 69825089 | 69826805 | 1717 | 6.58 | 1.03E-08 | 2.12E-08 |
| chrX | 2754591 | 2757427 | 2837 | -6.5 | 1.03E-08 | 2.12E-08 |
| chr8 | 74355597 | 74358422 | 2826 | -7.03 | 1.03E-08 | 2.12E-08 |
| chr17 | 9021686 | 9022469 | 784 | -6.49 | 1.03E-08 | 2.12E-08 |
| chr12 | 86878766 | 86883589 | 4824 | -6.48 | 1.03E-08 | 2.12E-08 |
| chr2 | 20950488 | 20952369 | 1882 | -6.48 | 1.03E-08 | 2.12E-08 |
| chr10 | 6788101 | 6793343 | 5243 | -6.65 | 1.04E-08 | 2.13E-08 |
| chr3 | 46571690 | 46573875 | 2186 | -6.62 | 1.04E-08 | 2.13E-08 |
| chrX | 13480488 | 13483233 | 2746 | -6.77 | 1.04E-08 | 2.13E-08 |
| chr3 | 123375616 | 123376956 | 1341 | -6.61 | 1.04E-08 | 2.13E-08 |
| chr11 | 3738558 | 3740768 | 2211 | -6.67 | 1.04E-08 | 2.14E-08 |
| chr9 | 12869650 | 12872408 | 2759 | -6.49 | 1.04E-08 | 2.14E-08 |
| chr11 | 36656771 | 36661901 | 5131 | -6.6 | 1.04E-08 | 2.14E-08 |
| chr12 | 1809193 | 1810696 | 1504 | -6.48 | 1.04E-08 | 2.14E-08 |
| chr6 | 84068757 | 84070137 | 1381 | -6.48 | 1.04E-08 | 2.14E-08 |
| chr15 | 56549120 | 56636952 | 87833 | -3.2 | 1.04E-08 | 2.14E-08 |
| chr5 | 42930789 | 42932120 | 1332 | -6.6 | 1.04E-08 | 2.14E-08 |
| chr9 | 87876066 | 88034035 | 157970 | 0.95 | 1.04E-08 | 2.14E-08 |
| chr3 | 155611682 | 155613084 | 1403 | -6.52 | 1.04E-08 | 2.14E-08 |
| chr13 | 112133911 | 112161450 | 27540 | -2.02 | 1.04E-08 | 2.15E-08 |
| chr2 | 33146789 | 33148011 | 1223 | -6.49 | 1.04E-08 | 2.15E-08 |
| chr5 | 93072809 | 93074440 | 1632 | -6.49 | 1.04E-08 | 2.15E-08 |
| chr1 | 120306932 | 120308286 | 1355 | -6.59 | 1.05E-08 | 2.15E-08 |
| chr11 | 114391160 | 114399016 | 7857 | -1.75 | 1.05E-08 | 2.15E-08 |
| chr8 | 8956368 | 8958351 | 1984 | -6.79 | 1.05E-08 | 2.15E-08 |
| chr2 | 150124947 | 150127010 | 2064 | -2.71 | 1.05E-08 | 2.15E-08 |
| chr4 | 96463790 | 96465038 | 1249 | -6.54 | 1.05E-08 | 2.16E-08 |
| chrX | 154204190 | 154206429 | 2240 | -6.54 | 1.05E-08 | 2.16E-08 |
| chr3 | 131494326 | 131499237 | 4912 | -6.69 | 1.05E-08 | 2.16E-08 |
| chr11 | 46254411 | 46258147 | 3737 | -6.48 | 1.05E-08 | 2.16E-08 |
| chr4 | 113080215 | 113082352 | 2138 | -6.48 | 1.05E-08 | 2.16E-08 |
| chr1 | 76220031 | 76224130 | 4100 | -6.52 | 1.05E-08 | 2.16E-08 |
| chr20 | 18000678 | 18002512 | 1835 | -6.52 | 1.05E-08 | 2.16E-08 |
| chr20 | 17578095 | 17579764 | 1670 | -6.88 | 1.05E-08 | 2.17E-08 |
| chrX | 116649594 | 116689786 | 40193 | 2.61 | 1.06E-08 | 2.17E-08 |
| chrX | 109475789 | 109481814 | 6026 | -6.63 | 1.06E-08 | 2.17E-08 |
| chr17 | 56768706 | 56769805 | 1100 | -6.5 | 1.06E-08 | 2.17E-08 |
| chr2 | 192535646 | 192539913 | 4268 | -6.5 | 1.06E-08 | 2.17E-08 |
| chr4 | 129888510 | 129891262 | 2753 | -6.65 | 1.06E-08 | 2.17E-08 |
| chrX | 45549934 | 45560659 | 10726 | -6.71 | 1.06E-08 | 2.17E-08 |
| chr15 | 28405739 | 28407308 | 1570 | -6.66 | 1.06E-08 | 2.17E-08 |
| chr11 | 126540078 | 126542071 | 1994 | -4.13 | 1.06E-08 | 2.17E-08 |
| chr4 | 57511672 | 57513691 | 2020 | -6.66 | 1.06E-08 | 2.17E-08 |
| chr3 | 62194433 | 62206844 | 12412 | 5.65 | 1.06E-08 | 2.17E-08 |
| chr5 | 31206895 | 31209130 | 2236 | 4.93 | 1.06E-08 | 2.18E-08 |
| chr5 | 41706807 | 41718645 | 11839 | -2.2 | 1.06E-08 | 2.18E-08 |
| chr11 | 101907859 | 101909645 | 1787 | -6.53 | 1.07E-08 | 2.19E-08 |
| chr22 | 31983609 | 31984527 | 919 | -6.68 | 1.07E-08 | 2.19E-08 |
| chr1 | 226782560 | 226784230 | 1671 | -6.48 | 1.07E-08 | 2.19E-08 |
| chr8 | 43227745 | 43231260 | 3516 | -6.48 | 1.07E-08 | 2.19E-08 |
| chr4 | 83388510 | 83389855 | 1346 | -6.49 | 1.07E-08 | 2.19E-08 |
| chr3 | 146244444 | 146257815 | 13372 | -6.56 | 1.07E-08 | 2.20E-08 |
| chr7 | 94077428 | 94085009 | 7582 | -7.31 | 1.07E-08 | 2.20E-08 |
| chr18 | 34565463 | 34580493 | 15031 | -6.73 | 1.08E-08 | 2.21E-08 |
| chr9 | 92226197 | 92227677 | 1481 | -6.73 | 1.08E-08 | 2.21E-08 |
| chr5 | 158155200 | 158167406 | 12207 | 7.36 | 1.08E-08 | 2.21E-08 |
| chr15 | 56001668 | 56002813 | 1146 | -6.5 | 1.08E-08 | 2.21E-08 |

| | | | | | | |
|---|---|---|---|---|---|---|
| chr4 | 178967162 | 178968627 | 1466 | -6.5 | 1.08E-08 | 2.21E-08 |
| chr7 | 75644990 | 75651752 | 6763 | -6.53 | 1.08E-08 | 2.21E-08 |
| chr9 | 15613236 | 15614459 | 1224 | -6.56 | 1.08E-08 | 2.21E-08 |
| chr10 | 17042857 | 17047752 | 4896 | -5.1 | 1.08E-08 | 2.22E-08 |
| chr14 | 81687980 | 81689056 | 1077 | -6.5 | 1.08E-08 | 2.22E-08 |
| chr2 | 215827000 | 215829699 | 2700 | -6.74 | 1.08E-08 | 2.22E-08 |
| chr5 | 168194840 | 168196229 | 1390 | -6.55 | 1.09E-08 | 2.22E-08 |
| chr14 | 66545818 | 66800194 | 254377 | 1.28 | 1.09E-08 | 2.23E-08 |
| chr3 | 158215720 | 158218551 | 2832 | -6.5 | 1.09E-08 | 2.23E-08 |
| chr3 | 130840941 | 130861118 | 20178 | -6.52 | 1.09E-08 | 2.23E-08 |
| chr15 | 72770087 | 72777523 | 7437 | -6.72 | 1.09E-08 | 2.23E-08 |
| chr4 | 90065696 | 90067767 | 2072 | -6.83 | 1.09E-08 | 2.23E-08 |
| chr1 | 87512784 | 87514674 | 1891 | -6.48 | 1.09E-08 | 2.23E-08 |
| chr18 | 67746706 | 67748273 | 1568 | -6.56 | 1.09E-08 | 2.24E-08 |
| chr5 | 130930250 | 130932929 | 2680 | -6.91 | 1.09E-08 | 2.24E-08 |
| chr8 | 30468273 | 30469694 | 1422 | -6.72 | 1.09E-08 | 2.24E-08 |
| chr17 | 30029568 | 30038392 | 8825 | -7.48 | 1.09E-08 | 2.24E-08 |
| chr1 | 194231712 | 194234647 | 2936 | -6.48 | 1.09E-08 | 2.24E-08 |
| chr7 | 132671101 | 132672644 | 1544 | -6.48 | 1.09E-08 | 2.24E-08 |
| chr1 | 100509328 | 100511076 | 1749 | -6.52 | 1.09E-08 | 2.24E-08 |
| chr10 | 26838480 | 26840437 | 1958 | -6.47 | 1.10E-08 | 2.25E-08 |
| chr22 | 26247508 | 26257876 | 10369 | -1.57 | 1.10E-08 | 2.26E-08 |
| chr9 | 104253556 | 104255562 | 2007 | -6.64 | 1.10E-08 | 2.26E-08 |
| chr8 | 103409625 | 103420456 | 10832 | -6.77 | 1.10E-08 | 2.26E-08 |
| chr10 | 91408484 | 91409859 | 1376 | -6.48 | 1.11E-08 | 2.27E-08 |
| chrX | 56189129 | 56193133 | 4005 | -6.47 | 1.11E-08 | 2.27E-08 |
| chr10 | 76817078 | 76818227 | 1150 | -6.59 | 1.11E-08 | 2.27E-08 |
| chr4 | 136667684 | 136672838 | 5155 | -6.77 | 1.11E-08 | 2.27E-08 |
| chr13 | 35435395 | 35445128 | 9734 | -6.59 | 1.11E-08 | 2.27E-08 |
| chr2 | 226498103 | 226511335 | 13233 | -2.61 | 1.11E-08 | 2.28E-08 |
| chr7 | 128940043 | 128941999 | 1957 | -6.58 | 1.11E-08 | 2.28E-08 |
| chr21 | 18715030 | 18724877 | 9848 | 2.09 | 1.11E-08 | 2.28E-08 |
| chrX | 82205341 | 82209308 | 3968 | -6.47 | 1.12E-08 | 2.28E-08 |
| chr20 | 56652127 | 56653381 | 1255 | -6.47 | 1.12E-08 | 2.28E-08 |
| chr1 | 115682633 | 115685178 | 2546 | -6.54 | 1.12E-08 | 2.28E-08 |
| chr7 | 156670212 | 156673732 | 3521 | -6.54 | 1.12E-08 | 2.28E-08 |
| chr12 | 843312 | 844701 | 1390 | -6.85 | 1.12E-08 | 2.29E-08 |
| chr2 | 109135646 | 109136624 | 979 | -6.48 | 1.12E-08 | 2.30E-08 |
| chr15 | 26071281 | 26073300 | 2020 | -6.54 | 1.12E-08 | 2.30E-08 |
| chr8 | 90294950 | 90297312 | 2363 | -6.79 | 1.12E-08 | 2.30E-08 |
| chr3 | 4662494 | 4663654 | 1161 | -6.52 | 1.13E-08 | 2.30E-08 |
| chr1 | 32549219 | 32550736 | 1518 | -6.7 | 1.13E-08 | 2.30E-08 |
| chr14 | 71213724 | 71226204 | 12481 | -7.17 | 1.13E-08 | 2.30E-08 |
| chr5 | 79589327 | 79611498 | 22172 | -7.54 | 1.13E-08 | 2.31E-08 |
| chr1 | 198342979 | 198344270 | 1292 | -6.49 | 1.13E-08 | 2.31E-08 |
| chr4 | 147969047 | 147973134 | 4088 | -6.86 | 1.13E-08 | 2.31E-08 |
| chr20 | 3102732 | 3103806 | 1075 | -6.56 | 1.13E-08 | 2.31E-08 |
| chr10 | 72597019 | 72598219 | 1201 | -6.79 | 1.13E-08 | 2.31E-08 |
| chr3 | 195331564 | 195335464 | 3901 | -5.95 | 1.13E-08 | 2.32E-08 |
| chr13 | 21908663 | 21909951 | 1289 | -6.56 | 1.13E-08 | 2.32E-08 |
| chr1 | 84822038 | 84834494 | 12457 | -6.87 | 1.14E-08 | 2.32E-08 |
| chr18 | 18639700 | 18642240 | 2541 | -6.71 | 1.14E-08 | 2.32E-08 |
| chr2 | 8250868 | 8253270 | 2403 | -4.85 | 1.14E-08 | 2.33E-08 |
| chr9 | 125289831 | 125536739 | 246909 | -2.38 | 1.14E-08 | 2.33E-08 |
| chr11 | 68858626 | 68860714 | 2089 | -6.53 | 1.14E-08 | 2.33E-08 |
| chr4 | 101921468 | 101923437 | 1970 | -6.47 | 1.14E-08 | 2.33E-08 |
| chr7 | 69211069 | 69212357 | 1289 | -6.47 | 1.14E-08 | 2.33E-08 |
| chr11 | 108242879 | 108244470 | 1592 | -6.46 | 1.14E-08 | 2.34E-08 |
| chr22 | 44690032 | 44691684 | 1653 | -6.46 | 1.14E-08 | 2.34E-08 |
| chr5 | 176782124 | 176783104 | 981 | -6.46 | 1.14E-08 | 2.34E-08 |
| chr14 | 92080391 | 92081967 | 1577 | -6.47 | 1.14E-08 | 2.34E-08 |

| | | | | | | |
|---|---|---|---|---|---|---|
| chr4 | 125539014 | 125540694 | 1681 | -6.47 | 1.14E-08 | 2.34E-08 |
| chr4 | 68980869 | 68982796 | 1928 | -3.17 | 1.14E-08 | 2.34E-08 |
| chr2 | 136046681 | 136064572 | 17892 | -7 | 1.14E-08 | 2.34E-08 |
| chr19 | 17174534 | 17176056 | 1523 | -6.78 | 1.15E-08 | 2.34E-08 |
| chr18 | 13107848 | 13109187 | 1340 | -6.6 | 1.15E-08 | 2.34E-08 |
| chr3 | 121493377 | 121494742 | 1366 | -6.65 | 1.15E-08 | 2.34E-08 |
| chr1 | 76521940 | 76538489 | 16550 | -7.12 | 1.15E-08 | 2.34E-08 |
| chr12 | 26367841 | 26369487 | 1647 | -1.97 | 1.15E-08 | 2.34E-08 |
| chr10 | 115476961 | 115478462 | 1502 | -6.47 | 1.15E-08 | 2.35E-08 |
| chr6 | 37384323 | 37386048 | 1726 | -6.66 | 1.15E-08 | 2.35E-08 |
| chr6 | 55568481 | 55571175 | 2695 | -6.52 | 1.15E-08 | 2.35E-08 |
| chr8 | 37926838 | 37928072 | 1235 | -6.51 | 1.15E-08 | 2.35E-08 |
| chrX | 12339702 | 12340860 | 1159 | -6.51 | 1.15E-08 | 2.35E-08 |
| chr1 | 12323870 | 12325628 | 1759 | -6.67 | 1.15E-08 | 2.36E-08 |
| chr8 | 41770625 | 41781019 | 10395 | -2.61 | 1.16E-08 | 2.36E-08 |
| chr4 | 3381208 | 3383564 | 2357 | -6.74 | 1.16E-08 | 2.37E-08 |
| chr4 | 99318148 | 99321479 | 3332 | -6.74 | 1.16E-08 | 2.37E-08 |
| chr14 | 58549539 | 58551139 | 1601 | -6.49 | 1.16E-08 | 2.37E-08 |
| chr5 | 77029575 | 77031095 | 1521 | -6.47 | 1.16E-08 | 2.37E-08 |
| chr2 | 9266449 | 9267836 | 1388 | -6.52 | 1.16E-08 | 2.37E-08 |
| chr15 | 35604956 | 35609410 | 4455 | -2.82 | 1.16E-08 | 2.37E-08 |
| chr1 | 193846305 | 193848261 | 1957 | -6.51 | 1.17E-08 | 2.38E-08 |
| chrX | 65086765 | 65088479 | 1715 | -5.09 | 1.17E-08 | 2.38E-08 |
| chr1 | 149192985 | 149194240 | 1256 | -6.46 | 1.17E-08 | 2.39E-08 |
| chr4 | 130381839 | 130385791 | 3953 | -5.32 | 1.17E-08 | 2.39E-08 |
| chr3 | 120327423 | 120329932 | 2510 | -6.83 | 1.17E-08 | 2.39E-08 |
| chr2 | 68541499 | 68542083 | 585 | -6.51 | 1.17E-08 | 2.39E-08 |
| chr16 | 24614513 | 24616252 | 1740 | -6.62 | 1.17E-08 | 2.39E-08 |
| chr18 | 33088635 | 33090275 | 1641 | -6.62 | 1.17E-08 | 2.39E-08 |
| chr3 | 41112483 | 41114682 | 2200 | 6.56 | 1.17E-08 | 2.39E-08 |
| chr15 | 35100215 | 35107574 | 7360 | -6.48 | 1.17E-08 | 2.40E-08 |
| chr8 | 127394550 | 127429919 | 35370 | -6.44 | 1.18E-08 | 2.40E-08 |
| chr1 | 180547139 | 180558653 | 11515 | -7.42 | 1.18E-08 | 2.41E-08 |
| chr1 | 225774426 | 225790148 | 15723 | -6.49 | 1.18E-08 | 2.41E-08 |
| chr17 | 36038283 | 36039552 | 1270 | -6.58 | 1.18E-08 | 2.41E-08 |
| chr2 | 55122290 | 55123372 | 1083 | -5.19 | 1.18E-08 | 2.41E-08 |
| chr4 | 85345953 | 85346976 | 1024 | -6.5 | 1.18E-08 | 2.41E-08 |
| chr4 | 98858978 | 98861620 | 2643 | -6.5 | 1.18E-08 | 2.41E-08 |
| chr12 | 132907109 | 132913950 | 6842 | -7.09 | 1.18E-08 | 2.41E-08 |
| chr2 | 205151341 | 205153750 | 2410 | -3.88 | 1.19E-08 | 2.42E-08 |
| chr12 | 96025722 | 96027090 | 1369 | -6.47 | 1.19E-08 | 2.43E-08 |
| chr12 | 125505909 | 125519934 | 14026 | -6.47 | 1.19E-08 | 2.43E-08 |
| chr2 | 189475577 | 189484843 | 9267 | -6.67 | 1.19E-08 | 2.43E-08 |
| chr4 | 126343973 | 126345720 | 1748 | -6.46 | 1.19E-08 | 2.43E-08 |
| chr1 | 155612837 | 155614147 | 1311 | -6.53 | 1.20E-08 | 2.44E-08 |
| chr8 | 29221256 | 29232663 | 11408 | -6.49 | 1.20E-08 | 2.44E-08 |
| chr12 | 88019938 | 88022438 | 2501 | -6.48 | 1.20E-08 | 2.44E-08 |
| chr15 | 102162923 | 102164523 | 1601 | -6.48 | 1.20E-08 | 2.44E-08 |
| chr19 | 11293571 | 11294592 | 1022 | -6.46 | 1.20E-08 | 2.44E-08 |
| chr8 | 123778289 | 123779978 | 1690 | -6.46 | 1.20E-08 | 2.44E-08 |
| chr17 | 9054842 | 9104826 | 49985 | -8.44 | 1.20E-08 | 2.45E-08 |
| chr5 | 68147588 | 68161066 | 13479 | -1.24 | 1.20E-08 | 2.45E-08 |
| chr5 | 96435004 | 96438019 | 3016 | -6.53 | 1.21E-08 | 2.46E-08 |
| chr11 | 110201820 | 110203277 | 1458 | -6.46 | 1.21E-08 | 2.46E-08 |
| chr14 | 74918085 | 74921608 | 3524 | -3.72 | 1.21E-08 | 2.46E-08 |
| chr2 | 17718144 | 17723663 | 5520 | -2.81 | 1.21E-08 | 2.46E-08 |
| chr20 | 13410486 | 13421654 | 11169 | -6.47 | 1.21E-08 | 2.47E-08 |
| chr4 | 129909010 | 129911238 | 2229 | -6.47 | 1.21E-08 | 2.47E-08 |
| chr11 | 129492698 | 129499421 | 6724 | -3.22 | 1.21E-08 | 2.47E-08 |
| chr18 | 2100221 | 2101763 | 1543 | -6.51 | 1.21E-08 | 2.47E-08 |
| chr2 | 11366 | 23279 | 11914 | -6.86 | 1.22E-08 | 2.47E-08 |

| | | | | | | |
|---|---|---|---|---|---|---|
| chr15 | 63677538 | 63678801 | 1264 | -6.71 | 1.22E-08 | 2.47E-08 |
| chr1 | 76012412 | 76014597 | 2186 | 6.53 | 1.22E-08 | 2.48E-08 |
| chr16 | 67101459 | 67103560 | 2102 | -6.48 | 1.22E-08 | 2.48E-08 |
| chrX | 4817236 | 4821508 | 4273 | -6.93 | 1.22E-08 | 2.48E-08 |
| chr5 | 132880980 | 132884730 | 3751 | -2.2 | 1.22E-08 | 2.48E-08 |
| chrX | 117317642 | 117318876 | 1235 | -6.45 | 1.22E-08 | 2.49E-08 |
| chr20 | 4457105 | 4459280 | 2176 | -6.72 | 1.22E-08 | 2.49E-08 |
| chr2 | 33491193 | 33492705 | 1513 | -6.46 | 1.23E-08 | 2.50E-08 |
| chr2 | 31528741 | 31534745 | 6005 | -2.31 | 1.23E-08 | 2.50E-08 |
| chr4 | 105349070 | 105350890 | 1821 | -6.47 | 1.23E-08 | 2.50E-08 |
| chr1 | 31514408 | 31516060 | 1653 | -6.5 | 1.23E-08 | 2.50E-08 |
| chr1 | 1881413 | 1883085 | 1673 | -6.72 | 1.23E-08 | 2.50E-08 |
| chr4 | 126870489 | 126874393 | 3905 | 6.74 | 1.23E-08 | 2.50E-08 |
| chr1 | 81573801 | 81576720 | 2920 | -6.45 | 1.23E-08 | 2.50E-08 |
| chr2 | 32461374 | 32462552 | 1179 | -6.45 | 1.23E-08 | 2.50E-08 |
| chr3 | 112853835 | 112857480 | 3646 | -6.53 | 1.23E-08 | 2.51E-08 |
| chr11 | 45409017 | 45410016 | 1000 | -6.46 | 1.24E-08 | 2.51E-08 |
| chr19 | 6747404 | 6748341 | 938 | -6.48 | 1.24E-08 | 2.52E-08 |
| chr8 | 22821830 | 22823417 | 1588 | -6.48 | 1.24E-08 | 2.52E-08 |
| chr13 | 21450856 | 21454764 | 3909 | -6.46 | 1.24E-08 | 2.52E-08 |
| chr3 | 158302646 | 158304031 | 1386 | -6.46 | 1.24E-08 | 2.52E-08 |
| chr5 | 124553231 | 124555785 | 2555 | -7.13 | 1.25E-08 | 2.54E-08 |
| chr12 | 87153246 | 87155569 | 2324 | -6.6 | 1.25E-08 | 2.54E-08 |
| chr19 | 38976995 | 38983244 | 6250 | -2.32 | 1.25E-08 | 2.54E-08 |
| chr18 | 18611626 | 18613150 | 1525 | -6.5 | 1.25E-08 | 2.55E-08 |
| chr2 | 135066040 | 135067115 | 1076 | -6.5 | 1.25E-08 | 2.55E-08 |
| chr5 | 153801861 | 153816717 | 14857 | -6.47 | 1.25E-08 | 2.55E-08 |
| chr3 | 16419254 | 16421040 | 1787 | -6.51 | 1.25E-08 | 2.55E-08 |
| chr2 | 166597933 | 166600957 | 3025 | 6.54 | 1.25E-08 | 2.55E-08 |
| chr4 | 162990609 | 162992395 | 1787 | -6.56 | 1.26E-08 | 2.55E-08 |
| chr5 | 43425903 | 43427636 | 1734 | -6.75 | 1.26E-08 | 2.56E-08 |
| chr10 | 25332790 | 25340892 | 8103 | -7.14 | 1.26E-08 | 2.56E-08 |
| chr2 | 43815590 | 43816230 | 641 | -6.47 | 1.26E-08 | 2.56E-08 |
| chr7 | 35834092 | 35836009 | 1918 | -7.02 | 1.26E-08 | 2.56E-08 |
| chr21 | 30596591 | 30598981 | 2391 | -6.96 | 1.26E-08 | 2.56E-08 |
| chr2 | 145492269 | 145520176 | 27908 | -5.81 | 1.26E-08 | 2.56E-08 |
| chr3 | 17434354 | 17435895 | 1542 | -6.46 | 1.26E-08 | 2.56E-08 |
| chr11 | 92026777 | 92030294 | 3518 | -6.79 | 1.26E-08 | 2.56E-08 |
| chr2 | 58198964 | 58200230 | 1267 | -6.57 | 1.26E-08 | 2.56E-08 |
| chrX | 152559336 | 152560240 | 905 | -4.86 | 1.26E-08 | 2.56E-08 |
| chr18 | 19841285 | 19870160 | 28876 | -7.08 | 1.26E-08 | 2.57E-08 |
| chr1 | 154495815 | 154509275 | 13461 | -7.43 | 1.26E-08 | 2.57E-08 |
| chr3 | 70884866 | 70898744 | 13879 | -6.65 | 1.26E-08 | 2.57E-08 |
| chr10 | 27131254 | 27132716 | 1463 | -6.55 | 1.26E-08 | 2.57E-08 |
| chr14 | 54868560 | 54869646 | 1087 | -6.5 | 1.26E-08 | 2.57E-08 |
| chr20 | 46081734 | 46083330 | 1597 | -6.5 | 1.26E-08 | 2.57E-08 |
| chr6 | 126511781 | 126513506 | 1726 | -6.66 | 1.27E-08 | 2.57E-08 |
| chr7 | 131835784 | 131838018 | 2235 | -6.66 | 1.27E-08 | 2.58E-08 |
| chr16 | 8687093 | 8704897 | 17805 | -3.21 | 1.27E-08 | 2.58E-08 |
| chr11 | 104352549 | 104354344 | 1796 | -6.47 | 1.27E-08 | 2.58E-08 |
| chr12 | 41208130 | 41212104 | 3975 | -6.84 | 1.27E-08 | 2.58E-08 |
| chr14 | 61134653 | 61150520 | 15868 | -6.84 | 1.27E-08 | 2.58E-08 |
| chr4 | 55993189 | 55994999 | 1811 | -4.65 | 1.27E-08 | 2.59E-08 |
| chr1 | 98353159 | 98365204 | 12046 | -6.55 | 1.28E-08 | 2.59E-08 |
| chr3 | 33153212 | 33154283 | 1072 | -6.45 | 1.28E-08 | 2.59E-08 |
| chr2 | 169161724 | 169188460 | 26737 | -2.81 | 1.28E-08 | 2.59E-08 |
| chr3 | 4832556 | 4833819 | 1264 | -6.44 | 1.28E-08 | 2.60E-08 |
| chr3 | 113805716 | 113807257 | 1542 | -6.44 | 1.28E-08 | 2.60E-08 |
| chr4 | 162661999 | 162664871 | 2873 | -6.44 | 1.28E-08 | 2.60E-08 |
| chr6 | 5550211 | 5551695 | 1485 | -6.44 | 1.28E-08 | 2.60E-08 |
| chr7 | 27955483 | 27956744 | 1262 | -6.46 | 1.28E-08 | 2.60E-08 |

| | | | | | | |
|---|---|---|---|---|---|---|
| chr14 | 35312652 | 35313614 | 963 | -6.46 | 1.28E-08 | 2.60E-08 |
| chr17 | 65844929 | 65846382 | 1454 | -6.69 | 1.29E-08 | 2.61E-08 |
| chr6 | 79810133 | 79812332 | 2200 | -6.45 | 1.29E-08 | 2.61E-08 |
| chr11 | 14247736 | 14248944 | 1209 | -6.45 | 1.29E-08 | 2.61E-08 |
| chr7 | 124435553 | 124437093 | 1541 | -4.78 | 1.29E-08 | 2.62E-08 |
| chr17 | 2487154 | 2488243 | 1090 | -6.53 | 1.29E-08 | 2.62E-08 |
| chr12 | 17808503 | 17812708 | 4206 | -6.78 | 1.29E-08 | 2.63E-08 |
| chr9 | 107776842 | 107778244 | 1403 | -6.52 | 1.29E-08 | 2.63E-08 |
| chr4 | 53505516 | 53509983 | 4468 | -6.7 | 1.29E-08 | 2.63E-08 |
| chr11 | 56990896 | 56993591 | 2696 | -6.5 | 1.30E-08 | 2.63E-08 |
| chr3 | 45602781 | 45613454 | 10674 | -6.44 | 1.30E-08 | 2.63E-08 |
| chrX | 117400815 | 117403034 | 2220 | -6.46 | 1.30E-08 | 2.64E-08 |
| chr9 | 17536850 | 17789531 | 252682 | 3.28 | 1.30E-08 | 2.64E-08 |
| chr13 | 101268644 | 101270733 | 2090 | -6.79 | 1.31E-08 | 2.65E-08 |
| chr6 | 78952890 | 78955790 | 2901 | -6.44 | 1.31E-08 | 2.66E-08 |
| chr9 | 110450691 | 110473919 | 23229 | -6.93 | 1.31E-08 | 2.66E-08 |
| chr16 | 28831054 | 28832232 | 1179 | -6.51 | 1.31E-08 | 2.66E-08 |
| chr7 | 93685282 | 93687460 | 2179 | -6.46 | 1.31E-08 | 2.66E-08 |
| chr11 | 75139696 | 75141021 | 1326 | -6.52 | 1.32E-08 | 2.67E-08 |
| chr6 | 52066244 | 52072263 | 6020 | -2.62 | 1.32E-08 | 2.67E-08 |
| chr20 | 30287947 | 30295619 | 7673 | -6.51 | 1.32E-08 | 2.68E-08 |
| chr2 | 230997312 | 231017452 | 20141 | -7.12 | 1.32E-08 | 2.68E-08 |
| chr6 | 160409399 | 160411027 | 1629 | -6.45 | 1.33E-08 | 2.69E-08 |
| chrX | 109378004 | 109379228 | 1225 | -6.45 | 1.33E-08 | 2.69E-08 |
| chr12 | 34258348 | 34261379 | 3032 | -6.46 | 1.33E-08 | 2.70E-08 |
| chr17 | 45301105 | 45302301 | 1197 | -6.44 | 1.33E-08 | 2.70E-08 |
| chr5 | 159795608 | 159796674 | 1067 | -6.59 | 1.33E-08 | 2.70E-08 |
| chr8 | 109208936 | 109217514 | 8579 | -7.09 | 1.34E-08 | 2.71E-08 |
| chr3 | 17243645 | 17245737 | 2093 | -6.63 | 1.34E-08 | 2.71E-08 |
| chr10 | 93298688 | 93301736 | 3049 | -6.95 | 1.34E-08 | 2.71E-08 |
| chr20 | 3816243 | 3817476 | 1234 | -6.6 | 1.34E-08 | 2.71E-08 |
| chr3 | 108221495 | 108223477 | 1983 | -6.6 | 1.34E-08 | 2.71E-08 |
| chrX | 148878530 | 148882437 | 3908 | -1.96 | 1.34E-08 | 2.72E-08 |
| chr12 | 66806205 | 66808636 | 2432 | -6.82 | 1.34E-08 | 2.72E-08 |
| chrX | 133117573 | 133121285 | 3713 | -4.83 | 1.34E-08 | 2.72E-08 |
| chr9 | 132773558 | 132790515 | 16958 | -7.2 | 1.34E-08 | 2.72E-08 |
| chr10 | 120270272 | 120321214 | 50943 | -1.5 | 1.35E-08 | 2.73E-08 |
| chr13 | 90653416 | 90654884 | 1469 | -5.09 | 1.35E-08 | 2.73E-08 |
| chr17 | 19829252 | 19830598 | 1347 | -6.55 | 1.35E-08 | 2.73E-08 |
| chr14 | 50511378 | 50513479 | 2102 | -6.64 | 1.35E-08 | 2.74E-08 |
| chr15 | 50894551 | 50895340 | 790 | -6.49 | 1.35E-08 | 2.74E-08 |
| chr2 | 214116756 | 214118095 | 1340 | -6.49 | 1.35E-08 | 2.74E-08 |
| chr2 | 161835046 | 161836232 | 1187 | -6.44 | 1.35E-08 | 2.74E-08 |
| chr3 | 108261983 | 108263445 | 1463 | -6.55 | 1.35E-08 | 2.74E-08 |
| chr5 | 139639797 | 139640488 | 692 | -6.55 | 1.35E-08 | 2.74E-08 |
| chr6 | 168655661 | 168657473 | 1813 | -6.55 | 1.35E-08 | 2.74E-08 |
| chr7 | 63248910 | 63250389 | 1480 | -6.54 | 1.35E-08 | 2.74E-08 |
| chr3 | 160330903 | 160332317 | 1415 | 6.52 | 1.36E-08 | 2.74E-08 |
| chr2 | 122223470 | 122224396 | 927 | -6.44 | 1.36E-08 | 2.75E-08 |
| chr8 | 48418124 | 48430084 | 11961 | -7.09 | 1.36E-08 | 2.75E-08 |
| chr8 | 126083910 | 126085531 | 1622 | -6.54 | 1.36E-08 | 2.76E-08 |
| chr21 | 18653109 | 18655812 | 2704 | 4.97 | 1.37E-08 | 2.76E-08 |
| chr4 | 88309366 | 88310271 | 906 | -6.44 | 1.37E-08 | 2.77E-08 |
| chr3 | 169454376 | 169455952 | 1577 | -6.49 | 1.37E-08 | 2.77E-08 |
| chrX | 99786269 | 99787399 | 1131 | -6.54 | 1.37E-08 | 2.77E-08 |
| chr7 | 4199997 | 4201674 | 1678 | -6.67 | 1.37E-08 | 2.77E-08 |
| chr13 | 91303161 | 91305556 | 2396 | -4.78 | 1.37E-08 | 2.77E-08 |
| chr5 | 149036186 | 149037310 | 1125 | -6.84 | 1.37E-08 | 2.77E-08 |
| chr1 | 203766623 | 203768343 | 1721 | -6.46 | 1.37E-08 | 2.77E-08 |
| chr4 | 103512124 | 103513310 | 1187 | -6.46 | 1.37E-08 | 2.77E-08 |
| chr22 | 44181145 | 44197828 | 16684 | -7.03 | 1.37E-08 | 2.77E-08 |

| | | | | | | |
|---|---|---|---|---|---|---|
| chr21 | 46482649 | 46484171 | 1523 | -6.43 | 1.37E-08 | 2.78E-08 |
| chr6 | 85854489 | 85855572 | 1084 | -6.53 | 1.37E-08 | 2.78E-08 |
| chr8 | 68045790 | 68047000 | 1211 | -6.45 | 1.38E-08 | 2.79E-08 |
| chr2 | 216923875 | 216924648 | 774 | -6.52 | 1.38E-08 | 2.80E-08 |
| chr7 | 28241405 | 28260013 | 18609 | -5.08 | 1.39E-08 | 2.81E-08 |
| chr6 | 87525626 | 87549645 | 24020 | 3.59 | 1.39E-08 | 2.82E-08 |
| chr1 | 91948441 | 91953567 | 5127 | -6.7 | 1.40E-08 | 2.82E-08 |
| chr17 | 59536916 | 59539634 | 2719 | -7.25 | 1.40E-08 | 2.83E-08 |
| chr1 | 36494217 | 36495134 | 918 | -6.48 | 1.40E-08 | 2.83E-08 |
| chr7 | 150492180 | 150531594 | 39415 | -2.4 | 1.40E-08 | 2.83E-08 |
| chr9 | 117409176 | 117410604 | 1429 | -6.81 | 1.40E-08 | 2.83E-08 |
| chr12 | 8950973 | 8953046 | 2074 | -6.45 | 1.40E-08 | 2.84E-08 |
| chr2 | 150667558 | 150669724 | 2167 | -6.45 | 1.40E-08 | 2.84E-08 |
| chr15 | 53982324 | 53983379 | 1056 | -6.44 | 1.40E-08 | 2.84E-08 |
| chr1 | 20745815 | 20752904 | 7090 | -6.42 | 1.41E-08 | 2.85E-08 |
| chr4 | 79062513 | 79064946 | 2434 | -6.46 | 1.41E-08 | 2.86E-08 |
| chr5 | 67728095 | 67731350 | 3256 | -6.46 | 1.41E-08 | 2.86E-08 |
| chr2 | 29302034 | 29313066 | 11033 | -6.8 | 1.41E-08 | 2.86E-08 |
| chr12 | 122626595 | 122627943 | 1349 | -6.43 | 1.42E-08 | 2.86E-08 |
| chr5 | 17755763 | 17757930 | 2168 | -6.5 | 1.42E-08 | 2.86E-08 |
| chr8 | 59888651 | 59890221 | 1571 | -6.47 | 1.42E-08 | 2.87E-08 |
| chr10 | 98397499 | 98400168 | 2670 | -7.13 | 1.42E-08 | 2.87E-08 |
| chr7 | 107552246 | 107553213 | 968 | -6.43 | 1.42E-08 | 2.87E-08 |
| chr7 | 133639322 | 133640854 | 1533 | -6.43 | 1.42E-08 | 2.87E-08 |
| chrX | 28416808 | 28428896 | 12089 | -7.26 | 1.42E-08 | 2.87E-08 |
| chr5 | 110067461 | 110069913 | 2453 | -6.57 | 1.42E-08 | 2.87E-08 |
| chr20 | 42915880 | 42917398 | 1519 | -6.6 | 1.42E-08 | 2.87E-08 |
| chr14 | 74745251 | 74747491 | 2241 | -6.57 | 1.42E-08 | 2.88E-08 |
| chr4 | 6321733 | 6336713 | 14981 | -6.94 | 1.42E-08 | 2.88E-08 |
| chr17 | 10663160 | 10666072 | 2913 | 6.49 | 1.43E-08 | 2.89E-08 |
| chr6 | 34363851 | 34364904 | 1054 | -6.43 | 1.43E-08 | 2.89E-08 |
| chr14 | 67569527 | 67573420 | 3894 | -6.63 | 1.43E-08 | 2.89E-08 |
| chr5 | 162846561 | 162848663 | 2103 | -6.55 | 1.43E-08 | 2.89E-08 |
| chr15 | 68535801 | 68541862 | 6062 | -7.08 | 1.44E-08 | 2.90E-08 |
| chr3 | 131759571 | 131760979 | 1409 | -6.43 | 1.44E-08 | 2.91E-08 |
| chr2 | 187093324 | 187095141 | 1818 | -4.89 | 1.44E-08 | 2.91E-08 |
| chr1 | 96920210 | 96923282 | 3073 | -6.49 | 1.44E-08 | 2.91E-08 |
| chrX | 13732830 | 13734167 | 1338 | -6.53 | 1.45E-08 | 2.92E-08 |
| chr7 | 142984071 | 142985166 | 1096 | -6.49 | 1.45E-08 | 2.92E-08 |
| chr6 | 41263543 | 41279502 | 15960 | -1.89 | 1.45E-08 | 2.93E-08 |
| chr19 | 34251425 | 34276535 | 25111 | -7.15 | 1.45E-08 | 2.93E-08 |
| chr9 | 94850459 | 94851861 | 1403 | -6.58 | 1.45E-08 | 2.93E-08 |
| chr2 | 198514932 | 198516318 | 1387 | -6.59 | 1.45E-08 | 2.93E-08 |
| chr11 | 110371175 | 110372837 | 1663 | -3.4 | 1.45E-08 | 2.93E-08 |
| chr19 | 36446826 | 36447785 | 960 | -6.42 | 1.45E-08 | 2.93E-08 |
| chr3 | 105514418 | 105516335 | 1918 | -6.96 | 1.45E-08 | 2.94E-08 |
| chr12 | 125314191 | 125315333 | 1143 | -6.44 | 1.46E-08 | 2.94E-08 |
| chr14 | 32170849 | 32173983 | 3135 | -6.62 | 1.46E-08 | 2.94E-08 |
| chr7 | 116177753 | 116179066 | 1314 | -6.48 | 1.46E-08 | 2.95E-08 |
| chr15 | 56338129 | 56339628 | 1500 | -6.54 | 1.46E-08 | 2.95E-08 |
| chr2 | 26973857 | 26974971 | 1115 | -6.54 | 1.46E-08 | 2.95E-08 |
| chr20 | 8008132 | 8030270 | 22139 | -7.39 | 1.47E-08 | 2.96E-08 |
| chr8 | 39421827 | 39422745 | 919 | -6.54 | 1.47E-08 | 2.96E-08 |
| chr12 | 1975081 | 2017079 | 41999 | -2.37 | 1.47E-08 | 2.96E-08 |
| chr8 | 6667996 | 6680769 | 12774 | -6.97 | 1.47E-08 | 2.97E-08 |
| chr7 | 39000749 | 39015302 | 14554 | -2.14 | 1.47E-08 | 2.97E-08 |
| chr15 | 32977234 | 32979529 | 2296 | -2.91 | 1.47E-08 | 2.97E-08 |
| chr8 | 92108997 | 92126006 | 17010 | -7.49 | 1.47E-08 | 2.97E-08 |
| chr4 | 40593366 | 40594896 | 1531 | -6.42 | 1.47E-08 | 2.97E-08 |
| chr4 | 189185338 | 189187150 | 1813 | -6.64 | 1.48E-08 | 2.98E-08 |
| chrX | 139145332 | 139155483 | 10152 | 3.97 | 1.48E-08 | 2.98E-08 |

| | | | | | | |
|---|---|---|---|---|---|---|
| chr3 | 29892919 | 29894530 | 1612 | -6.47 | 1.48E-08 | 2.98E-08 |
| chr2 | 144774302 | 144780490 | 6189 | -6.97 | 1.48E-08 | 2.98E-08 |
| chr7 | 150423318 | 150437536 | 14219 | -2.32 | 1.48E-08 | 2.98E-08 |
| chr5 | 83810419 | 83813893 | 3475 | 6.4 | 1.48E-08 | 2.98E-08 |
| chr15 | 38581719 | 38584044 | 2326 | -6.53 | 1.48E-08 | 2.98E-08 |
| chr4 | 126696932 | 126698897 | 1966 | -6.53 | 1.48E-08 | 2.98E-08 |
| chr11 | 93990170 | 93998595 | 8426 | -1.81 | 1.48E-08 | 2.99E-08 |
| chr4 | 110366832 | 110367802 | 971 | -6.42 | 1.48E-08 | 2.99E-08 |
| chr10 | 111622552 | 111623905 | 1354 | -6.66 | 1.49E-08 | 2.99E-08 |
| chr4 | 101749086 | 101751486 | 2401 | -6.69 | 1.49E-08 | 3.00E-08 |
| chr10 | 119825538 | 119826739 | 1202 | -6.42 | 1.49E-08 | 3.01E-08 |
| chr2 | 64657462 | 64658665 | 1204 | -6.47 | 1.50E-08 | 3.01E-08 |
| chr6 | 101450452 | 101451832 | 1381 | -3.03 | 1.50E-08 | 3.01E-08 |
| chr4 | 94123679 | 94125599 | 1921 | -6.49 | 1.50E-08 | 3.02E-08 |
| chr4 | 134818127 | 134831984 | 13858 | 7.43 | 1.50E-08 | 3.02E-08 |
| chr3 | 130918770 | 130920331 | 1562 | -6.43 | 1.50E-08 | 3.02E-08 |
| chr1 | 33125002 | 33127447 | 2446 | -6.47 | 1.50E-08 | 3.03E-08 |
| chr12 | 127584949 | 127587024 | 2076 | -6.47 | 1.50E-08 | 3.03E-08 |
| chrX | 127908762 | 127910632 | 1871 | 5.59 | 1.51E-08 | 3.03E-08 |
| chr12 | 117022150 | 117025988 | 3839 | -3.51 | 1.51E-08 | 3.04E-08 |
| chr5 | 54853476 | 54854851 | 1376 | -6.45 | 1.51E-08 | 3.04E-08 |
| chr1 | 38438361 | 38439033 | 673 | -5.16 | 1.51E-08 | 3.05E-08 |
| chr5 | 126391169 | 126392681 | 1513 | -3.51 | 1.52E-08 | 3.05E-08 |
| chr16 | 50270812 | 50272286 | 1475 | -6.58 | 1.52E-08 | 3.05E-08 |
| chr3 | 9972213 | 9973616 | 1404 | -6.57 | 1.52E-08 | 3.05E-08 |
| chr22 | 29529274 | 29530511 | 1238 | -6.56 | 1.52E-08 | 3.06E-08 |
| chr6 | 37411719 | 37412617 | 899 | -6.56 | 1.52E-08 | 3.06E-08 |
| chr6 | 145896676 | 145898530 | 1855 | -6.46 | 1.52E-08 | 3.06E-08 |
| chr7 | 15453899 | 15461644 | 7746 | -6.72 | 1.52E-08 | 3.06E-08 |
| chr1 | 40163945 | 40165233 | 1289 | -6.42 | 1.52E-08 | 3.06E-08 |
| chr14 | 77885100 | 77886834 | 1735 | -6.42 | 1.52E-08 | 3.06E-08 |
| chr8 | 71250967 | 71253432 | 2466 | -6.42 | 1.52E-08 | 3.06E-08 |
| chr1 | 212453082 | 212454279 | 1198 | -6.49 | 1.52E-08 | 3.06E-08 |
| chr4 | 67776577 | 67793096 | 16520 | 2.54 | 1.52E-08 | 3.07E-08 |
| chr10 | 95577929 | 95594000 | 16072 | -7.45 | 1.52E-08 | 3.07E-08 |
| chr12 | 104270160 | 104271486 | 1327 | -6.54 | 1.53E-08 | 3.07E-08 |
| chr5 | 99775772 | 99778343 | 2572 | 6.48 | 1.53E-08 | 3.09E-08 |
| chr3 | 18438689 | 18440883 | 2195 | -6.43 | 1.53E-08 | 3.09E-08 |
| chr9 | 128860005 | 128906762 | 46758 | -4.07 | 1.53E-08 | 3.09E-08 |
| chrX | 83396474 | 83399319 | 2846 | -6.41 | 1.53E-08 | 3.09E-08 |
| chr8 | 131403237 | 131406056 | 2820 | -6.63 | 1.54E-08 | 3.10E-08 |
| chr12 | 58764810 | 58766498 | 1689 | -6.45 | 1.54E-08 | 3.10E-08 |
| chr9 | 38767558 | 38774754 | 7197 | -6.44 | 1.54E-08 | 3.10E-08 |
| chr15 | 63331230 | 63332394 | 1165 | -6.47 | 1.54E-08 | 3.10E-08 |
| chr3 | 71817977 | 71820234 | 2258 | -4.91 | 1.54E-08 | 3.11E-08 |
| chr15 | 70137120 | 70142382 | 5263 | -2.27 | 1.54E-08 | 3.11E-08 |
| chr4 | 63872537 | 63878933 | 6397 | -6.88 | 1.55E-08 | 3.11E-08 |
| chr12 | 3233030 | 3234783 | 1754 | -6.74 | 1.55E-08 | 3.12E-08 |
| chr12 | 95017358 | 95020465 | 3108 | -6.51 | 1.55E-08 | 3.12E-08 |
| chr8 | 37952071 | 37953843 | 1773 | -6.51 | 1.55E-08 | 3.12E-08 |
| chr4 | 90248851 | 90250049 | 1199 | -5.74 | 1.55E-08 | 3.12E-08 |
| chr22 | 45439480 | 45443503 | 4024 | -3.02 | 1.55E-08 | 3.12E-08 |
| chr18 | 28632791 | 28634339 | 1549 | -6.48 | 1.55E-08 | 3.12E-08 |
| chr4 | 188753184 | 188755019 | 1836 | -6.48 | 1.55E-08 | 3.12E-08 |
| chr7 | 111873999 | 111875483 | 1485 | -6.45 | 1.56E-08 | 3.13E-08 |
| chr13 | 85385750 | 85393815 | 8066 | -6.95 | 1.56E-08 | 3.13E-08 |
| chrX | 21027813 | 21030560 | 2748 | -6.95 | 1.56E-08 | 3.13E-08 |
| chr17 | 17435361 | 17436389 | 1029 | -6.41 | 1.56E-08 | 3.13E-08 |
| chr10 | 5910583 | 5911619 | 1037 | -6.41 | 1.56E-08 | 3.13E-08 |
| chr4 | 133368924 | 133371053 | 2130 | -6.41 | 1.56E-08 | 3.13E-08 |
| chr18 | 61566195 | 61611637 | 45443 | 3.35 | 1.56E-08 | 3.14E-08 |

| | | | | | | |
|---|---|---|---|---|---|---|
| chr20 | 32215353 | 32216069 | 717 | -6.41 | 1.56E-08 | 3.14E-08 |
| chr6 | 111089034 | 111090269 | 1236 | -6.41 | 1.56E-08 | 3.14E-08 |
| chr7 | 110255389 | 110256397 | 1009 | -6.41 | 1.56E-08 | 3.14E-08 |
| chr22 | 35866381 | 35867810 | 1430 | -6.66 | 1.56E-08 | 3.14E-08 |
| chr17 | 15358354 | 15366167 | 7814 | -7.03 | 1.56E-08 | 3.14E-08 |
| chr2 | 97295647 | 97296637 | 991 | -6.47 | 1.57E-08 | 3.15E-08 |
| chr14 | 106897016 | 106977448 | 80433 | 1.88 | 1.57E-08 | 3.15E-08 |
| chr15 | 61420909 | 61421985 | 1077 | -3.83 | 1.57E-08 | 3.15E-08 |
| chr2 | 205912866 | 205924354 | 11489 | -6.41 | 1.57E-08 | 3.15E-08 |
| chr15 | 43563712 | 43564985 | 1274 | -6.42 | 1.57E-08 | 3.15E-08 |
| chr3 | 37440783 | 37442413 | 1631 | -6.42 | 1.57E-08 | 3.15E-08 |
| chr16 | 46603234 | 46606255 | 3022 | -6.43 | 1.57E-08 | 3.16E-08 |
| chr4 | 137389027 | 137390737 | 1711 | -6.67 | 1.57E-08 | 3.16E-08 |
| chr4 | 111389600 | 111391168 | 1569 | -6.56 | 1.58E-08 | 3.16E-08 |
| chr10 | 28958059 | 28959115 | 1057 | -6.49 | 1.58E-08 | 3.17E-08 |
| chr5 | 124276028 | 124277616 | 1589 | -6.84 | 1.58E-08 | 3.17E-08 |
| chr3 | 142038423 | 142039845 | 1423 | -6.68 | 1.58E-08 | 3.17E-08 |
| chr20 | 15078923 | 15080623 | 1701 | -6.41 | 1.58E-08 | 3.17E-08 |
| chr2 | 189393846 | 189424945 | 31100 | -6.77 | 1.58E-08 | 3.17E-08 |
| chr18 | 46085108 | 46086581 | 1474 | -6.54 | 1.58E-08 | 3.18E-08 |
| chr13 | 113581325 | 113582595 | 1271 | -6.6 | 1.58E-08 | 3.18E-08 |
| chr10 | 78991887 | 78993692 | 1806 | -6.42 | 1.58E-08 | 3.18E-08 |
| chr1 | 64228179 | 64230349 | 2171 | -6.44 | 1.59E-08 | 3.18E-08 |
| chr14 | 93781673 | 93782568 | 896 | -6.75 | 1.59E-08 | 3.19E-08 |
| chr15 | 76254171 | 76271444 | 17274 | -7.24 | 1.59E-08 | 3.19E-08 |
| chr2 | 182677538 | 182678961 | 1424 | -6.53 | 1.59E-08 | 3.19E-08 |
| chr2 | 18782168 | 18785478 | 3311 | 6.96 | 1.59E-08 | 3.19E-08 |
| chr14 | 69206103 | 69208667 | 2565 | -6.69 | 1.59E-08 | 3.20E-08 |
| chr2 | 718358 | 719916 | 1559 | -6.52 | 1.59E-08 | 3.20E-08 |
| chr2 | 24541584 | 24542988 | 1405 | -6.62 | 1.59E-08 | 3.20E-08 |
| chrX | 35981607 | 35983470 | 1864 | -6.62 | 1.59E-08 | 3.20E-08 |
| chrX | 53172896 | 53174394 | 1499 | -6.41 | 1.59E-08 | 3.20E-08 |
| chr2 | 31421972 | 31424441 | 2470 | 6.56 | 1.60E-08 | 3.20E-08 |
| chr5 | 151014113 | 151021348 | 7236 | -6.78 | 1.60E-08 | 3.20E-08 |
| chr4 | 130920791 | 130922631 | 1841 | -6.41 | 1.60E-08 | 3.22E-08 |
| chr7 | 44992868 | 44994396 | 1529 | -6.43 | 1.60E-08 | 3.22E-08 |
| chr1 | 147133771 | 147134926 | 1156 | -6.42 | 1.61E-08 | 3.22E-08 |
| chr1 | 181239807 | 181275774 | 35968 | -1.51 | 1.61E-08 | 3.23E-08 |
| chr20 | 43751422 | 43753392 | 1971 | -2.87 | 1.61E-08 | 3.23E-08 |
| chr12 | 89945980 | 89947725 | 1746 | -6.71 | 1.61E-08 | 3.24E-08 |
| chr6 | 64241732 | 64243663 | 1932 | -6.43 | 1.61E-08 | 3.24E-08 |
| chr13 | 31510162 | 31537514 | 27353 | -2.88 | 1.62E-08 | 3.24E-08 |
| chr12 | 18963530 | 18965723 | 2194 | -6.4 | 1.62E-08 | 3.24E-08 |
| chr1 | 196665167 | 196671499 | 6333 | -6.43 | 1.62E-08 | 3.24E-08 |
| chr1 | 97413848 | 97416727 | 2880 | -6.41 | 1.62E-08 | 3.24E-08 |
| chr14 | 24828873 | 24830094 | 1222 | -6.41 | 1.62E-08 | 3.24E-08 |
| chr22 | 35924774 | 35926240 | 1467 | -6.56 | 1.62E-08 | 3.24E-08 |
| chr5 | 140402605 | 140404336 | 1732 | -6.55 | 1.62E-08 | 3.25E-08 |
| chr6 | 11817873 | 11820647 | 2775 | -6.58 | 1.62E-08 | 3.25E-08 |
| chr11 | 92941026 | 92943099 | 2074 | -6.55 | 1.62E-08 | 3.25E-08 |
| chr3 | 196075873 | 196076881 | 1009 | -6.5 | 1.62E-08 | 3.25E-08 |
| chr1 | 117692658 | 117708489 | 15832 | -6.45 | 1.62E-08 | 3.26E-08 |
| chr4 | 124454076 | 124455935 | 1860 | -7 | 1.62E-08 | 3.26E-08 |
| chr18 | 76744762 | 76746151 | 1390 | -6.6 | 1.63E-08 | 3.26E-08 |
| chr19 | 13857369 | 13858511 | 1143 | -6.42 | 1.63E-08 | 3.26E-08 |
| chr12 | 86748475 | 86750401 | 1927 | -6.41 | 1.63E-08 | 3.26E-08 |
| chr16 | 27596130 | 27601386 | 5257 | -6.41 | 1.63E-08 | 3.26E-08 |
| chr4 | 173115391 | 173117785 | 2395 | -6.41 | 1.63E-08 | 3.26E-08 |
| chr12 | 68604997 | 68608379 | 3383 | -1.87 | 1.64E-08 | 3.28E-08 |
| chr20 | 61207727 | 61208759 | 1033 | -6.42 | 1.64E-08 | 3.29E-08 |
| chr5 | 107384805 | 107386615 | 1811 | -6.43 | 1.65E-08 | 3.30E-08 |

| | | | | | | |
|---|---|---|---|---|---|---|
| chr10 | 26859937 | 26861282 | 1346 | -6.4 | 1.65E-08 | 3.30E-08 |
| chr20 | 44382829 | 44385601 | 2773 | -5.44 | 1.65E-08 | 3.31E-08 |
| chr4 | 99658458 | 99660068 | 1611 | -6.41 | 1.66E-08 | 3.32E-08 |
| chr21 | 38321256 | 38322388 | 1133 | -6.44 | 1.66E-08 | 3.32E-08 |
| chr6 | 153808847 | 153821253 | 12407 | -6.74 | 1.66E-08 | 3.32E-08 |
| chr1 | 156134558 | 156136107 | 1550 | -6.5 | 1.66E-08 | 3.33E-08 |
| chr18 | 10907425 | 10908621 | 1197 | -4.77 | 1.67E-08 | 3.34E-08 |
| chr5 | 37518217 | 37522709 | 4493 | -6.4 | 1.67E-08 | 3.35E-08 |
| chr22 | 37343394 | 37353811 | 10418 | -6.66 | 1.67E-08 | 3.35E-08 |
| chr9 | 86878548 | 86879657 | 1110 | -6.66 | 1.67E-08 | 3.35E-08 |
| chr13 | 42873231 | 42875479 | 2249 | -6.4 | 1.68E-08 | 3.35E-08 |
| chr1 | 224747795 | 224749837 | 2043 | -4.99 | 1.68E-08 | 3.36E-08 |
| chr11 | 76950673 | 76952200 | 1528 | -6.66 | 1.68E-08 | 3.36E-08 |
| chr4 | 170698164 | 170699258 | 1095 | -6.4 | 1.68E-08 | 3.36E-08 |
| chrX | 51205270 | 51206901 | 1632 | -4.84 | 1.68E-08 | 3.37E-08 |
| chr10 | 88798416 | 88800116 | 1701 | -6.48 | 1.68E-08 | 3.37E-08 |
| chr7 | 130015556 | 130017398 | 1843 | -6.48 | 1.68E-08 | 3.37E-08 |
| chr11 | 122490340 | 122491507 | 1168 | -6.67 | 1.69E-08 | 3.37E-08 |
| chr4 | 28423287 | 28425676 | 2390 | -6.45 | 1.69E-08 | 3.37E-08 |
| chrX | 85141971 | 85144010 | 2040 | -6.45 | 1.69E-08 | 3.37E-08 |
| chr7 | 96818695 | 96828849 | 10155 | -5.89 | 1.69E-08 | 3.38E-08 |
| chr4 | 139028781 | 139043301 | 14521 | -6.9 | 1.69E-08 | 3.38E-08 |
| chr1 | 241013258 | 241015265 | 2008 | -6.46 | 1.69E-08 | 3.39E-08 |
| chr13 | 89912522 | 89915163 | 2642 | -6.42 | 1.69E-08 | 3.39E-08 |
| chr1 | 38020075 | 38020990 | 916 | -6.4 | 1.70E-08 | 3.40E-08 |
| chr14 | 78462478 | 78490333 | 27856 | -1.7 | 1.70E-08 | 3.40E-08 |
| chr15 | 25791940 | 25854834 | 62895 | -1.07 | 1.70E-08 | 3.40E-08 |
| chr10 | 3343733 | 3352296 | 8564 | -4.01 | 1.70E-08 | 3.41E-08 |
| chr17 | 44866438 | 44867600 | 1163 | -6.47 | 1.71E-08 | 3.42E-08 |
| chr4 | 187840608 | 187842574 | 1967 | -6.47 | 1.71E-08 | 3.42E-08 |
| chr12 | 124147202 | 124148505 | 1304 | -6.54 | 1.71E-08 | 3.42E-08 |
| chr4 | 126231231 | 126232318 | 1088 | -6.54 | 1.71E-08 | 3.42E-08 |
| chr8 | 1864945 | 1867000 | 2056 | -6.69 | 1.71E-08 | 3.43E-08 |
| chr10 | 90779027 | 90780835 | 1809 | -6.53 | 1.71E-08 | 3.43E-08 |
| chr9 | 80754175 | 80756709 | 2535 | -6.41 | 1.72E-08 | 3.43E-08 |
| chr12 | 122895048 | 122896368 | 1321 | -6.6 | 1.72E-08 | 3.44E-08 |
| chr16 | 46875703 | 46876907 | 1205 | -6.45 | 1.72E-08 | 3.44E-08 |
| chr15 | 99655515 | 99657046 | 1532 | -6.41 | 1.72E-08 | 3.45E-08 |
| chr2 | 177068692 | 177070077 | 1386 | -6.46 | 1.72E-08 | 3.45E-08 |
| chr12 | 125175795 | 125177595 | 1801 | -6.42 | 1.73E-08 | 3.45E-08 |
| chr4 | 14907887 | 14914990 | 7104 | -6.78 | 1.73E-08 | 3.45E-08 |
| chr2 | 224545315 | 224547028 | 1714 | -6.54 | 1.73E-08 | 3.46E-08 |
| chr10 | 127498700 | 127500047 | 1348 | -6.44 | 1.73E-08 | 3.46E-08 |
| chr9 | 29245476 | 29249516 | 4041 | -2.55 | 1.73E-08 | 3.46E-08 |
| chr17 | 72052347 | 72168777 | 116431 | -3.46 | 1.73E-08 | 3.47E-08 |
| chr21 | 45695470 | 45696451 | 982 | -6.45 | 1.74E-08 | 3.47E-08 |
| chr1 | 93899225 | 93900803 | 1579 | -6.46 | 1.74E-08 | 3.48E-08 |
| chrX | 83177473 | 83263915 | 86443 | 4.32 | 1.74E-08 | 3.48E-08 |
| chr17 | 1316342 | 1317601 | 1260 | -6.51 | 1.75E-08 | 3.49E-08 |
| chr4 | 160339154 | 160342093 | 2940 | -6.4 | 1.75E-08 | 3.49E-08 |
| chr11 | 44858107 | 44869696 | 11590 | -6.61 | 1.75E-08 | 3.49E-08 |
| chr4 | 155968388 | 155971729 | 3342 | -6.39 | 1.75E-08 | 3.49E-08 |
| chr2 | 122679242 | 122681374 | 2133 | -6.5 | 1.75E-08 | 3.50E-08 |
| chr11 | 63940872 | 63947655 | 6784 | -6.72 | 1.75E-08 | 3.50E-08 |
| chr9 | 128623620 | 128624547 | 928 | -6.43 | 1.76E-08 | 3.51E-08 |
| chr4 | 131513307 | 131516463 | 3157 | -6.39 | 1.76E-08 | 3.51E-08 |
| chr3 | 36547133 | 36547944 | 812 | -6.45 | 1.76E-08 | 3.51E-08 |
| chr2 | 151291196 | 151293890 | 2695 | -6.4 | 1.77E-08 | 3.53E-08 |
| chr19 | 20447203 | 20450198 | 2996 | -6.64 | 1.77E-08 | 3.54E-08 |
| chr15 | 62418422 | 62420254 | 1833 | -6.44 | 1.77E-08 | 3.54E-08 |
| chr1 | 118364229 | 118373637 | 9409 | -6.64 | 1.78E-08 | 3.55E-08 |

| | | | | | | |
|---|---|---|---|---|---|---|
| chr2 | 173887658 | 173888935 | 1278 | -6.64 | 1.78E-08 | 3.55E-08 |
| chr2 | 181920226 | 181922628 | 2403 | -6.44 | 1.78E-08 | 3.55E-08 |
| chr6 | 63741349 | 63744131 | 2783 | 2.72 | 1.78E-08 | 3.56E-08 |
| chr9 | 94881743 | 94893303 | 11561 | -7.08 | 1.78E-08 | 3.56E-08 |
| chr15 | 79150603 | 79151709 | 1107 | -6.42 | 1.78E-08 | 3.56E-08 |
| chr3 | 190853430 | 190854717 | 1288 | 5.64 | 1.79E-08 | 3.57E-08 |
| chr14 | 37337454 | 37339010 | 1557 | -6.4 | 1.79E-08 | 3.57E-08 |
| chr8 | 122452774 | 122458975 | 6202 | -6.89 | 1.79E-08 | 3.58E-08 |
| chr12 | 13469345 | 13470475 | 1131 | -6.39 | 1.79E-08 | 3.58E-08 |
| chr3 | 120172722 | 120173742 | 1021 | -6.43 | 1.79E-08 | 3.58E-08 |
| chr17 | 46771825 | 46772661 | 837 | -6.4 | 1.80E-08 | 3.58E-08 |
| chr4 | 149114661 | 149116704 | 2044 | -6.41 | 1.80E-08 | 3.59E-08 |
| chr4 | 101843483 | 101845827 | 2345 | -6.47 | 1.80E-08 | 3.59E-08 |
| chr4 | 181455595 | 181462520 | 6926 | 2.26 | 1.81E-08 | 3.61E-08 |
| chr1 | 223318629 | 223325076 | 6448 | -1.63 | 1.81E-08 | 3.61E-08 |
| chr2 | 19409634 | 19412709 | 3076 | -6.42 | 1.81E-08 | 3.62E-08 |
| chr5 | 33985924 | 33988342 | 2419 | -6.46 | 1.82E-08 | 3.63E-08 |
| chr7 | 148142353 | 148144252 | 1900 | -6.4 | 1.82E-08 | 3.63E-08 |
| chr12 | 91783632 | 91800685 | 17054 | -6.43 | 1.83E-08 | 3.64E-08 |
| chr9 | 17406983 | 17411883 | 4901 | -6.46 | 1.83E-08 | 3.64E-08 |
| chr5 | 106406233 | 106407970 | 1738 | -4.9 | 1.83E-08 | 3.65E-08 |
| chr5 | 144955931 | 144958317 | 2387 | -6.4 | 1.84E-08 | 3.67E-08 |
| chrX | 116560062 | 116634278 | 74217 | 2.02 | 1.84E-08 | 3.67E-08 |
| chr1 | 4393732 | 4395747 | 2016 | -3.48 | 1.84E-08 | 3.68E-08 |
| chr9 | 90697415 | 90698882 | 1468 | -6.55 | 1.84E-08 | 3.68E-08 |
| chr10 | 42856722 | 42914035 | 57314 | -2.17 | 1.85E-08 | 3.68E-08 |
| chr10 | 79081156 | 79082573 | 1418 | -5.01 | 1.85E-08 | 3.68E-08 |
| chr5 | 91979713 | 91985029 | 5317 | -7.19 | 1.85E-08 | 3.68E-08 |
| chr11 | 99561030 | 99563708 | 2679 | -6.57 | 1.85E-08 | 3.68E-08 |
| chr5 | 180579723 | 180582206 | 2484 | -6.52 | 1.85E-08 | 3.68E-08 |
| chr7 | 43909969 | 43912227 | 2259 | -6.58 | 1.85E-08 | 3.69E-08 |
| chr10 | 100088137 | 100089038 | 902 | -6.38 | 1.85E-08 | 3.69E-08 |
| chr2 | 45603658 | 45605339 | 1682 | -6.38 | 1.85E-08 | 3.69E-08 |
| chr20 | 8760216 | 8763289 | 3074 | -6.38 | 1.85E-08 | 3.69E-08 |
| chr4 | 142014136 | 142016383 | 2248 | -6.56 | 1.86E-08 | 3.70E-08 |
| chr1 | 189508719 | 189510647 | 1929 | -6.39 | 1.86E-08 | 3.70E-08 |
| chr2 | 231339618 | 231341105 | 1488 | -6.4 | 1.86E-08 | 3.70E-08 |
| chr10 | 7414605 | 7419683 | 5079 | -6.38 | 1.86E-08 | 3.71E-08 |
| chrX | 53154830 | 53156090 | 1261 | -6.38 | 1.86E-08 | 3.71E-08 |
| chr5 | 89877530 | 89884853 | 7324 | 3.97 | 1.86E-08 | 3.71E-08 |
| chr10 | 129547165 | 129571647 | 24483 | 2.21 | 1.86E-08 | 3.71E-08 |
| chr2 | 99534547 | 99540834 | 6288 | -6.57 | 1.86E-08 | 3.71E-08 |
| chr4 | 132111672 | 132114021 | 2350 | -6.38 | 1.86E-08 | 3.71E-08 |
| chr5 | 37028183 | 37029425 | 1243 | -6.52 | 1.86E-08 | 3.71E-08 |
| chr3 | 16965557 | 16966787 | 1231 | -6.5 | 1.86E-08 | 3.71E-08 |
| chr5 | 180094137 | 180201953 | 107817 | -1.62 | 1.86E-08 | 3.72E-08 |
| chr8 | 81176338 | 81177717 | 1380 | -6.51 | 1.87E-08 | 3.72E-08 |
| chr17 | 7652879 | 7654127 | 1249 | -6.6 | 1.87E-08 | 3.72E-08 |
| chr20 | 11200242 | 11203360 | 3119 | -6.59 | 1.87E-08 | 3.73E-08 |
| chr8 | 30444009 | 30444814 | 806 | -6.38 | 1.88E-08 | 3.74E-08 |
| chr14 | 104744104 | 104745121 | 1018 | -6.61 | 1.88E-08 | 3.74E-08 |
| chr16 | 84915629 | 84916662 | 1034 | -6.39 | 1.88E-08 | 3.75E-08 |
| chr4 | 137157848 | 137159808 | 1961 | -6.44 | 1.88E-08 | 3.75E-08 |
| chr4 | 91191926 | 91193834 | 1909 | -4.81 | 1.88E-08 | 3.75E-08 |
| chr6 | 46543952 | 46546919 | 2968 | 5 | 1.88E-08 | 3.75E-08 |
| chr11 | 98958363 | 98962410 | 4048 | 6.71 | 1.89E-08 | 3.76E-08 |
| chr2 | 151193706 | 151196261 | 2556 | -6.48 | 1.89E-08 | 3.76E-08 |
| chr3 | 64310786 | 64312276 | 1491 | -6.48 | 1.89E-08 | 3.76E-08 |
| chr13 | 90868500 | 90870740 | 2241 | -6.38 | 1.89E-08 | 3.77E-08 |
| chr16 | 87341644 | 87343306 | 1663 | -6.38 | 1.89E-08 | 3.77E-08 |
| chr7 | 36212744 | 36213886 | 1143 | -6.38 | 1.89E-08 | 3.77E-08 |

| | | | | | | |
|---|---|---|---|---|---|---|
| chr15 | 70866046 | 70867421 | 1376 | -6.4 | 1.89E-08 | 3.77E-08 |
| chr4 | 62310119 | 62313903 | 3785 | -6.39 | 1.89E-08 | 3.77E-08 |
| chr3 | 176699347 | 176700840 | 1494 | -6.47 | 1.90E-08 | 3.78E-08 |
| chr2 | 42375536 | 42377311 | 1776 | -6.94 | 1.90E-08 | 3.79E-08 |
| chr6 | 129049056 | 129050653 | 1598 | -6.43 | 1.90E-08 | 3.79E-08 |
| chr1 | 246107539 | 246109313 | 1775 | -6.38 | 1.90E-08 | 3.79E-08 |
| chr3 | 81613300 | 81614523 | 1224 | -6.38 | 1.90E-08 | 3.79E-08 |
| chr8 | 131641716 | 131649644 | 7929 | -7.08 | 1.91E-08 | 3.80E-08 |
| chr8 | 28677973 | 28679215 | 1243 | -6.42 | 1.92E-08 | 3.82E-08 |
| chr15 | 24694300 | 24726244 | 31945 | 2.03 | 1.92E-08 | 3.82E-08 |
| chr12 | 594064 | 603775 | 9712 | -3.89 | 1.92E-08 | 3.82E-08 |
| chr2 | 222432595 | 222435728 | 3134 | -6.46 | 1.92E-08 | 3.83E-08 |
| chr2 | 47193056 | 47194334 | 1279 | -6.39 | 1.92E-08 | 3.83E-08 |
| chr5 | 100659706 | 100662322 | 2617 | -6.39 | 1.92E-08 | 3.83E-08 |
| chr6 | 114212389 | 114214294 | 1906 | -6.4 | 1.92E-08 | 3.83E-08 |
| chr7 | 129088142 | 129090227 | 2086 | -6.4 | 1.92E-08 | 3.83E-08 |
| chr2 | 131566002 | 131567002 | 1001 | -6.43 | 1.93E-08 | 3.83E-08 |
| chr14 | 62058702 | 62073015 | 14314 | -7.59 | 1.93E-08 | 3.84E-08 |
| chr3 | 133248476 | 133258503 | 10028 | -6.66 | 1.93E-08 | 3.85E-08 |
| chr16 | 8837727 | 8839140 | 1414 | -6.45 | 1.94E-08 | 3.85E-08 |
| chr4 | 91717898 | 91719701 | 1804 | -6.4 | 1.94E-08 | 3.86E-08 |
| chr1 | 38718885 | 38726062 | 7178 | -6.82 | 1.94E-08 | 3.86E-08 |
| chrX | 79591278 | 79592499 | 1222 | -6.67 | 1.94E-08 | 3.86E-08 |
| chr4 | 76507046 | 76509587 | 2542 | -6.45 | 1.94E-08 | 3.87E-08 |
| chr7 | 91750809 | 91752218 | 1410 | -6.45 | 1.94E-08 | 3.87E-08 |
| chr20 | 37292123 | 37293603 | 1481 | -6.45 | 1.95E-08 | 3.88E-08 |
| chr8 | 127935570 | 127936887 | 1318 | -5.51 | 1.96E-08 | 3.89E-08 |
| chr7 | 75871072 | 75872251 | 1180 | -6.37 | 1.96E-08 | 3.90E-08 |
| chr11 | 113282141 | 113284856 | 2716 | 6.5 | 1.96E-08 | 3.90E-08 |
| chr7 | 50832517 | 50833377 | 861 | -6.41 | 1.96E-08 | 3.90E-08 |
| chr8 | 95359851 | 95360862 | 1012 | -6.41 | 1.96E-08 | 3.90E-08 |
| chr1 | 94119735 | 94121724 | 1990 | -6.38 | 1.97E-08 | 3.91E-08 |
| chrX | 37082554 | 37097824 | 15271 | 1.46 | 1.97E-08 | 3.91E-08 |
| chr9 | 131994733 | 131996808 | 2076 | -6.39 | 1.97E-08 | 3.91E-08 |
| chrX | 141091349 | 141093671 | 2323 | -6.52 | 1.97E-08 | 3.92E-08 |
| chr16 | 31110309 | 31111943 | 1635 | -6.39 | 1.97E-08 | 3.92E-08 |
| chr13 | 37471230 | 37474465 | 3236 | -3.46 | 1.97E-08 | 3.92E-08 |
| chrX | 21828888 | 21830895 | 2008 | -6.51 | 1.98E-08 | 3.93E-08 |
| chr4 | 146542623 | 146543760 | 1138 | -6.4 | 1.98E-08 | 3.93E-08 |
| chr14 | 95360313 | 95389529 | 29217 | -2.6 | 1.98E-08 | 3.93E-08 |
| chr6 | 153507201 | 153508767 | 1567 | -6.5 | 1.98E-08 | 3.94E-08 |
| chr9 | 2009455 | 2012292 | 2838 | -4.03 | 1.98E-08 | 3.94E-08 |
| chr6 | 5401166 | 5402928 | 1763 | -6.57 | 1.98E-08 | 3.94E-08 |
| chr9 | 119117999 | 119119138 | 1140 | -6.37 | 1.99E-08 | 3.95E-08 |
| chrX | 53804831 | 53809986 | 5156 | -2.39 | 2.00E-08 | 3.96E-08 |
| chr8 | 39702914 | 39705159 | 2246 | 6.69 | 2.00E-08 | 3.97E-08 |
| chr5 | 39987456 | 40025215 | 37760 | -3.1 | 2.00E-08 | 3.97E-08 |
| chr1 | 93873595 | 93882651 | 9057 | -6.7 | 2.01E-08 | 3.98E-08 |
| chr1 | 95736734 | 95739530 | 2797 | -2.94 | 2.01E-08 | 3.99E-08 |
| chr2 | 63391365 | 63415633 | 24269 | -7.4 | 2.01E-08 | 3.99E-08 |
| chr1 | 61974756 | 61976472 | 1717 | -6.4 | 2.01E-08 | 3.99E-08 |
| chr6 | 26225051 | 26226416 | 1366 | -6.41 | 2.01E-08 | 4.00E-08 |
| chr7 | 105780812 | 105783305 | 2494 | -6.41 | 2.01E-08 | 4.00E-08 |
| chr2 | 111703554 | 111732359 | 28806 | -2.44 | 2.02E-08 | 4.00E-08 |
| chr18 | 71257217 | 71259929 | 2713 | -6.39 | 2.02E-08 | 4.01E-08 |
| chr9 | 88554303 | 88555520 | 1218 | -6.39 | 2.02E-08 | 4.01E-08 |
| chr2 | 61495353 | 61496434 | 1082 | -6.55 | 2.02E-08 | 4.01E-08 |
| chr4 | 68730286 | 68739728 | 9443 | 7.3 | 2.03E-08 | 4.02E-08 |
| chr7 | 94832325 | 94857581 | 25257 | -6.62 | 2.03E-08 | 4.02E-08 |
| chr15 | 73275732 | 73279487 | 3756 | -1.84 | 2.03E-08 | 4.03E-08 |
| chr2 | 160522107 | 160524434 | 2328 | -6.42 | 2.03E-08 | 4.04E-08 |

| | | | | | | |
|---|---|---|---|---|---|---|
| chr9 | 77759164 | 77760497 | 1334 | -6.42 | 2.03E-08 | 4.04E-08 |
| chr10 | 128292901 | 128364913 | 72013 | 1.37 | 2.04E-08 | 4.04E-08 |
| chr19 | 2403952 | 2410320 | 6369 | -6.45 | 2.04E-08 | 4.05E-08 |
| chr2 | 45768006 | 45769489 | 1484 | -6.8 | 2.05E-08 | 4.07E-08 |
| chr2 | 129406099 | 129438440 | 32342 | -2.15 | 2.05E-08 | 4.07E-08 |
| chr12 | 120152425 | 120153768 | 1344 | -6.6 | 2.05E-08 | 4.07E-08 |
| chr14 | 76574747 | 76576067 | 1321 | -6.45 | 2.06E-08 | 4.08E-08 |
| chr13 | 73198113 | 73200096 | 1984 | 3.55 | 2.06E-08 | 4.08E-08 |
| chr20 | 3517569 | 3518132 | 564 | -6.47 | 2.06E-08 | 4.08E-08 |
| chr2 | 193995870 | 194008296 | 12427 | -7.02 | 2.06E-08 | 4.08E-08 |
| chr15 | 83575130 | 83579211 | 4082 | -5.79 | 2.06E-08 | 4.08E-08 |
| chr4 | 163902274 | 163905626 | 3353 | -6.41 | 2.06E-08 | 4.09E-08 |
| chr3 | 110544762 | 110546739 | 1978 | -6.37 | 2.06E-08 | 4.09E-08 |
| chrX | 38715407 | 38718313 | 2907 | 6.45 | 2.06E-08 | 4.09E-08 |
| chr2 | 228808773 | 228895769 | 86997 | 2.12 | 2.06E-08 | 4.09E-08 |
| chr1 | 223683206 | 223685602 | 2397 | -6.95 | 2.07E-08 | 4.10E-08 |
| chr9 | 115502004 | 115503749 | 1746 | -6.44 | 2.07E-08 | 4.11E-08 |
| chr2 | 46694234 | 46707201 | 12968 | -6.74 | 2.08E-08 | 4.12E-08 |
| chr2 | 154922654 | 154924520 | 1867 | -6.62 | 2.08E-08 | 4.12E-08 |
| chr10 | 30244705 | 30246250 | 1546 | -6.45 | 2.09E-08 | 4.13E-08 |
| chr11 | 13058042 | 13084972 | 26931 | -7.72 | 2.09E-08 | 4.14E-08 |
| chr4 | 14499780 | 14508448 | 8669 | -6.83 | 2.09E-08 | 4.15E-08 |
| chr12 | 90478008 | 90483652 | 5645 | -6.67 | 2.09E-08 | 4.15E-08 |
| chr1 | 5611027 | 5633250 | 22224 | -2.44 | 2.09E-08 | 4.15E-08 |
| chrX | 150303633 | 150316503 | 12871 | -2.1 | 2.12E-08 | 4.19E-08 |
| chr3 | 36326650 | 36327543 | 894 | -6.37 | 2.12E-08 | 4.20E-08 |
| chr11 | 35170840 | 35172229 | 1390 | -6.76 | 2.12E-08 | 4.20E-08 |
| chr14 | 68742776 | 68744930 | 2155 | -6.37 | 2.12E-08 | 4.20E-08 |
| chr2 | 136728888 | 136744297 | 15410 | -6.37 | 2.12E-08 | 4.20E-08 |
| chr5 | 122901481 | 122903405 | 1925 | -6.37 | 2.12E-08 | 4.20E-08 |
| chr4 | 111500849 | 111502305 | 1457 | -6.4 | 2.12E-08 | 4.20E-08 |
| chr2 | 73588987 | 73606786 | 17800 | -2.09 | 2.12E-08 | 4.21E-08 |
| chr18 | 3315540 | 3317385 | 1846 | -6.57 | 2.13E-08 | 4.22E-08 |
| chr19 | 51876845 | 51885492 | 8648 | -7.13 | 2.13E-08 | 4.22E-08 |
| chr5 | 63946723 | 63952316 | 5594 | -7.21 | 2.13E-08 | 4.22E-08 |
| chr12 | 132274675 | 132276039 | 1365 | -6.43 | 2.13E-08 | 4.23E-08 |
| chr12 | 38691649 | 38694261 | 2613 | -6.66 | 2.14E-08 | 4.23E-08 |
| chr3 | 177202072 | 177204040 | 1969 | -6.47 | 2.14E-08 | 4.23E-08 |
| chr1 | 114796576 | 114798903 | 2328 | -3.24 | 2.14E-08 | 4.24E-08 |
| chr3 | 17991243 | 17992417 | 1175 | -4.19 | 2.15E-08 | 4.25E-08 |
| chr6 | 50876986 | 50879930 | 2945 | 6.59 | 2.15E-08 | 4.25E-08 |
| chr1 | 91185185 | 91187083 | 1899 | -6.36 | 2.15E-08 | 4.26E-08 |
| chr2 | 136580151 | 136581530 | 1380 | -4.89 | 2.16E-08 | 4.27E-08 |
| chr2 | 237490121 | 237492764 | 2644 | 5.26 | 2.16E-08 | 4.29E-08 |
| chr19 | 46426743 | 46428230 | 1488 | -6.61 | 2.17E-08 | 4.29E-08 |
| chr16 | 54219963 | 54237761 | 17799 | -1.49 | 2.17E-08 | 4.29E-08 |
| chr1 | 25851374 | 25852542 | 1169 | -6.42 | 2.17E-08 | 4.29E-08 |
| chr6 | 170382081 | 170388678 | 6598 | -1.85 | 2.17E-08 | 4.29E-08 |
| chr2 | 43655186 | 43656655 | 1470 | -6.45 | 2.17E-08 | 4.29E-08 |
| chr4 | 104375955 | 104378105 | 2151 | -6.45 | 2.17E-08 | 4.29E-08 |
| chr10 | 80702513 | 80703864 | 1352 | -6.35 | 2.17E-08 | 4.30E-08 |
| chr17 | 819711 | 820681 | 971 | -6.35 | 2.17E-08 | 4.30E-08 |
| chr18 | 53455368 | 53457449 | 2082 | -6.39 | 2.17E-08 | 4.30E-08 |
| chr4 | 68280410 | 68281948 | 1539 | -6.36 | 2.18E-08 | 4.31E-08 |
| chr4 | 111423178 | 111424522 | 1345 | -6.36 | 2.18E-08 | 4.31E-08 |
| chr14 | 62023223 | 62032847 | 9625 | -6.41 | 2.18E-08 | 4.31E-08 |
| chr7 | 79064999 | 79070573 | 5575 | -7.08 | 2.18E-08 | 4.32E-08 |
| chr10 | 76402486 | 76403457 | 972 | -6.39 | 2.19E-08 | 4.33E-08 |
| chrX | 106189572 | 106190764 | 1193 | -6.39 | 2.19E-08 | 4.33E-08 |
| chr13 | 67322419 | 67324029 | 1611 | -6.36 | 2.19E-08 | 4.34E-08 |
| chr12 | 118693924 | 118694745 | 822 | -6.5 | 2.19E-08 | 4.34E-08 |

| | | | | | | |
|---|---|---|---|---|---|---|
| chr4 | 32131643 | 32133925 | 2283 | -6.5 | 2.19E-08 | 4.34E-08 |
| chr12 | 68693272 | 68694477 | 1206 | -6.49 | 2.20E-08 | 4.35E-08 |
| chr17 | 74850927 | 74851921 | 995 | -6.49 | 2.20E-08 | 4.35E-08 |
| chr7 | 100381069 | 100382667 | 1599 | -6.44 | 2.20E-08 | 4.35E-08 |
| chr11 | 129580469 | 129582750 | 2282 | -2.08 | 2.20E-08 | 4.35E-08 |
| chr1 | 191781482 | 191798954 | 17473 | -7.47 | 2.20E-08 | 4.36E-08 |
| chr11 | 62666927 | 62668994 | 2068 | -6.41 | 2.20E-08 | 4.36E-08 |
| chr4 | 188774933 | 188777935 | 3003 | -6.41 | 2.20E-08 | 4.36E-08 |
| chr9 | 117922260 | 117926001 | 3742 | -6.41 | 2.20E-08 | 4.36E-08 |
| chr6 | 132119643 | 132120951 | 1309 | -6.47 | 2.22E-08 | 4.38E-08 |
| chr11 | 12193186 | 12198629 | 5444 | -6.35 | 2.22E-08 | 4.39E-08 |
| chr7 | 46074288 | 46085174 | 10887 | -6.35 | 2.22E-08 | 4.39E-08 |
| chr6 | 111243640 | 111244944 | 1305 | -6.35 | 2.22E-08 | 4.40E-08 |
| chr7 | 72026533 | 72028056 | 1524 | -6.35 | 2.22E-08 | 4.40E-08 |
| chrX | 39708537 | 39710834 | 2298 | -6.35 | 2.22E-08 | 4.40E-08 |
| chr15 | 39527224 | 39529177 | 1954 | -5.21 | 2.23E-08 | 4.40E-08 |
| chr5 | 174917764 | 174918986 | 1223 | -6.58 | 2.23E-08 | 4.40E-08 |
| chr11 | 124677468 | 124679183 | 1716 | -6.38 | 2.23E-08 | 4.40E-08 |
| chr2 | 146927795 | 146929664 | 1870 | -6.38 | 2.23E-08 | 4.40E-08 |
| chr8 | 53813080 | 53814168 | 1089 | -6.38 | 2.23E-08 | 4.41E-08 |
| chr4 | 57533949 | 57535473 | 1525 | -6.46 | 2.23E-08 | 4.42E-08 |
| chr6 | 142102760 | 142104197 | 1438 | -6.42 | 2.23E-08 | 4.42E-08 |
| chrX | 134589840 | 134596894 | 7055 | -2.56 | 2.24E-08 | 4.42E-08 |
| chr1 | 90292136 | 90293500 | 1365 | -6.35 | 2.24E-08 | 4.43E-08 |
| chr8 | 99029962 | 99031655 | 1694 | -6.45 | 2.24E-08 | 4.43E-08 |
| chr6 | 107525182 | 107526367 | 1186 | -6.35 | 2.25E-08 | 4.45E-08 |
| chr8 | 94863367 | 94864529 | 1163 | -6.5 | 2.25E-08 | 4.45E-08 |
| chr13 | 20816520 | 20817662 | 1143 | -5.18 | 2.25E-08 | 4.45E-08 |
| chr15 | 101494555 | 101495803 | 1249 | -6.42 | 2.26E-08 | 4.45E-08 |
| chr9 | 84750457 | 84751773 | 1317 | -6.42 | 2.26E-08 | 4.45E-08 |
| chrX | 28573611 | 28579042 | 5432 | -5.17 | 2.26E-08 | 4.45E-08 |
| chr1 | 240888702 | 240890910 | 2209 | -6.49 | 2.26E-08 | 4.46E-08 |
| chrX | 70070123 | 70071276 | 1154 | -5.48 | 2.26E-08 | 4.46E-08 |
| chr4 | 148068235 | 148070028 | 1794 | -6.55 | 2.26E-08 | 4.46E-08 |
| chr5 | 143198916 | 143209622 | 10707 | -6.97 | 2.26E-08 | 4.47E-08 |
| chr12 | 51023764 | 51024808 | 1045 | -6.37 | 2.27E-08 | 4.47E-08 |
| chr1 | 18894234 | 18895399 | 1166 | -6.48 | 2.27E-08 | 4.47E-08 |
| chr20 | 31723937 | 31737614 | 13678 | -1.91 | 2.27E-08 | 4.47E-08 |
| chr2 | 33313204 | 33332038 | 18835 | -7.29 | 2.27E-08 | 4.48E-08 |
| chr10 | 82315970 | 82317042 | 1073 | -6.4 | 2.27E-08 | 4.48E-08 |
| chr12 | 80018795 | 80020360 | 1566 | -6.37 | 2.27E-08 | 4.48E-08 |
| chr1 | 194374722 | 194386253 | 11532 | -6.39 | 2.27E-08 | 4.48E-08 |
| chr15 | 73376766 | 73391354 | 14589 | -7.57 | 2.28E-08 | 4.50E-08 |
| chr4 | 133550615 | 133552451 | 1837 | -5.05 | 2.28E-08 | 4.50E-08 |
| chr16 | 51630188 | 51782604 | 152417 | -2.62 | 2.28E-08 | 4.51E-08 |
| chr17 | 19375950 | 19384314 | 8365 | -7.21 | 2.29E-08 | 4.51E-08 |
| chr2 | 158360973 | 158363908 | 2936 | -5.65 | 2.29E-08 | 4.51E-08 |
| chr6 | 133437556 | 133438760 | 1205 | -6.43 | 2.29E-08 | 4.51E-08 |
| chr11 | 4512362 | 4513457 | 1096 | -6.36 | 2.29E-08 | 4.51E-08 |
| chr7 | 105797622 | 105805819 | 8198 | -6.69 | 2.30E-08 | 4.53E-08 |
| chr4 | 85631440 | 85632545 | 1106 | -6.39 | 2.30E-08 | 4.53E-08 |
| chr3 | 148771200 | 148772315 | 1116 | -6.45 | 2.30E-08 | 4.53E-08 |
| chr7 | 149446234 | 149447829 | 1596 | -6.45 | 2.30E-08 | 4.53E-08 |
| chr19 | 714687 | 716019 | 1333 | -6.34 | 2.30E-08 | 4.54E-08 |
| chr6 | 158251512 | 158253949 | 2438 | -6.34 | 2.30E-08 | 4.54E-08 |
| chrX | 102461497 | 102463377 | 1881 | -6.34 | 2.30E-08 | 4.54E-08 |
| chr8 | 142825573 | 142846862 | 21290 | -4.32 | 2.30E-08 | 4.54E-08 |
| chr4 | 76658267 | 76659767 | 1501 | -6.4 | 2.30E-08 | 4.54E-08 |
| chr17 | 27510356 | 27511400 | 1045 | -6.44 | 2.31E-08 | 4.55E-08 |
| chr6 | 44550239 | 44551900 | 1662 | -5.13 | 2.31E-08 | 4.55E-08 |
| chr12 | 6912390 | 6914174 | 1785 | -6.35 | 2.31E-08 | 4.56E-08 |

| | | | | | | |
|---|---|---|---|---|---|---|
| chr4 | 66823030 | 66825422 | 2393 | -6.35 | 2.31E-08 | 4.56E-08 |
| chr15 | 35820931 | 35822246 | 1316 | -6.44 | 2.32E-08 | 4.58E-08 |
| chr4 | 77435074 | 77436250 | 1177 | -6.44 | 2.32E-08 | 4.58E-08 |
| chr2 | 114776497 | 114819606 | 43110 | -1.06 | 2.33E-08 | 4.59E-08 |
| chr2 | 33069201 | 33072488 | 3288 | -2.19 | 2.33E-08 | 4.59E-08 |
| chr14 | 36895917 | 36898577 | 2661 | 2.48 | 2.33E-08 | 4.59E-08 |
| chr20 | 10764727 | 10767012 | 2286 | -6.61 | 2.33E-08 | 4.59E-08 |
| chr7 | 75094726 | 75095730 | 1005 | -4.69 | 2.33E-08 | 4.60E-08 |
| chr11 | 122954263 | 122955612 | 1350 | -6.35 | 2.33E-08 | 4.60E-08 |
| chr12 | 57513357 | 57518143 | 4787 | -6.35 | 2.34E-08 | 4.60E-08 |
| chr6 | 22021482 | 22023254 | 1773 | -6.35 | 2.34E-08 | 4.60E-08 |
| chr10 | 71687977 | 71688757 | 781 | -6.34 | 2.34E-08 | 4.61E-08 |
| chr21 | 42663394 | 42688058 | 24665 | -1.33 | 2.34E-08 | 4.61E-08 |
| chrX | 43213108 | 43228675 | 15568 | 6.01 | 2.34E-08 | 4.61E-08 |
| chr2 | 160324561 | 160325923 | 1363 | -6.71 | 2.34E-08 | 4.62E-08 |
| chr18 | 12107292 | 12114057 | 6766 | -1.46 | 2.34E-08 | 4.62E-08 |
| chr3 | 127699534 | 127701006 | 1473 | -6.62 | 2.35E-08 | 4.63E-08 |
| chr8 | 100767565 | 100769391 | 1827 | -6.67 | 2.35E-08 | 4.63E-08 |
| chr4 | 152205189 | 152212341 | 7153 | -6.42 | 2.35E-08 | 4.63E-08 |
| chr1 | 182181862 | 182185244 | 3383 | 2.71 | 2.35E-08 | 4.63E-08 |
| chr5 | 153993444 | 153995443 | 2000 | -5.83 | 2.35E-08 | 4.64E-08 |
| chr7 | 72371078 | 72372681 | 1604 | -6.8 | 2.36E-08 | 4.64E-08 |
| chr17 | 71318737 | 71319703 | 967 | -6.63 | 2.36E-08 | 4.64E-08 |
| chr11 | 94024263 | 94026017 | 1755 | -6.4 | 2.37E-08 | 4.66E-08 |
| chr1 | 235277819 | 235280085 | 2267 | -6.67 | 2.37E-08 | 4.66E-08 |
| chr20 | 33665079 | 33666913 | 1835 | -6.35 | 2.37E-08 | 4.67E-08 |
| chr3 | 193194580 | 193195728 | 1149 | 5.15 | 2.37E-08 | 4.67E-08 |
| chr4 | 136849596 | 136852243 | 2648 | -6.73 | 2.38E-08 | 4.68E-08 |
| chr15 | 89275611 | 89339570 | 63960 | -2.11 | 2.38E-08 | 4.69E-08 |
| chr15 | 68898727 | 68900115 | 1389 | -6.34 | 2.38E-08 | 4.69E-08 |
| chr21 | 36760274 | 36761760 | 1487 | -6.34 | 2.38E-08 | 4.69E-08 |
| chr4 | 178670645 | 178672644 | 2000 | -6.34 | 2.38E-08 | 4.69E-08 |
| chr7 | 24822416 | 24823803 | 1388 | -6.5 | 2.38E-08 | 4.69E-08 |
| chr12 | 27422270 | 27423561 | 1292 | -6.37 | 2.38E-08 | 4.69E-08 |
| chr12 | 30814112 | 30815880 | 1769 | -6.52 | 2.39E-08 | 4.70E-08 |
| chr12 | 120202025 | 120203134 | 1110 | -6.34 | 2.39E-08 | 4.70E-08 |
| chrX | 117368584 | 117375950 | 7367 | -6.53 | 2.39E-08 | 4.70E-08 |
| chr18 | 70694410 | 70696583 | 2174 | -6.34 | 2.39E-08 | 4.70E-08 |
| chr4 | 160882427 | 160884596 | 2170 | -6.34 | 2.39E-08 | 4.70E-08 |
| chr20 | 35031261 | 35032586 | 1326 | -6.48 | 2.39E-08 | 4.70E-08 |
| chr8 | 33458283 | 33472284 | 14002 | -3.07 | 2.40E-08 | 4.72E-08 |
| chr6 | 4278509 | 4280319 | 1811 | -6.54 | 2.40E-08 | 4.72E-08 |
| chr17 | 14048266 | 14049880 | 1615 | -6.65 | 2.40E-08 | 4.72E-08 |
| chr4 | 53168351 | 53178463 | 10113 | 1.9 | 2.40E-08 | 4.72E-08 |
| chrX | 54917604 | 54919364 | 1761 | -6.51 | 2.41E-08 | 4.73E-08 |
| chr4 | 76106758 | 76108409 | 1652 | -6.34 | 2.41E-08 | 4.73E-08 |
| chr8 | 83663828 | 83671092 | 7265 | -6.34 | 2.41E-08 | 4.73E-08 |
| chr10 | 90636021 | 90637801 | 1781 | -6.48 | 2.41E-08 | 4.74E-08 |
| chr10 | 98962443 | 98963835 | 1393 | -6.48 | 2.41E-08 | 4.74E-08 |
| chr6 | 108206494 | 108208995 | 2502 | -6.48 | 2.41E-08 | 4.74E-08 |
| chr7 | 43852233 | 43853649 | 1417 | -6.48 | 2.41E-08 | 4.74E-08 |
| chr1 | 26416182 | 26418042 | 1861 | -6.53 | 2.41E-08 | 4.74E-08 |
| chr18 | 71966096 | 71980387 | 14292 | -6.48 | 2.41E-08 | 4.75E-08 |
| chr13 | 78164991 | 78166197 | 1207 | -6.37 | 2.41E-08 | 4.75E-08 |
| chr9 | 95159442 | 95160578 | 1137 | -6.37 | 2.41E-08 | 4.75E-08 |
| chr3 | 3206036 | 3208672 | 2637 | -6.89 | 2.41E-08 | 4.75E-08 |
| chr6 | 158736740 | 158738268 | 1529 | -6.56 | 2.41E-08 | 4.75E-08 |
| chr5 | 98097489 | 98098896 | 1408 | -3.59 | 2.42E-08 | 4.75E-08 |
| chr7 | 20464735 | 20466612 | 1878 | -6.47 | 2.42E-08 | 4.75E-08 |
| chr1 | 164578674 | 164592153 | 13480 | 7 | 2.42E-08 | 4.76E-08 |
| chr1 | 94182113 | 94183618 | 1506 | -6.46 | 2.42E-08 | 4.76E-08 |

| | | | | | | |
|---|---|---|---|---|---|---|
| chr11 | 119548310 | 119549572 | 1263 | -6.35 | 2.42E-08 | 4.76E-08 |
| chr6 | 170719285 | 170721024 | 1740 | -6.35 | 2.42E-08 | 4.76E-08 |
| chr11 | 14235826 | 14244651 | 8826 | 2.81 | 2.43E-08 | 4.77E-08 |
| chr3 | 177837880 | 177839892 | 2013 | -6.55 | 2.43E-08 | 4.78E-08 |
| chrX | 80002783 | 80005286 | 2504 | -6.55 | 2.43E-08 | 4.78E-08 |
| chr5 | 154392817 | 154409083 | 16267 | -5.63 | 2.43E-08 | 4.78E-08 |
| chr4 | 56374578 | 56375631 | 1054 | -6.34 | 2.43E-08 | 4.78E-08 |
| chr1 | 232448719 | 232454450 | 5732 | -6.36 | 2.43E-08 | 4.78E-08 |
| chr21 | 45802988 | 45810742 | 7755 | -6.34 | 2.44E-08 | 4.79E-08 |
| chr10 | 69373522 | 69392975 | 19454 | 1.82 | 2.44E-08 | 4.79E-08 |
| chr3 | 124817937 | 124824980 | 7044 | -6.4 | 2.44E-08 | 4.79E-08 |
| chr3 | 196135385 | 196137086 | 1702 | -6.4 | 2.44E-08 | 4.79E-08 |
| chr14 | 51195598 | 51196683 | 1086 | -6.71 | 2.44E-08 | 4.80E-08 |
| chr7 | 75820627 | 75821637 | 1011 | -6.33 | 2.44E-08 | 4.80E-08 |
| chr18 | 8387394 | 8388537 | 1144 | -6.67 | 2.44E-08 | 4.80E-08 |
| chr3 | 30068403 | 30092344 | 23942 | -7.76 | 2.45E-08 | 4.82E-08 |
| chr2 | 177991129 | 177992522 | 1394 | -5.01 | 2.45E-08 | 4.82E-08 |
| chr5 | 66487780 | 66498064 | 10285 | -6.44 | 2.46E-08 | 4.83E-08 |
| chr11 | 2898044 | 2900634 | 2591 | -6.69 | 2.47E-08 | 4.84E-08 |
| chr1 | 15299439 | 15301123 | 1685 | -6.39 | 2.47E-08 | 4.85E-08 |
| chr8 | 30327604 | 30328878 | 1275 | -4.78 | 2.47E-08 | 4.85E-08 |
| chr15 | 71089475 | 71092393 | 2919 | -6.84 | 2.47E-08 | 4.85E-08 |
| chr11 | 1285852 | 1287801 | 1950 | -6.35 | 2.47E-08 | 4.85E-08 |
| chr12 | 96443834 | 96451259 | 7426 | -6.37 | 2.47E-08 | 4.85E-08 |
| chr6 | 90625356 | 90627170 | 1815 | -6.33 | 2.47E-08 | 4.86E-08 |
| chr4 | 134976204 | 134979483 | 3280 | 6.59 | 2.47E-08 | 4.86E-08 |
| chr8 | 38593319 | 38598347 | 5029 | -6.77 | 2.48E-08 | 4.86E-08 |
| chr19 | 46758121 | 46759318 | 1198 | -6.42 | 2.48E-08 | 4.87E-08 |
| chr8 | 29382472 | 29385629 | 3158 | -6.6 | 2.48E-08 | 4.87E-08 |
| chr1 | 212485367 | 212486704 | 1338 | -6.33 | 2.48E-08 | 4.87E-08 |
| chr10 | 32274239 | 32277022 | 2784 | -6.43 | 2.48E-08 | 4.88E-08 |
| chr18 | 52869222 | 52870366 | 1145 | -6.33 | 2.49E-08 | 4.89E-08 |
| chr5 | 52944767 | 52945640 | 874 | -6.34 | 2.49E-08 | 4.89E-08 |
| chr6 | 11911902 | 11913727 | 1826 | -6.62 | 2.50E-08 | 4.91E-08 |
| chr2 | 221424377 | 221426384 | 2008 | -6.78 | 2.50E-08 | 4.91E-08 |
| chr2 | 236937959 | 236939704 | 1746 | -6.34 | 2.51E-08 | 4.92E-08 |
| chr13 | 105512465 | 105520150 | 7686 | -6.61 | 2.51E-08 | 4.93E-08 |
| chr11 | 108130541 | 108131448 | 908 | -6.36 | 2.51E-08 | 4.93E-08 |
| chr21 | 35508329 | 35518522 | 10194 | -6.37 | 2.52E-08 | 4.93E-08 |
| chr5 | 123629726 | 123631921 | 2196 | -5.75 | 2.53E-08 | 4.96E-08 |
| chr12 | 41257223 | 41258764 | 1542 | -6.4 | 2.53E-08 | 4.96E-08 |
| chr3 | 71209019 | 71210976 | 1958 | -6.38 | 2.53E-08 | 4.96E-08 |
| chr16 | 19617060 | 19620791 | 3732 | -6.41 | 2.54E-08 | 4.98E-08 |
| chr22 | 27483604 | 27484983 | 1380 | -6.41 | 2.54E-08 | 4.98E-08 |
| chrX | 154414310 | 154424309 | 10000 | -1.51 | 2.54E-08 | 4.98E-08 |
| chr13 | 68798101 | 68800738 | 2638 | -6.35 | 2.54E-08 | 4.98E-08 |
| chrX | 2769966 | 2771244 | 1279 | -6.35 | 2.54E-08 | 4.98E-08 |
| chr3 | 141860295 | 141862628 | 2334 | -6.63 | 2.54E-08 | 4.98E-08 |
| chr6 | 83213553 | 83215217 | 1665 | -6.4 | 2.55E-08 | 5.00E-08 |
| chr1 | 176187190 | 176188858 | 1669 | -5.24 | 2.55E-08 | 5.00E-08 |
| chr6 | 140492799 | 140494880 | 2082 | -6.64 | 2.55E-08 | 5.01E-08 |
| chr5 | 65425286 | 65427375 | 2090 | -6.33 | 2.56E-08 | 5.01E-08 |
| chr7 | 39982372 | 39983740 | 1369 | -6.33 | 2.56E-08 | 5.01E-08 |
| chr3 | 65918844 | 65921424 | 2581 | -6.33 | 2.57E-08 | 5.03E-08 |
| chr21 | 36136509 | 36138398 | 1890 | -6.37 | 2.57E-08 | 5.03E-08 |
| chr8 | 100419450 | 100422041 | 2592 | -6.49 | 2.57E-08 | 5.04E-08 |
| chr3 | 123731732 | 123741589 | 9858 | -3.35 | 2.58E-08 | 5.05E-08 |
| chr11 | 13470959 | 13475719 | 4761 | -6.36 | 2.58E-08 | 5.06E-08 |
| chr5 | 106726626 | 106728357 | 1732 | -4.9 | 2.58E-08 | 5.06E-08 |
| chr2 | 19880011 | 19882027 | 2017 | -4.77 | 2.58E-08 | 5.06E-08 |
| chr10 | 91040642 | 91060548 | 19907 | -6.89 | 2.59E-08 | 5.08E-08 |

| | | | | | | |
|---|---|---|---|---|---|---|
| chr13 | 46829346 | 46831238 | 1893 | -6.5 | 2.60E-08 | 5.08E-08 |
| chr6 | 21973993 | 21976160 | 2168 | -6.54 | 2.60E-08 | 5.08E-08 |
| chr1 | 66048170 | 66050356 | 2187 | -6.49 | 2.60E-08 | 5.08E-08 |
| chr2 | 39462849 | 39464832 | 1984 | -6.82 | 2.60E-08 | 5.08E-08 |
| chr7 | 156938509 | 156939995 | 1487 | -6.51 | 2.60E-08 | 5.09E-08 |
| chr7 | 107899633 | 107900781 | 1149 | -6.54 | 2.60E-08 | 5.09E-08 |
| chr9 | 130140612 | 130142974 | 2363 | -6.45 | 2.60E-08 | 5.09E-08 |
| chr20 | 42484999 | 42486414 | 1416 | -6.52 | 2.60E-08 | 5.09E-08 |
| chr20 | 21487547 | 21490876 | 3330 | -6.34 | 2.60E-08 | 5.09E-08 |
| chr8 | 64157799 | 64166790 | 8992 | -2.83 | 2.60E-08 | 5.10E-08 |
| chr11 | 134615443 | 134619669 | 4227 | -2.08 | 2.60E-08 | 5.10E-08 |
| chr7 | 47146009 | 47147458 | 1450 | -6.36 | 2.61E-08 | 5.11E-08 |
| chr11 | 111274373 | 111275742 | 1370 | -3.13 | 2.61E-08 | 5.12E-08 |
| chr5 | 107075960 | 107088877 | 12918 | -7.17 | 2.62E-08 | 5.14E-08 |
| chr22 | 27424412 | 27426878 | 2467 | -6.44 | 2.62E-08 | 5.14E-08 |
| chr4 | 174351338 | 174352902 | 1565 | -6.33 | 2.63E-08 | 5.14E-08 |
| chr4 | 67172590 | 67174907 | 2318 | -6.57 | 2.64E-08 | 5.16E-08 |
| chr5 | 30783489 | 31182472 | 398984 | 0.94 | 2.64E-08 | 5.17E-08 |
| chr1 | 110382671 | 110384374 | 1704 | -6.42 | 2.64E-08 | 5.18E-08 |
| chr15 | 33437051 | 33438450 | 1400 | -6.42 | 2.64E-08 | 5.18E-08 |
| chr5 | 108095244 | 108096437 | 1194 | -6.42 | 2.64E-08 | 5.18E-08 |
| chr3 | 69385653 | 69394289 | 8637 | -6.58 | 2.65E-08 | 5.18E-08 |
| chr21 | 45679177 | 45680761 | 1585 | -6.35 | 2.65E-08 | 5.19E-08 |
| chr12 | 60730500 | 60732525 | 2026 | -6.32 | 2.66E-08 | 5.20E-08 |
| chr4 | 57158489 | 57160226 | 1738 | -6.32 | 2.66E-08 | 5.21E-08 |
| chr7 | 137666880 | 137668148 | 1269 | -6.34 | 2.67E-08 | 5.22E-08 |
| chr21 | 44884331 | 44886522 | 2192 | -6.33 | 2.67E-08 | 5.22E-08 |
| chr4 | 110569600 | 110571029 | 1430 | -6.37 | 2.68E-08 | 5.24E-08 |
| chr6 | 119161077 | 119162834 | 1758 | -6.41 | 2.68E-08 | 5.24E-08 |
| chr6 | 40825126 | 40827300 | 2175 | -6.35 | 2.69E-08 | 5.27E-08 |
| chr3 | 156547440 | 156548966 | 1527 | -6.32 | 2.69E-08 | 5.27E-08 |
| chr4 | 20512572 | 20521637 | 9066 | 6.69 | 2.69E-08 | 5.27E-08 |
| chr1 | 218431874 | 218433424 | 1551 | -6.41 | 2.70E-08 | 5.27E-08 |
| chr8 | 16845759 | 16846758 | 1000 | -6.4 | 2.70E-08 | 5.28E-08 |
| chr5 | 10804371 | 10826840 | 22470 | -1.52 | 2.71E-08 | 5.31E-08 |
| chr6 | 48233525 | 48235856 | 2332 | -6.62 | 2.72E-08 | 5.32E-08 |
| chr1 | 18420266 | 18421908 | 1643 | -6.33 | 2.73E-08 | 5.33E-08 |
| chr1 | 84603763 | 84605433 | 1671 | -6.32 | 2.73E-08 | 5.35E-08 |
| chr10 | 27180499 | 27183507 | 3009 | -6.72 | 2.74E-08 | 5.36E-08 |
| chr15 | 75775207 | 75776026 | 820 | -6.39 | 2.74E-08 | 5.36E-08 |
| chr20 | 30691574 | 30693164 | 1591 | -5.17 | 2.74E-08 | 5.37E-08 |
| chrX | 79783897 | 79830310 | 46414 | 3.36 | 2.75E-08 | 5.38E-08 |
| chr2 | 240418062 | 240420179 | 2118 | -5.61 | 2.76E-08 | 5.39E-08 |
| chr7 | 65696584 | 65698310 | 1727 | -6.47 | 2.76E-08 | 5.39E-08 |
| chr13 | 46485992 | 46487339 | 1348 | -6.31 | 2.76E-08 | 5.39E-08 |
| chr16 | 53311415 | 53313469 | 2055 | -6.31 | 2.76E-08 | 5.40E-08 |
| chr4 | 96618448 | 96620772 | 2325 | -6.63 | 2.76E-08 | 5.40E-08 |
| chrX | 8472215 | 8488651 | 16437 | 5.17 | 2.76E-08 | 5.40E-08 |
| chr7 | 112057005 | 112057839 | 835 | -6.39 | 2.77E-08 | 5.41E-08 |
| chr4 | 83468919 | 83471271 | 2353 | -6.32 | 2.78E-08 | 5.43E-08 |
| chr5 | 176011449 | 176013155 | 1707 | -6.33 | 2.78E-08 | 5.44E-08 |
| chr7 | 138618747 | 138620493 | 1747 | -6.53 | 2.78E-08 | 5.44E-08 |
| chr2 | 187331245 | 187336729 | 5485 | -6.81 | 2.79E-08 | 5.44E-08 |
| chr5 | 124118297 | 124121208 | 2912 | -6.81 | 2.79E-08 | 5.44E-08 |
| chr6 | 14609393 | 14628376 | 18984 | -7.05 | 2.79E-08 | 5.44E-08 |
| chr10 | 4827826 | 4829482 | 1657 | -6.75 | 2.79E-08 | 5.45E-08 |
| chr2 | 122305924 | 122307444 | 1521 | -6.54 | 2.79E-08 | 5.45E-08 |
| chr2 | 239226272 | 239236902 | 10631 | -6.43 | 2.79E-08 | 5.46E-08 |
| chr5 | 157360858 | 157362943 | 2086 | -6.43 | 2.79E-08 | 5.46E-08 |
| chr6 | 160924287 | 160926724 | 2438 | -6.35 | 2.80E-08 | 5.46E-08 |
| chr3 | 177782620 | 177784267 | 1648 | -6.37 | 2.80E-08 | 5.47E-08 |

| | | | | | | |
|---|---|---|---|---|---|---|
| chr18 | 19365449 | 19366813 | 1365 | -6.42 | 2.80E-08 | 5.48E-08 |
| chr5 | 83190951 | 83201028 | 10078 | 7.36 | 2.81E-08 | 5.48E-08 |
| chrX | 20387756 | 20397888 | 10133 | -6.67 | 2.82E-08 | 5.51E-08 |
| chrX | 90970748 | 90992471 | 21724 | 1.86 | 2.82E-08 | 5.51E-08 |
| chr19 | 15000102 | 15001278 | 1177 | -6.48 | 2.83E-08 | 5.52E-08 |
| chr5 | 131029605 | 131030611 | 1007 | -6.48 | 2.83E-08 | 5.52E-08 |
| chr1 | 12006210 | 12009265 | 3056 | -6.47 | 2.83E-08 | 5.52E-08 |
| chr21 | 40320401 | 40335239 | 14839 | -7.06 | 2.83E-08 | 5.52E-08 |
| chr20 | 5797023 | 5798481 | 1459 | -6.57 | 2.83E-08 | 5.53E-08 |
| chr3 | 58205647 | 58207248 | 1602 | -6.57 | 2.83E-08 | 5.53E-08 |
| chr8 | 124080269 | 124081723 | 1455 | -6.57 | 2.83E-08 | 5.53E-08 |
| chr1 | 93535792 | 93550653 | 14862 | -6.5 | 2.83E-08 | 5.53E-08 |
| chr10 | 3273325 | 3275753 | 2429 | -6.41 | 2.83E-08 | 5.53E-08 |
| chr3 | 33412060 | 33413270 | 1211 | -6.41 | 2.83E-08 | 5.53E-08 |
| chr10 | 134569499 | 134571039 | 1541 | -6.33 | 2.83E-08 | 5.53E-08 |
| chr4 | 144187989 | 144199651 | 11663 | -6.33 | 2.83E-08 | 5.53E-08 |
| chr20 | 37097725 | 37098948 | 1224 | -6.37 | 2.83E-08 | 5.53E-08 |
| chr17 | 53585083 | 53597689 | 12607 | -8.01 | 2.84E-08 | 5.54E-08 |
| chr8 | 118048024 | 118049479 | 1456 | -4.73 | 2.84E-08 | 5.55E-08 |
| chr21 | 35316295 | 35317712 | 1418 | -6.34 | 2.84E-08 | 5.55E-08 |
| chr11 | 121184382 | 121186308 | 1927 | -6.4 | 2.85E-08 | 5.56E-08 |
| chr3 | 178888086 | 178890692 | 2607 | -6.4 | 2.85E-08 | 5.56E-08 |
| chr14 | 86011127 | 86013402 | 2276 | 6.38 | 2.85E-08 | 5.56E-08 |
| chr2 | 188692831 | 188694521 | 1691 | -6.32 | 2.86E-08 | 5.59E-08 |
| chr4 | 130078175 | 130080139 | 1965 | -6.32 | 2.86E-08 | 5.59E-08 |
| chr15 | 42662964 | 42664292 | 1329 | -6.36 | 2.87E-08 | 5.60E-08 |
| chr10 | 69950075 | 69951900 | 1826 | -6.31 | 2.87E-08 | 5.60E-08 |
| chr8 | 67651436 | 67652688 | 1253 | -6.34 | 2.87E-08 | 5.60E-08 |
| chr2 | 238838762 | 238856067 | 17306 | -6.84 | 2.87E-08 | 5.60E-08 |
| chrX | 140892039 | 140893403 | 1365 | -5.66 | 2.87E-08 | 5.60E-08 |
| chr2 | 72782629 | 72783708 | 1080 | -6.69 | 2.87E-08 | 5.61E-08 |
| chr17 | 65197012 | 65197709 | 698 | -6.42 | 2.88E-08 | 5.62E-08 |
| chr22 | 24241916 | 24243582 | 1667 | -6.42 | 2.88E-08 | 5.62E-08 |
| chr7 | 147966943 | 148027172 | 60230 | 2.44 | 2.88E-08 | 5.62E-08 |
| chr16 | 53406964 | 53409464 | 2501 | -6.6 | 2.88E-08 | 5.62E-08 |
| chr14 | 100392906 | 100393993 | 1088 | -6.55 | 2.89E-08 | 5.63E-08 |
| chr22 | 21332292 | 21333620 | 1329 | -6.34 | 2.89E-08 | 5.63E-08 |
| chr6 | 143524406 | 143525796 | 1391 | -6.41 | 2.89E-08 | 5.64E-08 |
| chr13 | 46807138 | 46814794 | 7657 | -3.1 | 2.89E-08 | 5.64E-08 |
| chr1 | 42733490 | 42734892 | 1403 | -6.31 | 2.90E-08 | 5.64E-08 |
| chr2 | 48356358 | 48357426 | 1069 | -6.31 | 2.90E-08 | 5.64E-08 |
| chr4 | 74462564 | 74463864 | 1301 | -6.31 | 2.90E-08 | 5.64E-08 |
| chr16 | 71716468 | 71717441 | 974 | -6.31 | 2.90E-08 | 5.66E-08 |
| chrX | 67761125 | 67762697 | 1573 | -6.61 | 2.91E-08 | 5.67E-08 |
| chr10 | 13817031 | 13818420 | 1390 | -6.38 | 2.91E-08 | 5.68E-08 |
| chr22 | 47806368 | 47916023 | 109656 | -2.5 | 2.92E-08 | 5.68E-08 |
| chr1 | 119800752 | 119812543 | 11792 | -6.71 | 2.92E-08 | 5.68E-08 |
| chr15 | 101598982 | 101600118 | 1137 | -6.57 | 2.92E-08 | 5.68E-08 |
| chr1 | 223912110 | 223913085 | 976 | -6.3 | 2.92E-08 | 5.69E-08 |
| chrX | 106634409 | 106666210 | 31802 | -1.73 | 2.92E-08 | 5.70E-08 |
| chr22 | 28457935 | 28459571 | 1637 | -6.31 | 2.94E-08 | 5.72E-08 |
| chr15 | 35242020 | 35243031 | 1012 | -6.32 | 2.94E-08 | 5.73E-08 |
| chr2 | 207544311 | 207545605 | 1295 | -6.33 | 2.95E-08 | 5.74E-08 |
| chr3 | 88070440 | 88071994 | 1555 | -6.33 | 2.95E-08 | 5.74E-08 |
| chr3 | 152893051 | 152894895 | 1845 | -6.63 | 2.95E-08 | 5.74E-08 |
| chr7 | 45133597 | 45135254 | 1658 | -6.39 | 2.95E-08 | 5.74E-08 |
| chr14 | 36933476 | 36943358 | 9883 | -1.81 | 2.95E-08 | 5.75E-08 |
| chr1 | 66391821 | 66392985 | 1165 | -6.3 | 2.97E-08 | 5.78E-08 |
| chr1 | 98331434 | 98333511 | 2078 | -6.3 | 2.97E-08 | 5.78E-08 |
| chr3 | 194036065 | 194038115 | 2051 | -6.6 | 2.97E-08 | 5.79E-08 |
| chr2 | 207510530 | 207531224 | 20695 | -2.71 | 2.97E-08 | 5.79E-08 |

| | | | | | | |
|---|---|---|---|---|---|---|
| chr12 | 25554320 | 25555931 | 1612 | -6.51 | 2.97E-08 | 5.79E-08 |
| chr10 | 118844428 | 118845813 | 1386 | -6.42 | 2.98E-08 | 5.81E-08 |
| chr13 | 48448136 | 48449511 | 1376 | -6.42 | 2.98E-08 | 5.81E-08 |
| chrX | 84442251 | 84444777 | 2527 | 6.56 | 3.00E-08 | 5.83E-08 |
| chr11 | 100651348 | 100653233 | 1886 | -6.34 | 3.00E-08 | 5.84E-08 |
| chr8 | 77279008 | 77287011 | 8004 | -6.31 | 3.00E-08 | 5.84E-08 |
| chr13 | 112584767 | 112589244 | 4478 | -2.01 | 3.00E-08 | 5.84E-08 |
| chr3 | 140817603 | 140832583 | 14981 | -6.54 | 3.00E-08 | 5.84E-08 |
| chr3 | 188134900 | 188136360 | 1461 | -6.36 | 3.00E-08 | 5.84E-08 |
| chr4 | 127816764 | 127818815 | 2052 | -6.36 | 3.00E-08 | 5.84E-08 |
| chr6 | 9059648 | 9061148 | 1501 | -6.36 | 3.00E-08 | 5.84E-08 |
| chr4 | 5519604 | 5526511 | 6908 | -6.41 | 3.00E-08 | 5.85E-08 |
| chr4 | 39766862 | 39768209 | 1348 | -6.31 | 3.01E-08 | 5.85E-08 |
| chr6 | 84270501 | 84283952 | 13452 | -5.47 | 3.01E-08 | 5.86E-08 |
| chr8 | 27823227 | 27824974 | 1748 | -3.39 | 3.02E-08 | 5.87E-08 |
| chr3 | 78672191 | 78674235 | 2045 | 6.37 | 3.03E-08 | 5.89E-08 |
| chr4 | 143998802 | 144000495 | 1694 | 6.42 | 3.03E-08 | 5.90E-08 |
| chr2 | 230689745 | 230691107 | 1363 | -6.75 | 3.05E-08 | 5.94E-08 |
| chr4 | 156089249 | 156091410 | 2162 | -6.36 | 3.05E-08 | 5.94E-08 |
| chr8 | 130354427 | 130363298 | 8872 | -2.79 | 3.06E-08 | 5.95E-08 |
| chr10 | 29038465 | 29050341 | 11877 | -7.39 | 3.06E-08 | 5.96E-08 |
| chr3 | 49631347 | 49632542 | 1196 | -6.3 | 3.07E-08 | 5.96E-08 |
| chr6 | 122040893 | 122052474 | 11582 | -6.84 | 3.07E-08 | 5.96E-08 |
| chr3 | 119261879 | 119262906 | 1028 | -4.92 | 3.07E-08 | 5.97E-08 |
| chr14 | 90477612 | 90480006 | 2395 | -4.87 | 3.07E-08 | 5.97E-08 |
| chr13 | 51218383 | 51220533 | 2151 | -6.31 | 3.07E-08 | 5.97E-08 |
| chr9 | 134600731 | 134602732 | 2002 | -6.35 | 3.08E-08 | 5.98E-08 |
| chr5 | 169803506 | 169804922 | 1417 | -6.31 | 3.08E-08 | 5.98E-08 |
| chr11 | 108445319 | 108449619 | 4301 | -6.29 | 3.08E-08 | 5.98E-08 |
| chr5 | 90750937 | 90751734 | 798 | -6.29 | 3.08E-08 | 5.98E-08 |
| chr7 | 39669694 | 39670634 | 941 | -6.29 | 3.08E-08 | 5.98E-08 |
| chr2 | 235714651 | 235716785 | 2135 | -7.08 | 3.08E-08 | 5.98E-08 |
| chr8 | 58523386 | 58580381 | 56996 | -6.81 | 3.08E-08 | 5.99E-08 |
| chr2 | 129453148 | 129490190 | 37043 | -2.58 | 3.08E-08 | 5.99E-08 |
| chr7 | 102707843 | 102711414 | 3572 | -6.29 | 3.08E-08 | 5.99E-08 |
| chr10 | 13104242 | 13106826 | 2585 | -6.44 | 3.09E-08 | 6.01E-08 |
| chr3 | 178826470 | 178827582 | 1113 | -6.44 | 3.09E-08 | 6.01E-08 |
| chr15 | 52192762 | 52193610 | 849 | -6.43 | 3.10E-08 | 6.02E-08 |
| chrX | 34972953 | 34974778 | 1826 | -6.43 | 3.10E-08 | 6.02E-08 |
| chr22 | 44455612 | 44456844 | 1233 | -6.32 | 3.10E-08 | 6.03E-08 |
| chr11 | 122162181 | 122178939 | 16759 | -1.83 | 3.10E-08 | 6.03E-08 |
| chr1 | 86057393 | 86067225 | 9833 | -7.18 | 3.10E-08 | 6.03E-08 |
| chr20 | 49499033 | 49500520 | 1488 | -6.35 | 3.11E-08 | 6.04E-08 |
| chr17 | 78576647 | 78578645 | 1999 | -6.51 | 3.11E-08 | 6.04E-08 |
| chr5 | 107447364 | 107449036 | 1673 | -6.42 | 3.12E-08 | 6.05E-08 |
| chr13 | 34413234 | 34414968 | 1735 | -6.34 | 3.12E-08 | 6.06E-08 |
| chr3 | 8905100 | 8926756 | 21657 | -6.53 | 3.13E-08 | 6.07E-08 |
| chr6 | 1845160 | 1846773 | 1614 | -6.53 | 3.13E-08 | 6.09E-08 |
| chr13 | 60638047 | 60644911 | 6865 | -6.35 | 3.14E-08 | 6.09E-08 |
| chr6 | 72149389 | 72150802 | 1414 | -6.35 | 3.14E-08 | 6.09E-08 |
| chr15 | 68300069 | 68301909 | 1841 | -6.54 | 3.14E-08 | 6.10E-08 |
| chr3 | 115421140 | 115422287 | 1148 | -6.54 | 3.14E-08 | 6.10E-08 |
| chr7 | 144341083 | 144343830 | 2748 | 3.24 | 3.15E-08 | 6.12E-08 |
| chr4 | 48930749 | 48955166 | 24418 | -7.31 | 3.16E-08 | 6.13E-08 |
| chr12 | 13302313 | 13304901 | 2589 | -6.32 | 3.16E-08 | 6.14E-08 |
| chr3 | 188689042 | 188690473 | 1432 | -6.32 | 3.16E-08 | 6.14E-08 |
| chr7 | 15435017 | 15437917 | 2901 | -6.62 | 3.17E-08 | 6.15E-08 |
| chr15 | 44937282 | 44939063 | 1782 | -6.56 | 3.17E-08 | 6.16E-08 |
| chr1 | 56544049 | 56554895 | 10847 | -6.5 | 3.18E-08 | 6.17E-08 |
| chr10 | 102516108 | 102518023 | 1916 | 6.71 | 3.18E-08 | 6.18E-08 |
| chr5 | 112941121 | 112943762 | 2642 | -6.42 | 3.19E-08 | 6.18E-08 |

| | | | | | | |
|---|---|---|---|---|---|---|
| chr3 | 47833802 | 47834873 | 1072 | -6.29 | 3.20E-08 | 6.20E-08 |
| chr5 | 100146739 | 100147664 | 926 | -6.29 | 3.20E-08 | 6.20E-08 |
| chr1 | 246052465 | 246054022 | 1558 | -6.33 | 3.21E-08 | 6.22E-08 |
| chr8 | 26034975 | 26055870 | 20896 | -6.33 | 3.21E-08 | 6.22E-08 |
| chr1 | 15612263 | 15625169 | 12907 | -6.64 | 3.21E-08 | 6.23E-08 |
| chr3 | 132291178 | 132292196 | 1019 | -6.29 | 3.21E-08 | 6.23E-08 |
| chr4 | 5796535 | 5797640 | 1106 | -6.29 | 3.21E-08 | 6.23E-08 |
| chr10 | 116244097 | 116245907 | 1811 | -6.37 | 3.22E-08 | 6.25E-08 |
| chr10 | 126174629 | 126176340 | 1712 | -6.37 | 3.22E-08 | 6.25E-08 |
| chr6 | 146270350 | 146270975 | 626 | -4.86 | 3.22E-08 | 6.26E-08 |
| chr1 | 150872077 | 150884809 | 12733 | -7.63 | 3.23E-08 | 6.26E-08 |
| chr16 | 47732830 | 47734488 | 1659 | -6.31 | 3.23E-08 | 6.26E-08 |
| chr14 | 54578914 | 54581042 | 2129 | -6.54 | 3.23E-08 | 6.27E-08 |
| chr18 | 47876723 | 47878662 | 1940 | -3.02 | 3.23E-08 | 6.27E-08 |
| chr15 | 102078922 | 102088472 | 9551 | -6.39 | 3.24E-08 | 6.28E-08 |
| chr15 | 102207041 | 102207873 | 833 | -6.39 | 3.24E-08 | 6.28E-08 |
| chrX | 139586452 | 139587675 | 1224 | -5.45 | 3.25E-08 | 6.31E-08 |
| chr17 | 66601078 | 66742777 | 141700 | 1.06 | 3.25E-08 | 6.31E-08 |
| chr2 | 99210437 | 99211089 | 653 | -6.34 | 3.26E-08 | 6.32E-08 |
| chr22 | 24619792 | 24621556 | 1765 | -6.34 | 3.26E-08 | 6.32E-08 |
| chr10 | 29627394 | 29633646 | 6253 | -2.04 | 3.26E-08 | 6.32E-08 |
| chr11 | 75643538 | 75644480 | 943 | -5.72 | 3.26E-08 | 6.32E-08 |
| chr18 | 56505976 | 56508120 | 2145 | -6.74 | 3.26E-08 | 6.33E-08 |
| chr19 | 30124698 | 30125536 | 839 | -6.61 | 3.27E-08 | 6.35E-08 |
| chr1 | 226526896 | 226530521 | 3626 | -2.68 | 3.28E-08 | 6.36E-08 |
| chr16 | 86742295 | 86746320 | 4026 | -6.91 | 3.28E-08 | 6.36E-08 |
| chr5 | 10382349 | 10383598 | 1250 | -6.32 | 3.29E-08 | 6.37E-08 |
| chr8 | 144982564 | 144984346 | 1783 | -6.72 | 3.29E-08 | 6.37E-08 |
| chr2 | 143050947 | 143069567 | 18621 | -2.82 | 3.29E-08 | 6.38E-08 |
| chr17 | 70578256 | 70580574 | 2319 | -6.3 | 3.30E-08 | 6.40E-08 |
| chr13 | 99924232 | 99925339 | 1108 | -6.28 | 3.30E-08 | 6.40E-08 |
| chr7 | 94794452 | 94796167 | 1716 | -6.29 | 3.31E-08 | 6.41E-08 |
| chr15 | 98453161 | 98505783 | 52623 | 1.19 | 3.31E-08 | 6.41E-08 |
| chr2 | 121091619 | 121092605 | 987 | -6.29 | 3.32E-08 | 6.43E-08 |
| chr4 | 147152599 | 147154311 | 1713 | -6.29 | 3.32E-08 | 6.43E-08 |
| chr9 | 6242233 | 6243825 | 1593 | -6.29 | 3.32E-08 | 6.43E-08 |
| chr5 | 179255116 | 179257961 | 2846 | -4.89 | 3.32E-08 | 6.44E-08 |
| chr6 | 20503109 | 20504807 | 1699 | -6.28 | 3.32E-08 | 6.44E-08 |
| chr5 | 100231652 | 100232359 | 708 | -6.28 | 3.32E-08 | 6.44E-08 |
| chr1 | 224052552 | 224064078 | 11527 | -3.77 | 3.33E-08 | 6.44E-08 |
| chr1 | 174637235 | 174639294 | 2060 | -6.32 | 3.33E-08 | 6.45E-08 |
| chr3 | 98710577 | 98712170 | 1594 | -6.32 | 3.33E-08 | 6.45E-08 |
| chr13 | 30849125 | 30850295 | 1171 | -6.63 | 3.33E-08 | 6.46E-08 |
| chr10 | 95335883 | 95340333 | 4451 | -6.84 | 3.33E-08 | 6.46E-08 |
| chr3 | 36456925 | 36458124 | 1200 | -6.33 | 3.35E-08 | 6.48E-08 |
| chrX | 46901573 | 46903357 | 1785 | -6.33 | 3.35E-08 | 6.48E-08 |
| chr18 | 8519295 | 8533585 | 14291 | -1.51 | 3.36E-08 | 6.50E-08 |
| chr5 | 156357614 | 156359245 | 1632 | -7.05 | 3.36E-08 | 6.51E-08 |
| chr2 | 106980434 | 106981969 | 1536 | -6.64 | 3.37E-08 | 6.53E-08 |
| chr3 | 140931437 | 140946251 | 14815 | -2.4 | 3.37E-08 | 6.53E-08 |
| chr1 | 240981763 | 240984163 | 2401 | -6.34 | 3.37E-08 | 6.53E-08 |
| chr22 | 46310010 | 46311093 | 1084 | -6.3 | 3.38E-08 | 6.54E-08 |
| chr10 | 89367460 | 89372976 | 5517 | -7.11 | 3.38E-08 | 6.55E-08 |
| chr7 | 35293799 | 35295684 | 1886 | -6.45 | 3.39E-08 | 6.55E-08 |
| chr1 | 159873332 | 159874403 | 1072 | -6.5 | 3.39E-08 | 6.56E-08 |
| chr2 | 31743551 | 31746127 | 2577 | -6.41 | 3.39E-08 | 6.56E-08 |
| chr8 | 70996379 | 70998588 | 2210 | -6.44 | 3.39E-08 | 6.56E-08 |
| chr3 | 115862231 | 115865985 | 3755 | -5.5 | 3.39E-08 | 6.56E-08 |
| chr13 | 109397586 | 109400040 | 2455 | -6.43 | 3.39E-08 | 6.56E-08 |
| chr3 | 151304452 | 151306118 | 1667 | -6.43 | 3.39E-08 | 6.56E-08 |
| chr9 | 99353348 | 99354374 | 1027 | -6.28 | 3.39E-08 | 6.56E-08 |

| | | | | | | |
|---|---|---|---|---|---|---|
| chr12 | 68065895 | 68068647 | 2753 | -6.61 | 3.39E-08 | 6.56E-08 |
| chr5 | 53384261 | 53386204 | 1944 | -6.47 | 3.39E-08 | 6.56E-08 |
| chr8 | 48327440 | 48329243 | 1804 | -6.47 | 3.39E-08 | 6.56E-08 |
| chr18 | 13063717 | 13064988 | 1272 | -6.32 | 3.40E-08 | 6.58E-08 |
| chr8 | 49573640 | 49580681 | 7042 | -5.18 | 3.40E-08 | 6.58E-08 |
| chr9 | 91163669 | 91164993 | 1325 | -6.41 | 3.41E-08 | 6.59E-08 |
| chr10 | 27396776 | 27398811 | 2036 | -6.66 | 3.41E-08 | 6.59E-08 |
| chr14 | 64794959 | 64808776 | 13818 | -7.22 | 3.41E-08 | 6.60E-08 |
| chr2 | 165736543 | 165738449 | 1907 | -6.5 | 3.42E-08 | 6.61E-08 |
| chr10 | 106271190 | 106344395 | 73206 | 2.64 | 3.43E-08 | 6.63E-08 |
| chr8 | 92068517 | 92069900 | 1384 | -6.4 | 3.43E-08 | 6.63E-08 |
| chr9 | 89425170 | 89426712 | 1543 | 5.11 | 3.44E-08 | 6.65E-08 |
| chr16 | 29951088 | 29962307 | 11220 | -6.88 | 3.44E-08 | 6.65E-08 |
| chr12 | 77152045 | 77153532 | 1488 | -6.27 | 3.44E-08 | 6.65E-08 |
| chr2 | 196607464 | 196608829 | 1366 | -6.27 | 3.44E-08 | 6.65E-08 |
| chr10 | 121623422 | 121624516 | 1095 | -6.29 | 3.45E-08 | 6.66E-08 |
| chr13 | 66455447 | 66457089 | 1643 | -6.38 | 3.45E-08 | 6.67E-08 |
| chr7 | 16120293 | 16123046 | 2754 | -6.52 | 3.45E-08 | 6.68E-08 |
| chr1 | 188161804 | 188169969 | 8166 | -6.64 | 3.45E-08 | 6.68E-08 |
| chr2 | 158136246 | 158138052 | 1807 | -6.31 | 3.46E-08 | 6.68E-08 |
| chr4 | 76163968 | 76164991 | 1024 | -4.66 | 3.46E-08 | 6.68E-08 |
| chr11 | 128560579 | 128561661 | 1083 | -6.27 | 3.46E-08 | 6.69E-08 |
| chr20 | 42870151 | 42871577 | 1427 | -6.29 | 3.46E-08 | 6.69E-08 |
| chr4 | 143242420 | 143243628 | 1209 | -6.3 | 3.47E-08 | 6.70E-08 |
| chr5 | 157273815 | 157274998 | 1184 | -6.3 | 3.47E-08 | 6.70E-08 |
| chr8 | 96292005 | 96294653 | 2649 | -4.08 | 3.48E-08 | 6.71E-08 |
| chr3 | 194349701 | 194352890 | 3190 | -6.3 | 3.48E-08 | 6.72E-08 |
| chr1 | 52295107 | 52296594 | 1488 | -6.32 | 3.48E-08 | 6.73E-08 |
| chr6 | 152770805 | 152772350 | 1546 | -6.32 | 3.48E-08 | 6.73E-08 |
| chr3 | 62225810 | 62228234 | 2425 | 6.59 | 3.48E-08 | 6.73E-08 |
| chr11 | 16204222 | 16205218 | 997 | -6.28 | 3.50E-08 | 6.75E-08 |
| chr3 | 18516595 | 18519360 | 2766 | 6.34 | 3.50E-08 | 6.75E-08 |
| chr12 | 28252040 | 28265572 | 13533 | -6.82 | 3.50E-08 | 6.76E-08 |
| chr3 | 105569174 | 105570097 | 924 | -6.27 | 3.50E-08 | 6.77E-08 |
| chr7 | 20131967 | 20135340 | 3374 | -6.55 | 3.51E-08 | 6.78E-08 |
| chr2 | 230535043 | 230587639 | 52597 | -6.16 | 3.51E-08 | 6.79E-08 |
| chr16 | 72521054 | 72522296 | 1243 | -6.3 | 3.52E-08 | 6.79E-08 |
| chr4 | 189691127 | 189693602 | 2476 | -6.3 | 3.52E-08 | 6.79E-08 |
| chr1 | 225978171 | 225979402 | 1232 | -6.28 | 3.52E-08 | 6.79E-08 |
| chr6 | 43174064 | 43175178 | 1115 | -6.28 | 3.52E-08 | 6.79E-08 |
| chrX | 20468463 | 20484202 | 15740 | -6.91 | 3.53E-08 | 6.82E-08 |
| chr1 | 48561500 | 48562819 | 1320 | -3.31 | 3.54E-08 | 6.83E-08 |
| chr6 | 36998145 | 36999175 | 1031 | -6.57 | 3.54E-08 | 6.83E-08 |
| chr13 | 73963996 | 73965596 | 1601 | -6.29 | 3.55E-08 | 6.85E-08 |
| chrX | 43580408 | 43587647 | 7240 | -4.8 | 3.55E-08 | 6.85E-08 |
| chrX | 9426553 | 9429614 | 3062 | -6.67 | 3.55E-08 | 6.86E-08 |
| chr21 | 45332407 | 45333759 | 1353 | -6.28 | 3.56E-08 | 6.87E-08 |
| chr18 | 53228934 | 53234763 | 5830 | -6.35 | 3.56E-08 | 6.88E-08 |
| chr2 | 204171169 | 204172549 | 1381 | -6.6 | 3.58E-08 | 6.90E-08 |
| chr9 | 72118675 | 72142059 | 23385 | -1.74 | 3.60E-08 | 6.95E-08 |
| chr6 | 22944032 | 22959343 | 15312 | -7.1 | 3.60E-08 | 6.96E-08 |
| chr6 | 157767782 | 157775760 | 7979 | -1.53 | 3.61E-08 | 6.97E-08 |
| chr14 | 89325265 | 89325866 | 602 | -6.3 | 3.61E-08 | 6.97E-08 |
| chr4 | 155323745 | 155325733 | 1989 | -6.3 | 3.61E-08 | 6.97E-08 |
| chr10 | 30061129 | 30090105 | 28977 | -7.35 | 3.63E-08 | 7.01E-08 |
| chr6 | 21409498 | 21411844 | 2347 | -6.46 | 3.64E-08 | 7.02E-08 |
| chrX | 102903164 | 102904459 | 1296 | -6.46 | 3.64E-08 | 7.02E-08 |
| chr4 | 167683323 | 167685349 | 2027 | -6.41 | 3.65E-08 | 7.03E-08 |
| chrX | 19209696 | 19210659 | 964 | -6.27 | 3.65E-08 | 7.04E-08 |
| chr4 | 66575465 | 66577699 | 2235 | -6.29 | 3.65E-08 | 7.04E-08 |
| chr18 | 43357890 | 43359435 | 1546 | -6.31 | 3.65E-08 | 7.04E-08 |

| | | | | | | |
|---|---|---|---|---|---|---|
| chr3 | 131779818 | 131782425 | 2608 | -6.27 | 3.65E-08 | 7.04E-08 |
| chr5 | 114627700 | 114629131 | 1432 | -6.27 | 3.65E-08 | 7.04E-08 |
| chr9 | 15195511 | 15196749 | 1239 | -6.27 | 3.65E-08 | 7.04E-08 |
| chr11 | 26625385 | 26626915 | 1531 | -6.3 | 3.66E-08 | 7.07E-08 |
| chr14 | 77862128 | 77863207 | 1080 | -6.3 | 3.66E-08 | 7.07E-08 |
| chr5 | 91465061 | 91466999 | 1939 | -6.3 | 3.66E-08 | 7.07E-08 |
| chr6 | 4167409 | 4168948 | 1540 | -6.43 | 3.67E-08 | 7.07E-08 |
| chr11 | 9101935 | 9103349 | 1415 | -4.73 | 3.67E-08 | 7.07E-08 |
| chr10 | 73830855 | 73832292 | 1438 | -6.27 | 3.67E-08 | 7.07E-08 |
| chr2 | 70286434 | 70287294 | 861 | -6.33 | 3.67E-08 | 7.07E-08 |
| chr13 | 42426951 | 42428499 | 1549 | -6.46 | 3.67E-08 | 7.08E-08 |
| chrX | 8971607 | 8972861 | 1255 | -6.41 | 3.67E-08 | 7.08E-08 |
| chr3 | 107810256 | 107811416 | 1161 | -6.39 | 3.67E-08 | 7.08E-08 |
| chr5 | 114658108 | 114659096 | 989 | -6.39 | 3.67E-08 | 7.08E-08 |
| chr12 | 67496775 | 67504260 | 7486 | -6.06 | 3.69E-08 | 7.12E-08 |
| chr4 | 66683657 | 66686537 | 2881 | -6.49 | 3.70E-08 | 7.13E-08 |
| chr6 | 117837366 | 117838735 | 1370 | -6.39 | 3.70E-08 | 7.13E-08 |
| chr15 | 77742884 | 77743886 | 1003 | -6.32 | 3.70E-08 | 7.14E-08 |
| chr7 | 45062807 | 45064466 | 1660 | -6.32 | 3.70E-08 | 7.14E-08 |
| chr10 | 27238124 | 27240336 | 2213 | -6.26 | 3.71E-08 | 7.14E-08 |
| chr4 | 63548214 | 63550409 | 2196 | -6.26 | 3.71E-08 | 7.15E-08 |
| chr6 | 107054037 | 107055525 | 1489 | -6.29 | 3.72E-08 | 7.16E-08 |
| chr20 | 33647990 | 33649317 | 1328 | -6.38 | 3.72E-08 | 7.17E-08 |
| chr10 | 122588974 | 122615842 | 26869 | -6.97 | 3.72E-08 | 7.17E-08 |
| chr4 | 168914938 | 168916477 | 1540 | -6.51 | 3.73E-08 | 7.18E-08 |
| chr5 | 79906341 | 79907839 | 1499 | -6.51 | 3.74E-08 | 7.20E-08 |
| chr3 | 182711043 | 182713116 | 2074 | -6.53 | 3.74E-08 | 7.20E-08 |
| chr14 | 78507238 | 80930754 | 2423517 | 3.73 | 3.74E-08 | 7.20E-08 |
| chr4 | 115134670 | 115136542 | 1873 | -6.31 | 3.74E-08 | 7.21E-08 |
| chr3 | 3171782 | 3173296 | 1515 | -6.76 | 3.74E-08 | 7.21E-08 |
| chr3 | 135665822 | 135680390 | 14569 | -3.53 | 3.76E-08 | 7.24E-08 |
| chr1 | 18577226 | 18578971 | 1746 | -3.51 | 3.76E-08 | 7.24E-08 |
| chr3 | 53665198 | 53667374 | 2177 | -5.23 | 3.76E-08 | 7.24E-08 |
| chr21 | 34845099 | 34846652 | 1554 | -6.26 | 3.76E-08 | 7.24E-08 |
| chr11 | 113206513 | 113208076 | 1564 | -6.53 | 3.76E-08 | 7.25E-08 |
| chr12 | 19632770 | 19634018 | 1249 | -6.35 | 3.77E-08 | 7.25E-08 |
| chr3 | 148539619 | 148541003 | 1385 | -6.35 | 3.77E-08 | 7.25E-08 |
| chr2 | 114455530 | 114457849 | 2320 | -6.29 | 3.77E-08 | 7.26E-08 |
| chr9 | 121752481 | 121760421 | 7941 | 3.76 | 3.77E-08 | 7.26E-08 |
| chr6 | 110741620 | 110803898 | 62279 | -3.84 | 3.78E-08 | 7.27E-08 |
| chr1 | 164722943 | 164733070 | 10128 | -3.42 | 3.78E-08 | 7.27E-08 |
| chrX | 103472010 | 103479794 | 7785 | -6.36 | 3.78E-08 | 7.28E-08 |
| chr1 | 111804073 | 111805899 | 1827 | -4.98 | 3.78E-08 | 7.28E-08 |
| chr11 | 109108074 | 109111171 | 3098 | -6.34 | 3.79E-08 | 7.30E-08 |
| chr20 | 34246810 | 34248706 | 1897 | -6.66 | 3.81E-08 | 7.32E-08 |
| chr3 | 38100932 | 38104739 | 3808 | 2.17 | 3.82E-08 | 7.34E-08 |
| chr4 | 1081796 | 1083526 | 1731 | -6.57 | 3.84E-08 | 7.39E-08 |
| chr10 | 89301153 | 89303786 | 2634 | -5.11 | 3.84E-08 | 7.39E-08 |
| chr14 | 96511274 | 96512695 | 1422 | -6.28 | 3.84E-08 | 7.40E-08 |
| chr11 | 128183428 | 128185408 | 1981 | -6.26 | 3.85E-08 | 7.40E-08 |
| chr9 | 33775743 | 33777481 | 1739 | -6.25 | 3.85E-08 | 7.41E-08 |
| chrX | 86752746 | 86754674 | 1929 | -6.25 | 3.85E-08 | 7.41E-08 |
| chr2 | 182193216 | 182195353 | 2138 | -6.58 | 3.86E-08 | 7.42E-08 |
| chr3 | 69266469 | 69268141 | 1673 | -6.58 | 3.86E-08 | 7.42E-08 |
| chr16 | 66499824 | 66500939 | 1116 | -6.3 | 3.86E-08 | 7.43E-08 |
| chr4 | 179803170 | 179805646 | 2477 | -6.3 | 3.86E-08 | 7.43E-08 |
| chr4 | 22081537 | 22093431 | 11895 | 5.5 | 3.86E-08 | 7.43E-08 |
| chr1 | 30973752 | 30975381 | 1630 | -6.34 | 3.87E-08 | 7.44E-08 |
| chr6 | 66394436 | 66397192 | 2757 | 3.91 | 3.87E-08 | 7.44E-08 |
| chr9 | 140973447 | 140990273 | 16827 | -2.43 | 3.88E-08 | 7.46E-08 |
| chrX | 129435184 | 129435927 | 744 | -6.26 | 3.88E-08 | 7.47E-08 |

| | | | | | | |
|---|---|---|---|---|---|---|
| chr1 | 173196125 | 173214884 | 18760 | -4.53 | 3.89E-08 | 7.47E-08 |
| chr12 | 111287572 | 111288892 | 1321 | -5.38 | 3.89E-08 | 7.48E-08 |
| chr13 | 110481656 | 110484366 | 2711 | -6.42 | 3.89E-08 | 7.49E-08 |
| chr19 | 13345663 | 13347394 | 1732 | -6.42 | 3.89E-08 | 7.49E-08 |
| chr1 | 15666272 | 15667299 | 1028 | -6.33 | 3.89E-08 | 7.49E-08 |
| chr13 | 28620608 | 28623481 | 2874 | -6.41 | 3.90E-08 | 7.49E-08 |
| chr2 | 109259039 | 109263541 | 4503 | -6.41 | 3.90E-08 | 7.49E-08 |
| chr1 | 162094342 | 162095638 | 1297 | -6.44 | 3.90E-08 | 7.49E-08 |
| chr6 | 48433583 | 48435348 | 1766 | -6.27 | 3.90E-08 | 7.49E-08 |
| chr7 | 152369630 | 152371256 | 1627 | -6.27 | 3.90E-08 | 7.49E-08 |
| chrX | 17207549 | 17257713 | 50165 | -1.14 | 3.90E-08 | 7.49E-08 |
| chr6 | 35760237 | 35761360 | 1124 | -6.45 | 3.90E-08 | 7.50E-08 |
| chr3 | 72502621 | 72504495 | 1875 | -6.25 | 3.90E-08 | 7.50E-08 |
| chr4 | 59860215 | 59863826 | 3612 | -6.25 | 3.90E-08 | 7.50E-08 |
| chr13 | 99026191 | 99027589 | 1399 | -6.27 | 3.91E-08 | 7.51E-08 |
| chr12 | 86243049 | 86245091 | 2043 | -6.94 | 3.91E-08 | 7.51E-08 |
| chr13 | 33849425 | 33851980 | 2556 | -5.61 | 3.91E-08 | 7.52E-08 |
| chr13 | 73732971 | 73735103 | 2133 | -6.46 | 3.91E-08 | 7.52E-08 |
| chr6 | 117033356 | 117035743 | 2388 | -6.46 | 3.91E-08 | 7.52E-08 |
| chr17 | 71499351 | 71500995 | 1645 | -6.39 | 3.92E-08 | 7.53E-08 |
| chr17 | 28202853 | 28203675 | 823 | -6.29 | 3.92E-08 | 7.54E-08 |
| chr10 | 88145680 | 88146784 | 1105 | -6.47 | 3.93E-08 | 7.54E-08 |
| chr17 | 38585146 | 38586619 | 1474 | -6.38 | 3.93E-08 | 7.55E-08 |
| chr17 | 80218835 | 80220467 | 1633 | -6.59 | 3.93E-08 | 7.56E-08 |
| chr12 | 19721948 | 19739323 | 17376 | -7.8 | 3.93E-08 | 7.56E-08 |
| chr8 | 105209767 | 105211662 | 1896 | 6.61 | 3.94E-08 | 7.56E-08 |
| chr6 | 128141606 | 128142707 | 1102 | -4.72 | 3.94E-08 | 7.57E-08 |
| chr18 | 3733083 | 3734385 | 1303 | -6.37 | 3.94E-08 | 7.57E-08 |
| chr3 | 36356802 | 36370189 | 13388 | -6.78 | 3.94E-08 | 7.57E-08 |
| chr11 | 93207670 | 93208946 | 1277 | -6.25 | 3.95E-08 | 7.58E-08 |
| chr17 | 67172400 | 67173271 | 872 | -6.25 | 3.95E-08 | 7.58E-08 |
| chr2 | 74059246 | 74060377 | 1132 | -6.32 | 3.96E-08 | 7.59E-08 |
| chr3 | 120306599 | 120310183 | 3585 | -5.7 | 3.96E-08 | 7.60E-08 |
| chr4 | 114367656 | 114368862 | 1207 | -6.37 | 3.96E-08 | 7.60E-08 |
| chr20 | 10402096 | 10404175 | 2080 | -6.72 | 3.96E-08 | 7.60E-08 |
| chr22 | 36108046 | 36109613 | 1568 | -6.36 | 3.97E-08 | 7.63E-08 |
| chr4 | 42241228 | 42248404 | 7177 | -6.71 | 3.98E-08 | 7.64E-08 |
| chr1 | 51084037 | 51086413 | 2377 | -6.31 | 3.98E-08 | 7.64E-08 |
| chr6 | 112668114 | 112672653 | 4540 | 2.45 | 3.98E-08 | 7.64E-08 |
| chr13 | 46312527 | 46314172 | 1646 | 6.36 | 3.99E-08 | 7.65E-08 |
| chr12 | 92252696 | 92260496 | 7801 | -6.29 | 3.99E-08 | 7.65E-08 |
| chr18 | 10799683 | 10800922 | 1240 | -6.29 | 3.99E-08 | 7.65E-08 |
| chr15 | 63184447 | 63186407 | 1961 | -6.35 | 3.99E-08 | 7.66E-08 |
| chr6 | 18403365 | 18418741 | 15377 | -6.42 | 4.00E-08 | 7.68E-08 |
| chr11 | 130835331 | 130941700 | 106370 | -3.5 | 4.00E-08 | 7.68E-08 |
| chr9 | 15465443 | 15466467 | 1025 | -6.41 | 4.00E-08 | 7.68E-08 |
| chr5 | 1324643 | 1326303 | 1661 | -6.44 | 4.01E-08 | 7.68E-08 |
| chr4 | 155434460 | 155436322 | 1863 | -4.83 | 4.01E-08 | 7.68E-08 |
| chr10 | 88061903 | 88062918 | 1016 | -6.26 | 4.01E-08 | 7.69E-08 |
| chr3 | 131379671 | 131381411 | 1741 | -6.26 | 4.01E-08 | 7.69E-08 |
| chr6 | 147329983 | 147331451 | 1469 | -6.26 | 4.01E-08 | 7.69E-08 |
| chr4 | 17699986 | 17701906 | 1921 | -2.31 | 4.01E-08 | 7.69E-08 |
| chr10 | 94150251 | 94160078 | 9828 | -6.25 | 4.01E-08 | 7.70E-08 |
| chr10 | 13936818 | 13952466 | 15649 | -7.17 | 4.02E-08 | 7.71E-08 |
| chr2 | 216111970 | 216114169 | 2200 | -6.31 | 4.03E-08 | 7.72E-08 |
| chr13 | 61206812 | 61208603 | 1792 | 6.37 | 4.03E-08 | 7.73E-08 |
| chr3 | 83530231 | 83609706 | 79476 | -1.13 | 4.03E-08 | 7.73E-08 |
| chr22 | 36631946 | 36632690 | 745 | -6.24 | 4.03E-08 | 7.73E-08 |
| chr9 | 101218979 | 101229502 | 10524 | -3.03 | 4.03E-08 | 7.73E-08 |
| chr13 | 31761623 | 31766574 | 4952 | -2.48 | 4.03E-08 | 7.73E-08 |
| chrX | 29138125 | 29139552 | 1428 | -6.34 | 4.04E-08 | 7.75E-08 |

| | | | | | | |
|---|---|---|---|---|---|---|
| chr9 | 21734892 | 21742505 | 7614 | -6.31 | 4.05E-08 | 7.76E-08 |
| chr19 | 11501116 | 11502440 | 1325 | -6.25 | 4.05E-08 | 7.77E-08 |
| chr6 | 129215132 | 129216189 | 1058 | -6.25 | 4.05E-08 | 7.77E-08 |
| chr12 | 118552251 | 118553712 | 1462 | -6.25 | 4.06E-08 | 7.79E-08 |
| chr4 | 56332203 | 56342218 | 10016 | -6.75 | 4.07E-08 | 7.80E-08 |
| chr12 | 16777910 | 16780946 | 3037 | 4.25 | 4.09E-08 | 7.83E-08 |
| chr18 | 69899451 | 69901694 | 2244 | -6.24 | 4.09E-08 | 7.83E-08 |
| chr6 | 17596813 | 17598328 | 1516 | -6.29 | 4.09E-08 | 7.83E-08 |
| chr7 | 113789199 | 113791533 | 2335 | 6.44 | 4.09E-08 | 7.84E-08 |
| chr12 | 42619760 | 42620827 | 1068 | -6.35 | 4.10E-08 | 7.86E-08 |
| chr20 | 49155806 | 49156915 | 1110 | -6.26 | 4.10E-08 | 7.86E-08 |
| chrX | 30354950 | 30356682 | 1733 | -6.26 | 4.10E-08 | 7.86E-08 |
| chr4 | 163615433 | 163616993 | 1561 | -6.3 | 4.11E-08 | 7.88E-08 |
| chr6 | 78457722 | 78534728 | 77007 | 1.6 | 4.12E-08 | 7.88E-08 |
| chr5 | 37741803 | 37743441 | 1639 | -6.35 | 4.12E-08 | 7.89E-08 |
| chr12 | 92175239 | 92177984 | 2746 | -6.25 | 4.12E-08 | 7.89E-08 |
| chr2 | 206008663 | 206009595 | 933 | -6.27 | 4.13E-08 | 7.92E-08 |
| chr4 | 173200400 | 173202909 | 2510 | -6.27 | 4.13E-08 | 7.92E-08 |
| chrX | 17106247 | 17112784 | 6538 | -5.41 | 4.14E-08 | 7.92E-08 |
| chr1 | 188054978 | 188056771 | 1794 | -6.52 | 4.14E-08 | 7.92E-08 |
| chr18 | 42286750 | 42288080 | 1331 | -6.34 | 4.14E-08 | 7.93E-08 |
| chr2 | 10843949 | 10845811 | 1863 | -6.92 | 4.14E-08 | 7.93E-08 |
| chr14 | 60996040 | 61017174 | 21135 | -2.9 | 4.14E-08 | 7.93E-08 |
| chr15 | 81574313 | 81576303 | 1991 | 6.36 | 4.14E-08 | 7.93E-08 |
| chr1 | 242063255 | 242064230 | 976 | -6.24 | 4.15E-08 | 7.95E-08 |
| chr18 | 43939138 | 43940424 | 1287 | -6.24 | 4.15E-08 | 7.95E-08 |
| chr4 | 93547234 | 93548480 | 1247 | -6.24 | 4.16E-08 | 7.96E-08 |
| chr6 | 109932504 | 109933462 | 959 | -6.24 | 4.16E-08 | 7.96E-08 |
| chr12 | 88941814 | 88943846 | 2033 | -6.25 | 4.16E-08 | 7.97E-08 |
| chr2 | 237867789 | 237870006 | 2218 | 6.42 | 4.18E-08 | 7.99E-08 |
| chr9 | 3616365 | 3620385 | 4021 | -3.18 | 4.18E-08 | 7.99E-08 |
| chr6 | 38853346 | 38862598 | 9253 | 3.53 | 4.18E-08 | 7.99E-08 |
| chr1 | 233054236 | 233055863 | 1628 | -6.42 | 4.18E-08 | 7.99E-08 |
| chr7 | 5380586 | 5383079 | 2494 | -6.78 | 4.18E-08 | 8.00E-08 |
| chr2 | 45706662 | 45707715 | 1054 | -6.24 | 4.18E-08 | 8.01E-08 |
| chr8 | 48042432 | 48045199 | 2768 | -5.72 | 4.20E-08 | 8.03E-08 |
| chr2 | 30673449 | 30674268 | 820 | -6.38 | 4.20E-08 | 8.03E-08 |
| chr10 | 115004046 | 115006794 | 2749 | 6.71 | 4.20E-08 | 8.04E-08 |
| chr4 | 106530062 | 106531126 | 1065 | -6.25 | 4.20E-08 | 8.04E-08 |
| chrX | 42017370 | 42021133 | 3764 | -1.69 | 4.21E-08 | 8.05E-08 |
| chr18 | 3783154 | 3786907 | 3754 | -6.38 | 4.21E-08 | 8.05E-08 |
| chr2 | 169460779 | 169463340 | 2562 | -6.59 | 4.21E-08 | 8.05E-08 |
| chr10 | 52057541 | 52059647 | 2107 | -6.32 | 4.21E-08 | 8.06E-08 |
| chr12 | 95065497 | 95066390 | 894 | -6.28 | 4.22E-08 | 8.06E-08 |
| chr18 | 20047712 | 20061818 | 14107 | -6.26 | 4.22E-08 | 8.07E-08 |
| chr11 | 126955127 | 126966599 | 11473 | -2.08 | 4.22E-08 | 8.07E-08 |
| chr4 | 166347932 | 166350551 | 2620 | -6.47 | 4.22E-08 | 8.08E-08 |
| chr5 | 37669519 | 37683988 | 14470 | -6.56 | 4.23E-08 | 8.09E-08 |
| chr10 | 88328354 | 88329652 | 1299 | -6.26 | 4.23E-08 | 8.09E-08 |
| chr14 | 88989620 | 88996767 | 7148 | -6.79 | 4.24E-08 | 8.10E-08 |
| chr16 | 3936116 | 3946167 | 10052 | -6.78 | 4.24E-08 | 8.11E-08 |
| chr9 | 15538062 | 15539571 | 1510 | -6.6 | 4.25E-08 | 8.12E-08 |
| chrX | 83640242 | 83705463 | 65222 | -1.14 | 4.25E-08 | 8.13E-08 |
| chr7 | 98683024 | 98698556 | 15533 | -6.49 | 4.26E-08 | 8.14E-08 |
| chr1 | 30788689 | 30789797 | 1109 | -6.35 | 4.27E-08 | 8.16E-08 |
| chr15 | 57398463 | 57399370 | 908 | -6.35 | 4.27E-08 | 8.16E-08 |
| chr2 | 101372424 | 101381613 | 9190 | -6.49 | 4.27E-08 | 8.16E-08 |
| chr5 | 78072507 | 78080746 | 8240 | -6.24 | 4.29E-08 | 8.19E-08 |
| chr2 | 235463639 | 235464788 | 1150 | -6.34 | 4.29E-08 | 8.20E-08 |
| chr6 | 125240651 | 125242305 | 1655 | -6.31 | 4.31E-08 | 8.23E-08 |
| chr1 | 20672574 | 20673639 | 1066 | -6.24 | 4.31E-08 | 8.23E-08 |

| | | | | | | |
|---|---|---|---|---|---|---|
| chr14 | 24341882 | 24359769 | 17888 | -2.74 | 4.31E-08 | 8.24E-08 |
| chr9 | 137773754 | 137775095 | 1342 | 5.43 | 4.31E-08 | 8.24E-08 |
| chr3 | 169682356 | 169683072 | 717 | -6.33 | 4.31E-08 | 8.24E-08 |
| chr13 | 30333744 | 30335613 | 1870 | -6.25 | 4.32E-08 | 8.25E-08 |
| chr19 | 55935882 | 55944296 | 8415 | -6.25 | 4.32E-08 | 8.25E-08 |
| chr16 | 56445462 | 56446143 | 682 | -6.24 | 4.33E-08 | 8.27E-08 |
| chr4 | 56801671 | 56806900 | 5230 | -1.69 | 4.33E-08 | 8.28E-08 |
| chr2 | 170161139 | 170162385 | 1247 | -6.29 | 4.33E-08 | 8.28E-08 |
| chr8 | 53897273 | 53906130 | 8858 | -2.3 | 4.35E-08 | 8.30E-08 |
| chr7 | 26304709 | 26307199 | 2491 | -6.63 | 4.35E-08 | 8.30E-08 |
| chr18 | 53307646 | 53310309 | 2664 | -4.82 | 4.35E-08 | 8.31E-08 |
| chr10 | 115732188 | 115735158 | 2971 | -5.32 | 4.36E-08 | 8.32E-08 |
| chr6 | 53028519 | 53029748 | 1230 | -6.32 | 4.38E-08 | 8.36E-08 |
| chr6 | 150726211 | 150727908 | 1698 | -2.63 | 4.38E-08 | 8.36E-08 |
| chr12 | 79307270 | 79309023 | 1754 | -6.23 | 4.38E-08 | 8.37E-08 |
| chr13 | 30718008 | 30718896 | 889 | -6.27 | 4.38E-08 | 8.37E-08 |
| chr15 | 32273873 | 32314818 | 40946 | 1.2 | 4.39E-08 | 8.38E-08 |
| chr8 | 115498908 | 115512424 | 13517 | -6.9 | 4.39E-08 | 8.38E-08 |
| chr11 | 70461093 | 70462607 | 1515 | -6.28 | 4.39E-08 | 8.39E-08 |
| chrX | 39296045 | 39297312 | 1268 | -2.93 | 4.40E-08 | 8.39E-08 |
| chr5 | 107870074 | 107876952 | 6879 | 3.35 | 4.40E-08 | 8.39E-08 |
| chr5 | 43695264 | 43697083 | 1820 | -6.31 | 4.41E-08 | 8.42E-08 |
| chr8 | 52726245 | 52728642 | 2398 | -6.31 | 4.41E-08 | 8.42E-08 |
| chr11 | 114231552 | 114232809 | 1258 | -6.25 | 4.42E-08 | 8.43E-08 |
| chr21 | 43514405 | 43515714 | 1310 | -6.25 | 4.42E-08 | 8.43E-08 |
| chr9 | 2863710 | 2864322 | 613 | -6.25 | 4.42E-08 | 8.43E-08 |
| chr7 | 23555790 | 23558577 | 2788 | -6.36 | 4.42E-08 | 8.43E-08 |
| chr22 | 26035296 | 26036333 | 1038 | -6.29 | 4.43E-08 | 8.45E-08 |
| chr6 | 161716159 | 161718139 | 1981 | -6.48 | 4.44E-08 | 8.48E-08 |
| chr13 | 67385189 | 67387319 | 2131 | -6.34 | 4.46E-08 | 8.51E-08 |
| chr10 | 91081935 | 91084718 | 2784 | -6.27 | 4.46E-08 | 8.51E-08 |
| chr5 | 75459922 | 75462164 | 2243 | -3.04 | 4.46E-08 | 8.51E-08 |
| chr8 | 116112948 | 116120296 | 7349 | -6.24 | 4.48E-08 | 8.54E-08 |
| chr20 | 1006958 | 1014717 | 7760 | -2.21 | 4.48E-08 | 8.54E-08 |
| chr10 | 119253195 | 119267429 | 14235 | -7.12 | 4.48E-08 | 8.54E-08 |
| chr1 | 96847884 | 96849130 | 1247 | -5.04 | 4.48E-08 | 8.55E-08 |
| chr2 | 182948545 | 182949952 | 1408 | -6.4 | 4.48E-08 | 8.55E-08 |
| chr13 | 89939203 | 89942279 | 3077 | -6.41 | 4.48E-08 | 8.55E-08 |
| chr4 | 114570115 | 114580536 | 10422 | -6.41 | 4.48E-08 | 8.55E-08 |
| chr6 | 158009993 | 158012488 | 2496 | -6.41 | 4.48E-08 | 8.55E-08 |
| chr16 | 27342438 | 27343444 | 1007 | -6.5 | 4.49E-08 | 8.56E-08 |
| chr21 | 43472669 | 43473937 | 1269 | -6.38 | 4.49E-08 | 8.56E-08 |
| chr1 | 67228034 | 67229266 | 1233 | -6.23 | 4.51E-08 | 8.59E-08 |
| chr6 | 155417657 | 155418896 | 1240 | -6.23 | 4.51E-08 | 8.59E-08 |
| chr6 | 25541178 | 25542034 | 857 | -6.23 | 4.51E-08 | 8.60E-08 |
| chr13 | 35409085 | 35411126 | 2042 | -6.23 | 4.51E-08 | 8.60E-08 |
| chr4 | 145021072 | 145037011 | 15940 | 1.85 | 4.51E-08 | 8.60E-08 |
| chr20 | 9083927 | 9085308 | 1382 | -6.51 | 4.51E-08 | 8.60E-08 |
| chr2 | 25899152 | 25900230 | 1079 | -6.25 | 4.51E-08 | 8.61E-08 |
| chr12 | 100284631 | 100286126 | 1496 | -6.33 | 4.52E-08 | 8.61E-08 |
| chr17 | 2407281 | 2408026 | 746 | -6.33 | 4.52E-08 | 8.61E-08 |
| chr2 | 63845754 | 63893100 | 47347 | -1.72 | 4.52E-08 | 8.62E-08 |
| chrX | 101317056 | 101319402 | 2347 | -6.28 | 4.52E-08 | 8.62E-08 |
| chr7 | 132547907 | 132549367 | 1461 | -6.24 | 4.53E-08 | 8.63E-08 |
| chr2 | 110453679 | 110454994 | 1316 | -6.29 | 4.55E-08 | 8.66E-08 |
| chr2 | 58853668 | 58855926 | 2259 | -6.22 | 4.56E-08 | 8.69E-08 |
| chr15 | 99079379 | 99084517 | 5139 | -1.73 | 4.57E-08 | 8.71E-08 |
| chr11 | 74516304 | 74517507 | 1204 | -6.27 | 4.58E-08 | 8.72E-08 |
| chr11 | 86736087 | 86738172 | 2086 | -6.27 | 4.58E-08 | 8.72E-08 |
| chr2 | 135309844 | 135415421 | 105578 | -3.17 | 4.58E-08 | 8.73E-08 |
| chr20 | 31963881 | 31966351 | 2471 | -5.37 | 4.58E-08 | 8.73E-08 |

| | | | | | | |
|---|---|---|---|---|---|---|
| chr1 | 232091551 | 232093422 | 1872 | -2.46 | 4.58E-08 | 8.73E-08 |
| chr12 | 66168093 | 66186897 | 18805 | -7.16 | 4.59E-08 | 8.74E-08 |
| chr12 | 42119342 | 42121298 | 1957 | -6.25 | 4.59E-08 | 8.75E-08 |
| chr17 | 25677960 | 25679864 | 1905 | -6.31 | 4.59E-08 | 8.75E-08 |
| chrX | 55477500 | 55480084 | 2585 | -6.49 | 4.59E-08 | 8.75E-08 |
| chr18 | 55408189 | 55409217 | 1029 | -6.28 | 4.60E-08 | 8.76E-08 |
| chr2 | 68629967 | 68632642 | 2676 | -1.87 | 4.60E-08 | 8.77E-08 |
| chr4 | 88444651 | 88446361 | 1711 | -6.26 | 4.61E-08 | 8.78E-08 |
| chr9 | 129557743 | 129559087 | 1345 | -6.26 | 4.61E-08 | 8.78E-08 |
| chr4 | 183345203 | 183362815 | 17613 | 2.85 | 4.62E-08 | 8.79E-08 |
| chr13 | 95433636 | 95435471 | 1836 | -6.23 | 4.62E-08 | 8.80E-08 |
| chr3 | 96679338 | 96682093 | 2756 | -6.22 | 4.64E-08 | 8.83E-08 |
| chr9 | 476237 | 478179 | 1943 | -6.51 | 4.64E-08 | 8.84E-08 |
| chr4 | 64488581 | 64491816 | 3236 | -6.23 | 4.65E-08 | 8.85E-08 |
| chrX | 76569865 | 76571146 | 1282 | -5.02 | 4.66E-08 | 8.87E-08 |
| chr12 | 45955863 | 45956645 | 783 | -6.31 | 4.67E-08 | 8.88E-08 |
| chrX | 44625530 | 44627301 | 1772 | -6.22 | 4.67E-08 | 8.88E-08 |
| chr10 | 120777810 | 120779243 | 1434 | -6.22 | 4.67E-08 | 8.88E-08 |
| chr12 | 83585420 | 83586854 | 1435 | -6.23 | 4.67E-08 | 8.89E-08 |
| chr8 | 103535506 | 103536673 | 1168 | -6.24 | 4.67E-08 | 8.89E-08 |
| chr1 | 221835621 | 221837131 | 1511 | -6.52 | 4.68E-08 | 8.90E-08 |
| chr2 | 120531208 | 120532588 | 1381 | -6.52 | 4.68E-08 | 8.90E-08 |
| chr3 | 125881957 | 125885209 | 3253 | -2.67 | 4.68E-08 | 8.91E-08 |
| chr13 | 66280925 | 66286141 | 5217 | 7.06 | 4.69E-08 | 8.92E-08 |
| chr11 | 21934716 | 21938401 | 3686 | -2.89 | 4.70E-08 | 8.94E-08 |
| chr4 | 41402468 | 41405045 | 2578 | -6.26 | 4.70E-08 | 8.94E-08 |
| chr5 | 36128719 | 36130082 | 1364 | -6.31 | 4.70E-08 | 8.95E-08 |
| chr4 | 159259799 | 159261665 | 1867 | -6.27 | 4.73E-08 | 8.99E-08 |
| chr3 | 107368382 | 107369509 | 1128 | -6.22 | 4.75E-08 | 9.03E-08 |
| chr6 | 110680977 | 110694106 | 13130 | -2.62 | 4.75E-08 | 9.04E-08 |
| chr17 | 33486726 | 33488566 | 1841 | -6.55 | 4.78E-08 | 9.08E-08 |
| chr4 | 100610618 | 100612640 | 2023 | -6.22 | 4.78E-08 | 9.09E-08 |
| chr12 | 118027843 | 118029198 | 1356 | -6.22 | 4.78E-08 | 9.09E-08 |
| chr18 | 29660513 | 29662059 | 1547 | -6.39 | 4.78E-08 | 9.09E-08 |
| chr3 | 149281494 | 149287383 | 5890 | -6.4 | 4.78E-08 | 9.10E-08 |
| chr5 | 65495590 | 65498047 | 2458 | -6.4 | 4.78E-08 | 9.10E-08 |
| chr11 | 36114753 | 36116030 | 1278 | -6.38 | 4.78E-08 | 9.10E-08 |
| chr4 | 140414736 | 140416227 | 1492 | -6.24 | 4.79E-08 | 9.10E-08 |
| chr16 | 24641580 | 24643512 | 1933 | -6.6 | 4.79E-08 | 9.11E-08 |
| chr2 | 242059987 | 242060697 | 711 | -6.42 | 4.79E-08 | 9.12E-08 |
| chr2 | 227935707 | 227966442 | 30736 | -7.54 | 4.80E-08 | 9.12E-08 |
| chr15 | 80437489 | 80438890 | 1402 | -6.26 | 4.80E-08 | 9.12E-08 |
| chr2 | 152176005 | 152178125 | 2121 | -3.75 | 4.82E-08 | 9.17E-08 |
| chr12 | 62731006 | 62733091 | 2086 | -6.36 | 4.83E-08 | 9.18E-08 |
| chr4 | 180006525 | 180009082 | 2558 | -6.36 | 4.83E-08 | 9.18E-08 |
| chr9 | 107524952 | 107525938 | 987 | -6.21 | 4.84E-08 | 9.20E-08 |
| chr14 | 91696807 | 91698071 | 1265 | -6.34 | 4.84E-08 | 9.20E-08 |
| chr3 | 117295009 | 117296192 | 1184 | -6.34 | 4.85E-08 | 9.23E-08 |
| chr3 | 120625622 | 120626983 | 1362 | -5.35 | 4.87E-08 | 9.25E-08 |
| chr1 | 21457399 | 21458704 | 1306 | -6.45 | 4.88E-08 | 9.27E-08 |
| chr4 | 79451160 | 79452053 | 894 | -6.23 | 4.88E-08 | 9.27E-08 |
| chr4 | 64446241 | 64449862 | 3622 | -6.21 | 4.88E-08 | 9.27E-08 |
| chr8 | 101458348 | 101459577 | 1230 | -6.33 | 4.89E-08 | 9.28E-08 |
| chr6 | 152018416 | 152020398 | 1983 | -3.52 | 4.90E-08 | 9.30E-08 |
| chr10 | 62768198 | 62769761 | 1564 | -5.44 | 4.90E-08 | 9.32E-08 |
| chr22 | 48867082 | 48868614 | 1533 | -6.47 | 4.92E-08 | 9.34E-08 |
| chr16 | 4330710 | 4332036 | 1327 | -6.22 | 4.92E-08 | 9.35E-08 |
| chr8 | 141391514 | 141392700 | 1187 | -6.22 | 4.92E-08 | 9.35E-08 |
| chr8 | 61061403 | 61065021 | 3619 | -2.08 | 4.92E-08 | 9.35E-08 |
| chr1 | 223172561 | 223174175 | 1615 | -6.21 | 4.93E-08 | 9.36E-08 |
| chr21 | 43801755 | 43812056 | 10302 | -6.87 | 4.94E-08 | 9.39E-08 |

| | | | | | | |
|---|---|---|---|---|---|---|
| chr3 | 87481659 | 87497170 | 15512 | -6.48 | 4.95E-08 | 9.40E-08 |
| chr18 | 46167990 | 46169177 | 1188 | -6.31 | 4.95E-08 | 9.40E-08 |
| chr8 | 124806625 | 124807596 | 972 | -6.5 | 4.95E-08 | 9.41E-08 |
| chr3 | 170495610 | 170517091 | 21482 | -5.67 | 4.96E-08 | 9.42E-08 |
| chr10 | 94473254 | 94474724 | 1471 | -6.7 | 4.96E-08 | 9.42E-08 |
| chr12 | 108747322 | 108749081 | 1760 | -6.51 | 4.97E-08 | 9.43E-08 |
| chr4 | 189523972 | 189526298 | 2327 | -6.31 | 4.98E-08 | 9.46E-08 |
| chr2 | 3511660 | 3512640 | 981 | -6.26 | 4.98E-08 | 9.46E-08 |
| chr5 | 92775566 | 92778048 | 2483 | -3.33 | 4.99E-08 | 9.47E-08 |
| chr9 | 30030955 | 30034631 | 3677 | -6.23 | 4.99E-08 | 9.47E-08 |
| chrX | 48683657 | 48684718 | 1062 | -6.23 | 4.99E-08 | 9.47E-08 |
| chr2 | 134340853 | 134343097 | 2245 | 6.29 | 5.00E-08 | 9.49E-08 |
| chr17 | 75559051 | 75564952 | 5902 | -6.8 | 5.01E-08 | 9.50E-08 |
| chr11 | 13849564 | 13850871 | 1308 | -6.22 | 5.01E-08 | 9.51E-08 |
| chr14 | 54819307 | 54822127 | 2821 | -6.22 | 5.01E-08 | 9.51E-08 |
| chr1 | 20542747 | 20552856 | 10110 | -6.26 | 5.01E-08 | 9.51E-08 |
| chr11 | 114276890 | 114277979 | 1090 | -6.26 | 5.01E-08 | 9.51E-08 |
| chr4 | 55040893 | 55042268 | 1376 | -6.26 | 5.01E-08 | 9.51E-08 |
| chr3 | 133328464 | 133329601 | 1138 | -6.52 | 5.01E-08 | 9.51E-08 |
| chr2 | 170239429 | 170240758 | 1330 | -5.44 | 5.01E-08 | 9.52E-08 |
| chrX | 105991324 | 105992143 | 820 | -4.72 | 5.02E-08 | 9.52E-08 |
| chr8 | 16475750 | 16480899 | 5150 | -1.85 | 5.02E-08 | 9.53E-08 |
| chr14 | 81971083 | 81972568 | 1486 | -6.21 | 5.02E-08 | 9.53E-08 |
| chr5 | 86696585 | 86697509 | 925 | -6.53 | 5.03E-08 | 9.54E-08 |
| chr1 | 207367848 | 207369642 | 1795 | -3.57 | 5.03E-08 | 9.55E-08 |
| chr2 | 102060108 | 102061785 | 1678 | -6.21 | 5.04E-08 | 9.55E-08 |
| chrX | 11063020 | 11064779 | 1760 | -6.25 | 5.04E-08 | 9.57E-08 |
| chrX | 77010564 | 77012580 | 2017 | -6.24 | 5.06E-08 | 9.59E-08 |
| chr5 | 171426844 | 171429060 | 2217 | -6.52 | 5.06E-08 | 9.60E-08 |
| chr3 | 71235474 | 71236810 | 1337 | -6.23 | 5.08E-08 | 9.63E-08 |
| chr10 | 21125961 | 21127772 | 1812 | -6.54 | 5.08E-08 | 9.63E-08 |
| chr5 | 41682714 | 41698648 | 15935 | 2.54 | 5.09E-08 | 9.66E-08 |
| chr19 | 6352231 | 6353152 | 922 | -6.29 | 5.10E-08 | 9.67E-08 |
| chr5 | 99852255 | 99869459 | 17205 | -6.01 | 5.10E-08 | 9.68E-08 |
| chr2 | 219805268 | 219807440 | 2173 | -4.03 | 5.10E-08 | 9.68E-08 |
| chr18 | 11875290 | 11876725 | 1436 | -6.28 | 5.12E-08 | 9.70E-08 |
| chr17 | 29523248 | 29524384 | 1137 | -6.22 | 5.12E-08 | 9.71E-08 |
| chr6 | 158145509 | 158146546 | 1038 | -4.53 | 5.14E-08 | 9.75E-08 |
| chr5 | 38390146 | 38411566 | 21421 | -2.24 | 5.14E-08 | 9.75E-08 |
| chr6 | 16195225 | 16197750 | 2526 | -6.21 | 5.14E-08 | 9.75E-08 |
| chr10 | 21253167 | 21254448 | 1282 | -6.28 | 5.15E-08 | 9.75E-08 |
| chr18 | 74407280 | 74408219 | 940 | -6.28 | 5.15E-08 | 9.75E-08 |
| chr2 | 210405990 | 210407419 | 1430 | -6.28 | 5.15E-08 | 9.75E-08 |
| chr4 | 124610508 | 124612187 | 1680 | -6.28 | 5.15E-08 | 9.75E-08 |
| chr4 | 137873363 | 137875872 | 2510 | -6.23 | 5.15E-08 | 9.77E-08 |
| chr9 | 21695117 | 21697096 | 1980 | -6.21 | 5.16E-08 | 9.77E-08 |
| chr2 | 121801025 | 121813014 | 11990 | -7.02 | 5.16E-08 | 9.79E-08 |
| chr5 | 143226775 | 143228121 | 1347 | -6.37 | 5.17E-08 | 9.80E-08 |
| chr12 | 86573688 | 86575407 | 1720 | -6.21 | 5.18E-08 | 9.82E-08 |
| chr13 | 74221050 | 74232044 | 10995 | -6.24 | 5.18E-08 | 9.82E-08 |
| chr13 | 99384282 | 99386341 | 2060 | -6.41 | 5.19E-08 | 9.83E-08 |
| chr3 | 55058248 | 55108700 | 50453 | -1 | 5.19E-08 | 9.84E-08 |
| chr7 | 134659000 | 134660946 | 1947 | -4.64 | 5.20E-08 | 9.84E-08 |
| chr15 | 81524875 | 81544021 | 19147 | -1.48 | 5.20E-08 | 9.86E-08 |
| chrX | 75661760 | 75664000 | 2241 | -4.83 | 5.20E-08 | 9.86E-08 |
| chr1 | 169082060 | 169084578 | 2519 | -6.42 | 5.20E-08 | 9.86E-08 |
| chr20 | 46544044 | 46567856 | 23813 | -1.93 | 5.21E-08 | 9.87E-08 |
| chrX | 39688032 | 39689510 | 1479 | -6.34 | 5.21E-08 | 9.88E-08 |
| chrX | 55190603 | 55192358 | 1756 | -6.34 | 5.21E-08 | 9.88E-08 |
| chr1 | 147300330 | 147302472 | 2143 | -3.54 | 5.21E-08 | 9.88E-08 |
| chr1 | 37647659 | 37697707 | 50049 | -2.55 | 5.23E-08 | 9.90E-08 |

| | | | | | | |
|---|---|---|---|---|---|---|
| chr2 | 142973910 | 142974875 | 966 | -6.39 | 5.23E-08 | 9.91E-08 |
| chrX | 123317403 | 123318755 | 1353 | -6.4 | 5.24E-08 | 9.92E-08 |
| chrX | 44108792 | 44111596 | 2805 | -6.76 | 5.24E-08 | 9.93E-08 |
| chr8 | 28406480 | 28407913 | 1434 | -6.32 | 5.25E-08 | 9.94E-08 |
| chr5 | 73897067 | 73899144 | 2078 | -6.24 | 5.25E-08 | 9.94E-08 |
| chr10 | 12759650 | 12761386 | 1737 | -6.2 | 5.25E-08 | 9.94E-08 |
| chr22 | 30321942 | 30322800 | 859 | -6.35 | 5.26E-08 | 9.95E-08 |
| chr5 | 16723580 | 16725118 | 1539 | -6.22 | 5.27E-08 | 9.97E-08 |
| chr6 | 54483564 | 54486248 | 2685 | -6.42 | 5.27E-08 | 9.97E-08 |
| chr16 | 3718195 | 3719951 | 1757 | -6.68 | 5.27E-08 | 9.97E-08 |
| chr3 | 105446215 | 105447445 | 1231 | -6.32 | 5.27E-08 | 9.98E-08 |
| chr1 | 5775438 | 5776363 | 926 | -4.71 | 5.29E-08 | 1.00E-07 |
| chr17 | 42796133 | 42797227 | 1095 | -6.44 | 5.30E-08 | 1.00E-07 |
| chr17 | 14114196 | 14115732 | 1537 | -6.2 | 5.31E-08 | 1.00E-07 |
| chr9 | 96247826 | 96249547 | 1722 | -6.59 | 5.31E-08 | 1.01E-07 |
| chr17 | 9706982 | 9709249 | 2268 | -6.3 | 5.32E-08 | 1.01E-07 |
| chr17 | 20048993 | 20050080 | 1088 | -6.31 | 5.34E-08 | 1.01E-07 |
| chr1 | 115734018 | 115735719 | 1702 | -6.23 | 5.34E-08 | 1.01E-07 |
| chr14 | 52578133 | 52580886 | 2754 | -6.23 | 5.34E-08 | 1.01E-07 |
| chr6 | 15105075 | 15106646 | 1572 | -6.29 | 5.36E-08 | 1.01E-07 |
| chr16 | 11427858 | 11429192 | 1335 | -6.6 | 5.36E-08 | 1.01E-07 |
| chr2 | 182829941 | 182831564 | 1624 | -6.25 | 5.36E-08 | 1.01E-07 |
| chr3 | 94683702 | 94688079 | 4378 | -6.25 | 5.36E-08 | 1.01E-07 |
| chrX | 153737527 | 153738573 | 1047 | -6.25 | 5.36E-08 | 1.01E-07 |
| chr14 | 103772028 | 103773903 | 1876 | -6.7 | 5.37E-08 | 1.02E-07 |
| chr11 | 69392374 | 69399799 | 7426 | -6.48 | 5.38E-08 | 1.02E-07 |
| chr4 | 90663218 | 90664494 | 1277 | -6.21 | 5.39E-08 | 1.02E-07 |
| chr7 | 69249653 | 69252029 | 2377 | -5.48 | 5.39E-08 | 1.02E-07 |
| chr1 | 86220360 | 86222241 | 1882 | -6.29 | 5.39E-08 | 1.02E-07 |
| chr18 | 60841733 | 60842635 | 903 | -4.89 | 5.40E-08 | 1.02E-07 |
| chr10 | 127992012 | 127998139 | 6128 | -4.99 | 5.42E-08 | 1.03E-07 |
| chr17 | 57706835 | 57708263 | 1429 | -6.22 | 5.42E-08 | 1.03E-07 |
| chr1 | 87719122 | 87726788 | 7667 | -1.44 | 5.42E-08 | 1.03E-07 |
| chr1 | 95214379 | 95215477 | 1099 | -6.19 | 5.43E-08 | 1.03E-07 |
| chr14 | 61470437 | 61471286 | 850 | -6.2 | 5.43E-08 | 1.03E-07 |
| chr9 | 3390080 | 3391627 | 1548 | -6.19 | 5.44E-08 | 1.03E-07 |
| chr13 | 44112950 | 44113955 | 1006 | -6.22 | 5.45E-08 | 1.03E-07 |
| chr5 | 140899732 | 140900754 | 1023 | -6.22 | 5.45E-08 | 1.03E-07 |
| chr4 | 156647858 | 156655669 | 7812 | 6.41 | 5.45E-08 | 1.03E-07 |
| chrX | 128467335 | 128468352 | 1018 | -6.24 | 5.47E-08 | 1.03E-07 |
| chr6 | 158159048 | 158175768 | 16721 | -7.01 | 5.48E-08 | 1.04E-07 |
| chr11 | 123268352 | 123269882 | 1531 | -6.2 | 5.48E-08 | 1.04E-07 |
| chr7 | 47042059 | 47043207 | 1149 | -6.2 | 5.48E-08 | 1.04E-07 |
| chr5 | 149829738 | 149830461 | 724 | -6.2 | 5.50E-08 | 1.04E-07 |
| chr8 | 62478266 | 62479848 | 1583 | -6.19 | 5.51E-08 | 1.04E-07 |
| chr8 | 42343781 | 42345466 | 1686 | -6.21 | 5.52E-08 | 1.04E-07 |
| chr8 | 18630588 | 18640614 | 10027 | -6.92 | 5.52E-08 | 1.04E-07 |
| chr2 | 170650374 | 170652295 | 1922 | -6.26 | 5.54E-08 | 1.05E-07 |
| chr6 | 164393556 | 164401736 | 8181 | -1.98 | 5.54E-08 | 1.05E-07 |
| chr2 | 157986229 | 157989660 | 3432 | -5.06 | 5.54E-08 | 1.05E-07 |
| chr7 | 117341196 | 117345935 | 4740 | -6.64 | 5.54E-08 | 1.05E-07 |
| chr4 | 28724010 | 28911930 | 187921 | 1.78 | 5.55E-08 | 1.05E-07 |
| chr2 | 187137872 | 187139471 | 1600 | -6.22 | 5.55E-08 | 1.05E-07 |
| chr4 | 148035929 | 148037093 | 1165 | -6.37 | 5.56E-08 | 1.05E-07 |
| chr3 | 61881613 | 61883246 | 1634 | -6.39 | 5.57E-08 | 1.05E-07 |
| chr3 | 113584265 | 113587151 | 2887 | -6.39 | 5.57E-08 | 1.05E-07 |
| chrX | 27909109 | 27910233 | 1125 | -6.19 | 5.57E-08 | 1.05E-07 |
| chr5 | 55960579 | 55963104 | 2526 | -6.55 | 5.57E-08 | 1.05E-07 |
| chr1 | 53885682 | 53888653 | 2972 | -4.25 | 5.57E-08 | 1.05E-07 |
| chr1 | 195969296 | 195973327 | 4032 | 2.99 | 5.58E-08 | 1.05E-07 |
| chr2 | 192409371 | 192410573 | 1203 | -6.4 | 5.58E-08 | 1.05E-07 |

| | | | | | | |
|---|---|---|---|---|---|---|
| chr10 | 101361429 | 101364860 | 3432 | -6.41 | 5.59E-08 | 1.06E-07 |
| chr6 | 144026865 | 144030922 | 4058 | -6.41 | 5.59E-08 | 1.06E-07 |
| chr6 | 84734541 | 84735574 | 1034 | -6.19 | 5.60E-08 | 1.06E-07 |
| chrX | 24351800 | 24354661 | 2862 | -1.97 | 5.60E-08 | 1.06E-07 |
| chr20 | 39634892 | 39636170 | 1279 | -6.21 | 5.62E-08 | 1.06E-07 |
| chr20 | 25049266 | 25050201 | 936 | -6.2 | 5.62E-08 | 1.06E-07 |
| chr7 | 133005828 | 133006569 | 742 | -6.2 | 5.62E-08 | 1.06E-07 |
| chr3 | 24736937 | 24748426 | 11490 | -6.85 | 5.63E-08 | 1.06E-07 |
| chr4 | 113337164 | 113338909 | 1746 | -6.22 | 5.64E-08 | 1.06E-07 |
| chr6 | 159117261 | 159118579 | 1319 | -6.31 | 5.65E-08 | 1.07E-07 |
| chr4 | 143315623 | 143317939 | 2317 | -6.25 | 5.66E-08 | 1.07E-07 |
| chr7 | 101275852 | 101276851 | 1000 | -6.19 | 5.67E-08 | 1.07E-07 |
| chr6 | 5250970 | 5252458 | 1489 | -6.21 | 5.68E-08 | 1.07E-07 |
| chr11 | 14472736 | 14474046 | 1311 | -6.25 | 5.68E-08 | 1.07E-07 |
| chr7 | 130369100 | 130406185 | 37086 | -2.55 | 5.68E-08 | 1.07E-07 |
| chr14 | 30306147 | 30325769 | 19623 | -1.83 | 5.68E-08 | 1.07E-07 |
| chr17 | 53385356 | 53387179 | 1824 | -6.86 | 5.69E-08 | 1.07E-07 |
| chr18 | 61522024 | 61523090 | 1067 | -4.83 | 5.70E-08 | 1.07E-07 |
| chr12 | 28035896 | 28037121 | 1226 | -6.3 | 5.70E-08 | 1.07E-07 |
| chr2 | 39839359 | 39844692 | 5334 | -1.85 | 5.71E-08 | 1.08E-07 |
| chr20 | 46966253 | 46995669 | 29417 | -7.49 | 5.71E-08 | 1.08E-07 |
| chr7 | 35536356 | 35537432 | 1077 | -5.19 | 5.72E-08 | 1.08E-07 |
| chr3 | 82119676 | 82121460 | 1785 | 6.34 | 5.72E-08 | 1.08E-07 |
| chr2 | 110315083 | 110317281 | 2199 | -6.19 | 5.73E-08 | 1.08E-07 |
| chr22 | 44479976 | 44481264 | 1289 | -6.19 | 5.73E-08 | 1.08E-07 |
| chr4 | 117870193 | 117878257 | 8065 | -6.24 | 5.73E-08 | 1.08E-07 |
| chr2 | 42408379 | 42409233 | 855 | -6.25 | 5.73E-08 | 1.08E-07 |
| chr19 | 43917335 | 43923749 | 6415 | -6.87 | 5.74E-08 | 1.08E-07 |
| chr4 | 177319751 | 177321929 | 2179 | -6.34 | 5.74E-08 | 1.08E-07 |
| chr10 | 126585838 | 126587492 | 1655 | -6.59 | 5.75E-08 | 1.08E-07 |
| chr4 | 69421072 | 69422658 | 1587 | -6.22 | 5.76E-08 | 1.09E-07 |
| chr1 | 147331745 | 147334136 | 2392 | -2.67 | 5.77E-08 | 1.09E-07 |
| chr15 | 70420544 | 70421463 | 920 | -6.18 | 5.78E-08 | 1.09E-07 |
| chr6 | 158546670 | 158547694 | 1025 | -6.18 | 5.78E-08 | 1.09E-07 |
| chr1 | 187579194 | 187580356 | 1163 | -3.81 | 5.81E-08 | 1.09E-07 |
| chr4 | 7319800 | 7321552 | 1753 | -6.31 | 5.82E-08 | 1.10E-07 |
| chr8 | 114457509 | 114459508 | 2000 | 6.7 | 5.83E-08 | 1.10E-07 |
| chr4 | 30996131 | 30998190 | 2060 | -6.19 | 5.83E-08 | 1.10E-07 |
| chr14 | 96065483 | 96117746 | 52264 | -1.16 | 5.84E-08 | 1.10E-07 |
| chr2 | 27750630 | 27752650 | 2021 | -3.29 | 5.84E-08 | 1.10E-07 |
| chr3 | 130252292 | 130272310 | 20019 | -3.27 | 5.85E-08 | 1.10E-07 |
| chr22 | 50045206 | 50047540 | 2335 | -3.63 | 5.86E-08 | 1.10E-07 |
| chr14 | 102060173 | 102065357 | 5185 | -6.8 | 5.86E-08 | 1.10E-07 |
| chr11 | 78819887 | 78822687 | 2801 | 6.25 | 5.87E-08 | 1.10E-07 |
| chr5 | 99836736 | 99838908 | 2173 | 6.25 | 5.87E-08 | 1.10E-07 |
| chr3 | 119926928 | 119927930 | 1003 | -4.67 | 5.87E-08 | 1.10E-07 |
| chr5 | 72709477 | 72711363 | 1887 | -3.14 | 5.87E-08 | 1.11E-07 |
| chr2 | 26216551 | 26219234 | 2684 | -6.18 | 5.87E-08 | 1.11E-07 |
| chr13 | 114035714 | 114043753 | 8040 | -5.84 | 5.87E-08 | 1.11E-07 |
| chr4 | 48613886 | 48615436 | 1551 | -6.46 | 5.88E-08 | 1.11E-07 |
| chr14 | 99895072 | 99896127 | 1056 | -6.18 | 5.89E-08 | 1.11E-07 |
| chr16 | 73034953 | 73036320 | 1368 | -6.18 | 5.89E-08 | 1.11E-07 |
| chr2 | 114083759 | 114084648 | 890 | -6.23 | 5.89E-08 | 1.11E-07 |
| chr20 | 55072262 | 55073142 | 881 | -6.21 | 5.90E-08 | 1.11E-07 |
| chr11 | 130043910 | 130045338 | 1429 | -6.26 | 5.90E-08 | 1.11E-07 |
| chr19 | 17213113 | 17214088 | 976 | -6.26 | 5.90E-08 | 1.11E-07 |
| chr1 | 15746692 | 15747771 | 1080 | -6.47 | 5.91E-08 | 1.11E-07 |
| chr17 | 71550183 | 71551164 | 982 | -6.23 | 5.92E-08 | 1.11E-07 |
| chr6 | 81766388 | 81873821 | 107434 | 1.73 | 5.95E-08 | 1.12E-07 |
| chr1 | 17588784 | 17589836 | 1053 | -6.25 | 5.96E-08 | 1.12E-07 |
| chr15 | 84805757 | 84806652 | 896 | -6.25 | 5.96E-08 | 1.12E-07 |

| | | | | | | |
|---|---|---|---|---|---|---|
| chr5 | 134175637 | 134176870 | 1234 | -6.19 | 5.97E-08 | 1.12E-07 |
| chr7 | 15919801 | 15921794 | 1994 | -6.35 | 5.98E-08 | 1.13E-07 |
| chr14 | 53730113 | 53733178 | 3066 | -6.83 | 5.98E-08 | 1.13E-07 |
| chr16 | 89780311 | 89781594 | 1284 | -6.2 | 5.98E-08 | 1.13E-07 |
| chr19 | 48193867 | 48194715 | 849 | -6.33 | 5.99E-08 | 1.13E-07 |
| chr1 | 167748529 | 167750112 | 1584 | -6.49 | 6.00E-08 | 1.13E-07 |
| chr5 | 2023304 | 2124273 | 100970 | -1.45 | 6.01E-08 | 1.13E-07 |
| chr8 | 19075414 | 19078156 | 2743 | -6.4 | 6.01E-08 | 1.13E-07 |
| chr14 | 69466909 | 69468215 | 1307 | -6.54 | 6.01E-08 | 1.13E-07 |
| chr2 | 139239774 | 139241082 | 1309 | -6.26 | 6.01E-08 | 1.13E-07 |
| chr12 | 60104649 | 60110097 | 5449 | -6.32 | 6.02E-08 | 1.13E-07 |
| chr1 | 200249158 | 200250371 | 1214 | -6.31 | 6.03E-08 | 1.13E-07 |
| chr12 | 17519013 | 17521630 | 2618 | -6.31 | 6.03E-08 | 1.13E-07 |
| chr1 | 55257381 | 55258702 | 1322 | -6.41 | 6.04E-08 | 1.14E-07 |
| chr1 | 191564277 | 191578851 | 14575 | -6.3 | 6.05E-08 | 1.14E-07 |
| chr18 | 23901985 | 23903369 | 1385 | -6.26 | 6.05E-08 | 1.14E-07 |
| chr2 | 72494673 | 72496044 | 1372 | -6.22 | 6.06E-08 | 1.14E-07 |
| chr7 | 116524207 | 116525201 | 995 | -6.22 | 6.06E-08 | 1.14E-07 |
| chrX | 21811823 | 21813948 | 2126 | -6.22 | 6.06E-08 | 1.14E-07 |
| chr5 | 90443217 | 90444733 | 1517 | -6.3 | 6.07E-08 | 1.14E-07 |
| chr20 | 34275432 | 34276466 | 1035 | -6.21 | 6.09E-08 | 1.15E-07 |
| chr6 | 101403349 | 101416020 | 12672 | 1.86 | 6.09E-08 | 1.15E-07 |
| chr1 | 192880645 | 192882519 | 1875 | -6.24 | 6.09E-08 | 1.15E-07 |
| chr13 | 89780324 | 89783161 | 2838 | -6.25 | 6.10E-08 | 1.15E-07 |
| chrX | 108771966 | 108773268 | 1303 | -6.25 | 6.10E-08 | 1.15E-07 |
| chr3 | 6039803 | 6042193 | 2391 | -3.13 | 6.10E-08 | 1.15E-07 |
| chr14 | 50375386 | 50376802 | 1417 | -6.18 | 6.11E-08 | 1.15E-07 |
| chr8 | 81030694 | 81031734 | 1041 | -6.18 | 6.11E-08 | 1.15E-07 |
| chr10 | 105533540 | 105542083 | 8544 | -6.19 | 6.11E-08 | 1.15E-07 |
| chr6 | 37682524 | 37684653 | 2130 | -6.44 | 6.12E-08 | 1.15E-07 |
| chr1 | 89648387 | 89649739 | 1353 | -6.18 | 6.12E-08 | 1.15E-07 |
| chr1 | 168767085 | 168768715 | 1631 | -6.18 | 6.12E-08 | 1.15E-07 |
| chr2 | 224875173 | 224876410 | 1238 | -6.19 | 6.12E-08 | 1.15E-07 |
| chr6 | 140222998 | 140237054 | 14057 | -6.67 | 6.12E-08 | 1.15E-07 |
| chr9 | 140922523 | 140929586 | 7064 | -2.4 | 6.12E-08 | 1.15E-07 |
| chr8 | 90413961 | 90415455 | 1495 | -6.17 | 6.13E-08 | 1.15E-07 |
| chr15 | 99772782 | 99773447 | 666 | -6.28 | 6.13E-08 | 1.15E-07 |
| chrX | 67253569 | 67256113 | 2545 | -2.45 | 6.13E-08 | 1.15E-07 |
| chr12 | 78439000 | 78441415 | 2416 | -6.21 | 6.15E-08 | 1.15E-07 |
| chr12 | 94295345 | 94296812 | 1468 | -6.17 | 6.17E-08 | 1.16E-07 |
| chr16 | 1472078 | 1473087 | 1010 | -6.17 | 6.17E-08 | 1.16E-07 |
| chr2 | 119147168 | 119187618 | 40451 | -2.52 | 6.17E-08 | 1.16E-07 |
| chr4 | 180310538 | 180314414 | 3877 | -6.18 | 6.19E-08 | 1.16E-07 |
| chr2 | 226535918 | 226538160 | 2243 | -6.2 | 6.21E-08 | 1.17E-07 |
| chr3 | 24306437 | 24307152 | 716 | -4.94 | 6.21E-08 | 1.17E-07 |
| chr11 | 33318620 | 33320289 | 1670 | -6.48 | 6.23E-08 | 1.17E-07 |
| chr8 | 121242235 | 121243937 | 1703 | 6.46 | 6.23E-08 | 1.17E-07 |
| chr9 | 92907147 | 92972496 | 65350 | -2.58 | 6.24E-08 | 1.17E-07 |
| chr2 | 183160598 | 183162340 | 1743 | 6.24 | 6.25E-08 | 1.17E-07 |
| chr4 | 32207622 | 32209635 | 2014 | -6.18 | 6.25E-08 | 1.17E-07 |
| chr12 | 125296984 | 125298165 | 1182 | -6.23 | 6.25E-08 | 1.17E-07 |
| chr11 | 107665120 | 107666912 | 1793 | -6.81 | 6.27E-08 | 1.18E-07 |
| chr4 | 168255809 | 168257438 | 1630 | -6.33 | 6.29E-08 | 1.18E-07 |
| chr19 | 2677999 | 2679826 | 1828 | -6.18 | 6.29E-08 | 1.18E-07 |
| chr9 | 86075793 | 86077408 | 1616 | -6.38 | 6.30E-08 | 1.18E-07 |
| chr3 | 182827107 | 182830464 | 3358 | -6.6 | 6.30E-08 | 1.18E-07 |
| chr5 | 14603241 | 14604344 | 1104 | -6.32 | 6.30E-08 | 1.18E-07 |
| chr13 | 22026863 | 22029198 | 2336 | -6.32 | 6.31E-08 | 1.18E-07 |
| chrX | 108430634 | 108432881 | 2248 | 5.34 | 6.32E-08 | 1.19E-07 |
| chr5 | 56295915 | 56296523 | 609 | -6.2 | 6.33E-08 | 1.19E-07 |
| chr4 | 123408338 | 123414112 | 5775 | -2.4 | 6.33E-08 | 1.19E-07 |

| | | | | | | |
|---|---|---|---|---|---|---|
| chr14 | 35541000 | 35543256 | 2257 | -6.3 | 6.34E-08 | 1.19E-07 |
| chr6 | 79987925 | 79989426 | 1502 | -6.4 | 6.34E-08 | 1.19E-07 |
| chr11 | 18645131 | 18646251 | 1121 | -6.16 | 6.35E-08 | 1.19E-07 |
| chr11 | 84279956 | 84281487 | 1532 | -6.21 | 6.35E-08 | 1.19E-07 |
| chr2 | 20309582 | 20310767 | 1186 | -6.51 | 6.35E-08 | 1.19E-07 |
| chr13 | 71863383 | 71866164 | 2782 | -6.3 | 6.35E-08 | 1.19E-07 |
| chr16 | 47163888 | 47165239 | 1352 | -6.16 | 6.39E-08 | 1.20E-07 |
| chr4 | 26775551 | 26776575 | 1025 | -6.16 | 6.39E-08 | 1.20E-07 |
| chr21 | 41102726 | 41104785 | 2060 | -4.2 | 6.39E-08 | 1.20E-07 |
| chr1 | 89200672 | 89202192 | 1521 | -6.17 | 6.39E-08 | 1.20E-07 |
| chr10 | 21538706 | 21540113 | 1408 | -6.17 | 6.39E-08 | 1.20E-07 |
| chr3 | 117327242 | 117328868 | 1627 | -6.17 | 6.39E-08 | 1.20E-07 |
| chr3 | 67883122 | 67885267 | 2146 | -3.14 | 6.41E-08 | 1.20E-07 |
| chr7 | 98613177 | 98614727 | 1551 | -6.28 | 6.42E-08 | 1.20E-07 |
| chr6 | 78242498 | 78293492 | 50995 | 6.58 | 6.43E-08 | 1.21E-07 |
| chr1 | 240928353 | 240936154 | 7802 | -6.44 | 6.43E-08 | 1.21E-07 |
| chr11 | 76416127 | 76417329 | 1203 | -6.22 | 6.45E-08 | 1.21E-07 |
| chr14 | 90357071 | 90358438 | 1368 | -6.22 | 6.45E-08 | 1.21E-07 |
| chr3 | 185733711 | 185735112 | 1402 | -6.22 | 6.45E-08 | 1.21E-07 |
| chr18 | 60553734 | 60555322 | 1589 | -6.21 | 6.45E-08 | 1.21E-07 |
| chr9 | 12957628 | 12959001 | 1374 | -6.38 | 6.46E-08 | 1.21E-07 |
| chr9 | 90804914 | 90807705 | 2792 | -6.38 | 6.46E-08 | 1.21E-07 |
| chr3 | 184621441 | 184622728 | 1288 | -6.45 | 6.47E-08 | 1.21E-07 |
| chr3 | 188119417 | 188120589 | 1173 | -6.31 | 6.47E-08 | 1.21E-07 |
| chrX | 40681859 | 40683800 | 1942 | -6.39 | 6.48E-08 | 1.21E-07 |
| chr10 | 133650744 | 133668762 | 18019 | -7.19 | 6.49E-08 | 1.22E-07 |
| chr5 | 148163106 | 148164451 | 1346 | 6.34 | 6.49E-08 | 1.22E-07 |
| chrX | 39196024 | 39203177 | 7154 | 5.61 | 6.50E-08 | 1.22E-07 |
| chr12 | 17737266 | 17740254 | 2989 | -6.4 | 6.51E-08 | 1.22E-07 |
| chr9 | 40770819 | 40771392 | 574 | -6.41 | 6.52E-08 | 1.22E-07 |
| chr7 | 158054596 | 158057897 | 3302 | -6.19 | 6.53E-08 | 1.22E-07 |
| chr2 | 122321151 | 122322498 | 1348 | -6.42 | 6.54E-08 | 1.23E-07 |
| chr13 | 42048098 | 42059282 | 11185 | -3.51 | 6.54E-08 | 1.23E-07 |
| chr3 | 93982418 | 93985324 | 2907 | 6.62 | 6.54E-08 | 1.23E-07 |
| chr4 | 187358002 | 187380850 | 22849 | -5.99 | 6.55E-08 | 1.23E-07 |
| chr10 | 30036997 | 30038282 | 1286 | -6.16 | 6.56E-08 | 1.23E-07 |
| chr5 | 168075908 | 168076973 | 1066 | -6.16 | 6.56E-08 | 1.23E-07 |
| chr5 | 138658668 | 138660365 | 1698 | -6.25 | 6.58E-08 | 1.23E-07 |
| chr2 | 61410297 | 61411142 | 846 | -6.16 | 6.58E-08 | 1.23E-07 |
| chr5 | 167081308 | 167082981 | 1674 | -6.16 | 6.58E-08 | 1.23E-07 |
| chr17 | 42730475 | 42731401 | 927 | -6.27 | 6.60E-08 | 1.24E-07 |
| chr3 | 98792033 | 98794245 | 2213 | -6.27 | 6.60E-08 | 1.24E-07 |
| chr11 | 130159695 | 130161863 | 2169 | -6.18 | 6.60E-08 | 1.24E-07 |
| chr3 | 184175105 | 184176650 | 1546 | -6.16 | 6.62E-08 | 1.24E-07 |
| chr8 | 19964315 | 19966792 | 2478 | 6.64 | 6.63E-08 | 1.24E-07 |
| chr17 | 41802645 | 41811574 | 8930 | -6.67 | 6.63E-08 | 1.24E-07 |
| chr10 | 74965704 | 74966873 | 1170 | -6.57 | 6.64E-08 | 1.24E-07 |
| chr6 | 21687317 | 21689236 | 1920 | -6.77 | 6.65E-08 | 1.24E-07 |
| chr8 | 123759098 | 123761164 | 2067 | -6.18 | 6.65E-08 | 1.25E-07 |
| chr9 | 290429 | 292021 | 1593 | 6.26 | 6.66E-08 | 1.25E-07 |
| chrX | 99712040 | 99713377 | 1338 | -3.12 | 6.66E-08 | 1.25E-07 |
| chr18 | 1209878 | 1275849 | 65972 | 1.14 | 6.66E-08 | 1.25E-07 |
| chr15 | 36845552 | 36854453 | 8902 | -6.5 | 6.67E-08 | 1.25E-07 |
| chr1 | 173764304 | 173765355 | 1052 | -6.15 | 6.68E-08 | 1.25E-07 |
| chr1 | 157020513 | 157034399 | 13887 | -6.38 | 6.71E-08 | 1.26E-07 |
| chr19 | 50806397 | 50807417 | 1021 | -6.51 | 6.71E-08 | 1.26E-07 |
| chr13 | 114132491 | 114133698 | 1208 | -6.16 | 6.74E-08 | 1.26E-07 |
| chr9 | 27072000 | 27073885 | 1886 | -6.24 | 6.78E-08 | 1.27E-07 |
| chr4 | 131164367 | 131167530 | 3164 | -6.19 | 6.80E-08 | 1.27E-07 |
| chr11 | 88330944 | 88333193 | 2250 | -6.16 | 6.80E-08 | 1.27E-07 |
| chr3 | 178978093 | 178978782 | 690 | -6.16 | 6.80E-08 | 1.27E-07 |

| | | | | | | |
|---|---|---|---|---|---|---|
| chr3 | 194861357 | 194862943 | 1587 | -6.16 | 6.80E-08 | 1.27E-07 |
| chrX | 57022788 | 57024645 | 1858 | -6.5 | 6.83E-08 | 1.28E-07 |
| chr3 | 153428811 | 153430648 | 1838 | -6.18 | 6.83E-08 | 1.28E-07 |
| chr3 | 129413989 | 129417645 | 3657 | -5.22 | 6.83E-08 | 1.28E-07 |
| chr3 | 43594689 | 43596029 | 1341 | -6.23 | 6.85E-08 | 1.28E-07 |
| chr1 | 240104075 | 240108677 | 4603 | -5.05 | 6.85E-08 | 1.28E-07 |
| chr12 | 9876939 | 9878418 | 1480 | -6.15 | 6.87E-08 | 1.29E-07 |
| chr21 | 42560437 | 42561700 | 1264 | -6.15 | 6.87E-08 | 1.29E-07 |
| chr6 | 5842066 | 5849220 | 7155 | -2.91 | 6.88E-08 | 1.29E-07 |
| chr14 | 106465449 | 106468052 | 2604 | -6.34 | 6.88E-08 | 1.29E-07 |
| chr12 | 99010479 | 99012528 | 2050 | -6.21 | 6.90E-08 | 1.29E-07 |
| chr11 | 21543507 | 21545627 | 2121 | -6.18 | 6.90E-08 | 1.29E-07 |
| chr4 | 7250451 | 7251295 | 845 | -6.18 | 6.90E-08 | 1.29E-07 |
| chr7 | 69191067 | 69192682 | 1616 | -6.36 | 6.91E-08 | 1.29E-07 |
| chr6 | 119400602 | 119406750 | 6149 | -3.03 | 6.91E-08 | 1.29E-07 |
| chr3 | 160913837 | 160914995 | 1159 | -6.15 | 6.91E-08 | 1.29E-07 |
| chr3 | 133365349 | 133367015 | 1667 | -6.17 | 6.92E-08 | 1.29E-07 |
| chr5 | 10716879 | 10718391 | 1513 | -6.17 | 6.92E-08 | 1.29E-07 |
| chr5 | 134782069 | 134789643 | 7575 | -6.72 | 6.92E-08 | 1.29E-07 |
| chr2 | 186910913 | 186913102 | 2190 | 5.65 | 6.94E-08 | 1.30E-07 |
| chr22 | 27442463 | 27444033 | 1571 | -6.29 | 6.94E-08 | 1.30E-07 |
| chr12 | 3207668 | 3208975 | 1308 | -6.16 | 6.94E-08 | 1.30E-07 |
| chr7 | 115161874 | 115162804 | 931 | -6.16 | 6.94E-08 | 1.30E-07 |
| chr8 | 62880886 | 62904701 | 23816 | -7.2 | 6.94E-08 | 1.30E-07 |
| chr2 | 102629427 | 102630951 | 1525 | -6.31 | 6.94E-08 | 1.30E-07 |
| chrX | 84491099 | 84493630 | 2532 | -5.07 | 6.95E-08 | 1.30E-07 |
| chr2 | 26246860 | 26248117 | 1258 | -6.3 | 6.96E-08 | 1.30E-07 |
| chr2 | 150960809 | 150963991 | 3183 | -6.37 | 6.96E-08 | 1.30E-07 |
| chr4 | 14544210 | 14546321 | 2112 | -6.17 | 6.97E-08 | 1.30E-07 |
| chr6 | 7358317 | 7359650 | 1334 | -6.17 | 6.97E-08 | 1.30E-07 |
| chr1 | 18645109 | 18646107 | 999 | -6.4 | 6.97E-08 | 1.30E-07 |
| chr3 | 162192209 | 162224237 | 32029 | -3.4 | 7.00E-08 | 1.31E-07 |
| chr14 | 61565053 | 61572450 | 7398 | -6.39 | 7.00E-08 | 1.31E-07 |
| chr4 | 2714850 | 2715815 | 966 | -6.18 | 7.01E-08 | 1.31E-07 |
| chr12 | 17717576 | 17718910 | 1335 | -6.28 | 7.01E-08 | 1.31E-07 |
| chr1 | 203215205 | 203224967 | 9763 | -2.48 | 7.03E-08 | 1.31E-07 |
| chr2 | 29196575 | 29197616 | 1042 | -6.42 | 7.03E-08 | 1.31E-07 |
| chr2 | 187596107 | 187603446 | 7340 | -6.41 | 7.04E-08 | 1.31E-07 |
| chr13 | 45281592 | 45282958 | 1367 | -6.27 | 7.04E-08 | 1.31E-07 |
| chr2 | 214017444 | 214018511 | 1068 | -6.42 | 7.04E-08 | 1.32E-07 |
| chr15 | 42313789 | 42324925 | 11137 | -7.25 | 7.06E-08 | 1.32E-07 |
| chrX | 130732710 | 130735721 | 3012 | -6.16 | 7.06E-08 | 1.32E-07 |
| chr3 | 61255686 | 61352574 | 96889 | 2.23 | 7.07E-08 | 1.32E-07 |
| chr2 | 138719243 | 138733785 | 14543 | -6.43 | 7.08E-08 | 1.32E-07 |
| chr8 | 33135092 | 33163212 | 28121 | -2.04 | 7.08E-08 | 1.32E-07 |
| chr3 | 142270298 | 142272084 | 1787 | -6.18 | 7.08E-08 | 1.32E-07 |
| chr4 | 2782251 | 2787058 | 4808 | -5.59 | 7.09E-08 | 1.32E-07 |
| chr10 | 62982295 | 63059185 | 76891 | -1.94 | 7.09E-08 | 1.32E-07 |
| chr13 | 77660726 | 77662218 | 1493 | -4.97 | 7.10E-08 | 1.33E-07 |
| chr22 | 39657097 | 39658582 | 1486 | -6.25 | 7.12E-08 | 1.33E-07 |
| chr9 | 5684408 | 5685319 | 912 | -6.2 | 7.13E-08 | 1.33E-07 |
| chr11 | 101843472 | 101846985 | 3514 | -6.15 | 7.13E-08 | 1.33E-07 |
| chr7 | 110818799 | 110820363 | 1565 | -6.17 | 7.13E-08 | 1.33E-07 |
| chrX | 40728015 | 40729495 | 1481 | -6.17 | 7.13E-08 | 1.33E-07 |
| chr12 | 104844462 | 104845181 | 720 | -6.16 | 7.14E-08 | 1.33E-07 |
| chr6 | 47353900 | 47355514 | 1615 | -6.16 | 7.14E-08 | 1.33E-07 |
| chr7 | 134487237 | 134488808 | 1572 | -6.45 | 7.14E-08 | 1.33E-07 |
| chr4 | 23243931 | 23246265 | 2335 | -6.14 | 7.14E-08 | 1.33E-07 |
| chr2 | 169086734 | 169087949 | 1216 | -6.25 | 7.15E-08 | 1.33E-07 |
| chr5 | 71341851 | 71359029 | 17179 | -1.55 | 7.16E-08 | 1.33E-07 |
| chr14 | 76056530 | 76082090 | 25561 | -6.38 | 7.16E-08 | 1.34E-07 |

| | | | | | | |
|---|---|---|---|---|---|---|
| chr6 | 168075119 | 168076306 | 1188 | -6.14 | 7.16E-08 | 1.34E-07 |
| chr4 | 6891889 | 6893230 | 1342 | -6.2 | 7.17E-08 | 1.34E-07 |
| chr10 | 4391722 | 4411221 | 19500 | -7.56 | 7.19E-08 | 1.34E-07 |
| chr12 | 41278227 | 41281186 | 2960 | -6.24 | 7.20E-08 | 1.34E-07 |
| chr4 | 131886152 | 131888798 | 2647 | -6.24 | 7.20E-08 | 1.34E-07 |
| chr17 | 53824035 | 53825148 | 1114 | -6.45 | 7.20E-08 | 1.34E-07 |
| chr15 | 101011183 | 101046324 | 35142 | -1.47 | 7.21E-08 | 1.34E-07 |
| chr21 | 18777664 | 18778743 | 1080 | -5.31 | 7.21E-08 | 1.34E-07 |
| chr3 | 105881250 | 105882319 | 1070 | -6.15 | 7.22E-08 | 1.35E-07 |
| chr4 | 114978407 | 114979976 | 1570 | -6.18 | 7.22E-08 | 1.35E-07 |
| chr7 | 19180094 | 19357846 | 177753 | 1.18 | 7.22E-08 | 1.35E-07 |
| chr16 | 1197016 | 1198138 | 1123 | -6.15 | 7.23E-08 | 1.35E-07 |
| chr2 | 48598113 | 48598830 | 718 | -6.15 | 7.23E-08 | 1.35E-07 |
| chrX | 52622965 | 52624909 | 1945 | -2.81 | 7.23E-08 | 1.35E-07 |
| chr7 | 87988041 | 87988742 | 702 | -6.14 | 7.24E-08 | 1.35E-07 |
| chr20 | 11259163 | 11260520 | 1358 | -6.17 | 7.25E-08 | 1.35E-07 |
| chr6 | 109896323 | 109914623 | 18301 | -2.57 | 7.25E-08 | 1.35E-07 |
| chr13 | 44915022 | 44916190 | 1169 | -6.23 | 7.25E-08 | 1.35E-07 |
| chr14 | 38521270 | 38522677 | 1408 | 6.21 | 7.27E-08 | 1.35E-07 |
| chr7 | 110374036 | 110375437 | 1402 | -6.58 | 7.27E-08 | 1.35E-07 |
| chr12 | 80121651 | 80122740 | 1090 | -6.19 | 7.27E-08 | 1.35E-07 |
| chr9 | 130760627 | 130771306 | 10680 | -7.01 | 7.28E-08 | 1.36E-07 |
| chrX | 20657797 | 20672353 | 14557 | -1.8 | 7.28E-08 | 1.36E-07 |
| chr18 | 46636681 | 46638183 | 1503 | -6.47 | 7.28E-08 | 1.36E-07 |
| chr7 | 97631585 | 97632638 | 1054 | -6.22 | 7.29E-08 | 1.36E-07 |
| chr3 | 162354960 | 162375413 | 20454 | -2.36 | 7.30E-08 | 1.36E-07 |
| chr1 | 185907146 | 185908389 | 1244 | -6.23 | 7.31E-08 | 1.36E-07 |
| chr17 | 17294386 | 17295550 | 1165 | -6.23 | 7.31E-08 | 1.36E-07 |
| chr15 | 40072936 | 40074100 | 1165 | -6.14 | 7.32E-08 | 1.36E-07 |
| chr6 | 45103885 | 45105224 | 1340 | -6.14 | 7.32E-08 | 1.36E-07 |
| chr10 | 90190618 | 90203303 | 12686 | -6.8 | 7.33E-08 | 1.37E-07 |
| chr6 | 141689340 | 141691805 | 2466 | -6.48 | 7.34E-08 | 1.37E-07 |
| chr2 | 147250867 | 147252387 | 1521 | -3.03 | 7.35E-08 | 1.37E-07 |
| chr7 | 72200552 | 72202137 | 1586 | -6.17 | 7.37E-08 | 1.37E-07 |
| chr19 | 55662886 | 55664798 | 1913 | -6.22 | 7.38E-08 | 1.37E-07 |
| chr4 | 40962344 | 40963340 | 997 | -6.22 | 7.38E-08 | 1.37E-07 |
| chrX | 135569561 | 135570237 | 677 | -5.36 | 7.38E-08 | 1.37E-07 |
| chr7 | 32865366 | 32866157 | 792 | -6.18 | 7.38E-08 | 1.37E-07 |
| chr15 | 37221200 | 37222607 | 1408 | -6.21 | 7.40E-08 | 1.38E-07 |
| chr1 | 62983769 | 62985524 | 1756 | -6.32 | 7.46E-08 | 1.39E-07 |
| chr11 | 47728669 | 47729817 | 1149 | -6.33 | 7.46E-08 | 1.39E-07 |
| chr3 | 101321670 | 101323429 | 1760 | -6.31 | 7.46E-08 | 1.39E-07 |
| chr4 | 169607280 | 169620497 | 13218 | -6.52 | 7.47E-08 | 1.39E-07 |
| chr4 | 57098036 | 57098840 | 805 | -4.63 | 7.49E-08 | 1.39E-07 |
| chr15 | 65858935 | 65860214 | 1280 | -6.15 | 7.50E-08 | 1.40E-07 |
| chr12 | 127533540 | 127536372 | 2833 | -6.15 | 7.53E-08 | 1.40E-07 |
| chr6 | 150153055 | 150153882 | 828 | -6.15 | 7.53E-08 | 1.40E-07 |
| chr22 | 33398684 | 33405375 | 6692 | -5.39 | 7.53E-08 | 1.40E-07 |
| chr2 | 191267563 | 191268649 | 1087 | -4.61 | 7.54E-08 | 1.40E-07 |
| chr8 | 57987702 | 57997520 | 9819 | 1.78 | 7.54E-08 | 1.40E-07 |
| chr2 | 65127645 | 65129252 | 1608 | -6.63 | 7.54E-08 | 1.40E-07 |
| chr2 | 24200552 | 24201341 | 790 | -6.3 | 7.57E-08 | 1.41E-07 |
| chr20 | 52832030 | 52833088 | 1059 | -6.3 | 7.57E-08 | 1.41E-07 |
| chr22 | 45208445 | 45209629 | 1185 | -6.14 | 7.58E-08 | 1.41E-07 |
| chr6 | 110459049 | 110460638 | 1590 | -6.14 | 7.58E-08 | 1.41E-07 |
| chr6 | 153437992 | 153438921 | 930 | -6.14 | 7.58E-08 | 1.41E-07 |
| chr12 | 69203354 | 69204212 | 859 | -6.29 | 7.58E-08 | 1.41E-07 |
| chr7 | 11957558 | 11959950 | 2393 | -6.29 | 7.58E-08 | 1.41E-07 |
| chrX | 122508636 | 122525111 | 16476 | -3.13 | 7.59E-08 | 1.41E-07 |
| chr11 | 46547814 | 46548955 | 1142 | -6.28 | 7.60E-08 | 1.41E-07 |
| chr5 | 64796293 | 64798085 | 1793 | -6.28 | 7.61E-08 | 1.42E-07 |

| | | | | | | |
|---|---|---|---|---|---|---|
| chr3 | 152083763 | 152096448 | 12686 | -6.64 | 7.61E-08 | 1.42E-07 |
| chr14 | 21410171 | 21412074 | 1904 | -2.68 | 7.63E-08 | 1.42E-07 |
| chr8 | 54176671 | 54216919 | 40249 | -4.63 | 7.63E-08 | 1.42E-07 |
| chr11 | 108865992 | 108867610 | 1619 | -6.16 | 7.65E-08 | 1.42E-07 |
| chr4 | 17727473 | 17732850 | 5378 | -2.14 | 7.65E-08 | 1.42E-07 |
| chr13 | 35198384 | 35212293 | 13910 | -6.86 | 7.65E-08 | 1.42E-07 |
| chr4 | 28256625 | 28265482 | 8858 | -6.55 | 7.67E-08 | 1.43E-07 |
| chr4 | 143484995 | 143486849 | 1855 | -4.69 | 7.69E-08 | 1.43E-07 |
| chr2 | 236352084 | 236353700 | 1617 | -6.43 | 7.70E-08 | 1.43E-07 |
| chr10 | 95395179 | 95396014 | 836 | -6.15 | 7.70E-08 | 1.43E-07 |
| chr1 | 86964389 | 86965756 | 1368 | -6.23 | 7.71E-08 | 1.43E-07 |
| chr13 | 82680480 | 82682315 | 1836 | -6.23 | 7.71E-08 | 1.43E-07 |
| chr3 | 138593031 | 138595460 | 2430 | -6.14 | 7.71E-08 | 1.43E-07 |
| chr1 | 20402884 | 20404385 | 1502 | -4.91 | 7.74E-08 | 1.44E-07 |
| chr8 | 29731148 | 29736040 | 4893 | -6.23 | 7.76E-08 | 1.44E-07 |
| chr2 | 70002709 | 70004021 | 1313 | -6.13 | 7.77E-08 | 1.44E-07 |
| chr4 | 90770120 | 90772811 | 2692 | -6.24 | 7.79E-08 | 1.45E-07 |
| chr5 | 38717696 | 38718811 | 1116 | -6.16 | 7.80E-08 | 1.45E-07 |
| chr5 | 14391152 | 14392347 | 1196 | -6.58 | 7.80E-08 | 1.45E-07 |
| chr3 | 111739998 | 111741273 | 1276 | -6.23 | 7.84E-08 | 1.46E-07 |
| chrX | 135102399 | 135103577 | 1179 | -6.23 | 7.84E-08 | 1.46E-07 |
| chr6 | 48856149 | 48858741 | 2593 | -6.44 | 7.84E-08 | 1.46E-07 |
| chr6 | 74291120 | 74292273 | 1154 | -6.13 | 7.86E-08 | 1.46E-07 |
| chr1 | 205703693 | 205704899 | 1207 | -6.44 | 7.86E-08 | 1.46E-07 |
| chr22 | 19901151 | 19902079 | 929 | -6.21 | 7.88E-08 | 1.46E-07 |
| chr3 | 9180828 | 9181571 | 744 | -6.21 | 7.88E-08 | 1.46E-07 |
| chr8 | 82714938 | 82716124 | 1187 | -6.15 | 7.90E-08 | 1.47E-07 |
| chr1 | 182371634 | 182376652 | 5019 | -4.07 | 7.90E-08 | 1.47E-07 |
| chr20 | 13866388 | 13875659 | 9272 | -2.96 | 7.91E-08 | 1.47E-07 |
| chr7 | 18994517 | 18996832 | 2316 | 6.2 | 7.92E-08 | 1.47E-07 |
| chr4 | 95520475 | 95521609 | 1135 | -6.22 | 7.93E-08 | 1.47E-07 |
| chr2 | 54959021 | 54960312 | 1292 | -6.2 | 7.96E-08 | 1.48E-07 |
| chr4 | 189719069 | 189720571 | 1503 | -6.2 | 7.96E-08 | 1.48E-07 |
| chr11 | 130310335 | 130311871 | 1537 | -6.17 | 7.97E-08 | 1.48E-07 |
| chr8 | 48959954 | 48961124 | 1171 | -6.21 | 7.98E-08 | 1.48E-07 |
| chr1 | 108484923 | 108490684 | 5762 | 2.79 | 7.98E-08 | 1.48E-07 |
| chr10 | 123671186 | 123679547 | 8362 | -6.32 | 8.04E-08 | 1.49E-07 |
| chr1 | 198616090 | 198617786 | 1697 | -6.13 | 8.04E-08 | 1.49E-07 |
| chrX | 140872007 | 140874597 | 2591 | -6.13 | 8.04E-08 | 1.49E-07 |
| chr15 | 41942276 | 41943569 | 1294 | -6.2 | 8.04E-08 | 1.49E-07 |
| chr21 | 42643629 | 42644724 | 1096 | -6.2 | 8.04E-08 | 1.49E-07 |
| chr4 | 110591912 | 110593052 | 1141 | -6.2 | 8.04E-08 | 1.49E-07 |
| chr3 | 22802129 | 22924786 | 122658 | 1.56 | 8.05E-08 | 1.49E-07 |
| chr2 | 103361428 | 103363536 | 2109 | -6.12 | 8.06E-08 | 1.50E-07 |
| chr2 | 121017682 | 121027542 | 9861 | -6.35 | 8.07E-08 | 1.50E-07 |
| chr19 | 43869856 | 43870985 | 1130 | -6.27 | 8.07E-08 | 1.50E-07 |
| chr4 | 146598456 | 146599753 | 1298 | -6.27 | 8.07E-08 | 1.50E-07 |
| chr7 | 83337903 | 83339640 | 1738 | -6.35 | 8.09E-08 | 1.50E-07 |
| chrX | 72996700 | 72997826 | 1127 | -5.29 | 8.09E-08 | 1.50E-07 |
| chr5 | 95411420 | 95412555 | 1136 | -6.12 | 8.10E-08 | 1.50E-07 |
| chr1 | 44803733 | 44804917 | 1185 | -6.16 | 8.11E-08 | 1.50E-07 |
| chr12 | 70102470 | 70103230 | 761 | -6.14 | 8.11E-08 | 1.50E-07 |
| chr3 | 66609104 | 66610646 | 1543 | -6.14 | 8.11E-08 | 1.50E-07 |
| chr1 | 89696409 | 89709226 | 12818 | 2.07 | 8.11E-08 | 1.50E-07 |
| chr11 | 17779763 | 17780959 | 1197 | -6.37 | 8.12E-08 | 1.51E-07 |
| chr4 | 151870364 | 151880798 | 10435 | -6.37 | 8.12E-08 | 1.51E-07 |
| chr5 | 60070504 | 60072510 | 2007 | -6.14 | 8.12E-08 | 1.51E-07 |
| chr5 | 120648915 | 120650448 | 1534 | -6.19 | 8.13E-08 | 1.51E-07 |
| chr7 | 12386425 | 12387489 | 1065 | -6.19 | 8.13E-08 | 1.51E-07 |
| chr10 | 102147944 | 102148977 | 1034 | -6.12 | 8.13E-08 | 1.51E-07 |
| chr19 | 39558971 | 39559882 | 912 | -6.12 | 8.13E-08 | 1.51E-07 |

| | | | | | | |
|---|---|---|---|---|---|---|
| chr14 | 73290447 | 73293494 | 3048 | 2.86 | 8.14E-08 | 1.51E-07 |
| chr6 | 2614133 | 2630027 | 15895 | -6.96 | 8.17E-08 | 1.52E-07 |
| chr1 | 113324553 | 113326343 | 1791 | -4.69 | 8.18E-08 | 1.52E-07 |
| chr11 | 110062270 | 110064142 | 1873 | -6.14 | 8.20E-08 | 1.52E-07 |
| chr1 | 37462142 | 37463545 | 1404 | 6.2 | 8.22E-08 | 1.52E-07 |
| chr7 | 81817805 | 81818553 | 749 | -6.18 | 8.23E-08 | 1.53E-07 |
| chr6 | 56986248 | 56988093 | 1846 | -6.19 | 8.25E-08 | 1.53E-07 |
| chr12 | 12498620 | 12500727 | 2108 | -6.41 | 8.26E-08 | 1.53E-07 |
| chr14 | 75148745 | 75150006 | 1262 | -6.23 | 8.26E-08 | 1.53E-07 |
| chr7 | 75704014 | 75705497 | 1484 | -6.3 | 8.30E-08 | 1.54E-07 |
| chr7 | 81459054 | 81463844 | 4791 | -5.85 | 8.30E-08 | 1.54E-07 |
| chr13 | 84353047 | 84355912 | 2866 | -6.22 | 8.31E-08 | 1.54E-07 |
| chr9 | 73045995 | 73047449 | 1455 | -6.22 | 8.31E-08 | 1.54E-07 |
| chr11 | 66086667 | 66087796 | 1130 | -6.14 | 8.32E-08 | 1.54E-07 |
| chr5 | 34662738 | 34664663 | 1926 | -6.14 | 8.32E-08 | 1.54E-07 |
| chr10 | 99770378 | 99773421 | 3044 | 6.97 | 8.32E-08 | 1.54E-07 |
| chr2 | 237889926 | 237962666 | 72741 | -5.05 | 8.33E-08 | 1.54E-07 |
| chr1 | 220768728 | 220770632 | 1905 | -6.18 | 8.33E-08 | 1.54E-07 |
| chr18 | 24070461 | 24072364 | 1904 | -6.18 | 8.33E-08 | 1.54E-07 |
| chr1 | 85100997 | 85102936 | 1940 | -6.13 | 8.34E-08 | 1.54E-07 |
| chr5 | 33336757 | 33338972 | 2216 | -6.43 | 8.34E-08 | 1.55E-07 |
| chr18 | 70378755 | 70380026 | 1272 | -6.17 | 8.35E-08 | 1.55E-07 |
| chr6 | 151314366 | 151315790 | 1425 | -6.17 | 8.35E-08 | 1.55E-07 |
| chr8 | 27475328 | 27484310 | 8983 | -6.17 | 8.35E-08 | 1.55E-07 |
| chr6 | 159560261 | 159584167 | 23907 | -2.66 | 8.35E-08 | 1.55E-07 |
| chrX | 55761364 | 55762489 | 1126 | -6.26 | 8.35E-08 | 1.55E-07 |
| chr8 | 140422817 | 140423454 | 638 | -6.26 | 8.37E-08 | 1.55E-07 |
| chr7 | 26211203 | 26212253 | 1051 | -6.13 | 8.37E-08 | 1.55E-07 |
| chr6 | 130285189 | 130286341 | 1153 | -6.11 | 8.39E-08 | 1.55E-07 |
| chr3 | 102082351 | 102083834 | 1484 | -4.98 | 8.39E-08 | 1.55E-07 |
| chr17 | 65596581 | 65597363 | 783 | -6.12 | 8.40E-08 | 1.56E-07 |
| chr1 | 171794043 | 171797908 | 3866 | -2.14 | 8.41E-08 | 1.56E-07 |
| chr18 | 44130037 | 44131380 | 1344 | -6.25 | 8.42E-08 | 1.56E-07 |
| chr8 | 75188670 | 75199494 | 10825 | 2.63 | 8.42E-08 | 1.56E-07 |
| chr6 | 131795278 | 131796942 | 1665 | -6.15 | 8.42E-08 | 1.56E-07 |
| chr7 | 91445700 | 91454086 | 8387 | -2.19 | 8.42E-08 | 1.56E-07 |
| chr10 | 75344820 | 75345473 | 654 | -6.11 | 8.43E-08 | 1.56E-07 |
| chr3 | 11703752 | 11705263 | 1512 | -6.21 | 8.43E-08 | 1.56E-07 |
| chr22 | 33443831 | 33445068 | 1238 | -6.12 | 8.45E-08 | 1.56E-07 |
| chr6 | 88075513 | 88083646 | 8134 | -7.19 | 8.46E-08 | 1.57E-07 |
| chr4 | 179147203 | 179149447 | 2245 | -6.16 | 8.47E-08 | 1.57E-07 |
| chr1 | 116890479 | 116891400 | 922 | -4.64 | 8.47E-08 | 1.57E-07 |
| chr3 | 17973146 | 17974502 | 1357 | -6.14 | 8.48E-08 | 1.57E-07 |
| chr4 | 105604454 | 105605893 | 1440 | -6.39 | 8.48E-08 | 1.57E-07 |
| chr13 | 106637680 | 106638713 | 1034 | -6.2 | 8.51E-08 | 1.57E-07 |
| chr11 | 116190722 | 116208798 | 18077 | 1.91 | 8.51E-08 | 1.58E-07 |
| chr6 | 66351214 | 66371847 | 20634 | 2.03 | 8.52E-08 | 1.58E-07 |
| chr8 | 124143616 | 124144286 | 671 | -6.41 | 8.53E-08 | 1.58E-07 |
| chr9 | 79056851 | 79058673 | 1823 | -6.69 | 8.53E-08 | 1.58E-07 |
| chr1 | 245002523 | 245003576 | 1054 | -6.13 | 8.53E-08 | 1.58E-07 |
| chr6 | 106850402 | 106852151 | 1750 | -6.22 | 8.55E-08 | 1.58E-07 |
| chr17 | 39119203 | 39127382 | 8180 | -6.69 | 8.57E-08 | 1.58E-07 |
| chr11 | 102889888 | 102892760 | 2873 | -2.79 | 8.58E-08 | 1.59E-07 |
| chr17 | 53299312 | 53300279 | 968 | 6.34 | 8.58E-08 | 1.59E-07 |
| chr12 | 108652348 | 108659934 | 7587 | -1.7 | 8.60E-08 | 1.59E-07 |
| chr7 | 16287689 | 16288800 | 1112 | -5.29 | 8.60E-08 | 1.59E-07 |
| chr3 | 40715158 | 40717912 | 2755 | -2.23 | 8.60E-08 | 1.59E-07 |
| chr6 | 125625849 | 125626943 | 1095 | -4.61 | 8.61E-08 | 1.59E-07 |
| chr12 | 106945549 | 106964809 | 19261 | -2.29 | 8.63E-08 | 1.60E-07 |
| chr1 | 68928444 | 68930109 | 1666 | -6.14 | 8.63E-08 | 1.60E-07 |
| chr6 | 135401369 | 135404626 | 3258 | -5.34 | 8.63E-08 | 1.60E-07 |

| | | | | | | |
|---|---|---|---|---|---|---|
| chr7 | 13486001 | 13492236 | 6236 | -6.29 | 8.66E-08 | 1.60E-07 |
| chr8 | 80823177 | 80824891 | 1715 | -6.29 | 8.66E-08 | 1.60E-07 |
| chr9 | 111013800 | 111014922 | 1123 | -6.18 | 8.68E-08 | 1.60E-07 |
| chr2 | 9849371 | 9850411 | 1041 | -6.11 | 8.69E-08 | 1.61E-07 |
| chr1 | 25502793 | 25504906 | 2114 | -6.34 | 8.71E-08 | 1.61E-07 |
| chr6 | 47733600 | 47734839 | 1240 | -6.15 | 8.73E-08 | 1.61E-07 |
| chrX | 134340957 | 134342154 | 1198 | -6.15 | 8.73E-08 | 1.61E-07 |
| chr4 | 38233636 | 38235577 | 1942 | -6.25 | 8.73E-08 | 1.61E-07 |
| chr7 | 43420147 | 43421084 | 938 | -6.35 | 8.73E-08 | 1.61E-07 |
| chr14 | 71486054 | 71486857 | 804 | -6.2 | 8.75E-08 | 1.62E-07 |
| chr1 | 110066304 | 110067365 | 1062 | -6.11 | 8.75E-08 | 1.62E-07 |
| chr3 | 174118019 | 174119140 | 1122 | -6.11 | 8.75E-08 | 1.62E-07 |
| chrX | 71737514 | 71750129 | 12616 | -6.25 | 8.75E-08 | 1.62E-07 |
| chr1 | 174688565 | 174689741 | 1177 | -6.12 | 8.77E-08 | 1.62E-07 |
| chr4 | 133056655 | 133058653 | 1999 | -6.13 | 8.78E-08 | 1.62E-07 |
| chr10 | 35586656 | 35587569 | 914 | -6.17 | 8.78E-08 | 1.62E-07 |
| chr7 | 83950935 | 83952376 | 1442 | -6.17 | 8.78E-08 | 1.62E-07 |
| chr17 | 35995633 | 35996456 | 824 | -6.13 | 8.78E-08 | 1.62E-07 |
| chr18 | 20691137 | 20693034 | 1898 | -6.13 | 8.78E-08 | 1.62E-07 |
| chr1 | 38654355 | 38666904 | 12550 | -6.24 | 8.78E-08 | 1.62E-07 |
| chr4 | 168204889 | 168206505 | 1617 | -6.1 | 8.83E-08 | 1.63E-07 |
| chr6 | 128648365 | 128649698 | 1334 | -6.1 | 8.83E-08 | 1.63E-07 |
| chr7 | 16558943 | 16560118 | 1176 | -6.38 | 8.84E-08 | 1.63E-07 |
| chr13 | 83128301 | 83129996 | 1696 | -6.15 | 8.84E-08 | 1.63E-07 |
| chr5 | 66472369 | 66473530 | 1162 | -6.15 | 8.84E-08 | 1.63E-07 |
| chr6 | 74844818 | 74846952 | 2135 | -4.63 | 8.85E-08 | 1.63E-07 |
| chr1 | 178640691 | 178645448 | 4758 | -6.63 | 8.87E-08 | 1.64E-07 |
| chr12 | 50964431 | 50965325 | 895 | -6.11 | 8.88E-08 | 1.64E-07 |
| chr16 | 69698086 | 69699245 | 1160 | -6.11 | 8.88E-08 | 1.64E-07 |
| chr9 | 138515269 | 138562057 | 46789 | -4.56 | 8.89E-08 | 1.64E-07 |
| chr10 | 17257226 | 17258083 | 858 | -6.11 | 8.90E-08 | 1.64E-07 |
| chr16 | 49575332 | 49576687 | 1356 | -6.21 | 8.91E-08 | 1.64E-07 |
| chr7 | 12171713 | 12172874 | 1162 | -6.12 | 8.95E-08 | 1.65E-07 |
| chr2 | 205328697 | 205329505 | 809 | -5.34 | 8.96E-08 | 1.65E-07 |
| chr22 | 36284251 | 36285230 | 980 | -6.1 | 8.96E-08 | 1.65E-07 |
| chr2 | 120141966 | 120143254 | 1289 | -6.11 | 8.97E-08 | 1.65E-07 |
| chr2 | 196654755 | 196656166 | 1412 | -6.65 | 8.98E-08 | 1.66E-07 |
| chr3 | 45946050 | 45948306 | 2257 | -6.42 | 8.99E-08 | 1.66E-07 |
| chr13 | 30796229 | 30797227 | 999 | -6.12 | 9.00E-08 | 1.66E-07 |
| chr6 | 34441588 | 34442936 | 1349 | -6.12 | 9.00E-08 | 1.66E-07 |
| chr20 | 12986400 | 13014299 | 27900 | -2.38 | 9.00E-08 | 1.66E-07 |
| chr2 | 97646223 | 97647401 | 1179 | -6.16 | 9.01E-08 | 1.66E-07 |
| chr9 | 36425840 | 36426850 | 1011 | -6.16 | 9.01E-08 | 1.66E-07 |
| chr4 | 136506239 | 136519693 | 13455 | -7.19 | 9.02E-08 | 1.66E-07 |
| chr4 | 96245388 | 96246829 | 1442 | -6.11 | 9.03E-08 | 1.66E-07 |
| chr9 | 92870693 | 92893557 | 22865 | -1.72 | 9.05E-08 | 1.67E-07 |
| chr11 | 95426568 | 95428568 | 2001 | -5.07 | 9.06E-08 | 1.67E-07 |
| chr21 | 19312588 | 19340688 | 28101 | 2.42 | 9.07E-08 | 1.67E-07 |
| chr2 | 199151922 | 199154513 | 2592 | 6.17 | 9.07E-08 | 1.67E-07 |
| chr7 | 106746851 | 106747849 | 999 | 6.17 | 9.07E-08 | 1.67E-07 |
| chr22 | 27797853 | 27799387 | 1535 | -6.1 | 9.10E-08 | 1.68E-07 |
| chr6 | 33524957 | 33526283 | 1327 | -6.1 | 9.10E-08 | 1.68E-07 |
| chr1 | 193896471 | 193897834 | 1364 | -6.1 | 9.10E-08 | 1.68E-07 |
| chr7 | 13127796 | 13129655 | 1860 | -6.1 | 9.10E-08 | 1.68E-07 |
| chr21 | 37679352 | 37680382 | 1031 | -6.13 | 9.11E-08 | 1.68E-07 |
| chr3 | 188659661 | 188662665 | 3005 | -6.29 | 9.12E-08 | 1.68E-07 |
| chr17 | 45940635 | 45941644 | 1010 | -6.12 | 9.13E-08 | 1.68E-07 |
| chr3 | 150332289 | 150334738 | 2450 | -6.12 | 9.13E-08 | 1.68E-07 |
| chr11 | 16378806 | 16380693 | 1888 | -6.26 | 9.15E-08 | 1.69E-07 |
| chr12 | 92044638 | 92045637 | 1000 | -6.15 | 9.15E-08 | 1.69E-07 |
| chr1 | 109021431 | 109022927 | 1497 | -6.32 | 9.15E-08 | 1.69E-07 |

| | | | | | | |
|---|---|---|---|---|---|---|
| chr12 | 40621204 | 40621877 | 674 | -6.1 | 9.15E-08 | 1.69E-07 |
| chr7 | 5512009 | 5513017 | 1009 | -6.1 | 9.15E-08 | 1.69E-07 |
| chr20 | 33868168 | 33870318 | 2151 | -6.26 | 9.16E-08 | 1.69E-07 |
| chr6 | 156880316 | 156946678 | 66363 | -3.32 | 9.18E-08 | 1.69E-07 |
| chr10 | 17759335 | 17760015 | 681 | -6.11 | 9.19E-08 | 1.69E-07 |
| chr4 | 132487983 | 132490250 | 2268 | -6.11 | 9.19E-08 | 1.69E-07 |
| chr10 | 86317427 | 86341087 | 23661 | -1.5 | 9.20E-08 | 1.69E-07 |
| chr4 | 102828018 | 102836471 | 8454 | -6.46 | 9.20E-08 | 1.70E-07 |
| chr1 | 214979398 | 214980287 | 890 | -3.95 | 9.23E-08 | 1.70E-07 |
| chr11 | 103387270 | 103388703 | 1434 | -6.11 | 9.23E-08 | 1.70E-07 |
| chr11 | 94186112 | 94187175 | 1064 | -6.13 | 9.26E-08 | 1.71E-07 |
| chr14 | 35601362 | 35602660 | 1299 | -6.13 | 9.26E-08 | 1.71E-07 |
| chr4 | 156849468 | 156851266 | 1799 | -6.13 | 9.26E-08 | 1.71E-07 |
| chr2 | 108880905 | 108882042 | 1138 | -6.23 | 9.27E-08 | 1.71E-07 |
| chr22 | 33015150 | 33016873 | 1724 | -2.64 | 9.28E-08 | 1.71E-07 |
| chrX | 36253911 | 36256943 | 3033 | -6.37 | 9.29E-08 | 1.71E-07 |
| chr8 | 119392742 | 119394274 | 1533 | -6.14 | 9.30E-08 | 1.71E-07 |
| chr13 | 81612066 | 81613902 | 1837 | -6.12 | 9.30E-08 | 1.71E-07 |
| chrX | 18024470 | 18026287 | 1818 | 6.2 | 9.32E-08 | 1.72E-07 |
| chr5 | 108992529 | 108994471 | 1943 | 6.25 | 9.34E-08 | 1.72E-07 |
| chr2 | 233046264 | 233060706 | 14443 | -6.28 | 9.34E-08 | 1.72E-07 |
| chr3 | 148804640 | 148805754 | 1115 | -6.27 | 9.35E-08 | 1.72E-07 |
| chr7 | 82873834 | 82875402 | 1569 | -6.27 | 9.35E-08 | 1.72E-07 |
| chr17 | 32981006 | 33140232 | 159227 | 2.37 | 9.35E-08 | 1.72E-07 |
| chr18 | 24254767 | 24256760 | 1994 | -6.48 | 9.36E-08 | 1.72E-07 |
| chr8 | 134355103 | 134357035 | 1933 | -6.31 | 9.36E-08 | 1.72E-07 |
| chr2 | 107140183 | 107141468 | 1286 | -6.26 | 9.36E-08 | 1.72E-07 |
| chr20 | 1043813 | 1084708 | 40896 | -2.63 | 9.36E-08 | 1.72E-07 |
| chrX | 73606379 | 73608053 | 1675 | -6.31 | 9.37E-08 | 1.72E-07 |
| chr12 | 52552306 | 52553779 | 1474 | -6.17 | 9.37E-08 | 1.72E-07 |
| chr15 | 91258583 | 91259764 | 1182 | -6.25 | 9.37E-08 | 1.72E-07 |
| chr8 | 143686376 | 143687529 | 1154 | -6.25 | 9.37E-08 | 1.72E-07 |
| chr15 | 32164841 | 32260476 | 95636 | -0.65 | 9.38E-08 | 1.73E-07 |
| chr18 | 60237323 | 60238510 | 1188 | -6.4 | 9.40E-08 | 1.73E-07 |
| chr12 | 68874005 | 68875261 | 1257 | -6.15 | 9.42E-08 | 1.73E-07 |
| chr5 | 73262251 | 73263915 | 1665 | -6.15 | 9.42E-08 | 1.73E-07 |
| chr2 | 84784981 | 84798774 | 13794 | -2.06 | 9.43E-08 | 1.73E-07 |
| chr11 | 45615210 | 45624327 | 9118 | -2.27 | 9.43E-08 | 1.73E-07 |
| chr5 | 158435581 | 158436979 | 1399 | -6.09 | 9.44E-08 | 1.74E-07 |
| chr20 | 55808120 | 55824033 | 15914 | -5.31 | 9.46E-08 | 1.74E-07 |
| chr4 | 121692084 | 121693189 | 1106 | -6.13 | 9.46E-08 | 1.74E-07 |
| chrX | 72351531 | 72360921 | 9391 | 2.73 | 9.47E-08 | 1.74E-07 |
| chr7 | 46731664 | 46732821 | 1158 | -4.89 | 9.47E-08 | 1.74E-07 |
| chr12 | 89508329 | 89509048 | 720 | -6.09 | 9.47E-08 | 1.74E-07 |
| chr18 | 9490713 | 9493027 | 2315 | -6.36 | 9.48E-08 | 1.74E-07 |
| chr3 | 69837538 | 69849878 | 12341 | -6.41 | 9.48E-08 | 1.74E-07 |
| chr22 | 36611134 | 36612276 | 1143 | -6.12 | 9.50E-08 | 1.75E-07 |
| chr17 | 80726488 | 80727421 | 934 | -6.09 | 9.50E-08 | 1.75E-07 |
| chrX | 7085657 | 7087884 | 2228 | -6.09 | 9.50E-08 | 1.75E-07 |
| chr7 | 5358384 | 5359636 | 1253 | -6.09 | 9.51E-08 | 1.75E-07 |
| chrX | 23979680 | 23981173 | 1494 | -6.22 | 9.52E-08 | 1.75E-07 |
| chr2 | 198706823 | 198710051 | 3229 | 6.67 | 9.53E-08 | 1.75E-07 |
| chr7 | 115087001 | 115088037 | 1037 | -6.1 | 9.54E-08 | 1.75E-07 |
| chr4 | 143805910 | 143808253 | 2344 | 3.61 | 9.59E-08 | 1.76E-07 |
| chr16 | 48274220 | 48275523 | 1304 | -6.18 | 9.59E-08 | 1.76E-07 |
| chrX | 37864191 | 37865294 | 1104 | -6.18 | 9.59E-08 | 1.76E-07 |
| chr8 | 102725935 | 102727467 | 1533 | -5.52 | 9.60E-08 | 1.76E-07 |
| chrX | 131254715 | 131255895 | 1181 | -6.15 | 9.60E-08 | 1.76E-07 |
| chr13 | 103433897 | 103434709 | 813 | -6.1 | 9.62E-08 | 1.77E-07 |
| chr4 | 127906231 | 127908281 | 2051 | -6.2 | 9.62E-08 | 1.77E-07 |
| chr1 | 223953950 | 223954879 | 930 | -6.44 | 9.63E-08 | 1.77E-07 |

| | | | | | | |
|---|---|---|---|---|---|---|
| chr8 | 71498636 | 71499818 | 1183 | -6.73 | 9.64E-08 | 1.77E-07 |
| chr6 | 4707496 | 4708720 | 1225 | -6.13 | 9.64E-08 | 1.77E-07 |
| chr16 | 20670228 | 20678669 | 8442 | -1.35 | 9.64E-08 | 1.77E-07 |
| chr13 | 95161868 | 95175177 | 13310 | -1.79 | 9.67E-08 | 1.78E-07 |
| chr3 | 107586989 | 107589398 | 2410 | -6.4 | 9.69E-08 | 1.78E-07 |
| chr6 | 9326585 | 9329269 | 2685 | -6.17 | 9.73E-08 | 1.79E-07 |
| chr8 | 71152571 | 71153619 | 1049 | -6.17 | 9.73E-08 | 1.79E-07 |
| chr7 | 133653670 | 133655647 | 1978 | -6.14 | 9.74E-08 | 1.79E-07 |
| chrX | 150105280 | 150106508 | 1229 | -6.19 | 9.75E-08 | 1.79E-07 |
| chr4 | 174785277 | 174786652 | 1376 | -6.42 | 9.76E-08 | 1.79E-07 |
| chr6 | 170482277 | 170483409 | 1133 | -6.1 | 9.77E-08 | 1.79E-07 |
| chr13 | 90846317 | 90848442 | 2126 | -6.1 | 9.77E-08 | 1.79E-07 |
| chr18 | 48521224 | 48522186 | 963 | -6.1 | 9.77E-08 | 1.79E-07 |
| chr4 | 131696815 | 131698426 | 1612 | -6.1 | 9.77E-08 | 1.79E-07 |
| chr9 | 30656465 | 30657817 | 1353 | 5.44 | 9.79E-08 | 1.80E-07 |
| chr13 | 112810137 | 112813552 | 3416 | -2.23 | 9.80E-08 | 1.80E-07 |
| chr9 | 77458016 | 77459439 | 1424 | -6.16 | 9.81E-08 | 1.80E-07 |
| chr3 | 42865524 | 42866673 | 1150 | -6.13 | 9.81E-08 | 1.80E-07 |
| chr19 | 45533601 | 45534660 | 1060 | -6.18 | 9.83E-08 | 1.80E-07 |
| chr1 | 211713412 | 211715195 | 1784 | -6.43 | 9.84E-08 | 1.81E-07 |
| chr1 | 39412581 | 39413762 | 1182 | -6.12 | 9.84E-08 | 1.81E-07 |
| chrX | 19767116 | 19768159 | 1044 | -6.12 | 9.84E-08 | 1.81E-07 |
| chr9 | 74110378 | 74116078 | 5701 | -1.8 | 9.84E-08 | 1.81E-07 |
| chr2 | 75009607 | 75036910 | 27304 | -6.24 | 9.85E-08 | 1.81E-07 |
| chr6 | 56909134 | 56910420 | 1287 | -6.08 | 9.87E-08 | 1.81E-07 |
| chr3 | 128748749 | 128749876 | 1128 | -6.09 | 9.87E-08 | 1.81E-07 |
| chrX | 62987527 | 63000871 | 13345 | -2.21 | 9.88E-08 | 1.81E-07 |
| chr19 | 3869018 | 3870256 | 1239 | -6.14 | 9.89E-08 | 1.81E-07 |
| chr2 | 100860954 | 100862403 | 1450 | -6.14 | 9.89E-08 | 1.81E-07 |
| chr6 | 1692039 | 1693185 | 1147 | -6.14 | 9.89E-08 | 1.81E-07 |
| chr2 | 177713311 | 177714878 | 1568 | 6.22 | 9.89E-08 | 1.81E-07 |
| chr7 | 106854059 | 106855434 | 1376 | -6.16 | 9.89E-08 | 1.82E-07 |
| chr16 | 56306740 | 56309872 | 3133 | -5.72 | 9.89E-08 | 1.82E-07 |
| chr3 | 16848264 | 16849188 | 925 | 4.69 | 9.91E-08 | 1.82E-07 |
| chr22 | 17804583 | 17806848 | 2266 | -6.17 | 9.91E-08 | 1.82E-07 |
| chr1 | 213312236 | 213313402 | 1167 | -6.09 | 9.94E-08 | 1.82E-07 |
| chr3 | 48295780 | 48297344 | 1565 | -6.09 | 9.94E-08 | 1.82E-07 |
| chr1 | 110262070 | 110264838 | 2769 | -6.08 | 9.94E-08 | 1.82E-07 |
| chr11 | 88240607 | 88242999 | 2393 | -5.25 | 9.96E-08 | 1.83E-07 |
| chr9 | 3860802 | 3875152 | 14351 | -5.72 | 9.99E-08 | 1.83E-07 |
| chr12 | 12429230 | 12431366 | 2137 | -6.8 | 1.00E-07 | 1.83E-07 |
| chr5 | 53761429 | 53775628 | 14200 | -6.16 | 1.00E-07 | 1.84E-07 |
| chr7 | 23344377 | 23345597 | 1221 | -6.28 | 1.00E-07 | 1.84E-07 |
| chr7 | 55270900 | 55272353 | 1454 | -6.28 | 1.00E-07 | 1.84E-07 |
| chr17 | 43168865 | 43169893 | 1029 | -6.3 | 1.00E-07 | 1.84E-07 |
| chr15 | 63564950 | 63566181 | 1232 | -6.31 | 1.01E-07 | 1.85E-07 |
| chr3 | 43504259 | 43505744 | 1486 | -6.27 | 1.01E-07 | 1.85E-07 |
| chr10 | 121403779 | 121405408 | 1630 | -6.26 | 1.01E-07 | 1.85E-07 |
| chr17 | 35681896 | 35683258 | 1363 | -6.26 | 1.01E-07 | 1.85E-07 |
| chr4 | 89353981 | 89355285 | 1305 | -6.29 | 1.01E-07 | 1.85E-07 |
| chr1 | 55577705 | 55578965 | 1261 | -6.22 | 1.01E-07 | 1.86E-07 |
| chr15 | 59561459 | 59562316 | 858 | -4.62 | 1.01E-07 | 1.86E-07 |
| chr16 | 20017652 | 20019187 | 1536 | 6.16 | 1.01E-07 | 1.86E-07 |
| chr6 | 140061186 | 140064123 | 2938 | -6.52 | 1.02E-07 | 1.86E-07 |
| chr2 | 227913631 | 227914624 | 994 | -6.33 | 1.02E-07 | 1.86E-07 |
| chr4 | 38877672 | 38892583 | 14912 | -6.35 | 1.02E-07 | 1.87E-07 |
| chr20 | 39544183 | 39545904 | 1722 | -6.14 | 1.02E-07 | 1.87E-07 |
| chr10 | 34546737 | 34549146 | 2410 | -6.49 | 1.02E-07 | 1.87E-07 |
| chr3 | 177390575 | 177392608 | 2034 | -6.34 | 1.02E-07 | 1.87E-07 |
| chr7 | 23228045 | 23229844 | 1800 | -6.53 | 1.02E-07 | 1.87E-07 |
| chr22 | 29380979 | 29382208 | 1230 | -6.21 | 1.02E-07 | 1.87E-07 |

| | | | | | | |
|---|---|---|---|---|---|---|
| chr3 | 183013898 | 183015005 | 1108 | -6.21 | 1.02E-07 | 1.87E-07 |
| chr4 | 111066941 | 111067998 | 1058 | -6.21 | 1.02E-07 | 1.87E-07 |
| chr4 | 150550319 | 150551837 | 1519 | -6.2 | 1.03E-07 | 1.88E-07 |
| chr15 | 62646186 | 62647237 | 1052 | -6.09 | 1.03E-07 | 1.88E-07 |
| chr2 | 165894773 | 165895716 | 944 | -6.09 | 1.03E-07 | 1.88E-07 |
| chr8 | 17587412 | 17588620 | 1209 | -6.1 | 1.03E-07 | 1.88E-07 |
| chr2 | 42563724 | 42564812 | 1089 | -6.08 | 1.03E-07 | 1.88E-07 |
| chr4 | 123658459 | 123659540 | 1082 | -6.08 | 1.03E-07 | 1.88E-07 |
| chr19 | 39424929 | 39426061 | 1133 | -6.2 | 1.03E-07 | 1.88E-07 |
| chr6 | 45624419 | 45625561 | 1143 | -6.08 | 1.03E-07 | 1.88E-07 |
| chr9 | 95118489 | 95120650 | 2162 | -6.08 | 1.03E-07 | 1.88E-07 |
| chr6 | 133197648 | 133198883 | 1236 | -6.11 | 1.03E-07 | 1.89E-07 |
| chr19 | 44115862 | 44117354 | 1493 | -6.36 | 1.03E-07 | 1.89E-07 |
| chrX | 41880573 | 41893594 | 13022 | -1.71 | 1.03E-07 | 1.89E-07 |
| chr2 | 42230940 | 42232397 | 1458 | -6.14 | 1.04E-07 | 1.90E-07 |
| chr7 | 98862971 | 98864344 | 1374 | -6.14 | 1.04E-07 | 1.90E-07 |
| chr21 | 39415447 | 39417092 | 1646 | -5.37 | 1.04E-07 | 1.90E-07 |
| chr11 | 12386258 | 12387721 | 1464 | -6.19 | 1.04E-07 | 1.90E-07 |
| chr17 | 79222375 | 79224226 | 1852 | -6.19 | 1.04E-07 | 1.90E-07 |
| chrX | 120575275 | 120576792 | 1518 | 6.54 | 1.04E-07 | 1.90E-07 |
| chr2 | 101297794 | 101317568 | 19775 | -6.19 | 1.04E-07 | 1.91E-07 |
| chr18 | 24481231 | 24504009 | 22779 | -1.99 | 1.04E-07 | 1.91E-07 |
| chrX | 70737143 | 70751138 | 13996 | -6.52 | 1.04E-07 | 1.91E-07 |
| chr11 | 31699888 | 31701015 | 1128 | -6.12 | 1.04E-07 | 1.91E-07 |
| chr6 | 166022687 | 166025101 | 2415 | -6.18 | 1.05E-07 | 1.91E-07 |
| chr10 | 44321887 | 44773566 | 451680 | 0.41 | 1.05E-07 | 1.92E-07 |
| chr20 | 30017432 | 30018265 | 834 | -6.13 | 1.05E-07 | 1.93E-07 |
| chr2 | 187552033 | 187553097 | 1065 | -6.17 | 1.05E-07 | 1.93E-07 |
| chr10 | 91242126 | 91243084 | 959 | -6.09 | 1.05E-07 | 1.93E-07 |
| chr3 | 90286573 | 90293194 | 6622 | -2.6 | 1.06E-07 | 1.93E-07 |
| chr3 | 58972057 | 58973196 | 1140 | -5.34 | 1.06E-07 | 1.94E-07 |
| chr7 | 95126436 | 95127821 | 1386 | -6.16 | 1.06E-07 | 1.94E-07 |
| chr8 | 16752114 | 16753914 | 1801 | 6.14 | 1.06E-07 | 1.94E-07 |
| chr12 | 12378373 | 12392129 | 13757 | -7.07 | 1.06E-07 | 1.95E-07 |
| chr11 | 86046280 | 86047184 | 905 | -6.1 | 1.06E-07 | 1.95E-07 |
| chrX | 48711393 | 48720860 | 9468 | -6.43 | 1.06E-07 | 1.95E-07 |
| chr10 | 87453442 | 87454651 | 1210 | -6.08 | 1.07E-07 | 1.95E-07 |
| chr5 | 157127659 | 157128944 | 1286 | -6.12 | 1.07E-07 | 1.95E-07 |
| chr2 | 57974150 | 57975048 | 899 | -6.45 | 1.07E-07 | 1.96E-07 |
| chr16 | 49904620 | 49905636 | 1017 | -6.07 | 1.07E-07 | 1.96E-07 |
| chr1 | 237114011 | 237165955 | 51945 | -2.05 | 1.07E-07 | 1.96E-07 |
| chr18 | 33013481 | 33015712 | 2232 | -6.6 | 1.07E-07 | 1.96E-07 |
| chr1 | 162011491 | 162013193 | 1703 | -6.07 | 1.08E-07 | 1.97E-07 |
| chr3 | 10519178 | 10520430 | 1253 | -6.07 | 1.08E-07 | 1.97E-07 |
| chr12 | 78294205 | 78304847 | 10643 | -6.57 | 1.08E-07 | 1.98E-07 |
| chr9 | 139161211 | 139163109 | 1899 | -6.45 | 1.08E-07 | 1.98E-07 |
| chr1 | 214448358 | 214450200 | 1843 | -6.15 | 1.08E-07 | 1.98E-07 |
| chr6 | 55437718 | 55438874 | 1157 | -6.15 | 1.08E-07 | 1.98E-07 |
| chr13 | 33970933 | 33972907 | 1975 | -6.08 | 1.08E-07 | 1.98E-07 |
| chr6 | 14351048 | 14352505 | 1458 | -6.08 | 1.08E-07 | 1.98E-07 |
| chr3 | 51525073 | 51525975 | 903 | -6.14 | 1.09E-07 | 1.98E-07 |
| chr1 | 210569617 | 210570422 | 806 | -6.46 | 1.09E-07 | 1.99E-07 |
| chr4 | 20180939 | 20183507 | 2569 | -6.26 | 1.09E-07 | 1.99E-07 |
| chr5 | 32321424 | 32322945 | 1522 | -6.26 | 1.09E-07 | 1.99E-07 |
| chr3 | 107399230 | 107400296 | 1067 | -6.09 | 1.09E-07 | 1.99E-07 |
| chr13 | 84604725 | 84608528 | 3804 | -6.29 | 1.09E-07 | 1.99E-07 |
| chr6 | 16142318 | 16146561 | 4244 | -6.47 | 1.09E-07 | 1.99E-07 |
| chr12 | 117890157 | 117891103 | 947 | -6.23 | 1.09E-07 | 2.00E-07 |
| chr6 | 137171608 | 137174792 | 3185 | -6.07 | 1.09E-07 | 2.00E-07 |
| chrX | 117663938 | 117665827 | 1890 | -6.22 | 1.09E-07 | 2.00E-07 |
| chr13 | 30872248 | 30873538 | 1291 | -6.14 | 1.10E-07 | 2.00E-07 |

| | | | | | | |
|---|---|---|---|---|---|---|
| chr14 | 32778227 | 32779447 | 1221 | -6.06 | 1.10E-07 | 2.00E-07 |
| chr11 | 118570584 | 118571910 | 1327 | -6.31 | 1.10E-07 | 2.00E-07 |
| chr3 | 40210390 | 40211324 | 935 | -3.18 | 1.10E-07 | 2.00E-07 |
| chr17 | 71955176 | 71980295 | 25120 | -2.57 | 1.10E-07 | 2.01E-07 |
| chr4 | 53950786 | 53952838 | 2053 | -6.27 | 1.11E-07 | 2.02E-07 |
| chr1 | 227054004 | 227054742 | 739 | -6.27 | 1.11E-07 | 2.02E-07 |
| chr12 | 26697877 | 26698961 | 1085 | -6.13 | 1.11E-07 | 2.02E-07 |
| chr15 | 71165448 | 71166202 | 755 | -6.13 | 1.11E-07 | 2.02E-07 |
| chr1 | 153530008 | 153531495 | 1488 | -6.24 | 1.11E-07 | 2.02E-07 |
| chr5 | 43989269 | 44014114 | 24846 | -1.48 | 1.11E-07 | 2.02E-07 |
| chr5 | 142804353 | 142805697 | 1345 | -6.1 | 1.11E-07 | 2.02E-07 |
| chr15 | 27125764 | 27128887 | 3124 | 4.74 | 1.11E-07 | 2.03E-07 |
| chr3 | 129493122 | 129499072 | 5951 | -6.19 | 1.11E-07 | 2.03E-07 |
| chr8 | 102451495 | 102455986 | 4492 | -6.31 | 1.11E-07 | 2.03E-07 |
| chr10 | 29748863 | 29750047 | 1185 | -6.09 | 1.11E-07 | 2.03E-07 |
| chr3 | 15634611 | 15647057 | 12447 | -6.09 | 1.11E-07 | 2.03E-07 |
| chr15 | 62844141 | 62845374 | 1234 | -6.31 | 1.11E-07 | 2.03E-07 |
| chr14 | 24076722 | 24096532 | 19811 | -3.44 | 1.11E-07 | 2.03E-07 |
| chr12 | 66780030 | 66781730 | 1701 | -6.21 | 1.11E-07 | 2.03E-07 |
| chr16 | 80198676 | 80201177 | 2502 | -6.21 | 1.11E-07 | 2.03E-07 |
| chr20 | 55442321 | 55451627 | 9307 | -7.31 | 1.12E-07 | 2.04E-07 |
| chr1 | 244150266 | 244159282 | 9017 | -6.18 | 1.12E-07 | 2.04E-07 |
| chr13 | 110825288 | 110826797 | 1510 | -6.2 | 1.12E-07 | 2.04E-07 |
| chr5 | 83906041 | 83917642 | 11602 | 1.77 | 1.12E-07 | 2.04E-07 |
| chr4 | 71441280 | 71442781 | 1502 | -5.51 | 1.12E-07 | 2.04E-07 |
| chr6 | 4928807 | 4930623 | 1817 | -6.06 | 1.12E-07 | 2.04E-07 |
| chr20 | 32036061 | 32037323 | 1263 | -6.12 | 1.12E-07 | 2.05E-07 |
| chr4 | 128920673 | 128921943 | 1271 | -6.17 | 1.12E-07 | 2.05E-07 |
| chr8 | 128844639 | 128847240 | 2602 | -6.19 | 1.13E-07 | 2.06E-07 |
| chr10 | 90374260 | 90375562 | 1303 | -6.12 | 1.13E-07 | 2.06E-07 |
| chrX | 140041426 | 140044929 | 3504 | -6.06 | 1.13E-07 | 2.06E-07 |
| chr22 | 30786058 | 30787716 | 1659 | -6.64 | 1.13E-07 | 2.06E-07 |
| chr4 | 106159985 | 106177941 | 17957 | -6.88 | 1.13E-07 | 2.06E-07 |
| chr13 | 35249937 | 35251987 | 2051 | -6.17 | 1.14E-07 | 2.07E-07 |
| chr18 | 52842848 | 52844387 | 1540 | -6.16 | 1.14E-07 | 2.08E-07 |
| chr12 | 52984157 | 52986127 | 1971 | -6.53 | 1.14E-07 | 2.08E-07 |
| chr4 | 87292926 | 87301710 | 8785 | -6.53 | 1.14E-07 | 2.08E-07 |
| chr8 | 120073352 | 120074163 | 812 | -6.09 | 1.15E-07 | 2.09E-07 |
| chrX | 29856261 | 29858017 | 1757 | -6.15 | 1.15E-07 | 2.09E-07 |
| chr8 | 28510843 | 28512154 | 1312 | -6.78 | 1.15E-07 | 2.10E-07 |
| chr20 | 49170635 | 49176255 | 5621 | -6.42 | 1.15E-07 | 2.10E-07 |
| chr11 | 74364000 | 74364979 | 980 | -6.15 | 1.16E-07 | 2.11E-07 |
| chr2 | 165448182 | 165449454 | 1273 | -6.15 | 1.16E-07 | 2.11E-07 |
| chr16 | 10826887 | 10828358 | 1472 | -6.11 | 1.16E-07 | 2.11E-07 |
| chr17 | 31268143 | 31269061 | 919 | -6.11 | 1.16E-07 | 2.11E-07 |
| chr2 | 152841698 | 152843683 | 1986 | 6.19 | 1.16E-07 | 2.11E-07 |
| chr7 | 14205548 | 14209198 | 3651 | -6.55 | 1.16E-07 | 2.11E-07 |
| chr6 | 143729312 | 143729912 | 601 | -6.07 | 1.16E-07 | 2.12E-07 |
| chr5 | 64195202 | 64196177 | 976 | -6.05 | 1.16E-07 | 2.12E-07 |
| chr5 | 67937076 | 67960398 | 23323 | -7.14 | 1.17E-07 | 2.12E-07 |
| chr15 | 79374553 | 79375377 | 825 | -6.13 | 1.17E-07 | 2.13E-07 |
| chr4 | 7347622 | 7349113 | 1492 | -6.13 | 1.17E-07 | 2.13E-07 |
| chr5 | 95140356 | 95141521 | 1166 | -6.45 | 1.17E-07 | 2.13E-07 |
| chr2 | 193672224 | 193674236 | 2013 | -6.14 | 1.17E-07 | 2.14E-07 |
| chr3 | 37252670 | 37253783 | 1114 | -6.07 | 1.17E-07 | 2.14E-07 |
| chr7 | 85471860 | 85473667 | 1808 | -6.07 | 1.17E-07 | 2.14E-07 |
| chrX | 30082792 | 30084534 | 1743 | -6.07 | 1.17E-07 | 2.14E-07 |
| chr4 | 151376591 | 151377897 | 1307 | -6.05 | 1.17E-07 | 2.14E-07 |
| chr2 | 138619059 | 138622245 | 3187 | -2.16 | 1.18E-07 | 2.14E-07 |
| chr10 | 73502070 | 73503509 | 1440 | -6.24 | 1.18E-07 | 2.14E-07 |
| chr8 | 126660589 | 126661616 | 1028 | -6.1 | 1.18E-07 | 2.14E-07 |

| | | | | | | |
|---|---|---|---|---|---|---|
| chrX | 44897931 | 44898863 | 933 | -6.1 | 1.18E-07 | 2.14E-07 |
| chr1 | 22606106 | 22607318 | 1213 | -6.1 | 1.18E-07 | 2.14E-07 |
| chr5 | 50067849 | 50069656 | 1808 | -6.1 | 1.18E-07 | 2.14E-07 |
| chrX | 44722460 | 44723749 | 1290 | -6.27 | 1.18E-07 | 2.14E-07 |
| chr2 | 197743342 | 197744435 | 1094 | -6.08 | 1.18E-07 | 2.15E-07 |
| chr6 | 148163725 | 148165124 | 1400 | -6.28 | 1.18E-07 | 2.15E-07 |
| chr17 | 59241642 | 59242646 | 1005 | -6.13 | 1.18E-07 | 2.15E-07 |
| chrX | 47234943 | 47236146 | 1204 | -6.29 | 1.18E-07 | 2.15E-07 |
| chr2 | 150313476 | 150319659 | 6184 | -1.91 | 1.18E-07 | 2.15E-07 |
| chr8 | 86034140 | 86035408 | 1269 | -6.21 | 1.18E-07 | 2.15E-07 |
| chr14 | 89247793 | 89251109 | 3317 | -2.31 | 1.18E-07 | 2.16E-07 |
| chrX | 14837353 | 14848777 | 11425 | -5.11 | 1.19E-07 | 2.16E-07 |
| chr14 | 96634953 | 96647193 | 12241 | -7.37 | 1.19E-07 | 2.16E-07 |
| chr4 | 98981695 | 98995933 | 14239 | -6.07 | 1.19E-07 | 2.16E-07 |
| chr5 | 173309381 | 173310737 | 1357 | -6.32 | 1.19E-07 | 2.17E-07 |
| chr9 | 100274246 | 100274982 | 737 | -6.05 | 1.19E-07 | 2.17E-07 |
| chr12 | 43619562 | 43621679 | 2118 | -6.33 | 1.20E-07 | 2.18E-07 |
| chr8 | 128253358 | 128255701 | 2344 | -6.33 | 1.20E-07 | 2.18E-07 |
| chr2 | 210331174 | 210332764 | 1591 | -6.09 | 1.20E-07 | 2.18E-07 |
| chr4 | 89638080 | 89639918 | 1839 | -6.17 | 1.20E-07 | 2.18E-07 |
| chr18 | 74439483 | 74442764 | 3282 | -4 | 1.20E-07 | 2.19E-07 |
| chr13 | 43299627 | 43300982 | 1356 | -6.12 | 1.20E-07 | 2.19E-07 |
| chr9 | 104080081 | 104093378 | 13298 | -6.86 | 1.20E-07 | 2.19E-07 |
| chr4 | 141692286 | 141693314 | 1029 | -6.07 | 1.21E-07 | 2.19E-07 |
| chr10 | 76183719 | 76184570 | 852 | -6.17 | 1.21E-07 | 2.20E-07 |
| chr2 | 7651716 | 7673367 | 21652 | -2 | 1.21E-07 | 2.20E-07 |
| chr6 | 168781882 | 168784052 | 2171 | -4.56 | 1.21E-07 | 2.20E-07 |
| chr1 | 93085059 | 93086085 | 1027 | -6.05 | 1.21E-07 | 2.21E-07 |
| chr7 | 23268330 | 23269182 | 853 | -6.05 | 1.21E-07 | 2.21E-07 |
| chr22 | 43492000 | 43493289 | 1290 | -6.06 | 1.21E-07 | 2.21E-07 |
| chr6 | 150513124 | 150515005 | 1882 | -6.06 | 1.21E-07 | 2.21E-07 |
| chr9 | 119517687 | 119518719 | 1033 | -6.11 | 1.21E-07 | 2.21E-07 |
| chr1 | 179624212 | 179649626 | 25415 | -6.17 | 1.21E-07 | 2.21E-07 |
| chrX | 45703722 | 45704751 | 1030 | -6.05 | 1.22E-07 | 2.21E-07 |
| chrX | 29046297 | 29048535 | 2239 | -6.37 | 1.22E-07 | 2.21E-07 |
| chr17 | 70854299 | 70855475 | 1177 | -6.23 | 1.22E-07 | 2.21E-07 |
| chr7 | 105214345 | 105215337 | 993 | -6.22 | 1.22E-07 | 2.22E-07 |
| chr1 | 149245493 | 149246581 | 1089 | -6.05 | 1.22E-07 | 2.22E-07 |
| chr11 | 44604060 | 44605260 | 1201 | -6.05 | 1.22E-07 | 2.22E-07 |
| chr15 | 53862174 | 53863391 | 1218 | -6.04 | 1.22E-07 | 2.22E-07 |
| chrX | 10447642 | 10449425 | 1784 | -6.28 | 1.22E-07 | 2.22E-07 |
| chr2 | 187302293 | 187304658 | 2366 | -6.37 | 1.22E-07 | 2.22E-07 |
| chr2 | 121150543 | 121152029 | 1487 | -6.08 | 1.22E-07 | 2.22E-07 |
| chr5 | 68605118 | 68606795 | 1678 | -6.08 | 1.22E-07 | 2.22E-07 |
| chr6 | 130199207 | 130200808 | 1602 | -3.43 | 1.22E-07 | 2.23E-07 |
| chr18 | 3608052 | 3617144 | 9093 | -6.29 | 1.23E-07 | 2.23E-07 |
| chr1 | 171310638 | 171319195 | 8558 | -6.05 | 1.23E-07 | 2.23E-07 |
| chr1 | 193014545 | 193016039 | 1495 | -6.65 | 1.23E-07 | 2.23E-07 |
| chr2 | 146658858 | 146660865 | 2008 | -6.39 | 1.23E-07 | 2.24E-07 |
| chr13 | 20530022 | 20530906 | 885 | -6.08 | 1.23E-07 | 2.24E-07 |
| chr20 | 48542407 | 48543267 | 861 | -6.18 | 1.23E-07 | 2.24E-07 |
| chr12 | 56184709 | 56185599 | 891 | -6.18 | 1.24E-07 | 2.25E-07 |
| chrX | 39442633 | 39443659 | 1027 | -6.06 | 1.24E-07 | 2.25E-07 |
| chr13 | 36615064 | 36616598 | 1535 | 6.53 | 1.24E-07 | 2.25E-07 |
| chrX | 42049564 | 42053658 | 4095 | -2.15 | 1.24E-07 | 2.25E-07 |
| chr14 | 94818348 | 94820098 | 1751 | -6.17 | 1.24E-07 | 2.25E-07 |
| chr14 | 52758430 | 52759139 | 710 | -6.06 | 1.24E-07 | 2.25E-07 |
| chr2 | 40442462 | 40446117 | 3656 | -2.39 | 1.24E-07 | 2.26E-07 |
| chrX | 18861499 | 18863181 | 1683 | -6.41 | 1.24E-07 | 2.26E-07 |
| chrX | 110430023 | 110431553 | 1531 | 4.76 | 1.24E-07 | 2.26E-07 |
| chr15 | 45743601 | 45744603 | 1003 | -6.05 | 1.25E-07 | 2.26E-07 |

| | | | | | | |
|---|---|---|---|---|---|---|
| chr3 | 37993813 | 37994875 | 1063 | -6.05 | 1.25E-07 | 2.26E-07 |
| chr4 | 148873289 | 148874888 | 1600 | -6.17 | 1.25E-07 | 2.26E-07 |
| chr1 | 164617037 | 164624299 | 7263 | 6.22 | 1.25E-07 | 2.27E-07 |
| chr10 | 59431752 | 59433775 | 2024 | -6.09 | 1.25E-07 | 2.27E-07 |
| chr19 | 49229613 | 49234922 | 5310 | -6.13 | 1.25E-07 | 2.27E-07 |
| chr6 | 141265448 | 141267161 | 1714 | -6.07 | 1.25E-07 | 2.28E-07 |
| chr7 | 14139749 | 14142527 | 2779 | -6.07 | 1.25E-07 | 2.28E-07 |
| chr9 | 95723358 | 95724769 | 1412 | -6.1 | 1.26E-07 | 2.28E-07 |
| chr12 | 27119756 | 27120469 | 714 | -6.04 | 1.26E-07 | 2.28E-07 |
| chr12 | 88116495 | 88117772 | 1278 | -6.04 | 1.26E-07 | 2.28E-07 |
| chr15 | 85572542 | 85573545 | 1004 | -6.04 | 1.26E-07 | 2.28E-07 |
| chr4 | 180351851 | 180354016 | 2166 | -6.04 | 1.26E-07 | 2.28E-07 |
| chr2 | 145474939 | 145476969 | 2031 | -5.1 | 1.26E-07 | 2.29E-07 |
| chr1 | 94315057 | 94316197 | 1141 | -6.12 | 1.27E-07 | 2.30E-07 |
| chr16 | 52628665 | 52637194 | 8530 | -6.81 | 1.27E-07 | 2.30E-07 |
| chr3 | 111346677 | 111363663 | 16987 | -6.97 | 1.27E-07 | 2.30E-07 |
| chr11 | 94658008 | 94659991 | 1984 | -6.05 | 1.27E-07 | 2.31E-07 |
| chr14 | 24149169 | 24185292 | 36124 | -1.35 | 1.27E-07 | 2.31E-07 |
| chr8 | 37565613 | 37566672 | 1060 | -6.24 | 1.27E-07 | 2.31E-07 |
| chr13 | 83601713 | 83603936 | 2224 | -6.22 | 1.27E-07 | 2.31E-07 |
| chr1 | 22145575 | 22147267 | 1693 | -6.09 | 1.27E-07 | 2.31E-07 |
| chr10 | 124148733 | 124150705 | 1973 | -6.09 | 1.27E-07 | 2.31E-07 |
| chr8 | 80687235 | 80688435 | 1201 | -6.25 | 1.27E-07 | 2.31E-07 |
| chr16 | 12310261 | 12311566 | 1306 | -6.21 | 1.27E-07 | 2.31E-07 |
| chr5 | 83593231 | 83594795 | 1565 | -6.21 | 1.27E-07 | 2.31E-07 |
| chr9 | 95830544 | 95831182 | 639 | -6.05 | 1.27E-07 | 2.31E-07 |
| chr1 | 199855940 | 199856887 | 948 | -6.08 | 1.27E-07 | 2.31E-07 |
| chr15 | 66705031 | 66716851 | 11821 | -6.27 | 1.27E-07 | 2.31E-07 |
| chr10 | 24853944 | 24861382 | 7439 | -6.2 | 1.28E-07 | 2.31E-07 |
| chr5 | 106353626 | 106355770 | 2145 | -6.2 | 1.28E-07 | 2.31E-07 |
| chr9 | 72015238 | 72017233 | 1996 | -6.19 | 1.28E-07 | 2.32E-07 |
| chr1 | 30548558 | 30550691 | 2134 | -6.13 | 1.28E-07 | 2.32E-07 |
| chr9 | 36679969 | 36713633 | 33665 | -6.5 | 1.28E-07 | 2.32E-07 |
| chr3 | 94136877 | 94145498 | 8622 | -6.46 | 1.29E-07 | 2.33E-07 |
| chr7 | 114126098 | 114128408 | 2311 | -6.3 | 1.29E-07 | 2.33E-07 |
| chr3 | 174766776 | 174769333 | 2558 | -6.04 | 1.29E-07 | 2.33E-07 |
| chr22 | 22949757 | 22950972 | 1216 | -6.08 | 1.29E-07 | 2.34E-07 |
| chr1 | 64453862 | 64469180 | 15319 | -6.59 | 1.29E-07 | 2.34E-07 |
| chr20 | 59608393 | 59609457 | 1065 | -6.04 | 1.29E-07 | 2.34E-07 |
| chrX | 41120750 | 41122806 | 2057 | -6.1 | 1.30E-07 | 2.35E-07 |
| chr7 | 117913558 | 117915408 | 1851 | 6.14 | 1.30E-07 | 2.35E-07 |
| chr13 | 90287901 | 90289292 | 1392 | -6.33 | 1.30E-07 | 2.35E-07 |
| chr8 | 119486775 | 119488035 | 1261 | -6.12 | 1.30E-07 | 2.35E-07 |
| chr7 | 81753518 | 81754327 | 810 | -6.16 | 1.30E-07 | 2.35E-07 |
| chrX | 54555163 | 54556053 | 891 | -6.06 | 1.30E-07 | 2.36E-07 |
| chr1 | 78551927 | 78553070 | 1144 | -4.58 | 1.30E-07 | 2.36E-07 |
| chr18 | 13384580 | 13386357 | 1778 | -6.15 | 1.31E-07 | 2.36E-07 |
| chr19 | 50162469 | 50164103 | 1635 | -6.15 | 1.31E-07 | 2.36E-07 |
| chr6 | 158202561 | 158203899 | 1339 | -6.05 | 1.31E-07 | 2.37E-07 |
| chr4 | 100246892 | 100249496 | 2605 | -6.04 | 1.31E-07 | 2.37E-07 |
| chr7 | 132577563 | 132579013 | 1451 | -6.14 | 1.31E-07 | 2.38E-07 |
| chr13 | 23395847 | 23397936 | 2090 | -6.09 | 1.31E-07 | 2.38E-07 |
| chr8 | 82312428 | 82327244 | 14817 | 1.88 | 1.32E-07 | 2.38E-07 |
| chr3 | 135831773 | 135832899 | 1127 | -6.11 | 1.32E-07 | 2.39E-07 |
| chr6 | 100307059 | 100308703 | 1645 | -3.45 | 1.32E-07 | 2.40E-07 |
| chr12 | 63368087 | 63368864 | 778 | -6.05 | 1.33E-07 | 2.40E-07 |
| chr3 | 96764841 | 96767187 | 2347 | 3.18 | 1.33E-07 | 2.41E-07 |
| chr9 | 88832655 | 88833854 | 1200 | -6.1 | 1.33E-07 | 2.41E-07 |
| chr12 | 89418282 | 89418876 | 595 | -6.08 | 1.33E-07 | 2.42E-07 |
| chr1 | 232497161 | 232498358 | 1198 | -6.04 | 1.34E-07 | 2.42E-07 |
| chr4 | 186159392 | 186160702 | 1311 | -6.04 | 1.34E-07 | 2.42E-07 |

| | | | | | | |
|---|---|---|---|---|---|---|
| chr18 | 33031043 | 33032957 | 1915 | -6.64 | 1.34E-07 | 2.42E-07 |
| chr10 | 97558873 | 97582678 | 23806 | -1.82 | 1.34E-07 | 2.42E-07 |
| chr3 | 119308772 | 119310196 | 1425 | -4.51 | 1.34E-07 | 2.42E-07 |
| chr6 | 129535105 | 129536390 | 1286 | 6.1 | 1.34E-07 | 2.42E-07 |
| chr6 | 114311046 | 114326588 | 15543 | -6.53 | 1.34E-07 | 2.43E-07 |
| chr10 | 134433166 | 134435700 | 2535 | -6.23 | 1.34E-07 | 2.43E-07 |
| chr19 | 33436892 | 33437623 | 732 | -6.23 | 1.34E-07 | 2.43E-07 |
| chr17 | 8199310 | 8200764 | 1455 | -6.12 | 1.34E-07 | 2.43E-07 |
| chr3 | 25690113 | 25692324 | 2212 | -6.2 | 1.35E-07 | 2.43E-07 |
| chr1 | 175878394 | 175879651 | 1258 | -6.2 | 1.35E-07 | 2.44E-07 |
| chr2 | 42082832 | 42086282 | 3451 | -5.45 | 1.35E-07 | 2.44E-07 |
| chr13 | 98704888 | 98706523 | 1636 | -6.06 | 1.35E-07 | 2.44E-07 |
| chr7 | 41141765 | 41144034 | 2270 | -6.06 | 1.35E-07 | 2.44E-07 |
| chr4 | 139202833 | 139203976 | 1144 | -6.29 | 1.35E-07 | 2.45E-07 |
| chr11 | 45339914 | 45350580 | 10667 | -6.21 | 1.35E-07 | 2.45E-07 |
| chr5 | 98425328 | 98426530 | 1203 | -6.08 | 1.36E-07 | 2.45E-07 |
| chr12 | 57359151 | 57359984 | 834 | -6.11 | 1.36E-07 | 2.45E-07 |
| chr3 | 45723223 | 45724452 | 1230 | -6.04 | 1.36E-07 | 2.46E-07 |
| chr2 | 230353765 | 230364920 | 11156 | -1.96 | 1.36E-07 | 2.46E-07 |
| chr11 | 119905766 | 119926364 | 20599 | -2.43 | 1.36E-07 | 2.46E-07 |
| chr5 | 92059131 | 92060906 | 1776 | -6.42 | 1.36E-07 | 2.47E-07 |
| chr2 | 216741777 | 216742791 | 1015 | -4.58 | 1.37E-07 | 2.47E-07 |
| chr5 | 102495295 | 102496632 | 1338 | -6.32 | 1.37E-07 | 2.48E-07 |
| chr6 | 3305706 | 3306969 | 1264 | -6.43 | 1.37E-07 | 2.48E-07 |
| chrX | 120244286 | 120245241 | 956 | -6.02 | 1.37E-07 | 2.48E-07 |
| chrX | 80018864 | 80020004 | 1141 | -6.03 | 1.37E-07 | 2.48E-07 |
| chr18 | 20441332 | 20444904 | 3573 | -2.46 | 1.37E-07 | 2.49E-07 |
| chr3 | 18372787 | 18375254 | 2468 | -6.19 | 1.38E-07 | 2.49E-07 |
| chr15 | 67082953 | 67084412 | 1460 | -6.02 | 1.38E-07 | 2.49E-07 |
| chr1 | 92015958 | 92030340 | 14383 | -6.07 | 1.38E-07 | 2.50E-07 |
| chr2 | 46354155 | 46355255 | 1101 | -6.04 | 1.38E-07 | 2.50E-07 |
| chr7 | 13856831 | 13860069 | 3239 | -6.03 | 1.38E-07 | 2.50E-07 |
| chr4 | 160154183 | 160155472 | 1290 | -6.14 | 1.38E-07 | 2.50E-07 |
| chr1 | 89560435 | 89567284 | 6850 | -3.39 | 1.39E-07 | 2.50E-07 |
| chr10 | 120336449 | 120338277 | 1829 | 6.33 | 1.39E-07 | 2.51E-07 |
| chr7 | 13802029 | 13814845 | 12817 | -6.84 | 1.39E-07 | 2.51E-07 |
| chr13 | 37589580 | 37590198 | 619 | -4.54 | 1.39E-07 | 2.51E-07 |
| chr5 | 88004220 | 88006819 | 2600 | 6.3 | 1.39E-07 | 2.51E-07 |
| chr6 | 82633171 | 82635117 | 1947 | -6.3 | 1.39E-07 | 2.52E-07 |
| chr8 | 39726146 | 39733853 | 7708 | -2.48 | 1.40E-07 | 2.52E-07 |
| chr16 | 53810743 | 53811635 | 893 | -6.12 | 1.40E-07 | 2.53E-07 |
| chr7 | 132711111 | 132712071 | 961 | -6.12 | 1.40E-07 | 2.53E-07 |
| chr4 | 152690839 | 152706328 | 15490 | -3 | 1.41E-07 | 2.54E-07 |
| chr20 | 52808357 | 52809945 | 1589 | -5.23 | 1.41E-07 | 2.55E-07 |
| chr6 | 112979704 | 112982179 | 2476 | 6.11 | 1.41E-07 | 2.55E-07 |
| chr16 | 86798802 | 86800696 | 1895 | -6.38 | 1.41E-07 | 2.55E-07 |
| chr8 | 27194972 | 27195874 | 903 | -6.14 | 1.41E-07 | 2.55E-07 |
| chr12 | 49098144 | 49099287 | 1144 | -6.02 | 1.42E-07 | 2.56E-07 |
| chr17 | 4867982 | 4869552 | 1571 | -6.02 | 1.42E-07 | 2.56E-07 |
| chr10 | 128602746 | 128610082 | 7337 | -6.11 | 1.42E-07 | 2.56E-07 |
| chr9 | 112775192 | 112776211 | 1020 | -6.11 | 1.42E-07 | 2.56E-07 |
| chr8 | 83228654 | 83234437 | 5784 | -6.48 | 1.42E-07 | 2.56E-07 |
| chr10 | 94400994 | 94401658 | 665 | -6.03 | 1.42E-07 | 2.57E-07 |
| chrX | 130263449 | 130264900 | 1452 | 6.14 | 1.43E-07 | 2.58E-07 |
| chr4 | 155570436 | 155583006 | 12571 | -6.1 | 1.43E-07 | 2.58E-07 |
| chr1 | 190335470 | 190337543 | 2074 | -6.36 | 1.43E-07 | 2.58E-07 |
| chr4 | 57912146 | 57914065 | 1920 | -6.06 | 1.43E-07 | 2.59E-07 |
| chr2 | 234133140 | 234134352 | 1213 | -6.01 | 1.44E-07 | 2.59E-07 |
| chr14 | 65700790 | 65702732 | 1943 | -6.05 | 1.44E-07 | 2.59E-07 |
| chr3 | 197013023 | 197014161 | 1139 | -6.05 | 1.44E-07 | 2.59E-07 |
| chr6 | 9144262 | 9145890 | 1629 | -6.02 | 1.44E-07 | 2.59E-07 |

| | | | | | | |
|---|---|---|---|---|---|---|
| chr1 | 20929283 | 20930572 | 1290 | -6.1 | 1.44E-07 | 2.60E-07 |
| chr13 | 33335351 | 33336067 | 717 | -6.1 | 1.44E-07 | 2.60E-07 |
| chr4 | 95629131 | 95674996 | 45866 | -8.16 | 1.44E-07 | 2.60E-07 |
| chr18 | 60116635 | 60117773 | 1139 | -6.11 | 1.44E-07 | 2.60E-07 |
| chr6 | 63302776 | 63304887 | 2112 | -6.11 | 1.44E-07 | 2.60E-07 |
| chr5 | 114383366 | 114384842 | 1477 | 6.13 | 1.44E-07 | 2.60E-07 |
| chr12 | 98878470 | 98879283 | 814 | -6.07 | 1.45E-07 | 2.61E-07 |
| chr14 | 36546661 | 36559097 | 12437 | -6.07 | 1.45E-07 | 2.61E-07 |
| chr5 | 37757031 | 37758462 | 1432 | -6.07 | 1.45E-07 | 2.61E-07 |
| chr18 | 3562904 | 3564844 | 1941 | -6.38 | 1.45E-07 | 2.61E-07 |
| chr1 | 64320901 | 64322089 | 1189 | -6.03 | 1.45E-07 | 2.62E-07 |
| chr14 | 45506889 | 45508400 | 1512 | -6.1 | 1.45E-07 | 2.62E-07 |
| chr2 | 72868213 | 72869488 | 1276 | -6.1 | 1.45E-07 | 2.62E-07 |
| chr1 | 44626355 | 44628650 | 2296 | -6.05 | 1.46E-07 | 2.63E-07 |
| chr13 | 63493630 | 63503847 | 10218 | -6.39 | 1.46E-07 | 2.63E-07 |
| chr5 | 36698980 | 36714939 | 15960 | -1.3 | 1.46E-07 | 2.64E-07 |
| chr3 | 191681608 | 191688593 | 6986 | 6.81 | 1.46E-07 | 2.64E-07 |
| chr5 | 95967219 | 95969081 | 1863 | -6.4 | 1.46E-07 | 2.64E-07 |
| chr9 | 100713294 | 100714413 | 1120 | -4.58 | 1.46E-07 | 2.64E-07 |
| chr1 | 78288491 | 78289777 | 1287 | -6.01 | 1.47E-07 | 2.64E-07 |
| chr17 | 1504725 | 1505984 | 1260 | -6.01 | 1.47E-07 | 2.64E-07 |
| chr12 | 109424971 | 109428243 | 3273 | -6.45 | 1.47E-07 | 2.64E-07 |
| chr17 | 71723232 | 71730386 | 7155 | -2.25 | 1.47E-07 | 2.65E-07 |
| chr5 | 102348346 | 102349438 | 1093 | -6.04 | 1.47E-07 | 2.65E-07 |
| chr12 | 95732642 | 95733545 | 904 | -6.02 | 1.47E-07 | 2.65E-07 |
| chr10 | 135154516 | 135155584 | 1069 | -6.06 | 1.47E-07 | 2.65E-07 |
| chr6 | 106440728 | 106442216 | 1489 | -6.06 | 1.47E-07 | 2.65E-07 |
| chr2 | 241252161 | 241257435 | 5275 | -3.93 | 1.47E-07 | 2.65E-07 |
| chr11 | 20676748 | 20677630 | 883 | 6.12 | 1.47E-07 | 2.66E-07 |
| chr16 | 29257230 | 29258766 | 1537 | -6.09 | 1.48E-07 | 2.68E-07 |
| chr5 | 173485493 | 173489122 | 3630 | -3.44 | 1.48E-07 | 2.68E-07 |
| chr2 | 130527861 | 130532770 | 4910 | -2.13 | 1.49E-07 | 2.68E-07 |
| chr12 | 63341788 | 63346053 | 4266 | -1.57 | 1.49E-07 | 2.68E-07 |
| chr6 | 148264996 | 148266918 | 1923 | -6.22 | 1.49E-07 | 2.68E-07 |
| chr12 | 80673322 | 80674784 | 1463 | -6.19 | 1.49E-07 | 2.68E-07 |
| chr1 | 240661729 | 240663763 | 2035 | -6.23 | 1.49E-07 | 2.68E-07 |
| chrX | 53353191 | 53353801 | 611 | -6.17 | 1.49E-07 | 2.68E-07 |
| chr12 | 108156351 | 108157260 | 910 | -6.07 | 1.50E-07 | 2.70E-07 |
| chr14 | 36497246 | 36498141 | 896 | -6.05 | 1.50E-07 | 2.70E-07 |
| chr5 | 91154617 | 91157216 | 2600 | -6.16 | 1.50E-07 | 2.70E-07 |
| chr6 | 160058711 | 160059718 | 1008 | -6.16 | 1.50E-07 | 2.70E-07 |
| chr10 | 126460165 | 126461658 | 1494 | -6.28 | 1.50E-07 | 2.70E-07 |
| chr5 | 127556240 | 127557320 | 1081 | -6.26 | 1.50E-07 | 2.70E-07 |
| chr7 | 128885266 | 128886050 | 785 | -6.01 | 1.50E-07 | 2.71E-07 |
| chr14 | 50221582 | 50222598 | 1017 | -6.08 | 1.50E-07 | 2.71E-07 |
| chr20 | 48738637 | 48739787 | 1151 | -6.08 | 1.50E-07 | 2.71E-07 |
| chr5 | 138507483 | 138509115 | 1633 | -6.44 | 1.51E-07 | 2.71E-07 |
| chr12 | 50181335 | 50181984 | 650 | -6.04 | 1.51E-07 | 2.71E-07 |
| chr16 | 11683011 | 11683786 | 776 | -6.04 | 1.51E-07 | 2.71E-07 |
| chr9 | 111338479 | 111353001 | 14523 | -1.21 | 1.51E-07 | 2.72E-07 |
| chr10 | 34180305 | 34188395 | 8091 | -6.86 | 1.51E-07 | 2.72E-07 |
| chr1 | 167878172 | 167879495 | 1324 | -6.29 | 1.51E-07 | 2.72E-07 |
| chr4 | 166931317 | 166932732 | 1416 | -6.14 | 1.51E-07 | 2.73E-07 |
| chr2 | 221772545 | 221857885 | 85341 | -6.72 | 1.52E-07 | 2.73E-07 |
| chr19 | 34766250 | 34766843 | 594 | -6.13 | 1.52E-07 | 2.73E-07 |
| chr6 | 22535420 | 22537599 | 2180 | -6 | 1.52E-07 | 2.73E-07 |
| chr4 | 141988830 | 141989738 | 909 | -6.02 | 1.52E-07 | 2.74E-07 |
| chr2 | 10944350 | 10946302 | 1953 | -6.01 | 1.52E-07 | 2.74E-07 |
| chr17 | 27355952 | 27364270 | 8319 | -6.59 | 1.52E-07 | 2.74E-07 |
| chr3 | 178115417 | 178119178 | 3762 | -4.95 | 1.52E-07 | 2.74E-07 |
| chr4 | 187691219 | 187701606 | 10388 | -6.73 | 1.52E-07 | 2.74E-07 |

| | | | | | | |
|---|---|---|---|---|---|---|
| chr5 | 168536373 | 168538243 | 1871 | -6.04 | 1.53E-07 | 2.75E-07 |
| chr19 | 39545163 | 39546033 | 871 | -6.12 | 1.53E-07 | 2.75E-07 |
| chr13 | 61125829 | 61127151 | 1323 | -6.06 | 1.53E-07 | 2.76E-07 |
| chr5 | 109127689 | 109128775 | 1087 | -6.03 | 1.53E-07 | 2.76E-07 |
| chr1 | 92259994 | 92260765 | 772 | -6.12 | 1.53E-07 | 2.76E-07 |
| chr6 | 52025034 | 52026372 | 1339 | -6.11 | 1.54E-07 | 2.77E-07 |
| chr19 | 14383336 | 14384613 | 1278 | -6.1 | 1.55E-07 | 2.79E-07 |
| chr7 | 20408449 | 20409275 | 827 | -6.1 | 1.55E-07 | 2.79E-07 |
| chr13 | 31613794 | 31614955 | 1162 | -6.05 | 1.55E-07 | 2.79E-07 |
| chr4 | 154426207 | 154427300 | 1094 | -6.05 | 1.55E-07 | 2.79E-07 |
| chr1 | 224375165 | 224376391 | 1227 | -6.54 | 1.55E-07 | 2.79E-07 |
| chr6 | 166595915 | 166602912 | 6998 | -4.61 | 1.55E-07 | 2.79E-07 |
| chr6 | 13287029 | 13289460 | 2432 | -6.36 | 1.56E-07 | 2.80E-07 |
| chr11 | 76336215 | 76342329 | 6115 | -6.37 | 1.56E-07 | 2.80E-07 |
| chr3 | 195931692 | 195932796 | 1105 | -6.02 | 1.56E-07 | 2.81E-07 |
| chr9 | 95910077 | 95911816 | 1740 | -6 | 1.56E-07 | 2.81E-07 |
| chr3 | 125608952 | 125610264 | 1313 | -4.47 | 1.56E-07 | 2.81E-07 |
| chr18 | 32691063 | 32693030 | 1968 | -6.05 | 1.57E-07 | 2.82E-07 |
| chr12 | 16389079 | 16415842 | 26764 | 1.61 | 1.57E-07 | 2.82E-07 |
| chr12 | 89634597 | 89637760 | 3164 | -1.9 | 1.57E-07 | 2.82E-07 |
| chr3 | 151238286 | 151240760 | 2475 | -2.62 | 1.57E-07 | 2.83E-07 |
| chr13 | 80253491 | 80258972 | 5482 | -5.05 | 1.57E-07 | 2.83E-07 |
| chr4 | 102458003 | 102460583 | 2581 | -6.38 | 1.57E-07 | 2.83E-07 |
| chr1 | 202005982 | 202008444 | 2463 | -6 | 1.58E-07 | 2.83E-07 |
| chr13 | 24167716 | 24169169 | 1454 | -6 | 1.58E-07 | 2.83E-07 |
| chr13 | 114955288 | 114956426 | 1139 | -6 | 1.58E-07 | 2.83E-07 |
| chr7 | 106430014 | 106430933 | 920 | -6 | 1.58E-07 | 2.83E-07 |
| chr4 | 154566017 | 154567187 | 1171 | -6.01 | 1.58E-07 | 2.84E-07 |
| chr14 | 22255063 | 22257513 | 2451 | -4.62 | 1.59E-07 | 2.85E-07 |
| chr6 | 141929586 | 141931642 | 2057 | -6 | 1.59E-07 | 2.85E-07 |
| chr14 | 85956759 | 85958219 | 1461 | -6.08 | 1.59E-07 | 2.86E-07 |
| chr15 | 63652406 | 63653569 | 1164 | -6.03 | 1.59E-07 | 2.86E-07 |
| chr7 | 103050307 | 103052782 | 2476 | -6.67 | 1.59E-07 | 2.86E-07 |
| chr4 | 111645289 | 111646940 | 1652 | 6.14 | 1.59E-07 | 2.87E-07 |
| chr13 | 95815994 | 95817949 | 1956 | -3.69 | 1.60E-07 | 2.87E-07 |
| chr2 | 215908518 | 215913365 | 4848 | -6.82 | 1.60E-07 | 2.87E-07 |
| chr6 | 126053500 | 126054763 | 1264 | -6.08 | 1.60E-07 | 2.87E-07 |
| chr7 | 77685548 | 77687017 | 1470 | -6.08 | 1.60E-07 | 2.87E-07 |
| chr2 | 96983501 | 96984742 | 1242 | -6.04 | 1.60E-07 | 2.87E-07 |
| chr5 | 75504185 | 75518053 | 13869 | -1.6 | 1.60E-07 | 2.87E-07 |
| chr6 | 113470893 | 113472751 | 1859 | -5.99 | 1.60E-07 | 2.87E-07 |
| chr11 | 10332955 | 10334131 | 1177 | -6.55 | 1.60E-07 | 2.88E-07 |
| chr17 | 47842983 | 47843965 | 983 | -5.69 | 1.61E-07 | 2.88E-07 |
| chr14 | 36281351 | 36282789 | 1439 | -6.07 | 1.61E-07 | 2.89E-07 |
| chr2 | 198153526 | 198154749 | 1224 | -6.2 | 1.61E-07 | 2.89E-07 |
| chr11 | 128871753 | 128873362 | 1610 | -6.19 | 1.61E-07 | 2.89E-07 |
| chr6 | 36716015 | 36717177 | 1163 | -6.21 | 1.61E-07 | 2.89E-07 |
| chr10 | 15902710 | 15914870 | 12161 | -1.59 | 1.61E-07 | 2.89E-07 |
| chr4 | 91991165 | 91998876 | 7712 | -6.17 | 1.61E-07 | 2.90E-07 |
| chr11 | 9716540 | 9717112 | 573 | -6.07 | 1.61E-07 | 2.90E-07 |
| chr13 | 67011131 | 67013411 | 2281 | -6.23 | 1.62E-07 | 2.90E-07 |
| chr15 | 42359140 | 42360579 | 1440 | -6.16 | 1.62E-07 | 2.90E-07 |
| chr15 | 66973719 | 66975516 | 1798 | -6.43 | 1.62E-07 | 2.91E-07 |
| chr13 | 66668491 | 66669868 | 1378 | -6.04 | 1.62E-07 | 2.91E-07 |
| chr14 | 102893287 | 102894748 | 1462 | -6.04 | 1.62E-07 | 2.91E-07 |
| chr4 | 103180349 | 103183145 | 2797 | -6.03 | 1.62E-07 | 2.92E-07 |
| chr9 | 90422756 | 90424692 | 1937 | -6.26 | 1.62E-07 | 2.92E-07 |
| chr1 | 6630421 | 6631862 | 1442 | -6.14 | 1.63E-07 | 2.92E-07 |
| chr10 | 4602765 | 4604178 | 1414 | -6.14 | 1.63E-07 | 2.92E-07 |
| chr3 | 27223573 | 27224454 | 882 | -6.01 | 1.63E-07 | 2.92E-07 |
| chr5 | 64283536 | 64284691 | 1156 | -6.01 | 1.63E-07 | 2.92E-07 |

| | | | | | | |
|---|---|---|---|---|---|---|
| chr7 | 75983069 | 75984636 | 1568 | -6.01 | 1.63E-07 | 2.92E-07 |
| chr1 | 80389354 | 80390956 | 1603 | -6.01 | 1.63E-07 | 2.92E-07 |
| chr20 | 3420374 | 3421228 | 855 | -6.01 | 1.63E-07 | 2.92E-07 |
| chr4 | 76779584 | 76780371 | 788 | -6.02 | 1.63E-07 | 2.92E-07 |
| chr7 | 54860282 | 54861770 | 1489 | -5.99 | 1.63E-07 | 2.93E-07 |
| chr9 | 115085780 | 115087452 | 1673 | -6.13 | 1.63E-07 | 2.93E-07 |
| chr16 | 12733108 | 12737293 | 4186 | -5.99 | 1.63E-07 | 2.93E-07 |
| chrX | 129102858 | 129103532 | 675 | -5.99 | 1.63E-07 | 2.93E-07 |
| chrX | 19258203 | 19305341 | 47139 | -3.05 | 1.64E-07 | 2.94E-07 |
| chr22 | 28111918 | 28112986 | 1069 | -5.99 | 1.64E-07 | 2.94E-07 |
| chr15 | 72401747 | 72402611 | 865 | -6 | 1.64E-07 | 2.94E-07 |
| chr12 | 109088437 | 109089449 | 1013 | -5.68 | 1.64E-07 | 2.94E-07 |
| chr2 | 65814484 | 65816135 | 1652 | -6.23 | 1.64E-07 | 2.94E-07 |
| chr2 | 198409181 | 198410202 | 1022 | -6.24 | 1.64E-07 | 2.95E-07 |
| chr6 | 128161588 | 128170980 | 9393 | -6.45 | 1.65E-07 | 2.95E-07 |
| chr19 | 42135811 | 42140594 | 4784 | -4.26 | 1.65E-07 | 2.96E-07 |
| chr16 | 68978045 | 68979098 | 1054 | -6.02 | 1.65E-07 | 2.96E-07 |
| chr3 | 112311506 | 112312626 | 1121 | -6.02 | 1.65E-07 | 2.96E-07 |
| chr4 | 93843362 | 93844447 | 1086 | -5.99 | 1.65E-07 | 2.96E-07 |
| chr7 | 98023351 | 98024301 | 951 | -5.99 | 1.65E-07 | 2.96E-07 |
| chr5 | 149074365 | 149090280 | 15916 | 6.36 | 1.65E-07 | 2.97E-07 |
| chrX | 43879259 | 43881778 | 2520 | -6.03 | 1.66E-07 | 2.97E-07 |
| chr8 | 135858186 | 135867760 | 9575 | -3.13 | 1.66E-07 | 2.97E-07 |
| chr5 | 174185119 | 174316832 | 131714 | -2.82 | 1.66E-07 | 2.97E-07 |
| chr10 | 63470364 | 63471730 | 1367 | -6.48 | 1.66E-07 | 2.98E-07 |
| chr6 | 35784409 | 35797323 | 12915 | -6.32 | 1.66E-07 | 2.98E-07 |
| chr3 | 183932703 | 183933942 | 1240 | -6.32 | 1.67E-07 | 2.99E-07 |
| chr3 | 115508439 | 115510532 | 2094 | -6.47 | 1.67E-07 | 2.99E-07 |
| chr16 | 81846495 | 81847523 | 1029 | -6.1 | 1.67E-07 | 2.99E-07 |
| chr12 | 31437543 | 31438924 | 1382 | -6.12 | 1.67E-07 | 2.99E-07 |
| chr5 | 75813644 | 75815161 | 1518 | -3.1 | 1.67E-07 | 2.99E-07 |
| chr3 | 115900613 | 115902631 | 2019 | 4.86 | 1.67E-07 | 3.00E-07 |
| chr1 | 199889684 | 199891350 | 1667 | -6.01 | 1.67E-07 | 3.00E-07 |
| chr16 | 63441537 | 63443389 | 1853 | -6.01 | 1.67E-07 | 3.00E-07 |
| chr18 | 12443358 | 12444505 | 1148 | -6.05 | 1.67E-07 | 3.00E-07 |
| chr10 | 32770580 | 32771536 | 957 | -6.49 | 1.68E-07 | 3.00E-07 |
| chr3 | 188637899 | 188639165 | 1267 | -6.09 | 1.68E-07 | 3.01E-07 |
| chr17 | 67521473 | 67522238 | 766 | -6 | 1.68E-07 | 3.01E-07 |
| chr2 | 38661460 | 38663080 | 1621 | -6.02 | 1.68E-07 | 3.01E-07 |
| chr22 | 23216221 | 23217434 | 1214 | -6.02 | 1.68E-07 | 3.01E-07 |
| chr3 | 12370780 | 12371640 | 861 | -6.02 | 1.68E-07 | 3.01E-07 |
| chr1 | 9318363 | 9319467 | 1105 | -6.04 | 1.68E-07 | 3.02E-07 |
| chr1 | 69153078 | 69154568 | 1491 | -6.04 | 1.68E-07 | 3.02E-07 |
| chr22 | 37687447 | 37688926 | 1480 | -5.99 | 1.69E-07 | 3.02E-07 |
| chr1 | 118248650 | 118272474 | 23825 | -5.45 | 1.69E-07 | 3.02E-07 |
| chr5 | 141621623 | 141622638 | 1016 | -5.99 | 1.69E-07 | 3.02E-07 |
| chr2 | 224142651 | 224144291 | 1641 | -6.1 | 1.69E-07 | 3.03E-07 |
| chr3 | 127875799 | 127876747 | 949 | -6.1 | 1.69E-07 | 3.03E-07 |
| chr18 | 54427667 | 54428594 | 928 | -6.02 | 1.69E-07 | 3.03E-07 |
| chr5 | 95432006 | 95434570 | 2565 | -6.36 | 1.70E-07 | 3.04E-07 |
| chr3 | 36754850 | 36755769 | 920 | -4.63 | 1.70E-07 | 3.04E-07 |
| chr12 | 56007822 | 56021876 | 14055 | -2.75 | 1.70E-07 | 3.04E-07 |
| chr1 | 32613125 | 32614829 | 1705 | -6.09 | 1.70E-07 | 3.04E-07 |
| chr4 | 187410264 | 187445782 | 35519 | -3.23 | 1.70E-07 | 3.04E-07 |
| chr9 | 123447683 | 123448489 | 807 | -6 | 1.70E-07 | 3.05E-07 |
| chr14 | 65133665 | 65134634 | 970 | -5.98 | 1.70E-07 | 3.05E-07 |
| chr6 | 11835913 | 11837934 | 2022 | -5.98 | 1.70E-07 | 3.05E-07 |
| chr10 | 69848683 | 69853841 | 5159 | -6.91 | 1.70E-07 | 3.05E-07 |
| chr2 | 235482060 | 235486941 | 4882 | -6.65 | 1.71E-07 | 3.05E-07 |
| chr17 | 76858723 | 76859457 | 735 | -6.04 | 1.71E-07 | 3.05E-07 |
| chr2 | 215427949 | 215429686 | 1738 | -6.04 | 1.71E-07 | 3.05E-07 |

| | | | | | | |
|---|---|---|---|---|---|---|
| chr6 | 37838599 | 37840015 | 1417 | -6.04 | 1.71E-07 | 3.05E-07 |
| chr9 | 133895384 | 133896728 | 1345 | -6.04 | 1.71E-07 | 3.05E-07 |
| chr2 | 143612246 | 143614073 | 1828 | -6.07 | 1.71E-07 | 3.06E-07 |
| chr22 | 34099369 | 34111229 | 11861 | -6.37 | 1.71E-07 | 3.06E-07 |
| chr1 | 116846894 | 116847928 | 1035 | -5.23 | 1.71E-07 | 3.06E-07 |
| chr4 | 137072196 | 137073964 | 1769 | -5.98 | 1.72E-07 | 3.08E-07 |
| chr7 | 133463658 | 133464690 | 1033 | -6 | 1.72E-07 | 3.08E-07 |
| chr8 | 110939536 | 110942347 | 2812 | -6.03 | 1.73E-07 | 3.10E-07 |
| chr5 | 86338759 | 86341005 | 2247 | -2.04 | 1.73E-07 | 3.10E-07 |
| chr6 | 128912157 | 128913527 | 1371 | -4.81 | 1.74E-07 | 3.11E-07 |
| chr10 | 15426235 | 15427634 | 1400 | -6.19 | 1.74E-07 | 3.12E-07 |
| chr5 | 141102670 | 141103770 | 1101 | -6.18 | 1.74E-07 | 3.12E-07 |
| chr2 | 56063529 | 56069194 | 5666 | -6.2 | 1.74E-07 | 3.12E-07 |
| chr1 | 113928965 | 113930980 | 2016 | -6.06 | 1.75E-07 | 3.12E-07 |
| chr16 | 53741530 | 53742766 | 1237 | -6.06 | 1.75E-07 | 3.12E-07 |
| chr15 | 35901958 | 35902873 | 916 | -6.07 | 1.75E-07 | 3.12E-07 |
| chr15 | 70441526 | 70442264 | 739 | -6 | 1.75E-07 | 3.12E-07 |
| chr14 | 64390855 | 64391612 | 758 | -5.98 | 1.75E-07 | 3.12E-07 |
| chr4 | 166274745 | 166275822 | 1078 | -5.98 | 1.75E-07 | 3.12E-07 |
| chr4 | 179547355 | 179549171 | 1817 | -5.98 | 1.75E-07 | 3.12E-07 |
| chr8 | 142622457 | 142623690 | 1234 | -5.98 | 1.75E-07 | 3.12E-07 |
| chr4 | 165688416 | 165689787 | 1372 | -5.22 | 1.75E-07 | 3.12E-07 |
| chr17 | 54819756 | 54823050 | 3295 | -6.16 | 1.75E-07 | 3.12E-07 |
| chr21 | 41074973 | 41077066 | 2094 | -6.16 | 1.75E-07 | 3.12E-07 |
| chr9 | 4750956 | 4751819 | 864 | -6.21 | 1.75E-07 | 3.12E-07 |
| chr4 | 100775542 | 100776750 | 1209 | -6.22 | 1.75E-07 | 3.13E-07 |
| chr15 | 90710265 | 90717935 | 7671 | -7.25 | 1.75E-07 | 3.13E-07 |
| chr8 | 62063565 | 62065180 | 1616 | 4.87 | 1.75E-07 | 3.13E-07 |
| chr4 | 108607221 | 108608246 | 1026 | -6.02 | 1.76E-07 | 3.14E-07 |
| chr11 | 85607645 | 85608604 | 960 | -6.13 | 1.76E-07 | 3.14E-07 |
| chr3 | 160207838 | 160208764 | 927 | -6.25 | 1.76E-07 | 3.15E-07 |
| chr17 | 39014362 | 39016210 | 1849 | -6.13 | 1.76E-07 | 3.15E-07 |
| chr19 | 45082685 | 45083825 | 1141 | -6.43 | 1.76E-07 | 3.15E-07 |
| chr2 | 114546837 | 114548756 | 1920 | -6.43 | 1.76E-07 | 3.15E-07 |
| chr17 | 62631442 | 62633249 | 1808 | -6.05 | 1.77E-07 | 3.16E-07 |
| chr6 | 145554385 | 145556686 | 2302 | -6.05 | 1.77E-07 | 3.16E-07 |
| chr2 | 150635272 | 150643338 | 8067 | -6.05 | 1.78E-07 | 3.17E-07 |
| chr9 | 138124201 | 138128424 | 4224 | -4.71 | 1.78E-07 | 3.17E-07 |
| chr17 | 60477520 | 60481297 | 3778 | -3.6 | 1.78E-07 | 3.17E-07 |
| chr4 | 166738136 | 166747388 | 9253 | -6.11 | 1.78E-07 | 3.17E-07 |
| chr10 | 133817971 | 133819363 | 1393 | -5.99 | 1.78E-07 | 3.19E-07 |
| chr20 | 9534462 | 9535556 | 1095 | -5.99 | 1.78E-07 | 3.19E-07 |
| chr1 | 52217244 | 52218520 | 1277 | -5.97 | 1.79E-07 | 3.19E-07 |
| chr6 | 14699540 | 14711576 | 12037 | -6.92 | 1.79E-07 | 3.20E-07 |
| chr13 | 85178387 | 85181050 | 2664 | -5.98 | 1.79E-07 | 3.20E-07 |
| chr18 | 44242110 | 44243732 | 1623 | -5.98 | 1.79E-07 | 3.20E-07 |
| chr20 | 32714238 | 32714998 | 761 | -5.98 | 1.79E-07 | 3.20E-07 |
| chr10 | 72139072 | 72139981 | 910 | -6.04 | 1.79E-07 | 3.20E-07 |
| chr19 | 5115698 | 5116838 | 1141 | -6.04 | 1.79E-07 | 3.20E-07 |
| chr2 | 162419269 | 162422028 | 2760 | -2.43 | 1.80E-07 | 3.21E-07 |
| chr14 | 59049055 | 59050765 | 1711 | -2.8 | 1.80E-07 | 3.21E-07 |
| chr11 | 126050039 | 126051111 | 1073 | -6.01 | 1.80E-07 | 3.21E-07 |
| chr5 | 141635419 | 141637063 | 1645 | -5.98 | 1.80E-07 | 3.21E-07 |
| chr18 | 57114045 | 57115739 | 1695 | -6.08 | 1.81E-07 | 3.23E-07 |
| chr3 | 94160547 | 94163238 | 2692 | -6.08 | 1.81E-07 | 3.23E-07 |
| chr17 | 17913675 | 17914327 | 653 | -5.97 | 1.81E-07 | 3.23E-07 |
| chr4 | 135256785 | 135263529 | 6745 | 6.91 | 1.81E-07 | 3.23E-07 |
| chr12 | 82813604 | 82815296 | 1693 | -6.32 | 1.81E-07 | 3.23E-07 |
| chr2 | 173976587 | 173978693 | 2107 | -6.32 | 1.81E-07 | 3.23E-07 |
| chr8 | 1689429 | 1692189 | 2761 | -6.32 | 1.81E-07 | 3.23E-07 |
| chr8 | 56943768 | 56944687 | 920 | -6.04 | 1.81E-07 | 3.23E-07 |

| | | | | | | |
|---|---|---|---|---|---|---|
| chr11 | 110114521 | 110115674 | 1154 | -6.16 | 1.81E-07 | 3.23E-07 |
| chr15 | 56279057 | 56280973 | 1917 | -6.08 | 1.82E-07 | 3.25E-07 |
| chr16 | 15861616 | 15862961 | 1346 | -6.08 | 1.82E-07 | 3.25E-07 |
| chr22 | 47042317 | 47043799 | 1483 | -6.08 | 1.82E-07 | 3.25E-07 |
| chr11 | 116263544 | 116266039 | 2496 | -2.27 | 1.82E-07 | 3.25E-07 |
| chr4 | 109578814 | 109580001 | 1188 | -5.99 | 1.82E-07 | 3.25E-07 |
| chrX | 45678456 | 45679312 | 857 | -5.99 | 1.82E-07 | 3.25E-07 |
| chr12 | 109248528 | 109257538 | 9011 | -6.13 | 1.83E-07 | 3.26E-07 |
| chr20 | 1468519 | 1551605 | 83087 | -2.46 | 1.83E-07 | 3.26E-07 |
| chr5 | 82562903 | 82565199 | 2297 | 6.58 | 1.83E-07 | 3.26E-07 |
| chr12 | 15989296 | 15990455 | 1160 | -5.98 | 1.83E-07 | 3.26E-07 |
| chr5 | 108359785 | 108360665 | 881 | -5.98 | 1.83E-07 | 3.26E-07 |
| chr2 | 188055239 | 188057256 | 2018 | 6.12 | 1.83E-07 | 3.26E-07 |
| chrX | 113731781 | 113755320 | 23540 | 1.79 | 1.83E-07 | 3.26E-07 |
| chr2 | 101460550 | 101462092 | 1543 | -6.25 | 1.83E-07 | 3.27E-07 |
| chr7 | 139878648 | 139880888 | 2241 | -4.99 | 1.83E-07 | 3.27E-07 |
| chr1 | 113796337 | 113797640 | 1304 | -6.07 | 1.84E-07 | 3.27E-07 |
| chr8 | 82549341 | 82550083 | 743 | -6 | 1.84E-07 | 3.28E-07 |
| chr6 | 130539789 | 130542254 | 2466 | -2.71 | 1.84E-07 | 3.28E-07 |
| chr5 | 113695021 | 113696398 | 1378 | -3.99 | 1.84E-07 | 3.28E-07 |
| chr3 | 151110501 | 151111371 | 871 | -5.16 | 1.84E-07 | 3.28E-07 |
| chr2 | 98659920 | 98661699 | 1780 | -6.11 | 1.84E-07 | 3.28E-07 |
| chr18 | 7185746 | 7186806 | 1061 | -5.97 | 1.85E-07 | 3.29E-07 |
| chr14 | 71336968 | 71339549 | 2582 | -6.02 | 1.85E-07 | 3.29E-07 |
| chr6 | 11792806 | 11794359 | 1554 | -6.02 | 1.85E-07 | 3.29E-07 |
| chr11 | 27424092 | 27425974 | 1883 | -6.1 | 1.85E-07 | 3.30E-07 |
| chr15 | 91109819 | 91110753 | 935 | -6.06 | 1.85E-07 | 3.30E-07 |
| chr1 | 60255564 | 60258548 | 2985 | -2.42 | 1.86E-07 | 3.31E-07 |
| chr1 | 221359615 | 221372799 | 13185 | -7.07 | 1.86E-07 | 3.31E-07 |
| chr1 | 45387955 | 45389200 | 1246 | -5.97 | 1.86E-07 | 3.31E-07 |
| chr7 | 109656590 | 109657404 | 815 | -5.97 | 1.86E-07 | 3.31E-07 |
| chr6 | 71950834 | 71952926 | 2093 | -4.52 | 1.86E-07 | 3.31E-07 |
| chr1 | 146542449 | 146583654 | 41206 | -2.37 | 1.86E-07 | 3.32E-07 |
| chr6 | 142852722 | 142857542 | 4821 | -5.24 | 1.86E-07 | 3.32E-07 |
| chr20 | 9821178 | 9893574 | 72397 | -0.83 | 1.86E-07 | 3.32E-07 |
| chr6 | 7949334 | 7950339 | 1006 | -4.42 | 1.87E-07 | 3.32E-07 |
| chr2 | 158259932 | 158261223 | 1292 | -6.09 | 1.87E-07 | 3.32E-07 |
| chr22 | 20993959 | 20994555 | 597 | -5.98 | 1.87E-07 | 3.33E-07 |
| chr4 | 180615166 | 180616858 | 1693 | -5.98 | 1.87E-07 | 3.33E-07 |
| chr11 | 126729539 | 126730985 | 1447 | -5.98 | 1.87E-07 | 3.33E-07 |
| chr13 | 45317226 | 45319196 | 1971 | -5.98 | 1.87E-07 | 3.33E-07 |
| chr4 | 175865812 | 175867212 | 1401 | -5.98 | 1.87E-07 | 3.33E-07 |
| chr8 | 37433309 | 37433995 | 687 | -5.98 | 1.87E-07 | 3.33E-07 |
| chrX | 18846415 | 18847728 | 1314 | -5.98 | 1.87E-07 | 3.33E-07 |
| chr12 | 748529 | 761574 | 13046 | -6.83 | 1.87E-07 | 3.33E-07 |
| chr4 | 129749628 | 129751116 | 1489 | -5.96 | 1.88E-07 | 3.34E-07 |
| chr2 | 89111028 | 89118247 | 7220 | -3.55 | 1.88E-07 | 3.34E-07 |
| chr15 | 76752751 | 76758716 | 5966 | -6.01 | 1.88E-07 | 3.34E-07 |
| chrX | 34519539 | 34527526 | 7988 | -2.42 | 1.88E-07 | 3.35E-07 |
| chr5 | 126140432 | 126141901 | 1470 | -6.32 | 1.88E-07 | 3.35E-07 |
| chr1 | 99878381 | 99881107 | 2727 | 3.02 | 1.89E-07 | 3.36E-07 |
| chr16 | 59313140 | 59324534 | 11395 | -2.33 | 1.89E-07 | 3.36E-07 |
| chr9 | 128737297 | 128745022 | 7726 | -6.01 | 1.89E-07 | 3.36E-07 |
| chr9 | 37203450 | 37205116 | 1667 | -6.18 | 1.89E-07 | 3.36E-07 |
| chrX | 114719554 | 114721338 | 1785 | -6.17 | 1.89E-07 | 3.36E-07 |
| chrX | 80688829 | 80738125 | 49297 | 2.39 | 1.89E-07 | 3.36E-07 |
| chr7 | 104851521 | 104855886 | 4366 | -5.97 | 1.89E-07 | 3.36E-07 |
| chr16 | 84189269 | 84190269 | 1001 | -6.16 | 1.89E-07 | 3.36E-07 |
| chr1 | 8371209 | 8372588 | 1380 | -6.15 | 1.89E-07 | 3.37E-07 |
| chr1 | 79740416 | 79741883 | 1468 | -6.04 | 1.89E-07 | 3.37E-07 |
| chr7 | 19111476 | 19115094 | 3619 | 6.65 | 1.89E-07 | 3.37E-07 |

| | | | | | | |
|---|---|---|---|---|---|---|
| chr8 | 128299935 | 128305222 | 5288 | -5.37 | 1.90E-07 | 3.37E-07 |
| chr11 | 32889668 | 32895418 | 5751 | -6.22 | 1.90E-07 | 3.38E-07 |
| chr2 | 201256061 | 201264373 | 8313 | -6.34 | 1.90E-07 | 3.38E-07 |
| chr2 | 23588597 | 23589184 | 588 | -6.13 | 1.90E-07 | 3.38E-07 |
| chr16 | 15834478 | 15835356 | 879 | -6.22 | 1.90E-07 | 3.38E-07 |
| chr7 | 33291217 | 33292296 | 1080 | -6.22 | 1.90E-07 | 3.38E-07 |
| chr12 | 118792093 | 118793372 | 1280 | -6.23 | 1.91E-07 | 3.39E-07 |
| chr10 | 49680713 | 49681855 | 1143 | -6.24 | 1.91E-07 | 3.40E-07 |
| chr11 | 72753325 | 72754288 | 964 | -6.06 | 1.91E-07 | 3.40E-07 |
| chr20 | 12953462 | 12977009 | 23548 | -7.84 | 1.91E-07 | 3.40E-07 |
| chr13 | 61045060 | 61045949 | 890 | -5.97 | 1.91E-07 | 3.41E-07 |
| chr7 | 152573177 | 152585742 | 12566 | -6.71 | 1.92E-07 | 3.41E-07 |
| chrX | 43611102 | 43612023 | 922 | -6.03 | 1.92E-07 | 3.41E-07 |
| chr12 | 16676370 | 16677993 | 1624 | -6.25 | 1.92E-07 | 3.41E-07 |
| chr12 | 110682944 | 110684073 | 1130 | -6.1 | 1.92E-07 | 3.41E-07 |
| chr4 | 104239586 | 104241095 | 1510 | 6.04 | 1.92E-07 | 3.42E-07 |
| chr3 | 57926386 | 57927587 | 1202 | -5.98 | 1.93E-07 | 3.42E-07 |
| chr18 | 22153416 | 22172163 | 18748 | -1.73 | 1.93E-07 | 3.43E-07 |
| chr11 | 8399440 | 8401214 | 1775 | -3.17 | 1.93E-07 | 3.43E-07 |
| chr12 | 2650075 | 2689792 | 39718 | 3.03 | 1.93E-07 | 3.43E-07 |
| chr1 | 81466044 | 81467625 | 1582 | -6.01 | 1.93E-07 | 3.43E-07 |
| chr13 | 48398063 | 48400249 | 2187 | -6.01 | 1.93E-07 | 3.43E-07 |
| chrX | 83375423 | 83376469 | 1047 | -6.01 | 1.93E-07 | 3.43E-07 |
| chr12 | 99354133 | 99359559 | 5427 | -1.56 | 1.93E-07 | 3.44E-07 |
| chr5 | 57729479 | 57841104 | 111626 | -1.84 | 1.93E-07 | 3.44E-07 |
| chr9 | 91562938 | 91569879 | 6942 | -6.28 | 1.93E-07 | 3.44E-07 |
| chr10 | 87528274 | 87529204 | 931 | -5.96 | 1.94E-07 | 3.44E-07 |
| chr6 | 117782291 | 117784195 | 1905 | -5.96 | 1.94E-07 | 3.45E-07 |
| chr12 | 93429931 | 93442153 | 12223 | -6.59 | 1.94E-07 | 3.46E-07 |
| chr6 | 168117018 | 168127452 | 10435 | -2.75 | 1.95E-07 | 3.46E-07 |
| chr1 | 213280431 | 213281249 | 819 | -6 | 1.95E-07 | 3.47E-07 |
| chr3 | 174358314 | 174360162 | 1849 | -6.07 | 1.96E-07 | 3.49E-07 |
| chr13 | 98058499 | 98059520 | 1022 | -6.4 | 1.96E-07 | 3.49E-07 |
| chr11 | 112357196 | 112363511 | 6316 | 1.71 | 1.97E-07 | 3.49E-07 |
| chr3 | 36694611 | 36696794 | 2184 | -5.97 | 1.97E-07 | 3.49E-07 |
| chr1 | 115140186 | 115141283 | 1098 | -6.31 | 1.97E-07 | 3.49E-07 |
| chr19 | 3591088 | 3592537 | 1450 | -4.87 | 1.98E-07 | 3.51E-07 |
| chr3 | 33980740 | 33981849 | 1110 | -5.98 | 1.98E-07 | 3.51E-07 |
| chrX | 77103054 | 77104220 | 1167 | -5.98 | 1.98E-07 | 3.51E-07 |
| chr14 | 62308454 | 62351430 | 42977 | -2.47 | 1.98E-07 | 3.51E-07 |
| chr7 | 20593653 | 20607485 | 13833 | -2.96 | 1.98E-07 | 3.52E-07 |
| chr12 | 87977443 | 87978625 | 1183 | -5.16 | 1.98E-07 | 3.52E-07 |
| chr2 | 107726439 | 108144132 | 417694 | 1.43 | 1.99E-07 | 3.53E-07 |
| chr2 | 233286703 | 233287761 | 1059 | -5.96 | 1.99E-07 | 3.53E-07 |
| chr4 | 171031499 | 171032780 | 1282 | -5.96 | 1.99E-07 | 3.53E-07 |
| chr5 | 156667127 | 156668051 | 925 | -5.96 | 1.99E-07 | 3.53E-07 |
| chr3 | 192959478 | 192968370 | 8893 | -2.73 | 1.99E-07 | 3.54E-07 |
| chr4 | 94895139 | 94896551 | 1413 | -6.02 | 2.00E-07 | 3.54E-07 |
| chr6 | 113037511 | 113038981 | 1471 | -3.97 | 2.00E-07 | 3.55E-07 |
| chr2 | 113243337 | 113244557 | 1221 | -5.97 | 2.00E-07 | 3.55E-07 |
| chr5 | 174169394 | 174172376 | 2983 | -3.51 | 2.00E-07 | 3.55E-07 |
| chr7 | 31705231 | 31707116 | 1886 | 6.39 | 2.00E-07 | 3.55E-07 |
| chr2 | 181506952 | 181508484 | 1533 | -6.17 | 2.00E-07 | 3.56E-07 |
| chr7 | 27712646 | 27727827 | 15182 | -6.18 | 2.00E-07 | 3.56E-07 |
| chr10 | 62396324 | 62398295 | 1972 | -6.15 | 2.00E-07 | 3.56E-07 |
| chr2 | 218322347 | 218325980 | 3634 | -2.88 | 2.01E-07 | 3.56E-07 |
| chr4 | 184105083 | 184105894 | 812 | -6.13 | 2.01E-07 | 3.56E-07 |
| chr2 | 221365404 | 221369513 | 4110 | -5.07 | 2.01E-07 | 3.57E-07 |
| chr5 | 167767138 | 167767856 | 719 | -6.12 | 2.01E-07 | 3.58E-07 |
| chr11 | 126184084 | 126186361 | 2278 | -6.22 | 2.02E-07 | 3.58E-07 |
| chr5 | 157197397 | 157198286 | 890 | -5.96 | 2.02E-07 | 3.58E-07 |

| | | | | | | |
|---|---|---|---|---|---|---|
| chr4 | 31407537 | 31410533 | 2997 | -5.95 | 2.02E-07 | 3.59E-07 |
| chr9 | 29968270 | 29970835 | 2566 | -6.23 | 2.02E-07 | 3.59E-07 |
| chr10 | 134990401 | 134991792 | 1392 | -5.98 | 2.03E-07 | 3.60E-07 |
| chr15 | 55779033 | 55780026 | 994 | -5.98 | 2.03E-07 | 3.60E-07 |
| chr3 | 94547184 | 94549881 | 2698 | -5.98 | 2.03E-07 | 3.60E-07 |
| chr8 | 32070162 | 32071039 | 878 | -5.98 | 2.03E-07 | 3.60E-07 |
| chr6 | 160472635 | 160473311 | 677 | -6.03 | 2.03E-07 | 3.61E-07 |
| chrX | 20425489 | 20426887 | 1399 | -5.95 | 2.04E-07 | 3.61E-07 |
| chr12 | 2054391 | 2069653 | 15263 | -6.46 | 2.04E-07 | 3.62E-07 |
| chr2 | 111885769 | 111887570 | 1802 | -6.26 | 2.04E-07 | 3.62E-07 |
| chr1 | 1301339 | 1302862 | 1524 | -6 | 2.04E-07 | 3.62E-07 |
| chr1 | 30441853 | 30451603 | 9751 | -5.98 | 2.04E-07 | 3.63E-07 |
| chr8 | 144280632 | 144282623 | 1992 | -5.98 | 2.04E-07 | 3.63E-07 |
| chr15 | 85834653 | 85835341 | 689 | -5.96 | 2.05E-07 | 3.63E-07 |
| chr7 | 150869504 | 150870982 | 1479 | -5.95 | 2.05E-07 | 3.64E-07 |
| chr19 | 1931331 | 1932322 | 992 | -6.15 | 2.05E-07 | 3.64E-07 |
| chr4 | 94625872 | 94627995 | 2124 | -4.65 | 2.05E-07 | 3.64E-07 |
| chr8 | 25150028 | 25152668 | 2641 | -6.14 | 2.05E-07 | 3.64E-07 |
| chrX | 124328283 | 124329323 | 1041 | 6.02 | 2.05E-07 | 3.64E-07 |
| chr14 | 38233723 | 38236026 | 2304 | -6.08 | 2.05E-07 | 3.64E-07 |
| chr16 | 57766551 | 57767835 | 1285 | -6.08 | 2.05E-07 | 3.64E-07 |
| chr3 | 23009271 | 23033009 | 23739 | -1.02 | 2.06E-07 | 3.65E-07 |
| chr17 | 66825138 | 66828140 | 3003 | -2.86 | 2.06E-07 | 3.65E-07 |
| chr2 | 113524571 | 113526096 | 1526 | -6.22 | 2.07E-07 | 3.67E-07 |
| chr6 | 149736025 | 149738272 | 2248 | -6.22 | 2.07E-07 | 3.67E-07 |
| chrX | 100951939 | 100953402 | 1464 | 6.02 | 2.07E-07 | 3.68E-07 |
| chrX | 40354270 | 40355790 | 1521 | -5.95 | 2.08E-07 | 3.69E-07 |
| chr4 | 187481119 | 187482276 | 1158 | -5.99 | 2.08E-07 | 3.69E-07 |
| chr11 | 113463248 | 113492067 | 28820 | -2.13 | 2.08E-07 | 3.69E-07 |
| chr8 | 143430943 | 143432704 | 1762 | -6.42 | 2.08E-07 | 3.69E-07 |
| chrX | 39772352 | 39773689 | 1338 | -5.95 | 2.09E-07 | 3.70E-07 |
| chrX | 120428728 | 120430471 | 1744 | 5.12 | 2.09E-07 | 3.71E-07 |
| chr8 | 103762481 | 103763733 | 1253 | -6.57 | 2.09E-07 | 3.71E-07 |
| chr10 | 112192318 | 112193579 | 1262 | -6.26 | 2.10E-07 | 3.71E-07 |
| chr19 | 17925549 | 17927375 | 1827 | -6.43 | 2.10E-07 | 3.72E-07 |
| chr3 | 57320137 | 57321052 | 916 | -5.96 | 2.10E-07 | 3.72E-07 |
| chr1 | 189764077 | 189766077 | 2001 | -6.44 | 2.11E-07 | 3.73E-07 |
| chr17 | 14721815 | 15113437 | 391623 | -0.44 | 2.11E-07 | 3.74E-07 |
| chrX | 34634807 | 34636128 | 1322 | -6.04 | 2.11E-07 | 3.74E-07 |
| chr1 | 165816862 | 165817857 | 996 | -6.01 | 2.11E-07 | 3.74E-07 |
| chr9 | 116383803 | 116384826 | 1024 | -6.06 | 2.12E-07 | 3.75E-07 |
| chr4 | 154858642 | 154874369 | 15728 | 5.78 | 2.12E-07 | 3.75E-07 |
| chrX | 130994157 | 130995389 | 1233 | -5.98 | 2.12E-07 | 3.76E-07 |
| chr18 | 2866145 | 2866578 | 434 | -4.04 | 2.12E-07 | 3.76E-07 |
| chr1 | 5759738 | 5761253 | 1516 | -4.67 | 2.12E-07 | 3.76E-07 |
| chr13 | 113501548 | 113502854 | 1307 | -5.99 | 2.13E-07 | 3.76E-07 |
| chr13 | 78060580 | 78062203 | 1624 | -6.05 | 2.13E-07 | 3.77E-07 |
| chr9 | 36533035 | 36534140 | 1106 | -6.05 | 2.13E-07 | 3.77E-07 |
| chr18 | 936662 | 937793 | 1132 | 6.02 | 2.13E-07 | 3.78E-07 |
| chr11 | 31798718 | 31799702 | 985 | -5.95 | 2.14E-07 | 3.78E-07 |
| chr8 | 24267708 | 24274393 | 6686 | -6.31 | 2.14E-07 | 3.78E-07 |
| chr1 | 181213806 | 181225575 | 11770 | -2.1 | 2.14E-07 | 3.79E-07 |
| chr17 | 67288355 | 67289468 | 1114 | -6.03 | 2.14E-07 | 3.79E-07 |
| chr3 | 96330339 | 96333140 | 2802 | -6.03 | 2.14E-07 | 3.79E-07 |
| chr18 | 70178345 | 70180312 | 1968 | -5.94 | 2.14E-07 | 3.79E-07 |
| chr19 | 15583130 | 15584021 | 892 | -5.96 | 2.15E-07 | 3.81E-07 |
| chr3 | 33223855 | 33224753 | 899 | -5.96 | 2.15E-07 | 3.81E-07 |
| chr9 | 36915185 | 36932779 | 17595 | -5.95 | 2.15E-07 | 3.81E-07 |
| chr12 | 19578955 | 19580257 | 1303 | -5.98 | 2.16E-07 | 3.82E-07 |
| chr11 | 59637285 | 59719782 | 82498 | -1.63 | 2.16E-07 | 3.82E-07 |
| chr8 | 39567271 | 39569725 | 2455 | 6.96 | 2.16E-07 | 3.82E-07 |

| | | | | | | |
|---|---|---|---|---|---|---|
| chr12 | 15645749 | 15646980 | 1232 | -5.94 | 2.16E-07 | 3.82E-07 |
| chr6 | 154611333 | 154613208 | 1876 | -6.04 | 2.16E-07 | 3.83E-07 |
| chr9 | 128608722 | 128609629 | 908 | -6.04 | 2.16E-07 | 3.83E-07 |
| chr11 | 35406010 | 35420358 | 14349 | -6.63 | 2.16E-07 | 3.83E-07 |
| chr4 | 17848584 | 17850228 | 1645 | -5.95 | 2.17E-07 | 3.84E-07 |
| chr3 | 8291873 | 8314642 | 22770 | -2.31 | 2.17E-07 | 3.84E-07 |
| chr15 | 99961123 | 99962424 | 1302 | -5.97 | 2.17E-07 | 3.84E-07 |
| chr7 | 111520367 | 111522043 | 1677 | -5.97 | 2.17E-07 | 3.84E-07 |
| chr3 | 109790460 | 109792480 | 2021 | -5.99 | 2.18E-07 | 3.86E-07 |
| chr11 | 96124048 | 96125574 | 1527 | -6.02 | 2.18E-07 | 3.86E-07 |
| chr15 | 48493142 | 48519734 | 26593 | 1.5 | 2.18E-07 | 3.86E-07 |
| chr2 | 151510317 | 151515627 | 5311 | -6.5 | 2.18E-07 | 3.86E-07 |
| chr11 | 78046061 | 78047345 | 1285 | -6.35 | 2.19E-07 | 3.86E-07 |
| chr11 | 70535503 | 70551856 | 16354 | -5.97 | 2.19E-07 | 3.88E-07 |
| chr5 | 815671 | 816804 | 1134 | -5.97 | 2.19E-07 | 3.88E-07 |
| chr3 | 61919255 | 61929247 | 9993 | -1.85 | 2.19E-07 | 3.88E-07 |
| chr11 | 101158858 | 101160216 | 1359 | -2.19 | 2.20E-07 | 3.88E-07 |
| chr2 | 85042916 | 85052953 | 10038 | 6.59 | 2.20E-07 | 3.88E-07 |
| chr4 | 146666489 | 146668260 | 1772 | -5.95 | 2.20E-07 | 3.88E-07 |
| chr2 | 232352633 | 232354835 | 2203 | -6.51 | 2.20E-07 | 3.89E-07 |
| chr15 | 31025932 | 31027517 | 1586 | -6.01 | 2.20E-07 | 3.89E-07 |
| chr2 | 152198326 | 152201490 | 3165 | -6.36 | 2.20E-07 | 3.89E-07 |
| chr14 | 66273169 | 66283309 | 10141 | -5.43 | 2.21E-07 | 3.90E-07 |
| chr7 | 142580729 | 142582398 | 1670 | 5.23 | 2.21E-07 | 3.90E-07 |
| chr11 | 104021093 | 104022098 | 1006 | -6.02 | 2.21E-07 | 3.90E-07 |
| chr15 | 59010488 | 59011776 | 1289 | -6.02 | 2.21E-07 | 3.90E-07 |
| chr2 | 19859830 | 19860987 | 1158 | -6.02 | 2.21E-07 | 3.90E-07 |
| chr5 | 123966319 | 123968399 | 2081 | -5.93 | 2.21E-07 | 3.90E-07 |
| chr4 | 95468082 | 95468814 | 733 | -5.94 | 2.21E-07 | 3.91E-07 |
| chr5 | 36931661 | 36933092 | 1432 | -5.94 | 2.21E-07 | 3.91E-07 |
| chr4 | 173228058 | 173229593 | 1536 | -5.94 | 2.21E-07 | 3.91E-07 |
| chr3 | 139151559 | 139152442 | 884 | -5.93 | 2.22E-07 | 3.92E-07 |
| chr7 | 135270639 | 135271657 | 1019 | -6.16 | 2.22E-07 | 3.92E-07 |
| chrX | 30140870 | 30142229 | 1360 | -6.16 | 2.22E-07 | 3.92E-07 |
| chr1 | 30733185 | 30734884 | 1700 | -6.14 | 2.22E-07 | 3.92E-07 |
| chr1 | 25369726 | 25370553 | 828 | -5.94 | 2.22E-07 | 3.92E-07 |
| chr4 | 23141251 | 23142925 | 1675 | -5.94 | 2.22E-07 | 3.92E-07 |
| chr6 | 99900948 | 99902157 | 1210 | -5.94 | 2.22E-07 | 3.92E-07 |
| chr1 | 154164079 | 154165738 | 1660 | 5.3 | 2.22E-07 | 3.93E-07 |
| chr16 | 50233315 | 50234093 | 779 | -6.15 | 2.22E-07 | 3.93E-07 |
| chr1 | 214307011 | 214307800 | 790 | -6.16 | 2.22E-07 | 3.93E-07 |
| chr2 | 27412482 | 27413436 | 955 | -5.96 | 2.23E-07 | 3.93E-07 |
| chr8 | 95039089 | 95039961 | 873 | -5.96 | 2.23E-07 | 3.93E-07 |
| chr13 | 106883640 | 106884776 | 1137 | -6 | 2.23E-07 | 3.93E-07 |
| chr7 | 134607796 | 134610261 | 2466 | -6.17 | 2.23E-07 | 3.93E-07 |
| chr14 | 55934317 | 55936027 | 1711 | -6.38 | 2.23E-07 | 3.93E-07 |
| chr4 | 47890562 | 47891829 | 1268 | -6.19 | 2.23E-07 | 3.94E-07 |
| chr17 | 27910106 | 27911111 | 1006 | -5.97 | 2.23E-07 | 3.94E-07 |
| chr22 | 31189699 | 31192187 | 2489 | -6.19 | 2.23E-07 | 3.94E-07 |
| chr12 | 105498644 | 105506847 | 8204 | -6.11 | 2.23E-07 | 3.94E-07 |
| chr2 | 43676828 | 43678079 | 1252 | -6.01 | 2.23E-07 | 3.94E-07 |
| chr6 | 168256671 | 168258149 | 1479 | -6.01 | 2.23E-07 | 3.94E-07 |
| chr4 | 115199300 | 115200828 | 1529 | -6.19 | 2.24E-07 | 3.95E-07 |
| chr7 | 93380921 | 93381743 | 823 | -6.19 | 2.24E-07 | 3.95E-07 |
| chr1 | 84408789 | 84410456 | 1668 | -6.39 | 2.24E-07 | 3.95E-07 |
| chr4 | 121503088 | 121518780 | 15693 | 4.24 | 2.24E-07 | 3.95E-07 |
| chr18 | 12913242 | 12914686 | 1445 | -6.1 | 2.24E-07 | 3.95E-07 |
| chr3 | 81537043 | 81537745 | 703 | -4.44 | 2.24E-07 | 3.95E-07 |
| chr9 | 139899340 | 139900393 | 1054 | -6.2 | 2.24E-07 | 3.95E-07 |
| chr9 | 97720441 | 97722020 | 1580 | -6.09 | 2.24E-07 | 3.95E-07 |
| chr11 | 44507278 | 44508954 | 1677 | -6.09 | 2.24E-07 | 3.96E-07 |

| | | | | | | |
|---|---|---|---|---|---|---|
| chr7 | 98834914 | 98835917 | 1004 | -6.09 | 2.24E-07 | 3.96E-07 |
| chr1 | 26064473 | 26065858 | 1386 | -5.94 | 2.24E-07 | 3.96E-07 |
| chr16 | 69590826 | 69591965 | 1140 | -6.55 | 2.25E-07 | 3.96E-07 |
| chr2 | 228486936 | 228487884 | 949 | -6.22 | 2.25E-07 | 3.97E-07 |
| chr9 | 118615172 | 118616708 | 1537 | -6.08 | 2.25E-07 | 3.97E-07 |
| chr12 | 118369064 | 118384947 | 15884 | -6.4 | 2.25E-07 | 3.98E-07 |
| chr6 | 54532064 | 54533725 | 1662 | -6.24 | 2.26E-07 | 3.98E-07 |
| chr5 | 18032798 | 18043156 | 10359 | -7.29 | 2.26E-07 | 3.98E-07 |
| chr3 | 171682317 | 171683402 | 1086 | -6.07 | 2.26E-07 | 3.99E-07 |
| chr5 | 100543982 | 100545617 | 1636 | -6.23 | 2.26E-07 | 3.99E-07 |
| chr13 | 82461468 | 82463325 | 1858 | -6.07 | 2.26E-07 | 3.99E-07 |
| chr14 | 31901120 | 31902040 | 921 | -5.97 | 2.26E-07 | 3.99E-07 |
| chr3 | 103194477 | 103196103 | 1627 | 4.54 | 2.26E-07 | 3.99E-07 |
| chr1 | 45644756 | 45645609 | 854 | -6 | 2.26E-07 | 3.99E-07 |
| chr1 | 56442133 | 56443894 | 1762 | -6 | 2.26E-07 | 3.99E-07 |
| chr19 | 54442578 | 54443257 | 680 | -6 | 2.26E-07 | 3.99E-07 |
| chr3 | 43674194 | 43675673 | 1480 | -6 | 2.26E-07 | 3.99E-07 |
| chr2 | 24081325 | 24082749 | 1425 | -6.42 | 2.27E-07 | 4.00E-07 |
| chr3 | 38385148 | 38386536 | 1389 | -6.06 | 2.27E-07 | 4.00E-07 |
| chr2 | 69107387 | 69110395 | 3009 | -2.25 | 2.27E-07 | 4.01E-07 |
| chr3 | 71390433 | 71391229 | 797 | -5.96 | 2.27E-07 | 4.01E-07 |
| chr8 | 133041457 | 133045677 | 4221 | 2.81 | 2.28E-07 | 4.01E-07 |
| chr7 | 37040873 | 37042708 | 1836 | -3.55 | 2.28E-07 | 4.02E-07 |
| chr9 | 72392341 | 72405524 | 13184 | -5.12 | 2.28E-07 | 4.02E-07 |
| chr14 | 63997789 | 63999225 | 1437 | -5.95 | 2.28E-07 | 4.02E-07 |
| chr19 | 19290588 | 19292444 | 1857 | -5.94 | 2.29E-07 | 4.03E-07 |
| chrX | 76522207 | 76554671 | 32465 | -1.4 | 2.29E-07 | 4.03E-07 |
| chr11 | 67410316 | 67424719 | 14404 | -6.26 | 2.29E-07 | 4.03E-07 |
| chr7 | 82213259 | 82214245 | 987 | -5.93 | 2.29E-07 | 4.03E-07 |
| chr1 | 210208626 | 210226056 | 17431 | -1.39 | 2.29E-07 | 4.04E-07 |
| chrX | 82919982 | 82994393 | 74412 | 1.79 | 2.29E-07 | 4.04E-07 |
| chr15 | 29917467 | 29961527 | 44061 | -8 | 2.29E-07 | 4.04E-07 |
| chr7 | 84807886 | 84809824 | 1939 | -4.62 | 2.30E-07 | 4.05E-07 |
| chr20 | 2461012 | 2462098 | 1087 | -6.04 | 2.30E-07 | 4.06E-07 |
| chrX | 154092362 | 154096987 | 4626 | -3.16 | 2.31E-07 | 4.06E-07 |
| chr18 | 69124656 | 69227902 | 103247 | 1.58 | 2.31E-07 | 4.06E-07 |
| chr15 | 90480904 | 90482022 | 1119 | -5.92 | 2.31E-07 | 4.07E-07 |
| chr7 | 46747202 | 46747971 | 770 | -5.92 | 2.31E-07 | 4.07E-07 |
| chr7 | 66494207 | 66495424 | 1218 | -5.92 | 2.31E-07 | 4.07E-07 |
| chr17 | 60941986 | 60944954 | 2969 | -2.65 | 2.32E-07 | 4.08E-07 |
| chr14 | 23476958 | 23478075 | 1118 | -5.92 | 2.33E-07 | 4.10E-07 |
| chr4 | 7926527 | 7936154 | 9628 | -5.92 | 2.33E-07 | 4.10E-07 |
| chr19 | 16847196 | 16848324 | 1129 | -5.93 | 2.33E-07 | 4.10E-07 |
| chr11 | 20683151 | 20693128 | 9978 | -1.76 | 2.33E-07 | 4.10E-07 |
| chr7 | 7716985 | 7717987 | 1003 | -5.98 | 2.33E-07 | 4.10E-07 |
| chr9 | 3443713 | 3444556 | 844 | -5.98 | 2.33E-07 | 4.10E-07 |
| chr17 | 80504808 | 80506229 | 1422 | -6.03 | 2.33E-07 | 4.10E-07 |
| chr2 | 191819830 | 191822162 | 2333 | -6.03 | 2.33E-07 | 4.10E-07 |
| chr10 | 34313804 | 34316410 | 2607 | -6.31 | 2.33E-07 | 4.11E-07 |
| chr7 | 152192399 | 152193787 | 1389 | -6.32 | 2.34E-07 | 4.12E-07 |
| chr6 | 21563318 | 21564888 | 1571 | -5.97 | 2.34E-07 | 4.12E-07 |
| chr6 | 13010548 | 13013225 | 2678 | 2.46 | 2.34E-07 | 4.13E-07 |
| chr4 | 178822676 | 178823996 | 1321 | -6.02 | 2.35E-07 | 4.13E-07 |
| chr17 | 32937646 | 32938634 | 989 | -6.02 | 2.35E-07 | 4.14E-07 |
| chr5 | 34035306 | 34038213 | 2908 | 6.04 | 2.36E-07 | 4.16E-07 |
| chr10 | 16656679 | 16658993 | 2315 | -5.95 | 2.36E-07 | 4.16E-07 |
| chr9 | 135342265 | 135349112 | 6848 | -5.95 | 2.36E-07 | 4.16E-07 |
| chr6 | 47335866 | 47338996 | 3131 | -5.92 | 2.37E-07 | 4.16E-07 |
| chr3 | 189750963 | 189753079 | 2117 | -5.97 | 2.37E-07 | 4.17E-07 |
| chr12 | 92885387 | 92887063 | 1677 | -6.55 | 2.37E-07 | 4.17E-07 |
| chr11 | 71838984 | 71840249 | 1266 | -6.01 | 2.38E-07 | 4.18E-07 |

| | | | | | | |
|---|---|---|---|---|---|---|
| chr13 | 23972947 | 23974684 | 1738 | -6.01 | 2.38E-07 | 4.18E-07 |
| chr6 | 159282243 | 159283895 | 1653 | -6.34 | 2.38E-07 | 4.18E-07 |
| chr14 | 52502906 | 52503883 | 978 | -5.97 | 2.38E-07 | 4.18E-07 |
| chr15 | 36165229 | 36167416 | 2188 | -5.97 | 2.38E-07 | 4.18E-07 |
| chr6 | 17509544 | 17510537 | 994 | -5.97 | 2.38E-07 | 4.18E-07 |
| chr6 | 22085268 | 22086810 | 1543 | -6.01 | 2.39E-07 | 4.20E-07 |
| chr8 | 39124270 | 39125815 | 1546 | -6.01 | 2.39E-07 | 4.20E-07 |
| chr5 | 132077859 | 132078596 | 738 | -6.36 | 2.39E-07 | 4.21E-07 |
| chr1 | 245728526 | 245729944 | 1419 | -4.41 | 2.40E-07 | 4.22E-07 |
| chr4 | 170757969 | 170759262 | 1294 | -6 | 2.41E-07 | 4.23E-07 |
| chr16 | 21192708 | 21214066 | 21359 | -7.68 | 2.41E-07 | 4.24E-07 |
| chr4 | 33314411 | 33318008 | 3598 | -3.55 | 2.41E-07 | 4.24E-07 |
| chr4 | 52894663 | 52895594 | 932 | -5.92 | 2.41E-07 | 4.24E-07 |
| chr3 | 101560701 | 101562776 | 2076 | -6.13 | 2.41E-07 | 4.24E-07 |
| chr9 | 15342035 | 15343658 | 1624 | -6.14 | 2.41E-07 | 4.24E-07 |
| chr14 | 21590408 | 21652579 | 62172 | -3.72 | 2.41E-07 | 4.24E-07 |
| chr9 | 75180312 | 75181290 | 979 | -6.12 | 2.41E-07 | 4.24E-07 |
| chr8 | 8448186 | 8450357 | 2172 | -5.96 | 2.41E-07 | 4.24E-07 |
| chr12 | 18475811 | 18477274 | 1464 | -5.94 | 2.41E-07 | 4.24E-07 |
| chr2 | 43954299 | 43955055 | 757 | -5.94 | 2.41E-07 | 4.24E-07 |
| chr4 | 102148857 | 102149479 | 623 | -5.93 | 2.41E-07 | 4.24E-07 |
| chr6 | 54041000 | 54042244 | 1245 | -5.93 | 2.41E-07 | 4.24E-07 |
| chr2 | 200672551 | 200673745 | 1195 | -6.15 | 2.42E-07 | 4.24E-07 |
| chr12 | 75852584 | 75858678 | 6095 | -6.7 | 2.42E-07 | 4.25E-07 |
| chr13 | 36250849 | 36252085 | 1237 | -6.1 | 2.42E-07 | 4.25E-07 |
| chr4 | 89200426 | 89201844 | 1419 | -6.1 | 2.42E-07 | 4.25E-07 |
| chr6 | 135859838 | 135861398 | 1561 | -6.1 | 2.42E-07 | 4.25E-07 |
| chr1 | 167670115 | 167672712 | 2598 | -2.69 | 2.42E-07 | 4.26E-07 |
| chr14 | 27383711 | 27385195 | 1485 | -6.17 | 2.42E-07 | 4.26E-07 |
| chr15 | 80305730 | 80308458 | 2729 | -5.92 | 2.42E-07 | 4.26E-07 |
| chr6 | 82918634 | 82919553 | 920 | -5.92 | 2.42E-07 | 4.26E-07 |
| chr11 | 95311455 | 95312920 | 1466 | 5.18 | 2.42E-07 | 4.26E-07 |
| chr5 | 76532715 | 76533643 | 929 | -5.92 | 2.42E-07 | 4.26E-07 |
| chr9 | 78990909 | 78992598 | 1690 | -5.92 | 2.42E-07 | 4.26E-07 |
| chr9 | 6520886 | 6522704 | 1819 | -6.18 | 2.43E-07 | 4.26E-07 |
| chr20 | 60422235 | 60423784 | 1550 | -5.92 | 2.43E-07 | 4.27E-07 |
| chr12 | 48924649 | 48931525 | 6877 | -3.11 | 2.43E-07 | 4.28E-07 |
| chr12 | 58249829 | 58251022 | 1194 | -6.07 | 2.44E-07 | 4.28E-07 |
| chr5 | 123863225 | 123875940 | 12716 | -7.01 | 2.44E-07 | 4.28E-07 |
| chr5 | 176157463 | 176158681 | 1219 | -6.21 | 2.45E-07 | 4.30E-07 |
| chr17 | 44993645 | 44994489 | 845 | -5.92 | 2.45E-07 | 4.30E-07 |
| chr5 | 111131157 | 111131985 | 829 | -5.91 | 2.45E-07 | 4.31E-07 |
| chrX | 39587654 | 39588693 | 1040 | -6.06 | 2.46E-07 | 4.31E-07 |
| chr1 | 235725617 | 235727207 | 1591 | -6.13 | 2.46E-07 | 4.32E-07 |
| chr13 | 90212218 | 90213719 | 1502 | -5.98 | 2.46E-07 | 4.32E-07 |
| chr8 | 110878972 | 110882756 | 3785 | -6.11 | 2.46E-07 | 4.33E-07 |
| chr9 | 103472356 | 103476185 | 3830 | -2 | 2.47E-07 | 4.33E-07 |
| chr15 | 62962597 | 62971887 | 9291 | -6.05 | 2.47E-07 | 4.33E-07 |
| chr5 | 50097297 | 50100225 | 2929 | -6.05 | 2.47E-07 | 4.33E-07 |
| chr7 | 12330923 | 12345035 | 14113 | -6.75 | 2.47E-07 | 4.33E-07 |
| chr6 | 127299086 | 127300278 | 1193 | -5.93 | 2.47E-07 | 4.34E-07 |
| chr7 | 109037034 | 109038384 | 1351 | -5.93 | 2.47E-07 | 4.34E-07 |
| chr18 | 46782523 | 46784386 | 1864 | -6.09 | 2.47E-07 | 4.34E-07 |
| chr2 | 201197024 | 201209263 | 12240 | -6.09 | 2.47E-07 | 4.34E-07 |
| chr7 | 132316701 | 132317412 | 712 | 5.34 | 2.47E-07 | 4.34E-07 |
| chr16 | 48066974 | 48067960 | 987 | -6.19 | 2.48E-07 | 4.34E-07 |
| chr2 | 214828782 | 214830497 | 1716 | -6.07 | 2.48E-07 | 4.36E-07 |
| chr5 | 90364181 | 90365321 | 1141 | -5.93 | 2.48E-07 | 4.36E-07 |
| chr10 | 106704176 | 106704960 | 785 | -5.98 | 2.48E-07 | 4.36E-07 |
| chr1 | 220898604 | 220906429 | 7826 | -6.27 | 2.50E-07 | 4.38E-07 |
| chrX | 80809195 | 80845741 | 36547 | 3.02 | 2.50E-07 | 4.38E-07 |

| | | | | | | |
|---|---|---|---|---|---|---|
| chrX | 49485928 | 49487031 | 1104 | -5.92 | 2.50E-07 | 4.39E-07 |
| chr4 | 159824164 | 159825223 | 1060 | -5.94 | 2.50E-07 | 4.39E-07 |
| chr6 | 146003006 | 146014711 | 11706 | -6.05 | 2.50E-07 | 4.39E-07 |
| chr3 | 23764924 | 23766022 | 1099 | -5.91 | 2.51E-07 | 4.40E-07 |
| chr3 | 118358416 | 118384884 | 26469 | 1.37 | 2.51E-07 | 4.41E-07 |
| chr13 | 108802262 | 108806761 | 4500 | -2.29 | 2.51E-07 | 4.41E-07 |
| chr2 | 160677287 | 160678661 | 1375 | -5.95 | 2.52E-07 | 4.41E-07 |
| chr2 | 21037767 | 21039333 | 1567 | -2.91 | 2.52E-07 | 4.43E-07 |
| chr6 | 119427600 | 119429071 | 1472 | -6.04 | 2.52E-07 | 4.43E-07 |
| chr14 | 57160055 | 57181551 | 21497 | 5.9 | 2.53E-07 | 4.43E-07 |
| chr1 | 229625875 | 229626660 | 786 | -5.91 | 2.53E-07 | 4.43E-07 |
| chr8 | 93912304 | 93913281 | 978 | -5.91 | 2.53E-07 | 4.43E-07 |
| chr3 | 161453628 | 161491486 | 37859 | 6.41 | 2.53E-07 | 4.44E-07 |
| chr20 | 5338956 | 5426316 | 87361 | -1.31 | 2.54E-07 | 4.45E-07 |
| chr1 | 167404994 | 167406025 | 1032 | -6.03 | 2.54E-07 | 4.45E-07 |
| chr3 | 126861980 | 126863253 | 1274 | -5.97 | 2.54E-07 | 4.45E-07 |
| chr5 | 60882977 | 60885408 | 2432 | -6.64 | 2.54E-07 | 4.46E-07 |
| chr12 | 123620258 | 123621648 | 1391 | -5.96 | 2.55E-07 | 4.47E-07 |
| chr2 | 189667865 | 189668843 | 979 | -6.03 | 2.55E-07 | 4.47E-07 |
| chr6 | 157989394 | 157990129 | 736 | -6.03 | 2.55E-07 | 4.47E-07 |
| chr5 | 93022360 | 93023633 | 1274 | -5.92 | 2.56E-07 | 4.48E-07 |
| chr9 | 93985293 | 93986528 | 1236 | -5.92 | 2.56E-07 | 4.48E-07 |
| chr16 | 16726023 | 16727287 | 1265 | 6.06 | 2.56E-07 | 4.48E-07 |
| chr7 | 36504430 | 36505360 | 931 | -6 | 2.56E-07 | 4.48E-07 |
| chr12 | 111993000 | 111993927 | 928 | -6.28 | 2.56E-07 | 4.49E-07 |
| chr1 | 76431048 | 76432363 | 1316 | -5.91 | 2.58E-07 | 4.52E-07 |
| chr17 | 59920872 | 59921951 | 1080 | -5.91 | 2.58E-07 | 4.52E-07 |
| chr5 | 156857142 | 156866246 | 9105 | -6.51 | 2.58E-07 | 4.52E-07 |
| chr13 | 49691192 | 49692381 | 1190 | -3.25 | 2.59E-07 | 4.54E-07 |
| chr10 | 127140808 | 127338341 | 197534 | -1.33 | 2.59E-07 | 4.54E-07 |
| chr15 | 81110079 | 81111266 | 1188 | -5.92 | 2.60E-07 | 4.55E-07 |
| chr1 | 188368454 | 188369668 | 1215 | -6.01 | 2.60E-07 | 4.55E-07 |
| chr2 | 134663662 | 134724484 | 60823 | -1.12 | 2.60E-07 | 4.56E-07 |
| chr7 | 73226884 | 73228237 | 1354 | -6.35 | 2.61E-07 | 4.56E-07 |
| chr5 | 84959349 | 85406895 | 447547 | 1.17 | 2.61E-07 | 4.57E-07 |
| chr2 | 45491445 | 45492697 | 1253 | -5.99 | 2.61E-07 | 4.57E-07 |
| chr6 | 129089065 | 129095281 | 6217 | -5.99 | 2.61E-07 | 4.57E-07 |
| chr13 | 112195981 | 112197623 | 1643 | -3.63 | 2.61E-07 | 4.58E-07 |
| chr8 | 143839962 | 143843884 | 3923 | -6.36 | 2.62E-07 | 4.58E-07 |
| chr4 | 133205921 | 133208103 | 2183 | -6 | 2.62E-07 | 4.58E-07 |
| chr17 | 53668981 | 53678670 | 9690 | -6.14 | 2.62E-07 | 4.59E-07 |
| chr16 | 19332047 | 19380203 | 48157 | -7.07 | 2.63E-07 | 4.60E-07 |
| chr4 | 143467178 | 143469699 | 2522 | -4.83 | 2.63E-07 | 4.60E-07 |
| chr2 | 127562607 | 127579188 | 16582 | -2.17 | 2.63E-07 | 4.60E-07 |
| chr6 | 22600852 | 22602657 | 1806 | -6.09 | 2.63E-07 | 4.60E-07 |
| chr20 | 9173650 | 9178901 | 5252 | -6.15 | 2.63E-07 | 4.60E-07 |
| chr9 | 75005199 | 75006612 | 1414 | -6.15 | 2.63E-07 | 4.60E-07 |
| chr17 | 64395186 | 64396172 | 987 | -5.95 | 2.63E-07 | 4.60E-07 |
| chr3 | 53014577 | 53015727 | 1151 | -5.95 | 2.63E-07 | 4.60E-07 |
| chr2 | 175585568 | 175586403 | 836 | -6.08 | 2.63E-07 | 4.60E-07 |
| chr6 | 54358591 | 54361873 | 3283 | -6.08 | 2.63E-07 | 4.60E-07 |
| chr8 | 99865491 | 99867293 | 1803 | -6.16 | 2.63E-07 | 4.61E-07 |
| chr7 | 103247005 | 103260598 | 13594 | -1.7 | 2.63E-07 | 4.61E-07 |
| chr5 | 138331693 | 138332922 | 1230 | -5.92 | 2.63E-07 | 4.61E-07 |
| chr1 | 87251459 | 87252762 | 1304 | -5.93 | 2.64E-07 | 4.62E-07 |
| chr19 | 50915141 | 50916201 | 1061 | -5.93 | 2.64E-07 | 4.62E-07 |
| chr2 | 236157809 | 236158707 | 899 | -6.07 | 2.64E-07 | 4.62E-07 |
| chr7 | 151388584 | 151389381 | 798 | -5.98 | 2.64E-07 | 4.63E-07 |
| chrX | 129449674 | 129459414 | 9741 | -2.07 | 2.65E-07 | 4.64E-07 |
| chr12 | 19014643 | 19016133 | 1491 | -5.9 | 2.65E-07 | 4.64E-07 |
| chr10 | 50833696 | 50835540 | 1845 | 6.21 | 2.66E-07 | 4.65E-07 |

| | | | | | | |
|---|---|---|---|---|---|---|
| chr6 | 141484264 | 141485547 | 1284 | -5.9 | 2.66E-07 | 4.65E-07 |
| chrX | 65935955 | 65938494 | 2540 | 2.74 | 2.66E-07 | 4.65E-07 |
| chr15 | 41174587 | 41175613 | 1027 | -5.94 | 2.66E-07 | 4.66E-07 |
| chr9 | 99294297 | 99295606 | 1310 | -5.94 | 2.66E-07 | 4.66E-07 |
| chr1 | 21816510 | 21826692 | 10183 | -6.39 | 2.67E-07 | 4.66E-07 |
| chr8 | 112113120 | 112114343 | 1224 | -6.04 | 2.67E-07 | 4.67E-07 |
| chr15 | 40020074 | 40022013 | 1940 | -6.4 | 2.68E-07 | 4.68E-07 |
| chr15 | 71658772 | 71659711 | 940 | -5.97 | 2.68E-07 | 4.68E-07 |
| chr2 | 56371866 | 56399721 | 27856 | 6.22 | 2.68E-07 | 4.69E-07 |
| chr21 | 45569078 | 45570012 | 935 | -5.94 | 2.68E-07 | 4.69E-07 |
| chr4 | 90084086 | 90085227 | 1142 | -5.94 | 2.68E-07 | 4.69E-07 |
| chrX | 140831683 | 140832855 | 1173 | -6.03 | 2.68E-07 | 4.69E-07 |
| chr11 | 18456084 | 18457432 | 1349 | -6.23 | 2.69E-07 | 4.70E-07 |
| chr3 | 101266271 | 101267190 | 920 | -5.98 | 2.69E-07 | 4.70E-07 |
| chr12 | 85864033 | 85865491 | 1459 | -5.9 | 2.69E-07 | 4.70E-07 |
| chr13 | 95144802 | 95146481 | 1680 | -2.3 | 2.69E-07 | 4.70E-07 |
| chr10 | 4668413 | 4682646 | 14234 | -7.49 | 2.71E-07 | 4.73E-07 |
| chr4 | 132441026 | 132442678 | 1653 | -5.92 | 2.71E-07 | 4.73E-07 |
| chr5 | 18063225 | 18064219 | 995 | -5.93 | 2.71E-07 | 4.73E-07 |
| chr1 | 108580735 | 108584934 | 4200 | 3.62 | 2.71E-07 | 4.74E-07 |
| chr16 | 54495074 | 54610187 | 115114 | -2.22 | 2.71E-07 | 4.74E-07 |
| chr9 | 115852806 | 115908595 | 55790 | -4.67 | 2.71E-07 | 4.74E-07 |
| chr10 | 116427066 | 116428153 | 1088 | -6.01 | 2.72E-07 | 4.75E-07 |
| chrX | 29316535 | 29318370 | 1836 | -5.91 | 2.72E-07 | 4.75E-07 |
| chr1 | 79774119 | 79776177 | 2059 | -5.96 | 2.72E-07 | 4.75E-07 |
| chr2 | 58347077 | 58348314 | 1238 | -5.96 | 2.72E-07 | 4.75E-07 |
| chr6 | 22360589 | 22362290 | 1702 | -5.96 | 2.72E-07 | 4.75E-07 |
| chr6 | 25445282 | 25446529 | 1248 | -5.96 | 2.72E-07 | 4.75E-07 |
| chr7 | 77637804 | 77639611 | 1808 | -6.26 | 2.72E-07 | 4.75E-07 |
| chr14 | 106873157 | 106874415 | 1259 | -3.26 | 2.72E-07 | 4.76E-07 |
| chr3 | 187189911 | 187217141 | 27231 | -2.67 | 2.72E-07 | 4.76E-07 |
| chr14 | 95800994 | 95822235 | 21242 | -7.74 | 2.73E-07 | 4.76E-07 |
| chr4 | 177393538 | 177396395 | 2858 | -6.12 | 2.73E-07 | 4.77E-07 |
| chr2 | 131841234 | 131842622 | 1389 | -6.11 | 2.73E-07 | 4.77E-07 |
| chr22 | 17421131 | 17422384 | 1254 | 5.97 | 2.73E-07 | 4.77E-07 |
| chr3 | 111812382 | 111813968 | 1587 | -6.07 | 2.74E-07 | 4.79E-07 |
| chr3 | 160694798 | 160699367 | 4570 | -2.45 | 2.75E-07 | 4.80E-07 |
| chr1 | 78405299 | 78406383 | 1085 | -6.07 | 2.75E-07 | 4.80E-07 |
| chr6 | 112344958 | 112346292 | 1335 | -6.07 | 2.75E-07 | 4.80E-07 |
| chr6 | 127810879 | 127812205 | 1327 | -6.07 | 2.75E-07 | 4.80E-07 |
| chrX | 84761417 | 84762865 | 1449 | 4.09 | 2.75E-07 | 4.80E-07 |
| chr8 | 143921331 | 143931804 | 10474 | -1.67 | 2.75E-07 | 4.81E-07 |
| chr2 | 63634995 | 63637197 | 2203 | -6.29 | 2.76E-07 | 4.81E-07 |
| chr4 | 114994681 | 114995768 | 1088 | -6.18 | 2.76E-07 | 4.81E-07 |
| chr2 | 164416758 | 164427955 | 11198 | -6.87 | 2.76E-07 | 4.81E-07 |
| chr17 | 76465007 | 76465924 | 918 | -5.93 | 2.76E-07 | 4.81E-07 |
| chr4 | 81100607 | 81102145 | 1539 | -5.93 | 2.76E-07 | 4.81E-07 |
| chr5 | 130214902 | 130216420 | 1519 | -5.93 | 2.76E-07 | 4.81E-07 |
| chr3 | 130593806 | 130605541 | 11736 | -5.56 | 2.76E-07 | 4.82E-07 |
| chr1 | 2094984 | 2096008 | 1025 | -5.95 | 2.76E-07 | 4.82E-07 |
| chr11 | 28515218 | 28517932 | 2715 | -5.95 | 2.76E-07 | 4.82E-07 |
| chr4 | 186691193 | 186692551 | 1359 | -5.95 | 2.76E-07 | 4.82E-07 |
| chr1 | 180948855 | 180950234 | 1380 | -5.89 | 2.76E-07 | 4.82E-07 |
| chr7 | 47548343 | 47549838 | 1496 | -5.89 | 2.76E-07 | 4.82E-07 |
| chr13 | 101904319 | 101905621 | 1303 | -6.05 | 2.77E-07 | 4.83E-07 |
| chr3 | 114719776 | 114721172 | 1397 | -6.05 | 2.77E-07 | 4.83E-07 |
| chr4 | 42057496 | 42058440 | 945 | -6.05 | 2.77E-07 | 4.83E-07 |
| chr5 | 6386184 | 6508613 | 122430 | -3.12 | 2.77E-07 | 4.83E-07 |
| chr18 | 52551966 | 52553233 | 1268 | -6.04 | 2.78E-07 | 4.84E-07 |
| chr18 | 74969551 | 75049674 | 80124 | -1.2 | 2.78E-07 | 4.86E-07 |
| chr4 | 137583085 | 137584328 | 1244 | -5.91 | 2.78E-07 | 4.86E-07 |

| | | | | | | |
|---|---|---|---|---|---|---|
| chr5 | 79978907 | 79980049 | 1143 | -6.22 | 2.79E-07 | 4.87E-07 |
| chrX | 117955942 | 117956813 | 872 | -5.89 | 2.79E-07 | 4.87E-07 |
| chr10 | 52577743 | 52592961 | 15219 | 1.37 | 2.79E-07 | 4.87E-07 |
| chr4 | 33867617 | 33934829 | 67213 | -1.53 | 2.80E-07 | 4.88E-07 |
| chrX | 13354935 | 13356064 | 1130 | -4.72 | 2.80E-07 | 4.89E-07 |
| chr12 | 21557724 | 21558942 | 1219 | -5.94 | 2.81E-07 | 4.90E-07 |
| chr19 | 14872893 | 14874277 | 1385 | -5.94 | 2.81E-07 | 4.90E-07 |
| chr1 | 46366866 | 46367915 | 1050 | -5.92 | 2.81E-07 | 4.90E-07 |
| chr11 | 9828786 | 9829504 | 719 | -6.02 | 2.81E-07 | 4.90E-07 |
| chr2 | 3223594 | 3224382 | 789 | -5.98 | 2.81E-07 | 4.90E-07 |
| chr1 | 178535217 | 178536590 | 1374 | -5.95 | 2.81E-07 | 4.91E-07 |
| chr2 | 61876046 | 61879468 | 3423 | -6.51 | 2.82E-07 | 4.92E-07 |
| chr2 | 180878194 | 180879721 | 1528 | -5.89 | 2.83E-07 | 4.93E-07 |
| chr8 | 24140099 | 24173523 | 33425 | -5.99 | 2.83E-07 | 4.94E-07 |
| chr1 | 213022371 | 213026316 | 3946 | -6.35 | 2.84E-07 | 4.95E-07 |
| chr3 | 176998542 | 177000004 | 1463 | -5.97 | 2.84E-07 | 4.96E-07 |
| chr3 | 81573741 | 81575404 | 1664 | -5.89 | 2.85E-07 | 4.96E-07 |
| chr6 | 144921474 | 144922865 | 1392 | -6.09 | 2.85E-07 | 4.97E-07 |
| chr7 | 17509798 | 17511124 | 1327 | -6.09 | 2.85E-07 | 4.97E-07 |
| chr13 | 50760768 | 50762295 | 1528 | -6.14 | 2.86E-07 | 4.98E-07 |
| chr16 | 81504461 | 81505194 | 734 | -6.28 | 2.86E-07 | 4.98E-07 |
| chr12 | 67625256 | 67629691 | 4436 | -1.56 | 2.86E-07 | 4.98E-07 |
| chr11 | 129885261 | 129886480 | 1220 | -5.93 | 2.86E-07 | 4.99E-07 |
| chr4 | 115043743 | 115045593 | 1851 | -5.93 | 2.86E-07 | 4.99E-07 |
| chr8 | 10638333 | 10639579 | 1247 | -5.91 | 2.87E-07 | 5.00E-07 |
| chr8 | 83041696 | 83044556 | 2861 | -5.91 | 2.87E-07 | 5.00E-07 |
| chr12 | 90714027 | 90715593 | 1567 | -6.16 | 2.87E-07 | 5.00E-07 |
| chr1 | 31004268 | 31015206 | 10939 | -6.05 | 2.87E-07 | 5.01E-07 |
| chr6 | 145013102 | 145014528 | 1427 | -6.05 | 2.87E-07 | 5.01E-07 |
| chr3 | 171627890 | 171638543 | 10654 | -6.54 | 2.87E-07 | 5.01E-07 |
| chr1 | 211208967 | 211210315 | 1349 | -5.99 | 2.88E-07 | 5.01E-07 |
| chr2 | 181680779 | 181682386 | 1608 | -5.99 | 2.88E-07 | 5.01E-07 |
| chr3 | 136848609 | 137373215 | 524607 | 9.64 | 2.89E-07 | 5.03E-07 |
| chr17 | 15269208 | 15270331 | 1124 | -6.18 | 2.89E-07 | 5.03E-07 |
| chr3 | 43786124 | 43788148 | 2025 | -6.18 | 2.89E-07 | 5.03E-07 |
| chr17 | 73020986 | 73021927 | 942 | -5.88 | 2.89E-07 | 5.03E-07 |
| chr7 | 43457135 | 43459521 | 2387 | -5.88 | 2.89E-07 | 5.03E-07 |
| chr2 | 184133275 | 184136130 | 2856 | -2.17 | 2.89E-07 | 5.04E-07 |
| chr11 | 108523255 | 108524414 | 1160 | -6.03 | 2.89E-07 | 5.04E-07 |
| chr2 | 144848761 | 144849711 | 951 | -4.46 | 2.89E-07 | 5.04E-07 |
| chr1 | 80227900 | 80229624 | 1725 | -5.98 | 2.90E-07 | 5.05E-07 |
| chr3 | 143508557 | 143513405 | 4849 | -2.64 | 2.90E-07 | 5.05E-07 |
| chr4 | 65391651 | 65393284 | 1634 | -5.88 | 2.90E-07 | 5.06E-07 |
| chr1 | 192688622 | 192689766 | 1145 | -6.03 | 2.91E-07 | 5.06E-07 |
| chr3 | 176160707 | 176172807 | 12101 | -6.03 | 2.91E-07 | 5.06E-07 |
| chr10 | 98146284 | 98148395 | 2112 | 6 | 2.91E-07 | 5.07E-07 |
| chr2 | 209015409 | 209018025 | 2617 | -3.97 | 2.91E-07 | 5.07E-07 |
| chr15 | 100421769 | 100422244 | 476 | -5.95 | 2.92E-07 | 5.08E-07 |
| chr6 | 154486931 | 154512199 | 25269 | -3.31 | 2.92E-07 | 5.08E-07 |
| chr1 | 217719927 | 217721487 | 1561 | -4.8 | 2.93E-07 | 5.10E-07 |
| chr1 | 243934824 | 243935356 | 533 | -5.89 | 2.93E-07 | 5.11E-07 |
| chr16 | 53235646 | 53236705 | 1060 | -5.89 | 2.93E-07 | 5.11E-07 |
| chr1 | 8287027 | 8288465 | 1439 | -6.01 | 2.94E-07 | 5.11E-07 |
| chr10 | 127351615 | 127367086 | 15472 | -2.38 | 2.94E-07 | 5.11E-07 |
| chr1 | 200375693 | 200387834 | 12142 | -6.24 | 2.94E-07 | 5.12E-07 |
| chr11 | 21778787 | 21783620 | 4834 | -3.13 | 2.95E-07 | 5.13E-07 |
| chr11 | 104003324 | 104004652 | 1329 | -5.88 | 2.95E-07 | 5.13E-07 |
| chr6 | 21180721 | 21181816 | 1096 | -5.88 | 2.95E-07 | 5.13E-07 |
| chr3 | 12239155 | 12263056 | 23902 | -6.6 | 2.95E-07 | 5.14E-07 |
| chr16 | 22810891 | 22816584 | 5694 | 1.68 | 2.95E-07 | 5.14E-07 |
| chr5 | 59296920 | 59346439 | 49520 | -1.98 | 2.95E-07 | 5.14E-07 |

| | | | | | | |
|---|---|---|---|---|---|---|
| chr12 | 114430223 | 114431562 | 1340 | -5.9 | 2.95E-07 | 5.14E-07 |
| chr4 | 63581258 | 63586028 | 4771 | -5.9 | 2.95E-07 | 5.14E-07 |
| chr7 | 1074997 | 1076452 | 1456 | -6.25 | 2.96E-07 | 5.15E-07 |
| chr8 | 125402050 | 125415063 | 13014 | -5.5 | 2.97E-07 | 5.16E-07 |
| chr17 | 39702800 | 39703591 | 792 | -5.89 | 2.97E-07 | 5.17E-07 |
| chr21 | 40739254 | 40740197 | 944 | -5.89 | 2.97E-07 | 5.17E-07 |
| chr2 | 85191941 | 85193569 | 1629 | -5.96 | 2.97E-07 | 5.17E-07 |
| chr16 | 19997287 | 20015239 | 17953 | -2.97 | 2.98E-07 | 5.18E-07 |
| chr1 | 239939405 | 239945248 | 5844 | -6.45 | 2.99E-07 | 5.20E-07 |
| chr2 | 62645915 | 62658939 | 13025 | -6.91 | 2.99E-07 | 5.20E-07 |
| chrX | 4884825 | 4886482 | 1658 | -6.28 | 2.99E-07 | 5.20E-07 |
| chr20 | 45199850 | 45200466 | 617 | -5.9 | 2.99E-07 | 5.20E-07 |
| chr5 | 114366845 | 114369568 | 2724 | -2.94 | 3.00E-07 | 5.22E-07 |
| chr2 | 136819411 | 136829916 | 10506 | -6.95 | 3.01E-07 | 5.22E-07 |
| chr18 | 32552713 | 32553558 | 846 | -5.88 | 3.02E-07 | 5.25E-07 |
| chr5 | 135102211 | 135103698 | 1488 | -5.88 | 3.02E-07 | 5.25E-07 |
| chrX | 66884839 | 66886135 | 1297 | -5.87 | 3.02E-07 | 5.26E-07 |
| chr12 | 15913477 | 15915527 | 2051 | -6.3 | 3.02E-07 | 5.26E-07 |
| chr18 | 64180071 | 64181448 | 1378 | -6.11 | 3.03E-07 | 5.27E-07 |
| chr17 | 33581669 | 33670429 | 88761 | -2.16 | 3.03E-07 | 5.27E-07 |
| chrX | 39056862 | 39111878 | 55017 | 1.29 | 3.04E-07 | 5.28E-07 |
| chr9 | 75076651 | 75077847 | 1197 | -5.88 | 3.04E-07 | 5.28E-07 |
| chr13 | 44206147 | 44207388 | 1242 | -6.06 | 3.04E-07 | 5.29E-07 |
| chrX | 44182524 | 44184295 | 1772 | -6.06 | 3.04E-07 | 5.29E-07 |
| chr12 | 9984863 | 9985568 | 706 | -6.05 | 3.05E-07 | 5.30E-07 |
| chr11 | 104136512 | 104137433 | 922 | -5.96 | 3.06E-07 | 5.32E-07 |
| chr14 | 85679512 | 85680445 | 934 | -5.96 | 3.06E-07 | 5.32E-07 |
| chr2 | 127742164 | 127747727 | 5564 | -3.12 | 3.06E-07 | 5.32E-07 |
| chr8 | 10292111 | 10333229 | 41119 | -2.38 | 3.06E-07 | 5.32E-07 |
| chr16 | 89522056 | 89523247 | 1192 | -6.04 | 3.06E-07 | 5.32E-07 |
| chr4 | 23261818 | 23263317 | 1500 | -5.92 | 3.06E-07 | 5.32E-07 |
| chr6 | 107282596 | 107284268 | 1673 | -5.92 | 3.06E-07 | 5.32E-07 |
| chr1 | 38903194 | 38909639 | 6446 | -5.49 | 3.07E-07 | 5.33E-07 |
| chr13 | 45100648 | 45101896 | 1249 | -6.02 | 3.08E-07 | 5.35E-07 |
| chr13 | 103070364 | 103173967 | 103604 | 5.04 | 3.08E-07 | 5.35E-07 |
| chr4 | 162291431 | 162294669 | 3239 | -4.71 | 3.08E-07 | 5.36E-07 |
| chr11 | 111465949 | 111467276 | 1328 | -4.85 | 3.09E-07 | 5.37E-07 |
| chr15 | 69598982 | 69599670 | 689 | -6.2 | 3.09E-07 | 5.37E-07 |
| chr13 | 83471642 | 83473903 | 2262 | -6.1 | 3.10E-07 | 5.38E-07 |
| chr12 | 101848724 | 101850528 | 1805 | -6.09 | 3.10E-07 | 5.38E-07 |
| chr2 | 234058264 | 234066913 | 8650 | -6.11 | 3.10E-07 | 5.39E-07 |
| chr6 | 167557983 | 167559197 | 1215 | -6.11 | 3.10E-07 | 5.39E-07 |
| chr12 | 683530 | 684387 | 858 | -5.87 | 3.11E-07 | 5.40E-07 |
| chr8 | 128452625 | 128456549 | 3925 | -6.14 | 3.11E-07 | 5.41E-07 |
| chr11 | 130071477 | 130072658 | 1182 | -5.87 | 3.12E-07 | 5.42E-07 |
| chr16 | 78163663 | 78164961 | 1299 | -5.87 | 3.12E-07 | 5.42E-07 |
| chr5 | 118335422 | 118336416 | 995 | -5.87 | 3.12E-07 | 5.42E-07 |
| chr3 | 18321378 | 18323251 | 1874 | -6.15 | 3.13E-07 | 5.43E-07 |
| chr8 | 102088437 | 102089567 | 1131 | -6.15 | 3.13E-07 | 5.43E-07 |
| chr2 | 231185466 | 231188025 | 2560 | -6.03 | 3.14E-07 | 5.44E-07 |
| chr3 | 111305269 | 111306865 | 1597 | -6.03 | 3.14E-07 | 5.44E-07 |
| chr11 | 114960763 | 114974850 | 14088 | 1.55 | 3.14E-07 | 5.45E-07 |
| chr7 | 203549 | 204809 | 1261 | -5.88 | 3.14E-07 | 5.45E-07 |
| chr19 | 2104937 | 2106778 | 1842 | -5.92 | 3.15E-07 | 5.47E-07 |
| chr5 | 53734042 | 53736725 | 2684 | 6.22 | 3.15E-07 | 5.47E-07 |
| chr8 | 55147037 | 55149622 | 2586 | 4.77 | 3.16E-07 | 5.48E-07 |
| chr12 | 105401265 | 105402949 | 1685 | -5.89 | 3.16E-07 | 5.49E-07 |
| chr5 | 40523071 | 40524016 | 946 | -5.87 | 3.17E-07 | 5.49E-07 |
| chrX | 46725136 | 46726919 | 1784 | -5.87 | 3.17E-07 | 5.50E-07 |
| chr4 | 118003264 | 118004706 | 1443 | -5.93 | 3.18E-07 | 5.51E-07 |
| chr9 | 36086495 | 36087419 | 925 | -5.93 | 3.18E-07 | 5.51E-07 |

| | | | | | | |
|---|---|---|---|---|---|---|
| chr13 | 52523852 | 52525355 | 1504 | -6 | 3.18E-07 | 5.51E-07 |
| chr13 | 41941677 | 41944760 | 3084 | -5.23 | 3.18E-07 | 5.52E-07 |
| chr22 | 38982814 | 38983550 | 737 | -5.86 | 3.18E-07 | 5.52E-07 |
| chr20 | 36211613 | 36242486 | 30874 | -6.1 | 3.19E-07 | 5.53E-07 |
| chr8 | 18996913 | 19000853 | 3941 | -5.9 | 3.19E-07 | 5.53E-07 |
| chr13 | 35794523 | 35795540 | 1018 | -4.36 | 3.19E-07 | 5.53E-07 |
| chr2 | 193388304 | 193389732 | 1429 | -5.87 | 3.19E-07 | 5.54E-07 |
| chr10 | 7142816 | 7194965 | 52150 | -1.79 | 3.20E-07 | 5.55E-07 |
| chr11 | 14937749 | 14956042 | 18294 | -1.36 | 3.20E-07 | 5.56E-07 |
| chr4 | 84927568 | 84928732 | 1165 | -5.87 | 3.21E-07 | 5.56E-07 |
| chr2 | 230152310 | 230155890 | 3581 | -2.42 | 3.21E-07 | 5.57E-07 |
| chr11 | 78786722 | 78787925 | 1204 | -5.98 | 3.22E-07 | 5.58E-07 |
| chr6 | 111932340 | 111934191 | 1852 | -5.88 | 3.23E-07 | 5.59E-07 |
| chr13 | 45386686 | 45388915 | 2230 | -5.87 | 3.23E-07 | 5.60E-07 |
| chr8 | 129896079 | 129899518 | 3440 | -5.42 | 3.24E-07 | 5.61E-07 |
| chr14 | 101112301 | 101113497 | 1197 | -6.25 | 3.24E-07 | 5.61E-07 |
| chr11 | 67442386 | 67466092 | 23707 | -7 | 3.24E-07 | 5.62E-07 |
| chr6 | 24943771 | 24954192 | 10422 | -1.91 | 3.24E-07 | 5.62E-07 |
| chr1 | 213232106 | 213232792 | 687 | -5.87 | 3.25E-07 | 5.63E-07 |
| chr13 | 35719669 | 35721040 | 1372 | -5.87 | 3.25E-07 | 5.63E-07 |
| chr2 | 152634144 | 152634968 | 825 | -5.87 | 3.25E-07 | 5.63E-07 |
| chr9 | 137169819 | 137170680 | 862 | -5.87 | 3.25E-07 | 5.63E-07 |
| chr3 | 64731009 | 64732575 | 1567 | 6.09 | 3.26E-07 | 5.65E-07 |
| chr1 | 80148622 | 80150060 | 1439 | -5.89 | 3.26E-07 | 5.65E-07 |
| chr4 | 115414182 | 115415110 | 929 | -5.89 | 3.26E-07 | 5.65E-07 |
| chr11 | 110224406 | 110225510 | 1105 | -5.92 | 3.28E-07 | 5.69E-07 |
| chr7 | 84717896 | 84719681 | 1786 | 5.96 | 3.29E-07 | 5.70E-07 |
| chr6 | 165693078 | 165705610 | 12533 | -1.79 | 3.30E-07 | 5.72E-07 |
| chr2 | 203787763 | 203788910 | 1148 | -5.96 | 3.30E-07 | 5.72E-07 |
| chr4 | 189671529 | 189673671 | 2143 | -5.96 | 3.30E-07 | 5.72E-07 |
| chr18 | 28757126 | 28758140 | 1015 | -5.88 | 3.30E-07 | 5.72E-07 |
| chr6 | 72026255 | 72027059 | 805 | -5.86 | 3.31E-07 | 5.74E-07 |
| chr1 | 38987351 | 38991329 | 3979 | -2.59 | 3.31E-07 | 5.74E-07 |
| chr1 | 152166658 | 152168021 | 1364 | -5.86 | 3.31E-07 | 5.74E-07 |
| chr11 | 111408948 | 111410029 | 1082 | -5.87 | 3.32E-07 | 5.76E-07 |
| chr11 | 118678458 | 118679357 | 900 | -5.86 | 3.33E-07 | 5.76E-07 |
| chr12 | 28666105 | 28667038 | 934 | -5.86 | 3.33E-07 | 5.76E-07 |
| chr2 | 19599649 | 19600273 | 625 | -5.86 | 3.33E-07 | 5.76E-07 |
| chr6 | 152795037 | 152796403 | 1367 | -5.86 | 3.33E-07 | 5.76E-07 |
| chr4 | 89717759 | 89718797 | 1039 | -5.95 | 3.34E-07 | 5.78E-07 |
| chrX | 134741515 | 134745797 | 4283 | 2.09 | 3.34E-07 | 5.78E-07 |
| chr4 | 15821985 | 15823133 | 1149 | -5.91 | 3.34E-07 | 5.79E-07 |
| chr9 | 104041790 | 104045527 | 3738 | -2.83 | 3.34E-07 | 5.79E-07 |
| chr5 | 129222494 | 129224287 | 1794 | 6.16 | 3.35E-07 | 5.79E-07 |
| chr10 | 81237957 | 81241704 | 3748 | -5.4 | 3.35E-07 | 5.80E-07 |
| chr3 | 43088078 | 43093891 | 5814 | -1.96 | 3.36E-07 | 5.81E-07 |
| chr13 | 84018416 | 84020494 | 2079 | -5.89 | 3.36E-07 | 5.81E-07 |
| chr20 | 30980742 | 30981710 | 969 | -5.89 | 3.36E-07 | 5.81E-07 |
| chr11 | 9655002 | 9655843 | 842 | -5.93 | 3.36E-07 | 5.82E-07 |
| chr1 | 211015715 | 211017572 | 1858 | 5.99 | 3.36E-07 | 5.82E-07 |
| chr11 | 18147088 | 18153103 | 6016 | -1.86 | 3.37E-07 | 5.82E-07 |
| chr8 | 70963468 | 70965851 | 2384 | -4.68 | 3.37E-07 | 5.83E-07 |
| chr10 | 69659171 | 69660328 | 1158 | -6.07 | 3.37E-07 | 5.84E-07 |
| chr7 | 16325261 | 16326307 | 1047 | -6.07 | 3.37E-07 | 5.84E-07 |
| chr8 | 120837136 | 120839768 | 2633 | -5.94 | 3.38E-07 | 5.84E-07 |
| chr6 | 125265416 | 125266819 | 1404 | -6.08 | 3.38E-07 | 5.84E-07 |
| chr6 | 147439760 | 147441729 | 1970 | -6.08 | 3.38E-07 | 5.84E-07 |
| chr5 | 124902047 | 124936126 | 34080 | -7.41 | 3.38E-07 | 5.84E-07 |
| chr17 | 9166984 | 9167991 | 1008 | -6.06 | 3.38E-07 | 5.84E-07 |
| chr10 | 123604765 | 123605634 | 870 | -5.87 | 3.38E-07 | 5.84E-07 |
| chr10 | 60998588 | 61004302 | 5715 | -6.1 | 3.38E-07 | 5.84E-07 |

| | | | | | | |
|---|---|---|---|---|---|---|
| chr15 | 90498361 | 90499788 | 1428 | -6.34 | 3.38E-07 | 5.84E-07 |
| chr5 | 127191054 | 127195814 | 4761 | -6.34 | 3.38E-07 | 5.84E-07 |
| chr21 | 27266209 | 27266944 | 736 | -6.12 | 3.38E-07 | 5.85E-07 |
| chr4 | 136956597 | 136958708 | 2112 | -6.03 | 3.40E-07 | 5.87E-07 |
| chr5 | 93095503 | 93097004 | 1502 | -6.03 | 3.40E-07 | 5.87E-07 |
| chr14 | 32069341 | 32070861 | 1521 | -6.03 | 3.40E-07 | 5.87E-07 |
| chr15 | 76546718 | 76547246 | 529 | -6.03 | 3.40E-07 | 5.87E-07 |
| chr2 | 241588950 | 241606587 | 17638 | -3.65 | 3.40E-07 | 5.88E-07 |
| chr10 | 12742036 | 12743357 | 1322 | -6.02 | 3.40E-07 | 5.89E-07 |
| chr13 | 41156843 | 41158115 | 1273 | -6.02 | 3.41E-07 | 5.89E-07 |
| chr18 | 54390689 | 54391751 | 1063 | -5.9 | 3.41E-07 | 5.89E-07 |
| chr11 | 87926040 | 87931420 | 5381 | -5.24 | 3.41E-07 | 5.90E-07 |
| chr4 | 37781862 | 37795103 | 13242 | -1.33 | 3.41E-07 | 5.90E-07 |
| chr22 | 39672671 | 39681554 | 8884 | -6.16 | 3.41E-07 | 5.90E-07 |
| chr5 | 142090565 | 142091573 | 1009 | -5.86 | 3.42E-07 | 5.91E-07 |
| chrX | 43246188 | 43315683 | 69496 | 1.36 | 3.43E-07 | 5.92E-07 |
| chr8 | 90758387 | 90760319 | 1933 | -6.17 | 3.43E-07 | 5.92E-07 |
| chr6 | 3147534 | 3148495 | 962 | -5.85 | 3.43E-07 | 5.93E-07 |
| chr6 | 162534286 | 162535621 | 1336 | -5.85 | 3.43E-07 | 5.93E-07 |
| chr15 | 64837746 | 64838244 | 499 | -5.86 | 3.43E-07 | 5.93E-07 |
| chr1 | 86169571 | 86170377 | 807 | -5.88 | 3.43E-07 | 5.93E-07 |
| chr17 | 17169045 | 17169706 | 662 | -6.18 | 3.43E-07 | 5.94E-07 |
| chrX | 114609784 | 114611306 | 1523 | -6 | 3.43E-07 | 5.94E-07 |
| chr11 | 87839681 | 87842645 | 2965 | -3.89 | 3.44E-07 | 5.94E-07 |
| chr3 | 108087913 | 108088478 | 566 | -4.44 | 3.44E-07 | 5.94E-07 |
| chr2 | 40663411 | 40664664 | 1254 | -5.99 | 3.45E-07 | 5.95E-07 |
| chr22 | 33474629 | 33500838 | 26210 | -1.89 | 3.45E-07 | 5.97E-07 |
| chr6 | 101489113 | 101491414 | 2302 | 5.96 | 3.45E-07 | 5.97E-07 |
| chr5 | 92702860 | 92713056 | 10197 | -5.48 | 3.46E-07 | 5.97E-07 |
| chr10 | 124240641 | 124241848 | 1208 | -5.99 | 3.46E-07 | 5.98E-07 |
| chr2 | 53353584 | 53355409 | 1826 | -5.99 | 3.46E-07 | 5.98E-07 |
| chr11 | 95817724 | 95819358 | 1635 | -5.98 | 3.46E-07 | 5.98E-07 |
| chr4 | 54601624 | 54603142 | 1519 | -5.98 | 3.46E-07 | 5.98E-07 |
| chrX | 109159901 | 109161375 | 1475 | -5.92 | 3.47E-07 | 5.99E-07 |
| chr19 | 50286475 | 50287518 | 1044 | -5.85 | 3.47E-07 | 5.99E-07 |
| chr2 | 174712543 | 174713525 | 983 | -5.85 | 3.47E-07 | 5.99E-07 |
| chr4 | 34354206 | 34355440 | 1235 | 4.64 | 3.48E-07 | 6.00E-07 |
| chr13 | 61412219 | 61473757 | 61539 | 1.49 | 3.48E-07 | 6.00E-07 |
| chr1 | 152590053 | 152621608 | 31556 | 2.11 | 3.48E-07 | 6.00E-07 |
| chrX | 74871367 | 74927015 | 55649 | 1.41 | 3.48E-07 | 6.01E-07 |
| chr11 | 103901975 | 103903111 | 1137 | -5.89 | 3.48E-07 | 6.01E-07 |
| chr13 | 113558891 | 113560139 | 1249 | -5.89 | 3.48E-07 | 6.01E-07 |
| chr20 | 35047619 | 35048683 | 1065 | -5.89 | 3.48E-07 | 6.01E-07 |
| chr17 | 57077375 | 57078048 | 674 | -5.98 | 3.48E-07 | 6.02E-07 |
| chr7 | 142115092 | 142115816 | 725 | -5.98 | 3.48E-07 | 6.02E-07 |
| chr8 | 128641802 | 128642595 | 794 | -5.98 | 3.48E-07 | 6.02E-07 |
| chr4 | 108210295 | 108218602 | 8308 | -2.06 | 3.48E-07 | 6.02E-07 |
| chr6 | 88394473 | 88396030 | 1558 | -5.91 | 3.49E-07 | 6.02E-07 |
| chr2 | 196870631 | 196883906 | 13276 | -6.6 | 3.49E-07 | 6.02E-07 |
| chr17 | 32901970 | 32902715 | 746 | -5.85 | 3.50E-07 | 6.03E-07 |
| chr2 | 120297343 | 120298380 | 1038 | -5.85 | 3.50E-07 | 6.03E-07 |
| chr16 | 80102971 | 80104327 | 1357 | -5.85 | 3.50E-07 | 6.04E-07 |
| chr13 | 76670276 | 76671288 | 1013 | 3.91 | 3.51E-07 | 6.06E-07 |
| chr2 | 8432860 | 8433861 | 1002 | -5.11 | 3.51E-07 | 6.06E-07 |
| chr15 | 83969781 | 83970687 | 907 | -5.96 | 3.52E-07 | 6.07E-07 |
| chrX | 150532725 | 150539337 | 6613 | -2.18 | 3.52E-07 | 6.07E-07 |
| chr4 | 91242900 | 91244065 | 1166 | -5.91 | 3.52E-07 | 6.07E-07 |
| chr9 | 76122037 | 76122981 | 945 | -6.25 | 3.52E-07 | 6.08E-07 |
| chr20 | 33256803 | 33257629 | 827 | -5.87 | 3.53E-07 | 6.08E-07 |
| chr4 | 176869626 | 176871292 | 1667 | -5.87 | 3.53E-07 | 6.08E-07 |
| chr6 | 41693401 | 41694314 | 914 | -5.87 | 3.53E-07 | 6.08E-07 |

| | | | | | | |
|---|---|---|---|---|---|---|
| chr11 | 27788678 | 27789990 | 1313 | -4.37 | 3.53E-07 | 6.08E-07 |
| chr17 | 61308486 | 61310041 | 1556 | -5.85 | 3.53E-07 | 6.08E-07 |
| chr6 | 48955574 | 48957428 | 1855 | -5.85 | 3.53E-07 | 6.08E-07 |
| chr6 | 141197001 | 141208157 | 11157 | -6.25 | 3.53E-07 | 6.10E-07 |
| chr1 | 55978374 | 55979979 | 1606 | -5.96 | 3.54E-07 | 6.10E-07 |
| chr14 | 55630382 | 55631156 | 775 | -5.96 | 3.54E-07 | 6.10E-07 |
| chr10 | 28770499 | 28773706 | 3208 | -2.27 | 3.54E-07 | 6.11E-07 |
| chr2 | 213420743 | 213422393 | 1651 | 5.16 | 3.54E-07 | 6.11E-07 |
| chr2 | 224656505 | 224666197 | 9693 | -5.86 | 3.56E-07 | 6.13E-07 |
| chr1 | 115035510 | 115037509 | 2000 | -5.84 | 3.56E-07 | 6.13E-07 |
| chr17 | 30926700 | 30927536 | 837 | -5.84 | 3.56E-07 | 6.13E-07 |
| chr1 | 98227067 | 98227970 | 904 | -5.88 | 3.56E-07 | 6.14E-07 |
| chr2 | 215785417 | 215786848 | 1432 | -5.88 | 3.56E-07 | 6.14E-07 |
| chr2 | 161891853 | 161892521 | 669 | -5.87 | 3.56E-07 | 6.14E-07 |
| chr11 | 27678585 | 27679786 | 1202 | -5.95 | 3.57E-07 | 6.15E-07 |
| chr1 | 12179938 | 12181042 | 1105 | -5.9 | 3.58E-07 | 6.17E-07 |
| chr7 | 155268230 | 155270229 | 2000 | -5.94 | 3.59E-07 | 6.19E-07 |
| chr21 | 30344623 | 30345555 | 933 | -5.94 | 3.60E-07 | 6.20E-07 |
| chr7 | 140862512 | 140863265 | 754 | -5.06 | 3.60E-07 | 6.21E-07 |
| chr18 | 47783949 | 47784930 | 982 | -5.44 | 3.63E-07 | 6.25E-07 |
| chr11 | 11100557 | 11156708 | 56152 | -1.34 | 3.63E-07 | 6.25E-07 |
| chr13 | 83207077 | 83208596 | 1520 | -5.84 | 3.63E-07 | 6.25E-07 |
| chr3 | 130152242 | 130153281 | 1040 | -5.84 | 3.63E-07 | 6.25E-07 |
| chr6 | 166150698 | 166152524 | 1827 | -5.86 | 3.63E-07 | 6.26E-07 |
| chr1 | 92901164 | 92902903 | 1740 | -4.35 | 3.64E-07 | 6.26E-07 |
| chr13 | 49667191 | 49668563 | 1373 | -5.93 | 3.64E-07 | 6.27E-07 |
| chr12 | 109999695 | 110000915 | 1221 | -5.84 | 3.64E-07 | 6.27E-07 |
| chr2 | 198592387 | 198593802 | 1416 | -5.84 | 3.64E-07 | 6.28E-07 |
| chr6 | 139058009 | 139060471 | 2463 | -5.84 | 3.64E-07 | 6.28E-07 |
| chr4 | 121592802 | 121594323 | 1522 | -5.84 | 3.65E-07 | 6.28E-07 |
| chr8 | 28978564 | 28979647 | 1084 | -5.89 | 3.65E-07 | 6.28E-07 |
| chr8 | 118673195 | 118675444 | 2250 | -5.89 | 3.65E-07 | 6.28E-07 |
| chr1 | 197154128 | 197156697 | 2570 | -5.43 | 3.65E-07 | 6.29E-07 |
| chr7 | 78948460 | 78949459 | 1000 | -5.85 | 3.66E-07 | 6.30E-07 |
| chr6 | 167842922 | 167851200 | 8279 | -3.02 | 3.67E-07 | 6.32E-07 |
| chr12 | 79445294 | 79453635 | 8342 | -6.07 | 3.68E-07 | 6.33E-07 |
| chr17 | 76096570 | 76098126 | 1557 | -6.07 | 3.68E-07 | 6.33E-07 |
| chr7 | 94285684 | 94287206 | 1523 | -6.07 | 3.68E-07 | 6.33E-07 |
| chr12 | 109500813 | 109502012 | 1200 | -6.06 | 3.68E-07 | 6.33E-07 |
| chr1 | 48921584 | 48923417 | 1834 | -5.92 | 3.68E-07 | 6.34E-07 |
| chr12 | 9796420 | 9798239 | 1820 | -5.92 | 3.68E-07 | 6.34E-07 |
| chr5 | 170252091 | 170253800 | 1710 | -6.04 | 3.68E-07 | 6.34E-07 |
| chr6 | 1828148 | 1829767 | 1620 | -5.84 | 3.68E-07 | 6.34E-07 |
| chr2 | 73065680 | 73066356 | 677 | -6.03 | 3.69E-07 | 6.35E-07 |
| chr13 | 84327580 | 84328924 | 1345 | -5.92 | 3.69E-07 | 6.35E-07 |
| chr19 | 11452023 | 11460695 | 8673 | -6.11 | 3.69E-07 | 6.35E-07 |
| chr17 | 77348291 | 77349379 | 1089 | -6.02 | 3.69E-07 | 6.36E-07 |
| chr11 | 115846722 | 116051731 | 205010 | 0.6 | 3.69E-07 | 6.36E-07 |
| chr3 | 149977111 | 149978099 | 989 | -6.01 | 3.70E-07 | 6.37E-07 |
| chr3 | 69333954 | 69335870 | 1917 | -6.12 | 3.70E-07 | 6.37E-07 |
| chr1 | 14968936 | 14973079 | 4144 | -4 | 3.71E-07 | 6.39E-07 |
| chr22 | 35954628 | 35955475 | 848 | -6 | 3.71E-07 | 6.39E-07 |
| chr2 | 174499431 | 174500889 | 1459 | -5.87 | 3.71E-07 | 6.39E-07 |
| chr1 | 113435540 | 113437105 | 1566 | -6.14 | 3.72E-07 | 6.40E-07 |
| chr2 | 84694214 | 84694904 | 691 | -5.88 | 3.72E-07 | 6.40E-07 |
| chr4 | 115560628 | 115562124 | 1497 | -4.64 | 3.73E-07 | 6.41E-07 |
| chr5 | 65097809 | 65098899 | 1091 | -5.86 | 3.75E-07 | 6.45E-07 |
| chrX | 77064057 | 77065733 | 1677 | -5.86 | 3.75E-07 | 6.45E-07 |
| chr3 | 132253903 | 132254950 | 1048 | -6.05 | 3.76E-07 | 6.46E-07 |
| chr3 | 23134834 | 23152718 | 17885 | -7.3 | 3.76E-07 | 6.47E-07 |
| chr13 | 83943145 | 83947963 | 4819 | -6.03 | 3.76E-07 | 6.47E-07 |

| | | | | | | |
|---|---|---|---|---|---|---|
| chr14 | 52411061 | 52431928 | 20868 | -3.46 | 3.76E-07 | 6.48E-07 |
| chr1 | 117584144 | 117585074 | 931 | -5.84 | 3.76E-07 | 6.48E-07 |
| chr2 | 231542445 | 231543576 | 1132 | -5.84 | 3.76E-07 | 6.48E-07 |
| chr7 | 12077644 | 12079223 | 1580 | -5.83 | 3.77E-07 | 6.49E-07 |
| chr12 | 76364790 | 76366748 | 1959 | -6.02 | 3.77E-07 | 6.49E-07 |
| chr8 | 74974439 | 75028808 | 54370 | -2.72 | 3.77E-07 | 6.49E-07 |
| chr11 | 33658680 | 33660686 | 2007 | -6.4 | 3.78E-07 | 6.50E-07 |
| chr12 | 110940152 | 110941110 | 959 | -6.01 | 3.78E-07 | 6.50E-07 |
| chr5 | 177539189 | 177539892 | 704 | -5.87 | 3.78E-07 | 6.50E-07 |
| chr6 | 111286093 | 111287190 | 1098 | -5.87 | 3.78E-07 | 6.50E-07 |
| chr6 | 122098522 | 122100157 | 1636 | -5.87 | 3.78E-07 | 6.50E-07 |
| chr6 | 3199061 | 3199852 | 792 | -5.9 | 3.78E-07 | 6.51E-07 |
| chr11 | 133948980 | 133951511 | 2532 | -2.16 | 3.79E-07 | 6.51E-07 |
| chr1 | 231079041 | 231079587 | 547 | -6.13 | 3.79E-07 | 6.51E-07 |
| chr6 | 169606953 | 169608872 | 1920 | -6.13 | 3.79E-07 | 6.51E-07 |
| chr1 | 86842926 | 86844301 | 1376 | -6.2 | 3.80E-07 | 6.53E-07 |
| chr10 | 99935457 | 99936445 | 989 | -5.87 | 3.80E-07 | 6.54E-07 |
| chrX | 9456477 | 9457790 | 1314 | -5.83 | 3.80E-07 | 6.54E-07 |
| chr11 | 62747551 | 62748577 | 1027 | 5.99 | 3.82E-07 | 6.56E-07 |
| chr1 | 197371232 | 197372104 | 873 | -5.85 | 3.82E-07 | 6.56E-07 |
| chr6 | 158785154 | 158786363 | 1210 | -5.85 | 3.82E-07 | 6.56E-07 |
| chr3 | 5089212 | 5090185 | 974 | -5.98 | 3.82E-07 | 6.56E-07 |
| chr2 | 163686682 | 163688151 | 1470 | 5.96 | 3.82E-07 | 6.57E-07 |
| chr1 | 98165956 | 98167476 | 1521 | -5.89 | 3.84E-07 | 6.60E-07 |
| chr1 | 59735526 | 59737070 | 1545 | -4.83 | 3.85E-07 | 6.61E-07 |
| chr7 | 150559229 | 150592367 | 33139 | -6.45 | 3.85E-07 | 6.62E-07 |
| chr14 | 69651948 | 69653099 | 1152 | -5.85 | 3.86E-07 | 6.62E-07 |
| chr1 | 150642791 | 150643766 | 976 | -5.86 | 3.86E-07 | 6.63E-07 |
| chr2 | 28819842 | 28820908 | 1067 | -5.89 | 3.86E-07 | 6.64E-07 |
| chr5 | 121329307 | 121330737 | 1431 | -6.21 | 3.88E-07 | 6.67E-07 |
| chrX | 151382615 | 151384181 | 1567 | -6.21 | 3.88E-07 | 6.67E-07 |
| chr4 | 130991890 | 130993178 | 1289 | -5.83 | 3.88E-07 | 6.67E-07 |
| chrX | 54641143 | 54641956 | 814 | -5.83 | 3.88E-07 | 6.67E-07 |
| chr22 | 48454334 | 48455162 | 829 | -5.84 | 3.89E-07 | 6.68E-07 |
| chr3 | 171781783 | 171782519 | 737 | -5.84 | 3.89E-07 | 6.68E-07 |
| chrX | 152996899 | 152997892 | 994 | -5.84 | 3.89E-07 | 6.68E-07 |
| chr17 | 28365535 | 28417769 | 52235 | -1.06 | 3.89E-07 | 6.68E-07 |
| chr8 | 11493198 | 11494441 | 1244 | -5.86 | 3.89E-07 | 6.69E-07 |
| chr11 | 29453075 | 29454047 | 973 | -5.83 | 3.91E-07 | 6.71E-07 |
| chr3 | 119055364 | 119056338 | 975 | -5.83 | 3.91E-07 | 6.71E-07 |
| chr7 | 140563343 | 140564873 | 1531 | -5.83 | 3.91E-07 | 6.71E-07 |
| chr10 | 15157930 | 15158944 | 1015 | -5.88 | 3.91E-07 | 6.71E-07 |
| chrX | 18181749 | 18183497 | 1749 | 5.97 | 3.91E-07 | 6.72E-07 |
| chr20 | 52507604 | 52508910 | 1307 | -5.88 | 3.92E-07 | 6.73E-07 |
| chr4 | 167631574 | 167634181 | 2608 | -5.88 | 3.92E-07 | 6.73E-07 |
| chr1 | 32977011 | 32977981 | 971 | -5.84 | 3.92E-07 | 6.73E-07 |
| chr8 | 125867502 | 125868873 | 1372 | -5.84 | 3.92E-07 | 6.73E-07 |
| chr12 | 2113958 | 2114984 | 1027 | -6.28 | 3.92E-07 | 6.73E-07 |
| chr7 | 16016525 | 16018848 | 2324 | -2.37 | 3.93E-07 | 6.74E-07 |
| chr5 | 105508242 | 105509542 | 1301 | -5.92 | 3.93E-07 | 6.74E-07 |
| chr14 | 52097707 | 52099536 | 1830 | -5.94 | 3.93E-07 | 6.75E-07 |
| chr4 | 128539455 | 128540108 | 654 | -5.93 | 3.96E-07 | 6.80E-07 |
| chr3 | 161307344 | 161439829 | 132486 | 3.44 | 3.97E-07 | 6.81E-07 |
| chr7 | 138091588 | 138092485 | 898 | -5.91 | 3.97E-07 | 6.81E-07 |
| chr15 | 37009359 | 37010551 | 1193 | -5.84 | 3.98E-07 | 6.82E-07 |
| chr1 | 188096778 | 188098224 | 1447 | -5.87 | 3.98E-07 | 6.82E-07 |
| chr16 | 47464896 | 47465740 | 845 | -5.82 | 3.98E-07 | 6.83E-07 |
| chr19 | 15963607 | 15964583 | 977 | -5.82 | 3.98E-07 | 6.83E-07 |
| chr8 | 122609132 | 122611265 | 2134 | -5.82 | 3.98E-07 | 6.83E-07 |
| chr20 | 48786132 | 48787026 | 895 | -5.82 | 3.99E-07 | 6.85E-07 |
| chr3 | 38057374 | 38058087 | 714 | -5.85 | 3.99E-07 | 6.85E-07 |

| | | | | | | |
|---|---|---|---|---|---|---|
| chr3 | 150939216 | 150941445 | 2230 | -5.85 | 3.99E-07 | 6.85E-07 |
| chr13 | 90257682 | 90259834 | 2153 | -6.53 | 4.00E-07 | 6.87E-07 |
| chr1 | 101215611 | 101217093 | 1483 | -5.82 | 4.00E-07 | 6.87E-07 |
| chr5 | 132501545 | 132507710 | 6166 | -4.97 | 4.01E-07 | 6.87E-07 |
| chr1 | 229028330 | 229029683 | 1354 | -6.07 | 4.01E-07 | 6.88E-07 |
| chr12 | 72329785 | 72330727 | 943 | -6.07 | 4.01E-07 | 6.88E-07 |
| chr3 | 33939470 | 33946579 | 7110 | -6.08 | 4.01E-07 | 6.88E-07 |
| chr2 | 73150154 | 73151065 | 912 | -6.02 | 4.01E-07 | 6.88E-07 |
| chr7 | 82129872 | 82131695 | 1824 | -6.08 | 4.02E-07 | 6.89E-07 |
| chr8 | 17196323 | 17211049 | 14727 | -6.91 | 4.02E-07 | 6.89E-07 |
| chr15 | 57285825 | 57286697 | 873 | -5.92 | 4.02E-07 | 6.89E-07 |
| chr2 | 232924339 | 232925389 | 1051 | -5.92 | 4.02E-07 | 6.89E-07 |
| chr11 | 123092363 | 123094085 | 1723 | -6.34 | 4.02E-07 | 6.90E-07 |
| chr7 | 30675910 | 30676689 | 780 | -5.84 | 4.02E-07 | 6.90E-07 |
| chr4 | 124684227 | 124685108 | 882 | -6.01 | 4.03E-07 | 6.90E-07 |
| chr8 | 63494969 | 63496155 | 1187 | 5.96 | 4.03E-07 | 6.91E-07 |
| chrX | 136086505 | 136087678 | 1174 | -5.83 | 4.03E-07 | 6.91E-07 |
| chr12 | 27598566 | 27599855 | 1290 | -5.83 | 4.03E-07 | 6.92E-07 |
| chr6 | 134674531 | 134676573 | 2043 | -6 | 4.04E-07 | 6.92E-07 |
| chr4 | 79514811 | 79515803 | 993 | -6.3 | 4.04E-07 | 6.92E-07 |
| chr22 | 42852100 | 42853583 | 1484 | -5.99 | 4.05E-07 | 6.94E-07 |
| chr3 | 113549782 | 113551253 | 1472 | -5.99 | 4.05E-07 | 6.94E-07 |
| chr5 | 60408581 | 60409759 | 1179 | -5.99 | 4.05E-07 | 6.94E-07 |
| chr13 | 85255893 | 85259107 | 3215 | -5.91 | 4.05E-07 | 6.95E-07 |
| chr13 | 34513374 | 34514988 | 1615 | -5.86 | 4.07E-07 | 6.97E-07 |
| chr15 | 79100914 | 79102153 | 1240 | -5.86 | 4.07E-07 | 6.97E-07 |
| chr3 | 72629391 | 72630861 | 1471 | -5.86 | 4.07E-07 | 6.97E-07 |
| chr3 | 114915534 | 114916062 | 529 | -5.86 | 4.07E-07 | 6.97E-07 |
| chr14 | 34858671 | 34860782 | 2112 | -6.14 | 4.07E-07 | 6.97E-07 |
| chr8 | 71470187 | 71477307 | 7121 | -6.14 | 4.07E-07 | 6.97E-07 |
| chr4 | 74162492 | 74165204 | 2713 | -4.85 | 4.07E-07 | 6.97E-07 |
| chr2 | 182512643 | 182513404 | 762 | -5.89 | 4.07E-07 | 6.98E-07 |
| chr4 | 166041995 | 166042852 | 858 | -5.97 | 4.08E-07 | 6.99E-07 |
| chr10 | 64315244 | 64316539 | 1296 | 4.6 | 4.08E-07 | 6.99E-07 |
| chr1 | 178559919 | 178561311 | 1393 | -6.16 | 4.09E-07 | 7.00E-07 |
| chr6 | 25784004 | 25785131 | 1128 | -5.9 | 4.09E-07 | 7.01E-07 |
| chr21 | 35201821 | 35203069 | 1249 | -6.17 | 4.10E-07 | 7.02E-07 |
| chr4 | 63621652 | 63624603 | 2952 | -5.84 | 4.10E-07 | 7.03E-07 |
| chr7 | 95276294 | 95277157 | 864 | -5.84 | 4.10E-07 | 7.03E-07 |
| chrX | 101006443 | 101007790 | 1348 | -4.42 | 4.10E-07 | 7.03E-07 |
| chr1 | 117988646 | 117989569 | 924 | -5.86 | 4.11E-07 | 7.04E-07 |
| chr4 | 166230086 | 166231063 | 978 | -5.83 | 4.11E-07 | 7.04E-07 |
| chr6 | 139649664 | 139650744 | 1081 | -5.83 | 4.11E-07 | 7.04E-07 |
| chrX | 34696692 | 34720322 | 23631 | -3.4 | 4.12E-07 | 7.05E-07 |
| chr7 | 111416834 | 111418054 | 1221 | -5.84 | 4.12E-07 | 7.05E-07 |
| chr9 | 116312750 | 116313687 | 938 | -5.84 | 4.12E-07 | 7.05E-07 |
| chr8 | 96333089 | 96348312 | 15224 | 1.86 | 4.12E-07 | 7.05E-07 |
| chr13 | 99221141 | 99222248 | 1108 | -5.88 | 4.13E-07 | 7.07E-07 |
| chr3 | 148308994 | 148326719 | 17726 | -7.22 | 4.13E-07 | 7.08E-07 |
| chr13 | 33669476 | 33673770 | 4295 | -5.23 | 4.14E-07 | 7.08E-07 |
| chr4 | 146877915 | 146878863 | 949 | -5.82 | 4.15E-07 | 7.10E-07 |
| chr5 | 83576147 | 83577663 | 1517 | -5.82 | 4.15E-07 | 7.10E-07 |
| chr6 | 6870417 | 6871327 | 911 | -5.82 | 4.15E-07 | 7.10E-07 |
| chr6 | 165839977 | 165840888 | 912 | -5.82 | 4.15E-07 | 7.10E-07 |
| chr8 | 117421321 | 117422850 | 1530 | -5.82 | 4.15E-07 | 7.10E-07 |
| chr2 | 85756414 | 85769703 | 13290 | -5.81 | 4.17E-07 | 7.14E-07 |
| chr4 | 7467182 | 7468214 | 1033 | -5.81 | 4.17E-07 | 7.14E-07 |
| chr10 | 17379547 | 17412318 | 32772 | -2.01 | 4.18E-07 | 7.14E-07 |
| chr13 | 20431327 | 20432599 | 1273 | -5.93 | 4.18E-07 | 7.14E-07 |
| chr1 | 99602524 | 99603621 | 1098 | -5.81 | 4.18E-07 | 7.15E-07 |
| chr16 | 49519854 | 49521258 | 1405 | -6.07 | 4.19E-07 | 7.17E-07 |

| | | | | | | |
|---|---|---|---|---|---|---|
| chrX | 71180112 | 71184562 | 4451 | -5.11 | 4.19E-07 | 7.17E-07 |
| chr11 | 66921844 | 66922743 | 900 | -6.09 | 4.20E-07 | 7.18E-07 |
| chr7 | 66227642 | 66228956 | 1315 | -5.82 | 4.20E-07 | 7.18E-07 |
| chr9 | 130470835 | 130471555 | 721 | -5.82 | 4.20E-07 | 7.18E-07 |
| chr1 | 41308043 | 41309118 | 1076 | -5.87 | 4.20E-07 | 7.18E-07 |
| chr3 | 71189422 | 71191942 | 2521 | -5.87 | 4.20E-07 | 7.18E-07 |
| chr6 | 162299026 | 162300515 | 1490 | -5.87 | 4.20E-07 | 7.18E-07 |
| chr20 | 16596690 | 16656127 | 59438 | -3.73 | 4.20E-07 | 7.18E-07 |
| chr2 | 56943586 | 57017740 | 74155 | -2.57 | 4.21E-07 | 7.20E-07 |
| chr15 | 42389857 | 42391259 | 1403 | -5.83 | 4.22E-07 | 7.22E-07 |
| chr13 | 27865263 | 27866540 | 1278 | -5.81 | 4.22E-07 | 7.22E-07 |
| chr16 | 57424880 | 57426869 | 1990 | -6.24 | 4.22E-07 | 7.22E-07 |
| chr1 | 94599064 | 94599959 | 896 | -5.88 | 4.23E-07 | 7.23E-07 |
| chr3 | 196697719 | 196711457 | 13739 | -6.73 | 4.25E-07 | 7.27E-07 |
| chr11 | 128543241 | 128546459 | 3219 | -3.6 | 4.26E-07 | 7.28E-07 |
| chr5 | 74379225 | 74380531 | 1307 | -5.84 | 4.26E-07 | 7.28E-07 |
| chr1 | 111330594 | 111336757 | 6164 | -1.84 | 4.26E-07 | 7.29E-07 |
| chr4 | 168417565 | 168419605 | 2041 | -5.84 | 4.27E-07 | 7.29E-07 |
| chr17 | 42568737 | 42570035 | 1299 | -5.96 | 4.27E-07 | 7.29E-07 |
| chr6 | 14799650 | 14800544 | 895 | -5.96 | 4.27E-07 | 7.29E-07 |
| chr3 | 157235922 | 157242229 | 6308 | -6.48 | 4.28E-07 | 7.31E-07 |
| chr4 | 39247205 | 39248029 | 825 | -5.81 | 4.28E-07 | 7.31E-07 |
| chr5 | 55861665 | 55862543 | 879 | -5.81 | 4.28E-07 | 7.31E-07 |
| chr8 | 281077 | 337451 | 56375 | -2.36 | 4.28E-07 | 7.31E-07 |
| chr3 | 196589706 | 196591111 | 1406 | -6.27 | 4.28E-07 | 7.31E-07 |
| chr21 | 36709894 | 36711559 | 1666 | -5.87 | 4.28E-07 | 7.32E-07 |
| chrX | 103057964 | 103058757 | 794 | -5.87 | 4.28E-07 | 7.32E-07 |
| chr17 | 18022519 | 18023879 | 1361 | -5.81 | 4.30E-07 | 7.35E-07 |
| chr2 | 145813936 | 145828612 | 14677 | -4.97 | 4.31E-07 | 7.37E-07 |
| chr1 | 164821214 | 164822629 | 1416 | 5.92 | 4.32E-07 | 7.38E-07 |
| chr2 | 174068750 | 174069525 | 776 | -5.82 | 4.33E-07 | 7.39E-07 |
| chr3 | 46002353 | 46003264 | 912 | -5.82 | 4.33E-07 | 7.39E-07 |
| chr7 | 134340528 | 134341392 | 865 | -5.89 | 4.34E-07 | 7.41E-07 |
| chr2 | 23901347 | 23902319 | 973 | -5.86 | 4.34E-07 | 7.42E-07 |
| chr5 | 108621601 | 108632290 | 10690 | -1.94 | 4.35E-07 | 7.43E-07 |
| chr12 | 69148697 | 69149358 | 662 | -5.82 | 4.35E-07 | 7.44E-07 |
| chr18 | 60077548 | 60079323 | 1776 | -5.85 | 4.35E-07 | 7.44E-07 |
| chr3 | 143187349 | 143188816 | 1468 | 5.88 | 4.36E-07 | 7.44E-07 |
| chr18 | 68374426 | 68381773 | 7348 | 2.03 | 4.36E-07 | 7.45E-07 |
| chr3 | 118966273 | 118967711 | 1439 | -6.04 | 4.37E-07 | 7.46E-07 |
| chr18 | 3124408 | 3125502 | 1095 | -5.8 | 4.37E-07 | 7.47E-07 |
| chr9 | 138828043 | 138829529 | 1487 | -5.8 | 4.37E-07 | 7.47E-07 |
| chr1 | 226316737 | 226318764 | 2028 | -6.06 | 4.37E-07 | 7.47E-07 |
| chr5 | 124197847 | 124200057 | 2211 | -6.06 | 4.37E-07 | 7.47E-07 |
| chr3 | 171750555 | 171752992 | 2438 | -6.01 | 4.38E-07 | 7.48E-07 |
| chr10 | 75154517 | 75155297 | 781 | -5.03 | 4.38E-07 | 7.48E-07 |
| chr2 | 223746442 | 223747442 | 1001 | -6 | 4.39E-07 | 7.49E-07 |
| chr12 | 65588988 | 65590846 | 1859 | -6.54 | 4.39E-07 | 7.49E-07 |
| chr6 | 140380177 | 140381945 | 1769 | -6.09 | 4.39E-07 | 7.49E-07 |
| chr6 | 131591249 | 131592358 | 1110 | -5.99 | 4.40E-07 | 7.50E-07 |
| chr6 | 85199192 | 85217629 | 18438 | -6.09 | 4.40E-07 | 7.51E-07 |
| chr4 | 68592876 | 68593792 | 917 | -5.85 | 4.41E-07 | 7.52E-07 |
| chrX | 43909553 | 43910623 | 1071 | -5.8 | 4.42E-07 | 7.54E-07 |
| chr8 | 133220352 | 133315799 | 95448 | 2.76 | 4.42E-07 | 7.55E-07 |
| chr17 | 6992795 | 6993804 | 1010 | -5.81 | 4.42E-07 | 7.55E-07 |
| chr6 | 10608181 | 10609773 | 1593 | -6.13 | 4.43E-07 | 7.56E-07 |
| chr2 | 173664905 | 173669704 | 4800 | -5.22 | 4.45E-07 | 7.59E-07 |
| chr15 | 56139770 | 56141506 | 1737 | -6.37 | 4.46E-07 | 7.60E-07 |
| chr2 | 46784798 | 46785481 | 684 | -5.85 | 4.48E-07 | 7.64E-07 |
| chr22 | 41143831 | 41144912 | 1082 | -5.85 | 4.48E-07 | 7.64E-07 |
| chr1 | 41716943 | 41721280 | 4338 | -5.14 | 4.48E-07 | 7.65E-07 |

| | | | | | | |
|---|---|---|---|---|---|---|
| chr1 | 47620130 | 47631736 | 11607 | -4.77 | 4.49E-07 | 7.66E-07 |
| chr3 | 29079597 | 29083861 | 4265 | -2.03 | 4.49E-07 | 7.66E-07 |
| chr12 | 1608701 | 1610086 | 1386 | -5.82 | 4.49E-07 | 7.66E-07 |
| chr3 | 4748606 | 4749324 | 719 | -5.82 | 4.49E-07 | 7.66E-07 |
| chr7 | 103206531 | 103222346 | 15816 | 6.15 | 4.49E-07 | 7.67E-07 |
| chr19 | 56071840 | 56081786 | 9947 | -5.93 | 4.50E-07 | 7.67E-07 |
| chr4 | 87360317 | 87365745 | 5429 | -5.81 | 4.50E-07 | 7.68E-07 |
| chr1 | 166860821 | 166911246 | 50426 | -2.05 | 4.51E-07 | 7.68E-07 |
| chr5 | 131447898 | 131506681 | 58784 | -2.95 | 4.51E-07 | 7.69E-07 |
| chr12 | 38916467 | 38927892 | 11426 | 2.26 | 4.51E-07 | 7.70E-07 |
| chr8 | 26300301 | 26301302 | 1002 | -5.93 | 4.53E-07 | 7.72E-07 |
| chr5 | 105608045 | 105610100 | 2056 | -6.3 | 4.53E-07 | 7.72E-07 |
| chr11 | 130456393 | 130457788 | 1396 | -6.2 | 4.54E-07 | 7.75E-07 |
| chr12 | 104873562 | 104874925 | 1364 | -5.83 | 4.55E-07 | 7.75E-07 |
| chr17 | 60805090 | 60805913 | 824 | -5.83 | 4.55E-07 | 7.75E-07 |
| chr22 | 49610962 | 49622038 | 11077 | -1.54 | 4.55E-07 | 7.76E-07 |
| chr6 | 113288683 | 113290193 | 1511 | -5.84 | 4.55E-07 | 7.76E-07 |
| chr1 | 95639787 | 95641117 | 1331 | -5.82 | 4.55E-07 | 7.76E-07 |
| chr18 | 20621660 | 20623207 | 1548 | -6.21 | 4.56E-07 | 7.77E-07 |
| chr4 | 134758572 | 134763226 | 4655 | -1.92 | 4.56E-07 | 7.77E-07 |
| chr7 | 116250855 | 116262095 | 11241 | -6.22 | 4.57E-07 | 7.79E-07 |
| chr10 | 4558951 | 4560200 | 1250 | -5.8 | 4.57E-07 | 7.79E-07 |
| chr7 | 127135558 | 127136293 | 736 | -5.06 | 4.58E-07 | 7.80E-07 |
| chr7 | 45231357 | 45233672 | 2316 | -2.43 | 4.58E-07 | 7.80E-07 |
| chr3 | 191849103 | 191850420 | 1318 | -5.8 | 4.58E-07 | 7.80E-07 |
| chr15 | 37132222 | 37143977 | 11756 | -6.44 | 4.58E-07 | 7.81E-07 |
| chr12 | 83028989 | 83063242 | 34254 | -7.42 | 4.59E-07 | 7.82E-07 |
| chr1 | 64889976 | 64890749 | 774 | -5.88 | 4.59E-07 | 7.83E-07 |
| chr18 | 57254479 | 57255916 | 1438 | -3.44 | 4.61E-07 | 7.85E-07 |
| chr1 | 223152364 | 223154703 | 2340 | -5.81 | 4.62E-07 | 7.87E-07 |
| chr8 | 9360210 | 9369892 | 9683 | -3.55 | 4.63E-07 | 7.88E-07 |
| chr5 | 82480915 | 82482323 | 1409 | -5.87 | 4.64E-07 | 7.91E-07 |
| chr9 | 119398393 | 119399635 | 1243 | -5.87 | 4.64E-07 | 7.91E-07 |
| chr8 | 124622911 | 124643965 | 21055 | -6.97 | 4.64E-07 | 7.91E-07 |
| chr1 | 115489201 | 115490487 | 1287 | -4.53 | 4.66E-07 | 7.93E-07 |
| chr21 | 17912061 | 17912934 | 874 | -5.82 | 4.66E-07 | 7.94E-07 |
| chr4 | 168538343 | 168539450 | 1108 | -5.82 | 4.66E-07 | 7.94E-07 |
| chr11 | 122391067 | 122417312 | 26246 | -2.76 | 4.67E-07 | 7.95E-07 |
| chr2 | 119891139 | 119892371 | 1233 | -5.81 | 4.67E-07 | 7.95E-07 |
| chr1 | 37729035 | 37729896 | 862 | -6.03 | 4.67E-07 | 7.95E-07 |
| chr5 | 139758501 | 139759214 | 714 | -6.04 | 4.67E-07 | 7.95E-07 |
| chr1 | 52002620 | 52004001 | 1382 | -6.02 | 4.67E-07 | 7.95E-07 |
| chr9 | 71326035 | 71370573 | 44539 | 3.38 | 4.67E-07 | 7.95E-07 |
| chr11 | 47056073 | 47057018 | 946 | -5.85 | 4.67E-07 | 7.96E-07 |
| chr20 | 56529335 | 56530950 | 1616 | -6.06 | 4.68E-07 | 7.97E-07 |
| chr17 | 55892121 | 55893925 | 1805 | -4.81 | 4.68E-07 | 7.97E-07 |
| chr18 | 21725559 | 21726755 | 1197 | -5.89 | 4.69E-07 | 7.98E-07 |
| chr4 | 164422064 | 164423459 | 1396 | -6.08 | 4.69E-07 | 7.98E-07 |
| chr1 | 191619191 | 191621958 | 2768 | -6.08 | 4.69E-07 | 7.99E-07 |
| chr3 | 104867279 | 104869683 | 2405 | -4.46 | 4.70E-07 | 8.00E-07 |
| chr4 | 2581268 | 2582575 | 1308 | -5.86 | 4.70E-07 | 8.00E-07 |
| chr4 | 140825592 | 140827372 | 1781 | -5.86 | 4.70E-07 | 8.00E-07 |
| chr4 | 140066463 | 140067455 | 993 | -5.97 | 4.70E-07 | 8.00E-07 |
| chr4 | 28507446 | 28508895 | 1450 | -5.04 | 4.71E-07 | 8.01E-07 |
| chrX | 54150964 | 54152448 | 1485 | -6.11 | 4.72E-07 | 8.03E-07 |
| chr12 | 85827972 | 85833344 | 5373 | -6.52 | 4.72E-07 | 8.04E-07 |
| chr1 | 188239250 | 188240566 | 1317 | -5.82 | 4.72E-07 | 8.04E-07 |
| chr18 | 72231944 | 72239535 | 7592 | -1.48 | 4.73E-07 | 8.04E-07 |
| chr20 | 31620775 | 31621922 | 1148 | -4.26 | 4.73E-07 | 8.05E-07 |
| chr1 | 113961916 | 113969020 | 7105 | -5.95 | 4.74E-07 | 8.07E-07 |
| chr11 | 27440734 | 27441562 | 829 | -5.94 | 4.76E-07 | 8.09E-07 |

| | | | | | | |
|---|---|---|---|---|---|---|
| chr4 | 85832304 | 85833549 | 1246 | -5.84 | 4.76E-07 | 8.11E-07 |
| chr1 | 81304643 | 81307332 | 2690 | -6.02 | 4.77E-07 | 8.11E-07 |
| chr2 | 217905836 | 217907541 | 1706 | -6.02 | 4.77E-07 | 8.11E-07 |
| chr14 | 101913649 | 101915520 | 1872 | -6.04 | 4.77E-07 | 8.12E-07 |
| chr11 | 13879848 | 13880910 | 1063 | -5.8 | 4.77E-07 | 8.12E-07 |
| chr22 | 37012104 | 37014049 | 1946 | -2.68 | 4.77E-07 | 8.12E-07 |
| chrX | 123065473 | 123066782 | 1310 | -6 | 4.78E-07 | 8.12E-07 |
| chr6 | 89486999 | 89488586 | 1588 | -5.8 | 4.78E-07 | 8.13E-07 |
| chr14 | 95901070 | 95902577 | 1508 | -5.99 | 4.78E-07 | 8.13E-07 |
| chr11 | 120166654 | 120167775 | 1122 | -5.81 | 4.78E-07 | 8.14E-07 |
| chr2 | 215770445 | 215772076 | 1632 | -5.81 | 4.78E-07 | 8.14E-07 |
| chr9 | 77526674 | 77527564 | 891 | -5.81 | 4.78E-07 | 8.14E-07 |
| chr7 | 158273595 | 158274555 | 961 | -4.46 | 4.79E-07 | 8.14E-07 |
| chr10 | 28475863 | 28493763 | 17901 | 2.92 | 4.79E-07 | 8.15E-07 |
| chr4 | 174155259 | 174156810 | 1552 | -6.08 | 4.80E-07 | 8.16E-07 |
| chr19 | 2023528 | 2036345 | 12818 | -5.97 | 4.80E-07 | 8.16E-07 |
| chr2 | 225870353 | 225871836 | 1484 | -5.78 | 4.80E-07 | 8.17E-07 |
| chr5 | 1288073 | 1289017 | 945 | -5.79 | 4.81E-07 | 8.18E-07 |
| chr15 | 56017583 | 56018419 | 837 | -6.1 | 4.81E-07 | 8.18E-07 |
| chr4 | 161319925 | 161321099 | 1175 | -5.81 | 4.82E-07 | 8.19E-07 |
| chr4 | 94172732 | 94174080 | 1349 | -5.96 | 4.82E-07 | 8.20E-07 |
| chr11 | 93702406 | 93732798 | 30393 | -2.48 | 4.83E-07 | 8.21E-07 |
| chr5 | 169709685 | 169724818 | 15134 | -2.26 | 4.83E-07 | 8.22E-07 |
| chr5 | 156766651 | 156767817 | 1167 | -6.19 | 4.84E-07 | 8.22E-07 |
| chr14 | 38670648 | 38672409 | 1762 | 6.15 | 4.84E-07 | 8.23E-07 |
| chr1 | 225643703 | 225644902 | 1200 | -5.91 | 4.84E-07 | 8.23E-07 |
| chr13 | 83149575 | 83151119 | 1545 | -5.79 | 4.85E-07 | 8.25E-07 |
| chr9 | 5552595 | 5554998 | 2404 | 6.19 | 4.85E-07 | 8.25E-07 |
| chr18 | 34547406 | 34551703 | 4298 | -5.94 | 4.86E-07 | 8.26E-07 |
| chr10 | 121357664 | 121358667 | 1004 | -5.83 | 4.87E-07 | 8.27E-07 |
| chr4 | 63762490 | 63764479 | 1990 | -5.83 | 4.87E-07 | 8.27E-07 |
| chr4 | 65924144 | 65936511 | 12368 | -4.9 | 4.87E-07 | 8.28E-07 |
| chr2 | 236112807 | 236119006 | 6200 | -6.37 | 4.87E-07 | 8.28E-07 |
| chr1 | 90346532 | 90348212 | 1681 | -5.79 | 4.87E-07 | 8.28E-07 |
| chr13 | 26816351 | 26818643 | 2293 | -5.08 | 4.88E-07 | 8.29E-07 |
| chr17 | 26258904 | 26265476 | 6573 | -1.92 | 4.88E-07 | 8.30E-07 |
| chr3 | 197628606 | 197630073 | 1468 | -5.79 | 4.91E-07 | 8.35E-07 |
| chr3 | 177371249 | 177373039 | 1791 | -5.81 | 4.93E-07 | 8.37E-07 |
| chr4 | 87593464 | 87594271 | 808 | -5.85 | 4.94E-07 | 8.39E-07 |
| chr3 | 96512512 | 96514644 | 2133 | -5.79 | 4.94E-07 | 8.39E-07 |
| chr8 | 19892802 | 19932527 | 39726 | -1.17 | 4.94E-07 | 8.39E-07 |
| chr12 | 86262067 | 86263225 | 1159 | -5.91 | 4.95E-07 | 8.40E-07 |
| chr17 | 26842259 | 26843042 | 784 | -5.91 | 4.95E-07 | 8.40E-07 |
| chr8 | 59800143 | 59800916 | 774 | -5.91 | 4.95E-07 | 8.40E-07 |
| chr10 | 59450396 | 59471433 | 21038 | -7.03 | 4.95E-07 | 8.40E-07 |
| chr8 | 25400639 | 25426528 | 25890 | -1.3 | 4.96E-07 | 8.42E-07 |
| chr13 | 27240593 | 27241955 | 1363 | -5.83 | 4.97E-07 | 8.43E-07 |
| chr14 | 74696317 | 74700238 | 3922 | -3.52 | 4.97E-07 | 8.44E-07 |
| chr6 | 153563112 | 153564752 | 1641 | -5.88 | 4.98E-07 | 8.45E-07 |
| chr18 | 45891237 | 45892365 | 1129 | -5.9 | 4.98E-07 | 8.46E-07 |
| chr6 | 36320428 | 36321881 | 1454 | -5.9 | 4.98E-07 | 8.46E-07 |
| chr3 | 28193999 | 28201812 | 7814 | -3.03 | 4.99E-07 | 8.47E-07 |
| chr15 | 83362263 | 83364527 | 2265 | -6.21 | 5.00E-07 | 8.48E-07 |
| chr18 | 75196292 | 75358565 | 162274 | 0.63 | 5.00E-07 | 8.49E-07 |
| chr20 | 986561 | 991175 | 4615 | -3.75 | 5.01E-07 | 8.50E-07 |
| chr5 | 111485866 | 111494203 | 8338 | -6.22 | 5.01E-07 | 8.51E-07 |
| chr7 | 110229040 | 110230261 | 1222 | -5.84 | 5.02E-07 | 8.52E-07 |
| chr1 | 4469993 | 4481382 | 11390 | -2.7 | 5.02E-07 | 8.52E-07 |
| chr12 | 88526799 | 88527640 | 842 | -5.78 | 5.03E-07 | 8.54E-07 |
| chr3 | 136265285 | 136266624 | 1340 | -5.8 | 5.04E-07 | 8.55E-07 |
| chr2 | 99244252 | 99245035 | 784 | -5.78 | 5.04E-07 | 8.56E-07 |

| | | | | | | |
|---|---|---|---|---|---|---|
| chr13 | 77531632 | 77537211 | 5580 | -5.3 | 5.04E-07 | 8.56E-07 |
| chr17 | 76269674 | 76270395 | 722 | -5.82 | 5.05E-07 | 8.57E-07 |
| chr4 | 94146032 | 94148196 | 2165 | -5.82 | 5.05E-07 | 8.57E-07 |
| chr9 | 81002423 | 81019236 | 16814 | -1.76 | 5.05E-07 | 8.57E-07 |
| chr21 | 41054394 | 41056016 | 1623 | -5.77 | 5.05E-07 | 8.57E-07 |
| chr6 | 168587091 | 168588510 | 1420 | -5.77 | 5.05E-07 | 8.57E-07 |
| chrX | 110028891 | 110031810 | 2920 | 4.76 | 5.06E-07 | 8.58E-07 |
| chr8 | 100208808 | 100209676 | 869 | -5.78 | 5.06E-07 | 8.58E-07 |
| chr9 | 134096285 | 134097320 | 1036 | -5.78 | 5.06E-07 | 8.58E-07 |
| chr11 | 115166390 | 115167714 | 1325 | -5.78 | 5.06E-07 | 8.58E-07 |
| chr6 | 23021530 | 23022599 | 1070 | -5.78 | 5.06E-07 | 8.58E-07 |
| chr20 | 9975181 | 9975912 | 732 | -5.87 | 5.06E-07 | 8.58E-07 |
| chr11 | 32258844 | 32264402 | 5559 | -1.43 | 5.06E-07 | 8.58E-07 |
| chr1 | 174363013 | 174363756 | 744 | -5.79 | 5.07E-07 | 8.59E-07 |
| chr4 | 59579189 | 59581212 | 2024 | -5.81 | 5.10E-07 | 8.65E-07 |
| chr8 | 18425344 | 18426268 | 925 | -5.81 | 5.10E-07 | 8.65E-07 |
| chrX | 122708028 | 122710415 | 2388 | -5.81 | 5.10E-07 | 8.65E-07 |
| chr13 | 29983748 | 30001878 | 18131 | -2.41 | 5.11E-07 | 8.67E-07 |
| chr6 | 161145904 | 161184647 | 38744 | -3.17 | 5.12E-07 | 8.68E-07 |
| chr1 | 209162802 | 209165045 | 2244 | -2.88 | 5.12E-07 | 8.68E-07 |
| chr5 | 72254259 | 72255906 | 1648 | -6.27 | 5.12E-07 | 8.69E-07 |
| chr1 | 203969428 | 203970580 | 1153 | -5.78 | 5.13E-07 | 8.69E-07 |
| chr4 | 137182743 | 137184565 | 1823 | -5.78 | 5.13E-07 | 8.69E-07 |
| chr1 | 10235771 | 10236540 | 770 | -5.81 | 5.14E-07 | 8.71E-07 |
| chr6 | 88792029 | 88793727 | 1699 | -2.51 | 5.15E-07 | 8.73E-07 |
| chrX | 69176892 | 69178125 | 1234 | 5.93 | 5.16E-07 | 8.74E-07 |
| chr15 | 24364610 | 24408492 | 43883 | 2.03 | 5.17E-07 | 8.77E-07 |
| chr11 | 103748890 | 103762747 | 13858 | -6.54 | 5.19E-07 | 8.79E-07 |
| chr15 | 100914751 | 100966222 | 51472 | -2.52 | 5.20E-07 | 8.82E-07 |
| chr3 | 109839573 | 109841452 | 1880 | -6.02 | 5.21E-07 | 8.83E-07 |
| chr17 | 39442110 | 39443306 | 1197 | -5.99 | 5.21E-07 | 8.83E-07 |
| chr6 | 145305194 | 145306508 | 1315 | -5.78 | 5.21E-07 | 8.83E-07 |
| chrX | 101371291 | 101373289 | 1999 | -2.26 | 5.22E-07 | 8.84E-07 |
| chr11 | 121376387 | 121377582 | 1196 | -5.99 | 5.22E-07 | 8.84E-07 |
| chr14 | 54142762 | 54144463 | 1702 | -5.99 | 5.22E-07 | 8.84E-07 |
| chr2 | 172754895 | 172755461 | 567 | -6.04 | 5.22E-07 | 8.85E-07 |
| chr2 | 102030964 | 102031933 | 970 | -6.05 | 5.22E-07 | 8.85E-07 |
| chr17 | 72418211 | 72419323 | 1113 | -5.77 | 5.23E-07 | 8.85E-07 |
| chr17 | 15889685 | 15890455 | 771 | -5.97 | 5.23E-07 | 8.86E-07 |
| chr5 | 80660796 | 80662034 | 1239 | -5.78 | 5.24E-07 | 8.87E-07 |
| chr7 | 97827762 | 97828644 | 883 | -5.78 | 5.24E-07 | 8.87E-07 |
| chr13 | 52181369 | 52182626 | 1258 | -6.06 | 5.24E-07 | 8.87E-07 |
| chr22 | 24475226 | 24475996 | 771 | -5.8 | 5.24E-07 | 8.87E-07 |
| chr12 | 104821782 | 104822956 | 1175 | -6.06 | 5.24E-07 | 8.87E-07 |
| chr13 | 90493638 | 90495140 | 1503 | -5.79 | 5.24E-07 | 8.88E-07 |
| chr7 | 39555223 | 39566989 | 11767 | -6.07 | 5.25E-07 | 8.88E-07 |
| chr15 | 25928956 | 25930107 | 1152 | -5.85 | 5.25E-07 | 8.89E-07 |
| chr14 | 38412941 | 38447736 | 34796 | 1.58 | 5.26E-07 | 8.91E-07 |
| chr13 | 102420895 | 102422022 | 1128 | -5.95 | 5.26E-07 | 8.91E-07 |
| chr8 | 12573533 | 12575867 | 2335 | -5.95 | 5.26E-07 | 8.91E-07 |
| chr21 | 16579485 | 16581433 | 1949 | -5.77 | 5.29E-07 | 8.95E-07 |
| chr4 | 110528577 | 110529807 | 1231 | -5.77 | 5.29E-07 | 8.95E-07 |
| chr4 | 144162789 | 144164000 | 1212 | -5.77 | 5.29E-07 | 8.95E-07 |
| chr6 | 143840716 | 143842089 | 1374 | -5.77 | 5.29E-07 | 8.95E-07 |
| chr8 | 42825439 | 42826505 | 1067 | -5.77 | 5.29E-07 | 8.95E-07 |
| chr3 | 187620911 | 187622178 | 1268 | -6.11 | 5.29E-07 | 8.95E-07 |
| chr5 | 158692232 | 158694133 | 1902 | -6.11 | 5.29E-07 | 8.95E-07 |
| chr12 | 74503850 | 74506041 | 2192 | -5.83 | 5.29E-07 | 8.95E-07 |
| chr16 | 81561090 | 81563090 | 2001 | -5.83 | 5.29E-07 | 8.95E-07 |
| chr4 | 110455300 | 110456166 | 867 | -5.83 | 5.29E-07 | 8.95E-07 |
| chr8 | 47985797 | 47987433 | 1637 | -5.83 | 5.29E-07 | 8.95E-07 |

| | | | | | | |
|---|---|---|---|---|---|---|
| chr21 | 43631682 | 43636866 | 5185 | -6.12 | 5.31E-07 | 8.99E-07 |
| chr3 | 194499312 | 194500852 | 1541 | 5.84 | 5.31E-07 | 8.99E-07 |
| chr14 | 92974005 | 92975877 | 1873 | -5.92 | 5.33E-07 | 9.01E-07 |
| chrX | 41152038 | 41153067 | 1030 | -5.92 | 5.33E-07 | 9.01E-07 |
| chr3 | 53895166 | 53895991 | 826 | -5.76 | 5.33E-07 | 9.02E-07 |
| chr9 | 138243464 | 138244166 | 703 | -5.76 | 5.33E-07 | 9.02E-07 |
| chr14 | 60698012 | 60700706 | 2695 | -5.28 | 5.34E-07 | 9.03E-07 |
| chr12 | 97079381 | 97081322 | 1942 | -4.72 | 5.34E-07 | 9.03E-07 |
| chr13 | 36312176 | 36331569 | 19394 | -1.64 | 5.34E-07 | 9.04E-07 |
| chr18 | 11498222 | 11500001 | 1780 | -5.8 | 5.34E-07 | 9.04E-07 |
| chr11 | 89656167 | 89657056 | 890 | 5.49 | 5.35E-07 | 9.05E-07 |
| chr1 | 171918007 | 171931111 | 13105 | -5.17 | 5.35E-07 | 9.05E-07 |
| chr3 | 23919729 | 23920635 | 907 | -6.15 | 5.35E-07 | 9.05E-07 |
| chr11 | 86031698 | 86032571 | 874 | -5.83 | 5.36E-07 | 9.06E-07 |
| chr6 | 5416543 | 5418446 | 1904 | -5.9 | 5.37E-07 | 9.09E-07 |
| chr13 | 84214122 | 84215526 | 1405 | -5.77 | 5.38E-07 | 9.10E-07 |
| chr6 | 15543689 | 15545163 | 1475 | -5.77 | 5.38E-07 | 9.10E-07 |
| chr7 | 15293917 | 15296395 | 2479 | -5.77 | 5.38E-07 | 9.10E-07 |
| chr3 | 59851777 | 59859162 | 7386 | -1.4 | 5.38E-07 | 9.10E-07 |
| chr6 | 53532310 | 53533235 | 926 | -4.56 | 5.38E-07 | 9.10E-07 |
| chr4 | 75773918 | 75788270 | 14353 | -2.31 | 5.39E-07 | 9.10E-07 |
| chr5 | 59464081 | 59497769 | 33689 | -2.01 | 5.40E-07 | 9.12E-07 |
| chr18 | 46315355 | 46316149 | 795 | -5.78 | 5.40E-07 | 9.12E-07 |
| chr10 | 105346687 | 105348131 | 1445 | -5.9 | 5.40E-07 | 9.13E-07 |
| chr22 | 46878430 | 46879434 | 1005 | -5.9 | 5.40E-07 | 9.13E-07 |
| chr3 | 115273549 | 115336930 | 63382 | 1.28 | 5.40E-07 | 9.13E-07 |
| chr14 | 57835899 | 57837440 | 1542 | -5.34 | 5.41E-07 | 9.14E-07 |
| chr10 | 97924818 | 97955876 | 31059 | -5.23 | 5.42E-07 | 9.15E-07 |
| chr4 | 124787856 | 124789696 | 1841 | -5.77 | 5.42E-07 | 9.16E-07 |
| chr3 | 65223984 | 65224806 | 823 | -5.76 | 5.42E-07 | 9.17E-07 |
| chr5 | 134891316 | 134892734 | 1419 | -5.89 | 5.43E-07 | 9.18E-07 |
| chr6 | 13308860 | 13309956 | 1097 | -5.89 | 5.43E-07 | 9.18E-07 |
| chr11 | 10975290 | 10985598 | 10309 | -2.11 | 5.43E-07 | 9.18E-07 |
| chr20 | 60302491 | 60303815 | 1325 | 6.06 | 5.43E-07 | 9.18E-07 |
| chr12 | 123574619 | 123575838 | 1220 | -6.19 | 5.44E-07 | 9.19E-07 |
| chr3 | 45276788 | 45278900 | 2113 | -5.79 | 5.46E-07 | 9.22E-07 |
| chr7 | 98009035 | 98009816 | 782 | -5.79 | 5.46E-07 | 9.22E-07 |
| chrX | 106121670 | 106131698 | 10029 | -5.05 | 5.47E-07 | 9.23E-07 |
| chr6 | 112272001 | 112273715 | 1715 | 5.9 | 5.47E-07 | 9.24E-07 |
| chr7 | 18139986 | 18145780 | 5795 | -6.21 | 5.48E-07 | 9.26E-07 |
| chr9 | 104320100 | 104321180 | 1081 | -4.66 | 5.49E-07 | 9.27E-07 |
| chr19 | 16583563 | 16584496 | 934 | -5.87 | 5.50E-07 | 9.29E-07 |
| chr12 | 79187697 | 79190487 | 2791 | -5.82 | 5.50E-07 | 9.29E-07 |
| chr7 | 98500615 | 98501740 | 1126 | -5.82 | 5.50E-07 | 9.29E-07 |
| chr16 | 63241158 | 63242911 | 1754 | -4.52 | 5.50E-07 | 9.29E-07 |
| chr15 | 53474877 | 53475851 | 975 | -4.54 | 5.50E-07 | 9.30E-07 |
| chr5 | 36750480 | 36790761 | 40282 | -1.6 | 5.53E-07 | 9.33E-07 |
| chr12 | 40363865 | 40364537 | 673 | -5.86 | 5.54E-07 | 9.36E-07 |
| chr1 | 39443545 | 39444973 | 1429 | -5.77 | 5.56E-07 | 9.39E-07 |
| chr11 | 119842401 | 119843600 | 1200 | -5.75 | 5.56E-07 | 9.40E-07 |
| chr6 | 83702577 | 83703806 | 1230 | -5.75 | 5.56E-07 | 9.40E-07 |
| chr17 | 20098826 | 20124335 | 25510 | -7.12 | 5.59E-07 | 9.44E-07 |
| chr16 | 48745246 | 48793076 | 47831 | 3.18 | 5.59E-07 | 9.44E-07 |
| chr19 | 8451623 | 8452272 | 650 | -5.77 | 5.59E-07 | 9.44E-07 |
| chr4 | 132386153 | 132387416 | 1264 | -5.77 | 5.59E-07 | 9.44E-07 |
| chrX | 56421314 | 56422797 | 1484 | 5.07 | 5.59E-07 | 9.44E-07 |
| chr4 | 174763050 | 174764275 | 1226 | -5.79 | 5.60E-07 | 9.44E-07 |
| chr12 | 78197889 | 78198824 | 936 | -4.51 | 5.60E-07 | 9.45E-07 |
| chr6 | 16793778 | 16794920 | 1143 | -5.81 | 5.60E-07 | 9.45E-07 |
| chr7 | 36292826 | 36293736 | 911 | -6.27 | 5.60E-07 | 9.45E-07 |
| chr12 | 13046427 | 13048615 | 2189 | -4.6 | 5.60E-07 | 9.46E-07 |

| | | | | | | |
|---|---|---|---|---|---|---|
| chr4 | 103215320 | 103216235 | 916 | -5.8 | 5.61E-07 | 9.46E-07 |
| chr10 | 73450649 | 73451892 | 1244 | -5.85 | 5.61E-07 | 9.46E-07 |
| chr6 | 39838829 | 39854808 | 15980 | 2.55 | 5.63E-07 | 9.49E-07 |
| chr1 | 245985991 | 245987966 | 1976 | -5.76 | 5.63E-07 | 9.50E-07 |
| chr5 | 114839023 | 114840254 | 1232 | -5.85 | 5.64E-07 | 9.51E-07 |
| chr15 | 76781672 | 76782465 | 794 | -5.75 | 5.64E-07 | 9.52E-07 |
| chr3 | 121307392 | 121308514 | 1123 | -5.75 | 5.64E-07 | 9.52E-07 |
| chr3 | 14779480 | 14793231 | 13752 | -3.99 | 5.64E-07 | 9.52E-07 |
| chr14 | 70892097 | 70920055 | 27959 | -4.99 | 5.64E-07 | 9.52E-07 |
| chr9 | 14697182 | 14710832 | 13651 | -2.12 | 5.65E-07 | 9.52E-07 |
| chr1 | 199373539 | 199375316 | 1778 | 3.15 | 5.65E-07 | 9.53E-07 |
| chr2 | 15909521 | 16070392 | 160872 | 3.3 | 5.65E-07 | 9.53E-07 |
| chr9 | 37713104 | 37713581 | 478 | -5.15 | 5.66E-07 | 9.54E-07 |
| chr6 | 10939405 | 10949795 | 10391 | -6.55 | 5.66E-07 | 9.55E-07 |
| chr3 | 46804285 | 46885762 | 81478 | -3.71 | 5.66E-07 | 9.55E-07 |
| chr1 | 164651596 | 164653445 | 1850 | 5.91 | 5.67E-07 | 9.56E-07 |
| chr1 | 17646104 | 17656814 | 10711 | -6.56 | 5.67E-07 | 9.56E-07 |
| chr8 | 64055378 | 64068259 | 12882 | -3.35 | 5.67E-07 | 9.57E-07 |
| chr2 | 144865911 | 144871279 | 5369 | -5.84 | 5.67E-07 | 9.57E-07 |
| chrX | 9268683 | 9272190 | 3508 | -6.3 | 5.67E-07 | 9.57E-07 |
| chr1 | 64111171 | 64112069 | 899 | -5.84 | 5.69E-07 | 9.60E-07 |
| chr1 | 162649132 | 162653895 | 4764 | -6 | 5.69E-07 | 9.60E-07 |
| chr6 | 6845383 | 6846994 | 1612 | -6 | 5.69E-07 | 9.60E-07 |
| chr3 | 174037243 | 174039804 | 2562 | -5.99 | 5.69E-07 | 9.60E-07 |
| chr9 | 79028752 | 79029943 | 1192 | -5.98 | 5.70E-07 | 9.60E-07 |
| chr12 | 27213651 | 27214958 | 1308 | -5.97 | 5.70E-07 | 9.61E-07 |
| chr1 | 95241707 | 95242620 | 914 | -5.79 | 5.71E-07 | 9.62E-07 |
| chr1 | 36248536 | 36249484 | 949 | -5.96 | 5.71E-07 | 9.63E-07 |
| chr3 | 57695579 | 57696579 | 1001 | -5.8 | 5.72E-07 | 9.63E-07 |
| chr1 | 224874445 | 224875816 | 1372 | -5.95 | 5.72E-07 | 9.64E-07 |
| chr1 | 185520507 | 185522666 | 2160 | -5.93 | 5.75E-07 | 9.69E-07 |
| chr18 | 34431745 | 34432915 | 1171 | -5.83 | 5.75E-07 | 9.69E-07 |
| chrX | 71231209 | 71232387 | 1179 | -5.83 | 5.75E-07 | 9.69E-07 |
| chr12 | 27699810 | 27700915 | 1106 | -5.78 | 5.75E-07 | 9.69E-07 |
| chr16 | 71620598 | 71621938 | 1341 | -4.87 | 5.75E-07 | 9.69E-07 |
| chr3 | 178660209 | 178665865 | 5657 | 2.18 | 5.76E-07 | 9.71E-07 |
| chr20 | 57278553 | 57280038 | 1486 | -5.76 | 5.76E-07 | 9.71E-07 |
| chr10 | 63592825 | 63601458 | 8634 | -5.75 | 5.77E-07 | 9.72E-07 |
| chr1 | 168904350 | 168905820 | 1471 | -4.56 | 5.78E-07 | 9.74E-07 |
| chr7 | 26171181 | 26175850 | 4670 | -6.35 | 5.78E-07 | 9.74E-07 |
| chr12 | 90283163 | 90285798 | 2636 | -6.1 | 5.79E-07 | 9.75E-07 |
| chr4 | 137093636 | 137094685 | 1050 | -5.91 | 5.79E-07 | 9.75E-07 |
| chrX | 82821628 | 82843144 | 21517 | 1.4 | 5.80E-07 | 9.77E-07 |
| chr13 | 61142355 | 61144733 | 2379 | -3.12 | 5.80E-07 | 9.77E-07 |
| chr1 | 67117980 | 67119844 | 1865 | -3.04 | 5.80E-07 | 9.77E-07 |
| chr19 | 5561602 | 5562579 | 978 | -5.79 | 5.81E-07 | 9.79E-07 |
| chr11 | 33923432 | 33955048 | 31617 | -6.14 | 5.81E-07 | 9.79E-07 |
| chr17 | 56597149 | 56598150 | 1002 | -5.82 | 5.81E-07 | 9.79E-07 |
| chr1 | 210650208 | 210651013 | 806 | -5.75 | 5.82E-07 | 9.79E-07 |
| chr4 | 168352734 | 168353715 | 982 | -5.75 | 5.82E-07 | 9.79E-07 |
| chr19 | 6037147 | 6038158 | 1012 | -5.98 | 5.83E-07 | 9.82E-07 |
| chr4 | 119926641 | 119927410 | 770 | -5.82 | 5.83E-07 | 9.82E-07 |
| chr10 | 80453968 | 80465462 | 11495 | -6.38 | 5.84E-07 | 9.83E-07 |
| chr10 | 75734727 | 75735947 | 1221 | -5.89 | 5.85E-07 | 9.84E-07 |
| chr10 | 71884476 | 71885612 | 1137 | -5.95 | 5.85E-07 | 9.85E-07 |
| chr10 | 120401599 | 120404069 | 2471 | 6.12 | 5.86E-07 | 9.86E-07 |
| chr3 | 15741690 | 15742441 | 752 | -5.75 | 5.86E-07 | 9.87E-07 |
| chr14 | 73085676 | 73088248 | 2573 | 6.09 | 5.86E-07 | 9.87E-07 |
| chr6 | 161824478 | 161825957 | 1480 | -5.76 | 5.86E-07 | 9.87E-07 |
| chr1 | 110963708 | 110967003 | 3296 | -4.56 | 5.87E-07 | 9.87E-07 |
| chr2 | 236027670 | 236028674 | 1005 | -5.94 | 5.87E-07 | 9.88E-07 |

| | | | | | | |
|---|---|---|---|---|---|---|
| chr6 | 149234911 | 149235745 | 835 | -5.74 | 5.88E-07 | 9.89E-07 |
| chr7 | 156740108 | 156740978 | 871 | -5.74 | 5.88E-07 | 9.89E-07 |
| chr2 | 69066501 | 69103420 | 36920 | 2.5 | 5.89E-07 | 9.91E-07 |
| chr15 | 24524640 | 24533445 | 8806 | 2.88 | 5.89E-07 | 9.92E-07 |
| chr14 | 70873402 | 70874522 | 1121 | -5.92 | 5.89E-07 | 9.92E-07 |
| chr10 | 116139999 | 116141182 | 1184 | -6.09 | 5.91E-07 | 9.95E-07 |
| chr1 | 67554904 | 67555842 | 939 | -5.87 | 5.92E-07 | 9.96E-07 |
| chr2 | 147161740 | 147163348 | 1609 | -5.77 | 5.92E-07 | 9.96E-07 |
| chr1 | 146666747 | 146668566 | 1820 | -4.46 | 5.92E-07 | 9.97E-07 |
| chr1 | 200110227 | 200110893 | 667 | -4.46 | 5.92E-07 | 9.97E-07 |
| chrX | 102502105 | 102503101 | 997 | -4.97 | 5.93E-07 | 9.97E-07 |
| chr1 | 68191625 | 68192854 | 1230 | -5.78 | 5.93E-07 | 9.97E-07 |
| chr12 | 105314804 | 105315715 | 912 | -5.78 | 5.93E-07 | 9.97E-07 |
| chr19 | 10033321 | 10034061 | 741 | -5.78 | 5.93E-07 | 9.97E-07 |
| chr2 | 65037561 | 65038333 | 773 | -5.78 | 5.93E-07 | 9.97E-07 |
| chr8 | 123643513 | 123644419 | 907 | -5.78 | 5.93E-07 | 9.97E-07 |
| chrX | 152202396 | 152219718 | 17323 | -3.33 | 5.94E-07 | 1.00E-06 |
| chr13 | 45426659 | 45427819 | 1161 | -5.75 | 5.95E-07 | 1.00E-06 |
| chr10 | 32441789 | 32444051 | 2263 | -5.81 | 5.96E-07 | 1.00E-06 |
| chr19 | 42504680 | 42507158 | 2479 | -6.19 | 5.96E-07 | 1.00E-06 |
| chr6 | 3519395 | 3520631 | 1237 | -6.19 | 5.96E-07 | 1.00E-06 |
| chr2 | 183719866 | 183722986 | 3121 | -4.87 | 5.97E-07 | 1.00E-06 |
| chr7 | 110841411 | 110842509 | 1099 | -5.86 | 5.97E-07 | 1.00E-06 |
| chr7 | 111388782 | 111389920 | 1139 | -5.78 | 5.98E-07 | 1.01E-06 |
| chr12 | 87217726 | 87219124 | 1399 | -5.74 | 5.99E-07 | 1.01E-06 |
| chrX | 47395041 | 47396425 | 1385 | -5.74 | 5.99E-07 | 1.01E-06 |
| chr20 | 60356 | 78309 | 17954 | -3.21 | 6.00E-07 | 1.01E-06 |
| chr21 | 38968639 | 38970415 | 1777 | -4.59 | 6.01E-07 | 1.01E-06 |
| chr12 | 132297638 | 132298485 | 848 | -5.8 | 6.03E-07 | 1.01E-06 |
| chr6 | 52119661 | 52121728 | 2068 | -5.8 | 6.03E-07 | 1.01E-06 |
| chr3 | 160625471 | 160629309 | 3839 | 2.1 | 6.03E-07 | 1.01E-06 |
| chr10 | 116819115 | 116821720 | 2606 | -2.17 | 6.05E-07 | 1.02E-06 |
| chr14 | 57662766 | 57663643 | 878 | -5.87 | 6.06E-07 | 1.02E-06 |
| chr8 | 38801433 | 38802369 | 937 | -5.87 | 6.06E-07 | 1.02E-06 |
| chr9 | 106826437 | 106843932 | 17496 | 5.4 | 6.08E-07 | 1.02E-06 |
| chr9 | 2463688 | 2465668 | 1981 | -4.78 | 6.09E-07 | 1.02E-06 |
| chr11 | 74665946 | 74666626 | 681 | -5.74 | 6.10E-07 | 1.02E-06 |
| chr15 | 42774192 | 42775202 | 1011 | -5.86 | 6.10E-07 | 1.02E-06 |
| chr11 | 65889294 | 65890445 | 1152 | -5.76 | 6.10E-07 | 1.03E-06 |
| chr13 | 111215832 | 111216548 | 717 | -5.74 | 6.13E-07 | 1.03E-06 |
| chr13 | 106731959 | 106733505 | 1547 | -5.79 | 6.14E-07 | 1.03E-06 |
| chr3 | 12747845 | 12748924 | 1080 | -5.79 | 6.14E-07 | 1.03E-06 |
| chr17 | 29567706 | 29568514 | 809 | -5.83 | 6.14E-07 | 1.03E-06 |
| chr5 | 23572601 | 23575305 | 2705 | -5.83 | 6.14E-07 | 1.03E-06 |
| chr8 | 106711685 | 106713536 | 1852 | 6.35 | 6.15E-07 | 1.03E-06 |
| chr16 | 49616003 | 49617742 | 1740 | 2.75 | 6.15E-07 | 1.03E-06 |
| chr5 | 124468258 | 124469378 | 1121 | -5.74 | 6.15E-07 | 1.03E-06 |
| chr15 | 31347898 | 31351570 | 3673 | -5.27 | 6.16E-07 | 1.03E-06 |
| chr15 | 98944228 | 99054356 | 110129 | -3.43 | 6.16E-07 | 1.03E-06 |
| chr1 | 75386683 | 75398150 | 11468 | 2.66 | 6.18E-07 | 1.04E-06 |
| chr20 | 18701203 | 18702021 | 819 | -5.85 | 6.19E-07 | 1.04E-06 |
| chr9 | 3744181 | 3764263 | 20083 | -7.34 | 6.19E-07 | 1.04E-06 |
| chr2 | 207680100 | 207680926 | 827 | -5.74 | 6.19E-07 | 1.04E-06 |
| chr2 | 232060814 | 232062009 | 1196 | -5.74 | 6.19E-07 | 1.04E-06 |
| chr7 | 6557769 | 6558900 | 1132 | -5.74 | 6.19E-07 | 1.04E-06 |
| chr18 | 25647193 | 25647944 | 752 | -5.76 | 6.19E-07 | 1.04E-06 |
| chr9 | 103271937 | 103273033 | 1097 | -5.76 | 6.19E-07 | 1.04E-06 |
| chr1 | 224655663 | 224656629 | 967 | -5.98 | 6.23E-07 | 1.05E-06 |
| chr5 | 143164115 | 143169508 | 5394 | -5.96 | 6.23E-07 | 1.05E-06 |
| chr6 | 74530294 | 74531255 | 962 | -5.84 | 6.24E-07 | 1.05E-06 |
| chr8 | 103583006 | 103584262 | 1257 | -5.95 | 6.24E-07 | 1.05E-06 |

| | | | | | | |
|---|---|---|---|---|---|---|
| chr7 | 29425568 | 29427039 | 1472 | -3.07 | 6.24E-07 | 1.05E-06 |
| chr5 | 81186571 | 81188382 | 1812 | -6.02 | 6.25E-07 | 1.05E-06 |
| chr18 | 55192114 | 55192898 | 785 | -5.31 | 6.25E-07 | 1.05E-06 |
| chr6 | 133493298 | 133500891 | 7594 | -6.32 | 6.25E-07 | 1.05E-06 |
| chr8 | 1985037 | 1986770 | 1734 | -6.04 | 6.26E-07 | 1.05E-06 |
| chr13 | 111820793 | 111821815 | 1023 | -5.78 | 6.27E-07 | 1.05E-06 |
| chr15 | 31366812 | 31367736 | 925 | -5.78 | 6.27E-07 | 1.05E-06 |
| chr8 | 110149122 | 110150160 | 1039 | -5.78 | 6.27E-07 | 1.05E-06 |
| chr11 | 126621670 | 126631648 | 9979 | -6.9 | 6.27E-07 | 1.05E-06 |
| chrX | 141043325 | 141054054 | 10730 | -5.01 | 6.28E-07 | 1.05E-06 |
| chr8 | 10810223 | 10812881 | 2659 | -5.37 | 6.30E-07 | 1.06E-06 |
| chr3 | 17287395 | 17288398 | 1004 | -5.76 | 6.31E-07 | 1.06E-06 |
| chr5 | 53423067 | 53424276 | 1210 | -5.76 | 6.31E-07 | 1.06E-06 |
| chr3 | 58587942 | 58591852 | 3911 | -5.17 | 6.32E-07 | 1.06E-06 |
| chr13 | 106167697 | 106168876 | 1180 | -5.75 | 6.34E-07 | 1.06E-06 |
| chr17 | 66302478 | 66304039 | 1562 | -5.75 | 6.34E-07 | 1.06E-06 |
| chr3 | 182172746 | 182174034 | 1289 | -5.75 | 6.34E-07 | 1.06E-06 |
| chr17 | 76590031 | 76591558 | 1528 | -5.78 | 6.34E-07 | 1.06E-06 |
| chr4 | 85062989 | 85063871 | 883 | -5.78 | 6.34E-07 | 1.06E-06 |
| chr3 | 56465310 | 56477968 | 12659 | -1.6 | 6.35E-07 | 1.06E-06 |
| chr13 | 42256941 | 42257635 | 695 | -5.89 | 6.37E-07 | 1.07E-06 |
| chr9 | 76010947 | 76015839 | 4893 | -4.97 | 6.37E-07 | 1.07E-06 |
| chr19 | 11208972 | 11209791 | 820 | -5.73 | 6.37E-07 | 1.07E-06 |
| chr2 | 189573323 | 189575786 | 2464 | -6.37 | 6.37E-07 | 1.07E-06 |
| chr1 | 167152148 | 167153386 | 1239 | 5.8 | 6.39E-07 | 1.07E-06 |
| chr2 | 71144547 | 71145514 | 968 | -5.8 | 6.39E-07 | 1.07E-06 |
| chr2 | 195760680 | 195762445 | 1766 | 5.92 | 6.41E-07 | 1.07E-06 |
| chr9 | 19329012 | 19329624 | 613 | -5.77 | 6.41E-07 | 1.07E-06 |
| chr19 | 4319145 | 4320449 | 1305 | -5.81 | 6.41E-07 | 1.07E-06 |
| chr14 | 103847147 | 103847962 | 816 | -6.13 | 6.42E-07 | 1.08E-06 |
| chr10 | 97134613 | 97135234 | 622 | -6.14 | 6.44E-07 | 1.08E-06 |
| chr2 | 225201593 | 225203250 | 1658 | -5.87 | 6.44E-07 | 1.08E-06 |
| chr11 | 113361086 | 113362157 | 1072 | 5.32 | 6.45E-07 | 1.08E-06 |
| chr1 | 245583588 | 245616285 | 32698 | -1.12 | 6.45E-07 | 1.08E-06 |
| chr8 | 82526856 | 82530514 | 3659 | -4.7 | 6.45E-07 | 1.08E-06 |
| chr9 | 104280484 | 104281741 | 1258 | -5.77 | 6.46E-07 | 1.08E-06 |
| chrX | 109461882 | 109462609 | 728 | -5.81 | 6.48E-07 | 1.09E-06 |
| chr4 | 55066817 | 55077989 | 11173 | -4.75 | 6.48E-07 | 1.09E-06 |
| chr17 | 77199790 | 77200534 | 745 | -5.72 | 6.48E-07 | 1.09E-06 |
| chr7 | 157128026 | 157129199 | 1174 | -5.72 | 6.48E-07 | 1.09E-06 |
| chr14 | 36956201 | 36957838 | 1638 | 6.08 | 6.48E-07 | 1.09E-06 |
| chr6 | 7756017 | 7757511 | 1495 | -5.86 | 6.49E-07 | 1.09E-06 |
| chr8 | 91913546 | 91914721 | 1176 | 5.81 | 6.49E-07 | 1.09E-06 |
| chr20 | 45444421 | 45454362 | 9942 | -6.43 | 6.50E-07 | 1.09E-06 |
| chr6 | 54822228 | 54823006 | 779 | -5.74 | 6.50E-07 | 1.09E-06 |
| chr4 | 170470220 | 170471764 | 1545 | -5.75 | 6.51E-07 | 1.09E-06 |
| chr9 | 75889358 | 75891097 | 1740 | -4.44 | 6.51E-07 | 1.09E-06 |
| chr4 | 147038925 | 147047475 | 8551 | -4.75 | 6.52E-07 | 1.09E-06 |
| chr2 | 159237963 | 159239113 | 1151 | -5.99 | 6.53E-07 | 1.09E-06 |
| chr17 | 55959772 | 55960471 | 700 | -5.95 | 6.53E-07 | 1.09E-06 |
| chrX | 120595046 | 120596302 | 1257 | 5.9 | 6.54E-07 | 1.09E-06 |
| chr15 | 62297922 | 62313480 | 15559 | -5.94 | 6.54E-07 | 1.09E-06 |
| chr6 | 4403419 | 4405456 | 2038 | -6.02 | 6.54E-07 | 1.10E-06 |
| chr12 | 114343577 | 114344485 | 909 | -6.02 | 6.55E-07 | 1.10E-06 |
| chr2 | 189819818 | 189820854 | 1037 | -5.73 | 6.55E-07 | 1.10E-06 |
| chr4 | 59897353 | 59899512 | 2160 | -5.73 | 6.55E-07 | 1.10E-06 |
| chr6 | 76271119 | 76278039 | 6921 | -2.47 | 6.55E-07 | 1.10E-06 |
| chr16 | 57217780 | 57218682 | 903 | -5.8 | 6.56E-07 | 1.10E-06 |
| chr4 | 52989177 | 53008998 | 19822 | -1.36 | 6.56E-07 | 1.10E-06 |
| chr7 | 132342204 | 132358153 | 15950 | -5.17 | 6.57E-07 | 1.10E-06 |
| chr10 | 71578143 | 71579386 | 1244 | -5.76 | 6.57E-07 | 1.10E-06 |

| | | | | | | |
|---|---|---|---|---|---|---|
| chr12 | 92363796 | 92365337 | 1542 | -5.76 | 6.57E-07 | 1.10E-06 |
| chr9 | 135428432 | 135435735 | 7304 | -5.32 | 6.58E-07 | 1.10E-06 |
| chr22 | 50866559 | 50867289 | 731 | -5.76 | 6.59E-07 | 1.10E-06 |
| chr11 | 67826804 | 67827411 | 608 | -5.91 | 6.60E-07 | 1.10E-06 |
| chr14 | 75785561 | 75786285 | 725 | -5.78 | 6.61E-07 | 1.11E-06 |
| chr6 | 121696152 | 121696927 | 776 | -5.78 | 6.61E-07 | 1.11E-06 |
| chr5 | 137745487 | 137746387 | 901 | -5.9 | 6.61E-07 | 1.11E-06 |
| chr4 | 54129666 | 54130908 | 1243 | -5.72 | 6.61E-07 | 1.11E-06 |
| chr14 | 60976387 | 60978942 | 2556 | -6.22 | 6.62E-07 | 1.11E-06 |
| chr5 | 72233515 | 72235143 | 1629 | -6.22 | 6.62E-07 | 1.11E-06 |
| chr8 | 117982682 | 117985674 | 2993 | 2.33 | 6.64E-07 | 1.11E-06 |
| chr16 | 14265930 | 14266628 | 699 | -5.79 | 6.64E-07 | 1.11E-06 |
| chr8 | 68271047 | 68272501 | 1455 | -5.83 | 6.66E-07 | 1.11E-06 |
| chr8 | 90202382 | 90203377 | 996 | -5.74 | 6.67E-07 | 1.12E-06 |
| chr1 | 206323143 | 206323878 | 736 | -5.88 | 6.68E-07 | 1.12E-06 |
| chr8 | 48819222 | 48820964 | 1743 | -5.88 | 6.68E-07 | 1.12E-06 |
| chr12 | 11526000 | 11542426 | 16427 | 1.66 | 6.69E-07 | 1.12E-06 |
| chr1 | 27602590 | 27604156 | 1567 | -5.74 | 6.71E-07 | 1.12E-06 |
| chr5 | 139296217 | 139296965 | 749 | -5.74 | 6.71E-07 | 1.12E-06 |
| chr20 | 52434771 | 52436178 | 1408 | -6.56 | 6.71E-07 | 1.12E-06 |
| chr15 | 44735149 | 44736236 | 1088 | -5.82 | 6.73E-07 | 1.13E-06 |
| chr2 | 145371715 | 145381616 | 9902 | -6.59 | 6.73E-07 | 1.13E-06 |
| chr6 | 142356463 | 142357905 | 1443 | -5.77 | 6.74E-07 | 1.13E-06 |
| chr7 | 56033577 | 56034969 | 1393 | -5.77 | 6.74E-07 | 1.13E-06 |
| chr2 | 19844322 | 19845412 | 1091 | -5.86 | 6.74E-07 | 1.13E-06 |
| chr11 | 36436598 | 36437122 | 525 | -5.72 | 6.77E-07 | 1.13E-06 |
| chr7 | 48141734 | 48142753 | 1020 | -5.72 | 6.77E-07 | 1.13E-06 |
| chr22 | 18079099 | 18080319 | 1221 | -5.85 | 6.78E-07 | 1.13E-06 |
| chr18 | 55171617 | 55176696 | 5080 | 1.6 | 6.79E-07 | 1.14E-06 |
| chr2 | 62682585 | 62683622 | 1038 | -5.72 | 6.81E-07 | 1.14E-06 |
| chr5 | 6523571 | 6588218 | 64648 | -2.59 | 6.81E-07 | 1.14E-06 |
| chr2 | 11287258 | 11288480 | 1223 | -5.72 | 6.81E-07 | 1.14E-06 |
| chr3 | 69407739 | 69409071 | 1333 | -5.72 | 6.81E-07 | 1.14E-06 |
| chr4 | 176008544 | 176010514 | 1971 | -5.81 | 6.82E-07 | 1.14E-06 |
| chr3 | 124695951 | 124696921 | 971 | -5.84 | 6.82E-07 | 1.14E-06 |
| chr20 | 14998524 | 15004871 | 6348 | -5.97 | 6.82E-07 | 1.14E-06 |
| chr4 | 141082649 | 141083860 | 1212 | -5.97 | 6.82E-07 | 1.14E-06 |
| chr1 | 45576757 | 45577901 | 1145 | -5.96 | 6.82E-07 | 1.14E-06 |
| chr17 | 46180151 | 46180960 | 810 | -5.77 | 6.83E-07 | 1.14E-06 |
| chr21 | 37844688 | 37845999 | 1312 | -5.77 | 6.83E-07 | 1.14E-06 |
| chr22 | 37135131 | 37136296 | 1166 | -5.77 | 6.83E-07 | 1.14E-06 |
| chr8 | 134659881 | 134660889 | 1009 | -5.77 | 6.83E-07 | 1.14E-06 |
| chrX | 36563443 | 36564950 | 1508 | -5.77 | 6.83E-07 | 1.14E-06 |
| chr6 | 147363630 | 147366377 | 2748 | -3.02 | 6.85E-07 | 1.14E-06 |
| chr2 | 69434199 | 69435077 | 879 | -5.92 | 6.86E-07 | 1.15E-06 |
| chr13 | 25237357 | 25238237 | 881 | -5.73 | 6.86E-07 | 1.15E-06 |
| chr8 | 101182090 | 101182960 | 871 | -5.84 | 6.86E-07 | 1.15E-06 |
| chr5 | 75993677 | 75994998 | 1322 | -6.03 | 6.86E-07 | 1.15E-06 |
| chr1 | 82108331 | 82109346 | 1016 | -5.71 | 6.88E-07 | 1.15E-06 |
| chr7 | 66429878 | 66431064 | 1187 | -5.71 | 6.88E-07 | 1.15E-06 |
| chrX | 76358764 | 76404952 | 46189 | 1.56 | 6.88E-07 | 1.15E-06 |
| chr13 | 102162012 | 102163718 | 1707 | -6.05 | 6.89E-07 | 1.15E-06 |
| chr2 | 128130512 | 128131764 | 1253 | -5.9 | 6.89E-07 | 1.15E-06 |
| chr14 | 59922217 | 59923728 | 1512 | -4.41 | 6.91E-07 | 1.15E-06 |
| chr4 | 140205585 | 140206397 | 813 | -5.89 | 6.91E-07 | 1.15E-06 |
| chr4 | 56060316 | 56072129 | 11814 | -3.33 | 6.92E-07 | 1.16E-06 |
| chr2 | 188651473 | 188674661 | 23189 | -6.08 | 6.93E-07 | 1.16E-06 |
| chr6 | 20352002 | 20364609 | 12608 | -2.1 | 6.94E-07 | 1.16E-06 |
| chr1 | 66803603 | 66804413 | 811 | -5.77 | 6.94E-07 | 1.16E-06 |
| chr11 | 123391040 | 123392075 | 1036 | -5.77 | 6.94E-07 | 1.16E-06 |
| chr1 | 12865134 | 12866642 | 1509 | -2.78 | 6.94E-07 | 1.16E-06 |

| | | | | | | |
|---|---|---|---|---|---|---|
| chr12 | 93647799 | 93648928 | 1130 | -5.88 | 6.94E-07 | 1.16E-06 |
| chr14 | 54277484 | 54278940 | 1457 | -5.88 | 6.94E-07 | 1.16E-06 |
| chr17 | 27684911 | 27685999 | 1089 | -5.88 | 6.94E-07 | 1.16E-06 |
| chr3 | 121646532 | 121647806 | 1275 | -5.88 | 6.94E-07 | 1.16E-06 |
| chr5 | 43266570 | 43268688 | 2119 | -3.38 | 6.95E-07 | 1.16E-06 |
| chr12 | 71069229 | 71069992 | 764 | -5.82 | 6.96E-07 | 1.16E-06 |
| chr1 | 64738173 | 64739288 | 1116 | -5.87 | 6.98E-07 | 1.16E-06 |
| chr13 | 43501731 | 43503957 | 2227 | -3.43 | 6.99E-07 | 1.17E-06 |
| chr20 | 13675346 | 13693708 | 18363 | -5.68 | 7.01E-07 | 1.17E-06 |
| chr1 | 45174485 | 45176698 | 2214 | -5.86 | 7.01E-07 | 1.17E-06 |
| chr11 | 69641726 | 69642614 | 889 | -5.86 | 7.01E-07 | 1.17E-06 |
| chr3 | 119091587 | 119093692 | 2106 | -6.12 | 7.02E-07 | 1.17E-06 |
| chr1 | 90585039 | 90606371 | 21333 | -1.32 | 7.02E-07 | 1.17E-06 |
| chr20 | 3676895 | 3677569 | 675 | -5.81 | 7.02E-07 | 1.17E-06 |
| chr1 | 166795975 | 166804441 | 8467 | -1.35 | 7.02E-07 | 1.17E-06 |
| chr8 | 141441409 | 141442559 | 1151 | -5.75 | 7.04E-07 | 1.17E-06 |
| chr1 | 189528963 | 189530809 | 1847 | -5.73 | 7.04E-07 | 1.17E-06 |
| chr22 | 22683536 | 22694068 | 10533 | -2.9 | 7.04E-07 | 1.18E-06 |
| chr3 | 96360782 | 96365851 | 5070 | -2.33 | 7.05E-07 | 1.18E-06 |
| chr1 | 189347748 | 189380102 | 32355 | -6.14 | 7.05E-07 | 1.18E-06 |
| chr8 | 28540258 | 28541109 | 852 | -5.85 | 7.06E-07 | 1.18E-06 |
| chr2 | 61460366 | 61461099 | 734 | -5.72 | 7.07E-07 | 1.18E-06 |
| chr8 | 10015203 | 10015917 | 715 | -5.71 | 7.08E-07 | 1.18E-06 |
| chr2 | 233031392 | 233032273 | 882 | -5.8 | 7.08E-07 | 1.18E-06 |
| chr7 | 27465501 | 27466842 | 1342 | -5.8 | 7.08E-07 | 1.18E-06 |
| chr6 | 138088163 | 138092188 | 4026 | -6.43 | 7.10E-07 | 1.18E-06 |
| chr5 | 62023458 | 62031452 | 7995 | -1.61 | 7.11E-07 | 1.18E-06 |
| chr5 | 134740439 | 134741276 | 838 | -5.84 | 7.11E-07 | 1.19E-06 |
| chr6 | 20281771 | 20285601 | 3831 | -1.57 | 7.11E-07 | 1.19E-06 |
| chr8 | 110516583 | 110517609 | 1027 | 6.12 | 7.11E-07 | 1.19E-06 |
| chr2 | 179546605 | 179547614 | 1010 | 6.06 | 7.12E-07 | 1.19E-06 |
| chr20 | 32095904 | 32098382 | 2479 | -5.77 | 7.13E-07 | 1.19E-06 |
| chr8 | 66943468 | 66944631 | 1164 | -5.77 | 7.13E-07 | 1.19E-06 |
| chr19 | 51551665 | 51579789 | 28125 | -2.37 | 7.14E-07 | 1.19E-06 |
| chr5 | 76876078 | 76902296 | 26219 | -2.61 | 7.16E-07 | 1.19E-06 |
| chr18 | 44803099 | 44803886 | 788 | -4.11 | 7.16E-07 | 1.19E-06 |
| chr13 | 67652300 | 67654181 | 1882 | -5.71 | 7.16E-07 | 1.19E-06 |
| chr22 | 29929947 | 29930688 | 742 | -5.71 | 7.16E-07 | 1.19E-06 |
| chr1 | 232350787 | 232352601 | 1815 | 5.8 | 7.17E-07 | 1.20E-06 |
| chr10 | 74868469 | 74869858 | 1390 | -5.75 | 7.19E-07 | 1.20E-06 |
| chr12 | 697579 | 698304 | 726 | -5.75 | 7.19E-07 | 1.20E-06 |
| chr4 | 163790889 | 163795687 | 4799 | -6.19 | 7.19E-07 | 1.20E-06 |
| chr3 | 32691461 | 32692056 | 596 | -5.71 | 7.21E-07 | 1.20E-06 |
| chr6 | 101027349 | 101028111 | 763 | -5.71 | 7.21E-07 | 1.20E-06 |
| chr13 | 82617693 | 82619151 | 1459 | -5.74 | 7.21E-07 | 1.20E-06 |
| chr3 | 11280501 | 11281282 | 782 | -5.74 | 7.21E-07 | 1.20E-06 |
| chr6 | 55905406 | 55906905 | 1500 | -5.72 | 7.22E-07 | 1.20E-06 |
| chrX | 116897233 | 116898551 | 1319 | 4.69 | 7.22E-07 | 1.20E-06 |
| chr15 | 57343279 | 57344347 | 1069 | -5.82 | 7.23E-07 | 1.20E-06 |
| chr18 | 74302660 | 74303243 | 584 | -5.82 | 7.23E-07 | 1.20E-06 |
| chr1 | 84738590 | 84739480 | 891 | -5.76 | 7.26E-07 | 1.21E-06 |
| chr17 | 36550093 | 36550699 | 607 | -5.76 | 7.26E-07 | 1.21E-06 |
| chr9 | 89066692 | 89068486 | 1795 | -2.46 | 7.28E-07 | 1.21E-06 |
| chr5 | 138567797 | 138573356 | 5560 | -6.56 | 7.30E-07 | 1.22E-06 |
| chr14 | 50677425 | 50678246 | 822 | -5.96 | 7.31E-07 | 1.22E-06 |
| chr16 | 9071013 | 9071738 | 726 | -5.95 | 7.31E-07 | 1.22E-06 |
| chr19 | 17764688 | 17765820 | 1133 | -5.7 | 7.31E-07 | 1.22E-06 |
| chr6 | 89404145 | 89405252 | 1108 | -5.78 | 7.32E-07 | 1.22E-06 |
| chr1 | 24787607 | 24788838 | 1232 | -4.92 | 7.32E-07 | 1.22E-06 |
| chr2 | 18035544 | 18038260 | 2717 | -5.94 | 7.32E-07 | 1.22E-06 |
| chr5 | 129142146 | 129162342 | 20197 | 2.04 | 7.33E-07 | 1.22E-06 |

| | | | | | | |
|---|---|---|---|---|---|---|
| chr14 | 37761053 | 37762760 | 1708 | -5.92 | 7.33E-07 | 1.22E-06 |
| chr3 | 156708370 | 156721807 | 13438 | -6.41 | 7.35E-07 | 1.22E-06 |
| chr3 | 61392473 | 61458778 | 66306 | 1.29 | 7.37E-07 | 1.23E-06 |
| chr10 | 63782594 | 63783794 | 1201 | -5.72 | 7.38E-07 | 1.23E-06 |
| chr5 | 41871277 | 41883293 | 12017 | -6.7 | 7.38E-07 | 1.23E-06 |
| chr18 | 55265250 | 55266121 | 872 | -5.77 | 7.41E-07 | 1.23E-06 |
| chr7 | 155478618 | 155479285 | 668 | -5.77 | 7.41E-07 | 1.23E-06 |
| chr22 | 25211381 | 25212313 | 933 | -5.72 | 7.41E-07 | 1.23E-06 |
| chr18 | 7051425 | 7052169 | 745 | -5.73 | 7.41E-07 | 1.23E-06 |
| chr2 | 192517886 | 192518832 | 947 | -5.73 | 7.41E-07 | 1.23E-06 |
| chrX | 109339753 | 109340505 | 753 | -5.73 | 7.41E-07 | 1.23E-06 |
| chrX | 41402748 | 41403513 | 766 | -5.7 | 7.42E-07 | 1.23E-06 |
| chr2 | 68768744 | 68769467 | 724 | -5.87 | 7.42E-07 | 1.24E-06 |
| chr13 | 90407788 | 90408911 | 1124 | -6.07 | 7.43E-07 | 1.24E-06 |
| chr18 | 71367481 | 71375604 | 8124 | -6.31 | 7.46E-07 | 1.24E-06 |
| chr3 | 175959343 | 175964471 | 5129 | 1.96 | 7.47E-07 | 1.24E-06 |
| chr19 | 46986572 | 46987636 | 1065 | -5.96 | 7.47E-07 | 1.24E-06 |
| chr5 | 78308120 | 78309974 | 1855 | -3 | 7.47E-07 | 1.24E-06 |
| chr10 | 22711994 | 22713639 | 1646 | -5.94 | 7.47E-07 | 1.24E-06 |
| chr14 | 55977361 | 55989797 | 12437 | -5.94 | 7.47E-07 | 1.24E-06 |
| chr10 | 29672084 | 29673188 | 1105 | -5.97 | 7.47E-07 | 1.24E-06 |
| chr1 | 52164608 | 52165484 | 877 | -5.7 | 7.48E-07 | 1.24E-06 |
| chr14 | 59121405 | 59122815 | 1411 | -5.7 | 7.48E-07 | 1.24E-06 |
| chr13 | 105901851 | 105903070 | 1220 | -5.93 | 7.48E-07 | 1.24E-06 |
| chr13 | 35010446 | 35011932 | 1487 | -5.73 | 7.48E-07 | 1.24E-06 |
| chr6 | 106683699 | 106684607 | 909 | -5.73 | 7.48E-07 | 1.24E-06 |
| chr9 | 103116368 | 103117199 | 832 | -5.73 | 7.48E-07 | 1.24E-06 |
| chr20 | 21458517 | 21467503 | 8987 | -1.63 | 7.48E-07 | 1.24E-06 |
| chr2 | 228705223 | 228716212 | 10990 | -5.99 | 7.49E-07 | 1.24E-06 |
| chrX | 13747523 | 13748100 | 578 | -5.99 | 7.49E-07 | 1.24E-06 |
| chr5 | 100626534 | 100627944 | 1411 | -5.92 | 7.49E-07 | 1.24E-06 |
| chr9 | 27024330 | 27025904 | 1575 | -5.92 | 7.49E-07 | 1.24E-06 |
| chr5 | 43218088 | 43219398 | 1311 | -4.41 | 7.49E-07 | 1.25E-06 |
| chr17 | 15327152 | 15330580 | 3429 | -6 | 7.49E-07 | 1.25E-06 |
| chr10 | 98121819 | 98122709 | 891 | -4.86 | 7.50E-07 | 1.25E-06 |
| chr11 | 103724024 | 103728793 | 4770 | -4.92 | 7.50E-07 | 1.25E-06 |
| chr11 | 106603585 | 106863636 | 260052 | 3.85 | 7.50E-07 | 1.25E-06 |
| chr15 | 82502863 | 82503598 | 736 | -5.7 | 7.50E-07 | 1.25E-06 |
| chrX | 19615279 | 19616293 | 1015 | -5.7 | 7.50E-07 | 1.25E-06 |
| chr4 | 159665365 | 159666257 | 893 | -5.76 | 7.51E-07 | 1.25E-06 |
| chr2 | 202983067 | 202985198 | 2132 | -6.02 | 7.51E-07 | 1.25E-06 |
| chr22 | 33810591 | 33818738 | 8148 | -6.02 | 7.51E-07 | 1.25E-06 |
| chr11 | 93646529 | 93665636 | 19108 | -3.25 | 7.52E-07 | 1.25E-06 |
| chrX | 67028461 | 67075384 | 46924 | -2.42 | 7.53E-07 | 1.25E-06 |
| chr19 | 44232293 | 44233249 | 957 | -5.7 | 7.54E-07 | 1.25E-06 |
| chr4 | 99471076 | 99472430 | 1355 | -5.7 | 7.54E-07 | 1.25E-06 |
| chr3 | 81828430 | 81833158 | 4729 | -2.86 | 7.55E-07 | 1.25E-06 |
| chr18 | 36503415 | 36506079 | 2665 | 2.12 | 7.56E-07 | 1.26E-06 |
| chr14 | 53931973 | 53933083 | 1111 | -5.88 | 7.56E-07 | 1.26E-06 |
| chrX | 149459668 | 149460902 | 1235 | -4.3 | 7.56E-07 | 1.26E-06 |
| chr12 | 75988612 | 75989931 | 1320 | -5.74 | 7.56E-07 | 1.26E-06 |
| chr8 | 76388424 | 76391756 | 3333 | -6.13 | 7.56E-07 | 1.26E-06 |
| chr2 | 180622771 | 180624197 | 1427 | -5.78 | 7.58E-07 | 1.26E-06 |
| chr7 | 129865851 | 129866611 | 761 | -5.78 | 7.58E-07 | 1.26E-06 |
| chr14 | 43016620 | 43150991 | 134372 | 1.97 | 7.60E-07 | 1.26E-06 |
| chr13 | 34177367 | 34179077 | 1711 | -5.69 | 7.60E-07 | 1.26E-06 |
| chr1 | 235627484 | 235628669 | 1186 | -5.71 | 7.61E-07 | 1.26E-06 |
| chr1 | 185296944 | 185298111 | 1168 | -5.72 | 7.64E-07 | 1.27E-06 |
| chr3 | 117228151 | 117229323 | 1173 | -5.72 | 7.64E-07 | 1.27E-06 |
| chr22 | 42597068 | 42598329 | 1262 | -5.85 | 7.65E-07 | 1.27E-06 |
| chr9 | 75754623 | 75755753 | 1131 | -5.85 | 7.65E-07 | 1.27E-06 |

| | | | | | | |
|---|---|---|---|---|---|---|
| chr2 | 158921350 | 158945548 | 24199 | -5.72 | 7.66E-07 | 1.27E-06 |
| chr2 | 208473671 | 208474380 | 710 | -5.82 | 7.68E-07 | 1.27E-06 |
| chr8 | 120020449 | 120026094 | 5646 | 1.74 | 7.68E-07 | 1.27E-06 |
| chr9 | 21343099 | 21357947 | 14849 | -1.27 | 7.69E-07 | 1.28E-06 |
| chr3 | 65844026 | 65845282 | 1257 | -5.77 | 7.70E-07 | 1.28E-06 |
| chr3 | 81502942 | 81504085 | 1144 | -5.77 | 7.70E-07 | 1.28E-06 |
| chr8 | 141698335 | 141699836 | 1502 | -6.41 | 7.71E-07 | 1.28E-06 |
| chr10 | 120011431 | 120027213 | 15783 | 2 | 7.73E-07 | 1.28E-06 |
| chr8 | 33309050 | 33310032 | 983 | -5.81 | 7.73E-07 | 1.28E-06 |
| chr1 | 221004581 | 221005688 | 1108 | -5.73 | 7.74E-07 | 1.28E-06 |
| chr11 | 27488307 | 27489173 | 867 | -5.73 | 7.74E-07 | 1.28E-06 |
| chr7 | 40254815 | 40258122 | 3308 | -5.73 | 7.74E-07 | 1.28E-06 |
| chr1 | 171640383 | 171641291 | 909 | -5.75 | 7.74E-07 | 1.28E-06 |
| chr9 | 132547825 | 132548844 | 1020 | -5.75 | 7.74E-07 | 1.28E-06 |
| chr14 | 65093338 | 65094909 | 1572 | -5.83 | 7.74E-07 | 1.28E-06 |
| chr13 | 40494469 | 40501502 | 7034 | -5.6 | 7.77E-07 | 1.29E-06 |
| chr19 | 51318032 | 51404178 | 86147 | -6.44 | 7.77E-07 | 1.29E-06 |
| chr1 | 101014650 | 101037442 | 22793 | -3.09 | 7.78E-07 | 1.29E-06 |
| chr7 | 95467079 | 95470103 | 3025 | -1.9 | 7.79E-07 | 1.29E-06 |
| chr2 | 38856682 | 38857639 | 958 | -5.69 | 7.79E-07 | 1.29E-06 |
| chr13 | 98775174 | 98776823 | 1650 | -5.82 | 7.80E-07 | 1.29E-06 |
| chr6 | 66566412 | 66567737 | 1326 | -4.77 | 7.80E-07 | 1.29E-06 |
| chr10 | 5644768 | 5645621 | 854 | -5.69 | 7.81E-07 | 1.29E-06 |
| chr7 | 91231071 | 91231802 | 732 | 5.77 | 7.81E-07 | 1.30E-06 |
| chr1 | 111262405 | 111263784 | 1380 | 6.06 | 7.81E-07 | 1.30E-06 |
| chr6 | 37233663 | 37234978 | 1316 | -5.71 | 7.82E-07 | 1.30E-06 |
| chr12 | 123908123 | 123909389 | 1267 | -5.69 | 7.83E-07 | 1.30E-06 |
| chr4 | 134026565 | 134028823 | 2259 | -4.53 | 7.85E-07 | 1.30E-06 |
| chr2 | 28140795 | 28141900 | 1106 | -6.26 | 7.86E-07 | 1.30E-06 |
| chr2 | 65019805 | 65020596 | 792 | -5.74 | 7.87E-07 | 1.30E-06 |
| chr4 | 105235041 | 105236781 | 1741 | -5.74 | 7.87E-07 | 1.30E-06 |
| chr8 | 40968687 | 40970186 | 1500 | -5.74 | 7.87E-07 | 1.30E-06 |
| chr10 | 59862204 | 59863506 | 1303 | -5.71 | 7.87E-07 | 1.31E-06 |
| chr16 | 47969870 | 47970998 | 1129 | -4.57 | 7.87E-07 | 1.31E-06 |
| chr2 | 213387009 | 213388714 | 1706 | 5.8 | 7.88E-07 | 1.31E-06 |
| chr10 | 115853147 | 115854578 | 1432 | -5.72 | 7.93E-07 | 1.31E-06 |
| chr3 | 170361617 | 170362944 | 1328 | -5.72 | 7.93E-07 | 1.31E-06 |
| chr4 | 171002603 | 171003468 | 866 | -5.72 | 7.93E-07 | 1.31E-06 |
| chr9 | 36404375 | 36405309 | 935 | -5.72 | 7.93E-07 | 1.31E-06 |
| chr1 | 234239522 | 234242869 | 3348 | -6.21 | 7.93E-07 | 1.32E-06 |
| chr15 | 70503184 | 70503985 | 802 | -5.8 | 7.94E-07 | 1.32E-06 |
| chr10 | 101587876 | 101588602 | 727 | -5.69 | 7.94E-07 | 1.32E-06 |
| chr11 | 45170787 | 45171608 | 822 | -5.74 | 7.97E-07 | 1.32E-06 |
| chr8 | 117097013 | 117100064 | 3052 | -5.78 | 8.00E-07 | 1.33E-06 |
| chr3 | 109906857 | 109926971 | 20115 | -7.24 | 8.00E-07 | 1.33E-06 |
| chr12 | 116965202 | 116966215 | 1014 | -5.73 | 8.01E-07 | 1.33E-06 |
| chr21 | 18147191 | 18148369 | 1179 | -5.73 | 8.01E-07 | 1.33E-06 |
| chr1 | 32006068 | 32009680 | 3613 | -5.12 | 8.04E-07 | 1.33E-06 |
| chr4 | 162496697 | 162498151 | 1455 | -4.56 | 8.08E-07 | 1.34E-06 |
| chr6 | 63393723 | 63395014 | 1292 | -5.68 | 8.10E-07 | 1.34E-06 |
| chrX | 101215232 | 101218237 | 3006 | 2.53 | 8.11E-07 | 1.34E-06 |
| chr2 | 157857755 | 157866143 | 8389 | -4.92 | 8.11E-07 | 1.34E-06 |
| chr4 | 25289541 | 25301257 | 11717 | -4.98 | 8.11E-07 | 1.34E-06 |
| chr7 | 47508434 | 47509671 | 1238 | -5.7 | 8.13E-07 | 1.35E-06 |
| chr8 | 67804835 | 67806110 | 1276 | -5.7 | 8.13E-07 | 1.35E-06 |
| chr10 | 15225837 | 15226880 | 1044 | -5.73 | 8.13E-07 | 1.35E-06 |
| chr4 | 92246988 | 92248753 | 1766 | -5.71 | 8.15E-07 | 1.35E-06 |
| chr8 | 8694995 | 8695892 | 898 | -5.71 | 8.15E-07 | 1.35E-06 |
| chr8 | 91002261 | 91003277 | 1017 | -5.71 | 8.15E-07 | 1.35E-06 |
| chr15 | 80582963 | 80662381 | 79419 | -1.68 | 8.16E-07 | 1.35E-06 |
| chr1 | 189637968 | 189638760 | 793 | -5.72 | 8.16E-07 | 1.35E-06 |

| | | | | | | |
|---|---|---|---|---|---|---|
| chr17 | 63667600 | 63668779 | 1180 | -5.72 | 8.16E-07 | 1.35E-06 |
| chr15 | 50671775 | 50672960 | 1186 | -6.3 | 8.16E-07 | 1.35E-06 |
| chr12 | 105851264 | 105874611 | 23348 | -5.93 | 8.19E-07 | 1.36E-06 |
| chr10 | 25029634 | 25030602 | 969 | -5.96 | 8.19E-07 | 1.36E-06 |
| chr7 | 100725022 | 100726476 | 1455 | -5.68 | 8.19E-07 | 1.36E-06 |
| chr1 | 246017297 | 246018423 | 1127 | -5.68 | 8.20E-07 | 1.36E-06 |
| chr12 | 32542588 | 32543825 | 1238 | -5.91 | 8.20E-07 | 1.36E-06 |
| chr14 | 74119659 | 74120599 | 941 | -5.93 | 8.20E-07 | 1.36E-06 |
| chr10 | 90057539 | 90059058 | 1520 | -5.95 | 8.21E-07 | 1.36E-06 |
| chr20 | 32388736 | 32389714 | 979 | -5.9 | 8.21E-07 | 1.36E-06 |
| chr3 | 182493257 | 182494805 | 1549 | -5.98 | 8.21E-07 | 1.36E-06 |
| chr4 | 184010216 | 184011541 | 1326 | -5.77 | 8.21E-07 | 1.36E-06 |
| chr4 | 120728937 | 120730040 | 1104 | -4.34 | 8.22E-07 | 1.36E-06 |
| chr8 | 56715030 | 56715598 | 569 | -5.91 | 8.22E-07 | 1.36E-06 |
| chr2 | 222918864 | 222919643 | 780 | -5.98 | 8.22E-07 | 1.36E-06 |
| chr12 | 87363007 | 87364437 | 1431 | -5.99 | 8.22E-07 | 1.36E-06 |
| chr1 | 81059369 | 81060793 | 1425 | -5.89 | 8.22E-07 | 1.36E-06 |
| chr13 | 45141837 | 45143129 | 1293 | -5.9 | 8.24E-07 | 1.36E-06 |
| chr3 | 192572162 | 192573106 | 945 | -5.88 | 8.24E-07 | 1.36E-06 |
| chr7 | 101430950 | 101432203 | 1254 | -5.88 | 8.24E-07 | 1.36E-06 |
| chr12 | 19221149 | 19222663 | 1515 | -6.02 | 8.26E-07 | 1.37E-06 |
| chr1 | 61900911 | 61901875 | 965 | -5.87 | 8.26E-07 | 1.37E-06 |
| chr11 | 63468404 | 63469388 | 985 | -5.68 | 8.27E-07 | 1.37E-06 |
| chr9 | 103331221 | 103332379 | 1159 | -5.68 | 8.27E-07 | 1.37E-06 |
| chr11 | 111875769 | 111877059 | 1291 | -6.03 | 8.28E-07 | 1.37E-06 |
| chrX | 123334850 | 123341189 | 6340 | -1.67 | 8.30E-07 | 1.37E-06 |
| chr1 | 210932051 | 210937681 | 5631 | -2.94 | 8.30E-07 | 1.37E-06 |
| chr10 | 95318311 | 95319853 | 1543 | -5.72 | 8.31E-07 | 1.37E-06 |
| chr17 | 55493606 | 55494465 | 860 | -5.72 | 8.31E-07 | 1.37E-06 |
| chr5 | 142126891 | 142128251 | 1361 | -5.72 | 8.31E-07 | 1.37E-06 |
| chr6 | 112141750 | 112142448 | 699 | -4.29 | 8.32E-07 | 1.38E-06 |
| chr8 | 99846954 | 99848754 | 1801 | -5.71 | 8.33E-07 | 1.38E-06 |
| chr4 | 75370116 | 75371333 | 1218 | -5.76 | 8.33E-07 | 1.38E-06 |
| chr12 | 12719944 | 12720538 | 595 | -5.86 | 8.33E-07 | 1.38E-06 |
| chr2 | 85590077 | 85591008 | 932 | -5.86 | 8.33E-07 | 1.38E-06 |
| chr2 | 194578353 | 194580374 | 2022 | -2.82 | 8.34E-07 | 1.38E-06 |
| chr5 | 142530444 | 142531045 | 602 | -5.85 | 8.35E-07 | 1.38E-06 |
| chr12 | 107960821 | 107961817 | 997 | -5.74 | 8.38E-07 | 1.39E-06 |
| chr18 | 19886193 | 19887504 | 1312 | -5.74 | 8.38E-07 | 1.39E-06 |
| chr3 | 70453709 | 70483560 | 29852 | -1.68 | 8.39E-07 | 1.39E-06 |
| chr1 | 22121119 | 22122379 | 1261 | -5.83 | 8.40E-07 | 1.39E-06 |
| chr1 | 85966415 | 85980904 | 14490 | -5.7 | 8.40E-07 | 1.39E-06 |
| chrX | 46708550 | 46709767 | 1218 | -5.7 | 8.40E-07 | 1.39E-06 |
| chr6 | 46675047 | 46676622 | 1576 | -4.28 | 8.40E-07 | 1.39E-06 |
| chrX | 116332345 | 116381726 | 49382 | 2.38 | 8.41E-07 | 1.39E-06 |
| chr8 | 142990942 | 142997850 | 6909 | -5.56 | 8.42E-07 | 1.39E-06 |
| chr4 | 118836035 | 118842929 | 6895 | 2.12 | 8.42E-07 | 1.39E-06 |
| chr1 | 67061584 | 67063493 | 1910 | -5.69 | 8.43E-07 | 1.39E-06 |
| chr17 | 75807383 | 75810438 | 3056 | 1.81 | 8.44E-07 | 1.39E-06 |
| chr7 | 109293474 | 109294234 | 761 | -5.82 | 8.46E-07 | 1.40E-06 |
| chr10 | 90893593 | 90894277 | 685 | -5.74 | 8.50E-07 | 1.41E-06 |
| chr10 | 120991075 | 120992439 | 1365 | -5.74 | 8.50E-07 | 1.41E-06 |
| chr3 | 120190900 | 120191917 | 1018 | -5.71 | 8.51E-07 | 1.41E-06 |
| chrX | 41790577 | 41792311 | 1735 | -5.7 | 8.51E-07 | 1.41E-06 |
| chr2 | 144636287 | 144645862 | 9576 | 2.3 | 8.52E-07 | 1.41E-06 |
| chr16 | 78372798 | 78373762 | 965 | -5.81 | 8.55E-07 | 1.41E-06 |
| chr18 | 3996779 | 3998137 | 1359 | -5.81 | 8.55E-07 | 1.41E-06 |
| chr7 | 31685525 | 31686809 | 1285 | 5.79 | 8.56E-07 | 1.41E-06 |
| chr2 | 168828070 | 168830046 | 1977 | -5.8 | 8.56E-07 | 1.41E-06 |
| chr12 | 22277389 | 22283299 | 5911 | -1.97 | 8.59E-07 | 1.42E-06 |
| chr6 | 80477493 | 80479827 | 2335 | -2.71 | 8.59E-07 | 1.42E-06 |

| | | | | | | |
|---|---|---|---|---|---|---|
| chr9 | 76142205 | 76142986 | 782 | -5.8 | 8.60E-07 | 1.42E-06 |
| chr5 | 57288208 | 57289955 | 1748 | -5.74 | 8.60E-07 | 1.42E-06 |
| chr7 | 27516490 | 27533342 | 16853 | -6.55 | 8.62E-07 | 1.42E-06 |
| chr1 | 186761802 | 186764418 | 2617 | -6.17 | 8.62E-07 | 1.42E-06 |
| chr15 | 23000402 | 23001104 | 703 | -5.73 | 8.63E-07 | 1.43E-06 |
| chr3 | 108916505 | 108918796 | 2292 | -5.67 | 8.64E-07 | 1.43E-06 |
| chr10 | 14741299 | 14742227 | 929 | -5.69 | 8.67E-07 | 1.43E-06 |
| chr17 | 38353762 | 38354324 | 563 | -5.69 | 8.67E-07 | 1.43E-06 |
| chr14 | 32908848 | 32909760 | 913 | -5.68 | 8.70E-07 | 1.44E-06 |
| chr1 | 227435165 | 227436057 | 893 | -5.69 | 8.71E-07 | 1.44E-06 |
| chr18 | 77460204 | 77461277 | 1074 | -5.69 | 8.71E-07 | 1.44E-06 |
| chr4 | 156316396 | 156318399 | 2004 | -5.69 | 8.71E-07 | 1.44E-06 |
| chr3 | 56515282 | 56516722 | 1441 | 6.15 | 8.71E-07 | 1.44E-06 |
| chr4 | 135044508 | 135045917 | 1410 | 6 | 8.71E-07 | 1.44E-06 |
| chr13 | 76628662 | 76643860 | 15199 | -3.36 | 8.72E-07 | 1.44E-06 |
| chr21 | 39373170 | 39392308 | 19139 | -5.49 | 8.73E-07 | 1.44E-06 |
| chr2 | 209476480 | 209477904 | 1425 | -4.69 | 8.75E-07 | 1.44E-06 |
| chr7 | 114916253 | 114917348 | 1096 | -6.23 | 8.76E-07 | 1.45E-06 |
| chrX | 35806805 | 35807850 | 1046 | -5.67 | 8.76E-07 | 1.45E-06 |
| chr15 | 82346542 | 82347628 | 1087 | -5.73 | 8.76E-07 | 1.45E-06 |
| chr6 | 2574824 | 2575708 | 885 | -5.67 | 8.77E-07 | 1.45E-06 |
| chr7 | 36704238 | 36705010 | 773 | -5.67 | 8.77E-07 | 1.45E-06 |
| chr10 | 101762610 | 101763232 | 623 | -5.72 | 8.78E-07 | 1.45E-06 |
| chr7 | 110050398 | 110051823 | 1426 | -6.23 | 8.78E-07 | 1.45E-06 |
| chrX | 75083932 | 75150342 | 66411 | 3.36 | 8.81E-07 | 1.45E-06 |
| chr3 | 65171620 | 65172788 | 1169 | -2.2 | 8.83E-07 | 1.46E-06 |
| chr2 | 218282428 | 218290473 | 8046 | -4.7 | 8.84E-07 | 1.46E-06 |
| chr7 | 35513695 | 35518843 | 5149 | -5.38 | 8.85E-07 | 1.46E-06 |
| chr8 | 47481705 | 47852639 | 370935 | -2.35 | 8.88E-07 | 1.46E-06 |
| chrX | 89439320 | 89440750 | 1431 | 3.11 | 8.88E-07 | 1.47E-06 |
| chr10 | 100239116 | 100258183 | 19068 | -1.79 | 8.90E-07 | 1.47E-06 |
| chr3 | 179338301 | 179338801 | 501 | -5.69 | 8.92E-07 | 1.47E-06 |
| chr4 | 110936689 | 110938474 | 1786 | -5.71 | 8.93E-07 | 1.47E-06 |
| chr2 | 171248065 | 171249909 | 1845 | -5.71 | 8.94E-07 | 1.48E-06 |
| chr8 | 119417007 | 119417871 | 865 | -5.71 | 8.94E-07 | 1.48E-06 |
| chr3 | 111517761 | 111518691 | 931 | -5.68 | 8.96E-07 | 1.48E-06 |
| chr12 | 55287851 | 55322151 | 34301 | 1.4 | 8.97E-07 | 1.48E-06 |
| chr16 | 9247444 | 9248186 | 743 | -5.69 | 8.98E-07 | 1.48E-06 |
| chr6 | 55161656 | 55162385 | 730 | -5.69 | 8.98E-07 | 1.48E-06 |
| chr10 | 4509145 | 4510170 | 1026 | -5.67 | 8.98E-07 | 1.48E-06 |
| chr18 | 21386351 | 21387383 | 1033 | -5.67 | 8.98E-07 | 1.48E-06 |
| chr2 | 146349087 | 146350041 | 955 | -5.67 | 8.98E-07 | 1.48E-06 |
| chr3 | 157872792 | 157873757 | 966 | -5.93 | 8.98E-07 | 1.48E-06 |
| chr4 | 177434639 | 177436464 | 1826 | -5.96 | 9.00E-07 | 1.48E-06 |
| chr8 | 126592775 | 126593955 | 1181 | -5.97 | 9.01E-07 | 1.48E-06 |
| chr10 | 70627527 | 70628514 | 988 | -5.87 | 9.03E-07 | 1.49E-06 |
| chr2 | 198224116 | 198225305 | 1190 | -5.87 | 9.03E-07 | 1.49E-06 |
| chr6 | 148127388 | 148128724 | 1337 | -5.75 | 9.03E-07 | 1.49E-06 |
| chr21 | 10872845 | 10895833 | 22989 | 1.4 | 9.04E-07 | 1.49E-06 |
| chr2 | 193924670 | 193926669 | 2000 | -6 | 9.04E-07 | 1.49E-06 |
| chr22 | 26679313 | 26679828 | 516 | -5.75 | 9.04E-07 | 1.49E-06 |
| chr11 | 70572907 | 70574248 | 1342 | -5.86 | 9.05E-07 | 1.49E-06 |
| chr11 | 32968623 | 32969824 | 1202 | -5.66 | 9.05E-07 | 1.49E-06 |
| chr4 | 60012548 | 60013774 | 1227 | -4.53 | 9.07E-07 | 1.49E-06 |
| chr12 | 54175998 | 54176813 | 816 | -5.7 | 9.10E-07 | 1.50E-06 |
| chrX | 152896404 | 152897311 | 908 | -5.84 | 9.10E-07 | 1.50E-06 |
| chr8 | 62736703 | 62748634 | 11932 | -6.37 | 9.11E-07 | 1.50E-06 |
| chr11 | 85455096 | 85456271 | 1176 | -6.39 | 9.12E-07 | 1.50E-06 |
| chr2 | 194652042 | 194653622 | 1581 | -6.04 | 9.12E-07 | 1.50E-06 |
| chr7 | 14484811 | 14488123 | 3313 | -6.04 | 9.12E-07 | 1.50E-06 |
| chr10 | 73714197 | 73715922 | 1726 | -5.83 | 9.13E-07 | 1.50E-06 |

| | | | | | | |
|---|---|---|---|---|---|---|
| chr7 | 47920188 | 47921545 | 1358 | -5.83 | 9.13E-07 | 1.50E-06 |
| chr7 | 102662882 | 102664478 | 1597 | -5.83 | 9.13E-07 | 1.50E-06 |
| chr16 | 84639031 | 84644611 | 5581 | -5.74 | 9.14E-07 | 1.51E-06 |
| chr9 | 115109578 | 115110705 | 1128 | -6.05 | 9.14E-07 | 1.51E-06 |
| chr1 | 188757069 | 188758133 | 1065 | -5.7 | 9.14E-07 | 1.51E-06 |
| chr15 | 90240964 | 90241666 | 703 | -5.7 | 9.14E-07 | 1.51E-06 |
| chr18 | 43691203 | 43692169 | 967 | -4.92 | 9.15E-07 | 1.51E-06 |
| chr2 | 189302039 | 189302705 | 667 | -5.66 | 9.15E-07 | 1.51E-06 |
| chr4 | 4467498 | 4468919 | 1422 | -5.68 | 9.16E-07 | 1.51E-06 |
| chr2 | 225322014 | 225323781 | 1768 | -6.06 | 9.16E-07 | 1.51E-06 |
| chr6 | 41156395 | 41158592 | 2198 | -5.02 | 9.18E-07 | 1.51E-06 |
| chr1 | 53511281 | 53512459 | 1179 | -5.81 | 9.22E-07 | 1.52E-06 |
| chr10 | 112208178 | 112209254 | 1077 | -5.81 | 9.22E-07 | 1.52E-06 |
| chr3 | 182157934 | 182158609 | 676 | -5.88 | 9.25E-07 | 1.52E-06 |
| chr12 | 49839613 | 49840711 | 1099 | -5.68 | 9.25E-07 | 1.52E-06 |
| chr12 | 101456851 | 101458824 | 1974 | -5.73 | 9.25E-07 | 1.52E-06 |
| chr12 | 124384113 | 124385280 | 1168 | -5.73 | 9.25E-07 | 1.52E-06 |
| chr2 | 181897117 | 181897761 | 645 | -5.73 | 9.25E-07 | 1.52E-06 |
| chr13 | 74322040 | 74322991 | 952 | -4.28 | 9.28E-07 | 1.53E-06 |
| chr5 | 120786067 | 120787478 | 1412 | -5.66 | 9.28E-07 | 1.53E-06 |
| chr9 | 96689617 | 96702649 | 13033 | -6.8 | 9.28E-07 | 1.53E-06 |
| chr19 | 16742202 | 16743247 | 1046 | -5.86 | 9.28E-07 | 1.53E-06 |
| chr1 | 207009251 | 207056933 | 47683 | -2.96 | 9.28E-07 | 1.53E-06 |
| chr12 | 70117828 | 70119165 | 1338 | -5.69 | 9.28E-07 | 1.53E-06 |
| chr17 | 73576307 | 73577599 | 1293 | -5.67 | 9.29E-07 | 1.53E-06 |
| chr13 | 60731329 | 60732254 | 926 | -5.85 | 9.30E-07 | 1.53E-06 |
| chr14 | 105810662 | 105812072 | 1411 | -5.85 | 9.30E-07 | 1.53E-06 |
| chr4 | 116004452 | 116007029 | 2578 | -5.85 | 9.30E-07 | 1.53E-06 |
| chr14 | 103740717 | 103741731 | 1015 | -5.73 | 9.30E-07 | 1.53E-06 |
| chr1 | 226645093 | 226657917 | 12825 | -2.16 | 9.32E-07 | 1.53E-06 |
| chr4 | 146704964 | 146705945 | 982 | -5.65 | 9.33E-07 | 1.53E-06 |
| chr14 | 68295118 | 68296262 | 1145 | -6.12 | 9.34E-07 | 1.54E-06 |
| chr6 | 112201501 | 112202499 | 999 | -5.79 | 9.34E-07 | 1.54E-06 |
| chr15 | 69441899 | 69443637 | 1739 | 4.92 | 9.34E-07 | 1.54E-06 |
| chr6 | 79589161 | 79590996 | 1836 | -5.84 | 9.35E-07 | 1.54E-06 |
| chr7 | 14735454 | 14744280 | 8827 | -6.04 | 9.36E-07 | 1.54E-06 |
| chr5 | 120286715 | 120289714 | 3000 | -1.99 | 9.36E-07 | 1.54E-06 |
| chr10 | 62162244 | 62174085 | 11842 | -6.13 | 9.36E-07 | 1.54E-06 |
| chr7 | 73850514 | 73867032 | 16519 | -7.04 | 9.36E-07 | 1.54E-06 |
| chr20 | 39356618 | 39378279 | 21662 | 5.3 | 9.37E-07 | 1.54E-06 |
| chr3 | 78879150 | 78880796 | 1647 | -5.69 | 9.37E-07 | 1.54E-06 |
| chr6 | 141023299 | 141025073 | 1775 | -5.72 | 9.37E-07 | 1.54E-06 |
| chr4 | 57128411 | 57144111 | 15701 | -6.05 | 9.39E-07 | 1.54E-06 |
| chr11 | 11247484 | 11249551 | 2068 | -2.47 | 9.39E-07 | 1.54E-06 |
| chr16 | 21590561 | 21591283 | 723 | -5.78 | 9.41E-07 | 1.55E-06 |
| chr3 | 31044740 | 31050052 | 5313 | -5.11 | 9.42E-07 | 1.55E-06 |
| chr12 | 85249403 | 85252789 | 3387 | 1.9 | 9.42E-07 | 1.55E-06 |
| chr1 | 150172669 | 150173823 | 1155 | -5.72 | 9.47E-07 | 1.56E-06 |
| chr2 | 120477947 | 120479249 | 1303 | -5.72 | 9.47E-07 | 1.56E-06 |
| chr9 | 121427933 | 121430184 | 2252 | -5.72 | 9.47E-07 | 1.56E-06 |
| chr15 | 63707086 | 63719583 | 12498 | -7.38 | 9.47E-07 | 1.56E-06 |
| chr3 | 63910462 | 63911244 | 783 | -5.68 | 9.48E-07 | 1.56E-06 |
| chr21 | 33672934 | 33673671 | 738 | -5.01 | 9.49E-07 | 1.56E-06 |
| chr1 | 215061553 | 215062241 | 689 | -5.77 | 9.49E-07 | 1.56E-06 |
| chr2 | 147128804 | 147131410 | 2607 | -5.77 | 9.49E-07 | 1.56E-06 |
| chr7 | 12596317 | 12597850 | 1534 | -5.65 | 9.51E-07 | 1.56E-06 |
| chr4 | 141971315 | 141974700 | 3386 | -1.83 | 9.52E-07 | 1.56E-06 |
| chr4 | 79266028 | 79267138 | 1111 | -4.6 | 9.54E-07 | 1.57E-06 |
| chr11 | 89341625 | 89347497 | 5873 | 5.57 | 9.56E-07 | 1.57E-06 |
| chr3 | 177287476 | 177289482 | 2007 | -5.65 | 9.57E-07 | 1.57E-06 |
| chr14 | 35637067 | 35637958 | 892 | -6.13 | 9.58E-07 | 1.57E-06 |

| | | | | | | |
|---|---|---|---|---|---|---|
| chr3 | 17733349 | 17734752 | 1404 | -5.76 | 9.59E-07 | 1.57E-06 |
| chr12 | 117262273 | 117263314 | 1042 | -6.14 | 9.62E-07 | 1.58E-06 |
| chr10 | 87651309 | 87652284 | 976 | -5.68 | 9.62E-07 | 1.58E-06 |
| chr2 | 171584823 | 171586903 | 2081 | -2.17 | 9.62E-07 | 1.58E-06 |
| chr12 | 46052983 | 46057640 | 4658 | -5.17 | 9.64E-07 | 1.58E-06 |
| chr15 | 44144764 | 44146247 | 1484 | -5.71 | 9.65E-07 | 1.58E-06 |
| chr9 | 29322306 | 29323235 | 930 | -5.71 | 9.65E-07 | 1.58E-06 |
| chr4 | 20332628 | 20334088 | 1461 | 5.84 | 9.67E-07 | 1.59E-06 |
| chr9 | 12923137 | 12924543 | 1407 | 5.84 | 9.67E-07 | 1.59E-06 |
| chr11 | 1559485 | 1560202 | 718 | -5.65 | 9.67E-07 | 1.59E-06 |
| chr13 | 106051402 | 106052568 | 1167 | -5.65 | 9.67E-07 | 1.59E-06 |
| chr19 | 47857178 | 47859156 | 1979 | -5.75 | 9.69E-07 | 1.59E-06 |
| chr2 | 230213636 | 230215196 | 1561 | -5.75 | 9.69E-07 | 1.59E-06 |
| chr12 | 116807708 | 116808924 | 1217 | -5.64 | 9.72E-07 | 1.60E-06 |
| chr12 | 44394707 | 44396116 | 1410 | 6.25 | 9.72E-07 | 1.60E-06 |
| chr17 | 39456815 | 39458519 | 1705 | -6.29 | 9.77E-07 | 1.60E-06 |
| chr16 | 70438649 | 70439622 | 974 | -5.76 | 9.78E-07 | 1.61E-06 |
| chr4 | 94803037 | 94804455 | 1419 | 2.96 | 9.78E-07 | 1.61E-06 |
| chr7 | 2099570 | 2100671 | 1102 | -5.7 | 9.81E-07 | 1.61E-06 |
| chr13 | 31591735 | 31592575 | 841 | -5.69 | 9.85E-07 | 1.62E-06 |
| chr19 | 46952579 | 46953437 | 859 | -5.69 | 9.85E-07 | 1.62E-06 |
| chrX | 44246589 | 44248147 | 1559 | -5.69 | 9.85E-07 | 1.62E-06 |
| chr14 | 71961921 | 71963352 | 1432 | -5.91 | 9.86E-07 | 1.62E-06 |
| chr9 | 124284528 | 124285273 | 746 | -5.91 | 9.86E-07 | 1.62E-06 |
| chr1 | 31188144 | 31189115 | 972 | -5.93 | 9.87E-07 | 1.62E-06 |
| chr1 | 56638309 | 56639176 | 868 | -5.93 | 9.87E-07 | 1.62E-06 |
| chr2 | 145299879 | 145301457 | 1579 | -5.89 | 9.87E-07 | 1.62E-06 |
| chr13 | 114575711 | 114576791 | 1081 | -5.94 | 9.87E-07 | 1.62E-06 |
| chr2 | 20294547 | 20295303 | 757 | -5.88 | 9.88E-07 | 1.62E-06 |
| chr2 | 236914348 | 236915248 | 901 | -5.88 | 9.88E-07 | 1.62E-06 |
| chr4 | 158793686 | 158794598 | 913 | 5.82 | 9.88E-07 | 1.62E-06 |
| chr4 | 159486439 | 159487538 | 1100 | -5.66 | 9.88E-07 | 1.62E-06 |
| chr20 | 62718039 | 62719554 | 1516 | -5.96 | 9.89E-07 | 1.62E-06 |
| chr10 | 133778258 | 133779306 | 1049 | -5.67 | 9.89E-07 | 1.62E-06 |
| chr7 | 32681148 | 32682410 | 1263 | -5.67 | 9.89E-07 | 1.62E-06 |
| chr18 | 10586027 | 10587760 | 1734 | -5.98 | 9.92E-07 | 1.63E-06 |
| chr14 | 91306570 | 91307606 | 1037 | -5.73 | 9.93E-07 | 1.63E-06 |
| chr14 | 78355144 | 78356722 | 1579 | -5.85 | 9.94E-07 | 1.63E-06 |
| chr5 | 93663360 | 93717585 | 54226 | 1.68 | 9.95E-07 | 1.63E-06 |
| chr4 | 4234690 | 4235628 | 939 | -5.64 | 9.96E-07 | 1.63E-06 |
| chr9 | 16962882 | 16964414 | 1533 | -5.64 | 9.96E-07 | 1.63E-06 |
| chr13 | 31972008 | 31991114 | 19107 | -1.57 | 9.97E-07 | 1.64E-06 |
| chr6 | 135755602 | 135756255 | 654 | -5.69 | 9.98E-07 | 1.64E-06 |
| chr15 | 78626704 | 78628102 | 1399 | -3.39 | 9.98E-07 | 1.64E-06 |
| chr5 | 148567710 | 148568621 | 912 | -6.02 | 9.98E-07 | 1.64E-06 |
| chr5 | 85765608 | 85766706 | 1099 | 5.73 | 1.00E-06 | 1.64E-06 |
| chrX | 85263143 | 85264982 | 1840 | -5.64 | 1.00E-06 | 1.64E-06 |
| chr2 | 145062900 | 145088072 | 25173 | -7.05 | 1.00E-06 | 1.64E-06 |
| chr8 | 120592212 | 120594630 | 2419 | -4.68 | 1.01E-06 | 1.65E-06 |
| chr2 | 205726769 | 205727413 | 645 | -5.68 | 1.01E-06 | 1.65E-06 |
| chr6 | 135074503 | 135075734 | 1232 | -5.72 | 1.01E-06 | 1.65E-06 |
| chrX | 18763348 | 18764234 | 887 | -5.72 | 1.01E-06 | 1.65E-06 |
| chr10 | 105416834 | 105417785 | 952 | -5.81 | 1.01E-06 | 1.65E-06 |
| chr15 | 34584876 | 34586149 | 1274 | -5.81 | 1.01E-06 | 1.65E-06 |
| chrX | 67345883 | 67346683 | 801 | -5.81 | 1.01E-06 | 1.65E-06 |
| chr6 | 152852537 | 152853454 | 918 | -5.64 | 1.01E-06 | 1.65E-06 |
| chr5 | 134110917 | 134111917 | 1001 | -5.73 | 1.01E-06 | 1.66E-06 |
| chr12 | 78712015 | 78713987 | 1973 | -5.8 | 1.01E-06 | 1.66E-06 |
| chr2 | 85966681 | 85975903 | 9223 | -6.49 | 1.02E-06 | 1.67E-06 |
| chr6 | 152415929 | 152417489 | 1561 | -5.66 | 1.02E-06 | 1.67E-06 |
| chr7 | 66058572 | 66060296 | 1725 | -5.78 | 1.02E-06 | 1.67E-06 |

| | | | | | | |
|---|---|---|---|---|---|---|
| chr7 | 66152725 | 66153846 | 1122 | -5.66 | 1.02E-06 | 1.67E-06 |
| chr6 | 14303194 | 14304192 | 999 | -5.71 | 1.02E-06 | 1.68E-06 |
| chr9 | 97282638 | 97283531 | 894 | -5.77 | 1.03E-06 | 1.68E-06 |
| chr11 | 87987778 | 87998193 | 10416 | -5.7 | 1.03E-06 | 1.68E-06 |
| chr2 | 105309881 | 105315684 | 5804 | 2.54 | 1.03E-06 | 1.69E-06 |
| chr3 | 177640720 | 177641525 | 806 | -5.64 | 1.03E-06 | 1.69E-06 |
| chrX | 36489335 | 36490543 | 1209 | -5.67 | 1.03E-06 | 1.69E-06 |
| chr13 | 43993577 | 43995351 | 1775 | -4.6 | 1.03E-06 | 1.69E-06 |
| chr4 | 89281358 | 89282384 | 1027 | -5.67 | 1.04E-06 | 1.70E-06 |
| chr6 | 127934579 | 127938270 | 3692 | -6.17 | 1.04E-06 | 1.70E-06 |
| chr11 | 93311655 | 93312823 | 1169 | -5.87 | 1.04E-06 | 1.70E-06 |
| chr10 | 30482140 | 30502713 | 20574 | -2.46 | 1.04E-06 | 1.70E-06 |
| chr14 | 90415045 | 90417095 | 2051 | -2.57 | 1.04E-06 | 1.70E-06 |
| chr8 | 107129774 | 107130419 | 646 | -4.41 | 1.04E-06 | 1.71E-06 |
| chr1 | 231913290 | 231914055 | 766 | -5.75 | 1.04E-06 | 1.71E-06 |
| chr8 | 82742951 | 82744646 | 1696 | -5.99 | 1.04E-06 | 1.71E-06 |
| chr4 | 127538202 | 127541194 | 2993 | -6.19 | 1.05E-06 | 1.71E-06 |
| chr4 | 156340134 | 156344709 | 4576 | -2.23 | 1.05E-06 | 1.71E-06 |
| chr4 | 21894472 | 21896897 | 2426 | 4.49 | 1.05E-06 | 1.72E-06 |
| chr7 | 12786516 | 12787754 | 1239 | -5.65 | 1.05E-06 | 1.72E-06 |
| chr22 | 20881894 | 20882996 | 1103 | -5.7 | 1.05E-06 | 1.72E-06 |
| chr11 | 4237201 | 4238839 | 1639 | 4.86 | 1.05E-06 | 1.72E-06 |
| chr1 | 186081821 | 186082432 | 612 | -5.63 | 1.05E-06 | 1.72E-06 |
| chr2 | 147453817 | 147462396 | 8580 | -2.8 | 1.05E-06 | 1.73E-06 |
| chr12 | 32372093 | 32372838 | 746 | -5.81 | 1.05E-06 | 1.73E-06 |
| chr2 | 9757305 | 9757985 | 681 | -5.81 | 1.05E-06 | 1.73E-06 |
| chr4 | 93584865 | 93586600 | 1736 | -5.81 | 1.05E-06 | 1.73E-06 |
| chr20 | 5502383 | 5504153 | 1771 | -5.03 | 1.05E-06 | 1.73E-06 |
| chr21 | 30725988 | 30727461 | 1474 | -5.65 | 1.05E-06 | 1.73E-06 |
| chr3 | 50345151 | 50345867 | 717 | -5.65 | 1.05E-06 | 1.73E-06 |
| chr7 | 133252209 | 133252850 | 642 | -5.65 | 1.05E-06 | 1.73E-06 |
| chr13 | 83011708 | 83025542 | 13835 | -6.23 | 1.06E-06 | 1.73E-06 |
| chr18 | 8836686 | 8837730 | 1045 | -5.63 | 1.06E-06 | 1.73E-06 |
| chr4 | 83649848 | 83650989 | 1142 | -5.63 | 1.06E-06 | 1.73E-06 |
| chrX | 68635567 | 68636470 | 904 | -5.63 | 1.06E-06 | 1.73E-06 |
| chr3 | 175329964 | 175332062 | 2099 | -5.69 | 1.06E-06 | 1.74E-06 |
| chr9 | 119883577 | 119884648 | 1072 | -5.7 | 1.06E-06 | 1.74E-06 |
| chr17 | 67505634 | 67506696 | 1063 | -5.63 | 1.07E-06 | 1.75E-06 |
| chr9 | 213349 | 214441 | 1093 | -5.63 | 1.07E-06 | 1.75E-06 |
| chr2 | 146100219 | 146101446 | 1228 | -5.78 | 1.07E-06 | 1.75E-06 |
| chr2 | 180166716 | 180167199 | 484 | -4.49 | 1.08E-06 | 1.76E-06 |
| chr1 | 168947804 | 168948736 | 933 | -5.62 | 1.08E-06 | 1.76E-06 |
| chr15 | 59328386 | 59329031 | 646 | -5.62 | 1.08E-06 | 1.76E-06 |
| chr4 | 64566321 | 64568012 | 1692 | -5.62 | 1.08E-06 | 1.76E-06 |
| chr2 | 24860292 | 24861839 | 1548 | -5.72 | 1.08E-06 | 1.76E-06 |
| chr7 | 20393243 | 20394082 | 840 | -5.72 | 1.08E-06 | 1.76E-06 |
| chr12 | 52507323 | 52508466 | 1144 | -5.64 | 1.08E-06 | 1.76E-06 |
| chr6 | 100942377 | 100944425 | 2049 | -5.64 | 1.08E-06 | 1.76E-06 |
| chr11 | 102049531 | 102051048 | 1518 | -6.13 | 1.08E-06 | 1.77E-06 |
| chr12 | 75779294 | 75780027 | 734 | -5.65 | 1.08E-06 | 1.77E-06 |
| chr4 | 162743385 | 162745019 | 1635 | -5.9 | 1.08E-06 | 1.77E-06 |
| chr6 | 135947382 | 135948491 | 1110 | -5.9 | 1.08E-06 | 1.77E-06 |
| chr1 | 189681807 | 189682888 | 1082 | -5.89 | 1.08E-06 | 1.77E-06 |
| chrX | 62934717 | 62935956 | 1240 | -5.89 | 1.08E-06 | 1.77E-06 |
| chr11 | 44528447 | 44529653 | 1207 | -5.92 | 1.08E-06 | 1.77E-06 |
| chr8 | 82991642 | 82993386 | 1745 | -5.92 | 1.08E-06 | 1.77E-06 |
| chr5 | 65148630 | 65149908 | 1279 | -5.87 | 1.09E-06 | 1.77E-06 |
| chr10 | 135134040 | 135135319 | 1280 | -5.67 | 1.09E-06 | 1.77E-06 |
| chr17 | 64713534 | 64714937 | 1404 | -5.94 | 1.09E-06 | 1.77E-06 |
| chrX | 10022126 | 10023481 | 1356 | -5.94 | 1.09E-06 | 1.78E-06 |
| chr2 | 182411067 | 182412365 | 1299 | -5.95 | 1.09E-06 | 1.78E-06 |

| | | | | | | |
|---|---|---|---|---|---|---|
| chr5 | 121109272 | 121111551 | 2280 | 3.64 | 1.09E-06 | 1.78E-06 |
| chr7 | 45907594 | 45919379 | 11786 | -5.84 | 1.09E-06 | 1.78E-06 |
| chr12 | 86197980 | 86199671 | 1692 | -3.1 | 1.09E-06 | 1.78E-06 |
| chr22 | 32920171 | 32921093 | 923 | -5.64 | 1.09E-06 | 1.78E-06 |
| chr5 | 80905420 | 80906402 | 983 | -5.64 | 1.09E-06 | 1.78E-06 |
| chr13 | 110539651 | 110562094 | 22444 | -6.73 | 1.10E-06 | 1.79E-06 |
| chr15 | 57850880 | 57851536 | 657 | -5.75 | 1.10E-06 | 1.79E-06 |
| chr13 | 105876412 | 105877826 | 1415 | -5.8 | 1.10E-06 | 1.80E-06 |
| chr2 | 27231404 | 27232057 | 654 | -5.8 | 1.10E-06 | 1.80E-06 |
| chr4 | 71956043 | 71965243 | 9201 | -6.65 | 1.11E-06 | 1.81E-06 |
| chr2 | 240236296 | 240237882 | 1587 | -5.62 | 1.11E-06 | 1.81E-06 |
| chr6 | 85395595 | 85396620 | 1026 | -5.62 | 1.11E-06 | 1.81E-06 |
| chrX | 28491531 | 28493305 | 1775 | -6.04 | 1.11E-06 | 1.81E-06 |
| chr2 | 193642495 | 193644334 | 1840 | -5.62 | 1.11E-06 | 1.81E-06 |
| chr11 | 101863675 | 101864365 | 691 | -5.64 | 1.11E-06 | 1.81E-06 |
| chr16 | 48154003 | 48159996 | 5994 | -5.64 | 1.11E-06 | 1.81E-06 |
| chr4 | 54331268 | 54333074 | 1807 | -3.67 | 1.11E-06 | 1.81E-06 |
| chr18 | 34672420 | 34674162 | 1743 | -5.7 | 1.11E-06 | 1.81E-06 |
| chr20 | 10205287 | 10213416 | 8130 | -1.96 | 1.11E-06 | 1.81E-06 |
| chr20 | 58199622 | 58240540 | 40919 | -2.33 | 1.11E-06 | 1.81E-06 |
| chr4 | 60260264 | 60261492 | 1229 | -3.49 | 1.11E-06 | 1.82E-06 |
| chr1 | 184865525 | 184869288 | 3764 | 2.13 | 1.11E-06 | 1.82E-06 |
| chr11 | 32604088 | 32605036 | 949 | -5.62 | 1.12E-06 | 1.82E-06 |
| chr19 | 39933917 | 39934480 | 564 | -5.62 | 1.12E-06 | 1.82E-06 |
| chr20 | 57446165 | 57447199 | 1035 | -5.62 | 1.12E-06 | 1.82E-06 |
| chr8 | 80904066 | 80904966 | 901 | -5.67 | 1.12E-06 | 1.82E-06 |
| chr11 | 94740631 | 94742659 | 2029 | -5.06 | 1.12E-06 | 1.82E-06 |
| chrX | 123466190 | 123467236 | 1047 | 5.71 | 1.12E-06 | 1.82E-06 |
| chr4 | 37520569 | 37523223 | 2655 | -2.03 | 1.12E-06 | 1.82E-06 |
| chrX | 88987840 | 88997355 | 9516 | 2.23 | 1.12E-06 | 1.82E-06 |
| chr18 | 72158948 | 72160044 | 1097 | -5.77 | 1.12E-06 | 1.83E-06 |
| chr4 | 179783706 | 179785104 | 1399 | -5.77 | 1.12E-06 | 1.83E-06 |
| chr12 | 49819667 | 49820368 | 702 | -5.61 | 1.12E-06 | 1.84E-06 |
| chr4 | 164088503 | 164089221 | 719 | -5.61 | 1.12E-06 | 1.84E-06 |
| chr6 | 20061427 | 20079617 | 18191 | -1.71 | 1.13E-06 | 1.84E-06 |
| chr6 | 141238165 | 141239522 | 1358 | -5.64 | 1.13E-06 | 1.84E-06 |
| chr1 | 160142772 | 160143743 | 972 | 5.77 | 1.13E-06 | 1.84E-06 |
| chr6 | 146589127 | 146590312 | 1186 | 5.77 | 1.13E-06 | 1.84E-06 |
| chr6 | 35202626 | 35204122 | 1497 | -5.69 | 1.13E-06 | 1.84E-06 |
| chr8 | 108186009 | 108189082 | 3074 | -6.13 | 1.13E-06 | 1.84E-06 |
| chr8 | 98347200 | 98435195 | 87996 | 2.4 | 1.13E-06 | 1.85E-06 |
| chr3 | 13114360 | 13115912 | 1553 | -6.14 | 1.13E-06 | 1.85E-06 |
| chr5 | 92249113 | 92250264 | 1152 | -5.62 | 1.13E-06 | 1.85E-06 |
| chr7 | 104962139 | 104962941 | 803 | -5.62 | 1.13E-06 | 1.85E-06 |
| chrX | 42658623 | 42659509 | 887 | -5.62 | 1.13E-06 | 1.85E-06 |
| chr2 | 219834665 | 219835477 | 813 | -5.71 | 1.14E-06 | 1.85E-06 |
| chr6 | 49010890 | 49012212 | 1323 | -5.71 | 1.14E-06 | 1.85E-06 |
| chr8 | 6403798 | 6404947 | 1150 | -5.71 | 1.14E-06 | 1.85E-06 |
| chr11 | 15991373 | 15992893 | 1521 | -5.64 | 1.14E-06 | 1.85E-06 |
| chr12 | 42529211 | 42529855 | 645 | -5.64 | 1.14E-06 | 1.85E-06 |
| chr22 | 35982334 | 35983596 | 1263 | -5.64 | 1.14E-06 | 1.85E-06 |
| chr10 | 51051603 | 51052476 | 874 | -5.74 | 1.14E-06 | 1.86E-06 |
| chr20 | 42945437 | 42946530 | 1094 | -6.16 | 1.14E-06 | 1.86E-06 |
| chr4 | 16086127 | 16087209 | 1083 | -5.62 | 1.15E-06 | 1.87E-06 |
| chr12 | 39109906 | 39110844 | 939 | -5.68 | 1.15E-06 | 1.87E-06 |
| chr3 | 11414939 | 11415968 | 1030 | -5.68 | 1.15E-06 | 1.87E-06 |
| chr3 | 33203911 | 33204669 | 759 | -5.68 | 1.15E-06 | 1.87E-06 |
| chr3 | 40962943 | 40964458 | 1516 | -5.68 | 1.15E-06 | 1.87E-06 |
| chr4 | 123672108 | 123673055 | 948 | -5.68 | 1.15E-06 | 1.87E-06 |
| chr12 | 120223215 | 120224410 | 1196 | -5.61 | 1.15E-06 | 1.87E-06 |
| chr4 | 27858560 | 27859665 | 1106 | -2.6 | 1.15E-06 | 1.88E-06 |

| | | | | | | |
|---|---|---|---|---|---|---|
| chr17 | 53698076 | 53787846 | 89771 | 1.25 | 1.15E-06 | 1.88E-06 |
| chr1 | 15570694 | 15572679 | 1986 | -6.22 | 1.16E-06 | 1.88E-06 |
| chr11 | 4218398 | 4219069 | 672 | -5.65 | 1.16E-06 | 1.89E-06 |
| chrX | 54626995 | 54627776 | 782 | -5.65 | 1.16E-06 | 1.89E-06 |
| chrX | 128513576 | 128514321 | 746 | -5.65 | 1.16E-06 | 1.89E-06 |
| chr1 | 230025250 | 230047084 | 21835 | -0.99 | 1.16E-06 | 1.90E-06 |
| chr7 | 80914638 | 80915425 | 788 | -5.63 | 1.17E-06 | 1.90E-06 |
| chr2 | 131887530 | 131888079 | 550 | -5.87 | 1.17E-06 | 1.90E-06 |
| chr6 | 212660 | 214834 | 2175 | -2.34 | 1.17E-06 | 1.90E-06 |
| chr1 | 113517125 | 113518223 | 1099 | -5.61 | 1.17E-06 | 1.90E-06 |
| chr12 | 80633567 | 80634119 | 553 | -5.63 | 1.17E-06 | 1.90E-06 |
| chr3 | 152114871 | 152115532 | 662 | -5.63 | 1.17E-06 | 1.90E-06 |
| chr6 | 150740756 | 150741985 | 1230 | -5.63 | 1.17E-06 | 1.90E-06 |
| chr6 | 25823264 | 25824145 | 882 | -5.86 | 1.17E-06 | 1.90E-06 |
| chr7 | 99351245 | 99353448 | 2204 | -5.85 | 1.17E-06 | 1.91E-06 |
| chr6 | 67845069 | 67847485 | 2417 | -5.64 | 1.17E-06 | 1.91E-06 |
| chr1 | 199426920 | 199428050 | 1131 | -5.67 | 1.17E-06 | 1.91E-06 |
| chr11 | 124510409 | 124511259 | 851 | -5.67 | 1.17E-06 | 1.91E-06 |
| chr14 | 74311580 | 74313489 | 1910 | -5.67 | 1.17E-06 | 1.91E-06 |
| chr16 | 84562781 | 84574258 | 11478 | -5.67 | 1.17E-06 | 1.91E-06 |
| chr18 | 61194289 | 61196062 | 1774 | -4.49 | 1.17E-06 | 1.91E-06 |
| chr5 | 135504572 | 135505637 | 1066 | -5.71 | 1.17E-06 | 1.91E-06 |
| chr2 | 160127712 | 160128168 | 457 | -5.83 | 1.18E-06 | 1.91E-06 |
| chr2 | 193208746 | 193209893 | 1148 | -5.61 | 1.18E-06 | 1.91E-06 |
| chr20 | 48039649 | 48040993 | 1345 | -5.61 | 1.18E-06 | 1.91E-06 |
| chr6 | 126341048 | 126342146 | 1099 | -5.61 | 1.18E-06 | 1.91E-06 |
| chr9 | 136564651 | 136565281 | 631 | -5.68 | 1.18E-06 | 1.92E-06 |
| chr5 | 132985855 | 132987393 | 1539 | -4.44 | 1.18E-06 | 1.92E-06 |
| chr13 | 105602952 | 105604798 | 1847 | -4.74 | 1.18E-06 | 1.92E-06 |
| chr5 | 156008545 | 156009821 | 1277 | -4.88 | 1.18E-06 | 1.92E-06 |
| chr13 | 67443699 | 67444658 | 960 | -5.61 | 1.18E-06 | 1.92E-06 |
| chr12 | 70376707 | 70387825 | 11119 | -6 | 1.18E-06 | 1.92E-06 |
| chr3 | 27872165 | 27878005 | 5841 | -5.33 | 1.18E-06 | 1.92E-06 |
| chr8 | 105182288 | 105186593 | 4306 | 2.73 | 1.19E-06 | 1.93E-06 |
| chr20 | 30995791 | 30996663 | 873 | -5.61 | 1.19E-06 | 1.93E-06 |
| chr4 | 144413831 | 144414818 | 988 | -5.61 | 1.19E-06 | 1.93E-06 |
| chr20 | 9602364 | 9611106 | 8743 | -6.02 | 1.19E-06 | 1.93E-06 |
| chr3 | 115839793 | 115845494 | 5702 | 1.76 | 1.19E-06 | 1.94E-06 |
| chrX | 4862929 | 4864826 | 1898 | -5.88 | 1.19E-06 | 1.94E-06 |
| chr6 | 167765790 | 167766900 | 1111 | -5.9 | 1.19E-06 | 1.94E-06 |
| chr1 | 205745288 | 205746109 | 822 | -5.68 | 1.19E-06 | 1.94E-06 |
| chr21 | 32983376 | 32984445 | 1070 | -5.68 | 1.19E-06 | 1.94E-06 |
| chr14 | 35893379 | 35894143 | 765 | -5.93 | 1.19E-06 | 1.94E-06 |
| chr4 | 49060389 | 49064066 | 3678 | 1.98 | 1.20E-06 | 1.95E-06 |
| chr2 | 29409343 | 29410965 | 1623 | -4.97 | 1.20E-06 | 1.95E-06 |
| chrX | 54676013 | 54724234 | 48222 | -2.73 | 1.20E-06 | 1.95E-06 |
| chr13 | 19450209 | 19458758 | 8550 | 1.81 | 1.20E-06 | 1.95E-06 |
| chr1 | 204729774 | 204731852 | 2079 | -5.82 | 1.20E-06 | 1.95E-06 |
| chr1 | 236939231 | 236945696 | 6466 | -3.49 | 1.20E-06 | 1.95E-06 |
| chr1 | 215023109 | 215024392 | 1284 | -2.53 | 1.20E-06 | 1.95E-06 |
| chr8 | 27117749 | 27119640 | 1892 | -4.65 | 1.20E-06 | 1.95E-06 |
| chr4 | 30255939 | 30290361 | 34423 | -1.36 | 1.20E-06 | 1.95E-06 |
| chrX | 69101762 | 69106218 | 4457 | 2.21 | 1.20E-06 | 1.95E-06 |
| chr8 | 23318951 | 23320964 | 2014 | -5.97 | 1.20E-06 | 1.95E-06 |
| chr2 | 121513619 | 121514413 | 795 | -5.69 | 1.21E-06 | 1.96E-06 |
| chr16 | 15974869 | 15977264 | 2396 | -5.63 | 1.21E-06 | 1.96E-06 |
| chr4 | 59938670 | 59940243 | 1574 | -5.63 | 1.21E-06 | 1.96E-06 |
| chr1 | 221065044 | 221074591 | 9548 | -6.52 | 1.21E-06 | 1.96E-06 |
| chr5 | 176094105 | 176095231 | 1127 | -5.63 | 1.21E-06 | 1.97E-06 |
| chr15 | 62332078 | 62332578 | 501 | -5.67 | 1.21E-06 | 1.97E-06 |
| chr7 | 25135260 | 25136039 | 780 | -5.67 | 1.21E-06 | 1.97E-06 |

| | | | | | | |
|---|---|---|---|---|---|---|
| chr15 | 75160135 | 75161058 | 924 | -5.75 | 1.21E-06 | 1.97E-06 |
| chr21 | 18419380 | 18421344 | 1965 | -2.48 | 1.21E-06 | 1.97E-06 |
| chr13 | 78017526 | 78020674 | 3149 | -2.63 | 1.21E-06 | 1.97E-06 |
| chr13 | 95563773 | 95568974 | 5202 | -1.63 | 1.21E-06 | 1.97E-06 |
| chr1 | 5640492 | 5646170 | 5679 | 2.32 | 1.22E-06 | 1.98E-06 |
| chr10 | 31515916 | 31542324 | 26409 | -6.07 | 1.22E-06 | 1.98E-06 |
| chr15 | 30188693 | 30204837 | 16145 | -6.92 | 1.22E-06 | 1.98E-06 |
| chrX | 34997999 | 35001835 | 3837 | -5.08 | 1.22E-06 | 1.98E-06 |
| chr8 | 125243618 | 125244715 | 1098 | -5.77 | 1.22E-06 | 1.99E-06 |
| chrX | 58148539 | 58283530 | 134992 | 0.75 | 1.22E-06 | 1.99E-06 |
| chr7 | 36613995 | 36621873 | 7879 | -5.84 | 1.22E-06 | 1.99E-06 |
| chr22 | 33938234 | 33939723 | 1490 | -5.64 | 1.23E-06 | 1.99E-06 |
| chr2 | 48180809 | 48302346 | 121538 | -1.97 | 1.23E-06 | 1.99E-06 |
| chr10 | 123869679 | 123870724 | 1046 | -6.06 | 1.23E-06 | 1.99E-06 |
| chrX | 38062669 | 38063640 | 972 | -5.66 | 1.23E-06 | 2.00E-06 |
| chr6 | 147989645 | 147991755 | 2111 | -3.23 | 1.23E-06 | 2.00E-06 |
| chr1 | 222390835 | 222403994 | 13160 | -1.41 | 1.23E-06 | 2.00E-06 |
| chr5 | 125960706 | 125961925 | 1220 | -5.6 | 1.23E-06 | 2.00E-06 |
| chr2 | 232899766 | 232900928 | 1163 | -5.6 | 1.23E-06 | 2.00E-06 |
| chr1 | 211393853 | 211395518 | 1666 | -4.89 | 1.23E-06 | 2.00E-06 |
| chr3 | 66658611 | 66659738 | 1128 | -5.6 | 1.24E-06 | 2.01E-06 |
| chr6 | 492129 | 492839 | 711 | -5.6 | 1.24E-06 | 2.01E-06 |
| chrX | 150969098 | 150971165 | 2068 | -5.75 | 1.24E-06 | 2.01E-06 |
| chr5 | 13789876 | 13791318 | 1443 | -6.1 | 1.24E-06 | 2.01E-06 |
| chr2 | 25070245 | 25071877 | 1633 | -5.62 | 1.24E-06 | 2.01E-06 |
| chr6 | 122236415 | 122237729 | 1315 | 5.76 | 1.24E-06 | 2.01E-06 |
| chr12 | 130916819 | 130919300 | 2482 | -6.12 | 1.24E-06 | 2.02E-06 |
| chr20 | 31789242 | 31791210 | 1969 | -4.67 | 1.24E-06 | 2.02E-06 |
| chr2 | 38267545 | 38268411 | 867 | -5.67 | 1.24E-06 | 2.02E-06 |
| chr13 | 90993121 | 90994094 | 974 | -5.59 | 1.25E-06 | 2.03E-06 |
| chr1 | 67648155 | 67648929 | 775 | -5.65 | 1.25E-06 | 2.03E-06 |
| chr6 | 52605565 | 52606655 | 1091 | -6.15 | 1.25E-06 | 2.03E-06 |
| chr8 | 99821650 | 99822321 | 672 | -5.71 | 1.25E-06 | 2.04E-06 |
| chr8 | 49696423 | 49697418 | 996 | -4.24 | 1.26E-06 | 2.04E-06 |
| chr22 | 25872956 | 25873808 | 853 | -5.63 | 1.26E-06 | 2.04E-06 |
| chr3 | 183504351 | 183505353 | 1003 | -5.63 | 1.26E-06 | 2.04E-06 |
| chr9 | 20888748 | 20890877 | 2130 | -4.65 | 1.26E-06 | 2.04E-06 |
| chr7 | 85171829 | 85173036 | 1208 | -6.29 | 1.26E-06 | 2.04E-06 |
| chrX | 49472913 | 49482049 | 9137 | 4.66 | 1.26E-06 | 2.04E-06 |
| chr4 | 78324544 | 78345922 | 21379 | 1.27 | 1.26E-06 | 2.05E-06 |
| chr9 | 4401255 | 4408158 | 6904 | -5.74 | 1.26E-06 | 2.05E-06 |
| chr2 | 23922387 | 23923446 | 1060 | -5.7 | 1.27E-06 | 2.05E-06 |
| chr7 | 32404977 | 32417561 | 12585 | -2.09 | 1.27E-06 | 2.05E-06 |
| chr1 | 155390358 | 155399062 | 8705 | -5.66 | 1.27E-06 | 2.06E-06 |
| chr18 | 74795486 | 74796088 | 603 | -5.61 | 1.27E-06 | 2.06E-06 |
| chr18 | 12577532 | 12578720 | 1189 | -5.64 | 1.27E-06 | 2.06E-06 |
| chr18 | 52591714 | 52595122 | 3409 | -3.46 | 1.27E-06 | 2.06E-06 |
| chr13 | 32324019 | 32393514 | 69496 | 1.06 | 1.28E-06 | 2.07E-06 |
| chr7 | 127571272 | 127572113 | 842 | -5.6 | 1.28E-06 | 2.07E-06 |
| chr7 | 91544782 | 91545982 | 1201 | -5.59 | 1.28E-06 | 2.08E-06 |
| chr7 | 93603178 | 93603922 | 745 | -5.59 | 1.28E-06 | 2.08E-06 |
| chr2 | 75213142 | 75214177 | 1036 | -4.46 | 1.28E-06 | 2.08E-06 |
| chr20 | 14235733 | 14236703 | 971 | -5.7 | 1.28E-06 | 2.08E-06 |
| chr7 | 95328770 | 95343769 | 15000 | -1.13 | 1.29E-06 | 2.09E-06 |
| chr4 | 1636306 | 1637315 | 1010 | -5.61 | 1.29E-06 | 2.09E-06 |
| chr3 | 143176691 | 143177394 | 704 | -4.18 | 1.29E-06 | 2.09E-06 |
| chr8 | 142023453 | 142024197 | 745 | -5.62 | 1.29E-06 | 2.10E-06 |
| chr6 | 122957681 | 122958336 | 656 | -5.65 | 1.29E-06 | 2.10E-06 |
| chr13 | 74535426 | 74536369 | 944 | -5.59 | 1.30E-06 | 2.10E-06 |
| chr17 | 1031854 | 1032604 | 751 | -5.69 | 1.30E-06 | 2.10E-06 |
| chr2 | 180829709 | 180830392 | 684 | -5.63 | 1.30E-06 | 2.10E-06 |

| | | | | | | |
|---|---|---|---|---|---|---|
| chr1 | 242530702 | 242535853 | 5152 | -6.32 | 1.30E-06 | 2.10E-06 |
| chr22 | 48254195 | 48269382 | 15188 | -3.29 | 1.30E-06 | 2.10E-06 |
| chr10 | 7081110 | 7124107 | 42998 | -1.02 | 1.30E-06 | 2.11E-06 |
| chr4 | 134318347 | 134321150 | 2804 | 4.67 | 1.30E-06 | 2.11E-06 |
| chr4 | 110951952 | 110952843 | 892 | -5.61 | 1.30E-06 | 2.12E-06 |
| chr20 | 8574170 | 8575704 | 1535 | -5.59 | 1.31E-06 | 2.12E-06 |
| chr1 | 113746399 | 113758867 | 12469 | -6.57 | 1.31E-06 | 2.13E-06 |
| chr1 | 205792080 | 205793376 | 1297 | -5.89 | 1.31E-06 | 2.13E-06 |
| chr22 | 38409518 | 38410684 | 1167 | -5.89 | 1.31E-06 | 2.13E-06 |
| chr4 | 79492608 | 79497254 | 4647 | -5.65 | 1.31E-06 | 2.13E-06 |
| chr11 | 134930342 | 134945789 | 15448 | -2.13 | 1.32E-06 | 2.13E-06 |
| chr3 | 30000420 | 30001922 | 1503 | -5.82 | 1.32E-06 | 2.13E-06 |
| chr8 | 99602330 | 99603469 | 1140 | -5.9 | 1.32E-06 | 2.13E-06 |
| chr1 | 36302036 | 36302845 | 810 | -5.84 | 1.32E-06 | 2.14E-06 |
| chr7 | 115599908 | 115600886 | 979 | -5.91 | 1.32E-06 | 2.14E-06 |
| chrX | 118203044 | 118203859 | 816 | -5.91 | 1.32E-06 | 2.14E-06 |
| chr2 | 73244393 | 73245327 | 935 | -5.93 | 1.32E-06 | 2.14E-06 |
| chr14 | 35771777 | 35772516 | 740 | -5.66 | 1.32E-06 | 2.14E-06 |
| chr2 | 219354397 | 219355926 | 1530 | -5.93 | 1.32E-06 | 2.14E-06 |
| chr8 | 62993376 | 62994527 | 1152 | -5.63 | 1.32E-06 | 2.14E-06 |
| chr13 | 102129927 | 102130948 | 1022 | -5.59 | 1.32E-06 | 2.14E-06 |
| chr19 | 4197179 | 4197841 | 663 | -5.62 | 1.32E-06 | 2.14E-06 |
| chr2 | 178446401 | 178447132 | 732 | -5.62 | 1.32E-06 | 2.14E-06 |
| chr2 | 193355869 | 193357317 | 1449 | -5.62 | 1.32E-06 | 2.14E-06 |
| chr9 | 15980332 | 15981345 | 1014 | -5.81 | 1.33E-06 | 2.15E-06 |
| chr4 | 154145527 | 154146760 | 1234 | -5.98 | 1.33E-06 | 2.15E-06 |
| chr2 | 215569215 | 215570388 | 1174 | -5.78 | 1.33E-06 | 2.15E-06 |
| chr9 | 110282457 | 110283449 | 993 | -5.78 | 1.33E-06 | 2.15E-06 |
| chr6 | 17642784 | 17644118 | 1335 | -5.98 | 1.33E-06 | 2.15E-06 |
| chrX | 11127275 | 11128390 | 1116 | -5.79 | 1.33E-06 | 2.15E-06 |
| chr1 | 193912423 | 193913655 | 1233 | -5.99 | 1.33E-06 | 2.16E-06 |
| chr5 | 107862195 | 107863552 | 1358 | -2.58 | 1.33E-06 | 2.16E-06 |
| chr2 | 120213138 | 120214401 | 1264 | -5.61 | 1.33E-06 | 2.16E-06 |
| chr15 | 66354148 | 66355215 | 1068 | -5.77 | 1.33E-06 | 2.16E-06 |
| chr19 | 4540471 | 4541149 | 679 | -5.77 | 1.33E-06 | 2.16E-06 |
| chr3 | 155439216 | 155440282 | 1067 | -5.77 | 1.33E-06 | 2.16E-06 |
| chr1 | 233784751 | 233785505 | 755 | -5.59 | 1.34E-06 | 2.16E-06 |
| chr4 | 103627695 | 103641198 | 13504 | -5.59 | 1.34E-06 | 2.16E-06 |
| chr4 | 176106211 | 176107125 | 915 | -5.59 | 1.34E-06 | 2.16E-06 |
| chr18 | 56624180 | 56626982 | 2803 | -5.66 | 1.34E-06 | 2.17E-06 |
| chr5 | 61387110 | 61511941 | 124832 | -1.39 | 1.34E-06 | 2.17E-06 |
| chr3 | 101689150 | 101691004 | 1855 | -5.76 | 1.34E-06 | 2.17E-06 |
| chr20 | 13915385 | 13939543 | 24159 | -4.87 | 1.34E-06 | 2.17E-06 |
| chr19 | 41276385 | 41277208 | 824 | -5.21 | 1.34E-06 | 2.17E-06 |
| chr1 | 154676297 | 154677221 | 925 | 5.65 | 1.34E-06 | 2.17E-06 |
| chr12 | 105979707 | 105980933 | 1227 | -5.75 | 1.35E-06 | 2.18E-06 |
| chr9 | 96813354 | 96814156 | 803 | -5.75 | 1.35E-06 | 2.18E-06 |
| chr2 | 19281779 | 19283318 | 1540 | -5.76 | 1.35E-06 | 2.18E-06 |
| chr1 | 56175275 | 56177564 | 2290 | -6.06 | 1.35E-06 | 2.18E-06 |
| chr12 | 17166226 | 17168561 | 2336 | -6.06 | 1.35E-06 | 2.18E-06 |
| chr14 | 50186930 | 50188066 | 1137 | -5.66 | 1.35E-06 | 2.18E-06 |
| chr21 | 17048864 | 17050207 | 1344 | -5.66 | 1.35E-06 | 2.18E-06 |
| chr4 | 2957428 | 2958684 | 1257 | -4.26 | 1.35E-06 | 2.19E-06 |
| chr4 | 113865666 | 113866432 | 767 | -5.75 | 1.35E-06 | 2.19E-06 |
| chr7 | 128013786 | 128015011 | 1226 | -5.75 | 1.35E-06 | 2.19E-06 |
| chr5 | 32167511 | 32168225 | 715 | -5.62 | 1.35E-06 | 2.19E-06 |
| chr8 | 40925469 | 40926633 | 1165 | -5.62 | 1.35E-06 | 2.19E-06 |
| chr5 | 112664692 | 112665637 | 946 | -5.65 | 1.36E-06 | 2.20E-06 |
| chr2 | 158448240 | 158449638 | 1399 | -6.08 | 1.36E-06 | 2.20E-06 |
| chr4 | 108273945 | 108275210 | 1266 | -4.82 | 1.36E-06 | 2.20E-06 |
| chr15 | 46033525 | 46034548 | 1024 | -5.58 | 1.36E-06 | 2.20E-06 |

| | | | | | | |
|---|---|---|---|---|---|---|
| chr6 | 112814541 | 112816180 | 1640 | 4.39 | 1.36E-06 | 2.21E-06 |
| chr1 | 235368947 | 235369520 | 574 | -6.12 | 1.37E-06 | 2.21E-06 |
| chr7 | 19445900 | 19447788 | 1889 | -5.72 | 1.37E-06 | 2.22E-06 |
| chr3 | 145988976 | 145989696 | 721 | -5.65 | 1.37E-06 | 2.22E-06 |
| chr8 | 77931902 | 77997908 | 66007 | 6.36 | 1.37E-06 | 2.22E-06 |
| chr13 | 51455892 | 51471491 | 15600 | -1.35 | 1.37E-06 | 2.22E-06 |
| chr12 | 110729901 | 110731003 | 1103 | -5.6 | 1.38E-06 | 2.23E-06 |
| chr7 | 43811893 | 43812717 | 825 | -5.6 | 1.38E-06 | 2.23E-06 |
| chr1 | 198887992 | 198889786 | 1795 | -5.64 | 1.38E-06 | 2.23E-06 |
| chr20 | 58692304 | 58704692 | 12389 | 2.5 | 1.38E-06 | 2.23E-06 |
| chr6 | 47642212 | 47645835 | 3624 | -3.06 | 1.38E-06 | 2.23E-06 |
| chr22 | 44842143 | 44843141 | 999 | -5.59 | 1.38E-06 | 2.24E-06 |
| chr5 | 137452024 | 137452667 | 644 | -5.71 | 1.39E-06 | 2.24E-06 |
| chr2 | 164272244 | 164272936 | 693 | -5.6 | 1.39E-06 | 2.24E-06 |
| chr5 | 55065224 | 55066149 | 926 | -5.6 | 1.39E-06 | 2.24E-06 |
| chr11 | 100571269 | 100572216 | 948 | -4.4 | 1.39E-06 | 2.24E-06 |
| chr8 | 73080108 | 73102554 | 22447 | -1.49 | 1.39E-06 | 2.25E-06 |
| chr10 | 33027540 | 33028921 | 1382 | -5.57 | 1.39E-06 | 2.25E-06 |
| chr15 | 22754909 | 22755415 | 507 | -5.57 | 1.39E-06 | 2.25E-06 |
| chr22 | 36981894 | 36983033 | 1140 | -5.58 | 1.39E-06 | 2.25E-06 |
| chr2 | 157079572 | 157101773 | 22202 | -2.46 | 1.40E-06 | 2.26E-06 |
| chr18 | 28503127 | 28504871 | 1745 | -4.61 | 1.40E-06 | 2.26E-06 |
| chr1 | 174833138 | 174834356 | 1219 | -5.64 | 1.40E-06 | 2.26E-06 |
| chr5 | 143259890 | 143262214 | 2325 | -5.64 | 1.40E-06 | 2.26E-06 |
| chrX | 15653640 | 15654504 | 865 | -5.64 | 1.40E-06 | 2.26E-06 |
| chr2 | 183933374 | 183934877 | 1504 | -5.7 | 1.40E-06 | 2.26E-06 |
| chr7 | 127080674 | 127102805 | 22132 | -2.23 | 1.40E-06 | 2.26E-06 |
| chr9 | 116825999 | 116828257 | 2259 | -6.28 | 1.41E-06 | 2.28E-06 |
| chrX | 13390197 | 13391900 | 1704 | -5.05 | 1.41E-06 | 2.28E-06 |
| chr6 | 160381560 | 160382433 | 874 | -5.68 | 1.41E-06 | 2.28E-06 |
| chr12 | 88308362 | 88309655 | 1294 | -5.57 | 1.43E-06 | 2.30E-06 |
| chr3 | 11638978 | 11639822 | 845 | -5.57 | 1.43E-06 | 2.30E-06 |
| chr3 | 190490128 | 190491045 | 918 | -5.57 | 1.43E-06 | 2.30E-06 |
| chr3 | 101788711 | 101790820 | 2110 | -6.36 | 1.43E-06 | 2.30E-06 |
| chr8 | 17895807 | 17896565 | 759 | -5.58 | 1.43E-06 | 2.30E-06 |
| chr5 | 6671985 | 6698437 | 26453 | -5.76 | 1.43E-06 | 2.30E-06 |
| chr8 | 42560428 | 42561282 | 855 | -4.41 | 1.43E-06 | 2.31E-06 |
| chr11 | 73326822 | 73327529 | 708 | -5.62 | 1.43E-06 | 2.31E-06 |
| chr2 | 54979598 | 54980271 | 674 | -5.62 | 1.43E-06 | 2.31E-06 |
| chr11 | 68412341 | 68416296 | 3956 | -5.75 | 1.43E-06 | 2.31E-06 |
| chr11 | 36503783 | 36505079 | 1297 | -5.67 | 1.43E-06 | 2.31E-06 |
| chr6 | 137471772 | 137479882 | 8111 | -2.37 | 1.43E-06 | 2.31E-06 |
| chr6 | 153200719 | 153204244 | 3526 | -2.08 | 1.43E-06 | 2.31E-06 |
| chr1 | 232140170 | 232143417 | 3248 | -2.19 | 1.43E-06 | 2.31E-06 |
| chr2 | 38504057 | 38504965 | 909 | -4.47 | 1.43E-06 | 2.31E-06 |
| chrX | 70174533 | 70187792 | 13260 | -5.03 | 1.44E-06 | 2.32E-06 |
| chr13 | 25110634 | 25111153 | 520 | -5.57 | 1.44E-06 | 2.32E-06 |
| chr3 | 119455157 | 119455938 | 782 | -5.57 | 1.44E-06 | 2.32E-06 |
| chr11 | 118602406 | 118603212 | 807 | -4.42 | 1.44E-06 | 2.33E-06 |
| chr1 | 201890785 | 201891802 | 1018 | -5.85 | 1.45E-06 | 2.33E-06 |
| chr22 | 29590228 | 29591139 | 912 | -5.85 | 1.45E-06 | 2.33E-06 |
| chr1 | 178589231 | 178590251 | 1021 | -5.86 | 1.45E-06 | 2.33E-06 |
| chr14 | 52323164 | 52324425 | 1262 | -5.84 | 1.45E-06 | 2.33E-06 |
| chr4 | 48221945 | 48223689 | 1745 | -5.84 | 1.45E-06 | 2.33E-06 |
| chr6 | 22165259 | 22166233 | 975 | -5.66 | 1.45E-06 | 2.34E-06 |
| chr14 | 56694687 | 56695618 | 932 | -5.57 | 1.45E-06 | 2.34E-06 |
| chr9 | 80982729 | 80983765 | 1037 | -5.57 | 1.45E-06 | 2.34E-06 |
| chr3 | 138418201 | 138418730 | 530 | -5.61 | 1.45E-06 | 2.35E-06 |
| chr3 | 132485431 | 132747414 | 261984 | 8.84 | 1.46E-06 | 2.35E-06 |
| chr5 | 95495880 | 95497286 | 1407 | -5.78 | 1.46E-06 | 2.35E-06 |
| chrX | 63602785 | 63605463 | 2679 | -2.56 | 1.46E-06 | 2.35E-06 |

| | | | | | | |
|---|---|---|---|---|---|---|
| chr22 | 37061120 | 37062055 | 936 | -5.58 | 1.46E-06 | 2.35E-06 |
| chr1 | 162170775 | 162171841 | 1067 | -5.77 | 1.46E-06 | 2.36E-06 |
| chr18 | 45231611 | 45232320 | 710 | -4.29 | 1.46E-06 | 2.36E-06 |
| chr10 | 8260446 | 8261295 | 850 | -5.76 | 1.47E-06 | 2.36E-06 |
| chr12 | 12290724 | 12291250 | 527 | -5.76 | 1.47E-06 | 2.36E-06 |
| chr5 | 39680898 | 39682126 | 1229 | -5.76 | 1.47E-06 | 2.36E-06 |
| chr11 | 116057867 | 116058733 | 867 | -5.65 | 1.47E-06 | 2.37E-06 |
| chr4 | 157569800 | 157573789 | 3990 | -4.67 | 1.47E-06 | 2.37E-06 |
| chr3 | 153134983 | 153142564 | 7582 | -2.06 | 1.47E-06 | 2.37E-06 |
| chr14 | 52132235 | 52133002 | 768 | -5.59 | 1.47E-06 | 2.38E-06 |
| chr14 | 91986970 | 91988046 | 1077 | -5.59 | 1.47E-06 | 2.38E-06 |
| chr18 | 46391886 | 46393030 | 1145 | -5.59 | 1.47E-06 | 2.38E-06 |
| chr5 | 16536790 | 16537637 | 848 | -5.59 | 1.47E-06 | 2.38E-06 |
| chr7 | 111734642 | 111736052 | 1411 | -5.74 | 1.48E-06 | 2.38E-06 |
| chr5 | 171231677 | 171260838 | 29162 | -2.42 | 1.48E-06 | 2.38E-06 |
| chr3 | 115684819 | 115687038 | 2220 | -3.58 | 1.48E-06 | 2.38E-06 |
| chr16 | 89141521 | 89143104 | 1584 | -5.57 | 1.48E-06 | 2.38E-06 |
| chr17 | 66542621 | 66543541 | 921 | -5.57 | 1.48E-06 | 2.38E-06 |
| chr4 | 168032888 | 168034299 | 1412 | -5.57 | 1.48E-06 | 2.38E-06 |
| chr8 | 55223883 | 55226305 | 2423 | -2.03 | 1.48E-06 | 2.39E-06 |
| chr1 | 244016161 | 244017088 | 928 | -5.56 | 1.48E-06 | 2.39E-06 |
| chr11 | 92077298 | 92079093 | 1796 | -2.08 | 1.48E-06 | 2.39E-06 |
| chr7 | 128604714 | 128605367 | 654 | -5.6 | 1.48E-06 | 2.39E-06 |
| chrX | 119365422 | 119366354 | 933 | -5.6 | 1.48E-06 | 2.39E-06 |
| chr15 | 85484345 | 85485231 | 887 | -5.73 | 1.49E-06 | 2.39E-06 |
| chr9 | 21000544 | 21001712 | 1169 | -5.64 | 1.49E-06 | 2.39E-06 |
| chr1 | 79945814 | 79947640 | 1827 | -6.05 | 1.49E-06 | 2.39E-06 |
| chr10 | 43766197 | 43767977 | 1781 | -6.05 | 1.49E-06 | 2.39E-06 |
| chr4 | 70386156 | 70387465 | 1310 | -4.94 | 1.49E-06 | 2.40E-06 |
| chr18 | 3851546 | 3852590 | 1045 | -5.64 | 1.49E-06 | 2.40E-06 |
| chr3 | 36576010 | 36591466 | 15457 | -6.07 | 1.49E-06 | 2.40E-06 |
| chr1 | 227397777 | 227398504 | 728 | -5.93 | 1.49E-06 | 2.41E-06 |
| chr4 | 3299043 | 3300201 | 1159 | -5.93 | 1.49E-06 | 2.41E-06 |
| chr4 | 31425054 | 31426967 | 1914 | -5.6 | 1.50E-06 | 2.41E-06 |
| chr5 | 127978933 | 127979537 | 605 | -5.6 | 1.50E-06 | 2.41E-06 |
| chr6 | 83331249 | 83332522 | 1274 | -5.57 | 1.50E-06 | 2.42E-06 |
| chr2 | 149896177 | 149905290 | 9114 | -5.4 | 1.50E-06 | 2.42E-06 |
| chr6 | 159892006 | 159900535 | 8530 | -1.54 | 1.50E-06 | 2.42E-06 |
| chr15 | 56353301 | 56354495 | 1195 | -5.71 | 1.50E-06 | 2.42E-06 |
| chr17 | 74584414 | 74585488 | 1075 | -5.71 | 1.50E-06 | 2.42E-06 |
| chr8 | 18521927 | 18522886 | 960 | -5.71 | 1.50E-06 | 2.42E-06 |
| chr12 | 80001879 | 80002453 | 575 | -5.64 | 1.51E-06 | 2.42E-06 |
| chr7 | 75627242 | 75627947 | 706 | -5.64 | 1.51E-06 | 2.42E-06 |
| chr2 | 230430488 | 230432018 | 1531 | 5.63 | 1.51E-06 | 2.43E-06 |
| chr4 | 101635296 | 101636101 | 806 | -5.76 | 1.51E-06 | 2.43E-06 |
| chr5 | 89789762 | 89790932 | 1171 | -5.76 | 1.51E-06 | 2.43E-06 |
| chr15 | 100587747 | 100878611 | 290865 | -0.93 | 1.51E-06 | 2.44E-06 |
| chr6 | 152721000 | 152721848 | 849 | -5.56 | 1.51E-06 | 2.44E-06 |
| chr10 | 63519063 | 63520665 | 1603 | -5.7 | 1.52E-06 | 2.44E-06 |
| chr4 | 24563076 | 24563806 | 731 | -5.59 | 1.52E-06 | 2.44E-06 |
| chr8 | 8678555 | 8679463 | 909 | -5.63 | 1.52E-06 | 2.44E-06 |
| chrX | 21690996 | 21691895 | 900 | -4.58 | 1.52E-06 | 2.45E-06 |
| chr1 | 189136877 | 189139097 | 2221 | -6.18 | 1.52E-06 | 2.45E-06 |
| chr12 | 53016753 | 53017750 | 998 | -5.57 | 1.52E-06 | 2.45E-06 |
| chr4 | 15842031 | 15843194 | 1164 | -5.57 | 1.52E-06 | 2.45E-06 |
| chr4 | 162472453 | 162473469 | 1017 | -5.57 | 1.52E-06 | 2.45E-06 |
| chr9 | 97452678 | 97454329 | 1652 | -6.19 | 1.53E-06 | 2.46E-06 |
| chr13 | 67769163 | 67770511 | 1349 | -5.63 | 1.53E-06 | 2.46E-06 |
| chr3 | 195153245 | 195154386 | 1142 | -5.63 | 1.53E-06 | 2.46E-06 |
| chr5 | 102628904 | 102630083 | 1180 | -5.63 | 1.53E-06 | 2.46E-06 |
| chr5 | 18323169 | 18334726 | 11558 | -1.99 | 1.53E-06 | 2.47E-06 |

| | | | | | | |
|---|---|---|---|---|---|---|
| chr4 | 20402600 | 20403434 | 835 | 5.68 | 1.54E-06 | 2.47E-06 |
| chr1 | 67692519 | 67721113 | 28595 | -2.55 | 1.54E-06 | 2.47E-06 |
| chr11 | 121992335 | 121992957 | 623 | -5.59 | 1.54E-06 | 2.47E-06 |
| chr15 | 29898626 | 29899382 | 757 | -5.59 | 1.54E-06 | 2.47E-06 |
| chr15 | 77839384 | 77840130 | 747 | -5.59 | 1.54E-06 | 2.47E-06 |
| chr18 | 34990587 | 34991568 | 982 | -5.59 | 1.54E-06 | 2.47E-06 |
| chr6 | 113872334 | 113873000 | 667 | -5.59 | 1.54E-06 | 2.47E-06 |
| chr7 | 129701800 | 129702514 | 715 | -5.59 | 1.54E-06 | 2.47E-06 |
| chr12 | 66294729 | 66295635 | 907 | -6.09 | 1.54E-06 | 2.47E-06 |
| chr17 | 47425183 | 47426022 | 840 | -5.71 | 1.54E-06 | 2.48E-06 |
| chr13 | 68012719 | 68282350 | 269632 | 8.91 | 1.54E-06 | 2.48E-06 |
| chr1 | 154888901 | 154889685 | 785 | -5.56 | 1.54E-06 | 2.48E-06 |
| chr9 | 133560024 | 133568214 | 8191 | -6.24 | 1.54E-06 | 2.48E-06 |
| chr2 | 61178357 | 61179037 | 681 | -5.67 | 1.54E-06 | 2.48E-06 |
| chr3 | 31929184 | 31930859 | 1676 | -5.67 | 1.54E-06 | 2.48E-06 |
| chr6 | 48044348 | 48053602 | 9255 | -5.67 | 1.54E-06 | 2.48E-06 |
| chr3 | 34310197 | 34315541 | 5345 | -2.17 | 1.54E-06 | 2.48E-06 |
| chr13 | 30473827 | 30475868 | 2042 | -2.87 | 1.55E-06 | 2.48E-06 |
| chr12 | 39080257 | 39081133 | 877 | -5.61 | 1.55E-06 | 2.49E-06 |
| chr17 | 70752168 | 70752895 | 728 | -5.61 | 1.55E-06 | 2.49E-06 |
| chr20 | 14528136 | 14528978 | 843 | -5.61 | 1.55E-06 | 2.49E-06 |
| chr12 | 122780532 | 122781032 | 501 | -5.7 | 1.55E-06 | 2.49E-06 |
| chr7 | 152273576 | 152275027 | 1452 | -5.7 | 1.55E-06 | 2.49E-06 |
| chr20 | 47885418 | 47905324 | 19907 | -6.21 | 1.55E-06 | 2.49E-06 |
| chr6 | 1542900 | 1543747 | 848 | -5.62 | 1.55E-06 | 2.49E-06 |
| chr21 | 18088847 | 18090371 | 1525 | -4.16 | 1.55E-06 | 2.49E-06 |
| chr1 | 111164465 | 111167250 | 2786 | -1.9 | 1.55E-06 | 2.50E-06 |
| chr3 | 97423917 | 97424409 | 493 | 5.63 | 1.56E-06 | 2.50E-06 |
| chr15 | 39133242 | 39134595 | 1354 | -5.66 | 1.56E-06 | 2.51E-06 |
| chr5 | 118493086 | 118502078 | 8993 | -6.4 | 1.56E-06 | 2.51E-06 |
| chr5 | 154600465 | 154940467 | 340003 | 0.89 | 1.56E-06 | 2.51E-06 |
| chr12 | 92589922 | 92591421 | 1500 | -4.74 | 1.57E-06 | 2.52E-06 |
| chr4 | 137690693 | 137691690 | 998 | -5.69 | 1.57E-06 | 2.52E-06 |
| chrX | 78587199 | 78616274 | 29076 | 1.57 | 1.57E-06 | 2.52E-06 |
| chr17 | 54808121 | 54810567 | 2447 | 5.65 | 1.57E-06 | 2.53E-06 |
| chr10 | 103780644 | 103781850 | 1207 | -5.61 | 1.57E-06 | 2.53E-06 |
| chr2 | 11249616 | 11250531 | 916 | -5.61 | 1.57E-06 | 2.53E-06 |
| chr3 | 135763244 | 135764188 | 945 | -5.61 | 1.57E-06 | 2.53E-06 |
| chr7 | 157970863 | 158041202 | 70340 | -2.61 | 1.58E-06 | 2.53E-06 |
| chrX | 36111115 | 36112489 | 1375 | -5.68 | 1.58E-06 | 2.54E-06 |
| chrX | 149890536 | 149891276 | 741 | -5.68 | 1.58E-06 | 2.54E-06 |
| chr1 | 188881322 | 188882335 | 1014 | -5.6 | 1.58E-06 | 2.54E-06 |
| chr10 | 87841191 | 87842088 | 898 | -5.6 | 1.58E-06 | 2.54E-06 |
| chr4 | 14357256 | 14358980 | 1725 | -5.56 | 1.58E-06 | 2.54E-06 |
| chrX | 75047273 | 75067997 | 20725 | 3.36 | 1.58E-06 | 2.54E-06 |
| chr17 | 64440920 | 64454911 | 13992 | -6.99 | 1.58E-06 | 2.54E-06 |
| chr18 | 2300929 | 2302165 | 1237 | -5.58 | 1.58E-06 | 2.54E-06 |
| chr2 | 138879094 | 138879841 | 748 | -5.58 | 1.58E-06 | 2.54E-06 |
| chr7 | 83542461 | 83543152 | 692 | -5.55 | 1.59E-06 | 2.55E-06 |
| chr19 | 14159116 | 14159666 | 551 | -5.57 | 1.59E-06 | 2.55E-06 |
| chr4 | 7111388 | 7112562 | 1175 | -5.57 | 1.59E-06 | 2.55E-06 |
| chr10 | 6886267 | 6904312 | 18046 | -6.53 | 1.59E-06 | 2.56E-06 |
| chr7 | 138510602 | 138511671 | 1070 | -5.84 | 1.59E-06 | 2.56E-06 |
| chr15 | 44517788 | 44518382 | 595 | -5.55 | 1.59E-06 | 2.56E-06 |
| chr10 | 126081206 | 126082792 | 1587 | -5.81 | 1.60E-06 | 2.56E-06 |
| chr2 | 20161205 | 20162295 | 1091 | -5.55 | 1.60E-06 | 2.56E-06 |
| chr4 | 109086666 | 109088261 | 1596 | -4.23 | 1.60E-06 | 2.56E-06 |
| chr6 | 15027663 | 15028864 | 1202 | -5.54 | 1.60E-06 | 2.57E-06 |
| chr5 | 89964468 | 89979359 | 14892 | 6.06 | 1.60E-06 | 2.57E-06 |
| chr12 | 82888507 | 82889512 | 1006 | -5.64 | 1.60E-06 | 2.57E-06 |
| chr17 | 40399479 | 40400090 | 612 | -5.6 | 1.60E-06 | 2.57E-06 |

| | | | | | | |
|---|---|---|---|---|---|---|
| chr2 | 70468761 | 70469408 | 648 | -5.6 | 1.60E-06 | 2.57E-06 |
| chr16 | 5219436 | 5248288 | 28853 | -1.14 | 1.61E-06 | 2.58E-06 |
| chr13 | 68726075 | 68727102 | 1028 | -5.76 | 1.61E-06 | 2.58E-06 |
| chr7 | 36341039 | 36341705 | 667 | -4.41 | 1.61E-06 | 2.58E-06 |
| chr16 | 46896089 | 46897272 | 1184 | -5.55 | 1.61E-06 | 2.58E-06 |
| chr6 | 155082362 | 155083607 | 1246 | -5.55 | 1.61E-06 | 2.58E-06 |
| chr10 | 1590049 | 1592176 | 2128 | -2.33 | 1.61E-06 | 2.58E-06 |
| chr2 | 164334791 | 164335837 | 1047 | -5.97 | 1.61E-06 | 2.59E-06 |
| chr3 | 133690673 | 133692583 | 1911 | -4.45 | 1.62E-06 | 2.59E-06 |
| chr3 | 177888503 | 177890312 | 1810 | -5.63 | 1.62E-06 | 2.60E-06 |
| chr6 | 165930846 | 165950860 | 20015 | -5.63 | 1.62E-06 | 2.60E-06 |
| chr2 | 169631748 | 169632871 | 1124 | -5.56 | 1.63E-06 | 2.62E-06 |
| chr2 | 170347237 | 170347703 | 467 | -5.56 | 1.63E-06 | 2.62E-06 |
| chr19 | 36186848 | 36187344 | 497 | -5.56 | 1.63E-06 | 2.62E-06 |
| chr4 | 152522526 | 152523356 | 831 | -5.56 | 1.63E-06 | 2.62E-06 |
| chrX | 118129820 | 118130996 | 1177 | -5.64 | 1.64E-06 | 2.62E-06 |
| chrX | 35243279 | 35244940 | 1662 | -6.05 | 1.64E-06 | 2.63E-06 |
| chr4 | 64170986 | 64173405 | 2420 | -5.55 | 1.64E-06 | 2.63E-06 |
| chr6 | 84002919 | 84003747 | 829 | -5.55 | 1.64E-06 | 2.63E-06 |
| chr14 | 106372060 | 106382708 | 10649 | -6.07 | 1.64E-06 | 2.64E-06 |
| chr4 | 130313113 | 130314757 | 1645 | -4.28 | 1.64E-06 | 2.64E-06 |
| chr5 | 110963558 | 110998130 | 34573 | -1.12 | 1.65E-06 | 2.64E-06 |
| chr11 | 101021670 | 101100396 | 78727 | 6.38 | 1.65E-06 | 2.65E-06 |
| chr8 | 97157772 | 97166388 | 8617 | 6.38 | 1.65E-06 | 2.65E-06 |
| chr10 | 90020775 | 90021683 | 909 | -5.54 | 1.65E-06 | 2.65E-06 |
| chr2 | 28881740 | 28882384 | 645 | -5.54 | 1.65E-06 | 2.65E-06 |
| chr7 | 127248162 | 127249302 | 1141 | -5.54 | 1.65E-06 | 2.65E-06 |
| chr4 | 114071695 | 114072821 | 1127 | -5.63 | 1.65E-06 | 2.65E-06 |
| chr12 | 28692291 | 28692913 | 623 | -5.58 | 1.66E-06 | 2.66E-06 |
| chr11 | 84982647 | 84986537 | 3891 | -2.41 | 1.66E-06 | 2.66E-06 |
| chr8 | 55332328 | 55333466 | 1139 | 5.82 | 1.66E-06 | 2.67E-06 |
| chr2 | 169341950 | 169342610 | 661 | -5.58 | 1.67E-06 | 2.67E-06 |
| chr4 | 65021601 | 65022879 | 1279 | -5.58 | 1.67E-06 | 2.67E-06 |
| chr6 | 16176716 | 16182422 | 5707 | -6.14 | 1.67E-06 | 2.67E-06 |
| chr6 | 11181094 | 11197882 | 16789 | -6.11 | 1.67E-06 | 2.67E-06 |
| chr3 | 86854910 | 86859626 | 4717 | -5.29 | 1.67E-06 | 2.67E-06 |
| chr8 | 110125852 | 110127584 | 1733 | -2.71 | 1.67E-06 | 2.67E-06 |
| chr2 | 206976672 | 206977710 | 1039 | -5.62 | 1.67E-06 | 2.68E-06 |
| chr15 | 102006275 | 102006934 | 660 | -5.68 | 1.68E-06 | 2.68E-06 |
| chr8 | 105464595 | 105466388 | 1794 | 2.86 | 1.68E-06 | 2.69E-06 |
| chr13 | 40337563 | 40338435 | 873 | -5.61 | 1.68E-06 | 2.69E-06 |
| chr3 | 12764245 | 12765379 | 1135 | -5.61 | 1.68E-06 | 2.69E-06 |
| chr3 | 62305195 | 62305979 | 785 | -5.55 | 1.68E-06 | 2.69E-06 |
| chr6 | 145569293 | 145570861 | 1569 | -5.86 | 1.68E-06 | 2.69E-06 |
| chr10 | 95486060 | 95500025 | 13966 | -1.24 | 1.68E-06 | 2.69E-06 |
| chr7 | 75151385 | 75151802 | 418 | -5.87 | 1.68E-06 | 2.69E-06 |
| chr4 | 177686182 | 177687360 | 1179 | -5.53 | 1.68E-06 | 2.69E-06 |
| chr7 | 80019899 | 80021148 | 1250 | -6.21 | 1.68E-06 | 2.70E-06 |
| chr4 | 15604762 | 15605384 | 623 | -5.79 | 1.69E-06 | 2.70E-06 |
| chr3 | 66054823 | 66056111 | 1289 | -5.67 | 1.69E-06 | 2.71E-06 |
| chr5 | 59405853 | 59406760 | 908 | -5.67 | 1.69E-06 | 2.71E-06 |
| chr1 | 186633254 | 186634125 | 872 | -5.55 | 1.69E-06 | 2.71E-06 |
| chr15 | 23189528 | 23190002 | 475 | -5.55 | 1.69E-06 | 2.71E-06 |
| chr4 | 107771326 | 107773625 | 2300 | 4.49 | 1.69E-06 | 2.71E-06 |
| chr1 | 81864245 | 81866012 | 1768 | -5.61 | 1.69E-06 | 2.71E-06 |
| chr12 | 34217121 | 34218092 | 972 | -5.76 | 1.69E-06 | 2.71E-06 |
| chr2 | 120071825 | 120072647 | 823 | -5.76 | 1.69E-06 | 2.71E-06 |
| chr1 | 99176904 | 99178384 | 1481 | -5.95 | 1.70E-06 | 2.72E-06 |
| chr1 | 25405240 | 25406304 | 1065 | -5.57 | 1.70E-06 | 2.73E-06 |
| chr6 | 790743 | 791551 | 809 | -5.57 | 1.70E-06 | 2.73E-06 |
| chr13 | 27944108 | 27945660 | 1553 | -6.3 | 1.70E-06 | 2.73E-06 |

| | | | | | | |
|---|---|---|---|---|---|---|
| chr4 | 1280188 | 1281028 | 841 | -5.54 | 1.71E-06 | 2.74E-06 |
| chr3 | 130155380 | 130159404 | 4025 | 2.58 | 1.71E-06 | 2.74E-06 |
| chr1 | 97975502 | 97976834 | 1333 | -5.59 | 1.71E-06 | 2.74E-06 |
| chr22 | 46687765 | 46688355 | 591 | -5.59 | 1.71E-06 | 2.74E-06 |
| chr6 | 100017984 | 100019317 | 1334 | -5.59 | 1.71E-06 | 2.74E-06 |
| chr8 | 70940997 | 70941829 | 833 | -5.59 | 1.71E-06 | 2.74E-06 |
| chr18 | 50048495 | 50050463 | 1969 | -4.23 | 1.71E-06 | 2.74E-06 |
| chr4 | 126430276 | 126436856 | 6581 | -5.73 | 1.71E-06 | 2.74E-06 |
| chr5 | 130790841 | 130791566 | 726 | -6 | 1.71E-06 | 2.74E-06 |
| chr10 | 17287715 | 17335418 | 47704 | -6.44 | 1.71E-06 | 2.74E-06 |
| chr8 | 62003302 | 62018603 | 15302 | -5.79 | 1.72E-06 | 2.74E-06 |
| chr20 | 56071219 | 56071987 | 769 | -5.6 | 1.72E-06 | 2.75E-06 |
| chr3 | 188030300 | 188031093 | 794 | -5.72 | 1.72E-06 | 2.75E-06 |
| chr6 | 139676127 | 139676972 | 846 | -4.57 | 1.72E-06 | 2.75E-06 |
| chr13 | 81245050 | 81248037 | 2988 | -2.88 | 1.72E-06 | 2.75E-06 |
| chr5 | 130702988 | 130703924 | 937 | -5.64 | 1.73E-06 | 2.76E-06 |
| chr2 | 11897964 | 11898680 | 717 | -5.53 | 1.73E-06 | 2.76E-06 |
| chr6 | 153597740 | 153598524 | 785 | -4.2 | 1.73E-06 | 2.76E-06 |
| chr5 | 121853429 | 122001067 | 147639 | 2.6 | 1.73E-06 | 2.77E-06 |
| chr6 | 139752452 | 139753642 | 1191 | -4.85 | 1.73E-06 | 2.77E-06 |
| chr11 | 12997118 | 12997884 | 767 | -4.77 | 1.73E-06 | 2.77E-06 |
| chrX | 57045630 | 57050538 | 4909 | -1.59 | 1.73E-06 | 2.77E-06 |
| chr1 | 220875697 | 220876510 | 814 | -5.56 | 1.74E-06 | 2.78E-06 |
| chr10 | 54396581 | 54397166 | 586 | -5.56 | 1.74E-06 | 2.78E-06 |
| chr10 | 8479931 | 8480664 | 734 | -5.6 | 1.74E-06 | 2.79E-06 |
| chr8 | 25181599 | 25182605 | 1007 | -5.6 | 1.74E-06 | 2.79E-06 |
| chr6 | 154954344 | 154955105 | 762 | -4.98 | 1.75E-06 | 2.79E-06 |
| chr17 | 59103195 | 59103976 | 782 | -5.63 | 1.75E-06 | 2.79E-06 |
| chr3 | 189783689 | 189784861 | 1173 | -5.63 | 1.75E-06 | 2.79E-06 |
| chr7 | 55125804 | 55126535 | 732 | -5.63 | 1.75E-06 | 2.79E-06 |
| chr12 | 38943650 | 38948405 | 4756 | 1.88 | 1.75E-06 | 2.79E-06 |
| chr17 | 17400550 | 17401213 | 664 | -5.58 | 1.75E-06 | 2.80E-06 |
| chr17 | 56735437 | 56736111 | 675 | -5.54 | 1.75E-06 | 2.80E-06 |
| chr2 | 203662581 | 203663273 | 693 | -5.54 | 1.75E-06 | 2.80E-06 |
| chr3 | 45981883 | 45982870 | 988 | -5.54 | 1.75E-06 | 2.80E-06 |
| chr4 | 8655967 | 8656714 | 748 | -5.54 | 1.75E-06 | 2.80E-06 |
| chr2 | 183433151 | 183434295 | 1145 | 5.74 | 1.76E-06 | 2.81E-06 |
| chr2 | 172386100 | 172386832 | 733 | -5.83 | 1.76E-06 | 2.81E-06 |
| chr4 | 163020572 | 163021714 | 1143 | -5.82 | 1.76E-06 | 2.81E-06 |
| chr7 | 65643048 | 65643795 | 748 | -5.86 | 1.76E-06 | 2.81E-06 |
| chrX | 77931649 | 77942335 | 10687 | -2.65 | 1.76E-06 | 2.81E-06 |
| chr8 | 131466540 | 131468514 | 1975 | -5.89 | 1.76E-06 | 2.82E-06 |
| chr1 | 47316505 | 47330234 | 13730 | -4.64 | 1.77E-06 | 2.82E-06 |
| chr6 | 118179605 | 118181316 | 1712 | 5.81 | 1.77E-06 | 2.83E-06 |
| chr13 | 83868105 | 83869585 | 1481 | -5.76 | 1.77E-06 | 2.83E-06 |
| chr6 | 153181348 | 153182190 | 843 | -5.76 | 1.77E-06 | 2.83E-06 |
| chr1 | 173785776 | 173786430 | 655 | -5.53 | 1.77E-06 | 2.83E-06 |
| chr11 | 48105471 | 48106146 | 676 | -5.52 | 1.78E-06 | 2.84E-06 |
| chr5 | 77463800 | 77464652 | 853 | -5.52 | 1.78E-06 | 2.84E-06 |
| chr6 | 114234920 | 114235779 | 860 | -5.52 | 1.78E-06 | 2.84E-06 |
| chr10 | 3773738 | 3774915 | 1178 | -5.74 | 1.78E-06 | 2.84E-06 |
| chr3 | 120099761 | 120101431 | 1671 | 2.78 | 1.78E-06 | 2.84E-06 |
| chr4 | 63802752 | 63806306 | 3555 | -5.95 | 1.78E-06 | 2.84E-06 |
| chr6 | 132530574 | 132532586 | 2013 | -5.95 | 1.78E-06 | 2.84E-06 |
| chr9 | 123860574 | 123861411 | 838 | -5.52 | 1.78E-06 | 2.84E-06 |
| chr18 | 54871416 | 54872505 | 1090 | -5.66 | 1.78E-06 | 2.85E-06 |
| chr20 | 35077924 | 35078729 | 806 | -5.66 | 1.78E-06 | 2.85E-06 |
| chr3 | 73076554 | 73077622 | 1069 | -5.55 | 1.78E-06 | 2.85E-06 |
| chr9 | 75054222 | 75055955 | 1734 | -5.72 | 1.79E-06 | 2.85E-06 |
| chr3 | 136213074 | 136214037 | 964 | -5.53 | 1.79E-06 | 2.86E-06 |
| chr5 | 141504597 | 141505521 | 925 | -5.57 | 1.79E-06 | 2.86E-06 |

| | | | | | | |
|---|---|---|---|---|---|---|
| chr9 | 15298018 | 15299325 | 1308 | -5.57 | 1.79E-06 | 2.86E-06 |
| chr1 | 183326332 | 183327688 | 1357 | -5.99 | 1.79E-06 | 2.86E-06 |
| chr9 | 80527627 | 80528857 | 1231 | -5.99 | 1.79E-06 | 2.86E-06 |
| chr4 | 93104200 | 93105366 | 1167 | -4.39 | 1.79E-06 | 2.86E-06 |
| chr17 | 53519697 | 53538260 | 18564 | -7.03 | 1.80E-06 | 2.87E-06 |
| chr11 | 120523710 | 120524413 | 704 | -5.61 | 1.80E-06 | 2.87E-06 |
| chr4 | 138596790 | 138597483 | 694 | -5.61 | 1.80E-06 | 2.87E-06 |
| chr8 | 23737167 | 23739505 | 2339 | -6.01 | 1.80E-06 | 2.87E-06 |
| chr1 | 62408351 | 62409016 | 666 | -5.65 | 1.80E-06 | 2.87E-06 |
| chr7 | 100315087 | 100317151 | 2065 | -6.03 | 1.80E-06 | 2.87E-06 |
| chr1 | 172406905 | 172407642 | 738 | -5.58 | 1.80E-06 | 2.88E-06 |
| chr13 | 76894923 | 76895598 | 676 | -5.58 | 1.80E-06 | 2.88E-06 |
| chr7 | 27894569 | 27895784 | 1216 | -5.58 | 1.80E-06 | 2.88E-06 |
| chr9 | 13255419 | 13256070 | 652 | -5.58 | 1.80E-06 | 2.88E-06 |
| chr4 | 123860567 | 123860911 | 345 | -4.45 | 1.80E-06 | 2.88E-06 |
| chr20 | 8793612 | 8794990 | 1379 | -5.52 | 1.80E-06 | 2.88E-06 |
| chr5 | 72502113 | 72503427 | 1315 | -5.52 | 1.80E-06 | 2.88E-06 |
| chr9 | 29428708 | 29429826 | 1119 | -5.52 | 1.80E-06 | 2.88E-06 |
| chr11 | 114058545 | 114063717 | 5173 | -4.37 | 1.81E-06 | 2.88E-06 |
| chr3 | 62243816 | 62245268 | 1453 | 5.62 | 1.81E-06 | 2.88E-06 |
| chr1 | 59411973 | 59413095 | 1123 | -5.54 | 1.81E-06 | 2.89E-06 |
| chrX | 96434724 | 96436416 | 1693 | -5.54 | 1.81E-06 | 2.89E-06 |
| chr3 | 171025548 | 171026824 | 1277 | -5.69 | 1.81E-06 | 2.89E-06 |
| chrX | 93646086 | 93741290 | 95205 | 0.99 | 1.82E-06 | 2.89E-06 |
| chr19 | 38787529 | 38788634 | 1106 | -6.08 | 1.82E-06 | 2.90E-06 |
| chr12 | 129264748 | 129266121 | 1374 | -5.53 | 1.82E-06 | 2.91E-06 |
| chr3 | 43693441 | 43694045 | 605 | -5.53 | 1.82E-06 | 2.91E-06 |
| chr1 | 210874690 | 210875626 | 937 | -5.68 | 1.82E-06 | 2.91E-06 |
| chr1 | 169166776 | 169167729 | 954 | -5.6 | 1.83E-06 | 2.91E-06 |
| chr22 | 31921378 | 31922143 | 766 | -5.6 | 1.83E-06 | 2.91E-06 |
| chr10 | 14682129 | 14685256 | 3128 | -2.71 | 1.83E-06 | 2.91E-06 |
| chr2 | 242816033 | 242817593 | 1561 | -5.63 | 1.83E-06 | 2.91E-06 |
| chr20 | 25607781 | 25608951 | 1171 | -6.13 | 1.83E-06 | 2.92E-06 |
| chr10 | 72853548 | 72884656 | 31109 | -3.34 | 1.83E-06 | 2.92E-06 |
| chr8 | 24391544 | 24425876 | 34333 | 1.5 | 1.83E-06 | 2.92E-06 |
| chr7 | 130714309 | 130715815 | 1507 | -5.54 | 1.83E-06 | 2.92E-06 |
| chr5 | 81947776 | 81950117 | 2342 | -4.24 | 1.83E-06 | 2.92E-06 |
| chr16 | 73151278 | 73169275 | 17998 | -6.56 | 1.84E-06 | 2.93E-06 |
| chr14 | 64371668 | 64372243 | 576 | -5.56 | 1.84E-06 | 2.93E-06 |
| chr9 | 84277682 | 84279103 | 1422 | -5.56 | 1.84E-06 | 2.93E-06 |
| chr4 | 108901509 | 108902503 | 995 | -5.52 | 1.84E-06 | 2.94E-06 |
| chr3 | 16839691 | 16841175 | 1485 | -6.18 | 1.85E-06 | 2.94E-06 |
| chr3 | 110345129 | 110346361 | 1233 | -5.62 | 1.85E-06 | 2.94E-06 |
| chr3 | 151261622 | 151262163 | 542 | -4.39 | 1.85E-06 | 2.94E-06 |
| chr17 | 58433721 | 58434650 | 930 | -6.22 | 1.85E-06 | 2.95E-06 |
| chr6 | 74509081 | 74510190 | 1110 | -5.66 | 1.85E-06 | 2.95E-06 |
| chr7 | 112208436 | 112383719 | 175284 | 4.24 | 1.86E-06 | 2.96E-06 |
| chr2 | 112293680 | 112294695 | 1016 | -5.58 | 1.86E-06 | 2.96E-06 |
| chr2 | 165824618 | 165826212 | 1595 | -5.58 | 1.86E-06 | 2.96E-06 |
| chrX | 23831688 | 23832637 | 950 | -5.58 | 1.86E-06 | 2.96E-06 |
| chr3 | 17823910 | 17830659 | 6750 | -6.27 | 1.86E-06 | 2.97E-06 |
| chr5 | 115189235 | 115201029 | 11795 | -6.32 | 1.87E-06 | 2.98E-06 |
| chr10 | 3472233 | 3485860 | 13628 | -6.38 | 1.87E-06 | 2.98E-06 |
| chr2 | 43208132 | 43209655 | 1524 | -6.35 | 1.87E-06 | 2.98E-06 |
| chr7 | 25955061 | 26022678 | 67618 | -1.55 | 1.87E-06 | 2.98E-06 |
| chr2 | 236804505 | 236805552 | 1048 | -5.56 | 1.87E-06 | 2.98E-06 |
| chr4 | 96054598 | 96055139 | 542 | -5.56 | 1.87E-06 | 2.98E-06 |
| chr6 | 90013125 | 90013915 | 791 | -5.51 | 1.87E-06 | 2.98E-06 |
| chr17 | 19541812 | 19556813 | 15002 | -5.51 | 1.88E-06 | 2.99E-06 |
| chr7 | 100412028 | 100412724 | 697 | -5.51 | 1.88E-06 | 2.99E-06 |
| chr8 | 49347283 | 49348375 | 1093 | -5.51 | 1.88E-06 | 2.99E-06 |

| | | | | | | |
|---|---|---|---|---|---|---|
| chr18 | 52810340 | 52813611 | 3272 | -2.27 | 1.88E-06 | 3.00E-06 |
| chr19 | 16649873 | 16650755 | 883 | -5.53 | 1.88E-06 | 3.00E-06 |
| chrX | 21431110 | 21438223 | 7114 | -5.09 | 1.88E-06 | 3.00E-06 |
| chr19 | 11270965 | 11271728 | 764 | -5.63 | 1.89E-06 | 3.01E-06 |
| chr4 | 112948651 | 112949783 | 1133 | -5.63 | 1.89E-06 | 3.01E-06 |
| chr22 | 18146348 | 18147761 | 1414 | -5.55 | 1.89E-06 | 3.01E-06 |
| chr10 | 134888183 | 134889227 | 1045 | -5.51 | 1.89E-06 | 3.01E-06 |
| chr22 | 22166050 | 22166779 | 730 | -5.51 | 1.89E-06 | 3.01E-06 |
| chr2 | 178670415 | 178672048 | 1634 | -2.96 | 1.89E-06 | 3.01E-06 |
| chr10 | 15676438 | 15754956 | 78519 | 2.52 | 1.89E-06 | 3.01E-06 |
| chr4 | 141630667 | 141631429 | 763 | -5.57 | 1.90E-06 | 3.02E-06 |
| chr1 | 60119451 | 60120337 | 887 | -4.37 | 1.90E-06 | 3.03E-06 |
| chr4 | 117965717 | 117967891 | 2175 | -5.79 | 1.91E-06 | 3.03E-06 |
| chr14 | 27523065 | 27524786 | 1722 | -5.62 | 1.91E-06 | 3.04E-06 |
| chr4 | 117657470 | 117660473 | 3004 | -5.62 | 1.91E-06 | 3.04E-06 |
| chr3 | 123700708 | 123719078 | 18371 | -2.04 | 1.91E-06 | 3.04E-06 |
| chr12 | 9914670 | 9915259 | 590 | -5.59 | 1.91E-06 | 3.04E-06 |
| chr5 | 93526031 | 93527548 | 1518 | 5.59 | 1.91E-06 | 3.04E-06 |
| chr18 | 45088767 | 45093271 | 4505 | -3.15 | 1.91E-06 | 3.04E-06 |
| chr1 | 116026817 | 116028986 | 2170 | -5.91 | 1.91E-06 | 3.05E-06 |
| chr2 | 119202417 | 119203738 | 1322 | -4.34 | 1.92E-06 | 3.05E-06 |
| chr1 | 232785514 | 232786364 | 851 | -5.93 | 1.92E-06 | 3.05E-06 |
| chr2 | 151923681 | 151935373 | 11693 | -5.51 | 1.93E-06 | 3.06E-06 |
| chr1 | 15388736 | 15389431 | 696 | -5.51 | 1.93E-06 | 3.06E-06 |
| chr11 | 15936964 | 15938618 | 1655 | -2.91 | 1.93E-06 | 3.06E-06 |
| chr12 | 102707414 | 102865448 | 158035 | 2.2 | 1.93E-06 | 3.07E-06 |
| chr4 | 176789179 | 176790321 | 1143 | -5.97 | 1.93E-06 | 3.07E-06 |
| chr1 | 171826484 | 171827463 | 980 | -5.72 | 1.93E-06 | 3.07E-06 |
| chr2 | 15831487 | 15832495 | 1009 | 5.59 | 1.93E-06 | 3.08E-06 |
| chr5 | 31719793 | 31721284 | 1492 | -4.3 | 1.94E-06 | 3.08E-06 |
| chr20 | 52962465 | 52963329 | 865 | -5.56 | 1.94E-06 | 3.08E-06 |
| chr6 | 163591083 | 163591994 | 912 | -5.56 | 1.94E-06 | 3.08E-06 |
| chr3 | 50856949 | 50858105 | 1157 | -5.71 | 1.94E-06 | 3.08E-06 |
| chr7 | 33310507 | 33311938 | 1432 | -5.82 | 1.94E-06 | 3.09E-06 |
| chr12 | 53333541 | 53334249 | 709 | -5.85 | 1.94E-06 | 3.09E-06 |
| chr4 | 132160072 | 132160980 | 909 | -5.87 | 1.95E-06 | 3.09E-06 |
| chr17 | 76241307 | 76242013 | 707 | -5.53 | 1.95E-06 | 3.09E-06 |
| chr7 | 77172890 | 77173415 | 526 | -5.53 | 1.95E-06 | 3.09E-06 |
| chr15 | 40770839 | 40772821 | 1983 | -4.18 | 1.95E-06 | 3.09E-06 |
| chr21 | 40301473 | 40302244 | 772 | -5.52 | 1.95E-06 | 3.10E-06 |
| chr6 | 142498737 | 142499579 | 843 | -5.52 | 1.95E-06 | 3.10E-06 |
| chr11 | 35994924 | 35996311 | 1388 | -5.88 | 1.95E-06 | 3.10E-06 |
| chr17 | 47058179 | 47058952 | 774 | -5.77 | 1.95E-06 | 3.10E-06 |
| chr2 | 56195325 | 56196122 | 798 | -5.76 | 1.95E-06 | 3.10E-06 |
| chr4 | 64126409 | 64128450 | 2042 | -5.76 | 1.95E-06 | 3.10E-06 |
| chr8 | 99586647 | 99587557 | 911 | -5.92 | 1.96E-06 | 3.11E-06 |
| chr16 | 69513393 | 69514118 | 726 | -5.74 | 1.96E-06 | 3.11E-06 |
| chr3 | 160595887 | 160606837 | 10951 | -5.74 | 1.96E-06 | 3.11E-06 |
| chr6 | 48584360 | 48585741 | 1382 | -5.6 | 1.96E-06 | 3.11E-06 |
| chr6 | 126273029 | 126274127 | 1099 | -5.6 | 1.96E-06 | 3.11E-06 |
| chr1 | 19601175 | 19602184 | 1010 | -5.68 | 1.96E-06 | 3.12E-06 |
| chr6 | 154631476 | 154632996 | 1521 | -5.73 | 1.96E-06 | 3.12E-06 |
| chr2 | 120979251 | 120980411 | 1161 | -5.57 | 1.97E-06 | 3.12E-06 |
| chr5 | 122045579 | 122046833 | 1255 | -5.57 | 1.97E-06 | 3.12E-06 |
| chr5 | 168517484 | 168518395 | 912 | -5.57 | 1.97E-06 | 3.12E-06 |
| chr9 | 84698993 | 84718456 | 19464 | -1.13 | 1.97E-06 | 3.13E-06 |
| chr3 | 54141104 | 54151162 | 10059 | -5.64 | 1.97E-06 | 3.13E-06 |
| chr18 | 77571044 | 77572651 | 1608 | -5.97 | 1.97E-06 | 3.14E-06 |
| chr4 | 7215518 | 7216474 | 957 | -5.5 | 1.98E-06 | 3.14E-06 |
| chr5 | 68646517 | 68646985 | 469 | -5.5 | 1.98E-06 | 3.14E-06 |
| chr12 | 107403035 | 107404270 | 1236 | -5.7 | 1.98E-06 | 3.14E-06 |

| | | | | | | |
|---|---|---|---|---|---|---|
| chr2 | 38927686 | 38928953 | 1268 | -5.7 | 1.98E-06 | 3.14E-06 |
| chrX | 18297442 | 18299017 | 1576 | -5.66 | 1.98E-06 | 3.14E-06 |
| chr18 | 57869303 | 57872197 | 2895 | -2.02 | 1.98E-06 | 3.15E-06 |
| chr3 | 172474517 | 172475370 | 854 | -6.16 | 1.98E-06 | 3.15E-06 |
| chr6 | 10733740 | 10734656 | 917 | -4.39 | 1.98E-06 | 3.15E-06 |
| chr12 | 113289361 | 113290192 | 832 | -5.5 | 1.98E-06 | 3.15E-06 |
| chr13 | 37609012 | 37609761 | 750 | -5.5 | 1.98E-06 | 3.15E-06 |
| chr17 | 26673735 | 26674433 | 699 | -5.5 | 1.98E-06 | 3.15E-06 |
| chr5 | 71571922 | 71573331 | 1410 | -5.5 | 1.98E-06 | 3.15E-06 |
| chr5 | 169467217 | 169468027 | 811 | -5.5 | 1.98E-06 | 3.15E-06 |
| chr10 | 98853100 | 98853666 | 567 | -5.5 | 1.98E-06 | 3.15E-06 |
| chr3 | 136293077 | 136293963 | 887 | -5.55 | 1.98E-06 | 3.15E-06 |
| chr7 | 4852256 | 4853017 | 762 | -5.55 | 1.98E-06 | 3.15E-06 |
| chr7 | 78832677 | 78834836 | 2160 | 6.55 | 1.99E-06 | 3.15E-06 |
| chr11 | 119654267 | 119655316 | 1050 | -5.59 | 1.99E-06 | 3.16E-06 |
| chr15 | 40274621 | 40275977 | 1357 | -5.59 | 1.99E-06 | 3.16E-06 |
| chr22 | 29890718 | 29891312 | 595 | -5.59 | 1.99E-06 | 3.16E-06 |
| chr4 | 118350022 | 118356349 | 6328 | -6.18 | 1.99E-06 | 3.16E-06 |
| chr10 | 16959874 | 16960558 | 685 | -5.5 | 1.99E-06 | 3.16E-06 |
| chr13 | 90126363 | 90127621 | 1259 | -5.66 | 1.99E-06 | 3.16E-06 |
| chr10 | 93899332 | 93900018 | 687 | -5.56 | 2.00E-06 | 3.17E-06 |
| chr19 | 4758367 | 4759198 | 832 | -5.56 | 2.00E-06 | 3.17E-06 |
| chr10 | 88547021 | 88547866 | 846 | -5.53 | 2.00E-06 | 3.18E-06 |
| chr5 | 73596683 | 73597682 | 1000 | -5.53 | 2.00E-06 | 3.18E-06 |
| chr6 | 40610115 | 40611105 | 991 | -5.53 | 2.00E-06 | 3.18E-06 |
| chr16 | 441790 | 442801 | 1012 | -5.67 | 2.01E-06 | 3.18E-06 |
| chr22 | 22766414 | 22775145 | 8732 | -1.35 | 2.01E-06 | 3.18E-06 |
| chr5 | 112678495 | 112702942 | 24448 | 1.21 | 2.01E-06 | 3.19E-06 |
| chr10 | 72505009 | 72506037 | 1029 | -5.52 | 2.01E-06 | 3.19E-06 |
| chr4 | 70485911 | 70487411 | 1501 | -5.52 | 2.01E-06 | 3.19E-06 |
| chr2 | 187027432 | 187028645 | 1214 | 5.57 | 2.01E-06 | 3.19E-06 |
| chr10 | 124632332 | 124632959 | 628 | -5.5 | 2.01E-06 | 3.19E-06 |
| chr5 | 115359098 | 115359749 | 652 | -5.5 | 2.01E-06 | 3.19E-06 |
| chr2 | 240603891 | 240607997 | 4107 | -2.06 | 2.01E-06 | 3.19E-06 |
| chr1 | 153292718 | 153305034 | 12317 | -2.63 | 2.02E-06 | 3.20E-06 |
| chr3 | 25910001 | 25912534 | 2534 | -2.97 | 2.02E-06 | 3.20E-06 |
| chr8 | 30356144 | 30357144 | 1001 | -5.64 | 2.02E-06 | 3.20E-06 |
| chr7 | 127038127 | 127038910 | 784 | -5.66 | 2.02E-06 | 3.20E-06 |
| chr16 | 80372056 | 80373842 | 1787 | 2.79 | 2.02E-06 | 3.20E-06 |
| chr3 | 104448381 | 104486093 | 37713 | 6.12 | 2.03E-06 | 3.22E-06 |
| chr4 | 185285845 | 185286860 | 1016 | -5.51 | 2.03E-06 | 3.22E-06 |
| chr11 | 130955157 | 130957652 | 2496 | -1.88 | 2.03E-06 | 3.22E-06 |
| chr3 | 182746101 | 182746854 | 754 | -5.55 | 2.03E-06 | 3.22E-06 |
| chr10 | 76841061 | 76841937 | 877 | -5.63 | 2.03E-06 | 3.22E-06 |
| chr2 | 15724914 | 15725583 | 670 | -5.63 | 2.03E-06 | 3.22E-06 |
| chr4 | 125150307 | 125163765 | 13459 | -5.65 | 2.03E-06 | 3.22E-06 |
| chr7 | 83378494 | 83384509 | 6016 | -6.19 | 2.03E-06 | 3.23E-06 |
| chr13 | 46733383 | 46734534 | 1152 | -5.54 | 2.04E-06 | 3.23E-06 |
| chr20 | 8496740 | 8499590 | 2851 | 4.69 | 2.04E-06 | 3.23E-06 |
| chr1 | 194404135 | 194411743 | 7609 | -6.22 | 2.04E-06 | 3.23E-06 |
| chr5 | 122386389 | 122417472 | 31084 | -3.63 | 2.04E-06 | 3.24E-06 |
| chr7 | 20799644 | 20800826 | 1183 | -5.64 | 2.05E-06 | 3.25E-06 |
| chr7 | 107956352 | 107957396 | 1045 | -5.64 | 2.05E-06 | 3.25E-06 |
| chr12 | 80880107 | 80880837 | 731 | 5.59 | 2.05E-06 | 3.25E-06 |
| chr15 | 64307577 | 64313804 | 6228 | -5.52 | 2.06E-06 | 3.26E-06 |
| chr13 | 40654649 | 40655374 | 726 | -4.38 | 2.06E-06 | 3.26E-06 |
| chr2 | 213755815 | 213757446 | 1632 | -4.12 | 2.06E-06 | 3.27E-06 |
| chr6 | 38431890 | 38432755 | 866 | -5.49 | 2.06E-06 | 3.27E-06 |
| chr9 | 72189687 | 72190311 | 625 | -4.35 | 2.07E-06 | 3.27E-06 |
| chr7 | 92212174 | 92212803 | 630 | -5.61 | 2.07E-06 | 3.27E-06 |
| chrX | 149271259 | 149272173 | 915 | -5.61 | 2.07E-06 | 3.27E-06 |

| | | | | | | |
|---|---|---|---|---|---|---|
| chr1 | 220378230 | 220378647 | 418 | -4.46 | 2.07E-06 | 3.27E-06 |
| chr13 | 28148405 | 28149336 | 932 | -5.5 | 2.07E-06 | 3.28E-06 |
| chr14 | 89137488 | 89138932 | 1445 | -5.5 | 2.07E-06 | 3.28E-06 |
| chr16 | 50298057 | 50298911 | 855 | -5.5 | 2.07E-06 | 3.28E-06 |
| chr3 | 66372889 | 66374170 | 1282 | -5.62 | 2.07E-06 | 3.28E-06 |
| chr3 | 172189421 | 172204611 | 15191 | -6.72 | 2.07E-06 | 3.29E-06 |
| chr5 | 102242549 | 102244190 | 1642 | -4.26 | 2.08E-06 | 3.29E-06 |
| chr2 | 198656091 | 198657165 | 1075 | -4.31 | 2.08E-06 | 3.29E-06 |
| chr1 | 235544612 | 235545111 | 500 | -5.51 | 2.08E-06 | 3.30E-06 |
| chr1 | 39768634 | 39769468 | 835 | -5.6 | 2.09E-06 | 3.30E-06 |
| chr8 | 23805651 | 23806246 | 596 | -5.49 | 2.09E-06 | 3.31E-06 |
| chr6 | 83348093 | 83348891 | 799 | -4.11 | 2.09E-06 | 3.31E-06 |
| chr12 | 41640731 | 41641886 | 1156 | -5.61 | 2.09E-06 | 3.31E-06 |
| chr13 | 83430478 | 83432003 | 1526 | -5.61 | 2.09E-06 | 3.31E-06 |
| chr5 | 126349202 | 126350520 | 1319 | -5.49 | 2.09E-06 | 3.31E-06 |
| chr3 | 112862425 | 112863606 | 1182 | 6 | 2.09E-06 | 3.31E-06 |
| chr9 | 92446061 | 92446921 | 861 | -5.52 | 2.10E-06 | 3.32E-06 |
| chr19 | 11256240 | 11256787 | 548 | -5.55 | 2.10E-06 | 3.33E-06 |
| chr12 | 100393836 | 100394720 | 885 | -5.54 | 2.11E-06 | 3.34E-06 |
| chr13 | 43550214 | 43551983 | 1770 | -5.54 | 2.11E-06 | 3.34E-06 |
| chr1 | 56418187 | 56419436 | 1250 | -5.49 | 2.11E-06 | 3.34E-06 |
| chr8 | 23346476 | 23347313 | 838 | -5.49 | 2.11E-06 | 3.34E-06 |
| chr7 | 47055910 | 47060670 | 4761 | -2.17 | 2.11E-06 | 3.35E-06 |
| chr6 | 157839917 | 157841044 | 1128 | -5.51 | 2.11E-06 | 3.35E-06 |
| chr21 | 47619263 | 47619907 | 645 | -5.58 | 2.13E-06 | 3.38E-06 |
| chr9 | 79302766 | 79306199 | 3434 | 2.35 | 2.14E-06 | 3.38E-06 |
| chrX | 151399775 | 151404361 | 4587 | -2.1 | 2.14E-06 | 3.39E-06 |
| chr14 | 85996389 | 85997797 | 1409 | -5.59 | 2.14E-06 | 3.39E-06 |
| chr5 | 177654027 | 177655231 | 1205 | -5.59 | 2.14E-06 | 3.39E-06 |
| chr7 | 139176562 | 139177702 | 1141 | -5.8 | 2.15E-06 | 3.40E-06 |
| chr9 | 94104206 | 94105105 | 900 | -5.8 | 2.15E-06 | 3.40E-06 |
| chr7 | 79529234 | 79530166 | 933 | -5.79 | 2.15E-06 | 3.40E-06 |
| chr16 | 10929679 | 10931034 | 1356 | -5.83 | 2.15E-06 | 3.40E-06 |
| chr17 | 237632 | 238556 | 925 | -5.78 | 2.15E-06 | 3.40E-06 |
| chr9 | 19020777 | 19022714 | 1938 | -2.48 | 2.15E-06 | 3.40E-06 |
| chr3 | 145970819 | 145972595 | 1777 | -5.77 | 2.15E-06 | 3.40E-06 |
| chr2 | 206089703 | 206090825 | 1123 | -4.19 | 2.15E-06 | 3.40E-06 |
| chr1 | 79285111 | 79286387 | 1277 | -5.48 | 2.15E-06 | 3.41E-06 |
| chr2 | 101989267 | 101989856 | 590 | -5.48 | 2.15E-06 | 3.41E-06 |
| chr7 | 101075130 | 101076735 | 1606 | -5.48 | 2.15E-06 | 3.41E-06 |
| chr1 | 61787217 | 61788376 | 1160 | -5.75 | 2.15E-06 | 3.41E-06 |
| chr16 | 12453711 | 12454937 | 1227 | -5.88 | 2.16E-06 | 3.41E-06 |
| chr9 | 29169710 | 29170483 | 774 | -5.88 | 2.16E-06 | 3.41E-06 |
| chr8 | 23781085 | 23787685 | 6601 | -5.74 | 2.16E-06 | 3.41E-06 |
| chr12 | 20052038 | 20053200 | 1163 | -5.57 | 2.16E-06 | 3.41E-06 |
| chr4 | 153999062 | 153999700 | 639 | -5.57 | 2.16E-06 | 3.41E-06 |
| chr7 | 19415090 | 19416225 | 1136 | -5.57 | 2.16E-06 | 3.41E-06 |
| chr8 | 81215366 | 81216301 | 936 | -5.57 | 2.16E-06 | 3.41E-06 |
| chr6 | 85330512 | 85332134 | 1623 | -5.83 | 2.16E-06 | 3.41E-06 |
| chr3 | 182563954 | 182564702 | 749 | -5.78 | 2.16E-06 | 3.41E-06 |
| chr5 | 76743093 | 76744049 | 957 | -5.86 | 2.16E-06 | 3.42E-06 |
| chr5 | 14717470 | 14719614 | 2145 | -5.9 | 2.16E-06 | 3.42E-06 |
| chr2 | 121726666 | 121727408 | 743 | -5.51 | 2.16E-06 | 3.42E-06 |
| chr7 | 2252159 | 2252949 | 791 | -5.51 | 2.16E-06 | 3.42E-06 |
| chr7 | 112502750 | 112510083 | 7334 | -5.91 | 2.16E-06 | 3.42E-06 |
| chr1 | 37587805 | 37588913 | 1109 | -5.76 | 2.16E-06 | 3.42E-06 |
| chr9 | 76336881 | 76337845 | 965 | -5.5 | 2.17E-06 | 3.42E-06 |
| chr12 | 106332162 | 106334003 | 1842 | 2.61 | 2.17E-06 | 3.42E-06 |
| chr13 | 36979193 | 36987497 | 8305 | 1.65 | 2.17E-06 | 3.42E-06 |
| chr5 | 33951393 | 33957345 | 5953 | -1.95 | 2.17E-06 | 3.43E-06 |
| chr1 | 88352711 | 88353983 | 1273 | -5.71 | 2.17E-06 | 3.43E-06 |

| | | | | | | |
|---|---|---|---|---|---|---|
| chr3 | 107570735 | 107571923 | 1189 | -5.71 | 2.17E-06 | 3.43E-06 |
| chr11 | 21659286 | 21660958 | 1673 | -5.72 | 2.18E-06 | 3.44E-06 |
| chr1 | 15222973 | 15223975 | 1003 | -5.5 | 2.18E-06 | 3.44E-06 |
| chr11 | 86541631 | 86542911 | 1281 | -5.5 | 2.18E-06 | 3.44E-06 |
| chr18 | 10493247 | 10518968 | 25722 | -2.04 | 2.18E-06 | 3.44E-06 |
| chr10 | 98057472 | 98061298 | 3827 | 1.73 | 2.18E-06 | 3.44E-06 |
| chr5 | 53544670 | 53546014 | 1345 | -5.69 | 2.18E-06 | 3.44E-06 |
| chr4 | 54696621 | 54699911 | 3291 | -3.32 | 2.19E-06 | 3.45E-06 |
| chr2 | 70116264 | 70116958 | 695 | -5.71 | 2.19E-06 | 3.45E-06 |
| chr13 | 25598743 | 25599842 | 1100 | -5.99 | 2.19E-06 | 3.45E-06 |
| chr7 | 79956075 | 79958298 | 2224 | -5.99 | 2.19E-06 | 3.45E-06 |
| chr11 | 12461246 | 12462681 | 1436 | -5.68 | 2.19E-06 | 3.46E-06 |
| chr13 | 24970046 | 24971339 | 1294 | -5.67 | 2.20E-06 | 3.47E-06 |
| chr20 | 58314038 | 58315164 | 1127 | -5.52 | 2.20E-06 | 3.48E-06 |
| chr12 | 59982946 | 59984719 | 1774 | -5.53 | 2.20E-06 | 3.48E-06 |
| chr13 | 43452645 | 43453566 | 922 | -5.53 | 2.20E-06 | 3.48E-06 |
| chr20 | 45791217 | 45792696 | 1480 | -5.53 | 2.20E-06 | 3.48E-06 |
| chr2 | 228596338 | 228597098 | 761 | -4.52 | 2.20E-06 | 3.48E-06 |
| chr3 | 33914997 | 33916128 | 1132 | -5.68 | 2.21E-06 | 3.48E-06 |
| chr10 | 114272858 | 114273799 | 942 | -5.57 | 2.21E-06 | 3.48E-06 |
| chr3 | 29803313 | 29804704 | 1392 | -5.66 | 2.21E-06 | 3.49E-06 |
| chr18 | 11539761 | 11562243 | 22483 | -2.18 | 2.21E-06 | 3.49E-06 |
| chr13 | 101758215 | 101759522 | 1308 | -5.47 | 2.21E-06 | 3.50E-06 |
| chr1 | 22961144 | 22961860 | 717 | -5.48 | 2.22E-06 | 3.50E-06 |
| chr18 | 44193083 | 44194043 | 961 | -5.55 | 2.22E-06 | 3.50E-06 |
| chr4 | 180572729 | 180574638 | 1910 | -5.55 | 2.22E-06 | 3.50E-06 |
| chr4 | 105775345 | 105784499 | 9155 | -5.65 | 2.22E-06 | 3.51E-06 |
| chr16 | 806343 | 807193 | 851 | -5.66 | 2.22E-06 | 3.51E-06 |
| chr15 | 30046174 | 30046693 | 520 | -5.65 | 2.23E-06 | 3.52E-06 |
| chr10 | 8077387 | 8078123 | 737 | -5.5 | 2.24E-06 | 3.53E-06 |
| chr1 | 220540171 | 220576014 | 35844 | -1.59 | 2.24E-06 | 3.54E-06 |
| chr11 | 89912801 | 89922227 | 9427 | -6.15 | 2.24E-06 | 3.54E-06 |
| chr8 | 101547405 | 101565578 | 18174 | -6.32 | 2.24E-06 | 3.54E-06 |
| chr4 | 56868419 | 56869147 | 729 | -5.63 | 2.25E-06 | 3.56E-06 |
| chr5 | 172364732 | 172365455 | 724 | -5.63 | 2.25E-06 | 3.56E-06 |
| chr21 | 33298717 | 33299290 | 574 | -5.51 | 2.25E-06 | 3.56E-06 |
| chr3 | 64629521 | 64630626 | 1106 | -5.51 | 2.25E-06 | 3.56E-06 |
| chr1 | 64908583 | 64910985 | 2403 | -2.53 | 2.26E-06 | 3.56E-06 |
| chr1 | 188778977 | 188780684 | 1708 | -5.48 | 2.26E-06 | 3.56E-06 |
| chr4 | 82204319 | 82265342 | 61024 | 2.91 | 2.26E-06 | 3.56E-06 |
| chr14 | 94942688 | 94972319 | 29632 | -1.31 | 2.26E-06 | 3.57E-06 |
| chr6 | 159852167 | 159867910 | 15744 | -1.17 | 2.26E-06 | 3.57E-06 |
| chr12 | 49788389 | 49789258 | 870 | -5.63 | 2.27E-06 | 3.58E-06 |
| chr8 | 121142922 | 121150459 | 7538 | -5.47 | 2.27E-06 | 3.58E-06 |
| chrX | 100955531 | 100956623 | 1093 | -2.38 | 2.28E-06 | 3.60E-06 |
| chr7 | 92007347 | 92010677 | 3331 | -5.04 | 2.28E-06 | 3.60E-06 |
| chr6 | 160331095 | 160367691 | 36597 | -5.82 | 2.28E-06 | 3.60E-06 |
| chr17 | 70971684 | 70972299 | 616 | -5.62 | 2.28E-06 | 3.60E-06 |
| chr1 | 189473870 | 189475389 | 1520 | -5.54 | 2.29E-06 | 3.61E-06 |
| chr12 | 113812878 | 113813704 | 827 | -5.54 | 2.29E-06 | 3.61E-06 |
| chr5 | 72289542 | 72290674 | 1133 | -5.54 | 2.29E-06 | 3.61E-06 |
| chr3 | 86781126 | 86784034 | 2909 | -4.44 | 2.29E-06 | 3.62E-06 |
| chrX | 75689832 | 75692993 | 3162 | 2.51 | 2.29E-06 | 3.62E-06 |
| chr20 | 2014720 | 2071485 | 56766 | -2.11 | 2.30E-06 | 3.62E-06 |
| chr16 | 14509940 | 14510675 | 736 | -5.61 | 2.30E-06 | 3.63E-06 |
| chr8 | 37251049 | 37328107 | 77059 | -1.9 | 2.30E-06 | 3.63E-06 |
| chr6 | 80454850 | 80455681 | 832 | -5.49 | 2.32E-06 | 3.65E-06 |
| chr2 | 29544180 | 29544784 | 605 | -5.59 | 2.32E-06 | 3.65E-06 |
| chr4 | 170185011 | 170186006 | 996 | -5.59 | 2.32E-06 | 3.65E-06 |
| chr7 | 857669 | 858394 | 726 | -5.48 | 2.32E-06 | 3.66E-06 |
| chr10 | 34101923 | 34102980 | 1058 | -5.47 | 2.33E-06 | 3.67E-06 |

| | | | | | | |
|---|---|---|---|---|---|---|
| chr20 | 42755673 | 42756400 | 728 | -5.47 | 2.33E-06 | 3.67E-06 |
| chr5 | 155433772 | 155434667 | 896 | -5.47 | 2.33E-06 | 3.67E-06 |
| chr17 | 17050811 | 17051818 | 1008 | -5.46 | 2.33E-06 | 3.67E-06 |
| chr13 | 98459762 | 98492186 | 32425 | -2.35 | 2.34E-06 | 3.69E-06 |
| chr2 | 135088539 | 135089626 | 1088 | -5.59 | 2.34E-06 | 3.69E-06 |
| chrX | 130804067 | 130805314 | 1248 | -5.59 | 2.34E-06 | 3.69E-06 |
| chr17 | 25531410 | 25552239 | 20830 | -5.58 | 2.35E-06 | 3.70E-06 |
| chr6 | 152299144 | 152300342 | 1199 | -2.06 | 2.35E-06 | 3.70E-06 |
| chr4 | 106800006 | 106815274 | 15269 | -3.95 | 2.35E-06 | 3.70E-06 |
| chr6 | 33471510 | 33472621 | 1112 | -5.47 | 2.36E-06 | 3.71E-06 |
| chr1 | 11326131 | 11326966 | 836 | -5.46 | 2.36E-06 | 3.71E-06 |
| chr7 | 64696241 | 64697706 | 1466 | -5.58 | 2.36E-06 | 3.72E-06 |
| chr11 | 109501023 | 109549279 | 48257 | 1 | 2.36E-06 | 3.72E-06 |
| chr13 | 27356034 | 27357933 | 1900 | -2.67 | 2.37E-06 | 3.74E-06 |
| chr5 | 159612927 | 159614200 | 1274 | -5.81 | 2.38E-06 | 3.74E-06 |
| chr11 | 119737740 | 119738810 | 1071 | -5.48 | 2.38E-06 | 3.74E-06 |
| chr12 | 76382432 | 76383878 | 1447 | -5.83 | 2.38E-06 | 3.74E-06 |
| chr1 | 95756364 | 95757820 | 1457 | -5.76 | 2.38E-06 | 3.74E-06 |
| chr20 | 46174049 | 46184325 | 10277 | -5.76 | 2.38E-06 | 3.74E-06 |
| chr4 | 105819390 | 105821095 | 1706 | -5.76 | 2.38E-06 | 3.74E-06 |
| chr7 | 17713724 | 17715803 | 2080 | -5.76 | 2.38E-06 | 3.74E-06 |
| chr6 | 138206518 | 138208076 | 1559 | -5.84 | 2.38E-06 | 3.75E-06 |
| chr3 | 143701050 | 143702220 | 1171 | -5.57 | 2.39E-06 | 3.76E-06 |
| chr11 | 33750530 | 33751402 | 873 | -5.52 | 2.39E-06 | 3.76E-06 |
| chr7 | 37493767 | 37906775 | 413009 | 1.86 | 2.39E-06 | 3.76E-06 |
| chr6 | 3412579 | 3413375 | 797 | -5.9 | 2.39E-06 | 3.77E-06 |
| chr14 | 74028423 | 74029995 | 1573 | -5.71 | 2.39E-06 | 3.77E-06 |
| chr3 | 159266702 | 159267784 | 1083 | -4.41 | 2.40E-06 | 3.78E-06 |
| chrX | 41228303 | 41229247 | 945 | -5.7 | 2.40E-06 | 3.78E-06 |
| chr4 | 174697488 | 174698563 | 1076 | -5.47 | 2.40E-06 | 3.78E-06 |
| chrX | 84249786 | 84250516 | 731 | -5.47 | 2.40E-06 | 3.78E-06 |
| chr11 | 20162221 | 20164137 | 1917 | -4.23 | 2.40E-06 | 3.78E-06 |
| chr12 | 107243302 | 107244076 | 775 | -5.49 | 2.40E-06 | 3.78E-06 |
| chr17 | 55672349 | 55673138 | 790 | -5.49 | 2.40E-06 | 3.78E-06 |
| chr20 | 18403480 | 18404024 | 545 | -5.49 | 2.40E-06 | 3.78E-06 |
| chrX | 100031222 | 100032228 | 1007 | -4.99 | 2.40E-06 | 3.78E-06 |
| chr18 | 30474569 | 30477906 | 3338 | -2.82 | 2.41E-06 | 3.79E-06 |
| chr15 | 64678724 | 64679459 | 736 | -5.47 | 2.41E-06 | 3.79E-06 |
| chr6 | 64445237 | 64446093 | 857 | -5.47 | 2.41E-06 | 3.79E-06 |
| chr7 | 62985420 | 62986060 | 641 | -5.47 | 2.41E-06 | 3.79E-06 |
| chr9 | 114933852 | 114935464 | 1613 | -5.47 | 2.41E-06 | 3.79E-06 |
| chr10 | 111676252 | 111676951 | 700 | -5.56 | 2.41E-06 | 3.80E-06 |
| chr22 | 19834577 | 19835478 | 902 | -5.67 | 2.42E-06 | 3.80E-06 |
| chr12 | 111257531 | 111259793 | 2263 | -6 | 2.42E-06 | 3.80E-06 |
| chrX | 22197698 | 22211906 | 14209 | -6.01 | 2.42E-06 | 3.81E-06 |
| chr6 | 137883891 | 137886046 | 2156 | -4.57 | 2.42E-06 | 3.81E-06 |
| chrX | 123144128 | 123145240 | 1113 | -5.5 | 2.43E-06 | 3.82E-06 |
| chrX | 30576115 | 30579202 | 3088 | 2.34 | 2.43E-06 | 3.82E-06 |
| chr4 | 63852455 | 63854014 | 1560 | -4.26 | 2.44E-06 | 3.84E-06 |
| chr18 | 29206339 | 29207053 | 715 | -5.55 | 2.44E-06 | 3.84E-06 |
| chr2 | 102646249 | 102647444 | 1196 | -6.1 | 2.44E-06 | 3.84E-06 |
| chr19 | 9953884 | 9954770 | 887 | -5.5 | 2.45E-06 | 3.85E-06 |
| chr19 | 15416182 | 15417034 | 853 | -5.5 | 2.45E-06 | 3.85E-06 |
| chr7 | 17336147 | 17338039 | 1893 | -3.37 | 2.45E-06 | 3.85E-06 |
| chr2 | 143945344 | 143947116 | 1773 | 6.13 | 2.45E-06 | 3.86E-06 |
| chr8 | 62041815 | 62044055 | 2241 | 6.37 | 2.45E-06 | 3.86E-06 |
| chr2 | 160417485 | 160431117 | 13633 | -6.16 | 2.45E-06 | 3.86E-06 |
| chr4 | 185600285 | 185601246 | 962 | -5.85 | 2.45E-06 | 3.86E-06 |
| chrX | 73899293 | 73900305 | 1013 | 4.45 | 2.45E-06 | 3.86E-06 |
| chrX | 73736389 | 73737568 | 1180 | -5.75 | 2.46E-06 | 3.86E-06 |
| chr12 | 32410699 | 32411410 | 712 | -5.74 | 2.46E-06 | 3.86E-06 |

| | | | | | | |
|---|---|---|---|---|---|---|
| chr9 | 130990654 | 130991462 | 809 | -5.45 | 2.46E-06 | 3.87E-06 |
| chr4 | 34376668 | 34377932 | 1265 | -5.63 | 2.46E-06 | 3.87E-06 |
| chr7 | 62121015 | 62146284 | 25270 | -2.13 | 2.47E-06 | 3.88E-06 |
| chr6 | 47011103 | 47031664 | 20562 | -6.68 | 2.47E-06 | 3.88E-06 |
| chr15 | 77616161 | 77616835 | 675 | -5.46 | 2.47E-06 | 3.89E-06 |
| chr18 | 44820587 | 44822665 | 2079 | 5.61 | 2.48E-06 | 3.89E-06 |
| chr12 | 67306524 | 67307314 | 791 | -5.45 | 2.48E-06 | 3.89E-06 |
| chr2 | 109218073 | 109219413 | 1341 | -5.7 | 2.48E-06 | 3.89E-06 |
| chr15 | 55733814 | 55735750 | 1937 | -5.49 | 2.48E-06 | 3.90E-06 |
| chr13 | 106422544 | 106423637 | 1094 | -5.69 | 2.48E-06 | 3.90E-06 |
| chr2 | 68077028 | 68152867 | 75840 | 1.08 | 2.49E-06 | 3.91E-06 |
| chr12 | 22730634 | 22731625 | 992 | -5.68 | 2.49E-06 | 3.91E-06 |
| chr4 | 123768391 | 123769288 | 898 | -5.68 | 2.49E-06 | 3.91E-06 |
| chr1 | 189443937 | 189445161 | 1225 | -5.53 | 2.49E-06 | 3.91E-06 |
| chr5 | 38675459 | 38676107 | 649 | -4.22 | 2.49E-06 | 3.92E-06 |
| chr10 | 105320449 | 105321329 | 881 | -5.61 | 2.50E-06 | 3.93E-06 |
| chr11 | 134148027 | 134148946 | 920 | -5.61 | 2.50E-06 | 3.93E-06 |
| chr12 | 31102618 | 31103963 | 1346 | -5.61 | 2.50E-06 | 3.93E-06 |
| chr19 | 10317707 | 10318696 | 990 | -5.53 | 2.51E-06 | 3.94E-06 |
| chr20 | 45632306 | 45633260 | 955 | -5.53 | 2.51E-06 | 3.94E-06 |
| chr6 | 153529941 | 153531046 | 1106 | -5.53 | 2.51E-06 | 3.94E-06 |
| chr4 | 166096004 | 166096880 | 877 | -5.49 | 2.52E-06 | 3.95E-06 |
| chr8 | 28732474 | 28733183 | 710 | -5.59 | 2.52E-06 | 3.96E-06 |
| chr13 | 111910930 | 111912415 | 1486 | -5.64 | 2.52E-06 | 3.96E-06 |
| chr7 | 148043974 | 148122244 | 78271 | 1.46 | 2.53E-06 | 3.98E-06 |
| chr3 | 36607026 | 36630553 | 23528 | -5.75 | 2.53E-06 | 3.98E-06 |
| chr5 | 172010182 | 172011506 | 1325 | 4.49 | 2.54E-06 | 3.99E-06 |
| chr21 | 46134761 | 46136383 | 1623 | -5.48 | 2.54E-06 | 3.99E-06 |
| chr8 | 123934990 | 123936181 | 1192 | -5.48 | 2.54E-06 | 3.99E-06 |
| chr11 | 107396451 | 107397278 | 828 | -5.52 | 2.55E-06 | 4.00E-06 |
| chr3 | 158411621 | 158412114 | 494 | -5.52 | 2.55E-06 | 4.00E-06 |
| chr14 | 80937972 | 80941684 | 3713 | -4.7 | 2.55E-06 | 4.00E-06 |
| chr22 | 31789613 | 31790628 | 1016 | -5.11 | 2.55E-06 | 4.01E-06 |
| chrX | 12613265 | 12639482 | 26218 | 5.64 | 2.55E-06 | 4.01E-06 |
| chr4 | 75932407 | 75934070 | 1664 | -2.4 | 2.55E-06 | 4.01E-06 |
| chr4 | 108754668 | 108755461 | 794 | -5.61 | 2.56E-06 | 4.02E-06 |
| chr4 | 152618158 | 152619034 | 877 | -5.61 | 2.56E-06 | 4.02E-06 |
| chrX | 3860070 | 3871785 | 11716 | -3.97 | 2.57E-06 | 4.03E-06 |
| chr3 | 124152548 | 124155190 | 2643 | -1.75 | 2.57E-06 | 4.03E-06 |
| chr2 | 153917241 | 153918166 | 926 | -5.57 | 2.57E-06 | 4.04E-06 |
| chr6 | 170539645 | 170540641 | 997 | -5.57 | 2.57E-06 | 4.04E-06 |
| chr1 | 230563189 | 230695734 | 132546 | -1.75 | 2.57E-06 | 4.04E-06 |
| chr16 | 72585440 | 72586448 | 1009 | -5.6 | 2.58E-06 | 4.04E-06 |
| chrX | 35842317 | 35843904 | 1588 | -3.74 | 2.58E-06 | 4.05E-06 |
| chr13 | 23601447 | 23603458 | 2012 | -4.22 | 2.58E-06 | 4.05E-06 |
| chr18 | 57039435 | 57044199 | 4765 | 2.12 | 2.59E-06 | 4.06E-06 |
| chr8 | 73894704 | 73904597 | 9894 | -2.08 | 2.59E-06 | 4.07E-06 |
| chr22 | 44338480 | 44339423 | 944 | -5.6 | 2.59E-06 | 4.07E-06 |
| chr3 | 109741052 | 109742731 | 1680 | -5.6 | 2.59E-06 | 4.07E-06 |
| chr15 | 77329328 | 77330487 | 1160 | -5.51 | 2.59E-06 | 4.07E-06 |
| chr7 | 79101893 | 79102720 | 828 | -5.44 | 2.60E-06 | 4.08E-06 |
| chr1 | 28459826 | 28461461 | 1636 | -5.56 | 2.60E-06 | 4.08E-06 |
| chr12 | 63136752 | 63150462 | 13711 | -5.56 | 2.60E-06 | 4.08E-06 |
| chr15 | 83987166 | 83988167 | 1002 | -5.47 | 2.61E-06 | 4.09E-06 |
| chr3 | 50294662 | 50295552 | 891 | -5.47 | 2.61E-06 | 4.09E-06 |
| chr5 | 129510307 | 129510938 | 632 | -5.47 | 2.61E-06 | 4.09E-06 |
| chr4 | 8139717 | 8140756 | 1040 | -5.44 | 2.61E-06 | 4.09E-06 |
| chr5 | 173120341 | 173121409 | 1069 | -5.45 | 2.61E-06 | 4.09E-06 |
| chr2 | 223023160 | 223028859 | 5700 | 4.99 | 2.61E-06 | 4.09E-06 |
| chr4 | 87711220 | 87712086 | 867 | -5.59 | 2.61E-06 | 4.09E-06 |
| chr7 | 95562590 | 95563592 | 1003 | 4.19 | 2.61E-06 | 4.10E-06 |

| | | | | | | |
|---|---|---|---|---|---|---|
| chr6 | 91133445 | 91142910 | 9466 | -4.59 | 2.62E-06 | 4.10E-06 |
| chr4 | 185141836 | 185158096 | 16261 | -1.48 | 2.62E-06 | 4.10E-06 |
| chr22 | 26645938 | 26647621 | 1684 | -2.55 | 2.62E-06 | 4.11E-06 |
| chrX | 131296123 | 131311968 | 15846 | -2.35 | 2.62E-06 | 4.11E-06 |
| chr2 | 18579643 | 18582457 | 2815 | 4.77 | 2.62E-06 | 4.11E-06 |
| chr2 | 179049440 | 179050806 | 1367 | -4.54 | 2.63E-06 | 4.12E-06 |
| chr1 | 218271939 | 218275144 | 3206 | -2.17 | 2.63E-06 | 4.12E-06 |
| chr1 | 62708748 | 62710558 | 1811 | -5.79 | 2.63E-06 | 4.12E-06 |
| chr6 | 89663663 | 89678153 | 14491 | -5.77 | 2.63E-06 | 4.12E-06 |
| chr12 | 105337648 | 105338826 | 1179 | -5.76 | 2.63E-06 | 4.12E-06 |
| chr3 | 24445257 | 24446147 | 891 | -5.75 | 2.63E-06 | 4.13E-06 |
| chrX | 151136400 | 151149455 | 13056 | -5.83 | 2.63E-06 | 4.13E-06 |
| chrX | 71255821 | 71257183 | 1363 | -5.74 | 2.63E-06 | 4.13E-06 |
| chr4 | 98539415 | 98541275 | 1861 | -5.5 | 2.63E-06 | 4.13E-06 |
| chrX | 75774818 | 75776389 | 1572 | -5.5 | 2.63E-06 | 4.13E-06 |
| chr7 | 109974159 | 109975349 | 1191 | -5.55 | 2.64E-06 | 4.13E-06 |
| chr11 | 102763767 | 102767282 | 3516 | -1.52 | 2.64E-06 | 4.14E-06 |
| chr11 | 17462337 | 17463430 | 1094 | -4.18 | 2.64E-06 | 4.14E-06 |
| chr1 | 172647797 | 172648991 | 1195 | -5.88 | 2.64E-06 | 4.14E-06 |
| chr10 | 64890092 | 64892849 | 2758 | -1.68 | 2.65E-06 | 4.16E-06 |
| chr11 | 47917188 | 47918516 | 1329 | -5.92 | 2.66E-06 | 4.16E-06 |
| chr4 | 108293200 | 108294970 | 1771 | -5.67 | 2.67E-06 | 4.18E-06 |
| chr2 | 42249804 | 42250594 | 791 | -4.72 | 2.67E-06 | 4.19E-06 |
| chr4 | 92876012 | 92877288 | 1277 | -5.46 | 2.68E-06 | 4.19E-06 |
| chr6 | 165813374 | 165814354 | 981 | -5.46 | 2.68E-06 | 4.19E-06 |
| chr8 | 118901095 | 118901720 | 626 | -5.46 | 2.68E-06 | 4.19E-06 |
| chr10 | 101471152 | 101472131 | 980 | -5.53 | 2.68E-06 | 4.19E-06 |
| chr13 | 100138348 | 100139364 | 1017 | -5.53 | 2.68E-06 | 4.19E-06 |
| chr9 | 6707491 | 6709273 | 1783 | -5.53 | 2.68E-06 | 4.19E-06 |
| chrX | 76941117 | 76942036 | 920 | -5.56 | 2.68E-06 | 4.20E-06 |
| chr19 | 56120844 | 56121791 | 948 | -5.65 | 2.68E-06 | 4.20E-06 |
| chr6 | 24431459 | 24432036 | 578 | -5.65 | 2.68E-06 | 4.20E-06 |
| chr12 | 58416990 | 58445883 | 28894 | -1.41 | 2.68E-06 | 4.20E-06 |
| chr8 | 103004533 | 103047880 | 43348 | -3.47 | 2.68E-06 | 4.20E-06 |
| chr1 | 68147503 | 68148679 | 1177 | -5.45 | 2.69E-06 | 4.21E-06 |
| chr4 | 188931717 | 188932479 | 763 | -5.45 | 2.69E-06 | 4.21E-06 |
| chr7 | 83096587 | 83097272 | 686 | -5.45 | 2.69E-06 | 4.21E-06 |
| chr9 | 115607363 | 115608523 | 1161 | -5.45 | 2.69E-06 | 4.21E-06 |
| chr6 | 169363341 | 169384331 | 20991 | -4.28 | 2.69E-06 | 4.21E-06 |
| chr7 | 151635433 | 151671400 | 35968 | -6.12 | 2.69E-06 | 4.21E-06 |
| chr1 | 243130611 | 243131108 | 498 | -4.17 | 2.69E-06 | 4.22E-06 |
| chr20 | 61379379 | 61389192 | 9814 | -6.09 | 2.70E-06 | 4.22E-06 |
| chr6 | 38893912 | 38898800 | 4889 | 4.86 | 2.70E-06 | 4.22E-06 |
| chr4 | 84947965 | 84948575 | 611 | -4.71 | 2.70E-06 | 4.23E-06 |
| chr4 | 63516590 | 63519178 | 2589 | -5.44 | 2.70E-06 | 4.23E-06 |
| chrX | 15053441 | 15098908 | 45468 | 1.92 | 2.71E-06 | 4.25E-06 |
| chr8 | 143062834 | 143068027 | 5194 | 1.82 | 2.72E-06 | 4.25E-06 |
| chr19 | 13890505 | 13891029 | 525 | -5.48 | 2.72E-06 | 4.26E-06 |
| chr4 | 129540642 | 129541952 | 1311 | -5.48 | 2.72E-06 | 4.26E-06 |
| chr8 | 117486142 | 117487491 | 1350 | -5.48 | 2.72E-06 | 4.26E-06 |
| chr6 | 112239213 | 112240385 | 1173 | 5.58 | 2.73E-06 | 4.28E-06 |
| chr4 | 103124565 | 103125996 | 1432 | -5.61 | 2.74E-06 | 4.28E-06 |
| chr3 | 118399232 | 118466554 | 67323 | 0.8 | 2.74E-06 | 4.29E-06 |
| chr4 | 37506863 | 37507675 | 813 | -3.15 | 2.75E-06 | 4.30E-06 |
| chr10 | 8382737 | 8383610 | 874 | -5.43 | 2.75E-06 | 4.30E-06 |
| chr4 | 141022847 | 141023780 | 934 | -5.43 | 2.75E-06 | 4.30E-06 |
| chr5 | 90644489 | 90645215 | 727 | -5.45 | 2.75E-06 | 4.31E-06 |
| chr15 | 59519588 | 59520358 | 771 | -5.43 | 2.76E-06 | 4.32E-06 |
| chr17 | 61163952 | 61164548 | 597 | -5.43 | 2.76E-06 | 4.32E-06 |
| chr3 | 129997030 | 129997990 | 961 | -5.24 | 2.76E-06 | 4.32E-06 |
| chr1 | 62385409 | 62386318 | 910 | -5.51 | 2.77E-06 | 4.33E-06 |

| | | | | | | |
|---|---|---|---|---|---|---|
| chr10 | 121056798 | 121057714 | 917 | -5.51 | 2.77E-06 | 4.33E-06 |
| chr9 | 36043580 | 36044164 | 585 | -5.43 | 2.77E-06 | 4.33E-06 |
| chr2 | 220004352 | 220005146 | 795 | -5.45 | 2.77E-06 | 4.33E-06 |
| chr6 | 105503606 | 105504339 | 734 | -5.58 | 2.78E-06 | 4.34E-06 |
| chr7 | 21999076 | 22000188 | 1113 | -5.58 | 2.78E-06 | 4.34E-06 |
| chr7 | 79602782 | 79603465 | 684 | -5.58 | 2.78E-06 | 4.34E-06 |
| chr22 | 49373787 | 49400253 | 26467 | -1.9 | 2.78E-06 | 4.34E-06 |
| chr13 | 51535550 | 51536100 | 551 | -5.77 | 2.79E-06 | 4.36E-06 |
| chr10 | 3176141 | 3176921 | 781 | -5.75 | 2.79E-06 | 4.36E-06 |
| chr2 | 141588625 | 141590364 | 1740 | -5.73 | 2.79E-06 | 4.37E-06 |
| chr13 | 105675007 | 105676125 | 1119 | -5.43 | 2.80E-06 | 4.38E-06 |
| chr3 | 9278369 | 9279006 | 638 | -5.43 | 2.80E-06 | 4.38E-06 |
| chr11 | 116524954 | 116556300 | 31347 | -4.06 | 2.80E-06 | 4.38E-06 |
| chr2 | 180676157 | 180677242 | 1086 | -5.47 | 2.80E-06 | 4.38E-06 |
| chr2 | 201316084 | 201317047 | 964 | -5.47 | 2.80E-06 | 4.38E-06 |
| chr18 | 37064598 | 37065485 | 888 | 5.51 | 2.80E-06 | 4.38E-06 |
| chr11 | 86185834 | 86197561 | 11728 | -6.39 | 2.80E-06 | 4.38E-06 |
| chr4 | 147987684 | 147997574 | 9891 | -5.44 | 2.81E-06 | 4.39E-06 |
| chr11 | 7701278 | 7702386 | 1109 | -5.68 | 2.82E-06 | 4.40E-06 |
| chr6 | 106335617 | 106348056 | 12440 | -5.29 | 2.82E-06 | 4.41E-06 |
| chr1 | 15797006 | 15797949 | 944 | -5.42 | 2.82E-06 | 4.41E-06 |
| chr6 | 123097085 | 123097707 | 623 | -5.42 | 2.82E-06 | 4.41E-06 |
| chr4 | 6740684 | 6741877 | 1194 | -5.5 | 2.82E-06 | 4.41E-06 |
| chr12 | 131370550 | 131371237 | 688 | -5.56 | 2.83E-06 | 4.42E-06 |
| chr6 | 144224721 | 144253532 | 28812 | -2.56 | 2.83E-06 | 4.43E-06 |
| chr13 | 68659050 | 68660277 | 1228 | -5.44 | 2.84E-06 | 4.44E-06 |
| chr2 | 207478221 | 207479099 | 879 | -5.44 | 2.84E-06 | 4.44E-06 |
| chr7 | 82114137 | 82115529 | 1393 | -5.44 | 2.84E-06 | 4.44E-06 |
| chr6 | 122895336 | 122896887 | 1552 | -3.66 | 2.84E-06 | 4.44E-06 |
| chr1 | 52635870 | 52636937 | 1068 | -5.51 | 2.84E-06 | 4.44E-06 |
| chr22 | 35614040 | 35614653 | 614 | -5.51 | 2.84E-06 | 4.44E-06 |
| chr12 | 99347768 | 99348487 | 720 | 5.66 | 2.84E-06 | 4.44E-06 |
| chr10 | 3452586 | 3454168 | 1583 | -5.64 | 2.85E-06 | 4.45E-06 |
| chr16 | 72267496 | 72268114 | 619 | -5.64 | 2.85E-06 | 4.45E-06 |
| chr4 | 3358738 | 3359909 | 1172 | -5.64 | 2.85E-06 | 4.45E-06 |
| chr5 | 137280007 | 137280547 | 541 | -5.46 | 2.87E-06 | 4.48E-06 |
| chr7 | 103143592 | 103146098 | 2507 | -1.95 | 2.87E-06 | 4.48E-06 |
| chr7 | 102790148 | 102790866 | 719 | -4.05 | 2.87E-06 | 4.48E-06 |
| chr1 | 229798167 | 229799046 | 880 | -5.44 | 2.88E-06 | 4.50E-06 |
| chr2 | 234249819 | 234250530 | 712 | -5.44 | 2.88E-06 | 4.50E-06 |
| chr10 | 33818574 | 33833465 | 14892 | -6.86 | 2.88E-06 | 4.50E-06 |
| chr5 | 153973531 | 153979235 | 5705 | -6.7 | 2.88E-06 | 4.50E-06 |
| chr3 | 105900713 | 105905511 | 4799 | -4.73 | 2.88E-06 | 4.50E-06 |
| chr7 | 129749451 | 129750409 | 959 | -5.5 | 2.88E-06 | 4.50E-06 |
| chr9 | 20388963 | 20389605 | 643 | -5.5 | 2.88E-06 | 4.50E-06 |
| chr1 | 1969626 | 1970845 | 1220 | -5.45 | 2.89E-06 | 4.51E-06 |
| chr2 | 203845457 | 203846182 | 726 | -5.45 | 2.89E-06 | 4.51E-06 |
| chr4 | 63783396 | 63785552 | 2157 | -5.48 | 2.89E-06 | 4.51E-06 |
| chr5 | 156169136 | 156170242 | 1107 | -5.48 | 2.89E-06 | 4.51E-06 |
| chr10 | 64259334 | 64261476 | 2143 | -5.6 | 2.90E-06 | 4.52E-06 |
| chr15 | 92608154 | 92611289 | 3136 | -6.41 | 2.91E-06 | 4.54E-06 |
| chr20 | 9549551 | 9550120 | 570 | -5.42 | 2.91E-06 | 4.54E-06 |
| chr20 | 37497706 | 37498410 | 705 | -5.42 | 2.91E-06 | 4.54E-06 |
| chr1 | 214901184 | 214903894 | 2711 | -4.55 | 2.91E-06 | 4.54E-06 |
| chr13 | 61010470 | 61011233 | 764 | -5.42 | 2.91E-06 | 4.54E-06 |
| chr3 | 190151484 | 190153088 | 1605 | -5.59 | 2.91E-06 | 4.55E-06 |
| chr6 | 148075117 | 148076805 | 1689 | -5.78 | 2.92E-06 | 4.55E-06 |
| chr1 | 62562585 | 62563661 | 1077 | -5.8 | 2.92E-06 | 4.55E-06 |
| chrX | 45517704 | 45518938 | 1235 | -5.75 | 2.92E-06 | 4.55E-06 |
| chr6 | 139951588 | 139953129 | 1542 | -5.81 | 2.92E-06 | 4.55E-06 |
| chr2 | 146492818 | 146493930 | 1113 | -5.82 | 2.92E-06 | 4.55E-06 |

| | | | | | | |
|---|---|---|---|---|---|---|
| chr6 | 45690140 | 45692056 | 1917 | -5.82 | 2.92E-06 | 4.55E-06 |
| chr13 | 50989089 | 50990346 | 1258 | -5.73 | 2.92E-06 | 4.55E-06 |
| chr1 | 79685551 | 79687846 | 2296 | -4.71 | 2.92E-06 | 4.56E-06 |
| chr14 | 51812178 | 51814174 | 1997 | -5.84 | 2.92E-06 | 4.56E-06 |
| chr10 | 111950992 | 111951943 | 952 | -5.71 | 2.93E-06 | 4.56E-06 |
| chr13 | 106598412 | 106606118 | 7707 | -5.71 | 2.93E-06 | 4.56E-06 |
| chr6 | 160947661 | 160951443 | 3783 | -2.85 | 2.93E-06 | 4.57E-06 |
| chr1 | 201556920 | 201557935 | 1016 | -5.7 | 2.93E-06 | 4.57E-06 |
| chr1 | 48862259 | 48863376 | 1118 | -5.49 | 2.93E-06 | 4.57E-06 |
| chr2 | 43063898 | 43065250 | 1353 | -5.88 | 2.93E-06 | 4.57E-06 |
| chr5 | 179237741 | 179238300 | 560 | -5.69 | 2.93E-06 | 4.58E-06 |
| chr1 | 81606078 | 81607520 | 1443 | -5.43 | 2.94E-06 | 4.58E-06 |
| chr20 | 36745014 | 36745875 | 862 | -5.43 | 2.94E-06 | 4.58E-06 |
| chr3 | 133277579 | 133278801 | 1223 | -5.68 | 2.94E-06 | 4.58E-06 |
| chr6 | 13534004 | 13535346 | 1343 | -5.68 | 2.94E-06 | 4.58E-06 |
| chr10 | 29166752 | 29175982 | 9231 | -5.94 | 2.94E-06 | 4.59E-06 |
| chr21 | 34359827 | 34386910 | 27084 | -3.49 | 2.95E-06 | 4.59E-06 |
| chr1 | 185862523 | 185863775 | 1253 | -5.67 | 2.95E-06 | 4.59E-06 |
| chr15 | 68141807 | 68142358 | 552 | -5.67 | 2.95E-06 | 4.59E-06 |
| chr5 | 55778670 | 55785018 | 6349 | -5.67 | 2.95E-06 | 4.59E-06 |
| chr2 | 211108684 | 211109678 | 995 | -5.57 | 2.95E-06 | 4.59E-06 |
| chr7 | 31764995 | 31765943 | 949 | -3.32 | 2.95E-06 | 4.59E-06 |
| chr6 | 13020449 | 13022365 | 1917 | -2.19 | 2.95E-06 | 4.60E-06 |
| chr12 | 89565656 | 89566754 | 1099 | -5.66 | 2.95E-06 | 4.61E-06 |
| chr14 | 70728339 | 70730054 | 1716 | -5.47 | 2.96E-06 | 4.61E-06 |
| chr18 | 20788052 | 20788979 | 928 | -5.47 | 2.96E-06 | 4.61E-06 |
| chr19 | 3139177 | 3140795 | 1619 | -5.47 | 2.96E-06 | 4.61E-06 |
| chr3 | 42043872 | 42044617 | 746 | -5.47 | 2.96E-06 | 4.61E-06 |
| chr6 | 155518360 | 155519182 | 823 | -5.47 | 2.96E-06 | 4.61E-06 |
| chr12 | 76023412 | 76025647 | 2236 | -6.31 | 2.96E-06 | 4.61E-06 |
| chr1 | 200184015 | 200185168 | 1154 | -5.41 | 2.96E-06 | 4.61E-06 |
| chr4 | 187749741 | 187750794 | 1054 | -5.41 | 2.96E-06 | 4.61E-06 |
| chr1 | 37619929 | 37634874 | 14946 | -3.14 | 2.96E-06 | 4.61E-06 |
| chr16 | 64203800 | 64274649 | 70850 | 1.06 | 2.96E-06 | 4.61E-06 |
| chr3 | 140760791 | 140761625 | 835 | -5.65 | 2.96E-06 | 4.62E-06 |
| chr7 | 79740522 | 79741766 | 1245 | -5.65 | 2.96E-06 | 4.62E-06 |
| chr6 | 88435391 | 88436333 | 943 | -5.56 | 2.97E-06 | 4.62E-06 |
| chrX | 81233047 | 81245534 | 12488 | 2.79 | 2.97E-06 | 4.63E-06 |
| chr3 | 181940857 | 181942029 | 1173 | -5.63 | 2.98E-06 | 4.64E-06 |
| chr1 | 233019269 | 233020228 | 960 | 4.29 | 2.98E-06 | 4.64E-06 |
| chr5 | 66172518 | 66180140 | 7623 | -6.22 | 2.98E-06 | 4.64E-06 |
| chr10 | 98104190 | 98120875 | 16686 | -1.57 | 2.98E-06 | 4.64E-06 |
| chr17 | 9841367 | 9842565 | 1199 | -6.1 | 2.98E-06 | 4.64E-06 |
| chr13 | 31863082 | 31864001 | 920 | -5.48 | 2.98E-06 | 4.64E-06 |
| chr4 | 99016199 | 99017359 | 1161 | -5.48 | 2.98E-06 | 4.64E-06 |
| chr5 | 106595988 | 106597465 | 1478 | -4.97 | 2.98E-06 | 4.65E-06 |
| chr2 | 147283697 | 147285168 | 1472 | -4.95 | 2.99E-06 | 4.65E-06 |
| chr14 | 57922885 | 57923760 | 876 | -5.62 | 2.99E-06 | 4.65E-06 |
| chr8 | 61794599 | 61805853 | 11255 | -2.44 | 3.00E-06 | 4.67E-06 |
| chr3 | 174692915 | 174693791 | 877 | -5.43 | 3.00E-06 | 4.67E-06 |
| chr5 | 81417860 | 81419283 | 1424 | -5.43 | 3.00E-06 | 4.67E-06 |
| chr9 | 133603394 | 133604557 | 1164 | -5.61 | 3.00E-06 | 4.68E-06 |
| chr1 | 104127401 | 104128804 | 1404 | 3.36 | 3.01E-06 | 4.68E-06 |
| chrX | 49917974 | 49919808 | 1835 | -2.72 | 3.01E-06 | 4.69E-06 |
| chr12 | 106921476 | 106925599 | 4124 | -4.9 | 3.01E-06 | 4.69E-06 |
| chr14 | 35869349 | 35870192 | 844 | -4.37 | 3.02E-06 | 4.69E-06 |
| chr6 | 167217080 | 167223267 | 6188 | -5.02 | 3.02E-06 | 4.70E-06 |
| chr1 | 15336288 | 15337237 | 950 | -5.5 | 3.03E-06 | 4.71E-06 |
| chr10 | 69434759 | 69455180 | 20422 | -1.67 | 3.03E-06 | 4.71E-06 |
| chr12 | 31482513 | 31483916 | 1404 | -5.46 | 3.03E-06 | 4.72E-06 |
| chr17 | 7020594 | 7021298 | 705 | -5.46 | 3.03E-06 | 4.72E-06 |

| | | | | | | |
|---|---|---|---|---|---|---|
| chr21 | 38811600 | 38812804 | 1205 | -5.41 | 3.03E-06 | 4.72E-06 |
| chr12 | 113481796 | 113482493 | 698 | -4.67 | 3.04E-06 | 4.73E-06 |
| chr4 | 96708953 | 96710892 | 1940 | -4.69 | 3.04E-06 | 4.73E-06 |
| chr17 | 2809819 | 2810625 | 807 | -5.53 | 3.04E-06 | 4.73E-06 |
| chr3 | 183788280 | 183789068 | 789 | -5.53 | 3.04E-06 | 4.73E-06 |
| chr8 | 134407302 | 134408078 | 777 | -5.57 | 3.06E-06 | 4.76E-06 |
| chr10 | 16936652 | 16946310 | 9659 | -3.11 | 3.07E-06 | 4.77E-06 |
| chr6 | 17434664 | 17435212 | 549 | -4.72 | 3.07E-06 | 4.77E-06 |
| chrX | 135803320 | 135804578 | 1259 | -2.23 | 3.07E-06 | 4.78E-06 |
| chr6 | 49034316 | 49035065 | 750 | -3.4 | 3.07E-06 | 4.78E-06 |
| chr20 | 24593402 | 24594633 | 1232 | 5.48 | 3.07E-06 | 4.78E-06 |
| chr8 | 131500062 | 131500872 | 811 | -5.52 | 3.07E-06 | 4.78E-06 |
| chr4 | 132128777 | 132143218 | 14442 | -5.49 | 3.08E-06 | 4.79E-06 |
| chr3 | 44924462 | 44925357 | 896 | -5.56 | 3.09E-06 | 4.80E-06 |
| chr6 | 38168445 | 38169419 | 975 | -5.56 | 3.09E-06 | 4.80E-06 |
| chr17 | 41523986 | 41524635 | 650 | -5.43 | 3.09E-06 | 4.81E-06 |
| chr3 | 42837532 | 42838213 | 682 | -5.46 | 3.10E-06 | 4.82E-06 |
| chr17 | 2235001 | 2235825 | 825 | -5.51 | 3.11E-06 | 4.83E-06 |
| chr6 | 109591882 | 109592737 | 856 | -5.51 | 3.11E-06 | 4.83E-06 |
| chr18 | 55462487 | 55463084 | 598 | -5.4 | 3.11E-06 | 4.83E-06 |
| chr6 | 152882493 | 152883164 | 672 | -5.4 | 3.11E-06 | 4.83E-06 |
| chr4 | 145896332 | 145897642 | 1311 | -5.55 | 3.11E-06 | 4.84E-06 |
| chr7 | 25150193 | 25150849 | 657 | -5.55 | 3.11E-06 | 4.84E-06 |
| chr20 | 35220562 | 35221508 | 947 | -5.44 | 3.12E-06 | 4.85E-06 |
| chr5 | 77899121 | 77900282 | 1162 | -5.44 | 3.12E-06 | 4.85E-06 |
| chr9 | 17371064 | 17372238 | 1175 | -4.11 | 3.12E-06 | 4.85E-06 |
| chr9 | 74076956 | 74078610 | 1655 | 2.67 | 3.13E-06 | 4.87E-06 |
| chr6 | 20381117 | 20382300 | 1184 | -4.21 | 3.13E-06 | 4.87E-06 |
| chrX | 114199847 | 114239926 | 40080 | 5.9 | 3.14E-06 | 4.88E-06 |
| chrX | 133006046 | 133008112 | 2067 | -2.2 | 3.14E-06 | 4.88E-06 |
| chr20 | 24573963 | 24585910 | 11948 | -3.38 | 3.14E-06 | 4.88E-06 |
| chr15 | 62371311 | 62372218 | 908 | -5.47 | 3.14E-06 | 4.89E-06 |
| chr15 | 75174460 | 75175550 | 1091 | -5.47 | 3.14E-06 | 4.89E-06 |
| chr19 | 7399553 | 7400731 | 1179 | -5.47 | 3.14E-06 | 4.89E-06 |
| chr4 | 97684451 | 97686629 | 2179 | -5.54 | 3.14E-06 | 4.89E-06 |
| chr19 | 30628858 | 30704810 | 75953 | -3.27 | 3.15E-06 | 4.89E-06 |
| chr13 | 106197420 | 106198577 | 1158 | -5.41 | 3.15E-06 | 4.90E-06 |
| chr5 | 83850069 | 83902292 | 52224 | 2.02 | 3.15E-06 | 4.90E-06 |
| chr3 | 161211697 | 161213631 | 1935 | -2.44 | 3.16E-06 | 4.91E-06 |
| chr20 | 3291432 | 3291909 | 478 | -5.4 | 3.17E-06 | 4.93E-06 |
| chr3 | 167418195 | 167419283 | 1089 | -5.81 | 3.17E-06 | 4.93E-06 |
| chr2 | 105212296 | 105235885 | 23590 | 1.18 | 3.18E-06 | 4.94E-06 |
| chr1 | 221664196 | 221680514 | 16319 | -6.95 | 3.18E-06 | 4.94E-06 |
| chr13 | 34079881 | 34081318 | 1438 | -5.53 | 3.18E-06 | 4.94E-06 |
| chr8 | 103917092 | 103918427 | 1336 | -4.71 | 3.18E-06 | 4.94E-06 |
| chr5 | 91082497 | 91083296 | 800 | -5.49 | 3.18E-06 | 4.94E-06 |
| chr1 | 30672904 | 30689873 | 16970 | -1.72 | 3.18E-06 | 4.94E-06 |
| chr4 | 52741612 | 52742382 | 771 | -5.87 | 3.18E-06 | 4.95E-06 |
| chr11 | 92169314 | 92170607 | 1294 | 5.49 | 3.18E-06 | 4.95E-06 |
| chr11 | 61653980 | 61654815 | 836 | -5.7 | 3.18E-06 | 4.95E-06 |
| chr11 | 85989921 | 85990787 | 867 | -5.7 | 3.18E-06 | 4.95E-06 |
| chr5 | 71035982 | 71038016 | 2035 | -4.48 | 3.19E-06 | 4.95E-06 |
| chrX | 132609093 | 132718202 | 109110 | 0.86 | 3.19E-06 | 4.96E-06 |
| chr7 | 147227 | 148766 | 1540 | -3 | 3.20E-06 | 4.96E-06 |
| chr11 | 16596172 | 16597152 | 981 | -5.67 | 3.20E-06 | 4.97E-06 |
| chr17 | 40132019 | 40133206 | 1188 | -5.97 | 3.20E-06 | 4.97E-06 |
| chr1 | 108883193 | 108884205 | 1013 | -5.42 | 3.20E-06 | 4.97E-06 |
| chr3 | 11588708 | 11589477 | 770 | -5.42 | 3.20E-06 | 4.97E-06 |
| chrX | 149828767 | 149829376 | 610 | -5.42 | 3.20E-06 | 4.97E-06 |
| chr3 | 142440269 | 142441511 | 1243 | -5.41 | 3.21E-06 | 4.99E-06 |
| chr9 | 123755217 | 123756200 | 984 | -5.41 | 3.21E-06 | 4.99E-06 |

| | | | | | | |
|---|---|---|---|---|---|---|
| chr4 | 72040480 | 72041191 | 712 | -5.43 | 3.22E-06 | 5.00E-06 |
| chr8 | 116541455 | 116542976 | 1522 | -5.51 | 3.22E-06 | 5.00E-06 |
| chr8 | 96054913 | 96055742 | 830 | -5.48 | 3.22E-06 | 5.00E-06 |
| chr11 | 45801226 | 45814753 | 13528 | -5.96 | 3.23E-06 | 5.01E-06 |
| chr6 | 149934195 | 149935363 | 1169 | -5.77 | 3.23E-06 | 5.02E-06 |
| chr4 | 57577613 | 57579026 | 1414 | -2.89 | 3.23E-06 | 5.02E-06 |
| chr9 | 116848198 | 116849851 | 1654 | -5.78 | 3.24E-06 | 5.02E-06 |
| chr1 | 187949327 | 187950577 | 1251 | -5.8 | 3.24E-06 | 5.02E-06 |
| chr6 | 165624659 | 165627078 | 2420 | -2.28 | 3.24E-06 | 5.02E-06 |
| chr6 | 139130284 | 139131852 | 1569 | -4.39 | 3.24E-06 | 5.03E-06 |
| chr12 | 30830013 | 30830897 | 885 | -5.44 | 3.24E-06 | 5.03E-06 |
| chr5 | 76950626 | 76951263 | 638 | -5.44 | 3.24E-06 | 5.03E-06 |
| chr1 | 192468686 | 192469708 | 1023 | -5.62 | 3.24E-06 | 5.03E-06 |
| chr3 | 27569621 | 27570267 | 647 | -5.69 | 3.25E-06 | 5.04E-06 |
| chr5 | 37553861 | 37555330 | 1470 | -5.69 | 3.25E-06 | 5.04E-06 |
| chr16 | 84834341 | 84848828 | 14488 | -5.89 | 3.25E-06 | 5.04E-06 |
| chr5 | 1215476 | 1216270 | 795 | -5.68 | 3.25E-06 | 5.05E-06 |
| chr11 | 101435230 | 101436291 | 1062 | -5.88 | 3.25E-06 | 5.05E-06 |
| chr12 | 106657524 | 106658613 | 1090 | -5.88 | 3.25E-06 | 5.05E-06 |
| chr5 | 14577777 | 14578683 | 907 | -5.4 | 3.26E-06 | 5.05E-06 |
| chr1 | 25909824 | 25910628 | 805 | -5.67 | 3.26E-06 | 5.06E-06 |
| chr19 | 34782685 | 34783458 | 774 | -5.94 | 3.26E-06 | 5.06E-06 |
| chr22 | 18460273 | 18461247 | 975 | -5.5 | 3.26E-06 | 5.06E-06 |
| chr4 | 26824357 | 26825435 | 1079 | -5.5 | 3.26E-06 | 5.06E-06 |
| chrX | 135876890 | 135878117 | 1228 | -5.5 | 3.26E-06 | 5.06E-06 |
| chr8 | 102437282 | 102438297 | 1016 | -5.96 | 3.27E-06 | 5.06E-06 |
| chrX | 140283242 | 140289353 | 6112 | -5.96 | 3.27E-06 | 5.06E-06 |
| chr9 | 75097586 | 75098441 | 856 | -5.66 | 3.27E-06 | 5.06E-06 |
| chr6 | 46733675 | 46735916 | 2242 | -4.28 | 3.27E-06 | 5.07E-06 |
| chr18 | 21221680 | 21222564 | 885 | -5.47 | 3.27E-06 | 5.07E-06 |
| chr12 | 89902568 | 89903798 | 1231 | -5.65 | 3.27E-06 | 5.08E-06 |
| chr16 | 72173711 | 72174533 | 823 | -5.65 | 3.27E-06 | 5.08E-06 |
| chr17 | 58605103 | 58606027 | 925 | -5.65 | 3.27E-06 | 5.08E-06 |
| chr8 | 27767504 | 27772187 | 4684 | -6.01 | 3.27E-06 | 5.08E-06 |
| chr11 | 72653022 | 72653986 | 965 | -5.4 | 3.27E-06 | 5.08E-06 |
| chr6 | 13764009 | 13764883 | 875 | -5.4 | 3.27E-06 | 5.08E-06 |
| chr11 | 96205941 | 96213160 | 7220 | -2.18 | 3.28E-06 | 5.08E-06 |
| chr13 | 52463540 | 52465281 | 1742 | -6.2 | 3.28E-06 | 5.08E-06 |
| chr5 | 53161040 | 53170384 | 9345 | -6.05 | 3.28E-06 | 5.08E-06 |
| chr1 | 32874696 | 32884849 | 10154 | -6.17 | 3.28E-06 | 5.09E-06 |
| chr21 | 17887416 | 17888668 | 1253 | -5.64 | 3.28E-06 | 5.09E-06 |
| chr19 | 16205950 | 16206718 | 769 | -5.4 | 3.28E-06 | 5.09E-06 |
| chr11 | 3966417 | 3967346 | 930 | -5.62 | 3.29E-06 | 5.11E-06 |
| chr9 | 100215568 | 100217070 | 1503 | -5.62 | 3.29E-06 | 5.11E-06 |
| chr20 | 33911731 | 33912576 | 846 | -5.58 | 3.30E-06 | 5.11E-06 |
| chr20 | 31564131 | 31565392 | 1262 | -5.61 | 3.31E-06 | 5.13E-06 |
| chr10 | 69600118 | 69601919 | 1802 | -5.41 | 3.31E-06 | 5.13E-06 |
| chr11 | 30366162 | 30367564 | 1403 | -5.57 | 3.32E-06 | 5.14E-06 |
| chr2 | 138818644 | 138839160 | 20517 | -6.02 | 3.32E-06 | 5.14E-06 |
| chrX | 54895032 | 54899517 | 4486 | -2.38 | 3.32E-06 | 5.14E-06 |
| chr2 | 56923852 | 56929637 | 5786 | -5.41 | 3.32E-06 | 5.15E-06 |
| chr7 | 84406184 | 84407038 | 855 | -5.6 | 3.32E-06 | 5.15E-06 |
| chr14 | 102180344 | 102181141 | 798 | -5.46 | 3.32E-06 | 5.15E-06 |
| chr18 | 46761251 | 46761954 | 704 | -5.46 | 3.32E-06 | 5.15E-06 |
| chr10 | 64592719 | 64700733 | 108015 | 1.58 | 3.33E-06 | 5.15E-06 |
| chr1 | 82736840 | 82738631 | 1792 | -5.42 | 3.33E-06 | 5.16E-06 |
| chr1 | 198991866 | 198992676 | 811 | -5.42 | 3.33E-06 | 5.16E-06 |
| chr1 | 18583289 | 18584168 | 880 | 5.52 | 3.34E-06 | 5.18E-06 |
| chr3 | 186183511 | 186184284 | 774 | 5.52 | 3.34E-06 | 5.18E-06 |
| chr9 | 134391878 | 134392402 | 525 | -4.91 | 3.35E-06 | 5.19E-06 |
| chr2 | 74466115 | 74466604 | 490 | -5.4 | 3.35E-06 | 5.19E-06 |

| | | | | | | |
|---|---|---|---|---|---|---|
| chr4 | 105880139 | 105880917 | 779 | -5.4 | 3.35E-06 | 5.19E-06 |
| chr11 | 40638554 | 40641772 | 3219 | 4.08 | 3.35E-06 | 5.19E-06 |
| chr14 | 28649243 | 28657712 | 8470 | 2.09 | 3.35E-06 | 5.19E-06 |
| chr4 | 64328856 | 64330874 | 2019 | -5.58 | 3.36E-06 | 5.20E-06 |
| chr9 | 125130910 | 125143246 | 12337 | -5.58 | 3.36E-06 | 5.20E-06 |
| chr1 | 112338544 | 112339346 | 803 | -4.19 | 3.37E-06 | 5.21E-06 |
| chr11 | 120618527 | 120620078 | 1552 | -4.12 | 3.37E-06 | 5.21E-06 |
| chr18 | 10064660 | 10066452 | 1793 | -2.33 | 3.37E-06 | 5.22E-06 |
| chr18 | 59832182 | 59833320 | 1139 | -5.48 | 3.37E-06 | 5.22E-06 |
| chr13 | 97559405 | 97561105 | 1701 | 5.48 | 3.37E-06 | 5.22E-06 |
| chr13 | 84466054 | 84467388 | 1335 | -5.54 | 3.38E-06 | 5.23E-06 |
| chr12 | 7585022 | 7586534 | 1513 | -5.45 | 3.38E-06 | 5.23E-06 |
| chr10 | 97693882 | 97722472 | 28591 | -2.58 | 3.39E-06 | 5.25E-06 |
| chr11 | 129391691 | 129408431 | 16741 | -2.02 | 3.40E-06 | 5.25E-06 |
| chrX | 41432595 | 41433519 | 925 | -5.53 | 3.40E-06 | 5.27E-06 |
| chr3 | 185756199 | 185756950 | 752 | -5.55 | 3.41E-06 | 5.27E-06 |
| chr4 | 53605741 | 53606541 | 801 | -5.55 | 3.41E-06 | 5.27E-06 |
| chr2 | 240758420 | 240783921 | 25502 | -6.02 | 3.41E-06 | 5.28E-06 |
| chr10 | 27067669 | 27069293 | 1625 | -5.41 | 3.41E-06 | 5.28E-06 |
| chr5 | 118426034 | 118427021 | 988 | -5.41 | 3.41E-06 | 5.28E-06 |
| chr16 | 59566613 | 59573959 | 7347 | -3.75 | 3.41E-06 | 5.28E-06 |
| chr10 | 106237943 | 106239192 | 1250 | 5.66 | 3.42E-06 | 5.29E-06 |
| chr4 | 5047135 | 5048673 | 1539 | -4.24 | 3.42E-06 | 5.29E-06 |
| chr17 | 55864888 | 55879325 | 14438 | -2.85 | 3.42E-06 | 5.29E-06 |
| chr20 | 26179652 | 26183437 | 3786 | -4.02 | 3.42E-06 | 5.29E-06 |
| chr15 | 95889847 | 95906449 | 16603 | -1.9 | 3.42E-06 | 5.29E-06 |
| chr7 | 43998732 | 43999515 | 784 | -5.4 | 3.42E-06 | 5.30E-06 |
| chr12 | 19763695 | 19765674 | 1980 | -4.77 | 3.43E-06 | 5.30E-06 |
| chr1 | 181415768 | 181435252 | 19485 | -1.6 | 3.43E-06 | 5.31E-06 |
| chr3 | 140642875 | 140643391 | 517 | -5.46 | 3.43E-06 | 5.31E-06 |
| chrX | 136174390 | 136183117 | 8728 | 4.91 | 3.43E-06 | 5.31E-06 |
| chr5 | 52754206 | 52755462 | 1257 | -5.44 | 3.44E-06 | 5.33E-06 |
| chr11 | 12572881 | 12595011 | 22131 | -5.84 | 3.45E-06 | 5.33E-06 |
| chr2 | 57829312 | 57833651 | 4340 | 3.98 | 3.45E-06 | 5.34E-06 |
| chr5 | 115723410 | 115723935 | 526 | -5.38 | 3.45E-06 | 5.34E-06 |
| chr3 | 94347837 | 94349490 | 1654 | -5.4 | 3.45E-06 | 5.34E-06 |
| chr4 | 187617852 | 187618738 | 887 | -5.4 | 3.45E-06 | 5.34E-06 |
| chr6 | 46909819 | 46911318 | 1500 | -5.38 | 3.46E-06 | 5.35E-06 |
| chr4 | 89084781 | 89086094 | 1314 | -5.53 | 3.47E-06 | 5.36E-06 |
| chr5 | 55875685 | 55881644 | 5960 | -1.88 | 3.47E-06 | 5.37E-06 |
| chr3 | 113832332 | 113833001 | 670 | 5.46 | 3.48E-06 | 5.37E-06 |
| chrX | 57540290 | 57541112 | 823 | 3.18 | 3.48E-06 | 5.38E-06 |
| chr1 | 87528505 | 87529273 | 769 | -5.42 | 3.48E-06 | 5.38E-06 |
| chr2 | 219231260 | 219232028 | 769 | -5.42 | 3.48E-06 | 5.38E-06 |
| chr3 | 48535561 | 48536299 | 739 | -5.42 | 3.48E-06 | 5.38E-06 |
| chr8 | 56901967 | 56902836 | 870 | -5.42 | 3.48E-06 | 5.38E-06 |
| chr5 | 114466620 | 114467615 | 996 | -5.38 | 3.49E-06 | 5.39E-06 |
| chr4 | 146913350 | 146914072 | 723 | -5.45 | 3.51E-06 | 5.42E-06 |
| chr2 | 129155625 | 129156597 | 973 | -5.39 | 3.51E-06 | 5.42E-06 |
| chr16 | 74996107 | 74997094 | 988 | -5.43 | 3.51E-06 | 5.43E-06 |
| chr7 | 129981017 | 129983665 | 2649 | -4.9 | 3.52E-06 | 5.44E-06 |
| chr6 | 1272069 | 1273604 | 1536 | -3.54 | 3.52E-06 | 5.44E-06 |
| chr5 | 167969738 | 167971155 | 1418 | -5.49 | 3.53E-06 | 5.46E-06 |
| chr12 | 99134550 | 99135325 | 776 | -5.5 | 3.54E-06 | 5.47E-06 |
| chr14 | 32015877 | 32019540 | 3664 | -5.29 | 3.54E-06 | 5.47E-06 |
| chr5 | 157409296 | 157410464 | 1169 | -5.38 | 3.55E-06 | 5.49E-06 |
| chr7 | 29978480 | 29979341 | 862 | -5.38 | 3.55E-06 | 5.49E-06 |
| chr4 | 161549443 | 161550565 | 1123 | -4.67 | 3.55E-06 | 5.49E-06 |
| chr3 | 43921281 | 43922359 | 1079 | -5.38 | 3.57E-06 | 5.51E-06 |
| chr12 | 19931426 | 19979845 | 48420 | -7.58 | 3.57E-06 | 5.52E-06 |
| chr20 | 5141243 | 5141778 | 536 | -5.48 | 3.57E-06 | 5.52E-06 |

| | | | | | | |
|---|---|---|---|---|---|---|
| chr16 | 69527382 | 69528451 | 1070 | -5.49 | 3.59E-06 | 5.54E-06 |
| chrX | 39875103 | 39876396 | 1294 | -5.49 | 3.59E-06 | 5.54E-06 |
| chr1 | 168088613 | 168090103 | 1491 | -5.42 | 3.59E-06 | 5.54E-06 |
| chr11 | 111667743 | 111668574 | 832 | -5.42 | 3.59E-06 | 5.54E-06 |
| chr11 | 2135898 | 2141438 | 5541 | -5.7 | 3.59E-06 | 5.54E-06 |
| chr4 | 134883005 | 134902126 | 19122 | 1.94 | 3.59E-06 | 5.55E-06 |
| chr8 | 125257366 | 125259678 | 2313 | -5.77 | 3.59E-06 | 5.55E-06 |
| chr7 | 5450876 | 5451854 | 979 | -5.39 | 3.59E-06 | 5.55E-06 |
| chr7 | 50518465 | 50519292 | 828 | -5.39 | 3.59E-06 | 5.55E-06 |
| chr2 | 101202044 | 101203313 | 1270 | -5.73 | 3.59E-06 | 5.55E-06 |
| chr8 | 134112930 | 134113813 | 884 | -5.72 | 3.60E-06 | 5.55E-06 |
| chr12 | 68256272 | 68257752 | 1481 | -5.71 | 3.60E-06 | 5.56E-06 |
| chr10 | 102165436 | 102166399 | 964 | -4.52 | 3.60E-06 | 5.56E-06 |
| chr3 | 72765430 | 72766992 | 1563 | -5.68 | 3.61E-06 | 5.57E-06 |
| chrX | 29027206 | 29028323 | 1118 | -5.91 | 3.61E-06 | 5.57E-06 |
| chr12 | 56882393 | 56883306 | 914 | -5.67 | 3.61E-06 | 5.57E-06 |
| chr10 | 33789177 | 33792126 | 2950 | -5.94 | 3.61E-06 | 5.58E-06 |
| chr4 | 174977278 | 174978430 | 1153 | -5.39 | 3.62E-06 | 5.58E-06 |
| chr16 | 57074391 | 57075832 | 1442 | -5.66 | 3.62E-06 | 5.58E-06 |
| chr10 | 116609332 | 116610038 | 707 | -5.71 | 3.62E-06 | 5.59E-06 |
| chr11 | 59915262 | 59947779 | 32518 | 1.37 | 3.63E-06 | 5.60E-06 |
| chr10 | 27759871 | 27792844 | 32974 | -1.61 | 3.63E-06 | 5.60E-06 |
| chr11 | 27770801 | 27772388 | 1588 | -2.46 | 3.63E-06 | 5.60E-06 |
| chr1 | 240242174 | 240243298 | 1125 | -5.64 | 3.63E-06 | 5.60E-06 |
| chr3 | 111791412 | 111792848 | 1437 | -5.64 | 3.63E-06 | 5.60E-06 |
| chr5 | 118375707 | 118376889 | 1183 | -5.64 | 3.63E-06 | 5.60E-06 |
| chr8 | 95202979 | 95204048 | 1070 | -5.64 | 3.63E-06 | 5.60E-06 |
| chr3 | 110564693 | 110566098 | 1406 | -5.68 | 3.63E-06 | 5.61E-06 |
| chr14 | 73308697 | 73312280 | 3584 | 2.16 | 3.63E-06 | 5.61E-06 |
| chr6 | 159918115 | 159920582 | 2468 | -4.61 | 3.64E-06 | 5.61E-06 |
| chr11 | 58337974 | 58338597 | 624 | -5.37 | 3.65E-06 | 5.64E-06 |
| chr11 | 20055041 | 20056054 | 1014 | -5.62 | 3.65E-06 | 5.64E-06 |
| chr14 | 36719112 | 36720439 | 1328 | -5.62 | 3.65E-06 | 5.64E-06 |
| chr6 | 80307242 | 80315275 | 8034 | -5.62 | 3.65E-06 | 5.64E-06 |
| chr9 | 136730199 | 136731379 | 1181 | -5.62 | 3.65E-06 | 5.64E-06 |
| chr1 | 159774364 | 159775982 | 1619 | 2.34 | 3.65E-06 | 5.64E-06 |
| chr6 | 150858184 | 150860933 | 2750 | -4.73 | 3.66E-06 | 5.64E-06 |
| chr1 | 59712490 | 59713602 | 1113 | -5.6 | 3.67E-06 | 5.65E-06 |
| chr5 | 174360305 | 174361658 | 1354 | -5.6 | 3.67E-06 | 5.65E-06 |
| chr13 | 27007098 | 27015194 | 8097 | -2.25 | 3.67E-06 | 5.66E-06 |
| chrX | 6980062 | 6981783 | 1722 | -5.41 | 3.67E-06 | 5.66E-06 |
| chr15 | 74184185 | 74185090 | 906 | -5.37 | 3.67E-06 | 5.66E-06 |
| chr4 | 44732404 | 44732965 | 562 | -5.37 | 3.67E-06 | 5.66E-06 |
| chr15 | 101827641 | 101828076 | 436 | -5.42 | 3.68E-06 | 5.67E-06 |
| chr4 | 178092614 | 178093272 | 659 | -5.59 | 3.68E-06 | 5.68E-06 |
| chr2 | 55933634 | 55934021 | 388 | -5.37 | 3.69E-06 | 5.69E-06 |
| chr5 | 118136791 | 118139153 | 2363 | 6.31 | 3.69E-06 | 5.69E-06 |
| chr18 | 72000264 | 72001168 | 905 | -5.61 | 3.69E-06 | 5.69E-06 |
| chr3 | 127150410 | 127154801 | 4392 | -1.84 | 3.69E-06 | 5.69E-06 |
| chr6 | 80827156 | 80828073 | 918 | -5.46 | 3.70E-06 | 5.70E-06 |
| chr7 | 115700347 | 115700864 | 518 | -5.46 | 3.70E-06 | 5.70E-06 |
| chrX | 117938738 | 117939788 | 1051 | -5.6 | 3.70E-06 | 5.71E-06 |
| chr2 | 106106706 | 106110794 | 4089 | -3.28 | 3.70E-06 | 5.71E-06 |
| chr7 | 17525147 | 17526310 | 1164 | -5.45 | 3.72E-06 | 5.73E-06 |
| chr8 | 18845930 | 18846909 | 980 | -5.57 | 3.72E-06 | 5.73E-06 |
| chr1 | 38927904 | 38945833 | 17930 | -4.8 | 3.72E-06 | 5.73E-06 |
| chr17 | 7207282 | 7208071 | 790 | -5.39 | 3.72E-06 | 5.73E-06 |
| chr22 | 50559696 | 50571137 | 11442 | -5.56 | 3.73E-06 | 5.75E-06 |
| chr9 | 124400932 | 124401770 | 839 | -5.38 | 3.74E-06 | 5.76E-06 |
| chr1 | 4172464 | 4183911 | 11448 | -4.33 | 3.75E-06 | 5.78E-06 |
| chr5 | 127734303 | 127735101 | 799 | -5.38 | 3.75E-06 | 5.78E-06 |

| | | | | | | |
|---|---|---|---|---|---|---|
| chr10 | 114303838 | 114304575 | 738 | -5.45 | 3.76E-06 | 5.79E-06 |
| chr20 | 4820280 | 4821167 | 888 | -5.45 | 3.76E-06 | 5.79E-06 |
| chr13 | 113810226 | 113823077 | 12852 | -5.4 | 3.76E-06 | 5.79E-06 |
| chr12 | 2801446 | 2826809 | 25364 | -1.37 | 3.76E-06 | 5.80E-06 |
| chr6 | 42250076 | 42251111 | 1036 | -5.56 | 3.76E-06 | 5.80E-06 |
| chr1 | 178089255 | 178090269 | 1015 | -5.44 | 3.78E-06 | 5.82E-06 |
| chrX | 73527311 | 73528284 | 974 | -5.44 | 3.78E-06 | 5.82E-06 |
| chr1 | 99985214 | 99986011 | 798 | -5.41 | 3.78E-06 | 5.82E-06 |
| chr9 | 116332922 | 116333967 | 1046 | -5.41 | 3.78E-06 | 5.82E-06 |
| chr11 | 1906335 | 1921569 | 15235 | 5.99 | 3.78E-06 | 5.82E-06 |
| chrX | 150272728 | 150278658 | 5931 | -1.86 | 3.79E-06 | 5.84E-06 |
| chr4 | 139401726 | 139402668 | 943 | -5.53 | 3.79E-06 | 5.84E-06 |
| chr3 | 72374392 | 72376049 | 1658 | -4.3 | 3.81E-06 | 5.87E-06 |
| chr10 | 59506760 | 59512316 | 5557 | -5 | 3.82E-06 | 5.88E-06 |
| chr2 | 205495595 | 205496358 | 764 | -5.52 | 3.83E-06 | 5.89E-06 |
| chr11 | 98205415 | 98220694 | 15280 | 1.69 | 3.84E-06 | 5.91E-06 |
| chr10 | 69470235 | 69476137 | 5903 | -2.34 | 3.84E-06 | 5.92E-06 |
| chr4 | 166575383 | 166576746 | 1364 | -5.36 | 3.84E-06 | 5.92E-06 |
| chr9 | 34737114 | 34888585 | 151472 | -1.18 | 3.85E-06 | 5.93E-06 |
| chrX | 43964799 | 43968385 | 3587 | -4.61 | 3.85E-06 | 5.93E-06 |
| chr6 | 48558258 | 48558985 | 728 | -5.51 | 3.86E-06 | 5.95E-06 |
| chr17 | 8104001 | 8104865 | 865 | -5.38 | 3.86E-06 | 5.95E-06 |
| chr17 | 20680297 | 20681918 | 1622 | -5.38 | 3.86E-06 | 5.95E-06 |
| chr2 | 143138834 | 143139445 | 612 | -5.38 | 3.86E-06 | 5.95E-06 |
| chr1 | 7598088 | 7598870 | 783 | -5.37 | 3.87E-06 | 5.95E-06 |
| chr5 | 171140091 | 171158240 | 18150 | -4.67 | 3.87E-06 | 5.95E-06 |
| chr5 | 52407779 | 52408448 | 670 | -5.36 | 3.88E-06 | 5.97E-06 |
| chr4 | 160817095 | 160818951 | 1857 | -5.51 | 3.88E-06 | 5.97E-06 |
| chr9 | 37381879 | 37382696 | 818 | -5.51 | 3.88E-06 | 5.97E-06 |
| chr8 | 133693488 | 133706347 | 12860 | 1.83 | 3.89E-06 | 5.99E-06 |
| chr1 | 192434990 | 192439114 | 4125 | -2.38 | 3.89E-06 | 5.99E-06 |
| chr2 | 217975287 | 217976861 | 1575 | -4.26 | 3.90E-06 | 6.00E-06 |
| chr1 | 1035333 | 1049286 | 13954 | -5.49 | 3.90E-06 | 6.01E-06 |
| chr2 | 217251801 | 217252776 | 976 | -5.49 | 3.90E-06 | 6.01E-06 |
| chr4 | 185476629 | 185477444 | 816 | -5.49 | 3.90E-06 | 6.01E-06 |
| chr5 | 77087770 | 77110931 | 23162 | -6.04 | 3.91E-06 | 6.01E-06 |
| chr6 | 3495299 | 3496067 | 769 | -5.42 | 3.91E-06 | 6.02E-06 |
| chr4 | 112516476 | 112517900 | 1425 | 5.53 | 3.91E-06 | 6.02E-06 |
| chr5 | 159261996 | 159262820 | 825 | -5.35 | 3.92E-06 | 6.03E-06 |
| chr17 | 36471984 | 36472597 | 614 | -5.36 | 3.93E-06 | 6.04E-06 |
| chr19 | 3402182 | 3402974 | 793 | -5.36 | 3.93E-06 | 6.04E-06 |
| chr8 | 67781830 | 67782596 | 767 | -5.36 | 3.93E-06 | 6.04E-06 |
| chr1 | 223700767 | 223701985 | 1219 | -4.5 | 3.94E-06 | 6.06E-06 |
| chr2 | 171891704 | 171892504 | 801 | -5.48 | 3.95E-06 | 6.07E-06 |
| chrX | 44341860 | 44342939 | 1080 | -5.48 | 3.95E-06 | 6.07E-06 |
| chr2 | 195786643 | 195787670 | 1028 | 4.1 | 3.95E-06 | 6.08E-06 |
| chr8 | 128712454 | 128734600 | 22147 | -7.3 | 3.95E-06 | 6.08E-06 |
| chr14 | 57131917 | 57148498 | 16582 | -1.89 | 3.96E-06 | 6.09E-06 |
| chr12 | 93054447 | 93057138 | 2692 | -4.71 | 3.97E-06 | 6.10E-06 |
| chr10 | 96561521 | 96562136 | 616 | -4.66 | 3.98E-06 | 6.11E-06 |
| chr3 | 136778241 | 136779792 | 1552 | 4.36 | 3.99E-06 | 6.13E-06 |
| chr10 | 101549884 | 101550822 | 939 | -5.41 | 3.99E-06 | 6.13E-06 |
| chr4 | 184091475 | 184092430 | 956 | -5.41 | 3.99E-06 | 6.13E-06 |
| chr5 | 127965209 | 127966206 | 998 | -5.41 | 3.99E-06 | 6.13E-06 |
| chrX | 135783112 | 135783839 | 728 | -5.41 | 3.99E-06 | 6.13E-06 |
| chr13 | 51423593 | 51438344 | 14752 | -1.33 | 3.99E-06 | 6.13E-06 |
| chr14 | 24483457 | 24485355 | 1899 | -5.76 | 3.99E-06 | 6.14E-06 |
| chr15 | 34880942 | 34881888 | 947 | -5.8 | 4.00E-06 | 6.14E-06 |
| chr5 | 118295086 | 118296340 | 1255 | -5.85 | 4.00E-06 | 6.15E-06 |
| chr7 | 3194182 | 3249353 | 55172 | -7.29 | 4.00E-06 | 6.15E-06 |
| chr8 | 107787453 | 107788813 | 1361 | -5.7 | 4.00E-06 | 6.15E-06 |

| | | | | | | |
|---|---|---|---|---|---|---|
| chr18 | 9018417 | 9019499 | 1083 | -5.47 | 4.00E-06 | 6.15E-06 |
| chr1 | 173803626 | 173804392 | 767 | -5.41 | 4.00E-06 | 6.15E-06 |
| chr2 | 190515131 | 190516223 | 1093 | -5.41 | 4.00E-06 | 6.15E-06 |
| chr8 | 110567101 | 110567714 | 614 | -5.41 | 4.00E-06 | 6.15E-06 |
| chr11 | 129663974 | 129679019 | 15046 | -4.95 | 4.01E-06 | 6.16E-06 |
| chr10 | 126864936 | 126865545 | 610 | -5.68 | 4.01E-06 | 6.16E-06 |
| chr3 | 178727495 | 178728132 | 638 | -5.36 | 4.01E-06 | 6.16E-06 |
| chr4 | 184125407 | 184126308 | 902 | -5.36 | 4.01E-06 | 6.16E-06 |
| chr4 | 103231814 | 103232598 | 785 | -5.35 | 4.01E-06 | 6.17E-06 |
| chr10 | 129314782 | 129318810 | 4029 | -1.94 | 4.02E-06 | 6.17E-06 |
| chr12 | 99149005 | 99150286 | 1282 | -5.38 | 4.02E-06 | 6.17E-06 |
| chr1 | 67983633 | 67985101 | 1469 | -5.65 | 4.02E-06 | 6.17E-06 |
| chr5 | 137310346 | 137310913 | 568 | -5.37 | 4.02E-06 | 6.18E-06 |
| chr10 | 101006763 | 101014184 | 7422 | -5.52 | 4.03E-06 | 6.19E-06 |
| chr9 | 95807390 | 95808675 | 1286 | -4.69 | 4.04E-06 | 6.21E-06 |
| chr17 | 45141856 | 45142697 | 842 | -5.62 | 4.04E-06 | 6.21E-06 |
| chr3 | 71713915 | 71716097 | 2183 | -2.23 | 4.05E-06 | 6.23E-06 |
| chr19 | 29998879 | 29999537 | 659 | -4.42 | 4.06E-06 | 6.23E-06 |
| chr12 | 49132467 | 49133734 | 1268 | -5.45 | 4.06E-06 | 6.23E-06 |
| chr1 | 8750179 | 8750890 | 712 | -5.4 | 4.07E-06 | 6.25E-06 |
| chr16 | 22139708 | 22140454 | 747 | -4.3 | 4.07E-06 | 6.26E-06 |
| chr22 | 23171905 | 23185568 | 13664 | -5.4 | 4.08E-06 | 6.27E-06 |
| chr6 | 151515200 | 151516584 | 1385 | -5.58 | 4.08E-06 | 6.27E-06 |
| chr6 | 89903331 | 89906092 | 2762 | -5.15 | 4.09E-06 | 6.28E-06 |
| chr2 | 39958397 | 39959308 | 912 | -5.37 | 4.09E-06 | 6.28E-06 |
| chrX | 102560663 | 102561149 | 487 | -5.37 | 4.09E-06 | 6.28E-06 |
| chr10 | 68840403 | 68842746 | 2344 | 4.75 | 4.10E-06 | 6.29E-06 |
| chr17 | 63247127 | 63284585 | 37459 | -2.22 | 4.10E-06 | 6.30E-06 |
| chr14 | 93216229 | 93217164 | 936 | -5.35 | 4.10E-06 | 6.30E-06 |
| chr9 | 20918692 | 20919212 | 521 | -5.35 | 4.10E-06 | 6.30E-06 |
| chr15 | 78711428 | 78712030 | 603 | -5.57 | 4.10E-06 | 6.30E-06 |
| chr2 | 45657588 | 45658938 | 1351 | -5.4 | 4.11E-06 | 6.30E-06 |
| chr8 | 71299678 | 71300715 | 1038 | -6.14 | 4.11E-06 | 6.31E-06 |
| chr3 | 170334164 | 170340026 | 5863 | -3.47 | 4.12E-06 | 6.32E-06 |
| chr3 | 106211292 | 106212025 | 734 | 4.37 | 4.12E-06 | 6.33E-06 |
| chr6 | 46891753 | 46892508 | 756 | -4.07 | 4.12E-06 | 6.33E-06 |
| chr4 | 81002459 | 81004152 | 1694 | -4.57 | 4.13E-06 | 6.34E-06 |
| chr4 | 41008934 | 41010188 | 1255 | -5.79 | 4.13E-06 | 6.34E-06 |
| chr1 | 226963923 | 226965698 | 1776 | -5.85 | 4.13E-06 | 6.34E-06 |
| chr6 | 138109899 | 138110972 | 1074 | -5.72 | 4.13E-06 | 6.34E-06 |
| chr2 | 183572844 | 183573827 | 984 | -5.95 | 4.13E-06 | 6.34E-06 |
| chr7 | 32265415 | 32282300 | 16886 | -3.2 | 4.14E-06 | 6.35E-06 |
| chr11 | 123313503 | 123314423 | 921 | -5.68 | 4.14E-06 | 6.36E-06 |
| chr7 | 25095947 | 25096515 | 569 | -5.68 | 4.14E-06 | 6.36E-06 |
| chr2 | 102868457 | 102913756 | 45300 | -0.93 | 4.15E-06 | 6.36E-06 |
| chr3 | 25629295 | 25629946 | 652 | -5.67 | 4.15E-06 | 6.36E-06 |
| chr22 | 20898046 | 20898782 | 737 | -5.65 | 4.16E-06 | 6.38E-06 |
| chr2 | 187853364 | 187856693 | 3330 | -1.87 | 4.16E-06 | 6.38E-06 |
| chr4 | 15862622 | 15863556 | 935 | -5.37 | 4.16E-06 | 6.39E-06 |
| chr5 | 111633455 | 111637256 | 3802 | -4.89 | 4.17E-06 | 6.39E-06 |
| chr13 | 82859452 | 82860546 | 1095 | -5.35 | 4.17E-06 | 6.39E-06 |
| chr6 | 133340016 | 133355328 | 15313 | -5.17 | 4.18E-06 | 6.42E-06 |
| chr3 | 181869354 | 181869962 | 609 | -4.74 | 4.19E-06 | 6.43E-06 |
| chr1 | 108541552 | 108557125 | 15574 | 4.36 | 4.19E-06 | 6.43E-06 |
| chr1 | 154195009 | 154195626 | 618 | -5.43 | 4.20E-06 | 6.44E-06 |
| chr11 | 73907012 | 73907696 | 685 | -5.43 | 4.20E-06 | 6.44E-06 |
| chr15 | 59308771 | 59309285 | 515 | -5.43 | 4.20E-06 | 6.44E-06 |
| chr4 | 37694479 | 37708210 | 13732 | -5.43 | 4.20E-06 | 6.44E-06 |
| chr5 | 162951627 | 162952772 | 1146 | -5.43 | 4.20E-06 | 6.44E-06 |
| chr1 | 196994504 | 196995771 | 1268 | -5.35 | 4.20E-06 | 6.44E-06 |
| chr3 | 196115361 | 196116595 | 1235 | -5.35 | 4.20E-06 | 6.44E-06 |

| | | | | | | |
|---|---|---|---|---|---|---|
| chr1 | 188992481 | 188993638 | 1158 | -5.34 | 4.20E-06 | 6.45E-06 |
| chr4 | 170169815 | 170170581 | 767 | -5.34 | 4.20E-06 | 6.45E-06 |
| chr12 | 11429130 | 11450805 | 21676 | 1.51 | 4.21E-06 | 6.46E-06 |
| chr11 | 27228612 | 27230137 | 1526 | -4.58 | 4.21E-06 | 6.46E-06 |
| chr8 | 127564134 | 127565916 | 1783 | -4.48 | 4.21E-06 | 6.46E-06 |
| chr4 | 56355960 | 56356875 | 916 | -5.38 | 4.22E-06 | 6.47E-06 |
| chr17 | 56212502 | 56249137 | 36636 | -1.76 | 4.23E-06 | 6.48E-06 |
| chr9 | 20194845 | 20197921 | 3077 | -5.12 | 4.23E-06 | 6.48E-06 |
| chr10 | 26751597 | 26753302 | 1706 | -5.51 | 4.23E-06 | 6.48E-06 |
| chr19 | 13330789 | 13331622 | 834 | -5.43 | 4.23E-06 | 6.49E-06 |
| chrX | 74304625 | 74305428 | 804 | -5.43 | 4.23E-06 | 6.49E-06 |
| chr6 | 152242207 | 152263605 | 21399 | -1.19 | 4.24E-06 | 6.50E-06 |
| chr19 | 36292792 | 36293497 | 706 | -5.38 | 4.27E-06 | 6.54E-06 |
| chr21 | 27555488 | 27556212 | 725 | -5.38 | 4.27E-06 | 6.54E-06 |
| chr1 | 91235881 | 91237035 | 1155 | 4.33 | 4.27E-06 | 6.54E-06 |
| chr11 | 95145134 | 95164784 | 19651 | 2.38 | 4.27E-06 | 6.54E-06 |
| chr5 | 157489615 | 157490652 | 1038 | -3.46 | 4.28E-06 | 6.55E-06 |
| chr22 | 29048210 | 29048989 | 780 | -5.41 | 4.28E-06 | 6.56E-06 |
| chr12 | 51968789 | 51979928 | 11140 | -6.3 | 4.28E-06 | 6.56E-06 |
| chr3 | 40438125 | 40439226 | 1102 | -5.55 | 4.28E-06 | 6.56E-06 |
| chrX | 84667296 | 84668281 | 986 | -5.42 | 4.30E-06 | 6.58E-06 |
| chr17 | 3156861 | 3343541 | 186681 | -0.89 | 4.30E-06 | 6.59E-06 |
| chr2 | 175191650 | 175192725 | 1076 | -5.48 | 4.31E-06 | 6.60E-06 |
| chr4 | 69036429 | 69038694 | 2266 | -2.15 | 4.31E-06 | 6.61E-06 |
| chr3 | 187229379 | 187230748 | 1370 | -2.75 | 4.32E-06 | 6.62E-06 |
| chr12 | 133302361 | 133303667 | 1307 | -5.53 | 4.32E-06 | 6.62E-06 |
| chr3 | 190134746 | 190135854 | 1109 | -5.35 | 4.32E-06 | 6.62E-06 |
| chr3 | 10402037 | 10413762 | 11726 | -6.24 | 4.33E-06 | 6.63E-06 |
| chr7 | 131525438 | 131526437 | 1000 | 5.42 | 4.33E-06 | 6.64E-06 |
| chr12 | 22701775 | 22702640 | 866 | -5.34 | 4.34E-06 | 6.64E-06 |
| chr6 | 33804752 | 33805419 | 668 | -5.34 | 4.34E-06 | 6.64E-06 |
| chr9 | 140440662 | 140441562 | 901 | -5.34 | 4.34E-06 | 6.64E-06 |
| chr8 | 98580780 | 98590104 | 9325 | -2.58 | 4.34E-06 | 6.65E-06 |
| chr14 | 96539569 | 96540512 | 944 | -5.52 | 4.35E-06 | 6.66E-06 |
| chr12 | 18390511 | 18391490 | 980 | -5.37 | 4.35E-06 | 6.66E-06 |
| chr6 | 111018586 | 111019318 | 733 | -5.37 | 4.35E-06 | 6.66E-06 |
| chr7 | 81163968 | 81164979 | 1012 | -5.37 | 4.35E-06 | 6.66E-06 |
| chr1 | 67760462 | 67761269 | 808 | -5.47 | 4.36E-06 | 6.67E-06 |
| chr12 | 74558792 | 74560083 | 1292 | -5.47 | 4.36E-06 | 6.67E-06 |
| chrX | 44755367 | 44756362 | 996 | -5.47 | 4.36E-06 | 6.67E-06 |
| chr9 | 3900617 | 3901492 | 876 | -5.33 | 4.36E-06 | 6.67E-06 |
| chr9 | 118968535 | 118970491 | 1957 | -5.33 | 4.36E-06 | 6.67E-06 |
| chr7 | 25041524 | 25049141 | 7618 | -6.2 | 4.36E-06 | 6.68E-06 |
| chr1 | 108244590 | 108245305 | 716 | -5.4 | 4.37E-06 | 6.69E-06 |
| chr12 | 67755899 | 67756625 | 727 | -5.51 | 4.37E-06 | 6.70E-06 |
| chr9 | 34711510 | 34714221 | 2712 | -4.93 | 4.38E-06 | 6.70E-06 |
| chr14 | 38391807 | 38394943 | 3137 | 4.29 | 4.39E-06 | 6.72E-06 |
| chr2 | 36453266 | 36496326 | 43061 | -2.34 | 4.40E-06 | 6.73E-06 |
| chr21 | 18521251 | 18634542 | 113292 | 6.43 | 4.40E-06 | 6.73E-06 |
| chr18 | 908366 | 909062 | 697 | -4.26 | 4.40E-06 | 6.73E-06 |
| chr14 | 103119579 | 103120277 | 699 | -5.5 | 4.40E-06 | 6.74E-06 |
| chr3 | 41165680 | 41188998 | 23319 | -5.65 | 4.41E-06 | 6.74E-06 |
| chr18 | 23688328 | 23704114 | 15787 | -2.14 | 4.41E-06 | 6.74E-06 |
| chr3 | 100259296 | 100259837 | 542 | -5.34 | 4.41E-06 | 6.74E-06 |
| chr14 | 105893204 | 105894017 | 814 | -5.46 | 4.41E-06 | 6.74E-06 |
| chr1 | 182951108 | 182951750 | 643 | -5.33 | 4.41E-06 | 6.75E-06 |
| chr14 | 57966933 | 57967561 | 629 | -5.33 | 4.41E-06 | 6.75E-06 |
| chr4 | 174581595 | 174582466 | 872 | -5.33 | 4.41E-06 | 6.75E-06 |
| chr8 | 128426375 | 128428207 | 1833 | -4.06 | 4.42E-06 | 6.77E-06 |
| chr4 | 72573548 | 72586406 | 12859 | -6.08 | 4.43E-06 | 6.77E-06 |
| chr4 | 187189911 | 187192957 | 3047 | -3.09 | 4.43E-06 | 6.77E-06 |

| | | | | | | |
|---|---|---|---|---|---|---|
| chr6 | 132362072 | 132363809 | 1738 | -5.83 | 4.44E-06 | 6.79E-06 |
| chr3 | 94704491 | 94707279 | 2789 | -5.87 | 4.44E-06 | 6.79E-06 |
| chr10 | 26782747 | 26784710 | 1964 | -5.71 | 4.44E-06 | 6.80E-06 |
| chr2 | 169396086 | 169397175 | 1090 | -5.39 | 4.45E-06 | 6.80E-06 |
| chr16 | 11471422 | 11473029 | 1608 | -5.69 | 4.45E-06 | 6.80E-06 |
| chr9 | 92022410 | 92026571 | 4162 | -5.69 | 4.45E-06 | 6.80E-06 |
| chr3 | 186234149 | 186235760 | 1612 | -4.13 | 4.45E-06 | 6.80E-06 |
| chr9 | 15413277 | 15414473 | 1197 | -5.68 | 4.45E-06 | 6.81E-06 |
| chr20 | 1620518 | 1636125 | 15608 | -1.36 | 4.46E-06 | 6.81E-06 |
| chr3 | 42528626 | 42529721 | 1096 | -3.64 | 4.46E-06 | 6.81E-06 |
| chr10 | 104563806 | 104564607 | 802 | -5.67 | 4.46E-06 | 6.81E-06 |
| chr6 | 121584662 | 121585445 | 784 | -5.67 | 4.46E-06 | 6.81E-06 |
| chr12 | 40813380 | 40814177 | 798 | -5.44 | 4.47E-06 | 6.83E-06 |
| chr17 | 28125572 | 28125980 | 409 | -5.48 | 4.47E-06 | 6.83E-06 |
| chr9 | 121547261 | 121548804 | 1544 | -5.64 | 4.47E-06 | 6.83E-06 |
| chr3 | 52239815 | 52240811 | 997 | -5.39 | 4.48E-06 | 6.84E-06 |
| chr19 | 48144614 | 48153246 | 8633 | -5.63 | 4.48E-06 | 6.84E-06 |
| chr2 | 159016954 | 159022521 | 5568 | -3.79 | 4.48E-06 | 6.84E-06 |
| chr1 | 164489419 | 164498993 | 9575 | -2.41 | 4.48E-06 | 6.85E-06 |
| chr5 | 71183199 | 71184524 | 1326 | -5.61 | 4.50E-06 | 6.87E-06 |
| chr1 | 82779281 | 82780755 | 1475 | -5.36 | 4.50E-06 | 6.87E-06 |
| chr13 | 81831069 | 81832666 | 1598 | -5.36 | 4.50E-06 | 6.87E-06 |
| chr6 | 154411161 | 154412488 | 1328 | 5.56 | 4.50E-06 | 6.88E-06 |
| chr14 | 22570110 | 22571268 | 1159 | -3.98 | 4.50E-06 | 6.88E-06 |
| chr13 | 100710700 | 100712143 | 1444 | -5.6 | 4.51E-06 | 6.89E-06 |
| chr6 | 107696663 | 107697465 | 803 | -5.58 | 4.52E-06 | 6.91E-06 |
| chr10 | 116124167 | 116125948 | 1782 | -5.33 | 4.53E-06 | 6.92E-06 |
| chr14 | 81782616 | 81783694 | 1079 | -5.33 | 4.53E-06 | 6.92E-06 |
| chr4 | 63993904 | 63995206 | 1303 | -5.33 | 4.53E-06 | 6.92E-06 |
| chr1 | 235018798 | 235019499 | 702 | -5.33 | 4.53E-06 | 6.92E-06 |
| chr8 | 40581177 | 40582001 | 825 | -5.33 | 4.53E-06 | 6.92E-06 |
| chr7 | 112021394 | 112021884 | 491 | -5.43 | 4.53E-06 | 6.93E-06 |
| chr17 | 45335651 | 45348009 | 12359 | -6.4 | 4.55E-06 | 6.94E-06 |
| chr20 | 58019707 | 58081518 | 61812 | -2.97 | 4.55E-06 | 6.95E-06 |
| chr2 | 167051019 | 167052428 | 1410 | 5.41 | 4.55E-06 | 6.95E-06 |
| chr3 | 162239161 | 162240274 | 1114 | -4.43 | 4.55E-06 | 6.95E-06 |
| chr1 | 171369639 | 171370968 | 1330 | -5.56 | 4.55E-06 | 6.96E-06 |
| chr13 | 98364827 | 98365623 | 797 | -5.56 | 4.55E-06 | 6.96E-06 |
| chr7 | 84360693 | 84361620 | 928 | -5.56 | 4.55E-06 | 6.96E-06 |
| chrX | 136196368 | 136204795 | 8428 | 5.25 | 4.56E-06 | 6.96E-06 |
| chr12 | 10232668 | 10238428 | 5761 | -2.44 | 4.57E-06 | 6.98E-06 |
| chr14 | 52549351 | 52553957 | 4607 | -5.3 | 4.57E-06 | 6.98E-06 |
| chr11 | 10513931 | 10515771 | 1841 | -5.55 | 4.58E-06 | 6.98E-06 |
| chr14 | 85932183 | 85933340 | 1158 | -5.55 | 4.58E-06 | 6.98E-06 |
| chr4 | 132950703 | 132952238 | 1536 | -5.55 | 4.58E-06 | 6.98E-06 |
| chrX | 4795187 | 4796173 | 987 | -5.55 | 4.58E-06 | 6.98E-06 |
| chr6 | 23055182 | 23076755 | 21574 | -1.17 | 4.58E-06 | 6.98E-06 |
| chr6 | 161947243 | 161955653 | 8411 | -5.13 | 4.58E-06 | 6.99E-06 |
| chr15 | 77466613 | 77467236 | 624 | -5.37 | 4.59E-06 | 7.01E-06 |
| chr16 | 57814903 | 57816488 | 1586 | -5.37 | 4.59E-06 | 7.01E-06 |
| chr19 | 18592362 | 18593109 | 748 | -5.37 | 4.59E-06 | 7.01E-06 |
| chr2 | 45007237 | 45007904 | 668 | -5.37 | 4.59E-06 | 7.01E-06 |
| chrX | 12586946 | 12588089 | 1144 | -5.37 | 4.59E-06 | 7.01E-06 |
| chr12 | 98211522 | 98225344 | 13823 | 5.95 | 4.61E-06 | 7.03E-06 |
| chr6 | 113205994 | 113206808 | 815 | -5.42 | 4.61E-06 | 7.03E-06 |
| chr7 | 626027 | 626749 | 723 | -5.42 | 4.61E-06 | 7.03E-06 |
| chr4 | 176578766 | 176579443 | 678 | -5.32 | 4.61E-06 | 7.04E-06 |
| chr6 | 15289702 | 15290636 | 935 | -5.37 | 4.63E-06 | 7.06E-06 |
| chr5 | 44958588 | 45028299 | 69712 | -1.05 | 4.63E-06 | 7.07E-06 |
| chr15 | 60180544 | 60182325 | 1782 | -2.5 | 4.63E-06 | 7.07E-06 |
| chr7 | 72976740 | 72977920 | 1181 | -5.32 | 4.63E-06 | 7.07E-06 |

| | | | | | | |
|---|---|---|---|---|---|---|
| chr3 | 15157310 | 15158108 | 799 | -5.34 | 4.64E-06 | 7.07E-06 |
| chr5 | 73028652 | 73043128 | 14477 | -5.44 | 4.65E-06 | 7.09E-06 |
| chr5 | 106428266 | 106428907 | 642 | -5.32 | 4.65E-06 | 7.09E-06 |
| chr20 | 62390043 | 62390879 | 837 | -5.51 | 4.65E-06 | 7.10E-06 |
| chr6 | 17025885 | 17045703 | 19819 | -2.7 | 4.67E-06 | 7.12E-06 |
| chr11 | 22302887 | 22317636 | 14750 | -1.49 | 4.67E-06 | 7.12E-06 |
| chrX | 55401095 | 55417113 | 16019 | 1.31 | 4.68E-06 | 7.14E-06 |
| chr2 | 20038176 | 20041006 | 2831 | 2.17 | 4.69E-06 | 7.15E-06 |
| chrX | 71097828 | 71106844 | 9017 | -2.31 | 4.70E-06 | 7.17E-06 |
| chr4 | 911679 | 912378 | 700 | -5.32 | 4.71E-06 | 7.18E-06 |
| chr5 | 74929379 | 74930403 | 1025 | -5.8 | 4.71E-06 | 7.19E-06 |
| chr6 | 17494390 | 17495347 | 958 | -5.33 | 4.72E-06 | 7.19E-06 |
| chr11 | 60676778 | 60677832 | 1055 | -5.49 | 4.72E-06 | 7.20E-06 |
| chr19 | 19266551 | 19267568 | 1018 | -5.49 | 4.72E-06 | 7.20E-06 |
| chr7 | 132620401 | 132621163 | 763 | -5.49 | 4.72E-06 | 7.20E-06 |
| chr14 | 103320232 | 103320911 | 680 | -5.7 | 4.73E-06 | 7.21E-06 |
| chr21 | 46383453 | 46384180 | 728 | -5.7 | 4.73E-06 | 7.21E-06 |
| chr12 | 25330663 | 25333842 | 3180 | -3.2 | 4.73E-06 | 7.21E-06 |
| chr16 | 86653570 | 86654308 | 739 | -5.36 | 4.73E-06 | 7.22E-06 |
| chr5 | 111792048 | 111803523 | 11476 | -1.6 | 4.73E-06 | 7.22E-06 |
| chr9 | 34951863 | 34952895 | 1033 | -4.27 | 4.74E-06 | 7.22E-06 |
| chr4 | 48865634 | 48866280 | 647 | -5.31 | 4.74E-06 | 7.23E-06 |
| chr10 | 127047642 | 127127611 | 79970 | -2.95 | 4.75E-06 | 7.25E-06 |
| chr1 | 113111577 | 113112217 | 641 | -5.41 | 4.76E-06 | 7.26E-06 |
| chr14 | 49268178 | 49287600 | 19423 | -1.74 | 4.78E-06 | 7.29E-06 |
| chr1 | 188467929 | 188468965 | 1037 | -5.39 | 4.79E-06 | 7.29E-06 |
| chr2 | 32384719 | 32385219 | 501 | -5.39 | 4.79E-06 | 7.29E-06 |
| chr11 | 6443943 | 6445367 | 1425 | 5.83 | 4.79E-06 | 7.30E-06 |
| chr10 | 28741330 | 28745014 | 3685 | -5.44 | 4.81E-06 | 7.32E-06 |
| chr14 | 75835401 | 75836523 | 1123 | -3.07 | 4.81E-06 | 7.33E-06 |
| chr13 | 34354641 | 34371280 | 16640 | -6.48 | 4.82E-06 | 7.34E-06 |
| chr3 | 153203208 | 153205132 | 1925 | 5.78 | 4.83E-06 | 7.36E-06 |
| chr17 | 34996877 | 34997498 | 622 | -5.56 | 4.83E-06 | 7.36E-06 |
| chrX | 135446154 | 135459212 | 13059 | -1.48 | 4.84E-06 | 7.38E-06 |
| chr10 | 93708568 | 93709321 | 754 | -5.35 | 4.85E-06 | 7.38E-06 |
| chr5 | 92083870 | 92085127 | 1258 | -5.35 | 4.85E-06 | 7.38E-06 |
| chr9 | 38340696 | 38342716 | 2021 | -5.33 | 4.85E-06 | 7.39E-06 |
| chrX | 83294838 | 83296179 | 1342 | -5.33 | 4.85E-06 | 7.39E-06 |
| chr13 | 109545348 | 109546763 | 1416 | -6.13 | 4.86E-06 | 7.40E-06 |
| chr4 | 67934230 | 67953989 | 19760 | 2.83 | 4.86E-06 | 7.40E-06 |
| chr12 | 108374817 | 108375620 | 804 | 5.48 | 4.86E-06 | 7.40E-06 |
| chr14 | 54535017 | 54542670 | 7654 | -2.06 | 4.87E-06 | 7.41E-06 |
| chr1 | 24388747 | 24389612 | 866 | -5.45 | 4.87E-06 | 7.41E-06 |
| chr11 | 72442455 | 72443280 | 826 | -5.45 | 4.87E-06 | 7.41E-06 |
| chr2 | 208924050 | 208924963 | 914 | -5.45 | 4.87E-06 | 7.41E-06 |
| chr3 | 122668582 | 122669904 | 1323 | -5.45 | 4.87E-06 | 7.41E-06 |
| chr2 | 128943951 | 128945516 | 1566 | -3.97 | 4.87E-06 | 7.41E-06 |
| chr12 | 71818433 | 71926706 | 108274 | 2.02 | 4.87E-06 | 7.42E-06 |
| chr18 | 566533 | 593459 | 26927 | -1.5 | 4.88E-06 | 7.43E-06 |
| chr10 | 12665824 | 12674826 | 9003 | -6.1 | 4.88E-06 | 7.43E-06 |
| chr13 | 63727627 | 63728965 | 1339 | -5.38 | 4.89E-06 | 7.45E-06 |
| chr3 | 130898788 | 130899771 | 984 | -5.31 | 4.90E-06 | 7.45E-06 |
| chr7 | 29740433 | 29740853 | 421 | -5.31 | 4.90E-06 | 7.45E-06 |
| chr1 | 173165397 | 173166401 | 1005 | -5.31 | 4.91E-06 | 7.47E-06 |
| chr14 | 85697221 | 85697795 | 575 | -5.31 | 4.91E-06 | 7.47E-06 |
| chr3 | 171923218 | 171923844 | 627 | -5.31 | 4.91E-06 | 7.47E-06 |
| chr12 | 9970396 | 9977522 | 7127 | 1.61 | 4.91E-06 | 7.48E-06 |
| chr2 | 182278801 | 182280766 | 1966 | -4.15 | 4.92E-06 | 7.49E-06 |
| chr6 | 146049950 | 146050727 | 778 | -5.43 | 4.93E-06 | 7.50E-06 |
| chr7 | 130254930 | 130255588 | 659 | -5.43 | 4.93E-06 | 7.50E-06 |
| chr12 | 41509705 | 41519114 | 9410 | -5.84 | 4.93E-06 | 7.51E-06 |

| | | | | | | |
|---|---|---|---|---|---|---|
| chr4 | 75959260 | 75967695 | 8436 | -5.84 | 4.93E-06 | 7.51E-06 |
| chr9 | 126155139 | 126155793 | 655 | -5.82 | 4.94E-06 | 7.51E-06 |
| chr4 | 148103931 | 148105818 | 1888 | -5.75 | 4.94E-06 | 7.52E-06 |
| chr12 | 70682520 | 70693006 | 10487 | -6.61 | 4.95E-06 | 7.53E-06 |
| chr20 | 48018761 | 48019519 | 759 | -5.71 | 4.95E-06 | 7.53E-06 |
| chr11 | 104968845 | 104969356 | 512 | 5.41 | 4.95E-06 | 7.53E-06 |
| chr14 | 75192891 | 75193368 | 478 | -5.32 | 4.95E-06 | 7.53E-06 |
| chr20 | 56389564 | 56390858 | 1295 | -5.32 | 4.95E-06 | 7.53E-06 |
| chr4 | 131608136 | 131609384 | 1249 | -5.32 | 4.95E-06 | 7.53E-06 |
| chr4 | 115235300 | 115236066 | 767 | -5.31 | 4.95E-06 | 7.53E-06 |
| chr1 | 100272632 | 100273264 | 633 | -5.69 | 4.95E-06 | 7.53E-06 |
| chr3 | 158112762 | 158113969 | 1208 | -5.69 | 4.95E-06 | 7.53E-06 |
| chr9 | 86963320 | 86964664 | 1345 | -5.69 | 4.95E-06 | 7.53E-06 |
| chr17 | 76677797 | 76685849 | 8053 | -7.05 | 4.95E-06 | 7.53E-06 |
| chr16 | 78496636 | 78499662 | 3027 | -3.61 | 4.95E-06 | 7.53E-06 |
| chr6 | 148033442 | 148034541 | 1100 | -5.68 | 4.96E-06 | 7.54E-06 |
| chr16 | 47203070 | 47204363 | 1294 | -5.67 | 4.96E-06 | 7.54E-06 |
| chr12 | 118451598 | 118452842 | 1245 | -5.65 | 4.97E-06 | 7.56E-06 |
| chr14 | 22555979 | 22556459 | 481 | -5.34 | 4.97E-06 | 7.56E-06 |
| chr8 | 116265840 | 116267362 | 1523 | -5.63 | 4.98E-06 | 7.57E-06 |
| chr2 | 169355777 | 169356521 | 745 | -5.3 | 4.98E-06 | 7.57E-06 |
| chrX | 91428004 | 91536681 | 108678 | 0.88 | 4.98E-06 | 7.58E-06 |
| chrX | 133087442 | 133089128 | 1687 | -2.88 | 4.98E-06 | 7.58E-06 |
| chr1 | 67735410 | 67741142 | 5733 | -2.31 | 4.99E-06 | 7.59E-06 |
| chr11 | 8446906 | 8447535 | 630 | -5.61 | 4.99E-06 | 7.59E-06 |
| chr22 | 24871671 | 24872869 | 1199 | -5.61 | 4.99E-06 | 7.59E-06 |
| chr6 | 137423891 | 137458462 | 34572 | -5.09 | 5.00E-06 | 7.61E-06 |
| chr20 | 8070295 | 8099969 | 29675 | 1.22 | 5.01E-06 | 7.62E-06 |
| chr2 | 214729880 | 214731110 | 1231 | -4.22 | 5.03E-06 | 7.64E-06 |
| chr1 | 3723388 | 3724578 | 1191 | -5.47 | 5.04E-06 | 7.66E-06 |
| chr18 | 8811854 | 8812718 | 865 | -5.47 | 5.04E-06 | 7.66E-06 |
| chr5 | 14454081 | 14454968 | 888 | -5.31 | 5.06E-06 | 7.69E-06 |
| chr8 | 8775552 | 8776179 | 628 | -5.31 | 5.06E-06 | 7.69E-06 |
| chr14 | 34716484 | 34717911 | 1428 | -4.34 | 5.07E-06 | 7.70E-06 |
| chr12 | 76865760 | 76867286 | 1527 | -5.37 | 5.07E-06 | 7.70E-06 |
| chr4 | 117230680 | 117231696 | 1017 | -5.37 | 5.07E-06 | 7.70E-06 |
| chr9 | 36256689 | 36257466 | 778 | -5.37 | 5.07E-06 | 7.70E-06 |
| chr17 | 76633594 | 76649059 | 15466 | -6.55 | 5.08E-06 | 7.72E-06 |
| chr2 | 222459408 | 222461121 | 1714 | -5.54 | 5.09E-06 | 7.73E-06 |
| chr1 | 198047547 | 198048299 | 753 | -5.53 | 5.11E-06 | 7.76E-06 |
| chr15 | 66135176 | 66136022 | 847 | -5.53 | 5.11E-06 | 7.76E-06 |
| chr18 | 29975016 | 29975866 | 851 | -5.45 | 5.11E-06 | 7.77E-06 |
| chr8 | 30386822 | 30387943 | 1122 | -5.3 | 5.12E-06 | 7.78E-06 |
| chr2 | 206439061 | 206442133 | 3073 | -4.76 | 5.12E-06 | 7.78E-06 |
| chr2 | 233400889 | 233401834 | 946 | -5.31 | 5.13E-06 | 7.80E-06 |
| chr5 | 56321508 | 56322135 | 628 | -5.31 | 5.13E-06 | 7.80E-06 |
| chr12 | 52617967 | 52618877 | 911 | -5.51 | 5.14E-06 | 7.80E-06 |
| chr3 | 193603671 | 193604822 | 1152 | -5.51 | 5.14E-06 | 7.80E-06 |
| chr4 | 84739608 | 84740914 | 1307 | -5.51 | 5.14E-06 | 7.80E-06 |
| chr4 | 103287236 | 103288396 | 1161 | -5.51 | 5.14E-06 | 7.80E-06 |
| chr6 | 118852006 | 118852766 | 761 | -5.51 | 5.14E-06 | 7.80E-06 |
| chr8 | 121084154 | 121085109 | 956 | -5.51 | 5.14E-06 | 7.80E-06 |
| chr19 | 46074870 | 46076331 | 1462 | -5.35 | 5.15E-06 | 7.82E-06 |
| chr9 | 17199169 | 17200353 | 1185 | -5.35 | 5.15E-06 | 7.82E-06 |
| chr6 | 38301063 | 38302043 | 981 | -5.39 | 5.16E-06 | 7.83E-06 |
| chr14 | 92373201 | 92382830 | 9630 | -6.52 | 5.16E-06 | 7.83E-06 |
| chr15 | 85951627 | 85952222 | 596 | -5.36 | 5.17E-06 | 7.84E-06 |
| chr4 | 161871401 | 161873022 | 1622 | -5.36 | 5.17E-06 | 7.84E-06 |
| chr1 | 30333128 | 30346762 | 13635 | -1.36 | 5.19E-06 | 7.88E-06 |
| chr3 | 170426849 | 170437105 | 10257 | -2.54 | 5.19E-06 | 7.88E-06 |
| chr16 | 20289274 | 20299318 | 10045 | -2.73 | 5.19E-06 | 7.89E-06 |

| | | | | | | |
|---|---|---|---|---|---|---|
| chr6 | 125835147 | 125839103 | 3957 | -3.3 | 5.20E-06 | 7.89E-06 |
| chr13 | 99980251 | 99980857 | 607 | -5.43 | 5.20E-06 | 7.90E-06 |
| chr10 | 129016530 | 129017312 | 783 | -5.3 | 5.21E-06 | 7.91E-06 |
| chr12 | 101933515 | 101942373 | 8859 | -5.18 | 5.21E-06 | 7.92E-06 |
| chr6 | 117132679 | 117133782 | 1104 | -4.44 | 5.22E-06 | 7.92E-06 |
| chr1 | 7494228 | 7495248 | 1021 | -5.32 | 5.23E-06 | 7.94E-06 |
| chr1 | 41137068 | 41137682 | 615 | -5.32 | 5.23E-06 | 7.94E-06 |
| chr1 | 186132576 | 186133647 | 1072 | -5.32 | 5.23E-06 | 7.94E-06 |
| chr2 | 148876185 | 148877014 | 830 | -5.32 | 5.23E-06 | 7.94E-06 |
| chr20 | 3893694 | 3894375 | 682 | -5.32 | 5.23E-06 | 7.94E-06 |
| chr3 | 189549805 | 189550587 | 783 | -5.32 | 5.23E-06 | 7.94E-06 |
| chr4 | 132979230 | 132980386 | 1157 | -5.29 | 5.24E-06 | 7.95E-06 |
| chr17 | 37324720 | 37325635 | 916 | -5.38 | 5.25E-06 | 7.97E-06 |
| chr5 | 112147020 | 112147588 | 569 | -5.38 | 5.25E-06 | 7.97E-06 |
| chrX | 114161086 | 114165213 | 4128 | 2.51 | 5.26E-06 | 7.98E-06 |
| chr1 | 211322491 | 211347683 | 25193 | -3.24 | 5.26E-06 | 7.98E-06 |
| chr4 | 135767781 | 135791904 | 24124 | 5.37 | 5.26E-06 | 7.99E-06 |
| chr16 | 84976168 | 84977148 | 981 | -5.32 | 5.26E-06 | 7.99E-06 |
| chr8 | 133866395 | 133874187 | 7793 | -4.03 | 5.27E-06 | 7.99E-06 |
| chr2 | 175949266 | 175949689 | 424 | -5.35 | 5.27E-06 | 8.00E-06 |
| chr3 | 122461152 | 122461868 | 717 | -5.35 | 5.27E-06 | 8.00E-06 |
| chr19 | 45653107 | 45653952 | 846 | -5.46 | 5.28E-06 | 8.01E-06 |
| chr2 | 148905250 | 148906459 | 1210 | -5.46 | 5.28E-06 | 8.01E-06 |
| chr7 | 128716570 | 128717552 | 983 | -5.46 | 5.28E-06 | 8.01E-06 |
| chr11 | 109351427 | 109352454 | 1028 | 5.56 | 5.28E-06 | 8.02E-06 |
| chr1 | 173664660 | 173666161 | 1502 | -2.07 | 5.30E-06 | 8.03E-06 |
| chr8 | 100089341 | 100090154 | 814 | -5.33 | 5.30E-06 | 8.03E-06 |
| chr1 | 97929875 | 97930394 | 520 | -4.64 | 5.30E-06 | 8.04E-06 |
| chr1 | 118461377 | 118462279 | 903 | -5.3 | 5.30E-06 | 8.04E-06 |
| chr18 | 44062021 | 44062655 | 635 | -5.41 | 5.31E-06 | 8.06E-06 |
| chr7 | 150388345 | 150392799 | 4455 | 4.95 | 5.31E-06 | 8.06E-06 |
| chr3 | 188255527 | 188256883 | 1357 | -5.45 | 5.33E-06 | 8.08E-06 |
| chr7 | 91615028 | 91616034 | 1007 | -5.45 | 5.33E-06 | 8.08E-06 |
| chr1 | 151903219 | 151903923 | 705 | -5.3 | 5.33E-06 | 8.08E-06 |
| chr1 | 244280246 | 244281054 | 809 | -5.3 | 5.33E-06 | 8.08E-06 |
| chr7 | 12714997 | 12715794 | 798 | -5.3 | 5.33E-06 | 8.08E-06 |
| chr6 | 159437426 | 159447277 | 9852 | -6.45 | 5.34E-06 | 8.10E-06 |
| chr13 | 74821892 | 74851147 | 29256 | -1.24 | 5.34E-06 | 8.10E-06 |
| chr12 | 122143145 | 122148874 | 5730 | -5.29 | 5.36E-06 | 8.12E-06 |
| chr14 | 95958460 | 95958969 | 510 | -5.29 | 5.36E-06 | 8.12E-06 |
| chr20 | 32886761 | 32887931 | 1171 | -5.29 | 5.36E-06 | 8.12E-06 |
| chr2 | 227601238 | 227612974 | 11737 | -5.93 | 5.36E-06 | 8.12E-06 |
| chrX | 18433968 | 18434945 | 978 | -5.37 | 5.36E-06 | 8.12E-06 |
| chr3 | 137803476 | 137812669 | 9194 | -5.9 | 5.37E-06 | 8.13E-06 |
| chr5 | 136911901 | 136929274 | 17374 | -1.68 | 5.37E-06 | 8.14E-06 |
| chr16 | 84321474 | 84322108 | 635 | -5.4 | 5.38E-06 | 8.15E-06 |
| chr17 | 47802993 | 47804025 | 1033 | -5.4 | 5.38E-06 | 8.15E-06 |
| chr12 | 22405514 | 22412242 | 6729 | 5.09 | 5.38E-06 | 8.15E-06 |
| chr1 | 212128804 | 212129609 | 806 | -5.83 | 5.38E-06 | 8.15E-06 |
| chrX | 52022259 | 52035013 | 12755 | -2.12 | 5.38E-06 | 8.15E-06 |
| chr3 | 124024259 | 124026690 | 2432 | -5.01 | 5.38E-06 | 8.16E-06 |
| chr4 | 5171515 | 5172009 | 495 | -5.34 | 5.39E-06 | 8.17E-06 |
| chr1 | 236297020 | 236298932 | 1913 | -4.54 | 5.41E-06 | 8.19E-06 |
| chr18 | 10155367 | 10157556 | 2190 | -4.04 | 5.42E-06 | 8.21E-06 |
| chr3 | 196548316 | 196549060 | 745 | -5.66 | 5.43E-06 | 8.22E-06 |
| chr2 | 169723605 | 169724022 | 418 | -5.31 | 5.43E-06 | 8.23E-06 |
| chrX | 132551176 | 132552907 | 1732 | -2.58 | 5.44E-06 | 8.24E-06 |
| chr6 | 90907275 | 90908460 | 1186 | -5.63 | 5.44E-06 | 8.25E-06 |
| chr14 | 74073525 | 74074501 | 977 | -5.42 | 5.45E-06 | 8.25E-06 |
| chrX | 49394268 | 49394775 | 508 | -5.42 | 5.45E-06 | 8.25E-06 |
| chr1 | 51547012 | 51549607 | 2596 | -4.55 | 5.46E-06 | 8.28E-06 |

| | | | | | | |
|---|---|---|---|---|---|---|
| chr1 | 18669140 | 18670096 | 957 | -5.32 | 5.47E-06 | 8.28E-06 |
| chr15 | 63610272 | 63610806 | 535 | -5.32 | 5.47E-06 | 8.28E-06 |
| chr6 | 160192803 | 160193652 | 850 | -5.32 | 5.47E-06 | 8.28E-06 |
| chr2 | 74254635 | 74255734 | 1100 | -5.6 | 5.47E-06 | 8.28E-06 |
| chr11 | 123328076 | 123329148 | 1073 | -5.59 | 5.48E-06 | 8.30E-06 |
| chr2 | 133382104 | 133382888 | 785 | -5.35 | 5.48E-06 | 8.30E-06 |
| chr22 | 26325891 | 26379924 | 54034 | 1.1 | 5.48E-06 | 8.30E-06 |
| chr18 | 13676668 | 13677901 | 1234 | -5.58 | 5.49E-06 | 8.31E-06 |
| chr6 | 16875654 | 16880911 | 5258 | -1.6 | 5.49E-06 | 8.31E-06 |
| chr19 | 39889512 | 39890192 | 681 | -5.75 | 5.51E-06 | 8.34E-06 |
| chr1 | 9763277 | 9764420 | 1144 | -5.72 | 5.51E-06 | 8.34E-06 |
| chr2 | 127653657 | 127684411 | 30755 | -3.14 | 5.51E-06 | 8.34E-06 |
| chr3 | 185381297 | 185381928 | 632 | -5.56 | 5.51E-06 | 8.35E-06 |
| chr22 | 43351724 | 43352530 | 807 | -5.28 | 5.51E-06 | 8.35E-06 |
| chr3 | 52745927 | 52746467 | 541 | -5.28 | 5.51E-06 | 8.35E-06 |
| chr7 | 16224513 | 16225260 | 748 | -5.28 | 5.51E-06 | 8.35E-06 |
| chr12 | 108177914 | 108230480 | 52567 | -2.14 | 5.52E-06 | 8.35E-06 |
| chr10 | 51103426 | 51104481 | 1056 | -5.33 | 5.52E-06 | 8.35E-06 |
| chr18 | 76874401 | 76875045 | 645 | -5.33 | 5.52E-06 | 8.35E-06 |
| chr5 | 133722621 | 133723216 | 596 | -5.33 | 5.52E-06 | 8.35E-06 |
| chr3 | 155229832 | 155231558 | 1727 | -5.66 | 5.53E-06 | 8.37E-06 |
| chr15 | 68736454 | 68753693 | 17240 | -1.81 | 5.54E-06 | 8.38E-06 |
| chr1 | 213491464 | 213492643 | 1180 | -4.12 | 5.55E-06 | 8.40E-06 |
| chr13 | 110057118 | 110058067 | 950 | -5.28 | 5.55E-06 | 8.40E-06 |
| chr7 | 107742050 | 107742699 | 650 | -5.28 | 5.55E-06 | 8.40E-06 |
| chr1 | 81637959 | 81639749 | 1791 | -5.61 | 5.56E-06 | 8.41E-06 |
| chr8 | 87635405 | 87638226 | 2822 | 2.59 | 5.56E-06 | 8.42E-06 |
| chrX | 125731064 | 125760460 | 29397 | 1.76 | 5.57E-06 | 8.42E-06 |
| chr3 | 159496728 | 159498767 | 2040 | -4.11 | 5.58E-06 | 8.44E-06 |
| chr1 | 111289368 | 111299347 | 9980 | -2.25 | 5.59E-06 | 8.46E-06 |
| chr14 | 77283298 | 77299107 | 15810 | -2.22 | 5.59E-06 | 8.46E-06 |
| chr2 | 178100234 | 178101376 | 1143 | -5.4 | 5.59E-06 | 8.46E-06 |
| chr5 | 141038911 | 141039624 | 714 | -5.37 | 5.60E-06 | 8.47E-06 |
| chr6 | 151636389 | 151637331 | 943 | -5.37 | 5.60E-06 | 8.47E-06 |
| chr5 | 105928662 | 105929498 | 837 | -4.63 | 5.61E-06 | 8.48E-06 |
| chr14 | 34701366 | 34702060 | 695 | -5.29 | 5.62E-06 | 8.49E-06 |
| chr6 | 22563002 | 22563909 | 908 | -5.29 | 5.62E-06 | 8.49E-06 |
| chr10 | 32303403 | 32304403 | 1001 | -5.34 | 5.62E-06 | 8.50E-06 |
| chr13 | 91012985 | 91013840 | 856 | -5.34 | 5.62E-06 | 8.50E-06 |
| chr17 | 17196064 | 17196701 | 638 | -5.34 | 5.62E-06 | 8.50E-06 |
| chr7 | 109991419 | 109992230 | 812 | -5.55 | 5.62E-06 | 8.50E-06 |
| chr10 | 126250237 | 126250888 | 652 | -5.28 | 5.62E-06 | 8.50E-06 |
| chr20 | 58140870 | 58141965 | 1096 | -5.28 | 5.62E-06 | 8.50E-06 |
| chr6 | 82196495 | 82222903 | 26409 | 3.44 | 5.64E-06 | 8.52E-06 |
| chr6 | 154334081 | 154334868 | 788 | 5.51 | 5.66E-06 | 8.55E-06 |
| chr13 | 20992148 | 20993406 | 1259 | -5.31 | 5.66E-06 | 8.56E-06 |
| chr18 | 11725417 | 11726157 | 741 | -5.31 | 5.66E-06 | 8.56E-06 |
| chr2 | 240655509 | 240656834 | 1326 | -5.31 | 5.66E-06 | 8.56E-06 |
| chr7 | 85412457 | 85427687 | 15231 | -5.21 | 5.67E-06 | 8.57E-06 |
| chr13 | 41918735 | 41919485 | 751 | -5.48 | 5.68E-06 | 8.58E-06 |
| chr18 | 13045546 | 13046293 | 748 | -5.38 | 5.68E-06 | 8.58E-06 |
| chr3 | 42014793 | 42015573 | 781 | -5.52 | 5.69E-06 | 8.59E-06 |
| chrX | 47146438 | 47155122 | 8685 | -5.19 | 5.69E-06 | 8.60E-06 |
| chr13 | 90904021 | 90905384 | 1364 | -5.29 | 5.69E-06 | 8.60E-06 |
| chr17 | 20665743 | 20666566 | 824 | -5.36 | 5.69E-06 | 8.60E-06 |
| chr19 | 1535581 | 1536143 | 563 | -5.36 | 5.69E-06 | 8.60E-06 |
| chr11 | 101378070 | 101379310 | 1241 | -5.47 | 5.71E-06 | 8.63E-06 |
| chr2 | 43993540 | 43994218 | 679 | -5.47 | 5.71E-06 | 8.63E-06 |
| chr8 | 99001637 | 99002913 | 1277 | -5.47 | 5.71E-06 | 8.63E-06 |
| chr8 | 120702735 | 120707800 | 5066 | -1.88 | 5.71E-06 | 8.63E-06 |
| chr1 | 82707181 | 82709437 | 2257 | -5.5 | 5.71E-06 | 8.63E-06 |

| | | | | | | |
|---|---|---|---|---|---|---|
| chr3 | 48823415 | 48823818 | 404 | -4.2 | 5.72E-06 | 8.65E-06 |
| chr1 | 95782186 | 95791110 | 8925 | 2.18 | 5.72E-06 | 8.65E-06 |
| chr1 | 78962076 | 79053812 | 91737 | 6.35 | 5.73E-06 | 8.65E-06 |
| chr13 | 32019161 | 32029030 | 9870 | 2.79 | 5.73E-06 | 8.66E-06 |
| chr16 | 58105609 | 58123554 | 17946 | -6.64 | 5.74E-06 | 8.67E-06 |
| chr17 | 26038004 | 26039898 | 1895 | -1.84 | 5.74E-06 | 8.67E-06 |
| chr12 | 40161175 | 40161859 | 685 | -5.49 | 5.74E-06 | 8.68E-06 |
| chr6 | 137751510 | 137752322 | 813 | 5.48 | 5.77E-06 | 8.71E-06 |
| chr4 | 86057308 | 86057967 | 660 | -4.93 | 5.77E-06 | 8.72E-06 |
| chr2 | 204374017 | 204375174 | 1158 | -5.32 | 5.77E-06 | 8.72E-06 |
| chr4 | 78716687 | 78717529 | 843 | -5.32 | 5.77E-06 | 8.72E-06 |
| chr15 | 63304617 | 63311088 | 6472 | -6.63 | 5.78E-06 | 8.73E-06 |
| chr13 | 27310010 | 27324150 | 14141 | -2.12 | 5.80E-06 | 8.75E-06 |
| chr17 | 858883 | 859882 | 1000 | -5.27 | 5.80E-06 | 8.76E-06 |
| chr5 | 130100243 | 130100813 | 571 | -5.3 | 5.82E-06 | 8.78E-06 |
| chr8 | 112063925 | 112065247 | 1323 | -5.3 | 5.82E-06 | 8.78E-06 |
| chr2 | 98556841 | 98557468 | 628 | -5.44 | 5.82E-06 | 8.79E-06 |
| chr5 | 156483209 | 156483904 | 696 | -5.44 | 5.82E-06 | 8.79E-06 |
| chr8 | 48760741 | 48761290 | 550 | -5.28 | 5.82E-06 | 8.79E-06 |
| chr7 | 8135079 | 8135955 | 877 | -3.96 | 5.85E-06 | 8.83E-06 |
| chr6 | 130299162 | 130300344 | 1183 | -5.27 | 5.86E-06 | 8.84E-06 |
| chr5 | 169439849 | 169441152 | 1304 | -4.17 | 5.86E-06 | 8.85E-06 |
| chr1 | 8969924 | 8970729 | 806 | -5.45 | 5.86E-06 | 8.85E-06 |
| chr3 | 133723105 | 133726617 | 3513 | -4.71 | 5.86E-06 | 8.85E-06 |
| chr1 | 38856762 | 38881826 | 25065 | -1.86 | 5.87E-06 | 8.86E-06 |
| chr5 | 75059747 | 75098885 | 39139 | -2.42 | 5.87E-06 | 8.87E-06 |
| chr2 | 72524435 | 72524985 | 551 | -5.29 | 5.88E-06 | 8.87E-06 |
| chr4 | 25379692 | 25380474 | 783 | -5.29 | 5.88E-06 | 8.87E-06 |
| chr10 | 103232263 | 103233733 | 1471 | -5.35 | 5.89E-06 | 8.88E-06 |
| chr6 | 42166900 | 42167706 | 807 | -5.35 | 5.89E-06 | 8.88E-06 |
| chr6 | 132572845 | 132580451 | 7607 | -4.67 | 5.89E-06 | 8.89E-06 |
| chr4 | 86087800 | 86088763 | 964 | -4.74 | 5.90E-06 | 8.91E-06 |
| chr10 | 124822555 | 124846046 | 23492 | -1.82 | 5.91E-06 | 8.91E-06 |
| chr1 | 111373172 | 111374215 | 1044 | -5.44 | 5.91E-06 | 8.92E-06 |
| chr1 | 174866397 | 174868044 | 1648 | -5.44 | 5.91E-06 | 8.92E-06 |
| chr20 | 17799668 | 17800532 | 865 | -5.44 | 5.91E-06 | 8.92E-06 |
| chr6 | 11743992 | 11758255 | 14264 | -6.12 | 5.92E-06 | 8.93E-06 |
| chr15 | 95835911 | 95870888 | 34978 | -3.03 | 5.93E-06 | 8.95E-06 |
| chr3 | 54044519 | 54064324 | 19806 | -1.16 | 5.94E-06 | 8.96E-06 |
| chr8 | 32003362 | 32012312 | 8951 | 1.43 | 5.95E-06 | 8.98E-06 |
| chr10 | 97380333 | 97380983 | 651 | -5.41 | 5.97E-06 | 9.00E-06 |
| chr14 | 65999542 | 66000821 | 1280 | -5.43 | 5.97E-06 | 9.00E-06 |
| chr4 | 91787048 | 91788188 | 1141 | -5.43 | 5.97E-06 | 9.00E-06 |
| chr11 | 50523672 | 50635318 | 111647 | -1.08 | 5.99E-06 | 9.03E-06 |
| chr3 | 172219010 | 172220389 | 1380 | -6.07 | 6.01E-06 | 9.06E-06 |
| chr16 | 90036957 | 90038183 | 1227 | -5.34 | 6.01E-06 | 9.06E-06 |
| chr8 | 130846588 | 130847064 | 477 | -5.34 | 6.01E-06 | 9.06E-06 |
| chr3 | 72409139 | 72412026 | 2888 | 1.87 | 6.01E-06 | 9.07E-06 |
| chr6 | 83284666 | 83287860 | 3195 | -4.99 | 6.02E-06 | 9.08E-06 |
| chr12 | 69091494 | 69092430 | 937 | -5.41 | 6.03E-06 | 9.09E-06 |
| chr5 | 56020586 | 56022262 | 1677 | -5.41 | 6.03E-06 | 9.09E-06 |
| chr16 | 79693514 | 79736233 | 42720 | -1.33 | 6.03E-06 | 9.10E-06 |
| chr12 | 124032507 | 124033104 | 598 | -5.27 | 6.05E-06 | 9.12E-06 |
| chr5 | 179785670 | 179786903 | 1234 | -5.27 | 6.05E-06 | 9.12E-06 |
| chr2 | 189547040 | 189548848 | 1809 | -5.93 | 6.06E-06 | 9.14E-06 |
| chr1 | 20855563 | 20856227 | 665 | -5.26 | 6.06E-06 | 9.14E-06 |
| chr13 | 82135713 | 82136907 | 1195 | -5.26 | 6.06E-06 | 9.14E-06 |
| chr5 | 133374610 | 133375257 | 648 | -5.26 | 6.06E-06 | 9.14E-06 |
| chr6 | 2401207 | 2401920 | 714 | -3.96 | 6.08E-06 | 9.16E-06 |
| chr5 | 56220920 | 56221579 | 660 | -5.39 | 6.09E-06 | 9.18E-06 |
| chr3 | 16559619 | 16568529 | 8911 | -3.05 | 6.09E-06 | 9.18E-06 |

| | | | | | | |
|---|---|---|---|---|---|---|
| chr3 | 108757647 | 108772587 | 14941 | -5.85 | 6.10E-06 | 9.19E-06 |
| chr1 | 193540873 | 193553413 | 12541 | -5.82 | 6.11E-06 | 9.20E-06 |
| chr2 | 141372405 | 141373635 | 1231 | -5.82 | 6.11E-06 | 9.20E-06 |
| chr13 | 115035028 | 115035668 | 641 | -4.04 | 6.13E-06 | 9.23E-06 |
| chr15 | 40116011 | 40117382 | 1372 | -5.75 | 6.13E-06 | 9.23E-06 |
| chr3 | 194579508 | 194580580 | 1073 | -5.75 | 6.13E-06 | 9.23E-06 |
| chr13 | 30633399 | 30634866 | 1468 | 5.72 | 6.13E-06 | 9.24E-06 |
| chr6 | 110657844 | 110659286 | 1443 | -5.74 | 6.14E-06 | 9.25E-06 |
| chrX | 40094801 | 40095825 | 1025 | -5.73 | 6.14E-06 | 9.25E-06 |
| chr22 | 39767689 | 39769000 | 1312 | -5.72 | 6.14E-06 | 9.25E-06 |
| chr1 | 204538995 | 204539873 | 879 | -5.29 | 6.14E-06 | 9.25E-06 |
| chr6 | 140314936 | 140315765 | 830 | -5.29 | 6.14E-06 | 9.25E-06 |
| chrX | 133305305 | 133306069 | 765 | -5.29 | 6.14E-06 | 9.25E-06 |
| chr11 | 95301329 | 95303307 | 1979 | -1.87 | 6.15E-06 | 9.26E-06 |
| chr2 | 39114759 | 39115692 | 934 | -5.33 | 6.15E-06 | 9.26E-06 |
| chr17 | 9279138 | 9279889 | 752 | -5.38 | 6.16E-06 | 9.28E-06 |
| chr8 | 27670945 | 27671647 | 703 | -5.38 | 6.16E-06 | 9.28E-06 |
| chr13 | 84515557 | 84516659 | 1103 | -5.66 | 6.16E-06 | 9.28E-06 |
| chr3 | 27323129 | 27323978 | 850 | -5.66 | 6.16E-06 | 9.28E-06 |
| chr3 | 70805361 | 70807286 | 1926 | -2.46 | 6.17E-06 | 9.29E-06 |
| chr2 | 63966309 | 63967101 | 793 | -5.28 | 6.18E-06 | 9.30E-06 |
| chr6 | 138445182 | 138455984 | 10803 | -1.63 | 6.18E-06 | 9.30E-06 |
| chr22 | 48287785 | 48386240 | 98456 | -2.25 | 6.19E-06 | 9.32E-06 |
| chr18 | 70260106 | 70261297 | 1192 | -4.01 | 6.19E-06 | 9.33E-06 |
| chrX | 41339501 | 41347327 | 7827 | -5.6 | 6.20E-06 | 9.33E-06 |
| chrX | 103734728 | 103740381 | 5654 | 4.9 | 6.21E-06 | 9.34E-06 |
| chr1 | 160527074 | 160533858 | 6785 | -2.57 | 6.21E-06 | 9.35E-06 |
| chr11 | 114439952 | 114441445 | 1494 | 4.34 | 6.21E-06 | 9.35E-06 |
| chr7 | 128053143 | 128054292 | 1150 | -5.59 | 6.21E-06 | 9.35E-06 |
| chr17 | 33505917 | 33506563 | 647 | -3.47 | 6.22E-06 | 9.36E-06 |
| chr11 | 43810455 | 43811078 | 624 | -5.26 | 6.22E-06 | 9.36E-06 |
| chr12 | 66899836 | 66900438 | 603 | -5.26 | 6.22E-06 | 9.36E-06 |
| chr12 | 78246484 | 78247175 | 692 | -5.26 | 6.22E-06 | 9.36E-06 |
| chr8 | 123614573 | 123615393 | 821 | -5.26 | 6.22E-06 | 9.36E-06 |
| chr8 | 59180360 | 59187439 | 7080 | -2.9 | 6.24E-06 | 9.39E-06 |
| chr6 | 163748281 | 163748971 | 691 | -5.57 | 6.24E-06 | 9.39E-06 |
| chr4 | 134353845 | 134354888 | 1044 | 5.34 | 6.24E-06 | 9.39E-06 |
| chr10 | 129730780 | 129731667 | 888 | -5.25 | 6.25E-06 | 9.41E-06 |
| chr21 | 46968708 | 46969286 | 579 | -5.25 | 6.25E-06 | 9.41E-06 |
| chr11 | 88150112 | 88151271 | 1160 | -5.56 | 6.25E-06 | 9.41E-06 |
| chr2 | 174336721 | 174337614 | 894 | -5.56 | 6.25E-06 | 9.41E-06 |
| chr18 | 74478410 | 74487413 | 9004 | -3.42 | 6.27E-06 | 9.43E-06 |
| chr3 | 182306775 | 182307368 | 594 | -5.26 | 6.28E-06 | 9.44E-06 |
| chr3 | 194773960 | 194774754 | 795 | -5.26 | 6.28E-06 | 9.44E-06 |
| chr6 | 76082645 | 76083120 | 476 | -5.26 | 6.28E-06 | 9.44E-06 |
| chr15 | 86178110 | 86178584 | 475 | -5.58 | 6.28E-06 | 9.45E-06 |
| chr12 | 13083236 | 13084066 | 831 | -5.53 | 6.29E-06 | 9.46E-06 |
| chr17 | 47009763 | 47010307 | 545 | -5.26 | 6.30E-06 | 9.48E-06 |
| chr1 | 69244973 | 69245963 | 991 | -5.31 | 6.31E-06 | 9.49E-06 |
| chr17 | 2686773 | 2687396 | 624 | -5.31 | 6.31E-06 | 9.49E-06 |
| chr4 | 178179965 | 178180623 | 659 | -5.31 | 6.31E-06 | 9.49E-06 |
| chr3 | 123346658 | 123347817 | 1160 | -5.52 | 6.31E-06 | 9.49E-06 |
| chr6 | 140507874 | 140509646 | 1773 | -5.52 | 6.31E-06 | 9.49E-06 |
| chr14 | 22039244 | 22064904 | 25661 | -2.93 | 6.32E-06 | 9.50E-06 |
| chr20 | 1297462 | 1298227 | 766 | -5.35 | 6.32E-06 | 9.50E-06 |
| chr2 | 135477782 | 135478969 | 1188 | -4.24 | 6.35E-06 | 9.55E-06 |
| chr1 | 32359951 | 32360899 | 949 | -5.28 | 6.36E-06 | 9.57E-06 |
| chr10 | 74154439 | 74155287 | 849 | -5.28 | 6.36E-06 | 9.57E-06 |
| chr10 | 89616103 | 89616779 | 677 | -5.49 | 6.37E-06 | 9.57E-06 |
| chr2 | 197086301 | 197087038 | 738 | -5.27 | 6.39E-06 | 9.60E-06 |
| chr3 | 109693424 | 109694358 | 935 | -5.27 | 6.39E-06 | 9.60E-06 |

| | | | | | | |
|---|---|---|---|---|---|---|
| chr7 | 2959475 | 2960005 | 531 | -5.27 | 6.39E-06 | 9.60E-06 |
| chrX | 7100347 | 7105800 | 5454 | 3.31 | 6.39E-06 | 9.61E-06 |
| chr2 | 223416565 | 223417414 | 850 | -3.94 | 6.40E-06 | 9.62E-06 |
| chr15 | 75482671 | 75483644 | 974 | -5.26 | 6.41E-06 | 9.63E-06 |
| chr14 | 65305519 | 65306239 | 721 | -5.34 | 6.41E-06 | 9.64E-06 |
| chr4 | 97454159 | 97456182 | 2024 | -4.06 | 6.44E-06 | 9.68E-06 |
| chr6 | 25073258 | 25073938 | 681 | -5.34 | 6.48E-06 | 9.74E-06 |
| chr14 | 56929944 | 56937068 | 7125 | -1.27 | 6.50E-06 | 9.78E-06 |
| chr12 | 65166814 | 65170617 | 3804 | -5.22 | 6.51E-06 | 9.78E-06 |
| chr1 | 226901781 | 226902467 | 687 | -5.44 | 6.52E-06 | 9.80E-06 |
| chr18 | 52521590 | 52522870 | 1281 | -5.44 | 6.52E-06 | 9.80E-06 |
| chr21 | 45253096 | 45254011 | 916 | -5.44 | 6.52E-06 | 9.80E-06 |
| chr20 | 8723240 | 8724907 | 1668 | -4.6 | 6.54E-06 | 9.83E-06 |
| chr6 | 13997778 | 14013904 | 16127 | -6.1 | 6.55E-06 | 9.84E-06 |
| chr7 | 86873677 | 86893409 | 19733 | -1.67 | 6.56E-06 | 9.85E-06 |
| chr5 | 79001577 | 79086867 | 85291 | -1.63 | 6.56E-06 | 9.86E-06 |
| chr5 | 139196477 | 139209742 | 13266 | -5.49 | 6.58E-06 | 9.89E-06 |
| chr18 | 43608586 | 43609602 | 1017 | -3.99 | 6.59E-06 | 9.90E-06 |
| chr7 | 91205694 | 91219225 | 13532 | 1.84 | 6.60E-06 | 9.92E-06 |
| chr16 | 89214994 | 89216351 | 1358 | -5.33 | 6.61E-06 | 9.93E-06 |
| chr1 | 196277351 | 196278258 | 908 | -5.26 | 6.62E-06 | 9.94E-06 |
| chr2 | 228081541 | 228082206 | 666 | -5.26 | 6.62E-06 | 9.94E-06 |
| chr3 | 189713748 | 189714427 | 680 | -5.26 | 6.62E-06 | 9.94E-06 |
| chr6 | 47524150 | 47524578 | 429 | -5.26 | 6.62E-06 | 9.94E-06 |
| chr7 | 47642704 | 47643242 | 539 | -5.26 | 6.62E-06 | 9.94E-06 |
| chr8 | 83401039 | 83402920 | 1882 | -5.26 | 6.62E-06 | 9.94E-06 |
| chr1 | 4927274 | 4970585 | 43312 | 0.93 | 6.62E-06 | 9.94E-06 |
| chr2 | 151630745 | 151631533 | 789 | -5.24 | 6.62E-06 | 9.95E-06 |
| chr2 | 223975578 | 223976629 | 1052 | -5.24 | 6.62E-06 | 9.95E-06 |
| chr7 | 100025347 | 100025855 | 509 | -5.24 | 6.62E-06 | 9.95E-06 |
| chr9 | 32502119 | 32503332 | 1214 | -5.24 | 6.62E-06 | 9.95E-06 |
| chr4 | 106734323 | 106735122 | 800 | -5.32 | 6.62E-06 | 9.95E-06 |
| chr20 | 36334901 | 36335587 | 687 | -5.28 | 6.63E-06 | 9.95E-06 |
| chr6 | 100443119 | 100443992 | 874 | -5.41 | 6.63E-06 | 9.95E-06 |
| chr11 | 19533547 | 19535144 | 1598 | 5.56 | 6.64E-06 | 9.97E-06 |
| chr7 | 42139507 | 42188903 | 49397 | -2.43 | 6.65E-06 | 9.98E-06 |
| chr5 | 90842905 | 90862903 | 19999 | -3.31 | 6.67E-06 | 1.00E-05 |
| chr20 | 52885630 | 52887454 | 1825 | -4.41 | 6.67E-06 | 1.00E-05 |
| chr7 | 69496401 | 69498290 | 1890 | -1.75 | 6.68E-06 | 1.00E-05 |
| chr17 | 28914647 | 28915392 | 746 | -5.28 | 6.68E-06 | 1.00E-05 |
| chr10 | 32662506 | 32663363 | 858 | -5.42 | 6.69E-06 | 1.00E-05 |
| chr1 | 53414494 | 53415192 | 699 | -5.24 | 6.70E-06 | 1.01E-05 |
| chr1 | 214186832 | 214188054 | 1223 | -5.24 | 6.70E-06 | 1.01E-05 |
| chr14 | 39311953 | 39358603 | 46651 | 5.85 | 6.71E-06 | 1.01E-05 |
| chr5 | 111054550 | 111055328 | 779 | 5.52 | 6.72E-06 | 1.01E-05 |
| chr13 | 98308256 | 98310721 | 2466 | -2.34 | 6.73E-06 | 1.01E-05 |
| chr4 | 85383763 | 85385415 | 1653 | -4.06 | 6.73E-06 | 1.01E-05 |
| chr1 | 66189022 | 66212271 | 23250 | -1.08 | 6.74E-06 | 1.01E-05 |
| chr8 | 81567807 | 81568614 | 808 | -5.31 | 6.75E-06 | 1.01E-05 |
| chrX | 39128020 | 39145516 | 17497 | -1.54 | 6.75E-06 | 1.01E-05 |
| chr11 | 60783869 | 60784579 | 711 | -5.31 | 6.75E-06 | 1.01E-05 |
| chr15 | 32956213 | 32957143 | 931 | -5.31 | 6.75E-06 | 1.01E-05 |
| chr11 | 27578932 | 27581026 | 2095 | 2.12 | 6.76E-06 | 1.01E-05 |
| chr6 | 5865465 | 5895077 | 29613 | -4.02 | 6.76E-06 | 1.01E-05 |
| chr3 | 39843846 | 39856284 | 12439 | -1.74 | 6.76E-06 | 1.01E-05 |
| chr5 | 92358990 | 92362882 | 3893 | -5.87 | 6.77E-06 | 1.01E-05 |
| chr10 | 54419474 | 54434198 | 14725 | -1.59 | 6.79E-06 | 1.02E-05 |
| chr11 | 71220149 | 71222373 | 2225 | -4.84 | 6.79E-06 | 1.02E-05 |
| chr4 | 166482132 | 166483111 | 980 | -5.26 | 6.82E-06 | 1.02E-05 |
| chr8 | 59990676 | 59991573 | 898 | -5.26 | 6.82E-06 | 1.02E-05 |
| chr5 | 175315448 | 175316628 | 1181 | -5.74 | 6.83E-06 | 1.03E-05 |

| | | | | | | |
|---|---|---|---|---|---|---|
| chr9 | 38648337 | 38649849 | 1513 | -5.72 | 6.84E-06 | 1.03E-05 |
| chr3 | 70268331 | 70269037 | 707 | -5.37 | 6.84E-06 | 1.03E-05 |
| chr8 | 121653398 | 121685011 | 31614 | -1.31 | 6.85E-06 | 1.03E-05 |
| chr22 | 26984055 | 26984507 | 453 | -5.38 | 6.85E-06 | 1.03E-05 |
| chr12 | 60390397 | 60393198 | 2802 | -5.68 | 6.87E-06 | 1.03E-05 |
| chr3 | 41910060 | 41911436 | 1377 | -5.66 | 6.88E-06 | 1.03E-05 |
| chr4 | 170777358 | 170778714 | 1357 | -5.66 | 6.88E-06 | 1.03E-05 |
| chr4 | 66259344 | 66260435 | 1092 | -5.23 | 6.88E-06 | 1.03E-05 |
| chr5 | 93047975 | 93048523 | 549 | -5.23 | 6.88E-06 | 1.03E-05 |
| chr21 | 36025456 | 36036435 | 10980 | -2.09 | 6.88E-06 | 1.03E-05 |
| chr5 | 142627495 | 142628461 | 967 | -4.03 | 6.88E-06 | 1.03E-05 |
| chrX | 43746936 | 43749933 | 2998 | -5.12 | 6.89E-06 | 1.03E-05 |
| chr4 | 40223726 | 40224669 | 944 | -5.63 | 6.90E-06 | 1.03E-05 |
| chr5 | 115045790 | 115061415 | 15626 | -2.29 | 6.90E-06 | 1.03E-05 |
| chr12 | 131697426 | 131698416 | 991 | -5.62 | 6.90E-06 | 1.04E-05 |
| chr1 | 56875531 | 56876210 | 680 | -5.3 | 6.91E-06 | 1.04E-05 |
| chr7 | 128797469 | 128798346 | 878 | -5.27 | 6.91E-06 | 1.04E-05 |
| chr9 | 83560743 | 83561809 | 1067 | -5.27 | 6.91E-06 | 1.04E-05 |
| chr12 | 57191395 | 57192306 | 912 | -5.59 | 6.92E-06 | 1.04E-05 |
| chr1 | 14162243 | 14163512 | 1270 | -4.09 | 6.92E-06 | 1.04E-05 |
| chr9 | 87161066 | 87294393 | 133328 | 0.7 | 6.96E-06 | 1.04E-05 |
| chr4 | 32463234 | 32465659 | 2426 | -4 | 6.98E-06 | 1.05E-05 |
| chr2 | 236307850 | 236334877 | 27028 | -6.72 | 6.99E-06 | 1.05E-05 |
| chr6 | 167083553 | 167084490 | 938 | -4.22 | 6.99E-06 | 1.05E-05 |
| chr12 | 39758984 | 39759521 | 538 | -5.36 | 6.99E-06 | 1.05E-05 |
| chr4 | 153672330 | 153673809 | 1480 | -5.36 | 6.99E-06 | 1.05E-05 |
| chr4 | 178855436 | 178856732 | 1297 | -5.23 | 7.03E-06 | 1.05E-05 |
| chr11 | 22330178 | 22330553 | 376 | -5.28 | 7.03E-06 | 1.05E-05 |
| chr13 | 50468364 | 50469137 | 774 | -5.25 | 7.04E-06 | 1.05E-05 |
| chr5 | 76718841 | 76719228 | 388 | -5.25 | 7.04E-06 | 1.05E-05 |
| chr16 | 66419107 | 66420010 | 904 | -5.35 | 7.04E-06 | 1.05E-05 |
| chr20 | 62246140 | 62247295 | 1156 | -5.35 | 7.04E-06 | 1.05E-05 |
| chr12 | 89544008 | 89545204 | 1197 | -5.81 | 7.04E-06 | 1.06E-05 |
| chr1 | 86253957 | 86254743 | 787 | -5.23 | 7.07E-06 | 1.06E-05 |
| chr21 | 43126571 | 43127523 | 953 | -5.23 | 7.07E-06 | 1.06E-05 |
| chr2 | 182298027 | 182300241 | 2215 | -4.21 | 7.07E-06 | 1.06E-05 |
| chr19 | 2614365 | 2615132 | 768 | -5.23 | 7.08E-06 | 1.06E-05 |
| chr4 | 189050030 | 189051104 | 1075 | -5.23 | 7.08E-06 | 1.06E-05 |
| chr2 | 205671386 | 205686205 | 14820 | -6.41 | 7.09E-06 | 1.06E-05 |
| chr18 | 59858854 | 59860022 | 1169 | -5.29 | 7.09E-06 | 1.06E-05 |
| chr8 | 62373615 | 62374347 | 733 | 3.02 | 7.10E-06 | 1.06E-05 |
| chr11 | 11819755 | 11820280 | 526 | -5.48 | 7.10E-06 | 1.06E-05 |
| chrX | 2788219 | 2789085 | 867 | -5.48 | 7.10E-06 | 1.06E-05 |
| chr13 | 48948912 | 48949461 | 550 | -5.7 | 7.11E-06 | 1.06E-05 |
| chrX | 42477193 | 42507093 | 29901 | 3.65 | 7.12E-06 | 1.07E-05 |
| chr20 | 57425919 | 57426988 | 1070 | -4.79 | 7.13E-06 | 1.07E-05 |
| chr5 | 130015671 | 130017148 | 1478 | -5.47 | 7.14E-06 | 1.07E-05 |
| chr10 | 129419530 | 129421570 | 2041 | 2.52 | 7.15E-06 | 1.07E-05 |
| chr6 | 101186350 | 101187982 | 1633 | -5.33 | 7.15E-06 | 1.07E-05 |
| chr11 | 28002482 | 28003202 | 721 | -4.65 | 7.15E-06 | 1.07E-05 |
| chr14 | 54679295 | 54680627 | 1333 | -5.23 | 7.16E-06 | 1.07E-05 |
| chr1 | 68638009 | 68638506 | 498 | -5.25 | 7.17E-06 | 1.07E-05 |
| chr8 | 135661873 | 135663047 | 1175 | -5.25 | 7.17E-06 | 1.07E-05 |
| chr1 | 207210403 | 207211363 | 961 | -4.21 | 7.18E-06 | 1.07E-05 |
| chr10 | 17110962 | 17111663 | 702 | -5.27 | 7.19E-06 | 1.08E-05 |
| chr12 | 59432123 | 59433291 | 1169 | -5.58 | 7.20E-06 | 1.08E-05 |
| chr6 | 116662697 | 116664031 | 1335 | -2.75 | 7.21E-06 | 1.08E-05 |
| chrX | 120401798 | 120411865 | 10068 | 2.01 | 7.21E-06 | 1.08E-05 |
| chr12 | 120424226 | 120424868 | 643 | -5.44 | 7.21E-06 | 1.08E-05 |
| chr3 | 122062013 | 122063250 | 1238 | -5.22 | 7.22E-06 | 1.08E-05 |
| chr3 | 131547450 | 131548651 | 1202 | -5.22 | 7.22E-06 | 1.08E-05 |

| | | | | | | |
|---|---|---|---|---|---|---|
| chr18 | 9111357 | 9112191 | 835 | -5.23 | 7.22E-06 | 1.08E-05 |
| chr1 | 189656397 | 189657644 | 1248 | -6.1 | 7.23E-06 | 1.08E-05 |
| chr4 | 98290679 | 98292309 | 1631 | -3.97 | 7.26E-06 | 1.09E-05 |
| chr17 | 65790228 | 65791330 | 1103 | -5.54 | 7.26E-06 | 1.09E-05 |
| chr11 | 45543180 | 45545536 | 2357 | -4.6 | 7.27E-06 | 1.09E-05 |
| chr22 | 28633970 | 28634911 | 942 | -5.24 | 7.27E-06 | 1.09E-05 |
| chr7 | 11032511 | 11034204 | 1694 | -5.53 | 7.28E-06 | 1.09E-05 |
| chr7 | 15111932 | 15113493 | 1562 | -5.32 | 7.28E-06 | 1.09E-05 |
| chr1 | 43089654 | 43090983 | 1330 | -3.91 | 7.29E-06 | 1.09E-05 |
| chr15 | 35715975 | 35716827 | 853 | -5.27 | 7.29E-06 | 1.09E-05 |
| chr21 | 30448309 | 30449068 | 760 | -5.27 | 7.29E-06 | 1.09E-05 |
| chr7 | 43792347 | 43793310 | 964 | -5.27 | 7.29E-06 | 1.09E-05 |
| chr8 | 22512759 | 22513542 | 784 | -3.94 | 7.30E-06 | 1.09E-05 |
| chr13 | 81257675 | 81260337 | 2663 | 4.19 | 7.31E-06 | 1.09E-05 |
| chr17 | 64154344 | 64154899 | 556 | -5.42 | 7.31E-06 | 1.09E-05 |
| chr18 | 74360856 | 74378649 | 17794 | -1.46 | 7.32E-06 | 1.09E-05 |
| chr4 | 22173385 | 22190453 | 17069 | 2.28 | 7.32E-06 | 1.09E-05 |
| chr1 | 225882397 | 225883197 | 801 | -5.4 | 7.37E-06 | 1.10E-05 |
| chr2 | 203040941 | 203042163 | 1223 | -5.4 | 7.37E-06 | 1.10E-05 |
| chr4 | 126789990 | 126790947 | 958 | -5.4 | 7.37E-06 | 1.10E-05 |
| chr3 | 136819054 | 136819679 | 626 | -5.26 | 7.37E-06 | 1.10E-05 |
| chr10 | 43373741 | 43429203 | 55463 | -6.04 | 7.38E-06 | 1.10E-05 |
| chr13 | 34050679 | 34060065 | 9387 | -5.32 | 7.39E-06 | 1.10E-05 |
| chr15 | 40962783 | 40963572 | 790 | -5.47 | 7.42E-06 | 1.11E-05 |
| chr6 | 34230794 | 34231911 | 1118 | -5.47 | 7.42E-06 | 1.11E-05 |
| chrX | 149746555 | 149747452 | 898 | -5.47 | 7.42E-06 | 1.11E-05 |
| chr12 | 109451863 | 109461739 | 9877 | -6.31 | 7.43E-06 | 1.11E-05 |
| chr4 | 180712529 | 180714208 | 1680 | -4.01 | 7.45E-06 | 1.11E-05 |
| chr6 | 112719855 | 112725386 | 5532 | -1.57 | 7.47E-06 | 1.12E-05 |
| chr1 | 20455584 | 20460176 | 4593 | -4.65 | 7.47E-06 | 1.12E-05 |
| chr4 | 138985880 | 138987525 | 1646 | -5.22 | 7.47E-06 | 1.12E-05 |
| chr9 | 29588894 | 29590152 | 1259 | -5.22 | 7.47E-06 | 1.12E-05 |
| chr4 | 102930613 | 102931930 | 1318 | -5.31 | 7.47E-06 | 1.12E-05 |
| chr7 | 107135674 | 107136222 | 549 | -5.31 | 7.47E-06 | 1.12E-05 |
| chr8 | 110809755 | 110812003 | 2249 | -5.45 | 7.48E-06 | 1.12E-05 |
| chrX | 134007419 | 134008338 | 920 | -5.45 | 7.48E-06 | 1.12E-05 |
| chr18 | 25769647 | 25770744 | 1098 | -5.21 | 7.49E-06 | 1.12E-05 |
| chr20 | 45809460 | 45810315 | 856 | -5.21 | 7.49E-06 | 1.12E-05 |
| chr5 | 129680926 | 129681616 | 691 | -5.21 | 7.49E-06 | 1.12E-05 |
| chr5 | 179759519 | 179760206 | 688 | -5.21 | 7.49E-06 | 1.12E-05 |
| chr3 | 93524959 | 93527509 | 2551 | -2.97 | 7.49E-06 | 1.12E-05 |
| chr22 | 28711127 | 28712495 | 1369 | -5.38 | 7.51E-06 | 1.12E-05 |
| chr8 | 104104232 | 104112139 | 7908 | -5.88 | 7.52E-06 | 1.12E-05 |
| chr3 | 131062096 | 131062923 | 828 | -5.26 | 7.52E-06 | 1.12E-05 |
| chr5 | 142059060 | 142060147 | 1088 | -5.26 | 7.52E-06 | 1.12E-05 |
| chr3 | 117866993 | 117890573 | 23581 | -1.76 | 7.53E-06 | 1.12E-05 |
| chr17 | 49212639 | 49213911 | 1273 | -5.82 | 7.57E-06 | 1.13E-05 |
| chr17 | 7819637 | 7820975 | 1339 | -5.25 | 7.57E-06 | 1.13E-05 |
| chr3 | 61647714 | 61648628 | 915 | -5.29 | 7.59E-06 | 1.13E-05 |
| chr7 | 42079333 | 42101749 | 22417 | 1.42 | 7.59E-06 | 1.13E-05 |
| chr13 | 60502554 | 60503353 | 800 | -5.22 | 7.59E-06 | 1.13E-05 |
| chr5 | 80995649 | 80996551 | 903 | -5.22 | 7.59E-06 | 1.13E-05 |
| chr6 | 46396465 | 46397704 | 1240 | 4.7 | 7.59E-06 | 1.13E-05 |
| chr2 | 113061024 | 113061572 | 549 | -5.36 | 7.60E-06 | 1.13E-05 |
| chr8 | 106870934 | 106871978 | 1045 | -5.21 | 7.60E-06 | 1.13E-05 |
| chr6 | 152313769 | 152317674 | 3906 | 2.36 | 7.61E-06 | 1.14E-05 |
| chr20 | 45547086 | 45548469 | 1384 | -5.75 | 7.61E-06 | 1.14E-05 |
| chr11 | 58682150 | 58689299 | 7150 | -2.07 | 7.65E-06 | 1.14E-05 |
| chr1 | 173843687 | 173844248 | 562 | -5.4 | 7.65E-06 | 1.14E-05 |
| chr13 | 29322588 | 29365200 | 42613 | -5.99 | 7.67E-06 | 1.15E-05 |
| chr1 | 201147681 | 201148406 | 726 | -2.88 | 7.68E-06 | 1.15E-05 |

| | | | | | | |
|---|---|---|---|---|---|---|
| chrX | 84593845 | 84594960 | 1116 | -5.35 | 7.69E-06 | 1.15E-05 |
| chr6 | 123227419 | 123231782 | 4364 | 4.7 | 7.69E-06 | 1.15E-05 |
| chr2 | 227865033 | 227865844 | 812 | -5.63 | 7.70E-06 | 1.15E-05 |
| chr11 | 124976025 | 124999239 | 23215 | -1.34 | 7.73E-06 | 1.15E-05 |
| chr15 | 100093845 | 100094646 | 802 | 5.72 | 7.73E-06 | 1.15E-05 |
| chr1 | 43638703 | 43639419 | 717 | -5.28 | 7.73E-06 | 1.15E-05 |
| chr2 | 171604596 | 171605733 | 1138 | -5.28 | 7.73E-06 | 1.15E-05 |
| chr6 | 24728416 | 24732899 | 4484 | -2.93 | 7.76E-06 | 1.16E-05 |
| chr17 | 71832817 | 71833764 | 948 | -5.27 | 7.77E-06 | 1.16E-05 |
| chr4 | 101191073 | 101192212 | 1140 | -5.27 | 7.77E-06 | 1.16E-05 |
| chr4 | 145999347 | 145999986 | 640 | -5.27 | 7.77E-06 | 1.16E-05 |
| chr12 | 52010690 | 52011355 | 666 | -5.22 | 7.79E-06 | 1.16E-05 |
| chr20 | 55628097 | 55628568 | 472 | -5.22 | 7.79E-06 | 1.16E-05 |
| chr7 | 67989924 | 67990908 | 985 | -5.22 | 7.79E-06 | 1.16E-05 |
| chr1 | 93413279 | 93413922 | 644 | -5.24 | 7.79E-06 | 1.16E-05 |
| chr13 | 23466809 | 23467804 | 996 | -5.24 | 7.79E-06 | 1.16E-05 |
| chrX | 29522670 | 29524243 | 1574 | -5.55 | 7.79E-06 | 1.16E-05 |
| chr14 | 55003419 | 55004011 | 593 | -5.24 | 7.79E-06 | 1.16E-05 |
| chr2 | 107494637 | 107495378 | 742 | -5.24 | 7.79E-06 | 1.16E-05 |
| chr18 | 32371782 | 32378573 | 6792 | -2.62 | 7.80E-06 | 1.16E-05 |
| chr10 | 92581103 | 92630902 | 49800 | -3.43 | 7.81E-06 | 1.16E-05 |
| chr13 | 84446335 | 84447155 | 821 | -5.21 | 7.82E-06 | 1.17E-05 |
| chr20 | 50184235 | 50191677 | 7443 | -5.37 | 7.82E-06 | 1.17E-05 |
| chr7 | 90061989 | 90069016 | 7028 | -2.55 | 7.82E-06 | 1.17E-05 |
| chr10 | 19198664 | 19204134 | 5471 | -1.75 | 7.82E-06 | 1.17E-05 |
| chr3 | 16484154 | 16484908 | 755 | -5.22 | 7.82E-06 | 1.17E-05 |
| chr11 | 93783367 | 93787527 | 4161 | -1.53 | 7.85E-06 | 1.17E-05 |
| chrX | 38095087 | 38098666 | 3580 | 2.32 | 7.86E-06 | 1.17E-05 |
| chr6 | 71813826 | 71814323 | 498 | 4.67 | 7.87E-06 | 1.17E-05 |
| chr11 | 44384924 | 44386070 | 1147 | -5.5 | 7.88E-06 | 1.17E-05 |
| chr18 | 29617520 | 29618484 | 965 | -3.98 | 7.88E-06 | 1.17E-05 |
| chr11 | 47812677 | 47813523 | 847 | -5.36 | 7.89E-06 | 1.18E-05 |
| chr2 | 30637922 | 30638480 | 559 | -5.36 | 7.89E-06 | 1.18E-05 |
| chr3 | 71609301 | 71610223 | 923 | -5.36 | 7.89E-06 | 1.18E-05 |
| chr20 | 25642474 | 25659691 | 17218 | -6.87 | 7.89E-06 | 1.18E-05 |
| chr3 | 108444257 | 108445475 | 1219 | 5.29 | 7.89E-06 | 1.18E-05 |
| chr10 | 52343299 | 52344294 | 996 | -5.49 | 7.90E-06 | 1.18E-05 |
| chr12 | 32226686 | 32227670 | 985 | -5.32 | 7.91E-06 | 1.18E-05 |
| chr7 | 24389629 | 24409458 | 19830 | -1.82 | 7.91E-06 | 1.18E-05 |
| chr6 | 108305668 | 108306489 | 822 | -5.47 | 7.94E-06 | 1.18E-05 |
| chr14 | 29136127 | 29137065 | 939 | 3.68 | 7.94E-06 | 1.18E-05 |
| chr3 | 52918924 | 52919565 | 642 | -5.2 | 7.96E-06 | 1.19E-05 |
| chr11 | 11653810 | 11720584 | 66775 | -1.19 | 7.96E-06 | 1.19E-05 |
| chr9 | 137033465 | 137034449 | 985 | -5.46 | 7.97E-06 | 1.19E-05 |
| chr16 | 89383163 | 89385431 | 2269 | -6.1 | 7.97E-06 | 1.19E-05 |
| chr15 | 37547269 | 37553986 | 6718 | 2.09 | 7.98E-06 | 1.19E-05 |
| chr20 | 34649461 | 34650359 | 899 | -5.26 | 7.98E-06 | 1.19E-05 |
| chr4 | 162639976 | 162641143 | 1168 | -5.26 | 7.98E-06 | 1.19E-05 |
| chr3 | 40849624 | 40855268 | 5645 | -2.87 | 7.99E-06 | 1.19E-05 |
| chr14 | 60374035 | 60375160 | 1126 | 5.6 | 8.00E-06 | 1.19E-05 |
| chr1 | 201948366 | 201949367 | 1002 | -5.2 | 8.01E-06 | 1.19E-05 |
| chr8 | 74707487 | 74708512 | 1026 | -5.2 | 8.01E-06 | 1.19E-05 |
| chr8 | 83755657 | 83756959 | 1303 | -5.2 | 8.01E-06 | 1.19E-05 |
| chrX | 152666951 | 152667626 | 676 | -5.2 | 8.02E-06 | 1.19E-05 |
| chr2 | 149561117 | 149679073 | 117957 | -3.13 | 8.03E-06 | 1.20E-05 |
| chr4 | 66841924 | 66842590 | 667 | -5.22 | 8.03E-06 | 1.20E-05 |
| chr12 | 64854010 | 64854653 | 644 | -5.34 | 8.04E-06 | 1.20E-05 |
| chr17 | 57395829 | 57396971 | 1143 | -5.31 | 8.04E-06 | 1.20E-05 |
| chr2 | 58620638 | 58624952 | 4315 | -1.46 | 8.05E-06 | 1.20E-05 |
| chr22 | 18394914 | 18395609 | 696 | -5.43 | 8.05E-06 | 1.20E-05 |
| chr4 | 139826664 | 139827284 | 621 | -5.43 | 8.05E-06 | 1.20E-05 |

| | | | | | | |
|---|---|---|---|---|---|---|
| chr11 | 104907889 | 104923544 | 15656 | -1.83 | 8.08E-06 | 1.20E-05 |
| chr7 | 50945782 | 50946199 | 418 | -5.23 | 8.08E-06 | 1.20E-05 |
| chr3 | 132102595 | 132103295 | 701 | -4.49 | 8.12E-06 | 1.21E-05 |
| chr4 | 67820485 | 67821565 | 1081 | 2.85 | 8.13E-06 | 1.21E-05 |
| chr8 | 54946308 | 54947258 | 951 | -5.2 | 8.15E-06 | 1.21E-05 |
| chr10 | 14844922 | 14845798 | 877 | -5.41 | 8.16E-06 | 1.21E-05 |
| chr3 | 149369191 | 149370040 | 850 | -5.41 | 8.16E-06 | 1.21E-05 |
| chrX | 152641795 | 152642813 | 1019 | -5.41 | 8.16E-06 | 1.21E-05 |
| chr8 | 8721518 | 8722193 | 676 | -5.21 | 8.16E-06 | 1.21E-05 |
| chr5 | 23827312 | 25360120 | 1532809 | 0.74 | 8.18E-06 | 1.22E-05 |
| chr11 | 10631041 | 10636248 | 5208 | -0.9 | 8.19E-06 | 1.22E-05 |
| chr4 | 135521205 | 135522713 | 1509 | 4 | 8.19E-06 | 1.22E-05 |
| chr1 | 56194720 | 56195895 | 1176 | -5.29 | 8.19E-06 | 1.22E-05 |
| chr14 | 51415592 | 51416277 | 686 | -5.29 | 8.19E-06 | 1.22E-05 |
| chr3 | 127904331 | 127904821 | 491 | -5.19 | 8.21E-06 | 1.22E-05 |
| chr4 | 135217148 | 135217750 | 603 | -5.19 | 8.21E-06 | 1.22E-05 |
| chr20 | 1599408 | 1600081 | 674 | -5.39 | 8.22E-06 | 1.22E-05 |
| chr4 | 138667460 | 138668217 | 758 | -5.39 | 8.22E-06 | 1.22E-05 |
| chr10 | 15818860 | 15819443 | 584 | -5.24 | 8.22E-06 | 1.22E-05 |
| chr22 | 35561212 | 35561892 | 681 | -5.25 | 8.24E-06 | 1.23E-05 |
| chr2 | 72923807 | 72924222 | 416 | -5.6 | 8.25E-06 | 1.23E-05 |
| chr8 | 100739017 | 100751332 | 12316 | -5.6 | 8.25E-06 | 1.23E-05 |
| chr2 | 152595751 | 152596426 | 676 | -4.17 | 8.25E-06 | 1.23E-05 |
| chr2 | 7602292 | 7635597 | 33306 | -1.11 | 8.26E-06 | 1.23E-05 |
| chr6 | 137269388 | 137270921 | 1534 | -4.01 | 8.26E-06 | 1.23E-05 |
| chrX | 95664280 | 95665765 | 1486 | -5.38 | 8.29E-06 | 1.23E-05 |
| chr19 | 5157972 | 5158709 | 738 | -4.56 | 8.29E-06 | 1.23E-05 |
| chrX | 41309829 | 41320270 | 10442 | -5.94 | 8.29E-06 | 1.23E-05 |
| chr7 | 117796572 | 117805180 | 8609 | -5.4 | 8.32E-06 | 1.24E-05 |
| chr11 | 45372043 | 45384443 | 12401 | -1.48 | 8.33E-06 | 1.24E-05 |
| chr5 | 138553555 | 138554730 | 1176 | -4.97 | 8.34E-06 | 1.24E-05 |
| chr16 | 19799545 | 19800319 | 775 | -5.3 | 8.34E-06 | 1.24E-05 |
| chr12 | 106151912 | 106153821 | 1910 | 3.89 | 8.35E-06 | 1.24E-05 |
| chr2 | 165433879 | 165434495 | 617 | -5.28 | 8.36E-06 | 1.24E-05 |
| chr5 | 38438321 | 38439281 | 961 | -5.28 | 8.36E-06 | 1.24E-05 |
| chr11 | 60508261 | 60525866 | 17606 | -5.16 | 8.37E-06 | 1.24E-05 |
| chr3 | 154802175 | 154828275 | 26101 | 1.01 | 8.37E-06 | 1.24E-05 |
| chr3 | 148126911 | 148129584 | 2674 | -2.82 | 8.37E-06 | 1.25E-05 |
| chr12 | 10256220 | 10264933 | 8714 | -2.09 | 8.39E-06 | 1.25E-05 |
| chr4 | 86769374 | 86770007 | 634 | -5.5 | 8.39E-06 | 1.25E-05 |
| chr4 | 70428100 | 70428658 | 559 | -5.23 | 8.44E-06 | 1.26E-05 |
| chr9 | 97678226 | 97678789 | 564 | -5.23 | 8.44E-06 | 1.26E-05 |
| chr14 | 90022362 | 90079577 | 57216 | -1.54 | 8.45E-06 | 1.26E-05 |
| chrX | 80643449 | 80644590 | 1142 | 4.27 | 8.45E-06 | 1.26E-05 |
| chr6 | 6690939 | 6691877 | 939 | -4.81 | 8.46E-06 | 1.26E-05 |
| chr12 | 62950255 | 62950920 | 666 | -5.29 | 8.46E-06 | 1.26E-05 |
| chr20 | 43286353 | 43287012 | 660 | -5.29 | 8.46E-06 | 1.26E-05 |
| chr3 | 43624959 | 43625781 | 823 | -5.29 | 8.46E-06 | 1.26E-05 |
| chrX | 48790097 | 48790731 | 635 | -5.29 | 8.46E-06 | 1.26E-05 |
| chr9 | 36954209 | 36955413 | 1205 | -5.47 | 8.46E-06 | 1.26E-05 |
| chr2 | 153735648 | 153736212 | 565 | 5.35 | 8.47E-06 | 1.26E-05 |
| chr11 | 40406004 | 40407046 | 1043 | -4.63 | 8.47E-06 | 1.26E-05 |
| chr4 | 4611431 | 4612699 | 1269 | -5.77 | 8.48E-06 | 1.26E-05 |
| chr3 | 24605509 | 24614199 | 8691 | -5.28 | 8.48E-06 | 1.26E-05 |
| chr13 | 105221884 | 105222825 | 942 | -5.23 | 8.48E-06 | 1.26E-05 |
| chr1 | 153758466 | 153758979 | 514 | -5.19 | 8.49E-06 | 1.26E-05 |
| chr11 | 111981363 | 111982014 | 652 | -5.18 | 8.49E-06 | 1.26E-05 |
| chr2 | 193251587 | 193257203 | 5617 | -5.18 | 8.49E-06 | 1.26E-05 |
| chr20 | 11102982 | 11103924 | 943 | -5.18 | 8.49E-06 | 1.26E-05 |
| chr4 | 162511449 | 162512280 | 832 | -5.18 | 8.49E-06 | 1.26E-05 |
| chr5 | 40483289 | 40483848 | 560 | -5.18 | 8.49E-06 | 1.26E-05 |

| | | | | | | |
|---|---|---|---|---|---|---|
| chr9 | 100984173 | 100984744 | 572 | -5.18 | 8.49E-06 | 1.26E-05 |
| chr19 | 34997837 | 35483760 | 485924 | -5.01 | 8.50E-06 | 1.26E-05 |
| chr7 | 23635873 | 23636671 | 799 | -5.19 | 8.50E-06 | 1.26E-05 |
| chr3 | 120250985 | 120263003 | 12019 | -2.63 | 8.52E-06 | 1.27E-05 |
| chr10 | 52043845 | 52044225 | 381 | 5.34 | 8.54E-06 | 1.27E-05 |
| chr9 | 29384248 | 29385033 | 786 | -5.34 | 8.54E-06 | 1.27E-05 |
| chr13 | 90600004 | 90608162 | 8159 | -5.69 | 8.55E-06 | 1.27E-05 |
| chr15 | 60034204 | 60040252 | 6049 | -1.29 | 8.55E-06 | 1.27E-05 |
| chr7 | 120436603 | 120438055 | 1453 | -2.06 | 8.58E-06 | 1.27E-05 |
| chr6 | 113732801 | 113734409 | 1609 | -5.43 | 8.59E-06 | 1.28E-05 |
| chr17 | 47192192 | 47193094 | 903 | -5.05 | 8.60E-06 | 1.28E-05 |
| chr1 | 77502748 | 77503835 | 1088 | -5.2 | 8.60E-06 | 1.28E-05 |
| chr11 | 2392392 | 2393341 | 950 | -5.2 | 8.60E-06 | 1.28E-05 |
| chr18 | 69715202 | 69716043 | 842 | -5.2 | 8.60E-06 | 1.28E-05 |
| chr4 | 143430594 | 143431221 | 628 | -5.18 | 8.63E-06 | 1.28E-05 |
| chr5 | 170828138 | 170829132 | 995 | -5.18 | 8.63E-06 | 1.28E-05 |
| chr11 | 109631012 | 109632338 | 1327 | -4.11 | 8.63E-06 | 1.28E-05 |
| chr19 | 41097636 | 41098236 | 601 | -5.32 | 8.65E-06 | 1.28E-05 |
| chr21 | 40843153 | 40843778 | 626 | -5.32 | 8.65E-06 | 1.28E-05 |
| chr4 | 131541337 | 131542202 | 866 | -5.32 | 8.65E-06 | 1.28E-05 |
| chr13 | 96207153 | 96208147 | 995 | -5.59 | 8.65E-06 | 1.28E-05 |
| chr2 | 180997231 | 180997831 | 601 | -5.58 | 8.67E-06 | 1.29E-05 |
| chr7 | 148995384 | 148996132 | 749 | -5.58 | 8.67E-06 | 1.29E-05 |
| chr4 | 105116489 | 105117326 | 838 | -5.22 | 8.67E-06 | 1.29E-05 |
| chr10 | 28496417 | 28507929 | 11513 | -2.81 | 8.69E-06 | 1.29E-05 |
| chr20 | 8487823 | 8492432 | 4610 | -2.2 | 8.71E-06 | 1.29E-05 |
| chr11 | 5300387 | 5301342 | 956 | -5.4 | 8.72E-06 | 1.29E-05 |
| chr19 | 47633347 | 47633898 | 552 | -5.4 | 8.72E-06 | 1.29E-05 |
| chr6 | 87843481 | 87844115 | 635 | -5.4 | 8.72E-06 | 1.29E-05 |
| chr10 | 97795806 | 97796449 | 644 | -5.27 | 8.73E-06 | 1.30E-05 |
| chr5 | 141975248 | 141976757 | 1510 | -5.27 | 8.73E-06 | 1.30E-05 |
| chr7 | 26036706 | 26039472 | 2767 | -2.06 | 8.74E-06 | 1.30E-05 |
| chr4 | 44753747 | 44755720 | 1974 | 3.19 | 8.74E-06 | 1.30E-05 |
| chr7 | 12500996 | 12501977 | 982 | -5.25 | 8.76E-06 | 1.30E-05 |
| chr14 | 43653916 | 43661293 | 7378 | -2.78 | 8.77E-06 | 1.30E-05 |
| chr18 | 21792704 | 21793421 | 718 | -5.31 | 8.77E-06 | 1.30E-05 |
| chr6 | 46198594 | 46200660 | 2067 | 1.8 | 8.77E-06 | 1.30E-05 |
| chrX | 23563199 | 23586523 | 23325 | -1.87 | 8.78E-06 | 1.30E-05 |
| chr4 | 7379820 | 7380583 | 764 | -5.19 | 8.80E-06 | 1.30E-05 |
| chr8 | 74011427 | 74012666 | 1240 | -5.19 | 8.80E-06 | 1.30E-05 |
| chr6 | 88817056 | 88837407 | 20352 | -1.78 | 8.81E-06 | 1.31E-05 |
| chr4 | 88335029 | 88335752 | 724 | -4.48 | 8.81E-06 | 1.31E-05 |
| chr4 | 107052791 | 107054365 | 1575 | -4.1 | 8.81E-06 | 1.31E-05 |
| chr7 | 11942173 | 11942892 | 720 | -5.49 | 8.81E-06 | 1.31E-05 |
| chr1 | 13899727 | 13980590 | 80864 | 0.71 | 8.81E-06 | 1.31E-05 |
| chr8 | 124735562 | 124736119 | 558 | -4.58 | 8.87E-06 | 1.31E-05 |
| chr20 | 20680610 | 20681006 | 397 | -5.25 | 8.89E-06 | 1.32E-05 |
| chrX | 13044315 | 13045334 | 1020 | -5.25 | 8.89E-06 | 1.32E-05 |
| chr11 | 11344233 | 11344808 | 576 | -5.18 | 8.90E-06 | 1.32E-05 |
| chr14 | 50814621 | 50816214 | 1594 | -5.18 | 8.90E-06 | 1.32E-05 |
| chr20 | 61169000 | 61169690 | 691 | -5.18 | 8.90E-06 | 1.32E-05 |
| chr10 | 22077838 | 22079793 | 1956 | -5.45 | 8.91E-06 | 1.32E-05 |
| chr1 | 209868742 | 209869355 | 614 | -5.21 | 8.92E-06 | 1.32E-05 |
| chr4 | 53067179 | 53068416 | 1238 | -5.21 | 8.92E-06 | 1.32E-05 |
| chr7 | 55662431 | 55731355 | 68925 | -6.12 | 8.93E-06 | 1.32E-05 |
| chr3 | 193562389 | 193563460 | 1072 | -5.19 | 8.93E-06 | 1.32E-05 |
| chr3 | 54853176 | 54883302 | 30127 | -3.47 | 8.93E-06 | 1.32E-05 |
| chr8 | 141863645 | 141863983 | 339 | -5.36 | 8.95E-06 | 1.33E-05 |
| chr13 | 40969366 | 40985324 | 15959 | -6.82 | 8.95E-06 | 1.33E-05 |
| chr6 | 52770525 | 52771403 | 879 | -3.38 | 8.95E-06 | 1.33E-05 |
| chr11 | 85647903 | 85664953 | 17051 | -6.47 | 8.98E-06 | 1.33E-05 |

| | | | | | | |
|---|---|---|---|---|---|---|
| chr4 | 149221345 | 149222419 | 1075 | -5.42 | 9.00E-06 | 1.33E-05 |
| chr5 | 125698817 | 125699531 | 715 | -5.42 | 9.00E-06 | 1.33E-05 |
| chrX | 101274069 | 101300182 | 26114 | 1.14 | 9.04E-06 | 1.34E-05 |
| chr5 | 7840940 | 7855169 | 14230 | -4.79 | 9.04E-06 | 1.34E-05 |
| chr1 | 242555215 | 242555659 | 445 | -5.18 | 9.07E-06 | 1.34E-05 |
| chr17 | 46514131 | 46515150 | 1020 | 4.36 | 9.07E-06 | 1.34E-05 |
| chr1 | 95395804 | 95419197 | 23394 | -6.58 | 9.09E-06 | 1.35E-05 |
| chr3 | 127347108 | 127347792 | 685 | -5.17 | 9.11E-06 | 1.35E-05 |
| chr17 | 63183807 | 63184653 | 847 | -5.4 | 9.11E-06 | 1.35E-05 |
| chrX | 103543171 | 103544609 | 1439 | -2.12 | 9.12E-06 | 1.35E-05 |
| chr11 | 72892685 | 72893270 | 586 | -5.2 | 9.13E-06 | 1.35E-05 |
| chr22 | 45653774 | 45654552 | 779 | -5.2 | 9.13E-06 | 1.35E-05 |
| chr14 | 95877133 | 95887962 | 10830 | -5.86 | 9.16E-06 | 1.36E-05 |
| chr4 | 181567251 | 181568260 | 1010 | 5.41 | 9.16E-06 | 1.36E-05 |
| chr2 | 173404047 | 173405200 | 1154 | -5.38 | 9.17E-06 | 1.36E-05 |
| chr6 | 110256463 | 110257612 | 1150 | -5.32 | 9.18E-06 | 1.36E-05 |
| chr3 | 148484565 | 148485857 | 1293 | 6.01 | 9.20E-06 | 1.36E-05 |
| chr5 | 14018185 | 14029098 | 10914 | -6.24 | 9.21E-06 | 1.36E-05 |
| chr10 | 89454825 | 89455591 | 767 | -5.18 | 9.23E-06 | 1.37E-05 |
| chr3 | 140659689 | 140660225 | 537 | -5.18 | 9.23E-06 | 1.37E-05 |
| chr4 | 126477503 | 126478418 | 916 | -5.18 | 9.23E-06 | 1.37E-05 |
| chr6 | 122083738 | 122084547 | 810 | -5.26 | 9.23E-06 | 1.37E-05 |
| chr13 | 49978672 | 49979649 | 978 | -5.37 | 9.24E-06 | 1.37E-05 |
| chr6 | 33833671 | 33834642 | 972 | -5.23 | 9.25E-06 | 1.37E-05 |
| chr11 | 72101372 | 72102107 | 736 | -5.22 | 9.27E-06 | 1.37E-05 |
| chr22 | 31508613 | 31509185 | 573 | -5.22 | 9.27E-06 | 1.37E-05 |
| chr12 | 25537105 | 25538118 | 1014 | -5.31 | 9.27E-06 | 1.37E-05 |
| chrX | 54936489 | 54938329 | 1841 | -5.9 | 9.28E-06 | 1.37E-05 |
| chr11 | 89138913 | 89140415 | 1503 | 4.25 | 9.31E-06 | 1.38E-05 |
| chrX | 36935155 | 36938379 | 3225 | 4.62 | 9.32E-06 | 1.38E-05 |
| chr2 | 225404413 | 225406274 | 1862 | -5.73 | 9.32E-06 | 1.38E-05 |
| chr2 | 152886255 | 152887539 | 1285 | -5.35 | 9.32E-06 | 1.38E-05 |
| chr2 | 108334519 | 108334998 | 480 | 4.55 | 9.38E-06 | 1.39E-05 |
| chr16 | 55717885 | 55729387 | 11503 | -1.56 | 9.39E-06 | 1.39E-05 |
| chr8 | 59089620 | 59090530 | 911 | -2.69 | 9.41E-06 | 1.39E-05 |
| chr17 | 9679372 | 9685631 | 6260 | -4.6 | 9.41E-06 | 1.39E-05 |
| chr12 | 132580110 | 132581051 | 942 | -5.34 | 9.41E-06 | 1.39E-05 |
| chr2 | 44496298 | 44496864 | 567 | -5.34 | 9.41E-06 | 1.39E-05 |
| chr2 | 74145800 | 74146304 | 505 | -5.34 | 9.41E-06 | 1.39E-05 |
| chr4 | 128708848 | 128710027 | 1180 | -5.34 | 9.41E-06 | 1.39E-05 |
| chr20 | 23507516 | 23570084 | 62569 | -3.85 | 9.42E-06 | 1.39E-05 |
| chr11 | 73672445 | 73673098 | 654 | -5.25 | 9.43E-06 | 1.39E-05 |
| chr14 | 62175364 | 62176062 | 699 | -5.25 | 9.43E-06 | 1.39E-05 |
| chr21 | 38106607 | 38107345 | 739 | -5.25 | 9.43E-06 | 1.39E-05 |
| chr4 | 157481345 | 157482326 | 982 | -5.25 | 9.43E-06 | 1.39E-05 |
| chrX | 39741812 | 39742833 | 1022 | -5.25 | 9.43E-06 | 1.39E-05 |
| chr12 | 48728588 | 48729438 | 851 | -4.49 | 9.46E-06 | 1.40E-05 |
| chr1 | 152320802 | 152321690 | 889 | -5.22 | 9.46E-06 | 1.40E-05 |
| chr6 | 55547116 | 55548246 | 1131 | -5.22 | 9.46E-06 | 1.40E-05 |
| chr5 | 90264369 | 90265855 | 1487 | -5.76 | 9.46E-06 | 1.40E-05 |
| chr8 | 125200478 | 125201785 | 1308 | -5.75 | 9.48E-06 | 1.40E-05 |
| chr17 | 38710810 | 38753302 | 42493 | -2.88 | 9.48E-06 | 1.40E-05 |
| chr2 | 160720758 | 160721418 | 661 | -5.29 | 9.49E-06 | 1.40E-05 |
| chr10 | 52307671 | 52308442 | 772 | -5.18 | 9.50E-06 | 1.40E-05 |
| chr20 | 36349747 | 36350556 | 810 | -5.18 | 9.50E-06 | 1.40E-05 |
| chr4 | 53722154 | 53727508 | 5355 | -5.23 | 9.51E-06 | 1.41E-05 |
| chr10 | 59585613 | 59586644 | 1032 | -5.33 | 9.51E-06 | 1.41E-05 |
| chr7 | 79293339 | 79294443 | 1105 | -5.18 | 9.52E-06 | 1.41E-05 |
| chr22 | 32848586 | 32849665 | 1080 | -3.94 | 9.52E-06 | 1.41E-05 |
| chr15 | 79460582 | 79461247 | 666 | -5.7 | 9.53E-06 | 1.41E-05 |
| chr1 | 13179342 | 13188521 | 9180 | 2.61 | 9.55E-06 | 1.41E-05 |

| | | | | | | |
|---|---|---|---|---|---|---|
| chr4 | 125393850 | 125394467 | 618 | -4.14 | 9.56E-06 | 1.41E-05 |
| chr4 | 52910047 | 52913754 | 3708 | -2.28 | 9.58E-06 | 1.42E-05 |
| chr1 | 30059678 | 30060235 | 558 | -5.2 | 9.58E-06 | 1.42E-05 |
| chr1 | 219450942 | 219452157 | 1216 | -5.2 | 9.58E-06 | 1.42E-05 |
| chr5 | 82569451 | 82569917 | 467 | -5.2 | 9.58E-06 | 1.42E-05 |
| chr7 | 106775506 | 106776580 | 1075 | -4.41 | 9.61E-06 | 1.42E-05 |
| chr1 | 208103666 | 208104380 | 715 | -5.28 | 9.61E-06 | 1.42E-05 |
| chrX | 101426146 | 101429265 | 3120 | 1.93 | 9.61E-06 | 1.42E-05 |
| chr17 | 4943400 | 4944550 | 1151 | -5.64 | 9.61E-06 | 1.42E-05 |
| chr1 | 80186573 | 80187997 | 1425 | -5.31 | 9.62E-06 | 1.42E-05 |
| chr17 | 72387993 | 72389546 | 1554 | -5.31 | 9.62E-06 | 1.42E-05 |
| chr7 | 14465165 | 14466925 | 1761 | -5.31 | 9.62E-06 | 1.42E-05 |
| chr10 | 60056168 | 60056592 | 425 | -5.16 | 9.63E-06 | 1.42E-05 |
| chr17 | 80617808 | 80619165 | 1358 | -5.23 | 9.64E-06 | 1.42E-05 |
| chr3 | 169909781 | 169910567 | 787 | -5.23 | 9.64E-06 | 1.42E-05 |
| chr4 | 41882424 | 41883290 | 867 | -5.23 | 9.64E-06 | 1.42E-05 |
| chr1 | 184046088 | 184046628 | 541 | -5.16 | 9.65E-06 | 1.43E-05 |
| chr19 | 50119629 | 50120421 | 793 | -5.16 | 9.65E-06 | 1.43E-05 |
| chr20 | 52901876 | 52902524 | 649 | -5.16 | 9.65E-06 | 1.43E-05 |
| chr13 | 62876415 | 62878204 | 1790 | -5.61 | 9.65E-06 | 1.43E-05 |
| chr2 | 55952655 | 55953100 | 446 | -5.21 | 9.69E-06 | 1.43E-05 |
| chr8 | 32257733 | 32258573 | 841 | -5.21 | 9.69E-06 | 1.43E-05 |
| chr10 | 87694883 | 87695458 | 576 | 5.24 | 9.70E-06 | 1.43E-05 |
| chr8 | 103562818 | 103563814 | 997 | 5.36 | 9.71E-06 | 1.43E-05 |
| chr4 | 16073805 | 16076454 | 2650 | 1.7 | 9.71E-06 | 1.43E-05 |
| chr4 | 59877859 | 59878661 | 803 | -5.57 | 9.71E-06 | 1.43E-05 |
| chr4 | 64615874 | 64617623 | 1750 | -5.57 | 9.71E-06 | 1.43E-05 |
| chr11 | 87603427 | 87666654 | 63228 | 3.59 | 9.74E-06 | 1.44E-05 |
| chr12 | 66863975 | 66865010 | 1036 | 4.24 | 9.74E-06 | 1.44E-05 |
| chr5 | 172870989 | 172871839 | 851 | -2.89 | 9.74E-06 | 1.44E-05 |
| chr12 | 57698997 | 57699350 | 354 | -5.26 | 9.75E-06 | 1.44E-05 |
| chr3 | 6065488 | 6095683 | 30196 | -2.02 | 9.76E-06 | 1.44E-05 |
| chr6 | 129407669 | 129408854 | 1186 | 5.72 | 9.79E-06 | 1.45E-05 |
| chr5 | 1040128 | 1041090 | 963 | -5.5 | 9.82E-06 | 1.45E-05 |
| chr19 | 47194653 | 47195555 | 903 | -5.44 | 9.82E-06 | 1.45E-05 |
| chr2 | 84021546 | 84035190 | 13645 | -4.98 | 9.82E-06 | 1.45E-05 |
| chrX | 118329853 | 118330663 | 811 | -5.15 | 9.83E-06 | 1.45E-05 |
| chr6 | 121355495 | 121356264 | 770 | -4.67 | 9.84E-06 | 1.45E-05 |
| chr6 | 150800634 | 150801779 | 1146 | -4.2 | 9.84E-06 | 1.45E-05 |
| chr20 | 31059535 | 31060489 | 955 | -5.49 | 9.85E-06 | 1.45E-05 |
| chr7 | 143729539 | 143808689 | 79151 | 1.7 | 9.85E-06 | 1.45E-05 |
| chr20 | 49719680 | 49720417 | 738 | -5.28 | 9.89E-06 | 1.46E-05 |
| chrX | 3489651 | 3490411 | 761 | -5.22 | 9.89E-06 | 1.46E-05 |
| chr3 | 192799663 | 192801187 | 1525 | -5.42 | 9.89E-06 | 1.46E-05 |
| chr5 | 178938415 | 178939540 | 1126 | -5.47 | 9.91E-06 | 1.46E-05 |
| chr4 | 142541087 | 142542020 | 934 | -5.18 | 9.93E-06 | 1.47E-05 |
| chr6 | 165648201 | 165676336 | 28136 | -1.43 | 9.93E-06 | 1.47E-05 |
| chr12 | 98238158 | 98278345 | 40188 | 5.63 | 9.95E-06 | 1.47E-05 |
| chr18 | 73993558 | 74020624 | 27067 | -1.01 | 9.97E-06 | 1.47E-05 |
| chr6 | 143320490 | 143321209 | 720 | -3.95 | 9.97E-06 | 1.47E-05 |
| chr18 | 32535341 | 32535958 | 618 | -4.46 | 1.00E-05 | 1.48E-05 |
| chr2 | 10863355 | 10864206 | 852 | -5.43 | 1.00E-05 | 1.48E-05 |
| chrX | 107352593 | 107353269 | 677 | -5.43 | 1.00E-05 | 1.48E-05 |
| chr6 | 133475968 | 133476960 | 993 | -3.93 | 1.00E-05 | 1.48E-05 |
| chr15 | 60645936 | 60657954 | 12019 | -5.27 | 1.00E-05 | 1.48E-05 |
| chr13 | 93944167 | 94126593 | 182427 | 3 | 1.00E-05 | 1.48E-05 |
| chr5 | 106896981 | 106897671 | 691 | -5.24 | 1.01E-05 | 1.48E-05 |
| chr5 | 168210299 | 168211049 | 751 | -5.24 | 1.01E-05 | 1.48E-05 |
| chrX | 46082368 | 46083223 | 856 | -5.41 | 1.01E-05 | 1.49E-05 |
| chr8 | 80564453 | 80565217 | 765 | -5.38 | 1.01E-05 | 1.49E-05 |
| chr1 | 5736912 | 5757031 | 20120 | 2.25 | 1.01E-05 | 1.49E-05 |

| | | | | | | |
|---|---|---|---|---|---|---|
| chr8 | 123470605 | 123544015 | 73411 | -0.96 | 1.01E-05 | 1.49E-05 |
| chr6 | 111462474 | 111464066 | 1593 | -4.62 | 1.01E-05 | 1.49E-05 |
| chr4 | 118451988 | 118455366 | 3379 | -5.36 | 1.01E-05 | 1.50E-05 |
| chr3 | 175395497 | 175396303 | 807 | -5.15 | 1.01E-05 | 1.50E-05 |
| chr4 | 86741159 | 86741676 | 518 | -5.15 | 1.01E-05 | 1.50E-05 |
| chr5 | 10405759 | 10406251 | 493 | -5.15 | 1.01E-05 | 1.50E-05 |
| chr7 | 5906930 | 5907682 | 753 | -5.15 | 1.01E-05 | 1.50E-05 |
| chr13 | 84048410 | 84050090 | 1681 | -5.2 | 1.02E-05 | 1.50E-05 |
| chr5 | 68477828 | 68478593 | 766 | -5.2 | 1.02E-05 | 1.50E-05 |
| chr9 | 6114557 | 6115349 | 793 | -5.2 | 1.02E-05 | 1.50E-05 |
| chr9 | 32893253 | 32909799 | 16547 | -1.76 | 1.02E-05 | 1.50E-05 |
| chr14 | 73338332 | 73342119 | 3788 | -2.55 | 1.02E-05 | 1.50E-05 |
| chr16 | 59736745 | 59769604 | 32860 | 1.94 | 1.02E-05 | 1.50E-05 |
| chr4 | 2046176 | 2046869 | 694 | -5.35 | 1.02E-05 | 1.50E-05 |
| chr4 | 60303946 | 60305324 | 1379 | -5.16 | 1.02E-05 | 1.50E-05 |
| chr4 | 83708631 | 83709309 | 679 | -5.16 | 1.02E-05 | 1.50E-05 |
| chr11 | 122126840 | 122131467 | 4628 | 2.5 | 1.02E-05 | 1.51E-05 |
| chr1 | 165623334 | 165623880 | 547 | -5.25 | 1.02E-05 | 1.51E-05 |
| chr11 | 82660308 | 82674924 | 14617 | -5.25 | 1.02E-05 | 1.51E-05 |
| chr6 | 72045396 | 72046289 | 894 | -5.25 | 1.02E-05 | 1.51E-05 |
| chr8 | 99712517 | 99712957 | 441 | -5.23 | 1.02E-05 | 1.51E-05 |
| chr2 | 215725242 | 215730828 | 5587 | -2.27 | 1.02E-05 | 1.51E-05 |
| chr13 | 78343687 | 78344200 | 514 | -5.14 | 1.03E-05 | 1.51E-05 |
| chr4 | 173546389 | 173547812 | 1424 | -5.14 | 1.03E-05 | 1.51E-05 |
| chr6 | 143819339 | 143819790 | 452 | -5.14 | 1.03E-05 | 1.51E-05 |
| chrX | 107281902 | 107282695 | 794 | -5.14 | 1.03E-05 | 1.51E-05 |
| chr7 | 116463973 | 116465818 | 1846 | -1.99 | 1.03E-05 | 1.51E-05 |
| chr11 | 47708923 | 47709347 | 425 | -5.15 | 1.03E-05 | 1.52E-05 |
| chr12 | 113867973 | 113868546 | 574 | -5.15 | 1.03E-05 | 1.52E-05 |
| chr15 | 31331241 | 31332078 | 838 | -5.15 | 1.03E-05 | 1.52E-05 |
| chr2 | 139511228 | 139512434 | 1207 | -5.93 | 1.03E-05 | 1.52E-05 |
| chr17 | 65552969 | 65553487 | 519 | -5.17 | 1.03E-05 | 1.52E-05 |
| chr6 | 1985373 | 1986235 | 863 | -5.17 | 1.03E-05 | 1.52E-05 |
| chr1 | 170649083 | 170650534 | 1452 | 2.36 | 1.03E-05 | 1.52E-05 |
| chr11 | 33696127 | 33697359 | 1233 | 4.49 | 1.04E-05 | 1.53E-05 |
| chr14 | 21891424 | 21892049 | 626 | -5.14 | 1.04E-05 | 1.53E-05 |
| chr5 | 72364683 | 72365145 | 463 | -5.14 | 1.04E-05 | 1.53E-05 |
| chr13 | 23780451 | 23789111 | 8661 | -1.51 | 1.04E-05 | 1.53E-05 |
| chrX | 111137872 | 111553318 | 415447 | 8.74 | 1.04E-05 | 1.53E-05 |
| chr1 | 172750132 | 172763816 | 13685 | -5.5 | 1.04E-05 | 1.53E-05 |
| chr7 | 26474261 | 26475311 | 1051 | -3.85 | 1.04E-05 | 1.53E-05 |
| chr4 | 127847791 | 127849220 | 1430 | -5.34 | 1.04E-05 | 1.53E-05 |
| chr6 | 145938949 | 145940218 | 1270 | -5.34 | 1.04E-05 | 1.53E-05 |
| chr18 | 850932 | 852920 | 1989 | -5.83 | 1.04E-05 | 1.54E-05 |
| chr17 | 57433845 | 57434457 | 613 | -5.22 | 1.04E-05 | 1.54E-05 |
| chr5 | 109080171 | 109080892 | 722 | -5.22 | 1.04E-05 | 1.54E-05 |
| chr11 | 103784620 | 103785268 | 649 | -3.86 | 1.04E-05 | 1.54E-05 |
| chr11 | 99095823 | 99097068 | 1246 | 5.52 | 1.05E-05 | 1.54E-05 |
| chr3 | 141421517 | 141423230 | 1714 | -5.33 | 1.05E-05 | 1.55E-05 |
| chrX | 62549995 | 62550899 | 905 | -5.33 | 1.05E-05 | 1.55E-05 |
| chr22 | 32715173 | 32715753 | 581 | -4.47 | 1.05E-05 | 1.55E-05 |
| chr5 | 142370893 | 142371539 | 647 | -5.31 | 1.05E-05 | 1.55E-05 |
| chr3 | 51758844 | 51808404 | 49561 | -2.33 | 1.06E-05 | 1.55E-05 |
| chr3 | 46405029 | 46407867 | 2839 | -1.53 | 1.06E-05 | 1.56E-05 |
| chr1 | 227040734 | 227041132 | 399 | -4.49 | 1.06E-05 | 1.56E-05 |
| chr12 | 60897616 | 60924368 | 26753 | 1.7 | 1.06E-05 | 1.56E-05 |
| chr6 | 44564457 | 44565100 | 644 | -5.31 | 1.06E-05 | 1.56E-05 |
| chr10 | 34208549 | 34209320 | 772 | -5.15 | 1.06E-05 | 1.56E-05 |
| chr5 | 72136417 | 72136972 | 556 | -5.2 | 1.06E-05 | 1.57E-05 |
| chr8 | 27209835 | 27210654 | 820 | -5.2 | 1.06E-05 | 1.57E-05 |
| chr5 | 50118061 | 50119721 | 1661 | -5.14 | 1.06E-05 | 1.57E-05 |

| | | | | | | |
|---|---|---|---|---|---|---|
| chr6 | 52794009 | 52795153 | 1145 | -5.14 | 1.06E-05 | 1.57E-05 |
| chr6 | 169171588 | 169237209 | 65622 | -1.65 | 1.07E-05 | 1.57E-05 |
| chr1 | 57197446 | 57216262 | 18817 | -1.6 | 1.07E-05 | 1.57E-05 |
| chr3 | 134272503 | 134272912 | 410 | -4.09 | 1.07E-05 | 1.57E-05 |
| chr21 | 28148451 | 28200081 | 51631 | 0.97 | 1.07E-05 | 1.57E-05 |
| chr11 | 17668957 | 17679541 | 10585 | -5.61 | 1.07E-05 | 1.58E-05 |
| chr1 | 55691358 | 55695564 | 4207 | -4.89 | 1.07E-05 | 1.58E-05 |
| chr7 | 43873540 | 43875075 | 1536 | -5.28 | 1.07E-05 | 1.58E-05 |
| chr1 | 235751757 | 235753022 | 1266 | -5.64 | 1.08E-05 | 1.58E-05 |
| chr1 | 54651689 | 54652129 | 441 | -5.15 | 1.08E-05 | 1.58E-05 |
| chr4 | 62382942 | 62384294 | 1353 | -5.15 | 1.08E-05 | 1.58E-05 |
| chr4 | 152251594 | 152252370 | 777 | -5.15 | 1.08E-05 | 1.58E-05 |
| chr16 | 63572760 | 63574103 | 1344 | -5.62 | 1.08E-05 | 1.59E-05 |
| chr3 | 106042582 | 106044323 | 1742 | -4.41 | 1.08E-05 | 1.59E-05 |
| chr18 | 30328797 | 30344272 | 15476 | 3.33 | 1.08E-05 | 1.59E-05 |
| chr2 | 214927206 | 214928292 | 1087 | -5.17 | 1.08E-05 | 1.59E-05 |
| chr4 | 37971512 | 37972374 | 863 | -5.17 | 1.08E-05 | 1.59E-05 |
| chr5 | 113656641 | 113680567 | 23927 | 3.75 | 1.08E-05 | 1.60E-05 |
| chr3 | 159292309 | 159321844 | 29536 | -4.48 | 1.09E-05 | 1.60E-05 |
| chr13 | 74503669 | 74504439 | 771 | -5.16 | 1.09E-05 | 1.60E-05 |
| chr14 | 93375053 | 93375873 | 821 | -5.16 | 1.09E-05 | 1.60E-05 |
| chr2 | 113329590 | 113330187 | 598 | -5.16 | 1.09E-05 | 1.60E-05 |
| chr1 | 12209520 | 12210251 | 732 | -5.21 | 1.09E-05 | 1.60E-05 |
| chr1 | 98310887 | 98311776 | 890 | -5.21 | 1.09E-05 | 1.60E-05 |
| chr9 | 74775292 | 74775794 | 503 | -5.21 | 1.09E-05 | 1.60E-05 |
| chr4 | 108649646 | 108650895 | 1250 | -3.94 | 1.09E-05 | 1.60E-05 |
| chr2 | 220181910 | 220182575 | 666 | -5.54 | 1.09E-05 | 1.60E-05 |
| chr7 | 47883152 | 47883979 | 828 | -5.54 | 1.09E-05 | 1.60E-05 |
| chr21 | 45294872 | 45295515 | 644 | -5.53 | 1.09E-05 | 1.61E-05 |
| chr6 | 39086862 | 39107190 | 20329 | -2 | 1.09E-05 | 1.61E-05 |
| chr1 | 153841904 | 153842716 | 813 | -5.13 | 1.09E-05 | 1.61E-05 |
| chr18 | 43623901 | 43624384 | 484 | -5.13 | 1.09E-05 | 1.61E-05 |
| chr2 | 174291449 | 174292081 | 633 | -5.13 | 1.09E-05 | 1.61E-05 |
| chr3 | 52064780 | 52066659 | 1880 | -6.04 | 1.10E-05 | 1.61E-05 |
| chr6 | 40487184 | 40487875 | 692 | -3.88 | 1.10E-05 | 1.61E-05 |
| chr4 | 123927676 | 123929526 | 1851 | -5.51 | 1.10E-05 | 1.61E-05 |
| chr7 | 155035563 | 155036718 | 1156 | -5.13 | 1.10E-05 | 1.62E-05 |
| chr6 | 122002735 | 122004177 | 1443 | -5.27 | 1.10E-05 | 1.62E-05 |
| chr3 | 174676695 | 174678551 | 1857 | -3.5 | 1.10E-05 | 1.62E-05 |
| chr20 | 39330777 | 39332231 | 1455 | -1.82 | 1.10E-05 | 1.62E-05 |
| chr15 | 57827558 | 57828738 | 1181 | -5.48 | 1.10E-05 | 1.62E-05 |
| chrX | 83760531 | 83787451 | 26921 | 1.58 | 1.11E-05 | 1.62E-05 |
| chr10 | 53934791 | 53937322 | 2532 | -2.5 | 1.11E-05 | 1.63E-05 |
| chr4 | 28540534 | 28556163 | 15630 | -1.78 | 1.11E-05 | 1.63E-05 |
| chr17 | 65085958 | 65086381 | 424 | -5.51 | 1.11E-05 | 1.63E-05 |
| chr5 | 83663453 | 83664244 | 792 | -4.04 | 1.11E-05 | 1.63E-05 |
| chr1 | 121244678 | 121252334 | 7657 | -4.89 | 1.11E-05 | 1.64E-05 |
| chr5 | 162769449 | 162770703 | 1255 | 5.21 | 1.11E-05 | 1.64E-05 |
| chr6 | 154285753 | 154286804 | 1052 | 5.21 | 1.11E-05 | 1.64E-05 |
| chr9 | 129308767 | 129309494 | 728 | -5.26 | 1.12E-05 | 1.64E-05 |
| chr15 | 24495759 | 24496104 | 346 | 4.27 | 1.12E-05 | 1.64E-05 |
| chr16 | 89580559 | 89580971 | 413 | -5.19 | 1.12E-05 | 1.64E-05 |
| chr2 | 29098718 | 29099300 | 583 | -5.43 | 1.12E-05 | 1.64E-05 |
| chr1 | 184947298 | 184947968 | 671 | -5.12 | 1.12E-05 | 1.65E-05 |
| chr7 | 12229325 | 12230130 | 806 | -5.12 | 1.12E-05 | 1.65E-05 |
| chr15 | 69621490 | 69622369 | 880 | -5.16 | 1.12E-05 | 1.65E-05 |
| chr6 | 141222240 | 141223071 | 832 | -5.16 | 1.12E-05 | 1.65E-05 |
| chrX | 8792545 | 8793485 | 941 | -5.16 | 1.12E-05 | 1.65E-05 |
| chr13 | 105947590 | 105948382 | 793 | -4.19 | 1.13E-05 | 1.65E-05 |
| chr2 | 165160463 | 165161054 | 592 | -5.14 | 1.13E-05 | 1.65E-05 |
| chr5 | 13857777 | 13859003 | 1227 | -5.14 | 1.13E-05 | 1.65E-05 |

| | | | | | | |
|---|---|---|---|---|---|---|
| chr4 | 107164112 | 107164513 | 402 | -5.14 | 1.13E-05 | 1.66E-05 |
| chr4 | 151486019 | 151486518 | 500 | -5.14 | 1.13E-05 | 1.66E-05 |
| chr1 | 5929736 | 5930568 | 833 | -5.4 | 1.13E-05 | 1.66E-05 |
| chr10 | 18421891 | 18422384 | 494 | -3.91 | 1.14E-05 | 1.67E-05 |
| chr16 | 70855487 | 70855837 | 351 | -4.06 | 1.14E-05 | 1.68E-05 |
| chr2 | 174138757 | 174139334 | 578 | -5.22 | 1.14E-05 | 1.68E-05 |
| chr22 | 22541210 | 22542521 | 1312 | -2.15 | 1.14E-05 | 1.68E-05 |
| chr8 | 37341156 | 37359112 | 17957 | -1.67 | 1.14E-05 | 1.68E-05 |
| chr5 | 20832323 | 20835282 | 2960 | 1.52 | 1.14E-05 | 1.68E-05 |
| chr14 | 99619550 | 99633919 | 14370 | -5.03 | 1.15E-05 | 1.68E-05 |
| chr5 | 40058922 | 40059796 | 875 | -5.39 | 1.15E-05 | 1.68E-05 |
| chr1 | 22505624 | 22506285 | 662 | -5.18 | 1.15E-05 | 1.69E-05 |
| chr7 | 50912333 | 50912929 | 597 | -5.18 | 1.15E-05 | 1.69E-05 |
| chr20 | 205238 | 208048 | 2811 | -2.32 | 1.15E-05 | 1.69E-05 |
| chr1 | 28492292 | 28493177 | 886 | -5.38 | 1.15E-05 | 1.69E-05 |
| chr1 | 84380549 | 84381085 | 537 | -5.38 | 1.15E-05 | 1.69E-05 |
| chr9 | 111750285 | 111751167 | 883 | -5.38 | 1.15E-05 | 1.69E-05 |
| chr1 | 187046999 | 187047674 | 676 | 5.26 | 1.15E-05 | 1.69E-05 |
| chr1 | 224978022 | 224978893 | 872 | 5.26 | 1.15E-05 | 1.69E-05 |
| chr14 | 94186346 | 94187799 | 1454 | 5.26 | 1.15E-05 | 1.69E-05 |
| chr12 | 39862242 | 39863510 | 1269 | -5.23 | 1.15E-05 | 1.69E-05 |
| chr13 | 46244761 | 46245668 | 908 | -5.23 | 1.15E-05 | 1.69E-05 |
| chr3 | 49668544 | 49670548 | 2005 | -4.75 | 1.16E-05 | 1.70E-05 |
| chr8 | 20137751 | 20189841 | 52091 | -1.59 | 1.16E-05 | 1.70E-05 |
| chr5 | 34848231 | 34849242 | 1012 | -5.33 | 1.17E-05 | 1.71E-05 |
| chrX | 44513727 | 44515117 | 1391 | -5.33 | 1.17E-05 | 1.71E-05 |
| chr1 | 186234960 | 186235661 | 702 | -5.11 | 1.17E-05 | 1.71E-05 |
| chr16 | 933728 | 934438 | 711 | -5.11 | 1.17E-05 | 1.71E-05 |
| chr13 | 88465707 | 88466447 | 741 | -5.13 | 1.17E-05 | 1.71E-05 |
| chr14 | 81395415 | 81396072 | 658 | -5.13 | 1.17E-05 | 1.71E-05 |
| chrX | 149873597 | 149874393 | 797 | -5.13 | 1.17E-05 | 1.71E-05 |
| chr9 | 4530675 | 4531324 | 650 | -5.12 | 1.17E-05 | 1.71E-05 |
| chrX | 71150375 | 71150786 | 412 | -4.13 | 1.17E-05 | 1.72E-05 |
| chr3 | 72890647 | 72891251 | 605 | -5.14 | 1.17E-05 | 1.72E-05 |
| chr2 | 157969376 | 157972619 | 3244 | -4.52 | 1.17E-05 | 1.72E-05 |
| chr2 | 108287281 | 108292938 | 5658 | -4.01 | 1.17E-05 | 1.72E-05 |
| chr12 | 3252804 | 3253775 | 972 | -5.34 | 1.18E-05 | 1.72E-05 |
| chr4 | 41913966 | 41923510 | 9545 | -5.77 | 1.18E-05 | 1.72E-05 |
| chr6 | 122023539 | 122024437 | 899 | -5.21 | 1.18E-05 | 1.73E-05 |
| chr1 | 207144081 | 207144591 | 511 | 5.36 | 1.18E-05 | 1.73E-05 |
| chr1 | 31626196 | 31627085 | 890 | -5.2 | 1.18E-05 | 1.73E-05 |
| chr7 | 99012088 | 99012961 | 874 | -5.2 | 1.18E-05 | 1.73E-05 |
| chr3 | 155326992 | 155344234 | 17243 | -5.74 | 1.18E-05 | 1.73E-05 |
| chrX | 108118004 | 108121621 | 3618 | 3.16 | 1.18E-05 | 1.73E-05 |
| chr19 | 40023938 | 40024514 | 577 | -5.11 | 1.18E-05 | 1.73E-05 |
| chr9 | 38322824 | 38323325 | 502 | -5.11 | 1.18E-05 | 1.73E-05 |
| chrX | 48636099 | 48637749 | 1651 | -2.79 | 1.18E-05 | 1.74E-05 |
| chr1 | 23789130 | 23789993 | 864 | -5.16 | 1.19E-05 | 1.74E-05 |
| chrX | 20442687 | 20443538 | 852 | -5.16 | 1.19E-05 | 1.74E-05 |
| chr18 | 11819671 | 11820405 | 735 | -5.12 | 1.19E-05 | 1.74E-05 |
| chr3 | 33453873 | 33454321 | 449 | -5.12 | 1.19E-05 | 1.74E-05 |
| chr2 | 158704886 | 158705505 | 620 | -3.48 | 1.19E-05 | 1.74E-05 |
| chr7 | 156016253 | 156166753 | 150501 | -1.16 | 1.19E-05 | 1.74E-05 |
| chr3 | 40941157 | 40942091 | 935 | -5.3 | 1.20E-05 | 1.76E-05 |
| chr17 | 60762588 | 60763555 | 968 | -5.19 | 1.20E-05 | 1.76E-05 |
| chr18 | 55810800 | 55811804 | 1005 | -5.19 | 1.20E-05 | 1.76E-05 |
| chr22 | 49566514 | 49596336 | 29823 | -1.46 | 1.21E-05 | 1.77E-05 |
| chrX | 119637329 | 119638237 | 909 | -5.29 | 1.21E-05 | 1.77E-05 |
| chr12 | 22848052 | 22848565 | 514 | -5.27 | 1.21E-05 | 1.77E-05 |
| chr15 | 51684362 | 51685278 | 917 | -5.58 | 1.21E-05 | 1.78E-05 |
| chr2 | 111924107 | 111924783 | 677 | -5.11 | 1.22E-05 | 1.78E-05 |

| | | | | | | |
|---|---|---|---|---|---|---|
| chr5 | 167729575 | 167730214 | 640 | -5.11 | 1.22E-05 | 1.78E-05 |
| chr6 | 170257529 | 170279816 | 22288 | -2.82 | 1.22E-05 | 1.78E-05 |
| chr18 | 25402833 | 25404797 | 1965 | -4.14 | 1.22E-05 | 1.78E-05 |
| chr4 | 152554245 | 152558166 | 3922 | -1.6 | 1.22E-05 | 1.79E-05 |
| chr15 | 53895900 | 53897920 | 2021 | -4.4 | 1.22E-05 | 1.79E-05 |
| chr6 | 37486860 | 37487525 | 666 | -5.12 | 1.22E-05 | 1.79E-05 |
| chr10 | 54107161 | 54108475 | 1315 | -5.55 | 1.22E-05 | 1.79E-05 |
| chr13 | 80450342 | 80451605 | 1264 | -5.55 | 1.22E-05 | 1.79E-05 |
| chr7 | 155344887 | 155347040 | 2154 | -5.53 | 1.23E-05 | 1.79E-05 |
| chr12 | 63543752 | 63544372 | 621 | -5.14 | 1.23E-05 | 1.80E-05 |
| chr7 | 90248910 | 90249825 | 916 | -5.14 | 1.23E-05 | 1.80E-05 |
| chr16 | 30931096 | 30932101 | 1006 | -5.17 | 1.23E-05 | 1.80E-05 |
| chr6 | 25640889 | 25641789 | 901 | -5.17 | 1.23E-05 | 1.80E-05 |
| chr9 | 20332447 | 20333111 | 665 | -5.51 | 1.23E-05 | 1.80E-05 |
| chr13 | 43399106 | 43400115 | 1010 | 3.25 | 1.23E-05 | 1.80E-05 |
| chr5 | 170611028 | 170611562 | 535 | -5.18 | 1.23E-05 | 1.80E-05 |
| chr8 | 120977204 | 120977955 | 752 | -5.18 | 1.23E-05 | 1.80E-05 |
| chr15 | 93331847 | 93332843 | 997 | -3.9 | 1.23E-05 | 1.81E-05 |
| chr11 | 51518586 | 51520807 | 2222 | 2.08 | 1.23E-05 | 1.81E-05 |
| chr2 | 149862316 | 149866030 | 3715 | -2.2 | 1.24E-05 | 1.81E-05 |
| chr7 | 158957572 | 158983328 | 25757 | -3.1 | 1.24E-05 | 1.81E-05 |
| chr8 | 29945820 | 29946794 | 975 | -3.9 | 1.24E-05 | 1.81E-05 |
| chr12 | 102087618 | 102088577 | 960 | -5.48 | 1.24E-05 | 1.81E-05 |
| chr2 | 153058207 | 153058976 | 770 | -5.48 | 1.24E-05 | 1.81E-05 |
| chr4 | 54707772 | 54708302 | 531 | 5.18 | 1.24E-05 | 1.81E-05 |
| chr1 | 241989044 | 241989986 | 943 | -5.47 | 1.24E-05 | 1.82E-05 |
| chr5 | 60693681 | 60694540 | 860 | -5.47 | 1.24E-05 | 1.82E-05 |
| chr19 | 6761260 | 6761713 | 454 | -5.26 | 1.24E-05 | 1.82E-05 |
| chr1 | 22103979 | 22104734 | 756 | -5.24 | 1.24E-05 | 1.82E-05 |
| chr1 | 65849807 | 65850624 | 818 | -5.24 | 1.24E-05 | 1.82E-05 |
| chr20 | 32760886 | 32761568 | 683 | -5.24 | 1.24E-05 | 1.82E-05 |
| chr1 | 41865848 | 41877493 | 11646 | -5.97 | 1.24E-05 | 1.82E-05 |
| chr3 | 105325669 | 105326658 | 990 | -5.1 | 1.25E-05 | 1.82E-05 |
| chr4 | 144035518 | 144036810 | 1293 | -5.1 | 1.25E-05 | 1.82E-05 |
| chr12 | 42143265 | 42166437 | 23173 | -1.15 | 1.25E-05 | 1.82E-05 |
| chr21 | 35601856 | 35703256 | 101401 | -0.61 | 1.25E-05 | 1.82E-05 |
| chr3 | 40886467 | 40888356 | 1890 | -4.08 | 1.25E-05 | 1.82E-05 |
| chr10 | 129426636 | 129437405 | 10770 | -1.76 | 1.25E-05 | 1.83E-05 |
| chr15 | 49297132 | 49297670 | 539 | -5.13 | 1.25E-05 | 1.83E-05 |
| chr3 | 187552145 | 187552733 | 589 | -5.13 | 1.25E-05 | 1.83E-05 |
| chr6 | 83085302 | 83086430 | 1129 | -3.71 | 1.25E-05 | 1.83E-05 |
| chr22 | 49707648 | 49754024 | 46377 | -2.14 | 1.25E-05 | 1.83E-05 |
| chr11 | 66508885 | 66509657 | 773 | -5.43 | 1.25E-05 | 1.83E-05 |
| chr2 | 70312746 | 70313371 | 626 | -5.1 | 1.26E-05 | 1.83E-05 |
| chr21 | 40378122 | 40378982 | 861 | -5.1 | 1.26E-05 | 1.83E-05 |
| chr5 | 162736114 | 162736889 | 776 | -4.27 | 1.26E-05 | 1.84E-05 |
| chr21 | 34475894 | 34477375 | 1482 | -4.4 | 1.26E-05 | 1.84E-05 |
| chr10 | 97034799 | 97035519 | 721 | -5.42 | 1.26E-05 | 1.84E-05 |
| chrX | 5068684 | 5071122 | 2439 | -2.46 | 1.26E-05 | 1.84E-05 |
| chr6 | 112717911 | 112719637 | 1727 | 2.25 | 1.26E-05 | 1.84E-05 |
| chr8 | 81715248 | 81716345 | 1098 | -5.23 | 1.26E-05 | 1.84E-05 |
| chr14 | 21979890 | 21980512 | 623 | -3.85 | 1.27E-05 | 1.85E-05 |
| chr10 | 92734616 | 92735509 | 894 | -5.57 | 1.27E-05 | 1.85E-05 |
| chr2 | 146299953 | 146301163 | 1211 | -5.39 | 1.27E-05 | 1.85E-05 |
| chr1 | 48792003 | 48792772 | 770 | -5.1 | 1.27E-05 | 1.85E-05 |
| chr10 | 93262554 | 93263081 | 528 | -5.1 | 1.27E-05 | 1.85E-05 |
| chr4 | 116317001 | 116318410 | 1410 | -5.1 | 1.27E-05 | 1.85E-05 |
| chr10 | 124482327 | 124534426 | 52100 | -1.68 | 1.27E-05 | 1.86E-05 |
| chr16 | 12496795 | 12497916 | 1122 | -5.38 | 1.27E-05 | 1.86E-05 |
| chr7 | 133184875 | 133185843 | 969 | -5.38 | 1.27E-05 | 1.86E-05 |
| chr12 | 57219310 | 57220558 | 1249 | -4.2 | 1.28E-05 | 1.87E-05 |

| | | | | | | |
|---|---|---|---|---|---|---|
| chr7 | 124224739 | 124266770 | 42032 | 1.47 | 1.28E-05 | 1.87E-05 |
| chr16 | 11728037 | 11728684 | 648 | -5.09 | 1.28E-05 | 1.87E-05 |
| chr7 | 48160126 | 48210177 | 50052 | -4.23 | 1.28E-05 | 1.87E-05 |
| chr2 | 162129054 | 162129755 | 702 | -5.36 | 1.28E-05 | 1.87E-05 |
| chr15 | 76466932 | 76467862 | 931 | -5.21 | 1.28E-05 | 1.87E-05 |
| chr1 | 9454838 | 9455604 | 767 | -5.11 | 1.28E-05 | 1.87E-05 |
| chr6 | 34755802 | 34756282 | 481 | -5.11 | 1.28E-05 | 1.87E-05 |
| chr6 | 153362375 | 153362838 | 464 | -5.11 | 1.28E-05 | 1.87E-05 |
| chr3 | 176560390 | 176561311 | 922 | 5.6 | 1.28E-05 | 1.88E-05 |
| chr3 | 197085278 | 197088850 | 3573 | -4.74 | 1.29E-05 | 1.88E-05 |
| chr6 | 161322511 | 161335997 | 13487 | -1.66 | 1.29E-05 | 1.88E-05 |
| chr14 | 70455073 | 70479396 | 24324 | -3.71 | 1.29E-05 | 1.89E-05 |
| chr11 | 33548196 | 33556902 | 8707 | -5.84 | 1.29E-05 | 1.89E-05 |
| chr11 | 121048908 | 121052972 | 4065 | -3.53 | 1.30E-05 | 1.89E-05 |
| chr12 | 305077 | 306748 | 1672 | 4.39 | 1.30E-05 | 1.89E-05 |
| chr4 | 188216162 | 188219672 | 3511 | -2.06 | 1.30E-05 | 1.90E-05 |
| chr6 | 4601041 | 4602519 | 1479 | -5.15 | 1.30E-05 | 1.90E-05 |
| chrX | 42526706 | 42530532 | 3827 | -3.17 | 1.30E-05 | 1.91E-05 |
| chr18 | 47467456 | 47468238 | 783 | -3.94 | 1.31E-05 | 1.91E-05 |
| chr18 | 28789704 | 28790319 | 616 | -5.2 | 1.31E-05 | 1.91E-05 |
| chr4 | 134273403 | 134274569 | 1167 | 5.21 | 1.31E-05 | 1.91E-05 |
| chr3 | 39544469 | 39568480 | 24012 | -1.95 | 1.31E-05 | 1.91E-05 |
| chr9 | 111174770 | 111194402 | 19633 | -6.1 | 1.31E-05 | 1.92E-05 |
| chr2 | 18574286 | 18575853 | 1568 | -2.87 | 1.32E-05 | 1.92E-05 |
| chr6 | 141948712 | 141949682 | 971 | -5.31 | 1.32E-05 | 1.92E-05 |
| chr5 | 132575457 | 132576226 | 770 | -5.14 | 1.32E-05 | 1.92E-05 |
| chr6 | 36277582 | 36278288 | 707 | -5.14 | 1.32E-05 | 1.92E-05 |
| chr3 | 131232903 | 131233879 | 977 | -4.29 | 1.32E-05 | 1.92E-05 |
| chr9 | 134220673 | 134221641 | 969 | -5.74 | 1.32E-05 | 1.93E-05 |
| chr21 | 43708906 | 43709577 | 672 | -5.39 | 1.32E-05 | 1.93E-05 |
| chr12 | 53286798 | 53287345 | 548 | -5.2 | 1.32E-05 | 1.93E-05 |
| chr5 | 65990105 | 66009690 | 19586 | -1.22 | 1.32E-05 | 1.93E-05 |
| chr8 | 16576309 | 16621207 | 44899 | -1.52 | 1.33E-05 | 1.93E-05 |
| chr5 | 179129574 | 179130603 | 1030 | -5.38 | 1.33E-05 | 1.93E-05 |
| chr4 | 186013556 | 186014077 | 522 | -5.09 | 1.33E-05 | 1.94E-05 |
| chr1 | 201648735 | 201649495 | 761 | -5.29 | 1.33E-05 | 1.94E-05 |
| chr17 | 53405735 | 53406423 | 689 | -5.29 | 1.33E-05 | 1.94E-05 |
| chr8 | 16507496 | 16524261 | 16766 | -1.43 | 1.33E-05 | 1.94E-05 |
| chr13 | 21937288 | 21938083 | 796 | -4.5 | 1.33E-05 | 1.94E-05 |
| chr11 | 117423930 | 117424625 | 696 | -5.11 | 1.33E-05 | 1.94E-05 |
| chr8 | 74244222 | 74259538 | 15317 | -7.36 | 1.33E-05 | 1.94E-05 |
| chr3 | 104377034 | 104407167 | 30134 | 1.44 | 1.33E-05 | 1.94E-05 |
| chr12 | 20537531 | 20538278 | 748 | -5.09 | 1.33E-05 | 1.94E-05 |
| chr12 | 42764965 | 42765768 | 804 | -5.09 | 1.33E-05 | 1.94E-05 |
| chrX | 42734417 | 42743015 | 8599 | 2.59 | 1.33E-05 | 1.95E-05 |
| chr18 | 74707075 | 74707703 | 629 | -5.28 | 1.34E-05 | 1.96E-05 |
| chr4 | 122966833 | 122967619 | 787 | -5.1 | 1.34E-05 | 1.96E-05 |
| chr2 | 105804997 | 105822272 | 17276 | -3.76 | 1.35E-05 | 1.96E-05 |
| chr3 | 72657734 | 72682094 | 24361 | -6.39 | 1.35E-05 | 1.96E-05 |
| chr2 | 128114045 | 128115019 | 975 | -5.62 | 1.35E-05 | 1.97E-05 |
| chr13 | 113691815 | 113692398 | 584 | -5.08 | 1.35E-05 | 1.97E-05 |
| chr21 | 17732774 | 17733328 | 555 | -5.08 | 1.35E-05 | 1.97E-05 |
| chr6 | 106933478 | 106933972 | 495 | -5.08 | 1.35E-05 | 1.97E-05 |
| chr6 | 84613847 | 84614483 | 637 | -5.12 | 1.35E-05 | 1.98E-05 |
| chr10 | 80748667 | 80749442 | 776 | -5.26 | 1.35E-05 | 1.98E-05 |
| chr11 | 82983005 | 82983443 | 439 | -5.26 | 1.35E-05 | 1.98E-05 |
| chr6 | 21318874 | 21319453 | 580 | -3.72 | 1.36E-05 | 1.98E-05 |
| chr1 | 212270273 | 212270894 | 622 | -5.17 | 1.36E-05 | 1.99E-05 |
| chr17 | 40969165 | 40969645 | 481 | -5.17 | 1.36E-05 | 1.99E-05 |
| chr2 | 177429208 | 177430084 | 877 | -5.17 | 1.36E-05 | 1.99E-05 |
| chr5 | 55576445 | 55577418 | 974 | -5.17 | 1.36E-05 | 1.99E-05 |

| | | | | | | |
|---|---|---|---|---|---|---|
| chr6 | 3078149 | 3079008 | 860 | -5.31 | 1.36E-05 | 1.99E-05 |
| chr1 | 51964861 | 51965908 | 1048 | -5.17 | 1.37E-05 | 1.99E-05 |
| chr18 | 34698005 | 34700002 | 1998 | -5.56 | 1.37E-05 | 1.99E-05 |
| chr8 | 59590077 | 59591849 | 1773 | 2.28 | 1.37E-05 | 2.00E-05 |
| chr2 | 10232515 | 10233105 | 591 | -5.25 | 1.37E-05 | 2.00E-05 |
| chr5 | 92128075 | 92139602 | 11528 | -6.37 | 1.37E-05 | 2.00E-05 |
| chr3 | 15330549 | 15331252 | 704 | -5.54 | 1.37E-05 | 2.00E-05 |
| chr8 | 122341776 | 122342399 | 624 | -5.54 | 1.37E-05 | 2.00E-05 |
| chr3 | 123809672 | 123852731 | 43060 | 1.16 | 1.38E-05 | 2.00E-05 |
| chr6 | 165208749 | 165315216 | 106468 | 4.27 | 1.38E-05 | 2.00E-05 |
| chr12 | 16360611 | 16364242 | 3632 | 3.9 | 1.38E-05 | 2.01E-05 |
| chr1 | 179342093 | 179342585 | 493 | -4.11 | 1.38E-05 | 2.02E-05 |
| chr10 | 8811589 | 8812153 | 565 | -5.5 | 1.38E-05 | 2.02E-05 |
| chr1 | 115509139 | 115546584 | 37446 | -2.94 | 1.39E-05 | 2.02E-05 |
| chr10 | 130070268 | 130070786 | 519 | -5.23 | 1.39E-05 | 2.02E-05 |
| chr22 | 24412470 | 24421137 | 8668 | -5.23 | 1.39E-05 | 2.02E-05 |
| chr22 | 40279901 | 40284397 | 4497 | -2.64 | 1.39E-05 | 2.02E-05 |
| chr1 | 111953407 | 111953973 | 567 | -5.49 | 1.39E-05 | 2.02E-05 |
| chr15 | 59231341 | 59232116 | 776 | -3.92 | 1.39E-05 | 2.02E-05 |
| chr2 | 177911889 | 177913317 | 1429 | -3.66 | 1.39E-05 | 2.02E-05 |
| chr18 | 12510591 | 12511676 | 1086 | -5.09 | 1.39E-05 | 2.02E-05 |
| chr4 | 90401131 | 90401619 | 489 | -5.09 | 1.39E-05 | 2.02E-05 |
| chr10 | 102961369 | 102969379 | 8011 | -5.47 | 1.39E-05 | 2.03E-05 |
| chr1 | 219751759 | 219761803 | 10045 | 1.68 | 1.39E-05 | 2.03E-05 |
| chr14 | 23937626 | 23938357 | 732 | -5.08 | 1.39E-05 | 2.03E-05 |
| chr14 | 101182359 | 101183070 | 712 | -5.08 | 1.39E-05 | 2.03E-05 |
| chr1 | 20573250 | 20573944 | 695 | -5.11 | 1.40E-05 | 2.03E-05 |
| chr11 | 18036984 | 18037603 | 620 | -5.11 | 1.40E-05 | 2.03E-05 |
| chr12 | 87286421 | 87289368 | 2948 | -5.46 | 1.40E-05 | 2.03E-05 |
| chr6 | 80098771 | 80099743 | 973 | -5.46 | 1.40E-05 | 2.03E-05 |
| chr1 | 183126922 | 183127983 | 1062 | -5.15 | 1.40E-05 | 2.03E-05 |
| chr1 | 186603047 | 186603605 | 559 | -5.15 | 1.40E-05 | 2.03E-05 |
| chr9 | 114208233 | 114208741 | 509 | -5.09 | 1.40E-05 | 2.03E-05 |
| chr3 | 69599 | 81437 | 11839 | 2.67 | 1.40E-05 | 2.03E-05 |
| chr21 | 42724888 | 42725539 | 652 | 4.02 | 1.40E-05 | 2.04E-05 |
| chr11 | 100607556 | 100608439 | 884 | -5.22 | 1.41E-05 | 2.05E-05 |
| chrX | 53857339 | 53858125 | 787 | -5.22 | 1.41E-05 | 2.05E-05 |
| chrX | 108178694 | 108182038 | 3345 | 2.59 | 1.41E-05 | 2.05E-05 |
| chrX | 80674419 | 80675505 | 1087 | 3.92 | 1.41E-05 | 2.05E-05 |
| chr3 | 188709074 | 188709987 | 914 | -4.49 | 1.41E-05 | 2.05E-05 |
| chr4 | 160456592 | 160519509 | 62918 | 5.93 | 1.41E-05 | 2.05E-05 |
| chr10 | 54476759 | 54478728 | 1970 | -4.5 | 1.41E-05 | 2.06E-05 |
| chr16 | 51369845 | 51371338 | 1494 | -5.41 | 1.41E-05 | 2.06E-05 |
| chr5 | 167668659 | 167669430 | 772 | -5.15 | 1.42E-05 | 2.07E-05 |
| chr15 | 43185242 | 43186344 | 1103 | -5.39 | 1.42E-05 | 2.07E-05 |
| chr12 | 7139460 | 7140220 | 761 | -5.07 | 1.42E-05 | 2.07E-05 |
| chr8 | 56622230 | 56622653 | 424 | -2.81 | 1.43E-05 | 2.07E-05 |
| chr11 | 12414632 | 12415288 | 657 | -5.2 | 1.43E-05 | 2.08E-05 |
| chr19 | 12873898 | 12874744 | 847 | -5.2 | 1.43E-05 | 2.08E-05 |
| chr6 | 7993735 | 7994144 | 410 | -5.07 | 1.43E-05 | 2.08E-05 |
| chr6 | 101117967 | 101118476 | 510 | -5.07 | 1.43E-05 | 2.08E-05 |
| chr13 | 42632561 | 42633109 | 549 | -5.07 | 1.43E-05 | 2.08E-05 |
| chr2 | 67731239 | 67732336 | 1098 | -5.07 | 1.43E-05 | 2.08E-05 |
| chr4 | 90570506 | 90571096 | 591 | -5.07 | 1.43E-05 | 2.08E-05 |
| chr5 | 36265767 | 36266609 | 843 | -5.37 | 1.43E-05 | 2.08E-05 |
| chr12 | 124060452 | 124061280 | 829 | -5.13 | 1.44E-05 | 2.09E-05 |
| chr3 | 195829511 | 195830092 | 582 | -5.13 | 1.44E-05 | 2.09E-05 |
| chr6 | 53961741 | 53962402 | 662 | -5.13 | 1.44E-05 | 2.09E-05 |
| chrX | 113901296 | 113934969 | 33674 | 5.52 | 1.44E-05 | 2.09E-05 |
| chr21 | 17094420 | 17095250 | 831 | -5.35 | 1.44E-05 | 2.10E-05 |
| chr14 | 105170376 | 105171233 | 858 | -5.23 | 1.44E-05 | 2.10E-05 |

| | | | | | | |
|---|---|---|---|---|---|---|
| chr6 | 140566509 | 140568857 | 2349 | -5.82 | 1.45E-05 | 2.11E-05 |
| chr1 | 27131818 | 27132607 | 790 | -5.1 | 1.45E-05 | 2.11E-05 |
| chr12 | 101514250 | 101515456 | 1207 | -5.1 | 1.45E-05 | 2.11E-05 |
| chr11 | 3942615 | 3943522 | 908 | -5.34 | 1.45E-05 | 2.11E-05 |
| chr10 | 18985227 | 18997719 | 12493 | -7 | 1.45E-05 | 2.11E-05 |
| chr3 | 136014987 | 136015405 | 419 | -5.13 | 1.45E-05 | 2.11E-05 |
| chr6 | 5461650 | 5462419 | 770 | -5.13 | 1.45E-05 | 2.11E-05 |
| chr2 | 167058336 | 167088938 | 30603 | 2.18 | 1.46E-05 | 2.12E-05 |
| chr5 | 92478647 | 92479709 | 1063 | -5.07 | 1.46E-05 | 2.12E-05 |
| chr13 | 35356722 | 35358454 | 1733 | -2.22 | 1.46E-05 | 2.12E-05 |
| chr12 | 110067973 | 110068266 | 294 | -5.08 | 1.46E-05 | 2.12E-05 |
| chrX | 48312770 | 48313288 | 519 | -5.08 | 1.46E-05 | 2.12E-05 |
| chr2 | 187795537 | 187796398 | 862 | 5.22 | 1.46E-05 | 2.13E-05 |
| chr13 | 25955767 | 25970093 | 14327 | -2.57 | 1.46E-05 | 2.13E-05 |
| chr10 | 100942830 | 100946812 | 3983 | -1.86 | 1.47E-05 | 2.13E-05 |
| chr6 | 148284586 | 148293297 | 8712 | -5.76 | 1.47E-05 | 2.13E-05 |
| chr3 | 34345906 | 34387170 | 41265 | -1.22 | 1.47E-05 | 2.14E-05 |
| chr12 | 57957304 | 57958374 | 1071 | 4.13 | 1.47E-05 | 2.14E-05 |
| chr9 | 124122219 | 124122990 | 772 | -5.21 | 1.47E-05 | 2.14E-05 |
| chr13 | 109854406 | 109855056 | 651 | -3.87 | 1.48E-05 | 2.14E-05 |
| chr4 | 188260670 | 188264565 | 3896 | -4.66 | 1.48E-05 | 2.15E-05 |
| chr10 | 114029269 | 114031172 | 1904 | 2.58 | 1.48E-05 | 2.15E-05 |
| chr3 | 182469739 | 182471147 | 1409 | -5.17 | 1.48E-05 | 2.15E-05 |
| chr16 | 23238518 | 23239164 | 647 | -5.12 | 1.48E-05 | 2.16E-05 |
| chr4 | 84178484 | 84179190 | 707 | -5.12 | 1.48E-05 | 2.16E-05 |
| chr6 | 9163463 | 9164055 | 593 | -5.12 | 1.48E-05 | 2.16E-05 |
| chr7 | 76898071 | 76898601 | 531 | -5.12 | 1.48E-05 | 2.16E-05 |
| chr11 | 104735291 | 104739805 | 4515 | -1.86 | 1.49E-05 | 2.16E-05 |
| chr1 | 80260324 | 80260906 | 583 | -4.56 | 1.49E-05 | 2.16E-05 |
| chr8 | 130589060 | 130590776 | 1717 | -1.49 | 1.49E-05 | 2.16E-05 |
| chr10 | 34244989 | 34259069 | 14081 | -5.7 | 1.49E-05 | 2.16E-05 |
| chr5 | 53496350 | 53498459 | 2110 | -5.69 | 1.49E-05 | 2.17E-05 |
| chr1 | 222408872 | 222409858 | 987 | 5.33 | 1.50E-05 | 2.17E-05 |
| chr6 | 87742745 | 87744093 | 1349 | -2.4 | 1.50E-05 | 2.18E-05 |
| chr19 | 55568385 | 55568962 | 578 | -5.44 | 1.50E-05 | 2.18E-05 |
| chr11 | 77228332 | 77229172 | 841 | -5.27 | 1.50E-05 | 2.18E-05 |
| chr12 | 74304028 | 74304560 | 533 | -5.27 | 1.50E-05 | 2.18E-05 |
| chr15 | 23651264 | 23651760 | 497 | -5.27 | 1.50E-05 | 2.18E-05 |
| chr5 | 36823373 | 36824313 | 941 | -2.84 | 1.51E-05 | 2.19E-05 |
| chr4 | 188797221 | 188807999 | 10779 | -6 | 1.51E-05 | 2.19E-05 |
| chr5 | 106445764 | 106446432 | 669 | -3.86 | 1.51E-05 | 2.19E-05 |
| chr6 | 49587853 | 49588828 | 976 | -5.08 | 1.51E-05 | 2.19E-05 |
| chr7 | 73725027 | 73725824 | 798 | -5.15 | 1.51E-05 | 2.19E-05 |
| chrX | 151606411 | 151607271 | 861 | -3.97 | 1.52E-05 | 2.20E-05 |
| chr3 | 127384517 | 127385267 | 751 | -5.41 | 1.52E-05 | 2.20E-05 |
| chr3 | 153035073 | 153044781 | 9709 | -5.61 | 1.52E-05 | 2.20E-05 |
| chr1 | 3024433 | 3025298 | 866 | -5.25 | 1.52E-05 | 2.20E-05 |
| chr9 | 103241856 | 103242359 | 504 | -5.06 | 1.52E-05 | 2.21E-05 |
| chr8 | 38869682 | 38870092 | 411 | -5.11 | 1.52E-05 | 2.21E-05 |
| chr1 | 149077385 | 149079481 | 2097 | 1.93 | 1.52E-05 | 2.21E-05 |
| chr4 | 23436723 | 23438665 | 1943 | -5.05 | 1.52E-05 | 2.21E-05 |
| chr9 | 33826780 | 33827447 | 668 | -5.05 | 1.52E-05 | 2.21E-05 |
| chr2 | 222308929 | 222310401 | 1473 | -5.06 | 1.52E-05 | 2.21E-05 |
| chr3 | 121460647 | 121461067 | 421 | -5.06 | 1.52E-05 | 2.21E-05 |
| chr6 | 34150466 | 34151739 | 1274 | -5.06 | 1.52E-05 | 2.21E-05 |
| chr2 | 160073059 | 160073902 | 844 | -5.1 | 1.53E-05 | 2.22E-05 |
| chr4 | 93743237 | 93744333 | 1097 | -5.1 | 1.53E-05 | 2.22E-05 |
| chr4 | 188355462 | 188360346 | 4885 | -3.54 | 1.53E-05 | 2.22E-05 |
| chr13 | 45450628 | 45451438 | 811 | -5.17 | 1.53E-05 | 2.23E-05 |
| chr2 | 231260097 | 231261216 | 1120 | -5.17 | 1.53E-05 | 2.23E-05 |
| chrX | 81036613 | 81047749 | 11137 | 1.98 | 1.53E-05 | 2.23E-05 |

| | | | | | | |
|---|---|---|---|---|---|---|
| chrX | 150124142 | 150124893 | 752 | -5.23 | 1.53E-05 | 2.23E-05 |
| chr11 | 15887483 | 15888754 | 1272 | -4.13 | 1.54E-05 | 2.23E-05 |
| chr6 | 162205112 | 162214183 | 9072 | -5.03 | 1.54E-05 | 2.23E-05 |
| chr15 | 66960253 | 66961195 | 943 | -5.54 | 1.54E-05 | 2.23E-05 |
| chr13 | 51925174 | 51925904 | 731 | -5.06 | 1.54E-05 | 2.23E-05 |
| chr15 | 77658400 | 77658729 | 330 | -5.06 | 1.54E-05 | 2.23E-05 |
| chr18 | 12845309 | 12846050 | 742 | -5.06 | 1.54E-05 | 2.23E-05 |
| chr2 | 201811826 | 201812325 | 500 | -5.06 | 1.54E-05 | 2.23E-05 |
| chr4 | 2857823 | 2858456 | 634 | -5.06 | 1.54E-05 | 2.23E-05 |
| chr5 | 110409733 | 110410586 | 854 | -5.06 | 1.54E-05 | 2.23E-05 |
| chr5 | 91544089 | 91546826 | 2738 | -4.45 | 1.54E-05 | 2.24E-05 |
| chr4 | 77208086 | 77222069 | 13984 | -5.34 | 1.54E-05 | 2.24E-05 |
| chr3 | 107718943 | 107719888 | 946 | -5.53 | 1.54E-05 | 2.24E-05 |
| chr12 | 15461372 | 15483839 | 22468 | 2.08 | 1.54E-05 | 2.24E-05 |
| chr1 | 4423562 | 4455919 | 32358 | -5.64 | 1.55E-05 | 2.24E-05 |
| chr19 | 51109795 | 51110517 | 723 | -5.07 | 1.55E-05 | 2.24E-05 |
| chr12 | 117211450 | 117211970 | 521 | -5.14 | 1.55E-05 | 2.24E-05 |
| chr4 | 48756179 | 48756805 | 627 | -5.14 | 1.55E-05 | 2.24E-05 |
| chr8 | 6463809 | 6464720 | 912 | -5.14 | 1.55E-05 | 2.24E-05 |
| chr18 | 54212492 | 54212913 | 422 | -5.05 | 1.55E-05 | 2.25E-05 |
| chr2 | 87314874 | 87315258 | 385 | -5.22 | 1.55E-05 | 2.25E-05 |
| chr2 | 218457438 | 218468662 | 11225 | 1.55 | 1.55E-05 | 2.25E-05 |
| chr2 | 180767802 | 180768760 | 959 | -4.03 | 1.56E-05 | 2.26E-05 |
| chr4 | 100097794 | 100120646 | 22853 | -2.71 | 1.56E-05 | 2.26E-05 |
| chr12 | 88719989 | 88721625 | 1637 | -5.48 | 1.56E-05 | 2.26E-05 |
| chr5 | 157011107 | 157012228 | 1122 | -4.01 | 1.56E-05 | 2.27E-05 |
| chr2 | 188258658 | 188259136 | 479 | -5.09 | 1.56E-05 | 2.27E-05 |
| chr7 | 47247932 | 47253484 | 5553 | -2.26 | 1.57E-05 | 2.27E-05 |
| chrX | 123433003 | 123433524 | 522 | -3.99 | 1.57E-05 | 2.27E-05 |
| chr5 | 17492798 | 17493493 | 696 | -5.31 | 1.57E-05 | 2.28E-05 |
| chr16 | 66522373 | 66523259 | 887 | -5.2 | 1.57E-05 | 2.28E-05 |
| chr2 | 120462510 | 120463493 | 984 | -5.2 | 1.57E-05 | 2.28E-05 |
| chr7 | 87897338 | 87914637 | 17300 | 2.21 | 1.58E-05 | 2.29E-05 |
| chr15 | 90358795 | 90359430 | 636 | -5.06 | 1.58E-05 | 2.29E-05 |
| chr22 | 28000466 | 28001297 | 832 | -5.06 | 1.58E-05 | 2.29E-05 |
| chr2 | 75839292 | 75840153 | 862 | 5.18 | 1.58E-05 | 2.29E-05 |
| chr1 | 66565023 | 66565952 | 930 | -5.14 | 1.58E-05 | 2.30E-05 |
| chr6 | 147072797 | 147073568 | 772 | -5.14 | 1.58E-05 | 2.30E-05 |
| chr1 | 62360977 | 62361660 | 684 | -5.08 | 1.58E-05 | 2.30E-05 |
| chr8 | 107380410 | 107381127 | 718 | -5.08 | 1.58E-05 | 2.30E-05 |
| chr4 | 7134579 | 7135219 | 641 | -5.12 | 1.59E-05 | 2.30E-05 |
| chr5 | 149245803 | 149246470 | 668 | -5.12 | 1.59E-05 | 2.30E-05 |
| chr6 | 36069241 | 36069746 | 506 | -5.12 | 1.59E-05 | 2.30E-05 |
| chr6 | 142145106 | 142145951 | 846 | -5.29 | 1.59E-05 | 2.30E-05 |
| chr4 | 175719668 | 175738340 | 18673 | -1.83 | 1.59E-05 | 2.30E-05 |
| chr2 | 131508512 | 131510536 | 2025 | -2.41 | 1.59E-05 | 2.31E-05 |
| chrX | 43388194 | 43394834 | 6641 | -1.45 | 1.60E-05 | 2.31E-05 |
| chr7 | 133328933 | 133329703 | 771 | -5.38 | 1.60E-05 | 2.32E-05 |
| chr3 | 142221117 | 142221520 | 404 | -5.05 | 1.60E-05 | 2.32E-05 |
| chr3 | 151954647 | 151955383 | 737 | -5.05 | 1.60E-05 | 2.32E-05 |
| chrX | 67173524 | 67180822 | 7299 | 2.19 | 1.60E-05 | 2.32E-05 |
| chr5 | 93561059 | 93586516 | 25458 | 1.94 | 1.60E-05 | 2.32E-05 |
| chr11 | 128007177 | 128010155 | 2979 | -1.64 | 1.60E-05 | 2.32E-05 |
| chr4 | 94767824 | 94779858 | 12035 | 1.48 | 1.60E-05 | 2.32E-05 |
| chrX | 5023212 | 5025171 | 1960 | -4.12 | 1.61E-05 | 2.33E-05 |
| chr12 | 88441188 | 88441809 | 622 | -5.26 | 1.61E-05 | 2.33E-05 |
| chr4 | 30455750 | 30457601 | 1852 | -5.08 | 1.61E-05 | 2.33E-05 |
| chr1 | 190540090 | 190647652 | 107563 | 1.36 | 1.61E-05 | 2.33E-05 |
| chr4 | 53276225 | 53279448 | 3224 | -1.47 | 1.61E-05 | 2.34E-05 |
| chr11 | 108227315 | 108227724 | 410 | -4.41 | 1.61E-05 | 2.34E-05 |
| chr3 | 40663931 | 40672408 | 8478 | -2.91 | 1.62E-05 | 2.34E-05 |

| | | | | | | |
|---|---|---|---|---|---|---|
| chr7 | 144360598 | 144362125 | 1528 | -4.11 | 1.62E-05 | 2.34E-05 |
| chr2 | 34736764 | 34803183 | 66420 | 0.95 | 1.62E-05 | 2.34E-05 |
| chrX | 134105158 | 134105812 | 655 | -4.13 | 1.62E-05 | 2.34E-05 |
| chr1 | 171724546 | 171727043 | 2498 | -2.26 | 1.62E-05 | 2.34E-05 |
| chr4 | 25342620 | 25343359 | 740 | -5.34 | 1.62E-05 | 2.35E-05 |
| chr6 | 13266475 | 13267381 | 907 | -2.72 | 1.62E-05 | 2.35E-05 |
| chr10 | 60705782 | 60706430 | 649 | -5.04 | 1.62E-05 | 2.35E-05 |
| chr5 | 108246374 | 108246823 | 450 | -5.04 | 1.62E-05 | 2.35E-05 |
| chr6 | 101264175 | 101264713 | 539 | -5.04 | 1.62E-05 | 2.35E-05 |
| chr4 | 186905117 | 186905911 | 795 | -5.33 | 1.63E-05 | 2.36E-05 |
| chr6 | 144462018 | 144462731 | 714 | -5.33 | 1.63E-05 | 2.36E-05 |
| chr2 | 109973707 | 109978094 | 4388 | -3.4 | 1.63E-05 | 2.36E-05 |
| chr13 | 28821528 | 28822196 | 669 | -5.04 | 1.63E-05 | 2.36E-05 |
| chr7 | 137611140 | 137611524 | 385 | -5.04 | 1.63E-05 | 2.36E-05 |
| chr4 | 87640032 | 87640564 | 533 | -5.24 | 1.63E-05 | 2.36E-05 |
| chr16 | 21699389 | 21712858 | 13470 | -5.77 | 1.64E-05 | 2.37E-05 |
| chr12 | 11463398 | 11504550 | 41153 | 2.03 | 1.64E-05 | 2.37E-05 |
| chr15 | 34805629 | 34806465 | 837 | -4.07 | 1.64E-05 | 2.37E-05 |
| chr6 | 114741514 | 114742574 | 1061 | 5.6 | 1.64E-05 | 2.37E-05 |
| chr17 | 41776517 | 41777001 | 485 | -5.31 | 1.64E-05 | 2.37E-05 |
| chr10 | 120036573 | 120040341 | 3769 | -4.84 | 1.64E-05 | 2.37E-05 |
| chr4 | 31524670 | 31526212 | 1543 | -4.09 | 1.64E-05 | 2.38E-05 |
| chr4 | 24358773 | 24366875 | 8103 | 4.83 | 1.65E-05 | 2.38E-05 |
| chr12 | 70204436 | 70205057 | 622 | -5.23 | 1.65E-05 | 2.38E-05 |
| chr2 | 239291827 | 239292471 | 645 | -5.16 | 1.65E-05 | 2.38E-05 |
| chr3 | 176132143 | 176133786 | 1644 | -5.07 | 1.65E-05 | 2.39E-05 |
| chr15 | 61505648 | 61507529 | 1882 | -2.27 | 1.65E-05 | 2.39E-05 |
| chr3 | 138927236 | 138932589 | 5354 | -5.12 | 1.66E-05 | 2.39E-05 |
| chr20 | 14250810 | 14251686 | 877 | -2.65 | 1.66E-05 | 2.40E-05 |
| chr11 | 9164643 | 9166469 | 1827 | -6.02 | 1.66E-05 | 2.40E-05 |
| chr8 | 793430 | 831861 | 38432 | -1.72 | 1.66E-05 | 2.40E-05 |
| chr3 | 142384622 | 142385800 | 1179 | -4.33 | 1.66E-05 | 2.40E-05 |
| chr18 | 48186592 | 48200571 | 13980 | -5.03 | 1.66E-05 | 2.40E-05 |
| chr1 | 3834360 | 3835719 | 1360 | -4.14 | 1.66E-05 | 2.40E-05 |
| chr1 | 40336893 | 40337524 | 632 | -5.07 | 1.66E-05 | 2.40E-05 |
| chr3 | 94461144 | 94461764 | 621 | -5.07 | 1.66E-05 | 2.40E-05 |
| chr6 | 168042717 | 168043636 | 920 | -2.95 | 1.66E-05 | 2.40E-05 |
| chr2 | 190444108 | 190444690 | 583 | -5.05 | 1.67E-05 | 2.41E-05 |
| chr10 | 31336822 | 31341999 | 5178 | -5.7 | 1.67E-05 | 2.41E-05 |
| chr6 | 44604996 | 44605854 | 859 | -2.7 | 1.67E-05 | 2.41E-05 |
| chr6 | 6667154 | 6674396 | 7243 | -1.57 | 1.67E-05 | 2.41E-05 |
| chr3 | 85107751 | 85110956 | 3206 | -2.79 | 1.67E-05 | 2.41E-05 |
| chr11 | 68112670 | 68129451 | 16782 | -6 | 1.67E-05 | 2.42E-05 |
| chr3 | 157128920 | 157130459 | 1540 | -5.68 | 1.67E-05 | 2.42E-05 |
| chr3 | 109658799 | 109659601 | 803 | -5.27 | 1.67E-05 | 2.42E-05 |
| chr6 | 55485613 | 55486235 | 623 | -5.27 | 1.67E-05 | 2.42E-05 |
| chr12 | 92844441 | 92845961 | 1521 | -1.7 | 1.68E-05 | 2.42E-05 |
| chr11 | 96001828 | 96002803 | 976 | -5.2 | 1.68E-05 | 2.43E-05 |
| chr17 | 55600451 | 55601672 | 1222 | -5.59 | 1.68E-05 | 2.43E-05 |
| chr4 | 2370553 | 2371155 | 603 | -5.14 | 1.68E-05 | 2.43E-05 |
| chr14 | 78386656 | 78387156 | 501 | -5.09 | 1.68E-05 | 2.43E-05 |
| chr21 | 40216297 | 40217693 | 1397 | -5.09 | 1.68E-05 | 2.43E-05 |
| chr3 | 111568595 | 111569553 | 959 | -5.09 | 1.68E-05 | 2.43E-05 |
| chr10 | 79979230 | 79980153 | 924 | -5.03 | 1.68E-05 | 2.43E-05 |
| chr16 | 53374620 | 53375898 | 1279 | -5.03 | 1.68E-05 | 2.43E-05 |
| chr5 | 79692964 | 79693789 | 826 | -5.03 | 1.68E-05 | 2.43E-05 |
| chr13 | 43340718 | 43363985 | 23268 | -1.21 | 1.69E-05 | 2.44E-05 |
| chrX | 105751165 | 105752949 | 1785 | 5.23 | 1.69E-05 | 2.44E-05 |
| chr2 | 19247092 | 19254057 | 6966 | -5.64 | 1.69E-05 | 2.44E-05 |
| chr3 | 15926937 | 15938899 | 11963 | -5.43 | 1.69E-05 | 2.44E-05 |
| chr4 | 48850230 | 48850963 | 734 | -5.56 | 1.69E-05 | 2.44E-05 |

| | | | | | | |
|---|---|---|---|---|---|---|
| chr4 | 22204411 | 22216968 | 12558 | 2.4 | 1.70E-05 | 2.45E-05 |
| chr7 | 138692995 | 138694633 | 1639 | -3.98 | 1.70E-05 | 2.45E-05 |
| chr13 | 41265043 | 41265663 | 621 | -4.44 | 1.70E-05 | 2.45E-05 |
| chr10 | 71457733 | 71481618 | 23886 | -5.57 | 1.70E-05 | 2.46E-05 |
| chr1 | 171237205 | 171238690 | 1486 | -3.31 | 1.70E-05 | 2.46E-05 |
| chr3 | 191631638 | 191632296 | 659 | -4.35 | 1.70E-05 | 2.46E-05 |
| chr4 | 11557991 | 11559163 | 1173 | -5.24 | 1.70E-05 | 2.46E-05 |
| chr7 | 73055377 | 73056466 | 1090 | -5.58 | 1.71E-05 | 2.47E-05 |
| chr12 | 91728776 | 91730884 | 2109 | -4.11 | 1.72E-05 | 2.48E-05 |
| chr11 | 2730968 | 2733238 | 2271 | -4.74 | 1.72E-05 | 2.48E-05 |
| chr1 | 230141866 | 230143736 | 1871 | -5.05 | 1.72E-05 | 2.48E-05 |
| chr10 | 75836934 | 75837363 | 430 | -5.05 | 1.72E-05 | 2.48E-05 |
| chr13 | 100798681 | 100799440 | 760 | -5.05 | 1.72E-05 | 2.49E-05 |
| chr2 | 242722014 | 242722766 | 753 | -5.05 | 1.72E-05 | 2.49E-05 |
| chr12 | 42183307 | 42243086 | 59780 | -1.51 | 1.72E-05 | 2.49E-05 |
| chr1 | 187029932 | 187044250 | 14319 | -6.3 | 1.73E-05 | 2.50E-05 |
| chr16 | 55003796 | 55004668 | 873 | -4.06 | 1.73E-05 | 2.50E-05 |
| chr4 | 100659624 | 100661824 | 2201 | -4.25 | 1.73E-05 | 2.50E-05 |
| chrX | 75453714 | 75458994 | 5281 | -1.83 | 1.74E-05 | 2.51E-05 |
| chr17 | 67228877 | 67230041 | 1165 | -5.52 | 1.74E-05 | 2.51E-05 |
| chr4 | 186301465 | 186302314 | 850 | -5.52 | 1.74E-05 | 2.51E-05 |
| chr22 | 28847090 | 28847764 | 675 | -5.03 | 1.74E-05 | 2.51E-05 |
| chr4 | 137440888 | 137441579 | 692 | -5.03 | 1.74E-05 | 2.51E-05 |
| chr20 | 37019207 | 37019836 | 630 | -5.07 | 1.74E-05 | 2.51E-05 |
| chr2 | 105460226 | 105461237 | 1012 | -5.21 | 1.74E-05 | 2.51E-05 |
| chr5 | 115476714 | 115477324 | 611 | -5.21 | 1.74E-05 | 2.51E-05 |
| chr5 | 73748487 | 73749081 | 595 | -5.02 | 1.75E-05 | 2.52E-05 |
| chrX | 44262655 | 44264236 | 1582 | -5.43 | 1.75E-05 | 2.52E-05 |
| chrX | 111114524 | 111115954 | 1431 | -3.98 | 1.75E-05 | 2.52E-05 |
| chr3 | 70640246 | 70650773 | 10528 | 1.86 | 1.75E-05 | 2.52E-05 |
| chr4 | 133769433 | 133772304 | 2872 | 6.01 | 1.75E-05 | 2.53E-05 |
| chr14 | 89701815 | 89702497 | 683 | -4.42 | 1.75E-05 | 2.53E-05 |
| chr1 | 101627084 | 101627525 | 442 | -5.03 | 1.75E-05 | 2.53E-05 |
| chr5 | 92986283 | 92987041 | 759 | -5.03 | 1.75E-05 | 2.53E-05 |
| chr3 | 4390160 | 4390641 | 482 | -5.11 | 1.76E-05 | 2.54E-05 |
| chr6 | 20182116 | 20182823 | 708 | -5.11 | 1.76E-05 | 2.54E-05 |
| chrX | 13214240 | 13214739 | 500 | -4.36 | 1.76E-05 | 2.54E-05 |
| chr10 | 12481454 | 12482460 | 1007 | -5.15 | 1.76E-05 | 2.54E-05 |
| chr6 | 146361353 | 146362035 | 683 | -5.15 | 1.76E-05 | 2.54E-05 |
| chr1 | 88610187 | 88611222 | 1036 | -5.03 | 1.76E-05 | 2.54E-05 |
| chr4 | 90756728 | 90757502 | 775 | -5.03 | 1.76E-05 | 2.54E-05 |
| chr9 | 86125449 | 86126086 | 638 | -5.03 | 1.76E-05 | 2.54E-05 |
| chr5 | 93782146 | 93782611 | 466 | 5.21 | 1.76E-05 | 2.54E-05 |
| chr11 | 116157222 | 116174958 | 17737 | 2.96 | 1.76E-05 | 2.54E-05 |
| chr10 | 6119142 | 6125115 | 5974 | -2.38 | 1.76E-05 | 2.54E-05 |
| chr1 | 182274739 | 182275444 | 706 | -5.08 | 1.76E-05 | 2.54E-05 |
| chr5 | 65313620 | 65314220 | 601 | -5.39 | 1.77E-05 | 2.55E-05 |
| chr1 | 190656903 | 190697074 | 40172 | 1.76 | 1.77E-05 | 2.55E-05 |
| chr13 | 49178876 | 49196278 | 17403 | -2.37 | 1.78E-05 | 2.56E-05 |
| chr13 | 49162316 | 49163002 | 687 | -5.02 | 1.78E-05 | 2.56E-05 |
| chr13 | 96441615 | 96442062 | 448 | -5.02 | 1.78E-05 | 2.56E-05 |
| chr8 | 142640270 | 142640949 | 680 | -5.02 | 1.78E-05 | 2.56E-05 |
| chrX | 9783667 | 9789387 | 5721 | -4.75 | 1.78E-05 | 2.56E-05 |
| chr6 | 77798159 | 77799655 | 1497 | -5.41 | 1.78E-05 | 2.57E-05 |
| chr8 | 90483114 | 90484020 | 907 | -5.41 | 1.78E-05 | 2.57E-05 |
| chr17 | 38161027 | 38162048 | 1022 | -5.14 | 1.78E-05 | 2.57E-05 |
| chr5 | 73361219 | 73361737 | 519 | -5.04 | 1.78E-05 | 2.57E-05 |
| chr12 | 50211101 | 50211702 | 602 | -5.18 | 1.79E-05 | 2.57E-05 |
| chr8 | 134024915 | 134028681 | 3767 | -4.38 | 1.79E-05 | 2.58E-05 |
| chr7 | 93847861 | 93848556 | 696 | -5.4 | 1.79E-05 | 2.58E-05 |
| chr10 | 134181204 | 134195988 | 14785 | -6.07 | 1.80E-05 | 2.59E-05 |

| | | | | | | |
|---|---|---|---|---|---|---|
| chr18 | 28597504 | 28598217 | 714 | -5.33 | 1.80E-05 | 2.59E-05 |
| chr18 | 73642657 | 73670118 | 27462 | -1.45 | 1.80E-05 | 2.59E-05 |
| chr8 | 62159626 | 62169460 | 9835 | -5.03 | 1.80E-05 | 2.60E-05 |
| chr6 | 168690088 | 168690800 | 713 | -5.37 | 1.80E-05 | 2.60E-05 |
| chr17 | 47156670 | 47169298 | 12629 | -6.37 | 1.80E-05 | 2.60E-05 |
| chr3 | 72397997 | 72398952 | 956 | -5.09 | 1.80E-05 | 2.60E-05 |
| chr6 | 38493736 | 38494472 | 737 | -5.09 | 1.80E-05 | 2.60E-05 |
| chr1 | 47942341 | 47943169 | 829 | -5.05 | 1.81E-05 | 2.60E-05 |
| chr17 | 75078707 | 75079544 | 838 | -5.05 | 1.81E-05 | 2.60E-05 |
| chr22 | 28215992 | 28216550 | 559 | -5.05 | 1.81E-05 | 2.60E-05 |
| chr22 | 28553271 | 28553961 | 691 | -5.03 | 1.81E-05 | 2.60E-05 |
| chr8 | 21599677 | 21600541 | 865 | -5.03 | 1.81E-05 | 2.60E-05 |
| chr5 | 140387220 | 140388581 | 1362 | -4.06 | 1.81E-05 | 2.61E-05 |
| chr6 | 88796384 | 88797456 | 1073 | 5.1 | 1.81E-05 | 2.61E-05 |
| chrX | 151106328 | 151107033 | 706 | -5.16 | 1.81E-05 | 2.61E-05 |
| chr1 | 64697064 | 64697763 | 700 | -5.13 | 1.81E-05 | 2.61E-05 |
| chr1 | 98188951 | 98189690 | 740 | -5.06 | 1.81E-05 | 2.61E-05 |
| chr11 | 1722799 | 1728058 | 5260 | -5.06 | 1.81E-05 | 2.61E-05 |
| chr12 | 10219006 | 10224276 | 5271 | 2.44 | 1.82E-05 | 2.62E-05 |
| chr1 | 86584730 | 86585644 | 915 | -2.87 | 1.82E-05 | 2.62E-05 |
| chr8 | 9688760 | 9748352 | 59593 | -1.81 | 1.82E-05 | 2.62E-05 |
| chr5 | 71970535 | 71988293 | 17759 | -3.65 | 1.83E-05 | 2.63E-05 |
| chr5 | 59606629 | 59607269 | 641 | 5.12 | 1.83E-05 | 2.63E-05 |
| chr15 | 27024664 | 27109571 | 84908 | -1.2 | 1.83E-05 | 2.63E-05 |
| chr1 | 196971972 | 196981356 | 9385 | -6.35 | 1.83E-05 | 2.64E-05 |
| chr6 | 141992266 | 141993290 | 1025 | -3.98 | 1.84E-05 | 2.64E-05 |
| chr3 | 53938773 | 54025962 | 87190 | -5.98 | 1.84E-05 | 2.64E-05 |
| chr4 | 23591811 | 23595412 | 3602 | -1.96 | 1.84E-05 | 2.64E-05 |
| chr2 | 213826354 | 213826996 | 643 | 5.36 | 1.84E-05 | 2.65E-05 |
| chr1 | 169471697 | 169472210 | 514 | -5.14 | 1.84E-05 | 2.65E-05 |
| chr19 | 1564611 | 1565498 | 888 | -5.14 | 1.84E-05 | 2.65E-05 |
| chr3 | 100045871 | 100046493 | 623 | -5.01 | 1.84E-05 | 2.65E-05 |
| chr7 | 111833857 | 111834398 | 542 | -5.01 | 1.84E-05 | 2.65E-05 |
| chr8 | 117147287 | 117147916 | 630 | -5.01 | 1.84E-05 | 2.65E-05 |
| chr2 | 196435676 | 196436820 | 1145 | -5.11 | 1.84E-05 | 2.65E-05 |
| chr2 | 226592103 | 226603668 | 11566 | -5.11 | 1.84E-05 | 2.65E-05 |
| chrX | 42115605 | 42118272 | 2668 | -2.06 | 1.84E-05 | 2.65E-05 |
| chr9 | 7103523 | 7104585 | 1063 | -5.77 | 1.84E-05 | 2.65E-05 |
| chr4 | 131778839 | 131780662 | 1824 | -5.3 | 1.85E-05 | 2.66E-05 |
| chr8 | 101682530 | 101683370 | 841 | -5.3 | 1.85E-05 | 2.66E-05 |
| chr12 | 117684322 | 117689355 | 5034 | -4.69 | 1.85E-05 | 2.67E-05 |
| chr21 | 17835151 | 17836327 | 1177 | -5.29 | 1.86E-05 | 2.68E-05 |
| chr10 | 65212261 | 65212707 | 447 | -5.05 | 1.87E-05 | 2.68E-05 |
| chr12 | 78467469 | 78467927 | 459 | -5.05 | 1.87E-05 | 2.68E-05 |
| chr4 | 180132581 | 180134011 | 1431 | -5.05 | 1.87E-05 | 2.68E-05 |
| chr12 | 33108591 | 33109179 | 589 | -5.01 | 1.87E-05 | 2.69E-05 |
| chr3 | 171288005 | 171288494 | 490 | -5.01 | 1.87E-05 | 2.69E-05 |
| chr6 | 90726358 | 90727284 | 927 | -3.92 | 1.87E-05 | 2.69E-05 |
| chr1 | 79905988 | 79907250 | 1263 | -5.13 | 1.87E-05 | 2.70E-05 |
| chr11 | 103883920 | 103884558 | 639 | -5.1 | 1.88E-05 | 2.70E-05 |
| chr1 | 38404025 | 38404390 | 366 | -5.24 | 1.88E-05 | 2.70E-05 |
| chr3 | 121792052 | 121799952 | 7901 | 1.92 | 1.88E-05 | 2.70E-05 |
| chr16 | 10697249 | 10698275 | 1027 | -5.04 | 1.88E-05 | 2.71E-05 |
| chr7 | 39454737 | 39455493 | 757 | -5.04 | 1.88E-05 | 2.71E-05 |
| chr3 | 30053308 | 30054185 | 878 | -5.26 | 1.89E-05 | 2.71E-05 |
| chr4 | 101829347 | 101830219 | 873 | -5.65 | 1.89E-05 | 2.71E-05 |
| chr3 | 141533949 | 141535514 | 1566 | -4.65 | 1.89E-05 | 2.72E-05 |
| chr4 | 1332841 | 1333648 | 808 | -4.38 | 1.89E-05 | 2.72E-05 |
| chr12 | 54871400 | 54872012 | 613 | 5.16 | 1.89E-05 | 2.72E-05 |
| chr5 | 54076344 | 54122575 | 46232 | 1.25 | 1.90E-05 | 2.73E-05 |
| chr21 | 16624613 | 16636313 | 11701 | -5.63 | 1.90E-05 | 2.73E-05 |

| | | | | | | |
|---|---|---|---|---|---|---|
| chr21 | 17432628 | 17433821 | 1194 | -5.63 | 1.90E-05 | 2.73E-05 |
| chr10 | 81219899 | 81220648 | 750 | -5.24 | 1.90E-05 | 2.73E-05 |
| chr19 | 8330185 | 8330979 | 795 | -5.24 | 1.90E-05 | 2.73E-05 |
| chr10 | 114479718 | 114480252 | 535 | -5.01 | 1.90E-05 | 2.74E-05 |
| chr2 | 231885207 | 231886025 | 819 | -5.01 | 1.90E-05 | 2.74E-05 |
| chr6 | 135623301 | 135623840 | 540 | -5.01 | 1.90E-05 | 2.74E-05 |
| chr21 | 40118469 | 40119254 | 786 | -5.21 | 1.91E-05 | 2.74E-05 |
| chr17 | 48489392 | 48490044 | 653 | -5.11 | 1.91E-05 | 2.75E-05 |
| chr4 | 155477203 | 155487131 | 9929 | -5.93 | 1.91E-05 | 2.75E-05 |
| chr12 | 79762576 | 79763278 | 703 | -5.09 | 1.91E-05 | 2.75E-05 |
| chr3 | 100223532 | 100244077 | 20546 | -1.53 | 1.92E-05 | 2.75E-05 |
| chr4 | 97375555 | 97376363 | 809 | -3.82 | 1.93E-05 | 2.77E-05 |
| chr3 | 46054590 | 46057419 | 2830 | 1.9 | 1.93E-05 | 2.77E-05 |
| chr11 | 133977403 | 133980018 | 2616 | -4.63 | 1.93E-05 | 2.77E-05 |
| chr5 | 89665222 | 89674987 | 9766 | -2.49 | 1.93E-05 | 2.77E-05 |
| chr20 | 9630470 | 9633326 | 2857 | 2.23 | 1.93E-05 | 2.78E-05 |
| chr10 | 112402387 | 112403271 | 885 | -5.21 | 1.94E-05 | 2.78E-05 |
| chr15 | 44531543 | 44532475 | 933 | -5.21 | 1.94E-05 | 2.78E-05 |
| chr6 | 136623803 | 136624713 | 911 | -5.21 | 1.94E-05 | 2.78E-05 |
| chr6 | 163991751 | 163992361 | 611 | -5 | 1.94E-05 | 2.78E-05 |
| chr1 | 11461563 | 11475976 | 14414 | -5.33 | 1.94E-05 | 2.79E-05 |
| chr10 | 119729015 | 119729718 | 704 | -2.7 | 1.95E-05 | 2.80E-05 |
| chr12 | 108528100 | 108528769 | 670 | -4.66 | 1.95E-05 | 2.80E-05 |
| chr18 | 13084528 | 13085228 | 701 | -4.66 | 1.95E-05 | 2.80E-05 |
| chr12 | 97754196 | 97766971 | 12776 | 5.37 | 1.95E-05 | 2.80E-05 |
| chr13 | 97606794 | 97644452 | 37659 | -2.33 | 1.95E-05 | 2.80E-05 |
| chr10 | 12856414 | 12857122 | 709 | -5.09 | 1.95E-05 | 2.81E-05 |
| chr9 | 668918 | 669777 | 860 | -5.09 | 1.95E-05 | 2.81E-05 |
| chr4 | 139275178 | 139275988 | 811 | -5.19 | 1.96E-05 | 2.81E-05 |
| chr4 | 56019225 | 56020288 | 1064 | -3.8 | 1.96E-05 | 2.81E-05 |
| chr18 | 33471344 | 33477731 | 6388 | -2.95 | 1.96E-05 | 2.82E-05 |
| chr15 | 62484502 | 62531751 | 47250 | -2.08 | 1.97E-05 | 2.82E-05 |
| chr22 | 25504549 | 25505073 | 525 | -5.02 | 1.97E-05 | 2.83E-05 |
| chr5 | 76293602 | 76294339 | 738 | -5.02 | 1.97E-05 | 2.83E-05 |
| chr6 | 37040946 | 37041723 | 778 | -5.02 | 1.97E-05 | 2.83E-05 |
| chr3 | 110419497 | 110424673 | 5177 | -4.77 | 1.97E-05 | 2.83E-05 |
| chr13 | 22378591 | 22382944 | 4354 | 1.7 | 1.97E-05 | 2.83E-05 |
| chr20 | 15359581 | 15423479 | 63899 | -1.86 | 1.97E-05 | 2.83E-05 |
| chr8 | 16814164 | 16828793 | 14630 | -1.08 | 1.97E-05 | 2.83E-05 |
| chr1 | 217830703 | 217831866 | 1164 | -5.52 | 1.98E-05 | 2.84E-05 |
| chr3 | 133158539 | 133159363 | 825 | -5.18 | 1.98E-05 | 2.84E-05 |
| chr4 | 66644092 | 66645125 | 1034 | -5.16 | 1.98E-05 | 2.84E-05 |
| chr16 | 89696159 | 89696819 | 661 | -5.04 | 1.98E-05 | 2.85E-05 |
| chr4 | 137205028 | 137205771 | 744 | -5.04 | 1.98E-05 | 2.85E-05 |
| chr12 | 25582340 | 25584080 | 1741 | -5.46 | 1.99E-05 | 2.85E-05 |
| chr12 | 17020317 | 17021971 | 1655 | 2.77 | 1.99E-05 | 2.85E-05 |
| chr4 | 137374778 | 137375576 | 799 | -5.44 | 1.99E-05 | 2.86E-05 |
| chr17 | 897645 | 898183 | 539 | -5.02 | 1.99E-05 | 2.86E-05 |
| chr18 | 56470701 | 56471349 | 649 | -5.02 | 1.99E-05 | 2.86E-05 |
| chr6 | 133628608 | 133629267 | 660 | -5.02 | 1.99E-05 | 2.86E-05 |
| chr10 | 89869255 | 89869973 | 719 | -4.99 | 1.99E-05 | 2.86E-05 |
| chr2 | 164236041 | 164236518 | 478 | -4.99 | 1.99E-05 | 2.86E-05 |
| chr4 | 155827322 | 155827912 | 591 | -4.99 | 1.99E-05 | 2.86E-05 |
| chr6 | 53736817 | 53737912 | 1096 | -5.43 | 2.00E-05 | 2.87E-05 |
| chr13 | 24570476 | 24571526 | 1051 | -5.08 | 2.00E-05 | 2.87E-05 |
| chr17 | 78496769 | 78497423 | 655 | -5.08 | 2.00E-05 | 2.87E-05 |
| chr12 | 74332231 | 74332923 | 693 | -5.16 | 2.00E-05 | 2.87E-05 |
| chr9 | 16882464 | 16883108 | 645 | -5.16 | 2.00E-05 | 2.87E-05 |
| chr5 | 61239785 | 61241464 | 1680 | 4.56 | 2.00E-05 | 2.87E-05 |
| chr1 | 197602139 | 197602914 | 776 | -4.99 | 2.00E-05 | 2.87E-05 |
| chr18 | 57579948 | 57580864 | 917 | -4.99 | 2.00E-05 | 2.87E-05 |

| | | | | | | |
|---|---|---|---|---|---|---|
| chr17 | 8744541 | 8775288 | 30748 | -7.16 | 2.01E-05 | 2.88E-05 |
| chr5 | 102369480 | 102371015 | 1536 | -3.8 | 2.01E-05 | 2.88E-05 |
| chr1 | 75220590 | 75223621 | 3032 | -4.65 | 2.01E-05 | 2.88E-05 |
| chr2 | 239139980 | 239141114 | 1135 | -4 | 2.01E-05 | 2.89E-05 |
| chr1 | 109927218 | 109927841 | 624 | -5 | 2.02E-05 | 2.89E-05 |
| chr1 | 234661252 | 234662028 | 777 | -5 | 2.02E-05 | 2.89E-05 |
| chr6 | 86261087 | 86261594 | 508 | -5 | 2.02E-05 | 2.89E-05 |
| chr11 | 76296981 | 76305969 | 8989 | -6.21 | 2.02E-05 | 2.89E-05 |
| chr13 | 63324543 | 63326405 | 1863 | -5.39 | 2.02E-05 | 2.90E-05 |
| chr7 | 107161297 | 107162059 | 763 | -5.39 | 2.02E-05 | 2.90E-05 |
| chr1 | 11941998 | 11942578 | 581 | -5.14 | 2.03E-05 | 2.90E-05 |
| chr5 | 102029975 | 102031918 | 1944 | 3.08 | 2.03E-05 | 2.90E-05 |
| chr4 | 5118375 | 5119333 | 959 | -5.37 | 2.03E-05 | 2.91E-05 |
| chr5 | 112513146 | 112513891 | 746 | 5.7 | 2.03E-05 | 2.91E-05 |
| chr22 | 37547157 | 37558012 | 10856 | -6.03 | 2.03E-05 | 2.91E-05 |
| chr13 | 49061728 | 49062467 | 740 | -5.15 | 2.03E-05 | 2.91E-05 |
| chr17 | 7685764 | 7686393 | 630 | -5.15 | 2.03E-05 | 2.91E-05 |
| chr2 | 47504983 | 47505629 | 647 | -5.15 | 2.03E-05 | 2.91E-05 |
| chr20 | 31659454 | 31660763 | 1310 | -5.15 | 2.03E-05 | 2.91E-05 |
| chr2 | 84726581 | 84750363 | 23783 | -5.51 | 2.03E-05 | 2.91E-05 |
| chr3 | 161614900 | 161617937 | 3038 | 2.37 | 2.04E-05 | 2.92E-05 |
| chr11 | 57185050 | 57185795 | 746 | -5.36 | 2.04E-05 | 2.92E-05 |
| chr17 | 72724213 | 72725197 | 985 | -5.36 | 2.04E-05 | 2.92E-05 |
| chr8 | 107970804 | 107971561 | 758 | -5.36 | 2.04E-05 | 2.92E-05 |
| chr13 | 114999427 | 114999994 | 568 | -5.35 | 2.04E-05 | 2.93E-05 |
| chr3 | 38721565 | 38963826 | 242262 | 1.57 | 2.04E-05 | 2.93E-05 |
| chr4 | 77148667 | 77182901 | 34235 | -5.62 | 2.05E-05 | 2.93E-05 |
| chr2 | 218905416 | 218910632 | 5217 | -4.7 | 2.05E-05 | 2.93E-05 |
| chr11 | 8817174 | 8817676 | 503 | -4.99 | 2.05E-05 | 2.93E-05 |
| chr20 | 269721 | 270314 | 594 | -4.99 | 2.05E-05 | 2.93E-05 |
| chr15 | 69183442 | 69274969 | 91528 | -2.13 | 2.05E-05 | 2.93E-05 |
| chr11 | 101537167 | 101537800 | 634 | -5.05 | 2.05E-05 | 2.94E-05 |
| chr9 | 139995799 | 139996215 | 417 | -5.06 | 2.06E-05 | 2.95E-05 |
| chr3 | 164480948 | 164481655 | 708 | 4.08 | 2.06E-05 | 2.95E-05 |
| chr19 | 29745710 | 29890755 | 145046 | 0.59 | 2.06E-05 | 2.95E-05 |
| chr12 | 86178741 | 86179577 | 837 | -5.13 | 2.06E-05 | 2.95E-05 |
| chr11 | 33183919 | 33185006 | 1088 | -5.02 | 2.06E-05 | 2.95E-05 |
| chr11 | 119106857 | 119107253 | 397 | -5.02 | 2.06E-05 | 2.95E-05 |
| chr9 | 93730430 | 93731100 | 671 | -5.02 | 2.06E-05 | 2.95E-05 |
| chr7 | 106574242 | 106575472 | 1231 | 5.12 | 2.07E-05 | 2.96E-05 |
| chr10 | 134867637 | 134869130 | 1494 | -1.96 | 2.07E-05 | 2.96E-05 |
| chr13 | 60158776 | 60159667 | 892 | 5.07 | 2.07E-05 | 2.97E-05 |
| chr4 | 96850980 | 96852257 | 1278 | -5.3 | 2.07E-05 | 2.97E-05 |
| chr3 | 162652655 | 162665929 | 13275 | 6.72 | 2.08E-05 | 2.97E-05 |
| chr6 | 63020375 | 63053122 | 32748 | 1.44 | 2.08E-05 | 2.97E-05 |
| chr5 | 34065624 | 34066259 | 636 | -3.83 | 2.08E-05 | 2.97E-05 |
| chr7 | 129584677 | 129585158 | 482 | -5.34 | 2.08E-05 | 2.97E-05 |
| chr9 | 94026135 | 94026998 | 864 | -5.29 | 2.09E-05 | 2.99E-05 |
| chr10 | 97976471 | 97980672 | 4202 | 1.47 | 2.09E-05 | 2.99E-05 |
| chr2 | 131157278 | 131157662 | 385 | -4.06 | 2.09E-05 | 2.99E-05 |
| chr6 | 28008504 | 28016672 | 8169 | 1.21 | 2.10E-05 | 3.00E-05 |
| chr16 | 69545463 | 69547358 | 1896 | -5.27 | 2.10E-05 | 3.00E-05 |
| chr2 | 108624249 | 108631844 | 7596 | -3.9 | 2.10E-05 | 3.00E-05 |
| chr18 | 68580601 | 68586952 | 6352 | -3.14 | 2.10E-05 | 3.01E-05 |
| chrX | 82677832 | 82678767 | 936 | -4.16 | 2.10E-05 | 3.01E-05 |
| chr14 | 39796428 | 39796892 | 465 | -4.33 | 2.10E-05 | 3.01E-05 |
| chr1 | 4703375 | 4704106 | 732 | -5.03 | 2.11E-05 | 3.01E-05 |
| chr9 | 123710493 | 123712618 | 2126 | -3.42 | 2.11E-05 | 3.02E-05 |
| chr4 | 142034325 | 142038878 | 4554 | -3.35 | 2.11E-05 | 3.02E-05 |
| chr22 | 26806882 | 26821192 | 14311 | -3.08 | 2.11E-05 | 3.02E-05 |
| chr22 | 43253529 | 43254071 | 543 | -5.1 | 2.11E-05 | 3.02E-05 |

| | | | | | | |
|---|---|---|---|---|---|---|
| chr1 | 79202634 | 79205141 | 2508 | -2.02 | 2.11E-05 | 3.02E-05 |
| chr16 | 31725946 | 31961232 | 235287 | -8.9 | 2.11E-05 | 3.03E-05 |
| chr4 | 66915592 | 66916469 | 878 | -5.04 | 2.12E-05 | 3.03E-05 |
| chr12 | 27055549 | 27055870 | 322 | -4.98 | 2.12E-05 | 3.04E-05 |
| chr5 | 171607893 | 171608451 | 559 | -4.98 | 2.12E-05 | 3.04E-05 |
| chr2 | 213789194 | 213789923 | 730 | -4.44 | 2.13E-05 | 3.05E-05 |
| chr2 | 194631341 | 194632453 | 1113 | -5.1 | 2.14E-05 | 3.05E-05 |
| chr2 | 197685665 | 197686292 | 628 | -4.97 | 2.14E-05 | 3.06E-05 |
| chr7 | 3842665 | 3858822 | 16158 | -3.31 | 2.14E-05 | 3.06E-05 |
| chr2 | 197568815 | 197569323 | 509 | -4.98 | 2.14E-05 | 3.07E-05 |
| chr6 | 17331262 | 17331783 | 522 | -4.98 | 2.14E-05 | 3.07E-05 |
| chr7 | 106395056 | 106395932 | 877 | -4.98 | 2.14E-05 | 3.07E-05 |
| chr9 | 35318383 | 35318934 | 552 | -5.26 | 2.14E-05 | 3.07E-05 |
| chr7 | 32361761 | 32389651 | 27891 | -2.35 | 2.14E-05 | 3.07E-05 |
| chr3 | 72950900 | 72951435 | 536 | -4.99 | 2.15E-05 | 3.07E-05 |
| chr1 | 200467906 | 200469239 | 1334 | -5.59 | 2.15E-05 | 3.07E-05 |
| chr11 | 101681702 | 101682722 | 1021 | -5 | 2.15E-05 | 3.08E-05 |
| chr7 | 72930773 | 72932369 | 1597 | -5.25 | 2.16E-05 | 3.08E-05 |
| chr12 | 18664008 | 18697836 | 33829 | 1.07 | 2.16E-05 | 3.08E-05 |
| chr18 | 10768452 | 10778791 | 10340 | -1.27 | 2.17E-05 | 3.10E-05 |
| chr6 | 129866756 | 129867293 | 538 | -4.97 | 2.17E-05 | 3.10E-05 |
| chr21 | 18173500 | 18180673 | 7174 | -5.08 | 2.18E-05 | 3.11E-05 |
| chr4 | 64472204 | 64473378 | 1175 | -5.08 | 2.18E-05 | 3.12E-05 |
| chr21 | 18471185 | 18493811 | 22627 | 4.01 | 2.18E-05 | 3.12E-05 |
| chr5 | 125680783 | 125683100 | 2318 | 1.61 | 2.18E-05 | 3.12E-05 |
| chr2 | 200693447 | 200694073 | 627 | -4.98 | 2.18E-05 | 3.12E-05 |
| chrX | 154125283 | 154175479 | 50197 | -0.94 | 2.19E-05 | 3.13E-05 |
| chr6 | 166912455 | 166913107 | 653 | -5.07 | 2.19E-05 | 3.13E-05 |
| chr3 | 139421359 | 139422797 | 1439 | -2.87 | 2.19E-05 | 3.13E-05 |
| chr15 | 101957072 | 101957563 | 492 | -5.03 | 2.19E-05 | 3.13E-05 |
| chr4 | 68001470 | 68002735 | 1266 | 3.54 | 2.19E-05 | 3.13E-05 |
| chr1 | 167050064 | 167051348 | 1285 | -3.07 | 2.20E-05 | 3.14E-05 |
| chr1 | 231615797 | 231616249 | 453 | -5.18 | 2.20E-05 | 3.15E-05 |
| chr2 | 45416931 | 45430020 | 13090 | -5.18 | 2.20E-05 | 3.15E-05 |
| chr7 | 67648692 | 67660489 | 11798 | -5.47 | 2.22E-05 | 3.18E-05 |
| chr2 | 155192640 | 155193203 | 564 | 5.16 | 2.23E-05 | 3.18E-05 |
| chr4 | 3455132 | 3458313 | 3182 | -4.62 | 2.23E-05 | 3.18E-05 |
| chr3 | 46472431 | 46473092 | 662 | -5.06 | 2.23E-05 | 3.19E-05 |
| chr7 | 117742507 | 117743055 | 549 | -5.06 | 2.23E-05 | 3.19E-05 |
| chr15 | 35684136 | 35684953 | 818 | -4.16 | 2.24E-05 | 3.19E-05 |
| chr11 | 63439343 | 63439895 | 553 | -5 | 2.24E-05 | 3.19E-05 |
| chr17 | 4655541 | 4656199 | 659 | -5 | 2.24E-05 | 3.19E-05 |
| chr4 | 151428734 | 151429155 | 422 | -4.98 | 2.24E-05 | 3.20E-05 |
| chr12 | 100415652 | 100416912 | 1261 | -5.43 | 2.25E-05 | 3.21E-05 |
| chrX | 16938881 | 16939848 | 968 | -3.53 | 2.25E-05 | 3.21E-05 |
| chr2 | 37011477 | 37012079 | 603 | -5.15 | 2.25E-05 | 3.22E-05 |
| chr3 | 117362938 | 117363850 | 913 | -5.15 | 2.25E-05 | 3.22E-05 |
| chr5 | 133417889 | 133418268 | 380 | -3.86 | 2.25E-05 | 3.22E-05 |
| chr1 | 79591105 | 79591790 | 686 | -4.99 | 2.25E-05 | 3.22E-05 |
| chr12 | 107611813 | 107612989 | 1177 | -4.99 | 2.25E-05 | 3.22E-05 |
| chr13 | 99068891 | 99070031 | 1141 | -4.99 | 2.25E-05 | 3.22E-05 |
| chr2 | 120876132 | 120876687 | 556 | -4.99 | 2.25E-05 | 3.22E-05 |
| chr6 | 155381872 | 155382787 | 916 | -4.99 | 2.25E-05 | 3.22E-05 |
| chr4 | 85282376 | 85291976 | 9601 | -1.98 | 2.26E-05 | 3.22E-05 |
| chr5 | 168460150 | 168471740 | 11591 | -6.02 | 2.26E-05 | 3.22E-05 |
| chr4 | 73824677 | 73838786 | 14110 | -4.42 | 2.26E-05 | 3.23E-05 |
| chr18 | 8461384 | 8462227 | 844 | -5.01 | 2.27E-05 | 3.24E-05 |
| chr2 | 138602630 | 138605081 | 2452 | -2.35 | 2.27E-05 | 3.25E-05 |
| chr3 | 185683099 | 185683644 | 546 | -4.29 | 2.28E-05 | 3.25E-05 |
| chr16 | 54262289 | 54263090 | 802 | -5.04 | 2.28E-05 | 3.25E-05 |
| chr6 | 79784360 | 79785050 | 691 | -5.04 | 2.28E-05 | 3.25E-05 |

| | | | | | | |
|---|---|---|---|---|---|---|
| chr20 | 57298641 | 57360030 | 61390 | -0.98 | 2.28E-05 | 3.26E-05 |
| chr7 | 85189399 | 85195963 | 6565 | -5.76 | 2.28E-05 | 3.26E-05 |
| chr12 | 108641895 | 108643883 | 1989 | 3.15 | 2.28E-05 | 3.26E-05 |
| chr7 | 29150205 | 29151963 | 1759 | -1.91 | 2.28E-05 | 3.26E-05 |
| chr20 | 44726574 | 44734357 | 7784 | -5.43 | 2.28E-05 | 3.26E-05 |
| chr3 | 93960072 | 93961492 | 1421 | 2.38 | 2.29E-05 | 3.26E-05 |
| chr3 | 109352586 | 109353129 | 544 | -5.15 | 2.29E-05 | 3.27E-05 |
| chr2 | 146170955 | 146181445 | 10491 | -5.18 | 2.29E-05 | 3.27E-05 |
| chrX | 17507460 | 17507992 | 533 | -4.96 | 2.29E-05 | 3.27E-05 |
| chr16 | 71215274 | 71338386 | 123113 | -1.91 | 2.29E-05 | 3.27E-05 |
| chr4 | 52819332 | 52865215 | 45884 | -2.25 | 2.30E-05 | 3.28E-05 |
| chr5 | 91096969 | 91107314 | 10346 | -4.87 | 2.30E-05 | 3.28E-05 |
| chr12 | 98102848 | 98115753 | 12906 | 2.07 | 2.30E-05 | 3.29E-05 |
| chr14 | 35334245 | 35335057 | 813 | -5.5 | 2.30E-05 | 3.29E-05 |
| chr6 | 62178790 | 62370387 | 191598 | 0.86 | 2.31E-05 | 3.29E-05 |
| chr10 | 116552449 | 116553397 | 949 | -4.96 | 2.31E-05 | 3.29E-05 |
| chr20 | 50362166 | 50362711 | 546 | -4.96 | 2.31E-05 | 3.29E-05 |
| chr9 | 111208483 | 111209204 | 722 | -4.96 | 2.31E-05 | 3.29E-05 |
| chr12 | 102556995 | 102557389 | 395 | -4.99 | 2.31E-05 | 3.30E-05 |
| chr2 | 193180308 | 193181010 | 703 | -4.99 | 2.31E-05 | 3.30E-05 |
| chr3 | 31730874 | 31731578 | 705 | -4.99 | 2.31E-05 | 3.30E-05 |
| chr2 | 2881255 | 3042190 | 160936 | -3.77 | 2.31E-05 | 3.30E-05 |
| chr14 | 90934028 | 90948656 | 14629 | -2.55 | 2.33E-05 | 3.32E-05 |
| chr6 | 21718993 | 21719911 | 919 | -5.31 | 2.33E-05 | 3.33E-05 |
| chr14 | 30258111 | 30260852 | 2742 | 4.12 | 2.33E-05 | 3.33E-05 |
| chr2 | 41965126 | 41977031 | 11906 | -4.99 | 2.34E-05 | 3.33E-05 |
| chr14 | 29181013 | 29187054 | 6042 | -3.34 | 2.34E-05 | 3.34E-05 |
| chr13 | 84640090 | 84641293 | 1204 | -4.96 | 2.34E-05 | 3.34E-05 |
| chr15 | 45556028 | 45556677 | 650 | -4.96 | 2.34E-05 | 3.34E-05 |
| chr14 | 54410656 | 54411270 | 615 | -5.29 | 2.35E-05 | 3.34E-05 |
| chr3 | 27785340 | 27785787 | 448 | -5.1 | 2.35E-05 | 3.35E-05 |
| chr6 | 164175662 | 164220710 | 45049 | -0.84 | 2.36E-05 | 3.36E-05 |
| chr7 | 28053239 | 28064986 | 11748 | -5.41 | 2.36E-05 | 3.36E-05 |
| chr3 | 124242691 | 124258590 | 15900 | -0.99 | 2.36E-05 | 3.36E-05 |
| chr10 | 121540316 | 121540684 | 369 | -4.99 | 2.36E-05 | 3.36E-05 |
| chr14 | 89840662 | 89841358 | 697 | -4.99 | 2.36E-05 | 3.36E-05 |
| chr22 | 24588825 | 24589298 | 474 | -4.99 | 2.36E-05 | 3.36E-05 |
| chr22 | 36270692 | 36271298 | 607 | -4.99 | 2.36E-05 | 3.36E-05 |
| chr16 | 23223355 | 23226722 | 3368 | 4.68 | 2.36E-05 | 3.36E-05 |
| chr5 | 42896227 | 42898102 | 1876 | -2.72 | 2.36E-05 | 3.37E-05 |
| chr8 | 59462154 | 59462601 | 448 | -4.97 | 2.37E-05 | 3.38E-05 |
| chr8 | 103239812 | 103240516 | 705 | -4.97 | 2.37E-05 | 3.38E-05 |
| chr9 | 140404987 | 140405384 | 398 | -4.97 | 2.37E-05 | 3.38E-05 |
| chr1 | 215279815 | 215281775 | 1961 | -2.42 | 2.37E-05 | 3.38E-05 |
| chr17 | 75936572 | 75957558 | 20987 | -1.78 | 2.38E-05 | 3.39E-05 |
| chr6 | 66326269 | 66327176 | 908 | 5.07 | 2.39E-05 | 3.40E-05 |
| chr15 | 72676449 | 72676949 | 501 | -5.01 | 2.39E-05 | 3.40E-05 |
| chrX | 17621611 | 17622085 | 475 | -5.01 | 2.39E-05 | 3.40E-05 |
| chr2 | 162971457 | 162972412 | 956 | -5.37 | 2.39E-05 | 3.41E-05 |
| chr12 | 103130068 | 103130658 | 591 | -5.08 | 2.39E-05 | 3.41E-05 |
| chr16 | 47025321 | 47026098 | 778 | -5.08 | 2.39E-05 | 3.41E-05 |
| chr4 | 129318666 | 129319325 | 660 | -5.08 | 2.39E-05 | 3.41E-05 |
| chr7 | 114779613 | 114781174 | 1562 | -3.88 | 2.39E-05 | 3.41E-05 |
| chr5 | 157295825 | 157296839 | 1015 | -4.97 | 2.40E-05 | 3.42E-05 |
| chr6 | 10200577 | 10201676 | 1100 | -4.97 | 2.40E-05 | 3.42E-05 |
| chr2 | 216276629 | 216277953 | 1325 | -5.36 | 2.40E-05 | 3.42E-05 |
| chrX | 100059632 | 100070772 | 11141 | -2.04 | 2.40E-05 | 3.42E-05 |
| chr15 | 72702371 | 72736525 | 34155 | -3.85 | 2.40E-05 | 3.42E-05 |
| chrX | 56947752 | 56948289 | 538 | 5.04 | 2.40E-05 | 3.42E-05 |
| chr4 | 36003098 | 36004551 | 1454 | -5.23 | 2.40E-05 | 3.42E-05 |
| chr1 | 89080617 | 89104236 | 23620 | -3.72 | 2.40E-05 | 3.42E-05 |

| | | | | | | |
|---|---|---|---|---|---|---|
| chr12 | 101073629 | 101091258 | 17630 | -1.5 | 2.41E-05 | 3.43E-05 |
| chr1 | 113624513 | 113625244 | 732 | -5.35 | 2.41E-05 | 3.43E-05 |
| chr2 | 96825914 | 96836931 | 11018 | -6.36 | 2.41E-05 | 3.43E-05 |
| chr6 | 116367833 | 116368324 | 492 | -3.93 | 2.41E-05 | 3.43E-05 |
| chr8 | 21664570 | 21665040 | 471 | -5.22 | 2.42E-05 | 3.45E-05 |
| chrX | 55848746 | 55853390 | 4645 | 1.97 | 2.42E-05 | 3.45E-05 |
| chr11 | 109700399 | 109701044 | 646 | -5.32 | 2.42E-05 | 3.45E-05 |
| chr6 | 84576648 | 84577436 | 789 | -5.32 | 2.42E-05 | 3.45E-05 |
| chr6 | 111704975 | 111726217 | 21243 | -5.78 | 2.43E-05 | 3.45E-05 |
| chr13 | 28176353 | 28189807 | 13455 | -5.31 | 2.43E-05 | 3.46E-05 |
| chr8 | 59373200 | 59374223 | 1024 | -5.31 | 2.43E-05 | 3.46E-05 |
| chrX | 81981110 | 81982236 | 1127 | -5.08 | 2.43E-05 | 3.47E-05 |
| chrX | 37400576 | 37400960 | 385 | 4.37 | 2.44E-05 | 3.47E-05 |
| chr2 | 157778906 | 157784149 | 5244 | -4.77 | 2.44E-05 | 3.47E-05 |
| chr4 | 141150860 | 141151476 | 617 | -4.35 | 2.44E-05 | 3.47E-05 |
| chr2 | 223558071 | 223558673 | 603 | -5.2 | 2.44E-05 | 3.47E-05 |
| chr3 | 105922765 | 105956019 | 33255 | -7.28 | 2.44E-05 | 3.48E-05 |
| chr1 | 184370069 | 184624493 | 254425 | 3.99 | 2.46E-05 | 3.50E-05 |
| chr5 | 1529372 | 1529996 | 625 | -5.19 | 2.46E-05 | 3.50E-05 |
| chr9 | 127201426 | 127202123 | 698 | -4.94 | 2.46E-05 | 3.51E-05 |
| chr4 | 97228700 | 97229919 | 1220 | -4.95 | 2.46E-05 | 3.51E-05 |
| chr7 | 1697455 | 1698110 | 656 | -4.95 | 2.46E-05 | 3.51E-05 |
| chr7 | 99367685 | 99378123 | 10439 | -1.51 | 2.47E-05 | 3.51E-05 |
| chr20 | 47599423 | 47599992 | 570 | -5.07 | 2.47E-05 | 3.52E-05 |
| chr8 | 116283641 | 116285706 | 2066 | -5.07 | 2.47E-05 | 3.52E-05 |
| chr4 | 22702049 | 22703429 | 1381 | -5.27 | 2.48E-05 | 3.52E-05 |
| chr11 | 127916600 | 127918960 | 2361 | 1.95 | 2.48E-05 | 3.52E-05 |
| chr1 | 217501701 | 217520523 | 18823 | -2.78 | 2.48E-05 | 3.53E-05 |
| chr11 | 64905951 | 64915924 | 9974 | -5.5 | 2.48E-05 | 3.53E-05 |
| chr20 | 48840851 | 48841470 | 620 | -5.17 | 2.48E-05 | 3.53E-05 |
| chrX | 150331313 | 150362558 | 31246 | -2.29 | 2.49E-05 | 3.54E-05 |
| chr17 | 26052705 | 26053727 | 1023 | -2.12 | 2.49E-05 | 3.55E-05 |
| chrX | 53282012 | 53282434 | 423 | -4.96 | 2.49E-05 | 3.55E-05 |
| chr5 | 115879252 | 115883946 | 4695 | -4.25 | 2.50E-05 | 3.55E-05 |
| chr12 | 132174069 | 132175021 | 953 | -2.65 | 2.50E-05 | 3.56E-05 |
| chr12 | 121099229 | 121099849 | 621 | 5.79 | 2.50E-05 | 3.56E-05 |
| chr1 | 168359893 | 168360541 | 649 | -4.94 | 2.50E-05 | 3.56E-05 |
| chr19 | 15503750 | 15504205 | 456 | -4.94 | 2.50E-05 | 3.56E-05 |
| chrX | 35909142 | 35910190 | 1049 | -4.94 | 2.50E-05 | 3.56E-05 |
| chr17 | 53182707 | 53183277 | 571 | -4.99 | 2.50E-05 | 3.56E-05 |
| chr20 | 43269277 | 43269820 | 544 | -4.99 | 2.50E-05 | 3.56E-05 |
| chr16 | 58272105 | 58272533 | 429 | -4.95 | 2.50E-05 | 3.56E-05 |
| chr6 | 9293986 | 9294430 | 445 | -4.95 | 2.50E-05 | 3.56E-05 |
| chr7 | 87361788 | 87363179 | 1392 | -5.15 | 2.51E-05 | 3.57E-05 |
| chr10 | 91261073 | 91265454 | 4382 | -5.46 | 2.51E-05 | 3.57E-05 |
| chr4 | 19616271 | 19617565 | 1295 | -5.46 | 2.51E-05 | 3.57E-05 |
| chr2 | 190752718 | 190753686 | 969 | -5.05 | 2.51E-05 | 3.57E-05 |
| chrX | 100250434 | 100251176 | 743 | -4.33 | 2.52E-05 | 3.58E-05 |
| chr5 | 44751798 | 44756008 | 4211 | 2.76 | 2.52E-05 | 3.58E-05 |
| chr3 | 127278835 | 127279314 | 480 | -4.29 | 2.52E-05 | 3.59E-05 |
| chr10 | 25052670 | 25055605 | 2936 | -4.17 | 2.52E-05 | 3.59E-05 |
| chr8 | 58369993 | 58371286 | 1294 | -4.99 | 2.52E-05 | 3.59E-05 |
| chr1 | 213534966 | 213536204 | 1239 | -3.77 | 2.53E-05 | 3.59E-05 |
| chrX | 79654125 | 79655985 | 1861 | -4.04 | 2.53E-05 | 3.60E-05 |
| chr7 | 11641331 | 11664750 | 23420 | -6.16 | 2.54E-05 | 3.61E-05 |
| chr11 | 10155369 | 10194902 | 39534 | -7.18 | 2.54E-05 | 3.61E-05 |
| chr11 | 45513536 | 45529002 | 15467 | -5.46 | 2.54E-05 | 3.61E-05 |
| chr10 | 13401923 | 13403762 | 1840 | -5.03 | 2.55E-05 | 3.62E-05 |
| chr11 | 107934571 | 107934966 | 396 | -5.03 | 2.55E-05 | 3.62E-05 |
| chr15 | 31517998 | 31518784 | 787 | -5.03 | 2.55E-05 | 3.62E-05 |
| chr11 | 95128919 | 95129749 | 831 | -5.41 | 2.55E-05 | 3.62E-05 |

| | | | | | | |
|---|---|---|---|---|---|---|
| chrX | 115358697 | 115359091 | 395 | 5.04 | 2.55E-05 | 3.62E-05 |
| chr2 | 242578736 | 242579248 | 513 | -5.04 | 2.56E-05 | 3.63E-05 |
| chr20 | 48514183 | 48514936 | 754 | -5.39 | 2.56E-05 | 3.64E-05 |
| chr1 | 59483893 | 59484960 | 1068 | -5.19 | 2.56E-05 | 3.64E-05 |
| chr11 | 21339656 | 21386810 | 47155 | 1.66 | 2.56E-05 | 3.64E-05 |
| chr7 | 100496929 | 100506724 | 9796 | -5.75 | 2.57E-05 | 3.65E-05 |
| chr8 | 101025074 | 101026777 | 1704 | -2.27 | 2.57E-05 | 3.65E-05 |
| chr22 | 30560709 | 30567955 | 7247 | -1.6 | 2.57E-05 | 3.65E-05 |
| chr6 | 140888677 | 140891166 | 2490 | -5.38 | 2.57E-05 | 3.65E-05 |
| chr2 | 35275543 | 35298955 | 23413 | -1.5 | 2.58E-05 | 3.66E-05 |
| chr6 | 13468372 | 13469717 | 1346 | -5.18 | 2.58E-05 | 3.66E-05 |
| chr2 | 209617313 | 209619440 | 2128 | -4.39 | 2.58E-05 | 3.67E-05 |
| chr13 | 34429512 | 34430066 | 555 | -4.95 | 2.59E-05 | 3.67E-05 |
| chr6 | 9128279 | 9129082 | 804 | -4.95 | 2.59E-05 | 3.67E-05 |
| chr1 | 214207261 | 214224100 | 16840 | -6.39 | 2.59E-05 | 3.67E-05 |
| chr10 | 34288724 | 34289508 | 785 | -4.93 | 2.59E-05 | 3.67E-05 |
| chr10 | 103973746 | 103974408 | 663 | -4.93 | 2.59E-05 | 3.67E-05 |
| chr13 | 31324876 | 31325624 | 749 | -4.93 | 2.59E-05 | 3.67E-05 |
| chr4 | 93804489 | 93805163 | 675 | -4.93 | 2.59E-05 | 3.67E-05 |
| chr2 | 145577279 | 145581791 | 4513 | -3.16 | 2.59E-05 | 3.67E-05 |
| chr4 | 117926027 | 117928417 | 2391 | -5.17 | 2.60E-05 | 3.69E-05 |
| chr1 | 242046391 | 242046897 | 507 | -4.95 | 2.60E-05 | 3.69E-05 |
| chr14 | 74150596 | 74151064 | 469 | -4.95 | 2.60E-05 | 3.69E-05 |
| chr6 | 227431 | 228413 | 983 | -4.98 | 2.60E-05 | 3.69E-05 |
| chrX | 28386126 | 28386874 | 749 | -4.98 | 2.60E-05 | 3.69E-05 |
| chr9 | 99196467 | 99209287 | 12821 | -6.26 | 2.60E-05 | 3.69E-05 |
| chr13 | 104160716 | 104161596 | 881 | -5.1 | 2.60E-05 | 3.70E-05 |
| chr21 | 39257761 | 39258562 | 802 | -5.1 | 2.60E-05 | 3.70E-05 |
| chr5 | 126043200 | 126044255 | 1056 | -5.1 | 2.60E-05 | 3.70E-05 |
| chr7 | 83281253 | 83281949 | 697 | -5.1 | 2.60E-05 | 3.70E-05 |
| chr19 | 40661177 | 40661702 | 526 | -5.34 | 2.60E-05 | 3.70E-05 |
| chr21 | 40534417 | 40543315 | 8899 | -4.75 | 2.61E-05 | 3.70E-05 |
| chr1 | 229343505 | 229356498 | 12994 | -6.93 | 2.62E-05 | 3.71E-05 |
| chr6 | 142620869 | 142621664 | 796 | -5.32 | 2.62E-05 | 3.71E-05 |
| chr17 | 38017482 | 38020521 | 3040 | -1.27 | 2.63E-05 | 3.73E-05 |
| chr6 | 153863354 | 153870863 | 7510 | -2.45 | 2.63E-05 | 3.73E-05 |
| chr3 | 67387974 | 67396838 | 8865 | -1.16 | 2.63E-05 | 3.73E-05 |
| chr1 | 202231428 | 202232579 | 1152 | -4.55 | 2.63E-05 | 3.74E-05 |
| chr10 | 28700434 | 28705771 | 5338 | -3.34 | 2.63E-05 | 3.74E-05 |
| chr7 | 31319638 | 31339465 | 19828 | -1.75 | 2.64E-05 | 3.74E-05 |
| chr6 | 143482374 | 143483106 | 733 | -5.14 | 2.64E-05 | 3.74E-05 |
| chr13 | 84112621 | 84113352 | 732 | -4.93 | 2.64E-05 | 3.75E-05 |
| chr3 | 133965709 | 133966376 | 668 | -4.93 | 2.64E-05 | 3.75E-05 |
| chr20 | 56050142 | 56050785 | 644 | -5.09 | 2.64E-05 | 3.75E-05 |
| chr1 | 244351552 | 244353650 | 2099 | -4.52 | 2.65E-05 | 3.76E-05 |
| chr2 | 219028338 | 219032546 | 4209 | 1.58 | 2.65E-05 | 3.76E-05 |
| chr11 | 8369564 | 8370530 | 967 | -5.28 | 2.66E-05 | 3.77E-05 |
| chr4 | 161400298 | 161401078 | 781 | -4.93 | 2.66E-05 | 3.77E-05 |
| chr5 | 70873735 | 70874508 | 774 | -4.93 | 2.66E-05 | 3.77E-05 |
| chr6 | 152066974 | 152067502 | 529 | -4.93 | 2.66E-05 | 3.77E-05 |
| chr1 | 63640706 | 63645810 | 5105 | -1.5 | 2.66E-05 | 3.77E-05 |
| chr7 | 2885230 | 2885847 | 618 | -5.13 | 2.66E-05 | 3.78E-05 |
| chr6 | 142636328 | 142636814 | 487 | -5.01 | 2.66E-05 | 3.78E-05 |
| chr1 | 59871847 | 59872492 | 646 | -4.92 | 2.66E-05 | 3.78E-05 |
| chr4 | 180183181 | 180186847 | 3667 | -4.52 | 2.67E-05 | 3.78E-05 |
| chr12 | 108345683 | 108369582 | 23900 | -2.61 | 2.67E-05 | 3.78E-05 |
| chr12 | 95092711 | 95093694 | 984 | -4.74 | 2.68E-05 | 3.79E-05 |
| chr16 | 10610255 | 10611013 | 759 | -4.74 | 2.68E-05 | 3.79E-05 |
| chr2 | 213268183 | 213278106 | 9924 | -2.39 | 2.68E-05 | 3.80E-05 |
| chr7 | 48308561 | 48326615 | 18055 | 1.6 | 2.68E-05 | 3.81E-05 |
| chr3 | 29874394 | 29875625 | 1232 | -5.07 | 2.68E-05 | 3.81E-05 |

| | | | | | | |
|---|---|---|---|---|---|---|
| chr11 | 63364006 | 63364678 | 673 | -5.12 | 2.69E-05 | 3.81E-05 |
| chrX | 30405975 | 30407157 | 1183 | 4.2 | 2.69E-05 | 3.81E-05 |
| chr16 | 20866037 | 20866718 | 682 | -5 | 2.69E-05 | 3.81E-05 |
| chr2 | 85564665 | 85565142 | 478 | -5 | 2.69E-05 | 3.81E-05 |
| chr14 | 20631238 | 20636470 | 5233 | -1.64 | 2.69E-05 | 3.82E-05 |
| chr11 | 36327367 | 36328214 | 848 | -3.79 | 2.70E-05 | 3.82E-05 |
| chr3 | 86935481 | 86935893 | 413 | -4.08 | 2.70E-05 | 3.82E-05 |
| chr1 | 20420674 | 20425283 | 4610 | -4.95 | 2.70E-05 | 3.83E-05 |
| chr5 | 18301889 | 18302735 | 847 | -5.61 | 2.70E-05 | 3.83E-05 |
| chr14 | 90001899 | 90005616 | 3718 | 4.71 | 2.71E-05 | 3.83E-05 |
| chr19 | 50449101 | 50449658 | 558 | -4.96 | 2.71E-05 | 3.84E-05 |
| chr3 | 66123918 | 66124423 | 506 | -4.96 | 2.71E-05 | 3.84E-05 |
| chr6 | 7873012 | 7873827 | 816 | -4.96 | 2.71E-05 | 3.84E-05 |
| chr8 | 106807383 | 106807922 | 540 | -3.77 | 2.71E-05 | 3.84E-05 |
| chr11 | 103434392 | 103435135 | 744 | -4.5 | 2.72E-05 | 3.85E-05 |
| chr20 | 46268251 | 46269984 | 1734 | -5.1 | 2.72E-05 | 3.85E-05 |
| chr6 | 142062133 | 142063613 | 1481 | -4.23 | 2.72E-05 | 3.85E-05 |
| chr2 | 108956682 | 108957384 | 703 | -5.22 | 2.72E-05 | 3.85E-05 |
| chr11 | 94505159 | 94505749 | 591 | -4.93 | 2.72E-05 | 3.86E-05 |
| chr18 | 25220265 | 25220853 | 589 | -4.93 | 2.72E-05 | 3.86E-05 |
| chr2 | 85947567 | 85948051 | 485 | -4.93 | 2.72E-05 | 3.86E-05 |
| chr8 | 83957424 | 83958460 | 1037 | -4.93 | 2.72E-05 | 3.86E-05 |
| chr9 | 34561915 | 34562559 | 645 | -4.93 | 2.72E-05 | 3.86E-05 |
| chr6 | 126541988 | 126542676 | 689 | -5 | 2.72E-05 | 3.86E-05 |
| chr9 | 3148164 | 3151785 | 3622 | 3.18 | 2.73E-05 | 3.86E-05 |
| chr10 | 111880261 | 111881191 | 931 | -4 | 2.73E-05 | 3.87E-05 |
| chr8 | 54613455 | 54617598 | 4144 | -4.82 | 2.73E-05 | 3.87E-05 |
| chr15 | 48861297 | 48861885 | 589 | -4.92 | 2.73E-05 | 3.87E-05 |
| chr7 | 36065573 | 36066512 | 940 | -4.92 | 2.73E-05 | 3.87E-05 |
| chr9 | 38679472 | 38680353 | 882 | -4.92 | 2.73E-05 | 3.87E-05 |
| chr6 | 79215829 | 79219246 | 3418 | -2.01 | 2.73E-05 | 3.87E-05 |
| chr5 | 78112172 | 78113052 | 881 | -5.2 | 2.74E-05 | 3.88E-05 |
| chrX | 105846802 | 105850346 | 3545 | -2.82 | 2.74E-05 | 3.88E-05 |
| chr2 | 9944326 | 9944871 | 546 | -5.09 | 2.75E-05 | 3.89E-05 |
| chr9 | 80667612 | 80739991 | 72380 | -2.41 | 2.75E-05 | 3.89E-05 |
| chr6 | 137903767 | 137932706 | 28940 | -3.82 | 2.75E-05 | 3.89E-05 |
| chr7 | 142710243 | 142857303 | 147061 | 2.52 | 2.76E-05 | 3.91E-05 |
| chr10 | 77242991 | 77243719 | 729 | -5.19 | 2.76E-05 | 3.91E-05 |
| chr2 | 232294755 | 232295757 | 1003 | -5.39 | 2.76E-05 | 3.91E-05 |
| chr22 | 28533102 | 28534101 | 1000 | -5.37 | 2.78E-05 | 3.94E-05 |
| chr5 | 150452945 | 150454503 | 1559 | -5.37 | 2.78E-05 | 3.94E-05 |
| chr3 | 124921149 | 124922409 | 1261 | -5.17 | 2.78E-05 | 3.94E-05 |
| chr11 | 86376941 | 86377529 | 589 | -3.81 | 2.79E-05 | 3.94E-05 |
| chr3 | 51840764 | 51895789 | 55026 | -1.49 | 2.79E-05 | 3.94E-05 |
| chr6 | 125480924 | 125481451 | 528 | -5.04 | 2.79E-05 | 3.95E-05 |
| chr9 | 95304970 | 95305813 | 844 | -5.04 | 2.79E-05 | 3.95E-05 |
| chr1 | 211415249 | 211415781 | 533 | -4.94 | 2.79E-05 | 3.95E-05 |
| chr13 | 101819139 | 101819808 | 670 | -4.98 | 2.79E-05 | 3.95E-05 |
| chr18 | 1963874 | 1986403 | 22530 | 1.2 | 2.79E-05 | 3.95E-05 |
| chr3 | 148504264 | 148505384 | 1121 | -2.98 | 2.80E-05 | 3.96E-05 |
| chr1 | 68240713 | 68243512 | 2800 | -4.54 | 2.80E-05 | 3.96E-05 |
| chrX | 12854849 | 12888386 | 33538 | -1.13 | 2.81E-05 | 3.97E-05 |
| chr1 | 69448019 | 69452401 | 4383 | 2.38 | 2.81E-05 | 3.98E-05 |
| chr9 | 129754089 | 129756082 | 1994 | -4.85 | 2.82E-05 | 3.98E-05 |
| chr6 | 166865268 | 166867675 | 2408 | -4.34 | 2.82E-05 | 3.98E-05 |
| chr15 | 31949226 | 31949729 | 504 | 5.04 | 2.82E-05 | 3.99E-05 |
| chr1 | 40067274 | 40067768 | 495 | -4.94 | 2.83E-05 | 4.00E-05 |
| chr4 | 20536053 | 20536599 | 547 | 4.45 | 2.83E-05 | 4.00E-05 |
| chrX | 73690294 | 73701074 | 10781 | -4.08 | 2.83E-05 | 4.01E-05 |
| chr12 | 68946538 | 68949065 | 2528 | -4.23 | 2.84E-05 | 4.01E-05 |
| chr1 | 1110651 | 1111069 | 419 | -4.91 | 2.84E-05 | 4.02E-05 |

| | | | | | | |
|---|---|---|---|---|---|---|
| chr14 | 96046990 | 96052568 | 5579 | -4.76 | 2.84E-05 | 4.02E-05 |
| chr1 | 48405166 | 48405841 | 676 | -4.91 | 2.84E-05 | 4.02E-05 |
| chr18 | 43161335 | 43164721 | 3387 | -2.11 | 2.85E-05 | 4.02E-05 |
| chr14 | 34477593 | 34480164 | 2572 | -4.3 | 2.85E-05 | 4.03E-05 |
| chr9 | 125950344 | 125951275 | 932 | -5.44 | 2.85E-05 | 4.03E-05 |
| chr6 | 22006870 | 22007388 | 519 | -4.92 | 2.85E-05 | 4.03E-05 |
| chr20 | 8669023 | 8672283 | 3261 | -4.75 | 2.85E-05 | 4.04E-05 |
| chr16 | 58478042 | 58478911 | 870 | -5.29 | 2.86E-05 | 4.04E-05 |
| chrX | 40883116 | 40884140 | 1025 | -5.05 | 2.86E-05 | 4.04E-05 |
| chr8 | 124671199 | 124672914 | 1716 | 2.25 | 2.86E-05 | 4.04E-05 |
| chrX | 81941533 | 81945416 | 3884 | -4.12 | 2.86E-05 | 4.04E-05 |
| chr12 | 41232111 | 41241987 | 9877 | -6.13 | 2.86E-05 | 4.05E-05 |
| chr6 | 154666954 | 154668515 | 1562 | -4.24 | 2.86E-05 | 4.05E-05 |
| chr11 | 48213217 | 48274600 | 61384 | -1.4 | 2.86E-05 | 4.05E-05 |
| chr13 | 97859057 | 97859760 | 704 | -4.97 | 2.87E-05 | 4.05E-05 |
| chr3 | 191779888 | 191781266 | 1379 | 5.01 | 2.87E-05 | 4.05E-05 |
| chr1 | 120235454 | 120248349 | 12896 | -1.71 | 2.87E-05 | 4.06E-05 |
| chr12 | 100263800 | 100264365 | 566 | -4.96 | 2.87E-05 | 4.06E-05 |
| chr17 | 64422708 | 64423425 | 718 | -4.96 | 2.87E-05 | 4.06E-05 |
| chr6 | 21336959 | 21337583 | 625 | -4.96 | 2.87E-05 | 4.06E-05 |
| chr7 | 82392279 | 82416135 | 23857 | -6.38 | 2.87E-05 | 4.06E-05 |
| chr3 | 46589981 | 46597466 | 7486 | -4.51 | 2.88E-05 | 4.07E-05 |
| chr2 | 20745443 | 20746605 | 1163 | -4.27 | 2.89E-05 | 4.08E-05 |
| chr21 | 29324584 | 29331284 | 6701 | 1.64 | 2.89E-05 | 4.09E-05 |
| chr2 | 192656850 | 192658287 | 1438 | -4.01 | 2.89E-05 | 4.09E-05 |
| chr9 | 3339392 | 3339948 | 557 | -4.9 | 2.90E-05 | 4.09E-05 |
| chr11 | 104123075 | 104123605 | 531 | -5.11 | 2.90E-05 | 4.10E-05 |
| chr8 | 87582770 | 87583619 | 850 | -5.11 | 2.90E-05 | 4.10E-05 |
| chr18 | 58042958 | 58044418 | 1461 | -3.87 | 2.90E-05 | 4.10E-05 |
| chr15 | 34395538 | 34396416 | 879 | -5.03 | 2.90E-05 | 4.10E-05 |
| chr12 | 63422857 | 63429559 | 6703 | -1.47 | 2.91E-05 | 4.11E-05 |
| chr16 | 69409231 | 69409697 | 467 | -5 | 2.92E-05 | 4.12E-05 |
| chr3 | 102933755 | 102934958 | 1204 | -5 | 2.92E-05 | 4.12E-05 |
| chr16 | 75759187 | 75785482 | 26296 | -2.73 | 2.92E-05 | 4.13E-05 |
| chr1 | 117511285 | 117511837 | 553 | -5.23 | 2.92E-05 | 4.13E-05 |
| chr1 | 194574951 | 194575676 | 726 | 5.12 | 2.92E-05 | 4.13E-05 |
| chr12 | 100910868 | 100911533 | 666 | -4.1 | 2.93E-05 | 4.13E-05 |
| chr12 | 113119455 | 113142409 | 22955 | -1.18 | 2.93E-05 | 4.14E-05 |
| chr18 | 34651590 | 34654402 | 2813 | -4.27 | 2.93E-05 | 4.14E-05 |
| chr5 | 73700939 | 73702930 | 1992 | -4.33 | 2.94E-05 | 4.15E-05 |
| chr6 | 84361876 | 84365350 | 3475 | 2.8 | 2.94E-05 | 4.15E-05 |
| chr12 | 92907153 | 92917389 | 10237 | -5.68 | 2.95E-05 | 4.16E-05 |
| chrX | 46297484 | 46298915 | 1432 | -5.68 | 2.95E-05 | 4.16E-05 |
| chr3 | 195573072 | 195574023 | 952 | -5.33 | 2.95E-05 | 4.17E-05 |
| chr6 | 63772781 | 63773721 | 941 | -3.85 | 2.96E-05 | 4.17E-05 |
| chr11 | 43521776 | 43568554 | 46779 | -2.38 | 2.96E-05 | 4.17E-05 |
| chr21 | 47375679 | 47385166 | 9488 | -2.84 | 2.96E-05 | 4.18E-05 |
| chr10 | 134254746 | 134255773 | 1028 | -5.31 | 2.97E-05 | 4.19E-05 |
| chr1 | 181087696 | 181088616 | 921 | -5.19 | 2.97E-05 | 4.20E-05 |
| chr11 | 108107904 | 108108467 | 564 | -4.92 | 2.97E-05 | 4.20E-05 |
| chr2 | 43139702 | 43140145 | 444 | -4.92 | 2.97E-05 | 4.20E-05 |
| chr2 | 106074194 | 106075249 | 1056 | -4.92 | 2.97E-05 | 4.20E-05 |
| chr9 | 123218398 | 123219272 | 875 | -5.07 | 2.98E-05 | 4.20E-05 |
| chr5 | 65521564 | 65533711 | 12148 | -5.89 | 2.98E-05 | 4.20E-05 |
| chr5 | 176260099 | 176260848 | 750 | -4.94 | 2.98E-05 | 4.21E-05 |
| chr6 | 147390003 | 147391251 | 1249 | -5.65 | 2.99E-05 | 4.21E-05 |
| chr9 | 3635459 | 3637355 | 1897 | 2.46 | 2.99E-05 | 4.22E-05 |
| chr6 | 42077842 | 42078666 | 825 | -5.18 | 2.99E-05 | 4.22E-05 |
| chr11 | 437209 | 437751 | 543 | -4.98 | 3.00E-05 | 4.23E-05 |
| chr3 | 43841799 | 43842438 | 640 | -4.98 | 3.00E-05 | 4.23E-05 |
| chr6 | 43291710 | 43292262 | 553 | -4.98 | 3.00E-05 | 4.23E-05 |

| | | | | | | |
|---|---|---|---|---|---|---|
| chrX | 40655513 | 40655998 | 486 | -4.98 | 3.00E-05 | 4.23E-05 |
| chr11 | 64785206 | 64785706 | 501 | -5.01 | 3.00E-05 | 4.23E-05 |
| chr14 | 58638576 | 58639042 | 467 | -5.01 | 3.00E-05 | 4.23E-05 |
| chr11 | 4629866 | 4649930 | 20065 | -5.16 | 3.00E-05 | 4.23E-05 |
| chr14 | 29195837 | 29197089 | 1253 | 5.35 | 3.01E-05 | 4.24E-05 |
| chrX | 57500512 | 57501303 | 792 | -4.3 | 3.01E-05 | 4.25E-05 |
| chr16 | 81810772 | 81811437 | 666 | -5.16 | 3.01E-05 | 4.25E-05 |
| chr5 | 173987883 | 173988877 | 995 | -2.62 | 3.01E-05 | 4.25E-05 |
| chr2 | 186438204 | 186450324 | 12121 | -4.6 | 3.01E-05 | 4.25E-05 |
| chr12 | 8001636 | 8002126 | 491 | -4.91 | 3.02E-05 | 4.27E-05 |
| chr2 | 134832988 | 134834643 | 1656 | -5.06 | 3.02E-05 | 4.27E-05 |
| chr7 | 6324027 | 6325477 | 1451 | -5.06 | 3.02E-05 | 4.27E-05 |
| chr2 | 64202958 | 64203308 | 351 | -3.92 | 3.03E-05 | 4.27E-05 |
| chr4 | 118428137 | 118429387 | 1251 | -5.25 | 3.03E-05 | 4.27E-05 |
| chr7 | 105700761 | 105701829 | 1069 | -5.25 | 3.03E-05 | 4.27E-05 |
| chr8 | 133922414 | 133923676 | 1263 | -3.79 | 3.03E-05 | 4.27E-05 |
| chr17 | 21370218 | 21383697 | 13480 | -1.56 | 3.04E-05 | 4.29E-05 |
| chr5 | 111967229 | 111968670 | 1442 | -4.2 | 3.05E-05 | 4.29E-05 |
| chr4 | 37610660 | 37611200 | 541 | -4.89 | 3.05E-05 | 4.30E-05 |
| chr6 | 138472420 | 138476137 | 3718 | -5.29 | 3.06E-05 | 4.31E-05 |
| chr16 | 59439934 | 59451231 | 11298 | -1.86 | 3.06E-05 | 4.31E-05 |
| chr16 | 66342120 | 66343001 | 882 | 5.01 | 3.06E-05 | 4.31E-05 |
| chr9 | 3578860 | 3595658 | 16799 | -5.09 | 3.07E-05 | 4.32E-05 |
| chr2 | 70428517 | 70428870 | 354 | -4.9 | 3.07E-05 | 4.32E-05 |
| chr2 | 228182347 | 228183009 | 663 | -4.9 | 3.07E-05 | 4.32E-05 |
| chrX | 140706572 | 140707313 | 742 | -4.9 | 3.07E-05 | 4.32E-05 |
| chr18 | 42434397 | 42436415 | 2019 | -2.58 | 3.07E-05 | 4.33E-05 |
| chr4 | 72351006 | 72351519 | 514 | -5.04 | 3.07E-05 | 4.33E-05 |
| chr9 | 4472658 | 4473395 | 738 | -4.89 | 3.08E-05 | 4.35E-05 |
| chr4 | 21700998 | 21703652 | 2655 | 2.25 | 3.08E-05 | 4.35E-05 |
| chr3 | 143573818 | 143600029 | 26212 | 1.62 | 3.08E-05 | 4.35E-05 |
| chr17 | 41632939 | 41633592 | 654 | -4.96 | 3.09E-05 | 4.35E-05 |
| chr5 | 1156545 | 1157148 | 604 | -4.96 | 3.09E-05 | 4.35E-05 |
| chr14 | 66882703 | 66884224 | 1522 | -2.64 | 3.09E-05 | 4.36E-05 |
| chr15 | 53708531 | 53711904 | 3374 | -2.71 | 3.10E-05 | 4.36E-05 |
| chr6 | 78897263 | 78909111 | 11849 | 2.46 | 3.10E-05 | 4.37E-05 |
| chr3 | 162841744 | 162850220 | 8477 | -2.19 | 3.10E-05 | 4.37E-05 |
| chr2 | 100447922 | 100483092 | 35171 | 0.97 | 3.10E-05 | 4.37E-05 |
| chrX | 135674608 | 135677047 | 2440 | -2.21 | 3.11E-05 | 4.39E-05 |
| chr16 | 55704342 | 55705133 | 792 | -4.62 | 3.12E-05 | 4.39E-05 |
| chr16 | 71991099 | 71991657 | 559 | -4.14 | 3.12E-05 | 4.39E-05 |
| chrX | 115333675 | 115334437 | 763 | 3.89 | 3.12E-05 | 4.40E-05 |
| chr17 | 10589658 | 10590081 | 424 | -4.98 | 3.12E-05 | 4.40E-05 |
| chr2 | 236988759 | 236989340 | 582 | -4.98 | 3.12E-05 | 4.40E-05 |
| chr19 | 34325307 | 34327575 | 2269 | -4.57 | 3.13E-05 | 4.41E-05 |
| chr4 | 130486296 | 130487260 | 965 | -5.18 | 3.13E-05 | 4.41E-05 |
| chr2 | 174699425 | 174700020 | 596 | -5.02 | 3.13E-05 | 4.41E-05 |
| chrX | 48237662 | 48238628 | 967 | -5.02 | 3.13E-05 | 4.41E-05 |
| chr5 | 154986254 | 154990721 | 4468 | -1.77 | 3.13E-05 | 4.42E-05 |
| chrX | 114060710 | 114081484 | 20775 | 1.63 | 3.14E-05 | 4.42E-05 |
| chr12 | 17668854 | 17669321 | 468 | -4.92 | 3.14E-05 | 4.43E-05 |
| chr5 | 33226013 | 33226535 | 523 | -4.92 | 3.14E-05 | 4.43E-05 |
| chr6 | 127046652 | 127047709 | 1058 | -3.88 | 3.15E-05 | 4.44E-05 |
| chr5 | 93498289 | 93500839 | 2551 | -4.61 | 3.15E-05 | 4.44E-05 |
| chr15 | 87109222 | 87207621 | 98400 | -2.8 | 3.16E-05 | 4.45E-05 |
| chr2 | 210521699 | 210522096 | 398 | -4.9 | 3.16E-05 | 4.45E-05 |
| chr7 | 114630213 | 114630734 | 522 | -4.9 | 3.16E-05 | 4.45E-05 |
| chr17 | 42291533 | 42292177 | 645 | -5.09 | 3.16E-05 | 4.45E-05 |
| chr2 | 69737424 | 69737841 | 418 | -5.09 | 3.16E-05 | 4.45E-05 |
| chr13 | 81768329 | 81768958 | 630 | -4.07 | 3.17E-05 | 4.46E-05 |
| chr14 | 90151819 | 90223716 | 71898 | -1.03 | 3.17E-05 | 4.47E-05 |

| | | | | | | |
|---|---|---|---|---|---|---|
| chr5 | 107945797 | 107946922 | 1126 | -2.18 | 3.18E-05 | 4.47E-05 |
| chr15 | 100473695 | 100575269 | 101575 | -1.64 | 3.18E-05 | 4.48E-05 |
| chr21 | 27853536 | 27871718 | 18183 | -5.16 | 3.18E-05 | 4.48E-05 |
| chr4 | 135316400 | 135321453 | 5054 | -1.98 | 3.19E-05 | 4.48E-05 |
| chr1 | 212887061 | 212887554 | 494 | -4.95 | 3.19E-05 | 4.49E-05 |
| chr14 | 92646106 | 92646759 | 654 | -4.95 | 3.19E-05 | 4.49E-05 |
| chr2 | 70845796 | 70846195 | 400 | -4.95 | 3.19E-05 | 4.49E-05 |
| chr7 | 79169950 | 79170393 | 444 | -4.96 | 3.19E-05 | 4.50E-05 |
| chr3 | 157115321 | 157115982 | 662 | -5 | 3.20E-05 | 4.50E-05 |
| chr3 | 161243662 | 161246663 | 3002 | 1.7 | 3.20E-05 | 4.50E-05 |
| chr3 | 150552210 | 150581882 | 29673 | -1.48 | 3.20E-05 | 4.51E-05 |
| chr19 | 36061830 | 36077383 | 15554 | -5.66 | 3.20E-05 | 4.51E-05 |
| chr1 | 31211032 | 31211701 | 670 | -5.13 | 3.21E-05 | 4.52E-05 |
| chr3 | 125115889 | 125117067 | 1179 | -4.02 | 3.22E-05 | 4.53E-05 |
| chr2 | 170283975 | 170303558 | 19584 | -5.29 | 3.22E-05 | 4.53E-05 |
| chr11 | 14621951 | 14628855 | 6905 | -5.01 | 3.22E-05 | 4.53E-05 |
| chr20 | 55324345 | 55341954 | 17610 | -6.82 | 3.22E-05 | 4.53E-05 |
| chrX | 152948867 | 152949531 | 665 | -5.06 | 3.23E-05 | 4.55E-05 |
| chr9 | 2590458 | 2593851 | 3394 | -1.62 | 3.24E-05 | 4.56E-05 |
| chr6 | 241873 | 252176 | 10304 | -1.4 | 3.24E-05 | 4.56E-05 |
| chr2 | 37225819 | 37226406 | 588 | -5.36 | 3.24E-05 | 4.56E-05 |
| chr2 | 175834366 | 175835479 | 1114 | -5.11 | 3.24E-05 | 4.56E-05 |
| chr7 | 18249089 | 18249707 | 619 | -4.23 | 3.25E-05 | 4.57E-05 |
| chr1 | 150547955 | 150550407 | 2453 | 2.03 | 3.25E-05 | 4.58E-05 |
| chr4 | 166246447 | 166247000 | 554 | -4.91 | 3.26E-05 | 4.58E-05 |
| chr7 | 134403019 | 134403465 | 447 | -4.91 | 3.26E-05 | 4.58E-05 |
| chr3 | 51276088 | 51279459 | 3372 | -4.31 | 3.26E-05 | 4.58E-05 |
| chr2 | 218391743 | 218411402 | 19660 | -1.51 | 3.26E-05 | 4.58E-05 |
| chr11 | 20189993 | 20208359 | 18367 | -4.22 | 3.26E-05 | 4.58E-05 |
| chr12 | 28723524 | 28724301 | 778 | -4.99 | 3.27E-05 | 4.60E-05 |
| chr21 | 36396441 | 36397406 | 966 | -4.99 | 3.27E-05 | 4.60E-05 |
| chr4 | 100021213 | 100021782 | 570 | -4.99 | 3.27E-05 | 4.60E-05 |
| chr6 | 13861139 | 13861623 | 485 | -4.99 | 3.27E-05 | 4.60E-05 |
| chrX | 114351032 | 114352435 | 1404 | -2.48 | 3.27E-05 | 4.60E-05 |
| chr1 | 230714228 | 230736980 | 22753 | -2.34 | 3.27E-05 | 4.60E-05 |
| chr5 | 17123643 | 17124717 | 1075 | -5.09 | 3.28E-05 | 4.61E-05 |
| chr10 | 94712297 | 94712675 | 379 | -4.88 | 3.28E-05 | 4.61E-05 |
| chr11 | 65603730 | 65619924 | 16195 | -4.88 | 3.28E-05 | 4.61E-05 |
| chr7 | 97890349 | 97891730 | 1382 | -5.37 | 3.28E-05 | 4.62E-05 |
| chr19 | 48463273 | 48604714 | 141442 | -3.36 | 3.29E-05 | 4.63E-05 |
| chr20 | 23745314 | 23745961 | 648 | -4.87 | 3.29E-05 | 4.63E-05 |
| chr2 | 38087577 | 38089858 | 2282 | 2.47 | 3.29E-05 | 4.63E-05 |
| chr16 | 51810457 | 51849393 | 38937 | -0.95 | 3.30E-05 | 4.64E-05 |
| chr3 | 23980096 | 23980585 | 490 | -4.93 | 3.30E-05 | 4.64E-05 |
| chr3 | 79014639 | 79015817 | 1179 | -4.93 | 3.30E-05 | 4.64E-05 |
| chr4 | 64033953 | 64034879 | 927 | -4.93 | 3.30E-05 | 4.64E-05 |
| chr4 | 71417282 | 71418218 | 937 | -2.8 | 3.30E-05 | 4.64E-05 |
| chr3 | 180419367 | 180422303 | 2937 | -1.9 | 3.32E-05 | 4.66E-05 |
| chr18 | 71719517 | 71720119 | 603 | -4.88 | 3.32E-05 | 4.66E-05 |
| chr3 | 194757763 | 194758662 | 900 | -4.88 | 3.32E-05 | 4.66E-05 |
| chr4 | 116051638 | 116052414 | 777 | -4.88 | 3.32E-05 | 4.66E-05 |
| chr13 | 95641720 | 95642484 | 765 | -4.88 | 3.33E-05 | 4.67E-05 |
| chr14 | 89198810 | 89199250 | 441 | -4.88 | 3.33E-05 | 4.67E-05 |
| chr5 | 49876289 | 49877356 | 1068 | -4.88 | 3.33E-05 | 4.67E-05 |
| chr8 | 101907311 | 101908104 | 794 | -5.33 | 3.33E-05 | 4.68E-05 |
| chr10 | 50863975 | 50923068 | 59094 | -1.17 | 3.34E-05 | 4.69E-05 |
| chrX | 88969766 | 88974206 | 4441 | 1.51 | 3.35E-05 | 4.70E-05 |
| chr14 | 65782652 | 65783570 | 919 | -4.97 | 3.35E-05 | 4.71E-05 |
| chr3 | 94063464 | 94064719 | 1256 | -4.97 | 3.35E-05 | 4.71E-05 |
| chr2 | 33233419 | 33233747 | 329 | -4.93 | 3.36E-05 | 4.72E-05 |
| chr3 | 98876140 | 98879081 | 2942 | -1.72 | 3.36E-05 | 4.72E-05 |

| | | | | | | |
|---|---|---|---|---|---|---|
| chr1 | 24942384 | 24943189 | 806 | -4.87 | 3.36E-05 | 4.72E-05 |
| chr16 | 23322038 | 23322672 | 635 | -4.87 | 3.36E-05 | 4.72E-05 |
| chr2 | 47632610 | 47633422 | 813 | -4.87 | 3.36E-05 | 4.72E-05 |
| chr14 | 24371586 | 24392720 | 21135 | -1.68 | 3.36E-05 | 4.72E-05 |
| chr2 | 234714927 | 234716092 | 1166 | -5.06 | 3.36E-05 | 4.72E-05 |
| chr4 | 22789567 | 22790435 | 869 | -3.26 | 3.37E-05 | 4.74E-05 |
| chr5 | 166216796 | 166259321 | 42526 | -0.83 | 3.38E-05 | 4.74E-05 |
| chr9 | 127940402 | 127940957 | 556 | -4.89 | 3.38E-05 | 4.75E-05 |
| chrX | 28900329 | 28900914 | 586 | -4.89 | 3.38E-05 | 4.75E-05 |
| chr3 | 55238087 | 55304107 | 66021 | -1.21 | 3.39E-05 | 4.76E-05 |
| chrX | 115280840 | 115283010 | 2171 | 6.09 | 3.39E-05 | 4.76E-05 |
| chr12 | 48618058 | 48681047 | 62990 | -1.71 | 3.39E-05 | 4.77E-05 |
| chr20 | 52150416 | 52150981 | 566 | -5 | 3.41E-05 | 4.79E-05 |
| chr18 | 25678151 | 25678935 | 785 | -5.04 | 3.41E-05 | 4.79E-05 |
| chr4 | 105199030 | 105199828 | 799 | -5.04 | 3.41E-05 | 4.79E-05 |
| chr1 | 214929781 | 214937215 | 7435 | -5 | 3.42E-05 | 4.80E-05 |
| chr19 | 55893685 | 55894435 | 751 | -5.26 | 3.42E-05 | 4.80E-05 |
| chr20 | 670636 | 670948 | 313 | -5.26 | 3.42E-05 | 4.80E-05 |
| chr14 | 51886036 | 51887595 | 1560 | -2.18 | 3.42E-05 | 4.81E-05 |
| chr14 | 59351656 | 59480217 | 128562 | 0.99 | 3.43E-05 | 4.81E-05 |
| chrX | 30766053 | 30804355 | 38303 | 1.48 | 3.43E-05 | 4.81E-05 |
| chr11 | 107459169 | 107460524 | 1356 | -4.42 | 3.43E-05 | 4.82E-05 |
| chr15 | 52927304 | 52927805 | 502 | -4.91 | 3.43E-05 | 4.82E-05 |
| chr2 | 19969059 | 19969725 | 667 | -4.91 | 3.43E-05 | 4.82E-05 |
| chr17 | 59360690 | 59361639 | 950 | -5.24 | 3.44E-05 | 4.82E-05 |
| chr1 | 192129876 | 192264600 | 134725 | 1.79 | 3.44E-05 | 4.82E-05 |
| chr1 | 248832856 | 248907751 | 74896 | -0.98 | 3.44E-05 | 4.83E-05 |
| chr14 | 77613510 | 77614094 | 585 | -4.95 | 3.44E-05 | 4.83E-05 |
| chr6 | 130491131 | 130491736 | 606 | -3.8 | 3.44E-05 | 4.83E-05 |
| chr3 | 10900100 | 10901786 | 1687 | -4.37 | 3.45E-05 | 4.83E-05 |
| chr1 | 63076642 | 63077084 | 443 | -4.92 | 3.45E-05 | 4.85E-05 |
| chr3 | 58187496 | 58187928 | 433 | -4.92 | 3.45E-05 | 4.85E-05 |
| chr14 | 105984215 | 105985265 | 1051 | -5.23 | 3.46E-05 | 4.85E-05 |
| chr9 | 6939036 | 6939813 | 778 | -5.03 | 3.47E-05 | 4.86E-05 |
| chr14 | 50169273 | 50169748 | 476 | -4.99 | 3.47E-05 | 4.87E-05 |
| chr3 | 52998282 | 52998894 | 613 | -4.99 | 3.47E-05 | 4.87E-05 |
| chr19 | 39470626 | 39476748 | 6123 | -5.8 | 3.48E-05 | 4.88E-05 |
| chr1 | 156334049 | 156335085 | 1037 | -5.21 | 3.48E-05 | 4.88E-05 |
| chr10 | 76662920 | 76663529 | 610 | -5.21 | 3.48E-05 | 4.88E-05 |
| chr18 | 76858398 | 76858871 | 474 | -5.21 | 3.48E-05 | 4.88E-05 |
| chr12 | 82866040 | 82867026 | 987 | -5.17 | 3.48E-05 | 4.88E-05 |
| chr6 | 2177972 | 2178669 | 698 | -5.17 | 3.48E-05 | 4.88E-05 |
| chr4 | 69988053 | 69989514 | 1462 | -4.87 | 3.49E-05 | 4.90E-05 |
| chr4 | 92333419 | 92334559 | 1141 | -4.87 | 3.49E-05 | 4.90E-05 |
| chr4 | 100406386 | 100407721 | 1336 | -2.61 | 3.50E-05 | 4.90E-05 |
| chr10 | 98901573 | 98903831 | 2259 | -4.05 | 3.50E-05 | 4.91E-05 |
| chr4 | 95367200 | 95368023 | 824 | -4.34 | 3.50E-05 | 4.91E-05 |
| chr12 | 50563295 | 50564067 | 773 | -5.2 | 3.50E-05 | 4.91E-05 |
| chr3 | 186039873 | 186041193 | 1321 | -4.01 | 3.51E-05 | 4.92E-05 |
| chr6 | 48724225 | 48725594 | 1370 | -5.15 | 3.53E-05 | 4.94E-05 |
| chr8 | 39014624 | 39015390 | 767 | -5.18 | 3.53E-05 | 4.94E-05 |
| chr21 | 42905082 | 42907313 | 2232 | -4.62 | 3.53E-05 | 4.94E-05 |
| chr13 | 24762767 | 24763455 | 689 | -4.97 | 3.53E-05 | 4.95E-05 |
| chr6 | 147870122 | 147870873 | 752 | -4.86 | 3.53E-05 | 4.95E-05 |
| chrX | 24277969 | 24278562 | 594 | -4.86 | 3.53E-05 | 4.95E-05 |
| chr15 | 26285288 | 26321531 | 36244 | -1.97 | 3.54E-05 | 4.96E-05 |
| chr4 | 147503046 | 147521703 | 18658 | -1.33 | 3.54E-05 | 4.96E-05 |
| chr3 | 187603557 | 187604167 | 611 | -4.86 | 3.55E-05 | 4.97E-05 |
| chr12 | 42988484 | 42989464 | 981 | -4.93 | 3.55E-05 | 4.97E-05 |
| chr7 | 107668973 | 107669976 | 1004 | -5.16 | 3.55E-05 | 4.98E-05 |
| chr6 | 142549218 | 142550564 | 1347 | -3.95 | 3.56E-05 | 4.99E-05 |

| | | | | | | |
|---|---|---|---|---|---|---|
| chr4 | 184693711 | 184694220 | 510 | -4.9 | 3.56E-05 | 4.99E-05 |
| chrX | 55352747 | 55363164 | 10418 | 4.99 | 3.57E-05 | 5.00E-05 |
| chr2 | 217289718 | 217290516 | 799 | -5.12 | 3.58E-05 | 5.01E-05 |
| chr2 | 57722138 | 57722684 | 547 | -4.85 | 3.58E-05 | 5.01E-05 |
| chr5 | 126324884 | 126325749 | 866 | -4.85 | 3.58E-05 | 5.01E-05 |
| chr2 | 192024647 | 192025479 | 833 | -5.15 | 3.58E-05 | 5.02E-05 |
| chr1 | 202941670 | 202942161 | 492 | -4.89 | 3.59E-05 | 5.02E-05 |
| chr3 | 135780826 | 135781753 | 928 | -4.89 | 3.59E-05 | 5.02E-05 |
| chr15 | 56304658 | 56309079 | 4422 | -4.61 | 3.59E-05 | 5.02E-05 |
| chr14 | 49357802 | 49359288 | 1487 | -3.82 | 3.59E-05 | 5.03E-05 |
| chr6 | 3666637 | 3680719 | 14083 | -4.04 | 3.59E-05 | 5.03E-05 |
| chr13 | 91845492 | 91848034 | 2543 | 1.96 | 3.59E-05 | 5.03E-05 |
| chr6 | 161093959 | 161097330 | 3372 | -2.14 | 3.59E-05 | 5.03E-05 |
| chrX | 107953463 | 107958617 | 5155 | -4.64 | 3.59E-05 | 5.03E-05 |
| chr2 | 62382175 | 62382911 | 737 | -4.96 | 3.60E-05 | 5.04E-05 |
| chr1 | 230496147 | 230497382 | 1236 | -5.13 | 3.61E-05 | 5.06E-05 |
| chr12 | 31017541 | 31022626 | 5086 | -5.26 | 3.61E-05 | 5.06E-05 |
| chr1 | 37142697 | 37143422 | 726 | -3.75 | 3.61E-05 | 5.06E-05 |
| chr3 | 75918045 | 75937283 | 19239 | 1.44 | 3.62E-05 | 5.06E-05 |
| chr14 | 106652889 | 106667502 | 14614 | 1.05 | 3.62E-05 | 5.07E-05 |
| chrX | 101350810 | 101352684 | 1875 | 2.28 | 3.62E-05 | 5.07E-05 |
| chr17 | 15629597 | 15630020 | 424 | -4.87 | 3.62E-05 | 5.07E-05 |
| chr4 | 73855774 | 73857702 | 1929 | -4.03 | 3.63E-05 | 5.08E-05 |
| chr1 | 29023560 | 29024286 | 727 | -5.1 | 3.63E-05 | 5.09E-05 |
| chr4 | 11435997 | 11436564 | 568 | -3.77 | 3.63E-05 | 5.09E-05 |
| chr1 | 97441924 | 97442793 | 870 | -3.19 | 3.63E-05 | 5.09E-05 |
| chr5 | 120863932 | 120876212 | 12281 | -5.43 | 3.64E-05 | 5.09E-05 |
| chr6 | 100465529 | 100467536 | 2008 | -4.19 | 3.64E-05 | 5.09E-05 |
| chrX | 51241449 | 51242565 | 1117 | -2.16 | 3.65E-05 | 5.11E-05 |
| chr13 | 93908313 | 93910143 | 1831 | 4.1 | 3.65E-05 | 5.11E-05 |
| chr18 | 52740887 | 52741680 | 794 | -3.9 | 3.66E-05 | 5.13E-05 |
| chr17 | 66807930 | 66808792 | 863 | -2.7 | 3.66E-05 | 5.13E-05 |
| chr6 | 167303932 | 167304710 | 779 | -2.4 | 3.66E-05 | 5.13E-05 |
| chr21 | 39839695 | 39840735 | 1041 | -5.08 | 3.66E-05 | 5.13E-05 |
| chr17 | 35327202 | 35327521 | 320 | -4.91 | 3.67E-05 | 5.13E-05 |
| chr8 | 10915641 | 10923358 | 7718 | -5.7 | 3.67E-05 | 5.13E-05 |
| chr11 | 69049257 | 69050366 | 1110 | -5.4 | 3.67E-05 | 5.14E-05 |
| chr5 | 167222084 | 167233274 | 11191 | 1.6 | 3.67E-05 | 5.14E-05 |
| chr15 | 100446686 | 100448399 | 1714 | -2.18 | 3.68E-05 | 5.15E-05 |
| chr14 | 30413267 | 30417888 | 4622 | -3.53 | 3.68E-05 | 5.15E-05 |
| chr5 | 156232186 | 156233163 | 978 | -5.1 | 3.68E-05 | 5.15E-05 |
| chr7 | 117995667 | 117996763 | 1097 | -5.1 | 3.68E-05 | 5.15E-05 |
| chr6 | 160843587 | 160843964 | 378 | -4.3 | 3.68E-05 | 5.16E-05 |
| chr8 | 123229570 | 123410985 | 181416 | -0.91 | 3.69E-05 | 5.16E-05 |
| chr14 | 37205688 | 37208694 | 3007 | 2.17 | 3.71E-05 | 5.19E-05 |
| chr14 | 96250426 | 96286471 | 36046 | 1.47 | 3.71E-05 | 5.19E-05 |
| chr18 | 26933097 | 26935939 | 2843 | -1.91 | 3.71E-05 | 5.20E-05 |
| chr2 | 184171181 | 184172156 | 976 | 4.95 | 3.72E-05 | 5.20E-05 |
| chr4 | 72273885 | 72274978 | 1094 | -4.03 | 3.72E-05 | 5.21E-05 |
| chrX | 84695459 | 84718081 | 22623 | 3.51 | 3.73E-05 | 5.22E-05 |
| chr22 | 47574324 | 47583452 | 9129 | -5.03 | 3.73E-05 | 5.22E-05 |
| chr9 | 136365117 | 136369884 | 4768 | -5.4 | 3.73E-05 | 5.22E-05 |
| chr14 | 56399911 | 56416500 | 16590 | -5.39 | 3.74E-05 | 5.23E-05 |
| chr9 | 137454555 | 137459508 | 4954 | -4.87 | 3.74E-05 | 5.23E-05 |
| chr17 | 46562501 | 46563109 | 609 | -5.35 | 3.74E-05 | 5.24E-05 |
| chr20 | 14488369 | 14497580 | 9212 | -6.16 | 3.76E-05 | 5.25E-05 |
| chr6 | 126067467 | 126068943 | 1477 | -4.95 | 3.76E-05 | 5.25E-05 |
| chr16 | 49818064 | 49818691 | 628 | -5.06 | 3.76E-05 | 5.26E-05 |
| chr4 | 152731627 | 152732307 | 681 | -5.06 | 3.76E-05 | 5.26E-05 |
| chr11 | 15813398 | 15816284 | 2887 | -4.67 | 3.77E-05 | 5.27E-05 |
| chr11 | 125518793 | 125519721 | 929 | -5.04 | 3.77E-05 | 5.28E-05 |

| | | | | | | |
|---|---|---|---|---|---|---|
| chr3 | 40770384 | 40772162 | 1779 | -4.11 | 3.79E-05 | 5.30E-05 |
| chr4 | 134798393 | 134800964 | 2572 | 2.31 | 3.80E-05 | 5.31E-05 |
| chr14 | 103755224 | 103755895 | 672 | -4.84 | 3.80E-05 | 5.31E-05 |
| chr4 | 131054597 | 131055081 | 485 | -4.84 | 3.80E-05 | 5.31E-05 |
| chr8 | 61976650 | 61977711 | 1062 | -3.73 | 3.80E-05 | 5.32E-05 |
| chr4 | 74505687 | 74506179 | 493 | -4.89 | 3.80E-05 | 5.32E-05 |
| chr4 | 85158881 | 85159244 | 364 | -4.89 | 3.80E-05 | 5.32E-05 |
| chr22 | 36047125 | 36047665 | 541 | -5.05 | 3.81E-05 | 5.32E-05 |
| chr1 | 188719058 | 188720325 | 1268 | -4.87 | 3.81E-05 | 5.33E-05 |
| chr5 | 72971495 | 72971879 | 385 | -3.89 | 3.82E-05 | 5.33E-05 |
| chr3 | 155579099 | 155579768 | 670 | -5.03 | 3.82E-05 | 5.33E-05 |
| chr2 | 194406715 | 194408242 | 1528 | -4.84 | 3.82E-05 | 5.34E-05 |
| chrX | 11749423 | 11750399 | 977 | -5.29 | 3.82E-05 | 5.34E-05 |
| chr2 | 224040622 | 224053067 | 12446 | -5.61 | 3.83E-05 | 5.35E-05 |
| chr3 | 15587569 | 15589222 | 1654 | -5.61 | 3.83E-05 | 5.35E-05 |
| chr12 | 25440298 | 25443932 | 3635 | -4.47 | 3.83E-05 | 5.35E-05 |
| chr7 | 23762804 | 23763443 | 640 | -5.33 | 3.83E-05 | 5.35E-05 |
| chr2 | 230123153 | 230128648 | 5496 | -4.98 | 3.83E-05 | 5.35E-05 |
| chr13 | 107313038 | 107340642 | 27605 | 1.07 | 3.84E-05 | 5.36E-05 |
| chr8 | 127307191 | 127308673 | 1483 | 1.85 | 3.84E-05 | 5.37E-05 |
| chr1 | 49133811 | 49134818 | 1008 | 3.96 | 3.84E-05 | 5.37E-05 |
| chr6 | 75606712 | 75612757 | 6046 | -4.37 | 3.84E-05 | 5.37E-05 |
| chr12 | 41031214 | 41031892 | 679 | -4.85 | 3.85E-05 | 5.38E-05 |
| chr8 | 121068497 | 121069070 | 574 | -4.85 | 3.85E-05 | 5.38E-05 |
| chr12 | 117108903 | 117109464 | 562 | -4.93 | 3.85E-05 | 5.38E-05 |
| chr15 | 41909849 | 41910643 | 795 | -4.93 | 3.85E-05 | 5.38E-05 |
| chr3 | 184371952 | 184372542 | 591 | -4.93 | 3.85E-05 | 5.38E-05 |
| chr6 | 159240854 | 159241393 | 540 | -4.93 | 3.85E-05 | 5.38E-05 |
| chr8 | 28488736 | 28489387 | 652 | -4.93 | 3.85E-05 | 5.38E-05 |
| chr12 | 80088419 | 80089736 | 1318 | -5.03 | 3.86E-05 | 5.39E-05 |
| chr16 | 49864412 | 49865415 | 1004 | -5.03 | 3.86E-05 | 5.39E-05 |
| chr3 | 41883715 | 41884686 | 972 | -5.03 | 3.86E-05 | 5.39E-05 |
| chr10 | 132906806 | 133006598 | 99793 | 0.55 | 3.87E-05 | 5.40E-05 |
| chr4 | 176134833 | 176135775 | 943 | -4.85 | 3.87E-05 | 5.40E-05 |
| chrX | 28815971 | 28816768 | 798 | -4.85 | 3.87E-05 | 5.40E-05 |
| chr3 | 38364635 | 38369992 | 5358 | -4.71 | 3.87E-05 | 5.40E-05 |
| chr8 | 120439788 | 120449147 | 9360 | 1.12 | 3.88E-05 | 5.42E-05 |
| chr20 | 50735022 | 50738842 | 3821 | -5.24 | 3.89E-05 | 5.43E-05 |
| chr20 | 25166220 | 25174438 | 8219 | -5.57 | 3.89E-05 | 5.43E-05 |
| chr12 | 15700738 | 15704280 | 3543 | -1.7 | 3.90E-05 | 5.45E-05 |
| chr15 | 63752065 | 63752714 | 650 | -4.83 | 3.91E-05 | 5.46E-05 |
| chr3 | 50945155 | 50945550 | 396 | -4.83 | 3.91E-05 | 5.46E-05 |
| chr1 | 5609453 | 5610996 | 1544 | 1.91 | 3.91E-05 | 5.46E-05 |
| chrX | 99804102 | 99805453 | 1352 | -2.23 | 3.92E-05 | 5.47E-05 |
| chr15 | 74136050 | 74136472 | 423 | -4.27 | 3.92E-05 | 5.47E-05 |
| chr3 | 175672752 | 175675111 | 2360 | -5.01 | 3.92E-05 | 5.47E-05 |
| chr5 | 34630527 | 34640801 | 10275 | -5.79 | 3.92E-05 | 5.47E-05 |
| chr15 | 79891109 | 79903886 | 12778 | 1.69 | 3.93E-05 | 5.49E-05 |
| chr1 | 45599756 | 45601751 | 1996 | -5.21 | 3.93E-05 | 5.49E-05 |
| chr1 | 79704565 | 79705859 | 1295 | -2.98 | 3.94E-05 | 5.50E-05 |
| chr2 | 218099450 | 218103522 | 4073 | -4.65 | 3.94E-05 | 5.50E-05 |
| chr7 | 144225396 | 144299794 | 74399 | 0.97 | 3.94E-05 | 5.50E-05 |
| chr11 | 69614581 | 69615438 | 858 | -4.9 | 3.95E-05 | 5.51E-05 |
| chr20 | 31248169 | 31249082 | 914 | -4.9 | 3.95E-05 | 5.51E-05 |
| chr14 | 73387489 | 73388463 | 975 | -3.83 | 3.95E-05 | 5.52E-05 |
| chr17 | 81065318 | 81066605 | 1288 | -5.2 | 3.96E-05 | 5.52E-05 |
| chr11 | 123050692 | 123051300 | 609 | -4.85 | 3.96E-05 | 5.53E-05 |
| chr17 | 20952839 | 20953248 | 410 | -4.85 | 3.96E-05 | 5.53E-05 |
| chrX | 77156962 | 77157432 | 471 | -4.85 | 3.96E-05 | 5.53E-05 |
| chr7 | 64517701 | 64518045 | 345 | -4.87 | 3.96E-05 | 5.53E-05 |
| chr11 | 134492979 | 134494449 | 1471 | 2.07 | 3.96E-05 | 5.53E-05 |

| | | | | | | |
|---|---|---|---|---|---|---|
| chr12 | 53245394 | 53246197 | 804 | -4.92 | 3.96E-05 | 5.53E-05 |
| chr4 | 14888732 | 14889714 | 983 | -4.92 | 3.96E-05 | 5.53E-05 |
| chr4 | 2304217 | 2309365 | 5149 | -5.77 | 3.97E-05 | 5.53E-05 |
| chr1 | 112086775 | 112089466 | 2692 | -4.16 | 3.97E-05 | 5.54E-05 |
| chr2 | 11747182 | 11747535 | 354 | -4.21 | 3.97E-05 | 5.54E-05 |
| chrX | 81835169 | 81837260 | 2092 | -3.48 | 3.98E-05 | 5.56E-05 |
| chr4 | 40694033 | 40694922 | 890 | -4.99 | 3.98E-05 | 5.56E-05 |
| chr4 | 167964556 | 167965218 | 663 | -4.99 | 3.98E-05 | 5.56E-05 |
| chr6 | 136911175 | 136911673 | 499 | -4.99 | 3.98E-05 | 5.56E-05 |
| chr8 | 39612901 | 39614533 | 1633 | 2.2 | 3.99E-05 | 5.56E-05 |
| chr13 | 110435239 | 110435866 | 628 | -4.19 | 3.99E-05 | 5.56E-05 |
| chr2 | 106238359 | 106247834 | 9476 | -1.57 | 4.02E-05 | 5.60E-05 |
| chr6 | 774098 | 774698 | 601 | 3.04 | 4.02E-05 | 5.61E-05 |
| chr4 | 26799218 | 26800340 | 1123 | -5.15 | 4.04E-05 | 5.63E-05 |
| chr2 | 40751380 | 40757203 | 5824 | -1.3 | 4.04E-05 | 5.64E-05 |
| chr17 | 39722576 | 39727438 | 4863 | 4.77 | 4.05E-05 | 5.65E-05 |
| chr17 | 74598680 | 74599482 | 803 | -4.98 | 4.06E-05 | 5.66E-05 |
| chr6 | 135650407 | 135650946 | 540 | -4.98 | 4.06E-05 | 5.66E-05 |
| chr7 | 12293202 | 12293658 | 457 | -4.98 | 4.06E-05 | 5.66E-05 |
| chr2 | 172611786 | 172612185 | 400 | -4.88 | 4.06E-05 | 5.66E-05 |
| chr2 | 239324240 | 239324857 | 618 | -5.13 | 4.07E-05 | 5.68E-05 |
| chr4 | 59679049 | 59680505 | 1457 | -5.13 | 4.07E-05 | 5.68E-05 |
| chrX | 107382810 | 107383236 | 427 | -4.83 | 4.07E-05 | 5.68E-05 |
| chr7 | 77499412 | 77499868 | 457 | -4.25 | 4.08E-05 | 5.68E-05 |
| chr1 | 152246588 | 152260392 | 13805 | -2.03 | 4.08E-05 | 5.69E-05 |
| chr17 | 16640376 | 16640871 | 496 | -4.9 | 4.09E-05 | 5.70E-05 |
| chr2 | 236844776 | 236845275 | 500 | -4.82 | 4.10E-05 | 5.71E-05 |
| chr3 | 18015804 | 18016360 | 557 | -4.82 | 4.10E-05 | 5.71E-05 |
| chr13 | 110160774 | 110161771 | 998 | -4.37 | 4.10E-05 | 5.71E-05 |
| chr9 | 72232062 | 72242835 | 10774 | 1.2 | 4.11E-05 | 5.72E-05 |
| chrX | 99753054 | 99755732 | 2679 | -4.17 | 4.11E-05 | 5.72E-05 |
| chrX | 38223805 | 38407773 | 183969 | 4.47 | 4.12E-05 | 5.74E-05 |
| chr17 | 2622918 | 2623605 | 688 | -4.96 | 4.14E-05 | 5.76E-05 |
| chr4 | 10424467 | 10425094 | 628 | -4.96 | 4.14E-05 | 5.76E-05 |
| chr8 | 66911077 | 66912103 | 1027 | -4.85 | 4.14E-05 | 5.77E-05 |
| chr16 | 58639062 | 58640267 | 1206 | -5.53 | 4.14E-05 | 5.77E-05 |
| chrX | 76630323 | 76631087 | 765 | -2.6 | 4.15E-05 | 5.78E-05 |
| chr5 | 169252905 | 169253582 | 678 | -4.82 | 4.16E-05 | 5.80E-05 |
| chr1 | 211283283 | 211295587 | 12305 | -6.28 | 4.17E-05 | 5.80E-05 |
| chr13 | 35899815 | 35900895 | 1081 | -5.12 | 4.17E-05 | 5.81E-05 |
| chr9 | 913986 | 1054295 | 140310 | -2.1 | 4.17E-05 | 5.81E-05 |
| chr4 | 134462580 | 134463152 | 573 | 4.91 | 4.17E-05 | 5.81E-05 |
| chr6 | 17216204 | 17217297 | 1094 | -5.08 | 4.18E-05 | 5.83E-05 |
| chr2 | 224754370 | 224754756 | 387 | -4.83 | 4.19E-05 | 5.84E-05 |
| chr6 | 163804262 | 163805056 | 795 | -4.83 | 4.19E-05 | 5.84E-05 |
| chr11 | 56568640 | 56577931 | 9292 | -2.76 | 4.20E-05 | 5.86E-05 |
| chrX | 74646029 | 74680986 | 34958 | 5.51 | 4.20E-05 | 5.86E-05 |
| chr6 | 116417274 | 116418161 | 888 | -4.94 | 4.23E-05 | 5.89E-05 |
| chr17 | 68778641 | 69279977 | 501337 | 0.61 | 4.23E-05 | 5.89E-05 |
| chr2 | 236532681 | 236533724 | 1044 | -5.35 | 4.23E-05 | 5.89E-05 |
| chr6 | 38416926 | 38417896 | 971 | -5.35 | 4.23E-05 | 5.89E-05 |
| chr13 | 90015343 | 90016170 | 828 | -4.88 | 4.23E-05 | 5.89E-05 |
| chr19 | 6651951 | 6652790 | 840 | -4.88 | 4.23E-05 | 5.89E-05 |
| chr2 | 188816067 | 188816739 | 673 | -4.88 | 4.23E-05 | 5.89E-05 |
| chr4 | 159850141 | 159850783 | 643 | -4.88 | 4.23E-05 | 5.89E-05 |
| chr17 | 71768869 | 71769902 | 1034 | 4.91 | 4.23E-05 | 5.90E-05 |
| chr2 | 7189545 | 7207848 | 18304 | -1.55 | 4.24E-05 | 5.90E-05 |
| chrX | 23475977 | 23511849 | 35873 | -1.98 | 4.24E-05 | 5.90E-05 |
| chrX | 3415376 | 3418450 | 3075 | 1.73 | 4.25E-05 | 5.92E-05 |
| chr2 | 56677619 | 56696331 | 18713 | -1.47 | 4.26E-05 | 5.93E-05 |
| chr11 | 6357585 | 6358016 | 432 | -5.05 | 4.27E-05 | 5.95E-05 |

| | | | | | | |
|---|---|---|---|---|---|---|
| chr9 | 138735673 | 138736510 | 838 | -5.05 | 4.27E-05 | 5.95E-05 |
| chr5 | 121185689 | 121188338 | 2650 | -1.79 | 4.28E-05 | 5.96E-05 |
| chr2 | 96406875 | 96407205 | 331 | -3.85 | 4.28E-05 | 5.96E-05 |
| chr5 | 108508512 | 108509179 | 668 | -4.63 | 4.29E-05 | 5.97E-05 |
| chr5 | 136888553 | 136889230 | 678 | -5.31 | 4.30E-05 | 5.98E-05 |
| chrX | 83469017 | 83469648 | 632 | -4.81 | 4.30E-05 | 5.99E-05 |
| chr15 | 79077525 | 79077886 | 362 | -4.83 | 4.32E-05 | 6.01E-05 |
| chr6 | 74706637 | 74707203 | 567 | -4.83 | 4.32E-05 | 6.01E-05 |
| chr8 | 116458576 | 116459526 | 951 | -4.83 | 4.32E-05 | 6.01E-05 |
| chr10 | 25418924 | 25443500 | 24577 | -1.32 | 4.32E-05 | 6.02E-05 |
| chr2 | 55667634 | 55669317 | 1684 | -4.85 | 4.32E-05 | 6.02E-05 |
| chr11 | 101795541 | 101796089 | 549 | -4.92 | 4.33E-05 | 6.02E-05 |
| chr4 | 75088596 | 75089162 | 567 | -4.92 | 4.33E-05 | 6.02E-05 |
| chr13 | 96582005 | 96582846 | 842 | -4.91 | 4.33E-05 | 6.03E-05 |
| chr21 | 30424138 | 30424938 | 801 | -4.91 | 4.33E-05 | 6.03E-05 |
| chr10 | 126047313 | 126060644 | 13332 | -5.8 | 4.35E-05 | 6.05E-05 |
| chr10 | 129050967 | 129051345 | 379 | -4.83 | 4.35E-05 | 6.05E-05 |
| chr3 | 97648996 | 97649662 | 667 | -4.83 | 4.35E-05 | 6.05E-05 |
| chr6 | 166705079 | 166705834 | 756 | -4.83 | 4.35E-05 | 6.05E-05 |
| chr1 | 64030035 | 64031135 | 1101 | -5.26 | 4.37E-05 | 6.08E-05 |
| chr1 | 197676033 | 197676621 | 589 | -5.02 | 4.38E-05 | 6.09E-05 |
| chr11 | 47231649 | 47232427 | 779 | -5.02 | 4.38E-05 | 6.09E-05 |
| chr17 | 75269784 | 75270422 | 639 | -5.02 | 4.38E-05 | 6.09E-05 |
| chr20 | 35537955 | 35538575 | 621 | -5.02 | 4.38E-05 | 6.09E-05 |
| chr5 | 107182120 | 107182624 | 505 | -5.02 | 4.38E-05 | 6.09E-05 |
| chr4 | 88741129 | 88741987 | 859 | -4.17 | 4.38E-05 | 6.10E-05 |
| chr9 | 21752994 | 21754272 | 1279 | 2.72 | 4.39E-05 | 6.10E-05 |
| chr20 | 4020711 | 4030864 | 10154 | -4.86 | 4.39E-05 | 6.11E-05 |
| chr8 | 99199387 | 99211081 | 11695 | -5.25 | 4.40E-05 | 6.12E-05 |
| chr4 | 150229065 | 150229818 | 754 | 3.82 | 4.41E-05 | 6.13E-05 |
| chr5 | 102120314 | 102121043 | 730 | -4.81 | 4.41E-05 | 6.14E-05 |
| chr1 | 85303484 | 85306259 | 2776 | -3.9 | 4.42E-05 | 6.14E-05 |
| chr5 | 154943934 | 154954811 | 10878 | -1.18 | 4.42E-05 | 6.15E-05 |
| chr8 | 67676845 | 67677484 | 640 | -5.23 | 4.42E-05 | 6.15E-05 |
| chr4 | 118141594 | 118142737 | 1144 | -3.69 | 4.43E-05 | 6.16E-05 |
| chrX | 100209363 | 100210026 | 664 | -4.9 | 4.43E-05 | 6.16E-05 |
| chr1 | 20056357 | 20066234 | 9878 | -5.54 | 4.44E-05 | 6.17E-05 |
| chr11 | 95192005 | 95195726 | 3722 | -2.32 | 4.44E-05 | 6.17E-05 |
| chr1 | 94375720 | 94376509 | 790 | -4.9 | 4.44E-05 | 6.18E-05 |
| chr16 | 69851443 | 69852031 | 589 | -4.9 | 4.44E-05 | 6.18E-05 |
| chr2 | 88962633 | 88963436 | 804 | -4.9 | 4.44E-05 | 6.18E-05 |
| chr10 | 89989383 | 89989910 | 528 | -4.8 | 4.45E-05 | 6.18E-05 |
| chr14 | 34377778 | 34378406 | 629 | -4.8 | 4.45E-05 | 6.18E-05 |
| chr16 | 86923621 | 86924384 | 764 | -4.8 | 4.45E-05 | 6.18E-05 |
| chr6 | 47178914 | 47180750 | 1837 | -4.1 | 4.46E-05 | 6.19E-05 |
| chr4 | 6655848 | 6656316 | 469 | -4.81 | 4.46E-05 | 6.20E-05 |
| chr13 | 25737940 | 25776425 | 38486 | -1.99 | 4.46E-05 | 6.20E-05 |
| chr1 | 177858780 | 177880228 | 21449 | -3.12 | 4.47E-05 | 6.21E-05 |
| chr19 | 51212452 | 51213496 | 1045 | -5.2 | 4.48E-05 | 6.22E-05 |
| chr2 | 193838309 | 193839785 | 1477 | -5.2 | 4.48E-05 | 6.22E-05 |
| chr3 | 57520618 | 57521930 | 1313 | -5.2 | 4.48E-05 | 6.22E-05 |
| chr13 | 84187719 | 84188705 | 987 | -3.84 | 4.48E-05 | 6.23E-05 |
| chr4 | 7606183 | 7607013 | 831 | -5.19 | 4.51E-05 | 6.26E-05 |
| chr18 | 11138478 | 11141523 | 3046 | 1.86 | 4.51E-05 | 6.26E-05 |
| chr12 | 89927404 | 89928137 | 734 | -4.98 | 4.51E-05 | 6.27E-05 |
| chr2 | 27733060 | 27733431 | 372 | -4.98 | 4.51E-05 | 6.27E-05 |
| chr3 | 190395524 | 190396208 | 685 | -4.98 | 4.51E-05 | 6.27E-05 |
| chr4 | 104823252 | 104824015 | 764 | -4.98 | 4.51E-05 | 6.27E-05 |
| chr5 | 172220656 | 172221608 | 953 | -4.81 | 4.53E-05 | 6.29E-05 |
| chr6 | 36664828 | 36665392 | 565 | -4.81 | 4.53E-05 | 6.29E-05 |
| chr18 | 44212861 | 44217432 | 4572 | -1.33 | 4.54E-05 | 6.30E-05 |

| | | | | | | |
|---|---|---|---|---|---|---|
| chr21 | 33333325 | 33334382 | 1058 | -3.89 | 4.55E-05 | 6.32E-05 |
| chr10 | 100993262 | 100993948 | 687 | -4.8 | 4.56E-05 | 6.34E-05 |
| chr14 | 51688297 | 51689927 | 1631 | -4.8 | 4.56E-05 | 6.34E-05 |
| chr2 | 191343872 | 191344230 | 359 | -4.8 | 4.56E-05 | 6.34E-05 |
| chr18 | 47163619 | 47164529 | 911 | -3.65 | 4.56E-05 | 6.34E-05 |
| chr11 | 43309680 | 43326634 | 16955 | -4 | 4.56E-05 | 6.34E-05 |
| chr1 | 230513333 | 230517241 | 3909 | -5.07 | 4.57E-05 | 6.34E-05 |
| chr6 | 16973164 | 17011776 | 38613 | -0.75 | 4.57E-05 | 6.35E-05 |
| chr1 | 199040661 | 199041212 | 552 | -4.88 | 4.57E-05 | 6.35E-05 |
| chr13 | 102259880 | 102260459 | 580 | -4.88 | 4.57E-05 | 6.35E-05 |
| chr18 | 68143902 | 68154920 | 11019 | -1.48 | 4.58E-05 | 6.36E-05 |
| chr2 | 129048052 | 129048679 | 628 | -4.84 | 4.58E-05 | 6.36E-05 |
| chr7 | 22835695 | 22836357 | 663 | -4.84 | 4.58E-05 | 6.36E-05 |
| chr2 | 64300497 | 64301164 | 668 | -4.96 | 4.59E-05 | 6.37E-05 |
| chr5 | 74083087 | 74083755 | 669 | -4.96 | 4.59E-05 | 6.37E-05 |
| chr1 | 20596170 | 20598232 | 2063 | -3.17 | 4.59E-05 | 6.37E-05 |
| chr20 | 11122350 | 11123391 | 1042 | -5.14 | 4.60E-05 | 6.39E-05 |
| chr20 | 50753834 | 50758274 | 4441 | -5.24 | 4.62E-05 | 6.41E-05 |
| chr6 | 101378749 | 101381855 | 3107 | -2.12 | 4.63E-05 | 6.42E-05 |
| chr10 | 37105753 | 37111345 | 5593 | -1.48 | 4.64E-05 | 6.44E-05 |
| chr1 | 190367928 | 190368590 | 663 | -4.86 | 4.66E-05 | 6.46E-05 |
| chr3 | 56059204 | 56079369 | 20166 | -1.5 | 4.66E-05 | 6.46E-05 |
| chr15 | 24640337 | 24642646 | 2310 | 2.89 | 4.66E-05 | 6.47E-05 |
| chrX | 67086882 | 67096822 | 9941 | 1.2 | 4.66E-05 | 6.47E-05 |
| chr2 | 154786004 | 154786560 | 557 | -5.22 | 4.67E-05 | 6.47E-05 |
| chr1 | 24019060 | 24019861 | 802 | -5.1 | 4.67E-05 | 6.49E-05 |
| chr13 | 81312287 | 81313283 | 997 | -5.1 | 4.67E-05 | 6.49E-05 |
| chr4 | 179592510 | 179594043 | 1534 | -5.1 | 4.67E-05 | 6.49E-05 |
| chr7 | 137781425 | 137782062 | 638 | -5.1 | 4.67E-05 | 6.49E-05 |
| chr20 | 49959956 | 49960790 | 835 | -4.94 | 4.68E-05 | 6.49E-05 |
| chr9 | 4347062 | 4348021 | 960 | -4.94 | 4.68E-05 | 6.49E-05 |
| chr4 | 94970995 | 94973132 | 2138 | -2.92 | 4.68E-05 | 6.50E-05 |
| chr1 | 101736784 | 101737579 | 796 | -3.78 | 4.68E-05 | 6.50E-05 |
| chr8 | 63955153 | 63967299 | 12147 | -3.66 | 4.69E-05 | 6.51E-05 |
| chrX | 23656950 | 23659626 | 2677 | -3.39 | 4.70E-05 | 6.52E-05 |
| chr4 | 56130504 | 56135152 | 4649 | 2.19 | 4.70E-05 | 6.52E-05 |
| chr7 | 132402897 | 132406518 | 3622 | -1.76 | 4.70E-05 | 6.53E-05 |
| chr14 | 97446022 | 97508566 | 62545 | 0.97 | 4.71E-05 | 6.53E-05 |
| chr5 | 93600133 | 93625473 | 25341 | 1.43 | 4.72E-05 | 6.54E-05 |
| chr3 | 148399991 | 148400438 | 448 | -4.86 | 4.72E-05 | 6.55E-05 |
| chr6 | 12177076 | 12177505 | 430 | -4.86 | 4.72E-05 | 6.55E-05 |
| chr11 | 93630607 | 93631211 | 605 | -3.95 | 4.73E-05 | 6.56E-05 |
| chr3 | 40010087 | 40010796 | 710 | -2.83 | 4.73E-05 | 6.56E-05 |
| chr8 | 52759826 | 52760458 | 633 | -5.18 | 4.74E-05 | 6.57E-05 |
| chr11 | 85376588 | 85385689 | 9102 | -5.63 | 4.76E-05 | 6.60E-05 |
| chr3 | 137688702 | 137689441 | 740 | -3.92 | 4.76E-05 | 6.60E-05 |
| chr8 | 60040210 | 60041639 | 1430 | -2.17 | 4.76E-05 | 6.61E-05 |
| chr15 | 43113466 | 43114238 | 773 | -4.92 | 4.77E-05 | 6.61E-05 |
| chr20 | 44795570 | 44796024 | 455 | -4.92 | 4.77E-05 | 6.61E-05 |
| chr1 | 9029189 | 9029777 | 589 | -4.78 | 4.77E-05 | 6.62E-05 |
| chr11 | 129857469 | 129858201 | 733 | -4.78 | 4.77E-05 | 6.62E-05 |
| chr15 | 30220839 | 30221509 | 671 | -4.78 | 4.77E-05 | 6.62E-05 |
| chr16 | 30141831 | 30142640 | 810 | -4.78 | 4.77E-05 | 6.62E-05 |
| chr8 | 31980472 | 31980868 | 397 | -4.78 | 4.77E-05 | 6.62E-05 |
| chr7 | 138492216 | 138492829 | 614 | -4.79 | 4.78E-05 | 6.62E-05 |
| chr14 | 49344306 | 49344649 | 344 | 4.88 | 4.79E-05 | 6.63E-05 |
| chr18 | 76504929 | 76507013 | 2085 | 2.12 | 4.79E-05 | 6.63E-05 |
| chr1 | 111818819 | 111822848 | 4030 | -2.95 | 4.79E-05 | 6.64E-05 |
| chr14 | 70435717 | 70436701 | 985 | -3.79 | 4.79E-05 | 6.64E-05 |
| chr13 | 113596599 | 113597090 | 492 | -4.82 | 4.79E-05 | 6.65E-05 |
| chr13 | 29926295 | 29929891 | 3597 | 1.9 | 4.80E-05 | 6.65E-05 |

| | | | | | | |
|---|---|---|---|---|---|---|
| chr16 | 58833314 | 58842403 | 9090 | -5.41 | 4.80E-05 | 6.66E-05 |
| chr10 | 34015755 | 34018278 | 2524 | -4.62 | 4.82E-05 | 6.69E-05 |
| chr3 | 108066889 | 108067997 | 1109 | -3.75 | 4.83E-05 | 6.69E-05 |
| chrX | 7895801 | 7896966 | 1166 | -4.15 | 4.83E-05 | 6.69E-05 |
| chr3 | 16622616 | 16623203 | 588 | -3.94 | 4.84E-05 | 6.70E-05 |
| chr3 | 40340709 | 40342390 | 1682 | -2.49 | 4.84E-05 | 6.71E-05 |
| chr8 | 39398207 | 39408368 | 10162 | -2.21 | 4.84E-05 | 6.71E-05 |
| chr16 | 86762295 | 86763012 | 718 | -4.8 | 4.85E-05 | 6.72E-05 |
| chr2 | 204027223 | 204027726 | 504 | -4.8 | 4.85E-05 | 6.72E-05 |
| chr10 | 12585616 | 12586205 | 590 | -4.78 | 4.86E-05 | 6.73E-05 |
| chr13 | 62901402 | 62901936 | 535 | -4.78 | 4.86E-05 | 6.73E-05 |
| chr14 | 45770049 | 45770526 | 478 | -5.04 | 4.86E-05 | 6.73E-05 |
| chr10 | 31698102 | 31698482 | 381 | -4.21 | 4.86E-05 | 6.73E-05 |
| chr6 | 10469321 | 10469893 | 573 | -4.79 | 4.87E-05 | 6.74E-05 |
| chr12 | 25454242 | 25458039 | 3798 | 1.65 | 4.87E-05 | 6.74E-05 |
| chr3 | 156212949 | 156223890 | 10942 | -4.8 | 4.87E-05 | 6.74E-05 |
| chr3 | 151971519 | 151972250 | 732 | -4.9 | 4.88E-05 | 6.76E-05 |
| chr6 | 153789129 | 153789713 | 585 | -4.9 | 4.88E-05 | 6.76E-05 |
| chr6 | 71845569 | 71857458 | 11890 | -1.27 | 4.89E-05 | 6.77E-05 |
| chr3 | 56386986 | 56430219 | 43234 | 1.31 | 4.89E-05 | 6.77E-05 |
| chr14 | 103628176 | 103628938 | 763 | -5.3 | 4.89E-05 | 6.77E-05 |
| chr1 | 203988060 | 203988872 | 813 | -4.84 | 4.89E-05 | 6.78E-05 |
| chr3 | 180397780 | 180398647 | 868 | -4.84 | 4.89E-05 | 6.78E-05 |
| chr14 | 59152997 | 59156992 | 3996 | -1.25 | 4.89E-05 | 6.78E-05 |
| chr13 | 48669668 | 48670245 | 578 | -5.28 | 4.92E-05 | 6.81E-05 |
| chr14 | 34454137 | 34455606 | 1470 | -5.28 | 4.92E-05 | 6.81E-05 |
| chr2 | 223722159 | 223723293 | 1135 | -5.28 | 4.92E-05 | 6.81E-05 |
| chr5 | 59231325 | 59231678 | 354 | -4.83 | 4.93E-05 | 6.83E-05 |
| chr8 | 90394780 | 90395603 | 824 | -4.83 | 4.93E-05 | 6.83E-05 |
| chr6 | 44516815 | 44518014 | 1200 | -3.31 | 4.94E-05 | 6.84E-05 |
| chr3 | 46161540 | 46217227 | 55688 | -2.64 | 4.96E-05 | 6.86E-05 |
| chr4 | 37446358 | 37460890 | 14533 | -3.14 | 4.97E-05 | 6.88E-05 |
| chr1 | 169706695 | 169707588 | 894 | -3.94 | 4.98E-05 | 6.89E-05 |
| chrX | 133319074 | 133346261 | 27188 | -3.65 | 4.98E-05 | 6.89E-05 |
| chr6 | 15958263 | 16007051 | 48789 | -5.31 | 4.98E-05 | 6.89E-05 |
| chr12 | 15091981 | 15093221 | 1241 | -5 | 4.98E-05 | 6.89E-05 |
| chr14 | 93392815 | 93393425 | 611 | -5 | 4.98E-05 | 6.89E-05 |
| chr3 | 158130390 | 158130935 | 546 | -5 | 4.98E-05 | 6.89E-05 |
| chr7 | 110910934 | 110911973 | 1040 | -5 | 4.98E-05 | 6.89E-05 |
| chrX | 101114012 | 101120540 | 6529 | 2.04 | 4.99E-05 | 6.90E-05 |
| chr4 | 82130125 | 82153866 | 23742 | -1.05 | 4.99E-05 | 6.90E-05 |
| chr2 | 84817416 | 84843153 | 25738 | -1.03 | 5.00E-05 | 6.93E-05 |
| chr16 | 85995031 | 86008764 | 13734 | -2.97 | 5.03E-05 | 6.96E-05 |
| chr10 | 65236067 | 65236476 | 410 | -4.78 | 5.03E-05 | 6.97E-05 |
| chr2 | 213773171 | 213774643 | 1473 | -5.22 | 5.04E-05 | 6.97E-05 |
| chr11 | 15002047 | 15004554 | 2508 | 4.29 | 5.04E-05 | 6.97E-05 |
| chr18 | 34023372 | 34024140 | 769 | -4.98 | 5.05E-05 | 6.98E-05 |
| chr13 | 39571796 | 39582985 | 11190 | -4.83 | 5.05E-05 | 6.99E-05 |
| chr22 | 29414524 | 29415103 | 580 | -5.2 | 5.07E-05 | 7.02E-05 |
| chr9 | 2523627 | 2539522 | 15896 | 5.23 | 5.10E-05 | 7.05E-05 |
| chr17 | 67396070 | 67407543 | 11474 | -5.72 | 5.10E-05 | 7.05E-05 |
| chr14 | 60800230 | 60800970 | 741 | -5.03 | 5.10E-05 | 7.06E-05 |
| chr9 | 72255505 | 72258805 | 3301 | -2.47 | 5.11E-05 | 7.06E-05 |
| chr12 | 15351486 | 15352107 | 622 | 3.94 | 5.11E-05 | 7.06E-05 |
| chr5 | 63916412 | 63920366 | 3955 | 1.47 | 5.11E-05 | 7.07E-05 |
| chr5 | 170929205 | 170930574 | 1370 | -5.38 | 5.11E-05 | 7.07E-05 |
| chr15 | 52900856 | 52902365 | 1510 | -3.98 | 5.12E-05 | 7.08E-05 |
| chr16 | 86872148 | 86872822 | 675 | -4.96 | 5.12E-05 | 7.08E-05 |
| chr5 | 172000377 | 172001053 | 677 | -4.96 | 5.12E-05 | 7.08E-05 |
| chr6 | 81980412 | 81983232 | 2821 | 1.86 | 5.12E-05 | 7.09E-05 |
| chrX | 12209236 | 12218546 | 9311 | -4.38 | 5.13E-05 | 7.10E-05 |

| | | | | | | |
|---|---|---|---|---|---|---|
| chr12 | 42690305 | 42690820 | 516 | -4.87 | 5.14E-05 | 7.10E-05 |
| chr2 | 174645146 | 174645862 | 717 | -4.87 | 5.14E-05 | 7.10E-05 |
| chr3 | 187258351 | 187258954 | 604 | -4.87 | 5.14E-05 | 7.10E-05 |
| chr6 | 143579870 | 143580553 | 684 | -4.87 | 5.14E-05 | 7.10E-05 |
| chrX | 76978548 | 76979044 | 497 | -4.87 | 5.14E-05 | 7.10E-05 |
| chr4 | 5846783 | 5879484 | 32702 | -2.6 | 5.16E-05 | 7.13E-05 |
| chr9 | 72854807 | 72855488 | 682 | -2.33 | 5.17E-05 | 7.14E-05 |
| chr4 | 147495985 | 147496513 | 529 | 4.91 | 5.18E-05 | 7.16E-05 |
| chr1 | 244418021 | 244418928 | 908 | -4.77 | 5.19E-05 | 7.17E-05 |
| chr22 | 38645538 | 38646338 | 801 | -4.77 | 5.19E-05 | 7.17E-05 |
| chr5 | 58547246 | 58547774 | 529 | -4.77 | 5.19E-05 | 7.17E-05 |
| chr7 | 105823286 | 105823737 | 452 | -4.77 | 5.19E-05 | 7.17E-05 |
| chr12 | 12776532 | 12777036 | 505 | -4.76 | 5.19E-05 | 7.17E-05 |
| chr8 | 30640647 | 30641311 | 665 | -4.76 | 5.19E-05 | 7.17E-05 |
| chr8 | 102009691 | 102010351 | 661 | -5.01 | 5.20E-05 | 7.18E-05 |
| chr7 | 94620064 | 94620513 | 450 | -4.95 | 5.21E-05 | 7.20E-05 |
| chr13 | 25934926 | 25935826 | 901 | -2.21 | 5.22E-05 | 7.21E-05 |
| chr5 | 131368068 | 131380671 | 12604 | -2.3 | 5.22E-05 | 7.22E-05 |
| chr3 | 70490055 | 70524083 | 34029 | -0.95 | 5.23E-05 | 7.23E-05 |
| chr6 | 169436268 | 169439754 | 3487 | -4.81 | 5.23E-05 | 7.23E-05 |
| chr2 | 130674400 | 130679184 | 4785 | -5.07 | 5.24E-05 | 7.24E-05 |
| chr3 | 58750421 | 58769736 | 19316 | -1.29 | 5.24E-05 | 7.25E-05 |
| chr1 | 5591026 | 5592089 | 1064 | 5.31 | 5.25E-05 | 7.25E-05 |
| chr4 | 184923265 | 184923854 | 590 | -4.99 | 5.25E-05 | 7.25E-05 |
| chr3 | 104715098 | 104718078 | 2981 | -1.56 | 5.27E-05 | 7.28E-05 |
| chr1 | 4334882 | 4357355 | 22474 | -1.07 | 5.28E-05 | 7.29E-05 |
| chr15 | 60335307 | 60411623 | 76317 | -0.88 | 5.31E-05 | 7.33E-05 |
| chr3 | 25734761 | 25735232 | 472 | -4.8 | 5.31E-05 | 7.34E-05 |
| chrX | 15935092 | 15943838 | 8747 | -4.45 | 5.31E-05 | 7.34E-05 |
| chr10 | 6971710 | 7020648 | 48939 | -1.07 | 5.31E-05 | 7.34E-05 |
| chr2 | 177735571 | 177738440 | 2870 | -4.59 | 5.31E-05 | 7.34E-05 |
| chr6 | 128889857 | 128890521 | 665 | -4.77 | 5.32E-05 | 7.34E-05 |
| chr1 | 211248445 | 211249160 | 716 | -3.69 | 5.32E-05 | 7.35E-05 |
| chr4 | 129735712 | 129736173 | 462 | -4.78 | 5.33E-05 | 7.36E-05 |
| chr10 | 61402084 | 61409318 | 7235 | -4.96 | 5.36E-05 | 7.40E-05 |
| chr14 | 91552337 | 91553223 | 887 | -4.84 | 5.37E-05 | 7.42E-05 |
| chr4 | 185963507 | 185983330 | 19824 | -4.86 | 5.38E-05 | 7.42E-05 |
| chr3 | 187475293 | 187476853 | 1561 | -4.13 | 5.38E-05 | 7.43E-05 |
| chr18 | 20893775 | 20894495 | 721 | -5.07 | 5.38E-05 | 7.43E-05 |
| chr12 | 66883568 | 66884141 | 574 | -3.63 | 5.40E-05 | 7.45E-05 |
| chr1 | 87946174 | 87946782 | 609 | -4.91 | 5.40E-05 | 7.46E-05 |
| chr13 | 21542523 | 21542987 | 465 | -4.91 | 5.40E-05 | 7.46E-05 |
| chr7 | 134563359 | 134563768 | 410 | -4.91 | 5.40E-05 | 7.46E-05 |
| chr1 | 110636805 | 110637594 | 790 | -5.36 | 5.40E-05 | 7.46E-05 |
| chr7 | 13570792 | 13572257 | 1466 | -5.36 | 5.40E-05 | 7.46E-05 |
| chr4 | 164140711 | 164144752 | 4042 | 2.15 | 5.42E-05 | 7.49E-05 |
| chr7 | 43967284 | 43968007 | 724 | -5.06 | 5.43E-05 | 7.50E-05 |
| chr2 | 198459967 | 198460518 | 552 | -4.95 | 5.43E-05 | 7.50E-05 |
| chr6 | 94550522 | 94756068 | 205547 | 1.01 | 5.44E-05 | 7.51E-05 |
| chr8 | 105781844 | 105783637 | 1794 | -4.38 | 5.45E-05 | 7.53E-05 |
| chr7 | 98179856 | 98220692 | 40837 | -5.06 | 5.46E-05 | 7.53E-05 |
| chr1 | 242214702 | 242215963 | 1262 | -5.33 | 5.47E-05 | 7.55E-05 |
| chr20 | 48385831 | 48386955 | 1125 | -5.33 | 5.47E-05 | 7.55E-05 |
| chr17 | 5773120 | 5776548 | 3429 | -2.06 | 5.47E-05 | 7.55E-05 |
| chr12 | 75917098 | 75917935 | 838 | -5.04 | 5.49E-05 | 7.57E-05 |
| chr7 | 12760021 | 12761430 | 1410 | -5.04 | 5.49E-05 | 7.57E-05 |
| chr9 | 43505019 | 43506212 | 1194 | -5.04 | 5.49E-05 | 7.57E-05 |
| chr10 | 120341226 | 120386213 | 44988 | -5.1 | 5.50E-05 | 7.59E-05 |
| chr11 | 134173055 | 134192875 | 19821 | -1.64 | 5.50E-05 | 7.59E-05 |
| chr11 | 63622586 | 63623209 | 624 | -4.93 | 5.51E-05 | 7.60E-05 |
| chrX | 129946426 | 129947101 | 676 | -4.83 | 5.51E-05 | 7.60E-05 |

| | | | | | | |
|---|---|---|---|---|---|---|
| chr7 | 118836222 | 119028160 | 191939 | 1.2 | 5.51E-05 | 7.61E-05 |
| chr13 | 50403796 | 50404501 | 706 | -4.89 | 5.51E-05 | 7.61E-05 |
| chr1 | 99449377 | 99451694 | 2318 | -5.79 | 5.54E-05 | 7.64E-05 |
| chr22 | 36750133 | 36750653 | 521 | -5.02 | 5.54E-05 | 7.65E-05 |
| chr5 | 10835026 | 10909787 | 74762 | -1.36 | 5.55E-05 | 7.66E-05 |
| chrX | 118971269 | 118971836 | 568 | -4.75 | 5.57E-05 | 7.68E-05 |
| chr13 | 84557563 | 84558356 | 794 | -4.78 | 5.57E-05 | 7.68E-05 |
| chr12 | 33064406 | 33065194 | 789 | -4.75 | 5.57E-05 | 7.69E-05 |
| chr15 | 75434076 | 75443124 | 9049 | -6.39 | 5.58E-05 | 7.70E-05 |
| chr7 | 109476505 | 109476932 | 428 | -4.92 | 5.58E-05 | 7.70E-05 |
| chrX | 132734511 | 132737364 | 2854 | 3.13 | 5.59E-05 | 7.71E-05 |
| chr13 | 101994106 | 101995037 | 932 | -5.16 | 5.59E-05 | 7.71E-05 |
| chr2 | 180446134 | 180446636 | 503 | -3.8 | 5.59E-05 | 7.71E-05 |
| chr12 | 14971395 | 14971887 | 493 | -4.75 | 5.59E-05 | 7.71E-05 |
| chr4 | 112766546 | 112766956 | 411 | -4.75 | 5.59E-05 | 7.71E-05 |
| chr1 | 104500015 | 104536654 | 36640 | 1.66 | 5.60E-05 | 7.72E-05 |
| chr1 | 244883512 | 244884342 | 831 | -5.27 | 5.60E-05 | 7.73E-05 |
| chr15 | 85531637 | 85532140 | 504 | -5 | 5.61E-05 | 7.73E-05 |
| chr7 | 100666841 | 100667476 | 636 | -5 | 5.61E-05 | 7.73E-05 |
| chr3 | 40141829 | 40143824 | 1996 | -4.43 | 5.62E-05 | 7.75E-05 |
| chr5 | 28965928 | 29428026 | 462099 | 1.18 | 5.63E-05 | 7.76E-05 |
| chr5 | 110426500 | 110427851 | 1352 | -4.14 | 5.64E-05 | 7.77E-05 |
| chr7 | 18467188 | 18468109 | 922 | -4.87 | 5.64E-05 | 7.78E-05 |
| chr2 | 23475897 | 23478932 | 3036 | -1.87 | 5.66E-05 | 7.80E-05 |
| chr19 | 12932315 | 12933201 | 887 | -4.76 | 5.66E-05 | 7.81E-05 |
| chr8 | 42731468 | 42732003 | 536 | -4.76 | 5.66E-05 | 7.81E-05 |
| chr1 | 248749503 | 248795172 | 45670 | 2.03 | 5.67E-05 | 7.82E-05 |
| chr14 | 71363088 | 71363700 | 613 | -4.81 | 5.67E-05 | 7.82E-05 |
| chr5 | 162922618 | 162923289 | 672 | -4.99 | 5.68E-05 | 7.82E-05 |
| chr1 | 235059759 | 235061343 | 1585 | -4.74 | 5.68E-05 | 7.83E-05 |
| chr11 | 73214601 | 73214959 | 359 | -4.74 | 5.68E-05 | 7.83E-05 |
| chr10 | 31028734 | 31030099 | 1366 | -5.22 | 5.71E-05 | 7.87E-05 |
| chr3 | 172328559 | 172329351 | 793 | -5.22 | 5.71E-05 | 7.87E-05 |
| chr9 | 102076942 | 102177812 | 100871 | 2.74 | 5.71E-05 | 7.87E-05 |
| chr17 | 43282274 | 43283160 | 887 | -5.5 | 5.71E-05 | 7.87E-05 |
| chr19 | 40174390 | 40281602 | 107213 | -1.26 | 5.73E-05 | 7.90E-05 |
| chr9 | 119854216 | 119855700 | 1485 | -4.97 | 5.75E-05 | 7.92E-05 |
| chr18 | 61624933 | 61627275 | 2343 | -4.43 | 5.76E-05 | 7.94E-05 |
| chr9 | 2573703 | 2574458 | 756 | 4.94 | 5.76E-05 | 7.94E-05 |
| chr5 | 85782204 | 85804591 | 22388 | 2.38 | 5.77E-05 | 7.95E-05 |
| chr2 | 200979434 | 200979941 | 508 | -2.96 | 5.78E-05 | 7.97E-05 |
| chr10 | 81256106 | 81256818 | 713 | -4.85 | 5.79E-05 | 7.97E-05 |
| chr12 | 32990865 | 32991309 | 445 | -4.85 | 5.79E-05 | 7.97E-05 |
| chr6 | 5574619 | 5575394 | 776 | -4.85 | 5.79E-05 | 7.97E-05 |
| chr10 | 45758579 | 45761810 | 3232 | -1.52 | 5.79E-05 | 7.98E-05 |
| chr10 | 64794677 | 64807546 | 12870 | -5.27 | 5.81E-05 | 8.00E-05 |
| chr2 | 163451236 | 163453396 | 2161 | -2.26 | 5.81E-05 | 8.00E-05 |
| chr14 | 30029140 | 30042114 | 12975 | -2.39 | 5.82E-05 | 8.02E-05 |
| chr6 | 84170430 | 84171376 | 947 | -3.89 | 5.84E-05 | 8.04E-05 |
| chr2 | 234226082 | 234226593 | 512 | -5.07 | 5.84E-05 | 8.04E-05 |
| chr15 | 45157892 | 45160440 | 2549 | -1.98 | 5.85E-05 | 8.05E-05 |
| chr10 | 58289047 | 58292046 | 3000 | -4.76 | 5.86E-05 | 8.07E-05 |
| chr16 | 46584635 | 46585808 | 1174 | -4.76 | 5.86E-05 | 8.07E-05 |
| chr5 | 68753844 | 68755585 | 1742 | -5.16 | 5.87E-05 | 8.08E-05 |
| chr1 | 22551486 | 22552060 | 575 | -5.06 | 5.88E-05 | 8.10E-05 |
| chr3 | 60172156 | 60173967 | 1812 | 2.54 | 5.88E-05 | 8.10E-05 |
| chr3 | 64691421 | 64692009 | 589 | -4.78 | 5.90E-05 | 8.13E-05 |
| chr3 | 24818031 | 24840431 | 22401 | -1.06 | 5.91E-05 | 8.13E-05 |
| chr8 | 99186150 | 99186771 | 622 | -4.74 | 5.91E-05 | 8.14E-05 |
| chr12 | 72342535 | 72655454 | 312920 | 8.6 | 5.92E-05 | 8.15E-05 |
| chr8 | 24121274 | 24124274 | 3001 | -1.65 | 5.92E-05 | 8.16E-05 |

| | | | | | | |
|---|---|---|---|---|---|---|
| chr2 | 226364044 | 226374634 | 10591 | 4.72 | 5.93E-05 | 8.16E-05 |
| chr4 | 108665330 | 108670854 | 5525 | -3.15 | 5.94E-05 | 8.18E-05 |
| chr3 | 172256321 | 172256894 | 574 | -4.83 | 5.95E-05 | 8.19E-05 |
| chr7 | 84622841 | 84623579 | 739 | -4.83 | 5.95E-05 | 8.19E-05 |
| chr10 | 127467113 | 127468191 | 1079 | -5.13 | 5.95E-05 | 8.19E-05 |
| chr3 | 138959589 | 138960423 | 835 | -4.13 | 5.96E-05 | 8.20E-05 |
| chr12 | 95814045 | 95814671 | 627 | -4.75 | 5.97E-05 | 8.22E-05 |
| chr4 | 105035653 | 105036263 | 611 | -3.77 | 5.97E-05 | 8.22E-05 |
| chr15 | 51727446 | 51738787 | 11342 | -2.81 | 5.97E-05 | 8.22E-05 |
| chr2 | 141631485 | 141632791 | 1307 | -3.99 | 5.97E-05 | 8.22E-05 |
| chr6 | 49766320 | 49793096 | 26777 | 5.43 | 5.98E-05 | 8.24E-05 |
| chr15 | 62435188 | 62455974 | 20787 | -4.1 | 5.99E-05 | 8.24E-05 |
| chr15 | 100363529 | 100365429 | 1901 | -2.78 | 6.00E-05 | 8.26E-05 |
| chr8 | 59592598 | 59596815 | 4218 | -1.93 | 6.01E-05 | 8.26E-05 |
| chr2 | 211041888 | 211042167 | 280 | -4.73 | 6.01E-05 | 8.26E-05 |
| chr4 | 3818736 | 3819265 | 530 | -4.73 | 6.01E-05 | 8.26E-05 |
| chr2 | 153750108 | 153752351 | 2244 | 2 | 6.01E-05 | 8.27E-05 |
| chr10 | 90273807 | 90274538 | 732 | -3.95 | 6.02E-05 | 8.28E-05 |
| chr10 | 94248292 | 94248884 | 593 | -4.91 | 6.02E-05 | 8.28E-05 |
| chr11 | 47011235 | 47011690 | 456 | -4.91 | 6.02E-05 | 8.28E-05 |
| chr11 | 117440002 | 117440371 | 370 | -4.91 | 6.02E-05 | 8.28E-05 |
| chr7 | 135353362 | 135354103 | 742 | -4.91 | 6.02E-05 | 8.28E-05 |
| chr17 | 37061062 | 37061340 | 279 | -4.78 | 6.03E-05 | 8.29E-05 |
| chr3 | 32379992 | 32380605 | 614 | -4.78 | 6.03E-05 | 8.29E-05 |
| chr18 | 69100123 | 69111100 | 10978 | 1.54 | 6.03E-05 | 8.30E-05 |
| chr17 | 27316991 | 27317764 | 774 | -5.39 | 6.05E-05 | 8.31E-05 |
| chr8 | 112087895 | 112088503 | 609 | -4.73 | 6.05E-05 | 8.32E-05 |
| chr22 | 34006709 | 34007322 | 614 | -4.72 | 6.08E-05 | 8.36E-05 |
| chrX | 121003624 | 121004133 | 510 | -4.72 | 6.08E-05 | 8.36E-05 |
| chr11 | 115457097 | 115471811 | 14715 | -1.47 | 6.09E-05 | 8.37E-05 |
| chr8 | 143033814 | 143043490 | 9677 | -5.21 | 6.11E-05 | 8.40E-05 |
| chr7 | 131382131 | 131396472 | 14342 | -5.08 | 6.11E-05 | 8.40E-05 |
| chr2 | 86492328 | 86501164 | 8837 | -5.56 | 6.12E-05 | 8.41E-05 |
| chrX | 18129692 | 18150791 | 21100 | -0.9 | 6.12E-05 | 8.42E-05 |
| chr10 | 14144229 | 14149474 | 5246 | -2.78 | 6.12E-05 | 8.42E-05 |
| chr7 | 118713650 | 118718901 | 5252 | -2.76 | 6.13E-05 | 8.42E-05 |
| chrX | 84649650 | 84650393 | 744 | -4.89 | 6.13E-05 | 8.43E-05 |
| chr4 | 136057706 | 136059797 | 2092 | -3.01 | 6.13E-05 | 8.43E-05 |
| chr13 | 52733908 | 52734438 | 531 | -4.81 | 6.14E-05 | 8.44E-05 |
| chr13 | 34538423 | 34539917 | 1495 | -3.95 | 6.14E-05 | 8.45E-05 |
| chr16 | 19676411 | 19676852 | 442 | -4.76 | 6.17E-05 | 8.48E-05 |
| chr5 | 80746329 | 80746932 | 604 | -4.76 | 6.17E-05 | 8.48E-05 |
| chr5 | 148270624 | 148275337 | 4714 | -1.34 | 6.20E-05 | 8.52E-05 |
| chr6 | 142448617 | 142453909 | 5293 | -3.99 | 6.20E-05 | 8.52E-05 |
| chr3 | 112213496 | 112232950 | 19455 | -1.99 | 6.22E-05 | 8.55E-05 |
| chr2 | 62354486 | 62355148 | 663 | -5.32 | 6.23E-05 | 8.56E-05 |
| chr4 | 31577356 | 31579281 | 1926 | -5.32 | 6.23E-05 | 8.56E-05 |
| chr7 | 141406136 | 141407336 | 1201 | -5.32 | 6.23E-05 | 8.56E-05 |
| chr1 | 54193155 | 54194066 | 912 | -3.81 | 6.23E-05 | 8.56E-05 |
| chr3 | 125857363 | 125859016 | 1654 | -4.09 | 6.24E-05 | 8.58E-05 |
| chr5 | 82954560 | 82955064 | 505 | -4.76 | 6.24E-05 | 8.58E-05 |
| chr6 | 150457336 | 150457708 | 373 | -4.76 | 6.24E-05 | 8.58E-05 |
| chr6 | 119217997 | 119218462 | 466 | -4.87 | 6.26E-05 | 8.60E-05 |
| chr10 | 102291641 | 102292165 | 525 | -4.73 | 6.26E-05 | 8.60E-05 |
| chr11 | 65874749 | 65875194 | 446 | -4.73 | 6.26E-05 | 8.60E-05 |
| chr18 | 47335751 | 47336372 | 622 | -4.73 | 6.26E-05 | 8.60E-05 |
| chr8 | 58206805 | 58254017 | 47213 | 0.86 | 6.27E-05 | 8.61E-05 |
| chr12 | 102209282 | 102209948 | 667 | -4.72 | 6.27E-05 | 8.61E-05 |
| chr17 | 63115049 | 63115505 | 457 | -4.72 | 6.27E-05 | 8.61E-05 |
| chr1 | 234702713 | 234703457 | 745 | -5.03 | 6.27E-05 | 8.61E-05 |
| chr18 | 57035478 | 57037830 | 2353 | -3.17 | 6.28E-05 | 8.63E-05 |

| | | | | | | |
|---|---|---|---|---|---|---|
| chr5 | 134994323 | 134996237 | 1915 | -4.48 | 6.31E-05 | 8.66E-05 |
| chr3 | 143677342 | 143682793 | 5452 | -2.47 | 6.32E-05 | 8.68E-05 |
| chr11 | 18913083 | 18924257 | 11175 | -1.55 | 6.32E-05 | 8.69E-05 |
| chr20 | 37416214 | 37427604 | 11391 | -5.73 | 6.33E-05 | 8.70E-05 |
| chr4 | 82393671 | 82394717 | 1047 | -3.08 | 6.34E-05 | 8.71E-05 |
| chr2 | 9144919 | 9145498 | 580 | -4.79 | 6.35E-05 | 8.72E-05 |
| chr2 | 237620690 | 237621583 | 894 | -4.79 | 6.35E-05 | 8.72E-05 |
| chr7 | 114411479 | 114412081 | 603 | -4.79 | 6.35E-05 | 8.72E-05 |
| chr5 | 100714134 | 100777975 | 63842 | 2.18 | 6.37E-05 | 8.74E-05 |
| chrX | 65971100 | 65971495 | 396 | 4.81 | 6.37E-05 | 8.74E-05 |
| chr1 | 13844132 | 13886169 | 42038 | 0.9 | 6.37E-05 | 8.75E-05 |
| chr15 | 51135853 | 51137973 | 2121 | -4.37 | 6.37E-05 | 8.75E-05 |
| chr1 | 239317213 | 239323512 | 6300 | 4.79 | 6.37E-05 | 8.75E-05 |
| chr4 | 156741785 | 156742616 | 832 | -3.55 | 6.39E-05 | 8.77E-05 |
| chrX | 13644083 | 13649464 | 5382 | -4.47 | 6.39E-05 | 8.77E-05 |
| chr15 | 36034963 | 36035517 | 555 | -4.85 | 6.39E-05 | 8.77E-05 |
| chr15 | 68870137 | 68870575 | 439 | -4.85 | 6.39E-05 | 8.77E-05 |
| chr4 | 58982557 | 59012172 | 29616 | 1.13 | 6.39E-05 | 8.78E-05 |
| chr13 | 104310858 | 104314274 | 3417 | 4.37 | 6.40E-05 | 8.79E-05 |
| chr6 | 112282884 | 112285712 | 2829 | -1.65 | 6.42E-05 | 8.81E-05 |
| chr8 | 121294388 | 121297696 | 3309 | 3.49 | 6.43E-05 | 8.82E-05 |
| chr18 | 14731369 | 14775161 | 43793 | 1.14 | 6.44E-05 | 8.84E-05 |
| chr2 | 152246667 | 152251471 | 4805 | -4.91 | 6.48E-05 | 8.89E-05 |
| chr3 | 24722407 | 24722932 | 526 | -4.97 | 6.48E-05 | 8.90E-05 |
| chr11 | 20386427 | 20387034 | 608 | -4.75 | 6.48E-05 | 8.90E-05 |
| chr9 | 74353485 | 74353897 | 413 | -4.75 | 6.48E-05 | 8.90E-05 |
| chr9 | 83528779 | 83544014 | 15236 | -5.07 | 6.48E-05 | 8.90E-05 |
| chr10 | 54339558 | 54349330 | 9773 | -3.97 | 6.49E-05 | 8.91E-05 |
| chrX | 113986456 | 114031587 | 45132 | 1.44 | 6.49E-05 | 8.91E-05 |
| chr1 | 52763905 | 52764475 | 571 | -4.71 | 6.50E-05 | 8.93E-05 |
| chr13 | 100107075 | 100107534 | 460 | -4.71 | 6.50E-05 | 8.93E-05 |
| chr14 | 31694090 | 31694559 | 470 | -4.71 | 6.52E-05 | 8.95E-05 |
| chr14 | 38331909 | 38333232 | 1324 | -4.71 | 6.52E-05 | 8.95E-05 |
| chr5 | 167510371 | 167523471 | 13101 | -4.71 | 6.52E-05 | 8.95E-05 |
| chr8 | 110704929 | 110708173 | 3245 | 2.45 | 6.54E-05 | 8.98E-05 |
| chr2 | 100153144 | 100165624 | 12481 | -1.94 | 6.56E-05 | 9.01E-05 |
| chrX | 128775072 | 128775765 | 694 | -4.9 | 6.57E-05 | 9.01E-05 |
| chr2 | 68605340 | 68617079 | 11740 | 3.2 | 6.57E-05 | 9.01E-05 |
| chr15 | 101093612 | 101094023 | 412 | -4.77 | 6.59E-05 | 9.04E-05 |
| chr6 | 130919389 | 130922019 | 2631 | 4.08 | 6.59E-05 | 9.05E-05 |
| chr4 | 188187652 | 188188658 | 1007 | -3.32 | 6.60E-05 | 9.05E-05 |
| chrX | 3335449 | 3348080 | 12632 | 1.82 | 6.60E-05 | 9.05E-05 |
| chr9 | 103169008 | 103175479 | 6472 | -5.97 | 6.60E-05 | 9.06E-05 |
| chr20 | 42357416 | 42357921 | 506 | -4.72 | 6.61E-05 | 9.06E-05 |
| chr4 | 98384680 | 98385732 | 1053 | -4.72 | 6.61E-05 | 9.06E-05 |
| chr13 | 27170219 | 27172734 | 2516 | -4.31 | 6.61E-05 | 9.06E-05 |
| chr16 | 58872394 | 58873039 | 646 | -5.18 | 6.61E-05 | 9.07E-05 |
| chr9 | 16265860 | 16266682 | 823 | -4.3 | 6.64E-05 | 9.11E-05 |
| chr1 | 219627019 | 219629098 | 2080 | -2.16 | 6.64E-05 | 9.11E-05 |
| chr6 | 149593185 | 149593816 | 632 | -2.82 | 6.65E-05 | 9.12E-05 |
| chr3 | 71562381 | 71563152 | 772 | -4.94 | 6.66E-05 | 9.13E-05 |
| chr18 | 49820250 | 49825380 | 5131 | 1.8 | 6.66E-05 | 9.13E-05 |
| chr10 | 104687986 | 104688478 | 493 | -4.81 | 6.72E-05 | 9.21E-05 |
| chr10 | 98193609 | 98200421 | 6813 | -4.86 | 6.72E-05 | 9.22E-05 |
| chr6 | 129328390 | 129335164 | 6775 | 4.04 | 6.72E-05 | 9.22E-05 |
| chr2 | 26674714 | 26675067 | 354 | -4.78 | 6.73E-05 | 9.22E-05 |
| chr22 | 25519378 | 25519870 | 493 | -4.78 | 6.73E-05 | 9.22E-05 |
| chr3 | 195987751 | 195988141 | 391 | -4.78 | 6.73E-05 | 9.22E-05 |
| chr7 | 12957452 | 12957907 | 456 | -4.78 | 6.73E-05 | 9.22E-05 |
| chr6 | 128117825 | 128125823 | 7999 | -1.62 | 6.74E-05 | 9.24E-05 |
| chr4 | 66036650 | 66037430 | 781 | -3.88 | 6.75E-05 | 9.25E-05 |

| | | | | | | |
|---|---|---|---|---|---|---|
| chrX | 129556660 | 129557161 | 502 | -3.88 | 6.75E-05 | 9.25E-05 |
| chr5 | 107409157 | 107409766 | 610 | -4.92 | 6.75E-05 | 9.26E-05 |
| chr1 | 97783037 | 97783650 | 614 | -4.87 | 6.76E-05 | 9.27E-05 |
| chr2 | 24237358 | 24237859 | 502 | -4.87 | 6.76E-05 | 9.27E-05 |
| chr14 | 106428188 | 106428810 | 623 | -2.21 | 6.77E-05 | 9.27E-05 |
| chr1 | 203061570 | 203084144 | 22575 | -5.55 | 6.78E-05 | 9.30E-05 |
| chr8 | 122291195 | 122292022 | 828 | -3.64 | 6.79E-05 | 9.30E-05 |
| chr14 | 89978297 | 89996859 | 18563 | -2.96 | 6.80E-05 | 9.32E-05 |
| chrX | 21783048 | 21785291 | 2244 | -2.51 | 6.83E-05 | 9.36E-05 |
| chr2 | 105615501 | 105646995 | 31495 | -3.12 | 6.83E-05 | 9.36E-05 |
| chr11 | 19913417 | 19914151 | 735 | -4.72 | 6.84E-05 | 9.37E-05 |
| chrX | 82762735 | 82764590 | 1856 | -2.53 | 6.84E-05 | 9.38E-05 |
| chr1 | 208123985 | 208129556 | 5572 | -2.04 | 6.86E-05 | 9.39E-05 |
| chr20 | 23031759 | 23050097 | 18339 | -6.74 | 6.86E-05 | 9.40E-05 |
| chr2 | 231463818 | 231464355 | 538 | -4.85 | 6.87E-05 | 9.41E-05 |
| chr6 | 160538564 | 160539364 | 801 | -4.85 | 6.87E-05 | 9.41E-05 |
| chr20 | 4050977 | 4051856 | 880 | -4.35 | 6.88E-05 | 9.42E-05 |
| chrX | 39315409 | 39318173 | 2765 | -3.12 | 6.90E-05 | 9.45E-05 |
| chr6 | 101453055 | 101455129 | 2075 | 3.99 | 6.91E-05 | 9.46E-05 |
| chr10 | 116926174 | 116926531 | 358 | -4.11 | 6.91E-05 | 9.46E-05 |
| chr1 | 230542354 | 230556972 | 14619 | -1.18 | 6.91E-05 | 9.47E-05 |
| chr1 | 43269423 | 43280384 | 10962 | -5.06 | 6.91E-05 | 9.47E-05 |
| chr9 | 122728678 | 123044554 | 315877 | 3.02 | 6.92E-05 | 9.48E-05 |
| chr10 | 29266380 | 29266854 | 475 | -5.08 | 6.92E-05 | 9.48E-05 |
| chr14 | 66225385 | 66236445 | 11061 | -2.4 | 6.92E-05 | 9.48E-05 |
| chrX | 109889301 | 109896397 | 7097 | 1.91 | 6.92E-05 | 9.48E-05 |
| chr16 | 49436132 | 49465288 | 29157 | -1.54 | 6.93E-05 | 9.49E-05 |
| chr2 | 61088944 | 61096070 | 7127 | -5.91 | 6.96E-05 | 9.53E-05 |
| chr16 | 1784630 | 1785136 | 507 | -4.7 | 6.96E-05 | 9.53E-05 |
| chr20 | 42093884 | 42094742 | 859 | -4.7 | 6.96E-05 | 9.53E-05 |
| chr3 | 33045743 | 33052735 | 6993 | -5.04 | 6.96E-05 | 9.53E-05 |
| chr1 | 19152747 | 19175489 | 22743 | -5.42 | 6.97E-05 | 9.54E-05 |
| chr7 | 133911839 | 133924064 | 12226 | -4.12 | 6.97E-05 | 9.55E-05 |
| chr2 | 233882063 | 233902872 | 20810 | -5.75 | 6.97E-05 | 9.55E-05 |
| chr11 | 134636788 | 134639835 | 3048 | -1.75 | 6.99E-05 | 9.56E-05 |
| chr13 | 96792811 | 96793329 | 519 | -4.84 | 6.99E-05 | 9.57E-05 |
| chr8 | 25474088 | 25512260 | 38173 | -2.12 | 7.01E-05 | 9.59E-05 |
| chr4 | 19471619 | 19472343 | 725 | -3.68 | 7.02E-05 | 9.61E-05 |
| chr17 | 6073210 | 6126520 | 53311 | -2.29 | 7.02E-05 | 9.61E-05 |
| chr9 | 6390502 | 6393721 | 3220 | -2.12 | 7.02E-05 | 9.61E-05 |
| chr3 | 42770316 | 42770767 | 452 | -4.69 | 7.04E-05 | 9.63E-05 |
| chr18 | 71463231 | 71486554 | 23324 | -1.22 | 7.05E-05 | 9.65E-05 |
| chr11 | 17603380 | 17617366 | 13987 | -2.64 | 7.06E-05 | 9.66E-05 |
| chr4 | 167753725 | 167754147 | 423 | -4.71 | 7.06E-05 | 9.66E-05 |
| chr19 | 40728147 | 40728973 | 827 | -5.01 | 7.07E-05 | 9.68E-05 |
| chr11 | 4523858 | 4552933 | 29076 | 2.87 | 7.07E-05 | 9.68E-05 |
| chr12 | 52166839 | 52168716 | 1878 | -4.46 | 7.08E-05 | 9.68E-05 |
| chr20 | 60217772 | 60218328 | 557 | -4.69 | 7.08E-05 | 9.68E-05 |
| chr4 | 31557097 | 31558010 | 914 | -4.69 | 7.08E-05 | 9.68E-05 |
| chr18 | 34799660 | 34800540 | 881 | -3.8 | 7.08E-05 | 9.68E-05 |
| chr1 | 76090062 | 76092362 | 2301 | 2.67 | 7.10E-05 | 9.72E-05 |
| chr12 | 41000616 | 41001311 | 696 | -4.86 | 7.11E-05 | 9.73E-05 |
| chr2 | 241840454 | 241841108 | 655 | -4.86 | 7.11E-05 | 9.73E-05 |
| chr7 | 28275563 | 28276714 | 1152 | 4.94 | 7.12E-05 | 9.74E-05 |
| chr21 | 43434644 | 43435200 | 557 | -4.82 | 7.12E-05 | 9.75E-05 |
| chr22 | 40134144 | 40159330 | 25187 | -1.86 | 7.13E-05 | 9.75E-05 |
| chr11 | 58963817 | 58964601 | 785 | -3.65 | 7.13E-05 | 9.75E-05 |
| chr10 | 120179277 | 120255866 | 76590 | -2.83 | 7.14E-05 | 9.76E-05 |
| chr12 | 125014257 | 125014684 | 428 | -4.69 | 7.14E-05 | 9.76E-05 |
| chr1 | 223476558 | 223511097 | 34540 | -1.65 | 7.14E-05 | 9.76E-05 |
| chr15 | 65225749 | 65226573 | 825 | -4.77 | 7.14E-05 | 9.76E-05 |

| | | | | | | |
|---|---|---|---|---|---|---|
| chr16 | 72984011 | 72984597 | 587 | -4.77 | 7.14E-05 | 9.76E-05 |
| chr17 | 72952428 | 72953092 | 665 | -4.77 | 7.14E-05 | 9.76E-05 |
| chr6 | 63625427 | 63626010 | 584 | 3.91 | 7.14E-05 | 9.77E-05 |
| chr11 | 45476581 | 45485840 | 9260 | -4.71 | 7.16E-05 | 9.80E-05 |
| chr18 | 32339942 | 32341298 | 1357 | 2.54 | 7.16E-05 | 9.80E-05 |
| chr18 | 33898960 | 33899348 | 389 | -4.16 | 7.17E-05 | 9.80E-05 |
| chr4 | 42375054 | 42385459 | 10406 | -1.66 | 7.18E-05 | 9.82E-05 |
| chr10 | 34562139 | 34576769 | 14631 | -5.71 | 7.18E-05 | 9.82E-05 |
| chr2 | 113844177 | 113865737 | 21561 | -2.19 | 7.19E-05 | 9.83E-05 |
| chr3 | 190613300 | 190622253 | 8954 | 2.39 | 7.19E-05 | 9.83E-05 |
| chr2 | 192457127 | 192457797 | 671 | -4.98 | 7.19E-05 | 9.83E-05 |
| chr10 | 73672415 | 73673203 | 789 | -4.73 | 7.19E-05 | 9.83E-05 |
| chr13 | 45851156 | 45851673 | 518 | -4.73 | 7.19E-05 | 9.83E-05 |
| chr5 | 138929874 | 138930215 | 342 | -4.73 | 7.19E-05 | 9.83E-05 |
| chr17 | 46595698 | 46596328 | 631 | -5.27 | 7.20E-05 | 9.84E-05 |
| chr8 | 16541317 | 16544639 | 3323 | 2.69 | 7.22E-05 | 9.86E-05 |
| chr4 | 151213022 | 151213469 | 448 | -3.75 | 7.25E-05 | 9.91E-05 |
| chr12 | 42048354 | 42048720 | 367 | -4.7 | 7.25E-05 | 9.91E-05 |
| chr13 | 39928785 | 39929556 | 772 | -4.7 | 7.25E-05 | 9.91E-05 |
| chr21 | 43605185 | 43618896 | 13712 | -4.7 | 7.25E-05 | 9.91E-05 |
| chr5 | 59714829 | 59717643 | 2815 | -1.45 | 7.25E-05 | 9.91E-05 |
| chr2 | 58918069 | 58918621 | 553 | -2.75 | 7.26E-05 | 9.92E-05 |
| chr4 | 163111104 | 163111848 | 745 | -4.97 | 7.26E-05 | 9.92E-05 |
| chr14 | 59241969 | 59242427 | 459 | -4.84 | 7.26E-05 | 9.93E-05 |
| chr18 | 61291191 | 61295563 | 4373 | -1.59 | 7.27E-05 | 9.93E-05 |
| chr17 | 72242584 | 72243352 | 769 | -4.8 | 7.27E-05 | 9.93E-05 |
| chr2 | 179892174 | 179893884 | 1711 | -4.16 | 7.28E-05 | 9.94E-05 |
| chr5 | 107982702 | 107983777 | 1076 | -3.93 | 7.28E-05 | 9.95E-05 |
| chr6 | 152376429 | 152377738 | 1310 | -2.23 | 7.29E-05 | 9.95E-05 |
| chr9 | 138452580 | 138472425 | 19846 | -2.37 | 7.32E-05 | 1.00E-04 |
| chr2 | 193725951 | 193726765 | 815 | -3.82 | 7.34E-05 | 1.00E-04 |
| chr1 | 224708417 | 224708890 | 474 | -5.23 | 7.35E-05 | 1.00E-04 |
| chr11 | 43769419 | 43769817 | 399 | -4.68 | 7.38E-05 | 0.000101 |
| chr3 | 46781017 | 46781555 | 539 | -4.68 | 7.38E-05 | 0.000101 |
| chr13 | 50185993 | 50186534 | 542 | -4.75 | 7.40E-05 | 0.000101 |
| chr13 | 99310891 | 99311612 | 722 | -4.75 | 7.40E-05 | 0.000101 |
| chr22 | 29363756 | 29364269 | 514 | -4.75 | 7.40E-05 | 0.000101 |
| chr1 | 35415649 | 35416864 | 1216 | -3.87 | 7.40E-05 | 0.000101 |
| chr2 | 203911299 | 203911784 | 486 | -4.69 | 7.40E-05 | 0.000101 |
| chr21 | 38429325 | 38429918 | 594 | -4.69 | 7.40E-05 | 0.000101 |
| chr10 | 125105798 | 125312220 | 206423 | 0.86 | 7.41E-05 | 0.000101 |
| chr10 | 96844633 | 96844951 | 319 | -4.72 | 7.41E-05 | 0.000101 |
| chr3 | 66036229 | 66036835 | 607 | -4.72 | 7.41E-05 | 0.000101 |
| chr14 | 54802593 | 54803315 | 723 | -4.96 | 7.42E-05 | 0.000101 |
| chr3 | 71062236 | 71062767 | 532 | -4.96 | 7.42E-05 | 0.000101 |
| chr1 | 32816120 | 32816593 | 474 | -4.82 | 7.43E-05 | 0.000101 |
| chr7 | 79769757 | 79770354 | 598 | -4.82 | 7.43E-05 | 0.000101 |
| chr3 | 60047555 | 60048900 | 1346 | 4.48 | 7.44E-05 | 0.000102 |
| chr7 | 110004986 | 110008216 | 3231 | -4.27 | 7.45E-05 | 0.000102 |
| chr20 | 56543748 | 56544309 | 562 | -4.16 | 7.45E-05 | 0.000102 |
| chr14 | 34418765 | 34419387 | 623 | -4.29 | 7.46E-05 | 0.000102 |
| chr19 | 50464523 | 50464954 | 432 | -4.93 | 7.48E-05 | 0.000102 |
| chr11 | 91513320 | 91519991 | 6672 | -2.36 | 7.50E-05 | 0.000102 |
| chr2 | 180144122 | 180144829 | 708 | -4.94 | 7.51E-05 | 0.000103 |
| chr8 | 86399592 | 86402639 | 3048 | -1.67 | 7.53E-05 | 0.000103 |
| chr12 | 76985709 | 76987010 | 1302 | 1.99 | 7.53E-05 | 0.000103 |
| chr10 | 28521997 | 28535574 | 13578 | -5.8 | 7.56E-05 | 0.000103 |
| chrX | 12747211 | 12749824 | 2614 | -2.63 | 7.56E-05 | 0.000103 |
| chr17 | 79348564 | 79349415 | 852 | -5.17 | 7.56E-05 | 0.000103 |
| chr10 | 91304976 | 91305729 | 754 | -4.7 | 7.56E-05 | 0.000103 |
| chr16 | 69434214 | 69434918 | 705 | -4.7 | 7.56E-05 | 0.000103 |

| | | | | | | |
|---|---|---|---|---|---|---|
| chrX | 107244447 | 107268775 | 24329 | -5.03 | 7.57E-05 | 0.000103 |
| chr10 | 51801409 | 51802685 | 1277 | -4.06 | 7.58E-05 | 0.000103 |
| chr2 | 183264870 | 183265341 | 472 | 4.8 | 7.61E-05 | 0.000104 |
| chr14 | 95239466 | 95240321 | 856 | -4.8 | 7.61E-05 | 0.000104 |
| chr8 | 53652379 | 53653370 | 992 | -4.8 | 7.61E-05 | 0.000104 |
| chr17 | 78887831 | 78888762 | 932 | -4.92 | 7.61E-05 | 0.000104 |
| chr12 | 83337684 | 83338184 | 501 | -4.67 | 7.63E-05 | 0.000104 |
| chr13 | 20611970 | 20612468 | 499 | -4.67 | 7.63E-05 | 0.000104 |
| chr1 | 56773641 | 56780586 | 6946 | -1.8 | 7.63E-05 | 0.000104 |
| chr2 | 204984600 | 205059101 | 74502 | -1.41 | 7.64E-05 | 0.000104 |
| chr5 | 118360514 | 118360954 | 441 | -4.67 | 7.67E-05 | 0.000105 |
| chr10 | 93114070 | 93140013 | 25944 | -1.43 | 7.67E-05 | 0.000105 |
| chr2 | 210147161 | 210147695 | 535 | -2.65 | 7.67E-05 | 0.000105 |
| chr2 | 75738589 | 75743232 | 4644 | -4.05 | 7.67E-05 | 0.000105 |
| chr2 | 155130992 | 155133644 | 2653 | -2.24 | 7.68E-05 | 0.000105 |
| chr7 | 112156901 | 112185086 | 28186 | -1.52 | 7.69E-05 | 0.000105 |
| chr1 | 178493494 | 178493974 | 481 | -4.04 | 7.71E-05 | 0.000105 |
| chr14 | 90828140 | 90828829 | 690 | -4.9 | 7.72E-05 | 0.000105 |
| chr15 | 23830864 | 23831799 | 936 | -4.68 | 7.74E-05 | 0.000105 |
| chr6 | 4185364 | 4185728 | 365 | -4.68 | 7.74E-05 | 0.000105 |
| chr7 | 44258454 | 44259266 | 813 | -5.12 | 7.74E-05 | 0.000106 |
| chr19 | 8820416 | 8842210 | 21795 | -1.25 | 7.77E-05 | 0.000106 |
| chr6 | 147112394 | 147118071 | 5678 | -4.62 | 7.78E-05 | 0.000106 |
| chr17 | 56702683 | 56703240 | 558 | -4.68 | 7.79E-05 | 0.000106 |
| chr19 | 14927792 | 14979711 | 51920 | -1.65 | 7.79E-05 | 0.000106 |
| chrX | 81489712 | 81494808 | 5097 | -3.47 | 7.80E-05 | 0.000106 |
| chr2 | 216717170 | 216719787 | 2618 | -4.47 | 7.80E-05 | 0.000106 |
| chr19 | 15055085 | 15107195 | 52111 | -3.37 | 7.81E-05 | 0.000106 |
| chrX | 134996852 | 134998533 | 1682 | -2.04 | 7.81E-05 | 0.000106 |
| chr20 | 4246110 | 4400355 | 154246 | -1.79 | 7.82E-05 | 0.000107 |
| chr1 | 63671934 | 63681439 | 9506 | -4.48 | 7.83E-05 | 0.000107 |
| chr13 | 47173055 | 47173740 | 686 | -4.88 | 7.84E-05 | 0.000107 |
| chr4 | 25550859 | 25551535 | 677 | -4.88 | 7.84E-05 | 0.000107 |
| chr5 | 122953565 | 123084593 | 131029 | -2.93 | 7.85E-05 | 0.000107 |
| chr4 | 31600806 | 31601504 | 699 | -4.66 | 7.86E-05 | 0.000107 |
| chr4 | 157701601 | 157702500 | 900 | -4.66 | 7.86E-05 | 0.000107 |
| chr6 | 105625500 | 105626184 | 685 | -4.66 | 7.86E-05 | 0.000107 |
| chr1 | 234257821 | 234258546 | 726 | -5.08 | 7.87E-05 | 0.000107 |
| chr5 | 167251979 | 167480898 | 228920 | 1.12 | 7.87E-05 | 0.000107 |
| chr2 | 107539220 | 107570045 | 30826 | -1.81 | 7.92E-05 | 0.000108 |
| chr11 | 20575554 | 20581816 | 6263 | -1.48 | 7.94E-05 | 0.000108 |
| chr8 | 98559952 | 98560481 | 530 | -5.07 | 7.94E-05 | 0.000108 |
| chrX | 76483946 | 76503077 | 19132 | -1.52 | 7.96E-05 | 0.000108 |
| chr1 | 200292917 | 200294180 | 1264 | -3.84 | 7.97E-05 | 0.000109 |
| chr19 | 14052976 | 14053792 | 817 | -4.86 | 7.97E-05 | 0.000109 |
| chr9 | 108086773 | 108087498 | 726 | -4.86 | 7.97E-05 | 0.000109 |
| chr6 | 129387596 | 129388188 | 593 | -3.66 | 7.98E-05 | 0.000109 |
| chr1 | 195792333 | 195793915 | 1583 | -4.07 | 7.98E-05 | 0.000109 |
| chr17 | 20529399 | 20529843 | 445 | -4.22 | 7.98E-05 | 0.000109 |
| chr6 | 155945807 | 156842508 | 896702 | 0.27 | 7.98E-05 | 0.000109 |
| chr14 | 62088246 | 62089695 | 1450 | -4 | 7.99E-05 | 0.000109 |
| chr14 | 55317426 | 55317880 | 455 | -4.68 | 7.99E-05 | 0.000109 |
| chr1 | 38609201 | 38623374 | 14174 | -1.44 | 8.00E-05 | 0.000109 |
| chr13 | 67536814 | 67537430 | 617 | -5.05 | 8.01E-05 | 0.000109 |
| chr15 | 51494912 | 51506366 | 11455 | 2.39 | 8.01E-05 | 0.000109 |
| chr14 | 37966335 | 37966660 | 326 | -4.71 | 8.02E-05 | 0.000109 |
| chr8 | 55075896 | 55076302 | 407 | -4.71 | 8.02E-05 | 0.000109 |
| chr7 | 37912695 | 37921703 | 9009 | -1.6 | 8.05E-05 | 0.00011 |
| chr14 | 93935300 | 94071692 | 136393 | -1.69 | 8.06E-05 | 0.00011 |
| chr2 | 55981675 | 55982351 | 677 | -5.07 | 8.09E-05 | 0.00011 |
| chr1 | 159649809 | 159654759 | 4951 | -2.01 | 8.11E-05 | 0.00011 |

| | | | | | | |
|---|---|---|---|---|---|---|
| chr1 | 100372104 | 100372589 | 486 | -4.84 | 8.12E-05 | 0.000111 |
| chr1 | 59096665 | 59097708 | 1044 | -2.37 | 8.13E-05 | 0.000111 |
| chr2 | 15819522 | 15820014 | 493 | -5.02 | 8.15E-05 | 0.000111 |
| chr2 | 150872023 | 150873110 | 1088 | -5.02 | 8.15E-05 | 0.000111 |
| chr8 | 61442587 | 61443321 | 735 | -5.02 | 8.15E-05 | 0.000111 |
| chr18 | 29173462 | 29174602 | 1141 | -3.9 | 8.16E-05 | 0.000111 |
| chr7 | 159018226 | 159081218 | 62993 | -0.84 | 8.16E-05 | 0.000111 |
| chr7 | 36567365 | 36568454 | 1090 | -3.29 | 8.16E-05 | 0.000111 |
| chr5 | 94767393 | 94769574 | 2182 | -4.69 | 8.19E-05 | 0.000111 |
| chr10 | 17667601 | 17668956 | 1356 | -5.26 | 8.22E-05 | 0.000112 |
| chr3 | 117496072 | 117497126 | 1055 | -3.8 | 8.22E-05 | 0.000112 |
| chr11 | 36271809 | 36278235 | 6427 | -5.68 | 8.23E-05 | 0.000112 |
| chr10 | 125654480 | 125695960 | 41481 | -5.67 | 8.24E-05 | 0.000112 |
| chr1 | 113832369 | 113832966 | 598 | -4.72 | 8.26E-05 | 0.000112 |
| chr6 | 159830973 | 159837460 | 6488 | -4.72 | 8.26E-05 | 0.000112 |
| chr1 | 94827892 | 94828436 | 545 | -4.65 | 8.27E-05 | 0.000113 |
| chr7 | 13178667 | 13179408 | 742 | -4.65 | 8.27E-05 | 0.000113 |
| chr9 | 30225412 | 30225897 | 486 | -4.65 | 8.27E-05 | 0.000113 |
| chr3 | 54968738 | 54970113 | 1376 | -3.51 | 8.28E-05 | 0.000113 |
| chr14 | 96346331 | 96348703 | 2373 | -4.08 | 8.28E-05 | 0.000113 |
| chr4 | 135238660 | 135241977 | 3318 | 2.83 | 8.29E-05 | 0.000113 |
| chr16 | 85386387 | 85386996 | 610 | -4.65 | 8.30E-05 | 0.000113 |
| chr6 | 136835700 | 136836221 | 522 | -4.65 | 8.30E-05 | 0.000113 |
| chr2 | 231567033 | 231567844 | 812 | -4.98 | 8.32E-05 | 0.000113 |
| chr8 | 130437586 | 130457130 | 19545 | -1.97 | 8.34E-05 | 0.000114 |
| chr18 | 52756840 | 52758001 | 1162 | -3.59 | 8.35E-05 | 0.000114 |
| chr4 | 120758615 | 120760773 | 2159 | -4.02 | 8.37E-05 | 0.000114 |
| chr6 | 898726 | 899383 | 658 | -4.67 | 8.37E-05 | 0.000114 |
| chr2 | 108691024 | 108734032 | 43009 | -1.32 | 8.37E-05 | 0.000114 |
| chr4 | 122502308 | 122503542 | 1235 | -3 | 8.38E-05 | 0.000114 |
| chr6 | 135923001 | 135923667 | 667 | -3.89 | 8.38E-05 | 0.000114 |
| chr10 | 63725679 | 63726192 | 514 | -4.69 | 8.41E-05 | 0.000114 |
| chr6 | 90103210 | 90103673 | 464 | -4.69 | 8.41E-05 | 0.000114 |
| chr21 | 45238412 | 45239033 | 622 | -4.64 | 8.41E-05 | 0.000114 |
| chr5 | 167687145 | 167687839 | 695 | -4.96 | 8.41E-05 | 0.000114 |
| chr6 | 64291494 | 64294335 | 2842 | -1.81 | 8.43E-05 | 0.000115 |
| chr6 | 87710536 | 87711884 | 1349 | -2.23 | 8.43E-05 | 0.000115 |
| chr7 | 566666 | 580844 | 14179 | -5.8 | 8.44E-05 | 0.000115 |
| chr22 | 22737486 | 22748921 | 11436 | -1.66 | 8.46E-05 | 0.000115 |
| chr2 | 150282155 | 150287741 | 5587 | -1.58 | 8.46E-05 | 0.000115 |
| chr3 | 82208887 | 82209236 | 350 | 5 | 8.47E-05 | 0.000115 |
| chr18 | 68419629 | 68432915 | 13287 | 4.89 | 8.48E-05 | 0.000115 |
| chr2 | 188884369 | 188885103 | 735 | -3.75 | 8.51E-05 | 0.000116 |
| chr8 | 54024136 | 54047756 | 23621 | 1.53 | 8.53E-05 | 0.000116 |
| chr12 | 100106489 | 100106905 | 417 | -4.7 | 8.53E-05 | 0.000116 |
| chr22 | 42746374 | 42746839 | 466 | -4.7 | 8.53E-05 | 0.000116 |
| chr21 | 33022757 | 33023241 | 485 | -4.78 | 8.53E-05 | 0.000116 |
| chr5 | 143561236 | 143562251 | 1016 | -4.78 | 8.53E-05 | 0.000116 |
| chr16 | 86243495 | 86318544 | 75050 | -1.14 | 8.59E-05 | 0.000117 |
| chr5 | 44732425 | 44733286 | 862 | -2.65 | 8.61E-05 | 0.000117 |
| chr17 | 61759458 | 61760107 | 650 | -4.92 | 8.62E-05 | 0.000117 |
| chr12 | 40948332 | 40950044 | 1713 | 2.23 | 8.62E-05 | 0.000117 |
| chr3 | 41772056 | 41772559 | 504 | -4.95 | 8.62E-05 | 0.000117 |
| chr15 | 75283654 | 75284367 | 714 | -4.71 | 8.63E-05 | 0.000117 |
| chr13 | 97465475 | 97468568 | 3094 | -2.18 | 8.65E-05 | 0.000118 |
| chr3 | 190794317 | 190795907 | 1591 | 2.71 | 8.65E-05 | 0.000118 |
| chr14 | 69782453 | 69782976 | 524 | -4.78 | 8.65E-05 | 0.000118 |
| chr3 | 129139695 | 129140412 | 718 | -4.78 | 8.65E-05 | 0.000118 |
| chr2 | 75312216 | 75312780 | 565 | -4.14 | 8.67E-05 | 0.000118 |
| chrX | 39382992 | 39411034 | 28043 | -1.81 | 8.68E-05 | 0.000118 |
| chr2 | 179214490 | 179215751 | 1262 | 5.2 | 8.69E-05 | 0.000118 |

| | | | | | | |
|---|---|---|---|---|---|---|
| chr14 | 59623559 | 59624926 | 1368 | -2.58 | 8.70E-05 | 0.000118 |
| chr15 | 39805455 | 39812343 | 6889 | -1.5 | 8.72E-05 | 0.000118 |
| chr3 | 176342737 | 176349398 | 6662 | 2.72 | 8.72E-05 | 0.000119 |
| chr9 | 112948408 | 112985411 | 37004 | -7.33 | 8.72E-05 | 0.000119 |
| chr2 | 85095372 | 85095696 | 325 | -4.64 | 8.72E-05 | 0.000119 |
| chr12 | 64989792 | 64990424 | 633 | -4.9 | 8.73E-05 | 0.000119 |
| chrX | 22254518 | 22255628 | 1111 | 3.13 | 8.73E-05 | 0.000119 |
| chr4 | 87540057 | 87540442 | 386 | -2.9 | 8.76E-05 | 0.000119 |
| chr17 | 6279456 | 6315205 | 35750 | -5.69 | 8.76E-05 | 0.000119 |
| chrX | 83736965 | 83748053 | 11089 | -1.16 | 8.76E-05 | 0.000119 |
| chr1 | 160676958 | 160679047 | 2090 | 3.33 | 8.76E-05 | 0.000119 |
| chr15 | 38665979 | 38666745 | 767 | 2.16 | 8.77E-05 | 0.000119 |
| chr14 | 102244152 | 102244572 | 421 | -4.65 | 8.78E-05 | 0.000119 |
| chr12 | 106590494 | 106614229 | 23736 | -5.48 | 8.80E-05 | 0.00012 |
| chr2 | 197232838 | 197233705 | 868 | -5.12 | 8.80E-05 | 0.00012 |
| chr17 | 65375545 | 65376133 | 589 | -4.68 | 8.83E-05 | 0.00012 |
| chr9 | 38731651 | 38732199 | 549 | -4.68 | 8.83E-05 | 0.00012 |
| chr19 | 2736397 | 2736989 | 593 | -4.89 | 8.85E-05 | 0.00012 |
| chr2 | 202295554 | 202296074 | 521 | -4.89 | 8.85E-05 | 0.00012 |
| chr3 | 31882056 | 31882897 | 842 | -4.89 | 8.85E-05 | 0.00012 |
| chr12 | 70801033 | 70802431 | 1399 | -2.28 | 8.85E-05 | 0.00012 |
| chr17 | 4325474 | 4326233 | 760 | -4.66 | 8.86E-05 | 0.00012 |
| chr2 | 206896924 | 206897383 | 460 | -4.66 | 8.86E-05 | 0.00012 |
| chr21 | 44571518 | 44572157 | 640 | -4.66 | 8.86E-05 | 0.00012 |
| chr4 | 95859430 | 95860121 | 692 | -4.66 | 8.86E-05 | 0.00012 |
| chr6 | 169723859 | 169724472 | 614 | -4.66 | 8.86E-05 | 0.00012 |
| chr2 | 11604178 | 11610396 | 6219 | -4.76 | 8.88E-05 | 0.00012 |
| chr1 | 248683113 | 248689988 | 6876 | 2.63 | 8.90E-05 | 0.000121 |
| chrX | 42110002 | 42112678 | 2677 | 2.18 | 8.94E-05 | 0.000121 |
| chr4 | 32728441 | 32734376 | 5936 | -1.63 | 8.94E-05 | 0.000121 |
| chr1 | 188806308 | 188806747 | 440 | -4.63 | 8.97E-05 | 0.000122 |
| chr1 | 146703199 | 146704009 | 811 | -4.87 | 8.98E-05 | 0.000122 |
| chr14 | 53023773 | 53039896 | 16124 | -5.35 | 9.01E-05 | 0.000122 |
| chr4 | 78733623 | 78734199 | 577 | -4.88 | 9.08E-05 | 0.000123 |
| chr13 | 44840045 | 44841170 | 1126 | -4.64 | 9.08E-05 | 0.000123 |
| chr19 | 11410824 | 11424670 | 13847 | -4.64 | 9.08E-05 | 0.000123 |
| chr15 | 63914124 | 63914647 | 524 | -3.81 | 9.09E-05 | 0.000123 |
| chr20 | 24799171 | 24803423 | 4253 | -2.11 | 9.10E-05 | 0.000123 |
| chr6 | 89016080 | 89018173 | 2094 | -2.88 | 9.10E-05 | 0.000123 |
| chr6 | 2872128 | 2872636 | 509 | -4.73 | 9.10E-05 | 0.000123 |
| chr6 | 141414537 | 141415020 | 484 | -4.73 | 9.10E-05 | 0.000123 |
| chr5 | 36190182 | 36190752 | 571 | -3.86 | 9.11E-05 | 0.000124 |
| chr21 | 44397044 | 44397466 | 423 | -5.05 | 9.11E-05 | 0.000124 |
| chr5 | 133139526 | 133140445 | 920 | -5.05 | 9.11E-05 | 0.000124 |
| chrX | 39802074 | 39810645 | 8572 | -5.05 | 9.11E-05 | 0.000124 |
| chr8 | 142351969 | 142352564 | 596 | -4.85 | 9.13E-05 | 0.000124 |
| chrX | 129881775 | 129882701 | 927 | -4.85 | 9.13E-05 | 0.000124 |
| chr7 | 106484789 | 106510663 | 25875 | 5.25 | 9.14E-05 | 0.000124 |
| chr5 | 32029479 | 32029989 | 511 | -4.67 | 9.16E-05 | 0.000124 |
| chr6 | 152107310 | 152157968 | 50659 | -1.4 | 9.19E-05 | 0.000125 |
| chr4 | 144626477 | 144706830 | 80354 | 5.79 | 9.21E-05 | 0.000125 |
| chr13 | 112866915 | 112867246 | 332 | -4.64 | 9.24E-05 | 0.000125 |
| chr6 | 131002885 | 131017295 | 14411 | 1.71 | 9.28E-05 | 0.000126 |
| chr2 | 67710125 | 67710787 | 663 | -3.58 | 9.28E-05 | 0.000126 |
| chr10 | 89723907 | 89724510 | 604 | -4.83 | 9.29E-05 | 0.000126 |
| chr17 | 14147666 | 14454284 | 306619 | -0.98 | 9.29E-05 | 0.000126 |
| chr20 | 30741086 | 30741874 | 789 | -4.84 | 9.31E-05 | 0.000126 |
| chr5 | 145305764 | 145306147 | 384 | -4.84 | 9.31E-05 | 0.000126 |
| chr14 | 28634820 | 28643537 | 8718 | -1.66 | 9.33E-05 | 0.000127 |
| chrX | 10670553 | 10683366 | 12814 | -5.81 | 9.35E-05 | 0.000127 |
| chr7 | 83402640 | 83403222 | 583 | -4.64 | 9.38E-05 | 0.000127 |

| | | | | | | |
|---|---|---|---|---|---|---|
| chr2 | 149044919 | 149046021 | 1103 | -4.67 | 9.39E-05 | 0.000127 |
| chr5 | 118787083 | 118787547 | 465 | -4.67 | 9.39E-05 | 0.000127 |
| chr9 | 5562153 | 5562473 | 321 | -4.67 | 9.39E-05 | 0.000127 |
| chr11 | 103854629 | 103855132 | 504 | -4.01 | 9.40E-05 | 0.000127 |
| chr21 | 38152378 | 38152946 | 569 | -5.25 | 9.40E-05 | 0.000127 |
| chr9 | 123074047 | 123102548 | 28502 | -2.37 | 9.41E-05 | 0.000127 |
| chr10 | 61311461 | 61350976 | 39516 | 1.52 | 9.42E-05 | 0.000128 |
| chr3 | 56121925 | 56123973 | 2049 | -2.73 | 9.43E-05 | 0.000128 |
| chr2 | 3059018 | 3103339 | 44322 | -2.23 | 9.43E-05 | 0.000128 |
| chr14 | 94724456 | 94739440 | 14985 | -1.47 | 9.45E-05 | 0.000128 |
| chr9 | 140789882 | 140860363 | 70482 | -4.33 | 9.49E-05 | 0.000129 |
| chr18 | 47895320 | 47899578 | 4259 | -4.87 | 9.51E-05 | 0.000129 |
| chr17 | 29397740 | 29398314 | 575 | -4.65 | 9.54E-05 | 0.000129 |
| chrX | 55332597 | 55335931 | 3335 | 2.46 | 9.55E-05 | 0.000129 |
| chr5 | 71719355 | 71722504 | 3150 | 1.67 | 9.56E-05 | 0.000129 |
| chr18 | 50110586 | 50112404 | 1819 | -3.87 | 9.56E-05 | 0.00013 |
| chr2 | 224013149 | 224014459 | 1311 | -3.31 | 9.57E-05 | 0.00013 |
| chr1 | 116733908 | 116734956 | 1049 | -2.14 | 9.57E-05 | 0.00013 |
| chr16 | 55688667 | 55689273 | 607 | -4.7 | 9.59E-05 | 0.00013 |
| chr3 | 119840715 | 119841327 | 613 | -4.7 | 9.59E-05 | 0.00013 |
| chr8 | 115475520 | 115476039 | 520 | -4.7 | 9.59E-05 | 0.00013 |
| chr7 | 104519725 | 104525760 | 6036 | -4.79 | 9.64E-05 | 0.000131 |
| chr1 | 35715058 | 35715514 | 457 | -4.79 | 9.65E-05 | 0.000131 |
| chr3 | 31567089 | 31567539 | 451 | -4.79 | 9.65E-05 | 0.000131 |
| chr1 | 209870726 | 209879053 | 8328 | 4.5 | 9.67E-05 | 0.000131 |
| chr8 | 104483171 | 104486238 | 3068 | -4.57 | 9.67E-05 | 0.000131 |
| chr2 | 239852361 | 239858049 | 5689 | -5.38 | 9.71E-05 | 0.000131 |
| chr7 | 137351961 | 137352381 | 421 | -4.61 | 9.76E-05 | 0.000132 |
| chr10 | 7661446 | 7718411 | 56966 | -1.06 | 9.77E-05 | 0.000132 |
| chr4 | 10545676 | 10546387 | 712 | -3.56 | 9.77E-05 | 0.000132 |
| chr6 | 138160899 | 138161524 | 626 | -4.93 | 9.78E-05 | 0.000132 |
| chr6 | 75469899 | 75491626 | 21728 | 1.77 | 9.79E-05 | 0.000133 |
| chr11 | 124773605 | 124781165 | 7561 | -4.18 | 9.81E-05 | 0.000133 |
| chr13 | 49256824 | 49299708 | 42885 | -0.87 | 9.83E-05 | 0.000133 |
| chr1 | 87056530 | 87059354 | 2825 | 2.13 | 9.85E-05 | 0.000133 |
| chr2 | 131002999 | 131003364 | 366 | -4.65 | 9.86E-05 | 0.000133 |
| chr9 | 76104799 | 76109340 | 4542 | -1.68 | 9.87E-05 | 0.000134 |
| chr9 | 108465337 | 108465694 | 358 | -4.76 | 9.87E-05 | 0.000134 |
| chr8 | 74017747 | 74057847 | 40101 | -1.21 | 9.87E-05 | 0.000134 |
| chr3 | 94615639 | 94616163 | 525 | -4.68 | 9.88E-05 | 0.000134 |
| chr2 | 222411509 | 222414969 | 3461 | -4.25 | 9.89E-05 | 0.000134 |
| chr10 | 121650087 | 121650624 | 538 | -4.91 | 9.90E-05 | 0.000134 |
| chr11 | 120712583 | 120713189 | 607 | -3.21 | 9.92E-05 | 0.000134 |
| chr10 | 18520058 | 18589312 | 69255 | 0.79 | 9.92E-05 | 0.000134 |
| chr2 | 53096203 | 53103304 | 7102 | -1.73 | 9.93E-05 | 0.000134 |
| chr22 | 29164804 | 29165349 | 546 | -4.6 | 9.94E-05 | 0.000135 |
| chrX | 70033890 | 70042180 | 8291 | -2.56 | 9.95E-05 | 0.000135 |
| chr2 | 182210252 | 182211167 | 916 | -3.83 | 9.97E-05 | 0.000135 |
| chr10 | 112111447 | 112112186 | 740 | -4.62 | 9.99E-05 | 0.000135 |
| chr6 | 63410052 | 63410421 | 370 | -4.62 | 9.99E-05 | 0.000135 |
| chrX | 43461588 | 43468211 | 6624 | -2.5 | 9.99E-05 | 0.000135 |
| chrX | 115734052 | 115736201 | 2150 | -5.37 | 1.00E-04 | 0.000136 |
| chr10 | 87635558 | 87636601 | 1044 | -3.87 | 1.00E-04 | 0.000136 |
| chr10 | 117732265 | 117890532 | 158268 | 2.09 | 0.000101 | 0.000136 |
| chrX | 65799432 | 65801009 | 1578 | 2.02 | 0.000101 | 0.000136 |
| chr8 | 133162172 | 133201512 | 39341 | 1.12 | 0.000101 | 0.000136 |
| chr14 | 37485536 | 37486005 | 470 | -4.67 | 0.000101 | 0.000137 |
| chr3 | 156931517 | 156932281 | 765 | -4.67 | 0.000101 | 0.000137 |
| chr4 | 138648484 | 138649348 | 865 | -4.67 | 0.000101 | 0.000137 |
| chr6 | 141045952 | 141047069 | 1118 | -4.87 | 0.000102 | 0.000138 |
| chr5 | 64010685 | 64011499 | 815 | -3.72 | 0.000102 | 0.000138 |

| | | | | | | |
|---|---|---|---|---|---|---|
| chr18 | 11594786 | 11596312 | 1527 | 1.47 | 0.000102 | 0.000138 |
| chr15 | 24551101 | 24551652 | 552 | 2.74 | 0.000102 | 0.000138 |
| chr18 | 61471413 | 61485114 | 13702 | 4.74 | 0.000102 | 0.000138 |
| chr2 | 53699835 | 53700267 | 433 | 4.83 | 0.000102 | 0.000138 |
| chr8 | 117776813 | 117777984 | 1172 | -3.78 | 0.000102 | 0.000138 |
| chr6 | 118041150 | 118042570 | 1421 | -3.68 | 0.000103 | 0.000139 |
| chr1 | 201462449 | 201463183 | 735 | -4.93 | 0.000103 | 0.000139 |
| chr4 | 159471620 | 159472674 | 1055 | -5.07 | 0.000103 | 0.000139 |
| chr1 | 223860707 | 223876871 | 16165 | -6.67 | 0.000103 | 0.000139 |
| chr4 | 115499606 | 115503420 | 3815 | 1.53 | 0.000103 | 0.000139 |
| chr5 | 173539755 | 173566932 | 27178 | -1.32 | 0.000103 | 0.00014 |
| chr10 | 87709214 | 87732041 | 22828 | -1.58 | 0.000103 | 0.00014 |
| chr4 | 13388409 | 13388847 | 439 | -4.6 | 0.000103 | 0.00014 |
| chr10 | 50849775 | 50852738 | 2964 | 4.18 | 0.000104 | 0.00014 |
| chr3 | 42384628 | 42413139 | 28512 | -1.99 | 0.000104 | 0.00014 |
| chr8 | 26252854 | 26253204 | 351 | -4.72 | 0.000104 | 0.00014 |
| chr8 | 143083195 | 143083725 | 531 | -3.99 | 0.000104 | 0.000141 |
| chr1 | 46229526 | 46230262 | 737 | -5.3 | 0.000104 | 0.000141 |
| chr2 | 166551835 | 166561811 | 9977 | 6.37 | 0.000104 | 0.000141 |
| chr19 | 43499836 | 43500301 | 466 | -4.61 | 0.000104 | 0.000141 |
| chr6 | 134451691 | 134452387 | 697 | -4.61 | 0.000104 | 0.000141 |
| chr6 | 137309102 | 137309903 | 802 | -3.84 | 0.000105 | 0.000141 |
| chr9 | 121780498 | 122711002 | 930505 | 8.87 | 0.000105 | 0.000141 |
| chr2 | 183072656 | 183073016 | 361 | 4.69 | 0.000105 | 0.000141 |
| chr13 | 47489640 | 47502021 | 12382 | 2 | 0.000105 | 0.000141 |
| chr13 | 89709406 | 89710139 | 734 | -5.04 | 0.000105 | 0.000142 |
| chr14 | 52201099 | 52201675 | 577 | -4.65 | 0.000105 | 0.000142 |
| chr14 | 105797656 | 105798154 | 499 | -4.64 | 0.000106 | 0.000143 |
| chr3 | 175166681 | 175167544 | 864 | -4.64 | 0.000106 | 0.000143 |
| chr10 | 14863223 | 14863784 | 562 | -4.59 | 0.000106 | 0.000143 |
| chr6 | 133260177 | 133260906 | 730 | -4.59 | 0.000106 | 0.000143 |
| chr6 | 8537459 | 8568904 | 31446 | -5.07 | 0.000106 | 0.000143 |
| chr14 | 93846668 | 93920551 | 73884 | -1.72 | 0.000106 | 0.000143 |
| chr6 | 156984218 | 156998076 | 13859 | -4.98 | 0.000106 | 0.000143 |
| chr9 | 5274919 | 5293085 | 18167 | -1.21 | 0.000106 | 0.000143 |
| chr1 | 31985294 | 31986087 | 794 | -4.81 | 0.000107 | 0.000144 |
| chr8 | 90876485 | 90876964 | 480 | -4.81 | 0.000107 | 0.000144 |
| chr4 | 39169343 | 39170470 | 1128 | -5 | 0.000107 | 0.000144 |
| chr4 | 94026675 | 94027150 | 476 | -4.71 | 0.000107 | 0.000144 |
| chr1 | 89165534 | 89165906 | 373 | -4.58 | 0.000107 | 0.000144 |
| chr1 | 95671892 | 95672506 | 615 | -4.58 | 0.000107 | 0.000144 |
| chr17 | 65252060 | 65252779 | 720 | -4.58 | 0.000107 | 0.000144 |
| chr2 | 231614620 | 231615057 | 438 | -4.87 | 0.000107 | 0.000144 |
| chr2 | 217510981 | 217511709 | 729 | -4.59 | 0.000107 | 0.000144 |
| chr4 | 77910795 | 77911283 | 489 | -4.59 | 0.000107 | 0.000144 |
| chr5 | 1194540 | 1194853 | 314 | -4.59 | 0.000107 | 0.000144 |
| chr2 | 220794861 | 220796657 | 1797 | -4.24 | 0.000107 | 0.000145 |
| chrX | 74992209 | 75001398 | 9190 | 4.79 | 0.000107 | 0.000145 |
| chr18 | 24743674 | 24749431 | 5758 | -1.32 | 0.000107 | 0.000145 |
| chr7 | 151525868 | 151526435 | 568 | -4 | 0.000107 | 0.000145 |
| chr17 | 72509103 | 72532919 | 23817 | -5.36 | 0.000107 | 0.000145 |
| chr20 | 20192435 | 20209938 | 17504 | -0.74 | 0.000107 | 0.000145 |
| chr3 | 191418724 | 191420549 | 1826 | -2.04 | 0.000108 | 0.000146 |
| chrX | 80783916 | 80795742 | 11827 | 1.35 | 0.000108 | 0.000146 |
| chr5 | 132708906 | 132716941 | 8036 | -5.16 | 0.000109 | 0.000147 |
| chr2 | 141418936 | 141419490 | 555 | -4.97 | 0.000109 | 0.000147 |
| chr13 | 34103975 | 34105855 | 1881 | -3.79 | 0.000109 | 0.000147 |
| chr7 | 135207609 | 135208410 | 802 | -4.69 | 0.000109 | 0.000148 |
| chrX | 71672679 | 71673126 | 448 | -4.69 | 0.000109 | 0.000148 |
| chr12 | 89615029 | 89615365 | 337 | -4.63 | 0.00011 | 0.000148 |
| chr7 | 2213849 | 2214744 | 896 | -4.63 | 0.00011 | 0.000148 |

| | | | | | | |
|---|---|---|---|---|---|---|
| chr10 | 21375516 | 21376126 | 611 | -4.59 | 0.00011 | 0.000148 |
| chr17 | 60377884 | 60446224 | 68341 | -2.4 | 0.00011 | 0.000149 |
| chr12 | 113668578 | 113669169 | 592 | -4.63 | 0.00011 | 0.000149 |
| chr8 | 110912097 | 110913146 | 1050 | -4.63 | 0.00011 | 0.000149 |
| chr4 | 7715699 | 7716293 | 595 | -4.01 | 0.00011 | 0.000149 |
| chr7 | 43220673 | 43223028 | 2356 | -3.19 | 0.00011 | 0.000149 |
| chr18 | 23550260 | 23553061 | 2802 | -2.23 | 0.000111 | 0.00015 |
| chr2 | 169977394 | 169979338 | 1945 | -4.07 | 0.000111 | 0.00015 |
| chr3 | 186162627 | 186165487 | 2861 | 4.34 | 0.000111 | 0.00015 |
| chr6 | 37314398 | 37314941 | 544 | -3.67 | 0.000111 | 0.00015 |
| chr19 | 51699293 | 51819624 | 120332 | -1.54 | 0.000112 | 0.000151 |
| chr20 | 44356162 | 44367244 | 11083 | -2.65 | 0.000112 | 0.000151 |
| chr2 | 173745056 | 173757296 | 12241 | 4.54 | 0.000112 | 0.000151 |
| chr3 | 193200374 | 193200890 | 517 | -2.97 | 0.000112 | 0.000151 |
| chr5 | 169636734 | 169637180 | 447 | -4.68 | 0.000112 | 0.000151 |
| chr12 | 14269084 | 14287921 | 18838 | -3.67 | 0.000112 | 0.000151 |
| chrX | 37460735 | 37462162 | 1428 | 4 | 0.000112 | 0.000151 |
| chr18 | 54749254 | 54779521 | 30268 | -5.07 | 0.000112 | 0.000151 |
| chr8 | 86902765 | 86905980 | 3216 | -1.56 | 0.000112 | 0.000151 |
| chr15 | 58341735 | 58342909 | 1175 | 2.04 | 0.000112 | 0.000151 |
| chr20 | 25063801 | 25065527 | 1727 | -5.39 | 0.000112 | 0.000152 |
| chr1 | 111542371 | 111553620 | 11250 | -1.16 | 0.000113 | 0.000152 |
| chr17 | 16242558 | 16243300 | 743 | -4.91 | 0.000113 | 0.000152 |
| chr6 | 63075239 | 63100814 | 25576 | 1.31 | 0.000113 | 0.000152 |
| chr17 | 38273 | 38934 | 662 | -4.75 | 0.000113 | 0.000152 |
| chr20 | 31091375 | 31091792 | 418 | -4.75 | 0.000113 | 0.000152 |
| chr5 | 91184699 | 91185139 | 441 | -4.75 | 0.000113 | 0.000152 |
| chr5 | 130673651 | 130674166 | 516 | -4.75 | 0.000113 | 0.000152 |
| chr14 | 81869376 | 81889452 | 20077 | -4.82 | 0.000113 | 0.000152 |
| chrX | 114140742 | 114144339 | 3598 | 1.64 | 0.000113 | 0.000152 |
| chrX | 57480294 | 57485655 | 5362 | -4.59 | 0.000113 | 0.000152 |
| chr5 | 54307187 | 54310184 | 2998 | -2.16 | 0.000114 | 0.000153 |
| chr15 | 45308096 | 45308740 | 645 | -3.03 | 0.000114 | 0.000153 |
| chr6 | 118761063 | 118777141 | 16079 | -1.21 | 0.000114 | 0.000153 |
| chr9 | 121617878 | 121627646 | 9769 | -1.62 | 0.000114 | 0.000154 |
| chr6 | 10437983 | 10438589 | 607 | -4.89 | 0.000114 | 0.000154 |
| chr4 | 118231951 | 118232721 | 771 | -4.61 | 0.000114 | 0.000154 |
| chr13 | 96479869 | 96480424 | 556 | -4.65 | 0.000114 | 0.000154 |
| chr2 | 64498084 | 64498682 | 599 | -3.74 | 0.000114 | 0.000154 |
| chr10 | 114026531 | 114028181 | 1651 | -4.2 | 0.000115 | 0.000155 |
| chr4 | 59693105 | 59694404 | 1300 | -3.57 | 0.000115 | 0.000155 |
| chr6 | 139822117 | 139822671 | 555 | -4.66 | 0.000115 | 0.000155 |
| chrX | 150836320 | 150836684 | 365 | -3.51 | 0.000115 | 0.000155 |
| chr6 | 119480640 | 119488111 | 7472 | 1.22 | 0.000115 | 0.000156 |
| chr7 | 110450022 | 110450632 | 611 | -4.56 | 0.000115 | 0.000156 |
| chrX | 65197065 | 65197355 | 291 | -4.56 | 0.000115 | 0.000156 |
| chr9 | 138485909 | 138493357 | 7449 | -2.48 | 0.000115 | 0.000156 |
| chr19 | 32727609 | 32796347 | 68739 | -2.98 | 0.000115 | 0.000156 |
| chr11 | 85316641 | 85317318 | 678 | -4.87 | 0.000116 | 0.000156 |
| chrX | 71376916 | 71379833 | 2918 | -4.41 | 0.000116 | 0.000156 |
| chr11 | 95641580 | 95642133 | 554 | -4.73 | 0.000116 | 0.000156 |
| chr10 | 90939383 | 90952598 | 13216 | -1.33 | 0.000116 | 0.000156 |
| chr16 | 20400326 | 20403081 | 2756 | 1.91 | 0.000116 | 0.000156 |
| chrX | 85340637 | 85365083 | 24447 | 5.04 | 0.000116 | 0.000156 |
| chr17 | 41068473 | 41069139 | 667 | -4.6 | 0.000116 | 0.000156 |
| chrX | 96293381 | 96294096 | 716 | -4.6 | 0.000116 | 0.000156 |
| chr1 | 52112174 | 52112603 | 430 | -4.57 | 0.000116 | 0.000156 |
| chr13 | 96128511 | 96128956 | 446 | -4.57 | 0.000116 | 0.000156 |
| chr4 | 97545228 | 97546267 | 1040 | -3.98 | 0.000116 | 0.000156 |
| chr1 | 166953195 | 166956447 | 3253 | -2.66 | 0.000116 | 0.000157 |
| chr11 | 127095445 | 127115689 | 20245 | 2.49 | 0.000116 | 0.000157 |

| | | | | | | |
|---|---|---|---|---|---|---|
| chr11 | 48519417 | 48519785 | 369 | 4.85 | 0.000117 | 0.000157 |
| chr15 | 76316769 | 76339516 | 22748 | -2.33 | 0.000117 | 0.000158 |
| chr14 | 35663254 | 35663666 | 413 | -3.5 | 0.000117 | 0.000158 |
| chr11 | 98846535 | 98881683 | 35149 | 2.36 | 0.000117 | 0.000158 |
| chr2 | 87337921 | 87338261 | 341 | -2.84 | 0.000118 | 0.000158 |
| chr10 | 72897919 | 72935999 | 38081 | 1.77 | 0.000118 | 0.000159 |
| chr8 | 116839063 | 116841100 | 2038 | -4.57 | 0.000118 | 0.000159 |
| chr2 | 133972911 | 133973803 | 893 | -2.77 | 0.000118 | 0.000159 |
| chr6 | 17939623 | 17940212 | 590 | -4.56 | 0.000118 | 0.000159 |
| chr13 | 82599734 | 82600774 | 1041 | -5.3 | 0.000118 | 0.000159 |
| chr6 | 136163784 | 136164987 | 1204 | -5.06 | 0.000118 | 0.000159 |
| chr1 | 88680502 | 88683426 | 2925 | -3.97 | 0.000118 | 0.000159 |
| chrX | 121347830 | 121351792 | 3963 | 3.64 | 0.000118 | 0.00016 |
| chr2 | 25640629 | 25641212 | 584 | -4.7 | 0.000118 | 0.00016 |
| chr8 | 32547578 | 32548198 | 621 | -4.7 | 0.000118 | 0.00016 |
| chr8 | 90031537 | 90046850 | 15314 | -2.23 | 0.000119 | 0.00016 |
| chr14 | 51923322 | 51923814 | 493 | -4.64 | 0.000119 | 0.00016 |
| chr2 | 26088726 | 26089311 | 586 | -4.64 | 0.000119 | 0.00016 |
| chr1 | 196487772 | 196488798 | 1027 | -4.63 | 0.000119 | 0.00016 |
| chr11 | 59335295 | 59336068 | 774 | -4.63 | 0.000119 | 0.00016 |
| chr3 | 182541350 | 182541821 | 472 | -4.63 | 0.000119 | 0.00016 |
| chr5 | 106822360 | 106822822 | 463 | -4.63 | 0.000119 | 0.00016 |
| chr6 | 22212369 | 22213106 | 738 | -4.63 | 0.000119 | 0.00016 |
| chr10 | 86284885 | 86285559 | 675 | -4.83 | 0.000119 | 0.00016 |
| chr12 | 7539162 | 7544357 | 5196 | -1.66 | 0.000119 | 0.00016 |
| chrX | 36436895 | 36437401 | 507 | -3.12 | 0.000119 | 0.00016 |
| chr4 | 125940296 | 125940871 | 576 | -4.74 | 0.000119 | 0.00016 |
| chr1 | 108343187 | 108344504 | 1318 | -3.86 | 0.00012 | 0.000161 |
| chr18 | 54073298 | 54074123 | 826 | -3.53 | 0.00012 | 0.000161 |
| chr16 | 66266544 | 66288912 | 22369 | -1.21 | 0.00012 | 0.000161 |
| chr7 | 13713831 | 13715331 | 1501 | -2.42 | 0.00012 | 0.000162 |
| chr6 | 150760323 | 150767382 | 7060 | -3.09 | 0.00012 | 0.000162 |
| chrX | 82776681 | 82788825 | 12145 | 1.73 | 0.00012 | 0.000162 |
| chr11 | 14309128 | 14309825 | 698 | -5.47 | 0.000121 | 0.000162 |
| chr7 | 30453268 | 30453914 | 647 | -4.81 | 0.000121 | 0.000162 |
| chr6 | 169622073 | 169623530 | 1458 | -3.81 | 0.000121 | 0.000162 |
| chr18 | 71754298 | 71780874 | 26577 | -5.25 | 0.000122 | 0.000164 |
| chr18 | 12234736 | 12242411 | 7676 | -4.68 | 0.000122 | 0.000164 |
| chr10 | 19432178 | 19541488 | 109311 | -5.42 | 0.000122 | 0.000164 |
| chrX | 76319969 | 76342281 | 22313 | 1.04 | 0.000122 | 0.000164 |
| chr3 | 162790763 | 162792103 | 1341 | -2.72 | 0.000122 | 0.000164 |
| chr4 | 153104425 | 153137561 | 33137 | -2.58 | 0.000122 | 0.000164 |
| chr3 | 160937592 | 160938404 | 813 | -5.24 | 0.000122 | 0.000165 |
| chr2 | 136795847 | 136797875 | 2029 | -4.18 | 0.000122 | 0.000165 |
| chr6 | 54991075 | 54993971 | 2897 | -2.08 | 0.000123 | 0.000165 |
| chr1 | 154726964 | 154727324 | 361 | -4.58 | 0.000123 | 0.000165 |
| chr6 | 110275381 | 110285210 | 9830 | -1.83 | 0.000123 | 0.000165 |
| chr1 | 187215844 | 187217321 | 1478 | -2.36 | 0.000123 | 0.000165 |
| chr5 | 130002016 | 130002974 | 959 | -4.99 | 0.000123 | 0.000165 |
| chr14 | 72144442 | 72145610 | 1169 | -3.69 | 0.000123 | 0.000166 |
| chr1 | 192453104 | 192453503 | 400 | -4.55 | 0.000123 | 0.000166 |
| chr2 | 8578050 | 8578503 | 454 | -4.55 | 0.000123 | 0.000166 |
| chr2 | 238496662 | 238497283 | 622 | -4.55 | 0.000123 | 0.000166 |
| chr2 | 163612298 | 163613191 | 894 | 4.66 | 0.000123 | 0.000166 |
| chr2 | 18986640 | 18986988 | 349 | -3.48 | 0.000123 | 0.000166 |
| chr19 | 29914460 | 29984951 | 70492 | -2.46 | 0.000124 | 0.000166 |
| chr12 | 87330474 | 87330906 | 433 | -4.61 | 0.000124 | 0.000166 |
| chr12 | 92610232 | 92610621 | 390 | -4.61 | 0.000124 | 0.000166 |
| chr18 | 26841080 | 26842366 | 1287 | -3.69 | 0.000124 | 0.000167 |
| chr12 | 108716191 | 108730977 | 14787 | -6.11 | 0.000124 | 0.000167 |
| chr6 | 85883587 | 85885659 | 2073 | -2.22 | 0.000124 | 0.000167 |

| | | | | | | |
|---|---|---|---|---|---|---|
| chr5 | 113465410 | 113465776 | 367 | 4.87 | 0.000125 | 0.000168 |
| chr22 | 20207013 | 20207496 | 484 | -4.57 | 0.000125 | 0.000168 |
| chr1 | 218305625 | 218315203 | 9579 | -2.18 | 0.000125 | 0.000168 |
| chr22 | 38240849 | 38241674 | 826 | -4.21 | 0.000125 | 0.000168 |
| chr13 | 36785148 | 36821484 | 36337 | -1.13 | 0.000125 | 0.000168 |
| chrX | 130548591 | 130549242 | 652 | -4.55 | 0.000125 | 0.000168 |
| chrX | 28674200 | 28674791 | 592 | -4.66 | 0.000125 | 0.000169 |
| chr3 | 67069604 | 67071282 | 1679 | -3.84 | 0.000125 | 0.000169 |
| chrX | 113783442 | 113825319 | 41878 | 1.13 | 0.000126 | 0.000169 |
| chr7 | 23909719 | 23911644 | 1926 | 2.68 | 0.000126 | 0.000169 |
| chr1 | 170639449 | 170643060 | 3612 | -2.78 | 0.000126 | 0.000169 |
| chr10 | 44218590 | 44297981 | 79392 | -0.94 | 0.000126 | 0.000169 |
| chr1 | 229154481 | 229155068 | 588 | -5.2 | 0.000126 | 0.000169 |
| chr4 | 75903215 | 75918682 | 15468 | -4.61 | 0.000126 | 0.000169 |
| chr16 | 51272830 | 51357603 | 84774 | -1.35 | 0.000126 | 0.000169 |
| chr7 | 94380241 | 94380984 | 744 | -4.69 | 0.000126 | 0.000169 |
| chr21 | 35914726 | 35927603 | 12878 | -5.72 | 0.000126 | 0.00017 |
| chr6 | 67829573 | 67831072 | 1500 | -3.85 | 0.000127 | 0.00017 |
| chr3 | 167844014 | 167846130 | 2117 | -1.98 | 0.000127 | 0.00017 |
| chr3 | 6044717 | 6046942 | 2226 | 4.22 | 0.000127 | 0.00017 |
| chr15 | 101941654 | 101942236 | 583 | -4.6 | 0.000127 | 0.00017 |
| chr13 | 66328045 | 66348053 | 20009 | -2.05 | 0.000127 | 0.000171 |
| chr9 | 109989164 | 110001843 | 12680 | -4.75 | 0.000127 | 0.000171 |
| chr20 | 57154150 | 57193241 | 39092 | -7.05 | 0.000127 | 0.000171 |
| chr1 | 60052332 | 60052729 | 398 | -3.95 | 0.000127 | 0.000171 |
| chr8 | 128022261 | 128043023 | 20763 | -3.59 | 0.000127 | 0.000171 |
| chr10 | 82107212 | 82118618 | 11407 | -5.97 | 0.000128 | 0.000172 |
| chr1 | 245714224 | 245714660 | 437 | 4.65 | 0.000128 | 0.000172 |
| chr10 | 28879276 | 28880183 | 908 | -2.52 | 0.000128 | 0.000172 |
| chr18 | 30296944 | 30300028 | 3085 | 1.62 | 0.000128 | 0.000172 |
| chr16 | 71202577 | 71213751 | 11175 | 2.17 | 0.000128 | 0.000172 |
| chrX | 131797452 | 131815763 | 18312 | 1.21 | 0.000128 | 0.000172 |
| chr11 | 27647915 | 27660939 | 13025 | -2.52 | 0.000128 | 0.000172 |
| chr22 | 18124805 | 18125384 | 580 | -4.92 | 0.000129 | 0.000172 |
| chr22 | 21425754 | 21427289 | 1536 | -4.92 | 0.000129 | 0.000172 |
| chr6 | 22276574 | 22277212 | 639 | -4.92 | 0.000129 | 0.000172 |
| chr6 | 167035955 | 167038619 | 2665 | -3.52 | 0.000129 | 0.000173 |
| chr4 | 77309756 | 77314968 | 5213 | -3.9 | 0.000129 | 0.000173 |
| chr6 | 137389215 | 137392548 | 3334 | -2.01 | 0.000129 | 0.000173 |
| chr11 | 112314342 | 112318415 | 4074 | -4.66 | 0.000129 | 0.000173 |
| chr4 | 112719899 | 112720981 | 1083 | 5.41 | 0.000129 | 0.000173 |
| chr12 | 32302238 | 32302715 | 478 | -4.67 | 0.000129 | 0.000173 |
| chr7 | 13516328 | 13517318 | 991 | -4.67 | 0.000129 | 0.000173 |
| chr14 | 42774421 | 42885205 | 110785 | 5.78 | 0.000129 | 0.000173 |
| chr5 | 73725302 | 73726590 | 1289 | -4.66 | 0.000129 | 0.000173 |
| chr15 | 80456071 | 80471604 | 15534 | -6.38 | 0.000129 | 0.000173 |
| chr8 | 30482845 | 30483582 | 738 | -4.84 | 0.000129 | 0.000173 |
| chr6 | 66694488 | 66699133 | 4646 | 3.74 | 0.000129 | 0.000174 |
| chr17 | 30446961 | 30447471 | 511 | -4.64 | 0.00013 | 0.000174 |
| chr3 | 176321310 | 176322512 | 1203 | 4.14 | 0.00013 | 0.000174 |
| chr7 | 7178751 | 7179288 | 538 | -4.73 | 0.00013 | 0.000174 |
| chr3 | 23390421 | 23390951 | 531 | -4.56 | 0.00013 | 0.000175 |
| chr8 | 54244839 | 54250525 | 5687 | -4.8 | 0.000131 | 0.000175 |
| chr17 | 6617282 | 6653374 | 36093 | -1.84 | 0.000131 | 0.000176 |
| chr22 | 30621318 | 30621979 | 662 | -4.58 | 0.000132 | 0.000176 |
| chr18 | 912825 | 921260 | 8436 | 1.79 | 0.000132 | 0.000177 |
| chr5 | 102177461 | 102178596 | 1136 | -3.76 | 0.000132 | 0.000177 |
| chr22 | 22366723 | 22369340 | 2618 | -4.27 | 0.000132 | 0.000177 |
| chr1 | 90528903 | 90530324 | 1422 | -3.93 | 0.000132 | 0.000177 |
| chr8 | 121107883 | 121120830 | 12948 | -1.26 | 0.000132 | 0.000177 |
| chr18 | 70439765 | 70444254 | 4490 | 1.26 | 0.000132 | 0.000177 |

| | | | | | | |
|---|---|---|---|---|---|---|
| chr5 | 85428904 | 85429338 | 435 | 4.69 | 0.000132 | 0.000177 |
| chr1 | 83241108 | 83241610 | 503 | -3.66 | 0.000132 | 0.000177 |
| chr15 | 81417769 | 81428492 | 10724 | -5.01 | 0.000132 | 0.000178 |
| chr3 | 161505545 | 161520660 | 15116 | 1.32 | 0.000132 | 0.000178 |
| chr2 | 170254951 | 170255996 | 1046 | -3.7 | 0.000133 | 0.000178 |
| chr4 | 128302665 | 128303656 | 992 | -2.95 | 0.000133 | 0.000178 |
| chr8 | 11230055 | 11244923 | 14869 | -1.49 | 0.000133 | 0.000179 |
| chr8 | 121567201 | 121571314 | 4114 | 1.63 | 0.000133 | 0.000179 |
| chr20 | 21505180 | 21505987 | 808 | 2.37 | 0.000134 | 0.000179 |
| chrX | 77400377 | 77407111 | 6735 | -1.93 | 0.000134 | 0.000179 |
| chr17 | 47755851 | 47756267 | 417 | -4.79 | 0.000134 | 0.00018 |
| chrX | 29918874 | 29929402 | 10529 | -5.79 | 0.000134 | 0.00018 |
| chr4 | 68018710 | 68037721 | 19012 | 1.61 | 0.000134 | 0.00018 |
| chr2 | 218623796 | 218649263 | 25468 | -5.42 | 0.000134 | 0.00018 |
| chr1 | 104768812 | 104803233 | 34422 | 1.42 | 0.000135 | 0.000181 |
| chr22 | 49310693 | 49313211 | 2519 | -1.97 | 0.000135 | 0.000181 |
| chr4 | 114492457 | 114492968 | 512 | -4.84 | 0.000135 | 0.000181 |
| chr1 | 249073694 | 249096949 | 23256 | -2.09 | 0.000136 | 0.000182 |
| chr2 | 230168421 | 230169199 | 779 | 3.17 | 0.000136 | 0.000183 |
| chr9 | 89840028 | 89992600 | 152573 | -0.93 | 0.000137 | 0.000183 |
| chrX | 53824744 | 53827363 | 2620 | -1.23 | 0.000137 | 0.000183 |
| chr17 | 514210 | 514788 | 579 | -4.53 | 0.000137 | 0.000183 |
| chr17 | 71266041 | 71266925 | 885 | -4.53 | 0.000137 | 0.000183 |
| chr22 | 42227671 | 42228119 | 449 | -4.53 | 0.000137 | 0.000183 |
| chr18 | 23513694 | 23514180 | 487 | -4.56 | 0.000137 | 0.000183 |
| chr7 | 56878667 | 56879051 | 385 | -4.56 | 0.000137 | 0.000183 |
| chrX | 130323300 | 130324271 | 972 | -3.23 | 0.000137 | 0.000183 |
| chr12 | 97167077 | 97167624 | 548 | -3.63 | 0.000137 | 0.000183 |
| chr6 | 166506656 | 166566546 | 59891 | -5.59 | 0.000137 | 0.000183 |
| chr2 | 181228530 | 181229038 | 509 | -4.82 | 0.000137 | 0.000184 |
| chr5 | 145157581 | 145158435 | 855 | -4.82 | 0.000137 | 0.000184 |
| chr9 | 79692006 | 79754366 | 62361 | -1.69 | 0.000137 | 0.000184 |
| chr2 | 210789634 | 210791652 | 2019 | -1.95 | 0.000138 | 0.000185 |
| chr6 | 100164158 | 100165634 | 1477 | -3.82 | 0.000139 | 0.000186 |
| chr4 | 38556905 | 38558776 | 1872 | -4.08 | 0.000139 | 0.000186 |
| chr6 | 125715682 | 125717896 | 2215 | -1.76 | 0.000139 | 0.000186 |
| chr9 | 88651914 | 88652336 | 423 | -4.53 | 0.000139 | 0.000186 |
| chr5 | 98408565 | 98408977 | 413 | -4.8 | 0.000139 | 0.000186 |
| chr4 | 20781368 | 20782374 | 1007 | -4.53 | 0.000139 | 0.000187 |
| chr7 | 25269045 | 25271754 | 2710 | -1.47 | 0.00014 | 0.000187 |
| chr17 | 61542220 | 61542753 | 534 | -4.59 | 0.00014 | 0.000187 |
| chr9 | 28877853 | 28878211 | 359 | -3.99 | 0.00014 | 0.000187 |
| chr5 | 99909198 | 99909972 | 775 | -3.74 | 0.00014 | 0.000188 |
| chr17 | 75853596 | 75857505 | 3910 | 1.92 | 0.00014 | 0.000188 |
| chr2 | 237840286 | 237841904 | 1619 | -2.51 | 0.00014 | 0.000188 |
| chr19 | 40414978 | 40415386 | 409 | -4.52 | 0.000141 | 0.000188 |
| chr2 | 172871666 | 172872037 | 372 | -4.52 | 0.000141 | 0.000188 |
| chr1 | 193577291 | 193578298 | 1008 | -5.42 | 0.000141 | 0.000189 |
| chr2 | 204705022 | 204707968 | 2947 | 2.45 | 0.000141 | 0.000189 |
| chr1 | 47459962 | 47460570 | 609 | -3.44 | 0.000141 | 0.000189 |
| chr12 | 50273016 | 50274490 | 1475 | -4.78 | 0.000141 | 0.000189 |
| chr6 | 19827701 | 19837954 | 10254 | -1.56 | 0.000142 | 0.000189 |
| chr3 | 171521212 | 171536842 | 15631 | -6.39 | 0.000142 | 0.00019 |
| chr7 | 120507679 | 120541143 | 33465 | 5.25 | 0.000142 | 0.00019 |
| chr8 | 28118889 | 28154496 | 35608 | -5.6 | 0.000143 | 0.000191 |
| chr4 | 16134691 | 16138719 | 4029 | -1.79 | 0.000143 | 0.000191 |
| chr10 | 72759862 | 72760396 | 535 | -4.54 | 0.000143 | 0.000191 |
| chr4 | 148932294 | 148932732 | 439 | -4.54 | 0.000143 | 0.000191 |
| chr6 | 55873761 | 55874196 | 436 | -4.54 | 0.000143 | 0.000191 |
| chr1 | 155593725 | 155594421 | 697 | -4.6 | 0.000143 | 0.000192 |
| chr18 | 8857991 | 8858586 | 596 | -4.6 | 0.000143 | 0.000192 |

| | | | | | | |
|---|---|---|---|---|---|---|
| chr2 | 112774177 | 112775277 | 1101 | -4.64 | 0.000143 | 0.000192 |
| chr3 | 39075001 | 39084875 | 9875 | -5.14 | 0.000143 | 0.000192 |
| chr16 | 75098261 | 75098930 | 670 | -4.96 | 0.000144 | 0.000192 |
| chr7 | 7990439 | 7991151 | 713 | -4.76 | 0.000144 | 0.000192 |
| chr9 | 36444783 | 36445382 | 600 | -4.7 | 0.000144 | 0.000193 |
| chr1 | 111349260 | 111349925 | 666 | -2.61 | 0.000144 | 0.000193 |
| chr14 | 31729077 | 31747025 | 17949 | -6.57 | 0.000144 | 0.000193 |
| chr4 | 169057829 | 169058870 | 1042 | -3.65 | 0.000145 | 0.000194 |
| chr11 | 103813584 | 103815023 | 1440 | -4.08 | 0.000145 | 0.000194 |
| chr4 | 133946675 | 133949000 | 2326 | -1.95 | 0.000145 | 0.000194 |
| chr14 | 35209897 | 35210656 | 760 | -4.94 | 0.000145 | 0.000194 |
| chr10 | 86344238 | 86360008 | 15771 | 1.58 | 0.000145 | 0.000194 |
| chr2 | 105576152 | 105602652 | 26501 | -1.25 | 0.000146 | 0.000195 |
| chr2 | 61446301 | 61447384 | 1084 | -4.04 | 0.000146 | 0.000195 |
| chr18 | 48268988 | 48310010 | 41023 | -1.9 | 0.000146 | 0.000195 |
| chr22 | 27293978 | 27295060 | 1083 | -3.6 | 0.000146 | 0.000195 |
| chr12 | 109009330 | 109010066 | 737 | -4.57 | 0.000146 | 0.000195 |
| chr3 | 94730142 | 94731390 | 1249 | -4.57 | 0.000146 | 0.000195 |
| chr9 | 121568649 | 121569213 | 565 | -4.57 | 0.000146 | 0.000195 |
| chr1 | 59077467 | 59078731 | 1265 | -4.11 | 0.000146 | 0.000196 |
| chr16 | 50883292 | 50886674 | 3383 | 1.47 | 0.000146 | 0.000196 |
| chr20 | 62055982 | 62056567 | 586 | -4.73 | 0.000146 | 0.000196 |
| chr7 | 123251396 | 123252321 | 926 | -3.73 | 0.000146 | 0.000196 |
| chr20 | 31521854 | 31547447 | 25594 | -5.51 | 0.000147 | 0.000196 |
| chr7 | 4501878 | 4614967 | 113090 | -7.41 | 0.000147 | 0.000196 |
| chr15 | 55881563 | 55882049 | 487 | -4.92 | 0.000147 | 0.000196 |
| chr1 | 15078823 | 15079462 | 640 | -4.51 | 0.000147 | 0.000197 |
| chr6 | 167480259 | 167501910 | 21652 | -2.04 | 0.000148 | 0.000198 |
| chr2 | 219787462 | 219789827 | 2366 | -4.17 | 0.000148 | 0.000198 |
| chr21 | 27976583 | 27992320 | 15738 | 1.2 | 0.000148 | 0.000198 |
| chr3 | 86357255 | 86363577 | 6323 | 2.41 | 0.000148 | 0.000198 |
| chr17 | 65624436 | 65624967 | 532 | -4.9 | 0.000149 | 0.000199 |
| chr4 | 113885556 | 113886090 | 535 | -4.5 | 0.000149 | 0.000199 |
| chr10 | 6501501 | 6501899 | 399 | -3.75 | 0.000149 | 0.000199 |
| chr2 | 107189260 | 107189946 | 687 | -3.45 | 0.000149 | 2.00E-04 |
| chr13 | 101565579 | 101568928 | 3350 | -1.39 | 0.00015 | 2.00E-04 |
| chr7 | 137371474 | 137380812 | 9339 | 3.28 | 0.00015 | 2.00E-04 |
| chr3 | 87708280 | 87708834 | 555 | -4.53 | 0.00015 | 2.00E-04 |
| chr7 | 148250058 | 148258021 | 7964 | -1.04 | 0.00015 | 2.00E-04 |
| chr5 | 74251360 | 74251784 | 425 | -4.51 | 0.00015 | 2.00E-04 |
| chr1 | 85539980 | 85540744 | 765 | -4.86 | 0.00015 | 2.00E-04 |
| chr1 | 147064180 | 147069511 | 5332 | -1.48 | 0.00015 | 0.000201 |
| chr10 | 23028577 | 23033426 | 4850 | -4.33 | 0.00015 | 0.000201 |
| chr2 | 84936036 | 85007305 | 71270 | -1.77 | 0.00015 | 0.000201 |
| chr18 | 73300446 | 73458219 | 157774 | -0.55 | 0.00015 | 0.000201 |
| chr5 | 63921778 | 63925022 | 3245 | -4.55 | 0.00015 | 0.000201 |
| chr5 | 7748191 | 7832024 | 83834 | 0.72 | 0.00015 | 0.000201 |
| chr7 | 82231402 | 82232196 | 795 | -4.88 | 0.00015 | 0.000201 |
| chr1 | 196706122 | 196711185 | 5064 | -4.62 | 0.000151 | 0.000201 |
| chr6 | 79431642 | 79432405 | 764 | -3.44 | 0.000151 | 0.000201 |
| chr9 | 96916767 | 96918536 | 1770 | -5.1 | 0.000151 | 0.000202 |
| chrX | 23631470 | 23637871 | 6402 | -4.07 | 0.000151 | 0.000202 |
| chr3 | 185442664 | 185443346 | 683 | -4.5 | 0.000151 | 0.000202 |
| chr6 | 118057043 | 118089157 | 32115 | -1.77 | 0.000151 | 0.000202 |
| chr4 | 111302703 | 111341507 | 38805 | -1.13 | 0.000152 | 0.000203 |
| chr21 | 36417764 | 36418397 | 634 | -4.86 | 0.000152 | 0.000203 |
| chr12 | 80522656 | 80566329 | 43674 | -5.14 | 0.000153 | 0.000204 |
| chr12 | 57048743 | 57049379 | 637 | -4.65 | 0.000153 | 0.000204 |
| chr6 | 143032889 | 143041724 | 8836 | -2.95 | 0.000153 | 0.000204 |
| chr11 | 74469449 | 74469886 | 438 | -4.56 | 0.000153 | 0.000204 |
| chr8 | 20905952 | 21428534 | 522583 | 0.64 | 0.000153 | 0.000204 |

| | | | | | | |
|---|---|---|---|---|---|---|
| chr4 | 41973514 | 41974140 | 627 | -4.54 | 0.000153 | 0.000205 |
| chr2 | 30272811 | 30326772 | 53962 | -1.98 | 0.000154 | 0.000205 |
| chr2 | 19765616 | 19766516 | 901 | -2.95 | 0.000154 | 0.000205 |
| chr14 | 50694582 | 50702945 | 8364 | -6.11 | 0.000154 | 0.000206 |
| chr9 | 92488327 | 92489903 | 1577 | -4.04 | 0.000154 | 0.000206 |
| chr4 | 134015336 | 134015820 | 485 | 3.66 | 0.000155 | 0.000207 |
| chr1 | 12786385 | 12834745 | 48361 | -3.47 | 0.000155 | 0.000207 |
| chr14 | 26838953 | 26841449 | 2497 | -5.28 | 0.000155 | 0.000207 |
| chr10 | 14725949 | 14726588 | 640 | -5.05 | 0.000156 | 0.000208 |
| chr9 | 28640964 | 28644510 | 3547 | -1.86 | 0.000156 | 0.000208 |
| chr3 | 45807678 | 45810087 | 2410 | -3.86 | 0.000156 | 0.000208 |
| chr19 | 55092984 | 55139346 | 46363 | -0.9 | 0.000156 | 0.000208 |
| chr2 | 222813322 | 222818040 | 4719 | 1.34 | 0.000156 | 0.000208 |
| chr2 | 45080307 | 45105018 | 24712 | -1.66 | 0.000156 | 0.000209 |
| chr16 | 79853748 | 79860129 | 6382 | 1.74 | 0.000156 | 0.000209 |
| chr3 | 4644079 | 4644592 | 514 | -3.9 | 0.000157 | 0.000209 |
| chr11 | 119797404 | 119800803 | 3400 | 1.52 | 0.000157 | 0.000209 |
| chr5 | 172916053 | 172917029 | 977 | -4.49 | 0.000157 | 0.000209 |
| chr7 | 72766832 | 72767602 | 771 | -4.49 | 0.000157 | 0.000209 |
| chr2 | 64028713 | 64029118 | 406 | -3.56 | 0.000157 | 0.00021 |
| chr12 | 367670 | 370362 | 2693 | 1.56 | 0.000157 | 0.00021 |
| chr17 | 25928142 | 25928768 | 627 | -5.03 | 0.000157 | 0.00021 |
| chr5 | 176989002 | 176989845 | 844 | -5.03 | 0.000157 | 0.00021 |
| chrX | 18197000 | 18222030 | 25031 | -1.73 | 0.000158 | 0.000211 |
| chr9 | 72528248 | 72529093 | 846 | -4.78 | 0.000159 | 0.000212 |
| chr20 | 13107720 | 13169477 | 61758 | -1.42 | 0.000159 | 0.000212 |
| chr14 | 101221323 | 101230271 | 8949 | -5.02 | 0.000159 | 0.000212 |
| chr13 | 63922364 | 63925471 | 3108 | 3.15 | 0.00016 | 0.000213 |
| chr16 | 46501419 | 46569359 | 67941 | 5.77 | 0.00016 | 0.000213 |
| chr13 | 95092604 | 95103465 | 10862 | -1.47 | 0.00016 | 0.000213 |
| chr1 | 13787910 | 13806129 | 18220 | -1.22 | 0.00016 | 0.000213 |
| chr13 | 31700603 | 31701167 | 565 | -4.65 | 0.00016 | 0.000213 |
| chr7 | 98234181 | 98237205 | 3025 | -3.02 | 0.000161 | 0.000215 |
| chr2 | 217429450 | 217451295 | 21846 | -4.98 | 0.000162 | 0.000216 |
| chrX | 23905089 | 23905636 | 548 | -3.44 | 0.000162 | 0.000216 |
| chr16 | 55832462 | 55832891 | 430 | -4.52 | 0.000162 | 0.000216 |
| chr3 | 177909973 | 177910648 | 676 | -4.52 | 0.000162 | 0.000216 |
| chrX | 118943061 | 118944717 | 1657 | -3.97 | 0.000162 | 0.000216 |
| chr7 | 148413490 | 148414089 | 600 | -4.98 | 0.000162 | 0.000216 |
| chr22 | 25989127 | 25989514 | 388 | -4.48 | 0.000163 | 0.000217 |
| chr4 | 26709843 | 26710307 | 465 | -4.48 | 0.000163 | 0.000217 |
| chr3 | 192851404 | 192928612 | 77209 | 3.06 | 0.000163 | 0.000217 |
| chr3 | 188369274 | 188370298 | 1025 | -4.75 | 0.000163 | 0.000217 |
| chr18 | 4116498 | 4124669 | 8172 | 4.85 | 0.000163 | 0.000218 |
| chrX | 51364892 | 51379971 | 15080 | 1.3 | 0.000163 | 0.000218 |
| chr9 | 128032285 | 128032702 | 418 | -4.47 | 0.000163 | 0.000218 |
| chr3 | 41087629 | 41090405 | 2777 | -4.13 | 0.000164 | 0.000218 |
| chr18 | 24525298 | 24527508 | 2211 | -4.01 | 0.000164 | 0.000218 |
| chr9 | 16398140 | 16403547 | 5408 | -4.62 | 0.000164 | 0.000219 |
| chr9 | 98317879 | 98318474 | 596 | -4.62 | 0.000164 | 0.000219 |
| chr4 | 83179398 | 83181062 | 1665 | -2.39 | 0.000164 | 0.000219 |
| chrX | 38635729 | 38639372 | 3644 | -2.78 | 0.000164 | 0.000219 |
| chr3 | 55381220 | 55405977 | 24758 | -0.91 | 0.000165 | 0.000219 |
| chr2 | 75577691 | 75593969 | 16279 | -1.1 | 0.000165 | 0.00022 |
| chr12 | 100247682 | 100248285 | 604 | -4.73 | 0.000165 | 0.00022 |
| chr5 | 64350828 | 64355722 | 4895 | -4.73 | 0.000165 | 0.00022 |
| chr1 | 77277270 | 77280625 | 3356 | 2.25 | 0.000165 | 0.00022 |
| chr5 | 56843367 | 56883619 | 40253 | -1.85 | 0.000165 | 0.000221 |
| chr11 | 102608597 | 102621083 | 12487 | -3.97 | 0.000166 | 0.000221 |
| chr5 | 54248275 | 54257028 | 8754 | -0.99 | 0.000167 | 0.000222 |
| chr1 | 84870732 | 84871230 | 499 | -4.48 | 0.000167 | 0.000222 |

| | | | | | | |
|---|---|---|---|---|---|---|
| chr16 | 1051298 | 1051898 | 601 | -4.48 | 0.000167 | 0.000222 |
| chr14 | 64870028 | 64871710 | 1683 | -3.79 | 0.000167 | 0.000222 |
| chr13 | 83387514 | 83388616 | 1103 | -4.72 | 0.000167 | 0.000223 |
| chr2 | 207788831 | 207819838 | 31008 | -5.23 | 0.000167 | 0.000223 |
| chr8 | 119642481 | 119662726 | 20246 | -0.92 | 0.000168 | 0.000224 |
| chr19 | 47659168 | 47659548 | 381 | -3.58 | 0.000168 | 0.000224 |
| chr3 | 112760069 | 112768011 | 7943 | -1.46 | 0.000168 | 0.000224 |
| chr16 | 57929054 | 57930281 | 1228 | -4.47 | 0.000168 | 0.000224 |
| chr3 | 111756596 | 111756985 | 390 | -4.47 | 0.000168 | 0.000224 |
| chr4 | 30557471 | 30559217 | 1747 | -4.47 | 0.000168 | 0.000224 |
| chr6 | 10176560 | 10177179 | 620 | -4.47 | 0.000168 | 0.000224 |
| chr5 | 126955898 | 126956475 | 578 | -3.49 | 0.000168 | 0.000224 |
| chr13 | 22390180 | 22390643 | 464 | -4.57 | 0.000168 | 0.000224 |
| chrX | 130338190 | 130352153 | 13964 | 2.13 | 0.000169 | 0.000225 |
| chrX | 39464497 | 39481967 | 17471 | -2.75 | 0.000169 | 0.000225 |
| chr8 | 49190764 | 49191477 | 714 | -4.6 | 0.000169 | 0.000225 |
| chr2 | 239062785 | 239063270 | 486 | -4.72 | 0.000169 | 0.000225 |
| chr4 | 33591577 | 33721531 | 129955 | -1.23 | 0.000169 | 0.000226 |
| chr21 | 39212816 | 39230503 | 17688 | -4.79 | 0.00017 | 0.000226 |
| chr9 | 119983398 | 119986520 | 3123 | 4.22 | 0.00017 | 0.000226 |
| chr12 | 105988657 | 105990356 | 1700 | 1.89 | 0.00017 | 0.000226 |
| chr2 | 147599157 | 147622018 | 22862 | 1.53 | 0.00017 | 0.000226 |
| chr2 | 133354908 | 133355773 | 866 | -4.49 | 0.000172 | 0.000229 |
| chr9 | 19915873 | 19916809 | 937 | -1.99 | 0.000172 | 0.000229 |
| chr11 | 120660718 | 120663707 | 2990 | -2.07 | 0.000172 | 0.000229 |
| chr8 | 86925318 | 86950829 | 25512 | 0.89 | 0.000172 | 0.000229 |
| chr14 | 32703143 | 32703798 | 656 | -4.7 | 0.000172 | 0.00023 |
| chr3 | 75355018 | 75361416 | 6399 | 3.72 | 0.000173 | 0.00023 |
| chr10 | 1214373 | 1215082 | 710 | -4.52 | 0.000173 | 0.00023 |
| chr18 | 50065691 | 50066324 | 634 | -4.52 | 0.000173 | 0.00023 |
| chr20 | 34359219 | 34359710 | 492 | -4.52 | 0.000173 | 0.00023 |
| chr3 | 123181481 | 123181828 | 348 | -4.52 | 0.000173 | 0.00023 |
| chr4 | 62226508 | 62227202 | 695 | -4.52 | 0.000173 | 0.00023 |
| chr5 | 169901975 | 169902557 | 583 | -4.52 | 0.000173 | 0.00023 |
| chr12 | 98079996 | 98093576 | 13581 | -1.74 | 0.000173 | 0.000231 |
| chr2 | 200006490 | 200019595 | 13106 | 2.15 | 0.000173 | 0.000231 |
| chr8 | 125079803 | 125082442 | 2640 | 3.24 | 0.000174 | 0.000231 |
| chr11 | 15772286 | 15791424 | 19139 | -1.48 | 0.000174 | 0.000231 |
| chr7 | 87393916 | 87426152 | 32237 | -4.96 | 0.000174 | 0.000232 |
| chr20 | 42382236 | 42388275 | 6040 | -5.3 | 0.000174 | 0.000232 |
| chr14 | 24573785 | 24574672 | 888 | -4.87 | 0.000174 | 0.000232 |
| chr4 | 189221940 | 189222497 | 558 | -4.87 | 0.000174 | 0.000232 |
| chr12 | 22420860 | 22462801 | 41942 | -2.74 | 0.000174 | 0.000232 |
| chr4 | 60486942 | 60487563 | 622 | -3.45 | 0.000174 | 0.000232 |
| chr18 | 42414136 | 42414968 | 833 | -4.57 | 0.000174 | 0.000232 |
| chr21 | 17419381 | 17419931 | 551 | -4.57 | 0.000174 | 0.000232 |
| chr7 | 12690858 | 12691651 | 794 | -4.57 | 0.000174 | 0.000232 |
| chr14 | 81546207 | 81546643 | 437 | 4.8 | 0.000175 | 0.000233 |
| chr12 | 67974572 | 67983616 | 9045 | 3.57 | 0.000175 | 0.000233 |
| chr17 | 63380615 | 63393166 | 12552 | 1.06 | 0.000175 | 0.000233 |
| chr18 | 32469936 | 32491257 | 21322 | -5.1 | 0.000175 | 0.000233 |
| chr14 | 105150846 | 105151360 | 515 | -4.66 | 0.000176 | 0.000234 |
| chr1 | 76332222 | 76333500 | 1279 | 3.79 | 0.000176 | 0.000234 |
| chr4 | 87171244 | 87172129 | 886 | -4.67 | 0.000176 | 0.000234 |
| chr4 | 189091873 | 189092377 | 505 | -4.67 | 0.000176 | 0.000234 |
| chr3 | 87724977 | 87725955 | 979 | -5.07 | 0.000176 | 0.000234 |
| chr7 | 87786528 | 87791155 | 4628 | -4.47 | 0.000176 | 0.000234 |
| chr14 | 22088043 | 22250693 | 162651 | 0.82 | 0.000176 | 0.000235 |
| chr6 | 106371713 | 106372164 | 452 | -4.46 | 0.000177 | 0.000235 |
| chr3 | 187355799 | 187373958 | 18160 | -1.13 | 0.000177 | 0.000235 |
| chr4 | 27037020 | 27047220 | 10201 | -2.35 | 0.000177 | 0.000236 |

| | | | | | | |
|---|---|---|---|---|---|---|
| chr19 | 42357179 | 42358197 | 1019 | 3.43 | 0.000177 | 0.000236 |
| chr7 | 43070452 | 43168066 | 97615 | -2.24 | 0.000177 | 0.000236 |
| chr12 | 66401798 | 66405500 | 3703 | -4.51 | 0.000178 | 0.000236 |
| chr1 | 230752388 | 230766269 | 13882 | -2.5 | 0.000178 | 0.000237 |
| chr7 | 120506133 | 120507516 | 1384 | -2.68 | 0.000178 | 0.000237 |
| chr7 | 117773582 | 117775441 | 1860 | -1.74 | 0.000178 | 0.000237 |
| chr12 | 27101367 | 27101788 | 422 | -4.46 | 0.000178 | 0.000237 |
| chr1 | 168316753 | 168317184 | 432 | -4.64 | 0.000179 | 0.000238 |
| chr6 | 131438347 | 131438669 | 323 | -4.64 | 0.000179 | 0.000238 |
| chr12 | 123399197 | 123400342 | 1146 | -4.53 | 0.000179 | 0.000238 |
| chr3 | 123899028 | 123911053 | 12026 | -1.69 | 0.00018 | 0.000239 |
| chr22 | 34131671 | 34132657 | 987 | -2.82 | 0.00018 | 0.000239 |
| chr2 | 230394473 | 230397377 | 2905 | -2.18 | 0.00018 | 0.000239 |
| chr19 | 43890400 | 43890905 | 506 | -3.43 | 0.00018 | 0.00024 |
| chr3 | 58672813 | 58688174 | 15362 | -4.02 | 0.00018 | 0.00024 |
| chr2 | 3801427 | 3907915 | 106489 | -1.08 | 0.000181 | 0.00024 |
| chr12 | 6293159 | 6293680 | 522 | -4.55 | 0.000181 | 0.00024 |
| chr9 | 103378962 | 103382201 | 3240 | -2.39 | 0.000181 | 0.00024 |
| chr1 | 2050283 | 2051108 | 826 | -4.81 | 0.000181 | 0.000241 |
| chr15 | 96606674 | 96608363 | 1690 | -3.87 | 0.000181 | 0.000241 |
| chr10 | 97505087 | 97508231 | 3145 | -3.13 | 0.000181 | 0.000241 |
| chr4 | 173360664 | 173361874 | 1211 | -3.69 | 0.000181 | 0.000241 |
| chr1 | 54909767 | 54910174 | 408 | -3.69 | 0.000181 | 0.000241 |
| chr11 | 102372478 | 102373045 | 568 | -4.5 | 0.000182 | 0.000242 |
| chr3 | 90311720 | 90312113 | 394 | -4.5 | 0.000182 | 0.000242 |
| chr4 | 38634114 | 38634982 | 869 | -4.5 | 0.000182 | 0.000242 |
| chr3 | 164022400 | 164029225 | 6826 | -3.92 | 0.000182 | 0.000242 |
| chr3 | 175834089 | 175837081 | 2993 | 1.68 | 0.000182 | 0.000242 |
| chrX | 112081899 | 112083437 | 1539 | -4.43 | 0.000183 | 0.000243 |
| chr7 | 56174573 | 56384813 | 210241 | -1.2 | 0.000183 | 0.000243 |
| chr11 | 63004235 | 63005431 | 1197 | -2.06 | 0.000183 | 0.000243 |
| chr16 | 76713341 | 76899345 | 186005 | -1.4 | 0.000184 | 0.000244 |
| chr1 | 155127671 | 155138104 | 10434 | -4.79 | 0.000184 | 0.000244 |
| chr9 | 117767904 | 117768731 | 828 | -5.55 | 0.000184 | 0.000244 |
| chr12 | 104901111 | 104901869 | 759 | -4.63 | 0.000184 | 0.000244 |
| chr16 | 67818035 | 67818452 | 418 | -4.63 | 0.000184 | 0.000244 |
| chr4 | 118770866 | 118771803 | 938 | -4.63 | 0.000184 | 0.000244 |
| chr12 | 64672337 | 64672722 | 386 | -4.81 | 0.000184 | 0.000245 |
| chr13 | 25976823 | 26002523 | 25701 | -0.9 | 0.000184 | 0.000245 |
| chr1 | 30130795 | 30136799 | 6005 | -5 | 0.000184 | 0.000245 |
| chr16 | 5180640 | 5181770 | 1131 | -2.01 | 0.000185 | 0.000245 |
| chr14 | 69627210 | 69641877 | 14668 | -4.88 | 0.000186 | 0.000246 |
| chr1 | 5913848 | 5914656 | 809 | -4.98 | 0.000186 | 0.000246 |
| chr22 | 27859443 | 27859936 | 494 | -3.87 | 0.000186 | 0.000247 |
| chrX | 102299689 | 102302480 | 2792 | -3.49 | 0.000187 | 0.000248 |
| chr2 | 152567144 | 152568046 | 903 | 3.91 | 0.000187 | 0.000249 |
| chr14 | 81563949 | 81610655 | 46707 | 0.83 | 0.000187 | 0.000249 |
| chr11 | 101266012 | 101272780 | 6769 | -2.14 | 0.000187 | 0.000249 |
| chr3 | 135619413 | 135622119 | 2707 | -2.78 | 0.000187 | 0.000249 |
| chr11 | 64094588 | 64095285 | 698 | -4.97 | 0.000188 | 0.000249 |
| chr11 | 101413924 | 101414405 | 482 | -4.53 | 0.000188 | 0.000249 |
| chr22 | 36256359 | 36256800 | 442 | -4.53 | 0.000188 | 0.000249 |
| chr5 | 106666936 | 106667417 | 482 | -4.53 | 0.000188 | 0.000249 |
| chr3 | 42305576 | 42337806 | 32231 | -3.22 | 0.000188 | 0.000249 |
| chr8 | 71602864 | 71617641 | 14778 | -4.78 | 0.000188 | 0.000249 |
| chr6 | 112907634 | 112932294 | 24661 | -0.88 | 0.000188 | 0.00025 |
| chr4 | 159777650 | 159778121 | 472 | -4.44 | 0.000188 | 0.00025 |
| chr6 | 7924752 | 7926393 | 1642 | -2.57 | 0.000189 | 0.00025 |
| chr18 | 53831451 | 53832895 | 1445 | -4.78 | 0.000189 | 0.00025 |
| chr17 | 60701252 | 60701923 | 672 | -4.6 | 0.000189 | 0.000251 |
| chr2 | 127597391 | 127636318 | 38928 | -3.07 | 0.000189 | 0.000251 |

| | | | | | | |
|---|---|---|---|---|---|---|
| chr3 | 189345227 | 189345944 | 718 | -5.17 | 0.000189 | 0.000251 |
| chr6 | 11956981 | 11957390 | 410 | -4.74 | 0.000189 | 0.000251 |
| chr4 | 160723873 | 160724873 | 1001 | -4.95 | 0.00019 | 0.000252 |
| chr7 | 73195672 | 73196418 | 747 | -4.95 | 0.00019 | 0.000252 |
| chr16 | 10719220 | 10735073 | 15854 | -6.73 | 0.00019 | 0.000252 |
| chr12 | 6207972 | 6208917 | 946 | -4.05 | 0.00019 | 0.000252 |
| chr5 | 4994396 | 5157029 | 162634 | -0.52 | 0.00019 | 0.000253 |
| chr13 | 46265365 | 46267028 | 1664 | -4.11 | 0.000191 | 0.000254 |
| chr12 | 69413861 | 69414383 | 523 | -4.49 | 0.000192 | 0.000255 |
| chr2 | 109195086 | 109195537 | 452 | -4.47 | 0.000192 | 0.000255 |
| chr15 | 71021715 | 71022167 | 453 | -4.72 | 0.000193 | 0.000255 |
| chr7 | 22198088 | 22482137 | 284050 | 7.61 | 0.000193 | 0.000256 |
| chr13 | 101370310 | 101399056 | 28747 | -4.95 | 0.000193 | 0.000257 |
| chr16 | 71354352 | 71371191 | 16840 | -5.17 | 0.000194 | 0.000257 |
| chr7 | 51481838 | 51482217 | 380 | -4.43 | 0.000194 | 0.000257 |
| chr11 | 102386330 | 102386906 | 577 | -4.58 | 0.000194 | 0.000257 |
| chr2 | 74912411 | 74912853 | 443 | -4.58 | 0.000194 | 0.000257 |
| chr6 | 157376902 | 157377456 | 555 | -4.58 | 0.000194 | 0.000257 |
| chr3 | 96605579 | 96611377 | 5799 | -1.94 | 0.000194 | 0.000258 |
| chr2 | 135280612 | 135283458 | 2847 | -1.76 | 0.000195 | 0.000258 |
| chr7 | 116351827 | 116352368 | 542 | -4.57 | 0.000195 | 0.000258 |
| chr13 | 27893047 | 27898722 | 5676 | -1.63 | 0.000195 | 0.000258 |
| chr8 | 140648733 | 140660402 | 11670 | -5.48 | 0.000196 | 0.000259 |
| chr9 | 140296361 | 140296815 | 455 | -4.5 | 0.000196 | 0.00026 |
| chr20 | 49195204 | 49195611 | 408 | -4.7 | 0.000196 | 0.00026 |
| chr14 | 76733877 | 76735345 | 1469 | -4.04 | 0.000196 | 0.00026 |
| chr3 | 13744113 | 13818571 | 74459 | -2.31 | 0.000196 | 0.00026 |
| chr2 | 187382220 | 187392555 | 10336 | -5.75 | 0.000196 | 0.00026 |
| chrX | 66546358 | 66576957 | 30600 | 1.06 | 0.000197 | 0.000261 |
| chr22 | 25234399 | 25235151 | 753 | -5.11 | 0.000197 | 0.000261 |
| chr19 | 36619723 | 36620158 | 436 | -4.44 | 0.000197 | 0.000261 |
| chr4 | 40494228 | 40494873 | 646 | -4.44 | 0.000197 | 0.000261 |
| chr7 | 106589451 | 106590573 | 1123 | -1.77 | 0.000197 | 0.000261 |
| chr17 | 639761 | 653534 | 13774 | -5.24 | 0.000197 | 0.000262 |
| chr14 | 42636498 | 42641933 | 5436 | -4.2 | 0.000198 | 0.000262 |
| chr10 | 128104196 | 128140143 | 35948 | -5.07 | 0.000198 | 0.000263 |
| chr11 | 63397956 | 63398489 | 534 | -4.71 | 0.000198 | 0.000263 |
| chr6 | 125924596 | 125926134 | 1539 | -4.12 | 0.000199 | 0.000263 |
| chr12 | 108393215 | 108494537 | 101323 | 0.88 | 0.000199 | 0.000264 |
| chr4 | 64795352 | 64797424 | 2073 | -4.87 | 0.000199 | 0.000264 |
| chr6 | 162032407 | 162034807 | 2401 | -2.29 | 2.00E-04 | 0.000265 |
| chr4 | 105065938 | 105066437 | 500 | -4.68 | 2.00E-04 | 0.000265 |
| chr13 | 23243743 | 23281546 | 37804 | -2.04 | 2.00E-04 | 0.000265 |
| chr4 | 128233983 | 128246238 | 12256 | -1.83 | 2.00E-04 | 0.000265 |
| chr5 | 148318814 | 148319274 | 461 | -4.44 | 0.000201 | 0.000266 |
| chr7 | 134789961 | 134819414 | 29454 | -5.01 | 0.000201 | 0.000267 |
| chr10 | 5671253 | 5671707 | 455 | -4.42 | 0.000202 | 0.000267 |
| chr4 | 137425372 | 137425854 | 483 | -4.42 | 0.000202 | 0.000267 |
| chr3 | 173111967 | 173112878 | 912 | -2.34 | 0.000202 | 0.000268 |
| chrX | 17997603 | 18002699 | 5097 | -1.9 | 0.000203 | 0.000269 |
| chr4 | 164308478 | 164308930 | 453 | 4.62 | 0.000203 | 0.000269 |
| chr16 | 11714043 | 11714784 | 742 | -4.65 | 0.000204 | 0.00027 |
| chr8 | 29262410 | 29274236 | 11827 | -6.27 | 0.000204 | 0.00027 |
| chr16 | 81291894 | 81292713 | 820 | -5.25 | 0.000204 | 0.00027 |
| chr12 | 15451585 | 15457714 | 6130 | -2.82 | 0.000204 | 0.000271 |
| chr7 | 15662614 | 15664314 | 1701 | -4.83 | 0.000205 | 0.000271 |
| chr8 | 9388818 | 9407554 | 18737 | -1.47 | 0.000205 | 0.000271 |
| chr4 | 97336305 | 97337721 | 1417 | -4.53 | 0.000205 | 0.000271 |
| chr3 | 61844942 | 61846085 | 1144 | -4.2 | 0.000205 | 0.000272 |
| chr4 | 73561563 | 73801035 | 239473 | 7.52 | 0.000206 | 0.000272 |
| chr16 | 83866092 | 83931893 | 65802 | -5.41 | 0.000206 | 0.000272 |

| | | | | | | |
|---|---|---|---|---|---|---|
| chr7 | 141800837 | 141808374 | 7538 | -1.05 | 0.000206 | 0.000272 |
| chr15 | 36344540 | 36345370 | 831 | -5.04 | 0.000206 | 0.000273 |
| chr3 | 160624012 | 160625222 | 1211 | -2.19 | 0.000206 | 0.000273 |
| chr11 | 115501997 | 115503241 | 1245 | 2.73 | 0.000206 | 0.000273 |
| chr1 | 64171758 | 64173332 | 1575 | -3.84 | 0.000207 | 0.000274 |
| chr11 | 123026713 | 123027359 | 647 | -4.53 | 0.000207 | 0.000274 |
| chr8 | 101356012 | 101359161 | 3150 | -3.4 | 0.000207 | 0.000274 |
| chr12 | 68428216 | 68429793 | 1578 | 1.91 | 0.000207 | 0.000275 |
| chr13 | 110283891 | 110296448 | 12558 | -4.95 | 0.000208 | 0.000275 |
| chr1 | 113882190 | 113883661 | 1472 | -4.09 | 0.000208 | 0.000275 |
| chr13 | 36973969 | 36974641 | 673 | -3.43 | 0.000208 | 0.000275 |
| chr5 | 107025511 | 107029385 | 3875 | -4.67 | 0.000208 | 0.000276 |
| chr10 | 81878933 | 81880962 | 2030 | -3.89 | 0.000209 | 0.000277 |
| chr4 | 123577796 | 123581904 | 4109 | -1.55 | 0.000209 | 0.000277 |
| chrX | 151598308 | 151599766 | 1459 | 3.76 | 0.00021 | 0.000277 |
| chr21 | 36903275 | 36953571 | 50297 | 0.97 | 0.00021 | 0.000278 |
| chr21 | 15494533 | 15506532 | 12000 | -1.39 | 0.00021 | 0.000278 |
| chr2 | 167014240 | 167014999 | 760 | 4.53 | 0.000211 | 0.000279 |
| chr9 | 88124201 | 88127485 | 3285 | 2.5 | 0.000211 | 0.000279 |
| chr8 | 27304615 | 27305073 | 459 | -4.51 | 0.000211 | 0.000279 |
| chr12 | 98297664 | 98298799 | 1136 | 2.25 | 0.000212 | 0.00028 |
| chr20 | 13888802 | 13889613 | 812 | -2.24 | 0.000212 | 0.00028 |
| chrX | 76283464 | 76283911 | 448 | 4.09 | 0.000212 | 0.00028 |
| chr7 | 15726405 | 15727186 | 782 | -3.33 | 0.000212 | 0.000281 |
| chr3 | 72083545 | 72087805 | 4261 | -1.6 | 0.000212 | 0.000281 |
| chrX | 38771068 | 38829616 | 58549 | 0.94 | 0.000212 | 0.000281 |
| chr16 | 75539745 | 75540382 | 638 | -4.41 | 0.000213 | 0.000281 |
| chr5 | 49681772 | 49682201 | 430 | -4.41 | 0.000213 | 0.000281 |
| chr5 | 79287158 | 79288911 | 1754 | -1.75 | 0.000213 | 0.000282 |
| chr1 | 192657724 | 192658101 | 378 | -4.41 | 0.000213 | 0.000282 |
| chrX | 76842749 | 76843144 | 396 | -4.41 | 0.000213 | 0.000282 |
| chr4 | 89997199 | 90015344 | 18146 | -5.02 | 0.000213 | 0.000282 |
| chr4 | 28207535 | 28208379 | 845 | -4.77 | 0.000214 | 0.000282 |
| chr12 | 20309161 | 20309581 | 421 | -4.51 | 0.000214 | 0.000284 |
| chr4 | 17673001 | 17675679 | 2679 | -4.14 | 0.000215 | 0.000284 |
| chr12 | 52840369 | 52883935 | 43567 | -0.79 | 0.000215 | 0.000284 |
| chr15 | 29779881 | 29808003 | 28123 | -5.17 | 0.000215 | 0.000284 |
| chrX | 84168613 | 84193494 | 24882 | 5.19 | 0.000215 | 0.000285 |
| chr20 | 7921157 | 7941126 | 19970 | -0.88 | 0.000216 | 0.000286 |
| chr4 | 123645131 | 123645437 | 307 | 4.74 | 0.000216 | 0.000286 |
| chr5 | 29957554 | 29973856 | 16303 | 2.09 | 0.000216 | 0.000286 |
| chrX | 107059200 | 107066118 | 6919 | -4.93 | 0.000217 | 0.000286 |
| chr6 | 39453675 | 39521868 | 68194 | 1.31 | 0.000217 | 0.000287 |
| chr2 | 32042676 | 32057960 | 15285 | -5.18 | 0.000217 | 0.000287 |
| chr10 | 98081620 | 98089435 | 7816 | 1.38 | 0.000217 | 0.000287 |
| chr7 | 101797996 | 101798365 | 370 | -4.4 | 0.000217 | 0.000287 |
| chr11 | 100709194 | 100710787 | 1594 | -3.71 | 0.000218 | 0.000288 |
| chr8 | 73922855 | 73923337 | 483 | -4.58 | 0.000218 | 0.000288 |
| chr6 | 46561445 | 46572005 | 10561 | -1.48 | 0.000218 | 0.000289 |
| chr1 | 2544662 | 2553301 | 8640 | -4.95 | 0.000218 | 0.000289 |
| chr5 | 38762377 | 38787250 | 24874 | -5.3 | 0.000219 | 0.000289 |
| chr9 | 73107960 | 73112404 | 4445 | 4.4 | 0.000219 | 0.000289 |
| chr10 | 80522094 | 80522664 | 571 | -4.42 | 0.000219 | 0.000289 |
| chr15 | 39369667 | 39370403 | 737 | -4.42 | 0.000219 | 0.000289 |
| chr7 | 15544163 | 15545167 | 1005 | -4.42 | 0.000219 | 0.000289 |
| chr11 | 122248351 | 122275792 | 27442 | -1.78 | 0.000219 | 0.000289 |
| chrX | 111079482 | 111110200 | 30719 | 2.56 | 0.000219 | 0.00029 |
| chr1 | 217310098 | 217323518 | 13421 | -1.4 | 0.000219 | 0.00029 |
| chr2 | 10156544 | 10166712 | 10169 | -6.31 | 0.00022 | 0.00029 |
| chrX | 82177533 | 82189706 | 12174 | -4.71 | 0.00022 | 0.000291 |
| chr18 | 70716465 | 70718295 | 1831 | -3.72 | 0.000221 | 0.000292 |

| | | | | | | |
|---|---|---|---|---|---|---|
| chr1 | 239474694 | 239541450 | 66757 | 0.72 | 0.000221 | 0.000292 |
| chr9 | 88982106 | 89004067 | 21962 | -5.41 | 0.000222 | 0.000293 |
| chr2 | 31402721 | 31409836 | 7116 | -1.96 | 0.000222 | 0.000293 |
| chr5 | 88144335 | 88147380 | 3046 | 4.34 | 0.000222 | 0.000294 |
| chr8 | 9023742 | 9024163 | 422 | -4.58 | 0.000223 | 0.000294 |
| chr7 | 109907542 | 109910430 | 2889 | -1.84 | 0.000223 | 0.000294 |
| chr6 | 113310036 | 113311010 | 975 | -4.91 | 0.000223 | 0.000294 |
| chr11 | 2358541 | 2372516 | 13976 | -4.74 | 0.000223 | 0.000295 |
| chr12 | 8311445 | 8312085 | 641 | -4.48 | 0.000223 | 0.000295 |
| chr13 | 100473090 | 100477041 | 3952 | -4.55 | 0.000224 | 0.000295 |
| chr7 | 47736307 | 47737692 | 1386 | -5.3 | 0.000224 | 0.000296 |
| chr6 | 75445321 | 75447522 | 2202 | -4.2 | 0.000224 | 0.000296 |
| chr2 | 162359560 | 162380586 | 21027 | -1.13 | 0.000225 | 0.000297 |
| chr1 | 80519535 | 80520093 | 559 | -4.47 | 0.000225 | 0.000298 |
| chr1 | 156475514 | 156476122 | 609 | -4.47 | 0.000225 | 0.000298 |
| chr1 | 160540062 | 160542905 | 2844 | 2.58 | 0.000226 | 0.000298 |
| chr11 | 85076482 | 85106678 | 30197 | -1.67 | 0.000226 | 0.000299 |
| chr21 | 15531277 | 15538846 | 7570 | -1.51 | 0.000226 | 0.000299 |
| chr9 | 119941694 | 119972455 | 30762 | -1.11 | 0.000226 | 0.000299 |
| chr20 | 9488232 | 9488648 | 417 | -4.4 | 0.000227 | 3.00E-04 |
| chr9 | 88456187 | 88456904 | 718 | -4.4 | 0.000227 | 3.00E-04 |
| chr14 | 73264475 | 73266622 | 2148 | 2.81 | 0.000227 | 3.00E-04 |
| chr4 | 15258600 | 15260767 | 2168 | 1.98 | 0.000228 | 0.000301 |
| chr10 | 79586695 | 79587240 | 546 | -3.37 | 0.000229 | 0.000302 |
| chr7 | 11334366 | 11372098 | 37733 | 0.66 | 0.000229 | 0.000302 |
| chr2 | 225132234 | 225132669 | 436 | -4.42 | 0.000229 | 0.000303 |
| chr8 | 110845214 | 110846018 | 805 | -4.42 | 0.000229 | 0.000303 |
| chr4 | 21965924 | 21979323 | 13400 | 3.23 | 0.000229 | 0.000303 |
| chr1 | 110681989 | 110688606 | 6618 | -1.92 | 0.00023 | 0.000303 |
| chrX | 132848873 | 132870810 | 21938 | -5.01 | 0.00023 | 0.000303 |
| chr7 | 103462049 | 103464721 | 2673 | -4.39 | 0.00023 | 0.000303 |
| chr12 | 26014621 | 26017212 | 2592 | -3.71 | 0.00023 | 0.000304 |
| chr2 | 102258818 | 102259482 | 665 | -4.41 | 0.00023 | 0.000304 |
| chr2 | 205453879 | 205465547 | 11669 | -6.58 | 0.00023 | 0.000304 |
| chr10 | 71731692 | 71793840 | 62149 | -3.31 | 0.00023 | 0.000304 |
| chr11 | 129317139 | 129373187 | 56049 | -1.8 | 0.000231 | 0.000305 |
| chr3 | 20280422 | 20301693 | 21272 | 2.71 | 0.000231 | 0.000305 |
| chrX | 109769511 | 109771984 | 2474 | -1.62 | 0.000231 | 0.000305 |
| chr9 | 28776729 | 28777547 | 819 | -4.85 | 0.000232 | 0.000306 |
| chrX | 14101014 | 14129109 | 28096 | -1.19 | 0.000232 | 0.000306 |
| chr20 | 34559724 | 34560229 | 506 | -4.38 | 0.000233 | 0.000307 |
| chr21 | 40471696 | 40472254 | 559 | -4.38 | 0.000233 | 0.000307 |
| chr22 | 28735740 | 28736332 | 593 | -4.38 | 0.000233 | 0.000307 |
| chr2 | 134790798 | 134816777 | 25980 | -1.84 | 0.000233 | 0.000307 |
| chr15 | 40041887 | 40047198 | 5312 | -4.41 | 0.000233 | 0.000307 |
| chr8 | 123424022 | 123453429 | 29408 | -0.95 | 0.000234 | 0.000308 |
| chr21 | 35366899 | 35414720 | 47822 | -5.26 | 0.000234 | 0.000309 |
| chr6 | 33896240 | 33896752 | 513 | -4.38 | 0.000235 | 0.00031 |
| chr9 | 117280457 | 117280834 | 378 | -4.38 | 0.000235 | 0.00031 |
| chrX | 101744540 | 101770929 | 26390 | -1.26 | 0.000236 | 0.000311 |
| chr12 | 103216149 | 103340416 | 124268 | 2.95 | 0.000236 | 0.000311 |
| chr2 | 154323219 | 154326673 | 3455 | 2.19 | 0.000237 | 0.000312 |
| chr6 | 1369581 | 1375222 | 5642 | -4.53 | 0.000238 | 0.000313 |
| chr4 | 73040553 | 73041771 | 1219 | -4.51 | 0.000238 | 0.000314 |
| chr9 | 29691956 | 29693227 | 1272 | -4.51 | 0.000238 | 0.000314 |
| chr10 | 106117168 | 106146912 | 29745 | -2.12 | 0.000238 | 0.000314 |
| chr21 | 37284599 | 37293720 | 9122 | -1.17 | 0.000238 | 0.000314 |
| chr10 | 80397078 | 80427165 | 30088 | -5.19 | 0.000238 | 0.000314 |
| chr3 | 28107559 | 28111063 | 3505 | -4.04 | 0.000238 | 0.000314 |
| chrX | 81137694 | 81140030 | 2337 | 4.24 | 0.000239 | 0.000315 |
| chr7 | 27480430 | 27480936 | 507 | -4.69 | 0.00024 | 0.000316 |

| | | | | | | |
|---|---|---|---|---|---|---|
| chr21 | 15817653 | 15844538 | 26886 | 0.8 | 0.00024 | 0.000317 |
| chr19 | 49385317 | 49386508 | 1192 | -5.01 | 0.00024 | 0.000317 |
| chr6 | 64466365 | 64478806 | 12442 | -5.79 | 0.000241 | 0.000317 |
| chr14 | 21102243 | 21105042 | 2800 | -4.22 | 0.000241 | 0.000317 |
| chr3 | 21453278 | 21458994 | 5717 | 1.5 | 0.000241 | 0.000318 |
| chr1 | 182424363 | 182452079 | 27717 | -0.86 | 0.000241 | 0.000318 |
| chr3 | 61770248 | 61774428 | 4181 | -3.91 | 0.000241 | 0.000318 |
| chr1 | 230158816 | 230181093 | 22278 | -6.34 | 0.000241 | 0.000318 |
| chr18 | 47124993 | 47133677 | 8685 | -3.31 | 0.000242 | 0.000318 |
| chr10 | 124884306 | 124889484 | 5179 | -4.5 | 0.000242 | 0.000319 |
| chr11 | 85196578 | 85300720 | 104143 | -1.09 | 0.000242 | 0.000319 |
| chrX | 148545319 | 148546190 | 872 | -3.58 | 0.000243 | 0.00032 |
| chr9 | 117834212 | 117834750 | 539 | -3.38 | 0.000243 | 0.00032 |
| chr6 | 43373750 | 43390861 | 17112 | -4.39 | 0.000243 | 0.00032 |
| chr2 | 97211961 | 97212467 | 507 | -4.37 | 0.000243 | 0.000321 |
| chr1 | 81327224 | 81327629 | 406 | -4.43 | 0.000243 | 0.000321 |
| chr2 | 232388494 | 232389892 | 1399 | -4.99 | 0.000244 | 0.000321 |
| chr9 | 134615845 | 134617325 | 1481 | -4.99 | 0.000244 | 0.000321 |
| chr12 | 6246114 | 6246641 | 528 | -4.4 | 0.000244 | 0.000322 |
| chr10 | 82021570 | 82023196 | 1627 | -4.02 | 0.000244 | 0.000322 |
| chr1 | 66926834 | 66942447 | 15614 | -1.64 | 0.000245 | 0.000322 |
| chr3 | 175418902 | 175419565 | 664 | -4.43 | 0.000245 | 0.000323 |
| chr3 | 139263421 | 139264759 | 1339 | -4.77 | 0.000245 | 0.000323 |
| chr15 | 38319756 | 38320588 | 833 | -2.89 | 0.000245 | 0.000323 |
| chr3 | 40091715 | 40092476 | 762 | -1.83 | 0.000245 | 0.000323 |
| chr4 | 87381343 | 87381850 | 508 | -3.99 | 0.000246 | 0.000324 |
| chr20 | 48661906 | 48662744 | 839 | -4.49 | 0.000246 | 0.000324 |
| chr4 | 92932646 | 92933138 | 493 | -4.49 | 0.000246 | 0.000324 |
| chr8 | 20194136 | 20203283 | 9148 | -1.26 | 0.000246 | 0.000324 |
| chr5 | 173293999 | 173295376 | 1378 | -5.35 | 0.000246 | 0.000324 |
| chr16 | 75818322 | 75819287 | 966 | -2.5 | 0.000247 | 0.000325 |
| chr3 | 133101299 | 133113784 | 12486 | -4.9 | 0.000247 | 0.000325 |
| chr18 | 71230504 | 71231038 | 535 | -4.59 | 0.000247 | 0.000325 |
| chr14 | 60543785 | 60544278 | 494 | -3.65 | 0.000247 | 0.000326 |
| chrX | 96897532 | 96905744 | 8213 | -4.46 | 0.000247 | 0.000326 |
| chr10 | 95366529 | 95367586 | 1058 | -3.48 | 0.000248 | 0.000326 |
| chr3 | 96727766 | 96746928 | 19163 | -4.32 | 0.000249 | 0.000328 |
| chr18 | 44867918 | 44872696 | 4779 | 2.09 | 0.00025 | 0.000329 |
| chr9 | 100464338 | 100464744 | 407 | -4.63 | 0.00025 | 0.00033 |
| chr5 | 94699269 | 94710643 | 11375 | 1.52 | 0.000251 | 0.00033 |
| chr4 | 10417133 | 10418248 | 1116 | 3.21 | 0.000251 | 0.00033 |
| chr8 | 140794475 | 140821563 | 27089 | -5.75 | 0.000251 | 0.00033 |
| chr13 | 106676279 | 106680596 | 4318 | -4.35 | 0.000251 | 0.000331 |
| chr16 | 49939570 | 49939940 | 371 | 3.57 | 0.000252 | 0.000332 |
| chr8 | 103464945 | 103465620 | 676 | -3.45 | 0.000252 | 0.000332 |
| chr18 | 10104581 | 10120981 | 16401 | -1.33 | 0.000253 | 0.000333 |
| chr1 | 90552054 | 90562359 | 10306 | -1 | 0.000253 | 0.000333 |
| chr4 | 24310811 | 24322328 | 11518 | 4.71 | 0.000254 | 0.000334 |
| chr9 | 27574535 | 27594520 | 19986 | -4.82 | 0.000254 | 0.000335 |
| chr3 | 116662378 | 116664337 | 1960 | -2.05 | 0.000255 | 0.000335 |
| chr19 | 41952716 | 42011774 | 59059 | -3.36 | 0.000255 | 0.000335 |
| chr15 | 62860607 | 62861440 | 834 | -4.05 | 0.000255 | 0.000335 |
| chr10 | 9055242 | 9061068 | 5827 | -4.4 | 0.000255 | 0.000336 |
| chr16 | 23363420 | 23364205 | 786 | -4.46 | 0.000255 | 0.000336 |
| chr19 | 5324963 | 5325361 | 399 | -4.46 | 0.000255 | 0.000336 |
| chr4 | 63894486 | 63894962 | 477 | -4.46 | 0.000255 | 0.000336 |
| chr5 | 142184612 | 142185298 | 687 | -4.46 | 0.000255 | 0.000336 |
| chr3 | 167874511 | 167876584 | 2074 | 3.93 | 0.000256 | 0.000336 |
| chr3 | 72519101 | 72519574 | 474 | -4.36 | 0.000256 | 0.000337 |
| chr1 | 65182479 | 65190143 | 7665 | -1.77 | 0.000257 | 0.000337 |
| chr6 | 75001696 | 75002942 | 1247 | -3.6 | 0.000257 | 0.000338 |

| | | | | | | |
|---|---|---|---|---|---|---|
| chr14 | 60665311 | 60677174 | 11864 | -1.49 | 0.000257 | 0.000338 |
| chr6 | 85810274 | 85812973 | 2700 | -2.05 | 0.000257 | 0.000338 |
| chr1 | 224259266 | 224262132 | 2867 | -4.45 | 0.000257 | 0.000338 |
| chrX | 110048366 | 110067381 | 19016 | 3.6 | 0.000257 | 0.000338 |
| chr8 | 118469274 | 118548265 | 78992 | -5.37 | 0.000257 | 0.000338 |
| chr3 | 191188666 | 191190052 | 1387 | 3.67 | 0.000257 | 0.000339 |
| chr3 | 104328003 | 104329204 | 1202 | 2.14 | 0.000257 | 0.000339 |
| chr6 | 15851264 | 15855169 | 3906 | -4.43 | 0.000258 | 0.000339 |
| chr9 | 5311126 | 5317635 | 6510 | -4.99 | 0.000258 | 0.000339 |
| chr19 | 10643157 | 10643585 | 429 | -4.37 | 0.000258 | 0.000339 |
| chr19 | 48661343 | 48661723 | 381 | -4.37 | 0.000258 | 0.000339 |
| chrX | 115641955 | 115643051 | 1097 | -4.37 | 0.000258 | 0.000339 |
| chr12 | 120842135 | 120842597 | 463 | -4.4 | 0.000258 | 0.00034 |
| chr4 | 8407924 | 8408344 | 421 | -4.4 | 0.000258 | 0.00034 |
| chr1 | 240121951 | 240158315 | 36365 | -2.96 | 0.000259 | 0.00034 |
| chrX | 130385789 | 130388912 | 3124 | 1.42 | 0.000261 | 0.000343 |
| chr8 | 39459493 | 39485132 | 25640 | -1.43 | 0.000261 | 0.000343 |
| chr2 | 16332340 | 17486923 | 1154584 | 6.35 | 0.000261 | 0.000343 |
| chr15 | 60921325 | 60977077 | 55753 | 1.87 | 0.000261 | 0.000344 |
| chr13 | 98344251 | 98344678 | 428 | -4.37 | 0.000262 | 0.000344 |
| chr18 | 2319623 | 2320055 | 433 | -4.37 | 0.000262 | 0.000344 |
| chr2 | 148945977 | 148946458 | 482 | -4.37 | 0.000262 | 0.000344 |
| chr7 | 102933744 | 102934634 | 891 | -4.37 | 0.000262 | 0.000344 |
| chr3 | 192292303 | 192292821 | 519 | -4.35 | 0.000262 | 0.000344 |
| chr21 | 15887707 | 15889374 | 1668 | -4.18 | 0.000262 | 0.000345 |
| chr8 | 129919864 | 129938320 | 18457 | -3.27 | 0.000263 | 0.000345 |
| chr12 | 9165066 | 9173089 | 8024 | -4.94 | 0.000263 | 0.000346 |
| chr1 | 99496548 | 99497056 | 509 | 4.52 | 0.000263 | 0.000346 |
| chr2 | 237314750 | 237392992 | 78243 | -3.11 | 0.000264 | 0.000347 |
| chr9 | 98281192 | 98295638 | 14447 | -2.89 | 0.000265 | 0.000348 |
| chr3 | 187265171 | 187269855 | 4685 | 4.54 | 0.000266 | 0.00035 |
| chrX | 114122636 | 114126838 | 4203 | 2.15 | 0.000266 | 0.00035 |
| chr15 | 29323450 | 29323944 | 495 | -4.52 | 0.000266 | 0.00035 |
| chr13 | 37388631 | 37389237 | 607 | -2.72 | 0.000266 | 0.00035 |
| chr2 | 188526184 | 188526848 | 665 | -3.19 | 0.000268 | 0.000352 |
| chr6 | 48327102 | 48337348 | 10247 | -4.03 | 0.000269 | 0.000353 |
| chr14 | 38614195 | 38615848 | 1654 | 4.11 | 0.000269 | 0.000353 |
| chr6 | 63911945 | 63925748 | 13804 | -3.71 | 0.000269 | 0.000354 |
| chr3 | 105867073 | 105867559 | 487 | -2.57 | 0.00027 | 0.000355 |
| chr1 | 217276159 | 217296395 | 20237 | 1.12 | 0.000272 | 0.000357 |
| chr4 | 121856987 | 121877445 | 20459 | -2.14 | 0.000272 | 0.000357 |
| chr2 | 151873629 | 151888727 | 15099 | -2.23 | 0.000272 | 0.000358 |
| chr8 | 19113226 | 19114194 | 969 | -4.82 | 0.000273 | 0.000359 |
| chr2 | 73535892 | 73574570 | 38679 | -1.68 | 0.000274 | 0.000359 |
| chr3 | 115801157 | 115801589 | 433 | -4.54 | 0.000274 | 0.000359 |
| chrX | 21007116 | 21009022 | 1907 | -2.01 | 0.000274 | 0.00036 |
| chr1 | 162464892 | 162467333 | 2442 | -2.45 | 0.000274 | 0.00036 |
| chr17 | 15939059 | 15939926 | 868 | -3.79 | 0.000274 | 0.00036 |
| chrX | 38594747 | 38619107 | 24361 | -1.1 | 0.000274 | 0.00036 |
| chr11 | 69941634 | 69942100 | 467 | -4.49 | 0.000274 | 0.00036 |
| chr14 | 73674497 | 73675054 | 558 | -4.49 | 0.000274 | 0.00036 |
| chr22 | 31577376 | 31577642 | 267 | -4.35 | 0.000275 | 0.000361 |
| chr3 | 178033595 | 178034069 | 475 | -4.35 | 0.000275 | 0.000361 |
| chr4 | 30494076 | 30494443 | 368 | -4.35 | 0.000275 | 0.000361 |
| chr11 | 112256743 | 112260444 | 3702 | -3.34 | 0.000276 | 0.000362 |
| chr15 | 85976589 | 85977164 | 576 | -5.01 | 0.000276 | 0.000362 |
| chr2 | 109940549 | 109945878 | 5330 | -4.74 | 0.000276 | 0.000363 |
| chr8 | 54010223 | 54010899 | 677 | -1.81 | 0.000277 | 0.000363 |
| chr5 | 66525869 | 66526829 | 961 | -2.38 | 0.000277 | 0.000363 |
| chrX | 51962817 | 51964829 | 2013 | -1.84 | 0.000277 | 0.000364 |
| chr6 | 58606837 | 58617116 | 10280 | 1.99 | 0.000277 | 0.000364 |

| | | | | | | |
|---|---|---|---|---|---|---|
| chr12 | 11670316 | 11731032 | 60717 | -5.1 | 0.000278 | 0.000364 |
| chr2 | 40737108 | 40737433 | 326 | -3.6 | 0.000278 | 0.000365 |
| chr2 | 217895387 | 217899762 | 4376 | 2.49 | 0.000279 | 0.000366 |
| chr13 | 83257562 | 83258210 | 649 | -4.41 | 0.000279 | 0.000366 |
| chr16 | 85447109 | 85447811 | 703 | -4.41 | 0.000279 | 0.000366 |
| chr20 | 30592574 | 30593081 | 508 | -4.41 | 0.000279 | 0.000366 |
| chr10 | 118901747 | 118920701 | 18955 | -4.11 | 0.00028 | 0.000367 |
| chr18 | 30011389 | 30012008 | 620 | -4.33 | 0.00028 | 0.000367 |
| chr2 | 3382614 | 3383193 | 580 | -4.36 | 0.00028 | 0.000368 |
| chr14 | 66862957 | 66863452 | 496 | -3.47 | 0.000281 | 0.000369 |
| chr4 | 185100230 | 185101375 | 1146 | -1.69 | 0.000281 | 0.000369 |
| chr12 | 99233612 | 99236421 | 2810 | -1.75 | 0.000281 | 0.000369 |
| chr12 | 66947248 | 66962880 | 15633 | -1.46 | 0.000282 | 0.00037 |
| chr18 | 72855823 | 72866912 | 11090 | -3.02 | 0.000282 | 0.00037 |
| chr6 | 159935554 | 159937853 | 2300 | -4.03 | 0.000282 | 0.00037 |
| chr6 | 78849425 | 78853561 | 4137 | 4.25 | 0.000282 | 0.00037 |
| chr6 | 116812815 | 116813628 | 814 | -2.03 | 0.000283 | 0.000371 |
| chr4 | 178541961 | 178542547 | 587 | -4.33 | 0.000284 | 0.000372 |
| chr10 | 78632281 | 78634804 | 2524 | 4.05 | 0.000284 | 0.000373 |
| chr13 | 78364172 | 78367741 | 3570 | -1.44 | 0.000284 | 0.000373 |
| chr2 | 173281035 | 173283457 | 2423 | -2.16 | 0.000284 | 0.000373 |
| chr1 | 210093006 | 210095959 | 2954 | -2.31 | 0.000285 | 0.000374 |
| chr7 | 22178207 | 22185186 | 6980 | 4.4 | 0.000285 | 0.000374 |
| chr3 | 66561875 | 66562742 | 868 | 1.98 | 0.000285 | 0.000374 |
| chr3 | 98193673 | 98195359 | 1687 | -4.11 | 0.000286 | 0.000375 |
| chr11 | 27511001 | 27511587 | 587 | -4.4 | 0.000286 | 0.000375 |
| chr11 | 11278116 | 11279265 | 1150 | 4.01 | 0.000286 | 0.000375 |
| chr4 | 106038550 | 106038909 | 360 | -4.1 | 0.000286 | 0.000375 |
| chr3 | 192694294 | 192697029 | 2736 | -4.42 | 0.000287 | 0.000376 |
| chr2 | 38069543 | 38072085 | 2543 | -2.85 | 0.000287 | 0.000376 |
| chr5 | 87697 | 88627 | 931 | -2.08 | 0.000288 | 0.000378 |
| chr1 | 227197511 | 227209207 | 11697 | -5.72 | 0.000289 | 0.000379 |
| chr8 | 74285246 | 74307056 | 21811 | -6.29 | 0.000289 | 0.000379 |
| chr8 | 77813956 | 77816943 | 2988 | 1.75 | 0.00029 | 0.00038 |
| chr20 | 57774853 | 57789708 | 14856 | 0.93 | 0.00029 | 0.00038 |
| chr1 | 172214102 | 172214857 | 756 | -4.73 | 0.00029 | 0.00038 |
| chr1 | 192637818 | 192638497 | 680 | -4.73 | 0.00029 | 0.00038 |
| chr12 | 20413944 | 20414709 | 766 | -4.73 | 0.00029 | 0.00038 |
| chr12 | 91066332 | 91068182 | 1851 | 3.8 | 0.00029 | 0.00038 |
| chr18 | 49915889 | 49917405 | 1517 | 1.84 | 0.00029 | 0.000381 |
| chr22 | 27948697 | 27949161 | 465 | -4.67 | 0.00029 | 0.000381 |
| chr20 | 10164609 | 10180995 | 16387 | -5 | 0.00029 | 0.000381 |
| chr4 | 134260802 | 134262663 | 1862 | -2.79 | 0.000291 | 0.000381 |
| chr6 | 114504375 | 114508480 | 4106 | 2.66 | 0.000293 | 0.000384 |
| chr4 | 145764393 | 145765462 | 1070 | -2.46 | 0.000293 | 0.000384 |
| chr2 | 236710645 | 236711080 | 436 | -4.35 | 0.000293 | 0.000384 |
| chr12 | 1443620 | 1444004 | 385 | -4.38 | 0.000293 | 0.000384 |
| chr17 | 15422760 | 15423259 | 500 | -4.38 | 0.000293 | 0.000384 |
| chr12 | 15846498 | 15846876 | 379 | -4.44 | 0.000294 | 0.000385 |
| chr7 | 80974204 | 80975012 | 809 | -4.44 | 0.000294 | 0.000385 |
| chr18 | 55926070 | 55926537 | 468 | -4.55 | 0.000294 | 0.000385 |
| chr19 | 20943620 | 20943937 | 318 | -4.32 | 0.000295 | 0.000386 |
| chr2 | 30406948 | 30413287 | 6340 | -1.89 | 0.000295 | 0.000387 |
| chr8 | 87089318 | 87089837 | 520 | -3.13 | 0.000296 | 0.000388 |
| chr2 | 222624977 | 222626005 | 1029 | -3.39 | 0.000296 | 0.000388 |
| chrX | 83625542 | 83627472 | 1931 | 2.11 | 0.000297 | 0.000389 |
| chr7 | 87112907 | 87114329 | 1423 | 2.79 | 0.000297 | 0.000389 |
| chr2 | 16070737 | 16071492 | 756 | -3.58 | 0.000297 | 0.000389 |
| chr8 | 25948020 | 25961586 | 13567 | -2.9 | 0.000298 | 0.00039 |
| chr12 | 60476970 | 60479324 | 2355 | -1.72 | 0.000298 | 0.00039 |
| chr16 | 5265268 | 5269906 | 4639 | 1.59 | 0.000298 | 0.00039 |

| | | | | | | |
|---|---|---|---|---|---|---|
| chrX | 153889002 | 153921486 | 32485 | -1.13 | 0.000298 | 0.00039 |
| chr3 | 133984744 | 133986699 | 1956 | -2.8 | 0.000298 | 0.00039 |
| chr10 | 24699671 | 24704763 | 5093 | -4.7 | 0.000298 | 0.00039 |
| chr12 | 41541680 | 41542447 | 768 | -4.88 | 0.000298 | 0.000391 |
| chrX | 66989853 | 66994583 | 4731 | -2.14 | 0.000298 | 0.000391 |
| chr4 | 123439697 | 123445442 | 5746 | -1.54 | 0.000299 | 0.000391 |
| chr6 | 162135197 | 162136113 | 917 | -2.42 | 0.000299 | 0.000391 |
| chr12 | 101368846 | 101373504 | 4659 | 1.26 | 0.000299 | 0.000392 |
| chr18 | 44308274 | 44322772 | 14499 | -4.96 | 0.000299 | 0.000392 |
| chr1 | 191592906 | 191593414 | 509 | -4.46 | 3.00E-04 | 0.000393 |
| chr10 | 11803399 | 11803776 | 378 | -4.46 | 3.00E-04 | 0.000393 |
| chr10 | 120688180 | 120707821 | 19642 | -4.93 | 3.00E-04 | 0.000393 |
| chr17 | 25755475 | 25755863 | 389 | -4.52 | 0.000301 | 0.000394 |
| chr11 | 10491076 | 10491625 | 550 | -3.72 | 0.000302 | 0.000396 |
| chr13 | 77077666 | 77081442 | 3777 | 3.51 | 0.000302 | 0.000396 |
| chr4 | 77359186 | 77360751 | 1566 | -2.17 | 0.000304 | 0.000398 |
| chr21 | 46174452 | 46175258 | 807 | -4.67 | 0.000304 | 0.000398 |
| chr8 | 80643307 | 80644838 | 1532 | -3.86 | 0.000305 | 0.000399 |
| chr2 | 169502241 | 169502556 | 316 | -4.41 | 0.000305 | 4.00E-04 |
| chr6 | 44626235 | 44626626 | 392 | -4.41 | 0.000305 | 4.00E-04 |
| chr4 | 60369250 | 60369773 | 524 | -4.16 | 0.000306 | 4.00E-04 |
| chr12 | 102694278 | 102696215 | 1938 | -2.84 | 0.000306 | 0.000401 |
| chr2 | 74925731 | 74929120 | 3390 | 2.49 | 0.000306 | 0.000401 |
| chr4 | 118968438 | 118969917 | 1480 | 2.34 | 0.000307 | 0.000402 |
| chr6 | 146329484 | 146331577 | 2094 | -4.13 | 0.000307 | 0.000402 |
| chr10 | 121222223 | 121234028 | 11806 | -5.9 | 0.000307 | 0.000402 |
| chr5 | 78397647 | 78399085 | 1439 | -1.69 | 0.000307 | 0.000402 |
| chrX | 113728099 | 113728835 | 737 | -2.3 | 0.000308 | 0.000402 |
| chr19 | 49330089 | 49330764 | 676 | -4.31 | 0.000308 | 0.000403 |
| chr8 | 25909153 | 25933735 | 24583 | -4.94 | 0.000309 | 0.000404 |
| chr22 | 25357171 | 25357644 | 474 | -4.5 | 0.000309 | 0.000404 |
| chr12 | 10899760 | 10921953 | 22194 | -1.73 | 0.000309 | 0.000405 |
| chr3 | 24858903 | 24870411 | 11509 | -1.96 | 0.00031 | 0.000405 |
| chr11 | 82439994 | 82467655 | 27662 | -2.21 | 0.00031 | 0.000405 |
| chr4 | 11579015 | 11580136 | 1122 | -4.65 | 0.00031 | 0.000405 |
| chr1 | 79451380 | 79451853 | 474 | -4.35 | 0.00031 | 0.000405 |
| chr7 | 95688293 | 95692112 | 3820 | -2.65 | 0.00031 | 0.000406 |
| chr2 | 139486380 | 139496587 | 10208 | -2.4 | 0.00031 | 0.000406 |
| chr3 | 102284560 | 102285590 | 1031 | 2.19 | 0.00031 | 0.000406 |
| chr8 | 107826460 | 107835951 | 9492 | -4.45 | 0.000311 | 0.000407 |
| chr3 | 184816504 | 184840867 | 24364 | -1.23 | 0.000311 | 0.000407 |
| chr7 | 83178582 | 83189141 | 10560 | -5.77 | 0.000312 | 0.000408 |
| chr11 | 94117374 | 94117831 | 458 | -2.2 | 0.000313 | 0.000409 |
| chr1 | 9694669 | 9695047 | 379 | -4.57 | 0.000313 | 0.00041 |
| chr3 | 135917138 | 135918272 | 1135 | -5.37 | 0.000313 | 0.00041 |
| chr6 | 70562289 | 70563659 | 1371 | -3.93 | 0.000314 | 0.00041 |
| chr18 | 51914146 | 51914918 | 773 | -4.32 | 0.000314 | 0.000411 |
| chr7 | 155111929 | 155115434 | 3506 | -4.15 | 0.000314 | 0.000411 |
| chr16 | 86340087 | 86341521 | 1435 | -3.44 | 0.000315 | 0.000412 |
| chr4 | 3327119 | 3327562 | 444 | -4.32 | 0.000315 | 0.000412 |
| chr5 | 156064304 | 156064747 | 444 | -4.32 | 0.000315 | 0.000412 |
| chr1 | 206974235 | 206987945 | 13711 | -1.41 | 0.000315 | 0.000412 |
| chr12 | 120825234 | 120825688 | 455 | -4.62 | 0.000315 | 0.000412 |
| chr2 | 176159613 | 176162465 | 2853 | -1.87 | 0.000315 | 0.000412 |
| chr1 | 242184693 | 242185571 | 879 | -5.02 | 0.000316 | 0.000413 |
| chr6 | 116829257 | 116831343 | 2087 | -3.6 | 0.000316 | 0.000413 |
| chr2 | 88653501 | 88699772 | 46272 | -2.63 | 0.000317 | 0.000414 |
| chr6 | 89468983 | 89469481 | 499 | -4.42 | 0.000317 | 0.000415 |
| chr18 | 11346131 | 11352900 | 6770 | -2.97 | 0.000317 | 0.000415 |
| chr1 | 151749132 | 151749617 | 486 | -3.3 | 0.000318 | 0.000416 |
| chr3 | 114706480 | 114706931 | 452 | -4.8 | 0.000319 | 0.000417 |

| | | | | | | |
|---|---|---|---|---|---|---|
| chr3 | 188289371 | 188289856 | 486 | -4.8 | 0.000319 | 0.000417 |
| chr1 | 118589495 | 118605896 | 16402 | 1.35 | 0.000319 | 0.000417 |
| chr7 | 141782665 | 141785371 | 2707 | 1.91 | 0.000321 | 0.00042 |
| chr1 | 235124210 | 235124750 | 541 | -4.6 | 0.000321 | 0.00042 |
| chr14 | 23729453 | 23731461 | 2009 | -4.21 | 0.000322 | 0.00042 |
| chr1 | 33991876 | 33992574 | 699 | 2.17 | 0.000322 | 0.00042 |
| chr8 | 112203330 | 112215332 | 12003 | -1.72 | 0.000322 | 0.000421 |
| chr3 | 93917656 | 93931183 | 13528 | 3.02 | 0.000323 | 0.000423 |
| chr9 | 28576288 | 28587057 | 10770 | -2.34 | 0.000324 | 0.000423 |
| chr11 | 93079082 | 93079768 | 687 | -3.46 | 0.000325 | 0.000424 |
| chr16 | 2852986 | 2857626 | 4641 | -4.81 | 0.000325 | 0.000424 |
| chr3 | 99835265 | 99835635 | 371 | -4.53 | 0.000325 | 0.000424 |
| chr3 | 109063283 | 109064014 | 732 | -4.53 | 0.000325 | 0.000424 |
| chr6 | 36542327 | 36542765 | 439 | -4.33 | 0.000326 | 0.000425 |
| chr2 | 8319973 | 8320654 | 682 | -3.35 | 0.000326 | 0.000426 |
| chr8 | 43126708 | 43127591 | 884 | -2.16 | 0.000327 | 0.000427 |
| chr1 | 56672434 | 56674072 | 1639 | -1.78 | 0.000327 | 0.000428 |
| chr4 | 90429360 | 90430819 | 1460 | -1.95 | 0.000328 | 0.000428 |
| chr12 | 27244098 | 27244897 | 800 | -3.48 | 0.000328 | 0.000428 |
| chr12 | 70959504 | 70959940 | 437 | -3.86 | 0.000329 | 0.00043 |
| chr1 | 199926091 | 199926683 | 593 | -4.32 | 0.00033 | 0.000431 |
| chrX | 106574491 | 106601836 | 27346 | -1.02 | 0.000332 | 0.000433 |
| chr13 | 29377919 | 29406718 | 28800 | -5.02 | 0.000332 | 0.000434 |
| chr1 | 19306073 | 19309554 | 3482 | -4.47 | 0.000332 | 0.000434 |
| chr6 | 35284021 | 35296818 | 12798 | -5.3 | 0.000333 | 0.000435 |
| chr10 | 14093738 | 14120823 | 27086 | -0.73 | 0.000333 | 0.000435 |
| chr1 | 241522843 | 241548041 | 25199 | -0.8 | 0.000333 | 0.000435 |
| chrX | 55983121 | 55992327 | 9207 | 1.67 | 0.000334 | 0.000436 |
| chr8 | 119767212 | 119769551 | 2340 | 1.75 | 0.000334 | 0.000436 |
| chr11 | 89889261 | 89898630 | 9370 | -1.3 | 0.000335 | 0.000437 |
| chr2 | 158465870 | 158492560 | 26691 | -2.9 | 0.000335 | 0.000437 |
| chrX | 34805722 | 34806279 | 558 | -4.29 | 0.000335 | 0.000437 |
| chr11 | 112407238 | 112419209 | 11972 | 4.35 | 0.000335 | 0.000438 |
| chr16 | 59463907 | 59495996 | 32090 | -4.78 | 0.000336 | 0.000439 |
| chr15 | 37607603 | 37764476 | 156874 | 0.71 | 0.000336 | 0.000439 |
| chr1 | 209689311 | 209694833 | 5523 | -1.3 | 0.000337 | 0.00044 |
| chr11 | 63059118 | 63063009 | 3892 | -1.37 | 0.000337 | 0.00044 |
| chr4 | 56179335 | 56180086 | 752 | -3.37 | 0.000338 | 0.000441 |
| chr18 | 71653863 | 71665582 | 11720 | -1.26 | 0.000338 | 0.000442 |
| chr15 | 86339491 | 86340146 | 656 | -4.28 | 0.000338 | 0.000442 |
| chr16 | 78572969 | 78573573 | 605 | -4.28 | 0.000338 | 0.000442 |
| chr2 | 24703173 | 24703595 | 423 | -4.28 | 0.000338 | 0.000442 |
| chr9 | 86650509 | 86652898 | 2390 | -3.93 | 0.000339 | 0.000442 |
| chr2 | 23504661 | 23546716 | 42056 | 1.07 | 0.000339 | 0.000442 |
| chr2 | 1230187 | 1517382 | 287196 | 0.49 | 0.000339 | 0.000443 |
| chr2 | 194146524 | 194147038 | 515 | -4.42 | 0.000339 | 0.000443 |
| chr19 | 40054737 | 40158218 | 103482 | -0.81 | 0.00034 | 0.000444 |
| chr9 | 79182391 | 79239307 | 56917 | 1.28 | 0.00034 | 0.000444 |
| chr6 | 129954590 | 129963983 | 9394 | -4.61 | 0.000341 | 0.000445 |
| chr3 | 186151837 | 186152524 | 688 | -3.46 | 0.000341 | 0.000445 |
| chr12 | 68448704 | 68584952 | 136249 | 1.92 | 0.000342 | 0.000446 |
| chr12 | 20261642 | 20262033 | 392 | -4.53 | 0.000343 | 0.000447 |
| chr8 | 57952109 | 57966650 | 14542 | 1.09 | 0.000343 | 0.000448 |
| chr12 | 38564626 | 38566969 | 2344 | -3.96 | 0.000344 | 0.000449 |
| chr7 | 27010632 | 27013638 | 3007 | -4.16 | 0.000344 | 0.000449 |
| chr16 | 78298208 | 78298649 | 442 | -4.28 | 0.000344 | 0.000449 |
| chr6 | 3848542 | 3849020 | 479 | -4.28 | 0.000344 | 0.000449 |
| chr2 | 57123361 | 57340374 | 217014 | 0.84 | 0.000344 | 0.000449 |
| chr8 | 57030022 | 57069346 | 39325 | -2.34 | 0.000345 | 0.00045 |
| chr3 | 164548489 | 164666868 | 118380 | 1.06 | 0.000345 | 0.00045 |
| chr1 | 56429351 | 56429763 | 413 | 4.48 | 0.000346 | 0.000452 |

| | | | | | | |
|---|---|---|---|---|---|---|
| chr2 | 211348035 | 211379245 | 31211 | -2.48 | 0.000346 | 0.000452 |
| chr6 | 140270546 | 140292947 | 22402 | -6.13 | 0.000347 | 0.000452 |
| chr2 | 230186772 | 230191355 | 4584 | -4.02 | 0.000347 | 0.000452 |
| chr8 | 62254876 | 62272648 | 17773 | 1.95 | 0.000347 | 0.000453 |
| chr6 | 71622908 | 71635295 | 12388 | -1.39 | 0.000348 | 0.000453 |
| chr1 | 95850135 | 95859895 | 9761 | 4.52 | 0.000348 | 0.000453 |
| chr5 | 148133380 | 148136032 | 2653 | -4.38 | 0.00035 | 0.000456 |
| chr20 | 24867982 | 24876017 | 8036 | 1.16 | 0.00035 | 0.000457 |
| chrX | 96839084 | 96840031 | 948 | -3.42 | 0.00035 | 0.000457 |
| chr3 | 110754869 | 110761556 | 6688 | -3.77 | 0.00035 | 0.000457 |
| chr5 | 5500581 | 6148091 | 647511 | 0.31 | 0.000351 | 0.000457 |
| chr2 | 84867017 | 84889525 | 22509 | -3.09 | 0.000351 | 0.000457 |
| chr4 | 38373733 | 38374636 | 904 | -3.75 | 0.000351 | 0.000457 |
| chr7 | 79655420 | 79655815 | 396 | -4.5 | 0.000351 | 0.000458 |
| chr11 | 16028767 | 16029434 | 668 | 2.08 | 0.000352 | 0.000459 |
| chr1 | 112066866 | 112073414 | 6549 | -4.45 | 0.000352 | 0.000459 |
| chr8 | 49375935 | 49376833 | 899 | -4.39 | 0.000352 | 0.000459 |
| chr9 | 100690268 | 100690616 | 349 | -4.39 | 0.000352 | 0.000459 |
| chr14 | 50433824 | 50434335 | 512 | -4.33 | 0.000352 | 0.000459 |
| chr2 | 197257403 | 197257816 | 414 | -4.33 | 0.000352 | 0.000459 |
| chr6 | 87434946 | 87441250 | 6305 | -1.51 | 0.000352 | 0.000459 |
| chr3 | 186997306 | 186998465 | 1160 | 2.97 | 0.000352 | 0.000459 |
| chr4 | 63648990 | 63649599 | 610 | -4.29 | 0.000353 | 0.00046 |
| chr6 | 27013777 | 27014164 | 388 | -4.29 | 0.000353 | 0.00046 |
| chr1 | 11641631 | 11653115 | 11485 | -3.14 | 0.000355 | 0.000463 |
| chr7 | 151435821 | 151436375 | 555 | -3.62 | 0.000355 | 0.000463 |
| chr4 | 97777943 | 97780457 | 2515 | 1.89 | 0.000355 | 0.000463 |
| chr9 | 26768248 | 26769727 | 1480 | 1.8 | 0.000356 | 0.000464 |
| chr4 | 34255929 | 34269374 | 13446 | 1.15 | 0.000356 | 0.000464 |
| chr5 | 56004092 | 56006794 | 2703 | -4.25 | 0.000356 | 0.000464 |
| chr9 | 25676387 | 26330357 | 653971 | -0.85 | 0.000356 | 0.000464 |
| chr2 | 111613096 | 111615469 | 2374 | -1.94 | 0.000356 | 0.000464 |
| chr10 | 14036734 | 14044778 | 8045 | -1.82 | 0.000357 | 0.000465 |
| chr21 | 27163121 | 27164517 | 1397 | -3.71 | 0.000359 | 0.000467 |
| chr1 | 81491575 | 81492271 | 697 | -4.27 | 0.000359 | 0.000468 |
| chr3 | 186904945 | 186926663 | 21719 | -1.58 | 0.000359 | 0.000468 |
| chr2 | 180893134 | 180895714 | 2581 | -2.33 | 0.000359 | 0.000468 |
| chrX | 130080811 | 130083695 | 2885 | -4.37 | 0.00036 | 0.000468 |
| chr9 | 107851039 | 107852078 | 1040 | -3.71 | 0.00036 | 0.000469 |
| chr17 | 16910024 | 16910602 | 579 | -4.48 | 0.00036 | 0.000469 |
| chr4 | 67859585 | 67861604 | 2020 | -2.73 | 0.00036 | 0.000469 |
| chr13 | 88915334 | 88917198 | 1865 | 2.36 | 0.00036 | 0.000469 |
| chr5 | 65558774 | 65559711 | 938 | -3.51 | 0.000361 | 0.000471 |
| chr2 | 147020201 | 147020837 | 637 | -4.83 | 0.000361 | 0.000471 |
| chr11 | 127953790 | 127963198 | 9409 | -1.53 | 0.000362 | 0.000471 |
| chr6 | 66430961 | 66431910 | 950 | -3.26 | 0.000362 | 0.000471 |
| chr10 | 3290028 | 3290476 | 449 | -4.29 | 0.000362 | 0.000471 |
| chr8 | 90570212 | 90570806 | 595 | -4.29 | 0.000362 | 0.000471 |
| chr9 | 110179253 | 110180496 | 1244 | -4.63 | 0.000363 | 0.000472 |
| chr20 | 31841885 | 31856139 | 14255 | -2.44 | 0.000363 | 0.000473 |
| chr4 | 153041480 | 153070911 | 29432 | -1.84 | 0.000363 | 0.000473 |
| chr7 | 34549599 | 34632799 | 83201 | -5.33 | 0.000364 | 0.000474 |
| chr19 | 55376542 | 55448740 | 72199 | -5.35 | 0.000364 | 0.000474 |
| chr11 | 8218075 | 8220764 | 2690 | 4.16 | 0.000364 | 0.000474 |
| chr8 | 9168770 | 9185968 | 17199 | 5.9 | 0.000364 | 0.000474 |
| chr7 | 108333764 | 108339425 | 5662 | 2.4 | 0.000366 | 0.000476 |
| chr1 | 39080830 | 39107193 | 26364 | -4.64 | 0.000367 | 0.000478 |
| chr11 | 33845325 | 33845939 | 615 | -2.55 | 0.000368 | 0.000479 |
| chr1 | 120068500 | 120074330 | 5831 | -1.26 | 0.000368 | 0.000479 |
| chr12 | 33154823 | 33194528 | 39706 | -1.46 | 0.000368 | 0.000479 |
| chr14 | 49382788 | 49387295 | 4508 | 1.79 | 0.000369 | 0.00048 |

| | | | | | | |
|---|---|---|---|---|---|---|
| chr10 | 60551145 | 60551576 | 432 | -4.45 | 0.00037 | 0.000482 |
| chr5 | 147746537 | 147758824 | 12288 | -4.21 | 0.000371 | 0.000483 |
| chr6 | 16031333 | 16041125 | 9793 | -4.35 | 0.000371 | 0.000483 |
| chr5 | 53667289 | 53669956 | 2668 | -4.14 | 0.000371 | 0.000483 |
| chr9 | 109901567 | 109932055 | 30489 | -1.19 | 0.000372 | 0.000484 |
| chr1 | 37293064 | 37293473 | 410 | -3.33 | 0.000372 | 0.000484 |
| chr8 | 98335721 | 98336748 | 1028 | -2.46 | 0.000372 | 0.000484 |
| chr3 | 181235406 | 181237455 | 2050 | 3.7 | 0.000372 | 0.000484 |
| chrX | 35417520 | 35418422 | 903 | -3.39 | 0.000373 | 0.000485 |
| chr2 | 20045879 | 20046703 | 825 | -4.26 | 0.000373 | 0.000485 |
| chr19 | 46721828 | 46722296 | 469 | -4.3 | 0.000373 | 0.000485 |
| chr10 | 80138251 | 80140845 | 2595 | -4.44 | 0.000374 | 0.000486 |
| chr17 | 79911534 | 79912046 | 513 | -4.25 | 0.000375 | 0.000488 |
| chr14 | 64738969 | 64741343 | 2375 | -4.16 | 0.000375 | 0.000488 |
| chr8 | 55392032 | 55449726 | 57695 | 0.96 | 0.000375 | 0.000488 |
| chrX | 34903812 | 34906019 | 2208 | -3.95 | 0.000375 | 0.000488 |
| chr5 | 120477165 | 120477573 | 409 | -4.58 | 0.000376 | 0.00049 |
| chr12 | 13447292 | 13460885 | 13594 | 1.47 | 0.000376 | 0.00049 |
| chr3 | 113963940 | 113995751 | 31812 | -2.18 | 0.000377 | 0.00049 |
| chr8 | 6743396 | 6744393 | 998 | -3.71 | 0.000377 | 0.000491 |
| chr6 | 18039312 | 18039897 | 586 | -4.76 | 0.000378 | 0.000492 |
| chr4 | 61214166 | 61216360 | 2195 | 2.51 | 0.000379 | 0.000493 |
| chrX | 74178008 | 74179805 | 1798 | 3.78 | 0.000379 | 0.000494 |
| chr12 | 99748448 | 99857218 | 108771 | -1.17 | 0.000379 | 0.000494 |
| chrX | 151652313 | 151664047 | 11735 | 4.26 | 0.00038 | 0.000494 |
| chr3 | 59066556 | 59083349 | 16794 | -1.35 | 0.000381 | 0.000495 |
| chr1 | 197022628 | 197023115 | 488 | -4.39 | 0.000381 | 0.000495 |
| chr12 | 117543410 | 117543946 | 537 | -4.42 | 0.000381 | 0.000496 |
| chr12 | 16243770 | 16339971 | 96202 | -0.45 | 0.000381 | 0.000496 |
| chr1 | 101709779 | 101710764 | 986 | -2.58 | 0.000382 | 0.000496 |
| chr1 | 219786594 | 219816102 | 29509 | 1.48 | 0.000382 | 0.000497 |
| chr9 | 16067729 | 16073312 | 5584 | -4.95 | 0.000382 | 0.000497 |
| chr5 | 31789040 | 31805202 | 16163 | -4.48 | 0.000382 | 0.000497 |
| chr12 | 43911509 | 43914621 | 3113 | 2.05 | 0.000383 | 0.000498 |
| chr16 | 10749476 | 10749952 | 477 | -4.33 | 0.000383 | 0.000498 |
| chr7 | 65574943 | 65575438 | 496 | -4.33 | 0.000383 | 0.000498 |
| chrX | 10975700 | 10976099 | 400 | -4.33 | 0.000383 | 0.000498 |
| chr7 | 68863494 | 68864204 | 711 | -4.74 | 0.000384 | 0.000499 |
| chr15 | 35451065 | 35454335 | 3271 | -1.34 | 0.000386 | 0.000502 |
| chr1 | 170082907 | 170479102 | 396196 | 7.88 | 0.000387 | 0.000503 |
| chr10 | 102341400 | 102350213 | 8814 | -1.85 | 0.000388 | 0.000504 |
| chrX | 108452386 | 108467388 | 15003 | 4.38 | 0.000388 | 0.000504 |
| chr6 | 151898257 | 151978291 | 80035 | -3.62 | 0.000388 | 0.000505 |
| chr1 | 89600310 | 89604730 | 4421 | -3.36 | 0.000389 | 0.000506 |
| chrX | 79882582 | 79893376 | 10795 | 1.06 | 0.000389 | 0.000506 |
| chrX | 67496840 | 67505026 | 8187 | -6.03 | 0.00039 | 0.000506 |
| chr3 | 192490587 | 192496314 | 5728 | -2.52 | 0.000391 | 0.000508 |
| chr8 | 6757113 | 6758299 | 1187 | -2.31 | 0.000391 | 0.000508 |
| chr2 | 19780968 | 19786286 | 5319 | -3.88 | 0.000391 | 0.000508 |
| chr10 | 28598365 | 28602212 | 3848 | -2.27 | 0.000393 | 0.00051 |
| chr1 | 112556732 | 112563125 | 6394 | -1.79 | 0.000393 | 0.00051 |
| chr3 | 161153079 | 161154850 | 1772 | -3.11 | 0.000393 | 0.000511 |
| chr17 | 41911594 | 41912016 | 423 | -4.4 | 0.000394 | 0.000511 |
| chr4 | 109045520 | 109045904 | 385 | -3.7 | 0.000394 | 0.000512 |
| chr2 | 214294480 | 214310350 | 15871 | -1.79 | 0.000394 | 0.000512 |
| chr6 | 46327986 | 46336549 | 8564 | 1.21 | 0.000395 | 0.000514 |
| chr13 | 65485580 | 65489422 | 3843 | -4.25 | 0.000396 | 0.000514 |
| chr15 | 51540169 | 51633199 | 93031 | 5.52 | 0.000396 | 0.000515 |
| chr10 | 134586551 | 134587226 | 676 | -4.7 | 0.000397 | 0.000515 |
| chr8 | 124257480 | 124258274 | 795 | -4.7 | 0.000397 | 0.000515 |
| chr5 | 78819780 | 78823932 | 4153 | -4.68 | 0.000397 | 0.000515 |

| | | | | | | |
|---|---|---|---|---|---|---|
| chr7 | 132209874 | 132243095 | 33222 | -0.75 | 0.000397 | 0.000516 |
| chr7 | 14885918 | 14929920 | 44003 | -4.58 | 0.000398 | 0.000517 |
| chr8 | 82091366 | 82098253 | 6888 | 1.83 | 0.000399 | 0.000518 |
| chr12 | 130971392 | 130975505 | 4114 | -1.77 | 0.000399 | 0.000519 |
| chr12 | 118043448 | 118090549 | 47102 | -5.17 | 4.00E-04 | 0.000519 |
| chr9 | 84687383 | 84687776 | 394 | -4.28 | 4.00E-04 | 0.00052 |
| chr1 | 187972162 | 187972808 | 647 | -4.51 | 0.000401 | 0.000521 |
| chr6 | 137865447 | 137866831 | 1385 | 2.05 | 0.000401 | 0.000521 |
| chr16 | 86229501 | 86230569 | 1069 | -2.22 | 0.000401 | 0.000521 |
| chr10 | 14002438 | 14009787 | 7350 | 2.75 | 0.000402 | 0.000522 |
| chr1 | 231743467 | 231759174 | 15708 | -1.64 | 0.000402 | 0.000523 |
| chr3 | 121680618 | 121704909 | 24292 | -3.66 | 0.000403 | 0.000523 |
| chrX | 116423680 | 116440489 | 16810 | 1.74 | 0.000404 | 0.000525 |
| chr17 | 77429963 | 77444626 | 14664 | -4.87 | 0.000405 | 0.000526 |
| chrX | 79447987 | 79448445 | 459 | -4.34 | 0.000405 | 0.000526 |
| chr4 | 74235805 | 74236795 | 991 | -2.27 | 0.000406 | 0.000527 |
| chr1 | 199462596 | 199473378 | 10783 | -4.72 | 0.000406 | 0.000527 |
| chrX | 12593261 | 12594629 | 1369 | 2.4 | 0.000407 | 0.000528 |
| chr18 | 49101325 | 49181344 | 80020 | 0.64 | 0.000407 | 0.000528 |
| chr4 | 13347880 | 13348807 | 928 | -3.2 | 0.000407 | 0.000528 |
| chr6 | 67802631 | 67803258 | 628 | -3.54 | 0.000407 | 0.000529 |
| chr1 | 99684749 | 99686591 | 1843 | -2.02 | 0.000409 | 0.000531 |
| chr16 | 72924228 | 72924674 | 447 | -4.24 | 0.00041 | 0.000531 |
| chr17 | 76986059 | 76986443 | 385 | -4.24 | 0.00041 | 0.000531 |
| chr5 | 29987665 | 29991626 | 3962 | 1.57 | 0.00041 | 0.000532 |
| chr13 | 22454376 | 22461730 | 7355 | 1.67 | 0.00041 | 0.000532 |
| chr22 | 40236174 | 40240729 | 4556 | -1.59 | 0.00041 | 0.000532 |
| chr12 | 121579779 | 121580362 | 584 | -4.48 | 0.00041 | 0.000533 |
| chr11 | 32140733 | 32189616 | 48884 | -0.81 | 0.000411 | 0.000533 |
| chr7 | 36252841 | 36253621 | 781 | -4.23 | 0.000411 | 0.000533 |
| chr8 | 80706707 | 80707107 | 401 | -4.23 | 0.000411 | 0.000533 |
| chr2 | 40790660 | 40806090 | 15431 | 1.18 | 0.000413 | 0.000535 |
| chr10 | 4147061 | 4149275 | 2215 | -3.85 | 0.000413 | 0.000535 |
| chr11 | 74382970 | 74383326 | 357 | -4.23 | 0.000414 | 0.000537 |
| chr16 | 54982463 | 54987633 | 5171 | -2.07 | 0.000414 | 0.000537 |
| chr11 | 110264440 | 110267068 | 2629 | -2.39 | 0.000414 | 0.000537 |
| chr16 | 64286488 | 64306788 | 20301 | 1.15 | 0.000415 | 0.000538 |
| chr8 | 132792749 | 132800275 | 7527 | -0.9 | 0.000415 | 0.000539 |
| chr13 | 97564719 | 97566160 | 1442 | -2.51 | 0.000416 | 0.000539 |
| chr2 | 187116715 | 187117321 | 607 | -3.6 | 0.000416 | 0.000539 |
| chr4 | 85112348 | 85114911 | 2564 | -3.7 | 0.000416 | 0.000539 |
| chr6 | 53247399 | 53247794 | 396 | -3.43 | 0.000416 | 0.00054 |
| chr2 | 69653062 | 69653556 | 495 | -4.83 | 0.000417 | 0.000541 |
| chr3 | 195084521 | 195084964 | 444 | -4.63 | 0.000418 | 0.000542 |
| chr12 | 91078885 | 91080247 | 1363 | -2.61 | 0.000418 | 0.000543 |
| chr8 | 33885895 | 33886303 | 409 | -4.22 | 0.000419 | 0.000544 |
| chr2 | 160784021 | 160786226 | 2206 | -4.03 | 0.000419 | 0.000544 |
| chr1 | 85230397 | 85232134 | 1738 | -2.99 | 0.00042 | 0.000545 |
| chrX | 111044372 | 111046032 | 1661 | -1.7 | 0.00042 | 0.000545 |
| chr3 | 94046899 | 94047873 | 975 | -5.01 | 0.00042 | 0.000545 |
| chr1 | 197721927 | 197722248 | 322 | -4.26 | 0.000421 | 0.000546 |
| chr6 | 169114689 | 169160713 | 46025 | -1.22 | 0.000421 | 0.000546 |
| chr17 | 6156175 | 6185143 | 28969 | -1 | 0.000422 | 0.000547 |
| chr10 | 73268957 | 73270122 | 1166 | -3.73 | 0.000422 | 0.000547 |
| chr2 | 222664010 | 222690871 | 26862 | 1.1 | 0.000422 | 0.000548 |
| chr3 | 115262178 | 115263326 | 1149 | -2.08 | 0.000423 | 0.000548 |
| chr3 | 27863419 | 27864599 | 1181 | 3.71 | 0.000423 | 0.000549 |
| chr1 | 172268648 | 172269159 | 512 | -4.34 | 0.000423 | 0.000549 |
| chr5 | 106175452 | 106175928 | 477 | -2.18 | 0.000425 | 0.00055 |
| chr13 | 40246504 | 40246882 | 379 | -4.28 | 0.000425 | 0.000551 |
| chr22 | 22797605 | 22821635 | 24031 | -5 | 0.000425 | 0.000551 |

| | | | | | | |
|---|---|---|---|---|---|---|
| chr6 | 17265857 | 17266453 | 597 | -4.61 | 0.000425 | 0.000551 |
| chr18 | 44159464 | 44175528 | 16065 | -0.81 | 0.000426 | 0.000552 |
| chr4 | 48316914 | 48317303 | 390 | -3.93 | 0.000427 | 0.000553 |
| chr2 | 137047342 | 137047805 | 464 | -4.24 | 0.000427 | 0.000553 |
| chr12 | 4442283 | 4450748 | 8466 | -5.01 | 0.000427 | 0.000554 |
| chr13 | 96110164 | 96112706 | 2543 | -3.87 | 0.000427 | 0.000554 |
| chr17 | 29445810 | 29446307 | 498 | -4.43 | 0.000428 | 0.000554 |
| chr2 | 238339319 | 238348823 | 9505 | -4.43 | 0.000432 | 0.00056 |
| chr10 | 129529962 | 129530871 | 910 | 5.11 | 0.000433 | 0.000561 |
| chr15 | 83700051 | 83707845 | 7795 | -5.13 | 0.000437 | 0.000566 |
| chr1 | 220042882 | 220043266 | 385 | -3.26 | 0.000438 | 0.000567 |
| chr11 | 77042881 | 77043162 | 282 | -4.6 | 0.000438 | 0.000568 |
| chr10 | 129944802 | 129945538 | 737 | -4.95 | 0.000439 | 0.000568 |
| chr9 | 83227906 | 83265877 | 37972 | -0.94 | 0.00044 | 0.00057 |
| chr2 | 68674115 | 68674756 | 642 | -3.36 | 0.00044 | 0.000571 |
| chr8 | 118041544 | 118045263 | 3720 | 2.1 | 0.00044 | 0.000571 |
| chr1 | 80131219 | 80132999 | 1781 | -3.54 | 0.000441 | 0.000571 |
| chr1 | 4139134 | 4159115 | 19982 | -4.83 | 0.000441 | 0.000571 |
| chr20 | 55598614 | 55614146 | 15533 | -4.53 | 0.000441 | 0.000571 |
| chrX | 108579264 | 108580204 | 941 | -1.72 | 0.000442 | 0.000573 |
| chr7 | 31306149 | 31306881 | 733 | -2.84 | 0.000443 | 0.000574 |
| chrX | 55917169 | 55922600 | 5432 | 1.32 | 0.000444 | 0.000575 |
| chr1 | 161252447 | 161253560 | 1114 | -2.91 | 0.000444 | 0.000575 |
| chr2 | 1837239 | 1859675 | 22437 | 1.41 | 0.000445 | 0.000576 |
| chr1 | 17628433 | 17629033 | 601 | -4.4 | 0.000445 | 0.000576 |
| chr2 | 218426769 | 218427395 | 627 | -4.4 | 0.000445 | 0.000576 |
| chr21 | 10935890 | 10936270 | 381 | 3.54 | 0.000445 | 0.000577 |
| chr11 | 95357147 | 95357886 | 740 | -1.87 | 0.000446 | 0.000577 |
| chr22 | 17484366 | 17484843 | 478 | -2.24 | 0.000448 | 0.00058 |
| chr1 | 157358620 | 157449522 | 90903 | 3.84 | 0.000449 | 0.000581 |
| chr11 | 74451125 | 74451570 | 446 | -4.38 | 0.000449 | 0.000581 |
| chr5 | 49747584 | 49748052 | 469 | -4.54 | 0.000451 | 0.000584 |
| chr9 | 115326167 | 115326780 | 614 | -4.54 | 0.000451 | 0.000584 |
| chr6 | 137827057 | 137855819 | 28763 | -1.33 | 0.000452 | 0.000585 |
| chr6 | 118015702 | 118018587 | 2886 | -3.97 | 0.000452 | 0.000585 |
| chr2 | 112009296 | 112010434 | 1139 | -2.78 | 0.000453 | 0.000586 |
| chr7 | 73906681 | 73907193 | 513 | -4.2 | 0.000453 | 0.000587 |
| chr20 | 42022011 | 42069235 | 47225 | -5.26 | 0.000455 | 0.000589 |
| chr7 | 103309832 | 103350059 | 40228 | 2.34 | 0.000455 | 0.000589 |
| chr5 | 60544289 | 60554767 | 10479 | -5.52 | 0.000456 | 0.00059 |
| chr18 | 60889930 | 60890907 | 978 | -2.24 | 0.000456 | 0.00059 |
| chr2 | 204537780 | 204538168 | 389 | -3.71 | 0.000456 | 0.000591 |
| chr21 | 38976038 | 38978649 | 2612 | 2.37 | 0.000457 | 0.000592 |
| chr2 | 130653353 | 130658703 | 5351 | -3.79 | 0.000457 | 0.000592 |
| chr1 | 211369743 | 211380784 | 11042 | -4.7 | 0.000457 | 0.000592 |
| chr2 | 204805983 | 204851914 | 45932 | -1.06 | 0.000457 | 0.000592 |
| chr11 | 34643950 | 34646331 | 2382 | -3.86 | 0.000458 | 0.000592 |
| chr9 | 121729977 | 121733530 | 3554 | 1.46 | 0.000458 | 0.000593 |
| chr7 | 94401955 | 94402636 | 682 | -4.54 | 0.000459 | 0.000594 |
| chr11 | 48437573 | 48437841 | 269 | 4.68 | 0.000461 | 0.000596 |
| chr14 | 75850105 | 75866642 | 16538 | -4.7 | 0.000462 | 0.000598 |
| chr6 | 125685817 | 125687380 | 1564 | -3.54 | 0.000462 | 0.000598 |
| chrX | 135161376 | 135162634 | 1259 | -3.56 | 0.000463 | 0.000598 |
| chr17 | 37977317 | 37996598 | 19282 | -5.19 | 0.000463 | 6.00E-04 |
| chr4 | 28568822 | 28570445 | 1624 | -2.11 | 0.000463 | 6.00E-04 |
| chr6 | 150602283 | 150620445 | 18163 | -4.84 | 0.000464 | 6.00E-04 |
| chr19 | 42151612 | 42198238 | 46627 | -1.12 | 0.000464 | 6.00E-04 |
| chr11 | 109081351 | 109083198 | 1848 | -3.94 | 0.000464 | 0.000601 |
| chr12 | 3337771 | 3338514 | 744 | -4.87 | 0.000465 | 0.000602 |
| chr6 | 146068208 | 146071517 | 3310 | -4.27 | 0.000466 | 0.000602 |
| chr3 | 61867828 | 61868101 | 274 | 3.52 | 0.000466 | 0.000602 |

| | | | | | | |
|---|---|---|---|---|---|---|
| chr13 | 32432116 | 32434043 | 1928 | 2.11 | 0.000467 | 0.000604 |
| chr16 | 10389382 | 10390765 | 1384 | -3.26 | 0.000467 | 0.000604 |
| chr11 | 119334427 | 119337651 | 3225 | -3.16 | 0.000467 | 0.000604 |
| chrX | 101332817 | 101335824 | 3008 | 4.01 | 0.000467 | 0.000604 |
| chr14 | 29168639 | 29170050 | 1412 | 4.12 | 0.000468 | 0.000605 |
| chr3 | 143621338 | 143659177 | 37840 | 5.05 | 0.000468 | 0.000605 |
| chrX | 69410558 | 69436906 | 26349 | -1.3 | 0.000469 | 0.000606 |
| chr1 | 90609095 | 90611205 | 2111 | 3.81 | 0.000469 | 0.000607 |
| chr11 | 115098344 | 115099840 | 1497 | -2.35 | 0.00047 | 0.000608 |
| chr15 | 31146669 | 31147370 | 702 | -2.11 | 0.000471 | 0.000608 |
| chr12 | 20558750 | 20559183 | 434 | -4.49 | 0.000471 | 0.000608 |
| chr3 | 178856026 | 178858439 | 2414 | -3.14 | 0.000471 | 0.000608 |
| chrX | 53732923 | 53734151 | 1229 | -3.41 | 0.000471 | 0.00061 |
| chr3 | 75956485 | 75979274 | 22790 | 0.93 | 0.000472 | 0.00061 |
| chr4 | 141929033 | 141931258 | 2226 | -2.86 | 0.000474 | 0.000613 |
| chr12 | 121917698 | 121918199 | 502 | -4.35 | 0.000474 | 0.000613 |
| chr3 | 104947845 | 104948276 | 432 | -4.35 | 0.000474 | 0.000613 |
| chr5 | 172758585 | 172759088 | 504 | -4.35 | 0.000474 | 0.000613 |
| chr1 | 153197155 | 153197607 | 453 | -3.22 | 0.000475 | 0.000614 |
| chr12 | 108498134 | 108514914 | 16781 | -1.99 | 0.000476 | 0.000615 |
| chr11 | 120809268 | 120845109 | 35842 | -1.22 | 0.000477 | 0.000617 |
| chrX | 51299262 | 51319956 | 20695 | 1.27 | 0.000477 | 0.000617 |
| chr6 | 54976460 | 54977877 | 1418 | -1.71 | 0.000478 | 0.000618 |
| chr12 | 18885190 | 18890977 | 5788 | -4.37 | 0.000478 | 0.000618 |
| chr1 | 184750456 | 184752544 | 2089 | -1.75 | 0.000479 | 0.000619 |
| chr5 | 83044578 | 83046888 | 2311 | -3.16 | 0.000479 | 0.000619 |
| chr8 | 49075339 | 49170198 | 94860 | -1.32 | 0.00048 | 0.00062 |
| chr3 | 158332415 | 158352342 | 19928 | -4.78 | 0.00048 | 0.00062 |
| chr2 | 206715425 | 206716328 | 904 | 4.99 | 0.00048 | 0.00062 |
| chr6 | 153684495 | 153685171 | 677 | -4.83 | 0.00048 | 0.00062 |
| chr5 | 73919711 | 73921098 | 1388 | -3.57 | 0.00048 | 0.00062 |
| chr17 | 4695854 | 4696317 | 464 | -4.46 | 0.000482 | 0.000622 |
| chrX | 5132174 | 5143569 | 11396 | 4.6 | 0.000482 | 0.000622 |
| chr6 | 81623333 | 81748086 | 124754 | 1.54 | 0.000482 | 0.000623 |
| chr2 | 172768632 | 172769045 | 414 | -4.22 | 0.000482 | 0.000623 |
| chrX | 70211236 | 70214738 | 3503 | -4.4 | 0.000484 | 0.000625 |
| chrX | 80758473 | 80763121 | 4649 | 3.32 | 0.000485 | 0.000627 |
| chr11 | 134699646 | 134702843 | 3198 | -1.58 | 0.000487 | 0.000629 |
| chr4 | 58028683 | 58029452 | 770 | -4.81 | 0.000488 | 0.00063 |
| chr4 | 9910295 | 9911698 | 1404 | -3.8 | 0.000488 | 0.000631 |
| chr12 | 12123778 | 12149724 | 25947 | -4.22 | 0.000489 | 0.000632 |
| chr2 | 100244138 | 100304608 | 60471 | -1.26 | 0.00049 | 0.000633 |
| chrX | 125777071 | 125801669 | 24599 | 2.21 | 0.000491 | 0.000634 |
| chr11 | 44838892 | 44839471 | 580 | -4.32 | 0.000492 | 0.000635 |
| chr5 | 145810619 | 145811220 | 602 | -3.23 | 0.000492 | 0.000636 |
| chr15 | 86130183 | 86130825 | 643 | -3.76 | 0.000493 | 0.000637 |
| chr13 | 29087619 | 29122774 | 35156 | -1.59 | 0.000493 | 0.000637 |
| chr10 | 77805828 | 77835688 | 29861 | 4.91 | 0.000494 | 0.000638 |
| chr9 | 92181026 | 92201491 | 20466 | -4.03 | 0.000496 | 0.000641 |
| chr16 | 67401153 | 67404735 | 3583 | -1.26 | 0.000497 | 0.000641 |
| chrX | 72740709 | 72755041 | 14333 | -0.96 | 0.000497 | 0.000642 |
| chr1 | 119920313 | 120018384 | 98072 | 5.39 | 0.000498 | 0.000643 |
| chr9 | 83278975 | 83380651 | 101677 | -1.4 | 0.000499 | 0.000645 |
| chr4 | 68173606 | 68177292 | 3687 | 2.04 | 0.000499 | 0.000645 |
| chr10 | 72547221 | 72563452 | 16232 | -1.14 | 0.000499 | 0.000645 |
| chr8 | 11711953 | 11712389 | 437 | -4.59 | 5.00E-04 | 0.000645 |
| chr1 | 210691489 | 210692047 | 559 | -3.32 | 0.000503 | 0.000649 |
| chr2 | 56041083 | 56041440 | 358 | -4.18 | 0.000504 | 0.00065 |
| chr6 | 48687087 | 48687506 | 420 | -4.18 | 0.000504 | 0.00065 |
| chr1 | 84500932 | 84503211 | 2280 | -4.08 | 0.000505 | 0.000651 |
| chr7 | 113710914 | 113724637 | 13724 | -5.28 | 0.000505 | 0.000651 |

| | | | | | | |
|---|---|---|---|---|---|---|
| chr12 | 77049761 | 77054166 | 4406 | -1.82 | 0.000505 | 0.000652 |
| chr9 | 94632184 | 94635155 | 2972 | 4.05 | 0.000506 | 0.000653 |
| chr5 | 41527848 | 41557819 | 29972 | -1.43 | 0.000506 | 0.000653 |
| chr21 | 19421232 | 19422652 | 1421 | 2.1 | 0.000506 | 0.000653 |
| chr9 | 91200188 | 91212105 | 11918 | -6.27 | 0.000506 | 0.000653 |
| chr8 | 133525619 | 133582709 | 57091 | 3.41 | 0.000507 | 0.000654 |
| chr4 | 22771226 | 22774770 | 3545 | -1.24 | 0.000507 | 0.000654 |
| chr8 | 117384330 | 117385967 | 1638 | -2.09 | 0.000509 | 0.000657 |
| chr6 | 18495494 | 18497567 | 2074 | -3.51 | 0.000509 | 0.000657 |
| chr5 | 59765970 | 59774306 | 8337 | -1.85 | 0.000509 | 0.000657 |
| chr19 | 55504242 | 55518409 | 14168 | -1.36 | 0.00051 | 0.000658 |
| chr15 | 49262233 | 49263308 | 1076 | -2.97 | 0.00051 | 0.000658 |
| chr6 | 135421028 | 135421363 | 336 | -4.29 | 0.000512 | 0.00066 |
| chr1 | 48205475 | 48207740 | 2266 | -3.88 | 0.000513 | 0.000662 |
| chr18 | 7122622 | 7143891 | 21270 | -2.78 | 0.000513 | 0.000662 |
| chr15 | 77946092 | 77947326 | 1235 | -4.1 | 0.000513 | 0.000662 |
| chr1 | 162119941 | 162120286 | 346 | -4.17 | 0.000513 | 0.000662 |
| chr4 | 69408100 | 69408483 | 384 | -4.22 | 0.000515 | 0.000664 |
| chr18 | 48815295 | 48819063 | 3769 | -2.13 | 0.000517 | 0.000667 |
| chr7 | 138421875 | 138422360 | 486 | -4.27 | 0.000521 | 0.000672 |
| chr11 | 44432424 | 44434792 | 2369 | -2.25 | 0.000521 | 0.000672 |
| chr5 | 41654155 | 41655413 | 1259 | -1.86 | 0.000521 | 0.000672 |
| chr2 | 11539578 | 11540137 | 560 | -4.38 | 0.000521 | 0.000672 |
| chr3 | 31754081 | 31754432 | 352 | -4.38 | 0.000521 | 0.000672 |
| chr12 | 15608114 | 15615683 | 7570 | -1.39 | 0.000522 | 0.000673 |
| chr21 | 27617426 | 27624576 | 7151 | -1.24 | 0.000522 | 0.000674 |
| chr5 | 95750276 | 95751713 | 1438 | 3.86 | 0.000523 | 0.000674 |
| chr17 | 53318982 | 53340328 | 21347 | -2.69 | 0.000524 | 0.000675 |
| chr6 | 47945932 | 47946770 | 839 | 2.16 | 0.000524 | 0.000676 |
| chr6 | 72257617 | 72271692 | 14076 | -2.55 | 0.000525 | 0.000678 |
| chrX | 148970700 | 149003769 | 33070 | -2.58 | 0.000527 | 0.00068 |
| chr4 | 53422275 | 53429305 | 7031 | -1.49 | 0.000527 | 0.00068 |
| chr13 | 112300060 | 112305048 | 4989 | 1.36 | 0.000528 | 0.000681 |
| chr15 | 66465888 | 66469005 | 3118 | 1.6 | 0.00053 | 0.000684 |
| chrX | 46669349 | 46670250 | 902 | -4.52 | 0.000531 | 0.000684 |
| chr10 | 27653905 | 27683678 | 29774 | -5.08 | 0.000532 | 0.000686 |
| chr8 | 62078866 | 62090890 | 12025 | 1.26 | 0.000532 | 0.000686 |
| chr3 | 157569391 | 157572032 | 2642 | -2.44 | 0.000532 | 0.000686 |
| chr5 | 14975505 | 14983984 | 8480 | -4.66 | 0.000533 | 0.000687 |
| chr4 | 134133587 | 134135430 | 1844 | -1.63 | 0.000533 | 0.000687 |
| chr4 | 89231009 | 89233596 | 2588 | -1.64 | 0.000534 | 0.000688 |
| chr1 | 75972327 | 75974072 | 1746 | 3.77 | 0.000534 | 0.000688 |
| chr10 | 87809783 | 87811827 | 2045 | -2.85 | 0.000534 | 0.000688 |
| chr11 | 114456997 | 114459417 | 2421 | -1.83 | 0.000534 | 0.000689 |
| chr2 | 188707063 | 188707393 | 331 | -4.26 | 0.000534 | 0.000689 |
| chr20 | 33204664 | 33205118 | 455 | -4.26 | 0.000534 | 0.000689 |
| chr11 | 19054497 | 19075575 | 21079 | -2 | 0.000535 | 0.000689 |
| chr17 | 21194883 | 21213262 | 18380 | -2.92 | 0.000536 | 0.000691 |
| chr15 | 81017400 | 81039242 | 21843 | -5.49 | 0.000538 | 0.000693 |
| chr6 | 128022788 | 128025252 | 2465 | -2.78 | 0.000538 | 0.000693 |
| chr14 | 30224566 | 30254793 | 30228 | 1.85 | 0.000539 | 0.000694 |
| chr2 | 100637567 | 100640364 | 2798 | 1.53 | 0.00054 | 0.000696 |
| chr12 | 38830942 | 38831769 | 828 | -3.32 | 0.00054 | 0.000696 |
| chr12 | 125852033 | 125866020 | 13988 | 1.59 | 0.00054 | 0.000696 |
| chr20 | 39488284 | 39492447 | 4164 | 4.18 | 0.000541 | 0.000697 |
| chr1 | 198446768 | 198448604 | 1837 | -3.42 | 0.000542 | 0.000699 |
| chr10 | 83027522 | 83063223 | 35702 | 1.11 | 0.000543 | 7.00E-04 |
| chr14 | 61891480 | 61899089 | 7610 | -4.06 | 0.000544 | 0.000701 |
| chr1 | 188583256 | 188589864 | 6609 | -4.86 | 0.000544 | 0.000701 |
| chr8 | 41222094 | 41248497 | 26404 | -4.97 | 0.000545 | 0.000702 |
| chr12 | 63169430 | 63177530 | 8101 | -5.73 | 0.000545 | 0.000702 |

| | | | | | | |
|---|---|---|---|---|---|---|
| chr3 | 174841820 | 174842879 | 1060 | -3.57 | 0.000546 | 0.000703 |
| chr3 | 152249117 | 152265424 | 16308 | -4.82 | 0.000546 | 0.000704 |
| chr11 | 85627157 | 85627974 | 818 | -3.43 | 0.000547 | 0.000704 |
| chr1 | 32904345 | 32905618 | 1274 | -2.18 | 0.000549 | 0.000708 |
| chr19 | 49654709 | 49655151 | 443 | -4.18 | 0.00055 | 0.000708 |
| chr5 | 174330055 | 174333855 | 3801 | -1.64 | 0.000551 | 0.000709 |
| chr8 | 76183252 | 76185018 | 1767 | -3.44 | 0.000551 | 0.00071 |
| chr3 | 162120943 | 162121372 | 430 | 4.4 | 0.000552 | 0.00071 |
| chr5 | 91385764 | 91386180 | 417 | -3.15 | 0.000552 | 0.000711 |
| chr15 | 96219957 | 96250950 | 30994 | -5.13 | 0.000552 | 0.000711 |
| chr5 | 27419287 | 27431764 | 12478 | -1.71 | 0.000552 | 0.000711 |
| chr8 | 83366625 | 83367485 | 861 | -4.47 | 0.000554 | 0.000713 |
| chr1 | 31302520 | 31302974 | 455 | -4.15 | 0.000554 | 0.000713 |
| chr4 | 117683082 | 117683860 | 779 | -4.32 | 0.000554 | 0.000714 |
| chr7 | 21822225 | 21919261 | 97037 | -2.5 | 0.000555 | 0.000715 |
| chrX | 17890304 | 17957505 | 67202 | -0.9 | 0.000556 | 0.000716 |
| chr3 | 180574787 | 180576137 | 1351 | -1.98 | 0.000557 | 0.000718 |
| chr3 | 113855994 | 113858372 | 2379 | -1.92 | 0.00056 | 0.00072 |
| chr9 | 139885101 | 139885731 | 631 | -4.23 | 0.000561 | 0.000722 |
| chr18 | 53756580 | 53773699 | 17120 | -0.96 | 0.000562 | 0.000724 |
| chr3 | 123942013 | 123952579 | 10567 | -0.87 | 0.000563 | 0.000724 |
| chr16 | 1380405 | 1380824 | 420 | -4.15 | 0.000563 | 0.000725 |
| chr12 | 48762921 | 48825096 | 62176 | 0.69 | 0.000565 | 0.000727 |
| chr5 | 78332489 | 78349819 | 17331 | -4.73 | 0.000566 | 0.000728 |
| chr4 | 33035718 | 33050387 | 14670 | 1.44 | 0.000566 | 0.000728 |
| chr4 | 152769195 | 152848599 | 79405 | -1.71 | 0.000566 | 0.000728 |
| chr16 | 64470788 | 64472021 | 1234 | -4.62 | 0.000567 | 0.00073 |
| chr18 | 13982223 | 13988547 | 6325 | 1.21 | 0.000568 | 0.000731 |
| chr2 | 210087308 | 210089268 | 1961 | 3.87 | 0.000569 | 0.000732 |
| chr2 | 222175901 | 222223370 | 47470 | 0.98 | 0.000569 | 0.000732 |
| chr14 | 96708217 | 96711929 | 3713 | -3.75 | 0.000569 | 0.000732 |
| chr6 | 91049067 | 91060542 | 11476 | -1.43 | 0.00057 | 0.000733 |
| chr3 | 40642764 | 40644578 | 1815 | -3.68 | 0.000571 | 0.000734 |
| chr5 | 42770140 | 42772412 | 2273 | -4.19 | 0.000571 | 0.000734 |
| chr5 | 134563821 | 134565337 | 1517 | -4.8 | 0.000571 | 0.000735 |
| chr8 | 117617998 | 117620448 | 2451 | 2.16 | 0.000571 | 0.000735 |
| chr4 | 106457991 | 106458826 | 836 | -3.99 | 0.000571 | 0.000735 |
| chr15 | 51491070 | 51492500 | 1431 | -1.96 | 0.000573 | 0.000737 |
| chrX | 132097396 | 132097768 | 373 | 4.24 | 0.000573 | 0.000737 |
| chr4 | 154726932 | 154735925 | 8994 | 1.25 | 0.000574 | 0.000738 |
| chr1 | 94759072 | 94759602 | 531 | -4.29 | 0.000574 | 0.000738 |
| chr2 | 31715092 | 31715588 | 497 | -4.29 | 0.000574 | 0.000738 |
| chr2 | 214553584 | 214561056 | 7473 | 1.39 | 0.000574 | 0.000739 |
| chrX | 74827010 | 74828946 | 1937 | 1.59 | 0.000575 | 0.000739 |
| chrX | 55050716 | 55051110 | 395 | 3.23 | 0.000575 | 0.000739 |
| chr9 | 29747375 | 29770450 | 23076 | 1.44 | 0.000577 | 0.000741 |
| chr2 | 186289974 | 186311706 | 21733 | 0.87 | 0.000577 | 0.000742 |
| chr5 | 14996935 | 15001326 | 4392 | -4.76 | 0.000578 | 0.000744 |
| chr8 | 61885996 | 61892375 | 6380 | -1.11 | 0.000578 | 0.000744 |
| chr2 | 222868604 | 222881769 | 13166 | 1.63 | 0.000578 | 0.000744 |
| chrX | 102535929 | 102545766 | 9838 | -2.38 | 0.000578 | 0.000744 |
| chr5 | 91306793 | 91309081 | 2289 | -3.88 | 0.000579 | 0.000745 |
| chr11 | 43971881 | 44000255 | 28375 | -5.09 | 0.00058 | 0.000746 |
| chr7 | 107285897 | 107286759 | 863 | -2.69 | 0.00058 | 0.000746 |
| chr21 | 33382322 | 33553680 | 171359 | -0.56 | 0.000581 | 0.000747 |
| chr8 | 119598562 | 119612082 | 13521 | -1.51 | 0.000582 | 0.000748 |
| chr3 | 97408561 | 97409714 | 1154 | -2.58 | 0.000582 | 0.000748 |
| chr14 | 81235969 | 81244036 | 8068 | -4.13 | 0.000582 | 0.000749 |
| chr2 | 33450107 | 33459478 | 9372 | -4.13 | 0.000582 | 0.000749 |
| chr6 | 91195559 | 91196060 | 502 | -4.13 | 0.000582 | 0.000749 |
| chr3 | 116625060 | 116630642 | 5583 | -1.57 | 0.000587 | 0.000755 |

| | | | | | | |
|---|---|---|---|---|---|---|
| chr7 | 129845810 | 129846301 | 492 | -4.16 | 0.000587 | 0.000755 |
| chr2 | 154359646 | 154708186 | 348541 | 7.51 | 0.000588 | 0.000756 |
| chr15 | 100000176 | 100006188 | 6013 | -4.85 | 0.000589 | 0.000758 |
| chr1 | 247015752 | 247016223 | 472 | -3.24 | 0.00059 | 0.000758 |
| chr14 | 89949908 | 89963724 | 13817 | 4.42 | 0.000591 | 0.00076 |
| chr8 | 133609482 | 133610877 | 1396 | 3.66 | 0.000593 | 0.000762 |
| chr5 | 79138793 | 79231101 | 92309 | -0.93 | 0.000593 | 0.000763 |
| chr12 | 15355059 | 15357886 | 2828 | -4.23 | 0.000594 | 0.000763 |
| chr10 | 122874387 | 122877063 | 2677 | -4.15 | 0.000594 | 0.000763 |
| chr8 | 87035522 | 87037668 | 2147 | -3.53 | 0.000595 | 0.000765 |
| chr2 | 44732093 | 44732436 | 344 | -4.39 | 0.000596 | 0.000766 |
| chr13 | 67959950 | 67961255 | 1306 | 2.68 | 0.000596 | 0.000766 |
| chr8 | 60980080 | 61026304 | 46225 | -0.73 | 0.000598 | 0.000769 |
| chr14 | 52789709 | 52790137 | 429 | -4.74 | 0.000601 | 0.000773 |
| chr12 | 12057391 | 12094155 | 36765 | -5.03 | 0.000602 | 0.000773 |
| chr2 | 30604576 | 30611157 | 6582 | -2.04 | 0.000602 | 0.000774 |
| chr5 | 157553028 | 157775474 | 222447 | 0.79 | 0.000602 | 0.000774 |
| chr16 | 20618763 | 20621913 | 3151 | 3.52 | 0.000604 | 0.000775 |
| chr11 | 27460303 | 27461127 | 825 | -4.44 | 0.000604 | 0.000775 |
| chr11 | 134789355 | 134807755 | 18401 | -1 | 0.000605 | 0.000777 |
| chr4 | 149382554 | 149387493 | 4940 | 3.3 | 0.000605 | 0.000777 |
| chr7 | 108310050 | 108330267 | 20218 | -1.49 | 0.000606 | 0.000778 |
| chr5 | 50155129 | 50217534 | 62406 | 0.95 | 0.000606 | 0.000778 |
| chr4 | 32424641 | 32427414 | 2774 | -3.61 | 0.000607 | 0.000779 |
| chrX | 76596439 | 76606879 | 10441 | -1.29 | 0.000613 | 0.000787 |
| chr1 | 199699973 | 199700437 | 465 | -4.36 | 0.000613 | 0.000787 |
| chrX | 51778747 | 51780840 | 2094 | 1.53 | 0.000613 | 0.000787 |
| chr12 | 101506029 | 101507776 | 1748 | 3.92 | 0.000613 | 0.000787 |
| chrX | 118569155 | 118569437 | 283 | -3.42 | 0.000613 | 0.000788 |
| chr7 | 31699203 | 31702534 | 3332 | -1.98 | 0.000615 | 0.000789 |
| chrX | 103157901 | 103167138 | 9238 | -4.43 | 0.000615 | 0.00079 |
| chr18 | 67558266 | 67558693 | 428 | -4.11 | 0.000615 | 0.00079 |
| chr2 | 17529718 | 17566450 | 36733 | -1.64 | 0.000616 | 0.000791 |
| chr7 | 148349956 | 148350571 | 616 | -4.12 | 0.000616 | 0.000791 |
| chr20 | 60030523 | 60035120 | 4598 | 1.96 | 0.000616 | 0.000791 |
| chr3 | 156128779 | 156129850 | 1072 | -2.98 | 0.000616 | 0.000792 |
| chr10 | 71514045 | 71515049 | 1005 | -3.44 | 0.000617 | 0.000792 |
| chr6 | 6646059 | 6648027 | 1969 | -2.64 | 0.000618 | 0.000794 |
| chr2 | 33865187 | 33894854 | 29668 | -1.05 | 0.000619 | 0.000795 |
| chr5 | 153588565 | 153588958 | 394 | -4.23 | 0.000621 | 0.000797 |
| chr8 | 6693916 | 6694569 | 654 | -4.23 | 0.000621 | 0.000797 |
| chr5 | 107157968 | 107158684 | 717 | -4.69 | 0.000621 | 0.000798 |
| chr5 | 137129322 | 137129789 | 468 | -4.13 | 0.000623 | 0.000799 |
| chr6 | 128717808 | 128718181 | 374 | -4.13 | 0.000623 | 0.000799 |
| chr4 | 71400165 | 71401975 | 1811 | 2.68 | 0.000625 | 0.000802 |
| chr10 | 25834907 | 25966366 | 131460 | 0.67 | 0.000625 | 0.000803 |
| chr6 | 155297532 | 155297997 | 466 | -4.5 | 0.000626 | 0.000804 |
| chr8 | 124560792 | 124561234 | 443 | -4.5 | 0.000626 | 0.000804 |
| chr2 | 190180807 | 190181157 | 351 | -4.17 | 0.000627 | 0.000804 |
| chr13 | 51347989 | 51354989 | 7001 | -1.93 | 0.000629 | 0.000807 |
| chr11 | 133706711 | 133727165 | 20455 | -2.17 | 0.000629 | 0.000807 |
| chr11 | 44295247 | 44297210 | 1964 | 2.52 | 0.00063 | 0.000808 |
| chr12 | 9265492 | 9267736 | 2245 | 3.87 | 0.00063 | 0.000808 |
| chr3 | 16294456 | 16294977 | 522 | -4.33 | 0.000631 | 0.000809 |
| chrX | 36134551 | 36141078 | 6528 | -4.32 | 0.000632 | 0.000811 |
| chr6 | 144494055 | 144494623 | 569 | -4.11 | 0.000632 | 0.000811 |
| chr6 | 154143345 | 154144531 | 1187 | 2.31 | 0.000634 | 0.000814 |
| chr3 | 36841832 | 36850601 | 8770 | -4.78 | 0.000634 | 0.000814 |
| chr2 | 238745555 | 238750194 | 4640 | -4.26 | 0.000637 | 0.000817 |
| chr6 | 146992209 | 146995650 | 3442 | 2.39 | 0.000641 | 0.000822 |
| chr6 | 129977979 | 129979313 | 1335 | -3.36 | 0.000641 | 0.000823 |

| | | | | | | |
|---|---|---|---|---|---|---|
| chr1 | 168515569 | 168515999 | 431 | -3.42 | 0.000642 | 0.000823 |
| chr21 | 37351251 | 37376518 | 25268 | -1.67 | 0.000644 | 0.000826 |
| chr2 | 194701202 | 194702037 | 836 | -3.05 | 0.000644 | 0.000826 |
| chr11 | 7040599 | 7045412 | 4814 | -3.65 | 0.000645 | 0.000827 |
| chr7 | 135032 | 136738 | 1707 | 2.12 | 0.000645 | 0.000828 |
| chr1 | 192381834 | 192382155 | 322 | -3.24 | 0.000646 | 0.000828 |
| chr5 | 170075126 | 170076298 | 1173 | -3.48 | 0.000647 | 0.000829 |
| chr1 | 154627194 | 154628809 | 1616 | -3.17 | 0.000647 | 0.00083 |
| chr22 | 19658692 | 19660937 | 2246 | -2.11 | 0.000648 | 0.000831 |
| chr1 | 59024845 | 59028190 | 3346 | -4.31 | 0.000648 | 0.000831 |
| chr14 | 56831518 | 56879011 | 47494 | -0.87 | 0.000648 | 0.000831 |
| chr6 | 85173101 | 85173710 | 610 | -4.3 | 0.000651 | 0.000834 |
| chr9 | 93526416 | 93566155 | 39740 | -2.41 | 0.000652 | 0.000836 |
| chr5 | 158462570 | 158463555 | 986 | -3.56 | 0.000655 | 0.000839 |
| chr10 | 115331861 | 115362909 | 31049 | -1.44 | 0.000655 | 0.00084 |
| chr3 | 104230102 | 104268670 | 38569 | 1.49 | 0.000657 | 0.000843 |
| chr2 | 162986758 | 162988722 | 1965 | -2.11 | 0.000659 | 0.000846 |
| chr9 | 91956526 | 91957824 | 1299 | -3.56 | 0.00066 | 0.000846 |
| chr1 | 160358368 | 160359170 | 803 | -3.92 | 0.000662 | 0.000849 |
| chr11 | 117874210 | 117899959 | 25750 | -4.93 | 0.000664 | 0.000852 |
| chr2 | 209743341 | 209750129 | 6789 | 4.28 | 0.00067 | 0.000859 |
| chr4 | 63456464 | 63467682 | 11219 | -5.4 | 0.000674 | 0.000864 |
| chr6 | 167867558 | 167871283 | 3726 | 1.36 | 0.000677 | 0.000867 |
| chr3 | 163074787 | 163083355 | 8569 | -3.91 | 0.000677 | 0.000867 |
| chr14 | 75803248 | 75812927 | 9680 | -1.32 | 0.000679 | 0.000871 |
| chr6 | 84254144 | 84256531 | 2388 | -1.57 | 0.000681 | 0.000873 |
| chr1 | 75557349 | 75557793 | 445 | -4.58 | 0.000682 | 0.000874 |
| chr14 | 57985923 | 57994219 | 8297 | -4.35 | 0.000684 | 0.000877 |
| chr1 | 116476382 | 116478103 | 1722 | -1.59 | 0.000684 | 0.000877 |
| chr3 | 50926100 | 50931860 | 5761 | -2.02 | 0.000685 | 0.000877 |
| chr17 | 58958265 | 58971672 | 13408 | -5.09 | 0.000686 | 0.000879 |
| chr15 | 76201955 | 76202236 | 282 | -4.3 | 0.000686 | 0.000879 |
| chr4 | 97190474 | 97191874 | 1401 | 3.7 | 0.000686 | 0.000879 |
| chr4 | 20275758 | 20276805 | 1048 | 3.62 | 0.000687 | 0.00088 |
| chr4 | 115478220 | 115481684 | 3465 | -4.12 | 0.000687 | 0.000881 |
| chr7 | 34505970 | 34511236 | 5267 | -1.92 | 0.000689 | 0.000883 |
| chr11 | 113381371 | 113383176 | 1806 | 3.82 | 0.00069 | 0.000884 |
| chrX | 3376166 | 3391439 | 15274 | 2.37 | 0.00069 | 0.000884 |
| chr12 | 68640570 | 68662662 | 22093 | 1.27 | 0.000691 | 0.000885 |
| chr3 | 78790060 | 78790791 | 732 | -3.47 | 0.000692 | 0.000886 |
| chr9 | 74044458 | 74048953 | 4496 | 2.23 | 0.000692 | 0.000886 |
| chr14 | 76986794 | 76989695 | 2902 | -4.51 | 0.000692 | 0.000887 |
| chr2 | 28734219 | 28742549 | 8331 | -1.69 | 0.000694 | 0.000889 |
| chr5 | 114418715 | 114421087 | 2373 | -2.05 | 0.000696 | 0.000892 |
| chr4 | 105930657 | 105932341 | 1685 | -3.65 | 0.000697 | 0.000893 |
| chr1 | 217421887 | 217440727 | 18841 | 1.31 | 0.000697 | 0.000893 |
| chr20 | 48855737 | 48856242 | 506 | -4.24 | 0.000698 | 0.000893 |
| chr4 | 23566831 | 23567368 | 538 | -4.24 | 0.000698 | 0.000893 |
| chr11 | 115405418 | 115410319 | 4902 | 1.45 | 0.000698 | 0.000894 |
| chr4 | 37356030 | 37359012 | 2983 | 3.97 | 0.000698 | 0.000894 |
| chr15 | 38387924 | 38390805 | 2882 | -4.04 | 0.000699 | 0.000895 |
| chr3 | 134420734 | 134421624 | 891 | -2.52 | 0.000699 | 0.000896 |
| chr4 | 121344227 | 121346573 | 2347 | 2.39 | 7.00E-04 | 0.000896 |
| chr22 | 45380644 | 45381006 | 363 | 3.19 | 7.00E-04 | 0.000896 |
| chr4 | 55148639 | 55170624 | 21986 | -0.87 | 7.00E-04 | 0.000897 |
| chr1 | 86297185 | 86297785 | 601 | -2.88 | 0.000703 | 9.00E-04 |
| chr13 | 110017372 | 110018485 | 1114 | -3.66 | 0.000705 | 0.000902 |
| chr7 | 97471300 | 97473287 | 1988 | -3.63 | 0.000706 | 0.000904 |
| chr1 | 5683570 | 5709172 | 25603 | -1.59 | 0.000708 | 0.000907 |
| chr5 | 110881240 | 110897533 | 16294 | -4.47 | 0.000708 | 0.000907 |
| chr11 | 124816885 | 124823284 | 6400 | -1.91 | 0.00071 | 0.000909 |

| | | | | | | |
|---|---|---|---|---|---|---|
| chr7 | 11258526 | 11259461 | 936 | -1.92 | 0.000712 | 0.000911 |
| chr5 | 85672930 | 85761994 | 89065 | 2.35 | 0.000712 | 0.000912 |
| chr8 | 119698391 | 119709786 | 11396 | 1.53 | 0.000713 | 0.000912 |
| chr4 | 145678915 | 145680234 | 1320 | -2.21 | 0.000714 | 0.000913 |
| chr1 | 199565317 | 199579055 | 13739 | -3.61 | 0.000716 | 0.000917 |
| chr20 | 22501076 | 22501745 | 670 | -4.7 | 0.00072 | 0.000922 |
| chr1 | 235108461 | 235109037 | 577 | -4.07 | 0.000721 | 0.000923 |
| chr2 | 180176 | 214326 | 34151 | -2.19 | 0.000721 | 0.000923 |
| chr15 | 52295096 | 52295514 | 419 | -4.14 | 0.000721 | 0.000923 |
| chrX | 70402434 | 70402926 | 493 | -4.14 | 0.000721 | 0.000923 |
| chr14 | 54109058 | 54109632 | 575 | -4.87 | 0.000723 | 0.000925 |
| chr9 | 124084209 | 124085590 | 1382 | -4.34 | 0.000723 | 0.000925 |
| chr18 | 56834589 | 56856976 | 22388 | -4.84 | 0.000724 | 0.000926 |
| chr3 | 176617843 | 176619349 | 1507 | 1.82 | 0.000724 | 0.000926 |
| chr1 | 224549170 | 224550048 | 879 | -4.5 | 0.000724 | 0.000927 |
| chr13 | 104074322 | 104077786 | 3465 | 1.43 | 0.000725 | 0.000928 |
| chr4 | 7739628 | 7740079 | 452 | -4.21 | 0.000726 | 0.000928 |
| chr1 | 108294816 | 108295449 | 634 | -3.27 | 0.000726 | 0.000929 |
| chrX | 10946352 | 10947153 | 802 | -3.29 | 0.000727 | 0.00093 |
| chr11 | 104156168 | 104157927 | 1760 | 2.28 | 0.000728 | 0.000931 |
| chr2 | 181626445 | 181629987 | 3543 | -4.43 | 0.000731 | 0.000935 |
| chr3 | 46439603 | 46441031 | 1429 | -2.05 | 0.000732 | 0.000936 |
| chr4 | 186387182 | 186390643 | 3462 | -3.98 | 0.000734 | 0.000938 |
| chr2 | 139612808 | 139615470 | 2663 | 2.8 | 0.000734 | 0.000938 |
| chr10 | 63072412 | 63072731 | 320 | 3.41 | 0.000735 | 0.00094 |
| chr8 | 63238149 | 63240918 | 2770 | -3.05 | 0.000736 | 0.000941 |
| chr5 | 71093989 | 71097135 | 3147 | -3.94 | 0.000736 | 0.000941 |
| chr11 | 19107709 | 19127439 | 19731 | -1.26 | 0.000736 | 0.000942 |
| chr8 | 9672495 | 9674525 | 2031 | -3.43 | 0.000737 | 0.000942 |
| chr13 | 112990289 | 112992479 | 2191 | -1.61 | 0.000739 | 0.000945 |
| chr17 | 29375131 | 29376308 | 1178 | 2 | 0.000739 | 0.000945 |
| chr15 | 78139892 | 78143762 | 3871 | -4.75 | 0.00074 | 0.000946 |
| chr3 | 100410334 | 100413640 | 3307 | 1.38 | 0.000741 | 0.000948 |
| chr4 | 176758159 | 176758820 | 662 | -3.09 | 0.000746 | 0.000953 |
| chr20 | 21437339 | 21445414 | 8076 | -1.32 | 0.000746 | 0.000954 |
| chr5 | 120224461 | 120224925 | 465 | -3.11 | 0.000747 | 0.000955 |
| chr11 | 59876013 | 59880876 | 4864 | 4.06 | 0.000747 | 0.000955 |
| chr9 | 107065285 | 107067668 | 2384 | -1.62 | 0.000748 | 0.000956 |
| chr11 | 116494284 | 116499648 | 5365 | -1.61 | 0.000748 | 0.000956 |
| chr5 | 169971303 | 169976492 | 5190 | -4.77 | 0.000748 | 0.000957 |
| chr10 | 105932470 | 105941090 | 8621 | -4.21 | 0.000748 | 0.000957 |
| chr4 | 21677313 | 21680125 | 2813 | 1.81 | 0.000749 | 0.000957 |
| chr10 | 9150231 | 9224980 | 74750 | -5.09 | 0.000749 | 0.000957 |
| chrX | 3690381 | 3713574 | 23194 | -1.24 | 0.00075 | 0.000959 |
| chr8 | 113759197 | 113760929 | 1733 | -3.43 | 0.000751 | 0.000959 |
| chr6 | 24902256 | 24908027 | 5772 | -1.07 | 0.000751 | 0.00096 |
| chrX | 148718330 | 148725665 | 7336 | -3.17 | 0.000752 | 0.000961 |
| chr9 | 20219777 | 20221488 | 1712 | 2.37 | 0.000754 | 0.000963 |
| chr16 | 55465519 | 55486221 | 20703 | 1.68 | 0.000754 | 0.000963 |
| chr2 | 132166961 | 132167735 | 775 | -2.66 | 0.000756 | 0.000966 |
| chr13 | 102403119 | 102404018 | 900 | -4.45 | 0.000757 | 0.000967 |
| chr6 | 44467028 | 44471527 | 4500 | -1.42 | 0.000757 | 0.000967 |
| chr7 | 28028418 | 28028688 | 271 | -4.06 | 0.000758 | 0.000968 |
| chrX | 124366381 | 124380056 | 13676 | 1.92 | 0.00076 | 0.000971 |
| chr3 | 50970187 | 50987634 | 17448 | -4.47 | 0.000762 | 0.000973 |
| chr6 | 75728650 | 75728967 | 318 | -2.97 | 0.000762 | 0.000974 |
| chr8 | 107240659 | 107245435 | 4777 | -1.84 | 0.000763 | 0.000976 |
| chr11 | 88486321 | 88516760 | 30440 | -1.37 | 0.000764 | 0.000977 |
| chr3 | 115921184 | 116463476 | 542293 | 7.75 | 0.000765 | 0.000978 |
| chr9 | 109587091 | 109598973 | 11883 | -2.64 | 0.000766 | 0.000979 |
| chr13 | 90145614 | 90146080 | 467 | -4.21 | 0.000766 | 0.000979 |

| | | | | | | |
|---|---|---|---|---|---|---|
| chr14 | 90327858 | 90335764 | 7907 | -4.21 | 0.000766 | 0.000979 |
| chr11 | 46035846 | 46036227 | 382 | -4.28 | 0.000767 | 0.00098 |
| chr1 | 34007549 | 34064130 | 56582 | -1.18 | 0.00077 | 0.000984 |
| chr12 | 9367914 | 9403674 | 35761 | 1.23 | 0.000771 | 0.000985 |
| chrX | 40606984 | 40613546 | 6563 | -4.56 | 0.000774 | 0.000989 |
| chr16 | 28084741 | 28085043 | 303 | -4.07 | 0.000774 | 0.000989 |
| chr10 | 128473224 | 128527318 | 54095 | -1.31 | 0.000775 | 0.00099 |
| chr12 | 44101654 | 44105766 | 4113 | 4.31 | 0.000776 | 0.000992 |
| chr8 | 104224591 | 104246158 | 21568 | -1.9 | 0.000777 | 0.000992 |
| chr18 | 70676010 | 70683716 | 7707 | 1.35 | 0.000778 | 0.000993 |
| chr11 | 27247280 | 27247739 | 460 | -4.07 | 0.000778 | 0.000994 |
| chr4 | 26001506 | 26057905 | 56400 | -2.08 | 0.00078 | 0.000996 |
| chr4 | 22640851 | 22655052 | 14202 | -4.17 | 0.000781 | 0.000998 |
| chr19 | 34606324 | 34607930 | 1607 | -2.25 | 0.000784 | 0.001 |
| chr3 | 171223665 | 171224146 | 482 | -3.09 | 0.000784 | 0.001 |
| chr2 | 222711385 | 222760117 | 48733 | 1.62 | 0.000786 | 0.001 |
| chr8 | 43187687 | 43191202 | 3516 | 2.34 | 0.000786 | 0.001 |
| chr12 | 46587602 | 46592683 | 5082 | -3.91 | 0.000787 | 0.001 |
| chr1 | 117258840 | 117260659 | 1820 | -2.37 | 0.00079 | 0.00101 |
| chr12 | 88328424 | 88332188 | 3765 | -4.03 | 0.000791 | 0.00101 |
| chr6 | 20131950 | 20132443 | 494 | -4.25 | 0.000792 | 0.00101 |
| chr8 | 27359942 | 27360248 | 307 | -4.25 | 0.000792 | 0.00101 |
| chr16 | 19901989 | 19902532 | 544 | -4.4 | 0.000793 | 0.00101 |
| chr5 | 78873913 | 78874453 | 541 | -4.4 | 0.000793 | 0.00101 |
| chr8 | 94702637 | 94702992 | 356 | -4.15 | 0.000794 | 0.00101 |
| chr18 | 57297220 | 57298145 | 926 | -3.15 | 0.000797 | 0.00102 |
| chr13 | 101642526 | 101646776 | 4251 | -4.22 | 0.000797 | 0.00102 |
| chr4 | 28616299 | 28618587 | 2289 | -2.55 | 0.000797 | 0.00102 |
| chr16 | 86379634 | 86410658 | 31025 | -4.95 | 0.000798 | 0.00102 |
| chr21 | 27889813 | 27891558 | 1746 | -2.75 | 0.000798 | 0.00102 |
| chr11 | 19439019 | 19439872 | 854 | -3.21 | 0.000799 | 0.00102 |
| chr8 | 119628855 | 119629794 | 940 | -3.69 | 8.00E-04 | 0.00102 |
| chr3 | 170222886 | 170226398 | 3513 | -1.93 | 0.000801 | 0.00102 |
| chr15 | 51390618 | 51397675 | 7058 | 1.04 | 0.000803 | 0.00102 |
| chr3 | 60157562 | 60159087 | 1526 | 3.2 | 0.000806 | 0.00103 |
| chr18 | 37078165 | 37079108 | 944 | -3.33 | 0.000807 | 0.00103 |
| chrX | 75166418 | 75178391 | 11974 | 2.25 | 0.000808 | 0.00103 |
| chr4 | 163979559 | 163997822 | 18264 | -4.22 | 0.00081 | 0.00103 |
| chr6 | 15771665 | 15772553 | 889 | 3 | 0.000812 | 0.00104 |
| chr3 | 50731402 | 50740286 | 8885 | -4.51 | 0.000813 | 0.00104 |
| chrX | 110113657 | 110133358 | 19702 | 1.9 | 0.000813 | 0.00104 |
| chr8 | 66476728 | 66492061 | 15334 | -4.42 | 0.000815 | 0.00104 |
| chr12 | 117833974 | 117842507 | 8534 | -4.45 | 0.000817 | 0.00104 |
| chr16 | 72193351 | 72224102 | 30752 | -1.07 | 0.000818 | 0.00104 |
| chr2 | 68798814 | 68799126 | 313 | -4.22 | 0.00082 | 0.00105 |
| chr3 | 115349550 | 115353316 | 3767 | 1.73 | 0.000821 | 0.00105 |
| chr3 | 140913523 | 140917215 | 3693 | -1.11 | 0.000822 | 0.00105 |
| chr14 | 34343469 | 34364979 | 21511 | -4.76 | 0.000822 | 0.00105 |
| chr12 | 47826650 | 47837357 | 10708 | -2.19 | 0.000824 | 0.00105 |
| chr11 | 49442472 | 49456289 | 13818 | 1.58 | 0.000827 | 0.00105 |
| chr6 | 49241946 | 49242964 | 1019 | -2.25 | 0.000827 | 0.00105 |
| chr10 | 118875176 | 118879954 | 4779 | -3.52 | 0.000829 | 0.00106 |
| chr10 | 64165784 | 64173196 | 7413 | -4.64 | 0.000829 | 0.00106 |
| chrX | 20304572 | 20307147 | 2576 | -3.17 | 0.00083 | 0.00106 |
| chr6 | 25038459 | 25038839 | 381 | -4.04 | 0.000831 | 0.00106 |
| chr1 | 29746280 | 29758802 | 12523 | -4.62 | 0.000834 | 0.00106 |
| chr4 | 186446678 | 186448515 | 1838 | -3.47 | 0.000834 | 0.00106 |
| chrX | 29394510 | 29398709 | 4200 | -4.14 | 0.000836 | 0.00107 |
| chr2 | 186674615 | 186684902 | 10288 | -1.35 | 0.000836 | 0.00107 |
| chr7 | 26053882 | 26059486 | 5605 | -1.77 | 0.000836 | 0.00107 |
| chr13 | 60830371 | 60837959 | 7589 | -1.98 | 0.000837 | 0.00107 |

| | | | | | | |
|---|---|---|---|---|---|---|
| chr7 | 30254461 | 30282035 | 27575 | -4.82 | 0.000837 | 0.00107 |
| chr9 | 87374918 | 87407441 | 32524 | -4.37 | 0.000838 | 0.00107 |
| chr12 | 48073184 | 48073614 | 431 | -3.32 | 0.000839 | 0.00107 |
| chr1 | 230292727 | 230303613 | 10887 | -4.51 | 0.00084 | 0.00107 |
| chrX | 35222009 | 35222426 | 418 | -4.03 | 0.00084 | 0.00107 |
| chr5 | 148455181 | 148456091 | 911 | -1.88 | 0.000843 | 0.00108 |
| chr4 | 121439612 | 121440213 | 602 | -3.07 | 0.000845 | 0.00108 |
| chr5 | 54167319 | 54167862 | 544 | 3.43 | 0.000845 | 0.00108 |
| chr6 | 130122537 | 130126704 | 4168 | -1.43 | 0.000846 | 0.00108 |
| chr15 | 50186882 | 50207571 | 20690 | -1.52 | 0.000847 | 0.00108 |
| chr1 | 190713364 | 190963655 | 250292 | 0.98 | 0.000847 | 0.00108 |
| chr10 | 26710627 | 26713812 | 3186 | -2.25 | 0.000848 | 0.00108 |
| chr5 | 173447401 | 173465648 | 18248 | -3.63 | 0.000848 | 0.00108 |
| chrX | 4427823 | 4438964 | 11142 | 4.23 | 0.000849 | 0.00108 |
| chr7 | 123532103 | 123534616 | 2514 | -1.71 | 0.000851 | 0.00108 |
| chr4 | 2433640 | 2434016 | 377 | -4.19 | 0.000852 | 0.00109 |
| chr22 | 22714034 | 22716626 | 2593 | -2.98 | 0.000853 | 0.00109 |
| chr18 | 13764290 | 13790124 | 25835 | -4.75 | 0.000853 | 0.00109 |
| chr5 | 130281930 | 130298876 | 16947 | -4.67 | 0.000854 | 0.00109 |
| chr5 | 95703413 | 95704752 | 1340 | -3.67 | 0.000854 | 0.00109 |
| chr2 | 230880698 | 230881226 | 529 | -4.02 | 0.000855 | 0.00109 |
| chr11 | 36211417 | 36211778 | 362 | -4.06 | 0.000855 | 0.00109 |
| chr5 | 149803465 | 149803893 | 429 | -4.06 | 0.000855 | 0.00109 |
| chr3 | 174408509 | 174409345 | 837 | -4.31 | 0.000856 | 0.00109 |
| chr4 | 56723696 | 56724043 | 348 | -4.31 | 0.000856 | 0.00109 |
| chr5 | 175709358 | 175709718 | 361 | -4.31 | 0.000856 | 0.00109 |
| chr9 | 132633597 | 132634132 | 536 | -4.48 | 0.000858 | 0.00109 |
| chr4 | 155300228 | 155301378 | 1151 | -2.25 | 0.000858 | 0.00109 |
| chr7 | 31394258 | 31397215 | 2958 | -3.69 | 0.000859 | 0.00109 |
| chr9 | 2295063 | 2389062 | 94000 | 0.96 | 0.00086 | 0.0011 |
| chr1 | 12906109 | 12911093 | 4985 | -1.47 | 0.00086 | 0.0011 |
| chr5 | 83776434 | 83779307 | 2874 | 4.05 | 0.000861 | 0.0011 |
| chr5 | 106067074 | 106071884 | 4811 | -2.15 | 0.000862 | 0.0011 |
| chr6 | 119446336 | 119465184 | 18849 | -3.24 | 0.000866 | 0.0011 |
| chr8 | 109185019 | 109186997 | 1979 | -3.83 | 0.000867 | 0.0011 |
| chrX | 124398846 | 124400752 | 1907 | -1.52 | 0.000868 | 0.00111 |
| chr10 | 117717917 | 117725898 | 7982 | 2.05 | 0.000869 | 0.00111 |
| chr16 | 63740030 | 63760184 | 20155 | 1.25 | 0.000869 | 0.00111 |
| chrX | 80149162 | 80151986 | 2825 | 2.29 | 0.000871 | 0.00111 |
| chr15 | 47368244 | 47381542 | 13299 | 1.92 | 0.000872 | 0.00111 |
| chr3 | 98076507 | 98079173 | 2667 | -1.58 | 0.000874 | 0.00111 |
| chr1 | 4985639 | 5016044 | 30406 | 0.69 | 0.000875 | 0.00111 |
| chr3 | 157496186 | 157522809 | 26624 | -4.36 | 0.000877 | 0.00112 |
| chr12 | 63401479 | 63406705 | 5227 | 2.93 | 0.000877 | 0.00112 |
| chr3 | 152983948 | 152984901 | 954 | -4.81 | 0.000879 | 0.00112 |
| chr9 | 4791310 | 4791789 | 480 | -4.28 | 0.000881 | 0.00112 |
| chr2 | 129326868 | 129347161 | 20294 | -4.59 | 0.000881 | 0.00112 |
| chr11 | 4582993 | 4586602 | 3610 | -1.22 | 0.000884 | 0.00112 |
| chr4 | 19989027 | 19989382 | 356 | -4.08 | 0.000885 | 0.00113 |
| chr1 | 153341347 | 153356104 | 14758 | -0.99 | 0.000887 | 0.00113 |
| chr14 | 73317470 | 73324284 | 6815 | -2.02 | 0.000889 | 0.00113 |
| chr7 | 87299133 | 87304943 | 5811 | 2.07 | 0.00089 | 0.00113 |
| chr1 | 18692696 | 18708122 | 15427 | -3.86 | 0.000892 | 0.00114 |
| chr9 | 94524181 | 94531419 | 7239 | -4.75 | 0.000894 | 0.00114 |
| chr6 | 85726649 | 85728028 | 1380 | -2.38 | 0.000895 | 0.00114 |
| chr18 | 2070495 | 2076176 | 5682 | 4.22 | 0.000897 | 0.00114 |
| chr1 | 62832427 | 62845724 | 13298 | -4.95 | 0.000897 | 0.00114 |
| chr4 | 29578839 | 29582861 | 4023 | 1.7 | 0.000898 | 0.00114 |
| chrX | 120497229 | 120497702 | 474 | 3.32 | 9.00E-04 | 0.00115 |
| chr1 | 191991705 | 192126472 | 134768 | 1.24 | 0.000903 | 0.00115 |
| chr3 | 153770366 | 153771294 | 929 | -1.61 | 0.000905 | 0.00115 |

| | | | | | | |
|---|---|---|---|---|---|---|
| chr2 | 26844276 | 26849265 | 4990 | -5.64 | 0.000906 | 0.00115 |
| chr6 | 144353111 | 144355572 | 2462 | -4.06 | 0.000906 | 0.00115 |
| chr19 | 6547223 | 6547633 | 411 | -4.25 | 0.000908 | 0.00115 |
| chr1 | 104824895 | 104826477 | 1583 | 2.29 | 0.000908 | 0.00115 |
| chr7 | 42048157 | 42061795 | 13639 | -4.49 | 0.000908 | 0.00116 |
| chr17 | 71437550 | 71482530 | 44981 | -7.52 | 0.000911 | 0.00116 |
| chr10 | 12384957 | 12385417 | 461 | -4.2 | 0.000913 | 0.00116 |
| chr8 | 132972133 | 132973317 | 1185 | -1.81 | 0.000914 | 0.00116 |
| chr8 | 82155378 | 82157371 | 1994 | -1.66 | 0.000915 | 0.00116 |
| chr6 | 87391475 | 87418444 | 26970 | 1.18 | 0.000916 | 0.00116 |
| chrX | 11044844 | 11045570 | 727 | -4.59 | 0.000918 | 0.00117 |
| chr11 | 64252687 | 64292183 | 39497 | -4.92 | 0.000919 | 0.00117 |
| chr8 | 98646766 | 98647202 | 437 | -4.4 | 0.000919 | 0.00117 |
| chr13 | 91732939 | 91734247 | 1309 | -2.39 | 0.000922 | 0.00117 |
| chr12 | 43878083 | 43881022 | 2940 | -1.57 | 0.000926 | 0.00118 |
| chr1 | 111365107 | 111366355 | 1249 | 3.52 | 0.000928 | 0.00118 |
| chr5 | 123220570 | 123230274 | 9705 | -2.57 | 0.00093 | 0.00118 |
| chr4 | 70343977 | 70344747 | 771 | -3.14 | 0.000931 | 0.00118 |
| chr2 | 150793606 | 150793955 | 350 | -4 | 0.000937 | 0.00119 |
| chr1 | 42021088 | 42022940 | 1853 | -3.83 | 0.000939 | 0.00119 |
| chrX | 48290112 | 48297873 | 7762 | -3.14 | 0.000941 | 0.0012 |
| chr4 | 177175003 | 177175446 | 444 | -3.99 | 0.000941 | 0.0012 |
| chr1 | 207319569 | 207320007 | 439 | 2.38 | 0.000942 | 0.0012 |
| chr5 | 54451477 | 54452335 | 859 | -2.4 | 0.000942 | 0.0012 |
| chr1 | 65375868 | 65376272 | 405 | -4.05 | 0.000943 | 0.0012 |
| chr5 | 31584425 | 31628686 | 44262 | -1.85 | 0.000947 | 0.0012 |
| chr22 | 49354213 | 49358498 | 4286 | -3.19 | 0.000947 | 0.0012 |
| chr4 | 146928151 | 146931399 | 3249 | -1.79 | 0.000948 | 0.00121 |
| chr10 | 90314940 | 90322911 | 7972 | -5.37 | 0.000949 | 0.00121 |
| chr3 | 122742582 | 122744477 | 1896 | -1.5 | 0.00095 | 0.00121 |
| chr4 | 65344379 | 65349199 | 4821 | -1.87 | 0.00095 | 0.00121 |
| chr1 | 103914152 | 103920050 | 5899 | 1.26 | 0.00095 | 0.00121 |
| chr12 | 78912728 | 78913547 | 820 | -2.04 | 0.00095 | 0.00121 |
| chr16 | 14460044 | 14460814 | 771 | -4.54 | 0.000955 | 0.00121 |
| chr2 | 188148000 | 188148610 | 611 | 2.22 | 0.000955 | 0.00121 |
| chr8 | 19760727 | 19778531 | 17805 | -1.99 | 0.000955 | 0.00121 |
| chr1 | 84249194 | 84250204 | 1011 | -1.88 | 0.000956 | 0.00121 |
| chr3 | 158571165 | 158576795 | 5631 | -4.07 | 0.000957 | 0.00122 |
| chr12 | 97394990 | 97729397 | 334408 | 7.61 | 0.000959 | 0.00122 |
| chr2 | 38397650 | 38398927 | 1278 | -2.2 | 0.00096 | 0.00122 |
| chr13 | 106905986 | 106908334 | 2349 | -1.7 | 0.00096 | 0.00122 |
| chr2 | 224457510 | 224457792 | 283 | -4 | 0.00096 | 0.00122 |
| chr5 | 135242316 | 135265229 | 22914 | -0.95 | 0.000961 | 0.00122 |
| chr6 | 81886529 | 81941849 | 55321 | 2.01 | 0.000963 | 0.00122 |
| chr9 | 85433594 | 85453844 | 20251 | -4.44 | 0.000963 | 0.00122 |
| chr2 | 136931126 | 136989139 | 58014 | -3.22 | 0.000965 | 0.00123 |
| chr5 | 38602883 | 38603350 | 468 | -4.35 | 0.000965 | 0.00123 |
| chr8 | 99699314 | 99699722 | 409 | -4.35 | 0.000965 | 0.00123 |
| chr16 | 64349495 | 64364284 | 14790 | 1.83 | 0.000966 | 0.00123 |
| chr20 | 22273473 | 22283083 | 9611 | -1.36 | 0.000968 | 0.00123 |
| chr2 | 217631560 | 217637495 | 5936 | -2.15 | 0.000969 | 0.00123 |
| chr21 | 19437146 | 19439123 | 1978 | 1.46 | 0.000974 | 0.00124 |
| chr11 | 123534015 | 123570955 | 36941 | -1.41 | 0.000974 | 0.00124 |
| chr15 | 81338314 | 81391152 | 52839 | -4.78 | 0.000978 | 0.00124 |
| chr5 | 63469740 | 63498848 | 29109 | 1.17 | 0.000979 | 0.00124 |
| chr2 | 195840588 | 195842337 | 1750 | 1.73 | 0.000979 | 0.00124 |
| chr7 | 106330659 | 106370629 | 39971 | -3.23 | 0.00098 | 0.00124 |
| chrX | 148772991 | 148785179 | 12189 | -2.59 | 0.00098 | 0.00125 |
| chrX | 114584595 | 114585211 | 617 | -4.68 | 0.000983 | 0.00125 |
| chr10 | 16004542 | 16426425 | 421884 | -1.03 | 0.000983 | 0.00125 |
| chr1 | 241380124 | 241387904 | 7781 | -1.15 | 0.000985 | 0.00125 |

| | | | | | | |
|---|---|---|---|---|---|---|
| chr17 | 4548388 | 4549550 | 1163 | -1.98 | 0.000989 | 0.00126 |
| chrX | 121461671 | 121476363 | 14693 | 1.62 | 0.000991 | 0.00126 |
| chr1 | 163246582 | 163248730 | 2149 | -2.93 | 0.000993 | 0.00126 |
| chr2 | 145611685 | 145613010 | 1326 | -3.68 | 0.000995 | 0.00126 |
| chr5 | 59798202 | 59799174 | 973 | -2.18 | 0.000996 | 0.00127 |
| chr2 | 143991794 | 143992426 | 633 | -3.19 | 0.000997 | 0.00127 |
| chr19 | 39291478 | 39292225 | 748 | 1.71 | 0.001 | 0.00127 |
| chr13 | 113059161 | 113095924 | 36764 | -4.3 | 0.001 | 0.00127 |
| chr7 | 69581862 | 69617938 | 36077 | -0.73 | 0.001 | 0.00127 |
| chr2 | 180562301 | 180562641 | 341 | -4.16 | 0.00101 | 0.00128 |
| chr16 | 13004692 | 14006955 | 1002264 | 0.2 | 0.00101 | 0.00128 |
| chr12 | 19427525 | 19428047 | 523 | -3.15 | 0.00101 | 0.00128 |
| chr4 | 35494848 | 35497322 | 2475 | -1.38 | 0.00101 | 0.00128 |
| chr11 | 13550062 | 13569846 | 19785 | -1.65 | 0.00101 | 0.00128 |
| chr1 | 178239222 | 178247255 | 8034 | -4.5 | 0.00101 | 0.00128 |
| chr15 | 61525472 | 61528679 | 3208 | -1.02 | 0.00101 | 0.00128 |
| chr11 | 64343776 | 64344081 | 306 | 4.12 | 0.00101 | 0.00128 |
| chr6 | 9711296 | 9713584 | 2289 | -3.68 | 0.00101 | 0.00128 |
| chr13 | 41007830 | 41008252 | 423 | -4.01 | 0.00101 | 0.00128 |
| chr4 | 31697541 | 31697950 | 410 | -4.01 | 0.00101 | 0.00128 |
| chr10 | 61012637 | 61033737 | 21101 | 1.67 | 0.00101 | 0.00129 |
| chr15 | 96678504 | 96699072 | 20569 | -4.7 | 0.00102 | 0.00129 |
| chrX | 72941382 | 72947833 | 6452 | 1.23 | 0.00102 | 0.00129 |
| chr2 | 7677547 | 7814521 | 136975 | -0.69 | 0.00102 | 0.00129 |
| chr4 | 100955576 | 100970564 | 14989 | -3.92 | 0.00102 | 0.00129 |
| chr1 | 99726063 | 99741321 | 15259 | -1.29 | 0.00102 | 0.00129 |
| chrX | 101958630 | 101958939 | 310 | -3.7 | 0.00102 | 0.00129 |
| chr10 | 54447536 | 54455577 | 8042 | -0.96 | 0.00102 | 0.0013 |
| chr1 | 56569288 | 56594069 | 24782 | -1.37 | 0.00102 | 0.0013 |
| chrX | 130786838 | 130797777 | 10940 | -0.8 | 0.00102 | 0.0013 |
| chr4 | 81352862 | 81356119 | 3258 | 3.96 | 0.00102 | 0.0013 |
| chr2 | 153889617 | 153896170 | 6554 | -1.47 | 0.00102 | 0.0013 |
| chr4 | 145305570 | 145312920 | 7351 | -2.94 | 0.00103 | 0.0013 |
| chr3 | 170467244 | 170468554 | 1311 | -2.95 | 0.00103 | 0.0013 |
| chr4 | 179637169 | 179637563 | 395 | -4.05 | 0.00103 | 0.0013 |
| chr21 | 33165837 | 33169518 | 3682 | 1.27 | 0.00103 | 0.0013 |
| chr11 | 89306358 | 89306799 | 442 | 2.96 | 0.00103 | 0.00131 |
| chr9 | 14431667 | 14451032 | 19366 | -4.58 | 0.00103 | 0.00131 |
| chr1 | 199417938 | 199419078 | 1141 | 3.38 | 0.00103 | 0.00131 |
| chr8 | 62048119 | 62049502 | 1384 | -1.88 | 0.00103 | 0.00131 |
| chr14 | 77969543 | 77969958 | 416 | -4.1 | 0.00104 | 0.00132 |
| chr14 | 34732234 | 34737994 | 5761 | -2.03 | 0.00104 | 0.00132 |
| chr13 | 28257980 | 28281930 | 23951 | -2.36 | 0.00104 | 0.00132 |
| chr8 | 107301178 | 107304190 | 3013 | -1.45 | 0.00104 | 0.00132 |
| chr2 | 7861091 | 7863123 | 2033 | -2.3 | 0.00104 | 0.00132 |
| chr12 | 39325787 | 39352043 | 26257 | -2.67 | 0.00104 | 0.00132 |
| chr7 | 156268849 | 156270489 | 1641 | -1.88 | 0.00104 | 0.00132 |
| chr15 | 67274727 | 67302060 | 27334 | -7.42 | 0.00105 | 0.00133 |
| chr15 | 93393109 | 93406042 | 12934 | -6.55 | 0.00105 | 0.00133 |
| chr4 | 14428207 | 14429390 | 1184 | -2.76 | 0.00105 | 0.00133 |
| chrX | 65912913 | 65914651 | 1739 | 1.78 | 0.00105 | 0.00133 |
| chrX | 150490637 | 150491728 | 1092 | 2.92 | 0.00105 | 0.00133 |
| chr1 | 199807978 | 199808370 | 393 | -3.97 | 0.00105 | 0.00133 |
| chr6 | 71068244 | 71068641 | 398 | -3.01 | 0.00106 | 0.00134 |
| chr2 | 166923330 | 166926499 | 3170 | 2.45 | 0.00106 | 0.00134 |
| chr2 | 84098458 | 84107976 | 9519 | -2.79 | 0.00106 | 0.00134 |
| chr7 | 141873222 | 141875878 | 2657 | -1.78 | 0.00106 | 0.00134 |
| chr2 | 74795882 | 74796423 | 542 | -3.14 | 0.00107 | 0.00135 |
| chr6 | 132893597 | 132903947 | 10351 | -2.3 | 0.00107 | 0.00135 |
| chr13 | 30185397 | 30208532 | 23136 | -0.9 | 0.00107 | 0.00136 |
| chr1 | 117865541 | 117868201 | 2661 | 2.43 | 0.00107 | 0.00136 |

| | | | | | | |
|---|---|---|---|---|---|---|
| chr3 | 117712183 | 117770379 | 58197 | -4.98 | 0.00107 | 0.00136 |
| chr6 | 8737796 | 8749180 | 11385 | -1.71 | 0.00107 | 0.00136 |
| chr7 | 35784550 | 35791805 | 7256 | 3.39 | 0.00107 | 0.00136 |
| chr1 | 184872640 | 184874294 | 1655 | -3.33 | 0.00107 | 0.00136 |
| chr12 | 40644445 | 40644807 | 363 | -3.07 | 0.00107 | 0.00136 |
| chr10 | 129998498 | 130002639 | 4142 | -4.16 | 0.00108 | 0.00136 |
| chr14 | 73031259 | 73061560 | 30302 | 0.97 | 0.00108 | 0.00136 |
| chr14 | 94660290 | 94701851 | 41562 | -4.99 | 0.00108 | 0.00136 |
| chr14 | 99851631 | 99852003 | 373 | -4.23 | 0.00108 | 0.00137 |
| chr1 | 66020971 | 66022370 | 1400 | -3.36 | 0.00108 | 0.00137 |
| chr10 | 77289067 | 77289575 | 509 | -3.05 | 0.00108 | 0.00137 |
| chr12 | 18779407 | 18790848 | 11442 | -3.82 | 0.00108 | 0.00137 |
| chr13 | 25621899 | 25640134 | 18236 | -2.66 | 0.00109 | 0.00137 |
| chr8 | 43252113 | 43266791 | 14679 | -1.91 | 0.00109 | 0.00138 |
| chr4 | 155239205 | 155247471 | 8267 | -1.77 | 0.00109 | 0.00138 |
| chr1 | 89670058 | 89677920 | 7863 | -1.17 | 0.00109 | 0.00138 |
| chr11 | 102145143 | 102145672 | 530 | -4.09 | 0.00109 | 0.00138 |
| chr19 | 47292286 | 47292657 | 372 | -4.02 | 0.00109 | 0.00138 |
| chr13 | 83966276 | 83968237 | 1962 | -3.35 | 0.00109 | 0.00138 |
| chr15 | 36126661 | 36127932 | 1272 | -3.58 | 0.00109 | 0.00138 |
| chr3 | 187015289 | 187022121 | 6833 | -1.12 | 0.00109 | 0.00138 |
| chr9 | 32936315 | 32939185 | 2871 | -3.2 | 0.00109 | 0.00138 |
| chr14 | 65350447 | 65351646 | 1200 | 2.36 | 0.00109 | 0.00139 |
| chr20 | 43771089 | 43799272 | 28184 | -1.26 | 0.00109 | 0.00139 |
| chr16 | 52300588 | 52314923 | 14336 | 1.08 | 0.0011 | 0.00139 |
| chrX | 148530912 | 148532203 | 1292 | -1.52 | 0.0011 | 0.00139 |
| chr6 | 169079764 | 169098403 | 18640 | -1.84 | 0.0011 | 0.00139 |
| chr1 | 66785655 | 66788434 | 2780 | -2.22 | 0.0011 | 0.00139 |
| chr8 | 52964441 | 52987468 | 23028 | -1.19 | 0.0011 | 0.0014 |
| chr4 | 35478931 | 35490688 | 11758 | 1.51 | 0.00111 | 0.0014 |
| chr5 | 59616196 | 59625021 | 8826 | -3.63 | 0.00111 | 0.0014 |
| chr10 | 102538880 | 102539596 | 717 | -4.54 | 0.00111 | 0.0014 |
| chr4 | 48973523 | 48993212 | 19690 | -4.34 | 0.00111 | 0.00141 |
| chr3 | 184194146 | 184196295 | 2150 | -4.1 | 0.00111 | 0.00141 |
| chr4 | 56094098 | 56106694 | 12597 | -1.16 | 0.00112 | 0.00141 |
| chr12 | 21811354 | 21811899 | 546 | -4.35 | 0.00112 | 0.00142 |
| chr15 | 62547718 | 62587534 | 39817 | -2.18 | 0.00112 | 0.00142 |
| chr8 | 63471053 | 63481826 | 10774 | 4.28 | 0.00112 | 0.00142 |
| chr2 | 84512320 | 84513051 | 732 | -3.79 | 0.00112 | 0.00142 |
| chrX | 30339804 | 30341301 | 1498 | -1.94 | 0.00112 | 0.00142 |
| chr4 | 19497614 | 19497930 | 317 | -3.95 | 0.00112 | 0.00142 |
| chr3 | 130310200 | 130347734 | 37535 | -0.79 | 0.00112 | 0.00142 |
| chr16 | 71638042 | 71638385 | 344 | -3.04 | 0.00113 | 0.00143 |
| chr16 | 73994761 | 74058126 | 63366 | 0.69 | 0.00113 | 0.00143 |
| chr10 | 9076944 | 9100383 | 23440 | -4.85 | 0.00113 | 0.00143 |
| chr9 | 17805505 | 17807065 | 1561 | 3.84 | 0.00113 | 0.00143 |
| chr11 | 62932235 | 62934701 | 2467 | 1.63 | 0.00113 | 0.00143 |
| chr9 | 87149645 | 87156335 | 6691 | -1.06 | 0.00113 | 0.00143 |
| chrX | 66619762 | 66652207 | 32446 | -1.24 | 0.00113 | 0.00143 |
| chr2 | 105248519 | 105287949 | 39431 | 2.88 | 0.00114 | 0.00144 |
| chr6 | 142244020 | 142244544 | 525 | -2.23 | 0.00114 | 0.00144 |
| chr4 | 20894957 | 20895371 | 415 | -3.15 | 0.00114 | 0.00144 |
| chr14 | 73139096 | 73141635 | 2540 | 2.16 | 0.00114 | 0.00144 |
| chr6 | 88321804 | 88322902 | 1099 | -2.39 | 0.00114 | 0.00144 |
| chr4 | 76005398 | 76024899 | 19502 | -4.35 | 0.00114 | 0.00144 |
| chr18 | 45017530 | 45040997 | 23468 | 2.21 | 0.00114 | 0.00144 |
| chr8 | 37094837 | 37111123 | 16287 | -2.61 | 0.00114 | 0.00144 |
| chr18 | 59573748 | 59582321 | 8574 | 4.21 | 0.00114 | 0.00145 |
| chr2 | 206504444 | 206505534 | 1091 | -3.57 | 0.00115 | 0.00145 |
| chr9 | 856943 | 897668 | 40726 | -4.63 | 0.00115 | 0.00146 |
| chr5 | 30017916 | 30018365 | 450 | -2.38 | 0.00115 | 0.00146 |

| | | | | | | |
|---|---|---|---|---|---|---|
| chr5 | 127571461 | 127572479 | 1019 | 2.41 | 0.00116 | 0.00146 |
| chr11 | 114105315 | 114106137 | 823 | -3.45 | 0.00116 | 0.00146 |
| chr1 | 91759832 | 91776037 | 16206 | -3.27 | 0.00116 | 0.00146 |
| chr11 | 121123787 | 121143835 | 20049 | -1.39 | 0.00116 | 0.00147 |
| chr13 | 79148927 | 79149672 | 746 | -2.22 | 0.00116 | 0.00147 |
| chr1 | 191811732 | 191817391 | 5660 | -3.7 | 0.00116 | 0.00147 |
| chr6 | 80561883 | 80564000 | 2118 | -1.7 | 0.00116 | 0.00147 |
| chr2 | 226209708 | 226217343 | 7636 | -1.15 | 0.00116 | 0.00147 |
| chr6 | 162059192 | 162085357 | 26166 | -3.95 | 0.00116 | 0.00147 |
| chr18 | 13829139 | 13866726 | 37588 | -3.81 | 0.00116 | 0.00147 |
| chr2 | 84412216 | 84412798 | 583 | 1.87 | 0.00117 | 0.00147 |
| chr18 | 54721663 | 54722003 | 341 | -3.93 | 0.00117 | 0.00147 |
| chr12 | 16466413 | 16468251 | 1839 | 3.88 | 0.00117 | 0.00147 |
| chr22 | 40326181 | 40327697 | 1517 | 3.8 | 0.00117 | 0.00147 |
| chr3 | 69442880 | 69455821 | 12942 | -4.51 | 0.00117 | 0.00147 |
| chr1 | 48401606 | 48402326 | 721 | 3.44 | 0.00117 | 0.00147 |
| chr9 | 85229814 | 85419439 | 189626 | -0.4 | 0.00117 | 0.00148 |
| chr9 | 37674876 | 37696043 | 21168 | -1.28 | 0.00117 | 0.00148 |
| chrX | 35265343 | 35268139 | 2797 | -3.17 | 0.00117 | 0.00148 |
| chr18 | 42007685 | 42092950 | 85266 | 0.9 | 0.00117 | 0.00148 |
| chr11 | 85117831 | 85118835 | 1005 | 3.34 | 0.00117 | 0.00148 |
| chr11 | 78933086 | 78936828 | 3743 | -2.22 | 0.00117 | 0.00148 |
| chr6 | 23171031 | 23172940 | 1910 | -3.38 | 0.00117 | 0.00148 |
| chr13 | 105690999 | 105696877 | 5879 | -3.36 | 0.00118 | 0.00149 |
| chrX | 72666583 | 72669781 | 3199 | 1.38 | 0.00118 | 0.00149 |
| chr3 | 192823888 | 192829079 | 5192 | 4.16 | 0.00118 | 0.00149 |
| chr16 | 59287560 | 59289386 | 1827 | -3.85 | 0.00118 | 0.00149 |
| chr13 | 82120131 | 82121872 | 1742 | -2.75 | 0.00118 | 0.00149 |
| chr12 | 11575017 | 11655453 | 80437 | 1.36 | 0.00118 | 0.00149 |
| chr8 | 125062565 | 125064706 | 2142 | -3.42 | 0.00118 | 0.00149 |
| chr18 | 60787486 | 60811909 | 24424 | -4.37 | 0.00118 | 0.00149 |
| chr16 | 72014306 | 72015297 | 992 | -3.25 | 0.00118 | 0.00149 |
| chr8 | 706424 | 777425 | 71002 | -1.68 | 0.00118 | 0.00149 |
| chrX | 67109695 | 67124971 | 15277 | -1.11 | 0.00119 | 0.0015 |
| chr21 | 16002557 | 16092780 | 90224 | -0.64 | 0.00119 | 0.0015 |
| chrX | 55700156 | 55701967 | 1812 | 1.98 | 0.00119 | 0.0015 |
| chr8 | 83618311 | 83619342 | 1032 | -4.47 | 0.00119 | 0.0015 |
| chr11 | 115555503 | 115556318 | 816 | -3.37 | 0.00119 | 0.0015 |
| chr5 | 134838705 | 134874314 | 35610 | -4.93 | 0.00119 | 0.00151 |
| chr4 | 26545452 | 26564485 | 19034 | -4.48 | 0.00119 | 0.00151 |
| chr11 | 63035501 | 63052309 | 16809 | 2.21 | 0.00119 | 0.00151 |
| chr6 | 154222147 | 154224192 | 2046 | 3.72 | 0.0012 | 0.00151 |
| chr5 | 1669322 | 1722321 | 53000 | -4.38 | 0.00121 | 0.00152 |
| chr2 | 100328893 | 100397592 | 68700 | -1.32 | 0.00121 | 0.00152 |
| chr8 | 58070181 | 58190229 | 120049 | -0.54 | 0.00121 | 0.00152 |
| chrX | 56536025 | 56536921 | 897 | -1.89 | 0.00121 | 0.00153 |
| chr4 | 22281425 | 22281987 | 563 | 2.15 | 0.00121 | 0.00153 |
| chr5 | 18081807 | 18090261 | 8455 | -1.79 | 0.00121 | 0.00153 |
| chr8 | 134233650 | 134234472 | 823 | 2.56 | 0.00121 | 0.00153 |
| chr16 | 52423750 | 52475359 | 51610 | -4.79 | 0.00121 | 0.00153 |
| chr4 | 26158922 | 26162194 | 3273 | -2.45 | 0.00122 | 0.00154 |
| chr11 | 108381133 | 108385627 | 4495 | -1.3 | 0.00122 | 0.00154 |
| chr4 | 60670401 | 60670733 | 333 | -3.92 | 0.00122 | 0.00154 |
| chr5 | 149700780 | 149701246 | 467 | -3.92 | 0.00122 | 0.00154 |
| chr5 | 27702283 | 27716543 | 14261 | -1.59 | 0.00122 | 0.00154 |
| chrX | 13914744 | 13921282 | 6539 | -1.39 | 0.00122 | 0.00154 |
| chr4 | 122939709 | 122946027 | 6319 | -4.11 | 0.00122 | 0.00154 |
| chr4 | 78114003 | 78147379 | 33377 | 1.36 | 0.00123 | 0.00155 |
| chr2 | 48640880 | 48641160 | 281 | -3.99 | 0.00123 | 0.00155 |
| chrX | 81394510 | 81396451 | 1942 | 1.89 | 0.00123 | 0.00155 |
| chr1 | 182659156 | 182688040 | 28885 | 4.62 | 0.00123 | 0.00155 |

| | | | | | | |
|---|---|---|---|---|---|---|
| chr8 | 140707440 | 140708719 | 1280 | 3.48 | 0.00123 | 0.00155 |
| chr12 | 28069118 | 28070128 | 1011 | -3.46 | 0.00124 | 0.00157 |
| chr4 | 22303738 | 22305810 | 2073 | -1.52 | 0.00124 | 0.00157 |
| chr3 | 193236950 | 193238213 | 1264 | 3.69 | 0.00125 | 0.00157 |
| chr12 | 12205233 | 12205793 | 561 | -3.3 | 0.00125 | 0.00157 |
| chrX | 90858441 | 90860048 | 1608 | 1.81 | 0.00125 | 0.00158 |
| chr11 | 18968763 | 18970396 | 1634 | -2.5 | 0.00125 | 0.00158 |
| chr6 | 81571704 | 81572775 | 1072 | 3.46 | 0.00125 | 0.00158 |
| chr21 | 27025793 | 27031129 | 5337 | -2.01 | 0.00125 | 0.00158 |
| chr7 | 50249324 | 50297364 | 48041 | -1.82 | 0.00126 | 0.00158 |
| chr4 | 66174494 | 66174870 | 377 | -3.41 | 0.00126 | 0.00159 |
| chr1 | 107996328 | 108015045 | 18718 | 2.67 | 0.00126 | 0.00159 |
| chr1 | 99251213 | 99252221 | 1009 | -2.64 | 0.00126 | 0.00159 |
| chr7 | 103167927 | 103187808 | 19882 | 1.86 | 0.00127 | 0.0016 |
| chr1 | 87023488 | 87025402 | 1915 | -1.8 | 0.00127 | 0.0016 |
| chr9 | 2666694 | 2667014 | 321 | -3.99 | 0.00127 | 0.0016 |
| chr18 | 10731775 | 10755374 | 23600 | 1.91 | 0.00127 | 0.00161 |
| chr4 | 181597490 | 181606836 | 9347 | 1.48 | 0.00128 | 0.00161 |
| chr1 | 91272722 | 91278516 | 5795 | 1.24 | 0.00128 | 0.00161 |
| chr2 | 99366561 | 99370042 | 3482 | -1.58 | 0.00128 | 0.00161 |
| chr18 | 69374330 | 69385735 | 11406 | 3.21 | 0.00128 | 0.00161 |
| chr4 | 22251867 | 22260578 | 8712 | 1.73 | 0.00128 | 0.00162 |
| chr3 | 177752709 | 177753176 | 468 | -3 | 0.00128 | 0.00162 |
| chr3 | 83904432 | 83914456 | 10025 | -1.67 | 0.00128 | 0.00162 |
| chr1 | 59554722 | 59567690 | 12969 | -5.33 | 0.00128 | 0.00162 |
| chr12 | 54161824 | 54162199 | 376 | -4.07 | 0.00129 | 0.00163 |
| chr19 | 51521593 | 51521960 | 368 | -4.07 | 0.00129 | 0.00163 |
| chr3 | 105838991 | 105839421 | 431 | -4.07 | 0.00129 | 0.00163 |
| chr16 | 53564581 | 53564949 | 369 | -3.96 | 0.00129 | 0.00163 |
| chr1 | 63344422 | 63347012 | 2591 | -4.24 | 0.00129 | 0.00163 |
| chr17 | 70005931 | 70048649 | 42719 | -0.59 | 0.00129 | 0.00163 |
| chr2 | 214611111 | 214611644 | 534 | 2.66 | 0.00129 | 0.00163 |
| chr13 | 105174238 | 105176203 | 1966 | -2.27 | 0.00129 | 0.00163 |
| chr2 | 35022510 | 35023941 | 1432 | -1.91 | 0.0013 | 0.00163 |
| chr5 | 1900512 | 1916009 | 15498 | -3.66 | 0.0013 | 0.00163 |
| chr8 | 64018762 | 64022390 | 3629 | -4.24 | 0.0013 | 0.00164 |
| chr16 | 80320839 | 80328626 | 7788 | -2.61 | 0.0013 | 0.00164 |
| chr8 | 67253720 | 67276877 | 23158 | -4.39 | 0.0013 | 0.00164 |
| chr12 | 28987355 | 28987741 | 387 | -3.11 | 0.0013 | 0.00164 |
| chr7 | 34350067 | 34472122 | 122056 | 1.45 | 0.00131 | 0.00165 |
| chr4 | 130940874 | 130942410 | 1537 | -2.61 | 0.00131 | 0.00166 |
| chr3 | 81413815 | 81416598 | 2784 | -2.1 | 0.00131 | 0.00166 |
| chrX | 21553310 | 21554619 | 1310 | -3.39 | 0.00131 | 0.00166 |
| chr3 | 82169545 | 82170551 | 1007 | 2.58 | 0.00132 | 0.00166 |
| chr11 | 125179470 | 125193012 | 13543 | -2.39 | 0.00132 | 0.00166 |
| chr10 | 1706778 | 1769570 | 62793 | -1.13 | 0.00132 | 0.00167 |
| chr4 | 92992346 | 92994170 | 1825 | -2.1 | 0.00132 | 0.00167 |
| chr11 | 94089771 | 94103650 | 13880 | -1.61 | 0.00132 | 0.00167 |
| chr19 | 15012656 | 15040561 | 27906 | -1.01 | 0.00132 | 0.00167 |
| chr5 | 73646960 | 73684845 | 37886 | -7.23 | 0.00133 | 0.00167 |
| chr4 | 148243574 | 148243983 | 410 | -4.17 | 0.00133 | 0.00167 |
| chrX | 115990536 | 115991007 | 472 | -4.17 | 0.00133 | 0.00167 |
| chr9 | 26734388 | 26747445 | 13058 | -2.95 | 0.00133 | 0.00167 |
| chr4 | 92534560 | 92534942 | 383 | -3.12 | 0.00133 | 0.00167 |
| chr9 | 89692554 | 89820410 | 127857 | 0.53 | 0.00133 | 0.00167 |
| chr1 | 215133973 | 215136953 | 2981 | -1.52 | 0.00133 | 0.00167 |
| chr10 | 83190955 | 83192781 | 1827 | 1.85 | 0.00133 | 0.00168 |
| chr15 | 96732705 | 96760322 | 27618 | -4.39 | 0.00133 | 0.00168 |
| chr11 | 91508443 | 91510570 | 2128 | 3.53 | 0.00133 | 0.00168 |
| chr12 | 40089683 | 40093013 | 3331 | 1.25 | 0.00133 | 0.00168 |
| chr5 | 93757838 | 93758168 | 331 | 4.15 | 0.00133 | 0.00168 |

| | | | | | | |
|---|---|---|---|---|---|---|
| chrX | 101044997 | 101053323 | 8327 | -2.98 | 0.00133 | 0.00168 |
| chr1 | 180777587 | 180809331 | 31745 | -6.01 | 0.00134 | 0.00168 |
| chr2 | 108747653 | 108749845 | 2193 | -3.08 | 0.00134 | 0.00169 |
| chrX | 72385464 | 72395824 | 10361 | 4.1 | 0.00134 | 0.00169 |
| chr18 | 30513315 | 30516044 | 2730 | -1.38 | 0.00134 | 0.00169 |
| chr6 | 47836683 | 47849398 | 12716 | 1.47 | 0.00134 | 0.00169 |
| chr19 | 43278826 | 43279377 | 552 | -1.94 | 0.00134 | 0.00169 |
| chr1 | 179530958 | 179540096 | 9139 | -4.04 | 0.00134 | 0.00169 |
| chr7 | 95234530 | 95254938 | 20409 | -4.42 | 0.00134 | 0.00169 |
| chr5 | 133168729 | 133174754 | 6026 | -1.1 | 0.00135 | 0.0017 |
| chr11 | 35823316 | 35823768 | 453 | -4.03 | 0.00135 | 0.0017 |
| chr4 | 165900151 | 165901620 | 1470 | 2.13 | 0.00135 | 0.0017 |
| chr9 | 93571147 | 93604548 | 33402 | -1.25 | 0.00135 | 0.0017 |
| chr10 | 75657693 | 75658275 | 583 | -4.5 | 0.00135 | 0.0017 |
| chr3 | 100426747 | 100427931 | 1185 | -3.64 | 0.00135 | 0.0017 |
| chr18 | 22940253 | 22941199 | 947 | -3.6 | 0.00135 | 0.0017 |
| chr8 | 24789147 | 24796239 | 7093 | -4.29 | 0.00135 | 0.0017 |
| chr5 | 105544613 | 105544960 | 348 | -3.95 | 0.00135 | 0.0017 |
| chr2 | 210169872 | 210172775 | 2904 | 3.8 | 0.00135 | 0.0017 |
| chr21 | 35784360 | 35785136 | 777 | -3.21 | 0.00136 | 0.00171 |
| chr3 | 27730587 | 27734305 | 3719 | -4.07 | 0.00136 | 0.00171 |
| chr8 | 946873 | 959235 | 12363 | -1.14 | 0.00136 | 0.00171 |
| chr12 | 18807412 | 18811242 | 3831 | 1.28 | 0.00136 | 0.00171 |
| chr10 | 61390447 | 61391891 | 1445 | 3.75 | 0.00136 | 0.00171 |
| chr7 | 104503904 | 104505039 | 1136 | -3.01 | 0.00136 | 0.00172 |
| chr9 | 106929875 | 106931277 | 1403 | -3.59 | 0.00136 | 0.00172 |
| chr6 | 168210413 | 168212019 | 1607 | -3.7 | 0.00136 | 0.00172 |
| chr5 | 41665386 | 41670142 | 4757 | 1.56 | 0.00137 | 0.00172 |
| chr12 | 59812735 | 59855065 | 42331 | -4.52 | 0.00137 | 0.00172 |
| chr1 | 183346595 | 183347486 | 892 | -3.29 | 0.00137 | 0.00172 |
| chr14 | 93348498 | 93362406 | 13909 | -4.7 | 0.00137 | 0.00172 |
| chrX | 108023582 | 108044492 | 20911 | 1.01 | 0.00137 | 0.00172 |
| chr4 | 132237832 | 132238172 | 341 | -4.3 | 0.00137 | 0.00173 |
| chrX | 77279544 | 77280046 | 503 | -4.3 | 0.00137 | 0.00173 |
| chr2 | 222586343 | 222610435 | 24093 | -1.66 | 0.00137 | 0.00173 |
| chr17 | 8148127 | 8148485 | 359 | -4.01 | 0.00137 | 0.00173 |
| chr3 | 129388017 | 129388681 | 665 | -3.11 | 0.00137 | 0.00173 |
| chrX | 130772692 | 130774264 | 1573 | 3.76 | 0.00138 | 0.00173 |
| chr7 | 154821096 | 154856736 | 35641 | -1.95 | 0.00138 | 0.00173 |
| chr1 | 230155469 | 230157324 | 1856 | 1.79 | 0.00138 | 0.00174 |
| chr7 | 132892028 | 132910846 | 18819 | -2.13 | 0.00139 | 0.00175 |
| chr3 | 122035088 | 122036652 | 1565 | 1.51 | 0.00139 | 0.00175 |
| chr13 | 28932958 | 28958793 | 25836 | 1.12 | 0.00139 | 0.00175 |
| chr8 | 126716348 | 126761041 | 44694 | -1.45 | 0.00139 | 0.00175 |
| chr2 | 75612338 | 75638962 | 26625 | -1.47 | 0.00139 | 0.00175 |
| chrX | 17844381 | 17845563 | 1183 | -3.1 | 0.00139 | 0.00175 |
| chr11 | 124289969 | 124301634 | 11666 | -0.89 | 0.0014 | 0.00176 |
| chr7 | 107376928 | 107377758 | 831 | -2.32 | 0.0014 | 0.00176 |
| chr5 | 151024339 | 151025593 | 1255 | 1.62 | 0.0014 | 0.00176 |
| chr1 | 222582770 | 222600233 | 17464 | -0.86 | 0.0014 | 0.00176 |
| chr5 | 129295435 | 129297999 | 2565 | 3.85 | 0.0014 | 0.00176 |
| chrX | 120363836 | 120364925 | 1090 | 3.65 | 0.0014 | 0.00176 |
| chr18 | 11292639 | 11302649 | 10011 | -0.8 | 0.0014 | 0.00176 |
| chr1 | 244518096 | 244529009 | 10914 | -5.24 | 0.00141 | 0.00177 |
| chr5 | 107800412 | 107810113 | 9702 | 4.3 | 0.00141 | 0.00177 |
| chr9 | 133387585 | 133388259 | 675 | -4.27 | 0.00141 | 0.00177 |
| chr6 | 154185088 | 154188491 | 3404 | 2.66 | 0.00141 | 0.00177 |
| chr10 | 25998738 | 26000918 | 2181 | -3.62 | 0.00141 | 0.00178 |
| chr1 | 37443201 | 37444568 | 1368 | -2.78 | 0.00141 | 0.00178 |
| chr2 | 147522672 | 147523757 | 1086 | -1.9 | 0.00141 | 0.00178 |
| chr1 | 65802178 | 65802667 | 490 | -4 | 0.00142 | 0.00178 |

| | | | | | | |
|---|---|---|---|---|---|---|
| chr13 | 91158609 | 91159119 | 511 | -4 | 0.00142 | 0.00178 |
| chr2 | 70364399 | 70364720 | 322 | -4 | 0.00142 | 0.00178 |
| chr8 | 83346726 | 83347061 | 336 | -4 | 0.00142 | 0.00178 |
| chr3 | 28062813 | 28081995 | 19183 | -1.73 | 0.00142 | 0.00178 |
| chr2 | 170205693 | 170207120 | 1428 | 1.49 | 0.00142 | 0.00178 |
| chr1 | 82899382 | 82899938 | 557 | -3.88 | 0.00142 | 0.00178 |
| chrX | 135467950 | 135468833 | 884 | 2.35 | 0.00142 | 0.00179 |
| chr11 | 110443337 | 110461052 | 17716 | 4.55 | 0.00143 | 0.00179 |
| chr2 | 13776813 | 13856826 | 80014 | -1.26 | 0.00143 | 0.0018 |
| chr6 | 140913858 | 140914391 | 534 | -2.98 | 0.00143 | 0.0018 |
| chr19 | 39501227 | 39532306 | 31080 | -7.17 | 0.00143 | 0.0018 |
| chr3 | 142908118 | 142914545 | 6428 | 1.28 | 0.00143 | 0.0018 |
| chr7 | 62825846 | 62842972 | 17127 | -2.15 | 0.00144 | 0.0018 |
| chr7 | 143827072 | 143875553 | 48482 | 4.89 | 0.00144 | 0.00181 |
| chr4 | 164209004 | 164209785 | 782 | 3.5 | 0.00144 | 0.00181 |
| chr14 | 55503917 | 55504777 | 861 | -3.1 | 0.00144 | 0.00181 |
| chr4 | 128393150 | 128393956 | 807 | -2.34 | 0.00144 | 0.00181 |
| chr2 | 159973319 | 159973737 | 419 | -4.42 | 0.00145 | 0.00182 |
| chr18 | 42196447 | 42199324 | 2878 | -2.23 | 0.00145 | 0.00182 |
| chr14 | 103492409 | 103492956 | 548 | -4.24 | 0.00145 | 0.00182 |
| chr11 | 109025936 | 109028094 | 2159 | -3.84 | 0.00145 | 0.00183 |
| chr11 | 21772782 | 21773988 | 1207 | 1.6 | 0.00145 | 0.00183 |
| chr6 | 66503042 | 66504408 | 1367 | -3.25 | 0.00145 | 0.00183 |
| chrX | 65262413 | 65299657 | 37245 | 4.7 | 0.00146 | 0.00183 |
| chr4 | 44449678 | 44450115 | 438 | -2.97 | 0.00146 | 0.00183 |
| chr4 | 104261747 | 104262942 | 1196 | -3.01 | 0.00146 | 0.00183 |
| chr2 | 183186587 | 183188268 | 1682 | -1.46 | 0.00146 | 0.00184 |
| chr5 | 5198129 | 5230200 | 32072 | -4.36 | 0.00146 | 0.00184 |
| chr14 | 29215234 | 29217897 | 2664 | -4.04 | 0.00147 | 0.00184 |
| chr3 | 170386129 | 170387226 | 1098 | -2.23 | 0.00147 | 0.00184 |
| chr17 | 51723750 | 52236298 | 512549 | 0.6 | 0.00147 | 0.00185 |
| chr2 | 103020521 | 103047661 | 27141 | -1.14 | 0.00147 | 0.00185 |
| chr2 | 240433640 | 240434255 | 616 | -4.58 | 0.00147 | 0.00185 |
| chr12 | 4366985 | 4381779 | 14795 | -1.4 | 0.00147 | 0.00185 |
| chr12 | 10093322 | 10094954 | 1633 | 3.88 | 0.00148 | 0.00185 |
| chr18 | 49873977 | 49900253 | 26277 | -2.92 | 0.00148 | 0.00185 |
| chrX | 13603820 | 13607178 | 3359 | -1.65 | 0.00148 | 0.00186 |
| chr17 | 63310937 | 63325972 | 15036 | 1.33 | 0.00148 | 0.00186 |
| chr1 | 199019815 | 199021009 | 1195 | -3.73 | 0.00149 | 0.00186 |
| chrX | 18017114 | 18018291 | 1178 | -2.07 | 0.00149 | 0.00187 |
| chr6 | 49130153 | 49130572 | 420 | -3.87 | 0.00149 | 0.00187 |
| chr13 | 111493006 | 111510809 | 17804 | -4.25 | 0.00149 | 0.00187 |
| chr11 | 112094965 | 112095273 | 309 | -3.96 | 0.00149 | 0.00187 |
| chr2 | 72373570 | 72374029 | 460 | -3.96 | 0.00149 | 0.00187 |
| chr3 | 25864480 | 25871933 | 7454 | -4.22 | 0.0015 | 0.00188 |
| chr10 | 25524557 | 25530830 | 6274 | -1.03 | 0.0015 | 0.00188 |
| chr13 | 28438314 | 28473065 | 34752 | -4.71 | 0.0015 | 0.00188 |
| chr2 | 27057900 | 27060036 | 2137 | -4.05 | 0.00151 | 0.00189 |
| chr8 | 26729524 | 26731655 | 2132 | -2.59 | 0.00151 | 0.00189 |
| chr3 | 159032163 | 159033832 | 1670 | -2 | 0.00151 | 0.00189 |
| chr14 | 54102011 | 54103303 | 1293 | 3.42 | 0.00152 | 0.00191 |
| chr5 | 40957052 | 40968597 | 11546 | -4.37 | 0.00152 | 0.00191 |
| chr13 | 107558455 | 107565496 | 7042 | 1.29 | 0.00152 | 0.00191 |
| chr9 | 129494874 | 129514088 | 19215 | -4.88 | 0.00152 | 0.00191 |
| chr13 | 93885486 | 93903077 | 17592 | -1.23 | 0.00152 | 0.00191 |
| chr6 | 99576853 | 99637891 | 61039 | 4.93 | 0.00152 | 0.00191 |
| chr8 | 88149622 | 88224071 | 74450 | 1.32 | 0.00153 | 0.00191 |
| chr18 | 52210190 | 52223878 | 13689 | -1.76 | 0.00153 | 0.00192 |
| chr10 | 71532620 | 71557445 | 24826 | -2.74 | 0.00153 | 0.00192 |
| chr2 | 173741635 | 173742579 | 945 | -2.16 | 0.00154 | 0.00193 |
| chr6 | 84532031 | 84552711 | 20681 | -4.36 | 0.00154 | 0.00194 |

| | | | | | | |
|---|---|---|---|---|---|---|
| chr18 | 25486282 | 25493655 | 7374 | -4.24 | 0.00155 | 0.00194 |
| chrX | 130298819 | 130302804 | 3986 | 1.49 | 0.00155 | 0.00194 |
| chr5 | 60210308 | 60211292 | 985 | -1.93 | 0.00155 | 0.00194 |
| chr7 | 18168006 | 18170913 | 2908 | 3.78 | 0.00155 | 0.00194 |
| chr3 | 138060243 | 138060804 | 562 | -2.99 | 0.00155 | 0.00195 |
| chr14 | 34760427 | 34765730 | 5304 | -1.35 | 0.00155 | 0.00195 |
| chr4 | 76294756 | 76295515 | 760 | -2.17 | 0.00156 | 0.00195 |
| chr9 | 131930894 | 131931607 | 714 | -4.34 | 0.00156 | 0.00196 |
| chr1 | 157479611 | 157833208 | 353598 | 7.21 | 0.00156 | 0.00196 |
| chr11 | 95371674 | 95372666 | 993 | -3.31 | 0.00157 | 0.00196 |
| chr4 | 65229941 | 65232687 | 2747 | -3.26 | 0.00157 | 0.00196 |
| chr3 | 181194512 | 181195736 | 1225 | 1.91 | 0.00157 | 0.00196 |
| chr4 | 109169270 | 109171251 | 1982 | -1.51 | 0.00157 | 0.00197 |
| chr6 | 165609431 | 165611320 | 1890 | -3.15 | 0.00157 | 0.00197 |
| chr13 | 41981557 | 41982585 | 1029 | -3.21 | 0.00157 | 0.00197 |
| chr1 | 106114648 | 106115230 | 583 | -2.91 | 0.00158 | 0.00198 |
| chr16 | 86130544 | 86144836 | 14293 | -4.19 | 0.00158 | 0.00198 |
| chr12 | 66152904 | 66154625 | 1722 | -1.63 | 0.00158 | 0.00198 |
| chr10 | 125443718 | 125616579 | 172862 | -0.54 | 0.00158 | 0.00199 |
| chr9 | 20269373 | 20302226 | 32854 | -4.45 | 0.00159 | 0.00199 |
| chr2 | 75704494 | 75715371 | 10878 | -1.85 | 0.00159 | 0.00199 |
| chr3 | 161592086 | 161604866 | 12781 | -1.98 | 0.00159 | 0.00199 |
| chr20 | 31501796 | 31508909 | 7114 | -4.51 | 0.0016 | 0.002 |
| chr4 | 30534880 | 30535448 | 569 | -4 | 0.0016 | 0.002 |
| chr5 | 115853335 | 115853705 | 371 | -4 | 0.0016 | 0.002 |
| chr5 | 154109255 | 154109649 | 395 | -4 | 0.0016 | 0.002 |
| chr8 | 118962197 | 118962713 | 517 | -4 | 0.0016 | 0.002 |
| chr4 | 110694228 | 110694838 | 611 | -2.28 | 0.0016 | 0.00201 |
| chr7 | 15419622 | 15420351 | 730 | -2.99 | 0.0016 | 0.00201 |
| chr1 | 153414651 | 153416950 | 2300 | -3.8 | 0.0016 | 0.00201 |
| chr3 | 155917927 | 155919216 | 1290 | -1.83 | 0.00161 | 0.00201 |
| chr6 | 53981329 | 53981900 | 572 | -2.27 | 0.00161 | 0.00201 |
| chr9 | 87613221 | 87653014 | 39794 | -4.79 | 0.00161 | 0.00202 |
| chr8 | 130323425 | 130336830 | 13406 | 0.78 | 0.00161 | 0.00202 |
| chr10 | 101240400 | 101271788 | 31389 | -4.42 | 0.00161 | 0.00202 |
| chr10 | 129483147 | 129510720 | 27574 | 1.53 | 0.00162 | 0.00203 |
| chr17 | 33871538 | 33878385 | 6848 | -3.93 | 0.00163 | 0.00204 |
| chr4 | 146296523 | 146309302 | 12780 | -0.84 | 0.00163 | 0.00204 |
| chr7 | 19072648 | 19093936 | 21289 | -2.25 | 0.00163 | 0.00204 |
| chr5 | 153507385 | 153507867 | 483 | -2.96 | 0.00163 | 0.00204 |
| chr4 | 180271312 | 180271717 | 406 | -3.82 | 0.00164 | 0.00205 |
| chr11 | 60724661 | 60737743 | 13083 | -4.55 | 0.00164 | 0.00206 |
| chr1 | 223721425 | 223722287 | 863 | -3.19 | 0.00164 | 0.00206 |
| chr11 | 14174395 | 14181798 | 7404 | -4.37 | 0.00165 | 0.00206 |
| chr3 | 10601732 | 10614769 | 13038 | -3.69 | 0.00165 | 0.00207 |
| chr9 | 72200978 | 72216102 | 15125 | 4.39 | 0.00165 | 0.00207 |
| chr4 | 34884634 | 34941827 | 57194 | 1.71 | 0.00165 | 0.00207 |
| chrX | 130771341 | 130771765 | 425 | -2.7 | 0.00165 | 0.00207 |
| chr1 | 179541081 | 179542572 | 1492 | 2.42 | 0.00166 | 0.00208 |
| chr7 | 87150235 | 87157574 | 7340 | -4.05 | 0.00166 | 0.00208 |
| chrX | 38735541 | 38747424 | 11884 | 3.63 | 0.00166 | 0.00208 |
| chr17 | 33287124 | 33287699 | 576 | 2.86 | 0.00166 | 0.00208 |
| chr2 | 205391000 | 205393087 | 2088 | -1.57 | 0.00166 | 0.00208 |
| chr4 | 15381158 | 15383443 | 2286 | -1.71 | 0.00166 | 0.00208 |
| chr3 | 140877679 | 140900669 | 22991 | -4.59 | 0.00166 | 0.00208 |
| chr3 | 152562892 | 152572383 | 9492 | -1.27 | 0.00167 | 0.00209 |
| chr11 | 125104770 | 125111200 | 6431 | -1.26 | 0.00168 | 0.0021 |
| chr8 | 26554262 | 26556113 | 1852 | 2.93 | 0.00168 | 0.0021 |
| chr6 | 70435210 | 70436084 | 875 | -3.28 | 0.00168 | 0.0021 |
| chr3 | 66941268 | 66967284 | 26017 | -0.89 | 0.00168 | 0.00211 |
| chrX | 36040017 | 36040432 | 416 | -3.08 | 0.00169 | 0.00212 |

| | | | | | | |
|---|---|---|---|---|---|---|
| chr1 | 114982411 | 114982697 | 287 | -3.81 | 0.00169 | 0.00212 |
| chr15 | 77269896 | 77273061 | 3166 | -2.72 | 0.00169 | 0.00212 |
| chr18 | 5277155 | 5280836 | 3682 | -2.36 | 0.00169 | 0.00212 |
| chr17 | 78688170 | 78697995 | 9826 | -5.81 | 0.0017 | 0.00212 |
| chr1 | 169066609 | 169066934 | 326 | -4.08 | 0.0017 | 0.00212 |
| chr11 | 120756500 | 120758024 | 1525 | -3.78 | 0.0017 | 0.00213 |
| chr4 | 65095262 | 65099403 | 4142 | -1.8 | 0.0017 | 0.00213 |
| chrX | 128389792 | 128422292 | 32501 | 4.67 | 0.0017 | 0.00213 |
| chr14 | 107188965 | 107192374 | 3410 | -3.64 | 0.0017 | 0.00213 |
| chr15 | 26197207 | 26238231 | 41025 | -1.1 | 0.00171 | 0.00213 |
| chr4 | 4207154 | 4208772 | 1619 | -2.11 | 0.00171 | 0.00214 |
| chr1 | 247511667 | 247639517 | 127851 | -3.34 | 0.00171 | 0.00214 |
| chr4 | 134663625 | 134666516 | 2892 | -1.56 | 0.00171 | 0.00214 |
| chr4 | 154763636 | 154842583 | 78948 | 4.97 | 0.00172 | 0.00215 |
| chrX | 108997758 | 109014966 | 17209 | -4.48 | 0.00172 | 0.00215 |
| chr2 | 152042292 | 152042840 | 549 | -2.57 | 0.00172 | 0.00215 |
| chr11 | 119869950 | 119878188 | 8239 | -1.2 | 0.00172 | 0.00215 |
| chr3 | 42353866 | 42363774 | 9909 | -4.43 | 0.00172 | 0.00216 |
| chr2 | 177768327 | 177769465 | 1139 | -3.2 | 0.00172 | 0.00216 |
| chr8 | 114426132 | 114427626 | 1495 | 2.03 | 0.00173 | 0.00216 |
| chr16 | 20322525 | 20339611 | 17087 | 1.09 | 0.00173 | 0.00217 |
| chr1 | 20153717 | 20165872 | 12156 | -3.48 | 0.00173 | 0.00217 |
| chr5 | 132978482 | 132984624 | 6143 | 1.43 | 0.00173 | 0.00217 |
| chr15 | 34751851 | 34752781 | 931 | 1.61 | 0.00174 | 0.00217 |
| chr1 | 187120499 | 187122302 | 1804 | -1.83 | 0.00174 | 0.00217 |
| chr15 | 29476464 | 29479554 | 3091 | -1.38 | 0.00174 | 0.00217 |
| chr8 | 54297287 | 54386214 | 88928 | -1.39 | 0.00174 | 0.00217 |
| chr8 | 121993262 | 121993941 | 680 | -2.09 | 0.00174 | 0.00217 |
| chr14 | 65846149 | 65846887 | 739 | -4.22 | 0.00174 | 0.00218 |
| chr4 | 100591104 | 100603366 | 12263 | -1.16 | 0.00174 | 0.00218 |
| chr10 | 87246008 | 87249748 | 3741 | 1.14 | 0.00174 | 0.00218 |
| chr3 | 163247235 | 163252525 | 5291 | 2.42 | 0.00174 | 0.00218 |
| chr13 | 94283994 | 94424493 | 140500 | 3.61 | 0.00175 | 0.00219 |
| chr3 | 67040383 | 67044103 | 3721 | -1.3 | 0.00175 | 0.00219 |
| chr10 | 122859903 | 122861294 | 1392 | 1.35 | 0.00175 | 0.00219 |
| chr14 | 38674573 | 38679680 | 5108 | -4.28 | 0.00175 | 0.00219 |
| chr16 | 55053469 | 55065259 | 11791 | -4.54 | 0.00175 | 0.00219 |
| chr7 | 123799952 | 124007890 | 207939 | 0.85 | 0.00175 | 0.00219 |
| chr8 | 58594934 | 58607893 | 12960 | -1.43 | 0.00176 | 0.0022 |
| chr14 | 22449710 | 22477466 | 27757 | -4.4 | 0.00177 | 0.00221 |
| chr4 | 111580031 | 111580939 | 909 | -3.08 | 0.00177 | 0.00221 |
| chrX | 15105315 | 15118921 | 13607 | 2.05 | 0.00177 | 0.00221 |
| chr6 | 159790364 | 159799387 | 9024 | -0.98 | 0.00177 | 0.00221 |
| chr3 | 175602930 | 175603472 | 543 | -3.03 | 0.00178 | 0.00222 |
| chr21 | 27995802 | 28014307 | 18506 | -1.53 | 0.00178 | 0.00222 |
| chr4 | 188408255 | 188409209 | 955 | -1.88 | 0.00178 | 0.00222 |
| chr4 | 147909578 | 147911320 | 1743 | -1.72 | 0.00178 | 0.00222 |
| chr8 | 74104962 | 74106645 | 1684 | -3.38 | 0.00179 | 0.00223 |
| chr1 | 209853372 | 209855069 | 1698 | -2.25 | 0.00179 | 0.00224 |
| chr6 | 74798774 | 74800996 | 2223 | -3.83 | 0.00179 | 0.00224 |
| chr16 | 56289361 | 56291528 | 2168 | -1.89 | 0.00179 | 0.00224 |
| chr3 | 16080242 | 16081845 | 1604 | -3.67 | 0.00179 | 0.00224 |
| chr10 | 81377122 | 81386905 | 9784 | -4.38 | 0.0018 | 0.00225 |
| chr9 | 5131045 | 5173137 | 42093 | -4.74 | 0.0018 | 0.00225 |
| chr2 | 213368467 | 213369440 | 974 | 2.03 | 0.0018 | 0.00225 |
| chr6 | 129250261 | 129264594 | 14334 | 4.33 | 0.0018 | 0.00225 |
| chr6 | 168057024 | 168059403 | 2380 | -1.4 | 0.0018 | 0.00225 |
| chr11 | 18999933 | 19002541 | 2609 | -1.98 | 0.0018 | 0.00225 |
| chr14 | 60946637 | 60947924 | 1288 | -3.76 | 0.00181 | 0.00226 |
| chr17 | 20849795 | 20883022 | 33228 | -6.98 | 0.00181 | 0.00226 |
| chr4 | 108112848 | 108113974 | 1127 | -3.45 | 0.00181 | 0.00226 |

| | | | | | | |
|---|---|---|---|---|---|---|
| chrX | 4919316 | 4924823 | 5508 | -3.91 | 0.00181 | 0.00227 |
| chr3 | 27132816 | 27134050 | 1235 | -2.65 | 0.00181 | 0.00227 |
| chr6 | 153842769 | 153846493 | 3725 | -1.35 | 0.00182 | 0.00227 |
| chrX | 75620442 | 75622103 | 1662 | 1.74 | 0.00182 | 0.00227 |
| chr2 | 102661878 | 102662869 | 992 | -3.52 | 0.00182 | 0.00228 |
| chr2 | 36263048 | 36315558 | 52511 | 0.94 | 0.00183 | 0.00228 |
| chr16 | 59582546 | 59583633 | 1088 | 1.96 | 0.00183 | 0.00228 |
| chrX | 11767937 | 11773969 | 6033 | -1.42 | 0.00183 | 0.00228 |
| chr10 | 53952691 | 53964142 | 11452 | -1.64 | 0.00183 | 0.00228 |
| chr18 | 13989837 | 14065121 | 75285 | -0.83 | 0.00183 | 0.00229 |
| chr12 | 76887226 | 76887659 | 434 | -4.01 | 0.00183 | 0.00229 |
| chr7 | 89735717 | 89736293 | 577 | -2.2 | 0.00183 | 0.00229 |
| chr18 | 11266413 | 11277710 | 11298 | 3.22 | 0.00183 | 0.00229 |
| chrX | 102517026 | 102518785 | 1760 | -3.75 | 0.00183 | 0.00229 |
| chr5 | 33552314 | 33553373 | 1060 | -2.59 | 0.00184 | 0.00229 |
| chr1 | 191695196 | 191695600 | 405 | -3.79 | 0.00184 | 0.0023 |
| chrX | 30518584 | 30519358 | 775 | -3.21 | 0.00184 | 0.0023 |
| chr15 | 89204926 | 89217311 | 12386 | -3.32 | 0.00184 | 0.0023 |
| chr10 | 26921649 | 26930645 | 8997 | -3.71 | 0.00184 | 0.0023 |
| chr9 | 111407605 | 111424784 | 17180 | -2.18 | 0.00184 | 0.0023 |
| chr11 | 58843547 | 58846100 | 2554 | 1.85 | 0.00185 | 0.00231 |
| chr10 | 7038446 | 7066336 | 27891 | -0.85 | 0.00185 | 0.00231 |
| chr15 | 37524790 | 37525223 | 434 | -2.92 | 0.00185 | 0.00231 |
| chr2 | 31002544 | 31017829 | 15286 | -1.06 | 0.00185 | 0.00231 |
| chr5 | 52169251 | 52173443 | 4193 | -1.45 | 0.00185 | 0.00231 |
| chr11 | 110597037 | 110597369 | 333 | -2 | 0.00185 | 0.00231 |
| chr4 | 35951759 | 35952752 | 994 | -3.35 | 0.00185 | 0.00231 |
| chr1 | 4088517 | 4100312 | 11796 | -1.01 | 0.00185 | 0.00231 |
| chr3 | 151327788 | 151330821 | 3034 | -3.97 | 0.00186 | 0.00232 |
| chr11 | 109199750 | 109200258 | 509 | -2.9 | 0.00186 | 0.00232 |
| chr2 | 213333055 | 213333588 | 534 | 2.33 | 0.00186 | 0.00232 |
| chr4 | 16096768 | 16098238 | 1471 | 2.89 | 0.00186 | 0.00232 |
| chr3 | 104282232 | 104301223 | 18992 | -4.37 | 0.00186 | 0.00232 |
| chr5 | 142785765 | 142789380 | 3616 | -3.79 | 0.00186 | 0.00232 |
| chr11 | 14131610 | 14132393 | 784 | -2.83 | 0.00186 | 0.00233 |
| chr1 | 39020392 | 39020985 | 594 | -2.58 | 0.00187 | 0.00233 |
| chr14 | 31315051 | 31320936 | 5886 | 0.94 | 0.00187 | 0.00234 |
| chr2 | 214755949 | 214788676 | 32728 | -2.15 | 0.00188 | 0.00234 |
| chr9 | 109685978 | 109688633 | 2656 | -1.41 | 0.00188 | 0.00235 |
| chrX | 134233761 | 134245873 | 12113 | -1.82 | 0.00188 | 0.00235 |
| chr6 | 126974711 | 126991848 | 17138 | -5.26 | 0.00188 | 0.00235 |
| chr1 | 187501645 | 187502122 | 478 | -2.57 | 0.00189 | 0.00235 |
| chr11 | 31751211 | 31759944 | 8734 | -4.18 | 0.00189 | 0.00235 |
| chr10 | 87874871 | 87902382 | 27512 | 4.45 | 0.00189 | 0.00236 |
| chr11 | 119683762 | 119717786 | 34025 | -4.55 | 0.00189 | 0.00236 |
| chr1 | 187465533 | 187466715 | 1183 | -1.48 | 0.00189 | 0.00236 |
| chr1 | 87075617 | 87081670 | 6054 | -1.44 | 0.0019 | 0.00237 |
| chr1 | 35183322 | 35184513 | 1192 | -3.55 | 0.0019 | 0.00238 |
| chrX | 81065832 | 81074642 | 8811 | -3.79 | 0.00191 | 0.00238 |
| chr3 | 137658329 | 137659236 | 908 | 1.98 | 0.00191 | 0.00238 |
| chr5 | 53947186 | 54075174 | 127989 | 0.56 | 0.00191 | 0.00239 |
| chr3 | 116643862 | 116647816 | 3955 | -3.12 | 0.00191 | 0.00239 |
| chr20 | 56921468 | 56922177 | 710 | -3.13 | 0.00191 | 0.00239 |
| chr3 | 54310806 | 54312067 | 1262 | -1.44 | 0.00192 | 0.00239 |
| chrX | 77473563 | 77476326 | 2764 | -2.82 | 0.00192 | 0.0024 |
| chr6 | 46161319 | 46162408 | 1090 | -2.19 | 0.00193 | 0.00241 |
| chr4 | 68803335 | 68804815 | 1481 | -2.24 | 0.00193 | 0.00241 |
| chr11 | 89011088 | 89012722 | 1635 | 3.41 | 0.00194 | 0.00241 |
| chr6 | 117695950 | 117699518 | 3569 | -3.95 | 0.00194 | 0.00242 |
| chr14 | 59268798 | 59294748 | 25951 | -6.92 | 0.00195 | 0.00243 |
| chr13 | 108986704 | 108987400 | 697 | -2.14 | 0.00195 | 0.00244 |

| | | | | | | |
|---|---|---|---|---|---|---|
| chr8 | 102532714 | 102556623 | 23910 | -6.91 | 0.00196 | 0.00244 |
| chr2 | 213850730 | 213853533 | 2804 | 3.67 | 0.00196 | 0.00245 |
| chrX | 140724975 | 140725478 | 504 | -4.2 | 0.00197 | 0.00245 |
| chr6 | 165555503 | 165556976 | 1474 | -1.81 | 0.00197 | 0.00245 |
| chr22 | 50124580 | 50142389 | 17810 | -4.69 | 0.00197 | 0.00246 |
| chr18 | 72191735 | 72196376 | 4642 | -4.21 | 0.00197 | 0.00246 |
| chr7 | 79544276 | 79544743 | 468 | -2.55 | 0.00197 | 0.00246 |
| chr12 | 70401074 | 70419924 | 18851 | -3.28 | 0.00198 | 0.00246 |
| chrX | 79309153 | 79368581 | 59429 | 1.84 | 0.00198 | 0.00247 |
| chr8 | 55262245 | 55276151 | 13907 | 1.44 | 0.00198 | 0.00247 |
| chr17 | 69382594 | 69385225 | 2632 | -3.59 | 0.00198 | 0.00247 |
| chr19 | 14705492 | 14713912 | 8421 | -1.27 | 0.00198 | 0.00247 |
| chr15 | 39835149 | 39840185 | 5037 | -4.2 | 0.00198 | 0.00247 |
| chr7 | 156622368 | 156622661 | 294 | -4 | 0.00198 | 0.00247 |
| chr2 | 166656099 | 166658892 | 2794 | -1.3 | 0.00198 | 0.00247 |
| chr4 | 67337534 | 67339216 | 1683 | -3.35 | 0.00198 | 0.00247 |
| chr11 | 76354896 | 76362080 | 7185 | -4.59 | 0.00199 | 0.00248 |
| chr8 | 142972769 | 142976918 | 4150 | -2.6 | 0.00199 | 0.00248 |
| chr18 | 43581143 | 43584182 | 3040 | -4.02 | 0.00199 | 0.00248 |
| chr17 | 49541099 | 49602987 | 61889 | -1.32 | 0.00199 | 0.00248 |
| chr3 | 117790205 | 117815023 | 24819 | -4.43 | 0.002 | 0.00249 |
| chr15 | 50288547 | 50290429 | 1883 | 1.53 | 0.002 | 0.00249 |
| chr7 | 119188600 | 119215769 | 27170 | -0.83 | 0.00202 | 0.00252 |
| chr7 | 62734963 | 62735327 | 365 | -2.88 | 0.00202 | 0.00252 |
| chr10 | 1182294 | 1194502 | 12209 | -4.69 | 0.00202 | 0.00252 |
| chr17 | 26155043 | 26160794 | 5752 | -3.71 | 0.00202 | 0.00252 |
| chrX | 106699594 | 106700898 | 1305 | -2.41 | 0.00203 | 0.00253 |
| chr12 | 97749062 | 97751567 | 2506 | -3.94 | 0.00203 | 0.00253 |
| chrX | 51654406 | 51695916 | 41511 | -0.99 | 0.00204 | 0.00253 |
| chr9 | 87677472 | 87686585 | 9114 | -3.97 | 0.00204 | 0.00254 |
| chr14 | 36793787 | 36828224 | 34438 | -2.74 | 0.00204 | 0.00254 |
| chr3 | 53419201 | 53421840 | 2640 | -1.95 | 0.00204 | 0.00254 |
| chrX | 97023537 | 97025277 | 1741 | 1.77 | 0.00205 | 0.00255 |
| chr4 | 144070933 | 144099154 | 28222 | -4.37 | 0.00205 | 0.00256 |
| chr2 | 193608327 | 193619607 | 11281 | -4.88 | 0.00205 | 0.00256 |
| chr3 | 17683520 | 17691101 | 7582 | -5.19 | 0.00205 | 0.00256 |
| chr18 | 73498415 | 73508574 | 10160 | -0.82 | 0.00205 | 0.00256 |
| chr11 | 95222846 | 95228941 | 6096 | -3.01 | 0.00205 | 0.00256 |
| chr11 | 130483653 | 130490424 | 6772 | -3.94 | 0.00206 | 0.00256 |
| chr4 | 68814328 | 68816164 | 1837 | 2.54 | 0.00206 | 0.00257 |
| chr6 | 63900268 | 63906624 | 6357 | 1.77 | 0.00207 | 0.00258 |
| chr9 | 137932902 | 137957955 | 25054 | -4.61 | 0.00208 | 0.00258 |
| chr12 | 62630354 | 62632365 | 2012 | -3.83 | 0.00208 | 0.00259 |
| chr11 | 116122380 | 116123626 | 1247 | -3.44 | 0.00208 | 0.00259 |
| chr11 | 77755666 | 77756268 | 603 | -3.75 | 0.00208 | 0.00259 |
| chr11 | 1693771 | 1701595 | 7825 | -5.17 | 0.00208 | 0.00259 |
| chr1 | 157468951 | 157469583 | 633 | -2.94 | 0.00208 | 0.00259 |
| chr17 | 65654607 | 65655596 | 990 | -4.71 | 0.00208 | 0.00259 |
| chr21 | 32755082 | 32755942 | 861 | -2.41 | 0.00209 | 0.0026 |
| chr5 | 32980647 | 32983052 | 2406 | -2.67 | 0.00209 | 0.0026 |
| chr16 | 80567204 | 80584996 | 17793 | -0.66 | 0.00209 | 0.0026 |
| chr8 | 120722806 | 120724185 | 1380 | -3.58 | 0.00209 | 0.0026 |
| chrX | 134799089 | 134799850 | 762 | -1.67 | 0.00209 | 0.0026 |
| chr1 | 228796337 | 228796776 | 440 | -2.89 | 0.00209 | 0.00261 |
| chr2 | 23422189 | 23475683 | 53495 | 1.46 | 0.0021 | 0.00261 |
| chr2 | 226639036 | 226640209 | 1174 | -3.24 | 0.0021 | 0.00261 |
| chr12 | 10423297 | 10435173 | 11877 | -4.06 | 0.0021 | 0.00262 |
| chrX | 139117776 | 139122544 | 4769 | -1.78 | 0.00211 | 0.00262 |
| chr12 | 43526552 | 43536487 | 9936 | -4.26 | 0.00211 | 0.00262 |
| chr17 | 77868235 | 77869118 | 884 | 3.4 | 0.00211 | 0.00262 |
| chr11 | 115356159 | 115358464 | 2306 | -3.7 | 0.00211 | 0.00262 |

| | | | | | | |
|---|---|---|---|---|---|---|
| chr20 | 22899968 | 22900430 | 463 | 3.36 | 0.00211 | 0.00263 |
| chr1 | 232048985 | 232049960 | 976 | 3.16 | 0.00211 | 0.00263 |
| chr12 | 60353336 | 60363810 | 10475 | -2.05 | 0.00211 | 0.00263 |
| chr11 | 115593034 | 115596793 | 3760 | -1.16 | 0.00211 | 0.00263 |
| chr9 | 140188895 | 140189249 | 355 | -4.02 | 0.00211 | 0.00263 |
| chr15 | 34917539 | 34986414 | 68876 | -0.91 | 0.00212 | 0.00263 |
| chr2 | 228791789 | 228793601 | 1813 | -3.92 | 0.00212 | 0.00263 |
| chr14 | 95529808 | 95542516 | 12709 | -6.19 | 0.00212 | 0.00263 |
| chr4 | 146509668 | 146511040 | 1373 | -3.04 | 0.00212 | 0.00263 |
| chr9 | 75530547 | 75532870 | 2324 | -3.58 | 0.00212 | 0.00263 |
| chr11 | 94055362 | 94057032 | 1671 | -1.78 | 0.00212 | 0.00264 |
| chr17 | 49515124 | 49526823 | 11700 | -2.12 | 0.00212 | 0.00264 |
| chr7 | 155858305 | 155862996 | 4692 | 4.23 | 0.00213 | 0.00264 |
| chr11 | 15021694 | 15028755 | 7062 | -1.61 | 0.00213 | 0.00265 |
| chr1 | 116146080 | 116183563 | 37484 | -4.6 | 0.00213 | 0.00265 |
| chr7 | 157533862 | 157535680 | 1819 | 1.61 | 0.00213 | 0.00265 |
| chr5 | 71313919 | 71324557 | 10639 | -5.78 | 0.00213 | 0.00265 |
| chr4 | 94728361 | 94740708 | 12348 | 0.89 | 0.00214 | 0.00266 |
| chrX | 11171462 | 11177564 | 6103 | 3.97 | 0.00214 | 0.00266 |
| chr10 | 18608109 | 18621114 | 13006 | 1.17 | 0.00214 | 0.00267 |
| chr1 | 82857574 | 82857997 | 424 | 3 | 0.00215 | 0.00268 |
| chr6 | 112363748 | 112364179 | 432 | -2.93 | 0.00216 | 0.00269 |
| chr20 | 49760569 | 49818529 | 57961 | -4.69 | 0.00216 | 0.00269 |
| chr9 | 34532399 | 34535935 | 3537 | -2.24 | 0.00217 | 0.0027 |
| chr3 | 65300846 | 65307017 | 6172 | -1.24 | 0.00217 | 0.0027 |
| chr3 | 176006064 | 176012415 | 6352 | 1.98 | 0.00217 | 0.0027 |
| chr7 | 96281079 | 96282467 | 1389 | -1.76 | 0.00218 | 0.0027 |
| chrX | 69777305 | 69822195 | 44891 | -4.63 | 0.00218 | 0.00271 |
| chr11 | 116273167 | 116274618 | 1452 | 1.7 | 0.00218 | 0.00271 |
| chr3 | 59751704 | 59755608 | 3905 | 3.79 | 0.00218 | 0.00271 |
| chr5 | 75170876 | 75276008 | 105133 | 0.53 | 0.00218 | 0.00271 |
| chr6 | 80664958 | 80681712 | 16755 | -3.88 | 0.00218 | 0.00272 |
| chr5 | 36839881 | 36842514 | 2634 | -4.12 | 0.00219 | 0.00272 |
| chr4 | 53023756 | 53048782 | 25027 | -1.17 | 0.00219 | 0.00272 |
| chr12 | 99477750 | 99520032 | 42283 | 1.3 | 0.00219 | 0.00272 |
| chr11 | 124897686 | 124898785 | 1100 | -2.46 | 0.0022 | 0.00273 |
| chr11 | 49828900 | 49829272 | 373 | 1.95 | 0.0022 | 0.00273 |
| chr3 | 133885926 | 133898496 | 12571 | -3.91 | 0.0022 | 0.00273 |
| chr11 | 128662303 | 128664479 | 2177 | 1.76 | 0.0022 | 0.00273 |
| chr20 | 23814145 | 23827557 | 13413 | -0.9 | 0.0022 | 0.00274 |
| chr7 | 29352521 | 29372896 | 20376 | 1.45 | 0.0022 | 0.00274 |
| chr6 | 155932445 | 155937570 | 5126 | -1.19 | 0.0022 | 0.00274 |
| chrX | 85404098 | 85406286 | 2189 | -2.61 | 0.0022 | 0.00274 |
| chrX | 42382543 | 42404865 | 22323 | 2.07 | 0.00221 | 0.00274 |
| chrX | 108097071 | 108098645 | 1575 | -1.75 | 0.00221 | 0.00274 |
| chr15 | 60057853 | 60063191 | 5339 | 1.26 | 0.00221 | 0.00275 |
| chr12 | 97800821 | 97815206 | 14386 | 1.92 | 0.00221 | 0.00275 |
| chr1 | 99542774 | 99543300 | 527 | -2.37 | 0.00221 | 0.00275 |
| chr1 | 222453112 | 222454932 | 1821 | -3.24 | 0.00222 | 0.00275 |
| chr7 | 124095306 | 124112675 | 17370 | -1.14 | 0.00223 | 0.00277 |
| chrX | 101162231 | 101166682 | 4452 | -1.31 | 0.00223 | 0.00277 |
| chr8 | 47962393 | 47973457 | 11065 | -0.92 | 0.00223 | 0.00277 |
| chr12 | 22092410 | 22115606 | 23197 | -4.48 | 0.00223 | 0.00277 |
| chr4 | 185430526 | 185447845 | 17320 | -4.3 | 0.00224 | 0.00278 |
| chr1 | 230851910 | 230875060 | 23151 | -4.39 | 0.00224 | 0.00278 |
| chr9 | 89655219 | 89682347 | 27129 | -0.67 | 0.00224 | 0.00278 |
| chr15 | 72315349 | 72330578 | 15230 | -6.8 | 0.00224 | 0.00279 |
| chr2 | 150500163 | 150521863 | 21701 | -5.25 | 0.00225 | 0.00279 |
| chr13 | 103349525 | 103405385 | 55861 | -2.99 | 0.00225 | 0.0028 |
| chr7 | 131433537 | 131448437 | 14901 | -4.14 | 0.00225 | 0.0028 |
| chr8 | 95618723 | 95621022 | 2300 | -3.37 | 0.00226 | 0.0028 |

| | | | | | | |
|---|---|---|---|---|---|---|
| chr2 | 206762985 | 206786389 | 23405 | -3.03 | 0.00226 | 0.00281 |
| chr6 | 12854202 | 12858557 | 4356 | -1.4 | 0.00226 | 0.00281 |
| chr12 | 3928515 | 3929695 | 1181 | 3.55 | 0.00227 | 0.00281 |
| chr11 | 59900507 | 59902111 | 1605 | -2.07 | 0.00227 | 0.00282 |
| chr13 | 101517952 | 101542017 | 24066 | -0.86 | 0.00227 | 0.00282 |
| chr22 | 49539586 | 49542465 | 2880 | -3.03 | 0.00228 | 0.00283 |
| chr4 | 144486679 | 144489046 | 2368 | -3.61 | 0.00228 | 0.00283 |
| chr8 | 140020001 | 140252521 | 232521 | 0.49 | 0.00228 | 0.00284 |
| chr11 | 49651139 | 49652121 | 983 | 1.8 | 0.00229 | 0.00284 |
| chr5 | 106119948 | 106120276 | 329 | -3.88 | 0.00229 | 0.00284 |
| chr17 | 79444005 | 79444356 | 352 | -3.95 | 0.00229 | 0.00284 |
| chrX | 85015070 | 85097232 | 82163 | 4.89 | 0.00229 | 0.00284 |
| chr4 | 151153688 | 151155385 | 1698 | -3.25 | 0.00229 | 0.00284 |
| chr18 | 59666336 | 59684692 | 18357 | -0.72 | 0.00229 | 0.00285 |
| chrX | 16314825 | 16328407 | 13583 | 1.19 | 0.00229 | 0.00285 |
| chr13 | 71833394 | 71835592 | 2199 | 2.47 | 0.00229 | 0.00285 |
| chr20 | 43819077 | 43866650 | 47574 | -0.63 | 0.0023 | 0.00286 |
| chr1 | 97713683 | 97714060 | 378 | -3.73 | 0.00231 | 0.00286 |
| chr11 | 88563771 | 88566176 | 2406 | -2.95 | 0.00231 | 0.00287 |
| chr21 | 10522237 | 10522709 | 473 | 1.93 | 0.00231 | 0.00287 |
| chr12 | 113304225 | 113306866 | 2642 | 2.75 | 0.00231 | 0.00287 |
| chr3 | 162780252 | 162782485 | 2234 | 3.46 | 0.00232 | 0.00287 |
| chr4 | 22048111 | 22050005 | 1895 | 1.48 | 0.00232 | 0.00288 |
| chr22 | 34031620 | 34032293 | 674 | -2.62 | 0.00232 | 0.00288 |
| chr4 | 9811584 | 9830351 | 18768 | 1.13 | 0.00232 | 0.00288 |
| chrX | 108848640 | 108849105 | 466 | -3.1 | 0.00232 | 0.00288 |
| chrX | 138294553 | 138777107 | 482555 | 7.16 | 0.00232 | 0.00288 |
| chr2 | 159695999 | 159696696 | 698 | -3.1 | 0.00232 | 0.00288 |
| chr15 | 101873325 | 101876591 | 3267 | -1.41 | 0.00232 | 0.00288 |
| chr3 | 31518360 | 31552793 | 34434 | -3.61 | 0.00232 | 0.00288 |
| chr3 | 164401444 | 164460133 | 58690 | -4.39 | 0.00232 | 0.00288 |
| chr12 | 95273054 | 95274532 | 1479 | 1.51 | 0.00233 | 0.00289 |
| chr3 | 164910523 | 164913705 | 3183 | -3.68 | 0.00233 | 0.00289 |
| chr15 | 74155260 | 74166560 | 11301 | -4.22 | 0.00233 | 0.0029 |
| chr1 | 38810477 | 38815097 | 4621 | -1.72 | 0.00234 | 0.0029 |
| chr2 | 195952081 | 195952837 | 757 | -1.8 | 0.00234 | 0.0029 |
| chr19 | 15850188 | 15874782 | 24595 | -1.99 | 0.00234 | 0.0029 |
| chr14 | 66348253 | 66350434 | 2182 | -3.9 | 0.00234 | 0.0029 |
| chr4 | 133961103 | 133973197 | 12095 | 0.85 | 0.00234 | 0.00291 |
| chr3 | 176027742 | 176049537 | 21796 | 1.36 | 0.00235 | 0.00291 |
| chr12 | 67996443 | 68000495 | 4053 | 1.17 | 0.00235 | 0.00291 |
| chr4 | 175954182 | 175954997 | 816 | -3.15 | 0.00235 | 0.00292 |
| chr4 | 107469173 | 107469490 | 318 | -3.24 | 0.00235 | 0.00292 |
| chr10 | 35935786 | 35950162 | 14377 | -1.77 | 0.00235 | 0.00292 |
| chr18 | 77302669 | 77303343 | 675 | 3.3 | 0.00236 | 0.00293 |
| chr14 | 95413744 | 95455763 | 42020 | -1.87 | 0.00237 | 0.00293 |
| chr5 | 4893492 | 4971074 | 77583 | 0.74 | 0.00237 | 0.00293 |
| chr12 | 48389217 | 48403545 | 14329 | 3.85 | 0.00237 | 0.00294 |
| chr10 | 9350715 | 9371421 | 20707 | 4.44 | 0.00237 | 0.00294 |
| chr18 | 54152834 | 54170592 | 17759 | -1.37 | 0.00237 | 0.00294 |
| chr5 | 45127008 | 45518685 | 391678 | -0.82 | 0.00237 | 0.00294 |
| chr2 | 193139073 | 193139650 | 578 | -4.22 | 0.00237 | 0.00294 |
| chr9 | 112591170 | 112597741 | 6572 | -1.63 | 0.00238 | 0.00295 |
| chr21 | 34229173 | 34302000 | 72828 | -0.96 | 0.00238 | 0.00295 |
| chr9 | 35973753 | 36015627 | 41875 | -1.26 | 0.00238 | 0.00295 |
| chr2 | 222082633 | 222087282 | 4650 | -3.59 | 0.00238 | 0.00295 |
| chr7 | 38876699 | 38877189 | 491 | -3.84 | 0.00239 | 0.00296 |
| chr17 | 3348649 | 3362645 | 13997 | -4.22 | 0.00239 | 0.00296 |
| chr1 | 5883518 | 5884632 | 1115 | -1.78 | 0.00239 | 0.00296 |
| chr11 | 82419045 | 82424853 | 5809 | 1.27 | 0.00239 | 0.00296 |
| chr13 | 27079929 | 27095551 | 15623 | -3.89 | 0.00239 | 0.00296 |

| | | | | | | |
|---|---|---|---|---|---|---|
| chr16 | 28992935 | 28999009 | 6075 | -4.57 | 0.00239 | 0.00296 |
| chr11 | 40624235 | 40629613 | 5379 | -2.21 | 0.00239 | 0.00297 |
| chr3 | 43158040 | 43242598 | 84559 | -2.75 | 0.0024 | 0.00297 |
| chr6 | 117612430 | 117614593 | 2164 | 1.75 | 0.0024 | 0.00298 |
| chr16 | 52195556 | 52224152 | 28597 | 0.76 | 0.0024 | 0.00298 |
| chr7 | 92554004 | 92637886 | 83883 | -6.44 | 0.0024 | 0.00298 |
| chrX | 95909550 | 95910780 | 1231 | 2.63 | 0.00241 | 0.00298 |
| chr2 | 33843665 | 33846393 | 2729 | -2.62 | 0.00241 | 0.00299 |
| chr12 | 77489823 | 77712292 | 222470 | 6.93 | 0.00242 | 0.00299 |
| chr18 | 69586728 | 69591333 | 4606 | 1.46 | 0.00242 | 0.00299 |
| chr7 | 31614682 | 31637076 | 22395 | -2.65 | 0.00242 | 0.003 |
| chr6 | 166622138 | 166626499 | 4362 | -4.25 | 0.00242 | 0.003 |
| chr21 | 47192059 | 47200010 | 7952 | -4.28 | 0.00243 | 0.003 |
| chr13 | 103586170 | 103618083 | 31914 | 0.93 | 0.00243 | 0.003 |
| chr3 | 155813149 | 155813938 | 790 | -3.4 | 0.00243 | 0.00301 |
| chr3 | 120621158 | 120623896 | 2739 | 1.87 | 0.00243 | 0.00301 |
| chr1 | 207445113 | 207449826 | 4714 | -3.9 | 0.00243 | 0.00302 |
| chr6 | 85066789 | 85098640 | 31852 | 1.09 | 0.00243 | 0.00302 |
| chr14 | 54187586 | 54202409 | 14824 | -6.73 | 0.00243 | 0.00302 |
| chr9 | 21071278 | 21088673 | 17396 | -6.73 | 0.00243 | 0.00302 |
| chr20 | 13191926 | 13227035 | 35110 | -3.34 | 0.00244 | 0.00302 |
| chr5 | 98092887 | 98093739 | 853 | 1.46 | 0.00245 | 0.00303 |
| chr18 | 52920951 | 52921885 | 935 | -3.24 | 0.00245 | 0.00303 |
| chr13 | 95190041 | 95190334 | 294 | 3.1 | 0.00245 | 0.00304 |
| chr19 | 56211988 | 56591181 | 379194 | -0.61 | 0.00246 | 0.00305 |
| chr8 | 87007672 | 87008955 | 1284 | -2.14 | 0.00247 | 0.00305 |
| chr6 | 121389227 | 121391588 | 2362 | -2.97 | 0.00247 | 0.00306 |
| chr1 | 56515235 | 56517473 | 2239 | -3.39 | 0.00247 | 0.00306 |
| chr7 | 107777385 | 107778261 | 877 | -3.04 | 0.00247 | 0.00306 |
| chr3 | 48426528 | 48434371 | 7844 | -4.17 | 0.00248 | 0.00307 |
| chr10 | 89927409 | 89937948 | 10540 | -4.49 | 0.00248 | 0.00307 |
| chr14 | 65312291 | 65316473 | 4183 | 1.55 | 0.00248 | 0.00307 |
| chr5 | 178964782 | 178965123 | 342 | -3.73 | 0.00248 | 0.00307 |
| chr9 | 103443598 | 103444672 | 1075 | 3.41 | 0.00248 | 0.00307 |
| chr12 | 131880718 | 131895068 | 14351 | -2.25 | 0.00248 | 0.00307 |
| chr17 | 60257683 | 60293764 | 36082 | -6.71 | 0.00248 | 0.00307 |
| chr16 | 55627269 | 55668288 | 41020 | -1.03 | 0.00248 | 0.00307 |
| chrX | 54857360 | 54863680 | 6321 | -2.03 | 0.00249 | 0.00308 |
| chr10 | 14057649 | 14072978 | 15330 | -0.94 | 0.00249 | 0.00308 |
| chr9 | 2402176 | 2461806 | 59631 | 0.8 | 0.00249 | 0.00308 |
| chr8 | 98288233 | 98288956 | 724 | -2.8 | 0.0025 | 0.00309 |
| chr11 | 45629795 | 45669369 | 39575 | -1.11 | 0.0025 | 0.0031 |
| chrX | 103254831 | 103256686 | 1856 | 3.44 | 0.00251 | 0.0031 |
| chr1 | 164611172 | 164613414 | 2243 | -1.49 | 0.00251 | 0.0031 |
| chr9 | 5368245 | 5368539 | 295 | -2.86 | 0.00251 | 0.00311 |
| chr7 | 31234756 | 31279728 | 44973 | -0.69 | 0.00252 | 0.00312 |
| chr18 | 45264036 | 45265682 | 1647 | -2.45 | 0.00252 | 0.00312 |
| chr10 | 54470708 | 54475899 | 5192 | 1.49 | 0.00253 | 0.00313 |
| chr1 | 183690244 | 183699072 | 8829 | -2.76 | 0.00253 | 0.00313 |
| chr1 | 7220167 | 7222715 | 2549 | -3.67 | 0.00254 | 0.00314 |
| chr3 | 84328378 | 84402888 | 74511 | -3.64 | 0.00254 | 0.00314 |
| chr15 | 86386163 | 86390769 | 4607 | -4.19 | 0.00254 | 0.00314 |
| chr7 | 521977 | 522632 | 656 | 1.67 | 0.00254 | 0.00314 |
| chrX | 13935901 | 13937158 | 1258 | -2.29 | 0.00254 | 0.00315 |
| chr4 | 127957295 | 127959999 | 2705 | -1.82 | 0.00254 | 0.00315 |
| chr15 | 84125399 | 84225181 | 99783 | -0.68 | 0.00255 | 0.00315 |
| chr4 | 104848625 | 104850219 | 1595 | -1.78 | 0.00255 | 0.00316 |
| chr10 | 91635175 | 91635579 | 405 | -2.98 | 0.00257 | 0.00317 |
| chr2 | 189723904 | 189733368 | 9465 | -4.13 | 0.00257 | 0.00318 |
| chr7 | 148340958 | 148343372 | 2415 | 1.18 | 0.00257 | 0.00318 |
| chr3 | 23612832 | 23638675 | 25844 | -6.68 | 0.00258 | 0.00319 |

| | | | | | | |
|---|---|---|---|---|---|---|
| chr1 | 41766257 | 41775297 | 9041 | -0.91 | 0.00258 | 0.00319 |
| chr6 | 38727811 | 38740914 | 13104 | -1.43 | 0.00258 | 0.00319 |
| chr6 | 146369119 | 146371413 | 2295 | 1.76 | 0.00259 | 0.0032 |
| chr4 | 20359938 | 20362562 | 2625 | 3.98 | 0.00259 | 0.00321 |
| chr6 | 63851900 | 63853203 | 1304 | -3.37 | 0.0026 | 0.00321 |
| chr1 | 75475979 | 75477054 | 1076 | 1.78 | 0.0026 | 0.00321 |
| chr6 | 55220702 | 55222199 | 1498 | -3.05 | 0.0026 | 0.00321 |
| chr20 | 37279530 | 37281546 | 2017 | 1.55 | 0.0026 | 0.00321 |
| chr10 | 69229860 | 69277429 | 47570 | -1.96 | 0.0026 | 0.00322 |
| chr3 | 39242836 | 39251469 | 8634 | -4.22 | 0.0026 | 0.00322 |
| chr4 | 114416257 | 114417352 | 1096 | -3.4 | 0.0026 | 0.00322 |
| chr7 | 129984389 | 129984761 | 373 | 2.96 | 0.0026 | 0.00322 |
| chrX | 42290061 | 42305163 | 15103 | 0.94 | 0.0026 | 0.00322 |
| chr11 | 133728901 | 133741304 | 12404 | 2.45 | 0.00261 | 0.00322 |
| chr13 | 104100299 | 104102489 | 2191 | -1.68 | 0.00261 | 0.00323 |
| chr13 | 63020231 | 63021288 | 1058 | -2.23 | 0.00262 | 0.00323 |
| chr9 | 79511992 | 79567065 | 55074 | -0.64 | 0.00262 | 0.00324 |
| chr9 | 112484274 | 112517451 | 33178 | 0.77 | 0.00263 | 0.00326 |
| chr9 | 91700686 | 91701703 | 1018 | 3.34 | 0.00264 | 0.00326 |
| chr1 | 112362795 | 112365075 | 2281 | -1.9 | 0.00264 | 0.00326 |
| chr7 | 106565421 | 106567701 | 2281 | -1.73 | 0.00264 | 0.00326 |
| chr9 | 28473572 | 28501649 | 28078 | 4.39 | 0.00264 | 0.00327 |
| chr5 | 83060788 | 83065876 | 5089 | -4.06 | 0.00265 | 0.00327 |
| chr1 | 39970134 | 39970400 | 267 | -3.67 | 0.00265 | 0.00328 |
| chr12 | 67515347 | 67523199 | 7853 | 2.2 | 0.00267 | 0.0033 |
| chr2 | 96374228 | 96375698 | 1471 | -1.42 | 0.00267 | 0.0033 |
| chr22 | 47932475 | 47956805 | 24331 | -2.81 | 0.00268 | 0.00331 |
| chr5 | 75019096 | 75040697 | 21602 | -4.26 | 0.00268 | 0.00331 |
| chr6 | 118946771 | 118947442 | 672 | -1.96 | 0.00268 | 0.00331 |
| chr14 | 107087917 | 107094136 | 6220 | 1.3 | 0.00268 | 0.00331 |
| chr13 | 39494595 | 39502125 | 7531 | 0.91 | 0.00268 | 0.00331 |
| chr7 | 92060339 | 92060770 | 432 | -3.22 | 0.00268 | 0.00331 |
| chr1 | 76463565 | 76465690 | 2126 | -3.63 | 0.00268 | 0.00331 |
| chr7 | 11390201 | 11400075 | 9875 | -1.4 | 0.00269 | 0.00332 |
| chr2 | 34962753 | 35021752 | 59000 | 0.74 | 0.00269 | 0.00332 |
| chr4 | 115572690 | 115574373 | 1684 | 3.21 | 0.00269 | 0.00333 |
| chr11 | 36534000 | 36547899 | 13900 | -4.38 | 0.00269 | 0.00333 |
| chr18 | 28705709 | 28706201 | 493 | -2.94 | 0.00269 | 0.00333 |
| chr16 | 27072630 | 27088271 | 15642 | -3.37 | 0.0027 | 0.00333 |
| chr2 | 71060192 | 71075925 | 15734 | -1.93 | 0.0027 | 0.00333 |
| chr4 | 104877497 | 104878525 | 1029 | -2.94 | 0.0027 | 0.00334 |
| chr11 | 115000333 | 115005595 | 5263 | 2.52 | 0.00271 | 0.00334 |
| chr12 | 95783865 | 95790287 | 6423 | -4.43 | 0.00271 | 0.00335 |
| chrX | 65318098 | 65322063 | 3966 | 3.75 | 0.00271 | 0.00335 |
| chr4 | 136075237 | 136077912 | 2676 | -1.81 | 0.00271 | 0.00335 |
| chr2 | 156940872 | 156953249 | 12378 | -1.15 | 0.00271 | 0.00335 |
| chr2 | 13552566 | 13553489 | 924 | 1.95 | 0.00271 | 0.00335 |
| chr9 | 89432939 | 89440633 | 7695 | -3.92 | 0.00271 | 0.00335 |
| chr2 | 119733308 | 119743753 | 10446 | -3.69 | 0.00272 | 0.00335 |
| chr12 | 60558647 | 60560253 | 1607 | -3.12 | 0.00272 | 0.00336 |
| chr6 | 61926715 | 61929013 | 2299 | -1.45 | 0.00272 | 0.00336 |
| chr11 | 62047823 | 62076927 | 29105 | -2.13 | 0.00272 | 0.00336 |
| chr12 | 92392133 | 92412900 | 20768 | -6.64 | 0.00272 | 0.00336 |
| chr13 | 60906454 | 60928374 | 21921 | -3.46 | 0.00272 | 0.00336 |
| chr2 | 57063894 | 57106005 | 42112 | -0.83 | 0.00272 | 0.00336 |
| chr11 | 106443160 | 106457531 | 14372 | -0.99 | 0.00273 | 0.00336 |
| chr7 | 25241765 | 25250636 | 8872 | -4.22 | 0.00273 | 0.00337 |
| chr12 | 106218973 | 106219787 | 815 | -3.21 | 0.00275 | 0.00339 |
| chr4 | 69025420 | 69026094 | 675 | 1.63 | 0.00275 | 0.00339 |
| chr15 | 60241174 | 60316256 | 75083 | -1.22 | 0.00275 | 0.00339 |
| chr13 | 23556149 | 23561198 | 5050 | -2.85 | 0.00275 | 0.0034 |

| | | | | | | |
|---|---|---|---|---|---|---|
| chr4 | 130642562 | 130643255 | 694 | -2.96 | 0.00276 | 0.0034 |
| chr7 | 84584308 | 84584749 | 442 | -2.96 | 0.00276 | 0.0034 |
| chr4 | 33956278 | 33974375 | 18098 | -1.11 | 0.00277 | 0.00341 |
| chr14 | 42730042 | 42755717 | 25676 | -2.17 | 0.00277 | 0.00342 |
| chr2 | 176110108 | 176114179 | 4072 | 3.88 | 0.00277 | 0.00342 |
| chr2 | 57867748 | 57883656 | 15909 | -0.98 | 0.00278 | 0.00343 |
| chr6 | 76102314 | 76102890 | 577 | -3.05 | 0.00279 | 0.00344 |
| chr9 | 121576334 | 121577464 | 1131 | 2.3 | 0.00279 | 0.00344 |
| chr14 | 36995767 | 37012922 | 17156 | -0.99 | 0.00279 | 0.00344 |
| chr1 | 4581287 | 4647030 | 65744 | -0.57 | 0.00279 | 0.00344 |
| chr3 | 86711024 | 86738174 | 27151 | 0.79 | 0.0028 | 0.00345 |
| chr1 | 203315396 | 203384256 | 68861 | -1.74 | 0.0028 | 0.00345 |
| chr11 | 85008224 | 85053091 | 44868 | -4.52 | 0.0028 | 0.00345 |
| chr16 | 55363481 | 55390131 | 26651 | 2.26 | 0.0028 | 0.00345 |
| chr2 | 199165076 | 199165773 | 698 | -3.17 | 0.0028 | 0.00346 |
| chr2 | 127699325 | 127726531 | 27207 | -4.52 | 0.0028 | 0.00346 |
| chr8 | 80585806 | 80594119 | 8314 | -3.1 | 0.0028 | 0.00346 |
| chr1 | 223216691 | 223229597 | 12907 | -1.72 | 0.00281 | 0.00347 |
| chr1 | 44861317 | 44863711 | 2395 | -1.31 | 0.00281 | 0.00347 |
| chr5 | 56766584 | 56772313 | 5730 | -2.41 | 0.00281 | 0.00347 |
| chr5 | 146620323 | 146659265 | 38943 | 4.56 | 0.00282 | 0.00348 |
| chr14 | 77178651 | 77210920 | 32270 | -3.85 | 0.00284 | 0.0035 |
| chr11 | 58445626 | 58517789 | 72164 | 4.79 | 0.00284 | 0.0035 |
| chr10 | 9322161 | 9330312 | 8152 | -3 | 0.00284 | 0.0035 |
| chr10 | 98025428 | 98027654 | 2227 | 1.54 | 0.00284 | 0.00351 |
| chr13 | 89243806 | 89248221 | 4416 | -2.03 | 0.00285 | 0.00351 |
| chr7 | 42855263 | 42863127 | 7865 | -2.4 | 0.00285 | 0.00351 |
| chr6 | 41948300 | 41972546 | 24247 | -6.59 | 0.00286 | 0.00352 |
| chr13 | 44289550 | 44292282 | 2733 | -1.42 | 0.00286 | 0.00352 |
| chr10 | 63089408 | 63149767 | 60360 | 0.87 | 0.00286 | 0.00352 |
| chr10 | 9125711 | 9131896 | 6186 | -1.36 | 0.00286 | 0.00353 |
| chr13 | 91235286 | 91244970 | 9685 | -4.55 | 0.00287 | 0.00353 |
| chrX | 114042687 | 114044667 | 1981 | 1.4 | 0.00287 | 0.00354 |
| chr9 | 101254892 | 101256628 | 1737 | -3.44 | 0.00287 | 0.00354 |
| chr5 | 100002741 | 100003533 | 793 | -4.03 | 0.00287 | 0.00354 |
| chr6 | 119666497 | 119668639 | 2143 | -3.37 | 0.00288 | 0.00355 |
| chr6 | 79523756 | 79527315 | 3560 | -1.03 | 0.00289 | 0.00356 |
| chr6 | 77958942 | 77961632 | 2691 | -3.58 | 0.00289 | 0.00356 |
| chr14 | 54760642 | 54762076 | 1435 | -3.49 | 0.00289 | 0.00356 |
| chrX | 99953057 | 99953816 | 760 | -2.92 | 0.00289 | 0.00356 |
| chr2 | 214531993 | 214548317 | 16325 | -1.36 | 0.00289 | 0.00357 |
| chr15 | 77298051 | 77299530 | 1480 | -1.82 | 0.00289 | 0.00357 |
| chr17 | 66582642 | 66588377 | 5736 | -4.01 | 0.0029 | 0.00357 |
| chr11 | 63075969 | 63105204 | 29236 | -0.95 | 0.0029 | 0.00358 |
| chrX | 68593969 | 68598287 | 4319 | -1.91 | 0.0029 | 0.00358 |
| chr6 | 45931627 | 45944586 | 12960 | -1.58 | 0.0029 | 0.00358 |
| chr6 | 100503548 | 100517467 | 13920 | -3.9 | 0.00291 | 0.00358 |
| chr1 | 43360039 | 43360632 | 594 | -2.1 | 0.00291 | 0.00358 |
| chr2 | 166594162 | 166596235 | 2074 | -1.86 | 0.00291 | 0.00358 |
| chr13 | 74176077 | 74187847 | 11771 | -4.69 | 0.00291 | 0.00359 |
| chr16 | 58885412 | 58998300 | 112889 | 0.79 | 0.00292 | 0.0036 |
| chr12 | 18618163 | 18653565 | 35403 | 1.08 | 0.00292 | 0.0036 |
| chr11 | 99102171 | 99103195 | 1025 | -2.49 | 0.00293 | 0.00361 |
| chr3 | 192228199 | 192231873 | 3675 | -1.7 | 0.00293 | 0.00361 |
| chr1 | 99347078 | 99348707 | 1630 | -2.45 | 0.00293 | 0.00361 |
| chr3 | 41091131 | 41092005 | 875 | 1.87 | 0.00293 | 0.00361 |
| chr12 | 125107844 | 125135092 | 27249 | -6.57 | 0.00294 | 0.00362 |
| chr5 | 31246040 | 31260439 | 14400 | 1.39 | 0.00294 | 0.00362 |
| chr7 | 93360661 | 93362027 | 1367 | -3.48 | 0.00295 | 0.00363 |
| chr9 | 73877695 | 73896959 | 19265 | 0.75 | 0.00295 | 0.00363 |
| chr6 | 84419603 | 84423263 | 3661 | -4.15 | 0.00295 | 0.00363 |

| | | | | | | |
|---|---|---|---|---|---|---|
| chr2 | 166447095 | 166449648 | 2554 | -1.97 | 0.00295 | 0.00364 |
| chr1 | 7936246 | 7940193 | 3948 | -4.19 | 0.00296 | 0.00364 |
| chr9 | 106983488 | 106983829 | 342 | -3.75 | 0.00296 | 0.00365 |
| chr3 | 130185643 | 130189017 | 3375 | 1.54 | 0.00296 | 0.00365 |
| chr13 | 32532189 | 32533819 | 1631 | -3.15 | 0.00297 | 0.00366 |
| chr4 | 141877370 | 141877970 | 601 | -3.1 | 0.00297 | 0.00366 |
| chr6 | 40667160 | 40676292 | 9133 | -4.27 | 0.00297 | 0.00366 |
| chr15 | 60128865 | 60175610 | 46746 | 1.18 | 0.00298 | 0.00367 |
| chr2 | 67377457 | 67380647 | 3191 | -1.39 | 0.00298 | 0.00367 |
| chr14 | 106985672 | 106997064 | 11393 | -1.68 | 0.00298 | 0.00367 |
| chr11 | 44399227 | 44419570 | 20344 | -4.06 | 0.003 | 0.00369 |
| chr13 | 68364583 | 68378300 | 13718 | 1.35 | 0.003 | 0.0037 |
| chr9 | 135881751 | 135894658 | 12908 | -4.56 | 0.00301 | 0.0037 |
| chr3 | 6169736 | 6172319 | 2584 | -2.64 | 0.00302 | 0.00372 |
| chr9 | 78894735 | 78906067 | 11333 | 1.15 | 0.00302 | 0.00372 |
| chr14 | 67370729 | 67371754 | 1026 | -3.25 | 0.00303 | 0.00373 |
| chr2 | 212932306 | 213072234 | 139929 | 0.9 | 0.00303 | 0.00373 |
| chr8 | 11840386 | 11856736 | 16351 | -2.2 | 0.00303 | 0.00373 |
| chr4 | 19294086 | 19299238 | 5153 | 1.04 | 0.00305 | 0.00375 |
| chr3 | 59959933 | 59960900 | 968 | 2.32 | 0.00305 | 0.00375 |
| chr4 | 54404035 | 54405803 | 1769 | 2.62 | 0.00305 | 0.00375 |
| chr16 | 77544506 | 77581265 | 36760 | -0.86 | 0.00305 | 0.00375 |
| chr12 | 131547463 | 131549152 | 1690 | 1.77 | 0.00305 | 0.00376 |
| chr3 | 61363528 | 61381000 | 17473 | 0.82 | 0.00305 | 0.00376 |
| chr14 | 67874242 | 67874679 | 438 | -3.62 | 0.00306 | 0.00376 |
| chr16 | 59355409 | 59412047 | 56639 | -2.29 | 0.00306 | 0.00376 |
| chr11 | 125054125 | 125070068 | 15944 | -4.52 | 0.00306 | 0.00377 |
| chr2 | 142880039 | 142880870 | 832 | 3.22 | 0.00306 | 0.00377 |
| chr16 | 12950326 | 12989684 | 39359 | -6.53 | 0.00306 | 0.00377 |
| chr8 | 22705844 | 22707598 | 1755 | -3.68 | 0.00307 | 0.00378 |
| chr5 | 125242708 | 125333776 | 91069 | -0.71 | 0.00307 | 0.00378 |
| chr7 | 67831209 | 67868335 | 37127 | -3.64 | 0.00307 | 0.00378 |
| chr14 | 39945527 | 39979901 | 34375 | -0.99 | 0.00307 | 0.00378 |
| chr2 | 186703874 | 186741870 | 37997 | -1.21 | 0.00308 | 0.00379 |
| chr1 | 164637070 | 164639238 | 2169 | -3.64 | 0.00308 | 0.00379 |
| chr3 | 80961823 | 81090749 | 128927 | 0.65 | 0.00308 | 0.0038 |
| chr13 | 72091576 | 72097482 | 5907 | 2.07 | 0.00309 | 0.0038 |
| chr12 | 89629985 | 89631692 | 1708 | 1.23 | 0.00309 | 0.0038 |
| chr2 | 215071006 | 215071375 | 370 | -2.76 | 0.00309 | 0.00381 |
| chr18 | 42666940 | 42676916 | 9977 | 1.98 | 0.0031 | 0.00382 |
| chr3 | 120548718 | 120550754 | 2037 | 3.65 | 0.0031 | 0.00382 |
| chr6 | 18582249 | 18583779 | 1531 | 3.26 | 0.00311 | 0.00382 |
| chr2 | 8062787 | 8072871 | 10085 | -3.68 | 0.00311 | 0.00382 |
| chr1 | 185167930 | 185194407 | 26478 | -6.52 | 0.00311 | 0.00382 |
| chr2 | 200075183 | 200088818 | 13636 | -1.29 | 0.00311 | 0.00383 |
| chrX | 83523011 | 83525831 | 2821 | 1.42 | 0.00312 | 0.00383 |
| chr2 | 105033202 | 105196301 | 163100 | -0.71 | 0.00312 | 0.00383 |
| chr4 | 76540326 | 76547881 | 7556 | -4.94 | 0.00312 | 0.00384 |
| chrX | 132931484 | 132934035 | 2552 | 1.5 | 0.00312 | 0.00384 |
| chr2 | 38128051 | 38129363 | 1313 | -3.3 | 0.00313 | 0.00385 |
| chr11 | 40566804 | 40578114 | 11311 | 2.65 | 0.00314 | 0.00386 |
| chr4 | 47819407 | 47820368 | 962 | 3.22 | 0.00314 | 0.00386 |
| chr7 | 116874071 | 116887656 | 13586 | -4.23 | 0.00315 | 0.00387 |
| chr1 | 101824426 | 101824833 | 408 | -2.86 | 0.00315 | 0.00387 |
| chr3 | 173984034 | 173986022 | 1989 | 1.51 | 0.00316 | 0.00389 |
| chr1 | 108446514 | 108448587 | 2074 | 2.17 | 0.00317 | 0.0039 |
| chr19 | 47811143 | 47825537 | 14395 | -6.5 | 0.00318 | 0.00391 |
| chr4 | 128216319 | 128218594 | 2276 | -1.82 | 0.00318 | 0.00391 |
| chrX | 17868951 | 17871400 | 2450 | 1.52 | 0.00318 | 0.00391 |
| chr6 | 79968692 | 79969534 | 843 | -2.82 | 0.00319 | 0.00392 |
| chr15 | 70777425 | 70777737 | 313 | -2.82 | 0.00319 | 0.00392 |

| | | | | | | |
|---|---|---|---|---|---|---|
| chr21 | 15617562 | 15621451 | 3890 | 1.65 | 0.00319 | 0.00392 |
| chrX | 43631713 | 43647435 | 15723 | -1.09 | 0.0032 | 0.00393 |
| chr4 | 33330669 | 33343608 | 12940 | -1.77 | 0.0032 | 0.00393 |
| chr1 | 93447669 | 93448967 | 1299 | -3.15 | 0.0032 | 0.00393 |
| chr7 | 86713358 | 86757353 | 43996 | 4.61 | 0.0032 | 0.00394 |
| chr7 | 79575763 | 79577842 | 2080 | -1.22 | 0.00321 | 0.00394 |
| chr7 | 82278672 | 82281574 | 2903 | -3.17 | 0.00321 | 0.00395 |
| chr8 | 113145863 | 113149359 | 3497 | -1.57 | 0.00321 | 0.00395 |
| chr14 | 106244351 | 106297745 | 53395 | -2.1 | 0.00321 | 0.00395 |
| chr2 | 98676604 | 98695064 | 18461 | -3.47 | 0.00322 | 0.00396 |
| chr14 | 94373905 | 94386883 | 12979 | -4.13 | 0.00322 | 0.00396 |
| chr11 | 93189206 | 93191668 | 2463 | -1.81 | 0.00323 | 0.00396 |
| chr2 | 222782602 | 222802575 | 19974 | 4.11 | 0.00323 | 0.00397 |
| chr15 | 96584967 | 96593290 | 8324 | -1.3 | 0.00323 | 0.00397 |
| chr5 | 142413212 | 142423061 | 9850 | -4.11 | 0.00324 | 0.00398 |
| chr3 | 81356489 | 81357387 | 899 | -2.91 | 0.00324 | 0.00399 |
| chr4 | 108231478 | 108232590 | 1113 | -2.13 | 0.00325 | 0.00399 |
| chr4 | 61290812 | 61300658 | 9847 | 1.16 | 0.00325 | 0.00399 |
| chr13 | 28327561 | 28330575 | 3015 | -3.88 | 0.00325 | 0.004 |
| chr14 | 81903798 | 81912427 | 8630 | -1.91 | 0.00327 | 0.00402 |
| chr6 | 47961121 | 47962732 | 1612 | 1.34 | 0.00327 | 0.00402 |
| chr4 | 39513814 | 39536985 | 23172 | -4.48 | 0.00328 | 0.00402 |
| chr18 | 68556505 | 68565067 | 8563 | 1.09 | 0.00328 | 0.00402 |
| chr4 | 145839081 | 145863034 | 23954 | 0.82 | 0.00328 | 0.00403 |
| chr4 | 157807326 | 157808033 | 708 | -4.58 | 0.00328 | 0.00403 |
| chr6 | 165403326 | 165405881 | 2556 | 1.52 | 0.00329 | 0.00404 |
| chrX | 73070087 | 73071748 | 1662 | -3.05 | 0.00329 | 0.00404 |
| chr1 | 112881170 | 112928171 | 47002 | -4.38 | 0.00329 | 0.00404 |
| chr21 | 30654249 | 30655605 | 1357 | -3.22 | 0.0033 | 0.00405 |
| chr9 | 80191128 | 80193563 | 2436 | 1.88 | 0.00331 | 0.00407 |
| chr12 | 119977970 | 119980694 | 2725 | 1.83 | 0.00331 | 0.00407 |
| chr4 | 95000764 | 95014935 | 14172 | -1.18 | 0.00332 | 0.00407 |
| chr6 | 152046729 | 152049169 | 2441 | -1.96 | 0.00332 | 0.00408 |
| chr1 | 233479391 | 233491334 | 11944 | -5.37 | 0.00333 | 0.00409 |
| chrX | 63052166 | 63058682 | 6517 | 0.98 | 0.00334 | 0.0041 |
| chr2 | 143233063 | 143234671 | 1609 | -3.38 | 0.00334 | 0.0041 |
| chr12 | 69349457 | 69349977 | 521 | -3.89 | 0.00334 | 0.0041 |
| chr13 | 43268981 | 43282525 | 13545 | 1.67 | 0.00334 | 0.0041 |
| chr11 | 18062644 | 18071131 | 8488 | -1.04 | 0.00336 | 0.00413 |
| chr6 | 1011932 | 1054882 | 42951 | -4.26 | 0.00337 | 0.00414 |
| chr1 | 66312947 | 66367306 | 54360 | -1.63 | 0.00337 | 0.00414 |
| chr1 | 38762639 | 38791145 | 28507 | -1.69 | 0.00338 | 0.00415 |
| chrX | 79730834 | 79763182 | 32349 | 1.05 | 0.00338 | 0.00415 |
| chr22 | 37275912 | 37315378 | 39467 | -4.32 | 0.0034 | 0.00417 |
| chr15 | 35930462 | 35936312 | 5851 | -0.87 | 0.0034 | 0.00418 |
| chrX | 57978469 | 58001094 | 22626 | 1.49 | 0.00341 | 0.00418 |
| chr13 | 36842952 | 36845030 | 2079 | -2.32 | 0.00341 | 0.00418 |
| chr12 | 65524785 | 65551768 | 26984 | -1.14 | 0.00341 | 0.00419 |
| chr20 | 45488047 | 45499972 | 11926 | -1.25 | 0.00341 | 0.00419 |
| chr4 | 29744272 | 29748756 | 4485 | 1.41 | 0.00342 | 0.00419 |
| chr6 | 160973344 | 161032058 | 58715 | -0.76 | 0.00342 | 0.00419 |
| chr1 | 4495388 | 4520188 | 24801 | -1.35 | 0.00342 | 0.00419 |
| chr12 | 97030504 | 97039187 | 8684 | -0.91 | 0.00342 | 0.0042 |
| chr6 | 122844349 | 122849274 | 4926 | -1.97 | 0.00342 | 0.0042 |
| chr3 | 108825386 | 108826915 | 1530 | 1.57 | 0.00343 | 0.00421 |
| chrX | 92156325 | 92157866 | 1542 | 1.5 | 0.00345 | 0.00423 |
| chr2 | 68251617 | 68253878 | 2262 | -1.07 | 0.00345 | 0.00424 |
| chr12 | 101463339 | 101476623 | 13285 | 5.95 | 0.00345 | 0.00424 |
| chr15 | 53497720 | 53498808 | 1089 | -1.77 | 0.00346 | 0.00424 |
| chr11 | 93940509 | 93942628 | 2120 | -1.26 | 0.00346 | 0.00425 |
| chr10 | 87468341 | 87474113 | 5773 | -4.02 | 0.00346 | 0.00425 |

| | | | | | | |
|---|---|---|---|---|---|---|
| chr6 | 164270482 | 164273190 | 2709 | -1.29 | 0.00347 | 0.00425 |
| chr2 | 127431305 | 127435142 | 3838 | -4.01 | 0.00347 | 0.00425 |
| chr6 | 105110699 | 105128537 | 17839 | -4.02 | 0.00347 | 0.00426 |
| chr6 | 128246205 | 128254742 | 8538 | -2.17 | 0.00347 | 0.00426 |
| chr10 | 95877081 | 95879456 | 2376 | -3.82 | 0.00347 | 0.00426 |
| chr6 | 148960186 | 148997274 | 37089 | -4.49 | 0.00348 | 0.00427 |
| chr6 | 79024180 | 79027310 | 3131 | 3.65 | 0.00348 | 0.00427 |
| chr5 | 38689373 | 38702903 | 13531 | -1.65 | 0.00348 | 0.00427 |
| chrX | 135359890 | 135362990 | 3101 | 1.4 | 0.00348 | 0.00427 |
| chr9 | 44194814 | 44195209 | 396 | -2.45 | 0.00349 | 0.00428 |
| chr14 | 21387903 | 21396896 | 8994 | -0.76 | 0.00349 | 0.00428 |
| chr13 | 30555525 | 30562831 | 7307 | -2.86 | 0.00349 | 0.00429 |
| chr16 | 56236253 | 56244780 | 8528 | -3.85 | 0.0035 | 0.00429 |
| chrX | 116704714 | 116741391 | 36678 | 1.06 | 0.0035 | 0.0043 |
| chr5 | 37835792 | 38374533 | 538742 | 0.33 | 0.00351 | 0.0043 |
| chr8 | 57999036 | 58000329 | 1294 | -2.04 | 0.00351 | 0.0043 |
| chr2 | 186886800 | 186889749 | 2950 | 1.94 | 0.00351 | 0.00431 |
| chrX | 66718158 | 66740297 | 22140 | -2.1 | 0.00352 | 0.00431 |
| chrX | 42317344 | 42365881 | 48538 | 4.55 | 0.00353 | 0.00433 |
| chr6 | 137608072 | 137609467 | 1396 | -3.43 | 0.00354 | 0.00434 |
| chr2 | 170097752 | 170105116 | 7365 | -1.06 | 0.00356 | 0.00436 |
| chr12 | 7647258 | 7699658 | 52401 | 0.67 | 0.00356 | 0.00436 |
| chr8 | 61844693 | 61869185 | 24493 | -2.92 | 0.00356 | 0.00437 |
| chr8 | 39681588 | 39684280 | 2693 | 1.63 | 0.00357 | 0.00437 |
| chr6 | 130975103 | 130976013 | 911 | 1.77 | 0.00357 | 0.00438 |
| chr10 | 28628045 | 28638890 | 10846 | -1.46 | 0.00357 | 0.00438 |
| chr3 | 70912657 | 70930008 | 17352 | -3.16 | 0.00358 | 0.00439 |
| chr15 | 34733151 | 34733511 | 361 | -3.6 | 0.00358 | 0.00439 |
| chr1 | 99273836 | 99276164 | 2329 | -1.63 | 0.00358 | 0.00439 |
| chrX | 34604654 | 34608784 | 4131 | -3.81 | 0.00359 | 0.0044 |
| chr1 | 157867078 | 157882890 | 15813 | -2.89 | 0.00359 | 0.0044 |
| chr12 | 15501751 | 15506962 | 5212 | 1.69 | 0.0036 | 0.00442 |
| chrX | 24469095 | 24472727 | 3633 | -3.93 | 0.00361 | 0.00442 |
| chr7 | 26576978 | 26611565 | 34588 | -1.13 | 0.00361 | 0.00442 |
| chrX | 58060472 | 58071662 | 11191 | 0.86 | 0.00361 | 0.00443 |
| chr1 | 213513056 | 213514821 | 1766 | 3.49 | 0.00362 | 0.00444 |
| chr4 | 156618941 | 156623280 | 4340 | 1.7 | 0.00362 | 0.00444 |
| chr6 | 90762103 | 90763963 | 1861 | -2.44 | 0.00363 | 0.00444 |
| chr6 | 28412217 | 28445317 | 33101 | -4.13 | 0.00363 | 0.00445 |
| chr1 | 31058246 | 31067738 | 9493 | -2.37 | 0.00363 | 0.00445 |
| chr2 | 157831371 | 157833157 | 1787 | -3.56 | 0.00364 | 0.00446 |
| chr4 | 23608346 | 23627691 | 19346 | -3.45 | 0.00364 | 0.00446 |
| chr3 | 24794784 | 24796197 | 1414 | -1.73 | 0.00366 | 0.00448 |
| chr11 | 104666415 | 104667607 | 1193 | -3.4 | 0.00366 | 0.00448 |
| chr4 | 65979438 | 65981199 | 1762 | 3.3 | 0.00366 | 0.00448 |
| chr18 | 26953181 | 26954162 | 982 | -2.69 | 0.00366 | 0.00448 |
| chr2 | 157070302 | 157072326 | 2025 | 2.81 | 0.00366 | 0.00448 |
| chr8 | 1033320 | 1111341 | 78022 | -0.66 | 0.00367 | 0.00449 |
| chr10 | 21403248 | 21418668 | 15421 | -6.38 | 0.00367 | 0.00449 |
| chrX | 81399734 | 81423793 | 24060 | -1.34 | 0.00367 | 0.0045 |
| chr4 | 49016107 | 49021065 | 4959 | -1.86 | 0.00367 | 0.0045 |
| chr3 | 61543118 | 61545120 | 2003 | -2.86 | 0.00367 | 0.0045 |
| chr14 | 54335137 | 54343804 | 8668 | -4.63 | 0.00368 | 0.0045 |
| chr8 | 18886531 | 18902517 | 15987 | -4.63 | 0.00368 | 0.0045 |
| chr10 | 57388735 | 57390844 | 2110 | -2.01 | 0.00368 | 0.00451 |
| chr4 | 155883661 | 155886082 | 2422 | -3.35 | 0.00368 | 0.00451 |
| chr1 | 167476960 | 167478751 | 1792 | 3.52 | 0.00368 | 0.00451 |
| chr6 | 13126754 | 13128333 | 1580 | -2.54 | 0.00369 | 0.00452 |
| chr9 | 93641059 | 93643374 | 2316 | 1.81 | 0.00369 | 0.00452 |
| chr6 | 110588424 | 110631460 | 43037 | -1.86 | 0.0037 | 0.00453 |
| chrX | 53061743 | 53062818 | 1076 | -2.17 | 0.0037 | 0.00453 |

| | | | | | | |
|---|---|---|---|---|---|---|
| chr11 | 62732798 | 62739412 | 6615 | -4.32 | 0.0037 | 0.00454 |
| chr11 | 110352268 | 110353128 | 861 | -2.58 | 0.00371 | 0.00454 |
| chr2 | 187745868 | 187747330 | 1463 | -3.49 | 0.00371 | 0.00454 |
| chr12 | 106027955 | 106047545 | 19591 | -4.4 | 0.00371 | 0.00454 |
| chr9 | 109600259 | 109602468 | 2210 | 1.59 | 0.00371 | 0.00454 |
| chrX | 72442298 | 72443103 | 806 | -3.02 | 0.00371 | 0.00454 |
| chr7 | 148188104 | 148191921 | 3818 | -4.01 | 0.00371 | 0.00455 |
| chr5 | 62046976 | 62055691 | 8716 | -3.61 | 0.00371 | 0.00455 |
| chr13 | 62979763 | 62982734 | 2972 | -1.45 | 0.00372 | 0.00455 |
| chr4 | 164148392 | 164149620 | 1229 | -1.61 | 0.00372 | 0.00455 |
| chrX | 141253211 | 141272782 | 19572 | 0.89 | 0.00372 | 0.00455 |
| chr6 | 169657703 | 169658679 | 977 | -3.01 | 0.00372 | 0.00456 |
| chr21 | 18126417 | 18128962 | 2546 | 1.8 | 0.00374 | 0.00457 |
| chr3 | 71784937 | 71798265 | 13329 | -4.45 | 0.00374 | 0.00457 |
| chrX | 129647482 | 129648680 | 1199 | 3.48 | 0.00374 | 0.00458 |
| chr2 | 146220007 | 146232568 | 12562 | -6.36 | 0.00376 | 0.0046 |
| chr7 | 23599314 | 23604115 | 4802 | -3.83 | 0.00376 | 0.0046 |
| chrX | 130174451 | 130174821 | 371 | -2.32 | 0.00376 | 0.0046 |
| chr2 | 102227399 | 102228252 | 854 | -2.9 | 0.00377 | 0.00462 |
| chr13 | 39721728 | 39757217 | 35490 | 0.68 | 0.00378 | 0.00462 |
| chr12 | 80910701 | 80912150 | 1450 | -3.32 | 0.00378 | 0.00463 |
| chr6 | 50931383 | 50932781 | 1399 | 1.92 | 0.00379 | 0.00463 |
| chr3 | 164907396 | 164907694 | 299 | 2.92 | 0.0038 | 0.00466 |
| chr2 | 98710490 | 98764996 | 54507 | -4.51 | 0.00381 | 0.00466 |
| chr4 | 135889269 | 135890492 | 1224 | -1.78 | 0.00381 | 0.00466 |
| chr5 | 59436293 | 59449802 | 13510 | -1.07 | 0.00381 | 0.00466 |
| chr4 | 143890239 | 143895716 | 5478 | 2.19 | 0.00381 | 0.00466 |
| chr10 | 27851027 | 27861630 | 10604 | -3.68 | 0.00382 | 0.00467 |
| chr9 | 101099001 | 101131242 | 32242 | -4.52 | 0.00382 | 0.00467 |
| chr3 | 162483335 | 162485069 | 1735 | -3.47 | 0.00382 | 0.00467 |
| chr7 | 132161219 | 132162929 | 1711 | -2.06 | 0.00382 | 0.00467 |
| chr6 | 75751902 | 75753294 | 1393 | -1.36 | 0.00382 | 0.00467 |
| chrX | 49460722 | 49462646 | 1925 | -1.29 | 0.00382 | 0.00468 |
| chrX | 48058843 | 48065860 | 7018 | -2.45 | 0.00383 | 0.00468 |
| chr3 | 55534526 | 55537319 | 2794 | 2.41 | 0.00383 | 0.00468 |
| chr11 | 106376882 | 106378499 | 1618 | 3.45 | 0.00383 | 0.00468 |
| chr12 | 10100845 | 10101378 | 534 | -1.9 | 0.00383 | 0.00469 |
| chr4 | 27690176 | 27694614 | 4439 | 4 | 0.00384 | 0.0047 |
| chr11 | 104469553 | 104479532 | 9980 | -2.16 | 0.00384 | 0.0047 |
| chr4 | 119087705 | 119089179 | 1475 | -3.17 | 0.00384 | 0.0047 |
| chr7 | 100677028 | 100705536 | 28509 | -1.91 | 0.00385 | 0.00471 |
| chr18 | 70565026 | 70586048 | 21023 | 4.35 | 0.00385 | 0.00471 |
| chrX | 47982112 | 47983329 | 1218 | -1.85 | 0.00387 | 0.00473 |
| chr16 | 49330048 | 49338441 | 8394 | 1.04 | 0.00387 | 0.00474 |
| chr12 | 53202081 | 53203644 | 1564 | 3.4 | 0.00389 | 0.00475 |
| chr8 | 31954844 | 31964628 | 9785 | -2.72 | 0.00389 | 0.00476 |
| chr3 | 99447113 | 99459510 | 12398 | 1.46 | 0.00389 | 0.00476 |
| chr5 | 143415953 | 143418015 | 2063 | 1.69 | 0.0039 | 0.00476 |
| chr1 | 88744141 | 88745301 | 1161 | -1.63 | 0.0039 | 0.00477 |
| chr2 | 163631732 | 163645920 | 14189 | 2.75 | 0.0039 | 0.00477 |
| chr2 | 158522196 | 158522882 | 687 | -2.85 | 0.0039 | 0.00477 |
| chr8 | 133507252 | 133508249 | 998 | 2.05 | 0.00391 | 0.00478 |
| chr14 | 53663458 | 53674613 | 11156 | -1.19 | 0.00391 | 0.00478 |
| chr5 | 22191567 | 22192016 | 450 | 1.73 | 0.00392 | 0.00479 |
| chr20 | 57673331 | 57760092 | 86762 | -0.84 | 0.00393 | 0.0048 |
| chr9 | 82819262 | 83215083 | 395822 | 0.28 | 0.00393 | 0.00481 |
| chr6 | 7699688 | 7712167 | 12480 | -5.06 | 0.00394 | 0.00482 |
| chr11 | 33868502 | 33870886 | 2385 | 2.04 | 0.00395 | 0.00482 |
| chr5 | 27889847 | 28946650 | 1056804 | 0.72 | 0.00396 | 0.00484 |
| chrX | 3440677 | 3457562 | 16886 | -1.11 | 0.00396 | 0.00484 |
| chr3 | 106131941 | 106153987 | 22047 | 4.18 | 0.00397 | 0.00485 |

| | | | | | | |
|---|---|---|---|---|---|---|
| chr1 | 115381128 | 115395688 | 14561 | 2.56 | 0.00397 | 0.00486 |
| chr4 | 4768089 | 4838505 | 70417 | -4.5 | 0.00397 | 0.00486 |
| chr6 | 25104340 | 25112553 | 8214 | -1.39 | 0.00398 | 0.00487 |
| chr22 | 25134272 | 25144315 | 10044 | -1.53 | 0.00399 | 0.00488 |
| chr15 | 77198225 | 77214140 | 15916 | 2.63 | 0.004 | 0.00488 |
| chrX | 135419235 | 135420468 | 1234 | -1.82 | 0.004 | 0.00489 |
| chr5 | 112556951 | 112559051 | 2101 | -1.88 | 0.00401 | 0.0049 |
| chr14 | 48614901 | 48708589 | 93689 | -1.6 | 0.00401 | 0.0049 |
| chr6 | 134392139 | 134392930 | 792 | -2.81 | 0.00402 | 0.00491 |
| chr8 | 123126538 | 123138426 | 11889 | -1.94 | 0.00402 | 0.00492 |
| chr11 | 104844301 | 104861786 | 17486 | -1.06 | 0.00403 | 0.00493 |
| chrX | 12907168 | 12909948 | 2781 | -1.67 | 0.00404 | 0.00493 |
| chr9 | 112394649 | 112395975 | 1327 | 2.01 | 0.00404 | 0.00494 |
| chr14 | 77034495 | 77156767 | 122273 | -6.28 | 0.00405 | 0.00494 |
| chr18 | 76675236 | 76679891 | 4656 | 1.03 | 0.00405 | 0.00495 |
| chr6 | 77488375 | 77493378 | 5004 | -1.59 | 0.00406 | 0.00496 |
| chrX | 75591975 | 75600132 | 8158 | 3.35 | 0.00408 | 0.00498 |
| chr9 | 86719729 | 86733731 | 14003 | -4.27 | 0.00408 | 0.00498 |
| chr7 | 151580634 | 151582900 | 2267 | 1.24 | 0.00408 | 0.00499 |
| chrX | 37152072 | 37156274 | 4203 | -1.02 | 0.00409 | 0.00499 |
| chr9 | 103406059 | 103422616 | 16558 | -0.95 | 0.00409 | 0.00499 |
| chrX | 8721225 | 8742978 | 21754 | -0.66 | 0.0041 | 0.00501 |
| chr2 | 67393442 | 67393982 | 541 | -2.69 | 0.0041 | 0.00501 |
| chr12 | 45371789 | 45507217 | 135429 | 0.7 | 0.00411 | 0.00502 |
| chr2 | 89137760 | 89145948 | 8189 | 1.09 | 0.00411 | 0.00502 |
| chr8 | 131922608 | 131923022 | 415 | 2.07 | 0.00411 | 0.00502 |
| chr13 | 27370744 | 27382946 | 12203 | -1.69 | 0.00412 | 0.00503 |
| chr5 | 40658171 | 40659315 | 1145 | -3.22 | 0.00412 | 0.00503 |
| chr13 | 36591900 | 36597410 | 5511 | -1.55 | 0.00412 | 0.00503 |
| chr7 | 124375680 | 124377283 | 1604 | -3.38 | 0.00413 | 0.00504 |
| chr1 | 159685343 | 159747609 | 62267 | -0.68 | 0.00413 | 0.00504 |
| chr4 | 106951003 | 106952858 | 1856 | -1.45 | 0.00413 | 0.00504 |
| chr2 | 84450405 | 84470878 | 20474 | -3.77 | 0.00414 | 0.00505 |
| chr14 | 97101503 | 97106409 | 4907 | 1.86 | 0.00414 | 0.00505 |
| chr2 | 222570149 | 222573123 | 2975 | 1.56 | 0.00414 | 0.00506 |
| chr6 | 47926364 | 47927282 | 919 | -3.2 | 0.00415 | 0.00506 |
| chr9 | 37181371 | 37185777 | 4407 | -4.08 | 0.00415 | 0.00507 |
| chr1 | 147015683 | 147016279 | 597 | -2.85 | 0.00415 | 0.00507 |
| chr10 | 3030574 | 3082533 | 51960 | -1.37 | 0.00416 | 0.00507 |
| chr8 | 80611305 | 80623520 | 12216 | -3.7 | 0.00416 | 0.00508 |
| chr14 | 106337881 | 106343392 | 5512 | 1.26 | 0.00417 | 0.00509 |
| chr3 | 167981644 | 168821642 | 839999 | 7.09 | 0.00418 | 0.0051 |
| chr2 | 84390386 | 84391589 | 1204 | 1.75 | 0.00419 | 0.00511 |
| chr3 | 130030606 | 130056120 | 25515 | -1.54 | 0.00419 | 0.00511 |
| chr18 | 4082680 | 4115258 | 32579 | -1.54 | 0.00419 | 0.00511 |
| chr3 | 112110609 | 112114692 | 4084 | -2.42 | 0.0042 | 0.00513 |
| chr6 | 8496513 | 8507946 | 11434 | -1.64 | 0.00423 | 0.00517 |
| chr6 | 100246096 | 100248582 | 2487 | 2 | 0.00424 | 0.00517 |
| chr7 | 50319229 | 50321155 | 1927 | -3.69 | 0.00424 | 0.00517 |
| chr9 | 117298830 | 117319755 | 20926 | -3.81 | 0.00424 | 0.00518 |
| chr1 | 77354549 | 77355923 | 1375 | -3.39 | 0.00425 | 0.00518 |
| chr9 | 104267104 | 104269065 | 1962 | 1.43 | 0.00426 | 0.0052 |
| chr5 | 82972516 | 82973547 | 1032 | -3.2 | 0.00426 | 0.0052 |
| chr12 | 17973059 | 17975693 | 2635 | -1.45 | 0.00427 | 0.00521 |
| chr22 | 48210241 | 48234914 | 24674 | -4.16 | 0.00427 | 0.00521 |
| chr8 | 79593990 | 79596214 | 2225 | -3.7 | 0.00428 | 0.00522 |
| chr8 | 71402426 | 71451429 | 49004 | -4.27 | 0.00428 | 0.00523 |
| chr6 | 14893780 | 14894868 | 1089 | -3.17 | 0.00428 | 0.00523 |
| chr7 | 36127469 | 36188725 | 61257 | -6.24 | 0.00429 | 0.00523 |
| chr6 | 131118295 | 131122984 | 4690 | -3.96 | 0.00429 | 0.00523 |
| chrX | 154915270 | 154926865 | 11596 | -3.66 | 0.00429 | 0.00524 |

| | | | | | | |
|---|---|---|---|---|---|---|
| chr9 | 101037915 | 101040388 | 2474 | 1.22 | 0.0043 | 0.00525 |
| chr11 | 32098174 | 32100259 | 2086 | 1.53 | 0.0043 | 0.00525 |
| chr15 | 92720652 | 92935774 | 215123 | 0.58 | 0.0043 | 0.00525 |
| chr12 | 63855344 | 63860370 | 5027 | -3.02 | 0.00431 | 0.00525 |
| chr18 | 58112490 | 58163782 | 51293 | 2.26 | 0.00432 | 0.00526 |
| chr15 | 29727590 | 29729061 | 1472 | 1.48 | 0.00432 | 0.00527 |
| chr6 | 122671314 | 122686099 | 14786 | 4.01 | 0.00432 | 0.00527 |
| chr7 | 157947036 | 157952762 | 5727 | -1.25 | 0.00433 | 0.00528 |
| chr6 | 162685880 | 162686718 | 839 | 2.43 | 0.00433 | 0.00529 |
| chr3 | 143220883 | 143381692 | 160810 | 6.25 | 0.00434 | 0.00529 |
| chr7 | 18972598 | 18973917 | 1320 | -1.54 | 0.00435 | 0.0053 |
| chr7 | 87942343 | 87942967 | 625 | -2.32 | 0.00435 | 0.0053 |
| chr2 | 211102740 | 211105988 | 3249 | 2.15 | 0.00436 | 0.00531 |
| chr6 | 147954761 | 147956252 | 1492 | -3.42 | 0.00436 | 0.00531 |
| chr7 | 81337364 | 81348220 | 10857 | -6.22 | 0.00436 | 0.00532 |
| chr1 | 168750395 | 168751095 | 701 | -1.97 | 0.00436 | 0.00532 |
| chr14 | 54500269 | 54505481 | 5213 | -4.19 | 0.00437 | 0.00533 |
| chr3 | 16020595 | 16056579 | 35985 | -1.99 | 0.00438 | 0.00534 |
| chr7 | 40318552 | 40364370 | 45819 | 1.46 | 0.00439 | 0.00535 |
| chr8 | 12624118 | 12627109 | 2992 | 1.79 | 0.00439 | 0.00536 |
| chr2 | 21122955 | 21150258 | 27304 | 1.29 | 0.0044 | 0.00537 |
| chr2 | 222529281 | 222531567 | 2287 | -2.25 | 0.00441 | 0.00538 |
| chr1 | 218052877 | 218064987 | 12111 | -4.02 | 0.00441 | 0.00538 |
| chr2 | 111443186 | 111474756 | 31571 | -1.48 | 0.00441 | 0.00538 |
| chr9 | 117902571 | 117908682 | 6112 | -3.8 | 0.00443 | 0.0054 |
| chr2 | 143113391 | 143114360 | 970 | -1.66 | 0.00444 | 0.00541 |
| chr13 | 96014577 | 96038858 | 24282 | -6.21 | 0.00444 | 0.00541 |
| chr9 | 85468515 | 85483261 | 14747 | -6.21 | 0.00444 | 0.00541 |
| chr4 | 35544492 | 35617192 | 72701 | 0.75 | 0.00446 | 0.00544 |
| chr11 | 49736421 | 49736829 | 409 | 1.64 | 0.00447 | 0.00544 |
| chr13 | 97507216 | 97536118 | 28903 | 4.33 | 0.00447 | 0.00545 |
| chr4 | 134732434 | 134734245 | 1812 | 1.94 | 0.00447 | 0.00545 |
| chr12 | 116746588 | 116790857 | 44270 | -6.2 | 0.00448 | 0.00545 |
| chr5 | 42702941 | 42706983 | 4043 | -3.75 | 0.00448 | 0.00546 |
| chr14 | 53989169 | 53989576 | 408 | -2.15 | 0.00448 | 0.00546 |
| chr4 | 85208170 | 85265954 | 57785 | -2.28 | 0.00448 | 0.00546 |
| chr16 | 58828058 | 58829464 | 1407 | 1.69 | 0.00449 | 0.00547 |
| chr15 | 48476405 | 48478001 | 1597 | -3.68 | 0.0045 | 0.00548 |
| chr6 | 16060489 | 16063561 | 3073 | -3.45 | 0.0045 | 0.00548 |
| chrX | 135708201 | 135710726 | 2526 | 1.56 | 0.0045 | 0.00548 |
| chr5 | 172031231 | 172043623 | 12393 | -4.45 | 0.0045 | 0.00549 |
| chr4 | 34315673 | 34318574 | 2902 | 1.38 | 0.00451 | 0.00549 |
| chr4 | 179764843 | 179765175 | 333 | -3.16 | 0.00451 | 0.00549 |
| chr2 | 135499710 | 135527271 | 27562 | -2.77 | 0.00451 | 0.00549 |
| chr5 | 180335430 | 180460321 | 124892 | -2.75 | 0.00451 | 0.0055 |
| chr7 | 29802590 | 29834867 | 32278 | -0.91 | 0.00451 | 0.0055 |
| chr10 | 12617199 | 12642633 | 25435 | -6.19 | 0.00452 | 0.0055 |
| chr13 | 89463266 | 89468023 | 4758 | -3.36 | 0.00452 | 0.00551 |
| chr2 | 213816969 | 213819514 | 2546 | -1.55 | 0.00453 | 0.00551 |
| chr1 | 112770763 | 112772752 | 1990 | -3.58 | 0.00453 | 0.00551 |
| chr4 | 20986819 | 20987862 | 1044 | 2.87 | 0.00453 | 0.00552 |
| chr1 | 207618479 | 207651533 | 33055 | 0.65 | 0.00456 | 0.00555 |
| chr1 | 30609331 | 30642314 | 32984 | -4.22 | 0.00456 | 0.00555 |
| chr2 | 218430851 | 218434488 | 3638 | 3.6 | 0.00456 | 0.00555 |
| chr8 | 9801468 | 9896973 | 95506 | -1.04 | 0.00457 | 0.00556 |
| chr7 | 24354841 | 24377738 | 22898 | 1.39 | 0.00457 | 0.00556 |
| chr17 | 72604043 | 72642619 | 38577 | -1.45 | 0.00458 | 0.00557 |
| chr11 | 16631585 | 16700073 | 68489 | -4.38 | 0.00458 | 0.00558 |
| chr5 | 76389402 | 76403289 | 13888 | -6.18 | 0.0046 | 0.0056 |
| chr11 | 109483040 | 109484486 | 1447 | 2.2 | 0.0046 | 0.0056 |
| chr7 | 12012032 | 12013370 | 1339 | -2.43 | 0.00461 | 0.00561 |

| | | | | | | |
|---|---|---|---|---|---|---|
| chr15 | 80499341 | 80521663 | 22323 | -1.52 | 0.00461 | 0.00561 |
| chr3 | 119405532 | 119407434 | 1903 | -2.95 | 0.00461 | 0.00561 |
| chr16 | 55809919 | 55811372 | 1454 | -3.13 | 0.00462 | 0.00562 |
| chr8 | 26522337 | 26536597 | 14261 | -3.8 | 0.00462 | 0.00562 |
| chr18 | 9998122 | 10001017 | 2896 | -3.55 | 0.00462 | 0.00563 |
| chrX | 151480247 | 151482111 | 1865 | -3.46 | 0.00462 | 0.00563 |
| chrX | 100841092 | 100843983 | 2892 | -1.78 | 0.00463 | 0.00564 |
| chr10 | 30296957 | 30298517 | 1561 | -3.57 | 0.00463 | 0.00564 |
| chrX | 51361959 | 51364094 | 2136 | -3.63 | 0.00463 | 0.00564 |
| chr1 | 169726546 | 169730425 | 3880 | -2 | 0.00464 | 0.00564 |
| chr6 | 9653078 | 9653831 | 754 | -2.83 | 0.00464 | 0.00565 |
| chr19 | 34382235 | 34584974 | 202740 | -3.32 | 0.00464 | 0.00565 |
| chr4 | 35224066 | 35285137 | 61072 | 1.43 | 0.00464 | 0.00565 |
| chr5 | 56745901 | 56746415 | 515 | 2.12 | 0.00465 | 0.00566 |
| chr4 | 143812744 | 143837929 | 25186 | -1.61 | 0.00465 | 0.00566 |
| chr4 | 144739575 | 144744050 | 4476 | 1.64 | 0.00466 | 0.00567 |
| chrX | 72211468 | 72212139 | 672 | -1.61 | 0.00466 | 0.00567 |
| chr14 | 81616325 | 81616674 | 350 | -2.7 | 0.00466 | 0.00567 |
| chr7 | 87654350 | 87662942 | 8593 | -3.35 | 0.00467 | 0.00568 |
| chr9 | 78946405 | 78959823 | 13419 | -6.16 | 0.00468 | 0.00569 |
| chr12 | 63561662 | 63599962 | 38301 | -4.31 | 0.0047 | 0.00571 |
| chr3 | 40274234 | 40275311 | 1078 | -3.12 | 0.0047 | 0.00572 |
| chr13 | 42834895 | 42836007 | 1113 | -2.62 | 0.0047 | 0.00572 |
| chr13 | 94530469 | 94531595 | 1127 | -1.69 | 0.00471 | 0.00573 |
| chr12 | 67795848 | 67806243 | 10396 | -2.06 | 0.00472 | 0.00574 |
| chr4 | 122471538 | 122472969 | 1432 | -1.37 | 0.00472 | 0.00575 |
| chr14 | 38634370 | 38649252 | 14883 | 3.22 | 0.00473 | 0.00575 |
| chr10 | 61189968 | 61288724 | 98757 | 0.56 | 0.00474 | 0.00577 |
| chr17 | 15218140 | 15219318 | 1179 | 3.32 | 0.00475 | 0.00578 |
| chr17 | 33777816 | 33818411 | 40596 | -1.42 | 0.00475 | 0.00578 |
| chrX | 57696256 | 57697734 | 1479 | 3.2 | 0.00476 | 0.00579 |
| chr3 | 56246195 | 56247362 | 1168 | -3.16 | 0.00476 | 0.00579 |
| chr17 | 64592399 | 64592710 | 312 | -3.58 | 0.00478 | 0.00581 |
| chr8 | 60973497 | 60978079 | 4583 | 1.25 | 0.00478 | 0.00582 |
| chr4 | 60090686 | 60091503 | 818 | -1.71 | 0.00479 | 0.00583 |
| chr2 | 100773982 | 100811707 | 37726 | -2.72 | 0.0048 | 0.00584 |
| chr21 | 17850961 | 17853371 | 2411 | -3.39 | 0.00481 | 0.00585 |
| chr2 | 73812879 | 73823146 | 10268 | -6.14 | 0.00481 | 0.00585 |
| chr3 | 105804483 | 105820492 | 16010 | -6.14 | 0.00481 | 0.00585 |
| chr11 | 33813806 | 33817657 | 3852 | -3.97 | 0.00481 | 0.00585 |
| chr4 | 107975480 | 108054525 | 79046 | 4.59 | 0.00482 | 0.00586 |
| chr4 | 15360213 | 15374669 | 14457 | 1.52 | 0.00485 | 0.0059 |
| chr9 | 431525 | 444941 | 13417 | -6.13 | 0.00485 | 0.0059 |
| chr2 | 186632522 | 186647797 | 15276 | -1.1 | 0.00487 | 0.00592 |
| chr2 | 45323261 | 45334748 | 11488 | -3.98 | 0.00487 | 0.00592 |
| chr17 | 21869883 | 21877835 | 7953 | 0.96 | 0.00487 | 0.00592 |
| chr6 | 150931844 | 150932986 | 1143 | 1.87 | 0.00487 | 0.00593 |
| chr5 | 39784908 | 39787950 | 3043 | -3.64 | 0.00487 | 0.00593 |
| chr10 | 23468217 | 23469602 | 1386 | -3.08 | 0.00489 | 0.00594 |
| chr4 | 37576577 | 37582305 | 5729 | -3.92 | 0.00489 | 0.00595 |
| chr1 | 101635032 | 101638255 | 3224 | 3.6 | 0.00489 | 0.00595 |
| chr20 | 2874875 | 2898933 | 24059 | -6.12 | 0.0049 | 0.00595 |
| chr2 | 237219077 | 237248688 | 29612 | -2.99 | 0.0049 | 0.00595 |
| chr18 | 32413713 | 32439972 | 26260 | -0.64 | 0.0049 | 0.00596 |
| chr11 | 110397575 | 110430804 | 33230 | -1.05 | 0.00491 | 0.00597 |
| chrX | 15130910 | 15131812 | 903 | -2.94 | 0.00491 | 0.00597 |
| chr5 | 32655168 | 32668232 | 13065 | -4.25 | 0.00491 | 0.00597 |
| chr4 | 135324707 | 135325588 | 882 | 3.68 | 0.00493 | 0.00598 |
| chr4 | 60064741 | 60067533 | 2793 | -2.94 | 0.00493 | 0.00599 |
| chr7 | 141901944 | 142038519 | 136576 | 0.53 | 0.00494 | 0.00601 |
| chr4 | 166077705 | 166079816 | 2112 | -3.6 | 0.00494 | 0.00601 |

| | | | | | | |
|---|---|---|---|---|---|---|
| chr18 | 71737125 | 71739057 | 1933 | -2.73 | 0.00494 | 0.00601 |
| chr13 | 26653309 | 26676760 | 23452 | -0.61 | 0.00495 | 0.00601 |
| chrX | 42701982 | 42716921 | 14940 | 0.93 | 0.00495 | 0.00601 |
| chrX | 102281383 | 102282917 | 1535 | -1.63 | 0.00496 | 0.00603 |
| chr13 | 98211280 | 98211668 | 389 | -1.67 | 0.00497 | 0.00603 |
| chr20 | 53282924 | 53283421 | 498 | -2.38 | 0.00497 | 0.00604 |
| chr7 | 14854035 | 14868682 | 14648 | -1 | 0.00497 | 0.00604 |
| chr6 | 90965777 | 90982018 | 16242 | -3.86 | 0.00498 | 0.00604 |
| chr8 | 10373140 | 10411553 | 38414 | -3.61 | 0.00498 | 0.00604 |
| chr22 | 35484577 | 35505762 | 21186 | -1.6 | 0.00498 | 0.00605 |
| chr15 | 49225690 | 49226932 | 1243 | -1.41 | 0.00498 | 0.00605 |
| chr16 | 79653155 | 79656586 | 3432 | -1.27 | 0.00498 | 0.00605 |
| chr3 | 118513680 | 118517525 | 3846 | 1.45 | 0.00498 | 0.00605 |
| chr2 | 113207138 | 113227201 | 20064 | -4.22 | 0.00499 | 0.00606 |
| chr12 | 72201818 | 72204021 | 2204 | 3.6 | 0.00499 | 0.00607 |
| chr4 | 145742459 | 145751865 | 9407 | 1.96 | 0.005 | 0.00607 |
| chr11 | 46200945 | 46213338 | 12394 | -4.24 | 0.005 | 0.00608 |
| chr19 | 43253652 | 43254217 | 566 | -1.52 | 0.005 | 0.00608 |
| chr3 | 58280714 | 58287981 | 7268 | -3.46 | 0.00503 | 0.00611 |
| chr1 | 167162808 | 167198152 | 35345 | -6.09 | 0.00503 | 0.00611 |
| chr10 | 92817244 | 92819074 | 1831 | 1.73 | 0.00505 | 0.00613 |
| chr3 | 9019273 | 9022063 | 2791 | -3.92 | 0.00506 | 0.00614 |
| chr9 | 136486077 | 136493460 | 7384 | -1.58 | 0.00506 | 0.00615 |
| chrX | 152369783 | 152371516 | 1734 | -1.74 | 0.00507 | 0.00615 |
| chr1 | 172993240 | 173002348 | 9109 | -3.82 | 0.00507 | 0.00615 |
| chr12 | 39035216 | 39037173 | 1958 | -1.65 | 0.00507 | 0.00615 |
| chr10 | 86006385 | 86019296 | 12912 | -5.7 | 0.00507 | 0.00616 |
| chr4 | 161572403 | 161574067 | 1665 | -1.83 | 0.00508 | 0.00617 |
| chr9 | 21239845 | 21242334 | 2490 | 1.3 | 0.00508 | 0.00617 |
| chr6 | 100485633 | 100487675 | 2043 | -1.25 | 0.00509 | 0.00618 |
| chr16 | 10953707 | 10955008 | 1302 | 3.22 | 0.00509 | 0.00618 |
| chr5 | 158289111 | 158290370 | 1260 | 3.34 | 0.00509 | 0.00618 |
| chr11 | 99633453 | 100295404 | 661952 | 5.94 | 0.00509 | 0.00618 |
| chr1 | 99047475 | 99048922 | 1448 | -1.71 | 0.0051 | 0.00618 |
| chr4 | 96551043 | 96552264 | 1222 | -1.72 | 0.0051 | 0.00619 |
| chr13 | 68436597 | 68442656 | 6060 | -1.41 | 0.0051 | 0.00619 |
| chr6 | 132594169 | 132596609 | 2441 | 3.54 | 0.00511 | 0.0062 |
| chrX | 134721992 | 134723744 | 1753 | 1.72 | 0.00511 | 0.0062 |
| chrX | 58013531 | 58047057 | 33527 | 0.76 | 0.00512 | 0.00621 |
| chr20 | 29898461 | 29933237 | 34777 | -0.95 | 0.00512 | 0.00622 |
| chr7 | 48906163 | 48909963 | 3801 | 1.66 | 0.00514 | 0.00623 |
| chrX | 41968035 | 41974680 | 6646 | -3.79 | 0.00515 | 0.00625 |
| chr2 | 180933671 | 180952917 | 19247 | -4.14 | 0.00515 | 0.00625 |
| chr2 | 95576256 | 95578372 | 2117 | 1.35 | 0.00515 | 0.00625 |
| chrX | 128432390 | 128432943 | 554 | -1.94 | 0.00516 | 0.00626 |
| chr4 | 190132412 | 190141270 | 8859 | -0.93 | 0.00516 | 0.00627 |
| chr12 | 131075713 | 131176195 | 100483 | -2.41 | 0.00519 | 0.0063 |
| chr18 | 49847299 | 49851626 | 4328 | 1.56 | 0.0052 | 0.00631 |
| chr3 | 127032259 | 127034227 | 1969 | -1.25 | 0.0052 | 0.00631 |
| chrX | 73580472 | 73582771 | 2300 | -3.67 | 0.00521 | 0.00632 |
| chr4 | 65496501 | 65500857 | 4357 | -1.39 | 0.00522 | 0.00633 |
| chr12 | 119427473 | 119539283 | 111811 | 6.32 | 0.00522 | 0.00633 |
| chr1 | 169600668 | 169607935 | 7268 | -4.01 | 0.00522 | 0.00633 |
| chr2 | 8813734 | 8829233 | 15500 | -6.06 | 0.00523 | 0.00634 |
| chr2 | 90003770 | 90004290 | 521 | 3.15 | 0.00523 | 0.00634 |
| chr2 | 222932814 | 222957861 | 25048 | 0.98 | 0.00523 | 0.00634 |
| chr4 | 30564861 | 30565657 | 797 | 1.89 | 0.00524 | 0.00636 |
| chr4 | 78445589 | 78491733 | 46145 | -4.36 | 0.00525 | 0.00637 |
| chrX | 108754554 | 108756078 | 1525 | -1.74 | 0.00525 | 0.00637 |
| chr4 | 134508775 | 134512221 | 3447 | 3.35 | 0.00525 | 0.00637 |
| chrX | 81007026 | 81011939 | 4914 | 1.46 | 0.00525 | 0.00637 |

| | | | | | | |
|---|---|---|---|---|---|---|
| chr1 | 239330989 | 239335959 | 4971 | -3.74 | 0.00526 | 0.00638 |
| chr2 | 159125301 | 159127370 | 2070 | 1.21 | 0.00526 | 0.00638 |
| chr1 | 89773387 | 89791025 | 17639 | -1 | 0.00526 | 0.00638 |
| chr7 | 144153601 | 144173215 | 19615 | 1.73 | 0.00526 | 0.00638 |
| chr4 | 138887619 | 138888949 | 1331 | -3.06 | 0.00528 | 0.0064 |
| chr1 | 39122530 | 39138032 | 15503 | -2.34 | 0.00528 | 0.0064 |
| chr12 | 19128651 | 19151264 | 22614 | -6.05 | 0.00528 | 0.0064 |
| chr15 | 58705492 | 58709590 | 4099 | -6.05 | 0.00528 | 0.0064 |
| chr12 | 102669048 | 102671052 | 2005 | -3.47 | 0.00528 | 0.0064 |
| chr9 | 28514502 | 28525450 | 10949 | 1.57 | 0.00529 | 0.00641 |
| chr20 | 20174703 | 20177987 | 3285 | 3.78 | 0.00531 | 0.00643 |
| chr11 | 121529884 | 121539523 | 9640 | -0.83 | 0.00532 | 0.00645 |
| chrX | 54900603 | 54903110 | 2508 | 1.8 | 0.00534 | 0.00647 |
| chr5 | 146091966 | 146092657 | 692 | 2.21 | 0.00535 | 0.00649 |
| chr7 | 99448329 | 99465818 | 17490 | -3.79 | 0.00535 | 0.00649 |
| chr14 | 95474134 | 95501457 | 27324 | -4.26 | 0.00536 | 0.00649 |
| chr6 | 39219854 | 39228948 | 9095 | -3.88 | 0.00536 | 0.0065 |
| chr10 | 81324771 | 81359324 | 34554 | -1.12 | 0.00536 | 0.0065 |
| chr6 | 130849861 | 130858883 | 9023 | 1.93 | 0.00536 | 0.0065 |
| chr3 | 28331880 | 28332755 | 876 | -2.94 | 0.00537 | 0.0065 |
| chr12 | 38991679 | 39000560 | 8882 | -1.28 | 0.00537 | 0.00651 |
| chr3 | 63767079 | 63767818 | 740 | -3.22 | 0.0054 | 0.00654 |
| chr11 | 99044036 | 99050885 | 6850 | 3.64 | 0.00542 | 0.00657 |
| chr3 | 106128960 | 106131182 | 2223 | -1.6 | 0.00543 | 0.00657 |
| chr1 | 56399829 | 56402177 | 2349 | -3.05 | 0.00543 | 0.00658 |
| chr14 | 81439962 | 81465801 | 25840 | -2.04 | 0.00543 | 0.00658 |
| chr3 | 29485008 | 29485722 | 715 | -3.07 | 0.00543 | 0.00658 |
| chr5 | 75334057 | 75344337 | 10281 | 0.84 | 0.00544 | 0.00659 |
| chr15 | 95956638 | 95987676 | 31039 | -0.78 | 0.00544 | 0.00659 |
| chrX | 73905559 | 73907001 | 1443 | -1.57 | 0.00545 | 0.0066 |
| chr8 | 31046100 | 31195273 | 149174 | -1.37 | 0.00545 | 0.0066 |
| chr5 | 75419468 | 75452071 | 32604 | 1.6 | 0.00545 | 0.00661 |
| chr7 | 140926954 | 140927578 | 625 | -2.27 | 0.00546 | 0.00661 |
| chr13 | 33990983 | 34004193 | 13211 | -1.93 | 0.00546 | 0.00661 |
| chr5 | 27744824 | 27753102 | 8279 | -2.13 | 0.00546 | 0.00661 |
| chr12 | 87997822 | 87999291 | 1470 | -1.44 | 0.00546 | 0.00662 |
| chr20 | 52992529 | 52992912 | 384 | -2.57 | 0.00547 | 0.00662 |
| chr10 | 16617111 | 16618161 | 1051 | -1.48 | 0.00547 | 0.00662 |
| chr20 | 9918740 | 9922145 | 3406 | -3.35 | 0.00547 | 0.00663 |
| chr15 | 50017331 | 50019101 | 1771 | -3.28 | 0.00548 | 0.00664 |
| chr11 | 31899821 | 31968859 | 69039 | 4.55 | 0.00548 | 0.00664 |
| chr10 | 23303144 | 23350109 | 46966 | -4.32 | 0.0055 | 0.00666 |
| chrX | 116455181 | 116494377 | 39197 | 2.66 | 0.0055 | 0.00666 |
| chr1 | 105013911 | 105066492 | 52582 | 1.04 | 0.0055 | 0.00666 |
| chr6 | 41205892 | 41209427 | 3536 | 2.2 | 0.0055 | 0.00666 |
| chr12 | 106304515 | 106313454 | 8940 | 0.73 | 0.00551 | 0.00667 |
| chr2 | 20030941 | 20032390 | 1450 | -3.27 | 0.00552 | 0.00668 |
| chr21 | 15869244 | 15887538 | 18295 | 0.98 | 0.00553 | 0.0067 |
| chr21 | 47593771 | 47598034 | 4264 | -3.94 | 0.00554 | 0.0067 |
| chr13 | 36832181 | 36837640 | 5460 | 2.12 | 0.00554 | 0.0067 |
| chr12 | 112990675 | 113004945 | 14271 | -6.01 | 0.00554 | 0.00671 |
| chr15 | 101634068 | 101649313 | 15246 | -6.01 | 0.00554 | 0.00671 |
| chr3 | 155487683 | 155515220 | 27538 | -6.01 | 0.00554 | 0.00671 |
| chrX | 21204644 | 21209368 | 4725 | -3.8 | 0.00555 | 0.00672 |
| chr3 | 159195357 | 159206113 | 10757 | -0.96 | 0.00556 | 0.00673 |
| chr13 | 35330590 | 35331079 | 490 | -2.68 | 0.00556 | 0.00673 |
| chr1 | 104307241 | 104319791 | 12551 | 2.68 | 0.00557 | 0.00674 |
| chr3 | 162337850 | 162341678 | 3829 | -3.56 | 0.00557 | 0.00674 |
| chr2 | 153970253 | 154032505 | 62253 | 4.49 | 0.00559 | 0.00677 |
| chr7 | 157356784 | 157384929 | 28146 | -6 | 0.00559 | 0.00677 |
| chr3 | 54317447 | 54334776 | 17330 | 1.07 | 0.00562 | 0.00681 |

| | | | | | | |
|---|---|---|---|---|---|---|
| chr9 | 137785798 | 137824235 | 38438 | -4.31 | 0.00562 | 0.00681 |
| chr10 | 61142038 | 61143373 | 1336 | 1.78 | 0.00563 | 0.00681 |
| chr2 | 8085898 | 8087029 | 1132 | -2.39 | 0.00564 | 0.00682 |
| chr11 | 122212470 | 122233388 | 20919 | -0.76 | 0.00564 | 0.00682 |
| chr22 | 23294161 | 23346381 | 52221 | -3.9 | 0.00564 | 0.00682 |
| chr1 | 63604121 | 63605549 | 1429 | -2.25 | 0.00564 | 0.00682 |
| chrX | 103767437 | 103770672 | 3236 | 1.42 | 0.00564 | 0.00683 |
| chr13 | 111983080 | 111988637 | 5558 | -0.94 | 0.00565 | 0.00683 |
| chr5 | 89983160 | 89984646 | 1487 | -2.13 | 0.00565 | 0.00684 |
| chr2 | 189039865 | 189040323 | 459 | -3.49 | 0.00565 | 0.00684 |
| chr6 | 52049756 | 52055718 | 5963 | 0.92 | 0.00566 | 0.00684 |
| chr13 | 97488922 | 97489785 | 864 | -3.16 | 0.00566 | 0.00685 |
| chr4 | 135407211 | 135412502 | 5292 | 2.3 | 0.00566 | 0.00685 |
| chr17 | 14467358 | 14655894 | 188537 | -1.12 | 0.00567 | 0.00686 |
| chr18 | 76814060 | 76828928 | 14869 | -3.72 | 0.00569 | 0.00688 |
| chr8 | 27426487 | 27431024 | 4538 | 3.83 | 0.00569 | 0.00689 |
| chr3 | 44163242 | 44429372 | 266131 | -4.21 | 0.00571 | 0.00691 |
| chrX | 65163432 | 65170973 | 7542 | 0.82 | 0.00571 | 0.00691 |
| chr3 | 187401836 | 187403826 | 1991 | -2.5 | 0.00572 | 0.00691 |
| chr1 | 104930222 | 104943396 | 13175 | 1.15 | 0.00572 | 0.00692 |
| chr7 | 95506594 | 95508954 | 2361 | -3.62 | 0.00572 | 0.00692 |
| chr8 | 43182989 | 43183488 | 500 | -2.85 | 0.00572 | 0.00692 |
| chr12 | 90928523 | 90940132 | 11610 | -3.47 | 0.00573 | 0.00693 |
| chr3 | 130279975 | 130290310 | 10336 | 0.88 | 0.00573 | 0.00693 |
| chr8 | 37042954 | 37081416 | 38463 | -0.91 | 0.00574 | 0.00694 |
| chr4 | 94822688 | 94823970 | 1283 | -1.59 | 0.00574 | 0.00694 |
| chr14 | 45345601 | 45366200 | 20600 | -2.83 | 0.00574 | 0.00695 |
| chr3 | 101630229 | 101635495 | 5267 | -3.57 | 0.00575 | 0.00695 |
| chr20 | 23467416 | 23486539 | 19124 | -5.97 | 0.00576 | 0.00697 |
| chr6 | 80601581 | 80603875 | 2295 | -1.35 | 0.00578 | 0.00698 |
| chr13 | 102482088 | 102484970 | 2883 | -1.49 | 0.00578 | 0.00699 |
| chr11 | 114343081 | 114345760 | 2680 | -1.37 | 0.0058 | 0.00701 |
| chr4 | 67869897 | 67873321 | 3425 | 2.44 | 0.0058 | 0.00701 |
| chr20 | 2163565 | 2171935 | 8371 | -4.23 | 0.0058 | 0.00701 |
| chr12 | 125761423 | 125762940 | 1518 | 1.87 | 0.00581 | 0.00703 |
| chr13 | 23575125 | 23580269 | 5145 | -3.05 | 0.00581 | 0.00703 |
| chr12 | 102682637 | 102684639 | 2003 | 1.24 | 0.00582 | 0.00704 |
| chr21 | 15902149 | 15902391 | 243 | -3.42 | 0.00584 | 0.00706 |
| chr13 | 94526546 | 94527827 | 1282 | 3.13 | 0.00584 | 0.00706 |
| chr5 | 122623150 | 122671815 | 48666 | -4.29 | 0.00586 | 0.00709 |
| chr5 | 168057447 | 168060630 | 3184 | -1.26 | 0.00587 | 0.00709 |
| chr2 | 84487823 | 84489781 | 1959 | -3.52 | 0.00587 | 0.00709 |
| chr7 | 111999106 | 112001051 | 1946 | -3.69 | 0.00588 | 0.0071 |
| chr7 | 99396400 | 99418829 | 22430 | -4.22 | 0.00588 | 0.00711 |
| chr11 | 19015568 | 19038717 | 23150 | 0.87 | 0.0059 | 0.00712 |
| chr1 | 181163812 | 181166218 | 2407 | -3.4 | 0.0059 | 0.00712 |
| chr22 | 17780454 | 17781273 | 820 | -1.84 | 0.0059 | 0.00712 |
| chr4 | 190239642 | 190466086 | 226445 | -0.43 | 0.0059 | 0.00713 |
| chr12 | 7434742 | 7450179 | 15438 | -1.12 | 0.00592 | 0.00715 |
| chrX | 114269035 | 114293529 | 24495 | 4.2 | 0.00593 | 0.00717 |
| chr21 | 32494520 | 32496485 | 1966 | -3.27 | 0.00594 | 0.00717 |
| chrX | 108136165 | 108147305 | 11141 | 1.3 | 0.00596 | 0.0072 |
| chr5 | 128010654 | 128425967 | 415314 | 6.61 | 0.00597 | 0.00722 |
| chr1 | 106027471 | 106042093 | 14623 | 0.81 | 0.00599 | 0.00724 |
| chr15 | 60760946 | 60768368 | 7423 | -3.71 | 0.00599 | 0.00724 |
| chrX | 81299892 | 81307572 | 7681 | 1.43 | 0.00602 | 0.00727 |
| chr1 | 169630120 | 169630534 | 415 | -1.74 | 0.00602 | 0.00727 |
| chr8 | 118005733 | 118007014 | 1282 | -2.72 | 0.00603 | 0.00729 |
| chr11 | 102723478 | 102727663 | 4186 | -3.67 | 0.00604 | 0.0073 |
| chr15 | 79491507 | 79565092 | 73586 | -1.03 | 0.00606 | 0.00732 |
| chr15 | 35127157 | 35127959 | 803 | -3.24 | 0.00607 | 0.00733 |

| | | | | | | |
|---|---|---|---|---|---|---|
| chr13 | 112926669 | 112929053 | 2385 | 2.6 | 0.00607 | 0.00733 |
| chr8 | 21444703 | 21450908 | 6206 | 1.24 | 0.00608 | 0.00734 |
| chr9 | 2875406 | 2887045 | 11640 | 1.81 | 0.00608 | 0.00734 |
| chr11 | 106322738 | 106325363 | 2626 | -3.33 | 0.00609 | 0.00735 |
| chr9 | 136086114 | 136095372 | 9259 | -3.96 | 0.00609 | 0.00735 |
| chr1 | 219844211 | 219844982 | 772 | -1.86 | 0.00609 | 0.00736 |
| chr1 | 84942482 | 84944844 | 2363 | -1 | 0.0061 | 0.00737 |
| chr3 | 121979917 | 121992075 | 12159 | -0.91 | 0.00611 | 0.00738 |
| chr15 | 58052262 | 58058489 | 6228 | 0.92 | 0.00612 | 0.00739 |
| chr10 | 30875146 | 30883387 | 8242 | -5.92 | 0.00613 | 0.0074 |
| chr2 | 42319103 | 42328267 | 9165 | -5.92 | 0.00613 | 0.0074 |
| chr1 | 184894932 | 184898117 | 3186 | 2.76 | 0.00613 | 0.0074 |
| chr18 | 69351180 | 69353459 | 2280 | 2.03 | 0.00613 | 0.0074 |
| chr3 | 66920585 | 66924119 | 3535 | 1.62 | 0.00614 | 0.00741 |
| chr11 | 21820328 | 21827286 | 6959 | -2.08 | 0.00614 | 0.00742 |
| chr22 | 35442653 | 35453567 | 10915 | -1.39 | 0.00615 | 0.00742 |
| chr12 | 15157213 | 15264019 | 106807 | 0.6 | 0.00615 | 0.00742 |
| chr20 | 1991458 | 2007428 | 15971 | -1.58 | 0.00615 | 0.00742 |
| chr3 | 84415396 | 84780416 | 365021 | -2 | 0.00615 | 0.00743 |
| chr7 | 13525988 | 13526320 | 333 | 2.61 | 0.00617 | 0.00744 |
| chr4 | 21632921 | 21648288 | 15368 | 1.57 | 0.00617 | 0.00745 |
| chr2 | 141493365 | 141494671 | 1307 | -3.29 | 0.00619 | 0.00747 |
| chr8 | 106982569 | 107009791 | 27223 | -5.91 | 0.00619 | 0.00747 |
| chr12 | 103088664 | 103091006 | 2343 | -1.54 | 0.00619 | 0.00747 |
| chr10 | 7720436 | 7772352 | 51917 | -2.88 | 0.00619 | 0.00747 |
| chr8 | 77545013 | 77548758 | 3746 | 1.07 | 0.00619 | 0.00748 |
| chr18 | 13458282 | 13480215 | 21934 | -4.47 | 0.0062 | 0.00748 |
| chr6 | 141598120 | 141610049 | 11930 | -4.47 | 0.0062 | 0.00748 |
| chrX | 67215776 | 67222239 | 6464 | -0.98 | 0.0062 | 0.00748 |
| chrX | 77836562 | 77850108 | 13547 | -2.21 | 0.0062 | 0.00749 |
| chrX | 130135586 | 130137185 | 1600 | -1.58 | 0.00621 | 0.00749 |
| chr2 | 84572823 | 84605058 | 32236 | 1.68 | 0.00621 | 0.00749 |
| chr13 | 97673420 | 97678841 | 5422 | -3.91 | 0.00621 | 0.0075 |
| chr11 | 63023715 | 63024375 | 661 | -2.54 | 0.00622 | 0.00751 |
| chr5 | 105669461 | 105669741 | 281 | -2.79 | 0.00623 | 0.00751 |
| chr2 | 84552008 | 84554292 | 2285 | 2.55 | 0.00625 | 0.00754 |
| chr2 | 210197667 | 210220785 | 23119 | -5.9 | 0.00626 | 0.00755 |
| chr10 | 18859892 | 18876320 | 16429 | -3.86 | 0.00626 | 0.00755 |
| chr6 | 8802381 | 8813010 | 10630 | -1.75 | 0.00626 | 0.00755 |
| chr12 | 92197046 | 92201804 | 4759 | -3.38 | 0.00627 | 0.00756 |
| chr18 | 35173603 | 35399724 | 226122 | 6.34 | 0.00627 | 0.00757 |
| chrX | 11560772 | 11562035 | 1264 | -1.63 | 0.00627 | 0.00757 |
| chr18 | 47543756 | 47553113 | 9358 | -0.74 | 0.00628 | 0.00758 |
| chr3 | 111997833 | 112001687 | 3855 | 3.75 | 0.00629 | 0.00759 |
| chr16 | 80794550 | 80926700 | 132151 | -0.84 | 0.0063 | 0.0076 |
| chr1 | 177210743 | 177212217 | 1475 | 1.62 | 0.0063 | 0.0076 |
| chr5 | 112641530 | 112646245 | 4716 | -1 | 0.0063 | 0.0076 |
| chr7 | 143117111 | 143172842 | 55732 | -2.01 | 0.00631 | 0.00761 |
| chr12 | 55372809 | 55442217 | 69409 | 4.49 | 0.00632 | 0.00762 |
| chr5 | 170226794 | 170233232 | 6439 | -2.8 | 0.00632 | 0.00762 |
| chr2 | 180194003 | 180210486 | 16484 | -5.89 | 0.00632 | 0.00762 |
| chrX | 140351826 | 140353358 | 1533 | -3.07 | 0.00632 | 0.00763 |
| chr9 | 111369414 | 111387047 | 17634 | 0.68 | 0.00635 | 0.00766 |
| chr2 | 167188797 | 167190080 | 1284 | 4.15 | 0.00635 | 0.00766 |
| chr6 | 46448811 | 46451309 | 2499 | 3.48 | 0.00636 | 0.00767 |
| chrX | 43186840 | 43191595 | 4756 | -1.16 | 0.00636 | 0.00767 |
| chr2 | 73521292 | 73521849 | 558 | -2.94 | 0.00637 | 0.00768 |
| chr11 | 64235501 | 64247139 | 11639 | -2.59 | 0.00638 | 0.00769 |
| chr1 | 179561416 | 179572398 | 10983 | -5.88 | 0.00639 | 0.0077 |
| chr15 | 63208604 | 63219397 | 10794 | -5.88 | 0.00639 | 0.0077 |
| chr18 | 60734673 | 60764115 | 29443 | -2.92 | 0.00639 | 0.00771 |

| | | | | | | |
|---|---|---|---|---|---|---|
| chr2 | 135458959 | 135462856 | 3898 | -1.69 | 0.0064 | 0.00771 |
| chr12 | 27659617 | 27660272 | 656 | -2.93 | 0.0064 | 0.00772 |
| chr1 | 220690964 | 220713857 | 22894 | -4.05 | 0.00641 | 0.00773 |
| chr7 | 31308692 | 31311329 | 2638 | 1.67 | 0.00641 | 0.00773 |
| chr5 | 89321975 | 89327043 | 5069 | 1.86 | 0.00641 | 0.00773 |
| chr13 | 75810370 | 75811971 | 1602 | -1.82 | 0.00642 | 0.00773 |
| chr13 | 31961361 | 31962711 | 1351 | 1.59 | 0.00643 | 0.00776 |
| chr8 | 1630482 | 1672161 | 41680 | -4.18 | 0.00644 | 0.00776 |
| chr3 | 172618947 | 172631457 | 12511 | 1.11 | 0.00644 | 0.00777 |
| chr2 | 187889305 | 187890205 | 901 | -2.08 | 0.00645 | 0.00778 |
| chr6 | 130814240 | 130817586 | 3347 | 3.47 | 0.00645 | 0.00778 |
| chr14 | 56002790 | 56020805 | 18016 | -5.87 | 0.00646 | 0.00778 |
| chr12 | 83732557 | 83739038 | 6482 | -2.12 | 0.00646 | 0.00779 |
| chr13 | 39556042 | 39558246 | 2205 | -3.48 | 0.00647 | 0.0078 |
| chr11 | 64363932 | 64364830 | 899 | 3.21 | 0.00647 | 0.0078 |
| chr2 | 157477846 | 157481672 | 3827 | -3.58 | 0.00648 | 0.0078 |
| chr20 | 21235467 | 21268003 | 32537 | -4.29 | 0.0065 | 0.00783 |
| chr2 | 150338893 | 150410134 | 71242 | -0.81 | 0.00652 | 0.00786 |
| chrX | 109995064 | 110012545 | 17482 | 0.8 | 0.00653 | 0.00787 |
| chr2 | 57694679 | 57695481 | 803 | -2.32 | 0.00653 | 0.00787 |
| chr11 | 26706488 | 26711392 | 4905 | 3.87 | 0.00654 | 0.00788 |
| chr4 | 33738279 | 33750282 | 12004 | -1.13 | 0.00654 | 0.00788 |
| chr1 | 205289204 | 205289926 | 723 | -2.09 | 0.00654 | 0.00788 |
| chr1 | 75918189 | 75919911 | 1723 | 2.8 | 0.00654 | 0.00788 |
| chr2 | 36510411 | 36512389 | 1979 | -1.68 | 0.00655 | 0.00788 |
| chr2 | 68205415 | 68234063 | 28649 | -0.66 | 0.00655 | 0.00789 |
| chr6 | 162706697 | 162710355 | 3659 | 1.87 | 0.00655 | 0.00789 |
| chr7 | 10933605 | 10952965 | 19361 | -0.7 | 0.00657 | 0.00792 |
| chr4 | 133163974 | 133174933 | 10960 | -3.36 | 0.00657 | 0.00792 |
| chr2 | 187239570 | 187264185 | 24616 | -5.85 | 0.0066 | 0.00795 |
| chr8 | 1282368 | 1286734 | 4367 | 1.45 | 0.0066 | 0.00795 |
| chr18 | 36968565 | 36970156 | 1592 | 2.14 | 0.0066 | 0.00795 |
| chr10 | 59995230 | 59995785 | 556 | -2.88 | 0.00661 | 0.00796 |
| chr12 | 38885737 | 38893023 | 7287 | -3.53 | 0.00662 | 0.00797 |
| chr13 | 109140960 | 109144312 | 3353 | -2.78 | 0.00662 | 0.00797 |
| chr8 | 62093826 | 62141766 | 47941 | -1.66 | 0.00662 | 0.00797 |
| chr8 | 120604601 | 120608901 | 4301 | 1.51 | 0.00662 | 0.00797 |
| chr2 | 157039780 | 157041506 | 1727 | 4.41 | 0.00663 | 0.00798 |
| chr11 | 100935637 | 100959739 | 24103 | 3.01 | 0.00664 | 0.00799 |
| chr3 | 67576209 | 67576711 | 503 | -2.68 | 0.00664 | 0.00799 |
| chr6 | 99422096 | 99422814 | 719 | 2.95 | 0.00664 | 0.00799 |
| chr16 | 51040605 | 51064657 | 24053 | -1.22 | 0.00664 | 0.008 |
| chr17 | 5450333 | 5453616 | 3284 | -0.76 | 0.00667 | 0.00802 |
| chr11 | 50324611 | 50505224 | 180614 | -0.53 | 0.00668 | 0.00804 |
| chr3 | 157620666 | 157624992 | 4327 | -0.95 | 0.00669 | 0.00805 |
| chr7 | 157487575 | 157491836 | 4262 | -1.36 | 0.00669 | 0.00805 |
| chr21 | 27812598 | 27815107 | 2510 | -3.61 | 0.00669 | 0.00806 |
| chr5 | 53082702 | 53084585 | 1884 | -1.16 | 0.0067 | 0.00806 |
| chr18 | 50224646 | 50226866 | 2221 | -1.12 | 0.00671 | 0.00808 |
| chr7 | 131474394 | 131475293 | 900 | -3.05 | 0.00672 | 0.00809 |
| chr13 | 76898172 | 76900233 | 2062 | 3.29 | 0.00673 | 0.0081 |
| chr16 | 23185833 | 23187601 | 1769 | 1.28 | 0.00673 | 0.0081 |
| chrX | 54747191 | 54771099 | 23909 | -3.86 | 0.00673 | 0.0081 |
| chrX | 109813086 | 109813898 | 813 | 2.95 | 0.00674 | 0.00811 |
| chr15 | 35481141 | 35499528 | 18388 | -4.11 | 0.00675 | 0.00812 |
| chr12 | 108542142 | 108544040 | 1899 | 1.34 | 0.00675 | 0.00812 |
| chr5 | 75886619 | 75888990 | 2372 | 3.35 | 0.00675 | 0.00813 |
| chr14 | 26815789 | 26823452 | 7664 | -1.48 | 0.00677 | 0.00815 |
| chr1 | 96749450 | 96750794 | 1345 | -2.66 | 0.00677 | 0.00815 |
| chr10 | 6677556 | 6705101 | 27546 | -0.72 | 0.00677 | 0.00815 |
| chr9 | 77803168 | 77805243 | 2076 | -1.24 | 0.00678 | 0.00816 |

| | | | | | | |
|---|---|---|---|---|---|---|
| chr8 | 74129507 | 74179230 | 49724 | -1.17 | 0.00678 | 0.00816 |
| chr3 | 51119043 | 51125456 | 6414 | -3.88 | 0.00679 | 0.00817 |
| chr4 | 183904197 | 183941595 | 37399 | -3.87 | 0.0068 | 0.00818 |
| chr9 | 110145950 | 110148173 | 2224 | 1.36 | 0.0068 | 0.00819 |
| chr11 | 56402898 | 56409922 | 7025 | 1.11 | 0.00681 | 0.00819 |
| chr12 | 20487315 | 20501929 | 14615 | -5.82 | 0.00682 | 0.0082 |
| chr3 | 176262060 | 176263564 | 1505 | 2.1 | 0.00684 | 0.00822 |
| chr6 | 137265316 | 137266630 | 1315 | 1.62 | 0.00685 | 0.00825 |
| chr6 | 131103362 | 131103839 | 478 | -2.57 | 0.00686 | 0.00826 |
| chr9 | 78878574 | 78880265 | 1692 | -3.65 | 0.00687 | 0.00826 |
| chr5 | 33958669 | 33964252 | 5584 | 2.19 | 0.00688 | 0.00827 |
| chr10 | 30376035 | 30378126 | 2092 | -2.71 | 0.00688 | 0.00828 |
| chrX | 101827857 | 101829494 | 1638 | -1.18 | 0.0069 | 0.00829 |
| chr7 | 38330946 | 38333044 | 2099 | -3.39 | 0.0069 | 0.0083 |
| chr15 | 50307080 | 50308174 | 1095 | -2.17 | 0.00691 | 0.00832 |
| chr11 | 44068828 | 44072223 | 3396 | 3.68 | 0.00691 | 0.00832 |
| chr4 | 141403154 | 141425327 | 22174 | -3.92 | 0.00693 | 0.00833 |
| chr12 | 108766216 | 108851101 | 84886 | -1.9 | 0.00693 | 0.00834 |
| chr10 | 25459708 | 25462960 | 3253 | 2.37 | 0.00694 | 0.00834 |
| chr6 | 8879658 | 8880548 | 891 | 1.48 | 0.00694 | 0.00834 |
| chr11 | 95167941 | 95177450 | 9510 | -1.12 | 0.00694 | 0.00834 |
| chr3 | 118552539 | 118559563 | 7025 | -3.91 | 0.00695 | 0.00836 |
| chr1 | 111775299 | 111780493 | 5195 | 3.67 | 0.00696 | 0.00837 |
| chr9 | 82520740 | 82528089 | 7350 | -2.81 | 0.00699 | 0.00841 |
| chr6 | 149392963 | 149393726 | 764 | -2.79 | 0.007 | 0.00841 |
| chr8 | 125872829 | 126009726 | 136898 | -6.27 | 0.007 | 0.00841 |
| chr6 | 146757920 | 146768080 | 10161 | 2.67 | 0.007 | 0.00842 |
| chr4 | 29120993 | 29137653 | 16661 | 0.83 | 0.007 | 0.00842 |
| chr13 | 67839560 | 67849118 | 9559 | -1.34 | 0.00701 | 0.00843 |
| chr1 | 2961202 | 2975884 | 14683 | -3.72 | 0.00702 | 0.00844 |
| chr21 | 37157730 | 37184031 | 26302 | 0.62 | 0.00703 | 0.00845 |
| chr9 | 90515494 | 90517416 | 1923 | -3.5 | 0.00703 | 0.00846 |
| chr16 | 73282836 | 73422777 | 139942 | -0.66 | 0.00704 | 0.00846 |
| chr8 | 62342019 | 62350472 | 8454 | -3.63 | 0.00704 | 0.00846 |
| chr4 | 78591097 | 78591840 | 744 | -1.99 | 0.00704 | 0.00846 |
| chr3 | 41469516 | 41482214 | 12699 | -5.79 | 0.00705 | 0.00847 |
| chr15 | 73236161 | 73271349 | 35189 | -2.12 | 0.00705 | 0.00848 |
| chr1 | 198019212 | 198019999 | 788 | -2.81 | 0.00705 | 0.00848 |
| chr18 | 8489358 | 8499829 | 10472 | -3.88 | 0.00706 | 0.00848 |
| chr10 | 61495573 | 61496729 | 1157 | -1.78 | 0.00706 | 0.00848 |
| chr2 | 16088957 | 16279451 | 190495 | 1.97 | 0.00707 | 0.00849 |
| chr8 | 17379854 | 17380790 | 937 | -1.64 | 0.00707 | 0.0085 |
| chr3 | 139444617 | 139450642 | 6026 | 3.76 | 0.00709 | 0.00852 |
| chr4 | 122022470 | 122033438 | 10969 | -2.2 | 0.00709 | 0.00852 |
| chr4 | 154887017 | 155219980 | 332964 | 6.27 | 0.0071 | 0.00853 |
| chr5 | 114745189 | 114760958 | 15770 | -5.12 | 0.0071 | 0.00854 |
| chr10 | 12903945 | 12937359 | 33415 | -4.06 | 0.00712 | 0.00855 |
| chr17 | 6009455 | 6058138 | 48684 | -0.8 | 0.00712 | 0.00855 |
| chr2 | 208943836 | 208955444 | 11609 | -3.7 | 0.00712 | 0.00855 |
| chr8 | 118423844 | 118459395 | 35552 | -1.57 | 0.00713 | 0.00856 |
| chr4 | 95031369 | 95032942 | 1574 | -3.48 | 0.00714 | 0.00857 |
| chr15 | 34895971 | 34897087 | 1117 | -2.02 | 0.00714 | 0.00857 |
| chr6 | 159660249 | 159662833 | 2585 | 1.78 | 0.00714 | 0.00858 |
| chr13 | 40580947 | 40583406 | 2460 | -1.58 | 0.00714 | 0.00858 |
| chr1 | 183665276 | 183672317 | 7042 | -3.53 | 0.00716 | 0.00861 |
| chr5 | 169744756 | 169780780 | 36025 | -2.59 | 0.00717 | 0.00861 |
| chr3 | 108966168 | 108969313 | 3146 | -1.88 | 0.00719 | 0.00864 |
| chr11 | 104080790 | 104082087 | 1298 | -1.44 | 0.0072 | 0.00864 |
| chr7 | 152153222 | 152169091 | 15870 | -5.77 | 0.00721 | 0.00866 |
| chr1 | 182185486 | 182190336 | 4851 | -1.73 | 0.00722 | 0.00867 |
| chr14 | 85813240 | 85821046 | 7807 | -3.49 | 0.00723 | 0.00868 |

| | | | | | | |
|---|---|---|---|---|---|---|
| chr22 | 40254256 | 40261352 | 7097 | 0.95 | 0.00723 | 0.00869 |
| chr6 | 169403060 | 169417798 | 14739 | -2.42 | 0.00724 | 0.00869 |
| chr13 | 31569067 | 31571661 | 2595 | -2.76 | 0.00724 | 0.00869 |
| chr7 | 142204735 | 142208034 | 3300 | 3.52 | 0.00724 | 0.00869 |
| chr2 | 53151989 | 53159193 | 7205 | -3.52 | 0.00724 | 0.0087 |
| chr11 | 22820084 | 22841555 | 21472 | -0.74 | 0.00725 | 0.00871 |
| chr14 | 43778945 | 43839755 | 60811 | 0.83 | 0.00726 | 0.00871 |
| chr9 | 112435405 | 112439854 | 4450 | 1.28 | 0.00726 | 0.00872 |
| chr14 | 90540540 | 90588291 | 47752 | -0.91 | 0.00726 | 0.00872 |
| chr3 | 24842338 | 24843645 | 1308 | -1.2 | 0.00727 | 0.00873 |
| chr12 | 94619806 | 94637399 | 17594 | 0.9 | 0.00727 | 0.00873 |
| chr4 | 68103254 | 68106867 | 3614 | 2.84 | 0.00727 | 0.00873 |
| chr18 | 47856405 | 47858225 | 1821 | -3.48 | 0.00728 | 0.00874 |
| chr2 | 7490868 | 7598951 | 108084 | -0.49 | 0.00728 | 0.00874 |
| chr4 | 164220097 | 164220517 | 421 | -1.87 | 0.00729 | 0.00875 |
| chr22 | 45407474 | 45419110 | 11637 | -5.76 | 0.00729 | 0.00875 |
| chr14 | 97573612 | 97576993 | 3382 | -3.66 | 0.00729 | 0.00875 |
| chr4 | 29192827 | 29195469 | 2643 | 3.34 | 0.0073 | 0.00876 |
| chr10 | 65484787 | 65486859 | 2073 | -0.87 | 0.00731 | 0.00877 |
| chr21 | 19084849 | 19086716 | 1868 | -1.21 | 0.00734 | 0.00881 |
| chr9 | 113339038 | 113361087 | 22050 | -4.02 | 0.00736 | 0.00883 |
| chr11 | 10441738 | 10454543 | 12806 | -2.44 | 0.00736 | 0.00883 |
| chr21 | 27039452 | 27054292 | 14841 | 0.88 | 0.00736 | 0.00883 |
| chrX | 54308970 | 54333699 | 24730 | -3.76 | 0.00738 | 0.00886 |
| chr3 | 19185323 | 19191141 | 5819 | -3.66 | 0.00739 | 0.00887 |
| chr12 | 109781500 | 109787021 | 5522 | -1.2 | 0.00741 | 0.00889 |
| chr4 | 35186707 | 35196103 | 9397 | 2.08 | 0.00742 | 0.0089 |
| chr20 | 22248499 | 22257727 | 9229 | 0.75 | 0.00742 | 0.0089 |
| chr10 | 113977658 | 114010863 | 33206 | -1.28 | 0.00743 | 0.00892 |
| chr10 | 24634304 | 24635263 | 960 | -2.94 | 0.00744 | 0.00893 |
| chr16 | 77596804 | 77612198 | 15395 | -3.8 | 0.00745 | 0.00894 |
| chr1 | 168459610 | 168472101 | 12492 | -5.73 | 0.00746 | 0.00895 |
| chr5 | 155316308 | 155317491 | 1184 | 1.44 | 0.00747 | 0.00896 |
| chrX | 57338851 | 57343895 | 5045 | -1.45 | 0.00747 | 0.00897 |
| chr6 | 88666142 | 88677291 | 11150 | -3.08 | 0.00747 | 0.00897 |
| chr3 | 162436410 | 162437335 | 926 | -1.86 | 0.00748 | 0.00897 |
| chr13 | 60765608 | 60766651 | 1044 | -1.43 | 0.00748 | 0.00898 |
| chr9 | 18394968 | 18515396 | 120429 | -5.72 | 0.00749 | 0.00898 |
| chrX | 51221549 | 51222109 | 561 | -2.94 | 0.0075 | 0.00899 |
| chr8 | 10856338 | 10868772 | 12435 | -0.77 | 0.0075 | 0.009 |
| chr4 | 188067561 | 188077294 | 9734 | -5.07 | 0.0075 | 0.009 |
| chr2 | 221311749 | 221318002 | 6254 | -2.29 | 0.00751 | 0.00901 |
| chr11 | 10742065 | 10754368 | 12304 | -5.72 | 0.00754 | 0.00905 |
| chr5 | 130315754 | 130318345 | 2592 | -1.38 | 0.00755 | 0.00906 |
| chr14 | 31986366 | 31988437 | 2072 | -2.1 | 0.00761 | 0.00913 |
| chr1 | 214132828 | 214135716 | 2889 | 1.88 | 0.00762 | 0.00913 |
| chr4 | 156577916 | 156580053 | 2138 | -3.62 | 0.00763 | 0.00915 |
| chr8 | 10541944 | 10542925 | 982 | -2.64 | 0.00765 | 0.00917 |
| chr2 | 149800141 | 149815524 | 15384 | -3.89 | 0.00765 | 0.00917 |
| chr7 | 54801057 | 54803274 | 2218 | -1.65 | 0.00768 | 0.00921 |
| chr1 | 213515751 | 213518308 | 2558 | -3.49 | 0.00769 | 0.00922 |
| chrX | 21336059 | 21383873 | 47815 | -4.09 | 0.0077 | 0.00923 |
| chr2 | 100646703 | 100647612 | 910 | -3.05 | 0.0077 | 0.00924 |
| chrX | 150286332 | 150289210 | 2879 | 1.65 | 0.00771 | 0.00925 |
| chr1 | 49230272 | 49231814 | 1543 | -2 | 0.00772 | 0.00926 |
| chr2 | 192681455 | 192681733 | 279 | -2.56 | 0.00772 | 0.00926 |
| chr10 | 7978732 | 7999547 | 20816 | -5.7 | 0.00772 | 0.00926 |
| chr3 | 46259940 | 46322378 | 62439 | -0.5 | 0.00773 | 0.00926 |
| chr1 | 85468605 | 85477514 | 8910 | -2.47 | 0.00773 | 0.00927 |
| chr8 | 39709689 | 39711750 | 2062 | -3.39 | 0.00776 | 0.0093 |
| chr12 | 113156369 | 113201582 | 45214 | -1.05 | 0.00776 | 0.0093 |

| | | | | | | |
|---|---|---|---|---|---|---|
| chr3 | 159833777 | 159838085 | 4309 | -3.38 | 0.00777 | 0.00931 |
| chr11 | 58982598 | 59190009 | 207412 | 6.12 | 0.00777 | 0.00932 |
| chr18 | 7373735 | 7398300 | 24566 | -2.98 | 0.00778 | 0.00932 |
| chr8 | 114398336 | 114408167 | 9832 | -1.9 | 0.00779 | 0.00934 |
| chr17 | 56152732 | 56156076 | 3345 | -2.08 | 0.00781 | 0.00936 |
| chr11 | 99518661 | 99519384 | 724 | -1.79 | 0.00782 | 0.00937 |
| chrX | 86725127 | 86727073 | 1947 | -3.25 | 0.00783 | 0.00938 |
| chr16 | 82217084 | 82220940 | 3857 | 1.35 | 0.00783 | 0.00939 |
| chr7 | 140823956 | 140848251 | 24296 | -1.96 | 0.00786 | 0.00942 |
| chr6 | 47865199 | 47878637 | 13439 | 1.86 | 0.00786 | 0.00942 |
| chr1 | 236774635 | 236798629 | 23995 | -4.21 | 0.00786 | 0.00942 |
| chr11 | 62920050 | 62921704 | 1655 | -1.96 | 0.00788 | 0.00945 |
| chr8 | 49597471 | 49600272 | 2802 | -3.17 | 0.00789 | 0.00945 |
| chr15 | 58077272 | 58086253 | 8982 | 0.9 | 0.00789 | 0.00946 |
| chr3 | 186059006 | 186069536 | 10531 | -5.68 | 0.00791 | 0.00947 |
| chr20 | 5438794 | 5451421 | 12628 | -1.44 | 0.00792 | 0.00949 |
| chr5 | 79853840 | 79856009 | 2170 | 3.42 | 0.00793 | 0.0095 |
| chr1 | 66131143 | 66137126 | 5984 | -1.91 | 0.00793 | 0.0095 |
| chr17 | 35053658 | 35056604 | 2947 | 2.62 | 0.00793 | 0.0095 |
| chrX | 82619776 | 82623116 | 3341 | 1.14 | 0.00794 | 0.00951 |
| chr10 | 125333547 | 125338543 | 4997 | -0.99 | 0.00797 | 0.00955 |
| chr6 | 70882580 | 70891205 | 8626 | 1.24 | 0.00799 | 0.00957 |
| chr4 | 139521869 | 139541590 | 19722 | -3.83 | 0.00799 | 0.00958 |
| chr7 | 28348966 | 28433558 | 84593 | 0.74 | 0.008 | 0.00958 |
| chr3 | 133760275 | 133773393 | 13119 | -2.73 | 0.00803 | 0.00961 |
| chrX | 116285864 | 116300128 | 14265 | 2.63 | 0.00804 | 0.00963 |
| chr9 | 88088307 | 88115747 | 27441 | -0.84 | 0.00804 | 0.00963 |
| chr2 | 56752067 | 56819060 | 66994 | 0.94 | 0.00806 | 0.00965 |
| chr9 | 113249781 | 113275291 | 25511 | 1.08 | 0.00807 | 0.00966 |
| chr2 | 108653167 | 108678110 | 24944 | -0.83 | 0.00808 | 0.00967 |
| chr1 | 224995906 | 225027280 | 31375 | -1.83 | 0.00808 | 0.00967 |
| chr10 | 65605997 | 65639499 | 33503 | -3.92 | 0.00808 | 0.00968 |
| chr4 | 105909248 | 105910508 | 1261 | 3.12 | 0.00809 | 0.00969 |
| chr15 | 89240849 | 89247995 | 7147 | -5.66 | 0.0081 | 0.0097 |
| chr4 | 141709712 | 141710985 | 1274 | -3.3 | 0.0081 | 0.0097 |
| chr8 | 52936473 | 52964105 | 27633 | 1.32 | 0.00812 | 0.00973 |
| chr3 | 28993045 | 29014057 | 21013 | -3.84 | 0.00813 | 0.00974 |
| chr8 | 142859425 | 142941112 | 81688 | -1.03 | 0.00815 | 0.00976 |
| chr3 | 79092146 | 79097986 | 5841 | 0.96 | 0.00817 | 0.00978 |
| chr16 | 74222744 | 74224048 | 1305 | 1.52 | 0.00818 | 0.00979 |
| chr4 | 142754344 | 142755003 | 660 | -1.64 | 0.00819 | 0.00981 |
| chrX | 67994886 | 67996082 | 1197 | 1.29 | 0.00821 | 0.00982 |
| chr12 | 18505155 | 18542418 | 37264 | -1.35 | 0.00821 | 0.00983 |
| chr9 | 80185107 | 80186591 | 1485 | -2.95 | 0.00822 | 0.00983 |
| chr21 | 19072193 | 19075367 | 3175 | 1.6 | 0.00822 | 0.00984 |
| chr7 | 106310702 | 106312410 | 1709 | 1.76 | 0.00822 | 0.00984 |
| chr4 | 141833678 | 141836834 | 3157 | 1.27 | 0.00822 | 0.00984 |
| chr2 | 218538601 | 218539535 | 935 | -1.53 | 0.00823 | 0.00985 |
| chr1 | 173290947 | 173291577 | 631 | -2.53 | 0.00825 | 0.00988 |
| chr1 | 42601375 | 42601986 | 612 | 1.75 | 0.00826 | 0.00988 |
| chr3 | 102038001 | 102042114 | 4114 | -3.55 | 0.00826 | 0.00989 |
| chr1 | 209339580 | 209678329 | 338750 | 0.57 | 0.00826 | 0.00989 |
| chr6 | 100066263 | 100087631 | 21369 | -1.13 | 0.00829 | 0.00992 |
| chr1 | 47523057 | 47524437 | 1381 | -3.08 | 0.00829 | 0.00992 |
| chr21 | 14944004 | 14944407 | 404 | 1.7 | 0.0083 | 0.00993 |
| chr11 | 20347069 | 20364228 | 17160 | 1.06 | 0.00831 | 0.00994 |
| chr16 | 79983624 | 80005720 | 22097 | -1.42 | 0.00831 | 0.00994 |
| chr5 | 98849173 | 98849515 | 343 | 2.81 | 0.00833 | 0.00996 |
| chr4 | 62791513 | 62793076 | 1564 | 1.75 | 0.00835 | 0.00999 |
| chr8 | 38494402 | 38496421 | 2020 | -1.92 | 0.00836 | 0.01 |
| chr5 | 83345137 | 83346374 | 1238 | -1.35 | 0.00838 | 0.01 |

| | | | | | | |
|---|---|---|---|---|---|---|
| chr12 | 76122132 | 76143922 | 21791 | -5.62 | 0.0084 | 0.0101 |
| chr2 | 113665705 | 113677778 | 12074 | -5.62 | 0.0084 | 0.0101 |
| chr12 | 41474029 | 41488190 | 14162 | -4.74 | 0.00841 | 0.0101 |
| chr2 | 204200989 | 204224536 | 23548 | -5.23 | 0.00843 | 0.0101 |
| chr7 | 40797066 | 40797829 | 764 | 2.23 | 0.00843 | 0.0101 |
| chr12 | 18365776 | 18366767 | 992 | -2.91 | 0.00843 | 0.0101 |
| chrX | 139058441 | 139059676 | 1236 | -3.12 | 0.00845 | 0.0101 |
| chrX | 105822304 | 105823685 | 1382 | -1.29 | 0.00845 | 0.0101 |
| chr8 | 75113677 | 75173733 | 60057 | 1.47 | 0.00848 | 0.0101 |
| chr21 | 27907899 | 27909186 | 1288 | -2.28 | 0.00848 | 0.0101 |
| chr1 | 112565270 | 112566903 | 1634 | 1.21 | 0.00849 | 0.0101 |
| chrX | 75339935 | 75351703 | 11769 | -1.99 | 0.00849 | 0.0101 |
| chr3 | 66992574 | 66994772 | 2199 | -2.04 | 0.00849 | 0.0101 |
| chr6 | 139525583 | 139531654 | 6072 | -1.58 | 0.00849 | 0.0102 |
| chr8 | 133411607 | 133448315 | 36709 | 4.22 | 0.0085 | 0.0102 |
| chrX | 30098727 | 30104498 | 5772 | -3.51 | 0.0085 | 0.0102 |
| chr2 | 108899171 | 108907604 | 8434 | -5.61 | 0.00851 | 0.0102 |
| chr5 | 155361875 | 155373511 | 11637 | -5.61 | 0.00851 | 0.0102 |
| chr1 | 209902456 | 209903968 | 1513 | -1.42 | 0.00851 | 0.0102 |
| chr20 | 57918462 | 57924193 | 5732 | -3.74 | 0.00851 | 0.0102 |
| chr6 | 127067855 | 127072386 | 4532 | -1.72 | 0.00853 | 0.0102 |
| chr2 | 458528 | 572846 | 114319 | -0.74 | 0.00854 | 0.0102 |
| chr10 | 37025903 | 37066692 | 40790 | 1.41 | 0.00854 | 0.0102 |
| chr19 | 14902174 | 14913188 | 11015 | -1.06 | 0.00854 | 0.0102 |
| chrX | 151524996 | 151528572 | 3577 | -1.1 | 0.00856 | 0.0102 |
| chr12 | 46409939 | 46510481 | 100543 | -5.91 | 0.00856 | 0.0102 |
| chr5 | 108322521 | 108323254 | 734 | -2.95 | 0.00857 | 0.0102 |
| chr7 | 31213584 | 31222177 | 8594 | -0.99 | 0.00858 | 0.0103 |
| chr4 | 123604335 | 123605639 | 1305 | -2.22 | 0.00858 | 0.0103 |
| chr18 | 25160666 | 25163403 | 2738 | -3.5 | 0.0086 | 0.0103 |
| chr6 | 57990396 | 58012346 | 21951 | 1.05 | 0.0086 | 0.0103 |
| chr8 | 16643081 | 16649156 | 6076 | -3.64 | 0.00862 | 0.0103 |
| chr8 | 40895780 | 40910874 | 15095 | -1.09 | 0.00863 | 0.0103 |
| chr9 | 91807481 | 91829259 | 21779 | -3.95 | 0.00864 | 0.0103 |
| chr2 | 136895273 | 136912076 | 16804 | -0.76 | 0.00865 | 0.0103 |
| chr1 | 171599139 | 171601856 | 2718 | -3.47 | 0.00867 | 0.0104 |
| chrX | 56368341 | 56371027 | 2687 | 1.35 | 0.00867 | 0.0104 |
| chr12 | 28807501 | 28914189 | 106689 | -0.76 | 0.00868 | 0.0104 |
| chr8 | 17725063 | 17768382 | 43320 | -5.62 | 0.00869 | 0.0104 |
| chr6 | 768249 | 769247 | 999 | -3.06 | 0.00871 | 0.0104 |
| chr14 | 37078196 | 37092267 | 14072 | -3.95 | 0.00873 | 0.0104 |
| chr4 | 145401483 | 145403805 | 2323 | -3.01 | 0.00874 | 0.0104 |
| chr4 | 139627368 | 139636073 | 8706 | 1.28 | 0.00874 | 0.0104 |
| chr7 | 124398883 | 124405714 | 6832 | -3.66 | 0.00874 | 0.0104 |
| chr18 | 76222141 | 76244532 | 22392 | 1.13 | 0.00875 | 0.0104 |
| chr10 | 15489841 | 15662493 | 172653 | 0.49 | 0.00875 | 0.0104 |
| chr6 | 749861 | 751327 | 1467 | -1.23 | 0.00875 | 0.0104 |
| chr1 | 48652849 | 48654923 | 2075 | -2.29 | 0.00876 | 0.0105 |
| chr1 | 245532945 | 245538697 | 5753 | -1.34 | 0.00876 | 0.0105 |
| chr16 | 63217805 | 63228609 | 10805 | -2.16 | 0.00877 | 0.0105 |
| chr14 | 36702213 | 36704300 | 2088 | 3.35 | 0.0088 | 0.0105 |
| chr7 | 24508272 | 24511047 | 2776 | -3.34 | 0.00881 | 0.0105 |
| chr4 | 78197080 | 78216001 | 18922 | 0.94 | 0.00881 | 0.0105 |
| chr9 | 111584931 | 111589155 | 4225 | -3.77 | 0.00881 | 0.0105 |
| chr12 | 53343846 | 53346737 | 2892 | 1.32 | 0.00883 | 0.0105 |
| chr10 | 22625243 | 22633936 | 8694 | -5.57 | 0.00883 | 0.0106 |
| chr3 | 83226193 | 83241552 | 15360 | -1.22 | 0.00884 | 0.0106 |
| chr11 | 134499371 | 134500951 | 1581 | -3.16 | 0.00885 | 0.0106 |
| chr12 | 131843370 | 131848350 | 4981 | -3.73 | 0.00886 | 0.0106 |
| chrX | 65793179 | 65793795 | 617 | -2.09 | 0.00886 | 0.0106 |
| chr4 | 146344234 | 146378986 | 34753 | -4.06 | 0.00887 | 0.0106 |

| | | | | | | |
|---|---|---|---|---|---|---|
| chr3 | 191750994 | 191751398 | 405 | 2.7 | 0.00888 | 0.0106 |
| chr1 | 219989273 | 220006287 | 17015 | -0.87 | 0.00889 | 0.0106 |
| chr7 | 106220736 | 106288589 | 67854 | -0.43 | 0.0089 | 0.0106 |
| chr3 | 119705344 | 119711537 | 6194 | -4.91 | 0.00894 | 0.0107 |
| chr3 | 16727116 | 16731282 | 4167 | -1 | 0.00895 | 0.0107 |
| chr10 | 121454275 | 121471060 | 16786 | -5.56 | 0.00895 | 0.0107 |
| chr15 | 57196076 | 57209491 | 13416 | -5.56 | 0.00895 | 0.0107 |
| chr2 | 19107779 | 19116706 | 8928 | -5.56 | 0.00895 | 0.0107 |
| chr2 | 19984400 | 19996935 | 12536 | -5.56 | 0.00895 | 0.0107 |
| chr6 | 70953937 | 70955130 | 1194 | 3.01 | 0.00896 | 0.0107 |
| chr15 | 78255728 | 78261621 | 5894 | -4.12 | 0.00896 | 0.0107 |
| chr10 | 116774736 | 116776462 | 1727 | 1.15 | 0.00897 | 0.0107 |
| chr5 | 41322701 | 41323420 | 720 | -2.89 | 0.00899 | 0.0107 |
| chr9 | 17826877 | 18178176 | 351300 | 6.44 | 0.009 | 0.0107 |
| chr5 | 41678682 | 41681370 | 2689 | -1.85 | 0.00901 | 0.0108 |
| chr8 | 105479527 | 105482719 | 3193 | 1.25 | 0.00902 | 0.0108 |
| chr4 | 118049067 | 118049678 | 612 | -2.65 | 0.00902 | 0.0108 |
| chr9 | 117589124 | 117668174 | 79051 | -5.66 | 0.00904 | 0.0108 |
| chr5 | 106560478 | 106562077 | 1600 | -3.2 | 0.00906 | 0.0108 |
| chr1 | 101294014 | 101305453 | 11440 | -5.55 | 0.00906 | 0.0108 |
| chr15 | 92436086 | 92446135 | 10050 | -5.55 | 0.00906 | 0.0108 |
| chr4 | 188720452 | 188727553 | 7102 | -3.54 | 0.00906 | 0.0108 |
| chr2 | 114735859 | 114763651 | 27793 | -1.99 | 0.00911 | 0.0109 |
| chr5 | 166180198 | 166197959 | 17762 | -1.73 | 0.00911 | 0.0109 |
| chr5 | 135171501 | 135195724 | 24224 | -3.95 | 0.00911 | 0.0109 |
| chr11 | 104592778 | 104595085 | 2308 | -1.73 | 0.00912 | 0.0109 |
| chr1 | 50839954 | 50856311 | 16358 | -3.56 | 0.00912 | 0.0109 |
| chr1 | 90763165 | 90836272 | 73108 | 4.34 | 0.00913 | 0.0109 |
| chr2 | 213142080 | 213146392 | 4313 | 2.91 | 0.00913 | 0.0109 |
| chr9 | 101339820 | 101349189 | 9370 | -2.38 | 0.00914 | 0.0109 |
| chr11 | 56959900 | 56962409 | 2510 | -1.12 | 0.00915 | 0.0109 |
| chr18 | 50185678 | 50190722 | 5045 | -3.4 | 0.00916 | 0.0109 |
| chr4 | 121912211 | 121917019 | 4809 | 3.58 | 0.00917 | 0.0109 |
| chr11 | 62704468 | 62713736 | 9269 | -5.54 | 0.00918 | 0.011 |
| chr8 | 142092013 | 142110998 | 18986 | -5.54 | 0.00918 | 0.011 |
| chr2 | 214417308 | 214420276 | 2969 | -1.28 | 0.00918 | 0.011 |
| chr10 | 87490086 | 87493776 | 3691 | 1.86 | 0.0092 | 0.011 |
| chr13 | 106252707 | 106253873 | 1167 | -2.77 | 0.00921 | 0.011 |
| chr5 | 122591539 | 122595736 | 4198 | -0.99 | 0.00922 | 0.011 |
| chrX | 55375890 | 55380364 | 4475 | 0.94 | 0.00925 | 0.011 |
| chr9 | 79127130 | 79169858 | 42729 | 1.77 | 0.00929 | 0.0111 |
| chr8 | 130026699 | 130037388 | 10690 | -3.61 | 0.00929 | 0.0111 |
| chr18 | 23527907 | 23529930 | 2024 | -1.91 | 0.00929 | 0.0111 |
| chr17 | 26415763 | 26438245 | 22483 | -5.53 | 0.0093 | 0.0111 |
| chr7 | 12979526 | 12992834 | 13309 | -5.53 | 0.0093 | 0.0111 |
| chr7 | 80065163 | 80081177 | 16015 | -5.53 | 0.0093 | 0.0111 |
| chr13 | 46692810 | 46693523 | 714 | -2.77 | 0.00931 | 0.0111 |
| chrX | 72907134 | 72922036 | 14903 | -2.78 | 0.00932 | 0.0111 |
| chr11 | 124385200 | 124429222 | 44023 | 0.87 | 0.00933 | 0.0111 |
| chr8 | 855631 | 929736 | 74106 | -0.97 | 0.00935 | 0.0111 |
| chr13 | 66942169 | 66944435 | 2267 | -3.2 | 0.00935 | 0.0112 |
| chr8 | 61199235 | 61217916 | 18682 | -0.75 | 0.00936 | 0.0112 |
| chr14 | 105072457 | 105097659 | 25203 | -3.57 | 0.0094 | 0.0112 |
| chr8 | 39209262 | 39220472 | 11211 | -0.77 | 0.0094 | 0.0112 |
| chr3 | 128224371 | 128263030 | 38660 | -4.1 | 0.00942 | 0.0112 |
| chr3 | 8766953 | 8781745 | 14793 | -5.51 | 0.00942 | 0.0112 |
| chr11 | 35140571 | 35143352 | 2782 | 1.69 | 0.00943 | 0.0112 |
| chrX | 118471118 | 118486097 | 14980 | -3.66 | 0.00944 | 0.0112 |
| chr1 | 182282429 | 182298040 | 15612 | 3.83 | 0.00947 | 0.0113 |
| chr6 | 130684398 | 130685032 | 635 | -2.75 | 0.00948 | 0.0113 |
| chrX | 5163730 | 5208617 | 44888 | 1.07 | 0.00948 | 0.0113 |

| | | | | | | |
|---|---|---|---|---|---|---|
| chr8 | 132894523 | 132896452 | 1930 | -1.24 | 0.0095 | 0.0113 |
| chrX | 63571882 | 63591121 | 19240 | 1.31 | 0.0095 | 0.0113 |
| chr3 | 16003888 | 16004485 | 598 | 1.61 | 0.00952 | 0.0113 |
| chr18 | 76292790 | 76396803 | 104014 | -0.55 | 0.00953 | 0.0114 |
| chr17 | 33523398 | 33563474 | 40077 | -4.18 | 0.00954 | 0.0114 |
| chr9 | 3199460 | 3213365 | 13906 | -1.48 | 0.00954 | 0.0114 |
| chr20 | 4602798 | 4629489 | 26692 | -4.18 | 0.00955 | 0.0114 |
| chrX | 123355845 | 123356745 | 901 | 3.11 | 0.00955 | 0.0114 |
| chr7 | 134745087 | 134751834 | 6748 | -0.75 | 0.00955 | 0.0114 |
| chr20 | 30032070 | 30047467 | 15398 | -5.5 | 0.00955 | 0.0114 |
| chr1 | 36160016 | 36160310 | 295 | -3.53 | 0.00957 | 0.0114 |
| chr4 | 112690566 | 112692106 | 1541 | 3.1 | 0.00957 | 0.0114 |
| chr13 | 37508817 | 37514304 | 5488 | -0.95 | 0.00959 | 0.0114 |
| chr2 | 103504046 | 105020396 | 1516351 | 0.32 | 0.0096 | 0.0114 |
| chr5 | 155099358 | 155102584 | 3227 | -1.58 | 0.00961 | 0.0115 |
| chr3 | 191831965 | 191832989 | 1025 | -1.49 | 0.00961 | 0.0115 |
| chr10 | 133127651 | 133169623 | 41973 | 0.81 | 0.00962 | 0.0115 |
| chrX | 110496532 | 110508489 | 11958 | 1.05 | 0.00962 | 0.0115 |
| chr11 | 116088693 | 116103539 | 14847 | -2.22 | 0.00962 | 0.0115 |
| chr13 | 89376399 | 89376847 | 449 | -2.77 | 0.00963 | 0.0115 |
| chr11 | 20279144 | 20293832 | 14689 | 1.01 | 0.00964 | 0.0115 |
| chr3 | 148193613 | 148205774 | 12162 | -3.65 | 0.00967 | 0.0115 |
| chr3 | 59979216 | 59999146 | 19931 | 2.4 | 0.00968 | 0.0115 |
| chrX | 141109678 | 141117426 | 7749 | -1.16 | 0.00968 | 0.0115 |
| chr9 | 72162209 | 72163554 | 1346 | -3.19 | 0.00968 | 0.0115 |
| chr8 | 24296772 | 24298636 | 1865 | 2.29 | 0.00971 | 0.0116 |
| chr9 | 106998600 | 106998867 | 268 | -3.68 | 0.00972 | 0.0116 |
| chr1 | 99473355 | 99475358 | 2004 | 1.31 | 0.00974 | 0.0116 |
| chr4 | 24996741 | 25000716 | 3976 | -1.56 | 0.00974 | 0.0116 |
| chr8 | 16730395 | 16748445 | 18051 | -0.7 | 0.00974 | 0.0116 |
| chr14 | 76766210 | 76768021 | 1812 | -3.37 | 0.00975 | 0.0116 |
| chr12 | 106337057 | 106343452 | 6396 | -1.76 | 0.00977 | 0.0116 |
| chr5 | 112560009 | 112560843 | 835 | 2.96 | 0.0098 | 0.0117 |
| chr2 | 46156028 | 46172831 | 16804 | -5.48 | 0.00981 | 0.0117 |
| chr5 | 77881250 | 77882093 | 844 | -2.91 | 0.00982 | 0.0117 |
| chr6 | 75356817 | 75397881 | 41065 | -0.83 | 0.00983 | 0.0117 |
| chr15 | 58336248 | 58338480 | 2233 | -1.07 | 0.00983 | 0.0117 |
| chr16 | 77040509 | 77058607 | 18099 | -3.08 | 0.00983 | 0.0117 |
| chr5 | 159923371 | 159925100 | 1730 | 3.4 | 0.00984 | 0.0117 |
| chr1 | 108460695 | 108461261 | 567 | -2.23 | 0.00985 | 0.0117 |
| chr4 | 135285668 | 135287737 | 2070 | -1.31 | 0.00985 | 0.0117 |
| chr1 | 245413013 | 245469809 | 56797 | -2.52 | 0.00988 | 0.0118 |
| chr13 | 88670797 | 88676043 | 5247 | 2.01 | 0.00988 | 0.0118 |
| chr4 | 37915749 | 37917796 | 2048 | 0.98 | 0.00988 | 0.0118 |
| chrX | 20977062 | 20978708 | 1647 | 1.31 | 0.0099 | 0.0118 |
| chrX | 81513118 | 81523755 | 10638 | -3.49 | 0.00991 | 0.0118 |
| chrX | 139925434 | 139926543 | 1110 | -2.79 | 0.00992 | 0.0118 |
| chr6 | 164375245 | 164376508 | 1264 | -1.66 | 0.00992 | 0.0118 |
| chr17 | 39146542 | 39157872 | 11331 | -5.46 | 0.00994 | 0.0118 |
| chrX | 20787958 | 20798732 | 10775 | -3.47 | 0.00996 | 0.0119 |
| chr13 | 109216701 | 109218369 | 1669 | -3.35 | 0.00997 | 0.0119 |
| chr2 | 115548 | 168910 | 53363 | 0.87 | 0.00997 | 0.0119 |
| chr2 | 217842152 | 217857119 | 14968 | -3.5 | 0.00997 | 0.0119 |
| chr2 | 95663689 | 95664668 | 980 | -1.73 | 0.00998 | 0.0119 |
| chr4 | 33195572 | 33210521 | 14950 | 1.02 | 0.00998 | 0.0119 |
| chr13 | 94630082 | 94631097 | 1016 | -1.4 | 0.00999 | 0.0119 |
| chr6 | 126195217 | 126196832 | 1616 | -3.21 | 0.01 | 0.0119 |
| chr2 | 88719684 | 88737761 | 18078 | -2.22 | 0.01 | 0.0119 |
| chr1 | 219831382 | 219831662 | 281 | 2.15 | 0.01 | 0.0119 |
| chr13 | 107099602 | 107101732 | 2131 | -3.21 | 0.01 | 0.0119 |
| chr4 | 67975060 | 67979684 | 4625 | 1.1 | 0.01 | 0.0119 |

| | | | | | | |
|---|---|---|---|---|---|---|
| chr10 | 105284642 | 105294666 | 10025 | -5.45 | 0.0101 | 0.012 |
| chr14 | 34778462 | 34788300 | 9839 | -1.15 | 0.0101 | 0.012 |
| chr9 | 121265884 | 121266947 | 1064 | 1.37 | 0.0101 | 0.012 |
| chr9 | 88060788 | 88065828 | 5041 | -1.41 | 0.0101 | 0.012 |
| chrX | 139195981 | 139196703 | 723 | -1.83 | 0.0101 | 0.012 |
| chr12 | 38761901 | 38763768 | 1868 | 1.18 | 0.0101 | 0.0121 |
| chr8 | 10141665 | 10144574 | 2910 | -3.25 | 0.0102 | 0.0121 |
| chr8 | 121608886 | 121609962 | 1077 | -2.06 | 0.0102 | 0.0121 |
| chr6 | 142877852 | 142879246 | 1395 | -2.54 | 0.0102 | 0.0121 |
| chrX | 20919327 | 20959881 | 40555 | -2.13 | 0.0102 | 0.0121 |
| chr16 | 32506670 | 32654435 | 147766 | -0.45 | 0.0102 | 0.0121 |
| chr6 | 142283023 | 142283338 | 316 | 2.96 | 0.0102 | 0.0121 |
| chr14 | 58243145 | 58276059 | 32915 | -1.61 | 0.0102 | 0.0121 |
| chr12 | 6353764 | 6367926 | 14163 | -5.44 | 0.0102 | 0.0121 |
| chr14 | 51597086 | 51598081 | 996 | 2.91 | 0.0102 | 0.0122 |
| chr6 | 8343988 | 8386796 | 42809 | 0.69 | 0.0102 | 0.0122 |
| chr4 | 135026178 | 135027012 | 835 | 4.41 | 0.0102 | 0.0122 |
| chr6 | 80438324 | 80442480 | 4157 | 1.74 | 0.0103 | 0.0122 |
| chr4 | 22866004 | 22879998 | 13995 | -3.55 | 0.0103 | 0.0122 |
| chrX | 12937852 | 12940485 | 2634 | 1.92 | 0.0103 | 0.0122 |
| chr8 | 58044399 | 58047476 | 3078 | -1.17 | 0.0103 | 0.0122 |
| chr9 | 101283489 | 101300278 | 16790 | -3.6 | 0.0103 | 0.0123 |
| chr2 | 214682227 | 214695263 | 13037 | -2.11 | 0.0103 | 0.0123 |
| chr4 | 66357890 | 66358459 | 570 | -2.49 | 0.0103 | 0.0123 |
| chr11 | 104387813 | 104389854 | 2042 | 1.87 | 0.0103 | 0.0123 |
| chr7 | 77735840 | 77737900 | 2061 | -3.46 | 0.0103 | 0.0123 |
| chr2 | 17746570 | 17747934 | 1365 | -3.38 | 0.0103 | 0.0123 |
| chrX | 68887724 | 68892250 | 4527 | -0.87 | 0.0104 | 0.0123 |
| chr11 | 36353737 | 36355882 | 2146 | -3.34 | 0.0104 | 0.0123 |
| chr5 | 155136371 | 155138357 | 1987 | -1.88 | 0.0104 | 0.0123 |
| chr8 | 16661949 | 16666120 | 4172 | -2.15 | 0.0104 | 0.0124 |
| chr12 | 10052835 | 10073792 | 20958 | -3.01 | 0.0104 | 0.0124 |
| chr13 | 101422274 | 101423500 | 1227 | -1.73 | 0.0104 | 0.0124 |
| chr8 | 82251341 | 82258158 | 6818 | 3.14 | 0.0104 | 0.0124 |
| chr7 | 15962470 | 15968392 | 5923 | -1.42 | 0.0104 | 0.0124 |
| chr22 | 32814108 | 32815001 | 894 | -2.22 | 0.0105 | 0.0124 |
| chr10 | 61095899 | 61108743 | 12845 | 0.92 | 0.0105 | 0.0124 |
| chrX | 141290646 | 141304680 | 14035 | 0.71 | 0.0105 | 0.0125 |
| chr12 | 124414299 | 124426979 | 12681 | -5.41 | 0.0105 | 0.0125 |
| chr14 | 89106593 | 89121766 | 15174 | -5.41 | 0.0105 | 0.0125 |
| chr10 | 63811968 | 63813721 | 1754 | 1.02 | 0.0105 | 0.0125 |
| chr3 | 71878449 | 71878944 | 496 | 2.55 | 0.0105 | 0.0125 |
| chr1 | 222203232 | 222210577 | 7346 | -3.07 | 0.0105 | 0.0125 |
| chrX | 39223753 | 39225112 | 1360 | -2 | 0.0106 | 0.0125 |
| chr1 | 20611703 | 20612732 | 1030 | -2.36 | 0.0106 | 0.0125 |
| chr10 | 33719482 | 33719836 | 355 | -1.73 | 0.0106 | 0.0126 |
| chr11 | 112304968 | 112310094 | 5127 | 1.98 | 0.0106 | 0.0126 |
| chr13 | 98545637 | 98561510 | 15874 | -3.65 | 0.0106 | 0.0126 |
| chr13 | 39786191 | 39817748 | 31558 | 1.07 | 0.0106 | 0.0126 |
| chr1 | 91592673 | 91604122 | 11450 | 2.47 | 0.0106 | 0.0126 |
| chr8 | 118013858 | 118019933 | 6076 | 3.63 | 0.0106 | 0.0126 |
| chr3 | 37833479 | 37834414 | 936 | 2.89 | 0.0106 | 0.0126 |
| chr1 | 104405461 | 104448870 | 43410 | 0.93 | 0.0107 | 0.0127 |
| chr15 | 72615505 | 72627908 | 12404 | -5.4 | 0.0107 | 0.0127 |
| chr8 | 18758306 | 18786189 | 27884 | -5.4 | 0.0107 | 0.0127 |
| chr5 | 14536693 | 14555604 | 18912 | -3.95 | 0.0107 | 0.0127 |
| chr2 | 150176556 | 150203842 | 27287 | -3.85 | 0.0107 | 0.0127 |
| chr12 | 91062870 | 91064637 | 1768 | -1.88 | 0.0107 | 0.0127 |
| chr8 | 119502280 | 119503908 | 1629 | -3.18 | 0.0107 | 0.0127 |
| chr11 | 124313570 | 124330944 | 17375 | 1.11 | 0.0107 | 0.0127 |
| chrX | 20577826 | 20578897 | 1072 | -2.98 | 0.0107 | 0.0127 |

| | | | | | | |
|---|---|---|---|---|---|---|
| chr8 | 61043278 | 61045357 | 2080 | -3.04 | 0.0107 | 0.0127 |
| chr5 | 113618405 | 113639325 | 20921 | 2.66 | 0.0107 | 0.0127 |
| chr22 | 17932704 | 17934537 | 1834 | -3.08 | 0.0107 | 0.0127 |
| chr1 | 80212063 | 80213847 | 1785 | -1.3 | 0.0107 | 0.0127 |
| chr4 | 145780550 | 145784220 | 3671 | -3.32 | 0.0107 | 0.0128 |
| chr5 | 45062036 | 45112968 | 50933 | -0.63 | 0.0108 | 0.0128 |
| chr4 | 24301815 | 24302712 | 898 | -1.46 | 0.0108 | 0.0128 |
| chr9 | 121714224 | 121716228 | 2005 | 1.02 | 0.0108 | 0.0128 |
| chrX | 110102777 | 110104262 | 1486 | -1.7 | 0.0108 | 0.0128 |
| chr8 | 124985576 | 125033404 | 47829 | -4.05 | 0.0108 | 0.0128 |
| chr6 | 148903524 | 148927018 | 23495 | -3.95 | 0.0108 | 0.0128 |
| chr4 | 23471584 | 23474696 | 3113 | -1.3 | 0.0108 | 0.0128 |
| chr5 | 167028160 | 167030329 | 2170 | -2.4 | 0.0108 | 0.0128 |
| chr10 | 80285405 | 80301503 | 16099 | -5.38 | 0.0108 | 0.0128 |
| chr8 | 94951250 | 94960797 | 9548 | -5.38 | 0.0108 | 0.0128 |
| chr4 | 183713761 | 183717274 | 3514 | -0.94 | 0.0108 | 0.0128 |
| chr10 | 94556214 | 94556725 | 512 | -1.47 | 0.0108 | 0.0128 |
| chr11 | 96150761 | 96175465 | 24705 | -3.84 | 0.0108 | 0.0128 |
| chr4 | 42344142 | 42344782 | 641 | -2.69 | 0.0108 | 0.0128 |
| chr14 | 53063244 | 53073462 | 10219 | -3.84 | 0.0108 | 0.0128 |
| chr7 | 124055333 | 124058331 | 2999 | 1.39 | 0.0108 | 0.0129 |
| chr1 | 29676562 | 29677233 | 672 | -2.61 | 0.0108 | 0.0129 |
| chr12 | 54195438 | 54196630 | 1193 | 1.77 | 0.0108 | 0.0129 |
| chrX | 51850030 | 51852123 | 2094 | 3.26 | 0.0109 | 0.0129 |
| chr2 | 206738371 | 206740067 | 1697 | -3.26 | 0.0109 | 0.0129 |
| chr3 | 31354633 | 31365963 | 11331 | -0.74 | 0.0109 | 0.0129 |
| chr3 | 62323905 | 62338990 | 15086 | 2.2 | 0.0109 | 0.0129 |
| chr3 | 161775528 | 161779245 | 3718 | -1.29 | 0.0109 | 0.013 |
| chr2 | 176074515 | 176076062 | 1548 | -3.04 | 0.0109 | 0.013 |
| chr5 | 138882207 | 138882656 | 450 | -2.24 | 0.011 | 0.013 |
| chr1 | 160021415 | 160025195 | 3781 | 1.74 | 0.011 | 0.013 |
| chr12 | 7702657 | 7703632 | 976 | -2.97 | 0.011 | 0.013 |
| chr2 | 58568648 | 58570336 | 1689 | -3.21 | 0.011 | 0.013 |
| chr2 | 67289519 | 67333994 | 44476 | 0.67 | 0.011 | 0.0131 |
| chr18 | 73691210 | 73755784 | 64575 | -0.4 | 0.011 | 0.0131 |
| chr3 | 168862181 | 168863773 | 1593 | -1.68 | 0.011 | 0.0131 |
| chr5 | 108603427 | 108605245 | 1819 | -1.67 | 0.011 | 0.0131 |
| chr2 | 132450815 | 132451139 | 325 | -2.34 | 0.011 | 0.0131 |
| chr8 | 58505688 | 58506119 | 432 | -3.73 | 0.0111 | 0.0131 |
| chr8 | 117887354 | 117918667 | 31314 | -1.72 | 0.0111 | 0.0131 |
| chr7 | 15492086 | 15492802 | 717 | -2.81 | 0.0111 | 0.0131 |
| chr6 | 154009098 | 154010127 | 1030 | -1.45 | 0.0111 | 0.0131 |
| chr13 | 109712977 | 109730825 | 17849 | -3.26 | 0.0111 | 0.0131 |
| chr13 | 26697324 | 26717131 | 19808 | 0.92 | 0.0111 | 0.0131 |
| chr5 | 129203199 | 129208295 | 5097 | -1.39 | 0.0111 | 0.0132 |
| chr16 | 73922126 | 73979482 | 57357 | 0.77 | 0.0111 | 0.0132 |
| chr12 | 7755461 | 7758145 | 2685 | 1.48 | 0.0111 | 0.0132 |
| chr5 | 90017527 | 90040106 | 22580 | 3.78 | 0.0111 | 0.0132 |
| chr10 | 35956982 | 35957987 | 1006 | 2.19 | 0.0111 | 0.0132 |
| chr1 | 4845448 | 4894571 | 49124 | -0.54 | 0.0111 | 0.0132 |
| chr5 | 123235352 | 123236761 | 1410 | 1.3 | 0.0111 | 0.0132 |
| chr3 | 65226912 | 65246934 | 20023 | 3.83 | 0.0111 | 0.0132 |
| chr4 | 108436548 | 108437468 | 921 | -2.51 | 0.0112 | 0.0132 |
| chr6 | 126002546 | 126033646 | 31101 | -2.21 | 0.0112 | 0.0132 |
| chr5 | 174566018 | 174809176 | 243159 | 0.81 | 0.0112 | 0.0133 |
| chr4 | 19234762 | 19236907 | 2146 | -1.5 | 0.0112 | 0.0133 |
| chr11 | 101142127 | 101143515 | 1389 | 3.18 | 0.0112 | 0.0133 |
| chr1 | 20256325 | 20298613 | 42289 | -3.57 | 0.0112 | 0.0133 |
| chr18 | 24423721 | 24437136 | 13416 | -3.62 | 0.0112 | 0.0133 |
| chr18 | 2886326 | 2896003 | 9678 | -3.05 | 0.0112 | 0.0133 |
| chr18 | 71668978 | 71679479 | 10502 | 0.83 | 0.0112 | 0.0133 |

| | | | | | | |
|---|---|---|---|---|---|---|
| chr16 | 70977802 | 70979834 | 2033 | 1.05 | 0.0112 | 0.0133 |
| chr6 | 129180447 | 129182851 | 2405 | -3.29 | 0.0112 | 0.0133 |
| chr3 | 173117061 | 173118299 | 1239 | 3.24 | 0.0112 | 0.0133 |
| chr9 | 110114610 | 110117728 | 3119 | -3.51 | 0.0113 | 0.0133 |
| chr2 | 136700152 | 136710807 | 10656 | -3.54 | 0.0113 | 0.0133 |
| chr6 | 14954273 | 14969886 | 15614 | -5.34 | 0.0113 | 0.0134 |
| chr1 | 88872607 | 88909243 | 36637 | -1.22 | 0.0113 | 0.0134 |
| chr18 | 13919266 | 13969081 | 49816 | -0.62 | 0.0113 | 0.0134 |
| chr2 | 56897001 | 56905974 | 8974 | -3.88 | 0.0113 | 0.0134 |
| chr3 | 155797335 | 155798585 | 1251 | -2.14 | 0.0113 | 0.0134 |
| chr1 | 179419003 | 179421947 | 2945 | -2.24 | 0.0113 | 0.0134 |
| chr11 | 98977309 | 98977689 | 381 | 2.72 | 0.0113 | 0.0134 |
| chr5 | 107564716 | 107565033 | 318 | -2.55 | 0.0114 | 0.0134 |
| chrX | 63071603 | 63148941 | 77339 | 0.65 | 0.0114 | 0.0135 |
| chr5 | 150920385 | 150926103 | 5719 | 0.95 | 0.0114 | 0.0135 |
| chr12 | 119676428 | 119679285 | 2858 | 3.49 | 0.0114 | 0.0135 |
| chr5 | 68267838 | 68284650 | 16813 | -2.68 | 0.0114 | 0.0135 |
| chr12 | 97347018 | 97369358 | 22341 | -4 | 0.0114 | 0.0135 |
| chr17 | 63331480 | 63366398 | 34919 | -4.13 | 0.0114 | 0.0135 |
| chr8 | 19154438 | 19156843 | 2406 | 1.88 | 0.0114 | 0.0135 |
| chr18 | 45907124 | 45908029 | 906 | -2.7 | 0.0114 | 0.0136 |
| chr15 | 38245137 | 38291971 | 46835 | -2.57 | 0.0115 | 0.0136 |
| chr10 | 85890901 | 85892242 | 1342 | 2.18 | 0.0115 | 0.0136 |
| chr17 | 71525967 | 71526978 | 1012 | -2.98 | 0.0115 | 0.0136 |
| chrX | 134735897 | 134736417 | 521 | -2.42 | 0.0115 | 0.0136 |
| chr11 | 88345833 | 88346537 | 705 | -1.54 | 0.0115 | 0.0136 |
| chr15 | 62470853 | 62474170 | 3318 | -2.62 | 0.0115 | 0.0136 |
| chr4 | 62126868 | 62138638 | 11771 | 1.07 | 0.0115 | 0.0136 |
| chr3 | 164511680 | 164532400 | 20721 | 0.77 | 0.0115 | 0.0137 |
| chr19 | 55195365 | 55228271 | 32907 | -1.01 | 0.0116 | 0.0137 |
| chr4 | 59026022 | 59028919 | 2898 | 1.16 | 0.0116 | 0.0137 |
| chr8 | 126786818 | 126814681 | 27864 | 1.25 | 0.0116 | 0.0137 |
| chr22 | 43889952 | 43894312 | 4361 | 1.18 | 0.0116 | 0.0137 |
| chr5 | 21447128 | 21449118 | 1991 | -1.36 | 0.0116 | 0.0137 |
| chr4 | 35846372 | 35854285 | 7914 | 1.22 | 0.0116 | 0.0137 |
| chr1 | 234730481 | 234754367 | 23887 | -5.31 | 0.0116 | 0.0138 |
| chr12 | 20146697 | 20163887 | 17191 | -5.31 | 0.0116 | 0.0138 |
| chr7 | 132063491 | 132087663 | 24173 | -5.31 | 0.0116 | 0.0138 |
| chr8 | 108557607 | 108565898 | 8292 | -5.31 | 0.0116 | 0.0138 |
| chr14 | 94991011 | 95046746 | 55736 | -4.08 | 0.0116 | 0.0138 |
| chr4 | 35012071 | 35036302 | 24232 | -1.03 | 0.0116 | 0.0138 |
| chr16 | 51224110 | 51262519 | 38410 | -1.08 | 0.0116 | 0.0138 |
| chr2 | 163941388 | 163941703 | 316 | 2.69 | 0.0117 | 0.0138 |
| chr3 | 190871402 | 190875471 | 4070 | -1.46 | 0.0117 | 0.0138 |
| chr2 | 195847542 | 195852263 | 4722 | -1.91 | 0.0117 | 0.0138 |
| chrX | 56497643 | 56501366 | 3724 | 2.24 | 0.0117 | 0.0139 |
| chr5 | 21405628 | 21407672 | 2045 | -1.18 | 0.0117 | 0.0139 |
| chr5 | 111915160 | 111921280 | 6121 | 3.6 | 0.0117 | 0.0139 |
| chr3 | 61559213 | 61560886 | 1674 | -1.45 | 0.0117 | 0.0139 |
| chr7 | 11282383 | 11330176 | 47794 | 4.17 | 0.0117 | 0.0139 |
| chr1 | 118522646 | 118539893 | 17248 | -0.9 | 0.0118 | 0.0139 |
| chrX | 103141309 | 103141933 | 625 | -2.84 | 0.0118 | 0.0139 |
| chrX | 93881243 | 93917251 | 36009 | 0.78 | 0.0118 | 0.0139 |
| chr6 | 77460781 | 77464488 | 3708 | 1.48 | 0.0118 | 0.0139 |
| chr15 | 57712572 | 57713752 | 1181 | -3.07 | 0.0118 | 0.0139 |
| chrX | 110867680 | 110868600 | 921 | 2.78 | 0.0118 | 0.0139 |
| chr17 | 6585813 | 6599020 | 13208 | 3.79 | 0.0118 | 0.014 |
| chr2 | 17963086 | 17965323 | 2238 | -1.69 | 0.0118 | 0.014 |
| chr6 | 6776620 | 6777561 | 942 | -1.37 | 0.0118 | 0.014 |
| chr12 | 108685135 | 108687632 | 2498 | 3.52 | 0.0118 | 0.014 |
| chr7 | 37209030 | 37214877 | 5848 | 1.23 | 0.0118 | 0.014 |

| | | | | | | |
|---|---|---|---|---|---|---|
| chrX | 132940174 | 132942059 | 1886 | -1.73 | 0.0119 | 0.014 |
| chr6 | 168986999 | 168988257 | 1259 | -1.42 | 0.0119 | 0.0141 |
| chr12 | 55099278 | 55102752 | 3475 | -3.37 | 0.0119 | 0.0141 |
| chr1 | 82968417 | 82985702 | 17286 | 0.88 | 0.0119 | 0.0141 |
| chr11 | 93746898 | 93760654 | 13757 | -1.62 | 0.0119 | 0.0141 |
| chrX | 134635469 | 134638163 | 2695 | 2.29 | 0.0119 | 0.0141 |
| chr12 | 25838355 | 25876621 | 38267 | -1.81 | 0.012 | 0.0141 |
| chr11 | 134867606 | 134870267 | 2662 | 3.32 | 0.012 | 0.0141 |
| chr20 | 18761178 | 18786417 | 25240 | -3.06 | 0.012 | 0.0141 |
| chr8 | 126167933 | 126173823 | 5891 | -5.14 | 0.012 | 0.0142 |
| chr18 | 3332495 | 3342331 | 9837 | -5.28 | 0.012 | 0.0142 |
| chr2 | 71746980 | 71759753 | 12774 | -5.28 | 0.012 | 0.0142 |
| chr1 | 204784693 | 204787340 | 2648 | -1.31 | 0.012 | 0.0142 |
| chr13 | 95529514 | 95530766 | 1253 | -2.82 | 0.012 | 0.0142 |
| chr18 | 345532 | 347419 | 1888 | -1.64 | 0.012 | 0.0142 |
| chr6 | 78997789 | 79000089 | 2301 | 3.12 | 0.0121 | 0.0143 |
| chr13 | 24946301 | 24951314 | 5014 | -1.09 | 0.0121 | 0.0143 |
| chr9 | 88533426 | 88534223 | 798 | -1.72 | 0.0121 | 0.0143 |
| chr4 | 104602940 | 104605759 | 2820 | 3.37 | 0.0121 | 0.0143 |
| chr12 | 26031453 | 26066549 | 35097 | -3.98 | 0.0121 | 0.0143 |
| chr3 | 66820000 | 66850852 | 30853 | -1.5 | 0.0121 | 0.0143 |
| chr20 | 60465918 | 60467908 | 1991 | -1.22 | 0.0121 | 0.0143 |
| chr11 | 106359102 | 106363501 | 4400 | -3.21 | 0.0121 | 0.0143 |
| chr22 | 35895496 | 35901048 | 5553 | -1.82 | 0.0121 | 0.0143 |
| chr4 | 21900321 | 21901983 | 1663 | -1.25 | 0.0121 | 0.0143 |
| chr4 | 27916892 | 27917841 | 950 | 1.64 | 0.0121 | 0.0143 |
| chr12 | 130878721 | 130906699 | 27979 | 1.18 | 0.0121 | 0.0144 |
| chr10 | 18776110 | 18780949 | 4840 | 3.52 | 0.0122 | 0.0144 |
| chr4 | 34465950 | 34469553 | 3604 | -1.36 | 0.0122 | 0.0144 |
| chr11 | 57014792 | 57026195 | 11404 | -5.27 | 0.0122 | 0.0144 |
| chr4 | 33546091 | 33549349 | 3259 | -1.43 | 0.0122 | 0.0144 |
| chr8 | 49893820 | 49895890 | 2071 | 1.64 | 0.0122 | 0.0144 |
| chr5 | 91342895 | 91362920 | 20026 | -3.62 | 0.0122 | 0.0144 |
| chr2 | 75663445 | 75688212 | 24768 | -1.43 | 0.0122 | 0.0144 |
| chr8 | 117468253 | 117470288 | 2036 | -2.62 | 0.0122 | 0.0144 |
| chr13 | 27409246 | 27426734 | 17489 | -1.45 | 0.0122 | 0.0144 |
| chr1 | 159114768 | 159128236 | 13469 | -1.16 | 0.0122 | 0.0144 |
| chr13 | 77524675 | 77526874 | 2200 | 2.45 | 0.0122 | 0.0144 |
| chr21 | 15921661 | 15952131 | 30471 | -0.58 | 0.0123 | 0.0145 |
| chr18 | 57980314 | 57991896 | 11583 | -1.07 | 0.0123 | 0.0145 |
| chr8 | 54446705 | 54449047 | 2343 | 2.57 | 0.0123 | 0.0145 |
| chr3 | 138864711 | 138871487 | 6777 | -1.22 | 0.0123 | 0.0145 |
| chr12 | 4698558 | 4699032 | 475 | -1.55 | 0.0123 | 0.0145 |
| chr3 | 18358116 | 18359021 | 906 | -2.81 | 0.0123 | 0.0145 |
| chr11 | 91190025 | 91223942 | 33918 | -3.22 | 0.0123 | 0.0145 |
| chr5 | 130337305 | 130352911 | 15607 | -1.94 | 0.0123 | 0.0145 |
| chr12 | 60412748 | 60413825 | 1078 | 1.51 | 0.0123 | 0.0145 |
| chr20 | 39577567 | 39583637 | 6071 | -1.82 | 0.0123 | 0.0146 |
| chr10 | 68888883 | 68895830 | 6948 | 3.6 | 0.0123 | 0.0146 |
| chr2 | 179825113 | 179827160 | 2048 | 1.45 | 0.0124 | 0.0146 |
| chr5 | 110830093 | 110831136 | 1044 | -1.15 | 0.0124 | 0.0146 |
| chr2 | 86619279 | 86679748 | 60470 | -5.55 | 0.0124 | 0.0146 |
| chr2 | 53813657 | 53814206 | 550 | 2.12 | 0.0124 | 0.0146 |
| chr18 | 43138719 | 43158923 | 20205 | 3.22 | 0.0124 | 0.0147 |
| chr3 | 83289160 | 83356514 | 67355 | -1.99 | 0.0125 | 0.0147 |
| chr1 | 242830765 | 242831956 | 1192 | -2.2 | 0.0125 | 0.0148 |
| chr15 | 62840624 | 62841493 | 870 | 3.06 | 0.0125 | 0.0148 |
| chr15 | 33881894 | 34008522 | 126629 | 0.61 | 0.0125 | 0.0148 |
| chr4 | 8923847 | 8939085 | 15239 | -3.9 | 0.0125 | 0.0148 |
| chr5 | 135051462 | 135052849 | 1388 | 2.78 | 0.0125 | 0.0148 |
| chr14 | 28674127 | 28699185 | 25059 | 3.85 | 0.0125 | 0.0148 |

| | | | | | | |
|---|---|---|---|---|---|---|
| chr4 | 135179943 | 135199080 | 19138 | 3 | 0.0125 | 0.0148 |
| chrX | 65212148 | 65212771 | 624 | -2.44 | 0.0125 | 0.0148 |
| chr5 | 71771506 | 71785824 | 14319 | -5.24 | 0.0125 | 0.0148 |
| chr5 | 111086448 | 111087469 | 1022 | -3.01 | 0.0125 | 0.0148 |
| chr3 | 162463751 | 162467222 | 3472 | -3.36 | 0.0126 | 0.0148 |
| chr4 | 100719635 | 100721449 | 1815 | -1.18 | 0.0126 | 0.0148 |
| chr19 | 51465000 | 51500407 | 35408 | -1.98 | 0.0126 | 0.0148 |
| chr7 | 142394919 | 142442476 | 47558 | 4.08 | 0.0126 | 0.0148 |
| chr4 | 61794889 | 61834245 | 39357 | 0.61 | 0.0126 | 0.0149 |
| chr5 | 154479166 | 154495694 | 16529 | -3.38 | 0.0126 | 0.0149 |
| chr14 | 54528675 | 54529953 | 1279 | 3 | 0.0126 | 0.0149 |
| chr4 | 15303360 | 15316394 | 13035 | -2.82 | 0.0126 | 0.0149 |
| chr1 | 64719607 | 64720614 | 1008 | -2.92 | 0.0126 | 0.0149 |
| chr17 | 37997941 | 37999198 | 1258 | 1.39 | 0.0126 | 0.0149 |
| chrX | 54995293 | 55011151 | 15859 | -1.93 | 0.0126 | 0.0149 |
| chrX | 93869932 | 93872181 | 2250 | -1.38 | 0.0127 | 0.015 |
| chr7 | 21324647 | 21376728 | 52082 | -2.45 | 0.0127 | 0.015 |
| chr1 | 105720293 | 105721402 | 1110 | 1.4 | 0.0127 | 0.015 |
| chr1 | 179439875 | 179504461 | 64587 | -1.58 | 0.0127 | 0.015 |
| chr7 | 97402589 | 97415157 | 12569 | -1.1 | 0.0127 | 0.015 |
| chr17 | 21311235 | 21314991 | 3757 | 1.72 | 0.0127 | 0.015 |
| chr5 | 94339783 | 94340798 | 1016 | -2.89 | 0.0127 | 0.015 |
| chr16 | 5099979 | 5102041 | 2063 | -3.41 | 0.0128 | 0.0151 |
| chr6 | 131022224 | 131024614 | 2391 | -1.2 | 0.0128 | 0.0151 |
| chr20 | 31808916 | 31829312 | 20397 | 3.72 | 0.0128 | 0.0151 |
| chr13 | 68915988 | 68918372 | 2385 | -1.28 | 0.0128 | 0.0151 |
| chr5 | 42773672 | 42776835 | 3164 | 2.25 | 0.0128 | 0.0151 |
| chr12 | 8714263 | 8719022 | 4760 | -0.94 | 0.0128 | 0.0151 |
| chr3 | 21425180 | 21448299 | 23120 | -3.52 | 0.0128 | 0.0151 |
| chr7 | 32467719 | 32511111 | 43393 | -3.94 | 0.0128 | 0.0151 |
| chr7 | 107918882 | 107928899 | 10018 | -5.07 | 0.0128 | 0.0151 |
| chr18 | 45009488 | 45011179 | 1692 | -1.23 | 0.0128 | 0.0151 |
| chr6 | 143633466 | 143641630 | 8165 | -3.57 | 0.0128 | 0.0151 |
| chr18 | 922521 | 922897 | 377 | -2.44 | 0.0128 | 0.0151 |
| chr10 | 128446984 | 128457207 | 10224 | -3.59 | 0.0128 | 0.0151 |
| chr2 | 205099054 | 205109373 | 10320 | -0.89 | 0.0129 | 0.0152 |
| chr12 | 67097488 | 67099346 | 1859 | -3.15 | 0.0129 | 0.0152 |
| chr20 | 41960148 | 42012011 | 51864 | -0.72 | 0.0129 | 0.0152 |
| chr3 | 85992671 | 85997219 | 4549 | -1.29 | 0.0129 | 0.0152 |
| chr13 | 43332612 | 43335954 | 3343 | 1.37 | 0.0129 | 0.0152 |
| chr7 | 10988678 | 11012728 | 24051 | -3.69 | 0.0129 | 0.0152 |
| chr2 | 122709719 | 122712599 | 2881 | -3.28 | 0.0129 | 0.0152 |
| chr3 | 18478217 | 18490833 | 12617 | 4.7 | 0.0129 | 0.0153 |
| chr5 | 96549554 | 96551509 | 1956 | -2.88 | 0.0129 | 0.0153 |
| chr22 | 25259892 | 25275379 | 15488 | -5.21 | 0.0129 | 0.0153 |
| chr5 | 41238342 | 41256324 | 17983 | -5.21 | 0.0129 | 0.0153 |
| chr9 | 92258952 | 92261026 | 2075 | -1.1 | 0.013 | 0.0153 |
| chr6 | 146779574 | 146807947 | 28374 | 0.99 | 0.013 | 0.0153 |
| chr2 | 15798021 | 15804549 | 6529 | -3.55 | 0.013 | 0.0153 |
| chr11 | 1620501 | 1623935 | 3435 | -1.74 | 0.013 | 0.0153 |
| chr9 | 28549552 | 28550626 | 1075 | -2.94 | 0.013 | 0.0153 |
| chr6 | 117227257 | 117229042 | 1786 | -2.55 | 0.013 | 0.0153 |
| chr11 | 13584510 | 13614758 | 30249 | -0.75 | 0.013 | 0.0153 |
| chr4 | 53622824 | 53625354 | 2531 | -1.28 | 0.013 | 0.0153 |
| chr3 | 124038519 | 124043571 | 5053 | 3.5 | 0.013 | 0.0153 |
| chr4 | 35156655 | 35171962 | 15308 | 2.77 | 0.013 | 0.0153 |
| chr1 | 167854772 | 167855344 | 573 | -2.76 | 0.013 | 0.0153 |
| chr3 | 12074644 | 12172665 | 98022 | -0.67 | 0.013 | 0.0154 |
| chr7 | 112450272 | 112451565 | 1294 | -3.07 | 0.013 | 0.0154 |
| chr18 | 61334073 | 61335020 | 948 | 3.04 | 0.013 | 0.0154 |
| chr5 | 2139101 | 2174104 | 35004 | -1.87 | 0.0131 | 0.0154 |

| | | | | | | |
|---|---|---|---|---|---|---|
| chr3 | 39739816 | 39740718 | 903 | -1.64 | 0.0131 | 0.0154 |
| chr12 | 20773730 | 20775402 | 1673 | -3.14 | 0.0131 | 0.0154 |
| chr8 | 31649500 | 31651060 | 1561 | 2.39 | 0.0131 | 0.0154 |
| chr4 | 21155752 | 21158739 | 2988 | 1.33 | 0.0131 | 0.0154 |
| chr12 | 52821392 | 52827587 | 6196 | 3.68 | 0.0131 | 0.0154 |
| chr11 | 40484201 | 40486498 | 2298 | 1.63 | 0.0131 | 0.0154 |
| chrX | 21725392 | 21726539 | 1148 | -1.81 | 0.0131 | 0.0155 |
| chr19 | 41040002 | 41053733 | 13732 | -5.19 | 0.0132 | 0.0155 |
| chr7 | 2494047 | 2504823 | 10777 | -5.19 | 0.0132 | 0.0155 |
| chrX | 106815453 | 106828214 | 12762 | -5.19 | 0.0132 | 0.0155 |
| chr12 | 118158817 | 118160131 | 1315 | 1.49 | 0.0132 | 0.0155 |
| chr12 | 102041946 | 102046082 | 4137 | 1.54 | 0.0132 | 0.0155 |
| chr17 | 26173769 | 26201780 | 28012 | -2.03 | 0.0132 | 0.0155 |
| chr5 | 102317915 | 102320167 | 2253 | -3.3 | 0.0133 | 0.0156 |
| chr18 | 76250464 | 76255570 | 5107 | -1.73 | 0.0133 | 0.0156 |
| chr11 | 134887989 | 134888832 | 844 | -1.62 | 0.0133 | 0.0156 |
| chr15 | 62390219 | 62392159 | 1941 | -3.41 | 0.0133 | 0.0156 |
| chr2 | 215235254 | 215235897 | 644 | 1.57 | 0.0133 | 0.0156 |
| chr13 | 83411637 | 83413742 | 2106 | -3.14 | 0.0133 | 0.0157 |
| chr3 | 191697548 | 191698734 | 1187 | -2.94 | 0.0133 | 0.0157 |
| chr4 | 27942310 | 27943958 | 1649 | -2.76 | 0.0133 | 0.0157 |
| chr13 | 46336859 | 46339330 | 2472 | 1.24 | 0.0133 | 0.0157 |
| chr14 | 43748371 | 43761636 | 13266 | 2.31 | 0.0133 | 0.0157 |
| chr16 | 20736299 | 20737517 | 1219 | -1.48 | 0.0133 | 0.0157 |
| chr10 | 45541988 | 45592577 | 50590 | -1.61 | 0.0133 | 0.0157 |
| chr1 | 56478870 | 56481919 | 3050 | -3.17 | 0.0133 | 0.0157 |
| chr5 | 135149168 | 135155641 | 6474 | -3.41 | 0.0133 | 0.0157 |
| chr4 | 90437371 | 90439149 | 1779 | 1.4 | 0.0133 | 0.0157 |
| chr12 | 103920347 | 103923365 | 3019 | -2.86 | 0.0133 | 0.0157 |
| chr5 | 150740617 | 150825360 | 84744 | -2.2 | 0.0133 | 0.0157 |
| chr9 | 20248156 | 20248991 | 836 | -2.55 | 0.0133 | 0.0157 |
| chr6 | 89171700 | 89172650 | 951 | -2.77 | 0.0133 | 0.0157 |
| chr7 | 149012449 | 149094094 | 81646 | -5.49 | 0.0133 | 0.0157 |
| chr17 | 21124742 | 21135858 | 11117 | -3.38 | 0.0134 | 0.0157 |
| chrX | 72269542 | 72271975 | 2434 | -1.52 | 0.0134 | 0.0157 |
| chr15 | 69434396 | 69439850 | 5455 | -5.17 | 0.0134 | 0.0157 |
| chr3 | 118659159 | 118675061 | 15903 | -5.17 | 0.0134 | 0.0157 |
| chr7 | 118672999 | 118695905 | 22907 | -1.47 | 0.0134 | 0.0158 |
| chr12 | 98771825 | 98775432 | 3608 | 1.11 | 0.0134 | 0.0158 |
| chr18 | 44344240 | 44346589 | 2350 | 3.14 | 0.0134 | 0.0158 |
| chr18 | 69270473 | 69284872 | 14400 | 1.1 | 0.0134 | 0.0158 |
| chr1 | 107968621 | 107969910 | 1290 | -1.84 | 0.0134 | 0.0158 |
| chr6 | 105691576 | 105692457 | 882 | -2.67 | 0.0135 | 0.0158 |
| chr4 | 65427480 | 65428545 | 1066 | -2.87 | 0.0135 | 0.0158 |
| chr9 | 116195212 | 116196851 | 1640 | -2.92 | 0.0135 | 0.0159 |
| chr1 | 166937225 | 166940664 | 3440 | 1.13 | 0.0135 | 0.0159 |
| chr14 | 97578645 | 97639486 | 60842 | 1.42 | 0.0135 | 0.0159 |
| chr11 | 14231890 | 14233214 | 1325 | -1.3 | 0.0135 | 0.0159 |
| chr4 | 34977705 | 34983953 | 6249 | 1.23 | 0.0135 | 0.0159 |
| chr20 | 47988922 | 47992214 | 3293 | 0.83 | 0.0135 | 0.0159 |
| chr2 | 45292713 | 45293863 | 1151 | -1.8 | 0.0136 | 0.016 |
| chr11 | 121658107 | 121798765 | 140659 | 0.35 | 0.0136 | 0.016 |
| chr20 | 44145824 | 44149470 | 3647 | -1.48 | 0.0136 | 0.016 |
| chr7 | 119270311 | 119307149 | 36839 | -1.65 | 0.0136 | 0.016 |
| chr20 | 2521346 | 2527620 | 6275 | -0.92 | 0.0136 | 0.016 |
| chr3 | 64826434 | 64829653 | 3220 | 2.37 | 0.0136 | 0.016 |
| chr3 | 61898723 | 61899215 | 493 | -2.5 | 0.0136 | 0.016 |
| chr2 | 73352825 | 73365835 | 13011 | -5.16 | 0.0136 | 0.016 |
| chr9 | 84381953 | 84398076 | 16124 | -5.16 | 0.0136 | 0.016 |
| chr1 | 57277893 | 57306537 | 28645 | -3.67 | 0.0136 | 0.016 |
| chrX | 74163087 | 74165423 | 2337 | -0.97 | 0.0136 | 0.016 |

| | | | | | | |
|---|---|---|---|---|---|---|
| chr12 | 107112465 | 107113111 | 647 | -1.48 | 0.0137 | 0.0161 |
| chr4 | 128007390 | 128010698 | 3309 | -1.52 | 0.0137 | 0.0161 |
| chr4 | 61942461 | 61980160 | 37700 | 0.91 | 0.0137 | 0.0161 |
| chr4 | 25035519 | 25060406 | 24888 | -0.74 | 0.0137 | 0.0161 |
| chr13 | 112888216 | 112889954 | 1739 | 1.37 | 0.0137 | 0.0161 |
| chr12 | 99283145 | 99339008 | 55864 | -0.6 | 0.0137 | 0.0161 |
| chr4 | 145079105 | 145084607 | 5503 | -1.43 | 0.0137 | 0.0161 |
| chr10 | 99701395 | 99758267 | 56873 | -0.67 | 0.0137 | 0.0161 |
| chr15 | 58356126 | 58357932 | 1807 | -3.36 | 0.0138 | 0.0162 |
| chr11 | 129249946 | 129251239 | 1294 | -1.81 | 0.0138 | 0.0162 |
| chr10 | 80480031 | 80498125 | 18095 | -3.84 | 0.0138 | 0.0162 |
| chrX | 57063937 | 57091543 | 27607 | -0.73 | 0.0138 | 0.0162 |
| chr12 | 101050498 | 101053047 | 2550 | -3.51 | 0.0138 | 0.0162 |
| chr10 | 32052790 | 32066409 | 13620 | -5.14 | 0.0138 | 0.0162 |
| chr12 | 117327649 | 117335717 | 8069 | -5.14 | 0.0138 | 0.0162 |
| chr6 | 100903793 | 100925345 | 21553 | -5.14 | 0.0138 | 0.0162 |
| chr10 | 133180975 | 133229629 | 48655 | 0.83 | 0.0138 | 0.0163 |
| chr4 | 89480793 | 89483781 | 2989 | -3.1 | 0.0138 | 0.0163 |
| chr2 | 66625949 | 66636758 | 10810 | -1.05 | 0.0139 | 0.0163 |
| chr6 | 154385557 | 154395117 | 9561 | 1.12 | 0.0139 | 0.0163 |
| chr16 | 33534927 | 33539102 | 4176 | 1.32 | 0.0139 | 0.0163 |
| chr11 | 121804166 | 121878176 | 74011 | 0.55 | 0.0139 | 0.0163 |
| chr6 | 97568248 | 97569412 | 1165 | -3.04 | 0.0139 | 0.0163 |
| chr16 | 80945029 | 80963572 | 18544 | -3.75 | 0.0139 | 0.0163 |
| chr1 | 104554192 | 104595216 | 41025 | 0.79 | 0.0139 | 0.0163 |
| chr9 | 18192890 | 18197795 | 4906 | 1.24 | 0.0139 | 0.0164 |
| chr12 | 2867403 | 2868450 | 1048 | -3.02 | 0.0139 | 0.0164 |
| chrX | 79381340 | 79402522 | 21183 | 3.83 | 0.014 | 0.0164 |
| chr14 | 21011293 | 21017354 | 6062 | 1.53 | 0.014 | 0.0164 |
| chr12 | 11355886 | 11376418 | 20533 | -3.82 | 0.014 | 0.0164 |
| chr18 | 70312229 | 70315364 | 3136 | 3.23 | 0.014 | 0.0165 |
| chr20 | 62813973 | 62823563 | 9591 | -3.59 | 0.014 | 0.0165 |
| chr18 | 71701626 | 71703179 | 1554 | -1.22 | 0.014 | 0.0165 |
| chr13 | 68330843 | 68344391 | 13549 | 3.71 | 0.014 | 0.0165 |
| chrX | 82654454 | 82656701 | 2248 | 1.99 | 0.0141 | 0.0165 |
| chr7 | 123494272 | 123495516 | 1245 | -3.05 | 0.0141 | 0.0165 |
| chr5 | 147859097 | 147859911 | 815 | -2.51 | 0.0141 | 0.0165 |
| chrX | 133408334 | 133474553 | 66220 | -3.98 | 0.0141 | 0.0165 |
| chr2 | 107172208 | 107173700 | 1493 | -3.17 | 0.0141 | 0.0166 |
| chr1 | 215258128 | 215264119 | 5992 | 0.83 | 0.0141 | 0.0166 |
| chr16 | 55914914 | 55997706 | 82793 | -0.78 | 0.0141 | 0.0166 |
| chr1 | 147273799 | 147291207 | 17409 | 3.77 | 0.0141 | 0.0166 |
| chr3 | 36787614 | 36826796 | 39183 | -1.15 | 0.0141 | 0.0166 |
| chr3 | 12282763 | 12292716 | 9954 | -3.85 | 0.0141 | 0.0166 |
| chr3 | 10677388 | 10699460 | 22073 | -2.76 | 0.0141 | 0.0166 |
| chrX | 102374548 | 102376081 | 1534 | -1.34 | 0.0142 | 0.0166 |
| chr4 | 147615243 | 147636512 | 21270 | 3.69 | 0.0142 | 0.0167 |
| chr6 | 106413976 | 106423413 | 9438 | -3.82 | 0.0142 | 0.0167 |
| chr2 | 226404310 | 226410619 | 6310 | -0.89 | 0.0142 | 0.0167 |
| chr6 | 106313810 | 106314759 | 950 | 2.54 | 0.0142 | 0.0167 |
| chr7 | 117629623 | 117632606 | 2984 | 3.27 | 0.0143 | 0.0167 |
| chr11 | 89547184 | 89547985 | 802 | 2.95 | 0.0143 | 0.0167 |
| chrX | 65889983 | 65890765 | 783 | 2.75 | 0.0143 | 0.0167 |
| chr17 | 70069791 | 70100082 | 30292 | -0.92 | 0.0143 | 0.0168 |
| chr20 | 93099 | 144164 | 51066 | -0.66 | 0.0143 | 0.0168 |
| chr2 | 58824266 | 58837789 | 13524 | 2.29 | 0.0143 | 0.0168 |
| chr6 | 9041764 | 9043264 | 1501 | -1.69 | 0.0143 | 0.0168 |
| chr4 | 28933205 | 28943657 | 10453 | 0.95 | 0.0143 | 0.0168 |
| chrX | 119978009 | 119979389 | 1381 | -1.13 | 0.0143 | 0.0168 |
| chr4 | 78394757 | 78420889 | 26133 | -0.54 | 0.0143 | 0.0168 |
| chrX | 109789201 | 109789846 | 646 | 1.55 | 0.0143 | 0.0168 |

| | | | | | | |
|---|---|---|---|---|---|---|
| chrX | 55111418 | 55114203 | 2786 | -1.33 | 0.0143 | 0.0168 |
| chr13 | 112633155 | 112679154 | 46000 | -0.45 | 0.0143 | 0.0168 |
| chr2 | 200364815 | 200366779 | 1965 | -1.23 | 0.0143 | 0.0168 |
| chr17 | 51657199 | 51706622 | 49424 | -3.75 | 0.0143 | 0.0168 |
| chr6 | 170214865 | 170237137 | 22273 | -3.84 | 0.0144 | 0.0169 |
| chrX | 21282366 | 21302843 | 20478 | -3.69 | 0.0144 | 0.0169 |
| chr12 | 127519721 | 127520844 | 1124 | 1.92 | 0.0144 | 0.0169 |
| chr10 | 26948343 | 26949525 | 1183 | -1.58 | 0.0144 | 0.0169 |
| chr3 | 130219247 | 130239443 | 20197 | 2.07 | 0.0144 | 0.0169 |
| chrX | 18055906 | 18077094 | 21189 | -1.18 | 0.0145 | 0.017 |
| chr11 | 121079999 | 121091495 | 11497 | 1.56 | 0.0145 | 0.017 |
| chrX | 80397676 | 80406825 | 9150 | -1.24 | 0.0145 | 0.017 |
| chrX | 150840916 | 150841890 | 975 | 2.82 | 0.0145 | 0.017 |
| chrX | 5100195 | 5107175 | 6981 | -3.37 | 0.0145 | 0.017 |
| chr6 | 38946124 | 38948160 | 2037 | 1.88 | 0.0145 | 0.017 |
| chr16 | 12534574 | 12548392 | 13819 | -5.09 | 0.0145 | 0.017 |
| chr17 | 75220872 | 75229981 | 9110 | -5.09 | 0.0145 | 0.017 |
| chrX | 41244065 | 41257910 | 13846 | -5.09 | 0.0145 | 0.017 |
| chr7 | 18020061 | 18021750 | 1690 | -3.14 | 0.0145 | 0.017 |
| chr5 | 108639175 | 108640024 | 850 | 1.26 | 0.0145 | 0.017 |
| chr3 | 135653495 | 135654160 | 666 | 1.49 | 0.0146 | 0.0171 |
| chr11 | 50053853 | 50072371 | 18519 | 0.52 | 0.0146 | 0.0171 |
| chr16 | 20517227 | 20533678 | 16452 | 0.68 | 0.0146 | 0.0171 |
| chr15 | 42267814 | 42268219 | 406 | -2.52 | 0.0146 | 0.0171 |
| chr13 | 88868173 | 88869896 | 1724 | 2.93 | 0.0146 | 0.0172 |
| chr18 | 53736862 | 53742819 | 5958 | -3.54 | 0.0147 | 0.0172 |
| chrX | 134207199 | 134212967 | 5769 | -0.86 | 0.0147 | 0.0172 |
| chrX | 73943452 | 73945555 | 2104 | -2.2 | 0.0147 | 0.0173 |
| chrX | 108075802 | 108094546 | 18745 | 3.16 | 0.0147 | 0.0173 |
| chr12 | 22869932 | 22870736 | 805 | -1.31 | 0.0148 | 0.0173 |
| chr3 | 7592809 | 7596648 | 3840 | -1.72 | 0.0148 | 0.0173 |
| chr12 | 93085592 | 93093096 | 7505 | -3.47 | 0.0148 | 0.0173 |
| chr4 | 93488787 | 93495724 | 6938 | -3.51 | 0.0148 | 0.0173 |
| chr8 | 103098234 | 103099051 | 818 | 1.63 | 0.0148 | 0.0173 |
| chr2 | 88767514 | 88779795 | 12282 | -1.26 | 0.0148 | 0.0173 |
| chr1 | 211066199 | 211068378 | 2180 | -3.26 | 0.0148 | 0.0174 |
| chr6 | 128988043 | 128988811 | 769 | -2.47 | 0.0148 | 0.0174 |
| chr11 | 121554371 | 121645685 | 91315 | 0.38 | 0.0148 | 0.0174 |
| chr2 | 186694211 | 186702293 | 8083 | 1.09 | 0.0148 | 0.0174 |
| chr5 | 106332589 | 106338551 | 5963 | -3.38 | 0.0148 | 0.0174 |
| chr18 | 73524185 | 73618918 | 94734 | 0.45 | 0.0149 | 0.0174 |
| chr10 | 116803156 | 116804034 | 879 | -2.83 | 0.0149 | 0.0174 |
| chr3 | 115199232 | 115199955 | 724 | 2.4 | 0.0149 | 0.0174 |
| chr8 | 54281438 | 54281858 | 421 | -1.43 | 0.0149 | 0.0175 |
| chr18 | 74926758 | 74961005 | 34248 | -3.85 | 0.0149 | 0.0175 |
| chr1 | 99711265 | 99712598 | 1334 | -1.79 | 0.0149 | 0.0175 |
| chr16 | 4276247 | 4278181 | 1935 | 1.03 | 0.0149 | 0.0175 |
| chrX | 30543188 | 30556049 | 12862 | -1.53 | 0.015 | 0.0176 |
| chr8 | 15607483 | 15609225 | 1743 | 3.04 | 0.015 | 0.0176 |
| chr2 | 182316223 | 182317213 | 991 | -1.51 | 0.015 | 0.0176 |
| chrX | 153948103 | 153951009 | 2907 | -2.12 | 0.015 | 0.0176 |
| chr20 | 42778474 | 42792353 | 13880 | -5.06 | 0.015 | 0.0176 |
| chr3 | 159347977 | 159358294 | 10318 | -5.06 | 0.015 | 0.0176 |
| chr8 | 27245721 | 27260408 | 14688 | -5.06 | 0.015 | 0.0176 |
| chr8 | 40800014 | 40832930 | 32917 | -3.75 | 0.015 | 0.0176 |
| chr5 | 113592736 | 113596542 | 3807 | 1.17 | 0.015 | 0.0176 |
| chrX | 67983946 | 67985436 | 1491 | -2.98 | 0.015 | 0.0176 |
| chr13 | 112909292 | 112913727 | 4436 | -1.31 | 0.015 | 0.0176 |
| chr11 | 26608311 | 26612376 | 4066 | -3.07 | 0.0151 | 0.0177 |
| chr4 | 104338549 | 104341003 | 2455 | -1.41 | 0.0151 | 0.0177 |
| chr3 | 53555364 | 53556513 | 1150 | -2.81 | 0.0151 | 0.0177 |

| | | | | | | |
|---|---|---|---|---|---|---|
| chr3 | 172579395 | 172581284 | 1890 | -1.51 | 0.0151 | 0.0177 |
| chrX | 79672038 | 79676464 | 4427 | 1.3 | 0.0151 | 0.0177 |
| chrX | 151550025 | 151553965 | 3941 | -1.36 | 0.0151 | 0.0177 |
| chr1 | 109976750 | 110004677 | 27928 | -3.64 | 0.0152 | 0.0178 |
| chr4 | 65687585 | 65696965 | 9381 | 1.02 | 0.0152 | 0.0178 |
| chr2 | 141160141 | 141166354 | 6214 | -3.35 | 0.0152 | 0.0178 |
| chrX | 131475728 | 131479815 | 4088 | -1.36 | 0.0152 | 0.0178 |
| chr4 | 145234234 | 145237978 | 3745 | -1.6 | 0.0152 | 0.0178 |
| chr12 | 16771805 | 16773421 | 1617 | -3.21 | 0.0152 | 0.0178 |
| chr8 | 63469417 | 63471014 | 1598 | -2.09 | 0.0152 | 0.0178 |
| chr1 | 116793263 | 116795004 | 1742 | -1.92 | 0.0152 | 0.0179 |
| chr5 | 124016799 | 124018652 | 1854 | -2.87 | 0.0153 | 0.0179 |
| chr2 | 108773128 | 108792292 | 19165 | -1.34 | 0.0153 | 0.0179 |
| chr20 | 37328716 | 37331811 | 3096 | 0.98 | 0.0153 | 0.0179 |
| chrX | 21406247 | 21410824 | 4578 | -3.34 | 0.0153 | 0.0179 |
| chr2 | 208138914 | 208147689 | 8776 | -5.04 | 0.0153 | 0.0179 |
| chr7 | 98343738 | 98374032 | 30295 | -5.04 | 0.0153 | 0.0179 |
| chr1 | 242857911 | 242906151 | 48241 | -0.8 | 0.0154 | 0.018 |
| chr11 | 85160258 | 85172564 | 12307 | 0.67 | 0.0154 | 0.018 |
| chr3 | 20245174 | 20257909 | 12736 | -3.49 | 0.0154 | 0.018 |
| chr12 | 111315100 | 111318353 | 3254 | -2.2 | 0.0154 | 0.018 |
| chr12 | 16383200 | 16387299 | 4100 | -1.55 | 0.0154 | 0.018 |
| chr13 | 23671416 | 23695988 | 24573 | 0.79 | 0.0154 | 0.018 |
| chr10 | 9267592 | 9306227 | 38636 | -0.65 | 0.0154 | 0.018 |
| chr5 | 54291178 | 54303618 | 12441 | 1.21 | 0.0154 | 0.018 |
| chr11 | 14101713 | 14102907 | 1195 | 1.47 | 0.0154 | 0.0181 |
| chr7 | 15682388 | 15683907 | 1520 | -2.84 | 0.0155 | 0.0181 |
| chr18 | 27743014 | 27756353 | 13340 | -1.52 | 0.0155 | 0.0181 |
| chr8 | 114307599 | 114308959 | 1361 | -2.33 | 0.0155 | 0.0182 |
| chr21 | 15760611 | 15809566 | 48956 | -3.43 | 0.0156 | 0.0182 |
| chr2 | 187202321 | 187219351 | 17031 | -5.02 | 0.0156 | 0.0182 |
| chr20 | 56009975 | 56033824 | 23850 | -5.02 | 0.0156 | 0.0182 |
| chr6 | 39162717 | 39163689 | 973 | -1.65 | 0.0156 | 0.0182 |
| chr20 | 14087566 | 14117351 | 29786 | -3.89 | 0.0156 | 0.0182 |
| chr9 | 91226734 | 91265175 | 38442 | -3.91 | 0.0156 | 0.0183 |
| chr13 | 60812201 | 60813785 | 1585 | -1.8 | 0.0156 | 0.0183 |
| chr11 | 18992911 | 18996832 | 3922 | 2.73 | 0.0157 | 0.0183 |
| chr1 | 5667187 | 5671200 | 4014 | -0.97 | 0.0157 | 0.0183 |
| chr18 | 76415548 | 76416660 | 1113 | -1.34 | 0.0157 | 0.0183 |
| chr17 | 6211998 | 6254213 | 42216 | -0.65 | 0.0157 | 0.0184 |
| chr15 | 37504634 | 37518131 | 13498 | 1.06 | 0.0157 | 0.0184 |
| chr3 | 37500975 | 37562023 | 61049 | -3.77 | 0.0157 | 0.0184 |
| chr6 | 142770763 | 142838814 | 68052 | -3.91 | 0.0157 | 0.0184 |
| chr1 | 116445145 | 116455202 | 10058 | 1.3 | 0.0157 | 0.0184 |
| chr3 | 67956173 | 67970837 | 14665 | 3.73 | 0.0157 | 0.0184 |
| chr5 | 6319332 | 6335884 | 16553 | -1.48 | 0.0157 | 0.0184 |
| chr3 | 93577004 | 93578585 | 1582 | -2.87 | 0.0158 | 0.0185 |
| chr8 | 121920855 | 121922349 | 1495 | -1.95 | 0.0158 | 0.0185 |
| chr8 | 41256789 | 41270421 | 13633 | -1.07 | 0.0158 | 0.0185 |
| chr1 | 232490297 | 232494018 | 3722 | 3.36 | 0.0158 | 0.0185 |
| chr3 | 98431714 | 98443493 | 11780 | -5.01 | 0.0158 | 0.0185 |
| chr16 | 51203382 | 51211186 | 7805 | -3.42 | 0.0159 | 0.0185 |
| chr4 | 57420321 | 57422393 | 2073 | -1.96 | 0.0159 | 0.0186 |
| chr7 | 29081706 | 29115975 | 34270 | 0.74 | 0.0159 | 0.0186 |
| chr17 | 69838758 | 69842052 | 3295 | -1.32 | 0.0159 | 0.0186 |
| chr15 | 60990817 | 61099083 | 108267 | 0.69 | 0.0159 | 0.0186 |
| chr15 | 101062057 | 101074559 | 12503 | -0.76 | 0.0159 | 0.0186 |
| chr11 | 122839120 | 122840734 | 1615 | -1.32 | 0.0159 | 0.0186 |
| chr6 | 46711739 | 46713943 | 2205 | 2.12 | 0.0159 | 0.0186 |
| chr2 | 13557765 | 13579794 | 22030 | 0.71 | 0.016 | 0.0187 |
| chr9 | 92517183 | 92526483 | 9301 | 2.18 | 0.016 | 0.0187 |

| | | | | | | |
|---|---|---|---|---|---|---|
| chrX | 121492533 | 121495946 | 3414 | 3.42 | 0.016 | 0.0187 |
| chr18 | 52280344 | 52310876 | 30533 | -3.61 | 0.016 | 0.0187 |
| chr12 | 46042967 | 46043937 | 971 | 1.29 | 0.016 | 0.0187 |
| chr3 | 40031271 | 40035615 | 4345 | -3.23 | 0.016 | 0.0187 |
| chr13 | 112736477 | 112751798 | 15322 | -0.82 | 0.016 | 0.0188 |
| chr9 | 15947146 | 15955140 | 7995 | -3.45 | 0.0161 | 0.0188 |
| chr6 | 46437532 | 46439260 | 1729 | -1.17 | 0.0161 | 0.0188 |
| chr8 | 22671300 | 22691095 | 19796 | -0.98 | 0.0161 | 0.0188 |
| chr4 | 11458924 | 11464516 | 5593 | -1.6 | 0.0161 | 0.0188 |
| chrX | 30024059 | 30030347 | 6289 | 1.5 | 0.0161 | 0.0188 |
| chr5 | 32061081 | 32062184 | 1104 | -2.75 | 0.0161 | 0.0188 |
| chr8 | 112304435 | 112333160 | 28726 | 2.18 | 0.0161 | 0.0189 |
| chr3 | 69492996 | 69504657 | 11662 | -4.99 | 0.0161 | 0.0189 |
| chr4 | 174649261 | 174665260 | 16000 | -4.99 | 0.0161 | 0.0189 |
| chr6 | 86486986 | 86539070 | 52085 | -0.85 | 0.0162 | 0.0189 |
| chr12 | 68095953 | 68097793 | 1841 | -3.06 | 0.0162 | 0.0189 |
| chr13 | 51406777 | 51407279 | 503 | -1.75 | 0.0162 | 0.0189 |
| chr18 | 59591633 | 59629543 | 37911 | 0.54 | 0.0162 | 0.0189 |
| chr12 | 25683858 | 25689747 | 5890 | -1.4 | 0.0162 | 0.0189 |
| chr18 | 25343139 | 25343929 | 791 | 2.79 | 0.0162 | 0.0189 |
| chr1 | 199764604 | 199765806 | 1203 | -3.92 | 0.0162 | 0.019 |
| chr8 | 105687206 | 105688015 | 810 | -2.89 | 0.0162 | 0.019 |
| chrX | 139163792 | 139166509 | 2718 | -1.11 | 0.0163 | 0.019 |
| chrX | 42235483 | 42236750 | 1268 | 1.57 | 0.0163 | 0.019 |
| chr3 | 152533032 | 152535408 | 2377 | -2.2 | 0.0163 | 0.019 |
| chr2 | 53504687 | 53522251 | 17565 | -1.57 | 0.0163 | 0.019 |
| chr22 | 33161695 | 33168842 | 7148 | -0.8 | 0.0163 | 0.019 |
| chr3 | 150602055 | 150606024 | 3970 | -0.81 | 0.0163 | 0.0191 |
| chr12 | 3484320 | 3590581 | 106262 | 0.62 | 0.0163 | 0.0191 |
| chr10 | 128222469 | 128242986 | 20518 | -1.13 | 0.0163 | 0.0191 |
| chr5 | 112807720 | 112849287 | 41568 | -3.96 | 0.0163 | 0.0191 |
| chr9 | 112704956 | 112706987 | 2032 | 1.81 | 0.0163 | 0.0191 |
| chr18 | 44951857 | 44963151 | 11295 | 2.44 | 0.0163 | 0.0191 |
| chr1 | 108364003 | 108380763 | 16761 | 0.72 | 0.0163 | 0.0191 |
| chr1 | 196007701 | 196009636 | 1936 | -3.16 | 0.0164 | 0.0191 |
| chr4 | 56616211 | 56616953 | 743 | -2.63 | 0.0164 | 0.0192 |
| chr20 | 10049446 | 10060325 | 10880 | -4.97 | 0.0164 | 0.0192 |
| chr6 | 75929951 | 75937981 | 8031 | -4.97 | 0.0164 | 0.0192 |
| chr3 | 15187849 | 15199984 | 12136 | -2.46 | 0.0164 | 0.0192 |
| chr6 | 99674712 | 99687418 | 12707 | 1.11 | 0.0165 | 0.0192 |
| chr5 | 82833501 | 82837388 | 3888 | -1.93 | 0.0165 | 0.0192 |
| chr3 | 163163898 | 163165930 | 2033 | -2.96 | 0.0165 | 0.0193 |
| chr7 | 90269494 | 90274401 | 4908 | -3.34 | 0.0166 | 0.0193 |
| chr11 | 113370166 | 113371028 | 863 | -2.07 | 0.0166 | 0.0193 |
| chr13 | 103216174 | 103217614 | 1441 | 1.43 | 0.0166 | 0.0193 |
| chr2 | 37654537 | 37670220 | 15684 | -3.58 | 0.0166 | 0.0194 |
| chr15 | 89999766 | 90000847 | 1082 | -1.48 | 0.0166 | 0.0194 |
| chr4 | 139673713 | 139677551 | 3839 | -3.32 | 0.0166 | 0.0194 |
| chr10 | 14254766 | 14479019 | 224254 | -0.47 | 0.0166 | 0.0194 |
| chrX | 150378997 | 150381524 | 2528 | 1.66 | 0.0166 | 0.0194 |
| chr6 | 120415381 | 120416942 | 1562 | 2.35 | 0.0166 | 0.0194 |
| chr9 | 85006434 | 85038756 | 32323 | 0.84 | 0.0166 | 0.0194 |
| chrX | 115398199 | 115409604 | 11406 | 2.32 | 0.0166 | 0.0194 |
| chr13 | 98325697 | 98328731 | 3035 | -0.88 | 0.0166 | 0.0194 |
| chr9 | 74065988 | 74067404 | 1417 | -3.24 | 0.0166 | 0.0194 |
| chr5 | 154440177 | 154441702 | 1526 | -3.06 | 0.0167 | 0.0194 |
| chr13 | 23726083 | 23731386 | 5304 | -3.3 | 0.0167 | 0.0194 |
| chr6 | 47908394 | 47909419 | 1026 | -2.88 | 0.0167 | 0.0195 |
| chr10 | 80348951 | 80353297 | 4347 | -3.49 | 0.0167 | 0.0195 |
| chr6 | 165331520 | 165334505 | 2986 | 0.95 | 0.0167 | 0.0195 |
| chr6 | 122816363 | 122819958 | 3596 | -1.59 | 0.0167 | 0.0195 |

| | | | | | | |
|---|---|---|---|---|---|---|
| chr12 | 12455517 | 12475582 | 20066 | -4.95 | 0.0167 | 0.0195 |
| chr2 | 95692683 | 95702245 | 9563 | -4.95 | 0.0167 | 0.0195 |
| chr20 | 5760301 | 5772760 | 12460 | -4.95 | 0.0167 | 0.0195 |
| chr20 | 52399254 | 52410572 | 11319 | -4.95 | 0.0167 | 0.0195 |
| chr15 | 35114353 | 35115562 | 1210 | 3.15 | 0.0168 | 0.0196 |
| chr17 | 67800199 | 67801701 | 1503 | 3.01 | 0.0168 | 0.0196 |
| chr19 | 14987633 | 14988420 | 788 | 1.68 | 0.0168 | 0.0196 |
| chr10 | 30539606 | 30564142 | 24537 | -3.88 | 0.0168 | 0.0197 |
| chr4 | 90989697 | 90992779 | 3083 | 1.75 | 0.0169 | 0.0197 |
| chr7 | 108353235 | 108855710 | 502476 | 5.98 | 0.0169 | 0.0197 |
| chr10 | 30181376 | 30229852 | 48477 | -1.28 | 0.0169 | 0.0197 |
| chr3 | 116704105 | 116704754 | 650 | -2.85 | 0.0169 | 0.0197 |
| chr6 | 112778179 | 112778753 | 575 | 2.57 | 0.0169 | 0.0198 |
| chr11 | 22043513 | 22047159 | 3647 | -3.41 | 0.017 | 0.0198 |
| chr13 | 95409846 | 95419687 | 9842 | -0.81 | 0.017 | 0.0198 |
| chr6 | 161296162 | 161296734 | 573 | -2.62 | 0.017 | 0.0199 |
| chr21 | 27589432 | 27593492 | 4061 | -3.26 | 0.017 | 0.0199 |
| chr3 | 180529711 | 180565560 | 35850 | 3.68 | 0.017 | 0.0199 |
| chr2 | 67423020 | 67424252 | 1233 | -1.28 | 0.0171 | 0.0199 |
| chr5 | 159309984 | 159322545 | 12562 | -4.93 | 0.0171 | 0.0199 |
| chr11 | 59270522 | 59271745 | 1224 | -2.95 | 0.0171 | 0.0199 |
| chr16 | 23308383 | 23309300 | 918 | -1.57 | 0.0171 | 0.02 |
| chrX | 84734270 | 84745654 | 11385 | 3.54 | 0.0171 | 0.02 |
| chr8 | 72774369 | 72875044 | 100676 | 4.03 | 0.0171 | 0.02 |
| chr6 | 5977947 | 5996367 | 18421 | -1.39 | 0.0171 | 0.02 |
| chr13 | 104332266 | 104341389 | 9124 | 1.65 | 0.0171 | 0.02 |
| chr6 | 108003713 | 108015870 | 12158 | -1.86 | 0.0172 | 0.02 |
| chr5 | 83109175 | 83122200 | 13026 | 1.72 | 0.0172 | 0.02 |
| chr3 | 7602071 | 8257654 | 655584 | 6.11 | 0.0172 | 0.02 |
| chr5 | 55799287 | 55800962 | 1676 | -3.12 | 0.0172 | 0.02 |
| chr11 | 116301767 | 116305182 | 3416 | 3.27 | 0.0172 | 0.0201 |
| chr1 | 112857017 | 112874816 | 17800 | 0.83 | 0.0172 | 0.0201 |
| chrX | 108703367 | 108708689 | 5323 | -1.33 | 0.0172 | 0.0201 |
| chr1 | 245392073 | 245394845 | 2773 | -2.98 | 0.0172 | 0.0201 |
| chr7 | 15066081 | 15078300 | 12220 | -3.02 | 0.0172 | 0.0201 |
| chr9 | 89040855 | 89054019 | 13165 | -3.7 | 0.0172 | 0.0201 |
| chr9 | 112612545 | 112639752 | 27208 | 0.65 | 0.0173 | 0.0201 |
| chr11 | 4595286 | 4611831 | 16546 | 1.24 | 0.0173 | 0.0201 |
| chr6 | 76693690 | 76807378 | 113689 | 5.37 | 0.0173 | 0.0201 |
| chr16 | 56175143 | 56188506 | 13364 | 3.65 | 0.0173 | 0.0201 |
| chr12 | 95167205 | 95259445 | 92241 | -5.23 | 0.0173 | 0.0202 |
| chr14 | 52250996 | 52276636 | 25641 | -3.77 | 0.0173 | 0.0202 |
| chr8 | 10201689 | 10202913 | 1225 | 1.84 | 0.0173 | 0.0202 |
| chr14 | 89529116 | 89529624 | 509 | 2.78 | 0.0173 | 0.0202 |
| chr10 | 81709410 | 81753833 | 44424 | -5.19 | 0.0173 | 0.0202 |
| chr3 | 56295719 | 56315582 | 19864 | -1.04 | 0.0174 | 0.0202 |
| chr21 | 39768109 | 39781374 | 13266 | -3.18 | 0.0174 | 0.0202 |
| chr1 | 170673542 | 170683549 | 10008 | -3.41 | 0.0174 | 0.0202 |
| chr2 | 225159200 | 225160226 | 1027 | -1.53 | 0.0174 | 0.0203 |
| chr6 | 16935023 | 16936174 | 1152 | 1.38 | 0.0174 | 0.0203 |
| chr8 | 81967023 | 81968475 | 1453 | -2.53 | 0.0174 | 0.0203 |
| chr1 | 88978952 | 88982712 | 3761 | 3.23 | 0.0175 | 0.0203 |
| chr11 | 134409257 | 134423378 | 14122 | -1.48 | 0.0175 | 0.0204 |
| chr12 | 99464773 | 99466535 | 1763 | -1.72 | 0.0175 | 0.0204 |
| chr17 | 49425433 | 49431366 | 5934 | 0.92 | 0.0175 | 0.0204 |
| chr2 | 27767951 | 27818784 | 50834 | -5.22 | 0.0175 | 0.0204 |
| chr18 | 54973823 | 54989468 | 15646 | -0.79 | 0.0175 | 0.0204 |
| chr1 | 116346083 | 116366253 | 20171 | -3.56 | 0.0175 | 0.0204 |
| chr7 | 119403755 | 119405010 | 1256 | 1.99 | 0.0175 | 0.0204 |
| chr3 | 175982146 | 175989440 | 7295 | 0.84 | 0.0176 | 0.0205 |
| chr6 | 129281394 | 129281885 | 492 | 1.68 | 0.0176 | 0.0205 |

| | | | | | | |
|---|---|---|---|---|---|---|
| chr8 | 117337123 | 117338735 | 1613 | 2.4 | 0.0176 | 0.0205 |
| chr4 | 23517896 | 23545169 | 27274 | 0.52 | 0.0176 | 0.0205 |
| chr5 | 1957258 | 1957712 | 455 | 2.82 | 0.0177 | 0.0206 |
| chr11 | 121056977 | 121062557 | 5581 | 1 | 0.0177 | 0.0206 |
| chr11 | 14843264 | 14848378 | 5115 | -0.85 | 0.0177 | 0.0206 |
| chr12 | 58530799 | 58539472 | 8674 | -1.16 | 0.0177 | 0.0206 |
| chr6 | 169580339 | 169580871 | 533 | -1.43 | 0.0177 | 0.0206 |
| chr11 | 106344470 | 106345879 | 1410 | -1.25 | 0.0177 | 0.0206 |
| chr22 | 33090377 | 33145117 | 54741 | -0.52 | 0.0177 | 0.0206 |
| chr6 | 130223184 | 130234773 | 11590 | -4.9 | 0.0177 | 0.0206 |
| chr16 | 55884578 | 55886033 | 1456 | 1.6 | 0.0177 | 0.0207 |
| chr17 | 46478298 | 46479064 | 767 | -2.88 | 0.0177 | 0.0207 |
| chr2 | 95607761 | 95642324 | 34564 | -1.47 | 0.0177 | 0.0207 |
| chr12 | 25734561 | 25791946 | 57386 | -0.69 | 0.0178 | 0.0207 |
| chr4 | 143927164 | 143953970 | 26807 | -0.95 | 0.0178 | 0.0207 |
| chr1 | 33652544 | 33693054 | 40511 | -3.85 | 0.0178 | 0.0207 |
| chr8 | 133461490 | 133484071 | 22582 | -0.47 | 0.0178 | 0.0208 |
| chr19 | 35897669 | 35906999 | 9331 | -1.85 | 0.0179 | 0.0208 |
| chr12 | 39595259 | 39664047 | 68789 | -1.68 | 0.0179 | 0.0208 |
| chr4 | 65816931 | 65820760 | 3830 | -0.97 | 0.018 | 0.0209 |
| chr3 | 111013375 | 111014849 | 1475 | -1.42 | 0.018 | 0.0209 |
| chr7 | 57294899 | 57544717 | 249819 | -5.58 | 0.018 | 0.0209 |
| chr11 | 27182283 | 27191166 | 8884 | -4.74 | 0.018 | 0.0209 |
| chr4 | 11791611 | 11794405 | 2795 | 1.57 | 0.018 | 0.021 |
| chr4 | 65824166 | 65827700 | 3535 | 0.96 | 0.018 | 0.021 |
| chr13 | 107292916 | 107297396 | 4481 | 1.24 | 0.018 | 0.021 |
| chr15 | 42979557 | 42994152 | 14596 | -4.88 | 0.0181 | 0.021 |
| chr8 | 62931250 | 62945826 | 14577 | -4.88 | 0.0181 | 0.021 |
| chrX | 80620936 | 80625261 | 4326 | 3.4 | 0.0181 | 0.021 |
| chr4 | 53191881 | 53201218 | 9338 | 0.76 | 0.0181 | 0.021 |
| chr11 | 102687537 | 102697940 | 10404 | -1.2 | 0.0181 | 0.0211 |
| chr8 | 88319473 | 88423216 | 103744 | -1.8 | 0.0181 | 0.0211 |
| chrX | 130154192 | 130160443 | 6252 | -0.64 | 0.0181 | 0.0211 |
| chrX | 25135898 | 25153974 | 18077 | -0.69 | 0.0182 | 0.0212 |
| chr2 | 58930357 | 58944945 | 14589 | 3.56 | 0.0182 | 0.0212 |
| chr8 | 83903181 | 83938446 | 35266 | -3.61 | 0.0182 | 0.0212 |
| chr2 | 18799164 | 18859816 | 60653 | 0.81 | 0.0183 | 0.0212 |
| chrX | 129595803 | 129605543 | 9741 | 1.21 | 0.0183 | 0.0213 |
| chr13 | 65339248 | 65343008 | 3761 | 2.94 | 0.0183 | 0.0213 |
| chr15 | 89232494 | 89236778 | 4285 | 1.6 | 0.0183 | 0.0213 |
| chrX | 119507403 | 119542665 | 35263 | -1.19 | 0.0183 | 0.0213 |
| chr1 | 106228925 | 106245964 | 17040 | 0.84 | 0.0183 | 0.0213 |
| chr8 | 144013356 | 144017618 | 4263 | -1.82 | 0.0183 | 0.0213 |
| chr19 | 34362058 | 34362575 | 518 | 1.28 | 0.0183 | 0.0213 |
| chrX | 17141662 | 17188423 | 46762 | -3.83 | 0.0184 | 0.0214 |
| chr1 | 114736045 | 114743129 | 7085 | -1.33 | 0.0184 | 0.0214 |
| chr13 | 46030615 | 46058211 | 27597 | -4.86 | 0.0184 | 0.0214 |
| chr4 | 154762802 | 154763480 | 679 | -2.77 | 0.0184 | 0.0214 |
| chr2 | 226233946 | 226235857 | 1912 | 0.96 | 0.0185 | 0.0215 |
| chr18 | 6894855 | 6898947 | 4093 | 1.52 | 0.0185 | 0.0215 |
| chr8 | 82225237 | 82227711 | 2475 | 1.51 | 0.0185 | 0.0215 |
| chr4 | 166880449 | 166882915 | 2467 | -2.94 | 0.0185 | 0.0215 |
| chrX | 73051101 | 73058675 | 7575 | 0.91 | 0.0185 | 0.0215 |
| chr3 | 190096034 | 190097580 | 1547 | -2.87 | 0.0185 | 0.0215 |
| chrX | 75708836 | 75713014 | 4179 | 1.74 | 0.0185 | 0.0215 |
| chr2 | 234997709 | 234998717 | 1009 | -1.39 | 0.0185 | 0.0215 |
| chr4 | 152746218 | 152749891 | 3674 | -3.25 | 0.0185 | 0.0216 |
| chr12 | 132025413 | 132037450 | 12038 | -1.08 | 0.0185 | 0.0216 |
| chr1 | 30710428 | 30711286 | 859 | -2.68 | 0.0186 | 0.0216 |
| chrX | 129570151 | 129589121 | 18971 | -1.89 | 0.0186 | 0.0216 |
| chr11 | 30375994 | 31009183 | 633190 | 6.2 | 0.0186 | 0.0216 |

| | | | | | | |
|---|---|---|---|---|---|---|
| chr13 | 81402726 | 81403817 | 1092 | 1.36 | 0.0186 | 0.0216 |
| chrX | 108397869 | 108413513 | 15645 | 1.21 | 0.0186 | 0.0216 |
| chr10 | 118347611 | 118349198 | 1588 | -1.53 | 0.0186 | 0.0216 |
| chr4 | 85130545 | 85131418 | 874 | -2.51 | 0.0186 | 0.0216 |
| chr1 | 67270027 | 67271156 | 1130 | -1.28 | 0.0186 | 0.0216 |
| chr17 | 9967363 | 10019604 | 52242 | -3.84 | 0.0186 | 0.0216 |
| chr21 | 40052412 | 40079215 | 26804 | -0.64 | 0.0186 | 0.0216 |
| chr12 | 2840195 | 2844053 | 3859 | 2.53 | 0.0186 | 0.0216 |
| chr2 | 107599296 | 107617042 | 17747 | -1.07 | 0.0186 | 0.0216 |
| chr3 | 131019168 | 131019883 | 716 | -2.62 | 0.0186 | 0.0217 |
| chr2 | 850285 | 851195 | 911 | -2.57 | 0.0187 | 0.0217 |
| chr5 | 27766503 | 27821320 | 54818 | -1.25 | 0.0187 | 0.0217 |
| chr16 | 66309808 | 66335270 | 25463 | -1.62 | 0.0187 | 0.0217 |
| chr6 | 108043590 | 108082457 | 38868 | 0.85 | 0.0187 | 0.0217 |
| chr3 | 42503323 | 42504437 | 1115 | 2.22 | 0.0187 | 0.0217 |
| chr11 | 40444797 | 40479598 | 34802 | -3.73 | 0.0187 | 0.0217 |
| chr9 | 26670116 | 26720262 | 50147 | 3.96 | 0.0187 | 0.0217 |
| chr6 | 130873432 | 130888785 | 15354 | 3.62 | 0.0187 | 0.0218 |
| chr6 | 159711760 | 159712396 | 637 | -1.59 | 0.0187 | 0.0218 |
| chr1 | 172740406 | 172743432 | 3027 | 1.21 | 0.0187 | 0.0218 |
| chr6 | 8591992 | 8607960 | 15969 | -2.41 | 0.0188 | 0.0218 |
| chr12 | 9225882 | 9239555 | 13674 | -1.12 | 0.0188 | 0.0218 |
| chr10 | 19227145 | 19229523 | 2379 | 0.98 | 0.0188 | 0.0218 |
| chr20 | 37252424 | 37264702 | 12279 | -4.84 | 0.0188 | 0.0218 |
| chr6 | 3033736 | 3048395 | 14660 | -4.84 | 0.0188 | 0.0218 |
| chr2 | 41994235 | 42064183 | 69949 | -5.43 | 0.0188 | 0.0219 |
| chr6 | 146892454 | 146897769 | 5316 | -2.48 | 0.0188 | 0.0219 |
| chr2 | 206718531 | 206721266 | 2736 | -1.13 | 0.0188 | 0.0219 |
| chr14 | 23859199 | 23864543 | 5345 | 1.51 | 0.0189 | 0.0219 |
| chr6 | 142204066 | 142212137 | 8072 | -2.69 | 0.0189 | 0.0219 |
| chr10 | 129969056 | 129984644 | 15589 | -3.71 | 0.0189 | 0.0219 |
| chr9 | 92679871 | 92709348 | 29478 | 3.72 | 0.0189 | 0.0219 |
| chr5 | 64634762 | 64643352 | 8591 | -3.56 | 0.0189 | 0.0219 |
| chr1 | 210138612 | 210139761 | 1150 | 1.49 | 0.0189 | 0.022 |
| chr21 | 39105791 | 39113288 | 7498 | 2.1 | 0.0189 | 0.022 |
| chr5 | 136575208 | 136576240 | 1033 | -1.12 | 0.0189 | 0.022 |
| chr1 | 211058091 | 211058635 | 545 | 2.42 | 0.019 | 0.022 |
| chr8 | 121831336 | 121835435 | 4100 | -1.13 | 0.019 | 0.022 |
| chr13 | 89191406 | 89194237 | 2832 | -2.82 | 0.019 | 0.022 |
| chr12 | 65290283 | 65293361 | 3079 | -3.22 | 0.019 | 0.0221 |
| chr14 | 66837504 | 66839395 | 1892 | 1.43 | 0.019 | 0.0221 |
| chr2 | 156744926 | 156807883 | 62958 | 3.93 | 0.019 | 0.0221 |
| chr2 | 217678005 | 217712090 | 34086 | 0.55 | 0.0191 | 0.0221 |
| chr2 | 66808612 | 67271653 | 463042 | 5.95 | 0.0191 | 0.0222 |
| chr3 | 54291169 | 54292707 | 1539 | 2.2 | 0.0191 | 0.0222 |
| chr20 | 59986484 | 59987796 | 1313 | 2.84 | 0.0192 | 0.0222 |
| chr14 | 105284118 | 105291661 | 7544 | -4.82 | 0.0192 | 0.0223 |
| chr11 | 59866297 | 59867611 | 1315 | -1.66 | 0.0192 | 0.0223 |
| chr14 | 59250374 | 59254474 | 4101 | 3.28 | 0.0192 | 0.0223 |
| chr4 | 22792366 | 22810907 | 18542 | 0.77 | 0.0192 | 0.0223 |
| chr5 | 30560826 | 30590954 | 30129 | -3.51 | 0.0192 | 0.0223 |
| chr5 | 75731319 | 75736690 | 5372 | -3.45 | 0.0192 | 0.0223 |
| chr4 | 30588331 | 30590308 | 1978 | -2.61 | 0.0193 | 0.0224 |
| chr14 | 21273537 | 21282248 | 8712 | 3.35 | 0.0193 | 0.0224 |
| chr12 | 15583196 | 15596143 | 12948 | 0.78 | 0.0193 | 0.0224 |
| chr12 | 21432596 | 21437449 | 4854 | 0.8 | 0.0193 | 0.0224 |
| chr7 | 124018276 | 124022093 | 3818 | 1.31 | 0.0193 | 0.0224 |
| chr10 | 122885909 | 122926237 | 40329 | 0.88 | 0.0193 | 0.0224 |
| chr13 | 85482152 | 85483945 | 1794 | 2.16 | 0.0193 | 0.0224 |
| chr13 | 58203530 | 58204384 | 855 | -2.14 | 0.0193 | 0.0224 |
| chr11 | 113499345 | 113503046 | 3702 | 2.18 | 0.0194 | 0.0225 |

| | | | | | | |
|---|---|---|---|---|---|---|
| chr19 | 42111976 | 42120725 | 8750 | 1.46 | 0.0194 | 0.0225 |
| chrX | 91880405 | 91882369 | 1965 | 1.4 | 0.0194 | 0.0225 |
| chr4 | 128198059 | 128199213 | 1155 | -2.96 | 0.0194 | 0.0226 |
| chr8 | 113780900 | 113800773 | 19874 | -2.35 | 0.0194 | 0.0226 |
| chr12 | 22286703 | 22330889 | 44187 | 1.24 | 0.0195 | 0.0226 |
| chr5 | 157506301 | 157539331 | 33031 | 0.63 | 0.0195 | 0.0226 |
| chr5 | 166701273 | 166701720 | 448 | -2.53 | 0.0195 | 0.0226 |
| chr13 | 108835704 | 108866092 | 30389 | -3.77 | 0.0195 | 0.0226 |
| chr9 | 101617176 | 101645660 | 28485 | -3.56 | 0.0195 | 0.0227 |
| chr13 | 30022952 | 30025449 | 2498 | -1.73 | 0.0195 | 0.0227 |
| chr6 | 19950500 | 19952733 | 2234 | -1.18 | 0.0195 | 0.0227 |
| chr2 | 136636414 | 136646843 | 10430 | -4.8 | 0.0196 | 0.0227 |
| chr15 | 84245039 | 84350719 | 105681 | 0.53 | 0.0196 | 0.0227 |
| chrX | 29119736 | 29121924 | 2189 | -2.97 | 0.0196 | 0.0228 |
| chr1 | 4027490 | 4033302 | 5813 | -1.46 | 0.0196 | 0.0228 |
| chr20 | 17820745 | 17831178 | 10434 | -4.65 | 0.0196 | 0.0228 |
| chr7 | 98250318 | 98327822 | 77505 | -5.36 | 0.0197 | 0.0228 |
| chr14 | 58050303 | 58085239 | 34937 | 1.89 | 0.0198 | 0.0229 |
| chr4 | 78362605 | 78368604 | 6000 | 1.31 | 0.0198 | 0.0229 |
| chr18 | 7570212 | 7582458 | 12247 | -4.41 | 0.0199 | 0.023 |
| chr9 | 82348508 | 82409635 | 61128 | -3.84 | 0.0199 | 0.0231 |
| chr4 | 152922730 | 152950353 | 27624 | -0.94 | 0.0199 | 0.0231 |
| chr15 | 89122504 | 89143080 | 20577 | -3.7 | 0.0199 | 0.0231 |
| chr19 | 16041532 | 16042281 | 750 | 2.84 | 0.02 | 0.0231 |
| chr2 | 25256774 | 25257607 | 834 | -2.79 | 0.02 | 0.0232 |
| chr20 | 1948515 | 1958236 | 9722 | -4.77 | 0.02 | 0.0232 |
| chr4 | 131412305 | 131414864 | 2560 | -3.06 | 0.02 | 0.0232 |
| chrX | 62822989 | 62825401 | 2413 | -1.22 | 0.02 | 0.0232 |
| chr3 | 111089524 | 111098489 | 8966 | -1.24 | 0.02 | 0.0232 |
| chr6 | 75675943 | 75707589 | 31647 | 1.81 | 0.02 | 0.0232 |
| chr12 | 70500238 | 70507777 | 7540 | -3.48 | 0.02 | 0.0232 |
| chr13 | 79872482 | 79874669 | 2188 | -0.87 | 0.0201 | 0.0233 |
| chr12 | 25292335 | 25294943 | 2609 | -3.08 | 0.0201 | 0.0233 |
| chr1 | 237188463 | 237238760 | 50298 | -1.55 | 0.0201 | 0.0233 |
| chr7 | 132126892 | 132156132 | 29241 | 1.64 | 0.0201 | 0.0234 |
| chr21 | 18227157 | 18229257 | 2101 | -3.02 | 0.0202 | 0.0234 |
| chr15 | 51431152 | 51478376 | 47225 | -1.78 | 0.0202 | 0.0234 |
| chrX | 108634677 | 108636229 | 1553 | -2.94 | 0.0202 | 0.0234 |
| chr13 | 88990877 | 88992084 | 1208 | -1.27 | 0.0202 | 0.0235 |
| chr2 | 128312544 | 128316320 | 3777 | -1.82 | 0.0202 | 0.0235 |
| chr2 | 41720514 | 41834931 | 114418 | 0.46 | 0.0202 | 0.0235 |
| chr6 | 162649335 | 162650602 | 1268 | 1.44 | 0.0203 | 0.0235 |
| chrX | 51331417 | 51332809 | 1393 | -1.11 | 0.0203 | 0.0235 |
| chr4 | 29979678 | 29982328 | 2651 | 1.09 | 0.0203 | 0.0235 |
| chr5 | 145929379 | 145969062 | 39684 | 1.86 | 0.0203 | 0.0235 |
| chr2 | 71860571 | 71861965 | 1395 | 1.14 | 0.0203 | 0.0235 |
| chr7 | 84829165 | 84832336 | 3172 | 1.05 | 0.0203 | 0.0236 |
| chr18 | 11123464 | 11127301 | 3838 | -1.34 | 0.0203 | 0.0236 |
| chr17 | 68740140 | 68746538 | 6399 | -3.39 | 0.0203 | 0.0236 |
| chr3 | 120527958 | 120540813 | 12856 | -3.35 | 0.0204 | 0.0236 |
| chr13 | 33985174 | 33987419 | 2246 | 1.56 | 0.0204 | 0.0236 |
| chr4 | 34282261 | 34297085 | 14825 | 2.03 | 0.0204 | 0.0236 |
| chr2 | 69963839 | 69973600 | 9762 | -4.75 | 0.0204 | 0.0236 |
| chr2 | 198690607 | 198692075 | 1469 | 2.99 | 0.0204 | 0.0236 |
| chr18 | 25200368 | 25213802 | 13435 | 2.74 | 0.0205 | 0.0237 |
| chr7 | 115428825 | 115431003 | 2179 | -1.74 | 0.0205 | 0.0237 |
| chr2 | 53549011 | 53551372 | 2362 | -1.1 | 0.0205 | 0.0237 |
| chr5 | 167193101 | 167198982 | 5882 | 0.63 | 0.0205 | 0.0237 |
| chr17 | 63206972 | 63210802 | 3831 | 3.42 | 0.0205 | 0.0238 |
| chr10 | 83095128 | 83097549 | 2422 | -1.74 | 0.0205 | 0.0238 |
| chrX | 51337399 | 51346150 | 8752 | 3.52 | 0.0205 | 0.0238 |

| | | | | | | |
|---|---|---|---|---|---|---|
| chr6 | 130859188 | 130861508 | 2321 | -1.89 | 0.0205 | 0.0238 |
| chr8 | 86872470 | 86893391 | 20922 | -0.91 | 0.0205 | 0.0238 |
| chr7 | 31071094 | 31083573 | 12480 | -1.87 | 0.0206 | 0.0238 |
| chr10 | 87756362 | 87758864 | 2503 | -3.33 | 0.0206 | 0.0239 |
| chrX | 99700975 | 99701981 | 1007 | 1.18 | 0.0206 | 0.0239 |
| chr13 | 91974434 | 91975695 | 1262 | -1.2 | 0.0206 | 0.0239 |
| chr12 | 2859097 | 2861267 | 2171 | 3.28 | 0.0206 | 0.0239 |
| chr9 | 121609767 | 121612096 | 2330 | 0.9 | 0.0206 | 0.0239 |
| chr3 | 54103684 | 54105773 | 2090 | -1.99 | 0.0207 | 0.0239 |
| chr14 | 29069379 | 29121276 | 51898 | -1.83 | 0.0207 | 0.024 |
| chr5 | 133105128 | 133109151 | 4024 | -3.44 | 0.0207 | 0.024 |
| chr15 | 39073051 | 39075120 | 2070 | -0.96 | 0.0207 | 0.024 |
| chr13 | 66235681 | 66245122 | 9442 | -1.63 | 0.0207 | 0.024 |
| chr2 | 171911619 | 171929481 | 17863 | -4.6 | 0.0207 | 0.024 |
| chr3 | 87955581 | 87990036 | 34456 | -3.8 | 0.0208 | 0.024 |
| chr3 | 138749424 | 138754690 | 5267 | -0.87 | 0.0208 | 0.024 |
| chr10 | 35962915 | 35972695 | 9781 | -2.36 | 0.0208 | 0.024 |
| chr5 | 56969468 | 56971432 | 1965 | -2.98 | 0.0208 | 0.024 |
| chr21 | 44810004 | 44845283 | 35280 | -3.61 | 0.0208 | 0.0241 |
| chr2 | 103468702 | 103482322 | 13621 | -0.82 | 0.0208 | 0.0241 |
| chr7 | 87697830 | 87699210 | 1381 | -1.51 | 0.0208 | 0.0241 |
| chr3 | 7479469 | 7480588 | 1120 | 1.37 | 0.0208 | 0.0241 |
| chr15 | 57681673 | 57695091 | 13419 | -4.73 | 0.0209 | 0.0241 |
| chr9 | 126639064 | 126651024 | 11961 | -4.73 | 0.0209 | 0.0241 |
| chr4 | 94301098 | 94302012 | 915 | -1.71 | 0.0209 | 0.0242 |
| chr8 | 59688165 | 59691895 | 3731 | -3.38 | 0.0209 | 0.0242 |
| chr13 | 21856148 | 21859055 | 2908 | -2.53 | 0.0209 | 0.0242 |
| chr5 | 99823305 | 99826300 | 2996 | -1.17 | 0.0209 | 0.0242 |
| chr15 | 35625752 | 35657637 | 31886 | -3.8 | 0.0209 | 0.0242 |
| chr4 | 21994058 | 22009746 | 15689 | 3.51 | 0.0209 | 0.0242 |
| chr15 | 39868372 | 39868996 | 625 | -1.69 | 0.0209 | 0.0242 |
| chr10 | 64517647 | 64545769 | 28123 | -3.38 | 0.021 | 0.0243 |
| chr1 | 207297174 | 207297709 | 536 | 1.6 | 0.021 | 0.0243 |
| chr9 | 108614021 | 108614848 | 828 | -1.56 | 0.021 | 0.0243 |
| chr2 | 162386122 | 162387734 | 1613 | 1.08 | 0.0211 | 0.0244 |
| chrX | 57823497 | 57825486 | 1990 | 1.09 | 0.0211 | 0.0244 |
| chrX | 67013402 | 67015670 | 2269 | -1.53 | 0.0211 | 0.0244 |
| chr1 | 115474262 | 115475324 | 1063 | -1.59 | 0.0211 | 0.0245 |
| chr10 | 92556826 | 92561018 | 4193 | -1.96 | 0.0212 | 0.0245 |
| chr14 | 56897232 | 56903898 | 6667 | -1.02 | 0.0212 | 0.0245 |
| chr1 | 79218916 | 79220439 | 1524 | -2.83 | 0.0212 | 0.0245 |
| chr5 | 2186912 | 2249043 | 62132 | -0.5 | 0.0212 | 0.0245 |
| chr2 | 153688003 | 153699253 | 11251 | -3.51 | 0.0212 | 0.0245 |
| chrX | 100628407 | 100629345 | 939 | -1.39 | 0.0212 | 0.0246 |
| chr11 | 129056011 | 129070725 | 14715 | -4.71 | 0.0213 | 0.0246 |
| chr15 | 74197666 | 74209443 | 11778 | -4.71 | 0.0213 | 0.0246 |
| chr4 | 89972182 | 89982407 | 10226 | -4.71 | 0.0213 | 0.0246 |
| chr3 | 150527073 | 150529241 | 2169 | 1.59 | 0.0213 | 0.0247 |
| chrX | 131083445 | 131084791 | 1347 | -1.9 | 0.0214 | 0.0247 |
| chr2 | 166527788 | 166529948 | 2161 | -2.84 | 0.0214 | 0.0247 |
| chr6 | 52006454 | 52008672 | 2219 | 1.76 | 0.0214 | 0.0247 |
| chr12 | 60183154 | 60190742 | 7589 | -3.27 | 0.0214 | 0.0247 |
| chr1 | 119848292 | 119849784 | 1493 | -2.74 | 0.0214 | 0.0247 |
| chr9 | 111447926 | 111500012 | 52087 | -5.11 | 0.0214 | 0.0248 |
| chr16 | 55852833 | 55854002 | 1170 | 1.28 | 0.0214 | 0.0248 |
| chr14 | 58484902 | 58485907 | 1006 | -2.81 | 0.0214 | 0.0248 |
| chr3 | 143554956 | 143557196 | 2241 | 2.06 | 0.0215 | 0.0248 |
| chr18 | 54792293 | 54795585 | 3293 | 1.51 | 0.0215 | 0.0249 |
| chr1 | 217460064 | 217463608 | 3545 | -3.28 | 0.0215 | 0.0249 |
| chr1 | 214948640 | 214955182 | 6543 | 3.15 | 0.0216 | 0.0249 |
| chr1 | 48614142 | 48615950 | 1809 | -1.17 | 0.0216 | 0.0249 |

| | | | | | | |
|---|---|---|---|---|---|---|
| chr2 | 222264868 | 222288647 | 23780 | -0.7 | 0.0217 | 0.025 |
| chr8 | 82470830 | 82495247 | 24418 | 2.89 | 0.0217 | 0.025 |
| chr9 | 21219600 | 21220244 | 645 | 1.28 | 0.0217 | 0.0251 |
| chr8 | 120563380 | 120565150 | 1771 | -2.98 | 0.0217 | 0.0251 |
| chr8 | 70753996 | 70820039 | 66044 | 0.64 | 0.0217 | 0.0251 |
| chr21 | 39175763 | 39186786 | 11024 | -3.17 | 0.0217 | 0.0251 |
| chr8 | 96806750 | 96812271 | 5522 | -1.59 | 0.0217 | 0.0251 |
| chr5 | 85989418 | 86254323 | 264906 | 5.72 | 0.0218 | 0.0252 |
| chr1 | 185740516 | 185743170 | 2655 | 3.28 | 0.0218 | 0.0252 |
| chr12 | 40574048 | 40584837 | 10790 | -4.69 | 0.0218 | 0.0252 |
| chr4 | 74781813 | 74793184 | 11372 | -4.69 | 0.0218 | 0.0252 |
| chr7 | 17066989 | 17081402 | 14414 | -4.69 | 0.0218 | 0.0252 |
| chr20 | 39562315 | 39565022 | 2708 | -1.26 | 0.0218 | 0.0252 |
| chr21 | 42753464 | 42788121 | 34658 | -2.3 | 0.0218 | 0.0252 |
| chrX | 11738930 | 11739757 | 828 | 1.7 | 0.0219 | 0.0253 |
| chr4 | 125052881 | 125055634 | 2754 | -3.1 | 0.0219 | 0.0253 |
| chr6 | 97791336 | 97792691 | 1356 | 2.89 | 0.0219 | 0.0253 |
| chr9 | 107461628 | 107466064 | 4437 | 3.21 | 0.0219 | 0.0253 |
| chr6 | 154348074 | 154350395 | 2322 | 3.1 | 0.022 | 0.0254 |
| chr5 | 156602418 | 156634000 | 31583 | -3.56 | 0.022 | 0.0254 |
| chr5 | 112470627 | 112472139 | 1513 | 1.24 | 0.022 | 0.0254 |
| chr11 | 8532304 | 8544186 | 11883 | -3.31 | 0.0221 | 0.0255 |
| chr8 | 26734164 | 26874469 | 140306 | 0.32 | 0.0221 | 0.0255 |
| chr1 | 86765437 | 86766865 | 1429 | -1.12 | 0.0221 | 0.0256 |
| chr21 | 48028829 | 48049097 | 20269 | -3.66 | 0.0222 | 0.0256 |
| chrX | 39342389 | 39342958 | 570 | -2.13 | 0.0222 | 0.0256 |
| chr6 | 18567153 | 18568838 | 1686 | 1.43 | 0.0222 | 0.0256 |
| chr2 | 139592197 | 139594219 | 2023 | 1.47 | 0.0222 | 0.0256 |
| chr5 | 94556124 | 94580342 | 24219 | 3.5 | 0.0222 | 0.0257 |
| chr12 | 91178060 | 91181946 | 3887 | -1.85 | 0.0223 | 0.0257 |
| chr4 | 84993812 | 84994691 | 880 | -1.56 | 0.0223 | 0.0257 |
| chr12 | 79151730 | 79163711 | 11982 | -4.67 | 0.0223 | 0.0257 |
| chr2 | 63299993 | 63307390 | 7398 | -4.67 | 0.0223 | 0.0257 |
| chr6 | 119555994 | 119557400 | 1407 | -1.09 | 0.0223 | 0.0257 |
| chr6 | 64589349 | 64597278 | 7930 | 3.4 | 0.0223 | 0.0258 |
| chr12 | 54908013 | 54927365 | 19353 | 1.29 | 0.0223 | 0.0258 |
| chr3 | 109025537 | 109029341 | 3805 | -1.84 | 0.0224 | 0.0258 |
| chrX | 56850432 | 56884558 | 34127 | -1.53 | 0.0224 | 0.0258 |
| chr3 | 162502656 | 162504902 | 2247 | -2.99 | 0.0224 | 0.0258 |
| chr11 | 59261807 | 59264901 | 3095 | 1.21 | 0.0224 | 0.0258 |
| chrX | 133030716 | 133031098 | 383 | 2.81 | 0.0224 | 0.0259 |
| chr9 | 27138402 | 27139592 | 1191 | -2.66 | 0.0224 | 0.0259 |
| chrX | 30369814 | 30372275 | 2462 | -3.08 | 0.0224 | 0.0259 |
| chr12 | 25818593 | 25823360 | 4768 | 0.95 | 0.0224 | 0.0259 |
| chr3 | 38634689 | 38636699 | 2011 | -3.13 | 0.0224 | 0.0259 |
| chr1 | 88773510 | 88776146 | 2637 | -3.14 | 0.0224 | 0.0259 |
| chr16 | 79922316 | 79925157 | 2842 | -2.88 | 0.0225 | 0.0259 |
| chr15 | 96056974 | 96117307 | 60334 | -2.08 | 0.0225 | 0.026 |
| chr3 | 5972898 | 5973932 | 1035 | 1.02 | 0.0225 | 0.026 |
| chr19 | 55481712 | 55483955 | 2244 | -2.86 | 0.0225 | 0.026 |
| chrX | 110778550 | 110779440 | 891 | -1.97 | 0.0225 | 0.026 |
| chr16 | 23822847 | 24018335 | 195489 | 0.52 | 0.0226 | 0.026 |
| chrX | 132746485 | 132755313 | 8829 | -0.71 | 0.0226 | 0.0261 |
| chr5 | 120952097 | 120952576 | 480 | -1.71 | 0.0226 | 0.0261 |
| chr14 | 77166701 | 77169423 | 2723 | 1.17 | 0.0227 | 0.0262 |
| chr4 | 173964389 | 173965218 | 830 | -2.78 | 0.0227 | 0.0262 |
| chr7 | 155844900 | 155845590 | 691 | 1.29 | 0.0227 | 0.0262 |
| chr1 | 37566961 | 37577612 | 10652 | 3.38 | 0.0227 | 0.0262 |
| chr14 | 21327544 | 21328654 | 1111 | -2.94 | 0.0227 | 0.0262 |
| chr20 | 52038612 | 52114746 | 76135 | -3.77 | 0.0227 | 0.0262 |
| chrX | 56032875 | 56033886 | 1012 | -1.71 | 0.0227 | 0.0262 |

| | | | | | | |
|---|---|---|---|---|---|---|
| chr20 | 9624760 | 9626341 | 1582 | -1.49 | 0.0227 | 0.0262 |
| chrX | 12900378 | 12905942 | 5565 | 0.79 | 0.0227 | 0.0262 |
| chr15 | 38179955 | 38181352 | 1398 | -1.85 | 0.0228 | 0.0263 |
| chr3 | 148936580 | 148946274 | 9695 | -4.64 | 0.0228 | 0.0263 |
| chr4 | 11249541 | 11294842 | 45302 | 0.58 | 0.0228 | 0.0263 |
| chr15 | 23778620 | 23783134 | 4515 | 0.68 | 0.0228 | 0.0263 |
| chr3 | 120608174 | 120609861 | 1688 | -2.44 | 0.0228 | 0.0263 |
| chrX | 91729863 | 91730245 | 383 | 1.35 | 0.0228 | 0.0264 |
| chr8 | 120679336 | 120697297 | 17962 | 3.55 | 0.0228 | 0.0264 |
| chr15 | 60075842 | 60095762 | 19921 | -3.74 | 0.0229 | 0.0264 |
| chr3 | 96643780 | 96664776 | 20997 | -3.32 | 0.0229 | 0.0264 |
| chr6 | 169753128 | 169757768 | 4641 | -0.98 | 0.023 | 0.0265 |
| chr10 | 87783726 | 87805536 | 21811 | 0.67 | 0.023 | 0.0265 |
| chr3 | 68958037 | 68960373 | 2337 | 1.33 | 0.023 | 0.0265 |
| chr5 | 107733244 | 107739648 | 6405 | -2.97 | 0.023 | 0.0265 |
| chrX | 150428472 | 150474508 | 46037 | 1.32 | 0.023 | 0.0266 |
| chr1 | 115326935 | 115335794 | 8860 | -3.28 | 0.023 | 0.0266 |
| chr11 | 20307375 | 20324276 | 16902 | -1.76 | 0.023 | 0.0266 |
| chr9 | 19483549 | 19484344 | 796 | -1.47 | 0.0231 | 0.0267 |
| chr2 | 108369680 | 108432402 | 62723 | 3.81 | 0.0231 | 0.0267 |
| chr8 | 25443774 | 25456969 | 13196 | 1.14 | 0.0232 | 0.0267 |
| chr15 | 26346735 | 27008590 | 661856 | 0.29 | 0.0233 | 0.0268 |
| chr7 | 157614349 | 157616310 | 1962 | 1.94 | 0.0233 | 0.0269 |
| chr11 | 110174147 | 110183476 | 9330 | -4.62 | 0.0233 | 0.0269 |
| chr9 | 92093908 | 92101934 | 8027 | -4.62 | 0.0233 | 0.0269 |
| chrX | 150766407 | 150784557 | 18151 | 3.57 | 0.0233 | 0.0269 |
| chr11 | 107604959 | 107611798 | 6840 | -2.54 | 0.0234 | 0.027 |
| chr13 | 112069710 | 112071791 | 2082 | 0.95 | 0.0234 | 0.027 |
| chr14 | 96290419 | 96334562 | 44144 | 0.54 | 0.0234 | 0.027 |
| chr19 | 32995707 | 33030215 | 34509 | -5.18 | 0.0234 | 0.027 |
| chr11 | 32021946 | 32022983 | 1038 | 1.48 | 0.0234 | 0.027 |
| chr9 | 111230716 | 111232097 | 1382 | -2.75 | 0.0234 | 0.027 |
| chr1 | 63489988 | 63510216 | 20229 | -3.4 | 0.0234 | 0.027 |
| chr8 | 97645696 | 98268407 | 622712 | 6.01 | 0.0235 | 0.0271 |
| chr6 | 80515608 | 80549153 | 33546 | -3.54 | 0.0235 | 0.0271 |
| chrX | 152465047 | 152471159 | 6113 | -0.77 | 0.0236 | 0.0272 |
| chrX | 65042686 | 65044195 | 1510 | -2.72 | 0.0236 | 0.0272 |
| chr4 | 100213550 | 100217917 | 4368 | 0.9 | 0.0236 | 0.0272 |
| chr1 | 37836832 | 37849109 | 12278 | -3.25 | 0.0236 | 0.0272 |
| chr4 | 83224639 | 83225975 | 1337 | -1.43 | 0.0236 | 0.0272 |
| chr5 | 131150011 | 131220632 | 70622 | -5.05 | 0.0237 | 0.0273 |
| chr2 | 75743326 | 75745662 | 2337 | 1.5 | 0.0237 | 0.0273 |
| chr2 | 186337280 | 186394328 | 57049 | -1.7 | 0.0237 | 0.0273 |
| chr1 | 152059181 | 152103951 | 44771 | -0.8 | 0.0237 | 0.0274 |
| chr16 | 9359360 | 9366490 | 7131 | -0.82 | 0.0237 | 0.0274 |
| chr1 | 49046583 | 49051188 | 4606 | 3.12 | 0.0238 | 0.0274 |
| chr7 | 108349358 | 108349929 | 572 | -1.28 | 0.0238 | 0.0274 |
| chr3 | 60085732 | 60088666 | 2935 | 2.39 | 0.0238 | 0.0274 |
| chr8 | 16790324 | 16801498 | 11175 | 0.75 | 0.0238 | 0.0274 |
| chr2 | 147541807 | 147546135 | 4329 | 2.06 | 0.0238 | 0.0274 |
| chr13 | 83563303 | 83574000 | 10698 | -4.06 | 0.0238 | 0.0275 |
| chr16 | 20033361 | 20051736 | 18376 | 0.56 | 0.0238 | 0.0275 |
| chrX | 129696440 | 129696980 | 541 | -2.59 | 0.0239 | 0.0275 |
| chr17 | 1140326 | 1162507 | 22182 | -4.6 | 0.0239 | 0.0275 |
| chr1 | 173256217 | 173262520 | 6304 | -0.79 | 0.0239 | 0.0276 |
| chr18 | 39522827 | 39528491 | 5665 | 0.7 | 0.024 | 0.0276 |
| chr2 | 177352557 | 177353673 | 1117 | -2.79 | 0.024 | 0.0276 |
| chr6 | 917091 | 943205 | 26115 | -3.45 | 0.024 | 0.0276 |
| chr8 | 140436099 | 140446412 | 10314 | 3.41 | 0.024 | 0.0276 |
| chr1 | 249193124 | 249196975 | 3852 | -1.74 | 0.024 | 0.0277 |
| chr4 | 161463780 | 161465550 | 1771 | -1.18 | 0.024 | 0.0277 |

| | | | | | | |
|---|---|---|---|---|---|---|
| chr12 | 91380176 | 91381751 | 1576 | -1.03 | 0.024 | 0.0277 |
| chr12 | 65345419 | 65464558 | 119140 | 0.5 | 0.0241 | 0.0277 |
| chr1 | 91796418 | 91797983 | 1566 | -1.12 | 0.0241 | 0.0277 |
| chr12 | 20073615 | 20089841 | 16227 | -3.41 | 0.0241 | 0.0277 |
| chr9 | 97358385 | 97358972 | 588 | -2.54 | 0.0241 | 0.0278 |
| chr11 | 115569445 | 115575755 | 6311 | -1.13 | 0.0241 | 0.0278 |
| chr14 | 70503921 | 70505440 | 1520 | 1.34 | 0.0241 | 0.0278 |
| chr3 | 104665844 | 104670284 | 4441 | 1.78 | 0.0241 | 0.0278 |
| chr11 | 19325277 | 19327524 | 2248 | -1.76 | 0.0242 | 0.0278 |
| chr1 | 173227845 | 173230692 | 2848 | -3.09 | 0.0242 | 0.0279 |
| chr4 | 136114949 | 136116059 | 1111 | -1.92 | 0.0242 | 0.0279 |
| chr21 | 15686641 | 15688412 | 1772 | 2.41 | 0.0242 | 0.0279 |
| chr2 | 7461426 | 7477534 | 16109 | 0.97 | 0.0242 | 0.0279 |
| chrX | 42415904 | 42416757 | 854 | -1.32 | 0.0243 | 0.0279 |
| chr4 | 41756951 | 41772320 | 15370 | -0.82 | 0.0243 | 0.0279 |
| chr13 | 112691864 | 112703815 | 11952 | -0.74 | 0.0243 | 0.028 |
| chr2 | 56709078 | 56718513 | 9436 | 0.8 | 0.0243 | 0.028 |
| chr9 | 43663486 | 43664524 | 1039 | -2.76 | 0.0243 | 0.028 |
| chr6 | 75525021 | 75531953 | 6933 | -1 | 0.0243 | 0.028 |
| chr11 | 120787068 | 120787743 | 676 | -1.79 | 0.0243 | 0.028 |
| chr7 | 150415364 | 150419849 | 4486 | 3.31 | 0.0243 | 0.028 |
| chr6 | 132547366 | 132547867 | 502 | -2.32 | 0.0243 | 0.028 |
| chrX | 68902964 | 68917076 | 14113 | 2.97 | 0.0244 | 0.028 |
| chr9 | 111513803 | 111517549 | 3747 | -1.73 | 0.0244 | 0.028 |
| chr11 | 88440598 | 88468028 | 27431 | -1.73 | 0.0244 | 0.0281 |
| chr12 | 78146363 | 78148986 | 2624 | -1.56 | 0.0244 | 0.0281 |
| chr10 | 50793643 | 50797927 | 4285 | -3.17 | 0.0244 | 0.0281 |
| chr12 | 4137800 | 4365246 | 227447 | 0.42 | 0.0244 | 0.0281 |
| chr1 | 232028587 | 232042097 | 13511 | -4.57 | 0.0244 | 0.0281 |
| chr19 | 43940403 | 43958157 | 17755 | -4.57 | 0.0244 | 0.0281 |
| chr2 | 153008124 | 153014725 | 6602 | -4.57 | 0.0244 | 0.0281 |
| chr2 | 236058981 | 236068328 | 9348 | -4.57 | 0.0244 | 0.0281 |
| chr7 | 79232645 | 79245917 | 13273 | -4.57 | 0.0244 | 0.0281 |
| chr20 | 59957209 | 59957888 | 680 | -2.55 | 0.0244 | 0.0281 |
| chr5 | 111995950 | 111997033 | 1084 | -1.5 | 0.0244 | 0.0281 |
| chr16 | 51031623 | 51033200 | 1578 | 2 | 0.0244 | 0.0281 |
| chr3 | 58710447 | 58710768 | 322 | -1.57 | 0.0245 | 0.0281 |
| chr8 | 62291011 | 62295364 | 4354 | 2.13 | 0.0245 | 0.0282 |
| chr6 | 153124285 | 153125172 | 888 | -2.85 | 0.0245 | 0.0282 |
| chr12 | 130990639 | 131004481 | 13843 | -0.91 | 0.0245 | 0.0282 |
| chr19 | 35981913 | 35983329 | 1417 | 1.32 | 0.0247 | 0.0284 |
| chr13 | 88694718 | 88695346 | 629 | 2.99 | 0.0247 | 0.0284 |
| chr13 | 103191314 | 103191727 | 414 | -2.53 | 0.0247 | 0.0284 |
| chr12 | 107506123 | 107507640 | 1518 | 2.46 | 0.0247 | 0.0284 |
| chrX | 16397869 | 16409604 | 11736 | -0.67 | 0.0247 | 0.0284 |
| chr9 | 13004657 | 13037758 | 33102 | -3.48 | 0.0248 | 0.0285 |
| chr3 | 115073341 | 115074563 | 1223 | -2.92 | 0.0248 | 0.0285 |
| chr8 | 26903002 | 26906187 | 3186 | 1.39 | 0.0248 | 0.0285 |
| chr12 | 55119913 | 55121135 | 1223 | -1.14 | 0.0248 | 0.0285 |
| chr18 | 4072380 | 4074447 | 2068 | 0.91 | 0.0248 | 0.0286 |
| chr6 | 145610454 | 145611956 | 1503 | -2.82 | 0.0249 | 0.0286 |
| chr14 | 59579569 | 59583500 | 3932 | -1.76 | 0.0249 | 0.0287 |
| chr4 | 79903906 | 79905129 | 1224 | 1.54 | 0.025 | 0.0287 |
| chrX | 85309283 | 85316933 | 7651 | -2.98 | 0.025 | 0.0287 |
| chr13 | 77974836 | 77976189 | 1354 | -2.39 | 0.025 | 0.0287 |
| chr3 | 98269953 | 98271329 | 1377 | -2.62 | 0.025 | 0.0288 |
| chr10 | 79499794 | 79511025 | 11232 | -4.55 | 0.025 | 0.0288 |
| chr9 | 74827423 | 74837980 | 10558 | -4.55 | 0.025 | 0.0288 |
| chr16 | 74073069 | 74137172 | 64104 | 0.44 | 0.025 | 0.0288 |
| chr7 | 87596629 | 87596954 | 326 | -2.23 | 0.025 | 0.0288 |
| chrX | 73783710 | 73784792 | 1083 | -2.24 | 0.0251 | 0.0288 |

| | | | | | | |
|---|---|---|---|---|---|---|
| chr7 | 56475849 | 56478532 | 2684 | -1.3 | 0.0251 | 0.0288 |
| chr7 | 109005490 | 109007960 | 2471 | -3.13 | 0.0252 | 0.0289 |
| chr1 | 75958318 | 75959966 | 1649 | -2.72 | 0.0252 | 0.029 |
| chr5 | 49653392 | 49655127 | 1736 | -2.95 | 0.0252 | 0.029 |
| chr18 | 71880449 | 71882829 | 2381 | 1.57 | 0.0252 | 0.029 |
| chr17 | 32613251 | 32618530 | 5280 | 0.83 | 0.0252 | 0.029 |
| chrX | 114321180 | 114331698 | 10519 | 3.38 | 0.0253 | 0.0291 |
| chr11 | 19359044 | 19382363 | 23320 | 1.97 | 0.0253 | 0.0291 |
| chr3 | 81395021 | 81395891 | 871 | 2.74 | 0.0253 | 0.0291 |
| chr12 | 60451729 | 60453917 | 2189 | 2.91 | 0.0253 | 0.0291 |
| chr1 | 187296776 | 187305885 | 9110 | 2.9 | 0.0254 | 0.0291 |
| chr17 | 10200435 | 10227478 | 27044 | -0.65 | 0.0254 | 0.0292 |
| chr3 | 120090222 | 120098719 | 8498 | -1.03 | 0.0254 | 0.0292 |
| chr12 | 91253944 | 91268355 | 14412 | -1.78 | 0.0254 | 0.0292 |
| chr2 | 157544265 | 157545249 | 985 | -1.78 | 0.0255 | 0.0293 |
| chr6 | 79045570 | 79046534 | 965 | 2.73 | 0.0255 | 0.0293 |
| chr2 | 171347839 | 171351853 | 4015 | 0.93 | 0.0255 | 0.0293 |
| chr5 | 56903284 | 56915402 | 12119 | -0.89 | 0.0255 | 0.0293 |
| chr15 | 61112557 | 61289061 | 176505 | 5.47 | 0.0255 | 0.0293 |
| chr19 | 8912541 | 9007026 | 94486 | -5.25 | 0.0255 | 0.0293 |
| chr10 | 3924327 | 3925295 | 969 | -2.54 | 0.0256 | 0.0294 |
| chr16 | 23339206 | 23350512 | 11307 | -4.52 | 0.0256 | 0.0295 |
| chr10 | 124660378 | 124664781 | 4404 | -1.26 | 0.0257 | 0.0295 |
| chr18 | 45041775 | 45043030 | 1256 | -2.85 | 0.0257 | 0.0295 |
| chr12 | 25987971 | 25991356 | 3386 | -3.13 | 0.0257 | 0.0295 |
| chr12 | 124345767 | 124370698 | 24932 | -0.75 | 0.0257 | 0.0295 |
| chr6 | 46963536 | 46964366 | 831 | -2.62 | 0.0257 | 0.0295 |
| chr12 | 86299691 | 86300109 | 419 | -2.32 | 0.0257 | 0.0295 |
| chr13 | 81219641 | 81221873 | 2233 | -1.55 | 0.0257 | 0.0295 |
| chr7 | 144391096 | 144402507 | 11412 | -3.21 | 0.0257 | 0.0296 |
| chr1 | 108050656 | 108053659 | 3004 | -1.51 | 0.0258 | 0.0296 |
| chr17 | 55845838 | 55850114 | 4277 | 3.52 | 0.0258 | 0.0296 |
| chr6 | 130708302 | 130780940 | 72639 | 1.19 | 0.0258 | 0.0296 |
| chr12 | 63112870 | 63114070 | 1201 | -1 | 0.0258 | 0.0296 |
| chrX | 135615792 | 135618315 | 2524 | 1.79 | 0.0259 | 0.0297 |
| chr1 | 18784027 | 18795283 | 11257 | -4.37 | 0.0259 | 0.0297 |
| chr18 | 76686230 | 76689991 | 3762 | -1.16 | 0.0259 | 0.0298 |
| chr10 | 27703464 | 27719589 | 16126 | -0.73 | 0.026 | 0.0298 |
| chr4 | 92628956 | 92632466 | 3511 | -2.66 | 0.026 | 0.0298 |
| chr5 | 150670063 | 150723831 | 53769 | -1.13 | 0.0261 | 0.0299 |
| chr17 | 25372415 | 25416707 | 44293 | 0.51 | 0.0261 | 0.0299 |
| chrX | 49624898 | 49625898 | 1001 | 1.23 | 0.0261 | 0.0299 |
| chrX | 150797534 | 150798124 | 591 | -1.22 | 0.0261 | 0.0299 |
| chr4 | 97797524 | 97798909 | 1386 | -2.55 | 0.0261 | 0.03 |
| chrX | 105968017 | 105968531 | 515 | 1.56 | 0.0261 | 0.03 |
| chr6 | 5970859 | 5971628 | 770 | 1.34 | 0.0261 | 0.03 |
| chr1 | 117284553 | 117340785 | 56233 | -3.63 | 0.0262 | 0.0301 |
| chr2 | 107508848 | 107526263 | 17416 | -1.21 | 0.0263 | 0.0301 |
| chr11 | 49630335 | 49630711 | 377 | 1.26 | 0.0263 | 0.0302 |
| chr3 | 134541186 | 135303596 | 762411 | 5.83 | 0.0263 | 0.0302 |
| chr7 | 85055308 | 85059698 | 4391 | 1.22 | 0.0263 | 0.0302 |
| chr11 | 113838418 | 113840017 | 1600 | -1.31 | 0.0264 | 0.0303 |
| chr6 | 1174001 | 1174661 | 661 | -1.66 | 0.0264 | 0.0303 |
| chr7 | 96662454 | 96715186 | 52733 | -0.87 | 0.0264 | 0.0303 |
| chr3 | 191478732 | 191496562 | 17831 | -0.82 | 0.0264 | 0.0303 |
| chr1 | 159176617 | 159177577 | 961 | 2.06 | 0.0264 | 0.0303 |
| chr5 | 159871840 | 159886852 | 15013 | -1.8 | 0.0264 | 0.0303 |
| chr2 | 157026954 | 157029238 | 2285 | -1.36 | 0.0264 | 0.0303 |
| chr7 | 131480925 | 131482447 | 1523 | 2.85 | 0.0264 | 0.0303 |
| chr12 | 19113106 | 19114968 | 1863 | -3.02 | 0.0264 | 0.0303 |
| chr10 | 64046936 | 64061684 | 14749 | -3.49 | 0.0264 | 0.0304 |

| | | | | | | |
|---|---|---|---|---|---|---|
| chr11 | 20328833 | 20329372 | 540 | 2.69 | 0.0264 | 0.0304 |
| chr17 | 5749125 | 5758162 | 9038 | -3.47 | 0.0265 | 0.0304 |
| chr10 | 128543986 | 128570536 | 26551 | -0.95 | 0.0265 | 0.0304 |
| chr10 | 117127940 | 117461362 | 333423 | 4.6 | 0.0265 | 0.0305 |
| chr4 | 149421967 | 149508138 | 86172 | 3.82 | 0.0266 | 0.0305 |
| chr6 | 89261211 | 89265904 | 4694 | 0.96 | 0.0267 | 0.0306 |
| chr17 | 16342999 | 16343603 | 605 | -1.93 | 0.0267 | 0.0306 |
| chr7 | 87282543 | 87286136 | 3594 | 3.22 | 0.0267 | 0.0306 |
| chr8 | 121709457 | 121712339 | 2883 | -1.11 | 0.0267 | 0.0307 |
| chr4 | 28592964 | 28608320 | 15357 | 1.17 | 0.0267 | 0.0307 |
| chr1 | 181453867 | 181476487 | 22621 | 1.63 | 0.0268 | 0.0307 |
| chr12 | 15555081 | 15557393 | 2313 | 1.8 | 0.0268 | 0.0307 |
| chr9 | 28612124 | 28625208 | 13085 | -3.2 | 0.0268 | 0.0307 |
| chr9 | 13343380 | 13343739 | 360 | -2.38 | 0.0268 | 0.0307 |
| chr6 | 152330385 | 152345629 | 15245 | -0.63 | 0.0268 | 0.0308 |
| chr7 | 116489291 | 116491613 | 2323 | -2.66 | 0.0269 | 0.0308 |
| chr18 | 54091760 | 54096290 | 4531 | -0.79 | 0.0269 | 0.0308 |
| chr11 | 67354446 | 67370919 | 16474 | -4.47 | 0.0269 | 0.0309 |
| chr5 | 30432621 | 30443230 | 10610 | -3.11 | 0.027 | 0.0309 |
| chr6 | 71682133 | 71684189 | 2057 | -0.99 | 0.027 | 0.031 |
| chr6 | 165477277 | 165479303 | 2027 | 1.46 | 0.027 | 0.031 |
| chr5 | 85511100 | 85512347 | 1248 | -1.68 | 0.027 | 0.031 |
| chrX | 107990614 | 108004284 | 13671 | -1.19 | 0.0271 | 0.0311 |
| chr11 | 44495784 | 44498610 | 2827 | 1.18 | 0.0271 | 0.0311 |
| chr4 | 135438285 | 135439521 | 1237 | 2.73 | 0.0271 | 0.0311 |
| chrX | 62251290 | 62280302 | 29013 | 0.7 | 0.0272 | 0.0311 |
| chr5 | 127009254 | 127020014 | 10761 | -3.13 | 0.0272 | 0.0312 |
| chr7 | 142269432 | 142271328 | 1897 | -1.41 | 0.0272 | 0.0312 |
| chr7 | 131985532 | 131988783 | 3252 | 3.15 | 0.0272 | 0.0312 |
| chr3 | 42382143 | 42383873 | 1731 | 1.07 | 0.0273 | 0.0313 |
| chr17 | 55815540 | 55824503 | 8964 | -1.81 | 0.0273 | 0.0313 |
| chr11 | 20650892 | 20652241 | 1350 | -1.27 | 0.0273 | 0.0313 |
| chrX | 151633932 | 151635252 | 1321 | -2.53 | 0.0273 | 0.0313 |
| chr3 | 186502307 | 186503549 | 1243 | 1.13 | 0.0273 | 0.0313 |
| chr3 | 68965877 | 68966712 | 836 | -2.34 | 0.0273 | 0.0313 |
| chr17 | 33165766 | 33208892 | 43127 | -0.63 | 0.0274 | 0.0314 |
| chrX | 79417200 | 79419567 | 2368 | -1.85 | 0.0274 | 0.0314 |
| chr13 | 26055587 | 26067451 | 11865 | -0.94 | 0.0274 | 0.0314 |
| chr7 | 151979410 | 151979899 | 490 | 1.2 | 0.0274 | 0.0314 |
| chr8 | 21710946 | 21711586 | 641 | -2.16 | 0.0274 | 0.0314 |
| chrX | 47518841 | 47653648 | 134808 | -5.07 | 0.0274 | 0.0314 |
| chr5 | 83836409 | 83838044 | 1636 | -2.02 | 0.0274 | 0.0314 |
| chr7 | 31208780 | 31211101 | 2322 | 1.48 | 0.0275 | 0.0315 |
| chr2 | 171063589 | 171064764 | 1176 | -1.65 | 0.0275 | 0.0316 |
| chr4 | 121527565 | 121528455 | 891 | -1.53 | 0.0276 | 0.0316 |
| chr7 | 105933669 | 105960888 | 27220 | -3.48 | 0.0276 | 0.0316 |
| chr16 | 9381997 | 9392781 | 10785 | -0.86 | 0.0276 | 0.0317 |
| chr16 | 85151482 | 85158924 | 7443 | -4.45 | 0.0276 | 0.0317 |
| chr17 | 65696832 | 65710323 | 13492 | -4.45 | 0.0276 | 0.0317 |
| chr8 | 54471805 | 54491729 | 19925 | -1.24 | 0.0277 | 0.0317 |
| chr18 | 59231417 | 59241765 | 10349 | -0.8 | 0.0277 | 0.0317 |
| chr4 | 34491435 | 34507068 | 15634 | 1.49 | 0.0277 | 0.0317 |
| chr2 | 207764987 | 207765843 | 857 | -2.47 | 0.0277 | 0.0317 |
| chr3 | 84855436 | 84986474 | 131039 | -1.14 | 0.0278 | 0.0318 |
| chr18 | 30062978 | 30067652 | 4675 | 1.39 | 0.0278 | 0.0319 |
| chr10 | 13910493 | 13910875 | 383 | -3.03 | 0.0278 | 0.0319 |
| chr1 | 111807656 | 111810219 | 2564 | 2.23 | 0.0278 | 0.0319 |
| chr4 | 24425028 | 24433698 | 8671 | -0.82 | 0.0279 | 0.0319 |
| chr8 | 1356153 | 1429638 | 73486 | -0.67 | 0.0279 | 0.0319 |
| chr4 | 165905771 | 165906529 | 759 | -2.52 | 0.0279 | 0.032 |
| chr10 | 118398451 | 118413489 | 15039 | -3.52 | 0.0279 | 0.032 |

| | | | | | | |
|---|---|---|---|---|---|---|
| chr11 | 106090738 | 106102334 | 11597 | -1.84 | 0.0279 | 0.032 |
| chr14 | 58225553 | 58232281 | 6729 | -0.75 | 0.028 | 0.0321 |
| chr6 | 97363238 | 97364594 | 1357 | -1.47 | 0.028 | 0.0321 |
| chr5 | 133703172 | 133703898 | 727 | -1.62 | 0.0281 | 0.0321 |
| chr21 | 19096132 | 19134958 | 38827 | 3.38 | 0.0281 | 0.0322 |
| chr10 | 69394732 | 69413141 | 18410 | -3.26 | 0.0281 | 0.0322 |
| chr1 | 171267172 | 171273664 | 6493 | -3.37 | 0.0282 | 0.0323 |
| chr2 | 71040438 | 71042428 | 1991 | 3.04 | 0.0282 | 0.0323 |
| chr2 | 230420854 | 230425725 | 4872 | -1.32 | 0.0282 | 0.0323 |
| chrX | 72257643 | 72258599 | 957 | 0.94 | 0.0282 | 0.0323 |
| chr15 | 23834115 | 23891010 | 56896 | -0.78 | 0.0282 | 0.0323 |
| chr12 | 1962069 | 1963389 | 1321 | 2.66 | 0.0283 | 0.0324 |
| chr10 | 100034418 | 100044087 | 9670 | -2.43 | 0.0283 | 0.0324 |
| chr2 | 41852408 | 41865260 | 12853 | 0.56 | 0.0283 | 0.0324 |
| chr4 | 21528272 | 21537489 | 9218 | 3.26 | 0.0283 | 0.0324 |
| chr4 | 108072015 | 108073877 | 1863 | 3.06 | 0.0283 | 0.0325 |
| chr4 | 145726200 | 145730741 | 4542 | -0.92 | 0.0284 | 0.0325 |
| chr6 | 125735494 | 125736936 | 1443 | -1.47 | 0.0284 | 0.0325 |
| chr9 | 29460036 | 29473442 | 13407 | -3.14 | 0.0284 | 0.0325 |
| chr20 | 1720643 | 1734779 | 14137 | -2.66 | 0.0285 | 0.0326 |
| chr10 | 23611177 | 23699998 | 88822 | 5 | 0.0285 | 0.0326 |
| chr6 | 80584356 | 80586378 | 2023 | 1.43 | 0.0285 | 0.0327 |
| chr4 | 128354661 | 128356147 | 1487 | -2.7 | 0.0286 | 0.0327 |
| chr14 | 96599220 | 96600534 | 1315 | 1.2 | 0.0286 | 0.0327 |
| chr14 | 57594329 | 57594980 | 652 | -2.41 | 0.0287 | 0.0328 |
| chr5 | 75558164 | 75576432 | 18269 | -1.49 | 0.0287 | 0.0329 |
| chr5 | 61098517 | 61099816 | 1300 | 1.18 | 0.0287 | 0.0329 |
| chr4 | 29939349 | 29941205 | 1857 | 1.05 | 0.0288 | 0.0329 |
| chr15 | 48586815 | 48589264 | 2450 | 1.06 | 0.0288 | 0.0329 |
| chr4 | 58243448 | 58245642 | 2195 | 2.87 | 0.0288 | 0.0329 |
| chr3 | 180286818 | 180288335 | 1518 | 2.12 | 0.0288 | 0.033 |
| chr7 | 155924803 | 155929367 | 4565 | -1.82 | 0.0289 | 0.033 |
| chr17 | 2979393 | 3132889 | 153497 | -0.42 | 0.0289 | 0.0331 |
| chr21 | 34332720 | 34334717 | 1998 | -1.9 | 0.029 | 0.0332 |
| chrX | 62068675 | 62073164 | 4490 | 1.15 | 0.029 | 0.0332 |
| chr2 | 48487815 | 48488516 | 702 | -1.97 | 0.0291 | 0.0332 |
| chr5 | 126681025 | 126683178 | 2154 | -2.24 | 0.0291 | 0.0333 |
| chr11 | 35655087 | 35668225 | 13139 | -4.39 | 0.0291 | 0.0333 |
| chr16 | 80506362 | 80552516 | 46155 | 0.61 | 0.0291 | 0.0333 |
| chr19 | 13585132 | 13613023 | 27892 | -3.51 | 0.0291 | 0.0333 |
| chr4 | 59611921 | 59612208 | 288 | -2.13 | 0.0292 | 0.0334 |
| chr3 | 50883168 | 50909332 | 26165 | -2.38 | 0.0292 | 0.0334 |
| chr4 | 143867401 | 143877226 | 9826 | -3.15 | 0.0293 | 0.0335 |
| chr22 | 49834044 | 49838668 | 4625 | -0.92 | 0.0293 | 0.0335 |
| chr18 | 77592894 | 77605743 | 12850 | -1.32 | 0.0293 | 0.0336 |
| chr5 | 155773618 | 155775817 | 2200 | -2.84 | 0.0293 | 0.0336 |
| chrX | 71955333 | 71956884 | 1552 | -1.41 | 0.0294 | 0.0336 |
| chr14 | 96407564 | 96417067 | 9504 | -1.08 | 0.0294 | 0.0336 |
| chrX | 101247928 | 101260552 | 12625 | 0.81 | 0.0294 | 0.0336 |
| chr14 | 49974670 | 50010117 | 35448 | -1.92 | 0.0294 | 0.0336 |
| chr11 | 104090165 | 104094725 | 4561 | 1.04 | 0.0294 | 0.0336 |
| chr3 | 159220585 | 159231177 | 10593 | -1.4 | 0.0294 | 0.0337 |
| chr20 | 55411346 | 55414398 | 3053 | 1.74 | 0.0295 | 0.0337 |
| chr2 | 68053665 | 68060397 | 6733 | 1.3 | 0.0295 | 0.0337 |
| chr3 | 30836376 | 30837754 | 1379 | -2.74 | 0.0295 | 0.0337 |
| chr21 | 27196871 | 27229894 | 33024 | -0.41 | 0.0295 | 0.0337 |
| chr9 | 21203653 | 21204659 | 1007 | -2.12 | 0.0295 | 0.0337 |
| chr6 | 78204652 | 78228169 | 23518 | 0.51 | 0.0295 | 0.0338 |
| chr8 | 88501096 | 88533916 | 32821 | 0.8 | 0.0295 | 0.0338 |
| chr20 | 52848558 | 52856661 | 8104 | -0.88 | 0.0295 | 0.0338 |
| chr10 | 120587086 | 120588117 | 1032 | -1.77 | 0.0296 | 0.0338 |

| | | | | | | |
|---|---|---|---|---|---|---|
| chr1 | 42453939 | 42455565 | 1627 | -2.91 | 0.0296 | 0.0339 |
| chr12 | 43832796 | 43835711 | 2916 | -2.8 | 0.0297 | 0.0339 |
| chrX | 65017841 | 65019037 | 1197 | 2.07 | 0.0297 | 0.0339 |
| chr2 | 230514397 | 230517858 | 3462 | -1.4 | 0.0297 | 0.0339 |
| chr1 | 220020722 | 220023986 | 3265 | -1.6 | 0.0297 | 0.034 |
| chr15 | 24661552 | 24677548 | 15997 | 1.36 | 0.0297 | 0.034 |
| chr11 | 51552529 | 51553333 | 805 | 1.21 | 0.0298 | 0.034 |
| chr3 | 113924446 | 113938296 | 13851 | 2.11 | 0.0298 | 0.034 |
| chr18 | 37380766 | 37412826 | 32061 | -0.6 | 0.0298 | 0.0341 |
| chr1 | 119898647 | 119900612 | 1966 | -1.97 | 0.0298 | 0.0341 |
| chr4 | 167005849 | 167007105 | 1257 | -2.58 | 0.0298 | 0.0341 |
| chr4 | 114277258 | 114278298 | 1041 | -1.86 | 0.0298 | 0.0341 |
| chr18 | 52008161 | 52012544 | 4384 | 1.99 | 0.0298 | 0.0341 |
| chr1 | 184962754 | 184979438 | 16685 | -3.37 | 0.0298 | 0.0341 |
| chr20 | 9693141 | 9786473 | 93333 | 0.43 | 0.0299 | 0.0342 |
| chr20 | 25091455 | 25119399 | 27945 | -4.37 | 0.0299 | 0.0342 |
| chr7 | 130892269 | 130903329 | 11061 | -4.37 | 0.0299 | 0.0342 |
| chr9 | 111249798 | 111260347 | 10550 | -4.37 | 0.0299 | 0.0342 |
| chr9 | 94203406 | 94236627 | 33222 | -3.61 | 0.0299 | 0.0342 |
| chr20 | 23760450 | 23762746 | 2297 | -1 | 0.0299 | 0.0342 |
| chr4 | 164237116 | 164237970 | 855 | 2.61 | 0.03 | 0.0343 |
| chr11 | 32031297 | 32032894 | 1598 | -2.04 | 0.03 | 0.0343 |
| chrX | 113168108 | 113168685 | 578 | -2.4 | 0.03 | 0.0343 |
| chr3 | 62377433 | 62599849 | 222417 | 5.32 | 0.03 | 0.0343 |
| chr12 | 67565394 | 67590336 | 24943 | 2.83 | 0.03 | 0.0343 |
| chr20 | 40281135 | 40283239 | 2105 | 2.92 | 0.0301 | 0.0344 |
| chr6 | 170243059 | 170243475 | 417 | 2.54 | 0.0301 | 0.0344 |
| chr11 | 134929878 | 134930314 | 437 | 2 | 0.0301 | 0.0344 |
| chr12 | 39366931 | 39393080 | 26150 | -1.29 | 0.0301 | 0.0344 |
| chr13 | 20859149 | 20861665 | 2517 | -2.89 | 0.0302 | 0.0345 |
| chr1 | 186988360 | 187007654 | 19295 | -3.4 | 0.0302 | 0.0345 |
| chr13 | 108534435 | 108791350 | 256916 | 5.35 | 0.0302 | 0.0345 |
| chr3 | 67940070 | 67942224 | 2155 | -1.25 | 0.0302 | 0.0345 |
| chr22 | 37116164 | 37117081 | 918 | 1.93 | 0.0303 | 0.0346 |
| chr3 | 75429173 | 75429788 | 616 | -1.3 | 0.0303 | 0.0346 |
| chr1 | 67783348 | 67785724 | 2377 | 0.87 | 0.0304 | 0.0347 |
| chr9 | 92773852 | 92785297 | 11446 | -3.03 | 0.0304 | 0.0347 |
| chr7 | 32089372 | 32090301 | 930 | -2.26 | 0.0304 | 0.0347 |
| chr15 | 65246027 | 65249632 | 3606 | 0.63 | 0.0304 | 0.0347 |
| chr6 | 164077632 | 164081674 | 4043 | -1.61 | 0.0305 | 0.0348 |
| chr12 | 41506135 | 41508419 | 2285 | 1 | 0.0305 | 0.0348 |
| chr9 | 94365752 | 94437299 | 71548 | -3.61 | 0.0306 | 0.0349 |
| chr2 | 209600403 | 209602888 | 2486 | 2.02 | 0.0306 | 0.0349 |
| chr6 | 97481289 | 97482968 | 1680 | -2.97 | 0.0306 | 0.0349 |
| chr10 | 85978406 | 85992115 | 13710 | -0.73 | 0.0306 | 0.0349 |
| chr3 | 25432683 | 25436232 | 3550 | -1.17 | 0.0306 | 0.035 |
| chr2 | 53270310 | 53272857 | 2548 | -2.74 | 0.0307 | 0.035 |
| chr11 | 113982098 | 114025456 | 43359 | 0.83 | 0.0307 | 0.035 |
| chr4 | 146212843 | 146213393 | 551 | -2.59 | 0.0307 | 0.035 |
| chr3 | 155938874 | 155940850 | 1977 | -1.76 | 0.0307 | 0.0351 |
| chrX | 72975507 | 72976923 | 1417 | -1.32 | 0.0307 | 0.0351 |
| chr3 | 46067334 | 46119231 | 51898 | -0.93 | 0.0308 | 0.0352 |
| chr9 | 89266143 | 89394821 | 128679 | 0.26 | 0.0308 | 0.0352 |
| chr4 | 136240721 | 136243812 | 3092 | -2.96 | 0.0308 | 0.0352 |
| chr2 | 132181495 | 132191100 | 9606 | -1.61 | 0.0308 | 0.0352 |
| chr7 | 77122348 | 77125321 | 2974 | -1.54 | 0.0309 | 0.0352 |
| chr1 | 112035066 | 112039728 | 4663 | -3.21 | 0.0309 | 0.0352 |
| chr6 | 75557352 | 75560915 | 3564 | 1.54 | 0.0309 | 0.0352 |
| chr4 | 79916952 | 79919119 | 2168 | -2.74 | 0.0309 | 0.0353 |
| chr2 | 152767143 | 152768014 | 872 | -1.29 | 0.0309 | 0.0353 |
| chr20 | 42403199 | 42443328 | 40130 | -4.93 | 0.0309 | 0.0353 |

| | | | | | | |
|---|---|---|---|---|---|---|
| chr12 | 17927199 | 17929786 | 2588 | 0.76 | 0.0309 | 0.0353 |
| chr8 | 25844794 | 25893181 | 48388 | -1.25 | 0.0309 | 0.0353 |
| chrX | 105796419 | 105798271 | 1853 | -0.93 | 0.031 | 0.0353 |
| chr4 | 158132386 | 158134090 | 1705 | 1.09 | 0.031 | 0.0353 |
| chr1 | 222431397 | 222442610 | 11214 | 3.32 | 0.031 | 0.0353 |
| chr2 | 34720862 | 34721671 | 810 | -2.45 | 0.0311 | 0.0354 |
| chrX | 102402009 | 102402570 | 562 | -2.64 | 0.0311 | 0.0354 |
| chr13 | 97481971 | 97485650 | 3680 | 2.84 | 0.0311 | 0.0355 |
| chr1 | 86443546 | 86444884 | 1339 | -2.67 | 0.0312 | 0.0356 |
| chr5 | 10160483 | 10161100 | 618 | 2.6 | 0.0313 | 0.0357 |
| chr5 | 41288963 | 41292196 | 3234 | -3.09 | 0.0313 | 0.0357 |
| chr13 | 29170384 | 29174809 | 4426 | -0.87 | 0.0313 | 0.0357 |
| chr9 | 81892577 | 81896606 | 4030 | -2.82 | 0.0314 | 0.0358 |
| chr9 | 109777539 | 109798707 | 21169 | -0.62 | 0.0314 | 0.0359 |
| chrX | 141189992 | 141195054 | 5063 | -1.03 | 0.0314 | 0.0359 |
| chr4 | 107965795 | 107968856 | 3062 | -1.12 | 0.0314 | 0.0359 |
| chr20 | 45212221 | 45212923 | 703 | 2.04 | 0.0315 | 0.0359 |
| chr10 | 65539579 | 65599914 | 60336 | -1.47 | 0.0315 | 0.036 |
| chr1 | 149160579 | 149177748 | 17170 | -0.74 | 0.0315 | 0.036 |
| chr4 | 47591535 | 47605229 | 13695 | -4.31 | 0.0316 | 0.036 |
| chr8 | 141190630 | 141218124 | 27495 | -4.31 | 0.0316 | 0.036 |
| chr4 | 8669560 | 8681492 | 11933 | -2.97 | 0.0316 | 0.0361 |
| chr2 | 64637547 | 64640782 | 3236 | 0.81 | 0.0316 | 0.0361 |
| chr10 | 82897091 | 82900118 | 3028 | -1.23 | 0.0316 | 0.0361 |
| chr7 | 7383167 | 7383901 | 735 | -2.51 | 0.0317 | 0.0361 |
| chr2 | 159078899 | 159088289 | 9391 | -0.8 | 0.0317 | 0.0361 |
| chr11 | 6296314 | 6299261 | 2948 | -2.52 | 0.0317 | 0.0361 |
| chr10 | 92708596 | 92709064 | 469 | -2.38 | 0.0317 | 0.0362 |
| chr9 | 136155927 | 136157741 | 1815 | 2.86 | 0.0317 | 0.0362 |
| chr7 | 103642733 | 103740165 | 97433 | 5.01 | 0.0317 | 0.0362 |
| chr3 | 153630753 | 153631893 | 1141 | 1.14 | 0.0318 | 0.0362 |
| chr1 | 182460755 | 182499079 | 38325 | -0.51 | 0.0318 | 0.0363 |
| chr20 | 23929670 | 23933221 | 3552 | 1.28 | 0.0319 | 0.0363 |
| chr5 | 55997972 | 55998754 | 783 | 1.08 | 0.0319 | 0.0363 |
| chr1 | 87025565 | 87026772 | 1208 | 1.49 | 0.0319 | 0.0363 |
| chr3 | 73163790 | 73165582 | 1793 | 1.88 | 0.0319 | 0.0363 |
| chrX | 56908624 | 56916713 | 8090 | 1.11 | 0.0319 | 0.0364 |
| chr4 | 33456950 | 33458169 | 1220 | -1.31 | 0.0319 | 0.0364 |
| chrX | 140738430 | 140739462 | 1033 | -2.54 | 0.032 | 0.0364 |
| chr18 | 53111124 | 53114048 | 2925 | -3.06 | 0.032 | 0.0365 |
| chr11 | 104453425 | 104456951 | 3527 | 0.87 | 0.032 | 0.0365 |
| chr9 | 111531512 | 111533493 | 1982 | -2.93 | 0.032 | 0.0365 |
| chr16 | 21244125 | 21244582 | 458 | -2.24 | 0.032 | 0.0365 |
| chr2 | 222039409 | 222044887 | 5479 | -3.15 | 0.032 | 0.0365 |
| chr7 | 132104782 | 132106310 | 1529 | -2.88 | 0.0321 | 0.0366 |
| chr5 | 27036231 | 27260281 | 224051 | 0.67 | 0.0321 | 0.0366 |
| chr1 | 222459349 | 222483141 | 23793 | 0.95 | 0.0322 | 0.0367 |
| chr11 | 116118327 | 116119092 | 766 | 2.72 | 0.0322 | 0.0367 |
| chr1 | 147035674 | 147037042 | 1369 | 1.73 | 0.0322 | 0.0367 |
| chr7 | 95511093 | 95515105 | 4013 | 0.91 | 0.0322 | 0.0367 |
| chr2 | 231135825 | 231144921 | 9097 | -3.3 | 0.0323 | 0.0368 |
| chr11 | 45784515 | 45788000 | 3486 | -1.33 | 0.0323 | 0.0368 |
| chr2 | 153897343 | 153898583 | 1241 | 1.21 | 0.0323 | 0.0368 |
| chr7 | 93051462 | 93055285 | 3824 | 3.18 | 0.0323 | 0.0368 |
| chr6 | 168919226 | 168923326 | 4101 | -2.96 | 0.0323 | 0.0368 |
| chr5 | 99786372 | 99787330 | 959 | -1.17 | 0.0323 | 0.0368 |
| chr16 | 81936370 | 81937162 | 793 | -2.5 | 0.0324 | 0.0369 |
| chr5 | 158244038 | 158245421 | 1384 | -2.06 | 0.0324 | 0.037 |
| chr18 | 8570657 | 8591427 | 20771 | -2.99 | 0.0324 | 0.037 |
| chr5 | 115943300 | 115943937 | 638 | -2.44 | 0.0325 | 0.037 |
| chr10 | 34450931 | 34460207 | 9277 | -4.28 | 0.0325 | 0.037 |

| | | | | | | |
|---|---|---|---|---|---|---|
| chr9 | 127407266 | 127419242 | 11977 | -4.28 | 0.0325 | 0.037 |
| chr5 | 148095929 | 148096530 | 602 | 2.26 | 0.0325 | 0.037 |
| chr10 | 6100056 | 6102146 | 2091 | 1.12 | 0.0325 | 0.0371 |
| chr7 | 18195498 | 18197919 | 2422 | 1.51 | 0.0326 | 0.0371 |
| chr5 | 94426405 | 94434262 | 7858 | -2.84 | 0.0326 | 0.0371 |
| chr3 | 157642517 | 157682164 | 39648 | 0.86 | 0.0327 | 0.0372 |
| chr6 | 119329303 | 119330985 | 1683 | -2.84 | 0.0327 | 0.0372 |
| chr14 | 59171124 | 59177997 | 6874 | 3.23 | 0.0327 | 0.0373 |
| chr2 | 186481667 | 186495566 | 13900 | -1 | 0.0327 | 0.0373 |
| chr10 | 130048587 | 130050297 | 1711 | -1.25 | 0.0327 | 0.0373 |
| chr3 | 124162863 | 124176769 | 13907 | 0.71 | 0.0327 | 0.0373 |
| chr5 | 86500078 | 86501194 | 1117 | 2.02 | 0.0328 | 0.0373 |
| chr12 | 43848233 | 43864886 | 16654 | 0.64 | 0.0328 | 0.0373 |
| chr7 | 154370894 | 154732748 | 361855 | 0.35 | 0.0328 | 0.0373 |
| chr8 | 82498610 | 82499999 | 1390 | -1.05 | 0.0328 | 0.0374 |
| chr9 | 101154973 | 101156571 | 1599 | -2.74 | 0.0329 | 0.0374 |
| chr13 | 104616209 | 104616762 | 554 | -1.55 | 0.0329 | 0.0374 |
| chr7 | 31654812 | 31659339 | 4528 | 1.2 | 0.0329 | 0.0375 |
| chr12 | 80854216 | 80855964 | 1749 | 1.41 | 0.033 | 0.0375 |
| chr11 | 27562632 | 27564165 | 1534 | -1.16 | 0.033 | 0.0376 |
| chr12 | 22391396 | 22392934 | 1539 | -3.07 | 0.033 | 0.0376 |
| chr4 | 33769916 | 33815258 | 45343 | -0.85 | 0.033 | 0.0376 |
| chr1 | 245653803 | 245691968 | 38166 | -0.84 | 0.033 | 0.0376 |
| chr2 | 8456380 | 8464112 | 7733 | -2.99 | 0.0331 | 0.0377 |
| chr2 | 200032813 | 200036701 | 3889 | -1.07 | 0.0331 | 0.0377 |
| chr3 | 115201572 | 115226751 | 25180 | -1.41 | 0.0331 | 0.0377 |
| chr2 | 143959770 | 143960274 | 505 | -1.24 | 0.0331 | 0.0377 |
| chr12 | 9277646 | 9298686 | 21041 | -1.42 | 0.0332 | 0.0377 |
| chr10 | 60904178 | 60906614 | 2437 | -1.52 | 0.0332 | 0.0378 |
| chr7 | 141853728 | 141857824 | 4097 | -2.11 | 0.0332 | 0.0378 |
| chr3 | 116522106 | 116530875 | 8770 | -3.13 | 0.0332 | 0.0378 |
| chr5 | 107823668 | 107838046 | 14379 | -0.64 | 0.0333 | 0.0379 |
| chr6 | 143697555 | 143704443 | 6889 | -1.88 | 0.0334 | 0.038 |
| chr8 | 127315477 | 127382687 | 67211 | -0.78 | 0.0334 | 0.038 |
| chr6 | 130571760 | 130578572 | 6813 | -0.82 | 0.0334 | 0.038 |
| chr4 | 1414386 | 1419498 | 5113 | -1.93 | 0.0334 | 0.038 |
| chr7 | 86695632 | 86698711 | 3080 | 2.94 | 0.0334 | 0.038 |
| chr12 | 46534119 | 46535144 | 1026 | 1.11 | 0.0334 | 0.038 |
| chr3 | 34402228 | 34420172 | 17945 | 0.86 | 0.0334 | 0.038 |
| chr3 | 64915310 | 64916808 | 1499 | 2.73 | 0.0335 | 0.0381 |
| chr22 | 23210887 | 23212537 | 1651 | 0.99 | 0.0335 | 0.0381 |
| chr1 | 146611903 | 146615171 | 3269 | -1.52 | 0.0335 | 0.0381 |
| chr8 | 1444238 | 1477856 | 33619 | -0.88 | 0.0335 | 0.0381 |
| chr5 | 90067578 | 90069056 | 1479 | 1.02 | 0.0336 | 0.0382 |
| chr1 | 18445535 | 18445913 | 379 | -1.41 | 0.0336 | 0.0382 |
| chr12 | 119566253 | 119582537 | 16285 | 3.4 | 0.0336 | 0.0382 |
| chrX | 118431133 | 118434133 | 3001 | -2.98 | 0.0336 | 0.0382 |
| chr1 | 210145489 | 210146590 | 1102 | -1.95 | 0.0337 | 0.0383 |
| chr1 | 37483252 | 37484275 | 1024 | -2.65 | 0.0337 | 0.0383 |
| chr7 | 142571485 | 142573643 | 2159 | -2.73 | 0.0338 | 0.0384 |
| chr8 | 531092 | 540993 | 9902 | -0.96 | 0.0338 | 0.0384 |
| chr9 | 87799978 | 87804123 | 4146 | 3.02 | 0.0338 | 0.0384 |
| chr8 | 113819161 | 113849115 | 29955 | -1.41 | 0.0338 | 0.0384 |
| chr9 | 121584303 | 121586925 | 2623 | -2.68 | 0.0338 | 0.0384 |
| chrX | 81285955 | 81288418 | 2464 | -2.69 | 0.0338 | 0.0384 |
| chr12 | 28978078 | 28979351 | 1274 | 2.23 | 0.0338 | 0.0385 |
| chr1 | 159191755 | 159248744 | 56990 | -0.59 | 0.0338 | 0.0385 |
| chr11 | 71085672 | 71105157 | 19486 | -0.6 | 0.0339 | 0.0385 |
| chr6 | 159742495 | 159762071 | 19577 | -0.51 | 0.0339 | 0.0386 |
| chr7 | 28460130 | 28460553 | 424 | -2.27 | 0.034 | 0.0387 |
| chr5 | 90987745 | 90993273 | 5529 | -2.59 | 0.034 | 0.0387 |

| | | | | | | |
|---|---|---|---|---|---|---|
| chr14 | 42971757 | 42988025 | 16269 | 3.45 | 0.034 | 0.0387 |
| chr12 | 63396041 | 63396415 | 375 | -2 | 0.034 | 0.0387 |
| chr8 | 140765753 | 140766826 | 1074 | 1.21 | 0.034 | 0.0387 |
| chr6 | 90670360 | 90673335 | 2976 | -1.14 | 0.034 | 0.0387 |
| chr14 | 70546492 | 70708714 | 162223 | 0.38 | 0.034 | 0.0387 |
| chr14 | 22947771 | 22967106 | 19336 | 1.63 | 0.0341 | 0.0388 |
| chr5 | 75296112 | 75316216 | 20105 | 0.87 | 0.0342 | 0.0389 |
| chr8 | 2116480 | 2178372 | 61893 | 0.67 | 0.0342 | 0.0389 |
| chr14 | 24203429 | 24261434 | 58006 | -0.81 | 0.0343 | 0.039 |
| chr2 | 222161478 | 222163518 | 2041 | -2.08 | 0.0343 | 0.039 |
| chrX | 62129695 | 62134347 | 4653 | -0.97 | 0.0343 | 0.039 |
| chr5 | 83423277 | 83424350 | 1074 | -1.99 | 0.0343 | 0.039 |
| chr10 | 89525661 | 89540191 | 14531 | -4.08 | 0.0344 | 0.0391 |
| chr16 | 2930007 | 2938791 | 8785 | -4.22 | 0.0344 | 0.0391 |
| chr2 | 135431126 | 135444956 | 13831 | -4.22 | 0.0344 | 0.0391 |
| chr2 | 231165685 | 231168266 | 2582 | -2.17 | 0.0345 | 0.0392 |
| chr5 | 123150878 | 123197854 | 46977 | -0.83 | 0.0345 | 0.0392 |
| chr8 | 54401242 | 54404048 | 2807 | -3.15 | 0.0345 | 0.0393 |
| chr3 | 173999951 | 174018285 | 18335 | 3.31 | 0.0346 | 0.0393 |
| chrX | 49507055 | 49509717 | 2663 | 0.86 | 0.0346 | 0.0393 |
| chr16 | 86088991 | 86092206 | 3216 | -0.87 | 0.0346 | 0.0393 |
| chr1 | 7665821 | 7666398 | 578 | -2.03 | 0.0346 | 0.0394 |
| chrX | 140794313 | 140795884 | 1572 | -2.05 | 0.0346 | 0.0394 |
| chr10 | 64393068 | 64487699 | 94632 | 0.41 | 0.0347 | 0.0395 |
| chr3 | 38135571 | 38136696 | 1126 | 2.74 | 0.0347 | 0.0395 |
| chr6 | 153234102 | 153240549 | 6448 | -3.14 | 0.0348 | 0.0395 |
| chr13 | 94153502 | 94197080 | 43579 | 3.5 | 0.0348 | 0.0396 |
| chr12 | 131019092 | 131062222 | 43131 | -0.58 | 0.0348 | 0.0396 |
| chr12 | 26900543 | 26935527 | 34985 | -3.53 | 0.0349 | 0.0397 |
| chr16 | 56194926 | 56219197 | 24272 | -3.42 | 0.0349 | 0.0397 |
| chr6 | 1248647 | 1250080 | 1434 | -1.26 | 0.0349 | 0.0397 |
| chr3 | 193173210 | 193173893 | 684 | -1.59 | 0.035 | 0.0397 |
| chr7 | 130425533 | 130433619 | 8087 | -0.88 | 0.035 | 0.0397 |
| chr10 | 6661878 | 6664787 | 2910 | 2.68 | 0.035 | 0.0397 |
| chr13 | 22935891 | 23042255 | 106365 | 0.33 | 0.035 | 0.0398 |
| chr7 | 15222010 | 15223565 | 1556 | 0.94 | 0.035 | 0.0398 |
| chr5 | 73319859 | 73320675 | 817 | -1.15 | 0.0351 | 0.0398 |
| chr11 | 27772639 | 27773244 | 606 | 1.39 | 0.0351 | 0.0398 |
| chr1 | 116755340 | 116756344 | 1005 | -1.44 | 0.0351 | 0.0399 |
| chrX | 68683634 | 68685261 | 1628 | -1.46 | 0.0351 | 0.0399 |
| chr5 | 50230683 | 50242911 | 12229 | 0.66 | 0.0352 | 0.0399 |
| chr1 | 147347747 | 147349061 | 1315 | 1.48 | 0.0352 | 0.04 |
| chr4 | 109642629 | 109650419 | 7791 | -3.26 | 0.0352 | 0.04 |
| chr4 | 136326623 | 136327675 | 1053 | -2.6 | 0.0353 | 0.0401 |
| chr5 | 81976313 | 81978856 | 2544 | -2.93 | 0.0353 | 0.0401 |
| chr2 | 17730363 | 17735699 | 5337 | 3.19 | 0.0354 | 0.0401 |
| chr10 | 117672177 | 117684944 | 12768 | -0.91 | 0.0354 | 0.0402 |
| chr8 | 39438131 | 39451968 | 13838 | -3.18 | 0.0354 | 0.0402 |
| chr6 | 12902535 | 12951080 | 48546 | -0.72 | 0.0355 | 0.0403 |
| chr7 | 117674712 | 117681808 | 7097 | 3.25 | 0.0355 | 0.0403 |
| chr12 | 67049338 | 67050123 | 786 | 1.28 | 0.0356 | 0.0404 |
| chr4 | 33359464 | 33369238 | 9775 | 1.33 | 0.0356 | 0.0404 |
| chrX | 36724340 | 36725073 | 734 | -1.21 | 0.0356 | 0.0404 |
| chr1 | 166992576 | 167010663 | 18088 | 0.86 | 0.0356 | 0.0404 |
| chr4 | 28587581 | 28588703 | 1123 | -1.41 | 0.0356 | 0.0405 |
| chr6 | 8858644 | 8859649 | 1006 | 2.69 | 0.0357 | 0.0405 |
| chr6 | 142120392 | 142121316 | 925 | -1.95 | 0.0357 | 0.0406 |
| chr15 | 61436899 | 61478652 | 41754 | -0.82 | 0.0357 | 0.0406 |
| chr11 | 122327425 | 122357221 | 29797 | -3.17 | 0.0357 | 0.0406 |
| chr13 | 40487465 | 40489554 | 2090 | 1.56 | 0.0358 | 0.0407 |
| chr2 | 217948949 | 217960368 | 11420 | -3.82 | 0.0359 | 0.0407 |

| | | | | | | |
|---|---|---|---|---|---|---|
| chr2 | 237775657 | 237778883 | 3227 | 2.89 | 0.0359 | 0.0408 |
| chr3 | 15981864 | 15986671 | 4808 | -1.47 | 0.0359 | 0.0408 |
| chr13 | 91719337 | 91725242 | 5906 | 2.14 | 0.036 | 0.0408 |
| chr4 | 88709898 | 88711324 | 1427 | 0.95 | 0.0361 | 0.0409 |
| chr14 | 96167050 | 96234703 | 67654 | 0.59 | 0.0361 | 0.041 |
| chrX | 42677160 | 42688694 | 11535 | -2.08 | 0.0361 | 0.041 |
| chr8 | 61906175 | 61926571 | 20397 | -3.13 | 0.0361 | 0.041 |
| chrX | 23447630 | 23459376 | 11747 | 3.23 | 0.0362 | 0.041 |
| chr12 | 97187547 | 97213978 | 26432 | -3.33 | 0.0362 | 0.0411 |
| chr13 | 76563055 | 76569198 | 6144 | -4.02 | 0.0362 | 0.0411 |
| chrX | 113572749 | 113576853 | 4105 | 0.8 | 0.0362 | 0.0411 |
| chr13 | 49212245 | 49214974 | 2730 | 1.19 | 0.0362 | 0.0411 |
| chr5 | 90041048 | 90042868 | 1821 | -2.81 | 0.0362 | 0.0411 |
| chr3 | 13911055 | 14144120 | 233066 | -4.4 | 0.0363 | 0.0412 |
| chr1 | 223338919 | 223340896 | 1978 | 1.79 | 0.0363 | 0.0412 |
| chr12 | 42290207 | 42292764 | 2558 | -3.07 | 0.0364 | 0.0413 |
| chr3 | 153108968 | 153113188 | 4221 | -3.13 | 0.0364 | 0.0413 |
| chr17 | 71648533 | 71704604 | 56072 | -4.76 | 0.0364 | 0.0413 |
| chr3 | 11989864 | 12060547 | 70684 | -0.6 | 0.0364 | 0.0413 |
| chr1 | 214234924 | 214293443 | 58520 | -3.41 | 0.0364 | 0.0413 |
| chr12 | 48947910 | 49023895 | 75986 | -0.48 | 0.0364 | 0.0413 |
| chr5 | 180315615 | 180317396 | 1782 | -1.19 | 0.0366 | 0.0415 |
| chr4 | 56005760 | 56009655 | 3896 | 1.01 | 0.0366 | 0.0415 |
| chr7 | 24554243 | 24565578 | 11336 | -0.56 | 0.0366 | 0.0415 |
| chr6 | 45916212 | 45917453 | 1242 | 1.15 | 0.0367 | 0.0416 |
| chr2 | 40486517 | 40490792 | 4276 | -0.79 | 0.0367 | 0.0417 |
| chr11 | 109741684 | 109798068 | 56385 | -3.46 | 0.0368 | 0.0417 |
| chr3 | 66881735 | 66900342 | 18608 | -3.07 | 0.0368 | 0.0417 |
| chr8 | 130008647 | 130009714 | 1068 | -2.74 | 0.0368 | 0.0418 |
| chr12 | 107631324 | 107653212 | 21889 | 0.48 | 0.0368 | 0.0418 |
| chr11 | 6731629 | 6926105 | 194477 | 5.12 | 0.0368 | 0.0418 |
| chr7 | 29182026 | 29224257 | 42232 | -0.89 | 0.0368 | 0.0418 |
| chr6 | 97291995 | 97307249 | 15255 | -3.28 | 0.0369 | 0.0418 |
| chr4 | 165714044 | 165714719 | 676 | -1.85 | 0.0369 | 0.0418 |
| chr2 | 30976074 | 30978652 | 2579 | -2.53 | 0.0369 | 0.0419 |
| chr12 | 113021326 | 113077077 | 55752 | -2.69 | 0.037 | 0.0419 |
| chr11 | 119785031 | 119788887 | 3857 | -1.75 | 0.037 | 0.0419 |
| chr4 | 85319126 | 85321538 | 2413 | 2.99 | 0.037 | 0.042 |
| chr3 | 55222118 | 55223377 | 1260 | -1.37 | 0.037 | 0.042 |
| chr4 | 29350282 | 29350692 | 411 | -2.06 | 0.0371 | 0.042 |
| chr4 | 62598632 | 62601379 | 2748 | -1.15 | 0.0371 | 0.0421 |
| chr9 | 71139232 | 71308193 | 168962 | 5.08 | 0.0371 | 0.0421 |
| chr14 | 54608037 | 54610220 | 2184 | 0.81 | 0.0372 | 0.0421 |
| chrX | 56750023 | 56751357 | 1335 | -2.67 | 0.0372 | 0.0422 |
| chr14 | 60509443 | 60527492 | 18050 | -3.28 | 0.0373 | 0.0422 |
| chr12 | 77372946 | 77412209 | 39264 | -3.43 | 0.0373 | 0.0423 |
| chr1 | 171187292 | 171188210 | 919 | -2.45 | 0.0374 | 0.0424 |
| chr16 | 64101277 | 64157709 | 56433 | 0.71 | 0.0374 | 0.0424 |
| chr14 | 24301536 | 24327391 | 25856 | -0.45 | 0.0375 | 0.0425 |
| chrX | 80170220 | 80210830 | 40611 | 2.34 | 0.0375 | 0.0425 |
| chr1 | 106457637 | 106480116 | 22480 | 0.72 | 0.0376 | 0.0426 |
| chr4 | 118797890 | 118799851 | 1962 | 3.01 | 0.0376 | 0.0426 |
| chr3 | 161661515 | 161664840 | 3326 | -2.95 | 0.0376 | 0.0426 |
| chr11 | 120869408 | 120881179 | 11772 | -0.79 | 0.0376 | 0.0427 |
| chr1 | 217878199 | 217893468 | 15270 | -4.12 | 0.0377 | 0.0427 |
| chr4 | 122623103 | 122637623 | 14521 | -4.12 | 0.0377 | 0.0427 |
| chr7 | 101010968 | 101025719 | 14752 | -4.12 | 0.0377 | 0.0427 |
| chr8 | 118717427 | 118727040 | 9614 | 0.83 | 0.0378 | 0.0429 |
| chr4 | 72663707 | 72665209 | 1503 | 1.49 | 0.0379 | 0.0429 |
| chr7 | 120924291 | 120925988 | 1698 | -1.53 | 0.0379 | 0.0429 |
| chr3 | 51385715 | 51386231 | 517 | -2.31 | 0.0379 | 0.043 |

| | | | | | | |
|---|---|---|---|---|---|---|
| chr11 | 127896589 | 127914985 | 18397 | -3.18 | 0.0379 | 0.043 |
| chrX | 100775136 | 100780121 | 4986 | -2.33 | 0.0379 | 0.043 |
| chr12 | 86466601 | 86469674 | 3074 | -0.92 | 0.038 | 0.0431 |
| chr13 | 27477424 | 27509327 | 31904 | -1.02 | 0.0381 | 0.0431 |
| chr14 | 42898557 | 42928029 | 29473 | 2.35 | 0.0381 | 0.0431 |
| chr13 | 51239999 | 51241334 | 1336 | -2.6 | 0.0381 | 0.0432 |
| chr5 | 46097160 | 46265259 | 168100 | -5.02 | 0.0382 | 0.0433 |
| chr14 | 96429957 | 96439014 | 9058 | -3.14 | 0.0382 | 0.0433 |
| chr8 | 74797925 | 74799203 | 1279 | -2.76 | 0.0382 | 0.0433 |
| chr14 | 66307651 | 66334836 | 27186 | 3.34 | 0.0383 | 0.0434 |
| chr10 | 92832336 | 92833935 | 1600 | -2.61 | 0.0383 | 0.0434 |
| chr4 | 94857023 | 94858117 | 1095 | -2.62 | 0.0384 | 0.0434 |
| chr4 | 4648688 | 4660851 | 12164 | -2.9 | 0.0384 | 0.0434 |
| chr18 | 51961366 | 51977580 | 16215 | -0.73 | 0.0384 | 0.0435 |
| chr13 | 42968326 | 42970141 | 1816 | 1.06 | 0.0384 | 0.0435 |
| chr9 | 89449343 | 89641968 | 192626 | 0.38 | 0.0385 | 0.0436 |
| chr11 | 122108412 | 122109657 | 1246 | -1.78 | 0.0386 | 0.0437 |
| chr4 | 57456248 | 57456864 | 617 | -2.48 | 0.0386 | 0.0437 |
| chr9 | 16110644 | 16115793 | 5150 | -1.37 | 0.0386 | 0.0437 |
| chrX | 119944078 | 119953525 | 9448 | 0.7 | 0.0386 | 0.0437 |
| chr5 | 135447925 | 135448195 | 271 | -3.27 | 0.0387 | 0.0438 |
| chr4 | 107759782 | 107760886 | 1105 | -2.44 | 0.0387 | 0.0438 |
| chr4 | 109175132 | 109424488 | 249357 | 5.14 | 0.0387 | 0.0438 |
| chr10 | 82675646 | 82689413 | 13768 | 0.79 | 0.0388 | 0.0439 |
| chr2 | 195973861 | 195979875 | 6015 | -2.94 | 0.0388 | 0.0439 |
| chr7 | 35559041 | 35566371 | 7331 | -2.84 | 0.0388 | 0.0439 |
| chr7 | 133760291 | 133783645 | 23355 | -3.47 | 0.0388 | 0.0439 |
| chr3 | 86394566 | 86402370 | 7805 | 0.77 | 0.0389 | 0.044 |
| chr12 | 100502401 | 100526669 | 24269 | -4.09 | 0.0389 | 0.0441 |
| chr5 | 125774983 | 125788302 | 13320 | -4.09 | 0.0389 | 0.0441 |
| chr4 | 156442354 | 156443975 | 1622 | -1.7 | 0.039 | 0.0441 |
| chr6 | 8788374 | 8789214 | 841 | -2.39 | 0.039 | 0.0441 |
| chr18 | 44710432 | 44726748 | 16317 | -3.18 | 0.039 | 0.0441 |
| chr13 | 88704061 | 88705198 | 1138 | -1.45 | 0.039 | 0.0441 |
| chr21 | 39886807 | 39888872 | 2066 | -1.3 | 0.039 | 0.0442 |
| chr13 | 31649983 | 31650914 | 932 | 1.6 | 0.0391 | 0.0443 |
| chr8 | 38474334 | 38477037 | 2704 | -1.27 | 0.0392 | 0.0443 |
| chr4 | 76077329 | 76081499 | 4171 | -1.28 | 0.0392 | 0.0443 |
| chr18 | 28462030 | 28464642 | 2613 | 0.91 | 0.0392 | 0.0444 |
| chr4 | 188034731 | 188035663 | 933 | 2.54 | 0.0393 | 0.0444 |
| chr8 | 126575399 | 126576431 | 1033 | -2.63 | 0.0393 | 0.0445 |
| chrX | 105744258 | 105745049 | 792 | -1.12 | 0.0394 | 0.0446 |
| chr4 | 146984560 | 146987809 | 3250 | -1.58 | 0.0394 | 0.0446 |
| chr13 | 89157624 | 89164303 | 6680 | -2.97 | 0.0394 | 0.0446 |
| chr4 | 143741041 | 143742389 | 1349 | -2.66 | 0.0395 | 0.0447 |
| chr10 | 92573025 | 92575831 | 2807 | 2.94 | 0.0395 | 0.0447 |
| chr1 | 105891026 | 105891475 | 450 | 1.85 | 0.0395 | 0.0447 |
| chrX | 53751556 | 53780368 | 28813 | -3.18 | 0.0395 | 0.0447 |
| chr7 | 56840173 | 56841994 | 1822 | -2.74 | 0.0395 | 0.0447 |
| chr3 | 153783808 | 153785178 | 1371 | 2.64 | 0.0395 | 0.0447 |
| chr10 | 16906246 | 16930321 | 24076 | -1.06 | 0.0396 | 0.0448 |
| chrX | 118391659 | 118393607 | 1949 | -1.42 | 0.0396 | 0.0448 |
| chr16 | 56004146 | 56149231 | 145086 | 4.94 | 0.0398 | 0.045 |
| chr11 | 96252762 | 96254007 | 1246 | -1.93 | 0.0398 | 0.045 |
| chr6 | 75425961 | 75432356 | 6396 | -3.11 | 0.0398 | 0.045 |
| chr2 | 176184669 | 176191752 | 7084 | -1.91 | 0.0398 | 0.045 |
| chr3 | 162247664 | 162248449 | 786 | 1.6 | 0.0398 | 0.0451 |
| chr6 | 167894054 | 167895998 | 1945 | 1.54 | 0.0399 | 0.0451 |
| chr9 | 98551867 | 98567134 | 15268 | -2 | 0.0399 | 0.0451 |
| chr6 | 119567849 | 119616096 | 48248 | -0.57 | 0.0399 | 0.0452 |
| chr13 | 91989451 | 91999155 | 9705 | -1.07 | 0.04 | 0.0452 |

| | | | | | | |
|---|---|---|---|---|---|---|
| chr6 | 147408383 | 147411896 | 3514 | -3.04 | 0.04 | 0.0452 |
| chr11 | 21400039 | 21401099 | 1061 | 1.38 | 0.04 | 0.0453 |
| chr4 | 4895279 | 4937975 | 42697 | -0.7 | 0.0401 | 0.0454 |
| chr14 | 81705866 | 81710385 | 4520 | -3.1 | 0.0401 | 0.0454 |
| chr11 | 84849443 | 84917637 | 68195 | -4.67 | 0.0402 | 0.0454 |
| chr3 | 186402159 | 186409115 | 6957 | -2.04 | 0.0402 | 0.0455 |
| chr8 | 101118317 | 101133515 | 15199 | -3.23 | 0.0402 | 0.0455 |
| chr5 | 93764062 | 93765422 | 1361 | -1.78 | 0.0403 | 0.0455 |
| chr14 | 24959277 | 24976568 | 17292 | 0.73 | 0.0403 | 0.0456 |
| chr13 | 76712918 | 76716349 | 3432 | 1.09 | 0.0403 | 0.0456 |
| chr12 | 108547843 | 108583838 | 35996 | -2.88 | 0.0403 | 0.0456 |
| chr9 | 15800054 | 15817318 | 17265 | -3.27 | 0.0404 | 0.0456 |
| chr7 | 62162978 | 62361349 | 198372 | -5.08 | 0.0404 | 0.0456 |
| chr15 | 51973209 | 51998036 | 24828 | 0.98 | 0.0404 | 0.0457 |
| chr18 | 64209844 | 64213057 | 3214 | 2.86 | 0.0405 | 0.0458 |
| chr1 | 82999725 | 83001966 | 2242 | 1.25 | 0.0405 | 0.0458 |
| chrX | 25104107 | 25114037 | 9931 | -3.13 | 0.0406 | 0.0459 |
| chr9 | 121246621 | 121247825 | 1205 | 1.29 | 0.0406 | 0.0459 |
| chr6 | 119367075 | 119380416 | 13342 | -0.75 | 0.0406 | 0.0459 |
| chr3 | 142930312 | 143126155 | 195844 | 4.99 | 0.0406 | 0.0459 |
| chr4 | 77495948 | 77496340 | 393 | -2.1 | 0.0407 | 0.046 |
| chrX | 7822765 | 7854182 | 31418 | 3.47 | 0.0407 | 0.046 |
| chr10 | 45881099 | 45908510 | 27412 | -3.28 | 0.0408 | 0.0461 |
| chr8 | 102258637 | 102260206 | 1570 | -2.81 | 0.0408 | 0.0461 |
| chr18 | 6882951 | 6884652 | 1702 | -2.81 | 0.0408 | 0.0461 |
| chr5 | 126802655 | 126833519 | 30865 | -3.15 | 0.0408 | 0.0461 |
| chr18 | 74041389 | 74066696 | 25308 | -3.29 | 0.0408 | 0.0461 |
| chr11 | 49802391 | 49802813 | 423 | 1.14 | 0.041 | 0.0464 |
| chr8 | 16702729 | 16711451 | 8723 | 0.71 | 0.0411 | 0.0464 |
| chr3 | 191253269 | 191255021 | 1753 | -2.41 | 0.0412 | 0.0465 |
| chr10 | 125711004 | 125714073 | 3070 | -0.79 | 0.0412 | 0.0466 |
| chr4 | 164481770 | 164482868 | 1099 | -1.27 | 0.0412 | 0.0466 |
| chr4 | 28622426 | 28632168 | 9743 | 0.73 | 0.0412 | 0.0466 |
| chr4 | 104194458 | 104194911 | 454 | -1.51 | 0.0413 | 0.0466 |
| chr13 | 60887021 | 60887525 | 505 | 1.81 | 0.0413 | 0.0467 |
| chr1 | 108525169 | 108528942 | 3774 | 1.33 | 0.0415 | 0.0468 |
| chr5 | 94449539 | 94542508 | 92970 | 0.46 | 0.0416 | 0.0469 |
| chr8 | 57913317 | 57936879 | 23563 | -3.15 | 0.0416 | 0.0469 |
| chr3 | 152377596 | 152381822 | 4227 | -2.56 | 0.0416 | 0.047 |
| chr15 | 30279693 | 30293432 | 13740 | -4.02 | 0.0416 | 0.047 |
| chr16 | 87642644 | 87643472 | 829 | 1.13 | 0.0416 | 0.047 |
| chr4 | 22025261 | 22027874 | 2614 | 2.76 | 0.0416 | 0.047 |
| chr8 | 108788780 | 108795644 | 6865 | -3.87 | 0.0416 | 0.047 |
| chr4 | 112737700 | 112740830 | 3131 | -1.74 | 0.0416 | 0.047 |
| chr5 | 91571865 | 91580281 | 8417 | -3.17 | 0.0417 | 0.0471 |
| chrX | 109807979 | 109809631 | 1653 | -2.38 | 0.0417 | 0.0471 |
| chr10 | 69291784 | 69293351 | 1568 | 1.17 | 0.0418 | 0.0472 |
| chr13 | 97597700 | 97598927 | 1228 | 1.28 | 0.0418 | 0.0472 |
| chr6 | 77521996 | 77539867 | 17872 | -3.23 | 0.0418 | 0.0472 |
| chr12 | 74288694 | 74290535 | 1842 | -2.63 | 0.0419 | 0.0473 |
| chrX | 132113978 | 132114520 | 543 | -1.03 | 0.0419 | 0.0473 |
| chr1 | 12697356 | 12766127 | 68772 | -4.61 | 0.0419 | 0.0473 |
| chr10 | 127959049 | 127960733 | 1685 | 2.71 | 0.042 | 0.0474 |
| chr1 | 56460826 | 56461495 | 670 | -1.19 | 0.042 | 0.0474 |
| chr16 | 50887171 | 51020966 | 133796 | -0.61 | 0.042 | 0.0474 |
| chr2 | 100582321 | 100612330 | 30010 | 0.48 | 0.042 | 0.0474 |
| chrX | 49431768 | 49437326 | 5559 | -2.8 | 0.042 | 0.0475 |
| chr8 | 91801746 | 91813679 | 11934 | -1.95 | 0.042 | 0.0475 |
| chr7 | 39940054 | 39941793 | 1740 | -2.73 | 0.0421 | 0.0475 |
| chr5 | 16638632 | 16639682 | 1051 | -1.19 | 0.0421 | 0.0475 |
| chr16 | 85938466 | 85940377 | 1912 | -2.92 | 0.0421 | 0.0475 |

| | | | | | | |
|---|---|---|---|---|---|---|
| chr11 | 134459539 | 134476369 | 16831 | -1.11 | 0.0422 | 0.0476 |
| chr20 | 2422207 | 2430719 | 8513 | -3.23 | 0.0422 | 0.0476 |
| chrX | 52887006 | 52887917 | 912 | -2.59 | 0.0422 | 0.0476 |
| chr8 | 133897865 | 133899439 | 1575 | -1.06 | 0.0422 | 0.0477 |
| chr4 | 35327622 | 35339522 | 11901 | 1.01 | 0.0422 | 0.0477 |
| chr8 | 102682603 | 102690319 | 7717 | -2.68 | 0.0423 | 0.0477 |
| chr16 | 27069400 | 27071187 | 1788 | 2.67 | 0.0423 | 0.0477 |
| chr3 | 100386583 | 100397419 | 10837 | -2.19 | 0.0423 | 0.0478 |
| chr2 | 65890662 | 65893718 | 3057 | 1.29 | 0.0424 | 0.0478 |
| chr12 | 124575986 | 124576841 | 856 | -2.22 | 0.0425 | 0.0479 |
| chr1 | 56294921 | 56304171 | 9251 | -2.82 | 0.0425 | 0.048 |
| chr7 | 154813551 | 154815685 | 2135 | 2.81 | 0.0425 | 0.048 |
| chr3 | 83629070 | 83685267 | 56198 | -1.95 | 0.0426 | 0.0481 |
| chr6 | 46411863 | 46430939 | 19077 | 0.93 | 0.0426 | 0.0481 |
| chr3 | 67107468 | 67111226 | 3759 | -0.73 | 0.0426 | 0.0481 |
| chr22 | 26292905 | 26313771 | 20867 | 0.66 | 0.0427 | 0.0481 |
| chr4 | 105913735 | 105914693 | 959 | -2.35 | 0.0427 | 0.0482 |
| chr1 | 157180579 | 157253278 | 72700 | 0.68 | 0.0428 | 0.0483 |
| chr3 | 162090354 | 162117709 | 27356 | -3.13 | 0.0428 | 0.0483 |
| chr4 | 145710472 | 145713045 | 2574 | -1.87 | 0.0429 | 0.0484 |
| chr15 | 51645205 | 51645707 | 503 | -2.35 | 0.0429 | 0.0484 |
| chr5 | 60581662 | 60590351 | 8690 | -3.84 | 0.0429 | 0.0484 |
| chr18 | 56896152 | 56924139 | 27988 | -3.11 | 0.0429 | 0.0484 |
| chr2 | 71419695 | 71503497 | 83803 | -4.56 | 0.0429 | 0.0484 |
| chr1 | 171384768 | 171399574 | 14807 | -3.98 | 0.043 | 0.0485 |
| chr2 | 237093027 | 237115321 | 22295 | -3.98 | 0.043 | 0.0485 |
| chr4 | 95420666 | 95434904 | 14239 | -3.98 | 0.043 | 0.0485 |
| chrX | 12573835 | 12575180 | 1346 | 1.49 | 0.0431 | 0.0486 |
| chr4 | 76361737 | 76367521 | 5785 | -2.16 | 0.0431 | 0.0486 |
| chr13 | 104712197 | 104715289 | 3093 | -1.41 | 0.0431 | 0.0487 |
| chr8 | 87648887 | 87649827 | 941 | -1.85 | 0.0432 | 0.0487 |
| chr14 | 56963528 | 57009558 | 46031 | 0.52 | 0.0432 | 0.0487 |
| chr1 | 232066724 | 232069367 | 2644 | -3.06 | 0.0433 | 0.0488 |
| chr19 | 35854675 | 35882375 | 27701 | -4.71 | 0.0435 | 0.049 |
| chr5 | 83390749 | 83394918 | 4170 | -2.92 | 0.0435 | 0.0491 |
| chr2 | 171208386 | 171215107 | 6722 | -2.93 | 0.0435 | 0.0491 |
| chr2 | 53753835 | 53779710 | 25876 | -3.14 | 0.0435 | 0.0491 |
| chr15 | 38226696 | 38240043 | 13348 | -0.71 | 0.0435 | 0.0491 |
| chr5 | 63527523 | 63896239 | 368717 | 5.3 | 0.0436 | 0.0491 |
| chr3 | 70103130 | 70106608 | 3479 | 0.94 | 0.0436 | 0.0492 |
| chr8 | 119542324 | 119543375 | 1052 | -0.88 | 0.0437 | 0.0492 |
| chr2 | 222120871 | 222136200 | 15330 | -0.75 | 0.0437 | 0.0493 |
| chr4 | 19953380 | 19954088 | 709 | 2.53 | 0.0437 | 0.0493 |
| chr20 | 4236942 | 4239678 | 2737 | 3 | 0.0438 | 0.0494 |
| chr1 | 226625255 | 226634413 | 9159 | -0.92 | 0.0439 | 0.0495 |
| chr15 | 34767370 | 34776985 | 9616 | 1.18 | 0.0439 | 0.0495 |
| chr18 | 27813048 | 27843348 | 30301 | 0.6 | 0.0439 | 0.0495 |
| chr4 | 123563757 | 123564320 | 564 | 2.44 | 0.0439 | 0.0495 |
| chr16 | 77630600 | 77798111 | 167512 | -0.79 | 0.0439 | 0.0495 |
| chrX | 11841731 | 11842711 | 981 | 1.53 | 0.0439 | 0.0495 |
| chr6 | 46512299 | 46513782 | 1484 | -1.09 | 0.0439 | 0.0495 |
| chr12 | 55356895 | 55358635 | 1741 | 1.28 | 0.044 | 0.0495 |
| chr8 | 31520911 | 31523194 | 2284 | -0.97 | 0.044 | 0.0495 |
| chr7 | 123773996 | 123775201 | 1206 | -1.04 | 0.044 | 0.0496 |
| chr11 | 48419004 | 48419326 | 323 | 2.28 | 0.044 | 0.0496 |
| chr1 | 226692809 | 226717016 | 24208 | -3.07 | 0.0441 | 0.0496 |
| chr5 | 59511987 | 59513209 | 1223 | -1.56 | 0.0441 | 0.0497 |
| chr7 | 19052198 | 19058411 | 6214 | -1.88 | 0.0441 | 0.0497 |
| chr10 | 129657697 | 129678337 | 20641 | -1.95 | 0.0442 | 0.0498 |
| chr3 | 163896592 | 163935582 | 38991 | -3.17 | 0.0442 | 0.0498 |
| chr15 | 58016591 | 58029760 | 13170 | -3.81 | 0.0442 | 0.0498 |

| | | | | | | |
|---|---|---|---|---|---|---|
| chr11 | 62290644 | 62300772 | 10129 | -2.62 | 0.0442 | 0.0498 |
| chr11 | 13103630 | 13105935 | 2306 | 1.08 | 0.0442 | 0.0498 |
| chr3 | 113920800 | 113923011 | 2212 | -2.72 | 0.0442 | 0.0498 |
| chr1 | 104880772 | 104905334 | 24563 | 0.82 | 0.0443 | 0.05 |
| chr5 | 86379910 | 86380386 | 477 | -2.28 | 0.0444 | 0.05 |
| chr4 | 34696996 | 34734073 | 37078 | -0.97 | 0.0444 | 0.05 |
| chr5 | 79867363 | 79873132 | 5770 | -1.06 | 0.0445 | 0.0501 |
| chr5 | 77596522 | 77611186 | 14665 | -3.95 | 0.0445 | 0.0501 |
| chr6 | 1495843 | 1510478 | 14636 | -3.95 | 0.0445 | 0.0501 |
| chr10 | 119340143 | 119367717 | 27575 | -0.42 | 0.0445 | 0.0501 |
| chr20 | 9802144 | 9807660 | 5517 | 1.13 | 0.0445 | 0.0502 |
| chr7 | 105645848 | 105649126 | 3279 | -2.78 | 0.0445 | 0.0502 |
| chr11 | 40600047 | 40607014 | 6968 | -0.89 | 0.0446 | 0.0502 |
| chr10 | 19635156 | 19636150 | 995 | 1.97 | 0.0446 | 0.0502 |
| chr14 | 76830997 | 76831769 | 773 | -2.36 | 0.0446 | 0.0502 |
| chr3 | 24791595 | 24792835 | 1241 | 1.05 | 0.0446 | 0.0503 |
| chr13 | 74777189 | 74778944 | 1756 | -1.21 | 0.0446 | 0.0503 |
| chr10 | 26865828 | 26866378 | 551 | 1.15 | 0.0448 | 0.0504 |
| chr6 | 159941381 | 159943440 | 2060 | 1.42 | 0.045 | 0.0506 |
| chr10 | 114155021 | 114155845 | 825 | -1.54 | 0.045 | 0.0507 |
| chrX | 131700484 | 131703184 | 2701 | 2.74 | 0.045 | 0.0507 |
| chr2 | 226248689 | 226262152 | 13464 | -0.84 | 0.0451 | 0.0507 |
| chr4 | 37477972 | 37479421 | 1450 | -1.08 | 0.0451 | 0.0508 |
| chr3 | 155966527 | 155967886 | 1360 | 2.7 | 0.0451 | 0.0508 |
| chr6 | 162742149 | 162756145 | 13997 | -2.99 | 0.0452 | 0.0508 |
| chr6 | 52112491 | 52114725 | 2235 | 1.18 | 0.0452 | 0.0509 |
| chr12 | 97059442 | 97061064 | 1623 | -0.89 | 0.0452 | 0.0509 |
| chr9 | 121456937 | 121461193 | 4257 | -2.71 | 0.0452 | 0.0509 |
| chr5 | 75389979 | 75391768 | 1790 | -1.69 | 0.0452 | 0.0509 |
| chr21 | 10989817 | 11000565 | 10749 | 0.75 | 0.0453 | 0.051 |
| chr3 | 71944399 | 72026460 | 82062 | -0.52 | 0.0453 | 0.051 |
| chr15 | 95995554 | 96009729 | 14176 | -1.76 | 0.0453 | 0.051 |
| chr2 | 41908868 | 41929764 | 20897 | -3.29 | 0.0453 | 0.051 |
| chr1 | 181181893 | 181200711 | 18819 | -1.51 | 0.0454 | 0.0511 |
| chr2 | 37027923 | 37030702 | 2780 | -1.58 | 0.0454 | 0.0511 |
| chr10 | 45627823 | 45628748 | 926 | 1.17 | 0.0454 | 0.0511 |
| chrX | 57746957 | 57748531 | 1575 | -1.19 | 0.0454 | 0.0512 |
| chr14 | 90092510 | 90134276 | 41767 | -0.37 | 0.0455 | 0.0512 |
| chr2 | 230504070 | 230505294 | 1225 | 1.13 | 0.0455 | 0.0512 |
| chr4 | 135658855 | 135673225 | 14371 | 1.24 | 0.0456 | 0.0513 |
| chr9 | 35300082 | 35305538 | 5457 | 0.7 | 0.0456 | 0.0513 |
| chr7 | 155128491 | 155129459 | 969 | -2.06 | 0.0456 | 0.0514 |
| chr4 | 42622988 | 42633208 | 10221 | 1.58 | 0.0457 | 0.0514 |
| chr6 | 63786155 | 63786923 | 769 | 1.51 | 0.0457 | 0.0514 |
| chr20 | 37358002 | 37365183 | 7182 | -2.74 | 0.0458 | 0.0515 |
| chr9 | 401184 | 402643 | 1460 | 2.02 | 0.0458 | 0.0515 |
| chr6 | 142255097 | 142276177 | 21081 | -0.64 | 0.0459 | 0.0516 |
| chr11 | 109266013 | 109268540 | 2528 | 2.45 | 0.0459 | 0.0516 |
| chr2 | 213192113 | 213193074 | 962 | 1.26 | 0.0459 | 0.0516 |
| chr2 | 186569919 | 186571101 | 1183 | 1.21 | 0.0459 | 0.0516 |
| chr1 | 18352785 | 18356168 | 3384 | 2.98 | 0.0459 | 0.0516 |
| chr20 | 23842204 | 23895092 | 52889 | -0.53 | 0.0459 | 0.0517 |
| chr2 | 122058345 | 122063889 | 5545 | -1.47 | 0.046 | 0.0517 |
| chr4 | 21055050 | 21057886 | 2837 | 2.75 | 0.046 | 0.0518 |
| chr12 | 96276613 | 96328376 | 51764 | -3.39 | 0.046 | 0.0518 |
| chr3 | 61574533 | 61589487 | 14955 | -3.91 | 0.0461 | 0.0519 |
| chr5 | 10667736 | 10688063 | 20328 | -3.91 | 0.0461 | 0.0519 |
| chr9 | 5507364 | 5521239 | 13876 | -3.91 | 0.0461 | 0.0519 |
| chr7 | 20832935 | 20847104 | 14170 | -3.14 | 0.0461 | 0.0519 |
| chr9 | 29723778 | 29734226 | 10449 | 0.75 | 0.0462 | 0.0519 |
| chr11 | 85131917 | 85148613 | 16697 | -0.65 | 0.0462 | 0.052 |

| | | | | | | |
|---|---|---|---|---|---|---|
| chr8 | 107571840 | 107573020 | 1181 | 1.86 | 0.0462 | 0.052 |
| chr19 | 15157445 | 15176908 | 19464 | -1.47 | 0.0464 | 0.0521 |
| chr4 | 42109725 | 42168437 | 58713 | -3.38 | 0.0464 | 0.0521 |
| chr3 | 67133037 | 67148247 | 15211 | 0.97 | 0.0464 | 0.0521 |
| chr10 | 79922518 | 79923305 | 788 | -2.52 | 0.0464 | 0.0522 |
| chr2 | 122539770 | 122637489 | 97720 | -4.53 | 0.0464 | 0.0522 |
| chr14 | 21344273 | 21368429 | 24157 | -0.66 | 0.0464 | 0.0522 |
| chr6 | 122909420 | 122909971 | 552 | -2.43 | 0.0465 | 0.0523 |
| chr2 | 157683362 | 157697153 | 13792 | 1.16 | 0.0465 | 0.0523 |
| chr2 | 209379577 | 209380613 | 1037 | -1.47 | 0.0465 | 0.0523 |
| chrX | 77864343 | 77865896 | 1554 | -1.02 | 0.0465 | 0.0523 |
| chr11 | 134657582 | 134658657 | 1076 | -1.36 | 0.0465 | 0.0523 |
| chr5 | 58880688 | 58881846 | 1159 | 1.17 | 0.0465 | 0.0523 |
| chr3 | 108372729 | 108374669 | 1941 | -2.93 | 0.0465 | 0.0523 |
| chr3 | 61486263 | 61503275 | 17013 | -0.64 | 0.0465 | 0.0523 |
| chrX | 131385645 | 131398724 | 13080 | -2.99 | 0.0465 | 0.0523 |
| chr12 | 132052486 | 132055547 | 3062 | 1.04 | 0.0466 | 0.0524 |
| chr12 | 114410211 | 114415178 | 4968 | -1.94 | 0.0467 | 0.0524 |
| chr2 | 176246621 | 176253834 | 7214 | -0.83 | 0.0467 | 0.0525 |
| chr7 | 87575816 | 87576241 | 426 | -2.34 | 0.0467 | 0.0525 |
| chr21 | 40021612 | 40022616 | 1005 | 1.14 | 0.0467 | 0.0525 |
| chr7 | 103888624 | 104462695 | 574072 | 5.35 | 0.0468 | 0.0526 |
| chr12 | 101897849 | 101915542 | 17694 | -3.11 | 0.0468 | 0.0526 |
| chr15 | 36008444 | 36012541 | 4098 | -2.68 | 0.0469 | 0.0527 |
| chr11 | 134905993 | 134909406 | 3414 | 2.32 | 0.047 | 0.0528 |
| chr1 | 217548805 | 217564822 | 16018 | -3.16 | 0.047 | 0.0528 |
| chr11 | 20594568 | 20622572 | 28005 | 2.24 | 0.047 | 0.0529 |
| chr13 | 76660099 | 76662348 | 2250 | -2.79 | 0.047 | 0.0529 |
| chr2 | 237268954 | 237292350 | 23397 | -3.19 | 0.0471 | 0.0529 |
| chr12 | 41996587 | 42027196 | 30610 | 0.59 | 0.0471 | 0.0529 |
| chr14 | 97233817 | 97237655 | 3839 | -1.21 | 0.0472 | 0.053 |
| chr12 | 95838659 | 95852429 | 13771 | -2.82 | 0.0472 | 0.053 |
| chr4 | 126837911 | 126856736 | 18826 | 3.15 | 0.0472 | 0.0531 |
| chr21 | 18334986 | 18340397 | 5412 | 0.72 | 0.0472 | 0.0531 |
| chrX | 10710602 | 10711832 | 1231 | 1.82 | 0.0473 | 0.0531 |
| chr3 | 118490819 | 118495921 | 5103 | 0.74 | 0.0473 | 0.0531 |
| chr5 | 27834866 | 27872082 | 37217 | -0.99 | 0.0473 | 0.0531 |
| chrX | 13620281 | 13621118 | 838 | -2.52 | 0.0473 | 0.0532 |
| chrX | 72404138 | 72424188 | 20051 | -0.69 | 0.0474 | 0.0532 |
| chr13 | 66291305 | 66294811 | 3507 | -1.54 | 0.0475 | 0.0533 |
| chr1 | 75137271 | 75138353 | 1083 | -1.04 | 0.0475 | 0.0534 |
| chr2 | 169245980 | 169273084 | 27105 | -0.93 | 0.0475 | 0.0534 |
| chrX | 150510864 | 150511587 | 724 | 2.51 | 0.0475 | 0.0534 |
| chr13 | 61217442 | 61217981 | 540 | -2.33 | 0.0476 | 0.0534 |
| chr9 | 87806671 | 87823785 | 17115 | -2.99 | 0.0476 | 0.0535 |
| chr4 | 134289716 | 134290182 | 467 | -1.49 | 0.0476 | 0.0535 |
| chrX | 129606243 | 129611451 | 5209 | -1.07 | 0.0477 | 0.0536 |
| chr3 | 8627643 | 8628264 | 622 | -2.37 | 0.0477 | 0.0536 |
| chr5 | 61974570 | 62018918 | 44349 | -1.96 | 0.0477 | 0.0536 |
| chr7 | 31129213 | 31133793 | 4581 | 1.06 | 0.0477 | 0.0536 |
| chr10 | 2979194 | 3008102 | 28909 | -3.13 | 0.0478 | 0.0536 |
| chr12 | 109131528 | 109141710 | 10183 | -3.87 | 0.0478 | 0.0537 |
| chr8 | 118063050 | 118064498 | 1449 | -2.44 | 0.0478 | 0.0537 |
| chr4 | 78517989 | 78521872 | 3884 | -2.78 | 0.0479 | 0.0537 |
| chr11 | 84286133 | 84288244 | 2112 | 1.13 | 0.0479 | 0.0537 |
| chr7 | 35450289 | 35455189 | 4901 | 1.27 | 0.0479 | 0.0538 |
| chr3 | 102347147 | 102348624 | 1478 | -2.43 | 0.048 | 0.0538 |
| chr11 | 51395341 | 51396277 | 937 | 1.37 | 0.048 | 0.0539 |
| chr20 | 13801171 | 13853836 | 52666 | -0.68 | 0.048 | 0.0539 |
| chr14 | 43331252 | 43332146 | 895 | 1.13 | 0.048 | 0.0539 |
| chr2 | 134516008 | 134528242 | 12235 | 0.75 | 0.0481 | 0.0541 |

| | | | | | | |
|---|---|---|---|---|---|---|
| chr11 | 133926969 | 133928112 | 1144 | -1.99 | 0.0482 | 0.0541 |
| chr16 | 24518980 | 24522874 | 3895 | -2.65 | 0.0482 | 0.0541 |
| chr11 | 104324588 | 104326220 | 1633 | -2.49 | 0.0482 | 0.0541 |
| chr5 | 157414783 | 157416742 | 1960 | 1.49 | 0.0483 | 0.0542 |
| chr4 | 24450552 | 24452112 | 1561 | -2.61 | 0.0483 | 0.0542 |
| chr1 | 110291501 | 110292048 | 548 | -1.1 | 0.0483 | 0.0542 |
| chrX | 84684008 | 84684949 | 942 | -1.01 | 0.0483 | 0.0542 |
| chr11 | 49864278 | 49864970 | 693 | 1.47 | 0.0483 | 0.0542 |
| chr2 | 60808027 | 60920174 | 112148 | -0.54 | 0.0483 | 0.0542 |
| chr10 | 71289227 | 71328594 | 39368 | -4.47 | 0.0484 | 0.0543 |
| chrX | 120139798 | 120140341 | 544 | -2.44 | 0.0485 | 0.0545 |
| chrX | 77949374 | 77950588 | 1215 | 1.17 | 0.0485 | 0.0545 |
| chr1 | 170064225 | 170066810 | 2586 | -1.66 | 0.0486 | 0.0545 |
| chr7 | 54638200 | 54639597 | 1398 | 2.68 | 0.0486 | 0.0546 |
| chr6 | 18537537 | 18547093 | 9557 | -2.99 | 0.0486 | 0.0546 |
| chr1 | 105630760 | 105642660 | 11901 | 0.64 | 0.0487 | 0.0547 |
| chr12 | 14835763 | 14837218 | 1456 | 1.26 | 0.0489 | 0.0548 |
| chr3 | 194740140 | 194741718 | 1579 | -2.66 | 0.0489 | 0.0549 |
| chr13 | 65527635 | 66172546 | 644912 | 5.25 | 0.0489 | 0.0549 |
| chr3 | 104348361 | 104360543 | 12183 | -0.53 | 0.0489 | 0.0549 |
| chr9 | 121263699 | 121264013 | 315 | -1.24 | 0.0489 | 0.0549 |
| chr1 | 182703260 | 182703993 | 734 | 1.6 | 0.049 | 0.055 |
| chrX | 79686947 | 79688457 | 1511 | -2.43 | 0.0491 | 0.0551 |
| chrX | 34510219 | 34511487 | 1269 | 2.54 | 0.0492 | 0.0552 |
| chr14 | 75746058 | 75747243 | 1186 | 2.14 | 0.0493 | 0.0553 |
| chr5 | 54176429 | 54176971 | 543 | -2.28 | 0.0494 | 0.0554 |
| chr2 | 217731275 | 217769540 | 38266 | -2.71 | 0.0494 | 0.0554 |
| chr5 | 173508175 | 173521655 | 13481 | -3.02 | 0.0494 | 0.0554 |
| chr9 | 121155574 | 121172329 | 16756 | 3.02 | 0.0494 | 0.0554 |
| chr17 | 70142588 | 70145113 | 2526 | -1.65 | 0.0494 | 0.0554 |
| chr12 | 67607768 | 67608634 | 867 | -1.46 | 0.0495 | 0.0555 |
| chr13 | 96757816 | 96762647 | 4832 | -2.14 | 0.0495 | 0.0556 |
| chr15 | 53658750 | 53675247 | 16498 | -3.23 | 0.0496 | 0.0556 |
| chr3 | 4920124 | 4934885 | 14762 | -3.83 | 0.0496 | 0.0556 |
| chr6 | 128076329 | 128093042 | 16714 | -3.83 | 0.0496 | 0.0556 |
| chrX | 93758551 | 93780803 | 22253 | 0.64 | 0.0496 | 0.0557 |
| chr8 | 1561725 | 1611100 | 49376 | -0.47 | 0.0497 | 0.0558 |
| chr11 | 16500587 | 16501219 | 633 | -2.44 | 0.0497 | 0.0558 |
| chr7 | 67280938 | 67388927 | 107990 | -4.59 | 0.0498 | 0.0558 |
| chr10 | 131721007 | 131722363 | 1357 | -2.58 | 0.0498 | 0.0559 |
| chr5 | 64574574 | 64576242 | 1669 | -1.93 | 0.0499 | 0.0559 |
| chr5 | 126462300 | 126463352 | 1053 | -2.55 | 0.0499 | 0.056 |
| chr1 | 199279177 | 199279724 | 548 | 1.22 | 0.05 | 0.0561 |
| chr15 | 96027259 | 96028873 | 1615 | -0.94 | 0.0501 | 0.0561 |
| chr10 | 82126377 | 82128371 | 1995 | 1.27 | 0.0501 | 0.0562 |
| chr11 | 127069417 | 127077179 | 7763 | -0.84 | 0.0501 | 0.0562 |
| chr17 | 15984946 | 15996916 | 11971 | -3.67 | 0.0501 | 0.0562 |
| chrX | 56344915 | 56345439 | 525 | 1.31 | 0.0501 | 0.0562 |
| chrX | 73845339 | 73846653 | 1315 | 2.58 | 0.0501 | 0.0562 |
| chr8 | 125095579 | 125097252 | 1674 | -1.31 | 0.0501 | 0.0562 |
| chr15 | 46113904 | 46128214 | 14311 | -0.84 | 0.0502 | 0.0563 |
| chrX | 55060648 | 55092917 | 32270 | -2.28 | 0.0502 | 0.0563 |
| chr6 | 45984168 | 45994307 | 10140 | -0.73 | 0.0503 | 0.0563 |
| chr8 | 53922338 | 53935789 | 13452 | -1.2 | 0.0503 | 0.0564 |
| chrX | 84420006 | 84420625 | 620 | 1.09 | 0.0504 | 0.0564 |
| chr10 | 30391538 | 30392226 | 689 | -2.28 | 0.0504 | 0.0564 |
| chr7 | 141732859 | 141740833 | 7975 | -1.11 | 0.0504 | 0.0565 |
| chr4 | 154698436 | 154701270 | 2835 | -1.02 | 0.0504 | 0.0565 |
| chr6 | 75778002 | 75780670 | 2669 | -2.85 | 0.0505 | 0.0566 |
| chr10 | 129464519 | 129468422 | 3904 | -2.86 | 0.0505 | 0.0566 |
| chr11 | 57155970 | 57157140 | 1171 | 2.61 | 0.0505 | 0.0566 |

| | | | | | | |
|---|---|---|---|---|---|---|
| chr6 | 152189418 | 152194356 | 4939 | -0.72 | 0.0505 | 0.0566 |
| chr6 | 63121673 | 63123985 | 2313 | 1.68 | 0.0506 | 0.0567 |
| chr11 | 127985397 | 127985831 | 435 | 1.64 | 0.0506 | 0.0567 |
| chr4 | 121979405 | 121981595 | 2191 | -0.98 | 0.0508 | 0.057 |
| chr15 | 69967957 | 70084747 | 116791 | -4.81 | 0.0509 | 0.057 |
| chr13 | 91945198 | 91947298 | 2101 | 2.8 | 0.0509 | 0.057 |
| chr3 | 148137835 | 148151190 | 13356 | 3.06 | 0.0509 | 0.057 |
| chr7 | 67249685 | 67267430 | 17746 | -1.56 | 0.0509 | 0.057 |
| chr8 | 26604031 | 26703209 | 99179 | 0.33 | 0.0509 | 0.057 |
| chr1 | 13781670 | 13785372 | 3703 | 1.07 | 0.051 | 0.0571 |
| chr8 | 68362454 | 68377497 | 15044 | -1.76 | 0.0511 | 0.0573 |
| chr3 | 162077521 | 162078407 | 887 | 2.2 | 0.0513 | 0.0575 |
| chr7 | 132820520 | 132840747 | 20228 | -2.01 | 0.0513 | 0.0575 |
| chr14 | 36972764 | 36979015 | 6252 | -1 | 0.0513 | 0.0575 |
| chr8 | 82027642 | 82030472 | 2831 | 2.93 | 0.0514 | 0.0575 |
| chr15 | 89360194 | 89367845 | 7652 | -2.41 | 0.0514 | 0.0576 |
| chr11 | 99014976 | 99015486 | 511 | -2.21 | 0.0515 | 0.0576 |
| chr15 | 73493794 | 73508337 | 14544 | -3.79 | 0.0515 | 0.0577 |
| chr22 | 27887559 | 27905830 | 18272 | -3.79 | 0.0515 | 0.0577 |
| chr6 | 148874713 | 148875424 | 712 | -2.25 | 0.0516 | 0.0578 |
| chr1 | 206190793 | 206194856 | 4064 | 0.74 | 0.0516 | 0.0578 |
| chr7 | 87952781 | 87973030 | 20250 | 3.1 | 0.0516 | 0.0578 |
| chr13 | 66303810 | 66312271 | 8462 | -0.59 | 0.0517 | 0.0579 |
| chr12 | 196143 | 211868 | 15726 | -3.26 | 0.0518 | 0.058 |
| chr9 | 102038933 | 102053246 | 14314 | -2.81 | 0.0518 | 0.0581 |
| chr7 | 22058243 | 22084378 | 26136 | -1.34 | 0.0518 | 0.0581 |
| chr6 | 149469970 | 149471726 | 1757 | 2.57 | 0.0519 | 0.0581 |
| chr22 | 32482399 | 32483560 | 1162 | 1.27 | 0.0519 | 0.0581 |
| chr1 | 160525381 | 160527069 | 1689 | 1.03 | 0.0519 | 0.0581 |
| chr1 | 99894428 | 99895124 | 697 | -1.06 | 0.0519 | 0.0581 |
| chr14 | 51334811 | 51335330 | 520 | 1.15 | 0.052 | 0.0582 |
| chr15 | 42298884 | 42300070 | 1187 | -2.49 | 0.052 | 0.0582 |
| chr1 | 91854393 | 91856116 | 1724 | -1.88 | 0.052 | 0.0583 |
| chr7 | 41006033 | 41022664 | 16632 | -0.47 | 0.052 | 0.0583 |
| chr9 | 83394168 | 83456547 | 62380 | -2.07 | 0.0521 | 0.0583 |
| chr21 | 15967206 | 15970082 | 2877 | -2.83 | 0.0521 | 0.0583 |
| chrX | 42130180 | 42135131 | 4952 | 0.7 | 0.0521 | 0.0583 |
| chr7 | 143614885 | 143660506 | 45622 | 0.61 | 0.0521 | 0.0583 |
| chr10 | 12994322 | 13010648 | 16327 | -3.06 | 0.0521 | 0.0583 |
| chr5 | 169976641 | 169977903 | 1263 | 1.88 | 0.0521 | 0.0583 |
| chr8 | 133072700 | 133153128 | 80429 | 0.45 | 0.0521 | 0.0583 |
| chr4 | 32680769 | 32683054 | 2286 | -0.96 | 0.0522 | 0.0584 |
| chr2 | 139568388 | 139570910 | 2523 | -1.01 | 0.0522 | 0.0584 |
| chr12 | 119995371 | 119996492 | 1122 | -2.52 | 0.0522 | 0.0584 |
| chrX | 90923429 | 90923874 | 446 | 1.47 | 0.0522 | 0.0584 |
| chr4 | 122439174 | 122440657 | 1484 | -2.6 | 0.0522 | 0.0584 |
| chr7 | 132030436 | 132043628 | 13193 | -0.66 | 0.0522 | 0.0585 |
| chr5 | 112527962 | 112543328 | 15367 | 2.03 | 0.0523 | 0.0585 |
| chr16 | 48670294 | 48715907 | 45614 | -4.4 | 0.0523 | 0.0585 |
| chr3 | 170205322 | 170206482 | 1161 | -2.35 | 0.0523 | 0.0585 |
| chr11 | 134560602 | 134562590 | 1989 | -1.31 | 0.0526 | 0.0589 |
| chr3 | 148081654 | 148082200 | 547 | -1.79 | 0.0526 | 0.0589 |
| chr1 | 206950824 | 206951244 | 421 | 1.6 | 0.0526 | 0.0589 |
| chr4 | 62077100 | 62087035 | 9936 | 1.28 | 0.0527 | 0.0589 |
| chr7 | 135450347 | 135452149 | 1803 | 2.04 | 0.0527 | 0.0589 |
| chr6 | 129357547 | 129360265 | 2719 | 2.64 | 0.0527 | 0.059 |
| chr12 | 22052061 | 22052476 | 416 | -2.03 | 0.0528 | 0.0591 |
| chr3 | 108641075 | 108652759 | 11685 | -1 | 0.0528 | 0.0591 |
| chr11 | 104628214 | 104640609 | 12396 | 1.09 | 0.0529 | 0.0592 |
| chr7 | 22089157 | 22141933 | 52777 | -0.6 | 0.0529 | 0.0592 |
| chr6 | 147765170 | 147765576 | 407 | 1.99 | 0.0529 | 0.0592 |

| | | | | | | |
|---|---|---|---|---|---|---|
| chr2 | 3994972 | 3999614 | 4643 | -1.08 | 0.0529 | 0.0592 |
| chr3 | 112195648 | 112196556 | 909 | -2.52 | 0.053 | 0.0592 |
| chr9 | 97767948 | 97770454 | 2507 | -0.97 | 0.053 | 0.0593 |
| chr4 | 60504406 | 60505613 | 1208 | -1.58 | 0.0532 | 0.0595 |
| chr7 | 21933229 | 21934650 | 1422 | -1.42 | 0.0532 | 0.0595 |
| chrX | 114241668 | 114243024 | 1357 | -1.24 | 0.0532 | 0.0595 |
| chr11 | 94786708 | 94787573 | 866 | -2.39 | 0.0533 | 0.0597 |
| chr15 | 35018645 | 35086337 | 67693 | 0.7 | 0.0534 | 0.0597 |
| chr8 | 63509913 | 63823589 | 313677 | 4.99 | 0.0535 | 0.0598 |
| chr6 | 125731285 | 125732519 | 1235 | 0.97 | 0.0535 | 0.0598 |
| chr22 | 26620584 | 26620981 | 398 | -1.19 | 0.0535 | 0.0598 |
| chr1 | 146989299 | 146992554 | 3256 | -2.17 | 0.0535 | 0.0599 |
| chr2 | 174533394 | 174559718 | 26325 | -3.75 | 0.0535 | 0.0599 |
| chr3 | 4358888 | 4372421 | 13534 | -3.75 | 0.0535 | 0.0599 |
| chr3 | 185051640 | 185069081 | 17442 | -3.75 | 0.0535 | 0.0599 |
| chr13 | 76994087 | 76996189 | 2103 | -2.54 | 0.0536 | 0.0599 |
| chr4 | 27065236 | 27085521 | 20286 | -0.98 | 0.0536 | 0.0599 |
| chr11 | 102564688 | 102565663 | 976 | 2.15 | 0.0536 | 0.06 |
| chr13 | 24490166 | 24492980 | 2815 | -0.83 | 0.0537 | 0.06 |
| chr2 | 220200830 | 220204959 | 4130 | -2.7 | 0.0537 | 0.06 |
| chr1 | 182334197 | 182336085 | 1889 | 0.99 | 0.0537 | 0.06 |
| chr14 | 81412755 | 81436599 | 23845 | -0.74 | 0.0537 | 0.0601 |
| chr3 | 106196804 | 106199334 | 2531 | -1.47 | 0.0538 | 0.0602 |
| chrX | 42521209 | 42521885 | 677 | 1.01 | 0.0538 | 0.0602 |
| chr9 | 101431393 | 101441783 | 10391 | -2.36 | 0.0538 | 0.0602 |
| chrX | 116260314 | 116260924 | 611 | -1.58 | 0.0538 | 0.0602 |
| chr4 | 134611987 | 134612623 | 637 | 1.13 | 0.0539 | 0.0603 |
| chr9 | 118984498 | 119008370 | 23873 | -3.22 | 0.054 | 0.0603 |
| chrX | 114166657 | 114167600 | 944 | -1.45 | 0.054 | 0.0604 |
| chr3 | 39405860 | 39414732 | 8873 | -2.8 | 0.054 | 0.0604 |
| chr3 | 190515091 | 190519506 | 4416 | -2.24 | 0.0541 | 0.0604 |
| chr3 | 133038407 | 133039770 | 1364 | -0.8 | 0.0541 | 0.0605 |
| chr2 | 13532338 | 13533206 | 869 | 1.39 | 0.0542 | 0.0605 |
| chr7 | 121116910 | 121119025 | 2116 | -2.43 | 0.0542 | 0.0606 |
| chr3 | 22941216 | 22956501 | 15286 | -0.58 | 0.0542 | 0.0606 |
| chrX | 118222291 | 118246400 | 24110 | 0.77 | 0.0542 | 0.0606 |
| chr10 | 80314219 | 80325745 | 11527 | -1.99 | 0.0543 | 0.0607 |
| chr1 | 30812139 | 30813175 | 1037 | 1.69 | 0.0543 | 0.0607 |
| chr2 | 147501338 | 147503673 | 2336 | 2.68 | 0.0544 | 0.0608 |
| chr4 | 63370766 | 63404560 | 33795 | -0.75 | 0.0544 | 0.0608 |
| chr2 | 3723842 | 3795291 | 71450 | -1.25 | 0.0545 | 0.0609 |
| chrX | 130750377 | 130752798 | 2422 | -2.53 | 0.0545 | 0.0609 |
| chrX | 81352339 | 81372036 | 19698 | 0.81 | 0.0545 | 0.0609 |
| chr10 | 123234439 | 123249997 | 15559 | -2.79 | 0.0545 | 0.0609 |
| chr10 | 117553009 | 117634898 | 81890 | 3.34 | 0.0546 | 0.061 |
| chr18 | 33384285 | 33385197 | 913 | 2.6 | 0.0546 | 0.061 |
| chr14 | 81820744 | 81822112 | 1369 | -2.6 | 0.0547 | 0.0611 |
| chr20 | 30006140 | 30007073 | 934 | 1.75 | 0.0547 | 0.0611 |
| chr17 | 20783600 | 20785137 | 1538 | -1.91 | 0.0547 | 0.0611 |
| chr12 | 68342593 | 68354407 | 11815 | -2.86 | 0.0547 | 0.0611 |
| chr6 | 45873020 | 45878638 | 5619 | -1.27 | 0.0547 | 0.0611 |
| chr2 | 109824429 | 109836642 | 12214 | -2.79 | 0.0547 | 0.0611 |
| chr4 | 61192511 | 61195175 | 2665 | 1.95 | 0.0548 | 0.0612 |
| chr9 | 30706171 | 30777536 | 71366 | 0.89 | 0.0548 | 0.0612 |
| chr15 | 25867124 | 25876205 | 9082 | -0.68 | 0.0548 | 0.0612 |
| chr12 | 111643328 | 111644713 | 1386 | 1.72 | 0.0548 | 0.0612 |
| chr4 | 27809450 | 27820200 | 10751 | -2.9 | 0.0548 | 0.0612 |
| chr18 | 48831737 | 48835665 | 3929 | -1.03 | 0.0549 | 0.0613 |
| chr17 | 28562107 | 28567570 | 5464 | -2.47 | 0.0549 | 0.0614 |
| chr13 | 64343874 | 64349333 | 5460 | 2.99 | 0.055 | 0.0615 |
| chr5 | 121257876 | 121258334 | 459 | -1.7 | 0.0551 | 0.0615 |

| | | | | | | |
|---|---|---|---|---|---|---|
| chr4 | 107618402 | 107620412 | 2011 | 1.59 | 0.0551 | 0.0615 |
| chr13 | 94990058 | 94992865 | 2808 | 2.53 | 0.0551 | 0.0615 |
| chr21 | 37929219 | 37957879 | 28661 | -2.99 | 0.0551 | 0.0615 |
| chr7 | 150458532 | 150472512 | 13981 | -2.99 | 0.0552 | 0.0616 |
| chr14 | 76978065 | 76979709 | 1645 | 0.8 | 0.0552 | 0.0617 |
| chr1 | 224277216 | 224281543 | 4328 | -2.83 | 0.0552 | 0.0617 |
| chr21 | 18698689 | 18699797 | 1109 | 2.44 | 0.0552 | 0.0617 |
| chr17 | 38871980 | 38923287 | 51308 | -4.58 | 0.0553 | 0.0617 |
| chr15 | 96274830 | 96309031 | 34202 | -1.81 | 0.0554 | 0.0618 |
| chr1 | 157281817 | 157285323 | 3507 | 2.88 | 0.0554 | 0.0618 |
| chr17 | 72687899 | 72690878 | 2980 | -2.77 | 0.0554 | 0.0619 |
| chr3 | 167972408 | 167974180 | 1773 | -1.15 | 0.0555 | 0.0619 |
| chrX | 42205493 | 42210133 | 4641 | 0.79 | 0.0556 | 0.062 |
| chr4 | 62029674 | 62031908 | 2235 | 1.51 | 0.0556 | 0.0621 |
| chr3 | 106167332 | 106168077 | 746 | -1.18 | 0.0557 | 0.0622 |
| chr8 | 82079725 | 82081228 | 1504 | -2.6 | 0.0557 | 0.0622 |
| chr15 | 33182805 | 33183850 | 1046 | -2.53 | 0.0557 | 0.0622 |
| chr3 | 82768496 | 82769046 | 551 | -1.57 | 0.0557 | 0.0622 |
| chr12 | 89651542 | 89665096 | 13555 | -3.71 | 0.0557 | 0.0622 |
| chr13 | 43097476 | 43112319 | 14844 | -1.07 | 0.0557 | 0.0622 |
| chr18 | 67477155 | 67490598 | 13444 | -0.91 | 0.0557 | 0.0622 |
| chrX | 150411261 | 150413517 | 2257 | -2.11 | 0.0558 | 0.0623 |
| chr1 | 91170469 | 91171497 | 1029 | 1.31 | 0.0558 | 0.0623 |
| chr19 | 8784825 | 8792173 | 7349 | -1.03 | 0.0559 | 0.0624 |
| chr9 | 14990711 | 14998708 | 7998 | -0.8 | 0.0559 | 0.0624 |
| chr9 | 74084681 | 74087166 | 2486 | -1.01 | 0.0559 | 0.0624 |
| chr8 | 95643531 | 95644370 | 840 | -1.35 | 0.056 | 0.0624 |
| chr8 | 121848073 | 121855486 | 7414 | 1.35 | 0.0561 | 0.0626 |
| chr4 | 29156262 | 29168513 | 12252 | 0.69 | 0.0561 | 0.0626 |
| chr11 | 88267417 | 88268657 | 1241 | -2.27 | 0.0561 | 0.0626 |
| chr18 | 58071303 | 58074898 | 3596 | 0.9 | 0.0561 | 0.0626 |
| chr3 | 97758879 | 97776596 | 17718 | -2.92 | 0.0561 | 0.0626 |
| chr12 | 14294469 | 14304348 | 9880 | 3.06 | 0.0561 | 0.0626 |
| chr8 | 59130341 | 59131218 | 878 | 1.5 | 0.0562 | 0.0627 |
| chr4 | 29270966 | 29324298 | 53333 | 0.54 | 0.0562 | 0.0627 |
| chr3 | 55367252 | 55370930 | 3679 | 1.15 | 0.0563 | 0.0628 |
| chrX | 80959293 | 80987649 | 28357 | 1.12 | 0.0564 | 0.0629 |
| chr6 | 129339793 | 129340428 | 636 | -2.4 | 0.0565 | 0.063 |
| chr17 | 71847877 | 71851676 | 3800 | -2.01 | 0.0565 | 0.0631 |
| chrX | 76302400 | 76305789 | 3390 | 0.89 | 0.0566 | 0.0631 |
| chr5 | 16626871 | 16628655 | 1785 | 1.5 | 0.0566 | 0.0631 |
| chr11 | 109313244 | 109315399 | 2156 | 1.3 | 0.0566 | 0.0631 |
| chr5 | 93747330 | 93748638 | 1309 | -1.24 | 0.0566 | 0.0631 |
| chrX | 85380037 | 85382230 | 2194 | 1.91 | 0.0566 | 0.0632 |
| chr4 | 136581685 | 136584007 | 2323 | -2.31 | 0.0567 | 0.0632 |
| chr5 | 33076836 | 33078886 | 2051 | -0.8 | 0.0567 | 0.0633 |
| chr3 | 124126674 | 124127861 | 1188 | -2.41 | 0.0568 | 0.0634 |
| chr1 | 4375781 | 4377393 | 1613 | 0.87 | 0.0568 | 0.0634 |
| chr10 | 118135710 | 118268205 | 132496 | 4.53 | 0.0569 | 0.0634 |
| chr5 | 6352342 | 6354002 | 1661 | -2.45 | 0.0569 | 0.0635 |
| chr15 | 53565115 | 53585025 | 19911 | 3.17 | 0.057 | 0.0635 |
| chr20 | 22856666 | 22883776 | 27111 | -3.15 | 0.057 | 0.0635 |
| chr1 | 15111502 | 15112569 | 1068 | -2.35 | 0.057 | 0.0635 |
| chr8 | 130392339 | 130419026 | 26688 | -0.92 | 0.057 | 0.0636 |
| chr3 | 24887523 | 25328902 | 441380 | 5.02 | 0.057 | 0.0636 |
| chr6 | 63818933 | 63822190 | 3258 | 2.68 | 0.0571 | 0.0637 |
| chr8 | 53448374 | 53449983 | 1610 | 1.13 | 0.0572 | 0.0637 |
| chrX | 43656877 | 43658231 | 1355 | 1 | 0.0572 | 0.0638 |
| chr6 | 129134273 | 129145770 | 11498 | 1.03 | 0.0573 | 0.0639 |
| chr13 | 22464026 | 22466499 | 2474 | -1.04 | 0.0573 | 0.0639 |
| chrX | 151513961 | 151514532 | 572 | 1.45 | 0.0573 | 0.0639 |

| | | | | | | |
|---|---|---|---|---|---|---|
| chr12 | 39963347 | 39964272 | 926 | 1.15 | 0.0574 | 0.0639 |
| chr3 | 51168102 | 51191404 | 23303 | -0.7 | 0.0574 | 0.064 |
| chr21 | 37223588 | 37228413 | 4826 | 2.91 | 0.0575 | 0.0641 |
| chr22 | 22908092 | 22908501 | 410 | -2.02 | 0.0575 | 0.0641 |
| chr11 | 27173369 | 27174734 | 1366 | 0.93 | 0.0575 | 0.0641 |
| chr12 | 60765323 | 60787418 | 22096 | -3.03 | 0.0576 | 0.0642 |
| chr12 | 7617719 | 7624471 | 6753 | -3 | 0.0577 | 0.0643 |
| chr12 | 77325267 | 77326828 | 1562 | -2.33 | 0.0577 | 0.0643 |
| chr3 | 181058427 | 181059067 | 641 | -1.18 | 0.0578 | 0.0644 |
| chr6 | 101680518 | 101833267 | 152750 | 4.63 | 0.0578 | 0.0644 |
| chr18 | 11708561 | 11711929 | 3369 | -2.84 | 0.0578 | 0.0644 |
| chr12 | 102644836 | 102645638 | 803 | 2.4 | 0.0579 | 0.0645 |
| chr1 | 104668238 | 104669295 | 1058 | 1.6 | 0.0579 | 0.0645 |
| chr4 | 135908243 | 135909715 | 1473 | -1.08 | 0.058 | 0.0646 |
| chr3 | 72926360 | 72933540 | 7181 | -2.98 | 0.058 | 0.0646 |
| chrX | 12767641 | 12791960 | 24320 | -1.02 | 0.058 | 0.0647 |
| chr8 | 6885048 | 6885729 | 682 | -1.32 | 0.0581 | 0.0647 |
| chr3 | 6005559 | 6017794 | 12236 | -0.82 | 0.0581 | 0.0647 |
| chr6 | 121339952 | 121346767 | 6816 | 2.99 | 0.0581 | 0.0647 |
| chr20 | 44090858 | 44108445 | 17588 | -2.87 | 0.0581 | 0.0647 |
| chrX | 83604781 | 83607309 | 2529 | 2.6 | 0.0581 | 0.0647 |
| chr5 | 145760090 | 145772233 | 12144 | -2.96 | 0.0582 | 0.0648 |
| chr17 | 52251399 | 52338065 | 86667 | -3.23 | 0.0582 | 0.0648 |
| chr4 | 122212347 | 122213709 | 1363 | -2.39 | 0.0582 | 0.0649 |
| chr2 | 150247105 | 150251970 | 4866 | 1.29 | 0.0583 | 0.0649 |
| chr14 | 31327862 | 31332203 | 4342 | -2.73 | 0.0583 | 0.0649 |
| chr15 | 49694154 | 49716542 | 22389 | -3.05 | 0.0584 | 0.065 |
| chr1 | 66978908 | 66979535 | 628 | -2.29 | 0.0584 | 0.0651 |
| chr7 | 152600885 | 152605166 | 4282 | 1.36 | 0.0585 | 0.0651 |
| chrX | 107976712 | 107977553 | 842 | -1.85 | 0.0585 | 0.0651 |
| chr12 | 60231430 | 60238766 | 7337 | -2.09 | 0.0585 | 0.0652 |
| chr6 | 55016749 | 55019367 | 2619 | -2.79 | 0.0586 | 0.0652 |
| chr12 | 108285069 | 108300199 | 15131 | -2.95 | 0.0586 | 0.0652 |
| chr8 | 127584712 | 127828358 | 243647 | -4.82 | 0.0586 | 0.0653 |
| chr3 | 29914156 | 29916171 | 2016 | -2.63 | 0.0586 | 0.0653 |
| chr4 | 15330790 | 15335589 | 4800 | 0.6 | 0.0587 | 0.0653 |
| chr9 | 138195534 | 138227210 | 31677 | -4.31 | 0.0587 | 0.0654 |
| chr10 | 115124779 | 115137892 | 13114 | 0.8 | 0.0589 | 0.0656 |
| chr10 | 80305537 | 80308609 | 3073 | 1.31 | 0.059 | 0.0657 |
| chr1 | 83708138 | 83708681 | 544 | 1.03 | 0.059 | 0.0657 |
| chr2 | 53420100 | 53421329 | 1230 | -1.49 | 0.059 | 0.0657 |
| chr15 | 87065180 | 87092396 | 27217 | -1.01 | 0.0591 | 0.0658 |
| chr5 | 42785031 | 42799079 | 14049 | -3.11 | 0.0591 | 0.0658 |
| chr13 | 112514598 | 112558736 | 44139 | -0.61 | 0.0591 | 0.0658 |
| chr8 | 54510546 | 54592296 | 81751 | -3.24 | 0.0593 | 0.066 |
| chr12 | 16666180 | 16667416 | 1237 | 1.68 | 0.0593 | 0.066 |
| chr10 | 45808010 | 45834373 | 26364 | -3.12 | 0.0593 | 0.066 |
| chr4 | 60554122 | 60555360 | 1239 | 1.37 | 0.0593 | 0.066 |
| chr4 | 134685827 | 134687213 | 1387 | 2.39 | 0.0595 | 0.0662 |
| chr12 | 119653292 | 119654393 | 1102 | 2.56 | 0.0595 | 0.0662 |
| chr12 | 7477588 | 7478131 | 544 | 1.48 | 0.0595 | 0.0662 |
| chr2 | 204865195 | 204913754 | 48560 | -0.54 | 0.0595 | 0.0662 |
| chr12 | 109348262 | 109350776 | 2515 | 0.79 | 0.0596 | 0.0663 |
| chr13 | 27123049 | 27153714 | 30666 | -2.42 | 0.0597 | 0.0664 |
| chr10 | 124551401 | 124553157 | 1757 | -1.34 | 0.0597 | 0.0664 |
| chr11 | 106171851 | 106174713 | 2863 | -1.88 | 0.0597 | 0.0664 |
| chr11 | 61929018 | 61957654 | 28637 | -1.35 | 0.0597 | 0.0664 |
| chrX | 83819546 | 84098725 | 279180 | 4.8 | 0.0597 | 0.0665 |
| chr5 | 54334782 | 54356536 | 21755 | 0.81 | 0.0598 | 0.0666 |
| chr1 | 170684944 | 170891825 | 206882 | 4.8 | 0.0599 | 0.0666 |
| chr3 | 10905146 | 10909267 | 4122 | 1.43 | 0.0599 | 0.0667 |

| | | | | | | |
|---|---|---|---|---|---|---|
| chrX | 113706926 | 113707710 | 785 | 2.6 | 0.0599 | 0.0667 |
| chr3 | 85938006 | 85964605 | 26600 | -1.78 | 0.06 | 0.0667 |
| chrX | 103032465 | 103047917 | 15453 | 3.03 | 0.06 | 0.0667 |
| chr9 | 134963615 | 135000926 | 37312 | -3.08 | 0.06 | 0.0668 |
| chr6 | 38982962 | 38984286 | 1325 | -1.65 | 0.0601 | 0.0668 |
| chr16 | 78048189 | 78129255 | 81067 | 1.11 | 0.0601 | 0.0668 |
| chr7 | 18949861 | 18959732 | 9872 | -1.04 | 0.0601 | 0.0669 |
| chr16 | 78431366 | 78445468 | 14103 | 0.99 | 0.0603 | 0.0671 |
| chr1 | 83011326 | 83026514 | 15189 | 0.51 | 0.0603 | 0.0671 |
| chr1 | 79317044 | 79317716 | 673 | -2.13 | 0.0603 | 0.0671 |
| chr1 | 198381907 | 198383029 | 1123 | 1.01 | 0.0604 | 0.0672 |
| chr16 | 52347425 | 52362023 | 14599 | 0.62 | 0.0605 | 0.0673 |
| chr13 | 77016746 | 77028641 | 11896 | -2.92 | 0.0605 | 0.0673 |
| chr6 | 74393431 | 74403489 | 10059 | -3.62 | 0.0605 | 0.0673 |
| chr8 | 10039206 | 10050239 | 11034 | -3.62 | 0.0605 | 0.0673 |
| chr10 | 83532880 | 83537845 | 4966 | -0.85 | 0.0605 | 0.0673 |
| chr4 | 62974062 | 62998229 | 24168 | 0.55 | 0.0606 | 0.0674 |
| chr17 | 75696209 | 75758152 | 61944 | -4.64 | 0.0606 | 0.0674 |
| chr3 | 98058759 | 98066882 | 8124 | 1.08 | 0.0607 | 0.0675 |
| chrX | 105930480 | 105931297 | 818 | -1.91 | 0.0608 | 0.0676 |
| chr2 | 231094622 | 231112373 | 17752 | -1.47 | 0.061 | 0.0678 |
| chr3 | 191158084 | 191158765 | 682 | -2.26 | 0.061 | 0.0679 |
| chr14 | 88549603 | 88649273 | 99671 | -0.38 | 0.0611 | 0.0679 |
| chr1 | 18231836 | 18232413 | 578 | -1.18 | 0.0611 | 0.0679 |
| chr10 | 99859534 | 99910091 | 50558 | -3.19 | 0.0612 | 0.0681 |
| chr1 | 190512281 | 190515855 | 3575 | 2.77 | 0.0612 | 0.0681 |
| chr6 | 154025927 | 154027587 | 1661 | -1.47 | 0.0613 | 0.0681 |
| chr2 | 200534079 | 200537685 | 3607 | 1.05 | 0.0615 | 0.0684 |
| chr1 | 116799802 | 116803284 | 3483 | 2.8 | 0.0616 | 0.0685 |
| chr6 | 164117797 | 164119729 | 1933 | 0.87 | 0.0616 | 0.0685 |
| chr3 | 17910118 | 17911690 | 1573 | 1.6 | 0.0617 | 0.0685 |
| chr16 | 55087754 | 55345629 | 257876 | 0.26 | 0.0617 | 0.0686 |
| chr14 | 70927869 | 70930082 | 2214 | 1.44 | 0.0618 | 0.0687 |
| chr12 | 39416672 | 39418310 | 1639 | -1.84 | 0.0618 | 0.0687 |
| chr12 | 99278323 | 99281187 | 2865 | -2.4 | 0.0618 | 0.0687 |
| chr7 | 37191126 | 37193025 | 1900 | 1.31 | 0.0619 | 0.0687 |
| chr3 | 93913183 | 93914331 | 1149 | -2.38 | 0.0619 | 0.0688 |
| chr3 | 111117400 | 111118493 | 1094 | -2.08 | 0.0619 | 0.0688 |
| chr5 | 95944643 | 95952348 | 7706 | -3.44 | 0.0619 | 0.0688 |
| chr15 | 54106782 | 54114690 | 7909 | 1.53 | 0.0619 | 0.0688 |
| chr1 | 223697123 | 223699113 | 1991 | 2.62 | 0.062 | 0.0689 |
| chr6 | 44713347 | 44714953 | 1607 | -1.21 | 0.0621 | 0.069 |
| chr3 | 163972169 | 163974909 | 2741 | -2.57 | 0.0622 | 0.0691 |
| chr22 | 27358720 | 27360522 | 1803 | 0.98 | 0.0623 | 0.0692 |
| chrX | 102242373 | 102249116 | 6744 | 2.93 | 0.0625 | 0.0694 |
| chr5 | 33563729 | 33889888 | 326160 | 4.87 | 0.0626 | 0.0695 |
| chr20 | 10123183 | 10151132 | 27950 | -0.84 | 0.0628 | 0.0698 |
| chr10 | 84248359 | 84322989 | 74631 | -3.13 | 0.0629 | 0.0699 |
| chr1 | 201201791 | 201214486 | 12696 | -1.28 | 0.063 | 0.07 |
| chr12 | 68263891 | 68265768 | 1878 | 1.65 | 0.063 | 0.07 |
| chrX | 72525457 | 72526399 | 943 | 1.28 | 0.0631 | 0.0701 |
| chrX | 72536931 | 72537907 | 977 | -2.23 | 0.0631 | 0.0701 |
| chr6 | 13139769 | 13141858 | 2090 | 1.04 | 0.0632 | 0.0702 |
| chr8 | 143884013 | 143907786 | 23774 | -3.57 | 0.0632 | 0.0702 |
| chrX | 19049385 | 19070584 | 21200 | -3.57 | 0.0632 | 0.0702 |
| chrX | 134691914 | 134694645 | 2732 | -2.57 | 0.0633 | 0.0704 |
| chr16 | 55006919 | 55008080 | 1162 | 0.88 | 0.0634 | 0.0704 |
| chr1 | 76175059 | 76181727 | 6669 | -2.1 | 0.0635 | 0.0705 |
| chr8 | 109818866 | 109839435 | 20570 | 3.12 | 0.0635 | 0.0706 |
| chr8 | 115529847 | 115530837 | 991 | -2.29 | 0.0636 | 0.0706 |
| chr3 | 70822408 | 70824729 | 2322 | 0.99 | 0.0636 | 0.0706 |

| | | | | | | |
|---|---|---|---|---|---|---|
| chr12 | 22133182 | 22134390 | 1209 | -1.53 | 0.0636 | 0.0707 |
| chr15 | 73942478 | 73965412 | 22935 | -3.02 | 0.0638 | 0.0708 |
| chr5 | 122070201 | 122072163 | 1963 | -2.29 | 0.0638 | 0.0708 |
| chr15 | 35414691 | 35436781 | 22091 | -2.91 | 0.0638 | 0.0709 |
| chr8 | 20208400 | 20225301 | 16902 | 0.52 | 0.0638 | 0.0709 |
| chr2 | 141088319 | 141090447 | 2129 | -2.29 | 0.0639 | 0.071 |
| chrX | 82693727 | 82695030 | 1304 | -1.3 | 0.0639 | 0.071 |
| chr3 | 117655977 | 117657843 | 1867 | -2.46 | 0.064 | 0.0711 |
| chr6 | 159668436 | 159679970 | 11535 | -0.54 | 0.064 | 0.0711 |
| chr9 | 120997415 | 121039258 | 41844 | -0.69 | 0.064 | 0.0711 |
| chr3 | 123872895 | 123890521 | 17627 | 0.81 | 0.064 | 0.0711 |
| chr8 | 48012503 | 48013710 | 1208 | -1.86 | 0.0641 | 0.0711 |
| chr4 | 61709684 | 61763537 | 53854 | 0.59 | 0.0641 | 0.0712 |
| chr3 | 109721103 | 109722055 | 953 | -2.28 | 0.0642 | 0.0712 |
| chr20 | 23908649 | 23921875 | 13227 | -2.82 | 0.0642 | 0.0713 |
| chr7 | 48223661 | 48298167 | 74507 | -0.44 | 0.0642 | 0.0713 |
| chr13 | 65409468 | 65476234 | 66767 | 3.26 | 0.0643 | 0.0714 |
| chr18 | 61662812 | 61673981 | 11170 | -1.01 | 0.0644 | 0.0715 |
| chr18 | 2125869 | 2127227 | 1359 | -0.83 | 0.0644 | 0.0715 |
| chr5 | 173858161 | 173944910 | 86750 | -4.45 | 0.0644 | 0.0715 |
| chrX | 68935305 | 68939023 | 3719 | -1.54 | 0.0645 | 0.0716 |
| chr2 | 58528088 | 58541398 | 13311 | -1.12 | 0.0645 | 0.0716 |
| chrX | 109835923 | 109837930 | 2008 | 2.54 | 0.0646 | 0.0717 |
| chrX | 82484429 | 82487350 | 2922 | -1.05 | 0.0646 | 0.0717 |
| chr1 | 106174431 | 106219138 | 44708 | 0.58 | 0.0646 | 0.0717 |
| chr16 | 75749148 | 75750744 | 1597 | 0.93 | 0.0647 | 0.0718 |
| chrX | 124445722 | 124447287 | 1566 | -2.47 | 0.0647 | 0.0718 |
| chr8 | 115065160 | 115092609 | 27450 | -2.88 | 0.0648 | 0.072 |
| chr16 | 24044379 | 24318982 | 274604 | 0.29 | 0.0649 | 0.072 |
| chr14 | 38478045 | 38496167 | 18123 | 0.7 | 0.0649 | 0.072 |
| chr5 | 88150809 | 88153228 | 2420 | -1.4 | 0.065 | 0.0721 |
| chrX | 69132179 | 69133413 | 1235 | -1.18 | 0.0651 | 0.0722 |
| chr5 | 114437776 | 114449646 | 11871 | -2.12 | 0.0651 | 0.0722 |
| chr6 | 51928568 | 51929883 | 1316 | -0.89 | 0.0651 | 0.0723 |
| chr3 | 153587712 | 153589969 | 2258 | 0.83 | 0.0651 | 0.0723 |
| chr17 | 79475535 | 79478944 | 3410 | -0.93 | 0.0653 | 0.0725 |
| chr18 | 74906466 | 74907648 | 1183 | -2.46 | 0.0653 | 0.0725 |
| chr12 | 83628401 | 83637125 | 8725 | -2.54 | 0.0654 | 0.0725 |
| chr4 | 62530880 | 62531665 | 786 | 1.23 | 0.0655 | 0.0726 |
| chr18 | 11165381 | 11173434 | 8054 | -0.49 | 0.0655 | 0.0727 |
| chr18 | 69255604 | 69256694 | 1091 | 0.94 | 0.0655 | 0.0727 |
| chr6 | 118617322 | 118742134 | 124813 | 4.42 | 0.0655 | 0.0727 |
| chr1 | 184075052 | 184347648 | 272597 | 4.8 | 0.0656 | 0.0727 |
| chr12 | 96998609 | 97004443 | 5835 | 0.5 | 0.0656 | 0.0727 |
| chr2 | 57777642 | 57779761 | 2120 | -2.6 | 0.0656 | 0.0728 |
| chr11 | 15033857 | 15082256 | 48400 | 3.26 | 0.0656 | 0.0728 |
| chr4 | 10501014 | 10534339 | 33326 | 0.5 | 0.0657 | 0.0728 |
| chr15 | 79840565 | 79842337 | 1773 | 2.67 | 0.0658 | 0.073 |
| chr13 | 61536793 | 61538462 | 1670 | 1.56 | 0.0658 | 0.073 |
| chr9 | 81183125 | 81184308 | 1184 | -2.07 | 0.0659 | 0.073 |
| chr9 | 77110568 | 77111226 | 659 | -2.13 | 0.0659 | 0.0731 |
| chr22 | 40163175 | 40200093 | 36919 | -0.52 | 0.066 | 0.0732 |
| chr1 | 93776403 | 93786555 | 10153 | -3.52 | 0.0661 | 0.0733 |
| chr14 | 92112467 | 92117063 | 4597 | -3.52 | 0.0661 | 0.0733 |
| chr2 | 152009035 | 152015184 | 6150 | -0.58 | 0.0661 | 0.0733 |
| chr20 | 53275622 | 53282364 | 6743 | 2.92 | 0.0661 | 0.0733 |
| chr10 | 103083442 | 103087440 | 3999 | -1.71 | 0.0661 | 0.0733 |
| chr13 | 78562025 | 78691051 | 129027 | 4.48 | 0.0661 | 0.0733 |
| chr5 | 65600524 | 65641479 | 40956 | -0.88 | 0.0661 | 0.0733 |
| chr20 | 22299451 | 22311354 | 11904 | -0.43 | 0.0662 | 0.0734 |
| chrX | 81442917 | 81445688 | 2772 | 1.03 | 0.0663 | 0.0735 |

| | | | | | | |
|---|---|---|---|---|---|---|
| chrX | 29752764 | 29754225 | 1462 | -2.49 | 0.0664 | 0.0736 |
| chr3 | 37604082 | 37738504 | 134423 | -0.54 | 0.0665 | 0.0737 |
| chr1 | 104724384 | 104741948 | 17565 | 1.57 | 0.0665 | 0.0737 |
| chr9 | 73063763 | 73064332 | 570 | -2.15 | 0.0665 | 0.0737 |
| chr4 | 21089792 | 21095164 | 5373 | -0.7 | 0.0665 | 0.0737 |
| chr3 | 27990201 | 28001621 | 11421 | -1.4 | 0.0665 | 0.0738 |
| chr4 | 155421335 | 155421887 | 553 | -2.13 | 0.0667 | 0.0739 |
| chr1 | 87611207 | 87611832 | 626 | -0.9 | 0.0667 | 0.0739 |
| chr7 | 19010824 | 19031611 | 20788 | 0.66 | 0.0668 | 0.074 |
| chr17 | 39595778 | 39608476 | 12699 | -0.99 | 0.0668 | 0.0741 |
| chr13 | 31831851 | 31845542 | 13692 | -3.35 | 0.0669 | 0.0742 |
| chr3 | 103286744 | 103289242 | 2499 | 1.27 | 0.067 | 0.0743 |
| chrX | 95915304 | 95916662 | 1359 | -1.49 | 0.0672 | 0.0744 |
| chr8 | 112951825 | 113074686 | 122862 | -0.96 | 0.0672 | 0.0745 |
| chr18 | 54119146 | 54121652 | 2507 | 0.98 | 0.0673 | 0.0746 |
| chrX | 134771088 | 134776690 | 5603 | 0.81 | 0.0673 | 0.0746 |
| chr1 | 227595050 | 227599774 | 4725 | -0.66 | 0.0674 | 0.0747 |
| chr7 | 121232072 | 121233300 | 1229 | -1.07 | 0.0674 | 0.0747 |
| chr8 | 7003754 | 7004971 | 1218 | 1 | 0.0674 | 0.0747 |
| chr7 | 37260317 | 37479746 | 219430 | 4.65 | 0.0675 | 0.0748 |
| chr4 | 30603726 | 30605727 | 2002 | -1.85 | 0.0676 | 0.0748 |
| chr14 | 49915146 | 49955896 | 40751 | -3.01 | 0.0676 | 0.0749 |
| chr14 | 54613485 | 54632256 | 18772 | -1.2 | 0.0676 | 0.0749 |
| chr15 | 24509340 | 24509855 | 516 | 1.27 | 0.0677 | 0.075 |
| chrX | 102265811 | 102268809 | 2999 | 0.68 | 0.0678 | 0.0751 |
| chr18 | 76455569 | 76471633 | 16065 | -0.55 | 0.0679 | 0.0752 |
| chr11 | 84774480 | 84817545 | 43066 | -0.91 | 0.0679 | 0.0752 |
| chr9 | 21242756 | 21249522 | 6767 | -2.56 | 0.0679 | 0.0752 |
| chr4 | 15426686 | 15428552 | 1867 | 2.59 | 0.068 | 0.0754 |
| chr1 | 77239177 | 77251169 | 11993 | 0.65 | 0.0681 | 0.0754 |
| chrX | 116109294 | 116110345 | 1052 | -1.79 | 0.0681 | 0.0754 |
| chr11 | 99583535 | 99592895 | 9361 | -1.4 | 0.0683 | 0.0757 |
| chrX | 108070516 | 108071259 | 744 | -0.98 | 0.0684 | 0.0757 |
| chr12 | 45963718 | 46001817 | 38100 | -3.08 | 0.0685 | 0.0759 |
| chr16 | 59198419 | 59210674 | 12256 | -0.7 | 0.0686 | 0.0759 |
| chr12 | 34248849 | 34251378 | 2530 | 2.59 | 0.0686 | 0.076 |
| chr1 | 20790362 | 20791421 | 1060 | 1.33 | 0.0686 | 0.076 |
| chr10 | 23712933 | 23713784 | 852 | 1.4 | 0.0686 | 0.076 |
| chr1 | 105213889 | 105218443 | 4555 | 0.77 | 0.0688 | 0.0762 |
| chr13 | 36554580 | 36556205 | 1626 | -2.45 | 0.0688 | 0.0762 |
| chrX | 70109466 | 70112708 | 3243 | -1.87 | 0.0689 | 0.0763 |
| chr1 | 78644248 | 78662122 | 17875 | 0.91 | 0.0689 | 0.0763 |
| chr11 | 84923643 | 84924038 | 396 | 1.21 | 0.069 | 0.0764 |
| chr1 | 248624266 | 248625385 | 1120 | 2.32 | 0.069 | 0.0764 |
| chr1 | 244378184 | 244388329 | 10146 | -3.47 | 0.0692 | 0.0766 |
| chr13 | 105633936 | 105635281 | 1346 | -1.54 | 0.0692 | 0.0766 |
| chr10 | 61159654 | 61179163 | 19510 | -0.79 | 0.0692 | 0.0766 |
| chr8 | 40645570 | 40660747 | 15178 | -2.8 | 0.0692 | 0.0766 |
| chr3 | 22985901 | 22997521 | 11621 | -0.81 | 0.0693 | 0.0767 |
| chr12 | 53123594 | 53166465 | 42872 | 3.18 | 0.0693 | 0.0768 |
| chr14 | 28959762 | 29056006 | 96245 | -0.86 | 0.0693 | 0.0768 |
| chr4 | 25970816 | 25971366 | 551 | -1.1 | 0.0694 | 0.0768 |
| chrX | 79905510 | 79906376 | 867 | -0.8 | 0.0694 | 0.0768 |
| chr1 | 88936777 | 88938472 | 1696 | 1.36 | 0.0694 | 0.0768 |
| chr6 | 88619389 | 88620895 | 1507 | -1.65 | 0.0694 | 0.0769 |
| chrX | 55723648 | 55728828 | 5181 | 0.98 | 0.0695 | 0.0769 |
| chr6 | 129364441 | 129365220 | 780 | -2.22 | 0.0696 | 0.0771 |
| chrX | 125714887 | 125715380 | 494 | -1.23 | 0.0697 | 0.0771 |
| chr1 | 182157875 | 182168998 | 11124 | 0.57 | 0.0697 | 0.0772 |
| chr7 | 135574702 | 135576321 | 1620 | -1.06 | 0.0697 | 0.0772 |
| chr3 | 73174281 | 73177265 | 2985 | -0.87 | 0.0697 | 0.0772 |

| | | | | | | |
|---|---|---|---|---|---|---|
| chr5 | 72003769 | 72004765 | 997 | -0.85 | 0.0698 | 0.0772 |
| chr2 | 184163858 | 184164211 | 354 | -1.98 | 0.0698 | 0.0773 |
| chr10 | 72002794 | 72006724 | 3931 | 2.7 | 0.0699 | 0.0773 |
| chr15 | 102341913 | 102381306 | 39394 | 0.55 | 0.07 | 0.0774 |
| chr2 | 145396434 | 145397349 | 916 | -2.1 | 0.07 | 0.0774 |
| chr4 | 22815458 | 22816410 | 953 | -2.08 | 0.07 | 0.0775 |
| chr4 | 32981638 | 32999114 | 17477 | 0.84 | 0.0702 | 0.0776 |
| chr14 | 30276365 | 30285037 | 8673 | 2.8 | 0.0702 | 0.0777 |
| chr5 | 43982057 | 43983578 | 1522 | 1.08 | 0.0705 | 0.078 |
| chr15 | 79391814 | 79406229 | 14416 | -0.64 | 0.0705 | 0.078 |
| chr3 | 186047406 | 186053697 | 6292 | 2.47 | 0.0706 | 0.0781 |
| chr10 | 120045675 | 120048909 | 3235 | 1.23 | 0.0706 | 0.0781 |
| chrX | 66583349 | 66606848 | 23500 | -0.78 | 0.0707 | 0.0782 |
| chr18 | 57077116 | 57079443 | 2328 | -1.8 | 0.0708 | 0.0783 |
| chr3 | 86910326 | 86913519 | 3194 | -2.17 | 0.0711 | 0.0787 |
| chrX | 12655545 | 12707027 | 51483 | 0.36 | 0.0711 | 0.0787 |
| chr2 | 183332748 | 183338793 | 6046 | -1.18 | 0.0712 | 0.0787 |
| chr5 | 170090571 | 170091839 | 1269 | -2.09 | 0.0712 | 0.0787 |
| chr12 | 8604311 | 8641026 | 36716 | 0.78 | 0.0714 | 0.079 |
| chrX | 68107016 | 68109584 | 2569 | 0.87 | 0.0714 | 0.079 |
| chr5 | 59115424 | 59129311 | 13888 | 1.41 | 0.0715 | 0.0791 |
| chr5 | 113459323 | 113464527 | 5205 | -0.97 | 0.0715 | 0.0791 |
| chr13 | 77489576 | 77491104 | 1529 | 1.27 | 0.0715 | 0.0791 |
| chr3 | 85551900 | 85785226 | 233327 | -1.58 | 0.0717 | 0.0793 |
| chrX | 151464130 | 151466067 | 1938 | -2.2 | 0.0717 | 0.0793 |
| chr3 | 36036036 | 36131379 | 95344 | 0.29 | 0.0717 | 0.0793 |
| chr4 | 42653177 | 42654562 | 1386 | -2.51 | 0.0718 | 0.0794 |
| chr12 | 59567957 | 59570117 | 2161 | -2.42 | 0.0718 | 0.0794 |
| chr3 | 72512879 | 72515815 | 2937 | 1.28 | 0.072 | 0.0796 |
| chr7 | 141756126 | 141758882 | 2757 | 0.99 | 0.072 | 0.0796 |
| chr7 | 87810184 | 87812385 | 2202 | -1.28 | 0.072 | 0.0796 |
| chr11 | 112269630 | 112297660 | 28031 | 0.56 | 0.0722 | 0.0798 |
| chr8 | 61952655 | 61959847 | 7193 | -0.79 | 0.0722 | 0.0798 |
| chr2 | 159655812 | 159669913 | 14102 | 2.58 | 0.0722 | 0.0799 |
| chr18 | 68612709 | 68613208 | 500 | -1.03 | 0.0723 | 0.0799 |
| chr5 | 108002331 | 108004567 | 2237 | -0.72 | 0.0723 | 0.0799 |
| chr10 | 38504066 | 38518490 | 14425 | -2.11 | 0.0724 | 0.08 |
| chr5 | 59143103 | 59157154 | 14052 | 1.09 | 0.0725 | 0.0801 |
| chr8 | 53309930 | 53315075 | 5146 | -2.79 | 0.0725 | 0.0801 |
| chr2 | 61062719 | 61064141 | 1423 | 2.51 | 0.0725 | 0.0801 |
| chr16 | 71652807 | 71655203 | 2397 | -1.21 | 0.0725 | 0.0801 |
| chr1 | 104687036 | 104704225 | 17190 | 0.85 | 0.0725 | 0.0801 |
| chr8 | 62639262 | 62652478 | 13217 | -3.41 | 0.0725 | 0.0802 |
| chr18 | 71594088 | 71599652 | 5565 | -0.63 | 0.0725 | 0.0802 |
| chr7 | 21389417 | 21459244 | 69828 | -4.2 | 0.0727 | 0.0803 |
| chr16 | 77416604 | 77419920 | 3317 | -1.21 | 0.0728 | 0.0805 |
| chr9 | 132157278 | 132159491 | 2214 | 0.75 | 0.0729 | 0.0806 |
| chrX | 16352725 | 16353395 | 671 | 1.22 | 0.0732 | 0.0809 |
| chr11 | 134301035 | 134340463 | 39429 | -2.92 | 0.0732 | 0.0809 |
| chr2 | 18878509 | 18896880 | 18372 | -2.81 | 0.0732 | 0.0809 |
| chr12 | 119586863 | 119591621 | 4759 | -2.87 | 0.0733 | 0.081 |
| chrX | 23802265 | 23804239 | 1975 | -1.04 | 0.0733 | 0.081 |
| chr11 | 94874602 | 94875151 | 550 | -1.31 | 0.0734 | 0.0811 |
| chr3 | 42897850 | 42902075 | 4226 | -2.55 | 0.0734 | 0.0811 |
| chr6 | 168143842 | 168145738 | 1897 | -2.61 | 0.0734 | 0.0811 |
| chr4 | 78162959 | 78164671 | 1713 | 1.35 | 0.0734 | 0.0811 |
| chr5 | 94647766 | 94653080 | 5315 | -0.69 | 0.0735 | 0.0812 |
| chr13 | 40398198 | 40409123 | 10926 | -0.46 | 0.0735 | 0.0812 |
| chr4 | 34954938 | 34956546 | 1609 | 2.35 | 0.0736 | 0.0813 |
| chr4 | 122302692 | 122304588 | 1897 | -1.01 | 0.0736 | 0.0813 |
| chr2 | 64524701 | 64528246 | 3546 | 1.81 | 0.0737 | 0.0814 |

| | | | | | | |
|---|---|---|---|---|---|---|
| chr1 | 47296813 | 47298367 | 1555 | -1.53 | 0.0737 | 0.0815 |
| chr7 | 150380206 | 150381280 | 1075 | -1.1 | 0.0738 | 0.0815 |
| chr20 | 58338407 | 58340555 | 2149 | 0.83 | 0.0738 | 0.0815 |
| chr9 | 121129276 | 121130472 | 1197 | 1.16 | 0.0738 | 0.0815 |
| chr10 | 83125468 | 83147922 | 22455 | 0.62 | 0.0739 | 0.0817 |
| chr7 | 92970851 | 92971382 | 532 | -2.21 | 0.074 | 0.0817 |
| chr18 | 44826875 | 44827687 | 813 | -1.77 | 0.074 | 0.0818 |
| chr13 | 104735944 | 104736367 | 424 | -1.89 | 0.074 | 0.0818 |
| chr8 | 114081412 | 114150683 | 69272 | -0.86 | 0.0743 | 0.082 |
| chr4 | 15433290 | 15448813 | 15524 | -2.65 | 0.0743 | 0.082 |
| chr11 | 84731552 | 84761595 | 30044 | 0.84 | 0.0743 | 0.0821 |
| chr8 | 10926202 | 10929383 | 3182 | 1.59 | 0.0743 | 0.0821 |
| chr13 | 78501419 | 78502241 | 823 | 2.27 | 0.0744 | 0.0821 |
| chr8 | 113567931 | 113582586 | 14656 | -1.09 | 0.0744 | 0.0822 |
| chr4 | 75839936 | 75853190 | 13255 | -1.05 | 0.0744 | 0.0822 |
| chr11 | 31876019 | 31877667 | 1649 | -1.19 | 0.0745 | 0.0823 |
| chr11 | 119781758 | 119782348 | 591 | 1.76 | 0.0745 | 0.0823 |
| chr8 | 30697800 | 30706746 | 8947 | 1.79 | 0.0746 | 0.0824 |
| chr3 | 108860561 | 108893478 | 32918 | -1.78 | 0.0746 | 0.0824 |
| chr4 | 76193964 | 76195844 | 1881 | -2.38 | 0.0748 | 0.0826 |
| chr8 | 105717822 | 105778754 | 60933 | 3.11 | 0.0748 | 0.0826 |
| chr12 | 104052297 | 104058335 | 6039 | -0.75 | 0.075 | 0.0828 |
| chr1 | 87626957 | 87631898 | 4942 | 0.86 | 0.0751 | 0.0829 |
| chr6 | 78874777 | 78876590 | 1814 | 2.32 | 0.0751 | 0.0829 |
| chr10 | 115152161 | 115159349 | 7189 | 2.88 | 0.0751 | 0.0829 |
| chr7 | 46220527 | 46222572 | 2046 | -2.53 | 0.0751 | 0.0829 |
| chr12 | 107536541 | 107558707 | 22167 | -1.37 | 0.0752 | 0.083 |
| chr2 | 106177 | 108450 | 2274 | -1.36 | 0.0752 | 0.083 |
| chr2 | 195750043 | 195750816 | 774 | -2.29 | 0.0752 | 0.083 |
| chr1 | 239796067 | 239865010 | 68944 | 3.17 | 0.0754 | 0.0832 |
| chr11 | 95281914 | 95283998 | 2085 | 2.54 | 0.0754 | 0.0832 |
| chr11 | 128508535 | 128513025 | 4491 | -1.57 | 0.0755 | 0.0833 |
| chr9 | 13062706 | 13066306 | 3601 | -0.76 | 0.0755 | 0.0834 |
| chr10 | 117649412 | 117658241 | 8830 | 2.8 | 0.0755 | 0.0834 |
| chr8 | 59176708 | 59177777 | 1070 | 0.86 | 0.0757 | 0.0835 |
| chr4 | 60899806 | 60908988 | 9183 | -1.18 | 0.0757 | 0.0835 |
| chr6 | 66443835 | 66459653 | 15819 | 0.94 | 0.0757 | 0.0836 |
| chr10 | 90603220 | 90604141 | 922 | -1.61 | 0.0758 | 0.0836 |
| chr4 | 32628160 | 32630522 | 2363 | -2.25 | 0.0758 | 0.0836 |
| chr5 | 79110174 | 79120059 | 9886 | -0.81 | 0.0759 | 0.0837 |
| chrX | 154390912 | 154393070 | 2159 | -1.43 | 0.0759 | 0.0838 |
| chr2 | 13597240 | 13762880 | 165641 | -0.48 | 0.076 | 0.0839 |
| chr11 | 49887535 | 49888027 | 493 | 1.78 | 0.0761 | 0.0839 |
| chr13 | 76821591 | 76828994 | 7404 | 0.88 | 0.0762 | 0.0841 |
| chr20 | 9909841 | 9911841 | 2001 | 0.98 | 0.0764 | 0.0843 |
| chr12 | 131584005 | 131585854 | 1850 | 0.88 | 0.0764 | 0.0843 |
| chr20 | 59747824 | 59820112 | 72289 | -4.09 | 0.0765 | 0.0844 |
| chr10 | 6032026 | 6088684 | 56659 | -4.25 | 0.0765 | 0.0844 |
| chr14 | 94221261 | 94222767 | 1507 | -0.91 | 0.0765 | 0.0844 |
| chrX | 51261013 | 51282716 | 21704 | 0.62 | 0.0766 | 0.0845 |
| chr2 | 171655691 | 171660402 | 4712 | -2.61 | 0.0767 | 0.0846 |
| chr11 | 104436145 | 104446828 | 10684 | -2.6 | 0.0768 | 0.0847 |
| chr1 | 239344844 | 239368456 | 23613 | 0.76 | 0.0769 | 0.0848 |
| chr2 | 177706196 | 177707124 | 929 | -2.32 | 0.0769 | 0.0848 |
| chr1 | 104053608 | 104054052 | 445 | -1.29 | 0.0769 | 0.0848 |
| chr11 | 106425338 | 106427077 | 1740 | -1.23 | 0.077 | 0.0849 |
| chr4 | 104258588 | 104259695 | 1108 | 2.42 | 0.077 | 0.0849 |
| chr4 | 94870626 | 94875123 | 4498 | 0.9 | 0.0771 | 0.0851 |
| chr4 | 145351413 | 145365236 | 13824 | -1.15 | 0.0772 | 0.0851 |
| chr1 | 35116409 | 35117651 | 1243 | -2.27 | 0.0773 | 0.0852 |
| chr13 | 23523629 | 23524413 | 785 | -1.57 | 0.0773 | 0.0853 |

| | | | | | | |
|---|---|---|---|---|---|---|
| chr1 | 242724144 | 242742437 | 18294 | -1.34 | 0.0773 | 0.0853 |
| chr5 | 89341285 | 89384840 | 43556 | 0.51 | 0.0774 | 0.0854 |
| chr9 | 113122741 | 113123274 | 534 | -2.18 | 0.0775 | 0.0855 |
| chr10 | 129257933 | 129273055 | 15123 | -1.5 | 0.0776 | 0.0856 |
| chrX | 150396111 | 150400301 | 4191 | 1.43 | 0.0777 | 0.0856 |
| chr8 | 118756746 | 118757559 | 814 | -1.55 | 0.0778 | 0.0858 |
| chr21 | 33184473 | 33217996 | 33524 | -0.52 | 0.0778 | 0.0858 |
| chr11 | 99164788 | 99171949 | 7162 | -0.82 | 0.0779 | 0.0858 |
| chr18 | 70343317 | 70344855 | 1539 | -2.36 | 0.078 | 0.086 |
| chr11 | 72241035 | 72257353 | 16319 | 0.65 | 0.078 | 0.086 |
| chr9 | 85053002 | 85225383 | 172382 | -0.22 | 0.078 | 0.086 |
| chr2 | 207586750 | 207598390 | 11641 | -2.83 | 0.0781 | 0.0861 |
| chr20 | 5875678 | 5878526 | 2849 | 1.25 | 0.0781 | 0.0861 |
| chr16 | 59665100 | 59687055 | 21956 | -2.92 | 0.0781 | 0.0861 |
| chr3 | 162925747 | 162927732 | 1986 | -2.33 | 0.0782 | 0.0862 |
| chrX | 114658529 | 114659909 | 1381 | -0.88 | 0.0782 | 0.0862 |
| chrX | 22417711 | 22421210 | 3500 | -0.86 | 0.0782 | 0.0862 |
| chrX | 16357596 | 16361542 | 3947 | -2.54 | 0.0783 | 0.0863 |
| chrX | 131424367 | 131449023 | 24657 | -1.71 | 0.0783 | 0.0863 |
| chr10 | 4374582 | 4377768 | 3187 | -2.65 | 0.0784 | 0.0864 |
| chr10 | 12979259 | 12986381 | 7123 | 2.88 | 0.0784 | 0.0864 |
| chr1 | 106166776 | 106167987 | 1212 | -0.93 | 0.0785 | 0.0865 |
| chrX | 132814290 | 132814739 | 450 | -2.05 | 0.0786 | 0.0866 |
| chr5 | 166785281 | 166832261 | 46981 | 0.44 | 0.0786 | 0.0867 |
| chr11 | 113899041 | 113901370 | 2330 | -0.68 | 0.0786 | 0.0867 |
| chr20 | 998082 | 1001970 | 3889 | 0.95 | 0.0787 | 0.0868 |
| chrX | 3005894 | 3097531 | 91638 | -0.44 | 0.0789 | 0.0869 |
| chr20 | 23689754 | 23698219 | 8466 | -0.68 | 0.0789 | 0.0869 |
| chr12 | 25915214 | 25916787 | 1574 | -2.52 | 0.0789 | 0.0869 |
| chr7 | 39426817 | 39440173 | 13357 | 2.86 | 0.079 | 0.087 |
| chr18 | 51930424 | 51932898 | 2475 | -0.75 | 0.0791 | 0.0871 |
| chr9 | 102544433 | 102545441 | 1009 | -2.24 | 0.0791 | 0.0871 |
| chr21 | 10967688 | 10969524 | 1837 | 0.84 | 0.0791 | 0.0871 |
| chr2 | 170170247 | 170173277 | 3031 | 1.23 | 0.0791 | 0.0872 |
| chr10 | 128263336 | 128286292 | 22957 | -0.59 | 0.0791 | 0.0872 |
| chr3 | 85808555 | 85873847 | 65293 | -0.85 | 0.0792 | 0.0872 |
| chrX | 73913471 | 73927883 | 14413 | 2.8 | 0.0792 | 0.0872 |
| chr11 | 131551926 | 131552756 | 831 | 0.95 | 0.0792 | 0.0872 |
| chr6 | 70991194 | 70991838 | 645 | -2.12 | 0.0792 | 0.0872 |
| chr6 | 46587990 | 46591015 | 3026 | -2.51 | 0.0796 | 0.0876 |
| chr1 | 156789909 | 156792834 | 2926 | -2.57 | 0.0796 | 0.0876 |
| chr22 | 49517430 | 49526336 | 8907 | -0.88 | 0.0796 | 0.0877 |
| chr13 | 40812343 | 40825051 | 12709 | -1.26 | 0.0797 | 0.0877 |
| chr16 | 49594629 | 49607939 | 13311 | -2.72 | 0.0797 | 0.0878 |
| chr1 | 146496394 | 146505988 | 9595 | 0.82 | 0.0797 | 0.0878 |
| chr13 | 68918495 | 69759008 | 840514 | 4.89 | 0.0799 | 0.0879 |
| chrX | 72322807 | 72326092 | 3286 | 2.54 | 0.0799 | 0.088 |
| chr14 | 92824981 | 92829365 | 4385 | -2.37 | 0.0801 | 0.0882 |
| chrX | 154881926 | 154892750 | 10825 | -1.71 | 0.0801 | 0.0882 |
| chr4 | 20160387 | 20162225 | 1839 | -0.98 | 0.0801 | 0.0882 |
| chr13 | 82960636 | 82961455 | 820 | -1.94 | 0.0802 | 0.0883 |
| chr20 | 52950468 | 52951767 | 1300 | 2.35 | 0.0802 | 0.0883 |
| chr1 | 160719659 | 160722125 | 2467 | 0.79 | 0.0802 | 0.0883 |
| chr8 | 133054213 | 133054983 | 771 | -0.83 | 0.0803 | 0.0884 |
| chr2 | 84902776 | 84905296 | 2521 | 2.55 | 0.0804 | 0.0885 |
| chr11 | 6185232 | 6201864 | 16633 | -1.01 | 0.0805 | 0.0886 |
| chr16 | 50757591 | 50769640 | 12050 | -2.19 | 0.0805 | 0.0886 |
| chr3 | 70660134 | 70774629 | 114496 | -0.26 | 0.0805 | 0.0886 |
| chr9 | 97811406 | 97815633 | 4228 | -0.78 | 0.0806 | 0.0887 |
| chrX | 42082810 | 42089412 | 6603 | -0.51 | 0.0806 | 0.0888 |
| chr5 | 166265669 | 166268212 | 2544 | 1.88 | 0.0807 | 0.0888 |

| | | | | | | |
|---|---|---|---|---|---|---|
| chr16 | 49353082 | 49362918 | 9837 | 0.44 | 0.0807 | 0.0888 |
| chr3 | 81106499 | 81209864 | 103366 | 0.54 | 0.0809 | 0.0891 |
| chr2 | 134321218 | 134322792 | 1575 | -1.48 | 0.081 | 0.0892 |
| chr14 | 66889866 | 66891944 | 2079 | 2.08 | 0.081 | 0.0892 |
| chr18 | 45964821 | 45966892 | 2072 | 0.93 | 0.081 | 0.0892 |
| chr7 | 117471154 | 117484742 | 13589 | -0.56 | 0.0811 | 0.0893 |
| chr9 | 129328035 | 129335356 | 7322 | -2.52 | 0.0811 | 0.0893 |
| chr10 | 83545028 | 83571660 | 26633 | 0.68 | 0.0812 | 0.0893 |
| chr19 | 55002721 | 55017214 | 14494 | -0.78 | 0.0812 | 0.0894 |
| chr15 | 46208809 | 46221726 | 12918 | -2.68 | 0.0812 | 0.0894 |
| chr13 | 20773810 | 20799149 | 25340 | -2.84 | 0.0812 | 0.0894 |
| chr18 | 11044424 | 11045641 | 1218 | -2.35 | 0.0813 | 0.0895 |
| chr8 | 144135809 | 144207883 | 72075 | -4.33 | 0.0813 | 0.0895 |
| chr8 | 43008756 | 43038809 | 30054 | -2.96 | 0.0813 | 0.0895 |
| chr12 | 102998437 | 102999435 | 999 | -2.28 | 0.0815 | 0.0896 |
| chr1 | 106130223 | 106158452 | 28230 | 0.51 | 0.0816 | 0.0897 |
| chr11 | 45540657 | 45541815 | 1159 | 2.28 | 0.0816 | 0.0898 |
| chrX | 120195744 | 120197863 | 2120 | -0.88 | 0.0816 | 0.0898 |
| chr8 | 30680658 | 30683807 | 3150 | 0.92 | 0.0818 | 0.09 |
| chr4 | 30404805 | 30410037 | 5233 | -1.8 | 0.0818 | 0.09 |
| chr8 | 17233435 | 17234501 | 1067 | -1.27 | 0.0819 | 0.0901 |
| chr12 | 4561519 | 4591189 | 29671 | -0.41 | 0.0819 | 0.0901 |
| chr12 | 4004060 | 4117025 | 112966 | -0.33 | 0.082 | 0.0902 |
| chr1 | 236818873 | 236861621 | 42749 | -2.97 | 0.082 | 0.0902 |
| chr3 | 113903292 | 113907567 | 4276 | -0.72 | 0.082 | 0.0902 |
| chr5 | 173614739 | 173616295 | 1557 | 1.41 | 0.0821 | 0.0903 |
| chr2 | 215332975 | 215336961 | 3987 | 2.54 | 0.0822 | 0.0904 |
| chr5 | 59397337 | 59399776 | 2440 | 1.07 | 0.0822 | 0.0904 |
| chrX | 36984891 | 36986137 | 1247 | -2.34 | 0.0822 | 0.0904 |
| chr8 | 83872428 | 83887005 | 14578 | -2.68 | 0.0822 | 0.0904 |
| chr1 | 69290703 | 69293240 | 2538 | 2.6 | 0.0823 | 0.0905 |
| chr1 | 90842651 | 90843718 | 1068 | -1.74 | 0.0823 | 0.0905 |
| chr11 | 40516964 | 40539394 | 22431 | -0.75 | 0.0824 | 0.0906 |
| chr14 | 96038910 | 96040525 | 1616 | 1.63 | 0.0824 | 0.0906 |
| chr3 | 163011722 | 163026462 | 14741 | -1.83 | 0.0825 | 0.0907 |
| chr16 | 51081758 | 51155308 | 73551 | -0.45 | 0.0825 | 0.0907 |
| chrX | 151429333 | 151442168 | 12836 | -1.1 | 0.0825 | 0.0907 |
| chrX | 69992936 | 70008979 | 16044 | -2.75 | 0.0826 | 0.0909 |
| chr15 | 60108706 | 60112436 | 3731 | 1.01 | 0.0827 | 0.0909 |
| chr14 | 22667438 | 22739101 | 71664 | -2.22 | 0.0828 | 0.0911 |
| chr18 | 51639206 | 51640367 | 1162 | 1.57 | 0.083 | 0.0913 |
| chr5 | 107163563 | 107165444 | 1882 | 2.5 | 0.0832 | 0.0914 |
| chr4 | 29098300 | 29106683 | 8384 | 0.98 | 0.0832 | 0.0915 |
| chr14 | 23895389 | 23903319 | 7931 | 2.79 | 0.0833 | 0.0915 |
| chr13 | 44150547 | 44151127 | 581 | -2.05 | 0.0834 | 0.0916 |
| chr11 | 125264198 | 125265965 | 1768 | -2.37 | 0.0834 | 0.0917 |
| chr11 | 29419435 | 29420093 | 659 | 0.99 | 0.0834 | 0.0917 |
| chr11 | 88877929 | 88878259 | 331 | 1.49 | 0.0834 | 0.0917 |
| chr9 | 123056307 | 123058855 | 2549 | -1.24 | 0.0834 | 0.0917 |
| chr11 | 43580578 | 43586903 | 6326 | 0.68 | 0.0835 | 0.0918 |
| chr6 | 46841141 | 46841819 | 679 | 1.08 | 0.0835 | 0.0918 |
| chr7 | 69433236 | 69455494 | 22259 | -0.44 | 0.0835 | 0.0918 |
| chr8 | 121313342 | 121321292 | 7951 | -0.49 | 0.0835 | 0.0918 |
| chr12 | 77472519 | 77474730 | 2212 | 2.09 | 0.0836 | 0.0919 |
| chr16 | 49621867 | 49623224 | 1358 | -2.41 | 0.0836 | 0.0919 |
| chr5 | 166996122 | 167005952 | 9831 | 1 | 0.0836 | 0.0919 |
| chr3 | 70046990 | 70088408 | 41419 | -0.46 | 0.0837 | 0.092 |
| chr2 | 105385139 | 105387782 | 2644 | -1.79 | 0.0837 | 0.092 |
| chr4 | 67876790 | 67882593 | 5804 | -1.22 | 0.0837 | 0.092 |
| chr11 | 129519519 | 129558150 | 38632 | -2.97 | 0.0838 | 0.0921 |
| chr16 | 9287305 | 9337865 | 50561 | -2.42 | 0.0838 | 0.0921 |

| | | | | | | |
|---|---|---|---|---|---|---|
| chr17 | 75972866 | 75981045 | 8180 | -2.83 | 0.0842 | 0.0926 |
| chr3 | 190897552 | 190900296 | 2745 | 2.47 | 0.0843 | 0.0926 |
| chrX | 150389038 | 150390411 | 1374 | -1.43 | 0.0843 | 0.0926 |
| chr7 | 50898138 | 50899595 | 1458 | -2.33 | 0.0844 | 0.0927 |
| chr2 | 74341002 | 74360996 | 19995 | -3.24 | 0.0844 | 0.0927 |
| chr2 | 102310212 | 102320126 | 9915 | -3.24 | 0.0844 | 0.0927 |
| chr4 | 114815345 | 114829256 | 13912 | -3.24 | 0.0844 | 0.0927 |
| chr5 | 115071542 | 115082260 | 10719 | -0.6 | 0.0846 | 0.0929 |
| chrX | 52864845 | 52867169 | 2325 | -2.32 | 0.0846 | 0.0929 |
| chr4 | 135135975 | 135136920 | 946 | -2.17 | 0.0848 | 0.0932 |
| chr4 | 154637351 | 154639335 | 1985 | -2.37 | 0.0848 | 0.0932 |
| chr14 | 58114040 | 58155654 | 41615 | 0.45 | 0.085 | 0.0933 |
| chr18 | 2168482 | 2195304 | 26823 | 0.46 | 0.085 | 0.0934 |
| chr2 | 34874707 | 34918089 | 43383 | 0.59 | 0.0851 | 0.0935 |
| chr8 | 113862617 | 113922987 | 60371 | -0.6 | 0.0851 | 0.0935 |
| chr3 | 148110595 | 148111786 | 1192 | 2.45 | 0.0851 | 0.0935 |
| chr3 | 175850138 | 175877655 | 27518 | 1.01 | 0.0851 | 0.0935 |
| chr3 | 142849115 | 142859546 | 10432 | -1.32 | 0.0852 | 0.0935 |
| chr3 | 64855401 | 64856467 | 1067 | -1.44 | 0.0852 | 0.0936 |
| chrX | 30444831 | 30447619 | 2789 | -2.41 | 0.0852 | 0.0936 |
| chr17 | 21891015 | 21891432 | 418 | 1.56 | 0.0853 | 0.0936 |
| chr11 | 78841543 | 78928508 | 86966 | 0.8 | 0.0854 | 0.0938 |
| chr21 | 15550767 | 15554539 | 3773 | 0.72 | 0.0854 | 0.0938 |
| chr13 | 35088230 | 35095996 | 7767 | -2.54 | 0.0855 | 0.0939 |
| chr11 | 134621813 | 134622684 | 872 | 0.86 | 0.0855 | 0.0939 |
| chr14 | 49138725 | 49249183 | 110459 | -2.99 | 0.0856 | 0.094 |
| chr9 | 140875380 | 140887942 | 12563 | -2.71 | 0.0856 | 0.094 |
| chr8 | 75040432 | 75047175 | 6744 | 0.76 | 0.0857 | 0.0941 |
| chrX | 131115138 | 131123399 | 8262 | 2.71 | 0.0857 | 0.0941 |
| chr2 | 53444077 | 53445647 | 1571 | -1.03 | 0.0857 | 0.0941 |
| chr3 | 144859831 | 144892847 | 33017 | 2.95 | 0.0858 | 0.0942 |
| chr8 | 113340867 | 113544748 | 203882 | -4.06 | 0.0858 | 0.0942 |
| chr4 | 147456516 | 147457855 | 1340 | -1.78 | 0.0858 | 0.0942 |
| chr6 | 130794225 | 130796735 | 2511 | 1 | 0.0858 | 0.0942 |
| chr6 | 119504115 | 119506219 | 2105 | 1.43 | 0.0861 | 0.0945 |
| chr4 | 86281947 | 86283816 | 1870 | -2.44 | 0.0861 | 0.0945 |
| chr1 | 231847964 | 231849685 | 1722 | -1.23 | 0.0861 | 0.0945 |
| chr13 | 28416646 | 28423722 | 7077 | -2.68 | 0.0862 | 0.0946 |
| chr11 | 18973210 | 18978533 | 5324 | 2.74 | 0.0863 | 0.0947 |
| chr12 | 92644293 | 92644692 | 400 | -1.97 | 0.0863 | 0.0948 |
| chr11 | 21902048 | 21906072 | 4025 | -2.55 | 0.0863 | 0.0948 |
| chr21 | 39050738 | 39051812 | 1075 | -2.19 | 0.0865 | 0.0949 |
| chr3 | 191802517 | 191803250 | 734 | -2.12 | 0.0866 | 0.095 |
| chr1 | 77312141 | 77331878 | 19738 | -0.94 | 0.0866 | 0.095 |
| chr18 | 33407583 | 33434969 | 27387 | -0.56 | 0.0867 | 0.0951 |
| chr10 | 115974119 | 115989299 | 15181 | -0.75 | 0.0867 | 0.0952 |
| chr3 | 90227158 | 90228489 | 1332 | 1.3 | 0.0868 | 0.0952 |
| chr7 | 131954531 | 131956108 | 1578 | 0.78 | 0.0869 | 0.0953 |
| chr8 | 141341034 | 141350356 | 9323 | -3.06 | 0.0869 | 0.0953 |
| chr2 | 11694565 | 11695972 | 1408 | 2.56 | 0.0871 | 0.0955 |
| chr4 | 34552461 | 34554659 | 2199 | 2.36 | 0.0871 | 0.0956 |
| chr14 | 106881095 | 106883162 | 2068 | 1.04 | 0.0872 | 0.0956 |
| chrX | 75540152 | 75571782 | 31631 | 0.89 | 0.0872 | 0.0957 |
| chrX | 75634250 | 75635507 | 1258 | -0.95 | 0.0872 | 0.0957 |
| chr12 | 120066914 | 120069901 | 2988 | 0.94 | 0.0873 | 0.0957 |
| chr8 | 47939039 | 47953400 | 14362 | 2.57 | 0.0875 | 0.0959 |
| chr15 | 35002553 | 35005341 | 2789 | 2.37 | 0.0875 | 0.096 |
| chr1 | 95889887 | 95905941 | 16055 | 0.74 | 0.0877 | 0.0962 |
| chr3 | 124073447 | 124110792 | 37346 | -0.49 | 0.0878 | 0.0963 |
| chr3 | 162857357 | 162858043 | 687 | 2.08 | 0.0878 | 0.0963 |
| chr5 | 111153624 | 111164052 | 10429 | -2.58 | 0.0878 | 0.0963 |

| | | | | | | |
|---|---|---|---|---|---|---|
| chr5 | 43814298 | 43817388 | 3091 | -0.83 | 0.0879 | 0.0964 |
| chr8 | 113663717 | 113692622 | 28906 | -0.88 | 0.0879 | 0.0964 |
| chr11 | 59488638 | 59491164 | 2527 | -2.42 | 0.088 | 0.0965 |
| chr1 | 222495504 | 222559323 | 63820 | 0.38 | 0.088 | 0.0965 |
| chr2 | 85338287 | 85341348 | 3062 | 0.92 | 0.0881 | 0.0966 |
| chr13 | 25274004 | 25276264 | 2261 | -0.98 | 0.0882 | 0.0967 |
| chr4 | 123451360 | 123451906 | 547 | 2.09 | 0.0883 | 0.0968 |
| chr2 | 53481972 | 53489777 | 7806 | -2.56 | 0.0883 | 0.0968 |
| chr2 | 214326849 | 214342767 | 15919 | -0.63 | 0.0884 | 0.097 |
| chrX | 13868097 | 13868899 | 803 | -1.65 | 0.0885 | 0.097 |
| chr5 | 81753074 | 81761385 | 8312 | -2.8 | 0.0886 | 0.0971 |
| chr11 | 120136259 | 120138263 | 2005 | -1.61 | 0.0887 | 0.0973 |
| chr3 | 28413826 | 28953210 | 539385 | 4.67 | 0.0888 | 0.0974 |
| chrX | 134334580 | 134336121 | 1542 | 0.89 | 0.0889 | 0.0974 |
| chr14 | 36747095 | 36748030 | 936 | -2.3 | 0.089 | 0.0976 |
| chr16 | 54116239 | 54130987 | 14749 | -3.18 | 0.0891 | 0.0977 |
| chr6 | 89928544 | 89941225 | 12682 | -2.56 | 0.0891 | 0.0977 |
| chr1 | 56614279 | 56615335 | 1057 | -1.71 | 0.0892 | 0.0978 |
| chr12 | 20210802 | 20211993 | 1192 | -2.15 | 0.0893 | 0.0979 |
| chr3 | 184858979 | 184877210 | 18232 | -1.06 | 0.0893 | 0.0979 |
| chr5 | 1973430 | 2011547 | 38118 | -0.38 | 0.0894 | 0.098 |
| chr10 | 59900140 | 59905463 | 5324 | -2.62 | 0.0894 | 0.098 |
| chr4 | 134622116 | 134630264 | 8149 | 0.66 | 0.0894 | 0.098 |
| chr8 | 25541488 | 25551862 | 10375 | 0.78 | 0.0895 | 0.0981 |
| chr4 | 78177311 | 78183270 | 5960 | 0.58 | 0.0895 | 0.0981 |
| chr11 | 55030781 | 55031120 | 340 | 1.19 | 0.0898 | 0.0984 |
| chr13 | 76685199 | 76698907 | 13709 | -0.56 | 0.0898 | 0.0985 |
| chrX | 136122431 | 136128571 | 6141 | 0.74 | 0.0899 | 0.0986 |
| chr17 | 64840880 | 64889688 | 48809 | -3.97 | 0.09 | 0.0986 |
| chr15 | 80545232 | 80580383 | 35152 | -0.67 | 0.0901 | 0.0987 |
| chr13 | 104838215 | 104839232 | 1018 | 1.9 | 0.0902 | 0.0988 |
| chr4 | 10559081 | 10569863 | 10783 | 0.89 | 0.0902 | 0.0988 |
| chr1 | 104364849 | 104366684 | 1836 | 2.35 | 0.0902 | 0.0988 |
| chr4 | 134424131 | 134426795 | 2665 | -1.44 | 0.0902 | 0.0988 |
| chr4 | 30350684 | 30363978 | 13295 | -1.25 | 0.0902 | 0.0989 |
| chr5 | 61263517 | 61327519 | 64003 | 0.29 | 0.0903 | 0.0989 |
| chrX | 71200069 | 71202204 | 2136 | -0.64 | 0.0903 | 0.0989 |
| chr3 | 61980230 | 62117217 | 136988 | 4.18 | 0.0903 | 0.099 |
| chr4 | 63421368 | 63424037 | 2670 | -2.27 | 0.0904 | 0.099 |
| chr11 | 49229610 | 49230462 | 853 | 2.23 | 0.0904 | 0.0991 |
| chr20 | 24863320 | 24865189 | 1870 | -1.26 | 0.0905 | 0.0992 |
| chr5 | 129113383 | 129114724 | 1342 | -1.48 | 0.0906 | 0.0993 |
| chr12 | 60160855 | 60161580 | 726 | -2.01 | 0.0906 | 0.0993 |
| chr1 | 40611184 | 40613373 | 2190 | -0.77 | 0.0906 | 0.0993 |
| chr6 | 100407017 | 100408288 | 1272 | -1.52 | 0.0906 | 0.0993 |
| chr10 | 116770099 | 116772020 | 1922 | -1.15 | 0.0906 | 0.0993 |
| chr11 | 118218959 | 118220225 | 1267 | 1.01 | 0.0907 | 0.0993 |
| chr1 | 168537027 | 168540239 | 3213 | -0.72 | 0.0907 | 0.0994 |
| chr7 | 131501948 | 131512626 | 10679 | -0.87 | 0.0908 | 0.0994 |
| chr18 | 58007164 | 58011681 | 4518 | 0.75 | 0.0908 | 0.0995 |
| chr6 | 165357041 | 165358596 | 1556 | -2.25 | 0.0911 | 0.0998 |
| chr12 | 412293 | 422060 | 9768 | -3 | 0.0912 | 0.0998 |
| chr8 | 130208165 | 130319172 | 111008 | -4.09 | 0.0912 | 0.0999 |
| chr7 | 119522883 | 119564880 | 41998 | 0.77 | 0.0912 | 0.0999 |
| chr18 | 10041331 | 10043210 | 1880 | -2.11 | 0.0913 | 0.0999 |
| chr3 | 137673927 | 137674936 | 1010 | -2.17 | 0.0913 | 0.1 |
| chr9 | 19505424 | 19526551 | 21128 | 1.24 | 0.0914 | 0.1 |
| chr3 | 175906598 | 175924650 | 18053 | 0.87 | 0.0915 | 0.1 |
| chr2 | 134361685 | 134413227 | 51543 | 0.55 | 0.0915 | 0.1 |
| chr11 | 92137296 | 92138527 | 1232 | -2.2 | 0.0915 | 0.1 |
| chr1 | 56725780 | 56736683 | 10904 | 2.59 | 0.0917 | 0.1 |

| | | | | | | |
|---|---|---|---|---|---|---|
| chr2 | 68562132 | 68577256 | 15125 | -0.62 | 0.0918 | 0.1 |
| chr4 | 183392580 | 183409525 | 16946 | -2.03 | 0.0919 | 0.101 |
| chr5 | 146070619 | 146073569 | 2951 | -1.2 | 0.0919 | 0.101 |
| chr4 | 37715754 | 37716854 | 1101 | 0.76 | 0.0919 | 0.101 |
| chr6 | 125851403 | 125856575 | 5173 | 2.68 | 0.0919 | 0.101 |
| chr11 | 54834766 | 54876370 | 41605 | 0.41 | 0.092 | 0.101 |
| chr4 | 13216441 | 13235764 | 19324 | 0.51 | 0.0921 | 0.101 |
| chr1 | 179680529 | 179722424 | 41896 | -3.99 | 0.0921 | 0.101 |
| chr5 | 134999849 | 135001697 | 1849 | 0.85 | 0.0921 | 0.101 |
| chr6 | 25044933 | 25047554 | 2622 | 1.4 | 0.0921 | 0.101 |
| chr5 | 100051182 | 100051726 | 545 | -1.66 | 0.0921 | 0.101 |
| chr2 | 166342036 | 166342669 | 634 | -2.13 | 0.0921 | 0.101 |
| chr2 | 100825627 | 100841650 | 16024 | -0.62 | 0.0922 | 0.101 |
| chr7 | 32421089 | 32422600 | 1512 | 0.98 | 0.0923 | 0.101 |
| chr2 | 106668094 | 106677706 | 9613 | -1.31 | 0.0923 | 0.101 |
| chr11 | 128577194 | 128582957 | 5764 | -1.04 | 0.0923 | 0.101 |
| chr17 | 48938319 | 48940203 | 1885 | -0.67 | 0.0925 | 0.101 |
| chr1 | 146978744 | 146980157 | 1414 | 0.84 | 0.0926 | 0.101 |
| chr4 | 24341320 | 24342581 | 1262 | 1.19 | 0.0927 | 0.101 |
| chr3 | 167823743 | 167825307 | 1565 | -2.39 | 0.0927 | 0.101 |
| chr13 | 95533056 | 95534411 | 1356 | 0.94 | 0.0928 | 0.102 |
| chr1 | 182765354 | 182765982 | 629 | -2.08 | 0.0928 | 0.102 |
| chr6 | 100222856 | 100223839 | 984 | -1.49 | 0.0928 | 0.102 |
| chr13 | 27036647 | 27039371 | 2725 | 0.97 | 0.0928 | 0.102 |
| chr5 | 76825225 | 76846879 | 21655 | -2.7 | 0.093 | 0.102 |
| chr17 | 21908708 | 21912359 | 3652 | 0.56 | 0.0931 | 0.102 |
| chr3 | 40072204 | 40073169 | 966 | -0.77 | 0.0932 | 0.102 |
| chrX | 106540137 | 106549496 | 9360 | -0.82 | 0.0936 | 0.102 |
| chr4 | 147546299 | 147547012 | 714 | 1.1 | 0.0936 | 0.102 |
| chr3 | 20310474 | 20310989 | 516 | -1.16 | 0.0937 | 0.102 |
| chr13 | 29166749 | 29168129 | 1381 | 0.77 | 0.0937 | 0.103 |
| chr22 | 17462227 | 17475731 | 13505 | -0.55 | 0.0939 | 0.103 |
| chr11 | 114356532 | 114359255 | 2724 | 1.22 | 0.0939 | 0.103 |
| chr11 | 93350514 | 93351284 | 771 | -1.59 | 0.0939 | 0.103 |
| chr2 | 41883534 | 41884188 | 655 | -1.31 | 0.094 | 0.103 |
| chr7 | 15515715 | 15518030 | 2316 | 2.49 | 0.094 | 0.103 |
| chr5 | 157539456 | 157540863 | 1408 | -1.44 | 0.094 | 0.103 |
| chr10 | 117475178 | 117479149 | 3972 | 1.99 | 0.0941 | 0.103 |
| chr11 | 114253299 | 114255471 | 2173 | -2.32 | 0.0942 | 0.103 |
| chr13 | 107301500 | 107303783 | 2284 | -2.39 | 0.0942 | 0.103 |
| chr3 | 51051381 | 51053137 | 1757 | -1.35 | 0.0942 | 0.103 |
| chr4 | 186845026 | 186845453 | 428 | -1.91 | 0.0942 | 0.103 |
| chr1 | 244925872 | 244939210 | 13339 | -3.12 | 0.0943 | 0.103 |
| chr9 | 35281708 | 35287526 | 5819 | -3.12 | 0.0943 | 0.103 |
| chr1 | 100726571 | 100727067 | 497 | -1.64 | 0.0943 | 0.103 |
| chr4 | 145243957 | 145245880 | 1924 | 1.84 | 0.0943 | 0.103 |
| chr8 | 62211548 | 62215214 | 3667 | -0.72 | 0.0944 | 0.103 |
| chr4 | 85367063 | 85369566 | 2504 | 1.77 | 0.0944 | 0.103 |
| chr4 | 60222165 | 60225867 | 3703 | -1.62 | 0.0944 | 0.103 |
| chr4 | 112746563 | 112748567 | 2005 | 2.39 | 0.0944 | 0.103 |
| chrX | 140473112 | 140473612 | 501 | -1.84 | 0.0946 | 0.103 |
| chr14 | 20997485 | 20998556 | 1072 | 1.28 | 0.0946 | 0.103 |
| chrX | 139771180 | 139777633 | 6454 | 1.39 | 0.0947 | 0.104 |
| chr1 | 236911321 | 236913950 | 2630 | -0.89 | 0.0947 | 0.104 |
| chr3 | 83697899 | 83748486 | 50588 | -1.5 | 0.0947 | 0.104 |
| chr9 | 110137886 | 110139483 | 1598 | -2.13 | 0.0947 | 0.104 |
| chr12 | 71134298 | 71144025 | 9728 | 0.55 | 0.0948 | 0.104 |
| chr12 | 131991050 | 132008284 | 17235 | -2.72 | 0.0948 | 0.104 |
| chr8 | 107995664 | 107996130 | 467 | -1.16 | 0.0949 | 0.104 |
| chr7 | 120552743 | 120554502 | 1760 | -1.15 | 0.0949 | 0.104 |
| chrX | 131238152 | 131239517 | 1366 | -1.58 | 0.0949 | 0.104 |

| | | | | | | |
|---|---|---|---|---|---|---|
| chr11 | 49023980 | 49025999 | 2020 | 0.83 | 0.0951 | 0.104 |
| chr9 | 103800612 | 103817293 | 16682 | 2.76 | 0.0951 | 0.104 |
| chr13 | 51393013 | 51395536 | 2524 | 0.82 | 0.0951 | 0.104 |
| chr2 | 88521468 | 88526863 | 5396 | 2.6 | 0.0953 | 0.104 |
| chr16 | 56246270 | 56256641 | 10372 | 1.97 | 0.0954 | 0.104 |
| chr6 | 45818290 | 45844033 | 25744 | 0.67 | 0.0954 | 0.104 |
| chr9 | 79394400 | 79415938 | 21539 | 0.68 | 0.0954 | 0.104 |
| chr6 | 84237798 | 84239891 | 2094 | -0.68 | 0.0954 | 0.104 |
| chr21 | 18362075 | 18363681 | 1607 | 1.03 | 0.0955 | 0.104 |
| chr8 | 62186485 | 62186903 | 419 | 1.81 | 0.0955 | 0.104 |
| chr4 | 76095477 | 76096464 | 988 | 0.94 | 0.0956 | 0.104 |
| chr1 | 104404486 | 104405447 | 962 | -0.88 | 0.0956 | 0.104 |
| chr3 | 134439758 | 134441332 | 1575 | 1.41 | 0.0957 | 0.105 |
| chr6 | 86562692 | 86563289 | 598 | 1.51 | 0.0957 | 0.105 |
| chr5 | 40317198 | 40317698 | 501 | -0.9 | 0.0957 | 0.105 |
| chr17 | 41945176 | 41946393 | 1218 | -0.99 | 0.0958 | 0.105 |
| chr4 | 134738260 | 134739578 | 1319 | -1.36 | 0.0959 | 0.105 |
| chr5 | 83296034 | 83299367 | 3334 | -2.39 | 0.096 | 0.105 |
| chr18 | 75099978 | 75176765 | 76788 | 0.27 | 0.096 | 0.105 |
| chr13 | 22418982 | 22420603 | 1622 | 0.71 | 0.096 | 0.105 |
| chr5 | 149467226 | 149475993 | 8768 | -1.38 | 0.0963 | 0.105 |
| chr11 | 18835433 | 18890863 | 55431 | -2.86 | 0.0967 | 0.106 |
| chr12 | 117856347 | 117863773 | 7427 | -2.35 | 0.0969 | 0.106 |
| chr20 | 44294890 | 44340990 | 46101 | -0.41 | 0.0969 | 0.106 |
| chr5 | 150939457 | 150948766 | 9310 | 0.49 | 0.0969 | 0.106 |
| chr18 | 52034882 | 52049899 | 15018 | 0.95 | 0.0969 | 0.106 |
| chr22 | 47778946 | 47780863 | 1918 | 1.25 | 0.097 | 0.106 |
| chr1 | 219730510 | 219731831 | 1322 | 0.84 | 0.097 | 0.106 |
| chr17 | 34280918 | 34281742 | 825 | 2.25 | 0.0971 | 0.106 |
| chr8 | 112273686 | 112275134 | 1449 | -1.13 | 0.0972 | 0.106 |
| chr8 | 125129344 | 125147420 | 18077 | -2.65 | 0.0972 | 0.106 |
| chr7 | 144201234 | 144203678 | 2445 | 2.38 | 0.0972 | 0.106 |
| chr1 | 182031537 | 182047128 | 15592 | -2.69 | 0.0972 | 0.106 |
| chr1 | 115360065 | 115362289 | 2225 | 0.93 | 0.0973 | 0.106 |
| chr11 | 51500746 | 51501397 | 652 | 2.23 | 0.0974 | 0.106 |
| chr15 | 38054878 | 38070052 | 15175 | -0.65 | 0.0975 | 0.106 |
| chr10 | 62817312 | 62856813 | 39502 | -2.77 | 0.0975 | 0.106 |
| chr12 | 3459416 | 3461295 | 1880 | -1.64 | 0.0976 | 0.107 |
| chr2 | 134587785 | 134629236 | 41452 | -0.79 | 0.0976 | 0.107 |
| chr20 | 58104777 | 58126086 | 21310 | -0.37 | 0.0977 | 0.107 |
| chr15 | 96787968 | 96790891 | 2924 | -0.83 | 0.0977 | 0.107 |
| chr5 | 171131302 | 171131979 | 678 | 0.95 | 0.0977 | 0.107 |
| chr7 | 156180969 | 156243328 | 62360 | -0.49 | 0.0978 | 0.107 |
| chr15 | 95939452 | 95942804 | 3353 | 0.85 | 0.0978 | 0.107 |
| chr7 | 103029873 | 103031767 | 1895 | -0.85 | 0.0978 | 0.107 |
| chr3 | 45385571 | 45425911 | 40341 | -2.73 | 0.0978 | 0.107 |
| chr5 | 167541046 | 167599380 | 58335 | 0.78 | 0.098 | 0.107 |
| chr5 | 132473126 | 132473710 | 585 | -1.97 | 0.0981 | 0.107 |
| chr6 | 66482014 | 66483865 | 1852 | 1.23 | 0.0982 | 0.107 |
| chr3 | 169658931 | 169660523 | 1593 | -2.36 | 0.0983 | 0.107 |
| chr3 | 152339670 | 152340866 | 1197 | 1.4 | 0.0984 | 0.107 |
| chr3 | 83369028 | 83511414 | 142387 | -1.12 | 0.0984 | 0.107 |
| chrX | 108174084 | 108176113 | 2030 | -1.1 | 0.0985 | 0.107 |
| chr4 | 34515122 | 34517893 | 2772 | -1.22 | 0.0985 | 0.108 |
| chr4 | 108381161 | 108381706 | 546 | -1.45 | 0.0985 | 0.108 |
| chr4 | 61096110 | 61098104 | 1995 | 0.96 | 0.0986 | 0.108 |
| chrX | 3320120 | 3321498 | 1379 | -0.75 | 0.0986 | 0.108 |
| chr12 | 59768609 | 59785169 | 16561 | -1.93 | 0.0986 | 0.108 |
| chrX | 151963010 | 151963418 | 409 | -1.86 | 0.0986 | 0.108 |
| chr13 | 42078068 | 42079088 | 1021 | -2.25 | 0.0987 | 0.108 |
| chr11 | 14154057 | 14154381 | 325 | 1.98 | 0.0987 | 0.108 |

| | | | | | | |
|---|---|---|---|---|---|---|
| chr4 | 147006495 | 147008770 | 2276 | -1.66 | 0.0987 | 0.108 |
| chr10 | 87264695 | 87272554 | 7860 | 0.54 | 0.0988 | 0.108 |
| chr1 | 229268292 | 229272409 | 4118 | -2.51 | 0.0989 | 0.108 |
| chr12 | 26890853 | 26893262 | 2410 | -2.55 | 0.099 | 0.108 |
| chr11 | 104488680 | 104489101 | 422 | 1.36 | 0.0991 | 0.108 |
| chr16 | 20091856 | 20259789 | 167934 | -0.32 | 0.0993 | 0.108 |
| chr7 | 87119915 | 87124204 | 4290 | -2.43 | 0.0993 | 0.108 |
| chr5 | 129344151 | 129352241 | 8091 | -0.77 | 0.0994 | 0.108 |
| chr3 | 67909702 | 67910252 | 551 | -1.97 | 0.0994 | 0.108 |
| chr1 | 187598850 | 187609454 | 10605 | -2.6 | 0.0995 | 0.108 |
| chr4 | 135092289 | 135112177 | 19889 | 0.53 | 0.0996 | 0.109 |
| chr5 | 120148318 | 120155164 | 6847 | -0.96 | 0.0997 | 0.109 |
| chrX | 3290711 | 3297525 | 6815 | -0.75 | 0.0997 | 0.109 |
| chr14 | 96364503 | 96401965 | 37463 | 0.51 | 0.0998 | 0.109 |
| chr10 | 112381207 | 112388465 | 7259 | -3.05 | 0.1 | 0.109 |
| chr4 | 65730866 | 65803887 | 73022 | 0.58 | 0.1 | 0.109 |
| chr1 | 116725834 | 116733192 | 7359 | 1.11 | 0.1 | 0.109 |
| chr1 | 147387094 | 147389952 | 2859 | 2.28 | 0.1 | 0.109 |
| chr9 | 17818559 | 17821073 | 2515 | -0.82 | 0.1 | 0.109 |
| chr4 | 32851644 | 32858910 | 7267 | 0.55 | 0.1 | 0.109 |
| chr3 | 84798257 | 84843917 | 45661 | -1.05 | 0.1 | 0.109 |
| chr7 | 108969881 | 108970500 | 620 | -1.2 | 0.1 | 0.109 |
| chr8 | 61247979 | 61248706 | 728 | 0.89 | 0.1 | 0.109 |
| chr10 | 61831427 | 61835836 | 4410 | 0.91 | 0.1 | 0.109 |
| chr8 | 114518268 | 114564151 | 45884 | -0.91 | 0.1 | 0.109 |
| chr13 | 113625816 | 113626427 | 612 | 1.34 | 0.1 | 0.109 |
| chr12 | 38970070 | 38972256 | 2187 | -0.66 | 0.1 | 0.109 |
| chr8 | 26538872 | 26539906 | 1035 | 1.13 | 0.101 | 0.11 |
| chr3 | 121925425 | 121927940 | 2516 | 0.87 | 0.101 | 0.11 |
| chr3 | 37750179 | 37776661 | 26483 | -2.74 | 0.101 | 0.11 |
| chr1 | 187733214 | 187734467 | 1254 | -0.83 | 0.101 | 0.11 |
| chr11 | 61967888 | 61969138 | 1251 | 1.2 | 0.101 | 0.11 |
| chr17 | 49626380 | 49626914 | 535 | -2.11 | 0.101 | 0.11 |
| chr12 | 4502705 | 4548156 | 45452 | -0.54 | 0.101 | 0.11 |
| chrX | 101107285 | 101108234 | 950 | -1.77 | 0.101 | 0.11 |
| chrX | 82711972 | 82713934 | 1963 | 2.45 | 0.101 | 0.11 |
| chr5 | 73489837 | 73490265 | 429 | -0.97 | 0.101 | 0.11 |
| chr5 | 134931469 | 134932619 | 1151 | -1.61 | 0.101 | 0.11 |
| chr2 | 235291353 | 235294989 | 3637 | -2.31 | 0.101 | 0.11 |
| chr4 | 129245150 | 129246748 | 1599 | -0.8 | 0.101 | 0.11 |
| chr4 | 16100622 | 16103181 | 2560 | -1.5 | 0.101 | 0.11 |
| chr9 | 21422939 | 21445048 | 22110 | -2.73 | 0.101 | 0.11 |
| chr2 | 177572983 | 177574920 | 1938 | 1.84 | 0.101 | 0.11 |
| chr9 | 14916272 | 14917549 | 1278 | -1.1 | 0.101 | 0.11 |
| chr11 | 122358314 | 122371606 | 13293 | 2.02 | 0.101 | 0.11 |
| chrX | 3130803 | 3143294 | 12492 | 0.59 | 0.101 | 0.111 |
| chr7 | 40293360 | 40297481 | 4122 | 1.66 | 0.101 | 0.111 |
| chr6 | 144368213 | 144370620 | 2408 | -2.19 | 0.102 | 0.111 |
| chr6 | 46086485 | 46087472 | 988 | -1.94 | 0.102 | 0.111 |
| chr5 | 89896342 | 89897626 | 1285 | -2.22 | 0.102 | 0.111 |
| chr14 | 60862827 | 60895272 | 32446 | -0.59 | 0.102 | 0.111 |
| chr6 | 167296115 | 167298313 | 2199 | 0.63 | 0.102 | 0.111 |
| chr12 | 76421237 | 76423725 | 2489 | -1.16 | 0.102 | 0.111 |
| chr3 | 17141466 | 17142738 | 1273 | -2.18 | 0.102 | 0.111 |
| chr2 | 188010933 | 188021903 | 10971 | -1.43 | 0.102 | 0.111 |
| chr14 | 43356192 | 43357507 | 1316 | -2 | 0.102 | 0.111 |
| chrX | 68784043 | 68784321 | 279 | -1.71 | 0.102 | 0.111 |
| chr3 | 106000317 | 106004050 | 3734 | 0.86 | 0.102 | 0.111 |
| chr1 | 166927203 | 166928311 | 1109 | -1.06 | 0.103 | 0.112 |
| chrX | 42164912 | 42189182 | 24271 | -0.44 | 0.103 | 0.112 |
| chr8 | 62386498 | 62394034 | 7537 | -1.11 | 0.103 | 0.112 |

| | | | | | | |
|---|---|---|---|---|---|---|
| chr13 | 79821756 | 79823299 | 1544 | -0.99 | 0.103 | 0.112 |
| chr7 | 57286090 | 57288249 | 2160 | 0.84 | 0.103 | 0.112 |
| chr7 | 92689714 | 92765745 | 76032 | -3.84 | 0.103 | 0.112 |
| chr8 | 82244346 | 82245368 | 1023 | -2.11 | 0.103 | 0.112 |
| chr2 | 15850007 | 15852493 | 2487 | 1.16 | 0.103 | 0.112 |
| chr6 | 130911801 | 130913591 | 1791 | -2.36 | 0.103 | 0.112 |
| chr1 | 170568431 | 170609274 | 40844 | -0.74 | 0.103 | 0.112 |
| chr17 | 56300206 | 56339954 | 39749 | -1.75 | 0.103 | 0.112 |
| chr4 | 22833504 | 22841302 | 7799 | -2.5 | 0.103 | 0.112 |
| chr3 | 55329384 | 55345144 | 15761 | -0.5 | 0.103 | 0.112 |
| chr12 | 110073891 | 110074491 | 601 | 1.93 | 0.103 | 0.113 |
| chr15 | 79849431 | 79878523 | 29093 | -0.65 | 0.103 | 0.113 |
| chr4 | 108723896 | 108725424 | 1529 | -2.11 | 0.104 | 0.113 |
| chr18 | 48241814 | 48242525 | 712 | -2 | 0.104 | 0.113 |
| chr20 | 13247684 | 13252961 | 5278 | 0.48 | 0.104 | 0.113 |
| chr5 | 89715413 | 89715946 | 534 | -2.02 | 0.104 | 0.113 |
| chr7 | 19125639 | 19135757 | 10119 | -2.52 | 0.104 | 0.113 |
| chr4 | 63065926 | 63111051 | 45126 | 0.54 | 0.104 | 0.113 |
| chrX | 115676346 | 115681599 | 5254 | -2.32 | 0.104 | 0.113 |
| chr17 | 9667751 | 9668202 | 452 | 2.11 | 0.104 | 0.113 |
| chr8 | 113234733 | 113325642 | 90910 | -2.15 | 0.104 | 0.114 |
| chr17 | 62344213 | 62349961 | 5749 | -2.41 | 0.104 | 0.114 |
| chr13 | 25698786 | 25723122 | 24337 | -0.68 | 0.104 | 0.114 |
| chr3 | 86087588 | 86201953 | 114366 | -0.72 | 0.105 | 0.114 |
| chr7 | 119149915 | 119175869 | 25955 | -0.97 | 0.105 | 0.114 |
| chr8 | 91568350 | 91570126 | 1777 | 1.3 | 0.105 | 0.114 |
| chr3 | 69518508 | 69524360 | 5853 | 0.55 | 0.105 | 0.114 |
| chr2 | 177620233 | 177649185 | 28953 | 0.65 | 0.105 | 0.114 |
| chr7 | 40206099 | 40207052 | 954 | 0.92 | 0.105 | 0.114 |
| chr4 | 55138426 | 55140252 | 1827 | 1.26 | 0.105 | 0.114 |
| chrX | 16376138 | 16378242 | 2105 | -1.47 | 0.105 | 0.115 |
| chr18 | 3939329 | 3951657 | 12329 | -1.47 | 0.105 | 0.115 |
| chr4 | 25002188 | 25004336 | 2149 | 2.31 | 0.105 | 0.115 |
| chr15 | 81483151 | 81513050 | 29900 | -0.45 | 0.106 | 0.115 |
| chr4 | 76359341 | 76360522 | 1182 | 1.65 | 0.106 | 0.115 |
| chr2 | 176271839 | 176346346 | 74508 | 3.89 | 0.106 | 0.115 |
| chr10 | 16588078 | 16604402 | 16325 | -1.38 | 0.106 | 0.115 |
| chr3 | 142922554 | 142924256 | 1703 | -0.97 | 0.106 | 0.115 |
| chr6 | 112733054 | 112748336 | 15283 | -0.52 | 0.106 | 0.116 |
| chr12 | 100128272 | 100128695 | 424 | 1.53 | 0.106 | 0.116 |
| chr1 | 51575166 | 51599862 | 24697 | -2.98 | 0.106 | 0.116 |
| chr12 | 120453760 | 120462815 | 9056 | -2.98 | 0.106 | 0.116 |
| chr9 | 133424524 | 133438696 | 14173 | -2.98 | 0.106 | 0.116 |
| chr18 | 48790278 | 48791207 | 930 | -1.67 | 0.106 | 0.116 |
| chr15 | 29734721 | 29736006 | 1286 | -0.77 | 0.106 | 0.116 |
| chr9 | 15746121 | 15763905 | 17785 | -2.24 | 0.107 | 0.116 |
| chr4 | 33300132 | 33301561 | 1430 | -1.18 | 0.107 | 0.116 |
| chr18 | 70621121 | 70628438 | 7318 | 0.92 | 0.107 | 0.116 |
| chr5 | 126438065 | 126438617 | 553 | -1.91 | 0.107 | 0.116 |
| chr14 | 22479421 | 22503962 | 24542 | 2.77 | 0.107 | 0.116 |
| chr2 | 177856183 | 177857504 | 1322 | -1.66 | 0.107 | 0.116 |
| chrX | 62159589 | 62221559 | 61971 | 0.42 | 0.107 | 0.116 |
| chr18 | 47609227 | 47610710 | 1484 | 1.05 | 0.107 | 0.116 |
| chr7 | 70062972 | 70119804 | 56833 | -0.47 | 0.107 | 0.117 |
| chr15 | 69403900 | 69405716 | 1817 | 2.3 | 0.107 | 0.117 |
| chr4 | 101587865 | 101588208 | 344 | -1.82 | 0.108 | 0.117 |
| chr13 | 88937736 | 88939773 | 2038 | 0.86 | 0.108 | 0.117 |
| chr4 | 16110848 | 16111341 | 494 | 1.04 | 0.108 | 0.117 |
| chrX | 119996681 | 119999589 | 2909 | 1.4 | 0.108 | 0.117 |
| chr18 | 64222777 | 64224795 | 2019 | -2.25 | 0.108 | 0.117 |
| chr5 | 6153536 | 6157607 | 4072 | -1.05 | 0.108 | 0.117 |

| | | | | | | |
|---|---|---|---|---|---|---|
| chr10 | 134606623 | 134615987 | 9365 | -2.28 | 0.108 | 0.117 |
| chr21 | 14392434 | 14392815 | 382 | -1.14 | 0.108 | 0.118 |
| chr17 | 20822015 | 20823026 | 1012 | -0.93 | 0.108 | 0.118 |
| chr4 | 188235637 | 188247939 | 12303 | 1.48 | 0.108 | 0.118 |
| chr7 | 68961847 | 68967670 | 5824 | -2.54 | 0.108 | 0.118 |
| chr3 | 37789366 | 37796861 | 7496 | 0.55 | 0.108 | 0.118 |
| chr5 | 139432797 | 139442861 | 10065 | -0.55 | 0.109 | 0.118 |
| chr11 | 106462657 | 106463690 | 1034 | 1.65 | 0.109 | 0.118 |
| chr6 | 85911419 | 85911961 | 543 | -1.11 | 0.109 | 0.118 |
| chr18 | 47590846 | 47593396 | 2551 | 0.73 | 0.109 | 0.118 |
| chr9 | 93462081 | 93503766 | 41686 | -0.59 | 0.109 | 0.118 |
| chr14 | 106483787 | 106529708 | 45922 | -0.42 | 0.109 | 0.118 |
| chr10 | 3737796 | 3741305 | 3510 | 1.46 | 0.109 | 0.119 |
| chr9 | 88144683 | 88145434 | 752 | 1.74 | 0.109 | 0.119 |
| chr4 | 35445298 | 35465464 | 20167 | 0.65 | 0.109 | 0.119 |
| chr6 | 52078807 | 52082466 | 3660 | 0.9 | 0.109 | 0.119 |
| chr7 | 127066432 | 127066810 | 379 | -0.99 | 0.109 | 0.119 |
| chr9 | 14484616 | 14485153 | 538 | -2.02 | 0.109 | 0.119 |
| chr5 | 168730292 | 168732608 | 2317 | -2.27 | 0.109 | 0.119 |
| chr18 | 73768488 | 73929429 | 160942 | 0.28 | 0.109 | 0.119 |
| chr8 | 114811428 | 114812598 | 1171 | 0.84 | 0.11 | 0.119 |
| chr11 | 19488668 | 19512820 | 24153 | 2.74 | 0.11 | 0.119 |
| chr9 | 13363595 | 13408724 | 45130 | 0.39 | 0.11 | 0.119 |
| chr5 | 99799775 | 99801708 | 1934 | 1.21 | 0.11 | 0.119 |
| chr7 | 132855275 | 132870710 | 15436 | -2.67 | 0.11 | 0.119 |
| chr2 | 119553928 | 119555790 | 1863 | 0.68 | 0.11 | 0.119 |
| chr11 | 129464476 | 129471927 | 7452 | -0.54 | 0.11 | 0.119 |
| chr13 | 112327591 | 112353770 | 26180 | 0.55 | 0.11 | 0.119 |
| chr1 | 89008709 | 89009751 | 1043 | 2.04 | 0.11 | 0.12 |
| chr9 | 109843048 | 109853710 | 10663 | -1.55 | 0.11 | 0.12 |
| chr4 | 181627418 | 181630468 | 3051 | -0.72 | 0.11 | 0.12 |
| chr14 | 60346214 | 60349357 | 3144 | 2.48 | 0.11 | 0.12 |
| chr5 | 170108600 | 170110719 | 2120 | 1.06 | 0.11 | 0.12 |
| chr2 | 206531234 | 206535155 | 3922 | -2.41 | 0.111 | 0.12 |
| chr7 | 37110458 | 37111802 | 1345 | 2.3 | 0.111 | 0.12 |
| chr3 | 49203971 | 49204928 | 958 | -0.95 | 0.111 | 0.12 |
| chr1 | 88747232 | 88758496 | 11265 | 0.87 | 0.111 | 0.12 |
| chr4 | 6006711 | 6007438 | 728 | 1.6 | 0.111 | 0.12 |
| chr15 | 38118749 | 38119599 | 851 | 1.02 | 0.111 | 0.12 |
| chr12 | 54091209 | 54092453 | 1245 | 0.89 | 0.111 | 0.12 |
| chrX | 69194427 | 69202587 | 8161 | -1.48 | 0.111 | 0.121 |
| chr7 | 112735678 | 112740059 | 4382 | 2.51 | 0.111 | 0.121 |
| chr4 | 88812662 | 88813160 | 499 | -1.97 | 0.111 | 0.121 |
| chrX | 124509728 | 124519076 | 9349 | 0.6 | 0.111 | 0.121 |
| chr1 | 218080686 | 218082410 | 1725 | -2.22 | 0.111 | 0.121 |
| chr21 | 19020527 | 19044943 | 24417 | -0.54 | 0.111 | 0.121 |
| chr6 | 112322591 | 112323737 | 1147 | -2.08 | 0.111 | 0.121 |
| chr6 | 112830109 | 112831695 | 1587 | -2.12 | 0.112 | 0.121 |
| chr12 | 22535669 | 22557760 | 22092 | -1.73 | 0.112 | 0.121 |
| chr7 | 86675251 | 86680261 | 5011 | -2.21 | 0.112 | 0.121 |
| chrX | 36732353 | 36733800 | 1448 | 1.27 | 0.112 | 0.121 |
| chr13 | 48774522 | 48775212 | 691 | -2.03 | 0.112 | 0.122 |
| chr11 | 101226589 | 101235192 | 8604 | -0.51 | 0.112 | 0.122 |
| chr7 | 106122495 | 106132112 | 9618 | -1.33 | 0.112 | 0.122 |
| chr1 | 239902808 | 239904643 | 1836 | -0.64 | 0.112 | 0.122 |
| chr12 | 91026520 | 91042119 | 15600 | -0.95 | 0.112 | 0.122 |
| chr7 | 24430397 | 24432073 | 1677 | 2.28 | 0.112 | 0.122 |
| chr13 | 88789098 | 88813634 | 24537 | 0.66 | 0.112 | 0.122 |
| chr5 | 95387593 | 95388771 | 1179 | -2.19 | 0.113 | 0.122 |
| chr4 | 81963824 | 81967621 | 3798 | 1.69 | 0.113 | 0.122 |
| chr6 | 139411226 | 139412353 | 1128 | -0.84 | 0.113 | 0.122 |

| | | | | | | |
|---|---|---|---|---|---|---|
| chr3 | 176595870 | 176599767 | 3898 | 0.61 | 0.113 | 0.122 |
| chr1 | 158098224 | 158176284 | 78061 | -1.1 | 0.113 | 0.122 |
| chr2 | 75555294 | 75556300 | 1007 | -0.9 | 0.113 | 0.122 |
| chr7 | 106515343 | 106559257 | 43915 | 2.78 | 0.113 | 0.123 |
| chrX | 148536541 | 148539025 | 2485 | 0.69 | 0.113 | 0.123 |
| chr12 | 60244809 | 60245609 | 801 | 1.42 | 0.113 | 0.123 |
| chr7 | 120645033 | 120645432 | 400 | -1.05 | 0.113 | 0.123 |
| chr4 | 56512435 | 56514031 | 1597 | 1.33 | 0.113 | 0.123 |
| chr12 | 29518068 | 29531091 | 13024 | -2.9 | 0.113 | 0.123 |
| chr15 | 68770021 | 68784420 | 14400 | -2.9 | 0.113 | 0.123 |
| chr17 | 39033167 | 39057636 | 24470 | -2.9 | 0.113 | 0.123 |
| chr20 | 17461377 | 17466118 | 4742 | -2.9 | 0.113 | 0.123 |
| chr22 | 31893454 | 31903837 | 10384 | -2.9 | 0.113 | 0.123 |
| chr22 | 40551010 | 40561945 | 10936 | -2.9 | 0.113 | 0.123 |
| chr6 | 89632630 | 89644110 | 11481 | -2.9 | 0.113 | 0.123 |
| chr8 | 121942126 | 121952530 | 10405 | -2.9 | 0.113 | 0.123 |
| chr7 | 30798297 | 30800488 | 2192 | -2.26 | 0.113 | 0.123 |
| chr7 | 37119814 | 37121325 | 1512 | -2.13 | 0.114 | 0.123 |
| chr11 | 111291154 | 111291630 | 477 | -1.57 | 0.114 | 0.123 |
| chrX | 135762102 | 135767984 | 5883 | 1.6 | 0.114 | 0.124 |
| chr16 | 10975565 | 10976059 | 495 | -1.87 | 0.114 | 0.124 |
| chr3 | 121955592 | 121959532 | 3941 | 1.04 | 0.114 | 0.124 |
| chr1 | 39214528 | 39234094 | 19567 | -0.43 | 0.114 | 0.124 |
| chr17 | 39414217 | 39425537 | 11321 | 1.25 | 0.114 | 0.124 |
| chr21 | 39891187 | 40013896 | 122710 | 0.26 | 0.115 | 0.124 |
| chr1 | 241177110 | 241371558 | 194449 | 4.04 | 0.115 | 0.124 |
| chr1 | 79163131 | 79165844 | 2714 | -2.25 | 0.115 | 0.124 |
| chr11 | 88368889 | 88379800 | 10912 | -0.49 | 0.115 | 0.124 |
| chr8 | 115598503 | 115599198 | 696 | -1.92 | 0.115 | 0.125 |
| chr4 | 130298826 | 130300397 | 1572 | -1.39 | 0.115 | 0.125 |
| chr12 | 104113600 | 104130127 | 16528 | -2.62 | 0.115 | 0.125 |
| chr17 | 70287066 | 70291787 | 4722 | 0.96 | 0.115 | 0.125 |
| chrX | 57789472 | 57791200 | 1729 | 2.29 | 0.115 | 0.125 |
| chr4 | 111288218 | 111289110 | 893 | -1.99 | 0.115 | 0.125 |
| chrX | 56318040 | 56318927 | 888 | 2.17 | 0.115 | 0.125 |
| chr11 | 6205367 | 6206618 | 1252 | 1.59 | 0.115 | 0.125 |
| chr15 | 39022042 | 39026573 | 4532 | -0.5 | 0.115 | 0.125 |
| chr8 | 76247011 | 76266603 | 19593 | -2.49 | 0.115 | 0.125 |
| chr19 | 43465058 | 43465454 | 397 | -1.26 | 0.115 | 0.125 |
| chr1 | 105371787 | 105373864 | 2078 | 0.72 | 0.115 | 0.125 |
| chr2 | 36936614 | 36994155 | 57542 | -3.76 | 0.115 | 0.125 |
| chr5 | 27530225 | 27545119 | 14895 | -2.05 | 0.116 | 0.125 |
| chr3 | 108603992 | 108604816 | 825 | 2.13 | 0.116 | 0.125 |
| chr9 | 18532166 | 18556715 | 24550 | -0.6 | 0.116 | 0.125 |
| chr9 | 98476841 | 98487963 | 11123 | -0.88 | 0.116 | 0.126 |
| chr4 | 123432590 | 123434034 | 1445 | 0.92 | 0.116 | 0.126 |
| chr10 | 97608208 | 97616479 | 8272 | -0.63 | 0.116 | 0.126 |
| chr1 | 104971816 | 104972985 | 1170 | 1.03 | 0.116 | 0.126 |
| chr4 | 30013718 | 30014730 | 1013 | -0.87 | 0.116 | 0.126 |
| chrX | 34681974 | 34684050 | 2077 | 1.15 | 0.116 | 0.126 |
| chr4 | 130546650 | 130548404 | 1755 | -0.8 | 0.116 | 0.126 |
| chr11 | 32267869 | 32271044 | 3176 | 2.38 | 0.116 | 0.126 |
| chr1 | 105823372 | 105869930 | 46559 | 0.48 | 0.116 | 0.126 |
| chr4 | 34009829 | 34041751 | 31923 | -0.43 | 0.116 | 0.126 |
| chr4 | 32563370 | 32563934 | 565 | -1.29 | 0.116 | 0.126 |
| chr7 | 91400420 | 91405787 | 5368 | 2.57 | 0.116 | 0.126 |
| chr4 | 15387747 | 15392577 | 4831 | 0.67 | 0.117 | 0.126 |
| chr18 | 49196679 | 49222480 | 25802 | -0.32 | 0.117 | 0.126 |
| chrX | 119493319 | 119494561 | 1243 | 2.18 | 0.117 | 0.127 |
| chr1 | 152375924 | 152377130 | 1207 | 1.32 | 0.117 | 0.127 |
| chr2 | 56631289 | 56664634 | 33346 | -0.49 | 0.117 | 0.127 |

| | | | | | | |
|---|---|---|---|---|---|---|
| chr10 | 61050736 | 61073304 | 22569 | 2.65 | 0.117 | 0.127 |
| chr2 | 135267285 | 135267719 | 435 | 1.66 | 0.117 | 0.127 |
| chr1 | 229174777 | 229189921 | 15145 | -1.17 | 0.117 | 0.127 |
| chr21 | 35434321 | 35438526 | 4206 | 0.66 | 0.117 | 0.127 |
| chr4 | 101327780 | 101329636 | 1857 | -2.27 | 0.118 | 0.127 |
| chr8 | 55182300 | 55185566 | 3267 | 2.29 | 0.118 | 0.127 |
| chr11 | 130805141 | 130817721 | 12581 | -0.52 | 0.118 | 0.128 |
| chr1 | 105326480 | 105347812 | 21333 | 0.4 | 0.118 | 0.128 |
| chr11 | 13820592 | 13833392 | 12801 | -0.66 | 0.118 | 0.128 |
| chr5 | 121242340 | 121243963 | 1624 | -0.82 | 0.118 | 0.128 |
| chr4 | 190690516 | 190704356 | 13841 | -0.58 | 0.118 | 0.128 |
| chr10 | 64768922 | 64785554 | 16633 | 0.37 | 0.118 | 0.128 |
| chr7 | 23951862 | 23953051 | 1190 | -0.92 | 0.118 | 0.128 |
| chr1 | 104637961 | 104640827 | 2867 | 1.12 | 0.118 | 0.128 |
| chr11 | 19608761 | 19610190 | 1430 | -0.98 | 0.118 | 0.128 |
| chr8 | 16630925 | 16632792 | 1868 | 0.63 | 0.118 | 0.128 |
| chr4 | 19383989 | 19395745 | 11757 | 0.76 | 0.119 | 0.128 |
| chr10 | 17350085 | 17363559 | 13475 | 1.69 | 0.119 | 0.129 |
| chr6 | 12790967 | 12800167 | 9201 | 0.5 | 0.119 | 0.129 |
| chr15 | 26245714 | 26248212 | 2499 | 0.88 | 0.119 | 0.129 |
| chrX | 109696590 | 109699564 | 2975 | 1.58 | 0.119 | 0.129 |
| chr2 | 73964885 | 73966377 | 1493 | -1.12 | 0.119 | 0.129 |
| chr3 | 139650711 | 139652660 | 1950 | -1.08 | 0.12 | 0.129 |
| chr7 | 124063983 | 124065371 | 1389 | -1.36 | 0.12 | 0.129 |
| chr5 | 32517115 | 32519981 | 2867 | -1.03 | 0.12 | 0.13 |
| chr7 | 97426113 | 97435770 | 9658 | 2.55 | 0.12 | 0.13 |
| chrX | 93794624 | 93849380 | 54757 | 0.47 | 0.12 | 0.13 |
| chr8 | 16773043 | 16775670 | 2628 | 1.33 | 0.12 | 0.13 |
| chr4 | 63037352 | 63048175 | 10824 | 0.53 | 0.12 | 0.13 |
| chrX | 51405243 | 51405646 | 404 | 1.96 | 0.12 | 0.13 |
| chr11 | 124461044 | 124469993 | 8950 | -2.34 | 0.12 | 0.13 |
| chr18 | 26588235 | 26589134 | 900 | 2.11 | 0.12 | 0.13 |
| chr2 | 154720352 | 154721053 | 702 | -2.05 | 0.12 | 0.13 |
| chr4 | 33372710 | 33415466 | 42757 | -0.64 | 0.121 | 0.13 |
| chrX | 113861418 | 113866584 | 5167 | 0.83 | 0.121 | 0.131 |
| chr1 | 49291895 | 49302272 | 10378 | 0.83 | 0.121 | 0.131 |
| chr10 | 120139882 | 120163271 | 23390 | -0.34 | 0.121 | 0.131 |
| chr5 | 121137590 | 121139946 | 2357 | 2.18 | 0.121 | 0.131 |
| chr8 | 143951761 | 143989984 | 38224 | -0.52 | 0.121 | 0.131 |
| chr6 | 137623663 | 137626078 | 2416 | -2.13 | 0.121 | 0.131 |
| chr14 | 38375742 | 38376391 | 650 | -2.03 | 0.121 | 0.131 |
| chr16 | 52387439 | 52408749 | 21311 | -0.61 | 0.121 | 0.131 |
| chrX | 38961386 | 38981565 | 20180 | 0.48 | 0.121 | 0.131 |
| chr7 | 156844548 | 156911037 | 66490 | -3.77 | 0.121 | 0.131 |
| chr8 | 112339420 | 112352209 | 12790 | 0.5 | 0.121 | 0.131 |
| chr8 | 121163963 | 121166423 | 2461 | 2.31 | 0.122 | 0.131 |
| chr15 | 96385268 | 96415959 | 30692 | -1.72 | 0.122 | 0.132 |
| chr13 | 47459018 | 47460193 | 1176 | 1.52 | 0.122 | 0.132 |
| chr12 | 2697215 | 2730604 | 33390 | -0.29 | 0.122 | 0.132 |
| chr6 | 24142661 | 24143920 | 1260 | -2.12 | 0.122 | 0.132 |
| chr14 | 43719380 | 43731390 | 12011 | 1.46 | 0.122 | 0.132 |
| chr21 | 37233665 | 37249388 | 15724 | -0.92 | 0.122 | 0.132 |
| chr18 | 22315266 | 22315987 | 722 | -1.05 | 0.122 | 0.132 |
| chr6 | 38759225 | 38760314 | 1090 | -1.22 | 0.122 | 0.132 |
| chr5 | 110502962 | 110517633 | 14672 | 0.98 | 0.122 | 0.132 |
| chr6 | 46054028 | 46064081 | 10054 | -0.84 | 0.122 | 0.132 |
| chr20 | 54899542 | 54918855 | 19314 | -0.62 | 0.122 | 0.132 |
| chr4 | 118606201 | 118606970 | 770 | 1.14 | 0.123 | 0.133 |
| chr18 | 33448777 | 33453498 | 4722 | -2.38 | 0.123 | 0.133 |
| chr18 | 11198000 | 11252985 | 54986 | 0.35 | 0.123 | 0.133 |
| chr5 | 74231823 | 74235967 | 4145 | -1.06 | 0.123 | 0.133 |

| | | | | | | |
|---|---|---|---|---|---|---|
| chr3 | 51144335 | 51153937 | 9603 | -0.78 | 0.123 | 0.134 |
| chr12 | 66646058 | 66656437 | 10380 | 2.56 | 0.123 | 0.134 |
| chr15 | 58303133 | 58304954 | 1822 | 0.99 | 0.123 | 0.134 |
| chr10 | 60069105 | 60070183 | 1079 | 2.06 | 0.123 | 0.134 |
| chr3 | 69543558 | 69546601 | 3044 | -0.61 | 0.124 | 0.134 |
| chr6 | 47792657 | 47794057 | 1401 | 1.47 | 0.124 | 0.134 |
| chr11 | 32039711 | 32090458 | 50748 | -0.41 | 0.124 | 0.134 |
| chr6 | 46313460 | 46315062 | 1603 | -1.5 | 0.124 | 0.134 |
| chr17 | 69341367 | 69359617 | 18251 | -0.66 | 0.124 | 0.134 |
| chr5 | 18731786 | 18734391 | 2606 | -2.27 | 0.124 | 0.134 |
| chr2 | 207565703 | 207566941 | 1239 | -1.1 | 0.124 | 0.134 |
| chr4 | 34653443 | 34681366 | 27924 | 1.21 | 0.124 | 0.134 |
| chr10 | 120553992 | 120556203 | 2212 | -1.06 | 0.124 | 0.134 |
| chr4 | 53298191 | 53303896 | 5706 | 2.37 | 0.124 | 0.134 |
| chr5 | 158880886 | 158881974 | 1089 | -1.04 | 0.124 | 0.134 |
| chrX | 54981245 | 54984920 | 3676 | 2.23 | 0.125 | 0.135 |
| chr7 | 15272822 | 15273740 | 919 | -1.27 | 0.125 | 0.135 |
| chr2 | 196008911 | 196009634 | 724 | -1.63 | 0.125 | 0.135 |
| chr10 | 120411732 | 120413565 | 1834 | -1.28 | 0.125 | 0.135 |
| chr7 | 124192295 | 124205117 | 12823 | 1.35 | 0.125 | 0.135 |
| chr4 | 37490605 | 37491703 | 1099 | 1.91 | 0.125 | 0.135 |
| chr8 | 91414573 | 91515366 | 100794 | 3.73 | 0.126 | 0.136 |
| chr4 | 147573914 | 147574720 | 807 | -1.15 | 0.126 | 0.136 |
| chr10 | 91574906 | 91604524 | 29619 | -2.14 | 0.126 | 0.136 |
| chr18 | 27730367 | 27738260 | 7894 | 1.01 | 0.126 | 0.136 |
| chr2 | 67919959 | 67948841 | 28883 | -0.55 | 0.126 | 0.137 |
| chr9 | 91344643 | 91352171 | 7529 | 2.5 | 0.126 | 0.137 |
| chr1 | 91207903 | 91221305 | 13403 | 1.09 | 0.126 | 0.137 |
| chr7 | 38419045 | 38445484 | 26440 | -2.63 | 0.126 | 0.137 |
| chr2 | 1872360 | 1901038 | 28679 | 0.44 | 0.127 | 0.137 |
| chr1 | 105950608 | 105952722 | 2115 | 0.8 | 0.127 | 0.137 |
| chr18 | 10816603 | 10884455 | 67853 | -0.25 | 0.127 | 0.137 |
| chr7 | 22028975 | 22035417 | 6443 | -2.45 | 0.127 | 0.137 |
| chr3 | 161966129 | 161979945 | 13817 | 2.08 | 0.127 | 0.137 |
| chr9 | 112575262 | 112577512 | 2251 | 2.32 | 0.127 | 0.137 |
| chr3 | 172685459 | 172686313 | 855 | -1.33 | 0.127 | 0.137 |
| chr9 | 14495812 | 14506469 | 10658 | 1.17 | 0.127 | 0.137 |
| chr15 | 24739936 | 24742290 | 2355 | 1.86 | 0.127 | 0.137 |
| chr1 | 152326277 | 152326874 | 598 | 1.69 | 0.127 | 0.137 |
| chr17 | 3391291 | 3412504 | 21214 | -0.69 | 0.127 | 0.138 |
| chr7 | 23954399 | 23986957 | 32559 | 0.75 | 0.127 | 0.138 |
| chr1 | 120220571 | 120221639 | 1069 | -1.07 | 0.128 | 0.138 |
| chr1 | 175018325 | 175020454 | 2130 | 1.16 | 0.128 | 0.138 |
| chr18 | 22106295 | 22115110 | 8816 | -0.51 | 0.128 | 0.138 |
| chr4 | 61356468 | 61454855 | 98388 | 0.53 | 0.128 | 0.138 |
| chr1 | 245866966 | 245867920 | 955 | 1.16 | 0.128 | 0.138 |
| chr10 | 52001005 | 52002365 | 1361 | -2.12 | 0.128 | 0.138 |
| chr4 | 35732811 | 35759895 | 27085 | 0.71 | 0.128 | 0.139 |
| chr15 | 84078800 | 84097844 | 19045 | -1.77 | 0.128 | 0.139 |
| chr10 | 91697213 | 91708490 | 11278 | -1.16 | 0.128 | 0.139 |
| chr13 | 25447912 | 25449078 | 1167 | -0.94 | 0.128 | 0.139 |
| chr22 | 49867839 | 49870269 | 2431 | 0.86 | 0.129 | 0.139 |
| chr14 | 34704457 | 34705018 | 562 | 2.04 | 0.129 | 0.139 |
| chr8 | 86013540 | 86018708 | 5169 | -2.38 | 0.129 | 0.139 |
| chr2 | 214388549 | 214390456 | 1908 | -2.11 | 0.129 | 0.14 |
| chr2 | 213911113 | 213911517 | 405 | 1.11 | 0.129 | 0.14 |
| chr5 | 126973820 | 126975554 | 1735 | 1.8 | 0.13 | 0.14 |
| chr6 | 99269909 | 99270571 | 663 | 0.83 | 0.13 | 0.14 |
| chr18 | 25171464 | 25174201 | 2738 | 2.3 | 0.13 | 0.14 |
| chr12 | 7565397 | 7572261 | 6865 | -1.11 | 0.13 | 0.14 |
| chr1 | 91039115 | 91040596 | 1482 | -1.07 | 0.13 | 0.14 |

| | | | | | | |
|---|---|---|---|---|---|---|
| chrX | 135495539 | 135507421 | 11883 | -1.38 | 0.13 | 0.14 |
| chr1 | 104373321 | 104379203 | 5883 | -0.65 | 0.13 | 0.14 |
| chr7 | 119349369 | 119374682 | 25314 | -2.39 | 0.13 | 0.141 |
| chr17 | 71098815 | 71111847 | 13033 | -2.74 | 0.13 | 0.141 |
| chr4 | 136814880 | 136827992 | 13113 | -2.74 | 0.13 | 0.141 |
| chr1 | 48835289 | 48836834 | 1546 | -1.17 | 0.13 | 0.141 |
| chr18 | 11056988 | 11096280 | 39293 | 2.65 | 0.131 | 0.141 |
| chr1 | 194503588 | 194504683 | 1096 | -1.93 | 0.131 | 0.141 |
| chr5 | 111764198 | 111771202 | 7005 | 0.59 | 0.131 | 0.141 |
| chr8 | 105706595 | 105708512 | 1918 | -1.45 | 0.131 | 0.141 |
| chr7 | 26618752 | 26623554 | 4803 | 1.42 | 0.131 | 0.141 |
| chr13 | 43525074 | 43526306 | 1233 | -1.53 | 0.131 | 0.141 |
| chr14 | 59324693 | 59337382 | 12690 | 0.91 | 0.131 | 0.141 |
| chr1 | 81087425 | 81087783 | 359 | -1.7 | 0.131 | 0.142 |
| chr3 | 82931517 | 83045602 | 114086 | -1.85 | 0.131 | 0.142 |
| chr3 | 161532587 | 161557286 | 24700 | 0.49 | 0.131 | 0.142 |
| chr4 | 29239674 | 29254336 | 14663 | 0.75 | 0.132 | 0.142 |
| chr8 | 124844098 | 124847900 | 3803 | 1.35 | 0.132 | 0.142 |
| chr4 | 128106986 | 128121225 | 14240 | -0.77 | 0.132 | 0.142 |
| chr14 | 29150187 | 29159350 | 9164 | -1.15 | 0.132 | 0.142 |
| chr3 | 150695060 | 150743388 | 48329 | -0.34 | 0.132 | 0.142 |
| chr7 | 45271363 | 45273192 | 1830 | 0.75 | 0.132 | 0.143 |
| chr4 | 22284264 | 22285697 | 1434 | -1 | 0.132 | 0.143 |
| chr9 | 17105878 | 17111765 | 5888 | -2.41 | 0.132 | 0.143 |
| chr4 | 18063770 | 18078572 | 14803 | -1.22 | 0.132 | 0.143 |
| chr6 | 66469967 | 66472757 | 2791 | -1.47 | 0.132 | 0.143 |
| chr4 | 115361300 | 115362524 | 1225 | 2.11 | 0.132 | 0.143 |
| chr15 | 30310802 | 30312604 | 1803 | 0.7 | 0.132 | 0.143 |
| chr5 | 83228258 | 83229741 | 1484 | 1.14 | 0.133 | 0.143 |
| chr8 | 117926656 | 117927710 | 1055 | -2.06 | 0.133 | 0.143 |
| chr2 | 150266842 | 150268345 | 1504 | 1.24 | 0.133 | 0.143 |
| chr16 | 33592963 | 33626367 | 33405 | -0.44 | 0.133 | 0.143 |
| chr4 | 35698980 | 35716697 | 17718 | 0.6 | 0.133 | 0.144 |
| chr13 | 94563065 | 94571034 | 7970 | 2.42 | 0.133 | 0.144 |
| chr3 | 81975894 | 81976648 | 755 | 2.05 | 0.133 | 0.144 |
| chr6 | 159686431 | 159693979 | 7549 | 2.42 | 0.134 | 0.144 |
| chr3 | 40386633 | 40420058 | 33426 | -2.44 | 0.134 | 0.144 |
| chr3 | 86219991 | 86271296 | 51306 | -0.55 | 0.134 | 0.145 |
| chr2 | 209705452 | 209727577 | 22126 | 2.58 | 0.134 | 0.145 |
| chr4 | 65834938 | 65836070 | 1133 | -0.75 | 0.134 | 0.145 |
| chr5 | 99948833 | 99949748 | 916 | -2.05 | 0.134 | 0.145 |
| chr15 | 91656093 | 91659107 | 3015 | 0.92 | 0.135 | 0.145 |
| chr3 | 16690399 | 16704806 | 14408 | -0.72 | 0.135 | 0.145 |
| chr5 | 87933196 | 87933656 | 461 | -0.98 | 0.135 | 0.145 |
| chr3 | 121733352 | 121736395 | 3044 | -0.78 | 0.135 | 0.146 |
| chr6 | 8820137 | 8848481 | 28345 | -0.34 | 0.135 | 0.146 |
| chr13 | 112259953 | 112261553 | 1601 | -1.06 | 0.135 | 0.146 |
| chr10 | 82778005 | 82778637 | 633 | -1.97 | 0.135 | 0.146 |
| chr3 | 85893879 | 85920636 | 26758 | -2.4 | 0.135 | 0.146 |
| chr4 | 29816478 | 29831716 | 15239 | 0.62 | 0.135 | 0.146 |
| chrX | 127898429 | 127900270 | 1842 | -1.15 | 0.135 | 0.146 |
| chr12 | 130656402 | 130863442 | 207041 | -0.32 | 0.136 | 0.146 |
| chr18 | 32356833 | 32358552 | 1720 | 0.7 | 0.136 | 0.146 |
| chr7 | 139980433 | 139982469 | 2037 | 2.26 | 0.136 | 0.146 |
| chr18 | 43261514 | 43262035 | 522 | 0.83 | 0.136 | 0.146 |
| chr17 | 36070648 | 36079060 | 8413 | -2.53 | 0.136 | 0.147 |
| chr1 | 152013546 | 152039856 | 26311 | -0.59 | 0.136 | 0.147 |
| chr5 | 125621716 | 125665543 | 43828 | 0.43 | 0.136 | 0.147 |
| chr20 | 58183831 | 58186075 | 2245 | -2.09 | 0.136 | 0.147 |
| chr6 | 118167983 | 118169595 | 1613 | -0.79 | 0.137 | 0.147 |
| chr1 | 175082062 | 175083020 | 959 | -2.11 | 0.137 | 0.147 |

| | | | | | | |
|---|---|---|---|---|---|---|
| chr5 | 68308858 | 68318993 | 10136 | 0.41 | 0.137 | 0.147 |
| chr11 | 34551691 | 34564342 | 12652 | -0.81 | 0.137 | 0.148 |
| chr16 | 77086540 | 77163316 | 76777 | 0.47 | 0.137 | 0.148 |
| chr5 | 132901140 | 132919018 | 17879 | -0.66 | 0.137 | 0.148 |
| chrX | 72509233 | 72510428 | 1196 | 1.12 | 0.137 | 0.148 |
| chr3 | 32860546 | 32861455 | 910 | -1.78 | 0.137 | 0.148 |
| chr2 | 230467611 | 230468809 | 1199 | -1.02 | 0.137 | 0.148 |
| chr1 | 230898748 | 230917672 | 18925 | -2.38 | 0.137 | 0.148 |
| chr6 | 137715708 | 137732262 | 16555 | -1.04 | 0.138 | 0.149 |
| chrX | 51871251 | 51883588 | 12338 | 0.59 | 0.138 | 0.149 |
| chr1 | 204930341 | 204933553 | 3213 | 2.18 | 0.138 | 0.149 |
| chr3 | 153241542 | 153245734 | 4193 | 0.84 | 0.138 | 0.149 |
| chr6 | 100148187 | 100150385 | 2199 | 0.56 | 0.138 | 0.149 |
| chr12 | 42666189 | 42666966 | 778 | -1.89 | 0.138 | 0.149 |
| chr9 | 21271102 | 21308727 | 37626 | 0.55 | 0.138 | 0.149 |
| chr1 | 14180740 | 14183667 | 2928 | -1.2 | 0.139 | 0.149 |
| chrX | 20816826 | 20906206 | 89381 | -3.61 | 0.139 | 0.15 |
| chr5 | 27345233 | 27348121 | 2889 | 1.64 | 0.139 | 0.15 |
| chr10 | 17456061 | 17459804 | 3744 | -1.24 | 0.139 | 0.15 |
| chr11 | 26505876 | 26512016 | 6141 | -2.21 | 0.14 | 0.151 |
| chr12 | 4731472 | 4750685 | 19214 | -2.35 | 0.14 | 0.151 |
| chrX | 57701273 | 57702334 | 1062 | -1.54 | 0.14 | 0.151 |
| chrX | 30061913 | 30063848 | 1936 | -2.13 | 0.14 | 0.151 |
| chr4 | 60864861 | 60866392 | 1532 | -0.92 | 0.14 | 0.151 |
| chr2 | 213124458 | 213126262 | 1805 | -0.74 | 0.14 | 0.151 |
| chr9 | 13951235 | 13965868 | 14634 | 2.45 | 0.14 | 0.151 |
| chr11 | 76710564 | 76730414 | 19851 | -2.65 | 0.14 | 0.151 |
| chr13 | 43663390 | 43676187 | 12798 | -2.65 | 0.14 | 0.151 |
| chr15 | 26255606 | 26256221 | 616 | -1.46 | 0.14 | 0.151 |
| chr10 | 100952052 | 100966788 | 14737 | 2.5 | 0.141 | 0.152 |
| chr1 | 100268386 | 100270858 | 2473 | 1.69 | 0.141 | 0.152 |
| chr6 | 101519928 | 101550218 | 30291 | -2.45 | 0.141 | 0.152 |
| chr6 | 79440136 | 79441079 | 944 | 1.07 | 0.142 | 0.152 |
| chrX | 131837060 | 131839034 | 1975 | 1.7 | 0.142 | 0.152 |
| chr2 | 154956197 | 154979828 | 23632 | -2.41 | 0.142 | 0.152 |
| chr1 | 104840138 | 104868163 | 28026 | 0.77 | 0.142 | 0.153 |
| chr4 | 30205005 | 30208645 | 3641 | -0.9 | 0.142 | 0.153 |
| chr18 | 378836 | 399051 | 20216 | 0.49 | 0.142 | 0.153 |
| chrX | 125635879 | 125637146 | 1268 | -0.99 | 0.142 | 0.153 |
| chr5 | 121824743 | 121827081 | 2339 | 1.46 | 0.143 | 0.153 |
| chr8 | 37126496 | 37210046 | 83551 | -0.58 | 0.143 | 0.154 |
| chr12 | 90993279 | 91014491 | 21213 | 2.53 | 0.143 | 0.154 |
| chr4 | 190158507 | 190188921 | 30415 | -0.47 | 0.143 | 0.154 |
| chr7 | 34755232 | 34757151 | 1920 | -1.35 | 0.143 | 0.154 |
| chr13 | 103630887 | 103632285 | 1399 | 1.03 | 0.143 | 0.154 |
| chr10 | 69306461 | 69318484 | 12024 | 1.31 | 0.143 | 0.154 |
| chr15 | 61541491 | 61542024 | 534 | -1.44 | 0.143 | 0.154 |
| chr2 | 84163799 | 84166188 | 2390 | 1.11 | 0.143 | 0.154 |
| chr7 | 119328321 | 119336765 | 8445 | -2.12 | 0.143 | 0.154 |
| chr4 | 143908973 | 143912013 | 3041 | 0.85 | 0.143 | 0.154 |
| chr12 | 106320407 | 106321945 | 1539 | -1.03 | 0.143 | 0.154 |
| chr4 | 90448391 | 90448944 | 554 | -1.88 | 0.143 | 0.154 |
| chr8 | 40784465 | 40787292 | 2828 | 0.66 | 0.143 | 0.154 |
| chr3 | 86038257 | 86040065 | 1809 | -1.21 | 0.143 | 0.154 |
| chr6 | 154263594 | 154264196 | 603 | -1.12 | 0.143 | 0.154 |
| chr10 | 84109824 | 84237223 | 127400 | -0.56 | 0.143 | 0.154 |
| chr17 | 63406202 | 63473879 | 67678 | -2.58 | 0.144 | 0.154 |
| chr1 | 47509301 | 47510799 | 1499 | 1.51 | 0.144 | 0.155 |
| chr12 | 103146199 | 103149847 | 3649 | 2.35 | 0.144 | 0.155 |
| chr2 | 222647164 | 222648496 | 1333 | -0.88 | 0.144 | 0.155 |
| chr3 | 70611584 | 70630010 | 18427 | -0.41 | 0.144 | 0.155 |

| | | | | | | |
|---|---|---|---|---|---|---|
| chr5 | 33893439 | 33939817 | 46379 | 0.29 | 0.144 | 0.155 |
| chr11 | 49366102 | 49370005 | 3904 | 1.09 | 0.144 | 0.155 |
| chr13 | 23074266 | 23077673 | 3408 | 0.95 | 0.144 | 0.155 |
| chr4 | 108134976 | 108137751 | 2776 | 1.17 | 0.144 | 0.155 |
| chr3 | 15525714 | 15528346 | 2633 | -0.71 | 0.144 | 0.155 |
| chr12 | 44084305 | 44086509 | 2205 | 1.01 | 0.144 | 0.155 |
| chr8 | 80537575 | 80539101 | 1527 | -1.1 | 0.144 | 0.155 |
| chr11 | 18208332 | 18221991 | 13660 | -2.33 | 0.144 | 0.155 |
| chr8 | 120660884 | 120662655 | 1772 | -2.09 | 0.144 | 0.155 |
| chr10 | 69479773 | 69482132 | 2360 | 0.62 | 0.144 | 0.155 |
| chr12 | 106112777 | 106125280 | 12504 | -2.37 | 0.145 | 0.155 |
| chr6 | 125700840 | 125702777 | 1938 | 1.07 | 0.145 | 0.156 |
| chr2 | 215124326 | 215125659 | 1334 | -1.34 | 0.145 | 0.156 |
| chr3 | 72115818 | 72130699 | 14882 | -0.45 | 0.145 | 0.156 |
| chr7 | 50452340 | 50454106 | 1767 | 0.8 | 0.145 | 0.156 |
| chr1 | 162210963 | 162216190 | 5228 | 2.39 | 0.146 | 0.157 |
| chr6 | 143362316 | 143363653 | 1338 | 2.15 | 0.146 | 0.157 |
| chr6 | 101502511 | 101503220 | 710 | -1.46 | 0.146 | 0.157 |
| chr15 | 58376383 | 58432061 | 55679 | -0.35 | 0.146 | 0.157 |
| chr4 | 76242871 | 76259004 | 16134 | -0.78 | 0.146 | 0.157 |
| chr4 | 54542996 | 54544478 | 1483 | -2.13 | 0.146 | 0.157 |
| chr3 | 39870681 | 39913268 | 42588 | 0.5 | 0.146 | 0.157 |
| chr7 | 22492913 | 22510145 | 17233 | -2.33 | 0.147 | 0.158 |
| chr4 | 145444239 | 145444818 | 580 | -1.92 | 0.147 | 0.158 |
| chr3 | 81763836 | 81770123 | 6288 | -2.33 | 0.147 | 0.158 |
| chr12 | 78870076 | 78873430 | 3355 | -2.31 | 0.147 | 0.158 |
| chr5 | 59201463 | 59208985 | 7523 | -0.39 | 0.147 | 0.158 |
| chr12 | 91130007 | 91175711 | 45705 | 2.59 | 0.147 | 0.158 |
| chr1 | 95807591 | 95808259 | 669 | 1.35 | 0.147 | 0.158 |
| chrX | 21705502 | 21707134 | 1633 | 2.08 | 0.147 | 0.158 |
| chr1 | 120417565 | 120431592 | 14028 | -1.24 | 0.147 | 0.158 |
| chrX | 141142072 | 141162119 | 20048 | 0.42 | 0.147 | 0.158 |
| chr2 | 21098584 | 21101600 | 3017 | 2.1 | 0.147 | 0.158 |
| chrX | 25171345 | 25181956 | 10612 | 0.42 | 0.147 | 0.158 |
| chr12 | 10125570 | 10131745 | 6176 | 0.42 | 0.147 | 0.158 |
| chr3 | 24748829 | 24749138 | 310 | 0.95 | 0.148 | 0.159 |
| chr6 | 50913094 | 50914970 | 1877 | 0.69 | 0.148 | 0.159 |
| chrX | 21054339 | 21055141 | 803 | -0.98 | 0.148 | 0.159 |
| chr5 | 122528662 | 122546281 | 17620 | -0.64 | 0.148 | 0.159 |
| chr12 | 39491257 | 39492677 | 1421 | 0.75 | 0.148 | 0.159 |
| chr3 | 112783989 | 112786177 | 2189 | -2.23 | 0.148 | 0.159 |
| chr13 | 68402144 | 68433233 | 31090 | 0.75 | 0.148 | 0.16 |
| chr14 | 63759384 | 63815983 | 56600 | 0.56 | 0.148 | 0.16 |
| chr18 | 52242200 | 52245367 | 3168 | -0.99 | 0.149 | 0.16 |
| chr13 | 76938837 | 76940576 | 1740 | -1.15 | 0.149 | 0.16 |
| chr1 | 191937215 | 191966632 | 29418 | 0.93 | 0.149 | 0.16 |
| chr8 | 62304654 | 62314673 | 10020 | -2.31 | 0.149 | 0.16 |
| chr13 | 23088986 | 23090223 | 1238 | -0.91 | 0.149 | 0.16 |
| chr6 | 153925831 | 153927133 | 1303 | 1.48 | 0.149 | 0.16 |
| chr2 | 186772505 | 186773662 | 1158 | 1.86 | 0.15 | 0.161 |
| chr2 | 186929053 | 186931235 | 2183 | 2.09 | 0.15 | 0.161 |
| chr4 | 20166574 | 20169054 | 2481 | 0.71 | 0.15 | 0.161 |
| chr12 | 101495845 | 101500098 | 4254 | -2.25 | 0.15 | 0.161 |
| chr4 | 34630787 | 34643773 | 12987 | -0.78 | 0.15 | 0.161 |
| chr7 | 144454241 | 144457077 | 2837 | -2.07 | 0.15 | 0.162 |
| chr1 | 86717135 | 86718572 | 1438 | -2.09 | 0.15 | 0.162 |
| chr20 | 40263284 | 40268172 | 4889 | 0.65 | 0.15 | 0.162 |
| chrX | 51708601 | 51722636 | 14036 | 2.38 | 0.151 | 0.162 |
| chr16 | 19927466 | 19981173 | 53708 | -2.5 | 0.151 | 0.162 |
| chr4 | 47640373 | 47642514 | 2142 | 1.24 | 0.151 | 0.162 |
| chr1 | 89037905 | 89049339 | 11435 | 2.37 | 0.151 | 0.162 |

| | | | | | | |
|---|---|---|---|---|---|---|
| chr7 | 85052285 | 85053249 | 965 | -0.87 | 0.151 | 0.162 |
| chr17 | 8658301 | 8696475 | 38175 | -3.41 | 0.151 | 0.162 |
| chr1 | 116282003 | 116282782 | 780 | 2.03 | 0.151 | 0.162 |
| chr8 | 118580871 | 118586261 | 5391 | -0.4 | 0.151 | 0.162 |
| chr4 | 119704392 | 119704854 | 463 | -1.73 | 0.151 | 0.163 |
| chr9 | 90758250 | 90766710 | 8461 | -0.46 | 0.151 | 0.163 |
| chr1 | 209224066 | 209231536 | 7471 | -2.36 | 0.151 | 0.163 |
| chrX | 13571059 | 13571679 | 621 | -1.42 | 0.151 | 0.163 |
| chr2 | 215192004 | 215205422 | 13419 | 1.29 | 0.151 | 0.163 |
| chr12 | 77467942 | 77468314 | 373 | -1.01 | 0.151 | 0.163 |
| chr5 | 27385912 | 27386546 | 635 | 1.4 | 0.152 | 0.163 |
| chr13 | 43126832 | 43263208 | 136377 | -0.32 | 0.152 | 0.163 |
| chr4 | 142081849 | 142122000 | 40152 | -1.85 | 0.152 | 0.163 |
| chr14 | 81559388 | 81562301 | 2914 | -0.85 | 0.152 | 0.163 |
| chr1 | 239433448 | 239449229 | 15782 | 0.47 | 0.152 | 0.163 |
| chr4 | 185078005 | 185079368 | 1364 | 0.89 | 0.152 | 0.163 |
| chrX | 132958782 | 132965886 | 7105 | -2.27 | 0.152 | 0.163 |
| chr10 | 117697678 | 117709384 | 11707 | -2.34 | 0.152 | 0.163 |
| chr4 | 33070423 | 33071597 | 1175 | -1.92 | 0.152 | 0.163 |
| chr2 | 238277227 | 238316395 | 39169 | -0.33 | 0.152 | 0.163 |
| chr1 | 182145930 | 182148387 | 2458 | -1.65 | 0.152 | 0.163 |
| chr9 | 119533431 | 119535446 | 2016 | -1.28 | 0.152 | 0.164 |
| chr2 | 56830004 | 56839653 | 9650 | 0.45 | 0.153 | 0.164 |
| chr15 | 22366710 | 22386677 | 19968 | 0.31 | 0.153 | 0.164 |
| chr1 | 78678528 | 78679418 | 891 | -1.92 | 0.153 | 0.164 |
| chr2 | 105743281 | 105744239 | 959 | -1.79 | 0.153 | 0.164 |
| chr9 | 71310929 | 71312079 | 1151 | -0.74 | 0.153 | 0.164 |
| chr6 | 71776640 | 71779113 | 2474 | 2.12 | 0.153 | 0.165 |
| chr18 | 26632952 | 26635713 | 2762 | 0.68 | 0.153 | 0.165 |
| chr7 | 39409659 | 39412319 | 2661 | -0.7 | 0.153 | 0.165 |
| chr14 | 43745767 | 43747357 | 1591 | -1.92 | 0.154 | 0.165 |
| chr5 | 27361935 | 27363732 | 1798 | -1.92 | 0.154 | 0.165 |
| chr8 | 82307166 | 82309874 | 2709 | -2.17 | 0.154 | 0.165 |
| chr16 | 10285997 | 10303966 | 17970 | -2.45 | 0.154 | 0.165 |
| chr4 | 61854264 | 61877588 | 23325 | 0.72 | 0.154 | 0.165 |
| chr1 | 184643112 | 184643705 | 594 | 2.05 | 0.154 | 0.166 |
| chr1 | 159155765 | 159163369 | 7605 | -0.7 | 0.154 | 0.166 |
| chr4 | 34330716 | 34340681 | 9966 | -1.6 | 0.155 | 0.166 |
| chr2 | 95438730 | 95439503 | 774 | -1.79 | 0.155 | 0.166 |
| chr1 | 242748833 | 242750829 | 1997 | 1.13 | 0.155 | 0.167 |
| chr12 | 60252326 | 60300557 | 48232 | -2.42 | 0.155 | 0.167 |
| chr4 | 95072333 | 95075385 | 3053 | -0.85 | 0.155 | 0.167 |
| chr2 | 186859181 | 186860880 | 1700 | 2.07 | 0.156 | 0.167 |
| chr9 | 30387007 | 30427126 | 40120 | -2.4 | 0.156 | 0.167 |
| chr4 | 58127244 | 58129325 | 2082 | 0.78 | 0.156 | 0.167 |
| chr9 | 112290237 | 112353268 | 63032 | -3.41 | 0.156 | 0.167 |
| chr12 | 94876576 | 94914866 | 38291 | -1.81 | 0.156 | 0.167 |
| chrX | 57561021 | 57579821 | 18801 | 0.5 | 0.156 | 0.167 |
| chr11 | 36337509 | 36340678 | 3170 | 2.25 | 0.156 | 0.168 |
| chr3 | 112069903 | 112076265 | 6363 | 0.71 | 0.156 | 0.168 |
| chr13 | 30254358 | 30272331 | 17974 | -1.33 | 0.156 | 0.168 |
| chr9 | 82489361 | 82507263 | 17903 | -0.85 | 0.157 | 0.168 |
| chr8 | 17235505 | 17239607 | 4103 | 2.23 | 0.157 | 0.168 |
| chr4 | 161655717 | 161656405 | 689 | -0.84 | 0.157 | 0.168 |
| chr17 | 15257928 | 15258674 | 747 | 0.89 | 0.157 | 0.168 |
| chr10 | 131698444 | 131704243 | 5800 | 0.64 | 0.157 | 0.169 |
| chr3 | 115874679 | 115877256 | 2578 | 1.3 | 0.157 | 0.169 |
| chr5 | 30536801 | 30541705 | 4905 | -2.14 | 0.157 | 0.169 |
| chr18 | 10249876 | 10254313 | 4438 | -0.69 | 0.158 | 0.169 |
| chr1 | 114879106 | 114881095 | 1990 | 0.85 | 0.158 | 0.169 |
| chr13 | 95486794 | 95488400 | 1607 | 0.88 | 0.158 | 0.169 |

| | | | | | | |
|---|---|---|---|---|---|---|
| chrX | 56018062 | 56019000 | 939 | -1.19 | 0.158 | 0.169 |
| chr9 | 137198425 | 137199261 | 837 | 1.16 | 0.158 | 0.169 |
| chrX | 129665917 | 129667278 | 1362 | -1.93 | 0.158 | 0.17 |
| chr4 | 77814447 | 77816660 | 2214 | 2.08 | 0.158 | 0.17 |
| chr5 | 144754484 | 144762212 | 7729 | -0.59 | 0.158 | 0.17 |
| chr1 | 105182299 | 105184831 | 2533 | -0.84 | 0.159 | 0.17 |
| chr11 | 100460719 | 100462559 | 1841 | -0.61 | 0.159 | 0.17 |
| chr5 | 30606694 | 30622254 | 15561 | -1.07 | 0.159 | 0.17 |
| chr12 | 60414775 | 60447943 | 33169 | -0.8 | 0.159 | 0.17 |
| chr18 | 7494328 | 7514546 | 20219 | 0.66 | 0.159 | 0.17 |
| chr4 | 67887018 | 67888236 | 1219 | 1.92 | 0.159 | 0.17 |
| chr10 | 30373447 | 30375929 | 2483 | 0.71 | 0.159 | 0.17 |
| chr7 | 135441194 | 135443469 | 2276 | -2.25 | 0.159 | 0.17 |
| chr12 | 45357781 | 45361447 | 3667 | -2.2 | 0.159 | 0.171 |
| chr6 | 120377287 | 120383022 | 5736 | -0.51 | 0.16 | 0.171 |
| chr7 | 29390791 | 29403123 | 12333 | 0.99 | 0.16 | 0.171 |
| chr15 | 101123618 | 101123981 | 364 | 1.84 | 0.16 | 0.172 |
| chr15 | 60688053 | 60689345 | 1293 | -1.15 | 0.16 | 0.172 |
| chr1 | 63557602 | 63560967 | 3366 | 0.64 | 0.16 | 0.172 |
| chr6 | 55272364 | 55276338 | 3975 | -0.52 | 0.16 | 0.172 |
| chr6 | 129313428 | 129314251 | 824 | -1.54 | 0.16 | 0.172 |
| chr18 | 48008294 | 48009969 | 1676 | 0.62 | 0.161 | 0.172 |
| chr1 | 245821890 | 245823145 | 1256 | 0.94 | 0.161 | 0.172 |
| chr14 | 97068361 | 97078783 | 10423 | -1.59 | 0.161 | 0.172 |
| chr19 | 34159042 | 34160623 | 1582 | -0.86 | 0.161 | 0.173 |
| chr4 | 21189070 | 21191717 | 2648 | 0.64 | 0.161 | 0.173 |
| chrX | 51831267 | 51835623 | 4357 | 0.53 | 0.161 | 0.173 |
| chr10 | 59488938 | 59490988 | 2051 | -1.56 | 0.161 | 0.173 |
| chrX | 123354127 | 123354931 | 805 | -1.89 | 0.161 | 0.173 |
| chr6 | 55041029 | 55042359 | 1331 | 0.78 | 0.161 | 0.173 |
| chr2 | 145973828 | 145974604 | 777 | -1.8 | 0.162 | 0.173 |
| chr13 | 60137587 | 60139961 | 2375 | 1.47 | 0.162 | 0.173 |
| chr10 | 84469932 | 84550172 | 80241 | -2.46 | 0.162 | 0.174 |
| chr18 | 7046843 | 7049492 | 2650 | 0.92 | 0.162 | 0.174 |
| chr11 | 104492836 | 104493942 | 1107 | -0.79 | 0.162 | 0.174 |
| chrX | 103712595 | 103713443 | 849 | -0.99 | 0.162 | 0.174 |
| chr9 | 91435778 | 91535252 | 99475 | -3.41 | 0.162 | 0.174 |
| chr1 | 91252522 | 91268943 | 16422 | 0.56 | 0.163 | 0.174 |
| chrX | 51408881 | 51410184 | 1304 | -0.96 | 0.163 | 0.175 |
| chr2 | 187905943 | 187913838 | 7896 | -0.7 | 0.163 | 0.175 |
| chrX | 12734395 | 12736164 | 1770 | 0.93 | 0.163 | 0.175 |
| chr8 | 97218087 | 97233971 | 15885 | -0.91 | 0.163 | 0.175 |
| chr2 | 36172735 | 36174939 | 2205 | 0.93 | 0.164 | 0.175 |
| chr8 | 85999411 | 86002391 | 2981 | 1.12 | 0.164 | 0.175 |
| chr5 | 66090015 | 66092139 | 2125 | 0.79 | 0.164 | 0.176 |
| chr4 | 133758632 | 133761829 | 3198 | -1.81 | 0.164 | 0.176 |
| chr6 | 146928027 | 146957209 | 29183 | 0.39 | 0.164 | 0.176 |
| chr12 | 39010149 | 39031437 | 21289 | 0.6 | 0.165 | 0.176 |
| chr5 | 91046473 | 91049632 | 3160 | 1.45 | 0.165 | 0.177 |
| chr22 | 17341546 | 17342200 | 655 | -0.72 | 0.165 | 0.177 |
| chr10 | 131766843 | 131815087 | 48245 | -3.31 | 0.165 | 0.177 |
| chr8 | 133708179 | 133787236 | 79058 | -2.47 | 0.165 | 0.177 |
| chrX | 81602283 | 81606176 | 3894 | -2.06 | 0.166 | 0.177 |
| chr10 | 79456960 | 79477841 | 20882 | -2.46 | 0.166 | 0.177 |
| chr10 | 90115836 | 90128109 | 12274 | -2.46 | 0.166 | 0.177 |
| chr8 | 88552997 | 88556873 | 3877 | -0.75 | 0.166 | 0.177 |
| chr4 | 29059373 | 29078216 | 18844 | 0.59 | 0.166 | 0.177 |
| chr8 | 9063061 | 9096374 | 33314 | -1.07 | 0.166 | 0.177 |
| chr8 | 82219034 | 82219987 | 954 | -0.91 | 0.166 | 0.178 |
| chr3 | 81272958 | 81283116 | 10159 | -0.97 | 0.166 | 0.178 |
| chr1 | 194615784 | 194617084 | 1301 | 1.69 | 0.166 | 0.178 |

| | | | | | | |
|---|---|---|---|---|---|---|
| chr8 | 19992818 | 20050259 | 57442 | -0.47 | 0.166 | 0.178 |
| chr5 | 86318589 | 86327462 | 8874 | 0.69 | 0.166 | 0.178 |
| chr1 | 170490686 | 170492514 | 1829 | -0.65 | 0.167 | 0.178 |
| chr1 | 63351942 | 63354074 | 2133 | 0.84 | 0.167 | 0.178 |
| chr11 | 84440572 | 84712598 | 272027 | 3.67 | 0.167 | 0.178 |
| chr10 | 127939653 | 127941230 | 1578 | 0.86 | 0.167 | 0.178 |
| chr5 | 167621359 | 167642962 | 21604 | 2.46 | 0.167 | 0.179 |
| chr11 | 15687748 | 15760373 | 72626 | -0.34 | 0.167 | 0.179 |
| chr1 | 105089637 | 105092696 | 3060 | 0.56 | 0.167 | 0.179 |
| chr19 | 51617947 | 51619958 | 2012 | -1.28 | 0.167 | 0.179 |
| chr1 | 230432237 | 230433431 | 1195 | -0.79 | 0.167 | 0.179 |
| chr3 | 70825610 | 70826653 | 1044 | -1.63 | 0.168 | 0.179 |
| chr17 | 14671352 | 14706805 | 35454 | -0.85 | 0.168 | 0.179 |
| chr7 | 97439175 | 97439911 | 737 | -0.73 | 0.168 | 0.179 |
| chr14 | 28923370 | 28925414 | 2045 | 1.96 | 0.168 | 0.18 |
| chrX | 66690009 | 66703022 | 13014 | -1.59 | 0.168 | 0.18 |
| chr7 | 3873197 | 3874864 | 1668 | -2.01 | 0.168 | 0.18 |
| chr15 | 81472310 | 81473206 | 897 | -1.8 | 0.168 | 0.18 |
| chr11 | 78990812 | 79141484 | 150673 | 3.61 | 0.169 | 0.18 |
| chr2 | 129564447 | 129581160 | 16714 | 0.34 | 0.169 | 0.181 |
| chr12 | 74094999 | 74125427 | 30429 | -2.34 | 0.169 | 0.181 |
| chr12 | 2891564 | 2892390 | 827 | 0.62 | 0.169 | 0.181 |
| chr2 | 172353099 | 172361232 | 8134 | -2.27 | 0.169 | 0.181 |
| chr10 | 98219603 | 98258704 | 39102 | -2.36 | 0.169 | 0.181 |
| chr15 | 23899985 | 23935292 | 35308 | 0.32 | 0.17 | 0.181 |
| chr4 | 65294522 | 65306153 | 11632 | -0.65 | 0.17 | 0.182 |
| chr2 | 292610 | 431966 | 139357 | -0.38 | 0.17 | 0.182 |
| chr18 | 437160 | 462817 | 25658 | -2.25 | 0.17 | 0.182 |
| chr8 | 121885443 | 121912945 | 27503 | 0.73 | 0.17 | 0.182 |
| chr1 | 220653154 | 220654681 | 1528 | 1.49 | 0.17 | 0.182 |
| chr21 | 28029202 | 28075899 | 46698 | 0.38 | 0.171 | 0.183 |
| chr21 | 15586296 | 15602556 | 16261 | -0.67 | 0.171 | 0.183 |
| chr3 | 106074675 | 106077461 | 2787 | -0.88 | 0.171 | 0.183 |
| chr3 | 130517784 | 130523606 | 5823 | -0.65 | 0.171 | 0.183 |
| chr10 | 19650237 | 19653794 | 3558 | 2.14 | 0.171 | 0.183 |
| chr22 | 30860549 | 30861231 | 683 | 1.66 | 0.171 | 0.183 |
| chr18 | 29762850 | 29775876 | 13027 | -0.44 | 0.172 | 0.184 |
| chr19 | 30611459 | 30614201 | 2743 | 0.68 | 0.172 | 0.184 |
| chr4 | 107357481 | 107358968 | 1488 | -1.84 | 0.172 | 0.184 |
| chr18 | 22092706 | 22093767 | 1062 | 0.81 | 0.172 | 0.184 |
| chr6 | 8629253 | 8646216 | 16964 | 0.48 | 0.172 | 0.184 |
| chr12 | 99457650 | 99460780 | 3131 | 1.7 | 0.172 | 0.184 |
| chr12 | 67838806 | 67839470 | 665 | -1.73 | 0.172 | 0.184 |
| chrX | 62320200 | 62321043 | 844 | 0.86 | 0.173 | 0.185 |
| chrX | 153935656 | 153938091 | 2436 | 0.89 | 0.173 | 0.185 |
| chr14 | 65407817 | 65408894 | 1078 | 1.88 | 0.173 | 0.185 |
| chr5 | 110483057 | 110487072 | 4016 | 2.16 | 0.173 | 0.185 |
| chrX | 30014875 | 30016331 | 1457 | -1 | 0.174 | 0.186 |
| chr9 | 112357305 | 112387136 | 29832 | -0.43 | 0.174 | 0.186 |
| chr11 | 92529331 | 92533382 | 4052 | 0.87 | 0.174 | 0.186 |
| chr2 | 57770089 | 57773966 | 3878 | 1.37 | 0.174 | 0.186 |
| chr8 | 9199768 | 9201927 | 2160 | 1.05 | 0.174 | 0.186 |
| chr14 | 43165851 | 43195384 | 29534 | 0.78 | 0.174 | 0.187 |
| chr3 | 116565217 | 116570158 | 4942 | -0.69 | 0.175 | 0.187 |
| chr14 | 75839450 | 75842668 | 3219 | 2.09 | 0.175 | 0.187 |
| chr7 | 15807754 | 15809646 | 1893 | -1.87 | 0.175 | 0.187 |
| chr11 | 134563582 | 134565030 | 1449 | 1.12 | 0.175 | 0.187 |
| chr10 | 129747806 | 129748902 | 1097 | 0.8 | 0.175 | 0.187 |
| chr6 | 46399893 | 46401698 | 1806 | -0.85 | 0.175 | 0.187 |
| chr1 | 159629871 | 159641376 | 11506 | -0.81 | 0.176 | 0.188 |
| chr2 | 180900322 | 180908467 | 8146 | 2.23 | 0.176 | 0.188 |

| | | | | | | |
|---|---|---|---|---|---|---|
| chr14 | 70520789 | 70523917 | 3129 | 0.87 | 0.176 | 0.188 |
| chr1 | 218123475 | 218125664 | 2190 | 1.05 | 0.176 | 0.188 |
| chr5 | 132958956 | 132965059 | 6104 | -2.14 | 0.176 | 0.188 |
| chr18 | 56771274 | 56803409 | 32136 | -0.49 | 0.176 | 0.188 |
| chr11 | 106383412 | 106385287 | 1876 | -0.65 | 0.176 | 0.188 |
| chr10 | 80237276 | 80239824 | 2549 | 2.14 | 0.176 | 0.188 |
| chr4 | 35877954 | 35881090 | 3137 | 0.89 | 0.176 | 0.188 |
| chr13 | 28991145 | 29047722 | 56578 | -0.29 | 0.176 | 0.188 |
| chr21 | 10475110 | 10475645 | 536 | 0.77 | 0.177 | 0.189 |
| chr4 | 60719234 | 60738930 | 19697 | 0.46 | 0.177 | 0.189 |
| chr15 | 46097973 | 46101865 | 3893 | 0.5 | 0.177 | 0.189 |
| chr5 | 54377057 | 54429855 | 52799 | -0.42 | 0.177 | 0.189 |
| chr8 | 107322904 | 107346557 | 23654 | -1.05 | 0.177 | 0.189 |
| chr13 | 22492549 | 22492929 | 381 | 0.92 | 0.177 | 0.189 |
| chrX | 121574408 | 121575765 | 1358 | -0.99 | 0.177 | 0.189 |
| chr4 | 105957510 | 105974508 | 16999 | -0.51 | 0.178 | 0.19 |
| chr3 | 82160004 | 82164593 | 4590 | -0.53 | 0.178 | 0.19 |
| chr18 | 22004301 | 22006365 | 2065 | 0.79 | 0.178 | 0.19 |
| chr3 | 68990037 | 69007938 | 17902 | 1.08 | 0.178 | 0.19 |
| chr14 | 57547806 | 57574322 | 26517 | -0.4 | 0.178 | 0.19 |
| chr9 | 79599320 | 79599700 | 381 | 1.7 | 0.178 | 0.19 |
| chr2 | 168788057 | 168791882 | 3826 | 2.09 | 0.178 | 0.19 |
| chrX | 100014419 | 100015655 | 1237 | -1.85 | 0.179 | 0.191 |
| chr6 | 94533938 | 94534919 | 982 | 1.58 | 0.179 | 0.191 |
| chr17 | 63297874 | 63310652 | 12779 | -2.23 | 0.179 | 0.192 |
| chr3 | 157175929 | 157176904 | 976 | 0.92 | 0.18 | 0.192 |
| chr4 | 134709992 | 134710257 | 266 | 1.15 | 0.18 | 0.192 |
| chr15 | 60199720 | 60222240 | 22521 | 0.31 | 0.18 | 0.192 |
| chr11 | 133755709 | 133760776 | 5068 | 0.54 | 0.18 | 0.192 |
| chr21 | 29292797 | 29293187 | 391 | 1.65 | 0.18 | 0.192 |
| chr1 | 209293426 | 209325657 | 32232 | 0.62 | 0.18 | 0.192 |
| chr2 | 214566048 | 214568421 | 2374 | -1 | 0.18 | 0.192 |
| chr18 | 73248725 | 73253044 | 4320 | 0.69 | 0.18 | 0.192 |
| chrX | 101825326 | 101825886 | 561 | 1.05 | 0.18 | 0.192 |
| chr3 | 28029080 | 28031232 | 2153 | 0.97 | 0.18 | 0.193 |
| chr2 | 58484087 | 58489730 | 5644 | -2.08 | 0.18 | 0.193 |
| chr17 | 34239206 | 34243825 | 4620 | -1.43 | 0.181 | 0.193 |
| chr11 | 95248113 | 95250055 | 1943 | 0.58 | 0.181 | 0.193 |
| chr1 | 56747554 | 56748713 | 1160 | -1.08 | 0.181 | 0.193 |
| chr13 | 107068704 | 107069405 | 702 | -1.74 | 0.181 | 0.193 |
| chr10 | 98187629 | 98189168 | 1540 | 0.66 | 0.181 | 0.193 |
| chr7 | 50099521 | 50179773 | 80253 | -2.39 | 0.181 | 0.193 |
| chr3 | 109274986 | 109277711 | 2726 | -1.18 | 0.181 | 0.193 |
| chr5 | 175985608 | 175994329 | 8722 | -2.35 | 0.181 | 0.194 |
| chr1 | 157906585 | 157921648 | 15064 | 2.29 | 0.182 | 0.194 |
| chr11 | 26382831 | 26402152 | 19322 | 0.42 | 0.182 | 0.194 |
| chr3 | 64844512 | 64849312 | 4801 | 2.1 | 0.182 | 0.194 |
| chr9 | 113168345 | 113174712 | 6368 | 0.81 | 0.182 | 0.194 |
| chr1 | 187791333 | 187803709 | 12377 | -2.19 | 0.182 | 0.194 |
| chr18 | 55006479 | 55089516 | 83038 | -0.43 | 0.182 | 0.194 |
| chr11 | 27742264 | 27743223 | 960 | -1.37 | 0.182 | 0.194 |
| chr1 | 75647581 | 75649030 | 1450 | 0.73 | 0.182 | 0.194 |
| chr4 | 32699408 | 32701894 | 2487 | -1.07 | 0.182 | 0.195 |
| chr6 | 91016432 | 91018897 | 2466 | 0.55 | 0.182 | 0.195 |
| chr1 | 184838387 | 184842993 | 4607 | 2.18 | 0.182 | 0.195 |
| chr9 | 18339586 | 18343946 | 4361 | -2.12 | 0.182 | 0.195 |
| chr4 | 133860438 | 133865237 | 4800 | 0.68 | 0.183 | 0.195 |
| chr8 | 117519593 | 117520812 | 1220 | 2.07 | 0.183 | 0.195 |
| chr9 | 87853265 | 87856771 | 3507 | -0.96 | 0.183 | 0.195 |
| chr13 | 82413513 | 82414544 | 1032 | -1.7 | 0.183 | 0.195 |
| chr5 | 118150456 | 118151737 | 1282 | -1.09 | 0.183 | 0.196 |

| | | | | | | |
|---|---|---|---|---|---|---|
| chr8 | 29190501 | 29194288 | 3788 | -2.07 | 0.184 | 0.196 |
| chr1 | 204736676 | 204737672 | 997 | 1.97 | 0.184 | 0.196 |
| chr16 | 59520042 | 59549071 | 29030 | -2.3 | 0.184 | 0.197 |
| chr18 | 5269674 | 5270077 | 404 | 0.92 | 0.184 | 0.197 |
| chr1 | 226742578 | 226745639 | 3062 | -0.44 | 0.184 | 0.197 |
| chr16 | 20639285 | 20655766 | 16482 | 0.48 | 0.184 | 0.197 |
| chr4 | 12734496 | 12735540 | 1045 | 1.19 | 0.185 | 0.197 |
| chr4 | 59116701 | 59118879 | 2179 | -0.54 | 0.185 | 0.197 |
| chr8 | 24717016 | 24731901 | 14886 | 0.85 | 0.185 | 0.197 |
| chr7 | 159002106 | 159003008 | 903 | 1.32 | 0.185 | 0.197 |
| chr4 | 53398145 | 53402550 | 4406 | -0.76 | 0.185 | 0.198 |
| chr17 | 49488337 | 49490679 | 2343 | -0.82 | 0.185 | 0.198 |
| chr11 | 56481142 | 56482632 | 1491 | 1.97 | 0.186 | 0.198 |
| chr2 | 194723905 | 194748654 | 24750 | -0.64 | 0.186 | 0.198 |
| chr11 | 27265011 | 27279498 | 14488 | -2.51 | 0.186 | 0.198 |
| chrX | 81107847 | 81109003 | 1157 | -1.58 | 0.186 | 0.198 |
| chr12 | 19082803 | 19083958 | 1156 | -0.64 | 0.186 | 0.198 |
| chr1 | 147021730 | 147022722 | 993 | 1.75 | 0.186 | 0.198 |
| chr8 | 91109728 | 91113905 | 4178 | 2.02 | 0.186 | 0.198 |
| chr8 | 77360212 | 77436765 | 76554 | -2.33 | 0.186 | 0.198 |
| chr2 | 134500801 | 134502169 | 1369 | 1.85 | 0.186 | 0.199 |
| chrX | 6938849 | 6939880 | 1032 | 1.46 | 0.187 | 0.199 |
| chr14 | 58290995 | 58320550 | 29556 | -0.53 | 0.187 | 0.199 |
| chr8 | 114465131 | 114508137 | 43007 | -0.73 | 0.187 | 0.199 |
| chr13 | 93924121 | 93928256 | 4136 | 2.12 | 0.187 | 0.199 |
| chr6 | 162841250 | 162841785 | 536 | -1.68 | 0.187 | 0.199 |
| chr5 | 66076681 | 66078525 | 1845 | -0.79 | 0.187 | 0.2 |
| chr13 | 88781097 | 88782274 | 1178 | -0.81 | 0.187 | 0.2 |
| chr13 | 97538862 | 97540227 | 1366 | -0.69 | 0.188 | 0.2 |
| chr9 | 72102475 | 72104581 | 2107 | -0.56 | 0.188 | 0.2 |
| chr4 | 22307733 | 22308337 | 605 | 0.83 | 0.188 | 0.2 |
| chr14 | 27564918 | 27565340 | 423 | 1.86 | 0.188 | 0.2 |
| chr8 | 31259695 | 31307320 | 47626 | -0.24 | 0.188 | 0.201 |
| chr5 | 30204567 | 30214041 | 9475 | -0.97 | 0.188 | 0.201 |
| chr6 | 55239566 | 55241148 | 1583 | -0.79 | 0.189 | 0.201 |
| chr18 | 30149859 | 30168502 | 18644 | 0.34 | 0.189 | 0.201 |
| chr1 | 18223670 | 18229969 | 6300 | 0.71 | 0.189 | 0.201 |
| chr2 | 156903920 | 156904800 | 881 | 1.86 | 0.189 | 0.201 |
| chr11 | 49127502 | 49127976 | 475 | 0.86 | 0.189 | 0.201 |
| chr5 | 71022845 | 71027905 | 5061 | 2.16 | 0.189 | 0.202 |
| chr4 | 88712888 | 88716246 | 3359 | -0.56 | 0.19 | 0.202 |
| chr11 | 51363119 | 51377748 | 14630 | -0.31 | 0.19 | 0.202 |
| chr4 | 9991236 | 9992770 | 1535 | 0.98 | 0.19 | 0.202 |
| chr6 | 123315641 | 123316626 | 986 | -0.69 | 0.19 | 0.202 |
| chr8 | 49848435 | 49874997 | 26563 | -0.39 | 0.19 | 0.202 |
| chr18 | 10302785 | 10395761 | 92977 | -0.32 | 0.19 | 0.203 |
| chr17 | 25458061 | 25487917 | 29857 | -2.21 | 0.19 | 0.203 |
| chr7 | 18897516 | 18898663 | 1148 | -0.88 | 0.19 | 0.203 |
| chr1 | 175023755 | 175028501 | 4747 | -0.69 | 0.19 | 0.203 |
| chr1 | 217477931 | 217479458 | 1528 | 1.02 | 0.191 | 0.203 |
| chr20 | 43887611 | 43905483 | 17873 | 0.4 | 0.191 | 0.203 |
| chr4 | 58964296 | 58970028 | 5733 | 0.61 | 0.191 | 0.203 |
| chr11 | 89052205 | 89052506 | 302 | 0.91 | 0.191 | 0.203 |
| chr4 | 60139483 | 60140537 | 1055 | -1.23 | 0.191 | 0.204 |
| chr4 | 176767928 | 176768344 | 417 | 0.9 | 0.191 | 0.204 |
| chr1 | 228695304 | 228743497 | 48194 | 0.74 | 0.191 | 0.204 |
| chr14 | 37035864 | 37042922 | 7059 | 0.42 | 0.192 | 0.204 |
| chr5 | 74393779 | 74400084 | 6306 | -1.88 | 0.192 | 0.204 |
| chr4 | 34753038 | 34753726 | 689 | -1.7 | 0.192 | 0.204 |
| chr9 | 84755263 | 84976634 | 221372 | -0.25 | 0.192 | 0.204 |
| chr7 | 50212447 | 50237063 | 24617 | -0.56 | 0.192 | 0.204 |

| | | | | | | |
|---|---|---|---|---|---|---|
| chr14 | 94863402 | 94865059 | 1658 | 1.11 | 0.192 | 0.204 |
| chr9 | 117683832 | 117685302 | 1471 | 2.03 | 0.192 | 0.205 |
| chr19 | 43077061 | 43105642 | 28582 | -0.68 | 0.192 | 0.205 |
| chr10 | 124569021 | 124580583 | 11563 | 1.02 | 0.192 | 0.205 |
| chr2 | 213314683 | 213315509 | 827 | 1.26 | 0.193 | 0.205 |
| chr2 | 41902903 | 41904960 | 2058 | 1.24 | 0.193 | 0.205 |
| chr4 | 60182579 | 60183728 | 1150 | 0.84 | 0.193 | 0.205 |
| chr3 | 39701588 | 39734760 | 33173 | -0.66 | 0.193 | 0.206 |
| chr2 | 67949929 | 67958997 | 9069 | 2.2 | 0.193 | 0.206 |
| chr12 | 54203006 | 54204795 | 1790 | -1.09 | 0.194 | 0.206 |
| chr13 | 63941233 | 63954779 | 13547 | 0.79 | 0.194 | 0.206 |
| chr10 | 117691639 | 117696663 | 5025 | 2.03 | 0.194 | 0.207 |
| chr18 | 49832848 | 49844201 | 11354 | -0.72 | 0.194 | 0.207 |
| chr5 | 112410559 | 112425348 | 14790 | -1.74 | 0.194 | 0.207 |
| chr4 | 61455212 | 61506672 | 51461 | 0.35 | 0.195 | 0.208 |
| chr12 | 25322468 | 25323869 | 1402 | 1.98 | 0.196 | 0.208 |
| chr1 | 187138646 | 187153481 | 14836 | -0.48 | 0.196 | 0.209 |
| chr3 | 192223537 | 192224389 | 853 | 0.99 | 0.196 | 0.209 |
| chr5 | 31385061 | 31389297 | 4237 | -0.9 | 0.196 | 0.209 |
| chr3 | 14336263 | 14337066 | 804 | -1.72 | 0.197 | 0.209 |
| chr3 | 50806915 | 50822640 | 15726 | -2.08 | 0.197 | 0.209 |
| chr4 | 59080447 | 59107768 | 27322 | 0.3 | 0.197 | 0.21 |
| chr5 | 94610792 | 94645611 | 34820 | 1.15 | 0.197 | 0.21 |
| chr14 | 92804968 | 92806194 | 1227 | 0.83 | 0.197 | 0.21 |
| chr3 | 153193503 | 153194264 | 762 | -0.66 | 0.197 | 0.21 |
| chr3 | 39983892 | 39995289 | 11398 | 0.48 | 0.197 | 0.21 |
| chr10 | 114169703 | 114170782 | 1080 | 1.47 | 0.197 | 0.21 |
| chr4 | 174471258 | 174473268 | 2011 | -0.8 | 0.198 | 0.21 |
| chr7 | 144493271 | 144495063 | 1793 | 1.19 | 0.198 | 0.21 |
| chr4 | 62047050 | 62049232 | 2183 | 0.97 | 0.198 | 0.21 |
| chr3 | 98830490 | 98831266 | 777 | 1.91 | 0.198 | 0.21 |
| chrX | 62300346 | 62305820 | 5475 | 0.71 | 0.198 | 0.21 |
| chr5 | 44784026 | 44799665 | 15640 | -0.82 | 0.198 | 0.211 |
| chr8 | 31197161 | 31255707 | 58547 | -0.28 | 0.199 | 0.212 |
| chr3 | 82142192 | 82143238 | 1047 | -1.23 | 0.199 | 0.212 |
| chr5 | 93629998 | 93634007 | 4010 | -0.67 | 0.199 | 0.212 |
| chr4 | 161708832 | 161711057 | 2226 | -0.57 | 0.199 | 0.212 |
| chr1 | 105443725 | 105446955 | 3231 | 0.66 | 0.199 | 0.212 |
| chr4 | 112388288 | 112388779 | 492 | -1.37 | 0.199 | 0.212 |
| chr2 | 194782980 | 194786235 | 3256 | -0.51 | 0.2 | 0.212 |
| chr20 | 55537828 | 55538483 | 656 | -1.69 | 0.2 | 0.213 |
| chr7 | 95692671 | 95695108 | 2438 | 0.84 | 0.2 | 0.213 |
| chr15 | 56717265 | 56734903 | 17639 | -2.23 | 0.2 | 0.213 |
| chr5 | 64126394 | 64142716 | 16323 | -2.23 | 0.2 | 0.213 |
| chr5 | 59977792 | 59991945 | 14154 | -2.07 | 0.2 | 0.213 |
| chr12 | 92707283 | 92707853 | 571 | -1.23 | 0.2 | 0.213 |
| chrX | 101232507 | 101233197 | 691 | 0.96 | 0.201 | 0.214 |
| chr15 | 83388744 | 83391689 | 2946 | -0.86 | 0.201 | 0.214 |
| chr13 | 112085744 | 112086983 | 1240 | 0.78 | 0.201 | 0.214 |
| chr19 | 49469179 | 49470369 | 1191 | -0.54 | 0.202 | 0.215 |
| chr11 | 98222935 | 98225023 | 2089 | -0.61 | 0.202 | 0.215 |
| chr8 | 112817772 | 112886496 | 68725 | -0.67 | 0.203 | 0.215 |
| chr3 | 78644906 | 78645525 | 620 | 1.01 | 0.203 | 0.216 |
| chr3 | 65311595 | 65326184 | 14590 | 0.42 | 0.203 | 0.216 |
| chr7 | 158870987 | 158871928 | 942 | 0.87 | 0.203 | 0.216 |
| chr6 | 154429913 | 154444639 | 14727 | 0.4 | 0.203 | 0.216 |
| chr20 | 57801681 | 57899334 | 97654 | 0.32 | 0.203 | 0.216 |
| chr6 | 161068741 | 161079034 | 10294 | -0.45 | 0.203 | 0.216 |
| chr16 | 84304874 | 84307386 | 2513 | 0.56 | 0.203 | 0.216 |
| chr7 | 142385064 | 142387776 | 2713 | -1.5 | 0.203 | 0.216 |
| chrX | 83551729 | 83556996 | 5268 | 0.74 | 0.204 | 0.217 |

| | | | | | | |
|---|---|---|---|---|---|---|
| chr4 | 104133343 | 104135194 | 1852 | 0.6 | 0.204 | 0.217 |
| chr4 | 54742609 | 54743182 | 574 | -0.83 | 0.204 | 0.217 |
| chr10 | 82323833 | 82348712 | 24880 | -2.1 | 0.204 | 0.217 |
| chr7 | 143969304 | 143969698 | 395 | 0.93 | 0.204 | 0.217 |
| chr5 | 59420196 | 59423032 | 2837 | 2 | 0.204 | 0.217 |
| chr4 | 34269998 | 34282155 | 12158 | -1.05 | 0.205 | 0.218 |
| chr5 | 129290925 | 129292807 | 1883 | -0.51 | 0.205 | 0.218 |
| chr1 | 157314957 | 157319980 | 5024 | -0.44 | 0.205 | 0.218 |
| chr15 | 43554993 | 43556841 | 1849 | 0.76 | 0.205 | 0.218 |
| chr4 | 36410601 | 36412738 | 2138 | 1.93 | 0.205 | 0.218 |
| chr13 | 109185788 | 109190169 | 4382 | -0.58 | 0.205 | 0.218 |
| chr13 | 107275501 | 107278836 | 3336 | 0.53 | 0.206 | 0.218 |
| chr4 | 100710446 | 100711456 | 1011 | 1.69 | 0.206 | 0.218 |
| chr4 | 65612583 | 65622540 | 9958 | -0.61 | 0.206 | 0.219 |
| chr6 | 116166412 | 116176287 | 9876 | -0.87 | 0.206 | 0.219 |
| chr13 | 94666886 | 94679563 | 12678 | 1.19 | 0.207 | 0.219 |
| chr4 | 135694008 | 135694596 | 589 | 1.76 | 0.207 | 0.22 |
| chr12 | 60832394 | 60840566 | 8173 | -0.64 | 0.207 | 0.22 |
| chr4 | 59399891 | 59403891 | 4001 | -1 | 0.207 | 0.22 |
| chr9 | 75544777 | 75545622 | 846 | 0.83 | 0.207 | 0.22 |
| chr13 | 22288049 | 22307255 | 19207 | 0.39 | 0.207 | 0.22 |
| chrX | 133209817 | 133213571 | 3755 | -0.45 | 0.207 | 0.22 |
| chr13 | 29052725 | 29055830 | 3106 | 1.99 | 0.207 | 0.22 |
| chr2 | 186511331 | 186514432 | 3102 | -0.8 | 0.207 | 0.22 |
| chr11 | 111121094 | 111132837 | 11744 | 0.4 | 0.208 | 0.221 |
| chrX | 17977636 | 17992445 | 14810 | 0.71 | 0.208 | 0.221 |
| chr3 | 83927412 | 84315651 | 388240 | -3.32 | 0.208 | 0.221 |
| chr16 | 80352470 | 80354424 | 1955 | 1.88 | 0.208 | 0.221 |
| chr7 | 45284745 | 45320303 | 35559 | -0.36 | 0.208 | 0.221 |
| chr11 | 118076046 | 118097611 | 21566 | -0.87 | 0.208 | 0.221 |
| chr16 | 77179312 | 77209601 | 30290 | -2.17 | 0.209 | 0.222 |
| chr4 | 60891111 | 60892176 | 1066 | 0.69 | 0.209 | 0.222 |
| chr13 | 108820627 | 108822268 | 1642 | 1.39 | 0.209 | 0.222 |
| chr8 | 107253118 | 107279255 | 26138 | 0.66 | 0.209 | 0.222 |
| chr3 | 36780162 | 36780537 | 376 | 1.09 | 0.209 | 0.222 |
| chr15 | 93135629 | 93138207 | 2579 | 0.8 | 0.209 | 0.222 |
| chr5 | 85613689 | 85615756 | 2068 | -0.87 | 0.209 | 0.222 |
| chr5 | 66028723 | 66046188 | 17466 | 0.63 | 0.209 | 0.222 |
| chr15 | 39096360 | 39099263 | 2904 | -0.82 | 0.209 | 0.222 |
| chr6 | 159975287 | 159977859 | 2573 | 1.93 | 0.21 | 0.223 |
| chr16 | 63266759 | 63298251 | 31493 | -2.15 | 0.21 | 0.223 |
| chr9 | 82573078 | 82598412 | 25335 | -0.43 | 0.21 | 0.223 |
| chr7 | 29430117 | 29432067 | 1951 | 0.84 | 0.21 | 0.223 |
| chr18 | 26965171 | 26977065 | 11895 | 0.62 | 0.21 | 0.223 |
| chr5 | 54316268 | 54317902 | 1635 | 1.86 | 0.212 | 0.225 |
| chr2 | 186805604 | 186837670 | 32067 | -0.78 | 0.212 | 0.225 |
| chr4 | 81291809 | 81292582 | 774 | -1.12 | 0.212 | 0.225 |
| chr7 | 158247476 | 158247792 | 317 | -0.69 | 0.212 | 0.225 |
| chr7 | 123465734 | 123473434 | 7701 | -1.22 | 0.212 | 0.225 |
| chrX | 36945544 | 36947223 | 1680 | -0.85 | 0.212 | 0.225 |
| chr22 | 49204574 | 49212091 | 7518 | -0.53 | 0.212 | 0.225 |
| chr21 | 36103735 | 36106562 | 2828 | 1.98 | 0.212 | 0.225 |
| chr22 | 39313635 | 39314773 | 1139 | 1.31 | 0.213 | 0.226 |
| chr5 | 61340870 | 61367717 | 26848 | -0.52 | 0.213 | 0.226 |
| chrX | 13586191 | 13587211 | 1021 | -0.66 | 0.213 | 0.226 |
| chr8 | 117402235 | 117403147 | 913 | -0.81 | 0.213 | 0.226 |
| chr11 | 59751113 | 59779694 | 28582 | 0.3 | 0.213 | 0.227 |
| chr9 | 35447356 | 35455210 | 7855 | -1.03 | 0.214 | 0.227 |
| chr14 | 22273275 | 22287155 | 13881 | -0.54 | 0.214 | 0.227 |
| chr9 | 87332140 | 87350458 | 18319 | -0.94 | 0.214 | 0.227 |
| chr8 | 118563346 | 118564024 | 679 | -1.77 | 0.214 | 0.227 |

| | | | | | | |
|---|---|---|---|---|---|---|
| chr5 | 29844997 | 29845361 | 365 | -0.75 | 0.214 | 0.227 |
| chr13 | 89041536 | 89045072 | 3537 | -0.71 | 0.215 | 0.228 |
| chr3 | 28098168 | 28099352 | 1185 | 1.27 | 0.215 | 0.228 |
| chr1 | 116500383 | 116501744 | 1362 | -0.77 | 0.215 | 0.228 |
| chr16 | 63737946 | 63739915 | 1970 | -1.79 | 0.215 | 0.228 |
| chr12 | 7410144 | 7412589 | 2446 | 1.94 | 0.215 | 0.228 |
| chr11 | 134587909 | 134589291 | 1383 | -0.69 | 0.215 | 0.228 |
| chr13 | 84779942 | 84781288 | 1347 | 0.66 | 0.215 | 0.228 |
| chr4 | 35396010 | 35425580 | 29571 | 0.8 | 0.216 | 0.229 |
| chr3 | 10831245 | 10852026 | 20782 | -2.1 | 0.216 | 0.229 |
| chr8 | 129970944 | 129972987 | 2044 | 0.66 | 0.216 | 0.229 |
| chr2 | 141557109 | 141557625 | 517 | -0.75 | 0.216 | 0.229 |
| chr13 | 101582210 | 101625246 | 43037 | 0.4 | 0.216 | 0.23 |
| chr11 | 76930644 | 76931996 | 1353 | -1.5 | 0.216 | 0.23 |
| chr6 | 106507481 | 106508597 | 1117 | 0.69 | 0.217 | 0.23 |
| chrX | 134299553 | 134306878 | 7326 | 0.5 | 0.217 | 0.231 |
| chr7 | 142134696 | 142135075 | 380 | -1 | 0.218 | 0.231 |
| chr7 | 13550800 | 13556763 | 5964 | 0.66 | 0.218 | 0.231 |
| chr2 | 194858907 | 194873717 | 14811 | 2.1 | 0.218 | 0.231 |
| chr3 | 104742073 | 104744452 | 2380 | 0.59 | 0.218 | 0.231 |
| chr3 | 83366170 | 83366799 | 630 | 1.16 | 0.218 | 0.232 |
| chr1 | 194661150 | 194662672 | 1523 | 0.68 | 0.219 | 0.232 |
| chr15 | 89269439 | 89270935 | 1497 | 0.76 | 0.219 | 0.232 |
| chr9 | 21124781 | 21171602 | 46822 | 0.25 | 0.219 | 0.232 |
| chr12 | 7718016 | 7740722 | 22707 | -0.41 | 0.219 | 0.233 |
| chr4 | 78289966 | 78296539 | 6574 | 0.41 | 0.22 | 0.233 |
| chr1 | 63582651 | 63584985 | 2335 | 1.89 | 0.22 | 0.233 |
| chr9 | 2263346 | 2272616 | 9271 | -1.86 | 0.22 | 0.233 |
| chr4 | 61577321 | 61587776 | 10456 | 0.7 | 0.22 | 0.233 |
| chr6 | 167109833 | 167125789 | 15957 | -0.99 | 0.22 | 0.233 |
| chr7 | 144180255 | 144191605 | 11351 | -0.63 | 0.22 | 0.233 |
| chr12 | 43509926 | 43513237 | 3312 | -1.82 | 0.22 | 0.233 |
| chr8 | 10343561 | 10359816 | 16256 | 0.71 | 0.22 | 0.233 |
| chr1 | 106060401 | 106061571 | 1171 | 0.8 | 0.22 | 0.234 |
| chrX | 55245167 | 55247324 | 2158 | 0.51 | 0.22 | 0.234 |
| chr13 | 107257083 | 107261121 | 4039 | 0.53 | 0.22 | 0.234 |
| chr6 | 110183177 | 110229088 | 45912 | -0.47 | 0.221 | 0.234 |
| chr8 | 82258201 | 82272203 | 14003 | 0.62 | 0.221 | 0.234 |
| chr2 | 84425093 | 84437377 | 12285 | -1.99 | 0.221 | 0.234 |
| chr6 | 47966855 | 47968361 | 1507 | -0.65 | 0.221 | 0.234 |
| chr2 | 195706755 | 195708706 | 1952 | -0.74 | 0.221 | 0.234 |
| chr7 | 131365682 | 131368627 | 2946 | 1.95 | 0.221 | 0.235 |
| chr18 | 23726436 | 23727606 | 1171 | 0.71 | 0.221 | 0.235 |
| chr6 | 72334924 | 72341853 | 6930 | -0.56 | 0.222 | 0.235 |
| chr15 | 74111240 | 74122096 | 10857 | -1.38 | 0.222 | 0.235 |
| chr13 | 39523002 | 39536118 | 13117 | -0.36 | 0.222 | 0.235 |
| chr4 | 133988019 | 133999479 | 11461 | -0.43 | 0.222 | 0.235 |
| chr3 | 71674470 | 71676011 | 1542 | 0.63 | 0.222 | 0.235 |
| chr6 | 142848389 | 142850465 | 2077 | 0.57 | 0.222 | 0.235 |
| chr2 | 238256734 | 238258004 | 1271 | 1.1 | 0.222 | 0.236 |
| chrX | 84429259 | 84429956 | 698 | -0.74 | 0.223 | 0.236 |
| chr1 | 106281201 | 106324811 | 43611 | 0.46 | 0.223 | 0.236 |
| chr3 | 136838438 | 136839989 | 1552 | -1.27 | 0.223 | 0.237 |
| chr14 | 36849098 | 36849878 | 781 | -1.22 | 0.223 | 0.237 |
| chr2 | 137148566 | 137178140 | 29575 | -0.49 | 0.224 | 0.237 |
| chr8 | 113086048 | 113129217 | 43170 | -2.14 | 0.224 | 0.237 |
| chr6 | 129101459 | 129104553 | 3095 | 0.54 | 0.224 | 0.237 |
| chr5 | 83218862 | 83219496 | 635 | -1.65 | 0.224 | 0.237 |
| chrX | 52579445 | 52579903 | 459 | -1.21 | 0.224 | 0.237 |
| chr1 | 101778174 | 101815826 | 37653 | 0.32 | 0.224 | 0.237 |
| chr10 | 67392509 | 67430560 | 38052 | -0.7 | 0.224 | 0.237 |

| | | | | | | |
|---|---|---|---|---|---|---|
| chr8 | 31348818 | 31350496 | 1679 | 1.82 | 0.224 | 0.237 |
| chr17 | 67147934 | 67148399 | 466 | -0.77 | 0.224 | 0.237 |
| chr17 | 32571911 | 32573834 | 1924 | -1.74 | 0.224 | 0.238 |
| chr20 | 31791340 | 31793513 | 2174 | 1.89 | 0.224 | 0.238 |
| chr16 | 79743768 | 79745123 | 1356 | -0.66 | 0.224 | 0.238 |
| chr7 | 127799027 | 127802849 | 3823 | 0.61 | 0.225 | 0.238 |
| chr5 | 112967930 | 113011582 | 43653 | -0.24 | 0.225 | 0.238 |
| chr6 | 24964741 | 24966109 | 1369 | 0.72 | 0.225 | 0.238 |
| chr2 | 70620423 | 70635429 | 15007 | -0.65 | 0.225 | 0.238 |
| chr6 | 6590556 | 6605617 | 15062 | -0.53 | 0.225 | 0.238 |
| chrX | 64687881 | 64689555 | 1675 | -0.69 | 0.225 | 0.239 |
| chr6 | 77506752 | 77508945 | 2194 | -0.86 | 0.226 | 0.239 |
| chr12 | 42256008 | 42266914 | 10907 | -0.47 | 0.226 | 0.239 |
| chr16 | 59870250 | 59871675 | 1426 | -0.88 | 0.226 | 0.24 |
| chr14 | 60960641 | 60963514 | 2874 | 0.72 | 0.227 | 0.24 |
| chr14 | 35942053 | 35943142 | 1090 | 1.66 | 0.227 | 0.24 |
| chr8 | 15439606 | 15453227 | 13622 | -1.31 | 0.227 | 0.24 |
| chr1 | 238046311 | 238058404 | 12094 | 0.35 | 0.227 | 0.241 |
| chr3 | 75404494 | 75404909 | 416 | 1.07 | 0.227 | 0.241 |
| chr3 | 50998747 | 50999414 | 668 | 1.69 | 0.227 | 0.241 |
| chr1 | 168528343 | 168532968 | 4626 | 0.69 | 0.227 | 0.241 |
| chr2 | 195719495 | 195720288 | 794 | 0.99 | 0.228 | 0.241 |
| chr18 | 27698949 | 27700702 | 1754 | -0.7 | 0.228 | 0.241 |
| chr2 | 108314546 | 108317308 | 2763 | 0.72 | 0.228 | 0.241 |
| chr22 | 22492825 | 22514819 | 21995 | -2.04 | 0.228 | 0.241 |
| chr1 | 50823133 | 50825248 | 2116 | 0.55 | 0.228 | 0.241 |
| chr2 | 217801971 | 217833731 | 31761 | -0.37 | 0.228 | 0.241 |
| chr1 | 239542393 | 239572236 | 29844 | -0.66 | 0.228 | 0.242 |
| chr7 | 124315196 | 124362030 | 46835 | 0.32 | 0.229 | 0.242 |
| chr11 | 116176031 | 116178276 | 2246 | -0.54 | 0.229 | 0.242 |
| chr3 | 51913704 | 51919327 | 5624 | -0.37 | 0.229 | 0.242 |
| chr14 | 22522113 | 22542139 | 20027 | -0.45 | 0.229 | 0.242 |
| chr1 | 84292015 | 84292882 | 868 | -1 | 0.229 | 0.242 |
| chr12 | 58946848 | 58947969 | 1122 | -1.69 | 0.229 | 0.243 |
| chr12 | 2764989 | 2784071 | 19083 | 0.3 | 0.229 | 0.243 |
| chr2 | 171452462 | 171453388 | 927 | 1.73 | 0.23 | 0.243 |
| chrX | 69107773 | 69109854 | 2082 | -1.76 | 0.23 | 0.243 |
| chr11 | 44452115 | 44459421 | 7307 | 0.67 | 0.23 | 0.243 |
| chr6 | 8676945 | 8699025 | 22081 | -0.27 | 0.23 | 0.243 |
| chr12 | 7763500 | 7786801 | 23302 | -2 | 0.23 | 0.243 |
| chr4 | 62608031 | 62608633 | 603 | 1.08 | 0.231 | 0.244 |
| chr1 | 194585304 | 194607776 | 22473 | 1.33 | 0.231 | 0.244 |
| chr2 | 156850274 | 156868146 | 17873 | -0.81 | 0.231 | 0.244 |
| chr1 | 164694223 | 164695754 | 1532 | -1.36 | 0.231 | 0.244 |
| chr3 | 37486366 | 37487123 | 758 | 0.64 | 0.231 | 0.244 |
| chr7 | 119109761 | 119130620 | 20860 | -2 | 0.231 | 0.244 |
| chr12 | 91196112 | 91198502 | 2391 | -0.66 | 0.231 | 0.244 |
| chr7 | 14936587 | 14955003 | 18417 | 0.62 | 0.231 | 0.245 |
| chr4 | 29045721 | 29049275 | 3555 | -0.52 | 0.232 | 0.245 |
| chr17 | 56832105 | 56832788 | 684 | -1.63 | 0.232 | 0.245 |
| chr1 | 160049348 | 160049767 | 420 | -1.54 | 0.232 | 0.245 |
| chr9 | 111572518 | 111573506 | 989 | 0.78 | 0.232 | 0.245 |
| chr16 | 22884193 | 22888614 | 4422 | 0.59 | 0.232 | 0.246 |
| chr13 | 39665420 | 39667563 | 2144 | 0.86 | 0.233 | 0.246 |
| chr2 | 153765695 | 153767602 | 1908 | 0.82 | 0.233 | 0.246 |
| chr1 | 245696603 | 245701090 | 4488 | 1.68 | 0.233 | 0.247 |
| chr6 | 128047321 | 128048427 | 1107 | 1.75 | 0.233 | 0.247 |
| chr8 | 120288343 | 120313896 | 25554 | -0.38 | 0.234 | 0.247 |
| chr3 | 70846028 | 70850352 | 4325 | 0.47 | 0.234 | 0.247 |
| chr4 | 135126376 | 135128893 | 2518 | 0.63 | 0.234 | 0.248 |
| chr16 | 34197975 | 35186582 | 988608 | -0.12 | 0.234 | 0.248 |

| | | | | | | |
|---|---|---|---|---|---|---|
| chr12 | 76562616 | 76564876 | 2261 | -0.48 | 0.234 | 0.248 |
| chr15 | 58452179 | 58486358 | 34180 | 0.4 | 0.234 | 0.248 |
| chr6 | 122259364 | 122267109 | 7746 | 0.47 | 0.235 | 0.248 |
| chr1 | 66171122 | 66173148 | 2027 | 0.53 | 0.235 | 0.249 |
| chr12 | 98042706 | 98066781 | 24076 | -2.01 | 0.236 | 0.249 |
| chrX | 69136147 | 69137507 | 1361 | 0.56 | 0.236 | 0.249 |
| chr11 | 48538644 | 48539936 | 1293 | 0.99 | 0.236 | 0.25 |
| chr6 | 110322150 | 110337725 | 15576 | 0.48 | 0.236 | 0.25 |
| chr3 | 149836929 | 149846551 | 9623 | -0.83 | 0.237 | 0.25 |
| chr2 | 161905133 | 161909066 | 3934 | 1.77 | 0.237 | 0.25 |
| chr18 | 14907985 | 14934393 | 26409 | -0.25 | 0.237 | 0.251 |
| chr11 | 57660622 | 57665749 | 5128 | -0.72 | 0.237 | 0.251 |
| chr6 | 154032843 | 154034308 | 1466 | 1.73 | 0.238 | 0.252 |
| chr5 | 29617706 | 29830051 | 212346 | -0.81 | 0.238 | 0.252 |
| chr6 | 72326800 | 72327221 | 422 | 0.86 | 0.239 | 0.252 |
| chr2 | 64518947 | 64520023 | 1077 | -0.74 | 0.239 | 0.253 |
| chr3 | 56498363 | 56501219 | 2857 | -0.48 | 0.239 | 0.253 |
| chr3 | 98267849 | 98269667 | 1819 | 0.87 | 0.239 | 0.253 |
| chr5 | 157799940 | 158115640 | 315701 | -0.16 | 0.239 | 0.253 |
| chr1 | 28833234 | 28834938 | 1705 | -0.63 | 0.24 | 0.253 |
| chr6 | 118104325 | 118111321 | 6997 | 0.78 | 0.24 | 0.254 |
| chr12 | 104675117 | 104687044 | 11928 | -1.89 | 0.24 | 0.254 |
| chr8 | 6660068 | 6661672 | 1605 | 1.86 | 0.24 | 0.254 |
| chr3 | 137636155 | 137640828 | 4674 | 0.76 | 0.241 | 0.254 |
| chr13 | 23090554 | 23092554 | 2001 | 0.48 | 0.241 | 0.254 |
| chr4 | 111248799 | 111258891 | 10093 | -1.3 | 0.241 | 0.255 |
| chr3 | 64917840 | 64920014 | 2175 | -0.53 | 0.242 | 0.255 |
| chr2 | 141601013 | 141601860 | 848 | 1.6 | 0.242 | 0.255 |
| chr12 | 15519920 | 15531169 | 11250 | 0.62 | 0.242 | 0.255 |
| chr5 | 72545478 | 72568484 | 23007 | -1.86 | 0.242 | 0.255 |
| chr18 | 68169536 | 68177913 | 8378 | -0.53 | 0.242 | 0.255 |
| chr9 | 80804857 | 80807085 | 2229 | -0.58 | 0.242 | 0.255 |
| chr12 | 13978241 | 13980925 | 2685 | -0.49 | 0.242 | 0.256 |
| chr3 | 27890679 | 27945777 | 55099 | -0.39 | 0.242 | 0.256 |
| chr4 | 22228974 | 22241806 | 12833 | -0.61 | 0.242 | 0.256 |
| chr9 | 82626092 | 82732462 | 106371 | -0.43 | 0.242 | 0.256 |
| chr2 | 135229684 | 135262306 | 32623 | -1.92 | 0.242 | 0.256 |
| chrX | 135726271 | 135748699 | 22429 | -0.27 | 0.242 | 0.256 |
| chr13 | 24089851 | 24098712 | 8862 | 0.99 | 0.243 | 0.256 |
| chr2 | 158988812 | 158989752 | 941 | -1.06 | 0.243 | 0.256 |
| chr11 | 134755390 | 134765763 | 10374 | 0.35 | 0.243 | 0.257 |
| chr12 | 98703482 | 98704882 | 1401 | 0.81 | 0.243 | 0.257 |
| chr7 | 117654939 | 117659150 | 4212 | -0.39 | 0.244 | 0.257 |
| chr14 | 102027701 | 102047032 | 19332 | -1.84 | 0.244 | 0.257 |
| chr2 | 1825410 | 1830112 | 4703 | 1.86 | 0.244 | 0.257 |
| chr12 | 40045635 | 40046447 | 813 | -1.31 | 0.244 | 0.257 |
| chr18 | 32282027 | 32305018 | 22992 | -1.18 | 0.244 | 0.258 |
| chr6 | 159646731 | 159647981 | 1251 | -0.73 | 0.244 | 0.258 |
| chr7 | 22584206 | 22585305 | 1100 | 1.71 | 0.244 | 0.258 |
| chr16 | 32489740 | 32490133 | 394 | -0.79 | 0.244 | 0.258 |
| chr4 | 134865469 | 134867339 | 1871 | 1.66 | 0.245 | 0.259 |
| chr1 | 223284673 | 223286655 | 1983 | 0.78 | 0.245 | 0.259 |
| chr7 | 14969548 | 14973064 | 3517 | 0.49 | 0.245 | 0.259 |
| chr17 | 21700419 | 21832243 | 131825 | 0.22 | 0.245 | 0.259 |
| chr9 | 120180304 | 120182255 | 1952 | -0.95 | 0.245 | 0.259 |
| chr2 | 67972052 | 68047545 | 75494 | 0.31 | 0.245 | 0.259 |
| chrX | 73034996 | 73036857 | 1862 | 1.81 | 0.245 | 0.259 |
| chr10 | 98440234 | 98440752 | 519 | -1.55 | 0.246 | 0.259 |
| chr14 | 85655610 | 85656894 | 1285 | 0.99 | 0.246 | 0.259 |
| chr11 | 109415301 | 109449359 | 34059 | -1.97 | 0.246 | 0.26 |
| chr7 | 157804965 | 157807670 | 2706 | -0.59 | 0.246 | 0.26 |

| | | | | | | |
|---|---|---|---|---|---|---|
| chr7 | 148336112 | 148337001 | 890 | -1.54 | 0.246 | 0.26 |
| chrX | 135023990 | 135024677 | 688 | -0.56 | 0.246 | 0.26 |
| chr3 | 156104470 | 156106244 | 1775 | 1.79 | 0.247 | 0.26 |
| chr1 | 75411031 | 75413717 | 2687 | 0.51 | 0.247 | 0.261 |
| chr4 | 63280489 | 63355553 | 75065 | -0.54 | 0.247 | 0.261 |
| chrX | 106751001 | 106762371 | 11371 | -1.81 | 0.248 | 0.262 |
| chr2 | 53522337 | 53524909 | 2573 | 1.83 | 0.248 | 0.262 |
| chr10 | 43204062 | 43205177 | 1116 | 0.62 | 0.248 | 0.262 |
| chr1 | 153206530 | 153209081 | 2552 | 0.45 | 0.248 | 0.262 |
| chr21 | 34344171 | 34346142 | 1972 | 1.77 | 0.248 | 0.262 |
| chr15 | 87210653 | 87212880 | 2228 | -1.08 | 0.248 | 0.262 |
| chr4 | 21173232 | 21174757 | 1526 | 0.57 | 0.248 | 0.262 |
| chr4 | 76374048 | 76385482 | 11435 | -1.56 | 0.249 | 0.262 |
| chr2 | 56882073 | 56883754 | 1682 | 0.54 | 0.249 | 0.262 |
| chr11 | 35442704 | 35489743 | 47040 | -0.31 | 0.249 | 0.263 |
| chrX | 57888557 | 57888935 | 379 | -0.72 | 0.249 | 0.263 |
| chr1 | 104385943 | 104388667 | 2725 | 0.81 | 0.249 | 0.263 |
| chr8 | 21677770 | 21682795 | 5026 | 0.82 | 0.249 | 0.263 |
| chr13 | 111789586 | 111794579 | 4994 | -0.9 | 0.25 | 0.263 |
| chr7 | 39516799 | 39538281 | 21483 | 0.66 | 0.25 | 0.263 |
| chr2 | 61980440 | 62026574 | 46135 | -1.96 | 0.25 | 0.264 |
| chr3 | 115121319 | 115122458 | 1140 | -1.58 | 0.25 | 0.264 |
| chr6 | 154075183 | 154077806 | 2624 | 1.75 | 0.25 | 0.264 |
| chr13 | 60211303 | 60212767 | 1465 | -1.64 | 0.25 | 0.264 |
| chr4 | 167299704 | 167300642 | 939 | -0.69 | 0.25 | 0.264 |
| chr7 | 23928942 | 23929989 | 1048 | -1.62 | 0.25 | 0.264 |
| chr14 | 96450163 | 96452024 | 1862 | 0.71 | 0.25 | 0.264 |
| chr2 | 171326134 | 171329438 | 3305 | -0.65 | 0.25 | 0.264 |
| chr4 | 32543261 | 32545099 | 1839 | -1.54 | 0.25 | 0.264 |
| chr3 | 104496495 | 104499124 | 2630 | -0.55 | 0.25 | 0.264 |
| chr15 | 70844316 | 70846949 | 2634 | -0.86 | 0.251 | 0.264 |
| chr4 | 97490693 | 97491559 | 867 | 0.63 | 0.251 | 0.264 |
| chr2 | 53492078 | 53500730 | 8653 | 1.96 | 0.251 | 0.265 |
| chr1 | 218247393 | 218248509 | 1117 | 1.2 | 0.251 | 0.265 |
| chr12 | 9421044 | 9423862 | 2819 | -0.71 | 0.251 | 0.265 |
| chr2 | 961493 | 1217036 | 255544 | 0.2 | 0.251 | 0.265 |
| chr20 | 4634459 | 4637472 | 3014 | 0.95 | 0.251 | 0.265 |
| chr6 | 101848668 | 101849395 | 728 | -0.77 | 0.252 | 0.265 |
| chr3 | 148478157 | 148478678 | 522 | -1.22 | 0.252 | 0.265 |
| chr12 | 80506976 | 80507592 | 617 | -1.54 | 0.253 | 0.267 |
| chr7 | 108951533 | 108953248 | 1716 | 0.58 | 0.253 | 0.267 |
| chr12 | 2129920 | 2155360 | 25441 | -0.42 | 0.253 | 0.267 |
| chr5 | 85935386 | 85936380 | 995 | -0.99 | 0.253 | 0.267 |
| chr4 | 47760267 | 47784733 | 24467 | -0.47 | 0.253 | 0.267 |
| chr4 | 33157736 | 33159433 | 1698 | -1.03 | 0.254 | 0.268 |
| chr1 | 111830984 | 111832911 | 1928 | 0.49 | 0.254 | 0.268 |
| chr3 | 86296770 | 86306364 | 9595 | -0.54 | 0.254 | 0.268 |
| chr10 | 1361086 | 1385998 | 24913 | -0.44 | 0.254 | 0.268 |
| chr12 | 22380489 | 22381316 | 828 | 1.27 | 0.254 | 0.268 |
| chr1 | 104993523 | 104998233 | 4711 | 0.44 | 0.254 | 0.268 |
| chr9 | 32744048 | 32863142 | 119095 | 0.21 | 0.254 | 0.268 |
| chr13 | 68386427 | 68387471 | 1045 | -0.72 | 0.255 | 0.269 |
| chrX | 43415595 | 43428635 | 13041 | -1.9 | 0.255 | 0.269 |
| chr9 | 71031615 | 71135462 | 103848 | 0.18 | 0.255 | 0.269 |
| chr12 | 55073087 | 55087782 | 14696 | -0.71 | 0.255 | 0.269 |
| chr14 | 101329127 | 101330344 | 1218 | 0.54 | 0.256 | 0.269 |
| chr17 | 49404097 | 49404513 | 417 | 0.79 | 0.256 | 0.27 |
| chr2 | 186618064 | 186620143 | 2080 | 1.7 | 0.256 | 0.27 |
| chr19 | 42216098 | 42246024 | 29927 | 0.79 | 0.256 | 0.27 |
| chrX | 49956522 | 49957974 | 1453 | -0.75 | 0.257 | 0.27 |
| chr17 | 37938195 | 37939832 | 1638 | 0.86 | 0.257 | 0.271 |

| | | | | | | |
|---|---|---|---|---|---|---|
| chr4 | 47796210 | 47797081 | 872 | 0.67 | 0.257 | 0.271 |
| chr3 | 85488291 | 85537382 | 49092 | -1.98 | 0.257 | 0.271 |
| chr5 | 30635476 | 30775980 | 140505 | 0.36 | 0.257 | 0.271 |
| chrX | 84388579 | 84389676 | 1098 | -0.63 | 0.257 | 0.271 |
| chr8 | 114818585 | 114833662 | 15078 | -0.73 | 0.257 | 0.271 |
| chr3 | 46346606 | 46398870 | 52265 | -0.23 | 0.257 | 0.271 |
| chr4 | 62759286 | 62760578 | 1293 | 1.4 | 0.258 | 0.272 |
| chr11 | 115380327 | 115401302 | 20976 | 0.37 | 0.258 | 0.272 |
| chr3 | 137491343 | 137492922 | 1580 | -0.57 | 0.258 | 0.272 |
| chr3 | 159251567 | 159255473 | 3907 | 1.87 | 0.258 | 0.272 |
| chr15 | 73120750 | 73136724 | 15975 | -1.47 | 0.258 | 0.272 |
| chr6 | 15742342 | 15744003 | 1662 | -1.02 | 0.258 | 0.272 |
| chr3 | 148159184 | 148160654 | 1471 | -1.11 | 0.258 | 0.272 |
| chr21 | 14925872 | 14926280 | 409 | 0.64 | 0.259 | 0.272 |
| chr21 | 18405311 | 18406791 | 1481 | -0.62 | 0.259 | 0.273 |
| chr14 | 30085650 | 30089290 | 3641 | -0.46 | 0.259 | 0.273 |
| chr12 | 125805722 | 125837881 | 32160 | -1.98 | 0.259 | 0.273 |
| chr3 | 175998110 | 175999382 | 1273 | -0.63 | 0.259 | 0.273 |
| chr11 | 123205664 | 123220547 | 14884 | -1.31 | 0.26 | 0.274 |
| chr6 | 101430600 | 101432865 | 2266 | 0.56 | 0.26 | 0.274 |
| chr10 | 83113703 | 83114465 | 763 | 0.6 | 0.26 | 0.274 |
| chr15 | 96196183 | 96206743 | 10561 | -1 | 0.26 | 0.274 |
| chr2 | 106949730 | 106961249 | 11520 | 1.86 | 0.26 | 0.274 |
| chr7 | 131464334 | 131466912 | 2579 | 0.77 | 0.26 | 0.274 |
| chrX | 13924748 | 13925107 | 360 | 1.47 | 0.261 | 0.274 |
| chr4 | 29685957 | 29692301 | 6345 | 0.98 | 0.261 | 0.275 |
| chr6 | 153259372 | 153260092 | 721 | 0.77 | 0.261 | 0.275 |
| chr2 | 150264540 | 150265382 | 843 | -0.84 | 0.261 | 0.275 |
| chr7 | 77659649 | 77663346 | 3698 | 1.85 | 0.261 | 0.275 |
| chr2 | 163601199 | 163601832 | 634 | -1.51 | 0.261 | 0.275 |
| chr4 | 61663610 | 61664147 | 538 | -1.09 | 0.262 | 0.276 |
| chr4 | 65212572 | 65214736 | 2165 | 0.57 | 0.262 | 0.276 |
| chrX | 82650821 | 82651378 | 558 | -1.45 | 0.262 | 0.276 |
| chr1 | 114809045 | 114810674 | 1630 | 0.56 | 0.262 | 0.276 |
| chrX | 113403996 | 113405791 | 1796 | -0.63 | 0.262 | 0.276 |
| chr2 | 217364707 | 217366387 | 1681 | -1.6 | 0.263 | 0.277 |
| chr4 | 90716726 | 90719450 | 2725 | -1.66 | 0.263 | 0.277 |
| chr4 | 22741699 | 22761766 | 20068 | -0.88 | 0.263 | 0.277 |
| chrX | 36827205 | 36829120 | 1916 | -0.7 | 0.263 | 0.277 |
| chr1 | 170622078 | 170623828 | 1751 | -0.5 | 0.263 | 0.277 |
| chr7 | 113895767 | 113897473 | 1707 | 0.84 | 0.264 | 0.278 |
| chr4 | 61074785 | 61076627 | 1843 | -1.58 | 0.264 | 0.278 |
| chr7 | 120947385 | 120952087 | 4703 | -0.49 | 0.265 | 0.279 |
| chr1 | 18387413 | 18393028 | 5616 | 0.37 | 0.265 | 0.279 |
| chr18 | 1909485 | 1945200 | 35716 | -0.46 | 0.265 | 0.279 |
| chr10 | 19557800 | 19558840 | 1041 | 0.93 | 0.265 | 0.279 |
| chr9 | 79103444 | 79111168 | 7725 | -0.62 | 0.265 | 0.279 |
| chrX | 96920995 | 96944892 | 23898 | -1.79 | 0.265 | 0.279 |
| chr8 | 6707496 | 6732106 | 24611 | -0.87 | 0.266 | 0.28 |
| chr10 | 81308368 | 81309199 | 832 | 1.37 | 0.266 | 0.28 |
| chr8 | 113940050 | 113983948 | 43899 | -1.92 | 0.266 | 0.28 |
| chr18 | 26626986 | 26628933 | 1948 | -0.54 | 0.266 | 0.28 |
| chr15 | 100986350 | 100991553 | 5204 | 0.58 | 0.267 | 0.281 |
| chr13 | 88831445 | 88832541 | 1097 | 1.61 | 0.267 | 0.281 |
| chr21 | 35594604 | 35596762 | 2159 | 0.55 | 0.267 | 0.281 |
| chr18 | 42661021 | 42662594 | 1574 | -0.6 | 0.267 | 0.281 |
| chr15 | 38089732 | 38092490 | 2759 | -0.36 | 0.268 | 0.282 |
| chr7 | 28333580 | 28335773 | 2194 | 0.56 | 0.268 | 0.282 |
| chr14 | 43286942 | 43299025 | 12084 | -0.51 | 0.268 | 0.282 |
| chr15 | 100431843 | 100434600 | 2758 | 0.86 | 0.269 | 0.283 |
| chr11 | 4417904 | 4418391 | 488 | -1.47 | 0.269 | 0.283 |

| | | | | | | |
|---|---|---|---|---|---|---|
| chr9 | 121553719 | 121554554 | 836 | 1.21 | 0.269 | 0.283 |
| chr3 | 39270999 | 39290915 | 19917 | -0.94 | 0.269 | 0.283 |
| chr9 | 89129240 | 89190970 | 61731 | -0.3 | 0.269 | 0.283 |
| chr6 | 91177106 | 91177467 | 362 | -1.4 | 0.269 | 0.283 |
| chr20 | 160569 | 179410 | 18842 | -0.41 | 0.27 | 0.284 |
| chr9 | 121306726 | 121310211 | 3486 | -0.49 | 0.27 | 0.284 |
| chr5 | 27590043 | 27673600 | 83558 | -1.02 | 0.27 | 0.284 |
| chr2 | 1778989 | 1812435 | 33447 | 0.43 | 0.27 | 0.284 |
| chr8 | 117948795 | 117966554 | 17760 | -1.84 | 0.27 | 0.284 |
| chr6 | 85957966 | 85972372 | 14407 | 0.51 | 0.27 | 0.284 |
| chr21 | 15992651 | 15993580 | 930 | -0.62 | 0.271 | 0.285 |
| chr4 | 147779195 | 147780057 | 863 | 1.61 | 0.271 | 0.285 |
| chr9 | 92500984 | 92503713 | 2730 | 1.76 | 0.271 | 0.285 |
| chr21 | 47066300 | 47066702 | 403 | 0.73 | 0.271 | 0.285 |
| chr5 | 65664275 | 65873599 | 209325 | 0.14 | 0.271 | 0.286 |
| chr7 | 142505155 | 142536850 | 31696 | -0.68 | 0.272 | 0.286 |
| chr4 | 139650350 | 139652004 | 1655 | 0.54 | 0.272 | 0.286 |
| chr1 | 199307012 | 199341778 | 34767 | 0.39 | 0.272 | 0.286 |
| chr12 | 99872373 | 99970816 | 98444 | -0.38 | 0.272 | 0.286 |
| chr6 | 46930016 | 46930897 | 882 | 0.73 | 0.272 | 0.286 |
| chrX | 102313658 | 102326611 | 12954 | -0.39 | 0.272 | 0.286 |
| chr12 | 80509213 | 80511234 | 2022 | 1.4 | 0.272 | 0.286 |
| chr2 | 197832019 | 197842545 | 10527 | -1.7 | 0.272 | 0.287 |
| chr6 | 152280647 | 152284669 | 4023 | -1.06 | 0.273 | 0.287 |
| chr8 | 104140734 | 104188131 | 47398 | -0.46 | 0.273 | 0.287 |
| chr18 | 45162275 | 45186919 | 24645 | -0.28 | 0.273 | 0.287 |
| chr12 | 7937755 | 7939066 | 1312 | 1.51 | 0.273 | 0.287 |
| chr11 | 48670916 | 48692457 | 21542 | 0.33 | 0.274 | 0.288 |
| chr7 | 157862359 | 157862841 | 483 | 0.64 | 0.274 | 0.288 |
| chr12 | 71199809 | 71200335 | 527 | 1.21 | 0.274 | 0.288 |
| chr3 | 103202914 | 103282545 | 79632 | -1.97 | 0.274 | 0.288 |
| chr14 | 43309719 | 43317565 | 7847 | -1.83 | 0.274 | 0.288 |
| chr10 | 31487976 | 31494556 | 6581 | -0.81 | 0.275 | 0.289 |
| chr12 | 59589585 | 59590825 | 1241 | -1.55 | 0.275 | 0.289 |
| chr10 | 82436128 | 82450758 | 14631 | -1.79 | 0.275 | 0.289 |
| chr1 | 76491590 | 76495878 | 4289 | 0.5 | 0.276 | 0.29 |
| chr13 | 23761021 | 23761633 | 613 | 1.46 | 0.276 | 0.29 |
| chr5 | 121125510 | 121127342 | 1833 | -1.61 | 0.276 | 0.29 |
| chr8 | 118301265 | 118312407 | 11143 | -0.34 | 0.276 | 0.29 |
| chr7 | 5568355 | 5568692 | 338 | -0.99 | 0.276 | 0.29 |
| chr17 | 70176645 | 70177941 | 1297 | -1.54 | 0.277 | 0.291 |
| chr17 | 67813384 | 68130112 | 316729 | 3.13 | 0.277 | 0.291 |
| chr1 | 111934844 | 111936693 | 1850 | -0.51 | 0.277 | 0.291 |
| chr16 | 73431793 | 73445780 | 13988 | -1.02 | 0.277 | 0.291 |
| chr3 | 83263102 | 83273994 | 10893 | -1.78 | 0.277 | 0.291 |
| chr3 | 85242124 | 85473750 | 231627 | -2.78 | 0.277 | 0.291 |
| chr13 | 112890152 | 112892870 | 2719 | -0.5 | 0.277 | 0.291 |
| chr3 | 111909228 | 111984740 | 75513 | -0.54 | 0.278 | 0.292 |
| chr11 | 115509359 | 115510672 | 1314 | -0.58 | 0.278 | 0.292 |
| chr2 | 88395707 | 88396614 | 908 | 0.96 | 0.278 | 0.292 |
| chr3 | 107032266 | 107032815 | 550 | -0.58 | 0.278 | 0.292 |
| chr5 | 154974131 | 154976230 | 2100 | 0.55 | 0.278 | 0.293 |
| chr3 | 62340174 | 62341336 | 1163 | -0.67 | 0.279 | 0.293 |
| chr16 | 49396147 | 49418744 | 22598 | 0.37 | 0.279 | 0.293 |
| chr9 | 89246443 | 89249992 | 3550 | -0.54 | 0.279 | 0.293 |
| chr8 | 113990267 | 113991058 | 792 | 1.61 | 0.279 | 0.293 |
| chr11 | 91288068 | 91494465 | 206398 | 2.98 | 0.279 | 0.294 |
| chr18 | 7231968 | 7241853 | 9886 | -1.74 | 0.279 | 0.294 |
| chr9 | 30670893 | 30692258 | 21366 | 0.41 | 0.28 | 0.294 |
| chrX | 124348060 | 124360330 | 12271 | 0.42 | 0.28 | 0.294 |
| chr7 | 7347906 | 7367304 | 19399 | -0.65 | 0.28 | 0.294 |

| | | | | | | |
|---|---|---|---|---|---|---|
| chr13 | 26024516 | 26033701 | 9186 | 0.4 | 0.28 | 0.294 |
| chr7 | 50308699 | 50310551 | 1853 | 1.58 | 0.28 | 0.294 |
| chr15 | 84783510 | 84784279 | 770 | -1.02 | 0.281 | 0.295 |
| chr13 | 68884828 | 68896916 | 12089 | 0.33 | 0.281 | 0.295 |
| chrX | 114104048 | 114105070 | 1023 | 1.21 | 0.281 | 0.295 |
| chr12 | 67880395 | 67905745 | 25351 | 0.28 | 0.281 | 0.296 |
| chr5 | 118893495 | 118894529 | 1035 | -1.07 | 0.282 | 0.296 |
| chr15 | 79583171 | 79596565 | 13395 | 0.36 | 0.282 | 0.296 |
| chr5 | 135599018 | 135624839 | 25822 | 1.92 | 0.282 | 0.296 |
| chr3 | 108562236 | 108563142 | 907 | -1.49 | 0.282 | 0.296 |
| chr1 | 159921066 | 159923312 | 2247 | 0.84 | 0.282 | 0.296 |
| chrX | 82860734 | 82862359 | 1626 | -1.37 | 0.282 | 0.296 |
| chr2 | 1757164 | 1763946 | 6783 | -0.38 | 0.282 | 0.296 |
| chr12 | 7634544 | 7635870 | 1327 | 0.47 | 0.283 | 0.297 |
| chr20 | 8120119 | 8180025 | 59907 | 0.32 | 0.283 | 0.297 |
| chr1 | 187255134 | 187256326 | 1193 | -1.38 | 0.283 | 0.297 |
| chr15 | 20172557 | 20173542 | 986 | 0.67 | 0.283 | 0.297 |
| chrX | 30006509 | 30009577 | 3069 | 1.73 | 0.283 | 0.297 |
| chr4 | 61118725 | 61172411 | 53687 | 0.36 | 0.284 | 0.298 |
| chr7 | 117926354 | 117927011 | 658 | -0.63 | 0.284 | 0.298 |
| chr4 | 29034510 | 29038558 | 4049 | 0.6 | 0.284 | 0.298 |
| chr15 | 40130465 | 40131420 | 956 | -1.51 | 0.284 | 0.298 |
| chr4 | 59284327 | 59299189 | 14863 | 0.54 | 0.285 | 0.299 |
| chr22 | 23083022 | 23085090 | 2069 | 0.56 | 0.285 | 0.299 |
| chr2 | 170057166 | 170064257 | 7092 | -0.35 | 0.285 | 0.299 |
| chr1 | 159261719 | 159616348 | 354630 | -0.15 | 0.285 | 0.299 |
| chr17 | 56210587 | 56212189 | 1603 | 0.61 | 0.285 | 0.299 |
| chr12 | 106236522 | 106265598 | 29077 | 1.62 | 0.285 | 0.3 |
| chr9 | 20166829 | 20189991 | 23163 | -0.31 | 0.285 | 0.3 |
| chr1 | 182392335 | 182408327 | 15993 | -0.4 | 0.286 | 0.3 |
| chr3 | 62365611 | 62376954 | 11344 | -0.54 | 0.286 | 0.3 |
| chr6 | 127985085 | 128008953 | 23869 | -1.44 | 0.286 | 0.3 |
| chr14 | 81479501 | 81539701 | 60201 | -0.19 | 0.286 | 0.3 |
| chr15 | 91747675 | 91748650 | 976 | 0.93 | 0.286 | 0.301 |
| chr5 | 4872919 | 4876409 | 3491 | 0.74 | 0.287 | 0.301 |
| chr8 | 27339823 | 27341322 | 1500 | -0.59 | 0.287 | 0.301 |
| chr4 | 11357257 | 11359061 | 1805 | -0.53 | 0.287 | 0.301 |
| chr11 | 15880394 | 15884430 | 4037 | 0.64 | 0.287 | 0.301 |
| chr3 | 67879531 | 67881145 | 1615 | 0.5 | 0.287 | 0.301 |
| chr8 | 972967 | 1012900 | 39934 | -0.38 | 0.287 | 0.301 |
| chr22 | 49283781 | 49284541 | 761 | 0.58 | 0.288 | 0.302 |
| chr3 | 117903974 | 117905281 | 1308 | 1.54 | 0.288 | 0.302 |
| chr19 | 38949694 | 38959477 | 9784 | -0.79 | 0.288 | 0.302 |
| chr17 | 5966777 | 5967249 | 473 | -0.77 | 0.288 | 0.302 |
| chr4 | 61602906 | 61612685 | 9780 | 0.54 | 0.289 | 0.303 |
| chr9 | 30494000 | 30577562 | 83563 | -0.47 | 0.289 | 0.303 |
| chr5 | 29848957 | 29884789 | 35833 | 0.52 | 0.289 | 0.303 |
| chr8 | 1483427 | 1484949 | 1523 | 1.09 | 0.289 | 0.303 |
| chr2 | 100658884 | 100695105 | 36222 | -0.31 | 0.289 | 0.304 |
| chr4 | 9871492 | 9886003 | 14512 | -1.32 | 0.289 | 0.304 |
| chr1 | 187777601 | 187778346 | 746 | -0.53 | 0.29 | 0.304 |
| chr12 | 54675242 | 54676104 | 863 | -0.48 | 0.29 | 0.304 |
| chr21 | 36838017 | 36874808 | 36792 | -0.31 | 0.29 | 0.304 |
| chr11 | 22205743 | 22210076 | 4334 | -0.37 | 0.29 | 0.304 |
| chr3 | 69557706 | 69565278 | 7573 | -0.29 | 0.292 | 0.306 |
| chr10 | 99814680 | 99828105 | 13426 | 1.82 | 0.292 | 0.306 |
| chr17 | 69978912 | 69989181 | 10270 | -1.67 | 0.292 | 0.306 |
| chr12 | 13421872 | 13438407 | 16536 | -0.83 | 0.293 | 0.307 |
| chr5 | 113426400 | 113429605 | 3206 | -0.42 | 0.293 | 0.307 |
| chr5 | 95767132 | 95786133 | 19002 | 1.88 | 0.293 | 0.307 |
| chr19 | 49572435 | 49573084 | 650 | -0.78 | 0.294 | 0.308 |

| | | | | | | |
|---|---|---|---|---|---|---|
| chr6 | 79114476 | 79117005 | 2530 | 0.79 | 0.294 | 0.308 |
| chr5 | 6181736 | 6316084 | 134349 | -0.17 | 0.294 | 0.308 |
| chr11 | 66186589 | 66190903 | 4315 | -0.73 | 0.294 | 0.309 |
| chr3 | 149091402 | 149093987 | 2586 | -0.39 | 0.294 | 0.309 |
| chr2 | 85020656 | 85021089 | 434 | -0.56 | 0.294 | 0.309 |
| chrX | 69463982 | 69486275 | 22294 | -1.78 | 0.295 | 0.309 |
| chr11 | 21580529 | 21581958 | 1430 | -1.48 | 0.295 | 0.309 |
| chr14 | 89539128 | 89541745 | 2618 | -1.58 | 0.295 | 0.309 |
| chr11 | 91547977 | 91899338 | 351362 | 2.96 | 0.295 | 0.309 |
| chr4 | 35902728 | 35906506 | 3779 | 0.38 | 0.295 | 0.309 |
| chr20 | 36103418 | 36121371 | 17954 | -0.48 | 0.295 | 0.31 |
| chr5 | 115951668 | 115955591 | 3924 | 1.74 | 0.296 | 0.31 |
| chr9 | 28347612 | 28348245 | 634 | -0.69 | 0.296 | 0.31 |
| chrX | 141243635 | 141247337 | 3703 | 0.48 | 0.296 | 0.31 |
| chrX | 152170129 | 152184137 | 14009 | -1.75 | 0.296 | 0.311 |
| chr15 | 70252426 | 70270012 | 17587 | -1.69 | 0.297 | 0.311 |
| chr8 | 11053124 | 11093185 | 40062 | -0.42 | 0.297 | 0.311 |
| chr3 | 83067636 | 83117801 | 50166 | -1.81 | 0.297 | 0.311 |
| chr3 | 108452811 | 108468374 | 15564 | -0.33 | 0.297 | 0.312 |
| chr3 | 191362651 | 191365155 | 2505 | -1.14 | 0.298 | 0.312 |
| chr14 | 103387087 | 103387918 | 832 | -0.77 | 0.298 | 0.312 |
| chr6 | 47807264 | 47817611 | 10348 | 0.29 | 0.298 | 0.312 |
| chr1 | 230477330 | 230480989 | 3660 | 1.7 | 0.299 | 0.313 |
| chr3 | 158910784 | 158936068 | 25285 | -0.24 | 0.299 | 0.313 |
| chr4 | 128088240 | 128089490 | 1251 | 1.54 | 0.3 | 0.314 |
| chr9 | 93702308 | 93711935 | 9628 | -0.44 | 0.3 | 0.314 |
| chr11 | 40313592 | 40314344 | 753 | -1.44 | 0.3 | 0.314 |
| chr10 | 48444342 | 48447155 | 2814 | -0.6 | 0.301 | 0.315 |
| chr7 | 137292878 | 137331726 | 38849 | 0.35 | 0.301 | 0.315 |
| chr3 | 156099758 | 156100565 | 808 | -1.42 | 0.301 | 0.315 |
| chrX | 64996268 | 64998030 | 1763 | 0.66 | 0.301 | 0.316 |
| chr3 | 191736212 | 191737713 | 1502 | -1.09 | 0.302 | 0.316 |
| chr11 | 120591317 | 120592879 | 1563 | 0.73 | 0.302 | 0.317 |
| chr4 | 63118331 | 63125625 | 7295 | 0.4 | 0.302 | 0.317 |
| chr3 | 127020070 | 127020753 | 684 | 0.55 | 0.302 | 0.317 |
| chr3 | 72039137 | 72071630 | 32494 | -0.32 | 0.303 | 0.317 |
| chr6 | 12673850 | 12768580 | 94731 | -0.3 | 0.303 | 0.317 |
| chr6 | 39693888 | 39704391 | 10504 | -1.72 | 0.303 | 0.317 |
| chr4 | 47631546 | 47638738 | 7193 | -0.45 | 0.303 | 0.318 |
| chrX | 62136524 | 62137426 | 903 | 0.99 | 0.304 | 0.318 |
| chrX | 75333960 | 75335947 | 1988 | 0.59 | 0.304 | 0.318 |
| chr8 | 10947798 | 10975829 | 28032 | 0.31 | 0.304 | 0.318 |
| chr12 | 83662488 | 83693293 | 30806 | -1.75 | 0.304 | 0.319 |
| chr5 | 36810253 | 36811256 | 1004 | 1.52 | 0.305 | 0.319 |
| chr5 | 29459333 | 29612113 | 152781 | 0.33 | 0.305 | 0.32 |
| chr1 | 105960929 | 106008774 | 47846 | 0.24 | 0.305 | 0.32 |
| chr11 | 124489766 | 124490765 | 1000 | 0.9 | 0.305 | 0.32 |
| chr6 | 160309268 | 160311763 | 2496 | -1.12 | 0.305 | 0.32 |
| chr3 | 161889153 | 161903672 | 14520 | -1.69 | 0.305 | 0.32 |
| chr18 | 76520502 | 76524319 | 3818 | -0.37 | 0.306 | 0.32 |
| chr2 | 136866645 | 136869565 | 2921 | 0.41 | 0.306 | 0.321 |
| chr8 | 114213156 | 114271610 | 58455 | -1.81 | 0.306 | 0.321 |
| chr9 | 98374236 | 98460060 | 85825 | -2.59 | 0.307 | 0.321 |
| chrX | 57729962 | 57731845 | 1884 | 0.52 | 0.307 | 0.321 |
| chr4 | 65560583 | 65570182 | 9600 | 0.55 | 0.307 | 0.322 |
| chr4 | 34571163 | 34587456 | 16294 | 1.26 | 0.307 | 0.322 |
| chrX | 57781635 | 57785010 | 3376 | -0.62 | 0.308 | 0.322 |
| chr8 | 16491251 | 16494520 | 3270 | 1.03 | 0.308 | 0.322 |
| chr18 | 70555114 | 70558063 | 2950 | -0.57 | 0.308 | 0.323 |
| chr13 | 46317088 | 46318393 | 1306 | -0.72 | 0.308 | 0.323 |
| chr3 | 162872964 | 162893586 | 20623 | -1.67 | 0.309 | 0.323 |

| | | | | | | |
|---|---|---|---|---|---|---|
| chr4 | 60962596 | 61006897 | 44302 | -0.44 | 0.31 | 0.324 |
| chr3 | 82878955 | 82913762 | 34808 | -0.73 | 0.31 | 0.324 |
| chr2 | 127525117 | 127526899 | 1783 | -0.49 | 0.31 | 0.324 |
| chrX | 151319388 | 151321149 | 1762 | 0.5 | 0.31 | 0.324 |
| chr8 | 53944736 | 54006263 | 61528 | 0.34 | 0.31 | 0.324 |
| chr5 | 57863011 | 58104934 | 241924 | 0.2 | 0.31 | 0.324 |
| chr17 | 75555741 | 75557094 | 1354 | 0.79 | 0.31 | 0.325 |
| chr6 | 13052346 | 13067123 | 14778 | 0.27 | 0.311 | 0.325 |
| chr4 | 61998783 | 62000310 | 1528 | 0.48 | 0.311 | 0.325 |
| chr22 | 26276054 | 26278116 | 2063 | 0.41 | 0.311 | 0.326 |
| chr9 | 138048816 | 138049565 | 750 | 0.67 | 0.312 | 0.326 |
| chr12 | 42037022 | 42038295 | 1274 | 0.46 | 0.312 | 0.326 |
| chr8 | 76071239 | 76166789 | 95551 | 0.29 | 0.312 | 0.327 |
| chr8 | 91746379 | 91747218 | 840 | 0.57 | 0.313 | 0.327 |
| chr2 | 35258825 | 35272588 | 13764 | 0.85 | 0.313 | 0.327 |
| chr16 | 74236167 | 74250068 | 13902 | -0.87 | 0.313 | 0.328 |
| chr2 | 170268623 | 170271225 | 2603 | 0.87 | 0.313 | 0.328 |
| chr17 | 59665519 | 59668838 | 3320 | 1.7 | 0.314 | 0.328 |
| chr2 | 38432196 | 38433239 | 1044 | 1.07 | 0.314 | 0.329 |
| chr14 | 22078196 | 22078687 | 492 | 1.48 | 0.314 | 0.329 |
| chrX | 24464039 | 24465617 | 1579 | 0.6 | 0.314 | 0.329 |
| chr4 | 33305470 | 33306354 | 885 | 0.64 | 0.315 | 0.329 |
| chr16 | 64319571 | 64344520 | 24950 | -0.41 | 0.315 | 0.329 |
| chr4 | 59193029 | 59195166 | 2138 | 0.62 | 0.315 | 0.329 |
| chr15 | 95929844 | 95938507 | 8664 | -1.71 | 0.315 | 0.329 |
| chr19 | 30727620 | 30861381 | 133762 | 0.24 | 0.315 | 0.33 |
| chr8 | 114849255 | 114927768 | 78514 | -0.45 | 0.315 | 0.33 |
| chr4 | 60931971 | 60947571 | 15601 | 0.41 | 0.315 | 0.33 |
| chr2 | 107266035 | 107415155 | 149121 | -0.12 | 0.316 | 0.33 |
| chr14 | 105329037 | 105329423 | 387 | 0.6 | 0.316 | 0.33 |
| chr8 | 118324934 | 118334521 | 9588 | -0.98 | 0.317 | 0.331 |
| chrX | 139950752 | 139964724 | 13973 | -0.47 | 0.317 | 0.332 |
| chr8 | 91760920 | 91789073 | 28154 | 0.4 | 0.317 | 0.332 |
| chr8 | 73865306 | 73867069 | 1764 | -0.56 | 0.317 | 0.332 |
| chr12 | 43918712 | 43922112 | 3401 | -0.44 | 0.318 | 0.332 |
| chr15 | 53801975 | 53802687 | 713 | -1.39 | 0.318 | 0.332 |
| chr11 | 62959145 | 62968091 | 8947 | 0.45 | 0.318 | 0.332 |
| chr18 | 10263650 | 10290323 | 26674 | 0.31 | 0.318 | 0.333 |
| chr1 | 204830235 | 204834271 | 4037 | -0.37 | 0.318 | 0.333 |
| chr5 | 68116675 | 68120980 | 4306 | 0.63 | 0.319 | 0.334 |
| chr1 | 105925636 | 105927654 | 2019 | 0.51 | 0.319 | 0.334 |
| chr4 | 61524272 | 61556794 | 32523 | 0.29 | 0.319 | 0.334 |
| chr5 | 34038870 | 34040289 | 1420 | -0.68 | 0.32 | 0.334 |
| chr8 | 26909429 | 27041474 | 132046 | -0.24 | 0.32 | 0.334 |
| chr6 | 46077305 | 46078444 | 1140 | 0.52 | 0.32 | 0.334 |
| chr8 | 73843801 | 73851032 | 7232 | 0.45 | 0.32 | 0.335 |
| chr1 | 218401557 | 218402546 | 990 | -0.67 | 0.32 | 0.335 |
| chr6 | 67929094 | 67933647 | 4554 | 0.36 | 0.321 | 0.336 |
| chr6 | 97780148 | 97790545 | 10398 | -0.78 | 0.321 | 0.336 |
| chr1 | 214333829 | 214336877 | 3049 | 0.38 | 0.321 | 0.336 |
| chr1 | 34104224 | 34120783 | 16560 | -0.24 | 0.322 | 0.336 |
| chr16 | 20699142 | 20715375 | 16234 | -1.64 | 0.322 | 0.336 |
| chr18 | 26908929 | 26911259 | 2331 | -1.48 | 0.322 | 0.336 |
| chr1 | 4194719 | 4195605 | 887 | 0.62 | 0.322 | 0.337 |
| chr4 | 135885944 | 135887781 | 1838 | 1.54 | 0.322 | 0.337 |
| chr6 | 50970779 | 50971406 | 628 | -1.14 | 0.323 | 0.337 |
| chr9 | 89203863 | 89213193 | 9331 | -0.25 | 0.323 | 0.337 |
| chr16 | 82018669 | 82019860 | 1192 | 1.57 | 0.323 | 0.338 |
| chrX | 110481358 | 110482495 | 1138 | -0.52 | 0.324 | 0.338 |
| chrX | 83792579 | 83798213 | 5635 | -0.48 | 0.325 | 0.339 |
| chr8 | 52903535 | 52905755 | 2221 | -0.87 | 0.326 | 0.34 |

| | | | | | | |
|---|---|---|---|---|---|---|
| chr2 | 7042334 | 7049599 | 7266 | -0.44 | 0.326 | 0.34 |
| chr12 | 67153551 | 67156140 | 2590 | -0.62 | 0.326 | 0.341 |
| chr8 | 88570403 | 88583478 | 13076 | -0.5 | 0.326 | 0.341 |
| chr20 | 36254811 | 36255951 | 1141 | 0.78 | 0.326 | 0.341 |
| chr3 | 78829051 | 78829581 | 531 | -1.18 | 0.326 | 0.341 |
| chr18 | 71592216 | 71592572 | 357 | 1.4 | 0.326 | 0.341 |
| chr5 | 31734180 | 31776455 | 42276 | -0.42 | 0.327 | 0.341 |
| chr7 | 3875646 | 3876938 | 1293 | 0.85 | 0.327 | 0.342 |
| chr4 | 33472089 | 33509398 | 37310 | -0.51 | 0.327 | 0.342 |
| chr10 | 11707820 | 11719406 | 11587 | -1.63 | 0.327 | 0.342 |
| chr10 | 52019574 | 52027166 | 7593 | -0.33 | 0.328 | 0.342 |
| chr16 | 49486996 | 49490432 | 3437 | -0.36 | 0.328 | 0.342 |
| chr1 | 232376017 | 232384423 | 8407 | -0.34 | 0.328 | 0.342 |
| chr4 | 112505372 | 112506002 | 631 | -0.86 | 0.328 | 0.343 |
| chr9 | 92339330 | 92341143 | 1814 | 1.56 | 0.328 | 0.343 |
| chr3 | 101996141 | 101999644 | 3504 | -0.43 | 0.328 | 0.343 |
| chr10 | 18842752 | 18845689 | 2938 | 0.81 | 0.328 | 0.343 |
| chr3 | 82788159 | 82865526 | 77368 | -0.41 | 0.329 | 0.343 |
| chr1 | 209711347 | 209716610 | 5264 | 1.28 | 0.329 | 0.344 |
| chr6 | 99651738 | 99660143 | 8406 | 1.19 | 0.33 | 0.344 |
| chr3 | 40797343 | 40809362 | 12020 | -0.3 | 0.33 | 0.344 |
| chr22 | 49467760 | 49470754 | 2995 | 0.37 | 0.33 | 0.345 |
| chr17 | 63502170 | 63516082 | 13913 | -1.57 | 0.331 | 0.345 |
| chr4 | 30051059 | 30052801 | 1743 | 1.26 | 0.331 | 0.345 |
| chr2 | 109806667 | 109807864 | 1198 | 1.42 | 0.331 | 0.346 |
| chr9 | 93347586 | 93448797 | 101212 | -0.23 | 0.332 | 0.346 |
| chr20 | 23952551 | 23955326 | 2776 | 0.82 | 0.332 | 0.347 |
| chr10 | 19212696 | 19214370 | 1675 | 0.43 | 0.332 | 0.347 |
| chr3 | 40729703 | 40748213 | 18511 | -0.32 | 0.332 | 0.347 |
| chr4 | 67705667 | 67708271 | 2605 | -0.54 | 0.332 | 0.347 |
| chr1 | 77232310 | 77232820 | 511 | -0.63 | 0.333 | 0.347 |
| chr2 | 147574619 | 147588546 | 13928 | 0.71 | 0.333 | 0.347 |
| chr18 | 22922255 | 22922866 | 612 | -0.63 | 0.333 | 0.347 |
| chr2 | 106926753 | 106927689 | 937 | 0.71 | 0.333 | 0.348 |
| chr4 | 62544657 | 62573573 | 28917 | 0.34 | 0.334 | 0.348 |
| chr11 | 18939043 | 18954869 | 15827 | 0.34 | 0.334 | 0.348 |
| chr8 | 53410804 | 53411804 | 1001 | 1.38 | 0.334 | 0.348 |
| chrX | 152507163 | 152510763 | 3601 | 0.46 | 0.334 | 0.349 |
| chr18 | 10420396 | 10420995 | 600 | -0.67 | 0.334 | 0.349 |
| chr4 | 49003324 | 49007342 | 4019 | 0.5 | 0.334 | 0.349 |
| chr1 | 38833469 | 38851509 | 18041 | 0.25 | 0.334 | 0.349 |
| chr7 | 132301044 | 132311440 | 10397 | -0.53 | 0.334 | 0.349 |
| chr5 | 112727756 | 112728312 | 557 | -0.82 | 0.335 | 0.35 |
| chr13 | 66386965 | 66387444 | 480 | -0.78 | 0.335 | 0.35 |
| chr4 | 134910041 | 134911410 | 1370 | -0.58 | 0.335 | 0.35 |
| chr8 | 82076518 | 82077095 | 578 | 0.6 | 0.336 | 0.351 |
| chrX | 18089358 | 18091898 | 2541 | 0.52 | 0.336 | 0.351 |
| chr2 | 226428630 | 226431516 | 2887 | 0.73 | 0.336 | 0.351 |
| chr4 | 35300906 | 35312812 | 11907 | 0.68 | 0.336 | 0.351 |
| chr7 | 97389090 | 97389780 | 691 | 1.46 | 0.336 | 0.351 |
| chr11 | 128458649 | 128468159 | 9511 | -0.34 | 0.337 | 0.351 |
| chr18 | 73096850 | 73097392 | 543 | 0.83 | 0.338 | 0.352 |
| chr3 | 155992689 | 155996338 | 3650 | -0.6 | 0.338 | 0.353 |
| chrX | 57867437 | 57869242 | 1806 | -0.5 | 0.338 | 0.353 |
| chr7 | 144121479 | 144131081 | 9603 | 0.42 | 0.338 | 0.353 |
| chr5 | 166138550 | 166164286 | 25737 | -1.63 | 0.338 | 0.353 |
| chr13 | 60782148 | 60783907 | 1760 | -1.11 | 0.338 | 0.353 |
| chr5 | 144775487 | 144820423 | 44937 | -0.39 | 0.338 | 0.353 |
| chr9 | 104023686 | 104024605 | 920 | 1.4 | 0.339 | 0.353 |
| chrX | 116205210 | 116252641 | 47432 | -0.21 | 0.339 | 0.354 |
| chr3 | 83763205 | 83890278 | 127074 | -2.41 | 0.339 | 0.354 |

| | | | | | | |
|---|---|---|---|---|---|---|
| chr7 | 124035643 | 124040103 | 4461 | -0.42 | 0.339 | 0.354 |
| chr5 | 96385077 | 96390485 | 5409 | -0.44 | 0.339 | 0.354 |
| chr10 | 33911920 | 33912654 | 735 | 0.56 | 0.34 | 0.354 |
| chr7 | 144315756 | 144317240 | 1485 | 0.52 | 0.341 | 0.355 |
| chr6 | 142410456 | 142421447 | 10992 | -0.52 | 0.341 | 0.355 |
| chr14 | 43677196 | 43703169 | 25974 | 0.76 | 0.341 | 0.356 |
| chr11 | 91238777 | 91275951 | 37175 | -0.39 | 0.341 | 0.356 |
| chr10 | 63638026 | 63638832 | 807 | 0.66 | 0.341 | 0.356 |
| chr10 | 14163592 | 14234508 | 70917 | -0.24 | 0.342 | 0.356 |
| chr1 | 181339851 | 181385119 | 45269 | -0.29 | 0.342 | 0.357 |
| chr4 | 63143776 | 63144297 | 522 | 0.58 | 0.342 | 0.357 |
| chr18 | 27679406 | 27680438 | 1033 | 0.58 | 0.342 | 0.357 |
| chr6 | 169061486 | 169066766 | 5281 | 0.28 | 0.342 | 0.357 |
| chr3 | 78765072 | 78765965 | 894 | -0.55 | 0.342 | 0.357 |
| chr7 | 158938705 | 158939295 | 591 | -1.32 | 0.342 | 0.357 |
| chr2 | 170042393 | 170044666 | 2274 | 0.65 | 0.342 | 0.357 |
| chr13 | 46648621 | 46649306 | 686 | -1.26 | 0.342 | 0.357 |
| chr10 | 19015387 | 19059513 | 44127 | -0.51 | 0.343 | 0.357 |
| chr8 | 31835233 | 31840809 | 5577 | -0.34 | 0.343 | 0.357 |
| chr5 | 76918174 | 76918867 | 694 | 0.98 | 0.343 | 0.357 |
| chr9 | 13985748 | 14015798 | 30051 | 0.45 | 0.343 | 0.358 |
| chrX | 62090798 | 62108552 | 17755 | 0.28 | 0.343 | 0.358 |
| chr8 | 104200177 | 104218138 | 17962 | 0.48 | 0.343 | 0.358 |
| chr9 | 34909805 | 34926471 | 16667 | -0.3 | 0.343 | 0.358 |
| chr11 | 71121496 | 71127249 | 5754 | 0.32 | 0.344 | 0.358 |
| chr4 | 59244544 | 59260522 | 15979 | 0.45 | 0.344 | 0.359 |
| chr7 | 37139635 | 37180984 | 41350 | 0.5 | 0.345 | 0.36 |
| chr11 | 27600055 | 27610472 | 10418 | -0.27 | 0.345 | 0.36 |
| chr7 | 155901870 | 155905232 | 3363 | 1.26 | 0.345 | 0.36 |
| chr1 | 86989747 | 86991509 | 1763 | -0.94 | 0.345 | 0.36 |
| chr6 | 142083067 | 142084746 | 1680 | 0.52 | 0.345 | 0.36 |
| chr15 | 57993182 | 57995366 | 2185 | 0.36 | 0.345 | 0.36 |
| chr14 | 21424134 | 21433974 | 9841 | 1.59 | 0.346 | 0.36 |
| chr8 | 113182031 | 113219196 | 37166 | -0.39 | 0.346 | 0.36 |
| chr4 | 60228774 | 60231522 | 2749 | 0.68 | 0.346 | 0.361 |
| chr4 | 62430121 | 62497648 | 67528 | -0.43 | 0.346 | 0.361 |
| chr2 | 220780154 | 220781307 | 1154 | -0.54 | 0.346 | 0.361 |
| chr15 | 37575006 | 37580441 | 5436 | 0.35 | 0.346 | 0.361 |
| chr2 | 103496515 | 103497314 | 800 | -1.31 | 0.346 | 0.361 |
| chr6 | 88765399 | 88767108 | 1710 | 0.45 | 0.347 | 0.361 |
| chr2 | 23576342 | 23578230 | 1889 | 0.82 | 0.347 | 0.361 |
| chr20 | 59617421 | 59684589 | 67169 | -0.3 | 0.347 | 0.361 |
| chr14 | 77225879 | 77226189 | 311 | -1.18 | 0.347 | 0.361 |
| chr1 | 210299042 | 210300255 | 1214 | -1.32 | 0.347 | 0.362 |
| chr7 | 86765924 | 86773495 | 7572 | -1.52 | 0.347 | 0.362 |
| chrX | 110864702 | 110867023 | 2322 | -0.47 | 0.347 | 0.362 |
| chr3 | 72296926 | 72307291 | 10366 | -0.43 | 0.348 | 0.362 |
| chr10 | 102239646 | 102241177 | 1532 | -0.52 | 0.348 | 0.363 |
| chrX | 101783497 | 101786242 | 2746 | -0.59 | 0.349 | 0.363 |
| chr2 | 213250370 | 213251348 | 979 | 0.82 | 0.349 | 0.364 |
| chr8 | 82419429 | 82436175 | 16747 | 1.57 | 0.349 | 0.364 |
| chr5 | 145982070 | 145996784 | 14715 | 0.3 | 0.349 | 0.364 |
| chr6 | 38793210 | 38795070 | 1861 | -0.92 | 0.35 | 0.364 |
| chr13 | 49229986 | 49231312 | 1327 | 0.63 | 0.35 | 0.365 |
| chr3 | 163040671 | 163054674 | 14004 | -0.89 | 0.35 | 0.365 |
| chr3 | 165573339 | 165574577 | 1239 | -1.03 | 0.351 | 0.365 |
| chr2 | 231867844 | 231869481 | 1638 | 0.86 | 0.351 | 0.365 |
| chr2 | 166086545 | 166088967 | 2423 | 0.92 | 0.351 | 0.366 |
| chr19 | 42025670 | 42057491 | 31822 | -1.72 | 0.351 | 0.366 |
| chr7 | 20865467 | 20867828 | 2362 | -0.48 | 0.351 | 0.366 |
| chr11 | 56534309 | 56535671 | 1363 | 0.81 | 0.352 | 0.366 |

| | | | | | | |
|---|---|---|---|---|---|---|
| chr4 | 30066556 | 30067068 | 513 | -0.61 | 0.352 | 0.366 |
| chr8 | 109433647 | 109446603 | 12957 | 1.6 | 0.352 | 0.366 |
| chr4 | 34627600 | 34628736 | 1137 | 0.5 | 0.352 | 0.366 |
| chr18 | 27607378 | 27618096 | 10719 | -0.63 | 0.352 | 0.367 |
| chr11 | 15824910 | 15831631 | 6722 | 0.46 | 0.352 | 0.367 |
| chr7 | 19377833 | 19391981 | 14149 | 0.58 | 0.352 | 0.367 |
| chr11 | 22168069 | 22173109 | 5041 | 0.43 | 0.352 | 0.367 |
| chr13 | 22405552 | 22407911 | 2360 | -0.46 | 0.353 | 0.367 |
| chr16 | 74269979 | 74289771 | 19793 | 0.34 | 0.353 | 0.367 |
| chr8 | 25556458 | 25821759 | 265302 | 0.15 | 0.353 | 0.368 |
| chr4 | 135748990 | 135750732 | 1743 | -0.6 | 0.353 | 0.368 |
| chr4 | 47719829 | 47753167 | 33339 | -0.21 | 0.353 | 0.368 |
| chrX | 151669795 | 151670134 | 340 | -0.58 | 0.354 | 0.368 |
| chr1 | 200344259 | 200353967 | 9709 | -1.39 | 0.354 | 0.369 |
| chr10 | 90924599 | 90926756 | 2158 | -0.57 | 0.354 | 0.369 |
| chr7 | 96632821 | 96643215 | 10395 | -1.03 | 0.354 | 0.369 |
| chr4 | 34058790 | 34076329 | 17540 | -0.4 | 0.355 | 0.369 |
| chr8 | 105150212 | 105152509 | 2298 | -0.84 | 0.355 | 0.37 |
| chr12 | 53295932 | 53297181 | 1250 | 0.59 | 0.355 | 0.37 |
| chr3 | 159054744 | 159070821 | 16078 | -0.69 | 0.355 | 0.37 |
| chr14 | 38449650 | 38450069 | 420 | -1.11 | 0.355 | 0.37 |
| chrX | 101409946 | 101410745 | 800 | -1.32 | 0.356 | 0.371 |
| chr4 | 35061471 | 35072021 | 10551 | 0.55 | 0.356 | 0.371 |
| chrX | 122106477 | 122120087 | 13611 | 1.14 | 0.357 | 0.371 |
| chr4 | 59055497 | 59057827 | 2331 | 0.42 | 0.357 | 0.371 |
| chr4 | 47419796 | 47420389 | 594 | -0.88 | 0.357 | 0.372 |
| chr3 | 30897763 | 30903790 | 6028 | -0.24 | 0.357 | 0.372 |
| chr3 | 159083629 | 159085087 | 1459 | -1.39 | 0.357 | 0.372 |
| chr3 | 11054968 | 11056702 | 1735 | -0.46 | 0.358 | 0.373 |
| chr8 | 112159249 | 112182486 | 23238 | -0.37 | 0.359 | 0.373 |
| chr18 | 35092911 | 35093488 | 578 | 0.72 | 0.359 | 0.373 |
| chr15 | 38956161 | 38958223 | 2063 | 0.7 | 0.359 | 0.374 |
| chr7 | 124151382 | 124171004 | 19623 | -0.48 | 0.359 | 0.374 |
| chr11 | 106397452 | 106402672 | 5221 | 1.55 | 0.361 | 0.375 |
| chrX | 39158259 | 39171789 | 13531 | 0.28 | 0.361 | 0.376 |
| chr8 | 88438335 | 88489767 | 51433 | 0.34 | 0.361 | 0.376 |
| chr1 | 165319453 | 165324820 | 5368 | -1.24 | 0.362 | 0.376 |
| chr6 | 159585145 | 159585889 | 745 | 0.69 | 0.362 | 0.377 |
| chr16 | 85856609 | 85858221 | 1613 | 1.42 | 0.362 | 0.377 |
| chr3 | 70561587 | 70566064 | 4478 | -0.38 | 0.363 | 0.377 |
| chr6 | 156867300 | 156869401 | 2102 | 0.46 | 0.363 | 0.377 |
| chr21 | 27008645 | 27010498 | 1854 | -0.43 | 0.363 | 0.377 |
| chr15 | 34045029 | 34113888 | 68860 | -0.3 | 0.363 | 0.377 |
| chr20 | 29993551 | 29998252 | 4702 | -0.45 | 0.363 | 0.378 |
| chr2 | 217600903 | 217621142 | 20240 | 0.26 | 0.363 | 0.378 |
| chr16 | 868431 | 869514 | 1084 | 0.52 | 0.364 | 0.379 |
| chr6 | 164236780 | 164256930 | 20151 | -0.31 | 0.364 | 0.379 |
| chr13 | 89274885 | 89277720 | 2836 | -1.36 | 0.364 | 0.379 |
| chr3 | 51064675 | 51079501 | 14827 | -0.26 | 0.364 | 0.379 |
| chr3 | 40103396 | 40105194 | 1799 | 1.05 | 0.364 | 0.379 |
| chr18 | 10422911 | 10425672 | 2762 | 0.39 | 0.364 | 0.379 |
| chr17 | 22132236 | 22155148 | 22913 | -0.51 | 0.365 | 0.379 |
| chr11 | 127125237 | 127232967 | 107731 | -0.15 | 0.365 | 0.379 |
| chr1 | 236880629 | 236881295 | 667 | -0.82 | 0.365 | 0.38 |
| chr2 | 180360168 | 180360553 | 386 | -1.08 | 0.365 | 0.38 |
| chrX | 120123523 | 120125223 | 1701 | 0.86 | 0.366 | 0.38 |
| chr13 | 22505966 | 22533926 | 27961 | 0.21 | 0.366 | 0.38 |
| chr4 | 107531415 | 107532292 | 878 | 0.48 | 0.366 | 0.381 |
| chr3 | 16071403 | 16073858 | 2456 | 1.52 | 0.366 | 0.381 |
| chr5 | 64732289 | 64734395 | 2107 | -0.75 | 0.367 | 0.381 |
| chr8 | 57327044 | 57341860 | 14817 | -0.49 | 0.367 | 0.381 |

| | | | | | | |
|---|---|---|---|---|---|---|
| chr8 | 59382880 | 59396019 | 13140 | 0.28 | 0.367 | 0.381 |
| chr9 | 92319889 | 92334817 | 14929 | -1.07 | 0.367 | 0.381 |
| chr3 | 54123305 | 54127044 | 3740 | -0.61 | 0.367 | 0.381 |
| chr3 | 144826732 | 144829216 | 2485 | 0.51 | 0.368 | 0.382 |
| chr4 | 19365310 | 19366263 | 954 | 0.68 | 0.368 | 0.383 |
| chr6 | 84205254 | 84205574 | 321 | -1.17 | 0.368 | 0.383 |
| chr10 | 19560210 | 19588031 | 27822 | -0.39 | 0.368 | 0.383 |
| chr5 | 112745704 | 112782405 | 36702 | 0.38 | 0.369 | 0.383 |
| chrX | 75303378 | 75304312 | 935 | -0.63 | 0.369 | 0.384 |
| chr2 | 166839004 | 166845773 | 6770 | -0.35 | 0.369 | 0.384 |
| chr19 | 13572942 | 13575047 | 2106 | 0.47 | 0.37 | 0.384 |
| chr3 | 111166122 | 111208732 | 42611 | -0.31 | 0.37 | 0.385 |
| chr8 | 31924992 | 31929537 | 4546 | -0.38 | 0.37 | 0.385 |
| chrX | 15539697 | 15541804 | 2108 | -0.46 | 0.37 | 0.385 |
| chr13 | 91862961 | 91863999 | 1039 | -0.79 | 0.37 | 0.385 |
| chr1 | 159935403 | 159955854 | 20452 | -0.31 | 0.37 | 0.385 |
| chr16 | 20067736 | 20070274 | 2539 | 1.43 | 0.37 | 0.385 |
| chr10 | 94576747 | 94577805 | 1059 | 0.51 | 0.37 | 0.385 |
| chr4 | 33846020 | 33846999 | 980 | -0.97 | 0.371 | 0.385 |
| chr11 | 92085395 | 92087658 | 2264 | 0.77 | 0.371 | 0.386 |
| chr4 | 35919297 | 35920947 | 1651 | 1.41 | 0.371 | 0.386 |
| chr17 | 33512442 | 33512811 | 370 | 0.65 | 0.372 | 0.386 |
| chr8 | 114001016 | 114078016 | 77001 | -0.83 | 0.372 | 0.386 |
| chr2 | 196086598 | 196087119 | 522 | -1.03 | 0.372 | 0.387 |
| chrX | 55264345 | 55265785 | 1441 | 0.68 | 0.372 | 0.387 |
| chr10 | 125351169 | 125420591 | 69423 | 0.23 | 0.372 | 0.387 |
| chr14 | 82004185 | 82006615 | 2431 | -1.34 | 0.372 | 0.387 |
| chr1 | 5509185 | 5538912 | 29728 | -0.27 | 0.372 | 0.387 |
| chr12 | 91115864 | 91116593 | 730 | -0.67 | 0.373 | 0.387 |
| chrX | 22297500 | 22301763 | 4264 | -0.55 | 0.373 | 0.388 |
| chr1 | 195757142 | 195780216 | 23075 | 0.71 | 0.374 | 0.388 |
| chr14 | 88663034 | 88796660 | 133627 | -0.17 | 0.374 | 0.388 |
| chrX | 74695151 | 74695660 | 510 | -0.5 | 0.374 | 0.388 |
| chr5 | 31633190 | 31685849 | 52660 | -0.33 | 0.374 | 0.389 |
| chr11 | 114493697 | 114494018 | 322 | 0.66 | 0.375 | 0.389 |
| chr1 | 106094685 | 106113033 | 18349 | 0.4 | 0.375 | 0.389 |
| chr6 | 70996421 | 70997440 | 1020 | 0.49 | 0.375 | 0.389 |
| chr7 | 7418501 | 7419892 | 1392 | 0.82 | 0.375 | 0.39 |
| chr5 | 173528678 | 173529402 | 725 | 0.51 | 0.376 | 0.39 |
| chr10 | 123261334 | 123265120 | 3787 | 0.51 | 0.376 | 0.391 |
| chr5 | 31702963 | 31706355 | 3393 | -0.38 | 0.376 | 0.391 |
| chr13 | 44515639 | 44516450 | 812 | 0.66 | 0.376 | 0.391 |
| chr17 | 67589485 | 67590421 | 937 | 1.38 | 0.378 | 0.392 |
| chr10 | 1423136 | 1424593 | 1458 | 1.41 | 0.378 | 0.392 |
| chr3 | 162683336 | 162685121 | 1786 | -0.54 | 0.379 | 0.394 |
| chr1 | 111906760 | 111920057 | 13298 | -0.29 | 0.379 | 0.394 |
| chr12 | 59244897 | 59245765 | 869 | 0.65 | 0.379 | 0.394 |
| chr12 | 18804424 | 18805525 | 1102 | -1.23 | 0.379 | 0.394 |
| chr3 | 150646561 | 150680914 | 34354 | 0.52 | 0.38 | 0.394 |
| chr12 | 39541312 | 39570851 | 29540 | -0.32 | 0.38 | 0.394 |
| chr2 | 222056725 | 222075730 | 19006 | 0.41 | 0.38 | 0.394 |
| chrX | 23605127 | 23607107 | 1981 | -0.48 | 0.381 | 0.396 |
| chr1 | 105244064 | 105279960 | 35897 | 0.35 | 0.381 | 0.396 |
| chr13 | 22746303 | 22920235 | 173933 | 0.14 | 0.382 | 0.397 |
| chr8 | 113612409 | 113643000 | 30592 | -1.55 | 0.382 | 0.397 |
| chr2 | 9178023 | 9178655 | 633 | -1.26 | 0.382 | 0.397 |
| chr2 | 89159321 | 89161871 | 2551 | -0.64 | 0.382 | 0.397 |
| chr14 | 58342659 | 58346433 | 3775 | -0.37 | 0.383 | 0.398 |
| chr8 | 10899408 | 10903468 | 4061 | -0.44 | 0.383 | 0.398 |
| chr14 | 92821300 | 92824256 | 2957 | 1.45 | 0.383 | 0.398 |
| chr1 | 114713240 | 114714488 | 1249 | 0.54 | 0.383 | 0.398 |

| | | | | | | |
|---|---|---|---|---|---|---|
| chr8 | 142687773 | 142688611 | 839 | 1.35 | 0.384 | 0.398 |
| chr14 | 34675402 | 34686663 | 11262 | -1.46 | 0.384 | 0.399 |
| chrX | 18315718 | 18328077 | 12360 | -1.44 | 0.384 | 0.399 |
| chr4 | 30079135 | 30088065 | 8931 | 0.57 | 0.384 | 0.399 |
| chr9 | 79624825 | 79671423 | 46599 | -1.01 | 0.385 | 0.399 |
| chr9 | 124351370 | 124351991 | 622 | 0.65 | 0.385 | 0.4 |
| chr6 | 168798813 | 168808180 | 9368 | -0.31 | 0.385 | 0.4 |
| chr9 | 72591000 | 72600567 | 9568 | 0.39 | 0.386 | 0.401 |
| chr10 | 101218658 | 101223032 | 4375 | -0.54 | 0.386 | 0.401 |
| chr18 | 71618184 | 71633463 | 15280 | 0.38 | 0.387 | 0.401 |
| chr8 | 32026772 | 32028989 | 2218 | -0.61 | 0.387 | 0.402 |
| chr4 | 59456325 | 59457599 | 1275 | -1.22 | 0.388 | 0.402 |
| chr6 | 50889465 | 50890197 | 733 | -0.64 | 0.388 | 0.403 |
| chr20 | 36055389 | 36060541 | 5153 | -0.34 | 0.388 | 0.403 |
| chr9 | 7254048 | 7269501 | 15454 | 0.36 | 0.389 | 0.404 |
| chrX | 79638674 | 79639350 | 677 | -0.41 | 0.389 | 0.404 |
| chr3 | 157590761 | 157597350 | 6590 | -0.35 | 0.389 | 0.404 |
| chr15 | 54016108 | 54017957 | 1850 | -0.42 | 0.39 | 0.405 |
| chr1 | 106259478 | 106265678 | 6201 | 0.6 | 0.391 | 0.406 |
| chrX | 69158133 | 69173271 | 15139 | -0.29 | 0.391 | 0.406 |
| chr2 | 215221705 | 215222171 | 467 | 0.6 | 0.391 | 0.406 |
| chr11 | 125587385 | 125592546 | 5162 | -0.64 | 0.392 | 0.407 |
| chr13 | 85011822 | 85013115 | 1294 | -0.66 | 0.392 | 0.407 |
| chr16 | 63174717 | 63176241 | 1525 | 1.27 | 0.392 | 0.407 |
| chrX | 132722036 | 132728692 | 6657 | 0.31 | 0.392 | 0.407 |
| chr3 | 112092577 | 112093913 | 1337 | 0.49 | 0.392 | 0.407 |
| chr1 | 111837289 | 111865594 | 28306 | -0.22 | 0.393 | 0.407 |
| chr10 | 24523600 | 24526044 | 2445 | -1.32 | 0.393 | 0.408 |
| chr5 | 73788445 | 73789144 | 700 | -1.1 | 0.393 | 0.408 |
| chr18 | 45723097 | 45738578 | 15482 | -1.29 | 0.394 | 0.408 |
| chr11 | 115561532 | 115562359 | 828 | 0.49 | 0.394 | 0.408 |
| chr4 | 136033334 | 136034091 | 758 | -0.43 | 0.394 | 0.409 |
| chr5 | 144732141 | 144739426 | 7286 | 1.4 | 0.394 | 0.409 |
| chr7 | 119436959 | 119484926 | 47968 | 0.38 | 0.394 | 0.409 |
| chr6 | 88961873 | 88963507 | 1635 | 0.39 | 0.395 | 0.409 |
| chr9 | 350979 | 351740 | 762 | 1.33 | 0.395 | 0.41 |
| chr3 | 42368540 | 42368868 | 329 | 0.58 | 0.395 | 0.41 |
| chr4 | 34560127 | 34567853 | 7727 | -1.06 | 0.396 | 0.41 |
| chr9 | 119480936 | 119484761 | 3826 | -0.3 | 0.396 | 0.41 |
| chrX | 55099751 | 55100041 | 291 | 0.53 | 0.396 | 0.41 |
| chr15 | 33661581 | 33868946 | 207366 | 0.13 | 0.396 | 0.411 |
| chr2 | 192595723 | 192596694 | 972 | 1.26 | 0.396 | 0.411 |
| chr1 | 218129643 | 218158052 | 28410 | -0.19 | 0.397 | 0.411 |
| chr8 | 112247748 | 112254255 | 6508 | -0.88 | 0.398 | 0.413 |
| chr20 | 47094402 | 47106818 | 12417 | -0.52 | 0.398 | 0.413 |
| chr6 | 159813567 | 159814828 | 1262 | -0.52 | 0.398 | 0.413 |
| chr11 | 32202833 | 32212566 | 9734 | 0.61 | 0.398 | 0.413 |
| chr13 | 22601096 | 22609212 | 8117 | 0.34 | 0.399 | 0.413 |
| chr11 | 101184012 | 101185318 | 1307 | -0.86 | 0.399 | 0.414 |
| chr8 | 19720028 | 19721416 | 1389 | 0.73 | 0.399 | 0.414 |
| chr5 | 79322342 | 79341920 | 19579 | -0.52 | 0.4 | 0.415 |
| chr3 | 29036778 | 29038358 | 1581 | -0.32 | 0.4 | 0.415 |
| chr1 | 194556800 | 194561705 | 4906 | -1.36 | 0.401 | 0.415 |
| chrX | 15174583 | 15175178 | 596 | 0.94 | 0.401 | 0.416 |
| chr3 | 108933320 | 108953307 | 19988 | -1.45 | 0.401 | 0.416 |
| chr11 | 20227418 | 20231316 | 3899 | -0.57 | 0.401 | 0.416 |
| chr3 | 86338345 | 86346348 | 8004 | -0.44 | 0.402 | 0.416 |
| chr8 | 114163907 | 114201330 | 37424 | -0.76 | 0.402 | 0.416 |
| chr7 | 18914919 | 18920268 | 5350 | -0.46 | 0.402 | 0.417 |
| chr1 | 105159490 | 105168426 | 8937 | -0.56 | 0.402 | 0.417 |
| chr1 | 96681224 | 96682548 | 1325 | 0.52 | 0.402 | 0.417 |

| | | | | | | |
|---|---|---|---|---|---|---|
| chr11 | 109684487 | 109684824 | 338 | 0.49 | 0.403 | 0.418 |
| chr11 | 124347394 | 124371112 | 23719 | -0.24 | 0.403 | 0.418 |
| chr5 | 155189500 | 155191727 | 2228 | 1.37 | 0.403 | 0.418 |
| chr16 | 63794140 | 63825407 | 31268 | -0.17 | 0.404 | 0.418 |
| chr10 | 118282955 | 118346578 | 63624 | 0.22 | 0.404 | 0.419 |
| chr4 | 29457714 | 29462381 | 4668 | -0.37 | 0.404 | 0.419 |
| chr13 | 37437420 | 37442342 | 4923 | -0.5 | 0.404 | 0.419 |
| chr14 | 36584150 | 36653846 | 69697 | 0.18 | 0.404 | 0.419 |
| chr20 | 48128245 | 48130718 | 2474 | -0.36 | 0.405 | 0.419 |
| chr5 | 88066462 | 88067829 | 1368 | 1.31 | 0.405 | 0.42 |
| chr10 | 49407637 | 49408691 | 1055 | -0.56 | 0.405 | 0.42 |
| chr9 | 84986966 | 84996531 | 9566 | -0.42 | 0.405 | 0.42 |
| chr6 | 125774650 | 125781520 | 6871 | -0.36 | 0.406 | 0.42 |
| chr4 | 32601711 | 32602957 | 1247 | 0.53 | 0.406 | 0.421 |
| chrX | 74977869 | 74979086 | 1218 | 0.76 | 0.407 | 0.421 |
| chr4 | 29440955 | 29441806 | 852 | -0.48 | 0.408 | 0.422 |
| chr8 | 106735519 | 106737647 | 2129 | -0.56 | 0.408 | 0.423 |
| chr8 | 57133954 | 57211112 | 77159 | -2.07 | 0.408 | 0.423 |
| chr4 | 54488588 | 54491430 | 2843 | 0.5 | 0.409 | 0.423 |
| chr4 | 60768618 | 60788329 | 19712 | 0.32 | 0.409 | 0.423 |
| chr11 | 18739314 | 18739657 | 344 | 0.86 | 0.409 | 0.423 |
| chr8 | 53257526 | 53301941 | 44416 | -0.21 | 0.409 | 0.424 |
| chr11 | 134291903 | 134292978 | 1076 | 0.47 | 0.409 | 0.424 |
| chr9 | 107095039 | 107098119 | 3081 | 0.87 | 0.41 | 0.424 |
| chr2 | 11658939 | 11675447 | 16509 | 1.03 | 0.41 | 0.424 |
| chr11 | 104511182 | 104522137 | 10956 | -1.28 | 0.41 | 0.424 |
| chr1 | 5493030 | 5494802 | 1773 | 0.38 | 0.41 | 0.425 |
| chrX | 79608626 | 79609198 | 573 | -1.11 | 0.411 | 0.425 |
| chr10 | 64811121 | 64827632 | 16512 | -0.34 | 0.412 | 0.426 |
| chr1 | 105541435 | 105617052 | 75618 | 0.37 | 0.413 | 0.427 |
| chr1 | 116824626 | 116825623 | 998 | -1.28 | 0.413 | 0.427 |
| chr7 | 137472152 | 137472668 | 517 | -1.12 | 0.413 | 0.428 |
| chr12 | 98335191 | 98351625 | 16435 | 0.29 | 0.413 | 0.428 |
| chr10 | 37087896 | 37092599 | 4704 | 1.4 | 0.413 | 0.428 |
| chr8 | 73914868 | 73915425 | 558 | 0.54 | 0.413 | 0.428 |
| chr8 | 11480136 | 11481191 | 1056 | 0.5 | 0.414 | 0.428 |
| chr1 | 241390291 | 241493637 | 103347 | 0.2 | 0.414 | 0.429 |
| chr2 | 186315038 | 186321655 | 6618 | -0.73 | 0.414 | 0.429 |
| chr4 | 29515780 | 29547007 | 31228 | 0.32 | 0.414 | 0.429 |
| chr9 | 227917 | 280319 | 52403 | -0.43 | 0.415 | 0.429 |
| chr2 | 222837914 | 222844975 | 7062 | 0.28 | 0.415 | 0.43 |
| chr5 | 123093595 | 123133712 | 40118 | -0.33 | 0.415 | 0.43 |
| chr12 | 110250541 | 110252430 | 1890 | 0.34 | 0.416 | 0.43 |
| chr1 | 115562753 | 115564886 | 2134 | -0.71 | 0.416 | 0.43 |
| chr3 | 36761086 | 36764131 | 3046 | 0.45 | 0.416 | 0.43 |
| chr3 | 124057773 | 124059864 | 2092 | 0.67 | 0.416 | 0.43 |
| chr1 | 106385886 | 106403686 | 17801 | 0.4 | 0.416 | 0.431 |
| chr4 | 134117002 | 134117697 | 696 | -1.13 | 0.416 | 0.431 |
| chr1 | 185667994 | 185671210 | 3217 | -0.74 | 0.416 | 0.431 |
| chr5 | 64385920 | 64387644 | 1725 | -1.23 | 0.417 | 0.432 |
| chr20 | 58255517 | 58299290 | 43774 | -0.18 | 0.418 | 0.432 |
| chr16 | 79900334 | 79903085 | 2752 | 0.43 | 0.418 | 0.433 |
| chr4 | 27968663 | 27970948 | 2286 | -0.43 | 0.418 | 0.433 |
| chr4 | 27119255 | 27120932 | 1678 | -1.2 | 0.418 | 0.433 |
| chr2 | 223242500 | 223244270 | 1771 | 0.51 | 0.418 | 0.433 |
| chr11 | 109229235 | 109253076 | 23842 | -0.21 | 0.418 | 0.433 |
| chr8 | 77451736 | 77463132 | 11397 | -0.45 | 0.419 | 0.433 |
| chr9 | 133055069 | 133061548 | 6480 | -0.37 | 0.419 | 0.434 |
| chr5 | 85633095 | 85634264 | 1170 | -0.75 | 0.419 | 0.434 |
| chr15 | 73702864 | 73705780 | 2917 | -0.35 | 0.419 | 0.434 |
| chr1 | 105923349 | 105923854 | 506 | 0.74 | 0.419 | 0.434 |

| | | | | | | |
|---|---|---|---|---|---|---|
| chr2 | 7393321 | 7441523 | 48203 | 0.27 | 0.42 | 0.434 |
| chr17 | 69853851 | 69957317 | 103467 | -0.26 | 0.42 | 0.434 |
| chr7 | 103067196 | 103138624 | 71429 | 0.26 | 0.42 | 0.434 |
| chr10 | 124851737 | 124853767 | 2031 | 0.51 | 0.42 | 0.435 |
| chr1 | 105781528 | 105786357 | 4830 | 0.59 | 0.421 | 0.435 |
| chr4 | 88533111 | 88534384 | 1274 | 0.67 | 0.421 | 0.435 |
| chr1 | 91526144 | 91531984 | 5841 | 0.33 | 0.421 | 0.436 |
| chr12 | 15122303 | 15124070 | 1768 | -0.44 | 0.422 | 0.437 |
| chr16 | 59017203 | 59130551 | 113349 | 0.2 | 0.422 | 0.437 |
| chr4 | 134555223 | 134558276 | 3054 | -0.34 | 0.422 | 0.437 |
| chr7 | 148216624 | 148220310 | 3687 | -0.42 | 0.423 | 0.437 |
| chr3 | 130097909 | 130098794 | 886 | -0.44 | 0.424 | 0.439 |
| chrX | 23429311 | 23431719 | 2409 | 0.43 | 0.424 | 0.439 |
| chr6 | 78964230 | 78966637 | 2408 | 1.2 | 0.425 | 0.44 |
| chr4 | 29852355 | 29904075 | 51721 | -0.4 | 0.425 | 0.44 |
| chr11 | 104796991 | 104806177 | 9187 | 0.34 | 0.426 | 0.441 |
| chr11 | 116354686 | 116356586 | 1901 | 0.4 | 0.426 | 0.441 |
| chr11 | 95205877 | 95207320 | 1444 | 1.29 | 0.426 | 0.441 |
| chr4 | 29810866 | 29811846 | 981 | -0.47 | 0.427 | 0.441 |
| chr10 | 124411326 | 124455691 | 44366 | 0.37 | 0.427 | 0.442 |
| chr7 | 159017153 | 159017450 | 298 | -0.71 | 0.427 | 0.442 |
| chr12 | 55051473 | 55052132 | 660 | -0.48 | 0.428 | 0.442 |
| chr17 | 8297897 | 8300133 | 2237 | 0.34 | 0.428 | 0.443 |
| chr2 | 240411592 | 240414028 | 2437 | 0.73 | 0.428 | 0.443 |
| chrX | 20723844 | 20743688 | 19845 | -1.32 | 0.429 | 0.443 |
| chr3 | 130096271 | 130096631 | 361 | 0.54 | 0.429 | 0.443 |
| chr8 | 6905685 | 6919564 | 13880 | 0.21 | 0.429 | 0.444 |
| chr5 | 23022218 | 23203849 | 181632 | 0.22 | 0.432 | 0.446 |
| chr4 | 127503956 | 127506547 | 2592 | -0.54 | 0.433 | 0.448 |
| chr22 | 22789335 | 22791002 | 1668 | 0.4 | 0.433 | 0.448 |
| chr12 | 15387395 | 15412131 | 24737 | 0.23 | 0.433 | 0.448 |
| chr7 | 34523915 | 34524274 | 360 | -0.8 | 0.434 | 0.449 |
| chr6 | 110391744 | 110392436 | 693 | 1.1 | 0.434 | 0.449 |
| chr4 | 21072480 | 21073015 | 536 | -1.05 | 0.435 | 0.449 |
| chr6 | 8755788 | 8768148 | 12361 | -0.29 | 0.435 | 0.449 |
| chr1 | 106359985 | 106362077 | 2093 | 0.38 | 0.435 | 0.45 |
| chr1 | 237981990 | 238020122 | 38133 | -0.25 | 0.435 | 0.45 |
| chr15 | 34134799 | 34136235 | 1437 | 1.24 | 0.436 | 0.45 |
| chr10 | 82914610 | 83000991 | 86382 | -1.96 | 0.436 | 0.451 |
| chrX | 58091210 | 58147214 | 56005 | 0.19 | 0.436 | 0.451 |
| chr2 | 100225988 | 100228466 | 2479 | -1.18 | 0.437 | 0.451 |
| chr2 | 215142966 | 215170895 | 27930 | -0.49 | 0.437 | 0.451 |
| chr12 | 93080186 | 93081443 | 1258 | 0.67 | 0.438 | 0.453 |
| chr5 | 85464854 | 85467193 | 2340 | -0.78 | 0.439 | 0.453 |
| chrX | 62148508 | 62156106 | 7599 | -0.3 | 0.439 | 0.453 |
| chr14 | 49300238 | 49333180 | 32943 | -0.4 | 0.439 | 0.454 |
| chr12 | 50240769 | 50250345 | 9577 | -1.32 | 0.439 | 0.454 |
| chr10 | 99842452 | 99846316 | 3865 | 1.28 | 0.44 | 0.454 |
| chr1 | 89757656 | 89759505 | 1850 | -0.37 | 0.44 | 0.455 |
| chr4 | 75804712 | 75813743 | 9032 | -0.28 | 0.44 | 0.455 |
| chr15 | 79818133 | 79826415 | 8283 | -0.21 | 0.44 | 0.455 |
| chrX | 36646396 | 36655696 | 9301 | -0.34 | 0.441 | 0.455 |
| chr14 | 60831238 | 60848486 | 17249 | -0.34 | 0.441 | 0.455 |
| chr11 | 69658749 | 69659679 | 931 | 0.47 | 0.441 | 0.455 |
| chr4 | 135708434 | 135717417 | 8984 | 0.42 | 0.441 | 0.456 |
| chr1 | 105470039 | 105524579 | 54541 | 0.27 | 0.442 | 0.456 |
| chr12 | 9338933 | 9340810 | 1878 | -0.34 | 0.442 | 0.456 |
| chr18 | 27491634 | 27513537 | 21904 | 0.39 | 0.442 | 0.456 |
| chr13 | 96778371 | 96779853 | 1483 | 0.79 | 0.442 | 0.457 |
| chr12 | 34348625 | 34351385 | 2761 | 0.54 | 0.442 | 0.457 |
| chr3 | 85052128 | 85094567 | 42440 | -1.38 | 0.443 | 0.457 |

| | | | | | | |
|---|---|---|---|---|---|---|
| chr10 | 25505976 | 25508016 | 2041 | -0.4 | 0.443 | 0.458 |
| chr12 | 119599784 | 119609857 | 10074 | -0.35 | 0.444 | 0.458 |
| chr12 | 18602857 | 18613319 | 10463 | 0.25 | 0.444 | 0.458 |
| chr9 | 123058909 | 123060023 | 1115 | 0.75 | 0.444 | 0.458 |
| chrX | 150685603 | 150714365 | 28763 | 0.26 | 0.445 | 0.459 |
| chr1 | 224760418 | 224762096 | 1679 | 0.71 | 0.445 | 0.46 |
| chr20 | 20223088 | 20335224 | 112137 | 0.2 | 0.445 | 0.46 |
| chr6 | 38829845 | 38830575 | 731 | 1.14 | 0.446 | 0.46 |
| chr3 | 107974116 | 107976593 | 2478 | 0.28 | 0.446 | 0.46 |
| chr12 | 41675454 | 41675725 | 272 | 0.56 | 0.446 | 0.461 |
| chr4 | 34394272 | 34418824 | 24553 | 0.38 | 0.447 | 0.461 |
| chr9 | 30598827 | 30640132 | 41306 | -1.35 | 0.447 | 0.462 |
| chr6 | 88569634 | 88593721 | 24088 | -0.33 | 0.447 | 0.462 |
| chr5 | 76228645 | 76231913 | 3269 | -0.39 | 0.448 | 0.462 |
| chr8 | 122045514 | 122046252 | 739 | 0.42 | 0.448 | 0.462 |
| chr16 | 82155480 | 82159294 | 3815 | -0.41 | 0.448 | 0.463 |
| chr4 | 135466859 | 135476730 | 9872 | 0.34 | 0.449 | 0.464 |
| chr2 | 238252967 | 238253517 | 551 | -0.64 | 0.449 | 0.464 |
| chr10 | 86375390 | 86376620 | 1231 | -0.91 | 0.45 | 0.464 |
| chr8 | 88227133 | 88229224 | 2092 | -0.38 | 0.45 | 0.464 |
| chr16 | 82079540 | 82142849 | 63310 | -0.25 | 0.45 | 0.465 |
| chr20 | 20085872 | 20087156 | 1285 | 0.44 | 0.451 | 0.465 |
| chr1 | 162714665 | 162716024 | 1360 | -0.32 | 0.451 | 0.465 |
| chr3 | 21384783 | 21406574 | 21792 | -1.32 | 0.452 | 0.466 |
| chr9 | 90401166 | 90402137 | 972 | -0.62 | 0.452 | 0.466 |
| chr9 | 121662356 | 121684777 | 22422 | -0.25 | 0.452 | 0.467 |
| chr13 | 112764659 | 112781661 | 17003 | 0.42 | 0.452 | 0.467 |
| chr7 | 39478007 | 39499357 | 21351 | -0.19 | 0.452 | 0.467 |
| chr16 | 32470743 | 32471178 | 436 | 0.41 | 0.453 | 0.467 |
| chr18 | 47993138 | 47995502 | 2365 | 0.54 | 0.453 | 0.467 |
| chr1 | 153217789 | 153237759 | 19971 | 0.57 | 0.453 | 0.468 |
| chr4 | 59225402 | 59230395 | 4994 | 0.35 | 0.454 | 0.468 |
| chr8 | 82451942 | 82453280 | 1339 | -0.41 | 0.454 | 0.469 |
| chr4 | 54675021 | 54675957 | 937 | -0.66 | 0.454 | 0.469 |
| chr2 | 183377781 | 183394754 | 16974 | 1.21 | 0.455 | 0.47 |
| chr4 | 54556040 | 54557293 | 1254 | 0.63 | 0.455 | 0.47 |
| chr8 | 26561172 | 26591161 | 29990 | -0.17 | 0.456 | 0.47 |
| chr2 | 186578310 | 186601182 | 22873 | -0.45 | 0.456 | 0.47 |
| chrX | 68555212 | 68570779 | 15568 | -0.31 | 0.456 | 0.47 |
| chr10 | 3010267 | 3010795 | 529 | 0.48 | 0.457 | 0.471 |
| chr3 | 16581474 | 16585785 | 4312 | 1.31 | 0.457 | 0.472 |
| chrX | 121950275 | 121952385 | 2111 | -0.33 | 0.458 | 0.472 |
| chr7 | 57786336 | 57786775 | 440 | 0.41 | 0.458 | 0.472 |
| chr17 | 37185989 | 37188059 | 2071 | 1.22 | 0.458 | 0.473 |
| chrX | 42067552 | 42069485 | 1934 | 0.35 | 0.458 | 0.473 |
| chr4 | 65592646 | 65593073 | 428 | 1.04 | 0.46 | 0.475 |
| chr8 | 114285617 | 114288428 | 2812 | -1.06 | 0.461 | 0.475 |
| chr17 | 21845644 | 21847773 | 2130 | -0.44 | 0.461 | 0.475 |
| chr18 | 74028787 | 74030405 | 1619 | 0.37 | 0.461 | 0.475 |
| chr13 | 31927433 | 31957767 | 30335 | -0.22 | 0.461 | 0.476 |
| chr15 | 73814805 | 73825055 | 10251 | -1.21 | 0.462 | 0.476 |
| chr16 | 15999964 | 16010008 | 10045 | -1.21 | 0.462 | 0.476 |
| chr5 | 143124885 | 143142101 | 17217 | -1.21 | 0.462 | 0.476 |
| chr7 | 36742122 | 36997686 | 255565 | 0.11 | 0.463 | 0.477 |
| chr15 | 48535772 | 48537678 | 1907 | -0.33 | 0.463 | 0.477 |
| chr19 | 55149568 | 55170644 | 21077 | -0.78 | 0.463 | 0.477 |
| chr4 | 60155332 | 60157124 | 1793 | -0.82 | 0.463 | 0.478 |
| chr5 | 75532464 | 75541345 | 8882 | -0.27 | 0.463 | 0.478 |
| chr1 | 207758249 | 207773083 | 14835 | 0.29 | 0.464 | 0.479 |
| chr4 | 155304000 | 155304280 | 281 | 0.92 | 0.465 | 0.48 |
| chr12 | 47319313 | 47321060 | 1748 | 0.82 | 0.465 | 0.48 |

| | | | | | | |
|---|---|---|---|---|---|---|
| chr7 | 32303755 | 32322352 | 18598 | -1.12 | 0.465 | 0.48 |
| chr3 | 68982532 | 68983558 | 1027 | -0.79 | 0.466 | 0.481 |
| chr14 | 58206977 | 58207692 | 716 | 0.42 | 0.468 | 0.483 |
| chr4 | 53358986 | 53383203 | 24218 | -0.4 | 0.468 | 0.483 |
| chr14 | 70370696 | 70372008 | 1313 | 0.33 | 0.469 | 0.483 |
| chr11 | 127929196 | 127934687 | 5492 | 0.29 | 0.469 | 0.484 |
| chr4 | 35803495 | 35822941 | 19447 | 0.27 | 0.47 | 0.484 |
| chrX | 74193634 | 74267780 | 74147 | 0.21 | 0.47 | 0.484 |
| chr5 | 27263466 | 27295459 | 31994 | 0.39 | 0.47 | 0.484 |
| chr12 | 30656411 | 30666337 | 9927 | 0.39 | 0.47 | 0.485 |
| chrX | 131039850 | 131050416 | 10567 | -0.25 | 0.47 | 0.485 |
| chr15 | 89396626 | 89414060 | 17435 | -0.65 | 0.47 | 0.485 |
| chr11 | 31375702 | 31377974 | 2273 | -0.46 | 0.471 | 0.486 |
| chr4 | 63010739 | 63015398 | 4660 | 0.28 | 0.471 | 0.486 |
| chr8 | 17950810 | 17964133 | 13324 | -0.31 | 0.472 | 0.486 |
| chr2 | 186796258 | 186797559 | 1302 | 0.51 | 0.472 | 0.486 |
| chr10 | 73553751 | 73554541 | 791 | 1.13 | 0.472 | 0.486 |
| chr8 | 86395353 | 86395982 | 630 | 0.38 | 0.472 | 0.486 |
| chr21 | 39442039 | 39447051 | 5013 | -0.24 | 0.473 | 0.487 |
| chr11 | 57681612 | 57702739 | 21128 | 0.19 | 0.473 | 0.488 |
| chr3 | 124199925 | 124201567 | 1643 | 0.51 | 0.473 | 0.488 |
| chr14 | 20977999 | 20988555 | 10557 | 1.18 | 0.473 | 0.488 |
| chr10 | 123215571 | 123224449 | 8879 | -0.42 | 0.473 | 0.488 |
| chr10 | 19626773 | 19629401 | 2629 | -1.03 | 0.474 | 0.488 |
| chr12 | 16223613 | 16224928 | 1316 | 0.41 | 0.474 | 0.489 |
| chr5 | 169672780 | 169679741 | 6962 | 0.45 | 0.475 | 0.49 |
| chr8 | 114629449 | 114630540 | 1092 | -0.47 | 0.475 | 0.49 |
| chr7 | 50341660 | 50370218 | 28559 | -0.38 | 0.476 | 0.491 |
| chr15 | 33591497 | 33647800 | 56304 | -0.21 | 0.477 | 0.491 |
| chrX | 73011562 | 73013703 | 2142 | 0.52 | 0.477 | 0.491 |
| chr18 | 69327873 | 69330828 | 2956 | 0.39 | 0.477 | 0.492 |
| chr6 | 81564787 | 81565690 | 904 | -0.77 | 0.477 | 0.492 |
| chr2 | 214497127 | 214502790 | 5664 | -0.55 | 0.478 | 0.492 |
| chr1 | 191839166 | 191920846 | 81681 | -0.35 | 0.478 | 0.493 |
| chr6 | 39255209 | 39257538 | 2330 | -0.36 | 0.478 | 0.493 |
| chr13 | 104856379 | 104857697 | 1319 | -0.35 | 0.479 | 0.493 |
| chr20 | 22518757 | 22535985 | 17229 | -1.26 | 0.479 | 0.494 |
| chr6 | 23104325 | 23126576 | 22252 | -0.33 | 0.479 | 0.494 |
| chr4 | 60851445 | 60852756 | 1312 | 0.64 | 0.479 | 0.494 |
| chr4 | 29597082 | 29674386 | 77305 | 0.28 | 0.479 | 0.494 |
| chr1 | 201981043 | 201983569 | 2527 | -0.33 | 0.48 | 0.494 |
| chr20 | 2547164 | 2555871 | 8708 | -0.26 | 0.48 | 0.495 |
| chr1 | 105425432 | 105429885 | 4454 | 0.69 | 0.481 | 0.495 |
| chr3 | 161783312 | 161783729 | 418 | 0.73 | 0.481 | 0.495 |
| chr6 | 162820419 | 162826582 | 6164 | -1.16 | 0.481 | 0.495 |
| chr4 | 62727074 | 62730033 | 2960 | -0.28 | 0.481 | 0.496 |
| chr4 | 90610040 | 90610984 | 945 | -0.39 | 0.483 | 0.497 |
| chr18 | 403073 | 413955 | 10883 | -0.65 | 0.484 | 0.498 |
| chr10 | 122951569 | 122998852 | 47284 | -0.32 | 0.484 | 0.498 |
| chr2 | 122702821 | 122704250 | 1430 | 0.39 | 0.484 | 0.498 |
| chr6 | 150372668 | 150376425 | 3758 | -0.29 | 0.484 | 0.498 |
| chr2 | 222695569 | 222697189 | 1621 | -0.46 | 0.484 | 0.499 |
| chr10 | 49423673 | 49431935 | 8263 | -0.39 | 0.485 | 0.5 |
| chr10 | 7510899 | 7512198 | 1300 | 0.36 | 0.485 | 0.5 |
| chr5 | 58119955 | 58158959 | 39005 | 0.32 | 0.486 | 0.5 |
| chr20 | 44153765 | 44207787 | 54023 | -0.22 | 0.486 | 0.501 |
| chr10 | 123446381 | 123448467 | 2087 | 1.17 | 0.486 | 0.501 |
| chr3 | 161907620 | 161909536 | 1917 | 0.43 | 0.486 | 0.501 |
| chr3 | 59804888 | 59833285 | 28398 | -0.3 | 0.487 | 0.501 |
| chr5 | 174880845 | 174883360 | 2516 | 1.19 | 0.487 | 0.501 |
| chr14 | 37107063 | 37111478 | 4416 | -0.24 | 0.487 | 0.502 |

| | | | | | | |
|---|---|---|---|---|---|---|
| chr8 | 19747596 | 19749999 | 2404 | 0.58 | 0.488 | 0.503 |
| chr8 | 67193866 | 67230182 | 36317 | -0.57 | 0.488 | 0.503 |
| chr1 | 160585791 | 160596902 | 11112 | 0.3 | 0.489 | 0.503 |
| chr6 | 133307095 | 133323441 | 16347 | 0.43 | 0.489 | 0.503 |
| chr3 | 90253332 | 90254231 | 900 | -0.3 | 0.489 | 0.503 |
| chr2 | 141064695 | 141064996 | 302 | 0.92 | 0.489 | 0.504 |
| chr2 | 204723430 | 204736407 | 12978 | -0.4 | 0.489 | 0.504 |
| chr15 | 73218592 | 73223121 | 4530 | 0.29 | 0.49 | 0.504 |
| chr2 | 34811096 | 34867580 | 56485 | -0.37 | 0.49 | 0.504 |
| chr11 | 129266569 | 129278492 | 11924 | 0.26 | 0.49 | 0.504 |
| chr10 | 22611047 | 22611329 | 283 | 0.52 | 0.491 | 0.505 |
| chr14 | 30430438 | 30431938 | 1501 | -0.32 | 0.491 | 0.505 |
| chr4 | 33440752 | 33447931 | 7180 | 0.28 | 0.491 | 0.506 |
| chr11 | 109711365 | 109711621 | 257 | 1 | 0.492 | 0.507 |
| chr4 | 62838527 | 62850554 | 12028 | -0.24 | 0.493 | 0.507 |
| chr6 | 146823298 | 146824469 | 1172 | -0.36 | 0.494 | 0.508 |
| chr11 | 92413381 | 92414267 | 887 | -0.68 | 0.494 | 0.508 |
| chr4 | 61673688 | 61691706 | 18019 | 0.31 | 0.494 | 0.509 |
| chr8 | 75215485 | 75249598 | 34114 | 0.25 | 0.495 | 0.509 |
| chr5 | 40833036 | 40834987 | 1952 | -0.52 | 0.495 | 0.51 |
| chr16 | 77083333 | 77085552 | 2220 | 0.45 | 0.495 | 0.51 |
| chr6 | 164298281 | 164353282 | 55002 | -0.18 | 0.496 | 0.51 |
| chr7 | 150612705 | 150617999 | 5295 | 1.2 | 0.496 | 0.511 |
| chrX | 133234040 | 133250253 | 16214 | -0.3 | 0.497 | 0.511 |
| chr19 | 15142407 | 15143542 | 1136 | -0.41 | 0.497 | 0.511 |
| chr7 | 119037346 | 119091831 | 54486 | -0.4 | 0.497 | 0.512 |
| chr10 | 69344560 | 69359023 | 14464 | -0.39 | 0.498 | 0.512 |
| chr10 | 115377334 | 115379016 | 1683 | 0.67 | 0.498 | 0.513 |
| chr6 | 85931234 | 85944895 | 13662 | 0.28 | 0.499 | 0.514 |
| chr11 | 40563019 | 40566789 | 3771 | -0.45 | 0.5 | 0.514 |
| chr11 | 21428762 | 21429924 | 1163 | -0.96 | 0.5 | 0.514 |
| chr1 | 112782089 | 112842610 | 60522 | -0.16 | 0.5 | 0.515 |
| chr8 | 53009069 | 53011184 | 2116 | 0.85 | 0.501 | 0.515 |
| chr10 | 45782366 | 45799453 | 17088 | -0.25 | 0.501 | 0.515 |
| chr2 | 31441470 | 31447181 | 5712 | 0.58 | 0.501 | 0.515 |
| chr11 | 51417622 | 51418134 | 513 | 0.34 | 0.501 | 0.515 |
| chr2 | 186602552 | 186603000 | 449 | 0.5 | 0.503 | 0.517 |
| chr8 | 114420717 | 114422178 | 1462 | -0.42 | 0.503 | 0.517 |
| chr4 | 62403060 | 62406911 | 3852 | -0.28 | 0.503 | 0.517 |
| chr15 | 53634978 | 53636914 | 1937 | -0.58 | 0.503 | 0.517 |
| chr13 | 22340691 | 22354816 | 14126 | -0.27 | 0.503 | 0.518 |
| chr11 | 61731744 | 61734431 | 2688 | -0.23 | 0.503 | 0.518 |
| chr3 | 83192196 | 83194072 | 1877 | -0.41 | 0.504 | 0.518 |
| chr12 | 68355182 | 68398718 | 43537 | 0.21 | 0.505 | 0.52 |
| chr7 | 152617504 | 152627637 | 10134 | -0.32 | 0.506 | 0.52 |
| chr21 | 18505898 | 18507252 | 1355 | -1.01 | 0.506 | 0.52 |
| chr4 | 63181228 | 63256089 | 74862 | 0.21 | 0.506 | 0.521 |
| chr1 | 147246575 | 147265147 | 18573 | 0.22 | 0.507 | 0.521 |
| chr2 | 134441720 | 134494250 | 52531 | -0.14 | 0.507 | 0.521 |
| chr20 | 11544268 | 11620454 | 76187 | 0.16 | 0.507 | 0.521 |
| chrX | 125647523 | 125650151 | 2629 | 0.46 | 0.507 | 0.521 |
| chr1 | 34066806 | 34068116 | 1311 | 0.4 | 0.508 | 0.522 |
| chr15 | 51349457 | 51350429 | 973 | 1.07 | 0.508 | 0.522 |
| chr1 | 104340478 | 104342283 | 1806 | -0.69 | 0.508 | 0.523 |
| chr14 | 66432613 | 66523197 | 90585 | -0.11 | 0.509 | 0.523 |
| chr9 | 3161770 | 3172576 | 10807 | 0.4 | 0.509 | 0.523 |
| chr10 | 17437146 | 17438007 | 862 | -0.49 | 0.509 | 0.523 |
| chr4 | 62926668 | 62940976 | 14309 | 0.37 | 0.509 | 0.524 |
| chr8 | 120151967 | 120153144 | 1178 | -0.43 | 0.509 | 0.524 |
| chr14 | 104407171 | 104408852 | 1682 | 0.76 | 0.51 | 0.525 |
| chr11 | 59208758 | 59229167 | 20410 | 0.3 | 0.511 | 0.525 |

| | | | | | | |
|---|---|---|---|---|---|---|
| chr2 | 56722748 | 56740419 | 17672 | -0.3 | 0.511 | 0.526 |
| chrX | 71054240 | 71076028 | 21789 | -0.29 | 0.511 | 0.526 |
| chr7 | 54687564 | 54753071 | 65508 | -0.24 | 0.512 | 0.526 |
| chr4 | 63156526 | 63168689 | 12164 | -0.39 | 0.512 | 0.526 |
| chrX | 72600331 | 72600951 | 621 | -0.45 | 0.512 | 0.526 |
| chr13 | 53447678 | 53466035 | 18358 | 0.18 | 0.512 | 0.527 |
| chr8 | 144009566 | 144011355 | 1790 | 0.97 | 0.513 | 0.527 |
| chr11 | 67635342 | 67636509 | 1168 | -0.72 | 0.513 | 0.527 |
| chr8 | 114847149 | 114849025 | 1877 | 0.35 | 0.513 | 0.527 |
| chr16 | 10939096 | 10940920 | 1825 | 0.37 | 0.513 | 0.527 |
| chr11 | 119822245 | 119825939 | 3695 | 0.22 | 0.514 | 0.528 |
| chr9 | 27305004 | 27326629 | 21626 | 0.25 | 0.514 | 0.528 |
| chr3 | 120582029 | 120591466 | 9438 | 0.51 | 0.514 | 0.529 |
| chr3 | 56328434 | 56334255 | 5822 | -0.22 | 0.515 | 0.529 |
| chr6 | 155883572 | 155920482 | 36911 | 0.24 | 0.515 | 0.529 |
| chr11 | 45498355 | 45509189 | 10835 | 0.97 | 0.516 | 0.53 |
| chr13 | 112839927 | 112841502 | 1576 | 0.78 | 0.516 | 0.53 |
| chr20 | 59949331 | 59950316 | 986 | 0.42 | 0.517 | 0.531 |
| chr13 | 22615642 | 22732132 | 116491 | -0.14 | 0.517 | 0.531 |
| chr8 | 87703248 | 87703640 | 393 | -0.93 | 0.517 | 0.531 |
| chr5 | 107751512 | 107766925 | 15414 | 0.56 | 0.517 | 0.531 |
| chr17 | 25344911 | 25358061 | 13151 | -0.17 | 0.517 | 0.532 |
| chr6 | 64557458 | 64558091 | 634 | 0.34 | 0.518 | 0.532 |
| chr18 | 43179331 | 43183747 | 4417 | -0.29 | 0.519 | 0.533 |
| chr4 | 35351379 | 35373601 | 22223 | -0.39 | 0.519 | 0.533 |
| chrX | 82634857 | 82635569 | 713 | -0.41 | 0.519 | 0.533 |
| chr7 | 47028198 | 47029389 | 1192 | -0.82 | 0.52 | 0.534 |
| chr2 | 186659846 | 186660763 | 918 | -0.91 | 0.52 | 0.534 |
| chr8 | 16654971 | 16656463 | 1493 | 0.31 | 0.521 | 0.535 |
| chr8 | 113709986 | 113735690 | 25705 | -0.66 | 0.522 | 0.536 |
| chr14 | 60015736 | 60036702 | 20967 | 0.67 | 0.522 | 0.536 |
| chr9 | 293207 | 332110 | 38904 | -0.21 | 0.522 | 0.536 |
| chr21 | 18381336 | 18382684 | 1349 | -0.34 | 0.522 | 0.536 |
| chr5 | 122553354 | 122579322 | 25969 | 0.39 | 0.523 | 0.537 |
| chr18 | 73983036 | 73983779 | 744 | 0.54 | 0.523 | 0.537 |
| chr8 | 130475403 | 130475979 | 577 | -0.4 | 0.523 | 0.537 |
| chr3 | 82207760 | 82208317 | 558 | -0.89 | 0.523 | 0.537 |
| chr12 | 54893856 | 54894264 | 409 | 0.63 | 0.523 | 0.537 |
| chr18 | 33368780 | 33370634 | 1855 | -0.36 | 0.523 | 0.537 |
| chr11 | 84839415 | 84840474 | 1060 | 0.5 | 0.524 | 0.538 |
| chr15 | 46048830 | 46049262 | 433 | -0.68 | 0.524 | 0.538 |
| chr2 | 213230259 | 213237599 | 7341 | 0.32 | 0.524 | 0.538 |
| chr1 | 218407402 | 218408849 | 1448 | 1.04 | 0.524 | 0.538 |
| chr1 | 4907279 | 4912743 | 5465 | -0.3 | 0.524 | 0.538 |
| chr4 | 53216742 | 53233953 | 17212 | 0.17 | 0.525 | 0.539 |
| chr10 | 1654360 | 1691922 | 37563 | -0.22 | 0.525 | 0.539 |
| chr12 | 55009182 | 55045788 | 36607 | -0.32 | 0.526 | 0.54 |
| chr4 | 134957913 | 134963530 | 5618 | 0.78 | 0.526 | 0.54 |
| chr4 | 145075006 | 145076773 | 1768 | 0.33 | 0.526 | 0.54 |
| chr9 | 79323581 | 79326517 | 2937 | 0.26 | 0.527 | 0.541 |
| chr11 | 36564991 | 36604540 | 39550 | -0.2 | 0.527 | 0.541 |
| chr6 | 168823048 | 168894725 | 71678 | -0.19 | 0.527 | 0.541 |
| chr10 | 2862843 | 2924283 | 61441 | -0.17 | 0.528 | 0.542 |
| chr12 | 51904868 | 51905698 | 831 | 0.37 | 0.528 | 0.542 |
| chr15 | 31925188 | 31925696 | 509 | -0.38 | 0.529 | 0.543 |
| chr18 | 68198414 | 68216552 | 18139 | -0.24 | 0.529 | 0.543 |
| chr10 | 15955312 | 15994849 | 39538 | -0.15 | 0.529 | 0.543 |
| chr4 | 19328991 | 19340272 | 11282 | 0.32 | 0.529 | 0.543 |
| chr1 | 120194894 | 120195669 | 776 | 0.51 | 0.53 | 0.544 |
| chr7 | 148282088 | 148284345 | 2258 | 0.39 | 0.53 | 0.544 |
| chr8 | 15533032 | 15561986 | 28955 | -0.3 | 0.53 | 0.544 |

| | | | | | | |
|---|---|---|---|---|---|---|
| chr8 | 9134784 | 9153941 | 19158 | 0.25 | 0.53 | 0.544 |
| chr4 | 47682329 | 47683562 | 1234 | 0.46 | 0.531 | 0.544 |
| chr12 | 108607592 | 108633854 | 26263 | -0.57 | 0.531 | 0.545 |
| chr16 | 64174648 | 64177823 | 3176 | -0.79 | 0.531 | 0.545 |
| chr13 | 88649529 | 88650592 | 1064 | 0.42 | 0.532 | 0.545 |
| chrX | 13969890 | 13974219 | 4330 | -0.31 | 0.532 | 0.546 |
| chr9 | 89080759 | 89120931 | 40173 | -0.18 | 0.532 | 0.546 |
| chr10 | 42926853 | 42928248 | 1396 | -0.58 | 0.532 | 0.546 |
| chr16 | 63194016 | 63204813 | 10798 | 0.2 | 0.533 | 0.547 |
| chr4 | 56640877 | 56641574 | 698 | 0.3 | 0.533 | 0.547 |
| chr12 | 97385719 | 97392078 | 6360 | -0.19 | 0.533 | 0.547 |
| chrX | 93864935 | 93867723 | 2789 | 0.61 | 0.534 | 0.547 |
| chr11 | 35939588 | 35940550 | 963 | 0.7 | 0.534 | 0.548 |
| chr4 | 33139832 | 33147689 | 7858 | 0.31 | 0.535 | 0.549 |
| chr12 | 48306512 | 48325451 | 18940 | 0.28 | 0.536 | 0.55 |
| chr18 | 2144262 | 2158015 | 13754 | 0.3 | 0.537 | 0.55 |
| chr10 | 133027583 | 133111738 | 84156 | -0.17 | 0.537 | 0.551 |
| chrX | 20769981 | 20771990 | 2010 | -0.76 | 0.538 | 0.551 |
| chrX | 82434438 | 82440078 | 5641 | -0.29 | 0.538 | 0.552 |
| chr6 | 114447337 | 114469819 | 22483 | 0.36 | 0.539 | 0.552 |
| chr6 | 40796727 | 40810120 | 13394 | -0.32 | 0.539 | 0.553 |
| chr3 | 111063392 | 111075636 | 12245 | -0.28 | 0.539 | 0.553 |
| chr4 | 133792434 | 133800679 | 8246 | 0.31 | 0.54 | 0.553 |
| chr4 | 57397885 | 57399127 | 1243 | 0.48 | 0.541 | 0.554 |
| chr10 | 97850243 | 97871421 | 21179 | -0.22 | 0.541 | 0.555 |
| chr14 | 43232632 | 43274350 | 41719 | 0.17 | 0.541 | 0.555 |
| chr10 | 12953585 | 12969425 | 15841 | -1.07 | 0.542 | 0.555 |
| chr1 | 164458075 | 164484398 | 26324 | 1.12 | 0.542 | 0.556 |
| chr5 | 70969466 | 70982173 | 12708 | -0.36 | 0.542 | 0.556 |
| chr10 | 14510385 | 14511874 | 1490 | 0.98 | 0.542 | 0.556 |
| chr5 | 71002705 | 71010537 | 7833 | -0.18 | 0.542 | 0.556 |
| chrX | 51898483 | 51903081 | 4599 | -0.26 | 0.543 | 0.556 |
| chr14 | 96142558 | 96151516 | 8959 | -0.36 | 0.543 | 0.557 |
| chr5 | 59263997 | 59280284 | 16288 | -0.42 | 0.543 | 0.557 |
| chr2 | 57751462 | 57756726 | 5265 | 0.37 | 0.544 | 0.557 |
| chr4 | 34592851 | 34605320 | 12470 | -0.24 | 0.544 | 0.557 |
| chr7 | 31343752 | 31374656 | 30905 | 0.31 | 0.544 | 0.557 |
| chr5 | 180473533 | 180475194 | 1662 | 0.42 | 0.544 | 0.557 |
| chr4 | 94440278 | 94440913 | 636 | -0.87 | 0.544 | 0.558 |
| chr13 | 23147442 | 23198406 | 50965 | -0.15 | 0.545 | 0.559 |
| chr4 | 82182013 | 82196788 | 14776 | 0.19 | 0.545 | 0.559 |
| chr10 | 93068885 | 93070153 | 1269 | 0.87 | 0.546 | 0.559 |
| chr9 | 2541743 | 2558400 | 16658 | -0.2 | 0.546 | 0.56 |
| chr1 | 224929407 | 224961838 | 32432 | 0.27 | 0.546 | 0.56 |
| chr2 | 56852637 | 56865682 | 13046 | 0.29 | 0.547 | 0.56 |
| chr4 | 56696136 | 56697113 | 978 | 0.57 | 0.547 | 0.56 |
| chr4 | 53324594 | 53325199 | 606 | 0.49 | 0.547 | 0.56 |
| chr6 | 16081290 | 16101772 | 20483 | 0.22 | 0.548 | 0.561 |
| chr10 | 124318662 | 124351732 | 33071 | 0.36 | 0.548 | 0.562 |
| chr13 | 88956525 | 88957885 | 1361 | 0.5 | 0.549 | 0.562 |
| chr10 | 2944182 | 2966668 | 22487 | -1.06 | 0.549 | 0.563 |
| chr10 | 124380183 | 124384371 | 4189 | 0.32 | 0.549 | 0.563 |
| chr1 | 167011509 | 167013045 | 1537 | -0.5 | 0.55 | 0.564 |
| chr1 | 105111261 | 105126507 | 15247 | -0.24 | 0.551 | 0.565 |
| chr5 | 154437737 | 154438408 | 672 | 0.44 | 0.552 | 0.565 |
| chr2 | 85076971 | 85081532 | 4562 | -0.24 | 0.552 | 0.565 |
| chr4 | 27063115 | 27064919 | 1805 | 0.35 | 0.552 | 0.565 |
| chr13 | 19750275 | 19753049 | 2775 | 0.29 | 0.552 | 0.566 |
| chr19 | 22564148 | 22564564 | 417 | 0.39 | 0.552 | 0.566 |
| chr18 | 73936653 | 73956011 | 19359 | 0.2 | 0.552 | 0.566 |
| chr9 | 112444873 | 112449613 | 4741 | -0.45 | 0.552 | 0.566 |

| | | | | | | |
|---|---|---|---|---|---|---|
| chrX | 13824188 | 13840227 | 16040 | -0.31 | 0.553 | 0.567 |
| chr12 | 108250140 | 108269395 | 19256 | 0.34 | 0.553 | 0.567 |
| chr16 | 78622257 | 78624677 | 2421 | 0.65 | 0.554 | 0.567 |
| chr4 | 29489965 | 29502153 | 12189 | 0.27 | 0.554 | 0.568 |
| chr1 | 153440522 | 153446001 | 5480 | -0.2 | 0.554 | 0.568 |
| chr2 | 53386248 | 53401171 | 14924 | -0.81 | 0.555 | 0.568 |
| chrX | 139799834 | 139809823 | 9990 | 0.23 | 0.557 | 0.57 |
| chr13 | 23210428 | 23226220 | 15793 | -0.17 | 0.558 | 0.571 |
| chr3 | 163143644 | 163145675 | 2032 | 0.43 | 0.558 | 0.572 |
| chrX | 57406081 | 57407454 | 1374 | -0.82 | 0.559 | 0.572 |
| chr3 | 144917170 | 144917757 | 588 | -0.54 | 0.56 | 0.573 |
| chr4 | 62189726 | 62193358 | 3633 | -0.3 | 0.56 | 0.574 |
| chr1 | 111059873 | 111061773 | 1901 | 0.24 | 0.561 | 0.574 |
| chr10 | 13974124 | 13997051 | 22928 | -0.25 | 0.561 | 0.574 |
| chr9 | 72282230 | 72283437 | 1208 | 0.32 | 0.561 | 0.575 |
| chr12 | 74029188 | 74082399 | 53212 | -0.34 | 0.561 | 0.575 |
| chr1 | 160403611 | 160406079 | 2469 | 0.27 | 0.562 | 0.575 |
| chr3 | 112131682 | 112134550 | 2869 | -0.21 | 0.562 | 0.575 |
| chrX | 100793166 | 100794749 | 1584 | 0.31 | 0.562 | 0.575 |
| chr8 | 114333972 | 114385080 | 51109 | 0.26 | 0.563 | 0.576 |
| chr21 | 18998914 | 19001329 | 2416 | -0.33 | 0.563 | 0.576 |
| chr9 | 118060929 | 118061616 | 688 | 0.38 | 0.564 | 0.577 |
| chr11 | 126974285 | 127042935 | 68651 | 0.17 | 0.564 | 0.578 |
| chr3 | 86046774 | 86070972 | 24199 | -0.31 | 0.564 | 0.578 |
| chr6 | 75661310 | 75663281 | 1972 | -0.28 | 0.564 | 0.578 |
| chr6 | 168542389 | 168546352 | 3964 | 0.31 | 0.566 | 0.58 |
| chr13 | 23108517 | 23145806 | 37290 | -0.16 | 0.567 | 0.581 |
| chr16 | 73468217 | 73906366 | 438150 | -0.07 | 0.567 | 0.581 |
| chr3 | 161681207 | 161701435 | 20229 | 0.28 | 0.568 | 0.581 |
| chr2 | 56285377 | 56319911 | 34535 | -0.24 | 0.569 | 0.583 |
| chr20 | 58188638 | 58189993 | 1356 | 0.44 | 0.57 | 0.583 |
| chr10 | 76152175 | 76164736 | 12562 | -0.93 | 0.571 | 0.584 |
| chr3 | 164482350 | 164496190 | 13841 | -0.36 | 0.573 | 0.586 |
| chr5 | 45586533 | 45931996 | 345464 | -0.12 | 0.573 | 0.587 |
| chr8 | 81250990 | 81252975 | 1986 | 0.38 | 0.574 | 0.587 |
| chr9 | 110984948 | 110992088 | 7141 | -0.93 | 0.574 | 0.587 |
| chr8 | 12238541 | 12239143 | 603 | -0.41 | 0.576 | 0.589 |
| chr15 | 60885277 | 60907297 | 22021 | 0.22 | 0.576 | 0.589 |
| chr3 | 159003945 | 159014531 | 10587 | -0.3 | 0.577 | 0.59 |
| chr11 | 14061455 | 14076095 | 14641 | -0.28 | 0.577 | 0.591 |
| chr2 | 222910045 | 222911986 | 1942 | 0.59 | 0.577 | 0.591 |
| chr2 | 214749627 | 214750954 | 1328 | 0.88 | 0.579 | 0.592 |
| chr9 | 91733413 | 91743434 | 10022 | -0.28 | 0.579 | 0.592 |
| chr10 | 115393592 | 115424913 | 31322 | -0.16 | 0.579 | 0.593 |
| chrX | 56448158 | 56450951 | 2794 | 0.34 | 0.581 | 0.594 |
| chr4 | 29729463 | 29730031 | 569 | -0.75 | 0.582 | 0.596 |
| chr1 | 15203218 | 15208424 | 5207 | -0.28 | 0.582 | 0.596 |
| chr10 | 44797827 | 44869083 | 71257 | -0.17 | 0.582 | 0.596 |
| chr14 | 89617726 | 89621025 | 3300 | -0.27 | 0.582 | 0.596 |
| chr2 | 214063842 | 214065244 | 1403 | -0.27 | 0.583 | 0.596 |
| chr14 | 24125314 | 24127130 | 1817 | -0.31 | 0.583 | 0.596 |
| chr2 | 100124667 | 100138741 | 14075 | -1.03 | 0.584 | 0.598 |
| chr14 | 73202338 | 73203157 | 820 | 0.45 | 0.584 | 0.598 |
| chr12 | 38905751 | 38906151 | 401 | -0.56 | 0.585 | 0.598 |
| chr4 | 29215045 | 29222717 | 7673 | -0.3 | 0.585 | 0.598 |
| chr11 | 134653788 | 134655646 | 1859 | 0.9 | 0.585 | 0.598 |
| chr6 | 45714291 | 45792666 | 78376 | -0.12 | 0.586 | 0.599 |
| chr4 | 19280098 | 19280573 | 476 | 0.36 | 0.588 | 0.601 |
| chr11 | 27294950 | 27298510 | 3561 | 0.27 | 0.588 | 0.601 |
| chr3 | 104324586 | 104326717 | 2132 | 0.4 | 0.589 | 0.602 |
| chr10 | 101274352 | 101276300 | 1949 | 0.89 | 0.589 | 0.602 |

| | | | | | | |
|---|---|---|---|---|---|---|
| chr10 | 63634250 | 63635704 | 1455 | -0.43 | 0.589 | 0.603 |
| chr5 | 75590169 | 75591257 | 1089 | -0.3 | 0.589 | 0.603 |
| chr3 | 55126762 | 55208231 | 81470 | -0.09 | 0.59 | 0.603 |
| chr15 | 31782530 | 31909737 | 127208 | -0.23 | 0.59 | 0.604 |
| chr4 | 22267761 | 22270111 | 2351 | -0.3 | 0.59 | 0.604 |
| chr9 | 94322650 | 94346948 | 24299 | 1.01 | 0.591 | 0.604 |
| chr1 | 153408247 | 153410863 | 2617 | 0.53 | 0.591 | 0.604 |
| chr20 | 9475608 | 9476999 | 1392 | 0.67 | 0.591 | 0.604 |
| chr7 | 41238825 | 41724645 | 485821 | 0.05 | 0.591 | 0.604 |
| chr7 | 47403049 | 47404450 | 1402 | -0.32 | 0.592 | 0.605 |
| chr17 | 39231435 | 39234797 | 3363 | -0.36 | 0.592 | 0.605 |
| chr4 | 23657456 | 23705855 | 48400 | -0.18 | 0.593 | 0.606 |
| chr5 | 7063804 | 7067240 | 3437 | -0.22 | 0.595 | 0.608 |
| chr2 | 108295009 | 108295869 | 861 | 0.79 | 0.595 | 0.609 |
| chr12 | 48840253 | 48903473 | 63221 | -0.13 | 0.595 | 0.609 |
| chrX | 18031764 | 18043432 | 11669 | -0.16 | 0.596 | 0.609 |
| chr8 | 30844870 | 30845387 | 518 | -0.8 | 0.596 | 0.609 |
| chr8 | 105176561 | 105178616 | 2056 | -0.82 | 0.597 | 0.61 |
| chr1 | 245509089 | 245518771 | 9683 | -0.24 | 0.597 | 0.61 |
| chr11 | 101000076 | 101001552 | 1477 | -0.26 | 0.597 | 0.61 |
| chr10 | 1263964 | 1267242 | 3279 | -0.22 | 0.597 | 0.61 |
| chr1 | 105699478 | 105709413 | 9936 | 0.22 | 0.597 | 0.611 |
| chr3 | 175945024 | 175948586 | 3563 | -0.34 | 0.598 | 0.611 |
| chrX | 151492458 | 151493791 | 1334 | 0.26 | 0.598 | 0.611 |
| chr13 | 22282030 | 22285508 | 3479 | -0.22 | 0.598 | 0.611 |
| chr6 | 13213842 | 13215508 | 1667 | 0.34 | 0.598 | 0.611 |
| chr3 | 106846208 | 106898708 | 52501 | -0.18 | 0.6 | 0.614 |
| chr9 | 21368354 | 21386976 | 18623 | -0.11 | 0.601 | 0.614 |
| chr10 | 1453774 | 1566256 | 112483 | 0.13 | 0.602 | 0.615 |
| chr6 | 142983446 | 143007012 | 23567 | -0.18 | 0.602 | 0.615 |
| chr16 | 59425181 | 59430381 | 5201 | 0.35 | 0.604 | 0.617 |
| chr20 | 8196795 | 8207960 | 11166 | 0.18 | 0.605 | 0.618 |
| chr3 | 59876897 | 59879978 | 3082 | -0.27 | 0.606 | 0.619 |
| chr3 | 39781775 | 39823647 | 41873 | 0.12 | 0.606 | 0.619 |
| chr3 | 176536879 | 176538128 | 1250 | -0.47 | 0.606 | 0.619 |
| chr1 | 218190575 | 218215884 | 25310 | -0.29 | 0.606 | 0.619 |
| chr20 | 44126094 | 44135435 | 9342 | 0.26 | 0.607 | 0.62 |
| chr14 | 34269998 | 34298219 | 28222 | 0.11 | 0.607 | 0.62 |
| chr7 | 108261905 | 108309727 | 47823 | -0.36 | 0.608 | 0.621 |
| chr11 | 40580424 | 40581334 | 911 | -0.36 | 0.608 | 0.621 |
| chr9 | 37053486 | 37054856 | 1371 | 0.53 | 0.608 | 0.621 |
| chr10 | 87494331 | 87495140 | 810 | -0.39 | 0.609 | 0.622 |
| chr2 | 34936728 | 34957748 | 21021 | -0.2 | 0.61 | 0.624 |
| chr22 | 33185779 | 33191518 | 5740 | 0.22 | 0.611 | 0.624 |
| chr9 | 119760235 | 119786363 | 26129 | -0.9 | 0.611 | 0.624 |
| chr3 | 149960929 | 149964267 | 3339 | -0.25 | 0.611 | 0.624 |
| chr13 | 104428738 | 104429587 | 850 | -0.76 | 0.611 | 0.624 |
| chr12 | 44100339 | 44101561 | 1223 | -0.78 | 0.611 | 0.624 |
| chr3 | 178211735 | 178214620 | 2886 | -0.31 | 0.612 | 0.625 |
| chr15 | 60431462 | 60608987 | 177526 | -0.11 | 0.612 | 0.625 |
| chr2 | 134740927 | 134773068 | 32142 | 0.55 | 0.612 | 0.625 |
| chr5 | 157447611 | 157476059 | 28449 | -0.13 | 0.613 | 0.627 |
| chr4 | 32910156 | 32912046 | 1891 | 0.23 | 0.615 | 0.628 |
| chr1 | 22664071 | 22664566 | 496 | 0.47 | 0.616 | 0.629 |
| chr13 | 89008015 | 89008361 | 347 | 0.31 | 0.616 | 0.63 |
| chr6 | 15663884 | 15678667 | 14784 | -0.71 | 0.618 | 0.631 |
| chr1 | 242771741 | 242785007 | 13267 | 0.17 | 0.619 | 0.632 |
| chr20 | 8045871 | 8049895 | 4025 | 0.19 | 0.62 | 0.633 |
| chr4 | 35629383 | 35633540 | 4158 | -0.61 | 0.62 | 0.634 |
| chr12 | 98139830 | 98140336 | 507 | 0.29 | 0.621 | 0.634 |
| chrX | 109672118 | 109673462 | 1345 | -0.34 | 0.623 | 0.636 |

| | | | | | | |
|---|---|---|---|---|---|---|
| chr4 | 56528520 | 56530313 | 1794 | 0.43 | 0.623 | 0.636 |
| chr7 | 142224654 | 142228893 | 4240 | -0.23 | 0.623 | 0.636 |
| chr5 | 60964241 | 60966570 | 2330 | -0.35 | 0.623 | 0.636 |
| chr10 | 25693808 | 25718634 | 24827 | 0.18 | 0.624 | 0.637 |
| chr11 | 31858109 | 31861404 | 3296 | -0.23 | 0.624 | 0.637 |
| chr7 | 131412810 | 131422524 | 9715 | 0.19 | 0.624 | 0.637 |
| chr20 | 59578028 | 59594768 | 16741 | -0.12 | 0.624 | 0.637 |
| chr9 | 30429639 | 30477843 | 48205 | -0.19 | 0.624 | 0.637 |
| chr2 | 218117111 | 218118205 | 1095 | 0.27 | 0.626 | 0.639 |
| chr3 | 43317518 | 43322698 | 5181 | 0.19 | 0.626 | 0.639 |
| chr6 | 167274299 | 167275460 | 1162 | 0.48 | 0.627 | 0.64 |
| chr1 | 105741689 | 105750393 | 8705 | 0.22 | 0.629 | 0.642 |
| chr11 | 113259934 | 113262041 | 2108 | -0.32 | 0.63 | 0.644 |
| chr18 | 7490041 | 7491822 | 1782 | -0.28 | 0.631 | 0.644 |
| chr4 | 112449653 | 112450888 | 1236 | 0.37 | 0.631 | 0.644 |
| chr3 | 162703616 | 162704991 | 1376 | -0.27 | 0.632 | 0.645 |
| chr7 | 14682044 | 14685648 | 3605 | 0.31 | 0.632 | 0.645 |
| chr13 | 88657223 | 88663575 | 6353 | -0.3 | 0.632 | 0.645 |
| chr2 | 99011465 | 99015553 | 4089 | -0.17 | 0.632 | 0.645 |
| chr12 | 14246417 | 14254666 | 8250 | 0.21 | 0.633 | 0.646 |
| chr1 | 212777589 | 212778433 | 845 | -0.71 | 0.633 | 0.646 |
| chr3 | 83145417 | 83146440 | 1024 | 0.27 | 0.634 | 0.647 |
| chr4 | 141510228 | 141514596 | 4369 | -0.24 | 0.634 | 0.647 |
| chr6 | 74229244 | 74229840 | 597 | -0.29 | 0.634 | 0.647 |
| chr11 | 45721298 | 45751086 | 29789 | 0.19 | 0.634 | 0.647 |
| chr5 | 43790525 | 43791961 | 1437 | 0.22 | 0.635 | 0.648 |
| chr8 | 40872122 | 40883849 | 11728 | 0.29 | 0.635 | 0.648 |
| chr10 | 1289474 | 1300358 | 10885 | 0.13 | 0.636 | 0.649 |
| chrX | 39237947 | 39249217 | 11271 | 0.18 | 0.636 | 0.649 |
| chr3 | 110774543 | 110775564 | 1022 | -0.34 | 0.636 | 0.649 |
| chrX | 3110378 | 3117360 | 6983 | -0.31 | 0.637 | 0.65 |
| chr2 | 107584549 | 107585378 | 830 | -0.62 | 0.638 | 0.65 |
| chr9 | 92834679 | 92836527 | 1849 | -0.31 | 0.638 | 0.651 |
| chr3 | 170318988 | 170320982 | 1995 | 0.25 | 0.639 | 0.652 |
| chr7 | 68132769 | 68134531 | 1763 | 0.54 | 0.639 | 0.652 |
| chr20 | 57378648 | 57379442 | 795 | 0.72 | 0.639 | 0.652 |
| chr7 | 85128259 | 85132458 | 4200 | -0.18 | 0.642 | 0.655 |
| chr4 | 62811944 | 62826726 | 14783 | -0.25 | 0.643 | 0.656 |
| chr6 | 27978170 | 27988652 | 10483 | -0.14 | 0.644 | 0.657 |
| chrX | 133035662 | 133051715 | 16054 | -0.29 | 0.645 | 0.657 |
| chr6 | 130895955 | 130896497 | 543 | -0.33 | 0.645 | 0.657 |
| chr3 | 70529764 | 70550900 | 21137 | 0.36 | 0.645 | 0.658 |
| chr20 | 22236487 | 22238353 | 1867 | -0.32 | 0.645 | 0.658 |
| chr13 | 53296218 | 53296781 | 564 | 0.25 | 0.646 | 0.659 |
| chr7 | 508294 | 509040 | 747 | 0.27 | 0.647 | 0.659 |
| chr18 | 73470723 | 73481640 | 10918 | 0.16 | 0.647 | 0.66 |
| chr1 | 105367193 | 105367893 | 701 | 0.29 | 0.647 | 0.66 |
| chr5 | 122006314 | 122018819 | 12506 | 0.17 | 0.647 | 0.66 |
| chr6 | 79505415 | 79507912 | 2498 | -0.26 | 0.648 | 0.661 |
| chr12 | 41745108 | 41745563 | 456 | -0.37 | 0.648 | 0.661 |
| chr16 | 24346920 | 24358467 | 11548 | -0.19 | 0.649 | 0.662 |
| chr4 | 10126834 | 10141822 | 14989 | -0.14 | 0.649 | 0.662 |
| chr7 | 109020621 | 109021375 | 755 | 0.24 | 0.649 | 0.662 |
| chr16 | 51923209 | 51976814 | 53606 | 0.09 | 0.649 | 0.662 |
| chr8 | 120474514 | 120481623 | 7110 | 0.2 | 0.649 | 0.662 |
| chr2 | 213194161 | 213210038 | 15878 | 0.15 | 0.649 | 0.662 |
| chr2 | 176205809 | 176224692 | 18884 | -0.23 | 0.65 | 0.662 |
| chr3 | 39614505 | 39628428 | 13924 | 0.14 | 0.65 | 0.663 |
| chr12 | 104396443 | 104397755 | 1313 | 0.26 | 0.651 | 0.663 |
| chr17 | 55786641 | 55795802 | 9162 | 0.14 | 0.651 | 0.664 |
| chrX | 49876519 | 49887500 | 10982 | 0.16 | 0.651 | 0.664 |

| | | | | | | |
|---|---|---|---|---|---|---|
| chr18 | 44091977 | 44103713 | 11737 | -0.29 | 0.651 | 0.664 |
| chr8 | 114946075 | 114995667 | 49593 | -0.2 | 0.651 | 0.664 |
| chr11 | 98993750 | 99001280 | 7531 | 0.24 | 0.652 | 0.664 |
| chr4 | 53107127 | 53153940 | 46814 | -0.1 | 0.652 | 0.664 |
| chr6 | 46772141 | 46782051 | 9911 | 0.14 | 0.653 | 0.665 |
| chr5 | 85537761 | 85538957 | 1197 | -0.48 | 0.653 | 0.666 |
| chr2 | 223127871 | 223177215 | 49345 | -0.17 | 0.654 | 0.666 |
| chr11 | 65267428 | 65274044 | 6617 | -0.18 | 0.654 | 0.667 |
| chr6 | 25840455 | 25841830 | 1376 | -0.22 | 0.655 | 0.668 |
| chr10 | 48612201 | 48661553 | 49353 | -0.15 | 0.656 | 0.668 |
| chr17 | 67601716 | 67717140 | 115425 | 0.2 | 0.657 | 0.669 |
| chr5 | 88187175 | 88188264 | 1090 | -0.3 | 0.657 | 0.669 |
| chr7 | 14989082 | 14989788 | 707 | -0.27 | 0.657 | 0.669 |
| chr4 | 61022224 | 61032397 | 10174 | 0.18 | 0.657 | 0.67 |
| chr12 | 106181968 | 106203720 | 21753 | 0.15 | 0.658 | 0.67 |
| chr5 | 112573505 | 112634200 | 60696 | 0.19 | 0.658 | 0.671 |
| chr12 | 66674388 | 66687595 | 13208 | -0.78 | 0.659 | 0.671 |
| chr11 | 102581282 | 102594509 | 13228 | -0.12 | 0.659 | 0.671 |
| chr1 | 152341084 | 152362845 | 21762 | -0.11 | 0.659 | 0.671 |
| chr17 | 68681413 | 68733953 | 52541 | -0.17 | 0.659 | 0.672 |
| chr7 | 92797789 | 92818820 | 21032 | 0.2 | 0.66 | 0.672 |
| chr8 | 12636678 | 12696134 | 59457 | 0.08 | 0.661 | 0.673 |
| chr14 | 38350714 | 38368859 | 18146 | -0.33 | 0.662 | 0.675 |
| chr1 | 218328492 | 218333531 | 5040 | 0.14 | 0.662 | 0.675 |
| chr4 | 90287525 | 90311330 | 23806 | 0.38 | 0.663 | 0.676 |
| chr19 | 43482770 | 43483354 | 585 | 0.38 | 0.664 | 0.676 |
| chr8 | 54086722 | 54160979 | 74258 | 0.07 | 0.664 | 0.677 |
| chr4 | 61893381 | 61927389 | 34009 | 0.1 | 0.665 | 0.677 |
| chr3 | 106225624 | 106271944 | 46321 | -0.26 | 0.666 | 0.678 |
| chr10 | 15933421 | 15941427 | 8007 | -0.3 | 0.666 | 0.678 |
| chr13 | 112001278 | 112018331 | 17054 | -0.14 | 0.666 | 0.678 |
| chr7 | 158353098 | 158354950 | 1853 | -0.18 | 0.666 | 0.678 |
| chr12 | 54294220 | 54300238 | 6019 | -0.14 | 0.667 | 0.679 |
| chr16 | 59009722 | 59010681 | 960 | 0.6 | 0.667 | 0.679 |
| chr2 | 57035210 | 57059374 | 24165 | -0.22 | 0.667 | 0.679 |
| chr3 | 86758148 | 86759019 | 872 | 0.22 | 0.667 | 0.68 |
| chr5 | 110562364 | 110563218 | 855 | -0.37 | 0.667 | 0.68 |
| chr10 | 129587277 | 129637696 | 50420 | -0.15 | 0.667 | 0.68 |
| chr5 | 135024342 | 135025356 | 1015 | 0.23 | 0.668 | 0.68 |
| chr7 | 142367051 | 142368437 | 1387 | -0.23 | 0.668 | 0.68 |
| chr16 | 80405828 | 80491708 | 85881 | 0.08 | 0.668 | 0.681 |
| chr20 | 29948082 | 29992763 | 44682 | -0.1 | 0.669 | 0.681 |
| chr4 | 35126551 | 35141626 | 15076 | 0.17 | 0.67 | 0.682 |
| chr12 | 10192128 | 10205112 | 12985 | -0.14 | 0.67 | 0.682 |
| chrX | 41855031 | 41863630 | 8600 | 0.14 | 0.67 | 0.683 |
| chr1 | 116395968 | 116401840 | 5873 | 0.16 | 0.671 | 0.683 |
| chr1 | 99498197 | 99504248 | 6052 | -0.22 | 0.672 | 0.684 |
| chr4 | 11386330 | 11389288 | 2959 | -0.18 | 0.672 | 0.684 |
| chr4 | 61308783 | 61341656 | 32874 | 0.12 | 0.672 | 0.684 |
| chr18 | 56945340 | 56957832 | 12493 | -0.17 | 0.673 | 0.685 |
| chr6 | 153158461 | 153168303 | 9843 | 0.26 | 0.673 | 0.685 |
| chr9 | 103956147 | 103957070 | 924 | 0.29 | 0.674 | 0.686 |
| chr2 | 237500500 | 237519815 | 19316 | 0.13 | 0.674 | 0.687 |
| chr9 | 34487007 | 34488829 | 1823 | 0.21 | 0.675 | 0.687 |
| chr8 | 63840018 | 63903553 | 63536 | 0.21 | 0.675 | 0.688 |
| chrX | 76418092 | 76419816 | 1725 | -0.29 | 0.676 | 0.688 |
| chr7 | 119413976 | 119416821 | 2846 | 0.2 | 0.676 | 0.688 |
| chr18 | 47443903 | 47445883 | 1981 | 0.26 | 0.676 | 0.688 |
| chr1 | 162412418 | 162425483 | 13066 | -0.15 | 0.676 | 0.688 |
| chr11 | 109214483 | 109216664 | 2182 | 0.19 | 0.677 | 0.689 |
| chr18 | 76453439 | 76454734 | 1296 | 0.23 | 0.678 | 0.69 |

| | | | | | | |
|---|---|---|---|---|---|---|
| chr9 | 94637031 | 94638677 | 1647 | -0.41 | 0.679 | 0.691 |
| chr3 | 150816159 | 150825206 | 9048 | -0.49 | 0.68 | 0.692 |
| chr10 | 67389371 | 67390429 | 1059 | 0.27 | 0.68 | 0.692 |
| chr4 | 61772507 | 61773807 | 1301 | 0.43 | 0.682 | 0.694 |
| chr4 | 67961023 | 67961857 | 835 | -0.32 | 0.683 | 0.695 |
| chr4 | 33111987 | 33120240 | 8254 | -0.26 | 0.683 | 0.695 |
| chr6 | 8249372 | 8328126 | 78755 | -0.07 | 0.683 | 0.695 |
| chr12 | 59687895 | 59709220 | 21326 | 0.28 | 0.684 | 0.696 |
| chr5 | 45550788 | 45574689 | 23902 | -0.17 | 0.684 | 0.696 |
| chr7 | 13697159 | 13698080 | 922 | 0.61 | 0.684 | 0.696 |
| chr3 | 190762260 | 190762731 | 472 | -0.59 | 0.685 | 0.697 |
| chr1 | 247653782 | 247660448 | 6667 | 0.16 | 0.686 | 0.698 |
| chr13 | 46284894 | 46287428 | 2535 | -0.16 | 0.687 | 0.699 |
| chrX | 150245954 | 150271907 | 25954 | -0.23 | 0.687 | 0.699 |
| chr4 | 29339970 | 29345352 | 5383 | -0.17 | 0.687 | 0.699 |
| chr7 | 92998477 | 93014018 | 15542 | -0.17 | 0.688 | 0.7 |
| chr3 | 175819194 | 175834062 | 14869 | 0.21 | 0.689 | 0.701 |
| chr10 | 64175964 | 64177107 | 1144 | 0.26 | 0.689 | 0.701 |
| chr4 | 29917017 | 29931450 | 14434 | 0.13 | 0.689 | 0.701 |
| chr14 | 37169121 | 37170136 | 1016 | -0.61 | 0.69 | 0.701 |
| chr4 | 32661164 | 32661968 | 805 | 0.37 | 0.69 | 0.701 |
| chr8 | 48054284 | 48070172 | 15889 | -0.17 | 0.69 | 0.702 |
| chr18 | 27760081 | 27804466 | 44386 | -0.14 | 0.69 | 0.702 |
| chr7 | 119240874 | 119248122 | 7249 | 0.15 | 0.691 | 0.702 |
| chr9 | 103480577 | 103483538 | 2962 | 0.17 | 0.692 | 0.704 |
| chr2 | 47805835 | 47830073 | 24239 | -0.18 | 0.694 | 0.706 |
| chr20 | 40295668 | 40317472 | 21805 | -0.5 | 0.694 | 0.706 |
| chr7 | 31578686 | 31598489 | 19804 | -0.18 | 0.695 | 0.706 |
| chr2 | 226302691 | 226308202 | 5512 | -0.18 | 0.695 | 0.706 |
| chr4 | 62872605 | 62889004 | 16400 | 0.15 | 0.695 | 0.706 |
| chr5 | 125356421 | 125599103 | 242683 | -0.1 | 0.695 | 0.707 |
| chr2 | 212905084 | 212919747 | 14664 | 0.16 | 0.695 | 0.707 |
| chr7 | 15218523 | 15219414 | 892 | -0.2 | 0.696 | 0.708 |
| chr15 | 96641454 | 96649558 | 8105 | 0.18 | 0.697 | 0.709 |
| chr2 | 215190968 | 215191353 | 386 | -0.31 | 0.698 | 0.71 |
| chr9 | 121118716 | 121128511 | 9796 | -0.25 | 0.698 | 0.71 |
| chr10 | 95550841 | 95563852 | 13012 | 0.15 | 0.699 | 0.71 |
| chr15 | 96910667 | 96920052 | 9386 | 0.27 | 0.699 | 0.71 |
| chr10 | 63166737 | 63185859 | 19123 | 0.17 | 0.699 | 0.711 |
| chr4 | 42668949 | 42670459 | 1511 | 0.26 | 0.7 | 0.711 |
| chr14 | 28708723 | 28739737 | 31015 | 0.16 | 0.7 | 0.712 |
| chr4 | 29420596 | 29424086 | 3491 | -0.2 | 0.701 | 0.713 |
| chr2 | 2714412 | 2730136 | 15725 | -0.14 | 0.702 | 0.714 |
| chr9 | 117538745 | 117559910 | 21166 | -0.13 | 0.702 | 0.714 |
| chr8 | 58014690 | 58033386 | 18697 | 0.1 | 0.703 | 0.715 |
| chr15 | 20188940 | 20197605 | 8666 | 0.12 | 0.703 | 0.715 |
| chr12 | 94536352 | 94549751 | 13400 | -0.8 | 0.704 | 0.716 |
| chr6 | 130836516 | 130838683 | 2168 | -0.43 | 0.704 | 0.716 |
| chr1 | 177134582 | 177136953 | 2372 | 0.21 | 0.705 | 0.716 |
| chr16 | 49540991 | 49541468 | 478 | -0.23 | 0.705 | 0.716 |
| chr2 | 166272653 | 166274956 | 2304 | -0.26 | 0.705 | 0.717 |
| chr10 | 19611456 | 19613407 | 1952 | -0.22 | 0.706 | 0.717 |
| chr14 | 92734507 | 92786565 | 52059 | -0.12 | 0.706 | 0.717 |
| chr8 | 10828527 | 10842460 | 13934 | -0.18 | 0.707 | 0.719 |
| chr2 | 222547710 | 222559364 | 11655 | -0.19 | 0.708 | 0.719 |
| chr12 | 70484695 | 70486816 | 2122 | -0.17 | 0.708 | 0.719 |
| chr2 | 865326 | 877643 | 12318 | -0.13 | 0.709 | 0.72 |
| chr9 | 113142488 | 113155818 | 13331 | -0.66 | 0.709 | 0.721 |
| chr14 | 39434645 | 39439079 | 4435 | 0.21 | 0.709 | 0.721 |
| chr4 | 33279802 | 33281285 | 1484 | -0.2 | 0.71 | 0.721 |
| chr4 | 61036017 | 61041424 | 5408 | -0.18 | 0.71 | 0.721 |

| | | | | | | |
|---|---|---|---|---|---|---|
| chr2 | 242912911 | 242931504 | 18594 | -0.37 | 0.71 | 0.722 |
| chr12 | 59741062 | 59752082 | 11021 | -0.37 | 0.711 | 0.722 |
| chr12 | 22242202 | 22269318 | 27117 | 0.1 | 0.711 | 0.723 |
| chr8 | 6770810 | 6829024 | 58215 | 0.09 | 0.714 | 0.725 |
| chrX | 30912457 | 30921236 | 8780 | -0.19 | 0.714 | 0.726 |
| chr7 | 16033621 | 16036706 | 3086 | -0.17 | 0.715 | 0.726 |
| chr13 | 68347328 | 68348319 | 992 | 0.21 | 0.715 | 0.726 |
| chr16 | 8599544 | 8611812 | 12269 | -0.12 | 0.715 | 0.726 |
| chrX | 81248442 | 81268117 | 19676 | -0.26 | 0.716 | 0.727 |
| chr16 | 79817100 | 79843920 | 26821 | 0.09 | 0.717 | 0.728 |
| chr5 | 42658700 | 42688273 | 29574 | 0.16 | 0.718 | 0.729 |
| chr9 | 16162311 | 16167539 | 5229 | -0.15 | 0.719 | 0.73 |
| chr13 | 78552465 | 78557496 | 5032 | -0.24 | 0.719 | 0.73 |
| chr3 | 34338422 | 34339204 | 783 | 0.24 | 0.72 | 0.731 |
| chr11 | 125144864 | 125146260 | 1397 | 0.17 | 0.72 | 0.731 |
| chr14 | 36865112 | 36881216 | 16105 | 0.15 | 0.72 | 0.731 |
| chrX | 131624844 | 131626320 | 1477 | 0.15 | 0.72 | 0.731 |
| chr2 | 220741461 | 220766214 | 24754 | -0.13 | 0.72 | 0.731 |
| chr3 | 180447660 | 180448689 | 1030 | -0.22 | 0.72 | 0.731 |
| chr10 | 129342322 | 129402821 | 60500 | -0.16 | 0.72 | 0.732 |
| chr11 | 56505449 | 56520110 | 14662 | 0.12 | 0.721 | 0.732 |
| chrX | 41920533 | 41927064 | 6532 | 0.11 | 0.722 | 0.733 |
| chr12 | 101137382 | 101201695 | 64314 | -0.14 | 0.722 | 0.733 |
| chr2 | 36345454 | 36354540 | 9087 | 0.1 | 0.722 | 0.733 |
| chr3 | 130483594 | 130486965 | 3372 | 0.14 | 0.723 | 0.734 |
| chrX | 56690975 | 56693066 | 2092 | 0.15 | 0.723 | 0.734 |
| chr8 | 9769991 | 9788052 | 18062 | -0.11 | 0.726 | 0.737 |
| chr7 | 15829628 | 15830500 | 873 | -0.55 | 0.726 | 0.737 |
| chr2 | 38214082 | 38228418 | 14337 | -0.6 | 0.726 | 0.737 |
| chr2 | 175878809 | 175890842 | 12034 | -0.6 | 0.726 | 0.737 |
| chr6 | 128193405 | 128208930 | 15526 | -0.6 | 0.726 | 0.737 |
| chr4 | 68951147 | 68952445 | 1299 | 0.21 | 0.727 | 0.738 |
| chr10 | 129297341 | 129301302 | 3962 | -0.24 | 0.727 | 0.738 |
| chr18 | 43214493 | 43228430 | 13938 | -0.18 | 0.728 | 0.739 |
| chr14 | 22305722 | 22445604 | 139883 | -0.1 | 0.728 | 0.74 |
| chr5 | 40302231 | 40303811 | 1581 | 0.22 | 0.729 | 0.74 |
| chr8 | 1530574 | 1543640 | 13067 | -0.26 | 0.731 | 0.742 |
| chr5 | 75111766 | 75162730 | 50965 | -0.13 | 0.731 | 0.742 |
| chr11 | 113162882 | 113167682 | 4801 | 0.19 | 0.731 | 0.742 |
| chr4 | 29778238 | 29791813 | 13576 | 0.12 | 0.734 | 0.745 |
| chr20 | 57989044 | 57995897 | 6854 | 0.13 | 0.734 | 0.745 |
| chr15 | 52037179 | 52053029 | 15851 | -0.78 | 0.734 | 0.745 |
| chr8 | 12620109 | 12621382 | 1274 | -0.19 | 0.734 | 0.745 |
| chr15 | 46180238 | 46192205 | 11968 | -0.11 | 0.734 | 0.745 |
| chr6 | 38920266 | 38925424 | 5159 | 0.11 | 0.735 | 0.745 |
| chr18 | 47672131 | 47674308 | 2178 | 0.16 | 0.735 | 0.746 |
| chr8 | 82776483 | 82779707 | 3225 | -0.14 | 0.735 | 0.746 |
| chr7 | 55366450 | 55367885 | 1436 | 0.42 | 0.736 | 0.747 |
| chr16 | 22831315 | 22842625 | 11311 | -0.19 | 0.737 | 0.748 |
| chr11 | 32514201 | 32515679 | 1479 | 0.17 | 0.738 | 0.748 |
| chr17 | 26317310 | 26322856 | 5547 | -0.11 | 0.738 | 0.749 |
| chrX | 37047198 | 37048185 | 988 | -0.21 | 0.74 | 0.751 |
| chr16 | 63177265 | 63179663 | 2399 | -0.16 | 0.741 | 0.751 |
| chr4 | 32801166 | 32805803 | 4638 | 0.18 | 0.741 | 0.751 |
| chr11 | 32293834 | 32295279 | 1446 | -0.26 | 0.741 | 0.752 |
| chr11 | 106115494 | 106123941 | 8448 | -0.15 | 0.741 | 0.752 |
| chr1 | 108590138 | 108614335 | 24198 | 0.1 | 0.741 | 0.752 |
| chr16 | 59143631 | 59183484 | 39854 | 0.16 | 0.742 | 0.752 |
| chr3 | 124007161 | 124019525 | 12365 | -0.16 | 0.742 | 0.753 |
| chr2 | 222506149 | 222511890 | 5742 | 0.15 | 0.743 | 0.754 |
| chr6 | 150873890 | 150893721 | 19832 | -0.59 | 0.744 | 0.754 |

| | | | | | | |
|---|---|---|---|---|---|---|
| chr3 | 60042589 | 60043811 | 1223 | -0.15 | 0.744 | 0.755 |
| chr7 | 155293496 | 155317846 | 24351 | -0.38 | 0.745 | 0.756 |
| chr3 | 150619165 | 150635120 | 15956 | -0.15 | 0.746 | 0.757 |
| chrX | 72553152 | 72567571 | 14420 | -0.12 | 0.747 | 0.758 |
| chr3 | 162953056 | 162954746 | 1691 | 0.16 | 0.747 | 0.758 |
| chr1 | 47430853 | 47438528 | 7676 | -0.33 | 0.748 | 0.759 |
| chr16 | 64181839 | 64190231 | 8393 | 0.16 | 0.748 | 0.759 |
| chr4 | 59405479 | 59406419 | 941 | 0.17 | 0.749 | 0.76 |
| chr4 | 61635818 | 61655879 | 20062 | 0.13 | 0.75 | 0.76 |
| chr2 | 64467479 | 64470711 | 3233 | -0.17 | 0.75 | 0.761 |
| chr4 | 60577604 | 60578674 | 1071 | -0.22 | 0.751 | 0.761 |
| chr13 | 88976297 | 88977523 | 1227 | -0.48 | 0.752 | 0.762 |
| chr13 | 29428768 | 29475652 | 46885 | 0.06 | 0.753 | 0.764 |
| chr1 | 169709905 | 169712084 | 2180 | 0.18 | 0.754 | 0.764 |
| chr1 | 20313727 | 20316097 | 2371 | 0.18 | 0.755 | 0.766 |
| chr10 | 127443348 | 127453352 | 10005 | -0.43 | 0.757 | 0.767 |
| chr13 | 25801943 | 25814730 | 12788 | -0.43 | 0.757 | 0.767 |
| chr10 | 135386300 | 135436064 | 49765 | -0.13 | 0.759 | 0.77 |
| chr1 | 209744983 | 209746957 | 1975 | 0.28 | 0.761 | 0.772 |
| chr15 | 49679475 | 49680679 | 1205 | 0.21 | 0.763 | 0.773 |
| chr13 | 31463119 | 31479832 | 16714 | -0.16 | 0.763 | 0.774 |
| chr8 | 16559767 | 16561604 | 1838 | -0.15 | 0.764 | 0.775 |
| chr4 | 29707372 | 29710337 | 2966 | -0.2 | 0.765 | 0.775 |
| chr3 | 27971910 | 27979156 | 7247 | 0.1 | 0.765 | 0.775 |
| chr9 | 85730833 | 85732661 | 1829 | 0.39 | 0.765 | 0.776 |
| chr1 | 105680433 | 105684181 | 3749 | -0.14 | 0.766 | 0.776 |
| chr13 | 23835373 | 23839851 | 4479 | -0.1 | 0.767 | 0.776 |
| chr4 | 139718681 | 139736163 | 17483 | -0.1 | 0.768 | 0.776 |
| chr1 | 15456290 | 15477333 | 21044 | 0.48 | 0.768 | 0.776 |
| chr1 | 179935587 | 179949397 | 13811 | 0.48 | 0.768 | 0.776 |
| chr1 | 183022730 | 183035647 | 12918 | 0.48 | 0.768 | 0.776 |
| chr10 | 3702943 | 3716483 | 13541 | 0.48 | 0.768 | 0.776 |
| chr10 | 5846832 | 5856961 | 10130 | 0.48 | 0.768 | 0.776 |
| chr10 | 22917030 | 22926298 | 9269 | 0.48 | 0.768 | 0.776 |
| chr10 | 27806530 | 27819033 | 12504 | 0.48 | 0.768 | 0.776 |
| chr10 | 34711095 | 34733060 | 21966 | 0.48 | 0.768 | 0.776 |
| chr10 | 81078476 | 81097924 | 19449 | 0.48 | 0.768 | 0.776 |
| chr10 | 94732519 | 94744806 | 12288 | 0.48 | 0.768 | 0.776 |
| chr10 | 99485802 | 99496547 | 10746 | 0.48 | 0.768 | 0.776 |
| chr10 | 106049900 | 106064078 | 14179 | 0.48 | 0.768 | 0.776 |
| chr11 | 20491900 | 20506321 | 14422 | 0.48 | 0.768 | 0.776 |
| chr11 | 31414239 | 31422106 | 7868 | 0.48 | 0.768 | 0.776 |
| chr11 | 32546064 | 32567247 | 21184 | 0.48 | 0.768 | 0.776 |
| chr11 | 46596007 | 46604781 | 8775 | 0.48 | 0.768 | 0.776 |
| chr11 | 64996996 | 65010409 | 13414 | 0.48 | 0.768 | 0.776 |
| chr11 | 70066497 | 70079618 | 13122 | 0.48 | 0.768 | 0.776 |
| chr11 | 75816149 | 75823886 | 7738 | 0.48 | 0.768 | 0.776 |
| chr11 | 108685942 | 108700929 | 14988 | 0.48 | 0.768 | 0.776 |
| chr12 | 9187736 | 9206828 | 19093 | 0.48 | 0.768 | 0.776 |
| chr12 | 15664304 | 15682005 | 17702 | 0.48 | 0.768 | 0.776 |
| chr12 | 27362270 | 27376762 | 14493 | 0.48 | 0.768 | 0.776 |
| chr12 | 44344691 | 44360053 | 15363 | 0.48 | 0.768 | 0.776 |
| chr12 | 79208805 | 79222914 | 14110 | 0.48 | 0.768 | 0.776 |
| chr12 | 95600570 | 95615081 | 14512 | 0.48 | 0.768 | 0.776 |
| chr12 | 104936223 | 104958591 | 22369 | 0.48 | 0.768 | 0.776 |
| chr12 | 106726029 | 106739691 | 13663 | 0.48 | 0.768 | 0.776 |
| chr12 | 107441195 | 107459386 | 18192 | 0.48 | 0.768 | 0.776 |
| chr12 | 120322708 | 120334253 | 11546 | 0.48 | 0.768 | 0.776 |
| chr12 | 120482020 | 120489947 | 7928 | 0.48 | 0.768 | 0.776 |
| chr13 | 45019792 | 45031334 | 11543 | 0.48 | 0.768 | 0.776 |
| chr14 | 32139071 | 32152869 | 13799 | 0.48 | 0.768 | 0.776 |

| | | | | | | |
|---|---|---|---|---|---|---|
| chr14 | 32511095 | 32524562 | 13468 | 0.48 | 0.768 | 0.776 |
| chr14 | 35120403 | 35143433 | 23031 | 0.48 | 0.768 | 0.776 |
| chr14 | 37902943 | 37914069 | 11127 | 0.48 | 0.768 | 0.776 |
| chr14 | 55084716 | 55099513 | 14798 | 0.48 | 0.768 | 0.776 |
| chr14 | 61995880 | 62009053 | 13174 | 0.48 | 0.768 | 0.776 |
| chr14 | 71919061 | 71931433 | 12373 | 0.48 | 0.768 | 0.776 |
| chr14 | 75391510 | 75405529 | 14020 | 0.48 | 0.768 | 0.776 |
| chr14 | 90884218 | 90895996 | 11779 | 0.48 | 0.768 | 0.776 |
| chr14 | 95751662 | 95762737 | 11076 | 0.48 | 0.768 | 0.776 |
| chr15 | 29345065 | 29362702 | 17638 | 0.48 | 0.768 | 0.776 |
| chr15 | 57639918 | 57653739 | 13822 | 0.48 | 0.768 | 0.776 |
| chr15 | 85444446 | 85457581 | 13136 | 0.48 | 0.768 | 0.776 |
| chr15 | 90952305 | 90961470 | 9166 | 0.48 | 0.768 | 0.776 |
| chr16 | 85735236 | 85746609 | 11374 | 0.48 | 0.768 | 0.776 |
| chr16 | 87286301 | 87295038 | 8738 | 0.48 | 0.768 | 0.776 |
| chr17 | 25881122 | 25891689 | 10568 | 0.48 | 0.768 | 0.776 |
| chr17 | 33386271 | 33399366 | 13096 | 0.48 | 0.768 | 0.776 |
| chr17 | 72290125 | 72318110 | 27986 | 0.48 | 0.768 | 0.776 |
| chr17 | 73108374 | 73117864 | 9491 | 0.48 | 0.768 | 0.776 |
| chr17 | 77228652 | 77242172 | 13521 | 0.48 | 0.768 | 0.776 |
| chr18 | 8307207 | 8319419 | 12213 | 0.48 | 0.768 | 0.776 |
| chr19 | 45179809 | 45191613 | 11805 | 0.48 | 0.768 | 0.776 |
| chr2 | 7109778 | 7131578 | 21801 | 0.48 | 0.768 | 0.776 |
| chr2 | 25199897 | 25214201 | 14305 | 0.48 | 0.768 | 0.776 |
| chr2 | 44596942 | 44612457 | 15516 | 0.48 | 0.768 | 0.776 |
| chr2 | 47943230 | 47968717 | 25488 | 0.48 | 0.768 | 0.776 |
| chr2 | 68319577 | 68333990 | 14414 | 0.48 | 0.768 | 0.776 |
| chr2 | 68721454 | 68735948 | 14495 | 0.48 | 0.768 | 0.776 |
| chr2 | 72392288 | 72396204 | 3917 | 0.48 | 0.768 | 0.776 |
| chr2 | 121595962 | 121607509 | 11548 | 0.48 | 0.768 | 0.776 |
| chr2 | 160842254 | 160880060 | 37807 | 0.48 | 0.768 | 0.776 |
| chr2 | 163190209 | 163200134 | 9926 | 0.48 | 0.768 | 0.776 |
| chr2 | 172787987 | 172798322 | 10336 | 0.48 | 0.768 | 0.776 |
| chr2 | 174781577 | 174793708 | 12132 | 0.48 | 0.768 | 0.776 |
| chr2 | 176046732 | 176057967 | 11236 | 0.48 | 0.768 | 0.776 |
| chr2 | 180115434 | 180126630 | 11197 | 0.48 | 0.768 | 0.776 |
| chr2 | 205870701 | 205884578 | 13878 | 0.48 | 0.768 | 0.776 |
| chr2 | 206187963 | 206201082 | 13120 | 0.48 | 0.768 | 0.776 |
| chr2 | 208372459 | 208384307 | 11849 | 0.48 | 0.768 | 0.776 |
| chr2 | 225217694 | 225227683 | 9990 | 0.48 | 0.768 | 0.776 |
| chr20 | 1819432 | 1828026 | 8595 | 0.48 | 0.768 | 0.776 |
| chr22 | 19138805 | 19152492 | 13688 | 0.48 | 0.768 | 0.776 |
| chr22 | 34291474 | 34301781 | 10308 | 0.48 | 0.768 | 0.776 |
| chr3 | 23497296 | 23510047 | 12752 | 0.48 | 0.768 | 0.776 |
| chr3 | 36653851 | 36668377 | 14527 | 0.48 | 0.768 | 0.776 |
| chr3 | 88219306 | 88231351 | 12046 | 0.48 | 0.768 | 0.776 |
| chr3 | 117529362 | 117539314 | 9953 | 0.48 | 0.768 | 0.776 |
| chr3 | 157043465 | 157058137 | 14673 | 0.48 | 0.768 | 0.776 |
| chr4 | 92027456 | 92037649 | 10194 | 0.48 | 0.768 | 0.776 |
| chr5 | 16976122 | 16986171 | 10050 | 0.48 | 0.768 | 0.776 |
| chr5 | 40681594 | 40696658 | 15065 | 0.48 | 0.768 | 0.776 |
| chr5 | 70923079 | 70934983 | 11905 | 0.48 | 0.768 | 0.776 |
| chr5 | 86601638 | 86616011 | 14374 | 0.48 | 0.768 | 0.776 |
| chr5 | 90715906 | 90726421 | 10516 | 0.48 | 0.768 | 0.776 |
| chr5 | 106262887 | 106297774 | 34888 | 0.48 | 0.768 | 0.776 |
| chr5 | 108199994 | 108217495 | 17502 | 0.48 | 0.768 | 0.776 |
| chr5 | 112177481 | 112191749 | 14269 | 0.48 | 0.768 | 0.776 |
| chr5 | 131738554 | 131749940 | 11387 | 0.48 | 0.768 | 0.776 |
| chr5 | 158387594 | 158400354 | 12761 | 0.48 | 0.768 | 0.776 |
| chr5 | 168685659 | 168694447 | 8789 | 0.48 | 0.768 | 0.776 |
| chr6 | 641258 | 666134 | 24877 | 0.48 | 0.768 | 0.776 |

| | | | | | | |
|---|---|---|---|---|---|---|
| chr6 | 10352765 | 10365132 | 12368 | 0.48 | 0.768 | 0.776 |
| chr6 | 71423601 | 71451848 | 28248 | 0.48 | 0.768 | 0.776 |
| chr6 | 134318464 | 134331346 | 12883 | 0.48 | 0.768 | 0.776 |
| chr7 | 2794664 | 2809936 | 15273 | 0.48 | 0.768 | 0.776 |
| chr7 | 2937771 | 2944948 | 7178 | 0.48 | 0.768 | 0.776 |
| chr7 | 82021602 | 82048804 | 27203 | 0.48 | 0.768 | 0.776 |
| chr7 | 83429257 | 83442737 | 13481 | 0.48 | 0.768 | 0.776 |
| chr7 | 83708474 | 83723040 | 14567 | 0.48 | 0.768 | 0.776 |
| chr7 | 91910126 | 91917921 | 7796 | 0.48 | 0.768 | 0.776 |
| chr7 | 106663687 | 106678197 | 14511 | 0.48 | 0.768 | 0.776 |
| chr7 | 108053056 | 108065244 | 12189 | 0.48 | 0.768 | 0.776 |
| chr7 | 111908701 | 111920059 | 11359 | 0.48 | 0.768 | 0.776 |
| chr7 | 133515702 | 133528152 | 12451 | 0.48 | 0.768 | 0.776 |
| chr7 | 158500182 | 158512502 | 12321 | 0.48 | 0.768 | 0.776 |
| chr8 | 8573463 | 8585469 | 12007 | 0.48 | 0.768 | 0.776 |
| chr8 | 18943379 | 18950820 | 7442 | 0.48 | 0.768 | 0.776 |
| chr8 | 27149009 | 27158599 | 9591 | 0.48 | 0.768 | 0.776 |
| chr8 | 28171835 | 28189445 | 17611 | 0.48 | 0.768 | 0.776 |
| chr8 | 90612549 | 90621056 | 8508 | 0.48 | 0.768 | 0.776 |
| chr8 | 100277908 | 100305530 | 27623 | 0.48 | 0.768 | 0.776 |
| chr8 | 103629887 | 103651427 | 21541 | 0.48 | 0.768 | 0.776 |
| chr9 | 4954463 | 4963800 | 9338 | 0.48 | 0.768 | 0.776 |
| chr9 | 5876321 | 5890403 | 14083 | 0.48 | 0.768 | 0.776 |
| chr9 | 14391185 | 14415896 | 24712 | 0.48 | 0.768 | 0.776 |
| chr9 | 35936727 | 35945633 | 8907 | 0.48 | 0.768 | 0.776 |
| chr9 | 90176772 | 90187680 | 10909 | 0.48 | 0.768 | 0.776 |
| chr9 | 101888817 | 101902483 | 13667 | 0.48 | 0.768 | 0.776 |
| chr9 | 128825096 | 128835669 | 10574 | 0.48 | 0.768 | 0.776 |
| chrX | 149552081 | 149559988 | 7908 | 0.48 | 0.768 | 0.776 |
| chrX | 149845529 | 149860351 | 14823 | 0.48 | 0.768 | 0.776 |
| chr12 | 6082644 | 6105313 | 22670 | 0.19 | 0.768 | 0.776 |
| chr19 | 16863523 | 16864399 | 877 | -0.18 | 0.768 | 0.776 |
| chr5 | 18347178 | 18348457 | 1280 | 0.19 | 0.768 | 0.776 |
| chr12 | 65478241 | 65507294 | 29054 | -0.19 | 0.768 | 0.776 |
| chr5 | 85805119 | 85806529 | 1411 | 0.23 | 0.769 | 0.777 |
| chr13 | 104095928 | 104097456 | 1529 | 0.16 | 0.771 | 0.778 |
| chr18 | 71532299 | 71544696 | 12398 | 0.14 | 0.771 | 0.779 |
| chr3 | 157381135 | 157492716 | 111582 | 0.11 | 0.771 | 0.779 |
| chr2 | 186527484 | 186554756 | 27273 | -0.38 | 0.772 | 0.779 |
| chr4 | 29363586 | 29398480 | 34895 | 0.12 | 0.772 | 0.779 |
| chr1 | 153424405 | 153425739 | 1335 | 0.17 | 0.773 | 0.78 |
| chr3 | 180495006 | 180496691 | 1686 | -0.12 | 0.773 | 0.781 |
| chrX | 23363782 | 23382401 | 18620 | 0.13 | 0.774 | 0.781 |
| chr3 | 129931571 | 129960199 | 28629 | 0.15 | 0.775 | 0.783 |
| chr3 | 152347744 | 152363444 | 15701 | -0.16 | 0.776 | 0.783 |
| chr20 | 5212964 | 5324667 | 111704 | -0.06 | 0.776 | 0.784 |
| chr14 | 73142526 | 73164268 | 21743 | 0.23 | 0.777 | 0.785 |
| chr12 | 106927614 | 106928431 | 818 | -0.2 | 0.777 | 0.785 |
| chr18 | 27637836 | 27666024 | 28189 | -0.1 | 0.777 | 0.785 |
| chr13 | 39684529 | 39700420 | 15892 | 0.15 | 0.778 | 0.786 |
| chr17 | 39845808 | 39847100 | 1293 | -0.14 | 0.778 | 0.786 |
| chr18 | 24674411 | 24688204 | 13794 | -0.1 | 0.779 | 0.786 |
| chr10 | 128382651 | 128435513 | 52863 | -0.06 | 0.78 | 0.787 |
| chr22 | 49643364 | 49651046 | 7683 | 0.12 | 0.78 | 0.788 |
| chr21 | 37202049 | 37203663 | 1615 | 0.16 | 0.781 | 0.789 |
| chr2 | 133943047 | 133959656 | 16610 | 0.13 | 0.782 | 0.789 |
| chr11 | 91908688 | 91918790 | 10103 | 0.21 | 0.783 | 0.79 |
| chr2 | 207327698 | 207359610 | 31913 | -0.51 | 0.783 | 0.79 |
| chr11 | 93802538 | 93814350 | 11813 | -0.1 | 0.783 | 0.79 |
| chr13 | 88764298 | 88765715 | 1418 | 0.29 | 0.783 | 0.79 |
| chrX | 109841572 | 109847558 | 5987 | 0.1 | 0.784 | 0.792 |

| | | | | | | |
|---|---|---|---|---|---|---|
| chr3 | 160556711 | 160573644 | 16934 | 0.58 | 0.785 | 0.792 |
| chr2 | 194602249 | 194603847 | 1599 | -0.18 | 0.785 | 0.793 |
| chr18 | 27562162 | 27592547 | 30386 | -0.1 | 0.786 | 0.793 |
| chr12 | 97237222 | 97288403 | 51182 | 0.09 | 0.786 | 0.794 |
| chr4 | 62144386 | 62145804 | 1419 | -0.18 | 0.787 | 0.795 |
| chr3 | 121751819 | 121778663 | 26845 | 0.11 | 0.789 | 0.796 |
| chr6 | 154003328 | 154005596 | 2269 | 0.14 | 0.79 | 0.798 |
| chr7 | 97374090 | 97374485 | 396 | -0.22 | 0.79 | 0.798 |
| chr8 | 127906677 | 127907418 | 742 | 0.18 | 0.791 | 0.798 |
| chr1 | 183905198 | 183921636 | 16439 | 0.08 | 0.792 | 0.799 |
| chr4 | 34231818 | 34247281 | 15464 | -0.14 | 0.792 | 0.799 |
| chr2 | 100521762 | 100563172 | 41411 | -0.06 | 0.793 | 0.8 |
| chr3 | 60057309 | 60057988 | 680 | -0.14 | 0.795 | 0.802 |
| chr14 | 95335902 | 95337308 | 1407 | 0.16 | 0.796 | 0.803 |
| chr7 | 142327887 | 142328657 | 771 | 0.14 | 0.797 | 0.804 |
| chr11 | 29436210 | 29437758 | 1549 | 0.16 | 0.797 | 0.805 |
| chr6 | 138464749 | 138466442 | 1694 | 0.14 | 0.798 | 0.805 |
| chr8 | 114586864 | 114621656 | 34793 | 0.07 | 0.799 | 0.806 |
| chr6 | 137800918 | 137807099 | 6182 | -0.09 | 0.801 | 0.808 |
| chr7 | 149700373 | 149700936 | 564 | 0.34 | 0.801 | 0.808 |
| chr20 | 23789234 | 23789711 | 478 | 0.13 | 0.801 | 0.808 |
| chr16 | 10307894 | 10321256 | 13363 | -0.47 | 0.801 | 0.808 |
| chr18 | 44911699 | 44924656 | 12958 | -0.1 | 0.802 | 0.809 |
| chr4 | 30015490 | 30037266 | 21777 | 0.08 | 0.802 | 0.81 |
| chr4 | 165851619 | 165864100 | 12482 | -0.21 | 0.803 | 0.81 |
| chr1 | 86654960 | 86657843 | 2884 | 0.13 | 0.803 | 0.81 |
| chr2 | 17527786 | 17528524 | 739 | 0.16 | 0.803 | 0.81 |
| chr1 | 105395931 | 105403222 | 7292 | -0.1 | 0.803 | 0.81 |
| chr1 | 206928571 | 206938007 | 9437 | -0.14 | 0.804 | 0.811 |
| chr14 | 52692492 | 52731590 | 39099 | -0.08 | 0.804 | 0.811 |
| chrX | 141214342 | 141236911 | 22570 | -0.07 | 0.804 | 0.811 |
| chr7 | 157876260 | 157883854 | 7595 | 0.16 | 0.804 | 0.811 |
| chr2 | 58615321 | 58616085 | 765 | -0.14 | 0.804 | 0.811 |
| chr4 | 20128053 | 20130448 | 2396 | 0.15 | 0.805 | 0.812 |
| chr15 | 87047098 | 87052403 | 5306 | -0.1 | 0.806 | 0.813 |
| chr1 | 82916754 | 82919542 | 2789 | -0.15 | 0.806 | 0.813 |
| chr7 | 106149718 | 106160312 | 10595 | -0.26 | 0.807 | 0.814 |
| chr4 | 62167953 | 62175976 | 8024 | -0.12 | 0.809 | 0.815 |
| chr3 | 123745922 | 123777382 | 31461 | 0.09 | 0.809 | 0.815 |
| chr18 | 44886635 | 44899203 | 12569 | 0.06 | 0.809 | 0.816 |
| chr16 | 82234253 | 82258855 | 24603 | 0.07 | 0.809 | 0.816 |
| chr7 | 95384037 | 95392601 | 8565 | 0.15 | 0.81 | 0.816 |
| chr1 | 228892527 | 228896112 | 3586 | -0.19 | 0.81 | 0.817 |
| chr11 | 57721275 | 57749563 | 28289 | 0.07 | 0.811 | 0.817 |
| chr4 | 33575126 | 33576877 | 1752 | 0.13 | 0.811 | 0.818 |
| chr3 | 171140364 | 171141014 | 651 | -0.13 | 0.812 | 0.819 |
| chrX | 111026404 | 111040345 | 13942 | -0.08 | 0.813 | 0.819 |
| chr21 | 36960507 | 36961038 | 532 | 0.15 | 0.813 | 0.819 |
| chrX | 73872192 | 73880543 | 8352 | -0.12 | 0.813 | 0.82 |
| chr18 | 26991975 | 27010931 | 18957 | -0.09 | 0.813 | 0.82 |
| chr3 | 104686214 | 104698348 | 12135 | 0.1 | 0.813 | 0.82 |
| chr4 | 128270438 | 128282949 | 12512 | -0.15 | 0.814 | 0.82 |
| chrX | 95597923 | 95611462 | 13540 | -0.07 | 0.814 | 0.821 |
| chr16 | 48735499 | 48743283 | 7785 | 0.39 | 0.814 | 0.821 |
| chr3 | 178166777 | 178167727 | 951 | -0.16 | 0.814 | 0.821 |
| chr12 | 22512885 | 22521678 | 8794 | 0.14 | 0.815 | 0.822 |
| chr17 | 55429064 | 55429519 | 456 | -0.16 | 0.816 | 0.822 |
| chr6 | 46023690 | 46035357 | 11668 | -0.07 | 0.816 | 0.822 |
| chrX | 89181400 | 89182095 | 696 | 0.11 | 0.817 | 0.823 |
| chr16 | 22906004 | 22952816 | 46813 | -0.1 | 0.818 | 0.824 |
| chr10 | 125070658 | 125094036 | 23379 | 0.05 | 0.819 | 0.825 |

| | | | | | | |
|---|---|---|---|---|---|---|
| chr5 | 43749338 | 43763659 | 14322 | 0.1 | 0.819 | 0.825 |
| chr2 | 68163840 | 68189467 | 25628 | -0.06 | 0.82 | 0.826 |
| chr2 | 213225841 | 213226540 | 700 | 0.28 | 0.82 | 0.827 |
| chr15 | 31962730 | 31964210 | 1481 | 0.1 | 0.821 | 0.827 |
| chr20 | 23656239 | 23661554 | 5316 | -0.1 | 0.821 | 0.827 |
| chr2 | 7349683 | 7380427 | 30745 | 0.06 | 0.821 | 0.827 |
| chr9 | 92818043 | 92818634 | 592 | 0.16 | 0.821 | 0.827 |
| chr3 | 6113971 | 6119302 | 5332 | -0.1 | 0.822 | 0.828 |
| chr6 | 81949972 | 81950774 | 803 | -0.19 | 0.823 | 0.829 |
| chr1 | 7421133 | 7422327 | 1195 | 0.14 | 0.824 | 0.83 |
| chr1 | 182513628 | 182536753 | 23126 | -0.07 | 0.824 | 0.83 |
| chr8 | 54068272 | 54069764 | 1493 | 0.16 | 0.824 | 0.831 |
| chr4 | 11306723 | 11344949 | 38227 | -0.07 | 0.825 | 0.831 |
| chr5 | 155339311 | 155341492 | 2182 | 0.11 | 0.826 | 0.832 |
| chr14 | 99401815 | 99405121 | 3307 | 0.12 | 0.826 | 0.832 |
| chr6 | 112698675 | 112699833 | 1159 | 0.11 | 0.826 | 0.833 |
| chr14 | 103546855 | 103548939 | 2085 | -0.1 | 0.827 | 0.833 |
| chr8 | 114643451 | 114805009 | 161559 | 0.09 | 0.828 | 0.834 |
| chr14 | 45320453 | 45323679 | 3227 | -0.12 | 0.829 | 0.835 |
| chrX | 62235185 | 62237644 | 2460 | 0.1 | 0.829 | 0.835 |
| chr13 | 79808442 | 79809584 | 1143 | -0.15 | 0.829 | 0.835 |
| chr8 | 121746539 | 121759726 | 13188 | -0.15 | 0.829 | 0.835 |
| chr10 | 133257647 | 133351752 | 94106 | 0.06 | 0.831 | 0.837 |
| chr7 | 45245149 | 45254341 | 9193 | 0.09 | 0.831 | 0.837 |
| chr3 | 93551380 | 93551863 | 484 | -0.12 | 0.832 | 0.838 |
| chr10 | 38458232 | 38476663 | 18432 | 0.07 | 0.832 | 0.838 |
| chr3 | 83282792 | 83284735 | 1944 | 0.14 | 0.833 | 0.839 |
| chr5 | 108996181 | 109013393 | 17213 | -0.15 | 0.834 | 0.84 |
| chr3 | 155886455 | 155888484 | 2030 | 0.15 | 0.834 | 0.84 |
| chr1 | 115218265 | 115230043 | 11779 | 0.24 | 0.835 | 0.84 |
| chr1 | 153309114 | 153309461 | 348 | -0.12 | 0.835 | 0.84 |
| chrX | 55947283 | 55947946 | 664 | -0.14 | 0.836 | 0.841 |
| chr16 | 22860491 | 22862453 | 1963 | -0.1 | 0.836 | 0.842 |
| chr13 | 94148834 | 94151106 | 2273 | 0.09 | 0.836 | 0.842 |
| chr14 | 22912565 | 22929058 | 16494 | -0.07 | 0.837 | 0.843 |
| chr11 | 82508764 | 82517293 | 8530 | 0.08 | 0.837 | 0.843 |
| chr6 | 84997958 | 85021046 | 23089 | 0.05 | 0.837 | 0.843 |
| chr14 | 92306033 | 92307918 | 1886 | -0.17 | 0.839 | 0.845 |
| chr14 | 62369238 | 62378672 | 9435 | 0.09 | 0.839 | 0.845 |
| chr3 | 104617966 | 104652912 | 34947 | -0.1 | 0.841 | 0.846 |
| chr3 | 104566075 | 104572311 | 6237 | -0.07 | 0.841 | 0.846 |
| chrX | 37025091 | 37025929 | 839 | -0.11 | 0.841 | 0.846 |
| chr2 | 122739776 | 122755625 | 15850 | 0.05 | 0.841 | 0.846 |
| chr4 | 53071488 | 53072820 | 1333 | -0.1 | 0.842 | 0.847 |
| chr3 | 156081233 | 156082402 | 1170 | -0.13 | 0.842 | 0.848 |
| chr1 | 207675914 | 207691384 | 15471 | -0.05 | 0.843 | 0.848 |
| chrX | 56456102 | 56456969 | 868 | -0.16 | 0.843 | 0.849 |
| chr8 | 112908926 | 112935079 | 26154 | -0.14 | 0.843 | 0.849 |
| chr15 | 38130907 | 38162542 | 31636 | 0.05 | 0.844 | 0.85 |
| chr8 | 133336451 | 133401605 | 65155 | 0.06 | 0.844 | 0.85 |
| chr16 | 59785789 | 59853369 | 67581 | -0.06 | 0.844 | 0.85 |
| chrX | 57525223 | 57526676 | 1454 | -0.09 | 0.844 | 0.85 |
| chr2 | 222630327 | 222630824 | 498 | 0.3 | 0.844 | 0.85 |
| chr14 | 42759393 | 42760627 | 1235 | 0.1 | 0.846 | 0.851 |
| chr8 | 19798020 | 19853507 | 55488 | -0.05 | 0.847 | 0.852 |
| chr3 | 81990660 | 81996345 | 5686 | 0.08 | 0.847 | 0.852 |
| chr11 | 11768372 | 11800914 | 32543 | -0.06 | 0.847 | 0.852 |
| chr2 | 2860289 | 2865233 | 4945 | -0.09 | 0.85 | 0.855 |
| chr6 | 83341499 | 83342502 | 1004 | -0.12 | 0.85 | 0.855 |
| chr18 | 32251304 | 32252855 | 1552 | 0.09 | 0.85 | 0.855 |
| chr7 | 57557751 | 57575559 | 17809 | -0.08 | 0.851 | 0.857 |

| | | | | | | |
|---|---|---|---|---|---|---|
| chr14 | 21174888 | 21248123 | 73236 | 0.06 | 0.851 | 0.857 |
| chr10 | 14019468 | 14020019 | 552 | -0.1 | 0.854 | 0.859 |
| chr21 | 41118182 | 41154705 | 36524 | -0.05 | 0.854 | 0.86 |
| chrX | 108803295 | 108814924 | 11630 | 0.07 | 0.855 | 0.86 |
| chr2 | 147563561 | 147565635 | 2075 | -0.12 | 0.855 | 0.861 |
| chr3 | 40824987 | 40832194 | 7208 | -0.06 | 0.856 | 0.861 |
| chr1 | 220612949 | 220619575 | 6627 | -0.06 | 0.857 | 0.862 |
| chr4 | 35643872 | 35667040 | 23169 | -0.06 | 0.857 | 0.862 |
| chr10 | 80508668 | 80510227 | 1560 | -0.08 | 0.858 | 0.863 |
| chr5 | 107901184 | 107903107 | 1924 | -0.09 | 0.859 | 0.864 |
| chr4 | 60756210 | 60757826 | 1617 | 0.13 | 0.863 | 0.868 |
| chr17 | 32758248 | 32761688 | 3441 | 0.09 | 0.864 | 0.869 |
| chr15 | 96010739 | 96012303 | 1565 | -0.19 | 0.864 | 0.869 |
| chr2 | 200110090 | 200111745 | 1656 | -0.15 | 0.866 | 0.871 |
| chr1 | 208456403 | 208457483 | 1081 | -0.13 | 0.867 | 0.872 |
| chr2 | 141052371 | 141053166 | 796 | -0.14 | 0.867 | 0.872 |
| chr3 | 104523289 | 104545931 | 22643 | -0.09 | 0.867 | 0.872 |
| chr4 | 161508271 | 161508930 | 660 | -0.1 | 0.868 | 0.873 |
| chr11 | 65191510 | 65194399 | 2890 | -0.06 | 0.868 | 0.873 |
| chr2 | 177687224 | 177689695 | 2472 | -0.09 | 0.868 | 0.873 |
| chr2 | 75404521 | 75407049 | 2529 | 0.09 | 0.869 | 0.874 |
| chr11 | 115793443 | 115830618 | 37176 | -0.04 | 0.87 | 0.874 |
| chr11 | 115641188 | 115782799 | 141612 | -0.03 | 0.871 | 0.876 |
| chr12 | 11407259 | 11415194 | 7936 | 0.05 | 0.872 | 0.877 |
| chr6 | 67901109 | 67915486 | 14378 | -0.06 | 0.872 | 0.877 |
| chr14 | 88658496 | 88661593 | 3098 | -0.11 | 0.873 | 0.878 |
| chr15 | 61390966 | 61392305 | 1340 | -0.12 | 0.873 | 0.878 |
| chr8 | 1514592 | 1516053 | 1462 | 0.14 | 0.873 | 0.878 |
| chr7 | 119391894 | 119392299 | 406 | -0.23 | 0.873 | 0.878 |
| chr13 | 22557668 | 22590152 | 32485 | -0.04 | 0.873 | 0.878 |
| chr5 | 6938952 | 6943347 | 4396 | 0.07 | 0.874 | 0.879 |
| chr4 | 62687968 | 62709813 | 21846 | 0.05 | 0.875 | 0.88 |
| chr5 | 29903110 | 29942676 | 39567 | -0.07 | 0.875 | 0.88 |
| chr6 | 143289563 | 143313224 | 23662 | 0.04 | 0.876 | 0.881 |
| chr15 | 25902453 | 25904523 | 2071 | -0.07 | 0.876 | 0.881 |
| chr8 | 127984532 | 128003371 | 18840 | 0.07 | 0.876 | 0.881 |
| chr10 | 7638102 | 7646703 | 8602 | -0.04 | 0.878 | 0.883 |
| chr8 | 10983496 | 11035484 | 51989 | 0.03 | 0.879 | 0.883 |
| chr4 | 121331898 | 121332562 | 665 | -0.14 | 0.879 | 0.884 |
| chr14 | 42066851 | 42078536 | 11686 | -0.08 | 0.879 | 0.884 |
| chr10 | 19151110 | 19161074 | 9965 | -0.06 | 0.879 | 0.884 |
| chr11 | 51200313 | 51244261 | 43949 | 0.04 | 0.88 | 0.885 |
| chr17 | 69294368 | 69306839 | 12472 | 0.05 | 0.881 | 0.886 |
| chr4 | 58939861 | 58943525 | 3665 | -0.1 | 0.882 | 0.887 |
| chr2 | 186872437 | 186878586 | 6150 | 0.1 | 0.883 | 0.887 |
| chr1 | 87085461 | 87086417 | 957 | 0.09 | 0.884 | 0.889 |
| chr18 | 10439170 | 10481328 | 42159 | -0.06 | 0.884 | 0.889 |
| chrX | 23398804 | 23399626 | 823 | 0.09 | 0.884 | 0.889 |
| chr21 | 15647028 | 15647442 | 415 | 0.08 | 0.885 | 0.889 |
| chr4 | 59034981 | 59038137 | 3157 | 0.12 | 0.885 | 0.889 |
| chr7 | 35627533 | 35628784 | 1252 | -0.06 | 0.885 | 0.89 |
| chr19 | 5367509 | 5368217 | 709 | 0.09 | 0.885 | 0.89 |
| chr2 | 214447346 | 214467477 | 20132 | 0.07 | 0.886 | 0.89 |
| chr16 | 51992387 | 52100110 | 107724 | 0.02 | 0.886 | 0.89 |
| chr15 | 58092493 | 58300407 | 207915 | -0.02 | 0.886 | 0.89 |
| chr8 | 71636398 | 71645043 | 8646 | 0.09 | 0.888 | 0.892 |
| chr10 | 64721865 | 64747755 | 25891 | 0.06 | 0.888 | 0.892 |
| chr9 | 109873370 | 109888437 | 15068 | 0.05 | 0.888 | 0.892 |
| chr8 | 75210194 | 75211893 | 1700 | 0.09 | 0.888 | 0.892 |
| chr7 | 50468659 | 50473869 | 5211 | 0.06 | 0.888 | 0.893 |
| chr5 | 89839405 | 89865911 | 26507 | 0.06 | 0.891 | 0.895 |

| | | | | | | |
|---|---|---|---|---|---|---|
| chr7 | 95419812 | 95432909 | 13098 | 0.07 | 0.891 | 0.895 |
| chr10 | 130027647 | 130047507 | 19861 | -0.06 | 0.891 | 0.896 |
| chr2 | 894153 | 948174 | 54022 | -0.03 | 0.893 | 0.897 |
| chr11 | 125606165 | 125636682 | 30518 | -0.05 | 0.893 | 0.897 |
| chr20 | 57394972 | 57410115 | 15144 | -0.13 | 0.894 | 0.898 |
| chr2 | 48808278 | 48810734 | 2457 | 0.06 | 0.894 | 0.898 |
| chr12 | 40872530 | 40877074 | 4545 | 0.04 | 0.894 | 0.898 |
| chr5 | 27302231 | 27332198 | 29968 | -0.07 | 0.895 | 0.898 |
| chr1 | 153371339 | 153377643 | 6305 | 0.05 | 0.895 | 0.899 |
| chr10 | 34071986 | 34074959 | 2974 | -0.07 | 0.897 | 0.901 |
| chr5 | 75477140 | 75486712 | 9573 | -0.05 | 0.901 | 0.905 |
| chr9 | 33643150 | 33657970 | 14821 | -0.05 | 0.901 | 0.905 |
| chr6 | 161941871 | 161943357 | 1487 | 0.05 | 0.901 | 0.905 |
| chr19 | 55275710 | 55276204 | 495 | -0.06 | 0.904 | 0.908 |
| chr13 | 78494417 | 78495578 | 1162 | -0.08 | 0.905 | 0.909 |
| chr1 | 104829390 | 104830819 | 1430 | 0.1 | 0.905 | 0.909 |
| chr9 | 30370041 | 30373984 | 3944 | 0.08 | 0.906 | 0.91 |
| chr6 | 39431269 | 39432565 | 1297 | -0.14 | 0.906 | 0.91 |
| chr15 | 92966473 | 92974010 | 7538 | -0.1 | 0.907 | 0.91 |
| chr12 | 54322580 | 54324976 | 2397 | -0.04 | 0.909 | 0.912 |
| chr1 | 104130186 | 104133314 | 3129 | 0.06 | 0.91 | 0.913 |
| chr8 | 113707117 | 113707530 | 414 | 0.08 | 0.911 | 0.914 |
| chr9 | 120942247 | 120974021 | 31775 | -0.05 | 0.911 | 0.915 |
| chr11 | 104707535 | 104713430 | 5896 | -0.04 | 0.911 | 0.915 |
| chr12 | 18331456 | 18332377 | 922 | 0.07 | 0.912 | 0.915 |
| chr12 | 6035487 | 6065054 | 29568 | -0.04 | 0.912 | 0.915 |
| chr5 | 137016410 | 137017390 | 981 | -0.11 | 0.913 | 0.917 |
| chr5 | 167645296 | 167646693 | 1398 | 0.07 | 0.914 | 0.917 |
| chr20 | 560703 | 562358 | 1656 | 0.05 | 0.914 | 0.918 |
| chr7 | 121050498 | 121052602 | 2105 | -0.05 | 0.914 | 0.918 |
| chr20 | 9656375 | 9678553 | 22179 | -0.04 | 0.915 | 0.918 |
| chr2 | 150221283 | 150225259 | 3977 | 0.05 | 0.915 | 0.919 |
| chr3 | 176112724 | 176114977 | 2254 | -0.08 | 0.917 | 0.92 |
| chr3 | 108482535 | 108534586 | 52052 | 0.03 | 0.918 | 0.922 |
| chr11 | 51289129 | 51340795 | 51667 | 0.02 | 0.919 | 0.922 |
| chr13 | 39648749 | 39652002 | 3254 | -0.04 | 0.919 | 0.923 |
| chr11 | 40647976 | 40650010 | 2035 | 0.08 | 0.92 | 0.923 |
| chr16 | 79760446 | 79786880 | 26435 | 0.03 | 0.921 | 0.924 |
| chr5 | 54210754 | 54236811 | 26058 | 0.03 | 0.921 | 0.924 |
| chr8 | 1123753 | 1268049 | 144297 | -0.02 | 0.921 | 0.925 |
| chr3 | 108664979 | 108666407 | 1429 | 0.05 | 0.921 | 0.925 |
| chr9 | 88036862 | 88048097 | 11236 | 0.04 | 0.922 | 0.925 |
| chr6 | 131060816 | 131063432 | 2617 | 0.06 | 0.923 | 0.926 |
| chr2 | 213085007 | 213106381 | 21375 | 0.05 | 0.924 | 0.927 |
| chr14 | 56509548 | 56511219 | 1672 | 0.04 | 0.924 | 0.927 |
| chr16 | 78706141 | 78706974 | 834 | 0.15 | 0.925 | 0.928 |
| chrX | 23540044 | 23548386 | 8343 | -0.04 | 0.926 | 0.929 |
| chr12 | 99533380 | 99735829 | 202450 | 0.02 | 0.926 | 0.929 |
| chr9 | 112532254 | 112541761 | 9508 | -0.05 | 0.927 | 0.93 |
| chrX | 131137303 | 131145132 | 7830 | -0.06 | 0.927 | 0.93 |
| chr7 | 25907444 | 25923927 | 16484 | 0.03 | 0.927 | 0.93 |
| chr1 | 210240144 | 210263829 | 23686 | 0.05 | 0.927 | 0.93 |
| chr4 | 34779703 | 34826932 | 47230 | -0.03 | 0.928 | 0.931 |
| chr20 | 39412932 | 39476063 | 63132 | 0.02 | 0.928 | 0.931 |
| chr16 | 77825844 | 78039704 | 213861 | 0.02 | 0.93 | 0.932 |
| chr1 | 69469626 | 69471572 | 1947 | -0.04 | 0.93 | 0.933 |
| chr3 | 162758602 | 162763414 | 4813 | -0.05 | 0.93 | 0.933 |
| chr16 | 20365385 | 20387494 | 22110 | -0.04 | 0.93 | 0.933 |
| chr18 | 73283115 | 73288211 | 5097 | -0.04 | 0.931 | 0.933 |
| chrX | 3240615 | 3273418 | 32804 | -0.02 | 0.933 | 0.936 |
| chr6 | 71786922 | 71803343 | 16422 | 0.07 | 0.934 | 0.937 |

| | | | | | | |
|---|---|---|---|---|---|---|
| chr4 | 61259987 | 61274619 | 14633 | 0.03 | 0.934 | 0.937 |
| chrX | 38684029 | 38698194 | 14166 | 0.03 | 0.934 | 0.937 |
| chr2 | 127443743 | 127501700 | 57958 | -0.01 | 0.935 | 0.938 |
| chr12 | 4393478 | 4413190 | 19713 | 0.05 | 0.937 | 0.939 |
| chr2 | 194834177 | 194835839 | 1663 | -0.04 | 0.938 | 0.941 |
| chrX | 134980205 | 134990571 | 10367 | 0.03 | 0.939 | 0.941 |
| chrX | 103273026 | 103305136 | 32111 | 0.02 | 0.939 | 0.941 |
| chr9 | 15076013 | 15123022 | 47010 | 0.02 | 0.94 | 0.942 |
| chr18 | 60919179 | 60948119 | 28941 | -0.04 | 0.942 | 0.945 |
| chr14 | 22590723 | 22625201 | 34479 | 0.02 | 0.942 | 0.945 |
| chr7 | 117538820 | 117540542 | 1723 | 0.05 | 0.943 | 0.946 |
| chr20 | 16590796 | 16592551 | 1756 | 0.09 | 0.943 | 0.946 |
| chr3 | 34433846 | 34446311 | 12466 | -0.03 | 0.944 | 0.946 |
| chr4 | 33230295 | 33248852 | 18558 | 0.04 | 0.944 | 0.947 |
| chr1 | 240050019 | 240071247 | 21229 | -0.15 | 0.945 | 0.947 |
| chr2 | 194163733 | 194174732 | 11000 | -0.15 | 0.945 | 0.947 |
| chr10 | 71426944 | 71441518 | 14575 | -0.03 | 0.946 | 0.949 |
| chr10 | 168361 | 169904 | 1544 | 0.03 | 0.948 | 0.95 |
| chrX | 16219709 | 16220867 | 1159 | 0.04 | 0.948 | 0.95 |
| chr16 | 10386508 | 10386970 | 463 | -0.04 | 0.948 | 0.95 |
| chr13 | 23056129 | 23066536 | 10408 | 0.02 | 0.948 | 0.95 |
| chr3 | 81220831 | 81240904 | 20074 | -0.02 | 0.948 | 0.951 |
| chr4 | 30151475 | 30170381 | 18907 | 0.03 | 0.949 | 0.951 |
| chr1 | 106435155 | 106443529 | 8375 | -0.04 | 0.949 | 0.951 |
| chr15 | 49969392 | 49970939 | 1548 | -0.04 | 0.949 | 0.952 |
| chr5 | 174386413 | 174539448 | 153036 | -0.02 | 0.95 | 0.952 |
| chr6 | 77426884 | 77436749 | 9866 | -0.03 | 0.95 | 0.953 |
| chr11 | 128707749 | 128714179 | 6431 | 0.02 | 0.951 | 0.953 |
| chr5 | 18103984 | 18105296 | 1313 | -0.05 | 0.951 | 0.953 |
| chr7 | 132270362 | 132296419 | 26058 | 0.02 | 0.952 | 0.954 |
| chr12 | 94521413 | 94524055 | 2643 | 0.03 | 0.952 | 0.954 |
| chr2 | 72032309 | 72061014 | 28706 | -0.02 | 0.952 | 0.954 |
| chr13 | 51759149 | 51788588 | 29440 | 0.02 | 0.953 | 0.955 |
| chr2 | 237824115 | 237825649 | 1535 | -0.03 | 0.955 | 0.957 |
| chr4 | 62923357 | 62925103 | 1747 | 0.04 | 0.956 | 0.958 |
| chr21 | 36719408 | 36720199 | 792 | -0.03 | 0.957 | 0.959 |
| chr9 | 103989445 | 104006664 | 17220 | 0.07 | 0.958 | 0.96 |
| chr20 | 57954375 | 57970655 | 16281 | -0.03 | 0.959 | 0.961 |
| chr11 | 133883980 | 133888248 | 4269 | 0.02 | 0.96 | 0.962 |
| chr19 | 9036897 | 9093283 | 56387 | 0.01 | 0.96 | 0.962 |
| chr13 | 53316642 | 53350700 | 34059 | 0.01 | 0.96 | 0.962 |
| chrX | 130220295 | 130241585 | 21291 | 0.01 | 0.961 | 0.963 |
| chr4 | 29445368 | 29449219 | 3852 | 0.02 | 0.961 | 0.963 |
| chr11 | 87076170 | 87117017 | 40848 | -0.01 | 0.963 | 0.964 |
| chr6 | 6041137 | 6064092 | 22956 | 0.01 | 0.963 | 0.965 |
| chr13 | 107576830 | 107578829 | 2000 | 0.03 | 0.965 | 0.966 |
| chr18 | 29340738 | 29342487 | 1750 | -0.03 | 0.965 | 0.967 |
| chr7 | 39894616 | 39894963 | 348 | 0.05 | 0.965 | 0.967 |
| chr16 | 77481119 | 77531792 | 50674 | 0.01 | 0.967 | 0.969 |
| chr2 | 118534096 | 118560835 | 26740 | 0.03 | 0.967 | 0.969 |
| chr14 | 28274701 | 28275285 | 585 | -0.02 | 0.967 | 0.969 |
| chr10 | 1622959 | 1650319 | 27361 | -0.02 | 0.969 | 0.97 |
| chr22 | 17034508 | 17035050 | 543 | 0.02 | 0.969 | 0.97 |
| chr4 | 60610387 | 60614964 | 4578 | -0.02 | 0.969 | 0.97 |
| chr2 | 29482155 | 29525721 | 43567 | -0.03 | 0.97 | 0.971 |
| chr3 | 113842095 | 113842541 | 447 | -0.04 | 0.97 | 0.971 |
| chr8 | 48016340 | 48017511 | 1172 | 0.02 | 0.97 | 0.972 |
| chr13 | 53375214 | 53388415 | 13202 | -0.02 | 0.97 | 0.972 |
| chr1 | 173682321 | 173683619 | 1299 | -0.03 | 0.971 | 0.973 |
| chr2 | 153870488 | 153873514 | 3027 | -0.03 | 0.972 | 0.973 |
| chr2 | 213149075 | 213169434 | 20360 | -0.02 | 0.972 | 0.974 |

| | | | | | | |
|---|---|---|---|---|---|---|
| chrX | 132888173 | 132899947 | 11775 | -0.01 | 0.973 | 0.974 |
| chr3 | 59932762 | 59935483 | 2722 | 0.02 | 0.973 | 0.974 |
| chr5 | 166754315 | 166765685 | 11371 | -0.01 | 0.975 | 0.976 |
| chr19 | 55026801 | 55082212 | 55412 | 0.01 | 0.975 | 0.976 |
| chr16 | 84456057 | 84458794 | 2738 | 0.01 | 0.976 | 0.977 |
| chr5 | 30601444 | 30603352 | 1909 | -0.03 | 0.976 | 0.977 |
| chr4 | 60804607 | 60851257 | 46651 | -0.02 | 0.976 | 0.977 |
| chr9 | 107125471 | 107126535 | 1065 | 0.02 | 0.976 | 0.977 |
| chr10 | 119505321 | 119582307 | 76987 | -0.01 | 0.976 | 0.978 |
| chr3 | 111124709 | 111153044 | 28336 | 0.01 | 0.977 | 0.978 |
| chr10 | 117499365 | 117511198 | 11834 | 0.01 | 0.977 | 0.978 |
| chr2 | 234812847 | 234817866 | 5020 | -0.01 | 0.978 | 0.979 |
| chr1 | 63580392 | 63582027 | 1636 | -0.02 | 0.979 | 0.979 |
| chr5 | 111013425 | 111014672 | 1248 | 0.01 | 0.979 | 0.98 |
| chr11 | 95263700 | 95267607 | 3908 | -0.01 | 0.979 | 0.98 |
| chr3 | 186085413 | 186113394 | 27982 | -0.01 | 0.98 | 0.981 |
| chr13 | 89223806 | 89228570 | 4765 | -0.02 | 0.98 | 0.981 |
| chr6 | 159615754 | 159631368 | 15615 | 0.01 | 0.98 | 0.981 |
| chr18 | 27527251 | 27535082 | 7832 | 0.01 | 0.981 | 0.982 |
| chr12 | 46752239 | 46765043 | 12805 | 0.01 | 0.981 | 0.982 |
| chr9 | 91774391 | 91776602 | 2212 | -0.01 | 0.982 | 0.983 |
| chr13 | 46186560 | 46188991 | 2432 | -0.01 | 0.984 | 0.984 |
| chr7 | 43183369 | 43186594 | 3226 | -0.01 | 0.984 | 0.984 |
| chr12 | 65311793 | 65335689 | 23897 | -0.01 | 0.984 | 0.984 |
| chr6 | 155876496 | 155880420 | 3925 | 0.01 | 0.984 | 0.984 |
| chr15 | 70112593 | 70114441 | 1849 | -0.02 | 0.984 | 0.985 |
| chr10 | 98090631 | 98091458 | 828 | -0.02 | 0.984 | 0.985 |
| chr20 | 56130677 | 56152372 | 21696 | 0.04 | 0.985 | 0.985 |
| chr14 | 58524486 | 58532268 | 7783 | 0.01 | 0.986 | 0.986 |
| chr19 | 10086063 | 10087631 | 1569 | -0.01 | 0.987 | 0.988 |
| chr1 | 152512974 | 152555305 | 42332 | 0.01 | 0.987 | 0.988 |
| chr4 | 28633945 | 28635122 | 1178 | -0.01 | 0.988 | 0.988 |
| chr18 | 6900290 | 6978306 | 78017 | 0 | 0.988 | 0.988 |
| chr5 | 115898420 | 115899526 | 1107 | 0 | 0.989 | 0.989 |
| chr6 | 13169837 | 13174642 | 4806 | 0 | 0.989 | 0.99 |
| chr11 | 121234475 | 121237754 | 3280 | 0.01 | 0.99 | 0.991 |
| chr4 | 62747075 | 62755657 | 8583 | 0 | 0.99 | 0.991 |
| chr21 | 34557585 | 34565646 | 8062 | 0 | 0.991 | 0.991 |
| chr6 | 166637563 | 166637956 | 394 | 0 | 0.992 | 0.993 |
| chr14 | 51600148 | 51600513 | 366 | 0.01 | 0.994 | 0.994 |
| chr7 | 35353899 | 35422196 | 68298 | 0 | 0.995 | 0.995 |
| chr6 | 77519044 | 77520392 | 1349 | 0 | 0.996 | 0.996 |
| chr5 | 155002961 | 155014342 | 11382 | 0 | 0.996 | 0.996 |
| chr6 | 159467618 | 159511184 | 43567 | 0 | 0.996 | 0.997 |
| chr1 | 164713063 | 164716002 | 2940 | 0 | 0.997 | 0.997 |
| chrX | 55879896 | 55884991 | 5096 | 0 | 0.998 | 0.998 |
| chr12 | 22344157 | 22367897 | 23741 | 0 | 0.999 | 0.999 |
| chr9 | 87074211 | 87135352 | 61142 | 0 | 0.999 | 0.999 |
| chr12 | 8657827 | 8691904 | 34078 | 0 | 1 | 1 |
| chrX | 42258637 | 42275270 | 16634 | 0 | 1 | 1 |

**Supplementary Table 4D, Differentially regulated super-enhancers, Related to Figure 3E and Supplementary Figure 3D**

| chrom | start | end | width | fold | pvalue | FDR |
|---|---|---|---|---|---|---|
| chr12 | 14508751 | 14629463 | 120713 | 3.22 | 0 | 0 |
| chr12 | 15835493 | 15887153 | 51661 | 2.02 | 0 | 0 |
| chr12 | 50594926 | 50721719 | 126794 | 2.66 | 0 | 0 |
| chr12 | 87805678 | 87836759 | 31082 | 5.25 | 0 | 0 |
| chr12 | 110974019 | 111024065 | 50047 | 3.06 | 0 | 0 |
| chr15 | 34423960 | 34502327 | 78368 | 3.04 | 0 | 0 |
| chr15 | 59824759 | 59950480 | 125722 | 1.25 | 0 | 0 |
| chr18 | 70912667 | 70964388 | 51722 | 3.65 | 0 | 0 |
| chr2 | 86916056 | 86934041 | 17986 | 3.34 | 0 | 0 |
| chr7 | 11624053 | 11711065 | 87013 | 2.12 | 0 | 0 |
| chr9 | 119026637 | 119162338 | 135702 | -3.71 | 0 | 0 |
| chr2 | 121515806 | 121752153 | 236348 | -1.33 | 2.17E-299 | 1.95E-297 |
| chr7 | 111124087 | 111203295 | 79209 | 1.72 | 1.58E-290 | 1.31E-288 |
| chr12 | 75337000 | 75360848 | 23849 | 2.5 | 1.83E-268 | 1.41E-266 |
| chr12 | 13015954 | 13083646 | 67693 | 1.24 | 3.22E-261 | 2.31E-259 |
| chr11 | 1050877 | 1078511 | 27635 | 1.95 | 2.46E-257 | 1.65E-255 |
| chr6 | 85443376 | 85487265 | 43890 | 2.01 | 5.69E-251 | 3.60E-249 |
| chr14 | 92198685 | 92247386 | 48702 | 3.62 | 1.43E-245 | 8.54E-244 |
| chr7 | 11723817 | 11874265 | 150449 | 1.23 | 4.41E-227 | 2.49E-225 |
| chr12 | 105702855 | 105752037 | 49183 | 2 | 1.64E-217 | 8.82E-216 |
| chr9 | 75635678 | 75710042 | 74365 | 1.07 | 6.17E-213 | 3.15E-211 |
| chr5 | 111541389 | 111576504 | 35116 | 2.11 | 2.64E-211 | 1.29E-209 |
| chr12 | 32108114 | 32163939 | 55826 | 1.93 | 1.76E-210 | 8.22E-209 |
| chr11 | 75022685 | 75141865 | 119181 | 1.27 | 6.28E-205 | 2.81E-203 |
| chr9 | 89984377 | 90041003 | 56627 | 1.35 | 2.41E-204 | 1.04E-202 |
| chr12 | 123346298 | 123383555 | 37258 | 1.93 | 1.18E-203 | 4.87E-202 |
| chr7 | 131185190 | 131242238 | 57049 | 2.15 | 3.36E-201 | 1.34E-199 |
| chr2 | 228189125 | 228281915 | 92791 | -1.41 | 3.41E-200 | 1.31E-198 |
| chr19 | 54023240 | 54083313 | 60074 | 2.05 | 9.16E-200 | 3.39E-198 |
| chr11 | 45273751 | 45311604 | 37854 | 2.32 | 1.27E-192 | 4.55E-191 |
| chr12 | 122177888 | 122278747 | 100860 | 1.11 | 2.84E-191 | 9.83E-190 |
| chr3 | 14443317 | 14532444 | 89128 | -1.33 | 1.25E-190 | 4.20E-189 |
| chr12 | 31856839 | 31933500 | 76662 | 1.39 | 9.96E-190 | 3.24E-188 |
| chr19 | 39105670 | 39227409 | 121740 | 0.74 | 6.75E-189 | 2.13E-187 |
| chr9 | 21466240 | 21617400 | 151161 | 0.98 | 2.51E-188 | 7.70E-187 |
| chr16 | 52579061 | 52613779 | 34719 | 2.73 | 5.33E-175 | 1.59E-173 |
| chr2 | 46858939 | 46883570 | 24632 | -1.79 | 2.34E-174 | 6.80E-173 |
| chr12 | 47463349 | 47500110 | 36762 | 1 | 3.56E-174 | 1.01E-172 |
| chr12 | 105338392 | 105354000 | 15609 | 2.02 | 4.58E-172 | 1.26E-170 |
| chr12 | 95461278 | 95734708 | 273431 | 0.73 | 2.38E-162 | 6.39E-161 |
| chr1 | 51424601 | 51451782 | 27182 | 1.47 | 5.28E-162 | 1.38E-160 |
| chr12 | 41082176 | 41123105 | 40930 | 1.48 | 7.12E-161 | 1.82E-159 |
| chr7 | 137973131 | 138039618 | 66488 | 2.65 | 1.36E-151 | 3.39E-150 |
| chr12 | 89884325 | 89920794 | 36470 | 1.51 | 1.06E-149 | 2.59E-148 |
| chr12 | 62972724 | 63060860 | 88137 | 1.07 | 4.95E-148 | 1.18E-146 |

| | | | | | | |
|---|---|---|---|---|---|---|
| chr11 | 100605436 | 100663607 | 58172 | 1.38 | 8.03E-140 | 1.88E-138 |
| chr2 | 26930734 | 27014403 | 83670 | -1.21 | 7.83E-139 | 1.79E-137 |
| chr9 | 90111828 | 90342920 | 231093 | 0.86 | 3.59E-138 | 8.03E-137 |
| chr8 | 30240017 | 30438997 | 198981 | 0.88 | 7.06E-136 | 1.55E-134 |
| chr5 | 175227100 | 175317570 | 90471 | -1.11 | 3.77E-135 | 8.10E-134 |
| chr1 | 150626581 | 150670983 | 44403 | 1.71 | 4.24E-135 | 8.92E-134 |
| chr2 | 234597302 | 234620659 | 23358 | -1.4 | 6.79E-135 | 1.40E-133 |
| chr12 | 112442970 | 112592297 | 149328 | 0.97 | 2.63E-133 | 5.32E-132 |
| chr13 | 67757035 | 67806157 | 49123 | 1.24 | 2.98E-132 | 5.93E-131 |
| chr12 | 95998692 | 96063505 | 64814 | 1.52 | 6.41E-132 | 1.25E-130 |
| chr1 | 84711788 | 84770013 | 58226 | 1.87 | 9.62E-131 | 1.84E-129 |
| chr5 | 142149059 | 142260991 | 111933 | 0.97 | 3.36E-128 | 6.33E-127 |
| chr12 | 79908021 | 79967015 | 58995 | 1.24 | 2.22E-125 | 4.11E-124 |
| chr12 | 12931954 | 13001053 | 69100 | 1.07 | 3.05E-125 | 5.56E-124 |
| chr3 | 30360118 | 30415850 | 55733 | 1.24 | 7.67E-125 | 1.37E-123 |
| chr20 | 47883574 | 47909864 | 26291 | 1.33 | 2.67E-123 | 4.70E-122 |
| chr2 | 189143981 | 189189323 | 45343 | 1.36 | 2.87E-121 | 4.97E-120 |
| chr9 | 21675040 | 21690822 | 15783 | 0.94 | 6.03E-121 | 1.03E-119 |
| chr20 | 36678045 | 36816165 | 138121 | 1.04 | 4.49E-120 | 7.54E-119 |
| chr17 | 73584907 | 73785345 | 200439 | 0.7 | 1.66E-119 | 2.74E-118 |
| chr6 | 128799446 | 128842876 | 43431 | 1.43 | 1.22E-116 | 1.98E-115 |
| chr2 | 47166410 | 47317390 | 150981 | -0.73 | 1.94E-116 | 3.12E-115 |
| chr5 | 42982060 | 43021416 | 39357 | 1.12 | 4.33E-116 | 6.84E-115 |
| chr10 | 95046712 | 95276079 | 229368 | 0.6 | 1.54E-115 | 2.40E-114 |
| chr8 | 23145602 | 23267977 | 122376 | -1.08 | 3.35E-115 | 5.14E-114 |
| chr20 | 60903810 | 60964836 | 61027 | 0.96 | 4.70E-111 | 7.10E-110 |
| chr10 | 126662301 | 126851531 | 189231 | 0.71 | 6.79E-111 | 1.01E-109 |
| chr15 | 69590281 | 69756337 | 166057 | -1.03 | 1.24E-108 | 1.82E-107 |
| chr2 | 235859835 | 235965263 | 105429 | -1.26 | 1.61E-106 | 2.33E-105 |
| chr14 | 95697852 | 95740338 | 42487 | 1.35 | 2.15E-106 | 3.08E-105 |
| chr1 | 207165964 | 207227223 | 61260 | 1.05 | 7.15E-106 | 1.01E-104 |
| chr12 | 12866397 | 12880189 | 13793 | 1.27 | 5.41E-104 | 7.55E-103 |
| chr17 | 15790006 | 15825723 | 35718 | 1.58 | 7.79E-103 | 1.07E-101 |
| chr14 | 55564101 | 55576900 | 12800 | 1.35 | 5.17E-102 | 7.03E-101 |
| chr3 | 146219041 | 146268352 | 49312 | 1.07 | 4.13E-101 | 5.55E-100 |
| chr5 | 71402400 | 71479301 | 76902 | -1.73 | 2.65E-100 | 3.52E-99 |
| chr14 | 103697992 | 103768594 | 70603 | 1.61 | 4.03E-100 | 5.28E-99 |
| chr16 | 66535966 | 66559183 | 23218 | 1.12 | 7.66E-100 | 9.91E-99 |
| chr10 | 93347301 | 93377615 | 30315 | 1.06 | 2.07E-99 | 2.65E-98 |
| chr12 | 63077029 | 63195151 | 118123 | 0.67 | 7.01E-99 | 8.86E-98 |
| chr2 | 161195310 | 161275877 | 80568 | -1.13 | 8.24E-99 | 1.03E-97 |
| chr12 | 56910033 | 57040676 | 130644 | 0.81 | 6.35E-97 | 7.84E-96 |
| chr18 | 67953891 | 68054695 | 100805 | 0.76 | 8.60E-97 | 1.05E-95 |
| chr8 | 128210581 | 128238341 | 27761 | 1.42 | 1.32E-96 | 1.59E-95 |
| chr9 | 97738129 | 97883827 | 145699 | 0.63 | 1.93E-95 | 2.31E-94 |
| chr4 | 77608902 | 77646566 | 37665 | 1.23 | 2.06E-95 | 2.43E-94 |
| chr21 | 30539992 | 30591090 | 51099 | 1.72 | 2.67E-94 | 3.12E-93 |
| chr10 | 105643785 | 105679366 | 35582 | 1.13 | 1.00E-93 | 1.16E-92 |
| chr15 | 93340059 | 93562582 | 222524 | 0.65 | 4.71E-93 | 5.38E-92 |
| chr12 | 51155737 | 51185976 | 30240 | 1.19 | 6.81E-93 | 7.70E-92 |
| chr9 | 75440898 | 75576601 | 135704 | 0.68 | 4.01E-92 | 4.47E-91 |
| chr7 | 36426276 | 36489460 | 63185 | 1.21 | 4.04E-92 | 4.47E-91 |
| chr9 | 75591811 | 75622667 | 30857 | 0.72 | 4.11E-92 | 4.50E-91 |
| chr15 | 72020752 | 72107256 | 86505 | -0.89 | 3.58E-89 | 3.88E-88 |
| chr3 | 129547421 | 129655807 | 108387 | 1.06 | 1.44E-88 | 1.55E-87 |
| chr22 | 37892896 | 37960821 | 67926 | 1.1 | 1.94E-87 | 2.06E-86 |
| chr12 | 98883763 | 98942922 | 59160 | 1.06 | 6.06E-87 | 6.38E-86 |
| chr13 | 67642518 | 67725357 | 82840 | 0.77 | 1.09E-86 | 1.14E-85 |
| chr15 | 52541693 | 52588933 | 47241 | 0.99 | 2.84E-86 | 2.93E-85 |

| | | | | | | |
|---|---|---|---|---|---|---|
| chr12 | 53921038 | 53970820 | 49783 | 1.43 | 7.27E-86 | 7.44E-85 |
| chr12 | 110537659 | 110603049 | 65391 | 0.95 | 1.54E-83 | 1.56E-82 |
| chr17 | 70382128 | 70478237 | 96110 | 0.6 | 5.73E-83 | 5.75E-82 |
| chr8 | 101469509 | 101525364 | 55856 | 1.11 | 3.83E-82 | 3.81E-81 |
| chr12 | 6469686 | 6542439 | 72754 | 0.68 | 4.20E-82 | 4.14E-81 |
| chr12 | 110630326 | 110842616 | 212291 | 0.64 | 6.80E-82 | 6.64E-81 |
| chr10 | 46965284 | 47151348 | 186065 | -0.55 | 1.90E-81 | 1.84E-80 |
| chr9 | 75723959 | 75800214 | 76256 | 0.65 | 2.44E-80 | 2.34E-79 |
| chr5 | 158584015 | 158639457 | 55443 | 0.67 | 3.30E-80 | 3.13E-79 |
| chr3 | 193829031 | 193860958 | 31928 | 2.15 | 6.84E-80 | 6.45E-79 |
| chr4 | 15653186 | 15704984 | 51799 | 1 | 3.68E-79 | 3.44E-78 |
| chr2 | 208528452 | 208616026 | 87575 | -1.32 | 1.74E-78 | 1.61E-77 |
| chr17 | 53471855 | 53537517 | 65663 | -1.04 | 1.91E-78 | 1.75E-77 |
| chr1 | 98490955 | 98532942 | 41988 | 1.23 | 3.63E-78 | 3.30E-77 |
| chr1 | 200040121 | 200099883 | 59763 | 1.89 | 5.58E-78 | 5.04E-77 |
| chr12 | 125185140 | 125254348 | 69209 | 1.53 | 1.23E-76 | 1.10E-75 |
| chr8 | 32402709 | 32628986 | 226278 | -0.59 | 2.43E-76 | 2.16E-75 |
| chr15 | 101458403 | 101553117 | 94715 | -0.97 | 5.71E-76 | 5.03E-75 |
| chr12 | 104642571 | 104772290 | 129720 | 0.65 | 2.73E-75 | 2.39E-74 |
| chr3 | 160108471 | 160169403 | 60933 | 0.94 | 7.17E-75 | 6.21E-74 |
| chr12 | 109007185 | 109130187 | 123003 | 0.65 | 4.69E-74 | 4.03E-73 |
| chr12 | 6386823 | 6455832 | 69010 | 0.84 | 6.28E-74 | 5.36E-73 |
| chr10 | 43799635 | 43858813 | 59179 | 1.01 | 9.21E-74 | 7.78E-73 |
| chr10 | 80826359 | 81010399 | 184041 | -0.57 | 5.33E-73 | 4.48E-72 |
| chr7 | 16830768 | 16866344 | 35577 | 0.91 | 5.68E-73 | 4.73E-72 |
| chr12 | 64236575 | 64384203 | 147629 | 0.59 | 8.43E-73 | 6.96E-72 |
| chr12 | 113611039 | 113710430 | 99392 | 0.93 | 5.93E-72 | 4.86E-71 |
| chr2 | 234258991 | 234406019 | 147029 | -0.69 | 9.35E-72 | 7.60E-71 |
| chr8 | 101312051 | 101364739 | 52689 | 1.16 | 2.06E-70 | 1.67E-69 |
| chr15 | 55650269 | 55702920 | 52652 | 0.99 | 2.01E-69 | 1.61E-68 |
| chr12 | 10514208 | 10550887 | 36680 | 3.15 | 5.02E-69 | 4.00E-68 |
| chr9 | 131625489 | 131690762 | 65274 | 0.83 | 1.50E-68 | 1.19E-67 |
| chr14 | 68972619 | 69075959 | 103341 | 0.89 | 2.42E-68 | 1.90E-67 |
| chr20 | 30248315 | 30352470 | 104156 | 0.54 | 2.85E-68 | 2.22E-67 |
| chr11 | 12064947 | 12112490 | 47544 | -0.97 | 4.94E-68 | 3.81E-67 |
| chr12 | 47575437 | 47676690 | 101254 | 0.58 | 1.90E-67 | 1.46E-66 |
| chr10 | 33176590 | 33306187 | 129598 | 0.54 | 2.43E-67 | 1.85E-66 |
| chr3 | 70297437 | 70333377 | 35941 | 1.06 | 3.53E-67 | 2.67E-66 |
| chr12 | 53758559 | 53903271 | 144713 | 0.7 | 6.74E-67 | 5.06E-66 |
| chr1 | 39849861 | 39877795 | 27935 | 1 | 7.10E-67 | 5.29E-66 |
| chr19 | 16177156 | 16223984 | 46829 | 0.94 | 6.17E-66 | 4.57E-65 |
| chr12 | 120612882 | 120715470 | 102589 | 0.7 | 1.17E-65 | 8.62E-65 |
| chr12 | 50897998 | 50992076 | 94079 | 0.89 | 2.16E-65 | 1.58E-64 |
| chr12 | 21787004 | 21828953 | 41950 | 1.01 | 2.84E-65 | 2.06E-64 |
| chr4 | 57894036 | 57985896 | 91861 | 1.05 | 3.86E-65 | 2.78E-64 |
| chr19 | 44143474 | 44210002 | 66529 | 0.94 | 4.05E-65 | 2.90E-64 |
| chr14 | 64319054 | 64391226 | 72173 | 1.27 | 9.90E-65 | 7.04E-64 |
| chr2 | 233699954 | 233767742 | 67789 | -1.27 | 1.74E-64 | 1.23E-63 |
| chr2 | 73247476 | 73341011 | 93536 | -1.07 | 1.57E-63 | 1.10E-62 |
| chr7 | 16920268 | 16976073 | 55806 | 1.27 | 5.82E-63 | 4.06E-62 |
| chr9 | 78979315 | 79039500 | 60186 | 0.77 | 1.05E-62 | 7.29E-62 |
| chrX | 106160283 | 106203160 | 42878 | -1.92 | 1.57E-61 | 1.08E-60 |
| chr15 | 52121436 | 52201158 | 79723 | 0.53 | 1.61E-61 | 1.10E-60 |
| chr1 | 201974746 | 202092396 | 117651 | 0.61 | 1.68E-61 | 1.14E-60 |
| chr12 | 122580659 | 122615989 | 35331 | 0.97 | 3.96E-61 | 2.68E-60 |
| chr12 | 76302643 | 76388956 | 86314 | 0.66 | 4.22E-61 | 2.82E-60 |
| chr17 | 76123719 | 76195956 | 72238 | 1.08 | 4.23E-61 | 2.82E-60 |
| chr16 | 19395521 | 19458247 | 62727 | 0.87 | 2.00E-60 | 1.33E-59 |
| chr5 | 179659187 | 179780996 | 121810 | 2.89 | 2.90E-60 | 1.91E-59 |
| chr3 | 101874488 | 101915671 | 41184 | 0.97 | 3.01E-60 | 1.97E-59 |
| chr3 | 171509381 | 171620305 | 110925 | 0.75 | 1.99E-59 | 1.30E-58 |

| | | | | | | |
|---|---|---|---|---|---|---|
| chr18 | 8698939 | 8831316 | 132378 | 0.62 | 9.54E-59 | 6.17E-58 |
| chr4 | 79472126 | 79493255 | 21130 | 0.94 | 2.10E-58 | 1.35E-57 |
| chr11 | 12131388 | 12205276 | 73889 | -1.61 | 4.17E-58 | 2.67E-57 |
| chr7 | 134198960 | 134250761 | 51802 | -0.77 | 8.55E-58 | 5.44E-57 |
| chr1 | 234733610 | 234795850 | 62241 | 0.62 | 1.03E-56 | 6.51E-56 |
| chr20 | 22355521 | 22411991 | 56471 | -0.97 | 1.27E-56 | 7.96E-56 |
| chr17 | 72977465 | 73209597 | 232133 | 0.47 | 2.68E-56 | 1.68E-55 |
| chr12 | 46823300 | 46889621 | 66322 | 0.78 | 8.10E-56 | 5.03E-55 |
| chr8 | 29444955 | 29633784 | 188830 | 0.52 | 1.62E-55 | 1.00E-54 |
| chr11 | 33710594 | 33759481 | 48888 | 0.85 | 1.76E-55 | 1.08E-54 |
| chr12 | 65001248 | 65101354 | 100107 | 0.6 | 5.61E-55 | 3.43E-54 |
| chr5 | 72132747 | 72204211 | 71465 | -0.77 | 6.91E-55 | 4.20E-54 |
| chr7 | 130816412 | 130883504 | 67093 | -0.71 | 2.90E-54 | 1.75E-53 |
| chr5 | 127243576 | 127330400 | 86825 | 0.81 | 4.32E-54 | 2.59E-53 |
| chr22 | 46447984 | 46512822 | 64839 | 0.61 | 4.76E-54 | 2.84E-53 |
| chr5 | 39071961 | 39139700 | 67740 | -0.79 | 5.64E-54 | 3.35E-53 |
| chr2 | 234725951 | 234777560 | 51610 | -1.04 | 7.47E-54 | 4.41E-53 |
| chr9 | 118909921 | 119014091 | 104171 | -2.08 | 2.07E-52 | 1.21E-51 |
| chr17 | 39042400 | 39079763 | 37364 | 0.77 | 4.55E-52 | 2.65E-51 |
| chr7 | 134103319 | 134168615 | 65297 | -0.78 | 8.15E-52 | 4.73E-51 |
| chr2 | 228314245 | 228433813 | 119569 | -0.7 | 7.82E-51 | 4.51E-50 |
| chr4 | 169707682 | 169787432 | 79751 | -1.16 | 1.40E-50 | 8.02E-50 |
| chr16 | 14568441 | 14624579 | 56139 | -0.84 | 1.57E-50 | 8.97E-50 |
| chr11 | 61447335 | 61498475 | 51141 | 0.95 | 1.17E-49 | 6.62E-49 |
| chr16 | 1727478 | 1780302 | 52825 | 0.79 | 1.68E-49 | 9.49E-49 |
| chr9 | 91002064 | 91076336 | 74273 | 1.04 | 2.99E-49 | 1.68E-48 |
| chr22 | 30581480 | 30659737 | 78258 | 0.56 | 3.61E-49 | 2.02E-48 |
| chr3 | 30267279 | 30306070 | 38792 | 0.88 | 5.71E-48 | 3.18E-47 |
| chr14 | 50318041 | 50387729 | 69689 | 0.8 | 4.55E-47 | 2.52E-46 |
| chr1 | 231512030 | 231561586 | 49557 | 0.72 | 1.51E-46 | 8.32E-46 |
| chr16 | 53765785 | 53815992 | 50208 | 0.92 | 3.35E-46 | 1.84E-45 |
| chr22 | 36717807 | 36802108 | 84302 | 0.56 | 6.17E-46 | 3.36E-45 |
| chr12 | 106471472 | 106534409 | 62938 | -0.73 | 1.44E-45 | 7.80E-45 |
| chr10 | 4692190 | 4747184 | 54995 | 0.69 | 2.97E-45 | 1.60E-44 |
| chr20 | 52257475 | 52278624 | 21150 | 0.74 | 3.03E-45 | 1.62E-44 |
| chr6 | 39152499 | 39197240 | 44742 | 1.14 | 3.79E-45 | 2.03E-44 |
| chr8 | 142237532 | 142303213 | 65682 | -0.95 | 4.79E-45 | 2.55E-44 |
| chr17 | 76296118 | 76357397 | 61280 | -0.86 | 5.53E-45 | 2.92E-44 |
| chr4 | 139078799 | 139198012 | 119214 | 0.45 | 7.70E-45 | 4.05E-44 |
| chr10 | 95750674 | 95803681 | 53008 | -1.19 | 1.19E-44 | 6.23E-44 |
| chr2 | 47073555 | 47117040 | 43486 | -0.96 | 1.22E-44 | 6.35E-44 |
| chr15 | 60632600 | 60744218 | 111619 | 0.44 | 1.78E-44 | 9.23E-44 |
| chr21 | 38251886 | 38302194 | 50309 | -0.78 | 6.84E-44 | 3.53E-43 |
| chr16 | 53075418 | 53182611 | 107194 | 0.64 | 3.96E-43 | 2.03E-42 |
| chr11 | 67873245 | 67918265 | 45021 | 1.03 | 3.98E-43 | 2.04E-42 |
| chr12 | 52533841 | 52686148 | 152308 | 0.45 | 4.46E-43 | 2.27E-42 |
| chr20 | 49031146 | 49068559 | 37414 | 0.88 | 7.55E-43 | 3.83E-42 |
| chr12 | 96176095 | 96203764 | 27670 | 0.7 | 8.02E-43 | 4.04E-42 |
| chr4 | 38664385 | 38698431 | 34047 | 0.82 | 8.98E-43 | 4.51E-42 |
| chr6 | 142600288 | 142719699 | 119412 | 0.56 | 2.39E-42 | 1.19E-41 |
| chr12 | 53253080 | 53455825 | 202746 | 0.92 | 2.53E-42 | 1.26E-41 |
| chr18 | 55394941 | 55488181 | 93241 | 0.69 | 3.15E-42 | 1.56E-41 |
| chr19 | 23683883 | 23755672 | 71790 | 13.89 | 3.71E-42 | 1.83E-41 |
| chrX | 118601851 | 118646624 | 44774 | 0.84 | 2.51E-41 | 1.23E-40 |
| chr5 | 95949215 | 96127482 | 178268 | 0.54 | 2.76E-41 | 1.35E-40 |
| chr16 | 24795053 | 24867902 | 72850 | -0.68 | 5.86E-41 | 2.85E-40 |
| chr3 | 172463398 | 172532214 | 68817 | 0.56 | 8.09E-41 | 3.91E-40 |
| chr3 | 105082759 | 105122613 | 39855 | 0.64 | 1.77E-40 | 8.53E-40 |
| chr6 | 10399675 | 10465031 | 65357 | 0.7 | 1.89E-40 | 9.08E-40 |
| chrX | 114791412 | 114860761 | 69350 | 0.68 | 6.34E-40 | 3.03E-39 |
| chr2 | 47371914 | 47422607 | 50694 | -0.69 | 8.43E-40 | 4.01E-39 |

| | | | | | | |
|---|---|---|---|---|---|---|
| chr15 | 71476445 | 71765585 | 289141 | -0.38 | 8.59E-40 | 4.06E-39 |
| chr21 | 44815591 | 44941960 | 126370 | 0.48 | 9.14E-40 | 4.31E-39 |
| chr2 | 43202440 | 43284285 | 81846 | 2.06 | 9.29E-40 | 4.36E-39 |
| chr14 | 102720545 | 102808221 | 87677 | 0.61 | 9.62E-40 | 4.49E-39 |
| chr8 | 48419378 | 48457360 | 37983 | 0.64 | 1.35E-39 | 6.29E-39 |
| chr3 | 185033211 | 185081564 | 48354 | 0.78 | 1.65E-39 | 7.64E-39 |
| chr7 | 27120369 | 27224521 | 104153 | 0.88 | 2.00E-39 | 9.24E-39 |
| chr15 | 89157737 | 89193308 | 35572 | 0.78 | 2.01E-39 | 9.24E-39 |
| chr19 | 33488334 | 33610090 | 121757 | 0.43 | 6.26E-39 | 2.86E-38 |
| chr1 | 178985887 | 179068060 | 82174 | -0.81 | 1.38E-38 | 6.27E-38 |
| chr11 | 123300952 | 123376784 | 75833 | -0.71 | 1.46E-38 | 6.62E-38 |
| chr4 | 95560208 | 95627092 | 66885 | 0.5 | 2.47E-38 | 1.12E-37 |
| chr4 | 95671133 | 95711939 | 40807 | 0.65 | 3.94E-38 | 1.77E-37 |
| chr4 | 39024069 | 39056794 | 32726 | 0.61 | 4.35E-38 | 1.95E-37 |
| chr17 | 60855231 | 60902909 | 47679 | 0.7 | 5.63E-38 | 2.51E-37 |
| chr16 | 24917329 | 25006871 | 89543 | -0.98 | 6.79E-38 | 3.01E-37 |
| chr11 | 102175105 | 102195002 | 19898 | 0.68 | 1.02E-37 | 4.51E-37 |
| chr15 | 41522129 | 41578186 | 56058 | 0.66 | 1.45E-37 | 6.37E-37 |
| chr11 | 86406638 | 86456745 | 50108 | 0.67 | 1.49E-37 | 6.54E-37 |
| chr17 | 19960386 | 20014376 | 53991 | -0.64 | 2.99E-37 | 1.30E-36 |
| chr7 | 107818727 | 107950135 | 131409 | -13.02 | 3.06E-37 | 1.33E-36 |
| chr19 | 23518964 | 23621534 | 102571 | 13.01 | 3.42E-37 | 1.48E-36 |
| chr12 | 110462825 | 110498824 | 36000 | 0.88 | 6.52E-37 | 2.81E-36 |
| chr9 | 118683537 | 118741604 | 58068 | -0.6 | 9.57E-37 | 4.11E-36 |
| chr11 | 122499916 | 122613410 | 113495 | -0.63 | 1.83E-36 | 7.83E-36 |
| chr19 | 37957055 | 38025237 | 68183 | 12.87 | 1.99E-36 | 8.48E-36 |
| chr7 | 93515035 | 93554104 | 39070 | 0.72 | 2.43E-36 | 1.03E-35 |
| chr8 | 8455999 | 8522831 | 66833 | 0.75 | 3.45E-36 | 1.46E-35 |
| chr19 | 56866492 | 56916403 | 49912 | 12.83 | 3.46E-36 | 1.46E-35 |
| chr5 | 159324748 | 159356111 | 31364 | 0.7 | 4.09E-36 | 1.72E-35 |
| chr14 | 64853309 | 64933043 | 79735 | 0.65 | 7.27E-36 | 3.04E-35 |
| chr17 | 19606533 | 19670207 | 63675 | -0.57 | 1.76E-35 | 7.35E-35 |
| chr15 | 67315787 | 67491898 | 176112 | 0.31 | 2.16E-35 | 8.98E-35 |
| chr17 | 39652470 | 39706532 | 54063 | 0.78 | 3.15E-35 | 1.30E-34 |
| chr8 | 74204149 | 74277107 | 72959 | 0.72 | 5.41E-35 | 2.23E-34 |
| chr1 | 27851810 | 27929344 | 77535 | 0.64 | 6.50E-35 | 2.66E-34 |
| chr6 | 36482209 | 36585298 | 103090 | 0.64 | 8.86E-35 | 3.62E-34 |
| chr19 | 23904555 | 24008573 | 104019 | 12.55 | 1.09E-34 | 4.44E-34 |
| chr3 | 187869983 | 188015032 | 145050 | 0.46 | 1.54E-34 | 6.24E-34 |
| chr6 | 56486609 | 56598453 | 111845 | 0.63 | 1.70E-34 | 6.88E-34 |
| chr8 | 22831700 | 22964330 | 132631 | 0.59 | 1.87E-34 | 7.52E-34 |
| chrX | 115399382 | 115417652 | 18271 | 12.48 | 2.53E-34 | 1.01E-33 |
| chr16 | 70703816 | 70816669 | 112854 | -0.6 | 2.54E-34 | 1.02E-33 |
| chr2 | 26161352 | 26368855 | 207504 | -0.42 | 2.87E-34 | 1.14E-33 |
| chr15 | 70784452 | 70802923 | 18472 | 0.64 | 5.53E-34 | 2.19E-33 |
| chr6 | 106038916 | 106059933 | 21018 | -0.69 | 8.10E-34 | 3.20E-33 |
| chr10 | 123792387 | 123831089 | 38703 | 0.67 | 1.02E-33 | 4.01E-33 |
| chr12 | 109162085 | 109252253 | 90169 | 0.46 | 1.33E-33 | 5.20E-33 |
| chr1 | 16447117 | 16516816 | 69700 | 0.52 | 2.81E-33 | 1.10E-32 |
| chr9 | 132489153 | 132545011 | 55859 | 0.63 | 3.15E-33 | 1.23E-32 |
| chr19 | 38117740 | 38162413 | 44674 | 12.28 | 3.32E-33 | 1.29E-32 |
| chr3 | 112464808 | 112543464 | 78657 | 0.61 | 3.66E-33 | 1.41E-32 |
| chr19 | 37281841 | 37342653 | 60813 | 12.26 | 4.40E-33 | 1.69E-32 |
| chr3 | 30636004 | 30739590 | 103587 | 0.47 | 8.90E-33 | 3.41E-32 |
| chr7 | 31962672 | 32002573 | 39902 | -0.65 | 1.07E-32 | 4.07E-32 |
| chr12 | 80028728 | 80087613 | 58886 | 0.63 | 1.52E-32 | 5.78E-32 |
| chr3 | 12327958 | 12361501 | 33544 | 0.66 | 1.52E-32 | 5.78E-32 |
| chr16 | 74760049 | 74809463 | 49415 | 0.63 | 1.81E-32 | 6.86E-32 |
| chr7 | 158575373 | 158623720 | 48348 | 0.5 | 2.32E-32 | 8.74E-32 |
| chr19 | 53424937 | 53450826 | 25890 | 12.11 | 2.51E-32 | 9.42E-32 |
| chr8 | 32160403 | 32208843 | 48441 | -0.62 | 2.63E-32 | 9.83E-32 |

| | | | | | | |
|---|---|---|---|---|---|---|
| chr3 | 114425202 | 114494969 | 69768 | 0.93 | 3.71E-32 | 1.38E-31 |
| chr7 | 108038936 | 108097275 | 58340 | -12.07 | 3.74E-32 | 1.39E-31 |
| chr6 | 72084958 | 72131838 | 46881 | 0.58 | 6.80E-32 | 2.52E-31 |
| chr3 | 110650830 | 110715017 | 64188 | 12.02 | 8.17E-32 | 3.01E-31 |
| chr4 | 75228456 | 75250002 | 21547 | 0.72 | 9.61E-32 | 3.53E-31 |
| chr6 | 85591253 | 85621774 | 30522 | 12 | 1.10E-31 | 4.03E-31 |
| chr6 | 127420891 | 127515320 | 94430 | -11.98 | 1.17E-31 | 4.27E-31 |
| chr12 | 6975787 | 7090365 | 114579 | 0.49 | 1.38E-31 | 5.02E-31 |
| chr14 | 105423784 | 105512737 | 88954 | 0.55 | 2.10E-31 | 7.61E-31 |
| chr1 | 179090189 | 179145390 | 55202 | 0.78 | 2.44E-31 | 8.81E-31 |
| chr10 | 5584118 | 5602393 | 18276 | 0.75 | 2.52E-31 | 9.07E-31 |
| chrX | 115357993 | 115373936 | 15944 | 11.92 | 2.99E-31 | 1.07E-30 |
| chr1 | 8062189 | 8160939 | 98751 | 0.51 | 6.23E-31 | 2.23E-30 |
| chr12 | 124812002 | 124949101 | 137100 | 0.47 | 6.80E-31 | 2.42E-30 |
| chr15 | 86176810 | 86281309 | 104500 | 0.57 | 6.83E-31 | 2.43E-30 |
| chr3 | 118889587 | 118960613 | 71027 | 0.39 | 9.74E-31 | 3.45E-30 |
| chr5 | 43034676 | 43079524 | 44849 | 0.66 | 1.31E-30 | 4.64E-30 |
| chr5 | 52283101 | 52335092 | 51992 | -0.96 | 1.53E-30 | 5.40E-30 |
| chrX | 93834020 | 93861828 | 27809 | 11.77 | 2.15E-30 | 7.54E-30 |
| chr11 | 86278004 | 86319991 | 41988 | 0.71 | 2.31E-30 | 8.09E-30 |
| chr8 | 126271327 | 126322657 | 51331 | 0.81 | 2.36E-30 | 8.22E-30 |
| chr11 | 65183215 | 65278982 | 95768 | 0.36 | 2.43E-30 | 8.45E-30 |
| chr18 | 70982643 | 71008776 | 26134 | 11.75 | 3.05E-30 | 1.06E-29 |
| chr5 | 95827452 | 95875641 | 48190 | 2.68 | 3.06E-30 | 1.06E-29 |
| chr5 | 115208260 | 115277561 | 69302 | 11.75 | 3.18E-30 | 1.09E-29 |
| chr8 | 95416010 | 95488019 | 72010 | 0.55 | 4.31E-30 | 1.48E-29 |
| chr17 | 36056777 | 36131047 | 74271 | 0.47 | 4.59E-30 | 1.57E-29 |
| chr1 | 150509901 | 150553190 | 43290 | 0.64 | 4.68E-30 | 1.60E-29 |
| chr10 | 81906856 | 81969426 | 62571 | 0.54 | 4.71E-30 | 1.60E-29 |
| chr9 | 4114544 | 4180803 | 66260 | -0.57 | 6.97E-30 | 2.36E-29 |
| chr4 | 90164399 | 90238294 | 73896 | -0.57 | 9.57E-30 | 3.23E-29 |
| chr1 | 164704911 | 164775489 | 70579 | -0.63 | 1.20E-29 | 4.04E-29 |
| chr10 | 5110167 | 5152135 | 41969 | -0.5 | 1.22E-29 | 4.08E-29 |
| chr10 | 5022940 | 5084986 | 62047 | -0.6 | 1.39E-29 | 4.65E-29 |
| chr3 | 98525264 | 98644797 | 119534 | 0.44 | 1.50E-29 | 5.01E-29 |
| chr10 | 74357753 | 74515230 | 157478 | 0.39 | 1.74E-29 | 5.79E-29 |
| chr1 | 151940227 | 151974305 | 34079 | 0.48 | 1.84E-29 | 6.11E-29 |
| chr11 | 114150486 | 114189411 | 38926 | 0.58 | 1.86E-29 | 6.15E-29 |
| chr6 | 159184168 | 159241325 | 57158 | 0.69 | 2.27E-29 | 7.46E-29 |
| chr5 | 10549674 | 10646751 | 97078 | -0.51 | 3.98E-29 | 1.31E-28 |
| chr10 | 3892530 | 3938747 | 46218 | 0.76 | 6.70E-29 | 2.20E-28 |
| chrX | 30293778 | 30327525 | 33748 | -0.71 | 7.86E-29 | 2.56E-28 |
| chr8 | 102446282 | 102487789 | 41508 | 0.62 | 7.88E-29 | 2.57E-28 |
| chr3 | 149011970 | 149121630 | 109661 | 0.33 | 8.44E-29 | 2.74E-28 |
| chr11 | 7621056 | 7679177 | 58122 | -0.77 | 1.35E-28 | 4.36E-28 |
| chr7 | 116301678 | 116379078 | 77401 | 0.48 | 1.43E-28 | 4.63E-28 |
| chr5 | 58739734 | 58895990 | 156257 | 0.3 | 2.87E-28 | 9.22E-28 |
| chr7 | 101726230 | 101816387 | 90158 | 0.48 | 3.30E-28 | 1.06E-27 |
| chr9 | 22123380 | 22172831 | 49452 | 0.49 | 3.75E-28 | 1.20E-27 |
| chr12 | 56636679 | 56711624 | 74946 | 0.6 | 3.76E-28 | 1.20E-27 |
| chr1 | 66696369 | 66826207 | 129839 | 0.44 | 6.12E-28 | 1.95E-27 |
| chr17 | 38439702 | 38527874 | 88173 | -0.56 | 9.06E-28 | 2.87E-27 |
| chr6 | 47071543 | 47138405 | 66863 | 0.99 | 9.81E-28 | 3.10E-27 |
| chr7 | 116057234 | 116229140 | 171907 | 0.38 | 1.09E-27 | 3.43E-27 |
| chr3 | 57935309 | 58164001 | 228693 | 0.33 | 1.17E-27 | 3.67E-27 |
| chr10 | 33402929 | 33630600 | 227672 | 0.32 | 1.27E-27 | 3.98E-27 |
| chr5 | 95093176 | 95179819 | 86644 | 0.57 | 1.40E-27 | 4.36E-27 |
| chr7 | 98794151 | 98828962 | 34812 | 0.63 | 1.61E-27 | 5.03E-27 |
| chr8 | 11656602 | 11760560 | 103959 | 0.56 | 1.70E-27 | 5.29E-27 |
| chr12 | 14391934 | 14437620 | 45687 | 4.36 | 1.74E-27 | 5.38E-27 |
| chr15 | 86099979 | 86163037 | 63059 | 0.7 | 2.02E-27 | 6.23E-27 |

| | | | | | | |
|---|---|---|---|---|---|---|
| chr9 | 123440698 | 123478211 | 37514 | -0.57 | 2.14E-27 | 6.58E-27 |
| chr9 | 33225369 | 33352880 | 127512 | 0.4 | 4.57E-27 | 1.40E-26 |
| chr2 | 143607036 | 143656067 | 49032 | -0.5 | 7.25E-27 | 2.22E-26 |
| chr8 | 22017143 | 22086170 | 69028 | 0.71 | 1.15E-26 | 3.51E-26 |
| chr17 | 39801383 | 39874837 | 73455 | 0.54 | 1.37E-26 | 4.17E-26 |
| chr18 | 3245732 | 3265593 | 19862 | 0.69 | 1.48E-26 | 4.49E-26 |
| chr11 | 86191434 | 86244822 | 53389 | 0.6 | 2.12E-26 | 6.41E-26 |
| chr2 | 113943002 | 114007358 | 64357 | -0.63 | 5.12E-26 | 1.54E-25 |
| chr9 | 75128756 | 75208441 | 79686 | 0.47 | 7.69E-26 | 2.31E-25 |
| chr20 | 4877298 | 4957688 | 80391 | 0.58 | 1.07E-25 | 3.21E-25 |
| chr2 | 161060481 | 161179251 | 118771 | -0.38 | 1.24E-25 | 3.69E-25 |
| chr17 | 2853051 | 2915051 | 62001 | 0.67 | 1.41E-25 | 4.19E-25 |
| chr12 | 92449650 | 92502617 | 52968 | 0.59 | 4.63E-25 | 1.38E-24 |
| chr10 | 21805790 | 21856852 | 51063 | 0.49 | 6.86E-25 | 2.04E-24 |
| chr11 | 10876937 | 10924235 | 47299 | -0.67 | 7.21E-25 | 2.13E-24 |
| chr22 | 35746313 | 35827976 | 81664 | -0.63 | 1.17E-24 | 3.45E-24 |
| chr16 | 87859209 | 87906021 | 46813 | 0.57 | 1.79E-24 | 5.28E-24 |
| chr15 | 29960775 | 30150267 | 189493 | -0.38 | 2.07E-24 | 6.09E-24 |
| chr15 | 63334044 | 63428897 | 94854 | 0.38 | 3.73E-24 | 1.09E-23 |
| chr8 | 8144098 | 8245317 | 101220 | 0.36 | 5.10E-24 | 1.49E-23 |
| chr12 | 64455250 | 64515486 | 60237 | 0.62 | 6.73E-24 | 1.96E-23 |
| chr11 | 69041303 | 69089663 | 48361 | 0.49 | 7.00E-24 | 2.03E-23 |
| chr5 | 142711201 | 142785671 | 74471 | 0.45 | 8.20E-24 | 2.37E-23 |
| chr6 | 133887836 | 133969358 | 81523 | 0.48 | 1.11E-23 | 3.19E-23 |
| chr15 | 33110999 | 33175680 | 64682 | -0.53 | 1.17E-23 | 3.36E-23 |
| chr19 | 44235143 | 44290171 | 55029 | 0.51 | 1.22E-23 | 3.51E-23 |
| chr11 | 56643840 | 56702627 | 58788 | 0.51 | 1.26E-23 | 3.60E-23 |
| chr3 | 98679812 | 98705756 | 25945 | 0.49 | 1.44E-23 | 4.12E-23 |
| chr7 | 73887511 | 73959617 | 72107 | -0.71 | 2.92E-23 | 8.33E-23 |
| chr18 | 74765597 | 74845305 | 79709 | 0.43 | 4.57E-23 | 1.30E-22 |
| chr4 | 74962511 | 75000283 | 37773 | 0.53 | 8.31E-23 | 2.35E-22 |
| chr12 | 54641974 | 54700907 | 58934 | 0.59 | 1.10E-22 | 3.12E-22 |
| chr12 | 46750923 | 46785531 | 34609 | 0.58 | 1.27E-22 | 3.57E-22 |
| chr7 | 97905061 | 98053141 | 148081 | 0.34 | 1.29E-22 | 3.64E-22 |
| chr9 | 130253602 | 130347063 | 93462 | 0.42 | 1.39E-22 | 3.91E-22 |
| chr17 | 48922335 | 49073293 | 150959 | 0.41 | 1.41E-22 | 3.94E-22 |
| chr14 | 77488170 | 77526589 | 38420 | 0.52 | 1.47E-22 | 4.10E-22 |
| chr1 | 85740163 | 85839333 | 99171 | 0.67 | 2.02E-22 | 5.61E-22 |
| chr13 | 73591352 | 73685869 | 94518 | 0.51 | 2.19E-22 | 6.06E-22 |
| chr9 | 113672377 | 113715580 | 43204 | -0.63 | 2.85E-22 | 7.89E-22 |
| chr12 | 12289047 | 12385532 | 96486 | 0.46 | 3.34E-22 | 9.21E-22 |
| chr19 | 46170561 | 46235422 | 64862 | 0.48 | 4.80E-22 | 1.32E-21 |
| chr12 | 6601342 | 6724171 | 122830 | 0.44 | 5.07E-22 | 1.39E-21 |
| chr1 | 174921607 | 174998998 | 77392 | 0.76 | 7.69E-22 | 2.11E-21 |
| chr16 | 2008985 | 2107353 | 98369 | 0.42 | 1.42E-21 | 3.89E-21 |
| chr18 | 45524139 | 45554488 | 30350 | 0.66 | 1.60E-21 | 4.35E-21 |
| chr12 | 122843254 | 122916572 | 73319 | 0.38 | 2.32E-21 | 6.30E-21 |
| chr5 | 60571235 | 60630161 | 58927 | 0.48 | 2.34E-21 | 6.33E-21 |
| chr11 | 13114667 | 13165428 | 50762 | 0.61 | 2.50E-21 | 6.76E-21 |
| chr5 | 173364793 | 173400956 | 36164 | 0.68 | 2.63E-21 | 7.11E-21 |
| chr11 | 9585924 | 9610852 | 24929 | 0.53 | 4.79E-21 | 1.29E-20 |
| chr12 | 125288261 | 125394204 | 105944 | 0.48 | 5.68E-21 | 1.53E-20 |
| chr14 | 68582901 | 68677961 | 95061 | 0.34 | 8.54E-21 | 2.29E-20 |
| chr11 | 27368154 | 27495348 | 127195 | 0.31 | 1.49E-20 | 3.97E-20 |
| chr1 | 8197642 | 8300469 | 102828 | 0.44 | 1.68E-20 | 4.47E-20 |
| chr19 | 13253371 | 13292969 | 39599 | 0.47 | 2.24E-20 | 5.95E-20 |
| chr5 | 90663112 | 90680792 | 17681 | 0.53 | 2.32E-20 | 6.16E-20 |
| chr10 | 21561786 | 21664762 | 102977 | -0.33 | 2.38E-20 | 6.29E-20 |
| chrX | 24857100 | 24918964 | 61865 | 0.52 | 6.35E-20 | 1.68E-19 |
| chr10 | 60372440 | 60432848 | 60409 | -0.56 | 6.52E-20 | 1.72E-19 |
| chr17 | 57694159 | 58000044 | 305886 | 0.28 | 6.95E-20 | 1.82E-19 |

| | | | | | | |
|---|---|---|---|---|---|---|
| chr7 | 100724240 | 100798893 | 74654 | -0.42 | 8.01E-20 | 2.10E-19 |
| chr5 | 77786675 | 77945802 | 159128 | -0.35 | 8.47E-20 | 2.21E-19 |
| chr17 | 18751744 | 18836820 | 85077 | -0.52 | 8.79E-20 | 2.29E-19 |
| chr5 | 148851007 | 148944451 | 93445 | 0.46 | 9.07E-20 | 2.36E-19 |
| chr7 | 116397873 | 116454016 | 56144 | 0.52 | 9.70E-20 | 2.52E-19 |
| chr8 | 38852583 | 38922916 | 70334 | 0.36 | 1.02E-19 | 2.65E-19 |
| chr5 | 162863352 | 162911740 | 48389 | 0.63 | 1.27E-19 | 3.28E-19 |
| chr7 | 92310046 | 92466703 | 156658 | 0.32 | 1.78E-19 | 4.57E-19 |
| chr2 | 143671513 | 143757112 | 85600 | -0.67 | 1.84E-19 | 4.72E-19 |
| chr1 | 156415843 | 156479435 | 63593 | 0.45 | 3.37E-19 | 8.65E-19 |
| chr9 | 74964115 | 74982759 | 18645 | 0.57 | 3.39E-19 | 8.67E-19 |
| chr2 | 65282769 | 65358554 | 75786 | -0.58 | 3.44E-19 | 8.78E-19 |
| chr11 | 57498968 | 57570145 | 71178 | 0.41 | 4.27E-19 | 1.09E-18 |
| chr5 | 132341287 | 132423543 | 82257 | 0.5 | 4.61E-19 | 1.17E-18 |
| chr17 | 43114047 | 43276582 | 162536 | 0.31 | 5.40E-19 | 1.37E-18 |
| chr21 | 33783102 | 33854038 | 70937 | -0.47 | 6.12E-19 | 1.55E-18 |
| chr11 | 34168459 | 34303344 | 134886 | -0.46 | 6.68E-19 | 1.68E-18 |
| chr1 | 41404936 | 41492694 | 87759 | -0.49 | 7.34E-19 | 1.85E-18 |
| chr16 | 85005738 | 85071403 | 65666 | 0.5 | 1.10E-18 | 2.77E-18 |
| chr2 | 102678541 | 102740179 | 61639 | -0.66 | 1.13E-18 | 2.83E-18 |
| chr22 | 29168266 | 29220597 | 52332 | 0.76 | 1.17E-18 | 2.92E-18 |
| chr9 | 137217087 | 137310830 | 93744 | -0.4 | 1.24E-18 | 3.10E-18 |
| chr20 | 5614820 | 5744571 | 129752 | 0.34 | 2.19E-18 | 5.45E-18 |
| chr8 | 128746131 | 128780072 | 33942 | -0.5 | 2.60E-18 | 6.44E-18 |
| chr12 | 6304378 | 6339845 | 35468 | 0.5 | 4.62E-18 | 1.14E-17 |
| chrX | 12991858 | 13049376 | 57519 | 0.48 | 5.23E-18 | 1.29E-17 |
| chr14 | 61978061 | 62053347 | 75287 | 0.33 | 5.31E-18 | 1.31E-17 |
| chr1 | 17481446 | 17537341 | 55896 | 0.71 | 5.73E-18 | 1.41E-17 |
| chr22 | 31222128 | 31304162 | 82035 | 0.5 | 5.82E-18 | 1.43E-17 |
| chr16 | 81001614 | 81042625 | 41012 | 0.53 | 5.85E-18 | 1.43E-17 |
| chr15 | 52261836 | 52351065 | 89230 | 0.39 | 7.34E-18 | 1.79E-17 |
| chr19 | 45574188 | 45658099 | 83912 | -0.52 | 8.86E-18 | 2.16E-17 |
| chr15 | 71385364 | 71447714 | 62351 | -0.54 | 9.12E-18 | 2.22E-17 |
| chr9 | 36296127 | 36342688 | 46562 | 0.49 | 9.57E-18 | 2.32E-17 |
| chr2 | 86244864 | 86308173 | 63310 | -0.55 | 1.03E-17 | 2.50E-17 |
| chr7 | 43655242 | 43715646 | 60405 | 0.45 | 1.14E-17 | 2.76E-17 |
| chr15 | 58729614 | 58768639 | 39026 | -0.54 | 1.40E-17 | 3.38E-17 |
| chr8 | 26427536 | 26517602 | 90067 | -0.44 | 1.55E-17 | 3.72E-17 |
| chr7 | 6406741 | 6494113 | 87373 | 0.43 | 1.62E-17 | 3.88E-17 |
| chr17 | 33894576 | 33973521 | 78946 | 0.4 | 1.86E-17 | 4.46E-17 |
| chr9 | 4222885 | 4300773 | 77889 | 0.33 | 2.09E-17 | 4.98E-17 |
| chr17 | 46549575 | 46725574 | 176000 | 0.35 | 2.30E-17 | 5.47E-17 |
| chr19 | 39831612 | 39984584 | 152973 | 0.28 | 3.00E-17 | 7.12E-17 |
| chr16 | 15965417 | 15996705 | 31289 | -0.49 | 3.34E-17 | 7.91E-17 |
| chr16 | 69164900 | 69257015 | 92116 | 0.53 | 3.72E-17 | 8.80E-17 |
| chr8 | 19231137 | 19616507 | 385371 | -0.24 | 3.76E-17 | 8.88E-17 |
| chr1 | 64157183 | 64199687 | 42505 | -0.53 | 4.46E-17 | 1.05E-16 |
| chr17 | 43298324 | 43385055 | 86732 | -0.33 | 4.96E-17 | 1.17E-16 |
| chrX | 133662851 | 133713784 | 50934 | -0.68 | 5.07E-17 | 1.19E-16 |
| chr2 | 206540164 | 206637566 | 97403 | -0.35 | 5.75E-17 | 1.35E-16 |
| chr2 | 10410512 | 10565411 | 154900 | -0.32 | 6.07E-17 | 1.42E-16 |
| chr7 | 28059692 | 28117626 | 57935 | -0.45 | 6.49E-17 | 1.51E-16 |
| chr12 | 718954 | 756641 | 37688 | 0.49 | 6.65E-17 | 1.55E-16 |
| chr12 | 122971018 | 123024169 | 53152 | 0.5 | 6.82E-17 | 1.58E-16 |
| chr11 | 34653445 | 34708218 | 54774 | -0.4 | 7.15E-17 | 1.65E-16 |
| chr10 | 112147364 | 112225875 | 78512 | -0.37 | 1.09E-16 | 2.52E-16 |
| chr7 | 27774201 | 27834148 | 59948 | 0.55 | 1.13E-16 | 2.59E-16 |
| chr3 | 152847784 | 152884547 | 36764 | 0.53 | 1.14E-16 | 2.63E-16 |
| chr1 | 153503932 | 153546074 | 42143 | 0.36 | 1.36E-16 | 3.12E-16 |
| chr7 | 139300412 | 139373365 | 72954 | -0.45 | 1.44E-16 | 3.30E-16 |
| chr8 | 29403131 | 29431536 | 28406 | 0.42 | 1.86E-16 | 4.26E-16 |

| | | | | | | |
|---|---|---|---|---|---|---|
| chr20 | 5037513 | 5114202 | 76690 | 0.35 | 2.02E-16 | 4.60E-16 |
| chr4 | 187562793 | 187623075 | 60283 | 0.55 | 2.04E-16 | 4.63E-16 |
| chr17 | 19171880 | 19283062 | 111183 | -0.43 | 2.09E-16 | 4.74E-16 |
| chr14 | 105131634 | 105178709 | 47076 | 0.41 | 2.68E-16 | 6.08E-16 |
| chr20 | 46046226 | 46098049 | 51824 | 0.43 | 4.32E-16 | 9.78E-16 |
| chr10 | 6313232 | 6357796 | 44565 | -0.36 | 4.83E-16 | 1.09E-15 |
| chr11 | 67763851 | 67841935 | 78085 | 0.35 | 5.76E-16 | 1.30E-15 |
| chr10 | 134208955 | 134266665 | 57711 | 0.55 | 5.89E-16 | 1.32E-15 |
| chr20 | 25175533 | 25319601 | 144069 | 0.28 | 7.89E-16 | 1.77E-15 |
| chr5 | 34655522 | 34762912 | 107391 | -0.43 | 1.84E-15 | 4.12E-15 |
| chr19 | 1239774 | 1276852 | 37079 | 0.5 | 2.65E-15 | 5.92E-15 |
| chr8 | 17742896 | 17782981 | 40086 | 0.37 | 2.71E-15 | 6.05E-15 |
| chr8 | 29292313 | 29325199 | 32887 | 0.45 | 4.80E-15 | 1.07E-14 |
| chr18 | 2977902 | 3052848 | 74947 | 0.48 | 4.94E-15 | 1.10E-14 |
| chr9 | 116217029 | 116357842 | 140814 | -0.38 | 5.48E-15 | 1.21E-14 |
| chr10 | 27465559 | 27542413 | 76855 | 0.51 | 5.73E-15 | 1.27E-14 |
| chr7 | 130523385 | 130800703 | 277319 | 0.21 | 5.92E-15 | 1.30E-14 |
| chr14 | 55490660 | 55545130 | 54471 | 0.6 | 6.10E-15 | 1.34E-14 |
| chr20 | 48381153 | 48435797 | 54645 | 0.47 | 6.56E-15 | 1.44E-14 |
| chr2 | 9370760 | 9429450 | 58691 | -0.56 | 6.77E-15 | 1.48E-14 |
| chr15 | 26855524 | 26881571 | 26048 | -0.5 | 7.72E-15 | 1.69E-14 |
| chrX | 23791493 | 23841841 | 50349 | 0.44 | 9.03E-15 | 1.97E-14 |
| chr3 | 149229820 | 149380595 | 150776 | 0.22 | 1.43E-14 | 3.11E-14 |
| chr1 | 223886109 | 223965855 | 79747 | 0.29 | 1.44E-14 | 3.13E-14 |
| chr18 | 68078364 | 68118011 | 39648 | 0.44 | 1.48E-14 | 3.21E-14 |
| chr13 | 21555508 | 21645868 | 90361 | 0.44 | 1.54E-14 | 3.34E-14 |
| chr3 | 189010602 | 189076062 | 65461 | 0.42 | 1.56E-14 | 3.37E-14 |
| chr8 | 17612333 | 17679445 | 67113 | 0.37 | 1.87E-14 | 4.03E-14 |
| chr20 | 33837946 | 33874339 | 36394 | 0.41 | 2.64E-14 | 5.69E-14 |
| chr8 | 54852850 | 54986865 | 134016 | 0.31 | 3.07E-14 | 6.59E-14 |
| chr16 | 81428895 | 81813436 | 384542 | 0.21 | 3.18E-14 | 6.82E-14 |
| chr1 | 33168006 | 33255189 | 87184 | 0.41 | 3.49E-14 | 7.47E-14 |
| chr21 | 27486793 | 27543903 | 57111 | 0.43 | 3.94E-14 | 8.42E-14 |
| chr7 | 1542567 | 1611574 | 69008 | 0.35 | 4.44E-14 | 9.47E-14 |
| chr16 | 16013321 | 16204659 | 191339 | -0.24 | 4.78E-14 | 1.02E-13 |
| chr1 | 181041806 | 181111704 | 69899 | -0.36 | 5.22E-14 | 1.11E-13 |
| chr19 | 14605532 | 14640080 | 34549 | 0.49 | 5.33E-14 | 1.13E-13 |
| chr17 | 80138694 | 80261467 | 122774 | 0.29 | 7.97E-14 | 1.68E-13 |
| chr2 | 191702247 | 191814237 | 111991 | -0.39 | 9.14E-14 | 1.93E-13 |
| chr4 | 170145730 | 170204670 | 58941 | 0.37 | 1.01E-13 | 2.13E-13 |
| chr19 | 39295939 | 39484528 | 188590 | 0.25 | 1.05E-13 | 2.21E-13 |
| chr17 | 9720702 | 9747955 | 27254 | -0.44 | 1.18E-13 | 2.48E-13 |
| chr1 | 68149927 | 68237474 | 87548 | 0.4 | 1.64E-13 | 3.44E-13 |
| chr19 | 1150963 | 1214166 | 63204 | 0.44 | 2.42E-13 | 5.05E-13 |
| chr3 | 46955338 | 47030584 | 75247 | 0.37 | 2.74E-13 | 5.71E-13 |
| chr17 | 17584215 | 17760416 | 176202 | 0.22 | 2.95E-13 | 6.15E-13 |
| chr20 | 56229954 | 56294501 | 64548 | 0.4 | 4.78E-13 | 9.93E-13 |
| chr8 | 141991242 | 142052116 | 60875 | -0.42 | 7.01E-13 | 1.45E-12 |
| chr5 | 142939604 | 142980925 | 41322 | 0.44 | 7.05E-13 | 1.46E-12 |
| chr2 | 69952696 | 70057709 | 105014 | -0.26 | 7.18E-13 | 1.48E-12 |
| chr11 | 63375743 | 63451412 | 75670 | 0.48 | 7.39E-13 | 1.52E-12 |
| chr20 | 52185888 | 52243428 | 57541 | 0.31 | 7.64E-13 | 1.57E-12 |
| chr12 | 124955459 | 125058858 | 103400 | 0.39 | 7.83E-13 | 1.61E-12 |
| chr14 | 103557421 | 103623849 | 66429 | 0.34 | 9.79E-13 | 2.01E-12 |
| chr8 | 22406096 | 22463872 | 57777 | 0.4 | 1.23E-12 | 2.52E-12 |
| chr6 | 12279085 | 12385178 | 106094 | 0.25 | 1.35E-12 | 2.76E-12 |
| chr8 | 38198187 | 38270423 | 72237 | 0.41 | 1.48E-12 | 3.02E-12 |
| chr10 | 34529377 | 34829672 | 300296 | -0.2 | 1.57E-12 | 3.20E-12 |
| chr1 | 200859096 | 200886602 | 27507 | 0.36 | 1.98E-12 | 4.02E-12 |
| chr3 | 127448627 | 127542477 | 93851 | 0.32 | 2.20E-12 | 4.45E-12 |
| chr18 | 11848736 | 11861639 | 12904 | 0.49 | 2.92E-12 | 5.91E-12 |

| | | | | | | |
|---|---|---|---|---|---|---|
| chr12 | 57457573 | 57506935 | 49363 | 0.41 | 3.09E-12 | 6.24E-12 |
| chr12 | 107711526 | 107834329 | 122804 | -0.25 | 3.30E-12 | 6.66E-12 |
| chr8 | 130922923 | 131029768 | 106846 | -0.26 | 3.74E-12 | 7.53E-12 |
| chr8 | 101911851 | 101967475 | 55625 | 0.39 | 4.31E-12 | 8.66E-12 |
| chr20 | 5929417 | 5991295 | 61879 | 0.32 | 4.99E-12 | 1.00E-11 |
| chr12 | 122460523 | 122530545 | 70023 | 0.41 | 5.04E-12 | 1.01E-11 |
| chr14 | 92968466 | 93032440 | 63975 | -0.39 | 5.17E-12 | 1.03E-11 |
| chr10 | 104844186 | 104959166 | 114981 | 0.28 | 5.33E-12 | 1.06E-11 |
| chr17 | 39914999 | 39993131 | 78133 | 0.32 | 5.36E-12 | 1.07E-11 |
| chr6 | 64265208 | 64289885 | 24678 | 0.39 | 5.69E-12 | 1.13E-11 |
| chr10 | 3085343 | 3185242 | 99900 | 0.3 | 5.98E-12 | 1.19E-11 |
| chr15 | 89630522 | 89693425 | 62904 | 0.32 | 6.69E-12 | 1.32E-11 |
| chr9 | 134225422 | 134353322 | 127901 | 0.37 | 6.73E-12 | 1.33E-11 |
| chr10 | 94349665 | 94413966 | 64302 | 0.39 | 6.76E-12 | 1.33E-11 |
| chr14 | 31417443 | 31561725 | 144283 | 0.36 | 6.82E-12 | 1.34E-11 |
| chr20 | 34638472 | 34771697 | 133226 | 0.29 | 8.21E-12 | 1.61E-11 |
| chr17 | 46313555 | 46347216 | 33662 | 0.29 | 8.76E-12 | 1.72E-11 |
| chr2 | 187406512 | 187477710 | 71199 | -0.39 | 9.54E-12 | 1.87E-11 |
| chr3 | 141079357 | 141174399 | 95043 | -0.24 | 1.13E-11 | 2.20E-11 |
| chr12 | 111970390 | 112039334 | 68945 | 0.43 | 1.31E-11 | 2.55E-11 |
| chr2 | 153321712 | 153365034 | 43323 | -0.4 | 1.38E-11 | 2.68E-11 |
| chr15 | 76578626 | 76630004 | 51379 | -0.4 | 1.54E-11 | 3.00E-11 |
| chr17 | 27023834 | 27091219 | 67386 | 0.32 | 1.65E-11 | 3.21E-11 |
| chr17 | 170143 | 236848 | 66706 | 0.31 | 1.79E-11 | 3.47E-11 |
| chr8 | 71018870 | 71087587 | 68718 | -0.44 | 2.17E-11 | 4.18E-11 |
| chr5 | 59038941 | 59096025 | 57085 | 0.29 | 2.61E-11 | 5.03E-11 |
| chr1 | 21156492 | 21231267 | 74776 | -0.46 | 2.71E-11 | 5.22E-11 |
| chr17 | 35797520 | 35853643 | 56124 | 0.41 | 2.76E-11 | 5.29E-11 |
| chr17 | 79448157 | 79487399 | 39243 | 0.46 | 3.28E-11 | 6.29E-11 |
| chr20 | 52712213 | 52794566 | 82354 | 0.2 | 3.90E-11 | 7.47E-11 |
| chr10 | 101487227 | 101612868 | 125642 | -0.24 | 3.92E-11 | 7.50E-11 |
| chr16 | 19098020 | 19152586 | 54567 | 0.38 | 4.06E-11 | 7.75E-11 |
| chr1 | 218488618 | 218614092 | 125475 | -0.23 | 4.85E-11 | 9.23E-11 |
| chrX | 10502952 | 10604706 | 101755 | 0.25 | 5.83E-11 | 1.11E-10 |
| chr2 | 227597478 | 227666827 | 69350 | 0.4 | 6.37E-11 | 1.21E-10 |
| chr1 | 90285714 | 90424156 | 138443 | -0.25 | 6.56E-11 | 1.24E-10 |
| chr1 | 151137394 | 151240443 | 103050 | 0.31 | 6.81E-11 | 1.29E-10 |
| chr1 | 200320609 | 200387121 | 66513 | 0.35 | 8.53E-11 | 1.61E-10 |
| chr3 | 156498381 | 156545966 | 47586 | 0.26 | 8.73E-11 | 1.64E-10 |
| chr8 | 29342543 | 29389633 | 47091 | 0.33 | 1.05E-10 | 1.98E-10 |
| chr9 | 130660808 | 130806062 | 145255 | -0.23 | 1.09E-10 | 2.04E-10 |
| chr20 | 17917676 | 17957851 | 40176 | 0.37 | 1.20E-10 | 2.25E-10 |
| chr3 | 10206431 | 10270898 | 64468 | -0.34 | 1.31E-10 | 2.44E-10 |
| chr2 | 232465490 | 232552625 | 87136 | -0.3 | 1.39E-10 | 2.59E-10 |
| chr11 | 118777054 | 118801602 | 24549 | 0.32 | 2.00E-10 | 3.72E-10 |
| chr7 | 6258743 | 6371945 | 113203 | 0.29 | 2.15E-10 | 4.00E-10 |
| chr5 | 1469573 | 1568958 | 99386 | 0.53 | 2.46E-10 | 4.57E-10 |
| chr1 | 235085129 | 235154672 | 69544 | 0.26 | 2.93E-10 | 5.44E-10 |
| chr11 | 76455640 | 76517418 | 61779 | -0.28 | 3.01E-10 | 5.56E-10 |
| chr20 | 30101454 | 30222605 | 121152 | 0.25 | 3.05E-10 | 5.63E-10 |
| chr2 | 30454396 | 30525247 | 70852 | -0.35 | 3.22E-10 | 5.95E-10 |
| chr7 | 107530897 | 107669312 | 138416 | 0.29 | 3.32E-10 | 6.12E-10 |
| chr8 | 25027512 | 25349920 | 322409 | 0.17 | 3.69E-10 | 6.79E-10 |
| chr10 | 79567673 | 79687145 | 119473 | -0.35 | 3.95E-10 | 7.25E-10 |
| chr8 | 102339428 | 102399259 | 59832 | 0.29 | 4.16E-10 | 7.63E-10 |
| chr17 | 38593898 | 38718809 | 124912 | -0.22 | 4.47E-10 | 8.18E-10 |
| chr11 | 71638189 | 71760568 | 122380 | -0.28 | 4.50E-10 | 8.21E-10 |
| chr9 | 74906519 | 74941628 | 35110 | 0.32 | 6.34E-10 | 1.16E-09 |
| chr20 | 5531039 | 5594415 | 63377 | 0.35 | 6.46E-10 | 1.18E-09 |
| chr11 | 46678973 | 46742153 | 63181 | -0.44 | 6.90E-10 | 1.25E-09 |
| chr15 | 56138572 | 56211515 | 72944 | 0.23 | 7.07E-10 | 1.28E-09 |

| | | | | | | |
|---|---|---|---|---|---|---|
| chr20 | 48191961 | 48345161 | 153201 | 0.23 | 7.08E-10 | 1.28E-09 |
| chr5 | 118590991 | 118682324 | 91334 | -0.31 | 8.80E-10 | 1.59E-09 |
| chr1 | 230201715 | 230294719 | 93005 | 0.36 | 1.17E-09 | 2.11E-09 |
| chr8 | 131053663 | 131165025 | 111363 | -0.32 | 1.20E-09 | 2.16E-09 |
| chr15 | 31616995 | 31698788 | 81794 | -0.38 | 1.31E-09 | 2.35E-09 |
| chr1 | 169073964 | 169132966 | 59003 | 0.27 | 1.40E-09 | 2.52E-09 |
| chr20 | 49240789 | 49318619 | 77831 | 0.41 | 1.46E-09 | 2.61E-09 |
| chr7 | 75216989 | 75370988 | 154000 | 0.24 | 1.79E-09 | 3.20E-09 |
| chr4 | 119989168 | 120035208 | 46041 | 0.34 | 1.96E-09 | 3.50E-09 |
| chr9 | 99139348 | 99184692 | 45345 | 0.36 | 2.02E-09 | 3.60E-09 |
| chr17 | 48103087 | 48173349 | 70263 | 0.24 | 2.04E-09 | 3.63E-09 |
| chrX | 123074513 | 123106242 | 31730 | 0.44 | 2.04E-09 | 3.64E-09 |
| chr6 | 121800043 | 121844327 | 44285 | 0.35 | 3.26E-09 | 5.78E-09 |
| chr10 | 119081699 | 119136354 | 54656 | 0.29 | 3.43E-09 | 6.08E-09 |
| chr10 | 64966700 | 65035418 | 68719 | 0.31 | 3.47E-09 | 6.15E-09 |
| chr11 | 128360509 | 128394004 | 33496 | -0.33 | 3.69E-09 | 6.53E-09 |
| chr8 | 62452520 | 62705391 | 252872 | 0.16 | 3.77E-09 | 6.65E-09 |
| chr4 | 15001995 | 15050345 | 48351 | 0.35 | 4.33E-09 | 7.62E-09 |
| chr17 | 15845209 | 15966400 | 121192 | 0.27 | 5.39E-09 | 9.47E-09 |
| chr8 | 48223291 | 48300704 | 77414 | 0.3 | 5.84E-09 | 1.03E-08 |
| chr15 | 85522896 | 85686691 | 163796 | 0.19 | 6.62E-09 | 1.16E-08 |
| chr16 | 70408394 | 70473828 | 65435 | -0.39 | 6.95E-09 | 1.22E-08 |
| chr20 | 49334499 | 49412405 | 77907 | 0.23 | 7.17E-09 | 1.25E-08 |
| chr11 | 2913500 | 3014550 | 101051 | 0.31 | 7.22E-09 | 1.26E-08 |
| chr3 | 81765839 | 81811986 | 46148 | -0.32 | 7.30E-09 | 1.27E-08 |
| chr3 | 5007364 | 5069707 | 62344 | 0.25 | 7.72E-09 | 1.34E-08 |
| chr17 | 73839610 | 73909058 | 69449 | 0.33 | 8.34E-09 | 1.45E-08 |
| chr9 | 139775542 | 139845789 | 70248 | 0.28 | 8.44E-09 | 1.46E-08 |
| chrX | 46175720 | 46194270 | 18551 | 0.29 | 1.03E-08 | 1.77E-08 |
| chr1 | 245114629 | 245137595 | 22967 | 0.31 | 1.36E-08 | 2.34E-08 |
| chr10 | 102100744 | 102155751 | 55008 | 0.32 | 1.41E-08 | 2.43E-08 |
| chr7 | 101491399 | 101668736 | 177338 | -0.24 | 1.49E-08 | 2.56E-08 |
| chr17 | 55923177 | 56092893 | 169717 | 0.16 | 1.80E-08 | 3.10E-08 |
| chr1 | 241754096 | 241836686 | 82591 | 0.27 | 2.08E-08 | 3.57E-08 |
| chr7 | 151383006 | 151452560 | 69555 | -0.41 | 2.14E-08 | 3.67E-08 |
| chr8 | 144534191 | 144641552 | 107362 | -0.49 | 2.20E-08 | 3.77E-08 |
| chr9 | 132220231 | 132270648 | 50418 | 0.21 | 2.21E-08 | 3.77E-08 |
| chr11 | 65654699 | 65688052 | 33354 | -0.29 | 2.30E-08 | 3.92E-08 |
| chr17 | 25619978 | 25708345 | 88368 | 0.31 | 2.47E-08 | 4.20E-08 |
| chr7 | 73093519 | 73158527 | 65009 | 0.31 | 2.92E-08 | 4.96E-08 |
| chr10 | 88196710 | 88296098 | 99389 | 0.34 | 4.10E-08 | 6.95E-08 |
| chr20 | 4089064 | 4174404 | 85341 | -0.22 | 4.84E-08 | 8.20E-08 |
| chr14 | 74162339 | 74262580 | 100242 | 0.23 | 5.47E-08 | 9.25E-08 |
| chr13 | 47169279 | 47242786 | 73508 | 0.35 | 6.28E-08 | 1.06E-07 |
| chr19 | 36526944 | 36681454 | 154511 | 0.23 | 7.02E-08 | 1.18E-07 |
| chr4 | 140931986 | 141076646 | 144661 | 0.18 | 7.56E-08 | 1.27E-07 |
| chr9 | 35707575 | 35775627 | 68053 | 0.34 | 7.61E-08 | 1.28E-07 |
| chr4 | 95994029 | 96030808 | 36780 | 0.3 | 9.25E-08 | 1.55E-07 |
| chr8 | 90893625 | 90938824 | 45200 | -0.25 | 9.77E-08 | 1.64E-07 |
| chr8 | 118949429 | 119044281 | 94853 | -0.32 | 9.88E-08 | 1.65E-07 |
| chr19 | 48101292 | 48182602 | 81311 | 0.3 | 1.13E-07 | 1.89E-07 |
| chr14 | 69240470 | 69292184 | 51715 | 0.27 | 1.15E-07 | 1.92E-07 |
| chr6 | 74269995 | 74320310 | 50316 | 0.32 | 1.31E-07 | 2.18E-07 |
| chr6 | 160768351 | 160828280 | 59930 | 0.27 | 1.41E-07 | 2.35E-07 |
| chr18 | 9073454 | 9159059 | 85606 | 0.28 | 1.43E-07 | 2.37E-07 |
| chr17 | 63013429 | 63075204 | 61776 | -0.41 | 1.49E-07 | 2.47E-07 |
| chr19 | 45894364 | 46118474 | 224111 | 0.19 | 1.54E-07 | 2.55E-07 |
| chr2 | 188390021 | 188446506 | 56486 | 0.21 | 1.55E-07 | 2.56E-07 |
| chr21 | 45107714 | 45210444 | 102731 | 0.24 | 1.70E-07 | 2.81E-07 |
| chr17 | 2282555 | 2323058 | 40504 | 0.28 | 1.82E-07 | 3.00E-07 |
| chr15 | 66990466 | 67079056 | 88591 | 0.19 | 1.91E-07 | 3.13E-07 |

| | | | | | | |
|---|---|---|---|---|---|---|
| chr7 | 98669821 | 98761627 | 91807 | 0.37 | 2.11E-07 | 3.46E-07 |
| chr11 | 61714804 | 61749308 | 34505 | -0.27 | 2.16E-07 | 3.54E-07 |
| chr17 | 57151836 | 57264507 | 112672 | 0.25 | 2.28E-07 | 3.73E-07 |
| chr1 | 150569056 | 150603556 | 34501 | 0.3 | 2.39E-07 | 3.92E-07 |
| chr3 | 43892509 | 43936204 | 43696 | -0.29 | 2.48E-07 | 4.06E-07 |
| chr2 | 11507688 | 11558976 | 51289 | 0.29 | 2.61E-07 | 4.25E-07 |
| chr5 | 173172719 | 173230971 | 58253 | -0.35 | 2.66E-07 | 4.32E-07 |
| chr11 | 34066104 | 34128865 | 62762 | 0.34 | 3.17E-07 | 5.16E-07 |
| chr9 | 75048138 | 75107077 | 58940 | 0.28 | 3.19E-07 | 5.18E-07 |
| chr22 | 43314757 | 43371160 | 56404 | 0.3 | 3.91E-07 | 6.34E-07 |
| chr17 | 26638869 | 26708038 | 69170 | 0.28 | 4.03E-07 | 6.52E-07 |
| chr2 | 43356641 | 43479000 | 122360 | 0.2 | 4.22E-07 | 6.81E-07 |
| chr16 | 87791168 | 87841551 | 50384 | -0.34 | 4.78E-07 | 7.72E-07 |
| chr17 | 4577745 | 4643480 | 65736 | -0.34 | 5.65E-07 | 9.10E-07 |
| chr5 | 137849969 | 137912332 | 62364 | 0.27 | 5.75E-07 | 9.24E-07 |
| chr12 | 123546996 | 123650779 | 103784 | -0.33 | 6.68E-07 | 1.07E-06 |
| chr5 | 16778164 | 16937061 | 158898 | -0.19 | 8.73E-07 | 1.40E-06 |
| chr6 | 26019853 | 26071887 | 52035 | -0.23 | 9.05E-07 | 1.45E-06 |
| chr19 | 54480934 | 54539902 | 58969 | -0.22 | 1.05E-06 | 1.67E-06 |
| chr10 | 5312791 | 5337093 | 24303 | -0.32 | 1.06E-06 | 1.70E-06 |
| chr22 | 46065420 | 46204275 | 138856 | -0.18 | 1.16E-06 | 1.84E-06 |
| chr2 | 9652420 | 9706894 | 54475 | -0.25 | 1.21E-06 | 1.92E-06 |
| chr4 | 7805084 | 7943557 | 138474 | 0.2 | 1.33E-06 | 2.11E-06 |
| chr7 | 75534589 | 75623892 | 89304 | 0.24 | 1.35E-06 | 2.14E-06 |
| chr7 | 80389489 | 80554420 | 164932 | 0.18 | 1.46E-06 | 2.31E-06 |
| chr2 | 110857107 | 110874770 | 17664 | -0.32 | 1.53E-06 | 2.42E-06 |
| chr12 | 123826983 | 123890174 | 63192 | 0.3 | 1.63E-06 | 2.58E-06 |
| chr19 | 35476449 | 35521266 | 44818 | -0.24 | 1.66E-06 | 2.62E-06 |
| chr11 | 62305101 | 62391599 | 86499 | 0.22 | 1.74E-06 | 2.74E-06 |
| chr15 | 99384251 | 99501110 | 116860 | -0.14 | 1.95E-06 | 3.07E-06 |
| chr15 | 69068389 | 69114747 | 46359 | -0.31 | 2.35E-06 | 3.69E-06 |
| chr19 | 40898611 | 40972799 | 74189 | 0.21 | 2.43E-06 | 3.81E-06 |
| chr5 | 150419708 | 150475692 | 55985 | -0.33 | 2.63E-06 | 4.11E-06 |
| chr21 | 36153331 | 36262461 | 109131 | 0.22 | 2.97E-06 | 4.64E-06 |
| chr12 | 858332 | 938735 | 80404 | 0.29 | 3.05E-06 | 4.77E-06 |
| chr11 | 66589566 | 66689454 | 99889 | -0.22 | 3.32E-06 | 5.17E-06 |
| chr17 | 35626129 | 35726598 | 100470 | -0.27 | 3.42E-06 | 5.32E-06 |
| chr2 | 9504957 | 9575172 | 70216 | -0.3 | 3.45E-06 | 5.36E-06 |
| chr18 | 20678341 | 20740468 | 62128 | 0.27 | 3.94E-06 | 6.12E-06 |
| chr20 | 52441878 | 52489998 | 48121 | 0.23 | 3.97E-06 | 6.15E-06 |
| chr14 | 50401200 | 50513097 | 111898 | 0.41 | 4.06E-06 | 6.28E-06 |
| chr16 | 57787026 | 57933973 | 146948 | 0.16 | 4.28E-06 | 6.61E-06 |
| chr14 | 31575111 | 31679071 | 103961 | 0.23 | 4.39E-06 | 6.78E-06 |
| chr10 | 73472797 | 73535002 | 62206 | 0.24 | 4.63E-06 | 7.14E-06 |
| chr1 | 110452708 | 110548372 | 95665 | -0.23 | 4.71E-06 | 7.25E-06 |
| chr15 | 99189570 | 99367158 | 177589 | 0.16 | 5.55E-06 | 8.53E-06 |
| chr14 | 68907637 | 68937263 | 29627 | 0.24 | 5.92E-06 | 9.08E-06 |
| chr11 | 122024009 | 122073712 | 49704 | 0.27 | 6.07E-06 | 9.30E-06 |
| chr5 | 126111184 | 126183019 | 71836 | 0.26 | 7.01E-06 | 1.07E-05 |
| chr6 | 159268140 | 159292715 | 24576 | 0.24 | 7.02E-06 | 1.07E-05 |
| chr5 | 73090828 | 73126742 | 35915 | 0.29 | 7.94E-06 | 1.21E-05 |
| chr6 | 34622496 | 34691948 | 69453 | -0.29 | 1.01E-05 | 1.53E-05 |
| chr20 | 52351511 | 52427651 | 76141 | -0.17 | 1.35E-05 | 2.05E-05 |
| chr11 | 65046177 | 65159966 | 113790 | 0.23 | 1.44E-05 | 2.19E-05 |
| chr5 | 133837332 | 133928945 | 91614 | 0.2 | 1.53E-05 | 2.32E-05 |
| chr6 | 11179353 | 11234451 | 55099 | 0.17 | 1.64E-05 | 2.49E-05 |
| chr14 | 65170467 | 65231852 | 61386 | -0.25 | 1.64E-05 | 2.49E-05 |
| chr10 | 105505196 | 105595907 | 90712 | 0.24 | 1.68E-05 | 2.54E-05 |
| chr1 | 43388259 | 43435387 | 47129 | 0.21 | 1.70E-05 | 2.57E-05 |
| chr12 | 104606470 | 104618302 | 11833 | 0.24 | 1.91E-05 | 2.88E-05 |
| chr11 | 28064422 | 28145235 | 80814 | 0.22 | 1.94E-05 | 2.93E-05 |

| | | | | | | |
|---|---|---|---|---|---|---|
| chr17 | 6478154 | 6556085 | 77932 | 0.19 | 1.96E-05 | 2.94E-05 |
| chr19 | 48823609 | 48931419 | 107811 | 0.2 | 2.00E-05 | 3.00E-05 |
| chr2 | 54750073 | 54901085 | 151013 | 0.15 | 2.04E-05 | 3.05E-05 |
| chr8 | 142126888 | 142220511 | 93624 | 0.18 | 2.24E-05 | 3.35E-05 |
| chr17 | 19800350 | 19881985 | 81636 | 0.24 | 2.25E-05 | 3.36E-05 |
| chr1 | 150915080 | 150982086 | 67007 | -0.32 | 2.33E-05 | 3.47E-05 |
| chr2 | 27576469 | 27637413 | 60945 | -0.27 | 2.34E-05 | 3.49E-05 |
| chr7 | 4721176 | 4794036 | 72861 | 0.19 | 2.41E-05 | 3.58E-05 |
| chr3 | 151482797 | 151537111 | 54315 | -0.17 | 2.50E-05 | 3.71E-05 |
| chr19 | 18384744 | 18461211 | 76468 | 0.22 | 2.64E-05 | 3.92E-05 |
| chr19 | 45453791 | 45560695 | 106905 | -0.26 | 2.97E-05 | 4.40E-05 |
| chr9 | 131375540 | 131471264 | 95725 | 0.2 | 2.99E-05 | 4.42E-05 |
| chr19 | 36192944 | 36250409 | 57466 | -0.24 | 3.06E-05 | 4.52E-05 |
| chr1 | 15478598 | 15547488 | 68891 | -0.24 | 3.15E-05 | 4.65E-05 |
| chr7 | 5534608 | 5573105 | 38498 | 0.29 | 3.67E-05 | 5.40E-05 |
| chr9 | 132309277 | 132405270 | 95994 | -0.23 | 3.72E-05 | 5.47E-05 |
| chr1 | 9350040 | 9412327 | 62288 | -0.27 | 3.72E-05 | 5.47E-05 |
| chr18 | 7900497 | 7963936 | 63440 | 0.19 | 3.95E-05 | 5.80E-05 |
| chr16 | 89352493 | 89576340 | 223848 | -0.14 | 4.07E-05 | 5.96E-05 |
| chr14 | 32542587 | 32620310 | 77724 | 0.22 | 4.11E-05 | 6.02E-05 |
| chr5 | 17302757 | 17326969 | 24213 | 0.23 | 4.36E-05 | 6.38E-05 |
| chr20 | 55326825 | 55384077 | 57253 | 0.21 | 4.69E-05 | 6.84E-05 |
| chrX | 24161197 | 24209759 | 48563 | 0.25 | 5.19E-05 | 7.57E-05 |
| chr1 | 37917504 | 37965876 | 48373 | 0.28 | 5.33E-05 | 7.75E-05 |
| chr19 | 41723871 | 41917491 | 193621 | -0.13 | 5.49E-05 | 7.98E-05 |
| chr20 | 45881412 | 46032555 | 151144 | -0.15 | 5.88E-05 | 8.53E-05 |
| chr17 | 35439561 | 35518385 | 78825 | -0.28 | 5.91E-05 | 8.57E-05 |
| chr17 | 48696555 | 48846539 | 149985 | 0.13 | 6.15E-05 | 8.90E-05 |
| chr9 | 114873737 | 114939644 | 65908 | -0.25 | 6.33E-05 | 9.15E-05 |
| chr8 | 145007428 | 145065503 | 58076 | 0.21 | 6.37E-05 | 9.19E-05 |
| chr1 | 157963103 | 158009037 | 45935 | -0.2 | 6.90E-05 | 9.95E-05 |
| chr5 | 156452078 | 156500135 | 48058 | 0.2 | 7.18E-05 | 0.000103 |
| chr9 | 3502350 | 3528782 | 26433 | 0.21 | 7.28E-05 | 0.000105 |
| chr14 | 77362138 | 77432287 | 70150 | 0.18 | 7.33E-05 | 0.000105 |
| chr9 | 84127261 | 84306039 | 178779 | 0.15 | 7.60E-05 | 0.000109 |
| chr7 | 32602181 | 32659043 | 56863 | -0.23 | 7.95E-05 | 0.000114 |
| chr16 | 56833217 | 56903601 | 70385 | 0.23 | 8.10E-05 | 0.000116 |
| chr2 | 239171430 | 239209710 | 38281 | -0.2 | 8.45E-05 | 0.000121 |
| chr3 | 129295786 | 129409077 | 113292 | 0.16 | 8.94E-05 | 0.000128 |
| chr19 | 42363074 | 42411639 | 48566 | 0.23 | 9.96E-05 | 0.000142 |
| chr19 | 40402625 | 40488289 | 85665 | 0.15 | 0.000107 | 0.000152 |
| chr11 | 61514747 | 61634322 | 119576 | 0.18 | 0.000109 | 0.000155 |
| chr17 | 27852631 | 27921466 | 68836 | -0.23 | 0.000116 | 0.000164 |
| chr5 | 167715107 | 167883372 | 168266 | 0.13 | 0.000119 | 0.000169 |
| chr12 | 132335754 | 132417008 | 81255 | 0.19 | 0.00012 | 0.00017 |
| chr6 | 151325072 | 151414054 | 88983 | 0.19 | 0.000121 | 0.000172 |
| chrX | 23889199 | 23926884 | 37686 | 0.2 | 0.000149 | 0.000211 |
| chr8 | 37683779 | 37774308 | 90530 | 0.24 | 0.000164 | 0.000231 |
| chr5 | 153617122 | 153796732 | 179611 | -0.14 | 0.000167 | 0.000235 |
| chr12 | 96587328 | 96636778 | 49451 | 0.19 | 0.000172 | 0.000242 |
| chr14 | 32395998 | 32438030 | 42033 | -0.23 | 0.000177 | 0.000249 |
| chr19 | 59054467 | 59093038 | 38572 | -0.2 | 0.000185 | 0.000259 |
| chr6 | 136707828 | 136769823 | 61996 | 0.19 | 0.00019 | 0.000266 |
| chr1 | 32375512 | 32429825 | 54314 | 0.19 | 0.000197 | 0.000275 |
| chr6 | 12449985 | 12496774 | 46790 | -0.21 | 0.000199 | 0.000278 |
| chr3 | 152004886 | 152050496 | 45611 | 0.21 | 0.000221 | 0.000308 |
| chr22 | 45969414 | 46044957 | 75544 | 0.16 | 0.000225 | 0.000313 |
| chr1 | 153912331 | 153967025 | 54695 | 0.22 | 0.000246 | 0.000342 |
| chr6 | 126209645 | 126279784 | 70140 | 0.17 | 0.00027 | 0.000376 |
| chr7 | 47794000 | 47859600 | 65601 | 0.15 | 0.000275 | 0.000381 |
| chr1 | 156694876 | 156754449 | 59574 | 0.19 | 0.000345 | 0.000478 |

| | | | | | | |
|---|---|---|---|---|---|---|
| chr6 | 27759214 | 27808433 | 49220 | -0.2 | 0.000365 | 0.000505 |
| chr8 | 103779785 | 103823766 | 43982 | -0.17 | 0.00038 | 0.000526 |
| chr4 | 40443262 | 40529606 | 86345 | 0.18 | 0.000395 | 0.000545 |
| chr17 | 42607140 | 42640125 | 32986 | -0.18 | 0.000396 | 0.000546 |
| chr2 | 29328174 | 29389161 | 60988 | -0.14 | 0.00041 | 0.000565 |
| chr19 | 49457252 | 49503506 | 46255 | -0.24 | 0.000423 | 0.000581 |
| chr19 | 2523348 | 2553348 | 30001 | -0.25 | 0.000494 | 0.000679 |
| chr19 | 18473971 | 18548946 | 74976 | -0.19 | 0.000502 | 0.000689 |
| chr14 | 62160736 | 62230961 | 70226 | 0.19 | 0.000613 | 0.000839 |
| chr5 | 149821929 | 149904432 | 82504 | -0.2 | 0.000621 | 0.000849 |
| chr20 | 35898306 | 36028220 | 129915 | -0.13 | 0.000663 | 0.000905 |
| chr2 | 20365879 | 20432439 | 66561 | -0.19 | 0.000669 | 0.000913 |
| chr10 | 75756747 | 75883605 | 126859 | 0.16 | 0.000756 | 0.00103 |
| chr11 | 795501 | 864096 | 68596 | 0.18 | 0.000763 | 0.00104 |
| chr5 | 154133594 | 154218624 | 85031 | -0.18 | 0.000781 | 0.00106 |
| chr17 | 61817588 | 61926890 | 109303 | 0.15 | 0.000805 | 0.00109 |
| chr6 | 26148057 | 26208228 | 60172 | 0.17 | 0.000809 | 0.0011 |
| chr15 | 64752102 | 64996149 | 244048 | 0.14 | 0.000879 | 0.00119 |
| chr1 | 154897855 | 155037074 | 139220 | 0.14 | 0.000913 | 0.00123 |
| chr3 | 15779061 | 15858679 | 79619 | -0.17 | 0.000971 | 0.00131 |
| chr1 | 244995950 | 245029921 | 33972 | 0.17 | 0.001 | 0.00135 |
| chr20 | 48861732 | 48938981 | 77250 | -0.17 | 0.00101 | 0.00136 |
| chr2 | 55179791 | 55249970 | 70180 | -0.23 | 0.00111 | 0.00149 |
| chr4 | 74861871 | 74905734 | 43864 | 0.19 | 0.00112 | 0.0015 |
| chr12 | 31576532 | 31702094 | 125563 | 0.12 | 0.00117 | 0.00157 |
| chr5 | 106694897 | 106955801 | 260905 | 0.09 | 0.00124 | 0.00167 |
| chr1 | 207970176 | 208025273 | 55098 | 0.19 | 0.0013 | 0.00173 |
| chr17 | 33306694 | 33418025 | 111332 | 0.11 | 0.00142 | 0.0019 |
| chr7 | 72867383 | 72972720 | 105338 | 0.19 | 0.00148 | 0.00198 |
| chr10 | 101639760 | 101804592 | 164833 | 0.12 | 0.00156 | 0.00209 |
| chr2 | 36581757 | 36808125 | 226369 | -0.08 | 0.00166 | 0.00221 |
| chr14 | 102413874 | 102470072 | 56199 | 0.19 | 0.00174 | 0.00231 |
| chr15 | 72471787 | 72535823 | 64037 | -0.14 | 0.00185 | 0.00246 |
| chr17 | 70490748 | 70614576 | 123829 | 0.14 | 0.00187 | 0.00248 |
| chr6 | 136807197 | 136871653 | 64457 | -0.22 | 0.00189 | 0.0025 |
| chr11 | 47863882 | 47908269 | 44388 | -0.19 | 0.00197 | 0.00261 |
| chr19 | 42695418 | 42807756 | 112339 | -0.13 | 0.00206 | 0.00272 |
| chr17 | 8020753 | 8131648 | 110896 | 0.11 | 0.00208 | 0.00275 |
| chr19 | 56135734 | 56209328 | 73595 | -0.17 | 0.00217 | 0.00286 |
| chr17 | 46867942 | 47052921 | 184980 | -0.12 | 0.00219 | 0.00288 |
| chr11 | 67005992 | 67086202 | 80211 | -0.18 | 0.00223 | 0.00294 |
| chr5 | 133292474 | 133356491 | 64018 | 0.17 | 0.00281 | 0.00369 |
| chr16 | 84593406 | 84655886 | 62481 | 0.17 | 0.00285 | 0.00374 |
| chr20 | 3730152 | 3802985 | 72834 | 0.14 | 0.00289 | 0.00379 |
| chr1 | 233242403 | 233253235 | 10833 | 0.16 | 0.00298 | 0.00391 |
| chr15 | 67192111 | 67257210 | 65100 | -0.2 | 0.00319 | 0.00417 |
| chr11 | 35159900 | 35250679 | 90780 | -0.36 | 0.00327 | 0.00427 |
| chr10 | 112240994 | 112296416 | 55423 | -0.15 | 0.00331 | 0.00432 |
| chr9 | 33101330 | 33185549 | 84220 | 0.14 | 0.00359 | 0.00469 |
| chr11 | 47931388 | 47983300 | 51913 | -0.14 | 0.00365 | 0.00475 |
| chr16 | 73017394 | 73131308 | 113915 | -0.12 | 0.00367 | 0.00477 |
| chr4 | 84456364 | 84504088 | 47725 | -0.16 | 0.00385 | 0.005 |
| chr16 | 9161332 | 9224240 | 62909 | 0.17 | 0.00386 | 0.00501 |
| chr8 | 67024693 | 67045990 | 21298 | -0.17 | 0.00395 | 0.00511 |
| chr1 | 45240043 | 45286625 | 46583 | 0.16 | 0.00407 | 0.00526 |
| chr17 | 27423327 | 27508129 | 84803 | -0.14 | 0.00458 | 0.00592 |
| chr1 | 218632342 | 218674832 | 42491 | -0.13 | 0.00468 | 0.00604 |
| chr1 | 165796171 | 165881224 | 85054 | -0.15 | 0.0049 | 0.00632 |
| chr17 | 47067954 | 47134241 | 66288 | 0.12 | 0.00496 | 0.00639 |
| chr10 | 43878798 | 43917750 | 38953 | 0.16 | 0.0051 | 0.00656 |
| chr9 | 37900533 | 38071262 | 170730 | -0.09 | 0.00512 | 0.00658 |

| | | | | | | |
|---|---|---|---|---|---|---|
| chr17 | 71274972 | 71309297 | 34326 | -0.14 | 0.00538 | 0.0069 |
| chr7 | 75919518 | 76054916 | 135399 | 0.1 | 0.00543 | 0.00695 |
| chr5 | 14142197 | 14366987 | 224791 | 0.09 | 0.00546 | 0.00699 |
| chrX | 150000624 | 150068360 | 67737 | -0.14 | 0.00574 | 0.00734 |
| chr12 | 116784243 | 116895860 | 111618 | -0.14 | 0.00617 | 0.00788 |
| chr18 | 56515544 | 56681295 | 165752 | 0.1 | 0.00641 | 0.00817 |
| chr19 | 49042494 | 49149908 | 107415 | -0.12 | 0.00641 | 0.00817 |
| chr19 | 47571511 | 47777928 | 206418 | 0.09 | 0.00652 | 0.0083 |
| chr1 | 155895546 | 155953677 | 58132 | 0.15 | 0.00727 | 0.00924 |
| chr16 | 4303093 | 4443772 | 140680 | -0.14 | 0.00778 | 0.00988 |
| chr17 | 37000627 | 37079637 | 79011 | -0.13 | 0.00802 | 0.0102 |
| chr1 | 94019962 | 94170451 | 150490 | 0.1 | 0.00813 | 0.0103 |
| chr17 | 1321334 | 1395622 | 74289 | 0.15 | 0.00813 | 0.0103 |
| chr8 | 23376009 | 23428400 | 52392 | 0.14 | 0.00882 | 0.0112 |
| chr20 | 35431028 | 35493522 | 62495 | -0.22 | 0.00926 | 0.0117 |
| chr17 | 40457762 | 40584443 | 126682 | -0.11 | 0.00945 | 0.0119 |
| chr10 | 75639265 | 75700856 | 61592 | 0.15 | 0.00972 | 0.0122 |
| chr9 | 130823395 | 130866894 | 43500 | 0.15 | 0.0102 | 0.0128 |
| chrX | 45606755 | 45711661 | 104907 | 0.12 | 0.0103 | 0.0129 |
| chr2 | 85725522 | 85789548 | 64027 | -0.14 | 0.0104 | 0.013 |
| chr6 | 117803151 | 117877116 | 73966 | -0.14 | 0.0104 | 0.0131 |
| chr17 | 40020376 | 40087916 | 67541 | -0.14 | 0.0105 | 0.0132 |
| chr17 | 25853468 | 25926813 | 73346 | -0.13 | 0.011 | 0.0138 |
| chr1 | 249111478 | 249168428 | 56951 | 0.16 | 0.0116 | 0.0144 |
| chr5 | 90568678 | 90629242 | 60565 | -0.13 | 0.0117 | 0.0146 |
| chr2 | 42310981 | 42375595 | 64615 | 0.13 | 0.0117 | 0.0146 |
| chr14 | 105912308 | 105967433 | 55126 | -0.14 | 0.0119 | 0.0148 |
| chr9 | 114773470 | 114837417 | 63948 | -0.15 | 0.014 | 0.0174 |
| chr18 | 24033240 | 24072093 | 38854 | 0.12 | 0.014 | 0.0174 |
| chr6 | 56703474 | 56757557 | 54084 | -0.17 | 0.0143 | 0.0177 |
| chr2 | 85636926 | 85700513 | 63588 | 0.13 | 0.015 | 0.0186 |
| chr17 | 18084804 | 18197132 | 112329 | -0.13 | 0.015 | 0.0186 |
| chr16 | 84732780 | 84789088 | 56309 | 0.14 | 0.0158 | 0.0195 |
| chr17 | 36849946 | 36937868 | 87923 | -0.14 | 0.0177 | 0.0219 |
| chr19 | 39601292 | 39686518 | 85227 | -0.13 | 0.0183 | 0.0226 |
| chr9 | 127001988 | 127120592 | 118605 | 0.1 | 0.0203 | 0.025 |
| chr9 | 97540773 | 97596707 | 55935 | -0.11 | 0.0218 | 0.0268 |
| chr20 | 50302818 | 50384907 | 82090 | 0.1 | 0.0227 | 0.0279 |
| chr6 | 151177797 | 151287632 | 109836 | -0.12 | 0.024 | 0.0294 |
| chr8 | 48718272 | 48906086 | 187815 | 0.1 | 0.0241 | 0.0296 |
| chr10 | 104986600 | 105015778 | 29179 | 0.12 | 0.025 | 0.0306 |
| chr11 | 59303725 | 59348475 | 44751 | 0.12 | 0.0251 | 0.0307 |
| chr15 | 63614705 | 63674732 | 60028 | -0.09 | 0.0253 | 0.0309 |
| chr2 | 42395799 | 42525065 | 129267 | 0.11 | 0.0253 | 0.0309 |
| chr8 | 30532067 | 30602605 | 70539 | -0.14 | 0.0256 | 0.0312 |
| chr4 | 129472726 | 129524064 | 51339 | -0.13 | 0.0277 | 0.0337 |
| chr20 | 342602 | 427037 | 84436 | -0.11 | 0.0279 | 0.0339 |
| chr10 | 76944500 | 77006494 | 61995 | 0.12 | 0.0284 | 0.0346 |
| chr1 | 23850616 | 23925900 | 75285 | 0.11 | 0.0289 | 0.0351 |
| chr1 | 204366527 | 204393494 | 26968 | 0.12 | 0.0311 | 0.0377 |
| chr19 | 49946386 | 50002212 | 55827 | 0.12 | 0.0327 | 0.0396 |
| chr17 | 80476588 | 80568256 | 91669 | -0.12 | 0.0331 | 0.04 |
| chr19 | 48948002 | 49017836 | 69835 | -0.16 | 0.0334 | 0.0404 |
| chr11 | 64602690 | 64662488 | 59799 | -0.11 | 0.0348 | 0.042 |
| chr8 | 27713268 | 27806311 | 93044 | 0.1 | 0.0358 | 0.0432 |
| chr10 | 14548826 | 14659920 | 111095 | 0.1 | 0.0377 | 0.0453 |
| chr3 | 156391077 | 156440400 | 49324 | 0.11 | 0.0413 | 0.0497 |
| chr16 | 67866355 | 67928294 | 61940 | 0.12 | 0.0416 | 0.05 |
| chr10 | 123855671 | 124029255 | 173585 | 0.08 | 0.0434 | 0.0521 |
| chrX | 73458691 | 73513656 | 54966 | 0.12 | 0.0451 | 0.0541 |
| chr9 | 131872533 | 131941234 | 68702 | 0.1 | 0.0454 | 0.0544 |

| | | | | | | |
|---|---|---|---|---|---|---|
| chr18 | 45386655 | 45459820 | 73166 | 0.11 | 0.0465 | 0.0556 |
| chr18 | 46448227 | 46487919 | 39693 | -0.14 | 0.0469 | 0.0561 |
| chr22 | 46839832 | 46944075 | 104244 | -0.11 | 0.0474 | 0.0566 |
| chr17 | 16281168 | 16366266 | 85099 | -0.13 | 0.0487 | 0.0581 |
| chr1 | 156051817 | 156117036 | 65220 | 0.08 | 0.0522 | 0.0621 |
| chr19 | 50083243 | 50191486 | 108244 | 0.1 | 0.0522 | 0.0621 |
| chr17 | 42071935 | 42202121 | 130187 | -0.1 | 0.053 | 0.063 |
| chr10 | 114708463 | 114900505 | 192043 | -0.06 | 0.0535 | 0.0635 |
| chr16 | 69684792 | 69797673 | 112882 | -0.1 | 0.0543 | 0.0644 |
| chr22 | 46298860 | 46375077 | 76218 | -0.13 | 0.0544 | 0.0644 |
| chr11 | 47995837 | 48192801 | 196965 | 0.07 | 0.0563 | 0.0666 |
| chr7 | 130903909 | 130939507 | 35599 | 0.1 | 0.0599 | 0.0708 |
| chr19 | 38388558 | 38811081 | 422524 | 0.05 | 0.0613 | 0.0724 |
| chr20 | 46113543 | 46294672 | 181130 | 0.07 | 0.0619 | 0.073 |
| chr8 | 29187869 | 29210610 | 22742 | 0.21 | 0.0632 | 0.0744 |
| chr7 | 104612119 | 104692307 | 80189 | 0.11 | 0.0677 | 0.0796 |
| chr19 | 41168402 | 41285658 | 117257 | -0.07 | 0.0685 | 0.0805 |
| chr17 | 64649015 | 64804245 | 155231 | -0.09 | 0.0686 | 0.0806 |
| chr18 | 3581995 | 3723299 | 141305 | -0.07 | 0.0707 | 0.0829 |
| chr16 | 28138037 | 28224214 | 86178 | -0.16 | 0.0709 | 0.083 |
| chr2 | 86104355 | 86191144 | 86790 | 0.08 | 0.0729 | 0.0853 |
| chr19 | 54658059 | 54715756 | 57698 | 0.12 | 0.0733 | 0.0856 |
| chr8 | 128805964 | 129223734 | 417771 | 0.04 | 0.0746 | 0.0871 |
| chr10 | 32201077 | 32251608 | 50532 | 0.08 | 0.0765 | 0.0892 |
| chr1 | 201416938 | 201482461 | 65524 | 0.09 | 0.0766 | 0.0892 |
| chr3 | 124478798 | 124628936 | 150139 | -0.06 | 0.0767 | 0.0893 |
| chr12 | 120516938 | 120593525 | 76588 | 0.11 | 0.0778 | 0.0904 |
| chr21 | 44738989 | 44776219 | 37231 | 0.23 | 0.0797 | 0.0925 |
| chr17 | 62615059 | 62705974 | 90916 | -0.1 | 0.0829 | 0.0961 |
| chr2 | 17876850 | 17950216 | 73367 | -0.08 | 0.0829 | 0.0961 |
| chr15 | 91466000 | 91546977 | 80978 | 0.09 | 0.083 | 0.0961 |
| chr4 | 39497320 | 39561110 | 63791 | -0.08 | 0.0834 | 0.0964 |
| chr11 | 71774115 | 71824920 | 50806 | 0.1 | 0.0858 | 0.099 |
| chr17 | 75276731 | 75488547 | 211817 | 0.05 | 0.0858 | 0.099 |
| chr17 | 1613122 | 1640274 | 27153 | -0.11 | 0.0876 | 0.101 |
| chr10 | 69767482 | 69870584 | 103103 | 0.08 | 0.0963 | 0.111 |
| chr20 | 19830019 | 19928726 | 98708 | 0.06 | 0.0964 | 0.111 |
| chr1 | 39623189 | 39680146 | 56958 | 0.09 | 0.0981 | 0.113 |
| chr12 | 48146989 | 48223563 | 76575 | 0.08 | 0.0994 | 0.114 |
| chr15 | 85220029 | 85363117 | 143089 | -0.07 | 0.0998 | 0.114 |
| chr10 | 30707426 | 30754051 | 46626 | -0.12 | 0.1 | 0.115 |
| chr5 | 72900069 | 73006756 | 106688 | -0.09 | 0.101 | 0.115 |
| chr15 | 75912593 | 75965998 | 53406 | 0.09 | 0.101 | 0.115 |
| chr10 | 69539979 | 69615469 | 75491 | 0.08 | 0.101 | 0.115 |
| chr16 | 69407481 | 69516805 | 109325 | 0.06 | 0.105 | 0.12 |
| chr4 | 129730134 | 129794174 | 64041 | -0.09 | 0.11 | 0.125 |
| chr1 | 16084798 | 16128784 | 43987 | 0.11 | 0.11 | 0.125 |
| chr11 | 47502314 | 47601494 | 99181 | -0.09 | 0.111 | 0.127 |
| chr3 | 9424337 | 9484552 | 60216 | 0.09 | 0.113 | 0.128 |
| chr3 | 45634261 | 45732826 | 98566 | 0.08 | 0.115 | 0.13 |
| chr16 | 68288725 | 68345962 | 57238 | -0.11 | 0.117 | 0.133 |
| chr17 | 16912156 | 17094805 | 182650 | 0.05 | 0.118 | 0.133 |
| chr15 | 59396469 | 59681961 | 285493 | 0.04 | 0.12 | 0.135 |
| chr17 | 73422264 | 73567854 | 145591 | 0.07 | 0.123 | 0.139 |
| chr1 | 235529578 | 235576107 | 46530 | 0.09 | 0.124 | 0.14 |
| chr17 | 41131435 | 41208129 | 76695 | 0.08 | 0.126 | 0.142 |
| chr10 | 73922645 | 74125679 | 203035 | 0.05 | 0.131 | 0.148 |
| chr11 | 12694811 | 12945753 | 250943 | -0.04 | 0.136 | 0.153 |
| chr17 | 41580365 | 41637417 | 57053 | -0.1 | 0.136 | 0.153 |
| chr20 | 10385709 | 10429814 | 44106 | -0.09 | 0.137 | 0.154 |
| chr2 | 201979539 | 202023403 | 43865 | -0.07 | 0.138 | 0.155 |

| | | | | | | |
|---|---|---|---|---|---|---|
| chr10 | 121410047 | 121442317 | 32271 | -0.08 | 0.145 | 0.163 |
| chr7 | 47292993 | 47731428 | 438436 | -0.04 | 0.149 | 0.167 |
| chr2 | 127808846 | 127874092 | 65247 | 0.09 | 0.159 | 0.178 |
| chr19 | 47392967 | 47462506 | 69540 | -0.09 | 0.167 | 0.186 |
| chr6 | 2952004 | 3025157 | 73154 | -0.1 | 0.168 | 0.187 |
| chr2 | 70312343 | 70371309 | 58967 | -0.07 | 0.169 | 0.188 |
| chr18 | 56448387 | 56464013 | 15627 | 0.08 | 0.171 | 0.19 |
| chr10 | 123665593 | 123735541 | 69949 | 0.07 | 0.174 | 0.193 |
| chr3 | 8524327 | 8621237 | 96911 | -0.07 | 0.193 | 0.214 |
| chr1 | 23493658 | 23552834 | 59177 | 0.07 | 0.194 | 0.215 |
| chr12 | 117558198 | 117628272 | 70075 | 0.07 | 0.199 | 0.22 |
| chr10 | 62701962 | 62761774 | 59813 | -0.08 | 0.209 | 0.231 |
| chr9 | 35488707 | 35564039 | 75333 | -0.05 | 0.211 | 0.233 |
| chr10 | 73575729 | 73660462 | 84734 | -0.06 | 0.212 | 0.234 |
| chr19 | 13943498 | 13992005 | 48508 | 0.06 | 0.227 | 0.251 |
| chr17 | 42214865 | 42300421 | 85557 | -0.06 | 0.235 | 0.259 |
| chr15 | 90686588 | 90793592 | 107005 | 0.05 | 0.241 | 0.265 |
| chr11 | 65867485 | 65925765 | 58281 | -0.06 | 0.242 | 0.266 |
| chr2 | 178075335 | 178141000 | 65666 | -0.05 | 0.243 | 0.267 |
| chr9 | 111822348 | 111883037 | 60690 | 0.06 | 0.246 | 0.27 |
| chr16 | 11623277 | 11692648 | 69372 | 0.09 | 0.248 | 0.272 |
| chr22 | 40849413 | 40945632 | 96220 | -0.06 | 0.251 | 0.275 |
| chr6 | 53357963 | 53414685 | 56723 | -0.07 | 0.251 | 0.275 |
| chr11 | 73833121 | 73883685 | 50565 | 0.06 | 0.253 | 0.277 |
| chr2 | 28612208 | 28673659 | 61452 | -0.06 | 0.258 | 0.282 |
| chr20 | 19944154 | 20034036 | 89883 | -0.05 | 0.259 | 0.283 |
| chr15 | 66764627 | 66818498 | 53872 | -0.07 | 0.267 | 0.291 |
| chr15 | 90293549 | 90392899 | 99351 | -0.08 | 0.271 | 0.295 |
| chr5 | 135328992 | 135403253 | 74262 | 0.05 | 0.271 | 0.295 |
| chr3 | 171843601 | 171894913 | 51313 | -0.06 | 0.276 | 0.301 |
| chr19 | 749882 | 800131 | 50250 | 0.07 | 0.282 | 0.306 |
| chr4 | 75572602 | 75609114 | 36513 | -0.06 | 0.285 | 0.309 |
| chr9 | 74289211 | 74383992 | 94782 | 0.05 | 0.286 | 0.31 |
| chr10 | 104352680 | 104430274 | 77595 | -0.07 | 0.303 | 0.328 |
| chr17 | 76857284 | 76932989 | 75706 | -0.05 | 0.308 | 0.334 |
| chr15 | 63728626 | 63816139 | 87514 | -0.05 | 0.311 | 0.336 |
| chr2 | 238368394 | 238455753 | 87360 | -0.04 | 0.324 | 0.35 |
| chr10 | 96968789 | 97069738 | 100950 | -0.04 | 0.328 | 0.353 |
| chr19 | 46849470 | 46930537 | 81068 | 0.05 | 0.33 | 0.356 |
| chr14 | 23025463 | 23084608 | 59146 | -0.05 | 0.332 | 0.357 |
| chr11 | 18378094 | 18442537 | 64444 | -0.06 | 0.345 | 0.371 |
| chr17 | 40108658 | 40178185 | 69528 | 0.05 | 0.389 | 0.417 |
| chr13 | 95702259 | 95954571 | 252313 | -0.03 | 0.391 | 0.419 |
| chr19 | 54341489 | 54398960 | 57472 | 0.04 | 0.393 | 0.421 |
| chr20 | 31026900 | 31073804 | 46905 | 0.05 | 0.398 | 0.427 |
| chr6 | 129944247 | 130038874 | 94628 | -0.03 | 0.411 | 0.44 |
| chr1 | 36805454 | 36864154 | 58701 | -0.06 | 0.412 | 0.44 |
| chr11 | 33059917 | 33118010 | 58094 | 0.04 | 0.416 | 0.444 |
| chr15 | 90930479 | 90973880 | 43402 | 0.04 | 0.43 | 0.458 |
| chr17 | 7451698 | 7531222 | 79525 | -0.05 | 0.433 | 0.461 |
| chr5 | 159687195 | 159746475 | 59281 | -0.06 | 0.443 | 0.471 |
| chr14 | 35789556 | 35874847 | 85292 | -0.04 | 0.449 | 0.478 |
| chr6 | 26086856 | 26129119 | 42264 | -0.04 | 0.458 | 0.486 |
| chr5 | 179222607 | 179269140 | 46534 | 0.05 | 0.463 | 0.492 |
| chr20 | 48702247 | 48814133 | 111887 | -0.03 | 0.474 | 0.502 |
| chr8 | 141501464 | 141608748 | 107285 | 0.04 | 0.474 | 0.502 |
| chr20 | 49511902 | 49576022 | 64121 | -0.06 | 0.49 | 0.518 |
| chr19 | 45127988 | 45327576 | 199589 | 0.03 | 0.495 | 0.524 |
| chr10 | 34970874 | 35058664 | 87791 | 0.03 | 0.496 | 0.524 |
| chr17 | 7737019 | 7762551 | 25533 | -0.04 | 0.498 | 0.525 |
| chr16 | 30726123 | 30799382 | 73260 | -0.05 | 0.508 | 0.536 |

| | | | | | | |
|---|---|---|---|---|---|---|
| chr2 | 201217291 | 201289794 | 72504 | 0.04 | 0.516 | 0.543 |
| chr9 | 96212017 | 96264507 | 52491 | -0.04 | 0.519 | 0.546 |
| chr18 | 3401941 | 3469665 | 67725 | -0.03 | 0.529 | 0.555 |
| chr19 | 44022978 | 44124853 | 101876 | -0.04 | 0.531 | 0.557 |
| chr5 | 17212860 | 17285736 | 72877 | 0.04 | 0.547 | 0.574 |
| chr9 | 6645344 | 6716398 | 71055 | 0.04 | 0.553 | 0.58 |
| chr11 | 67111155 | 67170083 | 58929 | -0.02 | 0.572 | 0.598 |
| chr17 | 27684016 | 27787574 | 103559 | -0.03 | 0.573 | 0.599 |
| chr6 | 88374544 | 88443308 | 68765 | 0.03 | 0.586 | 0.612 |
| chr5 | 58262858 | 58697930 | 435073 | -0.01 | 0.599 | 0.626 |
| chr5 | 172158093 | 172237014 | 78922 | 0.02 | 0.609 | 0.635 |
| chr11 | 66713584 | 66774740 | 61157 | 0.04 | 0.613 | 0.638 |
| chr22 | 27136235 | 27174690 | 38456 | -0.03 | 0.654 | 0.681 |
| chr14 | 75720905 | 75777920 | 57016 | 0.05 | 0.663 | 0.69 |
| chr9 | 86534990 | 86619220 | 84231 | -0.06 | 0.674 | 0.7 |
| chr20 | 32943113 | 33035327 | 92215 | -0.02 | 0.701 | 0.728 |
| chr8 | 40009246 | 40074809 | 65564 | 0.01 | 0.708 | 0.734 |
| chr18 | 9594770 | 9649552 | 54783 | 0.02 | 0.712 | 0.737 |
| chr8 | 8691352 | 8751853 | 60502 | -0.02 | 0.716 | 0.741 |
| chr6 | 26272351 | 26366298 | 93948 | -0.02 | 0.723 | 0.747 |
| chr7 | 95024407 | 95073237 | 48831 | 0.02 | 0.728 | 0.752 |
| chr6 | 27082561 | 27129989 | 47429 | 0.02 | 0.74 | 0.763 |
| chr2 | 85131849 | 85294540 | 162692 | -0.01 | 0.741 | 0.763 |
| chr9 | 37414529 | 37509066 | 94538 | -0.02 | 0.741 | 0.763 |
| chr1 | 8916908 | 8973084 | 56177 | -0.02 | 0.757 | 0.779 |
| chr17 | 42966370 | 43025979 | 59610 | 0.02 | 0.758 | 0.779 |
| chr19 | 34649830 | 34725377 | 75548 | 0.01 | 0.766 | 0.786 |
| chr17 | 38254274 | 38337271 | 82998 | 0.02 | 0.766 | 0.786 |
| chr1 | 12585023 | 12680686 | 95664 | -0.01 | 0.775 | 0.794 |
| chr2 | 101434687 | 101777553 | 342867 | -0.01 | 0.785 | 0.804 |
| chr19 | 34842144 | 34985543 | 143400 | 0.01 | 0.8 | 0.818 |
| chr9 | 33813254 | 33889685 | 76432 | 0.01 | 0.804 | 0.821 |
| chr17 | 78215803 | 78269887 | 54085 | 0.01 | 0.805 | 0.822 |
| chr10 | 71183654 | 71268009 | 84356 | 0.01 | 0.815 | 0.832 |
| chr17 | 7107411 | 7168432 | 61022 | -0.01 | 0.816 | 0.832 |
| chr19 | 32834631 | 32897759 | 63129 | 0.01 | 0.818 | 0.833 |
| chr1 | 205227243 | 205294676 | 67434 | -0.01 | 0.836 | 0.85 |
| chr17 | 40598562 | 40742439 | 143878 | 0.01 | 0.836 | 0.85 |
| chr2 | 61607144 | 61766915 | 159772 | 0.01 | 0.838 | 0.851 |
| chr14 | 69329380 | 69455577 | 126198 | -0.01 | 0.853 | 0.865 |
| chr17 | 79779047 | 79896086 | 117040 | -0.01 | 0.86 | 0.871 |
| chr20 | 17534812 | 17663630 | 128819 | -0.01 | 0.885 | 0.896 |
| chr1 | 65347100 | 65407667 | 60568 | 0.01 | 0.886 | 0.896 |
| chr5 | 172259111 | 172387764 | 128654 | -0.01 | 0.887 | 0.896 |
| chr1 | 10433679 | 10491656 | 57978 | 0.01 | 0.89 | 0.898 |
| chr6 | 158395456 | 158502122 | 106667 | -0.01 | 0.89 | 0.898 |
| chr3 | 53249891 | 53319681 | 69791 | 0.01 | 0.9 | 0.907 |
| chr10 | 82188064 | 82296632 | 108569 | 0 | 0.905 | 0.911 |
| chr15 | 75134796 | 75232018 | 97223 | 0 | 0.925 | 0.93 |
| chr9 | 132077781 | 132125164 | 47384 | 0 | 0.965 | 0.969 |
| chr17 | 46100231 | 46186589 | 86359 | 0 | 0.966 | 0.97 |
| chr9 | 97604868 | 97715661 | 110794 | 0 | 0.967 | 0.97 |
| chr7 | 55086047 | 55277370 | 191324 | 0 | 0.97 | 0.971 |
| chr3 | 8483335 | 8500814 | 17480 | 0 | 0.976 | 0.977 |
| chr9 | 135978297 | 136025508 | 47212 | 0 | 0.993 | 0.993 |

**Supplementary Table 4E, CTCF differential binding analysis, Related to Supplementary Figure 5N, Q**

| chrom | start | end | Density Control | Density SETDB1_LOF | Fold change |
|---|---|---|---|---|---|
| chr2 | 115866592 | 115866884 | 0.875205 | 0.0239187 | -36.591 |
| chr2 | 180383687 | 180383889 | 1.33511 | 0.0687639 | -19.416 |
| chr2 | 113940853 | 113941067 | 1.02822 | 0.0604691 | -17.004 |
| chr8 | 116365125 | 116365314 | 1.62473 | 0.122083 | -13.308 |
| chr5 | 45417075 | 45417293 | 1.07651 | 0.103044 | -10.447 |
| chr17 | 15659564 | 15659747 | 1.09435 | 0.111625 | -9.804 |
| chr3 | 176187580 | 176187790 | 1.3559 | 0.150165 | -9.029 |
| chr2 | 144355443 | 144355634 | 1.60373 | 0.183306 | -8.749 |
| chr2 | 55893685 | 55893911 | 1.34768 | 0.161195 | -8.361 |
| chr17 | 75811436 | 75811652 | 1.21959 | 0.147363 | -8.276 |
| chr2 | 52801720 | 52801952 | 1.22347 | 0.148171 | -8.257 |
| chr2 | 23552623 | 23552815 | 1.3067 | 0.159958 | -8.169 |
| chr6 | 129219953 | 129220157 | 0.93916 | 0.115682 | -8.118 |
| chr2 | 142520509 | 142520744 | 1.06485 | 0.133806 | -7.958 |
| chr14 | 85212181 | 85212365 | 1.09977 | 0.138498 | -7.941 |
| chr7 | 74017838 | 74018052 | 1.11729 | 0.141688 | -7.886 |
| chr6 | 121637593 | 121637780 | 0.943092 | 0.121428 | -7.767 |
| chr5 | 53901601 | 53901792 | 1.10723 | 0.142695 | -7.759 |
| chr2 | 133542715 | 133542932 | 1.2083 | 0.156466 | -7.722 |
| chr2 | 107831007 | 107831337 | 1.18601 | 0.159684 | -7.427 |
| chr1 | 179597019 | 179597212 | 1.06623 | 0.148022 | -7.203 |
| chr5 | 128664220 | 128664411 | 0.999357 | 0.139296 | -7.174 |
| chr8 | 113945538 | 113945785 | 1.13725 | 0.158609 | -7.170 |
| chr2 | 84725251 | 84725455 | 1.15364 | 0.165943 | -6.952 |
| chr2 | 134641496 | 134641776 | 1.08693 | 0.157212 | -6.914 |
| chr20 | 24530115 | 24530425 | 1.57054 | 0.228272 | -6.880 |
| chr2 | 207782968 | 207783263 | 1.12719 | 0.16408 | -6.870 |
| chr2 | 82045511 | 82045709 | 1.4099 | 0.20991 | -6.717 |
| chr2 | 21779282 | 21779465 | 1.13123 | 0.171175 | -6.609 |
| chr2 | 184595357 | 184595549 | 1.1108 | 0.171055 | -6.494 |
| chr7 | 2296739 | 2297138 | 1.09424 | 0.168537 | -6.493 |
| chr2 | 229682846 | 229683200 | 0.893519 | 0.137752 | -6.486 |
| chr2 | 105267343 | 105267546 | 0.905224 | 0.140233 | -6.455 |
| chr7 | 49432566 | 49432778 | 1.60035 | 0.251404 | -6.366 |
| chr11 | 18281879 | 18282165 | 1.06579 | 0.169016 | -6.306 |
| chr7 | 12186210 | 12186399 | 1.11298 | 0.178052 | -6.251 |
| chr8 | 14286760 | 14286993 | 0.976733 | 0.156637 | -6.236 |
| chr2 | 139988846 | 139989102 | 1.23515 | 0.198697 | -6.216 |
| chr7 | 150232239 | 150232489 | 0.624237 | 0.101068 | -6.176 |
| chr2 | 100152903 | 100153360 | 1.07334 | 0.175997 | -6.099 |
| chr2 | 22679840 | 22680046 | 0.993009 | 0.164542 | -6.035 |
| chr5 | 75569611 | 75569797 | 1.02991 | 0.170918 | -6.026 |
| chr2 | 146644705 | 146645049 | 1.42249 | 0.236973 | -6.003 |
| chr2 | 134207595 | 134207795 | 0.954583 | 0.159172 | -5.997 |

| | | | | | |
|---|---|---|---|---|---|
| chrX | 131752409 | 131752611 | 1.14077 | 0.192205 | -5.935 |
| chr20 | 18151475 | 18151687 | 0.900201 | 0.152032 | -5.921 |
| chr5 | 119866947 | 119867165 | 1.24764 | 0.212975 | -5.858 |
| chr2 | 238139062 | 238139246 | 1.07907 | 0.184336 | -5.854 |
| chr10 | 61159662 | 61159854 | 0.814894 | 0.140625 | -5.795 |
| chr11 | 125572506 | 125572699 | 0.998672 | 0.172844 | -5.778 |
| chr9 | 101258695 | 101258880 | 1.27939 | 0.22187 | -5.766 |
| chr2 | 131071336 | 131071533 | 0.853679 | 0.148459 | -5.750 |
| chr16 | 61535800 | 61536005 | 1.04337 | 0.181469 | -5.750 |
| chr8 | 16719611 | 16719835 | 0.895125 | 0.155788 | -5.746 |
| chr20 | 44226530 | 44226737 | 1.11902 | 0.195177 | -5.733 |
| chr2 | 8364563 | 8364756 | 2.03584 | 0.362221 | -5.620 |
| chr8 | 91752784 | 91753041 | 1.3024 | 0.231749 | -5.620 |
| chr9 | 94822123 | 94822413 | 1.2376 | 0.220476 | -5.613 |
| chr5 | 177539850 | 177540057 | 0.841119 | 0.150767 | -5.579 |
| chr2 | 118131720 | 118131986 | 0.8438 | 0.152509 | -5.533 |
| chrX | 75933883 | 75934088 | 1.08665 | 0.196739 | -5.523 |
| chr14 | 29769923 | 29770167 | 1.00551 | 0.182458 | -5.511 |
| chr5 | 139274749 | 139274937 | 1.23743 | 0.224701 | -5.507 |
| chr2 | 166259770 | 166259967 | 1.27464 | 0.232369 | -5.485 |
| chr5 | 43232999 | 43233210 | 1.24123 | 0.227036 | -5.467 |
| chr5 | 65638549 | 65638731 | 1.30609 | 0.239414 | -5.455 |
| chr3 | 30002947 | 30003132 | 1.13596 | 0.208268 | -5.454 |
| chr2 | 140287205 | 140287609 | 1.07564 | 0.199243 | -5.399 |
| chr2 | 148308659 | 148308850 | 0.975965 | 0.181231 | -5.385 |
| chr11 | 82274986 | 82275188 | 0.98538 | 0.183179 | -5.379 |
| chr14 | 77885070 | 77885256 | 1.15029 | 0.214148 | -5.371 |
| chr2 | 212956893 | 212957187 | 0.800687 | 0.149548 | -5.354 |
| chr2 | 156555832 | 156556101 | 1.25893 | 0.236172 | -5.331 |
| chr8 | 60285893 | 60286100 | 1.00734 | 0.188996 | -5.330 |
| chr5 | 7563753 | 7563948 | 1.25191 | 0.235544 | -5.315 |
| chr2 | 194305454 | 194305668 | 1.28804 | 0.242866 | -5.303 |
| chr9 | 5389277 | 5389482 | 1.15416 | 0.219827 | -5.250 |
| chr3 | 3651930 | 3652198 | 1.29253 | 0.24648 | -5.244 |
| chr2 | 186067294 | 186067476 | 1.17766 | 0.225501 | -5.222 |
| chr2 | 133976865 | 133977068 | 1.32444 | 0.254282 | -5.209 |
| chr10 | 122147556 | 122147828 | 1.3068 | 0.251864 | -5.189 |
| chr17 | 59471746 | 59472006 | 1.33556 | 0.257703 | -5.183 |
| chr5 | 22323005 | 22323221 | 0.847735 | 0.163987 | -5.170 |
| chr2 | 82577100 | 82577284 | 1.22805 | 0.238252 | -5.154 |
| chr4 | 60919116 | 60919314 | 1.15642 | 0.225388 | -5.131 |
| chr5 | 5598317 | 5598514 | 1.09444 | 0.21333 | -5.130 |
| chr2 | 14045917 | 14046146 | 1.11631 | 0.217815 | -5.125 |
| chr1 | 245840288 | 245840622 | 0.926582 | 0.181282 | -5.111 |
| chr11 | 57039806 | 57040094 | 1.05784 | 0.207221 | -5.105 |
| chr2 | 130573141 | 130573594 | 1.19316 | 0.234182 | -5.095 |
| chr2 | 186391022 | 186391214 | 1.46522 | 0.28797 | -5.088 |
| chr2 | 31026091 | 31026342 | 1.32277 | 0.261187 | -5.064 |
| chr14 | 78897318 | 78897580 | 0.850621 | 0.168297 | -5.054 |
| chr4 | 152774742 | 152774938 | 0.974292 | 0.193913 | -5.024 |
| chr5 | 110606054 | 110606292 | 1.23647 | 0.24615 | -5.023 |
| chr2 | 115800082 | 115800300 | 0.97812 | 0.195331 | -5.008 |
| chr12 | 116369140 | 116369328 | 1.45675 | 0.290936 | -5.007 |
| chr2 | 122739967 | 122740169 | 1.34814 | 0.271271 | -4.970 |
| chrX | 44271834 | 44272017 | 0.987186 | 0.199006 | -4.961 |
| chr8 | 113338781 | 113339127 | 1.14831 | 0.231841 | -4.953 |
| chr3 | 24753273 | 24753490 | 1.0772 | 0.218998 | -4.919 |
| chr2 | 51769463 | 51769646 | 1.21208 | 0.246934 | -4.909 |
| chr4 | 68070942 | 68071165 | 0.862598 | 0.175777 | -4.907 |

| | | | | | |
|---|---|---|---|---|---|
| chr11 | 5430237 | 5430527 | 1.34766 | 0.274742 | -4.905 |
| chr4 | 148581561 | 148581774 | 1.30388 | 0.266339 | -4.896 |
| chr10 | 116752919 | 116753101 | 1.80908 | 0.370106 | -4.888 |
| chr7 | 2235840 | 2236060 | 1.02298 | 0.209406 | -4.885 |
| chr8 | 81696432 | 81696643 | 1.0867 | 0.222503 | -4.884 |
| chr9 | 30598829 | 30599011 | 1.29242 | 0.265321 | -4.871 |
| chr2 | 213445738 | 213445951 | 1.08682 | 0.223464 | -4.864 |
| chr2 | 225641741 | 225641933 | 1.03103 | 0.212119 | -4.861 |
| chr2 | 112770536 | 112770759 | 1.10151 | 0.226804 | -4.857 |
| chr3 | 102698914 | 102699222 | 1.30682 | 0.269109 | -4.856 |
| chr9 | 96189008 | 96189365 | 1.18254 | 0.244214 | -4.842 |
| chr2 | 119294065 | 119294344 | 1.12329 | 0.232485 | -4.832 |
| chr2 | 53862747 | 53862948 | 1.60246 | 0.331998 | -4.827 |
| chr5 | 27749985 | 27750341 | 1.14615 | 0.238173 | -4.812 |
| chr3 | 119994543 | 119994735 | 1.37651 | 0.286543 | -4.804 |
| chr5 | 61485148 | 61485353 | 1.05411 | 0.219447 | -4.803 |
| chr8 | 119975158 | 119975340 | 1.3057 | 0.271991 | -4.801 |
| chr5 | 8573143 | 8573341 | 1.0298 | 0.214629 | -4.798 |
| chr2 | 4298484 | 4298700 | 1.37488 | 0.28666 | -4.796 |
| chr2 | 18331010 | 18331227 | 2.71766 | 0.567281 | -4.791 |
| chr20 | 18008469 | 18008702 | 1.04565 | 0.218994 | -4.775 |
| chr4 | 182827795 | 182828021 | 1.1077 | 0.232834 | -4.757 |
| chr9 | 96489666 | 96489858 | 1.22761 | 0.258392 | -4.751 |
| chr5 | 85013397 | 85013645 | 0.989593 | 0.208749 | -4.741 |
| chr15 | 76457686 | 76457869 | 1.20869 | 0.25499 | -4.740 |
| chr2 | 62596465 | 62596665 | 1.10085 | 0.233031 | -4.724 |
| chr1 | 180312890 | 180313150 | 1.29853 | 0.275228 | -4.718 |
| chr16 | 13305654 | 13305852 | 0.915965 | 0.194466 | -4.710 |
| chr10 | 65866853 | 65867045 | 1.86635 | 0.396325 | -4.709 |
| chr9 | 92681704 | 92681905 | 1.53874 | 0.326785 | -4.709 |
| chr8 | 83891782 | 83892000 | 1.32592 | 0.283447 | -4.678 |
| chr14 | 84156582 | 84156778 | 0.957517 | 0.204745 | -4.677 |
| chr7 | 118491943 | 118492125 | 1.10467 | 0.236466 | -4.672 |
| chr2 | 166782878 | 166783091 | 0.898916 | 0.192551 | -4.668 |
| chr2 | 106605144 | 106605398 | 1.00798 | 0.215976 | -4.667 |
| chr2 | 211524839 | 211525057 | 1.02887 | 0.2205 | -4.666 |
| chr7 | 104157364 | 104157580 | 1.22388 | 0.262426 | -4.664 |
| chr2 | 23577918 | 23578388 | 1.07909 | 0.231592 | -4.659 |
| chr5 | 161861224 | 161861497 | 0.929527 | 0.199902 | -4.650 |
| chr5 | 89344535 | 89344721 | 1.21927 | 0.262784 | -4.640 |
| chr10 | 100399077 | 100399261 | 0.957144 | 0.206338 | -4.639 |
| chr1 | 233704900 | 233705158 | 0.997799 | 0.21524 | -4.636 |
| chr2 | 212575318 | 212575535 | 0.873484 | 0.188769 | -4.627 |
| chr2 | 123286463 | 123286677 | 1.35506 | 0.293041 | -4.624 |
| chr2 | 131690486 | 131690678 | 1.43268 | 0.310332 | -4.617 |
| chr2 | 236564066 | 236564384 | 0.976754 | 0.211755 | -4.613 |
| chr7 | 11964970 | 11965166 | 0.998888 | 0.216877 | -4.606 |
| chr3 | 95743684 | 95743919 | 0.809645 | 0.17643 | -4.589 |
| chr10 | 98905203 | 98905415 | 1.20215 | 0.26256 | -4.579 |
| chr1 | 177917141 | 177917360 | 0.996001 | 0.217677 | -4.576 |
| chr2 | 150595610 | 150595869 | 1.06467 | 0.232708 | -4.575 |
| chr11 | 77948902 | 77949084 | 1.04081 | 0.227556 | -4.574 |
| chr2 | 3254025 | 3254285 | 0.921762 | 0.201753 | -4.569 |
| chr10 | 91940230 | 91940427 | 1.37912 | 0.301941 | -4.568 |
| chr5 | 85767748 | 85767951 | 1.57046 | 0.344221 | -4.562 |
| chr16 | 23268840 | 23269030 | 0.986554 | 0.216626 | -4.554 |
| chr19 | 56492500 | 56492780 | 1.28897 | 0.28316 | -4.552 |
| chr2 | 9455462 | 9455671 | 1.09628 | 0.24087 | -4.551 |
| chr2 | 100451792 | 100452218 | 1.26946 | 0.279279 | -4.545 |
| chr7 | 142658167 | 142658368 | 1.04889 | 0.231144 | -4.538 |
| chr2 | 121240683 | 121240888 | 0.951497 | 0.209786 | -4.536 |

| | | | | | |
|---|---|---|---|---|---|
| chr2 | 123976898 | 123977123 | 1.31981 | 0.291501 | -4.528 |
| chr17 | 32114533 | 32114812 | 1.23528 | 0.273133 | -4.523 |
| chr2 | 125072562 | 125072776 | 1.24785 | 0.276113 | -4.519 |
| chr2 | 60445393 | 60445616 | 1.33809 | 0.29817 | -4.488 |
| chr2 | 205161043 | 205161228 | 1.4349 | 0.319947 | -4.485 |
| chr1 | 73486892 | 73487135 | 1.21402 | 0.270802 | -4.483 |
| chr10 | 113663237 | 113663446 | 1.40628 | 0.313881 | -4.480 |
| chr4 | 54193537 | 54193728 | 1.34296 | 0.300502 | -4.469 |
| chr8 | 72140355 | 72140552 | 1.5667 | 0.350651 | -4.468 |
| chr8 | 114684675 | 114684860 | 0.855129 | 0.191399 | -4.468 |
| chr2 | 225677940 | 225678184 | 0.80002 | 0.179228 | -4.464 |
| chr3 | 83642489 | 83642760 | 1.00088 | 0.224428 | -4.460 |
| chr14 | 40684954 | 40685148 | 1.07029 | 0.24019 | -4.456 |
| chr2 | 98827476 | 98827661 | 1.18741 | 0.266587 | -4.454 |
| chr2 | 51824988 | 51825318 | 1.07749 | 0.242042 | -4.452 |
| chr2 | 217420280 | 217420467 | 1.29838 | 0.292302 | -4.442 |
| chr10 | 59293329 | 59293533 | 0.791119 | 0.178234 | -4.439 |
| chr9 | 84528936 | 84529151 | 0.777382 | 0.175542 | -4.428 |
| chr9 | 26313947 | 26314132 | 0.832799 | 0.188085 | -4.428 |
| chr1 | 216951630 | 216952004 | 0.90886 | 0.205428 | -4.424 |
| chr3 | 36623928 | 36624123 | 1.04239 | 0.23584 | -4.420 |
| chr3 | 95015747 | 95015992 | 1.18886 | 0.269337 | -4.414 |
| chr2 | 125498340 | 125498576 | 1.3484 | 0.306117 | -4.405 |
| chr2 | 194390542 | 194390733 | 0.8875 | 0.201669 | -4.401 |
| chr5 | 123271476 | 123271737 | 0.943243 | 0.214956 | -4.388 |
| chr2 | 140840850 | 140841156 | 0.900692 | 0.205383 | -4.385 |
| chr2 | 50008027 | 50008357 | 0.879994 | 0.200802 | -4.382 |
| chr8 | 115875838 | 115876087 | 1.72301 | 0.394597 | -4.367 |
| chr2 | 235063866 | 235064057 | 1.2199 | 0.279381 | -4.366 |
| chr14 | 101651597 | 101651786 | 1.32073 | 0.302798 | -4.362 |
| chr16 | 55907835 | 55908035 | 0.814198 | 0.186745 | -4.360 |
| chr2 | 213322177 | 213322423 | 0.840922 | 0.193131 | -4.354 |
| chr2 | 80992956 | 80993152 | 1.30008 | 0.298618 | -4.354 |
| chr15 | 44769270 | 44769454 | 0.90217 | 0.207732 | -4.343 |
| chr18 | 62989075 | 62989273 | 0.995699 | 0.229336 | -4.342 |
| chr1 | 215184564 | 215184781 | 0.858058 | 0.197762 | -4.339 |
| chr11 | 4934325 | 4934620 | 1.0084 | 0.232639 | -4.335 |
| chr2 | 147614928 | 147615166 | 0.80401 | 0.185773 | -4.328 |
| chr20 | 61822107 | 61822319 | 0.803414 | 0.185757 | -4.325 |
| chr1 | 241541598 | 241541844 | 1.06041 | 0.245765 | -4.315 |
| chr8 | 50289321 | 50289624 | 0.950092 | 0.220266 | -4.313 |
| chr17 | 52009329 | 52009555 | 1.68533 | 0.391069 | -4.310 |
| chr5 | 50925951 | 50926262 | 1.07648 | 0.250052 | -4.305 |
| chr1 | 192851642 | 192851828 | 1.28371 | 0.298288 | -4.304 |
| chr2 | 127139051 | 127139277 | 1.17256 | 0.27261 | -4.301 |
| chr10 | 124232717 | 124232906 | 1.01583 | 0.236258 | -4.300 |
| chr8 | 83230937 | 83231146 | 1.01808 | 0.237511 | -4.286 |
| chr16 | 7727739 | 7727930 | 1.19326 | 0.278557 | -4.284 |
| chr3 | 88345428 | 88345615 | 1.02773 | 0.240063 | -4.281 |
| chr2 | 90108167 | 90108452 | 2.53001 | 0.591746 | -4.275 |
| chr14 | 106266739 | 106266940 | 0.985617 | 0.230669 | -4.273 |
| chr8 | 15740092 | 15740281 | 1.14499 | 0.268287 | -4.268 |
| chr2 | 213635126 | 213635353 | 1.17037 | 0.274242 | -4.268 |
| chr2 | 98375260 | 98375493 | 1.22803 | 0.287793 | -4.267 |
| chr1 | 234379913 | 234380109 | 0.938339 | 0.219936 | -4.266 |
| chr8 | 14572805 | 14573015 | 1.27079 | 0.297968 | -4.265 |
| chr9 | 36234569 | 36234841 | 1.06555 | 0.249867 | -4.264 |
| chr10 | 65959199 | 65959438 | 1.24446 | 0.291864 | -4.264 |
| chr2 | 27050338 | 27050566 | 1.66185 | 0.390288 | -4.258 |
| chr7 | 39957053 | 39957261 | 1.18746 | 0.279202 | -4.253 |
| chr11 | 11392814 | 11393000 | 1.28753 | 0.30311 | -4.248 |

| | | | | | |
|---|---|---|---|---|---|
| chr16 | 13343638 | 13343909 | 1.11689 | 0.26325 | -4.243 |
| chr17 | 65896942 | 65897125 | 1.63617 | 0.38603 | -4.238 |
| chr2 | 215021154 | 215021430 | 1.13343 | 0.26752 | -4.237 |
| chr8 | 5207490 | 5207674 | 1.0716 | 0.253009 | -4.235 |
| chr5 | 86675664 | 86675862 | 0.910719 | 0.215048 | -4.235 |
| chr2 | 8647553 | 8647748 | 1.23343 | 0.292251 | -4.220 |
| chr8 | 87950784 | 87950975 | 1.10865 | 0.263032 | -4.215 |
| chr9 | 136628629 | 136628857 | 1.34364 | 0.318949 | -4.213 |
| chr5 | 61098868 | 61099094 | 1.70718 | 0.40586 | -4.206 |
| chr3 | 164545772 | 164545964 | 1.18551 | 0.282074 | -4.203 |
| chr5 | 44722884 | 44723086 | 1.32214 | 0.314988 | -4.197 |
| chr2 | 219017730 | 219018012 | 0.895363 | 0.213553 | -4.193 |
| chr5 | 117079768 | 117080155 | 1.24927 | 0.298002 | -4.192 |
| chr4 | 33035609 | 33035796 | 1.18081 | 0.281936 | -4.188 |
| chr16 | 4750583 | 4750888 | 0.943329 | 0.226348 | -4.168 |
| chr15 | 33641011 | 33641197 | 0.984059 | 0.236291 | -4.165 |
| chr11 | 84665073 | 84665264 | 1.34703 | 0.323483 | -4.164 |
| chr17 | 16305304 | 16305537 | 1.06524 | 0.255889 | -4.163 |
| chr8 | 39164966 | 39165153 | 1.2622 | 0.303882 | -4.154 |
| chr2 | 38659897 | 38660098 | 1.16796 | 0.281409 | -4.150 |
| chr2 | 206417896 | 206418164 | 1.13101 | 0.272604 | -4.149 |
| chr2 | 242805763 | 242806062 | 1.15685 | 0.278854 | -4.149 |
| chr17 | 29490946 | 29491179 | 1.18028 | 0.284611 | -4.147 |
| chr2 | 133137699 | 133137935 | 1.18134 | 0.285251 | -4.141 |
| chr7 | 126955010 | 126955196 | 1.3456 | 0.325004 | -4.140 |
| chr5 | 140103158 | 140103349 | 0.868388 | 0.209772 | -4.140 |
| chr7 | 139983939 | 139984165 | 1.11664 | 0.269759 | -4.139 |
| chr3 | 82273742 | 82274010 | 0.870311 | 0.210255 | -4.139 |
| chr2 | 187919718 | 187919906 | 1.46148 | 0.354491 | -4.123 |
| chr2 | 21636703 | 21636932 | 1.13562 | 0.275455 | -4.123 |
| chr1 | 189041370 | 189041639 | 1.13796 | 0.2761 | -4.122 |
| chr5 | 51359368 | 51359574 | 1.14508 | 0.27784 | -4.121 |
| chr2 | 11152301 | 11152524 | 1.08482 | 0.263831 | -4.112 |
| chr9 | 92792592 | 92792796 | 1.66831 | 0.406273 | -4.106 |
| chr2 | 80867181 | 80867364 | 1.11182 | 0.271019 | -4.102 |
| chr9 | 120683699 | 120683922 | 1.01256 | 0.246933 | -4.101 |
| chr2 | 239684469 | 239684819 | 0.947127 | 0.231605 | -4.089 |
| chr10 | 89715285 | 89715482 | 1.70567 | 0.417238 | -4.088 |
| chr2 | 116433596 | 116433809 | 1.3649 | 0.334115 | -4.085 |
| chr2 | 51092155 | 51092376 | 1.29295 | 0.31658 | -4.084 |
| chr1 | 227774365 | 227774654 | 1.25813 | 0.308247 | -4.082 |
| chr20 | 38629666 | 38629888 | 1.35898 | 0.333811 | -4.071 |
| chr5 | 75341456 | 75341784 | 1.21865 | 0.299602 | -4.068 |
| chr2 | 211988638 | 211988836 | 1.34133 | 0.329987 | -4.065 |
| chr16 | 85957076 | 85957308 | 1.19298 | 0.294041 | -4.057 |
| chr5 | 103991313 | 103991540 | 1.23637 | 0.304786 | -4.057 |
| chr2 | 144270762 | 144271000 | 1.86404 | 0.459524 | -4.056 |
| chr1 | 242034273 | 242034536 | 1.25753 | 0.31036 | -4.052 |
| chr7 | 146417426 | 146417625 | 1.10005 | 0.271894 | -4.046 |
| chr7 | 113678690 | 113678915 | 1.12859 | 0.278995 | -4.045 |
| chr2 | 83897496 | 83897762 | 1.1556 | 0.285893 | -4.042 |
| chr14 | 79591875 | 79592231 | 1.2621 | 0.312255 | -4.042 |
| chr8 | 115334853 | 115335095 | 0.771088 | 0.191266 | -4.031 |
| chr3 | 25250242 | 25250510 | 1.17352 | 0.291276 | -4.029 |
| chr1 | 241039637 | 241040050 | 1.18882 | 0.29584 | -4.018 |
| chr1 | 8172932 | 8173180 | 1.06687 | 0.265676 | -4.016 |
| chr19 | 57245920 | 57246105 | 1.08228 | 0.270135 | -4.006 |
| chr11 | 72409048 | 72409324 | 0.98324 | 0.245718 | -4.002 |
| chr10 | 28203127 | 28203320 | 1.19349 | 0.298731 | -3.995 |
| chr8 | 115794846 | 115795028 | 1.23322 | 0.308939 | -3.992 |
| chr7 | 141063172 | 141063470 | 0.799727 | 0.200351 | -3.992 |

| | | | | | |
|---|---|---|---|---|---|
| chr2 | 128587522 | 128587819 | 1.36425 | 0.341813 | -3.991 |
| chr16 | 17623721 | 17623906 | 1.16832 | 0.292827 | -3.990 |
| chr8 | 84278151 | 84278358 | 1.28487 | 0.322162 | -3.988 |
| chr16 | 62720823 | 62721056 | 0.997375 | 0.250248 | -3.986 |
| chr3 | 12775791 | 12776061 | 1.11302 | 0.279705 | -3.979 |
| chr5 | 146157358 | 146157571 | 0.864979 | 0.217665 | -3.974 |
| chr7 | 91590587 | 91590772 | 1.144 | 0.288385 | -3.967 |
| chr1 | 244914841 | 244915086 | 1.02467 | 0.258362 | -3.966 |
| chr9 | 10979643 | 10979847 | 1.16626 | 0.294186 | -3.964 |
| chr9 | 105198646 | 105198845 | 0.827548 | 0.208818 | -3.963 |
| chr19 | 22283400 | 22283589 | 1.56235 | 0.394347 | -3.962 |
| chr2 | 123937256 | 123937538 | 0.788782 | 0.199274 | -3.958 |
| chr5 | 154668465 | 154668713 | 1.07752 | 0.272741 | -3.951 |
| chr2 | 166986326 | 166986518 | 1.33005 | 0.336841 | -3.949 |
| chr2 | 126985710 | 126986097 | 1.15122 | 0.29166 | -3.947 |
| chrX | 54377884 | 54378240 | 1.11152 | 0.282041 | -3.941 |
| chr2 | 212139475 | 212139671 | 1.10735 | 0.281277 | -3.937 |
| chr8 | 9203798 | 9203986 | 1.16166 | 0.295501 | -3.931 |
| chr2 | 66604373 | 66604620 | 1.11066 | 0.282639 | -3.930 |
| chr8 | 2393277 | 2393459 | 0.927674 | 0.236288 | -3.926 |
| chr1 | 212603740 | 212603922 | 1.35285 | 0.344737 | -3.924 |
| chr1 | 56395475 | 56395684 | 1.29429 | 0.330025 | -3.922 |
| chr2 | 167517411 | 167517711 | 0.990829 | 0.252655 | -3.922 |
| chr7 | 41754609 | 41754878 | 1.26554 | 0.322784 | -3.921 |
| chr2 | 110407780 | 110408035 | 1.15665 | 0.295262 | -3.917 |
| chr7 | 12571378 | 12571601 | 0.94632 | 0.241961 | -3.911 |
| chr5 | 13264535 | 13264725 | 1.39542 | 0.35703 | -3.908 |
| chr10 | 80235456 | 80235703 | 1.46352 | 0.374457 | -3.908 |
| chr2 | 193896878 | 193897079 | 1.03315 | 0.264505 | -3.906 |
| chr8 | 69919923 | 69920231 | 1.13185 | 0.289885 | -3.904 |
| chr8 | 61484594 | 61484895 | 1.13662 | 0.291558 | -3.898 |
| chr8 | 121515733 | 121515944 | 1.2489 | 0.320609 | -3.895 |
| chr9 | 19569450 | 19569812 | 1.26411 | 0.324734 | -3.893 |
| chr4 | 61749101 | 61749302 | 0.7483 | 0.192388 | -3.890 |
| chr2 | 231109707 | 231109917 | 1.05372 | 0.271123 | -3.886 |
| chr10 | 102080683 | 102080877 | 0.943796 | 0.242853 | -3.886 |
| chr2 | 51529576 | 51529795 | 1.54724 | 0.39855 | -3.882 |
| chr9 | 26686746 | 26686981 | 1.37375 | 0.353972 | -3.881 |
| chr2 | 24443478 | 24443822 | 1.20373 | 0.310631 | -3.875 |
| chr10 | 57351503 | 57351808 | 0.826598 | 0.213326 | -3.875 |
| chr2 | 123611550 | 123611860 | 1.00147 | 0.258698 | -3.871 |
| chr1 | 219646170 | 219646353 | 1.10146 | 0.284827 | -3.867 |
| chr10 | 132440071 | 132440310 | 1.24404 | 0.322062 | -3.863 |
| chr2 | 15785576 | 15785932 | 1.20043 | 0.310845 | -3.862 |
| chr1 | 229259025 | 229259207 | 1.37617 | 0.356487 | -3.860 |
| chr2 | 152137414 | 152137620 | 0.941629 | 0.244182 | -3.856 |
| chr5 | 171009124 | 171009435 | 0.980192 | 0.254195 | -3.856 |
| chr9 | 128161733 | 128162055 | 1.00637 | 0.260994 | -3.856 |
| chr2 | 231671193 | 231671404 | 1.07875 | 0.280094 | -3.851 |
| chr17 | 54173400 | 54173602 | 1.18083 | 0.307465 | -3.841 |
| chr9 | 87704504 | 87704769 | 1.11509 | 0.290439 | -3.839 |
| chr11 | 65159281 | 65159578 | 0.822735 | 0.214349 | -3.838 |
| chr3 | 906147 | 906362 | 0.869757 | 0.2268 | -3.835 |
| chr1 | 179747413 | 179747617 | 0.996168 | 0.260199 | -3.828 |
| chr2 | 225549146 | 225549344 | 1.02854 | 0.268676 | -3.828 |
| chr1 | 165304788 | 165304980 | 1.11182 | 0.290443 | -3.828 |
| chr10 | 130705340 | 130705613 | 1.2622 | 0.330846 | -3.815 |
| chr7 | 49456745 | 49456933 | 1.07584 | 0.282305 | -3.811 |
| chr10 | 57118441 | 57118978 | 1.24728 | 0.327477 | -3.809 |
| chr2 | 113872807 | 113873077 | 0.906276 | 0.237989 | -3.808 |
| chr2 | 123615152 | 123615381 | 1.14402 | 0.301258 | -3.797 |

| | | | | | |
|---|---|---|---|---|---|
| chr4 | 146203538 | 146203740 | 1.13309 | 0.298468 | -3.796 |
| chr2 | 13909268 | 13909460 | 1.26678 | 0.33398 | -3.793 |
| chr2 | 19296856 | 19297136 | 1.23121 | 0.324612 | -3.793 |
| chr3 | 22974364 | 22974547 | 0.968959 | 0.257009 | -3.770 |
| chr15 | 95352077 | 95352314 | 0.9308 | 0.247051 | -3.768 |
| chr8 | 116996872 | 116997067 | 0.867847 | 0.230527 | -3.765 |
| chr3 | 84129485 | 84129703 | 0.921723 | 0.244867 | -3.764 |
| chr11 | 62015824 | 62016012 | 1.2929 | 0.343491 | -3.764 |
| chr4 | 17326086 | 17326272 | 1.13085 | 0.300441 | -3.764 |
| chr7 | 113617212 | 113617426 | 1.12297 | 0.298417 | -3.763 |
| chr16 | 51383111 | 51383301 | 1.24556 | 0.331173 | -3.761 |
| chr20 | 14955167 | 14955400 | 1.01249 | 0.269245 | -3.760 |
| chr9 | 107118935 | 107119159 | 1.36809 | 0.364511 | -3.753 |
| chr2 | 76565480 | 76565807 | 1.04768 | 0.279358 | -3.750 |
| chr2 | 122371623 | 122371824 | 1.25634 | 0.335276 | -3.747 |
| chr2 | 240043451 | 240043724 | 0.771625 | 0.206151 | -3.743 |
| chr2 | 133254190 | 133254461 | 0.927217 | 0.247813 | -3.742 |
| chr3 | 72056923 | 72057115 | 1.46556 | 0.391738 | -3.741 |
| chr12 | 26100308 | 26100490 | 1.43168 | 0.382719 | -3.741 |
| chr16 | 82983902 | 82984148 | 0.947486 | 0.253432 | -3.739 |
| chr4 | 85057917 | 85058275 | 0.938581 | 0.251321 | -3.735 |
| chr9 | 80665138 | 80665340 | 1.09853 | 0.29424 | -3.733 |
| chr9 | 30190667 | 30190877 | 0.71706 | 0.192301 | -3.729 |
| chr2 | 56488852 | 56489082 | 1.64013 | 0.440003 | -3.728 |
| chr3 | 89225956 | 89226201 | 0.799665 | 0.214551 | -3.727 |
| chr20 | 54214013 | 54214240 | 1.13271 | 0.303942 | -3.727 |
| chr2 | 173630397 | 173630697 | 1.38555 | 0.37182 | -3.726 |
| chr2 | 145444751 | 145444961 | 1.15994 | 0.311629 | -3.722 |
| chr2 | 137778641 | 137778824 | 0.817224 | 0.219565 | -3.722 |
| chr18 | 29476760 | 29476953 | 1.21553 | 0.326587 | -3.722 |
| chr2 | 193550082 | 193550273 | 1.47948 | 0.397627 | -3.721 |
| chr10 | 68550953 | 68551163 | 1.14107 | 0.307003 | -3.717 |
| chr20 | 59334989 | 59335210 | 1.04441 | 0.281364 | -3.712 |
| chr14 | 46109953 | 46110156 | 0.846372 | 0.228186 | -3.709 |
| chr15 | 53526135 | 53526328 | 1.69943 | 0.458495 | -3.707 |
| chr16 | 17408131 | 17408324 | 1.04643 | 0.282807 | -3.700 |
| chr5 | 52456110 | 52456295 | 1.60929 | 0.43513 | -3.698 |
| chr7 | 749408 | 749639 | 1.03167 | 0.278956 | -3.698 |
| chr2 | 23127075 | 23127302 | 1.9835 | 0.536792 | -3.695 |
| chr8 | 70288746 | 70288961 | 1.41885 | 0.384228 | -3.693 |
| chr11 | 43089026 | 43089255 | 1.09073 | 0.295446 | -3.692 |
| chr2 | 83469434 | 83469625 | 1.59771 | 0.432875 | -3.691 |
| chr2 | 48519795 | 48520175 | 1.51312 | 0.409964 | -3.691 |
| chr2 | 200095538 | 200095754 | 1.22115 | 0.331514 | -3.684 |
| chr7 | 132055067 | 132055410 | 0.819291 | 0.222571 | -3.681 |
| chr10 | 111145960 | 111146269 | 1.24466 | 0.338149 | -3.681 |
| chr2 | 78879344 | 78879648 | 0.94196 | 0.255985 | -3.680 |
| chr8 | 142605408 | 142605721 | 1.14344 | 0.311046 | -3.676 |
| chr15 | 42424817 | 42425042 | 1.18315 | 0.321859 | -3.676 |
| chr2 | 203653481 | 203653680 | 1.07826 | 0.293422 | -3.675 |
| chr5 | 11665864 | 11666072 | 0.937478 | 0.255304 | -3.672 |
| chr3 | 97244540 | 97244824 | 1.37465 | 0.374558 | -3.670 |
| chr11 | 558652 | 558844 | 1.20636 | 0.328707 | -3.670 |
| chr2 | 56776954 | 56777297 | 1.1605 | 0.316488 | -3.667 |
| chr9 | 30237797 | 30238049 | 0.888178 | 0.242226 | -3.667 |
| chr11 | 702703 | 702984 | 1.44446 | 0.394074 | -3.665 |
| chr3 | 86450651 | 86450866 | 0.981213 | 0.267716 | -3.665 |
| chr11 | 37105755 | 37106071 | 1.07448 | 0.29319 | -3.665 |
| chrX | 120726500 | 120726686 | 1.20836 | 0.329783 | -3.664 |
| chr7 | 147031891 | 147032073 | 0.982437 | 0.268205 | -3.663 |
| chr2 | 140005684 | 140005941 | 1.52918 | 0.417652 | -3.661 |

| | | | | | |
|---|---|---|---|---|---|
| chr2 | 21295421 | 21295652 | 1.41738 | 0.387708 | -3.656 |
| chr2 | 116535540 | 116535777 | 1.08985 | 0.298176 | -3.655 |
| chr2 | 218618829 | 218619019 | 0.886248 | 0.24251 | -3.654 |
| chr2 | 145304760 | 145305020 | 1.06991 | 0.292878 | -3.653 |
| chr2 | 36420328 | 36420556 | 1.16265 | 0.318804 | -3.647 |
| chr2 | 23111217 | 23111697 | 0.944547 | 0.259207 | -3.644 |
| chr5 | 120229320 | 120229523 | 0.876643 | 0.240644 | -3.643 |
| chr2 | 51628250 | 51628509 | 1.1635 | 0.320254 | -3.633 |
| chr1 | 191373470 | 191373697 | 1.28294 | 0.35364 | -3.628 |
| chr8 | 113942442 | 113942656 | 0.934681 | 0.257719 | -3.627 |
| chr14 | 34291334 | 34291537 | 1.56785 | 0.432773 | -3.623 |
| chr5 | 161642473 | 161642672 | 1.22933 | 0.339375 | -3.622 |
| chr8 | 115749646 | 115749860 | 0.960158 | 0.265892 | -3.611 |
| chr2 | 115458151 | 115458387 | 1.17861 | 0.326753 | -3.607 |
| chr1 | 195961721 | 195961913 | 1.35671 | 0.376205 | -3.606 |
| chr10 | 66719063 | 66719309 | 1.04767 | 0.290719 | -3.604 |
| chr2 | 59433393 | 59433626 | 1.34362 | 0.373103 | -3.601 |
| chr2 | 70184822 | 70185109 | 1.25659 | 0.349236 | -3.598 |
| chr2 | 186311313 | 186311512 | 0.954672 | 0.265358 | -3.598 |
| chr2 | 53020899 | 53021086 | 0.749931 | 0.208658 | -3.594 |
| chr12 | 118672364 | 118672752 | 1.11264 | 0.309615 | -3.594 |
| chr10 | 8309925 | 8310122 | 1.02798 | 0.286081 | -3.593 |
| chr8 | 20488904 | 20489103 | 1.06488 | 0.296369 | -3.593 |
| chr8 | 78089666 | 78089909 | 1.23673 | 0.344354 | -3.591 |
| chr11 | 39924456 | 39924658 | 1.01947 | 0.283899 | -3.591 |
| chr10 | 53894300 | 53894496 | 1.41209 | 0.393376 | -3.590 |
| chr2 | 71158652 | 71158926 | 1.49758 | 0.41744 | -3.588 |
| chr7 | 43673961 | 43674173 | 1.36499 | 0.380556 | -3.587 |
| chr4 | 113789475 | 113789657 | 0.853099 | 0.23796 | -3.585 |
| chr2 | 211714194 | 211714406 | 1.51065 | 0.421376 | -3.585 |
| chr5 | 63059303 | 63059526 | 1.64621 | 0.459744 | -3.581 |
| chr5 | 83340548 | 83340732 | 1.24091 | 0.346675 | -3.579 |
| chr2 | 172441253 | 172441459 | 1.0273 | 0.287755 | -3.570 |
| chr3 | 102798595 | 102799173 | 1.14054 | 0.320204 | -3.562 |
| chr2 | 5916231 | 5916518 | 1.29931 | 0.364898 | -3.561 |
| chr7 | 122497221 | 122497561 | 0.951741 | 0.267753 | -3.555 |
| chr8 | 66793179 | 66793372 | 1.10316 | 0.310541 | -3.552 |
| chr2 | 138521218 | 138521677 | 1.05508 | 0.297042 | -3.552 |
| chr4 | 181449176 | 181449379 | 1.14087 | 0.321244 | -3.551 |
| chr20 | 60772923 | 60773155 | 0.986003 | 0.278397 | -3.542 |
| chr5 | 38621984 | 38622169 | 1.14075 | 0.322879 | -3.533 |
| chr14 | 78914870 | 78915131 | 2.46912 | 0.699023 | -3.532 |
| chr16 | 10978580 | 10978768 | 0.822645 | 0.232922 | -3.532 |
| chr4 | 30187754 | 30187945 | 0.836715 | 0.236969 | -3.531 |
| chr11 | 81429744 | 81429979 | 0.797019 | 0.225744 | -3.531 |
| chr2 | 156968294 | 156968504 | 0.909905 | 0.257988 | -3.527 |
| chr2 | 180262968 | 180263199 | 1.06304 | 0.301472 | -3.526 |
| chr11 | 23858384 | 23858660 | 1.31269 | 0.372481 | -3.524 |
| chr14 | 71306154 | 71306358 | 0.891991 | 0.253129 | -3.524 |
| chr9 | 30771160 | 30771354 | 1.30201 | 0.369493 | -3.524 |
| chr8 | 13228837 | 13229139 | 1.03093 | 0.292654 | -3.523 |
| chr1 | 175298984 | 175299493 | 0.982182 | 0.278874 | -3.522 |
| chr2 | 124921070 | 124921274 | 0.824369 | 0.23407 | -3.522 |
| chr2 | 143008703 | 143008960 | 1.20099 | 0.34107 | -3.521 |
| chr11 | 127713781 | 127714035 | 2.33087 | 0.662153 | -3.520 |
| chr2 | 160431638 | 160431855 | 1.33154 | 0.378729 | -3.516 |
| chr20 | 51089646 | 51089848 | 1.07296 | 0.305284 | -3.515 |
| chr1 | 166573000 | 166573192 | 0.890677 | 0.253451 | -3.514 |
| chr11 | 70351285 | 70351489 | 0.888147 | 0.252744 | -3.514 |
| chr8 | 20963877 | 20964134 | 0.819961 | 0.233548 | -3.511 |
| chr5 | 24869597 | 24869792 | 1.02117 | 0.291381 | -3.505 |

| | | | | | |
|---|---|---|---|---|---|
| chr2 | 213145473 | 213145757 | 1.11961 | 0.319678 | -3.502 |
| chr19 | 37320949 | 37321160 | 1.58031 | 0.451507 | -3.500 |
| chr5 | 8106776 | 8107041 | 0.895928 | 0.256009 | -3.500 |
| chr5 | 8052388 | 8052593 | 1.08768 | 0.311158 | -3.496 |
| chr16 | 9317091 | 9317296 | 0.847534 | 0.242932 | -3.489 |
| chr15 | 82213105 | 82213309 | 0.981199 | 0.281321 | -3.488 |
| chr2 | 140357885 | 140358073 | 0.973846 | 0.279214 | -3.488 |
| chr16 | 52574131 | 52574464 | 1.02571 | 0.294136 | -3.487 |
| chr1 | 195121981 | 195122191 | 1.0256 | 0.294142 | -3.487 |
| chr2 | 31335054 | 31335271 | 1.22856 | 0.352415 | -3.486 |
| chr3 | 116491061 | 116491299 | 1.21283 | 0.348258 | -3.483 |
| chr14 | 47619429 | 47619668 | 1.12968 | 0.324387 | -3.483 |
| chr1 | 231260669 | 231260940 | 1.04796 | 0.301073 | -3.481 |
| chr19 | 23545718 | 23545984 | 1.21353 | 0.348809 | -3.479 |
| chr5 | 119544761 | 119545062 | 1.10679 | 0.318143 | -3.479 |
| chr5 | 25146958 | 25147201 | 1.14937 | 0.330392 | -3.479 |
| chr7 | 108320034 | 108320230 | 1.4248 | 0.409593 | -3.479 |
| chr20 | 17330225 | 17330409 | 0.998657 | 0.287392 | -3.475 |
| chr2 | 104735388 | 104735581 | 1.31987 | 0.380026 | -3.473 |
| chr3 | 124077694 | 124077888 | 1.45396 | 0.418713 | -3.472 |
| chr16 | 24248044 | 24248262 | 1.658 | 0.477631 | -3.471 |
| chr8 | 20229924 | 20230109 | 1.17449 | 0.338372 | -3.471 |
| chr11 | 30922939 | 30923150 | 0.968167 | 0.279262 | -3.467 |
| chr1 | 233587379 | 233587562 | 1.20614 | 0.348015 | -3.466 |
| chr11 | 72623813 | 72624004 | 1.00421 | 0.289904 | -3.464 |
| chr2 | 12924195 | 12924378 | 1.19403 | 0.344732 | -3.464 |
| chr8 | 79282838 | 79283046 | 1.52031 | 0.439311 | -3.461 |
| chr11 | 40504058 | 40504245 | 1.51549 | 0.438123 | -3.459 |
| chr16 | 16247155 | 16247338 | 0.878097 | 0.2542 | -3.454 |
| chr13 | 84679636 | 84679836 | 1.00842 | 0.292042 | -3.453 |
| chr2 | 142698981 | 142699166 | 1.19166 | 0.345228 | -3.452 |
| chr11 | 64956882 | 64957076 | 1.02369 | 0.296965 | -3.447 |
| chr8 | 117174786 | 117174969 | 1.07803 | 0.312842 | -3.446 |
| chr5 | 51721703 | 51721915 | 1.02182 | 0.296962 | -3.441 |
| chr3 | 5471310 | 5471527 | 0.927936 | 0.269945 | -3.438 |
| chr3 | 86482316 | 86482545 | 1.01822 | 0.296322 | -3.436 |
| chr16 | 71888687 | 71888936 | 1.18553 | 0.345031 | -3.436 |
| chr7 | 92003484 | 92003702 | 1.26663 | 0.368814 | -3.434 |
| chr5 | 98972584 | 98972954 | 0.843326 | 0.245595 | -3.434 |
| chr2 | 34750483 | 34750782 | 1.27753 | 0.37208 | -3.433 |
| chr8 | 75593795 | 75594009 | 0.943774 | 0.275161 | -3.430 |
| chr1 | 195439620 | 195439802 | 1.42726 | 0.416159 | -3.430 |
| chr8 | 51930583 | 51930771 | 1.02078 | 0.297836 | -3.427 |
| chr2 | 123573063 | 123573450 | 1.10013 | 0.321132 | -3.426 |
| chr11 | 7861291 | 7861526 | 0.955821 | 0.27903 | -3.426 |
| chr10 | 13518275 | 13518462 | 1.22761 | 0.358413 | -3.425 |
| chr7 | 41928524 | 41928773 | 0.842551 | 0.246013 | -3.425 |
| chrX | 15606984 | 15607170 | 1.16538 | 0.340786 | -3.420 |
| chr9 | 26003148 | 26003345 | 1.08847 | 0.318332 | -3.419 |
| chr20 | 21144680 | 21145002 | 1.13943 | 0.333539 | -3.416 |
| chr2 | 166398568 | 166398769 | 1.0795 | 0.316014 | -3.416 |
| chr2 | 151843320 | 151843611 | 1.31216 | 0.384365 | -3.414 |
| chr12 | 5854495 | 5854762 | 2.99968 | 0.87927 | -3.412 |
| chr16 | 6936301 | 6936541 | 1.32758 | 0.38916 | -3.411 |
| chr7 | 145605383 | 145605595 | 0.806066 | 0.236415 | -3.410 |
| chr19 | 53513142 | 53513404 | 1.22037 | 0.357991 | -3.409 |
| chr17 | 13191525 | 13191711 | 1.474 | 0.432651 | -3.407 |
| chr11 | 61989551 | 61989794 | 1.32833 | 0.390004 | -3.406 |
| chr7 | 23800353 | 23800600 | 1.25649 | 0.369041 | -3.405 |
| chr2 | 116897437 | 116897623 | 1.49972 | 0.44049 | -3.405 |
| chr9 | 23724386 | 23724580 | 1.24348 | 0.365326 | -3.404 |

| | | | | | |
|---|---|---|---|---|---|
| chr2 | 186405127 | 186405423 | 1.17934 | 0.346792 | -3.401 |
| chr6 | 73142598 | 73142865 | 1.39657 | 0.410758 | -3.400 |
| chr8 | 114696393 | 114696583 | 1.07307 | 0.315957 | -3.396 |
| chr2 | 188034763 | 188035001 | 1.52227 | 0.448335 | -3.395 |
| chr9 | 73462304 | 73462504 | 1.03999 | 0.306306 | -3.395 |
| chr7 | 17292820 | 17293040 | 1.05941 | 0.312094 | -3.395 |
| chr9 | 121038379 | 121038644 | 1.35513 | 0.399383 | -3.393 |
| chr15 | 70575992 | 70576228 | 1.20783 | 0.3561 | -3.392 |
| chr5 | 15370594 | 15370825 | 1.11669 | 0.329392 | -3.390 |
| chr11 | 23096443 | 23096664 | 0.919652 | 0.271397 | -3.389 |
| chr16 | 20065373 | 20065556 | 1.23737 | 0.365182 | -3.388 |
| chr2 | 50261553 | 50261767 | 1.20198 | 0.354869 | -3.387 |
| chr5 | 166857633 | 166857829 | 1.00324 | 0.296319 | -3.386 |
| chr9 | 112205838 | 112206033 | 1.22659 | 0.362345 | -3.385 |
| chr9 | 29806265 | 29806511 | 1.05421 | 0.311474 | -3.385 |
| chr8 | 66914309 | 66914516 | 1.63069 | 0.48188 | -3.384 |
| chr14 | 24130908 | 24131124 | 1.03759 | 0.306631 | -3.384 |
| chr2 | 29188383 | 29188587 | 1.20363 | 0.355757 | -3.383 |
| chr3 | 161058317 | 161058507 | 0.99596 | 0.294452 | -3.382 |
| chr9 | 117117722 | 117118023 | 0.679797 | 0.201204 | -3.379 |
| chr7 | 88381467 | 88381652 | 1.25569 | 0.371689 | -3.378 |
| chr3 | 36544830 | 36545228 | 0.982542 | 0.290867 | -3.378 |
| chr3 | 96954065 | 96954361 | 0.964109 | 0.285682 | -3.375 |
| chr20 | 11610113 | 11610382 | 0.986709 | 0.292409 | -3.374 |
| chr2 | 117480387 | 117480605 | 1.14953 | 0.340662 | -3.374 |
| chr5 | 29591495 | 29591730 | 1.18603 | 0.351543 | -3.374 |
| chr15 | 94720558 | 94720745 | 1.00927 | 0.299176 | -3.374 |
| chr16 | 1799562 | 1799748 | 1.00791 | 0.298812 | -3.373 |
| chr2 | 116976370 | 116976574 | 0.996057 | 0.2954 | -3.372 |
| chr5 | 106342332 | 106342526 | 0.938768 | 0.278993 | -3.365 |
| chr4 | 134395708 | 134395894 | 1.03761 | 0.308377 | -3.365 |
| chr3 | 1425421 | 1425623 | 1.43854 | 0.427683 | -3.364 |
| chr2 | 25770352 | 25770550 | 1.24972 | 0.371567 | -3.363 |
| chr5 | 154396212 | 154396476 | 1.07073 | 0.318471 | -3.362 |
| chr7 | 105341262 | 105341449 | 1.22854 | 0.36554 | -3.361 |
| chr17 | 3006355 | 3006754 | 0.95423 | 0.283974 | -3.360 |
| chr7 | 134225037 | 134225253 | 1.01842 | 0.303102 | -3.360 |
| chr8 | 83054037 | 83054231 | 1.04614 | 0.311354 | -3.360 |
| chr8 | 144259292 | 144259485 | 1.56643 | 0.46621 | -3.360 |
| chr16 | 7933413 | 7933663 | 1.0407 | 0.309878 | -3.358 |
| chr2 | 88298307 | 88298505 | 0.718729 | 0.214018 | -3.358 |
| chr5 | 63315705 | 63315890 | 1.44678 | 0.430814 | -3.358 |
| chr7 | 145293755 | 145294014 | 1.04318 | 0.310663 | -3.358 |
| chr16 | 5540657 | 5540844 | 2.36382 | 0.704061 | -3.357 |
| chr1 | 218250544 | 218250765 | 1.11041 | 0.330858 | -3.356 |
| chr11 | 79203683 | 79204058 | 0.853899 | 0.2546 | -3.354 |
| chr2 | 212827094 | 212827283 | 1.08339 | 0.323156 | -3.353 |
| chr2 | 209923277 | 209923487 | 1.0648 | 0.317762 | -3.351 |
| chr5 | 2019653 | 2019858 | 0.939329 | 0.280318 | -3.351 |
| chr9 | 11185620 | 11185866 | 1.24301 | 0.371014 | -3.350 |
| chr8 | 79512631 | 79512823 | 0.920231 | 0.2749 | -3.348 |
| chr5 | 113761427 | 113761869 | 1.27369 | 0.380726 | -3.345 |
| chr1 | 223486018 | 223486343 | 0.852611 | 0.254949 | -3.344 |
| chr10 | 70931854 | 70932088 | 1.15648 | 0.345955 | -3.343 |
| chr10 | 72191393 | 72191601 | 1.31458 | 0.39326 | -3.343 |
| chr7 | 54352431 | 54352774 | 1.0663 | 0.319211 | -3.340 |
| chr19 | 24069662 | 24069875 | 1.22404 | 0.367029 | -3.335 |
| chr5 | 89153077 | 89153370 | 1.15776 | 0.347369 | -3.333 |
| chr2 | 124905211 | 124905394 | 1.12084 | 0.336381 | -3.332 |
| chr10 | 119424590 | 119424844 | 1.34521 | 0.404024 | -3.330 |
| chr2 | 19668760 | 19668991 | 1.12849 | 0.339236 | -3.327 |

| | | | | | |
|---|---|---|---|---|---|
| chr10 | 108019702 | 108020033 | 0.99948 | 0.300559 | -3.325 |
| chr2 | 52627779 | 52627988 | 1.15804 | 0.34824 | -3.325 |
| chr8 | 26925947 | 26926172 | 0.824691 | 0.248007 | -3.325 |
| chr11 | 16822667 | 16822886 | 1.10864 | 0.333426 | -3.325 |
| chr10 | 109997608 | 109997819 | 1.06426 | 0.320365 | -3.322 |
| chr7 | 143792402 | 143792698 | 1.06641 | 0.321098 | -3.321 |
| chr11 | 25152914 | 25153310 | 1.00653 | 0.303406 | -3.317 |
| chr3 | 97643416 | 97643620 | 1.301 | 0.392258 | -3.317 |
| chr5 | 27916918 | 27917146 | 0.900453 | 0.271568 | -3.316 |
| chr7 | 136266232 | 136266543 | 1.2601 | 0.380088 | -3.315 |
| chr2 | 23858208 | 23858431 | 1.0606 | 0.319982 | -3.315 |
| chr3 | 458717 | 458942 | 1.215 | 0.366822 | -3.312 |
| chr2 | 114454711 | 114454926 | 1.18414 | 0.357916 | -3.308 |
| chr2 | 194478036 | 194478290 | 0.966507 | 0.2922 | -3.308 |
| chrX | 117542632 | 117542850 | 1.40427 | 0.424714 | -3.306 |
| chr3 | 52892486 | 52892670 | 1.15998 | 0.350855 | -3.306 |
| chr2 | 80772360 | 80772719 | 1.02605 | 0.310347 | -3.306 |
| chr15 | 96890895 | 96891117 | 1.17058 | 0.354093 | -3.306 |
| chr11 | 7484177 | 7484389 | 1.05686 | 0.31978 | -3.305 |
| chr2 | 53137946 | 53138129 | 0.931421 | 0.28211 | -3.302 |
| chr2 | 14023823 | 14024022 | 1.31395 | 0.398349 | -3.298 |
| chr6 | 105602568 | 105602761 | 0.974116 | 0.295368 | -3.298 |
| chr7 | 27503959 | 27504184 | 1.08638 | 0.329724 | -3.295 |
| chr3 | 80406826 | 80407097 | 1.1514 | 0.349518 | -3.294 |
| chr10 | 104315524 | 104315776 | 0.919702 | 0.279235 | -3.294 |
| chr11 | 35811888 | 35812185 | 1.0034 | 0.304821 | -3.292 |
| chr7 | 77655015 | 77655223 | 0.875692 | 0.266119 | -3.291 |
| chr12 | 45045483 | 45045942 | 0.958819 | 0.291388 | -3.291 |
| chr8 | 108971351 | 108971547 | 0.923747 | 0.280899 | -3.289 |
| chr14 | 88227356 | 88227570 | 1.35695 | 0.412922 | -3.286 |
| chr2 | 15198249 | 15198469 | 2.24442 | 0.683486 | -3.284 |
| chr16 | 75634872 | 75635099 | 1.11005 | 0.338171 | -3.283 |
| chr20 | 16649827 | 16650065 | 1.02159 | 0.311278 | -3.282 |
| chr5 | 132023795 | 132023984 | 1.06158 | 0.323527 | -3.281 |
| chr11 | 41993041 | 41993237 | 1.0198 | 0.310852 | -3.281 |
| chr20 | 24996567 | 24996790 | 1.16976 | 0.356601 | -3.280 |
| chr15 | 77982547 | 77982882 | 1.23079 | 0.375235 | -3.280 |
| chr14 | 41244070 | 41244281 | 1.02276 | 0.311952 | -3.279 |
| chr10 | 56124658 | 56124910 | 1.16007 | 0.353942 | -3.278 |
| chr3 | 82391292 | 82391504 | 1.23678 | 0.377387 | -3.277 |
| chr5 | 6325442 | 6325750 | 0.891585 | 0.272123 | -3.276 |
| chr16 | 88693839 | 88694082 | 1.14059 | 0.348341 | -3.274 |
| chr16 | 80694699 | 80694941 | 1.43864 | 0.439577 | -3.273 |
| chr2 | 50091634 | 50092003 | 1.46152 | 0.446615 | -3.272 |
| chr19 | 24008530 | 24008714 | 0.904935 | 0.276557 | -3.272 |
| chr2 | 21785773 | 21785988 | 1.19562 | 0.365733 | -3.269 |
| chr2 | 81629115 | 81629385 | 1.19388 | 0.36525 | -3.269 |
| chr5 | 119258850 | 119259042 | 0.736221 | 0.225286 | -3.268 |
| chr2 | 176712291 | 176712570 | 1.12563 | 0.3445 | -3.267 |
| chr10 | 36506192 | 36506431 | 1.43253 | 0.438475 | -3.267 |
| chr2 | 99191442 | 99191721 | 1.06034 | 0.324636 | -3.266 |
| chr10 | 120119484 | 120119676 | 1.17071 | 0.358688 | -3.264 |
| chr5 | 19049594 | 19049828 | 0.954915 | 0.292682 | -3.263 |
| chr5 | 31101102 | 31101317 | 1.21012 | 0.371022 | -3.262 |
| chr3 | 95723145 | 95723338 | 1.17019 | 0.358904 | -3.260 |
| chr17 | 70212253 | 70212470 | 0.974189 | 0.298874 | -3.260 |
| chr20 | 12479878 | 12480176 | 0.889123 | 0.272824 | -3.259 |
| chr12 | 22273875 | 22274065 | 1.07702 | 0.330595 | -3.258 |
| chr16 | 64192512 | 64192702 | 1.1868 | 0.36434 | -3.257 |
| chr3 | 55834688 | 55834921 | 0.940134 | 0.288645 | -3.257 |
| chr5 | 129452457 | 129452891 | 1.08323 | 0.332676 | -3.256 |

| | | | | | |
|---|---|---|---|---|---|
| chr6 | 131590446 | 131590653 | 1.13934 | 0.349987 | -3.255 |
| chr14 | 103419289 | 103419552 | 1.18209 | 0.363154 | -3.255 |
| chr16 | 56093893 | 56094129 | 0.993797 | 0.305325 | -3.255 |
| chr10 | 85068518 | 85068971 | 1.07712 | 0.331059 | -3.254 |
| chr16 | 34927730 | 34927992 | 0.767058 | 0.235821 | -3.253 |
| chr2 | 52792020 | 52792325 | 1.10833 | 0.340907 | -3.251 |
| chr7 | 63917988 | 63918230 | 1.08064 | 0.332531 | -3.250 |
| chr9 | 80634063 | 80634268 | 1.04726 | 0.32226 | -3.250 |
| chr8 | 11015036 | 11015267 | 1.27731 | 0.393197 | -3.249 |
| chr5 | 170976410 | 170976595 | 1.1623 | 0.357861 | -3.248 |
| chr7 | 67735690 | 67735978 | 1.02364 | 0.315505 | -3.244 |
| chr20 | 20719306 | 20719518 | 1.18697 | 0.36606 | -3.243 |
| chr2 | 103759470 | 103759702 | 1.08884 | 0.335851 | -3.242 |
| chr2 | 81252694 | 81252908 | 1.08865 | 0.335801 | -3.242 |
| chr14 | 77989696 | 77989887 | 1.00672 | 0.310683 | -3.240 |
| chr7 | 88783536 | 88783815 | 0.935971 | 0.288904 | -3.240 |
| chr10 | 85353541 | 85353733 | 1.17246 | 0.361959 | -3.239 |
| chr9 | 26127530 | 26127803 | 0.904891 | 0.279565 | -3.237 |
| chr16 | 80880681 | 80880867 | 1.22351 | 0.378032 | -3.237 |
| chr16 | 78924342 | 78924638 | 1.2381 | 0.383026 | -3.232 |
| chr9 | 93496174 | 93496427 | 1.04868 | 0.324498 | -3.232 |
| chr8 | 117009653 | 117009859 | 1.28522 | 0.397785 | -3.231 |
| chr1 | 191160018 | 191160212 | 1.27156 | 0.393566 | -3.231 |
| chr9 | 23883908 | 23884248 | 0.951339 | 0.294576 | -3.230 |
| chr15 | 86986131 | 86986415 | 1.62012 | 0.501833 | -3.228 |
| chr2 | 118919081 | 118919266 | 1.30707 | 0.404902 | -3.228 |
| chr1 | 165164693 | 165164907 | 0.943283 | 0.292253 | -3.228 |
| chr2 | 212627114 | 212627487 | 1.06734 | 0.330704 | -3.227 |
| chr8 | 112327871 | 112328108 | 1.21606 | 0.376839 | -3.227 |
| chr20 | 17046275 | 17046514 | 1.23873 | 0.383917 | -3.227 |
| chr2 | 241749330 | 241749529 | 1.06386 | 0.32989 | -3.225 |
| chr16 | 27782996 | 27783209 | 0.912878 | 0.283114 | -3.224 |
| chr7 | 13522285 | 13522513 | 1.06477 | 0.330245 | -3.224 |
| chr2 | 78708754 | 78709067 | 1.45947 | 0.452754 | -3.224 |
| chr8 | 94023874 | 94024226 | 1.06181 | 0.329429 | -3.223 |
| chr2 | 212839788 | 212840014 | 1.11778 | 0.346903 | -3.222 |
| chr14 | 48628449 | 48628661 | 1.42538 | 0.442559 | -3.221 |
| chr3 | 95079750 | 95079936 | 1.26996 | 0.39436 | -3.220 |
| chr2 | 32803569 | 32803753 | 1.30091 | 0.40415 | -3.219 |
| chr8 | 51378176 | 51378438 | 1.32177 | 0.410728 | -3.218 |
| chr5 | 3743980 | 3744261 | 0.829068 | 0.257632 | -3.218 |
| chr15 | 32366958 | 32367149 | 0.986307 | 0.306823 | -3.215 |
| chr9 | 132009469 | 132009666 | 1.2358 | 0.38449 | -3.214 |
| chr5 | 95859185 | 95859382 | 1.17883 | 0.366941 | -3.213 |
| chr2 | 189627915 | 189628121 | 1.12182 | 0.349758 | -3.207 |
| chr11 | 60263451 | 60263650 | 0.739202 | 0.230497 | -3.207 |
| chr8 | 106151766 | 106151993 | 0.879447 | 0.274235 | -3.207 |
| chr2 | 57282316 | 57282623 | 0.882201 | 0.275246 | -3.205 |
| chr1 | 159416844 | 159417045 | 1.25766 | 0.392728 | -3.202 |
| chr5 | 21721226 | 21721455 | 0.794775 | 0.248355 | -3.200 |
| chr14 | 49882385 | 49882591 | 0.953304 | 0.29797 | -3.199 |
| chr10 | 135126951 | 135127187 | 1.00649 | 0.314653 | -3.199 |
| chr7 | 80084981 | 80085217 | 0.888468 | 0.277756 | -3.199 |
| chr2 | 48392359 | 48392623 | 1.5354 | 0.480004 | -3.199 |
| chr9 | 99129359 | 99129545 | 0.776995 | 0.24295 | -3.198 |
| chr10 | 20272914 | 20273113 | 1.36839 | 0.427895 | -3.198 |
| chr9 | 109145571 | 109145850 | 1.31439 | 0.411115 | -3.197 |
| chr10 | 91568015 | 91568230 | 0.998496 | 0.312387 | -3.196 |
| chr2 | 1317393 | 1317623 | 0.985014 | 0.308385 | -3.194 |
| chr20 | 2466262 | 2466465 | 1.11922 | 0.350437 | -3.194 |
| chr9 | 117221724 | 117221914 | 1.40793 | 0.440844 | -3.194 |

| | | | | | |
|---|---|---|---|---|---|
| chr5 | 161350580 | 161351062 | 0.951535 | 0.297981 | -3.193 |
| chr7 | 32266468 | 32266760 | 1.12193 | 0.35142 | -3.193 |
| chr16 | 3773832 | 3774041 | 0.863324 | 0.270424 | -3.192 |
| chr5 | 54057168 | 54057374 | 1.24703 | 0.39063 | -3.192 |
| chr2 | 81799791 | 81799978 | 1.015 | 0.318068 | -3.191 |
| chr9 | 17865056 | 17865325 | 0.945719 | 0.296662 | -3.188 |
| chr5 | 117005626 | 117005843 | 0.881638 | 0.276648 | -3.187 |
| chr17 | 8856193 | 8856391 | 1.34073 | 0.420709 | -3.187 |
| chr2 | 117529306 | 117529514 | 1.14434 | 0.359117 | -3.187 |
| chr3 | 77238332 | 77238532 | 1.19886 | 0.376461 | -3.185 |
| chr9 | 95596511 | 95596791 | 0.981182 | 0.308161 | -3.184 |
| chr8 | 89096370 | 89096564 | 1.01809 | 0.319867 | -3.183 |
| chr2 | 231225839 | 231226037 | 1.35894 | 0.426977 | -3.183 |
| chr4 | 21652615 | 21652890 | 0.860725 | 0.270633 | -3.180 |
| chr15 | 27603950 | 27604445 | 2.6024 | 0.818999 | -3.178 |
| chr7 | 158871607 | 158871834 | 1.41956 | 0.446758 | -3.177 |
| chr16 | 64714230 | 64714427 | 1.05269 | 0.331372 | -3.177 |
| chr14 | 98393936 | 98394355 | 1.13436 | 0.357168 | -3.176 |
| chr2 | 186635659 | 186635866 | 0.79283 | 0.249657 | -3.176 |
| chr5 | 116745125 | 116745314 | 1.09197 | 0.3439 | -3.175 |
| chr9 | 23982407 | 23982611 | 1.35763 | 0.42762 | -3.175 |
| chr9 | 130132235 | 130132464 | 1.01668 | 0.320229 | -3.175 |
| chr2 | 188103854 | 188104052 | 1.07738 | 0.339351 | -3.175 |
| chr16 | 47784743 | 47785085 | 1.27372 | 0.401218 | -3.175 |
| chr2 | 34961340 | 34961541 | 1.14576 | 0.361058 | -3.173 |
| chr12 | 97612717 | 97612912 | 1.30037 | 0.409977 | -3.172 |
| chr3 | 110605198 | 110605599 | 1.26557 | 0.39913 | -3.171 |
| chr8 | 85362545 | 85362763 | 1.05372 | 0.332417 | -3.170 |
| chr5 | 160208255 | 160208528 | 1.04774 | 0.330547 | -3.170 |
| chr2 | 137695621 | 137695828 | 1.0776 | 0.339988 | -3.170 |
| chr14 | 33347438 | 33347634 | 0.907833 | 0.286909 | -3.164 |
| chr7 | 121888874 | 121889104 | 1.03975 | 0.328833 | -3.162 |
| chr8 | 83747293 | 83747522 | 1.18556 | 0.375 | -3.161 |
| chr6 | 87784119 | 87784330 | 0.910581 | 0.288276 | -3.159 |
| chr16 | 6829221 | 6829406 | 1.13685 | 0.36005 | -3.157 |
| chr14 | 93822697 | 93822906 | 1.25004 | 0.396141 | -3.156 |
| chr2 | 217866423 | 217866716 | 0.906917 | 0.287442 | -3.155 |
| chr18 | 25049813 | 25050109 | 1.34429 | 0.42607 | -3.155 |
| chr10 | 132788387 | 132788574 | 0.816088 | 0.258664 | -3.155 |
| chr3 | 97026855 | 97027138 | 0.93211 | 0.295534 | -3.154 |
| chr20 | 2219117 | 2219311 | 1.08674 | 0.344666 | -3.153 |
| chr9 | 134492675 | 134492978 | 1.13298 | 0.35937 | -3.153 |
| chr5 | 115334424 | 115334685 | 1.00052 | 0.317513 | -3.151 |
| chr16 | 20284244 | 20284584 | 1.01705 | 0.322934 | -3.149 |
| chr5 | 9841678 | 9841979 | 1.11645 | 0.354557 | -3.149 |
| chr3 | 94947667 | 94948020 | 0.80469 | 0.25555 | -3.149 |
| chr2 | 104489185 | 104489385 | 0.802084 | 0.254892 | -3.147 |
| chr2 | 4497419 | 4497609 | 1.16697 | 0.371174 | -3.144 |
| chr2 | 51133468 | 51133800 | 0.898009 | 0.28563 | -3.144 |
| chr9 | 86975118 | 86975337 | 1.44726 | 0.460602 | -3.142 |
| chr14 | 103400034 | 103400219 | 1.40433 | 0.446988 | -3.142 |
| chr3 | 86022272 | 86022526 | 1.45032 | 0.461662 | -3.142 |
| chr2 | 52248739 | 52249014 | 1.52639 | 0.486064 | -3.140 |
| chr3 | 34463929 | 34464164 | 1.25358 | 0.399273 | -3.140 |
| chr6 | 12986332 | 12986517 | 1.15273 | 0.367243 | -3.139 |
| chr2 | 73420571 | 73420779 | 1.21606 | 0.387691 | -3.137 |
| chr10 | 87619618 | 87619800 | 0.795569 | 0.253648 | -3.137 |
| chr12 | 56004760 | 56005030 | 1.50826 | 0.481157 | -3.135 |
| chr2 | 151881337 | 151881602 | 1.04514 | 0.333686 | -3.132 |
| chr16 | 6479251 | 6479630 | 0.863043 | 0.275557 | -3.132 |
| chr3 | 11157435 | 11157672 | 1.0801 | 0.344864 | -3.132 |

| | | | | | |
|---|---|---|---|---|---|
| chr11 | 129969737 | 129969988 | 0.859039 | 0.274568 | -3.129 |
| chr8 | 88119383 | 88119614 | 1.0296 | 0.329096 | -3.129 |
| chr1 | 230521882 | 230522070 | 1.0346 | 0.330826 | -3.127 |
| chr10 | 107073619 | 107073803 | 1.15281 | 0.368633 | -3.127 |
| chr2 | 34876757 | 34876979 | 1.39104 | 0.444881 | -3.127 |
| chr17 | 5200059 | 5200374 | 1.13796 | 0.364095 | -3.125 |
| chr8 | 76793840 | 76794090 | 1.07424 | 0.343951 | -3.123 |
| chr12 | 101998617 | 101998903 | 1.31524 | 0.421973 | -3.117 |
| chr2 | 50423879 | 50424128 | 1.05979 | 0.340165 | -3.116 |
| chr11 | 99739244 | 99739426 | 1.10968 | 0.356234 | -3.115 |
| chr2 | 25163422 | 25163631 | 1.1727 | 0.376526 | -3.115 |
| chr10 | 81066724 | 81066934 | 0.8927 | 0.286632 | -3.114 |
| chr9 | 125567993 | 125568191 | 1.32032 | 0.423944 | -3.114 |
| chr8 | 135286563 | 135286888 | 1.24749 | 0.40057 | -3.114 |
| chr2 | 22579739 | 22580070 | 1.05761 | 0.339615 | -3.114 |
| chr10 | 18249473 | 18249680 | 1.02975 | 0.330762 | -3.113 |
| chr7 | 53843220 | 53843456 | 1.1753 | 0.377516 | -3.113 |
| chr1 | 199282371 | 199282584 | 1.07118 | 0.344102 | -3.113 |
| chr2 | 99397647 | 99397889 | 0.952903 | 0.306126 | -3.113 |
| chr3 | 147580025 | 147580260 | 1.06476 | 0.342185 | -3.112 |
| chr2 | 14177843 | 14178146 | 0.988249 | 0.317602 | -3.112 |
| chr5 | 54435142 | 54435342 | 0.970228 | 0.312083 | -3.109 |
| chr19 | 52869632 | 52869846 | 1.04006 | 0.334582 | -3.109 |
| chr2 | 201931420 | 201931608 | 1.22675 | 0.394672 | -3.108 |
| chr9 | 19856654 | 19856919 | 1.08387 | 0.348717 | -3.108 |
| chr2 | 137587119 | 137587336 | 0.936883 | 0.301476 | -3.108 |
| chr5 | 98271132 | 98271336 | 1.46928 | 0.472825 | -3.107 |
| chr2 | 237717143 | 237717339 | 1.10419 | 0.355394 | -3.107 |
| chr15 | 49302829 | 49303060 | 0.981191 | 0.315873 | -3.106 |
| chr8 | 83841831 | 83842060 | 1.09192 | 0.351657 | -3.105 |
| chr16 | 24398175 | 24398474 | 0.739749 | 0.238271 | -3.105 |
| chr19 | 58598571 | 58598758 | 1.27572 | 0.411005 | -3.104 |
| chr5 | 54438284 | 54438486 | 1.36226 | 0.438927 | -3.104 |
| chr1 | 187614992 | 187615178 | 1.13581 | 0.366075 | -3.103 |
| chr5 | 1432177 | 1432390 | 0.872307 | 0.281295 | -3.101 |
| chr11 | 13218555 | 13218763 | 1.35995 | 0.438604 | -3.101 |
| chr11 | 56598712 | 56598906 | 1.13774 | 0.367109 | -3.099 |
| chr5 | 38128222 | 38128410 | 0.970565 | 0.313283 | -3.098 |
| chr9 | 22923517 | 22924073 | 0.97785 | 0.315825 | -3.096 |
| chr8 | 114220180 | 114220459 | 0.937968 | 0.302949 | -3.096 |
| chr2 | 80429490 | 80429690 | 1.03547 | 0.334609 | -3.095 |
| chr7 | 27658625 | 27658809 | 1.27083 | 0.410722 | -3.094 |
| chr9 | 15317681 | 15317864 | 0.97176 | 0.314198 | -3.093 |
| chr3 | 94719670 | 94719863 | 1.11404 | 0.360274 | -3.092 |
| chr3 | 71681669 | 71681930 | 0.806441 | 0.260845 | -3.092 |
| chr2 | 76586242 | 76586456 | 1.27762 | 0.413279 | -3.091 |
| chr9 | 8175443 | 8175666 | 1.17085 | 0.378758 | -3.091 |
| chr8 | 14810750 | 14810932 | 1.21314 | 0.392685 | -3.089 |
| chr10 | 10709748 | 10709947 | 1.00795 | 0.326386 | -3.088 |
| chr2 | 185950398 | 185950695 | 0.987837 | 0.319982 | -3.087 |
| chr5 | 79613204 | 79613421 | 0.852164 | 0.276276 | -3.084 |
| chr2 | 172495780 | 172495979 | 1.22472 | 0.397274 | -3.083 |
| chr6 | 162463412 | 162463620 | 1.28543 | 0.417077 | -3.082 |
| chr10 | 44263653 | 44263852 | 0.785425 | 0.25486 | -3.082 |
| chr5 | 131401518 | 131401805 | 0.850413 | 0.275978 | -3.081 |
| chr11 | 60232300 | 60232495 | 1.2052 | 0.39152 | -3.078 |
| chr2 | 199992507 | 199992797 | 0.934688 | 0.303643 | -3.078 |
| chr3 | 33395080 | 33395353 | 1.20622 | 0.392049 | -3.077 |
| chr17 | 32731415 | 32731599 | 1.11767 | 0.363285 | -3.077 |
| chr2 | 73193579 | 73193782 | 1.20627 | 0.392551 | -3.073 |
| chr8 | 132090423 | 132090620 | 1.42192 | 0.462737 | -3.073 |

| | | | | | |
|---|---|---|---|---|---|
| chr12 | 5593118 | 5593326 | 1.61693 | 0.526323 | -3.072 |
| chr3 | 81088808 | 81089041 | 1.2089 | 0.393688 | -3.071 |
| chr8 | 97420755 | 97420977 | 1.41535 | 0.460975 | -3.070 |
| chr2 | 7469700 | 7470256 | 1.15166 | 0.375168 | -3.070 |
| chr12 | 107108991 | 107109295 | 1.07196 | 0.349245 | -3.069 |
| chr8 | 50274919 | 50275129 | 1.08257 | 0.352743 | -3.069 |
| chr5 | 53890208 | 53890473 | 1.05994 | 0.345384 | -3.069 |
| chr17 | 76905593 | 76905786 | 1.59588 | 0.520223 | -3.068 |
| chr10 | 1961690 | 1961946 | 1.07585 | 0.351083 | -3.064 |
| chr18 | 45736817 | 45737063 | 1.02273 | 0.333805 | -3.064 |
| chr7 | 122745142 | 122745369 | 1.57102 | 0.512931 | -3.063 |
| chr8 | 78100533 | 78100784 | 1.03258 | 0.337228 | -3.062 |
| chr3 | 59294599 | 59294811 | 1.14817 | 0.374983 | -3.062 |
| chr16 | 64117117 | 64117374 | 1.01261 | 0.330804 | -3.061 |
| chr4 | 186010288 | 186010542 | 0.799169 | 0.261112 | -3.061 |
| chr2 | 80597243 | 80597468 | 0.914343 | 0.298763 | -3.060 |
| chr7 | 38917532 | 38917755 | 1.57055 | 0.513337 | -3.059 |
| chr7 | 146864624 | 146864833 | 0.952948 | 0.311569 | -3.059 |
| chr11 | 60014296 | 60014649 | 1.07124 | 0.350392 | -3.057 |
| chr16 | 22178067 | 22178277 | 1.2565 | 0.411065 | -3.057 |
| chr2 | 214563434 | 214563660 | 1.0049 | 0.328922 | -3.055 |
| chr3 | 110627455 | 110627730 | 0.983335 | 0.322069 | -3.053 |
| chr16 | 62801969 | 62802162 | 1.16055 | 0.380121 | -3.053 |
| chr9 | 25239635 | 25239836 | 1.43937 | 0.471593 | -3.052 |
| chr5 | 22787942 | 22788139 | 0.90941 | 0.298028 | -3.051 |
| chr5 | 17133791 | 17134007 | 0.883546 | 0.289591 | -3.051 |
| chr4 | 101312188 | 101312476 | 0.960927 | 0.314955 | -3.051 |
| chr2 | 207797400 | 207797685 | 0.870456 | 0.285345 | -3.051 |
| chr15 | 93673029 | 93673326 | 0.973762 | 0.319222 | -3.050 |
| chr10 | 121801906 | 121802171 | 0.959326 | 0.314555 | -3.050 |
| chr19 | 29894592 | 29894784 | 0.948682 | 0.311086 | -3.050 |
| chr11 | 41467945 | 41468248 | 1.09519 | 0.359167 | -3.049 |
| chr3 | 6224313 | 6224527 | 1.25068 | 0.410414 | -3.047 |
| chr2 | 120374054 | 120374373 | 1.20504 | 0.395545 | -3.047 |
| chr6 | 156712411 | 156712605 | 0.715912 | 0.235049 | -3.046 |
| chr10 | 110841945 | 110842137 | 0.917521 | 0.301289 | -3.045 |
| chr2 | 212800210 | 212800442 | 1.14468 | 0.37592 | -3.045 |
| chr2 | 7932446 | 7932776 | 1.01999 | 0.335024 | -3.045 |
| chr2 | 126032392 | 126032603 | 1.03797 | 0.341001 | -3.044 |
| chr2 | 130000038 | 130000353 | 1.22765 | 0.403321 | -3.044 |
| chr2 | 233438745 | 233438953 | 1.04459 | 0.343244 | -3.043 |
| chr2 | 186063229 | 186063428 | 0.998778 | 0.3283 | -3.042 |
| chr19 | 23936316 | 23936534 | 1.30287 | 0.428292 | -3.042 |
| chr7 | 42747335 | 42747525 | 1.14495 | 0.3765 | -3.041 |
| chr8 | 78175412 | 78175644 | 1.07666 | 0.354044 | -3.041 |
| chr14 | 99634680 | 99634881 | 0.873939 | 0.287402 | -3.041 |
| chr8 | 90357354 | 90357544 | 1.05483 | 0.347086 | -3.039 |
| chr10 | 131904146 | 131904379 | 0.903242 | 0.297222 | -3.039 |
| chr17 | 75267712 | 75267928 | 1.14099 | 0.375508 | -3.039 |
| chr5 | 179205762 | 179206010 | 1.2868 | 0.423721 | -3.037 |
| chr10 | 82794846 | 82795242 | 0.98957 | 0.325939 | -3.036 |
| chr5 | 32997448 | 32997652 | 1.05301 | 0.346868 | -3.036 |
| chr14 | 89327190 | 89327424 | 1.01228 | 0.333557 | -3.035 |
| chr10 | 96873857 | 96874069 | 1.00265 | 0.330484 | -3.034 |
| chr11 | 31304620 | 31304873 | 1.16148 | 0.383036 | -3.032 |
| chr15 | 34045641 | 34045871 | 1.18416 | 0.390746 | -3.031 |
| chr11 | 90259884 | 90260148 | 1.46172 | 0.482442 | -3.030 |
| chr2 | 194166246 | 194166444 | 1.32021 | 0.435751 | -3.030 |
| chr14 | 33091839 | 33092117 | 1.45513 | 0.480357 | -3.029 |
| chr3 | 34329792 | 34330027 | 1.02683 | 0.339042 | -3.029 |
| chr2 | 12936808 | 12937023 | 1.34778 | 0.445097 | -3.028 |

| | | | | | |
|---|---|---|---|---|---|
| chr2 | 84032925 | 84033120 | 1.35782 | 0.448456 | -3.028 |
| chr10 | 107398530 | 107398754 | 1.10285 | 0.36425 | -3.028 |
| chr3 | 77540243 | 77540476 | 1.39512 | 0.460834 | -3.027 |
| chr2 | 210764364 | 210764745 | 0.966338 | 0.319251 | -3.027 |
| chr9 | 96522565 | 96522835 | 1.07863 | 0.356436 | -3.026 |
| chr5 | 23583182 | 23583405 | 1.55578 | 0.514265 | -3.025 |
| chr2 | 188230516 | 188230723 | 1.14834 | 0.379894 | -3.023 |
| chr11 | 117674777 | 117675003 | 1.03211 | 0.341484 | -3.022 |
| chr10 | 126176554 | 126176788 | 1.07573 | 0.355958 | -3.022 |
| chr9 | 11952832 | 11953085 | 1.10425 | 0.36544 | -3.022 |
| chr11 | 76683024 | 76683260 | 0.926034 | 0.30652 | -3.021 |
| chr2 | 117978015 | 117978339 | 1.11116 | 0.368053 | -3.019 |
| chr5 | 146510216 | 146510401 | 1.25475 | 0.415616 | -3.019 |
| chr8 | 80040321 | 80040625 | 0.922348 | 0.305541 | -3.019 |
| chr16 | 66570357 | 66570541 | 0.866513 | 0.287084 | -3.018 |
| chr7 | 88272993 | 88273246 | 1.00018 | 0.331451 | -3.018 |
| chr16 | 81975639 | 81976005 | 1.20218 | 0.398475 | -3.017 |
| chr2 | 115308999 | 115309313 | 1.42626 | 0.472766 | -3.017 |
| chr5 | 105362432 | 105362619 | 1.16754 | 0.38716 | -3.016 |
| chr10 | 105911902 | 105912103 | 0.926303 | 0.307342 | -3.014 |
| chr3 | 82188194 | 82188378 | 1.09925 | 0.364857 | -3.013 |
| chr12 | 77501935 | 77502149 | 0.936469 | 0.31098 | -3.011 |
| chr9 | 10397165 | 10397367 | 1.1638 | 0.386664 | -3.010 |
| chr8 | 83413506 | 83413766 | 0.793917 | 0.26381 | -3.009 |
| chr10 | 129469046 | 129469265 | 1.52968 | 0.508388 | -3.009 |
| chr5 | 75406782 | 75406975 | 1.11553 | 0.370811 | -3.008 |
| chr1 | 162827113 | 162827362 | 0.699461 | 0.232546 | -3.008 |
| chr16 | 8758732 | 8758930 | 0.92329 | 0.307063 | -3.007 |
| chr5 | 63969823 | 63970061 | 1.08452 | 0.360858 | -3.005 |
| chr5 | 30606999 | 30607182 | 0.812983 | 0.27067 | -3.004 |
| chr8 | 30540617 | 30540830 | 1.00425 | 0.334435 | -3.003 |
| chr2 | 7189988 | 7190302 | 1.11082 | 0.369933 | -3.003 |
| chr2 | 174196301 | 174196532 | 0.90327 | 0.300948 | -3.001 |
| chr5 | 163434469 | 163434701 | 1.04162 | 0.347066 | -3.001 |
| chr3 | 95831808 | 95832143 | 0.768529 | 0.256148 | -3.000 |
| chr20 | 44127772 | 44128015 | 1.24097 | 0.413653 | -3.000 |
| chr7 | 136182513 | 136182772 | 0.951503 | 0.317178 | -3.000 |
| chr2 | 60483141 | 60483375 | 1.28913 | 0.429745 | -3.000 |
| chr3 | 89954019 | 89954201 | 0.897881 | 0.299684 | -2.996 |
| chr2 | 139514904 | 139515277 | 0.827801 | 0.2763 | -2.996 |
| chr1 | 190654585 | 190654788 | 1.25156 | 0.417756 | -2.996 |
| chr2 | 103805854 | 103806181 | 1.01375 | 0.338402 | -2.996 |
| chr2 | 34973029 | 34973289 | 0.910716 | 0.304147 | -2.994 |
| chr3 | 75057807 | 75058232 | 0.82612 | 0.275934 | -2.994 |
| chr8 | 57707653 | 57707836 | 1.0352 | 0.34581 | -2.994 |
| chr7 | 106235348 | 106235620 | 1.17335 | 0.391976 | -2.993 |
| chr3 | 118630210 | 118630416 | 1.10486 | 0.36953 | -2.990 |
| chr2 | 77492454 | 77492639 | 1.28034 | 0.428243 | -2.990 |
| chr2 | 47037615 | 47037799 | 1.28884 | 0.431142 | -2.989 |
| chr20 | 36581723 | 36581976 | 1.05801 | 0.353957 | -2.989 |
| chr11 | 21631020 | 21631270 | 0.983315 | 0.329225 | -2.987 |
| chr10 | 129577221 | 129577437 | 0.876344 | 0.293533 | -2.986 |
| chr5 | 120873743 | 120873963 | 1.00847 | 0.337922 | -2.984 |
| chr2 | 53304370 | 53304757 | 1.47751 | 0.495193 | -2.984 |
| chr8 | 93845509 | 93845745 | 1.25013 | 0.418992 | -2.984 |
| chr8 | 89223330 | 89223633 | 0.912283 | 0.305968 | -2.982 |
| chr5 | 25703612 | 25703854 | 1.0378 | 0.348086 | -2.981 |
| chr20 | 49482667 | 49482863 | 1.04381 | 0.350138 | -2.981 |
| chr8 | 5185328 | 5185587 | 1.39276 | 0.467221 | -2.981 |
| chr2 | 14419443 | 14419642 | 1.34714 | 0.451922 | -2.981 |
| chr9 | 79128050 | 79128236 | 0.893075 | 0.299681 | -2.980 |

| | | | | | |
|---|---|---|---|---|---|
| chr9 | 126040584 | 126040775 | 1.07853 | 0.361914 | -2.980 |
| chr15 | 27445903 | 27446109 | 0.945434 | 0.31733 | -2.979 |
| chr3 | 10820958 | 10821265 | 0.901904 | 0.302815 | -2.978 |
| chr5 | 10370324 | 10370519 | 1.05429 | 0.354068 | -2.978 |
| chr2 | 30957219 | 30957415 | 1.01546 | 0.341227 | -2.976 |
| chr1 | 232604761 | 232604979 | 1.17414 | 0.394653 | -2.975 |
| chr2 | 115960844 | 115961061 | 1.02181 | 0.343494 | -2.975 |
| chr1 | 175839624 | 175839850 | 0.980722 | 0.329786 | -2.974 |
| chr7 | 14375420 | 14375632 | 1.09171 | 0.367155 | -2.973 |
| chr2 | 29485068 | 29485320 | 1.14802 | 0.386156 | -2.973 |
| chr5 | 96623124 | 96623368 | 1.42145 | 0.478414 | -2.971 |
| chr1 | 233919601 | 233919842 | 0.984621 | 0.331475 | -2.970 |
| chr16 | 61285645 | 61285957 | 1.22988 | 0.41435 | -2.968 |
| chr2 | 13035427 | 13035893 | 1.13996 | 0.384138 | -2.968 |
| chr2 | 83572912 | 83573265 | 1.01257 | 0.341252 | -2.967 |
| chr5 | 121264457 | 121264673 | 1.45637 | 0.490851 | -2.967 |
| chr3 | 173279857 | 173280107 | 1.01013 | 0.340518 | -2.966 |
| chr2 | 64271300 | 64271549 | 1.42524 | 0.480621 | -2.965 |
| chr10 | 35491145 | 35491375 | 0.842232 | 0.284119 | -2.964 |
| chr2 | 167265312 | 167265495 | 1.03792 | 0.350231 | -2.964 |
| chr9 | 131173138 | 131173352 | 0.798529 | 0.269543 | -2.963 |
| chr1 | 248555571 | 248555757 | 1.0711 | 0.361555 | -2.962 |
| chr9 | 110123122 | 110123310 | 1.11243 | 0.375811 | -2.960 |
| chr3 | 142833818 | 142834144 | 1.40832 | 0.475967 | -2.959 |
| chr5 | 12566397 | 12566797 | 1.09441 | 0.369877 | -2.959 |
| chr2 | 117284464 | 117284806 | 1.15209 | 0.389572 | -2.957 |
| chr5 | 65753984 | 65754293 | 1.1473 | 0.387982 | -2.957 |
| chr3 | 95928763 | 95929202 | 0.958554 | 0.324324 | -2.956 |
| chr9 | 6278536 | 6278737 | 1.36844 | 0.463109 | -2.955 |
| chr10 | 120392218 | 120392444 | 1.50492 | 0.509356 | -2.955 |
| chr2 | 138028546 | 138028741 | 1.03483 | 0.350387 | -2.953 |
| chr12 | 83705618 | 83706070 | 0.988804 | 0.334941 | -2.952 |
| chr14 | 56444156 | 56444439 | 0.915649 | 0.310306 | -2.951 |
| chr12 | 2572924 | 2573234 | 1.25393 | 0.424955 | -2.951 |
| chr20 | 17997167 | 17997452 | 1.3197 | 0.447398 | -2.950 |
| chr16 | 25670348 | 25670545 | 0.999415 | 0.338959 | -2.948 |
| chr2 | 5623177 | 5623377 | 1.22611 | 0.416058 | -2.947 |
| chr8 | 35352639 | 35352872 | 0.910964 | 0.309265 | -2.946 |
| chr2 | 108588993 | 108589180 | 0.919178 | 0.312101 | -2.945 |
| chr9 | 1390724 | 1390980 | 0.830591 | 0.282043 | -2.945 |
| chr5 | 25413661 | 25413957 | 1.23757 | 0.420252 | -2.945 |
| chr2 | 223612590 | 223612805 | 1.37072 | 0.465506 | -2.945 |
| chr2 | 116680033 | 116680233 | 0.966851 | 0.328407 | -2.944 |
| chr9 | 105653831 | 105654021 | 1.26211 | 0.428883 | -2.943 |
| chr2 | 238153128 | 238153339 | 0.817918 | 0.277954 | -2.943 |
| chr9 | 129724995 | 129725234 | 1.23758 | 0.420628 | -2.942 |
| chr14 | 86911381 | 86911671 | 1.1352 | 0.385958 | -2.941 |
| chr19 | 51066302 | 51066500 | 1.21048 | 0.411562 | -2.941 |
| chr2 | 34443905 | 34444231 | 1.07554 | 0.365721 | -2.941 |
| chr8 | 115889631 | 115889883 | 0.87894 | 0.298987 | -2.940 |
| chr9 | 17943635 | 17943914 | 0.915741 | 0.311584 | -2.939 |
| chr1 | 209656222 | 209656492 | 1.28243 | 0.436489 | -2.938 |
| chr10 | 86753850 | 86754033 | 1.13428 | 0.386102 | -2.938 |
| chr16 | 48929118 | 48929314 | 1.02442 | 0.348745 | -2.937 |
| chr1 | 96132360 | 96132565 | 1.01329 | 0.345008 | -2.937 |
| chr8 | 51759899 | 51760096 | 1.14089 | 0.388565 | -2.936 |
| chr3 | 95834327 | 95834522 | 1.00302 | 0.341663 | -2.936 |
| chr11 | 90181581 | 90181764 | 1.15178 | 0.392419 | -2.935 |
| chr2 | 195966465 | 195966685 | 1.15417 | 0.393338 | -2.934 |
| chr10 | 124952121 | 124952353 | 1.26006 | 0.429463 | -2.934 |
| chr7 | 23692586 | 23693000 | 1.2269 | 0.418343 | -2.933 |

| | | | | | |
|---|---|---|---|---|---|
| chr19 | 53233342 | 53233597 | 0.746584 | 0.254622 | -2.932 |
| chr5 | 146757907 | 146758203 | 1.10087 | 0.375463 | -2.932 |
| chr2 | 80182855 | 80183038 | 1.21865 | 0.415763 | -2.931 |
| chr9 | 98444374 | 98444564 | 1.25355 | 0.42814 | -2.928 |
| chr2 | 202800 | 203041 | 0.997803 | 0.341151 | -2.925 |
| chr1 | 237538516 | 237538703 | 0.712041 | 0.243451 | -2.925 |
| chr11 | 81369836 | 81370051 | 1.42235 | 0.486348 | -2.925 |
| chr2 | 209018076 | 209018375 | 0.800156 | 0.273748 | -2.923 |
| chr10 | 19674130 | 19674375 | 1.6391 | 0.56105 | -2.921 |
| chr8 | 76476772 | 76476973 | 0.961703 | 0.329271 | -2.921 |
| chr2 | 193756680 | 193756904 | 1.06404 | 0.364392 | -2.920 |
| chr2 | 8650812 | 8651002 | 1.07683 | 0.368895 | -2.919 |
| chr12 | 43845516 | 43845848 | 0.85674 | 0.293606 | -2.918 |
| chr11 | 7328252 | 7328443 | 0.893168 | 0.306096 | -2.918 |
| chr2 | 40872482 | 40872700 | 1.13671 | 0.389735 | -2.917 |
| chr2 | 229358392 | 229358622 | 0.813989 | 0.27912 | -2.916 |
| chr17 | 41215419 | 41215622 | 1.12492 | 0.385751 | -2.916 |
| chr7 | 113899687 | 113900007 | 0.824307 | 0.282953 | -2.913 |
| chr2 | 41518172 | 41518448 | 1.16367 | 0.399451 | -2.913 |
| chr11 | 44223889 | 44224128 | 1.31307 | 0.450872 | -2.912 |
| chr3 | 104448311 | 104448540 | 1.21902 | 0.418638 | -2.912 |
| chr16 | 68861588 | 68861783 | 1.36883 | 0.470353 | -2.910 |
| chr16 | 13275412 | 13275612 | 1.22307 | 0.420474 | -2.909 |
| chr8 | 120293905 | 120294103 | 0.979111 | 0.33669 | -2.908 |
| chr10 | 68939504 | 68939693 | 0.940507 | 0.323541 | -2.907 |
| chr5 | 87879286 | 87879486 | 1.36542 | 0.469815 | -2.906 |
| chr1 | 74312054 | 74312258 | 1.11992 | 0.385393 | -2.906 |
| chr20 | 23808945 | 23809172 | 0.882835 | 0.304226 | -2.902 |
| chr10 | 105802141 | 105802423 | 1.28446 | 0.442661 | -2.902 |
| chr20 | 19634808 | 19635049 | 1.04984 | 0.361823 | -2.902 |
| chr2 | 1949070 | 1949252 | 1.12644 | 0.388242 | -2.901 |
| chr16 | 62556189 | 62556461 | 0.743443 | 0.256292 | -2.901 |
| chr9 | 24075728 | 24075920 | 0.904836 | 0.311957 | -2.901 |
| chr3 | 84664836 | 84665087 | 1.5043 | 0.518777 | -2.900 |
| chr10 | 1079860 | 1080048 | 0.948727 | 0.327299 | -2.899 |
| chr4 | 190441646 | 190441835 | 0.970793 | 0.334956 | -2.898 |
| chr9 | 25633416 | 25633610 | 1.15566 | 0.398756 | -2.898 |
| chr10 | 10954412 | 10954639 | 0.856269 | 0.295538 | -2.897 |
| chr5 | 86940904 | 86941105 | 1.19607 | 0.412833 | -2.897 |
| chr2 | 95958924 | 95959191 | 0.95564 | 0.329867 | -2.897 |
| chr3 | 41329147 | 41329329 | 0.805333 | 0.277996 | -2.897 |
| chr2 | 228699102 | 228699472 | 1.03118 | 0.356033 | -2.896 |
| chr15 | 95236555 | 95236760 | 1.06593 | 0.368275 | -2.894 |
| chr2 | 126398342 | 126398617 | 1.0309 | 0.356219 | -2.894 |
| chr2 | 167615253 | 167615442 | 0.844628 | 0.291941 | -2.893 |
| chr3 | 86037054 | 86037407 | 1.03489 | 0.357835 | -2.892 |
| chr12 | 126761961 | 126762195 | 0.873979 | 0.302204 | -2.892 |
| chr8 | 116872255 | 116872516 | 1.07314 | 0.371073 | -2.892 |
| chr7 | 92298997 | 92299211 | 1.13961 | 0.394178 | -2.891 |
| chr19 | 47069987 | 47070207 | 1.53326 | 0.530373 | -2.891 |
| chr3 | 11047516 | 11047761 | 0.913395 | 0.315961 | -2.891 |
| chr10 | 132740229 | 132740506 | 1.62418 | 0.562006 | -2.890 |
| chr3 | 6393766 | 6394024 | 0.744618 | 0.25773 | -2.889 |
| chr5 | 44090006 | 44090349 | 1.15278 | 0.399009 | -2.889 |
| chr5 | 140233488 | 140233675 | 1.32426 | 0.458512 | -2.888 |
| chr12 | 133410482 | 133410746 | 1.27362 | 0.440992 | -2.888 |
| chr2 | 127806589 | 127806991 | 0.919151 | 0.318303 | -2.888 |
| chr3 | 19584812 | 19585008 | 2.96126 | 1.02564 | -2.887 |
| chr10 | 2267878 | 2268184 | 1.06743 | 0.369851 | -2.886 |
| chr5 | 84741579 | 84741878 | 1.07153 | 0.371565 | -2.884 |
| chr5 | 147310756 | 147310964 | 1.01267 | 0.351246 | -2.883 |

| | | | | | |
|---|---|---|---|---|---|
| chr15 | 27926271 | 27926496 | 1.55271 | 0.538662 | -2.883 |
| chr3 | 88928545 | 88928815 | 1.03348 | 0.358596 | -2.882 |
| chr14 | 48119653 | 48119867 | 1.15263 | 0.399979 | -2.882 |
| chr2 | 186120996 | 186121259 | 0.980214 | 0.340245 | -2.881 |
| chr8 | 92791621 | 92791866 | 1.12945 | 0.392059 | -2.881 |
| chr11 | 26005501 | 26005768 | 1.20828 | 0.419447 | -2.881 |
| chr2 | 122873225 | 122873524 | 1.08037 | 0.375065 | -2.880 |
| chr2 | 13717832 | 13718108 | 1.28218 | 0.44515 | -2.880 |
| chr5 | 16288973 | 16289155 | 0.947005 | 0.328848 | -2.880 |
| chr2 | 70399377 | 70399571 | 1.05397 | 0.366036 | -2.879 |
| chr2 | 117832165 | 117832524 | 0.985748 | 0.342351 | -2.879 |
| chr3 | 162052455 | 162052657 | 1.52038 | 0.528047 | -2.879 |
| chr14 | 88908526 | 88908710 | 0.892395 | 0.30998 | -2.879 |
| chr15 | 54018990 | 54019311 | 1.11484 | 0.38737 | -2.878 |
| chr1 | 223723576 | 223723874 | 1.54648 | 0.537409 | -2.878 |
| chr2 | 6273990 | 6274276 | 1.01318 | 0.352085 | -2.878 |
| chr16 | 83050175 | 83050369 | 1.01857 | 0.354065 | -2.877 |
| chr6 | 95120251 | 95120527 | 1.10446 | 0.384046 | -2.876 |
| chr2 | 75213331 | 75213526 | 1.05892 | 0.368276 | -2.875 |
| chr8 | 140008528 | 140008816 | 1.30894 | 0.45527 | -2.875 |
| chr2 | 53024272 | 53024544 | 1.05802 | 0.367998 | -2.875 |
| chr2 | 141636247 | 141636493 | 0.895548 | 0.311745 | -2.873 |
| chr19 | 52418756 | 52418940 | 1.13013 | 0.39345 | -2.872 |
| chr11 | 70139861 | 70140094 | 1.13261 | 0.394462 | -2.871 |
| chr15 | 98133630 | 98133847 | 1.12138 | 0.390554 | -2.871 |
| chr13 | 24581174 | 24581380 | 0.785721 | 0.273828 | -2.869 |
| chr9 | 88667595 | 88667804 | 0.918908 | 0.320251 | -2.869 |
| chr2 | 206101916 | 206102118 | 1.0992 | 0.383607 | -2.865 |
| chr5 | 115687624 | 115687811 | 0.983095 | 0.343176 | -2.865 |
| chr2 | 13660029 | 13660275 | 1.20321 | 0.420052 | -2.864 |
| chr2 | 104499925 | 104500126 | 0.930791 | 0.325066 | -2.863 |
| chr12 | 61031402 | 61031643 | 1.04098 | 0.363579 | -2.863 |
| chr6 | 76452200 | 76452499 | 0.800746 | 0.27971 | -2.863 |
| chr5 | 163535668 | 163535874 | 1.2521 | 0.437389 | -2.863 |
| chr2 | 215943766 | 215943955 | 1.0098 | 0.352755 | -2.863 |
| chr2 | 14158557 | 14158752 | 1.39352 | 0.486894 | -2.862 |
| chr5 | 62595021 | 62595207 | 1.37319 | 0.479913 | -2.861 |
| chr13 | 49904071 | 49904292 | 0.976087 | 0.341164 | -2.861 |
| chr8 | 73192212 | 73192417 | 0.983232 | 0.343676 | -2.861 |
| chr2 | 73201682 | 73201887 | 1.31562 | 0.459943 | -2.860 |
| chr11 | 81645529 | 81645819 | 1.30526 | 0.456427 | -2.860 |
| chr2 | 49604001 | 49604616 | 1.15044 | 0.402361 | -2.859 |
| chr2 | 156516536 | 156516741 | 0.905236 | 0.316649 | -2.859 |
| chr2 | 196865961 | 196866305 | 0.979098 | 0.34249 | -2.859 |
| chr9 | 79759022 | 79759360 | 1.00307 | 0.350935 | -2.858 |
| chr3 | 32248946 | 32249208 | 1.0347 | 0.362106 | -2.857 |
| chr14 | 87081965 | 87082192 | 0.978686 | 0.342557 | -2.857 |
| chr5 | 163667385 | 163667575 | 0.889882 | 0.311595 | -2.856 |
| chr9 | 26129838 | 26130151 | 0.851067 | 0.298006 | -2.856 |
| chr8 | 13510938 | 13511177 | 1.13405 | 0.397148 | -2.855 |
| gl000198_r | 1747 | 2050 | 1.09612 | 0.383975 | -2.855 |
| chr8 | 70544186 | 70544511 | 2.02023 | 0.707746 | -2.854 |
| chr16 | 77993806 | 77994048 | 1.29999 | 0.455435 | -2.854 |
| chr9 | 71460082 | 71460378 | 1.2802 | 0.448526 | -2.854 |
| chr12 | 8732355 | 8732606 | 1.01397 | 0.355271 | -2.854 |
| chr16 | 74406047 | 74406232 | 1.06969 | 0.37485 | -2.854 |
| chr5 | 82797418 | 82797603 | 0.926873 | 0.324878 | -2.853 |
| chr7 | 125016233 | 125016519 | 1.07905 | 0.378273 | -2.853 |
| chr10 | 124506262 | 124506462 | 0.941337 | 0.330034 | -2.852 |
| chr1 | 152176472 | 152176693 | 1.1691 | 0.409958 | -2.852 |
| chr16 | 69194058 | 69194261 | 1.00916 | 0.353919 | -2.851 |

| | | | | | |
|---|---|---|---|---|---|
| chr14 | 48081761 | 48081946 | 0.859849 | 0.301616 | -2.851 |
| chr5 | 62253650 | 62253938 | 1.47325 | 0.516846 | -2.850 |
| chr2 | 32311412 | 32311630 | 1.51881 | 0.532869 | -2.850 |
| chr11 | 41728070 | 41728297 | 1.16421 | 0.408463 | -2.850 |
| chr14 | 87599687 | 87599998 | 1.08332 | 0.380255 | -2.849 |
| chr2 | 72297818 | 72298009 | 1.4033 | 0.49269 | -2.848 |
| chr8 | 3034294 | 3034507 | 1.04225 | 0.366131 | -2.847 |
| chr5 | 103592941 | 103593142 | 1.45032 | 0.509675 | -2.846 |
| chr8 | 88708423 | 88708633 | 1.02751 | 0.361123 | -2.845 |
| chr7 | 24935018 | 24935299 | 1.04458 | 0.36718 | -2.845 |
| chr5 | 4872487 | 4872763 | 0.972495 | 0.34204 | -2.843 |
| chr16 | 23481411 | 23481607 | 0.830474 | 0.292134 | -2.843 |
| chr2 | 212860524 | 212860797 | 1.26983 | 0.447074 | -2.840 |
| chr2 | 1820472 | 1820735 | 1.0868 | 0.382657 | -2.840 |
| chr13 | 50844686 | 50844917 | 0.83232 | 0.293202 | -2.839 |
| chr1 | 152304666 | 152304917 | 0.971584 | 0.342349 | -2.838 |
| chr10 | 20632803 | 20633001 | 0.836149 | 0.294628 | -2.838 |
| chr19 | 57870378 | 57870714 | 1.27461 | 0.449179 | -2.838 |
| chr8 | 64296989 | 64297185 | 0.780298 | 0.275075 | -2.837 |
| chr16 | 13008736 | 13008966 | 1.19502 | 0.421413 | -2.836 |
| chr8 | 73770828 | 73771020 | 0.811407 | 0.286155 | -2.836 |
| chr5 | 100870325 | 100870653 | 1.01424 | 0.35773 | -2.835 |
| chr2 | 142059088 | 142059434 | 0.992203 | 0.350103 | -2.834 |
| chr8 | 138534832 | 138535041 | 1.18661 | 0.418726 | -2.834 |
| chr15 | 83336384 | 83336578 | 0.968791 | 0.341906 | -2.834 |
| chr2 | 171380068 | 171380443 | 1.10666 | 0.390606 | -2.833 |
| chr7 | 153651811 | 153651999 | 1.14311 | 0.403521 | -2.833 |
| chr2 | 6501179 | 6501460 | 1.08739 | 0.383871 | -2.833 |
| chr9 | 8675930 | 8676215 | 0.838519 | 0.296095 | -2.832 |
| chr2 | 198370904 | 198371089 | 0.966256 | 0.341364 | -2.831 |
| chr3 | 74405557 | 74405741 | 1.10285 | 0.389734 | -2.830 |
| chr8 | 133225619 | 133225896 | 1.11139 | 0.392782 | -2.830 |
| chr5 | 161528502 | 161528684 | 1.13308 | 0.400546 | -2.829 |
| chr11 | 38129314 | 38129564 | 1.03691 | 0.366585 | -2.829 |
| chr11 | 78873846 | 78874074 | 1.17426 | 0.415143 | -2.829 |
| chr2 | 219651933 | 219652212 | 0.884491 | 0.312741 | -2.828 |
| chr16 | 83340392 | 83340643 | 0.899095 | 0.317954 | -2.828 |
| chr10 | 54737709 | 54737893 | 1.28352 | 0.45393 | -2.828 |
| chr5 | 12960045 | 12960268 | 1.03348 | 0.365721 | -2.826 |
| chr2 | 79753504 | 79753742 | 1.05681 | 0.374242 | -2.824 |
| chr14 | 54593806 | 54594018 | 1.01927 | 0.361282 | -2.821 |
| chr8 | 110855182 | 110855411 | 1.2595 | 0.446465 | -2.821 |
| chr19 | 22088689 | 22088871 | 1.34823 | 0.478004 | -2.821 |
| chr3 | 39527098 | 39527300 | 1.17558 | 0.416797 | -2.821 |
| chr14 | 106720174 | 106720472 | 1.22422 | 0.434108 | -2.820 |
| chr2 | 185704920 | 185705115 | 0.914618 | 0.324326 | -2.820 |
| chr5 | 152472684 | 152473001 | 0.878075 | 0.311405 | -2.820 |
| chr2 | 205021783 | 205022026 | 1.33769 | 0.47443 | -2.820 |
| chr16 | 87632766 | 87633007 | 1.05971 | 0.375929 | -2.819 |
| chr19 | 57379357 | 57379632 | 1.23815 | 0.439232 | -2.819 |
| chr8 | 68868551 | 68868941 | 1.03364 | 0.366788 | -2.818 |
| chr14 | 66896083 | 66896266 | 0.949344 | 0.33708 | -2.816 |
| chr5 | 168943429 | 168943635 | 0.943645 | 0.335093 | -2.816 |
| chr7 | 78064749 | 78065031 | 1.12758 | 0.400435 | -2.816 |
| chr5 | 84007774 | 84008040 | 1.03412 | 0.367308 | -2.815 |
| chr11 | 38203893 | 38204086 | 0.879963 | 0.312563 | -2.815 |
| chr2 | 79520157 | 79520434 | 1.20245 | 0.427136 | -2.815 |
| chr12 | 118325240 | 118325584 | 1.26438 | 0.449162 | -2.815 |
| chr11 | 61954662 | 61954958 | 1.08701 | 0.38627 | -2.814 |
| chr2 | 199612022 | 199612249 | 1.10938 | 0.394241 | -2.814 |
| chr9 | 103711687 | 103711943 | 1.56501 | 0.556265 | -2.813 |

| | | | | | |
|---|---|---|---|---|---|
| chr7 | 122240191 | 122240413 | 1.05173 | 0.373841 | -2.813 |
| chr11 | 30707481 | 30707678 | 1.35914 | 0.483283 | -2.812 |
| chr9 | 32069411 | 32069752 | 1.56615 | 0.556905 | -2.812 |
| chr1 | 239206279 | 239206560 | 1.03746 | 0.36892 | -2.812 |
| chr3 | 82797622 | 82797869 | 1.26522 | 0.450004 | -2.812 |
| chr14 | 79214747 | 79214985 | 1.08394 | 0.38556 | -2.811 |
| chr9 | 82975595 | 82975880 | 0.993094 | 0.353339 | -2.811 |
| chr10 | 132800642 | 132801163 | 0.987225 | 0.351297 | -2.810 |
| chr2 | 117970211 | 117970445 | 0.933988 | 0.33236 | -2.810 |
| chr14 | 76378420 | 76378672 | 1.281 | 0.455916 | -2.810 |
| chr1 | 184322595 | 184322948 | 1.3056 | 0.464674 | -2.810 |
| chr2 | 165995981 | 165996170 | 0.957513 | 0.340863 | -2.809 |
| chr2 | 149847584 | 149847838 | 1.11045 | 0.395322 | -2.809 |
| chr8 | 117572706 | 117572896 | 1.38678 | 0.493968 | -2.807 |
| chr3 | 158663813 | 158664123 | 0.957946 | 0.341244 | -2.807 |
| chr5 | 13129612 | 13129837 | 0.919498 | 0.327563 | -2.807 |
| chr2 | 125321306 | 125321495 | 0.886121 | 0.31569 | -2.807 |
| chr15 | 98430172 | 98430484 | 1.15991 | 0.413233 | -2.807 |
| chr11 | 30877022 | 30877273 | 1.09682 | 0.390774 | -2.807 |
| chr16 | 62595862 | 62596073 | 1.15492 | 0.411505 | -2.807 |
| chr2 | 78395308 | 78395818 | 1.17963 | 0.420318 | -2.807 |
| chr10 | 85126063 | 85126258 | 0.895639 | 0.319167 | -2.806 |
| chr10 | 92380003 | 92380203 | 1.03123 | 0.367546 | -2.806 |
| chr1 | 244842366 | 244842619 | 1.28622 | 0.458638 | -2.804 |
| chr8 | 140511255 | 140511502 | 0.949703 | 0.338837 | -2.803 |
| chr20 | 48113036 | 48113274 | 1.2008 | 0.428468 | -2.803 |
| chr9 | 98752859 | 98753123 | 1.33863 | 0.477811 | -2.802 |
| chr9 | 71514846 | 71515046 | 1.12029 | 0.399909 | -2.801 |
| chr2 | 18871712 | 18872041 | 1.40813 | 0.502695 | -2.801 |
| chr2 | 150321097 | 150321359 | 1.31003 | 0.467925 | -2.800 |
| chr2 | 81807375 | 81807723 | 1.007 | 0.359771 | -2.799 |
| chr1 | 211192207 | 211192452 | 1.28441 | 0.458942 | -2.799 |
| chr20 | 49867884 | 49868121 | 1.25458 | 0.4483 | -2.799 |
| chr10 | 99838495 | 99838801 | 0.994086 | 0.355394 | -2.797 |
| chr17 | 31847503 | 31847827 | 2.98014 | 1.06567 | -2.796 |
| chr2 | 117857068 | 117857364 | 1.16156 | 0.4154 | -2.796 |
| chr10 | 127985980 | 127986204 | 0.965787 | 0.345778 | -2.793 |
| chr8 | 136142824 | 136143091 | 0.986878 | 0.353339 | -2.793 |
| chr2 | 222211038 | 222211224 | 0.888806 | 0.318316 | -2.792 |
| chr1 | 247112653 | 247112885 | 0.859684 | 0.307894 | -2.792 |
| chr19 | 33699074 | 33699262 | 1.5042 | 0.538777 | -2.792 |
| chr12 | 124493328 | 124493517 | 1.15797 | 0.414823 | -2.791 |
| chr1 | 120018552 | 120018750 | 2.03131 | 0.727995 | -2.790 |
| chr16 | 17553626 | 17553834 | 0.947566 | 0.339696 | -2.789 |
| chr2 | 117206815 | 117207045 | 0.968197 | 0.347175 | -2.789 |
| chr10 | 124416823 | 124417092 | 1.17371 | 0.420971 | -2.788 |
| chr16 | 60685126 | 60685385 | 0.893436 | 0.320453 | -2.788 |
| chr2 | 5375967 | 5376174 | 1.34011 | 0.480728 | -2.788 |
| chr13 | 83416119 | 83416369 | 1.19754 | 0.429808 | -2.786 |
| chr1 | 217086088 | 217086365 | 1.06245 | 0.381513 | -2.785 |
| chr7 | 41300716 | 41301087 | 1.05777 | 0.380015 | -2.783 |
| chr2 | 206268114 | 206268405 | 0.832391 | 0.299066 | -2.783 |
| chr12 | 14154478 | 14154807 | 1.13444 | 0.40761 | -2.783 |
| chr19 | 56371835 | 56372108 | 0.979681 | 0.352064 | -2.783 |
| chr7 | 52097042 | 52097379 | 1.09496 | 0.393707 | -2.781 |
| chr8 | 85972502 | 85972774 | 1.32102 | 0.475116 | -2.780 |
| chr10 | 131460827 | 131461098 | 0.832921 | 0.299665 | -2.780 |
| chr11 | 38002706 | 38002916 | 0.788093 | 0.283588 | -2.779 |
| chr7 | 51658545 | 51658761 | 1.3151 | 0.473329 | -2.778 |
| chr5 | 145117379 | 145117617 | 1.21541 | 0.43751 | -2.778 |
| chr10 | 9651871 | 9652121 | 0.954204 | 0.343526 | -2.778 |

| | | | | | |
|---|---|---|---|---|---|
| chr2 | 172588116 | 172588315 | 1.20431 | 0.433578 | -2.778 |
| chr10 | 1131191 | 1131565 | 1.06006 | 0.381681 | -2.777 |
| chr7 | 72892312 | 72892533 | 0.989707 | 0.356458 | -2.777 |
| chr5 | 135031866 | 135032049 | 0.994243 | 0.358124 | -2.776 |
| chr17 | 68835217 | 68835424 | 1.07347 | 0.386741 | -2.776 |
| chr2 | 193592731 | 193592965 | 1.09927 | 0.396161 | -2.775 |
| chr3 | 96150736 | 96150992 | 0.963641 | 0.347468 | -2.773 |
| chr17 | 53997260 | 53997559 | 1.23502 | 0.445416 | -2.773 |
| chr15 | 100602304 | 100602595 | 1.33759 | 0.48242 | -2.773 |
| chr1 | 35063548 | 35063798 | 2.45701 | 0.886257 | -2.772 |
| chr7 | 19113115 | 19113362 | 1.0248 | 0.369662 | -2.772 |
| chr3 | 162844971 | 162845173 | 1.04788 | 0.378125 | -2.771 |
| chr11 | 61980000 | 61980322 | 0.917515 | 0.331147 | -2.771 |
| chr3 | 16434456 | 16434644 | 0.765327 | 0.276233 | -2.771 |
| chr11 | 13822433 | 13822625 | 1.00021 | 0.361044 | -2.770 |
| chr5 | 143903977 | 143904277 | 1.27749 | 0.461144 | -2.770 |
| chrX | 53159547 | 53159740 | 0.839782 | 0.303165 | -2.770 |
| chr7 | 145561582 | 145561808 | 0.965028 | 0.34839 | -2.770 |
| chr14 | 47546035 | 47546329 | 1.03541 | 0.373858 | -2.770 |
| chr5 | 8327261 | 8327466 | 1.46352 | 0.528718 | -2.768 |
| chr3 | 109296525 | 109296865 | 1.2871 | 0.465164 | -2.767 |
| chr2 | 207871928 | 207872161 | 1.26059 | 0.455619 | -2.767 |
| chr7 | 51784703 | 51784885 | 0.96438 | 0.34862 | -2.766 |
| chr7 | 69768973 | 69769176 | 1.22411 | 0.442611 | -2.766 |
| chr1 | 191271810 | 191272244 | 1.12927 | 0.408376 | -2.765 |
| chr20 | 42114679 | 42114871 | 1.4133 | 0.511114 | -2.765 |
| chr15 | 32145574 | 32145773 | 1.17219 | 0.423927 | -2.765 |
| chr2 | 147123557 | 147123742 | 1.22948 | 0.444685 | -2.765 |
| chr2 | 22939742 | 22939975 | 1.09588 | 0.396485 | -2.764 |
| chr7 | 93279858 | 93280083 | 1.0015 | 0.362409 | -2.763 |
| chr17 | 3513024 | 3513237 | 1.06136 | 0.384106 | -2.763 |
| chr8 | 13778976 | 13779225 | 1.06173 | 0.384492 | -2.761 |
| chr2 | 22106803 | 22107048 | 1.21018 | 0.438259 | -2.761 |
| chr2 | 49164826 | 49165044 | 0.947751 | 0.343397 | -2.760 |
| chr2 | 131684874 | 131685123 | 1.2574 | 0.45567 | -2.759 |
| chr10 | 72304490 | 72304736 | 1.06742 | 0.38695 | -2.759 |
| chr14 | 82294120 | 82294316 | 1.0161 | 0.368417 | -2.758 |
| chr16 | 80157768 | 80157964 | 1.36658 | 0.495681 | -2.757 |
| chr16 | 8341026 | 8341211 | 1.02709 | 0.372667 | -2.756 |
| chr2 | 236032311 | 236032529 | 0.801106 | 0.290931 | -2.754 |
| chr8 | 31950285 | 31950569 | 1.06098 | 0.385315 | -2.754 |
| chr17 | 51168586 | 51168867 | 1.37273 | 0.49863 | -2.753 |
| chr1 | 209722971 | 209723179 | 1.63073 | 0.592411 | -2.753 |
| chr17 | 73927260 | 73927470 | 1.14683 | 0.416951 | -2.751 |
| chr5 | 51260475 | 51260840 | 1.11752 | 0.406375 | -2.750 |
| chr9 | 25033235 | 25033490 | 0.891983 | 0.324401 | -2.750 |
| chr10 | 98177987 | 98178205 | 1.16973 | 0.425619 | -2.748 |
| chr19 | 56556577 | 56556760 | 1.1155 | 0.405894 | -2.748 |
| chr20 | 41412141 | 41412406 | 1.14877 | 0.418068 | -2.748 |
| chr2 | 206415940 | 206416283 | 1.1552 | 0.420498 | -2.747 |
| chr15 | 93011642 | 93011937 | 1.16263 | 0.423416 | -2.746 |
| chr9 | 98204066 | 98204270 | 1.11433 | 0.405852 | -2.746 |
| chr2 | 57426817 | 57427002 | 1.09986 | 0.400669 | -2.745 |
| chr8 | 105025684 | 105025878 | 1.42102 | 0.517683 | -2.745 |
| chr2 | 77380290 | 77380487 | 1.56134 | 0.569135 | -2.743 |
| chr2 | 212090595 | 212090862 | 1.04697 | 0.381659 | -2.743 |
| chr8 | 6017014 | 6017197 | 0.839245 | 0.305967 | -2.743 |
| chr14 | 83891327 | 83891522 | 0.862695 | 0.314592 | -2.742 |
| chr2 | 51265242 | 51265443 | 1.17734 | 0.429415 | -2.742 |
| chr2 | 1785133 | 1785413 | 1.00144 | 0.36537 | -2.741 |
| chr17 | 79503212 | 79503401 | 0.984208 | 0.359113 | -2.741 |

| | | | | | |
|---|---|---|---|---|---|
| chr12 | 84813831 | 84814190 | 1.16285 | 0.424489 | -2.739 |
| chr10 | 17337600 | 17337828 | 1.06739 | 0.389663 | -2.739 |
| chr15 | 75486851 | 75487078 | 1.19132 | 0.43508 | -2.738 |
| chr15 | 87847295 | 87847504 | 1.15681 | 0.422625 | -2.737 |
| chr8 | 47435400 | 47435732 | 0.875727 | 0.319962 | -2.737 |
| chr8 | 43059550 | 43059806 | 1.07151 | 0.391582 | -2.736 |
| chr2 | 4886956 | 4887311 | 1.3669 | 0.4997 | -2.735 |
| chr6 | 73232565 | 73232819 | 0.871078 | 0.318456 | -2.735 |
| chr3 | 104741937 | 104742220 | 1.00047 | 0.365762 | -2.735 |
| chr11 | 7438649 | 7438895 | 1.15975 | 0.424389 | -2.733 |
| chr8 | 69766293 | 69766555 | 1.19989 | 0.439083 | -2.733 |
| chr2 | 184982836 | 184983201 | 1.19405 | 0.437154 | -2.731 |
| chr8 | 17205283 | 17205529 | 1.2075 | 0.442138 | -2.731 |
| chr9 | 106761949 | 106762141 | 1.38419 | 0.506896 | -2.731 |
| chr17 | 10055158 | 10055364 | 1.0635 | 0.38956 | -2.730 |
| chr20 | 21255776 | 21255979 | 1.09553 | 0.401299 | -2.730 |
| chr8 | 13589148 | 13589368 | 0.921322 | 0.337561 | -2.729 |
| chr14 | 78698149 | 78698396 | 1.1767 | 0.431135 | -2.729 |
| chr3 | 1097316 | 1097557 | 1.10099 | 0.403565 | -2.728 |
| chr8 | 47422291 | 47422505 | 1.06315 | 0.389753 | -2.728 |
| chr5 | 28837438 | 28837899 | 0.804797 | 0.295165 | -2.727 |
| chr2 | 16502218 | 16502408 | 1.12067 | 0.411068 | -2.726 |
| chr2 | 143022453 | 143022648 | 0.905595 | 0.33219 | -2.726 |
| chr7 | 113201319 | 113201610 | 0.97972 | 0.359505 | -2.725 |
| chr2 | 119009661 | 119009906 | 0.848237 | 0.311408 | -2.724 |
| chr17 | 11049187 | 11049392 | 1.32242 | 0.485582 | -2.723 |
| chrX | 19025294 | 19025560 | 0.915087 | 0.336154 | -2.722 |
| chr9 | 129319980 | 129320194 | 1.0872 | 0.399437 | -2.722 |
| chr2 | 50900836 | 50901059 | 0.893896 | 0.32847 | -2.721 |
| chr3 | 78598830 | 78599028 | 1.29719 | 0.476729 | -2.721 |
| chr2 | 30428925 | 30429190 | 1.05442 | 0.387517 | -2.721 |
| chr16 | 78134427 | 78134629 | 1.05654 | 0.388306 | -2.721 |
| chr2 | 242509281 | 242509509 | 1.03191 | 0.379354 | -2.720 |
| chr13 | 59752324 | 59752535 | 1.05565 | 0.388401 | -2.718 |
| chr5 | 161951980 | 161952266 | 1.52336 | 0.560496 | -2.718 |
| chr17 | 44985799 | 44986043 | 1.52496 | 0.56109 | -2.718 |
| chr15 | 77927797 | 77927990 | 1.27431 | 0.468978 | -2.717 |
| chr2 | 21997836 | 21998282 | 1.11149 | 0.409239 | -2.716 |
| chr9 | 117690449 | 117690695 | 1.02514 | 0.377613 | -2.715 |
| chr5 | 63524074 | 63524321 | 1.01834 | 0.375143 | -2.715 |
| chr1 | 157624953 | 157625222 | 1.00549 | 0.370428 | -2.714 |
| chr9 | 136034778 | 136035047 | 1.091 | 0.401931 | -2.714 |
| chr11 | 61570493 | 61570731 | 0.8736 | 0.322009 | -2.713 |
| chr10 | 55202270 | 55202509 | 0.91446 | 0.33711 | -2.713 |
| chr8 | 14021988 | 14022388 | 0.94929 | 0.350044 | -2.712 |
| chr3 | 8747384 | 8747657 | 1.45635 | 0.537172 | -2.711 |
| chr11 | 57941708 | 57941893 | 0.919176 | 0.339146 | -2.710 |
| chr7 | 76834105 | 76834323 | 1.26202 | 0.465664 | -2.710 |
| chr5 | 4760895 | 4761171 | 1.11333 | 0.410914 | -2.709 |
| chr10 | 42986771 | 42987085 | 0.977299 | 0.360714 | -2.709 |
| chr17 | 72381538 | 72381731 | 1.50083 | 0.553982 | -2.709 |
| chr2 | 2419185 | 2419390 | 0.932635 | 0.344496 | -2.707 |
| chr7 | 103705548 | 103705733 | 1.30598 | 0.482515 | -2.707 |
| chr5 | 83129109 | 83129331 | 1.07577 | 0.39765 | -2.705 |
| chr5 | 164721762 | 164722080 | 1.20761 | 0.446411 | -2.705 |
| chr2 | 87038291 | 87038482 | 1.19061 | 0.440165 | -2.705 |
| chr9 | 135008948 | 135009154 | 1.32443 | 0.489698 | -2.705 |
| chr11 | 73944607 | 73944972 | 1.01086 | 0.37384 | -2.704 |
| chr10 | 118885482 | 118885712 | 0.985455 | 0.364453 | -2.704 |
| chr11 | 2461760 | 2461959 | 0.929488 | 0.343787 | -2.704 |
| chr5 | 163394471 | 163394684 | 1.161 | 0.429517 | -2.703 |

| | | | | | |
|---|---|---|---|---|---|
| chr8 | 107109010 | 107109221 | 1.09718 | 0.405957 | -2.703 |
| chr11 | 14171441 | 14171727 | 0.890733 | 0.32961 | -2.702 |
| chr18 | 20498414 | 20498616 | 0.731538 | 0.270707 | -2.702 |
| chr20 | 3002374 | 3002624 | 1.36182 | 0.504039 | -2.702 |
| chr5 | 2782783 | 2783165 | 1.16437 | 0.431 | -2.702 |
| chr2 | 237516569 | 237516757 | 1.48894 | 0.551297 | -2.701 |
| chr3 | 5640940 | 5641156 | 0.906146 | 0.33557 | -2.700 |
| chr14 | 30028203 | 30028390 | 1.24513 | 0.461345 | -2.699 |
| chr5 | 22757991 | 22758212 | 1.11539 | 0.413467 | -2.698 |
| chr10 | 117095810 | 117095997 | 0.968421 | 0.359033 | -2.697 |
| chr5 | 152527557 | 152527785 | 1.02626 | 0.380691 | -2.696 |
| chr2 | 135391868 | 135392051 | 0.748654 | 0.277791 | -2.695 |
| chr2 | 212064574 | 212064802 | 1.02468 | 0.380238 | -2.695 |
| chr2 | 88684554 | 88684825 | 1.06656 | 0.395811 | -2.695 |
| chr12 | 103702565 | 103702753 | 0.984562 | 0.3654 | -2.694 |
| chr3 | 1889051 | 1889378 | 1.15244 | 0.427733 | -2.694 |
| chr8 | 97942283 | 97942470 | 0.957072 | 0.355324 | -2.694 |
| chr5 | 94600168 | 94600423 | 1.00945 | 0.374872 | -2.693 |
| chr2 | 14889135 | 14889348 | 1.37629 | 0.511171 | -2.692 |
| chr2 | 166488874 | 166489081 | 1.04141 | 0.38686 | -2.692 |
| chr5 | 110276112 | 110276317 | 1.18423 | 0.440027 | -2.691 |
| chr2 | 9331271 | 9331531 | 1.14141 | 0.424191 | -2.691 |
| chr3 | 156212190 | 156212377 | 0.945761 | 0.351537 | -2.690 |
| chr2 | 2425364 | 2425621 | 1.03132 | 0.383421 | -2.690 |
| chr2 | 137717551 | 137717749 | 0.849755 | 0.316099 | -2.688 |
| chr19 | 27997327 | 27997593 | 1.26553 | 0.470923 | -2.687 |
| chr14 | 95662618 | 95662843 | 1.6323 | 0.607458 | -2.687 |
| chr20 | 13679928 | 13680115 | 1.13087 | 0.420897 | -2.687 |
| chr10 | 62397772 | 62398012 | 0.960003 | 0.357407 | -2.686 |
| chr8 | 72564156 | 72564421 | 2.49711 | 0.929965 | -2.685 |
| chr15 | 98130868 | 98131137 | 1.10173 | 0.410439 | -2.684 |
| chr2 | 195492858 | 195493042 | 0.769014 | 0.286523 | -2.684 |
| chr20 | 59411825 | 59412073 | 0.967297 | 0.360581 | -2.683 |
| chr10 | 23129633 | 23129887 | 1.18447 | 0.441645 | -2.682 |
| chr8 | 126851397 | 126851644 | 1.11374 | 0.415276 | -2.682 |
| chr17 | 11224611 | 11224794 | 1.03354 | 0.385429 | -2.682 |
| chr2 | 53644734 | 53645058 | 1.23551 | 0.460839 | -2.681 |
| chr2 | 22829248 | 22829481 | 0.828824 | 0.309216 | -2.680 |
| chr12 | 44625213 | 44625584 | 1.1547 | 0.430817 | -2.680 |
| chr5 | 13959230 | 13959477 | 1.3888 | 0.518187 | -2.680 |
| chr5 | 15706436 | 15706661 | 1.02239 | 0.381618 | -2.679 |
| chr9 | 31489224 | 31489423 | 0.969896 | 0.362196 | -2.678 |
| chr3 | 38756424 | 38756645 | 1.02387 | 0.3824 | -2.677 |
| chr2 | 83978237 | 83978500 | 1.08727 | 0.406192 | -2.677 |
| chr2 | 98748577 | 98748867 | 0.824641 | 0.30809 | -2.677 |
| chr5 | 22633702 | 22633903 | 1.18082 | 0.44116 | -2.677 |
| chr8 | 42957321 | 42957590 | 1.172 | 0.437962 | -2.676 |
| chr15 | 95089285 | 95089510 | 1.34544 | 0.502872 | -2.676 |
| chr9 | 16927484 | 16927736 | 1.10067 | 0.411789 | -2.673 |
| chr5 | 43167851 | 43168148 | 1.13354 | 0.424097 | -2.673 |
| chr14 | 79368440 | 79368622 | 1.55008 | 0.580013 | -2.672 |
| chr19 | 29784853 | 29785054 | 1.49115 | 0.557989 | -2.672 |
| chr8 | 113826004 | 113826242 | 1.2067 | 0.451566 | -2.672 |
| chr10 | 104232583 | 104232801 | 0.995723 | 0.372666 | -2.672 |
| chr8 | 137189266 | 137189518 | 1.08143 | 0.404832 | -2.671 |
| chr2 | 83123950 | 83124198 | 1.20104 | 0.449625 | -2.671 |
| chr2 | 32456885 | 32457181 | 1.02304 | 0.383146 | -2.670 |
| chr7 | 122749565 | 122749864 | 1.0747 | 0.402589 | -2.669 |
| chr16 | 65435413 | 65435615 | 1.07587 | 0.403096 | -2.669 |
| chr4 | 98967919 | 98968165 | 1.21505 | 0.455306 | -2.669 |
| chr5 | 12025013 | 12025277 | 0.963689 | 0.361151 | -2.668 |

| | | | | | |
|---|---|---|---|---|---|
| chr2 | 162625420 | 162625636 | 1.09498 | 0.410357 | -2.668 |
| chr5 | 25719281 | 25719497 | 0.969073 | 0.363338 | -2.667 |
| chr10 | 77740899 | 77741119 | 0.922126 | 0.345928 | -2.666 |
| chr2 | 147668295 | 147668480 | 0.971126 | 0.36436 | -2.665 |
| chr19 | 56459459 | 56459693 | 1.40597 | 0.527576 | -2.665 |
| chr8 | 131410068 | 131410270 | 1.29807 | 0.487091 | -2.665 |
| chr7 | 88497792 | 88498009 | 1.09965 | 0.412669 | -2.665 |
| chr16 | 7605690 | 7605918 | 0.82678 | 0.310285 | -2.665 |
| chr8 | 55591539 | 55591810 | 1.00244 | 0.376215 | -2.665 |
| chr8 | 137362139 | 137362370 | 1.06709 | 0.40052 | -2.664 |
| chr2 | 76946157 | 76946388 | 1.01677 | 0.381682 | -2.664 |
| chr12 | 89380874 | 89381056 | 1.35281 | 0.508083 | -2.663 |
| chr8 | 100898224 | 100898478 | 1.36188 | 0.511615 | -2.662 |
| chr14 | 82578194 | 82578379 | 0.973447 | 0.365932 | -2.660 |
| chrX | 6131198 | 6131419 | 0.867065 | 0.326157 | -2.658 |
| chr3 | 121145801 | 121145984 | 1.20521 | 0.453356 | -2.658 |
| chr2 | 191301762 | 191301985 | 1.15695 | 0.435298 | -2.658 |
| chr16 | 6744684 | 6744878 | 1.05341 | 0.396348 | -2.658 |
| chr2 | 16739769 | 16740038 | 1.17465 | 0.442175 | -2.657 |
| chr14 | 57655682 | 57655889 | 1.09776 | 0.413284 | -2.656 |
| chr17 | 27805923 | 27806156 | 1.16485 | 0.438558 | -2.656 |
| chr11 | 84464823 | 84465046 | 1.08782 | 0.40988 | -2.654 |
| chr10 | 68076027 | 68076308 | 1.06099 | 0.399793 | -2.654 |
| chr20 | 19054281 | 19054563 | 1.65176 | 0.62243 | -2.654 |
| chr1 | 201183200 | 201183407 | 1.20317 | 0.453558 | -2.653 |
| chr15 | 58548017 | 58548219 | 1.18836 | 0.448073 | -2.652 |
| chr2 | 217754660 | 217754894 | 1.15973 | 0.437278 | -2.652 |
| chr16 | 87156915 | 87157153 | 0.862621 | 0.325352 | -2.651 |
| chr8 | 117101629 | 117101885 | 0.904622 | 0.341523 | -2.649 |
| chr2 | 51594173 | 51594365 | 0.873891 | 0.33012 | -2.647 |
| chr2 | 174557510 | 174557715 | 1.1343 | 0.428522 | -2.647 |
| chr19 | 23000285 | 23000480 | 1.58294 | 0.598045 | -2.647 |
| chr5 | 28873521 | 28873868 | 1.16157 | 0.438998 | -2.646 |
| chr15 | 32140183 | 32140437 | 1.37391 | 0.519306 | -2.646 |
| chr1 | 239260158 | 239260360 | 1.42484 | 0.538895 | -2.644 |
| chr2 | 170235630 | 170235946 | 0.936181 | 0.35412 | -2.644 |
| chr3 | 82197720 | 82197959 | 1.17911 | 0.446018 | -2.644 |
| chr2 | 66365379 | 66365834 | 1.22037 | 0.46173 | -2.643 |
| chr2 | 70903719 | 70903918 | 0.94015 | 0.355808 | -2.642 |
| chr9 | 112005933 | 112006286 | 1.24325 | 0.47054 | -2.642 |
| chr8 | 73981732 | 73981928 | 1.24799 | 0.47269 | -2.640 |
| chr8 | 65502766 | 65502958 | 1.25281 | 0.47452 | -2.640 |
| chr5 | 86054991 | 86055183 | 1.12211 | 0.425174 | -2.639 |
| chr19 | 55138175 | 55138496 | 1.34422 | 0.509384 | -2.639 |
| chr8 | 145530653 | 145530887 | 1.10752 | 0.419698 | -2.639 |
| chr15 | 88622238 | 88622694 | 1.34247 | 0.508755 | -2.639 |
| chr8 | 124576243 | 124576502 | 0.824274 | 0.312413 | -2.638 |
| chr5 | 42514802 | 42514989 | 1.03979 | 0.394212 | -2.638 |
| chr2 | 11092271 | 11092536 | 0.891689 | 0.338115 | -2.637 |
| chr9 | 19599620 | 19599811 | 1.33389 | 0.50607 | -2.636 |
| chr3 | 85309609 | 85309850 | 0.991534 | 0.376479 | -2.634 |
| chr16 | 27809662 | 27809850 | 1.22837 | 0.466424 | -2.634 |
| chr2 | 77510791 | 77511018 | 1.09554 | 0.416045 | -2.633 |
| chr17 | 3073561 | 3073762 | 1.06026 | 0.402689 | -2.633 |
| chr5 | 72810492 | 72810695 | 0.948628 | 0.36037 | -2.632 |
| chr10 | 9657229 | 9657453 | 1.36658 | 0.519483 | -2.631 |
| chr1 | 152722904 | 152723129 | 1.01622 | 0.386312 | -2.631 |
| chr9 | 76821713 | 76821963 | 0.878799 | 0.334158 | -2.630 |
| chr8 | 111455868 | 111456080 | 0.897684 | 0.341374 | -2.630 |
| chr16 | 25282182 | 25282399 | 0.991754 | 0.377213 | -2.629 |
| chr12 | 123974547 | 123974755 | 1.05964 | 0.40304 | -2.629 |

| | | | | | |
|---|---|---|---|---|---|
| chr16 | 62693807 | 62693997 | 1.10121 | 0.418876 | -2.629 |
| chr15 | 98193590 | 98193826 | 1.05303 | 0.400614 | -2.629 |
| chr7 | 54231229 | 54231415 | 1.36433 | 0.519071 | -2.628 |
| chr9 | 28270193 | 28270623 | 1.21725 | 0.463133 | -2.628 |
| chr17 | 25414052 | 25414265 | 1.39285 | 0.529979 | -2.628 |
| chr8 | 32650591 | 32650861 | 0.866506 | 0.329738 | -2.628 |
| chr8 | 2431798 | 2432013 | 1.21239 | 0.461365 | -2.628 |
| chr5 | 140487493 | 140487696 | 0.915907 | 0.348584 | -2.628 |
| chr8 | 20225175 | 20225413 | 0.973753 | 0.370743 | -2.626 |
| chr8 | 84107671 | 84107900 | 1.26907 | 0.483228 | -2.626 |
| chr2 | 158205628 | 158205810 | 0.978101 | 0.37244 | -2.626 |
| chr8 | 73441635 | 73441835 | 1.14753 | 0.437023 | -2.626 |
| chr3 | 103507613 | 103507827 | 0.774683 | 0.295107 | -2.625 |
| chr3 | 186797061 | 186797247 | 0.988293 | 0.376485 | -2.625 |
| chr5 | 117554519 | 117554704 | 0.863714 | 0.329045 | -2.625 |
| chr3 | 88707348 | 88707583 | 1.11269 | 0.423917 | -2.625 |
| chr19 | 57572170 | 57572448 | 1.23393 | 0.470123 | -2.625 |
| chr3 | 37686887 | 37687071 | 1.07917 | 0.411188 | -2.625 |
| chr19 | 53653044 | 53653558 | 1.02884 | 0.39202 | -2.624 |
| chr2 | 123237921 | 123238154 | 1.07929 | 0.411322 | -2.624 |
| chr3 | 193279714 | 193280000 | 0.890045 | 0.339231 | -2.624 |
| chrX | 19086168 | 19086552 | 1.09235 | 0.416339 | -2.624 |
| chr2 | 83280809 | 83281173 | 1.16651 | 0.444659 | -2.623 |
| chr10 | 97256774 | 97256989 | 0.810029 | 0.308874 | -2.623 |
| chr7 | 50904390 | 50904577 | 1.1162 | 0.425715 | -2.622 |
| chr11 | 55721142 | 55721362 | 0.812046 | 0.309809 | -2.621 |
| chr14 | 74682229 | 74682474 | 1.3061 | 0.49836 | -2.621 |
| chr5 | 56347266 | 56347473 | 1.52406 | 0.581627 | -2.620 |
| chr8 | 4261096 | 4261339 | 0.958928 | 0.36602 | -2.620 |
| chr8 | 1468421 | 1468619 | 1.11884 | 0.427262 | -2.619 |
| chrX | 28439348 | 28439644 | 0.973404 | 0.371866 | -2.618 |
| chr16 | 63724004 | 63724195 | 1.27897 | 0.488634 | -2.617 |
| chr2 | 125929910 | 125930118 | 1.17545 | 0.44939 | -2.616 |
| chr11 | 29499637 | 29499825 | 1.0468 | 0.400325 | -2.615 |
| chr3 | 76891015 | 76891262 | 1.45294 | 0.555664 | -2.615 |
| chr2 | 155785368 | 155785594 | 1.0469 | 0.400492 | -2.614 |
| chr2 | 117531684 | 117531975 | 1.11573 | 0.426962 | -2.613 |
| chr9 | 8312967 | 8313195 | 1.03568 | 0.396579 | -2.612 |
| chr17 | 52425429 | 52425831 | 1.06518 | 0.408222 | -2.609 |
| chr16 | 13527524 | 13527725 | 0.83029 | 0.318311 | -2.608 |
| chr5 | 103963945 | 103964205 | 1.40829 | 0.540051 | -2.608 |
| chr1 | 216960012 | 216960222 | 0.962453 | 0.36912 | -2.607 |
| chr2 | 36303167 | 36303360 | 1.029 | 0.394741 | -2.607 |
| chr8 | 5653116 | 5653314 | 1.16515 | 0.447024 | -2.606 |
| chr14 | 78050906 | 78051405 | 1.14399 | 0.438936 | -2.606 |
| chr15 | 94275390 | 94275577 | 0.975042 | 0.37412 | -2.606 |
| chr5 | 38053084 | 38053301 | 1.19262 | 0.457652 | -2.606 |
| chr8 | 2680483 | 2680690 | 1.11806 | 0.429188 | -2.605 |
| chr10 | 129249299 | 129249495 | 0.855864 | 0.328812 | -2.603 |
| chr10 | 55889932 | 55890133 | 1.00611 | 0.386619 | -2.602 |
| chr5 | 116020723 | 116021084 | 0.903602 | 0.347398 | -2.601 |
| chr9 | 135618077 | 135618311 | 0.956363 | 0.36771 | -2.601 |
| chr2 | 195063454 | 195063642 | 0.819638 | 0.315166 | -2.601 |
| chr9 | 12135759 | 12135960 | 0.807075 | 0.310339 | -2.601 |
| chr2 | 16458549 | 16458799 | 1.15709 | 0.44507 | -2.600 |
| chr8 | 77068736 | 77068988 | 1.1495 | 0.442232 | -2.599 |
| chr10 | 104579541 | 104579756 | 1.00014 | 0.384801 | -2.599 |
| chr14 | 44155292 | 44155508 | 1.11157 | 0.427819 | -2.598 |
| chr2 | 5538473 | 5538748 | 1.07962 | 0.415551 | -2.598 |
| chr15 | 75507948 | 75508190 | 0.886272 | 0.341134 | -2.598 |
| chr7 | 149078105 | 149078301 | 0.921137 | 0.354571 | -2.598 |

| | | | | | |
|---|---|---|---|---|---|
| chr19 | 31091515 | 31091730 | 1.60003 | 0.6159 | -2.598 |
| chr2 | 76718537 | 76718805 | 1.08162 | 0.416499 | -2.597 |
| chr5 | 167967344 | 167967629 | 1.05877 | 0.407904 | -2.596 |
| chr2 | 62187900 | 62188123 | 1.27627 | 0.491814 | -2.595 |
| chr7 | 54598332 | 54598588 | 0.949082 | 0.365794 | -2.595 |
| chr2 | 51617137 | 51617392 | 0.994638 | 0.383447 | -2.594 |
| chr16 | 70764850 | 70765040 | 1.10176 | 0.424873 | -2.593 |
| chr2 | 2182935 | 2183171 | 1.09343 | 0.421688 | -2.593 |
| chr1 | 190300716 | 190300919 | 0.953701 | 0.367823 | -2.593 |
| chr3 | 147306955 | 147307154 | 1.15549 | 0.445666 | -2.593 |
| chr3 | 27108558 | 27108743 | 1.1832 | 0.456386 | -2.593 |
| chr16 | 5950695 | 5950934 | 0.827147 | 0.319184 | -2.591 |
| chr17 | 51033658 | 51033974 | 1.15942 | 0.447415 | -2.591 |
| chr8 | 79369672 | 79369976 | 1.12996 | 0.436058 | -2.591 |
| chr19 | 32392747 | 32392984 | 1.4182 | 0.547309 | -2.591 |
| chr11 | 19060281 | 19060471 | 1.11932 | 0.432012 | -2.591 |
| chr2 | 126943910 | 126944178 | 1.19003 | 0.459324 | -2.591 |
| chr7 | 110577327 | 110577544 | 1.24754 | 0.481613 | -2.590 |
| chr15 | 57671016 | 57671214 | 1.66391 | 0.642406 | -2.590 |
| chr2 | 229483855 | 229484048 | 0.783161 | 0.302396 | -2.590 |
| chr16 | 17288147 | 17288651 | 1.57184 | 0.607176 | -2.589 |
| chr2 | 84830334 | 84830742 | 0.88016 | 0.340211 | -2.587 |
| chr8 | 108821393 | 108821577 | 0.893074 | 0.345251 | -2.587 |
| chrX | 75127837 | 75128034 | 1.13518 | 0.438899 | -2.586 |
| chr15 | 66375255 | 66375467 | 1.55536 | 0.601382 | -2.586 |
| chr8 | 99025273 | 99025482 | 1.21894 | 0.471315 | -2.586 |
| chr5 | 172602167 | 172602452 | 0.989902 | 0.382841 | -2.586 |
| chr17 | 50067990 | 50068198 | 1.2044 | 0.465864 | -2.585 |
| chr7 | 44608772 | 44608971 | 0.944185 | 0.365231 | -2.585 |
| chr7 | 85130836 | 85131121 | 1.02073 | 0.395019 | -2.584 |
| chr11 | 21428671 | 21428940 | 0.766336 | 0.296687 | -2.583 |
| chr19 | 39745231 | 39745573 | 1.36157 | 0.527243 | -2.582 |
| chr5 | 164464468 | 164464776 | 1.29485 | 0.501503 | -2.582 |
| chr1 | 162910276 | 162910493 | 1.22095 | 0.472944 | -2.582 |
| chr11 | 80193457 | 80193851 | 1.07764 | 0.417553 | -2.581 |
| chr5 | 110614561 | 110614867 | 2.54712 | 0.987089 | -2.580 |
| chr5 | 52136239 | 52136482 | 1.13776 | 0.440933 | -2.580 |
| chr10 | 113427511 | 113427771 | 0.901291 | 0.349356 | -2.580 |
| chr17 | 63515590 | 63515794 | 1.14357 | 0.44327 | -2.580 |
| chr2 | 26795441 | 26795661 | 1.1208 | 0.434465 | -2.580 |
| chr12 | 33164216 | 33164410 | 1.27081 | 0.49262 | -2.580 |
| chr2 | 70290384 | 70290646 | 1.16289 | 0.450839 | -2.579 |
| chr5 | 121952451 | 121952667 | 0.904311 | 0.350664 | -2.579 |
| chr16 | 23058751 | 23058940 | 1.21673 | 0.471878 | -2.578 |
| chr3 | 13432924 | 13433139 | 1.9905 | 0.77203 | -2.578 |
| chr3 | 89850427 | 89850635 | 1.04143 | 0.403968 | -2.578 |
| chr20 | 45148496 | 45148844 | 1.04689 | 0.406149 | -2.578 |
| chr17 | 12740808 | 12741005 | 1.36754 | 0.530655 | -2.577 |
| chr1 | 163352386 | 163352582 | 1.10162 | 0.427845 | -2.575 |
| chr8 | 73781338 | 73781543 | 1.10497 | 0.429147 | -2.575 |
| chr7 | 106387785 | 106388213 | 1.24107 | 0.482087 | -2.574 |
| chr3 | 95621512 | 95621710 | 1.1331 | 0.440208 | -2.574 |
| chr5 | 155125907 | 155126323 | 1.18625 | 0.460857 | -2.574 |
| chr8 | 52558781 | 52559079 | 1.60976 | 0.625455 | -2.574 |
| chr2 | 83069635 | 83069861 | 1.0131 | 0.393714 | -2.573 |
| chr10 | 57220665 | 57220867 | 1.29754 | 0.50428 | -2.573 |
| chr2 | 45391454 | 45391657 | 1.44668 | 0.562438 | -2.572 |
| chr2 | 182305171 | 182305490 | 1.22227 | 0.475195 | -2.572 |
| chr11 | 79982090 | 79982536 | 1.08103 | 0.420308 | -2.572 |
| chr5 | 142638913 | 142639112 | 1.00211 | 0.389678 | -2.572 |
| chr2 | 40233817 | 40234020 | 1.30389 | 0.507084 | -2.571 |

| | | | | | |
|---|---|---|---|---|---|
| chr17 | 13765808 | 13766035 | 1.28247 | 0.498919 | -2.570 |
| chr10 | 55167629 | 55167812 | 1.03713 | 0.403546 | -2.570 |
| chr8 | 68910437 | 68910660 | 1.27851 | 0.497551 | -2.570 |
| chr15 | 34648272 | 34648513 | 1.23047 | 0.478859 | -2.570 |
| chr2 | 146993535 | 146993842 | 1.0021 | 0.389994 | -2.570 |
| chr2 | 22965827 | 22966019 | 1.17422 | 0.457086 | -2.569 |
| chr2 | 179171175 | 179171434 | 1.44055 | 0.560821 | -2.569 |
| chr13 | 47158436 | 47158650 | 1.13642 | 0.442552 | -2.568 |
| chr11 | 74628062 | 74628253 | 1.19965 | 0.467261 | -2.567 |
| chr16 | 5092357 | 5092564 | 1.26176 | 0.491591 | -2.567 |
| chr10 | 113207740 | 113207935 | 1.05981 | 0.413082 | -2.566 |
| chr12 | 4721834 | 4722042 | 1.02306 | 0.398819 | -2.565 |
| chrX | 30833546 | 30833755 | 0.924368 | 0.360378 | -2.565 |
| chr2 | 187085175 | 187085457 | 1.07408 | 0.418769 | -2.565 |
| chr8 | 69362680 | 69362913 | 0.865696 | 0.337523 | -2.565 |
| chr16 | 52388204 | 52388434 | 1.38106 | 0.538461 | -2.565 |
| chr17 | 32926170 | 32926373 | 1.01933 | 0.397503 | -2.564 |
| chr2 | 137373833 | 137374082 | 0.892217 | 0.347937 | -2.564 |
| chr20 | 5312233 | 5312425 | 1.16694 | 0.455084 | -2.564 |
| chr11 | 61637331 | 61637672 | 0.987414 | 0.385114 | -2.564 |
| chr3 | 96098087 | 96098284 | 1.06608 | 0.415802 | -2.564 |
| chr17 | 68747252 | 68747469 | 1.20664 | 0.47076 | -2.563 |
| chr8 | 50164177 | 50164436 | 1.38932 | 0.542232 | -2.562 |
| chr5 | 109219815 | 109219999 | 1.06172 | 0.414434 | -2.562 |
| chr1 | 245582934 | 245583163 | 0.949131 | 0.370498 | -2.562 |
| chr2 | 117855872 | 117856068 | 1.15655 | 0.451584 | -2.561 |
| chr2 | 125567794 | 125568025 | 1.19845 | 0.467992 | -2.561 |
| chr17 | 52649761 | 52649960 | 1.21486 | 0.474646 | -2.560 |
| chr2 | 4142506 | 4142756 | 1.01835 | 0.397882 | -2.559 |
| chr11 | 73070442 | 73070692 | 0.985139 | 0.384917 | -2.559 |
| chr2 | 52655200 | 52655577 | 1.20853 | 0.472527 | -2.558 |
| chr3 | 7848376 | 7848635 | 0.930759 | 0.363922 | -2.558 |
| chr5 | 117858456 | 117858719 | 1.10511 | 0.432103 | -2.558 |
| chr5 | 86181778 | 86182117 | 1.45433 | 0.568652 | -2.558 |
| chr3 | 43582152 | 43582406 | 0.884213 | 0.345759 | -2.557 |
| chr11 | 7210214 | 7210419 | 0.94573 | 0.36987 | -2.557 |
| chr2 | 49699785 | 49700017 | 0.94702 | 0.370537 | -2.556 |
| chr12 | 32027714 | 32027932 | 0.818265 | 0.320209 | -2.555 |
| chr2 | 2649228 | 2649492 | 0.993698 | 0.388906 | -2.555 |
| chr12 | 44376899 | 44377093 | 1.09188 | 0.427484 | -2.554 |
| chr5 | 87897059 | 87897281 | 0.790067 | 0.309323 | -2.554 |
| chr2 | 140615775 | 140616074 | 1.20221 | 0.470778 | -2.554 |
| chr21 | 19895012 | 19895202 | 1.11646 | 0.437232 | -2.553 |
| chr12 | 83716540 | 83716741 | 1.09925 | 0.430496 | -2.553 |
| chr11 | 23464219 | 23464461 | 1.37639 | 0.53907 | -2.553 |
| chr2 | 85941158 | 85941446 | 1.3715 | 0.537157 | -2.553 |
| chr1 | 170737210 | 170737426 | 1.13169 | 0.443281 | -2.553 |
| chr2 | 184301255 | 184301516 | 0.97688 | 0.382667 | -2.553 |
| chr14 | 24714834 | 24715030 | 1.3652 | 0.534916 | -2.552 |
| chr19 | 37802967 | 37803235 | 0.996896 | 0.390622 | -2.552 |
| chr2 | 214496538 | 214496768 | 0.853784 | 0.334577 | -2.552 |
| chr17 | 53257427 | 53257619 | 1.13051 | 0.443124 | -2.551 |
| chr5 | 21737316 | 21737507 | 1.20649 | 0.472974 | -2.551 |
| chr5 | 85096518 | 85096700 | 1.23213 | 0.483191 | -2.550 |
| chr2 | 126850116 | 126850387 | 1.11694 | 0.438025 | -2.550 |
| chr10 | 66704569 | 66704842 | 0.858927 | 0.336969 | -2.549 |
| chr2 | 82038534 | 82039176 | 1.08146 | 0.424313 | -2.549 |
| chr10 | 98572859 | 98573132 | 1.07371 | 0.421508 | -2.547 |
| chr8 | 108879996 | 108880238 | 0.828189 | 0.325138 | -2.547 |
| chr2 | 57051575 | 57052107 | 1.22048 | 0.479187 | -2.547 |
| chr8 | 136330893 | 136331126 | 1.02885 | 0.403995 | -2.547 |

| | | | | | |
|---|---|---|---|---|---|
| chr17 | 52640640 | 52640831 | 1.08521 | 0.426208 | -2.546 |
| chr9 | 34954378 | 34954578 | 1.32733 | 0.521366 | -2.546 |
| chr13 | 58324646 | 58324857 | 0.961687 | 0.377807 | -2.545 |
| chr5 | 110201458 | 110201673 | 1.45543 | 0.571853 | -2.545 |
| chr3 | 74685541 | 74685743 | 1.1915 | 0.468249 | -2.545 |
| chr11 | 100820329 | 100820513 | 1.03224 | 0.405712 | -2.544 |
| chr10 | 84934679 | 84934871 | 0.806001 | 0.316895 | -2.543 |
| chr16 | 62127225 | 62127492 | 1.03806 | 0.408137 | -2.543 |
| chr19 | 51929141 | 51929340 | 0.9635 | 0.378882 | -2.543 |
| chr19 | 36920282 | 36920520 | 1.21251 | 0.476883 | -2.543 |
| chr3 | 175840676 | 175840916 | 1.33815 | 0.526312 | -2.543 |
| chr3 | 20875962 | 20876320 | 0.884813 | 0.348044 | -2.542 |
| chr8 | 36341884 | 36342074 | 1.10364 | 0.434215 | -2.542 |
| chr1 | 215883569 | 215883980 | 1.49059 | 0.586616 | -2.541 |
| chr8 | 139930781 | 139931020 | 1.22036 | 0.480308 | -2.541 |
| chr7 | 53006307 | 53006499 | 1.01547 | 0.399764 | -2.540 |
| chr7 | 15180720 | 15180915 | 0.938535 | 0.369483 | -2.540 |
| chr14 | 48485144 | 48485331 | 0.883532 | 0.347875 | -2.540 |
| chr20 | 17428823 | 17429108 | 1.06972 | 0.421197 | -2.540 |
| chr12 | 41451702 | 41451981 | 1.19224 | 0.469582 | -2.539 |
| chr3 | 115833049 | 115833237 | 1.13506 | 0.447263 | -2.538 |
| chr8 | 55648991 | 55649415 | 1.30639 | 0.514962 | -2.537 |
| chr1 | 165373234 | 165373496 | 2.39544 | 0.944282 | -2.537 |
| chr15 | 100965283 | 100965511 | 1.17953 | 0.465245 | -2.535 |
| chr5 | 136680502 | 136680770 | 2.69736 | 1.06405 | -2.535 |
| chr14 | 63384579 | 63384785 | 1.42668 | 0.562913 | -2.534 |
| chr10 | 50116248 | 50116436 | 1.19949 | 0.473289 | -2.534 |
| chr14 | 92058341 | 92058576 | 0.728796 | 0.287577 | -2.534 |
| chr5 | 28306723 | 28306998 | 1.12978 | 0.445901 | -2.534 |
| chr5 | 12256108 | 12256293 | 1.23447 | 0.487238 | -2.534 |
| chr10 | 132147853 | 132148038 | 0.973114 | 0.384086 | -2.534 |
| chr2 | 115616652 | 115616850 | 0.919709 | 0.363014 | -2.534 |
| chr8 | 116355584 | 116355847 | 1.28657 | 0.507978 | -2.533 |
| chr2 | 162486178 | 162486461 | 0.944692 | 0.373014 | -2.533 |
| chr11 | 36888821 | 36889013 | 1.01456 | 0.400609 | -2.533 |
| chr7 | 31001151 | 31001355 | 1.21682 | 0.4805 | -2.532 |
| chr11 | 48244111 | 48244331 | 1.47346 | 0.581891 | -2.532 |
| chr9 | 22607109 | 22607488 | 1.00036 | 0.39513 | -2.532 |
| chr11 | 81122325 | 81122579 | 0.934985 | 0.369353 | -2.531 |
| chr2 | 4581266 | 4581480 | 1.47037 | 0.580991 | -2.531 |
| chr1 | 244790590 | 244790851 | 1.16166 | 0.45905 | -2.531 |
| chr5 | 88949991 | 88950229 | 1.2373 | 0.489179 | -2.529 |
| chr7 | 8553214 | 8553450 | 1.25062 | 0.494556 | -2.529 |
| chr12 | 18759900 | 18760111 | 1.08085 | 0.42744 | -2.529 |
| chr5 | 165061509 | 165061761 | 0.980619 | 0.387877 | -2.528 |
| chr2 | 16114700 | 16114955 | 1.17739 | 0.465853 | -2.527 |
| chr2 | 163829 | 164175 | 1.02742 | 0.406522 | -2.527 |
| chr8 | 76611039 | 76611236 | 1.05868 | 0.418897 | -2.527 |
| chr2 | 212797857 | 212798237 | 0.902532 | 0.357124 | -2.527 |
| chr2 | 49796783 | 49797073 | 1.20387 | 0.476434 | -2.527 |
| chr5 | 23896422 | 23896616 | 0.894302 | 0.353959 | -2.527 |
| chr12 | 108617555 | 108617772 | 1.06212 | 0.420412 | -2.526 |
| chr2 | 125441575 | 125441822 | 0.820622 | 0.324887 | -2.526 |
| chr2 | 82931139 | 82931374 | 0.995947 | 0.394366 | -2.525 |
| chr8 | 95130130 | 95130327 | 1.0726 | 0.42477 | -2.525 |
| chr3 | 17832170 | 17832373 | 1.03929 | 0.411684 | -2.524 |
| chr3 | 2812945 | 2813209 | 1.14821 | 0.454853 | -2.524 |
| chr3 | 96779728 | 96780327 | 0.893468 | 0.353956 | -2.524 |
| chr11 | 21290082 | 21290479 | 1.13907 | 0.451318 | -2.524 |
| chr5 | 44575152 | 44575334 | 0.993368 | 0.39384 | -2.522 |
| chr3 | 83271879 | 83272149 | 1.08049 | 0.42844 | -2.522 |

| | | | | | |
|---|---|---|---|---|---|
| chr5 | 142635255 | 142635461 | 1.28012 | 0.507643 | -2.522 |
| chr8 | 21104333 | 21104553 | 1.97872 | 0.784766 | -2.521 |
| chr11 | 39685529 | 39685752 | 1.26108 | 0.500291 | -2.521 |
| chr7 | 135610576 | 135610758 | 1.19895 | 0.47566 | -2.521 |
| chr10 | 53182427 | 53182699 | 2.184 | 0.86671 | -2.520 |
| chr11 | 25682612 | 25682828 | 1.04764 | 0.415836 | -2.519 |
| chr7 | 118968136 | 118968338 | 1.0631 | 0.421994 | -2.519 |
| chr9 | 124960260 | 124960572 | 0.795839 | 0.316081 | -2.518 |
| chr2 | 79415379 | 79415609 | 1.42982 | 0.568022 | -2.517 |
| chr2 | 202955628 | 202955830 | 0.930454 | 0.369658 | -2.517 |
| chr3 | 107897882 | 107898067 | 1.33982 | 0.532451 | -2.516 |
| chr7 | 125713459 | 125713674 | 1.11118 | 0.44171 | -2.516 |
| chr14 | 51676282 | 51676465 | 1.10899 | 0.4409 | -2.515 |
| chr12 | 127382390 | 127382638 | 0.775827 | 0.30856 | -2.514 |
| chr2 | 1343819 | 1344033 | 0.921536 | 0.36665 | -2.513 |
| chr2 | 49405213 | 49405655 | 1.05382 | 0.419302 | -2.513 |
| chr2 | 4860168 | 4860391 | 1.11486 | 0.443609 | -2.513 |
| chr12 | 83745356 | 83745596 | 0.976996 | 0.38879 | -2.513 |
| chr9 | 106099816 | 106100044 | 0.813423 | 0.323724 | -2.513 |
| chr11 | 57912348 | 57912643 | 1.56046 | 0.621126 | -2.512 |
| chr2 | 41088282 | 41088506 | 1.30748 | 0.520458 | -2.512 |
| chr12 | 14587713 | 14587993 | 1.12149 | 0.446449 | -2.512 |
| chr2 | 2579792 | 2580103 | 1.23111 | 0.490112 | -2.512 |
| chr2 | 22555546 | 22555758 | 1.10646 | 0.440506 | -2.512 |
| chr2 | 215362503 | 215362932 | 0.885067 | 0.352367 | -2.512 |
| chr7 | 113296627 | 113296875 | 1.31518 | 0.523664 | -2.511 |
| chr8 | 6551396 | 6551587 | 1.16914 | 0.465564 | -2.511 |
| chr12 | 81769023 | 81769237 | 1.08908 | 0.433768 | -2.511 |
| chr2 | 59690731 | 59691003 | 0.907955 | 0.361632 | -2.511 |
| chr20 | 2302285 | 2302553 | 1.2839 | 0.511453 | -2.510 |
| chr5 | 12605459 | 12605643 | 1.05188 | 0.419102 | -2.510 |
| chr16 | 63092658 | 63092945 | 0.857188 | 0.341547 | -2.510 |
| chr10 | 84091610 | 84091825 | 1.03231 | 0.411374 | -2.509 |
| chr14 | 62493962 | 62494154 | 1.1128 | 0.443461 | -2.509 |
| chr3 | 187572908 | 187573096 | 0.910307 | 0.362862 | -2.509 |
| chr1 | 213743287 | 213743482 | 0.972023 | 0.387476 | -2.509 |
| chr2 | 52553976 | 52554178 | 1.15854 | 0.461853 | -2.508 |
| chr7 | 30830300 | 30830584 | 1.15587 | 0.460942 | -2.508 |
| chr8 | 86289717 | 86289989 | 0.964503 | 0.384716 | -2.507 |
| chr13 | 92998287 | 92998552 | 1.15498 | 0.46071 | -2.507 |
| chr3 | 80967673 | 80967856 | 1.08085 | 0.43114 | -2.507 |
| chr2 | 52979828 | 52980025 | 1.03233 | 0.4119 | -2.506 |
| chr2 | 2309420 | 2309925 | 1.21481 | 0.484856 | -2.506 |
| chr1 | 194002851 | 194003066 | 0.94958 | 0.378999 | -2.505 |
| chr5 | 12634703 | 12634983 | 1.06852 | 0.426747 | -2.504 |
| chr8 | 51881797 | 51882010 | 1.57546 | 0.629261 | -2.504 |
| chr5 | 144455550 | 144455763 | 0.847307 | 0.338572 | -2.503 |
| chr8 | 28411101 | 28411419 | 1.30082 | 0.519827 | -2.502 |
| chr11 | 70364594 | 70364779 | 1.04012 | 0.415731 | -2.502 |
| chr16 | 84373012 | 84373251 | 1.05246 | 0.420788 | -2.501 |
| chr5 | 45753841 | 45754278 | 0.95181 | 0.380779 | -2.500 |
| chr2 | 35609666 | 35609862 | 1.55648 | 0.622705 | -2.500 |
| chr5 | 117918030 | 117918286 | 1.00488 | 0.402037 | -2.499 |
| chr12 | 18338478 | 18338670 | 1.07846 | 0.431478 | -2.499 |
| chr8 | 125771698 | 125771919 | 0.991582 | 0.396776 | -2.499 |
| chr9 | 106598934 | 106599194 | 1.2016 | 0.480973 | -2.498 |
| chr8 | 135948097 | 135948299 | 1.06522 | 0.426396 | -2.498 |
| chr5 | 145045843 | 145046133 | 1.4358 | 0.574818 | -2.498 |
| chr3 | 6596393 | 6596595 | 1.06857 | 0.427805 | -2.498 |
| chr7 | 4854272 | 4854499 | 1.01544 | 0.406656 | -2.497 |
| chr2 | 139945325 | 139945514 | 0.906498 | 0.363069 | -2.497 |

| | | | | | |
|---|---|---|---|---|---|
| chr8 | 9905408 | 9905760 | 0.947152 | 0.379446 | -2.496 |
| chr5 | 27181765 | 27181956 | 1.35334 | 0.542342 | -2.495 |
| chr20 | 57333704 | 57333886 | 0.973154 | 0.390005 | -2.495 |
| chr19 | 45334327 | 45334512 | 1.13283 | 0.454009 | -2.495 |
| chr2 | 119111502 | 119111697 | 0.777117 | 0.311468 | -2.495 |
| chr5 | 28102114 | 28102418 | 1.09183 | 0.437675 | -2.495 |
| chr5 | 99236190 | 99236495 | 1.1445 | 0.458845 | -2.494 |
| chr10 | 132797715 | 132797925 | 0.91513 | 0.366898 | -2.494 |
| chr3 | 73557628 | 73557834 | 3.05211 | 1.22379 | -2.494 |
| chr3 | 115312212 | 115312531 | 1.12941 | 0.452867 | -2.494 |
| chr19 | 32707613 | 32707876 | 1.20094 | 0.481575 | -2.494 |
| chr8 | 66255588 | 66255787 | 1.29889 | 0.520924 | -2.493 |
| chr19 | 35137146 | 35137400 | 1.00074 | 0.401379 | -2.493 |
| chr2 | 13849593 | 13849839 | 1.25812 | 0.504638 | -2.493 |
| chr14 | 68257069 | 68257472 | 1.10177 | 0.442082 | -2.492 |
| chr2 | 141190698 | 141190888 | 1.08211 | 0.434258 | -2.492 |
| chr11 | 15311149 | 15311367 | 1.14884 | 0.461055 | -2.492 |
| chr18 | 58989537 | 58989796 | 2.10517 | 0.844877 | -2.492 |
| chr20 | 53221970 | 53222187 | 1.51207 | 0.606886 | -2.492 |
| chr5 | 178380634 | 178380961 | 1.15452 | 0.463413 | -2.491 |
| chr8 | 69312707 | 69312899 | 1.00458 | 0.403299 | -2.491 |
| chr2 | 198476912 | 198477162 | 1.00462 | 0.403339 | -2.491 |
| chr7 | 29175480 | 29175670 | 1.48323 | 0.595598 | -2.490 |
| chr7 | 88972883 | 88973073 | 1.06439 | 0.42758 | -2.489 |
| chr17 | 42268119 | 42268315 | 1.19618 | 0.480563 | -2.489 |
| chr2 | 5370908 | 5371096 | 1.12948 | 0.453798 | -2.489 |
| chr3 | 146891986 | 146892224 | 0.977275 | 0.392688 | -2.489 |
| chr2 | 34802138 | 34802320 | 1.02765 | 0.413073 | -2.488 |
| chr3 | 108860399 | 108860593 | 0.895635 | 0.360045 | -2.488 |
| chr8 | 14947929 | 14948223 | 1.49564 | 0.601255 | -2.488 |
| chr12 | 124557603 | 124557890 | 1.08905 | 0.43785 | -2.487 |
| chr20 | 40454591 | 40454811 | 1.04231 | 0.41909 | -2.487 |
| chr13 | 103033525 | 103033769 | 0.832165 | 0.33463 | -2.487 |
| chr2 | 50783009 | 50783279 | 1.017 | 0.40899 | -2.487 |
| chr17 | 12856187 | 12856504 | 1.00027 | 0.40227 | -2.487 |
| chr5 | 84607464 | 84607696 | 1.36562 | 0.549303 | -2.486 |
| chr7 | 41905354 | 41905606 | 1.00628 | 0.404799 | -2.486 |
| chr17 | 54396899 | 54397142 | 2.08467 | 0.838658 | -2.486 |
| chr2 | 127949607 | 127949791 | 0.858534 | 0.345388 | -2.486 |
| chr16 | 65549117 | 65549299 | 0.905635 | 0.364378 | -2.485 |
| chr4 | 65854115 | 65854329 | 1.16546 | 0.468917 | -2.485 |
| chr1 | 191779338 | 191779662 | 1.04514 | 0.420528 | -2.485 |
| chr5 | 178302149 | 178302515 | 1.05746 | 0.425512 | -2.485 |
| chr2 | 226394191 | 226394402 | 1.17705 | 0.473754 | -2.485 |
| chr1 | 33153750 | 33153936 | 0.911885 | 0.367133 | -2.484 |
| chr11 | 8693333 | 8693715 | 1.25241 | 0.504277 | -2.484 |
| chr2 | 163548580 | 163548767 | 1.1044 | 0.444815 | -2.483 |
| chr5 | 37787377 | 37787583 | 1.30447 | 0.525396 | -2.483 |
| chr15 | 98721258 | 98721502 | 2.28395 | 0.920134 | -2.482 |
| chr9 | 106411165 | 106411361 | 1.12283 | 0.452484 | -2.481 |
| chr1 | 167173671 | 167173853 | 0.85169 | 0.343334 | -2.481 |
| chr10 | 66424365 | 66424684 | 1.20269 | 0.484881 | -2.480 |
| chr17 | 30201123 | 30201312 | 0.886017 | 0.357303 | -2.480 |
| chr2 | 576805 | 577046 | 1.06398 | 0.429132 | -2.479 |
| chr1 | 213699645 | 213699828 | 1.26966 | 0.512228 | -2.479 |
| chr5 | 138414204 | 138414391 | 1.19461 | 0.482004 | -2.478 |
| chr2 | 65992871 | 65993053 | 0.999316 | 0.40333 | -2.478 |
| chr3 | 95510029 | 95510233 | 1.00606 | 0.406118 | -2.477 |
| chr5 | 11631678 | 11631877 | 0.904595 | 0.365213 | -2.477 |
| chr5 | 110018502 | 110018718 | 1.37155 | 0.553749 | -2.477 |
| chr20 | 6436262 | 6436458 | 0.985089 | 0.397779 | -2.476 |

| | | | | | |
|---|---|---|---|---|---|
| chr10 | 130961420 | 130961654 | 1.08473 | 0.438042 | -2.476 |
| chr11 | 26440100 | 26440284 | 1.06469 | 0.430173 | -2.475 |
| chr16 | 61271847 | 61272166 | 1.31318 | 0.53058 | -2.475 |
| chr7 | 90207304 | 90207534 | 0.871407 | 0.352095 | -2.475 |
| chr11 | 42577259 | 42577484 | 1.48472 | 0.599989 | -2.475 |
| chr2 | 68762889 | 68763199 | 1.16389 | 0.470373 | -2.474 |
| chr17 | 7640494 | 7640769 | 0.927049 | 0.374664 | -2.474 |
| chr5 | 128466962 | 128467290 | 0.997488 | 0.403137 | -2.474 |
| chr2 | 20974419 | 20974611 | 1.25931 | 0.50899 | -2.474 |
| chr17 | 8403041 | 8403355 | 1.0164 | 0.41086 | -2.474 |
| chr2 | 84177538 | 84177722 | 0.963846 | 0.3897 | -2.473 |
| chr1 | 194593960 | 194594195 | 1.01952 | 0.412221 | -2.473 |
| chr5 | 9781423 | 9781687 | 1.10865 | 0.448456 | -2.472 |
| chr3 | 104723876 | 104724072 | 1.08858 | 0.440356 | -2.472 |
| chr8 | 90509185 | 90509375 | 1.05278 | 0.425878 | -2.472 |
| chr3 | 103131981 | 103132209 | 1.26094 | 0.510091 | -2.472 |
| chr8 | 82358800 | 82358999 | 0.947942 | 0.383627 | -2.471 |
| chr4 | 165350887 | 165351082 | 0.961547 | 0.389348 | -2.470 |
| chr15 | 91942812 | 91943095 | 1.62594 | 0.658524 | -2.469 |
| chr2 | 135290134 | 135290341 | 0.759956 | 0.307803 | -2.469 |
| chr14 | 40929395 | 40929631 | 1.23995 | 0.502362 | -2.468 |
| chr7 | 121654721 | 121655119 | 1.09082 | 0.442117 | -2.467 |
| chr12 | 81082168 | 81082358 | 1.22078 | 0.494793 | -2.467 |
| chr9 | 112734625 | 112734937 | 1.18446 | 0.480169 | -2.467 |
| chr5 | 21402113 | 21402349 | 1.3137 | 0.532579 | -2.467 |
| chr2 | 35010223 | 35010414 | 1.31171 | 0.531874 | -2.466 |
| chr8 | 85184544 | 85184781 | 1.17664 | 0.477126 | -2.466 |
| chr2 | 153889515 | 153889750 | 0.727265 | 0.294925 | -2.466 |
| chr11 | 87101894 | 87102083 | 0.774879 | 0.314243 | -2.466 |
| chr10 | 64583927 | 64584390 | 0.945594 | 0.383479 | -2.466 |
| chr2 | 548782 | 549004 | 1.10899 | 0.449759 | -2.466 |
| chr3 | 94090115 | 94090438 | 1.16429 | 0.472193 | -2.466 |
| chr9 | 124282663 | 124282950 | 1.52355 | 0.617983 | -2.465 |
| chr17 | 18020494 | 18020741 | 1.10739 | 0.449203 | -2.465 |
| chr19 | 54649177 | 54649380 | 1.62601 | 0.659608 | -2.465 |
| chr1 | 181557104 | 181557308 | 1.44389 | 0.585767 | -2.465 |
| chr20 | 20446495 | 20446686 | 1.25146 | 0.507702 | -2.465 |
| chr19 | 24022122 | 24022407 | 1.13254 | 0.459482 | -2.465 |
| chr10 | 71537240 | 71537434 | 1.08452 | 0.440008 | -2.465 |
| chr10 | 80753577 | 80753810 | 1.01543 | 0.412041 | -2.464 |
| chr15 | 87459146 | 87459351 | 1.83482 | 0.744582 | -2.464 |
| chr20 | 50516254 | 50516546 | 0.96843 | 0.393018 | -2.464 |
| chr9 | 31220490 | 31221144 | 1.01749 | 0.413093 | -2.463 |
| chr2 | 17026076 | 17026261 | 1.20223 | 0.488187 | -2.463 |
| chr7 | 14678361 | 14678569 | 0.977174 | 0.396809 | -2.463 |
| chr8 | 137230938 | 137231154 | 1.43234 | 0.581887 | -2.462 |
| chr5 | 42249275 | 42249491 | 1.20074 | 0.487826 | -2.461 |
| chr9 | 12149050 | 12149330 | 1.27306 | 0.517209 | -2.461 |
| chr2 | 81348510 | 81348742 | 1.06179 | 0.431414 | -2.461 |
| chr2 | 37419029 | 37419227 | 1.27623 | 0.518586 | -2.461 |
| chr8 | 5835184 | 5835416 | 1.12569 | 0.45745 | -2.461 |
| chr19 | 31391920 | 31392195 | 1.34863 | 0.548077 | -2.461 |
| chr19 | 23443583 | 23443888 | 0.979123 | 0.398037 | -2.460 |
| chr2 | 128004537 | 128004735 | 1.10762 | 0.450342 | -2.460 |
| chr2 | 4660436 | 4660782 | 1.02092 | 0.415149 | -2.459 |
| chr16 | 4729677 | 4729969 | 0.968154 | 0.393898 | -2.458 |
| chr2 | 3311739 | 3311930 | 1.15713 | 0.4708 | -2.458 |
| chr10 | 129953191 | 129953376 | 0.736464 | 0.299665 | -2.458 |
| chr4 | 108472220 | 108472511 | 0.919858 | 0.374639 | -2.455 |
| chr7 | 122174973 | 122175241 | 1.15059 | 0.468654 | -2.455 |
| chr1 | 155562402 | 155562641 | 0.913972 | 0.372377 | -2.454 |

| | | | | | |
|---|---|---|---|---|---|
| chr9 | 76553665 | 76553930 | 1.01994 | 0.415557 | -2.454 |
| chr8 | 60331296 | 60331491 | 1.51395 | 0.616861 | -2.454 |
| chr8 | 136832614 | 136832881 | 1.04911 | 0.427462 | -2.454 |
| chr5 | 172472308 | 172472537 | 1.07631 | 0.438585 | -2.454 |
| chr5 | 83969231 | 83969667 | 1.26129 | 0.514181 | -2.453 |
| chr10 | 68217593 | 68217831 | 1.13846 | 0.464143 | -2.453 |
| chr2 | 186414731 | 186414960 | 1.03852 | 0.423529 | -2.452 |
| chr10 | 106190835 | 106191079 | 0.783871 | 0.319684 | -2.452 |
| chr2 | 13275855 | 13276292 | 1.14037 | 0.465171 | -2.452 |
| chr2 | 218752780 | 218753096 | 1.03161 | 0.420895 | -2.451 |
| chr5 | 141243234 | 141243500 | 0.997447 | 0.406957 | -2.451 |
| chr12 | 127962009 | 127962394 | 1.16883 | 0.476967 | -2.451 |
| chr2 | 76787116 | 76787340 | 1.19021 | 0.485817 | -2.450 |
| chr16 | 62208228 | 62208437 | 1.29219 | 0.527475 | -2.450 |
| chr8 | 93400617 | 93400882 | 1.09057 | 0.44518 | -2.450 |
| chr11 | 69705240 | 69705480 | 1.14965 | 0.469347 | -2.449 |
| chr10 | 95708109 | 95708305 | 1.04144 | 0.425182 | -2.449 |
| chr1 | 238386633 | 238387099 | 1.30164 | 0.531487 | -2.449 |
| chr3 | 159119011 | 159119362 | 0.878324 | 0.35864 | -2.449 |
| chr8 | 25697296 | 25697503 | 0.78748 | 0.321676 | -2.448 |
| chr17 | 12645681 | 12645875 | 2.16957 | 0.88633 | -2.448 |
| chr2 | 78754283 | 78754499 | 1.29597 | 0.529459 | -2.448 |
| chr15 | 61163184 | 61163402 | 1.25984 | 0.514765 | -2.447 |
| chr9 | 24956233 | 24956423 | 0.973055 | 0.397609 | -2.447 |
| chr2 | 29508995 | 29509190 | 1.14885 | 0.469516 | -2.447 |
| chr15 | 61717271 | 61717487 | 1.38305 | 0.565319 | -2.446 |
| chr5 | 145162558 | 145162938 | 0.987477 | 0.403637 | -2.446 |
| chr3 | 129956679 | 129956936 | 1.0364 | 0.423646 | -2.446 |
| chr2 | 241209012 | 241209282 | 0.903223 | 0.369227 | -2.446 |
| chr2 | 246999 | 247461 | 1.06661 | 0.436026 | -2.446 |
| chr10 | 68089076 | 68089283 | 0.917019 | 0.374916 | -2.446 |
| chr2 | 79882735 | 79882979 | 1.20845 | 0.494066 | -2.446 |
| chr10 | 126164883 | 126165163 | 0.83307 | 0.340626 | -2.446 |
| chr5 | 67107881 | 67108118 | 1.43355 | 0.586194 | -2.446 |
| chr8 | 137110922 | 137111142 | 0.959692 | 0.392451 | -2.445 |
| chr3 | 78365595 | 78365826 | 0.997357 | 0.407872 | -2.445 |
| chr15 | 89232479 | 89232852 | 0.959095 | 0.392239 | -2.445 |
| chr2 | 25967184 | 25967371 | 1.0872 | 0.44481 | -2.444 |
| chr11 | 37971866 | 37972114 | 1.35351 | 0.553777 | -2.444 |
| chr2 | 17319623 | 17320064 | 1.22267 | 0.500246 | -2.444 |
| chr2 | 96220144 | 96220326 | 1.33339 | 0.545641 | -2.444 |
| chr16 | 8545327 | 8545614 | 0.909175 | 0.372051 | -2.444 |
| chr3 | 23921262 | 23921477 | 1.00289 | 0.410427 | -2.444 |
| chr11 | 79535809 | 79536218 | 1.23033 | 0.503556 | -2.443 |
| chr20 | 61690128 | 61690340 | 0.921358 | 0.377159 | -2.443 |
| chr2 | 63493655 | 63493885 | 0.915861 | 0.375039 | -2.442 |
| chr2 | 74074645 | 74074882 | 1.28637 | 0.526929 | -2.441 |
| chr2 | 4726286 | 4726806 | 1.36325 | 0.558458 | -2.441 |
| chr8 | 138493176 | 138493445 | 1.2191 | 0.499539 | -2.440 |
| chr2 | 156061581 | 156061936 | 1.04402 | 0.427816 | -2.440 |
| chr5 | 117583601 | 117583824 | 0.884109 | 0.362323 | -2.440 |
| chr2 | 18298985 | 18299184 | 1.54737 | 0.634204 | -2.440 |
| chr17 | 50210386 | 50210627 | 1.2871 | 0.527538 | -2.440 |
| chr2 | 43291997 | 43292335 | 1.16349 | 0.477061 | -2.439 |
| chr20 | 21789604 | 21790084 | 3.41645 | 1.4013 | -2.438 |
| chr8 | 138740418 | 138740730 | 1.10739 | 0.454238 | -2.438 |
| chr5 | 1152265 | 1152458 | 1.35369 | 0.555369 | -2.437 |
| chr8 | 134877208 | 134877403 | 1.04721 | 0.429635 | -2.437 |
| chr5 | 4704237 | 4704512 | 1.02999 | 0.422671 | -2.437 |
| chr5 | 67481846 | 67482054 | 1.15088 | 0.472371 | -2.436 |
| chr5 | 53780197 | 53780473 | 1.09756 | 0.450492 | -2.436 |

| | | | | | |
|---|---|---|---|---|---|
| chr12 | 81914485 | 81914701 | 1.26374 | 0.518716 | -2.436 |
| chr14 | 49730836 | 49731024 | 1.0008 | 0.410789 | -2.436 |
| chr5 | 102937917 | 102938116 | 1.18601 | 0.486865 | -2.436 |
| chr10 | 55537927 | 55538124 | 1.05542 | 0.433267 | -2.436 |
| chr2 | 105860294 | 105860500 | 1.24592 | 0.511508 | -2.436 |
| chr10 | 27454340 | 27454609 | 1.05085 | 0.431431 | -2.436 |
| chr8 | 83834791 | 83835450 | 1.11928 | 0.45964 | -2.435 |
| chr8 | 74318902 | 74319108 | 1.17795 | 0.48379 | -2.435 |
| chr12 | 5551626 | 5551856 | 2.18186 | 0.896133 | -2.435 |
| chr7 | 44984581 | 44984765 | 1.07134 | 0.440146 | -2.434 |
| chr2 | 153699874 | 153700184 | 1.04405 | 0.428972 | -2.434 |
| chr2 | 207450604 | 207450909 | 1.35354 | 0.55617 | -2.434 |
| chr2 | 80926005 | 80926242 | 1.26276 | 0.518887 | -2.434 |
| chr1 | 179864480 | 179864716 | 1.02048 | 0.419502 | -2.433 |
| chr15 | 58780454 | 58780660 | 0.927621 | 0.38133 | -2.433 |
| chr5 | 4017300 | 4017536 | 1.01615 | 0.417732 | -2.433 |
| chr2 | 71165369 | 71165620 | 1.07567 | 0.442449 | -2.431 |
| chr14 | 36889714 | 36889902 | 1.29747 | 0.533759 | -2.431 |
| chr7 | 15565554 | 15565794 | 1.21027 | 0.497943 | -2.431 |
| chr5 | 113315409 | 113315692 | 1.03361 | 0.425352 | -2.430 |
| chr16 | 3251652 | 3251838 | 1.08337 | 0.445832 | -2.430 |
| chr7 | 137792031 | 137792261 | 0.91423 | 0.376266 | -2.430 |
| chr1 | 157808773 | 157809022 | 0.997271 | 0.410444 | -2.430 |
| chr8 | 81583071 | 81583424 | 0.973697 | 0.400885 | -2.429 |
| chr2 | 22989762 | 22989949 | 1.44723 | 0.595846 | -2.429 |
| chr11 | 17657442 | 17657631 | 1.11298 | 0.458235 | -2.429 |
| chr2 | 148208638 | 148208873 | 1.05216 | 0.433218 | -2.429 |
| chr1 | 191311272 | 191311511 | 1.20362 | 0.495635 | -2.428 |
| chr2 | 193272526 | 193272806 | 0.996662 | 0.410433 | -2.428 |
| chr16 | 55357558 | 55357996 | 2.19691 | 0.90505 | -2.427 |
| chr8 | 41814512 | 41814722 | 1.16648 | 0.480646 | -2.427 |
| chr2 | 115789096 | 115789302 | 1.03262 | 0.425567 | -2.426 |
| chr10 | 86742059 | 86742248 | 0.944047 | 0.389117 | -2.426 |
| chr8 | 13843984 | 13844290 | 1.10335 | 0.454816 | -2.426 |
| chr2 | 177534388 | 177534610 | 0.944758 | 0.389507 | -2.426 |
| chr8 | 68778760 | 68779066 | 1.68072 | 0.693095 | -2.425 |
| chr9 | 9270537 | 9270749 | 1.35253 | 0.557775 | -2.425 |
| chr2 | 41004323 | 41004644 | 1.10067 | 0.453923 | -2.425 |
| chr8 | 108817316 | 108817529 | 0.895586 | 0.369399 | -2.424 |
| chr10 | 53149129 | 53149378 | 2.40788 | 0.993427 | -2.424 |
| chr5 | 24960692 | 24960881 | 0.962867 | 0.397304 | -2.424 |
| chr11 | 80170744 | 80170984 | 1.17837 | 0.48634 | -2.423 |
| chr7 | 155206242 | 155206482 | 1.19354 | 0.492646 | -2.423 |
| chr8 | 110381096 | 110381516 | 0.885329 | 0.365461 | -2.422 |
| chrX | 138472331 | 138472578 | 0.843546 | 0.348225 | -2.422 |
| chr19 | 56981956 | 56982156 | 0.914096 | 0.377357 | -2.422 |
| chr10 | 134092664 | 134092863 | 1.0021 | 0.413731 | -2.422 |
| chr8 | 113325231 | 113325482 | 0.968351 | 0.3998 | -2.422 |
| chr2 | 147047517 | 147047713 | 1.10516 | 0.456423 | -2.421 |
| chr9 | 109587192 | 109587564 | 1.01319 | 0.418461 | -2.421 |
| chr14 | 67573704 | 67573901 | 1.00902 | 0.416834 | -2.421 |
| chr5 | 163996583 | 163996853 | 1.14519 | 0.473268 | -2.420 |
| chr5 | 98662802 | 98663060 | 1.04664 | 0.432544 | -2.420 |
| chr10 | 65808410 | 65808617 | 0.917742 | 0.379322 | -2.419 |
| chr5 | 10715036 | 10715300 | 0.801083 | 0.331146 | -2.419 |
| chr7 | 136838613 | 136838990 | 1.26428 | 0.522716 | -2.419 |
| chr7 | 119684988 | 119685256 | 1.26911 | 0.524742 | -2.419 |
| chr9 | 95713155 | 95713381 | 0.865371 | 0.357857 | -2.418 |
| chr12 | 82275961 | 82276228 | 1.11506 | 0.461225 | -2.418 |
| chr8 | 88405934 | 88406254 | 0.981782 | 0.406187 | -2.417 |
| chr2 | 229063280 | 229063502 | 0.902347 | 0.373373 | -2.417 |

| | | | | | |
|---|---|---|---|---|---|
| chr2 | 36892021 | 36892244 | 1.07004 | 0.442833 | -2.416 |
| chrX | 134721203 | 134721410 | 1.1709 | 0.484595 | -2.416 |
| chr2 | 166712573 | 166712859 | 0.994267 | 0.411544 | -2.416 |
| chr7 | 80606354 | 80606573 | 1.22954 | 0.508944 | -2.416 |
| chr5 | 86748780 | 86749177 | 1.33834 | 0.554011 | -2.416 |
| chr5 | 19923137 | 19923411 | 1.26452 | 0.523642 | -2.415 |
| chr2 | 211297450 | 211297641 | 0.952073 | 0.394277 | -2.415 |
| chr6 | 102001875 | 102002069 | 1.05484 | 0.436836 | -2.415 |
| chr13 | 79617469 | 79617660 | 1.2964 | 0.536911 | -2.415 |
| chr5 | 95246280 | 95246868 | 1.28303 | 0.531375 | -2.415 |
| chr9 | 121831401 | 121831644 | 0.864107 | 0.357891 | -2.414 |
| chr17 | 58117874 | 58118085 | 1.27411 | 0.527867 | -2.414 |
| chr2 | 80542667 | 80542949 | 0.92959 | 0.38537 | -2.412 |
| chr12 | 62889031 | 62889353 | 0.850059 | 0.352461 | -2.412 |
| chr1 | 167121422 | 167121658 | 0.898528 | 0.372641 | -2.411 |
| chr5 | 136155067 | 136155250 | 1.15649 | 0.479635 | -2.411 |
| chr8 | 12899765 | 12900079 | 2.04693 | 0.849046 | -2.411 |
| chr3 | 163887894 | 163888413 | 1.34561 | 0.558185 | -2.411 |
| chr5 | 25914701 | 25914894 | 1.08972 | 0.452055 | -2.411 |
| chr5 | 144579227 | 144579434 | 1.12623 | 0.467239 | -2.410 |
| chr8 | 111451100 | 111451288 | 0.881718 | 0.365878 | -2.410 |
| chr1 | 166350449 | 166350695 | 1.1157 | 0.46301 | -2.410 |
| chr12 | 29993330 | 29993598 | 1.20816 | 0.501402 | -2.410 |
| chr2 | 49917814 | 49918022 | 1.21541 | 0.504454 | -2.409 |
| chr14 | 56346148 | 56346335 | 1.70952 | 0.709597 | -2.409 |
| chr2 | 82928253 | 82928479 | 1.36367 | 0.566148 | -2.409 |
| chr8 | 118251581 | 118251833 | 0.922264 | 0.383018 | -2.408 |
| chr1 | 175343014 | 175343274 | 2.03259 | 0.844154 | -2.408 |
| chr9 | 121085714 | 121085903 | 1.00981 | 0.4194 | -2.408 |
| chr10 | 28227596 | 28227782 | 1.04613 | 0.434521 | -2.408 |
| chr2 | 174204700 | 174204888 | 0.85086 | 0.353521 | -2.407 |
| chr8 | 60973501 | 60973804 | 1.22536 | 0.509226 | -2.406 |
| chr10 | 9634755 | 9635037 | 0.82573 | 0.343206 | -2.406 |
| chr2 | 208926109 | 208926377 | 1.16675 | 0.485058 | -2.405 |
| chr17 | 52150745 | 52150939 | 1.27039 | 0.528408 | -2.404 |
| chr5 | 127664256 | 127664466 | 1.19112 | 0.495445 | -2.404 |
| chr19 | 28357409 | 28357660 | 1.32726 | 0.552077 | -2.404 |
| chr3 | 163664415 | 163664632 | 1.29964 | 0.540594 | -2.404 |
| chr14 | 84452482 | 84452681 | 1.2861 | 0.535114 | -2.403 |
| chr5 | 161699072 | 161699333 | 1.05703 | 0.439917 | -2.403 |
| chr9 | 122014381 | 122014567 | 1.06996 | 0.445308 | -2.403 |
| chr12 | 81992295 | 81992513 | 0.937166 | 0.390075 | -2.403 |
| chr2 | 7157054 | 7157278 | 1.05646 | 0.439777 | -2.402 |
| chr17 | 52159150 | 52159388 | 1.13712 | 0.47346 | -2.402 |
| chr14 | 27770387 | 27770646 | 1.0696 | 0.4454 | -2.401 |
| chr5 | 45088594 | 45088827 | 1.50618 | 0.627227 | -2.401 |
| chr2 | 48937671 | 48937872 | 1.11556 | 0.464818 | -2.400 |
| chr8 | 36203401 | 36203786 | 1.06884 | 0.445366 | -2.400 |
| chr2 | 101971042 | 101971224 | 1.20316 | 0.50136 | -2.400 |
| chr2 | 16921469 | 16921808 | 1.07211 | 0.446753 | -2.400 |
| chr11 | 87265878 | 87266342 | 1.2077 | 0.503261 | -2.400 |
| chr20 | 11818906 | 11819149 | 1.41803 | 0.591 | -2.399 |
| chr7 | 13613721 | 13613909 | 1.10066 | 0.458738 | -2.399 |
| chr16 | 65936757 | 65936972 | 0.941136 | 0.39233 | -2.399 |
| chr2 | 5222534 | 5222719 | 1.10438 | 0.460433 | -2.399 |
| chr14 | 66643499 | 66643696 | 1.00409 | 0.418673 | -2.398 |
| chr2 | 16737527 | 16737869 | 0.814999 | 0.339918 | -2.398 |
| chr16 | 6453953 | 6454155 | 1.03167 | 0.430311 | -2.398 |
| chr5 | 53630203 | 53630491 | 1.16172 | 0.484632 | -2.397 |
| chr17 | 10991874 | 10992140 | 1.15846 | 0.483499 | -2.396 |
| chr16 | 82746950 | 82747132 | 0.878693 | 0.36676 | -2.396 |

| | | | | | |
|---|---|---|---|---|---|
| chr10 | 66853241 | 66853461 | 1.09763 | 0.4582 | -2.396 |
| chr7 | 144243125 | 144243314 | 1.06115 | 0.443097 | -2.395 |
| chr5 | 146073636 | 146073883 | 1.26688 | 0.529013 | -2.395 |
| chr9 | 31456962 | 31457186 | 0.795816 | 0.332311 | -2.395 |
| chr8 | 13080066 | 13080262 | 1.17251 | 0.489715 | -2.394 |
| chr8 | 79289181 | 79289363 | 1.06695 | 0.445641 | -2.394 |
| chr2 | 137641966 | 137642177 | 1.10564 | 0.461927 | -2.394 |
| chr2 | 52002270 | 52002554 | 1.4148 | 0.59116 | -2.393 |
| chr3 | 84185134 | 84185380 | 1.19826 | 0.500779 | -2.393 |
| chr7 | 126044362 | 126044645 | 1.14813 | 0.479923 | -2.392 |
| chr7 | 9870104 | 9870455 | 1.15379 | 0.482399 | -2.392 |
| chr8 | 50201301 | 50201519 | 1.22809 | 0.513621 | -2.391 |
| chr9 | 30590106 | 30590372 | 0.948684 | 0.396797 | -2.391 |
| chr16 | 86423296 | 86423493 | 0.933652 | 0.390544 | -2.391 |
| chr6 | 46678100 | 46678316 | 0.907959 | 0.379902 | -2.390 |
| chr19 | 54124444 | 54124987 | 0.96177 | 0.402486 | -2.390 |
| chr8 | 20197600 | 20197836 | 1.20632 | 0.504828 | -2.390 |
| chr2 | 32982378 | 32982606 | 0.906603 | 0.379549 | -2.389 |
| chr5 | 118891280 | 118891511 | 1.19769 | 0.501541 | -2.388 |
| chr19 | 23349699 | 23349900 | 0.837654 | 0.350856 | -2.387 |
| chr9 | 30912535 | 30912785 | 1.01734 | 0.426122 | -2.387 |
| chr11 | 110317318 | 110317589 | 1.16181 | 0.486637 | -2.387 |
| chr10 | 34354947 | 34355151 | 0.855594 | 0.358517 | -2.386 |
| chr8 | 140152222 | 140152410 | 1.14918 | 0.481645 | -2.386 |
| chr3 | 103009226 | 103009453 | 0.803967 | 0.336974 | -2.386 |
| chr7 | 14253165 | 14253386 | 1.23125 | 0.5161 | -2.386 |
| chr13 | 24552779 | 24553061 | 0.858809 | 0.360001 | -2.386 |
| chr2 | 56786430 | 56786833 | 1.14357 | 0.479438 | -2.385 |
| chr3 | 147162004 | 147162391 | 1.01365 | 0.424996 | -2.385 |
| chr2 | 2486453 | 2486865 | 1.14846 | 0.481537 | -2.385 |
| chr8 | 141448543 | 141448754 | 1.2232 | 0.512935 | -2.385 |
| chr2 | 162678699 | 162678899 | 1.1123 | 0.466475 | -2.384 |
| chr14 | 45996529 | 45996771 | 1.57661 | 0.661268 | -2.384 |
| chr16 | 13118551 | 13118772 | 0.705946 | 0.296107 | -2.384 |
| chr5 | 3807487 | 3807683 | 1.06058 | 0.444868 | -2.384 |
| chr2 | 176673812 | 176673995 | 0.91688 | 0.38462 | -2.384 |
| chr1 | 188319399 | 188319584 | 0.903759 | 0.379125 | -2.384 |
| chr2 | 237237631 | 237237820 | 1.04466 | 0.438238 | -2.384 |
| chr15 | 86304788 | 86305007 | 1.19998 | 0.503526 | -2.383 |
| chr5 | 13011255 | 13011501 | 1.04801 | 0.439759 | -2.383 |
| chr8 | 57519256 | 57519483 | 0.797798 | 0.334772 | -2.383 |
| chr3 | 96083690 | 96083872 | 1.07011 | 0.449041 | -2.383 |
| chr10 | 2435316 | 2435525 | 1.24948 | 0.524309 | -2.383 |
| chr10 | 126042757 | 126042967 | 1.02797 | 0.431558 | -2.382 |
| chr2 | 83327052 | 83327274 | 1.10493 | 0.463905 | -2.382 |
| chr3 | 144699979 | 144700234 | 1.0496 | 0.440736 | -2.381 |
| chr3 | 144721376 | 144721723 | 1.05294 | 0.442145 | -2.381 |
| chr14 | 77573177 | 77573433 | 1.12894 | 0.474161 | -2.381 |
| chr7 | 97326809 | 97327067 | 1.13795 | 0.477963 | -2.381 |
| chr7 | 89080483 | 89080673 | 0.873604 | 0.367018 | -2.380 |
| chr8 | 85934188 | 85934417 | 1.12747 | 0.473672 | -2.380 |
| chr8 | 134882624 | 134882827 | 1.10676 | 0.465062 | -2.380 |
| chr2 | 35037330 | 35037589 | 1.13063 | 0.47527 | -2.379 |
| chr4 | 152506460 | 152506697 | 0.852081 | 0.35826 | -2.378 |
| chr2 | 73973746 | 73973994 | 1.21715 | 0.51189 | -2.378 |
| chr9 | 4521861 | 4522119 | 1.17332 | 0.493549 | -2.377 |
| chr2 | 4832314 | 4832501 | 1.05356 | 0.443243 | -2.377 |
| chr8 | 132638045 | 132638282 | 1.19562 | 0.503038 | -2.377 |
| chr2 | 59313440 | 59313821 | 1.32031 | 0.555539 | -2.377 |
| chr20 | 19229336 | 19229685 | 1.30589 | 0.549599 | -2.376 |
| chr2 | 10023980 | 10024248 | 1.43801 | 0.605267 | -2.376 |

| | | | | | |
|---|---|---|---|---|---|
| chr3 | 75079940 | 75080234 | 1.47285 | 0.620029 | -2.375 |
| chr2 | 6850506 | 6850804 | 1.05732 | 0.445129 | -2.375 |
| chr2 | 141799566 | 141799843 | 1.13782 | 0.479026 | -2.375 |
| chr15 | 64174814 | 64174998 | 1.18287 | 0.498059 | -2.375 |
| chr11 | 4880872 | 4881104 | 1.39152 | 0.586154 | -2.374 |
| chr5 | 178069303 | 178069577 | 0.98414 | 0.414603 | -2.374 |
| chr16 | 17518877 | 17519143 | 1.10456 | 0.465443 | -2.373 |
| chr2 | 128917043 | 128917240 | 1.13504 | 0.478389 | -2.373 |
| chr7 | 145382813 | 145383075 | 1.05635 | 0.445234 | -2.373 |
| chr5 | 107826800 | 107827028 | 0.89279 | 0.376303 | -2.373 |
| chr12 | 72863911 | 72864133 | 1.2235 | 0.515877 | -2.372 |
| chr3 | 113028045 | 113028229 | 0.883014 | 0.372376 | -2.371 |
| chr11 | 20859629 | 20859852 | 0.841828 | 0.355022 | -2.371 |
| chr14 | 69131698 | 69131915 | 1.29633 | 0.546967 | -2.370 |
| chr8 | 79222041 | 79222273 | 1.3386 | 0.564876 | -2.370 |
| chr8 | 121295113 | 121295320 | 1.01336 | 0.427679 | -2.369 |
| chr15 | 79505095 | 79505391 | 1.12683 | 0.475628 | -2.369 |
| chr1 | 248313112 | 248313311 | 1.16238 | 0.490778 | -2.368 |
| chr8 | 38340207 | 38340469 | 0.839164 | 0.354324 | -2.368 |
| chr19 | 28874611 | 28874937 | 1.39808 | 0.590368 | -2.368 |
| chr19 | 52962337 | 52962528 | 0.94969 | 0.401212 | -2.367 |
| chr3 | 155787809 | 155788017 | 1.11151 | 0.469591 | -2.367 |
| chr12 | 111961055 | 111961267 | 0.996817 | 0.421213 | -2.367 |
| chr7 | 136575124 | 136575316 | 1.08512 | 0.458542 | -2.366 |
| chr8 | 112424499 | 112424685 | 0.980237 | 0.414246 | -2.366 |
| chr14 | 26355775 | 26356001 | 1.23598 | 0.522332 | -2.366 |
| chr10 | 12040411 | 12040638 | 1.88723 | 0.797603 | -2.366 |
| chr15 | 73767363 | 73767663 | 1.27854 | 0.540524 | -2.365 |
| chr8 | 78232569 | 78232800 | 0.86859 | 0.367225 | -2.365 |
| chr8 | 51977412 | 51977652 | 1.35673 | 0.573637 | -2.365 |
| chr3 | 84318928 | 84319426 | 1.03838 | 0.439182 | -2.364 |
| chr9 | 18993380 | 18993600 | 1.02657 | 0.434466 | -2.363 |
| chr11 | 26207198 | 26207520 | 1.18226 | 0.50057 | -2.362 |
| chr7 | 154490460 | 154490687 | 1.84799 | 0.78273 | -2.361 |
| chr3 | 85938353 | 85938584 | 1.11303 | 0.471553 | -2.360 |
| chr17 | 4487116 | 4487313 | 2.74649 | 1.1638 | -2.360 |
| chr8 | 51510715 | 51510907 | 0.961722 | 0.407591 | -2.360 |
| chr7 | 113392427 | 113392642 | 1.13429 | 0.480729 | -2.360 |
| chr3 | 158695501 | 158695700 | 1.07154 | 0.454204 | -2.359 |
| chr19 | 53635171 | 53635501 | 1.28394 | 0.544241 | -2.359 |
| chr12 | 51597055 | 51597299 | 0.864194 | 0.366351 | -2.359 |
| chr5 | 117097024 | 117097298 | 1.53427 | 0.650486 | -2.359 |
| chr10 | 705709 | 705954 | 1.03313 | 0.438031 | -2.359 |
| chr2 | 51842177 | 51842433 | 0.923002 | 0.391353 | -2.358 |
| chr15 | 81758627 | 81758885 | 1.46148 | 0.619676 | -2.358 |
| chr17 | 45784822 | 45785022 | 1.12694 | 0.477892 | -2.358 |
| chr19 | 38052068 | 38052281 | 1.18335 | 0.502005 | -2.357 |
| chr9 | 91311120 | 91311310 | 0.972922 | 0.412747 | -2.357 |
| chr16 | 5364311 | 5364648 | 1.34059 | 0.568843 | -2.357 |
| chr7 | 13686898 | 13687146 | 1.06923 | 0.453727 | -2.357 |
| chr20 | 56456367 | 56456779 | 1.11449 | 0.472976 | -2.356 |
| chr5 | 63858668 | 63858896 | 1.18845 | 0.504381 | -2.356 |
| chr3 | 4162166 | 4162382 | 1.08233 | 0.459345 | -2.356 |
| chr2 | 194185183 | 194185627 | 0.959514 | 0.407242 | -2.356 |
| chr11 | 81281051 | 81281233 | 1.00534 | 0.426784 | -2.356 |
| chr3 | 31817961 | 31818290 | 0.901401 | 0.382701 | -2.355 |
| chr2 | 18235724 | 18235987 | 1.27889 | 0.542993 | -2.355 |
| chr11 | 33735348 | 33735629 | 1.06507 | 0.45222 | -2.355 |
| chr2 | 1417938 | 1418125 | 1.09899 | 0.466627 | -2.355 |
| chr3 | 158576262 | 158576446 | 1.50889 | 0.640696 | -2.355 |
| chr11 | 57894719 | 57894947 | 1.0455 | 0.444012 | -2.355 |

| | | | | | |
|---|---|---|---|---|---|
| chr11 | 90372985 | 90373231 | 1.01661 | 0.43175 | -2.355 |
| chr2 | 143191307 | 143191545 | 1.09026 | 0.46308 | -2.354 |
| chr2 | 214227123 | 214227324 | 0.726527 | 0.308587 | -2.354 |
| chr2 | 76566878 | 76567135 | 1.13656 | 0.48279 | -2.354 |
| chr2 | 84378335 | 84378692 | 1.07769 | 0.457818 | -2.354 |
| chr2 | 193980925 | 193981188 | 1.02642 | 0.43606 | -2.354 |
| chr5 | 44641131 | 44641323 | 1.19765 | 0.50898 | -2.353 |
| chr20 | 30357604 | 30357786 | 1.25559 | 0.533653 | -2.353 |
| chr8 | 54620463 | 54620758 | 1.20331 | 0.511541 | -2.352 |
| chr3 | 95856659 | 95856902 | 1.14682 | 0.487568 | -2.352 |
| chr11 | 2336995 | 2337347 | 0.923785 | 0.392785 | -2.352 |
| chr11 | 118041595 | 118041794 | 1.12974 | 0.480536 | -2.351 |
| chr7 | 123549867 | 123550102 | 1.04027 | 0.442491 | -2.351 |
| chr11 | 79777120 | 79777423 | 1.17101 | 0.498124 | -2.351 |
| chr7 | 118244182 | 118244456 | 1.13703 | 0.483789 | -2.350 |
| chr10 | 65875682 | 65875960 | 1.2514 | 0.532454 | -2.350 |
| chr2 | 82413290 | 82413522 | 0.944294 | 0.401893 | -2.350 |
| chr8 | 50904628 | 50904901 | 1.11269 | 0.473719 | -2.349 |
| chr5 | 2364225 | 2364413 | 1.28687 | 0.54789 | -2.349 |
| chr8 | 99797733 | 99797936 | 1.00951 | 0.429864 | -2.348 |
| chr3 | 105316731 | 105316969 | 1.04619 | 0.445507 | -2.348 |
| chr5 | 8372177 | 8372469 | 1.16599 | 0.496709 | -2.347 |
| chr4 | 13695396 | 13695647 | 1.00162 | 0.42672 | -2.347 |
| chr2 | 50367555 | 50367753 | 1.23841 | 0.527605 | -2.347 |
| chr9 | 102651534 | 102651781 | 0.988476 | 0.421124 | -2.347 |
| chr5 | 51264484 | 51264731 | 1.15113 | 0.490552 | -2.347 |
| chr9 | 106443735 | 106443941 | 0.8045 | 0.342848 | -2.347 |
| chr5 | 12598983 | 12599303 | 1.10448 | 0.470696 | -2.346 |
| chr16 | 25286543 | 25286768 | 1.09338 | 0.466095 | -2.346 |
| chr14 | 88271683 | 88271903 | 1.57414 | 0.671041 | -2.346 |
| chr17 | 40295079 | 40295324 | 0.812902 | 0.346551 | -2.346 |
| chr11 | 30950843 | 30951028 | 1.4124 | 0.602157 | -2.346 |
| chr12 | 98219867 | 98220219 | 1.13007 | 0.481795 | -2.346 |
| chr2 | 22043222 | 22043406 | 1.04179 | 0.444213 | -2.345 |
| chr5 | 113902688 | 113902931 | 1.11294 | 0.474662 | -2.345 |
| chr8 | 55867846 | 55868028 | 0.996196 | 0.42494 | -2.344 |
| chr8 | 107330517 | 107330897 | 0.904587 | 0.385879 | -2.344 |
| chr2 | 46812915 | 46813268 | 0.837925 | 0.357542 | -2.344 |
| chr2 | 78155031 | 78155219 | 1.80747 | 0.771324 | -2.343 |
| chr15 | 87368154 | 87368404 | 1.20788 | 0.515484 | -2.343 |
| chr10 | 59234076 | 59234266 | 1.11538 | 0.476029 | -2.343 |
| chr1 | 182411551 | 182411768 | 0.93527 | 0.399278 | -2.342 |
| chr16 | 86086800 | 86087034 | 1.67705 | 0.715985 | -2.342 |
| chr19 | 23626120 | 23626305 | 1.40691 | 0.600663 | -2.342 |
| chr5 | 15547741 | 15547979 | 1.05938 | 0.452293 | -2.342 |
| chr2 | 67027219 | 67027491 | 1.14806 | 0.490183 | -2.342 |
| chr11 | 24392898 | 24393092 | 1.11747 | 0.477291 | -2.341 |
| chr17 | 14579076 | 14579283 | 0.974102 | 0.416114 | -2.341 |
| chr2 | 34986521 | 34986729 | 1.18342 | 0.505716 | -2.340 |
| chr3 | 156899229 | 156899443 | 0.979937 | 0.418761 | -2.340 |
| chr19 | 21302913 | 21303148 | 0.943105 | 0.403071 | -2.340 |
| chrX | 147153022 | 147153229 | 0.969973 | 0.414586 | -2.340 |
| chr7 | 153916948 | 153917152 | 0.961515 | 0.411011 | -2.339 |
| chr8 | 79844861 | 79845138 | 1.04316 | 0.446046 | -2.339 |
| chr10 | 110093494 | 110093810 | 1.10917 | 0.474312 | -2.338 |
| chr8 | 116281458 | 116281733 | 1.15928 | 0.495789 | -2.338 |
| chr5 | 11836931 | 11837163 | 1.16053 | 0.496667 | -2.337 |
| chr7 | 136992511 | 136992694 | 0.905019 | 0.38735 | -2.336 |
| chr5 | 171560402 | 171560587 | 1.19182 | 0.510107 | -2.336 |
| chr2 | 118146462 | 118146675 | 1.0253 | 0.438858 | -2.336 |
| chr10 | 49856750 | 49857007 | 1.09012 | 0.466695 | -2.336 |

| | | | | | |
|---|---|---|---|---|---|
| chr5 | 54566288 | 54566487 | 0.917255 | 0.392697 | -2.336 |
| chr9 | 105300555 | 105300785 | 1.17173 | 0.501674 | -2.336 |
| chr9 | 89216379 | 89216589 | 0.718964 | 0.307825 | -2.336 |
| chr5 | 169171838 | 169172037 | 1.1522 | 0.493344 | -2.335 |
| chr15 | 52844267 | 52844542 | 1.16936 | 0.500737 | -2.335 |
| chr2 | 207490898 | 207491117 | 0.705151 | 0.301966 | -2.335 |
| chr16 | 64772729 | 64773073 | 0.921101 | 0.394444 | -2.335 |
| chr12 | 2511394 | 2511611 | 1.18423 | 0.50747 | -2.334 |
| chr5 | 51663941 | 51664130 | 1.00586 | 0.431075 | -2.333 |
| chr5 | 27142255 | 27142463 | 1.3273 | 0.568965 | -2.333 |
| chr5 | 106417207 | 106417417 | 1.07407 | 0.460584 | -2.332 |
| chr9 | 79538134 | 79538325 | 1.01495 | 0.435289 | -2.332 |
| chr17 | 75849678 | 75849986 | 1.2613 | 0.540975 | -2.332 |
| chr15 | 92561755 | 92562000 | 0.995005 | 0.426849 | -2.331 |
| chr7 | 115231395 | 115231602 | 1.06786 | 0.458139 | -2.331 |
| chr1 | 169913168 | 169913536 | 1.06903 | 0.458688 | -2.331 |
| chr10 | 23815654 | 23815918 | 1.12473 | 0.482608 | -2.331 |
| chr12 | 8757827 | 8758013 | 0.881239 | 0.378304 | -2.329 |
| chr5 | 129000145 | 129000438 | 1.8149 | 0.779111 | -2.329 |
| chr5 | 55475445 | 55475851 | 1.10766 | 0.475504 | -2.329 |
| chr2 | 104294452 | 104295043 | 1.10273 | 0.473484 | -2.329 |
| chr10 | 124659775 | 124660039 | 1.0402 | 0.446675 | -2.329 |
| chr2 | 203670665 | 203670866 | 0.764322 | 0.328211 | -2.329 |
| chr8 | 13436310 | 13436519 | 1.05997 | 0.455181 | -2.329 |
| chr5 | 105788795 | 105789036 | 0.950679 | 0.408265 | -2.329 |
| chr20 | 16636451 | 16636703 | 1.03119 | 0.442893 | -2.328 |
| chr3 | 180508819 | 180509002 | 0.946167 | 0.406418 | -2.328 |
| chr11 | 9326857 | 9327061 | 1.00845 | 0.433188 | -2.328 |
| chr14 | 29806457 | 29806724 | 1.11025 | 0.477056 | -2.327 |
| chr1 | 114079743 | 114080059 | 0.968839 | 0.41639 | -2.327 |
| chr1 | 191597911 | 191598168 | 1.64127 | 0.705818 | -2.325 |
| chr3 | 102802986 | 102803255 | 0.909812 | 0.391436 | -2.324 |
| chr15 | 63324112 | 63324320 | 1.19888 | 0.515896 | -2.324 |
| chr2 | 4279466 | 4279704 | 1.65877 | 0.713813 | -2.324 |
| chr5 | 28627867 | 28628059 | 0.852987 | 0.367148 | -2.323 |
| chr5 | 85801432 | 85801677 | 0.951389 | 0.40957 | -2.323 |
| chr3 | 154258943 | 154259138 | 1.28932 | 0.555102 | -2.323 |
| chr10 | 10636247 | 10636434 | 0.872802 | 0.375895 | -2.322 |
| chr17 | 81078289 | 81078483 | 1.12233 | 0.483524 | -2.321 |
| chr12 | 130351350 | 130351612 | 1.06582 | 0.459217 | -2.321 |
| chr20 | 44948644 | 44948832 | 0.975545 | 0.420348 | -2.321 |
| chr2 | 13455175 | 13455593 | 1.13071 | 0.487249 | -2.321 |
| chr8 | 64925489 | 64925701 | 0.951973 | 0.410237 | -2.321 |
| chrX | 95135666 | 95135867 | 1.31116 | 0.565083 | -2.320 |
| chr5 | 7984370 | 7984564 | 0.890891 | 0.383969 | -2.320 |
| chr8 | 120018743 | 120018991 | 0.794634 | 0.342505 | -2.320 |
| chr19 | 52760769 | 52760951 | 1.22872 | 0.529652 | -2.320 |
| chr2 | 114993825 | 114994085 | 1.10527 | 0.476725 | -2.318 |
| chr8 | 82797765 | 82798112 | 0.885176 | 0.381809 | -2.318 |
| chr8 | 109318424 | 109318692 | 1.30264 | 0.562134 | -2.317 |
| chr10 | 57813197 | 57813393 | 1.22211 | 0.527479 | -2.317 |
| chr14 | 88023909 | 88024131 | 0.818234 | 0.353246 | -2.316 |
| chr17 | 12299193 | 12299612 | 1.7754 | 0.766541 | -2.316 |
| chr6 | 91791293 | 91791543 | 1.10439 | 0.476845 | -2.316 |
| chr3 | 95530371 | 95530564 | 1.18802 | 0.513005 | -2.316 |
| chr2 | 102585815 | 102586033 | 0.992329 | 0.428551 | -2.316 |
| chr8 | 106173787 | 106174018 | 0.993506 | 0.429072 | -2.315 |
| chr8 | 27916830 | 27917012 | 0.807146 | 0.348661 | -2.315 |
| chr5 | 104499424 | 104499606 | 0.844847 | 0.364962 | -2.315 |
| chr3 | 96194880 | 96195130 | 0.847274 | 0.366028 | -2.315 |
| chr2 | 52769515 | 52769720 | 0.95876 | 0.414207 | -2.315 |

| | | | | | |
|---|---|---|---|---|---|
| chr15 | 92845601 | 92845805 | 1.04777 | 0.452684 | -2.315 |
| chr1 | 238982741 | 238982943 | 1.04769 | 0.452684 | -2.314 |
| chr5 | 97749311 | 97749506 | 0.876909 | 0.378947 | -2.314 |
| chr7 | 35448966 | 35449227 | 1.26125 | 0.545119 | -2.314 |
| chr8 | 72667991 | 72668190 | 1.58052 | 0.683148 | -2.314 |
| chr2 | 20084276 | 20084488 | 1.23971 | 0.535859 | -2.314 |
| chr2 | 139660171 | 139660435 | 1.08037 | 0.467084 | -2.313 |
| chr14 | 29661530 | 29661823 | 1.06103 | 0.45875 | -2.313 |
| chr5 | 116985957 | 116986163 | 1.17471 | 0.507951 | -2.313 |
| chr8 | 10122036 | 10122272 | 1.265 | 0.547115 | -2.312 |
| chr5 | 19494991 | 19495324 | 1.10533 | 0.478238 | -2.311 |
| chr15 | 61488512 | 61488694 | 1.13555 | 0.4917 | -2.309 |
| chr11 | 21660757 | 21660957 | 1.09618 | 0.474845 | -2.308 |
| chr14 | 83924427 | 83924727 | 0.944659 | 0.409228 | -2.308 |
| chr2 | 163286913 | 163287145 | 0.887528 | 0.384529 | -2.308 |
| chr2 | 80190556 | 80190752 | 1.08154 | 0.468693 | -2.308 |
| chr14 | 48396524 | 48396726 | 1.24847 | 0.5411 | -2.307 |
| chr7 | 131944275 | 131944526 | 1.22713 | 0.531871 | -2.307 |
| chr12 | 34760351 | 34760646 | 1.18978 | 0.51574 | -2.307 |
| chr9 | 22991405 | 22991701 | 0.918415 | 0.398139 | -2.307 |
| chr2 | 156711227 | 156711428 | 0.962021 | 0.417158 | -2.306 |
| chr9 | 122510341 | 122510570 | 1.12779 | 0.48909 | -2.306 |
| chr7 | 91119422 | 91119680 | 1.31532 | 0.570425 | -2.306 |
| chr2 | 215295547 | 215295858 | 0.779825 | 0.338307 | -2.305 |
| chr7 | 67948300 | 67948510 | 0.75209 | 0.326281 | -2.305 |
| chr7 | 141948786 | 141948990 | 0.774358 | 0.335999 | -2.305 |
| chr16 | 89929209 | 89929532 | 0.913379 | 0.39635 | -2.304 |
| chr10 | 28085392 | 28085644 | 0.952123 | 0.413292 | -2.304 |
| chr14 | 85009274 | 85009461 | 1.06418 | 0.461971 | -2.304 |
| chr10 | 83409955 | 83410145 | 1.39946 | 0.607683 | -2.303 |
| chr11 | 23188634 | 23188818 | 0.943234 | 0.40959 | -2.303 |
| chr1 | 195008396 | 195008713 | 0.880188 | 0.38226 | -2.303 |
| chr17 | 52712669 | 52712872 | 1.29466 | 0.5623 | -2.302 |
| chr17 | 21734372 | 21734571 | 1.00222 | 0.43532 | -2.302 |
| chr5 | 52833411 | 52833926 | 1.08852 | 0.472826 | -2.302 |
| chr1 | 163617236 | 163617467 | 0.985157 | 0.427929 | -2.302 |
| chr19 | 31282119 | 31282326 | 1.29508 | 0.562644 | -2.302 |
| chr2 | 71757116 | 71757315 | 1.11374 | 0.483925 | -2.301 |
| chr5 | 103297331 | 103297534 | 0.979441 | 0.425668 | -2.301 |
| chr5 | 104828035 | 104828262 | 0.911199 | 0.396092 | -2.300 |
| chr16 | 28066166 | 28066400 | 1.01931 | 0.443094 | -2.300 |
| chr5 | 85066235 | 85066439 | 1.07928 | 0.469202 | -2.300 |
| chr3 | 159221246 | 159221455 | 1.22395 | 0.532144 | -2.300 |
| chr10 | 76451919 | 76452146 | 0.987693 | 0.429525 | -2.300 |
| chr8 | 636569 | 636976 | 1.32382 | 0.575704 | -2.299 |
| chr5 | 167863851 | 167864156 | 1.28417 | 0.55875 | -2.298 |
| chr7 | 126746560 | 126746870 | 1.14488 | 0.498596 | -2.296 |
| chr8 | 37201777 | 37202006 | 1.03573 | 0.451172 | -2.296 |
| chr9 | 97996997 | 97997248 | 1.55554 | 0.677656 | -2.295 |
| chr14 | 41831323 | 41831538 | 1.03423 | 0.45069 | -2.295 |
| chr3 | 164125675 | 164125918 | 0.89832 | 0.391552 | -2.294 |
| chr7 | 73307974 | 73308185 | 0.795657 | 0.346831 | -2.294 |
| chr1 | 176447914 | 176448187 | 0.71094 | 0.309932 | -2.294 |
| chr12 | 24112724 | 24113103 | 1.28937 | 0.562132 | -2.294 |
| chr16 | 31947172 | 31947478 | 0.974446 | 0.424987 | -2.293 |
| chr12 | 15170838 | 15171099 | 1.10988 | 0.484089 | -2.293 |
| chr7 | 45052185 | 45052384 | 1.00953 | 0.44036 | -2.293 |
| chr8 | 118821709 | 118821899 | 0.895526 | 0.390639 | -2.292 |
| chr6 | 93206261 | 93206460 | 0.749614 | 0.327155 | -2.291 |
| chr7 | 100961760 | 100961983 | 1.14239 | 0.498579 | -2.291 |
| chr14 | 77331817 | 77332108 | 0.831493 | 0.362915 | -2.291 |

| | | | | | |
|---|---|---|---|---|---|
| chr8 | 73152194 | 73152381 | 1.10618 | 0.482919 | -2.291 |
| chr17 | 53791036 | 53791260 | 1.06984 | 0.467068 | -2.291 |
| chr3 | 79476389 | 79476591 | 0.641744 | 0.280218 | -2.290 |
| chr7 | 34221495 | 34221785 | 1.19346 | 0.521135 | -2.290 |
| chr10 | 107726274 | 107726475 | 1.17815 | 0.514471 | -2.290 |
| chr7 | 13666670 | 13666976 | 1.48971 | 0.650603 | -2.290 |
| chr19 | 28472451 | 28472653 | 1.27142 | 0.5553 | -2.290 |
| chr10 | 58393809 | 58394081 | 1.03696 | 0.452918 | -2.290 |
| chr5 | 144137166 | 144137379 | 1.10237 | 0.481678 | -2.289 |
| chr2 | 51189345 | 51189613 | 1.22068 | 0.533476 | -2.288 |
| chr2 | 83332030 | 83332228 | 1.02819 | 0.449369 | -2.288 |
| chr7 | 24321774 | 24321988 | 1.00311 | 0.438425 | -2.288 |
| chr1 | 154309741 | 154309951 | 1.12665 | 0.492486 | -2.288 |
| chr2 | 60656998 | 60657210 | 1.25714 | 0.54954 | -2.288 |
| chr19 | 58840063 | 58840271 | 1.37352 | 0.600597 | -2.287 |
| chr8 | 1857485 | 1857741 | 1.44595 | 0.632282 | -2.287 |
| chr7 | 38700076 | 38700422 | 2.44402 | 1.06902 | -2.286 |
| chr2 | 114898905 | 114899163 | 1.15822 | 0.50666 | -2.286 |
| chr19 | 31101841 | 31102044 | 1.23738 | 0.541298 | -2.286 |
| chr2 | 56946121 | 56946308 | 0.968229 | 0.423612 | -2.286 |
| chr1 | 181570937 | 181571119 | 1.35035 | 0.590949 | -2.285 |
| chr9 | 15918291 | 15918512 | 1.01758 | 0.445392 | -2.285 |
| chr10 | 55628607 | 55628808 | 1.25586 | 0.549787 | -2.284 |
| chr5 | 50358085 | 50358284 | 0.856183 | 0.374966 | -2.283 |
| chr3 | 36298275 | 36298493 | 1.10228 | 0.48285 | -2.283 |
| chr12 | 84174766 | 84174999 | 1.01245 | 0.443618 | -2.282 |
| chr8 | 58814462 | 58814868 | 1.38192 | 0.605547 | -2.282 |
| chr7 | 152754590 | 152754910 | 0.823006 | 0.360664 | -2.282 |
| chr5 | 26368076 | 26368334 | 1.34879 | 0.591088 | -2.282 |
| chr9 | 9387536 | 9387879 | 0.999308 | 0.437971 | -2.282 |
| chr20 | 17104098 | 17104297 | 1.14773 | 0.503087 | -2.281 |
| chr14 | 81989774 | 81990026 | 1.27772 | 0.560189 | -2.281 |
| chr8 | 14544057 | 14544343 | 1.05248 | 0.461472 | -2.281 |
| chr3 | 20922217 | 20922545 | 1.34188 | 0.588547 | -2.280 |
| chr10 | 56361361 | 56361794 | 1.20122 | 0.526877 | -2.280 |
| chr2 | 48960471 | 48960654 | 1.28833 | 0.565273 | -2.279 |
| chr10 | 28778148 | 28778343 | 1.30136 | 0.571115 | -2.279 |
| chr8 | 51971978 | 51972161 | 0.875182 | 0.384096 | -2.279 |
| chr1 | 247339787 | 247340076 | 1.11745 | 0.490696 | -2.277 |
| chr2 | 22956965 | 22957399 | 1.09589 | 0.48132 | -2.277 |
| chr10 | 132751483 | 132751756 | 0.787929 | 0.346068 | -2.277 |
| chr9 | 101739622 | 101739854 | 1.00286 | 0.440508 | -2.277 |
| chr16 | 33535734 | 33535976 | 1.02799 | 0.451622 | -2.276 |
| chr8 | 135102934 | 135103197 | 0.794075 | 0.348868 | -2.276 |
| chr2 | 151116568 | 151116791 | 1.03969 | 0.456781 | -2.276 |
| chr2 | 2353999 | 2354201 | 1.31445 | 0.57754 | -2.276 |
| chr16 | 14361501 | 14361723 | 1.04332 | 0.458454 | -2.276 |
| chr20 | 31665235 | 31665524 | 1.0913 | 0.479569 | -2.276 |
| chr9 | 131107336 | 131107870 | 1.02354 | 0.449889 | -2.275 |
| chr14 | 88144242 | 88144459 | 1.07018 | 0.470407 | -2.275 |
| chrX | 134677892 | 134678102 | 1.13602 | 0.499411 | -2.275 |
| chr17 | 50540607 | 50540971 | 1.07022 | 0.470524 | -2.275 |
| chr2 | 167341950 | 167342155 | 0.951936 | 0.418525 | -2.275 |
| chr8 | 124426836 | 124427018 | 0.877573 | 0.385921 | -2.274 |
| chr8 | 110405108 | 110405297 | 1.07415 | 0.472377 | -2.274 |
| chr12 | 577811 | 578069 | 1.18459 | 0.521036 | -2.274 |
| chr2 | 63564415 | 63564783 | 0.904248 | 0.397729 | -2.274 |
| chr2 | 215702159 | 215702377 | 0.796801 | 0.350477 | -2.273 |
| chr9 | 116576685 | 116576871 | 1.13293 | 0.49834 | -2.273 |
| chr8 | 89422650 | 89422876 | 0.965019 | 0.424504 | -2.273 |
| chr20 | 47926603 | 47926833 | 0.951668 | 0.41873 | -2.273 |

| | | | | | |
|---|---|---|---|---|---|
| chr5 | 164300848 | 164301094 | 1.09275 | 0.480933 | -2.272 |
| chr2 | 14046986 | 14047335 | 0.912607 | 0.401741 | -2.272 |
| chr10 | 54909399 | 54909599 | 0.731412 | 0.32198 | -2.272 |
| chr11 | 3022824 | 3023045 | 1.20405 | 0.530047 | -2.272 |
| chr19 | 58520958 | 58521165 | 1.0215 | 0.449701 | -2.272 |
| chr8 | 107404390 | 107404675 | 0.945979 | 0.416494 | -2.271 |
| chr12 | 70223663 | 70223854 | 0.986352 | 0.434437 | -2.270 |
| chr2 | 120439121 | 120439332 | 0.906512 | 0.399282 | -2.270 |
| chr12 | 5620785 | 5621136 | 1.82042 | 0.801887 | -2.270 |
| chr14 | 83544607 | 83544907 | 1.2102 | 0.533419 | -2.269 |
| chr7 | 133708567 | 133708776 | 1.21958 | 0.537572 | -2.269 |
| chr11 | 8484700 | 8485087 | 1.14179 | 0.503367 | -2.268 |
| chr1 | 153220672 | 153221135 | 1.26389 | 0.557231 | -2.268 |
| chr3 | 164971871 | 164972095 | 1.04287 | 0.459933 | -2.267 |
| chr17 | 75500401 | 75500694 | 1.32537 | 0.584528 | -2.267 |
| chr7 | 47366628 | 47366814 | 1.50308 | 0.663422 | -2.266 |
| chr17 | 76459080 | 76459315 | 1.13167 | 0.499524 | -2.265 |
| chr10 | 9939513 | 9939760 | 1.17841 | 0.520164 | -2.265 |
| chr2 | 128470589 | 128470796 | 0.841868 | 0.371625 | -2.265 |
| chr2 | 167388499 | 167388852 | 1.20874 | 0.53359 | -2.265 |
| chr2 | 135026767 | 135027046 | 1.04103 | 0.459741 | -2.264 |
| chr7 | 105814967 | 105815167 | 1.02888 | 0.454401 | -2.264 |
| chr7 | 7869834 | 7870096 | 1.04608 | 0.462033 | -2.264 |
| chr15 | 98315989 | 98316223 | 1.06374 | 0.469933 | -2.264 |
| chr4 | 61513757 | 61514051 | 0.959632 | 0.423953 | -2.264 |
| chr16 | 74127630 | 74127928 | 1.13768 | 0.502624 | -2.263 |
| chr20 | 56060013 | 56060205 | 1.07625 | 0.47555 | -2.263 |
| chr2 | 155480383 | 155480600 | 1.9875 | 0.878776 | -2.262 |
| chr9 | 106081011 | 106081209 | 0.910806 | 0.402716 | -2.262 |
| chr2 | 78387354 | 78387582 | 1.00304 | 0.443515 | -2.262 |
| chr8 | 72975350 | 72975617 | 0.892646 | 0.394712 | -2.262 |
| chr7 | 147290213 | 147290395 | 0.87329 | 0.38618 | -2.261 |
| chr5 | 43570125 | 43570377 | 0.979519 | 0.43316 | -2.261 |
| chr2 | 126247770 | 126248067 | 0.949517 | 0.419988 | -2.261 |
| chr9 | 29858167 | 29858471 | 1.29127 | 0.571264 | -2.260 |
| chr10 | 134251864 | 134252098 | 1.07891 | 0.477379 | -2.260 |
| chr2 | 191138647 | 191138858 | 0.940022 | 0.415943 | -2.260 |
| chr10 | 125936360 | 125936630 | 1.36421 | 0.603744 | -2.260 |
| chr4 | 70025773 | 70025955 | 0.862307 | 0.381676 | -2.259 |
| chr20 | 6869052 | 6869403 | 1.08462 | 0.480136 | -2.259 |
| chr10 | 10722232 | 10722581 | 0.935858 | 0.414335 | -2.259 |
| chr7 | 63852174 | 63852507 | 1.12655 | 0.49877 | -2.259 |
| chr11 | 26384057 | 26384253 | 0.9153 | 0.405267 | -2.259 |
| chr11 | 69393648 | 69393909 | 1.15784 | 0.512765 | -2.258 |
| chr2 | 51436416 | 51436805 | 0.972008 | 0.430556 | -2.258 |
| chr4 | 63527740 | 63528053 | 1.58343 | 0.701448 | -2.257 |
| chr20 | 3945361 | 3945568 | 1.30794 | 0.579439 | -2.257 |
| chr8 | 121150682 | 121150871 | 0.890839 | 0.394844 | -2.256 |
| chr17 | 79020668 | 79021003 | 0.877357 | 0.388892 | -2.256 |
| chr2 | 22660198 | 22660585 | 1.07765 | 0.47778 | -2.256 |
| chr15 | 92366726 | 92367159 | 0.959035 | 0.425313 | -2.255 |
| chr14 | 41785131 | 41785382 | 1.01227 | 0.449002 | -2.254 |
| chr2 | 89218642 | 89219148 | 1.68594 | 0.747829 | -2.254 |
| chr2 | 26027720 | 26027937 | 1.22457 | 0.54319 | -2.254 |
| chr8 | 83705396 | 83705599 | 1.09059 | 0.483765 | -2.254 |
| chr8 | 3229785 | 3229976 | 1.06317 | 0.471607 | -2.254 |
| chr8 | 49696349 | 49696531 | 1.05439 | 0.46772 | -2.254 |
| chr8 | 133648624 | 133648829 | 1.50744 | 0.668734 | -2.254 |
| chr2 | 114786254 | 114786517 | 1.04218 | 0.462359 | -2.254 |
| chr2 | 60120826 | 60121030 | 1.03641 | 0.459842 | -2.254 |
| chr2 | 77605755 | 77606014 | 1.20014 | 0.532569 | -2.253 |

| | | | | | |
|---|---|---|---|---|---|
| chr20 | 58256551 | 58256775 | 1.23765 | 0.549231 | -2.253 |
| chr3 | 103175977 | 103176206 | 1.03528 | 0.459506 | -2.253 |
| chr7 | 139032509 | 139032783 | 1.13865 | 0.505395 | -2.253 |
| chr7 | 14810057 | 14810263 | 1.03806 | 0.460892 | -2.252 |
| chr14 | 85476624 | 85476895 | 0.876286 | 0.389151 | -2.252 |
| chr12 | 126759373 | 126759622 | 0.916996 | 0.407259 | -2.252 |
| chr10 | 107051408 | 107051595 | 1.24668 | 0.553709 | -2.252 |
| chr5 | 5295599 | 5296009 | 1.06599 | 0.47362 | -2.251 |
| chr3 | 37742416 | 37742697 | 0.944006 | 0.419441 | -2.251 |
| chr2 | 209924548 | 209924761 | 0.935241 | 0.415577 | -2.250 |
| chr20 | 39490051 | 39490237 | 0.917483 | 0.407741 | -2.250 |
| chr5 | 164027989 | 164028282 | 0.927854 | 0.412464 | -2.250 |
| chr20 | 21915760 | 21916061 | 1.17798 | 0.523714 | -2.249 |
| chr17 | 80142074 | 80142322 | 1.17301 | 0.521654 | -2.249 |
| chr2 | 123796402 | 123796586 | 0.825916 | 0.367386 | -2.248 |
| chr17 | 31060093 | 31060292 | 1.10611 | 0.492293 | -2.247 |
| chr11 | 31980994 | 31981181 | 1.42945 | 0.636651 | -2.245 |
| chr2 | 214320225 | 214320553 | 1.02149 | 0.455099 | -2.245 |
| chr10 | 30332570 | 30332790 | 1.09044 | 0.485834 | -2.244 |
| chr11 | 4883950 | 4884144 | 0.973166 | 0.433589 | -2.244 |
| chr7 | 126965916 | 126966142 | 0.929008 | 0.413919 | -2.244 |
| chr16 | 16218552 | 16218735 | 1.50837 | 0.672413 | -2.243 |
| chr7 | 112539534 | 112539800 | 1.00833 | 0.44957 | -2.243 |
| chr5 | 22529442 | 22529673 | 1.11591 | 0.497538 | -2.243 |
| chr2 | 103887366 | 103887551 | 1.02364 | 0.456436 | -2.243 |
| chr17 | 56362236 | 56362421 | 1.12992 | 0.503903 | -2.242 |
| chr3 | 24767941 | 24768147 | 1.15352 | 0.514441 | -2.242 |
| chr2 | 39108787 | 39109090 | 1.11541 | 0.497466 | -2.242 |
| chr12 | 123853525 | 123853729 | 0.843788 | 0.376337 | -2.242 |
| chr3 | 161175625 | 161175817 | 0.856378 | 0.382097 | -2.241 |
| chr5 | 89045934 | 89046129 | 0.925921 | 0.413194 | -2.241 |
| chr2 | 154631126 | 154631333 | 1.07122 | 0.478187 | -2.240 |
| chr1 | 175511338 | 175511781 | 0.902602 | 0.403017 | -2.240 |
| chr15 | 32250821 | 32251026 | 1.35266 | 0.60398 | -2.240 |
| chr7 | 9565710 | 9566034 | 1.19178 | 0.532174 | -2.239 |
| chr2 | 223069969 | 223070162 | 0.978966 | 0.437169 | -2.239 |
| chr20 | 9167917 | 9168102 | 1.35724 | 0.606143 | -2.239 |
| chr16 | 22727779 | 22727980 | 0.967038 | 0.431968 | -2.239 |
| chr1 | 221747275 | 221747465 | 1.2476 | 0.557435 | -2.238 |
| chr12 | 10913389 | 10913693 | 0.990128 | 0.442497 | -2.238 |
| chr2 | 77761840 | 77762031 | 1.28409 | 0.573941 | -2.237 |
| chr7 | 98540161 | 98540354 | 1.11608 | 0.498875 | -2.237 |
| chr14 | 28361705 | 28361895 | 1.18994 | 0.531926 | -2.237 |
| chr3 | 164841133 | 164841379 | 0.829025 | 0.370721 | -2.236 |
| chr7 | 31109892 | 31110074 | 1.02008 | 0.456162 | -2.236 |
| chr15 | 32217698 | 32217900 | 1.35886 | 0.607669 | -2.236 |
| chr8 | 116735737 | 116735930 | 1.07976 | 0.482874 | -2.236 |
| chr9 | 122917848 | 122918041 | 0.780965 | 0.349284 | -2.236 |
| chr7 | 156460879 | 156461144 | 1.02087 | 0.456656 | -2.236 |
| chr2 | 184423993 | 184424231 | 1.00932 | 0.451507 | -2.235 |
| chr1 | 197986159 | 197986350 | 0.878475 | 0.392979 | -2.235 |
| chr3 | 144108248 | 144108655 | 0.904431 | 0.404645 | -2.235 |
| chr14 | 68209707 | 68209909 | 1.11047 | 0.497057 | -2.234 |
| chr1 | 188713600 | 188713818 | 0.93144 | 0.416985 | -2.234 |
| chr8 | 117324724 | 117325035 | 1.09629 | 0.49082 | -2.234 |
| chrX | 50887770 | 50888016 | 1.42968 | 0.640138 | -2.233 |
| chr5 | 10878707 | 10879004 | 0.906747 | 0.40601 | -2.233 |
| chr2 | 156118681 | 156119090 | 0.907136 | 0.406298 | -2.233 |
| chr2 | 183073314 | 183073505 | 1.19493 | 0.535223 | -2.233 |
| chr8 | 73169376 | 73169742 | 1.08024 | 0.483871 | -2.232 |
| chr3 | 19823828 | 19824152 | 1.12578 | 0.504339 | -2.232 |

| | | | | | |
|---|---|---|---|---|---|
| chr11 | 48298712 | 48298919 | 1.04168 | 0.466689 | -2.232 |
| chr2 | 229627022 | 229627237 | 1.00933 | 0.45228 | -2.232 |
| chr2 | 125308572 | 125308832 | 1.13032 | 0.506738 | -2.231 |
| chr8 | 120075524 | 120075757 | 1.27529 | 0.571792 | -2.230 |
| chr3 | 23952498 | 23952717 | 1.15343 | 0.517204 | -2.230 |
| chr3 | 165593418 | 165593648 | 0.861676 | 0.386437 | -2.230 |
| chr5 | 18893892 | 18894094 | 1.29787 | 0.582089 | -2.230 |
| chr10 | 7510014 | 7510220 | 1.11534 | 0.500266 | -2.229 |
| chr2 | 168955424 | 168955727 | 1.14543 | 0.513858 | -2.229 |
| chr10 | 57189931 | 57190167 | 1.04588 | 0.4692 | -2.229 |
| chr8 | 2693558 | 2693783 | 1.20578 | 0.540945 | -2.229 |
| chr2 | 186364769 | 186365018 | 1.32545 | 0.594845 | -2.228 |
| chr5 | 76662527 | 76662849 | 1.14385 | 0.513366 | -2.228 |
| chr6 | 124414033 | 124414255 | 0.982176 | 0.440818 | -2.228 |
| chr7 | 111263135 | 111263467 | 1.09311 | 0.490714 | -2.228 |
| chr14 | 46178823 | 46179062 | 1.1119 | 0.499178 | -2.227 |
| chr3 | 165336627 | 165337083 | 0.814138 | 0.36556 | -2.227 |
| chr12 | 117821026 | 117821213 | 1.13222 | 0.508459 | -2.227 |
| chr2 | 66359740 | 66360002 | 0.992732 | 0.445855 | -2.227 |
| chr2 | 101635351 | 101635579 | 1.15277 | 0.517866 | -2.226 |
| chr3 | 30431442 | 30431729 | 1.34313 | 0.60342 | -2.226 |
| chr9 | 76288884 | 76289322 | 0.992385 | 0.44587 | -2.226 |
| chr16 | 49034535 | 49034804 | 1.84088 | 0.827202 | -2.225 |
| chr14 | 72470535 | 72470782 | 1.14199 | 0.513208 | -2.225 |
| chr2 | 202227567 | 202227879 | 1.12976 | 0.507712 | -2.225 |
| chr2 | 52775049 | 52775353 | 1.45568 | 0.654203 | -2.225 |
| chr10 | 653008 | 653215 | 1.13497 | 0.510092 | -2.225 |
| chr16 | 60945828 | 60946126 | 1.32485 | 0.595461 | -2.225 |
| chr8 | 5302248 | 5302601 | 0.914127 | 0.410974 | -2.224 |
| chr19 | 20631152 | 20631336 | 0.901447 | 0.405313 | -2.224 |
| chr2 | 199312121 | 199312370 | 1.31042 | 0.589253 | -2.224 |
| chr16 | 89584189 | 89584597 | 0.806053 | 0.362475 | -2.224 |
| chr10 | 133008470 | 133008699 | 1.1353 | 0.510671 | -2.223 |
| chr17 | 51884746 | 51885307 | 0.985416 | 0.44332 | -2.223 |
| chr9 | 74877253 | 74877488 | 1.00152 | 0.45061 | -2.223 |
| chr8 | 21699287 | 21699489 | 0.95775 | 0.43096 | -2.222 |
| chr5 | 132523738 | 132523934 | 0.952092 | 0.428469 | -2.222 |
| chr2 | 77768384 | 77768611 | 0.767168 | 0.345375 | -2.221 |
| chr1 | 152824357 | 152824548 | 1.00908 | 0.454342 | -2.221 |
| chr14 | 99137335 | 99137581 | 0.943393 | 0.424835 | -2.221 |
| chr15 | 26958144 | 26958351 | 1.22281 | 0.550691 | -2.221 |
| chr2 | 183115552 | 183115775 | 0.849297 | 0.382496 | -2.220 |
| chr8 | 143843023 | 143843302 | 1.10159 | 0.496173 | -2.220 |
| chr12 | 4236855 | 4237044 | 0.930331 | 0.419077 | -2.220 |
| chr19 | 49696760 | 49696944 | 1.00959 | 0.455014 | -2.219 |
| chr7 | 120333802 | 120334046 | 0.935052 | 0.421581 | -2.218 |
| chr3 | 165096171 | 165096431 | 1.223 | 0.551418 | -2.218 |
| chr14 | 104909881 | 104910069 | 0.880225 | 0.396876 | -2.218 |
| chr10 | 121201615 | 121201811 | 1.00891 | 0.454966 | -2.218 |
| chr16 | 85693013 | 85693220 | 1.26112 | 0.568737 | -2.217 |
| chr7 | 48584541 | 48584964 | 1.16624 | 0.526033 | -2.217 |
| chr3 | 86309606 | 86309801 | 1.14707 | 0.517439 | -2.217 |
| chr19 | 22943091 | 22943365 | 0.980783 | 0.442507 | -2.216 |
| chr9 | 72863600 | 72863809 | 1.45414 | 0.656089 | -2.216 |
| chr15 | 98334559 | 98334914 | 1.21312 | 0.547409 | -2.216 |
| chr3 | 73376665 | 73376862 | 1.3301 | 0.600227 | -2.216 |
| chr8 | 33009048 | 33009335 | 1.19541 | 0.539509 | -2.216 |
| chr20 | 32346755 | 32346962 | 1.27682 | 0.576267 | -2.216 |
| chr8 | 111746159 | 111746486 | 1.29793 | 0.585873 | -2.215 |
| chr19 | 23078879 | 23079169 | 1.29978 | 0.586748 | -2.215 |
| chr12 | 125959334 | 125959635 | 1.19072 | 0.53753 | -2.215 |

| | | | | | |
|---|---|---|---|---|---|
| chr2 | 50232905 | 50233139 | 1.14341 | 0.516182 | -2.215 |
| chr1 | 203265659 | 203265882 | 1.02155 | 0.461191 | -2.215 |
| chr20 | 8040725 | 8040929 | 1.12589 | 0.508296 | -2.215 |
| chr2 | 22761969 | 22762260 | 1.63966 | 0.740271 | -2.215 |
| chr15 | 67679565 | 67679751 | 1.11884 | 0.505152 | -2.215 |
| chr7 | 112110166 | 112110513 | 0.820225 | 0.370381 | -2.215 |
| chr3 | 85914209 | 85914414 | 1.41072 | 0.637053 | -2.214 |
| chr9 | 108160185 | 108160478 | 1.18488 | 0.535077 | -2.214 |
| chr8 | 88836068 | 88836253 | 1.05137 | 0.474946 | -2.214 |
| chr11 | 79111830 | 79112104 | 2.91887 | 1.31858 | -2.214 |
| chr2 | 125520261 | 125520482 | 1.02955 | 0.465235 | -2.213 |
| chr2 | 74365065 | 74365288 | 1.03118 | 0.465984 | -2.213 |
| chr2 | 44995738 | 44995960 | 1.12349 | 0.507755 | -2.213 |
| chr2 | 29552857 | 29553221 | 1.26636 | 0.572325 | -2.213 |
| chr19 | 42290526 | 42290735 | 0.916667 | 0.414322 | -2.212 |
| chr2 | 34777188 | 34777617 | 0.883152 | 0.399178 | -2.212 |
| chr17 | 63326470 | 63326652 | 1.13203 | 0.511734 | -2.212 |
| chr15 | 27974610 | 27974888 | 0.994035 | 0.449358 | -2.212 |
| chr2 | 52938584 | 52938787 | 1.19692 | 0.541101 | -2.212 |
| chr15 | 82207451 | 82207917 | 0.834371 | 0.377248 | -2.212 |
| chr3 | 76972583 | 76972792 | 1.14067 | 0.515744 | -2.212 |
| chr7 | 22264218 | 22264420 | 1.50398 | 0.680185 | -2.211 |
| chr8 | 100234055 | 100234261 | 1.04166 | 0.471173 | -2.211 |
| chr2 | 79247304 | 79247615 | 1.31563 | 0.595216 | -2.210 |
| chr9 | 16361941 | 16362133 | 1.02838 | 0.465281 | -2.210 |
| chr2 | 81205116 | 81205351 | 1.06604 | 0.482416 | -2.210 |
| chr2 | 129590715 | 129590966 | 1.11614 | 0.505131 | -2.210 |
| chr9 | 31320037 | 31320235 | 1.2665 | 0.57318 | -2.210 |
| chr2 | 173353625 | 173353895 | 0.958019 | 0.433614 | -2.209 |
| chr15 | 51471363 | 51471653 | 1.12906 | 0.51119 | -2.209 |
| chr8 | 87096021 | 87096285 | 1.15838 | 0.524473 | -2.209 |
| chr19 | 53619900 | 53620247 | 1.03746 | 0.469763 | -2.208 |
| chr9 | 25233951 | 25234178 | 0.865344 | 0.391829 | -2.208 |
| chr19 | 29904013 | 29904221 | 1.45473 | 0.658706 | -2.208 |
| chr13 | 94282521 | 94282745 | 0.888299 | 0.402268 | -2.208 |
| chr9 | 30497805 | 30498038 | 1.29564 | 0.586757 | -2.208 |
| chr1 | 112592700 | 112592966 | 0.878043 | 0.397665 | -2.208 |
| chr20 | 40644961 | 40645188 | 0.905401 | 0.410109 | -2.208 |
| chr15 | 94573354 | 94573702 | 1.19068 | 0.53935 | -2.208 |
| chr5 | 165744718 | 165744931 | 1.29388 | 0.586217 | -2.207 |
| chr10 | 114027335 | 114027598 | 1.25311 | 0.567882 | -2.207 |
| chr5 | 158152390 | 158152577 | 1.09388 | 0.495765 | -2.206 |
| chr19 | 45434677 | 45434880 | 0.893107 | 0.404783 | -2.206 |
| chr19 | 53832231 | 53832455 | 1.08177 | 0.490321 | -2.206 |
| chr1 | 241374734 | 241374941 | 1.05613 | 0.478728 | -2.206 |
| chr20 | 24587339 | 24587761 | 0.960761 | 0.435631 | -2.205 |
| chr2 | 155791473 | 155791715 | 1.38836 | 0.629557 | -2.205 |
| chr3 | 74000482 | 74000669 | 1.10217 | 0.499828 | -2.205 |
| chr7 | 33886073 | 33886380 | 1.01889 | 0.462162 | -2.205 |
| chr20 | 51396415 | 51396733 | 0.921239 | 0.417916 | -2.204 |
| chr2 | 79286217 | 79286418 | 1.14351 | 0.518826 | -2.204 |
| chr2 | 59619873 | 59620078 | 1.21096 | 0.549597 | -2.203 |
| chr8 | 2220806 | 2221196 | 1.08435 | 0.492183 | -2.203 |
| chr17 | 69639899 | 69640229 | 1.24242 | 0.563955 | -2.203 |
| chr5 | 147838018 | 147838389 | 1.23703 | 0.561522 | -2.203 |
| chr2 | 180389800 | 180390142 | 0.734196 | 0.33338 | -2.202 |
| chr9 | 1156987 | 1157324 | 0.892074 | 0.405177 | -2.202 |
| chr9 | 100080802 | 100081044 | 0.898179 | 0.408053 | -2.201 |
| chr16 | 48957492 | 48957681 | 1.29558 | 0.588618 | -2.201 |
| chr20 | 19267814 | 19268009 | 1.64811 | 0.749029 | -2.200 |
| chr8 | 136632911 | 136633120 | 1.31271 | 0.596631 | -2.200 |

| | | | | | |
|---|---|---|---|---|---|
| chr2 | 14271343 | 14271572 | 0.968507 | 0.440278 | -2.200 |
| chr3 | 160788035 | 160788232 | 0.990495 | 0.450318 | -2.200 |
| chr16 | 55580191 | 55580418 | 0.750281 | 0.341117 | -2.199 |
| chr2 | 186863872 | 186864142 | 0.999473 | 0.454471 | -2.199 |
| chr20 | 43502138 | 43502336 | 0.842841 | 0.3833 | -2.199 |
| chr4 | 27846515 | 27846804 | 0.934437 | 0.424969 | -2.199 |
| chr15 | 31479275 | 31479506 | 1.3484 | 0.613326 | -2.199 |
| chr19 | 57193100 | 57193305 | 1.52543 | 0.693993 | -2.198 |
| chr2 | 60317249 | 60317504 | 1.05698 | 0.480943 | -2.198 |
| chr11 | 79172222 | 79172449 | 1.28807 | 0.586269 | -2.197 |
| chr2 | 83647912 | 83648112 | 1.09037 | 0.496298 | -2.197 |
| chr12 | 44481319 | 44481533 | 1.05647 | 0.481044 | -2.196 |
| chr2 | 159172139 | 159172433 | 1.07542 | 0.489716 | -2.196 |
| chr2 | 9029164 | 9029413 | 1.3313 | 0.606248 | -2.196 |
| chr11 | 84123669 | 84123910 | 1.20041 | 0.546664 | -2.196 |
| chr16 | 55338190 | 55338412 | 1.00801 | 0.459088 | -2.196 |
| chr2 | 81027627 | 81027913 | 1.48951 | 0.678519 | -2.195 |
| chr17 | 78195408 | 78195805 | 1.33138 | 0.606521 | -2.195 |
| chr7 | 84753993 | 84754354 | 1.15811 | 0.52764 | -2.195 |
| chr20 | 16325646 | 16325848 | 1.45463 | 0.662935 | -2.194 |
| chr7 | 8900896 | 8901127 | 1.33573 | 0.608915 | -2.194 |
| chr15 | 78724327 | 78724652 | 1.11324 | 0.507533 | -2.193 |
| chr8 | 77443460 | 77443674 | 0.705831 | 0.321812 | -2.193 |
| chr6 | 168351293 | 168351498 | 0.760907 | 0.347037 | -2.193 |
| chr10 | 97663033 | 97663245 | 1.11076 | 0.506637 | -2.192 |
| chr8 | 129786139 | 129786324 | 0.945948 | 0.431529 | -2.192 |
| chr8 | 79109572 | 79109754 | 0.787144 | 0.359146 | -2.192 |
| chr5 | 97494144 | 97494438 | 1.19261 | 0.544168 | -2.192 |
| chr2 | 184942443 | 184942629 | 0.96452 | 0.44014 | -2.191 |
| chr17 | 20466146 | 20466378 | 0.869005 | 0.396565 | -2.191 |
| chr2 | 167923825 | 167924058 | 1.01927 | 0.465194 | -2.191 |
| chr1 | 188731049 | 188731444 | 0.972255 | 0.443762 | -2.191 |
| chr8 | 121583665 | 121583900 | 0.899088 | 0.410424 | -2.191 |
| chr5 | 26870911 | 26871265 | 1.07399 | 0.490324 | -2.190 |
| chr5 | 6315766 | 6315952 | 0.894497 | 0.408383 | -2.190 |
| chr2 | 212934538 | 212934744 | 1.03852 | 0.474255 | -2.190 |
| chr19 | 30780351 | 30780533 | 1.51335 | 0.69112 | -2.190 |
| chr12 | 67855888 | 67856311 | 1.08554 | 0.495808 | -2.189 |
| chr9 | 107045289 | 107045479 | 0.826387 | 0.377443 | -2.189 |
| chr2 | 73715421 | 73715793 | 1.12826 | 0.515476 | -2.189 |
| chr10 | 2584119 | 2584384 | 0.809645 | 0.369967 | -2.188 |
| chr2 | 105276924 | 105277124 | 0.773891 | 0.353632 | -2.188 |
| chr7 | 130097489 | 130097674 | 0.747587 | 0.341938 | -2.186 |
| chr10 | 23249732 | 23250046 | 1.08428 | 0.495964 | -2.186 |
| gl000205_r | 115360 | 115601 | 1.58699 | 0.72604 | -2.186 |
| chr3 | 79881500 | 79881691 | 0.909141 | 0.415993 | -2.185 |
| chr17 | 32473390 | 32473851 | 1.99345 | 0.912341 | -2.185 |
| chr8 | 50804767 | 50804970 | 1.17255 | 0.536713 | -2.185 |
| chr15 | 76488132 | 76488339 | 0.867229 | 0.39699 | -2.185 |
| chr5 | 50431032 | 50431430 | 0.959768 | 0.439352 | -2.185 |
| chr10 | 54939351 | 54939539 | 0.977569 | 0.447504 | -2.184 |
| chr5 | 110644862 | 110645176 | 1.26644 | 0.579788 | -2.184 |
| chr8 | 132437012 | 132437238 | 1.10096 | 0.504032 | -2.184 |
| chr1 | 91118400 | 91118582 | 0.711334 | 0.32568 | -2.184 |
| chr11 | 22077241 | 22077671 | 1.14249 | 0.523167 | -2.184 |
| chr2 | 7179875 | 7180060 | 1.19537 | 0.547479 | -2.183 |
| chr14 | 22712761 | 22712982 | 1.2579 | 0.576143 | -2.183 |
| chr7 | 49281014 | 49281407 | 1.20815 | 0.553365 | -2.183 |
| chr7 | 32253117 | 32253312 | 1.10267 | 0.505091 | -2.183 |
| chr2 | 104579635 | 104579848 | 1.03184 | 0.472664 | -2.183 |
| chr9 | 90236185 | 90236404 | 1.30931 | 0.599785 | -2.183 |

| | | | | | |
|---|---|---|---|---|---|
| chr9 | 131704951 | 131705146 | 0.859165 | 0.393707 | -2.182 |
| chr12 | 125992896 | 125993101 | 1.35348 | 0.620235 | -2.182 |
| chr2 | 22254262 | 22254575 | 0.892275 | 0.408914 | -2.182 |
| chr7 | 113610892 | 113611137 | 1.10923 | 0.50842 | -2.182 |
| chr2 | 116990460 | 116990687 | 0.968652 | 0.444004 | -2.182 |
| chr2 | 82751945 | 82752202 | 0.906569 | 0.415587 | -2.181 |
| chr2 | 84425046 | 84425402 | 1.73917 | 0.797646 | -2.180 |
| chr1 | 25435905 | 25436090 | 0.893716 | 0.410007 | -2.180 |
| chr10 | 36116721 | 36116973 | 1.29563 | 0.594467 | -2.179 |
| chr17 | 69104634 | 69104891 | 1.32403 | 0.607509 | -2.179 |
| chr12 | 69988320 | 69988568 | 1.20387 | 0.552395 | -2.179 |
| chr19 | 52322290 | 52322501 | 0.953779 | 0.437677 | -2.179 |
| chr3 | 124630638 | 124630872 | 1.48296 | 0.680554 | -2.179 |
| chr8 | 82888830 | 82889070 | 1.15327 | 0.52927 | -2.179 |
| chr2 | 56758190 | 56758431 | 0.998825 | 0.458436 | -2.179 |
| chr2 | 35209498 | 35209758 | 1.36942 | 0.628544 | -2.179 |
| chr1 | 187742152 | 187742413 | 1.00444 | 0.461057 | -2.179 |
| chr8 | 56774937 | 56775138 | 0.983265 | 0.451686 | -2.177 |
| chr9 | 23557038 | 23557343 | 0.831579 | 0.382045 | -2.177 |
| chr12 | 60484227 | 60484411 | 1.24955 | 0.574228 | -2.176 |
| chr11 | 40708369 | 40708560 | 1.20906 | 0.555706 | -2.176 |
| chr12 | 43939294 | 43939523 | 1.29446 | 0.594988 | -2.176 |
| chr20 | 18327951 | 18328223 | 1.18015 | 0.542461 | -2.176 |
| chr8 | 42037286 | 42037636 | 0.98289 | 0.451946 | -2.175 |
| chr3 | 95429092 | 95429399 | 1.04248 | 0.479498 | -2.174 |
| chr1 | 167144722 | 167144914 | 0.949872 | 0.436943 | -2.174 |
| chr15 | 68076761 | 68076963 | 1.41139 | 0.649395 | -2.173 |
| chr10 | 21187969 | 21188411 | 1.10263 | 0.507738 | -2.172 |
| chr5 | 9705558 | 9705745 | 0.836447 | 0.385207 | -2.171 |
| chr3 | 162885214 | 162885434 | 0.913601 | 0.420944 | -2.170 |
| chr2 | 35754649 | 35754836 | 0.979721 | 0.451422 | -2.170 |
| chr17 | 71497508 | 71497727 | 1.05885 | 0.487907 | -2.170 |
| chr2 | 186130565 | 186130789 | 1.07018 | 0.493198 | -2.170 |
| chr6 | 58399492 | 58399814 | 1.05886 | 0.488043 | -2.170 |
| chr8 | 36522175 | 36522522 | 1.16739 | 0.538232 | -2.169 |
| chr15 | 95521993 | 95522346 | 1.1316 | 0.521895 | -2.168 |
| chr7 | 126828734 | 126828997 | 1.03161 | 0.475825 | -2.168 |
| chr2 | 66644005 | 66644223 | 1.47899 | 0.682299 | -2.168 |
| chr10 | 61364200 | 61364539 | 0.971301 | 0.44816 | -2.167 |
| chr9 | 27745058 | 27745289 | 0.849113 | 0.391805 | -2.167 |
| chr7 | 66916961 | 66917157 | 1.43033 | 0.66002 | -2.167 |
| chr5 | 22625967 | 22626207 | 0.983967 | 0.454089 | -2.167 |
| chr12 | 62257470 | 62257839 | 1.3327 | 0.61504 | -2.167 |
| chr2 | 8868173 | 8868414 | 1.07751 | 0.497298 | -2.167 |
| chr8 | 137123859 | 137124111 | 1.57808 | 0.728348 | -2.167 |
| chr9 | 80046151 | 80046459 | 0.866143 | 0.399773 | -2.167 |
| chr11 | 69514094 | 69514279 | 0.826839 | 0.381652 | -2.166 |
| chr15 | 56878954 | 56879169 | 1.30827 | 0.603933 | -2.166 |
| chr17 | 77780466 | 77780649 | 0.994205 | 0.459022 | -2.166 |
| chr16 | 8422205 | 8422424 | 1.17941 | 0.544634 | -2.166 |
| chr3 | 74606500 | 74606738 | 1.02781 | 0.474639 | -2.165 |
| chr16 | 56659398 | 56659643 | 1.331 | 0.614657 | -2.165 |
| chr16 | 67254275 | 67254531 | 0.816997 | 0.377341 | -2.165 |
| chr3 | 26522472 | 26522721 | 1.04266 | 0.481574 | -2.165 |
| chr10 | 9937527 | 9937723 | 1.07637 | 0.497258 | -2.165 |
| chr8 | 13728965 | 13729233 | 1.1185 | 0.516794 | -2.164 |
| chr12 | 23997922 | 23998240 | 1.04429 | 0.482557 | -2.164 |
| chr7 | 28980597 | 28980855 | 0.827716 | 0.38248 | -2.164 |
| chr3 | 89925319 | 89925564 | 0.976847 | 0.451462 | -2.164 |
| chr20 | 21748321 | 21748574 | 2.37124 | 1.09598 | -2.164 |
| chr5 | 83398070 | 83398496 | 0.967275 | 0.447102 | -2.163 |

| | | | | | |
|---|---|---|---|---|---|
| chr5 | 28941774 | 28942112 | 1.05036 | 0.485511 | -2.163 |
| chr3 | 21305389 | 21305653 | 1.11438 | 0.515139 | -2.163 |
| chr10 | 49953786 | 49954188 | 1.36152 | 0.629455 | -2.163 |
| chr5 | 129028189 | 129028487 | 1.20823 | 0.558603 | -2.163 |
| chr17 | 12343180 | 12343473 | 0.9748 | 0.450711 | -2.163 |
| chr3 | 82970526 | 82970725 | 0.835339 | 0.386266 | -2.163 |
| chr2 | 156215288 | 156215512 | 0.948753 | 0.438757 | -2.162 |
| chr11 | 85401571 | 85401859 | 0.871432 | 0.403009 | -2.162 |
| chr2 | 51471336 | 51471534 | 1.39046 | 0.643215 | -2.162 |
| chr2 | 38704009 | 38704386 | 1.15891 | 0.536153 | -2.162 |
| chr9 | 7292992 | 7293184 | 1.23922 | 0.573341 | -2.161 |
| chr10 | 68183427 | 68183660 | 0.916498 | 0.424059 | -2.161 |
| chr8 | 65133428 | 65133683 | 1.004 | 0.464566 | -2.161 |
| chr8 | 21608225 | 21608453 | 1.01753 | 0.470847 | -2.161 |
| chr5 | 115429138 | 115429325 | 1.33174 | 0.61633 | -2.161 |
| chr7 | 146069838 | 146070123 | 1.1068 | 0.512239 | -2.161 |
| chr9 | 113876174 | 113876369 | 1.0353 | 0.479202 | -2.160 |
| chr8 | 5729866 | 5730117 | 0.7852 | 0.363479 | -2.160 |
| chr3 | 56432883 | 56433146 | 0.885034 | 0.409772 | -2.160 |
| chr5 | 146126431 | 146126644 | 0.808798 | 0.374497 | -2.160 |
| chr7 | 71949437 | 71949640 | 1.12606 | 0.521401 | -2.160 |
| chr8 | 73388381 | 73388590 | 1.21704 | 0.563565 | -2.160 |
| chr3 | 163706258 | 163706531 | 0.788814 | 0.365451 | -2.158 |
| chr3 | 8157161 | 8157433 | 1.39606 | 0.646988 | -2.158 |
| chr1 | 181664693 | 181664919 | 1.3848 | 0.641795 | -2.158 |
| chr12 | 13607174 | 13607386 | 0.999117 | 0.46307 | -2.158 |
| chr11 | 27933170 | 27933381 | 1.15217 | 0.534042 | -2.157 |
| chr19 | 56175318 | 56175541 | 0.977792 | 0.453357 | -2.157 |
| chr3 | 180111818 | 180112096 | 0.71598 | 0.331985 | -2.157 |
| chr8 | 138195765 | 138196206 | 1.16841 | 0.541827 | -2.156 |
| chr14 | 94398970 | 94399185 | 0.867424 | 0.402337 | -2.156 |
| chr10 | 124533918 | 124534217 | 1.06121 | 0.492329 | -2.155 |
| chr3 | 115567270 | 115567490 | 1.14648 | 0.531895 | -2.155 |
| chr14 | 46968690 | 46968909 | 1.34748 | 0.625189 | -2.155 |
| chr16 | 59086544 | 59086753 | 1.34779 | 0.625433 | -2.155 |
| chr10 | 83663233 | 83663473 | 1.02211 | 0.474347 | -2.155 |
| chr2 | 59293934 | 59294426 | 1.11297 | 0.516522 | -2.155 |
| chr19 | 34157981 | 34158193 | 1.19745 | 0.555812 | -2.154 |
| chr17 | 50413998 | 50414263 | 1.2997 | 0.603311 | -2.154 |
| chr9 | 124925657 | 124925879 | 0.886232 | 0.411498 | -2.154 |
| chr2 | 67209708 | 67209902 | 0.95848 | 0.4452 | -2.153 |
| chr7 | 147579485 | 147579764 | 1.19442 | 0.554921 | -2.152 |
| chr9 | 8357419 | 8357673 | 1.0334 | 0.480178 | -2.152 |
| chr8 | 137007025 | 137007210 | 1.56911 | 0.729151 | -2.152 |
| chr7 | 78120484 | 78120730 | 0.899652 | 0.418158 | -2.151 |
| chr19 | 56552132 | 56552353 | 1.04093 | 0.483827 | -2.151 |
| chr19 | 58233619 | 58233854 | 0.935336 | 0.435018 | -2.150 |
| chr12 | 116760031 | 116760213 | 1.22075 | 0.567778 | -2.150 |
| chr8 | 51466470 | 51466837 | 0.994957 | 0.462826 | -2.150 |
| chr7 | 40804383 | 40804573 | 1.19078 | 0.553929 | -2.150 |
| chr11 | 41995243 | 41995434 | 1.29149 | 0.601006 | -2.149 |
| chr2 | 12897638 | 12897847 | 1.12928 | 0.525528 | -2.149 |
| chr10 | 109644527 | 109644822 | 0.986602 | 0.459154 | -2.149 |
| chr1 | 209361978 | 209362162 | 1.11142 | 0.517255 | -2.149 |
| chr1 | 195486369 | 195486628 | 1.15655 | 0.538281 | -2.149 |
| chr10 | 111045651 | 111045968 | 1.03408 | 0.481336 | -2.148 |
| chr9 | 11947620 | 11947886 | 1.20879 | 0.562688 | -2.148 |
| chr5 | 30407683 | 30407871 | 0.938863 | 0.437154 | -2.148 |
| chr2 | 195395877 | 195396107 | 0.894626 | 0.416599 | -2.147 |
| chr2 | 97044202 | 97044662 | 0.809899 | 0.377157 | -2.147 |
| chr20 | 62704112 | 62704326 | 0.937008 | 0.43635 | -2.147 |

| | | | | | |
|---|---|---|---|---|---|
| chr7 | 92726450 | 92726668 | 0.921585 | 0.429188 | -2.147 |
| chr5 | 93424079 | 93424319 | 1.16723 | 0.543593 | -2.147 |
| chr19 | 38019708 | 38019952 | 0.976043 | 0.454642 | -2.147 |
| chr8 | 140992026 | 140992241 | 0.957923 | 0.446202 | -2.147 |
| chr2 | 638054 | 638244 | 1.10036 | 0.512604 | -2.147 |
| chr3 | 115385523 | 115385712 | 1.01629 | 0.473444 | -2.147 |
| chr19 | 31480514 | 31480720 | 1.51361 | 0.705283 | -2.146 |
| chr16 | 7659211 | 7659404 | 1.35379 | 0.63084 | -2.146 |
| chr2 | 167897422 | 167897647 | 1.29419 | 0.603094 | -2.146 |
| chr14 | 42742117 | 42742359 | 0.923721 | 0.430463 | -2.146 |
| chrX | 136276322 | 136276622 | 1.22615 | 0.571427 | -2.146 |
| chr5 | 29651696 | 29651910 | 1.1092 | 0.516932 | -2.146 |
| chr15 | 72897638 | 72897986 | 1.04655 | 0.487741 | -2.146 |
| chr7 | 146398236 | 146398530 | 1.03167 | 0.480816 | -2.146 |
| chr8 | 21943529 | 21943752 | 0.956411 | 0.445747 | -2.146 |
| chr20 | 3211441 | 3211680 | 0.967833 | 0.451242 | -2.145 |
| chr19 | 23333329 | 23333585 | 1.00441 | 0.468306 | -2.145 |
| chr3 | 3011992 | 3012265 | 0.852004 | 0.397293 | -2.145 |
| chr4 | 44115925 | 44116146 | 0.811252 | 0.37837 | -2.144 |
| chr2 | 292499 | 292718 | 1.14917 | 0.536076 | -2.144 |
| chr10 | 10210378 | 10210569 | 1.13628 | 0.530186 | -2.143 |
| chr1 | 248481741 | 248481965 | 0.846288 | 0.39493 | -2.143 |
| chr4 | 18332003 | 18332206 | 1.12358 | 0.524371 | -2.143 |
| chr3 | 95861233 | 95861624 | 1.04883 | 0.489542 | -2.142 |
| chr16 | 84208974 | 84209312 | 2.01896 | 0.942402 | -2.142 |
| chr2 | 77401433 | 77401617 | 1.01184 | 0.472304 | -2.142 |
| chr3 | 83022963 | 83023236 | 1.00912 | 0.471074 | -2.142 |
| chr10 | 1082499 | 1082722 | 1.05471 | 0.492381 | -2.142 |
| chr16 | 77506634 | 77506858 | 1.08472 | 0.506461 | -2.142 |
| chr19 | 33999907 | 34000172 | 0.838396 | 0.391533 | -2.141 |
| chr2 | 116042515 | 116042799 | 0.890607 | 0.415968 | -2.141 |
| chr19 | 52353504 | 52353931 | 1.27912 | 0.597468 | -2.141 |
| chr2 | 74415435 | 74415740 | 1.58506 | 0.740538 | -2.140 |
| chr2 | 51932498 | 51932680 | 0.973962 | 0.455055 | -2.140 |
| chr12 | 67984091 | 67984327 | 1.01372 | 0.473664 | -2.140 |
| chr8 | 106523333 | 106523518 | 0.837475 | 0.391322 | -2.140 |
| chr17 | 45857578 | 45857768 | 1.16903 | 0.546433 | -2.139 |
| chr8 | 50054892 | 50055103 | 0.990525 | 0.463002 | -2.139 |
| chr2 | 85943549 | 85943767 | 1.418 | 0.662938 | -2.139 |
| chr7 | 142420179 | 142420772 | 1.64337 | 0.768409 | -2.139 |
| chr7 | 95378979 | 95379216 | 0.947957 | 0.443357 | -2.138 |
| chr14 | 44435865 | 44436048 | 0.97591 | 0.456478 | -2.138 |
| chr2 | 13130759 | 13130942 | 1.29647 | 0.606614 | -2.137 |
| chr17 | 61527738 | 61527947 | 1.12972 | 0.528629 | -2.137 |
| chr2 | 120264390 | 120264581 | 0.832982 | 0.389787 | -2.137 |
| chr12 | 96478176 | 96478361 | 1.06954 | 0.500562 | -2.137 |
| chr8 | 83041867 | 83042117 | 1.33601 | 0.625324 | -2.137 |
| chr8 | 65586808 | 65586996 | 1.02282 | 0.47885 | -2.136 |
| chr7 | 47501161 | 47501449 | 0.935161 | 0.437815 | -2.136 |
| chr9 | 24728317 | 24728533 | 0.97677 | 0.457368 | -2.136 |
| chr2 | 62516323 | 62516516 | 1.07209 | 0.502148 | -2.135 |
| chr17 | 63399274 | 63399502 | 1.31925 | 0.61792 | -2.135 |
| chr16 | 61532516 | 61532795 | 1.12062 | 0.52491 | -2.135 |
| chr5 | 83820822 | 83821186 | 1.15429 | 0.540691 | -2.135 |
| chr19 | 46479567 | 46479854 | 1.50998 | 0.70768 | -2.134 |
| chr8 | 39015959 | 39016168 | 1.36745 | 0.640906 | -2.134 |
| chr10 | 77832422 | 77832710 | 1.2261 | 0.574763 | -2.133 |
| chr7 | 41476027 | 41476221 | 1.17641 | 0.551498 | -2.133 |
| chr10 | 63898047 | 63898251 | 1.00095 | 0.469262 | -2.133 |
| chr12 | 118248992 | 118249186 | 1.02532 | 0.480716 | -2.133 |
| chr2 | 53484443 | 53484807 | 1.1008 | 0.516161 | -2.133 |

| | | | | | |
|---|---|---|---|---|---|
| chr2 | 79448256 | 79448456 | 1.06912 | 0.501421 | -2.132 |
| chr1 | 216235151 | 216235401 | 0.953743 | 0.447338 | -2.132 |
| chr1 | 222585897 | 222586162 | 0.88513 | 0.415211 | -2.132 |
| chr9 | 32119625 | 32119860 | 1.13974 | 0.534678 | -2.132 |
| chr9 | 106679745 | 106680112 | 1.38472 | 0.64962 | -2.132 |
| chr7 | 54206230 | 54206428 | 0.868145 | 0.407286 | -2.132 |
| chr16 | 48848900 | 48849100 | 0.941219 | 0.441694 | -2.131 |
| chr1 | 237689690 | 237689877 | 1.02858 | 0.482799 | -2.130 |
| chr7 | 118658202 | 118658465 | 1.01487 | 0.47638 | -2.130 |
| chr7 | 119430392 | 119430593 | 0.903008 | 0.42398 | -2.130 |
| chr7 | 80811304 | 80811660 | 0.912218 | 0.428365 | -2.130 |
| chr7 | 52902497 | 52902711 | 1.03292 | 0.485066 | -2.129 |
| chr15 | 64033636 | 64033825 | 1.19956 | 0.563354 | -2.129 |
| chr3 | 83429012 | 83429212 | 1.0159 | 0.477195 | -2.129 |
| chr19 | 34984366 | 34984567 | 1.18027 | 0.554456 | -2.129 |
| chr2 | 59383085 | 59383315 | 1.18899 | 0.558695 | -2.128 |
| chr3 | 110429184 | 110429442 | 1.23875 | 0.582103 | -2.128 |
| chr11 | 24594733 | 24594917 | 0.984895 | 0.462834 | -2.128 |
| chr8 | 137479743 | 137480017 | 1.44484 | 0.679081 | -2.128 |
| chr2 | 117540123 | 117540311 | 1.3491 | 0.63423 | -2.127 |
| chr7 | 118152444 | 118152793 | 1.00831 | 0.474084 | -2.127 |
| chr17 | 56540214 | 56540405 | 1.64807 | 0.775316 | -2.126 |
| chr5 | 161488446 | 161488655 | 0.764724 | 0.359764 | -2.126 |
| chr2 | 116021959 | 116022167 | 1.02317 | 0.481374 | -2.126 |
| chr2 | 218389690 | 218389953 | 0.91305 | 0.429597 | -2.125 |
| chr5 | 62430517 | 62430780 | 0.972924 | 0.45777 | -2.125 |
| chr16 | 84083284 | 84083484 | 0.975139 | 0.458815 | -2.125 |
| chr2 | 15044020 | 15044272 | 1.95615 | 0.920461 | -2.125 |
| chr10 | 59040337 | 59040599 | 0.989754 | 0.465749 | -2.125 |
| chr2 | 172595222 | 172595442 | 1.10236 | 0.518835 | -2.125 |
| chr10 | 128943566 | 128943753 | 0.91031 | 0.428454 | -2.125 |
| chr12 | 128293332 | 128293571 | 1.28633 | 0.605464 | -2.125 |
| chr7 | 70285854 | 70286146 | 0.90907 | 0.427969 | -2.124 |
| chr17 | 71242366 | 71242661 | 1.10614 | 0.520749 | -2.124 |
| chr20 | 18386488 | 18386690 | 1.09951 | 0.517667 | -2.124 |
| chr1 | 166726293 | 166726564 | 0.997467 | 0.469647 | -2.124 |
| chr17 | 8403735 | 8404116 | 1.10057 | 0.518347 | -2.123 |
| chr12 | 56253711 | 56253958 | 0.813841 | 0.383311 | -2.123 |
| chr5 | 50850217 | 50850462 | 1.19846 | 0.564784 | -2.122 |
| chr15 | 75890808 | 75891001 | 1.00039 | 0.471451 | -2.122 |
| chr20 | 43912710 | 43912930 | 0.955794 | 0.450488 | -2.122 |
| chr2 | 240328273 | 240328479 | 1.24639 | 0.58746 | -2.122 |
| chr19 | 37680382 | 37680578 | 1.03327 | 0.48703 | -2.122 |
| chr9 | 86849778 | 86849973 | 1.51735 | 0.715314 | -2.121 |
| chr5 | 135760545 | 135760835 | 1.72646 | 0.813924 | -2.121 |
| chr5 | 132678827 | 132679052 | 0.982931 | 0.463399 | -2.121 |
| chr8 | 96618608 | 96618875 | 1.04763 | 0.493975 | -2.121 |
| chr19 | 23491831 | 23492050 | 0.99489 | 0.469145 | -2.121 |
| chr15 | 60054068 | 60054271 | 1.31083 | 0.618148 | -2.121 |
| chr7 | 115492735 | 115492967 | 0.920668 | 0.434238 | -2.120 |
| chr19 | 22108257 | 22108466 | 1.07697 | 0.508056 | -2.120 |
| chr5 | 178673269 | 178673464 | 1.01661 | 0.479698 | -2.119 |
| chr8 | 35302067 | 35302269 | 1.01348 | 0.478291 | -2.119 |
| chr5 | 116526944 | 116527195 | 0.902407 | 0.425915 | -2.119 |
| chr16 | 696713 | 696897 | 1.19407 | 0.563759 | -2.118 |
| chr8 | 133593431 | 133593677 | 1.02441 | 0.483667 | -2.118 |
| chr20 | 11853366 | 11853561 | 1.04167 | 0.491911 | -2.118 |
| chr20 | 21876875 | 21877064 | 0.898908 | 0.424522 | -2.117 |
| chr20 | 9755445 | 9755668 | 1.28469 | 0.606738 | -2.117 |
| chr5 | 23473752 | 23473970 | 1.00428 | 0.474327 | -2.117 |
| chr19 | 37405995 | 37406181 | 1.53422 | 0.724947 | -2.116 |

| | | | | | |
|---|---|---|---|---|---|
| chr3 | 154916044 | 154916273 | 1.01191 | 0.478174 | -2.116 |
| chr19 | 52256436 | 52256635 | 1.44918 | 0.684808 | -2.116 |
| chr5 | 135144708 | 135144934 | 1.00499 | 0.474956 | -2.116 |
| chr3 | 6246322 | 6246622 | 0.895572 | 0.423278 | -2.116 |
| chr3 | 147548702 | 147548965 | 1.32051 | 0.624137 | -2.116 |
| chr2 | 53446978 | 53447237 | 0.989238 | 0.467755 | -2.115 |
| chr14 | 47097363 | 47097674 | 1.30644 | 0.617754 | -2.115 |
| chr2 | 199503419 | 199503641 | 1.15548 | 0.546436 | -2.115 |
| chr8 | 127799526 | 127799807 | 1.21263 | 0.573479 | -2.115 |
| chr19 | 54418697 | 54419166 | 1.05146 | 0.497301 | -2.114 |
| chr2 | 118076305 | 118076543 | 1.06056 | 0.501667 | -2.114 |
| chr14 | 74141367 | 74141571 | 1.33783 | 0.632918 | -2.114 |
| chr3 | 28584985 | 28585252 | 1.13621 | 0.537593 | -2.114 |
| chr3 | 144236113 | 144236307 | 1.07124 | 0.506881 | -2.113 |
| chr9 | 38526680 | 38526924 | 1.08465 | 0.513241 | -2.113 |
| chr5 | 133798730 | 133799060 | 1.10658 | 0.523672 | -2.113 |
| chr8 | 118332741 | 118333027 | 0.788748 | 0.373288 | -2.113 |
| chr11 | 38112426 | 38112623 | 1.09683 | 0.519112 | -2.113 |
| chr8 | 133152885 | 133153172 | 1.94265 | 0.919622 | -2.112 |
| chr5 | 165749463 | 165749665 | 1.31901 | 0.624401 | -2.112 |
| chr15 | 54016467 | 54016699 | 1.08333 | 0.512997 | -2.112 |
| chr2 | 56817856 | 56818040 | 1.12066 | 0.530683 | -2.112 |
| chr8 | 125079574 | 125079935 | 1.03418 | 0.489742 | -2.112 |
| chr20 | 40270620 | 40270816 | 0.951477 | 0.450597 | -2.112 |
| chr20 | 12640195 | 12640416 | 1.00605 | 0.476451 | -2.112 |
| chr11 | 41989291 | 41989707 | 1.14481 | 0.542183 | -2.111 |
| chr2 | 59549977 | 59550234 | 1.32478 | 0.62753 | -2.111 |
| chr5 | 33032190 | 33032394 | 0.937247 | 0.444035 | -2.111 |
| chr1 | 119990955 | 119991181 | 0.777154 | 0.36824 | -2.110 |
| chr2 | 194028488 | 194028696 | 1.1368 | 0.538665 | -2.110 |
| chr7 | 14541344 | 14541593 | 1.12948 | 0.535258 | -2.110 |
| chr5 | 21838948 | 21839143 | 0.941238 | 0.446061 | -2.110 |
| chr10 | 53352219 | 53352401 | 1.05816 | 0.501473 | -2.110 |
| chr19 | 45375213 | 45375448 | 1.44914 | 0.686803 | -2.110 |
| chr2 | 518368 | 518611 | 0.987216 | 0.467879 | -2.110 |
| chr7 | 5402520 | 5402856 | 0.899378 | 0.426329 | -2.110 |
| chr2 | 17597867 | 17598049 | 1.17467 | 0.556852 | -2.109 |
| chr17 | 80651090 | 80651405 | 1.23609 | 0.586086 | -2.109 |
| chr19 | 39392335 | 39392541 | 0.893642 | 0.423907 | -2.108 |
| chr2 | 43437560 | 43437742 | 0.992973 | 0.471065 | -2.108 |
| chr9 | 82042153 | 82042363 | 1.11679 | 0.529977 | -2.107 |
| chr2 | 60492649 | 60492932 | 0.80635 | 0.38266 | -2.107 |
| chr10 | 87971552 | 87971759 | 1.04668 | 0.496773 | -2.107 |
| chr5 | 166477657 | 166477860 | 1.15164 | 0.546607 | -2.107 |
| chr5 | 152330440 | 152330671 | 0.803012 | 0.381168 | -2.107 |
| chr2 | 22876938 | 22877231 | 0.940948 | 0.446742 | -2.106 |
| chr17 | 27350825 | 27351051 | 1.04971 | 0.498423 | -2.106 |
| chr19 | 57430480 | 57430938 | 1.15842 | 0.550265 | -2.105 |
| chr8 | 80149639 | 80149821 | 0.896699 | 0.425947 | -2.105 |
| chr1 | 186158545 | 186158754 | 0.745401 | 0.35409 | -2.105 |
| chr3 | 83306771 | 83306985 | 1.40333 | 0.66664 | -2.105 |
| chr2 | 52868487 | 52868670 | 0.935932 | 0.444643 | -2.105 |
| chr11 | 21634452 | 21634643 | 0.845391 | 0.40164 | -2.105 |
| chr3 | 101316173 | 101316372 | 0.931978 | 0.442803 | -2.105 |
| chr7 | 118918611 | 118918801 | 0.821051 | 0.390123 | -2.105 |
| chr2 | 230253710 | 230253892 | 1.0028 | 0.476508 | -2.104 |
| chr15 | 82128651 | 82128845 | 1.34902 | 0.64107 | -2.104 |
| chr7 | 16616416 | 16616742 | 0.924276 | 0.439272 | -2.104 |
| chr17 | 49866339 | 49866639 | 1.06525 | 0.506315 | -2.104 |
| chr14 | 74814109 | 74814411 | 1.04125 | 0.49493 | -2.104 |
| chr20 | 54510000 | 54510346 | 1.06476 | 0.506217 | -2.103 |

| | | | | | |
|---|---|---|---|---|---|
| chr7 | 42567042 | 42567294 | 1.15406 | 0.548737 | -2.103 |
| chr8 | 125077748 | 125078003 | 1.04409 | 0.496502 | -2.103 |
| chr8 | 85311996 | 85312206 | 1.11739 | 0.531601 | -2.102 |
| chr2 | 564966 | 565203 | 0.81314 | 0.386933 | -2.102 |
| chr2 | 241742718 | 241742951 | 0.833399 | 0.396635 | -2.101 |
| chr8 | 88813506 | 88814029 | 0.990977 | 0.471693 | -2.101 |
| chr14 | 85246905 | 85247105 | 1.1917 | 0.567303 | -2.101 |
| chr18 | 23412070 | 23412366 | 1.86612 | 0.888446 | -2.100 |
| chr10 | 56338756 | 56338982 | 1.40374 | 0.668382 | -2.100 |
| chr7 | 39961196 | 39961424 | 1.27085 | 0.605364 | -2.099 |
| chr3 | 32861528 | 32861734 | 0.95037 | 0.452713 | -2.099 |
| chr17 | 19346360 | 19346654 | 1.52985 | 0.728813 | -2.099 |
| chr10 | 88289471 | 88289690 | 0.979249 | 0.466558 | -2.099 |
| chr8 | 83155140 | 83155373 | 0.909843 | 0.433582 | -2.098 |
| chr9 | 85377004 | 85377241 | 0.884471 | 0.421507 | -2.098 |
| chr2 | 44253286 | 44253750 | 1.13267 | 0.539848 | -2.098 |
| chr3 | 80832633 | 80832851 | 0.941499 | 0.448841 | -2.098 |
| chr8 | 65220125 | 65220334 | 1.10211 | 0.525452 | -2.097 |
| chr6 | 12623385 | 12623587 | 0.878491 | 0.418924 | -2.097 |
| chr8 | 60572470 | 60572666 | 1.06659 | 0.508649 | -2.097 |
| chr1 | 195021036 | 195021251 | 0.983897 | 0.469228 | -2.097 |
| chr2 | 212987787 | 212988107 | 0.863938 | 0.412071 | -2.097 |
| chr15 | 93247430 | 93247636 | 1.36318 | 0.650251 | -2.096 |
| chr2 | 53655789 | 53656114 | 0.99783 | 0.476033 | -2.096 |
| chr2 | 138540733 | 138540922 | 1.05438 | 0.503193 | -2.095 |
| chr11 | 37779506 | 37779718 | 1.20264 | 0.574025 | -2.095 |
| chr7 | 1019762 | 1019964 | 0.869235 | 0.414908 | -2.095 |
| chr8 | 138951653 | 138951843 | 1.13562 | 0.542071 | -2.095 |
| chr22 | 39889318 | 39889540 | 1.3643 | 0.65125 | -2.095 |
| chr16 | 62266119 | 62266453 | 1.0599 | 0.505959 | -2.095 |
| chr5 | 164090954 | 164091166 | 0.825435 | 0.394111 | -2.094 |
| chr8 | 102909927 | 102910117 | 0.951057 | 0.454136 | -2.094 |
| chr2 | 140946689 | 140946879 | 0.95913 | 0.457997 | -2.094 |
| chr8 | 134756588 | 134756791 | 0.941603 | 0.449648 | -2.094 |
| chr9 | 84809874 | 84810126 | 0.921084 | 0.439923 | -2.094 |
| chr12 | 28793879 | 28794102 | 1.47419 | 0.704397 | -2.093 |
| chr19 | 35830029 | 35830334 | 1.10313 | 0.527131 | -2.093 |
| chr2 | 4271226 | 4271593 | 1.30618 | 0.624175 | -2.093 |
| chr2 | 236062101 | 236062286 | 1.22091 | 0.583522 | -2.092 |
| chr5 | 2227553 | 2227743 | 0.914432 | 0.437049 | -2.092 |
| chr2 | 7816819 | 7817056 | 1.0146 | 0.484949 | -2.092 |
| chr2 | 6738012 | 6738226 | 1.06249 | 0.507841 | -2.092 |
| chr2 | 76074190 | 76074488 | 1.11621 | 0.533597 | -2.092 |
| chr8 | 55412282 | 55412593 | 1.38875 | 0.664167 | -2.091 |
| chr2 | 116036864 | 116037120 | 0.955768 | 0.457191 | -2.091 |
| chr17 | 67237594 | 67237819 | 1.0762 | 0.514802 | -2.091 |
| chr14 | 50647958 | 50648158 | 1.11115 | 0.531545 | -2.090 |
| chr8 | 69898773 | 69899059 | 1.13482 | 0.54292 | -2.090 |
| chr14 | 55084930 | 55085133 | 1.06941 | 0.511686 | -2.090 |
| chr8 | 135974594 | 135974797 | 0.897043 | 0.429268 | -2.090 |
| chr2 | 203015565 | 203015771 | 0.807758 | 0.38657 | -2.090 |
| chr3 | 100397457 | 100397937 | 0.956356 | 0.457688 | -2.090 |
| chr9 | 123537376 | 123537639 | 1.2109 | 0.579521 | -2.089 |
| chr7 | 154971030 | 154971239 | 0.915538 | 0.438209 | -2.089 |
| chr3 | 35485676 | 35485902 | 1.26139 | 0.603764 | -2.089 |
| chr3 | 159260182 | 159260408 | 1.13251 | 0.542124 | -2.089 |
| chr14 | 50824053 | 50824236 | 0.86953 | 0.416243 | -2.089 |
| chr2 | 51403251 | 51403553 | 1.40221 | 0.671258 | -2.089 |
| chr8 | 89835852 | 89836114 | 0.855926 | 0.40977 | -2.089 |
| chr5 | 98025266 | 98025468 | 0.874383 | 0.41866 | -2.089 |
| chr11 | 18226670 | 18226999 | 1.12822 | 0.540229 | -2.088 |

| | | | | | |
|---|---|---|---|---|---|
| chr5 | 98456757 | 98456985 | 0.993507 | 0.475747 | -2.088 |
| chr2 | 18553508 | 18553691 | 1.08923 | 0.521598 | -2.088 |
| chr16 | 82446359 | 82446575 | 1.10937 | 0.531543 | -2.087 |
| chr15 | 50922939 | 50923145 | 1.3638 | 0.653515 | -2.087 |
| chr8 | 80155713 | 80155967 | 0.805669 | 0.386129 | -2.087 |
| chr1 | 165098879 | 165099064 | 2.04356 | 0.979408 | -2.087 |
| chr10 | 124670904 | 124671116 | 0.899948 | 0.431323 | -2.086 |
| chr19 | 24059950 | 24060347 | 1.12772 | 0.540538 | -2.086 |
| chr14 | 38723603 | 38723804 | 0.67044 | 0.321392 | -2.086 |
| chr3 | 39511580 | 39511830 | 1.40299 | 0.672588 | -2.086 |
| chr2 | 212206309 | 212206944 | 1.01934 | 0.488673 | -2.086 |
| chr5 | 51299512 | 51299929 | 1.38907 | 0.666024 | -2.086 |
| chr7 | 54397793 | 54398202 | 0.968732 | 0.4645 | -2.086 |
| chr19 | 31504348 | 31504576 | 1.48856 | 0.713817 | -2.085 |
| chr5 | 12387111 | 12387335 | 1.03316 | 0.49546 | -2.085 |
| chr15 | 24936516 | 24936830 | 1.07399 | 0.515102 | -2.085 |
| chr9 | 74108293 | 74108510 | 1.02863 | 0.493575 | -2.084 |
| chr9 | 128919731 | 128919940 | 1.09196 | 0.524 | -2.084 |
| chr20 | 21721611 | 21721886 | 0.971364 | 0.466317 | -2.083 |
| chr9 | 8871961 | 8872242 | 0.881558 | 0.423237 | -2.083 |
| chr3 | 73702709 | 73702934 | 0.727693 | 0.349405 | -2.083 |
| chr15 | 94246933 | 94247168 | 1.07184 | 0.514756 | -2.082 |
| chr11 | 24832397 | 24832977 | 0.983082 | 0.472143 | -2.082 |
| chr11 | 23359882 | 23360111 | 1.07261 | 0.51523 | -2.082 |
| chr2 | 27844297 | 27844599 | 1.24843 | 0.599728 | -2.082 |
| chr4 | 36968424 | 36968656 | 0.727846 | 0.34974 | -2.081 |
| chr15 | 47763666 | 47763956 | 1.36741 | 0.657093 | -2.081 |
| chr10 | 18522882 | 18523101 | 1.04671 | 0.503016 | -2.081 |
| chr9 | 15958753 | 15959036 | 0.944261 | 0.453803 | -2.081 |
| chr2 | 58838264 | 58838558 | 1.14168 | 0.548787 | -2.080 |
| chr17 | 68951833 | 68952221 | 1.2826 | 0.61658 | -2.080 |
| chr1 | 219496696 | 219496890 | 0.993491 | 0.477696 | -2.080 |
| chr8 | 115161608 | 115161815 | 1.02281 | 0.491805 | -2.080 |
| chr16 | 17638748 | 17638976 | 1.05915 | 0.509285 | -2.080 |
| chr1 | 176660768 | 176661094 | 2.10424 | 1.01183 | -2.080 |
| chr10 | 109587956 | 109588218 | 0.98036 | 0.471493 | -2.079 |
| chr7 | 124957843 | 124958125 | 0.765348 | 0.368102 | -2.079 |
| chr10 | 82055071 | 82055292 | 1.08771 | 0.523175 | -2.079 |
| chr10 | 75409814 | 75410140 | 0.814804 | 0.391912 | -2.079 |
| chr2 | 7445805 | 7446037 | 1.10216 | 0.530231 | -2.079 |
| chr2 | 126752419 | 126752619 | 0.89189 | 0.429145 | -2.078 |
| chr8 | 110389082 | 110389270 | 0.984192 | 0.473649 | -2.078 |
| chr15 | 100921898 | 100922084 | 1.12515 | 0.541515 | -2.078 |
| chr11 | 38581745 | 38581947 | 1.07707 | 0.518391 | -2.078 |
| chr5 | 111250792 | 111250986 | 1.08095 | 0.520268 | -2.078 |
| chr2 | 66480482 | 66480664 | 0.690003 | 0.332129 | -2.078 |
| chr2 | 193553669 | 193553929 | 0.886592 | 0.426786 | -2.077 |
| chr2 | 13454382 | 13454593 | 0.858591 | 0.413361 | -2.077 |
| chr9 | 140370721 | 140370917 | 1.02049 | 0.491414 | -2.077 |
| chr11 | 31299860 | 31300063 | 1.1536 | 0.555519 | -2.077 |
| chr7 | 22752007 | 22752230 | 1.31049 | 0.63109 | -2.077 |
| chr2 | 85084875 | 85085065 | 1.08389 | 0.522005 | -2.076 |
| chr8 | 78884299 | 78884497 | 1.05634 | 0.508746 | -2.076 |
| chr16 | 61921406 | 61921639 | 0.818955 | 0.394508 | -2.076 |
| chr8 | 140765952 | 140766186 | 1.23368 | 0.594316 | -2.076 |
| chr11 | 29743042 | 29743233 | 1.15052 | 0.554275 | -2.076 |
| chr2 | 81795922 | 81796125 | 1.24171 | 0.598221 | -2.076 |
| chr11 | 22059050 | 22059269 | 1.20853 | 0.58228 | -2.076 |
| chr5 | 165316104 | 165316308 | 1.18584 | 0.57149 | -2.075 |
| chr9 | 114083764 | 114084027 | 1.01023 | 0.486858 | -2.075 |
| chr8 | 10183719 | 10184011 | 0.896764 | 0.432264 | -2.075 |

| | | | | | |
|---|---|---|---|---|---|
| chr2 | 83384855 | 83385070 | 1.17514 | 0.566486 | -2.074 |
| chr15 | 96088112 | 96088501 | 1.47441 | 0.711276 | -2.073 |
| chr9 | 37013625 | 37013818 | 0.634789 | 0.306253 | -2.073 |
| chr8 | 62753205 | 62753419 | 0.940066 | 0.453697 | -2.072 |
| chr2 | 80445376 | 80445641 | 1.00858 | 0.486802 | -2.072 |
| chr3 | 81192919 | 81193185 | 0.736038 | 0.355388 | -2.071 |
| chr7 | 115224101 | 115224318 | 0.98786 | 0.477025 | -2.071 |
| chr5 | 73906460 | 73906754 | 1.0381 | 0.501323 | -2.071 |
| chr10 | 120018894 | 120019077 | 2.34554 | 1.13283 | -2.071 |
| chr7 | 153049871 | 153050075 | 1.08516 | 0.524179 | -2.070 |
| chr3 | 37483859 | 37484365 | 0.940394 | 0.454257 | -2.070 |
| chr2 | 141083022 | 141083223 | 0.704791 | 0.34049 | -2.070 |
| chr1 | 191845686 | 191845885 | 0.965319 | 0.466357 | -2.070 |
| chr8 | 111335365 | 111335626 | 1.37019 | 0.66199 | -2.070 |
| chr1 | 225639585 | 225639801 | 1.70746 | 0.825095 | -2.069 |
| chr2 | 7909228 | 7909474 | 1.21508 | 0.587212 | -2.069 |
| chr19 | 45494501 | 45494764 | 0.981063 | 0.474211 | -2.069 |
| chr10 | 24486820 | 24487092 | 1.03401 | 0.499804 | -2.069 |
| chr8 | 144982846 | 144983089 | 1.55348 | 0.750958 | -2.069 |
| chr9 | 130200756 | 130200943 | 1.18009 | 0.570527 | -2.068 |
| chr8 | 75213686 | 75213932 | 1.07246 | 0.518509 | -2.068 |
| chr15 | 94938984 | 94939330 | 1.28641 | 0.622014 | -2.068 |
| chr1 | 152595546 | 152595958 | 1.43212 | 0.692519 | -2.068 |
| chr5 | 12416086 | 12416337 | 0.909173 | 0.439727 | -2.068 |
| chr6 | 127508170 | 127508359 | 2.37441 | 1.1484 | -2.068 |
| chr4 | 68329092 | 68329276 | 1.20494 | 0.582812 | -2.067 |
| chr14 | 46685203 | 46685469 | 1.41313 | 0.68353 | -2.067 |
| chr9 | 7171015 | 7171201 | 1.03414 | 0.500214 | -2.067 |
| chr20 | 34144453 | 34144774 | 0.892501 | 0.431708 | -2.067 |
| chr10 | 31376756 | 31376977 | 0.915474 | 0.442832 | -2.067 |
| chr5 | 66639447 | 66639743 | 1.12396 | 0.543756 | -2.067 |
| chr3 | 13257866 | 13258048 | 0.85624 | 0.414345 | -2.066 |
| chr5 | 160491738 | 160492045 | 1.0282 | 0.497585 | -2.066 |
| chr8 | 113275831 | 113276063 | 1.31826 | 0.637991 | -2.066 |
| chr16 | 11360585 | 11360871 | 1.04957 | 0.508006 | -2.066 |
| chr3 | 83559242 | 83559555 | 0.943496 | 0.456751 | -2.066 |
| chr2 | 16680407 | 16680658 | 0.908429 | 0.439969 | -2.065 |
| chr8 | 111605746 | 111605959 | 0.786983 | 0.381157 | -2.065 |
| chr15 | 100803879 | 100804086 | 1.21583 | 0.589122 | -2.064 |
| chr9 | 11059180 | 11059452 | 0.938258 | 0.454631 | -2.064 |
| chr8 | 136812760 | 136813078 | 0.972669 | 0.471409 | -2.063 |
| chr12 | 126937366 | 126937563 | 1.30052 | 0.630307 | -2.063 |
| chr2 | 52259156 | 52259473 | 1.10076 | 0.5338 | -2.062 |
| chr2 | 13640041 | 13640376 | 1.14921 | 0.557328 | -2.062 |
| chr7 | 118960456 | 118960707 | 1.2899 | 0.625662 | -2.062 |
| chr8 | 14420589 | 14420778 | 1.11448 | 0.540612 | -2.062 |
| chr1 | 202653695 | 202654139 | 1.15861 | 0.56204 | -2.061 |
| chr19 | 37423962 | 37424208 | 0.974518 | 0.472823 | -2.061 |
| chr2 | 192359672 | 192359857 | 1.00305 | 0.486675 | -2.061 |
| chr17 | 54066706 | 54067226 | 1.11858 | 0.54284 | -2.061 |
| chr20 | 20618781 | 20619060 | 1.21004 | 0.587329 | -2.060 |
| chr3 | 143574060 | 143574462 | 1.09595 | 0.531986 | -2.060 |
| chr8 | 59355293 | 59355482 | 1.19032 | 0.577841 | -2.060 |
| chr9 | 106363596 | 106363949 | 0.896035 | 0.435146 | -2.059 |
| chr14 | 27656697 | 27656924 | 0.892212 | 0.433467 | -2.058 |
| chr16 | 55285683 | 55285871 | 0.69996 | 0.340098 | -2.058 |
| chr16 | 52692538 | 52692751 | 1.04663 | 0.508732 | -2.057 |
| chr3 | 78073157 | 78073531 | 0.929133 | 0.451664 | -2.057 |
| chr2 | 19412563 | 19412898 | 1.1502 | 0.55913 | -2.057 |
| chr2 | 84156287 | 84156560 | 0.99094 | 0.481726 | -2.057 |
| chr2 | 185116238 | 185116493 | 1.21948 | 0.592855 | -2.057 |

| | | | | | |
|---|---|---|---|---|---|
| chr9 | 96956496 | 96956786 | 0.800697 | 0.389264 | -2.057 |
| chr15 | 97471270 | 97471453 | 1.21502 | 0.590767 | -2.057 |
| chr7 | 149269305 | 149269487 | 0.940329 | 0.457395 | -2.056 |
| chr3 | 120920561 | 120920848 | 1.34482 | 0.654307 | -2.055 |
| chr5 | 8314889 | 8315329 | 1.13519 | 0.552542 | -2.054 |
| chr1 | 154744474 | 154744711 | 0.956132 | 0.465391 | -2.054 |
| chr4 | 172807688 | 172807871 | 0.861096 | 0.419152 | -2.054 |
| chr5 | 8319512 | 8319708 | 1.18078 | 0.574798 | -2.054 |
| chr11 | 25709579 | 25709787 | 1.53567 | 0.74763 | -2.054 |
| chr2 | 127399217 | 127399416 | 0.780843 | 0.380223 | -2.054 |
| chr2 | 35834553 | 35834945 | 1.05152 | 0.512171 | -2.053 |
| chr5 | 98446192 | 98446419 | 1.54958 | 0.754801 | -2.053 |
| chr8 | 129308560 | 129308746 | 1.10876 | 0.540097 | -2.053 |
| chr3 | 21436228 | 21436497 | 1.1659 | 0.567966 | -2.053 |
| chr12 | 83924635 | 83925161 | 1.03343 | 0.503475 | -2.053 |
| chr7 | 128024024 | 128024396 | 1.05207 | 0.512606 | -2.052 |
| chr8 | 64874562 | 64874771 | 0.799549 | 0.389574 | -2.052 |
| chr5 | 27183063 | 27183254 | 1.29256 | 0.629846 | -2.052 |
| chr7 | 86235350 | 86235594 | 0.813572 | 0.396468 | -2.052 |
| chr8 | 139268734 | 139268967 | 1.0223 | 0.498191 | -2.052 |
| chrX | 142904943 | 142905139 | 1.056 | 0.514625 | -2.052 |
| chr8 | 14327284 | 14327515 | 1.20283 | 0.586183 | -2.052 |
| chr19 | 29635350 | 29635548 | 1.48346 | 0.722954 | -2.052 |
| chr1 | 231884672 | 231884997 | 0.897448 | 0.437374 | -2.052 |
| chr3 | 85115669 | 85115916 | 1.04477 | 0.509181 | -2.052 |
| chr2 | 690284 | 690471 | 1.30737 | 0.637168 | -2.052 |
| chr6 | 110361110 | 110361332 | 1.06257 | 0.517867 | -2.052 |
| chr9 | 29608368 | 29608572 | 1.32952 | 0.64799 | -2.052 |
| chr3 | 3758309 | 3758519 | 1.52127 | 0.741485 | -2.052 |
| chr15 | 70366598 | 70366818 | 1.07029 | 0.521737 | -2.051 |
| chr3 | 1369626 | 1369863 | 0.888256 | 0.433069 | -2.051 |
| chr12 | 76719940 | 76720136 | 1.13908 | 0.555493 | -2.051 |
| chr20 | 12373793 | 12373979 | 1.07028 | 0.521982 | -2.050 |
| chr2 | 155660967 | 155661261 | 0.849509 | 0.414329 | -2.050 |
| chr14 | 58295736 | 58296007 | 1.11118 | 0.542032 | -2.050 |
| chr6 | 72926143 | 72926356 | 1.02629 | 0.50067 | -2.050 |
| chr5 | 123625484 | 123625703 | 0.857186 | 0.418227 | -2.050 |
| chr15 | 90049468 | 90049658 | 1.34355 | 0.655556 | -2.049 |
| chr7 | 131671550 | 131671784 | 1.26924 | 0.619341 | -2.049 |
| chr16 | 64983586 | 64983792 | 0.954078 | 0.465699 | -2.049 |
| chr5 | 17881301 | 17881489 | 0.86554 | 0.422489 | -2.049 |
| chr8 | 3331526 | 3331726 | 1.13825 | 0.555656 | -2.048 |
| chr8 | 4350911 | 4351258 | 1.08221 | 0.528341 | -2.048 |
| chr15 | 87235416 | 87235747 | 0.969613 | 0.473434 | -2.048 |
| chr10 | 116242177 | 116242363 | 1.14255 | 0.557885 | -2.048 |
| chr8 | 107791176 | 107791401 | 0.973949 | 0.475604 | -2.048 |
| chr8 | 135034461 | 135034656 | 1.23741 | 0.604298 | -2.048 |
| chr9 | 26592090 | 26592345 | 1.22634 | 0.59916 | -2.047 |
| chr2 | 4320378 | 4320603 | 1.18441 | 0.578808 | -2.046 |
| chr2 | 166870410 | 166870835 | 0.777799 | 0.380143 | -2.046 |
| chr4 | 29414706 | 29414924 | 0.967213 | 0.472754 | -2.046 |
| chr17 | 22026589 | 22026861 | 0.971687 | 0.474942 | -2.046 |
| chr8 | 109866776 | 109867246 | 0.999016 | 0.488317 | -2.046 |
| chr12 | 97491532 | 97491826 | 1.51848 | 0.742262 | -2.046 |
| chr8 | 10979538 | 10979775 | 0.857802 | 0.419332 | -2.046 |
| chr2 | 17333520 | 17333720 | 0.905396 | 0.442632 | -2.045 |
| chr3 | 94599289 | 94599504 | 1.01754 | 0.49751 | -2.045 |
| chr5 | 88152415 | 88152759 | 1.11926 | 0.547257 | -2.045 |
| chr14 | 84118613 | 84118822 | 1.06719 | 0.521856 | -2.045 |
| chr19 | 28412190 | 28412430 | 1.23082 | 0.601929 | -2.045 |
| chr2 | 490345 | 490550 | 1.01594 | 0.49687 | -2.045 |

| | | | | | |
|---|---|---|---|---|---|
| chr7 | 105756493 | 105756763 | 0.82546 | 0.403742 | -2.045 |
| chr2 | 76981654 | 76981850 | 0.960697 | 0.46989 | -2.045 |
| chr10 | 4300274 | 4300497 | 1.21366 | 0.593802 | -2.044 |
| chr8 | 113550447 | 113550640 | 0.858274 | 0.419936 | -2.044 |
| chr8 | 77012127 | 77012310 | 0.875728 | 0.428512 | -2.044 |
| chr20 | 49151478 | 49151868 | 1.30932 | 0.640781 | -2.043 |
| chr5 | 162176247 | 162176522 | 1.12245 | 0.549346 | -2.043 |
| chr19 | 32366497 | 32366811 | 1.27448 | 0.623759 | -2.043 |
| chr1 | 234466725 | 234466917 | 1.24626 | 0.609988 | -2.043 |
| chr2 | 59663552 | 59663755 | 0.930117 | 0.455274 | -2.043 |
| chr12 | 115301792 | 115302031 | 1.43017 | 0.700314 | -2.042 |
| chr1 | 196187834 | 196188063 | 0.883219 | 0.432601 | -2.042 |
| chr2 | 71285151 | 71285400 | 0.74723 | 0.366159 | -2.041 |
| chr14 | 83195559 | 83195786 | 1.11833 | 0.548014 | -2.041 |
| chr1 | 34925979 | 34926171 | 1.22501 | 0.600362 | -2.040 |
| chr11 | 70187804 | 70187989 | 1.12215 | 0.549976 | -2.040 |
| chr16 | 25388523 | 25388802 | 1.04857 | 0.513975 | -2.040 |
| chr12 | 113941592 | 113941827 | 0.848284 | 0.41586 | -2.040 |
| chr8 | 31096541 | 31096727 | 0.945878 | 0.463761 | -2.040 |
| chr2 | 7818612 | 7818846 | 1.22887 | 0.602621 | -2.039 |
| chr16 | 4388380 | 4388720 | 1.12538 | 0.551885 | -2.039 |
| chr17 | 52322269 | 52322512 | 1.23247 | 0.604823 | -2.038 |
| chr9 | 9328730 | 9328986 | 1.04485 | 0.512775 | -2.038 |
| chr5 | 18857221 | 18857578 | 0.968543 | 0.475357 | -2.038 |
| chr20 | 38927651 | 38927843 | 1.19108 | 0.584592 | -2.037 |
| chr5 | 122583136 | 122583329 | 1.01544 | 0.498388 | -2.037 |
| chr8 | 136707941 | 136708144 | 0.981768 | 0.481862 | -2.037 |
| chr7 | 15465135 | 15465335 | 0.924365 | 0.453693 | -2.037 |
| chr7 | 56596385 | 56596623 | 1.07037 | 0.525496 | -2.037 |
| chr12 | 25673587 | 25673789 | 0.963881 | 0.473269 | -2.037 |
| chr3 | 152423590 | 152423801 | 1.15666 | 0.568175 | -2.036 |
| chr5 | 163154260 | 163154445 | 1.01837 | 0.500304 | -2.036 |
| chr19 | 53122996 | 53123213 | 0.907613 | 0.446042 | -2.035 |
| chr16 | 35107112 | 35107378 | 1.17472 | 0.577361 | -2.035 |
| chr5 | 153711675 | 153711861 | 1.26284 | 0.620713 | -2.035 |
| chr17 | 66486184 | 66486399 | 1.09358 | 0.537552 | -2.034 |
| chr3 | 2529807 | 2530063 | 0.87254 | 0.428946 | -2.034 |
| chr12 | 74556411 | 74556596 | 1.09984 | 0.540689 | -2.034 |
| chr9 | 71616133 | 71616323 | 0.993139 | 0.48825 | -2.034 |
| chr19 | 23611361 | 23611773 | 0.995463 | 0.489395 | -2.034 |
| chr8 | 89486241 | 89486477 | 1.1397 | 0.560342 | -2.034 |
| chr2 | 16998556 | 16998747 | 1.06561 | 0.524033 | -2.033 |
| chr12 | 80831549 | 80831754 | 1.33294 | 0.655521 | -2.033 |
| chr8 | 66707480 | 66707828 | 1.00674 | 0.495126 | -2.033 |
| chr8 | 117063502 | 117063886 | 0.974362 | 0.47922 | -2.033 |
| chr17 | 50035109 | 50035366 | 1.33186 | 0.655233 | -2.033 |
| chr5 | 104382297 | 104382574 | 1.05169 | 0.517465 | -2.032 |
| chr2 | 4229138 | 4229418 | 0.916125 | 0.450856 | -2.032 |
| chr14 | 86817742 | 86818051 | 1.09734 | 0.540096 | -2.032 |
| chr8 | 115307524 | 115307718 | 0.768345 | 0.378208 | -2.032 |
| chr19 | 38023195 | 38023481 | 1.19052 | 0.586048 | -2.031 |
| chr19 | 51213568 | 51213928 | 1.16869 | 0.575438 | -2.031 |
| chr3 | 31542391 | 31542719 | 1.07617 | 0.529895 | -2.031 |
| chr5 | 43815795 | 43816042 | 0.941984 | 0.463955 | -2.030 |
| chr5 | 85854504 | 85854743 | 1.44114 | 0.710021 | -2.030 |
| chr8 | 133047137 | 133047328 | 1.38267 | 0.681248 | -2.030 |
| chr1 | 233838679 | 233838880 | 1.10551 | 0.544702 | -2.030 |
| chr2 | 77944535 | 77944756 | 1.25798 | 0.619956 | -2.029 |
| chr19 | 58611576 | 58611806 | 0.988798 | 0.487345 | -2.029 |
| chr3 | 100882304 | 100882530 | 0.782165 | 0.385527 | -2.029 |
| chr7 | 228271 | 228454 | 0.866918 | 0.427398 | -2.028 |

| | | | | | |
|---|---|---|---|---|---|
| chr5 | 119771844 | 119772033 | 1.2058 | 0.594574 | -2.028 |
| chr3 | 84369251 | 84369460 | 1.06304 | 0.524241 | -2.028 |
| chr1 | 50737691 | 50737961 | 1.94498 | 0.959383 | -2.027 |
| chr2 | 124382368 | 124382631 | 0.990197 | 0.488426 | -2.027 |
| chr3 | 154659436 | 154659733 | 0.977228 | 0.482173 | -2.027 |
| chr8 | 80478373 | 80478684 | 1.28584 | 0.634586 | -2.026 |
| chr1 | 235675137 | 235675381 | 1.41434 | 0.698375 | -2.025 |
| chr5 | 84645532 | 84645931 | 1.05962 | 0.523251 | -2.025 |
| chr1 | 170206887 | 170207277 | 0.969188 | 0.478611 | -2.025 |
| chr15 | 94724238 | 94724549 | 1.0962 | 0.541425 | -2.025 |
| chr2 | 88494221 | 88494457 | 1.231 | 0.608107 | -2.024 |
| chr12 | 121102405 | 121102605 | 0.854434 | 0.422136 | -2.024 |
| chr9 | 106439940 | 106440242 | 0.905973 | 0.447634 | -2.024 |
| chr7 | 131475631 | 131475825 | 0.998249 | 0.493335 | -2.023 |
| chr20 | 9396379 | 9396619 | 1.39935 | 0.691579 | -2.023 |
| chr20 | 36532872 | 36533067 | 0.841828 | 0.416043 | -2.023 |
| chr19 | 28003306 | 28003562 | 0.98565 | 0.487224 | -2.023 |
| chr2 | 68572745 | 68572987 | 1.32804 | 0.656478 | -2.023 |
| chr2 | 167729145 | 167729415 | 0.910835 | 0.450297 | -2.023 |
| chr2 | 185244996 | 185245279 | 0.983263 | 0.486169 | -2.022 |
| chr19 | 29244698 | 29245197 | 1.22544 | 0.605965 | -2.022 |
| chr5 | 103381792 | 103382080 | 1.07332 | 0.530767 | -2.022 |
| chr17 | 50146387 | 50146826 | 1.16891 | 0.578044 | -2.022 |
| chr8 | 118511626 | 118511830 | 0.958144 | 0.473872 | -2.022 |
| chr17 | 30751118 | 30751351 | 1.19727 | 0.592144 | -2.022 |
| chr3 | 76773318 | 76773553 | 1.19774 | 0.592397 | -2.022 |
| chr3 | 145642812 | 145643041 | 1.11896 | 0.553481 | -2.022 |
| chr9 | 129858568 | 129858786 | 1.08931 | 0.538821 | -2.022 |
| chr2 | 84110454 | 84110640 | 0.96941 | 0.479524 | -2.022 |
| chr5 | 103820165 | 103820437 | 0.91648 | 0.453349 | -2.022 |
| chr14 | 102571764 | 102572207 | 1.13728 | 0.562607 | -2.021 |
| chr10 | 9004203 | 9004436 | 1.23684 | 0.611908 | -2.021 |
| chr3 | 33801235 | 33801644 | 0.968468 | 0.47917 | -2.021 |
| chr16 | 62148122 | 62148391 | 1.28244 | 0.634701 | -2.021 |
| chr14 | 49002792 | 49003083 | 0.799996 | 0.395989 | -2.020 |
| chr2 | 115742917 | 115743213 | 0.800005 | 0.396005 | -2.020 |
| chr7 | 125157704 | 125157932 | 0.957997 | 0.474228 | -2.020 |
| chr20 | 24878134 | 24878536 | 1.18959 | 0.588907 | -2.020 |
| chr19 | 33038661 | 33038846 | 1.32769 | 0.657301 | -2.020 |
| chr4 | 131269734 | 131269995 | 0.997674 | 0.493946 | -2.020 |
| chr7 | 139503763 | 139503964 | 1.18531 | 0.586962 | -2.019 |
| chr3 | 97232999 | 97233184 | 1.11382 | 0.551679 | -2.019 |
| chr5 | 108593950 | 108594137 | 1.24675 | 0.617596 | -2.019 |
| chr2 | 85349144 | 85349385 | 0.925379 | 0.458452 | -2.018 |
| chr9 | 137542147 | 137542572 | 1.14086 | 0.565239 | -2.018 |
| chr2 | 162873996 | 162874242 | 1.22759 | 0.608213 | -2.018 |
| chr8 | 136689671 | 136689866 | 1.351 | 0.669604 | -2.018 |
| chr9 | 90571016 | 90571292 | 0.760861 | 0.37717 | -2.017 |
| chr5 | 151330404 | 151330586 | 1.27104 | 0.630096 | -2.017 |
| chr9 | 110463274 | 110463468 | 0.912868 | 0.452625 | -2.017 |
| chr11 | 40152102 | 40152346 | 1.12851 | 0.559616 | -2.017 |
| chr5 | 116080933 | 116081116 | 1.11928 | 0.55509 | -2.016 |
| chr8 | 137554224 | 137554421 | 1.21815 | 0.604183 | -2.016 |
| chr14 | 43750700 | 43751104 | 1.16553 | 0.578098 | -2.016 |
| chr2 | 50594910 | 50595193 | 0.888593 | 0.440749 | -2.016 |
| chr9 | 123122106 | 123122345 | 0.939424 | 0.466139 | -2.015 |
| chr2 | 190945443 | 190945676 | 1.05341 | 0.522834 | -2.015 |
| chr2 | 35396531 | 35396788 | 0.964438 | 0.478754 | -2.014 |
| chr2 | 50767780 | 50768013 | 1.35055 | 0.670525 | -2.014 |
| chr14 | 27723284 | 27723568 | 1.037 | 0.514858 | -2.014 |
| chr5 | 8184644 | 8184886 | 2.16466 | 1.07478 | -2.014 |

| | | | | | |
|---|---|---|---|---|---|
| chr2 | 6017564 | 6017829 | 1.12573 | 0.55897 | -2.014 |
| chr2 | 147580679 | 147580923 | 0.894809 | 0.444336 | -2.014 |
| chr8 | 85465415 | 85465692 | 1.17217 | 0.582182 | -2.013 |
| chr3 | 18621570 | 18621793 | 1.04667 | 0.519946 | -2.013 |
| chr5 | 63929685 | 63930062 | 1.07746 | 0.535249 | -2.013 |
| chr4 | 132385817 | 132386012 | 0.916571 | 0.455474 | -2.012 |
| chr16 | 82490238 | 82490531 | 2.70649 | 1.34512 | -2.012 |
| chr10 | 64043901 | 64044189 | 1.14982 | 0.571513 | -2.012 |
| chr10 | 108414858 | 108415062 | 0.802982 | 0.399147 | -2.012 |
| chr2 | 140905682 | 140905881 | 1.23635 | 0.61464 | -2.012 |
| chr17 | 52224412 | 52224600 | 1.31345 | 0.653084 | -2.011 |
| chr5 | 22950053 | 22950287 | 1.07688 | 0.535492 | -2.011 |
| chr17 | 58037051 | 58037247 | 0.730694 | 0.363357 | -2.011 |
| chr2 | 237180776 | 237180960 | 1.07887 | 0.536615 | -2.011 |
| chr7 | 54277947 | 54278264 | 0.996674 | 0.495795 | -2.010 |
| chr7 | 125646084 | 125646274 | 0.859509 | 0.427604 | -2.010 |
| chr20 | 12391581 | 12391766 | 1.11266 | 0.553622 | -2.010 |
| chr12 | 100686543 | 100686748 | 0.704801 | 0.350785 | -2.009 |
| chr9 | 106189218 | 106189404 | 1.01311 | 0.504253 | -2.009 |
| chr2 | 210543464 | 210543648 | 0.627503 | 0.312363 | -2.009 |
| chr8 | 111664000 | 111664353 | 0.921573 | 0.458755 | -2.009 |
| chr9 | 106405035 | 106405279 | 0.87269 | 0.434432 | -2.009 |
| chr9 | 120150170 | 120150463 | 0.983859 | 0.48979 | -2.009 |
| chr2 | 144415516 | 144415709 | 1.15028 | 0.572644 | -2.009 |
| chr3 | 154404906 | 154405187 | 0.992347 | 0.494052 | -2.009 |
| chr1 | 162121914 | 162122314 | 0.891806 | 0.44402 | -2.008 |
| chr11 | 57948704 | 57948935 | 1.41982 | 0.706926 | -2.008 |
| chr19 | 42268288 | 42268492 | 1.16876 | 0.581963 | -2.008 |
| chr12 | 117972386 | 117972635 | 1.11879 | 0.557092 | -2.008 |
| chr3 | 144302323 | 144302630 | 0.940879 | 0.468508 | -2.008 |
| chr5 | 119386176 | 119386419 | 0.660545 | 0.328932 | -2.008 |
| chr9 | 77334081 | 77334320 | 1.27462 | 0.63474 | -2.008 |
| chr14 | 69766075 | 69766281 | 2.00313 | 0.997766 | -2.008 |
| chr16 | 6116642 | 6116837 | 1.37349 | 0.68417 | -2.008 |
| chr16 | 25761693 | 25761987 | 0.901599 | 0.449244 | -2.007 |
| chr3 | 88826285 | 88826509 | 1.14366 | 0.569911 | -2.007 |
| chr4 | 81331585 | 81331770 | 1.06645 | 0.531452 | -2.007 |
| chr9 | 97861583 | 97861791 | 1.24465 | 0.620341 | -2.006 |
| chr2 | 80210545 | 80210741 | 1.12014 | 0.558366 | -2.006 |
| chr19 | 57144751 | 57144999 | 1.20167 | 0.599051 | -2.006 |
| chr20 | 52894030 | 52894227 | 0.772024 | 0.384917 | -2.006 |
| chr2 | 177896338 | 177896533 | 0.703063 | 0.350553 | -2.006 |
| chr5 | 24140975 | 24141297 | 0.905315 | 0.451441 | -2.005 |
| chr12 | 29750406 | 29750625 | 1.18349 | 0.590195 | -2.005 |
| chr8 | 116079851 | 116080053 | 1.05715 | 0.527225 | -2.005 |
| chr2 | 156653383 | 156653593 | 1.00145 | 0.499477 | -2.005 |
| chr12 | 92735295 | 92735573 | 0.966197 | 0.481953 | -2.005 |
| chr3 | 4828237 | 4828419 | 0.971849 | 0.484793 | -2.005 |
| chr2 | 31034267 | 31034630 | 1.19233 | 0.594801 | -2.005 |
| chr5 | 112154797 | 112155000 | 1.221 | 0.609106 | -2.005 |
| chr15 | 88778473 | 88778765 | 1.22843 | 0.612835 | -2.005 |
| chr5 | 166058481 | 166058684 | 1.30769 | 0.652567 | -2.004 |
| chr5 | 166986264 | 166986461 | 1.16441 | 0.581068 | -2.004 |
| chr5 | 146457040 | 146457433 | 1.00324 | 0.500679 | -2.004 |
| chr2 | 82806970 | 82807193 | 1.39696 | 0.697214 | -2.004 |
| chr20 | 59203181 | 59203424 | 1.59538 | 0.796258 | -2.004 |
| chr10 | 56880638 | 56880862 | 1.50272 | 0.750147 | -2.003 |
| chr9 | 23463615 | 23463899 | 1.16314 | 0.580667 | -2.003 |
| chr5 | 117942762 | 117942960 | 1.03533 | 0.516889 | -2.003 |
| chr10 | 24367313 | 24367498 | 1.15948 | 0.578902 | -2.003 |
| chr2 | 148013689 | 148013907 | 0.848593 | 0.423689 | -2.003 |

| | | | | | |
|---|---|---|---|---|---|
| chr15 | 88734832 | 88735040 | 0.986217 | 0.492512 | -2.002 |
| chr5 | 15029957 | 15030139 | 1.31693 | 0.657735 | -2.002 |
| chr5 | 63417690 | 63417922 | 0.941427 | 0.470203 | -2.002 |
| chr8 | 79484148 | 79484409 | 0.747049 | 0.373188 | -2.002 |
| chr2 | 137017516 | 137017763 | 1.13904 | 0.569052 | -2.002 |
| chr4 | 81330900 | 81331227 | 0.842901 | 0.421129 | -2.002 |
| chr7 | 49440435 | 49440727 | 0.951797 | 0.475585 | -2.001 |
| chr8 | 78681613 | 78681879 | 0.877472 | 0.438456 | -2.001 |
| chr5 | 9372399 | 9372656 | 0.923268 | 0.461388 | -2.001 |
| chr14 | 86321548 | 86321894 | 0.925285 | 0.462443 | -2.001 |
| chr14 | 66276020 | 66276206 | 1.26301 | 0.631252 | -2.001 |
| chr3 | 21476212 | 21476403 | 1.0994 | 0.549481 | -2.001 |
| chr17 | 71277896 | 71278278 | 1.2335 | 0.616589 | -2.001 |
| chr1 | 164912995 | 164913291 | 0.862301 | 0.431053 | -2.000 |
| chr17 | 72216674 | 72216875 | 1.03581 | 0.517826 | -2.000 |
| chr9 | 98402364 | 98402562 | 1.00599 | 0.502949 | -2.000 |
| chr14 | 98095508 | 98095849 | 0.936449 | 0.468218 | -2.000 |
| chr8 | 87850101 | 87850288 | 1.3647 | 0.682368 | -2.000 |
| chr14 | 47157616 | 47157808 | 0.965591 | 0.48292 | -1.999 |
| chr3 | 5972713 | 5973045 | 1.20281 | 0.601563 | -1.999 |
| chr18 | 5425094 | 5425279 | 1.31531 | 0.657839 | -1.999 |
| chr7 | 156569738 | 156569940 | 0.991352 | 0.495873 | -1.999 |
| chr14 | 28159341 | 28159660 | 1.24099 | 0.620758 | -1.999 |
| chr19 | 23421472 | 23421816 | 1.00317 | 0.501807 | -1.999 |
| chr2 | 212723568 | 212723820 | 1.10167 | 0.551137 | -1.999 |
| chr17 | 10301700 | 10301902 | 1.0614 | 0.530992 | -1.999 |
| chr3 | 7420841 | 7421075 | 0.991327 | 0.495942 | -1.999 |
| chr7 | 46715343 | 46715622 | 0.958591 | 0.479581 | -1.999 |
| chr2 | 220560638 | 220560843 | 0.909486 | 0.455039 | -1.999 |
| chr8 | 143982915 | 143983135 | 1.12173 | 0.561268 | -1.999 |
| chr2 | 162775455 | 162775639 | 1.23981 | 0.620516 | -1.998 |
| chr5 | 131730010 | 131730287 | 1.29535 | 0.648342 | -1.998 |
| chr17 | 71458890 | 71459175 | 1.33759 | 0.669492 | -1.998 |
| chr3 | 143842463 | 143842658 | 1.05918 | 0.530184 | -1.998 |
| chr8 | 80923448 | 80923747 | 1.02982 | 0.515503 | -1.998 |
| chr7 | 39237026 | 39237286 | 1.06141 | 0.531328 | -1.998 |
| chr19 | 29124641 | 29124885 | 2.2777 | 1.1405 | -1.997 |
| chr2 | 29327432 | 29327654 | 1.02967 | 0.515656 | -1.997 |
| chr20 | 54199227 | 54199509 | 1.10112 | 0.551512 | -1.997 |
| chr8 | 115801888 | 115802078 | 0.988549 | 0.495198 | -1.996 |
| chr2 | 156770375 | 156770582 | 1.01874 | 0.510387 | -1.996 |
| chr10 | 83722354 | 83722653 | 0.851782 | 0.426787 | -1.996 |
| chr11 | 10980093 | 10980295 | 1.17226 | 0.587371 | -1.996 |
| chr2 | 35023741 | 35023967 | 1.02376 | 0.513125 | -1.995 |
| chr3 | 163949881 | 163950111 | 1.16082 | 0.581918 | -1.995 |
| chr9 | 9929322 | 9929687 | 0.959066 | 0.480877 | -1.994 |
| chr5 | 164454885 | 164455268 | 0.94139 | 0.472056 | -1.994 |
| chr8 | 9749302 | 9749512 | 0.920125 | 0.461509 | -1.994 |
| chr16 | 60497936 | 60498133 | 0.886888 | 0.44486 | -1.994 |
| chr1 | 176509068 | 176509311 | 0.824134 | 0.41341 | -1.994 |
| chr20 | 7176346 | 7176581 | 0.980179 | 0.491697 | -1.993 |
| chr12 | 66940027 | 66940239 | 1.033 | 0.518221 | -1.993 |
| chr5 | 4038012 | 4038201 | 1.24311 | 0.623628 | -1.993 |
| chr7 | 88739284 | 88739532 | 1.02966 | 0.516652 | -1.993 |
| chr11 | 22790709 | 22790995 | 1.15482 | 0.579655 | -1.992 |
| chr1 | 175744221 | 175744422 | 1.11581 | 0.560076 | -1.992 |
| chr2 | 163769971 | 163770273 | 1.05893 | 0.531594 | -1.992 |
| chr15 | 97413601 | 97413969 | 1.10179 | 0.553154 | -1.992 |
| chr10 | 72492127 | 72492395 | 0.954965 | 0.479533 | -1.991 |
| chr2 | 197679747 | 197679965 | 1.16967 | 0.587462 | -1.991 |
| chr14 | 64259478 | 64259714 | 1.1403 | 0.572912 | -1.990 |

| | | | | | |
|---|---|---|---|---|---|
| chr3 | 164459796 | 164459999 | 1.06837 | 0.536823 | -1.990 |
| chr2 | 1485783 | 1486251 | 1.03343 | 0.51932 | -1.990 |
| chr3 | 74414529 | 74414764 | 1.34573 | 0.676419 | -1.989 |
| chr10 | 120386507 | 120386779 | 1.74413 | 0.876718 | -1.989 |
| chr7 | 122692977 | 122693393 | 0.732181 | 0.368176 | -1.989 |
| chr3 | 7890875 | 7891223 | 1.14552 | 0.576047 | -1.989 |
| chr2 | 13832290 | 13832735 | 0.957118 | 0.481428 | -1.988 |
| chr7 | 141478790 | 141479072 | 1.0327 | 0.519446 | -1.988 |
| chr7 | 103926757 | 103926942 | 0.832536 | 0.41877 | -1.988 |
| chr3 | 6480746 | 6481045 | 1.06747 | 0.536956 | -1.988 |
| chr20 | 55037631 | 55037895 | 0.959267 | 0.482564 | -1.988 |
| chr7 | 112799919 | 112800128 | 1.52188 | 0.765611 | -1.988 |
| chr16 | 19067782 | 19067995 | 1.1518 | 0.579459 | -1.988 |
| chr12 | 126525086 | 126525277 | 1.04276 | 0.524663 | -1.987 |
| chr2 | 240601865 | 240602054 | 1.25706 | 0.632556 | -1.987 |
| chr5 | 45291608 | 45291957 | 1.23909 | 0.623531 | -1.987 |
| chr14 | 72490723 | 72491083 | 1.22911 | 0.618524 | -1.987 |
| chr10 | 84404770 | 84404998 | 0.909487 | 0.457682 | -1.987 |
| chr8 | 40957839 | 40958053 | 1.13414 | 0.570751 | -1.987 |
| chr11 | 39239478 | 39239701 | 0.843439 | 0.424482 | -1.987 |
| chr8 | 75684195 | 75684447 | 0.872891 | 0.439412 | -1.986 |
| chr8 | 24819299 | 24819677 | 1.18184 | 0.595024 | -1.986 |
| chr9 | 5040741 | 5040955 | 0.956826 | 0.481813 | -1.986 |
| chr7 | 125803663 | 125803942 | 1.19524 | 0.60198 | -1.986 |
| chr13 | 108748364 | 108748577 | 0.876829 | 0.441651 | -1.985 |
| chr5 | 117420271 | 117420592 | 0.977226 | 0.492234 | -1.985 |
| chr17 | 21208876 | 21209422 | 1.43471 | 0.722684 | -1.985 |
| chr14 | 36619109 | 36619552 | 1.07975 | 0.543892 | -1.985 |
| chr19 | 53674406 | 53674601 | 1.06263 | 0.535326 | -1.985 |
| chr3 | 98996898 | 98997102 | 1.10182 | 0.555084 | -1.985 |
| chr19 | 40709252 | 40709550 | 1.15329 | 0.581069 | -1.985 |
| chr1 | 202093130 | 202093405 | 1.10422 | 0.556375 | -1.985 |
| chr2 | 4605157 | 4605359 | 1.2197 | 0.614566 | -1.985 |
| chr5 | 31051972 | 31052213 | 0.859233 | 0.432972 | -1.985 |
| chr16 | 15049298 | 15049504 | 1.03273 | 0.520407 | -1.984 |
| chr8 | 137557305 | 137557492 | 1.12491 | 0.566911 | -1.984 |
| chr8 | 85571466 | 85571710 | 0.906257 | 0.456798 | -1.984 |
| chr17 | 63529067 | 63529265 | 1.46341 | 0.737675 | -1.984 |
| chr5 | 23336785 | 23337034 | 0.985316 | 0.496695 | -1.984 |
| chr8 | 136748826 | 136749029 | 0.868575 | 0.437846 | -1.984 |
| chr5 | 139090622 | 139090852 | 1.20851 | 0.609216 | -1.984 |
| chr1 | 237733917 | 237734134 | 1.28523 | 0.647914 | -1.984 |
| chr16 | 7862277 | 7862639 | 1.03654 | 0.522545 | -1.984 |
| chr5 | 44649698 | 44649913 | 1.22126 | 0.615697 | -1.984 |
| chr2 | 125865185 | 125865371 | 1.01227 | 0.510357 | -1.983 |
| chr11 | 80865745 | 80866056 | 1.40448 | 0.708574 | -1.982 |
| chr8 | 84280837 | 84281037 | 1.41483 | 0.713798 | -1.982 |
| chr19 | 44464720 | 44465003 | 1.15982 | 0.58521 | -1.982 |
| chr2 | 76091369 | 76091823 | 0.949321 | 0.479063 | -1.982 |
| chr8 | 132623507 | 132623827 | 1.0906 | 0.550448 | -1.981 |
| chr1 | 194756854 | 194757159 | 0.968633 | 0.488923 | -1.981 |
| chr8 | 103438676 | 103438924 | 0.872537 | 0.440465 | -1.981 |
| chr7 | 10124884 | 10125102 | 1.37666 | 0.695039 | -1.981 |
| chr2 | 184473624 | 184474029 | 0.912616 | 0.460775 | -1.981 |
| chr20 | 15794822 | 15795221 | 1.14754 | 0.579433 | -1.980 |
| chr20 | 16085697 | 16086081 | 1.24963 | 0.631159 | -1.980 |
| chr2 | 31231619 | 31231816 | 1.0714 | 0.541153 | -1.980 |
| chr1 | 176848004 | 176848244 | 0.889441 | 0.449263 | -1.980 |
| chr9 | 12213007 | 12213194 | 1.11508 | 0.563262 | -1.980 |
| chr3 | 80602438 | 80602650 | 1.21779 | 0.615217 | -1.979 |
| chr14 | 86409974 | 86410243 | 1.06203 | 0.536557 | -1.979 |

| | | | | | |
|---|---|---|---|---|---|
| chr17 | 75602934 | 75603252 | 1.65 | 0.833643 | -1.979 |
| chr10 | 28582820 | 28583005 | 1.04587 | 0.528414 | -1.979 |
| chr2 | 31602111 | 31602302 | 1.10775 | 0.559701 | -1.979 |
| chr7 | 28663525 | 28663741 | 1.68363 | 0.850732 | -1.979 |
| chr17 | 7277850 | 7278047 | 1.10231 | 0.557023 | -1.979 |
| chr2 | 227157128 | 227157345 | 0.793802 | 0.401272 | -1.978 |
| chr11 | 79219034 | 79219328 | 1.37945 | 0.697435 | -1.978 |
| chr8 | 52146166 | 52146376 | 1.36349 | 0.689491 | -1.978 |
| chr11 | 66930288 | 66930483 | 0.895375 | 0.452799 | -1.977 |
| chr3 | 88268276 | 88268645 | 1.16551 | 0.589462 | -1.977 |
| chr18 | 66947038 | 66947239 | 1.25461 | 0.634702 | -1.977 |
| chr2 | 77514743 | 77515119 | 0.979032 | 0.495418 | -1.976 |
| chr5 | 27551639 | 27551889 | 1.26218 | 0.638744 | -1.976 |
| chr14 | 39937738 | 39937967 | 0.992785 | 0.502457 | -1.976 |
| chrX | 131874688 | 131874906 | 1.56778 | 0.793477 | -1.976 |
| chr17 | 74983851 | 74984075 | 1.08234 | 0.547793 | -1.976 |
| chr1 | 166151160 | 166151391 | 1.09269 | 0.553051 | -1.976 |
| chr12 | 62387616 | 62387798 | 2.04785 | 1.03653 | -1.976 |
| chr2 | 82739372 | 82739638 | 0.777862 | 0.393745 | -1.976 |
| chr16 | 52262574 | 52262806 | 0.926906 | 0.469227 | -1.975 |
| chr19 | 54068362 | 54068683 | 0.823442 | 0.416878 | -1.975 |
| chr8 | 111556150 | 111556434 | 1.07928 | 0.546653 | -1.974 |
| chr3 | 94145065 | 94145331 | 0.960484 | 0.486485 | -1.974 |
| chr16 | 162424 | 162706 | 1.24375 | 0.630005 | -1.974 |
| chr9 | 73358494 | 73358677 | 1.5057 | 0.762934 | -1.974 |
| chr2 | 211950386 | 211950639 | 0.996455 | 0.50497 | -1.973 |
| gl000195_r | 94970 | 95155 | 1.16826 | 0.592053 | -1.973 |
| chr5 | 131680427 | 131680689 | 1.22292 | 0.619819 | -1.973 |
| chr15 | 25819157 | 25819397 | 0.963092 | 0.488155 | -1.973 |
| chr14 | 40637525 | 40637757 | 1.08529 | 0.550104 | -1.973 |
| chr3 | 90030725 | 90030920 | 1.06937 | 0.542061 | -1.973 |
| chr10 | 110688689 | 110689058 | 0.895913 | 0.454165 | -1.973 |
| chr9 | 7903201 | 7903508 | 1.07964 | 0.547313 | -1.973 |
| chr5 | 19228080 | 19228278 | 1.15067 | 0.583368 | -1.972 |
| chr12 | 79903526 | 79903723 | 1.01093 | 0.512541 | -1.972 |
| chr8 | 106113660 | 106113903 | 0.939801 | 0.4765 | -1.972 |
| chr8 | 142339542 | 142339739 | 1.10176 | 0.558664 | -1.972 |
| chr9 | 122707688 | 122707898 | 0.872366 | 0.442403 | -1.972 |
| chr17 | 73062760 | 73062987 | 0.998049 | 0.506194 | -1.972 |
| chr20 | 45801378 | 45801666 | 1.25332 | 0.635814 | -1.971 |
| chr16 | 82705409 | 82705599 | 1.398 | 0.709291 | -1.971 |
| chr2 | 124858163 | 124858398 | 0.770146 | 0.39075 | -1.971 |
| chr5 | 135727054 | 135727463 | 0.952917 | 0.483503 | -1.971 |
| chr12 | 109169529 | 109169729 | 1.02358 | 0.519366 | -1.971 |
| chr2 | 4506781 | 4507111 | 0.95838 | 0.486301 | -1.971 |
| chr14 | 30160521 | 30160817 | 1.14562 | 0.581421 | -1.970 |
| chr7 | 129964141 | 129964332 | 1.26597 | 0.642511 | -1.970 |
| chr2 | 51501804 | 51502103 | 1.01636 | 0.515889 | -1.970 |
| chr10 | 55875291 | 55875591 | 0.897839 | 0.455746 | -1.970 |
| chr4 | 12138778 | 12138964 | 1.00656 | 0.510966 | -1.970 |
| chr1 | 166326665 | 166327063 | 1.0701 | 0.543227 | -1.970 |
| chr3 | 145357584 | 145357870 | 1.29035 | 0.655055 | -1.970 |
| chr5 | 9898581 | 9898925 | 1.39651 | 0.708979 | -1.970 |
| chr6 | 154289949 | 154290133 | 0.885058 | 0.449368 | -1.970 |
| chr13 | 78815585 | 78815791 | 1.18695 | 0.602682 | -1.969 |
| chr1 | 154856320 | 154856516 | 1.15967 | 0.58884 | -1.969 |
| chr1 | 210701511 | 210701745 | 1.38885 | 0.705243 | -1.969 |
| chr12 | 33472835 | 33473019 | 1.23911 | 0.629355 | -1.969 |
| chr8 | 83707662 | 83707967 | 0.948084 | 0.481553 | -1.969 |
| chr2 | 151799486 | 151799676 | 1.27311 | 0.646653 | -1.969 |
| chr1 | 212956388 | 212956736 | 1.21051 | 0.614889 | -1.969 |

| | | | | | |
|---|---|---|---|---|---|
| chr8 | 2762856 | 2763060 | 1.13381 | 0.576088 | -1.968 |
| chr11 | 40205294 | 40205544 | 0.850956 | 0.432417 | -1.968 |
| chr16 | 50700637 | 50700819 | 1.01216 | 0.514333 | -1.968 |
| chr19 | 29436251 | 29436454 | 1.10918 | 0.563666 | -1.968 |
| chr7 | 49619034 | 49619281 | 1.02034 | 0.51852 | -1.968 |
| chr3 | 98284984 | 98285194 | 1.1901 | 0.604824 | -1.968 |
| chr15 | 100858645 | 100858896 | 1.00908 | 0.51286 | -1.968 |
| chr14 | 41741243 | 41741458 | 1.0884 | 0.553198 | -1.967 |
| chr5 | 78746594 | 78746848 | 0.922481 | 0.468868 | -1.967 |
| chr5 | 115081090 | 115081272 | 1.0166 | 0.516736 | -1.967 |
| chr19 | 58776929 | 58777147 | 1.37072 | 0.696816 | -1.967 |
| chr8 | 70535931 | 70536170 | 1.06242 | 0.54009 | -1.967 |
| chr8 | 13831675 | 13831868 | 0.806758 | 0.410189 | -1.967 |
| chr2 | 109674022 | 109674213 | 1.06423 | 0.541121 | -1.967 |
| chr9 | 139268019 | 139268227 | 0.817514 | 0.415686 | -1.967 |
| chr17 | 6454171 | 6454504 | 1.13042 | 0.5748 | -1.967 |
| chrX | 50490717 | 50490926 | 1.48772 | 0.756507 | -1.967 |
| chr15 | 31478311 | 31478547 | 1.05831 | 0.538192 | -1.966 |
| chr19 | 58654118 | 58654395 | 1.22871 | 0.624884 | -1.966 |
| chr12 | 130794052 | 130794240 | 0.924894 | 0.470415 | -1.966 |
| chr10 | 55874826 | 55875030 | 0.844082 | 0.429335 | -1.966 |
| chr5 | 147225305 | 147225576 | 1.22314 | 0.622253 | -1.966 |
| chr9 | 8913013 | 8913226 | 0.778021 | 0.395813 | -1.966 |
| chr5 | 164544137 | 164544482 | 1.13498 | 0.577429 | -1.966 |
| chr15 | 101429883 | 101430180 | 1.05809 | 0.538373 | -1.965 |
| chr14 | 34199662 | 34199873 | 1.19389 | 0.607486 | -1.965 |
| chr5 | 147175443 | 147175633 | 1.11824 | 0.569021 | -1.965 |
| chr15 | 101632759 | 101633208 | 1.01002 | 0.513964 | -1.965 |
| chr20 | 41731158 | 41731375 | 0.925673 | 0.471221 | -1.964 |
| chr1 | 163979494 | 163979801 | 1.18893 | 0.605273 | -1.964 |
| chr2 | 18152130 | 18152512 | 1.23196 | 0.627408 | -1.964 |
| chr12 | 85605789 | 85606038 | 1.00629 | 0.512563 | -1.963 |
| chr11 | 25654526 | 25654977 | 0.894467 | 0.455651 | -1.963 |
| chr9 | 104658128 | 104658389 | 1.00366 | 0.511278 | -1.963 |
| chr12 | 88612222 | 88612404 | 1.14723 | 0.584478 | -1.963 |
| chr17 | 6044252 | 6044482 | 1.26593 | 0.644967 | -1.963 |
| chr1 | 241303299 | 241303504 | 0.975347 | 0.496945 | -1.963 |
| chr3 | 74782426 | 74782645 | 0.869228 | 0.442902 | -1.963 |
| chr2 | 153977528 | 153977742 | 0.919908 | 0.468747 | -1.962 |
| chr10 | 124106778 | 124107016 | 1.1901 | 0.606432 | -1.962 |
| chr7 | 22821760 | 22821952 | 1.04847 | 0.534279 | -1.962 |
| chr12 | 114650739 | 114650962 | 1.11471 | 0.568106 | -1.962 |
| chr2 | 51604067 | 51604377 | 0.811608 | 0.413648 | -1.962 |
| chr2 | 62337302 | 62337546 | 1.07077 | 0.545886 | -1.962 |
| chr8 | 5631614 | 5631896 | 1.59322 | 0.812249 | -1.961 |
| chr2 | 165124507 | 165124716 | 0.81681 | 0.416449 | -1.961 |
| chr5 | 178523962 | 178524148 | 0.761951 | 0.388492 | -1.961 |
| chr5 | 44565520 | 44565895 | 1.14222 | 0.582404 | -1.961 |
| chr10 | 9440509 | 9440771 | 1.19938 | 0.611683 | -1.961 |
| chr2 | 157096087 | 157096293 | 1.06827 | 0.544826 | -1.961 |
| chr9 | 25683705 | 25683950 | 1.23437 | 0.629553 | -1.961 |
| chr2 | 12916909 | 12917118 | 1.29297 | 0.659552 | -1.960 |
| chr17 | 72477656 | 72477918 | 0.921855 | 0.470262 | -1.960 |
| chr9 | 90666281 | 90666643 | 1.08093 | 0.551496 | -1.960 |
| chr3 | 120555366 | 120555574 | 1.29117 | 0.658813 | -1.960 |
| chr1 | 109810476 | 109810682 | 1.29728 | 0.662073 | -1.959 |
| chr3 | 85173232 | 85173469 | 0.861339 | 0.439604 | -1.959 |
| chr20 | 53605734 | 53605968 | 1.09061 | 0.556621 | -1.959 |
| chr10 | 97696120 | 97696368 | 1.21596 | 0.620674 | -1.959 |
| chr16 | 61502741 | 61503165 | 1.11581 | 0.569587 | -1.959 |
| chr10 | 34306267 | 34306471 | 1.33668 | 0.682428 | -1.959 |

| | | | | | |
|---|---|---|---|---|---|
| chr11 | 76826340 | 76826782 | 1.57847 | 0.805996 | -1.958 |
| chr2 | 14333614 | 14333962 | 0.929686 | 0.474756 | -1.958 |
| chr9 | 133735963 | 133736239 | 0.935436 | 0.477707 | -1.958 |
| chr2 | 225048668 | 225048972 | 0.914773 | 0.467221 | -1.958 |
| chr5 | 163890133 | 163890329 | 1.19691 | 0.611367 | -1.958 |
| chrX | 108205319 | 108205521 | 0.925262 | 0.472654 | -1.958 |
| chr3 | 163060182 | 163060470 | 1.04991 | 0.536337 | -1.958 |
| chr11 | 16925399 | 16925790 | 1.03099 | 0.526761 | -1.957 |
| chr3 | 149942517 | 149942721 | 1.12723 | 0.576008 | -1.957 |
| chr2 | 126076301 | 126076719 | 0.999457 | 0.510731 | -1.957 |
| chr8 | 13212841 | 13213036 | 1.01449 | 0.518614 | -1.956 |
| chr10 | 10646055 | 10646256 | 1.09517 | 0.559882 | -1.956 |
| chr2 | 213289027 | 213289216 | 0.935384 | 0.478227 | -1.956 |
| chr11 | 85985608 | 85985797 | 2.2912 | 1.17145 | -1.956 |
| chr8 | 134766191 | 134766475 | 1.11238 | 0.568786 | -1.956 |
| chr8 | 112222217 | 112222506 | 0.870463 | 0.445191 | -1.955 |
| chr7 | 118877495 | 118877738 | 1.3528 | 0.691885 | -1.955 |
| chr5 | 88915568 | 88915808 | 1.03116 | 0.527478 | -1.955 |
| chr9 | 113256282 | 113256540 | 1.03912 | 0.531708 | -1.954 |
| chr9 | 29845869 | 29846163 | 1.10508 | 0.565543 | -1.954 |
| chr10 | 45905251 | 45905523 | 1.10861 | 0.567468 | -1.954 |
| chr3 | 3750140 | 3750370 | 1.15688 | 0.592177 | -1.954 |
| chr19 | 22998808 | 22998994 | 1.34772 | 0.689944 | -1.953 |
| chr17 | 60937951 | 60938150 | 1.15158 | 0.589548 | -1.953 |
| chr2 | 53678238 | 53678584 | 1.27293 | 0.651674 | -1.953 |
| chr12 | 5665297 | 5665497 | 1.7736 | 0.908055 | -1.953 |
| chr20 | 33405568 | 33405752 | 1.2248 | 0.627104 | -1.953 |
| chr2 | 16956164 | 16956438 | 1.24104 | 0.63545 | -1.953 |
| chr19 | 40626018 | 40626216 | 1.09877 | 0.562636 | -1.953 |
| chr11 | 40085517 | 40085743 | 0.712012 | 0.364634 | -1.953 |
| chr8 | 59035667 | 59035890 | 1.27933 | 0.655172 | -1.953 |
| chr5 | 99486702 | 99486932 | 0.840253 | 0.43035 | -1.952 |
| chr3 | 102731311 | 102731581 | 0.902629 | 0.462314 | -1.952 |
| chr7 | 118763413 | 118763624 | 0.878803 | 0.450127 | -1.952 |
| chr9 | 37384547 | 37384929 | 0.96721 | 0.495429 | -1.952 |
| chr1 | 169558216 | 169558705 | 0.98384 | 0.503975 | -1.952 |
| chr1 | 155976803 | 155977010 | 0.643799 | 0.32979 | -1.952 |
| chr9 | 120916342 | 120916694 | 1.08175 | 0.554142 | -1.952 |
| chr2 | 1263605 | 1263806 | 1.30773 | 0.66991 | -1.952 |
| chr7 | 145090417 | 145090610 | 1.12109 | 0.574301 | -1.952 |
| chr8 | 69856898 | 69857127 | 1.03048 | 0.527984 | -1.952 |
| chr2 | 34467872 | 34468103 | 0.941637 | 0.482541 | -1.951 |
| chr5 | 62995467 | 62995715 | 1.08245 | 0.55471 | -1.951 |
| chr7 | 53727046 | 53727244 | 1.28845 | 0.660276 | -1.951 |
| chr15 | 101392865 | 101393075 | 1.38224 | 0.708372 | -1.951 |
| chr8 | 50322767 | 50323033 | 1.03186 | 0.528836 | -1.951 |
| chr11 | 29769547 | 29769732 | 0.82217 | 0.421373 | -1.951 |
| chr2 | 180425185 | 180425371 | 1.07185 | 0.549343 | -1.951 |
| chr10 | 4968028 | 4968224 | 0.760806 | 0.38999 | -1.951 |
| chr12 | 22877203 | 22877466 | 1.0682 | 0.547573 | -1.951 |
| chr2 | 80683348 | 80683552 | 1.01863 | 0.522222 | -1.951 |
| chr5 | 169821130 | 169821342 | 0.961357 | 0.492873 | -1.951 |
| chr19 | 54240822 | 54241104 | 1.05641 | 0.541679 | -1.950 |
| chr19 | 40711665 | 40711870 | 1.10883 | 0.568581 | -1.950 |
| chr2 | 117910307 | 117910524 | 1.12237 | 0.57572 | -1.950 |
| chr5 | 161559384 | 161559586 | 0.99638 | 0.511116 | -1.949 |
| chr1 | 240847059 | 240847393 | 1.04659 | 0.536878 | -1.949 |
| chr2 | 84567820 | 84568061 | 0.921804 | 0.472878 | -1.949 |
| chr7 | 24444021 | 24444266 | 0.869479 | 0.446052 | -1.949 |
| chr8 | 55170851 | 55171033 | 0.783109 | 0.401758 | -1.949 |
| chr7 | 102456523 | 102456741 | 1.02891 | 0.527873 | -1.949 |

| | | | | | |
|---|---|---|---|---|---|
| chr8 | 82701021 | 82701232 | 1.27464 | 0.653962 | -1.949 |
| chr1 | 245724064 | 245724374 | 0.891797 | 0.457594 | -1.949 |
| chr5 | 25815339 | 25815645 | 1.08757 | 0.558248 | -1.948 |
| chr3 | 152376136 | 152376358 | 0.904041 | 0.464082 | -1.948 |
| chr7 | 139666834 | 139667023 | 0.795507 | 0.408411 | -1.948 |
| chr7 | 52623640 | 52623889 | 1.04142 | 0.534716 | -1.948 |
| chr1 | 208233771 | 208234196 | 0.932124 | 0.478601 | -1.948 |
| chr9 | 31768701 | 31768898 | 1.10438 | 0.567062 | -1.948 |
| chr10 | 109114542 | 109114777 | 1.10112 | 0.565426 | -1.947 |
| chr3 | 82823893 | 82824116 | 1.29415 | 0.664548 | -1.947 |
| chr18 | 63775485 | 63775715 | 1.0581 | 0.543399 | -1.947 |
| chr9 | 121317433 | 121317629 | 1.15572 | 0.593534 | -1.947 |
| chr2 | 6523060 | 6523311 | 1.1146 | 0.5725 | -1.947 |
| chr2 | 163279674 | 163279879 | 1.36394 | 0.700575 | -1.947 |
| chr12 | 82965194 | 82965465 | 1.08203 | 0.555889 | -1.946 |
| chr5 | 164348876 | 164349068 | 1.25753 | 0.646132 | -1.946 |
| chr20 | 21984670 | 21984867 | 1.24293 | 0.638642 | -1.946 |
| chr8 | 98230105 | 98230310 | 0.861339 | 0.442609 | -1.946 |
| chr2 | 24389981 | 24390188 | 1.02817 | 0.528346 | -1.946 |
| chr5 | 177594893 | 177595167 | 1.01452 | 0.521437 | -1.946 |
| chr15 | 93252848 | 93253088 | 1.04836 | 0.538925 | -1.945 |
| chr8 | 142146150 | 142146468 | 1.3726 | 0.705651 | -1.945 |
| chr3 | 168788880 | 168789076 | 0.86237 | 0.443375 | -1.945 |
| chr7 | 88354806 | 88355046 | 1.04789 | 0.538788 | -1.945 |
| chr9 | 88084066 | 88084357 | 1.5205 | 0.781793 | -1.945 |
| chr20 | 12154996 | 12155251 | 1.40526 | 0.722611 | -1.945 |
| chr5 | 27935009 | 27935214 | 1.13281 | 0.582556 | -1.945 |
| chr2 | 68896657 | 68897045 | 1.18257 | 0.608193 | -1.944 |
| chr10 | 19637865 | 19638055 | 1.21203 | 0.623359 | -1.944 |
| chr20 | 41424125 | 41424390 | 1.42246 | 0.731703 | -1.944 |
| chr2 | 186918612 | 186918925 | 0.82285 | 0.423307 | -1.944 |
| chr15 | 29909944 | 29910176 | 0.946359 | 0.486847 | -1.944 |
| chr9 | 1683684 | 1683917 | 0.94723 | 0.487334 | -1.944 |
| chr19 | 28338111 | 28338305 | 1.15733 | 0.595435 | -1.944 |
| chr10 | 10388928 | 10389193 | 1.0518 | 0.541195 | -1.943 |
| chr3 | 186312626 | 186312853 | 0.871948 | 0.448661 | -1.943 |
| chr5 | 147202164 | 147202631 | 2.8277 | 1.45508 | -1.943 |
| chr8 | 107946075 | 107946265 | 1.29992 | 0.668954 | -1.943 |
| chr2 | 49869552 | 49869749 | 1.26451 | 0.650786 | -1.943 |
| chr17 | 10934189 | 10934500 | 0.96529 | 0.496835 | -1.943 |
| chr2 | 81494787 | 81495034 | 1.08259 | 0.557288 | -1.943 |
| chr10 | 125522386 | 125522685 | 0.868936 | 0.447365 | -1.942 |
| chr8 | 145505860 | 145506103 | 1.19879 | 0.617258 | -1.942 |
| chr12 | 83725514 | 83725741 | 1.4687 | 0.756282 | -1.942 |
| chr2 | 4656054 | 4656723 | 1.12229 | 0.578076 | -1.941 |
| chr2 | 78223480 | 78223670 | 1.35554 | 0.698296 | -1.941 |
| chr3 | 83710550 | 83710746 | 1.54378 | 0.79539 | -1.941 |
| chr11 | 5668911 | 5669094 | 0.991145 | 0.510717 | -1.941 |
| chr7 | 147209204 | 147209443 | 1.16312 | 0.59935 | -1.941 |
| chr11 | 38394616 | 38394913 | 0.849896 | 0.438007 | -1.940 |
| chr2 | 13122663 | 13123009 | 0.986292 | 0.508445 | -1.940 |
| chr8 | 110518435 | 110518629 | 1.09301 | 0.563473 | -1.940 |
| chr2 | 79172102 | 79172285 | 1.13215 | 0.583733 | -1.939 |
| chr20 | 30055263 | 30055484 | 1.00558 | 0.518641 | -1.939 |
| chr5 | 54307769 | 54308055 | 0.829694 | 0.427965 | -1.939 |
| chr4 | 119259239 | 119259437 | 0.750751 | 0.387253 | -1.939 |
| chr2 | 4965953 | 4966150 | 1.03001 | 0.531302 | -1.939 |
| chr2 | 176052194 | 176052457 | 1.31304 | 0.677346 | -1.939 |
| chr2 | 81411439 | 81411633 | 0.990503 | 0.510975 | -1.938 |
| chr5 | 29387151 | 29387424 | 0.771162 | 0.397864 | -1.938 |
| chr2 | 79363031 | 79363218 | 1.117 | 0.57636 | -1.938 |

| | | | | | |
|---|---|---|---|---|---|
| chr15 | 96506351 | 96506574 | 1.02692 | 0.529883 | -1.938 |
| chr19 | 23493484 | 23493743 | 0.78467 | 0.404934 | -1.938 |
| chr3 | 155854408 | 155854593 | 0.857202 | 0.442366 | -1.938 |
| chr1 | 213840197 | 213840390 | 0.996869 | 0.51451 | -1.938 |
| chr1 | 191239721 | 191239943 | 0.997387 | 0.514828 | -1.937 |
| chr2 | 34024039 | 34024337 | 0.907722 | 0.468596 | -1.937 |
| chr8 | 37033179 | 37033389 | 1.04757 | 0.540825 | -1.937 |
| chr19 | 57136434 | 57136694 | 1.16184 | 0.599949 | -1.937 |
| chr2 | 185732200 | 185732412 | 0.797044 | 0.411678 | -1.936 |
| chr3 | 78792636 | 78792903 | 1.0444 | 0.539458 | -1.936 |
| chr9 | 31699217 | 31699412 | 0.99046 | 0.51164 | -1.936 |
| chr10 | 114996242 | 114996557 | 0.91281 | 0.471618 | -1.935 |
| chr5 | 63518354 | 63518655 | 1.03711 | 0.535882 | -1.935 |
| chr8 | 136807200 | 136807421 | 1.62589 | 0.840276 | -1.935 |
| chr1 | 191890426 | 191890735 | 1.28128 | 0.662261 | -1.935 |
| chr19 | 31739262 | 31739516 | 1.12381 | 0.580944 | -1.934 |
| chr3 | 80628734 | 80628939 | 1.29382 | 0.668887 | -1.934 |
| chr19 | 28661091 | 28661420 | 1.2835 | 0.663569 | -1.934 |
| chr10 | 132429419 | 132429606 | 1.20332 | 0.622128 | -1.934 |
| chr3 | 96874527 | 96874775 | 1.26859 | 0.655874 | -1.934 |
| chr17 | 17236701 | 17236975 | 0.90302 | 0.46696 | -1.934 |
| chr19 | 37679783 | 37680164 | 0.908831 | 0.470057 | -1.933 |
| chr5 | 82738286 | 82738621 | 1.02414 | 0.529722 | -1.933 |
| chr9 | 1941180 | 1941407 | 1.04063 | 0.538362 | -1.933 |
| chr17 | 31975057 | 31975322 | 2.08157 | 1.07693 | -1.933 |
| chr2 | 108083417 | 108083617 | 1.03225 | 0.534094 | -1.933 |
| chr17 | 41929221 | 41929480 | 1.08926 | 0.56362 | -1.933 |
| chr17 | 65693628 | 65693845 | 0.905722 | 0.468956 | -1.931 |
| chr9 | 81693927 | 81694112 | 0.95705 | 0.495576 | -1.931 |
| chr2 | 128127807 | 128128005 | 0.955455 | 0.494771 | -1.931 |
| chr2 | 81181487 | 81181768 | 1.14742 | 0.594352 | -1.931 |
| chr14 | 28627749 | 28627937 | 0.960193 | 0.497382 | -1.930 |
| chr17 | 44994795 | 44995018 | 1.36892 | 0.709113 | -1.930 |
| chr5 | 63282347 | 63282570 | 1.16518 | 0.603574 | -1.930 |
| chr2 | 34090566 | 34090778 | 1.29956 | 0.673185 | -1.930 |
| chr2 | 50552410 | 50552608 | 0.99714 | 0.516696 | -1.930 |
| chr2 | 167448800 | 167449106 | 1.12818 | 0.584638 | -1.930 |
| chr2 | 23129774 | 23130005 | 0.934875 | 0.484473 | -1.930 |
| chr8 | 91901908 | 91902116 | 1.19682 | 0.620239 | -1.930 |
| chr8 | 4096855 | 4097194 | 1.19029 | 0.616931 | -1.929 |
| chr5 | 43711726 | 43711915 | 1.03058 | 0.53421 | -1.929 |
| chr3 | 1733066 | 1733255 | 1.14876 | 0.595498 | -1.929 |
| chr12 | 48916211 | 48916456 | 1.1269 | 0.584186 | -1.929 |
| chr5 | 85235943 | 85236181 | 1.14434 | 0.593238 | -1.929 |
| chr10 | 27734042 | 27734253 | 1.29458 | 0.671265 | -1.929 |
| chr11 | 19561547 | 19561824 | 1.19849 | 0.621553 | -1.928 |
| chr20 | 38384608 | 38385243 | 0.973574 | 0.504917 | -1.928 |
| chr10 | 98204960 | 98205153 | 0.796623 | 0.413171 | -1.928 |
| chr8 | 97526630 | 97526827 | 0.850376 | 0.441107 | -1.928 |
| chr16 | 26279471 | 26279831 | 0.894197 | 0.463844 | -1.928 |
| chr10 | 31597314 | 31597508 | 1.33339 | 0.691698 | -1.928 |
| chr3 | 11965438 | 11965666 | 0.949356 | 0.492656 | -1.927 |
| chr9 | 77100609 | 77100793 | 0.81063 | 0.420693 | -1.927 |
| chr2 | 139748356 | 139748600 | 0.780744 | 0.405196 | -1.927 |
| chr12 | 30246028 | 30246214 | 1.08166 | 0.56138 | -1.927 |
| chr2 | 105229391 | 105229660 | 1.18377 | 0.614443 | -1.927 |
| chr17 | 1059420 | 1059689 | 0.848561 | 0.440455 | -1.927 |
| chr16 | 8388100 | 8388352 | 0.940973 | 0.488483 | -1.926 |
| chr11 | 41393372 | 41393633 | 1.09986 | 0.570967 | -1.926 |
| chr12 | 42918318 | 42918563 | 0.915764 | 0.475425 | -1.926 |
| chr5 | 111403184 | 111403515 | 0.995145 | 0.516641 | -1.926 |

| | | | | | |
|---|---|---|---|---|---|
| chr14 | 63430138 | 63430397 | 0.933455 | 0.484914 | -1.925 |
| chr2 | 207304721 | 207305033 | 1.20527 | 0.626165 | -1.925 |
| chr8 | 144221297 | 144221487 | 0.947819 | 0.492525 | -1.924 |
| chr2 | 193849005 | 193849370 | 0.829123 | 0.430857 | -1.924 |
| chr9 | 109133248 | 109133441 | 1.02045 | 0.530338 | -1.924 |
| chr5 | 74459147 | 74459330 | 0.982977 | 0.510925 | -1.924 |
| chr5 | 163023088 | 163023338 | 1.2156 | 0.631991 | -1.923 |
| chr19 | 41615829 | 41616108 | 0.94879 | 0.493343 | -1.923 |
| chr7 | 108843548 | 108843733 | 1.51378 | 0.787185 | -1.923 |
| chr1 | 192242988 | 192243445 | 0.967633 | 0.503332 | -1.922 |
| chr8 | 3389178 | 3389738 | 0.97626 | 0.507823 | -1.922 |
| chr10 | 59315341 | 59315830 | 0.850758 | 0.442592 | -1.922 |
| chr7 | 119315199 | 119315397 | 0.966579 | 0.50301 | -1.922 |
| chr12 | 6745556 | 6745760 | 0.984439 | 0.512336 | -1.921 |
| chr2 | 22040397 | 22040599 | 0.960487 | 0.500009 | -1.921 |
| chr2 | 22174914 | 22175149 | 0.896471 | 0.46673 | -1.921 |
| chr5 | 3137491 | 3137737 | 1.20662 | 0.628254 | -1.921 |
| chr5 | 136500718 | 136500977 | 0.989396 | 0.515178 | -1.920 |
| chr6 | 112501428 | 112501620 | 1.47816 | 0.769763 | -1.920 |
| chr15 | 84661588 | 84661773 | 1.06339 | 0.553786 | -1.920 |
| chr9 | 106088409 | 106088694 | 0.800155 | 0.416706 | -1.920 |
| chr14 | 46956828 | 46957068 | 1.0633 | 0.553767 | -1.920 |
| chr12 | 118939373 | 118939699 | 0.944033 | 0.491757 | -1.920 |
| chr4 | 71410108 | 71410323 | 1.77181 | 0.922989 | -1.920 |
| chr14 | 51646269 | 51646494 | 0.786679 | 0.409936 | -1.919 |
| chr19 | 35600696 | 35600969 | 1.30014 | 0.677611 | -1.919 |
| chr11 | 61986214 | 61986502 | 0.794873 | 0.414296 | -1.919 |
| chr14 | 56888792 | 56889018 | 0.842106 | 0.438929 | -1.919 |
| chr17 | 75377176 | 75377372 | 1.0832 | 0.564694 | -1.918 |
| chr2 | 50950733 | 50950923 | 1.23117 | 0.641925 | -1.918 |
| chr19 | 38109134 | 38109412 | 1.18544 | 0.618112 | -1.918 |
| chr19 | 36978355 | 36978571 | 1.14439 | 0.596736 | -1.918 |
| chr5 | 134504770 | 134504963 | 1.16266 | 0.606294 | -1.918 |
| chr10 | 71694209 | 71694789 | 0.939773 | 0.490123 | -1.917 |
| chr2 | 81724804 | 81724986 | 1.14926 | 0.599477 | -1.917 |
| chr5 | 30557790 | 30557994 | 1.18231 | 0.616745 | -1.917 |
| chr17 | 58235624 | 58235807 | 1.33937 | 0.698814 | -1.917 |
| chr6 | 97160813 | 97161119 | 1.03665 | 0.540943 | -1.916 |
| chr1 | 183295045 | 183295229 | 0.919397 | 0.479772 | -1.916 |
| chr10 | 28416782 | 28417049 | 0.958856 | 0.500477 | -1.916 |
| chr7 | 44501394 | 44501696 | 1.0385 | 0.542131 | -1.916 |
| chr2 | 50494964 | 50495225 | 1.04498 | 0.545515 | -1.916 |
| chr2 | 177943221 | 177943522 | 0.952168 | 0.497064 | -1.916 |
| chr7 | 49158436 | 49158626 | 1.14983 | 0.600308 | -1.915 |
| chr3 | 76717336 | 76717581 | 0.936571 | 0.488985 | -1.915 |
| chr14 | 89815887 | 89816080 | 1.01138 | 0.528057 | -1.915 |
| chr5 | 168385494 | 168385743 | 0.787277 | 0.411259 | -1.914 |
| chr3 | 14651023 | 14651218 | 1.05381 | 0.550563 | -1.914 |
| chr19 | 52608074 | 52608284 | 1.05141 | 0.549396 | -1.914 |
| chr2 | 20866645 | 20866965 | 1.53559 | 0.802421 | -1.914 |
| chr8 | 52991800 | 52992025 | 1.16053 | 0.606445 | -1.914 |
| chr5 | 51819916 | 51820110 | 0.790213 | 0.413038 | -1.913 |
| chr10 | 86723479 | 86723916 | 1.51993 | 0.794506 | -1.913 |
| chr20 | 302737 | 302931 | 1.03208 | 0.53951 | -1.913 |
| chr7 | 112001419 | 112001601 | 1.04056 | 0.54398 | -1.913 |
| chr2 | 34805520 | 34805776 | 1.04408 | 0.545867 | -1.913 |
| chr2 | 146346419 | 146346741 | 1.13188 | 0.591783 | -1.913 |
| chr2 | 104672913 | 104673104 | 0.977946 | 0.511327 | -1.913 |
| chr8 | 73738520 | 73738764 | 1.07805 | 0.563717 | -1.912 |
| chr15 | 80847485 | 80847730 | 0.934719 | 0.488829 | -1.912 |
| chr14 | 90400212 | 90400451 | 1.20394 | 0.629679 | -1.912 |

| | | | | | |
|---|---|---|---|---|---|
| chr3 | 80561787 | 80562002 | 1.6157 | 0.845159 | -1.912 |
| chr3 | 20511158 | 20511358 | 1.1045 | 0.57776 | -1.912 |
| chr8 | 113663796 | 113663986 | 1.09123 | 0.570819 | -1.912 |
| chr5 | 170951429 | 170951720 | 1.36789 | 0.715651 | -1.911 |
| chr5 | 134110366 | 134110557 | 0.860392 | 0.450162 | -1.911 |
| chr15 | 24995166 | 24995361 | 1.05371 | 0.551365 | -1.911 |
| chr2 | 77682061 | 77682300 | 1.23782 | 0.647782 | -1.911 |
| chr3 | 162271603 | 162271787 | 1.04584 | 0.547324 | -1.911 |
| chr2 | 382585 | 382959 | 0.983992 | 0.515131 | -1.910 |
| chr5 | 72097892 | 72098076 | 1.09347 | 0.572447 | -1.910 |
| chr2 | 49556597 | 49556828 | 1.29326 | 0.677179 | -1.910 |
| chr19 | 32127198 | 32127458 | 1.04764 | 0.548569 | -1.910 |
| chr8 | 140620557 | 140620801 | 1.17665 | 0.616175 | -1.910 |
| chr2 | 123868929 | 123869308 | 0.910686 | 0.476996 | -1.909 |
| chr8 | 135115296 | 135115533 | 1.23542 | 0.647105 | -1.909 |
| chr2 | 21853389 | 21853620 | 1.22855 | 0.643583 | -1.909 |
| chr1 | 234343628 | 234343895 | 1.2472 | 0.653363 | -1.909 |
| chr5 | 13331742 | 13331978 | 1.35779 | 0.711315 | -1.909 |
| chr16 | 85504574 | 85504756 | 0.821801 | 0.430537 | -1.909 |
| chr19 | 35551293 | 35551570 | 1.3253 | 0.694356 | -1.909 |
| chr5 | 150489634 | 150489851 | 1.05107 | 0.550699 | -1.909 |
| chr7 | 1648853 | 1649046 | 1.06209 | 0.556609 | -1.908 |
| chr10 | 70422437 | 70422628 | 1.1972 | 0.627452 | -1.908 |
| chr2 | 59212462 | 59212807 | 1.033 | 0.541535 | -1.908 |
| chr15 | 57973452 | 57973729 | 1.04436 | 0.547564 | -1.907 |
| chr11 | 90337994 | 90338194 | 1.02554 | 0.537699 | -1.907 |
| chr9 | 8579734 | 8579956 | 1.16033 | 0.60839 | -1.907 |
| chr2 | 184802104 | 184802358 | 1.27118 | 0.666515 | -1.907 |
| chr3 | 12788043 | 12788258 | 0.966709 | 0.506906 | -1.907 |
| chr10 | 66291253 | 66291439 | 0.808798 | 0.424213 | -1.907 |
| chr9 | 22875330 | 22875645 | 1.09324 | 0.573612 | -1.906 |
| chr19 | 58504872 | 58505084 | 0.931633 | 0.488871 | -1.906 |
| chr19 | 56962830 | 56963017 | 1.17431 | 0.616315 | -1.905 |
| chr14 | 86728173 | 86728431 | 0.946023 | 0.49651 | -1.905 |
| chr10 | 65806389 | 65806623 | 0.971177 | 0.509725 | -1.905 |
| chr5 | 16666772 | 16666999 | 1.09926 | 0.576987 | -1.905 |
| chr10 | 19008635 | 19008868 | 0.846545 | 0.444362 | -1.905 |
| chr8 | 17287172 | 17287366 | 0.937855 | 0.492309 | -1.905 |
| chr5 | 87820713 | 87820921 | 1.29527 | 0.680031 | -1.905 |
| chr8 | 2271355 | 2271558 | 1.51287 | 0.794368 | -1.904 |
| chr8 | 52300023 | 52300217 | 0.890484 | 0.46757 | -1.904 |
| chr2 | 169892302 | 169892607 | 0.951724 | 0.499732 | -1.904 |
| chr2 | 142095674 | 142095880 | 0.976415 | 0.512735 | -1.904 |
| chr18 | 1467757 | 1467976 | 0.841319 | 0.441871 | -1.904 |
| chr10 | 3690906 | 3691240 | 1.26833 | 0.666156 | -1.904 |
| chr9 | 133477739 | 133477977 | 0.983027 | 0.516316 | -1.904 |
| chr16 | 46540483 | 46541023 | 1.27845 | 0.671539 | -1.904 |
| chr16 | 31656908 | 31657149 | 1.01732 | 0.534415 | -1.904 |
| chr3 | 85217056 | 85217519 | 1.22732 | 0.644862 | -1.903 |
| chr14 | 83434660 | 83435020 | 1.06707 | 0.560713 | -1.903 |
| chr15 | 72600070 | 72600257 | 1.34364 | 0.706056 | -1.903 |
| chr3 | 57838580 | 57838833 | 0.868586 | 0.456511 | -1.903 |
| chr12 | 112238103 | 112238515 | 0.959341 | 0.504232 | -1.903 |
| chr17 | 52401675 | 52401928 | 1.26081 | 0.662756 | -1.902 |
| chr10 | 55443739 | 55443944 | 0.883259 | 0.464308 | -1.902 |
| chr2 | 159513249 | 159513480 | 0.840637 | 0.441994 | -1.902 |
| chr7 | 36522609 | 36522832 | 1.19957 | 0.630804 | -1.902 |
| chr10 | 50899840 | 50900049 | 1.08965 | 0.573117 | -1.901 |
| chr15 | 97458688 | 97459007 | 1.01864 | 0.535854 | -1.901 |
| chr1 | 228275731 | 228275935 | 1.03282 | 0.543318 | -1.901 |
| chr16 | 46595806 | 46596044 | 0.882495 | 0.464249 | -1.901 |

| | | | | | |
|---|---|---|---|---|---|
| chr3 | 22208987 | 22209203 | 1.06644 | 0.56106 | -1.901 |
| chr5 | 97536757 | 97536999 | 1.00975 | 0.531248 | -1.901 |
| chr8 | 77179736 | 77180049 | 0.72148 | 0.379614 | -1.901 |
| chr16 | 64712178 | 64712396 | 0.79734 | 0.419579 | -1.900 |
| chr7 | 3666635 | 3666854 | 1.09494 | 0.576332 | -1.900 |
| chr4 | 172604029 | 172604217 | 0.684352 | 0.360222 | -1.900 |
| chr5 | 23552851 | 23553077 | 1.43377 | 0.754693 | -1.900 |
| chr14 | 83121491 | 83121686 | 0.869424 | 0.457783 | -1.899 |
| chr9 | 103181242 | 103181477 | 1.31556 | 0.692692 | -1.899 |
| chr1 | 183432648 | 183432870 | 0.861113 | 0.453425 | -1.899 |
| chr10 | 58856005 | 58856187 | 1.25816 | 0.662554 | -1.899 |
| chr15 | 31854396 | 31854706 | 1.49223 | 0.785843 | -1.899 |
| chr10 | 2249739 | 2249988 | 1.1286 | 0.59439 | -1.899 |
| chr8 | 24120226 | 24120582 | 0.950504 | 0.50066 | -1.899 |
| chr2 | 81066266 | 81066489 | 1.04145 | 0.548602 | -1.898 |
| chr1 | 175713925 | 175714358 | 1.20381 | 0.63414 | -1.898 |
| chr3 | 78080428 | 78080614 | 1.10868 | 0.584079 | -1.898 |
| chr8 | 136809293 | 136809513 | 1.23289 | 0.649517 | -1.898 |
| chr15 | 35234507 | 35234733 | 0.972937 | 0.512602 | -1.898 |
| chr7 | 52239271 | 52239482 | 0.712807 | 0.375568 | -1.898 |
| chr8 | 6813598 | 6813821 | 1.11813 | 0.589175 | -1.898 |
| chr1 | 152712570 | 152712761 | 1.11113 | 0.585542 | -1.898 |
| chr11 | 58797680 | 58797911 | 0.949674 | 0.500492 | -1.897 |
| chr8 | 78126476 | 78126703 | 1.10055 | 0.580013 | -1.897 |
| chr20 | 37266341 | 37266558 | 1.08897 | 0.57394 | -1.897 |
| chr20 | 8917746 | 8917975 | 1.41469 | 0.745713 | -1.897 |
| chr20 | 8179208 | 8179395 | 1.50873 | 0.795389 | -1.897 |
| chr5 | 6920675 | 6920873 | 0.866636 | 0.456958 | -1.897 |
| chr10 | 74201485 | 74201913 | 0.874682 | 0.461249 | -1.896 |
| chr12 | 11605474 | 11605700 | 1.14518 | 0.603919 | -1.896 |
| chr14 | 47544272 | 47544526 | 1.08238 | 0.570851 | -1.896 |
| chr16 | 18155432 | 18155667 | 0.72741 | 0.38371 | -1.896 |
| chr10 | 80784019 | 80784216 | 0.723677 | 0.381751 | -1.896 |
| chr12 | 5309794 | 5310030 | 1.48445 | 0.783084 | -1.896 |
| chr8 | 139601522 | 139601833 | 1.26164 | 0.665592 | -1.896 |
| chr19 | 32433230 | 32433445 | 1.02001 | 0.538128 | -1.895 |
| chr8 | 30158007 | 30158214 | 1.01464 | 0.535305 | -1.895 |
| chr7 | 10654930 | 10655114 | 1.09323 | 0.576772 | -1.895 |
| chr20 | 20884980 | 20885207 | 1.11139 | 0.586355 | -1.895 |
| chr5 | 24701022 | 24701226 | 0.97714 | 0.515527 | -1.895 |
| chr8 | 77890490 | 77890681 | 1.09437 | 0.577389 | -1.895 |
| chr2 | 60376339 | 60376560 | 1.10751 | 0.584376 | -1.895 |
| chr1 | 151491434 | 151491825 | 1.02004 | 0.538231 | -1.895 |
| chr7 | 40154932 | 40155375 | 0.946 | 0.499184 | -1.895 |
| chr19 | 24491098 | 24491292 | 0.946278 | 0.499337 | -1.895 |
| chr11 | 39527864 | 39528058 | 0.957514 | 0.505326 | -1.895 |
| chr19 | 40875879 | 40876278 | 1.17905 | 0.622284 | -1.895 |
| chr1 | 210346318 | 210346520 | 0.988074 | 0.521604 | -1.894 |
| chr8 | 138562120 | 138562385 | 1.30187 | 0.687271 | -1.894 |
| chrX | 80112465 | 80112710 | 0.935436 | 0.493846 | -1.894 |
| chr16 | 6072405 | 6072601 | 1.09689 | 0.579101 | -1.894 |
| chr2 | 34775069 | 34775272 | 1.05483 | 0.556956 | -1.894 |
| chr17 | 76016973 | 76017427 | 1.05054 | 0.554734 | -1.894 |
| chr11 | 55953989 | 55954202 | 1.23345 | 0.651396 | -1.894 |
| chr7 | 106398451 | 106398701 | 1.31296 | 0.693492 | -1.893 |
| chr2 | 119726807 | 119727059 | 1.45078 | 0.766389 | -1.893 |
| chr5 | 168014320 | 168014509 | 1.15345 | 0.609367 | -1.893 |
| chr7 | 11385098 | 11385342 | 0.926515 | 0.489498 | -1.893 |
| chr19 | 57621419 | 57621617 | 1.14271 | 0.60384 | -1.892 |
| chr2 | 50226997 | 50227327 | 1.30024 | 0.68714 | -1.892 |
| chr14 | 41172034 | 41172217 | 1.00646 | 0.5319 | -1.892 |

| | | | | | |
|---|---|---|---|---|---|
| chr13 | 66821130 | 66821323 | 0.8639 | 0.456585 | -1.892 |
| chr5 | 50748227 | 50748658 | 1.07902 | 0.570283 | -1.892 |
| chr2 | 51718089 | 51718452 | 1.03744 | 0.548475 | -1.891 |
| chr3 | 165671774 | 165672034 | 1.07036 | 0.565879 | -1.891 |
| chr5 | 158036915 | 158037116 | 1.10067 | 0.581977 | -1.891 |
| chr19 | 31702353 | 31702535 | 1.435 | 0.758765 | -1.891 |
| chr10 | 87898526 | 87899208 | 1.63195 | 0.863 | -1.891 |
| chr5 | 121558482 | 121558723 | 1.26213 | 0.667521 | -1.891 |
| chr7 | 4157261 | 4157484 | 1.24207 | 0.656961 | -1.891 |
| chr12 | 125367374 | 125367613 | 0.819059 | 0.433225 | -1.891 |
| chr3 | 13274947 | 13275223 | 0.798805 | 0.422652 | -1.890 |
| chr9 | 35902576 | 35902785 | 1.1039 | 0.584245 | -1.889 |
| chr10 | 122096511 | 122096853 | 1.80356 | 0.954685 | -1.889 |
| chr3 | 11300730 | 11300921 | 1.28352 | 0.679435 | -1.889 |
| chr15 | 80775841 | 80776035 | 1.05538 | 0.558684 | -1.889 |
| chr3 | 8112724 | 8112953 | 1.03699 | 0.549052 | -1.889 |
| chr1 | 178535295 | 178535497 | 0.8043 | 0.425855 | -1.889 |
| chr3 | 84805101 | 84805362 | 1.38309 | 0.732318 | -1.889 |
| chr3 | 169508459 | 169508645 | 1.33071 | 0.704594 | -1.889 |
| chr17 | 63865754 | 63866109 | 1.06376 | 0.563327 | -1.888 |
| chr7 | 49405465 | 49405698 | 1.04007 | 0.550807 | -1.888 |
| chr18 | 5774866 | 5775061 | 0.856487 | 0.453639 | -1.888 |
| chr20 | 15947070 | 15947354 | 1.23332 | 0.653245 | -1.888 |
| chr8 | 32359380 | 32359649 | 1.01911 | 0.53984 | -1.888 |
| chr7 | 112858046 | 112858416 | 1.17421 | 0.622074 | -1.888 |
| chr5 | 173817544 | 173817888 | 0.837295 | 0.443604 | -1.887 |
| chr12 | 119327232 | 119327472 | 1.22926 | 0.651295 | -1.887 |
| chr19 | 37423187 | 37423625 | 1.28473 | 0.680695 | -1.887 |
| chr17 | 32399132 | 32399333 | 1.40033 | 0.741988 | -1.887 |
| chr11 | 22510310 | 22510518 | 0.957289 | 0.507355 | -1.887 |
| chr12 | 24209083 | 24209301 | 1.01436 | 0.537703 | -1.886 |
| chr16 | 15154877 | 15155142 | 1.09332 | 0.579615 | -1.886 |
| chr14 | 46124632 | 46124955 | 0.99123 | 0.525538 | -1.886 |
| chr2 | 163338738 | 163338950 | 1.08349 | 0.574495 | -1.886 |
| chr10 | 107111106 | 107111388 | 1.03247 | 0.547453 | -1.886 |
| chr16 | 78587647 | 78587883 | 1.29445 | 0.68667 | -1.885 |
| chr9 | 134056712 | 134056956 | 1.41789 | 0.752349 | -1.885 |
| chr2 | 52412721 | 52412986 | 1.44776 | 0.768211 | -1.885 |
| chr2 | 71812227 | 71812413 | 1.05679 | 0.560788 | -1.884 |
| chr5 | 51017236 | 51017586 | 0.883031 | 0.468639 | -1.884 |
| chr8 | 14972591 | 14972883 | 1.14973 | 0.610285 | -1.884 |
| chr5 | 23058821 | 23059171 | 1.00419 | 0.533043 | -1.884 |
| chr5 | 146773616 | 146773838 | 1.27174 | 0.675067 | -1.884 |
| chr15 | 94921425 | 94921668 | 1.23529 | 0.655737 | -1.884 |
| chr2 | 6011416 | 6011639 | 1.29669 | 0.688348 | -1.884 |
| chr17 | 73293234 | 73293426 | 0.896539 | 0.47594 | -1.884 |
| chr7 | 39579596 | 39580002 | 1.05713 | 0.561264 | -1.883 |
| chr5 | 13506593 | 13506884 | 1.01392 | 0.538384 | -1.883 |
| chr20 | 56070870 | 56071304 | 0.914029 | 0.48538 | -1.883 |
| chr20 | 25200875 | 25201123 | 1.78611 | 0.948517 | -1.883 |
| chr5 | 119622899 | 119623104 | 0.861401 | 0.457466 | -1.883 |
| chr2 | 99440351 | 99440604 | 1.01625 | 0.539741 | -1.883 |
| chr2 | 57862852 | 57863043 | 1.22625 | 0.651278 | -1.883 |
| chr10 | 55691701 | 55691974 | 1.02933 | 0.546751 | -1.883 |
| chr14 | 51836588 | 51837170 | 1.27032 | 0.674825 | -1.882 |
| chr14 | 84225215 | 84225559 | 1.06905 | 0.567929 | -1.882 |
| chr8 | 127991090 | 127991288 | 1.42742 | 0.758545 | -1.882 |
| chr3 | 16825350 | 16825581 | 1.03465 | 0.549916 | -1.881 |
| chr5 | 68759698 | 68759918 | 1.00005 | 0.53156 | -1.881 |
| chr2 | 188614825 | 188615009 | 0.79915 | 0.42481 | -1.881 |
| chr12 | 62247545 | 62247768 | 1.02015 | 0.542317 | -1.881 |

| | | | | | |
|---|---|---|---|---|---|
| chr8 | 5579185 | 5579416 | 1.22068 | 0.648919 | -1.881 |
| chr19 | 32018485 | 32018771 | 1.48498 | 0.789458 | -1.881 |
| chr8 | 84334464 | 84334748 | 0.853494 | 0.453875 | -1.880 |
| chr11 | 25848914 | 25849098 | 1.25238 | 0.666028 | -1.880 |
| chr20 | 5160114 | 5160313 | 1.09114 | 0.580308 | -1.880 |
| chr7 | 148642435 | 148642745 | 0.938126 | 0.498951 | -1.880 |
| chr2 | 60912574 | 60912770 | 0.905616 | 0.481673 | -1.880 |
| chr17 | 68940402 | 68940590 | 1.09287 | 0.581372 | -1.880 |
| chr9 | 129117527 | 129117869 | 1.13693 | 0.605005 | -1.879 |
| chr10 | 68480548 | 68480951 | 1.1063 | 0.588741 | -1.879 |
| chr7 | 153078746 | 153078940 | 1.35991 | 0.723709 | -1.879 |
| chr7 | 90170361 | 90170708 | 0.720675 | 0.383634 | -1.879 |
| chr10 | 22425348 | 22425553 | 0.834822 | 0.444505 | -1.878 |
| chr5 | 121239780 | 121239994 | 1.11229 | 0.592312 | -1.878 |
| chr11 | 48250314 | 48250652 | 1.0552 | 0.561916 | -1.878 |
| chr17 | 63422808 | 63423191 | 1.34372 | 0.715694 | -1.878 |
| chr5 | 27590616 | 27590834 | 1.22473 | 0.652342 | -1.877 |
| chr11 | 24785803 | 24786012 | 1.00451 | 0.535084 | -1.877 |
| chr14 | 84214558 | 84214742 | 0.854979 | 0.455472 | -1.877 |
| chr16 | 13299679 | 13299882 | 0.854766 | 0.455379 | -1.877 |
| chr17 | 32166638 | 32166875 | 0.950427 | 0.506376 | -1.877 |
| chr5 | 107189433 | 107189632 | 1.03534 | 0.551753 | -1.876 |
| chr3 | 95717103 | 95717300 | 0.947368 | 0.504977 | -1.876 |
| chr7 | 26755159 | 26755364 | 0.730705 | 0.389502 | -1.876 |
| chr10 | 36844097 | 36844489 | 0.998151 | 0.532071 | -1.876 |
| chr2 | 3304357 | 3304545 | 1.1362 | 0.605697 | -1.876 |
| chr2 | 21635813 | 21636087 | 0.92262 | 0.491851 | -1.876 |
| chr7 | 28875989 | 28876339 | 0.902674 | 0.481218 | -1.876 |
| chr19 | 29171412 | 29171621 | 1.57368 | 0.839202 | -1.875 |
| chr12 | 84137765 | 84137958 | 1.07338 | 0.572455 | -1.875 |
| chr20 | 53796520 | 53796850 | 1.20478 | 0.642652 | -1.875 |
| chr2 | 32015271 | 32015507 | 1.22725 | 0.654736 | -1.874 |
| chr11 | 70852000 | 70852208 | 0.827054 | 0.441257 | -1.874 |
| chr15 | 98341010 | 98341251 | 0.94676 | 0.505158 | -1.874 |
| chr2 | 5913586 | 5913790 | 1.24852 | 0.666176 | -1.874 |
| chr5 | 15264050 | 15264315 | 1.12329 | 0.599413 | -1.874 |
| chr8 | 130333397 | 130333605 | 0.834286 | 0.44522 | -1.874 |
| chr2 | 66945033 | 66945369 | 0.897921 | 0.479201 | -1.874 |
| chr5 | 157712164 | 157712379 | 0.979579 | 0.52288 | -1.873 |
| chr7 | 88788054 | 88788299 | 1.09403 | 0.584096 | -1.873 |
| chr10 | 83790097 | 83790379 | 0.88486 | 0.472513 | -1.873 |
| chr7 | 85164441 | 85164670 | 1.16868 | 0.624145 | -1.872 |
| chr16 | 62647036 | 62647300 | 0.895155 | 0.47807 | -1.872 |
| chr5 | 111920941 | 111921176 | 1.06759 | 0.570235 | -1.872 |
| chr7 | 148800720 | 148800913 | 1.28181 | 0.684764 | -1.872 |
| chr1 | 194943837 | 194944268 | 1.03723 | 0.554111 | -1.872 |
| chr5 | 110772140 | 110772344 | 1.1404 | 0.609235 | -1.872 |
| chr5 | 27539893 | 27540138 | 0.944198 | 0.504556 | -1.871 |
| chr5 | 110493314 | 110493510 | 1.08369 | 0.57921 | -1.871 |
| chr20 | 55964541 | 55964753 | 1.56319 | 0.835581 | -1.871 |
| chr7 | 87931712 | 87931939 | 0.929341 | 0.496773 | -1.871 |
| chr2 | 118048187 | 118048385 | 1.0518 | 0.562324 | -1.870 |
| chr8 | 139736575 | 139736851 | 1.00851 | 0.539183 | -1.870 |
| chr14 | 70704728 | 70704925 | 0.831765 | 0.444802 | -1.870 |
| chr2 | 66240739 | 66240975 | 1.03245 | 0.552269 | -1.869 |
| chr8 | 51417081 | 51417270 | 0.955965 | 0.511362 | -1.869 |
| chr19 | 23905603 | 23905828 | 0.873311 | 0.46715 | -1.869 |
| chr3 | 76942094 | 76942408 | 1.02851 | 0.550183 | -1.869 |
| chr2 | 59297680 | 59298029 | 0.926563 | 0.495686 | -1.869 |
| chr2 | 226215183 | 226215393 | 1.13955 | 0.60967 | -1.869 |
| chr22 | 42928534 | 42928823 | 1.05529 | 0.564629 | -1.869 |

| | | | | | |
|---|---|---|---|---|---|
| chr8 | 52668661 | 52668847 | 1.31985 | 0.706195 | -1.869 |
| chr16 | 63134088 | 63134327 | 1.08388 | 0.579941 | -1.869 |
| chr3 | 79871051 | 79871380 | 0.995818 | 0.532829 | -1.869 |
| chr5 | 105092661 | 105092903 | 1.49392 | 0.799355 | -1.869 |
| chr5 | 31655807 | 31656020 | 0.982932 | 0.52598 | -1.869 |
| chr5 | 9964491 | 9964678 | 0.969095 | 0.518798 | -1.868 |
| chr11 | 38036457 | 38036642 | 0.865597 | 0.463418 | -1.868 |
| chr12 | 43986266 | 43986450 | 0.876986 | 0.46956 | -1.868 |
| chr5 | 76705914 | 76706097 | 0.87975 | 0.471058 | -1.868 |
| chr17 | 55638199 | 55638430 | 0.811319 | 0.434439 | -1.868 |
| chr13 | 53471705 | 53471887 | 1.51905 | 0.813759 | -1.867 |
| chr6 | 67450768 | 67450979 | 1.18843 | 0.636719 | -1.866 |
| chr5 | 135432839 | 135433035 | 1.03868 | 0.556553 | -1.866 |
| chr3 | 114097324 | 114097553 | 0.932598 | 0.499804 | -1.866 |
| chr9 | 27900818 | 27901148 | 1.17964 | 0.632248 | -1.866 |
| chr8 | 119462494 | 119462684 | 1.25693 | 0.67369 | -1.866 |
| chr1 | 150865989 | 150866206 | 0.920375 | 0.493344 | -1.866 |
| chr3 | 106653043 | 106653642 | 1.14381 | 0.613115 | -1.866 |
| chr10 | 52797008 | 52797206 | 0.967519 | 0.518622 | -1.866 |
| chr8 | 49934984 | 49935363 | 1.02309 | 0.548507 | -1.865 |
| chr8 | 105042791 | 105042991 | 0.950751 | 0.509798 | -1.865 |
| chr2 | 35513613 | 35513855 | 1.01895 | 0.546442 | -1.865 |
| chr2 | 151899866 | 151900118 | 1.06732 | 0.572438 | -1.865 |
| chr9 | 9960245 | 9960548 | 1.24293 | 0.666666 | -1.864 |
| chr7 | 133313430 | 133313630 | 1.02164 | 0.547976 | -1.864 |
| chr11 | 604062 | 604447 | 0.887874 | 0.476309 | -1.864 |
| chr17 | 79910770 | 79911091 | 1.27882 | 0.686225 | -1.864 |
| chr14 | 86223854 | 86224043 | 0.985205 | 0.528715 | -1.863 |
| chr11 | 78509613 | 78510030 | 2.04994 | 1.10019 | -1.863 |
| chr5 | 88255783 | 88255965 | 0.928973 | 0.498581 | -1.863 |
| chr9 | 114024339 | 114024755 | 0.848718 | 0.455587 | -1.863 |
| chr8 | 136900571 | 136901062 | 1.1281 | 0.605657 | -1.863 |
| chr9 | 31204402 | 31204587 | 1.12267 | 0.602762 | -1.863 |
| chr15 | 95017782 | 95017972 | 0.959017 | 0.514915 | -1.862 |
| chr2 | 4476812 | 4477013 | 1.08959 | 0.585024 | -1.862 |
| chr14 | 35441181 | 35441407 | 1.00754 | 0.540991 | -1.862 |
| chr5 | 163216321 | 163216590 | 0.741066 | 0.39797 | -1.862 |
| chr1 | 226391405 | 226391790 | 1.0533 | 0.565671 | -1.862 |
| chr8 | 99297562 | 99297752 | 1.22185 | 0.656194 | -1.862 |
| chr17 | 50406703 | 50406948 | 0.834331 | 0.448109 | -1.862 |
| chr2 | 64321981 | 64322190 | 1.06319 | 0.571054 | -1.862 |
| chr8 | 59647005 | 59647266 | 0.861309 | 0.462644 | -1.862 |
| chr15 | 66928347 | 66928595 | 0.878195 | 0.471789 | -1.861 |
| chr19 | 37581293 | 37581552 | 1.15617 | 0.62121 | -1.861 |
| chr6 | 23189784 | 23189974 | 0.951197 | 0.51115 | -1.861 |
| chr16 | 34956900 | 34957168 | 0.946493 | 0.508672 | -1.861 |
| chr10 | 100497084 | 100497329 | 1.33925 | 0.719927 | -1.860 |
| chr16 | 7903045 | 7903276 | 1.0055 | 0.540597 | -1.860 |
| chr7 | 41291446 | 41291648 | 1.06102 | 0.570475 | -1.860 |
| chr10 | 9892469 | 9892731 | 1.22174 | 0.656925 | -1.860 |
| chr12 | 59327215 | 59327401 | 0.92542 | 0.497597 | -1.860 |
| chr5 | 38077930 | 38078154 | 1.19993 | 0.645295 | -1.860 |
| chr14 | 53359719 | 53359958 | 1.18629 | 0.637962 | -1.859 |
| chr2 | 64551934 | 64552136 | 1.28305 | 0.690023 | -1.859 |
| chr5 | 75948135 | 75948402 | 1.15086 | 0.61908 | -1.859 |
| chr9 | 979205 | 979417 | 1.13999 | 0.613282 | -1.859 |
| chr2 | 88712640 | 88712829 | 1.2812 | 0.689309 | -1.859 |
| chr12 | 24546062 | 24546289 | 0.985185 | 0.530053 | -1.859 |
| chr9 | 126821397 | 126821731 | 0.9964 | 0.536269 | -1.858 |
| chr17 | 69635297 | 69635607 | 1.43713 | 0.77349 | -1.858 |
| chr3 | 186708104 | 186708327 | 1.08904 | 0.586185 | -1.858 |

| | | | | | |
|---|---|---|---|---|---|
| chr2 | 129839706 | 129839906 | 1.14014 | 0.613695 | -1.858 |
| chr9 | 12366366 | 12366629 | 1.1028 | 0.593606 | -1.858 |
| chr16 | 9360196 | 9360410 | 0.794135 | 0.427506 | -1.858 |
| chr7 | 125187476 | 125187749 | 1.03985 | 0.559878 | -1.857 |
| chr7 | 121168825 | 121169104 | 1.0586 | 0.570045 | -1.857 |
| chr8 | 113665116 | 113665342 | 0.988096 | 0.532115 | -1.857 |
| chr19 | 33066841 | 33067035 | 1.20891 | 0.651071 | -1.857 |
| chr5 | 84093624 | 84093850 | 1.1811 | 0.636121 | -1.857 |
| chr9 | 8656307 | 8656533 | 0.902125 | 0.485914 | -1.857 |
| chr2 | 40241892 | 40242163 | 0.962051 | 0.51821 | -1.856 |
| chr15 | 101223729 | 101223926 | 1.19384 | 0.643148 | -1.856 |
| chr16 | 64118806 | 64119136 | 1.00872 | 0.543488 | -1.856 |
| chr10 | 111208697 | 111209080 | 1.16529 | 0.627984 | -1.856 |
| chr19 | 34239207 | 34239395 | 0.955493 | 0.514932 | -1.856 |
| chr9 | 104968315 | 104968506 | 0.990565 | 0.533842 | -1.856 |
| chr2 | 16886678 | 16886941 | 1.24169 | 0.66921 | -1.855 |
| chr8 | 137318223 | 137318439 | 1.22973 | 0.662774 | -1.855 |
| chr7 | 15807753 | 15808025 | 1.33987 | 0.722335 | -1.855 |
| chr3 | 147200558 | 147200743 | 1.03514 | 0.558169 | -1.855 |
| chr2 | 118064365 | 118064572 | 1.01902 | 0.549661 | -1.854 |
| chr17 | 73279627 | 73279831 | 1.04529 | 0.563834 | -1.854 |
| chr1 | 226549260 | 226549487 | 1.07823 | 0.581667 | -1.854 |
| chr20 | 7652911 | 7653104 | 1.11583 | 0.601969 | -1.854 |
| chr19 | 34354231 | 34354545 | 1.31152 | 0.707564 | -1.854 |
| chr2 | 124866981 | 124867202 | 0.886132 | 0.478075 | -1.854 |
| chr8 | 49641653 | 49641880 | 1.45062 | 0.782789 | -1.853 |
| chr17 | 49922078 | 49922323 | 1.03781 | 0.560067 | -1.853 |
| chr17 | 12449328 | 12449594 | 2.16361 | 1.16763 | -1.853 |
| chr20 | 30040592 | 30040884 | 1.08337 | 0.584662 | -1.853 |
| chr3 | 153814752 | 153814947 | 1.13161 | 0.610709 | -1.853 |
| chr7 | 54076484 | 54076703 | 1.63256 | 0.881088 | -1.853 |
| chr20 | 1732021 | 1732262 | 1.45588 | 0.785847 | -1.853 |
| chr14 | 98818015 | 98818259 | 0.920537 | 0.497039 | -1.852 |
| chr5 | 62384947 | 62385204 | 0.886089 | 0.478453 | -1.852 |
| chr10 | 16653087 | 16653275 | 1.23693 | 0.667965 | -1.852 |
| chr10 | 133136794 | 133136998 | 0.95329 | 0.514837 | -1.852 |
| chr10 | 86434342 | 86434567 | 0.846665 | 0.457328 | -1.851 |
| chr4 | 165653959 | 165654152 | 0.953205 | 0.514887 | -1.851 |
| chr10 | 126936334 | 126936535 | 0.830975 | 0.448952 | -1.851 |
| chr2 | 142312477 | 142312713 | 1.16067 | 0.62708 | -1.851 |
| chr2 | 14035335 | 14035823 | 1.18684 | 0.641294 | -1.851 |
| chr17 | 50133280 | 50133554 | 1.00786 | 0.544638 | -1.851 |
| chr5 | 85933354 | 85933579 | 1.11244 | 0.601293 | -1.850 |
| chr15 | 101797084 | 101797291 | 2.54069 | 1.37337 | -1.850 |
| chr5 | 171228790 | 171228976 | 1.21064 | 0.654482 | -1.850 |
| chr8 | 58794650 | 58795110 | 0.984657 | 0.532317 | -1.850 |
| chr8 | 5691550 | 5691817 | 1.32143 | 0.714383 | -1.850 |
| chr8 | 137309167 | 137309446 | 1.02482 | 0.554101 | -1.850 |
| chr16 | 5250311 | 5250580 | 1.00952 | 0.545932 | -1.849 |
| chr3 | 6676312 | 6676605 | 0.856709 | 0.463338 | -1.849 |
| chr2 | 76258536 | 76258798 | 1.26345 | 0.683412 | -1.849 |
| chr21 | 28839579 | 28839797 | 0.774336 | 0.41885 | -1.849 |
| chr14 | 25761613 | 25761840 | 0.836971 | 0.452831 | -1.848 |
| chr8 | 77366975 | 77367192 | 0.95267 | 0.515441 | -1.848 |
| chr11 | 57317444 | 57317663 | 1.19548 | 0.646855 | -1.848 |
| chr1 | 202148837 | 202149179 | 1.09514 | 0.592599 | -1.848 |
| chr2 | 54739141 | 54739393 | 1.10645 | 0.59874 | -1.848 |
| chr7 | 28593869 | 28594086 | 1.19906 | 0.648938 | -1.848 |
| chr3 | 104334230 | 104334469 | 1.16719 | 0.631822 | -1.847 |
| chr2 | 35563848 | 35564139 | 0.971662 | 0.525989 | -1.847 |
| chr10 | 48485837 | 48486061 | 1.02118 | 0.552912 | -1.847 |

| | | | | | |
|---|---|---|---|---|---|
| chr7 | 122663595 | 122663807 | 0.835425 | 0.452339 | -1.847 |
| chr10 | 122582511 | 122582697 | 0.966828 | 0.523534 | -1.847 |
| chr3 | 163906366 | 163906597 | 0.732677 | 0.396786 | -1.847 |
| chr8 | 130116091 | 130116342 | 1.17175 | 0.634587 | -1.846 |
| chr17 | 65758395 | 65758590 | 0.808181 | 0.437882 | -1.846 |
| chr19 | 34032135 | 34032344 | 0.979982 | 0.530972 | -1.846 |
| chr9 | 108999041 | 108999284 | 0.897288 | 0.486205 | -1.845 |
| chr15 | 59115597 | 59115787 | 1.10454 | 0.59852 | -1.845 |
| chr5 | 22721205 | 22721401 | 1.13102 | 0.612975 | -1.845 |
| chr14 | 49662743 | 49662960 | 1.0769 | 0.583744 | -1.845 |
| chr2 | 104237265 | 104237563 | 1.01923 | 0.552568 | -1.845 |
| chr17 | 34291046 | 34291231 | 0.948495 | 0.514251 | -1.844 |
| chr2 | 82006698 | 82007018 | 0.897054 | 0.48638 | -1.844 |
| chr5 | 140237632 | 140237929 | 0.975177 | 0.528802 | -1.844 |
| chr8 | 13390178 | 13390369 | 1.37112 | 0.74361 | -1.844 |
| chr1 | 187145559 | 187145900 | 0.884013 | 0.479454 | -1.844 |
| chr5 | 7622717 | 7623123 | 1.05104 | 0.570057 | -1.844 |
| chr5 | 111764395 | 111764644 | 1.0941 | 0.593445 | -1.844 |
| chr3 | 168964129 | 168964359 | 0.789374 | 0.428183 | -1.844 |
| chr8 | 133556192 | 133556474 | 0.901212 | 0.488883 | -1.843 |
| chr7 | 69585886 | 69586073 | 0.908345 | 0.492764 | -1.843 |
| chr8 | 84172262 | 84172444 | 0.903839 | 0.490398 | -1.843 |
| chr3 | 147171635 | 147172133 | 0.910115 | 0.493806 | -1.843 |
| chr15 | 76657268 | 76657471 | 1.22448 | 0.664375 | -1.843 |
| chr10 | 100810984 | 100811357 | 1.02442 | 0.555879 | -1.843 |
| chr10 | 108877030 | 108877318 | 1.01879 | 0.55288 | -1.843 |
| chr1 | 162844265 | 162844462 | 1.16706 | 0.633406 | -1.843 |
| chr20 | 38894771 | 38895214 | 1.30761 | 0.709919 | -1.842 |
| chr3 | 144281360 | 144281631 | 0.814403 | 0.44219 | -1.842 |
| chr10 | 20508491 | 20508786 | 1.00035 | 0.543161 | -1.842 |
| chr2 | 63686751 | 63686986 | 0.961757 | 0.522395 | -1.841 |
| chr3 | 155415439 | 155415719 | 1.01751 | 0.552708 | -1.841 |
| chr15 | 55350018 | 55350258 | 1.07778 | 0.585455 | -1.841 |
| chr5 | 24112790 | 24113014 | 1.24822 | 0.678142 | -1.841 |
| chr11 | 33770752 | 33771047 | 0.916837 | 0.498126 | -1.841 |
| chr9 | 75887649 | 75887876 | 1.13246 | 0.615285 | -1.841 |
| chr19 | 28630149 | 28630373 | 1.14105 | 0.619958 | -1.841 |
| chr17 | 71030752 | 71031002 | 1.10297 | 0.59934 | -1.840 |
| chr7 | 83968597 | 83968807 | 1.17517 | 0.638586 | -1.840 |
| chr13 | 59764827 | 59765050 | 1.05739 | 0.574833 | -1.839 |
| chr5 | 128855072 | 128855260 | 1.34205 | 0.729604 | -1.839 |
| chr5 | 112365397 | 112365645 | 0.982929 | 0.534404 | -1.839 |
| chrX | 69057852 | 69058067 | 0.926073 | 0.503494 | -1.839 |
| chr7 | 88131113 | 88131384 | 1.09062 | 0.593042 | -1.839 |
| chr14 | 82457916 | 82458100 | 0.905107 | 0.49231 | -1.838 |
| chr16 | 65453932 | 65454230 | 0.952365 | 0.518082 | -1.838 |
| chr2 | 176703740 | 176703942 | 1.34938 | 0.73411 | -1.838 |
| chr10 | 78345888 | 78346073 | 1.01891 | 0.554327 | -1.838 |
| chr12 | 67156454 | 67156670 | 0.805654 | 0.438318 | -1.838 |
| chr8 | 121463847 | 121464105 | 1.06421 | 0.579038 | -1.838 |
| chr10 | 132979421 | 132979681 | 0.82421 | 0.448455 | -1.838 |
| chr15 | 70707729 | 70707969 | 0.964985 | 0.525197 | -1.837 |
| chr19 | 37964290 | 37964486 | 1.07749 | 0.58647 | -1.837 |
| chr2 | 52685146 | 52685349 | 1.47502 | 0.802853 | -1.837 |
| chr17 | 5083380 | 5083682 | 1.09491 | 0.596078 | -1.837 |
| chr14 | 49766125 | 49766383 | 1.22278 | 0.665929 | -1.836 |
| chr20 | 7342860 | 7343056 | 1.10111 | 0.599851 | -1.836 |
| chr3 | 79889918 | 79890112 | 1.33708 | 0.728458 | -1.835 |
| chr9 | 103559252 | 103559436 | 0.830869 | 0.452723 | -1.835 |
| chr10 | 57026188 | 57026598 | 0.953677 | 0.519679 | -1.835 |
| chr3 | 80677118 | 80677447 | 1.23074 | 0.670703 | -1.835 |

| | | | | | |
|---|---|---|---|---|---|
| chr5 | 105259124 | 105259531 | 0.832349 | 0.453661 | -1.835 |
| chr2 | 182096039 | 182096230 | 0.952736 | 0.519297 | -1.835 |
| chr2 | 11802189 | 11802394 | 1.43989 | 0.784911 | -1.834 |
| chr7 | 130333609 | 130333829 | 1.03679 | 0.565275 | -1.834 |
| chr5 | 145057310 | 145057497 | 1.78668 | 0.974374 | -1.834 |
| chr15 | 93825094 | 93825331 | 1.24567 | 0.679354 | -1.834 |
| chr17 | 79038065 | 79038285 | 1.066 | 0.581374 | -1.834 |
| chr10 | 91714512 | 91714864 | 1.35425 | 0.73863 | -1.833 |
| chr17 | 33007491 | 33007731 | 0.986928 | 0.538336 | -1.833 |
| chr2 | 49717924 | 49718324 | 1.6026 | 0.874192 | -1.833 |
| chr2 | 81476122 | 81476312 | 1.04517 | 0.570142 | -1.833 |
| chr1 | 170593606 | 170593833 | 1.13218 | 0.61761 | -1.833 |
| chr14 | 84439368 | 84439598 | 0.945062 | 0.51556 | -1.833 |
| chr7 | 98234980 | 98235327 | 1.1069 | 0.603869 | -1.833 |
| chr8 | 138822742 | 138822983 | 0.939911 | 0.512777 | -1.833 |
| chr17 | 33179638 | 33179872 | 1.2854 | 0.701267 | -1.833 |
| chr7 | 24179702 | 24179919 | 1.18125 | 0.644493 | -1.833 |
| chr9 | 80875775 | 80875958 | 0.971751 | 0.530241 | -1.833 |
| chr2 | 79091298 | 79091491 | 1.25589 | 0.685305 | -1.833 |
| chr9 | 26226749 | 26227017 | 1.30525 | 0.712253 | -1.833 |
| chr12 | 104367063 | 104367271 | 1.14941 | 0.627269 | -1.832 |
| chr2 | 18616732 | 18617006 | 1.93004 | 1.05343 | -1.832 |
| chr5 | 84758605 | 84758912 | 1.27823 | 0.697737 | -1.832 |
| chr1 | 157800214 | 157800501 | 1.21852 | 0.66517 | -1.832 |
| chr8 | 113311088 | 113311485 | 1.06507 | 0.581425 | -1.832 |
| chr5 | 54011412 | 54011599 | 0.857325 | 0.468093 | -1.832 |
| chr5 | 173861813 | 173862016 | 0.931272 | 0.508477 | -1.831 |
| chr5 | 36944307 | 36944513 | 1.02922 | 0.561976 | -1.831 |
| chr2 | 1412060 | 1412283 | 0.775479 | 0.423438 | -1.831 |
| chr14 | 38785324 | 38785522 | 0.767261 | 0.418997 | -1.831 |
| chr2 | 16174121 | 16174563 | 1.25282 | 0.684273 | -1.831 |
| chr7 | 88106959 | 88107181 | 1.039 | 0.567526 | -1.831 |
| chr5 | 85311510 | 85311703 | 1.08433 | 0.592346 | -1.831 |
| chr14 | 84361044 | 84361308 | 1.31593 | 0.718922 | -1.830 |
| chr11 | 87923770 | 87923978 | 0.738643 | 0.403551 | -1.830 |
| chr8 | 80245615 | 80245806 | 0.959204 | 0.524099 | -1.830 |
| chr9 | 76994786 | 76995119 | 1.28445 | 0.701817 | -1.830 |
| chr8 | 6054987 | 6055178 | 1.21422 | 0.663454 | -1.830 |
| chr14 | 98456911 | 98457137 | 0.788765 | 0.430997 | -1.830 |
| chr15 | 27967220 | 27967422 | 0.958049 | 0.523522 | -1.830 |
| chr2 | 61032582 | 61032784 | 1.73666 | 0.949137 | -1.830 |
| chr8 | 138187937 | 138188160 | 1.03009 | 0.563021 | -1.830 |
| chr8 | 137194627 | 137194910 | 1.39876 | 0.764557 | -1.830 |
| chr19 | 37295254 | 37295459 | 1.00336 | 0.548528 | -1.829 |
| chr1 | 157883675 | 157883904 | 1.09343 | 0.597873 | -1.829 |
| chr7 | 32841623 | 32841865 | 0.95712 | 0.523344 | -1.829 |
| chr17 | 37183584 | 37183970 | 1.03936 | 0.568324 | -1.829 |
| chr20 | 62246386 | 62246708 | 0.950619 | 0.519812 | -1.829 |
| chr10 | 19523719 | 19523969 | 1.16599 | 0.637594 | -1.829 |
| chr17 | 52371691 | 52371969 | 0.902609 | 0.493693 | -1.828 |
| chr16 | 61393967 | 61394161 | 1.77925 | 0.973196 | -1.828 |
| chr9 | 97006331 | 97006536 | 0.891888 | 0.487868 | -1.828 |
| chr12 | 45168576 | 45169066 | 1.79515 | 0.98205 | -1.828 |
| chr10 | 71055590 | 71055818 | 0.830134 | 0.454167 | -1.828 |
| chr5 | 144874792 | 144875033 | 1.1484 | 0.6284 | -1.827 |
| chr11 | 83710391 | 83710649 | 1.19603 | 0.654556 | -1.827 |
| chr6 | 100903435 | 100903617 | 0.902396 | 0.493869 | -1.827 |
| chr5 | 104392498 | 104392746 | 1.00405 | 0.549633 | -1.827 |
| chr3 | 80320110 | 80320412 | 0.997647 | 0.546242 | -1.826 |
| chr2 | 61591242 | 61591425 | 1.12982 | 0.618639 | -1.826 |
| chr5 | 161587775 | 161587958 | 0.822577 | 0.450423 | -1.826 |

| | | | | | |
|---|---|---|---|---|---|
| chr1 | 166936373 | 166936564 | 0.793927 | 0.434815 | -1.826 |
| chr5 | 143847343 | 143847604 | 1.47385 | 0.807202 | -1.826 |
| chr5 | 114400318 | 114400507 | 0.890319 | 0.487637 | -1.826 |
| chr5 | 22325671 | 22325921 | 0.96256 | 0.527278 | -1.826 |
| chr3 | 164928663 | 164928911 | 0.951127 | 0.521054 | -1.825 |
| chr17 | 68985599 | 68985796 | 1.14962 | 0.629833 | -1.825 |
| chr10 | 19852786 | 19853002 | 1.25339 | 0.686806 | -1.825 |
| chr8 | 51452031 | 51452625 | 1.11741 | 0.612307 | -1.825 |
| chr19 | 49769026 | 49769210 | 0.999883 | 0.547925 | -1.825 |
| chr15 | 100568206 | 100568436 | 0.962047 | 0.527258 | -1.825 |
| chr1 | 71151128 | 71151333 | 1.77576 | 0.973303 | -1.824 |
| chr10 | 106398676 | 106398890 | 1.62695 | 0.891799 | -1.824 |
| chr3 | 102871291 | 102871504 | 1.16622 | 0.639342 | -1.824 |
| chr10 | 12141988 | 12142252 | 0.960891 | 0.526825 | -1.824 |
| chr8 | 139236586 | 139236807 | 1.07507 | 0.589616 | -1.823 |
| chr4 | 121814201 | 121814411 | 1.04573 | 0.573557 | -1.823 |
| chr2 | 185764873 | 185765339 | 1.09065 | 0.598244 | -1.823 |
| chr12 | 41719904 | 41720099 | 1.36916 | 0.751076 | -1.823 |
| chr5 | 101977307 | 101977565 | 0.961392 | 0.527435 | -1.823 |
| chr2 | 194258451 | 194258761 | 1.08018 | 0.592609 | -1.823 |
| chr19 | 31502468 | 31502674 | 1.31405 | 0.72094 | -1.823 |
| chr15 | 69738888 | 69739148 | 1.15622 | 0.634366 | -1.823 |
| chr7 | 96418608 | 96418921 | 1.2834 | 0.704182 | -1.823 |
| chr3 | 190659662 | 190659920 | 0.943078 | 0.517474 | -1.822 |
| chr20 | 38555887 | 38556075 | 1.10483 | 0.606282 | -1.822 |
| chr3 | 164233234 | 164233458 | 0.953464 | 0.523248 | -1.822 |
| chr5 | 7391359 | 7391558 | 1.09119 | 0.598892 | -1.822 |
| chr8 | 126929374 | 126929634 | 1.16048 | 0.636988 | -1.822 |
| chr7 | 38603905 | 38604165 | 1.4221 | 0.780665 | -1.822 |
| chr2 | 76693228 | 76693414 | 1.05024 | 0.576612 | -1.821 |
| chr2 | 51600709 | 51600939 | 1.17609 | 0.645782 | -1.821 |
| chr2 | 86281898 | 86282094 | 1.04131 | 0.571835 | -1.821 |
| chr5 | 27781724 | 27781974 | 1.11847 | 0.61428 | -1.821 |
| chr2 | 76048909 | 76049169 | 1.2018 | 0.660072 | -1.821 |
| chr5 | 135044305 | 135044576 | 0.97405 | 0.535063 | -1.820 |
| chr1 | 247390970 | 247391279 | 0.852617 | 0.468458 | -1.820 |
| chr2 | 16456478 | 16456670 | 0.827852 | 0.454869 | -1.820 |
| chr5 | 84134339 | 84134593 | 1.00458 | 0.551983 | -1.820 |
| chr2 | 48154799 | 48154981 | 0.808535 | 0.444312 | -1.820 |
| chr1 | 245589973 | 245590276 | 0.946377 | 0.520093 | -1.820 |
| chr11 | 37437762 | 37437997 | 0.870784 | 0.478574 | -1.820 |
| chr19 | 30354707 | 30355022 | 1.29892 | 0.713878 | -1.820 |
| chr17 | 32408265 | 32408591 | 1.26904 | 0.697555 | -1.819 |
| chr8 | 100401350 | 100401565 | 0.832105 | 0.457413 | -1.819 |
| chr19 | 5243768 | 5243950 | 1.01406 | 0.557438 | -1.819 |
| chr15 | 75174345 | 75174613 | 1.00877 | 0.554545 | -1.819 |
| chr19 | 41060188 | 41060373 | 0.869265 | 0.47787 | -1.819 |
| chr16 | 85819199 | 85819421 | 1.04595 | 0.575124 | -1.819 |
| chr20 | 12899511 | 12899733 | 1.15362 | 0.634423 | -1.818 |
| chr15 | 102381322 | 102381536 | 1.27352 | 0.700376 | -1.818 |
| chr1 | 184632741 | 184633493 | 1.5641 | 0.860188 | -1.818 |
| chr17 | 26521483 | 26521699 | 0.956784 | 0.526222 | -1.818 |
| chr2 | 150834439 | 150834812 | 1.18196 | 0.65008 | -1.818 |
| chr20 | 31501714 | 31502002 | 1.01459 | 0.558033 | -1.818 |
| chr4 | 89999000 | 89999195 | 0.90993 | 0.500495 | -1.818 |
| chr20 | 25583922 | 25584118 | 0.926587 | 0.509694 | -1.818 |
| chr14 | 43228531 | 43228790 | 1.17073 | 0.644065 | -1.818 |
| chr9 | 71300812 | 71301106 | 0.910263 | 0.500807 | -1.818 |
| chr16 | 81116938 | 81117170 | 1.29072 | 0.710127 | -1.818 |
| chr11 | 26488018 | 26488320 | 0.991899 | 0.545742 | -1.818 |
| chr5 | 88678359 | 88678706 | 1.18989 | 0.654763 | -1.817 |

| | | | | | |
|---|---|---|---|---|---|
| chr9 | 10997035 | 10997263 | 1.14806 | 0.631903 | -1.817 |
| chr16 | 47465045 | 47465287 | 1.18467 | 0.652099 | -1.817 |
| chr12 | 67772481 | 67772783 | 1.28079 | 0.705031 | -1.817 |
| chr2 | 88437517 | 88437747 | 0.939228 | 0.517045 | -1.817 |
| chr17 | 50184061 | 50184331 | 1.77399 | 0.976684 | -1.816 |
| chr17 | 60743502 | 60743731 | 0.967599 | 0.532738 | -1.816 |
| chr19 | 41012060 | 41012245 | 1.55477 | 0.856142 | -1.816 |
| chr5 | 91554152 | 91554389 | 0.996118 | 0.548571 | -1.816 |
| chr12 | 32380450 | 32380690 | 1.07112 | 0.58988 | -1.816 |
| chr10 | 113700692 | 113700876 | 0.941871 | 0.518707 | -1.816 |
| chr8 | 136153577 | 136153850 | 1.08005 | 0.594825 | -1.816 |
| chr3 | 80310479 | 80310886 | 0.761248 | 0.419268 | -1.816 |
| chr8 | 138188397 | 138188603 | 1.13999 | 0.627874 | -1.816 |
| chr2 | 53494867 | 53495256 | 1.07257 | 0.59082 | -1.815 |
| chr8 | 112042737 | 112042969 | 0.887727 | 0.48902 | -1.815 |
| chr15 | 24826467 | 24826708 | 1.13164 | 0.623445 | -1.815 |
| chr5 | 22937293 | 22937714 | 0.987855 | 0.544271 | -1.815 |
| chr9 | 30554949 | 30555137 | 0.990609 | 0.54582 | -1.815 |
| chr5 | 157985352 | 157985552 | 1.26113 | 0.69488 | -1.815 |
| chr9 | 36164018 | 36164206 | 0.818434 | 0.451036 | -1.815 |
| chr8 | 62840386 | 62840572 | 1.14901 | 0.633295 | -1.814 |
| chr8 | 56276877 | 56277088 | 1.15168 | 0.634797 | -1.814 |
| chr8 | 114110853 | 114111104 | 0.874145 | 0.481881 | -1.814 |
| chr12 | 84954227 | 84954476 | 0.778245 | 0.429062 | -1.814 |
| chr7 | 154870713 | 154870913 | 1.1564 | 0.637597 | -1.814 |
| chr19 | 34090907 | 34091152 | 1.07551 | 0.593 | -1.814 |
| chr10 | 92418601 | 92418860 | 1.14228 | 0.62984 | -1.814 |
| chr5 | 104582863 | 104583079 | 0.923019 | 0.509049 | -1.813 |
| chr3 | 7493522 | 7493767 | 0.907299 | 0.500387 | -1.813 |
| chr14 | 88354491 | 88354786 | 1.62097 | 0.894222 | -1.813 |
| chr8 | 5922293 | 5922604 | 1.07782 | 0.594649 | -1.813 |
| chr10 | 76573194 | 76573442 | 0.805829 | 0.4446 | -1.812 |
| chr11 | 38406868 | 38407072 | 0.926271 | 0.51118 | -1.812 |
| chr14 | 46027196 | 46027646 | 0.979432 | 0.540621 | -1.812 |
| chr1 | 180326831 | 180327042 | 0.849878 | 0.469126 | -1.812 |
| chr2 | 20856488 | 20856718 | 1.31347 | 0.725027 | -1.812 |
| chr17 | 40077908 | 40078148 | 1.0728 | 0.592182 | -1.812 |
| chr5 | 161799296 | 161799650 | 1.29396 | 0.714342 | -1.811 |
| chr19 | 54271725 | 54271945 | 0.990287 | 0.546731 | -1.811 |
| chr16 | 61374643 | 61375108 | 1.35933 | 0.750588 | -1.811 |
| chr3 | 95894487 | 95894669 | 1.16232 | 0.641817 | -1.811 |
| chr2 | 186281808 | 186282131 | 1.03297 | 0.570455 | -1.811 |
| chr11 | 29618416 | 29618819 | 0.977474 | 0.53981 | -1.811 |
| chr8 | 68733084 | 68733378 | 1.20643 | 0.666259 | -1.811 |
| chr17 | 58869650 | 58869873 | 0.935113 | 0.516489 | -1.811 |
| chr2 | 116790790 | 116790995 | 0.732966 | 0.404883 | -1.810 |
| chr8 | 136332686 | 136333091 | 1.17233 | 0.64762 | -1.810 |
| chr2 | 134220455 | 134220717 | 1.04132 | 0.575289 | -1.810 |
| chr10 | 112901748 | 112902146 | 1.32931 | 0.734408 | -1.810 |
| chr12 | 130914577 | 130914784 | 1.04306 | 0.576285 | -1.810 |
| chr2 | 79824485 | 79824682 | 1.32017 | 0.729433 | -1.810 |
| chr3 | 162649683 | 162649883 | 1.01931 | 0.56333 | -1.809 |
| chr16 | 56602642 | 56602934 | 1.04379 | 0.576885 | -1.809 |
| chr2 | 162919698 | 162920028 | 1.00683 | 0.556464 | -1.809 |
| chr2 | 104301612 | 104301905 | 0.959452 | 0.530311 | -1.809 |
| chr15 | 80134509 | 80134789 | 1.09587 | 0.605774 | -1.809 |
| chr12 | 103805317 | 103805534 | 1.22487 | 0.677164 | -1.809 |
| chr1 | 191558928 | 191559115 | 0.826821 | 0.45712 | -1.809 |
| chr2 | 148028549 | 148028923 | 1.12329 | 0.62105 | -1.809 |
| chr16 | 60653399 | 60653834 | 0.987968 | 0.546388 | -1.808 |
| chr20 | 50901892 | 50902136 | 1.15525 | 0.638995 | -1.808 |

| | | | | | |
|---|---|---|---|---|---|
| chr14 | 84751854 | 84752087 | 1.05649 | 0.584373 | -1.808 |
| chr17 | 41925798 | 41926013 | 1.203 | 0.665467 | -1.808 |
| chr5 | 12720833 | 12721362 | 1.12749 | 0.623705 | -1.808 |
| chr14 | 103551467 | 103551718 | 0.980191 | 0.542233 | -1.808 |
| chr19 | 52425496 | 52425871 | 1.05624 | 0.584611 | -1.807 |
| chr2 | 184762286 | 184762501 | 0.957885 | 0.530173 | -1.807 |
| chr15 | 97712073 | 97712517 | 0.864551 | 0.478548 | -1.807 |
| chr2 | 126986793 | 126987125 | 1.06501 | 0.589532 | -1.807 |
| chr15 | 98580885 | 98581102 | 1.1699 | 0.64767 | -1.806 |
| chr15 | 73575760 | 73576034 | 1.50045 | 0.830672 | -1.806 |
| chr15 | 29842168 | 29842381 | 1.11905 | 0.619532 | -1.806 |
| chr2 | 23514882 | 23515130 | 1.28119 | 0.709533 | -1.806 |
| chr3 | 163089326 | 163089559 | 0.874928 | 0.4847 | -1.805 |
| chr5 | 129337814 | 129338095 | 1.19684 | 0.66304 | -1.805 |
| chr1 | 223154189 | 223154564 | 1.28795 | 0.713629 | -1.805 |
| chr8 | 55887951 | 55888163 | 1.20651 | 0.668545 | -1.805 |
| chr1 | 240144811 | 240145083 | 1.04488 | 0.579082 | -1.804 |
| chr3 | 70581144 | 70581443 | 1.11342 | 0.617113 | -1.804 |
| chr10 | 26428574 | 26428766 | 1.19933 | 0.664843 | -1.804 |
| chr12 | 49925189 | 49925390 | 0.936147 | 0.519 | -1.804 |
| chr15 | 61132459 | 61132683 | 1.2124 | 0.67224 | -1.804 |
| chr5 | 59823004 | 59823289 | 0.925817 | 0.513465 | -1.803 |
| chr20 | 17450382 | 17450580 | 1.03393 | 0.573456 | -1.803 |
| chr5 | 117558586 | 117558933 | 0.94743 | 0.525538 | -1.803 |
| chr7 | 41551471 | 41551665 | 0.774671 | 0.429758 | -1.803 |
| chr9 | 27111257 | 27111490 | 1.0652 | 0.591112 | -1.802 |
| chr3 | 153147093 | 153147275 | 1.07471 | 0.596413 | -1.802 |
| chr2 | 11662612 | 11662835 | 1.0904 | 0.605137 | -1.802 |
| chr19 | 50749513 | 50749751 | 0.819846 | 0.455043 | -1.802 |
| chr3 | 165398711 | 165399032 | 0.874052 | 0.485237 | -1.801 |
| chr2 | 198992472 | 198992726 | 2.30972 | 1.28233 | -1.801 |
| chr12 | 126207373 | 126207649 | 0.89738 | 0.49828 | -1.801 |
| chr8 | 137178119 | 137178310 | 0.950671 | 0.528002 | -1.801 |
| chr11 | 55570664 | 55571121 | 1.38048 | 0.766743 | -1.800 |
| chr16 | 83107590 | 83107781 | 0.791282 | 0.439536 | -1.800 |
| chr14 | 92913902 | 92914087 | 1.24464 | 0.691404 | -1.800 |
| chr2 | 24326729 | 24326947 | 0.864077 | 0.480101 | -1.800 |
| chr7 | 85495625 | 85496057 | 0.744521 | 0.41369 | -1.800 |
| chr7 | 71024287 | 71024472 | 0.768648 | 0.42718 | -1.799 |
| chr2 | 116689070 | 116689305 | 1.23829 | 0.688206 | -1.799 |
| chr7 | 73947722 | 73948003 | 1.04057 | 0.578337 | -1.799 |
| chr1 | 220735541 | 220735770 | 1.0506 | 0.584085 | -1.799 |
| chr11 | 83613750 | 83613932 | 1.12286 | 0.624302 | -1.799 |
| chr14 | 38711161 | 38711393 | 1.3563 | 0.754127 | -1.799 |
| chr2 | 115969874 | 115970109 | 0.967791 | 0.538113 | -1.798 |
| chr7 | 98384448 | 98384661 | 0.782047 | 0.434936 | -1.798 |
| chr12 | 16865214 | 16865498 | 1.22046 | 0.678817 | -1.798 |
| chr20 | 51127799 | 51128203 | 0.933848 | 0.519419 | -1.798 |
| chr17 | 4193896 | 4194207 | 0.932508 | 0.518834 | -1.797 |
| chr14 | 38940707 | 38941000 | 0.92107 | 0.512471 | -1.797 |
| chr7 | 87206673 | 87206866 | 1.21256 | 0.674701 | -1.797 |
| chr17 | 13458254 | 13458485 | 1.00963 | 0.561829 | -1.797 |
| chr5 | 13279132 | 13279371 | 1.15768 | 0.644419 | -1.796 |
| chr14 | 26478804 | 26479014 | 1.20392 | 0.670162 | -1.796 |
| chr2 | 4216044 | 4216313 | 1.05235 | 0.585816 | -1.796 |
| chr17 | 40248807 | 40249116 | 1.28581 | 0.715786 | -1.796 |
| chr15 | 94342965 | 94343223 | 1.3779 | 0.767168 | -1.796 |
| chr2 | 17676184 | 17676374 | 0.898046 | 0.500151 | -1.796 |
| chr8 | 52156977 | 52157189 | 1.45432 | 0.810062 | -1.795 |
| chr8 | 76027521 | 76027738 | 0.824793 | 0.45942 | -1.795 |
| chr10 | 26151060 | 26151409 | 1.01911 | 0.5677 | -1.795 |

| | | | | | |
|---|---|---|---|---|---|
| chr2 | 168626495 | 168626807 | 0.971483 | 0.541203 | -1.795 |
| chr2 | 34773220 | 34773408 | 0.840917 | 0.468471 | -1.795 |
| chr2 | 63379543 | 63379743 | 1.47952 | 0.824356 | -1.795 |
| chr8 | 59071937 | 59072234 | 0.929793 | 0.518127 | -1.795 |
| chr7 | 8568904 | 8569148 | 0.983166 | 0.547882 | -1.794 |
| chr2 | 3249533 | 3249778 | 0.951744 | 0.530503 | -1.794 |
| chr3 | 73278205 | 73278434 | 0.996886 | 0.555744 | -1.794 |
| chr7 | 106309455 | 106309655 | 0.721792 | 0.402417 | -1.794 |
| chr5 | 50699261 | 50699547 | 1.32042 | 0.736431 | -1.793 |
| chr20 | 5369526 | 5369792 | 1.06823 | 0.595794 | -1.793 |
| chr9 | 107249397 | 107249605 | 0.753095 | 0.420038 | -1.793 |
| chr11 | 79521406 | 79521645 | 0.912618 | 0.509131 | -1.793 |
| chr9 | 34471276 | 34471476 | 1.2034 | 0.671375 | -1.792 |
| chr15 | 98333975 | 98334260 | 1.33381 | 0.744211 | -1.792 |
| chr15 | 35570393 | 35570638 | 1.08698 | 0.606495 | -1.792 |
| chr2 | 6532558 | 6532824 | 1.06412 | 0.593795 | -1.792 |
| chr11 | 28235178 | 28235362 | 1.00464 | 0.560653 | -1.792 |
| chr7 | 140028154 | 140028361 | 0.943557 | 0.526598 | -1.792 |
| chr11 | 64953406 | 64953644 | 1.2724 | 0.710232 | -1.792 |
| chr15 | 69273417 | 69273657 | 0.764392 | 0.426673 | -1.792 |
| chr15 | 87365112 | 87365418 | 1.09493 | 0.611201 | -1.791 |
| chr15 | 95453055 | 95453261 | 1.12835 | 0.630012 | -1.791 |
| chr2 | 12742950 | 12743195 | 1.27611 | 0.712613 | -1.791 |
| chr19 | 28694845 | 28695239 | 1.19547 | 0.667646 | -1.791 |
| chr2 | 155881733 | 155881925 | 0.993504 | 0.554869 | -1.791 |
| chr16 | 78965191 | 78965421 | 1.0834 | 0.605088 | -1.790 |
| chr15 | 77803166 | 77803408 | 1.0676 | 0.59632 | -1.790 |
| chr8 | 98183004 | 98183288 | 1.15632 | 0.645942 | -1.790 |
| chr2 | 215083186 | 215083531 | 0.921264 | 0.514671 | -1.790 |
| chr2 | 81941690 | 81942041 | 0.916739 | 0.51216 | -1.790 |
| chr3 | 145724127 | 145724403 | 0.950874 | 0.531425 | -1.789 |
| chr9 | 106972379 | 106972736 | 0.788505 | 0.440709 | -1.789 |
| chr14 | 99081019 | 99081296 | 0.913987 | 0.510895 | -1.789 |
| chr9 | 32847738 | 32848144 | 0.962132 | 0.537819 | -1.789 |
| chr7 | 126117931 | 126118123 | 1.10107 | 0.615745 | -1.788 |
| chr2 | 77933445 | 77933680 | 1.03901 | 0.581083 | -1.788 |
| chr14 | 77013137 | 77013482 | 1.87114 | 1.0465 | -1.788 |
| chr4 | 123111359 | 123111559 | 0.996632 | 0.557442 | -1.788 |
| chr5 | 84186810 | 84187081 | 0.92667 | 0.518336 | -1.788 |
| chr11 | 79944035 | 79944299 | 1.04401 | 0.584136 | -1.787 |
| chr5 | 86128489 | 86128763 | 1.06419 | 0.595427 | -1.787 |
| chr5 | 165428163 | 165428388 | 1.0129 | 0.566778 | -1.787 |
| chr14 | 42494468 | 42494658 | 1.27139 | 0.711423 | -1.787 |
| chr3 | 21271020 | 21271208 | 0.928172 | 0.519434 | -1.787 |
| chr20 | 6810239 | 6810713 | 1.01848 | 0.570015 | -1.787 |
| chr8 | 73823731 | 73823937 | 0.942816 | 0.527673 | -1.787 |
| chr7 | 77745912 | 77746176 | 1.13657 | 0.636176 | -1.787 |
| chr7 | 153063004 | 153063215 | 0.951309 | 0.53254 | -1.786 |
| chr20 | 60266642 | 60266963 | 0.974816 | 0.545799 | -1.786 |
| chr19 | 57307818 | 57308057 | 1.52038 | 0.851262 | -1.786 |
| chr1 | 183932979 | 183933197 | 1.00654 | 0.563583 | -1.786 |
| chr11 | 14214462 | 14214647 | 0.991885 | 0.555382 | -1.786 |
| chr2 | 53175600 | 53175806 | 1.11009 | 0.621584 | -1.786 |
| chr3 | 161404281 | 161404475 | 1.3524 | 0.757274 | -1.786 |
| chr15 | 54255188 | 54255409 | 1.90814 | 1.06865 | -1.786 |
| chr7 | 88850547 | 88850784 | 1.1048 | 0.618765 | -1.785 |
| chr12 | 84649261 | 84649553 | 1.30459 | 0.730971 | -1.785 |
| chr5 | 82687979 | 82688195 | 1.17377 | 0.657677 | -1.785 |
| chr12 | 33265223 | 33265429 | 1.13931 | 0.63843 | -1.785 |
| chr14 | 48195373 | 48195610 | 1.19984 | 0.672469 | -1.784 |
| chr5 | 38731674 | 38731931 | 1.2347 | 0.692008 | -1.784 |

| | | | | | |
|---|---|---|---|---|---|
| chr10 | 66305906 | 66306202 | 0.966228 | 0.541554 | -1.784 |
| chr8 | 83713677 | 83713883 | 0.86351 | 0.484002 | -1.784 |
| chr16 | 9509304 | 9509503 | 0.80182 | 0.449432 | -1.784 |
| chr8 | 136716224 | 136716457 | 0.966064 | 0.541556 | -1.784 |
| chr20 | 59161350 | 59161535 | 1.14815 | 0.643726 | -1.784 |
| chr3 | 109414070 | 109414288 | 1.07484 | 0.602694 | -1.783 |
| chr11 | 24579631 | 24579821 | 1.14419 | 0.641593 | -1.783 |
| chr1 | 192463711 | 192463938 | 1.09945 | 0.61659 | -1.783 |
| chr14 | 59498205 | 59498414 | 1.33336 | 0.747837 | -1.783 |
| chr3 | 73949517 | 73949891 | 0.900339 | 0.504978 | -1.783 |
| chr8 | 111568580 | 111568808 | 1.26255 | 0.708195 | -1.783 |
| chr19 | 29034871 | 29035057 | 1.14424 | 0.641867 | -1.783 |
| chr2 | 30919637 | 30919837 | 1.20372 | 0.675367 | -1.782 |
| chr10 | 106374072 | 106374427 | 1.8551 | 1.04094 | -1.782 |
| chr10 | 57256997 | 57257184 | 1.05245 | 0.590576 | -1.782 |
| chr2 | 166385259 | 166385621 | 1.0993 | 0.616952 | -1.782 |
| chr3 | 79922281 | 79922741 | 0.973757 | 0.546534 | -1.782 |
| chr1 | 166992401 | 166992585 | 1.092 | 0.613026 | -1.781 |
| chr3 | 97114586 | 97114828 | 1.09394 | 0.614169 | -1.781 |
| chr19 | 46042766 | 46043039 | 1.04328 | 0.585761 | -1.781 |
| chr20 | 49998162 | 49998403 | 1.22866 | 0.689898 | -1.781 |
| chr14 | 48445473 | 48445733 | 1.12656 | 0.632626 | -1.781 |
| chr15 | 98212878 | 98213154 | 0.782428 | 0.439381 | -1.781 |
| chr8 | 93190775 | 93190959 | 1.04283 | 0.585613 | -1.781 |
| chr2 | 47977005 | 47977245 | 1.10643 | 0.621397 | -1.781 |
| chr16 | 73991967 | 73992276 | 0.960363 | 0.539667 | -1.780 |
| chr3 | 80120476 | 80120716 | 1.19274 | 0.670537 | -1.779 |
| chr10 | 57876511 | 57876727 | 1.04347 | 0.58678 | -1.778 |
| chr3 | 99533256 | 99533447 | 1.0575 | 0.594802 | -1.778 |
| chr8 | 4808222 | 4808406 | 0.927252 | 0.52171 | -1.777 |
| chr10 | 107049171 | 107049518 | 0.870747 | 0.489934 | -1.777 |
| chr8 | 36114018 | 36114202 | 0.710704 | 0.399893 | -1.777 |
| chr2 | 36018063 | 36018262 | 1.02679 | 0.577768 | -1.777 |
| chr11 | 41001485 | 41001764 | 0.850635 | 0.478652 | -1.777 |
| chr17 | 27523010 | 27523230 | 1.07216 | 0.603392 | -1.777 |
| chr20 | 31549217 | 31549527 | 1.00882 | 0.567778 | -1.777 |
| chr12 | 114183378 | 114183580 | 1.1152 | 0.627665 | -1.777 |
| chr3 | 113354098 | 113354319 | 0.930546 | 0.523808 | -1.777 |
| chr5 | 28602052 | 28602306 | 1.10052 | 0.619664 | -1.776 |
| chr10 | 69034264 | 69034639 | 1.22053 | 0.687291 | -1.776 |
| chr7 | 119159699 | 119159889 | 1.15379 | 0.64972 | -1.776 |
| chr8 | 55852114 | 55852298 | 1.24698 | 0.702236 | -1.776 |
| chr7 | 2512597 | 2512786 | 1.34013 | 0.754811 | -1.775 |
| chr3 | 156009019 | 156009206 | 0.73359 | 0.413272 | -1.775 |
| chr2 | 80546561 | 80546801 | 0.978022 | 0.551156 | -1.774 |
| chr20 | 59383044 | 59383272 | 0.841205 | 0.474057 | -1.774 |
| chr2 | 151101790 | 151102068 | 1.05621 | 0.59531 | -1.774 |
| chr11 | 66325403 | 66325608 | 1.27524 | 0.718775 | -1.774 |
| chr16 | 20444971 | 20445156 | 0.972337 | 0.548082 | -1.774 |
| chr1 | 154792066 | 154792350 | 1.67091 | 0.941858 | -1.774 |
| chr7 | 136605094 | 136605374 | 0.920145 | 0.518692 | -1.774 |
| chr12 | 13430381 | 13430566 | 1.12359 | 0.633435 | -1.774 |
| chr14 | 46555919 | 46556151 | 0.900006 | 0.507406 | -1.774 |
| chr16 | 81992376 | 81992589 | 0.90077 | 0.507853 | -1.774 |
| chr2 | 14296914 | 14297148 | 0.956083 | 0.539065 | -1.774 |
| chr12 | 67046345 | 67046567 | 1.04133 | 0.587171 | -1.773 |
| chr20 | 54558987 | 54559235 | 1.08872 | 0.613924 | -1.773 |
| chr20 | 53939359 | 53939688 | 0.877488 | 0.494838 | -1.773 |
| chr2 | 100206651 | 100207112 | 1.12186 | 0.632658 | -1.773 |
| chr8 | 109331097 | 109331285 | 1.10406 | 0.622627 | -1.773 |
| chr7 | 122145887 | 122146224 | 1.15007 | 0.648612 | -1.773 |

| | | | | | |
|---|---|---|---|---|---|
| chr3 | 87355891 | 87356107 | 0.989735 | 0.558192 | -1.773 |
| chr8 | 77322072 | 77322258 | 0.799308 | 0.450825 | -1.773 |
| chr2 | 60064028 | 60064215 | 1.19299 | 0.672878 | -1.773 |
| chr11 | 64596952 | 64597155 | 0.787406 | 0.44415 | -1.773 |
| chr2 | 38748495 | 38748806 | 0.810655 | 0.457303 | -1.773 |
| chr9 | 25496522 | 25496796 | 0.912698 | 0.514878 | -1.773 |
| chr15 | 94517983 | 94518241 | 1.15801 | 0.653301 | -1.773 |
| chr16 | 46610369 | 46610583 | 0.900634 | 0.508105 | -1.773 |
| chr10 | 87630297 | 87630536 | 0.866382 | 0.488823 | -1.772 |
| chr7 | 152884376 | 152884727 | 0.854985 | 0.48242 | -1.772 |
| chr11 | 115401593 | 115401777 | 1.63412 | 0.922084 | -1.772 |
| chr17 | 48550154 | 48550390 | 1.87538 | 1.05828 | -1.772 |
| chr8 | 126992860 | 126993096 | 1.34616 | 0.759672 | -1.772 |
| chr14 | 41087522 | 41087906 | 1.19307 | 0.673299 | -1.772 |
| chr8 | 97746468 | 97746656 | 0.996191 | 0.562228 | -1.772 |
| chr5 | 158839350 | 158839549 | 1.15555 | 0.652168 | -1.772 |
| chr20 | 57591413 | 57591681 | 1.01469 | 0.572705 | -1.772 |
| chr5 | 61362010 | 61362352 | 0.963952 | 0.544069 | -1.772 |
| chr7 | 9382030 | 9382223 | 1.0114 | 0.570856 | -1.772 |
| chr2 | 77139732 | 77139991 | 0.921605 | 0.520192 | -1.772 |
| chr19 | 35153641 | 35153824 | 1.34768 | 0.76069 | -1.772 |
| chr15 | 61641612 | 61641802 | 0.994139 | 0.561182 | -1.772 |
| chr9 | 102062082 | 102062316 | 1.06161 | 0.599341 | -1.771 |
| chr12 | 123241023 | 123241272 | 1.23931 | 0.699839 | -1.771 |
| chr2 | 167149138 | 167149412 | 1.08003 | 0.6099 | -1.771 |
| chr9 | 110111250 | 110111547 | 1.03173 | 0.582637 | -1.771 |
| chr10 | 98177520 | 98177764 | 0.976966 | 0.551718 | -1.771 |
| chr2 | 76791004 | 76791394 | 1.29105 | 0.72916 | -1.771 |
| chr1 | 187883603 | 187883786 | 0.865778 | 0.488985 | -1.771 |
| chr17 | 78607062 | 78607282 | 1.04221 | 0.58867 | -1.770 |
| chr2 | 51425102 | 51425459 | 1.00364 | 0.566887 | -1.770 |
| chr2 | 116779699 | 116779922 | 0.856015 | 0.483515 | -1.770 |
| chr10 | 70533271 | 70533463 | 0.948327 | 0.53568 | -1.770 |
| chr1 | 89361298 | 89361483 | 1.15004 | 0.649645 | -1.770 |
| chr5 | 109389406 | 109389607 | 1.16093 | 0.655797 | -1.770 |
| chr5 | 104113733 | 104113930 | 0.946066 | 0.534499 | -1.770 |
| chr2 | 2260572 | 2260799 | 0.927036 | 0.523766 | -1.770 |
| chr1 | 174791735 | 174791946 | 0.975458 | 0.551174 | -1.770 |
| chr5 | 62682358 | 62682709 | 1.11781 | 0.631639 | -1.770 |
| chr1 | 194860266 | 194860523 | 0.876251 | 0.495211 | -1.769 |
| chr8 | 126755485 | 126755691 | 1.38876 | 0.784931 | -1.769 |
| chr19 | 24318274 | 24318522 | 1.11733 | 0.631604 | -1.769 |
| chr2 | 13477017 | 13477233 | 1.06836 | 0.603938 | -1.769 |
| chr18 | 11777995 | 11778188 | 1.01556 | 0.574151 | -1.769 |
| chr7 | 123137775 | 123138234 | 1.25159 | 0.707686 | -1.769 |
| chr5 | 104089875 | 104090090 | 0.931213 | 0.526549 | -1.769 |
| chr2 | 57236996 | 57237195 | 0.703787 | 0.397981 | -1.768 |
| chr5 | 89522427 | 89522625 | 1.37384 | 0.776898 | -1.768 |
| chr7 | 55975130 | 55975324 | 1.14764 | 0.649125 | -1.768 |
| chr2 | 151726691 | 151726949 | 1.02974 | 0.582491 | -1.768 |
| chr15 | 86185819 | 86186072 | 0.97666 | 0.5525 | -1.768 |
| chr1 | 177249678 | 177249872 | 0.787469 | 0.445552 | -1.767 |
| chr3 | 85075006 | 85075215 | 1.4047 | 0.794796 | -1.767 |
| chr3 | 81522670 | 81522924 | 0.830722 | 0.470154 | -1.767 |
| chr8 | 137325111 | 137325571 | 1.33396 | 0.75501 | -1.767 |
| chr3 | 7564989 | 7565221 | 0.834241 | 0.47223 | -1.767 |
| chr8 | 139253090 | 139253462 | 0.860429 | 0.487081 | -1.767 |
| chr1 | 247004561 | 247004828 | 0.996654 | 0.564259 | -1.766 |
| chr12 | 81964815 | 81965047 | 1.03536 | 0.586206 | -1.766 |
| chr11 | 43908996 | 43909205 | 0.89785 | 0.508391 | -1.766 |
| chr12 | 93309057 | 93309284 | 1.1876 | 0.672543 | -1.766 |

| | | | | | |
|---|---|---|---|---|---|
| chr11 | 55304027 | 55304219 | 0.879499 | 0.498096 | -1.766 |
| chr2 | 76789098 | 76789366 | 0.975467 | 0.552598 | -1.765 |
| chr20 | 42590861 | 42591173 | 1.20367 | 0.681877 | -1.765 |
| chr4 | 139519979 | 139520175 | 0.967034 | 0.54788 | -1.765 |
| chr2 | 49938629 | 49939169 | 0.852652 | 0.483153 | -1.765 |
| chr17 | 54663856 | 54664087 | 0.983727 | 0.557446 | -1.765 |
| chr2 | 80493418 | 80493637 | 1.02183 | 0.579066 | -1.765 |
| chr9 | 11428988 | 11429188 | 1.09937 | 0.623015 | -1.765 |
| chr5 | 164880691 | 164880885 | 1.2347 | 0.699788 | -1.764 |
| chr19 | 23510932 | 23511576 | 0.944188 | 0.53517 | -1.764 |
| chr9 | 25157623 | 25157818 | 1.08196 | 0.61328 | -1.764 |
| chr3 | 97995094 | 97995333 | 1.12864 | 0.639776 | -1.764 |
| chr9 | 140658096 | 140658285 | 0.833569 | 0.472514 | -1.764 |
| chr9 | 105170155 | 105170357 | 0.744422 | 0.421983 | -1.764 |
| chr15 | 92215880 | 92216163 | 1.00965 | 0.572373 | -1.764 |
| chr11 | 79428262 | 79428465 | 1.0296 | 0.583697 | -1.764 |
| chr2 | 185187163 | 185187412 | 0.978364 | 0.554668 | -1.764 |
| chr8 | 4609125 | 4609348 | 0.897121 | 0.508633 | -1.764 |
| chr12 | 131347810 | 131348136 | 1.03793 | 0.588509 | -1.764 |
| chr8 | 74566244 | 74566506 | 1.24011 | 0.703173 | -1.764 |
| chr2 | 155746276 | 155746717 | 1.1211 | 0.63572 | -1.764 |
| chr8 | 27937122 | 27937431 | 0.991348 | 0.562252 | -1.763 |
| chr19 | 28954375 | 28954623 | 1.10305 | 0.625622 | -1.763 |
| chr2 | 154198107 | 154198349 | 0.809162 | 0.458954 | -1.763 |
| chr11 | 78426785 | 78427061 | 2.03466 | 1.15427 | -1.763 |
| chr10 | 33337094 | 33337282 | 1.14031 | 0.646939 | -1.763 |
| chr1 | 245373197 | 245373564 | 0.646498 | 0.36679 | -1.763 |
| chr8 | 133979946 | 133980154 | 1.20074 | 0.681297 | -1.762 |
| chr8 | 139638438 | 139638771 | 1.19007 | 0.675315 | -1.762 |
| chr3 | 165441645 | 165441845 | 0.965138 | 0.547712 | -1.762 |
| chr11 | 55563438 | 55563797 | 1.25875 | 0.714371 | -1.762 |
| chr7 | 92518454 | 92518913 | 0.856576 | 0.486202 | -1.762 |
| chr3 | 146650479 | 146650721 | 0.755756 | 0.428983 | -1.762 |
| chr5 | 162153565 | 162153749 | 1.03041 | 0.584995 | -1.761 |
| chr8 | 107465449 | 107465885 | 0.79996 | 0.454217 | -1.761 |
| chr7 | 113561754 | 113562098 | 0.815287 | 0.462925 | -1.761 |
| chr12 | 62469135 | 62469327 | 1.0294 | 0.58455 | -1.761 |
| chr2 | 76903058 | 76903400 | 1.20125 | 0.682196 | -1.761 |
| chr2 | 38808646 | 38808983 | 1.72376 | 0.978968 | -1.761 |
| chr11 | 39388704 | 39388987 | 1.16245 | 0.660199 | -1.761 |
| chr9 | 136210906 | 136211102 | 1.01988 | 0.579232 | -1.761 |
| chr2 | 214785293 | 214785549 | 0.888082 | 0.504443 | -1.761 |
| chr3 | 84206231 | 84206507 | 0.783554 | 0.445244 | -1.760 |
| chr5 | 28455300 | 28455549 | 1.13249 | 0.643637 | -1.760 |
| chr9 | 105595310 | 105595753 | 1.0301 | 0.585464 | -1.759 |
| chr17 | 32393350 | 32393588 | 1.00685 | 0.572255 | -1.759 |
| chr19 | 56729075 | 56729376 | 1.09173 | 0.620517 | -1.759 |
| chr17 | 214095 | 214330 | 1.20486 | 0.684846 | -1.759 |
| chr2 | 81644408 | 81644640 | 0.86745 | 0.493088 | -1.759 |
| chr15 | 88215643 | 88215945 | 1.69207 | 0.961949 | -1.759 |
| chr9 | 2979981 | 2980287 | 1.14861 | 0.653154 | -1.759 |
| chr17 | 74113534 | 74113838 | 1.03936 | 0.591037 | -1.759 |
| chr3 | 62608352 | 62608589 | 1.94045 | 1.10348 | -1.758 |
| chr3 | 169326081 | 169326294 | 1.37764 | 0.783484 | -1.758 |
| chr2 | 46443987 | 46444334 | 1.20934 | 0.687831 | -1.758 |
| chr14 | 94202836 | 94203030 | 2.48191 | 1.41178 | -1.758 |
| chr7 | 28607291 | 28607494 | 0.934365 | 0.531506 | -1.758 |
| chr7 | 52220627 | 52220821 | 1.1088 | 0.630735 | -1.758 |
| chr19 | 57580012 | 57580356 | 1.34033 | 0.762468 | -1.758 |
| chr2 | 12585672 | 12585895 | 1.03976 | 0.591524 | -1.758 |
| chr14 | 87987511 | 87987703 | 0.93505 | 0.53208 | -1.757 |

| | | | | | |
|---|---|---|---|---|---|
| chr4 | 65837603 | 65837803 | 1.25307 | 0.713063 | -1.757 |
| chr7 | 54593402 | 54593774 | 1.12297 | 0.639138 | -1.757 |
| chr11 | 42620035 | 42620233 | 1.17162 | 0.666861 | -1.757 |
| chr2 | 30012910 | 30013093 | 0.886802 | 0.504755 | -1.757 |
| chr12 | 33245333 | 33245567 | 1.1207 | 0.638208 | -1.756 |
| chr17 | 50366921 | 50367152 | 1.02243 | 0.58225 | -1.756 |
| chr7 | 51879724 | 51879933 | 1.36553 | 0.77764 | -1.756 |
| chr8 | 76648289 | 76648734 | 1.13841 | 0.648324 | -1.756 |
| chr2 | 81193941 | 81194153 | 1.07274 | 0.610999 | -1.756 |
| chr19 | 29594011 | 29594292 | 1.43303 | 0.816227 | -1.756 |
| chr8 | 43627719 | 43627917 | 1.07602 | 0.612955 | -1.755 |
| chr9 | 24122691 | 24123006 | 0.796285 | 0.453615 | -1.755 |
| chr5 | 88308260 | 88308491 | 0.978988 | 0.557739 | -1.755 |
| chr14 | 84893277 | 84893552 | 1.21833 | 0.694117 | -1.755 |
| chr7 | 156478141 | 156478373 | 1.28653 | 0.732973 | -1.755 |
| chr15 | 72882235 | 72882480 | 1.16885 | 0.66599 | -1.755 |
| chr7 | 9755333 | 9755547 | 0.978061 | 0.557288 | -1.755 |
| chr5 | 111212808 | 111213048 | 0.876769 | 0.499623 | -1.755 |
| chr2 | 80913184 | 80913402 | 1.23325 | 0.702843 | -1.755 |
| chr16 | 24238150 | 24238514 | 0.837064 | 0.477134 | -1.754 |
| chr2 | 10139483 | 10139677 | 1.05486 | 0.601356 | -1.754 |
| chr2 | 41158265 | 41158522 | 1.10622 | 0.63074 | -1.754 |
| chr19 | 44921675 | 44921897 | 0.91905 | 0.524033 | -1.754 |
| chr12 | 101663751 | 101663961 | 1.02248 | 0.583034 | -1.754 |
| chr12 | 127822889 | 127823085 | 0.963488 | 0.54946 | -1.754 |
| chr19 | 57674582 | 57674892 | 1.00287 | 0.571941 | -1.753 |
| chr5 | 18436505 | 18436732 | 0.99733 | 0.568911 | -1.753 |
| chr8 | 25245172 | 25245362 | 1.55568 | 0.887538 | -1.753 |
| chr3 | 77254101 | 77254284 | 1.12371 | 0.64118 | -1.753 |
| chr3 | 86806362 | 86806587 | 1.08764 | 0.620608 | -1.753 |
| chr7 | 125414826 | 125415061 | 1.07816 | 0.615225 | -1.752 |
| chr12 | 15415206 | 15415436 | 1.22361 | 0.698237 | -1.752 |
| chr19 | 384005 | 384301 | 1.8524 | 1.05712 | -1.752 |
| chr9 | 12515990 | 12516239 | 0.907571 | 0.518001 | -1.752 |
| chrX | 132306098 | 132306286 | 1.42944 | 0.815899 | -1.752 |
| chr12 | 127980760 | 127980942 | 0.917387 | 0.523685 | -1.752 |
| chr16 | 63035859 | 63036041 | 1.18817 | 0.678373 | -1.752 |
| chr14 | 46616877 | 46617188 | 1.07075 | 0.611345 | -1.751 |
| chr3 | 19131758 | 19132056 | 1.01649 | 0.580421 | -1.751 |
| chr1 | 229302786 | 229303040 | 0.97428 | 0.556548 | -1.751 |
| chr2 | 219117299 | 219117539 | 1.01545 | 0.580112 | -1.750 |
| chr15 | 81808454 | 81808746 | 0.982261 | 0.561214 | -1.750 |
| chr20 | 54502728 | 54502949 | 0.985262 | 0.562952 | -1.750 |
| chr10 | 107069039 | 107069335 | 1.09611 | 0.62641 | -1.750 |
| chr1 | 165043552 | 165043821 | 1.17309 | 0.670516 | -1.750 |
| chr15 | 77251659 | 77251859 | 1.20087 | 0.686565 | -1.749 |
| chr10 | 83684637 | 83684961 | 0.941742 | 0.538428 | -1.749 |
| chr2 | 16819622 | 16819884 | 1.0003 | 0.571911 | -1.749 |
| chr6 | 15648459 | 15648659 | 0.841314 | 0.481095 | -1.749 |
| chr7 | 117852455 | 117852770 | 0.999942 | 0.571863 | -1.749 |
| chr5 | 104546907 | 104547130 | 1.07974 | 0.617528 | -1.748 |
| chr10 | 9763923 | 9764155 | 0.991107 | 0.566953 | -1.748 |
| chr8 | 21378804 | 21379001 | 0.852394 | 0.487622 | -1.748 |
| chr5 | 6793346 | 6793538 | 1.02764 | 0.587875 | -1.748 |
| chr1 | 157585585 | 157585774 | 0.781574 | 0.447125 | -1.748 |
| chr10 | 58214904 | 58215107 | 1.00251 | 0.573679 | -1.748 |
| chr2 | 26149987 | 26150187 | 1.04935 | 0.600517 | -1.747 |
| chr8 | 109308030 | 109308282 | 0.939801 | 0.537936 | -1.747 |
| chr2 | 154082420 | 154082815 | 0.924664 | 0.529289 | -1.747 |
| chr11 | 23023186 | 23023476 | 0.806456 | 0.461631 | -1.747 |
| chr14 | 46276343 | 46276547 | 0.875195 | 0.501022 | -1.747 |

| | | | | | |
|---|---|---|---|---|---|
| chr1 | 237813130 | 237813376 | 0.982157 | 0.562325 | -1.747 |
| chr17 | 65704543 | 65704776 | 0.941195 | 0.538953 | -1.746 |
| chr12 | 126256989 | 126257189 | 1.2753 | 0.73028 | -1.746 |
| chr3 | 143943812 | 143944054 | 0.951139 | 0.544746 | -1.746 |
| chr9 | 124298216 | 124298406 | 0.89555 | 0.512928 | -1.746 |
| chr14 | 104351839 | 104352052 | 0.970133 | 0.555663 | -1.746 |
| chr17 | 4859912 | 4860194 | 1.21575 | 0.696399 | -1.746 |
| chr19 | 53104621 | 53104869 | 0.722853 | 0.414066 | -1.746 |
| chr2 | 16519502 | 16519781 | 1.02904 | 0.589517 | -1.746 |
| chr15 | 50694465 | 50694799 | 0.990916 | 0.56775 | -1.745 |
| chr17 | 51255043 | 51255228 | 0.981635 | 0.562545 | -1.745 |
| chr9 | 9363499 | 9363694 | 1.00271 | 0.574666 | -1.745 |
| chr9 | 20200898 | 20201113 | 1.07388 | 0.615459 | -1.745 |
| chr7 | 90683750 | 90684063 | 0.835235 | 0.478694 | -1.745 |
| chr19 | 52765345 | 52765537 | 1.28553 | 0.736782 | -1.745 |
| chr3 | 73932140 | 73932429 | 0.843044 | 0.483196 | -1.745 |
| chr10 | 57397204 | 57397386 | 1.04647 | 0.599851 | -1.745 |
| chr8 | 121898030 | 121898229 | 0.959698 | 0.550132 | -1.744 |
| chr8 | 92972411 | 92972596 | 1.14303 | 0.655255 | -1.744 |
| chr14 | 22106212 | 22106423 | 1.17464 | 0.673391 | -1.744 |
| chr10 | 4453270 | 4453807 | 0.896266 | 0.513864 | -1.744 |
| chr15 | 82018758 | 82019022 | 1.03148 | 0.591389 | -1.744 |
| chr2 | 40231653 | 40231862 | 1.09181 | 0.625984 | -1.744 |
| chr2 | 51446631 | 51446838 | 1.02694 | 0.588838 | -1.744 |
| chr14 | 58063886 | 58064127 | 1.05018 | 0.602311 | -1.744 |
| chr5 | 179714832 | 179715057 | 0.70623 | 0.405192 | -1.743 |
| chr20 | 21851115 | 21851372 | 1.24176 | 0.712461 | -1.743 |
| chr17 | 69892471 | 69892782 | 0.88085 | 0.505392 | -1.743 |
| chr7 | 116946531 | 116946738 | 0.944709 | 0.542038 | -1.743 |
| chr12 | 29225585 | 29225853 | 1.08232 | 0.621014 | -1.743 |
| chr9 | 113545138 | 113545352 | 1.49216 | 0.856206 | -1.743 |
| chr5 | 78726926 | 78727171 | 1.11465 | 0.639612 | -1.743 |
| chr15 | 98925513 | 98925748 | 1.05034 | 0.602863 | -1.742 |
| chr17 | 42820239 | 42820543 | 1.16551 | 0.669001 | -1.742 |
| chr2 | 82097242 | 82097447 | 0.930912 | 0.534343 | -1.742 |
| chr2 | 179843396 | 179843587 | 0.96863 | 0.556002 | -1.742 |
| chr11 | 31128599 | 31128811 | 1.85831 | 1.06677 | -1.742 |
| chr2 | 171649978 | 171650206 | 1.12388 | 0.645214 | -1.742 |
| chr15 | 61392672 | 61392888 | 1.0287 | 0.590632 | -1.742 |
| chr8 | 73831476 | 73831689 | 1.0095 | 0.579669 | -1.742 |
| chr7 | 106658336 | 106658604 | 1.08532 | 0.62325 | -1.741 |
| chr2 | 57082434 | 57082806 | 0.994014 | 0.570896 | -1.741 |
| chr8 | 144411335 | 144411705 | 1.0692 | 0.61426 | -1.741 |
| rUn_gl0002 | 9273 | 9455 | 0.929047 | 0.53377 | -1.741 |
| chr19 | 35359761 | 35359975 | 1.10616 | 0.635557 | -1.740 |
| chr9 | 7947711 | 7947899 | 0.94994 | 0.54587 | -1.740 |
| chr8 | 139517342 | 139517535 | 1.06798 | 0.613762 | -1.740 |
| chr2 | 13351350 | 13351541 | 1.06666 | 0.613019 | -1.740 |
| chr15 | 92056451 | 92056725 | 1.14934 | 0.660553 | -1.740 |
| chr2 | 5011736 | 5011994 | 0.956154 | 0.549606 | -1.740 |
| chr8 | 33747460 | 33747714 | 0.961971 | 0.553013 | -1.740 |
| chr10 | 54500697 | 54501135 | 2.45122 | 1.40922 | -1.739 |
| chr9 | 96401950 | 96402193 | 0.918079 | 0.527821 | -1.739 |
| chr11 | 32189329 | 32189604 | 0.96214 | 0.553174 | -1.739 |
| chr16 | 56348794 | 56349007 | 1.09343 | 0.628735 | -1.739 |
| chr12 | 59813529 | 59813889 | 1.25188 | 0.719907 | -1.739 |
| chr14 | 79224669 | 79225031 | 1.35036 | 0.776553 | -1.739 |
| chr17 | 54216764 | 54216999 | 1.23556 | 0.71056 | -1.739 |
| chr15 | 32102912 | 32103131 | 1.18386 | 0.680848 | -1.739 |
| chr2 | 222156510 | 222156743 | 0.819243 | 0.471206 | -1.739 |
| chr8 | 145658880 | 145659145 | 0.988129 | 0.568347 | -1.739 |

| | | | | | |
|---|---|---|---|---|---|
| chr17 | 14862377 | 14862565 | 1.25175 | 0.719986 | -1.739 |
| chr2 | 79985706 | 79985925 | 0.795458 | 0.457606 | -1.738 |
| chr11 | 79658969 | 79659208 | 1.05376 | 0.606326 | -1.738 |
| chr10 | 100510071 | 100510290 | 0.871137 | 0.501288 | -1.738 |
| chr10 | 110851365 | 110851572 | 1.10845 | 0.637857 | -1.738 |
| chr5 | 15930691 | 15930995 | 0.841422 | 0.484213 | -1.738 |
| chr7 | 8731511 | 8731729 | 1.0376 | 0.597142 | -1.738 |
| chr7 | 66956757 | 66957002 | 0.958947 | 0.5519 | -1.738 |
| chr9 | 78641837 | 78642104 | 1.21717 | 0.700541 | -1.737 |
| chr14 | 32973580 | 32973767 | 0.928509 | 0.534551 | -1.737 |
| chr2 | 54598761 | 54598947 | 1.25109 | 0.720276 | -1.737 |
| chr2 | 4833734 | 4833974 | 1.49741 | 0.862089 | -1.737 |
| chr9 | 106462749 | 106463065 | 0.925636 | 0.532914 | -1.737 |
| chr5 | 165286607 | 165286791 | 1.04394 | 0.601069 | -1.737 |
| chr7 | 142003478 | 142003864 | 1.46696 | 0.844677 | -1.737 |
| chr19 | 30998838 | 30999338 | 1.49692 | 0.861955 | -1.737 |
| chr19 | 57318932 | 57319133 | 1.42845 | 0.822542 | -1.737 |
| chr5 | 147266620 | 147266893 | 1.40519 | 0.809148 | -1.737 |
| chr2 | 7690623 | 7690822 | 1.17578 | 0.677049 | -1.737 |
| chr8 | 36950347 | 36950535 | 1.0651 | 0.613343 | -1.737 |
| chr2 | 185710248 | 185710549 | 1.09538 | 0.630869 | -1.736 |
| chr5 | 21257088 | 21257293 | 0.884655 | 0.509529 | -1.736 |
| chr2 | 133206411 | 133206653 | 2.22595 | 1.28235 | -1.736 |
| chr2 | 11584880 | 11585125 | 1.51234 | 0.87136 | -1.736 |
| chr11 | 38499455 | 38499701 | 1.22368 | 0.705059 | -1.736 |
| chr16 | 6038441 | 6038640 | 1.12902 | 0.650538 | -1.736 |
| chr19 | 30901640 | 30901823 | 1.12476 | 0.64812 | -1.735 |
| chr5 | 12428702 | 12428884 | 0.897759 | 0.517571 | -1.735 |
| chr1 | 222236128 | 222236478 | 0.850109 | 0.490109 | -1.735 |
| chr7 | 126140395 | 126140648 | 0.933744 | 0.538329 | -1.735 |
| chr17 | 69633347 | 69633562 | 1.09597 | 0.632012 | -1.734 |
| chr17 | 20709960 | 20710245 | 0.849265 | 0.489757 | -1.734 |
| chr19 | 35764682 | 35764897 | 1.37998 | 0.795867 | -1.734 |
| chr3 | 155161837 | 155162096 | 1.02694 | 0.592261 | -1.734 |
| chr5 | 101233641 | 101233840 | 0.835799 | 0.482086 | -1.734 |
| chr5 | 27789635 | 27789822 | 0.924983 | 0.533548 | -1.734 |
| chr7 | 146828985 | 146829183 | 1.11326 | 0.642194 | -1.734 |
| chr7 | 125517439 | 125517637 | 1.29903 | 0.749409 | -1.733 |
| chr5 | 165422368 | 165422620 | 1.05625 | 0.609394 | -1.733 |
| chr7 | 115161929 | 115162124 | 1.06211 | 0.612784 | -1.733 |
| chr16 | 3500212 | 3500477 | 0.875087 | 0.504885 | -1.733 |
| chr2 | 230231543 | 230231764 | 1.05775 | 0.610293 | -1.733 |
| chr17 | 16566613 | 16566841 | 1.1839 | 0.683188 | -1.733 |
| chr10 | 100171052 | 100171234 | 0.939455 | 0.542179 | -1.733 |
| chr2 | 124069820 | 124070110 | 0.885458 | 0.511053 | -1.733 |
| chr2 | 5798287 | 5798558 | 1.0832 | 0.625198 | -1.733 |
| chr4 | 56050840 | 56051046 | 1.64895 | 0.951787 | -1.732 |
| chr7 | 103892740 | 103893106 | 1.44232 | 0.832592 | -1.732 |
| chr12 | 44680720 | 44681047 | 0.872655 | 0.503786 | -1.732 |
| chr11 | 55434928 | 55435156 | 0.818742 | 0.472689 | -1.732 |
| chr8 | 113599504 | 113599703 | 0.902494 | 0.521046 | -1.732 |
| chr3 | 18570954 | 18571156 | 0.854376 | 0.493281 | -1.732 |
| chr6 | 123539753 | 123539966 | 0.898196 | 0.518674 | -1.732 |
| chr10 | 57783180 | 57783383 | 0.913853 | 0.52777 | -1.732 |
| chr14 | 86473239 | 86473555 | 1.28046 | 0.739549 | -1.731 |
| chr16 | 62394157 | 62394363 | 0.881124 | 0.508931 | -1.731 |
| chr8 | 52277712 | 52277895 | 1.18188 | 0.682776 | -1.731 |
| chr8 | 134905836 | 134906100 | 0.937124 | 0.541403 | -1.731 |
| chr2 | 82263481 | 82263749 | 1.21255 | 0.700553 | -1.731 |
| chr3 | 99218012 | 99218322 | 1.74752 | 1.00965 | -1.731 |
| chr16 | 77694351 | 77694606 | 2.09946 | 1.21299 | -1.731 |

| | | | | | |
|---|---|---|---|---|---|
| chr17 | 48903417 | 48903607 | 1.60922 | 0.929831 | -1.731 |
| chr2 | 45483959 | 45484146 | 0.991051 | 0.572668 | -1.731 |
| chr2 | 14945790 | 14946045 | 0.933466 | 0.539409 | -1.731 |
| chr2 | 16399959 | 16400469 | 1.6823 | 0.97213 | -1.731 |
| chr5 | 132464958 | 132465328 | 0.965865 | 0.558205 | -1.730 |
| chr9 | 104026639 | 104026847 | 0.852395 | 0.492659 | -1.730 |
| chr11 | 66630250 | 66630432 | 1.00469 | 0.580716 | -1.730 |
| chr2 | 67529137 | 67529335 | 0.977343 | 0.56493 | -1.730 |
| chr16 | 7686024 | 7686218 | 1.28705 | 0.743956 | -1.730 |
| chr17 | 52520124 | 52520325 | 0.793532 | 0.45869 | -1.730 |
| chr12 | 15267520 | 15267716 | 1.11382 | 0.644087 | -1.729 |
| chr14 | 83725166 | 83725367 | 1.37931 | 0.797685 | -1.729 |
| chr11 | 66016815 | 66017066 | 0.883997 | 0.511269 | -1.729 |
| chr7 | 77867907 | 77868235 | 0.993793 | 0.574815 | -1.729 |
| chr5 | 161470462 | 161470700 | 1.03918 | 0.6011 | -1.729 |
| chr11 | 37113297 | 37113649 | 0.888106 | 0.513738 | -1.729 |
| chr3 | 3282141 | 3282415 | 0.912487 | 0.527893 | -1.729 |
| chr10 | 56485566 | 56485813 | 1.23549 | 0.714827 | -1.728 |
| chr7 | 119056015 | 119056244 | 1.39749 | 0.808641 | -1.728 |
| chr5 | 89376214 | 89376590 | 0.987958 | 0.571695 | -1.728 |
| chr8 | 130390510 | 130390772 | 0.999466 | 0.578366 | -1.728 |
| chr12 | 88769563 | 88769757 | 1.00555 | 0.581993 | -1.728 |
| chr4 | 142402843 | 142403056 | 1.17902 | 0.682406 | -1.728 |
| chr10 | 1534677 | 1534961 | 0.970865 | 0.562033 | -1.727 |
| chr17 | 69181065 | 69181251 | 1.12814 | 0.653114 | -1.727 |
| chr1 | 164937280 | 164937524 | 0.83079 | 0.480988 | -1.727 |
| chr2 | 77348707 | 77348964 | 0.992063 | 0.574444 | -1.727 |
| chr15 | 94993516 | 94993772 | 1.37677 | 0.79739 | -1.727 |
| chr11 | 41216092 | 41216295 | 1.16881 | 0.676984 | -1.726 |
| chr19 | 42170013 | 42170473 | 1.30635 | 0.756651 | -1.726 |
| chr14 | 84259966 | 84260205 | 1.28257 | 0.743035 | -1.726 |
| chr20 | 40651944 | 40652205 | 0.916904 | 0.531201 | -1.726 |
| chr16 | 64161489 | 64161700 | 0.798064 | 0.462363 | -1.726 |
| chr16 | 17813386 | 17813587 | 1.04983 | 0.608271 | -1.726 |
| chr11 | 17586345 | 17586546 | 1.50925 | 0.874545 | -1.726 |
| chr7 | 146433242 | 146433536 | 0.9557 | 0.553794 | -1.726 |
| chr2 | 147045044 | 147045292 | 0.989587 | 0.573452 | -1.726 |
| chr15 | 29251635 | 29251995 | 0.941348 | 0.54555 | -1.726 |
| chr8 | 50105705 | 50105906 | 0.939932 | 0.544733 | -1.725 |
| chr16 | 25685936 | 25686171 | 1.21139 | 0.702093 | -1.725 |
| chr9 | 31730572 | 31730813 | 1.36663 | 0.79215 | -1.725 |
| chr2 | 10679839 | 10680049 | 1.11437 | 0.646019 | -1.725 |
| chr9 | 121741930 | 121742158 | 0.903157 | 0.52365 | -1.725 |
| chr5 | 164274879 | 164275063 | 0.900456 | 0.522096 | -1.725 |
| chr3 | 97161151 | 97161415 | 0.974808 | 0.565476 | -1.724 |
| chr2 | 199364314 | 199364570 | 0.916679 | 0.531851 | -1.724 |
| chr7 | 77620802 | 77621066 | 1.02481 | 0.594606 | -1.724 |
| chr9 | 32233080 | 32233280 | 0.907373 | 0.526601 | -1.723 |
| chr5 | 42005701 | 42005924 | 1.48051 | 0.859289 | -1.723 |
| chr3 | 61922071 | 61922292 | 1.58449 | 0.919744 | -1.723 |
| chr5 | 38049421 | 38049630 | 0.888635 | 0.515841 | -1.723 |
| chr7 | 136689063 | 136689307 | 1.12678 | 0.654245 | -1.722 |
| chr19 | 23817780 | 23818018 | 0.760828 | 0.441763 | -1.722 |
| chr11 | 59254947 | 59255254 | 0.867338 | 0.503702 | -1.722 |
| chr10 | 78448368 | 78448676 | 1.23436 | 0.716962 | -1.722 |
| chr8 | 117992684 | 117992914 | 1.00005 | 0.58088 | -1.722 |
| chr1 | 164050226 | 164050502 | 0.917717 | 0.533071 | -1.722 |
| chr5 | 7449771 | 7450136 | 1.09168 | 0.634161 | -1.721 |
| chr9 | 76396172 | 76396369 | 1.29256 | 0.750892 | -1.721 |
| chr5 | 77288372 | 77288584 | 1.39708 | 0.811695 | -1.721 |
| chr8 | 64723529 | 64723786 | 1.17674 | 0.683743 | -1.721 |

| | | | | | |
|---|---|---|---|---|---|
| chr2 | 221624822 | 221625100 | 0.934307 | 0.543035 | -1.721 |
| chr8 | 139609389 | 139609625 | 0.881331 | 0.512262 | -1.720 |
| chr1 | 226445618 | 226445861 | 1.02157 | 0.5938 | -1.720 |
| chr8 | 137349800 | 137350148 | 1.00932 | 0.586683 | -1.720 |
| chr15 | 24451207 | 24451470 | 1.12857 | 0.65606 | -1.720 |
| chr11 | 37269864 | 37270053 | 1.01206 | 0.588338 | -1.720 |
| chr10 | 10682270 | 10682482 | 1.38802 | 0.806993 | -1.720 |
| chr3 | 95179252 | 95179453 | 0.83547 | 0.485756 | -1.720 |
| chr2 | 77285428 | 77285635 | 1.13178 | 0.658064 | -1.720 |
| chr17 | 80420110 | 80420520 | 1.2351 | 0.718189 | -1.720 |
| chr9 | 30717223 | 30717452 | 0.790363 | 0.459628 | -1.720 |
| chr12 | 115836157 | 115836498 | 1.58365 | 0.921249 | -1.719 |
| chr14 | 54843422 | 54843675 | 0.969608 | 0.564085 | -1.719 |
| chr3 | 30843626 | 30843879 | 0.983202 | 0.572056 | -1.719 |
| chr2 | 50224920 | 50225169 | 1.11016 | 0.645949 | -1.719 |
| chr15 | 63100765 | 63101011 | 1.21434 | 0.706572 | -1.719 |
| chr19 | 28511545 | 28511740 | 1.18818 | 0.691404 | -1.719 |
| chr10 | 10904228 | 10904515 | 1.08993 | 0.634275 | -1.718 |
| chr11 | 59168901 | 59169139 | 1.08676 | 0.632526 | -1.718 |
| chr12 | 42656897 | 42657119 | 0.927413 | 0.539922 | -1.718 |
| chr2 | 105188658 | 105188865 | 1.1208 | 0.652609 | -1.717 |
| chr7 | 48623995 | 48624220 | 1.0979 | 0.639398 | -1.717 |
| chr5 | 109975524 | 109975719 | 1.18312 | 0.68904 | -1.717 |
| chr7 | 49039187 | 49039508 | 0.992865 | 0.578305 | -1.717 |
| chr3 | 156923333 | 156923528 | 1.19463 | 0.695863 | -1.717 |
| chr15 | 85210298 | 85210501 | 1.05156 | 0.612564 | -1.717 |
| chr8 | 138056684 | 138056992 | 1.03162 | 0.600956 | -1.717 |
| chr7 | 121860138 | 121860328 | 1.04793 | 0.610552 | -1.716 |
| chr1 | 223705817 | 223706031 | 1.02876 | 0.59945 | -1.716 |
| chr8 | 114982149 | 114982388 | 0.940892 | 0.548279 | -1.716 |
| chr3 | 83312072 | 83312292 | 1.03194 | 0.601342 | -1.716 |
| chr10 | 100952281 | 100952474 | 0.997206 | 0.581122 | -1.716 |
| chr8 | 127695328 | 127695519 | 1.07714 | 0.627719 | -1.716 |
| chr9 | 131223044 | 131223243 | 1.18888 | 0.693063 | -1.715 |
| chr2 | 207396129 | 207396345 | 0.982732 | 0.572893 | -1.715 |
| chr1 | 223683926 | 223684117 | 1.16163 | 0.677188 | -1.715 |
| chr2 | 49498371 | 49498610 | 1.02194 | 0.595805 | -1.715 |
| chr12 | 116748086 | 116748313 | 0.865279 | 0.504483 | -1.715 |
| chr5 | 153841075 | 153841262 | 1.04956 | 0.611968 | -1.715 |
| chr9 | 90651756 | 90651968 | 1.01211 | 0.590162 | -1.715 |
| chr3 | 115227082 | 115227324 | 1.11872 | 0.652338 | -1.715 |
| chr7 | 13623155 | 13623409 | 1.02285 | 0.596616 | -1.714 |
| chr8 | 103124417 | 103124605 | 1.18113 | 0.688988 | -1.714 |
| chr9 | 121303335 | 121303547 | 1.01926 | 0.59462 | -1.714 |
| chr19 | 29005050 | 29005268 | 1.33885 | 0.78114 | -1.714 |
| chr12 | 126897751 | 126898165 | 0.971164 | 0.56663 | -1.714 |
| chr1 | 160112749 | 160112931 | 1.00139 | 0.584275 | -1.714 |
| chr9 | 82796662 | 82796863 | 1.18656 | 0.69247 | -1.714 |
| chr17 | 77660858 | 77661103 | 1.03913 | 0.606453 | -1.713 |
| chr7 | 49363031 | 49363221 | 0.973255 | 0.568149 | -1.713 |
| chr17 | 17688291 | 17688510 | 1.21503 | 0.709331 | -1.713 |
| chr8 | 13917985 | 13918211 | 0.824756 | 0.481507 | -1.713 |
| chr11 | 32583397 | 32583587 | 1.32784 | 0.775248 | -1.713 |
| chr15 | 91657647 | 91657943 | 1.05813 | 0.617808 | -1.713 |
| chr2 | 50058899 | 50059169 | 1.1415 | 0.666526 | -1.713 |
| chr2 | 186147589 | 186147782 | 1.06235 | 0.620412 | -1.712 |
| chr3 | 75958002 | 75958235 | 0.969634 | 0.56631 | -1.712 |
| chr20 | 18072638 | 18072833 | 0.873095 | 0.509931 | -1.712 |
| chr9 | 13725722 | 13725929 | 1.27292 | 0.743565 | -1.712 |
| chr14 | 89351401 | 89351592 | 0.957688 | 0.559559 | -1.712 |
| chr2 | 205365 | 205600 | 0.925859 | 0.541054 | -1.711 |

| | | | | | |
|---|---|---|---|---|---|
| chr2 | 46054006 | 46054193 | 1.4416 | 0.842496 | -1.711 |
| chr8 | 144837082 | 144837270 | 1.1774 | 0.688106 | -1.711 |
| chr10 | 10055636 | 10055837 | 0.743089 | 0.434302 | -1.711 |
| chr7 | 118101558 | 118101771 | 0.898568 | 0.525182 | -1.711 |
| chr2 | 77574147 | 77574439 | 1.42895 | 0.835321 | -1.711 |
| chr15 | 62501642 | 62501827 | 1.15984 | 0.678074 | -1.710 |
| chr8 | 2671772 | 2672335 | 2.13059 | 1.24566 | -1.710 |
| chr12 | 123984194 | 123984396 | 1.14384 | 0.668882 | -1.710 |
| chr10 | 363318 | 363520 | 0.961752 | 0.562407 | -1.710 |
| chr19 | 52292863 | 52293098 | 1.23399 | 0.721653 | -1.710 |
| chr9 | 122204798 | 122204998 | 1.0957 | 0.640796 | -1.710 |
| chr9 | 108171774 | 108171972 | 1.20452 | 0.704581 | -1.710 |
| chr14 | 96533349 | 96533548 | 1.20147 | 0.702822 | -1.709 |
| chr4 | 32466541 | 32466789 | 0.75005 | 0.438788 | -1.709 |
| chr7 | 114949868 | 114950097 | 1.01351 | 0.592966 | -1.709 |
| chr19 | 33055494 | 33055692 | 1.45681 | 0.852347 | -1.709 |
| chr9 | 18303473 | 18303686 | 0.932198 | 0.545459 | -1.709 |
| chr9 | 102516299 | 102516488 | 1.13616 | 0.664816 | -1.709 |
| chr16 | 5552374 | 5552612 | 1.36276 | 0.797513 | -1.709 |
| chr5 | 59421098 | 59421435 | 0.967875 | 0.566621 | -1.708 |
| chr2 | 218030968 | 218031182 | 1.36894 | 0.801432 | -1.708 |
| chr11 | 5898844 | 5899150 | 0.995842 | 0.583027 | -1.708 |
| chr8 | 78008989 | 78009288 | 1.08184 | 0.633455 | -1.708 |
| chr21 | 20117522 | 20117835 | 0.953064 | 0.558069 | -1.708 |
| chr8 | 127109556 | 127109740 | 1.14658 | 0.671394 | -1.708 |
| chr17 | 52169182 | 52169438 | 0.824316 | 0.482709 | -1.708 |
| chr12 | 81880790 | 81881010 | 0.913041 | 0.534707 | -1.708 |
| chr8 | 116387750 | 116387946 | 1.196 | 0.700439 | -1.708 |
| chr7 | 53925126 | 53925338 | 0.932053 | 0.545914 | -1.707 |
| chr3 | 145366286 | 145366489 | 1.2953 | 0.758699 | -1.707 |
| chr9 | 119724088 | 119724283 | 1.03442 | 0.605936 | -1.707 |
| chr7 | 132751371 | 132751630 | 1.08505 | 0.635601 | -1.707 |
| chr8 | 144436165 | 144436404 | 1.1831 | 0.693059 | -1.707 |
| chr12 | 5555640 | 5555844 | 1.09731 | 0.642806 | -1.707 |
| chr11 | 26299541 | 26299820 | 0.867874 | 0.508589 | -1.706 |
| chr11 | 58789385 | 58789572 | 0.797729 | 0.467535 | -1.706 |
| chr17 | 61482540 | 61482745 | 1.13069 | 0.662679 | -1.706 |
| chr5 | 98611666 | 98611889 | 0.974939 | 0.571436 | -1.706 |
| chr2 | 133246336 | 133246608 | 0.876578 | 0.513817 | -1.706 |
| chr2 | 171092581 | 171092774 | 2.52343 | 1.47917 | -1.706 |
| chr2 | 124613179 | 124613363 | 1.19321 | 0.699468 | -1.706 |
| chr10 | 131226452 | 131226837 | 0.933467 | 0.547247 | -1.706 |
| chr10 | 78392479 | 78392763 | 1.60596 | 0.941557 | -1.706 |
| chr19 | 28026724 | 28026919 | 0.986323 | 0.578304 | -1.706 |
| chr11 | 24756832 | 24757077 | 0.833364 | 0.48867 | -1.705 |
| chr2 | 81088620 | 81088838 | 0.818106 | 0.479724 | -1.705 |
| chr2 | 40265618 | 40265920 | 1.20505 | 0.706702 | -1.705 |
| chr1 | 238085344 | 238085543 | 1.0325 | 0.605567 | -1.705 |
| chr5 | 83083818 | 83084003 | 0.873147 | 0.512164 | -1.705 |
| chr8 | 2238081 | 2238283 | 1.2033 | 0.705842 | -1.705 |
| chr3 | 89460228 | 89460493 | 1.00509 | 0.589655 | -1.705 |
| chr5 | 35960454 | 35960740 | 0.863547 | 0.506618 | -1.705 |
| chr5 | 163376436 | 163376720 | 1.13936 | 0.668464 | -1.704 |
| chr2 | 76508263 | 76508473 | 1.14029 | 0.669042 | -1.704 |
| chr3 | 113845201 | 113845411 | 0.958045 | 0.562144 | -1.704 |
| chr19 | 30751510 | 30751804 | 1.10793 | 0.650106 | -1.704 |
| chr2 | 21567203 | 21567477 | 0.987534 | 0.57957 | -1.704 |
| chr19 | 48696809 | 48697113 | 0.973571 | 0.571443 | -1.704 |
| chr12 | 24114465 | 24114776 | 1.00452 | 0.589711 | -1.703 |
| chr11 | 81329780 | 81329987 | 1.20409 | 0.706876 | -1.703 |
| chr9 | 6534502 | 6534738 | 1.12632 | 0.661227 | -1.703 |

| | | | | | |
|---|---|---|---|---|---|
| chr2 | 256032 | 256233 | 0.922381 | 0.541535 | -1.703 |
| chr3 | 3422555 | 3422818 | 1.19583 | 0.702086 | -1.703 |
| chr5 | 2572125 | 2572618 | 1.02905 | 0.604278 | -1.703 |
| chr1 | 49038399 | 49038631 | 2.05375 | 1.20612 | -1.703 |
| chr3 | 14820421 | 14820620 | 0.910688 | 0.534851 | -1.703 |
| chr11 | 44961309 | 44961594 | 1.50789 | 0.885599 | -1.703 |
| chr8 | 68698010 | 68698376 | 1.03015 | 0.605082 | -1.702 |
| chr9 | 132556791 | 132556995 | 0.83941 | 0.493088 | -1.702 |
| chr5 | 103977885 | 103978102 | 1.00508 | 0.590477 | -1.702 |
| chr10 | 115881551 | 115881865 | 1.21288 | 0.712603 | -1.702 |
| chr9 | 87790536 | 87790807 | 1.02822 | 0.604157 | -1.702 |
| chr12 | 70956008 | 70956231 | 0.871969 | 0.512405 | -1.702 |
| chr9 | 7649121 | 7649317 | 1.03693 | 0.609434 | -1.701 |
| chr8 | 16126148 | 16126375 | 0.792611 | 0.465858 | -1.701 |
| chr1 | 215672763 | 215672959 | 1.06202 | 0.624218 | -1.701 |
| chr2 | 61611502 | 61611704 | 1.00149 | 0.588707 | -1.701 |
| chr2 | 222695701 | 222696036 | 1.07516 | 0.632013 | -1.701 |
| chr3 | 81525973 | 81526177 | 0.937084 | 0.550851 | -1.701 |
| chr15 | 51418311 | 51418496 | 1.01143 | 0.594613 | -1.701 |
| chr11 | 41870922 | 41871172 | 1.04835 | 0.616385 | -1.701 |
| chr3 | 26533622 | 26533863 | 0.86613 | 0.509247 | -1.701 |
| chr15 | 91742868 | 91743055 | 1.1515 | 0.677057 | -1.701 |
| chr2 | 22284545 | 22284857 | 1.4464 | 0.850453 | -1.701 |
| chr2 | 47837825 | 47838039 | 1.08412 | 0.637528 | -1.701 |
| chr3 | 77370329 | 77370886 | 0.99899 | 0.587466 | -1.701 |
| chr15 | 31877597 | 31877790 | 1.26901 | 0.746265 | -1.700 |
| chr3 | 144658553 | 144658736 | 1.06683 | 0.627373 | -1.700 |
| chr9 | 109223764 | 109224037 | 1.20215 | 0.706984 | -1.700 |
| chr8 | 137753977 | 137754231 | 1.30492 | 0.767514 | -1.700 |
| chr15 | 49402372 | 49402573 | 1.17729 | 0.692554 | -1.700 |
| chr8 | 13312144 | 13312416 | 1.13899 | 0.67015 | -1.700 |
| chr10 | 129168033 | 129168376 | 1.11348 | 0.655148 | -1.700 |
| chr2 | 163704606 | 163704827 | 1.0728 | 0.631248 | -1.699 |
| chr3 | 97953388 | 97953572 | 0.941925 | 0.55424 | -1.699 |
| chr16 | 59697484 | 59697672 | 0.811843 | 0.477738 | -1.699 |
| chr10 | 67871003 | 67871201 | 1.11893 | 0.658505 | -1.699 |
| chr19 | 28367401 | 28367583 | 1.27892 | 0.753009 | -1.698 |
| chr9 | 23430076 | 23430283 | 0.827779 | 0.487482 | -1.698 |
| chr14 | 95004584 | 95004800 | 0.744094 | 0.438212 | -1.698 |
| chr1 | 191334206 | 191334585 | 0.841156 | 0.495407 | -1.698 |
| chr17 | 78283016 | 78283210 | 0.959357 | 0.565053 | -1.698 |
| chr20 | 11971664 | 11971880 | 1.17587 | 0.692629 | -1.698 |
| chr8 | 145717447 | 145717711 | 0.860836 | 0.507075 | -1.698 |
| chr8 | 50758466 | 50759083 | 0.9912 | 0.583996 | -1.697 |
| chr7 | 118650393 | 118650688 | 1.12575 | 0.663284 | -1.697 |
| chr5 | 91171406 | 91171615 | 0.817456 | 0.48166 | -1.697 |
| chr7 | 134340566 | 134340762 | 1.14795 | 0.676401 | -1.697 |
| chr14 | 106877425 | 106877619 | 2.96792 | 1.74913 | -1.697 |
| chr1 | 217171371 | 217171565 | 0.8707 | 0.513192 | -1.697 |
| chr2 | 63741251 | 63741589 | 0.967506 | 0.570289 | -1.697 |
| chr11 | 105709622 | 105709826 | 0.971177 | 0.572494 | -1.696 |
| chr11 | 29879065 | 29879317 | 1.02769 | 0.60586 | -1.696 |
| chr2 | 45029776 | 45029970 | 1.27864 | 0.753813 | -1.696 |
| chr7 | 82847247 | 82847505 | 0.749891 | 0.442198 | -1.696 |
| chr1 | 119910185 | 119910403 | 2.30082 | 1.35676 | -1.696 |
| chr11 | 42104095 | 42104307 | 1.0129 | 0.597424 | -1.695 |
| chr2 | 14265746 | 14266109 | 1.24916 | 0.736884 | -1.695 |
| chr3 | 25263355 | 25263700 | 0.921421 | 0.543766 | -1.695 |
| chr11 | 25534020 | 25534555 | 0.951055 | 0.56129 | -1.694 |
| chr8 | 127322966 | 127323241 | 1.01688 | 0.600169 | -1.694 |
| chr11 | 33229151 | 33229403 | 1.17714 | 0.694778 | -1.694 |

| | | | | | |
|---|---|---|---|---|---|
| chr5 | 15584107 | 15584405 | 0.81392 | 0.480412 | -1.694 |
| chr7 | 53411998 | 53412197 | 0.743105 | 0.438629 | -1.694 |
| chr20 | 15809138 | 15809358 | 1.00257 | 0.591783 | -1.694 |
| chr6 | 46975457 | 46975654 | 1.19841 | 0.70749 | -1.694 |
| chr10 | 115862763 | 115862947 | 1.03584 | 0.611529 | -1.694 |
| chr12 | 24803059 | 24803279 | 0.945092 | 0.557962 | -1.694 |
| chr11 | 8069133 | 8069450 | 0.975452 | 0.576046 | -1.693 |
| chr3 | 156214290 | 156214472 | 0.695474 | 0.410731 | -1.693 |
| chr2 | 83854368 | 83854614 | 0.986631 | 0.582708 | -1.693 |
| chr16 | 27651129 | 27651725 | 1.95553 | 1.15496 | -1.693 |
| chr7 | 46567542 | 46567762 | 1.00981 | 0.596408 | -1.693 |
| chr16 | 64607114 | 64607355 | 0.86414 | 0.510392 | -1.693 |
| chr5 | 84685233 | 84685500 | 1.16386 | 0.687731 | -1.692 |
| chr3 | 73711220 | 73711427 | 0.918326 | 0.542656 | -1.692 |
| chr12 | 110137419 | 110137636 | 0.839786 | 0.496289 | -1.692 |
| chr5 | 169838205 | 169838472 | 1.35616 | 0.801453 | -1.692 |
| chr2 | 4442674 | 4442893 | 1.40487 | 0.830279 | -1.692 |
| chr6 | 5950744 | 5951123 | 1.44288 | 0.85281 | -1.692 |
| chr9 | 130119246 | 130119515 | 1.00485 | 0.593914 | -1.692 |
| chr15 | 53015045 | 53015259 | 1.53991 | 0.91019 | -1.692 |
| chr11 | 24772968 | 24773206 | 0.954417 | 0.564135 | -1.692 |
| chr3 | 109448792 | 109448978 | 0.893704 | 0.528462 | -1.691 |
| chr15 | 98220729 | 98220917 | 1.13265 | 0.669765 | -1.691 |
| chr13 | 55783774 | 55783975 | 0.81889 | 0.484235 | -1.691 |
| chr7 | 36885376 | 36885576 | 1.12528 | 0.665492 | -1.691 |
| chr2 | 13944414 | 13944859 | 0.955342 | 0.565017 | -1.691 |
| chr2 | 1212340 | 1212606 | 0.891946 | 0.527583 | -1.691 |
| chr10 | 129325194 | 129325385 | 0.923175 | 0.546157 | -1.690 |
| chr5 | 23242340 | 23242588 | 0.974997 | 0.576815 | -1.690 |
| chr10 | 69232319 | 69232553 | 1.15145 | 0.681291 | -1.690 |
| chr3 | 107050404 | 107050598 | 0.841827 | 0.49816 | -1.690 |
| chr2 | 181376495 | 181376698 | 1.09383 | 0.647363 | -1.690 |
| chr8 | 25462396 | 25462597 | 1.29046 | 0.763819 | -1.689 |
| chr3 | 77863072 | 77863269 | 0.81302 | 0.481234 | -1.689 |
| chr14 | 84846572 | 84846807 | 1.41904 | 0.83996 | -1.689 |
| chr10 | 57670461 | 57670665 | 0.953764 | 0.564578 | -1.689 |
| chr7 | 152200564 | 152200847 | 1.01321 | 0.599861 | -1.689 |
| chr10 | 123562083 | 123562310 | 0.935633 | 0.553987 | -1.689 |
| chr3 | 4718392 | 4718703 | 1.08877 | 0.644712 | -1.689 |
| chr2 | 205058105 | 205058336 | 1.02686 | 0.608065 | -1.689 |
| chr14 | 46785037 | 46785243 | 1.18336 | 0.700784 | -1.689 |
| chr17 | 56886749 | 56886993 | 1.2913 | 0.764763 | -1.688 |
| chr20 | 12764348 | 12764631 | 1.25708 | 0.744641 | -1.688 |
| chr10 | 58482759 | 58483035 | 1.1713 | 0.693838 | -1.688 |
| chr3 | 133468208 | 133468460 | 0.748112 | 0.44317 | -1.688 |
| chr5 | 166911295 | 166911482 | 1.32221 | 0.7833 | -1.688 |
| chr10 | 116843501 | 116843701 | 1.01812 | 0.60321 | -1.688 |
| chr7 | 13395693 | 13395887 | 0.781739 | 0.463166 | -1.688 |
| chr5 | 145947577 | 145947762 | 0.715814 | 0.424227 | -1.687 |
| chr5 | 91970864 | 91971063 | 0.995654 | 0.590121 | -1.687 |
| chr21 | 40547015 | 40547341 | 1.02274 | 0.606259 | -1.687 |
| chr7 | 135800043 | 135800358 | 1.82999 | 1.08504 | -1.687 |
| chr10 | 65737189 | 65737413 | 1.10524 | 0.655338 | -1.687 |
| chr7 | 108334185 | 108334499 | 1.03845 | 0.615746 | -1.686 |
| chr8 | 2457369 | 2457565 | 1.28366 | 0.76118 | -1.686 |
| chr1 | 44506879 | 44507083 | 0.833483 | 0.494241 | -1.686 |
| chr11 | 8329165 | 8329500 | 1.11653 | 0.662164 | -1.686 |
| chr1 | 183935814 | 183936016 | 1.05019 | 0.622859 | -1.686 |
| chr9 | 104543919 | 104544105 | 1.18181 | 0.700939 | -1.686 |
| chr12 | 126593677 | 126593889 | 1.17289 | 0.695731 | -1.686 |
| chr8 | 14443760 | 14444033 | 1.44078 | 0.854748 | -1.686 |

| | | | | | |
|---|---|---|---|---|---|
| chr8 | 76417346 | 76417618 | 0.942868 | 0.559468 | -1.685 |
| chr11 | 104871656 | 104871858 | 1.61326 | 0.957383 | -1.685 |
| chr7 | 14378828 | 14379038 | 1.02891 | 0.610607 | -1.685 |
| chr1 | 26689837 | 26690025 | 0.965728 | 0.573169 | -1.685 |
| chr19 | 28918459 | 28918687 | 1.0121 | 0.600724 | -1.685 |
| chr17 | 2589137 | 2589335 | 1.12624 | 0.668504 | -1.685 |
| chr8 | 51968694 | 51969099 | 0.966682 | 0.57381 | -1.685 |
| chr7 | 121396061 | 121396355 | 1.13631 | 0.674507 | -1.685 |
| chr3 | 1633354 | 1633575 | 0.772713 | 0.458738 | -1.684 |
| chr3 | 80866641 | 80866849 | 0.934756 | 0.554952 | -1.684 |
| chr2 | 205042749 | 205043062 | 1.4486 | 0.86006 | -1.684 |
| chr2 | 103116494 | 103116744 | 0.864496 | 0.513272 | -1.684 |
| chr15 | 86698406 | 86698650 | 1.13069 | 0.671393 | -1.684 |
| chr3 | 171171058 | 171171251 | 1.17164 | 0.69576 | -1.684 |
| chr2 | 49502183 | 49502441 | 1.00732 | 0.598209 | -1.684 |
| chr1 | 216372982 | 216373249 | 1.04662 | 0.621657 | -1.684 |
| chr11 | 44317106 | 44317298 | 0.971729 | 0.577201 | -1.684 |
| chr1 | 165545000 | 165545187 | 1.20452 | 0.715506 | -1.683 |
| chrX | 150254485 | 150254667 | 0.87522 | 0.519927 | -1.683 |
| chr17 | 13309244 | 13309492 | 0.958587 | 0.569524 | -1.683 |
| chr2 | 60118692 | 60118984 | 1.2986 | 0.771557 | -1.683 |
| chr7 | 125584990 | 125585288 | 0.80926 | 0.480819 | -1.683 |
| chr16 | 13297114 | 13297328 | 1.18551 | 0.704523 | -1.683 |
| chr5 | 160774544 | 160775039 | 1.0571 | 0.628247 | -1.683 |
| chr12 | 33987789 | 33988017 | 0.994558 | 0.591092 | -1.683 |
| chr9 | 9070028 | 9070261 | 0.881266 | 0.523862 | -1.682 |
| chr5 | 42841328 | 42841610 | 0.964051 | 0.573075 | -1.682 |
| chr3 | 119948612 | 119948820 | 1.01458 | 0.603173 | -1.682 |
| chr5 | 27169501 | 27169709 | 1.04496 | 0.621311 | -1.682 |
| chr10 | 87584114 | 87584318 | 0.971156 | 0.577464 | -1.682 |
| chr2 | 88497292 | 88497495 | 0.952189 | 0.566244 | -1.682 |
| chr5 | 51403155 | 51403339 | 1.00196 | 0.595903 | -1.681 |
| chr9 | 30408262 | 30408502 | 0.977315 | 0.581268 | -1.681 |
| chr3 | 88361530 | 88361778 | 1.10679 | 0.658327 | -1.681 |
| chr16 | 64101230 | 64101500 | 1.07186 | 0.637583 | -1.681 |
| chr19 | 52929176 | 52929557 | 1.11406 | 0.66271 | -1.681 |
| chr9 | 111738843 | 111739154 | 1.00909 | 0.600324 | -1.681 |
| chr20 | 20069526 | 20069710 | 1.35127 | 0.803961 | -1.681 |
| chr9 | 78604064 | 78604279 | 0.992644 | 0.590691 | -1.680 |
| chr20 | 43250746 | 43250974 | 0.93598 | 0.556986 | -1.680 |
| chr5 | 149819812 | 149820153 | 0.85656 | 0.509752 | -1.680 |
| chr2 | 22429666 | 22429976 | 0.868114 | 0.516651 | -1.680 |
| chr8 | 115538507 | 115538734 | 0.890629 | 0.530066 | -1.680 |
| chr15 | 33661430 | 33661664 | 1.07294 | 0.638632 | -1.680 |
| chr7 | 85953537 | 85953724 | 1.20449 | 0.716934 | -1.680 |
| chr8 | 86982060 | 86982313 | 0.949654 | 0.56532 | -1.680 |
| chr2 | 59288244 | 59288484 | 1.23842 | 0.737341 | -1.680 |
| chr8 | 50059275 | 50059483 | 0.84511 | 0.503172 | -1.680 |
| chr7 | 109054415 | 109054600 | 0.838492 | 0.499242 | -1.680 |
| chr2 | 73890211 | 73890448 | 1.08774 | 0.647662 | -1.679 |
| chr1 | 219572687 | 219572894 | 0.976979 | 0.581737 | -1.679 |
| chr10 | 83454525 | 83454825 | 0.929831 | 0.553678 | -1.679 |
| chr16 | 65322948 | 65323163 | 0.89096 | 0.530537 | -1.679 |
| chr19 | 22924077 | 22924311 | 1.021 | 0.608036 | -1.679 |
| chr5 | 68522721 | 68522913 | 1.1333 | 0.675309 | -1.678 |
| chr15 | 94180770 | 94180964 | 0.976273 | 0.581771 | -1.678 |
| chr3 | 32976202 | 32976395 | 1.06181 | 0.632766 | -1.678 |
| chr10 | 44626699 | 44626976 | 0.833282 | 0.496601 | -1.678 |
| chr12 | 73484868 | 73485096 | 0.901927 | 0.537681 | -1.677 |
| chr2 | 232221888 | 232222090 | 0.684122 | 0.40794 | -1.677 |
| chr15 | 27703280 | 27703523 | 0.963688 | 0.574684 | -1.677 |

| | | | | | |
|---|---|---|---|---|---|
| chr7 | 112946163 | 112946345 | 0.960288 | 0.572713 | -1.677 |
| chr15 | 22937593 | 22937815 | 1.14516 | 0.682987 | -1.677 |
| chr9 | 131205636 | 131205918 | 1.13041 | 0.674191 | -1.677 |
| chr2 | 162776510 | 162776789 | 1.13481 | 0.676843 | -1.677 |
| chr5 | 12123789 | 12124126 | 0.936692 | 0.558681 | -1.677 |
| chr11 | 29918074 | 29918269 | 1.09551 | 0.653497 | -1.676 |
| chr17 | 12610335 | 12610520 | 1.0261 | 0.612134 | -1.676 |
| chr1 | 209903742 | 209903954 | 0.8296 | 0.49504 | -1.676 |
| chr8 | 136183195 | 136183457 | 0.955412 | 0.570129 | -1.676 |
| chr7 | 73562172 | 73562477 | 0.74173 | 0.442643 | -1.676 |
| chr3 | 96831987 | 96832527 | 0.971345 | 0.579706 | -1.676 |
| chr11 | 32489678 | 32489913 | 1.03122 | 0.615441 | -1.676 |
| chr7 | 28785330 | 28785570 | 0.847864 | 0.506023 | -1.676 |
| chr9 | 95161202 | 95161493 | 0.793548 | 0.473638 | -1.675 |
| chr3 | 9970011 | 9970193 | 1.08659 | 0.648555 | -1.675 |
| chr20 | 23048239 | 23048564 | 0.97251 | 0.580496 | -1.675 |
| chr1 | 195784829 | 195785047 | 1.08632 | 0.6485 | -1.675 |
| chr19 | 23831040 | 23831368 | 1.15873 | 0.691786 | -1.675 |
| chr2 | 103919188 | 103919457 | 0.895808 | 0.534824 | -1.675 |
| chr3 | 84042188 | 84042425 | 1.01313 | 0.604903 | -1.675 |
| chr8 | 80219226 | 80219457 | 1.01339 | 0.605173 | -1.675 |
| chr20 | 53782578 | 53782865 | 0.897201 | 0.535855 | -1.674 |
| chr1 | 176449165 | 176449423 | 1.12824 | 0.673876 | -1.674 |
| chr2 | 154670421 | 154670714 | 1.03219 | 0.616606 | -1.674 |
| chr9 | 75966982 | 75967454 | 1.0641 | 0.63569 | -1.674 |
| chr1 | 230862631 | 230862824 | 1.3671 | 0.816711 | -1.674 |
| chr1 | 247906595 | 247906793 | 1.28428 | 0.767299 | -1.674 |
| chr3 | 19466814 | 19467111 | 0.823438 | 0.491982 | -1.674 |
| chr2 | 126300191 | 126300395 | 0.959803 | 0.573507 | -1.674 |
| chr8 | 138709517 | 138709795 | 0.938474 | 0.560808 | -1.673 |
| chr15 | 79887751 | 79887947 | 1.22598 | 0.732645 | -1.673 |
| chr12 | 48430930 | 48431303 | 1.23823 | 0.74003 | -1.673 |
| chr10 | 57058456 | 57058691 | 1.19286 | 0.712962 | -1.673 |
| chr8 | 118149799 | 118150100 | 1.00722 | 0.602086 | -1.673 |
| chr7 | 9965025 | 9965418 | 0.776016 | 0.463997 | -1.672 |
| chr16 | 1130187 | 1130376 | 0.889303 | 0.531737 | -1.672 |
| chr15 | 33569192 | 33569401 | 1.071 | 0.640417 | -1.672 |
| chr5 | 180250423 | 180250614 | 0.7838 | 0.468696 | -1.672 |
| chr2 | 34988065 | 34988277 | 1.07738 | 0.644332 | -1.672 |
| chr20 | 38126894 | 38127317 | 0.924378 | 0.552861 | -1.672 |
| chr8 | 100871030 | 100871222 | 1.37767 | 0.824059 | -1.672 |
| chr19 | 57034400 | 57034669 | 0.935484 | 0.559619 | -1.672 |
| chr19 | 52260140 | 52260332 | 0.91549 | 0.547679 | -1.672 |
| chr3 | 13178486 | 13178669 | 0.890167 | 0.53253 | -1.672 |
| chr2 | 50286746 | 50286966 | 1.20361 | 0.720074 | -1.672 |
| chr16 | 48235688 | 48235882 | 1.26802 | 0.758704 | -1.671 |
| chr5 | 163383608 | 163383920 | 0.992827 | 0.59414 | -1.671 |
| chr15 | 101895566 | 101895985 | 1.12784 | 0.675054 | -1.671 |
| chr14 | 49464302 | 49464489 | 0.889338 | 0.532304 | -1.671 |
| chr2 | 1399877 | 1400085 | 0.814129 | 0.487318 | -1.671 |
| chr3 | 1463386 | 1463650 | 1.17842 | 0.705458 | -1.670 |
| chr5 | 102866777 | 102866976 | 1.03061 | 0.61702 | -1.670 |
| chr8 | 83734539 | 83734730 | 1.39579 | 0.835756 | -1.670 |
| chr16 | 20238485 | 20238743 | 0.72051 | 0.431453 | -1.670 |
| chr11 | 46181639 | 46181909 | 1.06055 | 0.635076 | -1.670 |
| chr8 | 79809394 | 79809748 | 1.06034 | 0.635031 | -1.670 |
| chr8 | 105928605 | 105928911 | 1.06955 | 0.640547 | -1.670 |
| chr1 | 222499541 | 222499813 | 1.26766 | 0.759271 | -1.670 |
| chr5 | 65027823 | 65028009 | 0.765873 | 0.458753 | -1.669 |
| chr20 | 15838009 | 15838281 | 1.47001 | 0.880698 | -1.669 |
| chr3 | 82678770 | 82678958 | 0.840711 | 0.503739 | -1.669 |

| | | | | | |
|---|---|---|---|---|---|
| chrX | 116133677 | 116133888 | 1.06501 | 0.63815 | -1.669 |
| chr5 | 8474989 | 8475212 | 0.842829 | 0.505054 | -1.669 |
| chr19 | 53962271 | 53962454 | 1.03522 | 0.620348 | -1.669 |
| chr2 | 68254075 | 68254274 | 1.03411 | 0.619699 | -1.669 |
| chr5 | 24046763 | 24047160 | 0.984993 | 0.59029 | -1.669 |
| chr11 | 21233966 | 21234404 | 1.1915 | 0.714063 | -1.669 |
| chr5 | 121076667 | 121077038 | 0.979091 | 0.586798 | -1.669 |
| chr7 | 152890262 | 152890454 | 1.87939 | 1.12667 | -1.668 |
| chr2 | 73070546 | 73070861 | 1.26098 | 0.756064 | -1.668 |
| chr10 | 115586656 | 115586957 | 1.09331 | 0.655599 | -1.668 |
| chr2 | 52431086 | 52431313 | 1.08874 | 0.652907 | -1.668 |
| chr8 | 4081885 | 4082089 | 0.72887 | 0.437101 | -1.668 |
| chr2 | 54074404 | 54074663 | 1.14084 | 0.684216 | -1.667 |
| chr2 | 59355591 | 59355871 | 0.944617 | 0.566673 | -1.667 |
| chr11 | 31377377 | 31377634 | 1.08138 | 0.648727 | -1.667 |
| chr11 | 64472446 | 64472633 | 1.19607 | 0.717772 | -1.666 |
| chr2 | 195119621 | 195119969 | 1.0692 | 0.641873 | -1.666 |
| chr8 | 132667771 | 132667957 | 1.19207 | 0.715753 | -1.665 |
| chr7 | 125880368 | 125880729 | 0.77922 | 0.46788 | -1.665 |
| chr5 | 22304403 | 22304677 | 1.03801 | 0.623297 | -1.665 |
| chr15 | 77152174 | 77152388 | 1.26605 | 0.760293 | -1.665 |
| chr7 | 103720841 | 103721127 | 0.917914 | 0.551426 | -1.665 |
| chr3 | 71966316 | 71966545 | 1.17107 | 0.703517 | -1.665 |
| chr19 | 33956112 | 33956341 | 1.09732 | 0.659362 | -1.664 |
| chr3 | 79856921 | 79857124 | 1.11634 | 0.671017 | -1.664 |
| chr1 | 175712678 | 175713074 | 2.09632 | 1.26025 | -1.663 |
| chr11 | 45693392 | 45693633 | 1.16349 | 0.69949 | -1.663 |
| chr5 | 28235811 | 28236104 | 0.948247 | 0.570196 | -1.663 |
| chr15 | 79398571 | 79398857 | 1.40297 | 0.843732 | -1.663 |
| chr5 | 117294356 | 117294545 | 0.962857 | 0.579055 | -1.663 |
| chr8 | 63925829 | 63926024 | 0.905798 | 0.544774 | -1.663 |
| chr9 | 122842524 | 122842751 | 0.981699 | 0.590441 | -1.663 |
| chr2 | 197673970 | 197674159 | 1.12191 | 0.674952 | -1.662 |
| chr9 | 122380721 | 122380944 | 1.05186 | 0.632854 | -1.662 |
| chr19 | 43081176 | 43081372 | 1.25136 | 0.752985 | -1.662 |
| chr2 | 76213839 | 76214210 | 0.903513 | 0.543678 | -1.662 |
| chr2 | 234232538 | 234232727 | 1.17287 | 0.705771 | -1.662 |
| chr3 | 38080966 | 38081194 | 1.21309 | 0.729996 | -1.662 |
| chr16 | 79719577 | 79719779 | 1.63139 | 0.981947 | -1.661 |
| chr2 | 239629278 | 239629493 | 1.17963 | 0.71017 | -1.661 |
| chr3 | 197529031 | 197529256 | 1.45902 | 0.878441 | -1.661 |
| chr3 | 164038904 | 164039179 | 0.915299 | 0.551091 | -1.661 |
| chr8 | 48993663 | 48993850 | 0.923448 | 0.556016 | -1.661 |
| chr9 | 24400187 | 24400441 | 1.10345 | 0.664407 | -1.661 |
| chr12 | 73946006 | 73946251 | 0.720011 | 0.433608 | -1.661 |
| chr3 | 26612718 | 26612936 | 1.44454 | 0.869965 | -1.660 |
| chr19 | 28636851 | 28637082 | 1.26472 | 0.761752 | -1.660 |
| chr2 | 80420405 | 80420764 | 0.986764 | 0.594397 | -1.660 |
| chr2 | 182295600 | 182295812 | 1.06906 | 0.644001 | -1.660 |
| chr5 | 108893277 | 108893656 | 1.10984 | 0.668633 | -1.660 |
| chr7 | 110303620 | 110303818 | 0.98497 | 0.593411 | -1.660 |
| chr5 | 4272470 | 4272685 | 1.10999 | 0.66874 | -1.660 |
| chr2 | 6721062 | 6721396 | 1.16351 | 0.701054 | -1.660 |
| chr3 | 22935443 | 22935717 | 1.15789 | 0.697673 | -1.660 |
| chr8 | 24121204 | 24121405 | 1.10009 | 0.662884 | -1.660 |
| chr16 | 25447766 | 25447963 | 0.714001 | 0.430249 | -1.660 |
| chr2 | 7749426 | 7749623 | 1.47152 | 0.886746 | -1.659 |
| chr1 | 220210831 | 220211044 | 0.777451 | 0.46852 | -1.659 |
| chr2 | 122569566 | 122569778 | 0.735491 | 0.443268 | -1.659 |
| chr19 | 52890467 | 52890721 | 1.04184 | 0.627926 | -1.659 |
| chr2 | 57109243 | 57109429 | 1.47744 | 0.890467 | -1.659 |

| | | | | | |
|---|---|---|---|---|---|
| chr3 | 121936044 | 121936269 | 1.28289 | 0.773235 | -1.659 |
| chr19 | 54327201 | 54327406 | 0.991401 | 0.597664 | -1.659 |
| chr19 | 22012654 | 22012872 | 0.968186 | 0.583673 | -1.659 |
| chr9 | 105741844 | 105742171 | 1.09342 | 0.659234 | -1.659 |
| chr17 | 6715958 | 6716342 | 1.08852 | 0.656331 | -1.658 |
| chr11 | 129805741 | 129805970 | 1.04475 | 0.630066 | -1.658 |
| chr5 | 165000594 | 165000789 | 1.13476 | 0.684374 | -1.658 |
| chr10 | 102988140 | 102988411 | 0.86204 | 0.519919 | -1.658 |
| chr3 | 150458466 | 150458712 | 1.02933 | 0.620842 | -1.658 |
| chr16 | 85449115 | 85449309 | 1.52007 | 0.917024 | -1.658 |
| chr5 | 89346513 | 89346782 | 0.664535 | 0.400902 | -1.658 |
| chr2 | 207657383 | 207657584 | 0.957011 | 0.577538 | -1.657 |
| chr15 | 70070586 | 70070876 | 0.984551 | 0.59429 | -1.657 |
| chr12 | 98492518 | 98492708 | 1.12258 | 0.677858 | -1.656 |
| chr11 | 78364698 | 78364897 | 1.00502 | 0.606892 | -1.656 |
| chr16 | 52394666 | 52394848 | 1.09683 | 0.662354 | -1.656 |
| chr19 | 41409342 | 41409524 | 1.0632 | 0.642091 | -1.656 |
| chr19 | 37170402 | 37170615 | 1.1065 | 0.668249 | -1.656 |
| chr7 | 96044337 | 96044532 | 2.19015 | 1.3229 | -1.656 |
| chr19 | 31752562 | 31752808 | 1.0103 | 0.61025 | -1.656 |
| chr12 | 129744613 | 129744974 | 1.2322 | 0.7447 | -1.655 |
| chr14 | 77341190 | 77341469 | 0.887641 | 0.536705 | -1.654 |
| chr2 | 34753825 | 34754578 | 0.935128 | 0.565538 | -1.654 |
| chr9 | 133362810 | 133363070 | 0.83991 | 0.507967 | -1.653 |
| chr19 | 35316079 | 35316353 | 1.27382 | 0.770406 | -1.653 |
| chr7 | 100166977 | 100167211 | 1.39976 | 0.846663 | -1.653 |
| chr15 | 80875543 | 80875735 | 1.07372 | 0.649458 | -1.653 |
| chr22 | 18586243 | 18586451 | 0.858328 | 0.519194 | -1.653 |
| chr9 | 76595030 | 76595343 | 1.00345 | 0.607016 | -1.653 |
| chr17 | 51154205 | 51154479 | 0.977459 | 0.591331 | -1.653 |
| chr19 | 52439064 | 52439292 | 1.42528 | 0.862264 | -1.653 |
| chr15 | 83749152 | 83749361 | 1.06303 | 0.643173 | -1.653 |
| chr19 | 30077346 | 30077590 | 1.59574 | 0.96589 | -1.652 |
| chr10 | 91861258 | 91861441 | 0.986967 | 0.597409 | -1.652 |
| chr17 | 38103157 | 38103418 | 0.852236 | 0.515901 | -1.652 |
| chr20 | 59290409 | 59290630 | 1.57022 | 0.95054 | -1.652 |
| chr15 | 97321583 | 97321820 | 1.05427 | 0.638216 | -1.652 |
| chr2 | 102417639 | 102417881 | 1.26843 | 0.767861 | -1.652 |
| chr19 | 58229084 | 58229337 | 1.27326 | 0.770841 | -1.652 |
| chr2 | 167972093 | 167972339 | 1.03418 | 0.626127 | -1.652 |
| chr17 | 67429095 | 67429280 | 0.958176 | 0.580133 | -1.652 |
| chr2 | 194136295 | 194136528 | 1.11692 | 0.676335 | -1.651 |
| chr20 | 5394375 | 5394769 | 1.29172 | 0.78231 | -1.651 |
| chr7 | 53519331 | 53519555 | 1.06805 | 0.646871 | -1.651 |
| chr19 | 35076089 | 35076591 | 1.15091 | 0.697087 | -1.651 |
| chr7 | 33515148 | 33515345 | 0.791544 | 0.479482 | -1.651 |
| chr9 | 18615250 | 18615459 | 0.991929 | 0.600894 | -1.651 |
| chr9 | 8288362 | 8288572 | 0.8083 | 0.489688 | -1.651 |
| chr10 | 90071642 | 90071998 | 0.983041 | 0.595555 | -1.651 |
| chr5 | 128271416 | 128271641 | 0.881439 | 0.534022 | -1.651 |
| chr2 | 13291720 | 13292193 | 1.11538 | 0.675798 | -1.650 |
| chr8 | 9844533 | 9844730 | 0.963482 | 0.583797 | -1.650 |
| chr11 | 6013852 | 6014113 | 1.32538 | 0.803082 | -1.650 |
| chr15 | 94994885 | 94995125 | 1.06844 | 0.647508 | -1.650 |
| chr2 | 217148211 | 217148544 | 1.99511 | 1.2091 | -1.650 |
| chr2 | 173138708 | 173138904 | 0.787213 | 0.477081 | -1.650 |
| chr7 | 33756280 | 33756487 | 1.82872 | 1.10836 | -1.650 |
| chr19 | 36278240 | 36278429 | 0.993821 | 0.602426 | -1.650 |
| chr9 | 132846749 | 132846978 | 0.835806 | 0.506645 | -1.650 |
| chr8 | 135113888 | 135114143 | 1.00111 | 0.606917 | -1.650 |
| chr14 | 84618930 | 84619187 | 1.0746 | 0.651566 | -1.649 |

| | | | | | |
|---|---|---|---|---|---|
| chr7 | 10588017 | 10588208 | 1.01068 | 0.612833 | -1.649 |
| chr16 | 1162059 | 1162274 | 0.969389 | 0.587801 | -1.649 |
| chr1 | 194977167 | 194977395 | 0.878741 | 0.532876 | -1.649 |
| chr12 | 93218757 | 93218993 | 0.72605 | 0.440303 | -1.649 |
| chr2 | 126411846 | 126412191 | 1.00043 | 0.606743 | -1.649 |
| chr15 | 94958172 | 94958365 | 1.196 | 0.725413 | -1.649 |
| chr19 | 23564233 | 23564585 | 0.738778 | 0.448122 | -1.649 |
| chr5 | 44904971 | 44905343 | 0.869635 | 0.527515 | -1.649 |
| chr19 | 57952644 | 57952888 | 1.12888 | 0.684818 | -1.648 |
| chr16 | 7227530 | 7227737 | 0.988155 | 0.599451 | -1.648 |
| chr20 | 31105098 | 31105295 | 0.955821 | 0.579843 | -1.648 |
| chr8 | 114956660 | 114957068 | 0.912373 | 0.553553 | -1.648 |
| chr11 | 65581703 | 65582186 | 0.796119 | 0.483038 | -1.648 |
| chr9 | 11137522 | 11137716 | 0.838024 | 0.508467 | -1.648 |
| chr19 | 40419620 | 40420000 | 1.14631 | 0.69565 | -1.648 |
| chr7 | 88528048 | 88528407 | 1.0465 | 0.635142 | -1.648 |
| chr9 | 104908297 | 104908522 | 0.982657 | 0.596486 | -1.647 |
| chr5 | 10060939 | 10061140 | 0.933971 | 0.566968 | -1.647 |
| chr1 | 169833836 | 169834022 | 0.866915 | 0.526369 | -1.647 |
| chr17 | 30833317 | 30833504 | 0.914266 | 0.555166 | -1.647 |
| chr2 | 53403466 | 53403735 | 0.919386 | 0.558336 | -1.647 |
| chr19 | 57025520 | 57025748 | 0.978976 | 0.594574 | -1.647 |
| chr2 | 79124783 | 79125005 | 1.1023 | 0.669535 | -1.646 |
| chr7 | 64441451 | 64441657 | 0.918294 | 0.557808 | -1.646 |
| chr8 | 109742406 | 109742601 | 1.18866 | 0.722136 | -1.646 |
| chr2 | 128952574 | 128952757 | 1.36416 | 0.828814 | -1.646 |
| chr3 | 147079708 | 147079898 | 1.70888 | 1.03865 | -1.645 |
| chr11 | 38986467 | 38986686 | 1.09432 | 0.665324 | -1.645 |
| chr19 | 42435468 | 42435783 | 1.79911 | 1.09411 | -1.644 |
| chr7 | 125008208 | 125008611 | 1.25521 | 0.763364 | -1.644 |
| chr16 | 61712373 | 61712584 | 0.886926 | 0.539424 | -1.644 |
| chr1 | 66038677 | 66038906 | 1.07299 | 0.652851 | -1.644 |
| chr2 | 182573098 | 182573291 | 1.24606 | 0.758161 | -1.644 |
| chr5 | 166893164 | 166893353 | 0.928827 | 0.565152 | -1.643 |
| chr2 | 78269770 | 78270093 | 1.02365 | 0.622892 | -1.643 |
| chr8 | 111987963 | 111988193 | 1.11518 | 0.67862 | -1.643 |
| chr10 | 123431359 | 123431631 | 0.829969 | 0.505082 | -1.643 |
| chr8 | 117892616 | 117893004 | 1.09772 | 0.668031 | -1.643 |
| chr2 | 67141311 | 67141637 | 1.07671 | 0.655295 | -1.643 |
| chr2 | 115124340 | 115124538 | 0.790915 | 0.481503 | -1.643 |
| chr8 | 82448858 | 82449104 | 0.898091 | 0.54683 | -1.642 |
| chr9 | 20264952 | 20265148 | 1.38421 | 0.84288 | -1.642 |
| chr19 | 37864546 | 37864766 | 1.3411 | 0.816697 | -1.642 |
| chr9 | 19056581 | 19056773 | 1.01674 | 0.61918 | -1.642 |
| chr8 | 145237004 | 145237211 | 1.16153 | 0.707362 | -1.642 |
| chr2 | 104756682 | 104756879 | 1.02265 | 0.622873 | -1.642 |
| chr7 | 143855001 | 143855224 | 1.00544 | 0.612492 | -1.642 |
| chr5 | 114308358 | 114308635 | 1.04264 | 0.63523 | -1.641 |
| chr12 | 61228619 | 61228824 | 1.54246 | 0.939773 | -1.641 |
| chr2 | 21933899 | 21934144 | 0.97239 | 0.592459 | -1.641 |
| chr9 | 71573165 | 71573384 | 0.706949 | 0.430755 | -1.641 |
| chr14 | 56971359 | 56971653 | 1.07569 | 0.655442 | -1.641 |
| chr20 | 53474629 | 53474885 | 0.952623 | 0.580664 | -1.641 |
| chr8 | 48936310 | 48936550 | 0.901306 | 0.549405 | -1.641 |
| chr11 | 82105952 | 82106188 | 1.07296 | 0.654065 | -1.640 |
| chr12 | 114036606 | 114036816 | 0.867288 | 0.528709 | -1.640 |
| chr5 | 88753251 | 88753435 | 1.08516 | 0.66164 | -1.640 |
| chr10 | 55058168 | 55058355 | 0.877489 | 0.535033 | -1.640 |
| chr14 | 44573871 | 44574230 | 1.01454 | 0.618597 | -1.640 |
| chr7 | 66292792 | 66292974 | 1.09675 | 0.668726 | -1.640 |
| chr9 | 9239711 | 9239909 | 1.48039 | 0.902646 | -1.640 |

| | | | | | |
|---|---|---|---|---|---|
| chr9 | 96089059 | 96089269 | 1.05208 | 0.641547 | -1.640 |
| chr5 | 132500760 | 132501002 | 0.849915 | 0.518287 | -1.640 |
| chr14 | 46185763 | 46185999 | 1.1509 | 0.701933 | -1.640 |
| chr5 | 75221006 | 75221408 | 0.882461 | 0.538226 | -1.640 |
| chr19 | 28825066 | 28825261 | 1.06128 | 0.647309 | -1.640 |
| chr20 | 43887144 | 43887327 | 0.955896 | 0.58306 | -1.639 |
| chr7 | 83550741 | 83551044 | 0.887566 | 0.54145 | -1.639 |
| chr5 | 170105555 | 170105739 | 0.964493 | 0.588433 | -1.639 |
| chr1 | 177465030 | 177465233 | 0.932544 | 0.568948 | -1.639 |
| chr9 | 12489991 | 12490238 | 1.03397 | 0.63092 | -1.639 |
| chr5 | 98708024 | 98708284 | 0.95838 | 0.584813 | -1.639 |
| chr20 | 46508442 | 46508658 | 1.16938 | 0.713597 | -1.639 |
| chr9 | 73618948 | 73619178 | 1.06635 | 0.650769 | -1.639 |
| chr19 | 44836235 | 44836461 | 1.41242 | 0.862072 | -1.638 |
| chr11 | 30135267 | 30135575 | 1.00741 | 0.614914 | -1.638 |
| chr2 | 36156096 | 36156331 | 1.13188 | 0.690945 | -1.638 |
| chr8 | 92355667 | 92355994 | 1.51627 | 0.925681 | -1.638 |
| chr9 | 38096463 | 38096736 | 1.06544 | 0.650474 | -1.638 |
| chr2 | 221821201 | 221821386 | 0.896881 | 0.547756 | -1.637 |
| chr14 | 62912519 | 62912723 | 1.06724 | 0.651925 | -1.637 |
| chr19 | 57435059 | 57435287 | 0.908093 | 0.554771 | -1.637 |
| chr12 | 128183522 | 128183763 | 1.04651 | 0.639396 | -1.637 |
| chr19 | 32114415 | 32114675 | 1.08117 | 0.660604 | -1.637 |
| chr16 | 2552558 | 2552821 | 0.950848 | 0.581006 | -1.637 |
| chr9 | 93213789 | 93214003 | 1.01055 | 0.617598 | -1.636 |
| chr8 | 78171554 | 78171787 | 1.07076 | 0.654464 | -1.636 |
| chr14 | 87193408 | 87193675 | 1.31557 | 0.804143 | -1.636 |
| chr10 | 113267595 | 113267922 | 0.990406 | 0.605393 | -1.636 |
| chr5 | 14562107 | 14562312 | 0.897265 | 0.548525 | -1.636 |
| chr8 | 1965363 | 1965575 | 0.982039 | 0.600369 | -1.636 |
| chr5 | 105252015 | 105252212 | 0.893606 | 0.546306 | -1.636 |
| chr1 | 207015848 | 207016100 | 0.948652 | 0.580046 | -1.635 |
| chr2 | 78098682 | 78098895 | 1.13094 | 0.69153 | -1.635 |
| chr11 | 25364324 | 25364634 | 1.14829 | 0.702141 | -1.635 |
| chr7 | 85650668 | 85650886 | 0.756531 | 0.462621 | -1.635 |
| chr2 | 61210484 | 61210711 | 1.07647 | 0.658266 | -1.635 |
| chr5 | 105437354 | 105437575 | 0.681065 | 0.416517 | -1.635 |
| chr3 | 86628262 | 86628568 | 0.85772 | 0.52456 | -1.635 |
| chr5 | 8444564 | 8444777 | 1.06445 | 0.651189 | -1.635 |
| chr19 | 29501693 | 29502044 | 1.12179 | 0.686314 | -1.635 |
| chr12 | 45224677 | 45224955 | 0.965241 | 0.590589 | -1.634 |
| chr8 | 139351892 | 139352286 | 0.939175 | 0.574645 | -1.634 |
| chr5 | 116349414 | 116349798 | 1.12672 | 0.689439 | -1.634 |
| chr4 | 74495547 | 74495824 | 0.993208 | 0.607792 | -1.634 |
| chr5 | 119636217 | 119636409 | 0.861833 | 0.52743 | -1.634 |
| chr10 | 84923433 | 84923742 | 1.12251 | 0.687052 | -1.634 |
| chr11 | 3631533 | 3631715 | 1.10526 | 0.676525 | -1.634 |
| chr10 | 54986262 | 54986480 | 1.37982 | 0.844616 | -1.634 |
| chr8 | 118609583 | 118609781 | 0.965649 | 0.591138 | -1.634 |
| chr12 | 100695633 | 100695907 | 1.01011 | 0.618363 | -1.634 |
| chr2 | 206117916 | 206118380 | 1.03397 | 0.632993 | -1.633 |
| chr5 | 119094325 | 119094533 | 0.811122 | 0.496588 | -1.633 |
| chr14 | 84702613 | 84702868 | 1.13026 | 0.69207 | -1.633 |
| chr5 | 98504159 | 98504359 | 0.878542 | 0.538004 | -1.633 |
| chr2 | 203025706 | 203025976 | 1.13967 | 0.697957 | -1.633 |
| chr5 | 27166589 | 27166820 | 0.985683 | 0.603655 | -1.633 |
| chr17 | 66655277 | 66655584 | 0.97113 | 0.594824 | -1.633 |
| chr3 | 79442330 | 79442527 | 0.976262 | 0.598021 | -1.632 |
| chr3 | 146883373 | 146883577 | 0.792365 | 0.485407 | -1.632 |
| chr11 | 23517868 | 23518104 | 0.994137 | 0.609035 | -1.632 |
| chr11 | 89884738 | 89884941 | 0.751865 | 0.460672 | -1.632 |

| | | | | | |
|---|---|---|---|---|---|
| chr10 | 9522969 | 9523223 | 0.942312 | 0.57744 | -1.632 |
| chr14 | 33004822 | 33005036 | 0.930122 | 0.569971 | -1.632 |
| chr2 | 16026129 | 16026387 | 1.16824 | 0.716025 | -1.632 |
| chr2 | 142030596 | 142030803 | 0.786869 | 0.482297 | -1.632 |
| chr9 | 19759049 | 19759402 | 1.08047 | 0.662265 | -1.631 |
| chr2 | 176173734 | 176173928 | 0.868094 | 0.532133 | -1.631 |
| chr3 | 95131495 | 95131766 | 0.919474 | 0.563728 | -1.631 |
| chr5 | 179655991 | 179656215 | 0.819056 | 0.502165 | -1.631 |
| chr1 | 200603582 | 200603907 | 1.44774 | 0.887662 | -1.631 |
| chr7 | 49887936 | 49888180 | 1.27358 | 0.780879 | -1.631 |
| chr2 | 81766711 | 81766969 | 0.994878 | 0.610011 | -1.631 |
| chr7 | 54237154 | 54237413 | 0.888825 | 0.545109 | -1.631 |
| chr15 | 28352798 | 28353008 | 1.28789 | 0.789858 | -1.631 |
| chr8 | 137461588 | 137461836 | 0.861167 | 0.528176 | -1.630 |
| chr1 | 160759377 | 160759568 | 0.799994 | 0.490738 | -1.630 |
| chr8 | 123459695 | 123459957 | 2.12927 | 1.3062 | -1.630 |
| chr15 | 66531965 | 66532165 | 1.14594 | 0.702991 | -1.630 |
| chr17 | 5229391 | 5229598 | 1.06968 | 0.656274 | -1.630 |
| chr3 | 14687491 | 14687679 | 0.932336 | 0.572064 | -1.630 |
| chr19 | 53835039 | 53835458 | 0.852252 | 0.522941 | -1.630 |
| chr8 | 85887971 | 85888191 | 0.846192 | 0.519227 | -1.630 |
| chr3 | 103264318 | 103264505 | 0.946627 | 0.580944 | -1.629 |
| chr7 | 132892575 | 132892784 | 1.27754 | 0.7841 | -1.629 |
| chr5 | 44534551 | 44534780 | 0.825141 | 0.506444 | -1.629 |
| chr16 | 47889407 | 47889652 | 0.934651 | 0.573699 | -1.629 |
| chr5 | 905392 | 905690 | 0.916444 | 0.562575 | -1.629 |
| chr15 | 82256176 | 82256407 | 1.06693 | 0.655062 | -1.629 |
| chr5 | 18731456 | 18731848 | 0.692887 | 0.425457 | -1.629 |
| chr7 | 98354575 | 98354772 | 0.980645 | 0.602253 | -1.628 |
| chr19 | 52858359 | 52858695 | 0.974549 | 0.598579 | -1.628 |
| chr2 | 211803080 | 211803296 | 1.04395 | 0.64125 | -1.628 |
| chr1 | 210838909 | 210839162 | 0.666925 | 0.409822 | -1.627 |
| chr7 | 104552396 | 104552865 | 0.857138 | 0.526766 | -1.627 |
| chr14 | 47864097 | 47864301 | 1.15436 | 0.709444 | -1.627 |
| chr5 | 163396447 | 163396668 | 0.832157 | 0.511438 | -1.627 |
| chr14 | 72221333 | 72221537 | 1.03058 | 0.63339 | -1.627 |
| chr1 | 177367878 | 177368093 | 0.784228 | 0.482019 | -1.627 |
| chr7 | 136560159 | 136560353 | 1.17376 | 0.721603 | -1.627 |
| chr8 | 51366245 | 51366532 | 0.936316 | 0.575691 | -1.626 |
| chr3 | 73697563 | 73697791 | 1.06969 | 0.657698 | -1.626 |
| chr2 | 201709142 | 201709483 | 0.959794 | 0.590179 | -1.626 |
| chr19 | 37313728 | 37314104 | 1.21233 | 0.745536 | -1.626 |
| chr10 | 55709249 | 55709487 | 1.29458 | 0.796131 | -1.626 |
| chr9 | 29990684 | 29990971 | 1.00489 | 0.617983 | -1.626 |
| chr1 | 246549151 | 246549496 | 0.859573 | 0.528622 | -1.626 |
| chr1 | 173517325 | 173517513 | 0.897866 | 0.552289 | -1.626 |
| chr15 | 54375159 | 54375409 | 1.99274 | 1.22578 | -1.626 |
| chr5 | 62275538 | 62275830 | 1.0257 | 0.630957 | -1.626 |
| chr19 | 31371560 | 31371802 | 1.14809 | 0.706265 | -1.626 |
| chr5 | 95189151 | 95189349 | 0.950609 | 0.584806 | -1.626 |
| chr15 | 50864213 | 50864473 | 1.16152 | 0.71477 | -1.625 |
| chr15 | 81697770 | 81697989 | 1.06968 | 0.658255 | -1.625 |
| chr14 | 60282328 | 60282531 | 0.994173 | 0.611837 | -1.625 |
| chr16 | 9783763 | 9784024 | 0.850549 | 0.523489 | -1.625 |
| chr1 | 246544031 | 246544445 | 1.08654 | 0.668784 | -1.625 |
| chr3 | 145554377 | 145554619 | 1.00201 | 0.616848 | -1.624 |
| chr5 | 12217776 | 12218123 | 0.986305 | 0.607186 | -1.624 |
| chr10 | 16820426 | 16820625 | 0.999391 | 0.615283 | -1.624 |
| chr14 | 82485216 | 82485512 | 0.800618 | 0.492959 | -1.624 |
| chr15 | 90446954 | 90447187 | 1.26198 | 0.777094 | -1.624 |
| chr14 | 35425214 | 35425608 | 1.02096 | 0.628746 | -1.624 |

| | | | | | |
|---|---|---|---|---|---|
| chr12 | 103281220 | 103281425 | 1.07003 | 0.659107 | -1.623 |
| chr2 | 11580040 | 11580495 | 1.26914 | 0.781871 | -1.623 |
| chr5 | 176035372 | 176035561 | 1.17446 | 0.723547 | -1.623 |
| chr2 | 51611879 | 51612084 | 1.22668 | 0.755727 | -1.623 |
| chr5 | 116790494 | 116790681 | 1.22249 | 0.753185 | -1.623 |
| chr5 | 11444964 | 11445181 | 0.979571 | 0.603561 | -1.623 |
| chr2 | 77691712 | 77691966 | 0.873653 | 0.538309 | -1.623 |
| chr5 | 24314286 | 24314557 | 1.0566 | 0.651077 | -1.623 |
| chr5 | 94522451 | 94522635 | 0.995116 | 0.613226 | -1.623 |
| chr2 | 123464486 | 123464833 | 1.0285 | 0.633889 | -1.623 |
| chr5 | 139145796 | 139146122 | 0.991092 | 0.610841 | -1.623 |
| chr11 | 37489075 | 37489359 | 1.02343 | 0.630792 | -1.622 |
| chr7 | 71074022 | 71074223 | 1.11625 | 0.688002 | -1.622 |
| chr3 | 88739483 | 88739700 | 1.09265 | 0.673489 | -1.622 |
| chr14 | 45381507 | 45381737 | 0.953656 | 0.58788 | -1.622 |
| chr2 | 79972263 | 79972596 | 1.02783 | 0.633614 | -1.622 |
| chr1 | 103561577 | 103561881 | 1.04203 | 0.642369 | -1.622 |
| chr3 | 83771873 | 83772094 | 0.86191 | 0.531344 | -1.622 |
| chr8 | 43342434 | 43342656 | 1.05248 | 0.648825 | -1.622 |
| chr2 | 147838061 | 147838363 | 0.735934 | 0.453708 | -1.622 |
| chr2 | 76704774 | 76705061 | 1.21711 | 0.75055 | -1.622 |
| chr10 | 80087291 | 80087494 | 0.89153 | 0.549786 | -1.622 |
| chr6 | 108408502 | 108408687 | 1.03056 | 0.635589 | -1.621 |
| chr12 | 24713845 | 24714103 | 0.970695 | 0.598677 | -1.621 |
| chr7 | 43210302 | 43210509 | 0.774661 | 0.477816 | -1.621 |
| chr19 | 35299866 | 35300104 | 1.05732 | 0.652255 | -1.621 |
| chr15 | 70941382 | 70941576 | 1.36755 | 0.843698 | -1.621 |
| chr20 | 9846436 | 9846825 | 0.986461 | 0.608674 | -1.621 |
| chr5 | 88480226 | 88480477 | 1.09082 | 0.673142 | -1.620 |
| chr10 | 114156928 | 114157263 | 0.948212 | 0.585163 | -1.620 |
| chr17 | 70427259 | 70427464 | 1.41575 | 0.873744 | -1.620 |
| chr6 | 129896499 | 129896718 | 2.25282 | 1.39051 | -1.620 |
| chr19 | 28508380 | 28508575 | 1.52567 | 0.941811 | -1.620 |
| chr7 | 48098907 | 48099399 | 1.09245 | 0.674406 | -1.620 |
| chr10 | 55665079 | 55665419 | 1.03807 | 0.640858 | -1.620 |
| chr19 | 39068439 | 39068759 | 1.28783 | 0.795072 | -1.620 |
| chr10 | 16091172 | 16091373 | 1.27004 | 0.784181 | -1.620 |
| chr8 | 13844802 | 13844992 | 0.874983 | 0.540299 | -1.619 |
| chr2 | 64059313 | 64059502 | 1.13586 | 0.701428 | -1.619 |
| chr9 | 106948722 | 106949083 | 0.846725 | 0.522911 | -1.619 |
| chr1 | 71515587 | 71515982 | 1.33674 | 0.825639 | -1.619 |
| chr11 | 21748736 | 21748938 | 1.06381 | 0.657076 | -1.619 |
| chr1 | 203476007 | 203476203 | 1.38615 | 0.856207 | -1.619 |
| chr2 | 47671798 | 47672080 | 0.985793 | 0.608993 | -1.619 |
| chr2 | 527593 | 527849 | 1.06463 | 0.657832 | -1.618 |
| chr7 | 147839760 | 147839989 | 1.00943 | 0.623738 | -1.618 |
| chr19 | 58792691 | 58792894 | 1.11596 | 0.689849 | -1.618 |
| chr12 | 763103 | 763287 | 0.900967 | 0.556981 | -1.618 |
| chr8 | 136135130 | 136135334 | 1.11337 | 0.688312 | -1.618 |
| chr10 | 2806259 | 2806455 | 0.89518 | 0.553436 | -1.617 |
| chr3 | 6777156 | 6777419 | 1.00398 | 0.620789 | -1.617 |
| chr5 | 173056854 | 173057088 | 1.46519 | 0.906011 | -1.617 |
| chr9 | 24545768 | 24546012 | 0.990629 | 0.612635 | -1.617 |
| chr5 | 1993812 | 1994059 | 0.806979 | 0.499104 | -1.617 |
| chr8 | 112882348 | 112882703 | 1.04963 | 0.649189 | -1.617 |
| chr3 | 28479547 | 28479789 | 0.808188 | 0.49986 | -1.617 |
| chr17 | 6477548 | 6477879 | 0.994234 | 0.614946 | -1.617 |
| chr2 | 4646462 | 4646655 | 0.726109 | 0.449152 | -1.617 |
| chr1 | 170592566 | 170592758 | 1.34386 | 0.831763 | -1.616 |
| chr11 | 90102308 | 90102529 | 0.921656 | 0.570514 | -1.615 |
| chr5 | 45355480 | 45355785 | 0.974353 | 0.603139 | -1.615 |

| | | | | | |
|---|---|---|---|---|---|
| chr5 | 103760255 | 103760500 | 1.25203 | 0.775087 | -1.615 |
| chr5 | 22823710 | 22823968 | 0.995124 | 0.61609 | -1.615 |
| chr3 | 12129024 | 12129242 | 1.26825 | 0.785253 | -1.615 |
| chr16 | 6654069 | 6654259 | 1.10281 | 0.682839 | -1.615 |
| chr5 | 97722075 | 97722351 | 0.857406 | 0.531076 | -1.614 |
| chr8 | 136040419 | 136040608 | 0.779269 | 0.482692 | -1.614 |
| chr8 | 92045832 | 92046255 | 0.916253 | 0.567568 | -1.614 |
| chr2 | 83752408 | 83752623 | 1.17436 | 0.727463 | -1.614 |
| chr16 | 82667865 | 82668049 | 1.2594 | 0.78019 | -1.614 |
| chr5 | 26451119 | 26451306 | 0.92309 | 0.571894 | -1.614 |
| chr2 | 239176709 | 239177052 | 0.92313 | 0.571941 | -1.614 |
| chr3 | 164352259 | 164352887 | 0.948556 | 0.587697 | -1.614 |
| chr9 | 18788515 | 18788732 | 1.15503 | 0.715629 | -1.614 |
| chr19 | 52690137 | 52690370 | 1.09745 | 0.679959 | -1.614 |
| chr19 | 34168541 | 34168738 | 0.98564 | 0.610689 | -1.614 |
| chr20 | 40536480 | 40536731 | 0.972782 | 0.602744 | -1.614 |
| chr5 | 110233354 | 110233536 | 0.973687 | 0.603311 | -1.614 |
| chr2 | 103713672 | 103714007 | 1.25126 | 0.775324 | -1.614 |
| chr8 | 113101773 | 113101983 | 1.04405 | 0.64696 | -1.614 |
| chr4 | 40827571 | 40827767 | 0.925271 | 0.573362 | -1.614 |
| chr2 | 167005587 | 167005814 | 0.979766 | 0.607194 | -1.614 |
| chr17 | 11240915 | 11241128 | 0.780684 | 0.483855 | -1.613 |
| chr5 | 12890699 | 12890906 | 1.1994 | 0.74357 | -1.613 |
| chr2 | 73955313 | 73955668 | 0.949252 | 0.588492 | -1.613 |
| chr2 | 13736835 | 13737018 | 1.36545 | 0.84652 | -1.613 |
| chr1 | 170068233 | 170068529 | 0.941803 | 0.583918 | -1.613 |
| chr5 | 178066655 | 178066853 | 1.00516 | 0.623303 | -1.613 |
| chr7 | 118875032 | 118875291 | 1.05871 | 0.656567 | -1.612 |
| chr3 | 77211724 | 77211968 | 1.0397 | 0.644797 | -1.612 |
| chr10 | 117332642 | 117332859 | 1.47965 | 0.917722 | -1.612 |
| chr15 | 29591551 | 29591765 | 1.10402 | 0.684838 | -1.612 |
| chr3 | 12014342 | 12014559 | 1.1265 | 0.698791 | -1.612 |
| chr10 | 122874434 | 122874652 | 1.11458 | 0.691399 | -1.612 |
| chr10 | 124969715 | 124969901 | 1.18194 | 0.733299 | -1.612 |
| chr2 | 176236620 | 176236829 | 1.25635 | 0.77952 | -1.612 |
| chr12 | 58217328 | 58217535 | 0.876445 | 0.543824 | -1.612 |
| chr15 | 65108700 | 65108887 | 0.94087 | 0.583848 | -1.611 |
| chr2 | 34237847 | 34238083 | 1.84269 | 1.14352 | -1.611 |
| chr1 | 203387350 | 203387605 | 0.874902 | 0.542996 | -1.611 |
| chr14 | 85410694 | 85410887 | 1.0988 | 0.68204 | -1.611 |
| chr20 | 54160766 | 54160956 | 0.943397 | 0.5856 | -1.611 |
| chr3 | 94890928 | 94891112 | 0.981567 | 0.609498 | -1.610 |
| chr3 | 85229369 | 85229552 | 1.21217 | 0.752786 | -1.610 |
| chr10 | 1132797 | 1133006 | 1.20842 | 0.750506 | -1.610 |
| chr8 | 4938887 | 4939139 | 0.846365 | 0.525718 | -1.610 |
| chr20 | 21029424 | 21029839 | 2.57701 | 1.60086 | -1.610 |
| chr12 | 84939441 | 84939772 | 1.01237 | 0.62894 | -1.610 |
| chr2 | 221787240 | 221787473 | 0.854261 | 0.530733 | -1.610 |
| chr16 | 72923577 | 72923771 | 0.936975 | 0.582155 | -1.609 |
| chr2 | 140901780 | 140901974 | 0.977781 | 0.607755 | -1.609 |
| chr9 | 73255229 | 73255612 | 1.27035 | 0.789609 | -1.609 |
| chr1 | 237279462 | 237279661 | 1.00825 | 0.626712 | -1.609 |
| chr15 | 69971869 | 69972190 | 0.961936 | 0.597935 | -1.609 |
| chr8 | 83755854 | 83756044 | 1.25117 | 0.777757 | -1.609 |
| chr11 | 69811096 | 69811299 | 0.879002 | 0.546424 | -1.609 |
| chr9 | 10339368 | 10339586 | 1.09539 | 0.681004 | -1.608 |
| chr3 | 16011865 | 16012094 | 1.11237 | 0.691606 | -1.608 |
| chr20 | 61532572 | 61533016 | 1.08543 | 0.675041 | -1.608 |
| chr11 | 59030872 | 59031121 | 1.00388 | 0.624471 | -1.608 |
| chr5 | 22738237 | 22738486 | 0.928788 | 0.577803 | -1.607 |
| chr8 | 85613363 | 85613559 | 1.10292 | 0.686155 | -1.607 |

| | | | | | |
|---|---|---|---|---|---|
| chr2 | 95865102 | 95865321 | 1.06298 | 0.661339 | -1.607 |
| chr12 | 82202660 | 82202875 | 1.08702 | 0.676392 | -1.607 |
| chr17 | 18043120 | 18043524 | 1.58707 | 0.987657 | -1.607 |
| chr8 | 139926672 | 139926986 | 2.6943 | 1.67676 | -1.607 |
| chr1 | 197867673 | 197867862 | 1.25091 | 0.778688 | -1.606 |
| chr5 | 51652238 | 51652428 | 1.87103 | 1.16476 | -1.606 |
| chr5 | 3828910 | 3829202 | 0.917368 | 0.571119 | -1.606 |
| chr7 | 105811308 | 105811502 | 0.958592 | 0.596863 | -1.606 |
| chr2 | 139995741 | 139996025 | 0.792635 | 0.493535 | -1.606 |
| chr8 | 85861456 | 85861708 | 0.825738 | 0.51415 | -1.606 |
| chr1 | 195820369 | 195820594 | 0.907739 | 0.565247 | -1.606 |
| chr7 | 92504919 | 92505139 | 1.01902 | 0.634635 | -1.606 |
| chr8 | 112004636 | 112004926 | 0.903815 | 0.562936 | -1.606 |
| chr5 | 24280423 | 24280683 | 0.726988 | 0.452801 | -1.606 |
| chr20 | 6401546 | 6401874 | 1.01323 | 0.631123 | -1.605 |
| chr15 | 70683990 | 70684188 | 1.10653 | 0.689322 | -1.605 |
| chrX | 139654911 | 139655206 | 1.9183 | 1.19521 | -1.605 |
| chr3 | 109550334 | 109550706 | 0.949002 | 0.591291 | -1.605 |
| chr15 | 97769277 | 97769563 | 1.44694 | 0.901594 | -1.605 |
| chr1 | 163453317 | 163453508 | 1.00365 | 0.625428 | -1.605 |
| chr10 | 85472466 | 85472696 | 0.663547 | 0.413497 | -1.605 |
| chr2 | 21346689 | 21346925 | 1.50683 | 0.939026 | -1.605 |
| chr3 | 37683230 | 37683427 | 1.24011 | 0.772875 | -1.605 |
| chr20 | 12023552 | 12024019 | 0.885001 | 0.551562 | -1.605 |
| chr15 | 34220852 | 34221047 | 1.15646 | 0.720774 | -1.604 |
| chr19 | 28919881 | 28920158 | 0.89375 | 0.557047 | -1.604 |
| chr19 | 29290643 | 29291000 | 1.03214 | 0.643312 | -1.604 |
| chr9 | 90144601 | 90144836 | 1.0979 | 0.684413 | -1.604 |
| chr3 | 85839740 | 85840083 | 1.12627 | 0.702142 | -1.604 |
| chr10 | 56881535 | 56881832 | 0.858766 | 0.535512 | -1.604 |
| chr1 | 183521091 | 183521302 | 1.24946 | 0.779149 | -1.604 |
| chr11 | 44781775 | 44781975 | 0.900985 | 0.561882 | -1.604 |
| chr12 | 5131639 | 5132033 | 2.14839 | 1.3399 | -1.603 |
| chr3 | 95790113 | 95790319 | 0.940018 | 0.586275 | -1.603 |
| chr16 | 66798424 | 66798691 | 0.901132 | 0.562094 | -1.603 |
| chr8 | 92941790 | 92941992 | 1.14424 | 0.713774 | -1.603 |
| chr14 | 88232121 | 88232383 | 0.971561 | 0.606078 | -1.603 |
| chr10 | 61501263 | 61501501 | 1.07539 | 0.67088 | -1.603 |
| chr5 | 111959528 | 111959750 | 0.919725 | 0.573778 | -1.603 |
| chr15 | 35121080 | 35121269 | 0.969291 | 0.604715 | -1.603 |
| chr8 | 146035425 | 146035793 | 0.971282 | 0.605985 | -1.603 |
| chr8 | 92901907 | 92902169 | 1.24487 | 0.77668 | -1.603 |
| chr8 | 65236090 | 65236369 | 0.995043 | 0.620832 | -1.603 |
| chr5 | 11890507 | 11890695 | 0.921034 | 0.574663 | -1.603 |
| chr6 | 161771138 | 161771367 | 1.18058 | 0.736621 | -1.603 |
| chr10 | 55386688 | 55387049 | 1.05838 | 0.660613 | -1.602 |
| chr2 | 13436321 | 13436650 | 0.945905 | 0.590498 | -1.602 |
| chr5 | 161433102 | 161433291 | 0.946562 | 0.591011 | -1.602 |
| chr2 | 77921853 | 77922117 | 1.02461 | 0.639824 | -1.601 |
| chr8 | 86417503 | 86417755 | 0.854933 | 0.533908 | -1.601 |
| chr14 | 62481094 | 62481416 | 1.03703 | 0.647637 | -1.601 |
| chr15 | 28025458 | 28025751 | 1.52086 | 0.949809 | -1.601 |
| chr2 | 14862290 | 14862578 | 0.778168 | 0.486026 | -1.601 |
| chr17 | 79107732 | 79108001 | 0.919141 | 0.574106 | -1.601 |
| chr1 | 156529330 | 156529520 | 0.814335 | 0.508688 | -1.601 |
| chr2 | 605995 | 606298 | 0.931513 | 0.581922 | -1.601 |
| chr17 | 8424612 | 8424831 | 0.986448 | 0.616284 | -1.601 |
| chr2 | 6163710 | 6164009 | 0.921416 | 0.575656 | -1.601 |
| chr14 | 38716506 | 38716756 | 1.08378 | 0.677193 | -1.600 |
| chr19 | 41479656 | 41479863 | 1.15496 | 0.721753 | -1.600 |
| chr9 | 135442309 | 135442546 | 1.14527 | 0.715799 | -1.600 |

| | | | | | |
|---|---|---|---|---|---|
| chr6 | 80140563 | 80140812 | 0.776799 | 0.485507 | -1.600 |
| chr2 | 17068450 | 17068714 | 0.756654 | 0.472936 | -1.600 |
| chr8 | 96628643 | 96628849 | 0.952333 | 0.595242 | -1.600 |
| chr9 | 130241059 | 130241350 | 0.80937 | 0.505951 | -1.600 |
| chr19 | 23580669 | 23580976 | 0.970701 | 0.606804 | -1.600 |
| chr11 | 26629100 | 26629307 | 1.02705 | 0.642067 | -1.600 |
| chr3 | 97775461 | 97775829 | 1.02106 | 0.638458 | -1.599 |
| chr11 | 10893269 | 10893466 | 1.12408 | 0.702943 | -1.599 |
| chr9 | 122192498 | 122192747 | 0.959429 | 0.600015 | -1.599 |
| chr10 | 132053181 | 132053517 | 1.17906 | 0.737428 | -1.599 |
| chr8 | 52482606 | 52482939 | 0.994741 | 0.622165 | -1.599 |
| chr2 | 65000936 | 65001132 | 1.14341 | 0.715221 | -1.599 |
| chr15 | 60092798 | 60093101 | 1.03031 | 0.644505 | -1.599 |
| chr8 | 112416369 | 112416705 | 1.17044 | 0.732178 | -1.599 |
| chr9 | 93307400 | 93307598 | 0.977824 | 0.611687 | -1.599 |
| chr15 | 98095723 | 98095943 | 1.15693 | 0.723752 | -1.599 |
| chr11 | 26127232 | 26127465 | 0.818903 | 0.512363 | -1.598 |
| chr2 | 77362360 | 77362544 | 1.49681 | 0.936578 | -1.598 |
| chr7 | 42307661 | 42307919 | 1.06137 | 0.664118 | -1.598 |
| chr2 | 86234963 | 86235234 | 1.13043 | 0.707359 | -1.598 |
| chr3 | 169213382 | 169213571 | 0.939796 | 0.588093 | -1.598 |
| chr9 | 11618242 | 11618980 | 1.14789 | 0.718359 | -1.598 |
| chr3 | 122640633 | 122640853 | 1.01955 | 0.63806 | -1.598 |
| chr2 | 81996412 | 81996612 | 0.953167 | 0.596518 | -1.598 |
| chr11 | 26121768 | 26122091 | 1.25286 | 0.784096 | -1.598 |
| chr3 | 79631836 | 79632051 | 0.811068 | 0.507709 | -1.598 |
| chr1 | 202905063 | 202905265 | 1.15105 | 0.720881 | -1.597 |
| chr19 | 22873624 | 22873870 | 0.975796 | 0.611277 | -1.596 |
| chr8 | 139452220 | 139452482 | 0.911182 | 0.570849 | -1.596 |
| chr19 | 28627184 | 28627588 | 1.28034 | 0.802127 | -1.596 |
| chr10 | 31997993 | 31998271 | 1.07476 | 0.67341 | -1.596 |
| chr10 | 20503639 | 20503866 | 1.19384 | 0.748233 | -1.596 |
| chr16 | 13749443 | 13749670 | 1.09192 | 0.684415 | -1.595 |
| chr7 | 116556515 | 116556729 | 1.09872 | 0.688743 | -1.595 |
| chr5 | 106035062 | 106035250 | 1.08393 | 0.679514 | -1.595 |
| chr2 | 201339815 | 201340041 | 0.825707 | 0.517769 | -1.595 |
| chr8 | 69765182 | 69765432 | 0.77359 | 0.485167 | -1.594 |
| chr19 | 30654775 | 30654988 | 1.04087 | 0.652852 | -1.594 |
| chr1 | 226625842 | 226626219 | 1.30355 | 0.817622 | -1.594 |
| chr8 | 51475694 | 51476106 | 0.939309 | 0.58923 | -1.594 |
| chr19 | 39432990 | 39433402 | 1.22333 | 0.767482 | -1.594 |
| chr16 | 61109250 | 61109560 | 0.941768 | 0.590839 | -1.594 |
| chr2 | 13232359 | 13232596 | 1.22976 | 0.771566 | -1.594 |
| chr8 | 4727868 | 4728184 | 1.07674 | 0.675566 | -1.594 |
| chr19 | 38176275 | 38176598 | 0.991678 | 0.622212 | -1.594 |
| chr3 | 144135889 | 144136164 | 1.1384 | 0.714301 | -1.594 |
| chr10 | 110561549 | 110561743 | 0.83221 | 0.522213 | -1.594 |
| chr7 | 117480534 | 117480748 | 0.732491 | 0.459782 | -1.593 |
| chr17 | 10519041 | 10519236 | 1.02797 | 0.645695 | -1.592 |
| chr17 | 35868130 | 35868566 | 1.34716 | 0.846261 | -1.592 |
| chr8 | 83391213 | 83391498 | 0.875655 | 0.550108 | -1.592 |
| chr10 | 89549213 | 89549460 | 1.16733 | 0.733407 | -1.592 |
| chr9 | 10409653 | 10409981 | 0.989843 | 0.621995 | -1.591 |
| chr15 | 35569589 | 35569827 | 1.20864 | 0.759663 | -1.591 |
| chr7 | 31199595 | 31199810 | 1.17084 | 0.735914 | -1.591 |
| chr8 | 88862402 | 88862656 | 0.898232 | 0.564579 | -1.591 |
| chr7 | 126798798 | 126799238 | 1.24326 | 0.781481 | -1.591 |
| chr7 | 76918989 | 76919199 | 0.877258 | 0.551485 | -1.591 |
| chr8 | 135093513 | 135093706 | 1.14611 | 0.720541 | -1.591 |
| chr7 | 125026235 | 125026476 | 1.33754 | 0.840912 | -1.591 |
| chr2 | 13771505 | 13771714 | 1.07338 | 0.674896 | -1.590 |

| | | | | | |
|---|---|---|---|---|---|
| chr5 | 20640139 | 20640464 | 1.18347 | 0.744174 | -1.590 |
| chr2 | 4863008 | 4863226 | 0.963882 | 0.606127 | -1.590 |
| chr8 | 15840765 | 15840962 | 1.41035 | 0.88691 | -1.590 |
| chr8 | 132041051 | 132041343 | 1.4088 | 0.885958 | -1.590 |
| chr5 | 147601116 | 147601381 | 1.19851 | 0.753713 | -1.590 |
| chr15 | 95517562 | 95517877 | 1.05165 | 0.661383 | -1.590 |
| chr5 | 61364468 | 61364671 | 0.9893 | 0.622172 | -1.590 |
| chr1 | 90965248 | 90965547 | 0.929445 | 0.584533 | -1.590 |
| chr9 | 92116435 | 92116678 | 1.09636 | 0.68959 | -1.590 |
| chr5 | 8489912 | 8490146 | 0.954724 | 0.600535 | -1.590 |
| chr15 | 95656941 | 95657470 | 1.00718 | 0.633605 | -1.590 |
| chr1 | 158484875 | 158485063 | 0.925872 | 0.582571 | -1.589 |
| chr10 | 79831727 | 79831943 | 0.965595 | 0.607611 | -1.589 |
| chr9 | 2523663 | 2524093 | 0.991031 | 0.623703 | -1.589 |
| chr8 | 112844385 | 112844611 | 0.98568 | 0.620338 | -1.589 |
| chr2 | 4681604 | 4681917 | 0.914563 | 0.575591 | -1.589 |
| chr3 | 98272033 | 98272350 | 0.596746 | 0.375589 | -1.589 |
| chr11 | 40166607 | 40166903 | 0.736016 | 0.463274 | -1.589 |
| chr1 | 196124879 | 196125072 | 0.938149 | 0.590548 | -1.589 |
| chr11 | 5735131 | 5735420 | 0.798298 | 0.50264 | -1.588 |
| chr5 | 11859142 | 11859337 | 0.978053 | 0.615821 | -1.588 |
| chr4 | 15418011 | 15418273 | 0.795482 | 0.500908 | -1.588 |
| chr8 | 96393770 | 96393973 | 0.855346 | 0.538711 | -1.588 |
| chr12 | 5333198 | 5333436 | 0.861214 | 0.542489 | -1.588 |
| chr15 | 94688193 | 94688523 | 1.16638 | 0.734725 | -1.588 |
| chr6 | 129059088 | 129059334 | 0.741278 | 0.467005 | -1.587 |
| chr9 | 31048521 | 31049111 | 1.05459 | 0.664414 | -1.587 |
| chr2 | 4948825 | 4949030 | 1.1403 | 0.718421 | -1.587 |
| chr1 | 247834663 | 247834925 | 1.25599 | 0.791319 | -1.587 |
| chr15 | 98223999 | 98224234 | 1.15253 | 0.726178 | -1.587 |
| chr12 | 29786089 | 29786361 | 0.938048 | 0.591051 | -1.587 |
| chr7 | 15435783 | 15435998 | 1.01424 | 0.639066 | -1.587 |
| chr12 | 100828394 | 100828764 | 0.843971 | 0.531788 | -1.587 |
| chr8 | 24740424 | 24740650 | 1.16893 | 0.7367 | -1.587 |
| chr8 | 130835943 | 130836210 | 1.16947 | 0.7372 | -1.586 |
| chr7 | 22723838 | 22724078 | 1.02357 | 0.64527 | -1.586 |
| chr20 | 21642303 | 21642598 | 1.17041 | 0.737877 | -1.586 |
| chr2 | 105350141 | 105350366 | 1.40634 | 0.886634 | -1.586 |
| chr11 | 59760958 | 59761262 | 1.27154 | 0.801665 | -1.586 |
| chr3 | 80813862 | 80814052 | 1.06367 | 0.670689 | -1.586 |
| chr8 | 126670413 | 126670642 | 1.0699 | 0.67466 | -1.586 |
| chr10 | 78364819 | 78365002 | 1.19717 | 0.754945 | -1.586 |
| chr2 | 69030957 | 69031175 | 1.313 | 0.828085 | -1.586 |
| chr11 | 23930357 | 23930596 | 0.830483 | 0.523792 | -1.586 |
| chr8 | 36198913 | 36199139 | 0.957798 | 0.604144 | -1.585 |
| chr3 | 162995221 | 162995501 | 1.11694 | 0.704546 | -1.585 |
| chr10 | 83503003 | 83503225 | 0.89488 | 0.564478 | -1.585 |
| chr15 | 83596742 | 83596963 | 1.48919 | 0.939701 | -1.585 |
| chr2 | 78638482 | 78638692 | 1.01013 | 0.637512 | -1.584 |
| chr15 | 94545327 | 94545511 | 1.13497 | 0.716408 | -1.584 |
| chr14 | 96362180 | 96362363 | 1.31587 | 0.83061 | -1.584 |
| chr16 | 83292639 | 83292876 | 1.10267 | 0.696232 | -1.584 |
| chr7 | 52584082 | 52584303 | 0.975587 | 0.616046 | -1.584 |
| chr11 | 18208476 | 18208684 | 1.18627 | 0.74918 | -1.583 |
| chr7 | 99195689 | 99195907 | 0.804951 | 0.508447 | -1.583 |
| chr1 | 242555860 | 242556092 | 1.13056 | 0.714194 | -1.583 |
| chr2 | 212962997 | 212963262 | 0.975489 | 0.616249 | -1.583 |
| chr15 | 24124723 | 24124913 | 0.937011 | 0.591993 | -1.583 |
| chr8 | 133083361 | 133083557 | 1.26994 | 0.802349 | -1.583 |
| chr11 | 78851060 | 78851286 | 0.948733 | 0.599489 | -1.583 |
| chr2 | 28860405 | 28860624 | 0.996872 | 0.630099 | -1.582 |

| | | | | | |
|---|---|---|---|---|---|
| chr10 | 86425741 | 86425959 | 1.91363 | 1.20957 | -1.582 |
| chr15 | 61350500 | 61350789 | 1.04588 | 0.661242 | -1.582 |
| chr8 | 57731449 | 57731707 | 1.25681 | 0.794609 | -1.582 |
| chr2 | 213110509 | 213110735 | 0.962435 | 0.608597 | -1.581 |
| chr7 | 118758145 | 118758335 | 1.14749 | 0.725692 | -1.581 |
| chr2 | 138587789 | 138587971 | 1.10849 | 0.701068 | -1.581 |
| chr7 | 136833053 | 136833316 | 0.978817 | 0.619083 | -1.581 |
| chr14 | 86476015 | 86476207 | 0.816203 | 0.516277 | -1.581 |
| chr7 | 25288455 | 25288664 | 0.827886 | 0.523776 | -1.581 |
| chr20 | 12142020 | 12142217 | 0.919301 | 0.581633 | -1.581 |
| chr20 | 51407217 | 51407428 | 0.920917 | 0.582669 | -1.581 |
| chr12 | 71114489 | 71114714 | 0.936537 | 0.592674 | -1.580 |
| chr12 | 40228778 | 40229021 | 1.0362 | 0.655898 | -1.580 |
| chr10 | 14242649 | 14242989 | 1.97826 | 1.25227 | -1.580 |
| chr5 | 32353694 | 32353906 | 1.07922 | 0.683166 | -1.580 |
| chr8 | 136139790 | 136140018 | 1.18958 | 0.753179 | -1.579 |
| chr15 | 33730454 | 33730662 | 1.03727 | 0.656824 | -1.579 |
| chr8 | 111842508 | 111842973 | 1.42608 | 0.903074 | -1.579 |
| chr12 | 9190692 | 9190999 | 1.13848 | 0.721222 | -1.579 |
| chr7 | 39397343 | 39397551 | 1.16672 | 0.739212 | -1.578 |
| chr10 | 113786400 | 113786722 | 0.924704 | 0.585886 | -1.578 |
| chr7 | 9621330 | 9621620 | 1.05198 | 0.666605 | -1.578 |
| chr7 | 124266696 | 124267053 | 0.925165 | 0.586257 | -1.578 |
| chr1 | 163018495 | 163018689 | 0.907892 | 0.575313 | -1.578 |
| chr14 | 85282241 | 85282446 | 1.05224 | 0.666817 | -1.578 |
| chr17 | 55515466 | 55515658 | 1.08706 | 0.689122 | -1.577 |
| chr14 | 65644849 | 65645071 | 0.90869 | 0.5761 | -1.577 |
| chr9 | 94117551 | 94117780 | 0.847337 | 0.537224 | -1.577 |
| chr11 | 81751580 | 81751876 | 0.968992 | 0.614358 | -1.577 |
| chr17 | 42722728 | 42723015 | 0.97335 | 0.617234 | -1.577 |
| chr8 | 138192772 | 138192979 | 0.922233 | 0.585088 | -1.576 |
| chr5 | 160163345 | 160163790 | 1.0643 | 0.675267 | -1.576 |
| chr8 | 84907335 | 84907517 | 0.995946 | 0.631997 | -1.576 |
| chr11 | 8076835 | 8077092 | 0.96535 | 0.612592 | -1.576 |
| chr11 | 66336722 | 66336906 | 0.794479 | 0.504174 | -1.576 |
| chr8 | 54712453 | 54712795 | 1.06775 | 0.677598 | -1.576 |
| chr9 | 93567461 | 93567838 | 0.996587 | 0.632593 | -1.575 |
| chr10 | 12206105 | 12206303 | 1.38621 | 0.879978 | -1.575 |
| chr7 | 119470412 | 119470648 | 0.806875 | 0.512279 | -1.575 |
| chr16 | 74694103 | 74694310 | 0.939302 | 0.596418 | -1.575 |
| chr11 | 112665959 | 112666241 | 2.02822 | 1.28787 | -1.575 |
| chr11 | 26914323 | 26914555 | 1.0428 | 0.662281 | -1.575 |
| chr7 | 53525022 | 53525338 | 1.10554 | 0.702133 | -1.575 |
| chr5 | 165812929 | 165813139 | 0.911365 | 0.578819 | -1.575 |
| chr5 | 161516100 | 161516352 | 0.839521 | 0.533216 | -1.574 |
| chr5 | 71357475 | 71357696 | 0.979075 | 0.621918 | -1.574 |
| chr10 | 59013742 | 59014011 | 0.906612 | 0.575909 | -1.574 |
| chr2 | 147596995 | 147597222 | 1.04524 | 0.664005 | -1.574 |
| chr2 | 82051266 | 82051602 | 1.17871 | 0.74902 | -1.574 |
| chr10 | 88108239 | 88108472 | 0.917522 | 0.583069 | -1.574 |
| chr15 | 94646797 | 94647081 | 1.11982 | 0.711755 | -1.573 |
| chr2 | 76698188 | 76698429 | 0.888058 | 0.564459 | -1.573 |
| chr11 | 31509545 | 31509802 | 1.12146 | 0.712895 | -1.573 |
| chr2 | 34763317 | 34763532 | 0.868517 | 0.552115 | -1.573 |
| chr3 | 76588941 | 76589379 | 0.862246 | 0.548166 | -1.573 |
| chr1 | 239030334 | 239030615 | 1.10568 | 0.702938 | -1.573 |
| chr5 | 136341446 | 136341653 | 0.860609 | 0.54716 | -1.573 |
| chr14 | 46617892 | 46618112 | 0.984722 | 0.626207 | -1.573 |
| chr8 | 145800183 | 145800375 | 0.935854 | 0.595166 | -1.572 |
| chr8 | 127557435 | 127557709 | 0.83498 | 0.531022 | -1.572 |
| chr7 | 142430192 | 142430377 | 1.15182 | 0.732696 | -1.572 |

| | | | | | |
|---|---|---|---|---|---|
| chr10 | 111193068 | 111193279 | 0.749399 | 0.476711 | -1.572 |
| chr1 | 192019384 | 192019583 | 0.776127 | 0.493729 | -1.572 |
| chr2 | 41592277 | 41592539 | 1.013 | 0.644435 | -1.572 |
| chr16 | 534987 | 535249 | 1.10654 | 0.703967 | -1.572 |
| chr19 | 30656169 | 30656492 | 1.03238 | 0.656798 | -1.572 |
| chr5 | 53684375 | 53684581 | 0.889612 | 0.566031 | -1.572 |
| chr12 | 54987686 | 54987993 | 0.949534 | 0.604183 | -1.572 |
| chr11 | 20362630 | 20362830 | 1.11113 | 0.70711 | -1.571 |
| chr10 | 1797921 | 1798200 | 1.14731 | 0.730165 | -1.571 |
| chr11 | 29831672 | 29831972 | 0.999142 | 0.63587 | -1.571 |
| chr2 | 76666829 | 76667164 | 1.11616 | 0.71039 | -1.571 |
| chr5 | 51755868 | 51756132 | 1.36286 | 0.867447 | -1.571 |
| chr1 | 191938538 | 191938788 | 0.91511 | 0.582589 | -1.571 |
| chr14 | 55844069 | 55844261 | 1.02124 | 0.650203 | -1.571 |
| chr9 | 105748424 | 105748608 | 1.0214 | 0.650361 | -1.571 |
| chr9 | 9584716 | 9585011 | 1.17035 | 0.745435 | -1.570 |
| chr5 | 97042471 | 97042653 | 0.824801 | 0.525344 | -1.570 |
| chr5 | 84527044 | 84527226 | 0.933043 | 0.594442 | -1.570 |
| chr1 | 170517251 | 170517503 | 0.973523 | 0.620341 | -1.569 |
| chr2 | 102896833 | 102897048 | 0.864398 | 0.550875 | -1.569 |
| chr19 | 38022342 | 38022692 | 1.01448 | 0.646803 | -1.568 |
| chr2 | 240318898 | 240319186 | 1.27256 | 0.81146 | -1.568 |
| chr7 | 9102383 | 9102631 | 0.662565 | 0.422616 | -1.568 |
| chr16 | 76038822 | 76039236 | 0.882961 | 0.563257 | -1.568 |
| chr12 | 17084048 | 17084231 | 1.01565 | 0.647907 | -1.568 |
| chr5 | 101406754 | 101406996 | 0.750319 | 0.478693 | -1.567 |
| chr16 | 35005158 | 35005362 | 1.04852 | 0.668996 | -1.567 |
| chr2 | 48211280 | 48211601 | 0.900252 | 0.57449 | -1.567 |
| chr11 | 15326569 | 15326780 | 1.05295 | 0.671992 | -1.567 |
| chr10 | 9995110 | 9995403 | 0.993289 | 0.634069 | -1.567 |
| chr3 | 96171461 | 96171676 | 1.04165 | 0.66498 | -1.566 |
| chr19 | 29411699 | 29411908 | 1.2485 | 0.79712 | -1.566 |
| chr15 | 51402351 | 51402559 | 1.06603 | 0.680765 | -1.566 |
| chr3 | 73566073 | 73566340 | 0.963418 | 0.615272 | -1.566 |
| chr2 | 117343973 | 117344329 | 0.90359 | 0.577068 | -1.566 |
| chr14 | 95696130 | 95696339 | 0.985549 | 0.629469 | -1.566 |
| chr2 | 79315027 | 79315454 | 0.878144 | 0.560874 | -1.566 |
| chr5 | 16274906 | 16275117 | 0.932828 | 0.595803 | -1.566 |
| chr8 | 99284902 | 99285092 | 0.936769 | 0.598403 | -1.565 |
| chr10 | 129368258 | 129368497 | 0.854702 | 0.546009 | -1.565 |
| chr2 | 79164525 | 79164936 | 1.00924 | 0.644753 | -1.565 |
| chr2 | 50419729 | 50419998 | 0.896931 | 0.573051 | -1.565 |
| chr9 | 116750252 | 116750465 | 0.952771 | 0.608837 | -1.565 |
| chr19 | 32636158 | 32636384 | 1.22149 | 0.780627 | -1.565 |
| chr2 | 71346859 | 71347118 | 0.969687 | 0.619775 | -1.565 |
| chr2 | 194901305 | 194901524 | 0.980506 | 0.626788 | -1.564 |
| chr8 | 31269627 | 31269889 | 1.14441 | 0.731723 | -1.564 |
| chr3 | 105961651 | 105961849 | 1.00949 | 0.645487 | -1.564 |
| chr3 | 116077025 | 116077346 | 0.906587 | 0.579716 | -1.564 |
| chr2 | 96930722 | 96931076 | 1.07948 | 0.690275 | -1.564 |
| chr11 | 32098736 | 32098943 | 0.947353 | 0.605877 | -1.564 |
| chr10 | 38114468 | 38114682 | 1.27425 | 0.814998 | -1.564 |
| chr17 | 38345696 | 38345916 | 1.21955 | 0.78012 | -1.563 |
| chr19 | 29884220 | 29884432 | 1.14576 | 0.733006 | -1.563 |
| chr12 | 90313390 | 90313672 | 1.17066 | 0.748955 | -1.563 |
| chr7 | 113058775 | 113059207 | 0.946193 | 0.605398 | -1.563 |
| chr14 | 43984122 | 43984344 | 0.816817 | 0.522625 | -1.563 |
| chr19 | 38114999 | 38115213 | 1.24056 | 0.793836 | -1.563 |
| chr20 | 16733185 | 16733423 | 1.26697 | 0.810771 | -1.563 |
| chr16 | 64573995 | 64574242 | 0.897181 | 0.574207 | -1.562 |
| chr5 | 28483494 | 28483689 | 1.20235 | 0.769537 | -1.562 |

| | | | | | |
|---|---|---|---|---|---|
| chr8 | 49652519 | 49652740 | 1.09614 | 0.701675 | -1.562 |
| chr11 | 29823236 | 29823449 | 0.87098 | 0.557575 | -1.562 |
| chr19 | 34058321 | 34058524 | 1.09694 | 0.702233 | -1.562 |
| chr14 | 84176285 | 84176476 | 0.826179 | 0.528912 | -1.562 |
| chr7 | 48707455 | 48707653 | 0.999256 | 0.639725 | -1.562 |
| chr17 | 32052805 | 32053157 | 1.72426 | 1.10388 | -1.562 |
| chr2 | 35680107 | 35680303 | 1.06586 | 0.682416 | -1.562 |
| chr10 | 1001833 | 1002106 | 1.43197 | 0.917021 | -1.562 |
| chr1 | 248496799 | 248497011 | 1.17934 | 0.755242 | -1.562 |
| chr11 | 57709045 | 57709298 | 0.871286 | 0.557992 | -1.561 |
| chr10 | 82815734 | 82815934 | 0.901926 | 0.577759 | -1.561 |
| chr3 | 77424238 | 77424460 | 1.04529 | 0.66963 | -1.561 |
| chr7 | 97309314 | 97309513 | 1.09575 | 0.702092 | -1.561 |
| chr3 | 156306959 | 156307182 | 1.26639 | 0.811448 | -1.561 |
| chr17 | 8733787 | 8734195 | 1.12598 | 0.721513 | -1.561 |
| chr2 | 45453444 | 45453648 | 0.98826 | 0.63328 | -1.561 |
| chr2 | 184234458 | 184234924 | 1.23523 | 0.791589 | -1.560 |
| chr7 | 120506808 | 120507025 | 0.753597 | 0.48294 | -1.560 |
| chr15 | 94539863 | 94540058 | 1.14224 | 0.732032 | -1.560 |
| chr8 | 24913537 | 24913782 | 1.06609 | 0.68333 | -1.560 |
| chr1 | 32149691 | 32149902 | 1.75916 | 1.12762 | -1.560 |
| chr20 | 53683951 | 53684249 | 0.971331 | 0.622638 | -1.560 |
| chr3 | 82253624 | 82253855 | 1.05605 | 0.677115 | -1.560 |
| chr17 | 6641538 | 6641783 | 1.24112 | 0.796178 | -1.559 |
| chr1 | 230007181 | 230007364 | 0.78697 | 0.504858 | -1.559 |
| chr1 | 216754071 | 216754309 | 1.09862 | 0.70483 | -1.559 |
| chr5 | 119208402 | 119208628 | 1.1504 | 0.738061 | -1.559 |
| chr7 | 90905134 | 90905399 | 0.968764 | 0.621533 | -1.559 |
| chr15 | 84408526 | 84408848 | 1.1035 | 0.707997 | -1.559 |
| chr1 | 48996465 | 48996697 | 2.11625 | 1.35783 | -1.559 |
| chr3 | 21608897 | 21609086 | 0.854608 | 0.548357 | -1.558 |
| chr7 | 69474061 | 69474451 | 0.990678 | 0.63569 | -1.558 |
| chr3 | 97010718 | 97010919 | 0.974333 | 0.625212 | -1.558 |
| chrX | 13623381 | 13623606 | 0.765487 | 0.491226 | -1.558 |
| chr8 | 82451841 | 82452113 | 1.12148 | 0.719995 | -1.558 |
| chr5 | 152652722 | 152652912 | 0.71533 | 0.459263 | -1.558 |
| chr10 | 31428665 | 31429090 | 1.03422 | 0.664018 | -1.558 |
| chr11 | 42868796 | 42868998 | 0.855154 | 0.549098 | -1.557 |
| chr2 | 147954472 | 147954666 | 1.00213 | 0.643487 | -1.557 |
| chr14 | 94071235 | 94071516 | 0.91652 | 0.588516 | -1.557 |
| chr20 | 9907329 | 9907511 | 1.04036 | 0.668072 | -1.557 |
| chr19 | 28677803 | 28678013 | 1.01816 | 0.653833 | -1.557 |
| chr3 | 141297253 | 141297450 | 1.02813 | 0.660283 | -1.557 |
| chr10 | 112962712 | 112962977 | 1.18485 | 0.760962 | -1.557 |
| chr2 | 14601740 | 14602030 | 1.01331 | 0.65081 | -1.557 |
| chr5 | 84180405 | 84180729 | 0.887669 | 0.570116 | -1.557 |
| chr12 | 33252338 | 33252561 | 0.969327 | 0.622654 | -1.557 |
| chr17 | 52143950 | 52144138 | 1.06615 | 0.684981 | -1.556 |
| chr9 | 11658742 | 11658940 | 0.896102 | 0.575734 | -1.556 |
| chr3 | 83210056 | 83210330 | 0.955456 | 0.613977 | -1.556 |
| chr11 | 36810781 | 36811018 | 0.74427 | 0.478359 | -1.556 |
| chr7 | 19620998 | 19621330 | 1.1857 | 0.762082 | -1.556 |
| chr2 | 78874546 | 78874933 | 1.02638 | 0.6597 | -1.556 |
| chr20 | 19000404 | 19000587 | 1.12895 | 0.725687 | -1.556 |
| chr19 | 35105823 | 35106137 | 1.35635 | 0.871883 | -1.556 |
| chr14 | 20714055 | 20714305 | 1.0052 | 0.646223 | -1.556 |
| chr11 | 5911447 | 5911679 | 0.95188 | 0.611964 | -1.555 |
| chr2 | 88547460 | 88547923 | 1.00831 | 0.648308 | -1.555 |
| chr17 | 18816253 | 18816599 | 1.08527 | 0.697943 | -1.555 |
| chr2 | 199812133 | 199812356 | 0.738074 | 0.474736 | -1.555 |
| chr10 | 31552980 | 31553287 | 1.02966 | 0.662333 | -1.555 |

| | | | | | |
|---|---|---|---|---|---|
| chr15 | 87017153 | 87017452 | 0.726056 | 0.467069 | -1.554 |
| chr16 | 62057803 | 62058038 | 1.1451 | 0.736789 | -1.554 |
| chr10 | 16705840 | 16706025 | 1.10454 | 0.710762 | -1.554 |
| chr3 | 79835954 | 79836210 | 0.878258 | 0.565263 | -1.554 |
| chr20 | 58972082 | 58972272 | 1.15963 | 0.74636 | -1.554 |
| chr9 | 9034619 | 9034806 | 0.89899 | 0.57865 | -1.554 |
| chr7 | 156663084 | 156663448 | 0.835761 | 0.53804 | -1.553 |
| chr1 | 147176593 | 147176799 | 0.873866 | 0.562581 | -1.553 |
| chr15 | 89394948 | 89395229 | 0.919663 | 0.592178 | -1.553 |
| chr15 | 32133152 | 32133365 | 0.95436 | 0.614559 | -1.553 |
| chr5 | 176253119 | 176253393 | 0.821766 | 0.529205 | -1.553 |
| chr17 | 63869084 | 63869300 | 1.14042 | 0.734435 | -1.553 |
| chr5 | 89909388 | 89909680 | 1.20803 | 0.777978 | -1.553 |
| chr2 | 202042632 | 202042893 | 0.854944 | 0.55066 | -1.553 |
| chr20 | 19193583 | 19193770 | 1.57868 | 1.01697 | -1.552 |
| chr5 | 9572635 | 9572904 | 1.01091 | 0.651226 | -1.552 |
| chr5 | 87764750 | 87765100 | 1.24875 | 0.804443 | -1.552 |
| chr1 | 189005869 | 189006056 | 1.07727 | 0.694063 | -1.552 |
| chr9 | 27978080 | 27978287 | 0.851204 | 0.548503 | -1.552 |
| chr3 | 1565185 | 1565495 | 0.761113 | 0.490476 | -1.552 |
| chr20 | 23356250 | 23356434 | 0.97189 | 0.626424 | -1.551 |
| chr19 | 42118931 | 42119235 | 1.2689 | 0.817897 | -1.551 |
| chr7 | 125719240 | 125719484 | 1.46617 | 0.945182 | -1.551 |
| chr15 | 32382125 | 32382411 | 1.14548 | 0.738478 | -1.551 |
| chr20 | 30703945 | 30704174 | 0.912374 | 0.588212 | -1.551 |
| chr14 | 66427845 | 66428090 | 1.74376 | 1.12425 | -1.551 |
| chr2 | 232794718 | 232794961 | 1.85953 | 1.19907 | -1.551 |
| chr7 | 150460666 | 150460867 | 1.15713 | 0.746192 | -1.551 |
| chr3 | 107372150 | 107372340 | 0.899014 | 0.579772 | -1.551 |
| chr1 | 227524038 | 227524333 | 1.07461 | 0.693052 | -1.551 |
| chr1 | 49283257 | 49283493 | 2.21724 | 1.43002 | -1.550 |
| chr10 | 106896584 | 106896792 | 0.980919 | 0.632661 | -1.550 |
| chr12 | 1619184 | 1619444 | 1.07046 | 0.690518 | -1.550 |
| chr3 | 14251393 | 14251746 | 1.07718 | 0.694876 | -1.550 |
| chr2 | 98911441 | 98911630 | 0.929321 | 0.59953 | -1.550 |
| chr3 | 154617685 | 154617914 | 1.09354 | 0.705535 | -1.550 |
| chr7 | 9533638 | 9533851 | 0.953702 | 0.615315 | -1.550 |
| chr19 | 32212748 | 32212962 | 1.54207 | 0.995006 | -1.550 |
| chr11 | 90087093 | 90087303 | 1.16641 | 0.752932 | -1.549 |
| chr17 | 20515843 | 20516116 | 1.03443 | 0.667763 | -1.549 |
| chr12 | 113852509 | 113852705 | 0.925859 | 0.597681 | -1.549 |
| chr20 | 47239066 | 47239344 | 1.12254 | 0.724829 | -1.549 |
| chr10 | 35587413 | 35587632 | 0.917242 | 0.592344 | -1.548 |
| chr7 | 93394120 | 93394369 | 0.767761 | 0.495886 | -1.548 |
| chr12 | 5112226 | 5112649 | 2.08599 | 1.34756 | -1.548 |
| chr11 | 24727900 | 24728125 | 0.894584 | 0.577958 | -1.548 |
| chr2 | 187650309 | 187650586 | 0.840676 | 0.543149 | -1.548 |
| chr9 | 24821756 | 24822050 | 1.06487 | 0.68804 | -1.548 |
| chr7 | 150257679 | 150257902 | 0.882387 | 0.570144 | -1.548 |
| chr19 | 37506113 | 37506524 | 1.24997 | 0.807654 | -1.548 |
| chr2 | 126828294 | 126828595 | 0.813151 | 0.525425 | -1.548 |
| chr2 | 151706182 | 151706364 | 1.05804 | 0.683675 | -1.548 |
| chr5 | 15073542 | 15073795 | 1.04596 | 0.675921 | -1.547 |
| chr7 | 90063529 | 90063727 | 1.26829 | 0.819619 | -1.547 |
| chr4 | 89199642 | 89199890 | 0.835983 | 0.54029 | -1.547 |
| chr8 | 69064978 | 69065383 | 0.900826 | 0.582267 | -1.547 |
| chr12 | 96626643 | 96626911 | 0.978932 | 0.632776 | -1.547 |
| chr2 | 63892768 | 63892981 | 1.1064 | 0.715197 | -1.547 |
| chr17 | 32846413 | 32846621 | 1.78855 | 1.15626 | -1.547 |
| chr10 | 7755197 | 7755384 | 0.884303 | 0.571715 | -1.547 |
| chr11 | 124065912 | 124066101 | 1.67576 | 1.08344 | -1.547 |

| | | | | | |
|---|---|---|---|---|---|
| chr20 | 47937074 | 47937541 | 1.52402 | 0.98544 | -1.547 |
| chr5 | 96791822 | 96792113 | 0.902434 | 0.583526 | -1.547 |
| chr12 | 17070133 | 17070366 | 1.16989 | 0.756487 | -1.546 |
| chr8 | 5690278 | 5690574 | 1.07523 | 0.69532 | -1.546 |
| chr5 | 13469784 | 13470051 | 1.02695 | 0.664135 | -1.546 |
| chr20 | 41635130 | 41635592 | 0.908108 | 0.58738 | -1.546 |
| chr11 | 62014291 | 62014595 | 1.10001 | 0.711551 | -1.546 |
| chr15 | 63301697 | 63301903 | 1.11695 | 0.722725 | -1.545 |
| chr2 | 197727494 | 197727718 | 0.849297 | 0.549559 | -1.545 |
| chr2 | 125129451 | 125129642 | 0.967348 | 0.626028 | -1.545 |
| chr10 | 98928836 | 98929056 | 1.27885 | 0.827646 | -1.545 |
| chr8 | 79139506 | 79139715 | 1.18572 | 0.767419 | -1.545 |
| chr14 | 82693133 | 82693419 | 0.986262 | 0.63835 | -1.545 |
| chr19 | 39765745 | 39765927 | 1.09469 | 0.70855 | -1.545 |
| chr20 | 24704577 | 24704870 | 1.02928 | 0.666301 | -1.545 |
| chr5 | 150532769 | 150533030 | 0.851957 | 0.551524 | -1.545 |
| chr11 | 78661263 | 78661450 | 0.99162 | 0.642033 | -1.544 |
| chr9 | 93219046 | 93219311 | 0.816373 | 0.528701 | -1.544 |
| chr6 | 11154591 | 11154806 | 1.0038 | 0.650101 | -1.544 |
| chr2 | 1298622 | 1298943 | 1.04145 | 0.674527 | -1.544 |
| chr11 | 78275529 | 78275769 | 2.40197 | 1.55623 | -1.543 |
| chr17 | 15686366 | 15686647 | 1.00701 | 0.652451 | -1.543 |
| chr20 | 54829351 | 54829628 | 0.930448 | 0.602849 | -1.543 |
| chr5 | 166453125 | 166453334 | 1.02705 | 0.665469 | -1.543 |
| chr2 | 50868762 | 50868998 | 1.37024 | 0.887916 | -1.543 |
| chr7 | 66833618 | 66833935 | 0.771499 | 0.499986 | -1.543 |
| chr12 | 83345203 | 83345417 | 1.24139 | 0.804613 | -1.543 |
| chr19 | 29145719 | 29146185 | 1.14243 | 0.740514 | -1.543 |
| chr1 | 164059834 | 164060038 | 0.968749 | 0.627946 | -1.543 |
| chr11 | 58614123 | 58614368 | 0.827156 | 0.536281 | -1.542 |
| chr8 | 36383099 | 36383283 | 1.02314 | 0.663366 | -1.542 |
| chr5 | 138374802 | 138375018 | 1.00419 | 0.651182 | -1.542 |
| chr17 | 11084479 | 11084721 | 0.806893 | 0.523277 | -1.542 |
| chr3 | 169220093 | 169220296 | 0.869226 | 0.563718 | -1.542 |
| chr2 | 80012306 | 80012524 | 1.01295 | 0.657083 | -1.542 |
| chr11 | 83439200 | 83439574 | 0.934737 | 0.606511 | -1.541 |
| chr17 | 78362058 | 78362310 | 1.14092 | 0.740347 | -1.541 |
| chr8 | 16238493 | 16238826 | 0.841601 | 0.546181 | -1.541 |
| chr7 | 112768318 | 112768508 | 0.916971 | 0.595145 | -1.541 |
| chr12 | 88347065 | 88347348 | 2.17963 | 1.41484 | -1.541 |
| chr5 | 105785007 | 105785223 | 1.29313 | 0.839451 | -1.540 |
| chr1 | 215696344 | 215696609 | 1.1087 | 0.719738 | -1.540 |
| chr5 | 81357727 | 81357935 | 0.882192 | 0.572721 | -1.540 |
| chr14 | 98512379 | 98512569 | 1.09212 | 0.709025 | -1.540 |
| chr5 | 89135148 | 89135390 | 1.17654 | 0.763844 | -1.540 |
| chr5 | 6430898 | 6431124 | 0.805549 | 0.523159 | -1.540 |
| chr15 | 27609291 | 27609483 | 0.803414 | 0.521804 | -1.540 |
| chr8 | 2943248 | 2943526 | 1.16266 | 0.755128 | -1.540 |
| chr7 | 11957852 | 11958203 | 1.10687 | 0.719025 | -1.539 |
| chr19 | 31518286 | 31518711 | 1.74482 | 1.13345 | -1.539 |
| chr5 | 44101414 | 44101717 | 0.940186 | 0.610943 | -1.539 |
| chr5 | 116454964 | 116455157 | 0.912957 | 0.59339 | -1.539 |
| chr14 | 86043958 | 86044187 | 1.00873 | 0.655686 | -1.538 |
| chr2 | 49767557 | 49767822 | 1.24798 | 0.811208 | -1.538 |
| chr19 | 28744174 | 28744535 | 1.04778 | 0.681119 | -1.538 |
| chr2 | 208865011 | 208865238 | 1.11497 | 0.724893 | -1.538 |
| chr5 | 156303102 | 156303284 | 1.10435 | 0.718013 | -1.538 |
| chr10 | 10732536 | 10732825 | 0.839199 | 0.545765 | -1.538 |
| chr10 | 19674927 | 19675112 | 1.08372 | 0.704839 | -1.538 |
| chr16 | 26532776 | 26533195 | 0.940911 | 0.612104 | -1.537 |
| chr16 | 34205826 | 34206333 | 1.03614 | 0.674085 | -1.537 |

| | | | | | |
|---|---|---|---|---|---|
| chr1 | 41537392 | 41537657 | 0.798059 | 0.519219 | -1.537 |
| chr15 | 27171054 | 27171299 | 1.07402 | 0.698764 | -1.537 |
| chr9 | 95553880 | 95554200 | 0.936037 | 0.609222 | -1.536 |
| chr7 | 119005295 | 119005544 | 0.90627 | 0.58987 | -1.536 |
| chr5 | 170164457 | 170164774 | 0.918217 | 0.59768 | -1.536 |
| chr7 | 55152920 | 55153102 | 0.985082 | 0.641208 | -1.536 |
| chr5 | 151849100 | 151849483 | 0.807318 | 0.525524 | -1.536 |
| chr17 | 9001656 | 9001908 | 1.0719 | 0.697756 | -1.536 |
| chr5 | 120255506 | 120255697 | 0.721795 | 0.469871 | -1.536 |
| chr8 | 50850694 | 50850877 | 1.02644 | 0.668193 | -1.536 |
| chr8 | 53822439 | 53822635 | 0.954384 | 0.621288 | -1.536 |
| chr17 | 78007721 | 78007956 | 1.00576 | 0.654798 | -1.536 |
| chr15 | 88227329 | 88227757 | 1.03892 | 0.676438 | -1.536 |
| chr5 | 12187634 | 12187994 | 1.06671 | 0.694593 | -1.536 |
| chr2 | 156692428 | 156692639 | 0.725382 | 0.472338 | -1.536 |
| chr9 | 116352282 | 116352466 | 2.72378 | 1.77388 | -1.535 |
| chr2 | 36997587 | 36998182 | 1.99393 | 1.29861 | -1.535 |
| chr8 | 70342692 | 70342960 | 1.03799 | 0.676161 | -1.535 |
| chr8 | 139948525 | 139948749 | 2.66117 | 1.7336 | -1.535 |
| chr2 | 50023423 | 50023680 | 0.928132 | 0.604736 | -1.535 |
| chr2 | 59693083 | 59693319 | 1.10571 | 0.720474 | -1.535 |
| chr17 | 80978346 | 80978557 | 1.41858 | 0.924474 | -1.534 |
| chr5 | 89104276 | 89104596 | 1.052 | 0.685579 | -1.534 |
| chr14 | 46850672 | 46851114 | 0.937411 | 0.61094 | -1.534 |
| chr2 | 141790577 | 141790867 | 1.03283 | 0.673256 | -1.534 |
| chr7 | 141964250 | 141964479 | 0.866635 | 0.564952 | -1.534 |
| chr5 | 31366048 | 31366388 | 1.06712 | 0.69573 | -1.534 |
| chr2 | 444846 | 445064 | 1.14494 | 0.746507 | -1.534 |
| chr9 | 76919080 | 76919294 | 1.18973 | 0.775779 | -1.534 |
| chr19 | 32010766 | 32011047 | 1.55301 | 1.01274 | -1.533 |
| chr12 | 59840340 | 59840548 | 1.04464 | 0.681306 | -1.533 |
| chr9 | 78676914 | 78677148 | 1.02056 | 0.665779 | -1.533 |
| chr8 | 137268538 | 137268723 | 1.03212 | 0.673405 | -1.533 |
| chr9 | 107238805 | 107238990 | 1.01795 | 0.664199 | -1.533 |
| chr17 | 27109839 | 27110032 | 1.23237 | 0.80421 | -1.532 |
| chr7 | 14588017 | 14588312 | 0.89478 | 0.583938 | -1.532 |
| chr11 | 25595828 | 25596032 | 0.881121 | 0.575045 | -1.532 |
| chr7 | 128983025 | 128983420 | 1.45371 | 0.948766 | -1.532 |
| chr12 | 124710970 | 124711159 | 0.858212 | 0.560243 | -1.532 |
| chr10 | 50189404 | 50189727 | 2.47061 | 1.61311 | -1.532 |
| chr14 | 28020197 | 28020415 | 0.807833 | 0.527464 | -1.532 |
| chr5 | 22388800 | 22389059 | 0.811253 | 0.529776 | -1.531 |
| chr2 | 57776773 | 57776987 | 2.05919 | 1.34495 | -1.531 |
| chr8 | 85468527 | 85468716 | 0.975238 | 0.637021 | -1.531 |
| chr2 | 231091900 | 231092090 | 0.799249 | 0.522078 | -1.531 |
| chr12 | 127993683 | 127993921 | 0.922883 | 0.602868 | -1.531 |
| chr8 | 101031538 | 101031746 | 1.21063 | 0.790883 | -1.531 |
| chr3 | 80670048 | 80670241 | 0.970106 | 0.633778 | -1.531 |
| chr19 | 38292938 | 38293220 | 1.29087 | 0.843342 | -1.531 |
| chr15 | 35400926 | 35401254 | 1.27738 | 0.834539 | -1.531 |
| chr5 | 135020112 | 135020409 | 0.817774 | 0.534327 | -1.530 |
| chr5 | 41600781 | 41601120 | 0.96998 | 0.633818 | -1.530 |
| chr1 | 198053422 | 198053629 | 0.835736 | 0.546112 | -1.530 |
| chr2 | 160230086 | 160230279 | 1.01169 | 0.661162 | -1.530 |
| chr11 | 81617030 | 81617347 | 0.783908 | 0.512353 | -1.530 |
| chr10 | 11668043 | 11668300 | 1.10136 | 0.719854 | -1.530 |
| chr4 | 184178067 | 184178256 | 2.9528 | 1.93068 | -1.529 |
| chr10 | 94199668 | 94200071 | 1.04516 | 0.68339 | -1.529 |
| chr7 | 70084816 | 70085037 | 1.19443 | 0.781087 | -1.529 |
| chr10 | 85963953 | 85964256 | 2.0193 | 1.32055 | -1.529 |
| chr2 | 118995044 | 118995379 | 1.04678 | 0.684568 | -1.529 |

| | | | | | |
|---|---|---|---|---|---|
| chr8 | 55759358 | 55759576 | 0.954554 | 0.624256 | -1.529 |
| chr1 | 225238621 | 225238803 | 0.946099 | 0.618758 | -1.529 |
| chr5 | 174279181 | 174279417 | 0.86247 | 0.56407 | -1.529 |
| chr5 | 97429780 | 97430360 | 1.02323 | 0.669244 | -1.529 |
| chr5 | 32703293 | 32703481 | 0.669519 | 0.437946 | -1.529 |
| chr8 | 68553060 | 68553369 | 1.15844 | 0.757877 | -1.529 |
| chr2 | 60122845 | 60123156 | 0.963974 | 0.630778 | -1.528 |
| chr3 | 6263404 | 6263603 | 0.884938 | 0.579064 | -1.528 |
| chr15 | 91985495 | 91985787 | 0.984698 | 0.644397 | -1.528 |
| chr5 | 16133319 | 16133635 | 1.09993 | 0.71997 | -1.528 |
| chr20 | 41200928 | 41201123 | 0.929513 | 0.608486 | -1.528 |
| chr12 | 119287972 | 119288365 | 1.66959 | 1.09301 | -1.528 |
| chr2 | 79532864 | 79533146 | 1.21218 | 0.793633 | -1.527 |
| chr17 | 5338471 | 5338663 | 1.00362 | 0.657106 | -1.527 |
| chr10 | 103870946 | 103871218 | 1.24826 | 0.817349 | -1.527 |
| chr8 | 13427264 | 13427488 | 0.985333 | 0.645196 | -1.527 |
| chr2 | 61876929 | 61877185 | 0.899486 | 0.589133 | -1.527 |
| chr2 | 170038701 | 170038922 | 1.04481 | 0.684378 | -1.527 |
| chr5 | 118421838 | 118422063 | 0.75935 | 0.497487 | -1.526 |
| chr11 | 74199269 | 74199632 | 0.850746 | 0.557513 | -1.526 |
| chr5 | 32998305 | 32998526 | 1.09679 | 0.718917 | -1.526 |
| chr19 | 28520181 | 28520428 | 0.952471 | 0.62437 | -1.525 |
| chr5 | 104702304 | 104702533 | 0.900878 | 0.590593 | -1.525 |
| chr10 | 52760913 | 52761146 | 1.15147 | 0.754948 | -1.525 |
| chr12 | 1440666 | 1440897 | 0.835623 | 0.548016 | -1.525 |
| chr8 | 5345354 | 5345746 | 1.12623 | 0.738752 | -1.525 |
| chr3 | 159616038 | 159616257 | 1.10147 | 0.722523 | -1.524 |
| chr19 | 34185318 | 34185507 | 1.12679 | 0.739229 | -1.524 |
| chr10 | 108117091 | 108117465 | 1.07879 | 0.70784 | -1.524 |
| chr1 | 216831125 | 216831325 | 0.665932 | 0.43695 | -1.524 |
| chr11 | 39402714 | 39402896 | 0.918404 | 0.602643 | -1.524 |
| chr9 | 78873922 | 78874177 | 0.917729 | 0.6023 | -1.524 |
| chr5 | 14815908 | 14816096 | 1.08734 | 0.713686 | -1.524 |
| chr2 | 179669236 | 179669782 | 1.24407 | 0.816604 | -1.523 |
| chr17 | 75678053 | 75678475 | 1.09106 | 0.716389 | -1.523 |
| chr2 | 75950347 | 75950551 | 1.02502 | 0.673043 | -1.523 |
| chr10 | 10830696 | 10830958 | 1.17566 | 0.772005 | -1.523 |
| chr2 | 12998212 | 12998422 | 1.06818 | 0.701815 | -1.522 |
| chr16 | 65322340 | 65322608 | 1.1746 | 0.771887 | -1.522 |
| chr14 | 85232012 | 85232256 | 1.19337 | 0.784259 | -1.522 |
| chr7 | 13415748 | 13415937 | 1.09088 | 0.716942 | -1.522 |
| chr20 | 8022470 | 8022710 | 1.04435 | 0.686474 | -1.521 |
| chr2 | 177407793 | 177408061 | 0.864567 | 0.568327 | -1.521 |
| chr5 | 51215011 | 51215503 | 0.937128 | 0.616047 | -1.521 |
| chr12 | 23041605 | 23041822 | 1.14988 | 0.755922 | -1.521 |
| chr9 | 127819812 | 127820003 | 1.1277 | 0.741358 | -1.521 |
| chr2 | 124893283 | 124893553 | 0.843444 | 0.554535 | -1.521 |
| chr12 | 84451699 | 84452113 | 1.0577 | 0.695411 | -1.521 |
| chr8 | 127094524 | 127094895 | 1.30458 | 0.857986 | -1.521 |
| chr17 | 11569298 | 11569605 | 1.42531 | 0.937478 | -1.520 |
| chr10 | 38217434 | 38217630 | 1.06132 | 0.698093 | -1.520 |
| chr2 | 129910729 | 129910921 | 0.939156 | 0.617801 | -1.520 |
| chr5 | 154130088 | 154130592 | 0.952902 | 0.626866 | -1.520 |
| chr3 | 77171871 | 77172155 | 0.976337 | 0.642308 | -1.520 |
| chr5 | 83903921 | 83904121 | 1.10353 | 0.726073 | -1.520 |
| chr3 | 84639089 | 84639375 | 0.801482 | 0.527366 | -1.520 |
| chr5 | 31666778 | 31667048 | 0.724889 | 0.477057 | -1.520 |
| chr5 | 11662968 | 11663207 | 0.914737 | 0.602059 | -1.519 |
| chr20 | 17426723 | 17426978 | 1.05694 | 0.695745 | -1.519 |
| chr1 | 10752432 | 10752741 | 1.93078 | 1.27102 | -1.519 |
| chr5 | 76912093 | 76912401 | 1.33737 | 0.880458 | -1.519 |

| | | | | | |
|---|---|---|---|---|---|
| chr5 | 4591756 | 4592036 | 1.04655 | 0.68911 | -1.519 |
| chr2 | 23037218 | 23037415 | 0.882575 | 0.581269 | -1.518 |
| chr12 | 84809075 | 84809272 | 1.01588 | 0.669076 | -1.518 |
| chr17 | 66750932 | 66751178 | 0.982946 | 0.64759 | -1.518 |
| chr8 | 49931052 | 49931236 | 0.90763 | 0.598037 | -1.518 |
| chr8 | 112228879 | 112229073 | 1.227 | 0.808496 | -1.518 |
| chr9 | 13689096 | 13689322 | 1.50527 | 0.99189 | -1.518 |
| chr3 | 163722082 | 163722335 | 1.04319 | 0.687456 | -1.517 |
| chr1 | 191453266 | 191453462 | 1.23545 | 0.814237 | -1.517 |
| chr6 | 103080436 | 103080644 | 1.02921 | 0.678365 | -1.517 |
| chr17 | 49794477 | 49794706 | 0.946559 | 0.623895 | -1.517 |
| chr17 | 54682248 | 54682698 | 1.02228 | 0.673832 | -1.517 |
| chr20 | 50692725 | 50692960 | 0.973724 | 0.641865 | -1.517 |
| chr20 | 35871378 | 35871870 | 1.062 | 0.700066 | -1.517 |
| chr19 | 30393345 | 30393555 | 1.1142 | 0.7345 | -1.517 |
| chr4 | 126683587 | 126683939 | 1.04995 | 0.692245 | -1.517 |
| chr14 | 25064865 | 25065085 | 1.03988 | 0.685643 | -1.517 |
| chr4 | 47034659 | 47034884 | 1.52121 | 1.00308 | -1.517 |
| chr8 | 2795292 | 2795574 | 0.979641 | 0.645978 | -1.517 |
| chr7 | 70938419 | 70938617 | 1.19364 | 0.787152 | -1.516 |
| chr9 | 104996130 | 104996373 | 0.961732 | 0.634324 | -1.516 |
| chr3 | 147793972 | 147794233 | 1.68355 | 1.11043 | -1.516 |
| chr2 | 22210375 | 22210863 | 0.8246 | 0.54394 | -1.516 |
| chr2 | 80302204 | 80302440 | 1.05662 | 0.696994 | -1.516 |
| chr17 | 11040499 | 11040925 | 1.04839 | 0.691607 | -1.516 |
| chr2 | 16868577 | 16868767 | 1.06385 | 0.701878 | -1.516 |
| chr2 | 78981273 | 78981507 | 1.0513 | 0.693694 | -1.516 |
| chr3 | 85437199 | 85437467 | 0.976345 | 0.644351 | -1.515 |
| chr2 | 13415703 | 13416019 | 1.26184 | 0.832794 | -1.515 |
| chr17 | 11072393 | 11072692 | 0.728037 | 0.480548 | -1.515 |
| chr17 | 60458978 | 60459386 | 0.995269 | 0.657031 | -1.515 |
| chr12 | 115298559 | 115298799 | 0.985077 | 0.650317 | -1.515 |
| chr3 | 118029154 | 118029378 | 1.14208 | 0.754045 | -1.515 |
| chr5 | 173142691 | 173143017 | 1.02562 | 0.677304 | -1.514 |
| chr2 | 234774598 | 234774830 | 1.38018 | 0.911454 | -1.514 |
| chr15 | 98222481 | 98222713 | 1.27815 | 0.844077 | -1.514 |
| chr17 | 43246800 | 43246992 | 1.11564 | 0.736763 | -1.514 |
| chr12 | 70036311 | 70036509 | 1.11281 | 0.73496 | -1.514 |
| chr7 | 152936372 | 152936707 | 1.23849 | 0.818119 | -1.514 |
| chr8 | 49359152 | 49359360 | 1.05117 | 0.694499 | -1.514 |
| chr9 | 22867210 | 22867509 | 1.44226 | 0.953047 | -1.513 |
| chr19 | 47498622 | 47498933 | 1.04779 | 0.692561 | -1.513 |
| chr5 | 13218381 | 13218593 | 1.04901 | 0.693402 | -1.513 |
| chr8 | 56239040 | 56239226 | 0.908287 | 0.600391 | -1.513 |
| chr8 | 119858626 | 119858927 | 1.04633 | 0.691754 | -1.513 |
| chr17 | 73571956 | 73572138 | 1.00543 | 0.664739 | -1.513 |
| chr15 | 87713191 | 87713478 | 1.13161 | 0.748237 | -1.512 |
| chr15 | 101090059 | 101090281 | 0.951738 | 0.629398 | -1.512 |
| chr2 | 50780888 | 50781086 | 0.877646 | 0.580404 | -1.512 |
| chr2 | 53341386 | 53341591 | 1.10249 | 0.72915 | -1.512 |
| chr9 | 24979716 | 24979911 | 1.0304 | 0.681483 | -1.512 |
| chr2 | 31136071 | 31136418 | 1.0321 | 0.682648 | -1.512 |
| chr20 | 30367672 | 30367957 | 1.00834 | 0.667085 | -1.512 |
| chr9 | 140948872 | 140949119 | 1.2421 | 0.82178 | -1.511 |
| chr5 | 74132414 | 74132655 | 0.933925 | 0.617922 | -1.511 |
| chr10 | 1445778 | 1445975 | 1.03115 | 0.682255 | -1.511 |
| chr14 | 49796886 | 49797197 | 1.03665 | 0.685974 | -1.511 |
| chr15 | 31897664 | 31897846 | 1.22161 | 0.80839 | -1.511 |
| chr12 | 45862673 | 45862894 | 0.921104 | 0.609562 | -1.511 |
| chr8 | 69875229 | 69875428 | 1.07981 | 0.71465 | -1.511 |
| chr17 | 51380008 | 51380203 | 1.22237 | 0.809172 | -1.511 |

| | | | | | |
|---|---|---|---|---|---|
| chr2 | 54365909 | 54366116 | 1.21154 | 0.802022 | -1.511 |
| chr10 | 106966621 | 106966846 | 0.970259 | 0.642316 | -1.511 |
| chr8 | 85805168 | 85805397 | 1.08681 | 0.719484 | -1.511 |
| chr5 | 51014548 | 51014754 | 0.776056 | 0.513832 | -1.510 |
| chr2 | 22578579 | 22579067 | 1.54086 | 1.02025 | -1.510 |
| chr7 | 89506469 | 89506730 | 1.24291 | 0.82299 | -1.510 |
| chr8 | 139164274 | 139164466 | 0.858345 | 0.568376 | -1.510 |
| chr2 | 184728044 | 184728272 | 0.865461 | 0.573098 | -1.510 |
| chr1 | 179638421 | 179638627 | 0.963751 | 0.63831 | -1.510 |
| chr9 | 125035427 | 125035618 | 1.03644 | 0.686467 | -1.510 |
| chr5 | 166476999 | 166477198 | 0.939381 | 0.622226 | -1.510 |
| chr2 | 59515561 | 59515917 | 1.05546 | 0.699237 | -1.509 |
| chr19 | 55687283 | 55687514 | 1.17661 | 0.779518 | -1.509 |
| chr11 | 41053387 | 41053598 | 0.978967 | 0.648603 | -1.509 |
| chr11 | 39530274 | 39530499 | 0.949245 | 0.628931 | -1.509 |
| chr12 | 73637204 | 73637443 | 0.893793 | 0.592345 | -1.509 |
| chr17 | 52684852 | 52685090 | 0.916667 | 0.607511 | -1.509 |
| chr21 | 22003032 | 22003221 | 0.914515 | 0.606113 | -1.509 |
| chr12 | 23568977 | 23569214 | 0.875867 | 0.580602 | -1.509 |
| chr10 | 67857659 | 67857850 | 0.886097 | 0.58741 | -1.508 |
| chr2 | 185527772 | 185528004 | 1.14082 | 0.756271 | -1.508 |
| chr2 | 13417364 | 13417634 | 1.38536 | 0.918414 | -1.508 |
| chr11 | 26448778 | 26449041 | 1.01009 | 0.669718 | -1.508 |
| chr2 | 36196691 | 36196931 | 0.833216 | 0.552468 | -1.508 |
| chr3 | 108627550 | 108627903 | 0.955156 | 0.633345 | -1.508 |
| chr3 | 80657968 | 80658252 | 1.39969 | 0.92816 | -1.508 |
| chr5 | 147628432 | 147628653 | 0.885268 | 0.587065 | -1.508 |
| chr2 | 5701035 | 5701286 | 1.03517 | 0.686488 | -1.508 |
| chr7 | 48835377 | 48835611 | 1.04253 | 0.691387 | -1.508 |
| chr17 | 51953383 | 51953730 | 1.20755 | 0.800849 | -1.508 |
| chr1 | 195531302 | 195531485 | 0.820301 | 0.544054 | -1.508 |
| chr5 | 57819439 | 57819667 | 0.830112 | 0.550645 | -1.508 |
| chr8 | 59210294 | 59210490 | 0.759252 | 0.503725 | -1.507 |
| chr2 | 237649450 | 237649708 | 0.735694 | 0.488098 | -1.507 |
| chr11 | 18535114 | 18535405 | 1.07543 | 0.713541 | -1.507 |
| chr9 | 103546882 | 103547234 | 1.00538 | 0.667111 | -1.507 |
| chr3 | 1121356 | 1121569 | 0.842764 | 0.559215 | -1.507 |
| chr7 | 78140015 | 78140285 | 1.4819 | 0.983377 | -1.507 |
| chr3 | 153320247 | 153320488 | 0.930077 | 0.61723 | -1.507 |
| chr2 | 1215831 | 1216037 | 1.32366 | 0.878444 | -1.507 |
| chr10 | 96265070 | 96265306 | 0.759687 | 0.504252 | -1.507 |
| chr9 | 72147335 | 72147521 | 1.25205 | 0.831112 | -1.506 |
| chr2 | 197218893 | 197219135 | 0.968465 | 0.642871 | -1.506 |
| chr2 | 52163512 | 52163794 | 0.972124 | 0.645321 | -1.506 |
| chr15 | 88838363 | 88838548 | 0.940921 | 0.624627 | -1.506 |
| chr3 | 80697135 | 80697341 | 1.09185 | 0.724843 | -1.506 |
| chr12 | 127765266 | 127765656 | 2.20783 | 1.4658 | -1.506 |
| chr5 | 85371081 | 85371303 | 1.04726 | 0.695329 | -1.506 |
| chr11 | 70625652 | 70625934 | 0.884396 | 0.587264 | -1.506 |
| chr8 | 111188825 | 111189030 | 0.719732 | 0.478021 | -1.506 |
| chr19 | 35991478 | 35991672 | 1.08706 | 0.722037 | -1.506 |
| chr2 | 144040419 | 144040610 | 1.35601 | 0.90087 | -1.505 |
| chr20 | 54173488 | 54173680 | 1.11733 | 0.742347 | -1.505 |
| chr3 | 86431508 | 86431727 | 0.939224 | 0.624075 | -1.505 |
| chr7 | 39076796 | 39077000 | 1.12303 | 0.746314 | -1.505 |
| chr15 | 79399919 | 79400125 | 1.16158 | 0.771958 | -1.505 |
| chr8 | 94468332 | 94468743 | 1.02933 | 0.684092 | -1.505 |
| chr2 | 186223248 | 186223435 | 0.906139 | 0.602296 | -1.504 |
| chr5 | 114005012 | 114005230 | 1.125 | 0.747796 | -1.504 |
| chr5 | 11079472 | 11079815 | 1.05588 | 0.701856 | -1.504 |
| chr9 | 22731489 | 22731791 | 1.04762 | 0.696472 | -1.504 |

| | | | | | |
|---|---|---|---|---|---|
| chr20 | 13252561 | 13252792 | 1.3001 | 0.864439 | -1.504 |
| chr19 | 46124507 | 46124894 | 0.830091 | 0.551935 | -1.504 |
| chr5 | 163276647 | 163276868 | 0.690938 | 0.459418 | -1.504 |
| chr12 | 81488776 | 81488983 | 1.35269 | 0.899435 | -1.504 |
| chr8 | 135204282 | 135204501 | 1.04299 | 0.693566 | -1.504 |
| chr2 | 13127798 | 13128205 | 1.08052 | 0.718562 | -1.504 |
| chr17 | 51413599 | 51413854 | 1.08458 | 0.721377 | -1.503 |
| chr8 | 21814583 | 21814820 | 0.896774 | 0.596473 | -1.503 |
| chr20 | 7275576 | 7275908 | 1.16021 | 0.771884 | -1.503 |
| chr11 | 60110712 | 60111042 | 1.60242 | 1.06619 | -1.503 |
| chr2 | 110292957 | 110293197 | 0.789446 | 0.525301 | -1.503 |
| chr2 | 34135705 | 34135922 | 0.921934 | 0.613524 | -1.503 |
| chrX | 43798524 | 43798717 | 1.0218 | 0.680111 | -1.502 |
| chr9 | 107244981 | 107245379 | 0.909674 | 0.605482 | -1.502 |
| chr15 | 53069026 | 53069220 | 1.08649 | 0.723269 | -1.502 |
| chr8 | 137916950 | 137917132 | 0.795313 | 0.529675 | -1.502 |
| chr8 | 89522089 | 89522364 | 0.869646 | 0.579228 | -1.501 |
| chr14 | 44314433 | 44314710 | 0.887927 | 0.591502 | -1.501 |
| chr14 | 40260224 | 40260537 | 1.07701 | 0.717568 | -1.501 |
| chr2 | 72254056 | 72254293 | 1.35767 | 0.904711 | -1.501 |
| chr8 | 110860424 | 110860846 | 1.14865 | 0.765629 | -1.500 |
| chr19 | 29549284 | 29549774 | 1.35123 | 0.90069 | -1.500 |
| chr10 | 20873485 | 20873692 | 0.744211 | 0.49608 | -1.500 |
| chr7 | 2554989 | 2555279 | 0.814877 | 0.5432 | -1.500 |
| chr12 | 23360613 | 23361109 | 0.96121 | 0.640816 | -1.500 |
| chr3 | 83540368 | 83540590 | 1.03633 | 0.691005 | -1.500 |
| chr7 | 96392437 | 96392654 | 1.02584 | 0.68402 | -1.500 |
| chr2 | 197940545 | 197940781 | 1.32074 | 0.880942 | -1.499 |
| chr19 | 32364046 | 32364267 | 1.16239 | 0.775442 | -1.499 |
| chr2 | 7035630 | 7035827 | 1.09092 | 0.72797 | -1.499 |
| chr5 | 144159633 | 144159838 | 1.46816 | 0.979781 | -1.498 |
| chr8 | 6893263 | 6893645 | 2.33946 | 1.56135 | -1.498 |
| chr14 | 20588591 | 20588791 | 0.892205 | 0.595501 | -1.498 |
| chr2 | 75220210 | 75220397 | 0.914268 | 0.610424 | -1.498 |
| chr2 | 211998603 | 211998827 | 1.20504 | 0.804602 | -1.498 |
| chr3 | 164208360 | 164208562 | 0.585386 | 0.390961 | -1.497 |
| chr8 | 93528191 | 93528463 | 0.999664 | 0.667704 | -1.497 |
| chr5 | 140896399 | 140896615 | 1.04104 | 0.695365 | -1.497 |
| chr7 | 14158175 | 14158374 | 1.0444 | 0.697794 | -1.497 |
| chr17 | 71323469 | 71323667 | 1.03936 | 0.694572 | -1.496 |
| chr8 | 41136959 | 41137298 | 0.940602 | 0.628617 | -1.496 |
| chr2 | 209977846 | 209978078 | 0.957268 | 0.639756 | -1.496 |
| chr9 | 128965237 | 128965447 | 0.854781 | 0.571268 | -1.496 |
| chr11 | 26109270 | 26109595 | 1.15392 | 0.771389 | -1.496 |
| chr8 | 145052067 | 145052452 | 0.881102 | 0.589029 | -1.496 |
| chr20 | 12834173 | 12834420 | 1.26125 | 0.84373 | -1.495 |
| chr5 | 96657107 | 96657450 | 1.17431 | 0.785741 | -1.495 |
| chr15 | 65993644 | 65993871 | 0.955181 | 0.63921 | -1.494 |
| chr14 | 62871227 | 62871614 | 0.903187 | 0.60443 | -1.494 |
| chr3 | 26004282 | 26004568 | 1.02791 | 0.687972 | -1.494 |
| chr5 | 165421170 | 165421466 | 1.14908 | 0.769272 | -1.494 |
| chr2 | 7633450 | 7633634 | 1.31997 | 0.883768 | -1.494 |
| chr10 | 3717234 | 3717428 | 0.612369 | 0.410004 | -1.494 |
| chr1 | 238875529 | 238875740 | 1.15497 | 0.773309 | -1.494 |
| chr5 | 119283912 | 119284108 | 1.06921 | 0.715907 | -1.494 |
| chr2 | 118212137 | 118212445 | 0.87467 | 0.585659 | -1.493 |
| chr2 | 1032659 | 1033016 | 1.33281 | 0.89259 | -1.493 |
| chr10 | 53720105 | 53720621 | 1.39855 | 0.936635 | -1.493 |
| chr11 | 23687685 | 23687921 | 0.986765 | 0.660996 | -1.493 |
| chr8 | 86927134 | 86927365 | 1.20092 | 0.804586 | -1.493 |
| chr5 | 103748563 | 103748888 | 1.33223 | 0.89267 | -1.492 |

| | | | | | |
|---|---|---|---|---|---|
| chr2 | 178770914 | 178771194 | 2.89252 | 1.93837 | -1.492 |
| chr3 | 165371521 | 165371818 | 1.00493 | 0.673446 | -1.492 |
| chr5 | 107625191 | 107625428 | 0.916476 | 0.61417 | -1.492 |
| chr1 | 193726172 | 193726369 | 1.10498 | 0.740593 | -1.492 |
| chr20 | 7071289 | 7071550 | 1.25939 | 0.844115 | -1.492 |
| chr10 | 82792253 | 82792438 | 0.961752 | 0.644716 | -1.492 |
| chr5 | 161229440 | 161229663 | 1.11556 | 0.747993 | -1.491 |
| chr16 | 709288 | 709496 | 1.95991 | 1.31416 | -1.491 |
| chr2 | 106536272 | 106536636 | 1.22522 | 0.821659 | -1.491 |
| chr5 | 152633263 | 152633482 | 1.38477 | 0.9287 | -1.491 |
| chr12 | 74167663 | 74167952 | 0.957519 | 0.642196 | -1.491 |
| chr1 | 212840030 | 212840236 | 1.01999 | 0.684137 | -1.491 |
| chr5 | 98435329 | 98435559 | 1.03066 | 0.691319 | -1.491 |
| chr10 | 96464370 | 96464671 | 0.844915 | 0.566783 | -1.491 |
| chr9 | 12693978 | 12694222 | 1.4975 | 1.00457 | -1.491 |
| chr8 | 69359540 | 69359834 | 1.10336 | 0.740198 | -1.491 |
| chr12 | 94769087 | 94769278 | 1.01317 | 0.67989 | -1.490 |
| chr1 | 79149837 | 79150021 | 0.884197 | 0.59337 | -1.490 |
| chr1 | 102705012 | 102705223 | 0.822191 | 0.551813 | -1.490 |
| chr9 | 29783567 | 29783761 | 1.05087 | 0.705323 | -1.490 |
| chr10 | 43692738 | 43693034 | 0.869202 | 0.583473 | -1.490 |
| chr1 | 215515642 | 215515875 | 0.988464 | 0.663658 | -1.489 |
| chr7 | 27526779 | 27527025 | 0.767859 | 0.515785 | -1.489 |
| chr19 | 34511019 | 34511272 | 1.09383 | 0.734904 | -1.488 |
| chr17 | 3286441 | 3286896 | 1.05683 | 0.710077 | -1.488 |
| chr1 | 231306618 | 231307074 | 1.34838 | 0.906043 | -1.488 |
| chr15 | 87791104 | 87791443 | 0.975687 | 0.655652 | -1.488 |
| chr17 | 51922274 | 51922552 | 0.958481 | 0.644091 | -1.488 |
| chr12 | 9494767 | 9495004 | 0.989283 | 0.664837 | -1.488 |
| chr3 | 163806672 | 163806865 | 0.945505 | 0.635466 | -1.488 |
| chr2 | 168526332 | 168526623 | 0.864614 | 0.581112 | -1.488 |
| chr9 | 23696261 | 23696443 | 1.0471 | 0.703781 | -1.488 |
| chr9 | 1217288 | 1217503 | 1.15303 | 0.775024 | -1.488 |
| chr2 | 46656020 | 46656305 | 1.12456 | 0.756004 | -1.488 |
| chr18 | 48910826 | 48911054 | 1.80752 | 1.21518 | -1.487 |
| chr7 | 119870461 | 119870650 | 1.02778 | 0.691011 | -1.487 |
| chr2 | 155936979 | 155937202 | 1.03387 | 0.695182 | -1.487 |
| chr2 | 240171843 | 240172140 | 0.843617 | 0.567267 | -1.487 |
| chr6 | 152445638 | 152445844 | 2.3795 | 1.60024 | -1.487 |
| chr10 | 6635430 | 6635667 | 1.04053 | 0.699784 | -1.487 |
| chr2 | 62006484 | 62006682 | 1.15932 | 0.779686 | -1.487 |
| chr2 | 2107176 | 2107493 | 0.854335 | 0.574609 | -1.487 |
| chr8 | 142431522 | 142431707 | 0.830243 | 0.558527 | -1.486 |
| chr20 | 53790398 | 53790583 | 1.0011 | 0.67353 | -1.486 |
| chr15 | 73794406 | 73794590 | 0.934777 | 0.628992 | -1.486 |
| chr2 | 35758929 | 35759219 | 0.758356 | 0.510283 | -1.486 |
| chr16 | 6621058 | 6621285 | 0.808342 | 0.543928 | -1.486 |
| chr16 | 26563111 | 26563420 | 0.987796 | 0.664691 | -1.486 |
| chr3 | 1560305 | 1560513 | 1.24513 | 0.837868 | -1.486 |
| chr8 | 94577767 | 94578050 | 1.74879 | 1.17688 | -1.486 |
| chr7 | 96879137 | 96879381 | 1.08293 | 0.728884 | -1.486 |
| chr2 | 82484538 | 82484726 | 1.01462 | 0.682974 | -1.486 |
| chr19 | 57530895 | 57531095 | 1.24385 | 0.837304 | -1.486 |
| chr7 | 89047691 | 89047895 | 1.03437 | 0.69634 | -1.485 |
| chr8 | 4692859 | 4693077 | 1.12771 | 0.759229 | -1.485 |
| chr9 | 23321058 | 23321270 | 0.92863 | 0.625199 | -1.485 |
| chr8 | 146174399 | 146174754 | 1.09122 | 0.734747 | -1.485 |
| chr5 | 114722853 | 114723316 | 0.976758 | 0.657832 | -1.485 |
| chr8 | 47880203 | 47880484 | 0.742414 | 0.500126 | -1.484 |
| chr11 | 15230230 | 15230478 | 2.80938 | 1.89269 | -1.484 |
| chr7 | 14272597 | 14272782 | 0.857467 | 0.577766 | -1.484 |

| | | | | | |
|---|---|---|---|---|---|
| chr11 | 20870519 | 20870834 | 0.94457 | 0.636582 | -1.484 |
| chr15 | 90206058 | 90206315 | 1.01004 | 0.680738 | -1.484 |
| chr12 | 104626008 | 104626322 | 0.91643 | 0.617809 | -1.483 |
| chr20 | 54700242 | 54700463 | 1.19233 | 0.803817 | -1.483 |
| chr2 | 81440826 | 81441054 | 1.07593 | 0.72539 | -1.483 |
| chr3 | 34234782 | 34235053 | 0.991563 | 0.668606 | -1.483 |
| chr2 | 41639363 | 41639546 | 1.48994 | 1.00474 | -1.483 |
| chr7 | 92047106 | 92047482 | 1.17765 | 0.794156 | -1.483 |
| chr10 | 82615204 | 82615493 | 1.3154 | 0.88707 | -1.483 |
| chr17 | 75754146 | 75754339 | 1.11165 | 0.749689 | -1.483 |
| chr2 | 911984 | 912203 | 1.18049 | 0.796138 | -1.483 |
| chr7 | 21370220 | 21370439 | 0.870153 | 0.586853 | -1.483 |
| chr7 | 52676986 | 52677357 | 1.10097 | 0.742631 | -1.483 |
| chr1 | 191765128 | 191765327 | 0.791988 | 0.534261 | -1.482 |
| chr5 | 87390457 | 87390681 | 0.953935 | 0.64353 | -1.482 |
| chr20 | 12083768 | 12084022 | 1.18986 | 0.802817 | -1.482 |
| chr10 | 106972068 | 106972326 | 0.985827 | 0.66516 | -1.482 |
| chr5 | 99022861 | 99023047 | 0.88279 | 0.595685 | -1.482 |
| chr1 | 191781480 | 191781699 | 1.0193 | 0.687843 | -1.482 |
| chr7 | 86526701 | 86526977 | 0.9021 | 0.608853 | -1.482 |
| chr14 | 25050027 | 25050262 | 0.949958 | 0.641194 | -1.482 |
| chr1 | 182748288 | 182748506 | 0.867956 | 0.585923 | -1.481 |
| chr5 | 85203634 | 85203900 | 0.901926 | 0.608872 | -1.481 |
| chr5 | 132169484 | 132169695 | 0.926811 | 0.625676 | -1.481 |
| chr16 | 52203943 | 52204138 | 1.25698 | 0.848582 | -1.481 |
| chr8 | 143722627 | 143723015 | 1.07807 | 0.727802 | -1.481 |
| chr9 | 18626271 | 18626475 | 1.11261 | 0.751149 | -1.481 |
| chr3 | 144806778 | 144807017 | 1.14515 | 0.773172 | -1.481 |
| chr7 | 78114450 | 78114684 | 1.01745 | 0.687064 | -1.481 |
| chr20 | 18139566 | 18139767 | 1.09738 | 0.741045 | -1.481 |
| chr3 | 991440 | 991753 | 0.975523 | 0.658763 | -1.481 |
| chr9 | 107188883 | 107189182 | 0.885217 | 0.597796 | -1.481 |
| chr6 | 68344278 | 68344518 | 0.927164 | 0.626245 | -1.481 |
| chr10 | 18391605 | 18391845 | 0.856433 | 0.578483 | -1.480 |
| chr17 | 51923302 | 51923577 | 0.952595 | 0.643496 | -1.480 |
| chr5 | 10775834 | 10776080 | 1.19994 | 0.810619 | -1.480 |
| chr20 | 9804962 | 9805188 | 0.986 | 0.666335 | -1.480 |
| chr5 | 160981512 | 160981710 | 0.952101 | 0.643489 | -1.480 |
| chr8 | 80143309 | 80143534 | 0.893339 | 0.603785 | -1.480 |
| chrX | 45602457 | 45602707 | 1.24469 | 0.841283 | -1.480 |
| chr5 | 150350979 | 150351168 | 1.00897 | 0.681977 | -1.479 |
| chr15 | 24886401 | 24886677 | 1.06174 | 0.717763 | -1.479 |
| chr2 | 231818805 | 231819031 | 0.943331 | 0.637793 | -1.479 |
| chr2 | 26160221 | 26160618 | 1.20034 | 0.81158 | -1.479 |
| chr10 | 26223876 | 26224093 | 1.79657 | 1.21472 | -1.479 |
| chr1 | 178424790 | 178425003 | 1.16509 | 0.788011 | -1.479 |
| chr10 | 92248818 | 92249174 | 0.84724 | 0.57304 | -1.479 |
| chr12 | 10715708 | 10716335 | 1.04938 | 0.709885 | -1.478 |
| chr14 | 82663219 | 82663528 | 0.866467 | 0.586242 | -1.478 |
| chr5 | 30944132 | 30944358 | 1.03546 | 0.700717 | -1.478 |
| chr15 | 82102969 | 82103175 | 1.37062 | 0.92757 | -1.478 |
| chr14 | 28103451 | 28103659 | 1.18603 | 0.802705 | -1.478 |
| chr2 | 126256872 | 126257092 | 0.87922 | 0.595073 | -1.477 |
| chr18 | 70545666 | 70545872 | 1.15035 | 0.778612 | -1.477 |
| chr5 | 20104683 | 20104900 | 1.35707 | 0.918617 | -1.477 |
| chr12 | 33692817 | 33693041 | 1.02248 | 0.692135 | -1.477 |
| chr15 | 95088844 | 95089102 | 0.920142 | 0.622863 | -1.477 |
| chr7 | 1723464 | 1723698 | 1.11187 | 0.752724 | -1.477 |
| chr17 | 1122641 | 1122828 | 0.751446 | 0.50882 | -1.477 |
| chr5 | 24835839 | 24836130 | 1.31884 | 0.893086 | -1.477 |
| chr3 | 70011840 | 70012043 | 0.927128 | 0.62785 | -1.477 |

| | | | | | |
|---|---|---|---|---|---|
| chr5 | 116829771 | 116829967 | 1.04617 | 0.708471 | -1.477 |
| chr2 | 76515276 | 76515566 | 1.00463 | 0.680458 | -1.476 |
| chr10 | 83429450 | 83429761 | 0.849802 | 0.575615 | -1.476 |
| chr2 | 191682716 | 191682902 | 1.10762 | 0.750339 | -1.476 |
| chr8 | 2542158 | 2542370 | 1.22394 | 0.829165 | -1.476 |
| chr2 | 41134162 | 41134381 | 1.13013 | 0.765763 | -1.476 |
| chr7 | 110765950 | 110766163 | 1.23663 | 0.837999 | -1.476 |
| chr5 | 128934527 | 128934720 | 0.912004 | 0.618053 | -1.476 |
| chr12 | 47157061 | 47157251 | 1.13917 | 0.772027 | -1.476 |
| chr2 | 17626802 | 17627138 | 0.965245 | 0.654171 | -1.476 |
| chr8 | 77969979 | 77970163 | 1.58552 | 1.07455 | -1.476 |
| chr11 | 15060543 | 15060734 | 1.28027 | 0.867755 | -1.475 |
| chr15 | 93028879 | 93029080 | 1.01998 | 0.691488 | -1.475 |
| chr10 | 83877335 | 83877631 | 0.974893 | 0.660972 | -1.475 |
| gl000205_r | 117328 | 117666 | 1.29326 | 0.87683 | -1.475 |
| chr8 | 136213795 | 136214046 | 0.923078 | 0.625935 | -1.475 |
| chr8 | 139288879 | 139289207 | 1.14721 | 0.77797 | -1.475 |
| chr5 | 42728661 | 42728945 | 0.702788 | 0.476637 | -1.474 |
| chr2 | 126639808 | 126640028 | 0.749894 | 0.508614 | -1.474 |
| chr10 | 110305816 | 110306030 | 0.972795 | 0.659896 | -1.474 |
| chr8 | 93161044 | 93161311 | 0.963867 | 0.653878 | -1.474 |
| chr15 | 29154807 | 29155019 | 1.08761 | 0.737829 | -1.474 |
| chr17 | 32869814 | 32870186 | 4.16482 | 2.82565 | -1.474 |
| chr10 | 129559807 | 129560005 | 1.01627 | 0.689567 | -1.474 |
| chr8 | 65024234 | 65024587 | 0.913076 | 0.619591 | -1.474 |
| chr8 | 127674286 | 127674565 | 0.985342 | 0.668755 | -1.473 |
| chr2 | 84029870 | 84030084 | 1.05535 | 0.716308 | -1.473 |
| chr19 | 46654944 | 46655162 | 1.04084 | 0.706564 | -1.473 |
| chr2 | 16938499 | 16938739 | 2.66772 | 1.81096 | -1.473 |
| chr3 | 118210246 | 118210512 | 0.878753 | 0.596835 | -1.472 |
| chr2 | 12514084 | 12514327 | 1.473 | 1.00076 | -1.472 |
| chr1 | 187347880 | 187348072 | 0.936673 | 0.636409 | -1.472 |
| chr10 | 68498954 | 68499138 | 0.79032 | 0.537013 | -1.472 |
| chr7 | 119906345 | 119906672 | 0.811521 | 0.551489 | -1.472 |
| chr17 | 69653647 | 69653861 | 1.26496 | 0.859669 | -1.471 |
| chr9 | 19888894 | 19889165 | 1.09497 | 0.744195 | -1.471 |
| chr10 | 59079481 | 59079784 | 1.03814 | 0.705574 | -1.471 |
| chr3 | 80511740 | 80511968 | 1.66148 | 1.12926 | -1.471 |
| chr12 | 60803295 | 60803638 | 0.896053 | 0.609088 | -1.471 |
| chr7 | 88654019 | 88654348 | 0.926403 | 0.62981 | -1.471 |
| chr15 | 91661850 | 91662166 | 0.919873 | 0.625428 | -1.471 |
| chr3 | 77676468 | 77676726 | 1.19495 | 0.812502 | -1.471 |
| chr12 | 82977527 | 82977771 | 0.941357 | 0.64012 | -1.471 |
| chr19 | 28488113 | 28488354 | 1.42868 | 0.971549 | -1.471 |
| chr10 | 56121553 | 56121792 | 1.0173 | 0.691843 | -1.470 |
| chr2 | 66077445 | 66077641 | 0.859858 | 0.584834 | -1.470 |
| chr2 | 81191260 | 81191627 | 1.19319 | 0.811744 | -1.470 |
| chr12 | 46437894 | 46438146 | 1.10197 | 0.749726 | -1.470 |
| chr2 | 23017229 | 23017429 | 1.26592 | 0.86136 | -1.470 |
| chr2 | 80110004 | 80110362 | 0.820895 | 0.558595 | -1.470 |
| chr17 | 31918320 | 31918610 | 1.15868 | 0.788564 | -1.469 |
| chr7 | 147948734 | 147948922 | 0.921561 | 0.627298 | -1.469 |
| chr3 | 102907313 | 102907515 | 1.00194 | 0.682116 | -1.469 |
| chr12 | 129407219 | 129407406 | 0.835071 | 0.568586 | -1.469 |
| chr11 | 71127460 | 71127668 | 0.98715 | 0.672143 | -1.469 |
| chr12 | 72875395 | 72875602 | 0.677973 | 0.461696 | -1.468 |
| chr7 | 53317445 | 53317709 | 1.10429 | 0.752022 | -1.468 |
| chr14 | 89125586 | 89125846 | 0.940984 | 0.641024 | -1.468 |
| chr7 | 152451590 | 152451774 | 0.820908 | 0.559299 | -1.468 |
| chr2 | 79614572 | 79614864 | 1.0914 | 0.743603 | -1.468 |
| chr10 | 1540074 | 1540414 | 0.732886 | 0.499423 | -1.467 |

| | | | | | |
|---|---|---|---|---|---|
| chr15 | 90008402 | 90008599 | 1.1919 | 0.812263 | -1.467 |
| chr7 | 91006697 | 91006999 | 0.939427 | 0.640288 | -1.467 |
| chr1 | 216435036 | 216435305 | 1.07251 | 0.730995 | -1.467 |
| chr8 | 133152446 | 133152628 | 1.13875 | 0.776228 | -1.467 |
| chr16 | 18973525 | 18973892 | 0.705609 | 0.480985 | -1.467 |
| chr9 | 105751968 | 105752183 | 0.820113 | 0.559101 | -1.467 |
| chr1 | 238402076 | 238402288 | 1.86006 | 1.26836 | -1.467 |
| chr5 | 149778337 | 149778599 | 1.39913 | 0.954068 | -1.466 |
| chr14 | 32353262 | 32353501 | 1.09155 | 0.744346 | -1.466 |
| chr17 | 25992728 | 25992971 | 0.977704 | 0.666734 | -1.466 |
| chr5 | 104818703 | 104818929 | 0.88246 | 0.601819 | -1.466 |
| chr11 | 81508191 | 81508409 | 1.03497 | 0.705935 | -1.466 |
| chr2 | 56582058 | 56582335 | 1.23362 | 0.84149 | -1.466 |
| chr8 | 133427480 | 133427709 | 1.12083 | 0.764599 | -1.466 |
| chr7 | 90611253 | 90611491 | 1.01656 | 0.69347 | -1.466 |
| chr11 | 87267228 | 87267415 | 0.741435 | 0.505832 | -1.466 |
| chr13 | 52913034 | 52913274 | 0.817079 | 0.557451 | -1.466 |
| chr17 | 11274467 | 11274748 | 1.49345 | 1.01922 | -1.465 |
| chr3 | 76343111 | 76343505 | 1.30614 | 0.891573 | -1.465 |
| chr2 | 78045852 | 78046074 | 1.10701 | 0.755687 | -1.465 |
| chr3 | 143407183 | 143407426 | 1.09011 | 0.744212 | -1.465 |
| chr8 | 100246299 | 100246711 | 1.08718 | 0.742249 | -1.465 |
| chr2 | 13467031 | 13467254 | 0.720965 | 0.492307 | -1.464 |
| chr3 | 17218288 | 17218521 | 1.27664 | 0.871748 | -1.464 |
| chr5 | 41021844 | 41022070 | 0.711523 | 0.485864 | -1.464 |
| chr14 | 82664998 | 82665378 | 1.09147 | 0.745314 | -1.464 |
| chrX | 15177208 | 15177407 | 0.860542 | 0.587984 | -1.464 |
| chr5 | 145950174 | 145950409 | 0.811898 | 0.554792 | -1.463 |
| chr15 | 53692051 | 53692260 | 0.971918 | 0.664381 | -1.463 |
| chr12 | 98303186 | 98303396 | 0.867042 | 0.592764 | -1.463 |
| chr2 | 119332966 | 119333195 | 0.652406 | 0.44604 | -1.463 |
| chr19 | 48297926 | 48298115 | 1.37043 | 0.937149 | -1.462 |
| chr7 | 27006608 | 27006875 | 0.757101 | 0.5178 | -1.462 |
| chr5 | 19222363 | 19222619 | 0.984587 | 0.673512 | -1.462 |
| chr14 | 51376391 | 51376615 | 0.918893 | 0.628589 | -1.462 |
| chr2 | 213218465 | 213218688 | 0.861735 | 0.589501 | -1.462 |
| chr2 | 68166080 | 68166279 | 0.826607 | 0.565634 | -1.461 |
| chr8 | 3673655 | 3673915 | 1.04578 | 0.715617 | -1.461 |
| chr20 | 2722740 | 2722992 | 0.939226 | 0.642728 | -1.461 |
| chr14 | 48833102 | 48833430 | 0.987752 | 0.676015 | -1.461 |
| chr19 | 57380559 | 57380914 | 1.0085 | 0.69031 | -1.461 |
| chr9 | 86069251 | 86069550 | 0.980429 | 0.671098 | -1.461 |
| chr8 | 132229339 | 132229844 | 0.908799 | 0.622105 | -1.461 |
| chr7 | 145998460 | 145998665 | 1.12484 | 0.770241 | -1.460 |
| chr18 | 4068660 | 4068917 | 1.93467 | 1.32509 | -1.460 |
| chr1 | 154464692 | 154464955 | 0.960927 | 0.658281 | -1.460 |
| chr5 | 40736406 | 40736689 | 2.03425 | 1.39379 | -1.460 |
| chr8 | 59835365 | 59835554 | 1.09019 | 0.746969 | -1.459 |
| chr12 | 18606918 | 18607274 | 0.865856 | 0.593269 | -1.459 |
| chr11 | 82188375 | 82188701 | 0.784668 | 0.537649 | -1.459 |
| chr20 | 45143373 | 45143683 | 1.1098 | 0.760446 | -1.459 |
| chr2 | 74944481 | 74944668 | 0.983806 | 0.674126 | -1.459 |
| chr19 | 37247603 | 37247833 | 1.18924 | 0.81497 | -1.459 |
| chr8 | 84257322 | 84257586 | 1.21485 | 0.832523 | -1.459 |
| chr15 | 92821027 | 92821338 | 1.78788 | 1.22538 | -1.459 |
| chr20 | 22160769 | 22161161 | 1.55589 | 1.06654 | -1.459 |
| chr17 | 59471115 | 59471335 | 1.90839 | 1.30823 | -1.459 |
| chr2 | 46412412 | 46412597 | 1.5992 | 1.09646 | -1.459 |
| chr20 | 25318021 | 25318288 | 1.01895 | 0.698735 | -1.458 |
| chr9 | 107023307 | 107023499 | 1.00687 | 0.690479 | -1.458 |
| chr8 | 134811999 | 134812387 | 0.939191 | 0.644082 | -1.458 |

| | | | | | |
|---|---|---|---|---|---|
| chr10 | 108392548 | 108392751 | 1.08121 | 0.741505 | -1.458 |
| chr5 | 99247712 | 99247904 | 1.03698 | 0.711205 | -1.458 |
| chr7 | 122815527 | 122815733 | 0.878153 | 0.602372 | -1.458 |
| chr7 | 44992669 | 44992913 | 0.809041 | 0.554994 | -1.458 |
| chr9 | 122871503 | 122871855 | 1.14252 | 0.783839 | -1.458 |
| chr14 | 79062052 | 79062327 | 1.00435 | 0.689063 | -1.458 |
| chr1 | 214578458 | 214578644 | 0.930934 | 0.638702 | -1.458 |
| chr2 | 17126149 | 17126339 | 1.22359 | 0.839574 | -1.457 |
| chr16 | 7225475 | 7225954 | 0.956679 | 0.656535 | -1.457 |
| chr14 | 47114850 | 47115090 | 0.865548 | 0.593998 | -1.457 |
| chr1 | 200551304 | 200551492 | 0.971906 | 0.667108 | -1.457 |
| chr20 | 36957102 | 36957346 | 1.0038 | 0.689016 | -1.457 |
| chr5 | 129580161 | 129580484 | 0.850008 | 0.583506 | -1.457 |
| chr8 | 61777790 | 61777984 | 1.02996 | 0.707052 | -1.457 |
| chr19 | 29218284 | 29218649 | 1.2252 | 0.841124 | -1.457 |
| chr17 | 50039115 | 50039426 | 1.2448 | 0.854633 | -1.457 |
| chr3 | 95437871 | 95438062 | 1.25619 | 0.862469 | -1.457 |
| chr16 | 21152882 | 21153111 | 0.944512 | 0.64849 | -1.456 |
| chr2 | 80483242 | 80483623 | 1.11265 | 0.763947 | -1.456 |
| chr9 | 31506150 | 31506344 | 0.829819 | 0.569755 | -1.456 |
| chr19 | 49983549 | 49983738 | 1.03375 | 0.709924 | -1.456 |
| chr3 | 96840444 | 96840626 | 0.755116 | 0.518641 | -1.456 |
| chr5 | 95782926 | 95783110 | 1.01744 | 0.698853 | -1.456 |
| chr2 | 78766655 | 78766910 | 0.671917 | 0.461534 | -1.456 |
| chr10 | 115984768 | 115984975 | 1.11874 | 0.768479 | -1.456 |
| chr8 | 126933645 | 126933899 | 1.2063 | 0.828662 | -1.456 |
| chr7 | 104020013 | 104020267 | 1.25933 | 0.865209 | -1.456 |
| chr1 | 241431684 | 241431872 | 0.806439 | 0.554161 | -1.455 |
| chr7 | 147281842 | 147282051 | 0.980634 | 0.674327 | -1.454 |
| chr10 | 4529461 | 4529695 | 1.00194 | 0.688985 | -1.454 |
| chr15 | 64223739 | 64223969 | 1.0869 | 0.747466 | -1.454 |
| chr2 | 52466096 | 52466341 | 1.1144 | 0.766421 | -1.454 |
| chr7 | 14504516 | 14504711 | 0.840356 | 0.577968 | -1.454 |
| chr17 | 58908331 | 58908520 | 1.07256 | 0.737722 | -1.454 |
| chr2 | 53230831 | 53231043 | 1.39035 | 0.956342 | -1.454 |
| chr19 | 37457064 | 37457269 | 1.1806 | 0.812088 | -1.454 |
| chr17 | 78222880 | 78223156 | 1.04006 | 0.715613 | -1.453 |
| chr11 | 31142356 | 31142616 | 1.27951 | 0.880399 | -1.453 |
| chr7 | 50651777 | 50652012 | 0.974054 | 0.670416 | -1.453 |
| chr10 | 26162842 | 26163073 | 0.671498 | 0.462228 | -1.453 |
| chr10 | 130328659 | 130328845 | 1.68655 | 1.16114 | -1.452 |
| chr9 | 103901496 | 103901724 | 1.02076 | 0.70279 | -1.452 |
| chr2 | 49808965 | 49809154 | 1.07318 | 0.738977 | -1.452 |
| chr5 | 24283704 | 24283892 | 1.23156 | 0.84804 | -1.452 |
| chr16 | 79247114 | 79247300 | 1.04437 | 0.719331 | -1.452 |
| chr2 | 239186346 | 239186610 | 1.17442 | 0.808967 | -1.452 |
| chr11 | 124272333 | 124272532 | 2.36494 | 1.62906 | -1.452 |
| chr14 | 63563593 | 63563890 | 1.10855 | 0.763638 | -1.452 |
| chr8 | 75926126 | 75926323 | 1.07942 | 0.743803 | -1.451 |
| chr8 | 136312016 | 136312434 | 1.00033 | 0.689369 | -1.451 |
| chr20 | 17015224 | 17015510 | 0.958303 | 0.660455 | -1.451 |
| chr12 | 42122576 | 42122792 | 1.15318 | 0.794808 | -1.451 |
| chr2 | 155631615 | 155631893 | 0.873802 | 0.602344 | -1.451 |
| chr8 | 10913524 | 10913717 | 1.65205 | 1.13934 | -1.450 |
| chr12 | 83733585 | 83733796 | 0.829025 | 0.571927 | -1.450 |
| chr7 | 123661338 | 123661646 | 1.08223 | 0.746636 | -1.449 |
| chr3 | 85873582 | 85873782 | 1.21005 | 0.835029 | -1.449 |
| chr19 | 39228832 | 39229097 | 1.32338 | 0.913366 | -1.449 |
| chr5 | 31141565 | 31141757 | 0.637448 | 0.439985 | -1.449 |
| chr7 | 126944618 | 126944911 | 0.718457 | 0.495915 | -1.449 |
| chr17 | 12989423 | 12989606 | 1.12304 | 0.775314 | -1.448 |

| | | | | | |
|---|---|---|---|---|---|
| chr7 | 54133837 | 54134086 | 0.942796 | 0.650913 | -1.448 |
| chr17 | 29921237 | 29921443 | 1.06465 | 0.735312 | -1.448 |
| chr2 | 70424749 | 70424960 | 1.05483 | 0.728576 | -1.448 |
| chr14 | 29639570 | 29639824 | 0.860111 | 0.594102 | -1.448 |
| gl000193_ra | 30743 | 31016 | 1.00457 | 0.693897 | -1.448 |
| chr8 | 51578020 | 51578217 | 1.01402 | 0.700425 | -1.448 |
| chr15 | 61693489 | 61693677 | 0.949556 | 0.656044 | -1.447 |
| chr10 | 10435824 | 10436028 | 0.940593 | 0.650213 | -1.447 |
| chr8 | 50214430 | 50214768 | 0.888221 | 0.614052 | -1.446 |
| chr8 | 136371754 | 136372039 | 1.27868 | 0.884113 | -1.446 |
| chr12 | 61110256 | 61110552 | 1.01067 | 0.698864 | -1.446 |
| chr2 | 34431864 | 34432189 | 1.35574 | 0.93749 | -1.446 |
| chr9 | 94925706 | 94925913 | 1.00994 | 0.698415 | -1.446 |
| chr16 | 62638563 | 62638898 | 1.03579 | 0.716296 | -1.446 |
| chr10 | 87870087 | 87870354 | 0.928397 | 0.642041 | -1.446 |
| chr3 | 29011753 | 29011953 | 1.03595 | 0.716439 | -1.446 |
| chr2 | 81704526 | 81704737 | 1.02749 | 0.7106 | -1.446 |
| chr2 | 238001948 | 238002133 | 1.7857 | 1.23504 | -1.446 |
| chr12 | 107833838 | 107834146 | 1.15723 | 0.800443 | -1.446 |
| chr19 | 38082008 | 38082197 | 1.19694 | 0.827976 | -1.446 |
| chr14 | 26057990 | 26058268 | 1.44985 | 1.00296 | -1.446 |
| chr12 | 101291747 | 101291945 | 0.908723 | 0.628762 | -1.445 |
| chr12 | 128589319 | 128589519 | 1.04714 | 0.724661 | -1.445 |
| chr7 | 56247213 | 56247502 | 0.734092 | 0.508026 | -1.445 |
| chr7 | 24005478 | 24005671 | 0.798929 | 0.552951 | -1.445 |
| chr7 | 118711867 | 118712096 | 1.08308 | 0.749651 | -1.445 |
| chr10 | 90775874 | 90776098 | 0.967865 | 0.670133 | -1.444 |
| chr15 | 45545532 | 45545821 | 1.17217 | 0.811598 | -1.444 |
| chr14 | 86451264 | 86451559 | 0.941301 | 0.651802 | -1.444 |
| chr3 | 106808170 | 106808420 | 1.05971 | 0.733822 | -1.444 |
| chr10 | 19907948 | 19908188 | 0.875654 | 0.60637 | -1.444 |
| chr15 | 27963552 | 27963884 | 1.30632 | 0.904648 | -1.444 |
| chr2 | 81087095 | 81087279 | 1.22018 | 0.845377 | -1.443 |
| chr8 | 88353357 | 88353836 | 0.87535 | 0.606552 | -1.443 |
| chr2 | 73263355 | 73263652 | 1.08105 | 0.749204 | -1.443 |
| chr3 | 88548051 | 88548238 | 0.832981 | 0.577446 | -1.443 |
| chr10 | 53509055 | 53509283 | 1.13581 | 0.787461 | -1.442 |
| chr3 | 143048464 | 143048682 | 1.3263 | 0.919589 | -1.442 |
| chr5 | 50951251 | 50951460 | 1.17502 | 0.814771 | -1.442 |
| chr8 | 52014756 | 52014973 | 0.868265 | 0.602081 | -1.442 |
| chr3 | 79990668 | 79990926 | 1.28235 | 0.889366 | -1.442 |
| chr7 | 18487355 | 18487759 | 1.07705 | 0.747044 | -1.442 |
| chr3 | 164477598 | 164477931 | 1.26517 | 0.87755 | -1.442 |
| chr5 | 27871022 | 27871205 | 1.17117 | 0.812365 | -1.442 |
| chr16 | 3620598 | 3620796 | 1.16068 | 0.805127 | -1.442 |
| chr5 | 108310024 | 108310211 | 1.07314 | 0.744427 | -1.442 |
| chr14 | 26682484 | 26682716 | 1.00174 | 0.694978 | -1.441 |
| chr12 | 49826905 | 49827288 | 0.946212 | 0.656507 | -1.441 |
| chr11 | 4667234 | 4667435 | 0.876292 | 0.608117 | -1.441 |
| chr8 | 69914461 | 69914649 | 0.888356 | 0.616524 | -1.441 |
| chr16 | 65159995 | 65160215 | 1.28005 | 0.888468 | -1.441 |
| chr7 | 139260949 | 139261144 | 1.06205 | 0.737218 | -1.441 |
| chr2 | 148905404 | 148905628 | 0.914127 | 0.634589 | -1.441 |
| chr2 | 25047930 | 25048190 | 3.2843 | 2.28005 | -1.440 |
| chr17 | 51781869 | 51782057 | 1.03708 | 0.720015 | -1.440 |
| chr11 | 3734741 | 3734967 | 1.18993 | 0.826159 | -1.440 |
| chr19 | 38663954 | 38664276 | 1.17813 | 0.81813 | -1.440 |
| chr5 | 23200342 | 23200589 | 0.910315 | 0.632231 | -1.440 |
| chr5 | 19958804 | 19959036 | 0.806293 | 0.560013 | -1.440 |
| chr14 | 75260507 | 75260981 | 1.02914 | 0.714798 | -1.440 |
| chr14 | 102349018 | 102349237 | 0.991797 | 0.689028 | -1.439 |

| | | | | | |
|---|---|---|---|---|---|
| chr7 | 22959660 | 22959886 | 0.88497 | 0.614835 | -1.439 |
| chr1 | 151009376 | 151009633 | 1.0172 | 0.706847 | -1.439 |
| chr8 | 87890706 | 87890956 | 1.04761 | 0.728025 | -1.439 |
| chr3 | 22834185 | 22834420 | 1.13992 | 0.792178 | -1.439 |
| chr19 | 28894862 | 28895045 | 1.0705 | 0.743984 | -1.439 |
| chr8 | 4691287 | 4691700 | 1.4579 | 1.01323 | -1.439 |
| chr2 | 200447094 | 200447278 | 1.16192 | 0.807693 | -1.439 |
| chr10 | 106364964 | 106365192 | 0.807815 | 0.561543 | -1.439 |
| chr3 | 145006801 | 145007086 | 0.929272 | 0.645972 | -1.439 |
| chr7 | 126418364 | 126418554 | 0.94375 | 0.656045 | -1.439 |
| chr14 | 47571884 | 47572097 | 1.04943 | 0.729538 | -1.438 |
| chr11 | 42044699 | 42044952 | 0.928125 | 0.645266 | -1.438 |
| chr8 | 85149321 | 85149596 | 1.03239 | 0.717911 | -1.438 |
| chr2 | 83778682 | 83779025 | 0.764896 | 0.531957 | -1.438 |
| chr1 | 210214375 | 210214608 | 1.29563 | 0.901266 | -1.438 |
| chr16 | 56433931 | 56434139 | 1.24376 | 0.865204 | -1.438 |
| chr10 | 11393116 | 11393341 | 0.828881 | 0.576652 | -1.437 |
| chr5 | 18253718 | 18253934 | 1.21481 | 0.84521 | -1.437 |
| chr3 | 156167730 | 156168194 | 1.72869 | 1.20292 | -1.437 |
| chr1 | 229121861 | 229122074 | 0.885554 | 0.616241 | -1.437 |
| chr11 | 31254878 | 31255317 | 0.933695 | 0.649775 | -1.437 |
| chr12 | 45340285 | 45340479 | 1.00957 | 0.702608 | -1.437 |
| chr16 | 63003950 | 63004228 | 1.06374 | 0.740312 | -1.437 |
| chr14 | 25302268 | 25302451 | 1.2656 | 0.880881 | -1.437 |
| chr9 | 81570983 | 81571180 | 1.03384 | 0.719641 | -1.437 |
| chr11 | 15238264 | 15238512 | 1.68411 | 1.17233 | -1.437 |
| chr2 | 13706790 | 13707114 | 1.16453 | 0.810701 | -1.436 |
| chr9 | 121889621 | 121889936 | 0.961646 | 0.669541 | -1.436 |
| chr2 | 140773166 | 140773386 | 1.23407 | 0.859282 | -1.436 |
| chr13 | 20599538 | 20599740 | 1.14551 | 0.797739 | -1.436 |
| chr9 | 15997551 | 15997743 | 0.777648 | 0.541638 | -1.436 |
| chr7 | 96304858 | 96305149 | 1.0001 | 0.696595 | -1.436 |
| chr17 | 62243552 | 62243826 | 0.95881 | 0.667857 | -1.436 |
| chr8 | 16724222 | 16724429 | 0.66428 | 0.462819 | -1.435 |
| chr5 | 50811705 | 50811930 | 1.03493 | 0.721093 | -1.435 |
| chr7 | 90917648 | 90918112 | 1.36542 | 0.951371 | -1.435 |
| chr7 | 108770722 | 108770953 | 0.863562 | 0.601814 | -1.435 |
| chr2 | 21409761 | 21409946 | 0.877224 | 0.611404 | -1.435 |
| chr14 | 98085380 | 98085571 | 0.785531 | 0.547501 | -1.435 |
| chr3 | 99423401 | 99423586 | 0.98455 | 0.686291 | -1.435 |
| chr5 | 25125348 | 25125547 | 1.08556 | 0.756704 | -1.435 |
| chr3 | 27598874 | 27599272 | 0.868181 | 0.605387 | -1.434 |
| chr3 | 172341728 | 172341964 | 2.00865 | 1.40077 | -1.434 |
| chr20 | 62813115 | 62813315 | 1.07322 | 0.748475 | -1.434 |
| chr1 | 248512230 | 248512497 | 0.952203 | 0.664092 | -1.434 |
| chr12 | 75462022 | 75462245 | 1.13491 | 0.791551 | -1.434 |
| chr16 | 57130936 | 57131128 | 1.30803 | 0.912944 | -1.433 |
| chr15 | 31896910 | 31897108 | 0.766776 | 0.535229 | -1.433 |
| chr7 | 147709008 | 147709332 | 0.897684 | 0.626611 | -1.433 |
| chr20 | 6157515 | 6157753 | 1.29988 | 0.907374 | -1.433 |
| chr5 | 18958378 | 18958566 | 1.00733 | 0.703234 | -1.432 |
| chr5 | 28542367 | 28542624 | 0.852672 | 0.595293 | -1.432 |
| chr10 | 8714131 | 8714323 | 1.03026 | 0.719344 | -1.432 |
| chr18 | 27502768 | 27502951 | 1.30148 | 0.908872 | -1.432 |
| chr4 | 140577796 | 140578024 | 1.2308 | 0.859825 | -1.431 |
| chr5 | 139953398 | 139953590 | 1.01532 | 0.709392 | -1.431 |
| chr7 | 54765403 | 54765707 | 0.951133 | 0.664574 | -1.431 |
| chr2 | 122893755 | 122894083 | 1.22541 | 0.856362 | -1.431 |
| chr11 | 44355620 | 44355818 | 0.944714 | 0.660221 | -1.431 |
| chr15 | 94346590 | 94346872 | 1.24481 | 0.870061 | -1.431 |
| chr16 | 64484869 | 64485135 | 0.757444 | 0.529423 | -1.431 |

| | | | | | |
|---|---|---|---|---|---|
| chr12 | 51060700 | 51060907 | 1.07036 | 0.748143 | -1.431 |
| chr3 | 84277230 | 84277423 | 0.81948 | 0.572865 | -1.430 |
| chr17 | 52324837 | 52325119 | 1.14061 | 0.797383 | -1.430 |
| chr16 | 64682971 | 64683214 | 0.677207 | 0.473647 | -1.430 |
| chr1 | 202124054 | 202124237 | 1.6069 | 1.12396 | -1.430 |
| chr7 | 24457940 | 24458123 | 1.10931 | 0.775947 | -1.430 |
| chr8 | 140155469 | 140155729 | 1.15052 | 0.804777 | -1.430 |
| chr10 | 107284834 | 107285351 | 1.03539 | 0.724249 | -1.430 |
| chr3 | 143915939 | 143916196 | 1.17542 | 0.822215 | -1.430 |
| chr7 | 116555081 | 116555300 | 1.23892 | 0.866665 | -1.430 |
| chr6 | 36750681 | 36750906 | 2.08979 | 1.46196 | -1.429 |
| chr3 | 140730393 | 140730668 | 2.72671 | 1.90783 | -1.429 |
| chr7 | 103364928 | 103365123 | 0.920172 | 0.643861 | -1.429 |
| chr20 | 12040644 | 12040866 | 1.25242 | 0.876447 | -1.429 |
| chr10 | 66854848 | 66855149 | 0.945562 | 0.661792 | -1.429 |
| chr3 | 84206818 | 84207001 | 0.960998 | 0.672691 | -1.429 |
| chr3 | 39710002 | 39710322 | 2.17989 | 1.52595 | -1.429 |
| chr5 | 94596987 | 94597307 | 1.40681 | 0.984869 | -1.428 |
| chr8 | 92994162 | 92994407 | 0.916839 | 0.641885 | -1.428 |
| chr11 | 87208330 | 87208529 | 0.929975 | 0.65109 | -1.428 |
| chr2 | 15285738 | 15285963 | 0.923961 | 0.647017 | -1.428 |
| chr8 | 72061041 | 72061250 | 1.13527 | 0.795004 | -1.428 |
| chr2 | 22673550 | 22673754 | 1.25851 | 0.88136 | -1.428 |
| chr8 | 48613003 | 48613258 | 0.96243 | 0.674111 | -1.428 |
| chr10 | 2366080 | 2366377 | 1.1143 | 0.780562 | -1.428 |
| chr7 | 141176431 | 141176881 | 1.12195 | 0.785998 | -1.427 |
| chr16 | 6603057 | 6603258 | 0.917495 | 0.642795 | -1.427 |
| chr8 | 132595192 | 132595387 | 1.22153 | 0.855839 | -1.427 |
| chr1 | 210757735 | 210758041 | 2.96461 | 2.07754 | -1.427 |
| chr5 | 44187328 | 44187549 | 0.961584 | 0.673949 | -1.427 |
| chr2 | 233476362 | 233476686 | 2.82501 | 1.98002 | -1.427 |
| chr1 | 190663431 | 190663752 | 1.03594 | 0.72611 | -1.427 |
| chr12 | 31039184 | 31039434 | 1.17601 | 0.824298 | -1.427 |
| chr17 | 27957314 | 27957512 | 1.05053 | 0.736525 | -1.426 |
| chr11 | 26470400 | 26470715 | 0.943972 | 0.661886 | -1.426 |
| chr12 | 56740498 | 56740735 | 1.3989 | 0.980907 | -1.426 |
| chr2 | 5551448 | 5551650 | 1.09424 | 0.767315 | -1.426 |
| chr2 | 40885534 | 40885731 | 1.11219 | 0.780161 | -1.426 |
| chr15 | 62551983 | 62552170 | 0.992173 | 0.696062 | -1.425 |
| chr12 | 131047090 | 131047320 | 0.929573 | 0.652175 | -1.425 |
| chr7 | 106702093 | 106702321 | 1.21889 | 0.855397 | -1.425 |
| chr5 | 35556625 | 35556818 | 1.29122 | 0.906239 | -1.425 |
| chr19 | 23945871 | 23946099 | 1.10993 | 0.779036 | -1.425 |
| chr14 | 73952988 | 73953170 | 1.01233 | 0.710624 | -1.425 |
| chr10 | 32492760 | 32492952 | 0.861273 | 0.604725 | -1.424 |
| chr10 | 49650204 | 49650398 | 0.865553 | 0.607765 | -1.424 |
| chr2 | 186013423 | 186013652 | 0.802594 | 0.563594 | -1.424 |
| chr3 | 99297516 | 99297795 | 0.851482 | 0.598016 | -1.424 |
| chr12 | 126389446 | 126389768 | 0.979196 | 0.68794 | -1.423 |
| chr17 | 41422109 | 41422329 | 1.18295 | 0.83119 | -1.423 |
| chr14 | 50522338 | 50522528 | 1.39244 | 0.978403 | -1.423 |
| chr7 | 42065564 | 42065772 | 1.09068 | 0.766454 | -1.423 |
| chr1 | 237529498 | 237529691 | 0.952614 | 0.669448 | -1.423 |
| chr7 | 13605630 | 13605823 | 1.09222 | 0.767675 | -1.423 |
| chr2 | 86623620 | 86623849 | 2.50763 | 1.76252 | -1.423 |
| chr12 | 97690588 | 97690804 | 1.17871 | 0.828532 | -1.423 |
| chr2 | 103838273 | 103838586 | 0.915848 | 0.643775 | -1.423 |
| chr11 | 60401699 | 60401959 | 1.16821 | 0.821353 | -1.422 |
| chr3 | 16016120 | 16016361 | 1.26122 | 0.886937 | -1.422 |
| chr9 | 80592210 | 80592604 | 1.47222 | 1.03533 | -1.422 |
| chr7 | 29860998 | 29861195 | 1.27806 | 0.899052 | -1.422 |

| | | | | | |
|---|---|---|---|---|---|
| chr10 | 12790915 | 12791103 | 0.754032 | 0.530426 | -1.422 |
| chr20 | 16650292 | 16650474 | 1.10491 | 0.777264 | -1.422 |
| chr3 | 11092374 | 11092624 | 0.916589 | 0.6449 | -1.421 |
| chr16 | 13310666 | 13310856 | 1.45382 | 1.0229 | -1.421 |
| chr1 | 218100044 | 218100294 | 1.01936 | 0.71724 | -1.421 |
| chr5 | 5584745 | 5584933 | 0.932015 | 0.655947 | -1.421 |
| chr14 | 84841510 | 84841707 | 0.754082 | 0.530777 | -1.421 |
| chr19 | 35459023 | 35459209 | 1.2939 | 0.910828 | -1.421 |
| chr15 | 79693772 | 79693981 | 1.27263 | 0.89607 | -1.420 |
| chr9 | 8364575 | 8364836 | 1.39618 | 0.983252 | -1.420 |
| chr12 | 82460496 | 82460703 | 1.10494 | 0.778224 | -1.420 |
| chr8 | 4356825 | 4357009 | 1.10666 | 0.779483 | -1.420 |
| chr3 | 806400 | 806745 | 0.889802 | 0.626789 | -1.420 |
| chr11 | 83846627 | 83846924 | 0.904092 | 0.637027 | -1.419 |
| chr10 | 28616664 | 28616947 | 0.800514 | 0.564094 | -1.419 |
| chr10 | 77853524 | 77853746 | 1.20159 | 0.84676 | -1.419 |
| chr4 | 76988925 | 76989120 | 2.02926 | 1.43016 | -1.419 |
| chr5 | 8676749 | 8676965 | 1.1202 | 0.789493 | -1.419 |
| chr9 | 1267881 | 1268117 | 1.06857 | 0.753201 | -1.419 |
| chr15 | 96052083 | 96052345 | 1.2412 | 0.874917 | -1.419 |
| chr14 | 49183311 | 49183577 | 0.966344 | 0.681218 | -1.419 |
| chr5 | 139970519 | 139970760 | 1.06294 | 0.74933 | -1.419 |
| chr16 | 50878256 | 50878441 | 0.993359 | 0.700294 | -1.418 |
| chr10 | 64230011 | 64230289 | 0.982469 | 0.692631 | -1.418 |
| chr7 | 71398253 | 71398457 | 1.0991 | 0.774867 | -1.418 |
| chr8 | 92476168 | 92476353 | 1.00934 | 0.711615 | -1.418 |
| chr9 | 25996419 | 25996822 | 0.79812 | 0.562826 | -1.418 |
| chr15 | 84501028 | 84501281 | 0.898755 | 0.633923 | -1.418 |
| chr10 | 66418241 | 66418457 | 0.98464 | 0.694506 | -1.418 |
| chr12 | 92590648 | 92590872 | 0.974811 | 0.687603 | -1.418 |
| chr8 | 111312506 | 111312801 | 0.909185 | 0.641326 | -1.418 |
| chr7 | 41860054 | 41860282 | 1.13688 | 0.802004 | -1.418 |
| chr20 | 6247183 | 6247483 | 0.977868 | 0.689864 | -1.417 |
| chr14 | 82900143 | 82900596 | 0.831766 | 0.58687 | -1.417 |
| chr11 | 22800788 | 22800987 | 1.15683 | 0.816304 | -1.417 |
| chr14 | 45359107 | 45359361 | 1.04339 | 0.736312 | -1.417 |
| chr7 | 147758561 | 147758758 | 0.884524 | 0.624249 | -1.417 |
| chr5 | 92110954 | 92111170 | 1.03682 | 0.731879 | -1.417 |
| chr9 | 140072146 | 140072380 | 0.764115 | 0.539413 | -1.417 |
| chr10 | 2627757 | 2628070 | 0.807489 | 0.570082 | -1.416 |
| chr14 | 72542594 | 72542873 | 0.894635 | 0.631618 | -1.416 |
| chr5 | 100019560 | 100019892 | 1.0294 | 0.726847 | -1.416 |
| chr8 | 106321969 | 106322182 | 1.13178 | 0.799271 | -1.416 |
| chr5 | 29461271 | 29461472 | 0.807002 | 0.569951 | -1.416 |
| chr7 | 151901076 | 151901278 | 0.933012 | 0.658952 | -1.416 |
| chr3 | 69666803 | 69666995 | 1.28524 | 0.90777 | -1.416 |
| chr3 | 79159725 | 79159960 | 1.17568 | 0.830432 | -1.416 |
| chr15 | 40564973 | 40565167 | 2.2988 | 1.62395 | -1.416 |
| chr8 | 36862831 | 36863066 | 0.942292 | 0.66567 | -1.416 |
| chr11 | 42838029 | 42838297 | 0.896442 | 0.63333 | -1.415 |
| chr2 | 180349430 | 180349647 | 0.906188 | 0.640282 | -1.415 |
| chr15 | 24997589 | 24997778 | 1.12286 | 0.793598 | -1.415 |
| chr19 | 22407593 | 22407788 | 1.30974 | 0.925765 | -1.415 |
| chr19 | 59022652 | 59023007 | 0.928188 | 0.656096 | -1.415 |
| chr19 | 34836161 | 34836442 | 1.07907 | 0.762785 | -1.415 |
| chr20 | 23455927 | 23456111 | 0.938726 | 0.663591 | -1.415 |
| chr2 | 219122461 | 219122651 | 2.16027 | 1.52714 | -1.415 |
| chr8 | 92452489 | 92452719 | 1.18373 | 0.836941 | -1.414 |
| chr10 | 8999028 | 8999260 | 1.15164 | 0.814277 | -1.414 |
| chr8 | 133156963 | 133157469 | 1.99115 | 1.40791 | -1.414 |
| chr10 | 82548654 | 82548945 | 0.969581 | 0.685619 | -1.414 |

| | | | | | |
|---|---|---|---|---|---|
| chr10 | 120517776 | 120517963 | 0.891621 | 0.630504 | -1.414 |
| chr6 | 154386309 | 154386563 | 0.897762 | 0.63491 | -1.414 |
| chr19 | 23980028 | 23980227 | 0.858267 | 0.606982 | -1.414 |
| chr9 | 92089821 | 92090160 | 0.754655 | 0.533743 | -1.414 |
| chr8 | 78142947 | 78143180 | 1.0633 | 0.752193 | -1.414 |
| chr7 | 53392383 | 53392571 | 1.094 | 0.773914 | -1.414 |
| chr10 | 103224506 | 103224691 | 0.899814 | 0.63658 | -1.414 |
| chr5 | 26875673 | 26875904 | 1.04278 | 0.737805 | -1.413 |
| chr9 | 122410956 | 122411298 | 1.2022 | 0.850747 | -1.413 |
| chr8 | 79751250 | 79751476 | 0.751979 | 0.532164 | -1.413 |
| chr12 | 53086193 | 53086412 | 1.11044 | 0.785857 | -1.413 |
| chr20 | 53824416 | 53824675 | 0.85788 | 0.607193 | -1.413 |
| chr2 | 77614465 | 77614723 | 1.12035 | 0.793228 | -1.412 |
| chr7 | 24539992 | 24540217 | 0.56542 | 0.400364 | -1.412 |
| chr14 | 105134465 | 105134761 | 0.832396 | 0.589432 | -1.412 |
| chr2 | 138394420 | 138394685 | 0.848016 | 0.600566 | -1.412 |
| chr8 | 135597807 | 135597997 | 0.846731 | 0.599725 | -1.412 |
| chr8 | 113653839 | 113654067 | 0.870419 | 0.616546 | -1.412 |
| chr7 | 96995753 | 96995946 | 0.853825 | 0.604793 | -1.412 |
| chr3 | 66304523 | 66304709 | 0.992341 | 0.703027 | -1.412 |
| chr18 | 44576842 | 44577250 | 1.24337 | 0.880904 | -1.411 |
| chr17 | 49765323 | 49765535 | 1.37977 | 0.977544 | -1.411 |
| chr3 | 3883332 | 3883537 | 0.822604 | 0.582841 | -1.411 |
| chr8 | 83976784 | 83977099 | 0.959243 | 0.679674 | -1.411 |
| chr2 | 18011806 | 18012219 | 3.20685 | 2.27239 | -1.411 |
| chr3 | 146737831 | 146738257 | 0.954232 | 0.676221 | -1.411 |
| chr7 | 44552528 | 44552786 | 0.891248 | 0.631686 | -1.411 |
| chr5 | 116333736 | 116334067 | 0.951522 | 0.674472 | -1.411 |
| chr5 | 67039541 | 67039723 | 1.24352 | 0.881498 | -1.411 |
| chr11 | 6060463 | 6060720 | 1.07725 | 0.763646 | -1.411 |
| chr17 | 57579600 | 57579841 | 0.955977 | 0.677789 | -1.410 |
| chr2 | 86031699 | 86031971 | 1.83446 | 1.30095 | -1.410 |
| chr9 | 12065031 | 12065324 | 1.07082 | 0.759431 | -1.410 |
| chr15 | 24139669 | 24139946 | 1.12816 | 0.800379 | -1.410 |
| chr5 | 130358531 | 130358797 | 1.09782 | 0.778903 | -1.409 |
| chr1 | 212988456 | 212988654 | 1.90764 | 1.35356 | -1.409 |
| chr12 | 71381622 | 71381826 | 1.06056 | 0.752647 | -1.409 |
| chr17 | 11130664 | 11130863 | 1.34049 | 0.951355 | -1.409 |
| chr19 | 53538522 | 53538738 | 1.32922 | 0.943482 | -1.409 |
| chr8 | 97665266 | 97665615 | 1.19538 | 0.848482 | -1.409 |
| chr9 | 77166881 | 77167087 | 1.23055 | 0.873482 | -1.409 |
| chr5 | 101945153 | 101945346 | 0.812388 | 0.576683 | -1.409 |
| chr10 | 113781864 | 113782082 | 0.921933 | 0.654473 | -1.409 |
| chr3 | 159238071 | 159238309 | 1.12361 | 0.79769 | -1.409 |
| chr12 | 12508485 | 12508699 | 1.1396 | 0.809091 | -1.408 |
| chr10 | 122900020 | 122900356 | 1.33545 | 0.948159 | -1.408 |
| chr10 | 129447979 | 129448266 | 1.6949 | 1.20343 | -1.408 |
| chr5 | 11214761 | 11215192 | 0.756455 | 0.537152 | -1.408 |
| chr8 | 140385354 | 140385629 | 1.07543 | 0.763739 | -1.408 |
| chr5 | 77540203 | 77540387 | 0.870287 | 0.618161 | -1.408 |
| chr5 | 21712183 | 21712383 | 1.03329 | 0.733966 | -1.408 |
| chr12 | 55015775 | 55015966 | 1.06568 | 0.757016 | -1.408 |
| chr12 | 86882961 | 86883188 | 0.831823 | 0.591015 | -1.407 |
| chr8 | 83536894 | 83537216 | 0.999173 | 0.709951 | -1.407 |
| chr7 | 68233957 | 68234239 | 0.931905 | 0.662164 | -1.407 |
| chr8 | 57470446 | 57470796 | 3.29791 | 2.34373 | -1.407 |
| chr2 | 54260158 | 54260355 | 0.855789 | 0.608205 | -1.407 |
| chr2 | 152904480 | 152904812 | 2.1638 | 1.53833 | -1.407 |
| chr9 | 6568157 | 6568355 | 1.41072 | 1.00294 | -1.407 |
| chr8 | 139989369 | 139989671 | 1.24558 | 0.885624 | -1.406 |
| chr5 | 26701986 | 26702239 | 0.911025 | 0.647761 | -1.406 |

| | | | | | |
|---|---|---|---|---|---|
| chr18 | 68459714 | 68459908 | 0.789959 | 0.561711 | -1.406 |
| chr2 | 123096403 | 123096608 | 0.921689 | 0.655408 | -1.406 |
| chr10 | 50931929 | 50932158 | 1.21897 | 0.866927 | -1.406 |
| chr2 | 1281413 | 1281743 | 1.22407 | 0.870576 | -1.406 |
| chr16 | 15790795 | 15791015 | 0.769442 | 0.547241 | -1.406 |
| chr8 | 89407674 | 89407916 | 0.755271 | 0.537162 | -1.406 |
| chr5 | 19828629 | 19828883 | 0.914069 | 0.650316 | -1.406 |
| chr10 | 113924921 | 113925181 | 1.18399 | 0.842355 | -1.406 |
| chr2 | 209862219 | 209862417 | 0.908032 | 0.646029 | -1.406 |
| chr7 | 100882197 | 100882517 | 0.807844 | 0.574827 | -1.405 |
| chr3 | 19467487 | 19467763 | 1.19753 | 0.852149 | -1.405 |
| chr3 | 1934809 | 1935022 | 0.839117 | 0.597179 | -1.405 |
| chr9 | 2605527 | 2605718 | 1.03112 | 0.733835 | -1.405 |
| chr1 | 218258255 | 218258652 | 0.96468 | 0.686623 | -1.405 |
| chr5 | 660749 | 661090 | 0.930281 | 0.662265 | -1.405 |
| chr14 | 80010300 | 80010544 | 1.57656 | 1.12237 | -1.405 |
| chr5 | 125894701 | 125894906 | 0.86352 | 0.614795 | -1.405 |
| chr3 | 146906541 | 146906737 | 0.805316 | 0.573415 | -1.404 |
| chr12 | 128449250 | 128449440 | 0.666587 | 0.474725 | -1.404 |
| chr19 | 31155714 | 31155950 | 1.61804 | 1.15243 | -1.404 |
| chr14 | 25856002 | 25856289 | 0.764148 | 0.544362 | -1.404 |
| chr2 | 7789819 | 7790029 | 0.909256 | 0.647833 | -1.404 |
| chr9 | 31830597 | 31830842 | 0.875197 | 0.623573 | -1.404 |
| chr8 | 54591931 | 54592126 | 1.03931 | 0.740603 | -1.403 |
| chr14 | 57915767 | 57915961 | 1.05542 | 0.752236 | -1.403 |
| chr8 | 128219292 | 128219517 | 2.8718 | 2.04706 | -1.403 |
| chr21 | 10874662 | 10874864 | 0.995528 | 0.709632 | -1.403 |
| chr5 | 117409962 | 117410301 | 1.03397 | 0.737207 | -1.403 |
| chr12 | 61417020 | 61417306 | 1.08172 | 0.771268 | -1.403 |
| chr8 | 69446749 | 69446962 | 1.42641 | 1.01726 | -1.402 |
| chr7 | 48801738 | 48801931 | 0.665912 | 0.474903 | -1.402 |
| chr16 | 27870514 | 27870869 | 1.0911 | 0.778193 | -1.402 |
| chr2 | 57518453 | 57518801 | 0.994861 | 0.709565 | -1.402 |
| chr14 | 46442923 | 46443121 | 0.876444 | 0.625251 | -1.402 |
| chr11 | 13549308 | 13549513 | 1.18592 | 0.846082 | -1.402 |
| chr14 | 100737636 | 100737829 | 1.91424 | 1.36607 | -1.401 |
| rUn_gl0002 | 35195 | 35401 | 0.960811 | 0.68571 | -1.401 |
| chr7 | 157032384 | 157032568 | 1.00315 | 0.715944 | -1.401 |
| chr5 | 11085848 | 11086037 | 0.801816 | 0.572369 | -1.401 |
| chr2 | 104037604 | 104037956 | 0.642382 | 0.458689 | -1.400 |
| chr16 | 70627491 | 70627699 | 0.901519 | 0.643757 | -1.400 |
| chr2 | 172560237 | 172560483 | 1.65864 | 1.18457 | -1.400 |
| chr9 | 10016610 | 10016863 | 1.24271 | 0.887521 | -1.400 |
| chr20 | 15923913 | 15924146 | 1.25314 | 0.895352 | -1.400 |
| chr1 | 181667444 | 181667707 | 0.974438 | 0.696253 | -1.400 |
| chr19 | 32327679 | 32327930 | 1.23347 | 0.881377 | -1.399 |
| chr2 | 147684889 | 147685156 | 1.19374 | 0.853024 | -1.399 |
| chr3 | 35118881 | 35119115 | 1.28756 | 0.920126 | -1.399 |
| chr8 | 136682041 | 136682266 | 0.849234 | 0.606994 | -1.399 |
| chr14 | 41721903 | 41722326 | 0.894848 | 0.639641 | -1.399 |
| chr9 | 106361169 | 106361449 | 1.45573 | 1.04085 | -1.399 |
| chr7 | 8978301 | 8978534 | 0.897391 | 0.641705 | -1.398 |
| chr2 | 6921614 | 6921849 | 0.773385 | 0.553061 | -1.398 |
| chr2 | 79225634 | 79225940 | 1.27078 | 0.908827 | -1.398 |
| chr2 | 56394131 | 56394545 | 0.954092 | 0.682453 | -1.398 |
| chr10 | 109533892 | 109534087 | 0.967535 | 0.692164 | -1.398 |
| chr19 | 49700019 | 49700226 | 1.04535 | 0.747912 | -1.398 |
| chr3 | 163675104 | 163675339 | 0.95729 | 0.685 | -1.398 |
| chr1 | 158341374 | 158341624 | 1.00813 | 0.72162 | -1.397 |
| chr2 | 30504991 | 30505191 | 0.987091 | 0.706697 | -1.397 |
| chr1 | 156918157 | 156918369 | 1.12419 | 0.804885 | -1.397 |

| | | | | | |
|---|---|---|---|---|---|
| chr12 | 93651207 | 93651461 | 1.05376 | 0.754504 | -1.397 |
| chr16 | 65353631 | 65353923 | 1.25817 | 0.900881 | -1.397 |
| chr12 | 127375814 | 127376089 | 1.19864 | 0.858287 | -1.397 |
| chr16 | 83452392 | 83452642 | 0.93776 | 0.671489 | -1.397 |
| chr9 | 24975138 | 24975338 | 1.03646 | 0.7422 | -1.396 |
| chr2 | 211785993 | 211786211 | 0.670694 | 0.480281 | -1.396 |
| chr11 | 566272 | 566694 | 0.994715 | 0.712357 | -1.396 |
| chr2 | 65248398 | 65248651 | 1.13678 | 0.814201 | -1.396 |
| chr8 | 2230466 | 2230743 | 1.66055 | 1.18936 | -1.396 |
| chr12 | 60005498 | 60005894 | 0.825944 | 0.591579 | -1.396 |
| chr10 | 49871790 | 49872145 | 1.06356 | 0.761957 | -1.396 |
| chr1 | 8169957 | 8170163 | 0.82384 | 0.590255 | -1.396 |
| chr1 | 205210865 | 205211109 | 0.842845 | 0.603927 | -1.396 |
| chr10 | 112922762 | 112922970 | 1.02771 | 0.736447 | -1.395 |
| chr2 | 82270015 | 82270239 | 1.20203 | 0.861389 | -1.395 |
| chr3 | 83509630 | 83509859 | 0.96694 | 0.692933 | -1.395 |
| chr7 | 54313791 | 54314010 | 0.97545 | 0.699159 | -1.395 |
| chr8 | 2830186 | 2830411 | 1.08231 | 0.77581 | -1.395 |
| chr2 | 209330848 | 209331351 | 1.44179 | 1.03356 | -1.395 |
| chr2 | 157749124 | 157749319 | 1.74497 | 1.25094 | -1.395 |
| chr17 | 65791039 | 65791323 | 1.30104 | 0.932759 | -1.395 |
| chr14 | 83429633 | 83429877 | 0.893334 | 0.640497 | -1.395 |
| chr12 | 124495281 | 124495480 | 1.15727 | 0.82978 | -1.395 |
| chr14 | 100365732 | 100365940 | 1.38814 | 0.995319 | -1.395 |
| chr9 | 13848228 | 13848447 | 1.03489 | 0.74213 | -1.394 |
| chr2 | 102851582 | 102852180 | 1.10558 | 0.792914 | -1.394 |
| chr3 | 78530158 | 78530464 | 1.08648 | 0.779275 | -1.394 |
| chr8 | 136445788 | 136446062 | 1.15483 | 0.828882 | -1.393 |
| chr8 | 83036859 | 83037126 | 1.1056 | 0.793556 | -1.393 |
| chr2 | 17658690 | 17659006 | 0.880191 | 0.631775 | -1.393 |
| chr2 | 142479780 | 142479982 | 1.15972 | 0.832466 | -1.393 |
| chr14 | 54659356 | 54659669 | 0.969526 | 0.695955 | -1.393 |
| chr11 | 41941972 | 41942313 | 0.823982 | 0.591495 | -1.393 |
| chr9 | 135896825 | 135897019 | 0.78318 | 0.56221 | -1.393 |
| chr2 | 53508447 | 53508702 | 1.28028 | 0.919349 | -1.393 |
| chr17 | 60186246 | 60186438 | 0.874842 | 0.62822 | -1.393 |
| chr7 | 54182806 | 54182994 | 0.947746 | 0.680591 | -1.393 |
| chr1 | 190561917 | 190562125 | 1.1688 | 0.839504 | -1.392 |
| chr8 | 4457558 | 4457754 | 1.06559 | 0.765377 | -1.392 |
| chr7 | 40543929 | 40544163 | 1.13636 | 0.816318 | -1.392 |
| chr3 | 36761086 | 36761474 | 1.01846 | 0.73189 | -1.392 |
| chr8 | 137536385 | 137536637 | 0.820467 | 0.589614 | -1.392 |
| chr8 | 88675051 | 88675239 | 1.1419 | 0.820804 | -1.391 |
| chr7 | 16283950 | 16284140 | 0.825013 | 0.593044 | -1.391 |
| chr7 | 144677683 | 144677892 | 1.17422 | 0.844065 | -1.391 |
| chr10 | 28929460 | 28929648 | 0.985421 | 0.708415 | -1.391 |
| chr2 | 35812205 | 35812448 | 1.18609 | 0.852729 | -1.391 |
| chr9 | 92729867 | 92730057 | 1.04558 | 0.751928 | -1.391 |
| chr17 | 31365770 | 31366006 | 2.37593 | 1.70869 | -1.390 |
| chr22 | 33219094 | 33219281 | 2.14858 | 1.54559 | -1.390 |
| chr3 | 83702017 | 83702272 | 0.950589 | 0.683893 | -1.390 |
| chr1 | 191721407 | 191721634 | 0.860533 | 0.619249 | -1.390 |
| chr9 | 88501704 | 88502003 | 0.983735 | 0.707955 | -1.390 |
| chr8 | 116060247 | 116060560 | 1.01161 | 0.728034 | -1.390 |
| chr5 | 174322380 | 174322600 | 1.13801 | 0.819005 | -1.390 |
| chr5 | 84281180 | 84281405 | 1.09679 | 0.789377 | -1.389 |
| chr19 | 32007397 | 32007582 | 1.03246 | 0.743132 | -1.389 |
| chr19 | 56436397 | 56436798 | 0.955731 | 0.687939 | -1.389 |
| chr3 | 4149303 | 4149497 | 1.17144 | 0.84322 | -1.389 |
| chr17 | 16504675 | 16505010 | 1.11494 | 0.802603 | -1.389 |
| chr5 | 157690247 | 157690433 | 1.18112 | 0.850295 | -1.389 |

| | | | | | |
|---|---|---|---|---|---|
| chr1 | 216132907 | 216133161 | 0.747081 | 0.537829 | -1.389 |
| chr12 | 10966998 | 10967279 | 0.784728 | 0.564968 | -1.389 |
| chr1 | 228016391 | 228016670 | 1.0158 | 0.731518 | -1.389 |
| chr12 | 108522712 | 108523083 | 0.899333 | 0.647675 | -1.389 |
| chr3 | 3376669 | 3376879 | 0.830487 | 0.598096 | -1.389 |
| chr17 | 69092478 | 69092662 | 1.17962 | 0.84954 | -1.389 |
| chr12 | 132248747 | 132248959 | 1.41279 | 1.01749 | -1.389 |
| chr12 | 82190924 | 82191181 | 1.19084 | 0.85771 | -1.388 |
| chr2 | 196009115 | 196009528 | 1.16671 | 0.840359 | -1.388 |
| chr16 | 68553543 | 68553732 | 0.858907 | 0.618692 | -1.388 |
| chr11 | 39078478 | 39078680 | 0.833178 | 0.6002 | -1.388 |
| chr18 | 35587971 | 35588182 | 0.816827 | 0.588445 | -1.388 |
| chr5 | 54211563 | 54211769 | 0.725466 | 0.522665 | -1.388 |
| chr17 | 21766038 | 21766275 | 1.0964 | 0.789924 | -1.388 |
| chr17 | 50023786 | 50023975 | 0.970602 | 0.699304 | -1.388 |
| chr3 | 165384294 | 165384527 | 1.09657 | 0.790242 | -1.388 |
| chr8 | 57598084 | 57598291 | 1.26734 | 0.91338 | -1.388 |
| chr9 | 87383544 | 87383764 | 0.764125 | 0.550742 | -1.387 |
| chr12 | 38924425 | 38924621 | 1.23306 | 0.8888 | -1.387 |
| chr3 | 601264 | 601607 | 1.17195 | 0.844914 | -1.387 |
| chr19 | 32514399 | 32514617 | 0.903938 | 0.65186 | -1.387 |
| chr14 | 66421656 | 66421968 | 1.07104 | 0.772414 | -1.387 |
| chr5 | 173998042 | 173998264 | 1.05658 | 0.762255 | -1.386 |
| chr7 | 94423676 | 94423884 | 1.06609 | 0.769138 | -1.386 |
| chr17 | 12343728 | 12343990 | 1.00663 | 0.726272 | -1.386 |
| chr17 | 50279398 | 50279640 | 1.1682 | 0.842856 | -1.386 |
| chr15 | 97908896 | 97909106 | 1.16831 | 0.842987 | -1.386 |
| chr2 | 76464873 | 76465166 | 1.21284 | 0.875132 | -1.386 |
| chr11 | 82387651 | 82387958 | 1.42108 | 1.02552 | -1.386 |
| chr2 | 81392083 | 81392287 | 0.96664 | 0.697609 | -1.386 |
| chr9 | 12465019 | 12465256 | 1.60851 | 1.16127 | -1.385 |
| chr17 | 21288817 | 21289111 | 1.0147 | 0.73262 | -1.385 |
| chr7 | 123144428 | 123144635 | 0.892009 | 0.644133 | -1.385 |
| chr7 | 96229904 | 96230086 | 0.925612 | 0.668417 | -1.385 |
| chr16 | 8161113 | 8161302 | 1.03731 | 0.749122 | -1.385 |
| chr8 | 14710309 | 14710555 | 0.792565 | 0.572382 | -1.385 |
| chr3 | 6279832 | 6280070 | 1.0095 | 0.72908 | -1.385 |
| chr10 | 9577561 | 9577941 | 0.978869 | 0.707134 | -1.384 |
| chr16 | 20371500 | 20371853 | 1.21816 | 0.880162 | -1.384 |
| chr16 | 6010928 | 6011111 | 0.988052 | 0.714026 | -1.384 |
| chr4 | 104350546 | 104350784 | 1.50303 | 1.08634 | -1.384 |
| chr14 | 70865866 | 70866172 | 0.808947 | 0.584683 | -1.384 |
| chr12 | 126712043 | 126712371 | 1.11179 | 0.803701 | -1.383 |
| chr14 | 100175238 | 100175475 | 0.752978 | 0.544322 | -1.383 |
| chr19 | 28594961 | 28595167 | 1.30062 | 0.940232 | -1.383 |
| chr15 | 61660193 | 61660409 | 1.12034 | 0.809998 | -1.383 |
| chr14 | 72758919 | 72759403 | 1.51269 | 1.09373 | -1.383 |
| chr5 | 7087924 | 7088285 | 1.07037 | 0.773984 | -1.383 |
| chr14 | 32026025 | 32026361 | 0.896174 | 0.648065 | -1.383 |
| chr3 | 3856805 | 3857141 | 0.896149 | 0.648263 | -1.382 |
| chr10 | 130789396 | 130789643 | 1.11976 | 0.810022 | -1.382 |
| chr17 | 52545797 | 52546055 | 0.910052 | 0.65844 | -1.382 |
| chr2 | 165594195 | 165594468 | 2.15904 | 1.56214 | -1.382 |
| chr8 | 137072152 | 137072378 | 1.3248 | 0.958544 | -1.382 |
| chr5 | 116188077 | 116188294 | 0.794096 | 0.574612 | -1.382 |
| chr15 | 86683045 | 86683441 | 1.88311 | 1.36295 | -1.382 |
| chr19 | 41998860 | 41999066 | 1.26848 | 0.918098 | -1.382 |
| chr11 | 7476630 | 7476820 | 0.692404 | 0.501213 | -1.381 |
| chr5 | 12230700 | 12230920 | 1.1594 | 0.839264 | -1.381 |
| chr3 | 155040914 | 155041141 | 2.32727 | 1.68515 | -1.381 |
| chr15 | 95032021 | 95032311 | 1.0376 | 0.751559 | -1.381 |

| | | | | | |
|---|---|---|---|---|---|
| chr1 | 158583295 | 158583524 | 0.933662 | 0.676364 | -1.380 |
| chr11 | 36643848 | 36644117 | 0.834687 | 0.604693 | -1.380 |
| chr7 | 37674871 | 37675083 | 1.01946 | 0.738674 | -1.380 |
| chr17 | 41063144 | 41063377 | 1.08283 | 0.784632 | -1.380 |
| chr2 | 5727495 | 5727733 | 1.07904 | 0.781891 | -1.380 |
| chr1 | 215486167 | 215486391 | 1.03506 | 0.750036 | -1.380 |
| chr8 | 14822478 | 14822705 | 0.813826 | 0.589737 | -1.380 |
| chr15 | 84487210 | 84487409 | 0.991321 | 0.718404 | -1.380 |
| chr12 | 82311870 | 82312136 | 0.921959 | 0.668148 | -1.380 |
| chr2 | 57494174 | 57494396 | 1.50431 | 1.09036 | -1.380 |
| chr2 | 12363767 | 12363975 | 1.01251 | 0.733934 | -1.380 |
| chr2 | 140028063 | 140028365 | 0.77169 | 0.559433 | -1.379 |
| chr5 | 36305942 | 36306224 | 1.0439 | 0.756994 | -1.379 |
| chr6 | 71611059 | 71611453 | 0.793023 | 0.575091 | -1.379 |
| chr19 | 44923280 | 44923539 | 0.975252 | 0.707273 | -1.379 |
| chr10 | 53743207 | 53743390 | 1.00952 | 0.732161 | -1.379 |
| chr5 | 9023434 | 9023650 | 0.706325 | 0.512284 | -1.379 |
| chr10 | 86475568 | 86475954 | 1.46542 | 1.06285 | -1.379 |
| chr8 | 53891964 | 53892147 | 0.943527 | 0.684443 | -1.379 |
| chr8 | 139397330 | 139397621 | 0.876121 | 0.635698 | -1.378 |
| chr15 | 72691081 | 72691320 | 0.979934 | 0.711046 | -1.378 |
| chr3 | 157108517 | 157108888 | 1.09814 | 0.797112 | -1.378 |
| chr3 | 58306838 | 58307217 | 1.16725 | 0.847279 | -1.378 |
| chr2 | 35822778 | 35822962 | 0.925157 | 0.671659 | -1.377 |
| chr2 | 126097326 | 126097517 | 0.801902 | 0.58218 | -1.377 |
| chr14 | 25074654 | 25075052 | 1.79849 | 1.30577 | -1.377 |
| chr2 | 35444364 | 35444568 | 1.13839 | 0.826612 | -1.377 |
| chr2 | 34465982 | 34466380 | 0.936926 | 0.680337 | -1.377 |
| chr2 | 172399667 | 172399904 | 1.0284 | 0.746807 | -1.377 |
| chr3 | 3848517 | 3848745 | 0.781242 | 0.56742 | -1.377 |
| chr20 | 24081333 | 24081598 | 2.33151 | 1.69396 | -1.376 |
| chr11 | 43008718 | 43008940 | 0.848089 | 0.616313 | -1.376 |
| chr14 | 85570184 | 85570686 | 0.979197 | 0.711642 | -1.376 |
| chr17 | 30943954 | 30944220 | 1.08026 | 0.785208 | -1.376 |
| chr3 | 89202631 | 89202854 | 0.849585 | 0.617619 | -1.376 |
| chr2 | 156217188 | 156217394 | 0.794893 | 0.577871 | -1.376 |
| chr1 | 246851425 | 246851627 | 1.0765 | 0.7826 | -1.376 |
| chr12 | 24615804 | 24616114 | 1.00267 | 0.729123 | -1.375 |
| chr1 | 214132752 | 214133174 | 0.9357 | 0.680447 | -1.375 |
| chr6 | 166446295 | 166446510 | 1.23723 | 0.899732 | -1.375 |
| chr15 | 75882221 | 75882469 | 1.14359 | 0.831915 | -1.375 |
| chr3 | 85949067 | 85949266 | 1.26892 | 0.923373 | -1.374 |
| chr10 | 37162352 | 37162574 | 0.891099 | 0.648464 | -1.374 |
| chr7 | 39523115 | 39523360 | 1.02561 | 0.746351 | -1.374 |
| chr7 | 132470769 | 132471011 | 1.07308 | 0.780897 | -1.374 |
| chr19 | 41561008 | 41561202 | 1.02314 | 0.744653 | -1.374 |
| chr2 | 176199066 | 176199264 | 0.843097 | 0.613647 | -1.374 |
| chr5 | 122729967 | 122730203 | 0.856031 | 0.623066 | -1.374 |
| chr2 | 13349336 | 13349561 | 0.630034 | 0.458689 | -1.374 |
| chr8 | 93085521 | 93085735 | 1.61593 | 1.17646 | -1.374 |
| chr5 | 84483568 | 84483793 | 0.774461 | 0.563963 | -1.373 |
| chr2 | 16800301 | 16800525 | 1.0279 | 0.748684 | -1.373 |
| chr11 | 38258625 | 38258884 | 0.958988 | 0.698531 | -1.373 |
| chr5 | 117627272 | 117627614 | 1.00532 | 0.732341 | -1.373 |
| chr2 | 22953500 | 22953803 | 1.01618 | 0.740472 | -1.372 |
| chr12 | 83803822 | 83804064 | 0.951773 | 0.693564 | -1.372 |
| chr5 | 112385019 | 112385231 | 1.05767 | 0.770745 | -1.372 |
| chr10 | 83720443 | 83720683 | 0.813879 | 0.593129 | -1.372 |
| chr1 | 150527794 | 150527992 | 1.11695 | 0.814019 | -1.372 |
| chr9 | 10513586 | 10513862 | 1.04296 | 0.760127 | -1.372 |
| chr16 | 76858414 | 76858681 | 0.950422 | 0.692708 | -1.372 |

| | | | | | |
|---|---|---|---|---|---|
| chr5 | 66059819 | 66060192 | 1.2525 | 0.912947 | -1.372 |
| chr7 | 147296624 | 147296837 | 1.01803 | 0.742106 | -1.372 |
| chr3 | 121811456 | 121811641 | 1.29938 | 0.947463 | -1.371 |
| chr16 | 61635385 | 61635678 | 0.797618 | 0.581613 | -1.371 |
| chr3 | 110615594 | 110615876 | 0.933837 | 0.681032 | -1.371 |
| chr8 | 83026870 | 83027232 | 1.17314 | 0.855647 | -1.371 |
| chr8 | 74927206 | 74927392 | 0.899887 | 0.656361 | -1.371 |
| chr16 | 72229467 | 72229761 | 1.54057 | 1.12394 | -1.371 |
| chr12 | 33872686 | 33872972 | 1.21723 | 0.888054 | -1.371 |
| chr12 | 61295296 | 61295616 | 0.666384 | 0.486224 | -1.371 |
| chr8 | 137695930 | 137696160 | 1.10067 | 0.803173 | -1.370 |
| chr10 | 54368881 | 54369081 | 1.00561 | 0.73395 | -1.370 |
| chr8 | 57775391 | 57775606 | 1.36434 | 0.995819 | -1.370 |
| chr3 | 103283614 | 103283834 | 0.734486 | 0.536164 | -1.370 |
| chr15 | 31198094 | 31198297 | 1.81542 | 1.32527 | -1.370 |
| chr1 | 164873023 | 164873217 | 1.14811 | 0.838159 | -1.370 |
| chr15 | 24141342 | 24141546 | 0.928307 | 0.677796 | -1.370 |
| chr6 | 23775507 | 23775840 | 0.676711 | 0.494246 | -1.369 |
| chr8 | 13120107 | 13120365 | 0.893122 | 0.652493 | -1.369 |
| chr19 | 29638924 | 29639273 | 1.05532 | 0.771129 | -1.369 |
| chr10 | 106685080 | 106685364 | 1.97367 | 1.4423 | -1.368 |
| chr8 | 93353497 | 93353694 | 1.15596 | 0.844801 | -1.368 |
| chr16 | 59173060 | 59173273 | 1.01113 | 0.73896 | -1.368 |
| chr15 | 27483408 | 27483605 | 1.2481 | 0.912515 | -1.368 |
| chr15 | 94200889 | 94201073 | 0.85591 | 0.625802 | -1.368 |
| chr2 | 14796833 | 14797157 | 0.922085 | 0.674443 | -1.367 |
| chr11 | 87464018 | 87464272 | 0.951643 | 0.696097 | -1.367 |
| chr5 | 84892512 | 84892759 | 1.09862 | 0.80388 | -1.367 |
| chr2 | 31490659 | 31490983 | 1.01408 | 0.74207 | -1.367 |
| chr10 | 95734495 | 95734705 | 0.782411 | 0.572629 | -1.366 |
| chr7 | 156912883 | 156913122 | 0.892939 | 0.653601 | -1.366 |
| chr7 | 21256975 | 21257236 | 1.05257 | 0.770515 | -1.366 |
| chr7 | 48147655 | 48147899 | 0.97985 | 0.717311 | -1.366 |
| chr5 | 4282883 | 4283070 | 1.05275 | 0.770695 | -1.366 |
| chr5 | 161820393 | 161820832 | 1.14837 | 0.84071 | -1.366 |
| chr9 | 19732206 | 19732453 | 0.85886 | 0.628835 | -1.366 |
| chr9 | 73279842 | 73280058 | 1.27398 | 0.933058 | -1.365 |
| chr2 | 36979631 | 36979896 | 0.957741 | 0.701456 | -1.365 |
| chr3 | 146939516 | 146939793 | 1.05495 | 0.772709 | -1.365 |
| chr11 | 81707058 | 81707374 | 1.09382 | 0.801233 | -1.365 |
| chr14 | 98960803 | 98961000 | 0.829853 | 0.60788 | -1.365 |
| chr9 | 95200931 | 95201250 | 0.748586 | 0.548353 | -1.365 |
| chr7 | 136212518 | 136212801 | 0.931363 | 0.682402 | -1.365 |
| chr5 | 164637090 | 164637452 | 1.07701 | 0.789172 | -1.365 |
| chr19 | 35145505 | 35145710 | 1.31244 | 0.961757 | -1.365 |
| chr7 | 15377283 | 15377514 | 0.946842 | 0.693857 | -1.365 |
| chr11 | 20304822 | 20305045 | 0.910156 | 0.667094 | -1.364 |
| chr20 | 24932601 | 24932803 | 0.865397 | 0.634359 | -1.364 |
| chr5 | 9714468 | 9714667 | 1.00135 | 0.734074 | -1.364 |
| chr10 | 84770215 | 84770423 | 0.99511 | 0.729552 | -1.364 |
| chr2 | 54892606 | 54892832 | 2.38855 | 1.75117 | -1.364 |
| chr7 | 41639545 | 41639753 | 1.15946 | 0.850093 | -1.364 |
| chr12 | 43337551 | 43337914 | 1.28825 | 0.944669 | -1.364 |
| chr7 | 145145861 | 145146067 | 0.767513 | 0.56284 | -1.364 |
| chr7 | 122561892 | 122562109 | 1.49044 | 1.0932 | -1.363 |
| chr2 | 49020438 | 49020710 | 1.01088 | 0.741527 | -1.363 |
| chr12 | 81541198 | 81541443 | 1.18278 | 0.867709 | -1.363 |
| chr5 | 60529374 | 60529596 | 0.948632 | 0.695976 | -1.363 |
| chr19 | 57825760 | 57826024 | 1.0732 | 0.787916 | -1.362 |
| chr16 | 26583677 | 26583929 | 0.830623 | 0.609902 | -1.362 |
| chr16 | 34659752 | 34659980 | 1.1364 | 0.83444 | -1.362 |

| | | | | | |
|---|---|---|---|---|---|
| chr20 | 6785561 | 6785756 | 1.1297 | 0.829659 | -1.362 |
| chr12 | 10206050 | 10206270 | 0.84486 | 0.620534 | -1.362 |
| chr14 | 83918181 | 83918505 | 1.12905 | 0.829294 | -1.361 |
| chr17 | 11031710 | 11031962 | 0.988823 | 0.726436 | -1.361 |
| chr2 | 41141798 | 41141993 | 1.03044 | 0.7571 | -1.361 |
| chr2 | 142613033 | 142613217 | 0.722296 | 0.530725 | -1.361 |
| chr19 | 33880570 | 33880756 | 1.31351 | 0.965399 | -1.361 |
| chr1 | 223253804 | 223254063 | 0.844296 | 0.620585 | -1.360 |
| chr3 | 26150474 | 26150668 | 1.12035 | 0.823642 | -1.360 |
| chr8 | 116379685 | 116380145 | 0.958549 | 0.704716 | -1.360 |
| chr8 | 55676759 | 55676997 | 0.841674 | 0.618846 | -1.360 |
| chr5 | 175837599 | 175837842 | 0.817312 | 0.601012 | -1.360 |
| chr9 | 9637601 | 9637915 | 0.933087 | 0.686224 | -1.360 |
| chr5 | 113200579 | 113200773 | 1.15856 | 0.852077 | -1.360 |
| chr1 | 247884137 | 247884349 | 0.778314 | 0.572449 | -1.360 |
| chr2 | 1586809 | 1587006 | 0.990009 | 0.728171 | -1.360 |
| chr14 | 65481136 | 65481367 | 0.846894 | 0.6231 | -1.359 |
| chr2 | 95623368 | 95623713 | 1.22464 | 0.901238 | -1.359 |
| chr5 | 101077294 | 101077655 | 1.2059 | 0.88745 | -1.359 |
| chr7 | 137426155 | 137426469 | 1.05094 | 0.773465 | -1.359 |
| chr5 | 96633112 | 96633301 | 0.814171 | 0.599472 | -1.358 |
| chr2 | 45722589 | 45722831 | 1.07948 | 0.794864 | -1.358 |
| chr2 | 123141504 | 123141709 | 0.738909 | 0.544312 | -1.358 |
| chr20 | 18183448 | 18183812 | 0.904009 | 0.665972 | -1.357 |
| chr1 | 228150340 | 228150523 | 1.04653 | 0.771025 | -1.357 |
| chr5 | 87898431 | 87898837 | 1.88279 | 1.38739 | -1.357 |
| chr8 | 136922207 | 136922408 | 1.07365 | 0.791222 | -1.357 |
| chr12 | 103765646 | 103765932 | 0.972305 | 0.716543 | -1.357 |
| chr8 | 53978161 | 53978373 | 0.946216 | 0.697381 | -1.357 |
| chr17 | 21219108 | 21219313 | 1.15968 | 0.854742 | -1.357 |
| chr9 | 10531066 | 10531369 | 0.961667 | 0.708888 | -1.357 |
| chr7 | 42530607 | 42530919 | 0.952006 | 0.701771 | -1.357 |
| chr8 | 65349059 | 65349267 | 0.871137 | 0.642172 | -1.357 |
| chr14 | 46379491 | 46379767 | 0.904026 | 0.66642 | -1.357 |
| chr3 | 146817244 | 146817460 | 1.16244 | 0.857023 | -1.356 |
| chr3 | 163297751 | 163298094 | 1.0418 | 0.768213 | -1.356 |
| chr7 | 96372747 | 96372942 | 1.0983 | 0.809938 | -1.356 |
| chr15 | 29227500 | 29227899 | 1.68598 | 1.24338 | -1.356 |
| chr3 | 89330407 | 89330633 | 1.00534 | 0.741513 | -1.356 |
| chr10 | 109872072 | 109872316 | 1.20079 | 0.88568 | -1.356 |
| chr2 | 99583731 | 99584185 | 0.791662 | 0.584026 | -1.356 |
| chr8 | 137627429 | 137627682 | 1.13861 | 0.839988 | -1.356 |
| chr2 | 2332734 | 2332962 | 1.37094 | 1.01144 | -1.355 |
| chr6 | 117293201 | 117293432 | 1.31494 | 0.970177 | -1.355 |
| chr6 | 40296495 | 40296687 | 2.18905 | 1.61531 | -1.355 |
| chr10 | 58979793 | 58980126 | 1.06395 | 0.785138 | -1.355 |
| chr9 | 30161968 | 30162171 | 0.703308 | 0.519032 | -1.355 |
| chr3 | 144391326 | 144391540 | 0.859229 | 0.63414 | -1.355 |
| chr2 | 2957985 | 2958180 | 1.13522 | 0.838049 | -1.355 |
| chr19 | 31609075 | 31609286 | 0.907511 | 0.669988 | -1.355 |
| chr11 | 82504038 | 82504246 | 1.00659 | 0.743305 | -1.354 |
| chr2 | 100999949 | 101000149 | 1.00646 | 0.743395 | -1.354 |
| chr2 | 119025241 | 119025430 | 0.747222 | 0.551923 | -1.354 |
| chr16 | 83285169 | 83285379 | 1.0804 | 0.798039 | -1.354 |
| chr20 | 16062826 | 16063040 | 1.14312 | 0.844432 | -1.354 |
| chr12 | 44399365 | 44399570 | 0.706627 | 0.522045 | -1.354 |
| chr3 | 103611930 | 103612186 | 0.905443 | 0.669109 | -1.353 |
| chr11 | 62138585 | 62138777 | 0.967324 | 0.714911 | -1.353 |
| chr5 | 4456921 | 4457111 | 0.948512 | 0.701106 | -1.353 |
| chr20 | 19939033 | 19939251 | 1.24047 | 0.916972 | -1.353 |
| chr2 | 74492199 | 74492406 | 0.98837 | 0.73063 | -1.353 |

| | | | | | |
|---|---|---|---|---|---|
| chr7 | 96572651 | 96572839 | 1.20733 | 0.89285 | -1.352 |
| chr1 | 182751507 | 182752025 | 1.43186 | 1.059 | -1.352 |
| chr9 | 12119344 | 12119551 | 0.83029 | 0.614263 | -1.352 |
| chr14 | 34793649 | 34793856 | 1.7594 | 1.30174 | -1.352 |
| chr19 | 28779012 | 28779210 | 1.35658 | 1.00381 | -1.351 |
| chr5 | 36459616 | 36459889 | 1.17117 | 0.866743 | -1.351 |
| chr20 | 19564488 | 19564737 | 1.49137 | 1.10376 | -1.351 |
| chr2 | 173612240 | 173612543 | 1.0261 | 0.759586 | -1.351 |
| chr2 | 78553257 | 78553642 | 1.09966 | 0.81418 | -1.351 |
| chr1 | 182775491 | 182775833 | 1.20114 | 0.889438 | -1.350 |
| chr5 | 165996576 | 165996918 | 1.12499 | 0.833132 | -1.350 |
| chr5 | 8409557 | 8409858 | 1.05811 | 0.783607 | -1.350 |
| chr9 | 79475815 | 79476157 | 1.22756 | 0.909105 | -1.350 |
| chr19 | 40827626 | 40827989 | 1.13112 | 0.83787 | -1.350 |
| chr2 | 125277057 | 125277243 | 0.728482 | 0.539628 | -1.350 |
| chr14 | 64477027 | 64477516 | 1.21057 | 0.896797 | -1.350 |
| chr5 | 104905821 | 104906025 | 0.839633 | 0.622127 | -1.350 |
| chr8 | 26295153 | 26295526 | 1.61609 | 1.19752 | -1.350 |
| chr11 | 55560646 | 55560847 | 1.23315 | 0.913907 | -1.349 |
| chr1 | 194624918 | 194625106 | 0.914995 | 0.678159 | -1.349 |
| chr2 | 78900268 | 78900458 | 1.44563 | 1.07145 | -1.349 |
| chr8 | 1214404 | 1214647 | 0.876894 | 0.650025 | -1.349 |
| chr9 | 73704857 | 73705073 | 1.00986 | 0.748692 | -1.349 |
| chr8 | 69040746 | 69040994 | 1.00765 | 0.747058 | -1.349 |
| chr2 | 155986281 | 155986524 | 0.967221 | 0.71711 | -1.349 |
| chr20 | 12819028 | 12819221 | 1.51806 | 1.12576 | -1.348 |
| chr10 | 24186256 | 24186515 | 1.11203 | 0.824666 | -1.348 |
| chr10 | 6988047 | 6988285 | 0.976417 | 0.724116 | -1.348 |
| chr3 | 102449467 | 102450046 | 1.02216 | 0.758066 | -1.348 |
| chr15 | 80079253 | 80079568 | 0.88718 | 0.6581 | -1.348 |
| chr8 | 66483191 | 66483388 | 0.755881 | 0.560836 | -1.348 |
| chr2 | 15399835 | 15400052 | 1.18283 | 0.87774 | -1.348 |
| chr2 | 83944234 | 83944480 | 1.00231 | 0.744019 | -1.347 |
| chr11 | 37825666 | 37825985 | 0.973001 | 0.722268 | -1.347 |
| chr5 | 109517275 | 109517636 | 0.980715 | 0.727999 | -1.347 |
| chr2 | 206580981 | 206581164 | 1.91563 | 1.42201 | -1.347 |
| chr15 | 94598443 | 94598675 | 1.04039 | 0.772304 | -1.347 |
| chr16 | 67899324 | 67899563 | 1.97837 | 1.46878 | -1.347 |
| chr5 | 116586830 | 116587059 | 0.989451 | 0.734597 | -1.347 |
| chr11 | 23628675 | 23628882 | 0.992704 | 0.737037 | -1.347 |
| chr20 | 13676975 | 13677177 | 0.882018 | 0.654923 | -1.347 |
| chr5 | 32724712 | 32724904 | 1.07692 | 0.799838 | -1.346 |
| chr8 | 92292684 | 92292877 | 0.927588 | 0.688928 | -1.346 |
| chr3 | 88824324 | 88824555 | 0.847853 | 0.629753 | -1.346 |
| chr10 | 105483842 | 105484027 | 0.946372 | 0.703039 | -1.346 |
| chr10 | 123975299 | 123975502 | 0.997354 | 0.740929 | -1.346 |
| chr7 | 113144718 | 113145145 | 0.954128 | 0.709091 | -1.346 |
| chr2 | 142955947 | 142956156 | 0.929053 | 0.69057 | -1.345 |
| chr17 | 50558683 | 50558971 | 0.99496 | 0.740059 | -1.344 |
| chr8 | 77966742 | 77967101 | 0.917245 | 0.682255 | -1.344 |
| chr20 | 16620467 | 16620772 | 1.10854 | 0.824614 | -1.344 |
| chr2 | 36221970 | 36222176 | 1.25463 | 0.933421 | -1.344 |
| chr14 | 79293850 | 79294165 | 1.12042 | 0.833581 | -1.344 |
| chr5 | 104190057 | 104190240 | 1.01456 | 0.754842 | -1.344 |
| chr7 | 125596670 | 125596869 | 0.989319 | 0.736118 | -1.344 |
| chr11 | 36847431 | 36847732 | 1.09533 | 0.815055 | -1.344 |
| chr12 | 27814129 | 27814318 | 1.23207 | 0.916932 | -1.344 |
| chr15 | 51522186 | 51522485 | 1.5077 | 1.12211 | -1.344 |
| chr2 | 146116249 | 146116536 | 0.953594 | 0.710189 | -1.343 |
| chr5 | 132105351 | 132105558 | 1.01552 | 0.756354 | -1.343 |
| chr2 | 76739482 | 76739892 | 0.959397 | 0.714623 | -1.343 |

| | | | | | |
|---|---|---|---|---|---|
| rUn_gl0002 | 67348 | 67556 | 0.594563 | 0.442879 | -1.342 |
| chr2 | 180305466 | 180305736 | 1.76904 | 1.31788 | -1.342 |
| chr8 | 39587613 | 39587832 | 1.0956 | 0.816276 | -1.342 |
| chr5 | 173858034 | 173858218 | 0.932907 | 0.695092 | -1.342 |
| chr12 | 90835001 | 90835476 | 1.00859 | 0.751529 | -1.342 |
| chr10 | 108908945 | 108909163 | 0.799277 | 0.595784 | -1.342 |
| chr2 | 127391892 | 127392274 | 1.54478 | 1.15154 | -1.341 |
| chr2 | 118723678 | 118723977 | 0.831176 | 0.619675 | -1.341 |
| chr3 | 25938553 | 25938777 | 1.82113 | 1.35773 | -1.341 |
| chr11 | 22404959 | 22405151 | 0.948521 | 0.707203 | -1.341 |
| chr19 | 22708143 | 22708325 | 1.23564 | 0.921296 | -1.341 |
| chr3 | 22641327 | 22641613 | 1.04866 | 0.781905 | -1.341 |
| chr2 | 74621648 | 74621878 | 0.987619 | 0.736395 | -1.341 |
| chr17 | 79642517 | 79642739 | 0.899691 | 0.670848 | -1.341 |
| chr2 | 199522322 | 199522549 | 1.02451 | 0.76393 | -1.341 |
| chr12 | 91502900 | 91503148 | 2.19289 | 1.63522 | -1.341 |
| chr19 | 31143475 | 31143682 | 1.22781 | 0.915601 | -1.341 |
| chr12 | 82409567 | 82409811 | 0.897478 | 0.66934 | -1.341 |
| chr2 | 65577973 | 65578287 | 1.46201 | 1.09071 | -1.340 |
| chr13 | 92299695 | 92300031 | 1.26816 | 0.946432 | -1.340 |
| chr2 | 34792717 | 34792937 | 0.806069 | 0.6016 | -1.340 |
| chr2 | 36528077 | 36528345 | 0.941938 | 0.703222 | -1.339 |
| chr8 | 96861425 | 96861697 | 1.24765 | 0.931538 | -1.339 |
| chr5 | 171272594 | 171273056 | 1.0161 | 0.758765 | -1.339 |
| chr5 | 28771216 | 28771461 | 0.904892 | 0.675977 | -1.339 |
| chr9 | 24432357 | 24432543 | 0.738096 | 0.551504 | -1.338 |
| chr2 | 4898670 | 4898877 | 1.43558 | 1.07282 | -1.338 |
| chr2 | 3247512 | 3247719 | 0.935309 | 0.698972 | -1.338 |
| chr8 | 141867553 | 141867752 | 0.866582 | 0.647645 | -1.338 |
| chr7 | 35808101 | 35808302 | 1.12101 | 0.837812 | -1.338 |
| chr10 | 66190406 | 66190648 | 0.917907 | 0.686114 | -1.338 |
| chr16 | 60943335 | 60943558 | 1.03736 | 0.77545 | -1.338 |
| chr7 | 64380753 | 64381032 | 0.832311 | 0.622221 | -1.338 |
| chr8 | 138505156 | 138505400 | 2.19549 | 1.64134 | -1.338 |
| chr16 | 12351976 | 12352175 | 1.65098 | 1.23444 | -1.337 |
| chr2 | 237328135 | 237328330 | 0.974013 | 0.728292 | -1.337 |
| chr10 | 64766951 | 64767146 | 1.12785 | 0.843324 | -1.337 |
| chr8 | 3729959 | 3730182 | 0.948035 | 0.708875 | -1.337 |
| chr19 | 58717030 | 58717246 | 1.34422 | 1.0052 | -1.337 |
| chr2 | 34470986 | 34471325 | 1.03543 | 0.774293 | -1.337 |
| chr17 | 47477085 | 47477268 | 0.955849 | 0.714967 | -1.337 |
| chr3 | 174429565 | 174429895 | 0.824266 | 0.616602 | -1.337 |
| chr16 | 58172474 | 58172692 | 0.866411 | 0.648143 | -1.337 |
| chr8 | 33532133 | 33532503 | 0.924448 | 0.691712 | -1.336 |
| chr5 | 173082892 | 173083195 | 1.0746 | 0.8041 | -1.336 |
| chr10 | 120472774 | 120472989 | 1.22039 | 0.913253 | -1.336 |
| chr8 | 26781371 | 26781568 | 1.08743 | 0.81377 | -1.336 |
| chr2 | 123505516 | 123505701 | 0.812458 | 0.608107 | -1.336 |
| chr5 | 3070108 | 3070436 | 0.995979 | 0.745604 | -1.336 |
| chr20 | 23454201 | 23454396 | 0.766209 | 0.573652 | -1.336 |
| chr10 | 58915081 | 58915326 | 0.788675 | 0.590516 | -1.336 |
| chr10 | 111374744 | 111374932 | 0.84678 | 0.634023 | -1.336 |
| chr5 | 161560233 | 161560522 | 0.820228 | 0.614462 | -1.335 |
| chr5 | 113195670 | 113196019 | 1.11862 | 0.838005 | -1.335 |
| chr7 | 126235965 | 126236293 | 0.900435 | 0.67461 | -1.335 |
| chr12 | 31585090 | 31585421 | 1.00046 | 0.749568 | -1.335 |
| chr5 | 104344902 | 104345096 | 0.969548 | 0.726516 | -1.335 |
| chr9 | 104554281 | 104554556 | 1.50832 | 1.13026 | -1.334 |
| chr17 | 26519719 | 26519925 | 1.06904 | 0.801201 | -1.334 |
| chr11 | 67908632 | 67908871 | 0.830368 | 0.622536 | -1.334 |
| chr15 | 98046940 | 98047135 | 0.989141 | 0.741596 | -1.334 |

| | | | | | |
|---|---|---|---|---|---|
| chr7 | 48453684 | 48453911 | 0.919393 | 0.689315 | -1.334 |
| chr19 | 28509889 | 28510199 | 1.22797 | 0.920675 | -1.334 |
| chr11 | 29968885 | 29969087 | 0.924342 | 0.693165 | -1.334 |
| chr19 | 23044316 | 23044598 | 1.31139 | 0.983432 | -1.333 |
| chr8 | 65054217 | 65054485 | 1.03666 | 0.777448 | -1.333 |
| chr20 | 21698822 | 21699027 | 0.649764 | 0.487379 | -1.333 |
| chr19 | 55150269 | 55150593 | 1.09097 | 0.81835 | -1.333 |
| chr8 | 35300782 | 35301039 | 1.12224 | 0.841863 | -1.333 |
| chr1 | 239547226 | 239547419 | 1.20603 | 0.905096 | -1.332 |
| chr19 | 38307377 | 38307625 | 1.07766 | 0.808912 | -1.332 |
| chr8 | 4466586 | 4466901 | 0.901996 | 0.677093 | -1.332 |
| chr2 | 59252352 | 59252658 | 1.76785 | 1.32728 | -1.332 |
| chr7 | 138647636 | 138648031 | 0.821793 | 0.617079 | -1.332 |
| chr2 | 22613433 | 22613631 | 1.22861 | 0.922651 | -1.332 |
| chr17 | 52460623 | 52460865 | 0.953604 | 0.716139 | -1.332 |
| chr15 | 87409772 | 87410155 | 1.18072 | 0.886744 | -1.332 |
| chr5 | 87357250 | 87357439 | 1.01279 | 0.7607 | -1.331 |
| chr3 | 102605589 | 102605859 | 1.27785 | 0.959787 | -1.331 |
| chr7 | 39434628 | 39434846 | 0.895709 | 0.672823 | -1.331 |
| chr10 | 27773565 | 27773756 | 0.934355 | 0.7019 | -1.331 |
| chr8 | 104883550 | 104883779 | 1.20802 | 0.907541 | -1.331 |
| chr8 | 96646973 | 96647169 | 0.937401 | 0.704449 | -1.331 |
| chr9 | 15416924 | 15417144 | 1.42236 | 1.06892 | -1.331 |
| chr17 | 10924748 | 10925079 | 0.985425 | 0.740639 | -1.331 |
| chr2 | 111611737 | 111612138 | 1.05906 | 0.796029 | -1.330 |
| chr3 | 115901861 | 115902070 | 1.15773 | 0.870258 | -1.330 |
| chr1 | 235373205 | 235373388 | 0.92878 | 0.698361 | -1.330 |
| chr3 | 93581119 | 93581365 | 1.02561 | 0.771182 | -1.330 |
| chr10 | 115578944 | 115579134 | 0.674713 | 0.507486 | -1.330 |
| chr14 | 39190964 | 39191353 | 1.63763 | 1.23188 | -1.329 |
| chr2 | 78862225 | 78862432 | 1.04108 | 0.783218 | -1.329 |
| chr16 | 65493801 | 65494016 | 0.894638 | 0.673103 | -1.329 |
| chr5 | 101582727 | 101582928 | 1.0323 | 0.77668 | -1.329 |
| chr7 | 9822019 | 9822264 | 0.8888 | 0.668735 | -1.329 |
| chr13 | 68084825 | 68085024 | 0.799005 | 0.601185 | -1.329 |
| chr19 | 40912284 | 40912482 | 1.04143 | 0.7836 | -1.329 |
| chr5 | 88542717 | 88542900 | 0.922316 | 0.694148 | -1.329 |
| chr11 | 93381378 | 93381564 | 1.18878 | 0.894703 | -1.329 |
| chr5 | 109530141 | 109530451 | 0.930259 | 0.70043 | -1.328 |
| chr17 | 5756887 | 5757171 | 1.37 | 1.03169 | -1.328 |
| chr19 | 56295667 | 56295907 | 1.30069 | 0.979519 | -1.328 |
| chr10 | 2805106 | 2805345 | 0.861035 | 0.648448 | -1.328 |
| chr2 | 80885518 | 80885779 | 1.04776 | 0.789172 | -1.328 |
| chr5 | 153336549 | 153336916 | 1.10353 | 0.831479 | -1.327 |
| chr15 | 24201323 | 24201518 | 0.93737 | 0.706445 | -1.327 |
| chr21 | 35724139 | 35724334 | 1.67842 | 1.26498 | -1.327 |
| chr5 | 136232708 | 136232966 | 1.07017 | 0.806571 | -1.327 |
| chr8 | 5849411 | 5849680 | 1.1969 | 0.902145 | -1.327 |
| chr3 | 116024063 | 116024292 | 0.701231 | 0.528606 | -1.327 |
| chr3 | 10473885 | 10474075 | 0.707741 | 0.53359 | -1.326 |
| chr11 | 21986893 | 21987111 | 0.942744 | 0.710768 | -1.326 |
| chr7 | 151088482 | 151088677 | 1.0392 | 0.783508 | -1.326 |
| chr2 | 41603422 | 41603721 | 1.04401 | 0.787381 | -1.326 |
| chr2 | 60934865 | 60935385 | 1.66652 | 1.25692 | -1.326 |
| chr7 | 94463587 | 94463811 | 0.982692 | 0.741216 | -1.326 |
| chr8 | 2308390 | 2308673 | 1.41907 | 1.07045 | -1.326 |
| chr15 | 91768059 | 91768364 | 2.29373 | 1.73031 | -1.326 |
| chr12 | 73846863 | 73847083 | 0.973984 | 0.734814 | -1.325 |
| chr12 | 119305086 | 119305345 | 0.977393 | 0.73739 | -1.325 |
| chr2 | 4241835 | 4242064 | 1.09769 | 0.828172 | -1.325 |
| chr3 | 13147476 | 13147749 | 1.13261 | 0.854518 | -1.325 |

| | | | | | |
|---|---|---|---|---|---|
| chr11 | 70642178 | 70642456 | 0.800234 | 0.60377 | -1.325 |
| chr2 | 150211149 | 150211384 | 0.663558 | 0.500655 | -1.325 |
| chr5 | 8289280 | 8289526 | 1.01876 | 0.768705 | -1.325 |
| chr2 | 181420075 | 181420581 | 0.897552 | 0.677284 | -1.325 |
| chr12 | 130117041 | 130117387 | 0.924799 | 0.697913 | -1.325 |
| chr8 | 5997566 | 5997775 | 2.24528 | 1.69444 | -1.325 |
| chr2 | 139904949 | 139905153 | 0.888474 | 0.670592 | -1.325 |
| chr19 | 28607634 | 28607865 | 1.34653 | 1.01634 | -1.325 |
| chr12 | 30196415 | 30196613 | 1.05019 | 0.7927 | -1.325 |
| chr5 | 2877786 | 2877982 | 0.939755 | 0.709344 | -1.325 |
| chr6 | 73839625 | 73839964 | 2.86789 | 2.16576 | -1.324 |
| chr11 | 69933532 | 69934121 | 1.11794 | 0.844446 | -1.324 |
| chr7 | 138342518 | 138342724 | 0.916529 | 0.692653 | -1.323 |
| chr5 | 92124578 | 92124864 | 1.01145 | 0.764401 | -1.323 |
| chr11 | 26132841 | 26133063 | 0.797413 | 0.60275 | -1.323 |
| chr7 | 120076673 | 120076942 | 1.06734 | 0.80679 | -1.323 |
| chr3 | 164927848 | 164928180 | 1.14418 | 0.864942 | -1.323 |
| chr14 | 24796614 | 24796837 | 0.874076 | 0.660765 | -1.323 |
| chr10 | 113516082 | 113516451 | 1.16542 | 0.881085 | -1.323 |
| chr15 | 55400977 | 55401395 | 2.40666 | 1.81989 | -1.322 |
| chr4 | 292327 | 292539 | 0.77166 | 0.583525 | -1.322 |
| chrX | 30552024 | 30552207 | 1.07003 | 0.809187 | -1.322 |
| chr9 | 14545284 | 14545517 | 1.0128 | 0.766063 | -1.322 |
| chr19 | 31128864 | 31129047 | 1.35056 | 1.02161 | -1.322 |
| chr10 | 13484126 | 13484368 | 1.09301 | 0.82683 | -1.322 |
| chr3 | 1873037 | 1873229 | 0.836868 | 0.633077 | -1.322 |
| chr8 | 98271315 | 98271698 | 1.38951 | 1.05136 | -1.322 |
| chr8 | 78393832 | 78394044 | 0.938581 | 0.710214 | -1.322 |
| chr2 | 76233264 | 76233750 | 0.896269 | 0.678208 | -1.322 |
| chr1 | 245301758 | 245301944 | 0.765689 | 0.579432 | -1.321 |
| chr3 | 24860082 | 24860371 | 1.20497 | 0.911975 | -1.321 |
| chr7 | 77974610 | 77974910 | 1.20955 | 0.91553 | -1.321 |
| chr2 | 4623207 | 4623402 | 0.824104 | 0.623811 | -1.321 |
| chr2 | 98317822 | 98318011 | 1.20891 | 0.915125 | -1.321 |
| chr8 | 106410362 | 106410734 | 0.770516 | 0.583296 | -1.321 |
| chr19 | 56985454 | 56985838 | 0.814645 | 0.616705 | -1.321 |
| chr2 | 16463268 | 16463593 | 1.2166 | 0.921056 | -1.321 |
| chr14 | 90351104 | 90351323 | 0.986588 | 0.746984 | -1.321 |
| chr14 | 93990364 | 93990566 | 1.02642 | 0.777238 | -1.321 |
| chr2 | 228612911 | 228613164 | 1.50053 | 1.13627 | -1.321 |
| chr14 | 76413165 | 76413381 | 1.00439 | 0.760771 | -1.320 |
| chr10 | 111278105 | 111278345 | 0.872055 | 0.660704 | -1.320 |
| chr2 | 50371416 | 50371614 | 1.06781 | 0.809069 | -1.320 |
| chr9 | 88891060 | 88891305 | 1.10245 | 0.835353 | -1.320 |
| chr2 | 35442069 | 35442349 | 0.989099 | 0.749723 | -1.319 |
| chr15 | 97801090 | 97801372 | 0.911066 | 0.690608 | -1.319 |
| chr10 | 100359126 | 100359338 | 1.32291 | 1.00305 | -1.319 |
| chr3 | 150452470 | 150452686 | 0.965198 | 0.73185 | -1.319 |
| chr3 | 6233015 | 6233226 | 0.994417 | 0.754076 | -1.319 |
| chr9 | 105685154 | 105685369 | 1.12028 | 0.849592 | -1.319 |
| chr2 | 42740479 | 42740686 | 2.28419 | 1.73243 | -1.318 |
| chr14 | 93440881 | 93441110 | 0.855269 | 0.64893 | -1.318 |
| chr7 | 123875521 | 123875777 | 0.806727 | 0.612146 | -1.318 |
| chr16 | 64555642 | 64555856 | 0.819093 | 0.621539 | -1.318 |
| chr10 | 9755231 | 9755645 | 1.00027 | 0.759127 | -1.318 |
| chr12 | 79711064 | 79711268 | 1.03571 | 0.786028 | -1.318 |
| chr2 | 1105618 | 1105870 | 0.594419 | 0.451264 | -1.317 |
| chr8 | 115704731 | 115705010 | 0.715423 | 0.543197 | -1.317 |
| chr5 | 104663431 | 104663633 | 1.01501 | 0.770839 | -1.317 |
| chr7 | 54697546 | 54697747 | 1.22788 | 0.932565 | -1.317 |
| chr3 | 154227813 | 154228008 | 1.08722 | 0.825788 | -1.317 |

| | | | | | |
|---|---|---|---|---|---|
| chr20 | 16623605 | 16623794 | 1.24837 | 0.948362 | -1.316 |
| chr14 | 91622159 | 91622355 | 1.59838 | 1.21427 | -1.316 |
| chr10 | 115580193 | 115580541 | 1.04655 | 0.795078 | -1.316 |
| chr15 | 28197447 | 28197700 | 1.14255 | 0.868078 | -1.316 |
| chr3 | 545001 | 545299 | 1.34776 | 1.02405 | -1.316 |
| chr2 | 198033382 | 198033742 | 1.72728 | 1.31272 | -1.316 |
| chr14 | 88066564 | 88066821 | 0.832671 | 0.632839 | -1.316 |
| chr4 | 63510005 | 63510194 | 0.815162 | 0.619701 | -1.315 |
| chr16 | 83389922 | 83390149 | 2.78235 | 2.11529 | -1.315 |
| chr3 | 82883549 | 82883866 | 0.852235 | 0.647917 | -1.315 |
| chr1 | 208166904 | 208167134 | 1.48545 | 1.12948 | -1.315 |
| chr15 | 29527171 | 29527389 | 1.04939 | 0.798058 | -1.315 |
| chr2 | 76421511 | 76421785 | 1.17994 | 0.897381 | -1.315 |
| chr2 | 4331262 | 4331549 | 0.942501 | 0.716951 | -1.315 |
| chr3 | 154196173 | 154196503 | 1.00131 | 0.761846 | -1.314 |
| chr8 | 127134674 | 127135044 | 1.03517 | 0.787704 | -1.314 |
| chr16 | 29239810 | 29239993 | 0.966472 | 0.735492 | -1.314 |
| chr10 | 55521005 | 55521247 | 0.829702 | 0.631497 | -1.314 |
| chr18 | 26172428 | 26172648 | 0.684622 | 0.521106 | -1.314 |
| chr2 | 168451274 | 168451622 | 0.936414 | 0.712952 | -1.313 |
| chr5 | 15817726 | 15817947 | 1.08363 | 0.825071 | -1.313 |
| chr10 | 14490326 | 14490542 | 0.890685 | 0.678187 | -1.313 |
| chr5 | 120821162 | 120821386 | 1.02683 | 0.781971 | -1.313 |
| chr10 | 84455657 | 84455897 | 0.979885 | 0.746249 | -1.313 |
| chr17 | 13129118 | 13129379 | 1.02887 | 0.783623 | -1.313 |
| chr8 | 140050037 | 140050262 | 2.18792 | 1.66707 | -1.312 |
| chr5 | 84613420 | 84613660 | 0.904887 | 0.689556 | -1.312 |
| chr8 | 105884158 | 105884349 | 0.901714 | 0.687227 | -1.312 |
| chr1 | 186200838 | 186201219 | 0.961458 | 0.732776 | -1.312 |
| chr19 | 31956642 | 31957069 | 1.24826 | 0.951618 | -1.312 |
| chr5 | 165693325 | 165693698 | 0.820747 | 0.625728 | -1.312 |
| chr3 | 21863099 | 21863432 | 1.18477 | 0.903258 | -1.312 |
| chr2 | 141568362 | 141568708 | 1.11284 | 0.848446 | -1.312 |
| chr11 | 36755498 | 36755759 | 0.841605 | 0.6417 | -1.312 |
| chr7 | 87681405 | 87681741 | 0.939029 | 0.716023 | -1.311 |
| chr5 | 117668764 | 117668973 | 0.834651 | 0.636457 | -1.311 |
| chr3 | 3957887 | 3958076 | 0.983986 | 0.750364 | -1.311 |
| chr5 | 146001956 | 146002171 | 1.11931 | 0.853635 | -1.311 |
| chr3 | 25074932 | 25075211 | 1.51475 | 1.15523 | -1.311 |
| chr16 | 10727962 | 10728172 | 0.755025 | 0.575854 | -1.311 |
| chr1 | 54535818 | 54536056 | 2.103 | 1.60395 | -1.311 |
| chr7 | 79862814 | 79862997 | 1.103 | 0.841288 | -1.311 |
| chr2 | 221877416 | 221877634 | 0.943874 | 0.719943 | -1.311 |
| chr3 | 10135909 | 10136339 | 0.990426 | 0.755525 | -1.311 |
| chr17 | 72536194 | 72536394 | 0.99395 | 0.758225 | -1.311 |
| chr17 | 51266437 | 51266700 | 0.962523 | 0.734268 | -1.311 |
| chr7 | 99066107 | 99066408 | 0.690449 | 0.526747 | -1.311 |
| chr10 | 67660742 | 67661043 | 0.985726 | 0.752065 | -1.311 |
| chr4 | 81525462 | 81525688 | 0.994926 | 0.759123 | -1.311 |
| chr10 | 57098242 | 57098506 | 0.860763 | 0.656775 | -1.311 |
| chr8 | 127693030 | 127693306 | 1.20951 | 0.922937 | -1.311 |
| chr10 | 52562882 | 52563102 | 0.814638 | 0.621779 | -1.310 |
| chr5 | 99963301 | 99963531 | 1.07427 | 0.820023 | -1.310 |
| chr17 | 50219298 | 50219523 | 1.36063 | 1.03863 | -1.310 |
| chr8 | 139947056 | 139947403 | 1.08809 | 0.830672 | -1.310 |
| chr2 | 17801596 | 17801858 | 0.822841 | 0.628255 | -1.310 |
| chr3 | 154526460 | 154526667 | 0.864565 | 0.660323 | -1.309 |
| chr3 | 143939488 | 143939697 | 0.744268 | 0.56845 | -1.309 |
| chr2 | 222507614 | 222507886 | 0.817873 | 0.624744 | -1.309 |
| chr2 | 102387009 | 102387422 | 1.36119 | 1.03983 | -1.309 |
| chr17 | 36104346 | 36104584 | 1.62965 | 1.24509 | -1.309 |

| | | | | | |
|---|---|---|---|---|---|
| chr17 | 52073073 | 52073271 | 1.17706 | 0.899677 | -1.308 |
| chr9 | 73616691 | 73616960 | 1.09483 | 0.836936 | -1.308 |
| chr1 | 177860969 | 177861324 | 1.08445 | 0.829052 | -1.308 |
| chr8 | 127283607 | 127283846 | 1.35596 | 1.03667 | -1.308 |
| chr8 | 137045673 | 137045894 | 1.17144 | 0.895607 | -1.308 |
| chr1 | 185565224 | 185565529 | 2.44167 | 1.86689 | -1.308 |
| chr2 | 21645055 | 21645239 | 1.61636 | 1.23621 | -1.308 |
| chr2 | 155428140 | 155428372 | 1.09946 | 0.84096 | -1.307 |
| chr15 | 62526184 | 62526400 | 0.964883 | 0.738067 | -1.307 |
| chr17 | 63313805 | 63314090 | 1.06718 | 0.816327 | -1.307 |
| chr2 | 80178360 | 80178559 | 0.875602 | 0.669807 | -1.307 |
| chr5 | 140014632 | 140014888 | 3.09839 | 2.37021 | -1.307 |
| chr8 | 138749960 | 138750193 | 0.719344 | 0.550327 | -1.307 |
| chr2 | 121444371 | 121444659 | 2.26328 | 1.73151 | -1.307 |
| chr7 | 71480448 | 71480642 | 0.724179 | 0.554105 | -1.307 |
| chr10 | 25770072 | 25770273 | 0.906416 | 0.693606 | -1.307 |
| chr10 | 105600294 | 105600512 | 0.802415 | 0.61413 | -1.307 |
| chrX | 78165807 | 78166100 | 0.998429 | 0.7642 | -1.307 |
| chr20 | 19014363 | 19014594 | 1.07373 | 0.821955 | -1.306 |
| chr14 | 92156599 | 92156827 | 0.979325 | 0.749848 | -1.306 |
| chr19 | 52153363 | 52153568 | 0.963758 | 0.737958 | -1.306 |
| chr1 | 239681717 | 239681905 | 0.956997 | 0.732791 | -1.306 |
| chr2 | 81671188 | 81671477 | 0.818088 | 0.626504 | -1.306 |
| chr3 | 164200579 | 164200761 | 0.920364 | 0.704877 | -1.306 |
| chr8 | 64790120 | 64790302 | 0.647822 | 0.496153 | -1.306 |
| chr5 | 2625420 | 2625637 | 0.899736 | 0.689175 | -1.306 |
| chr14 | 40266751 | 40266975 | 1.07613 | 0.824302 | -1.306 |
| chr2 | 135535838 | 135536217 | 1.43906 | 1.10248 | -1.305 |
| chr7 | 136090957 | 136091396 | 2.0274 | 1.55353 | -1.305 |
| chr5 | 126709765 | 126710002 | 1.21239 | 0.929051 | -1.305 |
| chr12 | 3542728 | 3542918 | 0.844856 | 0.647518 | -1.305 |
| chr12 | 93269954 | 93270143 | 0.923501 | 0.707817 | -1.305 |
| chr8 | 123879406 | 123879633 | 0.788127 | 0.60408 | -1.305 |
| chr11 | 62788101 | 62788444 | 2.22176 | 1.70298 | -1.305 |
| chr7 | 109279875 | 109280109 | 0.891867 | 0.683622 | -1.305 |
| chr2 | 6158070 | 6158442 | 1.12081 | 0.859134 | -1.305 |
| chr1 | 194758246 | 194758581 | 1.20972 | 0.927313 | -1.305 |
| chr14 | 84650302 | 84650669 | 1.00286 | 0.768754 | -1.305 |
| chr7 | 106967352 | 106967548 | 0.960064 | 0.73596 | -1.305 |
| chr11 | 64418498 | 64418896 | 1.42008 | 1.08872 | -1.304 |
| chr7 | 89596114 | 89596407 | 1.076 | 0.824934 | -1.304 |
| chr7 | 97250710 | 97250897 | 1.08453 | 0.831548 | -1.304 |
| chr9 | 24406008 | 24406219 | 1.40882 | 1.08037 | -1.304 |
| chr3 | 71720337 | 71720571 | 1.13963 | 0.874031 | -1.304 |
| chr19 | 28591459 | 28591649 | 1.35039 | 1.03571 | -1.304 |
| chr9 | 1691139 | 1691400 | 0.932661 | 0.715325 | -1.304 |
| chr14 | 81630551 | 81630750 | 1.83585 | 1.40829 | -1.304 |
| chr10 | 59111470 | 59111677 | 0.850033 | 0.652101 | -1.304 |
| chr3 | 26107362 | 26107573 | 0.983952 | 0.754954 | -1.303 |
| chr14 | 85285323 | 85285604 | 1.25343 | 0.96186 | -1.303 |
| chr5 | 23962593 | 23962788 | 1.40579 | 1.07882 | -1.303 |
| chr15 | 100069582 | 100069818 | 1.06595 | 0.818106 | -1.303 |
| chr7 | 124399216 | 124399454 | 1.06269 | 0.815626 | -1.303 |
| chr5 | 131669017 | 131669206 | 1.0315 | 0.791719 | -1.303 |
| chr2 | 95819890 | 95820081 | 0.939034 | 0.720765 | -1.303 |
| chr8 | 18640151 | 18640385 | 1.306 | 1.00268 | -1.303 |
| chr9 | 1837478 | 1837712 | 1.14982 | 0.882785 | -1.302 |
| chr12 | 61341685 | 61341879 | 1.11777 | 0.858608 | -1.302 |
| chr20 | 682511 | 682753 | 1.13152 | 0.869308 | -1.302 |
| chr1 | 189095059 | 189095264 | 0.984253 | 0.75618 | -1.302 |
| chr3 | 106616121 | 106616395 | 0.836182 | 0.642543 | -1.301 |

| | | | | | |
|---|---|---|---|---|---|
| chr5 | 172437647 | 172437885 | 1.30946 | 1.00634 | -1.301 |
| chr5 | 54399316 | 54399515 | 0.740691 | 0.5693 | -1.301 |
| chr17 | 72004372 | 72005096 | 1.34789 | 1.0361 | -1.301 |
| chr7 | 88294018 | 88294316 | 1.02203 | 0.785676 | -1.301 |
| chr1 | 165112764 | 165113116 | 1.92542 | 1.48017 | -1.301 |
| chr9 | 105405518 | 105405706 | 0.994134 | 0.764274 | -1.301 |
| chr1 | 173930960 | 173931173 | 1.07049 | 0.823148 | -1.300 |
| chr15 | 94651515 | 94651892 | 0.951149 | 0.731549 | -1.300 |
| chr2 | 220035942 | 220036431 | 1.86245 | 1.43256 | -1.300 |
| chr3 | 38885517 | 38885707 | 1.14793 | 0.883002 | -1.300 |
| chr3 | 102681624 | 102681834 | 1.14095 | 0.877651 | -1.300 |
| chr2 | 199436211 | 199436440 | 0.877695 | 0.675167 | -1.300 |
| chr12 | 106895797 | 106896109 | 1.08803 | 0.837008 | -1.300 |
| chr8 | 135932113 | 135932316 | 1.02031 | 0.784934 | -1.300 |
| chr9 | 124201069 | 124201278 | 0.880072 | 0.677447 | -1.299 |
| chr16 | 4600503 | 4600909 | 0.881001 | 0.678335 | -1.299 |
| chr4 | 155271129 | 155271355 | 1.50055 | 1.15568 | -1.298 |
| chr9 | 9377547 | 9377835 | 0.886179 | 0.682569 | -1.298 |
| chr2 | 49387627 | 49387851 | 1.05306 | 0.811231 | -1.298 |
| chr2 | 52749365 | 52749602 | 1.05674 | 0.814171 | -1.298 |
| chr3 | 20388406 | 20388664 | 1.00228 | 0.772267 | -1.298 |
| chr5 | 129814901 | 129815134 | 0.907988 | 0.699891 | -1.297 |
| chr16 | 22016204 | 22016401 | 0.841715 | 0.648819 | -1.297 |
| chr2 | 105557926 | 105558186 | 1.06444 | 0.820739 | -1.297 |
| chr11 | 36098898 | 36099174 | 1.07778 | 0.831087 | -1.297 |
| chr1 | 190053768 | 190053952 | 1.57538 | 1.21488 | -1.297 |
| chr7 | 117883038 | 117883225 | 1.05469 | 0.813398 | -1.297 |
| chr14 | 67934299 | 67934486 | 1.15435 | 0.890291 | -1.297 |
| chr14 | 84691635 | 84691869 | 1.11196 | 0.857618 | -1.297 |
| chr7 | 10019169 | 10019487 | 0.907876 | 0.700273 | -1.296 |
| chr7 | 117701459 | 117701713 | 0.796535 | 0.614414 | -1.296 |
| chr10 | 130585774 | 130585981 | 0.83126 | 0.641257 | -1.296 |
| rUn_gl0002 | 177036 | 177355 | 1.20879 | 0.932533 | -1.296 |
| chr10 | 10096408 | 10096717 | 0.916914 | 0.707403 | -1.296 |
| chr2 | 89203323 | 89203684 | 2.30552 | 1.77915 | -1.296 |
| chr2 | 153802762 | 153803003 | 0.985729 | 0.760777 | -1.296 |
| chr8 | 14843027 | 14843279 | 0.897512 | 0.692723 | -1.296 |
| chr7 | 44826753 | 44826936 | 1.18607 | 0.915507 | -1.296 |
| chr9 | 113950503 | 113950728 | 0.893772 | 0.689937 | -1.295 |
| chr3 | 164801837 | 164802093 | 0.829842 | 0.640682 | -1.295 |
| chr9 | 18129932 | 18130187 | 1.02035 | 0.78778 | -1.295 |
| chr8 | 132258059 | 132258361 | 1.46661 | 1.13239 | -1.295 |
| chr4 | 110926762 | 110927081 | 1.55246 | 1.19873 | -1.295 |
| chr2 | 28769207 | 28769510 | 0.977371 | 0.754677 | -1.295 |
| chr2 | 122115292 | 122115478 | 1.00655 | 0.77737 | -1.295 |
| chr21 | 41471558 | 41471782 | 0.98366 | 0.759738 | -1.295 |
| chr7 | 154807106 | 154807361 | 0.894418 | 0.690861 | -1.295 |
| chr2 | 168226812 | 168227046 | 0.665524 | 0.514078 | -1.295 |
| chr15 | 87817327 | 87817793 | 1.02802 | 0.794254 | -1.294 |
| chr7 | 141490162 | 141490376 | 1.04338 | 0.806252 | -1.294 |
| chr8 | 50752281 | 50752492 | 1.10117 | 0.850953 | -1.294 |
| chr19 | 35587742 | 35588039 | 1.02238 | 0.79017 | -1.294 |
| chr15 | 60397950 | 60398185 | 1.07113 | 0.827861 | -1.294 |
| chr11 | 18238527 | 18238788 | 1.14477 | 0.884994 | -1.294 |
| chr1 | 245365502 | 245365764 | 0.80174 | 0.619863 | -1.293 |
| chr20 | 37897618 | 37898152 | 2.08517 | 1.6123 | -1.293 |
| chr8 | 137935869 | 137936062 | 0.897202 | 0.693785 | -1.293 |
| chr10 | 59096659 | 59096841 | 1.259 | 0.973582 | -1.293 |
| chr10 | 57251044 | 57251326 | 0.9917 | 0.767047 | -1.293 |
| chr2 | 156025640 | 156025882 | 1.12251 | 0.868228 | -1.293 |
| chr8 | 54969187 | 54969446 | 0.781676 | 0.60468 | -1.293 |

| | | | | | |
|---|---|---|---|---|---|
| chr7 | 125874134 | 125874381 | 1.11996 | 0.866516 | -1.292 |
| chr2 | 6698171 | 6698409 | 0.951777 | 0.736459 | -1.292 |
| chr11 | 13663582 | 13663844 | 1.08989 | 0.843346 | -1.292 |
| chr19 | 50668722 | 50669071 | 1.29454 | 1.00172 | -1.292 |
| chr5 | 161209710 | 161209941 | 0.900705 | 0.697166 | -1.292 |
| chr17 | 55708290 | 55708491 | 0.898222 | 0.695346 | -1.292 |
| chr9 | 29926453 | 29926730 | 0.759918 | 0.588319 | -1.292 |
| chr11 | 25263460 | 25263704 | 0.966137 | 0.74798 | -1.292 |
| chr2 | 8944958 | 8945184 | 1.27291 | 0.985579 | -1.292 |
| chr7 | 29279299 | 29279524 | 1.53476 | 1.1884 | -1.291 |
| chr2 | 216319191 | 216319499 | 1.82836 | 1.41618 | -1.291 |
| chr8 | 22573470 | 22573866 | 1.05532 | 0.817637 | -1.291 |
| chr17 | 17905877 | 17906094 | 1.01636 | 0.787474 | -1.291 |
| chr8 | 137433594 | 137434266 | 0.950728 | 0.736708 | -1.291 |
| chr17 | 73841330 | 73841658 | 2.16418 | 1.67712 | -1.290 |
| chr8 | 4678507 | 4678868 | 0.993006 | 0.769812 | -1.290 |
| chr6 | 143144181 | 143144383 | 1.36806 | 1.06077 | -1.290 |
| chr20 | 38795867 | 38796073 | 0.96301 | 0.746804 | -1.290 |
| chr2 | 14758090 | 14758352 | 1.24844 | 0.968258 | -1.289 |
| chr5 | 97637491 | 97637759 | 0.90528 | 0.702118 | -1.289 |
| chr5 | 16664538 | 16664746 | 1.30271 | 1.01051 | -1.289 |
| chr8 | 76076532 | 76076844 | 1.07049 | 0.830437 | -1.289 |
| chr3 | 74800173 | 74800367 | 0.85687 | 0.664745 | -1.289 |
| chr17 | 8834758 | 8834950 | 0.948442 | 0.735824 | -1.289 |
| chr19 | 53622532 | 53622756 | 0.917757 | 0.71214 | -1.289 |
| chr11 | 80724873 | 80725058 | 1.12573 | 0.873769 | -1.288 |
| chr19 | 28683249 | 28683578 | 1.29678 | 1.00677 | -1.288 |
| chr15 | 53280785 | 53281003 | 1.24803 | 0.969026 | -1.288 |
| chr20 | 54379615 | 54379843 | 1.11775 | 0.867891 | -1.288 |
| chr1 | 100823774 | 100824015 | 1.47911 | 1.14865 | -1.288 |
| chr20 | 17824151 | 17824350 | 1.44337 | 1.12097 | -1.288 |
| chr7 | 134721465 | 134721753 | 0.814318 | 0.632578 | -1.287 |
| chr8 | 132611461 | 132611664 | 1.00397 | 0.78002 | -1.287 |
| chr5 | 13392528 | 13392736 | 0.84548 | 0.656948 | -1.287 |
| chr8 | 3549565 | 3549863 | 0.856754 | 0.66588 | -1.287 |
| chr5 | 128241290 | 128241555 | 3.09458 | 2.40518 | -1.287 |
| chr2 | 3343763 | 3343994 | 0.840275 | 0.653138 | -1.287 |
| chr10 | 54538400 | 54538747 | 1.58674 | 1.23337 | -1.287 |
| chr9 | 14967339 | 14967819 | 0.926592 | 0.72024 | -1.287 |
| chr8 | 50732832 | 50733047 | 0.823161 | 0.639986 | -1.286 |
| chr8 | 69176777 | 69176967 | 0.841758 | 0.654547 | -1.286 |
| chr8 | 136976335 | 136976664 | 1.14975 | 0.894095 | -1.286 |
| chr9 | 8278156 | 8278344 | 0.986989 | 0.767697 | -1.286 |
| chr18 | 51918959 | 51919217 | 1.67518 | 1.30356 | -1.285 |
| chr20 | 18074047 | 18074410 | 0.909493 | 0.707941 | -1.285 |
| chr7 | 103737210 | 103737402 | 1.32045 | 1.02795 | -1.285 |
| chr8 | 111542803 | 111543023 | 0.859332 | 0.66916 | -1.284 |
| chr7 | 143857206 | 143857408 | 0.977727 | 0.761606 | -1.284 |
| chr8 | 13353491 | 13353674 | 0.773879 | 0.602823 | -1.284 |
| chr8 | 66362629 | 66362865 | 1.85387 | 1.4445 | -1.283 |
| chr7 | 53430238 | 53430536 | 1.02832 | 0.801309 | -1.283 |
| chr15 | 97831844 | 97832040 | 0.93557 | 0.729054 | -1.283 |
| chr17 | 4455167 | 4455404 | 1.15043 | 0.896486 | -1.283 |
| chr8 | 129766352 | 129766692 | 0.932344 | 0.726553 | -1.283 |
| chr13 | 101374238 | 101374469 | 2.35815 | 1.83772 | -1.283 |
| chr2 | 189630361 | 189630637 | 1.12202 | 0.874427 | -1.283 |
| chr16 | 31861683 | 31861952 | 1.06644 | 0.831174 | -1.283 |
| chr9 | 71273756 | 71274096 | 0.741247 | 0.577766 | -1.283 |
| chr2 | 68873481 | 68873806 | 1.418 | 1.10541 | -1.283 |
| chr7 | 142681034 | 142681259 | 0.927896 | 0.723367 | -1.283 |
| chr8 | 115301579 | 115301805 | 0.953007 | 0.74302 | -1.283 |

| | | | | | |
|---|---|---|---|---|---|
| chr2 | 182869335 | 182869577 | 0.912965 | 0.711859 | -1.283 |
| chr11 | 42361818 | 42362026 | 0.938399 | 0.731702 | -1.282 |
| chr19 | 28590725 | 28590941 | 1.099 | 0.85706 | -1.282 |
| chr20 | 954449 | 954645 | 1.71347 | 1.33629 | -1.282 |
| chr2 | 77722652 | 77722888 | 0.966492 | 0.753773 | -1.282 |
| chr3 | 9464872 | 9465061 | 1.2905 | 1.0066 | -1.282 |
| chr2 | 5481889 | 5482107 | 1.25471 | 0.978972 | -1.282 |
| chr18 | 11761498 | 11761833 | 0.979911 | 0.764591 | -1.282 |
| chr3 | 33946308 | 33946582 | 0.842764 | 0.657612 | -1.282 |
| chr20 | 51577955 | 51578145 | 1.01467 | 0.791809 | -1.281 |
| chr2 | 51169405 | 51169608 | 1.14032 | 0.890431 | -1.281 |
| chr2 | 73094498 | 73094707 | 0.899168 | 0.702138 | -1.281 |
| chr5 | 7537700 | 7537980 | 1.27616 | 0.996606 | -1.281 |
| chr8 | 77302052 | 77302293 | 1.00859 | 0.78765 | -1.281 |
| chr2 | 178623537 | 178623793 | 1.09866 | 0.858 | -1.280 |
| chr2 | 67275565 | 67275772 | 1.05415 | 0.823356 | -1.280 |
| chr1 | 194580853 | 194581080 | 1.15935 | 0.905588 | -1.280 |
| chr12 | 74616390 | 74616626 | 1.27442 | 0.995494 | -1.280 |
| chr3 | 1642634 | 1642841 | 0.756692 | 0.591136 | -1.280 |
| chr1 | 197495898 | 197496559 | 1.15017 | 0.898715 | -1.280 |
| chr8 | 140659061 | 140659679 | 1.90708 | 1.49045 | -1.280 |
| chr10 | 9710703 | 9710891 | 0.85709 | 0.669897 | -1.279 |
| chr5 | 54191632 | 54192058 | 1.32851 | 1.03843 | -1.279 |
| chr9 | 102518628 | 102518850 | 0.887273 | 0.693571 | -1.279 |
| chr19 | 40630122 | 40630336 | 1.45472 | 1.13715 | -1.279 |
| chr7 | 145345405 | 145345629 | 0.919625 | 0.71893 | -1.279 |
| chr20 | 11905813 | 11906031 | 1.10275 | 0.862157 | -1.279 |
| chr17 | 13349718 | 13350079 | 1.02884 | 0.804557 | -1.279 |
| chr7 | 87879816 | 87880021 | 1.07499 | 0.84075 | -1.279 |
| chr5 | 170071287 | 170071532 | 1.0367 | 0.810981 | -1.278 |
| chr11 | 79707686 | 79707880 | 0.846456 | 0.662228 | -1.278 |
| chr17 | 54197695 | 54197936 | 0.978373 | 0.765436 | -1.278 |
| chr2 | 113808701 | 113808922 | 0.806762 | 0.631201 | -1.278 |
| chr7 | 46024303 | 46024585 | 0.974146 | 0.76217 | -1.278 |
| chr5 | 59591246 | 59591585 | 1.57351 | 1.23116 | -1.278 |
| chr16 | 13396398 | 13396679 | 0.838685 | 0.656247 | -1.278 |
| chr19 | 29132279 | 29132476 | 1.05913 | 0.828821 | -1.278 |
| chr2 | 199146700 | 199146939 | 0.699105 | 0.547116 | -1.278 |
| chr22 | 45588680 | 45588964 | 0.928173 | 0.726494 | -1.278 |
| chr17 | 15608033 | 15608256 | 0.947822 | 0.741985 | -1.277 |
| chr17 | 2646750 | 2646942 | 0.834686 | 0.653502 | -1.277 |
| chr11 | 37778616 | 37778846 | 0.804491 | 0.629965 | -1.277 |
| chr15 | 84642649 | 84642960 | 1.68355 | 1.31835 | -1.277 |
| chr5 | 119871221 | 119871509 | 1.05571 | 0.826715 | -1.277 |
| chr2 | 79910941 | 79911148 | 0.917349 | 0.718423 | -1.277 |
| chr14 | 89203671 | 89203858 | 1.29401 | 1.01364 | -1.277 |
| chr11 | 58470572 | 58470813 | 0.902098 | 0.706688 | -1.277 |
| chr7 | 10081810 | 10081998 | 0.836943 | 0.655722 | -1.276 |
| chr17 | 49938389 | 49938630 | 1.99054 | 1.55991 | -1.276 |
| chr14 | 49569539 | 49569757 | 0.793464 | 0.621832 | -1.276 |
| chr17 | 50682955 | 50683141 | 1.29911 | 1.01811 | -1.276 |
| chr1 | 176850143 | 176850496 | 1.98055 | 1.55226 | -1.276 |
| chr20 | 23457756 | 23458029 | 1.10975 | 0.869947 | -1.276 |
| chr8 | 80425184 | 80425472 | 1.06847 | 0.837753 | -1.275 |
| chr7 | 33699923 | 33700299 | 0.916289 | 0.71845 | -1.275 |
| chr14 | 47747531 | 47747760 | 1.29468 | 1.01525 | -1.275 |
| chr19 | 38916830 | 38917015 | 1.04554 | 0.819926 | -1.275 |
| chr1 | 190870071 | 190870376 | 1.09934 | 0.862201 | -1.275 |
| chr3 | 21917073 | 21917355 | 0.904251 | 0.709221 | -1.275 |
| chr5 | 163689213 | 163689438 | 1.00315 | 0.786817 | -1.275 |
| chr17 | 29338660 | 29338908 | 0.759327 | 0.595874 | -1.274 |

| | | | | | |
|---|---|---|---|---|---|
| chr10 | 68642839 | 68643078 | 1.14096 | 0.895456 | -1.274 |
| chr7 | 88570603 | 88570852 | 0.726002 | 0.569874 | -1.274 |
| chr2 | 6548038 | 6548245 | 1.06369 | 0.834959 | -1.274 |
| chr10 | 68013681 | 68013903 | 0.655054 | 0.514222 | -1.274 |
| chr7 | 124939623 | 124939984 | 1.04232 | 0.818282 | -1.274 |
| chr7 | 40657854 | 40658038 | 1.09784 | 0.862039 | -1.274 |
| chr8 | 91760935 | 91761154 | 0.908559 | 0.713428 | -1.274 |
| chr12 | 42649159 | 42649415 | 1.48272 | 1.16446 | -1.273 |
| chr5 | 127935634 | 127935829 | 1.02107 | 0.801949 | -1.273 |
| chr9 | 31616089 | 31616296 | 0.968739 | 0.760869 | -1.273 |
| chr20 | 38094324 | 38094538 | 1.14195 | 0.89696 | -1.273 |
| chr7 | 8944109 | 8944365 | 0.768214 | 0.603442 | -1.273 |
| chr7 | 88418826 | 88419114 | 1.72184 | 1.35284 | -1.273 |
| chr1 | 97709564 | 97709750 | 1.13271 | 0.890016 | -1.273 |
| chr2 | 216128729 | 216129013 | 1.20047 | 0.943289 | -1.273 |
| chr7 | 133866902 | 133867135 | 1.06553 | 0.837289 | -1.273 |
| chr14 | 88579944 | 88580146 | 1.06564 | 0.837413 | -1.273 |
| chr5 | 98477656 | 98477903 | 0.918035 | 0.72142 | -1.273 |
| chr6 | 143923673 | 143923895 | 0.697827 | 0.548451 | -1.272 |
| chr5 | 77534965 | 77535159 | 1.10795 | 0.871117 | -1.272 |
| chr19 | 23515192 | 23515835 | 0.894314 | 0.703193 | -1.272 |
| chr5 | 34791842 | 34792053 | 0.976541 | 0.76789 | -1.272 |
| chr2 | 18342390 | 18342693 | 1.13324 | 0.891238 | -1.272 |
| chr1 | 175726495 | 175726848 | 2.37811 | 1.87027 | -1.272 |
| chr1 | 232321793 | 232322075 | 0.834357 | 0.65632 | -1.271 |
| chr10 | 1468717 | 1468938 | 0.999645 | 0.78634 | -1.271 |
| chr8 | 141018505 | 141018719 | 1.0052 | 0.791144 | -1.271 |
| chr5 | 164336779 | 164337031 | 0.907512 | 0.714325 | -1.270 |
| chr14 | 85277834 | 85278026 | 0.92039 | 0.724578 | -1.270 |
| chr3 | 14317076 | 14317495 | 1.48081 | 1.16579 | -1.270 |
| chr9 | 100528466 | 100528723 | 1.23784 | 0.974687 | -1.270 |
| chr7 | 1923552 | 1923819 | 0.865099 | 0.681199 | -1.270 |
| chr17 | 17959657 | 17959839 | 1.50915 | 1.18835 | -1.270 |
| chr8 | 50956213 | 50956403 | 1.05111 | 0.827811 | -1.270 |
| chr8 | 105936686 | 105936906 | 0.813539 | 0.640798 | -1.270 |
| chr8 | 82177302 | 82177493 | 0.837429 | 0.65968 | -1.269 |
| chr14 | 85575822 | 85576127 | 0.904516 | 0.712623 | -1.269 |
| chr19 | 52166939 | 52167147 | 1.21721 | 0.959258 | -1.269 |
| chr3 | 136840766 | 136841054 | 1.08686 | 0.856548 | -1.269 |
| chr14 | 83104407 | 83104639 | 0.764321 | 0.602443 | -1.269 |
| chr3 | 7170265 | 7170463 | 1.05736 | 0.833419 | -1.269 |
| chr22 | 33192980 | 33193276 | 2.3156 | 1.82556 | -1.268 |
| chr7 | 143953225 | 143953496 | 1.09771 | 0.865466 | -1.268 |
| chr10 | 56885833 | 56886043 | 1.17281 | 0.924845 | -1.268 |
| chr5 | 104447112 | 104447371 | 0.882527 | 0.696031 | -1.268 |
| chr10 | 89376401 | 89376600 | 1.02411 | 0.807739 | -1.268 |
| chr17 | 71664863 | 71665067 | 1.15873 | 0.913944 | -1.268 |
| chr7 | 1585221 | 1585415 | 1.28563 | 1.01429 | -1.268 |
| chr7 | 143052481 | 143052671 | 1.07135 | 0.845389 | -1.267 |
| chr1 | 176528917 | 176529248 | 2.08761 | 1.64734 | -1.267 |
| chr7 | 10830340 | 10830541 | 0.836944 | 0.66057 | -1.267 |
| chr2 | 22007952 | 22008142 | 1.22651 | 0.968169 | -1.267 |
| chr17 | 52800010 | 52800537 | 0.979281 | 0.773032 | -1.267 |
| chr3 | 74329776 | 74330003 | 1.0342 | 0.816444 | -1.267 |
| chr12 | 104147024 | 104147243 | 1.23961 | 0.978756 | -1.267 |
| chr20 | 41295730 | 41296040 | 0.955729 | 0.75475 | -1.266 |
| chr16 | 83775767 | 83776001 | 0.997374 | 0.787833 | -1.266 |
| chr10 | 55460202 | 55460430 | 0.64935 | 0.513185 | -1.265 |
| chr3 | 140967889 | 140968119 | 1.02285 | 0.808492 | -1.265 |
| chr20 | 23594739 | 23594958 | 1.45706 | 1.15177 | -1.265 |
| chr15 | 76104007 | 76104209 | 1.08343 | 0.856509 | -1.265 |

| | | | | | |
|---|---|---|---|---|---|
| chr5 | 103420590 | 103420811 | 0.93992 | 0.743165 | -1.265 |
| chr5 | 68213908 | 68214305 | 0.762668 | 0.603062 | -1.265 |
| chr17 | 31866811 | 31866996 | 0.996526 | 0.787999 | -1.265 |
| chr8 | 132390612 | 132390842 | 1.10719 | 0.87552 | -1.265 |
| chr8 | 137845942 | 137846258 | 0.954547 | 0.754832 | -1.265 |
| chr8 | 79255816 | 79256012 | 1.12202 | 0.887286 | -1.265 |
| chr2 | 18046470 | 18046659 | 1.03983 | 0.822399 | -1.264 |
| chr15 | 69767295 | 69767551 | 1.2595 | 0.996139 | -1.264 |
| chr3 | 89727661 | 89727853 | 0.977653 | 0.773248 | -1.264 |
| chr12 | 23862138 | 23862332 | 0.946248 | 0.748624 | -1.264 |
| chr3 | 25362996 | 25363247 | 0.841709 | 0.66594 | -1.264 |
| chr5 | 35436574 | 35436819 | 1.40249 | 1.10962 | -1.264 |
| chr5 | 104353643 | 104353830 | 0.822076 | 0.650683 | -1.263 |
| chr14 | 20384650 | 20384918 | 1.39771 | 1.1064 | -1.263 |
| chr12 | 82141879 | 82142088 | 0.827335 | 0.654905 | -1.263 |
| chr19 | 52297116 | 52297428 | 0.967938 | 0.7665 | -1.263 |
| chr2 | 81483003 | 81483185 | 0.81265 | 0.643682 | -1.263 |
| chr3 | 77245052 | 77245362 | 0.953788 | 0.755482 | -1.262 |
| chr7 | 102701956 | 102702160 | 0.989181 | 0.783531 | -1.262 |
| chr19 | 32532666 | 32533029 | 1.37632 | 1.0903 | -1.262 |
| chr8 | 143584872 | 143585244 | 1.06043 | 0.840066 | -1.262 |
| chr8 | 65829120 | 65829366 | 0.659601 | 0.522572 | -1.262 |
| chr8 | 112741989 | 112742176 | 0.987118 | 0.78207 | -1.262 |
| chr10 | 61331228 | 61331438 | 0.908526 | 0.719834 | -1.262 |
| chr16 | 22910183 | 22910496 | 2.36397 | 1.87306 | -1.262 |
| chr9 | 89098860 | 89099152 | 0.980141 | 0.776618 | -1.262 |
| chr9 | 104249301 | 104249567 | 1.44676 | 1.14643 | -1.262 |
| chr14 | 48682330 | 48682514 | 0.983122 | 0.779058 | -1.262 |
| chr8 | 138997294 | 138997503 | 1.049 | 0.831307 | -1.262 |
| chr10 | 45651176 | 45651505 | 1.15573 | 0.915916 | -1.262 |
| chr9 | 126151047 | 126151262 | 0.914893 | 0.725106 | -1.262 |
| chr8 | 2858435 | 2858622 | 0.879698 | 0.697256 | -1.262 |
| chr8 | 60545865 | 60546065 | 1.00702 | 0.798218 | -1.262 |
| chr2 | 6036785 | 6037148 | 0.848194 | 0.672428 | -1.261 |
| chr2 | 34321544 | 34321925 | 1.68266 | 1.33419 | -1.261 |
| chr11 | 11983232 | 11983433 | 1.18437 | 0.93916 | -1.261 |
| chr16 | 25398902 | 25399122 | 0.833736 | 0.661155 | -1.261 |
| chr19 | 23309150 | 23309462 | 1.04481 | 0.828543 | -1.261 |
| chr5 | 68344109 | 68344314 | 1.45781 | 1.15634 | -1.261 |
| chr2 | 213280795 | 213280991 | 0.903588 | 0.716747 | -1.261 |
| chr2 | 66285920 | 66286134 | 1.05518 | 0.83701 | -1.261 |
| chr8 | 102998593 | 102998878 | 0.799708 | 0.634461 | -1.260 |
| chr20 | 21580864 | 21581244 | 1.28453 | 1.01922 | -1.260 |
| chr7 | 92906339 | 92906527 | 0.824052 | 0.653935 | -1.260 |
| chr7 | 49102429 | 49102782 | 0.94363 | 0.748899 | -1.260 |
| chr19 | 22685279 | 22685495 | 0.815846 | 0.647591 | -1.260 |
| chr1 | 152007046 | 152007291 | 1.89057 | 1.50071 | -1.260 |
| chr7 | 37144014 | 37144207 | 0.956613 | 0.75935 | -1.260 |
| chr9 | 30230057 | 30230321 | 1.00239 | 0.795713 | -1.260 |
| chr5 | 139309970 | 139310185 | 0.901122 | 0.715351 | -1.260 |
| chr11 | 83123715 | 83123953 | 1.08898 | 0.864522 | -1.260 |
| chr19 | 38773471 | 38773654 | 1.16755 | 0.927046 | -1.259 |
| chr10 | 67213269 | 67213570 | 0.95381 | 0.757472 | -1.259 |
| chr2 | 79162623 | 79162913 | 0.865366 | 0.687238 | -1.259 |
| chr12 | 127410576 | 127410927 | 0.909157 | 0.722067 | -1.259 |
| chr6 | 70876312 | 70876532 | 1.93141 | 1.53435 | -1.259 |
| chr1 | 195596951 | 195597171 | 0.997341 | 0.792361 | -1.259 |
| chr8 | 140501317 | 140501642 | 0.927685 | 0.737232 | -1.258 |
| chr11 | 80994506 | 80994704 | 0.77781 | 0.618176 | -1.258 |
| chr12 | 24308512 | 24308853 | 1.0031 | 0.797249 | -1.258 |
| chr1 | 220633926 | 220634212 | 0.952451 | 0.757042 | -1.258 |

| | | | | | |
|---|---|---|---|---|---|
| chr12 | 30476611 | 30476870 | 0.838904 | 0.666832 | -1.258 |
| chr19 | 28576928 | 28577338 | 1.4337 | 1.13986 | -1.258 |
| chr11 | 81289681 | 81289887 | 0.95625 | 0.760296 | -1.258 |
| chr10 | 132712918 | 132713143 | 1.01965 | 0.810772 | -1.258 |
| chr8 | 3660810 | 3661389 | 1.05859 | 0.841878 | -1.257 |
| chr20 | 56767690 | 56767990 | 2.8047 | 2.23077 | -1.257 |
| chr17 | 35850195 | 35850417 | 1.76196 | 1.40202 | -1.257 |
| chr8 | 92700357 | 92700825 | 0.877808 | 0.698499 | -1.257 |
| chr11 | 37069699 | 37070057 | 1.02702 | 0.817264 | -1.257 |
| chr8 | 83545029 | 83545217 | 1.01268 | 0.805946 | -1.257 |
| chr17 | 72570641 | 72570886 | 1.09668 | 0.873128 | -1.256 |
| chr2 | 6728213 | 6728401 | 0.827787 | 0.659137 | -1.256 |
| chr11 | 7557424 | 7557624 | 1.75172 | 1.39496 | -1.256 |
| chr4 | 25268755 | 25269093 | 0.867242 | 0.690849 | -1.255 |
| chr20 | 4761059 | 4761305 | 1.11598 | 0.889081 | -1.255 |
| chr2 | 5747848 | 5748108 | 0.873712 | 0.696083 | -1.255 |
| chr8 | 70303653 | 70303866 | 0.645779 | 0.514507 | -1.255 |
| chr5 | 28119642 | 28119828 | 0.983078 | 0.783266 | -1.255 |
| chr2 | 76785625 | 76785971 | 0.957979 | 0.763409 | -1.255 |
| chr13 | 95048057 | 95048273 | 0.696813 | 0.555288 | -1.255 |
| chr8 | 65191616 | 65191861 | 0.830012 | 0.661539 | -1.255 |
| chr9 | 101146107 | 101146299 | 1.15226 | 0.918387 | -1.255 |
| chr19 | 53149290 | 53149591 | 0.96574 | 0.769823 | -1.254 |
| chr11 | 5045206 | 5045450 | 0.893588 | 0.712707 | -1.254 |
| chr5 | 9847838 | 9848114 | 1.19458 | 0.952997 | -1.253 |
| chr2 | 4121162 | 4121355 | 1.13733 | 0.907569 | -1.253 |
| chr5 | 16223196 | 16223535 | 0.941657 | 0.751439 | -1.253 |
| chr3 | 80586999 | 80587216 | 1.12716 | 0.899479 | -1.253 |
| chr20 | 50261474 | 50261671 | 0.816892 | 0.651888 | -1.253 |
| chr1 | 40138353 | 40138589 | 1.34289 | 1.07167 | -1.253 |
| chr20 | 2694007 | 2694237 | 1.6102 | 1.285 | -1.253 |
| chr3 | 10514832 | 10515037 | 0.966154 | 0.771066 | -1.253 |
| chr17 | 52235911 | 52236235 | 0.972598 | 0.776273 | -1.253 |
| chr12 | 128274197 | 128274529 | 0.979103 | 0.781605 | -1.253 |
| chr12 | 8296171 | 8296373 | 0.880809 | 0.703159 | -1.253 |
| chr7 | 53417965 | 53418157 | 1.08447 | 0.865862 | -1.252 |
| chr5 | 56292217 | 56292437 | 1.84014 | 1.46933 | -1.252 |
| chr3 | 82439895 | 82440198 | 0.948973 | 0.757889 | -1.252 |
| chr8 | 113894483 | 113894848 | 0.990451 | 0.791079 | -1.252 |
| chr16 | 4716514 | 4716726 | 0.828527 | 0.66179 | -1.252 |
| chr5 | 26519298 | 26519489 | 1.00279 | 0.801093 | -1.252 |
| chr5 | 109894420 | 109894727 | 0.916226 | 0.732019 | -1.252 |
| chr10 | 54805092 | 54805275 | 0.928783 | 0.742071 | -1.252 |
| chr5 | 44664351 | 44664544 | 1.05275 | 0.841175 | -1.252 |
| chr1 | 49336303 | 49336555 | 1.14444 | 0.914495 | -1.251 |
| chr9 | 24184181 | 24184369 | 0.53851 | 0.430351 | -1.251 |
| chr14 | 66484360 | 66484554 | 0.922492 | 0.737351 | -1.251 |
| chr8 | 5214267 | 5214479 | 0.845644 | 0.675992 | -1.251 |
| chr3 | 195249058 | 195249303 | 0.952553 | 0.761507 | -1.251 |
| chr12 | 68648513 | 68648754 | 0.988039 | 0.789884 | -1.251 |
| chr5 | 86967867 | 86968074 | 1.35376 | 1.08236 | -1.251 |
| chr10 | 17074951 | 17075186 | 1.07683 | 0.860984 | -1.251 |
| chr8 | 3485824 | 3486222 | 1.012 | 0.809168 | -1.251 |
| chr16 | 77946664 | 77946958 | 1.00532 | 0.803941 | -1.250 |
| chr3 | 102833523 | 102833705 | 0.817965 | 0.654451 | -1.250 |
| chr9 | 31253720 | 31254012 | 0.849775 | 0.680016 | -1.250 |
| chr7 | 100186220 | 100186685 | 1.0798 | 0.864166 | -1.250 |
| chr11 | 20293815 | 20294029 | 0.985343 | 0.788581 | -1.250 |
| chr8 | 93952015 | 93952371 | 1.2648 | 1.01238 | -1.249 |
| chr3 | 144347082 | 144347422 | 1.04701 | 0.838134 | -1.249 |
| chr12 | 71184149 | 71184361 | 1.07531 | 0.860903 | -1.249 |

| | | | | | |
|---|---|---|---|---|---|
| chr10 | 100931942 | 100932217 | 1.9101 | 1.52933 | -1.249 |
| chr1 | 157661196 | 157661584 | 1.49577 | 1.19765 | -1.249 |
| chr8 | 43356583 | 43356772 | 1.08942 | 0.872558 | -1.249 |
| chr5 | 131945231 | 131945446 | 1.07928 | 0.864495 | -1.248 |
| chr1 | 175981147 | 175981446 | 1.19166 | 0.954659 | -1.248 |
| chr9 | 2747524 | 2747756 | 1.14558 | 0.917823 | -1.248 |
| chr19 | 32404084 | 32404584 | 1.34038 | 1.0743 | -1.248 |
| chr3 | 95158303 | 95158490 | 0.89343 | 0.716091 | -1.248 |
| chr18 | 2104942 | 2105196 | 2.63345 | 2.11077 | -1.248 |
| chr2 | 156113306 | 156113691 | 0.835705 | 0.669955 | -1.247 |
| chr11 | 41660555 | 41660770 | 0.984918 | 0.789574 | -1.247 |
| chr9 | 116725363 | 116725621 | 2.35408 | 1.88763 | -1.247 |
| chr12 | 104647264 | 104647663 | 1.31677 | 1.05591 | -1.247 |
| chr4 | 41965216 | 41965482 | 1.55822 | 1.24966 | -1.247 |
| chr20 | 659431 | 659616 | 1.23082 | 0.987106 | -1.247 |
| chr5 | 146777249 | 146777438 | 1.04396 | 0.837265 | -1.247 |
| chr9 | 101993992 | 101994265 | 2.28228 | 1.83043 | -1.247 |
| chr8 | 60970953 | 60971386 | 0.792923 | 0.63598 | -1.247 |
| chr17 | 68983948 | 68984378 | 0.982571 | 0.788133 | -1.247 |
| chr2 | 50296196 | 50296431 | 0.972047 | 0.779839 | -1.246 |
| chr11 | 104202948 | 104203168 | 1.25957 | 1.01066 | -1.246 |
| chr5 | 80698853 | 80699038 | 0.710903 | 0.570461 | -1.246 |
| chr8 | 134132701 | 134132887 | 2.0537 | 1.64801 | -1.246 |
| chr2 | 104646191 | 104646373 | 0.785292 | 0.630182 | -1.246 |
| chr15 | 98656259 | 98656510 | 1.08636 | 0.871785 | -1.246 |
| chr1 | 220900056 | 220900314 | 1.53191 | 1.22937 | -1.246 |
| chr7 | 86556081 | 86556301 | 2.305 | 1.84998 | -1.246 |
| chr7 | 2434284 | 2434476 | 1.00113 | 0.80364 | -1.246 |
| chr16 | 26776508 | 26776753 | 1.38353 | 1.1108 | -1.246 |
| chr1 | 238014573 | 238014878 | 2.37545 | 1.90726 | -1.245 |
| chr7 | 131985404 | 131985622 | 0.842633 | 0.676641 | -1.245 |
| chr1 | 149920806 | 149921041 | 0.788766 | 0.633406 | -1.245 |
| chr12 | 114932111 | 114932312 | 0.631926 | 0.507494 | -1.245 |
| chr5 | 133638714 | 133638933 | 0.720837 | 0.578922 | -1.245 |
| chr2 | 238209565 | 238209858 | 0.867088 | 0.696426 | -1.245 |
| chr10 | 85383769 | 85384019 | 0.869338 | 0.698329 | -1.245 |
| chr16 | 64128760 | 64128987 | 0.839875 | 0.674761 | -1.245 |
| chr17 | 51020587 | 51020867 | 0.790127 | 0.634819 | -1.245 |
| chr12 | 128521639 | 128521832 | 0.886305 | 0.712116 | -1.245 |
| chr3 | 16662647 | 16662992 | 0.965215 | 0.775545 | -1.245 |
| chr10 | 65661077 | 65661372 | 1.21332 | 0.975034 | -1.244 |
| chr8 | 34199784 | 34199977 | 1.02247 | 0.821675 | -1.244 |
| chr16 | 62218103 | 62218436 | 0.944202 | 0.758865 | -1.244 |
| chr4 | 66081274 | 66081530 | 0.898535 | 0.722185 | -1.244 |
| chr14 | 25277671 | 25277926 | 1.10755 | 0.890238 | -1.244 |
| chr1 | 157094757 | 157094988 | 1.18963 | 0.956219 | -1.244 |
| chr2 | 95665375 | 95665577 | 0.810932 | 0.651823 | -1.244 |
| chr14 | 85824301 | 85824485 | 0.936992 | 0.753153 | -1.244 |
| chr12 | 61925915 | 61926098 | 0.742498 | 0.59686 | -1.244 |
| chr14 | 86474826 | 86475039 | 0.815771 | 0.655859 | -1.244 |
| chr2 | 14958163 | 14958414 | 1.02219 | 0.82186 | -1.244 |
| chr5 | 36667267 | 36667450 | 0.829679 | 0.667087 | -1.244 |
| chr11 | 24328218 | 24328493 | 0.963882 | 0.775151 | -1.243 |
| chr11 | 24340421 | 24340658 | 1.02881 | 0.827437 | -1.243 |
| chr1 | 66902164 | 66902507 | 1.24049 | 0.997756 | -1.243 |
| chr5 | 84307986 | 84308171 | 0.864218 | 0.695157 | -1.243 |
| chr7 | 87942512 | 87942751 | 0.631282 | 0.507813 | -1.243 |
| chr2 | 151908180 | 151908531 | 2.09595 | 1.68604 | -1.243 |
| chr1 | 247830657 | 247830876 | 0.885972 | 0.712724 | -1.243 |
| chr14 | 26024004 | 26024202 | 0.859955 | 0.691858 | -1.243 |
| chr2 | 14200176 | 14200611 | 0.965025 | 0.776464 | -1.243 |

| | | | | | |
|---|---|---|---|---|---|
| chr3 | 21201026 | 21201230 | 0.721459 | 0.580533 | -1.243 |
| chr9 | 7570190 | 7570404 | 1.68038 | 1.35226 | -1.243 |
| chr12 | 105483615 | 105483798 | 0.946078 | 0.761507 | -1.242 |
| chr12 | 126896842 | 126897032 | 1.017 | 0.81883 | -1.242 |
| chr2 | 137729567 | 137729763 | 0.671382 | 0.540597 | -1.242 |
| chr9 | 93678111 | 93678322 | 1.46185 | 1.17713 | -1.242 |
| chr17 | 12934598 | 12935584 | 1.72451 | 1.38867 | -1.242 |
| chr19 | 23227236 | 23227479 | 0.801791 | 0.645712 | -1.242 |
| chr9 | 25490685 | 25490919 | 0.994779 | 0.801246 | -1.242 |
| chr16 | 86451097 | 86451284 | 0.884155 | 0.712325 | -1.241 |
| chr4 | 140337289 | 140337530 | 2.03169 | 1.63687 | -1.241 |
| chr2 | 184339137 | 184339363 | 1.27297 | 1.02586 | -1.241 |
| chr2 | 239068968 | 239069188 | 1.49284 | 1.20329 | -1.241 |
| chr8 | 145096109 | 145096626 | 0.974975 | 0.78593 | -1.241 |
| chr12 | 75757972 | 75758164 | 1.09582 | 0.883593 | -1.240 |
| chr15 | 97507602 | 97507880 | 1.21207 | 0.977485 | -1.240 |
| chr8 | 92727697 | 92727986 | 1.00886 | 0.813608 | -1.240 |
| chr10 | 106382424 | 106382732 | 1.76001 | 1.41964 | -1.240 |
| chr19 | 54676724 | 54676940 | 1.3869 | 1.11877 | -1.240 |
| chr5 | 134222104 | 134222384 | 1.00798 | 0.813109 | -1.240 |
| chr9 | 28302206 | 28302484 | 1.00606 | 0.811594 | -1.240 |
| chr7 | 139305625 | 139305869 | 2.60561 | 2.10232 | -1.239 |
| chr22 | 21138474 | 21138816 | 1.63863 | 1.32217 | -1.239 |
| chr8 | 89708277 | 89708696 | 1.14573 | 0.924555 | -1.239 |
| chr8 | 60882002 | 60882292 | 0.84795 | 0.684421 | -1.239 |
| chr1 | 231227649 | 231228010 | 2.53061 | 2.04371 | -1.238 |
| chr11 | 36153326 | 36153535 | 1.88136 | 1.51966 | -1.238 |
| chr16 | 60957339 | 60957588 | 1.15954 | 0.936657 | -1.238 |
| chr10 | 103203876 | 103204145 | 1.32572 | 1.07092 | -1.238 |
| chr8 | 36822835 | 36823028 | 0.910735 | 0.735749 | -1.238 |
| chr11 | 33652990 | 33653173 | 1.10295 | 0.891093 | -1.238 |
| chr2 | 232665294 | 232665597 | 2.49553 | 2.01633 | -1.238 |
| chr17 | 1062119 | 1062307 | 1.13604 | 0.917984 | -1.238 |
| chr15 | 91811868 | 91812076 | 1.8342 | 1.48221 | -1.237 |
| chr11 | 83663080 | 83663438 | 1.10035 | 0.889201 | -1.237 |
| chr5 | 26014371 | 26014590 | 1.22817 | 0.99282 | -1.237 |
| chr16 | 76626300 | 76626563 | 0.913564 | 0.738681 | -1.237 |
| chr3 | 107513457 | 107513702 | 1.34359 | 1.08656 | -1.237 |
| chr20 | 40490259 | 40490458 | 0.955338 | 0.772611 | -1.237 |
| chr21 | 21620701 | 21621037 | 0.789922 | 0.63898 | -1.236 |
| chr15 | 96261643 | 96261879 | 1.23113 | 0.996012 | -1.236 |
| chr5 | 19947727 | 19947915 | 0.918638 | 0.743199 | -1.236 |
| chr8 | 3178299 | 3178719 | 1.17991 | 0.954771 | -1.236 |
| chr8 | 73759235 | 73759701 | 1.51539 | 1.2264 | -1.236 |
| chr5 | 97198722 | 97198924 | 0.966016 | 0.781883 | -1.235 |
| chr8 | 120366422 | 120366649 | 1.76532 | 1.42886 | -1.235 |
| chr10 | 76002783 | 76003047 | 1.24244 | 1.00604 | -1.235 |
| chr14 | 56549148 | 56549333 | 1.04021 | 0.842305 | -1.235 |
| chr16 | 56800952 | 56801165 | 0.797116 | 0.64548 | -1.235 |
| chr15 | 87590336 | 87590603 | 0.970456 | 0.785879 | -1.235 |
| chr3 | 89688453 | 89688818 | 1.29319 | 1.04732 | -1.235 |
| chr8 | 35009751 | 35010017 | 2.41846 | 1.95888 | -1.235 |
| chr2 | 66614411 | 66614616 | 1.34221 | 1.08717 | -1.235 |
| chr11 | 986991 | 987200 | 0.708061 | 0.57352 | -1.235 |
| chr18 | 22094458 | 22094650 | 1.41908 | 1.14947 | -1.235 |
| chr14 | 75373846 | 75374028 | 1.46598 | 1.18759 | -1.234 |
| chr5 | 19945121 | 19945386 | 0.863934 | 0.700031 | -1.234 |
| chr10 | 56298839 | 56299195 | 0.898588 | 0.728221 | -1.234 |
| chr14 | 69876126 | 69876336 | 0.992237 | 0.804267 | -1.234 |
| chr19 | 58475655 | 58475953 | 1.15607 | 0.937132 | -1.234 |
| chr2 | 212610084 | 212610344 | 1.45643 | 1.18067 | -1.234 |

| | | | | | |
|---|---|---|---|---|---|
| chr8 | 52169279 | 52169508 | 0.83924 | 0.680345 | -1.234 |
| chr5 | 104208073 | 104208280 | 0.70177 | 0.568925 | -1.234 |
| chr10 | 57585156 | 57585416 | 0.719656 | 0.58344 | -1.233 |
| chr10 | 114067650 | 114068047 | 2.97581 | 2.41278 | -1.233 |
| chr2 | 171436161 | 171436351 | 0.95769 | 0.776557 | -1.233 |
| chr3 | 147177273 | 147177587 | 1.01336 | 0.821826 | -1.233 |
| chr7 | 80028401 | 80028708 | 1.01758 | 0.825276 | -1.233 |
| chr5 | 147283447 | 147283698 | 0.976605 | 0.79212 | -1.233 |
| chr16 | 7795906 | 7796170 | 0.818337 | 0.663871 | -1.233 |
| chr14 | 47693715 | 47693910 | 0.962964 | 0.781234 | -1.233 |
| chr2 | 6538371 | 6538568 | 0.849733 | 0.68945 | -1.232 |
| chr8 | 3094466 | 3094774 | 1.05919 | 0.859424 | -1.232 |
| chr3 | 95557790 | 95558020 | 1.05875 | 0.859169 | -1.232 |
| chr8 | 36610924 | 36611228 | 0.711575 | 0.577452 | -1.232 |
| chr12 | 3862224 | 3862458 | 1.3016 | 1.05674 | -1.232 |
| chr14 | 72398719 | 72399322 | 1.44494 | 1.17341 | -1.231 |
| chr20 | 99269 | 99453 | 1.14775 | 0.932166 | -1.231 |
| chr8 | 112428570 | 112428759 | 1.15822 | 0.940743 | -1.231 |
| chr20 | 24948994 | 24949188 | 1.00619 | 0.817341 | -1.231 |
| chr16 | 59487054 | 59487294 | 1.11981 | 0.909678 | -1.231 |
| chr10 | 50933904 | 50934086 | 0.91634 | 0.744484 | -1.231 |
| chr5 | 44638492 | 44638743 | 1.20757 | 0.981162 | -1.231 |
| chr8 | 64192888 | 64193222 | 0.874772 | 0.710762 | -1.231 |
| chr16 | 67966011 | 67966213 | 1.35128 | 1.09802 | -1.231 |
| chr5 | 19291368 | 19291575 | 0.768035 | 0.624184 | -1.230 |
| chr4 | 83123264 | 83123572 | 1.17671 | 0.956444 | -1.230 |
| chr8 | 111796814 | 111797084 | 1.14064 | 0.927167 | -1.230 |
| chr14 | 54218249 | 54218433 | 1.02718 | 0.835196 | -1.230 |
| chr11 | 83740333 | 83740805 | 0.854755 | 0.695063 | -1.230 |
| chr3 | 111126085 | 111126289 | 1.32643 | 1.07899 | -1.229 |
| chr7 | 125591574 | 125591959 | 0.925296 | 0.752844 | -1.229 |
| chr14 | 63529434 | 63529684 | 0.885061 | 0.720128 | -1.229 |
| chr5 | 4345267 | 4345508 | 1.14512 | 0.931725 | -1.229 |
| chr8 | 15903356 | 15903670 | 0.866788 | 0.70526 | -1.229 |
| chr8 | 72917505 | 72917736 | 1.76156 | 1.43329 | -1.229 |
| chr5 | 142002270 | 142002453 | 1.18502 | 0.964196 | -1.229 |
| chr7 | 123007653 | 123007837 | 1.83094 | 1.48981 | -1.229 |
| chr5 | 21248486 | 21248690 | 1.08136 | 0.879889 | -1.229 |
| chr3 | 159327302 | 159327497 | 1.17081 | 0.952724 | -1.229 |
| chr19 | 40710315 | 40710597 | 0.963781 | 0.784424 | -1.229 |
| chr15 | 98044026 | 98044222 | 0.922567 | 0.750964 | -1.229 |
| chr8 | 76521704 | 76521973 | 1.06331 | 0.86561 | -1.228 |
| chr19 | 4811725 | 4812007 | 3.50242 | 2.85198 | -1.228 |
| chr12 | 81159321 | 81159580 | 0.958164 | 0.780313 | -1.228 |
| chr9 | 121575368 | 121575571 | 0.817333 | 0.665699 | -1.228 |
| chr17 | 74071469 | 74071663 | 1.90516 | 1.55214 | -1.227 |
| chr2 | 198135432 | 198135638 | 0.954757 | 0.777875 | -1.227 |
| chr2 | 70910665 | 70910948 | 1.06608 | 0.86874 | -1.227 |
| chr11 | 25069562 | 25069802 | 1.23033 | 1.00267 | -1.227 |
| chr2 | 220602007 | 220602326 | 1.6536 | 1.3478 | -1.227 |
| chr15 | 97734526 | 97734733 | 0.999579 | 0.814919 | -1.227 |
| chr7 | 47932861 | 47933144 | 2.03921 | 1.6627 | -1.226 |
| chr12 | 65461991 | 65462247 | 0.719605 | 0.586747 | -1.226 |
| chr5 | 94632870 | 94633060 | 0.862768 | 0.703504 | -1.226 |
| chr20 | 47252981 | 47253185 | 0.623883 | 0.508754 | -1.226 |
| chr7 | 94647848 | 94648279 | 0.946406 | 0.771864 | -1.226 |
| chr8 | 102698003 | 102698193 | 0.782467 | 0.638203 | -1.226 |
| chr10 | 74166358 | 74166547 | 1.02231 | 0.833895 | -1.226 |
| chr11 | 40830553 | 40830816 | 0.9402 | 0.767035 | -1.226 |
| chr1 | 213337964 | 213338221 | 1.06331 | 0.867497 | -1.226 |
| chr6 | 159531044 | 159531360 | 1.29973 | 1.06047 | -1.226 |

| | | | | | |
|---|---|---|---|---|---|
| chr2 | 105911442 | 105911632 | 2.14662 | 1.75155 | -1.226 |
| chr8 | 16093624 | 16093855 | 0.907707 | 0.740755 | -1.225 |
| chr14 | 36454791 | 36455011 | 1.43109 | 1.16795 | -1.225 |
| chr14 | 79435558 | 79435753 | 1.78988 | 1.46107 | -1.225 |
| chr11 | 23232289 | 23232550 | 0.958806 | 0.782708 | -1.225 |
| chr11 | 5985654 | 5985844 | 1.07346 | 0.876348 | -1.225 |
| chr2 | 139729864 | 139730182 | 0.896329 | 0.731818 | -1.225 |
| chr12 | 29945728 | 29945920 | 0.845084 | 0.690086 | -1.225 |
| chr10 | 44477726 | 44477945 | 0.942388 | 0.769553 | -1.225 |
| chr2 | 4538751 | 4538987 | 1.02158 | 0.834253 | -1.225 |
| chr14 | 59485802 | 59486055 | 1.47023 | 1.20089 | -1.224 |
| chr19 | 52952899 | 52953229 | 0.874516 | 0.714322 | -1.224 |
| chr14 | 82166126 | 82166461 | 1.00009 | 0.816927 | -1.224 |
| chr17 | 18553533 | 18553752 | 1.26443 | 1.03293 | -1.224 |
| chr6 | 69348282 | 69348488 | 1.43294 | 1.17113 | -1.224 |
| chr8 | 2350673 | 2350936 | 0.757751 | 0.619351 | -1.223 |
| chr8 | 1541995 | 1542195 | 0.675179 | 0.551981 | -1.223 |
| chr12 | 97383651 | 97383865 | 1.1712 | 0.957518 | -1.223 |
| chr15 | 91685410 | 91685668 | 1.8115 | 1.48121 | -1.223 |
| chr14 | 43064399 | 43064633 | 0.922506 | 0.754463 | -1.223 |
| chr2 | 201413697 | 201413990 | 1.05373 | 0.861809 | -1.223 |
| chr11 | 7463904 | 7464187 | 2.74626 | 2.24706 | -1.222 |
| chr12 | 93931657 | 93931886 | 1.18186 | 0.967091 | -1.222 |
| chr2 | 167064211 | 167064424 | 1.01668 | 0.832016 | -1.222 |
| chr15 | 53550763 | 53550964 | 1.26495 | 1.03537 | -1.222 |
| chr14 | 87394355 | 87394606 | 0.914179 | 0.748363 | -1.222 |
| chr7 | 13340803 | 13341087 | 0.986351 | 0.807507 | -1.221 |
| chr8 | 67100426 | 67100611 | 0.91756 | 0.751408 | -1.221 |
| chr2 | 54730312 | 54730667 | 2.275 | 1.86324 | -1.221 |
| chr5 | 26115325 | 26115573 | 1.23157 | 1.00874 | -1.221 |
| chr2 | 53379203 | 53379450 | 1.04495 | 0.856133 | -1.221 |
| chr2 | 154143762 | 154144285 | 1.78474 | 1.46251 | -1.220 |
| chr1 | 194541969 | 194542162 | 1.04003 | 0.85226 | -1.220 |
| chr16 | 795683 | 796103 | 3.17444 | 2.60267 | -1.220 |
| chr19 | 7706251 | 7706470 | 1.3366 | 1.0959 | -1.220 |
| chr10 | 24131294 | 24131511 | 0.964147 | 0.790622 | -1.219 |
| chr12 | 71572440 | 71572760 | 0.917054 | 0.752038 | -1.219 |
| chr22 | 25198997 | 25199411 | 2.55776 | 2.09791 | -1.219 |
| chr2 | 80359190 | 80359397 | 0.837055 | 0.686693 | -1.219 |
| chr19 | 57884063 | 57884388 | 0.844584 | 0.692907 | -1.219 |
| chr20 | 20805602 | 20806068 | 1.78886 | 1.46763 | -1.219 |
| chr7 | 146585957 | 146586149 | 0.866494 | 0.711061 | -1.219 |
| chr7 | 132275700 | 132276074 | 0.90036 | 0.739146 | -1.218 |
| chr1 | 71595602 | 71595816 | 0.850349 | 0.698208 | -1.218 |
| chr7 | 46169537 | 46169752 | 2.0119 | 1.65228 | -1.218 |
| chr12 | 111706637 | 111706873 | 2.00091 | 1.64334 | -1.218 |
| chr3 | 11450877 | 11451084 | 0.792545 | 0.65106 | -1.217 |
| chr3 | 164786214 | 164786459 | 0.984403 | 0.808807 | -1.217 |
| chr20 | 48581463 | 48581718 | 2.99243 | 2.45884 | -1.217 |
| chr12 | 72590920 | 72591107 | 0.75563 | 0.620926 | -1.217 |
| chr9 | 134285592 | 134285815 | 0.790629 | 0.649747 | -1.217 |
| chr8 | 111296632 | 111296825 | 0.740548 | 0.608645 | -1.217 |
| chr8 | 56433503 | 56433762 | 2.68326 | 2.20558 | -1.217 |
| chr15 | 24068668 | 24069040 | 1.57439 | 1.29414 | -1.217 |
| chr8 | 77981110 | 77981321 | 0.966055 | 0.794155 | -1.216 |
| chr2 | 7184448 | 7184794 | 3.55686 | 2.92421 | -1.216 |
| chr16 | 87686779 | 87687132 | 1.5034 | 1.23602 | -1.216 |
| chr12 | 77750585 | 77750796 | 0.699551 | 0.575168 | -1.216 |
| chr17 | 66925058 | 66925240 | 1.00157 | 0.823546 | -1.216 |
| chr3 | 143374010 | 143374251 | 1.17276 | 0.964343 | -1.216 |
| chr10 | 58959307 | 58959544 | 0.843594 | 0.69395 | -1.216 |

| | | | | | |
|---|---|---|---|---|---|
| chrX | 139223757 | 139223996 | 1.16317 | 0.956867 | -1.216 |
| chr3 | 190265423 | 190265607 | 0.709424 | 0.583708 | -1.215 |
| chr2 | 51398278 | 51398549 | 0.841258 | 0.692237 | -1.215 |
| chr3 | 175365490 | 175365683 | 0.81964 | 0.674516 | -1.215 |
| chr1 | 227746193 | 227746480 | 0.907448 | 0.746803 | -1.215 |
| chr12 | 58068507 | 58068707 | 1.10366 | 0.908299 | -1.215 |
| chr14 | 105798786 | 105799161 | 0.937685 | 0.771733 | -1.215 |
| chr10 | 43870759 | 43870963 | 0.673746 | 0.554753 | -1.214 |
| chr17 | 13073816 | 13074055 | 1.27341 | 1.04877 | -1.214 |
| chr5 | 25984340 | 25984602 | 0.869527 | 0.716261 | -1.214 |
| chr8 | 64947538 | 64947993 | 1.16397 | 0.958855 | -1.214 |
| chr2 | 70305641 | 70305894 | 1.03717 | 0.854459 | -1.214 |
| chr11 | 1588167 | 1588465 | 1.28088 | 1.05538 | -1.214 |
| chr10 | 85805055 | 85805401 | 1.18937 | 0.980055 | -1.214 |
| chr19 | 36242900 | 36243131 | 1.03067 | 0.849698 | -1.213 |
| chr10 | 99964068 | 99964265 | 0.949851 | 0.78316 | -1.213 |
| chr3 | 100127176 | 100127449 | 1.18337 | 0.9759 | -1.213 |
| chr19 | 55042750 | 55042958 | 1.01585 | 0.837904 | -1.212 |
| chr8 | 84242567 | 84242766 | 0.780499 | 0.643786 | -1.212 |
| chr16 | 78487057 | 78487281 | 1.1184 | 0.922943 | -1.212 |
| chr8 | 115037326 | 115037675 | 1.08106 | 0.89217 | -1.212 |
| chr3 | 16446168 | 16446360 | 1.12338 | 0.927254 | -1.212 |
| chr2 | 153483970 | 153484160 | 2.00979 | 1.65983 | -1.211 |
| chr20 | 60244 | 60620 | 1.02157 | 0.843782 | -1.211 |
| chr10 | 87787751 | 87787945 | 0.834184 | 0.689062 | -1.211 |
| chr2 | 3875271 | 3875532 | 2.70948 | 2.23844 | -1.210 |
| chr7 | 153047516 | 153047804 | 1.20134 | 0.99264 | -1.210 |
| chr20 | 7011994 | 7012189 | 1.27974 | 1.05773 | -1.210 |
| chr1 | 179561150 | 179561416 | 1.42458 | 1.17759 | -1.210 |
| chr12 | 61836411 | 61836678 | 0.704815 | 0.582779 | -1.209 |
| chr11 | 14270828 | 14271216 | 2.84989 | 2.35701 | -1.209 |
| chr10 | 24074132 | 24074326 | 1.08113 | 0.894153 | -1.209 |
| chr1 | 163572387 | 163572724 | 0.926977 | 0.766804 | -1.209 |
| chr12 | 128843445 | 128843711 | 0.958039 | 0.792557 | -1.209 |
| chr9 | 120348666 | 120348910 | 1.00778 | 0.833833 | -1.209 |
| chr6 | 161089011 | 161089229 | 1.70468 | 1.41047 | -1.209 |
| chr15 | 98984804 | 98985041 | 0.944891 | 0.78195 | -1.208 |
| chr3 | 88495489 | 88495688 | 0.969113 | 0.802032 | -1.208 |
| chr9 | 107809131 | 107809321 | 1.07131 | 0.886647 | -1.208 |
| chr15 | 79149928 | 79150142 | 0.918383 | 0.760109 | -1.208 |
| chr10 | 55572077 | 55572296 | 1.06619 | 0.882874 | -1.208 |
| chr2 | 113103380 | 113103572 | 1.18361 | 0.98028 | -1.207 |
| chr5 | 131470216 | 131470543 | 0.979451 | 0.811237 | -1.207 |
| chr2 | 29531838 | 29532415 | 2.61439 | 2.16539 | -1.207 |
| chr2 | 28573649 | 28573844 | 0.876092 | 0.725855 | -1.207 |
| chr3 | 6174367 | 6174594 | 0.72408 | 0.599914 | -1.207 |
| chr5 | 153354010 | 153354228 | 2.24297 | 1.85838 | -1.207 |
| chr1 | 152626572 | 152626917 | 2.34413 | 1.94221 | -1.207 |
| chr5 | 22338919 | 22339173 | 0.916165 | 0.759118 | -1.207 |
| chr2 | 170965690 | 170965945 | 0.890244 | 0.737697 | -1.207 |
| chr12 | 33503859 | 33504260 | 1.17307 | 0.972554 | -1.206 |
| chr8 | 126841054 | 126841294 | 1.77554 | 1.47211 | -1.206 |
| chr5 | 31169507 | 31169710 | 0.740901 | 0.614358 | -1.206 |
| chr8 | 26942701 | 26942927 | 1.237 | 1.02587 | -1.206 |
| chr2 | 170623792 | 170624099 | 2.27476 | 1.88767 | -1.205 |
| chr10 | 3462474 | 3462676 | 0.878185 | 0.728791 | -1.205 |
| chr15 | 29921010 | 29921290 | 1.00071 | 0.830589 | -1.205 |
| chr12 | 133242993 | 133243226 | 0.875815 | 0.726931 | -1.205 |
| chr11 | 24146490 | 24146682 | 0.908552 | 0.754353 | -1.204 |
| chr2 | 232336989 | 232337273 | 0.975464 | 0.81004 | -1.204 |
| chr16 | 85475478 | 85475713 | 0.897585 | 0.745388 | -1.204 |

| | | | | | |
|---|---|---|---|---|---|
| chr20 | 62204362 | 62204548 | 1.31493 | 1.09198 | -1.204 |
| chr10 | 84682848 | 84683070 | 0.815886 | 0.677574 | -1.204 |
| chr10 | 133559373 | 133559653 | 1.03025 | 0.855662 | -1.204 |
| chr5 | 42719058 | 42719426 | 1.06505 | 0.884637 | -1.204 |
| chr11 | 133900949 | 133901131 | 0.938193 | 0.779366 | -1.204 |
| chr14 | 78982485 | 78982852 | 0.814356 | 0.676495 | -1.204 |
| chr2 | 79112443 | 79112658 | 0.890471 | 0.739806 | -1.204 |
| chr2 | 78415585 | 78415771 | 0.942075 | 0.782724 | -1.204 |
| chr8 | 87494933 | 87495246 | 1.45416 | 1.20882 | -1.203 |
| chr8 | 58094874 | 58095112 | 1.08896 | 0.905331 | -1.203 |
| chr1 | 216824111 | 216824397 | 0.836402 | 0.695442 | -1.203 |
| chr19 | 35190401 | 35190796 | 0.927757 | 0.771445 | -1.203 |
| chr12 | 61736349 | 61736554 | 0.90161 | 0.749728 | -1.203 |
| chr7 | 122073921 | 122074118 | 1.02355 | 0.851642 | -1.202 |
| chr8 | 136224026 | 136224369 | 0.981435 | 0.816643 | -1.202 |
| chr20 | 56148966 | 56149209 | 2.17429 | 1.80928 | -1.202 |
| chr2 | 115591138 | 115591419 | 0.875659 | 0.728679 | -1.202 |
| chr2 | 139603850 | 139604095 | 1.93274 | 1.60847 | -1.202 |
| chr19 | 32290411 | 32290646 | 1.10791 | 0.9221 | -1.202 |
| chr8 | 140005123 | 140005426 | 0.709009 | 0.590111 | -1.201 |
| chr8 | 50239816 | 50240007 | 0.8835 | 0.735402 | -1.201 |
| chr7 | 45903410 | 45903616 | 1.52152 | 1.26648 | -1.201 |
| chr7 | 118990338 | 118990534 | 0.746788 | 0.62164 | -1.201 |
| chr2 | 14932218 | 14932556 | 0.873374 | 0.727019 | -1.201 |
| chr20 | 11550006 | 11550196 | 1.21487 | 1.01139 | -1.201 |
| chr17 | 27899936 | 27900124 | 1.52465 | 1.26949 | -1.201 |
| chr1 | 215896608 | 215896811 | 0.822944 | 0.685362 | -1.201 |
| chr9 | 92707564 | 92707830 | 1.46567 | 1.22066 | -1.201 |
| chr1 | 162731022 | 162731228 | 0.899778 | 0.749367 | -1.201 |
| chr19 | 55647100 | 55647370 | 1.66447 | 1.38635 | -1.201 |
| chr3 | 86032374 | 86032597 | 1.12953 | 0.940838 | -1.201 |
| chr8 | 138638574 | 138638777 | 0.790731 | 0.658733 | -1.200 |
| chr7 | 113521945 | 113522154 | 0.972367 | 0.810088 | -1.200 |
| chr1 | 192614171 | 192614493 | 0.896903 | 0.747327 | -1.200 |
| chr8 | 84849788 | 84849974 | 1.01155 | 0.842959 | -1.200 |
| chr1 | 111097949 | 111098207 | 1.73579 | 1.4466 | -1.200 |
| chr8 | 52149669 | 52149958 | 0.777217 | 0.647813 | -1.200 |
| chr2 | 10612723 | 10612976 | 0.869312 | 0.72459 | -1.200 |
| chr2 | 220492671 | 220492893 | 0.972138 | 0.810328 | -1.200 |
| chr19 | 37050182 | 37050558 | 1.11772 | 0.93174 | -1.200 |
| chr19 | 16882395 | 16882707 | 2.55338 | 2.12853 | -1.200 |
| chr12 | 33835125 | 33835341 | 1.27242 | 1.0608 | -1.199 |
| chr15 | 92065903 | 92066370 | 1.47342 | 1.22838 | -1.199 |
| chr8 | 92093733 | 92094097 | 1.15261 | 0.961055 | -1.199 |
| chr11 | 80123579 | 80123874 | 0.917378 | 0.764948 | -1.199 |
| chr16 | 48076998 | 48077183 | 0.897093 | 0.748363 | -1.199 |
| chr2 | 181728134 | 181728426 | 0.914942 | 0.763262 | -1.199 |
| chr2 | 105735485 | 105735753 | 1.0041 | 0.837769 | -1.199 |
| chr3 | 21607652 | 21607918 | 0.809696 | 0.675817 | -1.198 |
| chr3 | 27678812 | 27679028 | 1.02262 | 0.853564 | -1.198 |
| chr2 | 232894883 | 232895263 | 1.47367 | 1.23035 | -1.198 |
| chr6 | 89991494 | 89991805 | 1.24388 | 1.0385 | -1.198 |
| chr8 | 84206263 | 84206633 | 0.966628 | 0.807091 | -1.198 |
| chr8 | 33909212 | 33909446 | 1.07226 | 0.895332 | -1.198 |
| chr2 | 62524752 | 62525032 | 0.953394 | 0.796081 | -1.198 |
| chr13 | 67990517 | 67990793 | 1.98344 | 1.65629 | -1.198 |
| chr2 | 68585120 | 68585400 | 1.66388 | 1.38951 | -1.197 |
| chr3 | 86204198 | 86204706 | 0.865811 | 0.723256 | -1.197 |
| chr17 | 50226477 | 50226671 | 1.05567 | 0.881908 | -1.197 |
| chr15 | 53331821 | 53332079 | 1.03647 | 0.865935 | -1.197 |
| chr19 | 22123700 | 22123886 | 0.911181 | 0.761285 | -1.197 |

| | | | | | |
|---|---|---|---|---|---|
| chr2 | 6375666 | 6375994 | 0.739647 | 0.618009 | -1.197 |
| chr7 | 24827733 | 24828004 | 1.92715 | 1.61052 | -1.197 |
| chr22 | 20925698 | 20926096 | 2.64206 | 2.208 | -1.197 |
| chr6 | 40276416 | 40276694 | 2.24739 | 1.87818 | -1.197 |
| chr2 | 22872828 | 22873090 | 0.984609 | 0.822877 | -1.197 |
| chr2 | 7889103 | 7889301 | 1.2385 | 1.03545 | -1.196 |
| chr4 | 156129578 | 156129805 | 1.93654 | 1.6192 | -1.196 |
| chr17 | 52677450 | 52677698 | 0.865108 | 0.723456 | -1.196 |
| chr2 | 2537069 | 2537406 | 0.97564 | 0.816042 | -1.196 |
| chr2 | 34420567 | 34420827 | 1.19526 | 0.999932 | -1.195 |
| chr19 | 34442082 | 34442285 | 1.29174 | 1.08074 | -1.195 |
| chr4 | 53881843 | 53882075 | 1.90615 | 1.59485 | -1.195 |
| chr16 | 87450464 | 87450789 | 1.56813 | 1.31209 | -1.195 |
| chr17 | 46139533 | 46139806 | 1.08263 | 0.906192 | -1.195 |
| chr3 | 99251030 | 99251444 | 1.76582 | 1.47811 | -1.195 |
| chr2 | 152861310 | 152861537 | 1.77309 | 1.48429 | -1.195 |
| chr15 | 27864511 | 27864704 | 1.0937 | 0.915569 | -1.195 |
| chr14 | 78070140 | 78070685 | 1.64379 | 1.37618 | -1.194 |
| chr10 | 117177232 | 117177593 | 1.52242 | 1.27505 | -1.194 |
| chr3 | 3583373 | 3583577 | 0.9611 | 0.804998 | -1.194 |
| chr16 | 84834529 | 84834778 | 0.838189 | 0.702297 | -1.193 |
| chr12 | 23714526 | 23715021 | 0.978958 | 0.820278 | -1.193 |
| chr17 | 17759395 | 17759580 | 1.00291 | 0.840405 | -1.193 |
| chr16 | 61429689 | 61429903 | 0.773781 | 0.648413 | -1.193 |
| chr19 | 30043408 | 30043889 | 1.13052 | 0.947434 | -1.193 |
| chr8 | 132331076 | 132331316 | 1.20115 | 1.00682 | -1.193 |
| chr2 | 61268174 | 61268384 | 0.921895 | 0.772792 | -1.193 |
| chr11 | 18264189 | 18264519 | 1.1153 | 0.934918 | -1.193 |
| chr17 | 73056590 | 73056850 | 0.822687 | 0.689693 | -1.193 |
| chr1 | 246645522 | 246645909 | 1.18894 | 0.99698 | -1.193 |
| chr3 | 16251082 | 16251362 | 0.846596 | 0.709977 | -1.192 |
| chr5 | 117000670 | 117000967 | 0.919855 | 0.771446 | -1.192 |
| chr5 | 119580107 | 119580369 | 0.808509 | 0.6781 | -1.192 |
| chr11 | 125303383 | 125303572 | 1.4696 | 1.23262 | -1.192 |
| chr7 | 70532740 | 70532928 | 1.07627 | 0.902722 | -1.192 |
| chr5 | 101090905 | 101091125 | 1.05645 | 0.886205 | -1.192 |
| chr2 | 13123921 | 13124237 | 1.07671 | 0.90329 | -1.192 |
| chr9 | 100149211 | 100149837 | 2.47778 | 2.07871 | -1.192 |
| chr16 | 35057195 | 35057450 | 0.829879 | 0.69622 | -1.192 |
| chr7 | 95470076 | 95470457 | 1.15482 | 0.969106 | -1.192 |
| chr7 | 113018511 | 113018699 | 0.826752 | 0.693902 | -1.191 |
| chr4 | 109704969 | 109705201 | 1.04876 | 0.880368 | -1.191 |
| chr17 | 17120053 | 17120417 | 1.72406 | 1.44729 | -1.191 |
| chr11 | 30597132 | 30597343 | 0.981237 | 0.824068 | -1.191 |
| chr3 | 76712507 | 76712755 | 1.10098 | 0.924787 | -1.191 |
| chr7 | 90054823 | 90055022 | 1.11161 | 0.93382 | -1.190 |
| chr11 | 25264818 | 25265049 | 1.12598 | 0.945906 | -1.190 |
| chr6 | 161171317 | 161171616 | 1.23023 | 1.03354 | -1.190 |
| chr3 | 143670265 | 143670527 | 1.58755 | 1.33398 | -1.190 |
| chr8 | 104350188 | 104350483 | 1.00661 | 0.846004 | -1.190 |
| chr12 | 45515113 | 45515412 | 1.20455 | 1.01267 | -1.189 |
| chr3 | 84156307 | 84156509 | 0.99157 | 0.833622 | -1.189 |
| chr9 | 126147652 | 126148087 | 3.22155 | 2.70872 | -1.189 |
| chr7 | 125809895 | 125810279 | 0.88047 | 0.740413 | -1.189 |
| chr5 | 61858234 | 61858417 | 1.03192 | 0.867914 | -1.189 |
| chr20 | 8024443 | 8024652 | 2.41104 | 2.02788 | -1.189 |
| chr6 | 158564842 | 158565081 | 1.1979 | 1.0076 | -1.189 |
| chr10 | 7074094 | 7074278 | 1.58931 | 1.33688 | -1.189 |
| chr14 | 40536771 | 40536961 | 0.805077 | 0.677322 | -1.189 |
| chr5 | 101742831 | 101743148 | 1.12696 | 0.948146 | -1.189 |
| chr2 | 176116296 | 176116520 | 1.86345 | 1.56812 | -1.188 |

| | | | | | |
|---|---|---|---|---|---|
| chr5 | 103144215 | 103144686 | 1.22169 | 1.02809 | -1.188 |
| chr4 | 172731571 | 172731796 | 0.835626 | 0.703251 | -1.188 |
| chr9 | 113312067 | 113312286 | 0.897874 | 0.75565 | -1.188 |
| chr11 | 30218085 | 30218500 | 2.15375 | 1.81275 | -1.188 |
| chr12 | 103114677 | 103114877 | 1.16416 | 0.979861 | -1.188 |
| chr19 | 37739632 | 37739925 | 1.24512 | 1.0481 | -1.188 |
| chr2 | 17347925 | 17348268 | 0.817479 | 0.688228 | -1.188 |
| chr7 | 8717920 | 8718153 | 1.06838 | 0.899635 | -1.188 |
| chr10 | 122182929 | 122183173 | 1.79969 | 1.5155 | -1.188 |
| chr2 | 191545081 | 191545345 | 2.21153 | 1.8625 | -1.187 |
| chr1 | 235466790 | 235467027 | 0.984077 | 0.828818 | -1.187 |
| chr2 | 66814462 | 66814676 | 1.70342 | 1.43504 | -1.187 |
| chr2 | 164901113 | 164901389 | 0.781631 | 0.65849 | -1.187 |
| chr9 | 78378663 | 78379024 | 0.835561 | 0.704064 | -1.187 |
| chr16 | 55291968 | 55292329 | 1.53079 | 1.29003 | -1.187 |
| chr9 | 31918357 | 31918607 | 0.614819 | 0.518158 | -1.187 |
| chr20 | 20217874 | 20218332 | 2.0009 | 1.68687 | -1.186 |
| chr9 | 136492394 | 136492615 | 0.819594 | 0.690994 | -1.186 |
| chr15 | 94758186 | 94758386 | 0.771892 | 0.650795 | -1.186 |
| chr5 | 44725756 | 44725939 | 1.11287 | 0.938492 | -1.186 |
| chr7 | 122291358 | 122291552 | 0.789964 | 0.666189 | -1.186 |
| chrX | 118279228 | 118279509 | 2.28447 | 1.92669 | -1.186 |
| chr11 | 41864471 | 41864684 | 1.11025 | 0.936567 | -1.185 |
| chrX | 110544403 | 110544616 | 0.868008 | 0.732606 | -1.185 |
| chr10 | 29509364 | 29509627 | 1.07837 | 0.910252 | -1.185 |
| chr8 | 139250817 | 139251065 | 1.21106 | 1.02229 | -1.185 |
| chr5 | 29889231 | 29889446 | 1.14559 | 0.96703 | -1.185 |
| chr12 | 73019987 | 73020439 | 0.824276 | 0.695866 | -1.185 |
| chr2 | 14410309 | 14410514 | 0.916959 | 0.774178 | -1.184 |
| chr3 | 165544594 | 165544819 | 1.13444 | 0.957826 | -1.184 |
| chr5 | 133190432 | 133190632 | 0.9977 | 0.84239 | -1.184 |
| chr19 | 35371982 | 35372213 | 1.04047 | 0.878608 | -1.184 |
| chr11 | 14170391 | 14170723 | 1.04436 | 0.88211 | -1.184 |
| chr9 | 129983372 | 129983572 | 0.781331 | 0.659945 | -1.184 |
| chr16 | 22332470 | 22332944 | 2.96509 | 2.50445 | -1.184 |
| chr8 | 72687134 | 72687529 | 1.50481 | 1.27109 | -1.184 |
| chr10 | 50528271 | 50528533 | 1.24193 | 1.04913 | -1.184 |
| chr8 | 108767804 | 108768109 | 1.1207 | 0.946954 | -1.183 |
| chr3 | 192163445 | 192163746 | 0.900728 | 0.761123 | -1.183 |
| chr2 | 97025185 | 97025399 | 1.70364 | 1.43965 | -1.183 |
| chr19 | 15952738 | 15952960 | 1.08264 | 0.915246 | -1.183 |
| chr14 | 46706323 | 46706563 | 0.827336 | 0.699633 | -1.183 |
| chr16 | 61447092 | 61447358 | 1.14973 | 0.972269 | -1.183 |
| chr2 | 179387681 | 179387945 | 3.62738 | 3.06784 | -1.182 |
| chr10 | 130658046 | 130658379 | 1.29433 | 1.09473 | -1.182 |
| chr7 | 53724722 | 53724967 | 1.44729 | 1.22439 | -1.182 |
| chr16 | 20331703 | 20331965 | 1.20914 | 1.023 | -1.182 |
| chr2 | 220751422 | 220751608 | 1.14243 | 0.966625 | -1.182 |
| chr2 | 76652833 | 76653152 | 0.872547 | 0.73828 | -1.182 |
| chr3 | 83433334 | 83433672 | 0.862943 | 0.730232 | -1.182 |
| chr3 | 172858845 | 172859083 | 0.93937 | 0.79493 | -1.182 |
| chr2 | 3057894 | 3058096 | 0.973605 | 0.823904 | -1.182 |
| chr20 | 19250676 | 19250878 | 0.861527 | 0.729217 | -1.181 |
| chr4 | 77101054 | 77101280 | 2.3302 | 1.97249 | -1.181 |
| chr7 | 123010495 | 123010746 | 0.9504 | 0.804518 | -1.181 |
| chr2 | 77893980 | 77894163 | 0.966831 | 0.818502 | -1.181 |
| chr3 | 18628338 | 18628622 | 1.00353 | 0.849624 | -1.181 |
| chr9 | 84679708 | 84679895 | 0.749707 | 0.634791 | -1.181 |
| chr2 | 8109918 | 8110135 | 0.855912 | 0.724789 | -1.181 |
| chr2 | 203438631 | 203438914 | 2.27146 | 1.9235 | -1.181 |
| chr3 | 4039797 | 4040053 | 1.10789 | 0.938215 | -1.181 |

| | | | | | |
|---|---|---|---|---|---|
| chr12 | 23793776 | 23794077 | 0.834827 | 0.707015 | -1.181 |
| chr5 | 77806262 | 77806580 | 1.16823 | 0.989514 | -1.181 |
| chr5 | 152333030 | 152333250 | 0.961711 | 0.814658 | -1.181 |
| chr17 | 69603615 | 69603833 | 0.818417 | 0.693369 | -1.180 |
| chr2 | 79447589 | 79447778 | 1.08549 | 0.919639 | -1.180 |
| chr7 | 116958431 | 116958736 | 0.744491 | 0.630749 | -1.180 |
| chr8 | 74720621 | 74720824 | 0.957377 | 0.811415 | -1.180 |
| chr2 | 177788390 | 177788577 | 0.873929 | 0.740723 | -1.180 |
| chr19 | 52379506 | 52379751 | 0.969957 | 0.822133 | -1.180 |
| chr2 | 79922549 | 79922835 | 0.873311 | 0.740237 | -1.180 |
| chr3 | 39329852 | 39330262 | 2.95944 | 2.5088 | -1.180 |
| chr5 | 83589280 | 83589502 | 0.79436 | 0.673459 | -1.180 |
| chr10 | 67488171 | 67488477 | 0.950667 | 0.806024 | -1.179 |
| chr9 | 77804365 | 77804590 | 0.681019 | 0.577446 | -1.179 |
| chr9 | 102586913 | 102587183 | 1.20376 | 1.02071 | -1.179 |
| chr5 | 25891140 | 25891441 | 0.99269 | 0.841933 | -1.179 |
| chr2 | 127117060 | 127117285 | 1.23868 | 1.05057 | -1.179 |
| chr5 | 66092728 | 66093013 | 1.35977 | 1.15329 | -1.179 |
| chr2 | 127931885 | 127932180 | 1.70864 | 1.44938 | -1.179 |
| chr19 | 23871792 | 23872188 | 0.806953 | 0.684519 | -1.179 |
| chr19 | 33795557 | 33795792 | 1.41402 | 1.19948 | -1.179 |
| chr1 | 184175844 | 184176178 | 0.80522 | 0.683052 | -1.179 |
| chr19 | 42205548 | 42205869 | 1.97675 | 1.677 | -1.179 |
| chr11 | 42764593 | 42764778 | 0.807403 | 0.684991 | -1.179 |
| chr9 | 27888762 | 27888980 | 2.16125 | 1.83381 | -1.179 |
| chr1 | 41779455 | 41779749 | 1.0168 | 0.862767 | -1.179 |
| chr12 | 128806539 | 128806897 | 0.878739 | 0.745693 | -1.178 |
| chr12 | 60908239 | 60908468 | 0.923853 | 0.784021 | -1.178 |
| chr2 | 76430577 | 76430896 | 1.0383 | 0.881182 | -1.178 |
| chr9 | 21455418 | 21455667 | 2.03089 | 1.72362 | -1.178 |
| chr7 | 132781105 | 132781352 | 0.952026 | 0.808069 | -1.178 |
| chr8 | 138014043 | 138014381 | 1.24371 | 1.05566 | -1.178 |
| chr1 | 161454519 | 161454714 | 1.32411 | 1.12401 | -1.178 |
| chr16 | 28488864 | 28489065 | 0.807344 | 0.685373 | -1.178 |
| chr17 | 26494575 | 26494792 | 1.09968 | 0.934058 | -1.177 |
| chr12 | 60434930 | 60435112 | 1.23614 | 1.05 | -1.177 |
| chr11 | 17604750 | 17604933 | 0.837796 | 0.711652 | -1.177 |
| chr3 | 98048406 | 98048717 | 2.37254 | 2.0158 | -1.177 |
| chr15 | 87595005 | 87595260 | 0.916512 | 0.77874 | -1.177 |
| chr12 | 95319558 | 95319741 | 1.18364 | 1.00578 | -1.177 |
| chr19 | 32175952 | 32176259 | 1.22483 | 1.0408 | -1.177 |
| chr12 | 68892413 | 68892808 | 1.43087 | 1.21601 | -1.177 |
| chr9 | 18122151 | 18122338 | 0.724867 | 0.616283 | -1.176 |
| chr3 | 88748450 | 88748675 | 1.03069 | 0.876408 | -1.176 |
| chr2 | 67464479 | 67464927 | 1.1291 | 0.960127 | -1.176 |
| chr10 | 116992740 | 116992942 | 1.07519 | 0.9143 | -1.176 |
| chr1 | 238379992 | 238380174 | 0.906379 | 0.770813 | -1.176 |
| chr2 | 6479375 | 6479564 | 0.89099 | 0.757744 | -1.176 |
| chr5 | 101084074 | 101084322 | 1.31807 | 1.1212 | -1.176 |
| chr5 | 63933854 | 63934189 | 0.96499 | 0.820869 | -1.176 |
| chr14 | 83718166 | 83718386 | 0.851259 | 0.724138 | -1.176 |
| chr9 | 34809659 | 34809906 | 1.35713 | 1.15458 | -1.175 |
| chr2 | 14406508 | 14406978 | 1.10328 | 0.938636 | -1.175 |
| chr14 | 30991222 | 30991869 | 1.49814 | 1.27471 | -1.175 |
| chr8 | 111250745 | 111251000 | 1.25115 | 1.06506 | -1.175 |
| chr1 | 30705156 | 30705460 | 1.9462 | 1.65713 | -1.174 |
| chr8 | 133109555 | 133110004 | 1.11422 | 0.948832 | -1.174 |
| chr17 | 45794926 | 45795278 | 1.19478 | 1.01769 | -1.174 |
| chr15 | 98643357 | 98643776 | 0.953482 | 0.812203 | -1.174 |
| chr17 | 17292109 | 17292334 | 0.915267 | 0.779654 | -1.174 |
| chr11 | 18561747 | 18562007 | 1.29489 | 1.10321 | -1.174 |

| | | | | | |
|---|---|---|---|---|---|
| chr1 | 195131753 | 195132113 | 1.13689 | 0.968633 | -1.174 |
| chr10 | 29890380 | 29890684 | 1.04995 | 0.894573 | -1.174 |
| chrX | 144283541 | 144283770 | 1.04316 | 0.88904 | -1.173 |
| chr10 | 123201678 | 123201904 | 1.07591 | 0.917001 | -1.173 |
| chr9 | 11859132 | 11859425 | 0.924598 | 0.788398 | -1.173 |
| chr9 | 9615135 | 9615464 | 1.3371 | 1.14042 | -1.172 |
| chr12 | 82458269 | 82458519 | 0.967088 | 0.824978 | -1.172 |
| chr3 | 2680827 | 2681054 | 0.878958 | 0.749862 | -1.172 |
| chr8 | 79334604 | 79334823 | 1.0489 | 0.894929 | -1.172 |
| chr17 | 66438840 | 66439026 | 0.983425 | 0.839291 | -1.172 |
| chr2 | 219984206 | 219984522 | 1.87965 | 1.60434 | -1.172 |
| chr17 | 60694587 | 60694889 | 0.852376 | 0.727579 | -1.172 |
| chr14 | 49207582 | 49207839 | 0.744302 | 0.63545 | -1.171 |
| chr7 | 88471624 | 88471861 | 1.03736 | 0.885658 | -1.171 |
| chr9 | 90277859 | 90278142 | 3.09957 | 2.6468 | -1.171 |
| chr14 | 79084599 | 79084991 | 1.0772 | 0.919895 | -1.171 |
| chr5 | 114027338 | 114027585 | 0.838856 | 0.716412 | -1.171 |
| chr8 | 135308078 | 135308367 | 0.805685 | 0.688162 | -1.171 |
| chr2 | 649106 | 649420 | 0.730862 | 0.624267 | -1.171 |
| chr20 | 60299598 | 60299975 | 1.69443 | 1.44734 | -1.171 |
| chr3 | 74485223 | 74485586 | 0.91169 | 0.778756 | -1.171 |
| chr2 | 241553788 | 241554025 | 0.652518 | 0.557394 | -1.171 |
| chr3 | 18612787 | 18613010 | 1.04082 | 0.889563 | -1.170 |
| chr2 | 16689552 | 16689876 | 3.0208 | 2.58184 | -1.170 |
| chr3 | 155327827 | 155328045 | 0.604438 | 0.516616 | -1.170 |
| chr3 | 56591657 | 56591889 | 1.77198 | 1.51465 | -1.170 |
| chr7 | 122606055 | 122606275 | 0.807465 | 0.690295 | -1.170 |
| chr9 | 77159429 | 77159639 | 0.853913 | 0.730005 | -1.170 |
| chr3 | 138253 | 138500 | 0.990585 | 0.846866 | -1.170 |
| chr3 | 103685652 | 103685957 | 1.06522 | 0.910926 | -1.169 |
| chr20 | 37960586 | 37960877 | 0.9166 | 0.78386 | -1.169 |
| chr2 | 219137364 | 219137753 | 2.93116 | 2.50717 | -1.169 |
| chr9 | 73938516 | 73938735 | 0.957875 | 0.819331 | -1.169 |
| chr7 | 126279507 | 126279699 | 0.93728 | 0.801728 | -1.169 |
| chr4 | 42521894 | 42522103 | 1.46962 | 1.2572 | -1.169 |
| chr2 | 210846914 | 210847245 | 0.859957 | 0.735822 | -1.169 |
| chr9 | 113214614 | 113214950 | 1.31324 | 1.12387 | -1.168 |
| chr2 | 154375877 | 154376088 | 1.73577 | 1.48597 | -1.168 |
| chr1 | 244532419 | 244532616 | 0.824801 | 0.706326 | -1.168 |
| chr11 | 126064082 | 126064274 | 1.2618 | 1.08058 | -1.168 |
| chr2 | 220862027 | 220862252 | 1.66071 | 1.42257 | -1.167 |
| chr4 | 17669413 | 17669614 | 0.731695 | 0.626785 | -1.167 |
| chr12 | 29705311 | 29705607 | 2.18355 | 1.87068 | -1.167 |
| chr14 | 87764425 | 87764857 | 0.906355 | 0.776489 | -1.167 |
| chr2 | 238683366 | 238683860 | 2.01996 | 1.73059 | -1.167 |
| chr15 | 32106377 | 32106737 | 0.945044 | 0.809704 | -1.167 |
| chr7 | 21235804 | 21235990 | 1.02853 | 0.881266 | -1.167 |
| chr9 | 73539867 | 73540191 | 1.26673 | 1.08549 | -1.167 |
| chr11 | 7947752 | 7947987 | 0.917511 | 0.786333 | -1.167 |
| chr12 | 67535236 | 67535551 | 0.875991 | 0.750805 | -1.167 |
| chr11 | 75156346 | 75156683 | 2.53936 | 2.17676 | -1.167 |
| chr10 | 5530728 | 5531006 | 1.12355 | 0.963176 | -1.167 |
| chr8 | 127010043 | 127010265 | 1.24664 | 1.06875 | -1.166 |
| chrX | 150674374 | 150674602 | 1.89276 | 1.62271 | -1.166 |
| chr8 | 105708291 | 105708534 | 1.12156 | 0.96172 | -1.166 |
| chr12 | 118178818 | 118179010 | 0.866339 | 0.743002 | -1.166 |
| chr15 | 56573543 | 56573805 | 1.18773 | 1.0187 | -1.166 |
| chr11 | 15036304 | 15036546 | 0.912393 | 0.782601 | -1.166 |
| chr10 | 34323071 | 34323253 | 0.961763 | 0.825064 | -1.166 |
| chr2 | 12856164 | 12856410 | 2.31645 | 1.98728 | -1.166 |
| chr3 | 5268512 | 5268773 | 1.42389 | 1.22192 | -1.165 |

| | | | | | |
|---|---|---|---|---|---|
| chr12 | 23929688 | 23929879 | 0.912742 | 0.78334 | -1.165 |
| chr8 | 130699717 | 130699974 | 2.06115 | 1.76919 | -1.165 |
| chr2 | 163752263 | 163752459 | 0.866161 | 0.743492 | -1.165 |
| chr2 | 2427067 | 2427269 | 0.904379 | 0.776323 | -1.165 |
| chr12 | 24537166 | 24537392 | 0.895277 | 0.768774 | -1.165 |
| chr8 | 138436769 | 138437191 | 1.0643 | 0.914049 | -1.164 |
| chr8 | 2368221 | 2368443 | 1.18645 | 1.01899 | -1.164 |
| chr11 | 14991443 | 14991827 | 1.39899 | 1.20163 | -1.164 |
| chr3 | 22919901 | 22920105 | 1.03102 | 0.885613 | -1.164 |
| chr5 | 37216406 | 37216625 | 0.913339 | 0.784537 | -1.164 |
| chr5 | 15105936 | 15106274 | 2.30215 | 1.97755 | -1.164 |
| chr2 | 232428080 | 232428409 | 2.29612 | 1.97247 | -1.164 |
| chr11 | 81448489 | 81448753 | 1.01881 | 0.875227 | -1.164 |
| chr2 | 105854506 | 105854799 | 3.20244 | 2.7517 | -1.164 |
| chr3 | 80616119 | 80616309 | 0.86913 | 0.747005 | -1.163 |
| chr17 | 28348739 | 28349440 | 1.91757 | 1.64813 | -1.163 |
| chr17 | 80134391 | 80134602 | 0.769201 | 0.661289 | -1.163 |
| chrX | 132139168 | 132139359 | 1.44464 | 1.2422 | -1.163 |
| chr8 | 71921756 | 71921947 | 0.834986 | 0.718091 | -1.163 |
| chr7 | 25185911 | 25186094 | 0.9303 | 0.800164 | -1.163 |
| chr8 | 52159703 | 52159924 | 0.971976 | 0.836038 | -1.163 |
| chr16 | 71345511 | 71345815 | 2.23632 | 1.92377 | -1.162 |
| chr11 | 59269949 | 59270146 | 0.680713 | 0.585581 | -1.162 |
| chrX | 38613321 | 38613658 | 1.34317 | 1.15561 | -1.162 |
| chr17 | 14419481 | 14419691 | 1.12008 | 0.963744 | -1.162 |
| chr14 | 63561784 | 63562043 | 0.791197 | 0.680793 | -1.162 |
| chr12 | 112339012 | 112339203 | 1.05574 | 0.908582 | -1.162 |
| chr22 | 22863093 | 22863296 | 1.83452 | 1.57889 | -1.162 |
| chr1 | 110172837 | 110173177 | 1.64806 | 1.41842 | -1.162 |
| chr3 | 187427090 | 187427274 | 1.35876 | 1.16951 | -1.162 |
| chr3 | 145218814 | 145219002 | 0.954515 | 0.821621 | -1.162 |
| chr5 | 13081251 | 13081460 | 0.968364 | 0.833623 | -1.162 |
| chr17 | 72497887 | 72498124 | 1.03926 | 0.894674 | -1.162 |
| chr15 | 93584914 | 93585156 | 1.50696 | 1.29736 | -1.162 |
| chr2 | 58821298 | 58821663 | 1.48511 | 1.27872 | -1.161 |
| chr18 | 66639335 | 66639605 | 0.904945 | 0.779263 | -1.161 |
| chr20 | 15790141 | 15790466 | 1.19794 | 1.0316 | -1.161 |
| chr2 | 4752131 | 4752324 | 0.792332 | 0.682376 | -1.161 |
| chr14 | 26155861 | 26156163 | 1.01541 | 0.874501 | -1.161 |
| chr7 | 85849422 | 85849729 | 0.835704 | 0.719852 | -1.161 |
| chr3 | 163325005 | 163325281 | 1.07437 | 0.925434 | -1.161 |
| chr16 | 90051688 | 90051932 | 0.823704 | 0.709541 | -1.161 |
| chr15 | 79356677 | 79356889 | 0.901795 | 0.776825 | -1.161 |
| chr1 | 191095246 | 191095467 | 0.784774 | 0.676108 | -1.161 |
| chr2 | 78578507 | 78578950 | 0.891542 | 0.768359 | -1.160 |
| chr12 | 130118628 | 130118862 | 1.07102 | 0.923087 | -1.160 |
| chr5 | 117130845 | 117131287 | 1.10266 | 0.950506 | -1.160 |
| chr5 | 145429084 | 145429327 | 2.60848 | 2.24866 | -1.160 |
| chr12 | 82377455 | 82377664 | 0.775932 | 0.668919 | -1.160 |
| chr5 | 12321009 | 12321230 | 0.96043 | 0.828144 | -1.160 |
| chr10 | 28429087 | 28429419 | 0.829046 | 0.714938 | -1.160 |
| chr8 | 4355122 | 4355332 | 0.990653 | 0.854422 | -1.159 |
| chr8 | 3763027 | 3763235 | 1.27401 | 1.09883 | -1.159 |
| chr3 | 159816295 | 159816631 | 0.907479 | 0.782928 | -1.159 |
| chr11 | 3830398 | 3830584 | 1.52501 | 1.31577 | -1.159 |
| chr16 | 68734295 | 68734556 | 1.0171 | 0.877751 | -1.159 |
| chr2 | 199500677 | 199500859 | 0.686122 | 0.592215 | -1.159 |
| chr2 | 75280727 | 75280981 | 1.94678 | 1.68037 | -1.159 |
| chr10 | 15627334 | 15627623 | 2.35801 | 2.03621 | -1.158 |
| chr15 | 98196013 | 98196311 | 1.14169 | 0.985886 | -1.158 |
| chr4 | 118248782 | 118248989 | 0.812656 | 0.701787 | -1.158 |

| | | | | | |
|---|---|---|---|---|---|
| chr19 | 22101953 | 22102655 | 0.846795 | 0.731322 | -1.158 |
| chrX | 152846587 | 152846814 | 1.68496 | 1.45525 | -1.158 |
| chr8 | 51350005 | 51350221 | 1.11098 | 0.959527 | -1.158 |
| chr20 | 18770190 | 18770421 | 0.873638 | 0.754636 | -1.158 |
| chr2 | 232765092 | 232765573 | 3.92328 | 3.38999 | -1.157 |
| chr7 | 105229904 | 105230226 | 2.44941 | 2.1171 | -1.157 |
| chr7 | 51814280 | 51814500 | 0.848117 | 0.733139 | -1.157 |
| chr5 | 114050728 | 114050973 | 0.981276 | 0.848261 | -1.157 |
| chr2 | 73161953 | 73162345 | 1.30857 | 1.13144 | -1.157 |
| chr17 | 940942 | 941153 | 0.830347 | 0.717954 | -1.157 |
| chr14 | 39261458 | 39261667 | 0.95742 | 0.827861 | -1.156 |
| chr12 | 105243335 | 105243530 | 0.928405 | 0.802783 | -1.156 |
| chr9 | 138865913 | 138866270 | 1.02697 | 0.888093 | -1.156 |
| chr5 | 119906775 | 119907099 | 0.898771 | 0.77731 | -1.156 |
| chr5 | 61306457 | 61306639 | 0.804594 | 0.695878 | -1.156 |
| chr2 | 178628262 | 178628781 | 2.73387 | 2.36452 | -1.156 |
| chr9 | 23842770 | 23843032 | 1.06194 | 0.918545 | -1.156 |
| chr3 | 94744658 | 94744909 | 0.84332 | 0.729683 | -1.156 |
| chr14 | 68195919 | 68196109 | 1.06064 | 0.917797 | -1.156 |
| chr10 | 55188361 | 55188624 | 0.704033 | 0.609248 | -1.156 |
| chr3 | 105022144 | 105022336 | 0.922436 | 0.798283 | -1.156 |
| chr1 | 239095350 | 239095599 | 0.941578 | 0.814919 | -1.155 |
| chr2 | 190888721 | 190888929 | 1.39291 | 1.20572 | -1.155 |
| chr5 | 84465850 | 84466070 | 0.883652 | 0.764935 | -1.155 |
| chr14 | 59978753 | 59979127 | 1.62167 | 1.40385 | -1.155 |
| chr7 | 88470478 | 88470691 | 1.02553 | 0.887827 | -1.155 |
| chr3 | 80768745 | 80768992 | 1.27754 | 1.10621 | -1.155 |
| chr12 | 66123031 | 66123293 | 1.07696 | 0.932657 | -1.155 |
| chr2 | 211063918 | 211064155 | 1.83592 | 1.59003 | -1.155 |
| chr7 | 119753639 | 119753862 | 0.942834 | 0.816657 | -1.155 |
| chr7 | 145853781 | 145854006 | 1.03755 | 0.898762 | -1.154 |
| chr2 | 178847984 | 178848241 | 2.13762 | 1.85172 | -1.154 |
| chr10 | 119866086 | 119866299 | 1.34816 | 1.16795 | -1.154 |
| chr9 | 13817017 | 13817349 | 0.75184 | 0.651581 | -1.154 |
| chr7 | 145103483 | 145103673 | 1.02637 | 0.88954 | -1.154 |
| chr18 | 75362855 | 75363128 | 2.28554 | 1.98103 | -1.154 |
| chr5 | 98397180 | 98397375 | 0.759283 | 0.658161 | -1.154 |
| chr8 | 50253375 | 50253566 | 0.804935 | 0.697856 | -1.153 |
| chr2 | 18636701 | 18636989 | 1.0581 | 0.917355 | -1.153 |
| chr15 | 94866121 | 94866442 | 0.986519 | 0.855396 | -1.153 |
| chr9 | 87246161 | 87246485 | 1.06311 | 0.921864 | -1.153 |
| chr10 | 62371025 | 62371231 | 0.951377 | 0.825072 | -1.153 |
| chr3 | 112698142 | 112698369 | 1.68576 | 1.46199 | -1.153 |
| chr17 | 66160567 | 66160751 | 0.886165 | 0.7686 | -1.153 |
| chr19 | 8035661 | 8036179 | 2.69234 | 2.33528 | -1.153 |
| chr10 | 95440269 | 95440458 | 1.70458 | 1.47859 | -1.153 |
| chr8 | 141210289 | 141210576 | 1.94291 | 1.68535 | -1.153 |
| chr2 | 34355323 | 34355796 | 0.869683 | 0.75446 | -1.153 |
| chr17 | 14869862 | 14870275 | 1.03911 | 0.90159 | -1.153 |
| chr3 | 38385270 | 38385498 | 0.78113 | 0.677763 | -1.153 |
| chr7 | 113628916 | 113629103 | 1.05657 | 0.916756 | -1.153 |
| chr2 | 71953142 | 71953493 | 1.0079 | 0.87476 | -1.152 |
| chr14 | 34878682 | 34879140 | 0.900854 | 0.781942 | -1.152 |
| chr8 | 13834672 | 13834960 | 0.906296 | 0.786826 | -1.152 |
| chr16 | 17273874 | 17274090 | 0.739901 | 0.642377 | -1.152 |
| chr5 | 141191087 | 141191445 | 2.54868 | 2.21289 | -1.152 |
| chr14 | 87513139 | 87513396 | 1.05752 | 0.918252 | -1.152 |
| chr10 | 110500572 | 110500778 | 0.887046 | 0.77047 | -1.151 |
| chr10 | 56462947 | 56463135 | 0.942753 | 0.81886 | -1.151 |
| chr5 | 8908232 | 8908450 | 1.39027 | 1.20772 | -1.151 |
| chr14 | 20369593 | 20369790 | 1.20127 | 1.04366 | -1.151 |

| | | | | | |
|---|---|---|---|---|---|
| chr5 | 157936347 | 157936629 | 0.844975 | 0.734276 | -1.151 |
| chr1 | 194886847 | 194887228 | 0.866143 | 0.752863 | -1.150 |
| chr6 | 1596531 | 1596802 | 3.81167 | 3.31317 | -1.150 |
| chr8 | 88015947 | 88016170 | 1.19005 | 1.03446 | -1.150 |
| chr9 | 27134993 | 27135195 | 0.675928 | 0.587569 | -1.150 |
| chr8 | 71661809 | 71662008 | 0.911436 | 0.792493 | -1.150 |
| chr9 | 104505360 | 104505548 | 0.978806 | 0.851221 | -1.150 |
| chr20 | 59520517 | 59520704 | 1.12089 | 0.974868 | -1.150 |
| chr5 | 165973093 | 165973292 | 1.10132 | 0.958146 | -1.149 |
| chr3 | 187397475 | 187397712 | 2.58087 | 2.24546 | -1.149 |
| chr2 | 193517273 | 193517477 | 0.996175 | 0.866804 | -1.149 |
| chr3 | 146435282 | 146435478 | 0.879147 | 0.765175 | -1.149 |
| chr8 | 65480266 | 65480507 | 1.98267 | 1.72569 | -1.149 |
| chr20 | 3690851 | 3691192 | 1.16873 | 1.01744 | -1.149 |
| chr10 | 121832849 | 121833046 | 0.900296 | 0.783772 | -1.149 |
| chr2 | 121058092 | 121058360 | 0.992369 | 0.863974 | -1.149 |
| chr20 | 24219359 | 24219782 | 1.12912 | 0.983036 | -1.149 |
| chr3 | 74620349 | 74620618 | 1.2574 | 1.09472 | -1.149 |
| chr19 | 52954614 | 52954853 | 1.09938 | 0.957256 | -1.148 |
| chr14 | 49487706 | 49488005 | 0.928705 | 0.808671 | -1.148 |
| chr16 | 65594116 | 65594349 | 1.08493 | 0.944732 | -1.148 |
| chr3 | 20794157 | 20794350 | 0.830409 | 0.723147 | -1.148 |
| chr10 | 78071529 | 78071784 | 1.44687 | 1.26021 | -1.148 |
| chr1 | 215437633 | 215437854 | 0.992785 | 0.864914 | -1.148 |
| chr20 | 13800907 | 13801106 | 2.0475 | 1.78379 | -1.148 |
| chr15 | 97125863 | 97126137 | 1.13144 | 0.985846 | -1.148 |
| chr9 | 117401471 | 117401741 | 1.04775 | 0.913096 | -1.147 |
| chr11 | 83539029 | 83539384 | 1.61747 | 1.4096 | -1.147 |
| chr17 | 13017887 | 13018175 | 2.63259 | 2.295 | -1.147 |
| chr1 | 218297356 | 218297659 | 1.02907 | 0.897284 | -1.147 |
| chr5 | 112385586 | 112385931 | 0.833914 | 0.727408 | -1.146 |
| chr9 | 22892162 | 22892378 | 0.896378 | 0.781959 | -1.146 |
| chr8 | 133559052 | 133559390 | 1.08138 | 0.943387 | -1.146 |
| chr17 | 13951083 | 13951640 | 1.0561 | 0.9214 | -1.146 |
| chr5 | 12883654 | 12883895 | 1.13823 | 0.993113 | -1.146 |
| chr2 | 2048781 | 2049205 | 1.04807 | 0.91457 | -1.146 |
| chr19 | 16296652 | 16296843 | 1.25165 | 1.09275 | -1.145 |
| chr1 | 195433718 | 195433977 | 1.04936 | 0.91621 | -1.145 |
| chr10 | 124591549 | 124591736 | 1.16609 | 1.01813 | -1.145 |
| chr11 | 38828557 | 38828873 | 0.968128 | 0.845316 | -1.145 |
| chr5 | 114369514 | 114369773 | 1.0473 | 0.914796 | -1.145 |
| chr10 | 9847849 | 9848044 | 0.962683 | 0.840898 | -1.145 |
| chr9 | 1940075 | 1940359 | 1.42821 | 1.24764 | -1.145 |
| chr5 | 179636150 | 179636426 | 2.98422 | 2.60704 | -1.145 |
| chr19 | 43253870 | 43254104 | 0.716118 | 0.625719 | -1.144 |
| chr5 | 15247172 | 15247406 | 1.12863 | 0.986201 | -1.144 |
| chr12 | 119997415 | 119997606 | 1.01664 | 0.888369 | -1.144 |
| chr14 | 41328222 | 41328564 | 0.783094 | 0.6843 | -1.144 |
| chr6 | 134936088 | 134936290 | 0.751895 | 0.65707 | -1.144 |
| chr2 | 200805087 | 200805401 | 0.959318 | 0.838371 | -1.144 |
| chr9 | 121986900 | 121987362 | 1.08802 | 0.950948 | -1.144 |
| chr2 | 59995672 | 59995940 | 0.962998 | 0.841857 | -1.144 |
| chr5 | 162339604 | 162339833 | 1.17443 | 1.02683 | -1.144 |
| chr8 | 133227820 | 133228072 | 1.87353 | 1.63833 | -1.144 |
| chr8 | 137411856 | 137412142 | 0.95742 | 0.83738 | -1.143 |
| chr11 | 21446596 | 21446790 | 0.952986 | 0.833633 | -1.143 |
| chr2 | 152095412 | 152095674 | 1.81084 | 1.58461 | -1.143 |
| chr1 | 234458810 | 234459037 | 1.99762 | 1.7482 | -1.143 |
| chr10 | 74708611 | 74708902 | 1.09201 | 0.955705 | -1.143 |
| chr12 | 122199432 | 122199668 | 0.917303 | 0.802935 | -1.142 |
| chr12 | 79748623 | 79748941 | 1.00394 | 0.878895 | -1.142 |

| | | | | | |
|---|---|---|---|---|---|
| chr2 | 77505851 | 77506053 | 0.962154 | 0.842701 | -1.142 |
| chr12 | 78105331 | 78105564 | 1.21939 | 1.06816 | -1.142 |
| chr5 | 142362524 | 142362746 | 1.13636 | 0.995431 | -1.142 |
| chr16 | 9741588 | 9742253 | 1.25714 | 1.10135 | -1.141 |
| chr14 | 63786888 | 63787152 | 1.08021 | 0.946481 | -1.141 |
| chr20 | 49836686 | 49837181 | 4.20161 | 3.68178 | -1.141 |
| chr5 | 25054241 | 25054549 | 0.956663 | 0.838433 | -1.141 |
| chr2 | 164026095 | 164026315 | 1.4148 | 1.24013 | -1.141 |
| chr2 | 68589152 | 68589420 | 2.85884 | 2.50663 | -1.141 |
| chr18 | 42792757 | 42793017 | 2.05375 | 1.80089 | -1.140 |
| chr8 | 65403629 | 65403821 | 1.14412 | 1.00338 | -1.140 |
| chr14 | 47483982 | 47484181 | 0.700113 | 0.613997 | -1.140 |
| chr16 | 29853288 | 29853508 | 0.908621 | 0.796926 | -1.140 |
| chr11 | 42450215 | 42450403 | 0.894195 | 0.784317 | -1.140 |
| chr9 | 37820377 | 37820566 | 0.915474 | 0.803084 | -1.140 |
| chr2 | 142601306 | 142601507 | 0.973282 | 0.854021 | -1.140 |
| chr20 | 16131260 | 16131544 | 1.25064 | 1.0975 | -1.140 |
| chr19 | 52181186 | 52181498 | 1.05046 | 0.921895 | -1.139 |
| chr1 | 242767856 | 242768060 | 1.09722 | 0.963158 | -1.139 |
| chr7 | 29432238 | 29432525 | 0.979443 | 0.859783 | -1.139 |
| chr4 | 74312142 | 74312329 | 2.27811 | 1.99988 | -1.139 |
| chr19 | 57343754 | 57344181 | 0.900206 | 0.790373 | -1.139 |
| chr17 | 79187582 | 79187768 | 0.896318 | 0.786962 | -1.139 |
| chr11 | 31979748 | 31980077 | 1.96748 | 1.72759 | -1.139 |
| chr14 | 63765363 | 63765590 | 0.877302 | 0.770557 | -1.139 |
| chr5 | 161119048 | 161119423 | 0.903228 | 0.793437 | -1.138 |
| chrX | 38487018 | 38487232 | 1.28266 | 1.12679 | -1.138 |
| chr1 | 43609345 | 43609565 | 1.20766 | 1.06091 | -1.138 |
| chr19 | 43819991 | 43820372 | 1.04963 | 0.922085 | -1.138 |
| chr2 | 47164627 | 47165151 | 1.27638 | 1.12148 | -1.138 |
| chr2 | 78459851 | 78460066 | 1.17636 | 1.03379 | -1.138 |
| chr3 | 22003839 | 22004043 | 0.75107 | 0.660048 | -1.138 |
| chr10 | 10266519 | 10266753 | 1.25638 | 1.10419 | -1.138 |
| chr9 | 11070192 | 11070482 | 1.31644 | 1.15702 | -1.138 |
| chr2 | 52611014 | 52611231 | 0.753752 | 0.662487 | -1.138 |
| chr1 | 159798948 | 159799345 | 1.35803 | 1.19369 | -1.138 |
| chr2 | 14322603 | 14322899 | 0.851674 | 0.748643 | -1.138 |
| chr3 | 26767560 | 26767860 | 0.800254 | 0.703523 | -1.137 |
| chr8 | 75740443 | 75740709 | 1.04716 | 0.920609 | -1.137 |
| chr2 | 204349032 | 204349214 | 1.68481 | 1.48197 | -1.137 |
| chr20 | 43740601 | 43740970 | 1.61129 | 1.41747 | -1.137 |
| chr7 | 101215152 | 101215412 | 0.987316 | 0.868574 | -1.137 |
| chr14 | 36818569 | 36818988 | 0.989355 | 0.870537 | -1.136 |
| chr5 | 102367436 | 102367623 | 1.66476 | 1.46485 | -1.136 |
| chr8 | 75765641 | 75765957 | 2.14785 | 1.89038 | -1.136 |
| chr15 | 27134504 | 27134803 | 1.87016 | 1.64671 | -1.136 |
| chr8 | 105029988 | 105030295 | 1.01623 | 0.894957 | -1.136 |
| chr8 | 136571913 | 136572186 | 0.754062 | 0.664168 | -1.135 |
| chr5 | 21601471 | 21601834 | 0.913849 | 0.804929 | -1.135 |
| chr1 | 202630368 | 202630564 | 2.02296 | 1.78205 | -1.135 |
| chr18 | 30156397 | 30156599 | 1.56609 | 1.37977 | -1.135 |
| chr2 | 138093950 | 138094258 | 1.0286 | 0.90637 | -1.135 |
| chr1 | 230673197 | 230673429 | 1.01089 | 0.89079 | -1.135 |
| chr2 | 34856773 | 34857000 | 1.21425 | 1.06999 | -1.135 |
| chr18 | 13558170 | 13558484 | 1.32703 | 1.16938 | -1.135 |
| chr2 | 217908754 | 217909084 | 1.66935 | 1.47118 | -1.135 |
| chr3 | 147383742 | 147383941 | 1.12357 | 0.990495 | -1.134 |
| chr5 | 126651718 | 126651924 | 1.07646 | 0.949098 | -1.134 |
| chr5 | 102376923 | 102377157 | 1.77262 | 1.56302 | -1.134 |
| chr2 | 242017818 | 242018176 | 1.9665 | 1.73414 | -1.134 |
| chr7 | 50827547 | 50827769 | 0.828337 | 0.730561 | -1.134 |

| | | | | | |
|---|---|---|---|---|---|
| chr16 | 16975866 | 16976052 | 0.991879 | 0.874801 | -1.134 |
| chr7 | 137324170 | 137324358 | 0.895996 | 0.790367 | -1.134 |
| chr11 | 32481498 | 32481717 | 0.895798 | 0.790423 | -1.133 |
| chr19 | 45121409 | 45121695 | 1.04182 | 0.919284 | -1.133 |
| chr11 | 23529285 | 23529630 | 1.01827 | 0.898526 | -1.133 |
| chr3 | 97834722 | 97834997 | 1.09048 | 0.962266 | -1.133 |
| chr1 | 159167808 | 159168163 | 2.37428 | 2.09551 | -1.133 |
| chr2 | 40891142 | 40891346 | 1.12377 | 0.9919 | -1.133 |
| chr7 | 28666316 | 28666564 | 1.49265 | 1.31764 | -1.133 |
| chr5 | 28795268 | 28795477 | 0.809001 | 0.71432 | -1.133 |
| chr10 | 69067791 | 69068067 | 0.994937 | 0.87866 | -1.132 |
| chr19 | 52378810 | 52379122 | 0.946697 | 0.836057 | -1.132 |
| chr8 | 71859058 | 71859292 | 1.49648 | 1.32166 | -1.132 |
| chr20 | 48000888 | 48001580 | 2.30231 | 2.03337 | -1.132 |
| chr3 | 1518121 | 1518380 | 0.839863 | 0.741792 | -1.132 |
| chr3 | 111340415 | 111340714 | 1.90989 | 1.68692 | -1.132 |
| chr7 | 104345237 | 104345420 | 1.01252 | 0.894481 | -1.132 |
| chr3 | 4833032 | 4833278 | 1.01459 | 0.896467 | -1.132 |
| chr16 | 81933951 | 81934268 | 0.827599 | 0.731261 | -1.132 |
| chr7 | 104284112 | 104284450 | 1.40522 | 1.24166 | -1.132 |
| chr20 | 18340751 | 18341014 | 1.19898 | 1.05972 | -1.131 |
| chr11 | 61596288 | 61596591 | 1.92381 | 1.70049 | -1.131 |
| chr2 | 50718689 | 50719103 | 1.92615 | 1.70262 | -1.131 |
| chr5 | 16585459 | 16585794 | 2.56542 | 2.26782 | -1.131 |
| chr11 | 32253482 | 32253771 | 2.29672 | 2.03086 | -1.131 |
| chr8 | 72700968 | 72701177 | 1.15138 | 1.01835 | -1.131 |
| chr17 | 12246031 | 12246410 | 1.28409 | 1.13584 | -1.131 |
| chr20 | 51079951 | 51080228 | 0.795714 | 0.70398 | -1.130 |
| chr10 | 64466446 | 64466647 | 0.867981 | 0.767972 | -1.130 |
| chr3 | 1482269 | 1482534 | 0.895408 | 0.792323 | -1.130 |
| chr11 | 37916258 | 37916443 | 1.12738 | 0.997775 | -1.130 |
| chr10 | 77611541 | 77611821 | 0.955032 | 0.845587 | -1.129 |
| chr11 | 122376534 | 122376759 | 2.08207 | 1.8437 | -1.129 |
| chrX | 23634559 | 23634753 | 2.1912 | 1.94036 | -1.129 |
| chr6 | 100729523 | 100729708 | 1.01982 | 0.903168 | -1.129 |
| chr1 | 203671262 | 203671537 | 2.00804 | 1.77849 | -1.129 |
| chr5 | 135683438 | 135683654 | 1.46996 | 1.30199 | -1.129 |
| chr9 | 12071955 | 12072252 | 1.02567 | 0.908499 | -1.129 |
| chr9 | 11458264 | 11458618 | 0.93739 | 0.830305 | -1.129 |
| chr5 | 9156615 | 9156835 | 0.970772 | 0.860442 | -1.128 |
| chr15 | 60843946 | 60844178 | 1.2863 | 1.1403 | -1.128 |
| chr2 | 53469228 | 53469527 | 1.12494 | 0.997308 | -1.128 |
| chr7 | 7403488 | 7403739 | 0.89251 | 0.791273 | -1.128 |
| chr2 | 61050831 | 61051045 | 0.698026 | 0.618884 | -1.128 |
| chr10 | 56847999 | 56848259 | 0.813693 | 0.721467 | -1.128 |
| chr8 | 64097455 | 64097644 | 0.76909 | 0.682079 | -1.128 |
| chr7 | 62315672 | 62315920 | 0.999503 | 0.886502 | -1.127 |
| chr20 | 55622432 | 55622621 | 1.84009 | 1.63216 | -1.127 |
| chr2 | 18586054 | 18586278 | 1.42561 | 1.26487 | -1.127 |
| chr20 | 20314563 | 20314791 | 1.03137 | 0.915241 | -1.127 |
| chr3 | 155083216 | 155083399 | 1.06526 | 0.945491 | -1.127 |
| chr5 | 147238210 | 147238440 | 0.898782 | 0.797751 | -1.127 |
| chr10 | 111170622 | 111170804 | 0.943682 | 0.837658 | -1.127 |
| chr12 | 4827015 | 4827367 | 0.960523 | 0.852993 | -1.126 |
| chr8 | 60610714 | 60610969 | 1.07177 | 0.952124 | -1.126 |
| chr9 | 90569028 | 90569300 | 0.870953 | 0.773768 | -1.126 |
| chr14 | 29725947 | 29726200 | 0.962654 | 0.855351 | -1.125 |
| chr5 | 157951699 | 157951947 | 0.966616 | 0.858891 | -1.125 |
| chr5 | 33530445 | 33530693 | 0.839997 | 0.746403 | -1.125 |
| chr5 | 94352327 | 94352520 | 1.05445 | 0.937122 | -1.125 |
| chr8 | 135237754 | 135237953 | 0.996955 | 0.88608 | -1.125 |

| | | | | | |
|---|---|---|---|---|---|
| chr8 | 14535010 | 14535253 | 0.9599 | 0.853212 | -1.125 |
| chr10 | 15442603 | 15442832 | 0.805349 | 0.715866 | -1.125 |
| chr9 | 130108905 | 130109454 | 1.26628 | 1.12567 | -1.125 |
| chr3 | 102383128 | 102383328 | 0.651466 | 0.579164 | -1.125 |
| chr8 | 138191684 | 138191877 | 0.975888 | 0.867657 | -1.125 |
| chr1 | 40544245 | 40544572 | 2.06629 | 1.83752 | -1.124 |
| chr1 | 178269118 | 178269391 | 2.72646 | 2.4251 | -1.124 |
| chr4 | 103352592 | 103352869 | 2.01748 | 1.79466 | -1.124 |
| chr14 | 100984897 | 100985118 | 0.754598 | 0.671344 | -1.124 |
| chr17 | 26753234 | 26753532 | 1.65325 | 1.47097 | -1.124 |
| chr3 | 47292205 | 47292487 | 1.3917 | 1.23836 | -1.124 |
| chr2 | 10407832 | 10408058 | 1.05431 | 0.938191 | -1.124 |
| chr1 | 166651258 | 166651462 | 1.10359 | 0.982223 | -1.124 |
| chr19 | 28733360 | 28733631 | 1.71627 | 1.52779 | -1.123 |
| chr17 | 71837792 | 71838179 | 3.89313 | 3.4659 | -1.123 |
| chr2 | 77109994 | 77110234 | 1.01771 | 0.906052 | -1.123 |
| chr1 | 168424645 | 168424829 | 0.763549 | 0.679885 | -1.123 |
| chr12 | 128195586 | 128195920 | 1.08739 | 0.968372 | -1.123 |
| chr14 | 41114430 | 41114730 | 1.25443 | 1.1173 | -1.123 |
| chr3 | 95742510 | 95742873 | 0.855878 | 0.762371 | -1.123 |
| chr8 | 28176171 | 28176416 | 2.25395 | 2.00772 | -1.123 |
| chr9 | 124225363 | 124225713 | 2.08383 | 1.85623 | -1.123 |
| chr19 | 28742641 | 28742883 | 1.16191 | 1.03504 | -1.123 |
| chr7 | 18855184 | 18855485 | 0.962654 | 0.857696 | -1.122 |
| chr5 | 161555287 | 161555558 | 1.25036 | 1.11412 | -1.122 |
| chr1 | 165897440 | 165897957 | 1.50907 | 1.34482 | -1.122 |
| chr9 | 8592357 | 8592579 | 1.00868 | 0.898939 | -1.122 |
| chr10 | 68932463 | 68932698 | 0.797586 | 0.710908 | -1.122 |
| chr20 | 15735022 | 15735209 | 0.980268 | 0.873776 | -1.122 |
| chr11 | 44478254 | 44478772 | 1.66228 | 1.48199 | -1.122 |
| chr10 | 9402913 | 9403127 | 0.882305 | 0.786712 | -1.122 |
| chr19 | 36445934 | 36446128 | 1.00263 | 0.894043 | -1.121 |
| chr10 | 5748228 | 5748442 | 0.943424 | 0.841413 | -1.121 |
| chr9 | 11763388 | 11763611 | 0.978217 | 0.872531 | -1.121 |
| chr12 | 9287894 | 9288196 | 1.44709 | 1.29077 | -1.121 |
| chr1 | 195091717 | 195091972 | 0.811594 | 0.724017 | -1.121 |
| chr10 | 87703647 | 87703871 | 1.04953 | 0.936444 | -1.121 |
| chr6 | 43037856 | 43038061 | 1.80279 | 1.60885 | -1.121 |
| chr1 | 154318518 | 154319067 | 0.884064 | 0.788963 | -1.121 |
| chr2 | 23445316 | 23445611 | 1.09546 | 0.97782 | -1.120 |
| chr8 | 81756580 | 81756783 | 0.851017 | 0.759738 | -1.120 |
| chr11 | 22675979 | 22676181 | 0.782667 | 0.698769 | -1.120 |
| chr2 | 195732135 | 195732325 | 1.24048 | 1.10752 | -1.120 |
| chr21 | 35720217 | 35720492 | 1.97337 | 1.76189 | -1.120 |
| chr3 | 21888099 | 21888341 | 1.1828 | 1.05617 | -1.120 |
| chr3 | 32865315 | 32865530 | 0.875186 | 0.781593 | -1.120 |
| chr4 | 42867655 | 42867946 | 1.27045 | 1.13474 | -1.120 |
| chr3 | 143654461 | 143654667 | 0.754717 | 0.674108 | -1.120 |
| chr5 | 77361646 | 77361911 | 1.23066 | 1.09923 | -1.120 |
| chr5 | 8904630 | 8904863 | 1.10838 | 0.990136 | -1.119 |
| chr8 | 78786276 | 78786477 | 1.06259 | 0.949266 | -1.119 |
| chr14 | 78630664 | 78630901 | 1.97807 | 1.76764 | -1.119 |
| chr15 | 97942296 | 97942661 | 1.16563 | 1.04185 | -1.119 |
| chr8 | 49065062 | 49065297 | 0.932683 | 0.833727 | -1.119 |
| chr8 | 104612466 | 104612703 | 0.666003 | 0.595351 | -1.119 |
| chr1 | 214502919 | 214503143 | 0.966221 | 0.863888 | -1.118 |
| chr15 | 102399865 | 102400147 | 0.927634 | 0.829422 | -1.118 |
| chr2 | 230645545 | 230645892 | 2.03385 | 1.81855 | -1.118 |
| chr19 | 31191536 | 31191813 | 0.953132 | 0.852296 | -1.118 |
| chr7 | 83320418 | 83320675 | 1.13528 | 1.01527 | -1.118 |
| chr2 | 51401773 | 51402128 | 1.28253 | 1.14698 | -1.118 |

| | | | | | |
|---|---|---|---|---|---|
| chr2 | 185778767 | 185778972 | 0.977194 | 0.87396 | -1.118 |
| chr19 | 32122175 | 32122896 | 1.35912 | 1.21563 | -1.118 |
| chr2 | 76521034 | 76521390 | 0.875025 | 0.782691 | -1.118 |
| chr20 | 19186566 | 19187003 | 3.16733 | 2.83392 | -1.118 |
| chr8 | 115770054 | 115770294 | 1.04784 | 0.937565 | -1.118 |
| chr12 | 62576158 | 62576400 | 0.916548 | 0.820092 | -1.118 |
| chr5 | 141365217 | 141365426 | 0.794696 | 0.711161 | -1.117 |
| chr10 | 117194786 | 117195036 | 1.29303 | 1.1572 | -1.117 |
| chr19 | 32622152 | 32622370 | 1.18985 | 1.06516 | -1.117 |
| chr7 | 9877854 | 9878088 | 0.925484 | 0.828873 | -1.117 |
| chr9 | 128736000 | 128736187 | 1.0987 | 0.984026 | -1.117 |
| chr16 | 34358138 | 34358325 | 1.17591 | 1.05341 | -1.116 |
| chr2 | 204261538 | 204261747 | 1.43823 | 1.28856 | -1.116 |
| chr3 | 116549092 | 116549342 | 1.13761 | 1.01924 | -1.116 |
| chr19 | 32626078 | 32626309 | 1.11223 | 0.996518 | -1.116 |
| chr16 | 11480830 | 11481146 | 2.22124 | 1.99023 | -1.116 |
| chr7 | 85987746 | 85987937 | 0.886489 | 0.794347 | -1.116 |
| chr3 | 154651949 | 154652149 | 1.1081 | 0.993063 | -1.116 |
| chr11 | 58975402 | 58975758 | 1.89595 | 1.69929 | -1.116 |
| chr15 | 91141481 | 91141669 | 1.05103 | 0.942084 | -1.116 |
| chr14 | 46035819 | 46036086 | 0.776758 | 0.696282 | -1.116 |
| chr2 | 74917116 | 74917349 | 1.01208 | 0.907233 | -1.116 |
| chr7 | 125599286 | 125599552 | 0.913952 | 0.819388 | -1.115 |
| chr8 | 49615762 | 49615982 | 1.09364 | 0.980551 | -1.115 |
| chr5 | 136915774 | 136915999 | 1.39584 | 1.2516 | -1.115 |
| chr3 | 81048536 | 81048741 | 0.966926 | 0.867051 | -1.115 |
| chr2 | 74426127 | 74426458 | 2.1817 | 1.9564 | -1.115 |
| chr5 | 105061591 | 105061967 | 0.85421 | 0.766086 | -1.115 |
| chr5 | 22294420 | 22294693 | 0.852391 | 0.764649 | -1.115 |
| chr19 | 29868753 | 29868949 | 0.911431 | 0.81762 | -1.115 |
| chr2 | 37515240 | 37515465 | 1.16126 | 1.04174 | -1.115 |
| chr8 | 144940449 | 144940710 | 1.06103 | 0.951858 | -1.115 |
| chr9 | 10280525 | 10280737 | 0.679843 | 0.610143 | -1.114 |
| chr1 | 209686689 | 209687031 | 1.49594 | 1.34285 | -1.114 |
| chr8 | 94373378 | 94373681 | 2.21218 | 1.98584 | -1.114 |
| chr8 | 23599913 | 23600232 | 1.12849 | 1.01307 | -1.114 |
| chrX | 88276113 | 88276317 | 1.19853 | 1.07619 | -1.114 |
| chr2 | 135480407 | 135480621 | 1.49536 | 1.34273 | -1.114 |
| chr12 | 41649525 | 41649786 | 1.15222 | 1.03462 | -1.114 |
| chr6 | 98719965 | 98720191 | 0.801993 | 0.720202 | -1.114 |
| chr14 | 60099053 | 60099372 | 1.55644 | 1.39791 | -1.113 |
| chr17 | 80788228 | 80788455 | 0.986566 | 0.886271 | -1.113 |
| chr1 | 225629324 | 225629509 | 2.04292 | 1.83524 | -1.113 |
| chr20 | 6538996 | 6539251 | 1.08605 | 0.975717 | -1.113 |
| chr17 | 13661349 | 13661673 | 0.839404 | 0.754172 | -1.113 |
| chr7 | 53759576 | 53759770 | 0.857308 | 0.770349 | -1.113 |
| chr10 | 65861386 | 65861573 | 0.757612 | 0.680782 | -1.113 |
| chr17 | 55710641 | 55710836 | 1.26558 | 1.13744 | -1.113 |
| chr15 | 71777469 | 71777787 | 2.40311 | 2.15994 | -1.113 |
| chr8 | 65814434 | 65814634 | 0.95135 | 0.855246 | -1.112 |
| chr11 | 68351230 | 68351924 | 2.04221 | 1.83602 | -1.112 |
| chr8 | 68751872 | 68752105 | 1.02151 | 0.918456 | -1.112 |
| chr14 | 44811130 | 44811345 | 0.812596 | 0.730622 | -1.112 |
| chr2 | 216693374 | 216693570 | 2.08573 | 1.8756 | -1.112 |
| chr2 | 171265597 | 171265925 | 1.88016 | 1.69082 | -1.112 |
| chr17 | 11160755 | 11161010 | 0.9464 | 0.851116 | -1.112 |
| chr14 | 94518356 | 94518565 | 1.15787 | 1.0413 | -1.112 |
| chr8 | 89267949 | 89268193 | 1.03424 | 0.930144 | -1.112 |
| chr8 | 135643801 | 135644092 | 1.61531 | 1.45276 | -1.112 |
| chr8 | 133664283 | 133664646 | 0.936465 | 0.842291 | -1.112 |
| chr2 | 12939253 | 12939664 | 1.07522 | 0.967327 | -1.112 |

| | | | | | |
|---|---|---|---|---|---|
| chr17 | 36151337 | 36151664 | 2.06458 | 1.85808 | -1.111 |
| chr2 | 233626109 | 233626361 | 1.89442 | 1.70508 | -1.111 |
| chr19 | 32139259 | 32139441 | 1.18257 | 1.06441 | -1.111 |
| chr14 | 95540138 | 95540365 | 0.736972 | 0.663344 | -1.111 |
| chr2 | 220386050 | 220386287 | 2.02387 | 1.82186 | -1.111 |
| chr8 | 112109939 | 112110151 | 0.857106 | 0.771568 | -1.111 |
| chr19 | 58993838 | 58994104 | 0.952607 | 0.857903 | -1.110 |
| chr9 | 103913820 | 103914043 | 0.686036 | 0.617903 | -1.110 |
| chr17 | 78747415 | 78747773 | 1.20335 | 1.08407 | -1.110 |
| chr10 | 55182685 | 55182879 | 0.918645 | 0.827655 | -1.110 |
| chr7 | 50359332 | 50359565 | 0.886386 | 0.798668 | -1.110 |
| chr2 | 148747446 | 148747643 | 1.17513 | 1.05904 | -1.110 |
| chr15 | 31347645 | 31347849 | 1.11403 | 1.00414 | -1.109 |
| chr16 | 79495357 | 79495583 | 1.00403 | 0.905044 | -1.109 |
| chr7 | 104533513 | 104533696 | 0.986087 | 0.889137 | -1.109 |
| chr2 | 1900840 | 1901218 | 1.55167 | 1.39913 | -1.109 |
| chr1 | 49142945 | 49143414 | 1.87575 | 1.69154 | -1.109 |
| chr19 | 40138834 | 40139047 | 1.10085 | 0.993051 | -1.109 |
| chr12 | 81026696 | 81026878 | 1.22815 | 1.10789 | -1.109 |
| chr3 | 137787429 | 137787766 | 2.91211 | 2.62712 | -1.108 |
| chr20 | 20056779 | 20057030 | 1.1065 | 0.998314 | -1.108 |
| chr16 | 89838383 | 89838614 | 1.05698 | 0.953721 | -1.108 |
| chr16 | 83276714 | 83276919 | 0.974086 | 0.87899 | -1.108 |
| chr17 | 30185844 | 30186073 | 0.972299 | 0.877443 | -1.108 |
| chr7 | 121892189 | 121892422 | 0.896782 | 0.809325 | -1.108 |
| chr10 | 2724166 | 2724438 | 1.12346 | 1.01391 | -1.108 |
| chr11 | 72394530 | 72395215 | 2.33632 | 2.1086 | -1.108 |
| chr12 | 123594737 | 123595004 | 0.855186 | 0.771852 | -1.108 |
| chr7 | 80362113 | 80362297 | 1.18123 | 1.06638 | -1.108 |
| chr10 | 44309433 | 44309729 | 0.981818 | 0.886368 | -1.108 |
| chr9 | 24027770 | 24028060 | 0.916757 | 0.827842 | -1.107 |
| chr5 | 19284091 | 19284344 | 0.724013 | 0.653838 | -1.107 |
| chr15 | 94270975 | 94271344 | 1.02784 | 0.928224 | -1.107 |
| chr2 | 59582246 | 59582440 | 1.17796 | 1.06388 | -1.107 |
| chr19 | 31095638 | 31095869 | 1.0049 | 0.907668 | -1.107 |
| chr19 | 49791936 | 49792151 | 0.802639 | 0.725054 | -1.107 |
| chr8 | 138968369 | 138968589 | 0.900442 | 0.813831 | -1.106 |
| chr2 | 167667903 | 167668107 | 0.783067 | 0.70776 | -1.106 |
| chr2 | 84615017 | 84615221 | 0.973814 | 0.880183 | -1.106 |
| chr12 | 117135124 | 117135398 | 1.58644 | 1.43425 | -1.106 |
| chr15 | 84308133 | 84308921 | 1.26677 | 1.14562 | -1.106 |
| chr16 | 53576260 | 53576491 | 1.00877 | 0.912395 | -1.106 |
| chr5 | 111702628 | 111703344 | 1.26531 | 1.14454 | -1.106 |
| chr11 | 30164628 | 30165049 | 1.89332 | 1.71296 | -1.105 |
| chr14 | 96468307 | 96468531 | 0.763794 | 0.691364 | -1.105 |
| chr3 | 7327332 | 7327521 | 0.985494 | 0.892113 | -1.105 |
| chr3 | 160883779 | 160883991 | 0.989892 | 0.896149 | -1.105 |
| chr19 | 28602896 | 28603098 | 1.09459 | 0.99102 | -1.105 |
| chr1 | 118619891 | 118620188 | 1.01549 | 0.919585 | -1.104 |
| chr1 | 159977401 | 159977720 | 1.91917 | 1.73812 | -1.104 |
| chr11 | 82575325 | 82575548 | 2.0196 | 1.82908 | -1.104 |
| chr8 | 135206114 | 135206325 | 0.985966 | 0.893004 | -1.104 |
| chr17 | 58883311 | 58883557 | 0.882035 | 0.7989 | -1.104 |
| chr2 | 20377557 | 20377768 | 1.35342 | 1.22595 | -1.104 |
| chr19 | 42801273 | 42801464 | 1.13389 | 1.0272 | -1.104 |
| chr15 | 30289979 | 30290188 | 1.29003 | 1.16873 | -1.104 |
| chr10 | 30614369 | 30614582 | 1.12115 | 1.01573 | -1.104 |
| chr11 | 134077373 | 134077698 | 1.60101 | 1.4506 | -1.104 |
| chr18 | 44789663 | 44789981 | 1.87785 | 1.70169 | -1.104 |
| chr10 | 14438749 | 14438978 | 1.10428 | 1.00078 | -1.103 |
| chr14 | 96763367 | 96763662 | 1.94477 | 1.76283 | -1.103 |

| | | | | | |
|---|---|---|---|---|---|
| chr10 | 17418036 | 17418254 | 2.09824 | 1.902 | -1.103 |
| chr7 | 146967639 | 146967845 | 0.905668 | 0.821171 | -1.103 |
| chr15 | 62010092 | 62010397 | 0.869006 | 0.788167 | -1.103 |
| chr2 | 113842017 | 113842336 | 1.16532 | 1.05723 | -1.102 |
| chr9 | 80246971 | 80247261 | 0.764883 | 0.693952 | -1.102 |
| chr15 | 79476679 | 79477099 | 3.44547 | 3.12641 | -1.102 |
| chr7 | 122649385 | 122649572 | 1.15846 | 1.05131 | -1.102 |
| chr11 | 43440995 | 43441227 | 2.23498 | 2.0284 | -1.102 |
| chr9 | 82359369 | 82359587 | 1.13935 | 1.03411 | -1.102 |
| chr5 | 11275404 | 11275614 | 0.798032 | 0.724382 | -1.102 |
| chr21 | 44882901 | 44883158 | 1.53166 | 1.39035 | -1.102 |
| chr9 | 11817329 | 11817600 | 0.962985 | 0.874163 | -1.102 |
| chr22 | 28282595 | 28282814 | 1.81113 | 1.64465 | -1.101 |
| chr15 | 24331251 | 24331846 | 0.910323 | 0.826727 | -1.101 |
| chr10 | 12697087 | 12697286 | 0.877341 | 0.796779 | -1.101 |
| chr14 | 22471019 | 22471329 | 1.7766 | 1.61388 | -1.101 |
| chr7 | 103407195 | 103407572 | 1.13156 | 1.02793 | -1.101 |
| chr17 | 79204268 | 79204572 | 2.76669 | 2.51357 | -1.101 |
| chr10 | 112821017 | 112821325 | 1.00785 | 0.915854 | -1.100 |
| chr1 | 206321855 | 206322112 | 1.05725 | 0.960843 | -1.100 |
| chr16 | 24057030 | 24057345 | 1.51013 | 1.37282 | -1.100 |
| chr2 | 233006754 | 233007078 | 1.77033 | 1.6097 | -1.100 |
| chr15 | 84506749 | 84507037 | 2.11005 | 1.91928 | -1.099 |
| chr19 | 30018308 | 30018493 | 1.51744 | 1.3804 | -1.099 |
| chr19 | 29181453 | 29181744 | 1.00739 | 0.916585 | -1.099 |
| chr7 | 102551322 | 102551541 | 0.739413 | 0.67278 | -1.099 |
| chr16 | 65048405 | 65048629 | 0.862727 | 0.784993 | -1.099 |
| chr5 | 117332751 | 117333032 | 1.10309 | 1.00417 | -1.099 |
| chr5 | 97178202 | 97178393 | 0.976415 | 0.888993 | -1.098 |
| chr13 | 85733522 | 85733828 | 0.863287 | 0.786086 | -1.098 |
| chr16 | 70507025 | 70507262 | 0.693278 | 0.631563 | -1.098 |
| chr11 | 42270154 | 42270410 | 0.839964 | 0.765204 | -1.098 |
| chr16 | 26381923 | 26382202 | 0.82355 | 0.750347 | -1.098 |
| chr5 | 41968048 | 41968629 | 0.889052 | 0.810119 | -1.097 |
| chr17 | 50399229 | 50399422 | 0.773662 | 0.705 | -1.097 |
| chr5 | 50353177 | 50353638 | 1.013 | 0.923397 | -1.097 |
| chr13 | 50181293 | 50181503 | 1.81402 | 1.65439 | -1.096 |
| chr3 | 122739982 | 122740185 | 0.993199 | 0.905982 | -1.096 |
| chr9 | 795974 | 796213 | 1.10813 | 1.01088 | -1.096 |
| chr7 | 106535820 | 106536025 | 0.858666 | 0.783327 | -1.096 |
| chr8 | 140964816 | 140965077 | 1.30894 | 1.19436 | -1.096 |
| chr14 | 26512003 | 26512277 | 1.16101 | 1.05942 | -1.096 |
| chr3 | 30109269 | 30109499 | 1.13561 | 1.03626 | -1.096 |
| chr8 | 144211664 | 144211948 | 0.787715 | 0.718862 | -1.096 |
| chr5 | 71540816 | 71541040 | 1.58349 | 1.44513 | -1.096 |
| chr7 | 34536765 | 34537078 | 1.51789 | 1.38531 | -1.096 |
| chr10 | 107769810 | 107770092 | 1.05749 | 0.965167 | -1.096 |
| chr2 | 84925572 | 84925848 | 1.01219 | 0.924064 | -1.095 |
| chr2 | 142923418 | 142923721 | 1.07051 | 0.977343 | -1.095 |
| chr19 | 31253451 | 31253695 | 1.04766 | 0.956565 | -1.095 |
| chr15 | 28346284 | 28346522 | 0.960821 | 0.877381 | -1.095 |
| chr5 | 22910602 | 22910808 | 1.30332 | 1.19018 | -1.095 |
| chr10 | 72040891 | 72041094 | 1.09517 | 1.00034 | -1.095 |
| chr8 | 114143518 | 114143744 | 0.951623 | 0.869251 | -1.095 |
| chr5 | 132572497 | 132572702 | 1.17384 | 1.07236 | -1.095 |
| chr1 | 238721746 | 238721932 | 1.05413 | 0.963131 | -1.094 |
| chr14 | 82770035 | 82770254 | 0.628993 | 0.574709 | -1.094 |
| chr19 | 40320484 | 40320724 | 1.2784 | 1.16823 | -1.094 |
| chr20 | 14580275 | 14580539 | 1.2778 | 1.16779 | -1.094 |
| chr17 | 32649932 | 32650128 | 0.84359 | 0.771053 | -1.094 |
| chr2 | 166143641 | 166143825 | 0.917074 | 0.838231 | -1.094 |

| | | | | | |
|---|---|---|---|---|---|
| chr17 | 62759461 | 62759781 | 1.06898 | 0.977153 | -1.094 |
| chr16 | 86724731 | 86724942 | 1.02108 | 0.933385 | -1.094 |
| chr3 | 19888807 | 19889008 | 1.12705 | 1.03031 | -1.094 |
| chr14 | 87521291 | 87521537 | 0.872215 | 0.797379 | -1.094 |
| chr5 | 15596833 | 15597032 | 1.13454 | 1.03766 | -1.093 |
| chr11 | 69376359 | 69376554 | 0.835727 | 0.764367 | -1.093 |
| chr11 | 42181631 | 42181879 | 1.06045 | 0.969937 | -1.093 |
| chr7 | 120365560 | 120366047 | 1.49745 | 1.36967 | -1.093 |
| chr15 | 98703383 | 98703601 | 1.04101 | 0.952385 | -1.093 |
| chr8 | 32139868 | 32140136 | 1.06627 | 0.975603 | -1.093 |
| chr20 | 48150864 | 48151083 | 2.45468 | 2.24608 | -1.093 |
| chr9 | 126069285 | 126069511 | 0.998192 | 0.91361 | -1.093 |
| chr15 | 94687764 | 94688069 | 1.06521 | 0.975034 | -1.092 |
| chr10 | 84626292 | 84626483 | 0.836751 | 0.766051 | -1.092 |
| chr8 | 6232189 | 6232408 | 1.06225 | 0.972554 | -1.092 |
| chr5 | 9183330 | 9183701 | 1.02765 | 0.941021 | -1.092 |
| chr20 | 22075319 | 22075563 | 1.19192 | 1.0915 | -1.092 |
| chr10 | 105648113 | 105648418 | 2.05308 | 1.88017 | -1.092 |
| chr8 | 109625444 | 109625642 | 0.994326 | 0.91069 | -1.092 |
| chr14 | 86619853 | 86620180 | 1.08227 | 0.991295 | -1.092 |
| chr1 | 45037622 | 45038171 | 1.21548 | 1.11331 | -1.092 |
| chr15 | 62499675 | 62499879 | 1.07695 | 0.986502 | -1.092 |
| chr7 | 158528652 | 158528834 | 1.05096 | 0.963009 | -1.091 |
| chr5 | 54014310 | 54014577 | 0.899297 | 0.824201 | -1.091 |
| chr7 | 22589761 | 22590259 | 2.16854 | 1.9878 | -1.091 |
| chr12 | 43423004 | 43423264 | 1.21041 | 1.10954 | -1.091 |
| chr8 | 134283390 | 134283603 | 1.09354 | 1.00243 | -1.091 |
| chr2 | 77413284 | 77413494 | 1.03411 | 0.94806 | -1.091 |
| chr12 | 127215683 | 127215967 | 0.918504 | 0.842092 | -1.091 |
| chr16 | 2107276 | 2107515 | 1.27981 | 1.17335 | -1.091 |
| chr10 | 27257186 | 27257460 | 1.20317 | 1.10315 | -1.091 |
| chrX | 30078962 | 30079152 | 1.58846 | 1.45653 | -1.091 |
| chr7 | 149508483 | 149508734 | 1.41914 | 1.3014 | -1.090 |
| chr10 | 134172964 | 134173276 | 0.703266 | 0.644953 | -1.090 |
| chr12 | 5282886 | 5283262 | 0.990264 | 0.908289 | -1.090 |
| chr9 | 136398333 | 136398563 | 1.44146 | 1.32218 | -1.090 |
| chr5 | 153089364 | 153089550 | 0.878735 | 0.806088 | -1.090 |
| chr15 | 28436981 | 28437195 | 1.07455 | 0.985758 | -1.090 |
| chr1 | 209609104 | 209609346 | 2.34355 | 2.15007 | -1.090 |
| chr3 | 114983050 | 114983276 | 0.838011 | 0.768854 | -1.090 |
| chr2 | 219835732 | 219836121 | 0.934112 | 0.857112 | -1.090 |
| chr2 | 152855364 | 152855913 | 2.41206 | 2.21342 | -1.090 |
| chr8 | 133574225 | 133574505 | 0.935027 | 0.858046 | -1.090 |
| chr20 | 21231931 | 21232130 | 1.21996 | 1.11973 | -1.090 |
| chr1 | 174967711 | 174967950 | 2.08061 | 1.90986 | -1.089 |
| chr14 | 74768764 | 74769014 | 1.33657 | 1.22696 | -1.089 |
| chr1 | 235693935 | 235694145 | 1.02109 | 0.937369 | -1.089 |
| chr15 | 29579197 | 29579471 | 1.12255 | 1.0307 | -1.089 |
| chr7 | 37642903 | 37643231 | 0.925863 | 0.850214 | -1.089 |
| chr2 | 194845369 | 194845552 | 0.842931 | 0.774088 | -1.089 |
| chr3 | 144178985 | 144179333 | 0.804668 | 0.739011 | -1.089 |
| chr9 | 74331147 | 74331368 | 1.57909 | 1.45027 | -1.089 |
| chr17 | 54362045 | 54362236 | 0.949397 | 0.871978 | -1.089 |
| chr5 | 65623058 | 65623243 | 1.09331 | 1.00434 | -1.089 |
| chr2 | 78035843 | 78036159 | 1.01835 | 0.935837 | -1.088 |
| chr6 | 87849127 | 87849315 | 1.23167 | 1.13207 | -1.088 |
| chr15 | 73373747 | 73373983 | 1.7635 | 1.6209 | -1.088 |
| chr8 | 98748672 | 98748860 | 2.52048 | 2.31684 | -1.088 |
| chrX | 130128760 | 130129108 | 1.32641 | 1.21947 | -1.088 |
| chr11 | 81958310 | 81958526 | 0.969898 | 0.891749 | -1.088 |
| chr6 | 143234897 | 143235104 | 0.829637 | 0.762816 | -1.088 |

| | | | | | |
|---|---|---|---|---|---|
| chr15 | 62424810 | 62425085 | 1.14953 | 1.05695 | -1.088 |
| chr11 | 7374321 | 7374575 | 2.00397 | 1.84268 | -1.088 |
| chr15 | 98018837 | 98019024 | 1.10291 | 1.0143 | -1.087 |
| chr3 | 79845306 | 79845630 | 1.12159 | 1.0315 | -1.087 |
| chr2 | 77085255 | 77085510 | 0.852952 | 0.784474 | -1.087 |
| chr17 | 5101775 | 5101973 | 0.839294 | 0.77193 | -1.087 |
| chr12 | 13357448 | 13357648 | 0.865511 | 0.796129 | -1.087 |
| chr10 | 67691626 | 67691835 | 0.755123 | 0.694631 | -1.087 |
| chr19 | 42783930 | 42784247 | 3.21906 | 2.96129 | -1.087 |
| chr16 | 85709211 | 85709509 | 1.54542 | 1.42239 | -1.086 |
| chr2 | 206812852 | 206813093 | 2.52656 | 2.32545 | -1.086 |
| chr10 | 98670379 | 98670583 | 2.15417 | 1.98297 | -1.086 |
| chr6 | 19943082 | 19943362 | 0.837328 | 0.770932 | -1.086 |
| chr7 | 125529579 | 125529791 | 0.6164 | 0.56756 | -1.086 |
| chr16 | 74151821 | 74152036 | 1.19129 | 1.09707 | -1.086 |
| chr3 | 123444660 | 123444858 | 1.05541 | 0.971952 | -1.086 |
| chr3 | 89536525 | 89536877 | 0.922764 | 0.850026 | -1.086 |
| chr14 | 37216796 | 37217058 | 2.0498 | 1.88841 | -1.085 |
| chr19 | 58524013 | 58524213 | 0.73988 | 0.681754 | -1.085 |
| chr12 | 12082147 | 12082387 | 0.817871 | 0.753741 | -1.085 |
| chr5 | 17190600 | 17190883 | 2.4641 | 2.27103 | -1.085 |
| chr9 | 129055801 | 129056082 | 1.20297 | 1.10894 | -1.085 |
| chr12 | 84367168 | 84367356 | 0.732456 | 0.675264 | -1.085 |
| chr1 | 247918874 | 247919079 | 0.826505 | 0.762089 | -1.085 |
| chr11 | 22942563 | 22942822 | 1.82633 | 1.68408 | -1.084 |
| chr11 | 43691082 | 43691265 | 0.74175 | 0.684048 | -1.084 |
| chr14 | 47608466 | 47608671 | 0.855316 | 0.788795 | -1.084 |
| chr8 | 39622035 | 39622281 | 0.994474 | 0.917348 | -1.084 |
| chr3 | 184941499 | 184941706 | 0.813932 | 0.750941 | -1.084 |
| chr19 | 50767644 | 50768069 | 1.35652 | 1.25154 | -1.084 |
| chr1 | 205554025 | 205554211 | 1.61085 | 1.48684 | -1.083 |
| chr8 | 127912333 | 127912542 | 0.993376 | 0.916902 | -1.083 |
| chr15 | 53083535 | 53083759 | 0.9269 | 0.855689 | -1.083 |
| chr3 | 110799220 | 110799411 | 1.47498 | 1.36179 | -1.083 |
| chr2 | 21795845 | 21796208 | 1.17778 | 1.0875 | -1.083 |
| chr8 | 41416642 | 41416837 | 0.926438 | 0.85558 | -1.083 |
| chr19 | 41667916 | 41668162 | 1.8992 | 1.75479 | -1.082 |
| chr5 | 4233829 | 4234144 | 1.32314 | 1.22264 | -1.082 |
| chr5 | 165386079 | 165386306 | 0.925798 | 0.855531 | -1.082 |
| chr2 | 82997774 | 82998096 | 1.15232 | 1.06488 | -1.082 |
| chr11 | 14564359 | 14564583 | 0.959755 | 0.886985 | -1.082 |
| chr9 | 3090653 | 3090894 | 0.884547 | 0.817534 | -1.082 |
| chr2 | 162864047 | 162864498 | 1.34194 | 1.24032 | -1.082 |
| chr2 | 71384581 | 71384988 | 2.37124 | 2.19177 | -1.082 |
| chr11 | 118223839 | 118224078 | 1.08639 | 1.00421 | -1.082 |
| chr16 | 3137212 | 3137424 | 1.79017 | 1.65492 | -1.082 |
| chr1 | 198025479 | 198025732 | 1.68651 | 1.55956 | -1.081 |
| chr17 | 73162607 | 73162804 | 0.818747 | 0.757146 | -1.081 |
| chr2 | 75063643 | 75063890 | 1.87237 | 1.73158 | -1.081 |
| chr8 | 130838491 | 130838758 | 3.05328 | 2.82386 | -1.081 |
| chr2 | 78099639 | 78099948 | 0.913559 | 0.844931 | -1.081 |
| chr5 | 176245074 | 176245405 | 1.71212 | 1.58355 | -1.081 |
| chr14 | 42079137 | 42079336 | 1.06252 | 0.982984 | -1.081 |
| chr8 | 144212199 | 144212805 | 2.93898 | 2.71922 | -1.081 |
| chr1 | 224238494 | 224238861 | 1.93344 | 1.78916 | -1.081 |
| chr10 | 30530113 | 30530428 | 1.82911 | 1.69297 | -1.080 |
| chr11 | 128160020 | 128160290 | 1.58014 | 1.46264 | -1.080 |
| chr7 | 65660411 | 65660783 | 0.92215 | 0.853642 | -1.080 |
| chr9 | 27679823 | 27680148 | 0.844819 | 0.782154 | -1.080 |
| chr14 | 47298656 | 47298950 | 1.18236 | 1.09497 | -1.080 |
| chr2 | 5801508 | 5801743 | 0.677527 | 0.627575 | -1.080 |

| | | | | | |
|---|---|---|---|---|---|
| chr3 | 73135240 | 73135602 | 1.71059 | 1.58465 | -1.079 |
| chr8 | 51170347 | 51170644 | 0.936914 | 0.868016 | -1.079 |
| chr9 | 7536384 | 7536566 | 1.79886 | 1.66696 | -1.079 |
| chr20 | 46657201 | 46657389 | 0.906096 | 0.839705 | -1.079 |
| chr3 | 156171899 | 156172258 | 0.951009 | 0.881355 | -1.079 |
| chr17 | 13078603 | 13078810 | 1.07206 | 0.993757 | -1.079 |
| chr2 | 170785005 | 170785232 | 1.51894 | 1.40814 | -1.079 |
| chr2 | 157484627 | 157484916 | 0.996475 | 0.923851 | -1.079 |
| chr15 | 90038066 | 90038540 | 2.21709 | 2.05552 | -1.079 |
| chr14 | 65358429 | 65358764 | 1.35347 | 1.25499 | -1.078 |
| chr15 | 24103107 | 24103376 | 0.989349 | 0.917624 | -1.078 |
| chr11 | 133004534 | 133004779 | 1.3666 | 1.26755 | -1.078 |
| chr20 | 25146268 | 25146499 | 0.814313 | 0.755314 | -1.078 |
| chr7 | 139229340 | 139229563 | 0.859394 | 0.797353 | -1.078 |
| chr20 | 48227206 | 48227393 | 1.63946 | 1.52111 | -1.078 |
| chr2 | 40440290 | 40440645 | 1.35388 | 1.25615 | -1.078 |
| chr1 | 47641032 | 47641349 | 3.23435 | 3.001 | -1.078 |
| chr5 | 62894242 | 62894529 | 1.09713 | 1.018 | -1.078 |
| chr14 | 38980169 | 38980439 | 1.07578 | 0.998391 | -1.078 |
| chr7 | 7455029 | 7455214 | 0.824159 | 0.764887 | -1.077 |
| chr7 | 91404654 | 91404840 | 1.05528 | 0.979415 | -1.077 |
| chr8 | 133408094 | 133408477 | 1.59494 | 1.48031 | -1.077 |
| chr8 | 65294625 | 65295037 | 2.23551 | 2.07491 | -1.077 |
| chr11 | 90244049 | 90244269 | 0.914111 | 0.848506 | -1.077 |
| chr14 | 54814819 | 54815008 | 2.06092 | 1.91302 | -1.077 |
| chr7 | 121261286 | 121261483 | 0.827844 | 0.768448 | -1.077 |
| chr17 | 72642440 | 72642668 | 1.55215 | 1.44093 | -1.077 |
| chr12 | 24032820 | 24033104 | 0.764715 | 0.710078 | -1.077 |
| chr1 | 56874793 | 56874997 | 1.48383 | 1.37785 | -1.077 |
| chr11 | 8359610 | 8360126 | 2.68607 | 2.49428 | -1.077 |
| chr17 | 28530456 | 28530682 | 1.02169 | 0.948752 | -1.077 |
| chr10 | 110409441 | 110409700 | 0.814202 | 0.756114 | -1.077 |
| chr10 | 85497836 | 85498104 | 0.632839 | 0.587744 | -1.077 |
| chr8 | 4695350 | 4695618 | 1.47369 | 1.36905 | -1.076 |
| chr2 | 4317453 | 4317681 | 1.07678 | 1.00056 | -1.076 |
| chr7 | 96054857 | 96055329 | 0.924416 | 0.85906 | -1.076 |
| chr9 | 76903682 | 76903923 | 0.915729 | 0.850988 | -1.076 |
| chr5 | 138125429 | 138125750 | 1.8184 | 1.68985 | -1.076 |
| chr14 | 84740902 | 84741123 | 0.921293 | 0.856214 | -1.076 |
| chr15 | 96238185 | 96238392 | 0.918093 | 0.8533 | -1.076 |
| chr17 | 61934467 | 61934687 | 1.82017 | 1.69209 | -1.076 |
| chr11 | 38754955 | 38755215 | 0.894756 | 0.831942 | -1.076 |
| chr19 | 54819132 | 54819379 | 0.93157 | 0.866183 | -1.075 |
| chr8 | 138759620 | 138759822 | 1.06823 | 0.993479 | -1.075 |
| chr12 | 98960675 | 98960903 | 1.66625 | 1.54968 | -1.075 |
| chr10 | 28768536 | 28768807 | 0.828122 | 0.770273 | -1.075 |
| chr19 | 37388793 | 37389058 | 1.62152 | 1.50837 | -1.075 |
| chr19 | 29143890 | 29144149 | 1.162 | 1.08114 | -1.075 |
| chr9 | 133928414 | 133928976 | 2.47572 | 2.30356 | -1.075 |
| chr2 | 120275068 | 120275378 | 2.64175 | 2.45815 | -1.075 |
| chr8 | 29173190 | 29173392 | 1.61072 | 1.49899 | -1.075 |
| chr12 | 54920713 | 54921183 | 1.48086 | 1.37831 | -1.074 |
| chr10 | 23814476 | 23814705 | 1.18372 | 1.10175 | -1.074 |
| chr17 | 42060761 | 42061081 | 1.66831 | 1.55279 | -1.074 |
| chr11 | 87433718 | 87433972 | 1.4311 | 1.33232 | -1.074 |
| chr2 | 22693859 | 22694139 | 0.943509 | 0.878445 | -1.074 |
| chr12 | 114881557 | 114881888 | 0.812306 | 0.75634 | -1.074 |
| chr14 | 45895908 | 45896280 | 1.76058 | 1.63948 | -1.074 |
| chr20 | 41566202 | 41566410 | 0.876613 | 0.816317 | -1.074 |
| chr3 | 115533713 | 115533935 | 2.41639 | 2.25021 | -1.074 |
| chr8 | 105337421 | 105338519 | 1.79167 | 1.6686 | -1.074 |

| | | | | | |
|---|---|---|---|---|---|
| chr9 | 104473558 | 104473984 | 2.26264 | 2.10726 | -1.074 |
| chr5 | 11267281 | 11267754 | 1.88432 | 1.75542 | -1.073 |
| chr17 | 26633909 | 26634227 | 1.17743 | 1.09692 | -1.073 |
| chr10 | 66059134 | 66059354 | 1.10662 | 1.0311 | -1.073 |
| chr5 | 90990657 | 90990879 | 0.775597 | 0.722849 | -1.073 |
| chr3 | 80810293 | 80810482 | 0.926385 | 0.863464 | -1.073 |
| chr7 | 142023561 | 142023787 | 1.37328 | 1.28001 | -1.073 |
| chrX | 119499279 | 119499543 | 1.52977 | 1.42591 | -1.073 |
| chr19 | 34372316 | 34372512 | 1.08972 | 1.01582 | -1.073 |
| chr8 | 22862058 | 22862326 | 1.4674 | 1.36808 | -1.073 |
| chr11 | 2237151 | 2237451 | 0.976208 | 0.910181 | -1.073 |
| chrX | 52603767 | 52604052 | 1.7669 | 1.64744 | -1.073 |
| chr5 | 26843375 | 26843594 | 1.10729 | 1.03251 | -1.072 |
| chr9 | 10891319 | 10891650 | 0.678421 | 0.632653 | -1.072 |
| chr12 | 101632711 | 101632993 | 0.932877 | 0.870158 | -1.072 |
| chr1 | 197490336 | 197490602 | 2.04616 | 1.90862 | -1.072 |
| chr7 | 154201113 | 154201330 | 1.62448 | 1.51534 | -1.072 |
| chr11 | 58053757 | 58053955 | 1.47129 | 1.37247 | -1.072 |
| chr2 | 21868108 | 21868295 | 1.17052 | 1.09193 | -1.072 |
| chr2 | 6493172 | 6493372 | 1.3195 | 1.23109 | -1.072 |
| chr12 | 78011020 | 78011250 | 0.832965 | 0.777184 | -1.072 |
| chr6 | 7994372 | 7994717 | 0.827135 | 0.771933 | -1.072 |
| chr2 | 137181002 | 137181375 | 2.75347 | 2.56998 | -1.071 |
| chr4 | 168808806 | 168809089 | 0.874367 | 0.816124 | -1.071 |
| chr8 | 105015687 | 105015944 | 0.901116 | 0.841095 | -1.071 |
| chr16 | 74896651 | 74897126 | 2.25556 | 2.10571 | -1.071 |
| chr9 | 139842495 | 139842769 | 1.99193 | 1.85969 | -1.071 |
| chr1 | 240115756 | 240115998 | 0.778936 | 0.727268 | -1.071 |
| chr5 | 13060669 | 13060918 | 1.01818 | 0.950743 | -1.071 |
| chr8 | 118158666 | 118158873 | 0.834501 | 0.779253 | -1.071 |
| chr20 | 6917646 | 6917842 | 1.09341 | 1.02104 | -1.071 |
| chr2 | 29589144 | 29589420 | 0.889341 | 0.830501 | -1.071 |
| chr2 | 113961532 | 113961802 | 3.03765 | 2.83674 | -1.071 |
| chr8 | 78507910 | 78508125 | 0.996033 | 0.930281 | -1.071 |
| chr7 | 128101485 | 128101726 | 2.02864 | 1.89504 | -1.070 |
| chr17 | 35014462 | 35014774 | 0.897868 | 0.838769 | -1.070 |
| chr17 | 63986595 | 63986968 | 1.34297 | 1.25477 | -1.070 |
| chr2 | 69602911 | 69603194 | 1.97376 | 1.84432 | -1.070 |
| chr14 | 23947549 | 23947794 | 1.70673 | 1.59531 | -1.070 |
| chr16 | 46538642 | 46538928 | 1.22947 | 1.14925 | -1.070 |
| chr20 | 7603131 | 7603381 | 1.16056 | 1.08485 | -1.070 |
| chr2 | 236674925 | 236675197 | 2.30096 | 2.15106 | -1.070 |
| chr12 | 4146846 | 4147321 | 1.84262 | 1.72263 | -1.070 |
| chr5 | 11396960 | 11397167 | 0.849209 | 0.794356 | -1.069 |
| chr11 | 73625485 | 73625668 | 1.04831 | 0.980642 | -1.069 |
| chr3 | 161376481 | 161376699 | 0.788808 | 0.738088 | -1.069 |
| chr5 | 44332520 | 44332746 | 1.20007 | 1.12294 | -1.069 |
| chr8 | 20042258 | 20042681 | 1.49274 | 1.39683 | -1.069 |
| chr12 | 71786380 | 71786623 | 2.43894 | 2.28246 | -1.069 |
| chr10 | 18393153 | 18393653 | 1.60448 | 1.50166 | -1.068 |
| chr8 | 42243016 | 42243242 | 2.425 | 2.26983 | -1.068 |
| chr1 | 182959995 | 182960204 | 1.0852 | 1.01584 | -1.068 |
| chr8 | 120781065 | 120781540 | 2.39144 | 2.23871 | -1.068 |
| chr3 | 77008084 | 77008301 | 1.09005 | 1.02065 | -1.068 |
| chr7 | 19635423 | 19635611 | 0.954009 | 0.893466 | -1.068 |
| chr2 | 135218179 | 135218512 | 2.18806 | 2.04937 | -1.068 |
| chr12 | 85521936 | 85522137 | 0.935446 | 0.876226 | -1.068 |
| chr2 | 103370119 | 103370332 | 2.00931 | 1.88229 | -1.067 |
| chr5 | 86150834 | 86151038 | 0.894128 | 0.837612 | -1.067 |
| chr2 | 49871180 | 49871418 | 0.906435 | 0.849385 | -1.067 |
| chr7 | 33725608 | 33725810 | 1.01395 | 0.9502 | -1.067 |

| | | | | | |
|---|---|---|---|---|---|
| chr6 | 105765818 | 105766110 | 1.4873 | 1.39384 | -1.067 |
| chr14 | 63595321 | 63595587 | 1.00138 | 0.938559 | -1.067 |
| chr1 | 232752254 | 232752498 | 0.897532 | 0.841505 | -1.067 |
| chr16 | 5683919 | 5684187 | 1.56291 | 1.46538 | -1.067 |
| chr7 | 131190899 | 131191123 | 0.76181 | 0.714348 | -1.066 |
| chr10 | 15304081 | 15304310 | 1.22507 | 1.14884 | -1.066 |
| chr10 | 86369448 | 86369633 | 0.609129 | 0.571241 | -1.066 |
| chr2 | 217381335 | 217381923 | 1.3913 | 1.30478 | -1.066 |
| chr1 | 112933502 | 112933774 | 1.68949 | 1.58448 | -1.066 |
| chr9 | 11986918 | 11987206 | 0.845436 | 0.793137 | -1.066 |
| chr16 | 60936328 | 60936530 | 0.825308 | 0.774276 | -1.066 |
| chr10 | 78796890 | 78797421 | 1.47372 | 1.38272 | -1.066 |
| chr8 | 111127843 | 111128090 | 1.08819 | 1.02118 | -1.066 |
| chr1 | 209182281 | 209182505 | 0.87265 | 0.818928 | -1.066 |
| chr14 | 81702286 | 81702486 | 0.759889 | 0.713126 | -1.066 |
| chr19 | 39103332 | 39103588 | 1.16665 | 1.09511 | -1.065 |
| chr7 | 32250420 | 32250653 | 2.18526 | 2.05147 | -1.065 |
| chr2 | 149897045 | 149897244 | 1.32867 | 1.24734 | -1.065 |
| chr9 | 89252430 | 89252713 | 1.80426 | 1.69387 | -1.065 |
| chr11 | 56370734 | 56370930 | 1.11175 | 1.04382 | -1.065 |
| chr3 | 3098400 | 3098588 | 0.695065 | 0.652622 | -1.065 |
| chr12 | 23003244 | 23003540 | 1.04563 | 0.982051 | -1.065 |
| chr5 | 90907443 | 90907731 | 1.22893 | 1.15442 | -1.065 |
| chr2 | 139081629 | 139081865 | 1.25207 | 1.17621 | -1.064 |
| chr12 | 12014288 | 12014501 | 0.970852 | 0.912069 | -1.064 |
| chr9 | 11034138 | 11034360 | 0.823667 | 0.77382 | -1.064 |
| chr12 | 77316031 | 77316294 | 0.986403 | 0.92677 | -1.064 |
| chr3 | 34301340 | 34301544 | 0.82463 | 0.774792 | -1.064 |
| chr2 | 225148902 | 225149304 | 2.26574 | 2.12898 | -1.064 |
| chr14 | 100856114 | 100856297 | 0.999174 | 0.938921 | -1.064 |
| chr1 | 238535217 | 238535459 | 0.979145 | 0.920539 | -1.064 |
| chr14 | 57995087 | 57995317 | 0.88957 | 0.83656 | -1.063 |
| chr16 | 13529770 | 13529990 | 0.983814 | 0.9252 | -1.063 |
| chr2 | 5235863 | 5236069 | 0.843063 | 0.792893 | -1.063 |
| chr2 | 241779945 | 241780326 | 1.85844 | 1.74791 | -1.063 |
| chr16 | 76562234 | 76562513 | 0.89063 | 0.837804 | -1.063 |
| chr14 | 100154152 | 100154559 | 2.88438 | 2.71357 | -1.063 |
| chr3 | 157583200 | 157583464 | 1.42265 | 1.33863 | -1.063 |
| chr3 | 38044867 | 38045355 | 2.69027 | 2.53147 | -1.063 |
| chr2 | 51604836 | 51605043 | 0.824241 | 0.77562 | -1.063 |
| chr12 | 7478921 | 7479176 | 1.20698 | 1.13592 | -1.063 |
| chrX | 17806492 | 17806930 | 2.72213 | 2.56189 | -1.063 |
| chrX | 118832899 | 118833142 | 1.58104 | 1.48806 | -1.062 |
| chr7 | 85131786 | 85132014 | 1.04279 | 0.981608 | -1.062 |
| chr20 | 36228713 | 36229001 | 0.75027 | 0.706405 | -1.062 |
| chr8 | 52256622 | 52256823 | 1.26248 | 1.18905 | -1.062 |
| chr20 | 25236085 | 25236293 | 1.55334 | 1.46333 | -1.062 |
| chr15 | 92860325 | 92860569 | 1.10322 | 1.03932 | -1.061 |
| chr11 | 64137550 | 64137912 | 0.894806 | 0.843152 | -1.061 |
| chr10 | 87860329 | 87860693 | 1.56261 | 1.47261 | -1.061 |
| chr16 | 17933134 | 17933323 | 0.79553 | 0.749784 | -1.061 |
| chr2 | 240931195 | 240931410 | 1.95524 | 1.84291 | -1.061 |
| chr20 | 19076469 | 19076747 | 1.2589 | 1.18675 | -1.061 |
| chr12 | 124993334 | 124993688 | 1.70913 | 1.61139 | -1.061 |
| chr19 | 29598715 | 29598981 | 1.09938 | 1.03655 | -1.061 |
| chr12 | 124883215 | 124883422 | 1.77258 | 1.67148 | -1.060 |
| chr5 | 163330210 | 163330416 | 0.711383 | 0.670978 | -1.060 |
| chr4 | 170192255 | 170192536 | 1.42827 | 1.34735 | -1.060 |
| chr5 | 162488288 | 162488513 | 0.817332 | 0.771118 | -1.060 |
| chr9 | 139014705 | 139015136 | 3.44701 | 3.25224 | -1.060 |
| chr5 | 161689454 | 161689686 | 0.778805 | 0.734954 | -1.060 |

| | | | | | |
|---|---|---|---|---|---|
| chr7 | 118403921 | 118404156 | 1.17024 | 1.10454 | -1.059 |
| chr7 | 142606668 | 142607004 | 1.64636 | 1.55453 | -1.059 |
| chr10 | 52770758 | 52771127 | 2.25575 | 2.12993 | -1.059 |
| chr7 | 153040281 | 153040572 | 0.883327 | 0.834151 | -1.059 |
| chr15 | 29536805 | 29537108 | 1.04086 | 0.982984 | -1.059 |
| chr7 | 120364465 | 120364719 | 1.19535 | 1.12897 | -1.059 |
| chr16 | 2944236 | 2944429 | 1.04266 | 0.984855 | -1.059 |
| chr2 | 72343258 | 72343558 | 1.92039 | 1.81425 | -1.059 |
| chr11 | 23289092 | 23289375 | 1.09576 | 1.03525 | -1.058 |
| chr9 | 95925559 | 95925793 | 0.837432 | 0.791232 | -1.058 |
| chr19 | 31099291 | 31099633 | 1.38712 | 1.31075 | -1.058 |
| chr8 | 2574579 | 2574888 | 0.895748 | 0.846437 | -1.058 |
| chr9 | 26639132 | 26639324 | 0.838661 | 0.792609 | -1.058 |
| chr16 | 80353149 | 80353335 | 1.03738 | 0.980489 | -1.058 |
| chr19 | 53820124 | 53820397 | 1.05963 | 1.00162 | -1.058 |
| chr10 | 20108488 | 20108719 | 1.63853 | 1.54896 | -1.058 |
| chr5 | 163539426 | 163539650 | 0.670554 | 0.633937 | -1.058 |
| chr5 | 91290783 | 91291002 | 0.723606 | 0.68415 | -1.058 |
| chr8 | 4746048 | 4746273 | 0.841092 | 0.795238 | -1.058 |
| chr5 | 176952593 | 176952972 | 0.839873 | 0.794265 | -1.057 |
| chr10 | 56090144 | 56090464 | 0.949983 | 0.898528 | -1.057 |
| chr5 | 97791893 | 97792119 | 1.0121 | 0.957305 | -1.057 |
| chr2 | 86263072 | 86263280 | 0.994754 | 0.941003 | -1.057 |
| chr1 | 182017794 | 182018003 | 0.881612 | 0.834085 | -1.057 |
| chr16 | 14394672 | 14394985 | 2.76362 | 2.61479 | -1.057 |
| chr15 | 27566213 | 27566398 | 1.1147 | 1.05494 | -1.057 |
| chr11 | 60093675 | 60093863 | 0.887916 | 0.840326 | -1.057 |
| chr19 | 55599964 | 55600480 | 3.45427 | 3.26913 | -1.057 |
| chr5 | 143769771 | 143769988 | 1.11458 | 1.05503 | -1.056 |
| chr3 | 6249810 | 6250066 | 1.1038 | 1.04497 | -1.056 |
| chr15 | 29158652 | 29158834 | 0.904298 | 0.85611 | -1.056 |
| chr8 | 70744521 | 70744935 | 1.71417 | 1.62285 | -1.056 |
| chr20 | 52836475 | 52836660 | 1.53199 | 1.45083 | -1.056 |
| chr15 | 66517397 | 66517592 | 0.838071 | 0.793686 | -1.056 |
| chr14 | 68343329 | 68343557 | 0.865351 | 0.819564 | -1.056 |
| chr3 | 1410686 | 1410892 | 0.876653 | 0.830542 | -1.056 |
| chr15 | 88874549 | 88874739 | 1.97487 | 1.87108 | -1.055 |
| chr3 | 22148101 | 22148384 | 0.820755 | 0.777658 | -1.055 |
| chr8 | 52271223 | 52271444 | 0.828219 | 0.78475 | -1.055 |
| chrX | 79046800 | 79047003 | 0.763422 | 0.723379 | -1.055 |
| chr19 | 37493480 | 37493869 | 0.946821 | 0.897162 | -1.055 |
| chr7 | 33847467 | 33847680 | 1.51273 | 1.43344 | -1.055 |
| chr2 | 237218320 | 237218594 | 1.39668 | 1.32348 | -1.055 |
| chr16 | 10759632 | 10759835 | 0.901168 | 0.853958 | -1.055 |
| chr2 | 121062915 | 121063247 | 1.39363 | 1.32072 | -1.055 |
| chr2 | 11674923 | 11675206 | 1.47949 | 1.40217 | -1.055 |
| chr9 | 116795687 | 116796052 | 1.64781 | 1.56181 | -1.055 |
| chr8 | 68705083 | 68705351 | 1.07351 | 1.01751 | -1.055 |
| chr8 | 96972141 | 96972361 | 0.962248 | 0.912253 | -1.055 |
| chr10 | 91613656 | 91613885 | 1.079 | 1.02301 | -1.055 |
| chr9 | 110531458 | 110531916 | 0.856514 | 0.812111 | -1.055 |
| chr17 | 54129588 | 54129871 | 0.934942 | 0.886623 | -1.054 |
| chr19 | 15202778 | 15202982 | 1.31275 | 1.245 | -1.054 |
| chr12 | 10951794 | 10952016 | 1.05328 | 0.998999 | -1.054 |
| chr11 | 25972813 | 25973032 | 1.07844 | 1.02288 | -1.054 |
| chr1 | 179090181 | 179090620 | 1.37127 | 1.30071 | -1.054 |
| chr6 | 166263769 | 166264261 | 2.11796 | 2.00905 | -1.054 |
| chr2 | 160824926 | 160825140 | 1.85237 | 1.75725 | -1.054 |
| chr14 | 25764563 | 25764962 | 0.750055 | 0.711783 | -1.054 |
| chr8 | 126369084 | 126369294 | 0.716901 | 0.680525 | -1.053 |
| chr17 | 62748314 | 62748718 | 0.835217 | 0.792847 | -1.053 |

| | | | | | |
|---|---|---|---|---|---|
| chr18 | 5023417 | 5023645 | 1.5354 | 1.45769 | -1.053 |
| chr11 | 101396622 | 101396828 | 0.925344 | 0.878717 | -1.053 |
| chr12 | 30291211 | 30291617 | 1.01493 | 0.963814 | -1.053 |
| chr2 | 74875058 | 74875461 | 1.23845 | 1.17617 | -1.053 |
| chr15 | 100842777 | 100843063 | 0.678439 | 0.644383 | -1.053 |
| chr7 | 142255801 | 142256097 | 0.812154 | 0.771471 | -1.053 |
| chr8 | 141109380 | 141109566 | 1.32969 | 1.26322 | -1.053 |
| chr10 | 121171786 | 121172076 | 0.852031 | 0.809542 | -1.052 |
| chr16 | 23946214 | 23946451 | 0.728857 | 0.692555 | -1.052 |
| chr5 | 54702993 | 54703214 | 2.20523 | 2.09551 | -1.052 |
| chr19 | 42847544 | 42847742 | 1.97907 | 1.8807 | -1.052 |
| chr14 | 74757133 | 74757441 | 1.91721 | 1.82199 | -1.052 |
| chr7 | 51041846 | 51042051 | 0.897482 | 0.85293 | -1.052 |
| chr13 | 59892472 | 59892885 | 1.80277 | 1.71342 | -1.052 |
| chr9 | 106057645 | 106057940 | 0.807313 | 0.767335 | -1.052 |
| chr1 | 162293593 | 162293861 | 0.823071 | 0.782323 | -1.052 |
| chr20 | 19372972 | 19373330 | 1.21116 | 1.15159 | -1.052 |
| chr9 | 1499461 | 1499712 | 1.20424 | 1.14512 | -1.052 |
| chr3 | 141852927 | 141853165 | 0.875958 | 0.833132 | -1.051 |
| chr2 | 45124631 | 45124880 | 1.292 | 1.22886 | -1.051 |
| chr2 | 128236870 | 128237094 | 1.66727 | 1.58584 | -1.051 |
| chr8 | 139211572 | 139211810 | 1.10372 | 1.0499 | -1.051 |
| chr6 | 107668159 | 107668347 | 0.761122 | 0.724074 | -1.051 |
| chr10 | 91129197 | 91129489 | 1.31657 | 1.25262 | -1.051 |
| chr5 | 133625318 | 133625531 | 1.86553 | 1.77493 | -1.051 |
| chr1 | 78011912 | 78012140 | 0.64293 | 0.611782 | -1.051 |
| chr13 | 41242499 | 41242699 | 2.17664 | 2.07143 | -1.051 |
| chr8 | 21402483 | 21402704 | 0.990765 | 0.942889 | -1.051 |
| chr7 | 135369589 | 135369793 | 0.684496 | 0.651503 | -1.051 |
| chr15 | 81400672 | 81401028 | 1.1927 | 1.13543 | -1.050 |
| chr2 | 44570073 | 44570308 | 1.59573 | 1.51928 | -1.050 |
| chr12 | 69305168 | 69305428 | 0.915609 | 0.871901 | -1.050 |
| chr8 | 145944030 | 145944213 | 0.954728 | 0.909211 | -1.050 |
| chr8 | 105083774 | 105084011 | 0.894014 | 0.851662 | -1.050 |
| chr16 | 89631940 | 89632349 | 2.91841 | 2.78042 | -1.050 |
| chr20 | 12318056 | 12318266 | 0.98935 | 0.94258 | -1.050 |
| chr16 | 4229147 | 4229349 | 1.06337 | 1.01325 | -1.049 |
| chr11 | 60531283 | 60531551 | 1.18393 | 1.12816 | -1.049 |
| chr15 | 50606562 | 50606869 | 2.78356 | 2.65252 | -1.049 |
| chr11 | 13624318 | 13624672 | 1.09657 | 1.04505 | -1.049 |
| chr8 | 71747135 | 71747453 | 1.72387 | 1.6429 | -1.049 |
| chr3 | 89592190 | 89592632 | 0.918226 | 0.875164 | -1.049 |
| chr3 | 25905100 | 25905302 | 1.10819 | 1.05627 | -1.049 |
| chr12 | 9353350 | 9353667 | 1.09373 | 1.04253 | -1.049 |
| chr10 | 97704678 | 97704886 | 0.916792 | 0.873885 | -1.049 |
| chr2 | 5993994 | 5994200 | 1.09239 | 1.04168 | -1.049 |
| chr5 | 51132271 | 51132674 | 0.965682 | 0.920931 | -1.049 |
| chr7 | 6204143 | 6204386 | 1.63071 | 1.55517 | -1.049 |
| chr2 | 53590546 | 53590807 | 0.927044 | 0.884138 | -1.049 |
| chrX | 3403837 | 3404045 | 0.844975 | 0.805946 | -1.048 |
| chr8 | 133067727 | 133068011 | 2.61296 | 2.49228 | -1.048 |
| chr3 | 85506348 | 85506560 | 0.877462 | 0.836945 | -1.048 |
| chr14 | 103860396 | 103860859 | 1.4532 | 1.38621 | -1.048 |
| chr12 | 130188942 | 130189201 | 0.846777 | 0.807765 | -1.048 |
| chr3 | 13268314 | 13268517 | 0.861756 | 0.822105 | -1.048 |
| chr11 | 86630755 | 86631024 | 1.9004 | 1.81335 | -1.048 |
| chr3 | 78840912 | 78841151 | 0.769895 | 0.73469 | -1.048 |
| chr17 | 55518189 | 55518470 | 2.15713 | 2.0586 | -1.048 |
| chr9 | 133884270 | 133884510 | 1.81663 | 1.73402 | -1.048 |
| chr3 | 20717071 | 20717289 | 0.848805 | 0.810211 | -1.048 |
| chr8 | 15035456 | 15035696 | 1.01788 | 0.97169 | -1.048 |

| | | | | | |
|---|---|---|---|---|---|
| chr1 | 108324904 | 108325144 | 1.61426 | 1.54109 | -1.047 |
| chr11 | 5402011 | 5402334 | 2.0324 | 1.94062 | -1.047 |
| chr10 | 65659205 | 65659521 | 0.901704 | 0.860991 | -1.047 |
| chr16 | 86794728 | 86795074 | 2.36482 | 2.25808 | -1.047 |
| chr2 | 65381287 | 65381492 | 1.22568 | 1.17037 | -1.047 |
| chr14 | 47117031 | 47117221 | 1.00596 | 0.96059 | -1.047 |
| chrX | 9842007 | 9842293 | 1.93281 | 1.84573 | -1.047 |
| chr9 | 71293456 | 71293747 | 0.831549 | 0.794127 | -1.047 |
| chr12 | 3123229 | 3123421 | 1.02904 | 0.982746 | -1.047 |
| chr8 | 104719506 | 104719890 | 1.45422 | 1.38886 | -1.047 |
| chr12 | 129761409 | 129761685 | 1.00276 | 0.95775 | -1.047 |
| chr15 | 94917153 | 94917393 | 0.746073 | 0.712627 | -1.047 |
| chr12 | 66942685 | 66943107 | 1.13197 | 1.08131 | -1.047 |
| chr19 | 38478017 | 38478395 | 1.93883 | 1.8522 | -1.047 |
| chr15 | 98622817 | 98623101 | 1.04366 | 0.99712 | -1.047 |
| chr14 | 25714028 | 25714216 | 0.874291 | 0.835314 | -1.047 |
| chr15 | 83341533 | 83341823 | 1.12455 | 1.07454 | -1.047 |
| chr20 | 3425049 | 3425316 | 1.25746 | 1.20161 | -1.046 |
| chr1 | 21543368 | 21543748 | 2.1696 | 2.07334 | -1.046 |
| chr2 | 118542953 | 118543588 | 1.30224 | 1.24449 | -1.046 |
| chr15 | 55330231 | 55330424 | 1.36129 | 1.30115 | -1.046 |
| chr14 | 46068958 | 46069240 | 0.838416 | 0.801499 | -1.046 |
| chr16 | 8202802 | 8203125 | 0.791557 | 0.756929 | -1.046 |
| chr2 | 51296648 | 51296848 | 1.14054 | 1.09066 | -1.046 |
| chr17 | 45820037 | 45820274 | 1.60412 | 1.5345 | -1.045 |
| chr8 | 76727693 | 76727919 | 0.824084 | 0.788323 | -1.045 |
| chr5 | 114725631 | 114725832 | 1.15024 | 1.10057 | -1.045 |
| chr21 | 33630719 | 33630976 | 2.07191 | 1.98257 | -1.045 |
| chr1 | 234290510 | 234290716 | 1.85417 | 1.77459 | -1.045 |
| chr9 | 80264441 | 80264624 | 1.28184 | 1.22697 | -1.045 |
| chr11 | 39965097 | 39965473 | 0.983135 | 0.941117 | -1.045 |
| chr3 | 150690310 | 150690659 | 0.901627 | 0.863103 | -1.045 |
| chr15 | 66297493 | 66297701 | 0.84247 | 0.806593 | -1.044 |
| chr17 | 34399824 | 34400224 | 2.38016 | 2.27883 | -1.044 |
| chr1 | 214935251 | 214935564 | 0.989495 | 0.947566 | -1.044 |
| chr10 | 23133984 | 23134205 | 0.879961 | 0.842739 | -1.044 |
| chr7 | 49279267 | 49279505 | 0.798607 | 0.764934 | -1.044 |
| chr9 | 78480428 | 78480819 | 0.998568 | 0.956484 | -1.044 |
| chr8 | 139453161 | 139453380 | 0.987641 | 0.946079 | -1.044 |
| chr5 | 163718404 | 163718609 | 1.03846 | 0.994835 | -1.044 |
| chr19 | 56587494 | 56587685 | 0.967802 | 0.927238 | -1.044 |
| chr17 | 33291833 | 33292101 | 1.10288 | 1.05672 | -1.044 |
| chr5 | 26161525 | 26161795 | 0.820524 | 0.786215 | -1.044 |
| chr20 | 44258159 | 44258440 | 2.20988 | 2.11751 | -1.044 |
| chr3 | 193687954 | 193688199 | 1.23897 | 1.18739 | -1.043 |
| chr1 | 180593173 | 180593372 | 2.00873 | 1.92529 | -1.043 |
| chr12 | 84869338 | 84869525 | 1.01073 | 0.968759 | -1.043 |
| chr2 | 18931273 | 18931506 | 2.75908 | 2.6446 | -1.043 |
| chr3 | 121820526 | 121820743 | 1.61029 | 1.54354 | -1.043 |
| chr2 | 14514779 | 14515143 | 1.11587 | 1.06996 | -1.043 |
| chr9 | 105477786 | 105478046 | 0.855288 | 0.820324 | -1.043 |
| chr2 | 51502381 | 51502668 | 0.800847 | 0.768184 | -1.043 |
| chr8 | 64134934 | 64135205 | 1.23196 | 1.18174 | -1.042 |
| chr2 | 164011822 | 164012077 | 1.26105 | 1.20972 | -1.042 |
| chr2 | 188160532 | 188160750 | 0.58509 | 0.561311 | -1.042 |
| chr15 | 94755651 | 94755848 | 1.03057 | 0.988732 | -1.042 |
| chr5 | 14510552 | 14511146 | 0.891184 | 0.855062 | -1.042 |
| chr18 | 55526576 | 55526786 | 1.47924 | 1.41949 | -1.042 |
| chr16 | 71493024 | 71493252 | 1.07975 | 1.03618 | -1.042 |
| chr5 | 116683020 | 116683246 | 1.25892 | 1.20813 | -1.042 |
| chr2 | 177005224 | 177005864 | 3.0582 | 2.93484 | -1.042 |

| | | | | | |
|---|---|---|---|---|---|
| chr2 | 20071391 | 20071760 | 0.983298 | 0.943698 | -1.042 |
| chr11 | 31418720 | 31419237 | 0.883781 | 0.848293 | -1.042 |
| chr12 | 62129782 | 62129987 | 0.964959 | 0.926251 | -1.042 |
| chr14 | 37886372 | 37886576 | 0.758816 | 0.728383 | -1.042 |
| chr1 | 157475463 | 157475836 | 2.07573 | 1.99252 | -1.042 |
| chr19 | 28891143 | 28891424 | 1.10586 | 1.06153 | -1.042 |
| chr3 | 79545098 | 79545291 | 0.830548 | 0.797268 | -1.042 |
| chr2 | 176952813 | 176953436 | 1.64422 | 1.57838 | -1.042 |
| chr8 | 86996213 | 86996430 | 1.00496 | 0.964777 | -1.042 |
| chrX | 110487570 | 110487783 | 1.03032 | 0.989174 | -1.042 |
| chr9 | 126824670 | 126824878 | 1.08649 | 1.04318 | -1.042 |
| chr11 | 32354802 | 32354994 | 1.99512 | 1.91568 | -1.041 |
| chr4 | 143048113 | 143048366 | 2.24782 | 2.15856 | -1.041 |
| chr19 | 47312097 | 47312306 | 0.917226 | 0.880957 | -1.041 |
| chr3 | 90094534 | 90094773 | 0.858218 | 0.824356 | -1.041 |
| chr1 | 192549641 | 192549839 | 2.04851 | 1.96777 | -1.041 |
| chr1 | 59250858 | 59251047 | 1.87795 | 1.80407 | -1.041 |
| chr10 | 58567890 | 58568179 | 1.07964 | 1.03722 | -1.041 |
| chr1 | 1555553 | 1555760 | 1.30825 | 1.25688 | -1.041 |
| chr17 | 59540986 | 59541310 | 1.372 | 1.31839 | -1.041 |
| chr19 | 18581075 | 18581343 | 1.40365 | 1.34885 | -1.041 |
| chr5 | 12845784 | 12846100 | 0.758113 | 0.728526 | -1.041 |
| chr15 | 52455833 | 52456075 | 1.53502 | 1.47514 | -1.041 |
| chr2 | 126463853 | 126464098 | 0.872904 | 0.838904 | -1.041 |
| chr11 | 114131752 | 114132108 | 3.2785 | 3.15114 | -1.040 |
| chr17 | 26248254 | 26248486 | 0.898076 | 0.863379 | -1.040 |
| chr18 | 56448855 | 56449052 | 1.48233 | 1.42507 | -1.040 |
| chr17 | 63259753 | 63260016 | 1.00114 | 0.962596 | -1.040 |
| chr12 | 84485593 | 84485801 | 1.02579 | 0.986337 | -1.040 |
| chr17 | 6049401 | 6049685 | 1.03667 | 0.996809 | -1.040 |
| chr2 | 27601012 | 27601258 | 1.4094 | 1.35541 | -1.040 |
| chr5 | 136105830 | 136106155 | 2.05314 | 1.97453 | -1.040 |
| chr19 | 29933646 | 29933890 | 0.883047 | 0.84937 | -1.040 |
| chr14 | 20393845 | 20394137 | 1.37243 | 1.32029 | -1.039 |
| chr2 | 126274119 | 126274381 | 1.08127 | 1.04023 | -1.039 |
| chr10 | 15296006 | 15296228 | 1.06874 | 1.02829 | -1.039 |
| chr14 | 78488213 | 78488638 | 1.69341 | 1.62941 | -1.039 |
| chr14 | 46705690 | 46705886 | 1.02045 | 0.982068 | -1.039 |
| chr8 | 132371111 | 132371318 | 0.997614 | 0.960224 | -1.039 |
| chr19 | 33896710 | 33897021 | 1.01062 | 0.97275 | -1.039 |
| chr19 | 30966450 | 30966637 | 0.977482 | 0.940855 | -1.039 |
| chr12 | 57051214 | 57051531 | 0.852924 | 0.820983 | -1.039 |
| chr2 | 71109771 | 71110001 | 0.826819 | 0.795936 | -1.039 |
| chr11 | 121065507 | 121065772 | 1.66529 | 1.60317 | -1.039 |
| chr7 | 49923548 | 49924059 | 1.12547 | 1.08355 | -1.039 |
| chr9 | 127155250 | 127155618 | 1.91306 | 1.84198 | -1.039 |
| chr1 | 231888737 | 231889033 | 0.897882 | 0.86471 | -1.038 |
| chr1 | 54181190 | 54181450 | 2.31835 | 2.23312 | -1.038 |
| chr17 | 48872703 | 48872898 | 0.729462 | 0.702657 | -1.038 |
| chr8 | 69796629 | 69796848 | 1.14546 | 1.10338 | -1.038 |
| chr22 | 42956908 | 42957250 | 1.28828 | 1.24097 | -1.038 |
| chr5 | 157724627 | 157724865 | 0.828893 | 0.798583 | -1.038 |
| chr3 | 103930187 | 103930389 | 0.69114 | 0.665907 | -1.038 |
| chr15 | 97348237 | 97348458 | 1.04102 | 1.00308 | -1.038 |
| chr20 | 44078873 | 44079284 | 1.50575 | 1.45134 | -1.037 |
| chr2 | 71087667 | 71087877 | 0.905653 | 0.873012 | -1.037 |
| chr15 | 87761269 | 87761549 | 0.937433 | 0.903686 | -1.037 |
| chr14 | 33467760 | 33468144 | 1.07478 | 1.0363 | -1.037 |
| chr11 | 60160090 | 60160292 | 0.784123 | 0.756055 | -1.037 |
| chr5 | 61414435 | 61414697 | 0.770881 | 0.743289 | -1.037 |
| chr5 | 70993688 | 70994170 | 1.85355 | 1.78734 | -1.037 |

| | | | | | |
|---|---|---|---|---|---|
| chr11 | 5747908 | 5748320 | 0.829694 | 0.800108 | -1.037 |
| chr9 | 139621162 | 139621459 | 1.7096 | 1.64889 | -1.037 |
| chr15 | 98784470 | 98784686 | 0.978249 | 0.943565 | -1.037 |
| chr1 | 145588498 | 145588817 | 2.06592 | 1.99318 | -1.036 |
| chr9 | 109175391 | 109175597 | 0.916703 | 0.884638 | -1.036 |
| chr4 | 15958594 | 15958776 | 0.752421 | 0.726197 | -1.036 |
| chr19 | 48366678 | 48367286 | 2.91652 | 2.81542 | -1.036 |
| chr7 | 50132111 | 50132783 | 1.27 | 1.22603 | -1.036 |
| chr17 | 64990437 | 64990788 | 1.12155 | 1.08286 | -1.036 |
| chr6 | 23446721 | 23446932 | 1.11238 | 1.07415 | -1.036 |
| chr15 | 94048821 | 94049122 | 0.812078 | 0.784192 | -1.036 |
| chr17 | 15964936 | 15965150 | 0.962177 | 0.929261 | -1.035 |
| chr2 | 4809483 | 4809753 | 1.17729 | 1.13725 | -1.035 |
| chr3 | 141295374 | 141295574 | 1.8371 | 1.77514 | -1.035 |
| chr5 | 9161860 | 9162061 | 0.809557 | 0.782355 | -1.035 |
| chr2 | 173326687 | 173327145 | 3.04646 | 2.94426 | -1.035 |
| chr7 | 52249337 | 52249633 | 0.948651 | 0.91684 | -1.035 |
| chr17 | 33123148 | 33123360 | 0.861304 | 0.832474 | -1.035 |
| chr8 | 83792757 | 83792954 | 0.813697 | 0.786476 | -1.035 |
| chr2 | 112459772 | 112459996 | 1.91648 | 1.85252 | -1.035 |
| chr12 | 73439859 | 73440171 | 0.916831 | 0.886247 | -1.035 |
| chr8 | 120234414 | 120234827 | 0.754517 | 0.729538 | -1.034 |
| chr14 | 21226263 | 21226513 | 1.32182 | 1.27823 | -1.034 |
| chr5 | 51436023 | 51436366 | 1.24005 | 1.19935 | -1.034 |
| chr11 | 77459826 | 77460064 | 0.718682 | 0.695392 | -1.033 |
| chr13 | 27933433 | 27933673 | 1.92052 | 1.85848 | -1.033 |
| chr15 | 35522156 | 35522351 | 1.02211 | 0.989181 | -1.033 |
| chr7 | 37535252 | 37535498 | 1.85242 | 1.79284 | -1.033 |
| chr19 | 57420848 | 57421090 | 1.0481 | 1.01441 | -1.033 |
| chr19 | 1795880 | 1796237 | 2.69338 | 2.60694 | -1.033 |
| chr18 | 30844563 | 30844796 | 1.03904 | 1.0058 | -1.033 |
| chr3 | 14244933 | 14245280 | 2.39303 | 2.31687 | -1.033 |
| chr9 | 22836813 | 22837060 | 0.801211 | 0.775842 | -1.033 |
| chr8 | 95284543 | 95284905 | 2.50874 | 2.42947 | -1.033 |
| chr17 | 21078813 | 21079174 | 0.876514 | 0.848949 | -1.032 |
| chr8 | 11286804 | 11287036 | 1.4135 | 1.36911 | -1.032 |
| chr8 | 96640306 | 96640505 | 0.8751 | 0.847659 | -1.032 |
| chr3 | 88461481 | 88461824 | 0.810732 | 0.78533 | -1.032 |
| chr12 | 85500086 | 85500370 | 0.787575 | 0.762985 | -1.032 |
| chr7 | 108300590 | 108301086 | 0.917764 | 0.889236 | -1.032 |
| chr8 | 96519432 | 96519635 | 0.884925 | 0.857684 | -1.032 |
| chr5 | 122089179 | 122089372 | 1.11151 | 1.07731 | -1.032 |
| chr11 | 8230877 | 8231087 | 1.26566 | 1.22679 | -1.032 |
| chr2 | 242229029 | 242229259 | 1.31923 | 1.27882 | -1.032 |
| chr16 | 68881163 | 68881478 | 1.91174 | 1.85326 | -1.032 |
| chr10 | 67908181 | 67908447 | 0.996757 | 0.966383 | -1.031 |
| chr2 | 17306023 | 17306239 | 0.823082 | 0.798025 | -1.031 |
| chr3 | 93680035 | 93680255 | 0.916 | 0.888123 | -1.031 |
| chr2 | 49210873 | 49211162 | 1.10496 | 1.07154 | -1.031 |
| chr12 | 5926860 | 5927311 | 0.842843 | 0.817405 | -1.031 |
| chr12 | 44711403 | 44711689 | 0.887809 | 0.861021 | -1.031 |
| chr2 | 51159166 | 51159377 | 0.79503 | 0.771101 | -1.031 |
| chr1 | 184191638 | 184192036 | 2.41042 | 2.33803 | -1.031 |
| chr7 | 103909793 | 103910001 | 0.879469 | 0.853237 | -1.031 |
| chr10 | 124408303 | 124408542 | 2.17047 | 2.10611 | -1.031 |
| chr22 | 25858542 | 25858884 | 3.05146 | 2.96099 | -1.031 |
| chr7 | 31162191 | 31162555 | 3.21297 | 3.11786 | -1.031 |
| chr19 | 48466502 | 48466775 | 2.1341 | 2.07097 | -1.030 |
| chr8 | 96317570 | 96317911 | 2.32268 | 2.25442 | -1.030 |
| chr11 | 23833593 | 23833873 | 1.04123 | 1.01069 | -1.030 |
| chr3 | 57993870 | 57994120 | 2.2544 | 2.18838 | -1.030 |

| | | | | | |
|---|---|---|---|---|---|
| chr1 | 187529973 | 187530202 | 0.761763 | 0.739569 | -1.030 |
| chr12 | 45354572 | 45354903 | 1.02279 | 0.993058 | -1.030 |
| chr3 | 73982822 | 73983073 | 0.898874 | 0.872755 | -1.030 |
| chr7 | 142599293 | 142599476 | 0.797929 | 0.774834 | -1.030 |
| chr17 | 52945241 | 52945426 | 0.997777 | 0.968937 | -1.030 |
| chr5 | 7887124 | 7887473 | 1.13684 | 1.10406 | -1.030 |
| chr5 | 165353033 | 165353224 | 0.879032 | 0.853731 | -1.030 |
| chr1 | 111067163 | 111067404 | 2.35871 | 2.29102 | -1.030 |
| chr16 | 85380935 | 85381139 | 1.33498 | 1.29672 | -1.030 |
| chr2 | 218797553 | 218797875 | 1.03858 | 1.00888 | -1.029 |
| chr17 | 14335975 | 14336219 | 0.872489 | 0.847742 | -1.029 |
| chr18 | 61705190 | 61705496 | 1.03674 | 1.00735 | -1.029 |
| chr20 | 37893220 | 37893539 | 2.56427 | 2.49159 | -1.029 |
| chr10 | 71548837 | 71549194 | 2.15006 | 2.08932 | -1.029 |
| chr1 | 44783928 | 44784287 | 2.24416 | 2.18077 | -1.029 |
| chr1 | 165578576 | 165578789 | 0.975142 | 0.947707 | -1.029 |
| chr8 | 127554670 | 127554895 | 0.763804 | 0.742511 | -1.029 |
| chr7 | 56323926 | 56324165 | 0.683862 | 0.664798 | -1.029 |
| chr20 | 24401563 | 24401786 | 0.816109 | 0.793359 | -1.029 |
| chr1 | 187516576 | 187516791 | 0.800126 | 0.77794 | -1.029 |
| chr7 | 86337530 | 86337856 | 1.03983 | 1.01101 | -1.029 |
| chr19 | 37336137 | 37336336 | 0.896037 | 0.871286 | -1.028 |
| chr7 | 119837530 | 119837751 | 0.855159 | 0.831552 | -1.028 |
| chr5 | 38551472 | 38551832 | 1.82529 | 1.7752 | -1.028 |
| chr11 | 81630083 | 81630473 | 0.906899 | 0.882082 | -1.028 |
| chr16 | 77382272 | 77382457 | 0.894445 | 0.870059 | -1.028 |
| chr2 | 5751667 | 5751920 | 0.729913 | 0.710025 | -1.028 |
| chr2 | 1842869 | 1843079 | 0.748535 | 0.728177 | -1.028 |
| chr15 | 76177530 | 76177816 | 1.40834 | 1.37008 | -1.028 |
| chr6 | 2196425 | 2196650 | 1.48526 | 1.44507 | -1.028 |
| chr7 | 24216330 | 24216605 | 1.14557 | 1.11479 | -1.028 |
| chr17 | 67196867 | 67197152 | 0.948992 | 0.923551 | -1.028 |
| chr8 | 91893886 | 91894241 | 0.831005 | 0.80879 | -1.027 |
| chr8 | 137935594 | 137935826 | 0.700415 | 0.681801 | -1.027 |
| chr2 | 73120753 | 73121236 | 2.1243 | 2.06789 | -1.027 |
| chr10 | 65976846 | 65977294 | 0.869167 | 0.846149 | -1.027 |
| chr15 | 88787216 | 88788023 | 1.81654 | 1.76845 | -1.027 |
| chr15 | 24975724 | 24976071 | 1.16741 | 1.13652 | -1.027 |
| chr12 | 59501672 | 59501898 | 0.88677 | 0.86335 | -1.027 |
| chr5 | 22621076 | 22621478 | 0.888691 | 0.865461 | -1.027 |
| chr5 | 18921060 | 18921246 | 0.815408 | 0.794104 | -1.027 |
| chr3 | 79885300 | 79885506 | 0.845885 | 0.823966 | -1.027 |
| chr12 | 62021162 | 62021536 | 1.03945 | 1.01288 | -1.026 |
| chr8 | 140007800 | 140008066 | 0.8489 | 0.827241 | -1.026 |
| chr6 | 160030062 | 160030249 | 1.00253 | 0.97701 | -1.026 |
| chr16 | 77257103 | 77257357 | 1.48055 | 1.44297 | -1.026 |
| chr8 | 56013592 | 56013813 | 1.27063 | 1.23844 | -1.026 |
| chr9 | 94533320 | 94533617 | 1.04315 | 1.01673 | -1.026 |
| chr11 | 63406895 | 63407085 | 0.983004 | 0.958247 | -1.026 |
| chr2 | 212022645 | 212022948 | 1.49256 | 1.45518 | -1.026 |
| chr15 | 69328020 | 69328328 | 3.23352 | 3.15259 | -1.026 |
| chr4 | 182449790 | 182450009 | 1.96376 | 1.91478 | -1.026 |
| chr9 | 10906305 | 10906601 | 0.819581 | 0.799259 | -1.025 |
| chr20 | 45142070 | 45142355 | 0.915557 | 0.892958 | -1.025 |
| chr11 | 4016817 | 4017021 | 0.858031 | 0.836894 | -1.025 |
| chr2 | 219141590 | 219141883 | 0.964678 | 0.940921 | -1.025 |
| chr8 | 52576267 | 52576456 | 1.30608 | 1.27394 | -1.025 |
| chr15 | 28034658 | 28034890 | 0.89766 | 0.875729 | -1.025 |
| chr7 | 84630379 | 84630578 | 0.959406 | 0.935986 | -1.025 |
| chr11 | 35856895 | 35857279 | 2.0461 | 1.99665 | -1.025 |
| chr1 | 177793822 | 177794019 | 1.02047 | 0.996131 | -1.024 |

| | | | | | |
|---|---|---|---|---|---|
| chr11 | 4813697 | 4813977 | 0.750125 | 0.732244 | -1.024 |
| chr5 | 6666993 | 6667221 | 1.02492 | 1.00049 | -1.024 |
| chr15 | 78689219 | 78689550 | 2.30901 | 2.25419 | -1.024 |
| chr5 | 26613096 | 26613476 | 0.997731 | 0.974216 | -1.024 |
| chr5 | 151355975 | 151356276 | 1.25786 | 1.22836 | -1.024 |
| chr5 | 88850705 | 88850895 | 0.743196 | 0.725799 | -1.024 |
| chr7 | 32246754 | 32246962 | 1.63094 | 1.59284 | -1.024 |
| chr16 | 944697 | 945022 | 0.925006 | 0.903439 | -1.024 |
| gl000205_r | 135015 | 135325 | 1.51971 | 1.48446 | -1.024 |
| chr8 | 112482654 | 112483013 | 1.04659 | 1.02255 | -1.024 |
| chr8 | 137183736 | 137183943 | 0.919191 | 0.898129 | -1.023 |
| chr1 | 156918556 | 156918786 | 1.41319 | 1.38101 | -1.023 |
| chr3 | 134077335 | 134077594 | 1.36049 | 1.32956 | -1.023 |
| chr11 | 41759903 | 41760187 | 0.837395 | 0.818371 | -1.023 |
| chr9 | 101841243 | 101841478 | 0.952228 | 0.930626 | -1.023 |
| chr11 | 17757350 | 17757709 | 1.46397 | 1.43076 | -1.023 |
| chr2 | 37232312 | 37232572 | 0.984839 | 0.962712 | -1.023 |
| chr19 | 49649076 | 49649643 | 2.9836 | 2.91657 | -1.023 |
| chr20 | 4876845 | 4877133 | 1.13694 | 1.11142 | -1.023 |
| chr19 | 49010342 | 49010544 | 1.01742 | 0.994623 | -1.023 |
| chr16 | 83990350 | 83990806 | 2.65923 | 2.59986 | -1.023 |
| chr9 | 34785724 | 34786158 | 0.733288 | 0.717156 | -1.022 |
| chr8 | 136249519 | 136249744 | 0.989564 | 0.967797 | -1.022 |
| chr9 | 91849571 | 91849777 | 1.48916 | 1.45692 | -1.022 |
| chr15 | 95223775 | 95224229 | 0.94512 | 0.924791 | -1.022 |
| chr2 | 76227144 | 76227353 | 0.960315 | 0.939664 | -1.022 |
| chr5 | 102017168 | 102017426 | 2.49426 | 2.44108 | -1.022 |
| chr10 | 102855387 | 102855772 | 2.83826 | 2.778 | -1.022 |
| chr5 | 135402700 | 135402932 | 0.789861 | 0.773113 | -1.022 |
| chr5 | 104256124 | 104256490 | 1.07221 | 1.04948 | -1.022 |
| chr12 | 55247734 | 55248290 | 2.24753 | 2.20011 | -1.022 |
| chr10 | 12591725 | 12591948 | 0.930137 | 0.910643 | -1.021 |
| chr2 | 80206233 | 80206435 | 0.835302 | 0.817941 | -1.021 |
| chrX | 154322728 | 154323054 | 1.91386 | 1.87432 | -1.021 |
| chr5 | 43419905 | 43420132 | 0.834403 | 0.817167 | -1.021 |
| chr6 | 37709755 | 37710108 | 1.54846 | 1.51658 | -1.021 |
| chr7 | 136704456 | 136704727 | 0.869928 | 0.85208 | -1.021 |
| chr5 | 123060388 | 123060621 | 0.791619 | 0.775434 | -1.021 |
| chr12 | 24989257 | 24989607 | 1.63931 | 1.60592 | -1.021 |
| chr7 | 98838437 | 98838620 | 0.912014 | 0.893459 | -1.021 |
| chr8 | 120114798 | 120115041 | 0.976076 | 0.956317 | -1.021 |
| chr17 | 26867835 | 26868041 | 2.96401 | 2.90465 | -1.020 |
| chr10 | 69916204 | 69916481 | 2.04074 | 1.99996 | -1.020 |
| chr8 | 28169146 | 28169401 | 0.892774 | 0.874971 | -1.020 |
| chr12 | 12889927 | 12890283 | 1.64794 | 1.61538 | -1.020 |
| chr1 | 207030018 | 207030454 | 1.33717 | 1.31122 | -1.020 |
| chr15 | 60426875 | 60427069 | 1.49403 | 1.46506 | -1.020 |
| chr12 | 31165187 | 31165467 | 1.79114 | 1.75711 | -1.019 |
| chr11 | 66040513 | 66040791 | 0.764 | 0.749608 | -1.019 |
| chr17 | 75624884 | 75625068 | 0.9502 | 0.932615 | -1.019 |
| chr2 | 59930136 | 59930341 | 0.721229 | 0.707952 | -1.019 |
| chr8 | 145099992 | 145100593 | 0.996897 | 0.978573 | -1.019 |
| chr7 | 113130670 | 113130884 | 0.929926 | 0.912869 | -1.019 |
| chr7 | 42512509 | 42512795 | 1.09862 | 1.07852 | -1.019 |
| chr19 | 28655418 | 28655830 | 1.0223 | 1.00389 | -1.018 |
| rUn_gl0002 | 139159 | 139526 | 1.23654 | 1.21433 | -1.018 |
| chr5 | 85139345 | 85139576 | 0.896453 | 0.880464 | -1.018 |
| chr2 | 232765826 | 232766011 | 1.12497 | 1.10507 | -1.018 |
| chr8 | 135686278 | 135686462 | 0.865497 | 0.850242 | -1.018 |
| chr10 | 29879604 | 29880196 | 1.55394 | 1.5267 | -1.018 |
| chr5 | 170589659 | 170589904 | 1.75964 | 1.72904 | -1.018 |

| | | | | | |
|---|---|---|---|---|---|
| chr9 | 17531697 | 17531955 | 2.43118 | 2.38892 | -1.018 |
| chr8 | 94339947 | 94340219 | 0.753057 | 0.740083 | -1.018 |
| chr1 | 184063416 | 184063629 | 0.843218 | 0.828798 | -1.017 |
| chr11 | 81531570 | 81531827 | 0.967944 | 0.951715 | -1.017 |
| chr2 | 121886426 | 121886952 | 1.6894 | 1.6611 | -1.017 |
| chr8 | 142937844 | 142938498 | 1.0098 | 0.992924 | -1.017 |
| chr15 | 102000265 | 102000601 | 0.963278 | 0.947219 | -1.017 |
| chr8 | 27182030 | 27182255 | 2.07894 | 2.04509 | -1.017 |
| chr12 | 107313722 | 107314233 | 1.75122 | 1.72274 | -1.017 |
| chr7 | 129007435 | 129007777 | 2.87216 | 2.82554 | -1.016 |
| chr12 | 73093309 | 73093539 | 0.999713 | 0.98354 | -1.016 |
| chr15 | 59924952 | 59925236 | 2.61085 | 2.56865 | -1.016 |
| chr8 | 27699726 | 27700030 | 0.760315 | 0.748034 | -1.016 |
| chr10 | 27512692 | 27513191 | 1.87454 | 1.84435 | -1.016 |
| chr2 | 214111485 | 214111729 | 2.04187 | 2.00917 | -1.016 |
| chr12 | 105047641 | 105047898 | 1.62768 | 1.60184 | -1.016 |
| chr20 | 21008352 | 21008559 | 1.2935 | 1.27303 | -1.016 |
| chr11 | 25548903 | 25549134 | 0.980362 | 0.96489 | -1.016 |
| chr9 | 30414839 | 30415058 | 0.937163 | 0.922417 | -1.016 |
| chr8 | 60615580 | 60615806 | 0.858187 | 0.844874 | -1.016 |
| chr17 | 52466726 | 52467104 | 1.0488 | 1.03256 | -1.016 |
| chr11 | 10472597 | 10472835 | 0.941807 | 0.927226 | -1.016 |
| chr12 | 71265138 | 71265353 | 0.883138 | 0.869477 | -1.016 |
| chr10 | 107648559 | 107648766 | 0.664714 | 0.654492 | -1.016 |
| chr2 | 170088283 | 170088481 | 0.784183 | 0.772206 | -1.016 |
| chr6 | 40772765 | 40772985 | 1.06125 | 1.0452 | -1.015 |
| chr5 | 114502723 | 114502910 | 0.703615 | 0.69318 | -1.015 |
| chr11 | 123511666 | 123512013 | 2.8712 | 2.82863 | -1.015 |
| chr1 | 210573925 | 210574243 | 2.04065 | 2.01057 | -1.015 |
| chr19 | 43065726 | 43065942 | 1.52025 | 1.49785 | -1.015 |
| chr9 | 18791137 | 18791368 | 2.25485 | 2.22172 | -1.015 |
| chr12 | 24749848 | 24750052 | 1.55214 | 1.52949 | -1.015 |
| chr7 | 16770965 | 16771268 | 0.960855 | 0.947236 | -1.014 |
| chr11 | 58953829 | 58954085 | 1.8234 | 1.79773 | -1.014 |
| chr5 | 156240648 | 156240947 | 0.937767 | 0.924811 | -1.014 |
| chr17 | 22012520 | 22012848 | 1.34073 | 1.32229 | -1.014 |
| chr10 | 56327912 | 56328277 | 0.821398 | 0.810108 | -1.014 |
| chr1 | 150802305 | 150802561 | 1.58502 | 1.5633 | -1.014 |
| chr19 | 39578760 | 39579041 | 1.09479 | 1.07983 | -1.014 |
| chr8 | 14284998 | 14285320 | 0.954401 | 0.941513 | -1.014 |
| chr12 | 3977596 | 3977832 | 0.793403 | 0.782718 | -1.014 |
| chr7 | 129247626 | 129247884 | 1.94451 | 1.91886 | -1.013 |
| chr9 | 132637667 | 132637942 | 1.01275 | 0.999464 | -1.013 |
| chr3 | 94004671 | 94004953 | 0.85639 | 0.845192 | -1.013 |
| chr9 | 79088655 | 79088927 | 0.790104 | 0.77978 | -1.013 |
| chr5 | 27828451 | 27828671 | 0.643814 | 0.635503 | -1.013 |
| chr1 | 216923434 | 216923636 | 1.23957 | 1.22363 | -1.013 |
| chr15 | 88817746 | 88818177 | 2.5671 | 2.53416 | -1.013 |
| chr1 | 209508453 | 209508763 | 2.02878 | 2.00281 | -1.013 |
| chr5 | 9291358 | 9291567 | 0.944083 | 0.932233 | -1.013 |
| chr7 | 42123803 | 42124100 | 0.914875 | 0.903574 | -1.013 |
| chr2 | 146510045 | 146510442 | 1.53857 | 1.51976 | -1.012 |
| chr16 | 46555414 | 46555679 | 0.818768 | 0.808805 | -1.012 |
| chr2 | 143941527 | 143941717 | 0.961699 | 0.950133 | -1.012 |
| chr10 | 64770991 | 64771200 | 0.765034 | 0.755861 | -1.012 |
| chr15 | 102031530 | 102031843 | 0.83351 | 0.823629 | -1.012 |
| chr7 | 132394745 | 132395199 | 1.73628 | 1.71598 | -1.012 |
| chr3 | 184445255 | 184445529 | 2.2097 | 2.18409 | -1.012 |
| chr9 | 131590485 | 131590800 | 0.945678 | 0.934757 | -1.012 |
| chr3 | 164766549 | 164766757 | 0.765771 | 0.75696 | -1.012 |
| chr6 | 126080315 | 126080526 | 1.22683 | 1.21282 | -1.012 |

| | | | | | |
|---|---|---|---|---|---|
| chr2 | 229050958 | 229051199 | 0.763349 | 0.754688 | -1.011 |
| chr5 | 52260147 | 52260356 | 1.1083 | 1.09598 | -1.011 |
| chr5 | 137618891 | 137619089 | 2.01307 | 1.99106 | -1.011 |
| chr8 | 36281579 | 36281843 | 0.89801 | 0.888215 | -1.011 |
| chr5 | 25851455 | 25851670 | 0.734716 | 0.726767 | -1.011 |
| chr1 | 168054019 | 168054369 | 3.49502 | 3.45779 | -1.011 |
| chr10 | 45949195 | 45949404 | 0.942988 | 0.933219 | -1.010 |
| chr9 | 7266922 | 7267339 | 0.843458 | 0.834807 | -1.010 |
| chr19 | 48817715 | 48817968 | 1.85078 | 1.83192 | -1.010 |
| chr8 | 123091277 | 123091556 | 1.28547 | 1.27238 | -1.010 |
| chr5 | 11839512 | 11839811 | 1.26387 | 1.25101 | -1.010 |
| chr5 | 24851308 | 24851574 | 0.926201 | 0.9168 | -1.010 |
| chr10 | 11193823 | 11194100 | 0.847296 | 0.838728 | -1.010 |
| chr12 | 84407756 | 84408008 | 0.948545 | 0.93912 | -1.010 |
| chr20 | 2668716 | 2669156 | 1.39439 | 1.38055 | -1.010 |
| chr5 | 104550269 | 104550651 | 0.982506 | 0.972849 | -1.010 |
| chr7 | 115347280 | 115347513 | 1.34578 | 1.33265 | -1.010 |
| chr9 | 104901301 | 104901506 | 1.15286 | 1.14164 | -1.010 |
| chr8 | 48775019 | 48775239 | 0.855687 | 0.847494 | -1.010 |
| chr2 | 37007436 | 37007679 | 2.4619 | 2.43837 | -1.010 |
| chr20 | 1140238 | 1140458 | 0.930819 | 0.922021 | -1.010 |
| chr17 | 7646848 | 7647248 | 3.14452 | 3.11507 | -1.009 |
| chr17 | 7463610 | 7463986 | 1.67126 | 1.6561 | -1.009 |
| chr7 | 113061820 | 113062198 | 1.7013 | 1.68619 | -1.009 |
| chr5 | 79174769 | 79174996 | 1.15952 | 1.14929 | -1.009 |
| chr7 | 125065700 | 125066048 | 1.01344 | 1.00451 | -1.009 |
| chr3 | 165572495 | 165572813 | 1.30375 | 1.29229 | -1.009 |
| chr6 | 13229382 | 13229565 | 1.3555 | 1.34361 | -1.009 |
| chr19 | 58414231 | 58414419 | 0.773644 | 0.766874 | -1.009 |
| chr10 | 26616923 | 26617259 | 0.971024 | 0.962573 | -1.009 |
| chr7 | 128512532 | 128512907 | 1.06667 | 1.05748 | -1.009 |
| chr9 | 130268168 | 130268504 | 1.79628 | 1.78117 | -1.008 |
| chr18 | 5879451 | 5879759 | 1.8989 | 1.88297 | -1.008 |
| chr19 | 48255869 | 48256114 | 0.91947 | 0.911827 | -1.008 |
| chr18 | 66327625 | 66327870 | 1.56076 | 1.54799 | -1.008 |
| chr9 | 136860204 | 136860482 | 1.57266 | 1.55987 | -1.008 |
| chr15 | 44061815 | 44062128 | 2.6034 | 2.58255 | -1.008 |
| chr10 | 45168853 | 45169233 | 0.872509 | 0.865603 | -1.008 |
| chr12 | 9013128 | 9013435 | 1.78385 | 1.76973 | -1.008 |
| chr9 | 80650376 | 80650623 | 2.82798 | 2.8056 | -1.008 |
| chr19 | 57664289 | 57664521 | 0.898427 | 0.891353 | -1.008 |
| chr12 | 103378072 | 103378409 | 0.872798 | 0.86595 | -1.008 |
| chr1 | 163595286 | 163595505 | 0.68984 | 0.684466 | -1.008 |
| chr2 | 129494201 | 129494581 | 3.17063 | 3.14609 | -1.008 |
| chr2 | 48828602 | 48828815 | 1.70451 | 1.69139 | -1.008 |
| chr10 | 68691154 | 68691423 | 2.06237 | 2.04651 | -1.008 |
| chr20 | 6518358 | 6518647 | 0.910436 | 0.90386 | -1.007 |
| chr10 | 102242236 | 102242832 | 2.88489 | 2.86417 | -1.007 |
| chr1 | 19814379 | 19814584 | 1.18803 | 1.17958 | -1.007 |
| chr2 | 97534595 | 97535101 | 2.18929 | 2.17394 | -1.007 |
| chr12 | 108376012 | 108376228 | 0.877237 | 0.871119 | -1.007 |
| chr19 | 13708563 | 13708751 | 2.03313 | 2.01899 | -1.007 |
| chr7 | 101011497 | 101011758 | 0.69172 | 0.68691 | -1.007 |
| chr9 | 21434434 | 21434751 | 1.691 | 1.67937 | -1.007 |
| chr5 | 42057411 | 42057814 | 1.78837 | 1.77617 | -1.007 |
| chr12 | 129677891 | 129678082 | 1.43057 | 1.42096 | -1.007 |
| chr10 | 28623965 | 28624193 | 0.96808 | 0.961825 | -1.007 |
| chr20 | 43919508 | 43919890 | 3.7729 | 3.7492 | -1.006 |
| chr3 | 94794351 | 94794592 | 0.722159 | 0.717793 | -1.006 |
| chr15 | 78912308 | 78912655 | 1.17703 | 1.16992 | -1.006 |
| chr4 | 25047304 | 25047596 | 1.78233 | 1.77157 | -1.006 |

| | | | | | |
|---|---|---|---|---|---|
| chr1 | 174227266 | 174227773 | 1.09413 | 1.08773 | -1.006 |
| chr1 | 205399923 | 205400281 | 4.44025 | 4.41488 | -1.006 |
| chr17 | 38267936 | 38268227 | 3.09689 | 3.0794 | -1.006 |
| chr2 | 219523328 | 219523572 | 1.70095 | 1.69139 | -1.006 |
| chr6 | 45468757 | 45468980 | 1.58254 | 1.5738 | -1.006 |
| chr2 | 190615894 | 190616153 | 1.89802 | 1.88767 | -1.005 |
| chr19 | 48271767 | 48272056 | 1.105 | 1.09912 | -1.005 |
| chr2 | 186836770 | 186836988 | 0.771644 | 0.76776 | -1.005 |
| chr5 | 32076819 | 32077114 | 1.89829 | 1.88882 | -1.005 |
| chr7 | 71360808 | 71361043 | 0.813563 | 0.809538 | -1.005 |
| chr8 | 124706036 | 124706241 | 1.59302 | 1.58516 | -1.005 |
| chr7 | 22149450 | 22150161 | 1.29903 | 1.29275 | -1.005 |
| chr9 | 102409993 | 102410233 | 0.767568 | 0.763928 | -1.005 |
| chr17 | 73923429 | 73923842 | 1.73814 | 1.72996 | -1.005 |
| chr12 | 923477 | 923877 | 1.17373 | 1.16822 | -1.005 |
| chr5 | 103228131 | 103228434 | 0.84494 | 0.840991 | -1.005 |
| chr3 | 39072352 | 39072805 | 2.07373 | 2.06409 | -1.005 |
| chr8 | 63317397 | 63317923 | 1.26285 | 1.25711 | -1.005 |
| chr20 | 23976903 | 23977280 | 1.85198 | 1.84357 | -1.005 |
| chr2 | 4113072 | 4113271 | 0.911786 | 0.907773 | -1.004 |
| chr17 | 73909775 | 73909995 | 0.979304 | 0.975045 | -1.004 |
| chr20 | 38788845 | 38789060 | 1.29349 | 1.28792 | -1.004 |
| chr19 | 31967949 | 31968166 | 1.13865 | 1.13381 | -1.004 |
| chr15 | 94711511 | 94711833 | 1.18252 | 1.17751 | -1.004 |
| chr4 | 120707044 | 120707384 | 1.19658 | 1.19162 | -1.004 |
| chr17 | 49883291 | 49883509 | 0.937566 | 0.933719 | -1.004 |
| chr12 | 115063037 | 115063352 | 2.36993 | 2.36025 | -1.004 |
| chr7 | 81345369 | 81345643 | 2.37726 | 2.36765 | -1.004 |
| chr2 | 219271270 | 219271569 | 3.49985 | 3.48573 | -1.004 |
| chr8 | 91720296 | 91720487 | 0.675287 | 0.672701 | -1.004 |
| chr19 | 32474652 | 32475003 | 1.37738 | 1.37211 | -1.004 |
| chr7 | 50244860 | 50245301 | 1.54918 | 1.54331 | -1.004 |
| chr8 | 62310846 | 62311045 | 0.980573 | 0.977013 | -1.004 |
| chr10 | 66371689 | 66371921 | 0.994295 | 0.990689 | -1.004 |
| chr19 | 33146140 | 33146342 | 1.47563 | 1.47032 | -1.004 |
| chr6 | 77165075 | 77165319 | 0.974198 | 0.970721 | -1.004 |
| chr12 | 109025788 | 109025996 | 0.848907 | 0.846007 | -1.003 |
| chr7 | 31515762 | 31516143 | 1.51712 | 1.51255 | -1.003 |
| chr14 | 43744684 | 43744924 | 1.11657 | 1.11336 | -1.003 |
| chr10 | 134607552 | 134607880 | 1.33918 | 1.33543 | -1.003 |
| chr10 | 64917877 | 64918219 | 1.84023 | 1.83542 | -1.003 |
| chr1 | 169519568 | 169519788 | 1.83111 | 1.82641 | -1.003 |
| chr12 | 88616164 | 88616646 | 0.873471 | 0.871261 | -1.003 |
| chr2 | 42009750 | 42010037 | 2.8012 | 2.79441 | -1.002 |
| chr2 | 217561178 | 217561748 | 1.12386 | 1.12115 | -1.002 |
| chr10 | 124551957 | 124552256 | 0.661444 | 0.659898 | -1.002 |
| chr10 | 86027345 | 86027607 | 1.48031 | 1.47751 | -1.002 |
| chr16 | 84310876 | 84311263 | 2.94993 | 2.94465 | -1.002 |
| chr12 | 45324909 | 45325164 | 1.213 | 1.21084 | -1.002 |
| chr5 | 115921808 | 115921998 | 0.666047 | 0.664892 | -1.002 |
| chr21 | 41884496 | 41884850 | 2.04925 | 2.04571 | -1.002 |
| chr10 | 118569083 | 118569398 | 2.26769 | 2.26398 | -1.002 |
| chr8 | 39847261 | 39847650 | 2.15708 | 2.15357 | -1.002 |
| chr2 | 160153550 | 160153771 | 1.03198 | 1.03032 | -1.002 |
| chr11 | 78357464 | 78357959 | 1.5315 | 1.52911 | -1.002 |
| chr18 | 45122404 | 45122744 | 1.05623 | 1.05473 | -1.001 |
| chr9 | 82724386 | 82724720 | 0.91589 | 0.914651 | -1.001 |
| chr2 | 47821022 | 47821280 | 0.780441 | 0.779464 | -1.001 |
| chr3 | 103027570 | 103028078 | 1.03942 | 1.03819 | -1.001 |
| chr1 | 248875066 | 248875390 | 0.80146 | 0.800516 | -1.001 |
| chr20 | 31253239 | 31253545 | 1.31708 | 1.3156 | -1.001 |

| | | | | | |
|---|---|---|---|---|---|
| chr3 | 108712589 | 108712851 | 0.896968 | 0.896102 | -1.001 |
| chr14 | 98674498 | 98674728 | 0.9713 | 0.970376 | -1.001 |
| chr20 | 56089324 | 56089741 | 1.96251 | 1.96092 | -1.001 |
| chr2 | 77156330 | 77156566 | 1.05445 | 1.05388 | -1.001 |
| chr2 | 227898085 | 227898488 | 2.71483 | 2.71354 | -1.000 |
| chr6 | 53844390 | 53844590 | 1.96608 | 1.96518 | -1.000 |
| chrX | 56790813 | 56791134 | 1.46652 | 1.46589 | -1.000 |
| chr2 | 63771181 | 63771480 | 2.63441 | 2.63339 | -1.000 |
| chr11 | 5740888 | 5741094 | 0.951683 | 0.951326 | -1.000 |
| chr2 | 188039421 | 188039628 | 0.956421 | 0.956125 | -1.000 |
| chr5 | 96499126 | 96499311 | 1.23099 | 1.23071 | -1.000 |
| chr17 | 71646300 | 71646624 | 1.84942 | 1.84906 | -1.000 |
| chr2 | 3691552 | 3691814 | 0.814374 | 0.814262 | -1.000 |
| chr2 | 80451869 | 80452150 | 0.844194 | 0.844153 | -1.000 |
| chr14 | 29220651 | 29221314 | 1.52523 | 1.5255 | 1.000 |
| chr14 | 22456171 | 22456426 | 1.92477 | 1.92513 | 1.000 |
| chr2 | 212320028 | 212320280 | 1.7759 | 1.77631 | 1.000 |
| chr19 | 29421632 | 29421869 | 0.909301 | 0.909581 | 1.000 |
| chr7 | 124937091 | 124937280 | 0.738556 | 0.738796 | 1.000 |
| chr10 | 130066290 | 130066727 | 1.37257 | 1.37318 | 1.000 |
| chr2 | 225222138 | 225222410 | 2.45513 | 2.45675 | 1.001 |
| chr16 | 66773752 | 66774169 | 1.91165 | 1.91316 | 1.001 |
| chr9 | 23803520 | 23803856 | 0.79942 | 0.800053 | 1.001 |
| chr14 | 94598716 | 94598919 | 2.24561 | 2.24842 | 1.001 |
| chr8 | 104123739 | 104123999 | 2.6418 | 2.64522 | 1.001 |
| chrX | 14715575 | 14715898 | 1.48671 | 1.48874 | 1.001 |
| chr2 | 38953100 | 38953443 | 1.65962 | 1.66215 | 1.002 |
| chr7 | 49466409 | 49466727 | 0.804057 | 0.805291 | 1.002 |
| chr8 | 140419041 | 140419329 | 0.904444 | 0.905929 | 1.002 |
| chr1 | 118693935 | 118694260 | 1.48486 | 1.48813 | 1.002 |
| chr8 | 88974342 | 88974650 | 0.887703 | 0.889692 | 1.002 |
| chr16 | 28102307 | 28102514 | 1.25415 | 1.25711 | 1.002 |
| chr3 | 77629093 | 77629308 | 1.02665 | 1.02927 | 1.003 |
| chr12 | 111482585 | 111482855 | 0.871013 | 0.873294 | 1.003 |
| chr14 | 62460773 | 62461415 | 0.78414 | 0.78621 | 1.003 |
| chr16 | 70286426 | 70286839 | 2.25853 | 2.26465 | 1.003 |
| chr9 | 72728398 | 72728726 | 2.57586 | 2.58287 | 1.003 |
| chr17 | 12381100 | 12381376 | 0.767619 | 0.769864 | 1.003 |
| chr17 | 30936184 | 30936438 | 1.70868 | 1.71372 | 1.003 |
| chr9 | 13768396 | 13768698 | 0.826871 | 0.829366 | 1.003 |
| chr2 | 239892138 | 239892560 | 3.58349 | 3.59523 | 1.003 |
| chr17 | 75524792 | 75524981 | 1.44752 | 1.45241 | 1.003 |
| chr16 | 85403496 | 85404040 | 2.58255 | 2.59144 | 1.003 |
| chr8 | 132013323 | 132013556 | 1.08894 | 1.09269 | 1.003 |
| chr20 | 44953709 | 44953991 | 1.62452 | 1.63025 | 1.004 |
| chr10 | 28364318 | 28364587 | 1.08157 | 1.08546 | 1.004 |
| chr11 | 18090022 | 18090342 | 0.943617 | 0.947198 | 1.004 |
| chr22 | 42158933 | 42159353 | 1.31463 | 1.31966 | 1.004 |
| chr5 | 24701674 | 24702029 | 0.923718 | 0.927299 | 1.004 |
| chr5 | 39950907 | 39951121 | 0.876793 | 0.880335 | 1.004 |
| chr1 | 150138233 | 150138462 | 1.8032 | 1.81108 | 1.004 |
| chr15 | 95181466 | 95181735 | 0.975942 | 0.980255 | 1.004 |
| chr2 | 77992123 | 77992392 | 0.93363 | 0.937784 | 1.004 |
| chr9 | 124465417 | 124465666 | 1.1807 | 1.18601 | 1.005 |
| chr11 | 79204559 | 79204934 | 2.38691 | 2.39822 | 1.005 |
| chr1 | 48575156 | 48575381 | 1.93863 | 1.94806 | 1.005 |
| chr19 | 37637499 | 37637774 | 1.17857 | 1.18439 | 1.005 |
| chr7 | 23983992 | 23984194 | 0.869454 | 0.87378 | 1.005 |
| chr16 | 61443016 | 61443230 | 0.997235 | 1.00231 | 1.005 |
| chr2 | 78758021 | 78758317 | 0.987857 | 0.993029 | 1.005 |
| chr15 | 81678805 | 81679019 | 1.1519 | 1.15802 | 1.005 |

| | | | | | |
|---|---|---|---|---|---|
| chr11 | 35509095 | 35509349 | 1.44853 | 1.45656 | 1.006 |
| chr17 | 27946350 | 27946582 | 1.06791 | 1.07391 | 1.006 |
| chr11 | 116603485 | 116603735 | 2.37303 | 2.3866 | 1.006 |
| chrX | 13676820 | 13677071 | 1.97935 | 1.9908 | 1.006 |
| chr14 | 57594329 | 57594516 | 0.858019 | 0.862994 | 1.006 |
| chr11 | 15229722 | 15229963 | 0.895997 | 0.901286 | 1.006 |
| chr1 | 8411556 | 8412080 | 1.78817 | 1.79881 | 1.006 |
| chr8 | 70014289 | 70014496 | 0.97415 | 0.980184 | 1.006 |
| chr7 | 105400695 | 105400982 | 1.80022 | 1.81138 | 1.006 |
| chr2 | 73220277 | 73220645 | 2.08148 | 2.09464 | 1.006 |
| chr5 | 91225091 | 91225349 | 1.33214 | 1.34083 | 1.007 |
| chr8 | 68840039 | 68840327 | 0.892462 | 0.898339 | 1.007 |
| chrX | 64748585 | 64748980 | 2.85894 | 2.87781 | 1.007 |
| chr5 | 71934780 | 71934973 | 0.971239 | 0.977755 | 1.007 |
| chr5 | 101965986 | 101966258 | 0.949247 | 0.955725 | 1.007 |
| chr7 | 12644894 | 12645078 | 1.04875 | 1.05594 | 1.007 |
| chr12 | 7798001 | 7798339 | 1.96379 | 1.97731 | 1.007 |
| chr16 | 56310757 | 56311213 | 1.14079 | 1.1487 | 1.007 |
| chr22 | 18020277 | 18020461 | 1.04158 | 1.0488 | 1.007 |
| chr20 | 49930057 | 49930358 | 0.618748 | 0.62304 | 1.007 |
| chr3 | 2385862 | 2386065 | 1.03503 | 1.04234 | 1.007 |
| chr2 | 78547017 | 78547327 | 0.924591 | 0.931125 | 1.007 |
| chr8 | 59951292 | 59951512 | 1.08337 | 1.09109 | 1.007 |
| chr18 | 49216635 | 49216884 | 0.643623 | 0.648227 | 1.007 |
| chr3 | 26532879 | 26533071 | 0.996367 | 1.00353 | 1.007 |
| chr8 | 113969718 | 113970084 | 0.887964 | 0.894455 | 1.007 |
| chr10 | 9680685 | 9680936 | 0.849254 | 0.855603 | 1.007 |
| chr8 | 4792655 | 4792946 | 1.00221 | 1.00976 | 1.008 |
| chr2 | 127808821 | 127809037 | 1.66438 | 1.67696 | 1.008 |
| chr17 | 59503423 | 59503645 | 1.333 | 1.3432 | 1.008 |
| chr2 | 54884825 | 54885232 | 1.4487 | 1.45979 | 1.008 |
| chr17 | 61064611 | 61064852 | 1.66391 | 1.6767 | 1.008 |
| chr18 | 11995440 | 11995724 | 1.75143 | 1.76494 | 1.008 |
| chr8 | 137986631 | 137986963 | 0.754377 | 0.760246 | 1.008 |
| chr5 | 4388940 | 4389174 | 1.00887 | 1.01674 | 1.008 |
| chr3 | 184072082 | 184072424 | 2.43729 | 2.45647 | 1.008 |
| chr20 | 2401918 | 2402163 | 1.08977 | 1.09845 | 1.008 |
| chr10 | 66495282 | 66495504 | 1.05998 | 1.06844 | 1.008 |
| chr1 | 175094478 | 175094702 | 2.03686 | 2.05358 | 1.008 |
| chr11 | 72320115 | 72320449 | 1.80846 | 1.8233 | 1.008 |
| chr16 | 61667267 | 61667473 | 0.708425 | 0.71436 | 1.008 |
| chr8 | 42938886 | 42939083 | 0.721953 | 0.72803 | 1.008 |
| chr2 | 76425091 | 76425362 | 1.392 | 1.40392 | 1.009 |
| chr16 | 62705081 | 62705317 | 0.821519 | 0.828562 | 1.009 |
| chr15 | 26884117 | 26884315 | 0.880888 | 0.888495 | 1.009 |
| chr2 | 174748921 | 174749434 | 1.30416 | 1.31552 | 1.009 |
| chr9 | 2228209 | 2228520 | 1.42277 | 1.43543 | 1.009 |
| chr7 | 28336586 | 28336796 | 2.02916 | 2.04737 | 1.009 |
| chr1 | 199444436 | 199444624 | 0.767356 | 0.774462 | 1.009 |
| chr7 | 145136902 | 145137124 | 0.898429 | 0.906964 | 1.010 |
| chr15 | 97424591 | 97424919 | 1.21887 | 1.23064 | 1.010 |
| chr1 | 203456409 | 203457218 | 2.27506 | 2.29713 | 1.010 |
| chr20 | 32032703 | 32032962 | 1.16023 | 1.17194 | 1.010 |
| chr2 | 77834162 | 77834440 | 0.878683 | 0.887559 | 1.010 |
| chr4 | 47124728 | 47125027 | 1.56256 | 1.57842 | 1.010 |
| chr10 | 111922779 | 111922993 | 0.876158 | 0.885057 | 1.010 |
| chr3 | 150329512 | 150329863 | 1.11313 | 1.12457 | 1.010 |
| chr1 | 237608437 | 237608658 | 0.837567 | 0.846218 | 1.010 |
| chr12 | 2710862 | 2711084 | 0.719263 | 0.726727 | 1.010 |
| chr17 | 32498169 | 32498806 | 1.60616 | 1.62292 | 1.010 |
| chr16 | 21012662 | 21012864 | 1.33769 | 1.35168 | 1.010 |

| | | | | | |
|---|---|---|---|---|---|
| chr11 | 76029777 | 76030205 | 3.02053 | 3.05236 | 1.011 |
| chr2 | 34605760 | 34605975 | 0.91544 | 0.925085 | 1.011 |
| chr12 | 3557252 | 3557472 | 0.690285 | 0.697599 | 1.011 |
| chr1 | 152863930 | 152864179 | 0.871437 | 0.880763 | 1.011 |
| chr1 | 249101330 | 249101740 | 1.93187 | 1.95285 | 1.011 |
| chr8 | 79703264 | 79703468 | 1.16374 | 1.17645 | 1.011 |
| chr1 | 17820709 | 17821088 | 2.39459 | 2.42111 | 1.011 |
| chr3 | 153916 | 154102 | 1.07864 | 1.09058 | 1.011 |
| chr5 | 4482302 | 4482726 | 1.21486 | 1.22838 | 1.011 |
| chr2 | 232466251 | 232466666 | 2.62455 | 2.65396 | 1.011 |
| chr20 | 16971820 | 16972142 | 0.938078 | 0.948696 | 1.011 |
| chr8 | 87987617 | 87987826 | 0.796733 | 0.805772 | 1.011 |
| chr2 | 33605129 | 33605414 | 2.38898 | 2.41627 | 1.011 |
| chr5 | 14358145 | 14358385 | 0.805815 | 0.815164 | 1.012 |
| chr2 | 101435052 | 101435291 | 2.3478 | 2.37561 | 1.012 |
| chr9 | 77392111 | 77392315 | 1.56198 | 1.58049 | 1.012 |
| chr14 | 99746667 | 99746897 | 0.731781 | 0.740462 | 1.012 |
| chr14 | 33944926 | 33945230 | 1.29526 | 1.31078 | 1.012 |
| chr12 | 54150973 | 54151279 | 2.90555 | 2.94042 | 1.012 |
| chr3 | 158563615 | 158563920 | 1.94725 | 1.97095 | 1.012 |
| chrX | 40126294 | 40126575 | 2.04088 | 2.06646 | 1.013 |
| chr8 | 125948761 | 125949024 | 0.87751 | 0.888523 | 1.013 |
| chr6 | 108301564 | 108301814 | 1.62664 | 1.64739 | 1.013 |
| chr16 | 70192572 | 70192795 | 1.15703 | 1.17209 | 1.013 |
| chr7 | 145545448 | 145545661 | 0.733335 | 0.742895 | 1.013 |
| chr7 | 97464795 | 97465158 | 1.39036 | 1.40863 | 1.013 |
| chr10 | 67817936 | 67818183 | 0.777196 | 0.787593 | 1.013 |
| chr20 | 4812218 | 4812418 | 0.919022 | 0.931347 | 1.013 |
| chr19 | 46282450 | 46282820 | 2.48716 | 2.52055 | 1.013 |
| chr6 | 99882814 | 99883209 | 1.14334 | 1.15893 | 1.014 |
| chr2 | 2233252 | 2233471 | 0.815471 | 0.826612 | 1.014 |
| chr20 | 40573111 | 40573397 | 0.942015 | 0.954895 | 1.014 |
| chr2 | 8784985 | 8785393 | 1.80149 | 1.82618 | 1.014 |
| chr8 | 137839612 | 137839849 | 0.965807 | 0.979108 | 1.014 |
| chr19 | 31275838 | 31276109 | 1.54742 | 1.56877 | 1.014 |
| chr8 | 63875677 | 63875985 | 0.689259 | 0.698844 | 1.014 |
| chr8 | 2372099 | 2372308 | 0.779626 | 0.790533 | 1.014 |
| chr2 | 182331387 | 182331738 | 2.7829 | 2.82291 | 1.014 |
| chr9 | 20624861 | 20625070 | 2.14016 | 2.17096 | 1.014 |
| chr2 | 14964084 | 14964294 | 1.13626 | 1.15278 | 1.015 |
| chr17 | 76070861 | 76071128 | 1.20133 | 1.21885 | 1.015 |
| chr11 | 58969043 | 58969402 | 0.910568 | 0.92408 | 1.015 |
| chr17 | 3811821 | 3812072 | 0.800308 | 0.812206 | 1.015 |
| chr3 | 101214503 | 101214739 | 1.42128 | 1.44258 | 1.015 |
| chr2 | 97429461 | 97429731 | 1.38886 | 1.40971 | 1.015 |
| chr19 | 8400546 | 8400745 | 2.23931 | 2.27296 | 1.015 |
| chr10 | 84915756 | 84916350 | 0.944816 | 0.959168 | 1.015 |
| chr2 | 45201243 | 45201710 | 1.5852 | 1.60948 | 1.015 |
| chr1 | 116524070 | 116524325 | 2.034 | 2.06523 | 1.015 |
| chr10 | 12889522 | 12889934 | 1.54749 | 1.57125 | 1.015 |
| chr2 | 124232268 | 124232492 | 0.702772 | 0.713564 | 1.015 |
| chr19 | 43193362 | 43193597 | 1.16077 | 1.17898 | 1.016 |
| chr10 | 67753535 | 67753781 | 0.801194 | 0.813871 | 1.016 |
| chr1 | 3640962 | 3641307 | 3.00538 | 3.0534 | 1.016 |
| chr19 | 28692131 | 28692457 | 1.27411 | 1.2946 | 1.016 |
| chr7 | 17746471 | 17746676 | 2.0871 | 2.12076 | 1.016 |
| chr8 | 125437436 | 125437845 | 2.00403 | 2.03636 | 1.016 |
| chr1 | 44020165 | 44020381 | 2.28332 | 2.32037 | 1.016 |
| chr16 | 53193804 | 53194046 | 2.03922 | 2.0724 | 1.016 |
| chr12 | 33861189 | 33861540 | 0.829043 | 0.842683 | 1.016 |
| chr1 | 23290714 | 23291029 | 2.98208 | 3.03152 | 1.017 |

| | | | | | |
|---|---|---|---|---|---|
| chr17 | 38269430 | 38269689 | 3.32967 | 3.38528 | 1.017 |
| chr5 | 3173085 | 3173330 | 1.08819 | 1.10636 | 1.017 |
| chr3 | 142945662 | 142945908 | 0.936798 | 0.952682 | 1.017 |
| chr1 | 114575883 | 114576194 | 1.95451 | 1.98779 | 1.017 |
| chr1 | 90081596 | 90081805 | 1.91913 | 1.9519 | 1.017 |
| chr14 | 24031941 | 24032327 | 2.8932 | 2.94264 | 1.017 |
| chr8 | 118762343 | 118762547 | 0.746485 | 0.759253 | 1.017 |
| chr7 | 86568675 | 86569050 | 1.13834 | 1.15789 | 1.017 |
| chr15 | 38975864 | 38976119 | 1.58327 | 1.61088 | 1.017 |
| chr10 | 99787923 | 99788224 | 2.1278 | 2.16499 | 1.017 |
| chr17 | 40971953 | 40972299 | 2.85118 | 2.90107 | 1.018 |
| chr15 | 81795704 | 81795925 | 0.906083 | 0.922054 | 1.018 |
| chr20 | 20511933 | 20512415 | 0.98033 | 0.997945 | 1.018 |
| chr7 | 118908935 | 118909282 | 0.868893 | 0.884514 | 1.018 |
| chr5 | 117728259 | 117728476 | 0.964817 | 0.982186 | 1.018 |
| chr14 | 73975409 | 73975706 | 1.55017 | 1.57809 | 1.018 |
| chr12 | 73466458 | 73466677 | 0.786047 | 0.800233 | 1.018 |
| chr1 | 178743389 | 178743603 | 0.771289 | 0.785334 | 1.018 |
| chr5 | 101180985 | 101181416 | 0.832793 | 0.848117 | 1.018 |
| chr19 | 35348335 | 35348590 | 0.972061 | 0.989982 | 1.018 |
| chr3 | 162760971 | 162761351 | 1.17819 | 1.20003 | 1.019 |
| chr11 | 101307293 | 101307630 | 1.74317 | 1.77556 | 1.019 |
| chr7 | 8316982 | 8317327 | 1.44898 | 1.47595 | 1.019 |
| chr14 | 65226876 | 65227221 | 2.66182 | 2.7114 | 1.019 |
| chr3 | 136829303 | 136829537 | 1.41155 | 1.43803 | 1.019 |
| chr5 | 33869179 | 33869588 | 0.902438 | 0.919506 | 1.019 |
| chr11 | 37540997 | 37541199 | 0.918156 | 0.93558 | 1.019 |
| chr15 | 93340344 | 93340699 | 1.47348 | 1.5015 | 1.019 |
| chr10 | 15691481 | 15691715 | 1.18274 | 1.20554 | 1.019 |
| chr5 | 13268522 | 13268768 | 0.961522 | 0.980171 | 1.019 |
| chr14 | 105956090 | 105956287 | 1.74199 | 1.77591 | 1.019 |
| chr2 | 152803687 | 152804237 | 1.66548 | 1.69799 | 1.020 |
| chr8 | 90134174 | 90134380 | 0.91135 | 0.929425 | 1.020 |
| chr9 | 130965388 | 130965759 | 3.2217 | 3.28607 | 1.020 |
| chr20 | 53263464 | 53263647 | 1.92662 | 1.96537 | 1.020 |
| chr3 | 22223241 | 22223530 | 0.858075 | 0.875337 | 1.020 |
| chr12 | 133252847 | 133253070 | 0.636962 | 0.649793 | 1.020 |
| chr5 | 53539400 | 53539583 | 1.90309 | 1.94149 | 1.020 |
| chr3 | 72987968 | 72988219 | 1.06441 | 1.08602 | 1.020 |
| chr6 | 135505080 | 135505279 | 2.11283 | 2.15616 | 1.021 |
| chr16 | 79439867 | 79440083 | 1.02079 | 1.04176 | 1.021 |
| chr5 | 31855493 | 31855754 | 1.23886 | 1.26431 | 1.021 |
| chr8 | 14403633 | 14403818 | 1.27624 | 1.30245 | 1.021 |
| chr16 | 17257018 | 17257353 | 0.989814 | 1.01019 | 1.021 |
| chr9 | 24377014 | 24377287 | 1.0448 | 1.06641 | 1.021 |
| chr12 | 92615929 | 92616142 | 0.807273 | 0.824117 | 1.021 |
| chr2 | 31138404 | 31138642 | 1.31465 | 1.3421 | 1.021 |
| chr5 | 134797924 | 134798356 | 1.65407 | 1.68894 | 1.021 |
| chr17 | 66610110 | 66610384 | 1.76434 | 1.8016 | 1.021 |
| chr5 | 179335782 | 179335990 | 1.32482 | 1.35289 | 1.021 |
| chr2 | 1890253 | 1890495 | 2.32349 | 2.37287 | 1.021 |
| chr11 | 112045777 | 112046082 | 2.57687 | 2.63239 | 1.022 |
| chr2 | 84250020 | 84250248 | 2.39042 | 2.44199 | 1.022 |
| chr9 | 133307761 | 133307982 | 1.32292 | 1.35155 | 1.022 |
| chr10 | 79365300 | 79365545 | 1.57374 | 1.60791 | 1.022 |
| chr20 | 1489984 | 1490213 | 1.60048 | 1.63533 | 1.022 |
| chr2 | 35653703 | 35653897 | 1.35188 | 1.38149 | 1.022 |
| chr4 | 84181790 | 84182020 | 1.71411 | 1.75182 | 1.022 |
| chr5 | 156262993 | 156263219 | 1.36268 | 1.39269 | 1.022 |
| chr7 | 55338792 | 55339028 | 1.33637 | 1.36599 | 1.022 |
| chr8 | 53490170 | 53490739 | 1.45004 | 1.4823 | 1.022 |

| | | | | | |
|---|---|---|---|---|---|
| chr12 | 118942831 | 118943215 | 0.872822 | 0.892541 | 1.023 |
| chr2 | 1322477 | 1322685 | 0.910029 | 0.930618 | 1.023 |
| chr10 | 109914772 | 109915015 | 0.834467 | 0.8535 | 1.023 |
| chr8 | 112043741 | 112044050 | 0.844896 | 0.864198 | 1.023 |
| chr16 | 30093993 | 30094290 | 1.25982 | 1.28864 | 1.023 |
| chr17 | 4086781 | 4087064 | 1.00883 | 1.03191 | 1.023 |
| chr7 | 53305945 | 53306198 | 0.966705 | 0.98899 | 1.023 |
| chr20 | 61024940 | 61025288 | 2.23519 | 2.28696 | 1.023 |
| chr2 | 216878168 | 216878482 | 2.23959 | 2.29178 | 1.023 |
| chr12 | 67961578 | 67961906 | 0.819062 | 0.838189 | 1.023 |
| chr11 | 14996373 | 14996705 | 2.22897 | 2.2811 | 1.023 |
| chr3 | 34056999 | 34057510 | 1.40125 | 1.43403 | 1.023 |
| chr2 | 24283821 | 24284095 | 1.93543 | 1.98111 | 1.024 |
| chr8 | 71716197 | 71716450 | 1.01657 | 1.04071 | 1.024 |
| chr10 | 58360007 | 58360248 | 0.871377 | 0.892283 | 1.024 |
| chr2 | 215867307 | 215867559 | 1.42705 | 1.4615 | 1.024 |
| chr20 | 61538744 | 61539098 | 1.67926 | 1.7198 | 1.024 |
| chr15 | 95118970 | 95119336 | 0.925599 | 0.948033 | 1.024 |
| chr3 | 22346956 | 22347138 | 0.915136 | 0.937473 | 1.024 |
| chr11 | 84095769 | 84095986 | 2.28324 | 2.33927 | 1.025 |
| chr2 | 76014742 | 76014963 | 1.3165 | 1.3489 | 1.025 |
| chr2 | 80669346 | 80669547 | 0.947519 | 0.970834 | 1.025 |
| chr19 | 29517233 | 29517679 | 1.14707 | 1.17544 | 1.025 |
| chr1 | 6245855 | 6246099 | 1.44906 | 1.48503 | 1.025 |
| chr12 | 127174235 | 127174455 | 0.855584 | 0.876857 | 1.025 |
| chr16 | 20311578 | 20312150 | 2.36605 | 2.42494 | 1.025 |
| chr12 | 43790705 | 43790962 | 0.958709 | 0.982636 | 1.025 |
| chr8 | 104004752 | 104004997 | 0.796976 | 0.816916 | 1.025 |
| chr1 | 145507714 | 145508001 | 1.96277 | 2.01241 | 1.025 |
| chr19 | 37720914 | 37721197 | 1.31759 | 1.35163 | 1.026 |
| chr7 | 85600327 | 85600509 | 0.887259 | 0.910238 | 1.026 |
| chr7 | 105934882 | 105935369 | 0.98614 | 1.01187 | 1.026 |
| chr7 | 86539509 | 86539752 | 0.703366 | 0.721822 | 1.026 |
| chr9 | 92717159 | 92717366 | 0.887668 | 0.911134 | 1.026 |
| chr10 | 119624830 | 119625061 | 0.753969 | 0.773962 | 1.027 |
| chr3 | 39041354 | 39041763 | 1.67142 | 1.71584 | 1.027 |
| chr9 | 71122428 | 71122782 | 1.43247 | 1.47054 | 1.027 |
| chr20 | 39868688 | 39868941 | 0.857874 | 0.880722 | 1.027 |
| chr21 | 33306098 | 33306361 | 1.53376 | 1.57468 | 1.027 |
| chr17 | 74121441 | 74121642 | 0.913144 | 0.937564 | 1.027 |
| chr8 | 26621558 | 26621986 | 1.21101 | 1.2436 | 1.027 |
| chr2 | 105466018 | 105466338 | 2.40006 | 2.46511 | 1.027 |
| chr7 | 101959077 | 101959301 | 0.972996 | 0.999416 | 1.027 |
| chr19 | 42527688 | 42528330 | 2.38643 | 2.45181 | 1.027 |
| chr8 | 42928681 | 42928994 | 2.42184 | 2.48841 | 1.027 |
| chr10 | 35525762 | 35526021 | 2.04942 | 2.10616 | 1.028 |
| chr15 | 77265741 | 77265942 | 1.91486 | 1.96805 | 1.028 |
| chr17 | 65555549 | 65555737 | 1.54473 | 1.58793 | 1.028 |
| chr2 | 17181372 | 17181639 | 0.971978 | 0.999224 | 1.028 |
| chr5 | 92556961 | 92557179 | 0.915448 | 0.941174 | 1.028 |
| chr10 | 24028367 | 24028790 | 0.931121 | 0.957476 | 1.028 |
| chr2 | 54990814 | 54991228 | 1.43773 | 1.4787 | 1.029 |
| chr8 | 5587874 | 5588268 | 0.989307 | 1.01754 | 1.029 |
| chr12 | 117992825 | 117993076 | 0.847566 | 0.871806 | 1.029 |
| chr7 | 31348246 | 31348506 | 0.779022 | 0.801333 | 1.029 |
| chr6 | 84245660 | 84246068 | 1.99945 | 2.05674 | 1.029 |
| chr9 | 100230889 | 100231244 | 1.91304 | 1.96851 | 1.029 |
| chr1 | 40498017 | 40498205 | 1.46724 | 1.50983 | 1.029 |
| chr7 | 26404791 | 26404995 | 2.06973 | 2.13016 | 1.029 |
| chr17 | 40315139 | 40315416 | 2.47093 | 2.54336 | 1.029 |
| chr8 | 136914402 | 136914627 | 0.936423 | 0.963944 | 1.029 |

| | | | | | |
|---|---|---|---|---|---|
| chr2 | 11919327 | 11919702 | 2.93669 | 3.02318 | 1.029 |
| chr8 | 57101792 | 57102014 | 1.94889 | 2.00667 | 1.030 |
| chr5 | 42971412 | 42971696 | 0.978436 | 1.00752 | 1.030 |
| chr19 | 36435466 | 36435786 | 1.03648 | 1.06734 | 1.030 |
| chr2 | 75693888 | 75694139 | 2.33038 | 2.39976 | 1.030 |
| chr20 | 38248067 | 38248350 | 0.879867 | 0.906221 | 1.030 |
| chr3 | 10363826 | 10364227 | 1.08306 | 1.11574 | 1.030 |
| chr8 | 85944183 | 85944403 | 1.01927 | 1.05013 | 1.030 |
| chr7 | 156700049 | 156700298 | 0.852435 | 0.878336 | 1.030 |
| chr1 | 171157882 | 171158089 | 2.21499 | 2.28242 | 1.030 |
| chr8 | 10405611 | 10405998 | 2.44292 | 2.51773 | 1.031 |
| chr9 | 129967454 | 129967866 | 0.965922 | 0.995503 | 1.031 |
| chr11 | 16273951 | 16274219 | 1.40392 | 1.44697 | 1.031 |
| chr1 | 174665158 | 174665365 | 0.775933 | 0.799911 | 1.031 |
| chr10 | 116063744 | 116063985 | 1.90661 | 1.96558 | 1.031 |
| chr2 | 107203833 | 107204159 | 2.26317 | 2.33351 | 1.031 |
| chr9 | 135603528 | 135603721 | 0.663854 | 0.684533 | 1.031 |
| chr6 | 13873934 | 13874404 | 2.4389 | 2.51498 | 1.031 |
| chr10 | 48377599 | 48378139 | 2.14627 | 2.21323 | 1.031 |
| chr14 | 97600719 | 97601053 | 0.735763 | 0.758799 | 1.031 |
| chr1 | 207038266 | 207038637 | 2.93001 | 3.02247 | 1.032 |
| chr3 | 119400190 | 119400374 | 0.961417 | 0.991799 | 1.032 |
| chr19 | 52286818 | 52287083 | 1.11368 | 1.14889 | 1.032 |
| chr5 | 111521620 | 111522440 | 1.30505 | 1.34636 | 1.032 |
| chr20 | 20645470 | 20645709 | 1.97959 | 2.04241 | 1.032 |
| chr14 | 40537727 | 40537981 | 1.11315 | 1.14855 | 1.032 |
| chr5 | 119991477 | 119991866 | 1.60544 | 1.65657 | 1.032 |
| chr2 | 78877900 | 78878093 | 1.17846 | 1.21605 | 1.032 |
| chr19 | 28845702 | 28846035 | 0.964057 | 0.994818 | 1.032 |
| chr20 | 12235260 | 12235482 | 1.10054 | 1.13586 | 1.032 |
| chr5 | 99015919 | 99016101 | 1.22416 | 1.26362 | 1.032 |
| chr1 | 171954064 | 171954297 | 1.8676 | 1.92784 | 1.032 |
| chr2 | 56304764 | 56305012 | 2.10322 | 2.1712 | 1.032 |
| chr16 | 632195 | 632411 | 0.808495 | 0.83464 | 1.032 |
| chrX | 21763952 | 21764237 | 2.13057 | 2.19957 | 1.032 |
| chr19 | 7406059 | 7406267 | 2.32928 | 2.40488 | 1.032 |
| chr20 | 20264352 | 20264727 | 0.878389 | 0.90729 | 1.033 |
| chr14 | 45899226 | 45899561 | 0.713273 | 0.736786 | 1.033 |
| chr4 | 122647902 | 122648139 | 2.47684 | 2.55876 | 1.033 |
| chr2 | 220507229 | 220507575 | 4.18563 | 4.32427 | 1.033 |
| chr9 | 113978978 | 113979162 | 1.00816 | 1.04155 | 1.033 |
| chr15 | 94721893 | 94722077 | 1.00067 | 1.03384 | 1.033 |
| chr10 | 105887267 | 105887548 | 1.66396 | 1.71917 | 1.033 |
| chr14 | 91821318 | 91821864 | 1.25591 | 1.29771 | 1.033 |
| chr2 | 6762410 | 6762613 | 0.901932 | 0.93203 | 1.033 |
| chr14 | 67584984 | 67585318 | 1.72785 | 1.78581 | 1.034 |
| chr19 | 49717195 | 49717437 | 1.39811 | 1.44503 | 1.034 |
| chr8 | 4028555 | 4028797 | 1.1031 | 1.1403 | 1.034 |
| chr5 | 135957621 | 135957824 | 0.685695 | 0.708925 | 1.034 |
| chr11 | 14253873 | 14254584 | 1.33995 | 1.38536 | 1.034 |
| chr2 | 128348992 | 128349391 | 4.26884 | 4.4135 | 1.034 |
| chr20 | 24145059 | 24145272 | 1.91477 | 1.97974 | 1.034 |
| chr20 | 58173700 | 58174214 | 2.3353 | 2.41461 | 1.034 |
| chr11 | 58162724 | 58163007 | 0.831635 | 0.859885 | 1.034 |
| chr16 | 67678588 | 67679019 | 2.85745 | 2.95482 | 1.034 |
| chr8 | 141300148 | 141300653 | 0.99788 | 1.03216 | 1.034 |
| chr2 | 153597306 | 153597550 | 1.82116 | 1.88396 | 1.034 |
| chr2 | 139131546 | 139131828 | 1.65566 | 1.71279 | 1.035 |
| chr11 | 48329392 | 48329666 | 2.73524 | 2.82986 | 1.035 |
| chr10 | 49732321 | 49732765 | 3.02533 | 3.13063 | 1.035 |
| chr14 | 67752540 | 67752818 | 1.17727 | 1.21831 | 1.035 |

| | | | | | |
|---|---|---|---|---|---|
| chr3 | 17195876 | 17196156 | 0.822027 | 0.850856 | 1.035 |
| chr20 | 30550553 | 30550783 | 1.00584 | 1.04113 | 1.035 |
| chr19 | 31540386 | 31540652 | 1.51365 | 1.56696 | 1.035 |
| chr17 | 38346815 | 38347042 | 1.28139 | 1.32654 | 1.035 |
| chr7 | 46932554 | 46932768 | 1.39624 | 1.44556 | 1.035 |
| chr15 | 92052489 | 92052935 | 1.0538 | 1.09118 | 1.035 |
| chr8 | 93063998 | 93064284 | 0.83155 | 0.861247 | 1.036 |
| chr8 | 135173407 | 135173615 | 0.830591 | 0.860284 | 1.036 |
| chr1 | 220738711 | 220739014 | 0.749844 | 0.776707 | 1.036 |
| chr10 | 91838391 | 91838646 | 0.974591 | 1.00984 | 1.036 |
| chr8 | 57659800 | 57660030 | 0.910748 | 0.943745 | 1.036 |
| chr2 | 45296268 | 45296470 | 0.769806 | 0.797737 | 1.036 |
| chr11 | 87931144 | 87931361 | 0.984863 | 1.0206 | 1.036 |
| chr8 | 137776303 | 137776500 | 0.751658 | 0.778934 | 1.036 |
| chr20 | 60055214 | 60055424 | 1.69562 | 1.75742 | 1.036 |
| chr11 | 118901612 | 118901895 | 1.82896 | 1.89572 | 1.037 |
| chr12 | 45806341 | 45806539 | 0.860321 | 0.891786 | 1.037 |
| chr3 | 185720875 | 185721152 | 1.0563 | 1.09494 | 1.037 |
| chr2 | 105851972 | 105852425 | 1.71639 | 1.77935 | 1.037 |
| chr3 | 42654987 | 42655222 | 1.21116 | 1.25559 | 1.037 |
| chr7 | 141527203 | 141527585 | 0.908046 | 0.941352 | 1.037 |
| chr8 | 78979083 | 78979348 | 0.759435 | 0.787489 | 1.037 |
| chr9 | 123201434 | 123201690 | 2.13854 | 2.21778 | 1.037 |
| chr2 | 34772443 | 34772775 | 1.05533 | 1.09479 | 1.037 |
| chr5 | 76113880 | 76114130 | 1.14826 | 1.1913 | 1.037 |
| chr15 | 90053167 | 90053453 | 0.89943 | 0.933151 | 1.037 |
| chr8 | 137344319 | 137344524 | 0.885793 | 0.919079 | 1.038 |
| chr12 | 77257023 | 77257304 | 0.87083 | 0.903921 | 1.038 |
| chr2 | 158485551 | 158485974 | 2.47344 | 2.56745 | 1.038 |
| chr12 | 71714424 | 71714621 | 0.854978 | 0.887518 | 1.038 |
| chr15 | 95581316 | 95581522 | 0.859019 | 0.891758 | 1.038 |
| chr2 | 179626473 | 179626888 | 1.86146 | 1.93271 | 1.038 |
| chr16 | 13967542 | 13968195 | 1.91238 | 1.98565 | 1.038 |
| chr9 | 123492212 | 123492399 | 1.46952 | 1.52591 | 1.038 |
| chr7 | 131438450 | 131438776 | 3.12838 | 3.24874 | 1.038 |
| chr12 | 123724776 | 123724990 | 1.16337 | 1.20832 | 1.039 |
| chr2 | 10188782 | 10189124 | 1.45616 | 1.51279 | 1.039 |
| chr5 | 36944768 | 36944980 | 1.40223 | 1.45684 | 1.039 |
| chr6 | 49377728 | 49377945 | 2.02501 | 2.10388 | 1.039 |
| chr5 | 136466220 | 136466449 | 0.780132 | 0.81053 | 1.039 |
| chr5 | 17620681 | 17621165 | 1.47592 | 1.53359 | 1.039 |
| chr10 | 65685168 | 65685375 | 0.91731 | 0.953313 | 1.039 |
| chr14 | 22552629 | 22553055 | 2.07715 | 2.15906 | 1.039 |
| chr5 | 4670787 | 4670982 | 0.701216 | 0.728978 | 1.040 |
| chrX | 78337469 | 78337688 | 1.32051 | 1.3728 | 1.040 |
| chr3 | 127192204 | 127192490 | 1.13099 | 1.1758 | 1.040 |
| chr20 | 9448657 | 9448980 | 2.69243 | 2.79931 | 1.040 |
| chr5 | 137643644 | 137643857 | 0.87915 | 0.914227 | 1.040 |
| chr19 | 12782781 | 12783114 | 2.20165 | 2.28961 | 1.040 |
| chrX | 19400939 | 19401237 | 2.25385 | 2.34397 | 1.040 |
| chr5 | 22349561 | 22350048 | 0.988969 | 1.02861 | 1.040 |
| chr1 | 163328713 | 163329233 | 1.49432 | 1.55433 | 1.040 |
| chr14 | 64945106 | 64945650 | 1.04719 | 1.08926 | 1.040 |
| chr5 | 115403817 | 115404040 | 1.78595 | 1.85773 | 1.040 |
| chr9 | 3467638 | 3467980 | 3.27618 | 3.40799 | 1.040 |
| chr4 | 187476619 | 187476818 | 1.59826 | 1.66258 | 1.040 |
| chr12 | 128872345 | 128872676 | 1.44276 | 1.5009 | 1.040 |
| chr8 | 64519338 | 64519739 | 0.96638 | 1.0054 | 1.040 |
| chr12 | 23441714 | 23442005 | 0.877691 | 0.913199 | 1.040 |
| chr15 | 52451764 | 52452060 | 0.782452 | 0.81411 | 1.040 |
| chr7 | 137335653 | 137335869 | 0.701033 | 0.729394 | 1.040 |

| | | | | | |
|---|---|---|---|---|---|
| chr7 | 1126519 | 1127034 | 2.56034 | 2.664 | 1.040 |
| chr2 | 71737293 | 71737747 | 3.38434 | 3.52145 | 1.041 |
| chr8 | 108043138 | 108043369 | 1.01312 | 1.05419 | 1.041 |
| chr14 | 77519631 | 77519996 | 2.48219 | 2.58301 | 1.041 |
| chr17 | 48602329 | 48602536 | 0.788954 | 0.821239 | 1.041 |
| chr5 | 134793167 | 134793561 | 3.14259 | 3.27171 | 1.041 |
| chr1 | 230860418 | 230860639 | 0.748141 | 0.778897 | 1.041 |
| chr5 | 8011914 | 8012212 | 0.989288 | 1.03001 | 1.041 |
| chr19 | 30314888 | 30315132 | 1.04833 | 1.09154 | 1.041 |
| chr5 | 113014702 | 113014923 | 1.68977 | 1.75969 | 1.041 |
| chr3 | 15161807 | 15162021 | 0.794897 | 0.827796 | 1.041 |
| chr7 | 22772180 | 22772707 | 1.10044 | 1.146 | 1.041 |
| chr2 | 227990233 | 227990548 | 1.3089 | 1.36325 | 1.042 |
| chr3 | 169547578 | 169547791 | 0.883575 | 0.920335 | 1.042 |
| chr3 | 1126558 | 1126835 | 0.989177 | 1.03042 | 1.042 |
| chr2 | 78753579 | 78753924 | 0.872679 | 0.909178 | 1.042 |
| chr16 | 34618626 | 34619153 | 0.878288 | 0.915073 | 1.042 |
| chr3 | 1852768 | 1853006 | 0.989422 | 1.03125 | 1.042 |
| chr16 | 72434270 | 72434526 | 1.9307 | 2.01245 | 1.042 |
| chr8 | 6884252 | 6884573 | 1.09022 | 1.13638 | 1.042 |
| chr5 | 68561864 | 68562166 | 1.84959 | 1.92813 | 1.042 |
| chr15 | 93129271 | 93129494 | 0.730991 | 0.762173 | 1.043 |
| chr5 | 168975427 | 168975637 | 1.81692 | 1.89474 | 1.043 |
| chr8 | 69306867 | 69307126 | 1.60953 | 1.67854 | 1.043 |
| chr16 | 83102085 | 83102319 | 0.843582 | 0.879761 | 1.043 |
| chr3 | 110728437 | 110728646 | 0.784023 | 0.817662 | 1.043 |
| chr8 | 114052460 | 114052702 | 0.98138 | 1.0235 | 1.043 |
| chr1 | 101754028 | 101754288 | 1.78803 | 1.86512 | 1.043 |
| chr10 | 61385823 | 61386066 | 1.77724 | 1.85406 | 1.043 |
| chr5 | 93548311 | 93548499 | 0.874177 | 0.912099 | 1.043 |
| chr15 | 58024748 | 58025067 | 3.00641 | 3.13705 | 1.043 |
| chr12 | 44986311 | 44986593 | 0.830958 | 0.867132 | 1.044 |
| chr8 | 41943321 | 41943596 | 1.02711 | 1.07202 | 1.044 |
| chr17 | 77898158 | 77898534 | 2.53287 | 2.64365 | 1.044 |
| chr12 | 1202414 | 1202635 | 2.12755 | 2.22084 | 1.044 |
| chr22 | 45664885 | 45665153 | 2.66497 | 2.78194 | 1.044 |
| chr17 | 1532365 | 1532596 | 1.87267 | 1.95499 | 1.044 |
| chr7 | 18103972 | 18104194 | 1.32628 | 1.38467 | 1.044 |
| chr2 | 9918143 | 9918332 | 1.71033 | 1.78578 | 1.044 |
| chr18 | 10490786 | 10491000 | 2.04706 | 2.13764 | 1.044 |
| chr2 | 232539535 | 232539882 | 2.12539 | 2.21974 | 1.044 |
| chr8 | 15047109 | 15047316 | 0.904193 | 0.94439 | 1.044 |
| chr1 | 55743272 | 55743494 | 2.54085 | 2.65389 | 1.044 |
| chr6 | 39833161 | 39833406 | 1.38062 | 1.4421 | 1.045 |
| chr14 | 63119660 | 63119868 | 0.855336 | 0.893442 | 1.045 |
| chr22 | 22326697 | 22327126 | 2.74319 | 2.86565 | 1.045 |
| chr10 | 100185397 | 100185779 | 2.49604 | 2.6076 | 1.045 |
| chr5 | 156781948 | 156782180 | 1.85631 | 1.93952 | 1.045 |
| chr2 | 40177922 | 40178189 | 1.04991 | 1.09709 | 1.045 |
| chr5 | 143392534 | 143392731 | 1.0063 | 1.05207 | 1.045 |
| chr10 | 68741756 | 68742011 | 0.915687 | 0.957427 | 1.046 |
| chr6 | 108675333 | 108675549 | 0.698214 | 0.730083 | 1.046 |
| chr1 | 202688783 | 202689026 | 1.98683 | 2.07754 | 1.046 |
| chr15 | 97370811 | 97371026 | 0.915681 | 0.95749 | 1.046 |
| chr19 | 40843672 | 40843920 | 1.18664 | 1.24083 | 1.046 |
| chr2 | 50520382 | 50520827 | 0.889409 | 0.930034 | 1.046 |
| chr3 | 121209640 | 121209939 | 0.978998 | 1.02375 | 1.046 |
| chr19 | 51409428 | 51409756 | 1.65975 | 1.73563 | 1.046 |
| chr2 | 13790682 | 13790926 | 1.28342 | 1.34245 | 1.046 |
| chr12 | 82021360 | 82021672 | 1.43336 | 1.49959 | 1.046 |
| chr15 | 98182017 | 98182295 | 0.979355 | 1.02461 | 1.046 |

| | | | | | |
|---|---|---|---|---|---|
| chr8 | 99395366 | 99395646 | 1.92368 | 2.01258 | 1.046 |
| chr11 | 5497531 | 5497743 | 1.83982 | 1.92491 | 1.046 |
| chr1 | 177537887 | 177538159 | 0.817469 | 0.855313 | 1.046 |
| chr2 | 63919244 | 63919438 | 1.56717 | 1.63976 | 1.046 |
| chr2 | 45834926 | 45835177 | 1.70709 | 1.78628 | 1.046 |
| chr8 | 49984632 | 49984880 | 0.787617 | 0.824633 | 1.047 |
| chr2 | 75325951 | 75326355 | 1.50182 | 1.5725 | 1.047 |
| chr9 | 10511266 | 10511485 | 0.957643 | 1.00277 | 1.047 |
| chr18 | 53082565 | 53082833 | 2.22321 | 2.32827 | 1.047 |
| chr1 | 195614749 | 195614965 | 0.677268 | 0.709431 | 1.047 |
| chr12 | 50156715 | 50157254 | 2.34055 | 2.4519 | 1.048 |
| chr6 | 8219675 | 8219904 | 1.98063 | 2.07491 | 1.048 |
| chr3 | 26609978 | 26610179 | 0.865528 | 0.906935 | 1.048 |
| chr14 | 70591131 | 70591353 | 0.899833 | 0.942889 | 1.048 |
| chr16 | 77235232 | 77235480 | 0.845851 | 0.886323 | 1.048 |
| chr11 | 7145219 | 7145518 | 0.825589 | 0.865111 | 1.048 |
| chr19 | 34309426 | 34309659 | 0.846845 | 0.887423 | 1.048 |
| chr19 | 42144393 | 42144631 | 1.12276 | 1.17657 | 1.048 |
| chr14 | 54823021 | 54823217 | 1.07575 | 1.12758 | 1.048 |
| chr15 | 93573934 | 93574260 | 2.84693 | 2.98461 | 1.048 |
| chr1 | 186299969 | 186300191 | 1.21767 | 1.27674 | 1.049 |
| chr7 | 2584480 | 2584708 | 0.847028 | 0.888187 | 1.049 |
| chrX | 149885339 | 149885613 | 2.50551 | 2.62756 | 1.049 |
| chr5 | 130388243 | 130388454 | 0.858 | 0.89982 | 1.049 |
| chr17 | 60793245 | 60793520 | 0.808527 | 0.847972 | 1.049 |
| chr10 | 92639975 | 92640246 | 1.50944 | 1.58346 | 1.049 |
| chr15 | 47488249 | 47488496 | 1.47111 | 1.54363 | 1.049 |
| chr1 | 192457250 | 192457481 | 1.20574 | 1.26524 | 1.049 |
| chr7 | 148553445 | 148553786 | 1.87272 | 1.96554 | 1.050 |
| chr15 | 89244893 | 89245120 | 2.1966 | 2.30554 | 1.050 |
| chr19 | 38713218 | 38713455 | 0.94283 | 0.989637 | 1.050 |
| chr16 | 30705318 | 30705561 | 2.39336 | 2.51253 | 1.050 |
| chr6 | 118151084 | 118151331 | 1.09633 | 1.15108 | 1.050 |
| chr9 | 73963812 | 73964037 | 0.94586 | 0.993131 | 1.050 |
| chr2 | 14755975 | 14756307 | 0.985914 | 1.03536 | 1.050 |
| chr5 | 153537018 | 153537293 | 2.01475 | 2.11598 | 1.050 |
| chrX | 103072552 | 103072766 | 0.953462 | 1.0014 | 1.050 |
| chrX | 105658617 | 105659002 | 1.61577 | 1.69729 | 1.050 |
| chr13 | 33684065 | 33684264 | 1.01817 | 1.06966 | 1.051 |
| chr7 | 34715424 | 34715648 | 1.85013 | 1.94369 | 1.051 |
| chr11 | 57568251 | 57568468 | 2.45271 | 2.57685 | 1.051 |
| chr19 | 48606359 | 48606609 | 2.41782 | 2.54028 | 1.051 |
| chr2 | 113482514 | 113482794 | 1.51733 | 1.59423 | 1.051 |
| chr2 | 180615971 | 180616184 | 1.7052 | 1.79163 | 1.051 |
| chr2 | 206544769 | 206545100 | 1.93565 | 2.0338 | 1.051 |
| chr5 | 112974036 | 112974329 | 0.995273 | 1.04574 | 1.051 |
| chr7 | 144088936 | 144089156 | 2.30526 | 2.42234 | 1.051 |
| chr16 | 1913636 | 1913884 | 1.58478 | 1.66541 | 1.051 |
| chr17 | 72953952 | 72954444 | 3.01845 | 3.17213 | 1.051 |
| chr5 | 43609960 | 43610162 | 1.11207 | 1.16873 | 1.051 |
| chr2 | 24644456 | 24644778 | 1.49529 | 1.57173 | 1.051 |
| chr14 | 92852029 | 92852245 | 0.634897 | 0.667491 | 1.051 |
| chr12 | 23806516 | 23806732 | 0.960358 | 1.00998 | 1.052 |
| chr1 | 154017442 | 154017758 | 0.766263 | 0.805877 | 1.052 |
| chr3 | 39748918 | 39749129 | 1.79453 | 1.88733 | 1.052 |
| chr1 | 206971509 | 206971797 | 2.28988 | 2.40834 | 1.052 |
| chr17 | 74143714 | 74144121 | 1.50948 | 1.5877 | 1.052 |
| chr1 | 152430938 | 152431211 | 2.74551 | 2.88789 | 1.052 |
| chr15 | 91236984 | 91237224 | 1.86891 | 1.96596 | 1.052 |
| chr2 | 225314152 | 225314555 | 1.71559 | 1.80505 | 1.052 |
| chr19 | 30833833 | 30834087 | 1.09383 | 1.15103 | 1.052 |

| | | | | | |
|---|---|---|---|---|---|
| chr7 | 5527424 | 5527903 | 2.00415 | 2.10907 | 1.052 |
| chr14 | 79032840 | 79033297 | 1.75627 | 1.84832 | 1.052 |
| chr5 | 164703340 | 164703606 | 0.848471 | 0.893091 | 1.053 |
| chr1 | 221508082 | 221508341 | 0.923423 | 0.972086 | 1.053 |
| chr12 | 127778090 | 127778369 | 1.84887 | 1.94717 | 1.053 |
| chr5 | 97903885 | 97904118 | 0.970918 | 1.02259 | 1.053 |
| chr10 | 116218808 | 116218999 | 1.65029 | 1.73821 | 1.053 |
| chr15 | 31498586 | 31498917 | 1.45823 | 1.53593 | 1.053 |
| chr17 | 38776153 | 38776574 | 3.13183 | 3.29889 | 1.053 |
| chr8 | 6921075 | 6921312 | 1.78674 | 1.88227 | 1.053 |
| chr8 | 4031497 | 4031787 | 0.880381 | 0.927531 | 1.054 |
| chr11 | 25949104 | 25949351 | 0.775819 | 0.817387 | 1.054 |
| chr19 | 17836626 | 17836930 | 1.74454 | 1.83837 | 1.054 |
| chr16 | 13547906 | 13548089 | 1.34939 | 1.4221 | 1.054 |
| chr2 | 44564897 | 44565099 | 1.45128 | 1.52951 | 1.054 |
| chr16 | 9349822 | 9350328 | 1.48868 | 1.56912 | 1.054 |
| chr8 | 131989240 | 131989449 | 0.96891 | 1.02132 | 1.054 |
| chr8 | 52633363 | 52633660 | 1.02731 | 1.08302 | 1.054 |
| chr2 | 41925868 | 41926342 | 2.35256 | 2.48022 | 1.054 |
| chr8 | 41815474 | 41815668 | 1.32908 | 1.40119 | 1.054 |
| chr21 | 35615957 | 35616159 | 1.72988 | 1.82396 | 1.054 |
| chr7 | 42016791 | 42017131 | 1.47393 | 1.55426 | 1.055 |
| chr3 | 152512429 | 152512738 | 0.859476 | 0.906347 | 1.055 |
| chr8 | 132816257 | 132816804 | 2.41919 | 2.55121 | 1.055 |
| chr9 | 91120870 | 91121079 | 0.815782 | 0.860311 | 1.055 |
| chr10 | 52968382 | 52968644 | 0.738065 | 0.778473 | 1.055 |
| chr2 | 57881966 | 57882430 | 0.927811 | 0.978622 | 1.055 |
| chr12 | 53462142 | 53462338 | 0.944571 | 0.996304 | 1.055 |
| chr10 | 84609792 | 84610088 | 0.816944 | 0.861726 | 1.055 |
| chr2 | 58647272 | 58647505 | 1.07146 | 1.13026 | 1.055 |
| chr22 | 29709118 | 29709490 | 1.93143 | 2.03754 | 1.055 |
| chr11 | 58555857 | 58556235 | 2.10827 | 2.22416 | 1.055 |
| chr13 | 26636999 | 26637309 | 1.35127 | 1.42567 | 1.055 |
| chr2 | 139586929 | 139587222 | 1.81605 | 1.91607 | 1.055 |
| chr19 | 36422553 | 36422840 | 3.7908 | 3.99975 | 1.055 |
| chr5 | 76267405 | 76267613 | 0.870194 | 0.918244 | 1.055 |
| chr9 | 117424446 | 117424783 | 2.2733 | 2.39892 | 1.055 |
| chr7 | 2491020 | 2491527 | 2.32204 | 2.45045 | 1.055 |
| chr20 | 23121752 | 23122233 | 1.04142 | 1.09904 | 1.055 |
| chr8 | 136978735 | 136978951 | 0.83914 | 0.885762 | 1.056 |
| chr7 | 141079009 | 141079254 | 0.790469 | 0.834397 | 1.056 |
| chr5 | 89729991 | 89730246 | 2.12541 | 2.24353 | 1.056 |
| chr7 | 2199708 | 2199960 | 1.37804 | 1.45487 | 1.056 |
| chr2 | 49173678 | 49173959 | 1.69688 | 1.79156 | 1.056 |
| chr15 | 101137052 | 101137252 | 0.891794 | 0.941657 | 1.056 |
| chr16 | 82913925 | 82914124 | 0.90041 | 0.950767 | 1.056 |
| chr14 | 73761937 | 73762169 | 1.38124 | 1.45859 | 1.056 |
| chr10 | 83967094 | 83967294 | 0.622778 | 0.657701 | 1.056 |
| chr8 | 140417833 | 140418414 | 1.65708 | 1.75032 | 1.056 |
| chr14 | 87834544 | 87834849 | 0.775981 | 0.81965 | 1.056 |
| chr3 | 81835611 | 81836162 | 1.00588 | 1.06251 | 1.056 |
| chr2 | 220554519 | 220554746 | 1.86152 | 1.96667 | 1.056 |
| chr19 | 33893340 | 33893687 | 1.43154 | 1.51265 | 1.057 |
| chr3 | 106783249 | 106783650 | 2.39695 | 2.53281 | 1.057 |
| chr3 | 20537380 | 20537598 | 0.9788 | 1.03437 | 1.057 |
| chr14 | 32395361 | 32395549 | 1.73126 | 1.82979 | 1.057 |
| chr8 | 106785068 | 106785461 | 1.35875 | 1.43611 | 1.057 |
| chr3 | 140956973 | 140957216 | 0.853993 | 0.902732 | 1.057 |
| chr8 | 101576241 | 101576532 | 3.06885 | 3.2444 | 1.057 |
| chr3 | 89689718 | 89690000 | 0.875342 | 0.925515 | 1.057 |
| chr20 | 38254056 | 38254388 | 1.07841 | 1.14024 | 1.057 |

| | | | | | |
|---|---|---|---|---|---|
| chr2 | 205556693 | 205556917 | 2.31889 | 2.45189 | 1.057 |
| chr1 | 207085920 | 207086345 | 1.44776 | 1.53083 | 1.057 |
| chr10 | 100041755 | 100042172 | 0.805194 | 0.851406 | 1.057 |
| chr7 | 142578744 | 142579007 | 1.89438 | 2.00312 | 1.057 |
| chr17 | 36577942 | 36578183 | 0.874796 | 0.925045 | 1.057 |
| chr7 | 126268117 | 126268421 | 0.703062 | 0.743455 | 1.057 |
| chr9 | 109499934 | 109500142 | 0.997426 | 1.05476 | 1.057 |
| chr16 | 4561538 | 4561792 | 2.53425 | 2.68045 | 1.058 |
| chr3 | 189462205 | 189462465 | 0.907044 | 0.959495 | 1.058 |
| chr12 | 23101473 | 23101716 | 1.05701 | 1.11823 | 1.058 |
| chr2 | 11049837 | 11050180 | 2.49491 | 2.64005 | 1.058 |
| chr17 | 31570941 | 31571155 | 1.04103 | 1.1016 | 1.058 |
| chr12 | 2400346 | 2400683 | 2.06956 | 2.19016 | 1.058 |
| chr12 | 16052260 | 16052523 | 1.99993 | 2.1167 | 1.058 |
| chr20 | 3230781 | 3230994 | 2.10636 | 2.22948 | 1.058 |
| chr20 | 38209670 | 38209893 | 0.901125 | 0.953943 | 1.059 |
| chr3 | 182627137 | 182627416 | 1.10821 | 1.17325 | 1.059 |
| chr10 | 30287246 | 30287745 | 3.51903 | 3.72562 | 1.059 |
| chr19 | 31187518 | 31187796 | 1.68324 | 1.78234 | 1.059 |
| chr8 | 50485743 | 50485978 | 1.04051 | 1.10192 | 1.059 |
| chr17 | 8638354 | 8638558 | 0.841674 | 0.891371 | 1.059 |
| chr10 | 105619034 | 105619248 | 1.69166 | 1.79179 | 1.059 |
| chr8 | 95092003 | 95092383 | 3.02663 | 3.20618 | 1.059 |
| chr1 | 154414957 | 154415165 | 1.33443 | 1.41367 | 1.059 |
| chr17 | 60797853 | 60798040 | 1.64823 | 1.74619 | 1.059 |
| chr2 | 85413338 | 85413577 | 2.27675 | 2.41205 | 1.059 |
| chrX | 115336251 | 115336459 | 1.5344 | 1.6265 | 1.060 |
| chr7 | 27235890 | 27236195 | 1.6423 | 1.7409 | 1.060 |
| chr2 | 30709500 | 30709737 | 1.13975 | 1.20827 | 1.060 |
| chr10 | 86394554 | 86394756 | 0.953185 | 1.01051 | 1.060 |
| chr10 | 26805765 | 26806042 | 0.8126 | 0.861543 | 1.060 |
| chr12 | 82270653 | 82270956 | 0.858384 | 0.910081 | 1.060 |
| chr7 | 30204131 | 30204445 | 1.9132 | 2.02883 | 1.060 |
| chr3 | 123599645 | 123600297 | 1.24123 | 1.31631 | 1.060 |
| chr2 | 158159081 | 158159321 | 1.82122 | 1.93175 | 1.061 |
| chr9 | 87997040 | 87997256 | 0.741059 | 0.786097 | 1.061 |
| chr2 | 214060545 | 214060833 | 0.841222 | 0.892371 | 1.061 |
| chr10 | 24792600 | 24792886 | 2.9959 | 3.1784 | 1.061 |
| chr11 | 114609750 | 114609955 | 2.0477 | 2.17247 | 1.061 |
| chr10 | 135132136 | 135132543 | 2.76657 | 2.93525 | 1.061 |
| chr1 | 47190916 | 47191192 | 2.40495 | 2.55217 | 1.061 |
| chr1 | 214505173 | 214505675 | 2.41354 | 2.5615 | 1.061 |
| chr20 | 7759703 | 7759898 | 1.05675 | 1.12166 | 1.061 |
| chr19 | 31805949 | 31806304 | 0.9516 | 1.01008 | 1.061 |
| chr15 | 27584254 | 27584526 | 0.886397 | 0.940937 | 1.062 |
| chr5 | 107596247 | 107596644 | 2.81938 | 2.99298 | 1.062 |
| chr15 | 27219280 | 27219588 | 1.95545 | 2.07587 | 1.062 |
| chr13 | 54356624 | 54356882 | 1.69221 | 1.79645 | 1.062 |
| chr3 | 82558237 | 82558443 | 0.72722 | 0.772024 | 1.062 |
| chr10 | 15017974 | 15018163 | 1.27743 | 1.35623 | 1.062 |
| chr16 | 3704243 | 3704844 | 1.99353 | 2.11659 | 1.062 |
| chr7 | 56012620 | 56012824 | 0.768619 | 0.816092 | 1.062 |
| chr12 | 113452780 | 113452985 | 0.859149 | 0.912389 | 1.062 |
| chr7 | 119847018 | 119847381 | 0.844166 | 0.896483 | 1.062 |
| chr15 | 79500571 | 79500810 | 1.3116 | 1.393 | 1.062 |
| chr9 | 32805970 | 32806191 | 0.774595 | 0.823063 | 1.063 |
| chr7 | 37188432 | 37188784 | 0.76276 | 0.810613 | 1.063 |
| chr2 | 71377378 | 71377647 | 0.959835 | 1.02013 | 1.063 |
| chr20 | 19673314 | 19673598 | 3.9692 | 4.21858 | 1.063 |
| chr2 | 88650504 | 88651062 | 2.54137 | 2.70155 | 1.063 |
| chr12 | 55526864 | 55527155 | 1.1248 | 1.19582 | 1.063 |

| | | | | | |
|---|---|---|---|---|---|
| chr3 | 120277796 | 120278130 | 2.47715 | 2.63386 | 1.063 |
| chr14 | 99793533 | 99793879 | 1.02262 | 1.08749 | 1.063 |
| chr17 | 16826371 | 16826654 | 0.833729 | 0.886652 | 1.063 |
| chr19 | 35626126 | 35626649 | 3.51118 | 3.73421 | 1.064 |
| chr2 | 218764303 | 218764741 | 1.60005 | 1.70188 | 1.064 |
| chr19 | 36861391 | 36861640 | 1.00538 | 1.06953 | 1.064 |
| chr17 | 47445850 | 47446097 | 1.56311 | 1.66313 | 1.064 |
| chr7 | 146721852 | 146722115 | 0.954675 | 1.01587 | 1.064 |
| chr8 | 115530535 | 115530732 | 0.886814 | 0.943734 | 1.064 |
| chr20 | 4475392 | 4475688 | 0.975463 | 1.03808 | 1.064 |
| chr2 | 14572117 | 14572326 | 1.21883 | 1.29717 | 1.064 |
| chr2 | 207846540 | 207846778 | 2.24317 | 2.3875 | 1.064 |
| chr10 | 50417203 | 50417401 | 0.703683 | 0.748982 | 1.064 |
| chr18 | 19024527 | 19024761 | 1.73524 | 1.84716 | 1.065 |
| chr1 | 216857110 | 216857346 | 0.704839 | 0.750336 | 1.065 |
| chr2 | 128936855 | 128937133 | 1.91676 | 2.04088 | 1.065 |
| chr7 | 28698119 | 28698352 | 0.833856 | 0.887921 | 1.065 |
| chr1 | 243507314 | 243507587 | 2.16536 | 2.3061 | 1.065 |
| chr12 | 126563046 | 126563369 | 0.879698 | 0.936888 | 1.065 |
| chr20 | 20421966 | 20422195 | 1.17928 | 1.25594 | 1.065 |
| chr9 | 87019139 | 87019351 | 1.2461 | 1.32716 | 1.065 |
| chr5 | 136572771 | 136573185 | 0.796902 | 0.848769 | 1.065 |
| chr10 | 66388743 | 66389131 | 0.778141 | 0.828962 | 1.065 |
| chr20 | 1455541 | 1455816 | 1.41626 | 1.50877 | 1.065 |
| chr8 | 74659208 | 74659486 | 2.52479 | 2.68975 | 1.065 |
| chr10 | 15767745 | 15768022 | 1.76799 | 1.88407 | 1.066 |
| chr7 | 89390028 | 89390250 | 0.75505 | 0.804651 | 1.066 |
| chr17 | 69168618 | 69168859 | 1.06489 | 1.13486 | 1.066 |
| chr2 | 217087266 | 217087560 | 0.917572 | 0.977878 | 1.066 |
| chr5 | 125556520 | 125556747 | 0.993228 | 1.05871 | 1.066 |
| chr2 | 154335231 | 154335647 | 2.99677 | 3.19471 | 1.066 |
| chr1 | 185702984 | 185703193 | 1.88782 | 2.01305 | 1.066 |
| chr19 | 50043818 | 50044064 | 0.716238 | 0.763761 | 1.066 |
| chr3 | 33186031 | 33186308 | 1.90364 | 2.02995 | 1.066 |
| chr14 | 79314647 | 79314873 | 2.28899 | 2.44145 | 1.067 |
| chr12 | 62513293 | 62513513 | 0.743664 | 0.793363 | 1.067 |
| chr14 | 63506559 | 63506786 | 1.46789 | 1.56631 | 1.067 |
| chr2 | 30064310 | 30064590 | 2.40461 | 2.56591 | 1.067 |
| chr2 | 78447585 | 78447828 | 0.867868 | 0.926137 | 1.067 |
| chr8 | 69922856 | 69923408 | 1.04967 | 1.12021 | 1.067 |
| chr2 | 8059674 | 8059893 | 0.855204 | 0.912699 | 1.067 |
| chr2 | 128112929 | 128113350 | 1.50372 | 1.60487 | 1.067 |
| chr10 | 27171004 | 27171291 | 1.90309 | 2.03113 | 1.067 |
| chr17 | 67096082 | 67096331 | 0.864957 | 0.92317 | 1.067 |
| chr5 | 103968674 | 103968893 | 0.79532 | 0.848881 | 1.067 |
| chr2 | 61653463 | 61653851 | 1.65048 | 1.76177 | 1.067 |
| chr9 | 133557973 | 133558227 | 2.50161 | 2.67035 | 1.067 |
| chr11 | 60033021 | 60033360 | 1.34528 | 1.43652 | 1.068 |
| chr9 | 140245879 | 140246089 | 1.03499 | 1.1053 | 1.068 |
| chr21 | 43757979 | 43758250 | 2.12774 | 2.27236 | 1.068 |
| chr14 | 51595978 | 51596169 | 1.13717 | 1.21463 | 1.068 |
| chr5 | 92245310 | 92245644 | 1.58407 | 1.69204 | 1.068 |
| chr8 | 80178200 | 80178409 | 1.01354 | 1.08266 | 1.068 |
| chr7 | 153374347 | 153374567 | 0.885482 | 0.945945 | 1.068 |
| chr9 | 97443776 | 97444016 | 1.67103 | 1.78526 | 1.068 |
| chr2 | 85875790 | 85876136 | 2.15943 | 2.30709 | 1.068 |
| chr10 | 66793686 | 66793990 | 0.930586 | 0.994241 | 1.068 |
| chr14 | 36994064 | 36995072 | 1.87591 | 2.00434 | 1.068 |
| chr17 | 25758004 | 25758248 | 2.00294 | 2.14014 | 1.069 |
| chr17 | 55910522 | 55911136 | 3.06412 | 3.27411 | 1.069 |
| chr9 | 19712466 | 19712739 | 0.848688 | 0.906853 | 1.069 |

| | | | | | |
|---|---|---|---|---|---|
| chr11 | 66258743 | 66259121 | 1.50665 | 1.60993 | 1.069 |
| chr8 | 54223819 | 54224023 | 0.897344 | 0.958855 | 1.069 |
| chr1 | 39074060 | 39074293 | 1.05411 | 1.12643 | 1.069 |
| chr17 | 72812315 | 72812656 | 1.00661 | 1.07568 | 1.069 |
| chr1 | 216787433 | 216787641 | 0.804776 | 0.860006 | 1.069 |
| chr1 | 167237096 | 167237363 | 2.44966 | 2.61784 | 1.069 |
| chr2 | 224592286 | 224592509 | 1.73876 | 1.85826 | 1.069 |
| chr10 | 120000808 | 120001110 | 2.77627 | 2.96715 | 1.069 |
| chr14 | 82946848 | 82947052 | 0.923647 | 0.987164 | 1.069 |
| chr1 | 153353232 | 153353444 | 1.11456 | 1.19125 | 1.069 |
| chr10 | 128259830 | 128260067 | 1.08214 | 1.15697 | 1.069 |
| chr3 | 47879132 | 47879673 | 2.23777 | 2.39278 | 1.069 |
| chr4 | 160113274 | 160113493 | 0.912231 | 0.975495 | 1.069 |
| chr1 | 154307536 | 154307865 | 3.0724 | 3.28572 | 1.069 |
| chr17 | 76819914 | 76820266 | 0.994097 | 1.06329 | 1.070 |
| chr8 | 134773160 | 134773411 | 0.709642 | 0.759048 | 1.070 |
| chr17 | 78338053 | 78338278 | 2.53716 | 2.71413 | 1.070 |
| chr1 | 38955510 | 38955755 | 2.17328 | 2.32498 | 1.070 |
| chr3 | 52048771 | 52048968 | 1.1651 | 1.2467 | 1.070 |
| chr8 | 144875964 | 144876215 | 1.62081 | 1.73451 | 1.070 |
| chr17 | 4152748 | 4153016 | 2.29287 | 2.4538 | 1.070 |
| chr2 | 184766733 | 184766972 | 0.850536 | 0.910246 | 1.070 |
| chr12 | 125961151 | 125961439 | 0.791509 | 0.847132 | 1.070 |
| chr12 | 24376432 | 24376635 | 0.676639 | 0.724489 | 1.071 |
| chr11 | 63597278 | 63597703 | 1.26353 | 1.35344 | 1.071 |
| chr14 | 88050967 | 88051302 | 0.851905 | 0.91259 | 1.071 |
| chr2 | 219757495 | 219757765 | 1.92294 | 2.05996 | 1.071 |
| chr16 | 86374232 | 86374596 | 2.1201 | 2.27121 | 1.071 |
| chr4 | 187590594 | 187590845 | 1.0529 | 1.12803 | 1.071 |
| chr3 | 76478992 | 76479203 | 0.993119 | 1.06405 | 1.071 |
| chr3 | 127689912 | 127690157 | 2.20163 | 2.35911 | 1.072 |
| chr15 | 98859100 | 98859352 | 0.873538 | 0.936028 | 1.072 |
| chr12 | 12769640 | 12769822 | 0.912329 | 0.977651 | 1.072 |
| chr12 | 60999323 | 60999517 | 0.752456 | 0.806334 | 1.072 |
| chr15 | 28266666 | 28267144 | 1.33585 | 1.4316 | 1.072 |
| chr2 | 128407350 | 128407632 | 2.28219 | 2.446 | 1.072 |
| chr11 | 71855088 | 71855326 | 2.17278 | 2.32927 | 1.072 |
| chr11 | 43526203 | 43526402 | 1.32391 | 1.41941 | 1.072 |
| chr10 | 111896873 | 111897180 | 1.42391 | 1.52675 | 1.072 |
| chr2 | 219575443 | 219575803 | 2.09292 | 2.24407 | 1.072 |
| chr7 | 21081444 | 21081688 | 0.804657 | 0.862768 | 1.072 |
| chr5 | 141339638 | 141340081 | 1.25094 | 1.34137 | 1.072 |
| chr4 | 112204272 | 112204510 | 2.07133 | 2.22158 | 1.073 |
| chr2 | 78360126 | 78360416 | 1.19426 | 1.28094 | 1.073 |
| chr8 | 52441236 | 52441419 | 1.16499 | 1.24972 | 1.073 |
| chr19 | 737743 | 738041 | 2.16875 | 2.32677 | 1.073 |
| chr2 | 3492935 | 3493154 | 1.05084 | 1.12742 | 1.073 |
| chr14 | 57440945 | 57441247 | 0.718016 | 0.770448 | 1.073 |
| chrX | 17025364 | 17025596 | 1.15649 | 1.24108 | 1.073 |
| chr9 | 92125663 | 92126125 | 0.689445 | 0.739883 | 1.073 |
| chr19 | 38286835 | 38287085 | 1.11054 | 1.19189 | 1.073 |
| chr2 | 240536948 | 240537133 | 0.710919 | 0.763006 | 1.073 |
| chr2 | 204647012 | 204647508 | 1.97208 | 2.11671 | 1.073 |
| chr2 | 2440223 | 2440525 | 0.761542 | 0.817401 | 1.073 |
| chr5 | 149047166 | 149047505 | 2.40964 | 2.58663 | 1.073 |
| chr19 | 5154242 | 5154477 | 2.02091 | 2.16942 | 1.073 |
| chr7 | 127660269 | 127660545 | 1.66201 | 1.7842 | 1.074 |
| chr12 | 106372947 | 106373245 | 0.906136 | 0.972778 | 1.074 |
| chr1 | 236701992 | 236702215 | 0.914789 | 0.982089 | 1.074 |
| chr12 | 33905898 | 33906110 | 0.975297 | 1.04709 | 1.074 |
| chr2 | 241922800 | 241923089 | 1.97567 | 2.12124 | 1.074 |

| | | | | | |
|---|---|---|---|---|---|
| chr5 | 42881178 | 42881390 | 1.07433 | 1.15349 | 1.074 |
| chr17 | 7545847 | 7546099 | 0.562695 | 0.604301 | 1.074 |
| chr10 | 123456401 | 123456664 | 1.60102 | 1.71953 | 1.074 |
| chr7 | 97492767 | 97492964 | 0.736715 | 0.791375 | 1.074 |
| chr15 | 92934909 | 92935162 | 1.68779 | 1.81309 | 1.074 |
| chr1 | 239061217 | 239061829 | 0.810943 | 0.871254 | 1.074 |
| chr11 | 92244002 | 92244193 | 0.876232 | 0.941577 | 1.075 |
| chr20 | 54919001 | 54919437 | 1.74934 | 1.87995 | 1.075 |
| chr6 | 47015437 | 47015626 | 1.20535 | 1.29551 | 1.075 |
| chr1 | 176998025 | 176998220 | 0.942133 | 1.01267 | 1.075 |
| chr2 | 62126040 | 62126251 | 1.87714 | 2.01769 | 1.075 |
| chr2 | 161126596 | 161126835 | 1.38104 | 1.48491 | 1.075 |
| chr4 | 89290410 | 89290690 | 1.56164 | 1.67913 | 1.075 |
| chr15 | 98835683 | 98836349 | 2.25522 | 2.42508 | 1.075 |
| chr6 | 48444013 | 48444209 | 0.959072 | 1.03156 | 1.076 |
| chr10 | 25756581 | 25756787 | 0.797126 | 0.857381 | 1.076 |
| chr5 | 16307997 | 16308206 | 0.893214 | 0.960769 | 1.076 |
| chr8 | 42029193 | 42029461 | 2.25746 | 2.4282 | 1.076 |
| chr10 | 135422372 | 135422639 | 1.39849 | 1.50533 | 1.076 |
| chr2 | 38766257 | 38766593 | 1.14085 | 1.22802 | 1.076 |
| chr16 | 67213388 | 67213825 | 1.12494 | 1.211 | 1.077 |
| chr17 | 13018428 | 13018837 | 0.900208 | 0.969168 | 1.077 |
| chr5 | 160766969 | 160767217 | 0.732636 | 0.788879 | 1.077 |
| chr10 | 125318880 | 125319185 | 2.89806 | 3.12092 | 1.077 |
| chr2 | 235459235 | 235459532 | 2.5196 | 2.71348 | 1.077 |
| chr19 | 43196186 | 43196407 | 1.12962 | 1.21667 | 1.077 |
| chr7 | 122794309 | 122794580 | 1.48401 | 1.59838 | 1.077 |
| chr14 | 54887260 | 54887540 | 1.62922 | 1.75489 | 1.077 |
| rUn_gl0002 | 124839 | 125121 | 1.08698 | 1.17083 | 1.077 |
| chr10 | 35250063 | 35250269 | 1.48419 | 1.59915 | 1.077 |
| chr8 | 133305281 | 133305598 | 0.830448 | 0.894864 | 1.078 |
| chr14 | 61028857 | 61029216 | 2.71961 | 2.93133 | 1.078 |
| chr19 | 47032815 | 47033177 | 2.97273 | 3.20415 | 1.078 |
| chr3 | 16778958 | 16779237 | 0.778594 | 0.839238 | 1.078 |
| chr20 | 343224 | 343446 | 2.08209 | 2.2443 | 1.078 |
| chr5 | 54698050 | 54698260 | 1.03345 | 1.11427 | 1.078 |
| chr10 | 64843573 | 64844061 | 1.17857 | 1.27079 | 1.078 |
| rUn_gl0002 | 145252 | 145637 | 1.16289 | 1.25404 | 1.078 |
| chr12 | 30175408 | 30175690 | 1.11787 | 1.20591 | 1.079 |
| chr1 | 9844651 | 9844887 | 1.91941 | 2.0707 | 1.079 |
| chr3 | 165399237 | 165399543 | 0.858485 | 0.926168 | 1.079 |
| chr12 | 49691906 | 49692165 | 1.16502 | 1.25709 | 1.079 |
| chr8 | 71437991 | 71438211 | 0.698189 | 0.753493 | 1.079 |
| chr3 | 169660117 | 169660345 | 0.743264 | 0.80238 | 1.080 |
| gl000195_r | 20826 | 21096 | 0.93756 | 1.01223 | 1.080 |
| rUn_gl0002 | 123336 | 123636 | 1.10998 | 1.19851 | 1.080 |
| chr8 | 20292815 | 20293289 | 1.95111 | 2.10693 | 1.080 |
| chr1 | 184282926 | 184283329 | 0.88607 | 0.956924 | 1.080 |
| chr6 | 24934157 | 24934378 | 1.91321 | 2.06619 | 1.080 |
| chr2 | 28636351 | 28636644 | 1.33305 | 1.43968 | 1.080 |
| chr4 | 40151555 | 40151812 | 1.24305 | 1.34269 | 1.080 |
| chr19 | 4495495 | 4495887 | 1.99646 | 2.15658 | 1.080 |
| chr17 | 51358094 | 51358316 | 0.94589 | 1.02188 | 1.080 |
| chr3 | 7208123 | 7208389 | 0.883379 | 0.954408 | 1.080 |
| chr12 | 91338894 | 91339102 | 1.75612 | 1.89745 | 1.080 |
| chr13 | 32484157 | 32484360 | 1.98012 | 2.13961 | 1.081 |
| rUn_gl0002 | 129508 | 129749 | 0.937373 | 1.01302 | 1.081 |
| chr10 | 92452136 | 92452368 | 0.765122 | 0.826956 | 1.081 |
| chr17 | 12906151 | 12906969 | 2.21524 | 2.39453 | 1.081 |
| chr5 | 76856565 | 76856785 | 1.63997 | 1.77292 | 1.081 |
| chr19 | 36183003 | 36183249 | 0.918641 | 0.993232 | 1.081 |

| | | | | | |
|---|---|---|---|---|---|
| chr20 | 36175904 | 36176108 | 0.82384 | 0.890733 | 1.081 |
| chr2 | 46782138 | 46782399 | 1.401 | 1.51484 | 1.081 |
| chr2 | 165698494 | 165698881 | 2.18428 | 2.36198 | 1.081 |
| chr19 | 4557705 | 4557907 | 1.59444 | 1.72423 | 1.081 |
| chr17 | 2255653 | 2255886 | 0.789661 | 0.853972 | 1.081 |
| chr7 | 122662645 | 122662860 | 1.25731 | 1.3599 | 1.082 |
| chr10 | 20077061 | 20077300 | 1.83642 | 1.98646 | 1.082 |
| chr2 | 4770990 | 4771177 | 0.898999 | 0.972665 | 1.082 |
| chr12 | 64667625 | 64667863 | 1.29842 | 1.40493 | 1.082 |
| chrX | 113952387 | 113952629 | 1.40383 | 1.519 | 1.082 |
| chr10 | 36766735 | 36766947 | 0.661098 | 0.71538 | 1.082 |
| chr19 | 10594296 | 10594536 | 1.69869 | 1.8386 | 1.082 |
| chr5 | 4261884 | 4262152 | 0.872367 | 0.944218 | 1.082 |
| chr5 | 52012240 | 52012591 | 0.762996 | 0.825855 | 1.082 |
| chr11 | 21220825 | 21221065 | 0.583504 | 0.631579 | 1.082 |
| chr4 | 11425680 | 11425961 | 1.38178 | 1.49562 | 1.082 |
| chr5 | 151894534 | 151894734 | 1.11943 | 1.2117 | 1.082 |
| chr2 | 50410854 | 50411057 | 0.854043 | 0.924451 | 1.082 |
| chr19 | 30159758 | 30160462 | 1.69857 | 1.83869 | 1.082 |
| chr4 | 143641356 | 143641584 | 1.73179 | 1.87527 | 1.083 |
| chr1 | 78799192 | 78799416 | 1.9144 | 2.07304 | 1.083 |
| chr14 | 98604876 | 98605238 | 1.11291 | 1.20525 | 1.083 |
| chr1 | 220132187 | 220132419 | 0.690767 | 0.748106 | 1.083 |
| chr7 | 106790936 | 106791149 | 1.60207 | 1.73511 | 1.083 |
| chr6 | 127587627 | 127587811 | 2.1853 | 2.3668 | 1.083 |
| chr8 | 139070915 | 139071179 | 0.760467 | 0.823642 | 1.083 |
| chr17 | 66007259 | 66007672 | 3.34939 | 3.62781 | 1.083 |
| chr8 | 75824161 | 75824562 | 0.882116 | 0.955471 | 1.083 |
| chr3 | 26399573 | 26400029 | 0.839911 | 0.909837 | 1.083 |
| chr17 | 49565385 | 49565629 | 1.22795 | 1.33025 | 1.083 |
| chr20 | 17323159 | 17323437 | 0.9386 | 1.01683 | 1.083 |
| chr3 | 24903212 | 24903567 | 0.993213 | 1.07604 | 1.083 |
| chr20 | 51833608 | 51834031 | 2.81747 | 3.05265 | 1.083 |
| chr20 | 24847861 | 24848177 | 1.11728 | 1.21062 | 1.084 |
| chr9 | 131103734 | 131104006 | 1.57785 | 1.70967 | 1.084 |
| chr7 | 28338534 | 28338806 | 1.43827 | 1.55863 | 1.084 |
| chr16 | 72170436 | 72170667 | 0.881233 | 0.954994 | 1.084 |
| chr17 | 63194860 | 63195163 | 1.63081 | 1.7675 | 1.084 |
| chr11 | 122659545 | 122659786 | 1.5095 | 1.63617 | 1.084 |
| chr2 | 122732645 | 122732929 | 1.60387 | 1.73846 | 1.084 |
| chr2 | 34682230 | 34682446 | 0.698282 | 0.756897 | 1.084 |
| chr2 | 51202794 | 51203013 | 0.896886 | 0.972267 | 1.084 |
| chr14 | 20955025 | 20955281 | 1.00479 | 1.08938 | 1.084 |
| chr5 | 84242158 | 84242441 | 0.986337 | 1.0695 | 1.084 |
| chr20 | 56771266 | 56771535 | 0.909937 | 0.98673 | 1.084 |
| chr8 | 143632469 | 143633139 | 1.73582 | 1.88239 | 1.084 |
| chr12 | 118911109 | 118911354 | 0.863198 | 0.936193 | 1.085 |
| chr3 | 104315447 | 104315813 | 1.30757 | 1.41835 | 1.085 |
| chr14 | 57953536 | 57953807 | 1.8146 | 1.96847 | 1.085 |
| chr12 | 131644388 | 131644608 | 0.981621 | 1.06492 | 1.085 |
| chr5 | 30854223 | 30854639 | 0.80972 | 0.878485 | 1.085 |
| chr16 | 31470307 | 31470536 | 1.78778 | 1.93962 | 1.085 |
| chr9 | 82790187 | 82790491 | 1.81308 | 1.96718 | 1.085 |
| chr7 | 145885285 | 145885500 | 0.823437 | 0.893453 | 1.085 |
| chr10 | 53131448 | 53131786 | 0.977017 | 1.06016 | 1.085 |
| chr1 | 167684725 | 167684981 | 2.72206 | 2.95428 | 1.085 |
| chr17 | 72002229 | 72002446 | 0.775802 | 0.842281 | 1.086 |
| chr12 | 1027623 | 1027882 | 1.03108 | 1.11945 | 1.086 |
| chr1 | 27429243 | 27429504 | 2.68269 | 2.91282 | 1.086 |
| chr22 | 34507283 | 34507539 | 1.67516 | 1.81915 | 1.086 |
| chr14 | 85494322 | 85494576 | 0.736259 | 0.799573 | 1.086 |

| | | | | | |
|---|---|---|---|---|---|
| chr15 | 94652513 | 94653046 | 1.02638 | 1.11469 | 1.086 |
| chr2 | 37763353 | 37763579 | 1.69275 | 1.83841 | 1.086 |
| chr20 | 55152613 | 55152972 | 2.25936 | 2.45409 | 1.086 |
| chr11 | 9628685 | 9629064 | 1.25467 | 1.36284 | 1.086 |
| chr4 | 88386970 | 88387272 | 1.15778 | 1.2577 | 1.086 |
| chr2 | 84858274 | 84858541 | 1.37939 | 1.49854 | 1.086 |
| chr8 | 68872527 | 68872729 | 0.946308 | 1.02826 | 1.087 |
| chr14 | 41147981 | 41148204 | 1.0526 | 1.14379 | 1.087 |
| chr15 | 24304654 | 24304961 | 1.04336 | 1.13393 | 1.087 |
| chr9 | 103377300 | 103377570 | 0.750326 | 0.815564 | 1.087 |
| chr2 | 97073044 | 97073397 | 2.41435 | 2.62431 | 1.087 |
| chr16 | 69964135 | 69964506 | 1.97096 | 2.14238 | 1.087 |
| chr6 | 46188972 | 46189250 | 1.98048 | 2.15281 | 1.087 |
| chr14 | 49772900 | 49773316 | 1.21196 | 1.31742 | 1.087 |
| chr9 | 127539523 | 127539782 | 1.63482 | 1.77732 | 1.087 |
| chr1 | 108019829 | 108020044 | 0.903125 | 0.981864 | 1.087 |
| chr7 | 132202979 | 132203259 | 1.09555 | 1.19109 | 1.087 |
| chr7 | 17474382 | 17474618 | 0.778903 | 0.846961 | 1.087 |
| chr8 | 51464395 | 51464594 | 0.961409 | 1.04558 | 1.088 |
| chr7 | 158439899 | 158440268 | 1.93765 | 2.10737 | 1.088 |
| chr20 | 62586611 | 62587292 | 0.959222 | 1.04336 | 1.088 |
| chr2 | 25790407 | 25790603 | 1.48214 | 1.61217 | 1.088 |
| chr14 | 90526603 | 90527072 | 1.4572 | 1.58529 | 1.088 |
| chr2 | 212507770 | 212508032 | 0.836872 | 0.910589 | 1.088 |
| chr5 | 165999986 | 166000252 | 0.687249 | 0.747793 | 1.088 |
| chr1 | 23215788 | 23216027 | 1.72247 | 1.87423 | 1.088 |
| chr7 | 145166703 | 145166958 | 0.8544 | 0.929826 | 1.088 |
| chr16 | 34456218 | 34456526 | 0.870956 | 0.947974 | 1.088 |
| chr11 | 57107774 | 57108056 | 1.207 | 1.31377 | 1.088 |
| chr14 | 22306027 | 22306220 | 0.909526 | 0.990304 | 1.089 |
| chr20 | 38931362 | 38931586 | 1.0539 | 1.14751 | 1.089 |
| chr5 | 118127527 | 118128041 | 1.32074 | 1.43805 | 1.089 |
| chr2 | 6612859 | 6613129 | 0.877276 | 0.95531 | 1.089 |
| chr13 | 20763435 | 20763698 | 0.929808 | 1.01254 | 1.089 |
| chr2 | 228103005 | 228103290 | 2.97153 | 3.23615 | 1.089 |
| chr8 | 104685616 | 104686189 | 0.6961 | 0.758147 | 1.089 |
| chr5 | 7755132 | 7755510 | 2.52387 | 2.74891 | 1.089 |
| chr3 | 81249725 | 81249929 | 0.60579 | 0.659908 | 1.089 |
| chr9 | 74105809 | 74106042 | 1.45272 | 1.58263 | 1.089 |
| chr17 | 7900536 | 7900747 | 2.18442 | 2.37996 | 1.090 |
| chr2 | 69664089 | 69664273 | 1.44756 | 1.57715 | 1.090 |
| chr7 | 2720532 | 2720825 | 2.90251 | 3.16313 | 1.090 |
| chr2 | 176174563 | 176174823 | 1.68932 | 1.84102 | 1.090 |
| chr2 | 119704939 | 119705149 | 0.903655 | 0.985007 | 1.090 |
| chr19 | 33929163 | 33929420 | 1.32809 | 1.44786 | 1.090 |
| chr22 | 26173600 | 26174168 | 1.26937 | 1.38388 | 1.090 |
| chr7 | 153263403 | 153263632 | 1.16289 | 1.26789 | 1.090 |
| chr7 | 156556804 | 156557025 | 1.58913 | 1.73289 | 1.090 |
| chr6 | 138029080 | 138029304 | 1.74349 | 1.90149 | 1.091 |
| chrX | 102966426 | 102966728 | 1.86233 | 2.03122 | 1.091 |
| chr16 | 34782056 | 34782251 | 0.99509 | 1.0854 | 1.091 |
| chr6 | 116538214 | 116538477 | 1.26924 | 1.38447 | 1.091 |
| chr10 | 85631274 | 85631640 | 2.72198 | 2.97009 | 1.091 |
| chr1 | 248785889 | 248786333 | 1.27172 | 1.38789 | 1.091 |
| chr17 | 68937055 | 68937418 | 0.895071 | 0.976882 | 1.091 |
| chr11 | 27068714 | 27069041 | 0.828032 | 0.903785 | 1.091 |
| chr1 | 247537236 | 247537449 | 1.38789 | 1.51497 | 1.092 |
| chr7 | 51495995 | 51496264 | 1.57814 | 1.72267 | 1.092 |
| chr8 | 145767584 | 145767776 | 1.08692 | 1.18673 | 1.092 |
| chr16 | 67918677 | 67918890 | 0.906892 | 0.990199 | 1.092 |
| chr7 | 54148717 | 54148942 | 0.699605 | 0.764026 | 1.092 |

| | | | | | |
|---|---|---|---|---|---|
| chr9 | 72037174 | 72037488 | 1.87803 | 2.05144 | 1.092 |
| chr14 | 87641046 | 87641298 | 0.850884 | 0.929545 | 1.092 |
| chr8 | 87045502 | 87045954 | 1.3263 | 1.44894 | 1.092 |
| chr12 | 114858939 | 114859329 | 0.907393 | 0.991341 | 1.093 |
| chr8 | 2537385 | 2537660 | 0.704881 | 0.770153 | 1.093 |
| chr3 | 34774973 | 34775164 | 0.60744 | 0.663829 | 1.093 |
| chr14 | 55604906 | 55605157 | 1.60067 | 1.74942 | 1.093 |
| chr14 | 25927545 | 25927769 | 0.835384 | 0.913037 | 1.093 |
| chr12 | 77943202 | 77943385 | 0.8201 | 0.896516 | 1.093 |
| chr14 | 71034624 | 71034907 | 2.05273 | 2.24433 | 1.093 |
| chrX | 153279045 | 153279513 | 2.54328 | 2.78089 | 1.093 |
| chr2 | 153046498 | 153046726 | 2.08853 | 2.28416 | 1.094 |
| chr10 | 54023408 | 54023596 | 1.02561 | 1.1217 | 1.094 |
| chr2 | 217268762 | 217269042 | 2.57199 | 2.81322 | 1.094 |
| chr3 | 38780188 | 38780493 | 1.77338 | 1.9397 | 1.094 |
| chr9 | 38418529 | 38418774 | 0.690955 | 0.755782 | 1.094 |
| chr7 | 27968763 | 27968945 | 1.09405 | 1.19676 | 1.094 |
| chr19 | 31186087 | 31186344 | 1.45053 | 1.58672 | 1.094 |
| chr7 | 82356255 | 82356529 | 1.98556 | 2.17217 | 1.094 |
| chr17 | 79504528 | 79505248 | 1.59119 | 1.74119 | 1.094 |
| chr5 | 170323726 | 170324034 | 2.02615 | 2.21716 | 1.094 |
| chr1 | 200416009 | 200416215 | 0.997818 | 1.09202 | 1.094 |
| chr5 | 133241350 | 133241645 | 0.779292 | 0.85287 | 1.094 |
| rUn_gl0002 | 142006 | 143285 | 1.13332 | 1.24036 | 1.094 |
| chr4 | 56733240 | 56733442 | 2.38497 | 2.61037 | 1.095 |
| chr14 | 84568913 | 84569120 | 0.555681 | 0.608218 | 1.095 |
| chr8 | 62156592 | 62156803 | 2.71694 | 2.97438 | 1.095 |
| chr20 | 61508501 | 61508971 | 0.705237 | 0.772193 | 1.095 |
| chr12 | 132536748 | 132537078 | 1.06514 | 1.16633 | 1.095 |
| chr6 | 116160070 | 116160320 | 1.18869 | 1.30165 | 1.095 |
| chr15 | 89683052 | 89683434 | 1.13496 | 1.24283 | 1.095 |
| chr6 | 74036472 | 74036727 | 1.12274 | 1.22946 | 1.095 |
| chr1 | 179675548 | 179675773 | 2.30199 | 2.52115 | 1.095 |
| chr14 | 106743366 | 106743649 | 1.25646 | 1.37612 | 1.095 |
| chr7 | 2130187 | 2130463 | 0.915511 | 1.00284 | 1.095 |
| chr1 | 120510620 | 120510903 | 2.08837 | 2.28797 | 1.096 |
| chr2 | 68120934 | 68121143 | 0.753071 | 0.825067 | 1.096 |
| chr12 | 54797996 | 54798557 | 2.48693 | 2.72478 | 1.096 |
| chr8 | 21763409 | 21763900 | 3.22996 | 3.53906 | 1.096 |
| chr2 | 42613739 | 42614142 | 1.48053 | 1.62231 | 1.096 |
| chr9 | 98984411 | 98984710 | 1.85085 | 2.02857 | 1.096 |
| chr15 | 92914116 | 92914321 | 0.834792 | 0.915027 | 1.096 |
| chr2 | 219033667 | 219034101 | 1.76096 | 1.93031 | 1.096 |
| chr14 | 46081409 | 46081681 | 0.781823 | 0.857062 | 1.096 |
| chr5 | 93860638 | 93860861 | 1.45072 | 1.5904 | 1.096 |
| chr3 | 30077626 | 30077920 | 0.880092 | 0.964944 | 1.096 |
| chr1 | 171195979 | 171196219 | 1.81822 | 1.9938 | 1.097 |
| chr2 | 163154530 | 163154830 | 1.37536 | 1.50827 | 1.097 |
| chr10 | 10363077 | 10363326 | 0.799815 | 0.877161 | 1.097 |
| chr20 | 6149975 | 6150203 | 1.35133 | 1.48213 | 1.097 |
| chr14 | 89584194 | 89584382 | 0.850802 | 0.933481 | 1.097 |
| chr8 | 105388064 | 105388280 | 0.980386 | 1.07577 | 1.097 |
| chr5 | 87794392 | 87794631 | 0.783752 | 0.86008 | 1.097 |
| chr8 | 84289556 | 84289769 | 0.791656 | 0.868771 | 1.097 |
| chr7 | 37893066 | 37893375 | 1.68562 | 1.84985 | 1.097 |
| chr12 | 3886458 | 3886671 | 1.68551 | 1.84982 | 1.097 |
| chr2 | 71772849 | 71773259 | 2.14709 | 2.35661 | 1.098 |
| chr1 | 167503029 | 167503461 | 2.31945 | 2.54588 | 1.098 |
| chr6 | 13625130 | 13625543 | 1.3563 | 1.48877 | 1.098 |
| chr11 | 61152577 | 61152785 | 0.687489 | 0.754673 | 1.098 |
| chr8 | 33630114 | 33630337 | 0.577639 | 0.634101 | 1.098 |

| | | | | | |
|---|---|---|---|---|---|
| chr15 | 42351900 | 42352241 | 2.2415 | 2.46068 | 1.098 |
| chr2 | 235964627 | 235964911 | 2.12201 | 2.32954 | 1.098 |
| chr1 | 245205977 | 245206297 | 3.00794 | 3.30222 | 1.098 |
| chr8 | 56595184 | 56595505 | 0.747435 | 0.820588 | 1.098 |
| chr2 | 128615731 | 128615952 | 2.17085 | 2.38374 | 1.098 |
| chr8 | 20695912 | 20696204 | 0.890011 | 0.977364 | 1.098 |
| chr1 | 216300939 | 216301148 | 0.614831 | 0.67523 | 1.098 |
| chr17 | 11267210 | 11267467 | 0.766404 | 0.841824 | 1.098 |
| chr8 | 10116171 | 10116399 | 0.827714 | 0.909247 | 1.099 |
| chr10 | 104271570 | 104271774 | 0.769419 | 0.845226 | 1.099 |
| chr2 | 53360380 | 53360700 | 0.891546 | 0.979544 | 1.099 |
| chr20 | 8370008 | 8370411 | 1.27075 | 1.39617 | 1.099 |
| chr2 | 175327081 | 175327859 | 1.07959 | 1.18618 | 1.099 |
| chr2 | 16212751 | 16212984 | 0.852327 | 0.936505 | 1.099 |
| chr14 | 23399616 | 23399887 | 1.75332 | 1.92651 | 1.099 |
| chr18 | 49814832 | 49815147 | 2.09079 | 2.29742 | 1.099 |
| chr5 | 33970033 | 33970357 | 2.03024 | 2.23095 | 1.099 |
| chr2 | 177319674 | 177319954 | 2.18887 | 2.40561 | 1.099 |
| chr20 | 19921395 | 19921611 | 0.95752 | 1.05243 | 1.099 |
| chr1 | 34078482 | 34078822 | 2.24433 | 2.46708 | 1.099 |
| chr10 | 22839759 | 22840046 | 2.30271 | 2.53132 | 1.099 |
| chr5 | 57792409 | 57792616 | 1.20205 | 1.32141 | 1.099 |
| chr2 | 86599989 | 86600581 | 1.42452 | 1.56612 | 1.099 |
| chr15 | 65020402 | 65020794 | 3.14486 | 3.45768 | 1.099 |
| chr15 | 75544620 | 75544994 | 1.10773 | 1.21795 | 1.100 |
| chr19 | 13323732 | 13323960 | 2.24033 | 2.46346 | 1.100 |
| chr9 | 38068796 | 38069021 | 0.992976 | 1.09194 | 1.100 |
| chr6 | 10478779 | 10479028 | 1.52208 | 1.6738 | 1.100 |
| chr8 | 98572164 | 98572677 | 2.19575 | 2.41472 | 1.100 |
| chr15 | 76094463 | 76094812 | 2.69648 | 2.96575 | 1.100 |
| chr7 | 75649489 | 75649762 | 2.76577 | 3.04214 | 1.100 |
| chr1 | 163461957 | 163462159 | 0.732569 | 0.805799 | 1.100 |
| chr9 | 103267872 | 103268181 | 2.34378 | 2.57806 | 1.100 |
| chr10 | 71674265 | 71674911 | 2.18069 | 2.39878 | 1.100 |
| chr8 | 56153830 | 56154092 | 0.822114 | 0.904401 | 1.100 |
| chr3 | 58477543 | 58477921 | 1.72279 | 1.89536 | 1.100 |
| chr10 | 99436490 | 99436789 | 2.79127 | 3.07094 | 1.100 |
| chr1 | 195346756 | 195346973 | 0.822321 | 0.904723 | 1.100 |
| chr16 | 1544402 | 1544644 | 1.55026 | 1.70593 | 1.100 |
| chr20 | 42882243 | 42882525 | 1.00682 | 1.10793 | 1.100 |
| chr6 | 56769439 | 56769660 | 1.4915 | 1.64135 | 1.100 |
| chr1 | 222451258 | 222451488 | 0.798537 | 0.878813 | 1.101 |
| chr7 | 21009385 | 21009594 | 0.782449 | 0.861174 | 1.101 |
| chr6 | 90077860 | 90078163 | 2.23291 | 2.45772 | 1.101 |
| chr22 | 40328960 | 40329224 | 1.19141 | 1.31145 | 1.101 |
| chr12 | 101916821 | 101917040 | 0.948364 | 1.04393 | 1.101 |
| chr5 | 84753999 | 84754209 | 1.15202 | 1.26812 | 1.101 |
| chr22 | 20144451 | 20144833 | 2.77305 | 3.05281 | 1.101 |
| chr9 | 24306822 | 24307057 | 1.01431 | 1.11676 | 1.101 |
| chr8 | 134985906 | 134986145 | 0.849741 | 0.935652 | 1.101 |
| chr10 | 28699797 | 28700037 | 1.45857 | 1.60609 | 1.101 |
| chr20 | 56821596 | 56821793 | 1.39982 | 1.5414 | 1.101 |
| chr7 | 29565197 | 29565438 | 0.78776 | 0.867504 | 1.101 |
| chr7 | 17022313 | 17022554 | 3.08133 | 3.39339 | 1.101 |
| chr15 | 29906026 | 29906506 | 2.70737 | 2.98199 | 1.101 |
| chr12 | 57618698 | 57618999 | 1.84549 | 2.03282 | 1.102 |
| chr9 | 96764803 | 96765210 | 1.73038 | 1.90612 | 1.102 |
| chr4 | 117044174 | 117044395 | 0.783771 | 0.863513 | 1.102 |
| chr2 | 2687606 | 2687874 | 0.796269 | 0.877319 | 1.102 |
| chr17 | 52584191 | 52584394 | 0.952361 | 1.04947 | 1.102 |
| chr11 | 113865200 | 113865424 | 1.42057 | 1.56545 | 1.102 |

| | | | | | |
|---|---|---|---|---|---|
| chr16 | 56341595 | 56341839 | 0.752641 | 0.829412 | 1.102 |
| chr19 | 29803610 | 29803832 | 1.05813 | 1.16613 | 1.102 |
| chr2 | 193059838 | 193060101 | 1.47531 | 1.62589 | 1.102 |
| chr20 | 58101916 | 58102166 | 1.22661 | 1.35182 | 1.102 |
| chr8 | 137199270 | 137199530 | 0.844161 | 0.930363 | 1.102 |
| chrX | 85315196 | 85315465 | 0.868014 | 0.956794 | 1.102 |
| chr7 | 118213090 | 118213320 | 0.82631 | 0.91085 | 1.102 |
| chr3 | 98062811 | 98063055 | 1.98566 | 2.18889 | 1.102 |
| chr22 | 38808772 | 38809148 | 2.82321 | 3.11238 | 1.102 |
| chr7 | 139303676 | 139303938 | 2.41189 | 2.65918 | 1.103 |
| chr5 | 125570294 | 125570609 | 2.4282 | 2.67729 | 1.103 |
| chr2 | 77697551 | 77697761 | 0.588899 | 0.649343 | 1.103 |
| chr8 | 133776761 | 133776991 | 0.861265 | 0.949703 | 1.103 |
| chr10 | 276728 | 276985 | 1.5289 | 1.6859 | 1.103 |
| chr17 | 33303570 | 33303904 | 1.44008 | 1.5881 | 1.103 |
| chr20 | 58425000 | 58425316 | 1.25273 | 1.3817 | 1.103 |
| chr10 | 2702464 | 2702715 | 0.69862 | 0.77055 | 1.103 |
| chr8 | 139567431 | 139567654 | 0.913055 | 1.00712 | 1.103 |
| chr5 | 101632010 | 101632201 | 1.44319 | 1.59194 | 1.103 |
| chr3 | 158977960 | 158978174 | 1.63954 | 1.80858 | 1.103 |
| chr16 | 80500241 | 80500679 | 2.11146 | 2.32944 | 1.103 |
| chr2 | 208490614 | 208490865 | 1.28324 | 1.41586 | 1.103 |
| chr2 | 171943218 | 171943608 | 1.62054 | 1.78813 | 1.103 |
| chr15 | 101840564 | 101840821 | 2.36023 | 2.60466 | 1.104 |
| chr12 | 6776785 | 6777201 | 1.38846 | 1.53233 | 1.104 |
| chr2 | 109744442 | 109744819 | 1.88169 | 2.07674 | 1.104 |
| chr12 | 120755122 | 120755371 | 1.44112 | 1.59053 | 1.104 |
| chr17 | 9632394 | 9632696 | 4.53294 | 5.00296 | 1.104 |
| chr7 | 116072390 | 116072665 | 1.01994 | 1.12573 | 1.104 |
| chr10 | 91323721 | 91323918 | 1.38628 | 1.53015 | 1.104 |
| chr8 | 89038936 | 89039162 | 0.766895 | 0.846494 | 1.104 |
| chr1 | 32914807 | 32915029 | 1.71832 | 1.8968 | 1.104 |
| chr7 | 100492427 | 100492827 | 2.70589 | 2.987 | 1.104 |
| chr2 | 23066117 | 23066359 | 1.07355 | 1.18513 | 1.104 |
| chr16 | 30103263 | 30103540 | 2.2321 | 2.46422 | 1.104 |
| chr8 | 3023757 | 3023982 | 0.741728 | 0.81887 | 1.104 |
| chr6 | 35705376 | 35705590 | 2.11991 | 2.34056 | 1.104 |
| chr7 | 73617548 | 73617764 | 1.74963 | 1.93192 | 1.104 |
| chr7 | 149400955 | 149401170 | 0.798727 | 0.882048 | 1.104 |
| chr6 | 63485483 | 63485671 | 1.18444 | 1.30815 | 1.104 |
| chr5 | 104312904 | 104313126 | 0.819299 | 0.904895 | 1.104 |
| chr5 | 146956491 | 146956823 | 1.8023 | 1.99076 | 1.105 |
| chr9 | 126889106 | 126889503 | 3.12042 | 3.44686 | 1.105 |
| chr12 | 126220623 | 126220841 | 0.867536 | 0.958307 | 1.105 |
| chr2 | 71187617 | 71187859 | 1.28096 | 1.41502 | 1.105 |
| chr1 | 202243761 | 202244000 | 1.53082 | 1.69106 | 1.105 |
| chr14 | 31125443 | 31125794 | 1.99368 | 2.20258 | 1.105 |
| chr10 | 14367248 | 14367604 | 1.14904 | 1.26953 | 1.105 |
| chr19 | 50831377 | 50831769 | 2.40479 | 2.65759 | 1.105 |
| chr3 | 103577889 | 103578122 | 1.01981 | 1.12709 | 1.105 |
| chr10 | 65837719 | 65838010 | 0.769385 | 0.850437 | 1.105 |
| chr16 | 18087979 | 18088203 | 1.20134 | 1.32793 | 1.105 |
| chr2 | 107161384 | 107161569 | 0.933661 | 1.03204 | 1.105 |
| chr3 | 6497842 | 6498026 | 0.785069 | 0.867856 | 1.105 |
| chr10 | 105425982 | 105426269 | 1.44654 | 1.59923 | 1.106 |
| chr2 | 128820938 | 128821375 | 2.15362 | 2.38122 | 1.106 |
| chr17 | 71952782 | 71953036 | 1.91772 | 2.12043 | 1.106 |
| chr2 | 61922045 | 61922368 | 2.35411 | 2.60293 | 1.106 |
| chr15 | 97778581 | 97778882 | 2.41271 | 2.66823 | 1.106 |
| chr17 | 20201595 | 20201822 | 0.751548 | 0.831169 | 1.106 |
| chr7 | 99056872 | 99057145 | 1.22572 | 1.35566 | 1.106 |

| | | | | | |
|---|---|---|---|---|---|
| chr17 | 56621102 | 56621367 | 0.631725 | 0.69872 | 1.106 |
| chr11 | 19092741 | 19093057 | 4.51456 | 4.994 | 1.106 |
| chr8 | 81594877 | 81595483 | 2.17019 | 2.4007 | 1.106 |
| chr9 | 107505783 | 107505990 | 1.9227 | 2.12701 | 1.106 |
| chr13 | 107863384 | 107863672 | 2.33017 | 2.57782 | 1.106 |
| chr5 | 152527851 | 152528177 | 0.927244 | 1.02599 | 1.106 |
| chr11 | 14140903 | 14141110 | 0.629607 | 0.696696 | 1.107 |
| chr12 | 61956309 | 61956492 | 0.666778 | 0.737836 | 1.107 |
| chr12 | 111259941 | 111260127 | 0.759745 | 0.840709 | 1.107 |
| chr16 | 82847063 | 82847319 | 0.657525 | 0.727603 | 1.107 |
| chr11 | 110247429 | 110247826 | 1.90381 | 2.10686 | 1.107 |
| chr1 | 61444695 | 61444939 | 2.07476 | 2.29609 | 1.107 |
| chr7 | 41042728 | 41043103 | 1.48166 | 1.63977 | 1.107 |
| chr7 | 122614468 | 122614710 | 0.93222 | 1.03183 | 1.107 |
| chr7 | 120432042 | 120432527 | 2.62807 | 2.90894 | 1.107 |
| chr17 | 54646323 | 54646527 | 1.00724 | 1.11492 | 1.107 |
| chr10 | 121981478 | 121981686 | 2.42602 | 2.6854 | 1.107 |
| chr11 | 58668376 | 58668582 | 1.99907 | 2.213 | 1.107 |
| chr10 | 98841434 | 98841766 | 2.94943 | 3.26525 | 1.107 |
| chr17 | 1513252 | 1513464 | 0.657452 | 0.727908 | 1.107 |
| chr17 | 43351393 | 43351958 | 0.737599 | 0.81667 | 1.107 |
| chr12 | 23009131 | 23009491 | 1.44312 | 1.5979 | 1.107 |
| chr16 | 79801564 | 79801801 | 1.43319 | 1.58708 | 1.107 |
| chr12 | 111479802 | 111480402 | 0.734998 | 0.813936 | 1.107 |
| chr3 | 22929255 | 22929461 | 2.03993 | 2.25904 | 1.107 |
| chr14 | 39837653 | 39837860 | 1.27359 | 1.41041 | 1.107 |
| chr5 | 8075841 | 8076077 | 0.853184 | 0.944857 | 1.107 |
| chr20 | 55904778 | 55905258 | 1.06383 | 1.17816 | 1.107 |
| chr7 | 123795518 | 123795737 | 1.1069 | 1.22628 | 1.108 |
| chr17 | 35404943 | 35405272 | 3.14631 | 3.48567 | 1.108 |
| chr2 | 22074947 | 22075143 | 0.888669 | 0.984564 | 1.108 |
| chr22 | 47076992 | 47077240 | 1.48658 | 1.647 | 1.108 |
| chr16 | 58054662 | 58055170 | 1.96363 | 2.17575 | 1.108 |
| chr9 | 7673060 | 7673276 | 0.801896 | 0.888527 | 1.108 |
| chr1 | 236615205 | 236615686 | 2.20136 | 2.4397 | 1.108 |
| chr15 | 78534370 | 78534854 | 1.36384 | 1.51154 | 1.108 |
| chr16 | 25850070 | 25850395 | 1.63128 | 1.8083 | 1.109 |
| chr9 | 95848004 | 95848312 | 0.718511 | 0.796514 | 1.109 |
| chr14 | 73750898 | 73751085 | 0.996287 | 1.10447 | 1.109 |
| chr7 | 231014 | 231320 | 1.28778 | 1.42768 | 1.109 |
| chr1 | 176918215 | 176918487 | 1.56477 | 1.73507 | 1.109 |
| chr10 | 105445957 | 105446271 | 3.60777 | 4.00039 | 1.109 |
| chr12 | 48264145 | 48264660 | 1.16577 | 1.29287 | 1.109 |
| chr11 | 66131337 | 66131962 | 2.32471 | 2.57819 | 1.109 |
| chr15 | 100752199 | 100752545 | 1.13822 | 1.26234 | 1.109 |
| chr20 | 30001360 | 30001723 | 1.37641 | 1.52677 | 1.109 |
| chr7 | 51564440 | 51564754 | 0.919724 | 1.0202 | 1.109 |
| chr3 | 988605 | 988866 | 0.672086 | 0.745638 | 1.109 |
| chr11 | 24581568 | 24581777 | 0.998993 | 1.10834 | 1.109 |
| chr10 | 58733202 | 58733746 | 0.78276 | 0.868583 | 1.110 |
| chr10 | 78871689 | 78872112 | 1.96654 | 2.1825 | 1.110 |
| chr13 | 41796421 | 41796685 | 1.59646 | 1.7718 | 1.110 |
| chr5 | 59583058 | 59583341 | 2.55337 | 2.83425 | 1.110 |
| chr1 | 153250341 | 153250613 | 1.33425 | 1.48104 | 1.110 |
| chr2 | 119992052 | 119992449 | 1.3489 | 1.49751 | 1.110 |
| chr19 | 5354586 | 5354840 | 1.09763 | 1.21863 | 1.110 |
| chr2 | 20877273 | 20877736 | 1.10876 | 1.23107 | 1.110 |
| chr19 | 55929523 | 55929842 | 2.13367 | 2.3691 | 1.110 |
| chr1 | 221104575 | 221104833 | 0.67507 | 0.74962 | 1.110 |
| chr18 | 57490941 | 57491143 | 1.87008 | 2.07685 | 1.111 |
| chr6 | 42466336 | 42466596 | 0.766362 | 0.851098 | 1.111 |

| | | | | | |
|---|---|---|---|---|---|
| chr12 | 55673191 | 55673430 | 1.032 | 1.14621 | 1.111 |
| chr11 | 41169299 | 41169553 | 0.74129 | 0.823383 | 1.111 |
| chr9 | 7657884 | 7658130 | 1.80236 | 2.00217 | 1.111 |
| chr5 | 168175297 | 168175729 | 2.36174 | 2.62365 | 1.111 |
| chr11 | 41245086 | 41245416 | 0.887531 | 0.985965 | 1.111 |
| chr14 | 47131073 | 47131290 | 1.04135 | 1.15694 | 1.111 |
| chr3 | 186232191 | 186232408 | 1.23179 | 1.36852 | 1.111 |
| chr3 | 157809404 | 157809687 | 1.28592 | 1.42872 | 1.111 |
| chr15 | 27062157 | 27062455 | 0.991885 | 1.10208 | 1.111 |
| chr10 | 77154453 | 77154679 | 0.684938 | 0.761099 | 1.111 |
| chr9 | 127209077 | 127209482 | 1.65688 | 1.84132 | 1.111 |
| chr3 | 125852963 | 125853333 | 2.71351 | 3.01629 | 1.112 |
| chr5 | 140254720 | 140255088 | 1.08492 | 1.20598 | 1.112 |
| chr8 | 79305715 | 79305959 | 1.82855 | 2.03263 | 1.112 |
| chr2 | 120022765 | 120023097 | 2.67955 | 2.97868 | 1.112 |
| chr2 | 39286637 | 39286819 | 1.36681 | 1.51952 | 1.112 |
| chr17 | 70059606 | 70059849 | 0.899602 | 1.00014 | 1.112 |
| chr12 | 55457049 | 55457289 | 1.07765 | 1.19818 | 1.112 |
| chr10 | 29228235 | 29228523 | 2.48517 | 2.76344 | 1.112 |
| chr4 | 138045487 | 138045728 | 1.02419 | 1.13889 | 1.112 |
| chr5 | 159681011 | 159681221 | 0.87545 | 0.973698 | 1.112 |
| chr20 | 32920361 | 32920718 | 0.88344 | 0.982605 | 1.112 |
| chr7 | 149496823 | 149497258 | 1.35142 | 1.5032 | 1.112 |
| chr17 | 72248600 | 72248807 | 0.937269 | 1.04261 | 1.112 |
| chr2 | 47146065 | 47146554 | 1.52369 | 1.69501 | 1.112 |
| chr14 | 79064399 | 79064932 | 2.69276 | 2.99574 | 1.113 |
| chr19 | 28781868 | 28782307 | 1.02084 | 1.13578 | 1.113 |
| chr7 | 101201857 | 101202060 | 0.980729 | 1.09133 | 1.113 |
| chr3 | 165894845 | 165895093 | 0.946751 | 1.05368 | 1.113 |
| chr15 | 62164167 | 62164391 | 1.53323 | 1.70658 | 1.113 |
| chr2 | 51651751 | 51651957 | 0.968151 | 1.07784 | 1.113 |
| chr17 | 35866840 | 35867084 | 1.84109 | 2.04982 | 1.113 |
| chr17 | 26542248 | 26542491 | 0.85568 | 0.952792 | 1.113 |
| chr15 | 51425064 | 51425449 | 2.34483 | 2.61123 | 1.114 |
| chr9 | 101405348 | 101405558 | 0.843887 | 0.939782 | 1.114 |
| chr20 | 31240735 | 31241356 | 2.13521 | 2.37824 | 1.114 |
| chr2 | 33495684 | 33495903 | 1.54226 | 1.71789 | 1.114 |
| chrX | 102205917 | 102206133 | 1.39634 | 1.55574 | 1.114 |
| chr9 | 91943960 | 91944169 | 0.904839 | 1.00815 | 1.114 |
| chr7 | 148702845 | 148703237 | 3.04334 | 3.39095 | 1.114 |
| chr12 | 129128601 | 129128835 | 0.677787 | 0.755229 | 1.114 |
| chr2 | 24047653 | 24047892 | 1.68471 | 1.87724 | 1.114 |
| chr13 | 102594486 | 102594690 | 1.61994 | 1.80525 | 1.114 |
| chr1 | 38385043 | 38385262 | 1.15562 | 1.28785 | 1.114 |
| chr19 | 45348842 | 45349111 | 2.25583 | 2.51404 | 1.114 |
| chr2 | 148668502 | 148668835 | 1.51378 | 1.68728 | 1.115 |
| chr2 | 158192708 | 158193071 | 1.76815 | 1.97117 | 1.115 |
| chr14 | 78895991 | 78896294 | 0.965103 | 1.07597 | 1.115 |
| chr11 | 75578854 | 75579112 | 2.15949 | 2.40784 | 1.115 |
| chr1 | 48221078 | 48221407 | 2.02581 | 2.25906 | 1.115 |
| chr15 | 32269979 | 32270387 | 3.56099 | 3.97133 | 1.115 |
| chr17 | 80871600 | 80871869 | 1.31165 | 1.46281 | 1.115 |
| chr18 | 55578094 | 55578303 | 1.81865 | 2.02848 | 1.115 |
| chr13 | 33802753 | 33803024 | 2.10783 | 2.35111 | 1.115 |
| chr3 | 119099697 | 119099912 | 1.27279 | 1.41979 | 1.115 |
| chr7 | 84029128 | 84029330 | 1.03864 | 1.15875 | 1.116 |
| chr7 | 130084996 | 130085407 | 0.992178 | 1.10706 | 1.116 |
| chr6 | 146362470 | 146362692 | 1.524 | 1.70074 | 1.116 |
| chr2 | 84499266 | 84499551 | 0.667813 | 0.745322 | 1.116 |
| chr2 | 172397797 | 172398071 | 1.85082 | 2.06562 | 1.116 |
| chr6 | 25720671 | 25720890 | 1.73567 | 1.93714 | 1.116 |

| | | | | | |
|---|---|---|---|---|---|
| chr15 | 79315785 | 79316049 | 0.978743 | 1.09267 | 1.116 |
| chr3 | 50473446 | 50473900 | 2.80962 | 3.13682 | 1.116 |
| chr17 | 66556263 | 66556511 | 0.88557 | 0.988751 | 1.117 |
| chr3 | 11102275 | 11102851 | 2.35259 | 2.62685 | 1.117 |
| chr5 | 5063912 | 5064184 | 0.985001 | 1.09985 | 1.117 |
| chrX | 134476576 | 134476991 | 1.63905 | 1.83019 | 1.117 |
| chr8 | 50764068 | 50764310 | 0.896014 | 1.00059 | 1.117 |
| chr9 | 103476843 | 103477272 | 1.7536 | 1.95838 | 1.117 |
| chr3 | 15447039 | 15447354 | 1.56168 | 1.74433 | 1.117 |
| chr17 | 10779256 | 10779733 | 1.89051 | 2.11167 | 1.117 |
| chr15 | 27996764 | 27997199 | 1.02523 | 1.14518 | 1.117 |
| chr2 | 95687801 | 95688457 | 2.13175 | 2.38119 | 1.117 |
| chr17 | 73671094 | 73671343 | 1.87535 | 2.09483 | 1.117 |
| chr11 | 9187421 | 9187697 | 1.80518 | 2.01667 | 1.117 |
| chr8 | 72565938 | 72566161 | 0.979446 | 1.0942 | 1.117 |
| chr10 | 118221304 | 118221529 | 0.93196 | 1.04125 | 1.117 |
| chr2 | 198774147 | 198774414 | 0.984782 | 1.10054 | 1.118 |
| chr8 | 60670623 | 60670851 | 1.20805 | 1.35008 | 1.118 |
| chr12 | 132990553 | 132990983 | 2.27026 | 2.53725 | 1.118 |
| chr19 | 32333581 | 32333823 | 1.30236 | 1.45554 | 1.118 |
| chr10 | 32293922 | 32294215 | 2.20356 | 2.46286 | 1.118 |
| chr2 | 116220326 | 116220549 | 0.8524 | 0.952732 | 1.118 |
| chr10 | 82639513 | 82639767 | 0.767479 | 0.857841 | 1.118 |
| chr12 | 104144940 | 104145163 | 1.254 | 1.40166 | 1.118 |
| chr2 | 74233676 | 74233932 | 0.964916 | 1.07855 | 1.118 |
| chr9 | 110184557 | 110184880 | 2.38546 | 2.66649 | 1.118 |
| chr17 | 74469741 | 74469994 | 1.8465 | 2.0647 | 1.118 |
| chr19 | 41140361 | 41140781 | 7.93466 | 8.87279 | 1.118 |
| chr2 | 34369440 | 34369748 | 0.798218 | 0.892622 | 1.118 |
| chr12 | 111343255 | 111343439 | 0.884041 | 0.988605 | 1.118 |
| chr5 | 91518032 | 91518564 | 1.02194 | 1.14281 | 1.118 |
| chr2 | 6434612 | 6434912 | 0.86102 | 0.962871 | 1.118 |
| chr2 | 11585150 | 11585367 | 0.72178 | 0.807156 | 1.118 |
| chr10 | 20082847 | 20083098 | 0.976273 | 1.09199 | 1.119 |
| chr14 | 64761522 | 64761942 | 3.43636 | 3.84398 | 1.119 |
| chr3 | 154566793 | 154566999 | 0.699611 | 0.78267 | 1.119 |
| chr1 | 155056659 | 155056896 | 1.1204 | 1.25354 | 1.119 |
| chr5 | 169005109 | 169005329 | 1.54453 | 1.7282 | 1.119 |
| chr19 | 3586114 | 3586443 | 1.96582 | 2.19967 | 1.119 |
| chr20 | 32728577 | 32728878 | 0.668135 | 0.747643 | 1.119 |
| chr1 | 170403507 | 170403731 | 0.791643 | 0.886027 | 1.119 |
| chr12 | 117868029 | 117868247 | 0.69533 | 0.778304 | 1.119 |
| chr10 | 61858903 | 61859137 | 1.98101 | 2.21751 | 1.119 |
| chr1 | 208412390 | 208412715 | 2.46488 | 2.75944 | 1.120 |
| chr5 | 25273100 | 25273296 | 0.824528 | 0.923193 | 1.120 |
| chr2 | 189156476 | 189156719 | 1.68379 | 1.88555 | 1.120 |
| chr10 | 119078427 | 119078691 | 1.10817 | 1.24099 | 1.120 |
| chr3 | 193290216 | 193290563 | 1.35508 | 1.51769 | 1.120 |
| chr17 | 72860859 | 72861425 | 2.7973 | 3.13321 | 1.120 |
| chr19 | 32115891 | 32116312 | 0.818361 | 0.916628 | 1.120 |
| chr10 | 11312556 | 11312933 | 2.5179 | 2.82028 | 1.120 |
| chr17 | 40611344 | 40611555 | 1.53983 | 1.72475 | 1.120 |
| chr11 | 94519792 | 94519982 | 1.61198 | 1.80569 | 1.120 |
| chr5 | 112999712 | 112999928 | 0.981736 | 1.09976 | 1.120 |
| chr21 | 45533956 | 45534228 | 1.89233 | 2.11997 | 1.120 |
| chr2 | 15888885 | 15889213 | 2.01124 | 2.25327 | 1.120 |
| chr15 | 35000577 | 35001105 | 1.95676 | 2.19227 | 1.120 |
| chr9 | 23379979 | 23380202 | 1.00061 | 1.12115 | 1.120 |
| chr5 | 156436044 | 156436280 | 0.87839 | 0.984246 | 1.121 |
| chr16 | 79081204 | 79081550 | 1.08091 | 1.21131 | 1.121 |
| chr9 | 115188241 | 115188595 | 0.702941 | 0.787863 | 1.121 |

| | | | | | |
|---|---|---|---|---|---|
| chr10 | 114000307 | 114000563 | 0.788111 | 0.883348 | 1.121 |
| chr16 | 46663238 | 46663457 | 1.24374 | 1.39404 | 1.121 |
| chr19 | 41858394 | 41858812 | 2.83831 | 3.18201 | 1.121 |
| chr7 | 138648867 | 138649082 | 0.748394 | 0.83908 | 1.121 |
| chr8 | 145609908 | 145610141 | 0.781578 | 0.876311 | 1.121 |
| chr5 | 11182867 | 11183077 | 1.44338 | 1.61862 | 1.121 |
| rUn_gl0002 | 123995 | 124458 | 1.12169 | 1.25788 | 1.121 |
| chr20 | 55205253 | 55205660 | 2.51935 | 2.82529 | 1.121 |
| chr2 | 23043709 | 23043955 | 0.933914 | 1.04739 | 1.122 |
| chr5 | 135969354 | 135969597 | 0.988921 | 1.10914 | 1.122 |
| chr11 | 62791496 | 62791921 | 2.96815 | 3.32924 | 1.122 |
| chr3 | 17300408 | 17300602 | 1.76971 | 1.98518 | 1.122 |
| chr3 | 114009686 | 114010046 | 3.02649 | 3.39496 | 1.122 |
| chr12 | 130281580 | 130282252 | 0.951791 | 1.06768 | 1.122 |
| chr2 | 209436168 | 209436446 | 1.27218 | 1.42716 | 1.122 |
| chr8 | 95514983 | 95515205 | 1.4169 | 1.58965 | 1.122 |
| chr9 | 136859477 | 136859906 | 3.32501 | 3.73043 | 1.122 |
| chr20 | 4089255 | 4089626 | 4.18385 | 4.69413 | 1.122 |
| chr10 | 2788679 | 2788919 | 0.787949 | 0.884082 | 1.122 |
| chr8 | 130760705 | 130760968 | 1.40288 | 1.57428 | 1.122 |
| chr8 | 71040041 | 71040461 | 0.964721 | 1.08264 | 1.122 |
| chr7 | 71941640 | 71941868 | 1.00898 | 1.13257 | 1.122 |
| chr16 | 57979816 | 57980061 | 0.912928 | 1.02483 | 1.123 |
| chr16 | 7874664 | 7874846 | 0.795174 | 0.892872 | 1.123 |
| chr2 | 25586677 | 25586900 | 0.774593 | 0.869831 | 1.123 |
| chr3 | 88627784 | 88628095 | 0.850941 | 0.955608 | 1.123 |
| chr9 | 108663793 | 108664145 | 2.48077 | 2.78625 | 1.123 |
| chr5 | 110807959 | 110808156 | 1.18372 | 1.32953 | 1.123 |
| chr3 | 164774289 | 164774659 | 0.973912 | 1.09391 | 1.123 |
| chr20 | 15983019 | 15983317 | 0.889367 | 0.999121 | 1.123 |
| chr7 | 104488833 | 104489278 | 1.88732 | 2.1204 | 1.124 |
| chr2 | 60869168 | 60869797 | 1.4609 | 1.64139 | 1.124 |
| chr7 | 10961495 | 10961806 | 0.849143 | 0.954131 | 1.124 |
| chr3 | 170886040 | 170886263 | 1.35902 | 1.52708 | 1.124 |
| chr1 | 6532509 | 6532714 | 1.55611 | 1.74857 | 1.124 |
| chr12 | 93058368 | 93058583 | 0.940692 | 1.05704 | 1.124 |
| chr16 | 24264372 | 24264824 | 1.59974 | 1.79759 | 1.124 |
| chr7 | 37549686 | 37549985 | 1.45073 | 1.63015 | 1.124 |
| chr3 | 113631875 | 113632144 | 1.91036 | 2.14672 | 1.124 |
| chr13 | 76444543 | 76444791 | 1.94384 | 2.18457 | 1.124 |
| chr10 | 95314608 | 95314795 | 2.41101 | 2.70997 | 1.124 |
| chr16 | 1202721 | 1203002 | 1.78596 | 2.00746 | 1.124 |
| chr8 | 142118674 | 142118908 | 0.905947 | 1.0183 | 1.124 |
| chr5 | 153989924 | 153990301 | 3.3569 | 3.7733 | 1.124 |
| chr1 | 41961367 | 41961571 | 1.29576 | 1.45658 | 1.124 |
| chr1 | 190371945 | 190372127 | 0.867125 | 0.974763 | 1.124 |
| chr1 | 27879522 | 27879731 | 0.832714 | 0.936106 | 1.124 |
| chr3 | 122539722 | 122540018 | 0.682629 | 0.767422 | 1.124 |
| chr19 | 14897124 | 14897476 | 2.57085 | 2.89041 | 1.124 |
| chr19 | 48372150 | 48372494 | 2.3101 | 2.59725 | 1.124 |
| chr8 | 30621796 | 30621979 | 0.654088 | 0.735403 | 1.124 |
| chr12 | 19708378 | 19708632 | 1.01909 | 1.14579 | 1.124 |
| chr17 | 3946796 | 3947066 | 0.98575 | 1.10831 | 1.124 |
| chr8 | 112939425 | 112939648 | 0.774605 | 0.871 | 1.124 |
| chr2 | 21724649 | 21724880 | 1.05683 | 1.18841 | 1.125 |
| chr20 | 8086649 | 8086843 | 1.11728 | 1.25643 | 1.125 |
| chr7 | 112120534 | 112120777 | 1.19113 | 1.3397 | 1.125 |
| chr9 | 110887947 | 110888150 | 1.62993 | 1.83328 | 1.125 |
| chr10 | 2154222 | 2154534 | 0.834501 | 0.938645 | 1.125 |
| chr8 | 12628845 | 12629055 | 1.02797 | 1.15635 | 1.125 |
| chr3 | 42577892 | 42578169 | 1.1802 | 1.32763 | 1.125 |

| | | | | | |
|---|---|---|---|---|---|
| chr7 | 46236880 | 46237112 | 1.0087 | 1.1348 | 1.125 |
| chr15 | 28096093 | 28096388 | 0.729793 | 0.821127 | 1.125 |
| chr15 | 91921070 | 91921273 | 0.80562 | 0.906461 | 1.125 |
| chr20 | 50050442 | 50050649 | 0.739533 | 0.832273 | 1.125 |
| chr5 | 3196168 | 3196373 | 0.839686 | 0.945013 | 1.125 |
| chr12 | 23667463 | 23667709 | 0.745658 | 0.839262 | 1.126 |
| rUn_gl0002 | 140234 | 140562 | 0.980377 | 1.10354 | 1.126 |
| chr16 | 7318277 | 7318528 | 1.19957 | 1.35036 | 1.126 |
| chr15 | 28128682 | 28128899 | 1.01045 | 1.13754 | 1.126 |
| chr14 | 100412992 | 100413618 | 3.3008 | 3.71629 | 1.126 |
| chr1 | 234529111 | 234529363 | 0.883865 | 0.995132 | 1.126 |
| chr10 | 16587216 | 16587456 | 1.6199 | 1.82392 | 1.126 |
| chr20 | 58014340 | 58014546 | 0.985169 | 1.10929 | 1.126 |
| chr1 | 175793279 | 175793552 | 0.952206 | 1.07218 | 1.126 |
| chr8 | 136797074 | 136797256 | 0.904464 | 1.01849 | 1.126 |
| chr5 | 36015737 | 36016037 | 0.845694 | 0.952321 | 1.126 |
| chr17 | 9793409 | 9793735 | 1.92129 | 2.16374 | 1.126 |
| chr5 | 64080312 | 64080560 | 2.09541 | 2.35987 | 1.126 |
| chr5 | 33436005 | 33436358 | 2.29816 | 2.58857 | 1.126 |
| chr5 | 75604970 | 75605172 | 2.00363 | 2.25687 | 1.126 |
| chr3 | 164965253 | 164965462 | 0.780527 | 0.879199 | 1.126 |
| chr9 | 26656248 | 26656854 | 1.28653 | 1.44921 | 1.126 |
| chr9 | 93799507 | 93799737 | 2.22268 | 2.50399 | 1.127 |
| chr5 | 34538055 | 34538396 | 1.44239 | 1.62505 | 1.127 |
| chr15 | 98652665 | 98653100 | 0.842035 | 0.948852 | 1.127 |
| chr3 | 13696812 | 13697200 | 1.86485 | 2.10149 | 1.127 |
| chr9 | 85110440 | 85110636 | 0.720939 | 0.812421 | 1.127 |
| chr1 | 29220224 | 29220437 | 0.835412 | 0.941596 | 1.127 |
| chr4 | 74122973 | 74123235 | 1.67775 | 1.89101 | 1.127 |
| chr19 | 33299334 | 33299550 | 1.3303 | 1.49947 | 1.127 |
| chr9 | 88055207 | 88055500 | 0.678523 | 0.764885 | 1.127 |
| chr15 | 89257265 | 89257514 | 0.908145 | 1.0239 | 1.127 |
| chr17 | 80152669 | 80152939 | 0.873787 | 0.985491 | 1.128 |
| chr1 | 156589194 | 156589471 | 1.91831 | 2.16367 | 1.128 |
| chr1 | 228125514 | 228125755 | 0.773432 | 0.872469 | 1.128 |
| chr11 | 72975498 | 72975783 | 0.853711 | 0.963034 | 1.128 |
| chr2 | 232770840 | 232771109 | 2.21411 | 2.4978 | 1.128 |
| chr3 | 128291449 | 128291738 | 1.84087 | 2.07698 | 1.128 |
| chr10 | 29718699 | 29719074 | 0.986633 | 1.11327 | 1.128 |
| chr15 | 81549397 | 81549728 | 0.900386 | 1.01616 | 1.129 |
| chr1 | 213014559 | 213014782 | 0.923647 | 1.0425 | 1.129 |
| chr5 | 136379859 | 136380190 | 0.684387 | 0.772495 | 1.129 |
| chr8 | 4433669 | 4433921 | 0.959538 | 1.08308 | 1.129 |
| chr10 | 66105774 | 66105981 | 0.844681 | 0.953445 | 1.129 |
| chr2 | 27414461 | 27414687 | 1.84591 | 2.0837 | 1.129 |
| chr17 | 68804747 | 68805108 | 1.55535 | 1.75573 | 1.129 |
| chr5 | 135165730 | 135166241 | 2.0231 | 2.28381 | 1.129 |
| chr6 | 80280413 | 80280795 | 1.51793 | 1.71374 | 1.129 |
| chr8 | 26960620 | 26960903 | 0.968383 | 1.0933 | 1.129 |
| chr9 | 34591368 | 34592079 | 2.05123 | 2.31593 | 1.129 |
| chr13 | 51699940 | 51700196 | 2.18585 | 2.46799 | 1.129 |
| chr2 | 249715 | 249917 | 0.857498 | 0.968376 | 1.129 |
| chr20 | 16868595 | 16868825 | 0.867567 | 0.979746 | 1.129 |
| chr11 | 68811632 | 68812012 | 2.41869 | 2.73157 | 1.129 |
| chr2 | 27977054 | 27977300 | 2.50507 | 2.82919 | 1.129 |
| chr2 | 211092352 | 211092596 | 0.639261 | 0.722067 | 1.130 |
| chr9 | 108764887 | 108765157 | 0.812694 | 0.917962 | 1.130 |
| chr8 | 135489056 | 135489443 | 1.87249 | 2.11506 | 1.130 |
| chr11 | 123578292 | 123578617 | 1.82019 | 2.05599 | 1.130 |
| chr7 | 145962722 | 145962927 | 0.851765 | 0.962165 | 1.130 |
| chr5 | 40288830 | 40289037 | 0.801614 | 0.905602 | 1.130 |

| | | | | | |
|---|---|---|---|---|---|
| chr5 | 167792542 | 167792903 | 1.5875 | 1.79354 | 1.130 |
| chr19 | 39964753 | 39965235 | 4.0275 | 4.55053 | 1.130 |
| chr14 | 87750499 | 87750732 | 0.671231 | 0.758411 | 1.130 |
| chr9 | 112322615 | 112323003 | 1.15015 | 1.29974 | 1.130 |
| chr5 | 162613167 | 162613419 | 0.686782 | 0.776164 | 1.130 |
| chr11 | 60130882 | 60131356 | 2.12847 | 2.40555 | 1.130 |
| chrX | 102206537 | 102206791 | 1.77973 | 2.01163 | 1.130 |
| chr6 | 46645067 | 46645288 | 1.64161 | 1.85558 | 1.130 |
| chr20 | 50767162 | 50767511 | 2.17681 | 2.46062 | 1.130 |
| chr19 | 13215277 | 13215723 | 1.60525 | 1.81456 | 1.130 |
| chr4 | 76916992 | 76917234 | 1.4034 | 1.5865 | 1.130 |
| chr8 | 144889280 | 144889500 | 1.02685 | 1.16091 | 1.131 |
| chr13 | 83984250 | 83984447 | 0.517489 | 0.585057 | 1.131 |
| chr5 | 7850025 | 7850281 | 1.2398 | 1.40174 | 1.131 |
| chr3 | 156855296 | 156855557 | 2.12272 | 2.40014 | 1.131 |
| chr1 | 186373650 | 186373936 | 2.06229 | 2.33212 | 1.131 |
| chr17 | 31619444 | 31619967 | 1.42361 | 1.61021 | 1.131 |
| chr10 | 58583652 | 58583907 | 1.05383 | 1.19217 | 1.131 |
| chr7 | 1513461 | 1514256 | 2.27342 | 2.57196 | 1.131 |
| chr1 | 149204265 | 149204593 | 2.64124 | 2.98837 | 1.131 |
| chr2 | 209238236 | 209238605 | 1.99807 | 2.26069 | 1.131 |
| chr2 | 73200676 | 73201107 | 3.07118 | 3.47576 | 1.132 |
| chr11 | 63438237 | 63438479 | 1.99792 | 2.26132 | 1.132 |
| chr3 | 71446303 | 71446508 | 1.628 | 1.84275 | 1.132 |
| chr12 | 98633452 | 98633664 | 0.930894 | 1.05376 | 1.132 |
| chr17 | 37139443 | 37139786 | 1.81002 | 2.04894 | 1.132 |
| chr1 | 212629166 | 212629552 | 3.14947 | 3.56541 | 1.132 |
| chr9 | 9942828 | 9943144 | 2.20444 | 2.49578 | 1.132 |
| chr17 | 55860615 | 55861013 | 3.19755 | 3.62042 | 1.132 |
| chr14 | 87369960 | 87370242 | 1.16003 | 1.31345 | 1.132 |
| chr1 | 237288076 | 237288482 | 1.74771 | 1.97894 | 1.132 |
| chr12 | 23521156 | 23521511 | 0.749643 | 0.848842 | 1.132 |
| chr11 | 46705513 | 46705794 | 0.794536 | 0.899698 | 1.132 |
| chr8 | 133655158 | 133655411 | 0.983296 | 1.11355 | 1.132 |
| chr2 | 163077529 | 163077830 | 1.45492 | 1.64769 | 1.133 |
| chr10 | 10845213 | 10845451 | 0.966361 | 1.09444 | 1.133 |
| chr2 | 171217839 | 171218246 | 1.731 | 1.96077 | 1.133 |
| chr9 | 77853244 | 77853574 | 1.24026 | 1.40489 | 1.133 |
| chr16 | 20394775 | 20395046 | 1.87519 | 2.12425 | 1.133 |
| chr17 | 78688614 | 78688863 | 0.995529 | 1.1279 | 1.133 |
| chr5 | 6683413 | 6683619 | 0.792892 | 0.898383 | 1.133 |
| chr15 | 66300184 | 66300398 | 1.26259 | 1.43063 | 1.133 |
| chr9 | 28527413 | 28527743 | 0.859425 | 0.973897 | 1.133 |
| chrX | 16713228 | 16713529 | 1.71505 | 1.94354 | 1.133 |
| chr1 | 225040742 | 225040990 | 1.44109 | 1.63322 | 1.133 |
| chr2 | 120417665 | 120417971 | 2.03861 | 2.31063 | 1.133 |
| chr7 | 14464348 | 14464559 | 0.870381 | 0.986601 | 1.134 |
| chr2 | 165336052 | 165336347 | 1.46228 | 1.65756 | 1.134 |
| chr10 | 35929550 | 35929738 | 1.89314 | 2.14599 | 1.134 |
| chr7 | 152444615 | 152444820 | 0.668218 | 0.757534 | 1.134 |
| chr7 | 98460779 | 98461033 | 0.733473 | 0.831513 | 1.134 |
| chr15 | 96012189 | 96012419 | 1.04067 | 1.17984 | 1.134 |
| chr4 | 154495578 | 154495793 | 1.45843 | 1.6535 | 1.134 |
| chr9 | 129327022 | 129327369 | 0.967656 | 1.09708 | 1.134 |
| chr3 | 49876679 | 49876984 | 2.25382 | 2.55551 | 1.134 |
| chr2 | 191646274 | 191646592 | 1.01197 | 1.14747 | 1.134 |
| chr10 | 28030837 | 28031338 | 1.41289 | 1.60236 | 1.134 |
| chr10 | 121454222 | 121454436 | 0.749285 | 0.84983 | 1.134 |
| chr18 | 30132493 | 30132684 | 1.50641 | 1.70865 | 1.134 |
| chr14 | 37026078 | 37026326 | 2.88623 | 3.27377 | 1.134 |
| chr5 | 36739839 | 36740079 | 1.59212 | 1.8061 | 1.134 |

| | | | | | |
|---|---|---|---|---|---|
| chr12 | 92198519 | 92198701 | 0.890847 | 1.01063 | 1.134 |
| chr8 | 137726435 | 137726721 | 1.40635 | 1.5957 | 1.135 |
| chrX | 118650234 | 118650511 | 2.44305 | 2.77213 | 1.135 |
| chr2 | 212709665 | 212710032 | 0.7277 | 0.825845 | 1.135 |
| chr2 | 241089599 | 241089980 | 2.2066 | 2.50475 | 1.135 |
| chr12 | 115074908 | 115075269 | 1.43367 | 1.62741 | 1.135 |
| chr8 | 47312983 | 47313200 | 0.719709 | 0.817012 | 1.135 |
| chr3 | 50230085 | 50230416 | 2.02602 | 2.29997 | 1.135 |
| chr1 | 115595195 | 115595461 | 2.21651 | 2.51625 | 1.135 |
| chr15 | 50967806 | 50968064 | 1.64219 | 1.86427 | 1.135 |
| chr12 | 96526625 | 96526825 | 1.09658 | 1.24488 | 1.135 |
| chr2 | 45182743 | 45183103 | 3.35489 | 3.80912 | 1.135 |
| chr17 | 47300799 | 47301247 | 2.42643 | 2.75498 | 1.135 |
| chr10 | 15944665 | 15944998 | 1.25699 | 1.42737 | 1.136 |
| chr1 | 48667876 | 48668186 | 1.74211 | 1.97836 | 1.136 |
| chr9 | 127905875 | 127906200 | 2.68906 | 3.05385 | 1.136 |
| chr19 | 1240347 | 1240587 | 2.18321 | 2.47944 | 1.136 |
| chr17 | 7464347 | 7464644 | 2.50964 | 2.85024 | 1.136 |
| chr10 | 102631970 | 102632345 | 1.90964 | 2.16889 | 1.136 |
| chr2 | 25050452 | 25051099 | 1.68102 | 1.90943 | 1.136 |
| chr17 | 25533617 | 25533822 | 1.60612 | 1.82445 | 1.136 |
| chr11 | 66263087 | 66263316 | 1.7849 | 2.02806 | 1.136 |
| chr1 | 226065179 | 226065752 | 1.55278 | 1.76436 | 1.136 |
| chr7 | 135471386 | 135471630 | 0.85477 | 0.971257 | 1.136 |
| chr16 | 78036862 | 78037325 | 2.25717 | 2.56483 | 1.136 |
| chr7 | 97696301 | 97696584 | 0.72016 | 0.818327 | 1.136 |
| chr15 | 87402262 | 87402584 | 0.865916 | 0.98402 | 1.136 |
| chr2 | 238581765 | 238582035 | 2.43816 | 2.77073 | 1.136 |
| chr6 | 159463658 | 159463995 | 2.33439 | 2.65298 | 1.136 |
| chr19 | 39711725 | 39712032 | 0.981909 | 1.11593 | 1.136 |
| chr12 | 33180403 | 33180590 | 0.896566 | 1.01903 | 1.137 |
| chr9 | 95004859 | 95005282 | 1.10302 | 1.25369 | 1.137 |
| chr9 | 136226188 | 136226385 | 1.01978 | 1.15908 | 1.137 |
| chr6 | 51885851 | 51886035 | 1.42908 | 1.6245 | 1.137 |
| chr1 | 154968205 | 154968427 | 1.75352 | 1.99348 | 1.137 |
| chr11 | 84416211 | 84416444 | 0.800976 | 0.910714 | 1.137 |
| chr8 | 142322492 | 142322755 | 1.54458 | 1.75631 | 1.137 |
| chr19 | 30713221 | 30713740 | 4.87836 | 5.54731 | 1.137 |
| chr12 | 62227959 | 62228160 | 0.929316 | 1.05676 | 1.137 |
| chr1 | 19988304 | 19988596 | 1.94213 | 2.20854 | 1.137 |
| chr20 | 8622122 | 8622345 | 0.720179 | 0.818968 | 1.137 |
| chr5 | 7805577 | 7806035 | 1.03978 | 1.18252 | 1.137 |
| chr17 | 48628061 | 48628503 | 2.24012 | 2.54788 | 1.137 |
| chr8 | 130832323 | 130832699 | 2.58592 | 2.94151 | 1.138 |
| chr5 | 122890229 | 122890501 | 0.94611 | 1.07624 | 1.138 |
| chr10 | 90976784 | 90977129 | 1.87578 | 2.13427 | 1.138 |
| chr14 | 79113639 | 79113904 | 0.924082 | 1.05147 | 1.138 |
| chr19 | 33593702 | 33593989 | 1.00588 | 1.14456 | 1.138 |
| chr20 | 19248594 | 19248898 | 0.93769 | 1.06702 | 1.138 |
| chr2 | 237421651 | 237422049 | 1.64517 | 1.87217 | 1.138 |
| chr15 | 100319083 | 100319315 | 0.675853 | 0.769201 | 1.138 |
| chr2 | 117126417 | 117126615 | 0.650804 | 0.740711 | 1.138 |
| chr7 | 42916045 | 42916318 | 1.76643 | 2.01068 | 1.138 |
| chr2 | 80166959 | 80167182 | 0.760644 | 0.865836 | 1.138 |
| chr1 | 155995746 | 155996104 | 2.30571 | 2.6247 | 1.138 |
| chr7 | 31232586 | 31232912 | 2.40494 | 2.73768 | 1.138 |
| chr22 | 26726985 | 26727314 | 1.91016 | 2.1746 | 1.138 |
| chr3 | 162987524 | 162987727 | 0.711163 | 0.809685 | 1.139 |
| chr9 | 105797990 | 105798289 | 1.02821 | 1.17068 | 1.139 |
| chr1 | 151224261 | 151224481 | 2.38131 | 2.71155 | 1.139 |
| chr1 | 205407988 | 205408235 | 1.37585 | 1.56665 | 1.139 |

| | | | | | |
|---|---|---|---|---|---|
| chr8 | 144129650 | 144129975 | 2.64534 | 3.01227 | 1.139 |
| chr19 | 39209514 | 39209782 | 1.78992 | 2.03823 | 1.139 |
| chr7 | 105805752 | 105805970 | 0.598317 | 0.681334 | 1.139 |
| chr2 | 178029317 | 178029628 | 4.08925 | 4.65745 | 1.139 |
| chr12 | 111369919 | 111370254 | 1.57246 | 1.79118 | 1.139 |
| chr10 | 68578580 | 68578798 | 1.01878 | 1.16051 | 1.139 |
| chr3 | 12217366 | 12217736 | 2.08232 | 2.37214 | 1.139 |
| chr8 | 136289966 | 136290273 | 1.05118 | 1.19755 | 1.139 |
| chr2 | 115710069 | 115710286 | 0.66525 | 0.758005 | 1.139 |
| chr2 | 242642036 | 242642462 | 1.42389 | 1.62251 | 1.139 |
| chr11 | 74640851 | 74641144 | 2.76556 | 3.15141 | 1.140 |
| chr8 | 124775707 | 124775946 | 1.92347 | 2.19183 | 1.140 |
| chr10 | 106490661 | 106490872 | 0.855863 | 0.975318 | 1.140 |
| chr19 | 32697014 | 32697216 | 0.972586 | 1.10833 | 1.140 |
| chr7 | 149322624 | 149322945 | 2.68066 | 3.05494 | 1.140 |
| chr2 | 220341003 | 220341331 | 3.5433 | 4.03807 | 1.140 |
| chr2 | 68668745 | 68669208 | 1.96168 | 2.23579 | 1.140 |
| chr5 | 127026456 | 127026644 | 0.815547 | 0.929579 | 1.140 |
| chr8 | 144388661 | 144389180 | 1.69069 | 1.92717 | 1.140 |
| chr17 | 74878157 | 74878408 | 0.984104 | 1.12188 | 1.140 |
| chr22 | 20072389 | 20072782 | 1.10694 | 1.26193 | 1.140 |
| chr3 | 61550281 | 61550614 | 1.50181 | 1.71209 | 1.140 |
| chr16 | 46761188 | 46761407 | 0.636485 | 0.725653 | 1.140 |
| chr7 | 33712778 | 33713092 | 0.792421 | 0.903435 | 1.140 |
| chr15 | 98288705 | 98288976 | 1.54168 | 1.75768 | 1.140 |
| chr20 | 32319527 | 32319883 | 2.96011 | 3.37497 | 1.140 |
| chr3 | 154591392 | 154591637 | 0.872822 | 0.995165 | 1.140 |
| chr10 | 44167187 | 44167393 | 1.56998 | 1.79017 | 1.140 |
| chr19 | 56433092 | 56433281 | 0.826804 | 0.942994 | 1.141 |
| chr2 | 10282433 | 10282985 | 1.99188 | 2.27194 | 1.141 |
| chr16 | 85649499 | 85649982 | 2.12948 | 2.42936 | 1.141 |
| chr5 | 110932536 | 110932774 | 0.698733 | 0.797226 | 1.141 |
| chr8 | 130648382 | 130648614 | 0.808643 | 0.922632 | 1.141 |
| chr10 | 21558836 | 21559219 | 3.04694 | 3.47671 | 1.141 |
| chr5 | 164712950 | 164713189 | 0.840499 | 0.959061 | 1.141 |
| chr12 | 16645204 | 16645413 | 0.880624 | 1.00491 | 1.141 |
| chr11 | 848285 | 848694 | 2.49457 | 2.84698 | 1.141 |
| chr19 | 57375907 | 57376361 | 1.06942 | 1.22058 | 1.141 |
| chr9 | 122919415 | 122919653 | 0.764096 | 0.872143 | 1.141 |
| chr1 | 10398127 | 10398413 | 2.13693 | 2.43932 | 1.142 |
| chr19 | 11047371 | 11047619 | 2.02716 | 2.31443 | 1.142 |
| chr2 | 136467504 | 136467811 | 2.24339 | 2.5616 | 1.142 |
| chr12 | 96371731 | 96371947 | 0.711757 | 0.812774 | 1.142 |
| chr14 | 77925574 | 77925819 | 1.72628 | 1.97133 | 1.142 |
| chr1 | 209728273 | 209728572 | 0.830774 | 0.948893 | 1.142 |
| chr8 | 133197563 | 133197798 | 0.71324 | 0.814648 | 1.142 |
| chr5 | 18466296 | 18466572 | 1.16276 | 1.32812 | 1.142 |
| chr12 | 8261443 | 8261732 | 3.79114 | 4.33048 | 1.142 |
| chr7 | 52385632 | 52385857 | 0.921661 | 1.05285 | 1.142 |
| chr7 | 141374600 | 141374788 | 1.80054 | 2.05726 | 1.143 |
| chr8 | 136478107 | 136478547 | 2.56407 | 2.92988 | 1.143 |
| chr9 | 94496646 | 94497157 | 2.5348 | 2.89727 | 1.143 |
| chr14 | 22320088 | 22320593 | 1.48516 | 1.69757 | 1.143 |
| chr5 | 26656901 | 26657225 | 0.814231 | 0.930789 | 1.143 |
| chr8 | 94070052 | 94070258 | 1.04436 | 1.19386 | 1.143 |
| chr19 | 6736613 | 6736846 | 1.569 | 1.79404 | 1.143 |
| chr16 | 9071522 | 9071734 | 0.729075 | 0.833719 | 1.144 |
| chr4 | 135165167 | 135165654 | 0.717451 | 0.820563 | 1.144 |
| chr17 | 45924979 | 45925491 | 1.65501 | 1.893 | 1.144 |
| chr9 | 122731374 | 122731936 | 2.49614 | 2.85516 | 1.144 |
| chr3 | 115555561 | 115556121 | 1.56922 | 1.79504 | 1.144 |

| | | | | | |
|---|---|---|---|---|---|
| chr15 | 33905478 | 33905664 | 0.83769 | 0.958382 | 1.144 |
| chrX | 133428695 | 133428881 | 1.59924 | 1.82969 | 1.144 |
| chr7 | 113338693 | 113338943 | 1.48493 | 1.69912 | 1.144 |
| chr3 | 120225882 | 120226354 | 1.49494 | 1.71068 | 1.144 |
| chr18 | 48311174 | 48311416 | 1.33176 | 1.52405 | 1.144 |
| chr1 | 195792314 | 195792557 | 0.851803 | 0.97487 | 1.144 |
| chr3 | 145936756 | 145937063 | 1.08532 | 1.24222 | 1.145 |
| chr3 | 104942716 | 104942940 | 0.703603 | 0.805339 | 1.145 |
| chr10 | 107887621 | 107887830 | 1.03235 | 1.18181 | 1.145 |
| chr11 | 116094983 | 116095439 | 1.78008 | 2.03787 | 1.145 |
| chr2 | 103065924 | 103066148 | 1.83033 | 2.09583 | 1.145 |
| chr5 | 133315452 | 133316037 | 1.01923 | 1.16714 | 1.145 |
| chr3 | 12087942 | 12088259 | 1.83742 | 2.10411 | 1.145 |
| chr15 | 90035252 | 90035628 | 1.91549 | 2.19364 | 1.145 |
| chr5 | 151729241 | 151729585 | 1.55792 | 1.78427 | 1.145 |
| chr20 | 37287588 | 37288157 | 1.80151 | 2.06338 | 1.145 |
| chr15 | 90823268 | 90823577 | 0.803296 | 0.920102 | 1.145 |
| chr7 | 33498979 | 33499363 | 1.29215 | 1.48031 | 1.146 |
| chr10 | 61382027 | 61382328 | 1.56735 | 1.79566 | 1.146 |
| chr20 | 24083007 | 24083296 | 1.56668 | 1.79499 | 1.146 |
| chr8 | 96351932 | 96352156 | 0.539111 | 0.617703 | 1.146 |
| chr10 | 124386324 | 124386691 | 0.880969 | 1.00946 | 1.146 |
| chr1 | 227673323 | 227673555 | 0.993971 | 1.13909 | 1.146 |
| chr1 | 153467377 | 153467782 | 2.61741 | 2.99966 | 1.146 |
| chr8 | 76737311 | 76737547 | 0.815383 | 0.934522 | 1.146 |
| chr8 | 73524707 | 73524935 | 0.739609 | 0.847683 | 1.146 |
| chr9 | 32362380 | 32362652 | 0.928645 | 1.06436 | 1.146 |
| chr20 | 46354122 | 46354544 | 2.75039 | 3.15258 | 1.146 |
| chr8 | 86999584 | 86999962 | 1.104 | 1.26559 | 1.146 |
| chr17 | 27451094 | 27451504 | 2.48338 | 2.84771 | 1.147 |
| chr10 | 15757820 | 15758028 | 1.16111 | 1.33159 | 1.147 |
| chr2 | 9953852 | 9954466 | 2.48791 | 2.85327 | 1.147 |
| chr2 | 219221378 | 219221650 | 1.06282 | 1.21901 | 1.147 |
| chr11 | 13406413 | 13406693 | 1.65147 | 1.89454 | 1.147 |
| chr6 | 127869106 | 127869309 | 1.07708 | 1.23565 | 1.147 |
| chr10 | 82684024 | 82684416 | 2.25479 | 2.58682 | 1.147 |
| chr17 | 27280540 | 27280849 | 1.85151 | 2.12425 | 1.147 |
| chr19 | 12936407 | 12936672 | 1.86045 | 2.13452 | 1.147 |
| chr8 | 145081824 | 145082028 | 1.09522 | 1.25658 | 1.147 |
| chr2 | 33428181 | 33428513 | 0.748719 | 0.859052 | 1.147 |
| chr15 | 61995082 | 61995313 | 0.867322 | 0.995227 | 1.147 |
| chr2 | 189890006 | 189890239 | 1.50435 | 1.72627 | 1.148 |
| chr7 | 18357725 | 18358053 | 0.72039 | 0.826701 | 1.148 |
| chr3 | 104706902 | 104707156 | 0.942855 | 1.08205 | 1.148 |
| chr2 | 231679390 | 231679617 | 1.2078 | 1.38646 | 1.148 |
| chr11 | 37318924 | 37319189 | 0.634802 | 0.728739 | 1.148 |
| chr20 | 7133734 | 7134012 | 0.930001 | 1.06776 | 1.148 |
| chr18 | 8609209 | 8609513 | 2.14438 | 2.46212 | 1.148 |
| chr7 | 105586486 | 105586697 | 0.815945 | 0.936886 | 1.148 |
| chr9 | 115593890 | 115594121 | 0.684377 | 0.785816 | 1.148 |
| chr3 | 134211359 | 134211603 | 1.16732 | 1.34039 | 1.148 |
| chr20 | 34149188 | 34149496 | 1.1663 | 1.33932 | 1.148 |
| chr2 | 127955455 | 127955681 | 2.94907 | 3.38675 | 1.148 |
| chr15 | 73469092 | 73469275 | 1.06958 | 1.22833 | 1.148 |
| chr20 | 22284131 | 22284524 | 1.35435 | 1.55538 | 1.148 |
| chr11 | 61536998 | 61537250 | 1.22518 | 1.40707 | 1.148 |
| chr19 | 4567605 | 4567955 | 2.98996 | 3.43401 | 1.149 |
| chr8 | 138346931 | 138347172 | 1.61088 | 1.85011 | 1.149 |
| chr5 | 26803292 | 26803487 | 0.935452 | 1.0744 | 1.149 |
| chr10 | 133483270 | 133483491 | 0.694589 | 0.797777 | 1.149 |
| chr13 | 30498387 | 30498666 | 2.50077 | 2.87229 | 1.149 |

| | | | | | |
|---|---|---|---|---|---|
| chr11 | 59197432 | 59197794 | 2.19291 | 2.51875 | 1.149 |
| chr16 | 84099265 | 84099469 | 1.59788 | 1.83554 | 1.149 |
| chr10 | 88659517 | 88659749 | 1.97986 | 2.2753 | 1.149 |
| chr2 | 26667499 | 26667915 | 0.871612 | 1.00181 | 1.149 |
| chr9 | 36103027 | 36103380 | 2.22383 | 2.55605 | 1.149 |
| chr5 | 9544633 | 9544914 | 3.65685 | 4.20375 | 1.150 |
| chr2 | 102490115 | 102490330 | 1.40199 | 1.6117 | 1.150 |
| chr9 | 34252836 | 34253304 | 2.60541 | 2.99565 | 1.150 |
| chr20 | 6252330 | 6252564 | 0.927181 | 1.06613 | 1.150 |
| chr16 | 63714056 | 63714242 | 0.932714 | 1.07269 | 1.150 |
| chr2 | 36198319 | 36198525 | 1.81865 | 2.09169 | 1.150 |
| chr10 | 31007892 | 31008202 | 0.602489 | 0.693045 | 1.150 |
| chr11 | 5405614 | 5405821 | 0.999783 | 1.15013 | 1.150 |
| chrX | 149000820 | 149001079 | 1.91817 | 2.20675 | 1.150 |
| chr5 | 161871390 | 161871634 | 0.66058 | 0.760098 | 1.151 |
| chr15 | 91790684 | 91791152 | 1.57334 | 1.81044 | 1.151 |
| chr15 | 98903368 | 98903590 | 0.908227 | 1.04523 | 1.151 |
| chr1 | 161011310 | 161011801 | 2.22563 | 2.56168 | 1.151 |
| chr7 | 12107108 | 12107458 | 2.02996 | 2.3367 | 1.151 |
| chr6 | 36164175 | 36164445 | 1.54066 | 1.77355 | 1.151 |
| chr17 | 54002775 | 54002990 | 0.803769 | 0.925443 | 1.151 |
| chr1 | 224326580 | 224326812 | 0.710596 | 0.8182 | 1.151 |
| chr14 | 52146778 | 52147007 | 0.770336 | 0.88701 | 1.151 |
| chr3 | 1854645 | 1854867 | 0.555068 | 0.639144 | 1.151 |
| chr10 | 73368784 | 73369307 | 3.09974 | 3.56971 | 1.152 |
| chr17 | 25381898 | 25382186 | 0.741799 | 0.854281 | 1.152 |
| chr5 | 92587967 | 92588246 | 1.17405 | 1.35212 | 1.152 |
| chr12 | 124423310 | 124423539 | 1.52752 | 1.75928 | 1.152 |
| chr12 | 54382963 | 54383207 | 2.21181 | 2.54744 | 1.152 |
| chr15 | 93754342 | 93754612 | 0.899223 | 1.03585 | 1.152 |
| chr2 | 30588209 | 30588525 | 0.642551 | 0.74039 | 1.152 |
| chr3 | 168936202 | 168936385 | 1.22013 | 1.40598 | 1.152 |
| chr8 | 89520706 | 89520935 | 0.788217 | 0.908341 | 1.152 |
| chr7 | 12498212 | 12498448 | 2.46549 | 2.84152 | 1.153 |
| chr2 | 68519287 | 68519590 | 1.76279 | 2.03174 | 1.153 |
| chr11 | 72284409 | 72284732 | 2.00125 | 2.3067 | 1.153 |
| chr7 | 122572396 | 122572673 | 0.699689 | 0.806554 | 1.153 |
| chr12 | 55218054 | 55218272 | 0.861427 | 0.993014 | 1.153 |
| chr5 | 65732617 | 65732855 | 1.39578 | 1.60902 | 1.153 |
| chr11 | 44638364 | 44638840 | 2.67169 | 3.08019 | 1.153 |
| chr14 | 21341840 | 21342157 | 2.57338 | 2.96684 | 1.153 |
| chr17 | 65796451 | 65796865 | 2.54365 | 2.93258 | 1.153 |
| chr7 | 37472916 | 37473163 | 0.939195 | 1.0828 | 1.153 |
| chr1 | 100173202 | 100173440 | 1.61573 | 1.863 | 1.153 |
| chr16 | 56610707 | 56611299 | 1.92628 | 2.22108 | 1.153 |
| chr10 | 85786414 | 85786905 | 1.8181 | 2.0966 | 1.153 |
| chr6 | 2449024 | 2449273 | 1.8825 | 2.17092 | 1.153 |
| chr7 | 73598077 | 73598334 | 1.75208 | 2.02059 | 1.153 |
| chr19 | 32000355 | 32000565 | 0.789222 | 0.910267 | 1.153 |
| chr17 | 53813269 | 53813490 | 1.96162 | 2.26304 | 1.154 |
| chr19 | 10500635 | 10500886 | 1.07686 | 1.2424 | 1.154 |
| chr7 | 150685454 | 150685916 | 2.33125 | 2.68975 | 1.154 |
| chr2 | 16466283 | 16466521 | 0.620728 | 0.716219 | 1.154 |
| chr9 | 139025151 | 139025428 | 1.89443 | 2.18587 | 1.154 |
| chr11 | 6759199 | 6759416 | 1.58605 | 1.83025 | 1.154 |
| chr2 | 219938096 | 219938495 | 1.79498 | 2.07136 | 1.154 |
| chr2 | 2702460 | 2702761 | 0.999717 | 1.15372 | 1.154 |
| chr11 | 4935302 | 4935507 | 0.754234 | 0.870431 | 1.154 |
| chr19 | 46061635 | 46062046 | 2.40435 | 2.77531 | 1.154 |
| chr9 | 131174722 | 131174937 | 1.5493 | 1.78837 | 1.154 |
| chr11 | 4572892 | 4573141 | 1.73998 | 2.00854 | 1.154 |

| | | | | | |
|---|---|---|---|---|---|
| chr13 | 72051929 | 72052193 | 0.813821 | 0.939683 | 1.155 |
| chrX | 154209193 | 154209435 | 2.17258 | 2.5088 | 1.155 |
| chr10 | 21964078 | 21964452 | 2.25831 | 2.60803 | 1.155 |
| chr7 | 78806241 | 78806481 | 1.85464 | 2.14195 | 1.155 |
| chr16 | 68295375 | 68295866 | 2.48832 | 2.87385 | 1.155 |
| chr15 | 27708734 | 27709050 | 0.945301 | 1.09191 | 1.155 |
| chr1 | 2929056 | 2929391 | 2.36541 | 2.73239 | 1.155 |
| chr2 | 234159989 | 234160341 | 3.11125 | 3.59409 | 1.155 |
| chr19 | 29443286 | 29443698 | 1.05999 | 1.22475 | 1.155 |
| chr5 | 76425314 | 76425533 | 1.65915 | 1.9172 | 1.156 |
| chr17 | 52776007 | 52776405 | 0.770361 | 0.890188 | 1.156 |
| chr16 | 8754869 | 8755185 | 1.74068 | 2.01151 | 1.156 |
| chr20 | 42018999 | 42019279 | 1.45281 | 1.67887 | 1.156 |
| chr19 | 6769340 | 6769525 | 1.33804 | 1.54635 | 1.156 |
| chr9 | 28331493 | 28331747 | 0.848558 | 0.980742 | 1.156 |
| chr4 | 159091721 | 159092053 | 1.5658 | 1.8098 | 1.156 |
| chr7 | 108732298 | 108732602 | 1.03806 | 1.19984 | 1.156 |
| chr9 | 26433648 | 26433850 | 0.811304 | 0.937814 | 1.156 |
| chrX | 139848268 | 139848502 | 2.01466 | 2.32885 | 1.156 |
| chr11 | 67273601 | 67273971 | 3.1428 | 3.63346 | 1.156 |
| chr3 | 108817057 | 108817373 | 0.924358 | 1.06871 | 1.156 |
| chr12 | 92964100 | 92964408 | 1.50905 | 1.74473 | 1.156 |
| chr7 | 32386824 | 32387274 | 1.68504 | 1.94824 | 1.156 |
| chr9 | 26744313 | 26744604 | 1.01136 | 1.16938 | 1.156 |
| chr7 | 124442325 | 124442549 | 0.968315 | 1.11969 | 1.156 |
| chr12 | 127176237 | 127176514 | 0.813598 | 0.940822 | 1.156 |
| chr1 | 223885961 | 223886183 | 1.85841 | 2.14907 | 1.156 |
| chr12 | 121661208 | 121661893 | 0.844767 | 0.976895 | 1.156 |
| chr2 | 77347078 | 77347300 | 0.906818 | 1.04865 | 1.156 |
| chr17 | 10519779 | 10520311 | 2.85556 | 3.30262 | 1.157 |
| chr17 | 54087947 | 54088197 | 0.680398 | 0.786965 | 1.157 |
| chr11 | 13977503 | 13977815 | 2.00719 | 2.32193 | 1.157 |
| chr1 | 246058017 | 246058339 | 0.738337 | 0.854154 | 1.157 |
| chr5 | 51258082 | 51258362 | 0.770579 | 0.891458 | 1.157 |
| chr2 | 104807039 | 104807271 | 0.783189 | 0.906083 | 1.157 |
| chr1 | 241390897 | 241391133 | 0.706537 | 0.817424 | 1.157 |
| chr17 | 61943547 | 61943774 | 0.759812 | 0.879115 | 1.157 |
| chr7 | 45257022 | 45257329 | 1.54615 | 1.78947 | 1.157 |
| chr10 | 90488211 | 90488467 | 1.12115 | 1.29784 | 1.158 |
| chr2 | 17165090 | 17165323 | 0.588536 | 0.681322 | 1.158 |
| chr2 | 63715227 | 63715595 | 2.18384 | 2.52822 | 1.158 |
| chr10 | 94985643 | 94985868 | 0.923692 | 1.06939 | 1.158 |
| chr17 | 19308300 | 19308522 | 0.713824 | 0.82648 | 1.158 |
| chr9 | 23322540 | 23322782 | 0.750153 | 0.868541 | 1.158 |
| chr8 | 139000303 | 139000573 | 0.751942 | 0.87062 | 1.158 |
| chr8 | 114059073 | 114059336 | 0.758652 | 0.878566 | 1.158 |
| chr19 | 28296688 | 28296918 | 0.971008 | 1.12452 | 1.158 |
| chr17 | 72292769 | 72293027 | 2.13583 | 2.47375 | 1.158 |
| chr3 | 2699890 | 2700105 | 0.741132 | 0.858404 | 1.158 |
| chr6 | 91186403 | 91186629 | 1.03631 | 1.20031 | 1.158 |
| chr17 | 76133458 | 76134407 | 1.68013 | 1.94626 | 1.158 |
| chr15 | 33985488 | 33985739 | 0.699106 | 0.809895 | 1.158 |
| chr3 | 85540745 | 85540967 | 0.578548 | 0.670342 | 1.159 |
| chr12 | 2508879 | 2509096 | 0.792987 | 0.91883 | 1.159 |
| chr2 | 30338616 | 30338893 | 2.51754 | 2.9172 | 1.159 |
| chr22 | 50936669 | 50937011 | 1.54808 | 1.79385 | 1.159 |
| chr2 | 182321874 | 182322196 | 2.62004 | 3.03625 | 1.159 |
| chr2 | 21926798 | 21927016 | 0.95637 | 1.10838 | 1.159 |
| chr11 | 29920236 | 29920555 | 0.820965 | 0.951605 | 1.159 |
| chr16 | 15798360 | 15798808 | 2.30768 | 2.67503 | 1.159 |
| chr9 | 12598359 | 12598649 | 0.931399 | 1.07994 | 1.159 |

| | | | | | |
|---|---|---|---|---|---|
| chr2 | 13277814 | 13278066 | 1.08014 | 1.25263 | 1.160 |
| chr9 | 135675580 | 135675811 | 1.82361 | 2.11496 | 1.160 |
| chr2 | 43054717 | 43055129 | 2.84593 | 3.30101 | 1.160 |
| chr5 | 6789148 | 6789400 | 0.996937 | 1.15637 | 1.160 |
| chr17 | 50127464 | 50127666 | 0.833739 | 0.967132 | 1.160 |
| chr22 | 26794656 | 26794940 | 1.27215 | 1.47573 | 1.160 |
| chr19 | 42297037 | 42297437 | 2.55281 | 2.96158 | 1.160 |
| chr6 | 155249527 | 155249758 | 2.29585 | 2.66383 | 1.160 |
| chr16 | 53455212 | 53455593 | 1.45987 | 1.69396 | 1.160 |
| chr1 | 24853645 | 24853971 | 1.33202 | 1.54586 | 1.161 |
| chr9 | 34379349 | 34379658 | 2.65682 | 3.08346 | 1.161 |
| chr2 | 38209278 | 38209668 | 2.22994 | 2.58804 | 1.161 |
| chr13 | 53383490 | 53383754 | 1.46364 | 1.69878 | 1.161 |
| chr16 | 65161110 | 65161325 | 0.692094 | 0.803312 | 1.161 |
| chr2 | 175442646 | 175442953 | 1.22093 | 1.41719 | 1.161 |
| chr7 | 24859858 | 24860187 | 2.16864 | 2.51729 | 1.161 |
| chr8 | 55586728 | 55586943 | 0.764112 | 0.886971 | 1.161 |
| chr11 | 132948475 | 132948780 | 2.12941 | 2.47197 | 1.161 |
| chr17 | 53638597 | 53639044 | 0.910474 | 1.05698 | 1.161 |
| chr8 | 61850332 | 61850592 | 0.906422 | 1.05239 | 1.161 |
| chr5 | 18704260 | 18704462 | 0.982894 | 1.14135 | 1.161 |
| chr14 | 96150112 | 96150487 | 1.96452 | 2.28131 | 1.161 |
| chr3 | 39854675 | 39855118 | 0.922969 | 1.07185 | 1.161 |
| chr13 | 102794072 | 102794339 | 1.84838 | 2.14713 | 1.162 |
| chrX | 4468078 | 4468311 | 0.91572 | 1.06374 | 1.162 |
| chr8 | 139011149 | 139011357 | 0.886016 | 1.02931 | 1.162 |
| chr17 | 79375721 | 79376026 | 1.86158 | 2.16267 | 1.162 |
| chr10 | 15786052 | 15786720 | 1.27271 | 1.47862 | 1.162 |
| chr14 | 79426091 | 79426430 | 2.16366 | 2.51391 | 1.162 |
| chr7 | 131266142 | 131266349 | 0.867413 | 1.00783 | 1.162 |
| chr2 | 225511035 | 225511273 | 2.28498 | 2.65514 | 1.162 |
| chr9 | 105209762 | 105209966 | 0.860806 | 1.00033 | 1.162 |
| chr5 | 146803381 | 146803670 | 0.750538 | 0.87221 | 1.162 |
| chr1 | 159101812 | 159102502 | 1.75667 | 2.04152 | 1.162 |
| chr13 | 21066843 | 21067116 | 1.90246 | 2.21112 | 1.162 |
| chr2 | 5531612 | 5531843 | 0.816087 | 0.9485 | 1.162 |
| chr15 | 32153480 | 32153698 | 0.741113 | 0.861415 | 1.162 |
| chr21 | 42542439 | 42542678 | 1.77672 | 2.06526 | 1.162 |
| chr4 | 47688351 | 47688607 | 1.39149 | 1.61774 | 1.163 |
| chr19 | 39220424 | 39220668 | 1.24393 | 1.44628 | 1.163 |
| chr8 | 10654509 | 10654836 | 0.855709 | 0.99492 | 1.163 |
| chr19 | 51207562 | 51207820 | 1.42878 | 1.66124 | 1.163 |
| chr11 | 15096265 | 15096871 | 1.76616 | 2.05356 | 1.163 |
| chr19 | 34430040 | 34430304 | 1.13746 | 1.3227 | 1.163 |
| chr8 | 139285699 | 139286119 | 0.924809 | 1.07547 | 1.163 |
| chrX | 51163523 | 51163830 | 1.85412 | 2.15629 | 1.163 |
| chr8 | 133785399 | 133785642 | 1.73868 | 2.02222 | 1.163 |
| chr9 | 92875603 | 92875848 | 0.632898 | 0.736137 | 1.163 |
| chr9 | 92218757 | 92219023 | 1.41558 | 1.64656 | 1.163 |
| chr5 | 33934877 | 33935104 | 1.82476 | 2.12261 | 1.163 |
| chr1 | 179544985 | 179545212 | 0.644057 | 0.749212 | 1.163 |
| chr22 | 28416025 | 28416513 | 2.16629 | 2.52013 | 1.163 |
| chr5 | 90079465 | 90079894 | 2.41908 | 2.81421 | 1.163 |
| chr19 | 34488567 | 34488799 | 1.26048 | 1.46643 | 1.163 |
| chr2 | 179343814 | 179344055 | 2.41967 | 2.81513 | 1.163 |
| chr1 | 145585361 | 145585574 | 1.67505 | 1.94885 | 1.163 |
| chr2 | 31361346 | 31361805 | 2.42633 | 2.82298 | 1.163 |
| chr2 | 39867353 | 39867550 | 0.968564 | 1.1269 | 1.163 |
| chr8 | 139115090 | 139115343 | 1.08044 | 1.25712 | 1.164 |
| chr1 | 179704881 | 179705086 | 0.679822 | 0.791002 | 1.164 |
| chr5 | 16791170 | 16791431 | 1.07132 | 1.24672 | 1.164 |

| | | | | | |
|---|---|---|---|---|---|
| chr5 | 118041813 | 118042154 | 0.836631 | 0.973606 | 1.164 |
| chr1 | 177225410 | 177225650 | 1.54936 | 1.80311 | 1.164 |
| chr7 | 43622195 | 43622712 | 4.059 | 4.72391 | 1.164 |
| chr4 | 139602269 | 139602474 | 2.10239 | 2.44681 | 1.164 |
| chr12 | 81353181 | 81353421 | 1.08373 | 1.26139 | 1.164 |
| chr2 | 97175160 | 97175539 | 2.97927 | 3.46768 | 1.164 |
| chr3 | 39250041 | 39250282 | 1.87855 | 2.18658 | 1.164 |
| chr11 | 46918554 | 46918803 | 1.36239 | 1.58629 | 1.164 |
| chr7 | 52424050 | 52424409 | 0.844781 | 0.98361 | 1.164 |
| chr8 | 53917523 | 53917732 | 1.2036 | 1.40143 | 1.164 |
| chr2 | 86306830 | 86307331 | 2.1061 | 2.4523 | 1.164 |
| chr17 | 78932632 | 78933054 | 1.27776 | 1.48812 | 1.165 |
| chr2 | 12467006 | 12467392 | 1.40986 | 1.64216 | 1.165 |
| chr21 | 36095048 | 36095345 | 1.18541 | 1.38113 | 1.165 |
| chr19 | 39902494 | 39902761 | 2.53843 | 2.95769 | 1.165 |
| chr17 | 19549265 | 19549521 | 2.45413 | 2.86006 | 1.165 |
| chr9 | 93657565 | 93657894 | 1.64879 | 1.92157 | 1.165 |
| chr7 | 132222830 | 132223016 | 0.897321 | 1.04579 | 1.165 |
| chr2 | 3775790 | 3776522 | 1.13401 | 1.32175 | 1.166 |
| chr11 | 107469351 | 107469613 | 1.40159 | 1.63373 | 1.166 |
| chr1 | 228496810 | 228497034 | 1.67969 | 1.95792 | 1.166 |
| chr11 | 111475019 | 111475485 | 1.27121 | 1.48184 | 1.166 |
| chr9 | 132483544 | 132483805 | 1.66503 | 1.94102 | 1.166 |
| chr20 | 5289577 | 5289798 | 0.837292 | 0.976129 | 1.166 |
| chr2 | 59571321 | 59571532 | 0.897442 | 1.04628 | 1.166 |
| chr16 | 62247723 | 62247955 | 0.781164 | 0.91078 | 1.166 |
| chr11 | 32028420 | 32028669 | 1.76194 | 2.05448 | 1.166 |
| chr1 | 164587128 | 164587330 | 1.7008 | 1.98404 | 1.167 |
| chr15 | 94766933 | 94767156 | 0.836093 | 0.975344 | 1.167 |
| chr8 | 92793643 | 92793846 | 1.18706 | 1.38484 | 1.167 |
| chr1 | 156879876 | 156880564 | 2.81632 | 3.28566 | 1.167 |
| chr6 | 17505350 | 17505635 | 1.02127 | 1.1915 | 1.167 |
| chr8 | 33282149 | 33282476 | 2.62763 | 3.06571 | 1.167 |
| chr8 | 80945085 | 80945289 | 1.06819 | 1.24641 | 1.167 |
| chr2 | 180083694 | 180084071 | 2.16907 | 2.53129 | 1.167 |
| chr17 | 19370290 | 19370887 | 1.1657 | 1.36041 | 1.167 |
| chr1 | 160133495 | 160133699 | 1.16969 | 1.36522 | 1.167 |
| chr9 | 107266602 | 107267054 | 0.997735 | 1.16453 | 1.167 |
| chr7 | 124496410 | 124496638 | 1.48926 | 1.73825 | 1.167 |
| chr9 | 87008570 | 87008784 | 1.08513 | 1.2666 | 1.167 |
| chr15 | 76527588 | 76527881 | 1.51904 | 1.77331 | 1.167 |
| chr19 | 42683383 | 42683670 | 2.11541 | 2.4697 | 1.167 |
| chr14 | 34197187 | 34197436 | 0.847484 | 0.989477 | 1.168 |
| chr17 | 27618233 | 27618478 | 1.82358 | 2.12913 | 1.168 |
| chr1 | 204367725 | 204368003 | 2.40711 | 2.81055 | 1.168 |
| chr20 | 3096225 | 3096494 | 2.25303 | 2.6309 | 1.168 |
| chr21 | 46312746 | 46313072 | 2.70318 | 3.15674 | 1.168 |
| chr12 | 29226694 | 29226962 | 1.55362 | 1.81434 | 1.168 |
| chr2 | 23535938 | 23536193 | 1.69209 | 1.97629 | 1.168 |
| chr19 | 30055578 | 30055860 | 0.998161 | 1.166 | 1.168 |
| chr7 | 147500351 | 147500616 | 0.835928 | 0.976516 | 1.168 |
| chr8 | 113477372 | 113477638 | 1.05867 | 1.2369 | 1.168 |
| chr3 | 124217824 | 124218059 | 1.02171 | 1.19387 | 1.169 |
| chr11 | 7299642 | 7299852 | 1.332 | 1.5565 | 1.169 |
| chr10 | 53652976 | 53653233 | 0.873223 | 1.02052 | 1.169 |
| chr1 | 167509501 | 167510075 | 2.04488 | 2.39006 | 1.169 |
| chr3 | 114911206 | 114911533 | 0.893253 | 1.04431 | 1.169 |
| chr9 | 72725907 | 72726215 | 2.24367 | 2.62318 | 1.169 |
| chr8 | 53415848 | 53416065 | 1.4763 | 1.72613 | 1.169 |
| chr2 | 113894500 | 113894924 | 1.82175 | 2.13028 | 1.169 |
| chr20 | 1190129 | 1190369 | 1.5198 | 1.77722 | 1.169 |

| | | | | | |
|---|---|---|---|---|---|
| chr14 | 101119594 | 101120037 | 2.02247 | 2.36523 | 1.169 |
| chr19 | 41777202 | 41777427 | 0.997728 | 1.16682 | 1.169 |
| chr19 | 29084717 | 29084943 | 0.800766 | 0.936689 | 1.170 |
| chr18 | 48577536 | 48577794 | 2.64812 | 3.09776 | 1.170 |
| chr12 | 50866718 | 50866996 | 0.698115 | 0.816666 | 1.170 |
| chr22 | 36942405 | 36942627 | 1.36325 | 1.59479 | 1.170 |
| chr1 | 215999498 | 215999779 | 1.03658 | 1.21265 | 1.170 |
| chr10 | 12943892 | 12944530 | 1.48832 | 1.74154 | 1.170 |
| chr17 | 49230553 | 49230854 | 2.8138 | 3.29261 | 1.170 |
| chr20 | 51327839 | 51328082 | 0.803244 | 0.939923 | 1.170 |
| chr16 | 64671057 | 64671327 | 0.786979 | 0.921009 | 1.170 |
| chr1 | 183197159 | 183197627 | 2.08004 | 2.43448 | 1.170 |
| chr1 | 163144594 | 163145117 | 1.91834 | 2.2455 | 1.171 |
| chr2 | 42722165 | 42722476 | 2.29765 | 2.68951 | 1.171 |
| chr11 | 130731811 | 130732211 | 3.25916 | 3.81508 | 1.171 |
| chr20 | 60585962 | 60586210 | 1.70839 | 2.00001 | 1.171 |
| chr12 | 61266948 | 61267187 | 0.897822 | 1.0511 | 1.171 |
| chr6 | 11618932 | 11619116 | 1.02003 | 1.19418 | 1.171 |
| chr10 | 119469644 | 119470082 | 1.27698 | 1.49507 | 1.171 |
| chr3 | 74316825 | 74317123 | 1.56482 | 1.8322 | 1.171 |
| chr7 | 26726046 | 26726308 | 2.12695 | 2.49038 | 1.171 |
| chr8 | 100921759 | 100921983 | 0.80209 | 0.939179 | 1.171 |
| chr20 | 44598369 | 44598746 | 2.05235 | 2.40314 | 1.171 |
| chr15 | 86818329 | 86818562 | 0.882573 | 1.03362 | 1.171 |
| chr16 | 73738389 | 73738686 | 0.80292 | 0.940423 | 1.171 |
| chr15 | 45751024 | 45751293 | 2.3034 | 2.69792 | 1.171 |
| chr22 | 33396663 | 33397031 | 2.39452 | 2.80486 | 1.171 |
| chr5 | 65602232 | 65602555 | 2.64555 | 3.09902 | 1.171 |
| chr5 | 113714662 | 113714969 | 0.788003 | 0.92313 | 1.171 |
| chr2 | 203453390 | 203453744 | 2.09618 | 2.45583 | 1.172 |
| chr21 | 32830002 | 32830305 | 0.910815 | 1.06712 | 1.172 |
| chr3 | 31766164 | 31766518 | 1.22705 | 1.43771 | 1.172 |
| chr15 | 89287779 | 89288167 | 0.962959 | 1.12834 | 1.172 |
| chr8 | 19981268 | 19981645 | 0.95586 | 1.12011 | 1.172 |
| chr2 | 107189075 | 107189412 | 1.92905 | 2.26072 | 1.172 |
| chr7 | 2387712 | 2387958 | 1.37731 | 1.61413 | 1.172 |
| chr8 | 70147393 | 70147757 | 0.626072 | 0.733851 | 1.172 |
| chr4 | 112727630 | 112727843 | 1.0139 | 1.1886 | 1.172 |
| chrX | 118142153 | 118142457 | 2.14979 | 2.52033 | 1.172 |
| chr2 | 136333692 | 136333901 | 0.87561 | 1.02661 | 1.172 |
| chr2 | 42096898 | 42097196 | 2.54556 | 2.98456 | 1.172 |
| chr3 | 50582445 | 50582679 | 0.854692 | 1.00216 | 1.173 |
| chr8 | 141590686 | 141590991 | 0.882096 | 1.03429 | 1.173 |
| chr14 | 102918745 | 102918990 | 0.837565 | 0.982093 | 1.173 |
| chr3 | 167097872 | 167098333 | 1.5586 | 1.8277 | 1.173 |
| chr17 | 55377241 | 55377479 | 1.02202 | 1.19853 | 1.173 |
| chr2 | 225224376 | 225224587 | 0.948038 | 1.11177 | 1.173 |
| chr11 | 132810966 | 132811488 | 1.34938 | 1.5825 | 1.173 |
| chr3 | 107656957 | 107657170 | 1.44924 | 1.69976 | 1.173 |
| chr19 | 47082875 | 47083216 | 1.80769 | 2.12027 | 1.173 |
| chr8 | 139667130 | 139667413 | 0.714054 | 0.837548 | 1.173 |
| chr15 | 74340748 | 74341067 | 2.11747 | 2.48371 | 1.173 |
| chr19 | 52018075 | 52018478 | 1.8856 | 2.21184 | 1.173 |
| chr2 | 82442379 | 82442659 | 1.05347 | 1.23586 | 1.173 |
| chr5 | 11662613 | 11662836 | 0.648704 | 0.761101 | 1.173 |
| chr19 | 15666740 | 15666942 | 2.18944 | 2.56912 | 1.173 |
| chr15 | 70730194 | 70730446 | 1.17554 | 1.37961 | 1.174 |
| chr7 | 123264715 | 123265066 | 1.49802 | 1.75811 | 1.174 |
| chr1 | 116249399 | 116249783 | 1.63031 | 1.91342 | 1.174 |
| chr9 | 1232170 | 1232606 | 0.902149 | 1.05882 | 1.174 |
| chr16 | 15140733 | 15140915 | 1.90863 | 2.24019 | 1.174 |

| | | | | | |
|---|---|---|---|---|---|
| chr15 | 93579000 | 93579643 | 1.88816 | 2.21636 | 1.174 |
| chr12 | 133215199 | 133215710 | 2.08472 | 2.44714 | 1.174 |
| chr3 | 21205496 | 21205795 | 1.24545 | 1.46201 | 1.174 |
| chr10 | 72124176 | 72124725 | 2.17365 | 2.55168 | 1.174 |
| chr15 | 95192982 | 95193264 | 0.81086 | 0.95188 | 1.174 |
| chr1 | 46943064 | 46943418 | 2.16854 | 2.54582 | 1.174 |
| chr7 | 66103423 | 66103640 | 0.710959 | 0.8347 | 1.174 |
| chr10 | 131843339 | 131843649 | 1.51199 | 1.77517 | 1.174 |
| chr1 | 154452370 | 154452601 | 2.28841 | 2.6868 | 1.174 |
| chr12 | 105446876 | 105447199 | 2.20179 | 2.5854 | 1.174 |
| chr3 | 158635776 | 158635978 | 1.73407 | 2.03627 | 1.174 |
| chr5 | 149777384 | 149777571 | 1.08665 | 1.2764 | 1.175 |
| chr21 | 33246339 | 33246626 | 2.50619 | 2.94394 | 1.175 |
| chr19 | 38910372 | 38910633 | 0.754933 | 0.886875 | 1.175 |
| chr10 | 133619149 | 133619683 | 1.42803 | 1.67773 | 1.175 |
| chr2 | 29181539 | 29181784 | 1.97565 | 2.32113 | 1.175 |
| chr5 | 161842847 | 161843174 | 0.784714 | 0.921954 | 1.175 |
| chr5 | 179297561 | 179297836 | 1.7551 | 2.06213 | 1.175 |
| chr9 | 4759278 | 4759739 | 2.95559 | 3.47294 | 1.175 |
| chr1 | 237496653 | 237496969 | 0.826429 | 0.971096 | 1.175 |
| chr6 | 24359923 | 24360254 | 2.61972 | 3.07864 | 1.175 |
| chr10 | 122595814 | 122596057 | 1.13034 | 1.32837 | 1.175 |
| chr11 | 125792939 | 125793243 | 1.65634 | 1.9469 | 1.175 |
| chr14 | 30683658 | 30683896 | 0.960482 | 1.129 | 1.175 |
| chr5 | 943318 | 943687 | 0.908367 | 1.06777 | 1.175 |
| chr17 | 3239211 | 3239424 | 1.14213 | 1.34265 | 1.176 |
| chr12 | 76625717 | 76626254 | 0.930772 | 1.0942 | 1.176 |
| chr15 | 66998534 | 66998881 | 2.96168 | 3.48243 | 1.176 |
| chr8 | 52544827 | 52545074 | 0.739374 | 0.869387 | 1.176 |
| chr8 | 117402983 | 117403283 | 0.624174 | 0.733933 | 1.176 |
| chr1 | 215355015 | 215355258 | 0.702708 | 0.8263 | 1.176 |
| chr1 | 230631167 | 230631410 | 0.84065 | 0.988621 | 1.176 |
| chr10 | 18883597 | 18884047 | 1.4321 | 1.68437 | 1.176 |
| chr9 | 26704060 | 26704292 | 0.645225 | 0.758969 | 1.176 |
| chr22 | 35627519 | 35627809 | 2.06756 | 2.43224 | 1.176 |
| chr15 | 80189000 | 80189203 | 1.43028 | 1.68263 | 1.176 |
| chr5 | 174728409 | 174728645 | 0.92489 | 1.08811 | 1.176 |
| chr7 | 107302245 | 107302467 | 0.979929 | 1.1529 | 1.177 |
| chr10 | 59006916 | 59007315 | 0.841489 | 0.990032 | 1.177 |
| chr17 | 30986143 | 30986677 | 2.20029 | 2.5887 | 1.177 |
| chr1 | 9327226 | 9327510 | 2.78957 | 3.2828 | 1.177 |
| chr10 | 52808631 | 52809142 | 0.956321 | 1.12553 | 1.177 |
| chr2 | 77354781 | 77355039 | 0.823472 | 0.969189 | 1.177 |
| chr8 | 88613687 | 88613922 | 0.996027 | 1.17229 | 1.177 |
| chr19 | 32623592 | 32623879 | 0.762407 | 0.897343 | 1.177 |
| chr10 | 101805840 | 101806151 | 2.25512 | 2.65472 | 1.177 |
| chr3 | 143399699 | 143399914 | 1.28696 | 1.51501 | 1.177 |
| chr1 | 185076869 | 185077088 | 0.928306 | 1.09282 | 1.177 |
| chr14 | 62893332 | 62893618 | 0.879067 | 1.03486 | 1.177 |
| chr5 | 130718107 | 130718397 | 1.77977 | 2.0955 | 1.177 |
| chr20 | 19178296 | 19178510 | 0.807068 | 0.95025 | 1.177 |
| chr7 | 112974912 | 112975144 | 0.556278 | 0.654973 | 1.177 |
| chr10 | 56020477 | 56020678 | 0.805598 | 0.94859 | 1.178 |
| chr16 | 76352892 | 76353108 | 0.803159 | 0.945747 | 1.178 |
| chr1 | 206912323 | 206912649 | 1.56372 | 1.84136 | 1.178 |
| chr3 | 54462322 | 54462613 | 2.10495 | 2.47876 | 1.178 |
| chr5 | 12292750 | 12292964 | 1.03815 | 1.22252 | 1.178 |
| chr19 | 41171370 | 41171583 | 0.848308 | 0.999037 | 1.178 |
| chr5 | 121528686 | 121528907 | 1.94917 | 2.29573 | 1.178 |
| chr5 | 163912958 | 163913180 | 0.97173 | 1.14451 | 1.178 |
| chr5 | 176432820 | 176433022 | 1.36198 | 1.60425 | 1.178 |

| | | | | | |
|---|---|---|---|---|---|
| chr12 | 7059569 | 7059804 | 1.27457 | 1.50133 | 1.178 |
| chr22 | 31432711 | 31432936 | 1.22974 | 1.44853 | 1.178 |
| chr8 | 144671114 | 144671662 | 0.931807 | 1.09761 | 1.178 |
| chr8 | 61523670 | 61523932 | 1.46109 | 1.72113 | 1.178 |
| chr7 | 137075095 | 137075332 | 0.87442 | 1.03009 | 1.178 |
| chr19 | 54926395 | 54926637 | 1.4889 | 1.75408 | 1.178 |
| chr10 | 120969045 | 120969297 | 3.3122 | 3.90246 | 1.178 |
| chr12 | 113261720 | 113261953 | 1.15756 | 1.36409 | 1.178 |
| chr3 | 23521540 | 23521745 | 0.874444 | 1.0305 | 1.178 |
| chr2 | 175201298 | 175201730 | 1.09235 | 1.28731 | 1.178 |
| chr2 | 178589436 | 178589800 | 1.82985 | 2.15646 | 1.178 |
| chr17 | 9323490 | 9323959 | 0.955885 | 1.12652 | 1.179 |
| chr7 | 24122066 | 24122314 | 0.759195 | 0.89478 | 1.179 |
| chr2 | 85864591 | 85865145 | 1.98381 | 2.33824 | 1.179 |
| chr16 | 63652463 | 63652666 | 0.864295 | 1.01887 | 1.179 |
| chr2 | 23422263 | 23422623 | 2.15425 | 2.53953 | 1.179 |
| chr1 | 173532121 | 173532319 | 0.986808 | 1.16332 | 1.179 |
| chr4 | 1586282 | 1586532 | 0.714846 | 0.842722 | 1.179 |
| chrX | 152084153 | 152084495 | 2.06859 | 2.43863 | 1.179 |
| chr7 | 120649295 | 120649549 | 0.844145 | 0.995169 | 1.179 |
| chr14 | 86608352 | 86608619 | 0.571293 | 0.673521 | 1.179 |
| chr17 | 48894582 | 48894965 | 3.00883 | 3.5473 | 1.179 |
| chr2 | 19036332 | 19036555 | 1.30725 | 1.5413 | 1.179 |
| chr17 | 75865941 | 75866150 | 0.754437 | 0.889545 | 1.179 |
| chr12 | 108064847 | 108065258 | 2.20189 | 2.59627 | 1.179 |
| chrX | 23336301 | 23336533 | 2.4041 | 2.83483 | 1.179 |
| chr20 | 54342397 | 54342664 | 0.900579 | 1.06194 | 1.179 |
| chr8 | 55464250 | 55464629 | 2.00711 | 2.36676 | 1.179 |
| chr16 | 11450519 | 11450907 | 2.14729 | 2.53212 | 1.179 |
| chr12 | 85197606 | 85197839 | 1.0355 | 1.22116 | 1.179 |
| chr7 | 132003540 | 132003878 | 2.48729 | 2.93329 | 1.179 |
| chr8 | 14738846 | 14739076 | 0.776642 | 0.915923 | 1.179 |
| chr9 | 71614659 | 71614876 | 1.92412 | 2.26952 | 1.180 |
| chr1 | 165338274 | 165338822 | 1.81651 | 2.14261 | 1.180 |
| chr16 | 78293874 | 78294100 | 0.608443 | 0.717705 | 1.180 |
| chr10 | 113358150 | 113358503 | 1.20122 | 1.41703 | 1.180 |
| chr1 | 225506746 | 225507010 | 2.48186 | 2.92791 | 1.180 |
| chr17 | 36853033 | 36853255 | 1.05465 | 1.24421 | 1.180 |
| chr8 | 140349140 | 140349411 | 0.891262 | 1.05149 | 1.180 |
| chr3 | 120993318 | 120993602 | 0.892119 | 1.05256 | 1.180 |
| chr17 | 76783506 | 76783802 | 2.37978 | 2.80778 | 1.180 |
| chr18 | 62507238 | 62507446 | 0.61953 | 0.731009 | 1.180 |
| chr7 | 144091198 | 144091414 | 0.600955 | 0.709135 | 1.180 |
| chr17 | 71475008 | 71475373 | 1.14051 | 1.34601 | 1.180 |
| chr19 | 42034137 | 42034352 | 0.984439 | 1.1619 | 1.180 |
| chr1 | 204593233 | 204593458 | 1.40249 | 1.65533 | 1.180 |
| chr9 | 4797413 | 4797720 | 2.14492 | 2.53175 | 1.180 |
| chr4 | 6021471 | 6021696 | 2.3647 | 2.79134 | 1.180 |
| chr17 | 67522202 | 67522425 | 2.32548 | 2.74513 | 1.180 |
| chr12 | 45518709 | 45518976 | 0.662439 | 0.782018 | 1.181 |
| chr12 | 91334693 | 91334975 | 2.18637 | 2.58104 | 1.181 |
| chr16 | 67875701 | 67875970 | 2.18629 | 2.58117 | 1.181 |
| chr7 | 42158508 | 42158694 | 0.734763 | 0.86748 | 1.181 |
| chr11 | 838450 | 838712 | 1.20739 | 1.42581 | 1.181 |
| chr9 | 93677232 | 93677515 | 1.91026 | 2.25619 | 1.181 |
| chrX | 69671931 | 69672199 | 2.01267 | 2.37744 | 1.181 |
| chr7 | 122101738 | 122101965 | 0.808484 | 0.955031 | 1.181 |
| chr11 | 22897469 | 22897765 | 2.03767 | 2.40703 | 1.181 |
| chr5 | 148200472 | 148200745 | 1.8714 | 2.21095 | 1.181 |
| chr19 | 35920542 | 35920906 | 0.984987 | 1.16373 | 1.181 |
| chr17 | 33200883 | 33201137 | 1.08858 | 1.28619 | 1.182 |

| | | | | | |
|---|---|---|---|---|---|
| chr20 | 39159166 | 39159420 | 0.634594 | 0.749798 | 1.182 |
| chr2 | 232579919 | 232580337 | 1.39288 | 1.6458 | 1.182 |
| chr8 | 42713657 | 42713891 | 1.44994 | 1.71328 | 1.182 |
| chr19 | 49438235 | 49438469 | 0.924171 | 1.09205 | 1.182 |
| chr17 | 31325329 | 31325668 | 3.27979 | 3.87588 | 1.182 |
| chr16 | 53056355 | 53056704 | 1.81947 | 2.15038 | 1.182 |
| chr17 | 79634907 | 79635106 | 1.51363 | 1.78891 | 1.182 |
| chr9 | 95527543 | 95527962 | 2.59145 | 3.0628 | 1.182 |
| chr17 | 7227480 | 7227889 | 4.07614 | 4.81833 | 1.182 |
| chr1 | 67230785 | 67230987 | 1.40999 | 1.66684 | 1.182 |
| chr3 | 193179494 | 193179816 | 2.60544 | 3.0801 | 1.182 |
| chr1 | 219926489 | 219926934 | 0.772699 | 0.913535 | 1.182 |
| chr1 | 62857156 | 62857376 | 1.01261 | 1.19719 | 1.182 |
| chr5 | 154029800 | 154030065 | 2.52181 | 2.9815 | 1.182 |
| chr15 | 78608065 | 78608401 | 1.33897 | 1.58309 | 1.182 |
| chr1 | 198723984 | 198724200 | 0.840025 | 0.993353 | 1.183 |
| chr6 | 14044426 | 14044661 | 1.53696 | 1.8175 | 1.183 |
| chr5 | 179265141 | 179265356 | 1.56012 | 1.84492 | 1.183 |
| chr16 | 1773474 | 1773921 | 1.49384 | 1.76666 | 1.183 |
| chr5 | 139222905 | 139223172 | 0.750687 | 0.88779 | 1.183 |
| chr9 | 103462926 | 103463207 | 1.68185 | 1.98916 | 1.183 |
| chr3 | 150556426 | 150556811 | 0.828436 | 0.979817 | 1.183 |
| chr15 | 86345938 | 86346175 | 1.5556 | 1.83995 | 1.183 |
| chr13 | 31553395 | 31553840 | 1.66438 | 1.96867 | 1.183 |
| chr17 | 5608402 | 5608750 | 1.43747 | 1.70042 | 1.183 |
| chr15 | 51132824 | 51133073 | 1.24543 | 1.47333 | 1.183 |
| chr17 | 75888295 | 75888750 | 1.24994 | 1.47876 | 1.183 |
| chr2 | 173836746 | 173837161 | 2.80768 | 3.32167 | 1.183 |
| chr5 | 148377806 | 148377991 | 1.4261 | 1.68724 | 1.183 |
| chr18 | 29302297 | 29302674 | 2.27038 | 2.68625 | 1.183 |
| chr1 | 158581077 | 158581297 | 0.468341 | 0.554162 | 1.183 |
| chr11 | 61322663 | 61323016 | 1.0256 | 1.21353 | 1.183 |
| chr2 | 188002983 | 188003341 | 1.55359 | 1.83832 | 1.183 |
| chr5 | 74239890 | 74240186 | 0.702497 | 0.83126 | 1.183 |
| chr20 | 15680858 | 15681213 | 0.887706 | 1.05051 | 1.183 |
| chr10 | 114074536 | 114075056 | 2.12646 | 2.51652 | 1.183 |
| chr2 | 153492143 | 153492448 | 1.62533 | 1.92346 | 1.183 |
| chr13 | 27281952 | 27282144 | 1.68673 | 1.99617 | 1.183 |
| chr15 | 72448379 | 72448714 | 2.49011 | 2.94702 | 1.183 |
| chr2 | 69137015 | 69137261 | 1.72787 | 2.04509 | 1.184 |
| chr14 | 80522566 | 80522859 | 1.4737 | 1.74438 | 1.184 |
| chr5 | 119659532 | 119659922 | 1.00661 | 1.19161 | 1.184 |
| chr6 | 19073098 | 19073351 | 0.77477 | 0.917238 | 1.184 |
| chr5 | 54240202 | 54240552 | 1.59759 | 1.89137 | 1.184 |
| chr1 | 229635282 | 229635482 | 0.864046 | 1.02298 | 1.184 |
| chr8 | 143410635 | 143410837 | 0.790355 | 0.935755 | 1.184 |
| chr17 | 20044143 | 20044386 | 0.900154 | 1.0658 | 1.184 |
| chr11 | 39020496 | 39020726 | 0.971287 | 1.15003 | 1.184 |
| chr16 | 55281235 | 55281491 | 0.562145 | 0.665686 | 1.184 |
| chr20 | 7373573 | 7373805 | 0.839047 | 0.993608 | 1.184 |
| chr9 | 37034666 | 37035055 | 1.42432 | 1.68675 | 1.184 |
| chr11 | 86387696 | 86387887 | 1.10446 | 1.30813 | 1.184 |
| chr17 | 49930681 | 49930890 | 1.09599 | 1.29811 | 1.184 |
| chr14 | 75372298 | 75372829 | 2.32052 | 2.74853 | 1.184 |
| chr2 | 136671377 | 136671612 | 1.1898 | 1.40936 | 1.185 |
| chr5 | 174378202 | 174378514 | 1.71728 | 2.03435 | 1.185 |
| chr19 | 35748633 | 35748835 | 0.90555 | 1.07277 | 1.185 |
| chr10 | 129867605 | 129868038 | 1.92714 | 2.28315 | 1.185 |
| chr1 | 165790576 | 165790782 | 0.730365 | 0.865348 | 1.185 |
| chr8 | 23841945 | 23842232 | 2.26999 | 2.68956 | 1.185 |
| chr9 | 93886054 | 93886355 | 0.91407 | 1.08303 | 1.185 |

| | | | | | |
|---|---|---|---|---|---|
| chr11 | 83547902 | 83548189 | 2.01543 | 2.38801 | 1.185 |
| chr11 | 89228250 | 89228508 | 0.712989 | 0.844838 | 1.185 |
| chr10 | 84210115 | 84210370 | 0.721484 | 0.854997 | 1.185 |
| chr1 | 182568556 | 182568923 | 1.48376 | 1.75835 | 1.185 |
| chr10 | 35021793 | 35022044 | 1.2475 | 1.47838 | 1.185 |
| chr20 | 14170525 | 14170739 | 1.81164 | 2.14692 | 1.185 |
| chr2 | 210290312 | 210290627 | 1.13374 | 1.34361 | 1.185 |
| chr1 | 209948832 | 209949048 | 0.998624 | 1.18351 | 1.185 |
| chr19 | 31210571 | 31210825 | 1.03292 | 1.22415 | 1.185 |
| chr6 | 9251112 | 9251338 | 1.90095 | 2.25296 | 1.185 |
| chr2 | 67704028 | 67704232 | 1.45707 | 1.72714 | 1.185 |
| chr1 | 160386129 | 160386492 | 1.53505 | 1.81979 | 1.185 |
| chr9 | 134703584 | 134703916 | 0.8991 | 1.06589 | 1.186 |
| chr8 | 92730096 | 92730315 | 0.78866 | 0.934986 | 1.186 |
| chr6 | 162169882 | 162170166 | 1.41928 | 1.68279 | 1.186 |
| chr5 | 86505639 | 86505851 | 0.638571 | 0.757189 | 1.186 |
| chr7 | 73864269 | 73864623 | 2.47258 | 2.93213 | 1.186 |
| chr3 | 163480150 | 163480358 | 0.777192 | 0.921687 | 1.186 |
| chr1 | 240256052 | 240256257 | 1.42076 | 1.68519 | 1.186 |
| chr8 | 938999 | 939269 | 0.648313 | 0.769018 | 1.186 |
| chr12 | 99232317 | 99232568 | 1.29092 | 1.53128 | 1.186 |
| chr9 | 109872519 | 109872742 | 0.901677 | 1.06958 | 1.186 |
| chr1 | 194311717 | 194312260 | 1.15584 | 1.37108 | 1.186 |
| chr15 | 66116476 | 66116774 | 2.12717 | 2.5234 | 1.186 |
| chr8 | 119755896 | 119756169 | 3.21907 | 3.81868 | 1.186 |
| chr14 | 85260181 | 85260449 | 0.615532 | 0.730211 | 1.186 |
| chr15 | 88864165 | 88864393 | 0.792168 | 0.939792 | 1.186 |
| chr2 | 175022754 | 175023033 | 1.6232 | 1.92572 | 1.186 |
| chr7 | 43550524 | 43550883 | 0.930367 | 1.10378 | 1.186 |
| chr16 | 76299497 | 76299707 | 1.07718 | 1.27798 | 1.186 |
| chr5 | 125171313 | 125171531 | 1.71504 | 2.03492 | 1.187 |
| chr17 | 5537920 | 5538141 | 0.671932 | 0.797298 | 1.187 |
| chr9 | 11273243 | 11273459 | 1.05801 | 1.25542 | 1.187 |
| chr1 | 208049501 | 208049741 | 1.30403 | 1.54752 | 1.187 |
| chr10 | 65579570 | 65579833 | 2.18192 | 2.58935 | 1.187 |
| chr8 | 103029734 | 103029941 | 0.705207 | 0.836924 | 1.187 |
| chr5 | 143562896 | 143563157 | 1.88247 | 2.23418 | 1.187 |
| chr1 | 117041693 | 117041953 | 1.63091 | 1.93605 | 1.187 |
| chr5 | 87986722 | 87987127 | 4.20719 | 4.99441 | 1.187 |
| chr3 | 35320037 | 35320241 | 0.700262 | 0.831378 | 1.187 |
| chr2 | 85779263 | 85779570 | 1.6829 | 1.99812 | 1.187 |
| chr11 | 47358160 | 47358343 | 0.827932 | 0.983048 | 1.187 |
| chr9 | 120318549 | 120318820 | 1.72442 | 2.04751 | 1.187 |
| chr9 | 83436624 | 83436927 | 0.835341 | 0.991884 | 1.187 |
| chr1 | 21694966 | 21695446 | 4.69346 | 5.57315 | 1.187 |
| chr3 | 8705221 | 8705500 | 2.2764 | 2.70313 | 1.187 |
| chr2 | 62671200 | 62671502 | 0.627616 | 0.745407 | 1.188 |
| chr6 | 13909177 | 13909448 | 1.74967 | 2.07816 | 1.188 |
| chr19 | 38543517 | 38543839 | 0.866908 | 1.02975 | 1.188 |
| chr5 | 68159823 | 68160030 | 0.973446 | 1.15638 | 1.188 |
| chr5 | 167503699 | 167504051 | 2.08549 | 2.4774 | 1.188 |
| chr5 | 143713122 | 143713454 | 0.924937 | 1.09876 | 1.188 |
| chr4 | 86291623 | 86291877 | 2.05136 | 2.43698 | 1.188 |
| chr8 | 137019658 | 137019980 | 0.892678 | 1.0605 | 1.188 |
| chr15 | 97320285 | 97320544 | 0.737489 | 0.876172 | 1.188 |
| chr9 | 120173909 | 120174145 | 2.06484 | 2.45332 | 1.188 |
| chr9 | 27353912 | 27354133 | 0.900786 | 1.07037 | 1.188 |
| chr5 | 138893196 | 138893458 | 2.11963 | 2.5187 | 1.188 |
| chr14 | 52243042 | 52243307 | 2.32296 | 2.76032 | 1.188 |
| chr19 | 57112945 | 57113219 | 0.824294 | 0.979765 | 1.189 |
| chr10 | 5204891 | 5205137 | 0.73579 | 0.874596 | 1.189 |

| | | | | | |
|---|---|---|---|---|---|
| chr12 | 130080159 | 130080398 | 0.717091 | 0.852482 | 1.189 |
| chr2 | 22135226 | 22135435 | 0.741103 | 0.881052 | 1.189 |
| chr9 | 11613905 | 11614371 | 0.788303 | 0.937221 | 1.189 |
| chrX | 57909488 | 57909672 | 0.712289 | 0.846849 | 1.189 |
| chr5 | 16210071 | 16210290 | 0.771838 | 0.917678 | 1.189 |
| chr1 | 87758683 | 87758923 | 1.39712 | 1.66117 | 1.189 |
| chr7 | 45962854 | 45963134 | 2.81382 | 3.34609 | 1.189 |
| chr5 | 68320104 | 68320349 | 1.65099 | 1.9633 | 1.189 |
| chr16 | 11082435 | 11082693 | 1.55925 | 1.85424 | 1.189 |
| chr7 | 2608144 | 2608382 | 0.804781 | 0.957068 | 1.189 |
| chr2 | 105489950 | 105490397 | 1.40679 | 1.67307 | 1.189 |
| chr11 | 70313603 | 70313945 | 2.77005 | 3.29446 | 1.189 |
| chr2 | 78342792 | 78343017 | 0.925146 | 1.10053 | 1.190 |
| chr15 | 90615171 | 90615393 | 0.672026 | 0.799541 | 1.190 |
| chr3 | 18704430 | 18704659 | 0.608344 | 0.723864 | 1.190 |
| chr5 | 118304251 | 118304523 | 0.731331 | 0.870264 | 1.190 |
| chr2 | 100859382 | 100859859 | 1.9337 | 2.30137 | 1.190 |
| chr7 | 105670483 | 105670750 | 1.01218 | 1.20472 | 1.190 |
| chr17 | 28405879 | 28406144 | 0.87225 | 1.0382 | 1.190 |
| chr8 | 136639281 | 136639688 | 0.86621 | 1.03112 | 1.190 |
| chr10 | 20014100 | 20014603 | 0.926165 | 1.1025 | 1.190 |
| chr1 | 174998917 | 174999132 | 3.0889 | 3.67702 | 1.190 |
| chr10 | 54577816 | 54578054 | 1.12878 | 1.34377 | 1.190 |
| chr9 | 109810688 | 109810902 | 0.625262 | 0.744377 | 1.191 |
| chr17 | 3057308 | 3057695 | 0.777906 | 0.926133 | 1.191 |
| chr16 | 85852765 | 85853320 | 1.63016 | 1.9409 | 1.191 |
| chr20 | 333925 | 334138 | 0.942798 | 1.12252 | 1.191 |
| chr8 | 105744321 | 105744535 | 0.704771 | 0.839203 | 1.191 |
| chr15 | 27142463 | 27142679 | 1.10826 | 1.31969 | 1.191 |
| chr1 | 205304729 | 205305108 | 4.69783 | 5.59416 | 1.191 |
| chr11 | 46413323 | 46413849 | 1.41525 | 1.6853 | 1.191 |
| chr18 | 77915679 | 77915997 | 2.5877 | 3.08166 | 1.191 |
| chr5 | 143656154 | 143656396 | 0.650162 | 0.774276 | 1.191 |
| chr6 | 101758728 | 101758935 | 0.758303 | 0.903069 | 1.191 |
| chr4 | 22637507 | 22637715 | 0.715987 | 0.852688 | 1.191 |
| chr12 | 7150238 | 7150469 | 1.46173 | 1.74085 | 1.191 |
| chr6 | 68150039 | 68150293 | 0.71943 | 0.856907 | 1.191 |
| chr16 | 58997233 | 58997480 | 0.807053 | 0.961338 | 1.191 |
| chr2 | 21970281 | 21970500 | 0.65533 | 0.78062 | 1.191 |
| chr10 | 14346559 | 14346781 | 0.952186 | 1.13427 | 1.191 |
| chr2 | 114006799 | 114007065 | 2.31664 | 2.75977 | 1.191 |
| chr7 | 149391650 | 149391853 | 0.895053 | 1.06648 | 1.192 |
| chr12 | 82288987 | 82289231 | 0.917327 | 1.09312 | 1.192 |
| chr7 | 132915435 | 132915719 | 1.59837 | 1.90468 | 1.192 |
| chr11 | 63713427 | 63713665 | 1.44945 | 1.72734 | 1.192 |
| chr13 | 78067105 | 78067419 | 1.17233 | 1.39714 | 1.192 |
| chr14 | 89489338 | 89489532 | 0.721549 | 0.859948 | 1.192 |
| chr21 | 17371920 | 17372124 | 0.732335 | 0.872819 | 1.192 |
| chr1 | 242059747 | 242059998 | 2.0956 | 2.49763 | 1.192 |
| chr1 | 199580294 | 199580522 | 0.886531 | 1.05664 | 1.192 |
| chr19 | 49501216 | 49501623 | 1.82706 | 2.17766 | 1.192 |
| chr2 | 80743252 | 80743477 | 1.00695 | 1.20017 | 1.192 |
| chr2 | 71222617 | 71222839 | 1.62685 | 1.93948 | 1.192 |
| chr15 | 88632043 | 88632351 | 0.729837 | 0.870214 | 1.192 |
| chr10 | 94276839 | 94277106 | 2.50976 | 2.99264 | 1.192 |
| chr2 | 20391177 | 20391415 | 1.37819 | 1.6434 | 1.192 |
| chr12 | 81157565 | 81157873 | 1.16195 | 1.38565 | 1.193 |
| chr7 | 87311047 | 87311264 | 0.854512 | 1.01904 | 1.193 |
| chr17 | 34097164 | 34097475 | 1.75223 | 2.08973 | 1.193 |
| chr1 | 10567407 | 10567811 | 2.01758 | 2.40632 | 1.193 |
| chr12 | 6464800 | 6465002 | 1.6161 | 1.92754 | 1.193 |

| | | | | | |
|---|---|---|---|---|---|
| chr14 | 47852744 | 47853041 | 0.851286 | 1.01537 | 1.193 |
| chr17 | 62936103 | 62936315 | 0.650579 | 0.77611 | 1.193 |
| chr10 | 130380336 | 130380540 | 0.777993 | 0.928157 | 1.193 |
| chr1 | 32405504 | 32405706 | 1.3893 | 1.65756 | 1.193 |
| chr9 | 111420790 | 111420985 | 0.802937 | 0.958194 | 1.193 |
| chr1 | 205662532 | 205662834 | 0.612786 | 0.731303 | 1.193 |
| chr9 | 35696579 | 35696922 | 2.38908 | 2.85152 | 1.194 |
| chr9 | 36873241 | 36873624 | 2.35126 | 2.8064 | 1.194 |
| chr1 | 154965458 | 154965843 | 0.906942 | 1.08257 | 1.194 |
| chr11 | 61984655 | 61985001 | 1.51848 | 1.81256 | 1.194 |
| chr9 | 37364805 | 37365077 | 0.966902 | 1.15422 | 1.194 |
| chr8 | 135918293 | 135918728 | 2.50515 | 2.99063 | 1.194 |
| chr9 | 89434126 | 89434356 | 0.686707 | 0.819809 | 1.194 |
| chr12 | 81289696 | 81290011 | 0.797819 | 0.952474 | 1.194 |
| chr2 | 11724401 | 11724671 | 2.89444 | 3.45559 | 1.194 |
| chr9 | 123748118 | 123748352 | 1.76773 | 2.11058 | 1.194 |
| chr22 | 25334789 | 25335116 | 1.71438 | 2.04718 | 1.194 |
| chr14 | 36993397 | 36993603 | 0.981631 | 1.17259 | 1.195 |
| chr16 | 81666845 | 81667040 | 1.29403 | 1.54578 | 1.195 |
| chr2 | 144041695 | 144041999 | 1.32585 | 1.5838 | 1.195 |
| chr11 | 92215860 | 92216155 | 0.776932 | 0.928099 | 1.195 |
| chr14 | 103966542 | 103966915 | 2.87425 | 3.43348 | 1.195 |
| chr1 | 155034840 | 155035500 | 1.53879 | 1.83832 | 1.195 |
| chr12 | 62831964 | 62832235 | 2.27933 | 2.72315 | 1.195 |
| chr2 | 168976104 | 168976466 | 1.30543 | 1.55962 | 1.195 |
| chr7 | 6593083 | 6593311 | 0.836137 | 0.998986 | 1.195 |
| chr11 | 23145305 | 23145546 | 0.711007 | 0.849528 | 1.195 |
| chr7 | 70294105 | 70294644 | 1.90057 | 2.27088 | 1.195 |
| chr7 | 8819330 | 8819583 | 0.693084 | 0.828139 | 1.195 |
| chr8 | 102658563 | 102658845 | 1.83941 | 2.19836 | 1.195 |
| chr12 | 82910861 | 82911147 | 1.80198 | 2.15364 | 1.195 |
| chr2 | 20425070 | 20425328 | 2.61964 | 3.13144 | 1.195 |
| chr1 | 2738611 | 2738932 | 2.38472 | 2.8509 | 1.195 |
| chrX | 109411134 | 109411461 | 2.09166 | 2.50076 | 1.196 |
| rUn_gl0002 | 122372 | 122918 | 1.19653 | 1.43091 | 1.196 |
| chr2 | 238421423 | 238421847 | 2.32806 | 2.7841 | 1.196 |
| chr19 | 43197694 | 43198292 | 2.21867 | 2.65342 | 1.196 |
| chr19 | 46425565 | 46425914 | 0.918016 | 1.098 | 1.196 |
| chr8 | 133490893 | 133491271 | 1.42565 | 1.70519 | 1.196 |
| chr8 | 122273170 | 122273409 | 1.93881 | 2.3191 | 1.196 |
| chr4 | 137752853 | 137753058 | 0.926968 | 1.10881 | 1.196 |
| chr9 | 14515917 | 14516196 | 0.919697 | 1.10012 | 1.196 |
| chr9 | 73217411 | 73217827 | 1.86954 | 2.23636 | 1.196 |
| chr17 | 26151495 | 26151731 | 0.896863 | 1.07294 | 1.196 |
| chr17 | 56654969 | 56655257 | 1.69156 | 2.0238 | 1.196 |
| chr10 | 19605335 | 19605590 | 0.912812 | 1.09217 | 1.196 |
| chr19 | 30919142 | 30919520 | 0.899634 | 1.07644 | 1.197 |
| chr20 | 50582895 | 50583108 | 1.64576 | 1.96925 | 1.197 |
| chr2 | 72017041 | 72017264 | 1.10881 | 1.32711 | 1.197 |
| chr17 | 63274431 | 63274762 | 1.88191 | 2.25272 | 1.197 |
| chr1 | 223858254 | 223858468 | 1.58956 | 1.90282 | 1.197 |
| chr2 | 42080785 | 42081210 | 1.72062 | 2.0597 | 1.197 |
| chr2 | 97626037 | 97626548 | 2.39256 | 2.86407 | 1.197 |
| chr7 | 105039755 | 105040122 | 1.18623 | 1.42001 | 1.197 |
| chr1 | 226814071 | 226814571 | 2.83876 | 3.39842 | 1.197 |
| chr16 | 48289174 | 48289382 | 0.855232 | 1.02395 | 1.197 |
| chr2 | 173143688 | 173143939 | 0.932315 | 1.11635 | 1.197 |
| chr13 | 100552661 | 100552890 | 1.70153 | 2.03762 | 1.198 |
| chr5 | 59945214 | 59945420 | 1.87934 | 2.25077 | 1.198 |
| chr8 | 85295597 | 85295802 | 0.836355 | 1.00171 | 1.198 |
| chr1 | 158405031 | 158405278 | 0.858462 | 1.0282 | 1.198 |

| | | | | | |
|---|---|---|---|---|---|
| chr3 | 34924175 | 34924459 | 0.64275 | 0.769837 | 1.198 |
| chr8 | 75015304 | 75015518 | 0.83273 | 0.997558 | 1.198 |
| chr11 | 27520746 | 27520973 | 2.23127 | 2.67308 | 1.198 |
| chr3 | 97698542 | 97698769 | 0.888334 | 1.06423 | 1.198 |
| chr2 | 64836387 | 64836816 | 1.83334 | 2.19688 | 1.198 |
| chr15 | 88284739 | 88284973 | 1.06616 | 1.27759 | 1.198 |
| chr7 | 126443053 | 126443434 | 1.39577 | 1.6726 | 1.198 |
| chr8 | 126954853 | 126955059 | 0.738343 | 0.884957 | 1.199 |
| chr1 | 76796637 | 76797032 | 1.32906 | 1.59299 | 1.199 |
| chr7 | 85501565 | 85501776 | 0.644057 | 0.772004 | 1.199 |
| chr15 | 87901025 | 87901241 | 1.02148 | 1.22448 | 1.199 |
| chr16 | 73748089 | 73748334 | 1.60557 | 1.92467 | 1.199 |
| chr20 | 17030793 | 17031185 | 0.784668 | 0.940939 | 1.199 |
| chr12 | 43523278 | 43523504 | 0.730623 | 0.876241 | 1.199 |
| chr7 | 86463567 | 86463878 | 0.768185 | 0.921381 | 1.199 |
| chr9 | 131944972 | 131945195 | 1.65844 | 1.98919 | 1.199 |
| chr7 | 28133245 | 28133635 | 1.93405 | 2.32004 | 1.200 |
| chr3 | 144736799 | 144737029 | 0.954336 | 1.14481 | 1.200 |
| chr15 | 66748878 | 66749084 | 0.773734 | 0.928234 | 1.200 |
| chr1 | 157782518 | 157782708 | 0.862859 | 1.0352 | 1.200 |
| chr7 | 117685800 | 117686173 | 0.972993 | 1.16751 | 1.200 |
| chr19 | 29067437 | 29067671 | 0.748236 | 0.897924 | 1.200 |
| chr17 | 2427475 | 2427689 | 1.04891 | 1.25875 | 1.200 |
| chr11 | 11988362 | 11988723 | 2.58191 | 3.09847 | 1.200 |
| chrX | 69253317 | 69253542 | 1.61494 | 1.93804 | 1.200 |
| chr10 | 85857633 | 85857884 | 1.47862 | 1.77453 | 1.200 |
| chr11 | 12985617 | 12985916 | 2.8511 | 3.4217 | 1.200 |
| chr4 | 38862484 | 38862740 | 2.52382 | 3.02895 | 1.200 |
| chr16 | 22152957 | 22153387 | 1.40717 | 1.68885 | 1.200 |
| chr7 | 127432773 | 127433107 | 2.14188 | 2.57139 | 1.201 |
| chr10 | 123494091 | 123494505 | 1.86894 | 2.24379 | 1.201 |
| chr9 | 90806033 | 90806278 | 0.87502 | 1.05056 | 1.201 |
| chr8 | 139368232 | 139368583 | 0.707568 | 0.849521 | 1.201 |
| chr16 | 81771647 | 81771981 | 2.08432 | 2.50254 | 1.201 |
| chr12 | 117669568 | 117669883 | 1.14876 | 1.37952 | 1.201 |
| chr11 | 121526734 | 121526999 | 2.7431 | 3.29438 | 1.201 |
| chr8 | 139603583 | 139603838 | 0.839618 | 1.00836 | 1.201 |
| chr11 | 39083313 | 39083527 | 0.974378 | 1.17024 | 1.201 |
| chr11 | 32542917 | 32543348 | 0.921666 | 1.10694 | 1.201 |
| chr3 | 184798354 | 184798592 | 0.581735 | 0.698808 | 1.201 |
| chr7 | 49865103 | 49865343 | 0.821011 | 0.986239 | 1.201 |
| chr7 | 100302900 | 100303446 | 2.54099 | 3.0524 | 1.201 |
| chr7 | 99650929 | 99651131 | 1.10803 | 1.33115 | 1.201 |
| chr15 | 34354129 | 34354495 | 3.07082 | 3.68944 | 1.201 |
| chr8 | 102217280 | 102217469 | 1.34223 | 1.61265 | 1.201 |
| chr8 | 135697863 | 135698148 | 2.08192 | 2.50136 | 1.201 |
| chr8 | 115318699 | 115318912 | 0.874281 | 1.05044 | 1.201 |
| chr15 | 52209756 | 52210078 | 2.78359 | 3.34481 | 1.202 |
| chr5 | 102664633 | 102664848 | 0.664919 | 0.79906 | 1.202 |
| chr11 | 8247088 | 8247506 | 2.73823 | 3.29078 | 1.202 |
| chr14 | 93153153 | 93153715 | 2.6518 | 3.18719 | 1.202 |
| chr12 | 2791745 | 2791950 | 1.19844 | 1.44058 | 1.202 |
| chr15 | 26505754 | 26505944 | 1.46324 | 1.75895 | 1.202 |
| chr7 | 21301398 | 21301648 | 1.96826 | 2.36609 | 1.202 |
| chr7 | 84650495 | 84650751 | 1.38193 | 1.6614 | 1.202 |
| chr22 | 22414980 | 22415372 | 2.24942 | 2.70443 | 1.202 |
| chr15 | 83383685 | 83383970 | 0.668028 | 0.803206 | 1.202 |
| chr12 | 53746448 | 53746696 | 2.29474 | 2.75929 | 1.202 |
| chr3 | 87676715 | 87677061 | 0.993414 | 1.19462 | 1.203 |
| chr19 | 40931179 | 40931629 | 3.74523 | 4.50393 | 1.203 |
| chr5 | 77930297 | 77930726 | 0.884339 | 1.0635 | 1.203 |

| | | | | | |
|---|---|---|---|---|---|
| chr2 | 49926854 | 49927098 | 0.80579 | 0.969061 | 1.203 |
| chr10 | 106494838 | 106495049 | 0.700204 | 0.842129 | 1.203 |
| chr1 | 220687739 | 220687958 | 0.679696 | 0.817519 | 1.203 |
| chr17 | 77632661 | 77633017 | 2.73276 | 3.28707 | 1.203 |
| chr2 | 75292042 | 75292484 | 2.7395 | 3.2954 | 1.203 |
| chr2 | 88359960 | 88360168 | 1.99519 | 2.40006 | 1.203 |
| chr12 | 79574497 | 79574785 | 0.902584 | 1.0858 | 1.203 |
| chr18 | 46705238 | 46705466 | 1.4422 | 1.73498 | 1.203 |
| chr20 | 31481974 | 31482276 | 1.69244 | 2.03605 | 1.203 |
| chr4 | 94712746 | 94713139 | 1.99682 | 2.40237 | 1.203 |
| chr12 | 91046798 | 91047031 | 0.943513 | 1.13519 | 1.203 |
| chr17 | 73891482 | 73891720 | 1.59392 | 1.91779 | 1.203 |
| chr20 | 17400291 | 17400587 | 0.858588 | 1.03305 | 1.203 |
| chr11 | 112583796 | 112584067 | 0.823194 | 0.990483 | 1.203 |
| chr17 | 20008872 | 20009097 | 1.0197 | 1.22697 | 1.203 |
| chr2 | 80819958 | 80820218 | 0.642657 | 0.773334 | 1.203 |
| chr20 | 2024366 | 2024573 | 0.733858 | 0.883106 | 1.203 |
| chr5 | 150695450 | 150695943 | 2.58069 | 3.10552 | 1.203 |
| chr8 | 139779856 | 139780493 | 2.47412 | 2.97821 | 1.204 |
| chr17 | 57401657 | 57401849 | 1.28006 | 1.54099 | 1.204 |
| chr5 | 10358053 | 10358284 | 1.74244 | 2.09774 | 1.204 |
| chr20 | 24118463 | 24118671 | 0.842048 | 1.01381 | 1.204 |
| chr2 | 3427612 | 3428123 | 3.27194 | 3.93946 | 1.204 |
| chr6 | 109235490 | 109235784 | 1.48079 | 1.78288 | 1.204 |
| chr12 | 25736544 | 25736750 | 0.878278 | 1.05748 | 1.204 |
| chr8 | 2810272 | 2810476 | 0.733069 | 0.882801 | 1.204 |
| chr19 | 3210761 | 3210982 | 1.53049 | 1.84322 | 1.204 |
| chr17 | 43209016 | 43209708 | 2.89421 | 3.48561 | 1.204 |
| chr8 | 92942761 | 92942969 | 0.764886 | 0.921362 | 1.205 |
| chr8 | 57841628 | 57842096 | 2.36665 | 2.85106 | 1.205 |
| chr14 | 88846114 | 88846581 | 0.811151 | 0.977198 | 1.205 |
| chr15 | 87369528 | 87369757 | 0.670217 | 0.807448 | 1.205 |
| chr2 | 232422666 | 232422900 | 1.5023 | 1.81013 | 1.205 |
| chr22 | 45019588 | 45019878 | 2.03875 | 2.45692 | 1.205 |
| chr7 | 38836576 | 38836806 | 1.68856 | 2.03501 | 1.205 |
| chr8 | 38283227 | 38283510 | 1.53391 | 1.84889 | 1.205 |
| chr19 | 28689307 | 28689613 | 1.19227 | 1.43713 | 1.205 |
| chr11 | 42934296 | 42934549 | 0.687415 | 0.82865 | 1.205 |
| chr12 | 29470702 | 29471077 | 1.63598 | 1.9724 | 1.206 |
| chr5 | 51759884 | 51760198 | 0.745677 | 0.899086 | 1.206 |
| chr14 | 35882531 | 35882910 | 1.48593 | 1.79164 | 1.206 |
| chr3 | 40621897 | 40622404 | 1.77416 | 2.13924 | 1.206 |
| chr5 | 145059816 | 145060030 | 1.26866 | 1.52974 | 1.206 |
| chr4 | 6265981 | 6266216 | 1.75655 | 2.11804 | 1.206 |
| chr11 | 32982213 | 32982437 | 1.31315 | 1.5836 | 1.206 |
| chr1 | 228415009 | 228415281 | 1.1264 | 1.35871 | 1.206 |
| chr15 | 86273591 | 86273890 | 1.47023 | 1.77372 | 1.206 |
| chr11 | 26298171 | 26298740 | 2.10556 | 2.54021 | 1.206 |
| chr5 | 131836857 | 131837394 | 2.34179 | 2.82552 | 1.207 |
| chr12 | 30990553 | 30990985 | 1.00234 | 1.2094 | 1.207 |
| chr12 | 115563708 | 115564006 | 0.757371 | 0.913887 | 1.207 |
| chr4 | 10583728 | 10584012 | 1.5736 | 1.8988 | 1.207 |
| chr7 | 129712063 | 129712294 | 1.8574 | 2.24126 | 1.207 |
| chr1 | 210022265 | 210022513 | 0.743627 | 0.897365 | 1.207 |
| chr3 | 144634190 | 144634524 | 0.799715 | 0.965136 | 1.207 |
| chr17 | 77105647 | 77106000 | 1.42903 | 1.72464 | 1.207 |
| chr20 | 5037405 | 5037680 | 1.84032 | 2.22128 | 1.207 |
| chr3 | 154235295 | 154235612 | 0.841947 | 1.01627 | 1.207 |
| chr5 | 8488587 | 8488800 | 0.914145 | 1.10344 | 1.207 |
| chr11 | 117068547 | 117068927 | 1.1452 | 1.38258 | 1.207 |
| chr20 | 25624686 | 25624929 | 1.03319 | 1.24742 | 1.207 |

| | | | | | |
|---|---|---|---|---|---|
| chr10 | 71476575 | 71477061 | 2.31009 | 2.78922 | 1.207 |
| chr20 | 1617235 | 1617707 | 0.904902 | 1.09269 | 1.208 |
| chr2 | 142687994 | 142688197 | 0.755846 | 0.912756 | 1.208 |
| chr14 | 24101158 | 24101659 | 2.56523 | 3.09781 | 1.208 |
| chr20 | 10393304 | 10393526 | 1.60786 | 1.94166 | 1.208 |
| chr17 | 8864470 | 8864721 | 0.902059 | 1.08935 | 1.208 |
| chr17 | 60484942 | 60485156 | 0.980602 | 1.18424 | 1.208 |
| chr2 | 11078359 | 11078589 | 0.857986 | 1.03636 | 1.208 |
| chr7 | 142148259 | 142148479 | 1.93646 | 2.33923 | 1.208 |
| chr17 | 3810820 | 3811146 | 1.75595 | 2.12127 | 1.208 |
| chr11 | 67426856 | 67427170 | 1.50704 | 1.82082 | 1.208 |
| chr6 | 13336764 | 13337029 | 1.7387 | 2.10127 | 1.209 |
| chr10 | 103649397 | 103649723 | 2.98996 | 3.61428 | 1.209 |
| chr20 | 45309733 | 45310328 | 1.6066 | 1.94226 | 1.209 |
| chr5 | 1479751 | 1480142 | 2.83658 | 3.42936 | 1.209 |
| chr12 | 76972049 | 76972341 | 0.913039 | 1.10388 | 1.209 |
| chr11 | 70037656 | 70038039 | 3.07817 | 3.72171 | 1.209 |
| chr7 | 124983139 | 124983441 | 0.578679 | 0.699666 | 1.209 |
| chr2 | 95745793 | 95746072 | 2.81199 | 3.39996 | 1.209 |
| chr17 | 55592852 | 55593287 | 0.801342 | 0.968909 | 1.209 |
| chr15 | 22836513 | 22837095 | 2.36735 | 2.86241 | 1.209 |
| chr5 | 117710134 | 117710474 | 0.763466 | 0.923119 | 1.209 |
| chr5 | 112942290 | 112942518 | 1.96628 | 2.37753 | 1.209 |
| chr5 | 137400882 | 137401160 | 1.50805 | 1.82356 | 1.209 |
| chr19 | 32039939 | 32040249 | 1.25074 | 1.51243 | 1.209 |
| chr7 | 150663146 | 150663365 | 0.924006 | 1.11744 | 1.209 |
| chrX | 123345287 | 123345503 | 1.6172 | 1.95612 | 1.210 |
| chr2 | 98313791 | 98314045 | 1.83581 | 2.22062 | 1.210 |
| chr12 | 100628558 | 100628800 | 2.04428 | 2.47292 | 1.210 |
| chr4 | 95327910 | 95328191 | 1.91239 | 2.31344 | 1.210 |
| chr19 | 30502026 | 30502322 | 1.15053 | 1.39191 | 1.210 |
| chr2 | 35568208 | 35568443 | 1.14002 | 1.3792 | 1.210 |
| chrX | 46545456 | 46545987 | 1.5139 | 1.83154 | 1.210 |
| chr2 | 10849437 | 10850015 | 1.76894 | 2.14024 | 1.210 |
| chr5 | 15902965 | 15903200 | 0.9703 | 1.17409 | 1.210 |
| chr7 | 155848139 | 155848436 | 0.814718 | 0.985895 | 1.210 |
| chr2 | 158453904 | 158454225 | 1.84276 | 2.23003 | 1.210 |
| chr7 | 33090367 | 33090615 | 1.87611 | 2.27049 | 1.210 |
| chr1 | 119551445 | 119551725 | 0.801585 | 0.970152 | 1.210 |
| chr7 | 12714571 | 12714850 | 1.30576 | 1.5805 | 1.210 |
| chr2 | 200353487 | 200353868 | 2.1827 | 2.64204 | 1.210 |
| chr1 | 201968817 | 201969312 | 1.85509 | 2.24581 | 1.211 |
| chr12 | 6147291 | 6147892 | 1.81526 | 2.19783 | 1.211 |
| chr14 | 30844575 | 30844921 | 1.8973 | 2.29726 | 1.211 |
| chr2 | 118789795 | 118790386 | 1.54581 | 1.87178 | 1.211 |
| chr6 | 116433141 | 116433368 | 1.48825 | 1.80213 | 1.211 |
| chr9 | 140317640 | 140318009 | 1.73157 | 2.09689 | 1.211 |
| chr14 | 53500674 | 53500901 | 0.587976 | 0.712152 | 1.211 |
| chr13 | 43169170 | 43169428 | 1.85805 | 2.25047 | 1.211 |
| chr2 | 45509516 | 45509743 | 0.730268 | 0.884563 | 1.211 |
| chr2 | 231476148 | 231476404 | 1.89617 | 2.29686 | 1.211 |
| chr9 | 129319165 | 129319448 | 0.760083 | 0.920921 | 1.212 |
| chr14 | 63618885 | 63619142 | 0.711472 | 0.862041 | 1.212 |
| chr2 | 60114 | 60352 | 0.788645 | 0.955554 | 1.212 |
| chr12 | 47377779 | 47378148 | 2.30912 | 2.79787 | 1.212 |
| chr16 | 762897 | 763193 | 2.33354 | 2.82779 | 1.212 |
| chr20 | 9439058 | 9439738 | 1.16181 | 1.40792 | 1.212 |
| chr5 | 7016004 | 7016239 | 1.21731 | 1.47534 | 1.212 |
| chrX | 8711257 | 8711486 | 2.14299 | 2.59723 | 1.212 |
| chr20 | 1094681 | 1094976 | 0.862022 | 1.0448 | 1.212 |
| chr15 | 62314306 | 62314577 | 1.41329 | 1.71303 | 1.212 |

| | | | | | |
|---|---|---|---|---|---|
| chr12 | 44237859 | 44238149 | 0.97983 | 1.18766 | 1.212 |
| chr10 | 123660087 | 123660380 | 1.85332 | 2.24678 | 1.212 |
| chr17 | 35249438 | 35249649 | 1.32426 | 1.60545 | 1.212 |
| chr21 | 27234869 | 27235135 | 1.47516 | 1.78866 | 1.213 |
| chr17 | 75396427 | 75397067 | 1.3077 | 1.58563 | 1.213 |
| chr2 | 186037756 | 186038048 | 1.1476 | 1.39153 | 1.213 |
| chr9 | 36334659 | 36334903 | 0.739438 | 0.896638 | 1.213 |
| chr3 | 147620802 | 147621012 | 0.909915 | 1.1034 | 1.213 |
| chr15 | 30032980 | 30033233 | 2.21412 | 2.68557 | 1.213 |
| chr3 | 40220445 | 40220674 | 1.30396 | 1.58169 | 1.213 |
| chr11 | 20719541 | 20719808 | 0.702251 | 0.851831 | 1.213 |
| chr10 | 129548743 | 129548969 | 0.807701 | 0.979807 | 1.213 |
| chr19 | 50779333 | 50779574 | 0.718404 | 0.871485 | 1.213 |
| chr9 | 5327083 | 5327296 | 0.65409 | 0.793468 | 1.213 |
| chr15 | 101168052 | 101168348 | 2.09567 | 2.54228 | 1.213 |
| chr10 | 130062248 | 130062604 | 1.69731 | 2.0591 | 1.213 |
| chr15 | 78907437 | 78907762 | 0.727099 | 0.882161 | 1.213 |
| chr12 | 45568184 | 45568600 | 0.951321 | 1.15426 | 1.213 |
| chr2 | 236502684 | 236502956 | 1.28279 | 1.55652 | 1.213 |
| chr5 | 54442410 | 54442741 | 3.28494 | 3.98599 | 1.213 |
| chr10 | 19918567 | 19918824 | 0.779764 | 0.946212 | 1.213 |
| chr8 | 108793839 | 108794106 | 1.59327 | 1.93344 | 1.214 |
| chr20 | 12359018 | 12359343 | 1.14734 | 1.39275 | 1.214 |
| chr22 | 35546518 | 35546817 | 2.06079 | 2.50159 | 1.214 |
| gl000203_r | 34399 | 34614 | 1.24129 | 1.50683 | 1.214 |
| chr18 | 1664058 | 1664398 | 1.48869 | 1.80718 | 1.214 |
| chr7 | 141931587 | 141931790 | 1.26069 | 1.53045 | 1.214 |
| chr4 | 184183638 | 184183862 | 1.38923 | 1.68663 | 1.214 |
| chr9 | 13731788 | 13732110 | 0.826075 | 1.00298 | 1.214 |
| chr14 | 22580104 | 22580487 | 1.1048 | 1.3414 | 1.214 |
| chr9 | 125082660 | 125082912 | 0.815547 | 0.990279 | 1.214 |
| chr15 | 102181396 | 102181715 | 2.32675 | 2.82536 | 1.214 |
| chr1 | 4712613 | 4713048 | 2.73412 | 3.32005 | 1.214 |
| chr16 | 3053756 | 3053972 | 2.31009 | 2.80554 | 1.214 |
| chr7 | 152556876 | 152557212 | 1.50104 | 1.82299 | 1.214 |
| chr10 | 45470376 | 45470766 | 3.29904 | 4.00666 | 1.214 |
| chr2 | 99185710 | 99186103 | 2.62659 | 3.19013 | 1.215 |
| chr3 | 97807810 | 97808030 | 1.02904 | 1.24986 | 1.215 |
| chr20 | 45510160 | 45510403 | 0.928305 | 1.12764 | 1.215 |
| chr1 | 235761411 | 235761772 | 1.40817 | 1.7106 | 1.215 |
| chr5 | 98132093 | 98132476 | 1.67813 | 2.03884 | 1.215 |
| chr13 | 20868693 | 20868895 | 1.01064 | 1.22802 | 1.215 |
| chr10 | 98760178 | 98760403 | 0.978375 | 1.18884 | 1.215 |
| chr11 | 104325904 | 104326127 | 0.656013 | 0.797261 | 1.215 |
| chr3 | 157065680 | 157065987 | 1.16461 | 1.41545 | 1.215 |
| chr3 | 28709136 | 28709656 | 1.22037 | 1.48336 | 1.216 |
| chr8 | 144512987 | 144513864 | 2.32508 | 2.82618 | 1.216 |
| chr3 | 8378238 | 8378442 | 1.87477 | 2.27883 | 1.216 |
| chr17 | 56841860 | 56842222 | 0.987211 | 1.2001 | 1.216 |
| chr20 | 17214063 | 17214245 | 0.81774 | 0.994082 | 1.216 |
| chr3 | 122177961 | 122178164 | 1.38723 | 1.68648 | 1.216 |
| chr15 | 81772938 | 81773366 | 1.58051 | 1.92153 | 1.216 |
| chr12 | 128741762 | 128742049 | 0.807887 | 0.982267 | 1.216 |
| chr11 | 58006688 | 58006942 | 0.708279 | 0.861172 | 1.216 |
| chr20 | 10677294 | 10677633 | 2.07809 | 2.52668 | 1.216 |
| chr17 | 70657261 | 70657502 | 1.66339 | 2.02274 | 1.216 |
| chr18 | 18775126 | 18775371 | 0.953199 | 1.15913 | 1.216 |
| chr12 | 80131983 | 80132207 | 1.88726 | 2.29511 | 1.216 |
| chr20 | 36020868 | 36021102 | 1.65233 | 2.00947 | 1.216 |
| chr7 | 78411600 | 78411906 | 0.893415 | 1.08661 | 1.216 |
| chr10 | 88457864 | 88458403 | 0.837543 | 1.01866 | 1.216 |

| | | | | | |
|---|---|---|---|---|---|
| chr14 | 58539395 | 58539647 | 1.89102 | 2.3 | 1.216 |
| chr2 | 17043844 | 17044142 | 0.668307 | 0.81285 | 1.216 |
| chr20 | 46958737 | 46958942 | 1.21802 | 1.48145 | 1.216 |
| chr2 | 23637923 | 23638194 | 1.09645 | 1.33361 | 1.216 |
| chr2 | 217531345 | 217531635 | 2.18087 | 2.65265 | 1.216 |
| chr10 | 18601684 | 18601925 | 1.31421 | 1.59855 | 1.216 |
| chr17 | 38446965 | 38447312 | 1.87633 | 2.28259 | 1.217 |
| chr2 | 198572446 | 198572630 | 1.45003 | 1.76429 | 1.217 |
| chr3 | 145905157 | 145905432 | 1.67784 | 2.04148 | 1.217 |
| chr9 | 75697057 | 75697242 | 2.28781 | 2.78371 | 1.217 |
| chr15 | 59793777 | 59794064 | 2.24525 | 2.73196 | 1.217 |
| chr14 | 50053488 | 50053925 | 2.51054 | 3.05481 | 1.217 |
| chr20 | 980047 | 980322 | 0.732868 | 0.891743 | 1.217 |
| chr1 | 36192310 | 36192748 | 2.03266 | 2.47369 | 1.217 |
| chr5 | 137946059 | 137946313 | 1.07142 | 1.30392 | 1.217 |
| chr7 | 89160852 | 89161153 | 0.74156 | 0.902494 | 1.217 |
| chr13 | 107570344 | 107570667 | 2.52485 | 3.07291 | 1.217 |
| chr11 | 44505611 | 44506035 | 1.05463 | 1.28359 | 1.217 |
| chr20 | 20446015 | 20446299 | 1.21642 | 1.4805 | 1.217 |
| chr17 | 64896245 | 64896534 | 1.62797 | 1.98153 | 1.217 |
| chr12 | 111768902 | 111769402 | 1.80897 | 2.20197 | 1.217 |
| chr14 | 105263602 | 105263888 | 0.586764 | 0.714374 | 1.217 |
| chr21 | 46558376 | 46558880 | 1.75633 | 2.13874 | 1.218 |
| chr1 | 154732811 | 154733345 | 2.63865 | 3.21321 | 1.218 |
| chr20 | 11657104 | 11657539 | 2.02327 | 2.46386 | 1.218 |
| chr1 | 211308747 | 211309003 | 1.65525 | 2.01571 | 1.218 |
| chr7 | 2904606 | 2905064 | 2.48375 | 3.0249 | 1.218 |
| chr2 | 42313674 | 42313863 | 1.38153 | 1.68281 | 1.218 |
| chr6 | 74564798 | 74565027 | 1.57696 | 1.92095 | 1.218 |
| chr8 | 39405022 | 39405288 | 0.884659 | 1.07767 | 1.218 |
| chr19 | 42130980 | 42131399 | 0.799264 | 0.9737 | 1.218 |
| chr11 | 63683831 | 63684216 | 1.35527 | 1.65109 | 1.218 |
| chr1 | 110596313 | 110596799 | 1.70521 | 2.07748 | 1.218 |
| chr10 | 80300040 | 80300293 | 0.617954 | 0.752916 | 1.218 |
| chr3 | 1061037 | 1061255 | 0.847867 | 1.03305 | 1.218 |
| chr20 | 25447527 | 25447804 | 0.690596 | 0.84149 | 1.219 |
| chr14 | 22788284 | 22788528 | 2.17316 | 2.64811 | 1.219 |
| chr15 | 91147223 | 91147462 | 2.15753 | 2.62906 | 1.219 |
| chr9 | 72586854 | 72587218 | 2.62722 | 3.2019 | 1.219 |
| chr8 | 54793212 | 54793410 | 1.53032 | 1.8651 | 1.219 |
| chr5 | 24564973 | 24565171 | 0.860033 | 1.04819 | 1.219 |
| chr2 | 43086106 | 43086513 | 2.32103 | 2.82897 | 1.219 |
| chr8 | 81300913 | 81301240 | 0.908488 | 1.10736 | 1.219 |
| chr18 | 61517327 | 61517539 | 1.56679 | 1.90978 | 1.219 |
| chr2 | 223846768 | 223847033 | 1.39596 | 1.70168 | 1.219 |
| chr2 | 233157351 | 233157733 | 1.75196 | 2.13573 | 1.219 |
| chr12 | 103471254 | 103471524 | 0.909144 | 1.10848 | 1.219 |
| chr7 | 46192360 | 46192580 | 1.40365 | 1.71145 | 1.219 |
| chr4 | 124582375 | 124582606 | 1.35056 | 1.64682 | 1.219 |
| chr11 | 55897969 | 55898224 | 0.565985 | 0.690145 | 1.219 |
| chr1 | 24711772 | 24712010 | 1.83026 | 2.23179 | 1.219 |
| chrX | 12401376 | 12401608 | 1.7474 | 2.1308 | 1.219 |
| chr2 | 113484209 | 113484674 | 2.80991 | 3.42652 | 1.219 |
| chr18 | 70339651 | 70339895 | 1.77621 | 2.16635 | 1.220 |
| chr8 | 12929202 | 12929418 | 0.734208 | 0.895477 | 1.220 |
| chr10 | 96892676 | 96893060 | 1.18786 | 1.44901 | 1.220 |
| chr7 | 39936013 | 39936346 | 2.4678 | 3.01037 | 1.220 |
| chr14 | 101129465 | 101130341 | 1.44357 | 1.761 | 1.220 |
| chr20 | 35911016 | 35911231 | 1.11293 | 1.35765 | 1.220 |
| chr7 | 22491794 | 22492204 | 2.37724 | 2.90014 | 1.220 |
| chr15 | 91899381 | 91899596 | 0.827958 | 1.01019 | 1.220 |

| | | | | | |
|---|---|---|---|---|---|
| chr9 | 136208929 | 136209144 | 1.4897 | 1.81758 | 1.220 |
| chr15 | 66273163 | 66273636 | 1.83242 | 2.23581 | 1.220 |
| chr7 | 45739859 | 45740181 | 0.849234 | 1.0362 | 1.220 |
| chr11 | 91587165 | 91587390 | 0.797649 | 0.973276 | 1.220 |
| chr20 | 7952211 | 7952427 | 0.996592 | 1.21602 | 1.220 |
| chr5 | 147007648 | 147007942 | 1.60042 | 1.95285 | 1.220 |
| chr8 | 26764908 | 26765152 | 0.880146 | 1.07397 | 1.220 |
| chr16 | 74309012 | 74309218 | 1.58907 | 1.93924 | 1.220 |
| chr8 | 132873260 | 132873600 | 1.77246 | 2.16305 | 1.220 |
| chr16 | 13750413 | 13750634 | 0.853899 | 1.04214 | 1.220 |
| chr12 | 99252209 | 99252494 | 1.0327 | 1.26041 | 1.221 |
| chr16 | 61018716 | 61019182 | 0.680628 | 0.830845 | 1.221 |
| chr5 | 111887211 | 111887476 | 1.64478 | 2.00779 | 1.221 |
| chr8 | 140260328 | 140260620 | 1.3451 | 1.64206 | 1.221 |
| chr15 | 38856084 | 38856611 | 2.05588 | 2.50984 | 1.221 |
| chr20 | 43639675 | 43640074 | 1.79615 | 2.19278 | 1.221 |
| chr5 | 145854870 | 145855226 | 1.29202 | 1.57739 | 1.221 |
| chr15 | 94325178 | 94325386 | 1.01189 | 1.23544 | 1.221 |
| chr8 | 133626517 | 133626767 | 1.04758 | 1.27911 | 1.221 |
| chr9 | 124920595 | 124920829 | 2.07399 | 2.53247 | 1.221 |
| chr16 | 54227580 | 54227913 | 1.86328 | 2.2752 | 1.221 |
| chr19 | 46784069 | 46784473 | 2.01088 | 2.45542 | 1.221 |
| chr15 | 75140674 | 75140904 | 0.628692 | 0.767699 | 1.221 |
| chrX | 110343485 | 110343706 | 1.43237 | 1.74909 | 1.221 |
| chr3 | 3991567 | 3991920 | 0.858824 | 1.04898 | 1.221 |
| chr1 | 21571582 | 21571930 | 1.34717 | 1.64546 | 1.221 |
| chr3 | 170527323 | 170527630 | 1.58544 | 1.93653 | 1.221 |
| chr7 | 37630955 | 37631175 | 0.663572 | 0.810531 | 1.221 |
| chr9 | 13317679 | 13317968 | 0.825744 | 1.00865 | 1.222 |
| chr1 | 39052752 | 39053022 | 2.48678 | 3.03839 | 1.222 |
| chr17 | 32106187 | 32106497 | 2.44029 | 2.98197 | 1.222 |
| chr1 | 46035627 | 46036089 | 2.47293 | 3.02204 | 1.222 |
| chr10 | 134032890 | 134033326 | 1.33783 | 1.63524 | 1.222 |
| chr1 | 54344031 | 54344527 | 1.10398 | 1.34943 | 1.222 |
| chr19 | 36177966 | 36178168 | 0.890525 | 1.08852 | 1.222 |
| chr15 | 29394387 | 29394993 | 2.79658 | 3.41843 | 1.222 |
| chr11 | 47182591 | 47183629 | 1.99785 | 2.44232 | 1.222 |
| chr7 | 32204127 | 32204564 | 1.78541 | 2.18283 | 1.223 |
| chr16 | 69913337 | 69913552 | 0.683542 | 0.835749 | 1.223 |
| chr17 | 18281226 | 18281500 | 1.35093 | 1.65186 | 1.223 |
| chr2 | 135169641 | 135169868 | 2.0228 | 2.47342 | 1.223 |
| chr1 | 165166523 | 165166849 | 0.615198 | 0.752265 | 1.223 |
| chr2 | 102003743 | 102004202 | 2.09268 | 2.55893 | 1.223 |
| chr8 | 109715871 | 109716314 | 1.30003 | 1.5898 | 1.223 |
| chr7 | 131734399 | 131734773 | 1.83503 | 2.24426 | 1.223 |
| chr1 | 55462080 | 55462427 | 2.23081 | 2.72838 | 1.223 |
| chr15 | 85291921 | 85292249 | 1.43285 | 1.75244 | 1.223 |
| chr15 | 101680565 | 101680866 | 0.827736 | 1.01236 | 1.223 |
| chr7 | 119487619 | 119487813 | 0.877612 | 1.07338 | 1.223 |
| chr12 | 21926225 | 21926630 | 1.5084 | 1.84496 | 1.223 |
| chr5 | 11834991 | 11835227 | 0.835694 | 1.0222 | 1.223 |
| chr11 | 15962547 | 15963175 | 1.56646 | 1.91607 | 1.223 |
| chr9 | 138508201 | 138508412 | 1.46195 | 1.78836 | 1.223 |
| chr5 | 172823417 | 172823686 | 0.920734 | 1.12632 | 1.223 |
| chr12 | 67036109 | 67036357 | 0.870383 | 1.06475 | 1.223 |
| chr17 | 3505869 | 3506207 | 2.07675 | 2.54051 | 1.223 |
| chr12 | 78956395 | 78956640 | 0.876306 | 1.07206 | 1.223 |
| chr1 | 28685949 | 28686405 | 1.0421 | 1.27492 | 1.223 |
| chr14 | 31449396 | 31449642 | 2.27423 | 2.78245 | 1.223 |
| chr17 | 63573071 | 63573306 | 1.01325 | 1.23972 | 1.224 |
| chr5 | 141824817 | 141825083 | 0.716006 | 0.876239 | 1.224 |

| | | | | | |
|---|---|---|---|---|---|
| chr8 | 145733864 | 145734344 | 2.90931 | 3.56042 | 1.224 |
| chr2 | 122080507 | 122080710 | 2.16267 | 2.64677 | 1.224 |
| chr1 | 41115813 | 41116055 | 1.2544 | 1.53521 | 1.224 |
| chr9 | 124442476 | 124442686 | 0.69124 | 0.846032 | 1.224 |
| chr16 | 29830984 | 29831194 | 2.02943 | 2.48408 | 1.224 |
| chr17 | 2705316 | 2705744 | 0.667419 | 0.817015 | 1.224 |
| chr3 | 19270858 | 19271156 | 0.853668 | 1.04509 | 1.224 |
| chr7 | 145869364 | 145869571 | 0.929135 | 1.13753 | 1.224 |
| chr22 | 38694364 | 38694641 | 1.67608 | 2.05205 | 1.224 |
| chr17 | 16843342 | 16843706 | 0.730394 | 0.89424 | 1.224 |
| chr3 | 80442384 | 80442637 | 0.703647 | 0.861654 | 1.225 |
| chr21 | 27080878 | 27081102 | 1.60785 | 1.96902 | 1.225 |
| chr11 | 121986321 | 121986627 | 1.31742 | 1.6134 | 1.225 |
| chr10 | 97757218 | 97757443 | 0.738373 | 0.904309 | 1.225 |
| chr11 | 12065840 | 12066094 | 2.24288 | 2.7471 | 1.225 |
| chr3 | 165063941 | 165064149 | 0.795346 | 0.974256 | 1.225 |
| chr17 | 78098199 | 78098422 | 1.84908 | 2.26516 | 1.225 |
| chr15 | 44955892 | 44956134 | 1.85511 | 2.27261 | 1.225 |
| chr19 | 41090030 | 41090271 | 1.44056 | 1.76482 | 1.225 |
| chr10 | 109754706 | 109754971 | 0.932944 | 1.1432 | 1.225 |
| chr13 | 27954169 | 27954395 | 1.55824 | 1.90949 | 1.225 |
| chr5 | 132143202 | 132143473 | 1.25137 | 1.53345 | 1.225 |
| chr12 | 29207700 | 29207977 | 0.890443 | 1.09124 | 1.226 |
| chr9 | 93004153 | 93004421 | 0.622118 | 0.76241 | 1.226 |
| chr12 | 129129398 | 129129618 | 1.07263 | 1.31455 | 1.226 |
| chr17 | 72029983 | 72030222 | 0.829189 | 1.01625 | 1.226 |
| chr17 | 31494 | 31737 | 0.509467 | 0.624409 | 1.226 |
| chr2 | 216887874 | 216888097 | 1.55649 | 1.90775 | 1.226 |
| chr1 | 155247043 | 155247412 | 2.32238 | 2.8465 | 1.226 |
| chr7 | 128021191 | 128021691 | 1.82522 | 2.23715 | 1.226 |
| chr17 | 48624260 | 48624595 | 3.72997 | 4.57181 | 1.226 |
| chr20 | 54325759 | 54326037 | 0.695514 | 0.852503 | 1.226 |
| chr2 | 22048460 | 22048711 | 0.742525 | 0.91026 | 1.226 |
| chr14 | 73120916 | 73121170 | 0.656829 | 0.805234 | 1.226 |
| chr5 | 180388805 | 180389066 | 0.87253 | 1.06968 | 1.226 |
| chr19 | 33732449 | 33733182 | 3.26817 | 4.00686 | 1.226 |
| chr16 | 8712626 | 8712868 | 0.960438 | 1.17772 | 1.226 |
| chr5 | 54467973 | 54468206 | 1.76824 | 2.16833 | 1.226 |
| chr14 | 23207906 | 23208116 | 0.891257 | 1.09293 | 1.226 |
| chr10 | 20038959 | 20039220 | 0.852452 | 1.04544 | 1.226 |
| chr2 | 241543776 | 241544024 | 1.63902 | 2.01026 | 1.227 |
| chrX | 69250326 | 69250632 | 2.16002 | 2.64932 | 1.227 |
| chr15 | 69794947 | 69795269 | 1.08892 | 1.33566 | 1.227 |
| chr8 | 6815561 | 6815965 | 1.20043 | 1.4725 | 1.227 |
| chr11 | 5391080 | 5391350 | 0.812546 | 0.996767 | 1.227 |
| chr7 | 105683251 | 105683659 | 3.50742 | 4.30268 | 1.227 |
| chr8 | 101553081 | 101553305 | 1.21945 | 1.49596 | 1.227 |
| chr19 | 28950126 | 28950337 | 0.943671 | 1.15766 | 1.227 |
| chr4 | 78016538 | 78016912 | 1.55021 | 1.9019 | 1.227 |
| chr14 | 79531730 | 79532034 | 0.862562 | 1.05826 | 1.227 |
| chr5 | 81161500 | 81161798 | 0.758408 | 0.930504 | 1.227 |
| chr10 | 2546110 | 2546400 | 0.623936 | 0.765544 | 1.227 |
| chr17 | 3211854 | 3212180 | 0.83553 | 1.02521 | 1.227 |
| chr11 | 81506953 | 81507224 | 0.853573 | 1.0474 | 1.227 |
| chr11 | 61975525 | 61975777 | 1.84592 | 2.26519 | 1.227 |
| chr5 | 123272969 | 123273187 | 1.36108 | 1.67022 | 1.227 |
| chr3 | 11755259 | 11755492 | 0.704838 | 0.864934 | 1.227 |
| chr5 | 161762085 | 161762293 | 0.866241 | 1.063 | 1.227 |
| chr16 | 56035524 | 56035742 | 0.768194 | 0.942713 | 1.227 |
| chr20 | 2153170 | 2153534 | 3.0373 | 3.72734 | 1.227 |
| chr17 | 901362 | 901635 | 0.984606 | 1.20837 | 1.227 |

| | | | | | |
|---|---|---|---|---|---|
| chr10 | 24451128 | 24451435 | 0.817767 | 1.00373 | 1.227 |
| chr10 | 72109762 | 72110028 | 0.772189 | 0.947832 | 1.227 |
| chr11 | 66839164 | 66839551 | 2.61223 | 3.20652 | 1.228 |
| chr10 | 15921066 | 15921304 | 1.56291 | 1.91866 | 1.228 |
| chr3 | 67914193 | 67914483 | 0.898426 | 1.10296 | 1.228 |
| chr14 | 42069799 | 42070072 | 1.49658 | 1.83733 | 1.228 |
| chr20 | 58030875 | 58031114 | 1.70596 | 2.09455 | 1.228 |
| chr3 | 43112803 | 43113048 | 1.61058 | 1.97781 | 1.228 |
| chr17 | 52705419 | 52705671 | 0.8356 | 1.02628 | 1.228 |
| chr14 | 59375469 | 59375791 | 0.823104 | 1.01095 | 1.228 |
| chr19 | 31063178 | 31063380 | 1.04092 | 1.27869 | 1.228 |
| chr14 | 94121528 | 94121722 | 1.39371 | 1.7122 | 1.229 |
| chr7 | 136009857 | 136010059 | 0.657209 | 0.807467 | 1.229 |
| chr3 | 128216641 | 128217028 | 2.83414 | 3.48237 | 1.229 |
| chr6 | 154268958 | 154269319 | 1.61968 | 1.99017 | 1.229 |
| chr2 | 54562923 | 54563228 | 0.631756 | 0.77644 | 1.229 |
| chr2 | 220163396 | 220163763 | 2.80421 | 3.44645 | 1.229 |
| chr15 | 82570413 | 82570727 | 0.648136 | 0.796617 | 1.229 |
| chr17 | 51283618 | 51283824 | 0.795851 | 0.978169 | 1.229 |
| chr1 | 23058371 | 23058655 | 1.52772 | 1.87773 | 1.229 |
| chr8 | 87005176 | 87005407 | 0.929807 | 1.14285 | 1.229 |
| chr16 | 19570682 | 19570994 | 1.67826 | 2.06281 | 1.229 |
| chr19 | 29305967 | 29306201 | 0.788175 | 0.968843 | 1.229 |
| chr19 | 39076931 | 39077656 | 0.920097 | 1.13108 | 1.229 |
| chr12 | 120204910 | 120205241 | 0.931896 | 1.14565 | 1.229 |
| chr10 | 52181074 | 52181348 | 2.88474 | 3.54647 | 1.229 |
| chr15 | 31079968 | 31080340 | 0.701034 | 0.861845 | 1.229 |
| chr10 | 134650335 | 134650875 | 2.6287 | 3.23203 | 1.230 |
| chr8 | 122842343 | 122842608 | 1.48249 | 1.82275 | 1.230 |
| chr19 | 30685411 | 30685816 | 0.965256 | 1.18683 | 1.230 |
| chr7 | 136805838 | 136806450 | 0.812794 | 0.999406 | 1.230 |
| chr5 | 114937630 | 114937878 | 1.34505 | 1.65388 | 1.230 |
| chr7 | 118573490 | 118573734 | 0.717384 | 0.882105 | 1.230 |
| chr5 | 4132848 | 4133307 | 0.793554 | 0.975807 | 1.230 |
| chr9 | 140301314 | 140301830 | 1.66237 | 2.04416 | 1.230 |
| chr16 | 6131390 | 6131648 | 1.18206 | 1.45356 | 1.230 |
| chr12 | 98602763 | 98603145 | 0.596828 | 0.734026 | 1.230 |
| chr5 | 33588739 | 33589171 | 1.76105 | 2.16588 | 1.230 |
| chr15 | 61095487 | 61095733 | 0.761174 | 0.936199 | 1.230 |
| chr8 | 2808607 | 2808851 | 0.902201 | 1.10972 | 1.230 |
| chr2 | 36395501 | 36395713 | 0.781998 | 0.962019 | 1.230 |
| chr17 | 48029772 | 48030176 | 2.73403 | 3.36363 | 1.230 |
| chr7 | 2758508 | 2759010 | 2.37094 | 2.91693 | 1.230 |
| chr3 | 65770837 | 65771052 | 1.86709 | 2.29731 | 1.230 |
| chr2 | 43158682 | 43158915 | 1.8105 | 2.22771 | 1.230 |
| chr10 | 98956309 | 98956636 | 2.12929 | 2.62018 | 1.231 |
| chr9 | 80736457 | 80736745 | 2.21098 | 2.72079 | 1.231 |
| chr18 | 52901894 | 52902143 | 1.72684 | 2.12505 | 1.231 |
| chr3 | 49208419 | 49208790 | 2.53998 | 3.12617 | 1.231 |
| chr1 | 222912088 | 222912330 | 1.48389 | 1.82638 | 1.231 |
| chr5 | 115106240 | 115106447 | 1.57761 | 1.94186 | 1.231 |
| chr2 | 201336035 | 201336403 | 1.60746 | 1.97885 | 1.231 |
| chr8 | 109508678 | 109509011 | 0.929427 | 1.14419 | 1.231 |
| chr6 | 40880187 | 40880479 | 1.58208 | 1.94797 | 1.231 |
| chr1 | 22290555 | 22290797 | 1.89697 | 2.33585 | 1.231 |
| chr5 | 61439229 | 61439918 | 1.11509 | 1.3731 | 1.231 |
| chr17 | 54631519 | 54631728 | 0.835038 | 1.02829 | 1.231 |
| chr8 | 106918246 | 106918440 | 1.48678 | 1.8309 | 1.231 |
| chr8 | 126242983 | 126243242 | 2.03728 | 2.50881 | 1.231 |
| chr3 | 10925136 | 10925508 | 1.60579 | 1.97751 | 1.231 |
| chr19 | 33740162 | 33740486 | 1.40579 | 1.73142 | 1.232 |

| | | | | | |
|---|---|---|---|---|---|
| chr5 | 66715482 | 66715747 | 0.673453 | 0.829444 | 1.232 |
| chr16 | 68479408 | 68479671 | 1.62581 | 2.00242 | 1.232 |
| chr9 | 85221737 | 85222153 | 1.05187 | 1.29557 | 1.232 |
| chr10 | 71720622 | 71721139 | 1.13415 | 1.39712 | 1.232 |
| chr12 | 124326286 | 124326805 | 1.35619 | 1.67085 | 1.232 |
| chr1 | 221376833 | 221377111 | 2.20658 | 2.7186 | 1.232 |
| chr17 | 65783718 | 65783940 | 0.852011 | 1.04973 | 1.232 |
| chr10 | 133757018 | 133757339 | 1.50994 | 1.86042 | 1.232 |
| chr7 | 1084313 | 1084728 | 1.80219 | 2.22074 | 1.232 |
| chr10 | 105236108 | 105236390 | 1.19276 | 1.46982 | 1.232 |
| chr11 | 63360532 | 63361013 | 2.22235 | 2.73919 | 1.233 |
| chr3 | 58293433 | 58293674 | 1.00681 | 1.24098 | 1.233 |
| chr15 | 89849460 | 89849693 | 1.80658 | 2.22732 | 1.233 |
| chr16 | 31836293 | 31836521 | 0.696244 | 0.858525 | 1.233 |
| chr5 | 80595406 | 80595613 | 1.45786 | 1.79789 | 1.233 |
| chr9 | 32581525 | 32581818 | 1.54263 | 1.90248 | 1.233 |
| chr9 | 91147407 | 91147627 | 1.77979 | 2.19496 | 1.233 |
| chr19 | 38634541 | 38634831 | 1.84577 | 2.27645 | 1.233 |
| chr5 | 50296424 | 50296859 | 0.729325 | 0.899726 | 1.234 |
| chr18 | 60251615 | 60251867 | 2.69159 | 3.32061 | 1.234 |
| chr11 | 61432990 | 61433222 | 0.711038 | 0.877266 | 1.234 |
| chr18 | 35358489 | 35358783 | 2.11247 | 2.60641 | 1.234 |
| chr8 | 139900404 | 139900697 | 0.910247 | 1.12312 | 1.234 |
| chr10 | 53157499 | 53157821 | 0.697885 | 0.861102 | 1.234 |
| chr2 | 169805205 | 169805468 | 1.79755 | 2.21812 | 1.234 |
| chr8 | 131931888 | 131932258 | 0.72622 | 0.896134 | 1.234 |
| chr5 | 154061959 | 154062200 | 1.68067 | 2.07402 | 1.234 |
| chr8 | 89332808 | 89333115 | 1.34075 | 1.65467 | 1.234 |
| chr15 | 32160037 | 32160419 | 2.75949 | 3.40598 | 1.234 |
| chr2 | 171573396 | 171573647 | 1.96702 | 2.42787 | 1.234 |
| chr10 | 49759922 | 49760154 | 1.6264 | 2.00747 | 1.234 |
| chr7 | 54133557 | 54133796 | 0.800573 | 0.988151 | 1.234 |
| chr3 | 146865233 | 146865530 | 0.610919 | 0.754112 | 1.234 |
| chr17 | 13522975 | 13523337 | 0.878785 | 1.08478 | 1.234 |
| chr15 | 95460313 | 95460521 | 0.790414 | 0.975783 | 1.235 |
| chr14 | 66662073 | 66662534 | 0.92672 | 1.14411 | 1.235 |
| chr5 | 8645915 | 8646156 | 0.811593 | 1.00207 | 1.235 |
| chr19 | 39366996 | 39367280 | 1.0102 | 1.2473 | 1.235 |
| chr10 | 90692465 | 90692740 | 2.35751 | 2.91087 | 1.235 |
| chr15 | 64361717 | 64361930 | 1.71703 | 2.12018 | 1.235 |
| chr2 | 15728516 | 15728797 | 2.26002 | 2.79077 | 1.235 |
| chr10 | 87165559 | 87165805 | 0.641026 | 0.791591 | 1.235 |
| chr3 | 80731589 | 80731789 | 0.899401 | 1.11081 | 1.235 |
| chr7 | 140143765 | 140144148 | 2.50804 | 3.09803 | 1.235 |
| chr20 | 62422055 | 62422520 | 0.557788 | 0.689016 | 1.235 |
| chr18 | 21085438 | 21085761 | 1.33183 | 1.64518 | 1.235 |
| chr8 | 103284765 | 103284971 | 1.3884 | 1.71518 | 1.235 |
| chr5 | 84563446 | 84563680 | 0.728541 | 0.900034 | 1.235 |
| chr2 | 97426677 | 97427012 | 1.54914 | 1.9139 | 1.235 |
| chr22 | 45973899 | 45974174 | 1.9094 | 2.35898 | 1.235 |
| chr10 | 15564699 | 15564930 | 0.633713 | 0.782946 | 1.235 |
| chr8 | 144262665 | 144262968 | 0.971644 | 1.20049 | 1.236 |
| chr14 | 37074187 | 37074444 | 1.45172 | 1.79381 | 1.236 |
| chr17 | 39562508 | 39562720 | 0.932265 | 1.15201 | 1.236 |
| chr10 | 3264239 | 3264448 | 1.35665 | 1.67649 | 1.236 |
| chr3 | 136732806 | 136732997 | 1.76142 | 2.17681 | 1.236 |
| chr2 | 76187056 | 76187289 | 0.860236 | 1.06314 | 1.236 |
| chr2 | 86462807 | 86463028 | 0.546988 | 0.676105 | 1.236 |
| chr8 | 102570335 | 102570640 | 0.742877 | 0.918231 | 1.236 |
| chr6 | 1336577 | 1336794 | 1.59295 | 1.96912 | 1.236 |
| chr7 | 85616073 | 85616326 | 0.852924 | 1.05437 | 1.236 |

| | | | | | |
|---|---|---|---|---|---|
| chr11 | 66214382 | 66214605 | 0.796392 | 0.984494 | 1.236 |
| chr22 | 40832867 | 40833300 | 0.935317 | 1.15626 | 1.236 |
| chr8 | 1840553 | 1840934 | 1.98896 | 2.45915 | 1.236 |
| chr15 | 43668353 | 43669020 | 1.28618 | 1.59024 | 1.236 |
| chr5 | 151841715 | 151841971 | 1.59725 | 1.97485 | 1.236 |
| chr19 | 10620924 | 10621404 | 1.66159 | 2.05443 | 1.236 |
| chr7 | 137138965 | 137139230 | 1.11448 | 1.37801 | 1.236 |
| chr14 | 46728822 | 46729065 | 1.08239 | 1.33834 | 1.236 |
| chr8 | 33497708 | 33498080 | 1.93087 | 2.38746 | 1.236 |
| chr7 | 86150309 | 86150513 | 0.851693 | 1.0531 | 1.236 |
| chr15 | 86369011 | 86369222 | 0.845936 | 1.04599 | 1.236 |
| chr5 | 28965829 | 28966050 | 0.701639 | 0.867569 | 1.236 |
| chr8 | 141368660 | 141369163 | 3.4161 | 4.22422 | 1.237 |
| chr14 | 98564806 | 98565055 | 0.760255 | 0.940122 | 1.237 |
| chr8 | 139698785 | 139699079 | 1.94407 | 2.40408 | 1.237 |
| chr7 | 122848900 | 122849197 | 0.890407 | 1.10118 | 1.237 |
| chr17 | 6746636 | 6746877 | 1.19141 | 1.47359 | 1.237 |
| chr8 | 143781642 | 143782057 | 2.27974 | 2.81974 | 1.237 |
| chr7 | 26409195 | 26409469 | 1.50443 | 1.86103 | 1.237 |
| chr1 | 201846902 | 201847377 | 1.63333 | 2.02054 | 1.237 |
| chr6 | 161791857 | 161792175 | 2.8691 | 3.54963 | 1.237 |
| chr2 | 16803298 | 16803523 | 0.73092 | 0.904325 | 1.237 |
| chr9 | 138981350 | 138981744 | 0.966945 | 1.19638 | 1.237 |
| chr19 | 13388856 | 13389155 | 1.88383 | 2.33141 | 1.238 |
| chr14 | 22236433 | 22236882 | 1.51846 | 1.87931 | 1.238 |
| chr19 | 59093226 | 59093473 | 0.932177 | 1.15379 | 1.238 |
| chr7 | 67253765 | 67253994 | 2.11608 | 2.61931 | 1.238 |
| chr5 | 171177264 | 171177515 | 1.29056 | 1.59761 | 1.238 |
| chr10 | 73574750 | 73574961 | 0.923273 | 1.14297 | 1.238 |
| chr2 | 239248514 | 239248827 | 1.23441 | 1.52824 | 1.238 |
| chr8 | 113525830 | 113526052 | 0.59246 | 0.733517 | 1.238 |
| chr9 | 135286472 | 135286832 | 2.09647 | 2.59593 | 1.238 |
| chr19 | 36410014 | 36410216 | 0.789742 | 0.978121 | 1.239 |
| chr15 | 23742891 | 23743145 | 1.82807 | 2.26422 | 1.239 |
| chr10 | 110899922 | 110900227 | 0.809739 | 1.003 | 1.239 |
| chr10 | 85814424 | 85814747 | 0.963301 | 1.19332 | 1.239 |
| chr16 | 88094281 | 88094821 | 0.915239 | 1.13382 | 1.239 |
| chr13 | 97469686 | 97469926 | 1.08109 | 1.33948 | 1.239 |
| chr12 | 124103033 | 124103439 | 1.80058 | 2.23096 | 1.239 |
| chr14 | 93644421 | 93644703 | 1.94098 | 2.40501 | 1.239 |
| chr17 | 40730308 | 40730759 | 1.68221 | 2.08453 | 1.239 |
| chr2 | 223478531 | 223478774 | 1.86877 | 2.31579 | 1.239 |
| chr3 | 34599303 | 34599533 | 0.633776 | 0.785385 | 1.239 |
| chr8 | 134183807 | 134184247 | 0.735528 | 0.911645 | 1.239 |
| chr8 | 32826416 | 32826624 | 0.908868 | 1.12679 | 1.240 |
| chr20 | 8012866 | 8013089 | 0.789379 | 0.978765 | 1.240 |
| chr21 | 38302029 | 38302262 | 0.982715 | 1.21852 | 1.240 |
| chr2 | 172272602 | 172272894 | 2.25061 | 2.79068 | 1.240 |
| chr15 | 94460037 | 94460374 | 0.927118 | 1.14962 | 1.240 |
| chr17 | 40873103 | 40873358 | 1.75733 | 2.17915 | 1.240 |
| chr11 | 64885230 | 64885637 | 2.45323 | 3.0421 | 1.240 |
| chr10 | 107388265 | 107388527 | 0.777331 | 0.964014 | 1.240 |
| chr15 | 99645978 | 99646335 | 0.872163 | 1.08163 | 1.240 |
| chr3 | 194392230 | 194392444 | 1.38351 | 1.71579 | 1.240 |
| chr10 | 105136626 | 105136918 | 0.777316 | 0.964104 | 1.240 |
| chr1 | 181448354 | 181448582 | 1.9799 | 2.45579 | 1.240 |
| chr19 | 41847619 | 41847850 | 1.18771 | 1.47323 | 1.240 |
| chrX | 69984913 | 69985118 | 0.463906 | 0.57546 | 1.240 |
| chr5 | 140821402 | 140821687 | 0.919314 | 1.14041 | 1.241 |
| chr9 | 112665895 | 112666125 | 2.21564 | 2.74851 | 1.241 |
| chr2 | 78156940 | 78157186 | 1.00284 | 1.24408 | 1.241 |

| | | | | | |
|---|---|---|---|---|---|
| chr15 | 35487939 | 35488166 | 1.13611 | 1.40944 | 1.241 |
| chr9 | 78463967 | 78464241 | 0.647296 | 0.803021 | 1.241 |
| chr10 | 111716457 | 111716744 | 1.94122 | 2.40832 | 1.241 |
| chr10 | 67575427 | 67575657 | 1.05008 | 1.30281 | 1.241 |
| chr6 | 43691635 | 43691870 | 1.34219 | 1.66546 | 1.241 |
| chr1 | 237415408 | 237415641 | 0.707588 | 0.878022 | 1.241 |
| chr5 | 10988007 | 10988360 | 1.7534 | 2.17592 | 1.241 |
| chr6 | 4156159 | 4156414 | 1.97744 | 2.45419 | 1.241 |
| chr1 | 40265345 | 40265681 | 2.44412 | 3.0335 | 1.241 |
| chr8 | 18160567 | 18160775 | 0.578006 | 0.71742 | 1.241 |
| chr5 | 82769386 | 82769767 | 2.73843 | 3.39916 | 1.241 |
| chr11 | 59051910 | 59052358 | 1.33928 | 1.66243 | 1.241 |
| chr8 | 65341544 | 65341828 | 1.62258 | 2.01418 | 1.241 |
| chr17 | 77020519 | 77020740 | 1.6508 | 2.04927 | 1.241 |
| chr13 | 114992266 | 114992496 | 2.78508 | 3.4575 | 1.241 |
| chr5 | 97563128 | 97563569 | 0.750393 | 0.931587 | 1.241 |
| chr5 | 50972232 | 50972529 | 0.643924 | 0.799467 | 1.242 |
| chr3 | 11101262 | 11101537 | 2.30266 | 2.85894 | 1.242 |
| chr14 | 23546088 | 23546434 | 1.72732 | 2.14471 | 1.242 |
| chr12 | 107916907 | 107917102 | 1.50973 | 1.87464 | 1.242 |
| chr1 | 242181026 | 242181208 | 2.46785 | 3.06462 | 1.242 |
| chr15 | 81890337 | 81890551 | 0.715077 | 0.888164 | 1.242 |
| chr10 | 122584361 | 122584651 | 1.55135 | 1.92716 | 1.242 |
| chr12 | 103555863 | 103556110 | 0.941256 | 1.16932 | 1.242 |
| chr3 | 16900415 | 16900619 | 0.754775 | 0.937694 | 1.242 |
| chr17 | 56494983 | 56495222 | 3.06768 | 3.81137 | 1.242 |
| chr1 | 177882879 | 177883126 | 1.36145 | 1.69154 | 1.242 |
| chr11 | 125011635 | 125011937 | 1.84743 | 2.29562 | 1.243 |
| chr6 | 110968257 | 110968463 | 1.22737 | 1.52532 | 1.243 |
| chr2 | 130939381 | 130939858 | 1.54082 | 1.91488 | 1.243 |
| chr8 | 26835162 | 26835436 | 0.924046 | 1.14839 | 1.243 |
| chr15 | 56889595 | 56889844 | 0.963097 | 1.19695 | 1.243 |
| chr5 | 11109065 | 11109313 | 0.691456 | 0.859378 | 1.243 |
| chr1 | 159004536 | 159004739 | 0.682002 | 0.847641 | 1.243 |
| chr11 | 82406598 | 82407140 | 1.93987 | 2.41111 | 1.243 |
| chr18 | 74110916 | 74111195 | 2.42016 | 3.00829 | 1.243 |
| chr10 | 83871675 | 83871938 | 0.955726 | 1.18801 | 1.243 |
| chr11 | 69484928 | 69485340 | 2.21045 | 2.74771 | 1.243 |
| chr9 | 95478793 | 95479189 | 2.8573 | 3.55182 | 1.243 |
| chr12 | 39666970 | 39667349 | 1.64643 | 2.04667 | 1.243 |
| chr17 | 45307140 | 45307457 | 3.09298 | 3.84558 | 1.243 |
| chr12 | 126034449 | 126034754 | 1.37623 | 1.71123 | 1.243 |
| chr20 | 53297447 | 53297924 | 1.24964 | 1.55384 | 1.243 |
| chr20 | 6029338 | 6029568 | 0.81326 | 1.01135 | 1.244 |
| chr10 | 24649457 | 24649810 | 1.6168 | 2.01067 | 1.244 |
| chr1 | 223500763 | 223500992 | 0.8311 | 1.0337 | 1.244 |
| chr10 | 59912535 | 59912753 | 1.69598 | 2.10947 | 1.244 |
| chr20 | 11853614 | 11853844 | 1.13729 | 1.41471 | 1.244 |
| chr5 | 50440942 | 50441202 | 0.797956 | 0.992598 | 1.244 |
| chr9 | 5469943 | 5470338 | 0.929694 | 1.15672 | 1.244 |
| chr5 | 107817158 | 107817404 | 1.33833 | 1.66525 | 1.244 |
| chr3 | 4675025 | 4675402 | 1.82987 | 2.27692 | 1.244 |
| chr12 | 94723105 | 94723360 | 0.811557 | 1.00986 | 1.244 |
| chr9 | 99089932 | 99090338 | 1.60965 | 2.00298 | 1.244 |
| chr12 | 113877287 | 113877770 | 2.28566 | 2.84431 | 1.244 |
| chr2 | 80114905 | 80115482 | 1.11005 | 1.3815 | 1.245 |
| chr11 | 66095735 | 66096138 | 2.63276 | 3.27665 | 1.245 |
| chr11 | 61395252 | 61395982 | 1.62778 | 2.026 | 1.245 |
| chr12 | 96353022 | 96353416 | 0.914438 | 1.13815 | 1.245 |
| chr17 | 487324 | 487547 | 0.813998 | 1.0132 | 1.245 |
| chr6 | 51824619 | 51824862 | 1.80334 | 2.24476 | 1.245 |

| | | | | | |
|---|---|---|---|---|---|
| chr20 | 50627180 | 50627393 | 0.768744 | 0.957053 | 1.245 |
| chr15 | 77947233 | 77947617 | 1.30937 | 1.63014 | 1.245 |
| chr5 | 42909079 | 42909308 | 2.53764 | 3.15942 | 1.245 |
| chr7 | 134917542 | 134917800 | 1.71109 | 2.13043 | 1.245 |
| chr19 | 5038947 | 5039237 | 1.62968 | 2.02912 | 1.245 |
| chr2 | 213551090 | 213551476 | 1.57679 | 1.96342 | 1.245 |
| chr2 | 6723519 | 6723736 | 0.790055 | 0.983826 | 1.245 |
| chr11 | 94237949 | 94238175 | 1.36265 | 1.69691 | 1.245 |
| chr8 | 3141812 | 3142443 | 1.47511 | 1.837 | 1.245 |
| chr6 | 99782503 | 99782743 | 1.5513 | 1.93206 | 1.245 |
| chr1 | 11097702 | 11097996 | 2.53855 | 3.1621 | 1.246 |
| chr19 | 18521489 | 18521742 | 1.71766 | 2.13961 | 1.246 |
| chr9 | 129622455 | 129622969 | 4.05028 | 5.04525 | 1.246 |
| chr9 | 34500625 | 34501025 | 3.11912 | 3.88551 | 1.246 |
| chr8 | 23614543 | 23614788 | 0.706601 | 0.880235 | 1.246 |
| chr2 | 152294025 | 152294279 | 1.17326 | 1.46159 | 1.246 |
| chr16 | 23446671 | 23447193 | 2.63469 | 3.28228 | 1.246 |
| chr7 | 73027293 | 73027541 | 1.44379 | 1.79885 | 1.246 |
| chr10 | 129915571 | 129915762 | 1.00255 | 1.24922 | 1.246 |
| chr8 | 140642742 | 140643773 | 2.50759 | 3.12469 | 1.246 |
| chr3 | 149489493 | 149489872 | 2.15511 | 2.68555 | 1.246 |
| rUn_gl0002 | 93495 | 93896 | 4.17022 | 5.19666 | 1.246 |
| chr1 | 93214629 | 93214915 | 2.1654 | 2.69858 | 1.246 |
| chr1 | 36181043 | 36181413 | 2.42808 | 3.0261 | 1.246 |
| chr20 | 24203233 | 24203537 | 0.719089 | 0.896195 | 1.246 |
| chr21 | 20574781 | 20575088 | 0.636596 | 0.793385 | 1.246 |
| chr20 | 30421270 | 30421544 | 2.45547 | 3.06026 | 1.246 |
| chr2 | 168821583 | 168821799 | 0.972269 | 1.21178 | 1.246 |
| chr19 | 43110666 | 43111520 | 1.94525 | 2.42454 | 1.246 |
| chr7 | 91615330 | 91615595 | 0.613646 | 0.764946 | 1.247 |
| chr18 | 5868266 | 5868478 | 1.50471 | 1.87586 | 1.247 |
| chr6 | 6557219 | 6557573 | 1.84252 | 2.29704 | 1.247 |
| chr9 | 79371271 | 79371481 | 1.09043 | 1.35943 | 1.247 |
| chr10 | 14238909 | 14239523 | 1.82402 | 2.27418 | 1.247 |
| chr19 | 59092472 | 59093055 | 3.15106 | 3.92919 | 1.247 |
| chr10 | 81160216 | 81160577 | 2.48601 | 3.10028 | 1.247 |
| chr17 | 53760285 | 53760611 | 0.802591 | 1.0009 | 1.247 |
| chr14 | 37761953 | 37762195 | 0.746488 | 0.93095 | 1.247 |
| chr8 | 113403703 | 113403933 | 0.56401 | 0.703471 | 1.247 |
| chr17 | 69826923 | 69827302 | 0.75013 | 0.935625 | 1.247 |
| chr11 | 41158559 | 41158758 | 0.844135 | 1.05289 | 1.247 |
| chr16 | 17570683 | 17570922 | 1.89893 | 2.36856 | 1.247 |
| chr7 | 155172821 | 155173196 | 1.77627 | 2.21577 | 1.247 |
| chr16 | 60365738 | 60365972 | 0.590812 | 0.737005 | 1.247 |
| chr10 | 29372235 | 29372498 | 0.653726 | 0.815513 | 1.247 |
| chr7 | 49159326 | 49159595 | 0.569556 | 0.710513 | 1.247 |
| chr8 | 134571802 | 134572023 | 0.9693 | 1.20942 | 1.248 |
| chr18 | 13823967 | 13824193 | 2.25393 | 2.81253 | 1.248 |
| chr5 | 42144399 | 42144594 | 0.648679 | 0.809443 | 1.248 |
| chr7 | 70561662 | 70561916 | 0.773821 | 0.965662 | 1.248 |
| chr3 | 37493213 | 37493726 | 1.76529 | 2.20306 | 1.248 |
| chr20 | 14122796 | 14123072 | 2.01825 | 2.51926 | 1.248 |
| chr11 | 80634826 | 80635015 | 1.04998 | 1.31068 | 1.248 |
| chr7 | 112872109 | 112872350 | 0.704007 | 0.878817 | 1.248 |
| chr12 | 119860949 | 119861463 | 0.959305 | 1.19779 | 1.249 |
| chr1 | 158968418 | 158968948 | 1.30949 | 1.63533 | 1.249 |
| chr17 | 30560470 | 30560725 | 0.901162 | 1.12541 | 1.249 |
| chr10 | 81368628 | 81368834 | 1.57503 | 1.96711 | 1.249 |
| chr12 | 58150507 | 58150955 | 3.57213 | 4.46148 | 1.249 |
| chr15 | 48644730 | 48644976 | 2.46801 | 3.08269 | 1.249 |
| chr1 | 239073042 | 239073262 | 0.840976 | 1.05051 | 1.249 |

| | | | | | |
|---|---|---|---|---|---|
| chr5 | 121794097 | 121794369 | 1.17346 | 1.46584 | 1.249 |
| chr3 | 77369855 | 77370046 | 1.14123 | 1.42561 | 1.249 |
| chr7 | 125565648 | 125565884 | 0.990707 | 1.23762 | 1.249 |
| chr15 | 31780854 | 31781155 | 2.06805 | 2.58354 | 1.249 |
| chr8 | 87137272 | 87137625 | 1.80764 | 2.25823 | 1.249 |
| chr22 | 27061493 | 27061763 | 1.89613 | 2.3688 | 1.249 |
| chr4 | 179749386 | 179749599 | 0.786031 | 0.981993 | 1.249 |
| chr9 | 133582682 | 133583237 | 1.45344 | 1.8158 | 1.249 |
| chr20 | 53265902 | 53266377 | 2.05704 | 2.5701 | 1.249 |
| chr7 | 100378923 | 100379210 | 0.8212 | 1.02613 | 1.250 |
| chr1 | 32879598 | 32879832 | 1.62159 | 2.02631 | 1.250 |
| chr3 | 118753624 | 118753984 | 1.97883 | 2.47279 | 1.250 |
| chr5 | 153847211 | 153847659 | 2.7895 | 3.48582 | 1.250 |
| chr5 | 668905 | 669617 | 2.28237 | 2.85266 | 1.250 |
| chr8 | 143122046 | 143122388 | 0.885738 | 1.10724 | 1.250 |
| chr2 | 9022339 | 9022586 | 1.12944 | 1.41192 | 1.250 |
| chr1 | 102033188 | 102033423 | 1.60486 | 2.00627 | 1.250 |
| chr8 | 112631373 | 112631585 | 0.692807 | 0.866156 | 1.250 |
| chr16 | 15915707 | 15916120 | 2.34347 | 2.93005 | 1.250 |
| chr8 | 4706096 | 4706367 | 0.863334 | 1.07944 | 1.250 |
| chr2 | 42033430 | 42033719 | 1.34376 | 1.68018 | 1.250 |
| chr1 | 20966555 | 20967117 | 1.48358 | 1.85522 | 1.251 |
| chr1 | 245960934 | 245961187 | 1.54912 | 1.9372 | 1.251 |
| chr1 | 163379841 | 163380325 | 1.08392 | 1.35552 | 1.251 |
| chr8 | 71446770 | 71446996 | 1.72305 | 2.1548 | 1.251 |
| chr2 | 17852274 | 17852636 | 1.20404 | 1.50577 | 1.251 |
| chr3 | 95423591 | 95423823 | 0.743808 | 0.930261 | 1.251 |
| chr4 | 144695248 | 144695477 | 1.11468 | 1.39427 | 1.251 |
| chr12 | 30552581 | 30552803 | 0.640887 | 0.801659 | 1.251 |
| chr7 | 23851189 | 23851441 | 1.0108 | 1.26447 | 1.251 |
| chr19 | 51039953 | 51040185 | 1.20249 | 1.50432 | 1.251 |
| chr1 | 61314349 | 61314602 | 1.54757 | 1.93617 | 1.251 |
| chr20 | 57225576 | 57226117 | 1.91654 | 2.3978 | 1.251 |
| chr1 | 204092483 | 204092844 | 1.08532 | 1.35798 | 1.251 |
| chr2 | 50158076 | 50158324 | 0.911833 | 1.14095 | 1.251 |
| chr1 | 209014760 | 209015002 | 0.662261 | 0.828744 | 1.251 |
| chr10 | 61696158 | 61696375 | 0.907933 | 1.13629 | 1.252 |
| chr12 | 51656529 | 51656819 | 1.92005 | 2.40304 | 1.252 |
| chr3 | 89662452 | 89662709 | 0.980746 | 1.22752 | 1.252 |
| chr15 | 81558174 | 81558515 | 2.72178 | 3.40666 | 1.252 |
| chr12 | 114769376 | 114769677 | 0.696979 | 0.872369 | 1.252 |
| chr11 | 44693369 | 44693615 | 0.951803 | 1.19142 | 1.252 |
| chrX | 132887499 | 132887703 | 0.693459 | 0.868062 | 1.252 |
| chr12 | 15380094 | 15380297 | 1.79387 | 2.24561 | 1.252 |
| chr18 | 21166840 | 21167159 | 2.57009 | 3.21733 | 1.252 |
| chr11 | 72141980 | 72142339 | 2.63804 | 3.30243 | 1.252 |
| chr15 | 28331184 | 28331633 | 2.02615 | 2.53674 | 1.252 |
| chr11 | 3403980 | 3404368 | 0.802297 | 1.00448 | 1.252 |
| chr10 | 60964660 | 60964914 | 0.72862 | 0.912264 | 1.252 |
| chr7 | 100494698 | 100495108 | 2.36328 | 2.95929 | 1.252 |
| chr9 | 101469347 | 101469807 | 1.48325 | 1.8574 | 1.252 |
| chr1 | 170679818 | 170680035 | 0.701198 | 0.878151 | 1.252 |
| chr7 | 92857638 | 92857966 | 1.54588 | 1.93616 | 1.252 |
| chr20 | 48890346 | 48890671 | 0.608918 | 0.762729 | 1.253 |
| chr16 | 11047775 | 11048195 | 2.94689 | 3.69132 | 1.253 |
| chr11 | 60101525 | 60101834 | 1.5403 | 1.92961 | 1.253 |
| chr5 | 10911335 | 10911604 | 0.823227 | 1.03141 | 1.253 |
| chr9 | 7475331 | 7475588 | 0.622272 | 0.779668 | 1.253 |
| chr1 | 3535609 | 3535886 | 2.102 | 2.63374 | 1.253 |
| chr10 | 71905282 | 71905815 | 1.47794 | 1.85181 | 1.253 |
| chr1 | 150444337 | 150444609 | 0.854123 | 1.07022 | 1.253 |

| | | | | | |
|---|---|---|---|---|---|
| chr5 | 85163771 | 85164095 | 0.792927 | 0.993536 | 1.253 |
| chr8 | 72723041 | 72723328 | 1.99662 | 2.50192 | 1.253 |
| chr1 | 226163733 | 226164001 | 2.51764 | 3.15491 | 1.253 |
| chr11 | 40634817 | 40635070 | 0.747413 | 0.936643 | 1.253 |
| chr1 | 153919074 | 153919463 | 3.0754 | 3.85443 | 1.253 |
| chr17 | 77061257 | 77061494 | 1.5428 | 1.93397 | 1.254 |
| chr12 | 41991728 | 41991987 | 0.871273 | 1.09237 | 1.254 |
| chr16 | 9144242 | 9144615 | 1.94795 | 2.44232 | 1.254 |
| chr6 | 121147328 | 121147637 | 1.00625 | 1.26164 | 1.254 |
| chr5 | 10689668 | 10690207 | 1.2841 | 1.61009 | 1.254 |
| chr6 | 135323431 | 135323647 | 1.46631 | 1.83881 | 1.254 |
| chr15 | 95303690 | 95303903 | 0.598486 | 0.750622 | 1.254 |
| chr2 | 229053534 | 229053903 | 0.687381 | 0.86213 | 1.254 |
| chr8 | 94227863 | 94228077 | 0.556788 | 0.69834 | 1.254 |
| chr7 | 156497976 | 156498179 | 0.697362 | 0.874676 | 1.254 |
| chr12 | 103600663 | 103600948 | 0.766614 | 0.961542 | 1.254 |
| chr19 | 35050103 | 35050441 | 0.740162 | 0.92841 | 1.254 |
| chr8 | 37708962 | 37709304 | 2.04623 | 2.56672 | 1.254 |
| chr15 | 95881763 | 95882020 | 0.634265 | 0.795618 | 1.254 |
| chr19 | 28571618 | 28571828 | 0.855151 | 1.07276 | 1.254 |
| chr3 | 134474025 | 134474267 | 0.950798 | 1.1928 | 1.255 |
| chr12 | 124521181 | 124521509 | 2.28062 | 2.86118 | 1.255 |
| chr5 | 162932739 | 162933029 | 1.61896 | 2.03108 | 1.255 |
| chr2 | 173458504 | 173458707 | 1.22244 | 1.53368 | 1.255 |
| chr11 | 6833673 | 6834326 | 1.55388 | 1.94954 | 1.255 |
| chr7 | 131373737 | 131374244 | 2.2285 | 2.79609 | 1.255 |
| chr19 | 36468465 | 36468816 | 1.04022 | 1.30523 | 1.255 |
| chr9 | 95468854 | 95469078 | 1.5131 | 1.89867 | 1.255 |
| chr7 | 23719597 | 23719874 | 1.62872 | 2.04388 | 1.255 |
| chr8 | 85145604 | 85145892 | 0.827533 | 1.0385 | 1.255 |
| chr15 | 41877945 | 41878205 | 0.948799 | 1.19074 | 1.255 |
| chr17 | 4883957 | 4884264 | 1.4972 | 1.87902 | 1.255 |
| chr12 | 5193438 | 5193783 | 1.43616 | 1.80253 | 1.255 |
| chr15 | 51573616 | 51573947 | 2.08207 | 2.61322 | 1.255 |
| chr15 | 31865845 | 31866081 | 0.640139 | 0.803459 | 1.255 |
| chr5 | 11968142 | 11968505 | 0.965121 | 1.21165 | 1.255 |
| chr2 | 208599063 | 208599299 | 1.8614 | 2.33708 | 1.256 |
| chr5 | 141081679 | 141082089 | 2.70783 | 3.39984 | 1.256 |
| chr12 | 25283688 | 25283938 | 1.16417 | 1.46186 | 1.256 |
| chr7 | 32519259 | 32519571 | 2.55762 | 3.21169 | 1.256 |
| chr7 | 147233791 | 147234098 | 0.795485 | 0.998985 | 1.256 |
| chr5 | 33585858 | 33586069 | 0.745165 | 0.935802 | 1.256 |
| chr1 | 222967946 | 222968159 | 1.28792 | 1.61743 | 1.256 |
| chr12 | 117047575 | 117047855 | 1.91184 | 2.40102 | 1.256 |
| chr3 | 74214075 | 74214278 | 0.885883 | 1.1126 | 1.256 |
| chr8 | 16451470 | 16451652 | 0.992191 | 1.24627 | 1.256 |
| chr4 | 86048824 | 86049096 | 1.185 | 1.48849 | 1.256 |
| chr17 | 79412685 | 79412978 | 1.37519 | 1.72741 | 1.256 |
| chr5 | 14247012 | 14247214 | 0.791367 | 0.994127 | 1.256 |
| chr19 | 47987018 | 47987335 | 4.31983 | 5.42726 | 1.256 |
| chr5 | 114582149 | 114582393 | 2.17076 | 2.72727 | 1.256 |
| chr1 | 165336624 | 165337201 | 2.00892 | 2.52401 | 1.256 |
| chr13 | 66212702 | 66212946 | 1.29973 | 1.633 | 1.256 |
| chr9 | 136364332 | 136364522 | 0.845993 | 1.06296 | 1.256 |
| chr1 | 205323462 | 205323757 | 2.52809 | 3.17662 | 1.257 |
| chr20 | 6895190 | 6895425 | 0.719741 | 0.904405 | 1.257 |
| chr12 | 124832524 | 124832888 | 2.13474 | 2.68257 | 1.257 |
| chr16 | 75007248 | 75007568 | 0.762307 | 0.957979 | 1.257 |
| chr20 | 58632287 | 58632880 | 1.87835 | 2.36072 | 1.257 |
| chr15 | 99813198 | 99813566 | 2.37817 | 2.98906 | 1.257 |
| chr3 | 113203829 | 113204057 | 0.692091 | 0.870011 | 1.257 |

| | | | | | |
|---|---|---|---|---|---|
| chr19 | 4525144 | 4525497 | 1.92805 | 2.42374 | 1.257 |
| chr8 | 28269509 | 28269773 | 1.36466 | 1.71551 | 1.257 |
| chr9 | 12582233 | 12582439 | 0.697324 | 0.876651 | 1.257 |
| chrX | 78780255 | 78780455 | 1.25761 | 1.58112 | 1.257 |
| chr10 | 113308126 | 113308359 | 0.667898 | 0.839729 | 1.257 |
| chr10 | 115193001 | 115193221 | 0.737256 | 0.926981 | 1.257 |
| chr2 | 16789894 | 16790719 | 2.17371 | 2.73312 | 1.257 |
| chr8 | 135490716 | 135490987 | 1.69855 | 2.13583 | 1.257 |
| chr1 | 235267475 | 235267855 | 3.26509 | 4.10605 | 1.258 |
| chr9 | 100788548 | 100788774 | 0.545612 | 0.686154 | 1.258 |
| chr11 | 74281631 | 74281934 | 1.82497 | 2.29514 | 1.258 |
| chr18 | 46166200 | 46166580 | 0.986671 | 1.2409 | 1.258 |
| chr10 | 16870235 | 16870440 | 1.33766 | 1.68234 | 1.258 |
| chr3 | 33818000 | 33818195 | 1.30415 | 1.64028 | 1.258 |
| chr17 | 76796692 | 76796955 | 0.716823 | 0.901605 | 1.258 |
| chr3 | 77073360 | 77073629 | 0.729787 | 0.917928 | 1.258 |
| chr17 | 6306604 | 6306809 | 0.836396 | 1.05204 | 1.258 |
| chr10 | 102397420 | 102397973 | 1.34587 | 1.69299 | 1.258 |
| chr14 | 69764835 | 69765074 | 0.714451 | 0.898778 | 1.258 |
| chr7 | 78482781 | 78482996 | 0.657982 | 0.827741 | 1.258 |
| chr6 | 87788729 | 87789267 | 2.26414 | 2.84843 | 1.258 |
| chr5 | 85766327 | 85766588 | 0.870121 | 1.09473 | 1.258 |
| chr8 | 9178177 | 9178421 | 1.73171 | 2.17883 | 1.258 |
| chr8 | 117800633 | 117800991 | 1.6677 | 2.09845 | 1.258 |
| chr1 | 196972219 | 196972465 | 0.938216 | 1.18056 | 1.258 |
| chr8 | 143554974 | 143555447 | 2.44352 | 3.07512 | 1.258 |
| chr14 | 98859800 | 98860064 | 0.810369 | 1.01991 | 1.259 |
| chr7 | 107365557 | 107365922 | 1.47146 | 1.85215 | 1.259 |
| chr2 | 51573437 | 51573720 | 0.853827 | 1.0749 | 1.259 |
| chr17 | 44824645 | 44824849 | 0.955677 | 1.20313 | 1.259 |
| chr3 | 13957650 | 13957868 | 0.705197 | 0.88781 | 1.259 |
| chr2 | 5976069 | 5976291 | 0.672094 | 0.846232 | 1.259 |
| chr3 | 79541230 | 79541473 | 0.594579 | 0.748665 | 1.259 |
| chr17 | 36762542 | 36763031 | 4.74882 | 5.97957 | 1.259 |
| chr22 | 22457513 | 22457872 | 1.04544 | 1.31645 | 1.259 |
| chr17 | 34176550 | 34176858 | 2.46622 | 3.10589 | 1.259 |
| chr3 | 49943782 | 49944044 | 1.72842 | 2.17673 | 1.259 |
| chr22 | 28508718 | 28509018 | 1.14863 | 1.44663 | 1.259 |
| chr11 | 5912465 | 5912741 | 1.01116 | 1.27354 | 1.259 |
| chr19 | 1618232 | 1618460 | 1.27695 | 1.60837 | 1.260 |
| chr5 | 109786402 | 109786643 | 0.643051 | 0.809947 | 1.260 |
| chr1 | 242866720 | 242866954 | 0.88708 | 1.11733 | 1.260 |
| chr11 | 2171466 | 2171839 | 1.37339 | 1.73006 | 1.260 |
| chr6 | 1364461 | 1364671 | 1.61539 | 2.03492 | 1.260 |
| chr1 | 168291203 | 168291406 | 1.39283 | 1.75458 | 1.260 |
| chr5 | 18839121 | 18839344 | 0.60483 | 0.761935 | 1.260 |
| chr1 | 160162887 | 160163250 | 3.29401 | 4.14977 | 1.260 |
| chr3 | 71760144 | 71760354 | 1.77097 | 2.23147 | 1.260 |
| chr15 | 94264620 | 94264836 | 0.763779 | 0.962394 | 1.260 |
| chr3 | 88411560 | 88412138 | 0.883699 | 1.1135 | 1.260 |
| chr20 | 11524785 | 11525035 | 1.60947 | 2.02811 | 1.260 |
| chr7 | 100386789 | 100386998 | 1.31128 | 1.65244 | 1.260 |
| chr15 | 28368629 | 28368931 | 1.88814 | 2.37947 | 1.260 |
| chr12 | 5635121 | 5635341 | 0.845114 | 1.06505 | 1.260 |
| chr7 | 30199469 | 30199836 | 0.947758 | 1.19444 | 1.260 |
| chr3 | 87194869 | 87195059 | 0.955399 | 1.2041 | 1.260 |
| chr4 | 148629347 | 148629644 | 2.23792 | 2.82062 | 1.260 |
| chr15 | 87581760 | 87581980 | 0.807313 | 1.01756 | 1.260 |
| chr2 | 7223962 | 7224330 | 1.70475 | 2.14882 | 1.260 |
| chr10 | 99223358 | 99223739 | 2.73282 | 3.44482 | 1.261 |
| chr2 | 59434122 | 59434387 | 1.02125 | 1.28748 | 1.261 |

| | | | | | |
|---|---|---|---|---|---|
| chr1 | 43212891 | 43213240 | 1.45853 | 1.83883 | 1.261 |
| chr5 | 148465701 | 148465916 | 0.782825 | 0.986979 | 1.261 |
| chr2 | 20063102 | 20063539 | 2.58162 | 3.25497 | 1.261 |
| chr7 | 125893644 | 125893934 | 0.979166 | 1.2346 | 1.261 |
| chr1 | 161711321 | 161711541 | 1.39326 | 1.75681 | 1.261 |
| chr15 | 42996613 | 42997166 | 1.59106 | 2.00627 | 1.261 |
| chr10 | 102577320 | 102577502 | 1.15565 | 1.45742 | 1.261 |
| chr19 | 40514434 | 40514647 | 0.952457 | 1.20121 | 1.261 |
| chr6 | 150155648 | 150156063 | 1.3392 | 1.68903 | 1.261 |
| chr1 | 166483848 | 166484136 | 0.679073 | 0.856578 | 1.261 |
| chr5 | 26606359 | 26606542 | 0.737277 | 0.929997 | 1.261 |
| chrX | 153218747 | 153219217 | 2.68805 | 3.3907 | 1.261 |
| chr9 | 137581458 | 137581791 | 2.69604 | 3.40093 | 1.261 |
| chr9 | 133005427 | 133005841 | 2.39791 | 3.0249 | 1.261 |
| chrX | 22154101 | 22154340 | 1.72759 | 2.1793 | 1.261 |
| chr7 | 48267833 | 48268143 | 0.804532 | 1.01495 | 1.262 |
| chr3 | 48774398 | 48774678 | 1.67323 | 2.11087 | 1.262 |
| chr5 | 145510119 | 145510391 | 1.21149 | 1.52835 | 1.262 |
| chr5 | 166457539 | 166458030 | 1.14986 | 1.45073 | 1.262 |
| chr3 | 124278728 | 124279028 | 2.45585 | 3.09847 | 1.262 |
| chr7 | 155608017 | 155608334 | 3.33869 | 4.21244 | 1.262 |
| chr8 | 139105204 | 139105699 | 0.903937 | 1.14054 | 1.262 |
| chr1 | 20301338 | 20301579 | 2.33004 | 2.93999 | 1.262 |
| chr17 | 7441005 | 7441213 | 0.745557 | 0.940789 | 1.262 |
| chr17 | 59494249 | 59494671 | 2.84521 | 3.59045 | 1.262 |
| chr10 | 73189759 | 73190076 | 1.05297 | 1.32884 | 1.262 |
| chr5 | 93361790 | 93362235 | 1.43154 | 1.80659 | 1.262 |
| chr7 | 142135024 | 142135355 | 1.74242 | 2.19903 | 1.262 |
| chr5 | 102475018 | 102475344 | 2.41233 | 3.04452 | 1.262 |
| chr10 | 24608428 | 24608740 | 1.64557 | 2.07692 | 1.262 |
| chr21 | 44527714 | 44527980 | 2.55891 | 3.22996 | 1.262 |
| chr2 | 1359110 | 1359344 | 0.798559 | 1.00807 | 1.262 |
| chr17 | 46801978 | 46802309 | 1.99233 | 2.51505 | 1.262 |
| chr6 | 121640310 | 121640630 | 1.20606 | 1.52251 | 1.262 |
| chr10 | 129093500 | 129093723 | 0.619502 | 0.782084 | 1.262 |
| chr17 | 39478523 | 39478915 | 2.403 | 3.03378 | 1.263 |
| chr7 | 1224956 | 1225165 | 0.830121 | 1.04803 | 1.263 |
| chr2 | 17491105 | 17491310 | 1.93184 | 2.43909 | 1.263 |
| chr2 | 3753343 | 3753634 | 0.689443 | 0.870493 | 1.263 |
| chr10 | 85768588 | 85769179 | 0.983668 | 1.24205 | 1.263 |
| chr1 | 207108857 | 207109213 | 2.38128 | 3.00698 | 1.263 |
| chr19 | 35818604 | 35819020 | 2.0024 | 2.52869 | 1.263 |
| chr2 | 165394812 | 165395059 | 1.76137 | 2.22432 | 1.263 |
| chr13 | 26148642 | 26148902 | 2.27681 | 2.87539 | 1.263 |
| chr4 | 122865205 | 122865432 | 1.0788 | 1.36249 | 1.263 |
| chr1 | 19439935 | 19440403 | 1.41773 | 1.79058 | 1.263 |
| chr1 | 48190814 | 48191130 | 0.742603 | 0.938029 | 1.263 |
| chr14 | 81930908 | 81931157 | 2.25515 | 2.84871 | 1.263 |
| chr6 | 152527612 | 152527978 | 1.76226 | 2.22622 | 1.263 |
| chr1 | 247886788 | 247887002 | 0.831768 | 1.05079 | 1.263 |
| chr16 | 86508345 | 86508589 | 0.767726 | 0.969948 | 1.263 |
| chr20 | 48153571 | 48153912 | 1.51641 | 1.91594 | 1.263 |
| chr7 | 1452420 | 1452775 | 1.98905 | 2.51313 | 1.263 |
| chr12 | 49750645 | 49751132 | 1.37822 | 1.74139 | 1.264 |
| chr7 | 98406804 | 98407030 | 0.650887 | 0.822403 | 1.264 |
| chr10 | 53801205 | 53801426 | 0.5889 | 0.744117 | 1.264 |
| chrX | 117761376 | 117761613 | 1.3875 | 1.75322 | 1.264 |
| chr3 | 80609005 | 80609190 | 1.16217 | 1.46861 | 1.264 |
| chr20 | 18170389 | 18170575 | 0.851997 | 1.07675 | 1.264 |
| chr5 | 85487700 | 85487940 | 0.728884 | 0.92116 | 1.264 |
| chr5 | 112672743 | 112673011 | 0.92191 | 1.16515 | 1.264 |

| | | | | | |
|---|---|---|---|---|---|
| chr14 | 30753039 | 30753439 | 1.93626 | 2.44727 | 1.264 |
| chr12 | 29672543 | 29672843 | 1.29099 | 1.63177 | 1.264 |
| chr5 | 179334793 | 179335170 | 1.83056 | 2.31377 | 1.264 |
| chr5 | 112714680 | 112714999 | 1.81778 | 2.29772 | 1.264 |
| chr15 | 95033752 | 95034074 | 0.771381 | 0.975257 | 1.264 |
| chr1 | 238356132 | 238356417 | 0.925551 | 1.17019 | 1.264 |
| chr2 | 70781302 | 70781676 | 1.9868 | 2.51204 | 1.264 |
| chr2 | 166294913 | 166295126 | 1.02353 | 1.29414 | 1.264 |
| chr20 | 37419125 | 37419427 | 1.76843 | 2.23642 | 1.265 |
| chr5 | 65619513 | 65619722 | 0.720079 | 0.910655 | 1.265 |
| chr9 | 38642640 | 38642950 | 1.623 | 2.05263 | 1.265 |
| chr5 | 117208722 | 117208965 | 0.622454 | 0.787263 | 1.265 |
| chr19 | 35581434 | 35582365 | 1.3652 | 1.7271 | 1.265 |
| chr11 | 10600508 | 10600806 | 0.770486 | 0.974772 | 1.265 |
| chr17 | 39573734 | 39574116 | 1.89262 | 2.39471 | 1.265 |
| chr7 | 144338605 | 144338971 | 0.748729 | 0.947417 | 1.265 |
| chr2 | 219867169 | 219868109 | 1.93896 | 2.45356 | 1.265 |
| chr2 | 239956968 | 239957431 | 2.22436 | 2.81477 | 1.265 |
| chr16 | 66304288 | 66304741 | 2.88999 | 3.65726 | 1.265 |
| chr9 | 16266710 | 16266998 | 1.93188 | 2.44546 | 1.266 |
| chr5 | 113835248 | 113835461 | 0.701835 | 0.888428 | 1.266 |
| gl000198_ra | 60941 | 61152 | 0.919323 | 1.16379 | 1.266 |
| chr9 | 129702918 | 129703312 | 2.20018 | 2.78539 | 1.266 |
| chr6 | 147106227 | 147106446 | 1.21302 | 1.53568 | 1.266 |
| chr17 | 27912078 | 27912301 | 1.68895 | 2.13866 | 1.266 |
| chr11 | 68615313 | 68615634 | 0.864444 | 1.09463 | 1.266 |
| chr1 | 95267658 | 95268066 | 1.11805 | 1.41578 | 1.266 |
| chr11 | 20265572 | 20265776 | 1.12677 | 1.42683 | 1.266 |
| chr4 | 870586 | 870812 | 0.981916 | 1.24353 | 1.266 |
| chr14 | 72980772 | 72981114 | 1.6806 | 2.1284 | 1.266 |
| chr20 | 52918488 | 52918760 | 0.587614 | 0.744238 | 1.267 |
| chr19 | 51400977 | 51401172 | 1.09918 | 1.3922 | 1.267 |
| chr1 | 152413300 | 152413722 | 1.99422 | 2.526 | 1.267 |
| chr1 | 209888242 | 209888447 | 0.979262 | 1.2405 | 1.267 |
| chr10 | 77744032 | 77744240 | 0.717092 | 0.908403 | 1.267 |
| chr1 | 166549500 | 166549779 | 0.672849 | 0.852393 | 1.267 |
| chr20 | 39574682 | 39575027 | 1.11979 | 1.4186 | 1.267 |
| chr10 | 98785345 | 98785733 | 1.67758 | 2.12536 | 1.267 |
| chr8 | 107533991 | 107534296 | 1.38763 | 1.75807 | 1.267 |
| chr17 | 3586202 | 3586831 | 3.27533 | 4.14995 | 1.267 |
| chr14 | 86523477 | 86523699 | 1.35015 | 1.71071 | 1.267 |
| chr2 | 1754628 | 1754867 | 2.47649 | 3.13787 | 1.267 |
| chr7 | 103800760 | 103801111 | 0.795585 | 1.00808 | 1.267 |
| chr15 | 85421829 | 85422041 | 1.69332 | 2.14561 | 1.267 |
| chr6 | 160493421 | 160493653 | 1.39207 | 1.76397 | 1.267 |
| chr20 | 41820688 | 41820921 | 1.88151 | 2.38426 | 1.267 |
| chr3 | 8281388 | 8281765 | 2.01441 | 2.55271 | 1.267 |
| chr14 | 56941222 | 56941447 | 2.17309 | 2.75389 | 1.267 |
| chr16 | 61480334 | 61480610 | 0.822128 | 1.0419 | 1.267 |
| chr14 | 104274805 | 104275147 | 2.02193 | 2.56246 | 1.267 |
| chr11 | 81001592 | 81001850 | 0.805994 | 1.0215 | 1.267 |
| chr5 | 38258811 | 38259101 | 0.893963 | 1.13302 | 1.267 |
| chr15 | 82231999 | 82232214 | 1.52934 | 1.93833 | 1.267 |
| chr19 | 34996603 | 34997180 | 0.851146 | 1.07881 | 1.267 |
| chr2 | 38603713 | 38604082 | 2.01123 | 2.54919 | 1.267 |
| chr1 | 229471395 | 229471604 | 0.672462 | 0.852364 | 1.268 |
| chr15 | 52623685 | 52623986 | 1.61714 | 2.04987 | 1.268 |
| chr11 | 64018567 | 64019144 | 1.68996 | 2.14225 | 1.268 |
| chr12 | 12503178 | 12503507 | 1.16378 | 1.4753 | 1.268 |
| chr12 | 49012235 | 49012664 | 2.93213 | 3.7173 | 1.268 |
| chr3 | 117074510 | 117074706 | 1.34162 | 1.70092 | 1.268 |

| | | | | | |
|---|---|---|---|---|---|
| chr13 | 21285745 | 21286047 | 1.02622 | 1.3011 | 1.268 |
| chr8 | 56991699 | 56991990 | 1.95306 | 2.47623 | 1.268 |
| chr2 | 209260100 | 209260344 | 1.32295 | 1.6774 | 1.268 |
| chr1 | 176715049 | 176715372 | 0.76999 | 0.97638 | 1.268 |
| chr16 | 88924216 | 88924416 | 1.59066 | 2.01718 | 1.268 |
| chr10 | 134448106 | 134448314 | 0.723177 | 0.917122 | 1.268 |
| chr4 | 71467214 | 71467490 | 2.18638 | 2.77274 | 1.268 |
| chr8 | 124702572 | 124702817 | 1.54072 | 1.95397 | 1.268 |
| chr16 | 18813005 | 18813283 | 3.51897 | 4.46325 | 1.268 |
| chr17 | 71407601 | 71407806 | 0.692268 | 0.878053 | 1.268 |
| chr10 | 7163143 | 7163511 | 2.31546 | 2.93738 | 1.269 |
| chr4 | 190714159 | 190714515 | 1.75858 | 2.23094 | 1.269 |
| chr2 | 10927720 | 10927993 | 1.73134 | 2.19652 | 1.269 |
| chr16 | 2977033 | 2977392 | 2.59861 | 3.29683 | 1.269 |
| chr8 | 88265487 | 88265854 | 0.898456 | 1.13987 | 1.269 |
| chr5 | 164134141 | 164134375 | 0.747868 | 0.948919 | 1.269 |
| chr9 | 18885587 | 18885822 | 0.748973 | 0.950341 | 1.269 |
| chr8 | 70576063 | 70576367 | 1.56619 | 1.98732 | 1.269 |
| chr10 | 120897505 | 120897708 | 0.747811 | 0.949023 | 1.269 |
| chr2 | 240028022 | 240028340 | 2.40132 | 3.04763 | 1.269 |
| chr3 | 33925019 | 33925256 | 1.33937 | 1.6999 | 1.269 |
| chr16 | 72275792 | 72276083 | 1.45403 | 1.84553 | 1.269 |
| chr3 | 151143906 | 151144268 | 1.14524 | 1.45362 | 1.269 |
| chr9 | 12329294 | 12329628 | 0.776133 | 0.985134 | 1.269 |
| chr6 | 44437524 | 44437944 | 2.28416 | 2.8995 | 1.269 |
| chr8 | 139199053 | 139199294 | 0.679805 | 0.863039 | 1.270 |
| chr16 | 23445597 | 23445971 | 1.77527 | 2.25381 | 1.270 |
| chr2 | 173671221 | 173671440 | 2.39157 | 3.03653 | 1.270 |
| chr7 | 53704344 | 53704695 | 0.742638 | 0.94294 | 1.270 |
| chr7 | 103555571 | 103555849 | 0.549135 | 0.697277 | 1.270 |
| chr10 | 107440133 | 107440400 | 0.785231 | 0.99711 | 1.270 |
| chr2 | 119769045 | 119769300 | 2.14388 | 2.72273 | 1.270 |
| chr7 | 86860601 | 86860803 | 0.66667 | 0.846719 | 1.270 |
| chr19 | 14459294 | 14459607 | 3.01354 | 3.8278 | 1.270 |
| chr10 | 27847480 | 27847759 | 1.50268 | 1.90884 | 1.270 |
| chr2 | 30370681 | 30370984 | 1.88369 | 2.39328 | 1.271 |
| chr2 | 230207196 | 230207465 | 1.87321 | 2.38011 | 1.271 |
| chr1 | 184160410 | 184160733 | 0.724511 | 0.920628 | 1.271 |
| chr1 | 204588945 | 204589183 | 0.953454 | 1.21157 | 1.271 |
| chr19 | 3308220 | 3308434 | 2.18972 | 2.78254 | 1.271 |
| chr8 | 21669533 | 21670105 | 2.19706 | 2.79187 | 1.271 |
| chr14 | 25204203 | 25204739 | 1.14111 | 1.45019 | 1.271 |
| chr3 | 140987600 | 140987862 | 0.724613 | 0.92089 | 1.271 |
| chr19 | 28441943 | 28442194 | 0.898791 | 1.14227 | 1.271 |
| chr5 | 98024690 | 98024958 | 0.772021 | 0.981224 | 1.271 |
| chr7 | 120880199 | 120880439 | 0.735606 | 0.934958 | 1.271 |
| chr8 | 105430254 | 105430576 | 2.79931 | 3.55855 | 1.271 |
| chr1 | 199521172 | 199521488 | 2.36179 | 3.00245 | 1.271 |
| chr3 | 89698701 | 89698957 | 0.748743 | 0.95186 | 1.271 |
| chr14 | 23734357 | 23734566 | 1.21699 | 1.54732 | 1.271 |
| chr5 | 18526087 | 18526310 | 0.772415 | 0.982168 | 1.272 |
| chr2 | 59629913 | 59630119 | 0.619437 | 0.78765 | 1.272 |
| chr14 | 93582195 | 93582468 | 2.03955 | 2.59343 | 1.272 |
| chr9 | 581928 | 582231 | 2.11575 | 2.69043 | 1.272 |
| chr7 | 34173715 | 34174185 | 1.65924 | 2.11003 | 1.272 |
| chr10 | 21114246 | 21114488 | 0.880792 | 1.1201 | 1.272 |
| chr22 | 42093128 | 42093668 | 2.49174 | 3.16884 | 1.272 |
| chr5 | 25293173 | 25293402 | 0.865553 | 1.10085 | 1.272 |
| chr3 | 106897142 | 106897362 | 0.597681 | 0.760177 | 1.272 |
| chr7 | 25299153 | 25299591 | 1.64831 | 2.09647 | 1.272 |
| chr12 | 100462667 | 100463050 | 0.652614 | 0.830066 | 1.272 |

| | | | | | |
|---|---|---|---|---|---|
| chr16 | 3144540 | 3144898 | 1.49637 | 1.90327 | 1.272 |
| chr14 | 44096446 | 44096839 | 0.897697 | 1.14184 | 1.272 |
| chr11 | 29549282 | 29549573 | 1.20147 | 1.52826 | 1.272 |
| chr16 | 54611698 | 54611982 | 0.650749 | 0.827745 | 1.272 |
| chr17 | 55087427 | 55087670 | 1.79165 | 2.27896 | 1.272 |
| chr22 | 26568421 | 26568858 | 1.56381 | 1.9892 | 1.272 |
| chr19 | 14597151 | 14597527 | 2.00267 | 2.54762 | 1.272 |
| chr7 | 137340740 | 137341327 | 1.6488 | 2.09746 | 1.272 |
| chr16 | 23812791 | 23812998 | 2.86016 | 3.63847 | 1.272 |
| chr1 | 207152781 | 207153058 | 1.69851 | 2.1608 | 1.272 |
| chr18 | 60689749 | 60690036 | 1.71847 | 2.18647 | 1.272 |
| chr9 | 128503872 | 128504091 | 0.711292 | 0.905038 | 1.272 |
| chr20 | 5383521 | 5384035 | 2.31719 | 2.94845 | 1.272 |
| chr5 | 179755556 | 179755989 | 2.47393 | 3.1479 | 1.272 |
| chr8 | 95834855 | 95835142 | 1.70714 | 2.17222 | 1.272 |
| chr11 | 37933805 | 37934043 | 0.875461 | 1.11404 | 1.273 |
| chr8 | 34767139 | 34767341 | 0.768907 | 0.978456 | 1.273 |
| chr1 | 240618560 | 240618994 | 1.03471 | 1.31671 | 1.273 |
| chr17 | 11733098 | 11733333 | 0.611253 | 0.777846 | 1.273 |
| chr1 | 147099245 | 147099628 | 2.5059 | 3.18927 | 1.273 |
| chr3 | 184089368 | 184089676 | 1.99847 | 2.54353 | 1.273 |
| chr4 | 38759685 | 38759913 | 1.11977 | 1.42518 | 1.273 |
| chr12 | 81436696 | 81436925 | 0.69263 | 0.881556 | 1.273 |
| chr13 | 51375493 | 51375734 | 1.41784 | 1.8046 | 1.273 |
| chr1 | 239222538 | 239222740 | 0.679447 | 0.864827 | 1.273 |
| chr11 | 134731488 | 134731708 | 1.55572 | 1.9802 | 1.273 |
| chr5 | 136965333 | 136965576 | 2.18839 | 2.78568 | 1.273 |
| chr5 | 149829498 | 149829758 | 2.88212 | 3.66882 | 1.273 |
| chr15 | 89738590 | 89738795 | 0.680343 | 0.866058 | 1.273 |
| chr2 | 78816480 | 78816754 | 0.772286 | 0.983215 | 1.273 |
| chr3 | 124000616 | 124000881 | 1.51128 | 1.92413 | 1.273 |
| chr2 | 176962316 | 176962631 | 2.8669 | 3.65029 | 1.273 |
| chr5 | 104813852 | 104814104 | 0.907749 | 1.15581 | 1.273 |
| chr3 | 121788362 | 121788573 | 0.787157 | 1.00227 | 1.273 |
| chr15 | 31539270 | 31539850 | 0.716307 | 0.912102 | 1.273 |
| chr7 | 101083583 | 101084063 | 1.9486 | 2.48132 | 1.273 |
| chr5 | 31186860 | 31187077 | 1.81014 | 2.30509 | 1.273 |
| chr9 | 109274428 | 109274642 | 0.720489 | 0.917626 | 1.274 |
| chr7 | 151012174 | 151012542 | 2.20668 | 2.81049 | 1.274 |
| chr10 | 85974426 | 85974787 | 1.24813 | 1.58971 | 1.274 |
| chr5 | 12861938 | 12862573 | 0.922262 | 1.17469 | 1.274 |
| chr1 | 177761565 | 177761789 | 0.668939 | 0.852053 | 1.274 |
| chr11 | 14330229 | 14330444 | 1.31468 | 1.67473 | 1.274 |
| chr19 | 14182909 | 14183187 | 1.7727 | 2.25835 | 1.274 |
| chr19 | 40023424 | 40023764 | 4.39079 | 5.59413 | 1.274 |
| chr11 | 1108681 | 1108967 | 2.68681 | 3.42364 | 1.274 |
| chr6 | 39157821 | 39158110 | 1.65619 | 2.1105 | 1.274 |
| chrX | 118400337 | 118400675 | 1.62541 | 2.07146 | 1.274 |
| chr2 | 95917252 | 95917489 | 0.782085 | 0.996712 | 1.274 |
| chr15 | 66444242 | 66444739 | 1.21511 | 1.54864 | 1.274 |
| chr12 | 71123853 | 71124111 | 1.29756 | 1.65374 | 1.275 |
| chr8 | 6776798 | 6777242 | 2.00763 | 2.55873 | 1.275 |
| chr4 | 8609243 | 8609719 | 2.35264 | 2.99847 | 1.275 |
| chr9 | 24714456 | 24714675 | 0.671853 | 0.856358 | 1.275 |
| chr13 | 42908359 | 42908562 | 1.25015 | 1.5936 | 1.275 |
| chr18 | 56724512 | 56724875 | 1.07809 | 1.37434 | 1.275 |
| chr7 | 24851982 | 24852315 | 1.70172 | 2.16955 | 1.275 |
| chr8 | 110359171 | 110359442 | 1.33418 | 1.7011 | 1.275 |
| chr19 | 55813210 | 55813670 | 2.64781 | 3.37663 | 1.275 |
| chr20 | 50002489 | 50002876 | 1.23422 | 1.57395 | 1.275 |
| chr17 | 53104434 | 53104693 | 0.893375 | 1.13929 | 1.275 |

| | | | | | |
|---|---|---|---|---|---|
| chr19 | 41513483 | 41513879 | 0.794406 | 1.01318 | 1.275 |
| chr3 | 128040968 | 128041245 | 1.30208 | 1.66072 | 1.275 |
| chr5 | 6765816 | 6766122 | 2.90522 | 3.7056 | 1.276 |
| chr10 | 73035270 | 73035472 | 1.29828 | 1.65611 | 1.276 |
| chr17 | 51239379 | 51239583 | 0.709242 | 0.904727 | 1.276 |
| chr7 | 97650805 | 97651201 | 2.72427 | 3.47519 | 1.276 |
| chr2 | 107159269 | 107159515 | 2.38615 | 3.04394 | 1.276 |
| chrX | 74823750 | 74824002 | 1.11511 | 1.42254 | 1.276 |
| chr8 | 19910002 | 19910242 | 0.592087 | 0.755423 | 1.276 |
| chr8 | 89081175 | 89081418 | 0.667973 | 0.85226 | 1.276 |
| chr1 | 43222489 | 43222955 | 1.5752 | 2.0098 | 1.276 |
| chr8 | 134581724 | 134582208 | 2.63347 | 3.36027 | 1.276 |
| chr11 | 69789260 | 69789526 | 1.4231 | 1.81589 | 1.276 |
| chr5 | 15382858 | 15383082 | 0.80921 | 1.03263 | 1.276 |
| chr7 | 52689093 | 52689449 | 0.703071 | 0.897236 | 1.276 |
| chr5 | 165057276 | 165057731 | 1.36431 | 1.74112 | 1.276 |
| chr19 | 55392529 | 55393084 | 1.23569 | 1.57702 | 1.276 |
| chr3 | 71979795 | 71980101 | 1.07426 | 1.37111 | 1.276 |
| chr2 | 11733008 | 11733216 | 0.893288 | 1.14016 | 1.276 |
| chr14 | 22202765 | 22203201 | 1.66334 | 2.12308 | 1.276 |
| chr13 | 29937183 | 29937518 | 2.40037 | 3.06425 | 1.277 |
| chr3 | 49486631 | 49486846 | 1.21754 | 1.55427 | 1.277 |
| chr20 | 44937133 | 44937482 | 1.15245 | 1.47133 | 1.277 |
| chr19 | 28450175 | 28450455 | 1.16868 | 1.49207 | 1.277 |
| chr4 | 111436006 | 111436357 | 0.720564 | 0.919978 | 1.277 |
| chr15 | 81475253 | 81475753 | 0.753808 | 0.962448 | 1.277 |
| chr2 | 9346317 | 9346701 | 3.02506 | 3.86278 | 1.277 |
| chr8 | 81937788 | 81937996 | 1.2764 | 1.62987 | 1.277 |
| chr1 | 176344163 | 176344472 | 1.78465 | 2.27891 | 1.277 |
| chr4 | 77228363 | 77228565 | 1.51535 | 1.93503 | 1.277 |
| chr13 | 107188794 | 107189137 | 3.04187 | 3.88449 | 1.277 |
| chr16 | 783642 | 784251 | 1.62074 | 2.0698 | 1.277 |
| chr4 | 154671458 | 154671727 | 1.51281 | 1.93228 | 1.277 |
| chr10 | 103595704 | 103596057 | 2.05295 | 2.62228 | 1.277 |
| chr5 | 37940310 | 37940602 | 0.691333 | 0.883077 | 1.277 |
| chr10 | 102754087 | 102754332 | 1.60705 | 2.05279 | 1.277 |
| chr5 | 19605885 | 19606173 | 1.02405 | 1.30812 | 1.277 |
| chr5 | 73968870 | 73969128 | 1.48548 | 1.89761 | 1.277 |
| chr9 | 108643967 | 108644480 | 2.0801 | 2.65727 | 1.277 |
| chr9 | 112460306 | 112460603 | 0.879859 | 1.12403 | 1.278 |
| chr11 | 12431587 | 12431996 | 0.7328 | 0.936178 | 1.278 |
| chr7 | 75702808 | 75703160 | 0.747943 | 0.955533 | 1.278 |
| chr6 | 157556671 | 157557045 | 1.70913 | 2.18362 | 1.278 |
| chr17 | 3640969 | 3641446 | 2.63183 | 3.36252 | 1.278 |
| chr7 | 71562926 | 71563195 | 0.782677 | 1.00007 | 1.278 |
| chrX | 24694616 | 24694814 | 1.72309 | 2.2018 | 1.278 |
| chr5 | 93718733 | 93719067 | 0.711321 | 0.908958 | 1.278 |
| chr7 | 6565117 | 6565458 | 0.830919 | 1.0618 | 1.278 |
| chr16 | 57754092 | 57754499 | 2.45504 | 3.13722 | 1.278 |
| chr1 | 226369763 | 226370037 | 1.63698 | 2.09186 | 1.278 |
| chr16 | 13051081 | 13051302 | 0.562614 | 0.718957 | 1.278 |
| chr7 | 51821762 | 51821967 | 0.746983 | 0.95464 | 1.278 |
| chr5 | 52466636 | 52466887 | 1.47443 | 1.88433 | 1.278 |
| chr9 | 35729727 | 35730031 | 1.57578 | 2.01388 | 1.278 |
| chr11 | 66848891 | 66849235 | 2.33973 | 2.99053 | 1.278 |
| chr7 | 49482852 | 49483250 | 0.741073 | 0.947238 | 1.278 |
| chr17 | 26847321 | 26847569 | 0.600366 | 0.767435 | 1.278 |
| chr5 | 41856364 | 41857056 | 1.43428 | 1.83341 | 1.278 |
| chr12 | 125021503 | 125021726 | 0.543422 | 0.694655 | 1.278 |
| chr16 | 53777033 | 53777335 | 1.82115 | 2.32807 | 1.278 |
| chr2 | 239371514 | 239371765 | 1.35602 | 1.7335 | 1.278 |

| | | | | | |
|---|---|---|---|---|---|
| chr5 | 164909716 | 164910143 | 0.664365 | 0.849338 | 1.278 |
| chr15 | 32110832 | 32111092 | 0.730747 | 0.934251 | 1.278 |
| chr1 | 110659008 | 110659368 | 2.10795 | 2.69509 | 1.279 |
| chr13 | 96216535 | 96216752 | 0.858696 | 1.09793 | 1.279 |
| chr7 | 102706154 | 102706415 | 1.57539 | 2.01441 | 1.279 |
| chr9 | 110658189 | 110658443 | 0.688428 | 0.880292 | 1.279 |
| chr12 | 25836772 | 25837034 | 0.681467 | 0.8714 | 1.279 |
| chr12 | 33996478 | 33996726 | 0.753653 | 0.96371 | 1.279 |
| chr2 | 170384779 | 170385093 | 2.48476 | 3.17746 | 1.279 |
| chr2 | 6100894 | 6101155 | 0.581701 | 0.743956 | 1.279 |
| chr3 | 39540055 | 39540423 | 2.04576 | 2.61639 | 1.279 |
| chr5 | 96894588 | 96894800 | 0.880318 | 1.12587 | 1.279 |
| chr2 | 227994621 | 227994887 | 1.52084 | 1.9453 | 1.279 |
| chr19 | 47894614 | 47895059 | 2.47262 | 3.16302 | 1.279 |
| chr17 | 56174300 | 56174562 | 1.15016 | 1.47136 | 1.279 |
| chr17 | 71641368 | 71641622 | 0.979933 | 1.25403 | 1.280 |
| chr2 | 79174686 | 79174895 | 0.647626 | 0.82888 | 1.280 |
| chr16 | 66016594 | 66016805 | 0.762119 | 0.975418 | 1.280 |
| chr17 | 40374243 | 40374566 | 2.44992 | 3.1357 | 1.280 |
| chr14 | 70690472 | 70690711 | 1.03136 | 1.32008 | 1.280 |
| chr1 | 182415246 | 182415550 | 1.56108 | 1.99831 | 1.280 |
| chr10 | 73246148 | 73246740 | 1.54281 | 1.97503 | 1.280 |
| chr15 | 33508603 | 33508812 | 1.95868 | 2.50747 | 1.280 |
| chr11 | 62102750 | 62103021 | 0.924401 | 1.18346 | 1.280 |
| chr2 | 3828487 | 3828724 | 1.89217 | 2.42249 | 1.280 |
| chr17 | 10554607 | 10554812 | 0.682331 | 0.873613 | 1.280 |
| chr1 | 7130461 | 7130839 | 2.27715 | 2.91556 | 1.280 |
| chr17 | 13349254 | 13349468 | 0.845383 | 1.08253 | 1.281 |
| chr1 | 235764183 | 235764704 | 1.32283 | 1.69397 | 1.281 |
| chr19 | 42289509 | 42289840 | 2.56527 | 3.28502 | 1.281 |
| chr20 | 21924886 | 21925114 | 0.871957 | 1.11661 | 1.281 |
| chr9 | 19085391 | 19085595 | 1.5373 | 1.96866 | 1.281 |
| chr5 | 98781146 | 98781354 | 0.73763 | 0.94462 | 1.281 |
| chr15 | 98442281 | 98442520 | 1.94 | 2.48446 | 1.281 |
| chr5 | 170174815 | 170175159 | 1.61466 | 2.06794 | 1.281 |
| chr2 | 20525923 | 20526227 | 2.80377 | 3.59096 | 1.281 |
| chr20 | 20818191 | 20818566 | 1.74408 | 2.23386 | 1.281 |
| chr11 | 36171597 | 36171830 | 1.60222 | 2.05219 | 1.281 |
| chr3 | 1253301 | 1253540 | 0.627505 | 0.803754 | 1.281 |
| chr20 | 59526914 | 59527186 | 0.610819 | 0.782459 | 1.281 |
| chr19 | 12948992 | 12949449 | 1.94471 | 2.49141 | 1.281 |
| chr11 | 77849821 | 77850227 | 1.06939 | 1.37003 | 1.281 |
| chr17 | 22194597 | 22194850 | 1.48317 | 1.90013 | 1.281 |
| chr8 | 87744518 | 87744731 | 0.856686 | 1.09764 | 1.281 |
| chr10 | 57642503 | 57642720 | 0.811363 | 1.03961 | 1.281 |
| chr4 | 16002713 | 16003338 | 1.24069 | 1.58973 | 1.281 |
| chr7 | 31358212 | 31358528 | 0.644417 | 0.825749 | 1.281 |
| chr16 | 30103785 | 30104128 | 3.19404 | 4.09284 | 1.281 |
| chr10 | 984917 | 985160 | 0.657168 | 0.842157 | 1.281 |
| chr3 | 169567421 | 169567704 | 0.806123 | 1.03306 | 1.282 |
| chr8 | 14724076 | 14724489 | 0.711811 | 0.912203 | 1.282 |
| chr3 | 99364124 | 99364470 | 1.33028 | 1.70483 | 1.282 |
| chr9 | 93405217 | 93405524 | 1.26818 | 1.62525 | 1.282 |
| chr2 | 169988450 | 169988716 | 1.1673 | 1.496 | 1.282 |
| chr19 | 29754332 | 29754917 | 1.90403 | 2.44029 | 1.282 |
| chr8 | 38688249 | 38688550 | 1.53781 | 1.97093 | 1.282 |
| chr20 | 62212086 | 62212580 | 0.995837 | 1.27647 | 1.282 |
| chr1 | 154320834 | 154321124 | 0.866152 | 1.11025 | 1.282 |
| chr7 | 48008405 | 48008788 | 2.20402 | 2.8253 | 1.282 |
| chr13 | 52371902 | 52372200 | 1.55356 | 1.99165 | 1.282 |
| chr8 | 143230732 | 143231054 | 1.59118 | 2.04007 | 1.282 |

| | | | | | |
|---|---|---|---|---|---|
| chr10 | 73044795 | 73045406 | 1.60009 | 2.0515 | 1.282 |
| chr1 | 886867 | 887069 | 1.29044 | 1.65463 | 1.282 |
| chr9 | 89102171 | 89102476 | 1.93011 | 2.47488 | 1.282 |
| chr15 | 66940031 | 66940331 | 0.56838 | 0.728833 | 1.282 |
| chr17 | 73168038 | 73168253 | 1.52501 | 1.95553 | 1.282 |
| chr7 | 148308832 | 148309091 | 0.814608 | 1.04459 | 1.282 |
| chr16 | 1156290 | 1156714 | 2.23235 | 2.86273 | 1.282 |
| chr3 | 13699354 | 13699681 | 2.58099 | 3.30983 | 1.282 |
| chr6 | 75920978 | 75921201 | 1.48982 | 1.91057 | 1.282 |
| chr3 | 97015898 | 97016117 | 0.632154 | 0.810725 | 1.282 |
| chr3 | 129035388 | 129035692 | 1.22055 | 1.56533 | 1.282 |
| chr17 | 7155691 | 7155901 | 0.992419 | 1.27282 | 1.283 |
| chr11 | 82638378 | 82638633 | 1.88837 | 2.42206 | 1.283 |
| chr8 | 138680089 | 138680303 | 0.992134 | 1.27263 | 1.283 |
| chr5 | 73936353 | 73936614 | 1.3753 | 1.76435 | 1.283 |
| chr10 | 99801192 | 99801531 | 1.86829 | 2.39682 | 1.283 |
| chr2 | 239213709 | 239214050 | 1.09961 | 1.41077 | 1.283 |
| chr7 | 92528205 | 92528439 | 0.85669 | 1.09913 | 1.283 |
| chr12 | 328971 | 329450 | 2.91994 | 3.74653 | 1.283 |
| chr8 | 100403948 | 100404331 | 1.18117 | 1.51559 | 1.283 |
| chr17 | 65626105 | 65626476 | 1.30186 | 1.67055 | 1.283 |
| chr5 | 36481436 | 36481693 | 0.900979 | 1.15616 | 1.283 |
| chr19 | 39283429 | 39283698 | 0.684454 | 0.87833 | 1.283 |
| chr17 | 37779534 | 37780051 | 2.31744 | 2.9739 | 1.283 |
| chr9 | 135978238 | 135978955 | 1.98475 | 2.54717 | 1.283 |
| chr22 | 37473819 | 37474219 | 3.72441 | 4.77991 | 1.283 |
| chr3 | 48673140 | 48673427 | 1.58157 | 2.02993 | 1.283 |
| chr3 | 155868260 | 155868571 | 1.9622 | 2.51856 | 1.284 |
| chr15 | 85197989 | 85198248 | 2.25138 | 2.88976 | 1.284 |
| chr20 | 58557897 | 58558129 | 0.80647 | 1.03517 | 1.284 |
| chr19 | 58207871 | 58208323 | 0.780192 | 1.00148 | 1.284 |
| chr16 | 28288992 | 28289369 | 2.43629 | 3.12732 | 1.284 |
| chr7 | 93072140 | 93072400 | 2.18831 | 2.80906 | 1.284 |
| chr19 | 41313667 | 41314113 | 1.57303 | 2.01929 | 1.284 |
| chr22 | 23624035 | 23624373 | 2.02294 | 2.59716 | 1.284 |
| chr15 | 99111621 | 99111826 | 1.56039 | 2.00365 | 1.284 |
| chr17 | 70723860 | 70724169 | 2.32259 | 2.98241 | 1.284 |
| chr20 | 61299925 | 61300231 | 2.20103 | 2.82639 | 1.284 |
| chr1 | 211871819 | 211872254 | 3.20646 | 4.11781 | 1.284 |
| chr1 | 237120435 | 237120805 | 0.939577 | 1.20665 | 1.284 |
| chr11 | 48194332 | 48194739 | 3.30129 | 4.24031 | 1.284 |
| chr1 | 223348425 | 223348693 | 1.47359 | 1.8929 | 1.285 |
| chr15 | 50437689 | 50437910 | 1.25459 | 1.6116 | 1.285 |
| chr11 | 60530422 | 60530882 | 0.793838 | 1.01976 | 1.285 |
| chr10 | 56513037 | 56513259 | 0.688791 | 0.885071 | 1.285 |
| chr11 | 38000246 | 38000573 | 0.743399 | 0.95524 | 1.285 |
| chr2 | 72377537 | 72378152 | 1.48164 | 1.90392 | 1.285 |
| chr5 | 12347534 | 12347790 | 0.670377 | 0.861454 | 1.285 |
| chr10 | 113875642 | 113875899 | 1.57799 | 2.02795 | 1.285 |
| chr1 | 158173601 | 158174213 | 1.26427 | 1.62514 | 1.285 |
| chr20 | 18568495 | 18568789 | 2.15167 | 2.76592 | 1.285 |
| chr5 | 156454107 | 156454357 | 2.14567 | 2.75839 | 1.286 |
| chr1 | 90511636 | 90511875 | 1.49281 | 1.91918 | 1.286 |
| chr5 | 172614154 | 172614796 | 1.18329 | 1.52126 | 1.286 |
| chr2 | 1906234 | 1906504 | 0.804068 | 1.03376 | 1.286 |
| chr3 | 124211686 | 124211935 | 1.76877 | 2.27414 | 1.286 |
| chr6 | 157073688 | 157074044 | 1.1641 | 1.49676 | 1.286 |
| chr1 | 209800878 | 209801407 | 2.17923 | 2.80214 | 1.286 |
| chr9 | 108480759 | 108481013 | 1.29181 | 1.6615 | 1.286 |
| chr2 | 240065500 | 240066124 | 1.34988 | 1.7363 | 1.286 |
| chr10 | 131433449 | 131433693 | 0.619846 | 0.797319 | 1.286 |

| | | | | | |
|---|---|---|---|---|---|
| chr10 | 115079948 | 115080168 | 0.734388 | 0.944816 | 1.287 |
| chr7 | 42546020 | 42546385 | 0.764137 | 0.983299 | 1.287 |
| chr8 | 35371747 | 35372002 | 0.673444 | 0.866888 | 1.287 |
| chrX | 106268622 | 106269141 | 1.34567 | 1.73221 | 1.287 |
| chr5 | 109338253 | 109338705 | 0.674411 | 0.868155 | 1.287 |
| chr8 | 25434786 | 25435010 | 1.86288 | 2.39809 | 1.287 |
| chr20 | 2733406 | 2733751 | 3.40554 | 4.38413 | 1.287 |
| chr14 | 55162038 | 55162394 | 0.640685 | 0.824798 | 1.287 |
| chr8 | 77280088 | 77280359 | 2.43286 | 3.13205 | 1.287 |
| chr2 | 179261847 | 179262112 | 1.1689 | 1.50488 | 1.287 |
| chr5 | 104446691 | 104447106 | 0.921153 | 1.18606 | 1.288 |
| chr21 | 42539696 | 42539999 | 1.39794 | 1.79998 | 1.288 |
| chr11 | 100552951 | 100553243 | 0.933591 | 1.20213 | 1.288 |
| chrX | 33452524 | 33452915 | 1.32359 | 1.70434 | 1.288 |
| chr14 | 82032340 | 82032539 | 1.53575 | 1.97754 | 1.288 |
| chr14 | 37699828 | 37700172 | 1.74548 | 2.24777 | 1.288 |
| chr6 | 116765372 | 116765577 | 1.26806 | 1.63318 | 1.288 |
| chr5 | 148645176 | 148645420 | 1.45866 | 1.87872 | 1.288 |
| chr5 | 138587059 | 138587542 | 0.960212 | 1.23682 | 1.288 |
| chr3 | 18592559 | 18592794 | 0.703742 | 0.906515 | 1.288 |
| chr8 | 137469279 | 137469500 | 0.653848 | 0.842251 | 1.288 |
| chr1 | 51655177 | 51655429 | 2.04236 | 2.63114 | 1.288 |
| chr16 | 80858673 | 80859011 | 1.63341 | 2.10429 | 1.288 |
| chr8 | 85681533 | 85681799 | 0.567948 | 0.731698 | 1.288 |
| chr2 | 86305972 | 86306185 | 2.17343 | 2.80009 | 1.288 |
| chr2 | 11722802 | 11723229 | 2.33511 | 3.00852 | 1.288 |
| chrX | 68251153 | 68251434 | 0.914781 | 1.17865 | 1.288 |
| chr20 | 14035864 | 14036082 | 2.07819 | 2.67774 | 1.289 |
| chr20 | 61146810 | 61147052 | 1.47894 | 1.90572 | 1.289 |
| chr1 | 226252545 | 226252808 | 1.54231 | 1.98755 | 1.289 |
| chr15 | 23807224 | 23807467 | 0.976438 | 1.2585 | 1.289 |
| chr22 | 23296929 | 23297154 | 1.66869 | 2.15074 | 1.289 |
| chr10 | 110338846 | 110339088 | 0.606555 | 0.781863 | 1.289 |
| chr9 | 114875315 | 114875547 | 2.20497 | 2.84237 | 1.289 |
| chr14 | 65203185 | 65203394 | 0.607452 | 0.783178 | 1.289 |
| chr2 | 25427078 | 25427669 | 0.880408 | 1.13509 | 1.289 |
| chr14 | 82656205 | 82656528 | 0.602455 | 0.776737 | 1.289 |
| chr20 | 21587428 | 21588030 | 0.688561 | 0.887862 | 1.289 |
| chr14 | 71284378 | 71284775 | 3.08481 | 3.97777 | 1.289 |
| chr6 | 6697179 | 6697492 | 2.14229 | 2.7625 | 1.290 |
| chr5 | 133710315 | 133710589 | 0.969851 | 1.25066 | 1.290 |
| chr18 | 21498412 | 21498778 | 1.81882 | 2.34557 | 1.290 |
| chr3 | 157575956 | 157576403 | 0.942437 | 1.21543 | 1.290 |
| chr9 | 80048560 | 80048954 | 3.83156 | 4.94249 | 1.290 |
| chr22 | 46527008 | 46527314 | 1.9075 | 2.46069 | 1.290 |
| chr5 | 165181540 | 165181783 | 0.933228 | 1.20388 | 1.290 |
| chr7 | 128803834 | 128804184 | 1.72369 | 2.22369 | 1.290 |
| chr20 | 47252235 | 47252815 | 2.3115 | 2.9822 | 1.290 |
| chr12 | 30154937 | 30155132 | 0.901507 | 1.1632 | 1.290 |
| chr16 | 81175096 | 81175310 | 1.21859 | 1.57232 | 1.290 |
| chr11 | 66743393 | 66743734 | 1.977 | 2.55094 | 1.290 |
| chr2 | 167038032 | 167038358 | 1.3186 | 1.7014 | 1.290 |
| chr12 | 107312565 | 107312897 | 2.85515 | 3.68427 | 1.290 |
| chr18 | 32966535 | 32966866 | 2.93747 | 3.79065 | 1.290 |
| chr14 | 66868231 | 66868527 | 1.22023 | 1.57479 | 1.291 |
| chr2 | 50237703 | 50237937 | 0.73569 | 0.949483 | 1.291 |
| chr8 | 41261623 | 41261940 | 2.42399 | 3.12842 | 1.291 |
| chr16 | 28884945 | 28885331 | 2.29282 | 2.95919 | 1.291 |
| chr12 | 65468674 | 65469059 | 1.13204 | 1.46109 | 1.291 |
| chr16 | 80351735 | 80351939 | 1.31955 | 1.7031 | 1.291 |
| chr3 | 118347590 | 118347988 | 2.2539 | 2.90903 | 1.291 |

| | | | | | |
|---|---|---|---|---|---|
| chr8 | 36362013 | 36362221 | 0.679852 | 0.877502 | 1.291 |
| chr1 | 216724228 | 216724458 | 0.98665 | 1.27352 | 1.291 |
| chr12 | 104165275 | 104165559 | 2.5985 | 3.35405 | 1.291 |
| chr14 | 42394734 | 42394977 | 1.2302 | 1.58791 | 1.291 |
| chr15 | 90193087 | 90193290 | 0.673578 | 0.869511 | 1.291 |
| chr2 | 33597806 | 33598334 | 1.38224 | 1.78432 | 1.291 |
| chr9 | 80745000 | 80745366 | 1.23353 | 1.59264 | 1.291 |
| chr22 | 25348001 | 25348428 | 2.29306 | 2.96064 | 1.291 |
| chr1 | 156554635 | 156554891 | 1.68296 | 2.17295 | 1.291 |
| chr8 | 563672 | 564004 | 0.803136 | 1.03701 | 1.291 |
| chr11 | 62379838 | 62380261 | 3.84291 | 4.96207 | 1.291 |
| chr11 | 63535607 | 63536053 | 2.98762 | 3.85772 | 1.291 |
| chr10 | 124325212 | 124325514 | 2.00456 | 2.5884 | 1.291 |
| chr7 | 84670962 | 84671214 | 1.66232 | 2.14652 | 1.291 |
| chr8 | 9606683 | 9606955 | 1.43288 | 1.85025 | 1.291 |
| chr8 | 128412946 | 128413308 | 2.40347 | 3.10369 | 1.291 |
| chrX | 47942716 | 47942922 | 1.88977 | 2.44051 | 1.291 |
| chr3 | 39314752 | 39314969 | 0.680994 | 0.879528 | 1.292 |
| chr21 | 34747060 | 34747287 | 1.35759 | 1.7534 | 1.292 |
| chr12 | 72492403 | 72492743 | 0.81607 | 1.054 | 1.292 |
| chr17 | 15074784 | 15075038 | 2.99536 | 3.8687 | 1.292 |
| chr12 | 104571457 | 104571721 | 1.18751 | 1.53377 | 1.292 |
| chr8 | 120276115 | 120276534 | 1.58248 | 2.0439 | 1.292 |
| chrX | 15489179 | 15489413 | 1.78027 | 2.29946 | 1.292 |
| chr7 | 96656896 | 96657320 | 2.05766 | 2.65778 | 1.292 |
| chr4 | 2221684 | 2222159 | 1.64517 | 2.12535 | 1.292 |
| chr19 | 31097675 | 31098018 | 0.951089 | 1.22897 | 1.292 |
| chr2 | 28715888 | 28716116 | 1.18147 | 1.52667 | 1.292 |
| chr2 | 210825819 | 210826045 | 1.69984 | 2.19652 | 1.292 |
| chr15 | 50344104 | 50344332 | 2.11743 | 2.73614 | 1.292 |
| chr16 | 30080233 | 30080434 | 1.52952 | 1.9766 | 1.292 |
| chr14 | 23991614 | 23991884 | 0.545193 | 0.704568 | 1.292 |
| chr10 | 57260784 | 57261017 | 0.931928 | 1.20439 | 1.292 |
| chr19 | 55803357 | 55803585 | 1.29856 | 1.67828 | 1.292 |
| chr8 | 72211191 | 72211414 | 0.651057 | 0.841441 | 1.292 |
| chr15 | 27707290 | 27707524 | 0.834788 | 1.07904 | 1.293 |
| chr1 | 154158634 | 154158860 | 1.8297 | 2.36508 | 1.293 |
| chr3 | 27509132 | 27509340 | 1.17548 | 1.51951 | 1.293 |
| chr8 | 80125912 | 80126146 | 0.858636 | 1.10995 | 1.293 |
| chr11 | 925495 | 925899 | 2.88182 | 3.72536 | 1.293 |
| chr9 | 113385801 | 113386017 | 1.81805 | 2.35025 | 1.293 |
| chr8 | 69012742 | 69013077 | 0.637029 | 0.823582 | 1.293 |
| chr14 | 91159854 | 91160317 | 0.724177 | 0.93626 | 1.293 |
| chr13 | 27521030 | 27521432 | 1.90604 | 2.46435 | 1.293 |
| chr19 | 49609846 | 49610420 | 3.22501 | 4.16991 | 1.293 |
| chr9 | 129540127 | 129540334 | 1.14554 | 1.48122 | 1.293 |
| chr3 | 69459454 | 69459835 | 1.87499 | 2.42448 | 1.293 |
| chr10 | 10138241 | 10138477 | 0.614422 | 0.794498 | 1.293 |
| chr6 | 16183407 | 16183613 | 1.28533 | 1.66208 | 1.293 |
| chr10 | 30928875 | 30929243 | 2.56226 | 3.31358 | 1.293 |
| chr8 | 21924191 | 21924756 | 2.50842 | 3.24396 | 1.293 |
| chr14 | 32013283 | 32013622 | 0.934673 | 1.20877 | 1.293 |
| chr3 | 64519315 | 64519626 | 1.56857 | 2.02859 | 1.293 |
| chrX | 110811706 | 110811918 | 1.00537 | 1.30022 | 1.293 |
| chr14 | 97934456 | 97934666 | 0.917414 | 1.18648 | 1.293 |
| chr1 | 191773417 | 191773673 | 0.947507 | 1.22541 | 1.293 |
| chr5 | 166468781 | 166469059 | 0.933349 | 1.20712 | 1.293 |
| chr1 | 44396243 | 44396595 | 1.19976 | 1.55177 | 1.293 |
| chr7 | 3976419 | 3976732 | 0.859673 | 1.11192 | 1.293 |
| chr6 | 13195171 | 13195454 | 1.55069 | 2.00587 | 1.294 |
| chr21 | 43537506 | 43537882 | 1.81558 | 2.34853 | 1.294 |

| | | | | | |
|---|---|---|---|---|---|
| chr10 | 49808898 | 49809145 | 0.784484 | 1.01477 | 1.294 |
| chr3 | 145231535 | 145231885 | 1.07649 | 1.39254 | 1.294 |
| chr9 | 94712341 | 94712910 | 1.5262 | 1.9744 | 1.294 |
| chr7 | 146933226 | 146933445 | 0.571601 | 0.739479 | 1.294 |
| chr16 | 10283954 | 10284176 | 0.604302 | 0.781798 | 1.294 |
| chr17 | 54352460 | 54352744 | 0.775257 | 1.00301 | 1.294 |
| chr19 | 917687 | 918014 | 1.53859 | 1.99063 | 1.294 |
| chr16 | 22329807 | 22330347 | 1.82694 | 2.36386 | 1.294 |
| chr21 | 15625977 | 15626185 | 1.12997 | 1.46206 | 1.294 |
| chrX | 49518922 | 49519147 | 2.98621 | 3.86419 | 1.294 |
| chr1 | 152319635 | 152319948 | 2.08055 | 2.69247 | 1.294 |
| chr2 | 50059661 | 50059942 | 0.590812 | 0.764577 | 1.294 |
| chr17 | 9128835 | 9129220 | 0.830795 | 1.07516 | 1.294 |
| chr12 | 63813975 | 63814222 | 1.7378 | 2.24903 | 1.294 |
| chr14 | 72939506 | 72939976 | 0.865054 | 1.11956 | 1.294 |
| chr9 | 30582364 | 30582646 | 0.86439 | 1.11881 | 1.294 |
| chr5 | 61423649 | 61423871 | 0.585548 | 0.757904 | 1.294 |
| chr1 | 203185432 | 203185621 | 1.80724 | 2.33933 | 1.294 |
| chr17 | 19417246 | 19417577 | 2.61687 | 3.38751 | 1.294 |
| chr1 | 203008364 | 203008858 | 1.95124 | 2.52607 | 1.295 |
| chr3 | 192534181 | 192534401 | 1.38644 | 1.79491 | 1.295 |
| chr15 | 62201430 | 62201728 | 0.842635 | 1.09098 | 1.295 |
| chr1 | 165113929 | 165114545 | 1.42791 | 1.84887 | 1.295 |
| chr1 | 226672993 | 226673240 | 0.958273 | 1.24085 | 1.295 |
| chr3 | 11394902 | 11395111 | 0.63453 | 0.821675 | 1.295 |
| chr8 | 51392099 | 51392386 | 0.64709 | 0.837939 | 1.295 |
| chr2 | 69947747 | 69948041 | 1.17393 | 1.5203 | 1.295 |
| chr17 | 80194148 | 80194654 | 2.75616 | 3.56959 | 1.295 |
| chr7 | 131491006 | 131491216 | 0.550343 | 0.712767 | 1.295 |
| chr8 | 140107939 | 140108157 | 0.686024 | 0.888487 | 1.295 |
| chr10 | 116652080 | 116652297 | 0.753397 | 0.975764 | 1.295 |
| chr10 | 132985991 | 132986254 | 0.788325 | 1.02103 | 1.295 |
| chr19 | 7419317 | 7419635 | 2.07695 | 2.69011 | 1.295 |
| chr7 | 44627753 | 44628053 | 1.67311 | 2.16705 | 1.295 |
| chr7 | 135020499 | 135020704 | 1.71629 | 2.22313 | 1.295 |
| chr8 | 107439593 | 107439843 | 1.78777 | 2.31589 | 1.295 |
| chr9 | 35756532 | 35756882 | 2.29846 | 2.97749 | 1.295 |
| chr1 | 26612303 | 26612631 | 1.48247 | 1.92049 | 1.295 |
| chr16 | 19692437 | 19692705 | 1.46713 | 1.90065 | 1.295 |
| chr11 | 27914276 | 27914505 | 0.637338 | 0.825681 | 1.296 |
| chr9 | 30715242 | 30715446 | 0.595097 | 0.770961 | 1.296 |
| chr2 | 50154053 | 50154255 | 0.968131 | 1.2543 | 1.296 |
| chr12 | 72496054 | 72496320 | 0.586727 | 0.76021 | 1.296 |
| chr17 | 69469234 | 69469685 | 1.72457 | 2.2347 | 1.296 |
| chr8 | 75000424 | 75000811 | 1.49074 | 1.93176 | 1.296 |
| chr9 | 94443907 | 94444336 | 2.63099 | 3.4094 | 1.296 |
| chr1 | 224246110 | 224246403 | 0.69567 | 0.901544 | 1.296 |
| chr1 | 110630742 | 110631045 | 1.78333 | 2.31111 | 1.296 |
| chr19 | 38393264 | 38393586 | 2.82928 | 3.66691 | 1.296 |
| chr10 | 108254856 | 108255138 | 0.724043 | 0.938471 | 1.296 |
| chr6 | 165462958 | 165463146 | 1.406 | 1.82251 | 1.296 |
| chr5 | 161701660 | 161701885 | 0.951753 | 1.23379 | 1.296 |
| chr10 | 98081413 | 98081752 | 0.854916 | 1.10828 | 1.296 |
| chr2 | 178976888 | 178977227 | 2.71898 | 3.52496 | 1.296 |
| chr2 | 105525937 | 105526169 | 0.755079 | 0.978922 | 1.296 |
| chr1 | 167099369 | 167099579 | 0.707737 | 0.917597 | 1.297 |
| chr1 | 200885383 | 200885705 | 1.74385 | 2.26102 | 1.297 |
| chr7 | 104621790 | 104621988 | 1.98746 | 2.57692 | 1.297 |
| chr2 | 109252734 | 109253184 | 2.60122 | 3.37302 | 1.297 |
| chr20 | 13934001 | 13934203 | 1.07117 | 1.38905 | 1.297 |
| chr20 | 47237007 | 47237925 | 2.47366 | 3.20827 | 1.297 |

| | | | | | |
|---|---|---|---|---|---|
| chr1 | 168312837 | 168313108 | 1.01338 | 1.3144 | 1.297 |
| chr2 | 72552872 | 72553317 | 1.78805 | 2.31924 | 1.297 |
| chr3 | 122977759 | 122977995 | 1.6861 | 2.18707 | 1.297 |
| chr12 | 22969751 | 22969955 | 0.903836 | 1.17241 | 1.297 |
| chr15 | 81921986 | 81922333 | 0.662031 | 0.858811 | 1.297 |
| chr14 | 36349048 | 36349378 | 0.858909 | 1.11423 | 1.297 |
| chr7 | 38147076 | 38147351 | 2.48146 | 3.2192 | 1.297 |
| chr1 | 240656104 | 240656312 | 1.2777 | 1.65759 | 1.297 |
| chr14 | 94646939 | 94647249 | 0.971182 | 1.25995 | 1.297 |
| chr12 | 67082696 | 67083142 | 1.64287 | 2.13175 | 1.298 |
| chr8 | 126924230 | 126924460 | 2.07212 | 2.68883 | 1.298 |
| chr15 | 51942144 | 51942448 | 1.57181 | 2.03964 | 1.298 |
| chr10 | 58815000 | 58815290 | 0.710904 | 0.922539 | 1.298 |
| chr1 | 94279576 | 94279946 | 3.3278 | 4.31862 | 1.298 |
| chr5 | 116201558 | 116201773 | 0.877268 | 1.1385 | 1.298 |
| chr3 | 56481077 | 56481428 | 1.41263 | 1.83345 | 1.298 |
| chr11 | 82266806 | 82267124 | 0.760279 | 0.986802 | 1.298 |
| chr3 | 164903551 | 164903838 | 1.81825 | 2.36002 | 1.298 |
| chr22 | 22777030 | 22777373 | 1.926 | 2.5 | 1.298 |
| chr7 | 107754620 | 107754899 | 1.78076 | 2.31165 | 1.298 |
| chr17 | 5633696 | 5634232 | 0.85105 | 1.10489 | 1.298 |
| chr15 | 62418023 | 62418253 | 1.48783 | 1.93163 | 1.298 |
| chr7 | 118598433 | 118598656 | 0.885414 | 1.14955 | 1.298 |
| chr5 | 12889520 | 12889826 | 0.688348 | 0.893704 | 1.298 |
| chr2 | 181923353 | 181923651 | 1.27245 | 1.65231 | 1.299 |
| chr15 | 26028792 | 26029221 | 1.11706 | 1.45056 | 1.299 |
| chr17 | 26265742 | 26265951 | 0.724548 | 0.940961 | 1.299 |
| chr5 | 37006133 | 37006354 | 1.65688 | 2.15181 | 1.299 |
| chr16 | 68400992 | 68401304 | 2.13121 | 2.76796 | 1.299 |
| chr2 | 18559378 | 18559666 | 1.96386 | 2.55108 | 1.299 |
| chr2 | 25809300 | 25809507 | 0.65017 | 0.844588 | 1.299 |
| chr2 | 7911967 | 7912183 | 0.684213 | 0.888825 | 1.299 |
| chr5 | 4902653 | 4902894 | 0.57985 | 0.75331 | 1.299 |
| chr12 | 63462745 | 63462935 | 1.57744 | 2.04962 | 1.299 |
| chr2 | 211654693 | 211654901 | 0.965807 | 1.2549 | 1.299 |
| chr8 | 142941447 | 142941805 | 0.625592 | 0.812848 | 1.299 |
| chr8 | 52831618 | 52831864 | 0.77948 | 1.01281 | 1.299 |
| chr1 | 63614851 | 63615205 | 1.11437 | 1.44801 | 1.299 |
| chr7 | 44573987 | 44574495 | 1.64241 | 2.13417 | 1.299 |
| chr8 | 92586120 | 92586348 | 0.743573 | 0.966226 | 1.299 |
| chr11 | 78356243 | 78356563 | 2.57687 | 3.34893 | 1.300 |
| chr3 | 39333208 | 39333712 | 2.23542 | 2.90522 | 1.300 |
| chr19 | 13204589 | 13204913 | 2.67796 | 3.48069 | 1.300 |
| chr10 | 81134286 | 81134717 | 2.81824 | 3.66311 | 1.300 |
| chr14 | 88149344 | 88149567 | 0.839232 | 1.09083 | 1.300 |
| chr8 | 84312601 | 84312897 | 0.950577 | 1.23567 | 1.300 |
| chr11 | 83739597 | 83739868 | 0.721839 | 0.93842 | 1.300 |
| chr1 | 204445782 | 204445991 | 1.08528 | 1.41093 | 1.300 |
| chr1 | 39568236 | 39568482 | 1.73634 | 2.25744 | 1.300 |
| chr16 | 8389947 | 8390151 | 0.626983 | 0.815214 | 1.300 |
| chr11 | 84258698 | 84258900 | 0.639902 | 0.832052 | 1.300 |
| chr3 | 146767586 | 146768037 | 0.713727 | 0.928069 | 1.300 |
| chr20 | 6588729 | 6589055 | 0.749105 | 0.974224 | 1.301 |
| chr7 | 30785871 | 30786170 | 1.0582 | 1.37626 | 1.301 |
| chr5 | 99019194 | 99019403 | 0.816392 | 1.06183 | 1.301 |
| chr12 | 678941 | 679230 | 2.74361 | 3.56878 | 1.301 |
| chr19 | 45841949 | 45842206 | 2.52267 | 3.28147 | 1.301 |
| chr17 | 33070811 | 33071255 | 0.782915 | 1.01845 | 1.301 |
| chr8 | 15042033 | 15042236 | 0.882867 | 1.14847 | 1.301 |
| chr14 | 96388793 | 96389025 | 0.764966 | 0.995114 | 1.301 |
| chr1 | 205493425 | 205493970 | 1.39159 | 1.81037 | 1.301 |

| | | | | | |
|---|---|---|---|---|---|
| chr3 | 183967410 | 183967692 | 1.12741 | 1.46671 | 1.301 |
| chr4 | 3445030 | 3445593 | 1.99125 | 2.59065 | 1.301 |
| chr19 | 58850578 | 58850767 | 1.07783 | 1.40251 | 1.301 |
| chrX | 129162162 | 129162551 | 2.67946 | 3.48663 | 1.301 |
| chr1 | 78580120 | 78580355 | 1.68631 | 2.19454 | 1.301 |
| chr5 | 32093254 | 32093655 | 2.02568 | 2.6365 | 1.302 |
| chr6 | 56460271 | 56460615 | 1.39079 | 1.8104 | 1.302 |
| chr11 | 82416837 | 82417237 | 0.928759 | 1.20901 | 1.302 |
| chr5 | 36507509 | 36507756 | 2.04058 | 2.65633 | 1.302 |
| chr11 | 60542889 | 60543225 | 2.15194 | 2.80134 | 1.302 |
| chr14 | 70553891 | 70554146 | 1.75229 | 2.28143 | 1.302 |
| chr15 | 35370742 | 35371026 | 1.91946 | 2.4991 | 1.302 |
| chr20 | 459668 | 459890 | 1.46385 | 1.90612 | 1.302 |
| chr1 | 111091079 | 111091357 | 1.02722 | 1.33759 | 1.302 |
| chr1 | 18246455 | 18246644 | 1.16874 | 1.5219 | 1.302 |
| chr17 | 33696273 | 33696515 | 0.848992 | 1.1056 | 1.302 |
| chr3 | 122102973 | 122103193 | 1.47765 | 1.92437 | 1.302 |
| chr19 | 32965272 | 32965543 | 0.907426 | 1.1818 | 1.302 |
| chr18 | 23476139 | 23476476 | 1.53506 | 1.99948 | 1.303 |
| chr22 | 38259526 | 38259779 | 1.86047 | 2.42333 | 1.303 |
| chr20 | 49410189 | 49410438 | 1.70734 | 2.22403 | 1.303 |
| chr1 | 6614536 | 6614823 | 2.30376 | 3.00098 | 1.303 |
| chr10 | 133836589 | 133837046 | 2.42314 | 3.15647 | 1.303 |
| chr3 | 35544318 | 35544545 | 0.570375 | 0.743128 | 1.303 |
| chr16 | 57496375 | 57496662 | 2.8222 | 3.67708 | 1.303 |
| chr7 | 102688195 | 102688449 | 0.726846 | 0.947142 | 1.303 |
| chr17 | 48164677 | 48165193 | 2.12212 | 2.76533 | 1.303 |
| chr19 | 41859840 | 41860276 | 4.00311 | 5.21656 | 1.303 |
| chr14 | 106686763 | 106686974 | 0.719952 | 0.938239 | 1.303 |
| chr12 | 50367828 | 50368332 | 1.17908 | 1.53675 | 1.303 |
| chr9 | 115582180 | 115582438 | 1.54845 | 2.01823 | 1.303 |
| chr17 | 17936900 | 17937209 | 1.54558 | 2.01452 | 1.303 |
| chr20 | 30619010 | 30619665 | 2.51212 | 3.27434 | 1.303 |
| chr20 | 9304311 | 9304651 | 1.39624 | 1.81991 | 1.303 |
| chr15 | 60621061 | 60621627 | 1.75522 | 2.28814 | 1.304 |
| chr8 | 106920549 | 106920851 | 1.63086 | 2.12603 | 1.304 |
| chr8 | 145613952 | 145614170 | 0.48024 | 0.626056 | 1.304 |
| chr17 | 76741490 | 76741775 | 1.44219 | 1.88016 | 1.304 |
| chr3 | 116277554 | 116277798 | 0.711574 | 0.927698 | 1.304 |
| chr20 | 31262169 | 31262484 | 2.47474 | 3.22644 | 1.304 |
| chr8 | 87715561 | 87715995 | 1.98921 | 2.5935 | 1.304 |
| chr19 | 50414852 | 50415229 | 0.677393 | 0.88321 | 1.304 |
| chr9 | 35291392 | 35291688 | 0.720622 | 0.939589 | 1.304 |
| chr8 | 51185089 | 51185359 | 0.620157 | 0.808605 | 1.304 |
| chr2 | 153782309 | 153782730 | 1.89718 | 2.47372 | 1.304 |
| chr6 | 157197463 | 157197745 | 1.24189 | 1.6193 | 1.304 |
| chr4 | 72095404 | 72095738 | 1.20229 | 1.56769 | 1.304 |
| chrX | 11742542 | 11742884 | 1.59715 | 2.0826 | 1.304 |
| chr8 | 23177499 | 23177780 | 1.57222 | 2.05017 | 1.304 |
| chr2 | 219922746 | 219923098 | 2.46999 | 3.2209 | 1.304 |
| chr7 | 27729274 | 27729501 | 1.3459 | 1.75521 | 1.304 |
| chr1 | 211623862 | 211624108 | 0.645512 | 0.84183 | 1.304 |
| chr3 | 10376324 | 10376562 | 1.86684 | 2.43479 | 1.304 |
| chr6 | 18990694 | 18990913 | 1.46705 | 1.91337 | 1.304 |
| chr20 | 16741839 | 16742225 | 2.41444 | 3.14951 | 1.304 |
| chr3 | 103129121 | 103129418 | 1.24856 | 1.62883 | 1.305 |
| chr1 | 110745683 | 110745928 | 1.76111 | 2.29772 | 1.305 |
| chr12 | 23358184 | 23358425 | 1.56117 | 2.03737 | 1.305 |
| chr8 | 137342799 | 137343205 | 0.984038 | 1.28446 | 1.305 |
| chr3 | 160342232 | 160342437 | 0.736418 | 0.961269 | 1.305 |
| chr10 | 67686236 | 67686529 | 0.491712 | 0.641874 | 1.305 |

| | | | | | |
|---|---|---|---|---|---|
| chr3 | 21016899 | 21017146 | 0.553749 | 0.72287 | 1.305 |
| chr2 | 114723337 | 114723555 | 1.25304 | 1.63582 | 1.305 |
| chr5 | 148113167 | 148113460 | 1.78626 | 2.33196 | 1.306 |
| chr14 | 97033671 | 97034080 | 1.68797 | 2.20369 | 1.306 |
| chr1 | 192230830 | 192231094 | 0.698148 | 0.911476 | 1.306 |
| chr18 | 29086427 | 29086674 | 1.18037 | 1.54108 | 1.306 |
| chr6 | 161351645 | 161352080 | 1.72751 | 2.25558 | 1.306 |
| chr5 | 124903602 | 124903795 | 0.634188 | 0.828091 | 1.306 |
| chr13 | 112081339 | 112081971 | 0.935621 | 1.22172 | 1.306 |
| chr12 | 61052114 | 61052297 | 0.726446 | 0.948661 | 1.306 |
| chr18 | 9971371 | 9971628 | 1.43838 | 1.87859 | 1.306 |
| chr7 | 18385342 | 18385589 | 1.82857 | 2.38838 | 1.306 |
| chr8 | 26558601 | 26559109 | 1.38008 | 1.8026 | 1.306 |
| chr2 | 3698612 | 3698845 | 2.35745 | 3.07941 | 1.306 |
| chr10 | 96479203 | 96479510 | 0.644366 | 0.841716 | 1.306 |
| chr10 | 102974495 | 102974868 | 3.23654 | 4.22789 | 1.306 |
| chr5 | 112947714 | 112947965 | 1.18029 | 1.54182 | 1.306 |
| chr20 | 44427826 | 44428186 | 1.76681 | 2.30804 | 1.306 |
| chr10 | 87755614 | 87755874 | 1.42025 | 1.85538 | 1.306 |
| chr12 | 24589766 | 24590022 | 0.854438 | 1.11628 | 1.306 |
| chr6 | 65122951 | 65123154 | 0.715059 | 0.934186 | 1.306 |
| chr15 | 34640456 | 34640752 | 0.749603 | 0.979357 | 1.307 |
| chr3 | 115632029 | 115632419 | 1.06682 | 1.39385 | 1.307 |
| chr17 | 39942479 | 39942942 | 2.96215 | 3.87034 | 1.307 |
| chr18 | 43490412 | 43490787 | 2.05061 | 2.67938 | 1.307 |
| chr2 | 142938587 | 142939084 | 1.46271 | 1.9113 | 1.307 |
| chr17 | 37331120 | 37331580 | 2.81488 | 3.67876 | 1.307 |
| chr1 | 170898935 | 170899432 | 1.71372 | 2.23992 | 1.307 |
| chr6 | 150716610 | 150716873 | 1.72514 | 2.25511 | 1.307 |
| chr5 | 144876181 | 144876385 | 0.764893 | 0.999881 | 1.307 |
| chr16 | 55069058 | 55069295 | 1.37888 | 1.80251 | 1.307 |
| chr2 | 207029072 | 207029309 | 1.63143 | 2.13291 | 1.307 |
| chr2 | 10442591 | 10442895 | 2.15349 | 2.81549 | 1.307 |
| chr10 | 133672445 | 133672695 | 0.54936 | 0.718366 | 1.308 |
| chr12 | 128759581 | 128759854 | 0.81711 | 1.06854 | 1.308 |
| chr7 | 23312738 | 23312998 | 2.25337 | 2.94681 | 1.308 |
| chr5 | 28345045 | 28345266 | 1.26283 | 1.65147 | 1.308 |
| chr9 | 97889726 | 97890284 | 2.53851 | 3.32005 | 1.308 |
| chr1 | 110185954 | 110186270 | 2.22444 | 2.9093 | 1.308 |
| chr14 | 30596073 | 30596275 | 0.751074 | 0.982322 | 1.308 |
| chr16 | 50503975 | 50504221 | 0.727164 | 0.951087 | 1.308 |
| chr8 | 144136148 | 144136424 | 1.13391 | 1.48319 | 1.308 |
| chr1 | 204661409 | 204661785 | 1.15956 | 1.51679 | 1.308 |
| chr19 | 30221408 | 30221683 | 1.96047 | 2.56465 | 1.308 |
| chr16 | 47670215 | 47670553 | 1.08126 | 1.41452 | 1.308 |
| chr6 | 36692484 | 36692697 | 2.18627 | 2.86013 | 1.308 |
| chr20 | 19573822 | 19574111 | 0.808705 | 1.05803 | 1.308 |
| chr18 | 3246087 | 3246315 | 2.59295 | 3.39254 | 1.308 |
| chr8 | 76169565 | 76169806 | 2.00022 | 2.61703 | 1.308 |
| chr19 | 16552569 | 16552898 | 1.13425 | 1.48411 | 1.308 |
| chr4 | 71425047 | 71425293 | 1.29538 | 1.69506 | 1.309 |
| chr11 | 119227010 | 119227679 | 2.44051 | 3.19356 | 1.309 |
| chr7 | 95545916 | 95546446 | 1.86939 | 2.44647 | 1.309 |
| chr1 | 174084075 | 174084307 | 1.6625 | 2.17586 | 1.309 |
| chr8 | 133106796 | 133107004 | 0.908711 | 1.18932 | 1.309 |
| chr5 | 67344365 | 67344621 | 0.681162 | 0.891607 | 1.309 |
| chr19 | 29751459 | 29751728 | 1.58276 | 2.07191 | 1.309 |
| chr7 | 142551607 | 142552039 | 1.6306 | 2.13455 | 1.309 |
| chr7 | 94819825 | 94820067 | 1.14835 | 1.50335 | 1.309 |
| chr14 | 65689099 | 65689446 | 1.96629 | 2.57438 | 1.309 |
| chr2 | 217547237 | 217547514 | 2.26567 | 2.96686 | 1.309 |

| | | | | | |
|---|---|---|---|---|---|
| chr17 | 36917654 | 36918169 | 1.09736 | 1.4371 | 1.310 |
| chr2 | 26685377 | 26685687 | 2.1447 | 2.80892 | 1.310 |
| chr3 | 78499909 | 78500138 | 0.875465 | 1.1466 | 1.310 |
| chr1 | 176435706 | 176436026 | 0.958797 | 1.25576 | 1.310 |
| chr12 | 2195122 | 2195324 | 0.760708 | 0.99645 | 1.310 |
| chr5 | 110886191 | 110886534 | 1.3484 | 1.76629 | 1.310 |
| chr15 | 74924293 | 74924761 | 2.03195 | 2.66175 | 1.310 |
| chr5 | 84365722 | 84365954 | 0.819692 | 1.07388 | 1.310 |
| chr7 | 42425601 | 42425816 | 0.922438 | 1.20851 | 1.310 |
| chr18 | 47730411 | 47730640 | 1.96342 | 2.57239 | 1.310 |
| chr11 | 23373267 | 23373550 | 0.800384 | 1.04869 | 1.310 |
| chr3 | 9029119 | 9030017 | 1.3597 | 1.78152 | 1.310 |
| chr12 | 116309477 | 116310039 | 1.13883 | 1.49226 | 1.310 |
| chr16 | 55498921 | 55499426 | 2.72976 | 3.57694 | 1.310 |
| chr18 | 20379100 | 20379321 | 1.20073 | 1.57366 | 1.311 |
| chr5 | 141258716 | 141259095 | 3.08486 | 4.04322 | 1.311 |
| chr7 | 137253063 | 137253374 | 0.844041 | 1.10629 | 1.311 |
| chr2 | 61408908 | 61409138 | 0.620119 | 0.81282 | 1.311 |
| chr10 | 95112442 | 95112660 | 1.47546 | 1.93401 | 1.311 |
| chr2 | 21273447 | 21273684 | 0.738851 | 0.968586 | 1.311 |
| chr7 | 147841979 | 147842215 | 0.732744 | 0.960659 | 1.311 |
| chr12 | 6304823 | 6305166 | 2.90124 | 3.80372 | 1.311 |
| chr7 | 50549659 | 50549864 | 1.06135 | 1.39153 | 1.311 |
| chr3 | 131513682 | 131513955 | 1.32763 | 1.74071 | 1.311 |
| chrX | 80451161 | 80451515 | 1.14539 | 1.50197 | 1.311 |
| chr7 | 140346769 | 140347201 | 2.30049 | 3.01682 | 1.311 |
| chr1 | 88527269 | 88527471 | 1.7463 | 2.29009 | 1.311 |
| chr2 | 89131293 | 89131590 | 1.38088 | 1.81099 | 1.311 |
| chr10 | 64583576 | 64583782 | 0.779019 | 1.02172 | 1.312 |
| chr3 | 128157306 | 128157610 | 1.18161 | 1.54974 | 1.312 |
| chr22 | 23301415 | 23301712 | 2.17256 | 2.84951 | 1.312 |
| chr1 | 110182862 | 110183254 | 2.28244 | 2.99365 | 1.312 |
| chr20 | 56158103 | 56158534 | 1.84132 | 2.41514 | 1.312 |
| chr13 | 88321686 | 88321935 | 2.32473 | 3.04923 | 1.312 |
| chr5 | 12373190 | 12373527 | 0.683748 | 0.896861 | 1.312 |
| chr7 | 45013412 | 45013775 | 1.24039 | 1.62723 | 1.312 |
| chr1 | 29203747 | 29204046 | 2.00616 | 2.63184 | 1.312 |
| chr15 | 79320718 | 79321078 | 2.14314 | 2.81166 | 1.312 |
| chr16 | 71446107 | 71446924 | 1.99229 | 2.61406 | 1.312 |
| chr5 | 176736652 | 176737102 | 2.82926 | 3.71226 | 1.312 |
| chr15 | 88094324 | 88094573 | 0.704687 | 0.924688 | 1.312 |
| chr19 | 43046864 | 43047304 | 1.06531 | 1.39806 | 1.312 |
| chr2 | 178128233 | 178128578 | 2.27606 | 2.9874 | 1.313 |
| chr3 | 142705847 | 142706258 | 1.39856 | 1.83567 | 1.313 |
| chr14 | 105748184 | 105748499 | 2.34766 | 3.08286 | 1.313 |
| chr3 | 160991439 | 160991779 | 1.25032 | 1.64195 | 1.313 |
| chr8 | 89478111 | 89478359 | 0.702346 | 0.922402 | 1.313 |
| chr15 | 79320023 | 79320302 | 0.681418 | 0.894924 | 1.313 |
| chr8 | 135189260 | 135189785 | 0.773941 | 1.01644 | 1.313 |
| chr15 | 29382523 | 29383120 | 2.06156 | 2.70835 | 1.314 |
| chr2 | 233232300 | 233232485 | 1.29335 | 1.69912 | 1.314 |
| chr20 | 62084265 | 62084599 | 0.891705 | 1.17149 | 1.314 |
| chr2 | 13025204 | 13025437 | 0.544213 | 0.714982 | 1.314 |
| chr7 | 39781892 | 39782197 | 1.24405 | 1.63445 | 1.314 |
| chr1 | 232951088 | 232951380 | 0.744415 | 0.978067 | 1.314 |
| chr15 | 79510959 | 79511303 | 1.18582 | 1.5581 | 1.314 |
| chr14 | 59974088 | 59974386 | 2.03342 | 2.67198 | 1.314 |
| chr15 | 59753527 | 59753798 | 0.787321 | 1.03465 | 1.314 |
| chr11 | 80230936 | 80231156 | 0.856736 | 1.12596 | 1.314 |
| chr9 | 2006468 | 2006710 | 0.739416 | 0.971797 | 1.314 |
| chrX | 135291453 | 135291852 | 1.55666 | 2.04593 | 1.314 |

| | | | | | |
|---|---|---|---|---|---|
| chr5 | 28670483 | 28670750 | 0.802011 | 1.05415 | 1.314 |
| chr9 | 130248470 | 130248920 | 0.791989 | 1.041 | 1.314 |
| chr1 | 157133347 | 157133848 | 2.75567 | 3.6222 | 1.314 |
| chr14 | 67891380 | 67891852 | 0.594479 | 0.781459 | 1.315 |
| chr5 | 90005248 | 90005475 | 1.75506 | 2.3071 | 1.315 |
| chr6 | 163591418 | 163591652 | 1.51861 | 1.9964 | 1.315 |
| chr7 | 27985009 | 27985225 | 0.628038 | 0.825666 | 1.315 |
| chr12 | 114340850 | 114341091 | 0.714029 | 0.938737 | 1.315 |
| chr18 | 34152214 | 34152466 | 1.09124 | 1.4347 | 1.315 |
| chr20 | 24761996 | 24762445 | 1.99352 | 2.62096 | 1.315 |
| chr12 | 100798574 | 100798880 | 1.50801 | 1.98266 | 1.315 |
| chr10 | 127702616 | 127702828 | 0.969488 | 1.27475 | 1.315 |
| chr10 | 98402768 | 98403042 | 1.77259 | 2.33082 | 1.315 |
| chr8 | 118875807 | 118876012 | 1.3684 | 1.79941 | 1.315 |
| chr11 | 72853332 | 72853623 | 2.03949 | 2.68193 | 1.315 |
| chr8 | 27849986 | 27850394 | 1.6813 | 2.21098 | 1.315 |
| chr10 | 109398198 | 109398407 | 0.727965 | 0.957363 | 1.315 |
| chr11 | 3207468 | 3207687 | 0.908344 | 1.19459 | 1.315 |
| chr2 | 43804491 | 43804747 | 2.03824 | 2.6806 | 1.315 |
| chr12 | 128328016 | 128328458 | 0.803711 | 1.05702 | 1.315 |
| chr19 | 42076806 | 42077395 | 3.18865 | 4.19402 | 1.315 |
| chr5 | 7593046 | 7593455 | 0.89341 | 1.1751 | 1.315 |
| chr10 | 13922610 | 13922858 | 0.586771 | 0.771787 | 1.315 |
| chr3 | 10875051 | 10875291 | 0.4863 | 0.639644 | 1.315 |
| chr20 | 32241032 | 32241341 | 2.49959 | 3.28791 | 1.315 |
| chr1 | 196287133 | 196287383 | 1.64109 | 2.15874 | 1.315 |
| chr7 | 19627069 | 19627319 | 0.639216 | 0.840853 | 1.315 |
| chr2 | 220374443 | 220374773 | 2.7253 | 3.58516 | 1.316 |
| chr16 | 3083747 | 3083978 | 1.01951 | 1.34119 | 1.316 |
| chr8 | 15501726 | 15501935 | 1.23969 | 1.63086 | 1.316 |
| chr17 | 75737218 | 75737535 | 1.59501 | 2.09837 | 1.316 |
| chr14 | 58167359 | 58167584 | 0.768269 | 1.01076 | 1.316 |
| chr19 | 29085251 | 29085948 | 0.902388 | 1.18721 | 1.316 |
| chr8 | 2254624 | 2254857 | 0.78891 | 1.03793 | 1.316 |
| chr7 | 44280573 | 44280923 | 2.32139 | 3.05432 | 1.316 |
| chr1 | 117360232 | 117360595 | 1.81237 | 2.38487 | 1.316 |
| chr5 | 31819392 | 31819635 | 0.619499 | 0.815243 | 1.316 |
| chr5 | 71819748 | 71820064 | 1.09378 | 1.43941 | 1.316 |
| chr3 | 154189009 | 154189236 | 0.647233 | 0.851799 | 1.316 |
| chr19 | 47808494 | 47808908 | 3.22826 | 4.24888 | 1.316 |
| chr8 | 66330672 | 66330881 | 0.749813 | 0.986872 | 1.316 |
| chr8 | 144804547 | 144805046 | 1.3918 | 1.83186 | 1.316 |
| chr20 | 22319807 | 22320049 | 1.54329 | 2.03133 | 1.316 |
| chr2 | 13074454 | 13074822 | 0.793943 | 1.04502 | 1.316 |
| chr16 | 67687014 | 67687318 | 1.34109 | 1.76548 | 1.316 |
| chr2 | 81019172 | 81019407 | 0.760258 | 1.00086 | 1.316 |
| chr3 | 196913904 | 196914175 | 1.60817 | 2.11716 | 1.317 |
| chr15 | 78162783 | 78163110 | 2.03658 | 2.68122 | 1.317 |
| chr16 | 53979901 | 53980125 | 1.71981 | 2.26419 | 1.317 |
| chr3 | 52157168 | 52157504 | 2.64536 | 3.4831 | 1.317 |
| chr12 | 128748026 | 128748310 | 0.689861 | 0.908339 | 1.317 |
| chr5 | 120966262 | 120966505 | 0.729663 | 0.96078 | 1.317 |
| chr3 | 95119389 | 95119599 | 0.911176 | 1.1998 | 1.317 |
| chr8 | 99717874 | 99718058 | 1.7466 | 2.29985 | 1.317 |
| chr3 | 142623912 | 142624139 | 0.803571 | 1.05812 | 1.317 |
| chr19 | 32955233 | 32955593 | 1.41464 | 1.86286 | 1.317 |
| chr11 | 69519353 | 69519747 | 2.26092 | 2.97741 | 1.317 |
| chr10 | 104813712 | 104813913 | 1.8398 | 2.42313 | 1.317 |
| chr10 | 53940439 | 53940686 | 0.855549 | 1.12683 | 1.317 |
| chr13 | 82337892 | 82338199 | 0.507852 | 0.668959 | 1.317 |
| chr16 | 22197893 | 22198143 | 1.56395 | 2.06012 | 1.317 |

| | | | | | |
|---|---|---|---|---|---|
| chr3 | 32975646 | 32975954 | 1.86509 | 2.45682 | 1.317 |
| chr10 | 78536979 | 78537241 | 0.780208 | 1.02775 | 1.317 |
| chr7 | 42730572 | 42730945 | 1.11673 | 1.47109 | 1.317 |
| chr8 | 3969811 | 3970068 | 0.66587 | 0.877206 | 1.317 |
| chr10 | 104438510 | 104438952 | 3.28303 | 4.32511 | 1.317 |
| chr1 | 216305815 | 216306044 | 0.794121 | 1.04619 | 1.317 |
| chr8 | 99049383 | 99049604 | 2.75227 | 3.62608 | 1.317 |
| chr11 | 86650132 | 86650404 | 1.34444 | 1.77165 | 1.318 |
| chr14 | 100004767 | 100004991 | 1.21817 | 1.60545 | 1.318 |
| chr7 | 41980275 | 41980794 | 1.77018 | 2.33295 | 1.318 |
| chr17 | 79152879 | 79153166 | 0.540792 | 0.712784 | 1.318 |
| chr9 | 134646241 | 134646447 | 0.669575 | 0.882611 | 1.318 |
| chr5 | 107857613 | 107858027 | 1.2509 | 1.64897 | 1.318 |
| chr11 | 44914070 | 44914276 | 0.669901 | 0.883111 | 1.318 |
| chr8 | 5882740 | 5882970 | 0.637873 | 0.840954 | 1.318 |
| chr7 | 7438435 | 7438656 | 0.633281 | 0.834906 | 1.318 |
| chr19 | 39501854 | 39502071 | 0.995225 | 1.3121 | 1.318 |
| chr1 | 179542235 | 179542478 | 0.721353 | 0.951051 | 1.318 |
| chr20 | 44452825 | 44453123 | 0.955724 | 1.26006 | 1.318 |
| chr1 | 224721415 | 224721764 | 0.784176 | 1.0339 | 1.318 |
| chr15 | 95177247 | 95177623 | 0.895959 | 1.18131 | 1.318 |
| chr7 | 5214186 | 5214450 | 1.89056 | 2.4928 | 1.319 |
| chr2 | 103059791 | 103060269 | 1.33642 | 1.76232 | 1.319 |
| chr2 | 46088183 | 46088481 | 0.630804 | 0.831958 | 1.319 |
| chr3 | 6763096 | 6763330 | 0.658612 | 0.868646 | 1.319 |
| chr4 | 19989082 | 19989323 | 0.621719 | 0.81999 | 1.319 |
| chr19 | 17316124 | 17316389 | 1.87702 | 2.47569 | 1.319 |
| chr2 | 190331420 | 190331716 | 1.70795 | 2.25274 | 1.319 |
| chr9 | 94338806 | 94339019 | 0.5913 | 0.779908 | 1.319 |
| chr19 | 51184890 | 51185192 | 0.632048 | 0.833681 | 1.319 |
| chr17 | 31376807 | 31377647 | 1.55373 | 2.04942 | 1.319 |
| chr19 | 33889027 | 33889454 | 1.18806 | 1.56711 | 1.319 |
| chr7 | 23201449 | 23201750 | 0.80266 | 1.05879 | 1.319 |
| chr3 | 113655595 | 113655857 | 0.786642 | 1.03767 | 1.319 |
| chr4 | 135683260 | 135683482 | 1.02233 | 1.34858 | 1.319 |
| chr15 | 52068330 | 52068783 | 1.69543 | 2.23652 | 1.319 |
| chr5 | 4879757 | 4880219 | 3.09207 | 4.07908 | 1.319 |
| chr8 | 131106827 | 131107211 | 1.25494 | 1.6556 | 1.319 |
| chr5 | 96204543 | 96204927 | 2.52583 | 3.33256 | 1.319 |
| chr10 | 29734422 | 29734728 | 1.24263 | 1.63958 | 1.319 |
| chr10 | 80359904 | 80360147 | 0.847125 | 1.11778 | 1.320 |
| chr15 | 72077028 | 72077376 | 2.00066 | 2.64008 | 1.320 |
| chr5 | 174710972 | 174711257 | 0.52212 | 0.689026 | 1.320 |
| chr4 | 156275106 | 156275426 | 2.46992 | 3.25977 | 1.320 |
| chr5 | 136834902 | 136835167 | 1.94365 | 2.56549 | 1.320 |
| chrX | 17963296 | 17963506 | 1.6831 | 2.22158 | 1.320 |
| chr8 | 33589013 | 33589216 | 0.688309 | 0.908632 | 1.320 |
| chr20 | 37262254 | 37262736 | 2.2301 | 2.94409 | 1.320 |
| chr2 | 54643356 | 54643595 | 0.807713 | 1.06636 | 1.320 |
| chr3 | 82182122 | 82182596 | 1.59402 | 2.10462 | 1.320 |
| chr3 | 143604713 | 143605003 | 2.03246 | 2.68359 | 1.320 |
| chr10 | 2238734 | 2238966 | 0.877636 | 1.15882 | 1.320 |
| chr7 | 96574103 | 96574308 | 0.575225 | 0.759564 | 1.320 |
| chr17 | 44795395 | 44795680 | 0.942957 | 1.24515 | 1.320 |
| chr15 | 33314005 | 33314276 | 1.37156 | 1.81113 | 1.320 |
| chr16 | 71401322 | 71401618 | 2.42733 | 3.20579 | 1.321 |
| chr2 | 30420474 | 30420691 | 0.915277 | 1.20892 | 1.321 |
| chr20 | 6537120 | 6537370 | 0.826743 | 1.09199 | 1.321 |
| chr9 | 104849747 | 104850014 | 0.51147 | 0.675595 | 1.321 |
| chr3 | 45034154 | 45034577 | 1.44799 | 1.91278 | 1.321 |
| chr8 | 114239770 | 114239966 | 0.896134 | 1.18382 | 1.321 |

| | | | | | |
|---|---|---|---|---|---|
| chr8 | 40962619 | 40962880 | 1.70555 | 2.25347 | 1.321 |
| chr11 | 75303161 | 75303372 | 0.8005 | 1.05768 | 1.321 |
| chr19 | 50187523 | 50188201 | 2.60368 | 3.44017 | 1.321 |
| chr17 | 5599514 | 5599901 | 0.781657 | 1.03287 | 1.321 |
| chr5 | 59473103 | 59473355 | 0.684321 | 0.904318 | 1.321 |
| chr10 | 66641942 | 66642183 | 0.648271 | 0.856727 | 1.322 |
| chr5 | 94244630 | 94244987 | 1.46149 | 1.93148 | 1.322 |
| chr14 | 96240557 | 96241040 | 1.70292 | 2.25059 | 1.322 |
| chr15 | 84692470 | 84692710 | 1.56156 | 2.06383 | 1.322 |
| chr5 | 139725551 | 139725940 | 1.50599 | 1.99043 | 1.322 |
| chr8 | 55836819 | 55837055 | 0.820105 | 1.08398 | 1.322 |
| chr12 | 9333488 | 9333697 | 0.910177 | 1.20306 | 1.322 |
| chr10 | 2398470 | 2398704 | 0.755441 | 0.998605 | 1.322 |
| chr12 | 84180269 | 84180483 | 0.795098 | 1.05112 | 1.322 |
| chr11 | 77259405 | 77259627 | 1.53148 | 2.02476 | 1.322 |
| chr5 | 43115537 | 43115812 | 0.634982 | 0.839501 | 1.322 |
| chr5 | 44552289 | 44552540 | 0.744231 | 0.983948 | 1.322 |
| chr7 | 119927725 | 119928044 | 0.720897 | 0.953181 | 1.322 |
| chr19 | 56143006 | 56143344 | 1.77605 | 2.34848 | 1.322 |
| chr8 | 135474148 | 135474382 | 0.847647 | 1.12092 | 1.322 |
| chr18 | 55847997 | 55848293 | 1.64653 | 2.17741 | 1.322 |
| chr7 | 31822074 | 31822370 | 1.98276 | 2.62223 | 1.323 |
| chr19 | 45491546 | 45491772 | 0.75299 | 0.995859 | 1.323 |
| chr8 | 136215339 | 136215596 | 0.793108 | 1.04893 | 1.323 |
| chr12 | 81844864 | 81845248 | 1.0249 | 1.35551 | 1.323 |
| chr2 | 15008951 | 15009222 | 0.874377 | 1.15655 | 1.323 |
| chr8 | 135860102 | 135860410 | 0.694877 | 0.919138 | 1.323 |
| chr9 | 8739101 | 8739379 | 0.735845 | 0.973332 | 1.323 |
| chr3 | 148579110 | 148579386 | 1.89157 | 2.50208 | 1.323 |
| chr20 | 11992993 | 11993299 | 1.71193 | 2.26464 | 1.323 |
| chr8 | 100370813 | 100371112 | 1.89335 | 2.50465 | 1.323 |
| chr3 | 17882797 | 17883075 | 2.7647 | 3.65741 | 1.323 |
| chr1 | 111161934 | 111162355 | 2.56159 | 3.38903 | 1.323 |
| chr6 | 5143919 | 5144290 | 2.36146 | 3.12432 | 1.323 |
| chr3 | 147757233 | 147757669 | 0.786305 | 1.04036 | 1.323 |
| chr17 | 48347374 | 48347932 | 1.85074 | 2.44877 | 1.323 |
| chr11 | 7825843 | 7826139 | 0.549235 | 0.726762 | 1.323 |
| chr19 | 4941295 | 4941507 | 1.22061 | 1.61518 | 1.323 |
| chr14 | 96608237 | 96608446 | 0.710896 | 0.940707 | 1.323 |
| chr9 | 11806082 | 11806380 | 0.694661 | 0.919247 | 1.323 |
| chr7 | 134176776 | 134176978 | 0.998326 | 1.32112 | 1.323 |
| chr3 | 26633668 | 26633891 | 0.651872 | 0.862668 | 1.323 |
| chr5 | 94185899 | 94186370 | 1.318 | 1.74431 | 1.323 |
| chr17 | 7426466 | 7426815 | 1.37259 | 1.81673 | 1.324 |
| chr8 | 70391109 | 70391317 | 0.730885 | 0.96744 | 1.324 |
| chr20 | 17756479 | 17756922 | 0.785342 | 1.03958 | 1.324 |
| chr9 | 71199368 | 71199866 | 3.53946 | 4.68565 | 1.324 |
| chrX | 49648888 | 49649207 | 2.38807 | 3.16157 | 1.324 |
| chr2 | 120581386 | 120581631 | 1.36139 | 1.80256 | 1.324 |
| chr12 | 121441586 | 121441863 | 1.6785 | 2.22266 | 1.324 |
| chr12 | 63713308 | 63713523 | 1.48203 | 1.96263 | 1.324 |
| chr4 | 38181330 | 38181639 | 2.4669 | 3.26706 | 1.324 |
| chr4 | 147164603 | 147164914 | 1.43064 | 1.89477 | 1.324 |
| chr2 | 192555724 | 192555961 | 1.6803 | 2.22582 | 1.325 |
| chr1 | 111020145 | 111020530 | 0.774256 | 1.02567 | 1.325 |
| chr2 | 113384160 | 113384477 | 2.70138 | 3.57865 | 1.325 |
| chr17 | 17334915 | 17335208 | 0.818878 | 1.08487 | 1.325 |
| chr15 | 86048374 | 86048623 | 1.92334 | 2.54817 | 1.325 |
| chr19 | 30756220 | 30756651 | 0.877999 | 1.16324 | 1.325 |
| chr5 | 112688648 | 112688860 | 0.631048 | 0.836075 | 1.325 |
| chr1 | 31894310 | 31894582 | 1.88491 | 2.49741 | 1.325 |

| | | | | | |
|---|---|---|---|---|---|
| chr6 | 168240406 | 168240629 | 0.794951 | 1.05332 | 1.325 |
| chr1 | 26137298 | 26137633 | 2.00521 | 2.657 | 1.325 |
| chr11 | 113561660 | 113562039 | 1.74216 | 2.30848 | 1.325 |
| chr12 | 47465113 | 47465321 | 1.67411 | 2.2184 | 1.325 |
| chr3 | 146716174 | 146716412 | 0.868997 | 1.15167 | 1.325 |
| chr8 | 127833135 | 127833354 | 1.62933 | 2.15949 | 1.325 |
| chr10 | 87372909 | 87373248 | 0.74345 | 0.985378 | 1.325 |
| chr10 | 92833202 | 92833480 | 0.823871 | 1.09197 | 1.325 |
| chr17 | 34449270 | 34449589 | 0.760577 | 1.00809 | 1.325 |
| chr15 | 57161798 | 57162097 | 1.33059 | 1.76379 | 1.326 |
| chr6 | 143834537 | 143834850 | 1.502 | 1.99103 | 1.326 |
| chr14 | 36538715 | 36539019 | 1.96916 | 2.61037 | 1.326 |
| chr13 | 25949961 | 25950204 | 1.91762 | 2.54215 | 1.326 |
| chrX | 73602524 | 73602867 | 2.7181 | 3.60354 | 1.326 |
| chr2 | 163157805 | 163157990 | 2.41076 | 3.19621 | 1.326 |
| chr14 | 69568069 | 69568291 | 1.73734 | 2.30341 | 1.326 |
| chr5 | 156793830 | 156794046 | 1.33302 | 1.76736 | 1.326 |
| chr9 | 135144149 | 135144562 | 2.11531 | 2.80473 | 1.326 |
| chr14 | 88700344 | 88700558 | 1.19717 | 1.58739 | 1.326 |
| chr7 | 96808829 | 96809065 | 2.34753 | 3.11291 | 1.326 |
| chr1 | 155980809 | 155981047 | 0.674655 | 0.894674 | 1.326 |
| chr7 | 16117776 | 16118012 | 1.56217 | 2.07166 | 1.326 |
| chr8 | 126714961 | 126715200 | 0.980944 | 1.30087 | 1.326 |
| chr3 | 20451711 | 20451944 | 0.765044 | 1.01457 | 1.326 |
| chr12 | 51931152 | 51931358 | 0.710227 | 0.941906 | 1.326 |
| chr1 | 153261169 | 153261516 | 1.81968 | 2.41346 | 1.326 |
| chr1 | 203053666 | 203053922 | 1.55426 | 2.06158 | 1.326 |
| chr4 | 141558885 | 141559313 | 1.14438 | 1.51796 | 1.326 |
| chr1 | 209527763 | 209527980 | 1.4981 | 1.98721 | 1.326 |
| chr17 | 56867671 | 56868080 | 0.694327 | 0.921026 | 1.327 |
| chr2 | 73517153 | 73517675 | 2.08293 | 2.76315 | 1.327 |
| chr2 | 242604894 | 242605144 | 1.10978 | 1.4722 | 1.327 |
| chr4 | 184183097 | 184183327 | 1.80817 | 2.39881 | 1.327 |
| chr11 | 36766200 | 36766461 | 1.16265 | 1.54244 | 1.327 |
| chrX | 47611184 | 47611402 | 1.96313 | 2.60449 | 1.327 |
| chr3 | 69355563 | 69355907 | 1.77529 | 2.35533 | 1.327 |
| chr9 | 33921937 | 33922262 | 2.01956 | 2.67943 | 1.327 |
| chr2 | 208095077 | 208095296 | 1.38647 | 1.83961 | 1.327 |
| chr14 | 69574166 | 69574403 | 1.37435 | 1.82363 | 1.327 |
| chr3 | 150548035 | 150548374 | 0.981086 | 1.3018 | 1.327 |
| chr2 | 56604126 | 56604356 | 0.764545 | 1.01453 | 1.327 |
| chr11 | 64879897 | 64880400 | 2.96132 | 3.92974 | 1.327 |
| chr3 | 108381062 | 108381342 | 1.27215 | 1.68838 | 1.327 |
| chr1 | 66969602 | 66969883 | 1.37016 | 1.81858 | 1.327 |
| chr5 | 20182406 | 20182788 | 0.645044 | 0.856188 | 1.327 |
| chr1 | 220906463 | 220906671 | 1.98355 | 2.63284 | 1.327 |
| chr1 | 160738194 | 160738436 | 1.22879 | 1.63116 | 1.327 |
| chr20 | 25277203 | 25277590 | 1.83415 | 2.43491 | 1.328 |
| chr5 | 146345523 | 146345911 | 1.30656 | 1.73457 | 1.328 |
| chr8 | 52194142 | 52194382 | 0.728467 | 0.967105 | 1.328 |
| chr9 | 27726182 | 27726525 | 1.72145 | 2.28547 | 1.328 |
| chr17 | 74554648 | 74554878 | 1.65257 | 2.19408 | 1.328 |
| chr17 | 76967340 | 76967696 | 2.77539 | 3.68486 | 1.328 |
| chr7 | 112645151 | 112645361 | 0.64479 | 0.856163 | 1.328 |
| chr6 | 126227645 | 126227902 | 2.13316 | 2.83258 | 1.328 |
| chr2 | 233470559 | 233471124 | 2.46984 | 3.27968 | 1.328 |
| chr20 | 16801979 | 16802479 | 2.08778 | 2.77237 | 1.328 |
| chr12 | 33239834 | 33240142 | 0.640929 | 0.851099 | 1.328 |
| chr3 | 149647690 | 149647896 | 0.689388 | 0.915488 | 1.328 |
| chr2 | 5899778 | 5899975 | 0.87622 | 1.16363 | 1.328 |
| chr12 | 120814406 | 120814627 | 0.8304 | 1.10285 | 1.328 |

| | | | | | |
|---|---|---|---|---|---|
| chr20 | 57427739 | 57428043 | 2.39848 | 3.18553 | 1.328 |
| chr1 | 94309868 | 94310109 | 1.62427 | 2.1573 | 1.328 |
| chr15 | 46604416 | 46604654 | 1.39999 | 1.85945 | 1.328 |
| chr15 | 73014478 | 73014886 | 1.96689 | 2.61256 | 1.328 |
| chr2 | 219552223 | 219552500 | 1.54322 | 2.04981 | 1.328 |
| chr16 | 69320943 | 69321156 | 1.81654 | 2.41291 | 1.328 |
| chr4 | 40918953 | 40919217 | 1.72177 | 2.28719 | 1.328 |
| chr7 | 28961494 | 28961699 | 1.70474 | 2.26462 | 1.328 |
| chr2 | 21227003 | 21227366 | 0.880854 | 1.17016 | 1.328 |
| chr7 | 38414127 | 38414376 | 1.11533 | 1.48167 | 1.328 |
| chr6 | 168602105 | 168602348 | 1.4566 | 1.93517 | 1.329 |
| chr15 | 73345296 | 73345539 | 1.06852 | 1.41963 | 1.329 |
| chr3 | 98320955 | 98321267 | 2.34809 | 3.11977 | 1.329 |
| chr19 | 30245662 | 30246011 | 0.807979 | 1.07363 | 1.329 |
| chr8 | 25624955 | 25625303 | 0.536191 | 0.712492 | 1.329 |
| chr4 | 25532971 | 25533204 | 1.53572 | 2.04073 | 1.329 |
| chr2 | 16083502 | 16083867 | 1.38748 | 1.84377 | 1.329 |
| chr10 | 2477062 | 2477326 | 0.726378 | 0.965289 | 1.329 |
| chr10 | 53478673 | 53478979 | 0.651662 | 0.866028 | 1.329 |
| chrX | 123094218 | 123094478 | 2.60814 | 3.46613 | 1.329 |
| chr11 | 20149822 | 20150178 | 2.09509 | 2.78442 | 1.329 |
| chr2 | 169966253 | 169966600 | 2.03692 | 2.70723 | 1.329 |
| chr14 | 29923576 | 29923820 | 0.885705 | 1.17732 | 1.329 |
| chr17 | 41987529 | 41987949 | 2.03089 | 2.69958 | 1.329 |
| chr6 | 122702730 | 122702985 | 1.69407 | 2.25196 | 1.329 |
| chr12 | 132405439 | 132405740 | 0.697449 | 0.927167 | 1.329 |
| chr6 | 156437571 | 156437816 | 0.588642 | 0.782525 | 1.329 |
| chr1 | 201012067 | 201012494 | 1.81115 | 2.40775 | 1.329 |
| chr2 | 176213723 | 176213943 | 0.628596 | 0.835689 | 1.329 |
| chr1 | 240168617 | 240168880 | 1.6989 | 2.25865 | 1.329 |
| chr1 | 111372506 | 111372887 | 1.51175 | 2.01032 | 1.330 |
| chr2 | 3465954 | 3466651 | 1.5644 | 2.08036 | 1.330 |
| chr2 | 21811177 | 21811494 | 0.648729 | 0.862725 | 1.330 |
| chr10 | 126736679 | 126737253 | 2.18088 | 2.90034 | 1.330 |
| chr8 | 13584114 | 13584330 | 0.970174 | 1.29049 | 1.330 |
| chr17 | 7665934 | 7666148 | 0.673493 | 0.895865 | 1.330 |
| chr17 | 79215204 | 79215820 | 2.00406 | 2.66585 | 1.330 |
| chr9 | 9102345 | 9102606 | 0.992159 | 1.31985 | 1.330 |
| chr9 | 90317951 | 90318156 | 0.669757 | 0.890976 | 1.330 |
| chr5 | 103545730 | 103545970 | 0.727219 | 0.967426 | 1.330 |
| chr9 | 107154716 | 107154961 | 0.782356 | 1.04087 | 1.330 |
| chr10 | 88720152 | 88720609 | 3.0777 | 4.09478 | 1.330 |
| chr1 | 245826950 | 245827184 | 0.767358 | 1.02097 | 1.331 |
| chr19 | 17371672 | 17372011 | 2.57498 | 3.42602 | 1.331 |
| chr12 | 102775135 | 102775429 | 0.62698 | 0.834236 | 1.331 |
| chr1 | 207155480 | 207155784 | 1.92579 | 2.56253 | 1.331 |
| chr2 | 290180 | 290512 | 2.37937 | 3.16615 | 1.331 |
| chr16 | 46963506 | 46964018 | 2.25351 | 2.99876 | 1.331 |
| chr10 | 111626354 | 111626650 | 1.02696 | 1.36664 | 1.331 |
| chr3 | 45082283 | 45082507 | 2.68514 | 3.57327 | 1.331 |
| chr2 | 231627086 | 231627353 | 1.79492 | 2.3888 | 1.331 |
| chr8 | 299621 | 299823 | 0.936401 | 1.24623 | 1.331 |
| chr12 | 129023821 | 129024258 | 0.720493 | 0.958983 | 1.331 |
| chr7 | 49729034 | 49729315 | 0.753607 | 1.00308 | 1.331 |
| chr6 | 88639586 | 88639823 | 2.23729 | 2.97794 | 1.331 |
| chr19 | 17496778 | 17497178 | 1.99857 | 2.66035 | 1.331 |
| chr10 | 45353202 | 45353630 | 2.46746 | 3.28455 | 1.331 |
| chr17 | 62067678 | 62068304 | 2.58059 | 3.43515 | 1.331 |
| chr16 | 58066131 | 58066517 | 1.65852 | 2.20785 | 1.331 |
| chr17 | 31626280 | 31626635 | 1.02047 | 1.35847 | 1.331 |
| chr8 | 23267898 | 23268173 | 2.76262 | 3.67773 | 1.331 |

| | | | | | |
|---|---|---|---|---|---|
| chr8 | 123651141 | 123651404 | 1.8171 | 2.41948 | 1.332 |
| chr8 | 67351164 | 67351507 | 3.15641 | 4.20282 | 1.332 |
| chr17 | 38459296 | 38459534 | 1.89405 | 2.52216 | 1.332 |
| chr17 | 54318735 | 54318987 | 0.797596 | 1.0621 | 1.332 |
| chr9 | 19028935 | 19029198 | 1.55729 | 2.07373 | 1.332 |
| chr19 | 45562365 | 45563099 | 2.4341 | 3.24135 | 1.332 |
| chr4 | 41890178 | 41890386 | 1.82546 | 2.43105 | 1.332 |
| chr22 | 20810594 | 20810870 | 2.01844 | 2.68823 | 1.332 |
| chrX | 95246852 | 95247101 | 0.511703 | 0.681552 | 1.332 |
| chr10 | 72843279 | 72843522 | 0.639131 | 0.851332 | 1.332 |
| chr10 | 97123094 | 97123367 | 1.48138 | 1.9733 | 1.332 |
| chr10 | 59339776 | 59340028 | 1.25807 | 1.67587 | 1.332 |
| chr2 | 137136404 | 137136688 | 1.83988 | 2.45095 | 1.332 |
| chr7 | 31168749 | 31169044 | 1.11057 | 1.47946 | 1.332 |
| chr11 | 82413015 | 82413217 | 1.34836 | 1.79632 | 1.332 |
| chr6 | 139690705 | 139690932 | 1.41967 | 1.89137 | 1.332 |
| chr3 | 16360921 | 16361108 | 1.59312 | 2.1227 | 1.332 |
| chr4 | 25769566 | 25769812 | 1.95132 | 2.60074 | 1.333 |
| chr4 | 124017174 | 124017514 | 1.22425 | 1.63187 | 1.333 |
| chr14 | 67802538 | 67802799 | 1.42767 | 1.90321 | 1.333 |
| chr2 | 172968602 | 172968909 | 2.21806 | 2.95696 | 1.333 |
| chr2 | 101655084 | 101655400 | 1.75839 | 2.3442 | 1.333 |
| chr6 | 16332350 | 16332671 | 2.18665 | 2.91515 | 1.333 |
| chr10 | 105324525 | 105324999 | 2.29336 | 3.05744 | 1.333 |
| chr17 | 26821538 | 26822240 | 2.28324 | 3.04397 | 1.333 |
| chr19 | 42257817 | 42258205 | 1.47755 | 1.96992 | 1.333 |
| chr1 | 235946533 | 235946818 | 1.35432 | 1.80568 | 1.333 |
| chr7 | 372613 | 372952 | 2.29718 | 3.06276 | 1.333 |
| chr19 | 18362630 | 18362928 | 1.56016 | 2.08019 | 1.333 |
| chr20 | 12731322 | 12731571 | 0.769665 | 1.02622 | 1.333 |
| chr13 | 114427752 | 114428124 | 2.21675 | 2.95573 | 1.333 |
| chr16 | 71512674 | 71513027 | 0.798537 | 1.06479 | 1.333 |
| chr15 | 27049385 | 27049763 | 1.0679 | 1.42413 | 1.334 |
| chr16 | 62290080 | 62290547 | 0.765054 | 1.02026 | 1.334 |
| chr8 | 52600233 | 52600468 | 0.66851 | 0.891519 | 1.334 |
| chr5 | 109314416 | 109314670 | 0.623131 | 0.831015 | 1.334 |
| chr19 | 47040566 | 47041153 | 2.17384 | 2.89915 | 1.334 |
| chr2 | 56410705 | 56411092 | 2.93145 | 3.90968 | 1.334 |
| chr7 | 77280954 | 77281289 | 1.93161 | 2.57645 | 1.334 |
| chr1 | 204925540 | 204925796 | 0.876346 | 1.16893 | 1.334 |
| chr1 | 197488957 | 197489159 | 1.05449 | 1.40662 | 1.334 |
| chr11 | 58217615 | 58217985 | 0.951383 | 1.2691 | 1.334 |
| chr2 | 231712636 | 231712951 | 3.2104 | 4.28251 | 1.334 |
| chr17 | 5152857 | 5153145 | 2.3125 | 3.08483 | 1.334 |
| chr15 | 46082657 | 46082876 | 1.19659 | 1.59632 | 1.334 |
| chr2 | 238062767 | 238062985 | 1.94193 | 2.59076 | 1.334 |
| chr14 | 82514372 | 82514595 | 0.722001 | 0.963273 | 1.334 |
| chr22 | 28276864 | 28277085 | 1.44108 | 1.92288 | 1.334 |
| chr2 | 3786081 | 3786357 | 1.62093 | 2.16288 | 1.334 |
| chr6 | 44561260 | 44561518 | 1.25543 | 1.67521 | 1.334 |
| chr5 | 78195731 | 78196050 | 1.88428 | 2.51435 | 1.334 |
| chr10 | 48631740 | 48631957 | 0.810546 | 1.08159 | 1.334 |
| chr3 | 107460213 | 107460415 | 1.35198 | 1.80419 | 1.334 |
| chr12 | 82463707 | 82463934 | 0.772021 | 1.03028 | 1.335 |
| chr6 | 160559322 | 160559534 | 1.6712 | 2.23032 | 1.335 |
| chr19 | 33670908 | 33671207 | 0.87187 | 1.16366 | 1.335 |
| chr16 | 19910680 | 19911241 | 1.75165 | 2.33789 | 1.335 |
| chr5 | 119950494 | 119950699 | 1.03767 | 1.38512 | 1.335 |
| chr15 | 87667403 | 87667863 | 0.967863 | 1.29203 | 1.335 |
| chr17 | 12703812 | 12704135 | 0.894608 | 1.19424 | 1.335 |
| chr5 | 104333811 | 104334143 | 0.731646 | 0.976718 | 1.335 |

| | | | | | |
|---|---|---|---|---|---|
| chrX | 71241911 | 71242139 | 1.50981 | 2.01566 | 1.335 |
| chr22 | 23632506 | 23633042 | 1.83442 | 2.44904 | 1.335 |
| chr2 | 5875590 | 5875848 | 0.727058 | 0.970753 | 1.335 |
| chr22 | 41993350 | 41993652 | 2.60352 | 3.47641 | 1.335 |
| chr9 | 101440283 | 101441025 | 1.18132 | 1.57739 | 1.335 |
| chr15 | 89720978 | 89721321 | 2.85488 | 3.81212 | 1.335 |
| chr9 | 129544090 | 129544404 | 1.9639 | 2.62271 | 1.335 |
| chr4 | 128868390 | 128868685 | 0.728543 | 0.972955 | 1.335 |
| chr2 | 141621390 | 141621635 | 1.28941 | 1.72213 | 1.336 |
| chr7 | 93251409 | 93251966 | 1.38358 | 1.84794 | 1.336 |
| chr18 | 12934506 | 12934784 | 2.2211 | 2.96657 | 1.336 |
| chr10 | 79516242 | 79516533 | 1.47457 | 1.96958 | 1.336 |
| chr12 | 31000192 | 31000500 | 1.01934 | 1.36154 | 1.336 |
| chr8 | 117899719 | 117900071 | 1.95338 | 2.6093 | 1.336 |
| chr12 | 72030343 | 72030689 | 1.86163 | 2.48677 | 1.336 |
| chr17 | 11773866 | 11774135 | 0.686383 | 0.91694 | 1.336 |
| chr20 | 17387533 | 17387887 | 0.835186 | 1.11576 | 1.336 |
| chr8 | 96572015 | 96572376 | 0.592944 | 0.792154 | 1.336 |
| chr17 | 72733791 | 72734084 | 1.41389 | 1.88893 | 1.336 |
| chr7 | 127144939 | 127145211 | 0.992371 | 1.32583 | 1.336 |
| chr13 | 110790085 | 110790399 | 1.01149 | 1.3515 | 1.336 |
| chr14 | 65321614 | 65321881 | 0.824207 | 1.10128 | 1.336 |
| chr4 | 37979437 | 37979690 | 1.72069 | 2.29921 | 1.336 |
| chr14 | 77561078 | 77561370 | 2.12954 | 2.84569 | 1.336 |
| chr6 | 34577401 | 34577610 | 1.51687 | 2.02703 | 1.336 |
| chr2 | 219513225 | 219513727 | 1.46546 | 1.95835 | 1.336 |
| chr12 | 54645513 | 54645860 | 1.68154 | 2.24738 | 1.337 |
| chr2 | 219968877 | 219969228 | 2.08955 | 2.79272 | 1.337 |
| chr2 | 218852998 | 218853553 | 1.67194 | 2.23471 | 1.337 |
| chr1 | 26718054 | 26718282 | 2.19675 | 2.93632 | 1.337 |
| chr6 | 153218974 | 153219369 | 1.67914 | 2.24466 | 1.337 |
| chr3 | 57029454 | 57029720 | 1.70516 | 2.27946 | 1.337 |
| chr10 | 20453729 | 20454048 | 0.905839 | 1.21105 | 1.337 |
| chr11 | 56286822 | 56287029 | 0.741044 | 0.990819 | 1.337 |
| chr17 | 19648184 | 19648537 | 2.73789 | 3.66093 | 1.337 |
| chr17 | 30017221 | 30017440 | 0.685857 | 0.91715 | 1.337 |
| chr1 | 240608682 | 240608871 | 0.763325 | 1.02088 | 1.337 |
| chr2 | 10219959 | 10220358 | 1.60133 | 2.14173 | 1.337 |
| chr12 | 102714328 | 102714556 | 0.898979 | 1.2025 | 1.338 |
| chr4 | 15459264 | 15459553 | 1.42955 | 1.91226 | 1.338 |
| chr16 | 78471181 | 78471618 | 2.5839 | 3.4565 | 1.338 |
| chr14 | 77679758 | 77680095 | 1.91451 | 2.5611 | 1.338 |
| chrX | 137842768 | 137842983 | 0.550956 | 0.737039 | 1.338 |
| chr2 | 113238266 | 113238907 | 1.83835 | 2.45929 | 1.338 |
| chr16 | 84518146 | 84518527 | 1.65286 | 2.21124 | 1.338 |
| chr2 | 75880861 | 75881372 | 0.742667 | 0.993612 | 1.338 |
| chr11 | 41800252 | 41800546 | 0.595806 | 0.797156 | 1.338 |
| chr5 | 164824792 | 164825001 | 0.878083 | 1.17489 | 1.338 |
| chr9 | 385872 | 386117 | 2.01169 | 2.69173 | 1.338 |
| chr12 | 113863085 | 113863542 | 3.02017 | 4.04117 | 1.338 |
| chr8 | 107775366 | 107775581 | 1.24016 | 1.6595 | 1.338 |
| chr8 | 135365837 | 135366030 | 0.738396 | 0.988078 | 1.338 |
| chr22 | 38496840 | 38497156 | 1.78359 | 2.38676 | 1.338 |
| chr3 | 156100280 | 156100701 | 1.49159 | 1.99628 | 1.338 |
| chr7 | 18391988 | 18392265 | 1.53294 | 2.05169 | 1.338 |
| chr12 | 6806980 | 6807291 | 1.79826 | 2.4069 | 1.338 |
| chr4 | 68566583 | 68566794 | 1.49513 | 2.00119 | 1.338 |
| chr12 | 61035675 | 61035920 | 0.823756 | 1.1026 | 1.339 |
| chr5 | 24633917 | 24634150 | 1.16824 | 1.56372 | 1.339 |
| chr6 | 16771215 | 16771487 | 1.91177 | 2.559 | 1.339 |
| chr7 | 154860594 | 154860786 | 0.619992 | 0.829923 | 1.339 |

| | | | | | |
|---|---|---|---|---|---|
| chr7 | 153206518 | 153207014 | 0.747183 | 1.00028 | 1.339 |
| chr5 | 41715848 | 41716069 | 0.709501 | 0.949869 | 1.339 |
| chr11 | 94701568 | 94701813 | 1.49388 | 2.00002 | 1.339 |
| chr11 | 40862902 | 40863144 | 0.78916 | 1.05656 | 1.339 |
| chr10 | 62095565 | 62095806 | 1.79229 | 2.39972 | 1.339 |
| chr9 | 21326594 | 21326877 | 2.19203 | 2.9352 | 1.339 |
| chr8 | 130951827 | 130952043 | 1.60991 | 2.15581 | 1.339 |
| chr7 | 24705291 | 24705580 | 1.46119 | 1.95677 | 1.339 |
| chr8 | 14686052 | 14686402 | 0.716102 | 0.959041 | 1.339 |
| chr20 | 51401396 | 51401604 | 0.809326 | 1.08391 | 1.339 |
| chr9 | 83174142 | 83174478 | 0.678121 | 0.908289 | 1.339 |
| chr2 | 231917578 | 231917932 | 2.81305 | 3.76799 | 1.339 |
| chr19 | 28505740 | 28506134 | 0.85024 | 1.13892 | 1.340 |
| chr5 | 130207229 | 130207546 | 1.04653 | 1.40191 | 1.340 |
| chr5 | 110092798 | 110093189 | 1.53563 | 2.0572 | 1.340 |
| chr1 | 206718171 | 206718484 | 2.4394 | 3.26814 | 1.340 |
| chr2 | 3211195 | 3211457 | 0.645057 | 0.864222 | 1.340 |
| chr20 | 16498989 | 16499234 | 1.57644 | 2.1121 | 1.340 |
| chr17 | 1933992 | 1934201 | 1.35075 | 1.80995 | 1.340 |
| chr5 | 123006701 | 123006979 | 1.88905 | 2.53135 | 1.340 |
| chr1 | 152843628 | 152844104 | 1.14243 | 1.53088 | 1.340 |
| chr15 | 56138341 | 56138735 | 1.89572 | 2.54048 | 1.340 |
| chr3 | 51716159 | 51716413 | 1.47319 | 1.97431 | 1.340 |
| chr2 | 30462186 | 30462495 | 2.9998 | 4.02026 | 1.340 |
| chr5 | 154285516 | 154285724 | 0.787598 | 1.05553 | 1.340 |
| chr20 | 36162785 | 36163243 | 1.49905 | 2.00907 | 1.340 |
| chr2 | 15299951 | 15300175 | 1.09044 | 1.4615 | 1.340 |
| chr22 | 18225278 | 18225610 | 2.02137 | 2.70936 | 1.340 |
| chr2 | 193027272 | 193027549 | 1.16976 | 1.56793 | 1.340 |
| chr1 | 238415392 | 238416139 | 0.727044 | 0.974674 | 1.341 |
| chr8 | 98464971 | 98465178 | 1.26618 | 1.69745 | 1.341 |
| chr10 | 47592145 | 47592357 | 0.872378 | 1.16955 | 1.341 |
| chr19 | 53759234 | 53759471 | 0.894481 | 1.19927 | 1.341 |
| chr10 | 86373206 | 86373651 | 1.28208 | 1.71896 | 1.341 |
| chrX | 117778324 | 117778643 | 0.805179 | 1.07955 | 1.341 |
| chr19 | 28910045 | 28910321 | 0.798433 | 1.07056 | 1.341 |
| chr4 | 48559560 | 48559793 | 1.17171 | 1.57107 | 1.341 |
| chr8 | 11622947 | 11623337 | 2.40132 | 3.21998 | 1.341 |
| chr9 | 130341928 | 130342133 | 1.44409 | 1.93659 | 1.341 |
| chr22 | 39930799 | 39931184 | 1.98493 | 2.66202 | 1.341 |
| chr7 | 5598569 | 5598838 | 2.19906 | 2.94921 | 1.341 |
| gl000202_r | 7132 | 8038 | 2.09971 | 2.81621 | 1.341 |
| chr19 | 55187723 | 55188088 | 2.55981 | 3.43338 | 1.341 |
| chr13 | 78711818 | 78712079 | 1.39344 | 1.86911 | 1.341 |
| chr12 | 132652139 | 132652351 | 0.890216 | 1.19415 | 1.341 |
| chr11 | 60524011 | 60524357 | 1.16013 | 1.55623 | 1.341 |
| chr16 | 58695191 | 58695408 | 2.14531 | 2.8779 | 1.341 |
| chr7 | 157847390 | 157847693 | 0.625562 | 0.839215 | 1.342 |
| chr10 | 27461297 | 27461613 | 0.706469 | 0.9478 | 1.342 |
| chr15 | 48542932 | 48543185 | 0.933084 | 1.25184 | 1.342 |
| chr5 | 176297012 | 176297505 | 1.85191 | 2.4846 | 1.342 |
| chr12 | 95976267 | 95976646 | 1.41327 | 1.89651 | 1.342 |
| chr2 | 233386671 | 233387150 | 2.16477 | 2.90504 | 1.342 |
| chr5 | 170737199 | 170737427 | 1.45024 | 1.94618 | 1.342 |
| chr14 | 22188234 | 22188614 | 0.714762 | 0.959211 | 1.342 |
| chr17 | 80087063 | 80087282 | 1.33559 | 1.79251 | 1.342 |
| chr2 | 85293948 | 85294249 | 2.03056 | 2.72523 | 1.342 |
| chr1 | 176866570 | 176867366 | 1.52662 | 2.0489 | 1.342 |
| chr8 | 65532518 | 65532743 | 0.703545 | 0.944255 | 1.342 |
| chr17 | 44928876 | 44929158 | 0.995589 | 1.33639 | 1.342 |
| chr12 | 65284547 | 65284999 | 1.59212 | 2.13714 | 1.342 |

| | | | | | |
|---|---|---|---|---|---|
| chr9 | 15403287 | 15403488 | 1.60323 | 2.15208 | 1.342 |
| chr12 | 6435588 | 6435812 | 1.23436 | 1.65702 | 1.342 |
| chr15 | 73214019 | 73214345 | 0.746032 | 1.00148 | 1.342 |
| chr20 | 44780473 | 44780908 | 1.64124 | 2.20339 | 1.343 |
| chr20 | 58508948 | 58509480 | 2.90127 | 3.89506 | 1.343 |
| chr2 | 42660578 | 42660825 | 0.698282 | 0.937498 | 1.343 |
| chr15 | 68846106 | 68846869 | 1.86416 | 2.50281 | 1.343 |
| chr20 | 39763729 | 39764236 | 2.46944 | 3.31574 | 1.343 |
| chr2 | 69170649 | 69171260 | 1.56378 | 2.09986 | 1.343 |
| chr9 | 112163653 | 112164069 | 2.92814 | 3.93202 | 1.343 |
| chr19 | 17895147 | 17895596 | 1.79399 | 2.40913 | 1.343 |
| chr3 | 172207973 | 172208263 | 1.74693 | 2.34603 | 1.343 |
| chr4 | 185960961 | 185961218 | 1.6625 | 2.23267 | 1.343 |
| chr15 | 53423266 | 53423505 | 0.749285 | 1.00627 | 1.343 |
| chr2 | 101223397 | 101223663 | 1.77281 | 2.3809 | 1.343 |
| chr7 | 53633001 | 53633237 | 0.691779 | 0.929092 | 1.343 |
| chr16 | 19083498 | 19084180 | 1.71505 | 2.30363 | 1.343 |
| chr1 | 28844487 | 28844793 | 1.88314 | 2.52949 | 1.343 |
| chr1 | 145578341 | 145578567 | 0.572391 | 0.768851 | 1.343 |
| chr4 | 42284096 | 42284467 | 1.19859 | 1.61001 | 1.343 |
| chr1 | 87207719 | 87207937 | 1.70888 | 2.29552 | 1.343 |
| chr11 | 43150421 | 43150763 | 0.542156 | 0.728327 | 1.343 |
| chr6 | 5030951 | 5031154 | 1.6565 | 2.22538 | 1.343 |
| chr14 | 22179341 | 22179975 | 1.51559 | 2.03615 | 1.343 |
| chr11 | 7570197 | 7570532 | 1.41321 | 1.89867 | 1.344 |
| chr7 | 33240446 | 33240660 | 0.928108 | 1.24695 | 1.344 |
| chr9 | 96827775 | 96828045 | 1.85712 | 2.49511 | 1.344 |
| chr5 | 91520345 | 91520579 | 0.762875 | 1.02504 | 1.344 |
| chr14 | 25275265 | 25275761 | 1.52567 | 2.0501 | 1.344 |
| chr7 | 36527390 | 36527673 | 1.9697 | 2.64685 | 1.344 |
| chr3 | 150643428 | 150643748 | 2.62478 | 3.52748 | 1.344 |
| chr4 | 3108353 | 3108674 | 2.75744 | 3.70577 | 1.344 |
| chr7 | 78057996 | 78058199 | 0.741543 | 0.996585 | 1.344 |
| chr10 | 113899873 | 113900303 | 2.52537 | 3.394 | 1.344 |
| chr2 | 14280189 | 14280672 | 0.775733 | 1.0426 | 1.344 |
| chr16 | 85350022 | 85350515 | 2.9458 | 3.96068 | 1.345 |
| chr17 | 27239415 | 27239679 | 1.58153 | 2.12641 | 1.345 |
| chr2 | 98885679 | 98886157 | 1.26959 | 1.70706 | 1.345 |
| chr9 | 139974026 | 139974258 | 0.69133 | 0.929553 | 1.345 |
| chr8 | 144815543 | 144815900 | 1.63037 | 2.19238 | 1.345 |
| chr21 | 45395375 | 45395785 | 2.49552 | 3.35596 | 1.345 |
| chr2 | 31392987 | 31393374 | 1.94485 | 2.61552 | 1.345 |
| chr7 | 16735428 | 16735683 | 1.14836 | 1.54439 | 1.345 |
| chr2 | 75146904 | 75147349 | 2.53299 | 3.40696 | 1.345 |
| chr17 | 38095850 | 38096091 | 1.52572 | 2.05223 | 1.345 |
| chr1 | 166604261 | 166604573 | 0.748146 | 1.00634 | 1.345 |
| chr2 | 23574385 | 23574794 | 2.57539 | 3.4642 | 1.345 |
| chr1 | 229885845 | 229886082 | 0.937228 | 1.26071 | 1.345 |
| chr19 | 54974006 | 54974729 | 0.869993 | 1.17029 | 1.345 |
| chr1 | 6535511 | 6535912 | 2.88743 | 3.88413 | 1.345 |
| chr19 | 50733767 | 50734125 | 0.86327 | 1.16126 | 1.345 |
| chr5 | 121836514 | 121836750 | 1.87237 | 2.51898 | 1.345 |
| chr22 | 26857150 | 26857563 | 1.41027 | 1.89739 | 1.345 |
| chr19 | 42538676 | 42539299 | 2.49926 | 3.36271 | 1.345 |
| chr17 | 39818821 | 39819361 | 6.58849 | 8.86576 | 1.346 |
| chr7 | 38787664 | 38787867 | 1.1395 | 1.53345 | 1.346 |
| chr17 | 66968228 | 66968565 | 1.27392 | 1.71439 | 1.346 |
| chr5 | 65667746 | 65668027 | 0.55504 | 0.747022 | 1.346 |
| chr14 | 33484033 | 33484374 | 0.754086 | 1.01496 | 1.346 |
| chr5 | 52796041 | 52796265 | 0.496169 | 0.667825 | 1.346 |
| chr7 | 27169657 | 27169934 | 0.874696 | 1.17731 | 1.346 |

| | | | | | |
|---|---|---|---|---|---|
| chr5 | 96566828 | 96567068 | 1.37471 | 1.85035 | 1.346 |
| chr11 | 78127929 | 78128191 | 1.46366 | 1.97029 | 1.346 |
| chr12 | 7033467 | 7033770 | 2.83013 | 3.80974 | 1.346 |
| chr10 | 10495558 | 10495802 | 0.850507 | 1.14497 | 1.346 |
| chr17 | 60749716 | 60750024 | 2.14284 | 2.88478 | 1.346 |
| chr1 | 225668586 | 225668987 | 2.55661 | 3.4422 | 1.346 |
| chr14 | 55821372 | 55821640 | 2.05697 | 2.76985 | 1.347 |
| chr22 | 22307196 | 22307458 | 2.25171 | 3.03209 | 1.347 |
| chr4 | 108939661 | 108940076 | 1.8232 | 2.45512 | 1.347 |
| chr9 | 77813402 | 77813619 | 1.28879 | 1.73548 | 1.347 |
| chr7 | 103087623 | 103087913 | 2.64202 | 3.55791 | 1.347 |
| chr5 | 33649596 | 33649831 | 0.654324 | 0.881178 | 1.347 |
| chr17 | 39735905 | 39736184 | 1.90564 | 2.56645 | 1.347 |
| chr10 | 5915424 | 5915734 | 1.4156 | 1.90651 | 1.347 |
| chr1 | 152432663 | 152433128 | 2.03971 | 2.74713 | 1.347 |
| chr15 | 92640867 | 92641151 | 2.03525 | 2.74118 | 1.347 |
| chr6 | 158402433 | 158402841 | 1.21196 | 1.63233 | 1.347 |
| chr5 | 163440198 | 163440540 | 0.734245 | 0.989057 | 1.347 |
| chr9 | 85054727 | 85054931 | 0.585221 | 0.788322 | 1.347 |
| chr7 | 129469911 | 129470144 | 1.91925 | 2.58538 | 1.347 |
| chr3 | 80635353 | 80635708 | 1.07659 | 1.45032 | 1.347 |
| chr19 | 38734471 | 38734859 | 2.73394 | 3.68348 | 1.347 |
| chr19 | 40970264 | 40970577 | 2.75259 | 3.70883 | 1.347 |
| chr19 | 50394671 | 50394877 | 0.659539 | 0.888667 | 1.347 |
| chr5 | 109804867 | 109805117 | 0.760978 | 1.02539 | 1.347 |
| chr11 | 10325980 | 10326405 | 1.99975 | 2.69474 | 1.348 |
| chr15 | 65393754 | 65394217 | 2.16761 | 2.92099 | 1.348 |
| chr17 | 41616539 | 41616812 | 0.836424 | 1.12713 | 1.348 |
| chr19 | 16358351 | 16358770 | 2.21353 | 2.98289 | 1.348 |
| chr15 | 89080769 | 89080985 | 0.915498 | 1.23371 | 1.348 |
| chr3 | 183945203 | 183945559 | 2.51859 | 3.39399 | 1.348 |
| chr1 | 214832279 | 214832489 | 1.26035 | 1.69846 | 1.348 |
| chr6 | 132344738 | 132344932 | 0.955794 | 1.28805 | 1.348 |
| chr8 | 6429837 | 6430073 | 1.63202 | 2.19935 | 1.348 |
| chr21 | 39724615 | 39724810 | 1.61034 | 2.17014 | 1.348 |
| chr7 | 47219807 | 47220082 | 1.58102 | 2.13082 | 1.348 |
| chr5 | 176000950 | 176001403 | 2.56575 | 3.45826 | 1.348 |
| chr17 | 38169918 | 38170376 | 2.36593 | 3.18901 | 1.348 |
| chr11 | 117817597 | 117817993 | 1.6165 | 2.17888 | 1.348 |
| chr13 | 99118904 | 99119236 | 1.709 | 2.30367 | 1.348 |
| chr2 | 121852907 | 121853202 | 2.22814 | 3.00362 | 1.348 |
| chr7 | 73620879 | 73621361 | 2.4948 | 3.36323 | 1.348 |
| chr1 | 192425523 | 192425740 | 0.698032 | 0.941059 | 1.348 |
| chr9 | 22946456 | 22946665 | 0.715347 | 0.964425 | 1.348 |
| chr12 | 131237286 | 131237791 | 0.90577 | 1.22116 | 1.348 |
| chr2 | 120048451 | 120048763 | 2.10059 | 2.83208 | 1.348 |
| chr18 | 69609599 | 69609860 | 0.850966 | 1.14735 | 1.348 |
| chr13 | 40190207 | 40190568 | 2.01896 | 2.72234 | 1.348 |
| chr1 | 67614909 | 67615175 | 1.74121 | 2.34793 | 1.348 |
| chr11 | 83286586 | 83286804 | 0.611241 | 0.824247 | 1.348 |
| chr7 | 31673583 | 31673857 | 1.84724 | 2.49103 | 1.349 |
| chr17 | 77184071 | 77184586 | 2.20335 | 2.97129 | 1.349 |
| chr8 | 90265533 | 90265830 | 1.57474 | 2.12363 | 1.349 |
| chr10 | 24417494 | 24417706 | 0.700011 | 0.944012 | 1.349 |
| chr20 | 3474821 | 3475190 | 1.01829 | 1.37324 | 1.349 |
| chr11 | 47198937 | 47199216 | 1.59633 | 2.15283 | 1.349 |
| chr8 | 14734342 | 14734666 | 0.863175 | 1.16412 | 1.349 |
| chr1 | 2718218 | 2718465 | 2.00486 | 2.70389 | 1.349 |
| chr2 | 152266462 | 152266691 | 1.27914 | 1.72522 | 1.349 |
| chr16 | 58131139 | 58131508 | 3.18314 | 4.29322 | 1.349 |
| chr11 | 14476416 | 14476933 | 1.66522 | 2.246 | 1.349 |

| | | | | | |
|---|---|---|---|---|---|
| chr12 | 122611794 | 122612024 | 2.1666 | 2.92225 | 1.349 |
| chr14 | 98664780 | 98665018 | 0.693139 | 0.934887 | 1.349 |
| chr8 | 35467161 | 35467383 | 0.771607 | 1.04075 | 1.349 |
| chr14 | 38379401 | 38379762 | 0.811204 | 1.09422 | 1.349 |
| chr7 | 33825419 | 33825664 | 0.791495 | 1.06764 | 1.349 |
| chr10 | 35902836 | 35903062 | 1.02119 | 1.3775 | 1.349 |
| chr11 | 18225308 | 18225825 | 1.15441 | 1.55725 | 1.349 |
| chr12 | 55073080 | 55073295 | 0.553947 | 0.747257 | 1.349 |
| chr7 | 146581241 | 146581626 | 0.790774 | 1.06677 | 1.349 |
| chr20 | 5392279 | 5392511 | 0.821352 | 1.10809 | 1.349 |
| chr11 | 102537152 | 102537366 | 1.64467 | 2.21895 | 1.349 |
| chr4 | 24421070 | 24421348 | 1.71826 | 2.31835 | 1.349 |
| chr9 | 17802341 | 17802546 | 1.49014 | 2.01073 | 1.349 |
| chrX | 99926100 | 99926442 | 1.30031 | 1.75461 | 1.349 |
| chr2 | 7006589 | 7006794 | 1.17542 | 1.58614 | 1.349 |
| chr9 | 138235691 | 138236225 | 2.53587 | 3.42219 | 1.350 |
| chr5 | 146749387 | 146749661 | 1.41018 | 1.9033 | 1.350 |
| chr11 | 3391908 | 3392155 | 0.739054 | 0.997505 | 1.350 |
| chr12 | 107093866 | 107094167 | 0.915004 | 1.23507 | 1.350 |
| chr12 | 40801497 | 40801720 | 1.00922 | 1.36228 | 1.350 |
| chr4 | 186055643 | 186055862 | 1.47434 | 1.99014 | 1.350 |
| chr10 | 120303315 | 120303967 | 1.51036 | 2.03883 | 1.350 |
| chr2 | 20981323 | 20981842 | 1.3366 | 1.80427 | 1.350 |
| chr3 | 114245962 | 114246289 | 1.87085 | 2.52563 | 1.350 |
| chr15 | 88943250 | 88943462 | 0.824845 | 1.11362 | 1.350 |
| chr10 | 102488796 | 102489509 | 1.72386 | 2.32777 | 1.350 |
| chr3 | 104502003 | 104502189 | 0.867345 | 1.17129 | 1.350 |
| chr12 | 44917006 | 44917389 | 0.790139 | 1.06714 | 1.351 |
| chr15 | 94749180 | 94749438 | 0.684381 | 0.924368 | 1.351 |
| chr3 | 113557427 | 113557842 | 2.95852 | 3.9969 | 1.351 |
| chr9 | 101762826 | 101763172 | 1.3591 | 1.83619 | 1.351 |
| chr1 | 88419535 | 88419813 | 1.70113 | 2.29845 | 1.351 |
| chr10 | 87632314 | 87632650 | 0.680246 | 0.919131 | 1.351 |
| chr5 | 9188969 | 9189271 | 0.529909 | 0.716008 | 1.351 |
| chr5 | 122040801 | 122041031 | 0.743829 | 1.00508 | 1.351 |
| chr17 | 64197979 | 64198185 | 1.66235 | 2.24639 | 1.351 |
| chr14 | 93598847 | 93599204 | 1.47105 | 1.98794 | 1.351 |
| chr3 | 14589670 | 14590174 | 1.92061 | 2.59572 | 1.352 |
| chr19 | 54415860 | 54416330 | 1.45913 | 1.97205 | 1.352 |
| chr2 | 134644258 | 134644499 | 1.67245 | 2.26038 | 1.352 |
| chr11 | 27741948 | 27742249 | 0.999776 | 1.35136 | 1.352 |
| chr12 | 44070081 | 44070604 | 1.04567 | 1.41342 | 1.352 |
| chr11 | 3630931 | 3631488 | 1.90403 | 2.5737 | 1.352 |
| chr22 | 22818420 | 22818668 | 1.26924 | 1.71568 | 1.352 |
| chr19 | 31160159 | 31160958 | 2.50833 | 3.39087 | 1.352 |
| chr8 | 793367 | 793595 | 1.06835 | 1.44424 | 1.352 |
| chr17 | 53947184 | 53947419 | 0.620355 | 0.838713 | 1.352 |
| chr17 | 47485293 | 47485696 | 3.20304 | 4.33053 | 1.352 |
| chr14 | 44529970 | 44530380 | 0.80131 | 1.08339 | 1.352 |
| chr10 | 67913905 | 67914172 | 0.543605 | 0.734977 | 1.352 |
| chr20 | 11696939 | 11697145 | 0.921583 | 1.24602 | 1.352 |
| chr15 | 51829142 | 51829486 | 1.5886 | 2.14788 | 1.352 |
| chr9 | 130683873 | 130684234 | 2.03568 | 2.75242 | 1.352 |
| chr4 | 8919710 | 8920093 | 1.94138 | 2.62499 | 1.352 |
| chr12 | 108731040 | 108731415 | 1.56993 | 2.12279 | 1.352 |
| chr5 | 125601674 | 125601896 | 0.824455 | 1.1149 | 1.352 |
| chr19 | 55657454 | 55657836 | 3.27439 | 4.42816 | 1.352 |
| chr2 | 119785992 | 119786248 | 1.66249 | 2.24835 | 1.352 |
| chr17 | 6157272 | 6157487 | 0.84647 | 1.1448 | 1.352 |
| chr1 | 9294504 | 9294728 | 1.47746 | 1.99827 | 1.353 |
| chr2 | 131554958 | 131555955 | 2.37303 | 3.20959 | 1.353 |

| | | | | | |
|---|---|---|---|---|---|
| chr1 | 180451678 | 180451886 | 1.23319 | 1.66809 | 1.353 |
| chr8 | 41278604 | 41278843 | 1.25336 | 1.69539 | 1.353 |
| chr15 | 61270955 | 61271214 | 0.656011 | 0.88744 | 1.353 |
| chr2 | 110873974 | 110874264 | 2.27194 | 3.07357 | 1.353 |
| chr20 | 118472 | 119136 | 1.15401 | 1.5612 | 1.353 |
| chr13 | 110760537 | 110760811 | 0.935484 | 1.26559 | 1.353 |
| chr10 | 99519101 | 99519577 | 2.38477 | 3.2263 | 1.353 |
| chr5 | 38420780 | 38421105 | 1.95471 | 2.64486 | 1.353 |
| chr14 | 29691875 | 29692119 | 0.746116 | 1.00957 | 1.353 |
| chr5 | 157602011 | 157602259 | 1.79559 | 2.42964 | 1.353 |
| chr9 | 115432656 | 115432953 | 0.680441 | 0.920753 | 1.353 |
| chr4 | 113558516 | 113558719 | 1.67337 | 2.26439 | 1.353 |
| chr10 | 861601 | 861967 | 0.930825 | 1.2596 | 1.353 |
| chr12 | 2591381 | 2591648 | 0.606795 | 0.821119 | 1.353 |
| chr16 | 4454402 | 4454863 | 2.79144 | 3.77754 | 1.353 |
| chr6 | 159673936 | 159674233 | 1.6193 | 2.19148 | 1.353 |
| chr8 | 141118361 | 141118714 | 0.731297 | 0.989728 | 1.353 |
| chr10 | 114812035 | 114812253 | 1.13403 | 1.53484 | 1.353 |
| chr7 | 100080757 | 100081350 | 3.55272 | 4.80859 | 1.354 |
| chr4 | 171059868 | 171060074 | 1.80034 | 2.43682 | 1.354 |
| chr1 | 226796459 | 226796771 | 2.37657 | 3.21707 | 1.354 |
| chr12 | 102715182 | 102715406 | 0.608982 | 0.824375 | 1.354 |
| chr11 | 6384264 | 6384708 | 2.7225 | 3.68571 | 1.354 |
| chr2 | 228726973 | 228727255 | 1.68817 | 2.28548 | 1.354 |
| chr19 | 28245565 | 28245783 | 0.95696 | 1.29561 | 1.354 |
| chr20 | 2859811 | 2860031 | 0.626982 | 0.848856 | 1.354 |
| chr13 | 74282628 | 74282818 | 1.80945 | 2.44983 | 1.354 |
| chr8 | 138623420 | 138623765 | 0.802911 | 1.08715 | 1.354 |
| chr8 | 22775974 | 22776373 | 1.84188 | 2.49395 | 1.354 |
| chr1 | 189398760 | 189398991 | 1.57932 | 2.13845 | 1.354 |
| chr21 | 37525367 | 37525728 | 1.70151 | 2.30396 | 1.354 |
| chr5 | 110696055 | 110696494 | 0.686231 | 0.929261 | 1.354 |
| chr8 | 138657578 | 138658072 | 0.858815 | 1.16308 | 1.354 |
| chr3 | 98891691 | 98892352 | 1.49801 | 2.02878 | 1.354 |
| chr18 | 12727007 | 12727363 | 1.40814 | 1.9071 | 1.354 |
| chr16 | 30964977 | 30965380 | 2.05052 | 2.77742 | 1.355 |
| chr8 | 101619332 | 101619559 | 0.713284 | 0.966161 | 1.355 |
| chr1 | 207223269 | 207223510 | 1.75555 | 2.37801 | 1.355 |
| chr5 | 172124555 | 172124808 | 2.48008 | 3.35979 | 1.355 |
| chr20 | 796643 | 797158 | 2.66355 | 3.60839 | 1.355 |
| chr19 | 55708001 | 55708205 | 0.649922 | 0.880475 | 1.355 |
| chr7 | 3735325 | 3735643 | 1.23414 | 1.67198 | 1.355 |
| chr2 | 97173804 | 97174225 | 2.29885 | 3.11445 | 1.355 |
| chr15 | 101555338 | 101555795 | 3.38083 | 4.58136 | 1.355 |
| chr1 | 151969361 | 151969615 | 1.37227 | 1.85961 | 1.355 |
| chr15 | 90102367 | 90102637 | 0.699991 | 0.948596 | 1.355 |
| chr9 | 35790326 | 35790732 | 2.28596 | 3.09801 | 1.355 |
| chr6 | 108542778 | 108543040 | 0.58807 | 0.79699 | 1.355 |
| chr17 | 32001677 | 32001910 | 1.71591 | 2.32562 | 1.355 |
| chr1 | 160261049 | 160261466 | 1.94408 | 2.63495 | 1.355 |
| chr3 | 32381668 | 32381872 | 1.68048 | 2.27776 | 1.355 |
| chr12 | 4966727 | 4967229 | 2.02608 | 2.74622 | 1.355 |
| chr5 | 148813964 | 148814344 | 2.51681 | 3.4115 | 1.355 |
| chr3 | 58163407 | 58163824 | 1.80331 | 2.44444 | 1.356 |
| chr17 | 46024044 | 46024668 | 1.951 | 2.64476 | 1.356 |
| chr17 | 45867444 | 45867766 | 1.7098 | 2.31786 | 1.356 |
| chr10 | 121692758 | 121692981 | 1.16294 | 1.5767 | 1.356 |
| chr14 | 105363032 | 105363396 | 2.78371 | 3.77415 | 1.356 |
| chr7 | 49698289 | 49698765 | 0.685483 | 0.929388 | 1.356 |
| chr2 | 220324129 | 220324522 | 1.98371 | 2.68965 | 1.356 |
| chr10 | 78764771 | 78765023 | 2.07004 | 2.8069 | 1.356 |

| | | | | | |
|---|---|---|---|---|---|
| chr10 | 120385081 | 120385295 | 0.660741 | 0.895971 | 1.356 |
| chr17 | 72172771 | 72173171 | 2.44212 | 3.31154 | 1.356 |
| chr17 | 45920423 | 45921039 | 1.98441 | 2.69091 | 1.356 |
| chr6 | 43231163 | 43231542 | 2.23634 | 3.03308 | 1.356 |
| chr19 | 50009788 | 50010170 | 2.56475 | 3.47858 | 1.356 |
| chr17 | 48217187 | 48217613 | 3.22281 | 4.37116 | 1.356 |
| chr8 | 117838416 | 117838629 | 0.620523 | 0.841645 | 1.356 |
| chr5 | 29724177 | 29724442 | 0.662352 | 0.898397 | 1.356 |
| chr6 | 34427735 | 34428001 | 1.53064 | 2.07612 | 1.356 |
| chr1 | 149933971 | 149934403 | 1.45422 | 1.97249 | 1.356 |
| chr14 | 72780034 | 72780385 | 0.676407 | 0.917507 | 1.356 |
| chr20 | 60473252 | 60473488 | 0.580238 | 0.787112 | 1.357 |
| chr17 | 46413348 | 46413586 | 1.61903 | 2.19631 | 1.357 |
| chr2 | 103457638 | 103458010 | 1.45812 | 1.97816 | 1.357 |
| chr7 | 107623295 | 107623542 | 1.01255 | 1.37378 | 1.357 |
| chr19 | 34489630 | 34489956 | 1.32736 | 1.80098 | 1.357 |
| chr4 | 15397786 | 15398135 | 1.4219 | 1.92926 | 1.357 |
| chr14 | 101372851 | 101373075 | 1.4898 | 2.02153 | 1.357 |
| chr1 | 193152248 | 193152477 | 1.9015 | 2.58019 | 1.357 |
| chr1 | 104441709 | 104441934 | 0.682186 | 0.925708 | 1.357 |
| chr13 | 20392281 | 20392751 | 1.91651 | 2.60069 | 1.357 |
| chr10 | 117357185 | 117357530 | 0.721617 | 0.979252 | 1.357 |
| chr6 | 106532583 | 106532863 | 2.15397 | 2.92297 | 1.357 |
| chr17 | 71187553 | 71187929 | 1.89836 | 2.57621 | 1.357 |
| chr9 | 73835610 | 73835953 | 0.827224 | 1.12262 | 1.357 |
| chr9 | 24805657 | 24805930 | 0.770624 | 1.04588 | 1.357 |
| chr15 | 27109894 | 27110181 | 1.69898 | 2.30587 | 1.357 |
| chr1 | 166791724 | 166791949 | 1.21419 | 1.64793 | 1.357 |
| chr19 | 35130366 | 35130835 | 2.48184 | 3.3686 | 1.357 |
| chr3 | 165275108 | 165275357 | 0.519259 | 0.704789 | 1.357 |
| chr1 | 153600043 | 153600328 | 2.23854 | 3.03858 | 1.357 |
| chr5 | 31117469 | 31117675 | 0.882831 | 1.19835 | 1.357 |
| chr10 | 98155689 | 98156137 | 2.24339 | 3.04524 | 1.357 |
| chr7 | 19091519 | 19091761 | 2.0896 | 2.83655 | 1.357 |
| chr18 | 54536458 | 54536716 | 1.58522 | 2.15196 | 1.358 |
| chr10 | 102430735 | 102431303 | 2.61385 | 3.54842 | 1.358 |
| chr2 | 40987215 | 40987692 | 1.86495 | 2.53177 | 1.358 |
| chr2 | 45324243 | 45324485 | 0.796893 | 1.08182 | 1.358 |
| chr20 | 48701990 | 48702303 | 2.53958 | 3.44761 | 1.358 |
| chr17 | 286557 | 286760 | 0.70626 | 0.958792 | 1.358 |
| chr4 | 76498631 | 76499118 | 3.08218 | 4.18427 | 1.358 |
| chr2 | 209120024 | 209120309 | 1.33547 | 1.81301 | 1.358 |
| chr1 | 55013450 | 55013669 | 1.20925 | 1.64179 | 1.358 |
| chr19 | 2199259 | 2199587 | 1.42608 | 1.93654 | 1.358 |
| chr1 | 95271341 | 95271732 | 2.22745 | 3.02487 | 1.358 |
| chr1 | 230389901 | 230390180 | 0.605357 | 0.822117 | 1.358 |
| chrX | 154146268 | 154146533 | 1.32137 | 1.79458 | 1.358 |
| chr18 | 44562687 | 44562963 | 1.29296 | 1.75614 | 1.358 |
| chr12 | 48911899 | 48912132 | 0.858836 | 1.16652 | 1.358 |
| chr11 | 59521375 | 59521761 | 2.56828 | 3.48842 | 1.358 |
| chr17 | 26207052 | 26207499 | 2.52391 | 3.42822 | 1.358 |
| chr11 | 24336373 | 24336648 | 0.791366 | 1.07493 | 1.358 |
| chr2 | 56359387 | 56359614 | 1.95062 | 2.64965 | 1.358 |
| chr1 | 74969868 | 74970135 | 1.79168 | 2.43377 | 1.358 |
| chr10 | 97989352 | 97989656 | 2.50893 | 3.40852 | 1.359 |
| chr10 | 95271136 | 95271378 | 1.07452 | 1.45984 | 1.359 |
| chr20 | 30440502 | 30440886 | 3.10704 | 4.22132 | 1.359 |
| chr9 | 83183711 | 83183927 | 0.726436 | 0.986956 | 1.359 |
| chr20 | 20072504 | 20072727 | 0.67469 | 0.916677 | 1.359 |
| chr5 | 141205646 | 141206022 | 1.15755 | 1.57272 | 1.359 |
| chr1 | 25830390 | 25830594 | 1.69211 | 2.29902 | 1.359 |

| | | | | | |
|---|---|---|---|---|---|
| chr1 | 208735049 | 208735275 | 0.69285 | 0.941397 | 1.359 |
| chr2 | 150417865 | 150418118 | 1.76259 | 2.39491 | 1.359 |
| chr2 | 218858553 | 218858869 | 2.39456 | 3.25387 | 1.359 |
| chr20 | 1987102 | 1987943 | 1.53471 | 2.08548 | 1.359 |
| chr2 | 4852973 | 4853213 | 0.59747 | 0.811941 | 1.359 |
| chr19 | 18107515 | 18108015 | 1.888 | 2.56577 | 1.359 |
| chr16 | 66623182 | 66623606 | 2.05849 | 2.79768 | 1.359 |
| chr8 | 14000199 | 14000479 | 0.722777 | 0.982324 | 1.359 |
| chr9 | 135134552 | 135134983 | 2.77246 | 3.76814 | 1.359 |
| chr1 | 38027448 | 38027846 | 1.2963 | 1.76186 | 1.359 |
| chr9 | 96589524 | 96590178 | 2.06621 | 2.80856 | 1.359 |
| chr3 | 40032127 | 40032363 | 1.64749 | 2.23949 | 1.359 |
| chr8 | 1972055 | 1972564 | 1.87194 | 2.54466 | 1.359 |
| chr10 | 53145156 | 53145458 | 0.678388 | 0.922224 | 1.359 |
| chr20 | 16149876 | 16150264 | 0.875081 | 1.18961 | 1.359 |
| chrX | 48369890 | 48370227 | 1.25874 | 1.71135 | 1.360 |
| chr19 | 5340073 | 5340283 | 1.05455 | 1.43387 | 1.360 |
| chr10 | 98386632 | 98386842 | 1.17553 | 1.59838 | 1.360 |
| chr8 | 60668146 | 60668369 | 0.747911 | 1.01695 | 1.360 |
| chr22 | 40921364 | 40921577 | 1.55098 | 2.10902 | 1.360 |
| chr5 | 114194802 | 114195133 | 0.805362 | 1.09513 | 1.360 |
| chr1 | 151861597 | 151861954 | 1.01463 | 1.37973 | 1.360 |
| chr4 | 104707273 | 104707551 | 1.54223 | 2.09735 | 1.360 |
| chr11 | 116087815 | 116088063 | 1.68011 | 2.28514 | 1.360 |
| chr2 | 180866330 | 180866657 | 0.727015 | 0.988817 | 1.360 |
| chr15 | 70124078 | 70124338 | 0.70968 | 0.965263 | 1.360 |
| chr9 | 27033904 | 27034146 | 1.42035 | 1.932 | 1.360 |
| chr4 | 119849141 | 119849365 | 0.791023 | 1.07599 | 1.360 |
| chr7 | 150011030 | 150011327 | 0.569415 | 0.774606 | 1.360 |
| chr5 | 78405827 | 78406033 | 1.30004 | 1.76857 | 1.360 |
| chr1 | 77613993 | 77614272 | 1.22266 | 1.66341 | 1.360 |
| chr16 | 55657234 | 55657540 | 1.17025 | 1.59213 | 1.361 |
| chr2 | 29857038 | 29857254 | 0.393308 | 0.5351 | 1.361 |
| chr3 | 124620177 | 124620563 | 2.94913 | 4.01246 | 1.361 |
| chr17 | 10529667 | 10530031 | 0.744499 | 1.01294 | 1.361 |
| chr1 | 203309256 | 203309527 | 1.98224 | 2.69706 | 1.361 |
| chr10 | 6689733 | 6690274 | 1.6788 | 2.2843 | 1.361 |
| gl000195_ra | 1259 | 1524 | 0.951522 | 1.29488 | 1.361 |
| chr8 | 67108135 | 67108384 | 0.731674 | 0.99575 | 1.361 |
| chr15 | 98462661 | 98462889 | 0.828313 | 1.12733 | 1.361 |
| chr17 | 52650909 | 52651125 | 0.846003 | 1.15146 | 1.361 |
| chr7 | 52082224 | 52082430 | 0.72791 | 0.990729 | 1.361 |
| chr12 | 125251489 | 125251886 | 3.201 | 4.35677 | 1.361 |
| chr2 | 46318182 | 46318490 | 1.14004 | 1.55178 | 1.361 |
| chr14 | 88082536 | 88082750 | 1.0772 | 1.46627 | 1.361 |
| chr1 | 197893711 | 197893956 | 2.3789 | 3.23836 | 1.361 |
| chr10 | 107638556 | 107638809 | 0.908288 | 1.23656 | 1.361 |
| chr5 | 71906996 | 71907280 | 2.38987 | 3.25361 | 1.361 |
| chr20 | 7000490 | 7000797 | 0.960288 | 1.30741 | 1.361 |
| chr14 | 65289929 | 65290311 | 2.79374 | 3.80376 | 1.362 |
| chr15 | 78259129 | 78259735 | 1.73186 | 2.35809 | 1.362 |
| chr17 | 19471038 | 19471279 | 0.985988 | 1.34257 | 1.362 |
| chr8 | 57511725 | 57511963 | 0.744516 | 1.01391 | 1.362 |
| chr19 | 36282056 | 36282266 | 1.1559 | 1.57416 | 1.362 |
| chr19 | 32271028 | 32271403 | 0.786714 | 1.07141 | 1.362 |
| chr5 | 16456367 | 16456760 | 1.56099 | 2.12609 | 1.362 |
| chr1 | 3369670 | 3370124 | 2.54284 | 3.46349 | 1.362 |
| chr16 | 81929560 | 81929799 | 1.37679 | 1.87572 | 1.362 |
| chr6 | 2738683 | 2738945 | 1.48525 | 2.02354 | 1.362 |
| chr10 | 61968517 | 61968714 | 1.80766 | 2.46283 | 1.362 |
| chr12 | 114630889 | 114631127 | 0.599925 | 0.817368 | 1.362 |

| | | | | | |
|---|---|---|---|---|---|
| chr2 | 200350756 | 200351032 | 2.13559 | 2.90966 | 1.362 |
| chr7 | 56142014 | 56142353 | 2.43678 | 3.32005 | 1.362 |
| chr2 | 113732686 | 113732927 | 0.983744 | 1.34036 | 1.363 |
| chr20 | 20048836 | 20049200 | 0.695377 | 0.947489 | 1.363 |
| chr8 | 38614244 | 38614558 | 2.12193 | 2.89125 | 1.363 |
| chr17 | 77948557 | 77949008 | 2.23026 | 3.03904 | 1.363 |
| chr1 | 165561999 | 165562523 | 1.50405 | 2.04949 | 1.363 |
| chrX | 102788381 | 102788684 | 1.44305 | 1.96641 | 1.363 |
| chr3 | 80818955 | 80819186 | 0.806713 | 1.09933 | 1.363 |
| chr10 | 9251187 | 9251478 | 2.24967 | 3.06583 | 1.363 |
| chr19 | 33790234 | 33790565 | 0.743698 | 1.01352 | 1.363 |
| chr8 | 26184982 | 26185195 | 1.73039 | 2.35828 | 1.363 |
| chr14 | 95215171 | 95215500 | 2.34134 | 3.19108 | 1.363 |
| chr20 | 13100186 | 13100561 | 2.73142 | 3.7229 | 1.363 |
| chr9 | 122723986 | 122724434 | 2.83105 | 3.85878 | 1.363 |
| chr12 | 96832575 | 96832793 | 0.746564 | 1.01759 | 1.363 |
| chr9 | 132372840 | 132373190 | 2.99217 | 4.07864 | 1.363 |
| chr5 | 72002894 | 72003144 | 0.781096 | 1.06472 | 1.363 |
| chr1 | 19567740 | 19568093 | 2.25174 | 3.06966 | 1.363 |
| chr17 | 74609288 | 74609571 | 0.815347 | 1.11153 | 1.363 |
| chr10 | 36813090 | 36813364 | 0.484345 | 0.660291 | 1.363 |
| chr9 | 101775225 | 101775544 | 0.872235 | 1.18919 | 1.363 |
| chr11 | 119350478 | 119350686 | 1.64628 | 2.2446 | 1.363 |
| chr12 | 109931007 | 109931233 | 0.623963 | 0.850762 | 1.363 |
| chr10 | 118986415 | 118986718 | 1.95448 | 2.66498 | 1.364 |
| chr10 | 97006291 | 97006668 | 2.08887 | 2.84824 | 1.364 |
| chr10 | 32656828 | 32657322 | 1.2203 | 1.66393 | 1.364 |
| chr3 | 120854585 | 120854790 | 0.644543 | 0.878873 | 1.364 |
| chr19 | 47082011 | 47082226 | 0.687278 | 0.937224 | 1.364 |
| chr17 | 50498288 | 50498605 | 0.708785 | 0.966563 | 1.364 |
| chr17 | 12085797 | 12086133 | 0.704561 | 0.960811 | 1.364 |
| chr10 | 1156051 | 1156499 | 1.59121 | 2.17002 | 1.364 |
| chr10 | 72663046 | 72663583 | 3.10687 | 4.237 | 1.364 |
| chr19 | 46512981 | 46513622 | 2.18064 | 2.97395 | 1.364 |
| chr20 | 33126452 | 33126891 | 2.59794 | 3.54332 | 1.364 |
| chr2 | 232477251 | 232477576 | 2.07088 | 2.82456 | 1.364 |
| chr14 | 88787259 | 88787476 | 0.552766 | 0.753944 | 1.364 |
| chr15 | 91576285 | 91576779 | 1.31166 | 1.78909 | 1.364 |
| chr7 | 98983100 | 98983419 | 0.587997 | 0.802042 | 1.364 |
| chr10 | 122495068 | 122495865 | 1.2062 | 1.64532 | 1.364 |
| chr1 | 206781450 | 206781854 | 2.74175 | 3.73993 | 1.364 |
| chr16 | 76888205 | 76888522 | 1.35507 | 1.84841 | 1.364 |
| chrX | 51357359 | 51357567 | 1.488 | 2.02976 | 1.364 |
| chr16 | 51852282 | 51852643 | 1.72827 | 2.35762 | 1.364 |
| chr4 | 6548367 | 6548717 | 2.80371 | 3.82491 | 1.364 |
| chr5 | 164911820 | 164912150 | 0.834018 | 1.13791 | 1.364 |
| chr11 | 64535041 | 64535622 | 1.98027 | 2.70188 | 1.364 |
| chr10 | 112114120 | 112114468 | 2.38964 | 3.26056 | 1.364 |
| chr20 | 20190533 | 20191018 | 0.760608 | 1.03786 | 1.365 |
| chr8 | 86209518 | 86210358 | 1.44652 | 1.9738 | 1.365 |
| chr12 | 71360581 | 71360949 | 0.705961 | 0.963322 | 1.365 |
| chr16 | 69184088 | 69184367 | 1.56546 | 2.13619 | 1.365 |
| chr9 | 136114446 | 136114872 | 3.63995 | 4.96707 | 1.365 |
| chr12 | 24238031 | 24238243 | 0.78068 | 1.06533 | 1.365 |
| chr14 | 20376956 | 20377203 | 1.2022 | 1.6406 | 1.365 |
| chr8 | 82023891 | 82024185 | 0.717788 | 0.979576 | 1.365 |
| chr2 | 43836513 | 43836768 | 1.69029 | 2.30677 | 1.365 |
| chr3 | 2501984 | 2502192 | 0.749388 | 1.02274 | 1.365 |
| chr5 | 170700415 | 170700642 | 1.33415 | 1.82084 | 1.365 |
| chr1 | 167772526 | 167772890 | 2.54264 | 3.47021 | 1.365 |
| chr12 | 44480135 | 44480536 | 0.799485 | 1.09117 | 1.365 |

| | | | | | |
|---|---|---|---|---|---|
| chr1 | 224832232 | 224832649 | 1.80475 | 2.46325 | 1.365 |
| chr17 | 27864012 | 27864300 | 1.23429 | 1.68467 | 1.365 |
| chr1 | 158005708 | 158006109 | 1.74623 | 2.38343 | 1.365 |
| chr8 | 37144976 | 37145208 | 0.7485 | 1.02166 | 1.365 |
| chr2 | 169777203 | 169777411 | 1.40122 | 1.91262 | 1.365 |
| chr6 | 97842081 | 97842352 | 1.64656 | 2.24755 | 1.365 |
| chr16 | 710489 | 711163 | 1.39842 | 1.90887 | 1.365 |
| chr17 | 53719039 | 53719450 | 1.33213 | 1.81844 | 1.365 |
| chr17 | 5783851 | 5784130 | 0.594873 | 0.812068 | 1.365 |
| chr18 | 57450294 | 57450635 | 1.07746 | 1.47087 | 1.365 |
| chrX | 130811674 | 130812074 | 1.53999 | 2.10234 | 1.365 |
| chr17 | 73996065 | 73997019 | 1.83012 | 2.49846 | 1.365 |
| chrX | 102090694 | 102091072 | 1.19875 | 1.63663 | 1.365 |
| chr8 | 136849398 | 136849609 | 0.767397 | 1.04779 | 1.365 |
| chr5 | 137474934 | 137475169 | 0.794851 | 1.08531 | 1.365 |
| chr7 | 122328487 | 122328699 | 0.603865 | 0.824536 | 1.365 |
| chr15 | 99710804 | 99711055 | 0.840441 | 1.14765 | 1.366 |
| chr6 | 143013913 | 143014291 | 1.55363 | 2.12165 | 1.366 |
| chr19 | 40040166 | 40040464 | 0.899365 | 1.22826 | 1.366 |
| chr10 | 26846036 | 26846357 | 2.3081 | 3.15222 | 1.366 |
| chr17 | 35930651 | 35930954 | 1.84281 | 2.5168 | 1.366 |
| chr20 | 42166722 | 42167149 | 2.05572 | 2.80781 | 1.366 |
| chr3 | 47891484 | 47891961 | 1.17327 | 1.60253 | 1.366 |
| chr8 | 57361382 | 57361738 | 1.28812 | 1.75941 | 1.366 |
| chr20 | 45376639 | 45376916 | 2.31242 | 3.15853 | 1.366 |
| chr14 | 106015927 | 106016171 | 0.952609 | 1.3012 | 1.366 |
| chr18 | 74092750 | 74093083 | 2.27581 | 3.10865 | 1.366 |
| chr16 | 48088852 | 48089312 | 2.44883 | 3.34508 | 1.366 |
| chr15 | 83995983 | 83996211 | 0.612416 | 0.836573 | 1.366 |
| chr2 | 25014004 | 25014208 | 1.47992 | 2.02166 | 1.366 |
| chr16 | 84985799 | 84986184 | 1.87249 | 2.55799 | 1.366 |
| chr12 | 123673654 | 123673893 | 0.747401 | 1.02111 | 1.366 |
| chr5 | 1174494 | 1174899 | 2.61384 | 3.57105 | 1.366 |
| chr11 | 67381627 | 67382171 | 1.75433 | 2.39681 | 1.366 |
| chr18 | 39486723 | 39486951 | 1.03446 | 1.41336 | 1.366 |
| chr8 | 64213288 | 64213674 | 1.65637 | 2.26309 | 1.366 |
| chr5 | 127830355 | 127830638 | 1.23664 | 1.6898 | 1.366 |
| chr15 | 75474814 | 75475327 | 1.12528 | 1.53784 | 1.367 |
| chr8 | 72732077 | 72732534 | 1.47668 | 2.01812 | 1.367 |
| chr15 | 77909882 | 77910228 | 2.22166 | 3.03633 | 1.367 |
| chr15 | 35498543 | 35498804 | 1.98913 | 2.71854 | 1.367 |
| chr5 | 12023834 | 12024033 | 1.41711 | 1.93709 | 1.367 |
| chr2 | 171678806 | 171679065 | 1.64785 | 2.2526 | 1.367 |
| chr3 | 172341109 | 172341381 | 1.35267 | 1.84913 | 1.367 |
| chr3 | 33481453 | 33481686 | 1.07457 | 1.46897 | 1.367 |
| chr11 | 59292239 | 59292514 | 0.767899 | 1.04982 | 1.367 |
| chr19 | 32715362 | 32716015 | 2.94191 | 4.02196 | 1.367 |
| chr3 | 193855624 | 193855971 | 1.73704 | 2.37496 | 1.367 |
| chr1 | 157550055 | 157550293 | 1.63134 | 2.23053 | 1.367 |
| chrX | 149756158 | 149756407 | 1.68057 | 2.29794 | 1.367 |
| chrX | 149940657 | 149941020 | 3.04047 | 4.15744 | 1.367 |
| chr2 | 130607254 | 130607492 | 1.48214 | 2.02672 | 1.367 |
| chr10 | 15637755 | 15638006 | 0.651201 | 0.890482 | 1.367 |
| chr6 | 147704171 | 147704374 | 1.14587 | 1.56696 | 1.367 |
| chr2 | 86322020 | 86322228 | 0.765678 | 1.04709 | 1.368 |
| chr5 | 66045020 | 66045240 | 0.637517 | 0.87183 | 1.368 |
| chr21 | 47300375 | 47300887 | 1.35252 | 1.84976 | 1.368 |
| chr1 | 164642924 | 164643185 | 1.80724 | 2.47168 | 1.368 |
| chr15 | 92769338 | 92769631 | 1.23767 | 1.69281 | 1.368 |
| chr15 | 86654306 | 86654598 | 0.642302 | 0.878524 | 1.368 |
| chr10 | 103989491 | 103989871 | 2.95288 | 4.03919 | 1.368 |

| | | | | | |
|---|---|---|---|---|---|
| chr2 | 128173500 | 128173745 | 1.6517 | 2.25944 | 1.368 |
| chr7 | 154535977 | 154536304 | 2.21065 | 3.02408 | 1.368 |
| chr3 | 64449261 | 64449547 | 1.28182 | 1.75371 | 1.368 |
| chr3 | 128880486 | 128880696 | 1.43041 | 1.95702 | 1.368 |
| chr2 | 79590272 | 79590503 | 0.830549 | 1.13643 | 1.368 |
| chr2 | 9126743 | 9127094 | 1.39289 | 1.9059 | 1.368 |
| chr2 | 71451123 | 71451401 | 2.24484 | 3.07164 | 1.368 |
| chr10 | 71559819 | 71560297 | 1.83276 | 2.50814 | 1.369 |
| chr19 | 29369861 | 29370456 | 1.20017 | 1.64251 | 1.369 |
| chr10 | 115682909 | 115683195 | 0.79157 | 1.08334 | 1.369 |
| chr20 | 62700830 | 62701571 | 2.53916 | 3.47544 | 1.369 |
| chr17 | 80512307 | 80512738 | 1.79092 | 2.45138 | 1.369 |
| chrX | 109399449 | 109399780 | 1.16324 | 1.59236 | 1.369 |
| chr12 | 26276190 | 26276422 | 1.49168 | 2.04212 | 1.369 |
| chr1 | 223705130 | 223705633 | 1.53352 | 2.09948 | 1.369 |
| chr18 | 74271201 | 74271425 | 0.608429 | 0.833018 | 1.369 |
| chr6 | 138328095 | 138328402 | 1.50056 | 2.05456 | 1.369 |
| chr17 | 27893028 | 27893364 | 1.22687 | 1.67995 | 1.369 |
| chr3 | 20381816 | 20382237 | 2.57336 | 3.52397 | 1.369 |
| chr2 | 71757717 | 71758137 | 1.25288 | 1.71572 | 1.369 |
| chr12 | 16977083 | 16977382 | 0.747334 | 1.02343 | 1.369 |
| chr8 | 3219719 | 3219960 | 0.676495 | 0.926421 | 1.369 |
| chr19 | 28503307 | 28503513 | 0.776482 | 1.06335 | 1.369 |
| chr19 | 7149172 | 7149444 | 2.55826 | 3.50347 | 1.369 |
| chr17 | 55817262 | 55817796 | 1.40071 | 1.91824 | 1.369 |
| chr19 | 39369536 | 39370158 | 0.779158 | 1.06704 | 1.369 |
| chr11 | 85537218 | 85537528 | 1.64716 | 2.25604 | 1.370 |
| chr22 | 23516108 | 23516442 | 2.02221 | 2.76974 | 1.370 |
| chr16 | 81238743 | 81238964 | 1.44383 | 1.97757 | 1.370 |
| chr1 | 177501255 | 177501501 | 0.606066 | 0.830114 | 1.370 |
| chr17 | 72732901 | 72733173 | 2.06944 | 2.83449 | 1.370 |
| chr5 | 71564983 | 71565367 | 2.00404 | 2.745 | 1.370 |
| chr20 | 20164726 | 20165131 | 0.852428 | 1.16765 | 1.370 |
| chr2 | 11924440 | 11925490 | 2.20481 | 3.02031 | 1.370 |
| chr14 | 24038034 | 24038264 | 1.59738 | 2.1884 | 1.370 |
| chr15 | 78167456 | 78167870 | 1.51629 | 2.07741 | 1.370 |
| chr7 | 47305876 | 47306208 | 2.41274 | 3.30562 | 1.370 |
| chr10 | 72024910 | 72025140 | 1.29471 | 1.77385 | 1.370 |
| chr19 | 58840419 | 58840709 | 1.18053 | 1.61742 | 1.370 |
| chr10 | 115559426 | 115559661 | 1.64191 | 2.24975 | 1.370 |
| chr18 | 64771985 | 64772243 | 1.53489 | 2.10324 | 1.370 |
| chr17 | 61773941 | 61774241 | 1.49584 | 2.04983 | 1.370 |
| chr17 | 68164067 | 68164488 | 2.12102 | 2.90654 | 1.370 |
| chr15 | 75976076 | 75976417 | 2.29548 | 3.14572 | 1.370 |
| chr1 | 49162659 | 49163109 | 1.62488 | 2.22685 | 1.370 |
| chr2 | 18458316 | 18458571 | 0.678216 | 0.929479 | 1.370 |
| chr2 | 141344041 | 141344319 | 1.25862 | 1.72496 | 1.371 |
| chr20 | 17919592 | 17920009 | 2.3238 | 3.18491 | 1.371 |
| chr10 | 102098242 | 102098560 | 2.71725 | 3.72424 | 1.371 |
| chr18 | 55637757 | 55638063 | 1.1838 | 1.62259 | 1.371 |
| chr17 | 14953976 | 14954286 | 0.787036 | 1.07885 | 1.371 |
| chr2 | 178011928 | 178012146 | 1.13844 | 1.56059 | 1.371 |
| chr19 | 32672522 | 32672755 | 0.809365 | 1.10955 | 1.371 |
| chr10 | 112835882 | 112836622 | 1.69575 | 2.32474 | 1.371 |
| chr11 | 61822065 | 61822269 | 1.50301 | 2.06051 | 1.371 |
| chr6 | 17804465 | 17804712 | 1.98899 | 2.72675 | 1.371 |
| chr6 | 41673649 | 41674077 | 2.1226 | 2.90992 | 1.371 |
| chr12 | 106620614 | 106620878 | 1.68594 | 2.31144 | 1.371 |
| chr7 | 132959799 | 132960014 | 0.974689 | 1.33636 | 1.371 |
| chr18 | 19494755 | 19494964 | 0.887576 | 1.21697 | 1.371 |
| chr17 | 71325544 | 71325750 | 0.637891 | 0.874651 | 1.371 |

| | | | | | |
|---|---|---|---|---|---|
| chr17 | 40845282 | 40845506 | 0.630982 | 0.865188 | 1.371 |
| chr7 | 107162689 | 107162921 | 0.590802 | 0.810142 | 1.371 |
| chr8 | 42100245 | 42100539 | 0.690348 | 0.946732 | 1.371 |
| chr20 | 47274261 | 47274696 | 2.61232 | 3.58251 | 1.371 |
| chr17 | 10220515 | 10220946 | 2.07716 | 2.84874 | 1.371 |
| chr4 | 152722764 | 152723143 | 2.15487 | 2.95564 | 1.372 |
| chr11 | 70962815 | 70963131 | 2.06063 | 2.82648 | 1.372 |
| chr7 | 91459252 | 91459655 | 1.65269 | 2.26698 | 1.372 |
| chr1 | 6101573 | 6101888 | 1.44007 | 1.97594 | 1.372 |
| chr17 | 77715680 | 77716187 | 3.7639 | 5.16452 | 1.372 |
| chr17 | 42317947 | 42318156 | 0.886258 | 1.21607 | 1.372 |
| chr5 | 51901551 | 51901833 | 0.655131 | 0.898961 | 1.372 |
| chr2 | 77965135 | 77965443 | 0.745211 | 1.02258 | 1.372 |
| chr12 | 81639621 | 81640070 | 0.85185 | 1.16896 | 1.372 |
| chr19 | 4589638 | 4589885 | 1.63473 | 2.2433 | 1.372 |
| chr1 | 214506165 | 214506792 | 2.35663 | 3.23413 | 1.372 |
| chr20 | 25171556 | 25171781 | 0.708077 | 0.971808 | 1.372 |
| chr1 | 19667915 | 19668376 | 1.86605 | 2.56113 | 1.372 |
| chr10 | 98270970 | 98271341 | 1.80554 | 2.47809 | 1.372 |
| chr9 | 115606970 | 115607332 | 2.43295 | 3.33919 | 1.372 |
| chr9 | 138494503 | 138494978 | 2.32565 | 3.19203 | 1.373 |
| chr16 | 81730797 | 81731218 | 2.61766 | 3.59288 | 1.373 |
| chr15 | 64978424 | 64978823 | 1.63039 | 2.2379 | 1.373 |
| chr8 | 107868797 | 107869029 | 1.82334 | 2.50295 | 1.373 |
| chr15 | 91436436 | 91436666 | 1.47402 | 2.02344 | 1.373 |
| chr10 | 57634510 | 57634764 | 0.855303 | 1.17417 | 1.373 |
| chr2 | 78545685 | 78545947 | 0.640987 | 0.879982 | 1.373 |
| chr2 | 239279461 | 239279665 | 1.44149 | 1.9791 | 1.373 |
| chr1 | 201118552 | 201118840 | 1.72311 | 2.36586 | 1.373 |
| chr1 | 63907400 | 63907625 | 1.69998 | 2.33415 | 1.373 |
| chr4 | 73192514 | 73192758 | 1.23332 | 1.69343 | 1.373 |
| chr7 | 28379111 | 28379565 | 0.682837 | 0.937586 | 1.373 |
| chr2 | 181969291 | 181969728 | 1.44066 | 1.97816 | 1.373 |
| chr11 | 1727908 | 1728192 | 1.75577 | 2.41091 | 1.373 |
| chr17 | 12421578 | 12421889 | 0.865337 | 1.18824 | 1.373 |
| chr22 | 38019570 | 38019987 | 1.94268 | 2.66776 | 1.373 |
| chr16 | 85206242 | 85206503 | 1.48794 | 2.04331 | 1.373 |
| chr17 | 71366056 | 71366444 | 1.72934 | 2.37482 | 1.373 |
| chr16 | 77203578 | 77203872 | 1.88691 | 2.59121 | 1.373 |
| chr12 | 111889499 | 111889788 | 1.96402 | 2.69715 | 1.373 |
| chr8 | 1273330 | 1273746 | 1.20457 | 1.6543 | 1.373 |
| chrX | 154314765 | 154315049 | 1.73941 | 2.38902 | 1.373 |
| chr20 | 44715821 | 44716069 | 1.23941 | 1.70241 | 1.374 |
| chr3 | 122279526 | 122279790 | 1.74096 | 2.39166 | 1.374 |
| chr3 | 196669237 | 196669503 | 1.51027 | 2.07496 | 1.374 |
| chr2 | 69792250 | 69792800 | 1.86785 | 2.56643 | 1.374 |
| chr17 | 61996659 | 61996884 | 2.83332 | 3.8935 | 1.374 |
| chr1 | 24098590 | 24098989 | 2.03952 | 2.80269 | 1.374 |
| chr17 | 28106205 | 28106610 | 2.10794 | 2.89673 | 1.374 |
| chr12 | 79767725 | 79768070 | 2.74612 | 3.77375 | 1.374 |
| chr1 | 6306813 | 6307169 | 2.49676 | 3.4311 | 1.374 |
| chr5 | 5692162 | 5692387 | 0.698763 | 0.960286 | 1.374 |
| chr16 | 59306484 | 59306691 | 0.899979 | 1.23686 | 1.374 |
| chr19 | 50181201 | 50181464 | 2.08038 | 2.85913 | 1.374 |
| chr16 | 22960150 | 22960430 | 1.93814 | 2.66366 | 1.374 |
| chr19 | 39467171 | 39467641 | 3.30415 | 4.54127 | 1.374 |
| chr7 | 29986460 | 29986718 | 1.26537 | 1.73923 | 1.374 |
| chr16 | 56698357 | 56698600 | 0.724474 | 0.995784 | 1.374 |
| chr7 | 93344573 | 93344865 | 1.91407 | 2.63101 | 1.375 |
| chr17 | 38689579 | 38689793 | 1.50784 | 2.07265 | 1.375 |
| chr5 | 131526000 | 131526242 | 1.77783 | 2.44382 | 1.375 |

| | | | | | |
|---|---|---|---|---|---|
| chr11 | 119883985 | 119884219 | 1.55475 | 2.1372 | 1.375 |
| chr15 | 75061170 | 75061715 | 1.75358 | 2.41068 | 1.375 |
| chr12 | 98816730 | 98816967 | 1.53601 | 2.11165 | 1.375 |
| chr1 | 119345602 | 119345811 | 0.605596 | 0.832579 | 1.375 |
| chr8 | 141562248 | 141562491 | 0.619091 | 0.85115 | 1.375 |
| chr5 | 62086913 | 62087169 | 1.28394 | 1.76531 | 1.375 |
| chr16 | 2473068 | 2473486 | 2.58145 | 3.54935 | 1.375 |
| chr6 | 90142621 | 90142964 | 2.54733 | 3.50242 | 1.375 |
| chr1 | 156859957 | 156860233 | 2.21854 | 3.05039 | 1.375 |
| chr20 | 18225889 | 18226127 | 0.871525 | 1.1983 | 1.375 |
| chr11 | 21533420 | 21533790 | 1.54187 | 2.12001 | 1.375 |
| chr11 | 19959819 | 19960179 | 2.37057 | 3.25951 | 1.375 |
| chr2 | 54725113 | 54725394 | 0.783576 | 1.07745 | 1.375 |
| chr14 | 69556952 | 69557226 | 2.11944 | 2.91463 | 1.375 |
| chr10 | 118047482 | 118047760 | 1.54919 | 2.13053 | 1.375 |
| chr19 | 4636253 | 4636593 | 2.17439 | 2.99037 | 1.375 |
| chr15 | 62637088 | 62637483 | 1.82741 | 2.5133 | 1.375 |
| chr17 | 49464697 | 49464912 | 0.850466 | 1.16985 | 1.376 |
| chr5 | 168374288 | 168374703 | 1.86792 | 2.5694 | 1.376 |
| chr2 | 22970552 | 22970783 | 0.742695 | 1.02164 | 1.376 |
| chr1 | 247654660 | 247654931 | 0.614699 | 0.8456 | 1.376 |
| chr17 | 46055703 | 46056052 | 1.65795 | 2.28082 | 1.376 |
| chr2 | 71098415 | 71099059 | 2.07346 | 2.85261 | 1.376 |
| chr11 | 65902091 | 65902287 | 1.47953 | 2.03578 | 1.376 |
| chr19 | 30598122 | 30598446 | 0.921513 | 1.268 | 1.376 |
| chr6 | 56435329 | 56435941 | 0.992394 | 1.36555 | 1.376 |
| chr22 | 32476975 | 32477212 | 1.56781 | 2.15735 | 1.376 |
| chr10 | 99445432 | 99445662 | 1.32922 | 1.82911 | 1.376 |
| chr1 | 69925242 | 69925448 | 0.650242 | 0.894899 | 1.376 |
| chr7 | 32472646 | 32472848 | 0.822205 | 1.1316 | 1.376 |
| chr1 | 57269246 | 57269470 | 1.08338 | 1.49112 | 1.376 |
| chr19 | 22770091 | 22770380 | 0.644159 | 0.886632 | 1.376 |
| chr5 | 164546585 | 164546857 | 0.667369 | 0.918604 | 1.376 |
| chr8 | 106386768 | 106386990 | 0.657988 | 0.905739 | 1.377 |
| chr8 | 110083312 | 110083560 | 0.828346 | 1.14025 | 1.377 |
| chr3 | 144521409 | 144521762 | 0.626145 | 0.861922 | 1.377 |
| chr6 | 10951954 | 10952210 | 1.1733 | 1.61515 | 1.377 |
| chr7 | 106842346 | 106842592 | 1.55632 | 2.14246 | 1.377 |
| chr20 | 21552580 | 21553087 | 2.07246 | 2.85302 | 1.377 |
| chr17 | 55320424 | 55321026 | 2.84835 | 3.92118 | 1.377 |
| chr3 | 182428416 | 182428669 | 1.32418 | 1.82296 | 1.377 |
| chr1 | 165019471 | 165019758 | 0.765165 | 1.05353 | 1.377 |
| chr8 | 2211755 | 2212615 | 0.933789 | 1.28574 | 1.377 |
| chr10 | 112039193 | 112039427 | 1.71717 | 2.36462 | 1.377 |
| chr7 | 31014424 | 31014868 | 1.79264 | 2.46859 | 1.377 |
| chr9 | 134466518 | 134466918 | 2.68489 | 3.69735 | 1.377 |
| chr11 | 32814402 | 32814747 | 1.74111 | 2.39778 | 1.377 |
| chr3 | 170679486 | 170679862 | 0.909009 | 1.25189 | 1.377 |
| chr19 | 10041616 | 10041900 | 1.72506 | 2.37581 | 1.377 |
| chr2 | 191895302 | 191895560 | 1.28091 | 1.76418 | 1.377 |
| chr10 | 81247393 | 81247671 | 1.57165 | 2.16481 | 1.377 |
| chr14 | 91122295 | 91122518 | 0.537538 | 0.740425 | 1.377 |
| chr22 | 45371577 | 45372065 | 1.85312 | 2.55262 | 1.377 |
| chr16 | 30662621 | 30662891 | 2.58376 | 3.55909 | 1.377 |
| chr1 | 78596519 | 78596853 | 1.10545 | 1.52279 | 1.378 |
| chr1 | 182045179 | 182045400 | 0.666633 | 0.91831 | 1.378 |
| chr6 | 66348009 | 66348217 | 0.962023 | 1.32523 | 1.378 |
| chr12 | 102311522 | 102311866 | 1.54201 | 2.12427 | 1.378 |
| chr7 | 155147894 | 155148160 | 0.740762 | 1.02049 | 1.378 |
| chr1 | 208195161 | 208195383 | 1.85882 | 2.56083 | 1.378 |
| chr12 | 117301038 | 117301359 | 2.02428 | 2.78884 | 1.378 |

| | | | | | |
|---|---|---|---|---|---|
| chr16 | 231168 | 231807 | 2.62485 | 3.61627 | 1.378 |
| chr11 | 65655783 | 65656245 | 2.75792 | 3.79976 | 1.378 |
| chr2 | 143044198 | 143044399 | 1.20938 | 1.66624 | 1.378 |
| chr7 | 41750956 | 41751573 | 1.04917 | 1.4456 | 1.378 |
| chr8 | 146238888 | 146239159 | 0.917823 | 1.26473 | 1.378 |
| chr10 | 76782087 | 76782406 | 1.92815 | 2.65713 | 1.378 |
| chr19 | 50393163 | 50393826 | 2.11665 | 2.91697 | 1.378 |
| chr7 | 48779514 | 48779772 | 0.824829 | 1.13689 | 1.378 |
| chr7 | 155165205 | 155165555 | 1.94454 | 2.6803 | 1.378 |
| chr6 | 43111292 | 43111557 | 2.10849 | 2.90631 | 1.378 |
| chr13 | 99492377 | 99492738 | 1.82014 | 2.50893 | 1.378 |
| chr22 | 45793738 | 45793958 | 0.8838 | 1.21834 | 1.379 |
| chr14 | 35192220 | 35192543 | 1.54953 | 2.13613 | 1.379 |
| chr8 | 141447843 | 141448063 | 1.43112 | 1.97301 | 1.379 |
| chr12 | 112111414 | 112111596 | 1.43945 | 1.98463 | 1.379 |
| chr16 | 24510607 | 24510859 | 1.71014 | 2.35784 | 1.379 |
| chr15 | 33411441 | 33411693 | 0.87684 | 1.2091 | 1.379 |
| chr17 | 63552294 | 63552905 | 1.8385 | 2.53522 | 1.379 |
| chr4 | 115307859 | 115308161 | 1.31348 | 1.81125 | 1.379 |
| chr13 | 99686173 | 99686387 | 1.51553 | 2.0899 | 1.379 |
| chr15 | 66483817 | 66484167 | 1.67198 | 2.30571 | 1.379 |
| chr9 | 85926023 | 85926362 | 0.594139 | 0.819364 | 1.379 |
| chr9 | 133769027 | 133769255 | 1.56751 | 2.16172 | 1.379 |
| chr14 | 103207177 | 103207442 | 0.659676 | 0.909788 | 1.379 |
| chr20 | 37522425 | 37522760 | 2.61047 | 3.60024 | 1.379 |
| chr7 | 2961464 | 2961797 | 1.05529 | 1.45547 | 1.379 |
| chrX | 44929737 | 44929946 | 1.40713 | 1.94109 | 1.379 |
| chr5 | 133332355 | 133332657 | 2.14995 | 2.96582 | 1.379 |
| chr2 | 37747561 | 37747773 | 0.667113 | 0.920321 | 1.380 |
| chr16 | 16277378 | 16277856 | 0.975078 | 1.34519 | 1.380 |
| chr17 | 28452329 | 28452846 | 1.53641 | 2.11972 | 1.380 |
| chr19 | 28523679 | 28523999 | 1.0635 | 1.46738 | 1.380 |
| chr15 | 97213030 | 97213283 | 0.696494 | 0.961124 | 1.380 |
| chr15 | 56578539 | 56578777 | 1.01815 | 1.40506 | 1.380 |
| chr9 | 79626822 | 79627173 | 3.13045 | 4.32018 | 1.380 |
| chr10 | 52548688 | 52549035 | 0.971524 | 1.34082 | 1.380 |
| chr19 | 55164674 | 55165019 | 1.79723 | 2.48042 | 1.380 |
| chr12 | 26919972 | 26920275 | 0.892218 | 1.23141 | 1.380 |
| chr5 | 178921070 | 178921325 | 1.22764 | 1.69448 | 1.380 |
| chr7 | 138571304 | 138571508 | 1.43426 | 1.97971 | 1.380 |
| chr2 | 220360015 | 220360377 | 1.66242 | 2.29472 | 1.380 |
| chr3 | 147754094 | 147754354 | 0.561406 | 0.774959 | 1.380 |
| chr9 | 113102696 | 113102927 | 1.65874 | 2.28975 | 1.380 |
| chr19 | 19601121 | 19601540 | 1.60092 | 2.20999 | 1.380 |
| chr4 | 75094473 | 75094752 | 1.90614 | 2.63158 | 1.381 |
| chr4 | 108658122 | 108658470 | 1.58425 | 2.18735 | 1.381 |
| chr17 | 10587226 | 10587415 | 1.29151 | 1.7834 | 1.381 |
| chr19 | 39670621 | 39670833 | 0.755456 | 1.04318 | 1.381 |
| chr7 | 100161464 | 100162038 | 2.23807 | 3.09086 | 1.381 |
| chr15 | 77327412 | 77327857 | 2.45217 | 3.38659 | 1.381 |
| chr2 | 18442758 | 18442982 | 0.95172 | 1.31443 | 1.381 |
| chr3 | 18595316 | 18595574 | 0.585613 | 0.808797 | 1.381 |
| chr2 | 10800154 | 10800460 | 1.99311 | 2.753 | 1.381 |
| chr9 | 79241469 | 79241672 | 0.709775 | 0.980488 | 1.381 |
| chr1 | 26690313 | 26690700 | 2.92981 | 4.04756 | 1.382 |
| chr17 | 7765941 | 7766310 | 2.35159 | 3.24898 | 1.382 |
| chr7 | 124059155 | 124059414 | 0.80321 | 1.10977 | 1.382 |
| chr5 | 10140507 | 10140924 | 1.13617 | 1.56992 | 1.382 |
| chr5 | 7068347 | 7068577 | 0.558761 | 0.7721 | 1.382 |
| chr2 | 75836194 | 75836573 | 2.71675 | 3.7541 | 1.382 |
| chr2 | 233918649 | 233919045 | 2.87081 | 3.96719 | 1.382 |

| | | | | | |
|---|---|---|---|---|---|
| chr6 | 130257763 | 130258039 | 1.63462 | 2.25897 | 1.382 |
| chr10 | 102503354 | 102503917 | 1.73415 | 2.39653 | 1.382 |
| chr3 | 48199629 | 48199907 | 2.50775 | 3.46563 | 1.382 |
| chr10 | 53324537 | 53324979 | 0.649546 | 0.897731 | 1.382 |
| chr8 | 77085050 | 77085417 | 0.682586 | 0.943392 | 1.382 |
| chr2 | 53079727 | 53080086 | 0.691528 | 0.955791 | 1.382 |
| chr5 | 153289871 | 153290145 | 0.512691 | 0.708617 | 1.382 |
| chr2 | 241496201 | 241496696 | 1.3669 | 1.88936 | 1.382 |
| chr2 | 127643861 | 127644249 | 1.92649 | 2.66299 | 1.382 |
| chr19 | 49728136 | 49728440 | 2.06062 | 2.84844 | 1.382 |
| chr12 | 51745677 | 51746117 | 2.11231 | 2.91991 | 1.382 |
| chr20 | 61422685 | 61423415 | 2.05034 | 2.83424 | 1.382 |
| chr16 | 68408966 | 68409203 | 0.514853 | 0.711702 | 1.382 |
| chr2 | 106192849 | 106193084 | 1.74846 | 2.41696 | 1.382 |
| chr8 | 140276421 | 140276739 | 0.82688 | 1.14304 | 1.382 |
| chr5 | 177409146 | 177409368 | 1.12368 | 1.55335 | 1.382 |
| chr17 | 76408477 | 76408787 | 1.36817 | 1.89137 | 1.382 |
| chr1 | 223263243 | 223263779 | 2.21924 | 3.06827 | 1.383 |
| chr15 | 101320111 | 101320329 | 0.826241 | 1.14235 | 1.383 |
| chr5 | 125992187 | 125992405 | 1.67262 | 2.3126 | 1.383 |
| chr2 | 10295561 | 10295975 | 0.827495 | 1.14424 | 1.383 |
| chr12 | 70379413 | 70379688 | 1.90259 | 2.63094 | 1.383 |
| chr15 | 69946445 | 69946811 | 0.781266 | 1.08039 | 1.383 |
| chr15 | 73127710 | 73128046 | 1.03973 | 1.43783 | 1.383 |
| chr2 | 31556396 | 31556790 | 2.06908 | 2.86135 | 1.383 |
| chr11 | 5067776 | 5068135 | 0.722097 | 0.998621 | 1.383 |
| chr9 | 5590046 | 5590351 | 2.17569 | 3.00902 | 1.383 |
| chr10 | 52737562 | 52737803 | 0.612008 | 0.846431 | 1.383 |
| chr17 | 27292689 | 27292954 | 1.82126 | 2.51894 | 1.383 |
| chr2 | 21586763 | 21586994 | 0.643535 | 0.890091 | 1.383 |
| chr16 | 56985240 | 56985518 | 1.96003 | 2.71099 | 1.383 |
| chr19 | 49178662 | 49179066 | 2.68214 | 3.71026 | 1.383 |
| chr3 | 31097169 | 31097603 | 0.691668 | 0.9568 | 1.383 |
| chr6 | 151691642 | 151691877 | 0.991352 | 1.37139 | 1.383 |
| chr4 | 13543627 | 13544014 | 1.07516 | 1.48738 | 1.383 |
| chr19 | 51250203 | 51250409 | 0.947505 | 1.31088 | 1.384 |
| chr12 | 67419315 | 67419578 | 2.13831 | 2.95859 | 1.384 |
| chr3 | 116105304 | 116105699 | 0.753693 | 1.04283 | 1.384 |
| chr4 | 164476093 | 164476485 | 1.28299 | 1.77522 | 1.384 |
| chr5 | 150035858 | 150036389 | 1.88903 | 2.6138 | 1.384 |
| chr1 | 40119096 | 40119542 | 2.06762 | 2.86104 | 1.384 |
| chr7 | 7463974 | 7464329 | 1.53298 | 2.12134 | 1.384 |
| chr20 | 41876939 | 41877190 | 0.673765 | 0.932396 | 1.384 |
| chr3 | 163033248 | 163033471 | 0.773262 | 1.07009 | 1.384 |
| chr2 | 194222222 | 194222434 | 0.699766 | 0.968417 | 1.384 |
| chr11 | 90981408 | 90981646 | 1.22223 | 1.6915 | 1.384 |
| chr15 | 40167126 | 40167453 | 1.29224 | 1.78849 | 1.384 |
| chr4 | 122291959 | 122292184 | 1.19887 | 1.65927 | 1.384 |
| chrX | 108728905 | 108729340 | 1.88121 | 2.60369 | 1.384 |
| chr19 | 47253367 | 47253863 | 1.79651 | 2.48666 | 1.384 |
| chr1 | 201417198 | 201417564 | 2.19096 | 3.03266 | 1.384 |
| chr12 | 56789760 | 56789967 | 0.746008 | 1.03264 | 1.384 |
| chr18 | 72143485 | 72143729 | 1.54461 | 2.1381 | 1.384 |
| chr19 | 30615772 | 30616256 | 1.1516 | 1.59415 | 1.384 |
| chr12 | 15429091 | 15429367 | 1.7624 | 2.43971 | 1.384 |
| chr5 | 60954365 | 60954790 | 2.06941 | 2.86492 | 1.384 |
| chr8 | 120779995 | 120780346 | 1.35317 | 1.87336 | 1.384 |
| chr8 | 40983702 | 40983994 | 1.68441 | 2.33198 | 1.384 |
| chr11 | 129793254 | 129793487 | 1.9257 | 2.66608 | 1.384 |
| chr9 | 111524117 | 111524407 | 2.05582 | 2.84625 | 1.384 |
| chr14 | 100991229 | 100991572 | 1.50108 | 2.07829 | 1.385 |

| | | | | | |
|---|---|---|---|---|---|
| chr2 | 24307196 | 24307681 | 2.23819 | 3.09887 | 1.385 |
| chr7 | 42376958 | 42377215 | 0.664454 | 0.919969 | 1.385 |
| chr20 | 23259532 | 23259761 | 0.973516 | 1.34789 | 1.385 |
| chr17 | 27077776 | 27078037 | 0.661098 | 0.915372 | 1.385 |
| chr11 | 127901513 | 127901759 | 1.51354 | 2.09571 | 1.385 |
| chr3 | 1424706 | 1424930 | 0.784042 | 1.08562 | 1.385 |
| chr7 | 149359586 | 149360042 | 1.80836 | 2.50399 | 1.385 |
| chr22 | 35934924 | 35935179 | 1.50146 | 2.07908 | 1.385 |
| chr7 | 47162141 | 47162343 | 0.873112 | 1.20904 | 1.385 |
| chr2 | 192109829 | 192110071 | 1.63989 | 2.27091 | 1.385 |
| chr10 | 67508164 | 67508500 | 0.76028 | 1.05284 | 1.385 |
| chr13 | 92058930 | 92059208 | 1.21647 | 1.68495 | 1.385 |
| chr3 | 116738018 | 116738270 | 1.81672 | 2.51644 | 1.385 |
| chr2 | 27165326 | 27165817 | 1.99035 | 2.7571 | 1.385 |
| chr14 | 32721332 | 32721590 | 1.71173 | 2.37122 | 1.385 |
| chr12 | 15421932 | 15422226 | 2.11126 | 2.92477 | 1.385 |
| chr1 | 241663309 | 241663519 | 1.29487 | 1.79389 | 1.385 |
| chr15 | 88245491 | 88245787 | 0.816065 | 1.13056 | 1.385 |
| chr1 | 11021114 | 11021350 | 2.22469 | 3.0822 | 1.385 |
| chr12 | 15737327 | 15737689 | 1.69378 | 2.34673 | 1.386 |
| chr11 | 4658187 | 4658398 | 1.6298 | 2.25815 | 1.386 |
| chr12 | 48583941 | 48584152 | 1.29114 | 1.78893 | 1.386 |
| chr8 | 65274760 | 65275008 | 0.699528 | 0.969223 | 1.386 |
| chr5 | 109212499 | 109212780 | 0.750889 | 1.04041 | 1.386 |
| chr16 | 85598091 | 85598382 | 2.12779 | 2.9485 | 1.386 |
| chr12 | 44394179 | 44394620 | 1.65456 | 2.29275 | 1.386 |
| chr2 | 232394992 | 232395406 | 2.5217 | 3.49461 | 1.386 |
| chr9 | 139637417 | 139637740 | 1.34393 | 1.86246 | 1.386 |
| chr10 | 102295393 | 102295729 | 2.80254 | 3.884 | 1.386 |
| chr11 | 78811193 | 78811578 | 1.68088 | 2.32958 | 1.386 |
| chr10 | 123583533 | 123583803 | 2.03869 | 2.82551 | 1.386 |
| chr5 | 28211568 | 28211872 | 0.666593 | 0.923858 | 1.386 |
| chr17 | 66196901 | 66197325 | 2.86976 | 3.97737 | 1.386 |
| chr2 | 10410655 | 10410940 | 1.44005 | 1.99587 | 1.386 |
| chr7 | 33876483 | 33876756 | 1.07877 | 1.49515 | 1.386 |
| chr1 | 207752714 | 207753067 | 1.20948 | 1.67641 | 1.386 |
| chr20 | 8702999 | 8703259 | 2.36498 | 3.27803 | 1.386 |
| chr6 | 135483741 | 135484128 | 1.34665 | 1.86657 | 1.386 |
| chr8 | 32020114 | 32020450 | 1.75315 | 2.43004 | 1.386 |
| chr1 | 200772057 | 200772378 | 2.28659 | 3.16947 | 1.386 |
| chr7 | 119710426 | 119710630 | 0.542296 | 0.751713 | 1.386 |
| chr16 | 11744778 | 11745054 | 0.612261 | 0.848721 | 1.386 |
| chr11 | 122982191 | 122982440 | 1.79377 | 2.48669 | 1.386 |
| chr14 | 100769504 | 100769812 | 1.63973 | 2.27322 | 1.386 |
| chr3 | 45266643 | 45266941 | 1.99221 | 2.76188 | 1.386 |
| chr13 | 100039777 | 100040028 | 1.67316 | 2.31964 | 1.386 |
| chr20 | 12695185 | 12695394 | 0.76001 | 1.05367 | 1.386 |
| chr11 | 17782015 | 17782240 | 1.2784 | 1.77239 | 1.386 |
| chr5 | 142473802 | 142474030 | 0.439749 | 0.609716 | 1.387 |
| chr16 | 87583968 | 87584539 | 2.60158 | 3.60722 | 1.387 |
| chr9 | 126323995 | 126324310 | 0.749799 | 1.03971 | 1.387 |
| chr20 | 2736680 | 2736999 | 1.75078 | 2.42787 | 1.387 |
| chr16 | 78512376 | 78512651 | 1.81646 | 2.51899 | 1.387 |
| chr11 | 27232981 | 27233223 | 1.28363 | 1.78019 | 1.387 |
| chr11 | 94252773 | 94253208 | 0.934302 | 1.29578 | 1.387 |
| chr2 | 219712370 | 219712823 | 3.34462 | 4.63864 | 1.387 |
| chr5 | 169497779 | 169498232 | 2.86215 | 3.9697 | 1.387 |
| chr2 | 174267169 | 174267447 | 1.82241 | 2.52797 | 1.387 |
| chr12 | 108001235 | 108001701 | 2.35889 | 3.27221 | 1.387 |
| chr6 | 50783868 | 50784108 | 1.61496 | 2.24036 | 1.387 |
| chr3 | 196950950 | 196951183 | 1.52246 | 2.11209 | 1.387 |

| | | | | | |
|---|---|---|---|---|---|
| chr5 | 137412836 | 137413098 | 1.76023 | 2.44205 | 1.387 |
| chr10 | 34375727 | 34376144 | 0.734307 | 1.01876 | 1.387 |
| chr5 | 66989921 | 66990195 | 0.437022 | 0.606342 | 1.387 |
| chr18 | 34902385 | 34902789 | 1.94087 | 2.69302 | 1.388 |
| chr7 | 90715628 | 90715837 | 0.525145 | 0.728672 | 1.388 |
| chr5 | 11464425 | 11464739 | 0.79037 | 1.0967 | 1.388 |
| chr9 | 15232018 | 15232355 | 1.8308 | 2.54053 | 1.388 |
| chr6 | 155569315 | 155569853 | 1.15658 | 1.60496 | 1.388 |
| chr19 | 31291048 | 31291469 | 1.55131 | 2.15276 | 1.388 |
| chr1 | 9927415 | 9927896 | 2.06531 | 2.8661 | 1.388 |
| chr5 | 121531541 | 121531902 | 1.46156 | 2.02825 | 1.388 |
| chr5 | 52558260 | 52558530 | 1.39933 | 1.94196 | 1.388 |
| chr8 | 120136144 | 120136422 | 0.763974 | 1.06032 | 1.388 |
| chr2 | 70568227 | 70568727 | 1.87753 | 2.60585 | 1.388 |
| chr18 | 13630154 | 13630510 | 1.44143 | 2.00067 | 1.388 |
| chrX | 52004686 | 52004924 | 2.10502 | 2.92175 | 1.388 |
| chr17 | 34982175 | 34982789 | 1.92487 | 2.67174 | 1.388 |
| chr10 | 29681381 | 29681657 | 2.67635 | 3.7151 | 1.388 |
| chr1 | 158476279 | 158476520 | 0.915661 | 1.27112 | 1.388 |
| chrX | 28273040 | 28273271 | 0.598536 | 0.830932 | 1.388 |
| chr3 | 54180749 | 54180993 | 2.29757 | 3.18971 | 1.388 |
| chr3 | 8098917 | 8099236 | 0.654277 | 0.908356 | 1.388 |
| chr3 | 14967184 | 14967696 | 1.60526 | 2.22864 | 1.388 |
| chr11 | 1616684 | 1617242 | 1.10012 | 1.52735 | 1.388 |
| chr9 | 96588568 | 96589129 | 2.10934 | 2.92858 | 1.388 |
| chr7 | 149487151 | 149487575 | 1.29176 | 1.79366 | 1.389 |
| chr3 | 143363408 | 143363691 | 1.55302 | 2.1565 | 1.389 |
| chr10 | 85107941 | 85108309 | 0.745019 | 1.03453 | 1.389 |
| chr7 | 92672794 | 92673080 | 0.731732 | 1.01609 | 1.389 |
| chr12 | 70132342 | 70132626 | 1.85918 | 2.58169 | 1.389 |
| chr5 | 179016741 | 179017072 | 0.841135 | 1.16803 | 1.389 |
| chr17 | 18011210 | 18011559 | 0.839511 | 1.16584 | 1.389 |
| chr3 | 43544666 | 43544919 | 0.989585 | 1.37425 | 1.389 |
| chr9 | 24648709 | 24648932 | 0.749048 | 1.04025 | 1.389 |
| chr2 | 218621196 | 218621685 | 2.13738 | 2.96842 | 1.389 |
| chr18 | 66647193 | 66647413 | 0.530738 | 0.73715 | 1.389 |
| chr12 | 119600317 | 119600658 | 0.644061 | 0.89457 | 1.389 |
| chr5 | 59105822 | 59106127 | 1.79053 | 2.48738 | 1.389 |
| chr5 | 12119842 | 12120133 | 0.973927 | 1.353 | 1.389 |
| chr16 | 34586693 | 34587219 | 0.674885 | 0.937576 | 1.389 |
| chr1 | 159059006 | 159059240 | 0.894947 | 1.24337 | 1.389 |
| chr13 | 28898395 | 28898710 | 2.49339 | 3.4644 | 1.389 |
| chr10 | 73699535 | 73699768 | 0.630673 | 0.876283 | 1.389 |
| chr12 | 78035661 | 78035900 | 1.00098 | 1.39099 | 1.390 |
| chr22 | 30881416 | 30881673 | 2.19226 | 3.04644 | 1.390 |
| chr1 | 161261281 | 161261581 | 1.92387 | 2.67355 | 1.390 |
| chr4 | 186104854 | 186105089 | 1.71813 | 2.38764 | 1.390 |
| chr1 | 27481558 | 27481834 | 2.80366 | 3.89646 | 1.390 |
| chr3 | 8190194 | 8190399 | 0.97533 | 1.3556 | 1.390 |
| chr8 | 15998128 | 15998354 | 0.739569 | 1.02795 | 1.390 |
| chr9 | 139413083 | 139413526 | 0.710445 | 0.987529 | 1.390 |
| chr1 | 21835573 | 21835937 | 1.62804 | 2.26302 | 1.390 |
| chr5 | 51121030 | 51121308 | 0.872938 | 1.21353 | 1.390 |
| chr9 | 99619180 | 99619468 | 1.89707 | 2.63727 | 1.390 |
| chr13 | 77477711 | 77478000 | 2.22103 | 3.08767 | 1.390 |
| chr11 | 55944434 | 55944814 | 0.962388 | 1.33796 | 1.390 |
| chr3 | 22657575 | 22657939 | 0.605615 | 0.841973 | 1.390 |
| chr19 | 40042007 | 40042559 | 2.07982 | 2.89158 | 1.390 |
| chr1 | 40123792 | 40124004 | 1.52046 | 2.11393 | 1.390 |
| chr8 | 60326606 | 60326833 | 0.675687 | 0.93943 | 1.390 |
| chr19 | 56291697 | 56291927 | 0.806995 | 1.12202 | 1.390 |

| | | | | | |
|---|---|---|---|---|---|
| chr20 | 1822060 | 1822292 | 0.72396 | 1.00658 | 1.390 |
| chrX | 107827997 | 107828221 | 0.920666 | 1.28019 | 1.391 |
| chr5 | 172141718 | 172142040 | 1.8274 | 2.54114 | 1.391 |
| chr10 | 87938166 | 87938476 | 1.2379 | 1.72144 | 1.391 |
| chr10 | 102638310 | 102638549 | 1.7768 | 2.47083 | 1.391 |
| chr17 | 79093735 | 79093952 | 1.38977 | 1.93279 | 1.391 |
| chr15 | 51480546 | 51480849 | 0.661602 | 0.920149 | 1.391 |
| chr3 | 144394691 | 144394976 | 0.630351 | 0.876768 | 1.391 |
| chr7 | 136659079 | 136659368 | 0.704434 | 0.979823 | 1.391 |
| chr16 | 52615478 | 52615736 | 1.60401 | 2.23117 | 1.391 |
| chr22 | 43742463 | 43742798 | 2.09739 | 2.91749 | 1.391 |
| chr8 | 52470092 | 52470294 | 0.771097 | 1.07263 | 1.391 |
| chr8 | 133501834 | 133502150 | 2.09022 | 2.9077 | 1.391 |
| chr7 | 139322699 | 139323067 | 0.872617 | 1.21396 | 1.391 |
| chr5 | 162733410 | 162733701 | 0.967423 | 1.34596 | 1.391 |
| chr5 | 165056577 | 165056854 | 0.618381 | 0.860343 | 1.391 |
| chr15 | 79673112 | 79673694 | 1.61153 | 2.2421 | 1.391 |
| chr10 | 108412074 | 108412290 | 0.642844 | 0.894436 | 1.391 |
| chr10 | 9436947 | 9437267 | 0.627235 | 0.872762 | 1.391 |
| chr20 | 60711848 | 60712235 | 1.92621 | 2.68021 | 1.391 |
| chr7 | 10648093 | 10648311 | 0.672077 | 0.935191 | 1.391 |
| chr9 | 38087943 | 38088333 | 2.83713 | 3.94786 | 1.392 |
| chr10 | 72318954 | 72319426 | 1.43224 | 1.99301 | 1.392 |
| chr6 | 116966771 | 116966992 | 0.996072 | 1.3861 | 1.392 |
| chr16 | 86570310 | 86570525 | 0.67139 | 0.934295 | 1.392 |
| chr15 | 65325361 | 65325834 | 3.23694 | 4.50465 | 1.392 |
| chr2 | 176991711 | 176992089 | 1.83724 | 2.55678 | 1.392 |
| chr17 | 17473529 | 17474128 | 1.92629 | 2.68087 | 1.392 |
| chr9 | 16133786 | 16134057 | 2.56977 | 3.57643 | 1.392 |
| chr1 | 211751079 | 211751498 | 2.53705 | 3.531 | 1.392 |
| chr5 | 107863143 | 107863506 | 0.68977 | 0.960002 | 1.392 |
| chr14 | 84884613 | 84884911 | 0.707833 | 0.985157 | 1.392 |
| chr9 | 117501149 | 117501463 | 1.50486 | 2.09446 | 1.392 |
| chr16 | 698090 | 698383 | 1.10978 | 1.5446 | 1.392 |
| chr2 | 210906 | 211171 | 0.779267 | 1.0846 | 1.392 |
| chr11 | 56351084 | 56351287 | 0.686175 | 0.955049 | 1.392 |
| chr1 | 110075513 | 110075767 | 1.72497 | 2.40098 | 1.392 |
| chr5 | 33380011 | 33380223 | 1.12054 | 1.55975 | 1.392 |
| chr7 | 28962324 | 28962638 | 2.34521 | 3.26452 | 1.392 |
| chr8 | 110249778 | 110250025 | 0.712962 | 0.992484 | 1.392 |
| chr15 | 46087759 | 46087983 | 1.67184 | 2.32731 | 1.392 |
| chr1 | 6408019 | 6408335 | 2.10279 | 2.92729 | 1.392 |
| chr15 | 73212626 | 73212829 | 0.949398 | 1.32166 | 1.392 |
| chr19 | 34921694 | 34921983 | 2.7541 | 3.83398 | 1.392 |
| chr14 | 64099716 | 64099990 | 1.38949 | 1.93435 | 1.392 |
| chr19 | 18599710 | 18599928 | 0.92073 | 1.28178 | 1.392 |
| chr5 | 170236632 | 170236904 | 1.20012 | 1.67073 | 1.392 |
| chr19 | 32975261 | 32975593 | 0.857481 | 1.19374 | 1.392 |
| chr7 | 50182027 | 50182480 | 1.29732 | 1.80606 | 1.392 |
| chr1 | 236772522 | 236772771 | 1.95483 | 2.72151 | 1.392 |
| chr12 | 132703597 | 132703814 | 1.3928 | 1.93905 | 1.392 |
| chr3 | 37595967 | 37596241 | 1.03618 | 1.44259 | 1.392 |
| chr22 | 44341692 | 44342402 | 1.4665 | 2.04173 | 1.392 |
| chr8 | 19670881 | 19671186 | 1.64853 | 2.29524 | 1.392 |
| chr8 | 136463046 | 136463250 | 0.765727 | 1.06622 | 1.392 |
| chr6 | 91103039 | 91103373 | 1.64723 | 2.29368 | 1.392 |
| chr7 | 19531498 | 19531892 | 1.0594 | 1.47521 | 1.393 |
| chr14 | 49814416 | 49814676 | 0.589344 | 0.820681 | 1.393 |
| chr10 | 104429390 | 104429897 | 1.60302 | 2.23239 | 1.393 |
| chr9 | 11795029 | 11795262 | 0.721816 | 1.00525 | 1.393 |
| chr2 | 62303978 | 62304245 | 0.411513 | 0.573104 | 1.393 |

| | | | | | |
|---|---|---|---|---|---|
| chr5 | 36729132 | 36729429 | 1.65985 | 2.31176 | 1.393 |
| chr17 | 74393917 | 74394474 | 1.81514 | 2.52807 | 1.393 |
| chr2 | 18632760 | 18632981 | 0.583343 | 0.812468 | 1.393 |
| chr5 | 110330482 | 110330867 | 0.666742 | 0.928676 | 1.393 |
| chr5 | 157189015 | 157189451 | 1.58368 | 2.20584 | 1.393 |
| chr17 | 8894886 | 8895208 | 2.46077 | 3.42768 | 1.393 |
| chr9 | 79830934 | 79831228 | 1.48155 | 2.06376 | 1.393 |
| chr5 | 35925100 | 35925357 | 0.984658 | 1.37165 | 1.393 |
| chr8 | 55660218 | 55660481 | 0.65351 | 0.910364 | 1.393 |
| chr20 | 20978885 | 20979181 | 0.61164 | 0.852057 | 1.393 |
| chr12 | 42712284 | 42712497 | 0.826822 | 1.15188 | 1.393 |
| chr15 | 79199872 | 79200287 | 1.25392 | 1.7469 | 1.393 |
| chr1 | 91483664 | 91483883 | 0.940638 | 1.31062 | 1.393 |
| chr11 | 62462073 | 62462525 | 2.12098 | 2.95576 | 1.394 |
| chr2 | 103456420 | 103456705 | 1.54646 | 2.15521 | 1.394 |
| chr5 | 139241933 | 139242261 | 0.773704 | 1.0783 | 1.394 |
| chr2 | 240745945 | 240746223 | 1.05268 | 1.46714 | 1.394 |
| chr7 | 115550666 | 115550914 | 1.8321 | 2.55355 | 1.394 |
| chr20 | 10407804 | 10408081 | 2.50758 | 3.49516 | 1.394 |
| chr19 | 35536722 | 35537111 | 1.43746 | 2.00364 | 1.394 |
| chr12 | 30610664 | 30610862 | 1.27781 | 1.78122 | 1.394 |
| chr14 | 34308861 | 34309195 | 2.01894 | 2.81449 | 1.394 |
| chr19 | 41108110 | 41108777 | 2.02601 | 2.82434 | 1.394 |
| chr2 | 21185771 | 21186401 | 1.62131 | 2.26022 | 1.394 |
| chr3 | 87019007 | 87019299 | 1.63467 | 2.27893 | 1.394 |
| chr5 | 75840054 | 75840410 | 1.04831 | 1.46158 | 1.394 |
| chr2 | 222922040 | 222922287 | 1.67808 | 2.33972 | 1.394 |
| chr12 | 10388355 | 10388961 | 1.35648 | 1.89134 | 1.394 |
| chr7 | 116443016 | 116443234 | 1.40205 | 1.9549 | 1.394 |
| chr15 | 45814724 | 45814933 | 1.26288 | 1.76104 | 1.394 |
| chr12 | 128406697 | 128406920 | 0.884414 | 1.23331 | 1.394 |
| chr6 | 146250284 | 146250511 | 1.2302 | 1.71553 | 1.395 |
| chr14 | 53272981 | 53273261 | 0.748025 | 1.04316 | 1.395 |
| chr5 | 4942257 | 4943114 | 1.94198 | 2.70827 | 1.395 |
| chr10 | 3281847 | 3282232 | 2.09334 | 2.91939 | 1.395 |
| chr20 | 25021995 | 25022260 | 0.720385 | 1.00471 | 1.395 |
| chr16 | 88803695 | 88804202 | 3.5827 | 4.99715 | 1.395 |
| chr2 | 159610534 | 159610861 | 1.97882 | 2.76009 | 1.395 |
| chr15 | 89200565 | 89201008 | 2.11044 | 2.94398 | 1.395 |
| chr10 | 80915498 | 80915708 | 0.815949 | 1.1383 | 1.395 |
| chr17 | 78335370 | 78335573 | 1.30239 | 1.81696 | 1.395 |
| chr6 | 37293510 | 37293842 | 1.91747 | 2.67521 | 1.395 |
| chr15 | 97894104 | 97894346 | 0.895359 | 1.24921 | 1.395 |
| chr10 | 34926387 | 34926762 | 2.47891 | 3.45873 | 1.395 |
| chr17 | 30789433 | 30789641 | 1.34729 | 1.87982 | 1.395 |
| chr5 | 148120827 | 148121213 | 1.36281 | 1.90153 | 1.395 |
| chr13 | 31253969 | 31254179 | 1.12435 | 1.56887 | 1.395 |
| chr2 | 219860673 | 219861090 | 2.69808 | 3.76498 | 1.395 |
| chrX | 110342140 | 110342401 | 1.87018 | 2.60988 | 1.396 |
| chr2 | 37109690 | 37109968 | 0.651028 | 0.908534 | 1.396 |
| chr6 | 65626926 | 65627134 | 0.507753 | 0.708604 | 1.396 |
| chr12 | 132235938 | 132236316 | 0.74871 | 1.0449 | 1.396 |
| chr17 | 49662930 | 49663237 | 2.28515 | 3.18916 | 1.396 |
| chr7 | 56636904 | 56637116 | 0.781971 | 1.09136 | 1.396 |
| chr4 | 134151583 | 134151812 | 0.752239 | 1.04987 | 1.396 |
| chr1 | 223276529 | 223276796 | 0.486656 | 0.679256 | 1.396 |
| chr17 | 12933689 | 12934229 | 1.4569 | 2.03351 | 1.396 |
| chr19 | 47910676 | 47910888 | 0.636995 | 0.889144 | 1.396 |
| chr17 | 14115166 | 14115428 | 1.21265 | 1.69269 | 1.396 |
| chr1 | 116563249 | 116563594 | 1.4567 | 2.03336 | 1.396 |
| chr2 | 78982151 | 78982406 | 0.861903 | 1.20311 | 1.396 |

| | | | | | |
|---|---|---|---|---|---|
| chr5 | 20132162 | 20132358 | 0.847838 | 1.18349 | 1.396 |
| chr5 | 171877827 | 171878576 | 1.48123 | 2.06764 | 1.396 |
| chr8 | 6658225 | 6658653 | 2.43754 | 3.40259 | 1.396 |
| chr1 | 213916792 | 213917030 | 0.622119 | 0.868457 | 1.396 |
| chr12 | 3351561 | 3351806 | 0.680828 | 0.950436 | 1.396 |
| chr8 | 64220165 | 64220392 | 0.75682 | 1.05655 | 1.396 |
| chr14 | 93605533 | 93605880 | 2.99264 | 4.17794 | 1.396 |
| chr11 | 66862389 | 66862690 | 1.68111 | 2.34719 | 1.396 |
| chr1 | 219581522 | 219581753 | 0.622175 | 0.868717 | 1.396 |
| chr6 | 160877038 | 160877351 | 1.96058 | 2.7375 | 1.396 |
| chr12 | 16132138 | 16132383 | 1.33911 | 1.86989 | 1.396 |
| chr7 | 36352292 | 36352494 | 1.42603 | 1.99134 | 1.396 |
| chr2 | 70044166 | 70044392 | 1.97579 | 2.75908 | 1.396 |
| chr9 | 109899834 | 109900214 | 1.94584 | 2.71726 | 1.396 |
| chr3 | 140641514 | 140641746 | 1.01993 | 1.42429 | 1.396 |
| chr17 | 74084044 | 74084373 | 0.950463 | 1.3273 | 1.396 |
| chr7 | 102052899 | 102053250 | 1.11625 | 1.55884 | 1.397 |
| chr5 | 128553275 | 128553549 | 0.699128 | 0.976349 | 1.397 |
| chr14 | 51568144 | 51568376 | 1.00541 | 1.4041 | 1.397 |
| chr3 | 113543989 | 113544292 | 0.786888 | 1.09892 | 1.397 |
| chr17 | 8203977 | 8204283 | 2.35894 | 3.29444 | 1.397 |
| chr11 | 31261949 | 31262156 | 1.66591 | 2.32659 | 1.397 |
| chr1 | 248782198 | 248782696 | 0.946083 | 1.32132 | 1.397 |
| chr1 | 208058469 | 208058767 | 2.90297 | 4.05448 | 1.397 |
| chr6 | 97762078 | 97762303 | 0.493471 | 0.689344 | 1.397 |
| chr8 | 132540852 | 132541093 | 0.709112 | 0.990595 | 1.397 |
| chr16 | 84341799 | 84342055 | 1.72379 | 2.40811 | 1.397 |
| chr2 | 218938737 | 218938974 | 1.83977 | 2.57024 | 1.397 |
| chr15 | 87371287 | 87371606 | 0.606897 | 0.847923 | 1.397 |
| chr10 | 75129816 | 75130112 | 0.709773 | 0.991691 | 1.397 |
| chr18 | 72531707 | 72531978 | 1.61069 | 2.25066 | 1.397 |
| chr10 | 77846783 | 77847040 | 2.15939 | 3.01763 | 1.397 |
| chr15 | 22912531 | 22912932 | 0.653201 | 0.912817 | 1.397 |
| chr2 | 27487931 | 27488224 | 1.9106 | 2.67 | 1.397 |
| chr1 | 214296589 | 214296792 | 1.65124 | 2.30769 | 1.398 |
| chr9 | 75637930 | 75638197 | 2.39542 | 3.34786 | 1.398 |
| chr17 | 73873871 | 73874234 | 1.36609 | 1.90934 | 1.398 |
| chr6 | 80523671 | 80523896 | 1.74985 | 2.44572 | 1.398 |
| chr9 | 72630107 | 72630319 | 0.492614 | 0.688519 | 1.398 |
| chr1 | 22746981 | 22747263 | 2.25885 | 3.15735 | 1.398 |
| chr17 | 26009026 | 26009246 | 0.878473 | 1.22791 | 1.398 |
| chr6 | 168756570 | 168757003 | 1.77882 | 2.48649 | 1.398 |
| chr3 | 143857383 | 143857640 | 0.586897 | 0.820411 | 1.398 |
| chr9 | 71946744 | 71947032 | 0.688441 | 0.962481 | 1.398 |
| chr5 | 176744905 | 176745198 | 2.27902 | 3.1863 | 1.398 |
| chr9 | 28193848 | 28194058 | 0.802231 | 1.12169 | 1.398 |
| chr14 | 94085538 | 94085836 | 1.44167 | 2.01581 | 1.398 |
| chr10 | 62642919 | 62643279 | 1.64316 | 2.29758 | 1.398 |
| chr2 | 192020408 | 192020676 | 1.76226 | 2.46412 | 1.398 |
| chr10 | 104238645 | 104239181 | 2.33361 | 3.26306 | 1.398 |
| chr2 | 100879151 | 100879651 | 1.73598 | 2.4275 | 1.398 |
| chr1 | 170952194 | 170952518 | 0.640002 | 0.895027 | 1.398 |
| chr8 | 78496051 | 78496283 | 0.6808 | 0.952111 | 1.399 |
| chr19 | 33610034 | 33610322 | 1.36293 | 1.90613 | 1.399 |
| chr2 | 48698019 | 48698386 | 1.34088 | 1.87537 | 1.399 |
| chr10 | 44381304 | 44381512 | 0.811197 | 1.13456 | 1.399 |
| chr9 | 131495462 | 131496003 | 2.79705 | 3.91204 | 1.399 |
| chr17 | 32117857 | 32118166 | 0.952823 | 1.33277 | 1.399 |
| chr6 | 109103533 | 109103813 | 1.60063 | 2.23892 | 1.399 |
| chr1 | 216368627 | 216368832 | 0.765007 | 1.07008 | 1.399 |
| chr2 | 108832190 | 108832423 | 1.61789 | 2.26311 | 1.399 |

| | | | | | |
|---|---|---|---|---|---|
| chr2 | 220159301 | 220159915 | 3.03821 | 4.25031 | 1.399 |
| chr9 | 128170219 | 128170714 | 1.99588 | 2.79214 | 1.399 |
| chr2 | 232717054 | 232717411 | 1.94453 | 2.72042 | 1.399 |
| chr12 | 2504451 | 2504891 | 2.89162 | 4.04546 | 1.399 |
| chr15 | 91191286 | 91191640 | 2.11217 | 2.95503 | 1.399 |
| chrX | 118887717 | 118887970 | 1.51901 | 2.12519 | 1.399 |
| chr1 | 171971986 | 171972206 | 2.12962 | 2.97955 | 1.399 |
| chr1 | 184383303 | 184383599 | 0.691589 | 0.96762 | 1.399 |
| chr11 | 17408237 | 17408491 | 2.18514 | 3.05732 | 1.399 |
| chr11 | 25187705 | 25187918 | 0.575671 | 0.805448 | 1.399 |
| chr12 | 108323390 | 108323863 | 0.856383 | 1.19822 | 1.399 |
| chr8 | 89697787 | 89697991 | 0.861647 | 1.20561 | 1.399 |
| chr7 | 90718743 | 90719145 | 2.04973 | 2.86816 | 1.399 |
| chr16 | 15859722 | 15860031 | 1.74614 | 2.44338 | 1.399 |
| chr20 | 25450754 | 25450943 | 1.89461 | 2.6512 | 1.399 |
| chr7 | 16417426 | 16417643 | 0.674781 | 0.944343 | 1.399 |
| chr15 | 45410238 | 45410638 | 2.58322 | 3.61521 | 1.400 |
| chr16 | 25066880 | 25067242 | 1.29616 | 1.81406 | 1.400 |
| chr5 | 19294822 | 19295031 | 0.713923 | 0.999209 | 1.400 |
| chr8 | 67334037 | 67334259 | 1.88204 | 2.63412 | 1.400 |
| chr17 | 882638 | 882896 | 1.13974 | 1.59529 | 1.400 |
| chr14 | 79758806 | 79759032 | 0.553363 | 0.774548 | 1.400 |
| chr2 | 36734511 | 36734830 | 2.3093 | 3.23236 | 1.400 |
| chr9 | 71395050 | 71395264 | 1.11618 | 1.56233 | 1.400 |
| chr5 | 25121671 | 25121904 | 0.638025 | 0.893061 | 1.400 |
| chr11 | 26630502 | 26630739 | 0.703327 | 0.984528 | 1.400 |
| chr3 | 1710601 | 1710822 | 0.739223 | 1.03487 | 1.400 |
| chr4 | 140973301 | 140973560 | 1.51564 | 2.12183 | 1.400 |
| chr12 | 8834718 | 8835022 | 1.43622 | 2.01068 | 1.400 |
| chr7 | 64042920 | 64043186 | 0.894061 | 1.2517 | 1.400 |
| chr17 | 68202704 | 68202916 | 0.717917 | 1.00515 | 1.400 |
| chr17 | 56032630 | 56032935 | 1.10672 | 1.54953 | 1.400 |
| chr3 | 165200459 | 165200915 | 0.626344 | 0.87697 | 1.400 |
| chr8 | 70204171 | 70204456 | 0.753316 | 1.05491 | 1.400 |
| chr17 | 57638445 | 57638682 | 1.0125 | 1.41787 | 1.400 |
| chr8 | 78557159 | 78557434 | 0.536348 | 0.751104 | 1.400 |
| chr5 | 42244237 | 42244443 | 0.658887 | 0.922762 | 1.400 |
| chr5 | 133082654 | 133082868 | 1.31246 | 1.83817 | 1.401 |
| chr2 | 129079315 | 129079798 | 2.43554 | 3.41123 | 1.401 |
| chr1 | 54286991 | 54287269 | 0.655899 | 0.918704 | 1.401 |
| chr5 | 38441826 | 38442165 | 2.50755 | 3.51233 | 1.401 |
| chr17 | 14427683 | 14427911 | 0.552884 | 0.774479 | 1.401 |
| chr6 | 5471222 | 5471532 | 1.72186 | 2.41199 | 1.401 |
| chr2 | 24605285 | 24605689 | 1.82349 | 2.55446 | 1.401 |
| chr11 | 11287541 | 11287963 | 1.99703 | 2.79768 | 1.401 |
| chr12 | 27297458 | 27297804 | 0.735876 | 1.03091 | 1.401 |
| chr3 | 21849976 | 21850474 | 0.823851 | 1.1542 | 1.401 |
| chr5 | 147823456 | 147823838 | 1.46936 | 2.05854 | 1.401 |
| chr3 | 185395051 | 185395285 | 0.950271 | 1.33133 | 1.401 |
| chr15 | 27308199 | 27308415 | 0.629815 | 0.882385 | 1.401 |
| chr14 | 74760958 | 74761367 | 1.7518 | 2.45454 | 1.401 |
| chr7 | 89345319 | 89345548 | 0.722364 | 1.01214 | 1.401 |
| chrX | 109693370 | 109693681 | 1.72205 | 2.41315 | 1.401 |
| chr9 | 36798259 | 36798613 | 1.31821 | 1.84729 | 1.401 |
| chr7 | 96076019 | 96076291 | 0.507455 | 0.711139 | 1.401 |
| chr4 | 153319241 | 153319477 | 1.02292 | 1.43352 | 1.401 |
| chr19 | 42596558 | 42596921 | 2.72816 | 3.82343 | 1.401 |
| chr4 | 16445252 | 16445503 | 1.10048 | 1.54237 | 1.402 |
| chr12 | 110133917 | 110134236 | 0.748419 | 1.04896 | 1.402 |
| chr11 | 31892522 | 31892994 | 1.68242 | 2.35804 | 1.402 |
| chr20 | 32450383 | 32450792 | 2.44713 | 3.42985 | 1.402 |

| | | | | | |
|---|---|---|---|---|---|
| chr8 | 107589872 | 107590142 | 1.7565 | 2.46207 | 1.402 |
| chr2 | 128453323 | 128453616 | 2.17189 | 3.0444 | 1.402 |
| chrX | 53340673 | 53340875 | 0.880356 | 1.23403 | 1.402 |
| chr7 | 67528236 | 67528691 | 1.23165 | 1.72655 | 1.402 |
| chr19 | 30277173 | 30277394 | 1.69734 | 2.3794 | 1.402 |
| chr9 | 123984801 | 123984987 | 1.89117 | 2.6512 | 1.402 |
| chr2 | 4790423 | 4790700 | 0.631578 | 0.885488 | 1.402 |
| chr10 | 30976613 | 30976961 | 2.15385 | 3.01995 | 1.402 |
| chr20 | 41759564 | 41759922 | 0.654306 | 0.917425 | 1.402 |
| chr7 | 4889118 | 4889392 | 1.28584 | 1.80292 | 1.402 |
| chr9 | 27442562 | 27442768 | 0.577839 | 0.810208 | 1.402 |
| chr11 | 45234642 | 45234962 | 1.271 | 1.78222 | 1.402 |
| chr10 | 71167424 | 71167830 | 1.32486 | 1.85797 | 1.402 |
| chr17 | 43176477 | 43176979 | 2.8827 | 4.04269 | 1.402 |
| chr17 | 69198554 | 69198797 | 0.712881 | 0.999743 | 1.402 |
| chr19 | 48993006 | 48993438 | 2.77973 | 3.89831 | 1.402 |
| chr14 | 92573037 | 92573270 | 3.33405 | 4.67595 | 1.402 |
| chr9 | 125753696 | 125753903 | 1.29511 | 1.81636 | 1.402 |
| chr7 | 137039191 | 137039468 | 0.484515 | 0.679543 | 1.403 |
| chr3 | 119278377 | 119278669 | 1.65278 | 2.31819 | 1.403 |
| chr10 | 78307197 | 78307424 | 1.56849 | 2.20003 | 1.403 |
| chr8 | 98424142 | 98424451 | 0.678581 | 0.951819 | 1.403 |
| chr14 | 94879378 | 94879642 | 1.21714 | 1.7074 | 1.403 |
| chr16 | 65097965 | 65098176 | 0.576117 | 0.808191 | 1.403 |
| chr16 | 835503 | 835988 | 2.5413 | 3.56511 | 1.403 |
| chr8 | 97173052 | 97173526 | 1.571 | 2.20391 | 1.403 |
| chr10 | 105479790 | 105480000 | 1.1021 | 1.54641 | 1.403 |
| chr20 | 37048564 | 37048803 | 0.681358 | 0.956048 | 1.403 |
| chr14 | 76466830 | 76467093 | 0.687592 | 0.964871 | 1.403 |
| chr22 | 38326510 | 38326962 | 0.99433 | 1.39544 | 1.403 |
| chr1 | 154263667 | 154263949 | 0.605175 | 0.849333 | 1.403 |
| chr20 | 30539800 | 30540165 | 2.35158 | 3.30036 | 1.403 |
| chr16 | 66762092 | 66762380 | 1.65511 | 2.3229 | 1.403 |
| chr9 | 12656900 | 12657120 | 0.536897 | 0.753553 | 1.404 |
| chr2 | 75426735 | 75427049 | 2.07584 | 2.91354 | 1.404 |
| chr12 | 99092725 | 99092983 | 2.1101 | 2.96179 | 1.404 |
| chr8 | 41470290 | 41470681 | 1.85672 | 2.6065 | 1.404 |
| chr11 | 2553950 | 2554374 | 3.7479 | 5.26157 | 1.404 |
| chr2 | 26311008 | 26311288 | 1.5116 | 2.12226 | 1.404 |
| chr1 | 11779795 | 11780349 | 1.95366 | 2.74295 | 1.404 |
| chr20 | 20408092 | 20408416 | 0.749759 | 1.05272 | 1.404 |
| chr5 | 4814471 | 4814730 | 0.628928 | 0.883067 | 1.404 |
| chr20 | 54201250 | 54201506 | 0.678436 | 0.952604 | 1.404 |
| chr9 | 36855459 | 36855745 | 1.90003 | 2.66792 | 1.404 |
| chr8 | 40110642 | 40110940 | 2.20568 | 3.09715 | 1.404 |
| chr17 | 26083742 | 26084048 | 1.23285 | 1.73116 | 1.404 |
| chr16 | 22166867 | 22167178 | 0.501401 | 0.704088 | 1.404 |
| chr7 | 97896223 | 97896611 | 1.18227 | 1.66023 | 1.404 |
| chr3 | 27175703 | 27176024 | 2.39391 | 3.3618 | 1.404 |
| chr7 | 144552013 | 144552280 | 1.70222 | 2.39047 | 1.404 |
| chr17 | 52027649 | 52028017 | 0.692196 | 0.972115 | 1.404 |
| chr20 | 40968221 | 40968456 | 0.659041 | 0.925558 | 1.404 |
| chr7 | 51498121 | 51498524 | 1.98778 | 2.79192 | 1.405 |
| chr16 | 50543462 | 50544011 | 1.62485 | 2.28227 | 1.405 |
| chr8 | 100716968 | 100717171 | 0.765493 | 1.07525 | 1.405 |
| chr10 | 35863785 | 35864014 | 1.03632 | 1.45571 | 1.405 |
| chr15 | 63340196 | 63340797 | 3.07135 | 4.31466 | 1.405 |
| chr3 | 96012905 | 96013301 | 0.850521 | 1.19484 | 1.405 |
| chr5 | 119662053 | 119662285 | 0.792402 | 1.11319 | 1.405 |
| chr9 | 124383269 | 124383620 | 1.65881 | 2.33036 | 1.405 |
| chrX | 75675063 | 75675291 | 1.14804 | 1.61288 | 1.405 |

| | | | | | |
|---|---|---|---|---|---|
| chr20 | 23422500 | 23422881 | 2.3533 | 3.30617 | 1.405 |
| chr16 | 55746256 | 55746647 | 1.66759 | 2.34314 | 1.405 |
| chr1 | 20756863 | 20757211 | 2.98394 | 4.19281 | 1.405 |
| chr10 | 118496947 | 118497362 | 1.19549 | 1.67988 | 1.405 |
| chrX | 152910126 | 152910329 | 0.8241 | 1.1581 | 1.405 |
| chr5 | 168683640 | 168683898 | 0.581264 | 0.81685 | 1.405 |
| chr15 | 84536906 | 84537112 | 0.682504 | 0.959214 | 1.405 |
| chr9 | 125227326 | 125227570 | 2.27514 | 3.19778 | 1.406 |
| chr8 | 5880753 | 5881106 | 0.587597 | 0.825909 | 1.406 |
| chr10 | 98040668 | 98040947 | 0.807787 | 1.13542 | 1.406 |
| chr16 | 85562567 | 85562786 | 1.35176 | 1.9002 | 1.406 |
| chr1 | 159862927 | 159863159 | 0.687932 | 0.967179 | 1.406 |
| chrX | 102155563 | 102155829 | 1.36111 | 1.91365 | 1.406 |
| chr19 | 46532083 | 46532569 | 2.49152 | 3.50304 | 1.406 |
| chr8 | 5518833 | 5519069 | 0.519151 | 0.729954 | 1.406 |
| chr10 | 11194836 | 11195047 | 0.688993 | 0.968772 | 1.406 |
| chr1 | 155698883 | 155699395 | 1.18757 | 1.66983 | 1.406 |
| chr10 | 110681764 | 110682159 | 0.707253 | 0.994499 | 1.406 |
| chr9 | 94902580 | 94903024 | 1.7789 | 2.50148 | 1.406 |
| chr11 | 111457093 | 111457315 | 1.43923 | 2.02393 | 1.406 |
| chr7 | 107526187 | 107526438 | 0.47789 | 0.672049 | 1.406 |
| chr17 | 21183265 | 21183708 | 3.42454 | 4.81607 | 1.406 |
| chr17 | 468893 | 469150 | 1.12045 | 1.57574 | 1.406 |
| chr17 | 67231649 | 67231991 | 2.48757 | 3.4984 | 1.406 |
| chr14 | 25602057 | 25602329 | 1.66554 | 2.3425 | 1.406 |
| chr8 | 59425848 | 59426076 | 1.63716 | 2.30262 | 1.406 |
| chr14 | 96565066 | 96565379 | 2.2307 | 3.13757 | 1.407 |
| chr2 | 232574987 | 232575866 | 1.72688 | 2.42893 | 1.407 |
| chr2 | 18033019 | 18033428 | 1.65337 | 2.32559 | 1.407 |
| chr3 | 153127103 | 153127306 | 0.596831 | 0.83949 | 1.407 |
| chr15 | 73969415 | 73969668 | 1.16411 | 1.63745 | 1.407 |
| chr6 | 149294960 | 149295264 | 1.11863 | 1.57349 | 1.407 |
| chr17 | 60817907 | 60818125 | 0.65341 | 0.919116 | 1.407 |
| chr15 | 89954548 | 89955075 | 2.42315 | 3.40861 | 1.407 |
| chr10 | 10146025 | 10146369 | 0.594855 | 0.83682 | 1.407 |
| chr8 | 112809467 | 112809927 | 1.5987 | 2.24898 | 1.407 |
| chr4 | 31472940 | 31473275 | 0.563284 | 0.792425 | 1.407 |
| chr8 | 124934668 | 124934998 | 2.32539 | 3.27148 | 1.407 |
| chr2 | 95872838 | 95873121 | 1.34259 | 1.88892 | 1.407 |
| chr22 | 37825058 | 37825388 | 1.93291 | 2.71946 | 1.407 |
| chr7 | 142886244 | 142886476 | 0.614803 | 0.865039 | 1.407 |
| chr10 | 23935962 | 23936182 | 0.662145 | 0.931659 | 1.407 |
| chr9 | 73125111 | 73125620 | 1.35006 | 1.89959 | 1.407 |
| chr8 | 96468067 | 96468345 | 0.600809 | 0.845373 | 1.407 |
| chr20 | 42843712 | 42844036 | 1.93103 | 2.71714 | 1.407 |
| chr1 | 202205453 | 202205891 | 2.19428 | 3.08774 | 1.407 |
| chr5 | 73508172 | 73508521 | 1.27618 | 1.79582 | 1.407 |
| chr5 | 83018209 | 83018455 | 1.70859 | 2.40431 | 1.407 |
| chr3 | 97874128 | 97874434 | 0.690693 | 0.971948 | 1.407 |
| chr15 | 92597225 | 92597569 | 1.53222 | 2.15628 | 1.407 |
| chr6 | 150392201 | 150392587 | 1.35039 | 1.90039 | 1.407 |
| chr9 | 119334440 | 119334796 | 2.26774 | 3.19165 | 1.407 |
| chr7 | 107949815 | 107950063 | 0.746198 | 1.05023 | 1.407 |
| chr3 | 169290862 | 169291153 | 0.752328 | 1.05888 | 1.407 |
| chr1 | 76457202 | 76457494 | 2.24944 | 3.16605 | 1.407 |
| chr5 | 77423717 | 77424002 | 1.35413 | 1.90605 | 1.408 |
| chr1 | 181007575 | 181007959 | 2.00655 | 2.82441 | 1.408 |
| chr19 | 10755433 | 10755787 | 3.16261 | 4.45192 | 1.408 |
| chr1 | 205294627 | 205294872 | 2.1534 | 3.03165 | 1.408 |
| chr2 | 16635887 | 16636172 | 0.731143 | 1.02933 | 1.408 |
| chr12 | 133449199 | 133449467 | 0.787001 | 1.108 | 1.408 |

| | | | | | |
|---|---|---|---|---|---|
| chrX | 23774308 | 23774666 | 1.91658 | 2.69837 | 1.408 |
| chr1 | 109820092 | 109820434 | 2.02434 | 2.85013 | 1.408 |
| chr11 | 87308663 | 87308887 | 0.745479 | 1.0496 | 1.408 |
| chr16 | 66378030 | 66378870 | 1.32536 | 1.8661 | 1.408 |
| chr12 | 49690255 | 49690738 | 2.20596 | 3.106 | 1.408 |
| chr15 | 32246261 | 32246521 | 1.09245 | 1.53819 | 1.408 |
| chr9 | 126885090 | 126885557 | 1.20262 | 1.69332 | 1.408 |
| chr4 | 85886857 | 85887128 | 0.573216 | 0.80714 | 1.408 |
| chr16 | 61004223 | 61004569 | 0.693506 | 0.97658 | 1.408 |
| chr14 | 74195447 | 74195793 | 2.45046 | 3.45072 | 1.408 |
| chr2 | 235430822 | 235431035 | 0.479156 | 0.674745 | 1.408 |
| chr19 | 13973091 | 13973379 | 2.08319 | 2.9337 | 1.408 |
| chr1 | 165893683 | 165894316 | 2.22705 | 3.13634 | 1.408 |
| chr1 | 194574908 | 194575178 | 0.756731 | 1.06573 | 1.408 |
| chr16 | 52587291 | 52587553 | 1.50032 | 2.11312 | 1.408 |
| chr14 | 75773701 | 75773906 | 1.24328 | 1.75114 | 1.408 |
| chr14 | 64896745 | 64897158 | 2.55045 | 3.59268 | 1.409 |
| chr17 | 45776195 | 45776901 | 2.42008 | 3.40907 | 1.409 |
| chr2 | 60045872 | 60046107 | 0.709164 | 0.999025 | 1.409 |
| chr18 | 68122019 | 68122230 | 1.66865 | 2.35075 | 1.409 |
| chr9 | 96382394 | 96382669 | 0.772237 | 1.08793 | 1.409 |
| chr5 | 18113890 | 18114118 | 1.92615 | 2.7136 | 1.409 |
| chr19 | 55610218 | 55610425 | 0.854479 | 1.20382 | 1.409 |
| chr5 | 98175540 | 98175742 | 0.99531 | 1.40225 | 1.409 |
| gl000198_r | 57233 | 57502 | 0.728371 | 1.02619 | 1.409 |
| chr5 | 180226205 | 180226612 | 1.44591 | 2.0372 | 1.409 |
| chr3 | 163350510 | 163350884 | 0.795655 | 1.12105 | 1.409 |
| chr3 | 51744724 | 51744984 | 0.89413 | 1.25983 | 1.409 |
| chrX | 71348805 | 71349036 | 1.31809 | 1.8573 | 1.409 |
| chr16 | 84077817 | 84078082 | 1.35785 | 1.9136 | 1.409 |
| chr1 | 211649568 | 211649900 | 2.53419 | 3.57143 | 1.409 |
| chr14 | 72887460 | 72887844 | 2.4047 | 3.38894 | 1.409 |
| chr8 | 127147097 | 127147460 | 0.720384 | 1.01524 | 1.409 |
| chr10 | 119501201 | 119501499 | 1.07281 | 1.51192 | 1.409 |
| chr6 | 136868551 | 136868798 | 2.031 | 2.8623 | 1.409 |
| chr12 | 108312635 | 108313054 | 0.600412 | 0.846198 | 1.409 |
| chr2 | 63673334 | 63673548 | 0.921671 | 1.29905 | 1.409 |
| chr5 | 98670703 | 98671009 | 0.811698 | 1.14405 | 1.409 |
| chr2 | 175636198 | 175636686 | 1.54211 | 2.17375 | 1.410 |
| chr16 | 67219235 | 67219659 | 2.33716 | 3.29451 | 1.410 |
| chr10 | 89610584 | 89610803 | 1.95082 | 2.75021 | 1.410 |
| chr12 | 31806167 | 31806400 | 1.65666 | 2.33562 | 1.410 |
| chr19 | 42612430 | 42613122 | 3.52658 | 4.97197 | 1.410 |
| chr8 | 19125193 | 19125522 | 2.00217 | 2.82304 | 1.410 |
| chr8 | 40794994 | 40795276 | 0.945107 | 1.33273 | 1.410 |
| chr5 | 132888635 | 132889011 | 2.316 | 3.26605 | 1.410 |
| chr8 | 136266741 | 136267282 | 0.71905 | 1.01402 | 1.410 |
| chr3 | 194323870 | 194324123 | 0.684123 | 0.964785 | 1.410 |
| chr17 | 33058100 | 33058317 | 0.402714 | 0.567947 | 1.410 |
| chr9 | 14846757 | 14847083 | 2.10287 | 2.96569 | 1.410 |
| chr14 | 21081582 | 21081859 | 1.85454 | 2.61553 | 1.410 |
| chr1 | 20901749 | 20902057 | 2.37577 | 3.35075 | 1.410 |
| chr10 | 76990405 | 76990671 | 1.91414 | 2.6999 | 1.411 |
| chr2 | 15291476 | 15291880 | 0.674835 | 0.951852 | 1.411 |
| chr5 | 155213898 | 155214101 | 0.711306 | 1.00334 | 1.411 |
| chr1 | 17563616 | 17563836 | 1.7896 | 2.52438 | 1.411 |
| chr3 | 142915542 | 142915953 | 0.661943 | 0.933733 | 1.411 |
| chr17 | 21812872 | 21813484 | 0.701427 | 0.989455 | 1.411 |
| chr9 | 136262156 | 136262411 | 1.29821 | 1.83133 | 1.411 |
| chr19 | 33599024 | 33599239 | 0.787305 | 1.11064 | 1.411 |
| chr6 | 105850975 | 105851624 | 1.36237 | 1.92201 | 1.411 |

| | | | | | |
|---|---|---|---|---|---|
| chr5 | 116759969 | 116760518 | 0.711679 | 1.00406 | 1.411 |
| chr1 | 234634811 | 234635169 | 2.64342 | 3.72952 | 1.411 |
| chr6 | 151186650 | 151186942 | 2.34835 | 3.31334 | 1.411 |
| chr15 | 100882138 | 100882610 | 2.33081 | 3.2886 | 1.411 |
| chr3 | 18430626 | 18431119 | 1.44002 | 2.0319 | 1.411 |
| chr8 | 143273482 | 143273716 | 0.577253 | 0.814529 | 1.411 |
| chr5 | 149501714 | 149502031 | 1.4005 | 1.97621 | 1.411 |
| chr19 | 39600758 | 39601382 | 2.06579 | 2.91504 | 1.411 |
| chr2 | 100436293 | 100436578 | 0.640338 | 0.903591 | 1.411 |
| chr8 | 17460160 | 17460477 | 1.62471 | 2.29273 | 1.411 |
| chr17 | 65374196 | 65374538 | 1.48635 | 2.0975 | 1.411 |
| chr2 | 64749933 | 64750215 | 1.85746 | 2.62131 | 1.411 |
| chr20 | 3544272 | 3544554 | 2.06227 | 2.91033 | 1.411 |
| chr11 | 57260817 | 57261247 | 2.27109 | 3.20509 | 1.411 |
| chr6 | 41335487 | 41335788 | 1.75694 | 2.47966 | 1.411 |
| chr7 | 29142188 | 29142506 | 1.23472 | 1.74266 | 1.411 |
| chr7 | 134671327 | 134671651 | 2.00228 | 2.826 | 1.411 |
| chr1 | 222638679 | 222639055 | 3.23371 | 4.56415 | 1.411 |
| chr20 | 1744892 | 1745490 | 1.74457 | 2.46236 | 1.411 |
| chr12 | 55117528 | 55118040 | 1.08452 | 1.53087 | 1.412 |
| chr2 | 211613413 | 211613629 | 0.822181 | 1.16056 | 1.412 |
| chr5 | 41270610 | 41270840 | 1.79301 | 2.53096 | 1.412 |
| chr9 | 12819874 | 12820232 | 1.48313 | 2.09354 | 1.412 |
| chr2 | 2415355 | 2415609 | 0.685459 | 0.967589 | 1.412 |
| chr19 | 47733671 | 47734205 | 3.3407 | 4.71572 | 1.412 |
| chr5 | 151912243 | 151912432 | 1.08923 | 1.53761 | 1.412 |
| chr5 | 173137053 | 173137279 | 0.54194 | 0.765031 | 1.412 |
| chr11 | 74851552 | 74852049 | 2.76145 | 3.89822 | 1.412 |
| chr1 | 231726399 | 231726706 | 1.96371 | 2.77212 | 1.412 |
| chr7 | 42892411 | 42892612 | 0.889964 | 1.25638 | 1.412 |
| chr6 | 43737094 | 43737532 | 2.89069 | 4.08092 | 1.412 |
| chr7 | 18360605 | 18360903 | 0.989046 | 1.39645 | 1.412 |
| chr13 | 27272829 | 27273162 | 3.11548 | 4.39889 | 1.412 |
| chr11 | 78297557 | 78297870 | 1.58877 | 2.24353 | 1.412 |
| chr2 | 153717903 | 153718237 | 1.98196 | 2.79876 | 1.412 |
| chr7 | 994102 | 994518 | 0.953412 | 1.34637 | 1.412 |
| chr13 | 101225324 | 101225624 | 1.78967 | 2.52757 | 1.412 |
| chr14 | 22668870 | 22669272 | 1.98254 | 2.79997 | 1.412 |
| chr10 | 61569138 | 61569469 | 2.13818 | 3.02 | 1.412 |
| chr12 | 129256933 | 129257146 | 0.645386 | 0.911601 | 1.412 |
| chr9 | 136218912 | 136219202 | 1.7218 | 2.4321 | 1.413 |
| chr15 | 42196703 | 42197077 | 2.39527 | 3.38341 | 1.413 |
| chr15 | 90614411 | 90614796 | 2.0906 | 2.95315 | 1.413 |
| chr1 | 65152480 | 65152712 | 1.77841 | 2.51218 | 1.413 |
| chr14 | 84725056 | 84725312 | 0.856143 | 1.20949 | 1.413 |
| chr2 | 43202106 | 43202569 | 2.4414 | 3.44904 | 1.413 |
| chr21 | 34572152 | 34572381 | 1.68981 | 2.38758 | 1.413 |
| chr17 | 48438069 | 48438373 | 0.85471 | 1.20768 | 1.413 |
| chr1 | 150532381 | 150532768 | 2.59518 | 3.66714 | 1.413 |
| chr3 | 17472394 | 17472742 | 0.913028 | 1.29018 | 1.413 |
| chr10 | 131908014 | 131908327 | 2.16933 | 3.06554 | 1.413 |
| chr16 | 8999091 | 8999384 | 1.27955 | 1.80819 | 1.413 |
| chr3 | 196264254 | 196264651 | 1.34633 | 1.90277 | 1.413 |
| chr11 | 69395370 | 69395592 | 0.936344 | 1.32335 | 1.413 |
| chr17 | 78954959 | 78955516 | 1.90068 | 2.68626 | 1.413 |
| chr10 | 127786110 | 127786393 | 1.09852 | 1.55257 | 1.413 |
| chr1 | 35331917 | 35332289 | 2.48475 | 3.5118 | 1.413 |
| chr3 | 107471581 | 107471838 | 1.20692 | 1.70596 | 1.413 |
| chr14 | 79606603 | 79606807 | 0.51465 | 0.727479 | 1.414 |
| chrX | 26667515 | 26668058 | 1.10417 | 1.56081 | 1.414 |
| chr9 | 72950092 | 72950301 | 0.752061 | 1.06309 | 1.414 |

| | | | | | |
|---|---|---|---|---|---|
| chr5 | 159923247 | 159923458 | 0.624535 | 0.882832 | 1.414 |
| chr2 | 110273940 | 110274309 | 2.53669 | 3.58612 | 1.414 |
| chr15 | 78714977 | 78715555 | 2.08337 | 2.94535 | 1.414 |
| chr22 | 45131670 | 45132100 | 2.7023 | 3.82035 | 1.414 |
| chr2 | 26691958 | 26692346 | 2.06156 | 2.91456 | 1.414 |
| chr15 | 33767624 | 33767879 | 0.759309 | 1.07349 | 1.414 |
| chr9 | 77079573 | 77079982 | 0.639821 | 0.904577 | 1.414 |
| chr3 | 196705675 | 196705963 | 1.16355 | 1.64504 | 1.414 |
| chr1 | 157163364 | 157163743 | 2.12326 | 3.0019 | 1.414 |
| chr10 | 64976926 | 64977227 | 1.41668 | 2.00301 | 1.414 |
| chr2 | 60009926 | 60010159 | 0.748342 | 1.05807 | 1.414 |
| chr6 | 42499851 | 42500300 | 1.01957 | 1.44156 | 1.414 |
| chr9 | 109934506 | 109934784 | 2.18194 | 3.08508 | 1.414 |
| chr11 | 76432669 | 76433145 | 1.84415 | 2.60764 | 1.414 |
| chr19 | 39078597 | 39078834 | 0.676071 | 0.955976 | 1.414 |
| chr3 | 150928667 | 150928888 | 0.770238 | 1.08914 | 1.414 |
| chr19 | 50740814 | 50741126 | 1.68281 | 2.37956 | 1.414 |
| chr8 | 137493641 | 137493888 | 0.85784 | 1.21306 | 1.414 |
| chr14 | 69727828 | 69728195 | 2.34183 | 3.31164 | 1.414 |
| chr16 | 27166343 | 27166838 | 2.35396 | 3.32893 | 1.414 |
| chr2 | 201770408 | 201770761 | 2.06209 | 2.91625 | 1.414 |
| chr9 | 37565735 | 37566003 | 0.581563 | 0.822459 | 1.414 |
| chr14 | 73141668 | 73141878 | 0.702526 | 0.993583 | 1.414 |
| chr4 | 169057795 | 169058001 | 0.687203 | 0.971912 | 1.414 |
| chr12 | 22837552 | 22837905 | 1.58825 | 2.24627 | 1.414 |
| chr10 | 128272435 | 128272646 | 0.760401 | 1.07545 | 1.414 |
| chr8 | 103503650 | 103503873 | 0.516757 | 0.730911 | 1.414 |
| chr10 | 57132251 | 57132533 | 0.647042 | 0.915195 | 1.414 |
| chr1 | 220454503 | 220454785 | 1.07024 | 1.51382 | 1.414 |
| chr10 | 83655387 | 83655640 | 0.581093 | 0.822 | 1.415 |
| chr5 | 38382993 | 38383270 | 2.28526 | 3.23276 | 1.415 |
| chr7 | 112509216 | 112509581 | 2.8782 | 4.07171 | 1.415 |
| chr9 | 79425349 | 79425655 | 1.92658 | 2.72547 | 1.415 |
| chr3 | 69488912 | 69489134 | 1.41124 | 1.99645 | 1.415 |
| chr8 | 3886888 | 3887125 | 0.775348 | 1.09692 | 1.415 |
| chr11 | 20408659 | 20409004 | 2.94241 | 4.1629 | 1.415 |
| chr13 | 20501352 | 20501592 | 1.03016 | 1.45746 | 1.415 |
| chr16 | 73990488 | 73990900 | 1.9179 | 2.71345 | 1.415 |
| chr17 | 14829844 | 14830218 | 1.96361 | 2.77811 | 1.415 |
| chr2 | 13216722 | 13216938 | 0.639889 | 0.905326 | 1.415 |
| chr15 | 64968046 | 64968290 | 0.662273 | 0.93702 | 1.415 |
| chr1 | 151072324 | 151072642 | 1.27516 | 1.80421 | 1.415 |
| chr7 | 66370731 | 66371008 | 1.12289 | 1.58879 | 1.415 |
| chr5 | 98390269 | 98390521 | 0.749994 | 1.06126 | 1.415 |
| chr19 | 41521905 | 41522116 | 0.778934 | 1.10226 | 1.415 |
| chr2 | 7650628 | 7650927 | 0.952999 | 1.34858 | 1.415 |
| chr19 | 10499585 | 10499884 | 2.10535 | 2.97928 | 1.415 |
| chr9 | 5600471 | 5600782 | 1.74951 | 2.47573 | 1.415 |
| chr19 | 4523096 | 4523690 | 1.70889 | 2.4183 | 1.415 |
| chr20 | 20341077 | 20341432 | 2.10047 | 2.97252 | 1.415 |
| chr2 | 65116148 | 65116380 | 1.77125 | 2.50671 | 1.415 |
| chr13 | 53542238 | 53542591 | 2.6188 | 3.70627 | 1.415 |
| chr2 | 207853940 | 207854458 | 1.52777 | 2.16224 | 1.415 |
| chr12 | 114350500 | 114350881 | 2.40419 | 3.40268 | 1.415 |
| chr7 | 12529818 | 12530022 | 0.973965 | 1.37846 | 1.415 |
| chr10 | 34229066 | 34229290 | 0.82723 | 1.17081 | 1.415 |
| chr3 | 943910 | 944175 | 0.957084 | 1.35461 | 1.415 |
| chr11 | 48328140 | 48328371 | 0.790721 | 1.11917 | 1.415 |
| chr9 | 105792463 | 105792703 | 0.659839 | 0.934016 | 1.416 |
| chr10 | 25927165 | 25927623 | 0.915323 | 1.29568 | 1.416 |
| chr10 | 10318450 | 10318703 | 0.742421 | 1.05117 | 1.416 |

| | | | | | |
|---|---|---|---|---|---|
| chr13 | 49306140 | 49306436 | 2.49938 | 3.53891 | 1.416 |
| chr3 | 17274918 | 17275158 | 1.61265 | 2.28348 | 1.416 |
| chr19 | 30239905 | 30240239 | 0.782043 | 1.10739 | 1.416 |
| chr5 | 25858154 | 25858442 | 1.01797 | 1.44149 | 1.416 |
| chr1 | 219589955 | 219590297 | 1.95206 | 2.76423 | 1.416 |
| chr9 | 21392186 | 21392567 | 0.972669 | 1.37738 | 1.416 |
| chr12 | 61199314 | 61199539 | 0.680547 | 0.963773 | 1.416 |
| chr1 | 203095546 | 203095916 | 1.26868 | 1.79673 | 1.416 |
| chr2 | 10176189 | 10176506 | 1.83969 | 2.60541 | 1.416 |
| chr2 | 200715962 | 200716224 | 1.35666 | 1.92137 | 1.416 |
| chr3 | 72924713 | 72924978 | 2.14784 | 3.04194 | 1.416 |
| chr12 | 46572183 | 46572593 | 0.750392 | 1.06282 | 1.416 |
| chr7 | 154100438 | 154100665 | 0.58541 | 0.829222 | 1.416 |
| chr3 | 23852358 | 23852608 | 1.51754 | 2.14961 | 1.417 |
| chr7 | 117835952 | 117836221 | 1.62506 | 2.30196 | 1.417 |
| chr16 | 89400102 | 89400371 | 1.35589 | 1.92071 | 1.417 |
| chr17 | 26768620 | 26768869 | 0.877659 | 1.24329 | 1.417 |
| chr4 | 10551387 | 10551600 | 1.56826 | 2.22162 | 1.417 |
| chr17 | 7452034 | 7452309 | 0.7915 | 1.12127 | 1.417 |
| chr5 | 146241978 | 146242239 | 0.923695 | 1.30861 | 1.417 |
| chr4 | 3049167 | 3049393 | 1.06144 | 1.50376 | 1.417 |
| chr11 | 32013997 | 32014296 | 0.550622 | 0.780138 | 1.417 |
| chr6 | 167403217 | 167403448 | 2.14202 | 3.03503 | 1.417 |
| chr8 | 145734800 | 145735489 | 2.56351 | 3.63224 | 1.417 |
| chr15 | 84728799 | 84729054 | 0.867154 | 1.22868 | 1.417 |
| chr3 | 72871584 | 72871823 | 1.77198 | 2.51074 | 1.417 |
| chr2 | 42316506 | 42316788 | 1.8185 | 2.57668 | 1.417 |
| chr3 | 123442460 | 123442713 | 2.02686 | 2.87206 | 1.417 |
| chr6 | 33995379 | 33995790 | 1.82192 | 2.58167 | 1.417 |
| chr14 | 76777917 | 76778206 | 0.64891 | 0.919545 | 1.417 |
| chr1 | 194858427 | 194858644 | 0.899256 | 1.27431 | 1.417 |
| chr1 | 195551917 | 195552165 | 0.662779 | 0.93924 | 1.417 |
| chr11 | 76378941 | 76379148 | 0.688683 | 0.975996 | 1.417 |
| chr7 | 12723322 | 12723525 | 1.25859 | 1.78374 | 1.417 |
| chr15 | 86775452 | 86775751 | 1.25571 | 1.77967 | 1.417 |
| chr12 | 130488450 | 130488697 | 0.543828 | 0.77076 | 1.417 |
| chr2 | 79746054 | 79746304 | 0.690072 | 0.978045 | 1.417 |
| chr19 | 22134679 | 22134952 | 0.658233 | 0.932934 | 1.417 |
| chr9 | 130808211 | 130808477 | 0.549688 | 0.779097 | 1.417 |
| chr8 | 59307785 | 59308087 | 2.04141 | 2.89344 | 1.417 |
| chr14 | 82759373 | 82759734 | 0.782796 | 1.10952 | 1.417 |
| chr1 | 39283655 | 39284185 | 1.81932 | 2.57882 | 1.417 |
| chr6 | 118297259 | 118297608 | 1.45069 | 2.05635 | 1.418 |
| chr5 | 16141235 | 16141486 | 0.666543 | 0.944863 | 1.418 |
| chr6 | 147223623 | 147223812 | 1.35686 | 1.92344 | 1.418 |
| chr1 | 179048405 | 179048649 | 1.40781 | 1.9957 | 1.418 |
| chr1 | 6140622 | 6140889 | 1.01236 | 1.43512 | 1.418 |
| chr14 | 38666126 | 38666732 | 0.959064 | 1.35958 | 1.418 |
| chr4 | 1312429 | 1312682 | 1.3403 | 1.90009 | 1.418 |
| chr2 | 20964218 | 20964431 | 0.728641 | 1.03299 | 1.418 |
| chr14 | 74578425 | 74578679 | 1.35291 | 1.91804 | 1.418 |
| chr10 | 118280938 | 118281283 | 1.77488 | 2.51641 | 1.418 |
| chr17 | 36693412 | 36693777 | 1.63221 | 2.31415 | 1.418 |
| chr1 | 94527044 | 94527306 | 1.87634 | 2.66032 | 1.418 |
| chr2 | 2427791 | 2428044 | 0.798086 | 1.13158 | 1.418 |
| chr8 | 145068798 | 145069000 | 0.846827 | 1.2007 | 1.418 |
| chr8 | 103714213 | 103714425 | 1.3543 | 1.92039 | 1.418 |
| chr2 | 171672256 | 171672661 | 1.59126 | 2.25651 | 1.418 |
| chr2 | 16079879 | 16080301 | 1.3204 | 1.87247 | 1.418 |
| chr2 | 241000502 | 241000708 | 1.40816 | 1.99693 | 1.418 |
| chr16 | 15644737 | 15645002 | 2.03211 | 2.88183 | 1.418 |

| | | | | | |
|---|---|---|---|---|---|
| chr21 | 41515853 | 41516080 | 1.39403 | 1.97695 | 1.418 |
| chr20 | 22132164 | 22132488 | 0.903329 | 1.28121 | 1.418 |
| chr5 | 180046029 | 180046493 | 2.53563 | 3.59634 | 1.418 |
| chr8 | 105342173 | 105342417 | 1.13461 | 1.60931 | 1.418 |
| chr6 | 36328323 | 36328570 | 1.54433 | 2.19051 | 1.418 |
| chr5 | 75470644 | 75471420 | 1.48158 | 2.10158 | 1.418 |
| chr1 | 27524483 | 27524734 | 1.72302 | 2.44408 | 1.418 |
| chr7 | 92239349 | 92239561 | 0.871721 | 1.23664 | 1.419 |
| chr15 | 98434863 | 98435157 | 0.773044 | 1.09668 | 1.419 |
| chr19 | 7989940 | 7990398 | 2.14627 | 3.04487 | 1.419 |
| chr20 | 3733683 | 3734007 | 2.34407 | 3.32549 | 1.419 |
| chr18 | 3129222 | 3129656 | 2.22127 | 3.15166 | 1.419 |
| chr20 | 4701389 | 4701642 | 2.32727 | 3.30205 | 1.419 |
| chr2 | 27672675 | 27672882 | 0.652881 | 0.926364 | 1.419 |
| chr8 | 56884754 | 56885087 | 1.71803 | 2.43769 | 1.419 |
| chr8 | 27181300 | 27181522 | 1.39382 | 1.97771 | 1.419 |
| chr3 | 89074360 | 89074571 | 0.724719 | 1.02832 | 1.419 |
| chr3 | 1877715 | 1877909 | 0.840284 | 1.19237 | 1.419 |
| chr9 | 2200122 | 2200384 | 0.872602 | 1.23829 | 1.419 |
| chr12 | 96989410 | 96989655 | 0.821964 | 1.16648 | 1.419 |
| chr11 | 124610402 | 124610709 | 1.35851 | 1.92793 | 1.419 |
| chr3 | 38692640 | 38693076 | 2.68838 | 3.81521 | 1.419 |
| chr11 | 8284721 | 8284944 | 0.777143 | 1.1029 | 1.419 |
| chr8 | 59230915 | 59231179 | 1.15967 | 1.64578 | 1.419 |
| chr3 | 18578794 | 18579083 | 0.571661 | 0.811369 | 1.419 |
| chr1 | 208797598 | 208797877 | 0.796104 | 1.12995 | 1.419 |
| chr11 | 76409327 | 76409691 | 1.57326 | 2.23318 | 1.419 |
| chr1 | 94505084 | 94505422 | 2.65082 | 3.76286 | 1.420 |
| chr9 | 7358832 | 7359059 | 0.618806 | 0.878454 | 1.420 |
| chr12 | 49331080 | 49331286 | 1.37691 | 1.95479 | 1.420 |
| chr17 | 47634605 | 47635098 | 1.65882 | 2.35505 | 1.420 |
| chr15 | 58366902 | 58367196 | 0.855746 | 1.21494 | 1.420 |
| chr8 | 145692118 | 145692714 | 1.84602 | 2.621 | 1.420 |
| chr11 | 18001255 | 18001918 | 1.64844 | 2.34057 | 1.420 |
| chr21 | 43237622 | 43237925 | 1.75298 | 2.48929 | 1.420 |
| chr17 | 12034640 | 12034900 | 0.954218 | 1.35505 | 1.420 |
| chr5 | 19995096 | 19995313 | 0.498252 | 0.707568 | 1.420 |
| chr17 | 70553046 | 70553416 | 2.65651 | 3.77261 | 1.420 |
| chr1 | 32230011 | 32230360 | 0.609037 | 0.865034 | 1.420 |
| chr19 | 35843684 | 35844007 | 2.44651 | 3.47488 | 1.420 |
| chr2 | 197296777 | 197296996 | 2.08921 | 2.96751 | 1.420 |
| chr15 | 46006763 | 46007052 | 1.71963 | 2.44273 | 1.421 |
| chr8 | 63897784 | 63898015 | 0.773537 | 1.09882 | 1.421 |
| chr3 | 186857452 | 186857828 | 1.89575 | 2.69302 | 1.421 |
| chr2 | 241172344 | 241172613 | 2.03732 | 2.89415 | 1.421 |
| chr2 | 39758467 | 39758707 | 0.723875 | 1.02837 | 1.421 |
| chr7 | 25633529 | 25633805 | 0.885865 | 1.25857 | 1.421 |
| chr9 | 102790420 | 102790729 | 2.40635 | 3.4188 | 1.421 |
| chr8 | 56837733 | 56837980 | 0.544499 | 0.773595 | 1.421 |
| chr12 | 91720251 | 91720534 | 2.25676 | 3.20636 | 1.421 |
| chr15 | 34029822 | 34030228 | 2.02936 | 2.88347 | 1.421 |
| chr17 | 54969169 | 54969416 | 1.35662 | 1.92768 | 1.421 |
| chr10 | 106743341 | 106743629 | 0.782326 | 1.11166 | 1.421 |
| chr8 | 59929512 | 59929732 | 0.920794 | 1.30843 | 1.421 |
| chr3 | 161226723 | 161226965 | 0.891915 | 1.26745 | 1.421 |
| chr5 | 156497019 | 156497223 | 1.10613 | 1.57192 | 1.421 |
| chr5 | 94433132 | 94433406 | 0.591702 | 0.840871 | 1.421 |
| chr14 | 31755998 | 31756318 | 1.5719 | 2.23402 | 1.421 |
| chr3 | 154173356 | 154173571 | 0.742175 | 1.05493 | 1.421 |
| chr3 | 42581378 | 42581821 | 2.19052 | 3.11369 | 1.421 |
| chr8 | 97295260 | 97295464 | 1.27538 | 1.81297 | 1.422 |

| | | | | | |
|---|---|---|---|---|---|
| chr15 | 51694035 | 51694408 | 1.97436 | 2.80671 | 1.422 |
| chr20 | 49598045 | 49598312 | 1.81229 | 2.57632 | 1.422 |
| chr9 | 33452739 | 33453131 | 2.11445 | 3.00585 | 1.422 |
| chr2 | 153572436 | 153572686 | 1.83963 | 2.61542 | 1.422 |
| chr6 | 152916777 | 152916998 | 1.16354 | 1.65428 | 1.422 |
| chr3 | 124869464 | 124869741 | 1.82666 | 2.59717 | 1.422 |
| chr19 | 6738511 | 6738773 | 1.63443 | 2.32393 | 1.422 |
| chr15 | 32260541 | 32260786 | 0.678837 | 0.965223 | 1.422 |
| chr3 | 123277955 | 123278233 | 1.57717 | 2.2426 | 1.422 |
| chr16 | 23382760 | 23383115 | 0.974823 | 1.38619 | 1.422 |
| chr17 | 43146030 | 43146327 | 0.687297 | 0.977425 | 1.422 |
| chr1 | 168606739 | 168606964 | 1.40677 | 2.00069 | 1.422 |
| chr19 | 42793143 | 42793540 | 1.01202 | 1.43938 | 1.422 |
| chr8 | 69448338 | 69448545 | 0.760155 | 1.08118 | 1.422 |
| chr16 | 48573186 | 48573419 | 0.735709 | 1.04644 | 1.422 |
| chr20 | 37676648 | 37677085 | 2.84327 | 4.04431 | 1.422 |
| chr19 | 54634743 | 54634985 | 0.610292 | 0.868101 | 1.422 |
| chr7 | 146814342 | 146814545 | 0.759133 | 1.07983 | 1.422 |
| chr19 | 40660235 | 40660596 | 0.778314 | 1.10718 | 1.423 |
| chr12 | 123548134 | 123548342 | 1.08081 | 1.53752 | 1.423 |
| chr2 | 173886695 | 173886950 | 0.9607 | 1.36671 | 1.423 |
| chr14 | 20320409 | 20320697 | 0.62711 | 0.892143 | 1.423 |
| chr1 | 116049885 | 116050126 | 1.22182 | 1.73825 | 1.423 |
| chr2 | 96079860 | 96080137 | 1.64066 | 2.33414 | 1.423 |
| chr5 | 15995541 | 15995842 | 1.03325 | 1.47025 | 1.423 |
| chr8 | 128190846 | 128191159 | 1.26474 | 1.79965 | 1.423 |
| chr17 | 72805657 | 72806053 | 0.566609 | 0.806256 | 1.423 |
| chr3 | 22225648 | 22225941 | 0.503482 | 0.716475 | 1.423 |
| chr11 | 39337968 | 39338170 | 0.637061 | 0.906601 | 1.423 |
| chr5 | 143923262 | 143923508 | 0.590122 | 0.83982 | 1.423 |
| chr1 | 208579902 | 208580118 | 0.728562 | 1.03685 | 1.423 |
| chr10 | 127210679 | 127211083 | 2.2035 | 3.13601 | 1.423 |
| chr15 | 48578002 | 48578207 | 1.47793 | 2.10342 | 1.423 |
| chr19 | 50486656 | 50487318 | 1.99205 | 2.83516 | 1.423 |
| chr1 | 200227541 | 200227957 | 1.23027 | 1.75098 | 1.423 |
| chr16 | 59226612 | 59226809 | 1.07279 | 1.52685 | 1.423 |
| chr20 | 40326434 | 40327000 | 1.37175 | 1.95241 | 1.423 |
| chr20 | 8732192 | 8732435 | 2.29237 | 3.26276 | 1.423 |
| chr13 | 113683711 | 113684134 | 2.20019 | 3.13158 | 1.423 |
| chr8 | 35998836 | 35999064 | 1.30887 | 1.86307 | 1.423 |
| chr15 | 49517904 | 49518131 | 1.13343 | 1.61339 | 1.423 |
| chr16 | 69157935 | 69158157 | 1.40278 | 1.99704 | 1.424 |
| chr8 | 4854498 | 4854813 | 1.4288 | 2.0342 | 1.424 |
| chr7 | 36513028 | 36513313 | 1.89764 | 2.70172 | 1.424 |
| chr8 | 52721727 | 52721970 | 0.880685 | 1.25387 | 1.424 |
| chr5 | 131862256 | 131862798 | 2.16578 | 3.0836 | 1.424 |
| chr3 | 119917235 | 119917441 | 1.10499 | 1.5733 | 1.424 |
| chr7 | 29145958 | 29146353 | 2.46107 | 3.50427 | 1.424 |
| chr8 | 130261312 | 130261886 | 0.85066 | 1.21126 | 1.424 |
| chr14 | 96487747 | 96488064 | 0.804287 | 1.14534 | 1.424 |
| chr17 | 50153121 | 50153359 | 0.753466 | 1.07299 | 1.424 |
| chr1 | 46993734 | 46994308 | 2.07634 | 2.95731 | 1.424 |
| chr12 | 114258916 | 114259129 | 0.715252 | 1.01883 | 1.424 |
| chr3 | 76154694 | 76154984 | 0.655034 | 0.933058 | 1.424 |
| chr2 | 74317547 | 74317766 | 0.709467 | 1.01073 | 1.425 |
| chr3 | 186882967 | 186883183 | 1.25116 | 1.78256 | 1.425 |
| chr5 | 176889616 | 176890012 | 0.617726 | 0.880112 | 1.425 |
| chr2 | 65914183 | 65914612 | 0.74385 | 1.05984 | 1.425 |
| chr2 | 128991161 | 128991442 | 2.3159 | 3.29969 | 1.425 |
| chr7 | 91265572 | 91265797 | 0.947213 | 1.34961 | 1.425 |
| chr22 | 37868437 | 37868726 | 1.86872 | 2.66263 | 1.425 |

| | | | | | |
|---|---|---|---|---|---|
| chr7 | 84660135 | 84660436 | 1.13247 | 1.61368 | 1.425 |
| chr12 | 72426584 | 72426822 | 0.755258 | 1.0762 | 1.425 |
| chr3 | 117195465 | 117195677 | 0.643862 | 0.917487 | 1.425 |
| chr8 | 80058496 | 80058795 | 0.862047 | 1.22842 | 1.425 |
| chr7 | 22900392 | 22900671 | 1.36888 | 1.95069 | 1.425 |
| chr19 | 51901398 | 51901742 | 1.55679 | 2.21855 | 1.425 |
| chr15 | 76214177 | 76214388 | 1.2628 | 1.79962 | 1.425 |
| chr17 | 43348152 | 43348623 | 2.74997 | 3.91903 | 1.425 |
| chr13 | 44362258 | 44362546 | 1.03225 | 1.47109 | 1.425 |
| chr1 | 23730491 | 23730725 | 2.6393 | 3.76145 | 1.425 |
| chr17 | 15917283 | 15917585 | 1.4928 | 2.12751 | 1.425 |
| chr7 | 131870953 | 131871239 | 0.62409 | 0.889489 | 1.425 |
| chr10 | 135121254 | 135121747 | 1.73962 | 2.47945 | 1.425 |
| chr10 | 92622741 | 92623013 | 1.85541 | 2.64458 | 1.425 |
| chr12 | 93896844 | 93897296 | 1.73382 | 2.47156 | 1.426 |
| chr12 | 50260922 | 50261312 | 2.77107 | 3.95042 | 1.426 |
| chr16 | 167688 | 168165 | 2.44624 | 3.48733 | 1.426 |
| chr10 | 69062977 | 69063199 | 0.730921 | 1.042 | 1.426 |
| chr12 | 110371827 | 110372079 | 1.07661 | 1.53483 | 1.426 |
| chrX | 13768410 | 13768645 | 1.45966 | 2.08107 | 1.426 |
| chr7 | 140344979 | 140345304 | 0.656042 | 0.93541 | 1.426 |
| chr17 | 16356659 | 16356891 | 3.40995 | 4.86251 | 1.426 |
| chr1 | 18935012 | 18935269 | 0.798449 | 1.13858 | 1.426 |
| chr10 | 462013 | 462269 | 1.32121 | 1.88408 | 1.426 |
| chr11 | 46148729 | 46148995 | 1.56403 | 2.2304 | 1.426 |
| chr17 | 5258372 | 5258606 | 0.639666 | 0.912241 | 1.426 |
| chr5 | 321511 | 321866 | 2.20757 | 3.14841 | 1.426 |
| chr10 | 24485409 | 24485819 | 3.08344 | 4.3976 | 1.426 |
| chr19 | 33920990 | 33921209 | 1.23886 | 1.76689 | 1.426 |
| chr7 | 4519065 | 4519538 | 1.66661 | 2.37702 | 1.426 |
| chr1 | 16399996 | 16400332 | 1.84939 | 2.63793 | 1.426 |
| chr12 | 86944825 | 86945177 | 0.960222 | 1.3697 | 1.426 |
| chr2 | 18580339 | 18580557 | 0.652272 | 0.930447 | 1.426 |
| chr11 | 50174539 | 50174826 | 0.62917 | 0.897524 | 1.427 |
| chr17 | 9862497 | 9863062 | 1.68189 | 2.39929 | 1.427 |
| chr17 | 69081413 | 69081655 | 1.19711 | 1.70777 | 1.427 |
| chr15 | 68607854 | 68608101 | 0.577603 | 0.824081 | 1.427 |
| chr5 | 133189391 | 133189594 | 0.474288 | 0.67668 | 1.427 |
| chr16 | 59878481 | 59878736 | 0.709065 | 1.01166 | 1.427 |
| chr17 | 41946106 | 41946319 | 0.636029 | 0.90755 | 1.427 |
| chr8 | 27311898 | 27312446 | 1.47498 | 2.10478 | 1.427 |
| chr14 | 21287055 | 21287319 | 1.13701 | 1.62256 | 1.427 |
| chr8 | 75878069 | 75878475 | 0.994403 | 1.41913 | 1.427 |
| chr17 | 9579510 | 9579725 | 1.28675 | 1.83648 | 1.427 |
| chr3 | 99472209 | 99472503 | 1.44896 | 2.06802 | 1.427 |
| chr9 | 126161966 | 126162377 | 1.38328 | 1.97439 | 1.427 |
| chr8 | 52053345 | 52053624 | 0.688084 | 0.982197 | 1.427 |
| chrX | 24517181 | 24517380 | 1.89115 | 2.6996 | 1.427 |
| chr7 | 128587384 | 128587733 | 1.45021 | 2.07022 | 1.428 |
| chr14 | 36291051 | 36291462 | 1.64855 | 2.35337 | 1.428 |
| chr5 | 156680544 | 156680901 | 2.04022 | 2.91252 | 1.428 |
| chr11 | 34731100 | 34731356 | 0.758785 | 1.08321 | 1.428 |
| chr20 | 20287155 | 20287431 | 0.66853 | 0.954419 | 1.428 |
| chr9 | 133739008 | 133739318 | 2.20861 | 3.15309 | 1.428 |
| chr1 | 156427437 | 156427653 | 1.55499 | 2.22005 | 1.428 |
| chr7 | 78302705 | 78303104 | 0.67376 | 0.961929 | 1.428 |
| chr8 | 121101447 | 121101865 | 1.92437 | 2.74752 | 1.428 |
| chr1 | 212803406 | 212803763 | 1.70624 | 2.43613 | 1.428 |
| chr7 | 131339518 | 131339864 | 3.49729 | 4.99339 | 1.428 |
| chr1 | 201330591 | 201331187 | 1.6936 | 2.41828 | 1.428 |
| chr2 | 68942567 | 68943225 | 1.97412 | 2.81886 | 1.428 |

| | | | | | |
|---|---|---|---|---|---|
| chr17 | 6904969 | 6905189 | 2.03632 | 2.90776 | 1.428 |
| chr17 | 1998285 | 1998499 | 1.04118 | 1.48677 | 1.428 |
| chr20 | 48125069 | 48125468 | 1.6517 | 2.35862 | 1.428 |
| chr2 | 202615395 | 202615648 | 0.826199 | 1.17984 | 1.428 |
| chr5 | 160038815 | 160039021 | 0.724512 | 1.03466 | 1.428 |
| chrX | 68723421 | 68723755 | 1.30089 | 1.85779 | 1.428 |
| chr20 | 44386733 | 44387034 | 0.718 | 1.0254 | 1.428 |
| chr10 | 118922569 | 118922853 | 1.64448 | 2.34856 | 1.428 |
| chr16 | 58790309 | 58790672 | 1.79389 | 2.56199 | 1.428 |
| chr6 | 7276180 | 7276554 | 2.34477 | 3.34883 | 1.428 |
| chr8 | 142639541 | 142639798 | 1.18726 | 1.69568 | 1.428 |
| chr8 | 132088582 | 132088886 | 0.719359 | 1.0275 | 1.428 |
| chr15 | 27772725 | 27773021 | 1.37256 | 1.96061 | 1.428 |
| chr9 | 21252979 | 21253468 | 1.18484 | 1.69248 | 1.428 |
| chr13 | 32316257 | 32316614 | 2.68732 | 3.83883 | 1.429 |
| chr11 | 18230731 | 18230976 | 0.762189 | 1.08885 | 1.429 |
| chr7 | 4034837 | 4035139 | 0.928518 | 1.32658 | 1.429 |
| chr18 | 57655393 | 57655687 | 1.79399 | 2.56323 | 1.429 |
| chr2 | 38782306 | 38782623 | 1.1594 | 1.65654 | 1.429 |
| chr3 | 16592328 | 16592570 | 1.69125 | 2.41656 | 1.429 |
| chr16 | 87735824 | 87736424 | 1.49071 | 2.13003 | 1.429 |
| chr8 | 91865277 | 91865479 | 0.8555 | 1.22244 | 1.429 |
| chr11 | 64146217 | 64146480 | 0.646734 | 0.924152 | 1.429 |
| chr8 | 123807220 | 123807607 | 3.19205 | 4.56128 | 1.429 |
| chr1 | 248406522 | 248406840 | 1.4828 | 2.11886 | 1.429 |
| chr11 | 128502100 | 128502410 | 1.40115 | 2.00219 | 1.429 |
| chr16 | 87883531 | 87884037 | 2.52619 | 3.60999 | 1.429 |
| chr11 | 118166519 | 118166915 | 1.2256 | 1.75153 | 1.429 |
| chr17 | 46385237 | 46385485 | 2.18174 | 3.11811 | 1.429 |
| chr7 | 207802 | 208007 | 0.968516 | 1.38422 | 1.429 |
| chr12 | 67555049 | 67555536 | 1.33875 | 1.91349 | 1.429 |
| chr7 | 21293661 | 21294036 | 2.17138 | 3.10369 | 1.429 |
| chr11 | 24088744 | 24088995 | 0.809585 | 1.1573 | 1.430 |
| chr3 | 152553130 | 152553353 | 1.58433 | 2.2649 | 1.430 |
| chr7 | 29961073 | 29961400 | 1.65903 | 2.37174 | 1.430 |
| chrX | 16141293 | 16141627 | 1.65481 | 2.36571 | 1.430 |
| chr19 | 33897410 | 33898243 | 2.2268 | 3.18352 | 1.430 |
| chr7 | 25891171 | 25891602 | 2.05091 | 2.93209 | 1.430 |
| chr9 | 127557195 | 127557620 | 0.932245 | 1.33279 | 1.430 |
| chr5 | 72747062 | 72747577 | 2.1002 | 3.00265 | 1.430 |
| chr16 | 2390626 | 2391248 | 4.8736 | 6.96816 | 1.430 |
| chr1 | 70994479 | 70994703 | 1.71022 | 2.44537 | 1.430 |
| chr5 | 104007425 | 104007812 | 0.880205 | 1.25859 | 1.430 |
| chr11 | 26831396 | 26831664 | 0.648128 | 0.926782 | 1.430 |
| chr2 | 76936075 | 76936356 | 0.752827 | 1.07651 | 1.430 |
| chr1 | 41748415 | 41748768 | 1.94043 | 2.77483 | 1.430 |
| chr1 | 36600224 | 36600651 | 1.60321 | 2.29264 | 1.430 |
| chr7 | 34130097 | 34130619 | 0.648926 | 0.928006 | 1.430 |
| chr8 | 58338103 | 58338307 | 0.633324 | 0.905693 | 1.430 |
| chr19 | 32659693 | 32660001 | 0.800504 | 1.14479 | 1.430 |
| chr3 | 123361144 | 123361620 | 2.32168 | 3.32021 | 1.430 |
| chr12 | 49725423 | 49725710 | 2.36525 | 3.38268 | 1.430 |
| chr10 | 69457503 | 69457723 | 1.37377 | 1.96478 | 1.430 |
| chr11 | 79419401 | 79419898 | 1.09948 | 1.57254 | 1.430 |
| chr10 | 74034561 | 74034951 | 1.21549 | 1.73851 | 1.430 |
| chr3 | 39142550 | 39142957 | 2.09141 | 2.99134 | 1.430 |
| chr3 | 165163879 | 165164083 | 0.834315 | 1.19332 | 1.430 |
| chr21 | 39542409 | 39542634 | 1.13763 | 1.62719 | 1.430 |
| chr19 | 4660418 | 4660769 | 1.37751 | 1.97036 | 1.430 |
| chr2 | 73390924 | 73391254 | 2.2407 | 3.20511 | 1.430 |
| chr2 | 109224247 | 109224526 | 1.29414 | 1.85121 | 1.430 |

| | | | | | |
|---|---|---|---|---|---|
| chr1 | 214758558 | 214758822 | 2.10705 | 3.01426 | 1.431 |
| chr11 | 123940161 | 123940345 | 2.31378 | 3.31006 | 1.431 |
| chr14 | 74201357 | 74201650 | 0.712142 | 1.01878 | 1.431 |
| chr9 | 8631401 | 8631606 | 0.61927 | 0.885947 | 1.431 |
| chr15 | 24887690 | 24887944 | 0.795321 | 1.13782 | 1.431 |
| chr11 | 67030144 | 67030503 | 2.36318 | 3.38088 | 1.431 |
| chr12 | 105814488 | 105814751 | 2.49041 | 3.56324 | 1.431 |
| chr4 | 92034355 | 92034589 | 1.32651 | 1.898 | 1.431 |
| chr4 | 154049109 | 154049343 | 1.74846 | 2.50198 | 1.431 |
| chr10 | 103051192 | 103051666 | 1.55467 | 2.22489 | 1.431 |
| chr10 | 6960343 | 6960673 | 1.47639 | 2.11291 | 1.431 |
| chr15 | 28119382 | 28119647 | 0.665068 | 0.951817 | 1.431 |
| chr1 | 25363741 | 25363962 | 1.38968 | 1.989 | 1.431 |
| chr8 | 145258258 | 145258565 | 2.54362 | 3.64086 | 1.431 |
| chr12 | 98394080 | 98394271 | 0.666125 | 0.953527 | 1.431 |
| chr20 | 58625149 | 58625603 | 1.07581 | 1.5401 | 1.432 |
| chr3 | 124480103 | 124480470 | 2.29043 | 3.27891 | 1.432 |
| chr9 | 116071551 | 116071850 | 1.68878 | 2.41767 | 1.432 |
| chr2 | 15717251 | 15717521 | 2.54343 | 3.64146 | 1.432 |
| chr19 | 44289382 | 44289831 | 3.22065 | 4.61113 | 1.432 |
| chr6 | 56112314 | 56112517 | 0.827655 | 1.18499 | 1.432 |
| chr17 | 11839038 | 11839464 | 3.34045 | 4.7828 | 1.432 |
| chr3 | 50359155 | 50359550 | 0.982672 | 1.40713 | 1.432 |
| chr2 | 101769312 | 101769593 | 1.67624 | 2.40033 | 1.432 |
| chr1 | 107664891 | 107665139 | 0.681156 | 0.975436 | 1.432 |
| chr5 | 113727447 | 113727732 | 0.683739 | 0.979232 | 1.432 |
| chr8 | 127522618 | 127522853 | 1.68748 | 2.41678 | 1.432 |
| chr11 | 107870565 | 107870839 | 0.826097 | 1.18314 | 1.432 |
| chr1 | 184569315 | 184569600 | 0.552612 | 0.79146 | 1.432 |
| chr13 | 60254607 | 60254864 | 1.58668 | 2.2725 | 1.432 |
| chr5 | 121495766 | 121495999 | 0.786547 | 1.12653 | 1.432 |
| chr12 | 73147833 | 73148089 | 0.582338 | 0.834105 | 1.432 |
| chr3 | 66124369 | 66124595 | 1.91922 | 2.74918 | 1.432 |
| chr3 | 127266712 | 127266961 | 1.63135 | 2.33682 | 1.432 |
| chr17 | 32806529 | 32806769 | 0.770658 | 1.10394 | 1.432 |
| chr8 | 104385477 | 104385897 | 1.63931 | 2.34837 | 1.433 |
| chr8 | 11116206 | 11116480 | 0.63777 | 0.913651 | 1.433 |
| chr10 | 57608237 | 57608575 | 0.893983 | 1.28089 | 1.433 |
| chr5 | 66110282 | 66110526 | 1.55442 | 2.22719 | 1.433 |
| chr2 | 10692760 | 10693158 | 0.675248 | 0.967524 | 1.433 |
| chr8 | 145539447 | 145540147 | 2.15218 | 3.08375 | 1.433 |
| chr21 | 30720871 | 30721160 | 2.03067 | 2.9098 | 1.433 |
| chr20 | 61603690 | 61603983 | 2.79204 | 4.00112 | 1.433 |
| chr17 | 67244846 | 67245132 | 0.907799 | 1.30109 | 1.433 |
| chr12 | 123507567 | 123507818 | 0.724327 | 1.03814 | 1.433 |
| chr19 | 16502678 | 16502911 | 0.551454 | 0.790409 | 1.433 |
| chr4 | 85180438 | 85180642 | 1.06169 | 1.52176 | 1.433 |
| chr2 | 101958610 | 101959022 | 1.66453 | 2.38589 | 1.433 |
| chr11 | 63403349 | 63403659 | 2.06618 | 2.96176 | 1.433 |
| chr20 | 34051570 | 34051882 | 1.86038 | 2.66688 | 1.434 |
| chr3 | 194408441 | 194408790 | 1.45733 | 2.08912 | 1.434 |
| chr8 | 131141862 | 131142140 | 2.6909 | 3.85762 | 1.434 |
| chr15 | 82204586 | 82204864 | 0.73661 | 1.05601 | 1.434 |
| chr19 | 38969054 | 38969652 | 1.37426 | 1.97017 | 1.434 |
| chr20 | 43013054 | 43013380 | 2.27757 | 3.26518 | 1.434 |
| chr7 | 31783955 | 31784242 | 1.83962 | 2.63753 | 1.434 |
| chr8 | 89034913 | 89035172 | 0.506069 | 0.725573 | 1.434 |
| chr2 | 232347391 | 232347777 | 2.88183 | 4.13184 | 1.434 |
| chr6 | 36648241 | 36648557 | 2.43002 | 3.48409 | 1.434 |
| chr5 | 170792428 | 170792753 | 1.19248 | 1.70976 | 1.434 |
| chr15 | 69758327 | 69758599 | 2.29621 | 3.29234 | 1.434 |

| | | | | | |
|---|---|---|---|---|---|
| chr14 | 104489788 | 104490084 | 0.659023 | 0.944927 | 1.434 |
| chr11 | 5536050 | 5536299 | 0.647469 | 0.928388 | 1.434 |
| chr9 | 91893902 | 91894230 | 1.47205 | 2.11073 | 1.434 |
| chr7 | 2926612 | 2926934 | 2.15266 | 3.0867 | 1.434 |
| chr16 | 87802884 | 87803244 | 1.15736 | 1.65956 | 1.434 |
| chr1 | 210547783 | 210548042 | 2.40774 | 3.45255 | 1.434 |
| chr17 | 19665830 | 19666179 | 2.42154 | 3.47243 | 1.434 |
| chr5 | 20833102 | 20833547 | 0.714574 | 1.0247 | 1.434 |
| chr16 | 75760949 | 75761338 | 0.733032 | 1.05118 | 1.434 |
| chr15 | 64183132 | 64183552 | 2.8496 | 4.08656 | 1.434 |
| chr3 | 72632494 | 72632714 | 0.687781 | 0.986423 | 1.434 |
| chr2 | 51458393 | 51458791 | 0.649506 | 0.931574 | 1.434 |
| chr12 | 49485964 | 49486242 | 1.58946 | 2.27982 | 1.434 |
| chr2 | 240821751 | 240822011 | 1.43699 | 2.06125 | 1.434 |
| chr8 | 63805588 | 63805839 | 0.65714 | 0.942647 | 1.434 |
| chr17 | 40555213 | 40555449 | 1.22357 | 1.75521 | 1.435 |
| chr1 | 162410951 | 162411172 | 0.857063 | 1.22949 | 1.435 |
| chr11 | 38812941 | 38813162 | 1.12988 | 1.62101 | 1.435 |
| chr7 | 45957188 | 45957580 | 2.62823 | 3.77093 | 1.435 |
| chr2 | 121278780 | 121279292 | 1.76143 | 2.52731 | 1.435 |
| chr9 | 127615043 | 127615725 | 2.78197 | 3.99194 | 1.435 |
| chr9 | 100978174 | 100978562 | 1.89051 | 2.71287 | 1.435 |
| chr8 | 19786392 | 19786640 | 0.950332 | 1.36374 | 1.435 |
| chr3 | 65327746 | 65328077 | 1.42449 | 2.04434 | 1.435 |
| chr9 | 101537073 | 101537446 | 1.92054 | 2.75644 | 1.435 |
| chr11 | 81335867 | 81336152 | 0.626268 | 0.898855 | 1.435 |
| chr7 | 105070197 | 105070460 | 1.63237 | 2.34294 | 1.435 |
| chr2 | 221551997 | 221552305 | 1.75333 | 2.51676 | 1.435 |
| chr5 | 75031053 | 75031284 | 1.45325 | 2.08606 | 1.435 |
| chr3 | 34831858 | 34832068 | 0.468351 | 0.672296 | 1.435 |
| chr10 | 61768619 | 61768871 | 1.43591 | 2.06122 | 1.435 |
| chr7 | 38409446 | 38409919 | 2.63421 | 3.78146 | 1.436 |
| chr5 | 78998779 | 78999085 | 1.63485 | 2.34691 | 1.436 |
| chr15 | 68128721 | 68129131 | 1.9784 | 2.84032 | 1.436 |
| chr1 | 206780311 | 206780522 | 1.39774 | 2.00674 | 1.436 |
| chr20 | 30434344 | 30434790 | 2.45349 | 3.52251 | 1.436 |
| chr5 | 10521643 | 10521955 | 1.40564 | 2.01814 | 1.436 |
| chr10 | 70108179 | 70108575 | 1.93105 | 2.77252 | 1.436 |
| chr16 | 4567845 | 4568332 | 1.37999 | 1.98135 | 1.436 |
| chr17 | 35798339 | 35798618 | 1.61044 | 2.31225 | 1.436 |
| chr7 | 29283871 | 29284236 | 1.47245 | 2.11423 | 1.436 |
| chr1 | 156790136 | 156790450 | 0.773905 | 1.11123 | 1.436 |
| chr20 | 45824996 | 45825320 | 2.4945 | 3.58182 | 1.436 |
| chr8 | 114214530 | 114214717 | 0.912256 | 1.31003 | 1.436 |
| chr13 | 45885251 | 45885500 | 2.19805 | 3.15653 | 1.436 |
| chrX | 67238424 | 67239247 | 0.996896 | 1.43164 | 1.436 |
| chr5 | 139592135 | 139592356 | 0.623568 | 0.895517 | 1.436 |
| chr9 | 14068795 | 14069031 | 1.25444 | 1.80156 | 1.436 |
| chr5 | 116176033 | 116176325 | 0.671949 | 0.965041 | 1.436 |
| chr5 | 176497589 | 176497826 | 0.729341 | 1.04748 | 1.436 |
| chr2 | 241083220 | 241083564 | 2.27621 | 3.26915 | 1.436 |
| chr12 | 117674254 | 117674721 | 1.8299 | 2.62821 | 1.436 |
| chr2 | 95768150 | 95768369 | 0.795342 | 1.14232 | 1.436 |
| chr6 | 166726848 | 166727065 | 1.41692 | 2.0351 | 1.436 |
| chr9 | 80984571 | 80984890 | 2.23021 | 3.2033 | 1.436 |
| chr12 | 30000612 | 30000823 | 0.673963 | 0.968078 | 1.436 |
| chr2 | 120584027 | 120584252 | 1.76429 | 2.53422 | 1.436 |
| chr16 | 82673357 | 82673662 | 1.80606 | 2.59437 | 1.436 |
| chr10 | 3509522 | 3509802 | 2.07841 | 2.98561 | 1.436 |
| chr15 | 31378524 | 31378757 | 0.808673 | 1.1617 | 1.437 |
| chr20 | 41690332 | 41690540 | 0.612991 | 0.880594 | 1.437 |

| | | | | | |
|---|---|---|---|---|---|
| chr20 | 11254754 | 11255012 | 1.91406 | 2.74999 | 1.437 |
| chr7 | 123220925 | 123221252 | 1.47911 | 2.12516 | 1.437 |
| chr17 | 7492610 | 7492986 | 2.24965 | 3.23255 | 1.437 |
| chr8 | 106125823 | 106126037 | 0.72729 | 1.0451 | 1.437 |
| chr9 | 87191835 | 87192079 | 0.796877 | 1.1451 | 1.437 |
| chr2 | 70814682 | 70815004 | 0.665088 | 0.955728 | 1.437 |
| chr1 | 204227461 | 204227890 | 3.1565 | 4.53614 | 1.437 |
| chr20 | 8778486 | 8778756 | 0.711962 | 1.02323 | 1.437 |
| chr12 | 84298164 | 84298371 | 0.682638 | 0.981091 | 1.437 |
| chr7 | 73062065 | 73062321 | 1.14665 | 1.64806 | 1.437 |
| chr19 | 46585536 | 46585760 | 1.57824 | 2.2684 | 1.437 |
| chr2 | 208838006 | 208838230 | 1.75153 | 2.51755 | 1.437 |
| chr17 | 7755063 | 7755738 | 2.39892 | 3.44809 | 1.437 |
| chr17 | 4468909 | 4469183 | 2.38285 | 3.42519 | 1.437 |
| chr14 | 78173741 | 78173999 | 0.933502 | 1.34189 | 1.437 |
| chr11 | 117678255 | 117678516 | 1.40253 | 2.01627 | 1.438 |
| chr14 | 99801180 | 99801388 | 0.527575 | 0.758434 | 1.438 |
| chr2 | 65092682 | 65093039 | 2.53792 | 3.64857 | 1.438 |
| chrX | 128494258 | 128494531 | 1.13464 | 1.63119 | 1.438 |
| chr7 | 148768976 | 148769189 | 2.33807 | 3.3614 | 1.438 |
| chr11 | 34380643 | 34381374 | 1.08466 | 1.55942 | 1.438 |
| chr5 | 160434914 | 160435238 | 0.698675 | 1.00451 | 1.438 |
| chr3 | 157290441 | 157290686 | 1.67591 | 2.40954 | 1.438 |
| chr10 | 2799705 | 2799983 | 0.67219 | 0.966464 | 1.438 |
| chr8 | 26579188 | 26579408 | 0.689819 | 0.991809 | 1.438 |
| chr20 | 34580682 | 34581015 | 2.73915 | 3.93846 | 1.438 |
| chr1 | 158497068 | 158497392 | 1.2603 | 1.81212 | 1.438 |
| chr8 | 63789865 | 63790085 | 0.635707 | 0.914074 | 1.438 |
| chr12 | 95967725 | 95967972 | 0.868548 | 1.24892 | 1.438 |
| chr16 | 66376093 | 66376622 | 2.28814 | 3.29023 | 1.438 |
| chr8 | 131550426 | 131550645 | 1.00695 | 1.44795 | 1.438 |
| chr20 | 41981596 | 41981874 | 0.571979 | 0.822491 | 1.438 |
| chr9 | 77801752 | 77802092 | 2.37081 | 3.4092 | 1.438 |
| chr9 | 86235694 | 86236097 | 1.58457 | 2.27863 | 1.438 |
| chr14 | 21159273 | 21159763 | 1.39766 | 2.00987 | 1.438 |
| chr10 | 47004281 | 47004601 | 1.32341 | 1.90314 | 1.438 |
| chr9 | 87312310 | 87312799 | 1.86337 | 2.6797 | 1.438 |
| chr14 | 88141440 | 88141672 | 0.616239 | 0.886298 | 1.438 |
| chr11 | 71736150 | 71736512 | 2.70206 | 3.8863 | 1.438 |
| chr1 | 234040280 | 234040552 | 2.41633 | 3.47542 | 1.438 |
| chr1 | 187703563 | 187703823 | 0.813675 | 1.17042 | 1.438 |
| chr2 | 231921149 | 231921441 | 2.47546 | 3.56081 | 1.438 |
| chr2 | 173892221 | 173892470 | 1.54782 | 2.22659 | 1.439 |
| chr7 | 105681612 | 105681972 | 1.99975 | 2.87672 | 1.439 |
| chr19 | 36067974 | 36068311 | 0.757918 | 1.09032 | 1.439 |
| chr9 | 129036402 | 129036670 | 0.519624 | 0.747517 | 1.439 |
| chr11 | 47926980 | 47927227 | 0.867226 | 1.24763 | 1.439 |
| chr2 | 96926463 | 96926865 | 1.5829 | 2.27726 | 1.439 |
| chr7 | 104781639 | 104781844 | 1.62541 | 2.33847 | 1.439 |
| chr1 | 156553854 | 156554120 | 2.03288 | 2.92497 | 1.439 |
| chr19 | 40898852 | 40899172 | 1.43233 | 2.06094 | 1.439 |
| chr2 | 139585069 | 139585312 | 1.37105 | 1.97282 | 1.439 |
| chr2 | 71173920 | 71174344 | 1.75857 | 2.53051 | 1.439 |
| chr15 | 94160564 | 94160813 | 0.811169 | 1.16728 | 1.439 |
| chr17 | 41299023 | 41299226 | 0.712834 | 1.02578 | 1.439 |
| chr15 | 84198723 | 84198926 | 0.775832 | 1.11649 | 1.439 |
| chr12 | 8683643 | 8683910 | 0.991717 | 1.42734 | 1.439 |
| chr8 | 56727638 | 56727943 | 1.1233 | 1.61672 | 1.439 |
| chr5 | 113785627 | 113786153 | 1.66966 | 2.4031 | 1.439 |
| chr20 | 37101890 | 37102114 | 0.734992 | 1.05788 | 1.439 |
| chr10 | 19716926 | 19717134 | 0.692423 | 0.996688 | 1.439 |

| | | | | | |
|---|---|---|---|---|---|
| chr8 | 137333748 | 137333989 | 0.721146 | 1.03815 | 1.440 |
| chr12 | 53135974 | 53136244 | 0.737049 | 1.06106 | 1.440 |
| chr12 | 115290961 | 115291262 | 0.605339 | 0.871458 | 1.440 |
| chr9 | 102028853 | 102029210 | 2.17631 | 3.13308 | 1.440 |
| chr17 | 40253421 | 40253981 | 2.41718 | 3.47997 | 1.440 |
| chr20 | 30200642 | 30201020 | 4.45174 | 6.40929 | 1.440 |
| chr2 | 11270097 | 11270423 | 1.28065 | 1.84382 | 1.440 |
| chr17 | 73126713 | 73127111 | 1.62199 | 2.3354 | 1.440 |
| chr7 | 94412304 | 94412520 | 0.64245 | 0.925023 | 1.440 |
| chr9 | 8284162 | 8284443 | 0.531811 | 0.76573 | 1.440 |
| chr1 | 36034485 | 36034892 | 2.0476 | 2.94825 | 1.440 |
| chr1 | 92951356 | 92951603 | 1.10554 | 1.59185 | 1.440 |
| chr14 | 102155291 | 102155511 | 0.550767 | 0.79305 | 1.440 |
| chr16 | 1419859 | 1420783 | 1.5054 | 2.16763 | 1.440 |
| chr15 | 75494266 | 75494738 | 2.75737 | 3.97056 | 1.440 |
| chr11 | 72524027 | 72524259 | 2.23963 | 3.22505 | 1.440 |
| chr17 | 80309487 | 80310089 | 2.16955 | 3.12417 | 1.440 |
| chr7 | 101087211 | 101087415 | 0.71666 | 1.03201 | 1.440 |
| chr3 | 2636968 | 2637433 | 0.657075 | 0.946229 | 1.440 |
| chr20 | 46306541 | 46306944 | 2.70981 | 3.90242 | 1.440 |
| chr12 | 118815744 | 118816088 | 1.92341 | 2.7703 | 1.440 |
| chr9 | 124446487 | 124447094 | 1.47588 | 2.12575 | 1.440 |
| chr3 | 102392399 | 102392764 | 2.19128 | 3.15622 | 1.440 |
| chr3 | 155183411 | 155183616 | 0.92981 | 1.33928 | 1.440 |
| chr11 | 60335840 | 60336115 | 0.593903 | 0.855471 | 1.440 |
| chr3 | 163466860 | 163467260 | 0.784649 | 1.13023 | 1.440 |
| chr4 | 144404017 | 144404275 | 1.36345 | 1.96396 | 1.440 |
| chr12 | 120273483 | 120273690 | 0.700573 | 1.00919 | 1.441 |
| chr19 | 18278896 | 18279262 | 1.42133 | 2.04756 | 1.441 |
| chr11 | 35433890 | 35434141 | 1.46194 | 2.10631 | 1.441 |
| chr2 | 231851956 | 231852328 | 2.84942 | 4.10539 | 1.441 |
| chr11 | 76130671 | 76131276 | 1.02374 | 1.47507 | 1.441 |
| chr2 | 168325047 | 168325291 | 0.702967 | 1.01288 | 1.441 |
| chr1 | 223815935 | 223816183 | 1.54753 | 2.22995 | 1.441 |
| chr17 | 42477530 | 42477947 | 1.4781 | 2.12997 | 1.441 |
| chr16 | 4700450 | 4700845 | 2.39183 | 3.44689 | 1.441 |
| chr11 | 56057923 | 56058131 | 0.591624 | 0.8526 | 1.441 |
| chr15 | 62127371 | 62128037 | 2.42227 | 3.49099 | 1.441 |
| chr20 | 708936 | 709491 | 1.69608 | 2.44456 | 1.441 |
| chr6 | 131193418 | 131193784 | 2.00195 | 2.88566 | 1.441 |
| chr17 | 66432087 | 66432948 | 1.81812 | 2.62069 | 1.441 |
| chr12 | 14169894 | 14170104 | 0.904986 | 1.30448 | 1.441 |
| chr5 | 19419772 | 19419989 | 0.512779 | 0.739144 | 1.441 |
| chr3 | 8755635 | 8756137 | 1.71254 | 2.4687 | 1.442 |
| chr1 | 203238154 | 203238534 | 2.50538 | 3.61167 | 1.442 |
| chr2 | 225643199 | 225643425 | 1.45843 | 2.10256 | 1.442 |
| chr10 | 129932266 | 129932488 | 0.700696 | 1.01019 | 1.442 |
| chr7 | 23584584 | 23584861 | 1.39907 | 2.0171 | 1.442 |
| chr1 | 156414165 | 156414499 | 1.30665 | 1.88387 | 1.442 |
| chr9 | 86787994 | 86788449 | 1.51462 | 2.18372 | 1.442 |
| chr1 | 109642577 | 109642794 | 1.99924 | 2.88258 | 1.442 |
| chr6 | 109156480 | 109156854 | 2.34118 | 3.37591 | 1.442 |
| chr20 | 5186059 | 5186490 | 2.25459 | 3.25115 | 1.442 |
| chr1 | 20427510 | 20427721 | 2.15604 | 3.10906 | 1.442 |
| chr17 | 74993165 | 74993368 | 0.82712 | 1.1928 | 1.442 |
| chr13 | 113585763 | 113586084 | 1.83235 | 2.64249 | 1.442 |
| chr7 | 107749803 | 107750015 | 1.09913 | 1.58526 | 1.442 |
| chr4 | 110574044 | 110574248 | 1.01413 | 1.4627 | 1.442 |
| chr18 | 2547345 | 2547748 | 0.790394 | 1.14002 | 1.442 |
| chr5 | 58470136 | 58470377 | 2.29214 | 3.30625 | 1.442 |
| chr12 | 85470424 | 85470678 | 1.47536 | 2.12817 | 1.442 |

| | | | | | |
|---|---|---|---|---|---|
| chr14 | 59251595 | 59251801 | 0.555495 | 0.801301 | 1.443 |
| chr4 | 91938051 | 91938370 | 0.627276 | 0.904881 | 1.443 |
| chr13 | 32593887 | 32594105 | 1.12199 | 1.61864 | 1.443 |
| chr7 | 148201095 | 148201302 | 1.20836 | 1.74334 | 1.443 |
| chr1 | 208171190 | 208171426 | 1.76236 | 2.54278 | 1.443 |
| chr5 | 97925049 | 97925315 | 0.592707 | 0.855192 | 1.443 |
| chr8 | 21731546 | 21731781 | 0.646746 | 0.933168 | 1.443 |
| chr2 | 71146929 | 71147279 | 2.62704 | 3.79067 | 1.443 |
| chr8 | 123854458 | 123854670 | 0.747407 | 1.0785 | 1.443 |
| chr2 | 190498032 | 190498300 | 0.877688 | 1.2665 | 1.443 |
| chr5 | 116423072 | 116423274 | 1.03591 | 1.49498 | 1.443 |
| chr4 | 130265181 | 130265446 | 1.28946 | 1.8611 | 1.443 |
| chr19 | 3324406 | 3324779 | 1.64522 | 2.37488 | 1.444 |
| chr16 | 9228663 | 9228979 | 2.19099 | 3.16282 | 1.444 |
| chr7 | 128784969 | 128785169 | 0.892173 | 1.28795 | 1.444 |
| chr3 | 75241211 | 75241798 | 1.6835 | 2.43048 | 1.444 |
| chr14 | 20395883 | 20396183 | 0.696435 | 1.00555 | 1.444 |
| chr18 | 11741558 | 11741834 | 1.77042 | 2.55624 | 1.444 |
| chr16 | 11749040 | 11749301 | 0.665078 | 0.960351 | 1.444 |
| chr2 | 50738383 | 50738716 | 0.729274 | 1.05307 | 1.444 |
| chr2 | 158880845 | 158881091 | 1.78388 | 2.57605 | 1.444 |
| chr6 | 85132121 | 85132432 | 1.97803 | 2.85656 | 1.444 |
| chr18 | 35024758 | 35024996 | 1.20571 | 1.74133 | 1.444 |
| chr7 | 117793674 | 117793916 | 1.43757 | 2.0762 | 1.444 |
| chr5 | 140180476 | 140180809 | 1.21937 | 1.7612 | 1.444 |
| chr2 | 216134604 | 216134840 | 1.66468 | 2.40457 | 1.444 |
| chr7 | 107103606 | 107103868 | 1.33484 | 1.92814 | 1.444 |
| chr15 | 34027890 | 34028162 | 1.75873 | 2.54058 | 1.445 |
| chr17 | 18224686 | 18224919 | 0.883231 | 1.27592 | 1.445 |
| chr2 | 26828234 | 26828829 | 1.28705 | 1.85928 | 1.445 |
| chr12 | 19880645 | 19880923 | 0.836532 | 1.2085 | 1.445 |
| chr17 | 13603837 | 13604157 | 0.658069 | 0.950743 | 1.445 |
| chr10 | 65819719 | 65819942 | 0.546121 | 0.789018 | 1.445 |
| chr2 | 149267039 | 149267304 | 2.6976 | 3.8975 | 1.445 |
| chr3 | 76600869 | 76601131 | 0.687088 | 0.992702 | 1.445 |
| chr2 | 113507292 | 113507610 | 3.03658 | 4.38727 | 1.445 |
| chr7 | 134434843 | 134435209 | 2.65988 | 3.84319 | 1.445 |
| chr15 | 88993418 | 88993744 | 2.06508 | 2.98382 | 1.445 |
| chr14 | 73639889 | 73640132 | 1.46427 | 2.11576 | 1.445 |
| chr10 | 45360280 | 45360626 | 2.49774 | 3.60905 | 1.445 |
| chr14 | 36283690 | 36283942 | 2.00518 | 2.89736 | 1.445 |
| chr16 | 11349422 | 11349965 | 2.05398 | 2.96787 | 1.445 |
| chr7 | 78378879 | 78379104 | 0.840974 | 1.21518 | 1.445 |
| chr1 | 36653840 | 36654083 | 1.80003 | 2.60113 | 1.445 |
| chr2 | 135761827 | 135762157 | 2.79599 | 4.04043 | 1.445 |
| chr10 | 11445081 | 11445299 | 0.56509 | 0.816622 | 1.445 |
| chr9 | 23676741 | 23677031 | 0.625052 | 0.903283 | 1.445 |
| chr1 | 1237285 | 1237732 | 1.44826 | 2.09317 | 1.445 |
| chr13 | 27070758 | 27071093 | 2.5046 | 3.61998 | 1.445 |
| chr2 | 114463889 | 114464096 | 0.632299 | 0.913933 | 1.445 |
| chr6 | 56622513 | 56622763 | 1.66113 | 2.40108 | 1.445 |
| chr11 | 45132229 | 45132446 | 1.61289 | 2.33143 | 1.446 |
| chr12 | 61422663 | 61422901 | 0.760737 | 1.09968 | 1.446 |
| chr8 | 140556201 | 140556534 | 0.587252 | 0.848904 | 1.446 |
| chr18 | 55228928 | 55229349 | 1.26985 | 1.83576 | 1.446 |
| chr15 | 75890001 | 75890369 | 1.02794 | 1.48612 | 1.446 |
| chr11 | 29470122 | 29470347 | 0.958255 | 1.3854 | 1.446 |
| chr7 | 137214443 | 137214659 | 0.692403 | 1.00104 | 1.446 |
| chr1 | 226645118 | 226645363 | 0.705032 | 1.01932 | 1.446 |
| chr12 | 9895730 | 9896078 | 1.46964 | 2.12491 | 1.446 |
| chr3 | 133931758 | 133932112 | 2.64191 | 3.82003 | 1.446 |

| | | | | | |
|---|---|---|---|---|---|
| chr2 | 11833079 | 11833405 | 1.96128 | 2.83599 | 1.446 |
| chr18 | 47721480 | 47721806 | 1.68814 | 2.44105 | 1.446 |
| chr19 | 16825565 | 16825862 | 2.26276 | 3.27215 | 1.446 |
| chr19 | 3275412 | 3275814 | 1.9798 | 2.86306 | 1.446 |
| chr10 | 55040091 | 55040363 | 0.681517 | 0.985626 | 1.446 |
| chr20 | 21553540 | 21553770 | 0.671077 | 0.970535 | 1.446 |
| chr1 | 58283060 | 58283316 | 0.581346 | 0.840808 | 1.446 |
| chr5 | 13070700 | 13070916 | 0.979091 | 1.41619 | 1.446 |
| chr7 | 87987608 | 87987832 | 1.78384 | 2.58025 | 1.446 |
| chr10 | 52938177 | 52938396 | 0.551969 | 0.798409 | 1.446 |
| chr3 | 156956889 | 156957524 | 1.86094 | 2.69185 | 1.447 |
| chr8 | 87100673 | 87101086 | 1.25269 | 1.81203 | 1.447 |
| chr9 | 133576439 | 133576985 | 2.61342 | 3.78058 | 1.447 |
| chr9 | 33852246 | 33852579 | 1.80044 | 2.60466 | 1.447 |
| chr16 | 6664996 | 6665432 | 0.66589 | 0.963344 | 1.447 |
| chr17 | 16905063 | 16905493 | 3.07199 | 4.44427 | 1.447 |
| chr3 | 49740908 | 49741254 | 2.10044 | 3.0388 | 1.447 |
| chr5 | 151122393 | 151122642 | 1.41198 | 2.04291 | 1.447 |
| chr8 | 109258080 | 109258276 | 1.5137 | 2.19013 | 1.447 |
| chr9 | 127082524 | 127083022 | 2.56921 | 3.71775 | 1.447 |
| chr2 | 33700765 | 33701031 | 2.04195 | 2.95515 | 1.447 |
| chr5 | 50751829 | 50752043 | 0.501593 | 0.725944 | 1.447 |
| chr11 | 2440936 | 2441204 | 1.82551 | 2.64205 | 1.447 |
| chr2 | 120867467 | 120867697 | 1.17281 | 1.69747 | 1.447 |
| chr14 | 29543573 | 29543895 | 0.716988 | 1.03777 | 1.447 |
| chr1 | 20688318 | 20688628 | 2.13299 | 3.08742 | 1.447 |
| chr20 | 719618 | 720063 | 2.41718 | 3.49881 | 1.447 |
| chr2 | 77017964 | 77018363 | 0.684168 | 0.990328 | 1.447 |
| chr20 | 50996249 | 50996614 | 2.34241 | 3.39081 | 1.448 |
| chr15 | 92883480 | 92883722 | 0.858534 | 1.24282 | 1.448 |
| chr6 | 123052008 | 123052265 | 2.52929 | 3.66163 | 1.448 |
| chr19 | 37468727 | 37469160 | 0.855769 | 1.23896 | 1.448 |
| chr19 | 17505759 | 17506027 | 1.16207 | 1.68246 | 1.448 |
| chr9 | 17771595 | 17771816 | 0.618524 | 0.895549 | 1.448 |
| chr11 | 108568645 | 108568951 | 2.15198 | 3.11592 | 1.448 |
| chr8 | 105391221 | 105391535 | 0.793293 | 1.14864 | 1.448 |
| chr3 | 25898399 | 25898977 | 1.42898 | 2.06909 | 1.448 |
| chr17 | 54897043 | 54897371 | 2.44941 | 3.54667 | 1.448 |
| chr18 | 49147554 | 49147849 | 1.10509 | 1.60023 | 1.448 |
| chr12 | 32518444 | 32518825 | 2.01494 | 2.91782 | 1.448 |
| chr19 | 42439045 | 42439444 | 3.79267 | 5.49225 | 1.448 |
| chr20 | 31886898 | 31887301 | 2.65492 | 3.84478 | 1.448 |
| chr11 | 36314010 | 36314196 | 0.963801 | 1.39579 | 1.448 |
| chr8 | 134508640 | 134509003 | 1.16548 | 1.68787 | 1.448 |
| chr17 | 15059476 | 15059697 | 1.54053 | 2.23116 | 1.448 |
| chr2 | 23789705 | 23789930 | 1.40506 | 2.035 | 1.448 |
| chr2 | 201450604 | 201450846 | 0.705886 | 1.02238 | 1.448 |
| chr5 | 106583796 | 106584169 | 1.87407 | 2.71448 | 1.448 |
| chr7 | 120672872 | 120673109 | 0.462306 | 0.669637 | 1.448 |
| chr5 | 153283932 | 153284160 | 0.619739 | 0.897682 | 1.448 |
| chr17 | 55191214 | 55191754 | 1.83546 | 2.65865 | 1.448 |
| chr1 | 1285090 | 1285354 | 1.02869 | 1.4901 | 1.449 |
| chr1 | 53743915 | 53744207 | 2.06885 | 2.99682 | 1.449 |
| chr17 | 49874485 | 49874706 | 0.598893 | 0.867527 | 1.449 |
| chr10 | 110623881 | 110624226 | 0.832178 | 1.20553 | 1.449 |
| chr10 | 73596945 | 73597239 | 1.63157 | 2.36359 | 1.449 |
| chr11 | 67314489 | 67314740 | 0.622858 | 0.902345 | 1.449 |
| chr2 | 148695338 | 148695627 | 1.3716 | 1.98712 | 1.449 |
| chr1 | 53647755 | 53647992 | 1.92997 | 2.79612 | 1.449 |
| chr1 | 152505865 | 152506116 | 2.40698 | 3.48723 | 1.449 |
| chr4 | 108511272 | 108511484 | 0.730799 | 1.05879 | 1.449 |

| | | | | | |
|---|---|---|---|---|---|
| chr16 | 24082222 | 24082805 | 0.86276 | 1.25 | 1.449 |
| chr5 | 131161084 | 131161399 | 0.578779 | 0.838575 | 1.449 |
| chr2 | 56498232 | 56498587 | 0.594232 | 0.860991 | 1.449 |
| chr14 | 71439095 | 71439513 | 2.64931 | 3.83869 | 1.449 |
| chr14 | 89133727 | 89134121 | 2.14022 | 3.10109 | 1.449 |
| chr6 | 138266060 | 138266353 | 2.08221 | 3.01706 | 1.449 |
| chr19 | 43045787 | 43046188 | 2.33507 | 3.38349 | 1.449 |
| chr17 | 32078092 | 32078523 | 2.41057 | 3.49291 | 1.449 |
| chr1 | 234348539 | 234348869 | 1.6117 | 2.33539 | 1.449 |
| chr3 | 116477124 | 116477328 | 1.78137 | 2.58126 | 1.449 |
| chr22 | 33382080 | 33382409 | 1.42955 | 2.0715 | 1.449 |
| chr10 | 55533190 | 55533442 | 0.61998 | 0.898403 | 1.449 |
| chr7 | 127268458 | 127268780 | 0.829801 | 1.20246 | 1.449 |
| chr1 | 63377770 | 63378035 | 1.50729 | 2.18431 | 1.449 |
| chr2 | 164047966 | 164048238 | 1.42248 | 2.06148 | 1.449 |
| chr11 | 60942924 | 60943255 | 0.657644 | 0.95312 | 1.449 |
| chr2 | 113716368 | 113716711 | 2.21028 | 3.20353 | 1.449 |
| chr15 | 79376779 | 79377151 | 0.687615 | 0.996632 | 1.449 |
| chr3 | 88472580 | 88472937 | 0.66455 | 0.963216 | 1.449 |
| chr9 | 32191343 | 32191592 | 0.743594 | 1.07779 | 1.449 |
| chr3 | 146413786 | 146414019 | 0.779233 | 1.12948 | 1.449 |
| chr19 | 14224967 | 14225255 | 1.11444 | 1.61538 | 1.450 |
| chr16 | 28229505 | 28229736 | 1.44951 | 2.10123 | 1.450 |
| chr3 | 73481575 | 73481921 | 1.45355 | 2.10709 | 1.450 |
| chr20 | 11612277 | 11612722 | 0.788893 | 1.14364 | 1.450 |
| chr3 | 122018823 | 122019033 | 1.91732 | 2.77967 | 1.450 |
| chr9 | 14616456 | 14616661 | 1.51037 | 2.18985 | 1.450 |
| chr14 | 94468104 | 94468750 | 2.41987 | 3.50879 | 1.450 |
| chr2 | 24027155 | 24027363 | 1.22327 | 1.77374 | 1.450 |
| chr11 | 11049552 | 11050029 | 2.04095 | 2.95948 | 1.450 |
| chr13 | 79552635 | 79552877 | 0.835225 | 1.21115 | 1.450 |
| chr9 | 133816406 | 133816883 | 2.22847 | 3.23148 | 1.450 |
| chr11 | 119210546 | 119211085 | 2.05511 | 2.98022 | 1.450 |
| chr20 | 20211951 | 20212288 | 0.827134 | 1.19951 | 1.450 |
| chr7 | 103977217 | 103977425 | 0.525061 | 0.761445 | 1.450 |
| chr7 | 157179762 | 157180239 | 2.32682 | 3.37471 | 1.450 |
| chr3 | 143960524 | 143960778 | 0.725089 | 1.05166 | 1.450 |
| chr15 | 82432161 | 82432422 | 1.74909 | 2.53693 | 1.450 |
| chr17 | 37821953 | 37822470 | 1.15091 | 1.66932 | 1.450 |
| chr17 | 72447146 | 72447476 | 0.622107 | 0.902335 | 1.450 |
| chr15 | 27816743 | 27816960 | 0.946666 | 1.37312 | 1.450 |
| chr1 | 216419937 | 216420149 | 0.89501 | 1.2982 | 1.450 |
| chr11 | 72300811 | 72301260 | 2.67522 | 3.88039 | 1.450 |
| chrX | 18408201 | 18408505 | 1.1474 | 1.66434 | 1.451 |
| chr14 | 22782726 | 22782963 | 1.51921 | 2.20374 | 1.451 |
| chr7 | 50472331 | 50472623 | 0.654856 | 0.949995 | 1.451 |
| chr15 | 66155031 | 66155271 | 1.62091 | 2.35145 | 1.451 |
| chr2 | 59423055 | 59423385 | 0.654649 | 0.949709 | 1.451 |
| chr11 | 116324799 | 116325050 | 1.17081 | 1.69853 | 1.451 |
| chr11 | 29454328 | 29454591 | 1.22895 | 1.78292 | 1.451 |
| chr12 | 114169892 | 114170136 | 1.36415 | 1.97928 | 1.451 |
| chr19 | 57559920 | 57560411 | 0.637044 | 0.924334 | 1.451 |
| chr3 | 18020887 | 18021093 | 1.14692 | 1.66416 | 1.451 |
| chr1 | 29529409 | 29529730 | 1.6712 | 2.42495 | 1.451 |
| chr16 | 74684910 | 74685339 | 1.34212 | 1.94751 | 1.451 |
| chr8 | 54318200 | 54318643 | 0.639242 | 0.927584 | 1.451 |
| chr10 | 66043638 | 66044000 | 0.64694 | 0.938773 | 1.451 |
| chr20 | 47521651 | 47521935 | 1.43811 | 2.08691 | 1.451 |
| chr5 | 15046105 | 15046349 | 0.634247 | 0.920386 | 1.451 |
| chr2 | 28030388 | 28030675 | 1.62519 | 2.35843 | 1.451 |
| chr20 | 23870429 | 23870790 | 1.42136 | 2.06274 | 1.451 |

| | | | | | |
|---|---|---|---|---|---|
| chr3 | 114759602 | 114759862 | 1.79618 | 2.60668 | 1.451 |
| chr7 | 101185832 | 101186450 | 1.75883 | 2.55267 | 1.451 |
| chr12 | 71346181 | 71346484 | 0.702516 | 1.01966 | 1.451 |
| chr17 | 74001330 | 74002020 | 1.51768 | 2.20284 | 1.451 |
| chr11 | 76673611 | 76673933 | 0.47774 | 0.693464 | 1.452 |
| chr5 | 18698559 | 18699047 | 0.861431 | 1.25042 | 1.452 |
| chr8 | 135379897 | 135380121 | 0.941274 | 1.36635 | 1.452 |
| chr17 | 73500312 | 73500896 | 2.21325 | 3.21287 | 1.452 |
| chr5 | 42025780 | 42025990 | 0.506724 | 0.735597 | 1.452 |
| chrX | 128739482 | 128739916 | 1.26429 | 1.83534 | 1.452 |
| chr15 | 88116780 | 88116989 | 0.719599 | 1.04469 | 1.452 |
| chr17 | 33488537 | 33488872 | 0.650151 | 0.943878 | 1.452 |
| chr1 | 26324663 | 26324927 | 2.46991 | 3.5858 | 1.452 |
| chr17 | 61851814 | 61852049 | 0.97889 | 1.42115 | 1.452 |
| chr12 | 17664281 | 17664561 | 1.01515 | 1.47383 | 1.452 |
| chr4 | 154064162 | 154064398 | 2.21119 | 3.21028 | 1.452 |
| chr21 | 42493798 | 42494070 | 2.87881 | 4.17957 | 1.452 |
| chr1 | 231686148 | 231686505 | 1.40479 | 2.03955 | 1.452 |
| chr5 | 148971998 | 148972608 | 0.617483 | 0.896517 | 1.452 |
| chr15 | 97540280 | 97540736 | 0.834218 | 1.21131 | 1.452 |
| chr10 | 134185176 | 134185563 | 1.6678 | 2.42177 | 1.452 |
| chr11 | 59570130 | 59570339 | 1.21332 | 1.76199 | 1.452 |
| chr7 | 143875520 | 143875755 | 0.903236 | 1.31169 | 1.452 |
| chr21 | 45031811 | 45032056 | 1.88839 | 2.74243 | 1.452 |
| chr12 | 54764032 | 54764492 | 2.5427 | 3.69269 | 1.452 |
| chr1 | 200528071 | 200528301 | 1.91747 | 2.78485 | 1.452 |
| chr1 | 111173374 | 111173702 | 2.04796 | 2.97448 | 1.452 |
| chr15 | 90844454 | 90844967 | 1.75771 | 2.55313 | 1.453 |
| chr12 | 79400210 | 79400419 | 1.05114 | 1.52693 | 1.453 |
| chr14 | 77767313 | 77767798 | 1.93356 | 2.80891 | 1.453 |
| chr5 | 169407401 | 169407939 | 2.10502 | 3.05799 | 1.453 |
| chr2 | 3005437 | 3005659 | 1.38817 | 2.01667 | 1.453 |
| chr7 | 149052113 | 149052350 | 0.674909 | 0.98048 | 1.453 |
| chr17 | 43096602 | 43096926 | 1.68785 | 2.45214 | 1.453 |
| chr19 | 41245350 | 41245609 | 0.844841 | 1.22743 | 1.453 |
| chr11 | 47280990 | 47281467 | 1.84485 | 2.68035 | 1.453 |
| chr17 | 27894849 | 27895062 | 1.606 | 2.33345 | 1.453 |
| chr2 | 200844810 | 200845063 | 1.84329 | 2.67828 | 1.453 |
| chr22 | 21271336 | 21271666 | 2.23437 | 3.24657 | 1.453 |
| chr2 | 169818233 | 169818474 | 2.18912 | 3.18113 | 1.453 |
| chr11 | 65341513 | 65341967 | 2.6373 | 3.83259 | 1.453 |
| chr4 | 47928649 | 47928970 | 1.69158 | 2.45837 | 1.453 |
| chr2 | 147427297 | 147427573 | 0.472133 | 0.686186 | 1.453 |
| chr8 | 137477457 | 137477749 | 0.776222 | 1.12818 | 1.453 |
| chr17 | 1696390 | 1696621 | 0.58703 | 0.853212 | 1.453 |
| chr3 | 70599952 | 70600421 | 1.58794 | 2.30812 | 1.454 |
| chr10 | 101120426 | 101120975 | 0.73586 | 1.06962 | 1.454 |
| chr6 | 42694400 | 42694650 | 2.17669 | 3.16398 | 1.454 |
| chr20 | 8150280 | 8150571 | 0.707937 | 1.02913 | 1.454 |
| chr16 | 85426981 | 85427684 | 2.09836 | 3.05058 | 1.454 |
| chr2 | 60839121 | 60839680 | 0.763072 | 1.10941 | 1.454 |
| chr19 | 18705088 | 18705530 | 2.13831 | 3.10889 | 1.454 |
| chr10 | 92617384 | 92617792 | 2.0262 | 2.94628 | 1.454 |
| chr17 | 27527727 | 27527974 | 0.632915 | 0.92034 | 1.454 |
| chr11 | 95438735 | 95439047 | 2.4273 | 3.52977 | 1.454 |
| chr8 | 41692209 | 41692597 | 2.0762 | 3.01934 | 1.454 |
| chr15 | 88024421 | 88024666 | 0.747307 | 1.0869 | 1.454 |
| chr7 | 28101782 | 28102114 | 1.95592 | 2.84473 | 1.454 |
| chr4 | 146877550 | 146877818 | 1.49077 | 2.1683 | 1.454 |
| chr7 | 47221146 | 47221401 | 0.669969 | 0.974454 | 1.454 |
| chr12 | 8699104 | 8699410 | 0.643956 | 0.936644 | 1.455 |

| | | | | | |
|---|---|---|---|---|---|
| chr14 | 23736443 | 23736663 | 0.733198 | 1.06648 | 1.455 |
| chr8 | 100298973 | 100299302 | 1.25606 | 1.82704 | 1.455 |
| chr8 | 138394051 | 138394300 | 0.755169 | 1.09852 | 1.455 |
| chr12 | 61282527 | 61282855 | 0.621289 | 0.903795 | 1.455 |
| chr5 | 112513847 | 112514090 | 0.939754 | 1.3672 | 1.455 |
| chr1 | 227085009 | 227085237 | 0.669455 | 0.973989 | 1.455 |
| chr5 | 63525058 | 63525283 | 1.06262 | 1.54609 | 1.455 |
| chr9 | 13427592 | 13427875 | 1.26616 | 1.84229 | 1.455 |
| chr9 | 133814525 | 133814754 | 1.5101 | 2.19729 | 1.455 |
| chr5 | 112093271 | 112093560 | 1.16 | 1.68789 | 1.455 |
| chr1 | 187234176 | 187234446 | 1.9993 | 2.90919 | 1.455 |
| chr19 | 12943288 | 12943832 | 2.29091 | 3.33354 | 1.455 |
| chr11 | 73189773 | 73190096 | 2.08347 | 3.03175 | 1.455 |
| chrX | 2793969 | 2794204 | 1.24642 | 1.81379 | 1.455 |
| chr11 | 30336193 | 30336439 | 0.864762 | 1.25843 | 1.455 |
| chr7 | 47868321 | 47868556 | 0.515234 | 0.749796 | 1.455 |
| chr9 | 77656472 | 77656714 | 0.578994 | 0.842604 | 1.455 |
| chr12 | 17432011 | 17432195 | 1.35054 | 1.96561 | 1.455 |
| chr3 | 19868486 | 19868736 | 0.679885 | 0.989522 | 1.455 |
| chr10 | 117042722 | 117042975 | 0.770775 | 1.12182 | 1.455 |
| chr17 | 73851989 | 73852333 | 1.14512 | 1.66667 | 1.455 |
| chr8 | 67904647 | 67904898 | 0.668575 | 0.973105 | 1.455 |
| chr2 | 35855400 | 35855719 | 0.847406 | 1.23344 | 1.456 |
| chr7 | 146536730 | 146537026 | 0.747218 | 1.08765 | 1.456 |
| chr5 | 62074830 | 62075096 | 1.84063 | 2.67927 | 1.456 |
| chrX | 49148382 | 49148727 | 1.65109 | 2.40353 | 1.456 |
| chr15 | 67258635 | 67258946 | 1.28739 | 1.87412 | 1.456 |
| chr2 | 2124574 | 2124973 | 2.73779 | 3.98607 | 1.456 |
| chr2 | 198650643 | 198651049 | 2.96747 | 4.32073 | 1.456 |
| chr15 | 78910635 | 78911060 | 1.75253 | 2.5519 | 1.456 |
| chr12 | 112628708 | 112628988 | 0.847915 | 1.23469 | 1.456 |
| chr19 | 42637181 | 42637636 | 3.35166 | 4.88056 | 1.456 |
| chr1 | 54326544 | 54327023 | 2.17465 | 3.16684 | 1.456 |
| chr1 | 10212997 | 10213318 | 1.82999 | 2.66513 | 1.456 |
| chr16 | 4409021 | 4409503 | 2.02349 | 2.94704 | 1.456 |
| chr12 | 116190997 | 116191219 | 1.57541 | 2.29448 | 1.456 |
| chr8 | 133728199 | 133728495 | 0.889007 | 1.29482 | 1.456 |
| chr19 | 49503058 | 49503403 | 2.24638 | 3.27195 | 1.457 |
| chr5 | 176087773 | 176088272 | 2.91633 | 4.24813 | 1.457 |
| chr2 | 102018358 | 102018664 | 0.800599 | 1.16624 | 1.457 |
| chr9 | 136608012 | 136608527 | 2.71901 | 3.96102 | 1.457 |
| chr9 | 116569724 | 116570059 | 2.11967 | 3.08793 | 1.457 |
| chr22 | 39531164 | 39531421 | 1.51374 | 2.20524 | 1.457 |
| chr7 | 143434209 | 143434535 | 1.19885 | 1.74657 | 1.457 |
| chr20 | 48227622 | 48228090 | 3.74778 | 5.46026 | 1.457 |
| chr17 | 47662952 | 47663277 | 1.05058 | 1.53066 | 1.457 |
| chr2 | 81139952 | 81140175 | 0.793973 | 1.15681 | 1.457 |
| chr15 | 65563343 | 65563635 | 2.12909 | 3.10218 | 1.457 |
| chr10 | 90789650 | 90789910 | 2.06711 | 3.01191 | 1.457 |
| chr6 | 8708164 | 8708437 | 1.10984 | 1.6172 | 1.457 |
| chr4 | 5814659 | 5814974 | 1.61619 | 2.35507 | 1.457 |
| chr8 | 143946270 | 143946769 | 1.61313 | 2.35063 | 1.457 |
| chr20 | 31533610 | 31533935 | 1.4214 | 2.07126 | 1.457 |
| chr4 | 122821578 | 122821867 | 1.43003 | 2.0839 | 1.457 |
| chr11 | 71291196 | 71291702 | 1.44636 | 2.10772 | 1.457 |
| chr8 | 101214094 | 101214298 | 0.482755 | 0.703523 | 1.457 |
| chr2 | 14774364 | 14774945 | 1.76215 | 2.56803 | 1.457 |
| chr1 | 197296929 | 197297252 | 1.36619 | 1.99106 | 1.457 |
| chr19 | 42473532 | 42473845 | 0.642367 | 0.936187 | 1.457 |
| chr3 | 187334179 | 187334409 | 1.51241 | 2.20425 | 1.457 |
| chr6 | 2744678 | 2744920 | 0.876326 | 1.27721 | 1.457 |

| | | | | | |
|---|---|---|---|---|---|
| chr19 | 39988541 | 39988762 | 1.12418 | 1.63854 | 1.458 |
| chr1 | 114825433 | 114825741 | 1.73025 | 2.52195 | 1.458 |
| chr8 | 67658017 | 67658334 | 0.597721 | 0.871285 | 1.458 |
| chr16 | 47689954 | 47690217 | 0.685959 | 1.00004 | 1.458 |
| chr9 | 139944120 | 139944512 | 1.8135 | 2.64407 | 1.458 |
| chr5 | 63151751 | 63152004 | 0.732683 | 1.06831 | 1.458 |
| chr8 | 67601214 | 67601489 | 2.17475 | 3.17116 | 1.458 |
| chr9 | 74818165 | 74818409 | 1.68943 | 2.46348 | 1.458 |
| chr9 | 27165755 | 27165991 | 1.19073 | 1.73633 | 1.458 |
| chr4 | 150920699 | 150920921 | 1.94155 | 2.83126 | 1.458 |
| chr19 | 28138929 | 28139519 | 0.755372 | 1.1016 | 1.458 |
| chr9 | 102978511 | 102978851 | 1.31317 | 1.91507 | 1.458 |
| chr8 | 19266002 | 19266261 | 1.79494 | 2.61777 | 1.458 |
| chr3 | 161277262 | 161277579 | 1.80461 | 2.63191 | 1.458 |
| chr5 | 152984525 | 152984835 | 0.599465 | 0.8743 | 1.458 |
| chr1 | 25258703 | 25259153 | 2.08529 | 3.04136 | 1.458 |
| chr10 | 80734501 | 80734920 | 2.18843 | 3.19188 | 1.459 |
| chr5 | 45090045 | 45090258 | 0.790904 | 1.15356 | 1.459 |
| chr3 | 159161593 | 159161911 | 0.574735 | 0.838298 | 1.459 |
| chr20 | 42320608 | 42320946 | 2.20972 | 3.22311 | 1.459 |
| chr3 | 2669915 | 2670124 | 0.693158 | 1.01105 | 1.459 |
| chr2 | 141454124 | 141454338 | 1.58347 | 2.30972 | 1.459 |
| chr5 | 131515918 | 131516170 | 1.91441 | 2.79253 | 1.459 |
| chr5 | 84571654 | 84571896 | 0.82965 | 1.21023 | 1.459 |
| chr3 | 10328665 | 10328947 | 1.11178 | 1.62187 | 1.459 |
| chr17 | 17472041 | 17472428 | 2.35043 | 3.42898 | 1.459 |
| chr7 | 118584760 | 118584945 | 0.682796 | 0.996167 | 1.459 |
| chr12 | 33420495 | 33420703 | 0.683108 | 0.996691 | 1.459 |
| chr9 | 112579949 | 112580417 | 0.996286 | 1.45384 | 1.459 |
| chr2 | 206790526 | 206790904 | 2.49126 | 3.63568 | 1.459 |
| chr9 | 139519567 | 139519826 | 1.31003 | 1.91183 | 1.459 |
| chr17 | 11778993 | 11779202 | 0.814328 | 1.18843 | 1.459 |
| chr3 | 8366497 | 8366995 | 1.40518 | 2.05072 | 1.459 |
| chr3 | 102189879 | 102190200 | 1.26074 | 1.83993 | 1.459 |
| chr10 | 104028254 | 104028507 | 0.699125 | 1.02031 | 1.459 |
| chr9 | 138859237 | 138859743 | 2.74714 | 4.00928 | 1.459 |
| chr19 | 41710161 | 41710509 | 2.38599 | 3.48227 | 1.459 |
| chr3 | 2339480 | 2339683 | 0.857367 | 1.25136 | 1.460 |
| chr10 | 128767555 | 128767783 | 0.91103 | 1.32974 | 1.460 |
| chr7 | 81794241 | 81794471 | 0.896984 | 1.30929 | 1.460 |
| chr8 | 119688399 | 119688653 | 0.761116 | 1.11102 | 1.460 |
| chr15 | 94898962 | 94899171 | 0.734854 | 1.07272 | 1.460 |
| chr7 | 3340431 | 3340782 | 2.7025 | 3.94503 | 1.460 |
| chr19 | 1194559 | 1194810 | 2.33846 | 3.41364 | 1.460 |
| chr3 | 147029751 | 147030016 | 0.689919 | 1.00713 | 1.460 |
| chr2 | 43055998 | 43056316 | 2.8542 | 4.16664 | 1.460 |
| chr1 | 157933415 | 157933746 | 2.30879 | 3.37062 | 1.460 |
| chr9 | 83051959 | 83052165 | 0.683391 | 0.997719 | 1.460 |
| chr10 | 99629254 | 99629541 | 1.46151 | 2.13376 | 1.460 |
| chr7 | 43428399 | 43428849 | 0.839136 | 1.22511 | 1.460 |
| chr7 | 25894226 | 25894476 | 1.46914 | 2.14492 | 1.460 |
| chr12 | 126017890 | 126018398 | 2.40539 | 3.51188 | 1.460 |
| chr14 | 100129201 | 100129445 | 0.602066 | 0.879028 | 1.460 |
| chr19 | 40175764 | 40176019 | 0.869132 | 1.26905 | 1.460 |
| chr15 | 45455238 | 45455625 | 1.66386 | 2.4295 | 1.460 |
| chr5 | 22034183 | 22034407 | 0.788865 | 1.15187 | 1.460 |
| chr12 | 132318327 | 132318768 | 1.11888 | 1.63376 | 1.460 |
| chr11 | 42662162 | 42662464 | 0.649503 | 0.948419 | 1.460 |
| chr12 | 73849620 | 73849833 | 0.775233 | 1.13213 | 1.460 |
| chr5 | 162774636 | 162774975 | 1.68266 | 2.45732 | 1.460 |
| chr8 | 59161516 | 59162120 | 1.34796 | 1.96854 | 1.460 |

| | | | | | |
|---|---|---|---|---|---|
| chr1 | 38276796 | 38277054 | 2.26298 | 3.30486 | 1.460 |
| chr11 | 8252134 | 8252354 | 0.569023 | 0.831011 | 1.460 |
| chr3 | 127370996 | 127371353 | 2.31374 | 3.37909 | 1.460 |
| chr9 | 89373703 | 89373911 | 0.997314 | 1.45653 | 1.460 |
| chr7 | 142937672 | 142938144 | 1.95011 | 2.84813 | 1.461 |
| chr1 | 97652020 | 97652232 | 1.46896 | 2.14549 | 1.461 |
| chr7 | 23316113 | 23316380 | 1.14887 | 1.67804 | 1.461 |
| chr2 | 36152887 | 36153190 | 0.724108 | 1.05765 | 1.461 |
| chr4 | 5792075 | 5792365 | 1.59337 | 2.32733 | 1.461 |
| chr2 | 47499372 | 47499901 | 2.82008 | 4.11938 | 1.461 |
| chr17 | 26070196 | 26070434 | 0.713937 | 1.04291 | 1.461 |
| chr10 | 131767195 | 131767531 | 1.56004 | 2.27894 | 1.461 |
| chr11 | 82015944 | 82016155 | 0.548229 | 0.80087 | 1.461 |
| chr16 | 81437195 | 81437494 | 3.04468 | 4.44781 | 1.461 |
| chr1 | 246917101 | 246917717 | 1.64048 | 2.39667 | 1.461 |
| chr10 | 73191528 | 73191870 | 0.748832 | 1.09403 | 1.461 |
| chr9 | 85081904 | 85082170 | 1.31326 | 1.91866 | 1.461 |
| chr7 | 123341595 | 123341906 | 1.9296 | 2.81918 | 1.461 |
| chr10 | 28468495 | 28468741 | 1.47031 | 2.14832 | 1.461 |
| chr4 | 145708468 | 145708786 | 0.886995 | 1.29607 | 1.461 |
| chr16 | 13990714 | 13990975 | 1.59723 | 2.3339 | 1.461 |
| chr8 | 2309986 | 2310346 | 0.846647 | 1.23714 | 1.461 |
| chr16 | 85120352 | 85120665 | 2.59565 | 3.79307 | 1.461 |
| chr11 | 68870324 | 68870774 | 1.08712 | 1.58864 | 1.461 |
| chr22 | 21948132 | 21948478 | 1.29144 | 1.88723 | 1.461 |
| chr8 | 114449953 | 114450320 | 2.04831 | 2.99346 | 1.461 |
| chr1 | 167895708 | 167895983 | 2.06426 | 3.01679 | 1.461 |
| chr9 | 139538820 | 139539335 | 2.3984 | 3.50529 | 1.462 |
| chr18 | 68157699 | 68157907 | 1.71827 | 2.51128 | 1.462 |
| chr5 | 102005923 | 102006306 | 1.39643 | 2.04091 | 1.462 |
| chr19 | 18572521 | 18572960 | 2.16448 | 3.16346 | 1.462 |
| chr8 | 126743552 | 126743833 | 0.762668 | 1.11475 | 1.462 |
| chr19 | 41034996 | 41035331 | 3.10787 | 4.54295 | 1.462 |
| chr3 | 157977405 | 157977689 | 0.752992 | 1.10069 | 1.462 |
| chr5 | 105642938 | 105643213 | 0.720884 | 1.05377 | 1.462 |
| chr5 | 133519733 | 133520290 | 2.02721 | 2.96333 | 1.462 |
| chr1 | 117671281 | 117671489 | 0.911423 | 1.33232 | 1.462 |
| chr15 | 97213490 | 97213996 | 0.91678 | 1.34024 | 1.462 |
| chr2 | 176554891 | 176555145 | 1.02762 | 1.5024 | 1.462 |
| chr15 | 74166274 | 74166555 | 1.17324 | 1.71538 | 1.462 |
| chr9 | 115205579 | 115205844 | 1.22756 | 1.79482 | 1.462 |
| chr8 | 82190038 | 82190247 | 1.2074 | 1.76535 | 1.462 |
| chr16 | 89233690 | 89234314 | 1.76639 | 2.58267 | 1.462 |
| chr8 | 133379884 | 133380236 | 0.681703 | 0.996767 | 1.462 |
| chr7 | 89260394 | 89260629 | 0.761125 | 1.11292 | 1.462 |
| chr17 | 18864109 | 18864518 | 2.44632 | 3.57748 | 1.462 |
| chr15 | 63074271 | 63074520 | 2.37467 | 3.47278 | 1.462 |
| chr7 | 25608445 | 25608897 | 2.3675 | 3.4624 | 1.462 |
| chr2 | 33102352 | 33102647 | 0.695399 | 1.01704 | 1.463 |
| chr3 | 89148169 | 89148477 | 2.26821 | 3.31763 | 1.463 |
| chr6 | 131206554 | 131206855 | 1.37448 | 2.01043 | 1.463 |
| chr11 | 11058406 | 11058676 | 1.64613 | 2.408 | 1.463 |
| chr5 | 72815597 | 72815818 | 2.10799 | 3.08366 | 1.463 |
| chr10 | 102261940 | 102262210 | 0.66897 | 0.978659 | 1.463 |
| chr2 | 176950935 | 176951383 | 1.30297 | 1.90618 | 1.463 |
| chr22 | 21995856 | 21996076 | 1.18815 | 1.7382 | 1.463 |
| chr9 | 25812147 | 25812487 | 0.583542 | 0.853725 | 1.463 |
| chr17 | 64149959 | 64150207 | 0.703678 | 1.02952 | 1.463 |
| chr5 | 99766019 | 99766480 | 2.51452 | 3.67907 | 1.463 |
| chr2 | 166095716 | 166096080 | 1.54529 | 2.26099 | 1.463 |
| chr9 | 133933063 | 133933577 | 2.23686 | 3.27294 | 1.463 |

| | | | | | |
|---|---|---|---|---|---|
| chr17 | 80574287 | 80574492 | 0.672635 | 0.984219 | 1.463 |
| chr12 | 120763308 | 120763638 | 2.52903 | 3.70066 | 1.463 |
| chr20 | 21614487 | 21614819 | 0.922583 | 1.35 | 1.463 |
| chr17 | 78055526 | 78055845 | 0.930795 | 1.36204 | 1.463 |
| chr19 | 41489816 | 41490125 | 0.801007 | 1.17224 | 1.463 |
| chr8 | 120521937 | 120522261 | 1.37509 | 2.01239 | 1.463 |
| chr5 | 109978609 | 109978918 | 0.863766 | 1.26412 | 1.464 |
| chr14 | 26645072 | 26645297 | 0.537776 | 0.787048 | 1.464 |
| chr1 | 174211998 | 174212286 | 0.756309 | 1.10688 | 1.464 |
| chr22 | 22958663 | 22958898 | 0.912768 | 1.33588 | 1.464 |
| chr5 | 22516690 | 22516951 | 0.729425 | 1.06755 | 1.464 |
| chr16 | 2771192 | 2771547 | 2.04172 | 2.98829 | 1.464 |
| chrX | 130202921 | 130203213 | 1.77548 | 2.59869 | 1.464 |
| chr3 | 29206874 | 29207143 | 0.588557 | 0.861507 | 1.464 |
| chr12 | 123350062 | 123350802 | 1.47004 | 2.15183 | 1.464 |
| chr7 | 73244210 | 73244462 | 0.758112 | 1.10972 | 1.464 |
| chr4 | 175077445 | 175077665 | 1.48355 | 2.17165 | 1.464 |
| chr21 | 44923291 | 44923602 | 1.86145 | 2.72512 | 1.464 |
| chr10 | 333828 | 334071 | 1.08858 | 1.59408 | 1.464 |
| chr1 | 155063850 | 155064317 | 2.91496 | 4.26882 | 1.464 |
| chr1 | 27204555 | 27204880 | 1.18594 | 1.73684 | 1.465 |
| chr19 | 51286870 | 51287245 | 1.69409 | 2.48143 | 1.465 |
| chr3 | 87820489 | 87821102 | 1.32908 | 1.94683 | 1.465 |
| chr15 | 81558720 | 81559125 | 2.14414 | 3.14082 | 1.465 |
| chr16 | 1104730 | 1105521 | 1.30795 | 1.91599 | 1.465 |
| chr10 | 73407367 | 73407611 | 1.59703 | 2.33949 | 1.465 |
| chr20 | 2073763 | 2074165 | 1.71656 | 2.51481 | 1.465 |
| chr6 | 107014434 | 107014790 | 1.59441 | 2.33588 | 1.465 |
| chr8 | 97345316 | 97345740 | 2.18155 | 3.19613 | 1.465 |
| chr20 | 15680118 | 15680355 | 0.97036 | 1.42174 | 1.465 |
| chr17 | 39477306 | 39477515 | 0.617654 | 0.904988 | 1.465 |
| chr15 | 80805096 | 80805386 | 1.6325 | 2.39199 | 1.465 |
| chr19 | 28779507 | 28779920 | 0.747582 | 1.09538 | 1.465 |
| chr8 | 36097864 | 36098094 | 0.636427 | 0.932527 | 1.465 |
| chr11 | 77330602 | 77330958 | 1.92054 | 2.81422 | 1.465 |
| chr2 | 192676915 | 192677152 | 1.63306 | 2.39305 | 1.465 |
| chr5 | 105752903 | 105753184 | 0.66007 | 0.967338 | 1.466 |
| chr4 | 160124208 | 160124450 | 1.69319 | 2.4814 | 1.466 |
| chr20 | 55082740 | 55082965 | 1.02945 | 1.50874 | 1.466 |
| chr5 | 147754320 | 147754642 | 2.03001 | 2.97524 | 1.466 |
| chr7 | 9220221 | 9220506 | 0.70916 | 1.03938 | 1.466 |
| chr20 | 30147865 | 30148223 | 2.40327 | 3.52247 | 1.466 |
| chr10 | 80956417 | 80957061 | 2.17587 | 3.18934 | 1.466 |
| chr12 | 100548570 | 100549103 | 1.65091 | 2.41988 | 1.466 |
| chr6 | 3279866 | 3280096 | 1.39247 | 2.04108 | 1.466 |
| chr5 | 163931276 | 163931484 | 1.45653 | 2.13516 | 1.466 |
| chr2 | 180275651 | 180275970 | 1.83479 | 2.68969 | 1.466 |
| chr11 | 14440411 | 14440677 | 1.6402 | 2.40451 | 1.466 |
| chr20 | 34389926 | 34390158 | 0.503668 | 0.738375 | 1.466 |
| chr3 | 195576563 | 195576930 | 1.93716 | 2.83996 | 1.466 |
| chr2 | 231766915 | 231767187 | 2.56547 | 3.7612 | 1.466 |
| chr9 | 4679302 | 4679848 | 1.05532 | 1.54719 | 1.466 |
| chr2 | 44104881 | 44105377 | 2.47434 | 3.62769 | 1.466 |
| chrX | 13524731 | 13525040 | 2.67616 | 3.92369 | 1.466 |
| chr8 | 145056619 | 145057072 | 2.29455 | 3.36427 | 1.466 |
| chr1 | 227107903 | 227108160 | 0.659162 | 0.966565 | 1.466 |
| chr8 | 133198305 | 133198529 | 0.659492 | 0.967047 | 1.466 |
| chr17 | 56254158 | 56254575 | 0.913714 | 1.33988 | 1.466 |
| chr5 | 176386208 | 176386440 | 0.858506 | 1.25897 | 1.466 |
| chr9 | 135526676 | 135526867 | 1.42079 | 2.08359 | 1.467 |
| chr15 | 38274119 | 38274348 | 0.937671 | 1.37511 | 1.467 |

| | | | | | |
|---|---|---|---|---|---|
| chr7 | 53976547 | 53976866 | 0.654849 | 0.960375 | 1.467 |
| chr17 | 37703921 | 37704724 | 1.84005 | 2.69873 | 1.467 |
| chr1 | 110750407 | 110750831 | 1.65312 | 2.42462 | 1.467 |
| chr16 | 83147052 | 83147261 | 1.0262 | 1.50512 | 1.467 |
| chr10 | 129833737 | 129834163 | 1.89653 | 2.78197 | 1.467 |
| chr1 | 199078784 | 199079111 | 0.816969 | 1.19841 | 1.467 |
| chr12 | 23093497 | 23093734 | 0.5804 | 0.851429 | 1.467 |
| chr15 | 44195999 | 44196298 | 1.53505 | 2.25192 | 1.467 |
| chr20 | 58524248 | 58524456 | 1.08398 | 1.59036 | 1.467 |
| chr9 | 130526284 | 130526600 | 0.85808 | 1.25896 | 1.467 |
| chr10 | 113793161 | 113793460 | 0.670023 | 0.983052 | 1.467 |
| chr16 | 68273817 | 68274237 | 2.73115 | 4.00726 | 1.467 |
| chr22 | 43624357 | 43624627 | 1.36363 | 2.00109 | 1.467 |
| chr11 | 91833499 | 91833714 | 0.509688 | 0.747969 | 1.468 |
| chr15 | 92006624 | 92006896 | 0.703634 | 1.0326 | 1.468 |
| chrX | 40510973 | 40511178 | 1.52748 | 2.2416 | 1.468 |
| chr2 | 172797435 | 172797759 | 0.969778 | 1.42326 | 1.468 |
| chr15 | 56066594 | 56066900 | 2.09607 | 3.07623 | 1.468 |
| chr5 | 111021761 | 111021981 | 1.48306 | 2.17657 | 1.468 |
| chr19 | 34515104 | 34515310 | 0.98238 | 1.44181 | 1.468 |
| chr8 | 78625762 | 78625984 | 0.740426 | 1.08671 | 1.468 |
| chr8 | 22871907 | 22872273 | 1.56756 | 2.30069 | 1.468 |
| chr9 | 29533706 | 29533919 | 0.668874 | 0.98174 | 1.468 |
| chr9 | 124397271 | 124397638 | 1.68357 | 2.47113 | 1.468 |
| chr14 | 78801954 | 78802207 | 0.678616 | 0.996121 | 1.468 |
| chr1 | 223297049 | 223297551 | 2.15336 | 3.16106 | 1.468 |
| chr17 | 26649007 | 26649244 | 1.2417 | 1.82278 | 1.468 |
| chr19 | 14258952 | 14259238 | 1.72502 | 2.53229 | 1.468 |
| chr1 | 91748287 | 91748604 | 1.76518 | 2.59127 | 1.468 |
| chr3 | 146409949 | 146410319 | 0.926417 | 1.35998 | 1.468 |
| chr17 | 43275146 | 43275564 | 3.77476 | 5.54147 | 1.468 |
| chr17 | 11890246 | 11890571 | 1.00723 | 1.47868 | 1.468 |
| chr11 | 61196847 | 61197058 | 1.12315 | 1.64892 | 1.468 |
| chr2 | 191044865 | 191045223 | 1.52745 | 2.2425 | 1.468 |
| chr11 | 17620359 | 17620848 | 2.51372 | 3.69054 | 1.468 |
| chr10 | 124781108 | 124781322 | 1.1005 | 1.61574 | 1.468 |
| chr15 | 98035293 | 98035514 | 1.24743 | 1.83148 | 1.468 |
| chr3 | 53972705 | 53972973 | 1.31486 | 1.93054 | 1.468 |
| chr19 | 31611288 | 31611493 | 0.635994 | 0.933807 | 1.468 |
| chr10 | 81928874 | 81929527 | 1.35854 | 1.99472 | 1.468 |
| chr12 | 55466930 | 55467159 | 1.96176 | 2.88049 | 1.468 |
| chr3 | 52516214 | 52516431 | 0.831022 | 1.22023 | 1.468 |
| chr16 | 76636549 | 76636753 | 0.747134 | 1.09722 | 1.469 |
| chr11 | 32706256 | 32706517 | 0.769338 | 1.12986 | 1.469 |
| chr16 | 5929089 | 5929365 | 0.685363 | 1.00653 | 1.469 |
| chr5 | 107428848 | 107429211 | 1.40786 | 2.06767 | 1.469 |
| chr15 | 96888810 | 96889064 | 1.5475 | 2.27283 | 1.469 |
| chr8 | 4219780 | 4220188 | 0.73639 | 1.08155 | 1.469 |
| chr17 | 42997416 | 42997752 | 2.27904 | 3.34748 | 1.469 |
| chr12 | 119024900 | 119025152 | 0.699553 | 1.02759 | 1.469 |
| chr20 | 34067017 | 34067258 | 0.846543 | 1.2435 | 1.469 |
| chr5 | 7381613 | 7381905 | 0.713601 | 1.04822 | 1.469 |
| chr15 | 57834445 | 57834733 | 2.65545 | 3.90079 | 1.469 |
| chr11 | 25774720 | 25775075 | 0.572821 | 0.841502 | 1.469 |
| chr19 | 46026915 | 46027268 | 3.0766 | 4.5197 | 1.469 |
| chr11 | 78143302 | 78143579 | 1.67063 | 2.45427 | 1.469 |
| chr11 | 19925112 | 19925495 | 1.70362 | 2.50289 | 1.469 |
| chr6 | 99689889 | 99690100 | 1.02533 | 1.50645 | 1.469 |
| chr4 | 41646329 | 41646580 | 1.31908 | 1.93818 | 1.469 |
| chr20 | 22254633 | 22254957 | 0.669937 | 0.984398 | 1.469 |
| chr2 | 68978896 | 68979290 | 2.26328 | 3.3257 | 1.469 |

| | | | | | |
|---|---|---|---|---|---|
| chr18 | 72956407 | 72956622 | 0.871909 | 1.28123 | 1.469 |
| chr9 | 36991603 | 36991906 | 1.81148 | 2.66195 | 1.469 |
| chr17 | 36478028 | 36478479 | 2.27309 | 3.34042 | 1.470 |
| chr7 | 139138067 | 139138310 | 0.841699 | 1.23694 | 1.470 |
| chr12 | 42726173 | 42726398 | 1.58836 | 2.33428 | 1.470 |
| chr3 | 113345625 | 113345994 | 2.60718 | 3.83156 | 1.470 |
| chrX | 107020493 | 107020754 | 0.994772 | 1.46194 | 1.470 |
| chr1 | 153325870 | 153326337 | 0.980322 | 1.44082 | 1.470 |
| chr9 | 21396073 | 21396518 | 2.86149 | 4.20579 | 1.470 |
| chr4 | 174427079 | 174427317 | 1.89199 | 2.7809 | 1.470 |
| chr5 | 173472505 | 173472737 | 2.29104 | 3.36756 | 1.470 |
| chr6 | 119399173 | 119399399 | 0.702338 | 1.03238 | 1.470 |
| chr15 | 73557888 | 73558237 | 2.40221 | 3.5311 | 1.470 |
| chr14 | 46868361 | 46868619 | 0.80592 | 1.18471 | 1.470 |
| chr8 | 38242518 | 38242803 | 1.76718 | 2.59802 | 1.470 |
| chr11 | 74164463 | 74164762 | 2.22302 | 3.26821 | 1.470 |
| chr20 | 3387979 | 3388278 | 1.2203 | 1.7941 | 1.470 |
| chrX | 30653818 | 30654026 | 1.47712 | 2.1717 | 1.470 |
| chr12 | 87936628 | 87936964 | 0.797144 | 1.17202 | 1.470 |
| chr2 | 231187932 | 231188306 | 1.58543 | 2.33114 | 1.470 |
| chr20 | 15604869 | 15605276 | 0.697433 | 1.02556 | 1.470 |
| chr11 | 3382924 | 3383198 | 0.677459 | 0.996219 | 1.471 |
| chr2 | 101629162 | 101629429 | 1.30984 | 1.92617 | 1.471 |
| chr19 | 42160598 | 42160959 | 0.834012 | 1.22651 | 1.471 |
| chr18 | 55225098 | 55225483 | 2.63457 | 3.87451 | 1.471 |
| chr3 | 1130931 | 1131172 | 1.10556 | 1.62589 | 1.471 |
| chr5 | 122946875 | 122947138 | 2.91107 | 4.28141 | 1.471 |
| chr9 | 95916608 | 95916996 | 1.66462 | 2.44826 | 1.471 |
| chr7 | 97870528 | 97871005 | 1.64493 | 2.41936 | 1.471 |
| chr3 | 49051527 | 49051834 | 1.86796 | 2.7475 | 1.471 |
| chr15 | 92926685 | 92927059 | 1.79983 | 2.64732 | 1.471 |
| chr8 | 131893931 | 131894178 | 1.58307 | 2.32864 | 1.471 |
| chr8 | 84462960 | 84463207 | 0.913137 | 1.3432 | 1.471 |
| chr5 | 113076970 | 113077190 | 0.563909 | 0.829518 | 1.471 |
| chr9 | 129961392 | 129961809 | 2.32875 | 3.42569 | 1.471 |
| chr9 | 19624016 | 19624250 | 0.940553 | 1.38362 | 1.471 |
| chr11 | 119345366 | 119345759 | 1.96212 | 2.88661 | 1.471 |
| chr22 | 24585594 | 24585926 | 2.37966 | 3.50088 | 1.471 |
| chr2 | 11264624 | 11264912 | 2.5592 | 3.7655 | 1.471 |
| chr6 | 16316967 | 16317263 | 1.72481 | 2.53783 | 1.471 |
| chr3 | 72282984 | 72283216 | 0.692849 | 1.01946 | 1.471 |
| chr17 | 69955240 | 69955606 | 1.47959 | 2.17714 | 1.471 |
| chr7 | 78869188 | 78869455 | 0.800135 | 1.17739 | 1.471 |
| chr3 | 56285448 | 56285670 | 1.52972 | 2.251 | 1.472 |
| chr20 | 44977644 | 44977957 | 2.66392 | 3.92005 | 1.472 |
| chr10 | 122702298 | 122702956 | 1.67263 | 2.46141 | 1.472 |
| chr3 | 26410723 | 26410940 | 0.890702 | 1.31074 | 1.472 |
| chr4 | 33091520 | 33091731 | 1.12858 | 1.66096 | 1.472 |
| chr1 | 165362049 | 165362492 | 2.01099 | 2.95971 | 1.472 |
| chr8 | 141152471 | 141152749 | 0.927341 | 1.36488 | 1.472 |
| chr10 | 35507316 | 35507763 | 1.5995 | 2.35427 | 1.472 |
| chr2 | 69039362 | 69039644 | 0.641271 | 0.943879 | 1.472 |
| chr2 | 73973298 | 73973603 | 0.887963 | 1.30701 | 1.472 |
| chr5 | 28202592 | 28202844 | 0.751954 | 1.10686 | 1.472 |
| chr19 | 17684734 | 17685264 | 2.03984 | 3.00284 | 1.472 |
| chr3 | 187458655 | 187458970 | 2.08452 | 3.06862 | 1.472 |
| chr15 | 94690361 | 94690645 | 0.678106 | 0.99825 | 1.472 |
| chr1 | 172714169 | 172714371 | 1.20024 | 1.76696 | 1.472 |
| chr9 | 105730558 | 105730788 | 0.645118 | 0.949763 | 1.472 |
| chr16 | 19513968 | 19514299 | 1.41158 | 2.07825 | 1.472 |
| chr9 | 137291372 | 137291725 | 1.90445 | 2.80395 | 1.472 |

| | | | | | |
|---|---|---|---|---|---|
| chr11 | 116512326 | 116512611 | 2.05039 | 3.01887 | 1.472 |
| chr2 | 38303518 | 38304065 | 1.70069 | 2.50414 | 1.472 |
| chr19 | 43117921 | 43118123 | 0.892203 | 1.31372 | 1.472 |
| chr1 | 31654352 | 31654744 | 1.83578 | 2.70315 | 1.472 |
| chr19 | 13088699 | 13089029 | 2.45703 | 3.61793 | 1.472 |
| chr4 | 47670900 | 47671165 | 1.19218 | 1.75548 | 1.473 |
| chr21 | 27543007 | 27543366 | 2.32231 | 3.41974 | 1.473 |
| chr1 | 19210593 | 19210965 | 1.73786 | 2.55912 | 1.473 |
| chr9 | 100663303 | 100663592 | 2.19194 | 3.22785 | 1.473 |
| chr9 | 92213100 | 92213370 | 1.71237 | 2.52169 | 1.473 |
| chr19 | 16222109 | 16222375 | 1.66806 | 2.45657 | 1.473 |
| chr8 | 8887393 | 8887660 | 1.62752 | 2.39688 | 1.473 |
| chr2 | 72025427 | 72025860 | 2.555 | 3.76293 | 1.473 |
| chrX | 65166855 | 65167080 | 0.722812 | 1.06461 | 1.473 |
| chr20 | 3297275 | 3297592 | 1.83247 | 2.69901 | 1.473 |
| chr7 | 95317051 | 95317392 | 1.12582 | 1.65828 | 1.473 |
| chr9 | 136964180 | 136964612 | 2.23962 | 3.29902 | 1.473 |
| chr6 | 88541784 | 88541988 | 0.957969 | 1.41123 | 1.473 |
| chr1 | 110334747 | 110334954 | 1.19602 | 1.76204 | 1.473 |
| chr15 | 69022836 | 69023122 | 1.25645 | 1.85117 | 1.473 |
| chr2 | 101601246 | 101601580 | 1.77474 | 2.61481 | 1.473 |
| chr15 | 25955245 | 25955617 | 1.83004 | 2.69641 | 1.473 |
| chr2 | 45768105 | 45768399 | 0.78046 | 1.14995 | 1.473 |
| chr15 | 72093530 | 72093793 | 2.07827 | 3.06224 | 1.473 |
| chr10 | 96204158 | 96204377 | 1.52836 | 2.252 | 1.473 |
| chr10 | 20493634 | 20493881 | 0.635776 | 0.93683 | 1.474 |
| chr11 | 37643655 | 37643867 | 0.846828 | 1.24782 | 1.474 |
| chr9 | 104278991 | 104279204 | 0.546047 | 0.804623 | 1.474 |
| chr12 | 122840444 | 122841037 | 1.32556 | 1.95333 | 1.474 |
| chr9 | 10815446 | 10815774 | 0.857364 | 1.26345 | 1.474 |
| chr22 | 49339630 | 49339974 | 1.99264 | 2.93655 | 1.474 |
| chr1 | 6549871 | 6550261 | 2.60038 | 3.83229 | 1.474 |
| chr15 | 68842848 | 68843423 | 2.11542 | 3.11758 | 1.474 |
| chr16 | 7638294 | 7638541 | 0.608958 | 0.897491 | 1.474 |
| chr12 | 52364627 | 52364928 | 2.01682 | 2.97262 | 1.474 |
| chr17 | 63227786 | 63228150 | 1.65915 | 2.44551 | 1.474 |
| chr16 | 4452329 | 4453210 | 2.6363 | 3.88589 | 1.474 |
| chr11 | 120800365 | 120800620 | 1.94776 | 2.87106 | 1.474 |
| chr10 | 69539696 | 69540043 | 1.56844 | 2.31194 | 1.474 |
| chr11 | 43873663 | 43873887 | 1.96704 | 2.89951 | 1.474 |
| chr7 | 101181954 | 101182477 | 2.52351 | 3.72009 | 1.474 |
| chr1 | 3481769 | 3482205 | 2.61831 | 3.86012 | 1.474 |
| chr17 | 71591682 | 71591977 | 1.18821 | 1.75177 | 1.474 |
| chr1 | 27627250 | 27627693 | 2.22279 | 3.27715 | 1.474 |
| chr7 | 53850213 | 53850425 | 0.689098 | 1.01597 | 1.474 |
| chr9 | 139999874 | 140000080 | 0.72033 | 1.06202 | 1.474 |
| chr11 | 116661967 | 116662325 | 2.22014 | 3.27339 | 1.474 |
| chr14 | 75075581 | 75076099 | 1.9716 | 2.90695 | 1.474 |
| chr6 | 147021302 | 147021648 | 1.26425 | 1.86404 | 1.474 |
| chr1 | 204183302 | 204183630 | 1.37519 | 2.02764 | 1.474 |
| chr17 | 54395730 | 54395996 | 0.70674 | 1.04207 | 1.474 |
| chr17 | 48559770 | 48560172 | 2.52699 | 3.72601 | 1.474 |
| chr1 | 86174083 | 86174340 | 0.948451 | 1.39849 | 1.475 |
| chr16 | 87331493 | 87331837 | 1.96741 | 2.90109 | 1.475 |
| chr8 | 2824835 | 2825210 | 0.714727 | 1.05392 | 1.475 |
| chr7 | 123672449 | 123672664 | 0.603833 | 0.890414 | 1.475 |
| chr10 | 120587424 | 120587647 | 1.24387 | 1.83426 | 1.475 |
| chr7 | 120159306 | 120159579 | 0.804157 | 1.18587 | 1.475 |
| chr21 | 43236887 | 43237246 | 1.68553 | 2.48566 | 1.475 |
| chr8 | 120875094 | 120875522 | 0.76242 | 1.12438 | 1.475 |
| chr21 | 18884850 | 18885120 | 1.97824 | 2.91749 | 1.475 |

| | | | | | |
|---|---|---|---|---|---|
| chr1 | 238619204 | 238619490 | 0.660738 | 0.974487 | 1.475 |
| chr2 | 241507549 | 241507844 | 0.837061 | 1.23457 | 1.475 |
| chr10 | 97144139 | 97144419 | 0.799954 | 1.17987 | 1.475 |
| chr10 | 98130201 | 98130761 | 1.76617 | 2.60499 | 1.475 |
| chr15 | 69075994 | 69076367 | 2.24365 | 3.30927 | 1.475 |
| chr9 | 1179284 | 1179536 | 0.50311 | 0.742076 | 1.475 |
| chr16 | 49619776 | 49619986 | 0.505932 | 0.746299 | 1.475 |
| chr17 | 6925258 | 6925649 | 3.28419 | 4.8446 | 1.475 |
| chr9 | 133537746 | 133538005 | 0.866196 | 1.27779 | 1.475 |
| chr1 | 30584343 | 30584672 | 1.46226 | 2.1571 | 1.475 |
| chr2 | 80469774 | 80470028 | 0.724048 | 1.06814 | 1.475 |
| chr9 | 106349116 | 106349319 | 0.772129 | 1.13909 | 1.475 |
| chr15 | 78077293 | 78077643 | 1.95663 | 2.88659 | 1.475 |
| chr19 | 13144829 | 13145213 | 2.69333 | 3.97356 | 1.475 |
| chr20 | 25227812 | 25228039 | 1.33362 | 1.96753 | 1.475 |
| chr4 | 5052211 | 5052467 | 1.75691 | 2.59202 | 1.475 |
| chr19 | 46417173 | 46417452 | 1.15338 | 1.70166 | 1.475 |
| chr10 | 88705384 | 88705643 | 1.0625 | 1.56759 | 1.475 |
| chr3 | 14822419 | 14822805 | 2.33219 | 3.44092 | 1.475 |
| chr9 | 21318081 | 21318791 | 1.24839 | 1.84187 | 1.475 |
| chr2 | 43456735 | 43456990 | 1.93418 | 2.85379 | 1.475 |
| chr2 | 22656076 | 22656291 | 0.821547 | 1.21221 | 1.476 |
| chr15 | 92249661 | 92250071 | 2.31519 | 3.41624 | 1.476 |
| chr7 | 85838343 | 85838620 | 0.682338 | 1.00695 | 1.476 |
| chr1 | 213547649 | 213548044 | 2.69743 | 3.98107 | 1.476 |
| chr3 | 51428387 | 51428663 | 2.43478 | 3.59348 | 1.476 |
| chr17 | 603790 | 604116 | 1.21094 | 1.78732 | 1.476 |
| chr3 | 9109066 | 9109429 | 2.16165 | 3.19087 | 1.476 |
| chr9 | 38363052 | 38363359 | 1.57769 | 2.32894 | 1.476 |
| chr12 | 99047742 | 99047984 | 1.30433 | 1.9255 | 1.476 |
| chr8 | 109735165 | 109735501 | 1.87489 | 2.7678 | 1.476 |
| chr1 | 201223599 | 201224061 | 3.33753 | 4.92735 | 1.476 |
| chr3 | 148665023 | 148665301 | 1.12895 | 1.66673 | 1.476 |
| chr19 | 33882831 | 33883149 | 0.683438 | 1.00902 | 1.476 |
| chr5 | 140464043 | 140464264 | 1.62414 | 2.39787 | 1.476 |
| chr8 | 9196730 | 9196940 | 0.712386 | 1.05183 | 1.476 |
| chr2 | 211442294 | 211442538 | 1.05 | 1.55043 | 1.477 |
| chr16 | 13836916 | 13837163 | 0.622764 | 0.919591 | 1.477 |
| chr7 | 43163201 | 43163403 | 0.66207 | 0.977645 | 1.477 |
| chr8 | 51745518 | 51745751 | 0.824635 | 1.21771 | 1.477 |
| chr5 | 169128889 | 169129099 | 1.10451 | 1.63101 | 1.477 |
| chr11 | 57250470 | 57250911 | 1.88472 | 2.78356 | 1.477 |
| chr3 | 125304518 | 125304746 | 2.05537 | 3.03569 | 1.477 |
| chr14 | 23025890 | 23026475 | 3.0086 | 4.44366 | 1.477 |
| chr15 | 92660520 | 92660933 | 0.79578 | 1.17536 | 1.477 |
| chr1 | 156352714 | 156353136 | 2.39877 | 3.54326 | 1.477 |
| chr11 | 91944843 | 91945059 | 1.71121 | 2.52766 | 1.477 |
| chr17 | 71556878 | 71557210 | 0.896695 | 1.32457 | 1.477 |
| chr11 | 32407571 | 32407910 | 1.22939 | 1.81605 | 1.477 |
| chr10 | 43248506 | 43248920 | 2.32121 | 3.42927 | 1.477 |
| chr12 | 56907592 | 56907779 | 1.41794 | 2.095 | 1.478 |
| chr1 | 171214642 | 171215006 | 1.86218 | 2.7514 | 1.478 |
| chr2 | 84240931 | 84241159 | 0.740856 | 1.09471 | 1.478 |
| chr1 | 238101903 | 238102441 | 0.950773 | 1.4049 | 1.478 |
| chr11 | 74160474 | 74160729 | 1.88122 | 2.77987 | 1.478 |
| chr5 | 155061824 | 155062051 | 0.571017 | 0.843823 | 1.478 |
| chr1 | 169515695 | 169516099 | 1.97089 | 2.91272 | 1.478 |
| chr10 | 124802916 | 124803207 | 1.84777 | 2.73083 | 1.478 |
| chr8 | 61338688 | 61338908 | 0.54295 | 0.802465 | 1.478 |
| chr1 | 224358433 | 224358771 | 1.62799 | 2.40622 | 1.478 |
| chr21 | 41492966 | 41493208 | 1.42261 | 2.10273 | 1.478 |

| | | | | | |
|---|---|---|---|---|---|
| chr1 | 218472788 | 218473063 | 1.34129 | 1.98259 | 1.478 |
| chr10 | 120353936 | 120354398 | 1.14737 | 1.69596 | 1.478 |
| chr10 | 93011268 | 93011519 | 1.73281 | 2.56143 | 1.478 |
| chr15 | 89330475 | 89330859 | 0.566006 | 0.836668 | 1.478 |
| chr11 | 62537029 | 62537478 | 1.0096 | 1.49243 | 1.478 |
| chr16 | 80746925 | 80747153 | 0.583202 | 0.862115 | 1.478 |
| chr5 | 174871594 | 174872084 | 2.67595 | 3.95582 | 1.478 |
| chr18 | 20787340 | 20787564 | 1.05569 | 1.56066 | 1.478 |
| chr7 | 100322886 | 100323265 | 1.91626 | 2.83307 | 1.478 |
| chr7 | 136164983 | 136165395 | 0.651506 | 0.963226 | 1.478 |
| chr10 | 1034138 | 1034350 | 1.13172 | 1.6733 | 1.479 |
| chr2 | 183903106 | 183903410 | 1.61318 | 2.38531 | 1.479 |
| chr12 | 25721552 | 25721790 | 0.663308 | 0.980811 | 1.479 |
| chr17 | 46231033 | 46231328 | 1.32082 | 1.95306 | 1.479 |
| chr21 | 37513833 | 37514063 | 1.5737 | 2.32698 | 1.479 |
| chr7 | 25605507 | 25605840 | 3.53574 | 5.22841 | 1.479 |
| chr15 | 29861160 | 29861459 | 1.35776 | 2.00785 | 1.479 |
| chr1 | 228367896 | 228368370 | 1.01842 | 1.50607 | 1.479 |
| chr15 | 62987473 | 62987712 | 0.66595 | 0.984828 | 1.479 |
| chr16 | 68389177 | 68389628 | 0.919165 | 1.35939 | 1.479 |
| chr17 | 48891876 | 48892171 | 0.769586 | 1.13818 | 1.479 |
| chr2 | 15777465 | 15777771 | 2.28849 | 3.38471 | 1.479 |
| chr1 | 186574906 | 186575161 | 1.80308 | 2.66679 | 1.479 |
| chr13 | 111719968 | 111720420 | 1.02155 | 1.51093 | 1.479 |
| chr17 | 39846041 | 39846427 | 1.64097 | 2.4271 | 1.479 |
| chr5 | 90099909 | 90100126 | 0.656506 | 0.97104 | 1.479 |
| chr2 | 74619079 | 74619441 | 3.05601 | 4.52036 | 1.479 |
| chr11 | 62361079 | 62361430 | 1.9621 | 2.9023 | 1.479 |
| chr19 | 42194933 | 42195858 | 1.95303 | 2.88893 | 1.479 |
| chr9 | 5869812 | 5870039 | 1.67785 | 2.48187 | 1.479 |
| chr15 | 38732506 | 38732797 | 2.01957 | 2.98737 | 1.479 |
| chr4 | 2765723 | 2766269 | 2.12014 | 3.13637 | 1.479 |
| chr5 | 122680545 | 122680839 | 1.54364 | 2.28354 | 1.479 |
| chr2 | 70995149 | 70995486 | 2.43325 | 3.59968 | 1.479 |
| chr20 | 3212035 | 3212298 | 0.628517 | 0.929948 | 1.480 |
| chr2 | 180942139 | 180942354 | 1.32619 | 1.96229 | 1.480 |
| chr5 | 118547864 | 118548090 | 0.754374 | 1.11625 | 1.480 |
| chr20 | 59426871 | 59427200 | 1.60215 | 2.37073 | 1.480 |
| chr1 | 208825645 | 208825961 | 2.06195 | 3.05115 | 1.480 |
| chr15 | 92110145 | 92110434 | 0.620185 | 0.917738 | 1.480 |
| chr20 | 58838147 | 58838361 | 1.95027 | 2.886 | 1.480 |
| chr8 | 91773356 | 91773578 | 0.683008 | 1.01074 | 1.480 |
| chr12 | 47899546 | 47899793 | 0.776083 | 1.1485 | 1.480 |
| chr2 | 57948016 | 57948336 | 2.16645 | 3.20632 | 1.480 |
| chr14 | 96518084 | 96518393 | 1.40054 | 2.07285 | 1.480 |
| chr17 | 54817182 | 54817532 | 0.93639 | 1.38589 | 1.480 |
| chr9 | 113800204 | 113800519 | 1.114 | 1.64883 | 1.480 |
| chr12 | 57861879 | 57862107 | 0.619424 | 0.916813 | 1.480 |
| chr3 | 114102646 | 114102932 | 2.60025 | 3.84897 | 1.480 |
| chr15 | 94026653 | 94026872 | 0.718735 | 1.0639 | 1.480 |
| chr16 | 30626639 | 30627028 | 2.37412 | 3.51476 | 1.480 |
| chr5 | 131398940 | 131399536 | 2.64904 | 3.92176 | 1.480 |
| chr8 | 28451447 | 28452154 | 1.73881 | 2.57427 | 1.480 |
| chr12 | 101062514 | 101063075 | 2.36285 | 3.49821 | 1.481 |
| chr17 | 61483072 | 61483461 | 1.53066 | 2.26615 | 1.481 |
| chr5 | 7261155 | 7261452 | 1.41294 | 2.09187 | 1.481 |
| chr12 | 54787874 | 54788375 | 1.94586 | 2.88092 | 1.481 |
| chr5 | 177624041 | 177624291 | 1.69775 | 2.51386 | 1.481 |
| chr14 | 89493572 | 89494079 | 2.10463 | 3.11644 | 1.481 |
| chr10 | 11182177 | 11182412 | 0.631697 | 0.935436 | 1.481 |
| chr2 | 131911606 | 131911934 | 1.78534 | 2.64382 | 1.481 |

| | | | | | |
|---|---|---|---|---|---|
| chr2 | 177502874 | 177503185 | 1.88547 | 2.79211 | 1.481 |
| chr5 | 137993216 | 137993498 | 0.650167 | 0.962807 | 1.481 |
| chr1 | 204096757 | 204097167 | 2.65351 | 3.92952 | 1.481 |
| chr6 | 105130947 | 105131409 | 1.47277 | 2.1811 | 1.481 |
| chr1 | 219794207 | 219794415 | 0.599591 | 0.888025 | 1.481 |
| chr7 | 29229307 | 29229779 | 2.46348 | 3.64854 | 1.481 |
| chr8 | 11815083 | 11815350 | 0.824499 | 1.22114 | 1.481 |
| chr14 | 31218703 | 31219004 | 0.821292 | 1.21642 | 1.481 |
| chr2 | 120233718 | 120234275 | 2.34965 | 3.48009 | 1.481 |
| chr22 | 29866367 | 29866838 | 2.64831 | 3.92261 | 1.481 |
| chr12 | 122287486 | 122287987 | 0.606724 | 0.898668 | 1.481 |
| chr7 | 49743083 | 49743313 | 0.859683 | 1.27336 | 1.481 |
| chr10 | 71173027 | 71173294 | 1.2921 | 1.91389 | 1.481 |
| chrX | 49605379 | 49605754 | 0.986146 | 1.46071 | 1.481 |
| chr2 | 45241169 | 45241579 | 3.27246 | 4.84741 | 1.481 |
| chr7 | 112628331 | 112628623 | 0.83615 | 1.2386 | 1.481 |
| chr12 | 34276050 | 34276314 | 2.21596 | 3.28267 | 1.481 |
| chr3 | 152184571 | 152184777 | 1.40329 | 2.07884 | 1.481 |
| chr5 | 36357459 | 36357669 | 0.866792 | 1.28414 | 1.481 |
| chr20 | 647994 | 648443 | 2.43618 | 3.6092 | 1.482 |
| chr20 | 36244321 | 36244636 | 1.86566 | 2.76398 | 1.482 |
| chr16 | 50402753 | 50403118 | 1.82299 | 2.7008 | 1.482 |
| chr11 | 2713675 | 2713906 | 1.52753 | 2.26328 | 1.482 |
| chr17 | 73621956 | 73622412 | 2.46659 | 3.65466 | 1.482 |
| chr17 | 48192221 | 48192572 | 1.88089 | 2.78701 | 1.482 |
| chr2 | 200569475 | 200569769 | 1.01422 | 1.50285 | 1.482 |
| chr8 | 103135975 | 103136293 | 2.07915 | 3.08105 | 1.482 |
| chr1 | 19201695 | 19202144 | 1.61764 | 2.39719 | 1.482 |
| chr11 | 22928311 | 22928522 | 0.777643 | 1.15245 | 1.482 |
| chr14 | 26311171 | 26311394 | 0.693035 | 1.02708 | 1.482 |
| chr20 | 23648950 | 23649523 | 1.86376 | 2.7623 | 1.482 |
| chr2 | 38702062 | 38702381 | 0.723703 | 1.07266 | 1.482 |
| chr9 | 133559416 | 133559772 | 2.26862 | 3.36253 | 1.482 |
| chr16 | 82297070 | 82297562 | 1.72005 | 2.54945 | 1.482 |
| chr14 | 86339050 | 86339532 | 0.618387 | 0.916607 | 1.482 |
| chr17 | 21022959 | 21023271 | 3.00568 | 4.45525 | 1.482 |
| chr16 | 10350665 | 10351029 | 1.97413 | 2.92626 | 1.482 |
| chr7 | 19753374 | 19753561 | 1.20479 | 1.78589 | 1.482 |
| chr2 | 9278746 | 9279173 | 1.63305 | 2.42081 | 1.482 |
| chrX | 48774372 | 48774881 | 2.9952 | 4.44007 | 1.482 |
| chr2 | 238041522 | 238041735 | 1.35412 | 2.00737 | 1.482 |
| chr5 | 136375193 | 136375578 | 0.91522 | 1.35683 | 1.483 |
| chr7 | 69658527 | 69658747 | 0.992155 | 1.4709 | 1.483 |
| chr5 | 67772873 | 67773104 | 0.998171 | 1.47983 | 1.483 |
| chr15 | 80711306 | 80711664 | 1.74856 | 2.59237 | 1.483 |
| chr3 | 2462075 | 2462377 | 0.630229 | 0.934357 | 1.483 |
| chr2 | 11969446 | 11969653 | 1.29611 | 1.92174 | 1.483 |
| chr19 | 36478208 | 36478526 | 0.804328 | 1.19264 | 1.483 |
| chr1 | 3341250 | 3341590 | 2.08034 | 3.08481 | 1.483 |
| chr6 | 51914309 | 51914660 | 1.35408 | 2.00804 | 1.483 |
| chr12 | 17962191 | 17962449 | 0.685175 | 1.01618 | 1.483 |
| chr20 | 23226362 | 23226712 | 2.46962 | 3.66284 | 1.483 |
| chr1 | 26221929 | 26222232 | 2.34178 | 3.47337 | 1.483 |
| chr6 | 74801153 | 74801430 | 1.31637 | 1.95247 | 1.483 |
| chr2 | 191861034 | 191861516 | 2.33578 | 3.46465 | 1.483 |
| chr2 | 168832159 | 168832365 | 0.572894 | 0.849862 | 1.483 |
| chr14 | 25179393 | 25180238 | 1.46834 | 2.17825 | 1.483 |
| chr14 | 21604099 | 21604404 | 1.44677 | 2.14641 | 1.484 |
| chr4 | 6875693 | 6876258 | 1.98785 | 2.94917 | 1.484 |
| chr9 | 124907146 | 124907571 | 1.41369 | 2.09741 | 1.484 |
| chr4 | 178302092 | 178302387 | 1.00639 | 1.49315 | 1.484 |

| | | | | | |
|---|---|---|---|---|---|
| chr12 | 45323250 | 45323461 | 0.777421 | 1.15345 | 1.484 |
| chr6 | 127927335 | 127927599 | 0.840712 | 1.24736 | 1.484 |
| chr9 | 112542151 | 112542497 | 1.0102 | 1.49885 | 1.484 |
| chr8 | 105899048 | 105899284 | 0.594784 | 0.882555 | 1.484 |
| chr11 | 81209407 | 81209741 | 0.730433 | 1.08389 | 1.484 |
| chr14 | 31551157 | 31551574 | 1.55416 | 2.30622 | 1.484 |
| chr2 | 37004946 | 37005215 | 0.782008 | 1.16043 | 1.484 |
| chr20 | 20167692 | 20167970 | 0.946165 | 1.40402 | 1.484 |
| chr19 | 16136453 | 16136794 | 1.11729 | 1.65837 | 1.484 |
| chr12 | 122106615 | 122106981 | 1.51618 | 2.25065 | 1.484 |
| chr8 | 132441704 | 132442030 | 0.630336 | 0.935706 | 1.484 |
| chr20 | 58800043 | 58800452 | 1.36187 | 2.02166 | 1.484 |
| chr5 | 140947094 | 140947456 | 2.65241 | 3.93743 | 1.484 |
| chr1 | 10490289 | 10490615 | 2.27581 | 3.37846 | 1.485 |
| chr22 | 50324640 | 50324845 | 1.77549 | 2.63574 | 1.485 |
| chr11 | 71131521 | 71131820 | 1.89724 | 2.8165 | 1.485 |
| chr2 | 242535119 | 242535353 | 0.947155 | 1.40608 | 1.485 |
| chr3 | 19972498 | 19972795 | 0.640804 | 0.951362 | 1.485 |
| chr7 | 141676055 | 141676412 | 2.78601 | 4.13659 | 1.485 |
| chr2 | 63867447 | 63867688 | 0.786422 | 1.16767 | 1.485 |
| chr8 | 75671312 | 75671522 | 0.730267 | 1.0843 | 1.485 |
| chr10 | 30673726 | 30673961 | 0.848953 | 1.26055 | 1.485 |
| chr3 | 146909737 | 146910286 | 0.719483 | 1.06833 | 1.485 |
| chr12 | 119850986 | 119851381 | 1.8605 | 2.76261 | 1.485 |
| chr9 | 132647991 | 132648293 | 2.38006 | 3.53418 | 1.485 |
| chr14 | 100230769 | 100231067 | 0.58347 | 0.866493 | 1.485 |
| chr10 | 103603103 | 103603605 | 1.96387 | 2.91653 | 1.485 |
| chr1 | 20244496 | 20244801 | 1.9093 | 2.83557 | 1.485 |
| chr17 | 19945688 | 19945903 | 0.648007 | 0.962453 | 1.485 |
| chr5 | 169689350 | 169689594 | 0.53787 | 0.798872 | 1.485 |
| chr12 | 4530695 | 4531028 | 1.79716 | 2.66937 | 1.485 |
| chr15 | 66790051 | 66790549 | 3.64003 | 5.40689 | 1.485 |
| chr1 | 210413224 | 210413436 | 1.20935 | 1.79641 | 1.485 |
| chr20 | 32044928 | 32045397 | 2.45953 | 3.6536 | 1.485 |
| chr11 | 117688483 | 117688778 | 2.11261 | 3.13833 | 1.486 |
| chrX | 135790324 | 135790867 | 1.09979 | 1.63382 | 1.486 |
| chr17 | 79979029 | 79979400 | 1.90111 | 2.82433 | 1.486 |
| chr9 | 36728936 | 36729329 | 2.26099 | 3.35933 | 1.486 |
| chr9 | 140699300 | 140699672 | 0.85097 | 1.26444 | 1.486 |
| chr7 | 1619145 | 1619482 | 2.22934 | 3.31255 | 1.486 |
| chr8 | 13519434 | 13520065 | 1.45858 | 2.1673 | 1.486 |
| chr2 | 59347602 | 59347861 | 0.585979 | 0.870714 | 1.486 |
| chr2 | 163009952 | 163010453 | 1.08716 | 1.61545 | 1.486 |
| chr1 | 10437492 | 10437762 | 1.42667 | 2.12001 | 1.486 |
| chr7 | 128850275 | 128850910 | 1.77016 | 2.63078 | 1.486 |
| chr10 | 16819497 | 16819737 | 1.40201 | 2.08366 | 1.486 |
| chr2 | 60379368 | 60379702 | 1.48997 | 2.21443 | 1.486 |
| chr6 | 2158846 | 2159119 | 2.05299 | 3.05127 | 1.486 |
| chr7 | 69396967 | 69397321 | 1.63792 | 2.43438 | 1.486 |
| chr6 | 217110 | 217296 | 1.46426 | 2.17633 | 1.486 |
| chr3 | 9842315 | 9842553 | 1.25395 | 1.86384 | 1.486 |
| chr15 | 60996403 | 60996657 | 0.816998 | 1.21448 | 1.487 |
| chr11 | 129201795 | 129202009 | 1.84509 | 2.7429 | 1.487 |
| chr9 | 107625578 | 107625793 | 1.56431 | 2.32563 | 1.487 |
| chr10 | 89878102 | 89878419 | 1.03013 | 1.5316 | 1.487 |
| chr14 | 20617608 | 20617896 | 0.87381 | 1.29924 | 1.487 |
| chr15 | 66373432 | 66373745 | 1.06907 | 1.58959 | 1.487 |
| chr2 | 72833046 | 72833268 | 1.52033 | 2.26073 | 1.487 |
| chr11 | 7857868 | 7858164 | 0.63998 | 0.951687 | 1.487 |
| chr10 | 15476037 | 15476410 | 1.30888 | 1.9464 | 1.487 |
| chr1 | 91977055 | 91977268 | 1.27186 | 1.89147 | 1.487 |

| | | | | | |
|---|---|---|---|---|---|
| chr2 | 30153478 | 30153704 | 0.809495 | 1.20387 | 1.487 |
| chr22 | 28447730 | 28448034 | 1.62648 | 2.41925 | 1.487 |
| chr21 | 41758248 | 41758648 | 1.47383 | 2.19226 | 1.487 |
| chr20 | 20298124 | 20298502 | 0.803431 | 1.1951 | 1.488 |
| chr19 | 1503464 | 1503832 | 2.34787 | 3.49264 | 1.488 |
| chr19 | 40671472 | 40671801 | 2.15731 | 3.20948 | 1.488 |
| chr1 | 155145675 | 155146046 | 3.17277 | 4.72022 | 1.488 |
| chr7 | 153356294 | 153356539 | 0.578343 | 0.86042 | 1.488 |
| chr1 | 244960102 | 244960445 | 1.39985 | 2.0827 | 1.488 |
| chr6 | 18122838 | 18123078 | 2.09429 | 3.11592 | 1.488 |
| chr5 | 94933112 | 94933347 | 0.694758 | 1.03373 | 1.488 |
| chr5 | 4741077 | 4741468 | 0.776255 | 1.15504 | 1.488 |
| chr8 | 101692845 | 101693426 | 1.82796 | 2.72014 | 1.488 |
| chr5 | 140904073 | 140904693 | 2.42656 | 3.61097 | 1.488 |
| chr2 | 170789685 | 170789869 | 1.34184 | 1.99681 | 1.488 |
| chr13 | 79832895 | 79833139 | 0.621665 | 0.925194 | 1.488 |
| chr5 | 139015336 | 139015581 | 1.33257 | 1.98323 | 1.488 |
| chr6 | 52264597 | 52264916 | 1.64498 | 2.44851 | 1.488 |
| chr16 | 88764314 | 88764852 | 1.23439 | 1.83738 | 1.488 |
| chr2 | 75691629 | 75692048 | 1.26688 | 1.88574 | 1.488 |
| chr8 | 145181295 | 145181695 | 1.44223 | 2.14676 | 1.489 |
| chr3 | 112238286 | 112238569 | 1.34651 | 2.0043 | 1.489 |
| chr5 | 179528688 | 179529014 | 0.685606 | 1.02054 | 1.489 |
| chr10 | 44185449 | 44186228 | 1.57475 | 2.34409 | 1.489 |
| chr19 | 42905357 | 42906157 | 1.77003 | 2.63493 | 1.489 |
| chr20 | 937058 | 937422 | 2.57541 | 3.83391 | 1.489 |
| chr19 | 44098848 | 44099187 | 2.00067 | 2.97838 | 1.489 |
| chr10 | 96580446 | 96580709 | 0.64479 | 0.959932 | 1.489 |
| chr11 | 75118924 | 75119250 | 1.45491 | 2.16624 | 1.489 |
| chr22 | 46512343 | 46512697 | 1.6029 | 2.38662 | 1.489 |
| chr3 | 25482734 | 25483121 | 2.00827 | 2.9904 | 1.489 |
| chr7 | 100887444 | 100887848 | 2.50338 | 3.72767 | 1.489 |
| chr2 | 132249939 | 132250430 | 1.10746 | 1.64907 | 1.489 |
| chr12 | 108851058 | 108851350 | 1.52647 | 2.27304 | 1.489 |
| chr17 | 8013434 | 8013953 | 2.75655 | 4.10502 | 1.489 |
| chr12 | 70013855 | 70014060 | 0.690309 | 1.02801 | 1.489 |
| chr8 | 58909578 | 58909853 | 2.04122 | 3.04002 | 1.489 |
| chrX | 84364295 | 84364499 | 1.31851 | 1.96391 | 1.489 |
| chr8 | 25027652 | 25028048 | 1.90256 | 2.83386 | 1.490 |
| chr3 | 145439743 | 145440113 | 0.721892 | 1.07527 | 1.490 |
| chr20 | 35710260 | 35710505 | 1.24481 | 1.85422 | 1.490 |
| chr19 | 7716933 | 7717309 | 1.76754 | 2.6329 | 1.490 |
| chrX | 57199288 | 57199526 | 0.634163 | 0.944674 | 1.490 |
| chr10 | 5671826 | 5672152 | 3.19911 | 4.76554 | 1.490 |
| chr11 | 34947628 | 34947916 | 1.1036 | 1.644 | 1.490 |
| chr12 | 27974213 | 27974467 | 1.33937 | 1.99522 | 1.490 |
| chr19 | 31665887 | 31666096 | 0.861962 | 1.28407 | 1.490 |
| chr7 | 113414326 | 113414592 | 1.42278 | 2.11955 | 1.490 |
| chr9 | 93662325 | 93662693 | 0.848537 | 1.2641 | 1.490 |
| chr12 | 130147858 | 130148162 | 0.609295 | 0.907705 | 1.490 |
| chr9 | 80103469 | 80103702 | 1.83413 | 2.73242 | 1.490 |
| chr9 | 19782037 | 19782287 | 0.577283 | 0.860101 | 1.490 |
| chr2 | 65414353 | 65414714 | 2.14503 | 3.19601 | 1.490 |
| chr7 | 83285298 | 83285633 | 2.78315 | 4.14712 | 1.490 |
| chr17 | 10199134 | 10199586 | 1.11182 | 1.65686 | 1.490 |
| chr12 | 66803079 | 66803347 | 1.30302 | 1.9418 | 1.490 |
| chrX | 132910223 | 132910433 | 1.18389 | 1.76431 | 1.490 |
| chr8 | 62475862 | 62476217 | 1.24751 | 1.85918 | 1.490 |
| chr19 | 30232835 | 30233052 | 0.949603 | 1.41528 | 1.490 |
| chr16 | 13274213 | 13274546 | 0.63586 | 0.947706 | 1.490 |
| chr13 | 96728166 | 96728609 | 1.42535 | 2.1245 | 1.491 |

| | | | | | |
|---|---|---|---|---|---|
| chr1 | 210181760 | 210182039 | 0.587418 | 0.875559 | 1.491 |
| chr3 | 10785855 | 10786075 | 0.671223 | 1.0005 | 1.491 |
| chr14 | 69813440 | 69813821 | 2.18513 | 3.25731 | 1.491 |
| chr5 | 37413088 | 37413427 | 2.75163 | 4.10187 | 1.491 |
| chrX | 152073096 | 152073510 | 2.34585 | 3.49717 | 1.491 |
| chr16 | 27709527 | 27709925 | 1.6217 | 2.41767 | 1.491 |
| chr16 | 74025167 | 74025384 | 0.831852 | 1.24019 | 1.491 |
| chr17 | 63382457 | 63382676 | 0.689726 | 1.02831 | 1.491 |
| chr8 | 103598079 | 103598282 | 1.43311 | 2.13662 | 1.491 |
| chr9 | 134353600 | 134354177 | 1.5769 | 2.35104 | 1.491 |
| chr11 | 10427734 | 10427954 | 1.537 | 2.29165 | 1.491 |
| chr7 | 53898974 | 53899317 | 0.613018 | 0.914018 | 1.491 |
| chr1 | 160696660 | 160696926 | 2.03586 | 3.03552 | 1.491 |
| chr16 | 545439 | 545762 | 1.11298 | 1.65957 | 1.491 |
| chr3 | 38570038 | 38570274 | 0.933309 | 1.39169 | 1.491 |
| chr16 | 11404083 | 11404292 | 0.6229 | 0.928838 | 1.491 |
| chr9 | 32378569 | 32378791 | 1.9561 | 2.91686 | 1.491 |
| chr3 | 151157405 | 151157651 | 1.44715 | 2.15795 | 1.491 |
| chr17 | 54526922 | 54527176 | 0.748095 | 1.11558 | 1.491 |
| chr10 | 81903459 | 81903780 | 1.25176 | 1.86671 | 1.491 |
| chr11 | 73115201 | 73115426 | 1.36967 | 2.04278 | 1.491 |
| chrX | 132926438 | 132926861 | 2.38236 | 3.55346 | 1.492 |
| chr19 | 32069219 | 32069517 | 0.777839 | 1.16022 | 1.492 |
| chr3 | 170905582 | 170905826 | 1.15565 | 1.72385 | 1.492 |
| chr12 | 68507410 | 68507751 | 0.886621 | 1.3226 | 1.492 |
| chr10 | 52374318 | 52374589 | 1.04633 | 1.56092 | 1.492 |
| chr17 | 17720242 | 17720467 | 0.903173 | 1.3474 | 1.492 |
| chr5 | 160772685 | 160772970 | 0.595179 | 0.887942 | 1.492 |
| chr12 | 49381804 | 49382250 | 2.45797 | 3.66731 | 1.492 |
| chr5 | 61756340 | 61756621 | 1.5079 | 2.24983 | 1.492 |
| chr12 | 120097122 | 120097468 | 2.20185 | 3.28525 | 1.492 |
| chr2 | 204889245 | 204889601 | 1.43264 | 2.13764 | 1.492 |
| chr14 | 24008847 | 24009191 | 1.78888 | 2.66923 | 1.492 |
| chr8 | 65607608 | 65607865 | 0.699059 | 1.04312 | 1.492 |
| chr15 | 87446969 | 87447298 | 0.696317 | 1.03907 | 1.492 |
| chr6 | 109394679 | 109394912 | 1.07854 | 1.60945 | 1.492 |
| chr1 | 14286898 | 14287229 | 2.40964 | 3.59585 | 1.492 |
| chr8 | 63117323 | 63117660 | 0.836309 | 1.24805 | 1.492 |
| chr15 | 98170278 | 98170610 | 0.714311 | 1.06601 | 1.492 |
| chr16 | 30953036 | 30953771 | 1.72559 | 2.57521 | 1.492 |
| chr17 | 33887192 | 33887527 | 0.632727 | 0.944266 | 1.492 |
| chr2 | 25198390 | 25198698 | 1.64814 | 2.45969 | 1.492 |
| chr11 | 14928967 | 14929606 | 1.19617 | 1.78518 | 1.492 |
| chr2 | 69941822 | 69942066 | 3.13303 | 4.67576 | 1.492 |
| chr15 | 92686955 | 92687347 | 0.688851 | 1.02806 | 1.492 |
| chr2 | 14940358 | 14940568 | 0.745309 | 1.11237 | 1.493 |
| chr19 | 35558020 | 35558416 | 2.80196 | 4.18199 | 1.493 |
| chr7 | 120747911 | 120748274 | 1.98065 | 2.95616 | 1.493 |
| chr11 | 76838103 | 76838726 | 3.69867 | 5.52039 | 1.493 |
| chr2 | 220406277 | 220406884 | 4.12756 | 6.16058 | 1.493 |
| chr7 | 1216653 | 1216978 | 1.83428 | 2.73778 | 1.493 |
| chr1 | 89343145 | 89343435 | 1.04437 | 1.5589 | 1.493 |
| chr8 | 145781110 | 145781423 | 0.759271 | 1.1334 | 1.493 |
| chr19 | 3472986 | 3473334 | 1.57074 | 2.34479 | 1.493 |
| chr1 | 177582125 | 177582594 | 0.637128 | 0.951128 | 1.493 |
| chr10 | 29806137 | 29806690 | 1.73008 | 2.58275 | 1.493 |
| chr7 | 139703763 | 139704044 | 1.48751 | 2.22063 | 1.493 |
| chr11 | 72296742 | 72297159 | 2.71027 | 4.04616 | 1.493 |
| chr14 | 55904858 | 55905183 | 1.22838 | 1.83389 | 1.493 |
| chr14 | 74967952 | 74968331 | 1.41709 | 2.11574 | 1.493 |
| chr9 | 112980005 | 112980246 | 1.50619 | 2.24877 | 1.493 |

| | | | | | |
|---|---|---|---|---|---|
| chr5 | 99175925 | 99176303 | 0.769507 | 1.1489 | 1.493 |
| chr19 | 54984790 | 54985054 | 0.636808 | 0.95086 | 1.493 |
| chr17 | 17715489 | 17716081 | 2.63643 | 3.93679 | 1.493 |
| chr19 | 29483703 | 29483915 | 0.728748 | 1.0882 | 1.493 |
| chr19 | 31520909 | 31521152 | 0.799258 | 1.19355 | 1.493 |
| chr2 | 218578229 | 218578534 | 3.51406 | 5.24763 | 1.493 |
| chr8 | 14917345 | 14917568 | 0.513192 | 0.766369 | 1.493 |
| chr11 | 4093997 | 4094206 | 1.68413 | 2.51507 | 1.493 |
| chr20 | 24898965 | 24899565 | 1.44884 | 2.16374 | 1.493 |
| chr20 | 23181617 | 23181929 | 1.39381 | 2.08165 | 1.494 |
| chr5 | 10019149 | 10019387 | 0.685757 | 1.0242 | 1.494 |
| chr2 | 210842450 | 210842694 | 0.965043 | 1.44135 | 1.494 |
| chr3 | 5630273 | 5630499 | 0.70133 | 1.04755 | 1.494 |
| chr17 | 50496776 | 50497055 | 0.829536 | 1.23907 | 1.494 |
| chr5 | 133863284 | 133863606 | 1.75288 | 2.61827 | 1.494 |
| chr15 | 101173225 | 101173500 | 1.9253 | 2.87585 | 1.494 |
| chr1 | 86483954 | 86484315 | 1.50339 | 2.24566 | 1.494 |
| chr20 | 5429150 | 5429464 | 1.90354 | 2.8435 | 1.494 |
| chr12 | 3143337 | 3143681 | 1.77674 | 2.65413 | 1.494 |
| chr19 | 45931778 | 45932376 | 1.20105 | 1.79415 | 1.494 |
| chr17 | 45741682 | 45742042 | 2.40101 | 3.58679 | 1.494 |
| chr2 | 21355894 | 21356196 | 0.692927 | 1.03514 | 1.494 |
| chr16 | 59933909 | 59934110 | 1.39233 | 2.08002 | 1.494 |
| chr19 | 8067200 | 8067676 | 2.57801 | 3.85138 | 1.494 |
| chr8 | 66557224 | 66557528 | 1.91914 | 2.86717 | 1.494 |
| chr9 | 100745421 | 100745816 | 2.37529 | 3.54874 | 1.494 |
| chr16 | 6732229 | 6732706 | 1.07933 | 1.61261 | 1.494 |
| chr17 | 33734509 | 33734787 | 0.931704 | 1.39223 | 1.494 |
| chr8 | 62238944 | 62239203 | 0.702936 | 1.05041 | 1.494 |
| chr12 | 109823383 | 109823593 | 1.07383 | 1.6048 | 1.494 |
| chr12 | 102063920 | 102064172 | 1.30975 | 1.9574 | 1.494 |
| chr13 | 47405798 | 47406146 | 1.23554 | 1.8465 | 1.494 |
| chr17 | 1179462 | 1179870 | 2.64603 | 3.95449 | 1.495 |
| chr1 | 159789613 | 159789986 | 0.488636 | 0.730291 | 1.495 |
| chr19 | 2019542 | 2019904 | 1.98104 | 2.96081 | 1.495 |
| chr15 | 94625191 | 94625418 | 0.633526 | 0.946882 | 1.495 |
| chrX | 17064649 | 17064969 | 1.74513 | 2.60834 | 1.495 |
| chr16 | 30905419 | 30905717 | 1.96091 | 2.93088 | 1.495 |
| chr1 | 41958049 | 41958298 | 1.89951 | 2.83914 | 1.495 |
| chr15 | 90216323 | 90216723 | 1.75409 | 2.62181 | 1.495 |
| chr2 | 70882390 | 70882668 | 1.70637 | 2.55055 | 1.495 |
| chr2 | 97215693 | 97216148 | 2.23604 | 3.34248 | 1.495 |
| chr15 | 24233377 | 24233616 | 0.650113 | 0.971854 | 1.495 |
| chr2 | 197318344 | 197318588 | 1.19878 | 1.79206 | 1.495 |
| chr2 | 75904428 | 75904815 | 2.96275 | 4.4291 | 1.495 |
| chr7 | 6595842 | 6596118 | 0.892907 | 1.33484 | 1.495 |
| chr8 | 140605350 | 140605576 | 0.761812 | 1.13886 | 1.495 |
| chr6 | 153552199 | 153552556 | 2.14348 | 3.20454 | 1.495 |
| chr8 | 13220554 | 13220979 | 0.500695 | 0.74857 | 1.495 |
| chr11 | 45124656 | 45124991 | 2.55089 | 3.81397 | 1.495 |
| chr14 | 52729282 | 52729591 | 0.604512 | 0.903908 | 1.495 |
| chr10 | 12306169 | 12306527 | 2.68463 | 4.01448 | 1.495 |
| chr11 | 73087124 | 73087651 | 1.90586 | 2.85002 | 1.495 |
| chr7 | 55580413 | 55580733 | 0.600869 | 0.89857 | 1.495 |
| chr1 | 185650243 | 185650526 | 1.66587 | 2.49129 | 1.495 |
| chr7 | 157030928 | 157031135 | 1.2305 | 1.84036 | 1.496 |
| chr2 | 17050515 | 17051248 | 0.711579 | 1.06426 | 1.496 |
| chr9 | 130073508 | 130073747 | 1.49376 | 2.23423 | 1.496 |
| chr12 | 40510264 | 40510484 | 0.693046 | 1.03667 | 1.496 |
| chr19 | 49601056 | 49601572 | 2.16632 | 3.24046 | 1.496 |
| chr2 | 25500107 | 25500511 | 3.0978 | 4.63391 | 1.496 |

| | | | | | |
|---|---|---|---|---|---|
| chr7 | 26051895 | 26052132 | 0.624404 | 0.934025 | 1.496 |
| chr22 | 21978297 | 21978717 | 1.93089 | 2.88838 | 1.496 |
| chr9 | 78481324 | 78481579 | 1.17613 | 1.75938 | 1.496 |
| chr2 | 88459203 | 88459898 | 1.54827 | 2.31615 | 1.496 |
| chr8 | 117460216 | 117460469 | 1.49257 | 2.23294 | 1.496 |
| chr5 | 153436531 | 153436752 | 0.587238 | 0.878589 | 1.496 |
| chr14 | 59427831 | 59428060 | 0.563242 | 0.842731 | 1.496 |
| chr5 | 164456587 | 164456812 | 0.732705 | 1.09629 | 1.496 |
| chr1 | 234918977 | 234919208 | 0.666901 | 0.997854 | 1.496 |
| chr21 | 40813521 | 40813775 | 1.4774 | 2.21068 | 1.496 |
| chr2 | 43021534 | 43021897 | 1.05477 | 1.57833 | 1.496 |
| chr14 | 81421101 | 81421450 | 2.46811 | 3.69325 | 1.496 |
| chr20 | 20840818 | 20841172 | 2.43137 | 3.63839 | 1.496 |
| chr8 | 73232082 | 73232314 | 0.991106 | 1.48315 | 1.496 |
| chr1 | 172688773 | 172689160 | 1.9472 | 2.91393 | 1.496 |
| chr1 | 46806316 | 46806543 | 2.09221 | 3.13128 | 1.497 |
| chr5 | 43631453 | 43631724 | 1.46375 | 2.19073 | 1.497 |
| chr8 | 6687557 | 6688011 | 2.03423 | 3.04458 | 1.497 |
| chr15 | 97982641 | 97983159 | 0.657293 | 0.983764 | 1.497 |
| chr2 | 15010312 | 15010547 | 0.681569 | 1.0201 | 1.497 |
| chr14 | 23039252 | 23039587 | 1.5414 | 2.30701 | 1.497 |
| chr16 | 23814910 | 23815193 | 1.40712 | 2.1061 | 1.497 |
| chr5 | 78182516 | 78182857 | 1.3518 | 2.02331 | 1.497 |
| chr21 | 44596768 | 44597060 | 2.22994 | 3.33773 | 1.497 |
| chr14 | 36915703 | 36915934 | 0.534881 | 0.800616 | 1.497 |
| chr18 | 42258939 | 42259207 | 1.37548 | 2.05883 | 1.497 |
| chr5 | 134301030 | 134301344 | 0.657147 | 0.983749 | 1.497 |
| chr10 | 129301033 | 129301293 | 0.606345 | 0.907733 | 1.497 |
| chr1 | 208159628 | 208159874 | 0.966957 | 1.44763 | 1.497 |
| chr2 | 118979647 | 118979880 | 1.24281 | 1.86071 | 1.497 |
| chr7 | 154380746 | 154380949 | 0.593982 | 0.8893 | 1.497 |
| chr20 | 21246665 | 21246873 | 0.614066 | 0.9194 | 1.497 |
| chr20 | 38144614 | 38144880 | 0.642699 | 0.962308 | 1.497 |
| chr5 | 152703075 | 152703286 | 0.537946 | 0.805472 | 1.497 |
| chr2 | 11586652 | 11587237 | 1.9857 | 2.97322 | 1.497 |
| chr1 | 147350915 | 147351181 | 0.689083 | 1.03189 | 1.497 |
| chr14 | 103482385 | 103482645 | 0.571378 | 0.855699 | 1.498 |
| chr20 | 62013188 | 62013581 | 2.6163 | 3.91819 | 1.498 |
| chr15 | 51695321 | 51695898 | 1.27666 | 1.91207 | 1.498 |
| chr7 | 54608450 | 54608696 | 0.679469 | 1.01767 | 1.498 |
| chr2 | 147070571 | 147070813 | 0.660836 | 0.989765 | 1.498 |
| chr19 | 47045969 | 47046383 | 1.40832 | 2.10935 | 1.498 |
| chr6 | 133284187 | 133284423 | 1.96797 | 2.94758 | 1.498 |
| chr10 | 63656771 | 63657148 | 1.47909 | 2.21539 | 1.498 |
| chrX | 153536587 | 153537022 | 1.67561 | 2.50978 | 1.498 |
| chr19 | 42502560 | 42503406 | 1.97306 | 2.9554 | 1.498 |
| chr2 | 238318748 | 238318972 | 1.15088 | 1.7239 | 1.498 |
| chr21 | 35815853 | 35816331 | 1.68162 | 2.51891 | 1.498 |
| chr16 | 63036061 | 63036269 | 0.732518 | 1.09731 | 1.498 |
| chr2 | 176983710 | 176984051 | 2.05664 | 3.081 | 1.498 |
| chr7 | 39785769 | 39786112 | 2.32378 | 3.4812 | 1.498 |
| chr6 | 144537204 | 144537658 | 1.52527 | 2.28501 | 1.498 |
| chr19 | 35075180 | 35075646 | 0.949682 | 1.42277 | 1.498 |
| chr7 | 1546624 | 1546886 | 1.09591 | 1.64186 | 1.498 |
| chr5 | 98432921 | 98433195 | 0.851657 | 1.27599 | 1.498 |
| chr2 | 199174465 | 199174715 | 2.53164 | 3.7931 | 1.498 |
| chr15 | 69168258 | 69168504 | 2.90702 | 4.35564 | 1.498 |
| chr2 | 36001365 | 36001667 | 0.552577 | 0.827957 | 1.498 |
| chr12 | 76980613 | 76980926 | 1.44651 | 2.1674 | 1.498 |
| chr8 | 140225505 | 140225985 | 2.84375 | 4.2613 | 1.498 |
| chr10 | 130011154 | 130011399 | 0.620454 | 0.92989 | 1.499 |

| | | | | | |
|---|---|---|---|---|---|
| chr22 | 50118448 | 50118819 | 1.22604 | 1.83757 | 1.499 |
| chr5 | 122707346 | 122707579 | 1.63005 | 2.44311 | 1.499 |
| chr17 | 49731334 | 49731569 | 0.758803 | 1.1373 | 1.499 |
| chr1 | 204186664 | 204186870 | 0.754093 | 1.13031 | 1.499 |
| chr11 | 4923515 | 4923812 | 0.524239 | 0.785827 | 1.499 |
| chr17 | 10517809 | 10518226 | 1.86762 | 2.79957 | 1.499 |
| chr19 | 5805915 | 5806214 | 1.69334 | 2.53834 | 1.499 |
| chr3 | 182555758 | 182556001 | 0.830704 | 1.24525 | 1.499 |
| chr5 | 5598931 | 5599224 | 0.459515 | 0.688892 | 1.499 |
| chr14 | 25881229 | 25881735 | 0.76717 | 1.15015 | 1.499 |
| chr18 | 56737720 | 56737980 | 1.50618 | 2.25808 | 1.499 |
| chr3 | 1529383 | 1529763 | 0.782606 | 1.17333 | 1.499 |
| chr15 | 93290284 | 93290548 | 1.06151 | 1.5915 | 1.499 |
| chr17 | 76246899 | 76247490 | 4.07731 | 6.11302 | 1.499 |
| chr17 | 47970933 | 47971139 | 0.944929 | 1.41685 | 1.499 |
| chr12 | 62198246 | 62198492 | 0.666823 | 0.999887 | 1.499 |
| chr12 | 112071527 | 112071873 | 0.573129 | 0.859408 | 1.500 |
| chr20 | 54823040 | 54823441 | 1.73376 | 2.59987 | 1.500 |
| chr12 | 4320636 | 4320868 | 0.771722 | 1.15728 | 1.500 |
| chr8 | 99370315 | 99370708 | 2.54701 | 3.81952 | 1.500 |
| chr14 | 62629761 | 62630027 | 1.47728 | 2.21553 | 1.500 |
| chr1 | 32053415 | 32053680 | 1.75939 | 2.63869 | 1.500 |
| chr19 | 8808421 | 8808824 | 1.38272 | 2.07394 | 1.500 |
| chr1 | 40870609 | 40870871 | 1.6099 | 2.41477 | 1.500 |
| chr12 | 49189653 | 49189977 | 1.75141 | 2.62726 | 1.500 |
| chr5 | 1008655 | 1009080 | 1.51296 | 2.26966 | 1.500 |
| chr22 | 38213618 | 38214053 | 2.72997 | 4.0958 | 1.500 |
| chr11 | 75095550 | 75095872 | 2.81695 | 4.22637 | 1.500 |
| chr20 | 20845100 | 20845395 | 1.24888 | 1.87374 | 1.500 |
| chrX | 12098460 | 12098664 | 0.981046 | 1.47192 | 1.500 |
| chr11 | 12910330 | 12910622 | 1.51183 | 2.2683 | 1.500 |
| chr5 | 119342966 | 119343249 | 0.543844 | 0.815992 | 1.500 |
| chr1 | 46998494 | 46998801 | 2.13342 | 3.20117 | 1.500 |
| chr17 | 54825055 | 54825268 | 0.590154 | 0.885568 | 1.501 |
| chr10 | 10099457 | 10099731 | 0.494278 | 0.741706 | 1.501 |
| chr10 | 73467885 | 73468330 | 1.9798 | 2.97095 | 1.501 |
| chr6 | 151007079 | 151007348 | 1.68053 | 2.5219 | 1.501 |
| chr6 | 141998795 | 141999000 | 1.2172 | 1.82662 | 1.501 |
| chr2 | 74702084 | 74702409 | 1.28384 | 1.92663 | 1.501 |
| chr6 | 103911611 | 103911925 | 0.52376 | 0.785998 | 1.501 |
| chr15 | 45507739 | 45507943 | 0.846703 | 1.27069 | 1.501 |
| chr18 | 64227313 | 64227639 | 0.878288 | 1.31814 | 1.501 |
| chr3 | 24833736 | 24834130 | 0.893268 | 1.34063 | 1.501 |
| chr1 | 216194240 | 216194537 | 0.632099 | 0.948703 | 1.501 |
| chr9 | 31412215 | 31412454 | 0.489283 | 0.734362 | 1.501 |
| chr15 | 90195927 | 90196129 | 1.04003 | 1.56101 | 1.501 |
| chr9 | 82859760 | 82860007 | 0.664333 | 0.99713 | 1.501 |
| chr2 | 95989049 | 95989409 | 1.40432 | 2.10783 | 1.501 |
| chr15 | 59147868 | 59148077 | 0.487827 | 0.732246 | 1.501 |
| chr19 | 34611984 | 34612324 | 2.31655 | 3.47735 | 1.501 |
| chr12 | 118558750 | 118559006 | 2.29894 | 3.45097 | 1.501 |
| chr5 | 59109598 | 59110126 | 1.48913 | 2.23536 | 1.501 |
| chr17 | 41427904 | 41428187 | 0.775329 | 1.16387 | 1.501 |
| chr3 | 108769503 | 108769781 | 1.79409 | 2.69322 | 1.501 |
| chr8 | 103209712 | 103209973 | 2.38552 | 3.58107 | 1.501 |
| chr6 | 113627800 | 113628040 | 1.36216 | 2.04486 | 1.501 |
| chr2 | 214018484 | 214018709 | 1.57829 | 2.36936 | 1.501 |
| chr7 | 100289199 | 100289690 | 2.54398 | 3.81909 | 1.501 |
| chr10 | 69280650 | 69281001 | 1.22207 | 1.83467 | 1.501 |
| chr12 | 30287320 | 30287544 | 0.900887 | 1.35254 | 1.501 |
| chr17 | 28134630 | 28135157 | 1.62788 | 2.44406 | 1.501 |

| | | | | | |
|---|---|---|---|---|---|
| chr7 | 85715652 | 85715878 | 0.482748 | 0.724831 | 1.501 |
| chr11 | 64828614 | 64828911 | 1.15369 | 1.73229 | 1.502 |
| chr16 | 50839793 | 50841034 | 1.59484 | 2.39469 | 1.502 |
| chr5 | 8099470 | 8099780 | 0.515995 | 0.774809 | 1.502 |
| chr15 | 27243539 | 27243746 | 0.824742 | 1.23844 | 1.502 |
| chrX | 10936171 | 10936383 | 0.590755 | 0.887108 | 1.502 |
| chr15 | 98452592 | 98452854 | 1.77615 | 2.66725 | 1.502 |
| chr3 | 32060719 | 32060912 | 1.12927 | 1.69582 | 1.502 |
| chr2 | 161920222 | 161920523 | 1.7232 | 2.58781 | 1.502 |
| chr19 | 48339603 | 48339849 | 0.793197 | 1.1912 | 1.502 |
| chr14 | 20346248 | 20346518 | 1.33489 | 2.00475 | 1.502 |
| chr12 | 89804231 | 89804500 | 0.452769 | 0.680026 | 1.502 |
| chr11 | 63913486 | 63914073 | 2.29051 | 3.44021 | 1.502 |
| chr8 | 25333092 | 25333564 | 0.961316 | 1.44386 | 1.502 |
| chr9 | 93039187 | 93039430 | 0.739865 | 1.11129 | 1.502 |
| chr12 | 124292827 | 124293037 | 1.30626 | 1.96213 | 1.502 |
| chr20 | 58135678 | 58136183 | 1.18812 | 1.7847 | 1.502 |
| chr2 | 103089441 | 103089699 | 1.53571 | 2.30724 | 1.502 |
| chr5 | 111907932 | 111908147 | 0.56655 | 0.851187 | 1.502 |
| chr1 | 18249421 | 18249617 | 1.63174 | 2.45158 | 1.502 |
| chr1 | 45484384 | 45484672 | 1.79251 | 2.69318 | 1.502 |
| chr2 | 185177833 | 185178042 | 0.531812 | 0.799091 | 1.503 |
| chr16 | 69029169 | 69029395 | 1.00871 | 1.51568 | 1.503 |
| chr8 | 38268124 | 38268478 | 1.79857 | 2.70259 | 1.503 |
| chr2 | 20842142 | 20842580 | 2.8351 | 4.26023 | 1.503 |
| chr8 | 42269860 | 42270182 | 2.25763 | 3.39249 | 1.503 |
| chr10 | 121302817 | 121303227 | 1.64525 | 2.47252 | 1.503 |
| chr19 | 18561771 | 18562146 | 2.58096 | 3.87871 | 1.503 |
| chr5 | 156693242 | 156693489 | 0.821308 | 1.2343 | 1.503 |
| chr10 | 118927463 | 118928068 | 2.18548 | 3.28456 | 1.503 |
| chr19 | 46201856 | 46202109 | 0.707785 | 1.0638 | 1.503 |
| chr20 | 9883581 | 9884210 | 1.61617 | 2.42937 | 1.503 |
| chr2 | 8833314 | 8833793 | 2.57036 | 3.86368 | 1.503 |
| chr5 | 79070544 | 79070822 | 0.574001 | 0.862829 | 1.503 |
| chrX | 119014787 | 119015153 | 1.4411 | 2.16623 | 1.503 |
| chr17 | 20614750 | 20614993 | 0.65406 | 0.983326 | 1.503 |
| chr2 | 121490750 | 121490974 | 0.53596 | 0.805781 | 1.503 |
| chr11 | 9168397 | 9168681 | 0.683701 | 1.02795 | 1.504 |
| chr22 | 24191656 | 24192120 | 2.80877 | 4.2231 | 1.504 |
| chr6 | 33668269 | 33668570 | 2.01611 | 3.03133 | 1.504 |
| chr19 | 506691 | 507068 | 2.27871 | 3.42627 | 1.504 |
| chr2 | 119639197 | 119639464 | 1.68575 | 2.5347 | 1.504 |
| chr6 | 37019186 | 37019665 | 2.07655 | 3.12232 | 1.504 |
| chr8 | 132871887 | 132872126 | 1.15174 | 1.73177 | 1.504 |
| chr9 | 113430071 | 113430363 | 0.609253 | 0.916078 | 1.504 |
| chr15 | 83663964 | 83664235 | 0.75715 | 1.13847 | 1.504 |
| chr7 | 73280428 | 73280741 | 1.22358 | 1.83981 | 1.504 |
| chr1 | 209866294 | 209866602 | 1.61281 | 2.42512 | 1.504 |
| chr1 | 175857059 | 175857777 | 1.86636 | 2.8064 | 1.504 |
| chr1 | 54147785 | 54148126 | 1.0362 | 1.55818 | 1.504 |
| chr1 | 206234403 | 206234727 | 1.84608 | 2.77609 | 1.504 |
| chr5 | 171057132 | 171057389 | 1.27214 | 1.91305 | 1.504 |
| chr8 | 85461776 | 85462062 | 0.821801 | 1.2359 | 1.504 |
| chr5 | 36282824 | 36283059 | 0.87777 | 1.32009 | 1.504 |
| chr12 | 29970480 | 29970761 | 0.567707 | 0.853814 | 1.504 |
| chr14 | 52445835 | 52446262 | 2.3022 | 3.46246 | 1.504 |
| chr15 | 51628015 | 51628448 | 0.763563 | 1.14839 | 1.504 |
| chr19 | 32100306 | 32100644 | 0.691733 | 1.04039 | 1.504 |
| chr3 | 38045915 | 38046304 | 2.07295 | 3.1178 | 1.504 |
| chr8 | 36115956 | 36116173 | 0.705274 | 1.06077 | 1.504 |
| chrX | 153943213 | 153943559 | 1.28762 | 1.93666 | 1.504 |

| | | | | | |
|---|---|---|---|---|---|
| chr12 | 124229379 | 124229631 | 1.23234 | 1.85352 | 1.504 |
| chr12 | 81947246 | 81947510 | 0.613572 | 0.922878 | 1.504 |
| chr19 | 48014170 | 48014399 | 0.964328 | 1.45055 | 1.504 |
| chr17 | 46026143 | 46026968 | 1.74978 | 2.63206 | 1.504 |
| chr6 | 33670188 | 33670413 | 1.00173 | 1.50684 | 1.504 |
| chr1 | 208267129 | 208267478 | 2.91872 | 4.39057 | 1.504 |
| chr6 | 138912084 | 138912317 | 1.66105 | 2.49869 | 1.504 |
| chr14 | 91064493 | 91064732 | 0.666844 | 1.00315 | 1.504 |
| chr11 | 126180383 | 126180725 | 1.83266 | 2.75695 | 1.504 |
| chr17 | 69080365 | 69080627 | 1.16177 | 1.74781 | 1.504 |
| chr17 | 62398451 | 62398833 | 1.72703 | 2.59829 | 1.504 |
| chr3 | 28932755 | 28933095 | 0.565172 | 0.850288 | 1.504 |
| chr17 | 12084065 | 12084278 | 1.66858 | 2.51056 | 1.505 |
| chr3 | 142856602 | 142856812 | 0.4828 | 0.726432 | 1.505 |
| chr3 | 158257303 | 158257516 | 1.29017 | 1.94128 | 1.505 |
| chr10 | 129362272 | 129362559 | 2.05183 | 3.08772 | 1.505 |
| chr15 | 94082247 | 94082677 | 0.764302 | 1.15018 | 1.505 |
| chr1 | 239624916 | 239625132 | 0.686551 | 1.03319 | 1.505 |
| chr9 | 134615426 | 134615795 | 7.11834 | 10.7125 | 1.505 |
| chr2 | 35056108 | 35056369 | 0.702697 | 1.05763 | 1.505 |
| chr19 | 32118689 | 32118937 | 0.703148 | 1.05835 | 1.505 |
| chr6 | 36062995 | 36063364 | 1.65028 | 2.48394 | 1.505 |
| chr8 | 11271211 | 11271547 | 1.9522 | 2.93839 | 1.505 |
| chr2 | 129512936 | 129513255 | 2.4993 | 3.76197 | 1.505 |
| chr2 | 58133343 | 58133595 | 2.22088 | 3.34291 | 1.505 |
| chr6 | 86632688 | 86633087 | 1.76894 | 2.66269 | 1.505 |
| chr19 | 3126020 | 3126372 | 2.23424 | 3.36313 | 1.505 |
| chr5 | 80007847 | 80008087 | 1.5686 | 2.3613 | 1.505 |
| chr16 | 46942804 | 46943141 | 1.76188 | 2.65236 | 1.505 |
| chr2 | 3736290 | 3736522 | 0.740411 | 1.11463 | 1.505 |
| chr11 | 17640384 | 17640871 | 0.52027 | 0.783245 | 1.505 |
| chr2 | 219741348 | 219741802 | 2.30442 | 3.4695 | 1.506 |
| chr10 | 6130538 | 6131059 | 4.22749 | 6.3652 | 1.506 |
| chr1 | 30176504 | 30176711 | 1.81655 | 2.73516 | 1.506 |
| chr16 | 19421934 | 19422145 | 1.95959 | 2.95053 | 1.506 |
| chr19 | 49273235 | 49273563 | 0.657983 | 0.990766 | 1.506 |
| chr5 | 143347637 | 143347957 | 1.15096 | 1.73309 | 1.506 |
| chr4 | 99666550 | 99666843 | 1.38444 | 2.08469 | 1.506 |
| chr2 | 18568647 | 18569312 | 1.4602 | 2.19878 | 1.506 |
| chr20 | 6746192 | 6746428 | 1.34817 | 2.03011 | 1.506 |
| chr14 | 60220346 | 60220630 | 0.554589 | 0.835156 | 1.506 |
| chr14 | 86504735 | 86504994 | 0.983199 | 1.48064 | 1.506 |
| chr2 | 192029905 | 192030146 | 1.05361 | 1.58667 | 1.506 |
| chr22 | 46390748 | 46391064 | 2.05948 | 3.10146 | 1.506 |
| chr7 | 33282919 | 33283207 | 1.31193 | 1.97569 | 1.506 |
| chr8 | 41003935 | 41004209 | 1.64244 | 2.47341 | 1.506 |
| chr7 | 127197746 | 127198100 | 0.62184 | 0.936499 | 1.506 |
| chr6 | 97945723 | 97945936 | 1.1818 | 1.7799 | 1.506 |
| chr2 | 238653614 | 238654011 | 2.24445 | 3.38046 | 1.506 |
| chr9 | 76543282 | 76543631 | 0.689289 | 1.0382 | 1.506 |
| chr2 | 23083594 | 23083834 | 0.512614 | 0.772101 | 1.506 |
| chr8 | 26647512 | 26647760 | 0.642682 | 0.968018 | 1.506 |
| chr1 | 212872291 | 212872901 | 1.15896 | 1.74577 | 1.506 |
| chr9 | 87011973 | 87012300 | 1.84137 | 2.77377 | 1.506 |
| chr10 | 69615382 | 69615734 | 1.91034 | 2.87774 | 1.506 |
| chr2 | 188364737 | 188364977 | 1.68444 | 2.53747 | 1.506 |
| chr12 | 25021937 | 25022265 | 0.892318 | 1.34425 | 1.506 |
| chr17 | 2729458 | 2729730 | 1.60661 | 2.42041 | 1.507 |
| chr5 | 117527343 | 117527576 | 0.595405 | 0.896995 | 1.507 |
| chr16 | 27931889 | 27932141 | 1.07693 | 1.62247 | 1.507 |
| chr11 | 78822666 | 78822874 | 0.578111 | 0.87097 | 1.507 |

| | | | | | |
|---|---|---|---|---|---|
| chr5 | 152563442 | 152563733 | 0.454231 | 0.684346 | 1.507 |
| chr5 | 159701480 | 159701789 | 0.812431 | 1.22403 | 1.507 |
| chr19 | 7934073 | 7934416 | 1.50897 | 2.27348 | 1.507 |
| chr17 | 4524276 | 4524650 | 0.718689 | 1.08284 | 1.507 |
| chr21 | 24952349 | 24952582 | 0.591273 | 0.890867 | 1.507 |
| chr12 | 97160100 | 97160517 | 0.613419 | 0.924251 | 1.507 |
| chr10 | 32458426 | 32458817 | 0.739628 | 1.11444 | 1.507 |
| chr2 | 5901232 | 5901435 | 0.554094 | 0.834908 | 1.507 |
| chrX | 130961991 | 130962213 | 1.09451 | 1.64924 | 1.507 |
| chr2 | 127534929 | 127535341 | 2.03389 | 3.06493 | 1.507 |
| chr10 | 5136907 | 5137119 | 1.01676 | 1.53226 | 1.507 |
| chr2 | 50944862 | 50945111 | 0.682995 | 1.0293 | 1.507 |
| chr10 | 15721215 | 15721501 | 0.517106 | 0.779319 | 1.507 |
| chr5 | 51635497 | 51635714 | 0.613341 | 0.924357 | 1.507 |
| chr4 | 187255332 | 187255593 | 0.696947 | 1.05036 | 1.507 |
| chr1 | 201317306 | 201317565 | 1.91572 | 2.88733 | 1.507 |
| chr17 | 14591581 | 14591786 | 1.19566 | 1.80212 | 1.507 |
| chr9 | 128321678 | 128321936 | 0.678772 | 1.02306 | 1.507 |
| chr20 | 13189383 | 13189670 | 0.944077 | 1.423 | 1.507 |
| chr11 | 18137079 | 18137594 | 1.8204 | 2.74393 | 1.507 |
| chr2 | 68509999 | 68510280 | 1.95676 | 2.94951 | 1.507 |
| chr15 | 86438796 | 86439122 | 0.665874 | 1.00372 | 1.507 |
| chr22 | 35746685 | 35747019 | 1.42732 | 2.15158 | 1.507 |
| chr17 | 45967317 | 45967718 | 3.19314 | 4.81371 | 1.508 |
| chr12 | 49483582 | 49483945 | 1.75011 | 2.63836 | 1.508 |
| chrX | 70349063 | 70349347 | 1.78222 | 2.68683 | 1.508 |
| chr13 | 25201291 | 25201568 | 2.40845 | 3.63103 | 1.508 |
| chr17 | 1358636 | 1359109 | 1.09835 | 1.65591 | 1.508 |
| chr3 | 35038566 | 35038782 | 0.633163 | 0.954573 | 1.508 |
| chr10 | 67940558 | 67940806 | 1.131 | 1.70518 | 1.508 |
| chr17 | 43226498 | 43226903 | 2.74019 | 4.13135 | 1.508 |
| chr2 | 240960926 | 240961265 | 2.76884 | 4.1746 | 1.508 |
| chr3 | 15511932 | 15512150 | 0.971634 | 1.46495 | 1.508 |
| chr10 | 78744567 | 78744912 | 2.55957 | 3.85914 | 1.508 |
| chr10 | 131687357 | 131687822 | 1.56257 | 2.35594 | 1.508 |
| chr20 | 34207484 | 34207793 | 1.6203 | 2.44314 | 1.508 |
| chr10 | 103991446 | 103991901 | 2.62755 | 3.96229 | 1.508 |
| chr15 | 73626460 | 73626733 | 0.748667 | 1.12907 | 1.508 |
| chr1 | 238027618 | 238028004 | 2.28985 | 3.45351 | 1.508 |
| chr15 | 75335164 | 75335526 | 2.0508 | 3.09303 | 1.508 |
| chr2 | 232578551 | 232578933 | 2.42547 | 3.65844 | 1.508 |
| chr2 | 85954957 | 85955217 | 0.725313 | 1.09406 | 1.508 |
| chr19 | 31173578 | 31174213 | 1.78427 | 2.69141 | 1.508 |
| chr7 | 65670219 | 65670463 | 1.39651 | 2.10652 | 1.508 |
| chr10 | 74777826 | 74778371 | 0.620135 | 0.935448 | 1.508 |
| chr2 | 62022102 | 62022413 | 0.604736 | 0.912275 | 1.509 |
| chr9 | 35736094 | 35736463 | 1.96726 | 2.96783 | 1.509 |
| chr2 | 113837631 | 113838029 | 1.73577 | 2.61871 | 1.509 |
| chr5 | 105457659 | 105458132 | 1.66433 | 2.51114 | 1.509 |
| chrX | 16844372 | 16844606 | 0.927974 | 1.40025 | 1.509 |
| chr2 | 235197632 | 235197915 | 1.45528 | 2.19598 | 1.509 |
| chr1 | 65486583 | 65486805 | 1.33977 | 2.02181 | 1.509 |
| chr2 | 201827901 | 201828239 | 1.44697 | 2.18363 | 1.509 |
| chr8 | 50396594 | 50396798 | 0.544872 | 0.822301 | 1.509 |
| chr3 | 34645288 | 34645513 | 0.739412 | 1.11593 | 1.509 |
| chr5 | 142268149 | 142268399 | 1.49764 | 2.26026 | 1.509 |
| chr2 | 33808358 | 33808651 | 2.58845 | 3.90676 | 1.509 |
| chr5 | 53785275 | 53785695 | 1.11956 | 1.68982 | 1.509 |
| chr7 | 51293059 | 51293326 | 1.5961 | 2.4092 | 1.509 |
| chr1 | 156767128 | 156767453 | 1.67514 | 2.52862 | 1.510 |
| chr12 | 23307723 | 23307982 | 0.748691 | 1.13023 | 1.510 |

| | | | | | |
|---|---|---|---|---|---|
| chr5 | 29543436 | 29543644 | 0.684837 | 1.03386 | 1.510 |
| chr5 | 128031561 | 128031878 | 1.28742 | 1.9436 | 1.510 |
| chr4 | 37391663 | 37391944 | 1.34163 | 2.02549 | 1.510 |
| chr7 | 22041045 | 22041339 | 1.60433 | 2.4221 | 1.510 |
| chr19 | 42585784 | 42586039 | 1.83738 | 2.77407 | 1.510 |
| chr7 | 90981217 | 90981587 | 0.627212 | 0.946975 | 1.510 |
| chr16 | 67011640 | 67011929 | 2.3645 | 3.57 | 1.510 |
| chr5 | 166659886 | 166660101 | 0.529286 | 0.799144 | 1.510 |
| chr21 | 28088283 | 28088580 | 2.32896 | 3.5164 | 1.510 |
| chr8 | 96309723 | 96309978 | 1.55651 | 2.35015 | 1.510 |
| chr19 | 4267081 | 4267350 | 1.3655 | 2.06175 | 1.510 |
| chr15 | 89672762 | 89673068 | 2.79803 | 4.22474 | 1.510 |
| chr21 | 46898009 | 46898336 | 1.3678 | 2.06531 | 1.510 |
| chr12 | 117320573 | 117320818 | 1.25748 | 1.89876 | 1.510 |
| chr1 | 205457215 | 205457743 | 2.08419 | 3.14711 | 1.510 |
| chr12 | 23469720 | 23469958 | 0.834358 | 1.25988 | 1.510 |
| chr2 | 160966549 | 160966784 | 1.59137 | 2.40314 | 1.510 |
| chr2 | 218650235 | 218650547 | 2.59855 | 3.92449 | 1.510 |
| chr12 | 129750123 | 129750341 | 0.697781 | 1.05384 | 1.510 |
| chr1 | 10270141 | 10270613 | 2.15469 | 3.25429 | 1.510 |
| chr14 | 96046987 | 96047207 | 0.584668 | 0.883053 | 1.510 |
| chr12 | 26375061 | 26375383 | 0.703366 | 1.06236 | 1.510 |
| chr3 | 85870353 | 85870740 | 1.31517 | 1.98645 | 1.510 |
| chr14 | 22217080 | 22217583 | 1.82339 | 2.75408 | 1.510 |
| chr2 | 202545519 | 202545853 | 1.56783 | 2.36815 | 1.510 |
| chrX | 102154963 | 102155302 | 0.899986 | 1.35939 | 1.510 |
| chr19 | 42828385 | 42828662 | 0.776604 | 1.17307 | 1.511 |
| chr12 | 48080516 | 48080735 | 1.77448 | 2.68044 | 1.511 |
| chr1 | 97311121 | 97311329 | 1.12162 | 1.69429 | 1.511 |
| chr1 | 203473606 | 203474346 | 1.34151 | 2.02644 | 1.511 |
| chr5 | 115174602 | 115174985 | 1.19947 | 1.81192 | 1.511 |
| chr15 | 33534866 | 33535477 | 1.35452 | 2.04623 | 1.511 |
| chr8 | 109852347 | 109852628 | 0.690657 | 1.0434 | 1.511 |
| chr5 | 168067990 | 168068256 | 0.549619 | 0.830339 | 1.511 |
| chr7 | 43479346 | 43479808 | 2.47996 | 3.74663 | 1.511 |
| chr8 | 101170640 | 101170878 | 1.99419 | 3.01277 | 1.511 |
| chr5 | 36738213 | 36738421 | 0.498399 | 0.753027 | 1.511 |
| chr15 | 81854527 | 81854744 | 0.737601 | 1.1145 | 1.511 |
| chr7 | 142000617 | 142001117 | 0.854761 | 1.29153 | 1.511 |
| chr8 | 49438611 | 49438824 | 0.544279 | 0.822447 | 1.511 |
| chr8 | 34294829 | 34295085 | 0.5293 | 0.799878 | 1.511 |
| chr7 | 92090726 | 92091187 | 1.17164 | 1.7707 | 1.511 |
| chr17 | 36010532 | 36010757 | 1.16965 | 1.76782 | 1.511 |
| chr19 | 7917782 | 7918548 | 1.67266 | 2.5281 | 1.511 |
| chr15 | 71168593 | 71168843 | 0.71511 | 1.08084 | 1.511 |
| chr9 | 36866853 | 36867161 | 1.26408 | 1.91062 | 1.511 |
| chr5 | 145671152 | 145671373 | 1.37552 | 2.07919 | 1.512 |
| chr11 | 42627333 | 42627578 | 0.67797 | 1.02481 | 1.512 |
| chr22 | 32029746 | 32030131 | 2.32495 | 3.51438 | 1.512 |
| chr1 | 5805783 | 5806489 | 2.02567 | 3.06215 | 1.512 |
| chr1 | 167411304 | 167411738 | 2.25807 | 3.41354 | 1.512 |
| chr2 | 41085795 | 41086033 | 0.643053 | 0.97213 | 1.512 |
| chr1 | 245906062 | 245906311 | 0.632588 | 0.956313 | 1.512 |
| chrX | 68607404 | 68607774 | 1.63017 | 2.46442 | 1.512 |
| chr7 | 99177747 | 99178001 | 0.823559 | 1.24512 | 1.512 |
| chr20 | 24205891 | 24206112 | 0.589554 | 0.891393 | 1.512 |
| chr15 | 87557958 | 87558171 | 0.760256 | 1.14951 | 1.512 |
| chr12 | 4465010 | 4465447 | 1.24579 | 1.88365 | 1.512 |
| chr16 | 19594867 | 19595422 | 1.87881 | 2.84087 | 1.512 |
| chr22 | 30031831 | 30032068 | 1.52831 | 2.31113 | 1.512 |
| chr20 | 4930532 | 4930833 | 1.33904 | 2.02493 | 1.512 |

| | | | | | |
|---|---|---|---|---|---|
| chr19 | 54902547 | 54903162 | 1.63534 | 2.47307 | 1.512 |
| chr5 | 129774287 | 129774564 | 0.834396 | 1.26185 | 1.512 |
| chr12 | 122084367 | 122084591 | 0.546233 | 0.826114 | 1.512 |
| chr20 | 29900966 | 29901485 | 1.77543 | 2.68512 | 1.512 |
| chr15 | 33518865 | 33519110 | 1.86032 | 2.81353 | 1.512 |
| chr20 | 19457675 | 19457887 | 0.619017 | 0.936214 | 1.512 |
| chr10 | 73078741 | 73079051 | 2.03758 | 3.08169 | 1.512 |
| chr2 | 233198122 | 233198731 | 1.61906 | 2.44875 | 1.512 |
| chr12 | 1254050 | 1254275 | 0.663136 | 1.00301 | 1.513 |
| chr2 | 178119168 | 178119555 | 1.78998 | 2.70768 | 1.513 |
| chr8 | 144110699 | 144111148 | 3.14331 | 4.75485 | 1.513 |
| chr3 | 133916573 | 133916823 | 1.66974 | 2.52582 | 1.513 |
| chr1 | 109655307 | 109655617 | 2.37473 | 3.5923 | 1.513 |
| chr1 | 19239501 | 19240007 | 3.13946 | 4.74932 | 1.513 |
| chr10 | 30960039 | 30960292 | 1.43445 | 2.17003 | 1.513 |
| chr10 | 120563150 | 120563352 | 1.16099 | 1.75647 | 1.513 |
| chr16 | 48664343 | 48664792 | 2.53754 | 3.83932 | 1.513 |
| chr5 | 129188426 | 129188838 | 2.33197 | 3.52838 | 1.513 |
| chr5 | 28385059 | 28385338 | 0.698523 | 1.05692 | 1.513 |
| chr8 | 53408740 | 53409060 | 2.28104 | 3.45149 | 1.513 |
| chr7 | 40847828 | 40848061 | 0.778908 | 1.17877 | 1.513 |
| chr11 | 74154252 | 74154454 | 1.00781 | 1.52523 | 1.513 |
| chr14 | 72983653 | 72983907 | 2.40869 | 3.64541 | 1.513 |
| chr20 | 42738609 | 42738823 | 0.915257 | 1.38531 | 1.514 |
| chr3 | 9343925 | 9344591 | 1.63 | 2.46734 | 1.514 |
| chr7 | 41409222 | 41409586 | 0.492715 | 0.745859 | 1.514 |
| chr18 | 24048862 | 24049124 | 1.41978 | 2.14923 | 1.514 |
| chr1 | 51240490 | 51240753 | 1.13163 | 1.71307 | 1.514 |
| chr20 | 799578 | 799894 | 1.83431 | 2.77682 | 1.514 |
| chr9 | 123518143 | 123518430 | 2.0814 | 3.15089 | 1.514 |
| chr7 | 120065529 | 120065826 | 0.650012 | 0.984086 | 1.514 |
| chr20 | 44642253 | 44643144 | 1.96742 | 2.97879 | 1.514 |
| chr17 | 53070372 | 53070717 | 1.4334 | 2.17032 | 1.514 |
| chr8 | 121276933 | 121277155 | 0.765314 | 1.1588 | 1.514 |
| chr5 | 11364765 | 11364983 | 0.800597 | 1.21227 | 1.514 |
| chr9 | 9640163 | 9640458 | 0.563092 | 0.852656 | 1.514 |
| chr17 | 8633346 | 8633593 | 0.745726 | 1.12924 | 1.514 |
| chr19 | 44698364 | 44698642 | 0.891701 | 1.3504 | 1.514 |
| chr7 | 150773783 | 150774309 | 1.00751 | 1.5258 | 1.514 |
| chr15 | 24210276 | 24210584 | 0.56105 | 0.849682 | 1.514 |
| chr16 | 8974095 | 8974470 | 1.71012 | 2.58991 | 1.514 |
| chr8 | 143706626 | 143706832 | 0.723851 | 1.09627 | 1.515 |
| chr1 | 30321999 | 30322286 | 1.12258 | 1.70018 | 1.515 |
| chr1 | 22968550 | 22968912 | 2.6597 | 4.02835 | 1.515 |
| chr4 | 101293693 | 101293912 | 1.21691 | 1.84313 | 1.515 |
| chr12 | 94518521 | 94518725 | 0.657381 | 0.995677 | 1.515 |
| chr20 | 60985193 | 60985625 | 1.21419 | 1.83902 | 1.515 |
| chr8 | 36200854 | 36201172 | 0.660529 | 1.00045 | 1.515 |
| chr4 | 77008323 | 77008649 | 1.25742 | 1.90453 | 1.515 |
| chr7 | 157132846 | 157133114 | 1.88473 | 2.8547 | 1.515 |
| chr19 | 29394430 | 29394641 | 0.680946 | 1.03142 | 1.515 |
| chr10 | 121362063 | 121362321 | 1.73852 | 2.63334 | 1.515 |
| chr19 | 5052609 | 5052812 | 0.783365 | 1.18661 | 1.515 |
| chr16 | 66988061 | 66988304 | 1.55909 | 2.36173 | 1.515 |
| chr17 | 61912148 | 61912597 | 2.64623 | 4.00864 | 1.515 |
| chr5 | 169179383 | 169179667 | 0.500702 | 0.758489 | 1.515 |
| chr15 | 57598828 | 57599200 | 2.93473 | 4.4458 | 1.515 |
| chr12 | 2692156 | 2692551 | 2.47409 | 3.74822 | 1.515 |
| chr8 | 62380023 | 62380417 | 2.28085 | 3.45547 | 1.515 |
| chr17 | 8022043 | 8022397 | 2.62591 | 3.97832 | 1.515 |
| chr1 | 7962867 | 7963055 | 3.13439 | 4.74888 | 1.515 |

| | | | | | |
|---|---|---|---|---|---|
| chr17 | 42281042 | 42281369 | 1.69177 | 2.56337 | 1.515 |
| chr6 | 166795293 | 166795665 | 2.16353 | 3.27827 | 1.515 |
| chr2 | 25599447 | 25599973 | 3.17801 | 4.81569 | 1.515 |
| chr1 | 180385243 | 180385453 | 1.54896 | 2.3472 | 1.515 |
| chrX | 17708295 | 17708526 | 1.76595 | 2.67604 | 1.515 |
| chr1 | 112879601 | 112879932 | 1.56886 | 2.37763 | 1.516 |
| chr19 | 50969115 | 50969579 | 0.842869 | 1.27739 | 1.516 |
| chr17 | 71251809 | 71252178 | 2.60077 | 3.94157 | 1.516 |
| chr2 | 37012268 | 37012696 | 3.10729 | 4.70942 | 1.516 |
| chr17 | 75485137 | 75485443 | 0.749337 | 1.13572 | 1.516 |
| chr20 | 23427312 | 23427523 | 1.43562 | 2.17603 | 1.516 |
| chr7 | 123505667 | 123505869 | 1.51932 | 2.30308 | 1.516 |
| chr7 | 95530413 | 95530665 | 0.938819 | 1.42313 | 1.516 |
| chr1 | 21985614 | 21986006 | 3.60334 | 5.4623 | 1.516 |
| chr11 | 43588879 | 43589275 | 1.60025 | 2.42583 | 1.516 |
| chr17 | 77763530 | 77764070 | 2.01514 | 3.05485 | 1.516 |
| chr12 | 88332972 | 88333190 | 1.01741 | 1.54241 | 1.516 |
| chr8 | 13593635 | 13593900 | 0.606666 | 0.919743 | 1.516 |
| chr3 | 120280129 | 120280419 | 1.56264 | 2.36908 | 1.516 |
| chr17 | 73127405 | 73127835 | 1.99869 | 3.03055 | 1.516 |
| chr12 | 40384277 | 40384825 | 2.2311 | 3.38313 | 1.516 |
| chr19 | 29676226 | 29676579 | 0.709707 | 1.07618 | 1.516 |
| chr17 | 27375618 | 27375847 | 0.63937 | 0.969526 | 1.516 |
| chr12 | 103469529 | 103469811 | 0.572465 | 0.868088 | 1.516 |
| chr17 | 70114738 | 70115049 | 2.31873 | 3.51636 | 1.517 |
| chr10 | 12028649 | 12028931 | 1.62064 | 2.45772 | 1.517 |
| chr1 | 5976005 | 5976379 | 1.05566 | 1.60094 | 1.517 |
| chr11 | 93385095 | 93385579 | 1.19473 | 1.81198 | 1.517 |
| chr12 | 81104359 | 81104636 | 0.734569 | 1.11412 | 1.517 |
| chr20 | 36933108 | 36933563 | 1.04507 | 1.58506 | 1.517 |
| chr12 | 54137369 | 54137708 | 1.90421 | 2.88818 | 1.517 |
| chr21 | 34288978 | 34289305 | 1.12623 | 1.70838 | 1.517 |
| chr1 | 156577913 | 156578171 | 2.54879 | 3.86629 | 1.517 |
| chrX | 122911714 | 122911940 | 2.13874 | 3.24429 | 1.517 |
| chr1 | 10554745 | 10555303 | 1.6474 | 2.49901 | 1.517 |
| chr10 | 72297257 | 72297794 | 1.35663 | 2.05793 | 1.517 |
| chr16 | 11465629 | 11465988 | 1.82207 | 2.76402 | 1.517 |
| chr14 | 39336788 | 39337104 | 1.6446 | 2.4949 | 1.517 |
| chr2 | 242598738 | 242599036 | 1.64141 | 2.49017 | 1.517 |
| chr3 | 163564974 | 163565178 | 0.629905 | 0.955638 | 1.517 |
| chr8 | 70310495 | 70310748 | 0.691657 | 1.04933 | 1.517 |
| chr2 | 237089568 | 237089799 | 1.2114 | 1.83787 | 1.517 |
| chr17 | 10758007 | 10758307 | 0.713401 | 1.08235 | 1.517 |
| chr3 | 42922470 | 42922789 | 2.15588 | 3.27084 | 1.517 |
| chr9 | 23757438 | 23757654 | 0.59549 | 0.903458 | 1.517 |
| chr14 | 76922653 | 76922887 | 1.10496 | 1.6765 | 1.517 |
| chr20 | 62669702 | 62670704 | 1.43929 | 2.18381 | 1.517 |
| chr15 | 73533263 | 73533603 | 2.99574 | 4.54576 | 1.517 |
| chr17 | 1552694 | 1553033 | 0.661026 | 1.00309 | 1.517 |
| chr17 | 43384120 | 43384376 | 0.925088 | 1.40379 | 1.517 |
| chr7 | 140931295 | 140931624 | 1.33767 | 2.02987 | 1.517 |
| chr9 | 86552853 | 86553095 | 1.20606 | 1.8303 | 1.518 |
| chr12 | 57576731 | 57577064 | 2.17245 | 3.29696 | 1.518 |
| chr5 | 114193494 | 114193832 | 0.637695 | 0.967791 | 1.518 |
| chr15 | 27562252 | 27562513 | 0.751938 | 1.14121 | 1.518 |
| chr4 | 7721210 | 7721699 | 1.71563 | 2.60407 | 1.518 |
| chr3 | 114546419 | 114546653 | 1.17532 | 1.78407 | 1.518 |
| chr5 | 132701036 | 132701334 | 1.28397 | 1.94912 | 1.518 |
| chr16 | 16885257 | 16885461 | 0.837324 | 1.27112 | 1.518 |
| chr8 | 143750715 | 143751457 | 2.20335 | 3.34502 | 1.518 |
| chr9 | 111612584 | 111612789 | 0.84184 | 1.27804 | 1.518 |

| | | | | | |
|---|---|---|---|---|---|
| chr8 | 100010104 | 100010477 | 1.786 | 2.71146 | 1.518 |
| chr11 | 74915824 | 74916407 | 1.76766 | 2.68368 | 1.518 |
| chr5 | 36310388 | 36310631 | 0.785333 | 1.19233 | 1.518 |
| chr1 | 48204283 | 48204559 | 1.86515 | 2.83186 | 1.518 |
| chr5 | 146937599 | 146937923 | 0.672467 | 1.02101 | 1.518 |
| chr8 | 137387068 | 137387276 | 0.758963 | 1.15234 | 1.518 |
| chr14 | 22855862 | 22856119 | 1.85855 | 2.82193 | 1.518 |
| chr17 | 81077429 | 81077786 | 0.828011 | 1.25721 | 1.518 |
| chr16 | 60908217 | 60908427 | 0.583633 | 0.886238 | 1.518 |
| chr1 | 228688666 | 228688990 | 2.7635 | 4.19661 | 1.519 |
| chr15 | 42281703 | 42282040 | 1.49763 | 2.27436 | 1.519 |
| chr15 | 81503491 | 81503761 | 0.463806 | 0.704353 | 1.519 |
| chr22 | 37888923 | 37889130 | 1.18424 | 1.79844 | 1.519 |
| chr2 | 77967068 | 77967300 | 0.628272 | 0.954139 | 1.519 |
| chr14 | 39576775 | 39577014 | 0.91716 | 1.39287 | 1.519 |
| chr15 | 81269935 | 81270266 | 1.62693 | 2.47089 | 1.519 |
| chr19 | 46640743 | 46640946 | 0.74601 | 1.13303 | 1.519 |
| chr1 | 68908844 | 68909049 | 1.05964 | 1.60944 | 1.519 |
| chr3 | 184007210 | 184007491 | 2.15303 | 3.27017 | 1.519 |
| chr2 | 121828510 | 121828972 | 2.81413 | 4.27435 | 1.519 |
| chrX | 102744766 | 102744985 | 1.47006 | 2.23291 | 1.519 |
| chr4 | 8412403 | 8412662 | 1.15248 | 1.75058 | 1.519 |
| chr9 | 132546058 | 132546346 | 2.10348 | 3.19542 | 1.519 |
| chr15 | 89377500 | 89377962 | 1.82458 | 2.77187 | 1.519 |
| chr1 | 28574972 | 28575316 | 1.872 | 2.84397 | 1.519 |
| chr8 | 145051186 | 145051643 | 2.18153 | 3.31436 | 1.519 |
| chr10 | 100754167 | 100754394 | 0.912566 | 1.38646 | 1.519 |
| chr4 | 1514929 | 1515298 | 1.94497 | 2.95514 | 1.519 |
| chr19 | 33513813 | 33514089 | 0.834682 | 1.26827 | 1.519 |
| chr1 | 153744510 | 153744907 | 1.62587 | 2.47051 | 1.520 |
| chr9 | 127618234 | 127618526 | 1.1471 | 1.74302 | 1.520 |
| chr11 | 67033652 | 67033909 | 2.71026 | 4.11834 | 1.520 |
| chr13 | 105870853 | 105871065 | 0.557259 | 0.846901 | 1.520 |
| chr1 | 171856221 | 171856439 | 0.751305 | 1.14185 | 1.520 |
| chr5 | 150944096 | 150944312 | 0.568484 | 0.864008 | 1.520 |
| chr7 | 20212366 | 20212679 | 1.69198 | 2.57171 | 1.520 |
| chr20 | 16615357 | 16615583 | 1.09168 | 1.65931 | 1.520 |
| chr12 | 101221624 | 101221976 | 1.78646 | 2.71555 | 1.520 |
| chr3 | 30969568 | 30970078 | 1.03308 | 1.57054 | 1.520 |
| chr11 | 64614590 | 64615021 | 2.54362 | 3.86712 | 1.520 |
| chr19 | 47783493 | 47783808 | 1.82689 | 2.77749 | 1.520 |
| chr3 | 88533363 | 88533582 | 0.54513 | 0.828814 | 1.520 |
| chr11 | 94356380 | 94356659 | 1.47054 | 2.23584 | 1.520 |
| chr17 | 48611669 | 48612123 | 1.0919 | 1.66027 | 1.521 |
| chr14 | 63855582 | 63855875 | 2.7465 | 4.17621 | 1.521 |
| chr20 | 12950982 | 12951259 | 1.43537 | 2.18262 | 1.521 |
| chr8 | 52936978 | 52937180 | 0.681005 | 1.03569 | 1.521 |
| chr13 | 103568390 | 103568664 | 2.04761 | 3.1144 | 1.521 |
| chr1 | 154506709 | 154507055 | 1.30074 | 1.97844 | 1.521 |
| chr7 | 78136593 | 78136809 | 0.614728 | 0.935015 | 1.521 |
| chr2 | 135208432 | 135208950 | 2.54121 | 3.86532 | 1.521 |
| chr10 | 94640130 | 94640333 | 1.14029 | 1.73452 | 1.521 |
| chr5 | 66688824 | 66689063 | 0.691136 | 1.05131 | 1.521 |
| chr4 | 102090414 | 102090674 | 1.49387 | 2.27248 | 1.521 |
| chr5 | 103960715 | 103960997 | 0.738548 | 1.12352 | 1.521 |
| chr10 | 104594724 | 104595066 | 1.5881 | 2.41598 | 1.521 |
| chrX | 52963921 | 52964221 | 1.80289 | 2.74283 | 1.521 |
| chr8 | 29177933 | 29178145 | 1.60161 | 2.43667 | 1.521 |
| chr11 | 8285365 | 8285907 | 2.2945 | 3.49088 | 1.521 |
| chr15 | 95057031 | 95057245 | 0.643155 | 0.978513 | 1.521 |
| chr7 | 52086131 | 52086376 | 0.75241 | 1.14479 | 1.522 |

| | | | | | |
|---|---|---|---|---|---|
| chr19 | 10602807 | 10603081 | 1.95608 | 2.97623 | 1.522 |
| chr19 | 47117376 | 47117712 | 2.23938 | 3.4075 | 1.522 |
| chr1 | 10927204 | 10927512 | 1.34574 | 2.04799 | 1.522 |
| chr14 | 51887596 | 51887875 | 0.655632 | 0.997795 | 1.522 |
| chr7 | 44890156 | 44890512 | 0.739236 | 1.12503 | 1.522 |
| chrX | 15367241 | 15367517 | 1.3983 | 2.12826 | 1.522 |
| chr8 | 135843981 | 135844305 | 1.45435 | 2.21389 | 1.522 |
| chr14 | 92818912 | 92819250 | 1.63647 | 2.49114 | 1.522 |
| chr2 | 201677094 | 201677386 | 2.13442 | 3.24915 | 1.522 |
| chr5 | 147862452 | 147862791 | 2.05965 | 3.13543 | 1.522 |
| chr20 | 24436411 | 24436618 | 0.528013 | 0.803809 | 1.522 |
| chr2 | 74594912 | 74595119 | 0.612146 | 0.931901 | 1.522 |
| chr1 | 160346345 | 160346695 | 2.82033 | 4.29357 | 1.522 |
| chr8 | 144130191 | 144130606 | 1.34175 | 2.04263 | 1.522 |
| chr20 | 48252822 | 48253053 | 0.526071 | 0.800881 | 1.522 |
| chr22 | 27065012 | 27065215 | 1.30219 | 1.98244 | 1.522 |
| chr14 | 101112091 | 101112365 | 2.62985 | 4.00369 | 1.522 |
| chr5 | 162228070 | 162228279 | 0.67892 | 1.03361 | 1.522 |
| chr11 | 82061702 | 82061923 | 0.487427 | 0.742087 | 1.522 |
| chr17 | 16819314 | 16819680 | 2.27719 | 3.46695 | 1.522 |
| chr15 | 99579243 | 99579550 | 0.799797 | 1.21774 | 1.523 |
| chr9 | 34664277 | 34664682 | 3.26924 | 4.97762 | 1.523 |
| chr3 | 24815353 | 24815599 | 2.08086 | 3.16833 | 1.523 |
| chr1 | 11001967 | 11002332 | 2.09556 | 3.19081 | 1.523 |
| chr19 | 5001457 | 5001908 | 1.70919 | 2.60252 | 1.523 |
| chr2 | 22676162 | 22676379 | 0.712023 | 1.08421 | 1.523 |
| chr12 | 101065843 | 101066286 | 1.63088 | 2.48356 | 1.523 |
| chr15 | 60736427 | 60736666 | 1.85544 | 2.82569 | 1.523 |
| chr7 | 43251058 | 43251462 | 1.07623 | 1.63902 | 1.523 |
| chr3 | 38173126 | 38173473 | 1.50429 | 2.29102 | 1.523 |
| chrX | 106961272 | 106961608 | 1.64866 | 2.51092 | 1.523 |
| chr2 | 113642532 | 113642833 | 2.01345 | 3.06665 | 1.523 |
| chr3 | 99405841 | 99406346 | 1.60151 | 2.43933 | 1.523 |
| chr5 | 86901608 | 86901872 | 0.624795 | 0.9517 | 1.523 |
| chr8 | 41219020 | 41219326 | 0.763806 | 1.16345 | 1.523 |
| chr12 | 55310930 | 55311183 | 0.7937 | 1.209 | 1.523 |
| chr3 | 22971106 | 22971494 | 0.561894 | 0.855906 | 1.523 |
| chr2 | 21265729 | 21266135 | 3.35061 | 5.10399 | 1.523 |
| chr7 | 150788463 | 150788808 | 1.76535 | 2.68923 | 1.523 |
| chr2 | 85525944 | 85526282 | 1.4206 | 2.16419 | 1.523 |
| chr9 | 36555657 | 36556168 | 1.70947 | 2.60431 | 1.523 |
| chr1 | 40804392 | 40804721 | 2.77873 | 4.2334 | 1.524 |
| chr9 | 129160157 | 129161111 | 1.87153 | 2.85131 | 1.524 |
| chr19 | 4576099 | 4576366 | 1.44798 | 2.20604 | 1.524 |
| chr19 | 15667249 | 15667671 | 1.39034 | 2.11832 | 1.524 |
| chr2 | 5502766 | 5503018 | 0.663075 | 1.01028 | 1.524 |
| chr12 | 27536958 | 27537298 | 2.4343 | 3.70901 | 1.524 |
| chrX | 2953765 | 2954135 | 1.49325 | 2.27519 | 1.524 |
| chr3 | 153813659 | 153813883 | 1.0044 | 1.53048 | 1.524 |
| chr12 | 69751285 | 69751593 | 2.91981 | 4.44927 | 1.524 |
| chr14 | 81453817 | 81454176 | 2.61671 | 3.98746 | 1.524 |
| chr7 | 116767708 | 116767957 | 1.36597 | 2.08157 | 1.524 |
| chr10 | 54998246 | 54998461 | 0.592402 | 0.902803 | 1.524 |
| chr17 | 72489228 | 72489844 | 1.47263 | 2.24428 | 1.524 |
| chr12 | 101527957 | 101528205 | 0.661239 | 1.00773 | 1.524 |
| chr7 | 108573800 | 108574021 | 0.648334 | 0.988061 | 1.524 |
| chr9 | 73007130 | 73007397 | 0.885079 | 1.34886 | 1.524 |
| chr5 | 217956 | 218220 | 1.05288 | 1.60461 | 1.524 |
| chr1 | 180223502 | 180223751 | 0.614006 | 0.935776 | 1.524 |
| chr11 | 64240113 | 64240347 | 0.543096 | 0.827708 | 1.524 |
| chr2 | 231689219 | 231689486 | 1.50676 | 2.2964 | 1.524 |

| | | | | | |
|---|---|---|---|---|---|
| chr20 | 13305633 | 13305960 | 1.97161 | 3.00491 | 1.524 |
| chr11 | 72001926 | 72002142 | 1.32447 | 2.01869 | 1.524 |
| chr5 | 52774209 | 52774420 | 1.33461 | 2.03423 | 1.524 |
| chr7 | 86863973 | 86864183 | 0.583039 | 0.888691 | 1.524 |
| chr1 | 33603447 | 33603899 | 1.35389 | 2.06368 | 1.524 |
| chr16 | 10673804 | 10674139 | 2.40203 | 3.66132 | 1.524 |
| chr2 | 71016638 | 71017198 | 1.65605 | 2.52428 | 1.524 |
| chr8 | 31330674 | 31331262 | 0.688588 | 1.04962 | 1.524 |
| chr14 | 21074409 | 21074699 | 1.1863 | 1.80833 | 1.524 |
| chr9 | 104489982 | 104490713 | 1.0354 | 1.57832 | 1.524 |
| chr1 | 205366199 | 205366562 | 1.57426 | 2.39981 | 1.524 |
| chr5 | 31825662 | 31825906 | 1.36922 | 2.08727 | 1.524 |
| chr4 | 101017253 | 101017470 | 1.82091 | 2.77588 | 1.524 |
| chr5 | 32690304 | 32690550 | 1.17796 | 1.79586 | 1.525 |
| chr1 | 32129460 | 32129729 | 2.98919 | 4.55726 | 1.525 |
| chr3 | 124283865 | 124284078 | 2.00526 | 3.05727 | 1.525 |
| chr5 | 104163087 | 104163362 | 0.55335 | 0.843752 | 1.525 |
| chr7 | 5490289 | 5490663 | 1.88226 | 2.87021 | 1.525 |
| chr3 | 15149548 | 15150056 | 1.9742 | 3.01042 | 1.525 |
| chr2 | 85690119 | 85690385 | 0.577462 | 0.88057 | 1.525 |
| chr2 | 36096757 | 36096961 | 0.711309 | 1.08468 | 1.525 |
| chr11 | 2996910 | 2997179 | 1.40161 | 2.13746 | 1.525 |
| chr17 | 17908283 | 17908548 | 1.10514 | 1.68534 | 1.525 |
| chr5 | 31741708 | 31741912 | 0.718636 | 1.09592 | 1.525 |
| chr3 | 76157224 | 76157441 | 1.11591 | 1.70179 | 1.525 |
| chr15 | 64359739 | 64360058 | 2.1724 | 3.31299 | 1.525 |
| chr20 | 56070335 | 56070706 | 1.77778 | 2.71119 | 1.525 |
| chr3 | 164601855 | 164602132 | 0.803635 | 1.22569 | 1.525 |
| chr20 | 8326162 | 8326365 | 0.821542 | 1.25304 | 1.525 |
| chr2 | 95742474 | 95742944 | 2.09145 | 3.19003 | 1.525 |
| chr13 | 82504146 | 82504380 | 0.937239 | 1.42955 | 1.525 |
| chr2 | 55847348 | 55847617 | 1.37777 | 2.10157 | 1.525 |
| chr22 | 19957980 | 19958429 | 2.20904 | 3.36957 | 1.525 |
| chr5 | 171710405 | 171710724 | 1.53185 | 2.33669 | 1.525 |
| chr16 | 88450149 | 88450490 | 1.49225 | 2.27635 | 1.525 |
| chr11 | 82578595 | 82578884 | 1.97688 | 3.01596 | 1.526 |
| chr19 | 46146362 | 46146744 | 3.37487 | 5.14889 | 1.526 |
| chrX | 133387917 | 133388324 | 1.18395 | 1.8063 | 1.526 |
| chr10 | 81227224 | 81227602 | 1.92828 | 2.94192 | 1.526 |
| chr19 | 22955340 | 22955570 | 0.529851 | 0.808416 | 1.526 |
| chrX | 115110258 | 115110461 | 0.932867 | 1.42334 | 1.526 |
| chr1 | 154989971 | 154990385 | 3.60766 | 5.50458 | 1.526 |
| chr20 | 40336482 | 40336687 | 0.46174 | 0.704536 | 1.526 |
| chr19 | 31613571 | 31613996 | 0.518655 | 0.791409 | 1.526 |
| chr17 | 51546194 | 51546412 | 0.787809 | 1.20217 | 1.526 |
| chr5 | 135280716 | 135281029 | 1.44055 | 2.19831 | 1.526 |
| chr17 | 35887392 | 35887703 | 1.87287 | 2.85825 | 1.526 |
| chr5 | 103093342 | 103093545 | 0.647809 | 0.98868 | 1.526 |
| chr7 | 155391464 | 155391776 | 2.23102 | 3.40498 | 1.526 |
| chr9 | 36937430 | 36937673 | 0.525562 | 0.80216 | 1.526 |
| chr2 | 172970994 | 172971236 | 1.29507 | 1.97671 | 1.526 |
| chr7 | 147355655 | 147355902 | 0.659621 | 1.00681 | 1.526 |
| chr13 | 39848133 | 39848366 | 0.903464 | 1.37908 | 1.526 |
| chr9 | 89048947 | 89049168 | 1.46005 | 2.22873 | 1.526 |
| chr16 | 77709674 | 77709960 | 1.61392 | 2.46365 | 1.527 |
| chr1 | 150968234 | 150968708 | 1.40984 | 2.15213 | 1.527 |
| chr3 | 129653378 | 129653644 | 1.58848 | 2.42488 | 1.527 |
| chr2 | 224904231 | 224904560 | 1.59392 | 2.43323 | 1.527 |
| chr10 | 16186020 | 16186387 | 0.660726 | 1.00871 | 1.527 |
| chr1 | 47854453 | 47854755 | 1.5283 | 2.33335 | 1.527 |
| chr4 | 1747980 | 1748359 | 2.40009 | 3.66445 | 1.527 |

| | | | | | |
|---|---|---|---|---|---|
| chr19 | 48697654 | 48698138 | 1.21045 | 1.84817 | 1.527 |
| chr1 | 45121281 | 45121593 | 1.86194 | 2.843 | 1.527 |
| chr12 | 127101012 | 127101323 | 0.618579 | 0.944511 | 1.527 |
| chr15 | 29386055 | 29386445 | 0.750173 | 1.14544 | 1.527 |
| chr3 | 9768729 | 9769258 | 1.95095 | 2.97896 | 1.527 |
| chr5 | 38922226 | 38922597 | 2.58219 | 3.94282 | 1.527 |
| chr4 | 124485960 | 124486241 | 2.17924 | 3.32758 | 1.527 |
| chr19 | 18958020 | 18958391 | 1.8747 | 2.86265 | 1.527 |
| chr4 | 47566452 | 47566662 | 1.05167 | 1.60589 | 1.527 |
| chr10 | 58824763 | 58824975 | 0.682712 | 1.0425 | 1.527 |
| chr2 | 234885445 | 234885649 | 1.08603 | 1.65837 | 1.527 |
| chrX | 79479717 | 79479967 | 0.927139 | 1.41584 | 1.527 |
| chr7 | 86219359 | 86219620 | 0.684109 | 1.04473 | 1.527 |
| chr15 | 99884122 | 99884343 | 1.17654 | 1.79684 | 1.527 |
| chr7 | 29448425 | 29448841 | 1.12838 | 1.72331 | 1.527 |
| chr12 | 69224056 | 69224347 | 2.09783 | 3.20406 | 1.527 |
| chr10 | 70981943 | 70982277 | 1.66228 | 2.53885 | 1.527 |
| chr12 | 13447980 | 13448426 | 0.63453 | 0.969152 | 1.527 |
| chr9 | 88313328 | 88313616 | 1.39093 | 2.12445 | 1.527 |
| chr19 | 50978098 | 50978383 | 0.821732 | 1.2551 | 1.527 |
| chr1 | 19670843 | 19671193 | 1.96696 | 3.00433 | 1.527 |
| chr6 | 3411236 | 3411446 | 1.62492 | 2.48193 | 1.527 |
| chr13 | 75932073 | 75932330 | 1.00634 | 1.53714 | 1.527 |
| chr5 | 35651335 | 35651608 | 1.18313 | 1.80729 | 1.528 |
| chr7 | 157092540 | 157092764 | 1.42507 | 2.17704 | 1.528 |
| chr9 | 72534456 | 72534663 | 1.71654 | 2.62249 | 1.528 |
| chr16 | 87350474 | 87350718 | 1.78349 | 2.7249 | 1.528 |
| chr2 | 58675195 | 58675498 | 2.03387 | 3.10746 | 1.528 |
| chr5 | 128188466 | 128188680 | 0.469894 | 0.717941 | 1.528 |
| chr10 | 80511618 | 80511879 | 0.755825 | 1.15482 | 1.528 |
| chr7 | 99058906 | 99059341 | 0.817105 | 1.24847 | 1.528 |
| chrX | 135418845 | 135419166 | 0.757736 | 1.15777 | 1.528 |
| chr10 | 24003095 | 24003407 | 0.587971 | 0.898395 | 1.528 |
| chr1 | 241130402 | 241130625 | 0.905364 | 1.38342 | 1.528 |
| chr19 | 42250582 | 42250886 | 2.19311 | 3.3512 | 1.528 |
| chr10 | 28965973 | 28966383 | 1.98769 | 3.03738 | 1.528 |
| chr12 | 114843351 | 114843578 | 0.459926 | 0.702896 | 1.528 |
| chr1 | 3407803 | 3408139 | 2.46734 | 3.77113 | 1.528 |
| chr2 | 86315129 | 86315761 | 1.02909 | 1.57293 | 1.528 |
| chr9 | 137590284 | 137590911 | 2.36973 | 3.6221 | 1.528 |
| chr15 | 100923972 | 100924305 | 1.37215 | 2.09735 | 1.529 |
| chr4 | 16918365 | 16918727 | 1.41748 | 2.16672 | 1.529 |
| chr3 | 19908297 | 19908536 | 1.45827 | 2.22913 | 1.529 |
| chr12 | 128071129 | 128071530 | 0.696196 | 1.06427 | 1.529 |
| chr7 | 36060932 | 36061413 | 1.15006 | 1.75815 | 1.529 |
| chr7 | 108137588 | 108137958 | 1.80175 | 2.75446 | 1.529 |
| chr19 | 1767428 | 1767800 | 2.59043 | 3.9602 | 1.529 |
| chr11 | 61151432 | 61151663 | 0.5615 | 0.858514 | 1.529 |
| chr8 | 27918543 | 27918902 | 2.00745 | 3.06952 | 1.529 |
| chr5 | 108432168 | 108432431 | 0.677825 | 1.03651 | 1.529 |
| chr5 | 141371942 | 141372263 | 1.44773 | 2.21383 | 1.529 |
| chr3 | 52506011 | 52506325 | 1.75366 | 2.68166 | 1.529 |
| chr20 | 25275053 | 25275494 | 2.86224 | 4.37697 | 1.529 |
| chr7 | 100171407 | 100171740 | 1.45286 | 2.22182 | 1.529 |
| chr8 | 141169734 | 141170099 | 1.96471 | 3.00463 | 1.529 |
| chr14 | 20883995 | 20884198 | 1.22076 | 1.86693 | 1.529 |
| chr12 | 22236212 | 22236494 | 0.9356 | 1.43099 | 1.529 |
| chr17 | 9497064 | 9497308 | 0.671853 | 1.02759 | 1.529 |
| chr12 | 94157609 | 94157827 | 1.25967 | 1.9267 | 1.530 |
| chr1 | 166448213 | 166448600 | 1.58412 | 2.42301 | 1.530 |
| chr5 | 87508375 | 87508624 | 0.610819 | 0.934298 | 1.530 |

| | | | | | |
|---|---|---|---|---|---|
| chr14 | 20375642 | 20375889 | 0.796577 | 1.21844 | 1.530 |
| chr15 | 57852070 | 57852385 | 3.07054 | 4.69675 | 1.530 |
| chr17 | 4890415 | 4890975 | 2.4915 | 3.81113 | 1.530 |
| chr14 | 73442587 | 73442792 | 1.21388 | 1.85696 | 1.530 |
| chr4 | 187394095 | 187394583 | 0.939217 | 1.4368 | 1.530 |
| chr5 | 21581681 | 21582094 | 0.658156 | 1.00695 | 1.530 |
| chr21 | 44847190 | 44847556 | 1.6211 | 2.48025 | 1.530 |
| chr3 | 138198211 | 138198679 | 1.55473 | 2.3788 | 1.530 |
| chr5 | 156361979 | 156362181 | 1.39539 | 2.13505 | 1.530 |
| gl000195_ra | 17 | 726 | 1.54565 | 2.36496 | 1.530 |
| chr13 | 42790263 | 42790593 | 1.55532 | 2.37989 | 1.530 |
| chr1 | 225082812 | 225083004 | 1.00154 | 1.53254 | 1.530 |
| chr2 | 131090083 | 131090526 | 1.49492 | 2.2875 | 1.530 |
| chr2 | 39062178 | 39062564 | 1.23545 | 1.89061 | 1.530 |
| chr4 | 8596103 | 8596529 | 1.72148 | 2.63452 | 1.530 |
| chr5 | 98368221 | 98368505 | 1.98471 | 3.03748 | 1.530 |
| chr11 | 17752192 | 17752966 | 1.80181 | 2.75758 | 1.530 |
| chr12 | 81465453 | 81465655 | 0.651228 | 0.996681 | 1.530 |
| chr7 | 152062278 | 152062522 | 0.638772 | 0.977627 | 1.530 |
| chr10 | 105245424 | 105245825 | 2.31349 | 3.54089 | 1.531 |
| chr19 | 47787607 | 47788151 | 1.6517 | 2.52801 | 1.531 |
| chr10 | 114868237 | 114868588 | 1.88935 | 2.89201 | 1.531 |
| chr8 | 55545416 | 55545688 | 1.63553 | 2.50357 | 1.531 |
| chr17 | 18120703 | 18121159 | 2.34839 | 3.59484 | 1.531 |
| chr4 | 158125746 | 158126022 | 1.67277 | 2.56068 | 1.531 |
| chr15 | 35307529 | 35307764 | 0.714575 | 1.09389 | 1.531 |
| chr9 | 38691033 | 38691241 | 0.675959 | 1.03487 | 1.531 |
| chr11 | 67124314 | 67124655 | 2.31537 | 3.54481 | 1.531 |
| chr17 | 74835806 | 74836240 | 1.55169 | 2.37578 | 1.531 |
| chr16 | 61084346 | 61084883 | 0.865037 | 1.3245 | 1.531 |
| chr16 | 61796664 | 61797036 | 0.632209 | 0.968072 | 1.531 |
| chr17 | 34135926 | 34136194 | 1.95315 | 2.99076 | 1.531 |
| chr14 | 51244664 | 51244902 | 1.70872 | 2.61657 | 1.531 |
| chr5 | 12563626 | 12563844 | 0.611417 | 0.936267 | 1.531 |
| chr2 | 4510179 | 4510453 | 0.679687 | 1.04085 | 1.531 |
| chr10 | 71503004 | 71503240 | 1.70561 | 2.61195 | 1.531 |
| chr7 | 6388635 | 6388879 | 2.59617 | 3.97617 | 1.532 |
| chr1 | 233926776 | 233927190 | 2.16571 | 3.31696 | 1.532 |
| chr9 | 90240281 | 90240506 | 0.66206 | 1.01401 | 1.532 |
| chr12 | 53370098 | 53370442 | 2.24275 | 3.43505 | 1.532 |
| chr2 | 74378522 | 74378921 | 1.83342 | 2.80812 | 1.532 |
| chr9 | 11421337 | 11421543 | 0.504555 | 0.7728 | 1.532 |
| chr9 | 130548964 | 130549338 | 0.867406 | 1.32872 | 1.532 |
| chr12 | 29235959 | 29236180 | 0.67219 | 1.02969 | 1.532 |
| chr8 | 73199272 | 73199844 | 1.34308 | 2.05751 | 1.532 |
| chr5 | 171339226 | 171339449 | 1.36733 | 2.09481 | 1.532 |
| chr14 | 87718631 | 87718886 | 0.58363 | 0.89415 | 1.532 |
| chr19 | 2236219 | 2236613 | 2.82803 | 4.33274 | 1.532 |
| chr17 | 50637529 | 50637852 | 0.844588 | 1.29398 | 1.532 |
| chr5 | 77488395 | 77488731 | 0.614712 | 0.941885 | 1.532 |
| chr11 | 64510220 | 64511414 | 1.61346 | 2.47234 | 1.532 |
| chr5 | 75937410 | 75937685 | 1.42225 | 2.17941 | 1.532 |
| chr12 | 93221687 | 93221956 | 2.05712 | 3.15238 | 1.532 |
| chr17 | 56340979 | 56341387 | 0.766688 | 1.17489 | 1.532 |
| chr11 | 134047063 | 134047367 | 0.546772 | 0.83789 | 1.532 |
| chr17 | 37910180 | 37910805 | 2.11088 | 3.23487 | 1.532 |
| chr16 | 8024953 | 8025224 | 0.683963 | 1.04817 | 1.533 |
| chr9 | 86211825 | 86212152 | 1.63146 | 2.50037 | 1.533 |
| chrX | 101323047 | 101323249 | 1.08338 | 1.66042 | 1.533 |
| chr4 | 88072061 | 88072259 | 1.62981 | 2.49793 | 1.533 |
| chr2 | 11228899 | 11229157 | 0.686292 | 1.05188 | 1.533 |

| | | | | | |
|---|---|---|---|---|---|
| chr15 | 69244492 | 69244728 | 0.832851 | 1.27653 | 1.533 |
| chr5 | 161394942 | 161395150 | 0.786249 | 1.2051 | 1.533 |
| chr19 | 33725755 | 33726659 | 1.86763 | 2.86269 | 1.533 |
| chr9 | 104421101 | 104421319 | 0.809744 | 1.24122 | 1.533 |
| chr20 | 17592021 | 17592325 | 1.48167 | 2.2712 | 1.533 |
| chr14 | 79075178 | 79075399 | 0.529633 | 0.811856 | 1.533 |
| chr8 | 106793988 | 106794338 | 2.19802 | 3.36935 | 1.533 |
| chr4 | 145239928 | 145240230 | 0.707734 | 1.08496 | 1.533 |
| chr6 | 169295186 | 169295412 | 1.89074 | 2.89866 | 1.533 |
| chr20 | 2432718 | 2433128 | 2.66226 | 4.08154 | 1.533 |
| chr3 | 164177529 | 164177761 | 0.653163 | 1.00138 | 1.533 |
| chr7 | 100227048 | 100227287 | 1.2059 | 1.84891 | 1.533 |
| chr11 | 111293699 | 111294074 | 2.83601 | 4.34844 | 1.533 |
| chr12 | 120591547 | 120591999 | 0.709725 | 1.08825 | 1.533 |
| chr2 | 69534079 | 69534833 | 3.1984 | 4.90423 | 1.533 |
| chr7 | 27762659 | 27762940 | 1.14687 | 1.75868 | 1.533 |
| chr2 | 134601905 | 134602132 | 1.62461 | 2.4913 | 1.533 |
| chr10 | 83875213 | 83875478 | 0.615982 | 0.944603 | 1.533 |
| chr5 | 171483297 | 171483865 | 0.67341 | 1.03269 | 1.534 |
| chr19 | 3547417 | 3547714 | 1.19036 | 1.82546 | 1.534 |
| chr19 | 50762073 | 50762581 | 2.39307 | 3.66987 | 1.534 |
| chr20 | 43934993 | 43935545 | 3.14573 | 4.82414 | 1.534 |
| chr2 | 23726471 | 23726725 | 0.651522 | 0.999161 | 1.534 |
| chr2 | 89173594 | 89173933 | 1.46446 | 2.24588 | 1.534 |
| chr4 | 169506609 | 169506969 | 1.30351 | 1.9991 | 1.534 |
| chr5 | 92733624 | 92733892 | 0.64853 | 0.994611 | 1.534 |
| chr17 | 56596036 | 56596316 | 1.70766 | 2.61896 | 1.534 |
| chr14 | 102930172 | 102930421 | 0.682563 | 1.04682 | 1.534 |
| chr7 | 32244063 | 32244315 | 1.41647 | 2.17248 | 1.534 |
| chr10 | 75717112 | 75717392 | 1.77318 | 2.7198 | 1.534 |
| chr9 | 123743915 | 123744343 | 1.57509 | 2.41609 | 1.534 |
| chr9 | 11583976 | 11584261 | 0.527672 | 0.80942 | 1.534 |
| chr7 | 96225217 | 96225455 | 0.806153 | 1.23665 | 1.534 |
| chr10 | 28670797 | 28671178 | 1.89283 | 2.90365 | 1.534 |
| chr2 | 170681231 | 170681771 | 2.94467 | 4.51747 | 1.534 |
| chr22 | 50587527 | 50587969 | 2.07067 | 3.17667 | 1.534 |
| chr15 | 70759312 | 70759685 | 2.07499 | 3.18334 | 1.534 |
| chr5 | 137914982 | 137915325 | 1.86931 | 2.8678 | 1.534 |
| chr1 | 100921073 | 100921335 | 0.770949 | 1.18276 | 1.534 |
| chr16 | 4790558 | 4790796 | 0.690669 | 1.0596 | 1.534 |
| chr10 | 102193526 | 102193922 | 2.48158 | 3.80758 | 1.534 |
| chr8 | 59614353 | 59614615 | 1.33469 | 2.04809 | 1.535 |
| chr3 | 14832856 | 14833191 | 2.16962 | 3.32933 | 1.535 |
| chr11 | 8240809 | 8241320 | 1.68704 | 2.58881 | 1.535 |
| chr3 | 121977674 | 121977937 | 1.61248 | 2.47441 | 1.535 |
| chr3 | 55933883 | 55934100 | 1.11616 | 1.71283 | 1.535 |
| chr17 | 62211005 | 62211264 | 1.57846 | 2.42227 | 1.535 |
| chr10 | 97751516 | 97751719 | 0.69309 | 1.06361 | 1.535 |
| chr16 | 26818479 | 26818692 | 0.687242 | 1.05466 | 1.535 |
| chr4 | 88894792 | 88895075 | 1.79252 | 2.75094 | 1.535 |
| chr9 | 139686098 | 139686721 | 3.73565 | 5.73322 | 1.535 |
| chrX | 149582495 | 149582851 | 1.11147 | 1.7059 | 1.535 |
| chr5 | 6468994 | 6469219 | 0.623557 | 0.957057 | 1.535 |
| chr4 | 115299519 | 115299729 | 1.40786 | 2.16112 | 1.535 |
| chr7 | 2508518 | 2509031 | 1.88743 | 2.89741 | 1.535 |
| chr5 | 168414598 | 168414902 | 2.28391 | 3.50651 | 1.535 |
| chr3 | 45970590 | 45970979 | 1.85655 | 2.85041 | 1.535 |
| chr12 | 112856841 | 112857268 | 1.58099 | 2.42739 | 1.535 |
| chr11 | 41155669 | 41156033 | 0.788561 | 1.21074 | 1.535 |
| chr14 | 53684235 | 53684576 | 1.50754 | 2.31468 | 1.535 |
| chr3 | 14941559 | 14941777 | 0.60584 | 0.930272 | 1.536 |

| | | | | | |
|---|---|---|---|---|---|
| chr5 | 151120174 | 151120705 | 2.00179 | 3.07384 | 1.536 |
| chr17 | 76513466 | 76513710 | 0.622752 | 0.956273 | 1.536 |
| chr1 | 108072240 | 108072514 | 1.43737 | 2.20718 | 1.536 |
| chr7 | 31824763 | 31824980 | 0.801134 | 1.23025 | 1.536 |
| chr8 | 68956729 | 68956991 | 0.575287 | 0.883432 | 1.536 |
| chr3 | 155470942 | 155471339 | 1.18963 | 1.82685 | 1.536 |
| chr12 | 91748989 | 91749554 | 1.70806 | 2.62302 | 1.536 |
| chr17 | 33716906 | 33717480 | 0.635286 | 0.975596 | 1.536 |
| chr7 | 54544943 | 54545270 | 0.693003 | 1.06426 | 1.536 |
| chr12 | 12487327 | 12487756 | 2.13136 | 3.27341 | 1.536 |
| chr8 | 113819172 | 113819504 | 0.804776 | 1.23602 | 1.536 |
| chr11 | 91307146 | 91307457 | 0.590891 | 0.907529 | 1.536 |
| chrX | 109389841 | 109390098 | 2.07151 | 3.18156 | 1.536 |
| chr11 | 64132827 | 64133062 | 0.618547 | 0.95007 | 1.536 |
| chr20 | 18756533 | 18756764 | 0.677286 | 1.0403 | 1.536 |
| chr4 | 38784539 | 38784754 | 1.1971 | 1.83886 | 1.536 |
| chr1 | 205454565 | 205455055 | 3.04352 | 4.67522 | 1.536 |
| chr18 | 55803072 | 55803405 | 1.83164 | 2.81366 | 1.536 |
| chr8 | 78359741 | 78359987 | 0.534837 | 0.82159 | 1.536 |
| chr8 | 99766648 | 99766871 | 0.794115 | 1.21996 | 1.536 |
| chr7 | 45175767 | 45176025 | 1.65021 | 2.53526 | 1.536 |
| chr17 | 72855886 | 72856283 | 2.04911 | 3.14824 | 1.536 |
| chr12 | 108888780 | 108889085 | 1.42671 | 2.19203 | 1.536 |
| chr8 | 145550223 | 145550570 | 2.57494 | 3.95621 | 1.536 |
| chr1 | 158065531 | 158065987 | 2.0155 | 3.09675 | 1.536 |
| chr1 | 249080408 | 249080652 | 0.562542 | 0.864362 | 1.537 |
| chr10 | 75608739 | 75609287 | 2.37722 | 3.65268 | 1.537 |
| chr16 | 63384839 | 63385071 | 0.813794 | 1.25043 | 1.537 |
| chr9 | 81216920 | 81217199 | 1.47002 | 2.25882 | 1.537 |
| chr8 | 30404789 | 30405086 | 1.51368 | 2.32592 | 1.537 |
| chr1 | 19233475 | 19233897 | 1.52117 | 2.33744 | 1.537 |
| chr12 | 127025467 | 127025723 | 0.874084 | 1.34313 | 1.537 |
| chr7 | 26579227 | 26579447 | 0.80483 | 1.23676 | 1.537 |
| chr16 | 74068894 | 74069176 | 1.23369 | 1.89579 | 1.537 |
| chr22 | 38003032 | 38003425 | 3.18268 | 4.89154 | 1.537 |
| chr1 | 55276192 | 55276479 | 1.90012 | 2.92039 | 1.537 |
| chr8 | 117840557 | 117840792 | 0.68255 | 1.04905 | 1.537 |
| chr12 | 17592297 | 17592723 | 0.781313 | 1.20086 | 1.537 |
| chr19 | 11052765 | 11053028 | 2.00261 | 3.07802 | 1.537 |
| chr17 | 1961617 | 1962280 | 2.76152 | 4.24456 | 1.537 |
| chr2 | 213432401 | 213432664 | 1.66295 | 2.55602 | 1.537 |
| chr1 | 241790810 | 241791062 | 1.41827 | 2.17996 | 1.537 |
| chr10 | 102478215 | 102478434 | 0.759646 | 1.16763 | 1.537 |
| chr7 | 144565781 | 144565996 | 0.600381 | 0.92283 | 1.537 |
| chr5 | 170171318 | 170171822 | 1.91331 | 2.94112 | 1.537 |
| chr12 | 109278208 | 109278698 | 1.66846 | 2.56478 | 1.537 |
| chr1 | 110041013 | 110041221 | 2.05119 | 3.15323 | 1.537 |
| chr2 | 6913158 | 6913716 | 2.40973 | 3.70444 | 1.537 |
| chr14 | 59422041 | 59422257 | 0.722934 | 1.11136 | 1.537 |
| chr2 | 74366010 | 74366486 | 1.98617 | 3.05338 | 1.537 |
| chr18 | 29004428 | 29004681 | 1.83128 | 2.8154 | 1.537 |
| chr1 | 29658796 | 29659083 | 1.48189 | 2.27832 | 1.537 |
| chr19 | 40895655 | 40895947 | 0.770578 | 1.18475 | 1.537 |
| chr4 | 110354558 | 110354947 | 1.8989 | 2.91961 | 1.538 |
| chr10 | 134166224 | 134166649 | 2.54139 | 3.90776 | 1.538 |
| chr16 | 80487386 | 80487624 | 0.85975 | 1.32203 | 1.538 |
| chr12 | 97702402 | 97702681 | 1.44276 | 2.21871 | 1.538 |
| chr11 | 133323386 | 133323610 | 0.666617 | 1.02515 | 1.538 |
| chr3 | 194403351 | 194403623 | 1.17135 | 1.80137 | 1.538 |
| chr8 | 37557774 | 37558241 | 2.25819 | 3.47285 | 1.538 |
| chr10 | 118204095 | 118204412 | 1.71598 | 2.63911 | 1.538 |

| | | | | | |
|---|---|---|---|---|---|
| chr8 | 139506443 | 139506674 | 0.868045 | 1.33502 | 1.538 |
| chr5 | 89532353 | 89532602 | 0.798451 | 1.22802 | 1.538 |
| chr5 | 9203338 | 9203552 | 0.544239 | 0.837069 | 1.538 |
| chr1 | 46877692 | 46878029 | 1.16503 | 1.79191 | 1.538 |
| chr11 | 3042514 | 3042747 | 1.60411 | 2.46741 | 1.538 |
| chrX | 17553430 | 17553746 | 1.01947 | 1.56815 | 1.538 |
| chr2 | 127825599 | 127826230 | 1.37878 | 2.12105 | 1.538 |
| chr22 | 44244671 | 44244938 | 1.51638 | 2.33285 | 1.538 |
| chr12 | 93467918 | 93468137 | 0.607446 | 0.934519 | 1.538 |
| chr19 | 13700054 | 13700387 | 2.19087 | 3.37062 | 1.538 |
| chr12 | 75069911 | 75070175 | 0.592848 | 0.912106 | 1.539 |
| chr19 | 14184071 | 14184583 | 1.76471 | 2.71515 | 1.539 |
| chr8 | 88937014 | 88937234 | 0.769495 | 1.18393 | 1.539 |
| chr16 | 83019869 | 83020181 | 0.610822 | 0.939832 | 1.539 |
| chr3 | 58626355 | 58626692 | 1.99157 | 3.0643 | 1.539 |
| chr6 | 41869986 | 41870194 | 1.0182 | 1.56666 | 1.539 |
| chr12 | 83808820 | 83809037 | 0.799683 | 1.2305 | 1.539 |
| chr1 | 147113648 | 147114188 | 1.56856 | 2.41382 | 1.539 |
| chr9 | 116533180 | 116533431 | 0.672057 | 1.03424 | 1.539 |
| chr9 | 98498587 | 98498983 | 1.36635 | 2.10273 | 1.539 |
| chr1 | 98933723 | 98934055 | 1.86008 | 2.86274 | 1.539 |
| chr1 | 192415080 | 192415299 | 1.48837 | 2.29066 | 1.539 |
| chr14 | 23212879 | 23213151 | 0.511065 | 0.786609 | 1.539 |
| chr1 | 162817581 | 162818006 | 1.90187 | 2.92733 | 1.539 |
| chrX | 109661478 | 109661724 | 1.23869 | 1.90661 | 1.539 |
| chr3 | 161023201 | 161023515 | 1.70872 | 2.63012 | 1.539 |
| chr5 | 8913514 | 8913739 | 0.518272 | 0.79777 | 1.539 |
| chr6 | 44202566 | 44202911 | 0.991841 | 1.52684 | 1.539 |
| chr5 | 52726459 | 52726805 | 1.53031 | 2.35578 | 1.539 |
| chr15 | 23874183 | 23874686 | 0.824431 | 1.26916 | 1.539 |
| chr9 | 127540141 | 127540356 | 0.617957 | 0.951335 | 1.539 |
| chr17 | 48350401 | 48350925 | 3.08487 | 4.74948 | 1.540 |
| chr1 | 152161070 | 152161397 | 2.00547 | 3.0877 | 1.540 |
| chr17 | 75079529 | 75079953 | 0.773817 | 1.1914 | 1.540 |
| chr19 | 31224232 | 31224522 | 0.782234 | 1.20447 | 1.540 |
| chr8 | 96902858 | 96903254 | 1.13062 | 1.74091 | 1.540 |
| chr8 | 97246965 | 97247214 | 1.33226 | 2.0515 | 1.540 |
| chr7 | 138056228 | 138056475 | 1.04928 | 1.61575 | 1.540 |
| chr16 | 76856979 | 76857219 | 1.05164 | 1.61942 | 1.540 |
| chr8 | 137728757 | 137729067 | 0.891095 | 1.37224 | 1.540 |
| chr11 | 44972446 | 44972818 | 3.28396 | 5.05728 | 1.540 |
| chr6 | 152744825 | 152745055 | 0.496609 | 0.764786 | 1.540 |
| chr3 | 159750027 | 159750448 | 1.92764 | 2.96864 | 1.540 |
| chr20 | 43724229 | 43724797 | 1.41022 | 2.17182 | 1.540 |
| chr3 | 32440541 | 32440856 | 1.43833 | 2.21513 | 1.540 |
| chr8 | 3603141 | 3603367 | 0.76386 | 1.17641 | 1.540 |
| chr5 | 103341515 | 103341744 | 0.62666 | 0.965116 | 1.540 |
| chr1 | 223175572 | 223175866 | 0.984256 | 1.516 | 1.540 |
| chr12 | 103204996 | 103205239 | 2.22023 | 3.41987 | 1.540 |
| chr2 | 237086239 | 237087068 | 2.48941 | 3.83475 | 1.540 |
| chr11 | 32195005 | 32195300 | 1.49778 | 2.30725 | 1.540 |
| chr15 | 68503094 | 68503574 | 1.80029 | 2.77338 | 1.541 |
| chr12 | 29151678 | 29152041 | 0.963362 | 1.48416 | 1.541 |
| chr8 | 139739621 | 139740218 | 2.80183 | 4.31655 | 1.541 |
| chr15 | 52548502 | 52548775 | 2.56703 | 3.9549 | 1.541 |
| chr15 | 72825390 | 72825715 | 1.60079 | 2.46626 | 1.541 |
| chr12 | 109871641 | 109871874 | 1.07431 | 1.6553 | 1.541 |
| chr7 | 100202112 | 100202521 | 0.991563 | 1.52786 | 1.541 |
| chr1 | 6094488 | 6094858 | 1.25989 | 1.94133 | 1.541 |
| chrX | 16454629 | 16455085 | 1.66575 | 2.56684 | 1.541 |
| chr12 | 58376911 | 58377250 | 1.57263 | 2.42336 | 1.541 |

| | | | | | |
|---|---|---|---|---|---|
| chr8 | 132886190 | 132886616 | 0.63719 | 0.981888 | 1.541 |
| chr2 | 65276192 | 65276598 | 0.656067 | 1.01103 | 1.541 |
| chr9 | 76298797 | 76299061 | 1.27089 | 1.95853 | 1.541 |
| chr2 | 190007906 | 190008220 | 1.60157 | 2.46819 | 1.541 |
| chr3 | 98315786 | 98316075 | 1.11137 | 1.71277 | 1.541 |
| chr15 | 70647749 | 70647952 | 0.864141 | 1.33176 | 1.541 |
| chr3 | 113874326 | 113874603 | 1.64049 | 2.52845 | 1.541 |
| chr1 | 165883487 | 165883690 | 1.58855 | 2.44845 | 1.541 |
| chr16 | 1200797 | 1201055 | 0.970742 | 1.49621 | 1.541 |
| chr15 | 78195198 | 78195573 | 2.39566 | 3.69256 | 1.541 |
| chr12 | 91247851 | 91248134 | 1.64368 | 2.53355 | 1.541 |
| chr5 | 81937259 | 81937490 | 1.57171 | 2.42267 | 1.541 |
| chr12 | 48232138 | 48232745 | 1.56365 | 2.41028 | 1.541 |
| chr1 | 231860896 | 231861259 | 1.69406 | 2.61133 | 1.541 |
| chr12 | 52822026 | 52822670 | 0.662118 | 1.02063 | 1.541 |
| chr14 | 100706611 | 100706948 | 2.01329 | 3.10341 | 1.541 |
| chr11 | 66175840 | 66176448 | 3.1192 | 4.80841 | 1.542 |
| chr22 | 31159827 | 31160044 | 1.16582 | 1.7977 | 1.542 |
| chr7 | 140683629 | 140684147 | 1.05167 | 1.62169 | 1.542 |
| chr20 | 49844211 | 49844539 | 0.778978 | 1.2012 | 1.542 |
| chrX | 136134567 | 136134904 | 2.35148 | 3.62615 | 1.542 |
| chr3 | 106837109 | 106837366 | 0.787821 | 1.21495 | 1.542 |
| chr8 | 30239715 | 30240119 | 2.10175 | 3.24135 | 1.542 |
| chr1 | 228134306 | 228134666 | 1.78182 | 2.74804 | 1.542 |
| chr2 | 74561986 | 74562404 | 2.45165 | 3.7812 | 1.542 |
| chr1 | 179836862 | 179837243 | 2.05525 | 3.16987 | 1.542 |
| chr1 | 235580501 | 235580744 | 1.38114 | 2.13019 | 1.542 |
| chr8 | 36580516 | 36580851 | 0.694569 | 1.07127 | 1.542 |
| chr17 | 40825918 | 40826493 | 1.58567 | 2.44583 | 1.542 |
| chr10 | 18574027 | 18574452 | 0.61711 | 0.951895 | 1.543 |
| chr8 | 39853255 | 39853555 | 1.87436 | 2.89132 | 1.543 |
| chr10 | 9343235 | 9343447 | 1.12241 | 1.73148 | 1.543 |
| chr10 | 52823496 | 52823763 | 1.1959 | 1.84484 | 1.543 |
| chr2 | 190538961 | 190539388 | 2.31054 | 3.56441 | 1.543 |
| chr19 | 49009386 | 49009785 | 1.30481 | 2.0129 | 1.543 |
| chr2 | 97220060 | 97220496 | 2.57814 | 3.97733 | 1.543 |
| chr7 | 25934987 | 25935267 | 1.94191 | 2.99581 | 1.543 |
| chr13 | 36527872 | 36528084 | 1.537 | 2.3712 | 1.543 |
| chr19 | 4558356 | 4558718 | 3.06004 | 4.7209 | 1.543 |
| chr14 | 22745157 | 22745401 | 1.3003 | 2.00606 | 1.543 |
| chr3 | 38084575 | 38084836 | 0.428448 | 0.661007 | 1.543 |
| chr7 | 44111878 | 44112292 | 1.66753 | 2.57278 | 1.543 |
| chr1 | 202162023 | 202162313 | 2.11176 | 3.25824 | 1.543 |
| chr3 | 97758144 | 97758791 | 2.17059 | 3.34903 | 1.543 |
| chr12 | 122062476 | 122062827 | 2.53777 | 3.91575 | 1.543 |
| chrX | 67416743 | 67417053 | 1.27067 | 1.96066 | 1.543 |
| chr14 | 73276071 | 73276336 | 0.519162 | 0.801089 | 1.543 |
| chr16 | 68430429 | 68430743 | 1.51244 | 2.33383 | 1.543 |
| chr11 | 40261922 | 40262145 | 0.663193 | 1.02338 | 1.543 |
| chr17 | 27405549 | 27405782 | 0.65113 | 1.00477 | 1.543 |
| chr19 | 39715118 | 39715622 | 0.758421 | 1.17035 | 1.543 |
| chr17 | 53679177 | 53679454 | 2.18163 | 3.36666 | 1.543 |
| chr11 | 6374563 | 6375012 | 2.7935 | 4.31114 | 1.543 |
| chr6 | 36270290 | 36270629 | 2.20498 | 3.40303 | 1.543 |
| chr7 | 31174076 | 31174423 | 1.70284 | 2.62821 | 1.543 |
| chr9 | 99056681 | 99056943 | 0.60697 | 0.936823 | 1.543 |
| chr11 | 16957447 | 16957685 | 1.91234 | 2.95195 | 1.544 |
| chr15 | 75108788 | 75109083 | 0.737951 | 1.13918 | 1.544 |
| chr2 | 235409133 | 235409370 | 1.8527 | 2.86005 | 1.544 |
| chr17 | 11048058 | 11048406 | 1.13075 | 1.74557 | 1.544 |
| chr8 | 125605190 | 125605574 | 2.21623 | 3.42138 | 1.544 |

| | | | | | |
|---|---|---|---|---|---|
| chr4 | 62607677 | 62607890 | 0.77682 | 1.19928 | 1.544 |
| chr2 | 72364500 | 72365036 | 3.08233 | 4.75884 | 1.544 |
| chr14 | 105505755 | 105506031 | 0.61749 | 0.953365 | 1.544 |
| chr1 | 222347657 | 222347938 | 0.820642 | 1.26709 | 1.544 |
| chr3 | 13395494 | 13395938 | 1.12604 | 1.73869 | 1.544 |
| chr2 | 128929780 | 128930027 | 0.890214 | 1.37459 | 1.544 |
| chr16 | 71576111 | 71576556 | 1.84795 | 2.85348 | 1.544 |
| chr3 | 49839946 | 49840367 | 2.19867 | 3.39503 | 1.544 |
| chr3 | 182339806 | 182340032 | 1.94535 | 3.0041 | 1.544 |
| chr12 | 94671188 | 94671436 | 0.625408 | 0.96579 | 1.544 |
| chr1 | 47013627 | 47013912 | 0.993169 | 1.53372 | 1.544 |
| chr17 | 74361691 | 74361893 | 1.42924 | 2.20716 | 1.544 |
| chr15 | 85369544 | 85370010 | 1.98055 | 3.05875 | 1.544 |
| chr1 | 174695624 | 174695925 | 1.13196 | 1.74822 | 1.544 |
| chr10 | 79896145 | 79896446 | 0.709048 | 1.09507 | 1.544 |
| chr9 | 105865259 | 105865541 | 0.794513 | 1.22711 | 1.544 |
| chr12 | 98129743 | 98130058 | 0.656054 | 1.01332 | 1.545 |
| chr13 | 46734630 | 46734922 | 1.45785 | 2.25176 | 1.545 |
| chr19 | 34587269 | 34587544 | 1.30917 | 2.02216 | 1.545 |
| chr15 | 78487903 | 78488788 | 1.54412 | 2.38508 | 1.545 |
| chr22 | 39892422 | 39892767 | 1.59887 | 2.46964 | 1.545 |
| chr1 | 156594214 | 156594433 | 1.28084 | 1.9787 | 1.545 |
| chr14 | 56953838 | 56954092 | 1.53099 | 2.36516 | 1.545 |
| chr9 | 111034779 | 111035331 | 1.32682 | 2.04979 | 1.545 |
| chr5 | 137785630 | 137785981 | 2.72307 | 4.20692 | 1.545 |
| chr13 | 29393966 | 29394170 | 1.61736 | 2.49876 | 1.545 |
| chr1 | 155196467 | 155196672 | 1.16871 | 1.80569 | 1.545 |
| chr20 | 22664692 | 22665015 | 1.81741 | 2.80815 | 1.545 |
| chr21 | 27254029 | 27254426 | 1.76352 | 2.7249 | 1.545 |
| chrX | 79903341 | 79903877 | 0.979027 | 1.51287 | 1.545 |
| chr10 | 118976113 | 118976699 | 2.73244 | 4.22274 | 1.545 |
| chr9 | 125112814 | 125113222 | 2.24988 | 3.47703 | 1.545 |
| chr7 | 149450695 | 149451108 | 2.26738 | 3.50436 | 1.546 |
| chr6 | 26596073 | 26596328 | 1.87345 | 2.8958 | 1.546 |
| chr18 | 44731355 | 44731590 | 1.85455 | 2.86668 | 1.546 |
| chr10 | 99805549 | 99805950 | 2.61436 | 4.0412 | 1.546 |
| chr13 | 27845032 | 27845418 | 1.84343 | 2.84954 | 1.546 |
| chr2 | 68661960 | 68662218 | 0.790969 | 1.22268 | 1.546 |
| chr9 | 9667814 | 9668024 | 0.661198 | 1.0222 | 1.546 |
| chr5 | 59287784 | 59288009 | 0.848087 | 1.31118 | 1.546 |
| chr8 | 124031224 | 124031448 | 1.19504 | 1.84758 | 1.546 |
| chr10 | 123463853 | 123464061 | 0.578903 | 0.89503 | 1.546 |
| chr19 | 40219797 | 40220152 | 0.665045 | 1.02822 | 1.546 |
| chr5 | 127244871 | 127245148 | 1.89466 | 2.92937 | 1.546 |
| chr2 | 85442396 | 85442612 | 0.681108 | 1.05315 | 1.546 |
| chr14 | 60902379 | 60902588 | 0.591559 | 0.914697 | 1.546 |
| chr10 | 52708199 | 52708449 | 0.658704 | 1.01854 | 1.546 |
| chr9 | 30732102 | 30732431 | 0.920166 | 1.42287 | 1.546 |
| chr8 | 142837999 | 142838285 | 0.709856 | 1.09769 | 1.546 |
| chr1 | 62759965 | 62760175 | 0.951917 | 1.47203 | 1.546 |
| chr15 | 80285702 | 80286114 | 2.1673 | 3.35166 | 1.546 |
| chr17 | 20811693 | 20811941 | 1.40546 | 2.1736 | 1.547 |
| chr12 | 124221309 | 124221701 | 0.78637 | 1.21619 | 1.547 |
| chr15 | 74220067 | 74220496 | 1.24502 | 1.92553 | 1.547 |
| chr2 | 150571806 | 150572016 | 2.16463 | 3.34781 | 1.547 |
| chr5 | 43389833 | 43390228 | 1.88773 | 2.91957 | 1.547 |
| chr5 | 139154654 | 139155228 | 2.45503 | 3.79702 | 1.547 |
| chr14 | 34218843 | 34219088 | 0.737303 | 1.14036 | 1.547 |
| chr17 | 961633 | 962008 | 2.15383 | 3.33125 | 1.547 |
| chr17 | 61159729 | 61159979 | 1.23351 | 1.90782 | 1.547 |
| chr1 | 9515809 | 9516211 | 1.38975 | 2.14977 | 1.547 |

| | | | | | |
|---|---|---|---|---|---|
| chr17 | 72421164 | 72421382 | 0.643316 | 0.995132 | 1.547 |
| chrX | 10157743 | 10157956 | 1.28048 | 1.98079 | 1.547 |
| chr16 | 18957523 | 18957786 | 1.44961 | 2.2425 | 1.547 |
| chr12 | 31834831 | 31835382 | 1.70156 | 2.63228 | 1.547 |
| chr8 | 1643914 | 1644346 | 1.56925 | 2.4277 | 1.547 |
| chr2 | 45208788 | 45209059 | 1.87868 | 2.90656 | 1.547 |
| chr16 | 14486733 | 14486947 | 0.498257 | 0.770871 | 1.547 |
| chr11 | 112160641 | 112161071 | 1.9897 | 3.0784 | 1.547 |
| chr4 | 100298598 | 100298855 | 1.16363 | 1.80033 | 1.547 |
| chr12 | 124860124 | 124860497 | 0.823071 | 1.2735 | 1.547 |
| chr19 | 1853303 | 1853631 | 2.30853 | 3.57195 | 1.547 |
| chr2 | 182602783 | 182602995 | 0.915596 | 1.41671 | 1.547 |
| chr20 | 40363002 | 40363614 | 1.76864 | 2.73667 | 1.547 |
| chr11 | 12352885 | 12353130 | 1.29214 | 1.99944 | 1.547 |
| chr12 | 26272429 | 26272664 | 1.16771 | 1.80692 | 1.547 |
| chr5 | 132848660 | 132848884 | 0.549258 | 0.849986 | 1.548 |
| chr12 | 88389453 | 88389693 | 1.84691 | 2.85818 | 1.548 |
| chr11 | 116613539 | 116613848 | 1.66295 | 2.57362 | 1.548 |
| chr11 | 16513742 | 16514118 | 0.636112 | 0.984481 | 1.548 |
| chr5 | 179095477 | 179095734 | 2.24318 | 3.47173 | 1.548 |
| chr8 | 6673747 | 6674068 | 1.73019 | 2.67778 | 1.548 |
| chr8 | 18054749 | 18055137 | 1.46309 | 2.2644 | 1.548 |
| chr1 | 2838675 | 2838912 | 1.59279 | 2.4652 | 1.548 |
| chr9 | 19776316 | 19776600 | 0.704238 | 1.09006 | 1.548 |
| chr1 | 240116277 | 240116578 | 1.17524 | 1.81918 | 1.548 |
| chr1 | 62548429 | 62548870 | 1.50539 | 2.33038 | 1.548 |
| chr20 | 43399764 | 43399969 | 0.787337 | 1.21882 | 1.548 |
| chr15 | 79103816 | 79104249 | 2.86974 | 4.44265 | 1.548 |
| chr2 | 27145576 | 27145988 | 1.52631 | 2.36291 | 1.548 |
| chr14 | 49601768 | 49602085 | 0.753256 | 1.16614 | 1.548 |
| chr5 | 146286377 | 146286621 | 0.771701 | 1.19472 | 1.548 |
| chr19 | 3706013 | 3706462 | 2.57055 | 3.97968 | 1.548 |
| chr10 | 99046951 | 99047269 | 0.594783 | 0.92096 | 1.548 |
| chr4 | 80838393 | 80838642 | 1.28962 | 1.99685 | 1.548 |
| chr2 | 46688535 | 46688887 | 2.92782 | 4.53358 | 1.548 |
| chr1 | 18923876 | 18924248 | 2.78333 | 4.30994 | 1.548 |
| chr8 | 11828057 | 11828680 | 1.28463 | 1.98932 | 1.549 |
| chr3 | 193505990 | 193506263 | 1.45204 | 2.24873 | 1.549 |
| chr10 | 71906180 | 71906573 | 1.81732 | 2.81445 | 1.549 |
| chr12 | 120080669 | 120081090 | 0.898004 | 1.39077 | 1.549 |
| chr13 | 21498938 | 21499223 | 1.83927 | 2.84854 | 1.549 |
| chr9 | 93796005 | 93796388 | 1.52305 | 2.35884 | 1.549 |
| chr4 | 160647851 | 160648204 | 0.647275 | 1.00249 | 1.549 |
| chr10 | 105176593 | 105176994 | 1.34743 | 2.08696 | 1.549 |
| chr18 | 46303463 | 46303828 | 2.27676 | 3.52642 | 1.549 |
| chr15 | 80748027 | 80748561 | 2.16318 | 3.35068 | 1.549 |
| chr20 | 48802850 | 48803131 | 2.40331 | 3.72264 | 1.549 |
| chr12 | 87668405 | 87668659 | 1.78412 | 2.76354 | 1.549 |
| chr5 | 105376527 | 105376733 | 0.715138 | 1.10784 | 1.549 |
| chr2 | 36347143 | 36347413 | 0.690579 | 1.06981 | 1.549 |
| chr8 | 137663217 | 137663458 | 0.756947 | 1.17263 | 1.549 |
| chr12 | 132505890 | 132506125 | 0.844524 | 1.30832 | 1.549 |
| chr9 | 73976948 | 73977192 | 0.577914 | 0.895348 | 1.549 |
| chr10 | 82356510 | 82356746 | 1.63163 | 2.52794 | 1.549 |
| chr6 | 158294769 | 158295164 | 1.70724 | 2.64511 | 1.549 |
| chr11 | 43333341 | 43333595 | 1.7284 | 2.67794 | 1.549 |
| chr3 | 154736335 | 154736664 | 0.545335 | 0.844942 | 1.549 |
| chr6 | 148899342 | 148899626 | 2.55415 | 3.95774 | 1.550 |
| chr17 | 62024088 | 62024732 | 1.86537 | 2.89062 | 1.550 |
| chr17 | 76285890 | 76286224 | 2.80474 | 4.34629 | 1.550 |
| chr20 | 6054798 | 6055131 | 1.06207 | 1.64587 | 1.550 |

| | | | | | |
|---|---|---|---|---|---|
| chr14 | 36837139 | 36837397 | 0.652706 | 1.01153 | 1.550 |
| chr21 | 32953100 | 32953403 | 1.62323 | 2.5156 | 1.550 |
| chrX | 11776416 | 11776806 | 2.11462 | 3.27756 | 1.550 |
| chr9 | 1009092 | 1009486 | 2.78448 | 4.3159 | 1.550 |
| chr16 | 69373383 | 69373590 | 1.30694 | 2.02575 | 1.550 |
| chr3 | 37896873 | 37897279 | 2.09013 | 3.2397 | 1.550 |
| chr10 | 28530737 | 28531051 | 1.98307 | 3.07383 | 1.550 |
| chr11 | 120206591 | 120206849 | 1.37931 | 2.13799 | 1.550 |
| chr4 | 6332628 | 6333078 | 1.50243 | 2.32888 | 1.550 |
| chr14 | 29990807 | 29991207 | 1.14865 | 1.78051 | 1.550 |
| chr9 | 99328982 | 99329344 | 1.35811 | 2.10527 | 1.550 |
| chr2 | 4776892 | 4777105 | 0.582391 | 0.902801 | 1.550 |
| chr5 | 175605896 | 175606153 | 0.921984 | 1.42924 | 1.550 |
| chr10 | 79471173 | 79471554 | 3.12537 | 4.84512 | 1.550 |
| chr7 | 97924150 | 97924493 | 2.49058 | 3.86103 | 1.550 |
| chr8 | 73765242 | 73765731 | 0.807072 | 1.25116 | 1.550 |
| chr2 | 47748273 | 47748790 | 0.992571 | 1.53874 | 1.550 |
| chr10 | 96560596 | 96560825 | 0.646652 | 1.00249 | 1.550 |
| chr15 | 31802269 | 31802500 | 1.08334 | 1.6795 | 1.550 |
| chr20 | 43929703 | 43930025 | 1.31013 | 2.03116 | 1.550 |
| chr16 | 26464643 | 26464892 | 0.605228 | 0.938383 | 1.550 |
| chr16 | 70772341 | 70772636 | 1.86841 | 2.89691 | 1.550 |
| chr7 | 154241798 | 154242043 | 0.733744 | 1.1377 | 1.551 |
| chr9 | 133873665 | 133874034 | 1.88305 | 2.91977 | 1.551 |
| chr9 | 132482870 | 132483195 | 1.30614 | 2.02528 | 1.551 |
| chr8 | 22722240 | 22722702 | 2.17715 | 3.37599 | 1.551 |
| chr9 | 120407007 | 120407263 | 1.78782 | 2.7723 | 1.551 |
| chr16 | 66463470 | 66463954 | 0.772509 | 1.19794 | 1.551 |
| chr3 | 12933530 | 12933810 | 1.8194 | 2.82141 | 1.551 |
| chr11 | 63920535 | 63920937 | 3.67486 | 5.69887 | 1.551 |
| chr1 | 168115006 | 168115328 | 3.43206 | 5.32237 | 1.551 |
| chr17 | 63499777 | 63500107 | 0.789233 | 1.224 | 1.551 |
| chr7 | 121634681 | 121635312 | 1.21746 | 1.88817 | 1.551 |
| chr11 | 65083051 | 65083430 | 2.8202 | 4.37402 | 1.551 |
| chr14 | 51560452 | 51560888 | 1.82479 | 2.83021 | 1.551 |
| chr8 | 110643674 | 110643912 | 0.687887 | 1.06692 | 1.551 |
| chr16 | 28886496 | 28886835 | 0.996416 | 1.54548 | 1.551 |
| chr5 | 12530373 | 12530797 | 0.740284 | 1.14822 | 1.551 |
| chr10 | 101492435 | 101492659 | 0.930328 | 1.44308 | 1.551 |
| chr17 | 79451295 | 79451727 | 3.09807 | 4.80558 | 1.551 |
| chr20 | 16054408 | 16054625 | 0.862212 | 1.3375 | 1.551 |
| chr9 | 111945498 | 111946142 | 1.55409 | 2.41076 | 1.551 |
| chr16 | 73477343 | 73477760 | 0.545038 | 0.845513 | 1.551 |
| chr19 | 29201304 | 29201531 | 1.01357 | 1.57239 | 1.551 |
| chr11 | 44915335 | 44916027 | 2.14259 | 3.32408 | 1.551 |
| chr7 | 53080174 | 53080402 | 0.715061 | 1.10937 | 1.551 |
| chr7 | 33981851 | 33982108 | 0.7745 | 1.2016 | 1.551 |
| chr7 | 126946132 | 126946492 | 0.871691 | 1.3524 | 1.551 |
| chr16 | 1123953 | 1124228 | 1.39137 | 2.15868 | 1.551 |
| chr21 | 46548435 | 46548866 | 1.7394 | 2.69883 | 1.552 |
| chr5 | 171741514 | 171741718 | 0.5982 | 0.928174 | 1.552 |
| chr19 | 45665333 | 45665633 | 1.65149 | 2.56262 | 1.552 |
| chr2 | 115674872 | 115675161 | 0.587531 | 0.911676 | 1.552 |
| chr5 | 68258757 | 68259061 | 1.99144 | 3.09022 | 1.552 |
| chr13 | 79739421 | 79739652 | 0.588084 | 0.912574 | 1.552 |
| chr8 | 22774913 | 22775145 | 1.39528 | 2.16526 | 1.552 |
| chr12 | 53044105 | 53044446 | 0.595752 | 0.924537 | 1.552 |
| chr6 | 33819758 | 33819992 | 1.633 | 2.53447 | 1.552 |
| chr3 | 20349087 | 20349407 | 0.673669 | 1.0456 | 1.552 |
| chr16 | 65892044 | 65892252 | 0.716765 | 1.11256 | 1.552 |
| chr14 | 50319574 | 50319963 | 1.55882 | 2.41961 | 1.552 |

| | | | | | |
|---|---|---|---|---|---|
| chr21 | 15776121 | 15776338 | 1.59183 | 2.47087 | 1.552 |
| chr19 | 40255967 | 40256627 | 2.13143 | 3.30861 | 1.552 |
| chr7 | 143187889 | 143188113 | 1.46445 | 2.27336 | 1.552 |
| chr10 | 130454717 | 130454967 | 0.501964 | 0.779241 | 1.552 |
| chr10 | 21309583 | 21309832 | 1.34001 | 2.08038 | 1.553 |
| chr8 | 74045459 | 74045681 | 0.544361 | 0.845194 | 1.553 |
| chr16 | 4311556 | 4311993 | 2.03006 | 3.15202 | 1.553 |
| chr1 | 204910894 | 204911203 | 2.93205 | 4.55258 | 1.553 |
| chr11 | 33992653 | 33992895 | 1.51269 | 2.34879 | 1.553 |
| chr17 | 33046477 | 33046808 | 0.669444 | 1.0395 | 1.553 |
| chr19 | 32409241 | 32409460 | 0.809469 | 1.25697 | 1.553 |
| chr17 | 6067923 | 6068218 | 0.896272 | 1.39177 | 1.553 |
| chr18 | 12092749 | 12093083 | 1.55862 | 2.42028 | 1.553 |
| chr5 | 63908160 | 63908510 | 1.90618 | 2.96007 | 1.553 |
| chr22 | 45949661 | 45949935 | 2.29605 | 3.56559 | 1.553 |
| chr19 | 39125210 | 39125665 | 2.52869 | 3.92689 | 1.553 |
| chr10 | 134018342 | 134018560 | 0.882797 | 1.37094 | 1.553 |
| chr3 | 161569776 | 161570121 | 1.87187 | 2.907 | 1.553 |
| chr14 | 50833567 | 50833947 | 0.6328 | 0.982747 | 1.553 |
| chr19 | 19106571 | 19106890 | 1.24539 | 1.93411 | 1.553 |
| chr1 | 154721598 | 154721829 | 1.14505 | 1.77838 | 1.553 |
| chr3 | 149111213 | 149111468 | 2.3516 | 3.65236 | 1.553 |
| chr9 | 116870184 | 116870563 | 3.32137 | 5.15864 | 1.553 |
| chr7 | 93003760 | 93003968 | 1.77461 | 2.75658 | 1.553 |
| chr1 | 171624593 | 171625000 | 1.3997 | 2.17424 | 1.553 |
| chr19 | 3326976 | 3327241 | 1.47606 | 2.29288 | 1.553 |
| chr1 | 7812840 | 7813241 | 1.27569 | 1.98181 | 1.554 |
| chr11 | 34587080 | 34587287 | 1.60835 | 2.49862 | 1.554 |
| chr11 | 32073735 | 32074089 | 1.54974 | 2.40764 | 1.554 |
| chr5 | 74561531 | 74561882 | 1.27184 | 1.97598 | 1.554 |
| chr2 | 88487590 | 88487800 | 0.871358 | 1.35379 | 1.554 |
| chr19 | 1266893 | 1267278 | 1.79833 | 2.79426 | 1.554 |
| chr2 | 19562747 | 19563099 | 1.48178 | 2.30247 | 1.554 |
| chr14 | 82756939 | 82757279 | 0.663446 | 1.03091 | 1.554 |
| chr2 | 85956076 | 85956471 | 3.12053 | 4.84903 | 1.554 |
| chr3 | 37814480 | 37814734 | 0.539613 | 0.838537 | 1.554 |
| chr1 | 161041664 | 161041931 | 2.0008 | 3.10928 | 1.554 |
| chr16 | 56634364 | 56634714 | 1.78467 | 2.77348 | 1.554 |
| chr19 | 31059594 | 31059845 | 0.807732 | 1.25527 | 1.554 |
| chr3 | 52035496 | 52035862 | 1.84393 | 2.86568 | 1.554 |
| chr15 | 85201397 | 85201838 | 2.13257 | 3.3143 | 1.554 |
| chr19 | 50981783 | 50982016 | 0.564914 | 0.877991 | 1.554 |
| chr1 | 24476972 | 24477319 | 1.49264 | 2.31993 | 1.554 |
| chr8 | 97891990 | 97892255 | 0.664658 | 1.03308 | 1.554 |
| chr1 | 53370550 | 53370765 | 0.979437 | 1.52238 | 1.554 |
| chr10 | 65772656 | 65772979 | 0.58149 | 0.903844 | 1.554 |
| chr7 | 135357649 | 135357896 | 1.92091 | 2.98583 | 1.554 |
| chr18 | 56751455 | 56751727 | 1.61006 | 2.5027 | 1.554 |
| chr20 | 22672126 | 22672595 | 1.91194 | 2.97196 | 1.554 |
| chr7 | 104137950 | 104138204 | 0.655531 | 1.01919 | 1.555 |
| chr16 | 3224274 | 3224650 | 1.48669 | 2.31148 | 1.555 |
| chrX | 41340550 | 41340788 | 1.04442 | 1.62384 | 1.555 |
| chrX | 49020051 | 49020336 | 2.12998 | 3.31164 | 1.555 |
| chr11 | 113906809 | 113907197 | 2.21967 | 3.45111 | 1.555 |
| chr2 | 181260127 | 181260336 | 0.521217 | 0.8104 | 1.555 |
| chr11 | 74099814 | 74100030 | 1.40068 | 2.17783 | 1.555 |
| chr5 | 139243655 | 139244027 | 1.88256 | 2.92719 | 1.555 |
| chr9 | 79782377 | 79782564 | 1.31314 | 2.0418 | 1.555 |
| chr17 | 38511178 | 38511667 | 2.91699 | 4.53569 | 1.555 |
| chr5 | 85519761 | 85519981 | 0.586649 | 0.912197 | 1.555 |
| chr5 | 122435272 | 122435732 | 1.29973 | 2.02107 | 1.555 |

| | | | | | |
|---|---|---|---|---|---|
| chr10 | 22278495 | 22278820 | 1.33849 | 2.08138 | 1.555 |
| chr2 | 55945421 | 55945627 | 1.49205 | 2.32019 | 1.555 |
| chr3 | 173296957 | 173297192 | 0.525114 | 0.816582 | 1.555 |
| chr7 | 50850014 | 50850511 | 1.04846 | 1.63042 | 1.555 |
| chr7 | 101245586 | 101246085 | 2.14423 | 3.3344 | 1.555 |
| chr12 | 65707570 | 65707777 | 1.44234 | 2.24301 | 1.555 |
| chr2 | 75855058 | 75855341 | 2.72775 | 4.24199 | 1.555 |
| chr9 | 6579816 | 6580187 | 0.687151 | 1.06875 | 1.555 |
| chr9 | 97316632 | 97316962 | 1.87177 | 2.9113 | 1.555 |
| chr15 | 101870016 | 101870241 | 0.712089 | 1.10757 | 1.555 |
| chr9 | 126785031 | 126785394 | 0.549646 | 0.854919 | 1.555 |
| chr14 | 31919241 | 31919530 | 1.12674 | 1.75257 | 1.555 |
| chr8 | 53513975 | 53514271 | 1.00293 | 1.55999 | 1.555 |
| chr2 | 179756184 | 179756461 | 1.71267 | 2.66395 | 1.555 |
| chr1 | 22141033 | 22141264 | 0.89071 | 1.3855 | 1.556 |
| chr6 | 125632293 | 125632535 | 1.62843 | 2.53303 | 1.556 |
| chr22 | 22549704 | 22550110 | 2.11839 | 3.29528 | 1.556 |
| chr16 | 90085915 | 90086233 | 1.82493 | 2.83884 | 1.556 |
| chr1 | 212980583 | 212980893 | 0.59219 | 0.921224 | 1.556 |
| chr18 | 12246916 | 12247488 | 1.13242 | 1.76167 | 1.556 |
| chr7 | 24329992 | 24330207 | 1.30147 | 2.02469 | 1.556 |
| chr22 | 25168546 | 25168863 | 1.4268 | 2.21971 | 1.556 |
| chr20 | 36306521 | 36306881 | 0.536376 | 0.834542 | 1.556 |
| chr5 | 161930799 | 161931064 | 0.534206 | 0.831169 | 1.556 |
| chr20 | 32503478 | 32503829 | 1.92823 | 3.00018 | 1.556 |
| chr12 | 130444203 | 130444433 | 0.693257 | 1.07867 | 1.556 |
| chr20 | 50722203 | 50722480 | 1.8842 | 2.93191 | 1.556 |
| chr11 | 65149745 | 65150010 | 1.33201 | 2.07291 | 1.556 |
| chr1 | 209749917 | 209750452 | 2.1489 | 3.34424 | 1.556 |
| chr20 | 21032878 | 21033094 | 0.634674 | 0.987728 | 1.556 |
| chr5 | 176944350 | 176944632 | 0.536156 | 0.834415 | 1.556 |
| chr10 | 54067992 | 54068284 | 2.079 | 3.23603 | 1.557 |
| chr9 | 32008745 | 32009023 | 0.60382 | 0.939877 | 1.557 |
| chr9 | 105453368 | 105453610 | 0.687159 | 1.06965 | 1.557 |
| chr2 | 80380145 | 80380395 | 0.785092 | 1.22212 | 1.557 |
| chr7 | 93532157 | 93532427 | 1.93698 | 3.01522 | 1.557 |
| chr1 | 206899670 | 206899954 | 1.73205 | 2.69623 | 1.557 |
| chr14 | 26863328 | 26863555 | 1.24152 | 1.93264 | 1.557 |
| chr16 | 82343881 | 82344122 | 0.593994 | 0.924662 | 1.557 |
| chr5 | 108903135 | 108903366 | 0.6204 | 0.965791 | 1.557 |
| chr5 | 146958026 | 146958238 | 1.07096 | 1.66727 | 1.557 |
| chr3 | 160295387 | 160295674 | 1.41656 | 2.2054 | 1.557 |
| chr3 | 74382419 | 74382733 | 0.765927 | 1.1925 | 1.557 |
| chr7 | 39593630 | 39593848 | 0.69291 | 1.07883 | 1.557 |
| chr10 | 2970407 | 2970802 | 1.52587 | 2.3758 | 1.557 |
| chr17 | 58220267 | 58220584 | 2.17418 | 3.38534 | 1.557 |
| chr17 | 72810658 | 72810863 | 1.191 | 1.85452 | 1.557 |
| chr9 | 9495866 | 9496079 | 0.694035 | 1.08069 | 1.557 |
| chr11 | 75099836 | 75100137 | 1.63386 | 2.54414 | 1.557 |
| chr17 | 78997593 | 78998009 | 2.34822 | 3.6565 | 1.557 |
| rUn_gl0002 | 144156 | 144622 | 1.53607 | 2.39195 | 1.557 |
| chr8 | 25041138 | 25041390 | 1.88609 | 2.93715 | 1.557 |
| chr4 | 15957920 | 15958233 | 1.91063 | 2.97543 | 1.557 |
| chr14 | 77339158 | 77339697 | 1.17471 | 1.82941 | 1.557 |
| chr14 | 106501390 | 106501779 | 1.67755 | 2.61251 | 1.557 |
| chr16 | 82208254 | 82208486 | 0.806455 | 1.25599 | 1.557 |
| chr21 | 37961717 | 37962135 | 1.38234 | 2.15288 | 1.557 |
| chr1 | 155931890 | 155932136 | 1.09562 | 1.70646 | 1.558 |
| chr16 | 75032315 | 75032729 | 1.75766 | 2.73766 | 1.558 |
| chr17 | 30771153 | 30771526 | 3.63662 | 5.66432 | 1.558 |
| chr6 | 70818163 | 70818404 | 0.77572 | 1.20831 | 1.558 |

| | | | | | |
|---|---|---|---|---|---|
| chr1 | 12166951 | 12167229 | 1.42049 | 2.21282 | 1.558 |
| chr2 | 27502496 | 27503406 | 1.16593 | 1.81631 | 1.558 |
| chr10 | 128436653 | 128436925 | 0.568415 | 0.885585 | 1.558 |
| chr3 | 22598648 | 22599097 | 0.61511 | 0.958343 | 1.558 |
| chr17 | 13036722 | 13036951 | 0.721527 | 1.12415 | 1.558 |
| chr9 | 32136816 | 32137010 | 0.83141 | 1.29536 | 1.558 |
| chr7 | 147232684 | 147232934 | 0.622939 | 0.970564 | 1.558 |
| chr8 | 18143649 | 18143927 | 0.657281 | 1.02411 | 1.558 |
| chr7 | 123744751 | 123745033 | 0.592354 | 0.922958 | 1.558 |
| chr7 | 106013583 | 106013793 | 0.535919 | 0.835044 | 1.558 |
| chr2 | 151467240 | 151467506 | 1.66619 | 2.59619 | 1.558 |
| chr2 | 43839631 | 43839869 | 1.4673 | 2.28636 | 1.558 |
| chr17 | 18151336 | 18151869 | 1.35379 | 2.10969 | 1.558 |
| chr1 | 32228239 | 32228515 | 1.90729 | 2.97234 | 1.558 |
| chr8 | 81599396 | 81599781 | 2.00706 | 3.12795 | 1.558 |
| chr8 | 63924443 | 63924756 | 0.865596 | 1.34904 | 1.559 |
| chr7 | 89111101 | 89111348 | 0.657146 | 1.02418 | 1.559 |
| chr8 | 120879412 | 120879654 | 1.57513 | 2.45488 | 1.559 |
| chr22 | 41318826 | 41319047 | 1.2983 | 2.02358 | 1.559 |
| chr2 | 57986957 | 57987292 | 2.31095 | 3.60202 | 1.559 |
| chr14 | 57533652 | 57533956 | 2.0329 | 3.16867 | 1.559 |
| chr16 | 86868114 | 86868538 | 0.980053 | 1.52763 | 1.559 |
| chr19 | 15374839 | 15375177 | 1.408 | 2.19469 | 1.559 |
| chr11 | 87063618 | 87064205 | 2.42698 | 3.78303 | 1.559 |
| chr18 | 43271315 | 43271536 | 2.09559 | 3.26656 | 1.559 |
| chr20 | 12188568 | 12188777 | 1.12671 | 1.7563 | 1.559 |
| chr16 | 8728950 | 8729683 | 2.26902 | 3.53731 | 1.559 |
| chr8 | 12724405 | 12724674 | 1.58684 | 2.47395 | 1.559 |
| chr5 | 72599507 | 72599894 | 0.922528 | 1.43827 | 1.559 |
| chr11 | 21255826 | 21256036 | 0.505138 | 0.787538 | 1.559 |
| chr5 | 173738896 | 173739395 | 2.22783 | 3.47333 | 1.559 |
| chr5 | 140857917 | 140858228 | 1.08346 | 1.68921 | 1.559 |
| chr9 | 38334294 | 38334563 | 1.20769 | 1.88297 | 1.559 |
| chr9 | 114860043 | 114860366 | 2.14992 | 3.35213 | 1.559 |
| chr7 | 40226207 | 40226497 | 1.02893 | 1.60444 | 1.559 |
| chr12 | 113486534 | 113486749 | 2.39825 | 3.73971 | 1.559 |
| chrX | 19390354 | 19390791 | 1.69882 | 2.64919 | 1.559 |
| chr16 | 30685645 | 30685850 | 0.582211 | 0.907927 | 1.559 |
| chr11 | 78787598 | 78787887 | 1.45546 | 2.26975 | 1.559 |
| chr16 | 19998218 | 19998454 | 0.505824 | 0.788887 | 1.560 |
| chr10 | 120142600 | 120142867 | 0.669369 | 1.044 | 1.560 |
| chr12 | 119275878 | 119276257 | 0.537867 | 0.838913 | 1.560 |
| chr17 | 32965138 | 32965692 | 2.18161 | 3.40266 | 1.560 |
| chr14 | 102024456 | 102024700 | 0.97702 | 1.5239 | 1.560 |
| chr2 | 103232310 | 103232712 | 2.85084 | 4.4472 | 1.560 |
| chr22 | 37249567 | 37249889 | 1.55284 | 2.42248 | 1.560 |
| chr12 | 66724870 | 66725158 | 0.849019 | 1.32453 | 1.560 |
| chr8 | 70488120 | 70488331 | 0.78473 | 1.22424 | 1.560 |
| chr15 | 99599967 | 99600404 | 0.853501 | 1.3316 | 1.560 |
| chr8 | 97918182 | 97918426 | 0.532274 | 0.830434 | 1.560 |
| chr18 | 34888921 | 34889174 | 0.525575 | 0.820005 | 1.560 |
| chr12 | 113612068 | 113612596 | 3.13805 | 4.89608 | 1.560 |
| chr19 | 55944489 | 55944699 | 0.358858 | 0.559902 | 1.560 |
| chr22 | 24110499 | 24110812 | 2.04213 | 3.18626 | 1.560 |
| chr14 | 42299805 | 42300074 | 0.755335 | 1.17859 | 1.560 |
| chr16 | 66785639 | 66785935 | 1.86297 | 2.90697 | 1.560 |
| chr5 | 68760229 | 68760533 | 0.692978 | 1.08132 | 1.560 |
| chr2 | 155309468 | 155309657 | 0.866563 | 1.3522 | 1.560 |
| chr6 | 112658423 | 112658686 | 1.47118 | 2.29566 | 1.560 |
| chr1 | 217507262 | 217507565 | 1.4742 | 2.30049 | 1.561 |
| chr14 | 79107732 | 79108127 | 0.762975 | 1.19063 | 1.561 |

| | | | | | |
|---|---|---|---|---|---|
| chr17 | 4529245 | 4529587 | 1.4195 | 2.21527 | 1.561 |
| chr15 | 89454354 | 89454556 | 1.18224 | 1.84505 | 1.561 |
| chr8 | 95199655 | 95200219 | 0.827922 | 1.2921 | 1.561 |
| chr2 | 4772487 | 4772825 | 0.673294 | 1.05079 | 1.561 |
| chr5 | 154813868 | 154814097 | 0.497234 | 0.776034 | 1.561 |
| chr10 | 91553433 | 91553744 | 2.018 | 3.14955 | 1.561 |
| chr1 | 158072823 | 158073103 | 0.654826 | 1.02202 | 1.561 |
| chr15 | 29214205 | 29214645 | 0.831069 | 1.2971 | 1.561 |
| chr3 | 158984666 | 158984966 | 1.19748 | 1.86904 | 1.561 |
| chr12 | 60994897 | 60995167 | 0.71806 | 1.12087 | 1.561 |
| chr20 | 51551222 | 51551500 | 0.688454 | 1.07469 | 1.561 |
| chr2 | 159202860 | 159203143 | 0.604975 | 0.944391 | 1.561 |
| chr4 | 8470513 | 8470880 | 2.16313 | 3.37673 | 1.561 |
| chr20 | 9130326 | 9130744 | 0.566979 | 0.885085 | 1.561 |
| chr13 | 21286712 | 21287073 | 1.96005 | 3.06033 | 1.561 |
| chr11 | 1890782 | 1891683 | 1.18396 | 1.84862 | 1.561 |
| chr11 | 6760018 | 6760281 | 0.502899 | 0.785274 | 1.561 |
| chr10 | 124027482 | 124028131 | 1.87866 | 2.93359 | 1.562 |
| chr12 | 61153175 | 61153435 | 0.824011 | 1.28672 | 1.562 |
| chr5 | 61064549 | 61064904 | 2.43738 | 3.80615 | 1.562 |
| chr2 | 60777533 | 60777905 | 1.61917 | 2.52848 | 1.562 |
| chr15 | 80534531 | 80534834 | 1.76696 | 2.75935 | 1.562 |
| chr5 | 127537044 | 127537379 | 2.65638 | 4.14833 | 1.562 |
| chr6 | 117926468 | 117926696 | 1.37784 | 2.1517 | 1.562 |
| chr1 | 245988549 | 245988861 | 0.593766 | 0.927282 | 1.562 |
| chr7 | 33934837 | 33935254 | 2.38568 | 3.72573 | 1.562 |
| chr17 | 45879717 | 45880161 | 1.7764 | 2.77425 | 1.562 |
| chr16 | 88662199 | 88662445 | 0.527489 | 0.823815 | 1.562 |
| chr2 | 86902757 | 86903080 | 0.998211 | 1.55902 | 1.562 |
| chr14 | 53762715 | 53763139 | 1.36222 | 2.12761 | 1.562 |
| chr17 | 73521047 | 73521280 | 1.38008 | 2.1555 | 1.562 |
| chr12 | 4713833 | 4714182 | 2.64597 | 4.13269 | 1.562 |
| chr3 | 182401404 | 182401785 | 1.61687 | 2.52549 | 1.562 |
| chr8 | 22322458 | 22322839 | 0.57067 | 0.8914 | 1.562 |
| chr18 | 74061092 | 74061357 | 1.6407 | 2.56288 | 1.562 |
| chr3 | 143410412 | 143410661 | 0.514175 | 0.8032 | 1.562 |
| chr14 | 66537414 | 66537834 | 1.95838 | 3.05925 | 1.562 |
| chr6 | 36945075 | 36945524 | 1.35185 | 2.11181 | 1.562 |
| chr9 | 34760473 | 34760983 | 1.87675 | 2.93183 | 1.562 |
| chr12 | 82410546 | 82410845 | 0.830716 | 1.29774 | 1.562 |
| chr16 | 72866455 | 72866829 | 3.22282 | 5.03506 | 1.562 |
| chr12 | 56053369 | 56053889 | 1.16562 | 1.82111 | 1.562 |
| chr3 | 70157313 | 70157523 | 1.24678 | 1.94791 | 1.562 |
| chr11 | 114266500 | 114266739 | 0.943836 | 1.47466 | 1.562 |
| chr12 | 12714686 | 12714948 | 0.770542 | 1.2039 | 1.562 |
| chr2 | 39956023 | 39956401 | 2.16285 | 3.37949 | 1.563 |
| chrX | 111885652 | 111885979 | 1.61315 | 2.52064 | 1.563 |
| chr15 | 68943330 | 68943779 | 2.83008 | 4.42224 | 1.563 |
| chr18 | 56164177 | 56164445 | 1.09197 | 1.70632 | 1.563 |
| chr11 | 116562476 | 116562767 | 1.47758 | 2.30892 | 1.563 |
| chr12 | 125687787 | 125688015 | 1.84879 | 2.88899 | 1.563 |
| chr7 | 12428794 | 12429047 | 2.11901 | 3.31127 | 1.563 |
| chr16 | 13959921 | 13960325 | 1.6744 | 2.61653 | 1.563 |
| chr11 | 68611448 | 68611711 | 1.50255 | 2.34806 | 1.563 |
| chr6 | 33729951 | 33730345 | 1.83761 | 2.87169 | 1.563 |
| chr2 | 121070757 | 121071116 | 2.36976 | 3.70338 | 1.563 |
| chr12 | 44915684 | 44915970 | 0.747301 | 1.16788 | 1.563 |
| chr2 | 67120435 | 67120650 | 0.674524 | 1.05416 | 1.563 |
| chr10 | 125341610 | 125342022 | 1.83708 | 2.87108 | 1.563 |
| chr2 | 206640826 | 206641169 | 2.71112 | 4.23721 | 1.563 |
| chrX | 49049235 | 49049557 | 1.53578 | 2.40035 | 1.563 |

| | | | | | |
|---|---|---|---|---|---|
| chr5 | 54201794 | 54202106 | 1.56605 | 2.4478 | 1.563 |
| chr17 | 73844492 | 73844830 | 2.0102 | 3.14206 | 1.563 |
| chr5 | 99780920 | 99781282 | 0.608694 | 0.95144 | 1.563 |
| chr17 | 45393085 | 45393347 | 1.43837 | 2.24831 | 1.563 |
| chr12 | 109784816 | 109785085 | 0.884816 | 1.38307 | 1.563 |
| chr5 | 122609715 | 122610100 | 1.03442 | 1.61693 | 1.563 |
| chr12 | 30593811 | 30594069 | 1.39227 | 2.17634 | 1.563 |
| chr10 | 67860823 | 67861078 | 0.764708 | 1.19538 | 1.563 |
| chr12 | 43644841 | 43645084 | 0.862114 | 1.34764 | 1.563 |
| chr2 | 8667887 | 8668105 | 0.656328 | 1.02596 | 1.563 |
| chr8 | 35558741 | 35558996 | 0.776982 | 1.21458 | 1.563 |
| chr10 | 101619455 | 101619718 | 2.16138 | 3.37885 | 1.563 |
| chr6 | 25556723 | 25556952 | 1.63365 | 2.5539 | 1.563 |
| chr20 | 2502215 | 2502801 | 2.19042 | 3.42486 | 1.564 |
| chr5 | 173000106 | 173000440 | 2.53124 | 3.95802 | 1.564 |
| chr1 | 188751406 | 188751657 | 1.60062 | 2.50289 | 1.564 |
| chr20 | 36531846 | 36532158 | 1.78432 | 2.79016 | 1.564 |
| chr11 | 72947910 | 72948269 | 1.59895 | 2.50037 | 1.564 |
| chr12 | 83891953 | 83892171 | 0.740821 | 1.15847 | 1.564 |
| chr8 | 125901041 | 125901308 | 1.1718 | 1.83244 | 1.564 |
| chr17 | 28875057 | 28875326 | 0.664042 | 1.03842 | 1.564 |
| chr17 | 79988302 | 79988832 | 1.97875 | 3.09437 | 1.564 |
| chr8 | 17058204 | 17058461 | 0.949452 | 1.4848 | 1.564 |
| chr9 | 132379591 | 132379871 | 2.05752 | 3.21772 | 1.564 |
| chr2 | 74007161 | 74007452 | 2.57838 | 4.03238 | 1.564 |
| chr1 | 203086823 | 203087381 | 2.00716 | 3.13907 | 1.564 |
| chr12 | 30334380 | 30334599 | 0.819468 | 1.28161 | 1.564 |
| chr8 | 1175062 | 1175306 | 0.554668 | 0.867541 | 1.564 |
| chr17 | 74109245 | 74109468 | 0.552748 | 0.864542 | 1.564 |
| chr3 | 77402957 | 77403182 | 0.546613 | 0.854946 | 1.564 |
| chr5 | 142393611 | 142393974 | 1.10199 | 1.72364 | 1.564 |
| chr9 | 78917918 | 78918205 | 1.13564 | 1.77633 | 1.564 |
| chrX | 49603094 | 49603364 | 1.49203 | 2.33379 | 1.564 |
| chr11 | 67156849 | 67157438 | 1.16791 | 1.82682 | 1.564 |
| chr15 | 90190216 | 90190991 | 2.30096 | 3.59926 | 1.564 |
| chr1 | 147266698 | 147266932 | 0.688894 | 1.07768 | 1.564 |
| chr19 | 4208845 | 4209122 | 1.56516 | 2.44848 | 1.564 |
| chr2 | 234312964 | 234313198 | 1.88803 | 2.95357 | 1.564 |
| chr20 | 18393915 | 18394202 | 1.33338 | 2.08597 | 1.564 |
| chr1 | 112289918 | 112290186 | 1.83483 | 2.87052 | 1.564 |
| chr6 | 11496529 | 11496820 | 1.49227 | 2.33472 | 1.565 |
| chr3 | 27934341 | 27934571 | 0.501724 | 0.785007 | 1.565 |
| chr2 | 223572965 | 223573366 | 2.19177 | 3.4293 | 1.565 |
| chr15 | 100455553 | 100455796 | 0.565576 | 0.884949 | 1.565 |
| chr6 | 142310475 | 142310767 | 2.19855 | 3.44007 | 1.565 |
| chr3 | 111387853 | 111388059 | 1.23656 | 1.93499 | 1.565 |
| chr18 | 7164642 | 7164930 | 1.30081 | 2.03572 | 1.565 |
| chr16 | 56903493 | 56903701 | 1.63315 | 2.55583 | 1.565 |
| chrX | 36885314 | 36885526 | 0.958881 | 1.50067 | 1.565 |
| chr14 | 93500749 | 93501077 | 1.54682 | 2.42082 | 1.565 |
| chr1 | 211124246 | 211124599 | 0.675685 | 1.0575 | 1.565 |
| chr9 | 127379817 | 127380095 | 1.87861 | 2.9403 | 1.565 |
| chr14 | 22336655 | 22337029 | 1.66092 | 2.59963 | 1.565 |
| chr10 | 73571700 | 73572087 | 2.48059 | 3.88267 | 1.565 |
| chr12 | 19035692 | 19035948 | 1.02538 | 1.60499 | 1.565 |
| chr12 | 61383029 | 61383281 | 0.787216 | 1.23223 | 1.565 |
| chr3 | 38687934 | 38688393 | 1.58154 | 2.47585 | 1.565 |
| chr12 | 91302858 | 91303259 | 2.02701 | 3.17325 | 1.565 |
| chr17 | 73615990 | 73616469 | 1.452 | 2.27307 | 1.565 |
| chr9 | 129276510 | 129276884 | 2.2815 | 3.57166 | 1.565 |
| chr11 | 59961117 | 59961507 | 1.42306 | 2.22789 | 1.566 |

| | | | | | |
|---|---|---|---|---|---|
| chr2 | 50719639 | 50719965 | 1.42202 | 2.22631 | 1.566 |
| chr3 | 127787009 | 127787385 | 1.36194 | 2.13225 | 1.566 |
| chr1 | 62596840 | 62597129 | 2.16652 | 3.39195 | 1.566 |
| chr2 | 88301415 | 88301703 | 2.69377 | 4.21758 | 1.566 |
| chr8 | 29732547 | 29732817 | 1.25236 | 1.9608 | 1.566 |
| chr20 | 15839017 | 15839373 | 0.728369 | 1.1404 | 1.566 |
| chr3 | 149008981 | 149009232 | 1.28876 | 2.0178 | 1.566 |
| chr10 | 126429837 | 126430247 | 1.71101 | 2.67897 | 1.566 |
| chr14 | 24398985 | 24399327 | 1.81446 | 2.84097 | 1.566 |
| chr16 | 19234605 | 19234916 | 1.97922 | 3.09901 | 1.566 |
| chr9 | 25609166 | 25609411 | 0.577464 | 0.904173 | 1.566 |
| chr3 | 88162661 | 88163032 | 1.53913 | 2.40997 | 1.566 |
| chr2 | 204090193 | 204090502 | 1.53341 | 2.40108 | 1.566 |
| chr8 | 23550160 | 23550583 | 2.15881 | 3.38061 | 1.566 |
| chr20 | 22149044 | 22149294 | 0.724076 | 1.13392 | 1.566 |
| chr1 | 230446369 | 230446622 | 1.52272 | 2.38467 | 1.566 |
| chr9 | 88515426 | 88515759 | 1.77713 | 2.78316 | 1.566 |
| chr5 | 132795814 | 132796189 | 1.85971 | 2.91259 | 1.566 |
| chr5 | 98107216 | 98107568 | 1.99513 | 3.12475 | 1.566 |
| chr2 | 89179883 | 89180423 | 1.58586 | 2.48378 | 1.566 |
| chr5 | 139091041 | 139091284 | 0.967056 | 1.5146 | 1.566 |
| chr16 | 3081531 | 3082107 | 1.36115 | 2.13187 | 1.566 |
| chr1 | 3326772 | 3327049 | 1.18793 | 1.86058 | 1.566 |
| chr21 | 43912350 | 43912606 | 0.705667 | 1.10526 | 1.566 |
| chr21 | 42832447 | 42832852 | 1.50579 | 2.35852 | 1.566 |
| chr19 | 3471938 | 3472367 | 1.78964 | 2.80319 | 1.566 |
| chr16 | 765594 | 765986 | 0.963368 | 1.50901 | 1.566 |
| chr10 | 115376114 | 115376342 | 1.27243 | 1.99318 | 1.566 |
| chr12 | 89864573 | 89864863 | 1.77107 | 2.77428 | 1.566 |
| chr1 | 117215192 | 117215511 | 1.3474 | 2.11065 | 1.566 |
| chr8 | 3177572 | 3177826 | 0.682664 | 1.06949 | 1.567 |
| chr3 | 1123805 | 1124135 | 1.07771 | 1.68872 | 1.567 |
| chr20 | 60957765 | 60958187 | 2.31147 | 3.62217 | 1.567 |
| chr9 | 17838105 | 17838460 | 0.531376 | 0.832709 | 1.567 |
| chr3 | 120137806 | 120138135 | 1.65447 | 2.59279 | 1.567 |
| chr20 | 43948240 | 43948527 | 1.61763 | 2.53508 | 1.567 |
| chr12 | 53721318 | 53721564 | 1.07576 | 1.68601 | 1.567 |
| chr2 | 21086584 | 21086838 | 0.680195 | 1.06608 | 1.567 |
| chr1 | 172578791 | 172578995 | 1.25948 | 1.97406 | 1.567 |
| chr5 | 135866062 | 135866273 | 0.808453 | 1.26715 | 1.567 |
| chr1 | 179261541 | 179261819 | 1.90003 | 2.97823 | 1.567 |
| chr1 | 180779465 | 180779714 | 0.800327 | 1.25453 | 1.568 |
| chr11 | 61276759 | 61276977 | 1.27785 | 2.00305 | 1.568 |
| chr5 | 50676600 | 50676809 | 0.842656 | 1.32092 | 1.568 |
| chr16 | 68027077 | 68027509 | 2.88481 | 4.52226 | 1.568 |
| chrX | 40403197 | 40403439 | 1.81805 | 2.84999 | 1.568 |
| chr19 | 39005706 | 39006074 | 3.14336 | 4.92758 | 1.568 |
| chr7 | 122910182 | 122910464 | 0.688834 | 1.07986 | 1.568 |
| chr7 | 28117860 | 28118274 | 2.59496 | 4.06828 | 1.568 |
| chr19 | 4370035 | 4370260 | 1.27686 | 2.00195 | 1.568 |
| chr7 | 37274539 | 37274746 | 1.52898 | 2.39724 | 1.568 |
| chr1 | 228646949 | 228647219 | 1.8297 | 2.86881 | 1.568 |
| chr5 | 8129415 | 8129692 | 0.78037 | 1.22355 | 1.568 |
| chr7 | 14731285 | 14731573 | 1.67432 | 2.62521 | 1.568 |
| chr7 | 40369158 | 40369447 | 1.3402 | 2.10144 | 1.568 |
| chr1 | 200859900 | 200860270 | 1.71258 | 2.68554 | 1.568 |
| chr2 | 150860022 | 150860228 | 0.428125 | 0.671364 | 1.568 |
| chr9 | 84122147 | 84122385 | 0.682975 | 1.07103 | 1.568 |
| chr10 | 6962030 | 6962329 | 1.87839 | 2.94586 | 1.568 |
| chr14 | 53521674 | 53522001 | 1.46569 | 2.29866 | 1.568 |
| chr4 | 11767551 | 11767903 | 1.59664 | 2.50403 | 1.568 |

| | | | | | |
|---|---|---|---|---|---|
| chr7 | 143824825 | 143825164 | 1.91861 | 3.00902 | 1.568 |
| chr12 | 25363954 | 25364278 | 0.835449 | 1.31029 | 1.568 |
| chr3 | 11024992 | 11025469 | 1.94825 | 3.05562 | 1.568 |
| chr13 | 50550832 | 50551173 | 1.79032 | 2.80802 | 1.568 |
| chr8 | 9929411 | 9929635 | 0.712726 | 1.11788 | 1.568 |
| chr1 | 173398694 | 173398978 | 1.58376 | 2.48409 | 1.568 |
| chr7 | 36331128 | 36331413 | 2.30772 | 3.61967 | 1.569 |
| chr14 | 102968112 | 102968414 | 1.40943 | 2.21073 | 1.569 |
| chr1 | 179778894 | 179779562 | 1.66088 | 2.60525 | 1.569 |
| chr8 | 81904190 | 81904546 | 1.98335 | 3.11159 | 1.569 |
| chr2 | 152476104 | 152476374 | 0.540791 | 0.848429 | 1.569 |
| chr17 | 56385122 | 56385384 | 0.740849 | 1.16234 | 1.569 |
| chr7 | 8872386 | 8872670 | 0.482732 | 0.757507 | 1.569 |
| chrX | 134618655 | 134618913 | 1.03901 | 1.63042 | 1.569 |
| chr7 | 103797306 | 103797539 | 1.31564 | 2.06455 | 1.569 |
| chr8 | 78066458 | 78066766 | 0.624428 | 0.979882 | 1.569 |
| chr2 | 7343091 | 7343467 | 2.02301 | 3.17479 | 1.569 |
| chr1 | 37116087 | 37116419 | 2.31222 | 3.6288 | 1.569 |
| chr9 | 83965492 | 83965727 | 0.679695 | 1.06672 | 1.569 |
| chr7 | 130581709 | 130582037 | 1.50467 | 2.36149 | 1.569 |
| chr2 | 6001783 | 6002010 | 0.729908 | 1.14558 | 1.569 |
| chr22 | 23520644 | 23520975 | 2.46026 | 3.86146 | 1.570 |
| chr1 | 8042434 | 8042706 | 1.46436 | 2.29837 | 1.570 |
| chr5 | 179402269 | 179402516 | 1.59217 | 2.49917 | 1.570 |
| chr5 | 64396849 | 64397213 | 1.3872 | 2.17747 | 1.570 |
| chr3 | 50579805 | 50580242 | 2.57699 | 4.04514 | 1.570 |
| chr16 | 69528862 | 69529116 | 1.12911 | 1.77249 | 1.570 |
| chr14 | 74945712 | 74946055 | 2.16271 | 3.39506 | 1.570 |
| chr11 | 12303408 | 12303718 | 2.04355 | 3.2082 | 1.570 |
| chr11 | 13185091 | 13185305 | 0.474809 | 0.745458 | 1.570 |
| chrX | 109158423 | 109158699 | 0.964915 | 1.51501 | 1.570 |
| chr15 | 38682217 | 38682577 | 1.50659 | 2.36554 | 1.570 |
| chr5 | 14871942 | 14872499 | 1.49228 | 2.34308 | 1.570 |
| chr20 | 55907622 | 55907858 | 1.10718 | 1.73858 | 1.570 |
| chr19 | 49356843 | 49357060 | 0.633243 | 0.994372 | 1.570 |
| chr19 | 51298162 | 51298395 | 1.12662 | 1.76913 | 1.570 |
| chr2 | 135223813 | 135224028 | 0.961733 | 1.51023 | 1.570 |
| chr3 | 136824201 | 136824449 | 1.21735 | 1.91163 | 1.570 |
| chr2 | 129145092 | 129145333 | 1.00563 | 1.57917 | 1.570 |
| chr12 | 28065970 | 28066240 | 0.614764 | 0.965442 | 1.570 |
| chr7 | 142996893 | 142997278 | 1.97097 | 3.09531 | 1.570 |
| chr8 | 21894177 | 21894764 | 2.25135 | 3.53565 | 1.570 |
| chr15 | 89497999 | 89498368 | 0.637501 | 1.00126 | 1.571 |
| chr11 | 18806494 | 18806714 | 0.663539 | 1.04227 | 1.571 |
| chr2 | 121842533 | 121842855 | 1.7514 | 2.75138 | 1.571 |
| chr8 | 33013605 | 33013834 | 0.681808 | 1.07114 | 1.571 |
| chr22 | 46259821 | 46260170 | 1.67217 | 2.62709 | 1.571 |
| chr12 | 34523002 | 34523401 | 0.492172 | 0.773268 | 1.571 |
| chr17 | 48578501 | 48578833 | 1.64277 | 2.58101 | 1.571 |
| chr17 | 62384174 | 62385022 | 1.40061 | 2.20061 | 1.571 |
| chr16 | 81069056 | 81069512 | 3.70846 | 5.82675 | 1.571 |
| chr1 | 15753562 | 15753780 | 1.18308 | 1.85888 | 1.571 |
| chr17 | 46259051 | 46259309 | 1.68163 | 2.64227 | 1.571 |
| chr17 | 56332192 | 56332541 | 0.917138 | 1.44107 | 1.571 |
| chr8 | 144256501 | 144256773 | 0.644558 | 1.01278 | 1.571 |
| chr9 | 124623625 | 124624081 | 1.29375 | 2.03289 | 1.571 |
| chr2 | 22479777 | 22480103 | 0.734309 | 1.15384 | 1.571 |
| chr3 | 49603811 | 49604042 | 0.966286 | 1.51843 | 1.571 |
| chr5 | 170065593 | 170066104 | 2.6688 | 4.19384 | 1.571 |
| chr9 | 36980153 | 36980429 | 0.822499 | 1.29253 | 1.571 |
| chr3 | 95477701 | 95477988 | 0.781278 | 1.22777 | 1.571 |

| | | | | | |
|---|---|---|---|---|---|
| chr11 | 46691447 | 46691910 | 0.844144 | 1.32658 | 1.572 |
| chr5 | 59973604 | 59973808 | 1.48413 | 2.33237 | 1.572 |
| chr19 | 4547522 | 4547790 | 1.73744 | 2.73053 | 1.572 |
| chr14 | 76853319 | 76853896 | 2.20518 | 3.46564 | 1.572 |
| chr17 | 19896130 | 19896376 | 1.97275 | 3.10035 | 1.572 |
| chr12 | 129599189 | 129599476 | 0.587103 | 0.922801 | 1.572 |
| chr17 | 10400845 | 10401171 | 0.600575 | 0.943977 | 1.572 |
| chr8 | 121140849 | 121141052 | 0.682936 | 1.07343 | 1.572 |
| chr9 | 132386400 | 132386894 | 1.64619 | 2.58753 | 1.572 |
| chr1 | 208418649 | 208418906 | 1.53221 | 2.40839 | 1.572 |
| chr11 | 16809931 | 16810289 | 3.56794 | 5.60829 | 1.572 |
| chr3 | 160638850 | 160639137 | 1.7815 | 2.80032 | 1.572 |
| chr1 | 219905266 | 219905786 | 0.560857 | 0.881618 | 1.572 |
| chr16 | 53316238 | 53316558 | 0.751556 | 1.18138 | 1.572 |
| chr5 | 19670559 | 19670804 | 0.523611 | 0.823163 | 1.572 |
| chr10 | 18505408 | 18505718 | 1.06006 | 1.66655 | 1.572 |
| chr3 | 135614024 | 135614367 | 1.84661 | 2.90323 | 1.572 |
| chr15 | 82364710 | 82364954 | 0.653316 | 1.02718 | 1.572 |
| chr4 | 21790932 | 21791160 | 1.13904 | 1.7909 | 1.572 |
| chr6 | 159590160 | 159590510 | 1.45672 | 2.29038 | 1.572 |
| chr8 | 131476956 | 131477197 | 0.728172 | 1.1449 | 1.572 |
| chr3 | 14675365 | 14675773 | 1.88191 | 2.95899 | 1.572 |
| chr13 | 33224909 | 33225235 | 1.80216 | 2.8336 | 1.572 |
| chr18 | 3481848 | 3482063 | 1.04337 | 1.64063 | 1.572 |
| chr15 | 61209067 | 61209448 | 1.64222 | 2.58231 | 1.572 |
| chr17 | 53604970 | 53605211 | 1.70406 | 2.6796 | 1.572 |
| chr1 | 184044625 | 184044902 | 1.53372 | 2.41177 | 1.573 |
| chr17 | 11014103 | 11014323 | 0.487986 | 0.767382 | 1.573 |
| chr1 | 227100424 | 227100715 | 1.26005 | 1.98152 | 1.573 |
| chr15 | 74674749 | 74675089 | 2.01961 | 3.176 | 1.573 |
| chr5 | 132759482 | 132760288 | 2.04984 | 3.22389 | 1.573 |
| chr7 | 100005276 | 100005653 | 1.32273 | 2.08032 | 1.573 |
| chr8 | 81402209 | 81402493 | 1.33998 | 2.1075 | 1.573 |
| chr1 | 204958410 | 204958987 | 2.70824 | 4.25961 | 1.573 |
| chr1 | 63428910 | 63429187 | 1.25583 | 1.97526 | 1.573 |
| chr19 | 3821863 | 3822257 | 2.73221 | 4.29765 | 1.573 |
| chrX | 18395925 | 18396296 | 1.58171 | 2.48797 | 1.573 |
| chr7 | 132584728 | 132584933 | 1.15552 | 1.81765 | 1.573 |
| chr12 | 120703585 | 120703923 | 2.35702 | 3.70771 | 1.573 |
| chr7 | 148272659 | 148273043 | 1.78853 | 2.81345 | 1.573 |
| chr17 | 69760710 | 69760968 | 1.24816 | 1.96351 | 1.573 |
| chr2 | 17307415 | 17307636 | 0.815172 | 1.28236 | 1.573 |
| chr14 | 99971991 | 99972216 | 0.502505 | 0.790507 | 1.573 |
| chr17 | 79492769 | 79493071 | 0.529234 | 0.83256 | 1.573 |
| chr15 | 60799999 | 60800321 | 2.34528 | 3.68951 | 1.573 |
| chr15 | 52519621 | 52520009 | 1.93411 | 3.04278 | 1.573 |
| chr3 | 674257 | 674555 | 1.09045 | 1.71564 | 1.573 |
| chr8 | 31268183 | 31268455 | 0.851484 | 1.33967 | 1.573 |
| chr19 | 1546208 | 1546524 | 2.14068 | 3.36829 | 1.573 |
| chr20 | 542323 | 542802 | 2.35365 | 3.70349 | 1.574 |
| chr7 | 132341685 | 132342019 | 0.997657 | 1.56988 | 1.574 |
| chr17 | 6918113 | 6918334 | 1.59632 | 2.51206 | 1.574 |
| chr9 | 34836948 | 34837172 | 0.846354 | 1.3319 | 1.574 |
| chr12 | 41820189 | 41820398 | 0.462887 | 0.72846 | 1.574 |
| chr3 | 141363482 | 141364063 | 1.52739 | 2.40373 | 1.574 |
| chr10 | 134467293 | 134467738 | 0.77223 | 1.21533 | 1.574 |
| chr19 | 39542821 | 39543406 | 3.25376 | 5.12078 | 1.574 |
| chr11 | 62690501 | 62690798 | 1.67596 | 2.63787 | 1.574 |
| chr10 | 54369448 | 54369717 | 0.539895 | 0.8498 | 1.574 |
| chr15 | 74329304 | 74329506 | 1.87903 | 2.95767 | 1.574 |
| chr4 | 114873639 | 114873901 | 1.22609 | 1.92995 | 1.574 |

| | | | | | |
|---|---|---|---|---|---|
| chrX | 128735161 | 128735406 | 1.19363 | 1.87886 | 1.574 |
| chr19 | 18512595 | 18512886 | 1.36476 | 2.14827 | 1.574 |
| chr7 | 53739873 | 53740199 | 0.827891 | 1.30319 | 1.574 |
| chr10 | 2275979 | 2276278 | 0.616021 | 0.969694 | 1.574 |
| chr11 | 8350286 | 8350797 | 2.16852 | 3.41355 | 1.574 |
| chr3 | 184187144 | 184187473 | 1.63027 | 2.56635 | 1.574 |
| chr6 | 106787239 | 106787531 | 1.76968 | 2.78585 | 1.574 |
| chr11 | 67368529 | 67368786 | 0.649488 | 1.02248 | 1.574 |
| chr17 | 40189745 | 40190199 | 0.718075 | 1.13052 | 1.574 |
| chr22 | 36590961 | 36591234 | 0.681808 | 1.07345 | 1.574 |
| chr5 | 56433951 | 56434225 | 1.8844 | 2.96716 | 1.575 |
| chr15 | 84115609 | 84115900 | 1.28877 | 2.02932 | 1.575 |
| chr20 | 62066261 | 62066562 | 1.97216 | 3.1055 | 1.575 |
| chr9 | 19493075 | 19493554 | 1.36128 | 2.14365 | 1.575 |
| chr19 | 30992326 | 30992684 | 0.707723 | 1.11459 | 1.575 |
| chr14 | 105191460 | 105192320 | 1.00297 | 1.57961 | 1.575 |
| chr6 | 46387324 | 46387594 | 1.26591 | 1.99372 | 1.575 |
| chr12 | 48231435 | 48231743 | 2.51728 | 3.96465 | 1.575 |
| chr11 | 123382018 | 123382422 | 1.64877 | 2.59687 | 1.575 |
| chr2 | 57724660 | 57724972 | 1.87235 | 2.949 | 1.575 |
| chr7 | 155752814 | 155753104 | 0.742394 | 1.16931 | 1.575 |
| chr11 | 36786114 | 36786339 | 0.559821 | 0.881826 | 1.575 |
| chr12 | 21401213 | 21401547 | 0.67204 | 1.0587 | 1.575 |
| chr20 | 21594970 | 21595424 | 0.777672 | 1.22517 | 1.575 |
| chr19 | 45981622 | 45981876 | 2.34597 | 3.696 | 1.575 |
| chr1 | 232505839 | 232506129 | 0.605612 | 0.954184 | 1.576 |
| chr5 | 121944159 | 121944364 | 0.490004 | 0.772037 | 1.576 |
| chr5 | 178932704 | 178932935 | 0.62958 | 0.991958 | 1.576 |
| chr2 | 219051014 | 219051352 | 1.18849 | 1.87258 | 1.576 |
| chr2 | 99453534 | 99453841 | 1.74323 | 2.74666 | 1.576 |
| chr12 | 98818661 | 98818922 | 1.53127 | 2.41273 | 1.576 |
| chr7 | 42784993 | 42785360 | 1.56063 | 2.459 | 1.576 |
| chr19 | 35010637 | 35010936 | 0.766988 | 1.20854 | 1.576 |
| chr10 | 58936493 | 58936747 | 0.561538 | 0.884833 | 1.576 |
| chr19 | 57666234 | 57666501 | 0.821208 | 1.294 | 1.576 |
| chr1 | 11724056 | 11724378 | 1.49066 | 2.3489 | 1.576 |
| chr7 | 155655306 | 155655624 | 2.02916 | 3.19769 | 1.576 |
| chr11 | 47433617 | 47434118 | 2.6275 | 4.14081 | 1.576 |
| chr12 | 21218795 | 21219060 | 1.31302 | 2.06927 | 1.576 |
| chrX | 53508150 | 53508450 | 1.23153 | 1.94086 | 1.576 |
| chr6 | 125191802 | 125192042 | 0.824168 | 1.2989 | 1.576 |
| chr7 | 151402735 | 151402994 | 1.6808 | 2.64904 | 1.576 |
| chr16 | 24592050 | 24592287 | 1.15074 | 1.81365 | 1.576 |
| chr10 | 30870667 | 30871135 | 1.61608 | 2.54709 | 1.576 |
| chr7 | 134143141 | 134143497 | 1.22293 | 1.92746 | 1.576 |
| chr7 | 100958532 | 100959037 | 1.50958 | 2.37927 | 1.576 |
| chr14 | 88161711 | 88161944 | 0.552198 | 0.870337 | 1.576 |
| chr11 | 94802893 | 94803156 | 1.8445 | 2.90743 | 1.576 |
| chr1 | 157847933 | 157848202 | 1.2705 | 2.00266 | 1.576 |
| chr8 | 119724674 | 119724940 | 0.744921 | 1.17422 | 1.576 |
| chr5 | 13250897 | 13251136 | 0.918144 | 1.44733 | 1.576 |
| chr8 | 22419352 | 22419705 | 2.04709 | 3.22702 | 1.576 |
| chr7 | 40644430 | 40644729 | 1.27354 | 2.00761 | 1.576 |
| chr9 | 97367029 | 97367251 | 0.546663 | 0.861768 | 1.576 |
| chr6 | 64017776 | 64018021 | 1.14493 | 1.80498 | 1.577 |
| chr10 | 107364043 | 107364362 | 0.617096 | 0.972869 | 1.577 |
| chr6 | 76006985 | 76007252 | 0.926864 | 1.46124 | 1.577 |
| chr17 | 37736113 | 37736337 | 0.720733 | 1.13628 | 1.577 |
| chr15 | 65370175 | 65370623 | 1.94647 | 3.06893 | 1.577 |
| chr16 | 77618135 | 77618419 | 1.30278 | 2.05404 | 1.577 |
| chr2 | 207646275 | 207646616 | 1.49868 | 2.36292 | 1.577 |

| | | | | | |
|---|---|---|---|---|---|
| chr6 | 39315656 | 39315916 | 1.98456 | 3.12901 | 1.577 |
| chr14 | 94451289 | 94451526 | 1.1063 | 1.74439 | 1.577 |
| chr2 | 202101178 | 202101419 | 1.28534 | 2.02685 | 1.577 |
| chr1 | 17828808 | 17829169 | 2.1039 | 3.31787 | 1.577 |
| chr1 | 162067924 | 162068153 | 0.543667 | 0.857387 | 1.577 |
| chr9 | 93551709 | 93552081 | 0.695381 | 1.09665 | 1.577 |
| chr5 | 162991359 | 162991602 | 1.7447 | 2.75149 | 1.577 |
| chr13 | 36392771 | 36393159 | 1.12074 | 1.76756 | 1.577 |
| chr3 | 71635756 | 71636033 | 1.91568 | 3.02148 | 1.577 |
| chr4 | 48447700 | 48447947 | 1.08863 | 1.71712 | 1.577 |
| chr2 | 42465904 | 42466231 | 1.56631 | 2.47062 | 1.577 |
| chr11 | 16967507 | 16967849 | 2.72359 | 4.29608 | 1.577 |
| chr10 | 43788744 | 43788949 | 1.87871 | 2.96352 | 1.577 |
| chr3 | 14509558 | 14509891 | 2.53283 | 3.99534 | 1.577 |
| chr3 | 9061215 | 9061708 | 0.603382 | 0.951817 | 1.577 |
| chr6 | 43920537 | 43920739 | 1.16915 | 1.84437 | 1.578 |
| chr20 | 48806784 | 48807145 | 2.0875 | 3.29323 | 1.578 |
| chr20 | 35830644 | 35831087 | 1.55144 | 2.44768 | 1.578 |
| chr8 | 88998502 | 88998842 | 0.758348 | 1.19647 | 1.578 |
| chr13 | 35731384 | 35731594 | 1.30131 | 2.05327 | 1.578 |
| chr7 | 154581168 | 154581623 | 0.841609 | 1.32794 | 1.578 |
| chr1 | 117827291 | 117827535 | 1.41009 | 2.22497 | 1.578 |
| chr12 | 103493967 | 103494380 | 0.773978 | 1.22135 | 1.578 |
| chr17 | 56381551 | 56382015 | 2.02084 | 3.18892 | 1.578 |
| chr11 | 107670026 | 107670304 | 1.91352 | 3.01959 | 1.578 |
| chr19 | 10971879 | 10972137 | 1.17079 | 1.84755 | 1.578 |
| chr15 | 89157761 | 89158107 | 2.95258 | 4.65934 | 1.578 |
| chr1 | 16531241 | 16531855 | 0.969696 | 1.53026 | 1.578 |
| chr12 | 113392046 | 113392357 | 0.501644 | 0.791653 | 1.578 |
| chr11 | 121978955 | 121979271 | 1.80247 | 2.84454 | 1.578 |
| chr7 | 75903316 | 75903563 | 0.531608 | 0.838971 | 1.578 |
| chr12 | 50038248 | 50038736 | 2.55476 | 4.0319 | 1.578 |
| chrX | 153188336 | 153188621 | 1.59221 | 2.51282 | 1.578 |
| chr16 | 15776004 | 15776243 | 1.42622 | 2.2511 | 1.578 |
| chr12 | 29863083 | 29863296 | 0.696293 | 1.09902 | 1.578 |
| chr5 | 126625605 | 126625937 | 1.87593 | 2.96094 | 1.578 |
| chrX | 25041136 | 25041380 | 1.40486 | 2.21749 | 1.578 |
| chr2 | 152882212 | 152882736 | 2.99493 | 4.72739 | 1.578 |
| chr6 | 119121660 | 119121981 | 1.6322 | 2.57639 | 1.578 |
| chr7 | 31132320 | 31132590 | 0.619009 | 0.977143 | 1.579 |
| chr1 | 187520301 | 187520558 | 1.18228 | 1.86637 | 1.579 |
| chr5 | 1645101 | 1645360 | 2.34968 | 3.7094 | 1.579 |
| chr1 | 156808633 | 156809087 | 2.12831 | 3.35996 | 1.579 |
| chr16 | 1850044 | 1850319 | 0.695209 | 1.09755 | 1.579 |
| chr1 | 165797199 | 165797488 | 0.906723 | 1.43154 | 1.579 |
| chr10 | 135071096 | 135071517 | 1.57144 | 2.481 | 1.579 |
| chr3 | 81388311 | 81388540 | 1.72976 | 2.73104 | 1.579 |
| chr6 | 150045009 | 150045306 | 2.03597 | 3.21455 | 1.579 |
| chr2 | 103136560 | 103136769 | 1.15911 | 1.83014 | 1.579 |
| chr2 | 210937752 | 210937965 | 1.37873 | 2.17691 | 1.579 |
| chr2 | 232789969 | 232790350 | 1.91237 | 3.01949 | 1.579 |
| chr11 | 73676284 | 73676602 | 0.581354 | 0.917922 | 1.579 |
| chr22 | 19072644 | 19072930 | 1.14117 | 1.80188 | 1.579 |
| chr8 | 104807786 | 104807998 | 1.25478 | 1.98138 | 1.579 |
| chr16 | 80118580 | 80118815 | 1.41264 | 2.23071 | 1.579 |
| chr2 | 27473045 | 27473486 | 2.29867 | 3.62994 | 1.579 |
| chr5 | 9012485 | 9012689 | 0.569323 | 0.899062 | 1.579 |
| chr22 | 39712419 | 39712846 | 1.6379 | 2.58659 | 1.579 |
| chr9 | 129468764 | 129469108 | 2.23829 | 3.53499 | 1.579 |
| chr16 | 66943678 | 66944051 | 1.33129 | 2.1026 | 1.579 |
| chr15 | 32267780 | 32268449 | 0.753178 | 1.18963 | 1.579 |

| | | | | | |
|---|---|---|---|---|---|
| chr16 | 79476565 | 79476807 | 0.541799 | 0.855765 | 1.579 |
| chr17 | 36665408 | 36665916 | 2.01156 | 3.17725 | 1.580 |
| chr3 | 137770957 | 137771412 | 1.59563 | 2.52035 | 1.580 |
| chr4 | 187508605 | 187508888 | 1.51266 | 2.38937 | 1.580 |
| chr10 | 102063533 | 102063800 | 1.68109 | 2.65543 | 1.580 |
| chr17 | 47921167 | 47921596 | 2.09246 | 3.30523 | 1.580 |
| chr1 | 161456832 | 161457279 | 1.2989 | 2.05183 | 1.580 |
| chr13 | 113295103 | 113295424 | 2.16185 | 3.4151 | 1.580 |
| chr21 | 35763099 | 35763519 | 1.67499 | 2.64619 | 1.580 |
| chr10 | 27649568 | 27649846 | 2.08977 | 3.30161 | 1.580 |
| chr1 | 216825399 | 216825698 | 0.815592 | 1.28856 | 1.580 |
| chr9 | 30304116 | 30304329 | 0.533842 | 0.843478 | 1.580 |
| chr10 | 87484148 | 87484409 | 0.647871 | 1.02367 | 1.580 |
| chr14 | 97643389 | 97643625 | 1.57706 | 2.49186 | 1.580 |
| chr22 | 23583168 | 23583390 | 0.94041 | 1.48595 | 1.580 |
| chrX | 130050827 | 130051141 | 1.39413 | 2.20303 | 1.580 |
| chr3 | 13133629 | 13134009 | 2.08075 | 3.28807 | 1.580 |
| chr1 | 16118477 | 16118946 | 2.08672 | 3.29757 | 1.580 |
| chr17 | 42175613 | 42175849 | 1.23429 | 1.95055 | 1.580 |
| chr7 | 122411776 | 122412016 | 0.826674 | 1.30642 | 1.580 |
| chr10 | 53613391 | 53614130 | 0.478932 | 0.756879 | 1.580 |
| chr7 | 876127 | 876405 | 1.81364 | 2.86619 | 1.580 |
| chr11 | 72559994 | 72560280 | 1.60291 | 2.53324 | 1.580 |
| chr17 | 19715772 | 19716028 | 1.44331 | 2.28106 | 1.580 |
| chr17 | 1670438 | 1670832 | 0.796325 | 1.2586 | 1.581 |
| chr2 | 99218022 | 99218347 | 1.33468 | 2.10951 | 1.581 |
| chr15 | 60878041 | 60878323 | 2.35158 | 3.7168 | 1.581 |
| chr12 | 127835293 | 127835499 | 0.802868 | 1.269 | 1.581 |
| chr15 | 41151310 | 41151658 | 1.77527 | 2.80597 | 1.581 |
| chr5 | 55566382 | 55566694 | 3.27097 | 5.1703 | 1.581 |
| chr16 | 735996 | 736348 | 2.20726 | 3.48914 | 1.581 |
| chr12 | 2392842 | 2393074 | 0.709171 | 1.12106 | 1.581 |
| chr7 | 151000441 | 151000982 | 2.35347 | 3.72037 | 1.581 |
| chr5 | 139742807 | 139743147 | 2.5107 | 3.96904 | 1.581 |
| chr1 | 244374445 | 244374710 | 1.26184 | 1.99479 | 1.581 |
| chr11 | 6400542 | 6400818 | 1.677 | 2.65111 | 1.581 |
| chr18 | 5057755 | 5058048 | 1.51867 | 2.40089 | 1.581 |
| chr8 | 104768420 | 104768749 | 0.61746 | 0.976216 | 1.581 |
| chr17 | 4792292 | 4792951 | 1.97474 | 3.12213 | 1.581 |
| chr5 | 121877707 | 121877997 | 0.826209 | 1.30638 | 1.581 |
| chr3 | 69062744 | 69062954 | 2.05569 | 3.25042 | 1.581 |
| chr7 | 150595525 | 150595951 | 2.5074 | 3.96468 | 1.581 |
| chr11 | 66346490 | 66346780 | 2.77569 | 4.38903 | 1.581 |
| chr5 | 170218433 | 170218687 | 1.85717 | 2.93663 | 1.581 |
| chr21 | 40105856 | 40106162 | 1.4223 | 2.24914 | 1.581 |
| chr6 | 158493200 | 158493509 | 1.68629 | 2.66661 | 1.581 |
| chr10 | 12824409 | 12824803 | 2.05434 | 3.24869 | 1.581 |
| chr6 | 124655792 | 124656016 | 0.613249 | 0.969789 | 1.581 |
| chr16 | 70301374 | 70301742 | 1.2664 | 2.0027 | 1.581 |
| chr17 | 70475293 | 70475621 | 0.968812 | 1.5321 | 1.581 |
| chr5 | 109307614 | 109307834 | 0.628773 | 0.994352 | 1.581 |
| chr20 | 23232976 | 23233290 | 0.726914 | 1.14965 | 1.582 |
| chr3 | 159220789 | 159221083 | 0.770364 | 1.21842 | 1.582 |
| chr17 | 16318246 | 16318790 | 1.46475 | 2.31671 | 1.582 |
| chr2 | 208905471 | 208905783 | 1.43658 | 2.27216 | 1.582 |
| chrX | 130396724 | 130397061 | 1.89038 | 2.98992 | 1.582 |
| chr12 | 62051410 | 62051707 | 0.696851 | 1.10221 | 1.582 |
| chr5 | 180217022 | 180217237 | 1.0112 | 1.59947 | 1.582 |
| chr7 | 92617496 | 92617717 | 1.4045 | 2.22167 | 1.582 |
| chrX | 133530170 | 133530533 | 1.11706 | 1.76701 | 1.582 |
| chr7 | 145972814 | 145973095 | 0.681433 | 1.07794 | 1.582 |

| | | | | | |
|---|---|---|---|---|---|
| chr8 | 116088998 | 116089353 | 1.54004 | 2.43614 | 1.582 |
| chr2 | 39502207 | 39502445 | 0.56909 | 0.900279 | 1.582 |
| chr8 | 76254986 | 76255360 | 1.27094 | 2.01065 | 1.582 |
| chr1 | 109440808 | 109441042 | 1.33043 | 2.10482 | 1.582 |
| chr5 | 125564606 | 125564890 | 0.605449 | 0.957859 | 1.582 |
| chr12 | 8696576 | 8697215 | 1.74054 | 2.75371 | 1.582 |
| chr2 | 89221971 | 89222317 | 0.687744 | 1.0881 | 1.582 |
| chr22 | 23226186 | 23226446 | 1.73527 | 2.74547 | 1.582 |
| chr5 | 146568438 | 146568775 | 1.59677 | 2.52647 | 1.582 |
| chr17 | 29764224 | 29764698 | 2.19747 | 3.47695 | 1.582 |
| chr17 | 77038326 | 77038980 | 1.93821 | 3.06674 | 1.582 |
| chr19 | 36661612 | 36661959 | 2.35543 | 3.72697 | 1.582 |
| chr5 | 126898000 | 126898362 | 1.34219 | 2.12375 | 1.582 |
| chr18 | 9575461 | 9575790 | 1.50072 | 2.3746 | 1.582 |
| chr12 | 113866987 | 113867208 | 0.727371 | 1.15094 | 1.582 |
| chr5 | 116324730 | 116325125 | 0.582107 | 0.921111 | 1.582 |
| chr22 | 48970898 | 48971248 | 2.31514 | 3.66365 | 1.582 |
| chr2 | 210199176 | 210199524 | 2.08859 | 3.30515 | 1.582 |
| chr1 | 40071899 | 40072210 | 1.35294 | 2.14101 | 1.582 |
| chr8 | 49891531 | 49891927 | 1.50094 | 2.37524 | 1.583 |
| chr11 | 44932922 | 44933297 | 1.75351 | 2.77495 | 1.583 |
| chr11 | 32914835 | 32915408 | 1.46262 | 2.31469 | 1.583 |
| chr7 | 22893471 | 22893981 | 1.74968 | 2.76897 | 1.583 |
| chr5 | 79220858 | 79221217 | 0.696761 | 1.10269 | 1.583 |
| chr11 | 17967216 | 17967493 | 0.617317 | 0.977005 | 1.583 |
| chr19 | 4803871 | 4804403 | 2.21141 | 3.49997 | 1.583 |
| chr2 | 27239755 | 27240110 | 1.71326 | 2.71167 | 1.583 |
| chr11 | 73006262 | 73006964 | 1.86247 | 2.94807 | 1.583 |
| chr10 | 95301759 | 95302121 | 2.51956 | 3.98835 | 1.583 |
| chr12 | 45342523 | 45342879 | 4.70832 | 7.45304 | 1.583 |
| chr19 | 45905583 | 45905874 | 2.36065 | 3.7368 | 1.583 |
| chr2 | 166420390 | 166420613 | 0.896612 | 1.41934 | 1.583 |
| chr22 | 31709278 | 31709674 | 1.75854 | 2.78399 | 1.583 |
| chr2 | 200328112 | 200328430 | 2.00803 | 3.17899 | 1.583 |
| chr1 | 236238816 | 236239285 | 0.641615 | 1.01579 | 1.583 |
| chr19 | 36083200 | 36083519 | 1.30221 | 2.06163 | 1.583 |
| chr6 | 149867372 | 149867630 | 1.73361 | 2.74475 | 1.583 |
| chr7 | 96206091 | 96206509 | 0.606915 | 0.960904 | 1.583 |
| chr6 | 3742685 | 3742951 | 1.86654 | 2.95525 | 1.583 |
| chr10 | 89376184 | 89376387 | 0.651309 | 1.03123 | 1.583 |
| chr14 | 25287025 | 25287295 | 0.828691 | 1.31212 | 1.583 |
| chr17 | 4785221 | 4785694 | 2.76946 | 4.38517 | 1.583 |
| chr3 | 100991278 | 100991601 | 0.632967 | 1.00224 | 1.583 |
| chr11 | 69616212 | 69616702 | 1.72568 | 2.73247 | 1.583 |
| chr11 | 44654395 | 44654602 | 1.16898 | 1.85101 | 1.583 |
| chr1 | 183121061 | 183121552 | 0.690375 | 1.09318 | 1.583 |
| chr4 | 183727428 | 183727872 | 2.70806 | 4.28837 | 1.584 |
| chr16 | 67499724 | 67500169 | 2.30149 | 3.64459 | 1.584 |
| chr9 | 14541630 | 14541850 | 0.606221 | 0.960009 | 1.584 |
| chr3 | 34876528 | 34876758 | 0.693396 | 1.09811 | 1.584 |
| chr7 | 151329420 | 151329784 | 1.73254 | 2.74389 | 1.584 |
| chr5 | 137977509 | 137977937 | 1.94053 | 3.07331 | 1.584 |
| chr12 | 104099775 | 104100183 | 0.953197 | 1.50971 | 1.584 |
| chr8 | 126766564 | 126766836 | 0.639563 | 1.01303 | 1.584 |
| chr11 | 44307879 | 44308277 | 1.92768 | 3.05355 | 1.584 |
| chr5 | 151885052 | 151885267 | 1.15616 | 1.83145 | 1.584 |
| chr19 | 10968862 | 10969284 | 1.91287 | 3.03017 | 1.584 |
| chr6 | 33739268 | 33739661 | 1.99712 | 3.16367 | 1.584 |
| chr7 | 21806972 | 21807264 | 2.58482 | 4.09475 | 1.584 |
| chr20 | 61814625 | 61815251 | 1.8976 | 3.0063 | 1.584 |
| chr20 | 52532339 | 52532584 | 1.96848 | 3.11863 | 1.584 |

| | | | | | |
|---|---|---|---|---|---|
| chr15 | 94707576 | 94707785 | 1.34072 | 2.12416 | 1.584 |
| chr19 | 23076764 | 23077194 | 1.04206 | 1.65098 | 1.584 |
| chr7 | 158235708 | 158235915 | 0.779254 | 1.23465 | 1.584 |
| chr14 | 103607533 | 103607930 | 2.38799 | 3.78371 | 1.584 |
| chr10 | 116064327 | 116064652 | 2.16966 | 3.43786 | 1.585 |
| chr9 | 117415769 | 117416112 | 2.11922 | 3.35811 | 1.585 |
| chr11 | 8835119 | 8835459 | 2.5107 | 3.97855 | 1.585 |
| chr5 | 75633724 | 75633990 | 1.35476 | 2.14686 | 1.585 |
| chr19 | 56182883 | 56183287 | 2.75848 | 4.37182 | 1.585 |
| chr12 | 5249829 | 5250131 | 0.869406 | 1.37791 | 1.585 |
| chr15 | 34533530 | 34533846 | 1.38463 | 2.19454 | 1.585 |
| chr4 | 122173477 | 122173717 | 1.4817 | 2.34844 | 1.585 |
| chr2 | 201346184 | 201346440 | 1.4176 | 2.24695 | 1.585 |
| chr6 | 152085462 | 152085763 | 1.76306 | 2.79465 | 1.585 |
| chr1 | 212541288 | 212541644 | 1.61182 | 2.55497 | 1.585 |
| chr14 | 24834874 | 24835169 | 1.17768 | 1.8669 | 1.585 |
| chr7 | 76982951 | 76983300 | 0.518634 | 0.822164 | 1.585 |
| chr16 | 81763860 | 81764181 | 1.95285 | 3.09584 | 1.585 |
| chr17 | 65268773 | 65269069 | 1.38394 | 2.19409 | 1.585 |
| chr7 | 118910091 | 118910318 | 2.0898 | 3.31315 | 1.585 |
| chr2 | 1322183 | 1322403 | 0.623337 | 0.988244 | 1.585 |
| chr6 | 24389573 | 24389791 | 1.09547 | 1.73685 | 1.585 |
| chr5 | 150544182 | 150544739 | 1.54729 | 2.45326 | 1.586 |
| chr12 | 42766051 | 42766299 | 1.86359 | 2.95483 | 1.586 |
| chr17 | 61045821 | 61046153 | 2.15732 | 3.42061 | 1.586 |
| chr16 | 61519413 | 61519624 | 0.610419 | 0.967882 | 1.586 |
| chr7 | 6641723 | 6642353 | 1.99709 | 3.1666 | 1.586 |
| chr5 | 39714295 | 39714504 | 1.61986 | 2.56849 | 1.586 |
| chr7 | 47140483 | 47140772 | 1.9004 | 3.01366 | 1.586 |
| chr1 | 145491017 | 145491228 | 0.516786 | 0.819528 | 1.586 |
| chr19 | 31514208 | 31514628 | 0.699665 | 1.10956 | 1.586 |
| chr12 | 114392765 | 114393312 | 2.16786 | 3.43797 | 1.586 |
| chr15 | 91477361 | 91477882 | 1.69906 | 2.69452 | 1.586 |
| chr15 | 73105763 | 73106275 | 0.750915 | 1.19094 | 1.586 |
| chr5 | 149818730 | 149819073 | 1.64544 | 2.60982 | 1.586 |
| chr11 | 80286620 | 80286914 | 0.607098 | 0.96295 | 1.586 |
| chr9 | 89563462 | 89563686 | 1.66492 | 2.64089 | 1.586 |
| chr19 | 36287496 | 36287838 | 0.608348 | 0.96499 | 1.586 |
| chr11 | 124456242 | 124456460 | 1.59241 | 2.52597 | 1.586 |
| chr7 | 3105420 | 3105676 | 1.82441 | 2.89403 | 1.586 |
| chr1 | 155025001 | 155025225 | 1.54694 | 2.45394 | 1.586 |
| chr1 | 205030652 | 205031109 | 2.15628 | 3.4206 | 1.586 |
| chr11 | 72182303 | 72182701 | 1.87771 | 2.97873 | 1.586 |
| chr1 | 150533334 | 150533760 | 1.96051 | 3.11036 | 1.587 |
| chr19 | 34607743 | 34607963 | 0.627151 | 0.994991 | 1.587 |
| chr2 | 80897689 | 80898009 | 0.623075 | 0.988532 | 1.587 |
| chr18 | 13680694 | 13680905 | 1.26358 | 2.00479 | 1.587 |
| chr4 | 38732815 | 38733127 | 2.08597 | 3.30961 | 1.587 |
| chr22 | 22918732 | 22919235 | 1.73095 | 2.74638 | 1.587 |
| chr1 | 220059701 | 220059987 | 0.641108 | 1.01722 | 1.587 |
| chr1 | 1337392 | 1337661 | 1.42624 | 2.26301 | 1.587 |
| chr1 | 201315658 | 201316023 | 1.29337 | 2.05223 | 1.587 |
| chr12 | 26804549 | 26804855 | 1.6437 | 2.60813 | 1.587 |
| chr20 | 43100152 | 43100467 | 1.65872 | 2.63199 | 1.587 |
| chr12 | 33538534 | 33538786 | 0.64282 | 1.02002 | 1.587 |
| chr10 | 72906728 | 72906931 | 0.568024 | 0.901381 | 1.587 |
| chr8 | 101271134 | 101271429 | 1.68195 | 2.6692 | 1.587 |
| chr1 | 171614093 | 171614365 | 1.97611 | 3.13607 | 1.587 |
| chr4 | 119145464 | 119145729 | 1.50949 | 2.39566 | 1.587 |
| chr19 | 35880274 | 35880478 | 0.876776 | 1.39151 | 1.587 |
| chr12 | 53937116 | 53937558 | 1.73817 | 2.75864 | 1.587 |

| | | | | | |
|---|---|---|---|---|---|
| chr14 | 90581041 | 90581303 | 0.694154 | 1.10171 | 1.587 |
| chr9 | 80897726 | 80897936 | 0.51937 | 0.824315 | 1.587 |
| chr17 | 5495771 | 5496182 | 1.30173 | 2.06606 | 1.587 |
| chr12 | 92205506 | 92205770 | 2.06628 | 3.27956 | 1.587 |
| chr10 | 5665151 | 5665492 | 2.19871 | 3.48979 | 1.587 |
| chr11 | 33851093 | 33851419 | 1.79333 | 2.8464 | 1.587 |
| chr10 | 125754055 | 125754494 | 2.11003 | 3.34929 | 1.587 |
| chr2 | 219381351 | 219381666 | 1.4009 | 2.2237 | 1.587 |
| chr19 | 36039909 | 36040495 | 2.1667 | 3.43937 | 1.587 |
| chr8 | 103694782 | 103695158 | 1.15601 | 1.83505 | 1.587 |
| chr16 | 66684868 | 66685248 | 2.33493 | 3.70653 | 1.587 |
| chr20 | 24478388 | 24478632 | 0.600813 | 0.953776 | 1.587 |
| chr5 | 79125147 | 79125492 | 0.875228 | 1.38942 | 1.587 |
| chr20 | 21878007 | 21878242 | 0.709857 | 1.12694 | 1.588 |
| chr14 | 62327093 | 62327310 | 1.58634 | 2.51856 | 1.588 |
| chr17 | 72956688 | 72957150 | 1.92101 | 3.0499 | 1.588 |
| chr19 | 39993380 | 39993872 | 3.06472 | 4.86583 | 1.588 |
| chr2 | 177517107 | 177517363 | 1.55792 | 2.47363 | 1.588 |
| chr12 | 31118708 | 31118937 | 1.4037 | 2.22882 | 1.588 |
| chr5 | 9394193 | 9394521 | 0.7344 | 1.16612 | 1.588 |
| chr8 | 126823072 | 126823552 | 0.752424 | 1.19477 | 1.588 |
| chr16 | 20786026 | 20786581 | 1.7646 | 2.80217 | 1.588 |
| chr3 | 147152523 | 147152868 | 0.63207 | 1.00372 | 1.588 |
| chr2 | 26673043 | 26673329 | 0.7479 | 1.18768 | 1.588 |
| chr11 | 69656841 | 69657224 | 1.42518 | 2.26326 | 1.588 |
| chr22 | 23285044 | 23285367 | 1.15289 | 1.83087 | 1.588 |
| chr9 | 77880391 | 77880827 | 1.67774 | 2.6644 | 1.588 |
| chr17 | 27175733 | 27176093 | 1.93334 | 3.07047 | 1.588 |
| chr7 | 25436733 | 25436970 | 0.700558 | 1.11273 | 1.588 |
| chr12 | 133138471 | 133138799 | 1.13268 | 1.79915 | 1.588 |
| chr1 | 159103991 | 159104219 | 0.50286 | 0.798751 | 1.588 |
| chr14 | 51086355 | 51086596 | 1.4023 | 2.22745 | 1.588 |
| chr17 | 41176462 | 41176758 | 1.97343 | 3.13481 | 1.589 |
| chr6 | 166814223 | 166814599 | 1.47812 | 2.34805 | 1.589 |
| chr8 | 43135959 | 43136292 | 2.72713 | 4.33223 | 1.589 |
| chr8 | 142441372 | 142441981 | 2.28506 | 3.6301 | 1.589 |
| chr5 | 126649518 | 126649937 | 1.7049 | 2.70858 | 1.589 |
| chr13 | 97610611 | 97610863 | 1.35267 | 2.14906 | 1.589 |
| chr12 | 128618232 | 128618770 | 0.804307 | 1.27788 | 1.589 |
| chr16 | 2515696 | 2515990 | 1.12976 | 1.79497 | 1.589 |
| chr3 | 69039557 | 69039767 | 0.724116 | 1.15048 | 1.589 |
| chr14 | 93685553 | 93685760 | 1.4537 | 2.3097 | 1.589 |
| chr15 | 78623761 | 78623973 | 0.712438 | 1.132 | 1.589 |
| chr4 | 79513608 | 79513820 | 1.27822 | 2.03107 | 1.589 |
| chr3 | 588346 | 588581 | 0.511748 | 0.813165 | 1.589 |
| chr9 | 77274461 | 77274667 | 0.588372 | 0.934925 | 1.589 |
| chr10 | 102758816 | 102759273 | 2.13923 | 3.39925 | 1.589 |
| chr11 | 91689976 | 91690180 | 0.613173 | 0.97436 | 1.589 |
| chr8 | 101847732 | 101847989 | 1.74114 | 2.76688 | 1.589 |
| chr11 | 85565910 | 85566258 | 1.88071 | 2.98875 | 1.589 |
| chr16 | 86077757 | 86078427 | 0.954462 | 1.51692 | 1.589 |
| chr10 | 111612184 | 111612627 | 2.30719 | 3.66704 | 1.589 |
| chr15 | 99322843 | 99323216 | 2.90703 | 4.62046 | 1.589 |
| chr8 | 104524347 | 104524579 | 1.40334 | 2.23064 | 1.590 |
| chr19 | 55570584 | 55570847 | 0.83104 | 1.32107 | 1.590 |
| chr7 | 28428928 | 28429351 | 1.23904 | 1.96966 | 1.590 |
| chr8 | 121464976 | 121465192 | 0.88069 | 1.40001 | 1.590 |
| chr8 | 144483329 | 144483667 | 4.41943 | 7.02542 | 1.590 |
| chr17 | 48277835 | 48278453 | 3.30992 | 5.26171 | 1.590 |
| chr3 | 39002498 | 39002838 | 2.06379 | 3.28077 | 1.590 |
| chr8 | 37232860 | 37233093 | 0.711874 | 1.13165 | 1.590 |

| | | | | | |
|---|---|---|---|---|---|
| chr2 | 113357599 | 113357941 | 2.6262 | 4.17485 | 1.590 |
| chrX | 15583430 | 15583740 | 1.50708 | 2.39593 | 1.590 |
| chr8 | 2117879 | 2118110 | 0.70872 | 1.12676 | 1.590 |
| chr5 | 85664680 | 85665293 | 0.702451 | 1.11684 | 1.590 |
| chr7 | 123490401 | 123490715 | 0.515315 | 0.819311 | 1.590 |
| chr15 | 41849781 | 41850213 | 2.38167 | 3.78668 | 1.590 |
| chr9 | 90721382 | 90721644 | 1.17113 | 1.86202 | 1.590 |
| chr11 | 84269176 | 84269406 | 1.00073 | 1.59114 | 1.590 |
| chr12 | 20791310 | 20791597 | 0.838361 | 1.33299 | 1.590 |
| chr1 | 230767958 | 230768365 | 1.61143 | 2.56221 | 1.590 |
| chr1 | 27179592 | 27179923 | 1.86064 | 2.95867 | 1.590 |
| chr1 | 15631281 | 15631583 | 1.88878 | 3.00344 | 1.590 |
| chr15 | 71865194 | 71865437 | 1.30411 | 2.07377 | 1.590 |
| chr16 | 1244455 | 1245051 | 2.50888 | 3.98966 | 1.590 |
| chr20 | 22314868 | 22315124 | 1.49421 | 2.37616 | 1.590 |
| chr16 | 17386697 | 17387039 | 0.549724 | 0.874212 | 1.590 |
| chr1 | 76374384 | 76374566 | 1.26984 | 2.0194 | 1.590 |
| chr19 | 7633291 | 7633588 | 1.9109 | 3.03891 | 1.590 |
| chr3 | 73181359 | 73181742 | 0.555564 | 0.883532 | 1.590 |
| chr10 | 114211226 | 114211530 | 0.72576 | 1.15421 | 1.590 |
| chr10 | 126362498 | 126362742 | 0.461634 | 0.734231 | 1.591 |
| chr11 | 82100137 | 82100353 | 0.723671 | 1.151 | 1.591 |
| chr18 | 60683489 | 60683777 | 1.40032 | 2.22721 | 1.591 |
| chr1 | 161979477 | 161979763 | 0.427492 | 0.679951 | 1.591 |
| chr12 | 7372138 | 7372590 | 1.87754 | 2.98634 | 1.591 |
| chr15 | 79272027 | 79272455 | 1.81738 | 2.89074 | 1.591 |
| chr5 | 101014265 | 101014695 | 1.2511 | 1.99001 | 1.591 |
| chr16 | 75145144 | 75145461 | 1.84609 | 2.93658 | 1.591 |
| chr17 | 20900469 | 20900674 | 0.587386 | 0.934395 | 1.591 |
| chr2 | 47562173 | 47562539 | 1.69912 | 2.70291 | 1.591 |
| chr2 | 47238571 | 47238799 | 0.532909 | 0.847764 | 1.591 |
| chr1 | 2986279 | 2986555 | 0.699718 | 1.11322 | 1.591 |
| chr4 | 7989188 | 7989437 | 1.56479 | 2.4897 | 1.591 |
| chr4 | 11625406 | 11625626 | 1.25082 | 1.99016 | 1.591 |
| chr18 | 49194532 | 49194744 | 1.62652 | 2.58804 | 1.591 |
| chr8 | 118896542 | 118896759 | 1.23771 | 1.9694 | 1.591 |
| chr10 | 105805234 | 105805571 | 2.03147 | 3.23245 | 1.591 |
| chr14 | 59986355 | 59986708 | 0.679431 | 1.08111 | 1.591 |
| chr7 | 139802208 | 139802442 | 0.618849 | 0.984716 | 1.591 |
| chr5 | 86727497 | 86727992 | 0.576689 | 0.917637 | 1.591 |
| chr14 | 29223491 | 29223788 | 1.53242 | 2.43851 | 1.591 |
| chr7 | 47293992 | 47294366 | 2.64977 | 4.21659 | 1.591 |
| chr1 | 221631662 | 221631990 | 0.865091 | 1.37665 | 1.591 |
| chr1 | 18436837 | 18437046 | 1.01937 | 1.62219 | 1.591 |
| chr9 | 73761861 | 73762094 | 0.805927 | 1.28255 | 1.591 |
| chr2 | 6057521 | 6057803 | 0.518037 | 0.824484 | 1.592 |
| chr5 | 79379388 | 79379718 | 1.5445 | 2.45828 | 1.592 |
| chr15 | 53489516 | 53489885 | 3.13358 | 4.98766 | 1.592 |
| chr1 | 153482205 | 153482582 | 2.36772 | 3.76868 | 1.592 |
| chr2 | 223426509 | 223426856 | 1.4578 | 2.32039 | 1.592 |
| chr19 | 35546596 | 35547049 | 1.90383 | 3.0305 | 1.592 |
| chr5 | 144480919 | 144481123 | 0.788786 | 1.2556 | 1.592 |
| chr5 | 20075097 | 20075552 | 0.734084 | 1.16858 | 1.592 |
| chr10 | 117114699 | 117115052 | 1.93082 | 3.07376 | 1.592 |
| chr8 | 5934279 | 5934540 | 1.42791 | 2.27337 | 1.592 |
| chr2 | 51419617 | 51419834 | 0.503836 | 0.802171 | 1.592 |
| chr12 | 54687819 | 54688477 | 1.4713 | 2.34258 | 1.592 |
| chr3 | 123208129 | 123208556 | 1.74299 | 2.77516 | 1.592 |
| chr10 | 87195723 | 87195973 | 0.680078 | 1.08281 | 1.592 |
| chr1 | 31252375 | 31252746 | 1.5385 | 2.44967 | 1.592 |
| chr2 | 66017113 | 66017390 | 0.788051 | 1.25478 | 1.592 |

| | | | | | |
|---|---|---|---|---|---|
| chr6 | 22828754 | 22829034 | 2.18228 | 3.47483 | 1.592 |
| chr10 | 111965855 | 111966254 | 1.23578 | 1.96785 | 1.592 |
| chr1 | 174739070 | 174739425 | 1.28438 | 2.04527 | 1.592 |
| chr3 | 100488671 | 100488951 | 0.865886 | 1.37888 | 1.592 |
| chr5 | 149502432 | 149502744 | 0.732161 | 1.16593 | 1.592 |
| chr8 | 23153456 | 23153958 | 1.50105 | 2.39036 | 1.592 |
| chr12 | 49318383 | 49318598 | 1.19031 | 1.89554 | 1.592 |
| chr15 | 71200187 | 71200444 | 0.944432 | 1.50401 | 1.593 |
| chr10 | 80424766 | 80425044 | 1.33123 | 2.12007 | 1.593 |
| chr12 | 41623332 | 41623618 | 0.746766 | 1.18928 | 1.593 |
| chr10 | 10971120 | 10971345 | 0.413942 | 0.659235 | 1.593 |
| chr3 | 184135615 | 184135961 | 1.91474 | 3.0494 | 1.593 |
| chr4 | 141782592 | 141782856 | 1.45723 | 2.32083 | 1.593 |
| chr7 | 90942557 | 90942770 | 0.66564 | 1.06013 | 1.593 |
| chr20 | 3050179 | 3050540 | 1.06308 | 1.69312 | 1.593 |
| chr9 | 90698849 | 90699266 | 0.50519 | 0.804597 | 1.593 |
| chr2 | 111874428 | 111874884 | 1.66228 | 2.64754 | 1.593 |
| chr10 | 2404613 | 2404867 | 0.920383 | 1.46598 | 1.593 |
| chr19 | 48893926 | 48894162 | 1.30201 | 2.07385 | 1.593 |
| chr10 | 92671371 | 92671688 | 1.99333 | 3.17519 | 1.593 |
| chr5 | 76440384 | 76440721 | 2.62041 | 4.17411 | 1.593 |
| chr9 | 91615948 | 91616269 | 1.32427 | 2.10949 | 1.593 |
| chr7 | 37660644 | 37660869 | 0.542086 | 0.863536 | 1.593 |
| chr8 | 144481339 | 144481822 | 0.640122 | 1.01974 | 1.593 |
| chr10 | 99740381 | 99740610 | 0.499088 | 0.79509 | 1.593 |
| chr22 | 46041527 | 46041764 | 1.5653 | 2.49375 | 1.593 |
| chr5 | 141340299 | 141340606 | 1.79656 | 2.86223 | 1.593 |
| chr5 | 61739680 | 61739890 | 1.07831 | 1.71798 | 1.593 |
| chr1 | 155030329 | 155030760 | 0.887002 | 1.41323 | 1.593 |
| chr8 | 124835679 | 124835915 | 1.13406 | 1.80688 | 1.593 |
| chr2 | 112645366 | 112645625 | 1.76729 | 2.81588 | 1.593 |
| chr1 | 80683880 | 80684088 | 1.29704 | 2.06671 | 1.593 |
| chr17 | 3575505 | 3575714 | 0.646911 | 1.03081 | 1.593 |
| chr16 | 46835306 | 46835640 | 0.972165 | 1.5491 | 1.593 |
| chr10 | 87489836 | 87490082 | 0.455812 | 0.726341 | 1.594 |
| chr19 | 50037502 | 50037763 | 0.845759 | 1.3478 | 1.594 |
| chr1 | 51070263 | 51070483 | 0.682977 | 1.08843 | 1.594 |
| chr19 | 45302053 | 45302260 | 0.451053 | 0.71884 | 1.594 |
| chr12 | 44491988 | 44492294 | 0.658597 | 1.04961 | 1.594 |
| chr16 | 47071658 | 47072049 | 2.47459 | 3.94381 | 1.594 |
| chr7 | 108770352 | 108770614 | 0.591594 | 0.942843 | 1.594 |
| chr12 | 114256536 | 114257097 | 1.46432 | 2.33387 | 1.594 |
| chr15 | 91632380 | 91632595 | 0.604379 | 0.963304 | 1.594 |
| chr2 | 86115802 | 86116449 | 1.87189 | 2.98357 | 1.594 |
| chr3 | 13537706 | 13538136 | 1.82499 | 2.90881 | 1.594 |
| chr1 | 153116019 | 153116276 | 0.570257 | 0.908938 | 1.594 |
| chr12 | 95795916 | 95796213 | 0.965802 | 1.53946 | 1.594 |
| chr9 | 124449102 | 124449498 | 2.47047 | 3.93791 | 1.594 |
| chr19 | 39423403 | 39423743 | 1.53102 | 2.44048 | 1.594 |
| chr3 | 109346500 | 109346787 | 0.677286 | 1.07964 | 1.594 |
| chr18 | 47883603 | 47883968 | 1.49545 | 2.38394 | 1.594 |
| chr2 | 30951329 | 30951540 | 0.683809 | 1.09022 | 1.594 |
| chr2 | 5918629 | 5918873 | 0.587773 | 0.937118 | 1.594 |
| chr5 | 156243065 | 156243335 | 1.93149 | 3.07953 | 1.594 |
| chr5 | 144035927 | 144036156 | 0.5001 | 0.797399 | 1.594 |
| chr7 | 76145336 | 76145555 | 2.11425 | 3.37124 | 1.595 |
| chr8 | 123745759 | 123746284 | 2.03753 | 3.24913 | 1.595 |
| chr7 | 142647347 | 142647846 | 0.741591 | 1.18258 | 1.595 |
| chr3 | 8459085 | 8459430 | 1.76792 | 2.81925 | 1.595 |
| chr9 | 130285780 | 130286139 | 1.88568 | 3.00716 | 1.595 |
| chr14 | 20341244 | 20341637 | 0.776659 | 1.23858 | 1.595 |

| | | | | | |
|---|---|---|---|---|---|
| chr14 | 67213055 | 67213306 | 1.35073 | 2.15413 | 1.595 |
| chr12 | 7370152 | 7370697 | 0.694463 | 1.10755 | 1.595 |
| chr6 | 72546856 | 72547060 | 0.541324 | 0.863357 | 1.595 |
| chr13 | 25115070 | 25115308 | 1.40786 | 2.24547 | 1.595 |
| chr1 | 24741949 | 24742176 | 1.83617 | 2.92865 | 1.595 |
| chr20 | 48795768 | 48796147 | 1.96904 | 3.14063 | 1.595 |
| chr13 | 37432373 | 37432609 | 0.91273 | 1.45589 | 1.595 |
| chr18 | 68101205 | 68101470 | 1.6724 | 2.66764 | 1.595 |
| chr2 | 138623549 | 138623869 | 1.46961 | 2.34426 | 1.595 |
| chrX | 152933364 | 152933882 | 1.4653 | 2.33744 | 1.595 |
| chr5 | 52490592 | 52490817 | 0.570994 | 0.9109 | 1.595 |
| chr11 | 119611909 | 119612371 | 2.01233 | 3.21033 | 1.595 |
| chr7 | 116311822 | 116312368 | 1.36813 | 2.18262 | 1.595 |
| chr11 | 19579221 | 19579557 | 2.03613 | 3.24843 | 1.595 |
| chr12 | 102845409 | 102845744 | 0.626839 | 1.00008 | 1.595 |
| chr2 | 231039250 | 231039454 | 0.72417 | 1.15539 | 1.595 |
| chr9 | 11901689 | 11902062 | 0.542394 | 0.865385 | 1.595 |
| chr10 | 72027510 | 72027721 | 0.952846 | 1.52031 | 1.596 |
| chr22 | 31151460 | 31151663 | 1.53421 | 2.44801 | 1.596 |
| chrX | 68758079 | 68758448 | 1.69031 | 2.69713 | 1.596 |
| chr2 | 75636860 | 75637173 | 0.578812 | 0.923585 | 1.596 |
| chr1 | 93770241 | 93770514 | 1.48313 | 2.36667 | 1.596 |
| chr2 | 219874951 | 219875207 | 1.97102 | 3.14533 | 1.596 |
| chrX | 108718371 | 108718989 | 1.42812 | 2.279 | 1.596 |
| chr10 | 71560730 | 71561240 | 1.92174 | 3.06691 | 1.596 |
| chr14 | 64783953 | 64784205 | 0.51866 | 0.827781 | 1.596 |
| chr19 | 32778777 | 32779019 | 1.92407 | 3.07087 | 1.596 |
| chr1 | 11953933 | 11954348 | 2.10295 | 3.35656 | 1.596 |
| chr8 | 92478456 | 92478718 | 0.603855 | 0.963836 | 1.596 |
| chr1 | 9349098 | 9349433 | 2.32033 | 3.70372 | 1.596 |
| chr3 | 157717050 | 157717319 | 0.433289 | 0.691705 | 1.596 |
| chr14 | 24559750 | 24560102 | 2.47496 | 3.95107 | 1.596 |
| chr10 | 129214463 | 129214746 | 0.628388 | 1.00318 | 1.596 |
| chr3 | 140718962 | 140719199 | 1.14575 | 1.82916 | 1.596 |
| chr17 | 4525252 | 4525505 | 0.813752 | 1.29915 | 1.596 |
| chr9 | 139565884 | 139566613 | 2.20721 | 3.5238 | 1.597 |
| chr16 | 24811996 | 24812367 | 2.10321 | 3.35793 | 1.597 |
| chr12 | 121443137 | 121443532 | 2.46804 | 3.94069 | 1.597 |
| chr12 | 121447618 | 121448157 | 1.3469 | 2.1506 | 1.597 |
| chr1 | 168062307 | 168062532 | 1.38011 | 2.20363 | 1.597 |
| chr16 | 22419828 | 22420058 | 0.78938 | 1.2605 | 1.597 |
| chr8 | 119674623 | 119675022 | 1.48964 | 2.37878 | 1.597 |
| chr17 | 36600272 | 36600769 | 2.45023 | 3.9128 | 1.597 |
| chr11 | 6272329 | 6272730 | 1.20375 | 1.9223 | 1.597 |
| chr17 | 80732814 | 80733428 | 2.21045 | 3.52994 | 1.597 |
| chr18 | 712597 | 713070 | 1.78928 | 2.85738 | 1.597 |
| chr15 | 65931000 | 65931268 | 1.70974 | 2.73058 | 1.597 |
| chrX | 151989768 | 151990018 | 1.83776 | 2.93507 | 1.597 |
| chr3 | 101324268 | 101324473 | 0.463477 | 0.740245 | 1.597 |
| chr10 | 103362515 | 103362920 | 2.84427 | 4.54289 | 1.597 |
| chr19 | 54992386 | 54992847 | 1.45689 | 2.32696 | 1.597 |
| chr19 | 3665904 | 3666363 | 2.17367 | 3.47197 | 1.597 |
| chr20 | 61071293 | 61071901 | 1.43364 | 2.29 | 1.597 |
| chr8 | 134867781 | 134867986 | 0.905351 | 1.44619 | 1.597 |
| chr11 | 80124557 | 80124830 | 0.523451 | 0.836153 | 1.597 |
| chr2 | 73312739 | 73313160 | 2.52431 | 4.0326 | 1.598 |
| chr7 | 134783205 | 134783437 | 0.558841 | 0.892753 | 1.598 |
| chr12 | 72332569 | 72332891 | 1.46888 | 2.34662 | 1.598 |
| chr1 | 183602820 | 183603135 | 1.29045 | 2.06187 | 1.598 |
| chr3 | 150004437 | 150004666 | 1.40045 | 2.2377 | 1.598 |
| chr18 | 59249116 | 59249492 | 2.64199 | 4.22152 | 1.598 |

| | | | | | |
|---|---|---|---|---|---|
| chr17 | 73754721 | 73755150 | 1.15626 | 1.84756 | 1.598 |
| chr5 | 53712295 | 53712555 | 0.675385 | 1.0792 | 1.598 |
| chr10 | 64926616 | 64926960 | 1.48625 | 2.37491 | 1.598 |
| chr1 | 8409037 | 8409406 | 1.763 | 2.81715 | 1.598 |
| chr5 | 106907741 | 106908058 | 1.61934 | 2.58759 | 1.598 |
| chr2 | 20442005 | 20442363 | 2.3159 | 3.70081 | 1.598 |
| chr1 | 1597134 | 1597384 | 1.33605 | 2.13507 | 1.598 |
| chr14 | 69215126 | 69215402 | 0.582461 | 0.930804 | 1.598 |
| chr14 | 97622982 | 97623258 | 0.662408 | 1.05858 | 1.598 |
| chr9 | 113189849 | 113190174 | 1.87373 | 2.99442 | 1.598 |
| chr17 | 49923540 | 49923743 | 0.613816 | 0.98099 | 1.598 |
| chr7 | 50269658 | 50269874 | 1.3706 | 2.19046 | 1.598 |
| chr14 | 81678277 | 81678503 | 1.3163 | 2.10381 | 1.598 |
| chr14 | 101317412 | 101317920 | 2.29661 | 3.67063 | 1.598 |
| chr17 | 61651152 | 61651438 | 1.0216 | 1.63285 | 1.598 |
| chr4 | 55099482 | 55099813 | 1.75111 | 2.79893 | 1.598 |
| chr12 | 58120680 | 58120888 | 1.09078 | 1.74358 | 1.598 |
| chr7 | 105807219 | 105807539 | 2.13048 | 3.40568 | 1.599 |
| chr10 | 101195866 | 101196132 | 1.18764 | 1.89864 | 1.599 |
| chr9 | 24456653 | 24457008 | 0.592154 | 0.946666 | 1.599 |
| chr18 | 71863494 | 71863899 | 1.19475 | 1.9101 | 1.599 |
| chr7 | 98434404 | 98434791 | 1.72665 | 2.76048 | 1.599 |
| chr14 | 54468320 | 54468560 | 1.07886 | 1.72484 | 1.599 |
| chr1 | 54220300 | 54220720 | 1.08841 | 1.74018 | 1.599 |
| chr12 | 34079844 | 34080179 | 0.80941 | 1.2942 | 1.599 |
| chr16 | 11735219 | 11735555 | 2.03873 | 3.25985 | 1.599 |
| chr3 | 80177393 | 80177684 | 0.633638 | 1.01318 | 1.599 |
| chr19 | 5504241 | 5504448 | 1.31579 | 2.10398 | 1.599 |
| chr2 | 238832260 | 238832615 | 3.24918 | 5.19561 | 1.599 |
| chr9 | 86535680 | 86536270 | 0.838926 | 1.3415 | 1.599 |
| chr1 | 42277615 | 42278022 | 1.60905 | 2.57307 | 1.599 |
| chr20 | 58003582 | 58003893 | 1.20119 | 1.92094 | 1.599 |
| chr17 | 77814332 | 77814544 | 0.899222 | 1.43807 | 1.599 |
| chr12 | 49628975 | 49629276 | 2.20448 | 3.52558 | 1.599 |
| chr9 | 95719469 | 95719813 | 2.11009 | 3.37467 | 1.599 |
| chr19 | 45520242 | 45520508 | 1.89694 | 3.03379 | 1.599 |
| chr19 | 10828083 | 10828290 | 1.39208 | 2.22638 | 1.599 |
| chr7 | 134409393 | 134409774 | 1.41539 | 2.26369 | 1.599 |
| chr19 | 48058659 | 48059078 | 0.708949 | 1.13386 | 1.599 |
| chr3 | 4031197 | 4031418 | 0.715892 | 1.14511 | 1.600 |
| chr1 | 57297561 | 57297916 | 2.20598 | 3.52868 | 1.600 |
| chr3 | 172384665 | 172384968 | 1.56632 | 2.50549 | 1.600 |
| chr5 | 56594132 | 56594364 | 1.73511 | 2.77559 | 1.600 |
| chr1 | 68936108 | 68936390 | 1.25717 | 2.01108 | 1.600 |
| chr14 | 104597228 | 104597688 | 1.72575 | 2.76085 | 1.600 |
| chr9 | 119257298 | 119257676 | 1.75851 | 2.81329 | 1.600 |
| chr11 | 62553950 | 62554970 | 2.26098 | 3.61717 | 1.600 |
| chr4 | 116858350 | 116858573 | 0.650743 | 1.04107 | 1.600 |
| chr19 | 14143090 | 14143294 | 1.58711 | 2.53914 | 1.600 |
| chr19 | 5567555 | 5567838 | 0.939927 | 1.50376 | 1.600 |
| chr5 | 9714175 | 9714393 | 0.635734 | 1.0171 | 1.600 |
| chrX | 77851552 | 77851795 | 0.564219 | 0.902745 | 1.600 |
| chr3 | 100181841 | 100182067 | 1.60504 | 2.56807 | 1.600 |
| chr12 | 101714023 | 101714294 | 0.815448 | 1.30474 | 1.600 |
| chr2 | 81354604 | 81354953 | 0.967858 | 1.54861 | 1.600 |
| chr14 | 105293907 | 105294290 | 1.31557 | 2.10502 | 1.600 |
| chr10 | 2964582 | 2964851 | 1.19412 | 1.9107 | 1.600 |
| chr11 | 7914380 | 7914582 | 1.20901 | 1.93456 | 1.600 |
| chr14 | 60816524 | 60817089 | 1.53435 | 2.45518 | 1.600 |
| chr16 | 57696171 | 57696391 | 0.469021 | 0.750507 | 1.600 |
| chr19 | 31544298 | 31544515 | 0.818953 | 1.31048 | 1.600 |

| | | | | | |
|---|---|---|---|---|---|
| chr19 | 10975149 | 10975407 | 2.35459 | 3.76781 | 1.600 |
| chr1 | 182374044 | 182374257 | 0.596099 | 0.953887 | 1.600 |
| chr14 | 70540575 | 70541049 | 2.41827 | 3.86977 | 1.600 |
| chr3 | 117934486 | 117934710 | 0.600766 | 0.961359 | 1.600 |
| chr2 | 167395949 | 167396192 | 1.34168 | 2.147 | 1.600 |
| chr9 | 117068102 | 117068476 | 2.1718 | 3.47558 | 1.600 |
| chr3 | 106328625 | 106328856 | 1.55358 | 2.48627 | 1.600 |
| chr1 | 32986033 | 32986422 | 2.84828 | 4.55857 | 1.600 |
| chr2 | 44059184 | 44059402 | 0.550334 | 0.880823 | 1.601 |
| chr15 | 90757873 | 90758083 | 1.40659 | 2.2515 | 1.601 |
| chr4 | 90359340 | 90359608 | 0.896047 | 1.4343 | 1.601 |
| chr8 | 27023389 | 27023658 | 0.593087 | 0.949385 | 1.601 |
| chr11 | 12068864 | 12069237 | 1.72837 | 2.76711 | 1.601 |
| chrX | 9281287 | 9281526 | 1.6037 | 2.56751 | 1.601 |
| chr2 | 51321463 | 51321672 | 0.543781 | 0.870593 | 1.601 |
| chr8 | 141841270 | 141841487 | 1.40553 | 2.25031 | 1.601 |
| chr8 | 84075392 | 84075768 | 0.975047 | 1.56124 | 1.601 |
| chr17 | 42852106 | 42852599 | 2.85952 | 4.5787 | 1.601 |
| chr20 | 48545251 | 48545532 | 2.08954 | 3.34591 | 1.601 |
| chr12 | 115122138 | 115122601 | 1.27754 | 2.04571 | 1.601 |
| chr5 | 80404303 | 80404509 | 1.19727 | 1.91722 | 1.601 |
| chr11 | 669883 | 670155 | 0.459298 | 0.735529 | 1.601 |
| chr15 | 99619984 | 99620277 | 1.4735 | 2.35982 | 1.602 |
| chr1 | 170942708 | 170943009 | 0.577112 | 0.92431 | 1.602 |
| chr1 | 176706678 | 176707315 | 0.581405 | 0.931188 | 1.602 |
| chr14 | 72322164 | 72322464 | 1.96867 | 3.15314 | 1.602 |
| chr9 | 78889672 | 78889934 | 0.542321 | 0.868634 | 1.602 |
| chr17 | 74723020 | 74723499 | 1.49785 | 2.39919 | 1.602 |
| chr5 | 119671399 | 119671700 | 0.510843 | 0.818262 | 1.602 |
| chr10 | 105899079 | 105899290 | 1.2723 | 2.0381 | 1.602 |
| chr17 | 71630869 | 71631082 | 0.459634 | 0.736291 | 1.602 |
| chr5 | 43400240 | 43400538 | 1.3808 | 2.21194 | 1.602 |
| chr1 | 24868524 | 24868787 | 1.79421 | 2.87429 | 1.602 |
| chr3 | 192284544 | 192284808 | 1.50482 | 2.41076 | 1.602 |
| chr20 | 557624 | 558405 | 0.882537 | 1.41391 | 1.602 |
| chr8 | 118295189 | 118295445 | 0.640951 | 1.02691 | 1.602 |
| chr19 | 30207416 | 30207827 | 2.95507 | 4.73469 | 1.602 |
| chr3 | 104857252 | 104857514 | 0.59861 | 0.959148 | 1.602 |
| chrX | 19757863 | 19758124 | 2.37053 | 3.79835 | 1.602 |
| chr3 | 25689583 | 25689918 | 1.22873 | 1.96886 | 1.602 |
| chr11 | 118042260 | 118042680 | 2.898 | 4.64372 | 1.602 |
| chr12 | 102489503 | 102489729 | 0.781221 | 1.25184 | 1.602 |
| chr17 | 71592229 | 71592446 | 0.625576 | 1.00243 | 1.602 |
| chr8 | 66674102 | 66674509 | 1.89166 | 3.03121 | 1.602 |
| chr17 | 10230925 | 10231405 | 1.84971 | 2.96404 | 1.602 |
| chr3 | 140824690 | 140824962 | 1.40149 | 2.2459 | 1.603 |
| chr7 | 38972215 | 38972533 | 2.14859 | 3.44315 | 1.603 |
| chr19 | 55677257 | 55677762 | 2.17782 | 3.4901 | 1.603 |
| chr16 | 67439074 | 67439410 | 0.500076 | 0.801499 | 1.603 |
| chr2 | 102637067 | 102637502 | 2.89618 | 4.64188 | 1.603 |
| chr6 | 15691488 | 15691938 | 3.54704 | 5.68504 | 1.603 |
| chr16 | 24021810 | 24022130 | 1.48964 | 2.38763 | 1.603 |
| chr20 | 42846024 | 42846341 | 2.8639 | 4.59037 | 1.603 |
| chr20 | 42896664 | 42896903 | 0.507576 | 0.81358 | 1.603 |
| chr22 | 46959241 | 46959911 | 1.66735 | 2.67263 | 1.603 |
| chr18 | 34389601 | 34389823 | 1.35443 | 2.17106 | 1.603 |
| chr19 | 32167213 | 32167438 | 0.86345 | 1.38406 | 1.603 |
| chr15 | 73532231 | 73532504 | 0.910221 | 1.45907 | 1.603 |
| chr3 | 149597686 | 149598019 | 1.46469 | 2.34793 | 1.603 |
| chr20 | 1283634 | 1283878 | 0.631905 | 1.01296 | 1.603 |
| chr11 | 765132 | 765658 | 1.79062 | 2.87048 | 1.603 |

| | | | | | |
|---|---|---|---|---|---|
| chr1 | 96701803 | 96702021 | 1.02548 | 1.64393 | 1.603 |
| chr16 | 31310379 | 31310590 | 0.987006 | 1.58226 | 1.603 |
| chr5 | 45020801 | 45021009 | 0.428311 | 0.686644 | 1.603 |
| chr3 | 25155038 | 25155461 | 0.71702 | 1.1495 | 1.603 |
| chr2 | 150169955 | 150170248 | 1.41997 | 2.2765 | 1.603 |
| chr1 | 169078231 | 169078683 | 0.756927 | 1.21351 | 1.603 |
| chr3 | 144483080 | 144483325 | 0.817811 | 1.31115 | 1.603 |
| chr10 | 13628697 | 13629017 | 2.97196 | 4.76479 | 1.603 |
| chr5 | 29042372 | 29042583 | 0.762376 | 1.22229 | 1.603 |
| chrX | 119115687 | 119115932 | 2.17984 | 3.4949 | 1.603 |
| chr20 | 44757153 | 44757559 | 1.6192 | 2.59613 | 1.603 |
| chr17 | 25572345 | 25572640 | 1.86973 | 2.99795 | 1.603 |
| chr12 | 53109282 | 53109717 | 0.839394 | 1.34596 | 1.603 |
| chr19 | 12907351 | 12907784 | 1.71102 | 2.74362 | 1.604 |
| chr7 | 48178943 | 48179294 | 1.32581 | 2.12593 | 1.604 |
| chr22 | 32602923 | 32603182 | 1.65665 | 2.65648 | 1.604 |
| chr7 | 101458683 | 101459190 | 1.98629 | 3.18511 | 1.604 |
| chr20 | 9619031 | 9619254 | 0.66262 | 1.06255 | 1.604 |
| chr7 | 37324370 | 37324671 | 1.45878 | 2.33926 | 1.604 |
| chr11 | 26960986 | 26961280 | 0.527213 | 0.845437 | 1.604 |
| chr1 | 64602829 | 64603141 | 1.99981 | 3.20695 | 1.604 |
| chr3 | 35965657 | 35965880 | 1.52668 | 2.44827 | 1.604 |
| chr2 | 171729738 | 171730164 | 2.22116 | 3.56209 | 1.604 |
| chr20 | 36858227 | 36858600 | 1.51471 | 2.42929 | 1.604 |
| chr2 | 73089620 | 73089991 | 3.39675 | 5.44774 | 1.604 |
| chr8 | 68535829 | 68536074 | 0.740738 | 1.18802 | 1.604 |
| chr17 | 2624118 | 2624401 | 0.603309 | 0.967613 | 1.604 |
| chr16 | 85394135 | 85394341 | 0.44594 | 0.715243 | 1.604 |
| chr19 | 32142448 | 32142999 | 0.767409 | 1.2309 | 1.604 |
| chr4 | 7158502 | 7158840 | 1.46536 | 2.3504 | 1.604 |
| chr1 | 29562722 | 29563093 | 1.76686 | 2.83426 | 1.604 |
| chr16 | 20397074 | 20397462 | 1.45912 | 2.34064 | 1.604 |
| chr10 | 26524154 | 26524427 | 0.631625 | 1.01322 | 1.604 |
| chr6 | 27763647 | 27763937 | 1.15784 | 1.8574 | 1.604 |
| chr11 | 12621060 | 12621390 | 2.83015 | 4.54026 | 1.604 |
| chr12 | 108472182 | 108472454 | 0.541886 | 0.869318 | 1.604 |
| chr14 | 77617686 | 77617944 | 2.14962 | 3.44864 | 1.604 |
| chr14 | 43231389 | 43231598 | 0.597982 | 0.959362 | 1.604 |
| chr2 | 68349538 | 68349841 | 2.98161 | 4.78375 | 1.604 |
| chr8 | 64303884 | 64304321 | 1.64226 | 2.63496 | 1.604 |
| chr7 | 30779982 | 30780435 | 2.91174 | 4.67188 | 1.605 |
| chr5 | 153344629 | 153344903 | 1.96622 | 3.15501 | 1.605 |
| chr1 | 206907920 | 206908461 | 1.38435 | 2.22138 | 1.605 |
| chr11 | 86689921 | 86690134 | 1.29499 | 2.07806 | 1.605 |
| chr15 | 90316349 | 90316561 | 0.958882 | 1.53874 | 1.605 |
| chr1 | 183885153 | 183885440 | 1.88042 | 3.01759 | 1.605 |
| chr7 | 146610293 | 146610509 | 0.666437 | 1.06946 | 1.605 |
| chr8 | 15969229 | 15969629 | 0.70731 | 1.1351 | 1.605 |
| chrX | 20460786 | 20461063 | 1.2308 | 1.97522 | 1.605 |
| chr9 | 77357725 | 77358152 | 1.52299 | 2.44421 | 1.605 |
| chr9 | 33813720 | 33813982 | 1.18736 | 1.90562 | 1.605 |
| chr19 | 33239449 | 33239702 | 1.35158 | 2.16936 | 1.605 |
| chr8 | 143534032 | 143534473 | 1.72024 | 2.76113 | 1.605 |
| chr4 | 81123315 | 81123671 | 1.7816 | 2.85963 | 1.605 |
| chr11 | 58421424 | 58421856 | 2.1458 | 3.44422 | 1.605 |
| chr12 | 46536812 | 46537088 | 0.527811 | 0.847191 | 1.605 |
| chr16 | 3151308 | 3151719 | 2.09312 | 3.35968 | 1.605 |
| chr12 | 99780689 | 99781121 | 1.39557 | 2.24017 | 1.605 |
| chr17 | 45078459 | 45078872 | 1.69034 | 2.71336 | 1.605 |
| chr10 | 29675381 | 29675682 | 1.46862 | 2.35751 | 1.605 |
| chr5 | 177608735 | 177609001 | 0.567552 | 0.911067 | 1.605 |

| | | | | | |
|---|---|---|---|---|---|
| chr10 | 11466417 | 11466912 | 2.10326 | 3.3765 | 1.605 |
| chr1 | 57286703 | 57287089 | 2.15237 | 3.4554 | 1.605 |
| chr7 | 105649914 | 105650295 | 2.85349 | 4.581 | 1.605 |
| chr17 | 74864167 | 74864613 | 2.19772 | 3.52827 | 1.605 |
| chr3 | 38140014 | 38140237 | 0.597835 | 0.95978 | 1.605 |
| chr17 | 7726060 | 7726415 | 2.35723 | 3.78442 | 1.605 |
| chr12 | 78658470 | 78658663 | 1.04899 | 1.68413 | 1.605 |
| chr7 | 123842849 | 123843078 | 0.518209 | 0.831974 | 1.605 |
| chr3 | 103940392 | 103940716 | 0.637401 | 1.02334 | 1.605 |
| chr6 | 42071934 | 42072192 | 1.10556 | 1.77505 | 1.606 |
| chr5 | 140937440 | 140937759 | 1.86209 | 2.98979 | 1.606 |
| chr14 | 78328487 | 78329037 | 2.65228 | 4.25855 | 1.606 |
| chr10 | 126254395 | 126254690 | 1.84393 | 2.96067 | 1.606 |
| chr11 | 44948415 | 44949214 | 1.92587 | 3.09246 | 1.606 |
| chr17 | 72364203 | 72364885 | 2.43316 | 3.90715 | 1.606 |
| chr3 | 44370695 | 44371019 | 0.388186 | 0.623381 | 1.606 |
| chr3 | 66458144 | 66458579 | 1.88269 | 3.02347 | 1.606 |
| chr7 | 48028472 | 48028701 | 1.6239 | 2.60796 | 1.606 |
| chr8 | 86500368 | 86500860 | 0.570502 | 0.916247 | 1.606 |
| chr14 | 21402330 | 21402683 | 2.31075 | 3.71128 | 1.606 |
| chr14 | 45315508 | 45315734 | 1.65202 | 2.65339 | 1.606 |
| chr7 | 12362000 | 12362193 | 1.63744 | 2.62997 | 1.606 |
| chr10 | 49743884 | 49744089 | 0.864857 | 1.38915 | 1.606 |
| chr8 | 6914038 | 6914262 | 0.724361 | 1.16357 | 1.606 |
| chr16 | 57443839 | 57444050 | 0.771799 | 1.23982 | 1.606 |
| chr20 | 62127358 | 62127795 | 2.45611 | 3.94552 | 1.606 |
| chr17 | 42443582 | 42443796 | 0.747789 | 1.20126 | 1.606 |
| chr4 | 40058323 | 40058651 | 1.61758 | 2.59878 | 1.607 |
| chr2 | 20595705 | 20596058 | 2.18468 | 3.50991 | 1.607 |
| chr1 | 21029009 | 21029299 | 1.51455 | 2.43335 | 1.607 |
| chr12 | 116107282 | 116107665 | 0.537431 | 0.863469 | 1.607 |
| chr2 | 121834506 | 121834986 | 2.69325 | 4.32726 | 1.607 |
| chr16 | 70687644 | 70687922 | 1.77616 | 2.85387 | 1.607 |
| chr3 | 93838268 | 93838495 | 0.673952 | 1.08289 | 1.607 |
| chr2 | 240873894 | 240874292 | 1.31296 | 2.10967 | 1.607 |
| chr8 | 36104286 | 36104625 | 0.634492 | 1.01954 | 1.607 |
| chr14 | 40872588 | 40872800 | 0.803458 | 1.29106 | 1.607 |
| chr2 | 242892878 | 242893338 | 2.26828 | 3.64508 | 1.607 |
| chr5 | 114111936 | 114112239 | 0.494509 | 0.794689 | 1.607 |
| chr11 | 16841808 | 16842243 | 1.65535 | 2.66037 | 1.607 |
| chr12 | 72476668 | 72476943 | 0.602907 | 0.96897 | 1.607 |
| chr8 | 26754773 | 26755118 | 0.770556 | 1.23841 | 1.607 |
| chr8 | 9435564 | 9435793 | 0.625041 | 1.00465 | 1.607 |
| chr1 | 203409975 | 203410267 | 0.824049 | 1.32453 | 1.607 |
| chr17 | 65938929 | 65939239 | 1.28919 | 2.07219 | 1.607 |
| chr17 | 76356845 | 76357233 | 2.74731 | 4.41603 | 1.607 |
| chr19 | 48794296 | 48794926 | 2.61624 | 4.20542 | 1.607 |
| chr5 | 155133116 | 155133524 | 0.558744 | 0.898155 | 1.607 |
| chr11 | 64111297 | 64111615 | 1.19722 | 1.92451 | 1.607 |
| chr12 | 16512816 | 16513158 | 2.75416 | 4.42749 | 1.608 |
| chr9 | 102899066 | 102899315 | 0.586207 | 0.942389 | 1.608 |
| chr2 | 27543908 | 27544135 | 1.11682 | 1.79546 | 1.608 |
| chr7 | 78901530 | 78901715 | 1.39795 | 2.24742 | 1.608 |
| chr1 | 116307272 | 116307482 | 1.27273 | 2.04613 | 1.608 |
| chr1 | 180798059 | 180798261 | 0.557001 | 0.895497 | 1.608 |
| chr19 | 51600608 | 51600914 | 2.83039 | 4.55052 | 1.608 |
| chr14 | 79636876 | 79637446 | 0.531745 | 0.854977 | 1.608 |
| chr8 | 41801722 | 41802016 | 1.51861 | 2.44182 | 1.608 |
| chr11 | 116370164 | 116370440 | 1.36478 | 2.1945 | 1.608 |
| chr4 | 165689795 | 165690232 | 0.66343 | 1.06676 | 1.608 |
| chr16 | 12499221 | 12499540 | 0.634811 | 1.02084 | 1.608 |

| | | | | | |
|---|---|---|---|---|---|
| chr16 | 28295550 | 28295759 | 0.876616 | 1.40969 | 1.608 |
| chr19 | 43905611 | 43906011 | 0.593385 | 0.954229 | 1.608 |
| chr1 | 23991804 | 23992040 | 0.932103 | 1.49896 | 1.608 |
| chr6 | 39134160 | 39134526 | 2.25996 | 3.63436 | 1.608 |
| chr2 | 209040121 | 209040338 | 2.0467 | 3.29143 | 1.608 |
| chr5 | 133090338 | 133090580 | 1.30668 | 2.10153 | 1.608 |
| chr12 | 12798484 | 12798761 | 1.36298 | 2.19209 | 1.608 |
| chr9 | 97893603 | 97893990 | 1.70846 | 2.74802 | 1.608 |
| chr5 | 132013048 | 132013303 | 1.28579 | 2.06821 | 1.609 |
| chr6 | 37659592 | 37660124 | 1.42684 | 2.2951 | 1.609 |
| chr7 | 948390 | 948746 | 2.6044 | 4.18928 | 1.609 |
| chr19 | 28780746 | 28781005 | 0.848538 | 1.36492 | 1.609 |
| chrX | 73770073 | 73770556 | 3.76635 | 6.05835 | 1.609 |
| chr11 | 65097450 | 65097811 | 1.61988 | 2.60567 | 1.609 |
| chrX | 131426880 | 131427288 | 1.30879 | 2.1053 | 1.609 |
| chr5 | 162835365 | 162835936 | 1.16124 | 1.86799 | 1.609 |
| chr14 | 96113795 | 96114045 | 0.585046 | 0.941131 | 1.609 |
| chr20 | 54233023 | 54233263 | 0.68082 | 1.09521 | 1.609 |
| chr2 | 55808690 | 55808892 | 1.28636 | 2.06936 | 1.609 |
| chr1 | 42501252 | 42501557 | 1.28609 | 2.06901 | 1.609 |
| chr6 | 84937717 | 84937965 | 1.14623 | 1.84403 | 1.609 |
| chr17 | 74010251 | 74010709 | 2.84163 | 4.57171 | 1.609 |
| chr3 | 141193111 | 141193411 | 1.58993 | 2.55801 | 1.609 |
| chr20 | 11892887 | 11893305 | 1.67481 | 2.69471 | 1.609 |
| chr20 | 13251126 | 13251373 | 0.709231 | 1.14117 | 1.609 |
| chr17 | 48707933 | 48708290 | 2.70903 | 4.35894 | 1.609 |
| chr2 | 191209020 | 191209280 | 0.714582 | 1.14986 | 1.609 |
| chr16 | 57893294 | 57893589 | 1.4267 | 2.29578 | 1.609 |
| chr6 | 114429400 | 114429886 | 1.29538 | 2.08451 | 1.609 |
| chr3 | 107565094 | 107565465 | 1.75729 | 2.82784 | 1.609 |
| chr1 | 195280927 | 195281278 | 0.657795 | 1.05855 | 1.609 |
| chr1 | 173883389 | 173883979 | 1.28454 | 2.06717 | 1.609 |
| chr8 | 133124381 | 133124584 | 0.480866 | 0.77385 | 1.609 |
| chr17 | 4060142 | 4060589 | 0.895837 | 1.44166 | 1.609 |
| chr10 | 73151290 | 73151501 | 0.635912 | 1.02338 | 1.609 |
| chr8 | 2241468 | 2242113 | 0.990872 | 1.59463 | 1.609 |
| chr12 | 67782639 | 67782845 | 1.13888 | 1.8329 | 1.609 |
| chr7 | 44961701 | 44962119 | 2.32832 | 3.74725 | 1.609 |
| chr5 | 163122582 | 163122909 | 0.483396 | 0.778005 | 1.609 |
| chr17 | 61072759 | 61073132 | 1.56692 | 2.52193 | 1.609 |
| chr10 | 36310052 | 36310277 | 0.524236 | 0.843756 | 1.610 |
| chr18 | 9771484 | 9771853 | 1.97886 | 3.18503 | 1.610 |
| chr8 | 57006181 | 57006623 | 1.62776 | 2.62003 | 1.610 |
| chr7 | 1471689 | 1472703 | 2.05937 | 3.31486 | 1.610 |
| chr11 | 37130561 | 37130766 | 0.534782 | 0.860836 | 1.610 |
| chr7 | 42882848 | 42883163 | 1.14164 | 1.83781 | 1.610 |
| chr4 | 152541427 | 152541643 | 0.695132 | 1.11904 | 1.610 |
| chr14 | 105635506 | 105635809 | 1.37688 | 2.21655 | 1.610 |
| chr4 | 15731610 | 15731814 | 1.03822 | 1.67139 | 1.610 |
| chr2 | 241790593 | 241791013 | 2.65739 | 4.27806 | 1.610 |
| chr8 | 73220547 | 73220760 | 0.503465 | 0.810523 | 1.610 |
| chr17 | 41610976 | 41611278 | 0.824554 | 1.32747 | 1.610 |
| chr19 | 34891443 | 34891657 | 0.798042 | 1.28479 | 1.610 |
| chr3 | 154435079 | 154435352 | 0.538313 | 0.866704 | 1.610 |
| chr7 | 6199746 | 6200132 | 1.89775 | 3.0556 | 1.610 |
| chr9 | 33623958 | 33624324 | 1.64348 | 2.64626 | 1.610 |
| chr17 | 18044584 | 18044981 | 1.20337 | 1.93768 | 1.610 |
| chr1 | 163141663 | 163141917 | 0.561617 | 0.904374 | 1.610 |
| chr2 | 222715042 | 222715255 | 0.515314 | 0.829832 | 1.610 |
| chr8 | 79836894 | 79837252 | 0.619837 | 0.998161 | 1.610 |
| chr3 | 185661289 | 185661629 | 2.66488 | 4.29143 | 1.610 |

| | | | | | |
|---|---|---|---|---|---|
| chr14 | 72219765 | 72220173 | 2.44074 | 3.93051 | 1.610 |
| chr18 | 71168024 | 71168270 | 0.639042 | 1.02913 | 1.610 |
| chr3 | 48595656 | 48595919 | 2.03706 | 3.28061 | 1.610 |
| chr5 | 169527447 | 169527699 | 1.56817 | 2.52547 | 1.610 |
| chr8 | 73907198 | 73907404 | 1.56721 | 2.52397 | 1.610 |
| chr1 | 212769232 | 212769487 | 1.25904 | 2.02771 | 1.611 |
| chr8 | 135296984 | 135297232 | 0.905981 | 1.45914 | 1.611 |
| chr16 | 70690720 | 70691019 | 0.572246 | 0.921653 | 1.611 |
| chr7 | 75831048 | 75831367 | 2.90717 | 4.68244 | 1.611 |
| chr14 | 21131235 | 21131460 | 1.23341 | 1.98661 | 1.611 |
| chr11 | 67029178 | 67029618 | 0.906479 | 1.46005 | 1.611 |
| chr3 | 38562943 | 38563229 | 2.05601 | 3.31158 | 1.611 |
| chr7 | 150118146 | 150118511 | 2.1637 | 3.48516 | 1.611 |
| chr17 | 76839280 | 76839818 | 1.95143 | 3.14349 | 1.611 |
| chr19 | 5480589 | 5480895 | 1.79142 | 2.88573 | 1.611 |
| chr5 | 76535032 | 76535258 | 1.27743 | 2.05779 | 1.611 |
| chr12 | 80254881 | 80255109 | 0.584518 | 0.941628 | 1.611 |
| chr12 | 119673274 | 119673679 | 0.797421 | 1.28462 | 1.611 |
| chr5 | 161748934 | 161749188 | 0.560203 | 0.902471 | 1.611 |
| chr14 | 23122156 | 23122625 | 1.52889 | 2.46301 | 1.611 |
| chr16 | 29898945 | 29899200 | 0.786735 | 1.2675 | 1.611 |
| chr4 | 37568420 | 37568668 | 1.02016 | 1.64359 | 1.611 |
| chr5 | 131805321 | 131805544 | 0.836513 | 1.34773 | 1.611 |
| chr3 | 153735916 | 153736191 | 1.7524 | 2.82348 | 1.611 |
| chr8 | 78812120 | 78812385 | 0.61544 | 0.991649 | 1.611 |
| chr8 | 136125672 | 136125953 | 0.622945 | 1.00374 | 1.611 |
| chr9 | 34547909 | 34548429 | 2.14824 | 3.46149 | 1.611 |
| chr11 | 61875517 | 61875889 | 1.51473 | 2.44083 | 1.611 |
| chr1 | 185639537 | 185639775 | 1.33809 | 2.15642 | 1.612 |
| chr1 | 100475995 | 100476219 | 0.515076 | 0.830128 | 1.612 |
| chr9 | 35481454 | 35481668 | 1.66183 | 2.67846 | 1.612 |
| chr17 | 64196792 | 64197053 | 2.69963 | 4.35134 | 1.612 |
| chr9 | 1077802 | 1078201 | 0.472391 | 0.761465 | 1.612 |
| chr8 | 53878458 | 53878660 | 1.09125 | 1.7591 | 1.612 |
| chr17 | 12190051 | 12190388 | 0.873053 | 1.40738 | 1.612 |
| chr15 | 60967115 | 60967395 | 0.593521 | 0.95678 | 1.612 |
| chr1 | 111574964 | 111575236 | 1.1623 | 1.87377 | 1.612 |
| chr17 | 67502675 | 67502922 | 1.17136 | 1.88841 | 1.612 |
| chr8 | 19727922 | 19728137 | 1.48832 | 2.39948 | 1.612 |
| chr2 | 85109300 | 85109556 | 2.00187 | 3.22746 | 1.612 |
| chr6 | 33640635 | 33640875 | 1.58184 | 2.55031 | 1.612 |
| chr7 | 29095142 | 29095368 | 0.6724 | 1.08412 | 1.612 |
| chr5 | 84421874 | 84422173 | 0.610225 | 0.983942 | 1.612 |
| chr5 | 62866614 | 62866827 | 0.564097 | 0.909604 | 1.613 |
| chr11 | 130304429 | 130304683 | 1.86494 | 3.00742 | 1.613 |
| chrX | 67731162 | 67731423 | 1.563 | 2.52056 | 1.613 |
| chr2 | 72151054 | 72151491 | 1.42849 | 2.30366 | 1.613 |
| chr8 | 20463000 | 20463230 | 0.611084 | 0.98547 | 1.613 |
| chr2 | 166060486 | 166060764 | 1.80338 | 2.90833 | 1.613 |
| chr1 | 206757013 | 206757480 | 2.07379 | 3.34447 | 1.613 |
| chr12 | 121931919 | 121932531 | 2.47387 | 3.98971 | 1.613 |
| chr14 | 97207761 | 97208142 | 2.26175 | 3.64763 | 1.613 |
| chr20 | 21683424 | 21684029 | 1.56386 | 2.52229 | 1.613 |
| chr15 | 84240797 | 84241244 | 1.37305 | 2.21477 | 1.613 |
| chr22 | 43627269 | 43627625 | 1.93435 | 3.12026 | 1.613 |
| chr15 | 63385850 | 63386109 | 2.54766 | 4.11004 | 1.613 |
| chr1 | 156115812 | 156116114 | 1.58302 | 2.55393 | 1.613 |
| chr19 | 49474127 | 49474549 | 2.30248 | 3.71466 | 1.613 |
| chr5 | 109940412 | 109940625 | 0.716254 | 1.15556 | 1.613 |
| chr12 | 55428116 | 55428491 | 0.789973 | 1.27455 | 1.613 |
| chr1 | 185139307 | 185139951 | 1.75796 | 2.83656 | 1.614 |

| | | | | | |
|---|---|---|---|---|---|
| chr19 | 6048814 | 6049187 | 2.11327 | 3.40986 | 1.614 |
| chr8 | 133206110 | 133206324 | 0.944457 | 1.52398 | 1.614 |
| chr5 | 84092236 | 84092620 | 0.665317 | 1.07358 | 1.614 |
| chr4 | 26331877 | 26332131 | 1.27007 | 2.04948 | 1.614 |
| chr19 | 49977150 | 49977634 | 3.30942 | 5.34043 | 1.614 |
| chr17 | 73234699 | 73235189 | 2.10256 | 3.39296 | 1.614 |
| chr14 | 52643703 | 52643953 | 0.650197 | 1.04926 | 1.614 |
| chr2 | 39820925 | 39821185 | 1.39125 | 2.24517 | 1.614 |
| chr1 | 33342167 | 33342638 | 1.21601 | 1.96241 | 1.614 |
| chr14 | 105287138 | 105287381 | 1.28892 | 2.08008 | 1.614 |
| chr3 | 147560778 | 147561037 | 0.648299 | 1.04626 | 1.614 |
| chr1 | 111984256 | 111984552 | 1.42347 | 2.29732 | 1.614 |
| chr12 | 126679959 | 126680500 | 0.726926 | 1.17318 | 1.614 |
| chr5 | 148337148 | 148337509 | 1.65698 | 2.67421 | 1.614 |
| chr1 | 182205550 | 182205826 | 0.724616 | 1.16948 | 1.614 |
| chr18 | 77140789 | 77141196 | 2.18344 | 3.52397 | 1.614 |
| chr3 | 139656563 | 139656886 | 1.67499 | 2.70336 | 1.614 |
| chr17 | 38191370 | 38191801 | 1.68905 | 2.72607 | 1.614 |
| chr12 | 59783683 | 59783992 | 0.736679 | 1.18906 | 1.614 |
| chr7 | 98246910 | 98247097 | 0.672571 | 1.08559 | 1.614 |
| chr11 | 32735122 | 32735349 | 1.47632 | 2.38296 | 1.614 |
| chr14 | 20400547 | 20400823 | 0.705275 | 1.1384 | 1.614 |
| chr8 | 110742611 | 110742824 | 0.467153 | 0.754046 | 1.614 |
| chr8 | 11421767 | 11422382 | 1.49063 | 2.40609 | 1.614 |
| chr7 | 38110302 | 38110689 | 1.28388 | 2.07239 | 1.614 |
| chr12 | 62191096 | 62191311 | 0.56277 | 0.908433 | 1.614 |
| chr2 | 207423918 | 207424248 | 1.11155 | 1.79429 | 1.614 |
| chr3 | 39772446 | 39772706 | 0.593987 | 0.958913 | 1.614 |
| chr14 | 78013943 | 78014175 | 0.707859 | 1.14277 | 1.614 |
| chr11 | 18013390 | 18013661 | 2.99129 | 4.82916 | 1.614 |
| chr11 | 63246352 | 63246575 | 2.01395 | 3.25138 | 1.614 |
| chr16 | 87715878 | 87716174 | 1.01783 | 1.64322 | 1.614 |
| chr9 | 77617150 | 77617385 | 1.62871 | 2.62964 | 1.615 |
| chr2 | 103209124 | 103209334 | 0.851188 | 1.37429 | 1.615 |
| chr9 | 99007325 | 99007575 | 0.527726 | 0.852137 | 1.615 |
| chr17 | 2610965 | 2611462 | 1.52504 | 2.46256 | 1.615 |
| chr16 | 11490051 | 11490458 | 2.84063 | 4.587 | 1.615 |
| chr9 | 130221158 | 130221427 | 0.559688 | 0.903771 | 1.615 |
| chr3 | 169386844 | 169387231 | 2.08925 | 3.37371 | 1.615 |
| chr3 | 7054289 | 7054548 | 0.604215 | 0.975708 | 1.615 |
| chr16 | 49384332 | 49384748 | 2.46845 | 3.98631 | 1.615 |
| chr12 | 109571948 | 109572364 | 1.96805 | 3.17828 | 1.615 |
| chr2 | 714306 | 714624 | 1.75126 | 2.82819 | 1.615 |
| chr3 | 20332630 | 20332872 | 0.552027 | 0.891496 | 1.615 |
| chr5 | 171821911 | 171822369 | 0.718201 | 1.15992 | 1.615 |
| chr17 | 20847355 | 20847998 | 1.30802 | 2.11252 | 1.615 |
| chr9 | 98489095 | 98489397 | 1.32627 | 2.14201 | 1.615 |
| chr5 | 94619733 | 94620221 | 1.13814 | 1.83825 | 1.615 |
| chr1 | 120156508 | 120156770 | 1.31829 | 2.12927 | 1.615 |
| chr10 | 71695978 | 71696296 | 0.684333 | 1.10533 | 1.615 |
| chr14 | 76175593 | 76175920 | 1.5939 | 2.57447 | 1.615 |
| chr2 | 226147610 | 226147969 | 1.49263 | 2.41103 | 1.615 |
| chr16 | 48387525 | 48387822 | 2.09477 | 3.38389 | 1.615 |
| chr9 | 32915736 | 32915947 | 0.73137 | 1.1816 | 1.616 |
| chr15 | 94649035 | 94649248 | 0.567713 | 0.917207 | 1.616 |
| chr16 | 84788032 | 84788309 | 1.23352 | 1.99298 | 1.616 |
| chr19 | 18580414 | 18580811 | 2.17428 | 3.51296 | 1.616 |
| chr5 | 150431216 | 150431556 | 1.69187 | 2.73354 | 1.616 |
| chr19 | 35046357 | 35046753 | 0.999699 | 1.61528 | 1.616 |
| chr19 | 3429474 | 3429861 | 2.61067 | 4.21839 | 1.616 |
| chr2 | 226530098 | 226530417 | 1.34971 | 2.18097 | 1.616 |

| | | | | | |
|---|---|---|---|---|---|
| chr17 | 49839561 | 49839849 | 0.645038 | 1.04235 | 1.616 |
| chr2 | 11822098 | 11822515 | 2.3888 | 3.86017 | 1.616 |
| chr17 | 6955380 | 6955781 | 1.9134 | 3.09199 | 1.616 |
| chr8 | 118056975 | 118057289 | 2.16328 | 3.49592 | 1.616 |
| chr11 | 17405393 | 17405906 | 2.03694 | 3.29183 | 1.616 |
| chr2 | 175411681 | 175411911 | 1.3372 | 2.16108 | 1.616 |
| chr9 | 18442655 | 18443032 | 2.12298 | 3.4313 | 1.616 |
| chr1 | 204176077 | 204176772 | 2.17612 | 3.51725 | 1.616 |
| chr11 | 64532838 | 64533095 | 1.07833 | 1.74289 | 1.616 |
| chr22 | 44388941 | 44389474 | 1.38171 | 2.23332 | 1.616 |
| chr3 | 14643628 | 14643966 | 1.11196 | 1.79744 | 1.616 |
| chr17 | 20020018 | 20020307 | 2.00596 | 3.24271 | 1.617 |
| chr5 | 99814877 | 99815133 | 1.41186 | 2.28238 | 1.617 |
| chr14 | 26523018 | 26523319 | 0.682805 | 1.10383 | 1.617 |
| chr2 | 44895474 | 44895824 | 0.928217 | 1.50063 | 1.617 |
| chr15 | 85832984 | 85833321 | 1.5878 | 2.56712 | 1.617 |
| chr16 | 75498586 | 75498954 | 3.22584 | 5.21555 | 1.617 |
| chr17 | 18873859 | 18874117 | 0.896908 | 1.45022 | 1.617 |
| chr10 | 101320359 | 101320657 | 1.60634 | 2.59735 | 1.617 |
| chr22 | 39077723 | 39078020 | 2.02867 | 3.28024 | 1.617 |
| chr8 | 54721392 | 54721707 | 1.1563 | 1.86967 | 1.617 |
| chr14 | 41331770 | 41332075 | 0.616168 | 0.996344 | 1.617 |
| chr7 | 75892233 | 75892604 | 2.40443 | 3.88799 | 1.617 |
| chr5 | 39080768 | 39081001 | 2.26997 | 3.67062 | 1.617 |
| chr1 | 159183940 | 159184297 | 2.37673 | 3.84329 | 1.617 |
| chr2 | 96840194 | 96840633 | 2.22039 | 3.59067 | 1.617 |
| chr12 | 22612876 | 22613123 | 1.53943 | 2.48948 | 1.617 |
| chr17 | 62103106 | 62103606 | 2.6139 | 4.22719 | 1.617 |
| chr2 | 238363612 | 238363834 | 0.658841 | 1.06549 | 1.617 |
| chr11 | 76170276 | 76170688 | 1.42872 | 2.31069 | 1.617 |
| chr3 | 46491338 | 46491590 | 1.20056 | 1.94168 | 1.617 |
| chr8 | 6345659 | 6345955 | 0.987697 | 1.59744 | 1.617 |
| chr10 | 71455920 | 71456261 | 1.51021 | 2.44254 | 1.617 |
| chr19 | 33634823 | 33635155 | 2.96506 | 4.79602 | 1.618 |
| chrX | 130975457 | 130975728 | 2.17455 | 3.51741 | 1.618 |
| chr5 | 116073296 | 116073500 | 0.850871 | 1.37632 | 1.618 |
| chr15 | 98532577 | 98532816 | 0.738334 | 1.19431 | 1.618 |
| chr6 | 159555810 | 159556088 | 2.08602 | 3.37434 | 1.618 |
| chr8 | 22553236 | 22553550 | 0.941709 | 1.52336 | 1.618 |
| chr7 | 127848349 | 127848837 | 1.4178 | 2.29353 | 1.618 |
| chr5 | 32136042 | 32136291 | 0.740252 | 1.1975 | 1.618 |
| chr15 | 90443384 | 90443620 | 0.566461 | 0.916365 | 1.618 |
| chr1 | 163285588 | 163285883 | 1.38467 | 2.23999 | 1.618 |
| chr2 | 38968111 | 38968594 | 1.03599 | 1.67594 | 1.618 |
| chr4 | 104128933 | 104129167 | 0.989747 | 1.60114 | 1.618 |
| chr4 | 181585995 | 181586227 | 1.47221 | 2.38178 | 1.618 |
| chr12 | 120602165 | 120602418 | 0.800468 | 1.29507 | 1.618 |
| chr8 | 25905333 | 25905586 | 0.931051 | 1.50638 | 1.618 |
| chr2 | 86070683 | 86070986 | 1.78441 | 2.88709 | 1.618 |
| chrX | 57355558 | 57355964 | 1.28998 | 2.08716 | 1.618 |
| chr8 | 110664730 | 110665089 | 1.11155 | 1.79849 | 1.618 |
| chr2 | 231714759 | 231715202 | 2.46347 | 3.98608 | 1.618 |
| chr6 | 85056158 | 85056363 | 0.916296 | 1.48264 | 1.618 |
| chr19 | 42800025 | 42800637 | 2.81571 | 4.55651 | 1.618 |
| chr12 | 120907453 | 120907893 | 2.46365 | 3.98682 | 1.618 |
| chr7 | 131192643 | 131192974 | 2.55333 | 4.13204 | 1.618 |
| chr3 | 186539270 | 186539503 | 1.09422 | 1.77079 | 1.618 |
| chr4 | 1345595 | 1346050 | 1.95721 | 3.16744 | 1.618 |
| chr10 | 49866339 | 49867219 | 1.28985 | 2.08744 | 1.618 |
| chr2 | 13407261 | 13407520 | 0.673002 | 1.08916 | 1.618 |
| chr9 | 98098641 | 98098916 | 1.8766 | 3.03704 | 1.618 |

| | | | | | |
|---|---|---|---|---|---|
| chr19 | 4872670 | 4872967 | 1.8743 | 3.03335 | 1.618 |
| chr12 | 31270026 | 31270409 | 1.57805 | 2.55396 | 1.618 |
| chr17 | 2654926 | 2655131 | 0.82521 | 1.33556 | 1.618 |
| chr1 | 216667326 | 216667606 | 0.596399 | 0.965271 | 1.619 |
| chr20 | 44651457 | 44651833 | 1.97838 | 3.20208 | 1.619 |
| chr11 | 125274683 | 125275259 | 2.08604 | 3.37642 | 1.619 |
| chr2 | 235453611 | 235453892 | 2.00673 | 3.24805 | 1.619 |
| chr19 | 38801292 | 38801587 | 0.818915 | 1.32555 | 1.619 |
| chr21 | 47393599 | 47393865 | 1.40328 | 2.2716 | 1.619 |
| chr19 | 35203623 | 35203907 | 1.43082 | 2.31621 | 1.619 |
| chr22 | 24921354 | 24921812 | 1.26774 | 2.05222 | 1.619 |
| chr17 | 78944321 | 78944603 | 1.02365 | 1.65709 | 1.619 |
| chr5 | 170832606 | 170832862 | 1.36477 | 2.20934 | 1.619 |
| chr2 | 75631118 | 75631402 | 1.80873 | 2.92821 | 1.619 |
| chr14 | 49964439 | 49964792 | 1.97843 | 3.20296 | 1.619 |
| chr2 | 81686276 | 81686527 | 0.699512 | 1.13247 | 1.619 |
| chr5 | 34486335 | 34486593 | 1.98194 | 3.20872 | 1.619 |
| chr5 | 79603458 | 79603691 | 1.26135 | 2.04211 | 1.619 |
| chr17 | 76009856 | 76010313 | 1.17671 | 1.90518 | 1.619 |
| chr22 | 46710186 | 46710538 | 1.73917 | 2.81584 | 1.619 |
| chr19 | 14582990 | 14583372 | 2.40607 | 3.89569 | 1.619 |
| chr15 | 88299878 | 88300130 | 0.518241 | 0.83914 | 1.619 |
| chr1 | 205760690 | 205761079 | 2.12943 | 3.44802 | 1.619 |
| chr17 | 45056262 | 45056505 | 0.655311 | 1.06109 | 1.619 |
| chr10 | 104829404 | 104829694 | 1.25369 | 2.03015 | 1.619 |
| chr1 | 201058275 | 201058488 | 0.626123 | 1.01395 | 1.619 |
| chr16 | 83350014 | 83350286 | 0.545476 | 0.883366 | 1.619 |
| chr20 | 7917242 | 7917646 | 1.59145 | 2.57725 | 1.619 |
| chr6 | 111580224 | 111580509 | 2.22413 | 3.60198 | 1.620 |
| chr2 | 203143762 | 203143971 | 1.2984 | 2.10283 | 1.620 |
| chr14 | 70651977 | 70652418 | 0.552289 | 0.894466 | 1.620 |
| chr17 | 55198770 | 55199024 | 1.50454 | 2.43671 | 1.620 |
| chr2 | 218751411 | 218751781 | 1.92175 | 3.11242 | 1.620 |
| chr4 | 181587441 | 181587662 | 1.27907 | 2.07179 | 1.620 |
| chr12 | 10343401 | 10343759 | 1.77342 | 2.87261 | 1.620 |
| chr11 | 11820273 | 11820705 | 2.31673 | 3.75272 | 1.620 |
| chr3 | 66988826 | 66989096 | 1.49622 | 2.42368 | 1.620 |
| chr16 | 56901177 | 56901575 | 1.52208 | 2.46571 | 1.620 |
| chr14 | 95156797 | 95157030 | 0.57391 | 0.929735 | 1.620 |
| chr22 | 33414241 | 33414711 | 1.52697 | 2.47381 | 1.620 |
| chr12 | 118502358 | 118502655 | 0.888486 | 1.43943 | 1.620 |
| chr20 | 43926810 | 43927140 | 0.650178 | 1.05336 | 1.620 |
| chr12 | 64150338 | 64150548 | 0.677941 | 1.09836 | 1.620 |
| chr3 | 74267589 | 74268030 | 0.716511 | 1.16086 | 1.620 |
| chr10 | 82020379 | 82020952 | 2.01804 | 3.26976 | 1.620 |
| chr17 | 6605064 | 6605411 | 1.58175 | 2.5629 | 1.620 |
| chr8 | 28861827 | 28862256 | 1.56889 | 2.54215 | 1.620 |
| chr17 | 46102473 | 46102759 | 2.21861 | 3.5953 | 1.621 |
| chr10 | 122086653 | 122086968 | 0.583967 | 0.946406 | 1.621 |
| chr8 | 34026135 | 34026415 | 0.665724 | 1.07898 | 1.621 |
| chr15 | 75824811 | 75825130 | 1.38304 | 2.24161 | 1.621 |
| chrX | 129254613 | 129255138 | 1.51977 | 2.46323 | 1.621 |
| chr2 | 28817170 | 28817526 | 2.04188 | 3.30951 | 1.621 |
| chr1 | 176558948 | 176559156 | 0.487203 | 0.789712 | 1.621 |
| chr2 | 9282834 | 9283203 | 2.52291 | 4.08975 | 1.621 |
| chr1 | 94929359 | 94929573 | 1.21789 | 1.97432 | 1.621 |
| chr11 | 17885811 | 17886350 | 1.66671 | 2.70192 | 1.621 |
| chr2 | 179884958 | 179885194 | 0.912934 | 1.48004 | 1.621 |
| chr18 | 13216133 | 13216406 | 1.15272 | 1.86879 | 1.621 |
| chr18 | 71784147 | 71784524 | 1.12107 | 1.81749 | 1.621 |
| chr16 | 21695515 | 21695880 | 1.6791 | 2.72229 | 1.621 |

| | | | | | |
|---|---|---|---|---|---|
| chr11 | 13992540 | 13992869 | 1.38253 | 2.24153 | 1.621 |
| chr12 | 6679453 | 6679821 | 1.69399 | 2.74655 | 1.621 |
| chr12 | 81797482 | 81797746 | 0.956491 | 1.55081 | 1.621 |
| chr11 | 66079262 | 66079635 | 1.96691 | 3.1891 | 1.621 |
| chr7 | 118961244 | 118961806 | 0.687516 | 1.11474 | 1.621 |
| chr7 | 51567561 | 51567842 | 0.598819 | 0.970971 | 1.621 |
| chr14 | 69255694 | 69255937 | 2.82605 | 4.58243 | 1.622 |
| chr1 | 236653290 | 236653525 | 0.956477 | 1.55099 | 1.622 |
| chr19 | 36109786 | 36110059 | 1.17134 | 1.89955 | 1.622 |
| chr1 | 2314111 | 2314430 | 1.07569 | 1.74448 | 1.622 |
| chr3 | 19055641 | 19055990 | 0.599429 | 0.972118 | 1.622 |
| chrX | 148552450 | 148552778 | 1.78308 | 2.89169 | 1.622 |
| chr2 | 46220446 | 46220879 | 1.38478 | 2.24594 | 1.622 |
| chr19 | 18340278 | 18340542 | 1.47615 | 2.39429 | 1.622 |
| chr6 | 21523296 | 21523566 | 0.800702 | 1.29877 | 1.622 |
| chr1 | 235891894 | 235892119 | 0.638798 | 1.03618 | 1.622 |
| chr6 | 10801478 | 10801805 | 1.92577 | 3.12376 | 1.622 |
| chr17 | 34987683 | 34988076 | 0.68182 | 1.10603 | 1.622 |
| chr10 | 71661347 | 71661717 | 1.39752 | 2.26706 | 1.622 |
| chr1 | 182564833 | 182565104 | 2.05931 | 3.34065 | 1.622 |
| chr1 | 23188527 | 23188923 | 1.84884 | 2.9993 | 1.622 |
| chr6 | 167669473 | 167669714 | 0.897217 | 1.45554 | 1.622 |
| chr1 | 152114422 | 152114843 | 1.98505 | 3.22044 | 1.622 |
| chr16 | 62822251 | 62822475 | 0.702824 | 1.14024 | 1.622 |
| chr5 | 146575572 | 146575802 | 0.63551 | 1.03104 | 1.622 |
| chr3 | 149198243 | 149198529 | 2.30762 | 3.74387 | 1.622 |
| chr14 | 101006860 | 101007167 | 1.65179 | 2.68002 | 1.622 |
| chr4 | 54774866 | 54775089 | 1.02921 | 1.66991 | 1.623 |
| chr20 | 3199029 | 3199575 | 2.81164 | 4.56198 | 1.623 |
| chr2 | 6987112 | 6987331 | 0.824329 | 1.33751 | 1.623 |
| chr9 | 140705849 | 140706321 | 0.721954 | 1.17141 | 1.623 |
| chr1 | 237977693 | 237978014 | 1.85543 | 3.01064 | 1.623 |
| chr9 | 103364624 | 103365030 | 1.31046 | 2.12637 | 1.623 |
| chr4 | 174209447 | 174209747 | 1.16313 | 1.88735 | 1.623 |
| chr1 | 6583672 | 6583995 | 1.62201 | 2.63199 | 1.623 |
| chr2 | 56418816 | 56419085 | 2.03084 | 3.29551 | 1.623 |
| chr10 | 100069218 | 100069695 | 1.95987 | 3.18035 | 1.623 |
| chr14 | 45737766 | 45738054 | 2.197 | 3.56536 | 1.623 |
| chr2 | 79075996 | 79076214 | 0.831515 | 1.34942 | 1.623 |
| chr11 | 102665269 | 102665489 | 1.07425 | 1.74337 | 1.623 |
| chr3 | 81055134 | 81055398 | 0.717013 | 1.16362 | 1.623 |
| chr4 | 6089481 | 6089726 | 1.47548 | 2.39451 | 1.623 |
| chr21 | 17198570 | 17198800 | 1.24407 | 2.01903 | 1.623 |
| chr3 | 64516193 | 64516399 | 1.09164 | 1.77164 | 1.623 |
| chr21 | 40948642 | 40948868 | 0.560318 | 0.909381 | 1.623 |
| chr13 | 23981086 | 23981349 | 1.82331 | 2.95922 | 1.623 |
| chrX | 152110189 | 152110439 | 2.15524 | 3.49813 | 1.623 |
| chr11 | 38676442 | 38676713 | 0.515684 | 0.837038 | 1.623 |
| chr6 | 159595296 | 159595663 | 2.39205 | 3.88297 | 1.623 |
| chr1 | 12139627 | 12139899 | 1.48426 | 2.40943 | 1.623 |
| chr17 | 61502674 | 61503199 | 2.68524 | 4.35922 | 1.623 |
| chr19 | 57515947 | 57516173 | 0.755705 | 1.22682 | 1.623 |
| chrX | 114877282 | 114877606 | 1.98124 | 3.21638 | 1.623 |
| chr10 | 82142733 | 82143134 | 1.57763 | 2.56128 | 1.624 |
| chr1 | 230450926 | 230451158 | 0.744867 | 1.20944 | 1.624 |
| chr9 | 140071093 | 140071562 | 1.32017 | 2.14359 | 1.624 |
| chr10 | 9445259 | 9445494 | 0.616415 | 1.00089 | 1.624 |
| chr8 | 3412342 | 3412731 | 0.773983 | 1.2568 | 1.624 |
| chr16 | 59872507 | 59872718 | 0.670128 | 1.08823 | 1.624 |
| chr11 | 34171412 | 34171623 | 1.6499 | 2.67944 | 1.624 |
| chr11 | 35637777 | 35638115 | 1.61481 | 2.62247 | 1.624 |

| | | | | | |
|---|---|---|---|---|---|
| chr8 | 2498425 | 2498709 | 0.404645 | 0.657155 | 1.624 |
| chr8 | 9688442 | 9688802 | 2.08525 | 3.38653 | 1.624 |
| chrX | 40017416 | 40017816 | 1.09535 | 1.77893 | 1.624 |
| chr5 | 117068622 | 117068851 | 0.699548 | 1.13619 | 1.624 |
| chr3 | 155869646 | 155870008 | 1.87227 | 3.04119 | 1.624 |
| chr8 | 142517891 | 142518631 | 2.12158 | 3.44616 | 1.624 |
| chr17 | 59049131 | 59049424 | 1.57004 | 2.5503 | 1.624 |
| chr16 | 28333016 | 28333639 | 1.94314 | 3.15648 | 1.624 |
| chr5 | 167014851 | 167015084 | 1.57467 | 2.55797 | 1.624 |
| chr10 | 92048249 | 92048469 | 0.738339 | 1.19941 | 1.624 |
| chr12 | 123850208 | 123850552 | 1.818 | 2.95335 | 1.625 |
| chr16 | 2892506 | 2892816 | 1.56965 | 2.54996 | 1.625 |
| chr2 | 233893583 | 233893896 | 1.42243 | 2.31097 | 1.625 |
| chr1 | 152433948 | 152434353 | 1.46867 | 2.3861 | 1.625 |
| chr3 | 76619524 | 76619935 | 0.920988 | 1.49642 | 1.625 |
| chr15 | 65023440 | 65023832 | 2.32516 | 3.77799 | 1.625 |
| chr6 | 35262167 | 35262478 | 2.07608 | 3.3734 | 1.625 |
| chr3 | 58615857 | 58616197 | 1.90514 | 3.09568 | 1.625 |
| chr7 | 73021344 | 73021811 | 2.58735 | 4.20462 | 1.625 |
| chr2 | 99001512 | 99001974 | 2.74685 | 4.46384 | 1.625 |
| chr17 | 72297339 | 72297783 | 2.06122 | 3.34983 | 1.625 |
| chr20 | 6381092 | 6381776 | 0.713439 | 1.15948 | 1.625 |
| chr19 | 17966435 | 17966755 | 1.72865 | 2.8096 | 1.625 |
| chr11 | 10229710 | 10230013 | 1.21565 | 1.97593 | 1.625 |
| chr5 | 109888883 | 109889109 | 0.640688 | 1.04138 | 1.625 |
| chr9 | 126112077 | 126112525 | 2.69966 | 4.38816 | 1.625 |
| chr9 | 77567152 | 77567532 | 2.65369 | 4.31351 | 1.625 |
| chr9 | 127601955 | 127602261 | 1.40832 | 2.28921 | 1.625 |
| chr3 | 98932065 | 98932423 | 0.525331 | 0.853958 | 1.626 |
| chr20 | 47950424 | 47950888 | 3.41922 | 5.55818 | 1.626 |
| chr9 | 117101382 | 117101771 | 2.62167 | 4.26179 | 1.626 |
| chr11 | 24203031 | 24203310 | 0.517617 | 0.841466 | 1.626 |
| chr1 | 239472058 | 239472348 | 1.71591 | 2.78952 | 1.626 |
| chr8 | 22158516 | 22158730 | 0.433575 | 0.704862 | 1.626 |
| chr11 | 43300277 | 43300536 | 1.29857 | 2.11113 | 1.626 |
| chr4 | 140578167 | 140578459 | 2.19331 | 3.56584 | 1.626 |
| chr9 | 135776962 | 135777615 | 1.43999 | 2.34116 | 1.626 |
| chr15 | 40971312 | 40971617 | 2.13856 | 3.47693 | 1.626 |
| chr5 | 132155823 | 132156283 | 1.86175 | 3.02726 | 1.626 |
| chr3 | 33876957 | 33877162 | 0.801665 | 1.30356 | 1.626 |
| chr19 | 41955206 | 41955702 | 3.67343 | 5.9734 | 1.626 |
| chr3 | 81056839 | 81057113 | 0.63741 | 1.03651 | 1.626 |
| chr11 | 11808361 | 11808632 | 1.19123 | 1.93712 | 1.626 |
| chr17 | 30580040 | 30580641 | 2.5381 | 4.12737 | 1.626 |
| chr19 | 42210546 | 42211180 | 4.42601 | 7.19768 | 1.626 |
| chr11 | 122734484 | 122734718 | 1.35812 | 2.20865 | 1.626 |
| chr8 | 24369087 | 24369424 | 1.617 | 2.62967 | 1.626 |
| chr14 | 77336558 | 77336841 | 1.10638 | 1.79927 | 1.626 |
| chr4 | 169864150 | 169864519 | 1.13739 | 1.84972 | 1.626 |
| chr8 | 145150899 | 145151338 | 1.42919 | 2.32433 | 1.626 |
| chr1 | 54414750 | 54415074 | 1.31057 | 2.13148 | 1.626 |
| chr17 | 41081354 | 41081557 | 1.103 | 1.79391 | 1.626 |
| chr19 | 36207565 | 36208207 | 2.43243 | 3.95608 | 1.626 |
| chr5 | 11261310 | 11261772 | 0.660788 | 1.07473 | 1.626 |
| chr2 | 80158941 | 80159249 | 0.625172 | 1.01683 | 1.626 |
| chr10 | 129557013 | 129557228 | 0.535504 | 0.871015 | 1.627 |
| chr2 | 50998259 | 50998646 | 0.711168 | 1.15676 | 1.627 |
| chr8 | 38042028 | 38042271 | 1.61854 | 2.63276 | 1.627 |
| chr8 | 28627172 | 28627575 | 2.04325 | 3.32423 | 1.627 |
| chr12 | 19441679 | 19441930 | 1.11262 | 1.81017 | 1.627 |
| chr7 | 42069756 | 42070014 | 0.614425 | 0.999631 | 1.627 |

| | | | | | |
|---|---|---|---|---|---|
| chr5 | 51422389 | 51422631 | 0.840768 | 1.3679 | 1.627 |
| chr12 | 130064361 | 130064668 | 0.745473 | 1.21286 | 1.627 |
| chr1 | 217920574 | 217920847 | 1.50193 | 2.44368 | 1.627 |
| chr3 | 76833324 | 76833646 | 0.713919 | 1.16157 | 1.627 |
| chr11 | 6337074 | 6337426 | 2.01306 | 3.27555 | 1.627 |
| chr12 | 9483550 | 9484148 | 2.37836 | 3.86996 | 1.627 |
| chr5 | 6401831 | 6402041 | 0.416359 | 0.677478 | 1.627 |
| chr17 | 27438771 | 27439347 | 2.62561 | 4.27251 | 1.627 |
| chr1 | 151148385 | 151148744 | 1.43474 | 2.3347 | 1.627 |
| chr12 | 29959803 | 29960010 | 0.626398 | 1.01938 | 1.627 |
| chr15 | 84376760 | 84377031 | 0.595315 | 0.968803 | 1.627 |
| chr12 | 27016535 | 27016803 | 1.11751 | 1.81868 | 1.627 |
| chr15 | 40509369 | 40509673 | 1.50601 | 2.45103 | 1.628 |
| chr8 | 92866961 | 92867183 | 0.636836 | 1.03651 | 1.628 |
| chr8 | 19991202 | 19991570 | 1.52008 | 2.47408 | 1.628 |
| chr15 | 81262411 | 81262718 | 2.60325 | 4.23732 | 1.628 |
| chr3 | 113464034 | 113464250 | 1.24867 | 2.03246 | 1.628 |
| chr17 | 76108645 | 76109198 | 2.8485 | 4.63663 | 1.628 |
| chr22 | 50964756 | 50965067 | 2.02555 | 3.29708 | 1.628 |
| chr5 | 34958276 | 34958529 | 1.42456 | 2.31893 | 1.628 |
| chr8 | 94433206 | 94433426 | 0.581951 | 0.94731 | 1.628 |
| chr10 | 81806632 | 81807148 | 2.20225 | 3.5852 | 1.628 |
| chr1 | 150951871 | 150952327 | 3.28203 | 5.34314 | 1.628 |
| chr3 | 82747188 | 82747405 | 0.599215 | 0.975526 | 1.628 |
| chr19 | 54963863 | 54964125 | 0.744671 | 1.21239 | 1.628 |
| chr17 | 32526871 | 32527316 | 4.99421 | 8.13105 | 1.628 |
| chr19 | 42155487 | 42155693 | 0.619577 | 1.00875 | 1.628 |
| chr19 | 42317819 | 42318303 | 0.738078 | 1.2017 | 1.628 |
| chr9 | 92217872 | 92218253 | 2.18619 | 3.55948 | 1.628 |
| chr11 | 13856109 | 13856333 | 1.25255 | 2.03949 | 1.628 |
| chr1 | 87550001 | 87550215 | 1.09794 | 1.78776 | 1.628 |
| chr4 | 57141293 | 57141597 | 0.494143 | 0.804611 | 1.628 |
| chr9 | 92163986 | 92164323 | 2.53762 | 4.13215 | 1.628 |
| chr8 | 26417532 | 26418048 | 0.893579 | 1.45509 | 1.628 |
| chr10 | 129477971 | 129478363 | 0.893021 | 1.45421 | 1.628 |
| chr20 | 40367292 | 40367629 | 2.01005 | 3.27327 | 1.628 |
| chr12 | 109125304 | 109125606 | 2.15872 | 3.51546 | 1.628 |
| chr17 | 30788731 | 30789211 | 1.42775 | 2.32509 | 1.629 |
| chr2 | 68936633 | 68936842 | 1.36366 | 2.22072 | 1.629 |
| chr7 | 35662231 | 35662481 | 0.656819 | 1.06964 | 1.629 |
| chr7 | 25878645 | 25878962 | 0.964794 | 1.57123 | 1.629 |
| chr12 | 132314864 | 132315178 | 2.03861 | 3.32005 | 1.629 |
| chr12 | 51735171 | 51735505 | 1.96602 | 3.20198 | 1.629 |
| chr12 | 113748420 | 113748736 | 0.894661 | 1.45715 | 1.629 |
| chr12 | 53459721 | 53460087 | 2.55218 | 4.15682 | 1.629 |
| chr17 | 66347700 | 66347919 | 0.628999 | 1.02449 | 1.629 |
| chr1 | 220550004 | 220550588 | 0.569848 | 0.928155 | 1.629 |
| chr10 | 2181910 | 2182146 | 0.684251 | 1.11455 | 1.629 |
| chr12 | 50030665 | 50031007 | 1.45737 | 2.37392 | 1.629 |
| chr10 | 99331800 | 99332380 | 1.93628 | 3.15436 | 1.629 |
| chr11 | 7709431 | 7709839 | 1.77442 | 2.8909 | 1.629 |
| chr16 | 68362544 | 68362870 | 1.94671 | 3.17173 | 1.629 |
| chr19 | 4633217 | 4633554 | 1.83191 | 2.9847 | 1.629 |
| chr8 | 130548010 | 130548361 | 2.11773 | 3.45048 | 1.629 |
| chr3 | 52325514 | 52325992 | 1.03885 | 1.69267 | 1.629 |
| chr17 | 980337 | 980945 | 0.615471 | 1.00286 | 1.629 |
| chrX | 99702899 | 99703212 | 1.56838 | 2.55556 | 1.629 |
| chr8 | 27062444 | 27062844 | 1.77847 | 2.89797 | 1.629 |
| chr8 | 131027135 | 131027725 | 1.38918 | 2.26363 | 1.629 |
| chr5 | 93234766 | 93235095 | 1.02271 | 1.66651 | 1.630 |
| chr5 | 54410294 | 54410497 | 0.717225 | 1.16877 | 1.630 |

| | | | | | |
|---|---|---|---|---|---|
| chr16 | 26841868 | 26842073 | 0.621572 | 1.01291 | 1.630 |
| chr17 | 42942521 | 42942852 | 1.87797 | 3.0604 | 1.630 |
| chr2 | 70844736 | 70845091 | 2.52612 | 4.11677 | 1.630 |
| chr17 | 17545154 | 17545477 | 2.21917 | 3.61673 | 1.630 |
| chr13 | 36363577 | 36363787 | 0.998191 | 1.62685 | 1.630 |
| chr1 | 191686436 | 191686669 | 0.610704 | 0.995359 | 1.630 |
| chr17 | 37809965 | 37811025 | 2.10411 | 3.42942 | 1.630 |
| chr2 | 4006940 | 4007231 | 1.84809 | 3.01219 | 1.630 |
| chr16 | 12990477 | 12990727 | 1.73221 | 2.82347 | 1.630 |
| chr2 | 65100557 | 65101002 | 3.34219 | 5.44795 | 1.630 |
| chr9 | 136351171 | 136351787 | 1.80823 | 2.94765 | 1.630 |
| chr15 | 73928775 | 73929252 | 3.33749 | 5.44061 | 1.630 |
| chr5 | 86180654 | 86180956 | 0.552312 | 0.90038 | 1.630 |
| chr1 | 150997045 | 150997378 | 1.88624 | 3.07499 | 1.630 |
| chr12 | 6944821 | 6945202 | 2.96358 | 4.83128 | 1.630 |
| chr17 | 40456924 | 40457229 | 0.542433 | 0.884343 | 1.630 |
| chr17 | 21360314 | 21360685 | 2.97738 | 4.85422 | 1.630 |
| chr8 | 38642617 | 38642857 | 1.28636 | 2.09731 | 1.630 |
| chr11 | 73041720 | 73042054 | 1.01237 | 1.65077 | 1.631 |
| chr2 | 48647902 | 48648172 | 1.87877 | 3.06367 | 1.631 |
| chr3 | 139281026 | 139281249 | 1.42817 | 2.32889 | 1.631 |
| chr17 | 65887907 | 65888251 | 1.50484 | 2.45396 | 1.631 |
| chr4 | 47838666 | 47838884 | 1.5051 | 2.45438 | 1.631 |
| chr10 | 32545620 | 32545966 | 2.20653 | 3.59831 | 1.631 |
| chr10 | 62194265 | 62194477 | 1.58926 | 2.59172 | 1.631 |
| chr7 | 30978242 | 30978717 | 2.73033 | 4.45257 | 1.631 |
| chr14 | 26066862 | 26067258 | 0.508239 | 0.828841 | 1.631 |
| chr7 | 4991752 | 4992073 | 0.579918 | 0.94576 | 1.631 |
| chr11 | 61244992 | 61245536 | 2.21373 | 3.61038 | 1.631 |
| chr16 | 87096358 | 87096590 | 1.56924 | 2.55927 | 1.631 |
| chr21 | 46265941 | 46266186 | 1.26814 | 2.06823 | 1.631 |
| chr12 | 81085947 | 81086158 | 0.745671 | 1.21614 | 1.631 |
| chr10 | 102694109 | 102694319 | 1.59737 | 2.60523 | 1.631 |
| chr1 | 229393861 | 229394159 | 1.61386 | 2.63226 | 1.631 |
| chr3 | 154518979 | 154519307 | 0.780223 | 1.27257 | 1.631 |
| chr2 | 234620147 | 234620380 | 2.48267 | 4.04937 | 1.631 |
| chr14 | 102245360 | 102245690 | 0.466983 | 0.761738 | 1.631 |
| chr6 | 154822061 | 154822303 | 1.36328 | 2.22386 | 1.631 |
| chr1 | 159760136 | 159760432 | 1.15449 | 1.88333 | 1.631 |
| chr8 | 3404468 | 3404677 | 0.685385 | 1.1181 | 1.631 |
| chr7 | 121732951 | 121733354 | 1.51364 | 2.46939 | 1.631 |
| chr6 | 134385059 | 134385312 | 2.14752 | 3.50366 | 1.631 |
| chr3 | 192520829 | 192521044 | 1.01325 | 1.65319 | 1.632 |
| chr3 | 143494252 | 143494495 | 1.37151 | 2.23776 | 1.632 |
| chr2 | 50258735 | 50258941 | 0.683778 | 1.11569 | 1.632 |
| chr15 | 78297467 | 78297682 | 0.771064 | 1.25816 | 1.632 |
| chr7 | 141349696 | 141350007 | 1.38035 | 2.25234 | 1.632 |
| chr8 | 10331922 | 10332165 | 1.2365 | 2.01772 | 1.632 |
| chr1 | 9202356 | 9202653 | 1.62213 | 2.64705 | 1.632 |
| chr15 | 77926795 | 77927378 | 1.68044 | 2.74221 | 1.632 |
| chr3 | 97742145 | 97742407 | 1.94576 | 3.17532 | 1.632 |
| chr7 | 30101497 | 30101730 | 2.05326 | 3.35075 | 1.632 |
| chr8 | 99176865 | 99177072 | 1.08913 | 1.77737 | 1.632 |
| chr1 | 12719060 | 12719291 | 1.9359 | 3.15926 | 1.632 |
| chrX | 77165909 | 77166147 | 1.52627 | 2.4908 | 1.632 |
| chr20 | 19472061 | 19472561 | 0.582857 | 0.951217 | 1.632 |
| chr11 | 76758087 | 76758273 | 1.21639 | 1.98515 | 1.632 |
| chr14 | 34568011 | 34568203 | 1.31582 | 2.14755 | 1.632 |
| chr2 | 231024895 | 231025255 | 1.84694 | 3.01448 | 1.632 |
| chr13 | 29379370 | 29379692 | 1.90716 | 3.11279 | 1.632 |
| chr16 | 81923902 | 81924297 | 2.01233 | 3.28445 | 1.632 |

| | | | | | |
|---|---|---|---|---|---|
| chr15 | 29960185 | 29960745 | 2.37711 | 3.87994 | 1.632 |
| chr2 | 50803974 | 50804193 | 0.687987 | 1.12295 | 1.632 |
| chr5 | 671667 | 672095 | 2.04585 | 3.33929 | 1.632 |
| chr11 | 68214596 | 68215102 | 2.65949 | 4.34093 | 1.632 |
| chr3 | 45706736 | 45706945 | 1.41183 | 2.30448 | 1.632 |
| chr8 | 20806533 | 20806760 | 0.611798 | 0.998701 | 1.632 |
| chr11 | 75919801 | 75920330 | 2.20305 | 3.59647 | 1.633 |
| chr15 | 89447724 | 89448336 | 1.7895 | 2.92142 | 1.633 |
| chr2 | 196524962 | 196525341 | 0.925605 | 1.51108 | 1.633 |
| chr4 | 48580458 | 48580698 | 1.12044 | 1.8292 | 1.633 |
| chr10 | 134271996 | 134272452 | 2.29442 | 3.74585 | 1.633 |
| chr10 | 79667660 | 79667917 | 1.51629 | 2.47554 | 1.633 |
| chr9 | 134829646 | 134829981 | 1.16436 | 1.90097 | 1.633 |
| chr14 | 104690768 | 104691208 | 1.84995 | 3.02036 | 1.633 |
| chr3 | 46936766 | 46937065 | 2.07134 | 3.38188 | 1.633 |
| chr3 | 4277513 | 4277759 | 0.505664 | 0.825606 | 1.633 |
| chr22 | 19718143 | 19718522 | 1.69038 | 2.76006 | 1.633 |
| chr2 | 241929211 | 241929610 | 1.77893 | 2.90492 | 1.633 |
| chr2 | 25565241 | 25565674 | 2.41152 | 3.93794 | 1.633 |
| chr11 | 1649867 | 1650476 | 2.16025 | 3.52774 | 1.633 |
| chr20 | 31044653 | 31045170 | 2.25587 | 3.68397 | 1.633 |
| chr4 | 48242454 | 48242680 | 0.461255 | 0.753269 | 1.633 |
| chr2 | 102857331 | 102857711 | 1.99531 | 3.25858 | 1.633 |
| chr10 | 76995635 | 76996082 | 5.73493 | 9.36618 | 1.633 |
| chr1 | 220863375 | 220863931 | 2.52947 | 4.13124 | 1.633 |
| chr19 | 47231891 | 47232771 | 2.38206 | 3.89048 | 1.633 |
| chr16 | 31221262 | 31221662 | 0.601323 | 0.982117 | 1.633 |
| chr11 | 975837 | 976094 | 1.64923 | 2.69365 | 1.633 |
| chr19 | 46110577 | 46110845 | 2.20013 | 3.59343 | 1.633 |
| chr17 | 21447028 | 21447235 | 0.602937 | 0.984802 | 1.633 |
| chr15 | 90762738 | 90763157 | 2.22861 | 3.64043 | 1.634 |
| chr17 | 27696803 | 27697431 | 1.5016 | 2.45286 | 1.634 |
| chr10 | 94674278 | 94674571 | 1.75734 | 2.87076 | 1.634 |
| chr14 | 29485565 | 29485915 | 1.2684 | 2.07209 | 1.634 |
| chr9 | 129235489 | 129235707 | 0.509722 | 0.832709 | 1.634 |
| chr7 | 31413004 | 31413272 | 1.7395 | 2.84183 | 1.634 |
| chr17 | 75768526 | 75768818 | 1.88387 | 3.07771 | 1.634 |
| chr12 | 88260345 | 88260805 | 1.21467 | 1.98446 | 1.634 |
| chr8 | 95104140 | 95104349 | 0.687598 | 1.12339 | 1.634 |
| chr5 | 54059861 | 54060169 | 0.730013 | 1.19273 | 1.634 |
| chr5 | 121706669 | 121707159 | 0.549247 | 0.897395 | 1.634 |
| chr13 | 23733404 | 23733736 | 2.06021 | 3.36613 | 1.634 |
| chrX | 153316635 | 153317030 | 1.67998 | 2.74494 | 1.634 |
| chr4 | 88296882 | 88297153 | 1.072 | 1.75158 | 1.634 |
| chr5 | 132689543 | 132689753 | 0.83723 | 1.36801 | 1.634 |
| chr10 | 99674495 | 99674892 | 2.40201 | 3.92491 | 1.634 |
| chr2 | 97466685 | 97467085 | 1.91201 | 3.12425 | 1.634 |
| chr18 | 77288811 | 77289156 | 1.583 | 2.58675 | 1.634 |
| chr7 | 144645700 | 144645937 | 0.604 | 0.986999 | 1.634 |
| chr9 | 101025974 | 101026416 | 2.03881 | 3.33161 | 1.634 |
| chr13 | 25302203 | 25302467 | 2.16104 | 3.53138 | 1.634 |
| chr19 | 48770302 | 48770634 | 1.67617 | 2.73919 | 1.634 |
| chr8 | 94648128 | 94648395 | 1.23631 | 2.02044 | 1.634 |
| chr8 | 52594055 | 52594265 | 0.60034 | 0.981148 | 1.634 |
| chr3 | 124269356 | 124269630 | 1.9949 | 3.26044 | 1.634 |
| chr6 | 151139027 | 151139318 | 1.57238 | 2.56989 | 1.634 |
| chr2 | 20335405 | 20335732 | 2.59342 | 4.23874 | 1.634 |
| chr11 | 82268438 | 82268651 | 0.634435 | 1.03696 | 1.634 |
| chr9 | 71895731 | 71895948 | 0.549748 | 0.898543 | 1.634 |
| chr3 | 80088247 | 80088469 | 0.450568 | 0.736439 | 1.634 |
| chr12 | 114127805 | 114128083 | 0.517841 | 0.846428 | 1.635 |

| | | | | | |
|---|---|---|---|---|---|
| chr16 | 1231324 | 1231616 | 1.42413 | 2.32791 | 1.635 |
| chrX | 30531466 | 30531679 | 1.47954 | 2.41858 | 1.635 |
| chr22 | 41799826 | 41800223 | 1.79184 | 2.92913 | 1.635 |
| chr1 | 36834579 | 36834969 | 2.13432 | 3.48909 | 1.635 |
| chr1 | 204776103 | 204776835 | 3.1955 | 5.22397 | 1.635 |
| chr3 | 58159639 | 58159854 | 1.19967 | 1.96121 | 1.635 |
| chr8 | 145689406 | 145689639 | 2.18036 | 3.56448 | 1.635 |
| chr3 | 4070614 | 4071070 | 0.595234 | 0.97316 | 1.635 |
| chr17 | 15847924 | 15848668 | 1.66045 | 2.71472 | 1.635 |
| chr9 | 123294058 | 123294285 | 1.58391 | 2.58969 | 1.635 |
| chr13 | 109567542 | 109567820 | 0.897661 | 1.4677 | 1.635 |
| chr8 | 79248523 | 79248763 | 0.62208 | 1.01712 | 1.635 |
| chr5 | 90680685 | 90681019 | 1.53896 | 2.51631 | 1.635 |
| chr12 | 124806761 | 124807000 | 0.603437 | 0.986673 | 1.635 |
| chr3 | 141594354 | 141594640 | 2.02979 | 3.3189 | 1.635 |
| chr16 | 65766617 | 65766906 | 0.683502 | 1.1176 | 1.635 |
| chr19 | 47799323 | 47799719 | 1.80151 | 2.94567 | 1.635 |
| chr8 | 87606081 | 87606827 | 0.718256 | 1.17443 | 1.635 |
| chr15 | 97921527 | 97922165 | 0.781255 | 1.27746 | 1.635 |
| chr8 | 21952709 | 21953334 | 1.3922 | 2.27649 | 1.635 |
| chr17 | 27940221 | 27940844 | 2.16461 | 3.53957 | 1.635 |
| chr1 | 32469194 | 32469425 | 0.822112 | 1.34433 | 1.635 |
| chr17 | 6908158 | 6908506 | 1.23503 | 2.01961 | 1.635 |
| chr12 | 68728058 | 68728282 | 1.04477 | 1.70852 | 1.635 |
| chr17 | 3977952 | 3978498 | 1.23642 | 2.02193 | 1.635 |
| chr17 | 58237002 | 58237231 | 0.524876 | 0.858387 | 1.635 |
| chr1 | 85773808 | 85774235 | 2.72299 | 4.45324 | 1.635 |
| chr2 | 73487377 | 73487871 | 1.54197 | 2.52193 | 1.636 |
| chr1 | 2703007 | 2703540 | 1.16574 | 1.90661 | 1.636 |
| chr7 | 744104 | 744387 | 0.511247 | 0.83618 | 1.636 |
| chr10 | 6245325 | 6245657 | 1.34706 | 2.20336 | 1.636 |
| chr11 | 75442294 | 75442526 | 0.544374 | 0.890477 | 1.636 |
| chr9 | 38400518 | 38400974 | 1.30605 | 2.13662 | 1.636 |
| chr7 | 146231699 | 146231905 | 0.653669 | 1.0694 | 1.636 |
| chr3 | 27105249 | 27105520 | 0.636053 | 1.0406 | 1.636 |
| chr17 | 48246784 | 48247076 | 2.30562 | 3.77213 | 1.636 |
| chr1 | 40609741 | 40609966 | 1.33242 | 2.17999 | 1.636 |
| chr1 | 213168859 | 213169114 | 1.33958 | 2.19171 | 1.636 |
| chr3 | 163933197 | 163933490 | 0.609315 | 0.996964 | 1.636 |
| chr17 | 56293188 | 56293400 | 0.997208 | 1.63168 | 1.636 |
| chr14 | 24370031 | 24370273 | 0.74046 | 1.21161 | 1.636 |
| chr10 | 124463388 | 124463840 | 2.50235 | 4.0946 | 1.636 |
| chr20 | 32778430 | 32778814 | 1.92602 | 3.15154 | 1.636 |
| chr9 | 89334655 | 89334893 | 0.521533 | 0.853439 | 1.636 |
| chr2 | 28843999 | 28844273 | 1.61685 | 2.64583 | 1.636 |
| chr7 | 93852790 | 93853025 | 1.00665 | 1.6473 | 1.636 |
| chr1 | 220948607 | 220949095 | 1.97499 | 3.23203 | 1.636 |
| chr12 | 88242064 | 88242325 | 1.5929 | 2.60695 | 1.637 |
| chr11 | 30122970 | 30123181 | 0.394516 | 0.645678 | 1.637 |
| chr2 | 121706977 | 121707251 | 1.39073 | 2.27624 | 1.637 |
| chr6 | 132919694 | 132919916 | 1.02822 | 1.68291 | 1.637 |
| chr14 | 41324431 | 41324898 | 0.576831 | 0.944167 | 1.637 |
| chr6 | 13332083 | 13332363 | 1.37325 | 2.24777 | 1.637 |
| chr7 | 131936820 | 131937051 | 1.18798 | 1.94454 | 1.637 |
| chr5 | 153877309 | 153877647 | 0.637916 | 1.04422 | 1.637 |
| chr3 | 50653330 | 50653740 | 2.7614 | 4.52032 | 1.637 |
| chr10 | 24846050 | 24846461 | 1.50787 | 2.46835 | 1.637 |
| chr16 | 50918046 | 50918254 | 1.62472 | 2.65967 | 1.637 |
| chr17 | 79347254 | 79347553 | 0.711287 | 1.16439 | 1.637 |
| chr5 | 71074535 | 71074827 | 2.02196 | 3.31002 | 1.637 |
| chr16 | 50058754 | 50059119 | 3.17345 | 5.19554 | 1.637 |

| | | | | | |
|---|---|---|---|---|---|
| chr7 | 132265370 | 132265895 | 1.35996 | 2.22657 | 1.637 |
| chr2 | 170336055 | 170336295 | 1.38133 | 2.26161 | 1.637 |
| chr10 | 27868990 | 27869194 | 0.624893 | 1.02319 | 1.637 |
| chr5 | 25581418 | 25581797 | 0.654543 | 1.07179 | 1.637 |
| chr7 | 137693625 | 137693884 | 1.88942 | 3.09396 | 1.638 |
| chr9 | 2824613 | 2824842 | 1.43835 | 2.35534 | 1.638 |
| chr2 | 10575076 | 10575484 | 2.94807 | 4.82783 | 1.638 |
| chr5 | 166744671 | 166745037 | 0.939027 | 1.53777 | 1.638 |
| chr2 | 74917628 | 74918002 | 1.42916 | 2.34053 | 1.638 |
| chr13 | 49875302 | 49875586 | 1.48617 | 2.43393 | 1.638 |
| chr5 | 4868294 | 4868566 | 2.53196 | 4.14674 | 1.638 |
| chr1 | 21596526 | 21597066 | 2.01193 | 3.29513 | 1.638 |
| chr13 | 94764608 | 94764821 | 0.398318 | 0.652379 | 1.638 |
| chr14 | 100806247 | 100806456 | 0.547362 | 0.896501 | 1.638 |
| chr9 | 112336401 | 112336886 | 0.938115 | 1.53654 | 1.638 |
| chr15 | 52263587 | 52263921 | 1.09668 | 1.79626 | 1.638 |
| chr1 | 160221767 | 160222077 | 1.27951 | 2.09578 | 1.638 |
| chr15 | 87688018 | 87688255 | 0.744893 | 1.22014 | 1.638 |
| chr5 | 180033005 | 180033547 | 1.56793 | 2.56829 | 1.638 |
| chr10 | 82265275 | 82265693 | 2.03629 | 3.33558 | 1.638 |
| chr17 | 38324657 | 38324931 | 1.8136 | 2.97087 | 1.638 |
| chr3 | 31810344 | 31810628 | 1.29533 | 2.12195 | 1.638 |
| chr15 | 94996472 | 94996720 | 0.828053 | 1.35656 | 1.638 |
| chr12 | 98495009 | 98495193 | 0.746272 | 1.22263 | 1.638 |
| chr3 | 114018318 | 114018578 | 1.71053 | 2.8024 | 1.638 |
| chr4 | 84565856 | 84566153 | 1.51415 | 2.48066 | 1.638 |
| chr16 | 3179396 | 3179869 | 2.29081 | 3.7532 | 1.638 |
| chr9 | 13044602 | 13044991 | 0.651092 | 1.06673 | 1.638 |
| chr6 | 56716647 | 56716870 | 1.14797 | 1.88081 | 1.638 |
| chr4 | 82327034 | 82327433 | 1.212 | 1.98581 | 1.638 |
| chr3 | 48646866 | 48647260 | 0.831469 | 1.36238 | 1.639 |
| chr5 | 55231987 | 55232278 | 1.9013 | 3.11542 | 1.639 |
| chr15 | 97331225 | 97331606 | 0.610053 | 0.999633 | 1.639 |
| chr2 | 174265601 | 174265867 | 1.12709 | 1.84687 | 1.639 |
| chr16 | 67203564 | 67203886 | 1.5479 | 2.53646 | 1.639 |
| chr1 | 240167366 | 240167950 | 1.69124 | 2.77145 | 1.639 |
| chr1 | 153163503 | 153163710 | 0.448105 | 0.734361 | 1.639 |
| chr16 | 58521581 | 58521950 | 1.78751 | 2.92953 | 1.639 |
| chr9 | 99079655 | 99080039 | 1.89352 | 3.10336 | 1.639 |
| chr14 | 77379600 | 77380355 | 2.23975 | 3.67085 | 1.639 |
| chr17 | 18855131 | 18855568 | 0.985114 | 1.6146 | 1.639 |
| chr19 | 17559419 | 17559740 | 1.68604 | 2.76342 | 1.639 |
| chr12 | 119340325 | 119340594 | 0.538478 | 0.882571 | 1.639 |
| chr10 | 1869253 | 1869519 | 0.524082 | 0.859012 | 1.639 |
| chr16 | 30803274 | 30803606 | 0.67985 | 1.11434 | 1.639 |
| chr6 | 26572885 | 26573154 | 1.6145 | 2.64638 | 1.639 |
| chr11 | 43911811 | 43912193 | 1.91682 | 3.14195 | 1.639 |
| chr19 | 42069675 | 42070199 | 2.57592 | 4.22251 | 1.639 |
| chr22 | 46374416 | 46375241 | 2.05498 | 3.36883 | 1.639 |
| chr5 | 150494835 | 150495237 | 2.90029 | 4.75464 | 1.639 |
| chr21 | 26758264 | 26758520 | 0.856983 | 1.40493 | 1.639 |
| chr10 | 8565717 | 8565956 | 0.384993 | 0.631163 | 1.639 |
| chr7 | 22122186 | 22122908 | 2.02397 | 3.31813 | 1.639 |
| chr19 | 31549722 | 31550018 | 0.810299 | 1.32843 | 1.639 |
| chr15 | 84705633 | 84705874 | 0.781563 | 1.2814 | 1.640 |
| chr17 | 14708760 | 14708988 | 1.23411 | 2.02339 | 1.640 |
| chr3 | 108633839 | 108634287 | 1.7126 | 2.80792 | 1.640 |
| chr2 | 65867718 | 65867959 | 2.4446 | 4.00819 | 1.640 |
| chr3 | 40369193 | 40369431 | 2.10103 | 3.44493 | 1.640 |
| chr1 | 154934324 | 154934629 | 1.54661 | 2.53593 | 1.640 |
| chr7 | 112051295 | 112051583 | 1.79842 | 2.94888 | 1.640 |

| | | | | | |
|---|---|---|---|---|---|
| chr7 | 73819595 | 73820016 | 1.16849 | 1.91601 | 1.640 |
| chrX | 16910885 | 16911246 | 1.35858 | 2.22774 | 1.640 |
| chr1 | 174231805 | 174232294 | 1.89791 | 3.1123 | 1.640 |
| chr2 | 147491723 | 147491956 | 0.921895 | 1.51188 | 1.640 |
| chrX | 132109361 | 132109612 | 1.18955 | 1.95091 | 1.640 |
| chr8 | 52438617 | 52438962 | 1.65469 | 2.71377 | 1.640 |
| chr8 | 112370889 | 112371252 | 0.661968 | 1.08566 | 1.640 |
| chr9 | 80071814 | 80072069 | 1.86721 | 3.06236 | 1.640 |
| chr14 | 33769697 | 33770177 | 0.571322 | 0.937033 | 1.640 |
| chr5 | 116283693 | 116283902 | 1.49671 | 2.45486 | 1.640 |
| chr10 | 89265082 | 89265425 | 1.63499 | 2.68172 | 1.640 |
| chr8 | 64624308 | 64624530 | 0.629259 | 1.03212 | 1.640 |
| chr8 | 131606781 | 131606983 | 0.607839 | 0.99703 | 1.640 |
| chr20 | 42612045 | 42612334 | 1.64096 | 2.69185 | 1.640 |
| chr14 | 99984600 | 99985194 | 2.1425 | 3.51482 | 1.641 |
| chr10 | 11740042 | 11740437 | 1.2044 | 1.97607 | 1.641 |
| chr3 | 30083818 | 30084085 | 1.89582 | 3.11059 | 1.641 |
| chr16 | 53972556 | 53973108 | 0.482141 | 0.791095 | 1.641 |
| chr9 | 34546971 | 34547184 | 1.48527 | 2.43708 | 1.641 |
| chr12 | 25710827 | 25711095 | 1.52792 | 2.50745 | 1.641 |
| chr7 | 55323263 | 55323565 | 2.70229 | 4.43475 | 1.641 |
| chr5 | 139624030 | 139624390 | 2.86508 | 4.70252 | 1.641 |
| chr15 | 32321625 | 32321856 | 1.89833 | 3.11582 | 1.641 |
| chr22 | 18640003 | 18640218 | 1.54363 | 2.53366 | 1.641 |
| chr11 | 118965582 | 118965901 | 1.28251 | 2.10507 | 1.641 |
| chr17 | 38050843 | 38051089 | 0.589465 | 0.967583 | 1.641 |
| chr20 | 57657138 | 57657477 | 0.854019 | 1.40186 | 1.641 |
| chr20 | 58070865 | 58071402 | 2.19517 | 3.6034 | 1.642 |
| chr7 | 94285371 | 94285658 | 2.34856 | 3.85522 | 1.642 |
| chr11 | 88218015 | 88218337 | 1.44459 | 2.37142 | 1.642 |
| chr6 | 43576059 | 43576354 | 1.6786 | 2.75582 | 1.642 |
| chr19 | 50748792 | 50749041 | 0.482435 | 0.79206 | 1.642 |
| chr14 | 70175413 | 70175834 | 1.06266 | 1.74469 | 1.642 |
| chr12 | 72356557 | 72356882 | 0.682816 | 1.12108 | 1.642 |
| chr5 | 119388109 | 119388412 | 0.686674 | 1.12751 | 1.642 |
| chr1 | 39357197 | 39357560 | 0.509424 | 0.836478 | 1.642 |
| chr1 | 186955118 | 186955320 | 1.1157 | 1.83209 | 1.642 |
| chr11 | 4063015 | 4063281 | 0.564439 | 0.926878 | 1.642 |
| chr8 | 40057260 | 40057468 | 1.27757 | 2.09797 | 1.642 |
| chr9 | 95217123 | 95217846 | 1.04604 | 1.7178 | 1.642 |
| chr5 | 125436316 | 125436889 | 1.21782 | 1.9999 | 1.642 |
| chr8 | 127579185 | 127579587 | 1.41631 | 2.32594 | 1.642 |
| chr16 | 85752190 | 85752738 | 1.71347 | 2.81415 | 1.642 |
| chr9 | 108177658 | 108178039 | 1.78425 | 2.93039 | 1.642 |
| chr6 | 127786086 | 127786357 | 1.52764 | 2.50907 | 1.642 |
| chr2 | 152664778 | 152665114 | 1.04753 | 1.7206 | 1.643 |
| chr5 | 145921868 | 145922138 | 1.60795 | 2.64111 | 1.643 |
| chr7 | 86099064 | 86099283 | 1.11123 | 1.82524 | 1.643 |
| chr17 | 9891071 | 9891556 | 2.5247 | 4.14698 | 1.643 |
| chr3 | 141022221 | 141022541 | 1.47841 | 2.42842 | 1.643 |
| chr11 | 1938839 | 1939333 | 1.06849 | 1.75512 | 1.643 |
| chr18 | 53447521 | 53447776 | 0.696218 | 1.14364 | 1.643 |
| chr8 | 94239749 | 94239998 | 0.651135 | 1.06968 | 1.643 |
| chr3 | 107212067 | 107212302 | 0.818894 | 1.34531 | 1.643 |
| chr17 | 40336118 | 40336554 | 2.88003 | 4.73197 | 1.643 |
| chr8 | 24779348 | 24779559 | 0.518639 | 0.852154 | 1.643 |
| chr17 | 7535819 | 7536150 | 1.44541 | 2.37492 | 1.643 |
| chr20 | 56793789 | 56794043 | 0.625491 | 1.02773 | 1.643 |
| chr17 | 31559876 | 31560155 | 0.728607 | 1.19728 | 1.643 |
| chr2 | 1549776 | 1550091 | 1.54736 | 2.5429 | 1.643 |
| chr1 | 15599067 | 15599326 | 1.22977 | 2.02101 | 1.643 |

| | | | | | |
|---|---|---|---|---|---|
| chr15 | 79341862 | 79342105 | 0.633701 | 1.04144 | 1.643 |
| chr6 | 17167681 | 17167889 | 1.73104 | 2.84498 | 1.644 |
| chr5 | 38265255 | 38266161 | 1.46517 | 2.40803 | 1.644 |
| chr15 | 69917155 | 69917667 | 0.528709 | 0.868958 | 1.644 |
| chr15 | 32085617 | 32085998 | 0.593959 | 0.976229 | 1.644 |
| chr2 | 72098748 | 72099166 | 1.62658 | 2.67359 | 1.644 |
| chr3 | 116489745 | 116490074 | 2.12617 | 3.49478 | 1.644 |
| chr1 | 231052981 | 231053384 | 1.50211 | 2.46905 | 1.644 |
| chr9 | 88099961 | 88100250 | 0.496648 | 0.816381 | 1.644 |
| chr4 | 146538140 | 146538400 | 1.25687 | 2.06608 | 1.644 |
| chr19 | 47861135 | 47861665 | 1.35425 | 2.22621 | 1.644 |
| chr16 | 89681983 | 89682411 | 3.52135 | 5.78871 | 1.644 |
| chr7 | 42565886 | 42566183 | 0.426287 | 0.700768 | 1.644 |
| chr16 | 48151288 | 48151790 | 2.59094 | 4.25925 | 1.644 |
| chr2 | 220298635 | 220298854 | 1.5714 | 2.58337 | 1.644 |
| chr8 | 103826845 | 103827196 | 2.41951 | 3.9779 | 1.644 |
| chr19 | 34636585 | 34637040 | 1.87944 | 3.09006 | 1.644 |
| chr10 | 106034411 | 106034855 | 2.7222 | 4.47577 | 1.644 |
| chr7 | 40518839 | 40519064 | 0.528837 | 0.869512 | 1.644 |
| chr2 | 25532246 | 25532578 | 1.31058 | 2.15498 | 1.644 |
| chr1 | 160048368 | 160048784 | 2.20713 | 3.62926 | 1.644 |
| chr16 | 57757102 | 57757428 | 1.17243 | 1.92789 | 1.644 |
| chr2 | 102447802 | 102448055 | 1.1562 | 1.9012 | 1.644 |
| chr20 | 14929128 | 14929340 | 1.05534 | 1.73543 | 1.644 |
| chrX | 71219008 | 71219309 | 1.81574 | 2.98604 | 1.645 |
| chr1 | 157059946 | 157060250 | 2.09361 | 3.44306 | 1.645 |
| chr15 | 30322731 | 30323024 | 0.720303 | 1.18461 | 1.645 |
| chr11 | 69702512 | 69702756 | 0.744126 | 1.2238 | 1.645 |
| chr1 | 151906527 | 151906836 | 2.3683 | 3.89499 | 1.645 |
| chr9 | 2132579 | 2132801 | 1.25614 | 2.06591 | 1.645 |
| chr7 | 38438323 | 38438687 | 2.47045 | 4.06307 | 1.645 |
| chr16 | 54400188 | 54400434 | 0.566022 | 0.930925 | 1.645 |
| chr7 | 1215774 | 1216149 | 2.54557 | 4.1868 | 1.645 |
| chr2 | 46238081 | 46238285 | 1.30069 | 2.13936 | 1.645 |
| chr8 | 112864550 | 112864771 | 0.661441 | 1.08795 | 1.645 |
| chr12 | 102153208 | 102153542 | 1.3919 | 2.28945 | 1.645 |
| chr14 | 37619752 | 37619987 | 1.43129 | 2.35426 | 1.645 |
| chr12 | 79941679 | 79941956 | 1.54955 | 2.54883 | 1.645 |
| chr12 | 2531884 | 2532210 | 0.675948 | 1.11188 | 1.645 |
| chr2 | 239481968 | 239482308 | 1.62225 | 2.66847 | 1.645 |
| chr12 | 108306072 | 108306768 | 2.33826 | 3.8464 | 1.645 |
| chr7 | 130406807 | 130407057 | 1.3021 | 2.14211 | 1.645 |
| chr17 | 3747278 | 3747749 | 1.64164 | 2.70077 | 1.645 |
| chr2 | 127807722 | 127808250 | 1.8095 | 2.97695 | 1.645 |
| chr20 | 10028103 | 10028394 | 1.4313 | 2.35489 | 1.645 |
| chr9 | 76562185 | 76562407 | 0.635926 | 1.04628 | 1.645 |
| chr14 | 85994965 | 85995235 | 1.53268 | 2.52172 | 1.645 |
| chr11 | 61202994 | 61203338 | 2.02169 | 3.32634 | 1.645 |
| chr1 | 8922546 | 8923077 | 1.09605 | 1.80339 | 1.645 |
| chr5 | 36546589 | 36547060 | 0.562705 | 0.925889 | 1.645 |
| chr11 | 122086172 | 122086506 | 1.91771 | 3.1556 | 1.646 |
| chr1 | 201191921 | 201192405 | 2.21043 | 3.6374 | 1.646 |
| chr7 | 16080095 | 16080341 | 0.838705 | 1.38018 | 1.646 |
| chr12 | 47426239 | 47426620 | 0.724699 | 1.19261 | 1.646 |
| chr20 | 8286125 | 8286441 | 1.28478 | 2.11437 | 1.646 |
| chr7 | 120195251 | 120195493 | 0.761949 | 1.25401 | 1.646 |
| chr7 | 1747737 | 1747963 | 1.26653 | 2.08455 | 1.646 |
| chr19 | 37149736 | 37150216 | 0.6931 | 1.14079 | 1.646 |
| chr17 | 63112049 | 63112343 | 0.834911 | 1.37422 | 1.646 |
| chr7 | 134943473 | 134943744 | 0.501976 | 0.826253 | 1.646 |
| chr16 | 66929122 | 66929495 | 2.21846 | 3.65167 | 1.646 |

| | | | | | |
|---|---|---|---|---|---|
| chr2 | 3740046 | 3740262 | 1.30362 | 2.14588 | 1.646 |
| chr7 | 8458359 | 8458656 | 1.2173 | 2.00379 | 1.646 |
| chr16 | 24917618 | 24917876 | 1.09056 | 1.79517 | 1.646 |
| chr5 | 66283889 | 66284092 | 1.86359 | 3.06769 | 1.646 |
| chr12 | 109004363 | 109004607 | 0.580311 | 0.955268 | 1.646 |
| chr1 | 5681156 | 5681399 | 1.35528 | 2.23099 | 1.646 |
| chr2 | 9279743 | 9280291 | 2.23764 | 3.68387 | 1.646 |
| chr14 | 28763961 | 28764362 | 0.858306 | 1.41307 | 1.646 |
| chr1 | 206831809 | 206832179 | 2.15658 | 3.55051 | 1.646 |
| chr2 | 120460415 | 120460792 | 3.15662 | 5.19717 | 1.646 |
| chr7 | 158645858 | 158646273 | 0.607766 | 1.00065 | 1.646 |
| chr9 | 103200181 | 103200479 | 2.50247 | 4.12016 | 1.646 |
| chr5 | 59020398 | 59020683 | 1.54185 | 2.53861 | 1.646 |
| chr1 | 12091232 | 12091615 | 1.28442 | 2.11487 | 1.647 |
| chr14 | 24777130 | 24777698 | 1.89167 | 3.11481 | 1.647 |
| chr10 | 122759280 | 122759654 | 0.73628 | 1.21237 | 1.647 |
| chr9 | 129261397 | 129262233 | 1.76592 | 2.90781 | 1.647 |
| chr17 | 5210529 | 5210950 | 1.02284 | 1.68431 | 1.647 |
| chr16 | 73230718 | 73230954 | 1.39172 | 2.29182 | 1.647 |
| chr19 | 47269903 | 47270502 | 2.01129 | 3.31209 | 1.647 |
| chr1 | 28417775 | 28418021 | 1.76091 | 2.89985 | 1.647 |
| chr17 | 65356077 | 65356325 | 0.971758 | 1.60035 | 1.647 |
| chr17 | 12927723 | 12928138 | 4.655 | 7.66627 | 1.647 |
| chr2 | 40781911 | 40782161 | 1.92795 | 3.17512 | 1.647 |
| chr6 | 143600633 | 143600893 | 0.750105 | 1.23534 | 1.647 |
| chr3 | 119901675 | 119902004 | 0.69196 | 1.13961 | 1.647 |
| chr6 | 137113561 | 137114433 | 1.50837 | 2.4842 | 1.647 |
| chr3 | 47463977 | 47464204 | 1.12908 | 1.85961 | 1.647 |
| chr3 | 14550342 | 14551086 | 2.03627 | 3.35395 | 1.647 |
| chr22 | 46751344 | 46751631 | 1.79151 | 2.95081 | 1.647 |
| chr7 | 5125045 | 5125461 | 2.69514 | 4.43941 | 1.647 |
| chr20 | 8701778 | 8702075 | 1.74448 | 2.87351 | 1.647 |
| chr1 | 107973187 | 107973681 | 1.2944 | 2.13216 | 1.647 |
| chrX | 99767536 | 99767768 | 1.22041 | 2.0103 | 1.647 |
| chr17 | 40669320 | 40669827 | 1.99184 | 3.28123 | 1.647 |
| chr2 | 236655064 | 236655285 | 1.73284 | 2.85459 | 1.647 |
| chr15 | 29832148 | 29832371 | 0.582542 | 0.959669 | 1.647 |
| chr17 | 11081816 | 11082068 | 0.580522 | 0.956357 | 1.647 |
| chr17 | 63096709 | 63096974 | 3.45835 | 5.69734 | 1.647 |
| chr1 | 165687155 | 165687458 | 1.66579 | 2.74434 | 1.647 |
| chr22 | 33975463 | 33975727 | 1.58965 | 2.6189 | 1.647 |
| chr14 | 73980885 | 73981133 | 0.84259 | 1.38815 | 1.647 |
| chr6 | 152964176 | 152964443 | 1.41327 | 2.32846 | 1.648 |
| chr10 | 104805576 | 104805867 | 0.637692 | 1.05065 | 1.648 |
| chr1 | 45118287 | 45118926 | 1.71004 | 2.81755 | 1.648 |
| chr17 | 80509705 | 80510073 | 1.98994 | 3.27875 | 1.648 |
| chr4 | 110624212 | 110624414 | 1.17368 | 1.93391 | 1.648 |
| chr9 | 17063622 | 17063843 | 1.55295 | 2.55888 | 1.648 |
| chr18 | 43861173 | 43861482 | 2.0909 | 3.44532 | 1.648 |
| chr17 | 51008335 | 51008649 | 0.653044 | 1.07609 | 1.648 |
| chr11 | 120381789 | 120382530 | 2.06813 | 3.4079 | 1.648 |
| chr9 | 129253623 | 129253901 | 2.13443 | 3.51717 | 1.648 |
| chr18 | 53528560 | 53528768 | 1.35985 | 2.24081 | 1.648 |
| chr18 | 9226117 | 9226349 | 0.473987 | 0.781075 | 1.648 |
| chr5 | 79968013 | 79968301 | 1.93071 | 3.18185 | 1.648 |
| chr14 | 61005185 | 61005758 | 0.524804 | 0.864902 | 1.648 |
| chr1 | 32574941 | 32575229 | 1.76426 | 2.90762 | 1.648 |
| chr5 | 120342765 | 120342975 | 0.60141 | 0.991167 | 1.648 |
| chr5 | 141660599 | 141660975 | 0.548185 | 0.90345 | 1.648 |
| chr7 | 117390783 | 117391010 | 1.32351 | 2.18125 | 1.648 |
| chr17 | 7039586 | 7039915 | 1.86263 | 3.06987 | 1.648 |

| | | | | | |
|---|---|---|---|---|---|
| chr2 | 11828574 | 11828926 | 2.70991 | 4.46631 | 1.648 |
| chr20 | 8316255 | 8316561 | 0.698866 | 1.1519 | 1.648 |
| chr16 | 22988621 | 22988955 | 0.830121 | 1.36827 | 1.648 |
| chr2 | 241453297 | 241453608 | 2.30675 | 3.80218 | 1.648 |
| chr1 | 177727020 | 177727331 | 0.532791 | 0.878223 | 1.648 |
| chr18 | 39878789 | 39879024 | 1.34072 | 2.21001 | 1.648 |
| chr14 | 60937389 | 60937636 | 0.538228 | 0.887214 | 1.648 |
| chr17 | 50598858 | 50599123 | 0.844374 | 1.39191 | 1.648 |
| chr12 | 71654532 | 71654758 | 0.6467 | 1.06609 | 1.649 |
| chr15 | 90015985 | 90016559 | 2.01783 | 3.32643 | 1.649 |
| chr5 | 35003443 | 35003869 | 1.59399 | 2.62783 | 1.649 |
| chr22 | 36576265 | 36576737 | 1.75113 | 2.88716 | 1.649 |
| chr1 | 146635052 | 146635270 | 0.717258 | 1.1826 | 1.649 |
| chr5 | 140305903 | 140306633 | 1.95077 | 3.2169 | 1.649 |
| chr20 | 20549316 | 20549585 | 2.03935 | 3.3632 | 1.649 |
| chr12 | 70983736 | 70984106 | 1.90546 | 3.14245 | 1.649 |
| chr5 | 167541460 | 167541769 | 1.92405 | 3.17317 | 1.649 |
| chr5 | 180075672 | 180076291 | 1.37643 | 2.27003 | 1.649 |
| chr10 | 64197448 | 64197765 | 1.65678 | 2.73245 | 1.649 |
| chr16 | 15850365 | 15850770 | 2.27599 | 3.75371 | 1.649 |
| chr11 | 2172616 | 2173542 | 1.4656 | 2.41728 | 1.649 |
| chr7 | 32849962 | 32850220 | 0.505939 | 0.834479 | 1.649 |
| chr1 | 2792549 | 2792796 | 1.86019 | 3.0682 | 1.649 |
| chr1 | 152025090 | 152025372 | 1.79646 | 2.96316 | 1.649 |
| chr11 | 122503957 | 122504311 | 1.01527 | 1.67463 | 1.649 |
| chr14 | 24802745 | 24803195 | 2.28116 | 3.76264 | 1.649 |
| chr8 | 98144520 | 98144791 | 0.604889 | 0.997744 | 1.649 |
| chr17 | 39065605 | 39065844 | 1.76111 | 2.905 | 1.650 |
| chr3 | 77582537 | 77582822 | 0.864138 | 1.42543 | 1.650 |
| chr19 | 51597983 | 51598651 | 1.39829 | 2.30659 | 1.650 |
| chr11 | 85198959 | 85199167 | 0.503045 | 0.829832 | 1.650 |
| chr11 | 64124046 | 64124289 | 1.39888 | 2.30763 | 1.650 |
| chr3 | 119430682 | 119430948 | 0.654836 | 1.08026 | 1.650 |
| chr3 | 159099476 | 159099747 | 0.643783 | 1.06207 | 1.650 |
| chr11 | 6552554 | 6552864 | 2.31369 | 3.81711 | 1.650 |
| chr1 | 232763511 | 232763721 | 0.981237 | 1.61885 | 1.650 |
| chr20 | 38506216 | 38506641 | 0.574568 | 0.947943 | 1.650 |
| chr14 | 92116361 | 92116563 | 1.65377 | 2.72849 | 1.650 |
| chr15 | 93497884 | 93498130 | 1.1946 | 1.97093 | 1.650 |
| chr12 | 52414416 | 52414949 | 1.69613 | 2.79841 | 1.650 |
| chr8 | 135292315 | 135292560 | 0.504957 | 0.833124 | 1.650 |
| chr12 | 110243058 | 110243524 | 2.87034 | 4.73594 | 1.650 |
| chr16 | 49891560 | 49892182 | 1.9781 | 3.26383 | 1.650 |
| chr19 | 7431923 | 7432153 | 0.869128 | 1.43408 | 1.650 |
| chr11 | 13793981 | 13794292 | 1.04031 | 1.71658 | 1.650 |
| chr21 | 33784651 | 33784912 | 0.706416 | 1.16565 | 1.650 |
| chr8 | 80439331 | 80439708 | 0.715103 | 1.18 | 1.650 |
| chr7 | 85936204 | 85936409 | 0.490163 | 0.808871 | 1.650 |
| chr16 | 55737310 | 55737662 | 0.997925 | 1.64684 | 1.650 |
| chr5 | 34905742 | 34905967 | 1.38281 | 2.28204 | 1.650 |
| chr9 | 115665493 | 115665698 | 1.32466 | 2.18613 | 1.650 |
| chr1 | 7102834 | 7103041 | 0.925064 | 1.52667 | 1.650 |
| chr14 | 50320574 | 50320920 | 2.50362 | 4.13191 | 1.650 |
| chr2 | 45007412 | 45007709 | 0.732428 | 1.2088 | 1.650 |
| chr6 | 116832600 | 116832815 | 0.854436 | 1.41016 | 1.650 |
| chr8 | 54647128 | 54647365 | 2.08423 | 3.43983 | 1.650 |
| chr3 | 147254736 | 147255037 | 0.529456 | 0.873958 | 1.651 |
| chr8 | 138447791 | 138448008 | 0.561486 | 0.926854 | 1.651 |
| chr3 | 63947590 | 63947840 | 1.15583 | 1.90796 | 1.651 |
| chr14 | 54908076 | 54908361 | 2.63263 | 4.34582 | 1.651 |
| chr11 | 66789242 | 66789464 | 1.34573 | 2.22148 | 1.651 |

| | | | | | |
|---|---|---|---|---|---|
| chr11 | 1324517 | 1324957 | 1.46744 | 2.42245 | 1.651 |
| chr1 | 45251781 | 45252162 | 2.59617 | 4.28589 | 1.651 |
| chr6 | 49499769 | 49500172 | 1.60269 | 2.64611 | 1.651 |
| chr1 | 109104402 | 109104720 | 1.74742 | 2.88509 | 1.651 |
| chr13 | 23594456 | 23594697 | 1.39492 | 2.30316 | 1.651 |
| chr17 | 56394729 | 56395132 | 2.54814 | 4.20738 | 1.651 |
| chr1 | 37019462 | 37019727 | 1.69764 | 2.80309 | 1.651 |
| chr7 | 5219992 | 5220261 | 0.645179 | 1.06543 | 1.651 |
| chr16 | 88882373 | 88882821 | 2.60136 | 4.29611 | 1.651 |
| chr8 | 67243038 | 67243383 | 0.629998 | 1.04049 | 1.652 |
| chr15 | 31489524 | 31490008 | 2.16346 | 3.57324 | 1.652 |
| chrX | 68505809 | 68506198 | 2.48545 | 4.10505 | 1.652 |
| chr7 | 98963142 | 98963569 | 3.13168 | 5.1725 | 1.652 |
| chr2 | 70660558 | 70660807 | 0.470286 | 0.77678 | 1.652 |
| chr19 | 9190504 | 9190777 | 2.69839 | 4.45727 | 1.652 |
| chr19 | 30348892 | 30349491 | 1.90435 | 3.14567 | 1.652 |
| chr3 | 141681397 | 141681720 | 2.25832 | 3.7304 | 1.652 |
| chr1 | 154297094 | 154297524 | 2.28713 | 3.77804 | 1.652 |
| chr14 | 83321071 | 83321551 | 0.571083 | 0.9434 | 1.652 |
| chr13 | 60252424 | 60252630 | 1.40969 | 2.32878 | 1.652 |
| chr4 | 166264971 | 166265190 | 0.939275 | 1.55166 | 1.652 |
| chr11 | 71096705 | 71096982 | 0.539552 | 0.891378 | 1.652 |
| chr12 | 67645607 | 67645900 | 0.509909 | 0.842418 | 1.652 |
| chr10 | 85965397 | 85965660 | 2.10665 | 3.48046 | 1.652 |
| chr1 | 15426847 | 15427159 | 1.92471 | 3.18003 | 1.652 |
| chr8 | 67682827 | 67683066 | 0.645718 | 1.06686 | 1.652 |
| chr17 | 12116876 | 12117125 | 0.494317 | 0.816731 | 1.652 |
| chr8 | 104086069 | 104086502 | 1.58903 | 2.62552 | 1.652 |
| chr7 | 140426226 | 140426429 | 1.23314 | 2.0375 | 1.652 |
| chr17 | 56524879 | 56525278 | 1.62844 | 2.69068 | 1.652 |
| chr19 | 47535416 | 47535891 | 2.741 | 4.52899 | 1.652 |
| chr17 | 57407005 | 57407246 | 1.69992 | 2.80882 | 1.652 |
| chr22 | 43263718 | 43264113 | 2.91451 | 4.81575 | 1.652 |
| chr5 | 158560218 | 158560509 | 1.46626 | 2.4228 | 1.652 |
| chr6 | 107814389 | 107814643 | 0.488853 | 0.80778 | 1.652 |
| chr22 | 34319760 | 34320001 | 1.39552 | 2.30602 | 1.652 |
| chr12 | 55508000 | 55508203 | 0.611491 | 1.01049 | 1.653 |
| chr20 | 60462201 | 60462691 | 2.33784 | 3.86327 | 1.653 |
| chr22 | 50648153 | 50648587 | 1.73223 | 2.86272 | 1.653 |
| chr12 | 27168621 | 27168927 | 1.11348 | 1.8402 | 1.653 |
| chr12 | 55095919 | 55096286 | 1.17609 | 1.9437 | 1.653 |
| chr8 | 11723841 | 11724082 | 1.06023 | 1.75225 | 1.653 |
| chr5 | 172139998 | 172140441 | 2.04582 | 3.38116 | 1.653 |
| chr2 | 64873497 | 64873832 | 2.77707 | 4.58986 | 1.653 |
| chr12 | 4803907 | 4804264 | 0.562846 | 0.930259 | 1.653 |
| chr5 | 126935626 | 126935852 | 0.692432 | 1.14451 | 1.653 |
| chr17 | 8024899 | 8025326 | 2.87502 | 4.75219 | 1.653 |
| chr22 | 27193958 | 27194162 | 1.10284 | 1.82292 | 1.653 |
| chr5 | 172198968 | 172199285 | 2.09281 | 3.45929 | 1.653 |
| chr9 | 77692133 | 77692423 | 1.59384 | 2.6346 | 1.653 |
| chr9 | 117444366 | 117444696 | 1.56971 | 2.59508 | 1.653 |
| chr19 | 51059633 | 51060380 | 2.70235 | 4.46764 | 1.653 |
| chr20 | 19975976 | 19976295 | 2.08328 | 3.44416 | 1.653 |
| chr5 | 121761020 | 121761247 | 0.706922 | 1.16871 | 1.653 |
| chr1 | 3200577 | 3200846 | 1.66374 | 2.75058 | 1.653 |
| chr8 | 65003883 | 65004100 | 0.540506 | 0.89359 | 1.653 |
| chr1 | 5574626 | 5574991 | 3.05546 | 5.05151 | 1.653 |
| chr12 | 50708493 | 50708763 | 1.59483 | 2.63693 | 1.653 |
| chr3 | 142712876 | 142713116 | 1.67867 | 2.77555 | 1.653 |
| chr3 | 13691858 | 13692391 | 3.8722 | 6.40252 | 1.653 |
| chr6 | 73326238 | 73326445 | 0.922115 | 1.52469 | 1.653 |

| | | | | | |
|---|---|---|---|---|---|
| chr2 | 77739082 | 77739290 | 0.559015 | 0.92432 | 1.653 |
| chr10 | 54539830 | 54540309 | 2.15941 | 3.57059 | 1.654 |
| chr17 | 78169832 | 78170261 | 3.34958 | 5.53871 | 1.654 |
| chr11 | 60064286 | 60064697 | 0.559374 | 0.924963 | 1.654 |
| chr10 | 109983967 | 109984397 | 0.534812 | 0.884381 | 1.654 |
| chr7 | 96190654 | 96190903 | 0.675852 | 1.11761 | 1.654 |
| chr1 | 214801459 | 214801936 | 1.86243 | 3.07979 | 1.654 |
| chr12 | 48172966 | 48173401 | 2.16363 | 3.57787 | 1.654 |
| chr9 | 138392248 | 138392704 | 2.46859 | 4.08241 | 1.654 |
| chr7 | 72866059 | 72866493 | 1.99026 | 3.29141 | 1.654 |
| chr19 | 36449915 | 36450370 | 1.16572 | 1.92787 | 1.654 |
| chr9 | 139358172 | 139358456 | 0.548564 | 0.90722 | 1.654 |
| chr15 | 91683166 | 91683372 | 0.854095 | 1.41265 | 1.654 |
| chr5 | 10701650 | 10701907 | 2.08603 | 3.45033 | 1.654 |
| chr9 | 33466613 | 33467021 | 2.81484 | 4.65585 | 1.654 |
| chr16 | 57610273 | 57610719 | 2.11096 | 3.49167 | 1.654 |
| chr7 | 23317038 | 23317331 | 1.43428 | 2.37247 | 1.654 |
| chr16 | 47634823 | 47635144 | 0.760402 | 1.25781 | 1.654 |
| chr3 | 53147959 | 53148373 | 1.91694 | 3.17089 | 1.654 |
| chr5 | 75699525 | 75699849 | 0.816842 | 1.35119 | 1.654 |
| chr7 | 52105636 | 52105908 | 0.701203 | 1.15996 | 1.654 |
| chr11 | 118827292 | 118827694 | 1.88499 | 3.11836 | 1.654 |
| chr7 | 24921346 | 24921548 | 1.05634 | 1.74753 | 1.654 |
| chr8 | 137906847 | 137907083 | 0.796072 | 1.317 | 1.654 |
| chr16 | 2954195 | 2954550 | 2.73421 | 4.52345 | 1.654 |
| chr4 | 156717249 | 156717487 | 1.33546 | 2.20937 | 1.654 |
| chr10 | 15251830 | 15252231 | 2.1016 | 3.47695 | 1.654 |
| chr3 | 11985526 | 11985927 | 1.91003 | 3.16004 | 1.654 |
| chr1 | 220950072 | 220950493 | 2.25049 | 3.72341 | 1.654 |
| chr2 | 30929857 | 30930473 | 1.86308 | 3.0825 | 1.655 |
| chr5 | 145205314 | 145205557 | 1.53655 | 2.54225 | 1.655 |
| chr3 | 107646358 | 107646676 | 1.98723 | 3.28795 | 1.655 |
| chrX | 82913762 | 82913985 | 0.666178 | 1.10222 | 1.655 |
| chr20 | 49203178 | 49203412 | 0.763346 | 1.26301 | 1.655 |
| chr19 | 1113212 | 1113602 | 2.04504 | 3.38378 | 1.655 |
| chr11 | 25851082 | 25851316 | 0.61257 | 1.01359 | 1.655 |
| chr8 | 146012757 | 146013038 | 2.04225 | 3.3792 | 1.655 |
| chr8 | 56830538 | 56831004 | 0.505662 | 0.836707 | 1.655 |
| chr10 | 12926115 | 12926502 | 0.632757 | 1.04702 | 1.655 |
| chr13 | 45883593 | 45883910 | 2.02048 | 3.34366 | 1.655 |
| chr9 | 3817789 | 3818000 | 0.653171 | 1.08092 | 1.655 |
| chr6 | 125474666 | 125475036 | 1.86585 | 3.08779 | 1.655 |
| chr3 | 52854770 | 52855127 | 1.58995 | 2.63124 | 1.655 |
| chr14 | 60798487 | 60798910 | 1.40967 | 2.33296 | 1.655 |
| chr22 | 18312783 | 18313106 | 2.69031 | 4.45258 | 1.655 |
| chr12 | 132397606 | 132397905 | 1.208 | 1.99931 | 1.655 |
| chr19 | 30181900 | 30182191 | 1.44452 | 2.39082 | 1.655 |
| chr11 | 106888163 | 106888707 | 1.62378 | 2.68754 | 1.655 |
| chr17 | 54468774 | 54469315 | 0.63479 | 1.05065 | 1.655 |
| chr6 | 137167237 | 137167517 | 1.46942 | 2.43208 | 1.655 |
| chr1 | 226309002 | 226309508 | 1.84856 | 3.05971 | 1.655 |
| chr17 | 2628077 | 2628408 | 0.891204 | 1.47511 | 1.655 |
| chr15 | 34650574 | 34650857 | 1.08677 | 1.79882 | 1.655 |
| chr8 | 111096264 | 111096495 | 0.713534 | 1.18109 | 1.655 |
| chr1 | 215249755 | 215250132 | 0.97066 | 1.60676 | 1.655 |
| chr16 | 72991952 | 72992182 | 0.557293 | 0.922586 | 1.655 |
| chr20 | 1946290 | 1946952 | 2.10717 | 3.48843 | 1.656 |
| chr8 | 93703734 | 93703971 | 0.583134 | 0.965377 | 1.656 |
| chr4 | 123523845 | 123524235 | 0.753413 | 1.24728 | 1.656 |
| chr16 | 84930598 | 84930849 | 0.82839 | 1.37142 | 1.656 |
| chr15 | 68608731 | 68609207 | 1.53612 | 2.54317 | 1.656 |

| | | | | | |
|---|---|---|---|---|---|
| chr20 | 934253 | 934759 | 1.10026 | 1.82158 | 1.656 |
| chr11 | 108741922 | 108742185 | 1.12541 | 1.86323 | 1.656 |
| chr17 | 27443974 | 27444561 | 2.63563 | 4.36374 | 1.656 |
| chr11 | 61213649 | 61214063 | 2.34105 | 3.87611 | 1.656 |
| chr11 | 78992896 | 78993167 | 0.535603 | 0.886822 | 1.656 |
| chr1 | 1840324 | 1840675 | 1.81232 | 3.00097 | 1.656 |
| chr6 | 69989828 | 69990086 | 0.51833 | 0.858291 | 1.656 |
| chr8 | 37773735 | 37774085 | 1.4713 | 2.43634 | 1.656 |
| chr8 | 142471563 | 142471877 | 0.726496 | 1.20303 | 1.656 |
| chr12 | 14964075 | 14964339 | 1.54806 | 2.56397 | 1.656 |
| chr17 | 47938538 | 47938992 | 1.86022 | 3.08102 | 1.656 |
| chr12 | 63192950 | 63193230 | 1.91343 | 3.16918 | 1.656 |
| chr9 | 100716811 | 100717183 | 1.43133 | 2.3708 | 1.656 |
| chr15 | 45418831 | 45419057 | 1.48664 | 2.46245 | 1.656 |
| chr10 | 84137424 | 84137612 | 1.06754 | 1.76828 | 1.656 |
| chr17 | 63790709 | 63790950 | 1.40037 | 2.31959 | 1.656 |
| chr14 | 81350600 | 81350805 | 0.677343 | 1.122 | 1.656 |
| chr3 | 22441944 | 22442191 | 0.637971 | 1.05678 | 1.656 |
| chr15 | 78112665 | 78113184 | 0.984072 | 1.63017 | 1.657 |
| chr2 | 110303660 | 110304025 | 2.37417 | 3.93299 | 1.657 |
| chr15 | 44069577 | 44069919 | 1.66056 | 2.75108 | 1.657 |
| chr1 | 194880978 | 194881233 | 0.686546 | 1.13747 | 1.657 |
| chr7 | 120178721 | 120178977 | 0.652837 | 1.08166 | 1.657 |
| chr9 | 140130819 | 140131365 | 2.64549 | 4.38325 | 1.657 |
| chr1 | 205626664 | 205626973 | 1.79537 | 2.97473 | 1.657 |
| chr14 | 60043370 | 60043697 | 2.03918 | 3.37885 | 1.657 |
| chr17 | 42956024 | 42956423 | 1.30162 | 2.15677 | 1.657 |
| chr22 | 42833449 | 42833843 | 2.11982 | 3.51251 | 1.657 |
| chr3 | 116333728 | 116334247 | 0.826871 | 1.37012 | 1.657 |
| chr10 | 48398410 | 48398821 | 3.21084 | 5.32037 | 1.657 |
| chrX | 62495087 | 62495348 | 1.33688 | 2.21522 | 1.657 |
| chr12 | 104451532 | 104451818 | 1.32018 | 2.18762 | 1.657 |
| chr5 | 58663633 | 58663930 | 1.59045 | 2.63554 | 1.657 |
| chr7 | 118269821 | 118270026 | 0.832638 | 1.37977 | 1.657 |
| chr14 | 22568449 | 22568818 | 1.26143 | 2.09034 | 1.657 |
| chr3 | 78694949 | 78695247 | 1.70024 | 2.81752 | 1.657 |
| chr17 | 73743852 | 73744180 | 1.53229 | 2.53923 | 1.657 |
| chr4 | 77679487 | 77679784 | 1.34564 | 2.23018 | 1.657 |
| chr8 | 23019358 | 23019651 | 1.05627 | 1.7506 | 1.657 |
| chr13 | 108807882 | 108808215 | 1.67051 | 2.76869 | 1.657 |
| chr7 | 50736215 | 50736523 | 0.607368 | 1.00665 | 1.657 |
| chr13 | 101298618 | 101298928 | 1.60735 | 2.66406 | 1.657 |
| chr22 | 38909119 | 38909347 | 0.754093 | 1.24985 | 1.657 |
| chr3 | 153715245 | 153715429 | 0.713933 | 1.18329 | 1.657 |
| chr5 | 149652177 | 149652370 | 1.11891 | 1.85462 | 1.658 |
| chr17 | 28420495 | 28420856 | 1.96653 | 3.25959 | 1.658 |
| chr15 | 32941625 | 32941850 | 1.2984 | 2.15216 | 1.658 |
| chrX | 15161471 | 15161716 | 1.38884 | 2.30207 | 1.658 |
| chr6 | 44201919 | 44202192 | 2.02227 | 3.35219 | 1.658 |
| chr10 | 104111853 | 104112401 | 1.79489 | 2.97536 | 1.658 |
| chr12 | 120752025 | 120752367 | 1.19856 | 1.98689 | 1.658 |
| chr3 | 73150205 | 73150656 | 1.32113 | 2.19011 | 1.658 |
| chr9 | 100968019 | 100968322 | 2.0991 | 3.47994 | 1.658 |
| chr9 | 139524701 | 139525170 | 1.50732 | 2.49887 | 1.658 |
| chr2 | 85408517 | 85409012 | 0.719553 | 1.19291 | 1.658 |
| chr4 | 166277159 | 166277390 | 0.951131 | 1.57684 | 1.658 |
| chr6 | 84324766 | 84325006 | 1.61619 | 2.67944 | 1.658 |
| chr11 | 20650676 | 20650959 | 0.654506 | 1.0851 | 1.658 |
| chrX | 30588692 | 30588985 | 0.677474 | 1.1232 | 1.658 |
| chr14 | 83516694 | 83516924 | 0.649608 | 1.07702 | 1.658 |
| chr17 | 47918846 | 47919183 | 0.893597 | 1.48159 | 1.658 |

| | | | | | |
|---|---|---|---|---|---|
| chr16 | 8232723 | 8233021 | 0.688479 | 1.14154 | 1.658 |
| chr8 | 89443396 | 89443604 | 0.510656 | 0.846697 | 1.658 |
| chr10 | 100856605 | 100856832 | 0.744452 | 1.23435 | 1.658 |
| chr6 | 41007048 | 41007414 | 1.29492 | 2.14707 | 1.658 |
| chr19 | 44658005 | 44658222 | 0.548876 | 0.910112 | 1.658 |
| chr2 | 218874904 | 218875212 | 1.12973 | 1.87331 | 1.658 |
| chr15 | 32267035 | 32267487 | 1.21316 | 2.01166 | 1.658 |
| chr17 | 2256314 | 2256663 | 3.29908 | 5.47062 | 1.658 |
| chr7 | 150475847 | 150476159 | 2.26911 | 3.76291 | 1.658 |
| chr10 | 72972230 | 72972489 | 0.722572 | 1.19829 | 1.658 |
| chr17 | 78400615 | 78401232 | 2.36744 | 3.92614 | 1.658 |
| chr2 | 78511212 | 78511553 | 0.568722 | 0.943161 | 1.658 |
| chr17 | 79700765 | 79701090 | 2.49819 | 4.143 | 1.658 |
| chr10 | 118691790 | 118692043 | 0.771495 | 1.27953 | 1.659 |
| chr6 | 36583016 | 36583235 | 2.0221 | 3.35382 | 1.659 |
| chr7 | 135347061 | 135347264 | 1.54817 | 2.56786 | 1.659 |
| chr1 | 247844650 | 247844885 | 1.61009 | 2.67067 | 1.659 |
| chr12 | 30476176 | 30476590 | 0.666774 | 1.10599 | 1.659 |
| chr10 | 24041898 | 24042144 | 0.554602 | 0.919972 | 1.659 |
| chr11 | 75288261 | 75288566 | 1.57655 | 2.6153 | 1.659 |
| chr14 | 64061599 | 64061867 | 0.61385 | 1.01832 | 1.659 |
| chr8 | 129965401 | 129965621 | 1.00863 | 1.67339 | 1.659 |
| chr7 | 133538590 | 133538997 | 2.00374 | 3.32459 | 1.659 |
| chr2 | 180286851 | 180287098 | 1.53099 | 2.54025 | 1.659 |
| chr7 | 100157365 | 100157850 | 2.39566 | 3.97522 | 1.659 |
| chr1 | 234205602 | 234205856 | 1.12046 | 1.85927 | 1.659 |
| chr3 | 63804299 | 63804530 | 1.60323 | 2.66037 | 1.659 |
| chr9 | 131690228 | 131690538 | 2.4064 | 3.99321 | 1.659 |
| chr9 | 15142888 | 15143235 | 2.12018 | 3.51841 | 1.659 |
| chr2 | 200336187 | 200336523 | 1.97799 | 3.28247 | 1.660 |
| chr11 | 2909571 | 2909970 | 2.66043 | 4.41508 | 1.660 |
| chrX | 154277291 | 154277683 | 1.81735 | 3.01606 | 1.660 |
| chr12 | 15459153 | 15459698 | 0.622488 | 1.03309 | 1.660 |
| chr17 | 12163079 | 12163346 | 0.717544 | 1.19089 | 1.660 |
| chr11 | 44154947 | 44155170 | 1.30951 | 2.17352 | 1.660 |
| chr3 | 50712723 | 50713078 | 1.38715 | 2.30245 | 1.660 |
| chr21 | 36007763 | 36008023 | 0.774566 | 1.28569 | 1.660 |
| chr12 | 94528050 | 94528424 | 1.34843 | 2.23829 | 1.660 |
| chr20 | 4270422 | 4270880 | 1.69349 | 2.81107 | 1.660 |
| chr7 | 140175602 | 140175869 | 1.43523 | 2.38238 | 1.660 |
| chr7 | 39328707 | 39328911 | 0.644809 | 1.0704 | 1.660 |
| chr17 | 1928948 | 1929602 | 1.35421 | 2.24815 | 1.660 |
| chr13 | 27778141 | 27778350 | 0.787548 | 1.30743 | 1.660 |
| chr18 | 10787375 | 10787609 | 1.86259 | 3.09215 | 1.660 |
| chr6 | 132271671 | 132271936 | 1.49563 | 2.48296 | 1.660 |
| chr16 | 8617639 | 8618037 | 2.51268 | 4.17146 | 1.660 |
| chr7 | 125040520 | 125040766 | 0.522441 | 0.867335 | 1.660 |
| chr6 | 71722603 | 71722930 | 1.36155 | 2.26045 | 1.660 |
| chr2 | 13354472 | 13354742 | 0.458912 | 0.761918 | 1.660 |
| chr16 | 68014381 | 68014659 | 0.88192 | 1.46429 | 1.660 |
| chr10 | 48426816 | 48427206 | 2.33207 | 3.87217 | 1.660 |
| chr10 | 79807019 | 79807448 | 1.69081 | 2.80754 | 1.660 |
| chr15 | 29602519 | 29602729 | 0.596626 | 0.990706 | 1.661 |
| chr1 | 179568835 | 179569205 | 1.127 | 1.87148 | 1.661 |
| chr10 | 33395782 | 33396035 | 0.825299 | 1.37051 | 1.661 |
| chr5 | 11721834 | 11722095 | 0.515252 | 0.855657 | 1.661 |
| chr6 | 38839719 | 38840264 | 1.12754 | 1.87247 | 1.661 |
| chr17 | 46961811 | 46962577 | 1.75935 | 2.92172 | 1.661 |
| chr20 | 6563793 | 6564104 | 1.64059 | 2.72453 | 1.661 |
| chr12 | 9786822 | 9787234 | 1.62512 | 2.69885 | 1.661 |
| chr8 | 139473203 | 139473517 | 0.52443 | 0.870954 | 1.661 |

| | | | | | |
|---|---|---|---|---|---|
| chr8 | 145288517 | 145288918 | 2.01505 | 3.34653 | 1.661 |
| chr9 | 115555572 | 115555915 | 1.69566 | 2.8161 | 1.661 |
| chr11 | 76426154 | 76426396 | 0.60591 | 1.00631 | 1.661 |
| chr22 | 37655630 | 37655895 | 1.43909 | 2.39018 | 1.661 |
| chr5 | 149288695 | 149288946 | 0.632489 | 1.05056 | 1.661 |
| chr14 | 88176553 | 88176797 | 0.554447 | 0.920952 | 1.661 |
| chr15 | 75135431 | 75135764 | 1.5167 | 2.51933 | 1.661 |
| chr22 | 24551564 | 24551927 | 1.32521 | 2.20126 | 1.661 |
| chr7 | 149321763 | 149322135 | 2.07619 | 3.44875 | 1.661 |
| chr5 | 167150325 | 167150720 | 2.04956 | 3.40455 | 1.661 |
| chr12 | 26611758 | 26612035 | 0.520799 | 0.865148 | 1.661 |
| chr10 | 22242504 | 22242769 | 1.01801 | 1.69116 | 1.661 |
| chr14 | 104560922 | 104561434 | 2.31844 | 3.85159 | 1.661 |
| chr2 | 217642452 | 217642720 | 1.63599 | 2.71798 | 1.661 |
| chrX | 100851165 | 100851450 | 1.23298 | 2.04844 | 1.661 |
| chr17 | 59555447 | 59555663 | 0.631502 | 1.04918 | 1.661 |
| chr1 | 182365907 | 182366223 | 2.12888 | 3.53697 | 1.661 |
| chr15 | 25985079 | 25985292 | 1.01803 | 1.69139 | 1.661 |
| chr10 | 111466315 | 111466620 | 0.49768 | 0.826876 | 1.661 |
| chr3 | 126398081 | 126398682 | 1.52676 | 2.53665 | 1.661 |
| chr14 | 31825263 | 31825483 | 1.5016 | 2.49487 | 1.661 |
| chr3 | 185486374 | 185486610 | 1.63899 | 2.72328 | 1.662 |
| chr22 | 30658836 | 30659242 | 1.99706 | 3.31845 | 1.662 |
| chr2 | 26968599 | 26968945 | 1.9694 | 3.27252 | 1.662 |
| chr20 | 62541019 | 62541264 | 1.46286 | 2.43088 | 1.662 |
| chr2 | 31749034 | 31749259 | 1.53911 | 2.55766 | 1.662 |
| chr19 | 39127486 | 39127919 | 3.3907 | 5.63481 | 1.662 |
| chr5 | 115473544 | 115473833 | 1.57627 | 2.61966 | 1.662 |
| chr16 | 70287721 | 70288358 | 1.0071 | 1.67376 | 1.662 |
| chr16 | 24528407 | 24528662 | 1.94471 | 3.23215 | 1.662 |
| chr10 | 125785355 | 125785632 | 0.886707 | 1.47382 | 1.662 |
| chr18 | 33519136 | 33519381 | 0.579762 | 0.96364 | 1.662 |
| chr7 | 123512335 | 123512614 | 1.54562 | 2.56916 | 1.662 |
| chr2 | 233533858 | 233534073 | 1.06903 | 1.77703 | 1.662 |
| chr9 | 138377762 | 138377970 | 0.65482 | 1.0885 | 1.662 |
| chr2 | 201983242 | 201983490 | 1.05666 | 1.75649 | 1.662 |
| chr3 | 149211977 | 149212351 | 1.8926 | 3.14614 | 1.662 |
| chr8 | 63555930 | 63556276 | 0.537024 | 0.892717 | 1.662 |
| chr2 | 96054887 | 96055343 | 3.28248 | 5.45669 | 1.662 |
| chr5 | 70897084 | 70897330 | 1.10847 | 1.84271 | 1.662 |
| chr12 | 9481534 | 9481749 | 0.642343 | 1.06785 | 1.662 |
| chr12 | 67809347 | 67809549 | 1.09977 | 1.82834 | 1.662 |
| chr1 | 237549832 | 237550240 | 1.07942 | 1.79457 | 1.663 |
| chr5 | 75484654 | 75484865 | 0.625154 | 1.0394 | 1.663 |
| chr19 | 41029183 | 41029594 | 3.54277 | 5.89049 | 1.663 |
| chr1 | 162824083 | 162824363 | 1.68714 | 2.80521 | 1.663 |
| chr11 | 120173667 | 120174130 | 1.42754 | 2.37361 | 1.663 |
| chr12 | 121916739 | 121917105 | 1.57965 | 2.62654 | 1.663 |
| chr7 | 38006369 | 38006683 | 0.553875 | 0.920948 | 1.663 |
| chr7 | 34816435 | 34816640 | 0.82075 | 1.36472 | 1.663 |
| chr10 | 15209868 | 15210177 | 0.987457 | 1.64197 | 1.663 |
| chr8 | 90425717 | 90425927 | 0.788232 | 1.31074 | 1.663 |
| chr12 | 4317096 | 4317300 | 0.846955 | 1.4084 | 1.663 |
| chr17 | 63937764 | 63938096 | 0.687067 | 1.14258 | 1.663 |
| chr10 | 52314756 | 52314962 | 1.17414 | 1.95261 | 1.663 |
| chr7 | 120627628 | 120628068 | 2.29114 | 3.8103 | 1.663 |
| chr5 | 141966965 | 141967360 | 1.68644 | 2.80468 | 1.663 |
| chr2 | 75692684 | 75693023 | 0.792808 | 1.31851 | 1.663 |
| chr1 | 11898684 | 11899096 | 2.40234 | 3.99549 | 1.663 |
| chr1 | 100929312 | 100929563 | 1.6419 | 2.73077 | 1.663 |
| chr5 | 171466976 | 171467592 | 2.08769 | 3.47226 | 1.663 |

| | | | | | |
|---|---|---|---|---|---|
| chr11 | 44131689 | 44132145 | 1.62664 | 2.70549 | 1.663 |
| chr17 | 37388224 | 37388685 | 2.69974 | 4.49045 | 1.663 |
| chr22 | 25154109 | 25154484 | 1.35962 | 2.26148 | 1.663 |
| chrX | 109224246 | 109224466 | 0.58126 | 0.966822 | 1.663 |
| chr9 | 128018358 | 128018600 | 1.11152 | 1.84887 | 1.663 |
| chr5 | 136656211 | 136656470 | 1.75482 | 2.91894 | 1.663 |
| chr1 | 3400265 | 3400826 | 1.33662 | 2.22343 | 1.663 |
| chr12 | 22890147 | 22890446 | 0.619272 | 1.03014 | 1.663 |
| chr17 | 35062577 | 35062984 | 1.95648 | 3.25468 | 1.664 |
| chr12 | 113899396 | 113899868 | 1.56558 | 2.60444 | 1.664 |
| chr2 | 151887229 | 151887644 | 0.868678 | 1.4451 | 1.664 |
| chr14 | 40201041 | 40201400 | 0.653132 | 1.08653 | 1.664 |
| chr3 | 975094 | 975331 | 0.819797 | 1.36381 | 1.664 |
| chr9 | 15104873 | 15105171 | 1.50219 | 2.4991 | 1.664 |
| chr7 | 125330712 | 125330937 | 0.623207 | 1.03681 | 1.664 |
| chr16 | 23568697 | 23569001 | 2.12076 | 3.52845 | 1.664 |
| chr8 | 85540940 | 85541145 | 0.458415 | 0.762714 | 1.664 |
| chr11 | 44449056 | 44449530 | 0.874005 | 1.4542 | 1.664 |
| chr5 | 174891710 | 174891985 | 1.63262 | 2.71642 | 1.664 |
| chr16 | 24241545 | 24241843 | 0.620813 | 1.03301 | 1.664 |
| chr10 | 118932924 | 118933153 | 0.712101 | 1.18496 | 1.664 |
| chr1 | 167093289 | 167093790 | 1.34364 | 2.236 | 1.664 |
| chr14 | 100196283 | 100196732 | 1.68804 | 2.80917 | 1.664 |
| chr11 | 75136559 | 75136888 | 1.19479 | 1.98833 | 1.664 |
| chr19 | 43159969 | 43160273 | 0.818742 | 1.36253 | 1.664 |
| chr22 | 22555551 | 22555969 | 1.76808 | 2.94246 | 1.664 |
| chr7 | 134291105 | 134291390 | 2.39552 | 3.98674 | 1.664 |
| chr17 | 72858317 | 72858942 | 1.95314 | 3.2506 | 1.664 |
| chr15 | 80602130 | 80602416 | 1.66869 | 2.77729 | 1.664 |
| chr12 | 94495099 | 94495524 | 2.21855 | 3.69262 | 1.664 |
| chr11 | 62803208 | 62803782 | 1.21656 | 2.02503 | 1.665 |
| chr10 | 12172369 | 12172607 | 1.32696 | 2.2089 | 1.665 |
| chr7 | 23711093 | 23711502 | 0.628938 | 1.047 | 1.665 |
| chr6 | 105932550 | 105932866 | 1.75132 | 2.9155 | 1.665 |
| chr1 | 155220236 | 155220887 | 3.80159 | 6.32902 | 1.665 |
| chr4 | 10462952 | 10463223 | 1.11272 | 1.85251 | 1.665 |
| chr8 | 143715852 | 143716893 | 2.29931 | 3.828 | 1.665 |
| chr5 | 153921324 | 153921587 | 1.61037 | 2.6811 | 1.665 |
| chr2 | 24246995 | 24247274 | 1.66754 | 2.77633 | 1.665 |
| chr3 | 156140045 | 156140296 | 0.516076 | 0.859274 | 1.665 |
| chr17 | 41411594 | 41411877 | 1.13838 | 1.89546 | 1.665 |
| chr19 | 45982228 | 45982561 | 2.30776 | 3.84255 | 1.665 |
| chr14 | 23341820 | 23342155 | 2.28698 | 3.80808 | 1.665 |
| chr9 | 36023294 | 36023876 | 2.01501 | 3.35539 | 1.665 |
| chr5 | 33997491 | 33997828 | 1.50174 | 2.50092 | 1.665 |
| chr16 | 74657152 | 74657477 | 1.27826 | 2.12876 | 1.665 |
| chr17 | 48374719 | 48375144 | 2.32917 | 3.87893 | 1.665 |
| chr2 | 144177630 | 144177978 | 0.667217 | 1.11117 | 1.665 |
| chr5 | 22387564 | 22387847 | 0.622312 | 1.03639 | 1.665 |
| chr20 | 12947208 | 12947440 | 1.2672 | 2.11041 | 1.665 |
| chr11 | 102785462 | 102785776 | 0.852492 | 1.41983 | 1.666 |
| chr3 | 98531770 | 98531988 | 0.975508 | 1.62478 | 1.666 |
| chrX | 69317284 | 69317562 | 1.52956 | 2.54759 | 1.666 |
| chrX | 117941291 | 117941694 | 1.42882 | 2.37981 | 1.666 |
| chr16 | 7627334 | 7627596 | 0.557261 | 0.92819 | 1.666 |
| chr2 | 7037406 | 7037911 | 0.860109 | 1.43265 | 1.666 |
| chr12 | 101265224 | 101265541 | 0.541268 | 0.901579 | 1.666 |
| chr6 | 169111146 | 169111579 | 0.693108 | 1.15455 | 1.666 |
| chr5 | 172571291 | 172571601 | 2.7932 | 4.65283 | 1.666 |
| chr18 | 48033560 | 48033879 | 1.59885 | 2.66333 | 1.666 |
| chr2 | 158348790 | 158349070 | 2.16324 | 3.6037 | 1.666 |

| | | | | | |
|---|---|---|---|---|---|
| chr4 | 155473309 | 155473603 | 1.25501 | 2.09074 | 1.666 |
| chr1 | 219834460 | 219834732 | 1.69764 | 2.82817 | 1.666 |
| chr4 | 84412050 | 84412346 | 1.8875 | 3.14456 | 1.666 |
| chr18 | 59568604 | 59569017 | 1.39257 | 2.32015 | 1.666 |
| chr15 | 43517471 | 43517993 | 1.11152 | 1.85196 | 1.666 |
| chr17 | 74619961 | 74620301 | 3.25041 | 5.4159 | 1.666 |
| chr4 | 153422043 | 153422278 | 1.22657 | 2.0438 | 1.666 |
| chr9 | 80072959 | 80073315 | 2.40512 | 4.00757 | 1.666 |
| chr11 | 18127557 | 18127897 | 2.75671 | 4.59349 | 1.666 |
| chr13 | 113097664 | 113098072 | 1.06545 | 1.77552 | 1.666 |
| chr7 | 6556061 | 6556280 | 1.27476 | 2.12435 | 1.666 |
| chr10 | 118435848 | 118436051 | 0.758422 | 1.26393 | 1.667 |
| chr13 | 22212326 | 22212694 | 1.11259 | 1.85421 | 1.667 |
| chr20 | 59539920 | 59540209 | 0.550269 | 0.917065 | 1.667 |
| chr2 | 159974958 | 159975444 | 1.51081 | 2.51793 | 1.667 |
| chr7 | 1704228 | 1704711 | 2.02039 | 3.36721 | 1.667 |
| chr10 | 111242962 | 111243224 | 0.517517 | 0.862684 | 1.667 |
| chr22 | 23096118 | 23096364 | 1.49486 | 2.49189 | 1.667 |
| chr2 | 218515501 | 218515811 | 2.24874 | 3.74863 | 1.667 |
| chr3 | 57204213 | 57204529 | 2.25216 | 3.75438 | 1.667 |
| chr9 | 32251504 | 32251845 | 1.26586 | 2.11022 | 1.667 |
| chr7 | 14360646 | 14360851 | 0.819714 | 1.36651 | 1.667 |
| chr5 | 82089731 | 82089968 | 1.38056 | 2.30151 | 1.667 |
| chr2 | 236579357 | 236579568 | 1.73062 | 2.88519 | 1.667 |
| chrX | 94851296 | 94851506 | 0.649337 | 1.08254 | 1.667 |
| chr15 | 50408744 | 50408959 | 0.989352 | 1.64944 | 1.667 |
| chr2 | 26207553 | 26207896 | 1.36418 | 2.27439 | 1.667 |
| chr9 | 21154013 | 21154237 | 0.923046 | 1.53893 | 1.667 |
| chr21 | 34569228 | 34569608 | 0.905311 | 1.50937 | 1.667 |
| chr2 | 69152442 | 69152804 | 2.33029 | 3.8852 | 1.667 |
| chr2 | 151928020 | 151928244 | 1.12005 | 1.86745 | 1.667 |
| chr19 | 39903251 | 39903696 | 3.10857 | 5.18291 | 1.667 |
| chr5 | 127548996 | 127549286 | 1.3322 | 2.22118 | 1.667 |
| chr17 | 72598113 | 72598491 | 2.38072 | 3.96965 | 1.667 |
| chr3 | 121975685 | 121976094 | 0.630723 | 1.05171 | 1.667 |
| chr18 | 77929096 | 77929456 | 2.19953 | 3.66767 | 1.667 |
| chr7 | 82110844 | 82111219 | 1.50103 | 2.50294 | 1.667 |
| chr11 | 64332623 | 64333154 | 0.61731 | 1.02936 | 1.667 |
| chr14 | 78624309 | 78624513 | 0.480113 | 0.800592 | 1.668 |
| chr14 | 88655687 | 88656000 | 1.73548 | 2.89394 | 1.668 |
| chr9 | 130160257 | 130160667 | 1.60777 | 2.6811 | 1.668 |
| chr15 | 38646619 | 38646849 | 1.07914 | 1.79961 | 1.668 |
| chr19 | 36228462 | 36228668 | 0.785667 | 1.31022 | 1.668 |
| chr15 | 101008546 | 101008840 | 0.762244 | 1.27127 | 1.668 |
| chr7 | 10842371 | 10842624 | 0.450653 | 0.751629 | 1.668 |
| chr8 | 111188059 | 111188275 | 0.412766 | 0.688466 | 1.668 |
| chr8 | 96770991 | 96771236 | 0.820973 | 1.36933 | 1.668 |
| chr17 | 42248161 | 42248379 | 0.779974 | 1.30102 | 1.668 |
| chr18 | 61178558 | 61178840 | 1.57432 | 2.62606 | 1.668 |
| chr1 | 245248872 | 245249264 | 0.549 | 0.915831 | 1.668 |
| chr1 | 12661345 | 12661537 | 1.10109 | 1.83684 | 1.668 |
| chr17 | 39643001 | 39643611 | 1.1216 | 1.87108 | 1.668 |
| chr2 | 68822255 | 68822705 | 1.54844 | 2.58321 | 1.668 |
| chr11 | 75839515 | 75839812 | 0.70788 | 1.18095 | 1.668 |
| chr14 | 41862579 | 41862819 | 0.592011 | 0.987649 | 1.668 |
| chr11 | 123447763 | 123448082 | 1.92463 | 3.21087 | 1.668 |
| chr7 | 36011050 | 36011281 | 0.954949 | 1.59319 | 1.668 |
| chr1 | 237613571 | 237613805 | 0.59485 | 0.992465 | 1.668 |
| chr7 | 101922667 | 101923302 | 1.68411 | 2.80984 | 1.668 |
| chr14 | 103572108 | 103572325 | 1.16427 | 1.94257 | 1.668 |
| chr15 | 101043816 | 101044054 | 0.860682 | 1.43605 | 1.669 |

| | | | | | |
|---|---|---|---|---|---|
| chr2 | 119612911 | 119613620 | 2.27026 | 3.78792 | 1.669 |
| chr8 | 54654708 | 54654970 | 1.37698 | 2.29753 | 1.669 |
| chr15 | 56788769 | 56789072 | 1.39281 | 2.32413 | 1.669 |
| chr1 | 95320008 | 95320239 | 1.51869 | 2.53424 | 1.669 |
| chr8 | 131814782 | 131815266 | 1.59909 | 2.66847 | 1.669 |
| chr22 | 19619751 | 19620028 | 1.37363 | 2.29235 | 1.669 |
| chr1 | 229290172 | 229290402 | 0.493875 | 0.824199 | 1.669 |
| chr6 | 155054320 | 155054633 | 1.93796 | 3.23427 | 1.669 |
| chr19 | 30228465 | 30228668 | 0.666465 | 1.11229 | 1.669 |
| chr7 | 66949305 | 66949568 | 2.08645 | 3.48217 | 1.669 |
| chr12 | 101808399 | 101808601 | 1.11016 | 1.8528 | 1.669 |
| chr6 | 143896580 | 143896787 | 1.51116 | 2.52209 | 1.669 |
| chr5 | 104075757 | 104075976 | 0.751049 | 1.25351 | 1.669 |
| chr8 | 106305375 | 106305812 | 0.628327 | 1.04869 | 1.669 |
| chr17 | 26255399 | 26255681 | 0.612914 | 1.02298 | 1.669 |
| chr3 | 37833060 | 37833277 | 0.506779 | 0.845872 | 1.669 |
| chr1 | 198668415 | 198668942 | 1.99865 | 3.33601 | 1.669 |
| chr13 | 39591032 | 39591329 | 1.08065 | 1.80375 | 1.669 |
| chr5 | 135216054 | 135216297 | 1.48809 | 2.48391 | 1.669 |
| chr12 | 130511125 | 130511535 | 1.98907 | 3.32031 | 1.669 |
| chr1 | 60442491 | 60442795 | 1.62173 | 2.70716 | 1.669 |
| chr17 | 26881495 | 26881802 | 1.20044 | 2.00389 | 1.669 |
| chr1 | 239089655 | 239089972 | 0.780109 | 1.30225 | 1.669 |
| chr7 | 30722302 | 30722542 | 0.645475 | 1.07753 | 1.669 |
| chr1 | 151974185 | 151974511 | 2.38625 | 3.98353 | 1.669 |
| chr5 | 83838854 | 83839079 | 0.87187 | 1.45553 | 1.669 |
| chr11 | 120105697 | 120106094 | 2.21283 | 3.69419 | 1.669 |
| chr15 | 51366611 | 51366839 | 0.598591 | 0.999321 | 1.669 |
| chr16 | 1387940 | 1388568 | 1.99325 | 3.32772 | 1.669 |
| chr2 | 220142368 | 220142854 | 2.52009 | 4.20728 | 1.670 |
| chr19 | 58045072 | 58045354 | 0.61972 | 1.03463 | 1.670 |
| chr1 | 145457160 | 145457519 | 1.56463 | 2.6122 | 1.670 |
| chr8 | 22445266 | 22445707 | 1.39917 | 2.33618 | 1.670 |
| chr8 | 17908834 | 17909082 | 0.695706 | 1.16163 | 1.670 |
| chr17 | 10654742 | 10654974 | 0.534049 | 0.891713 | 1.670 |
| chr20 | 36849519 | 36849860 | 2.42743 | 4.05322 | 1.670 |
| chr17 | 74543396 | 74543720 | 2.9274 | 4.88816 | 1.670 |
| chr4 | 5823175 | 5823795 | 1.2353 | 2.06271 | 1.670 |
| chr20 | 13439523 | 13439985 | 1.8934 | 3.16167 | 1.670 |
| chr2 | 135002981 | 135003293 | 2.77183 | 4.62855 | 1.670 |
| chr18 | 33864710 | 33865023 | 1.46784 | 2.45109 | 1.670 |
| chr7 | 100738184 | 100738472 | 2.11608 | 3.53361 | 1.670 |
| chr20 | 8655236 | 8655458 | 0.680494 | 1.13635 | 1.670 |
| chr17 | 73745833 | 73746377 | 2.6448 | 4.41656 | 1.670 |
| chr15 | 65128007 | 65128643 | 1.55622 | 2.59884 | 1.670 |
| chr5 | 111830292 | 111830576 | 0.552154 | 0.922092 | 1.670 |
| chr11 | 8112297 | 8112646 | 1.71835 | 2.86968 | 1.670 |
| chr2 | 30313746 | 30314065 | 2.04234 | 3.4109 | 1.670 |
| chr11 | 46390861 | 46391609 | 1.52304 | 2.54363 | 1.670 |
| chr1 | 79142161 | 79142371 | 1.37669 | 2.29926 | 1.670 |
| chr4 | 152197962 | 152198283 | 2.13949 | 3.57331 | 1.670 |
| chr1 | 36912034 | 36912236 | 1.24774 | 2.08396 | 1.670 |
| chr8 | 42285787 | 42286113 | 0.690106 | 1.15261 | 1.670 |
| chr1 | 17732609 | 17733000 | 1.70877 | 2.85411 | 1.670 |
| chr12 | 128162165 | 128162394 | 1.35864 | 2.26929 | 1.670 |
| chr8 | 3241396 | 3241607 | 0.722033 | 1.20602 | 1.670 |
| chr3 | 2074805 | 2075135 | 0.547774 | 0.914972 | 1.670 |
| chr5 | 106440423 | 106440662 | 1.50611 | 2.51594 | 1.670 |
| chr19 | 10616310 | 10616546 | 1.44707 | 2.41737 | 1.671 |
| chr11 | 128987565 | 128987850 | 1.76121 | 2.94227 | 1.671 |
| chr3 | 58517607 | 58517913 | 0.669361 | 1.11826 | 1.671 |

| | | | | | |
|---|---|---|---|---|---|
| chr12 | 55977454 | 55977728 | 0.557315 | 0.931145 | 1.671 |
| chr2 | 237969360 | 237969838 | 0.864097 | 1.44375 | 1.671 |
| chr19 | 2391314 | 2391552 | 1.36443 | 2.27975 | 1.671 |
| chr7 | 16471813 | 16472059 | 1.63416 | 2.7305 | 1.671 |
| chr17 | 40111713 | 40111987 | 1.78444 | 2.98164 | 1.671 |
| chr2 | 71175583 | 71176035 | 2.21146 | 3.69515 | 1.671 |
| chr11 | 64416886 | 64417273 | 1.05178 | 1.75751 | 1.671 |
| chr5 | 44594573 | 44594809 | 0.545539 | 0.911599 | 1.671 |
| chr7 | 142115670 | 142115882 | 1.88673 | 3.15287 | 1.671 |
| chr3 | 78790328 | 78790540 | 1.25253 | 2.09314 | 1.671 |
| chr17 | 46157657 | 46158016 | 1.28708 | 2.15089 | 1.671 |
| chr10 | 17649570 | 17649846 | 1.63891 | 2.73895 | 1.671 |
| chr2 | 7264748 | 7264959 | 0.543158 | 0.907777 | 1.671 |
| chr16 | 66962617 | 66962978 | 1.0464 | 1.74889 | 1.671 |
| chr16 | 48291626 | 48291886 | 1.30272 | 2.1773 | 1.671 |
| chr9 | 112214013 | 112214232 | 1.22667 | 2.05025 | 1.671 |
| chr2 | 162107420 | 162107679 | 1.59319 | 2.66299 | 1.671 |
| chr14 | 53282830 | 53283144 | 0.713356 | 1.19239 | 1.672 |
| chr2 | 29276719 | 29277055 | 1.08715 | 1.81726 | 1.672 |
| chr19 | 30774967 | 30775540 | 0.628256 | 1.05019 | 1.672 |
| chr3 | 129062698 | 129063129 | 1.28157 | 2.14233 | 1.672 |
| chr4 | 24981806 | 24982082 | 1.60137 | 2.67691 | 1.672 |
| chr12 | 71467443 | 71467691 | 0.567064 | 0.947946 | 1.672 |
| chr14 | 47559884 | 47560089 | 0.627811 | 1.04949 | 1.672 |
| chr16 | 82065087 | 82065297 | 1.25661 | 2.1008 | 1.672 |
| chr12 | 130143643 | 130143852 | 0.635844 | 1.06301 | 1.672 |
| chr11 | 84487382 | 84487592 | 0.675103 | 1.12866 | 1.672 |
| chr14 | 21966716 | 21967053 | 1.28987 | 2.15644 | 1.672 |
| chr5 | 98405070 | 98405388 | 0.576563 | 0.963917 | 1.672 |
| chr19 | 48352304 | 48352712 | 2.30401 | 3.85196 | 1.672 |
| chr1 | 19700830 | 19701295 | 1.30477 | 2.18146 | 1.672 |
| chr2 | 46163580 | 46163833 | 1.66585 | 2.7852 | 1.672 |
| chr15 | 66380622 | 66381029 | 0.568347 | 0.950263 | 1.672 |
| chr10 | 102268505 | 102268768 | 1.63664 | 2.7365 | 1.672 |
| chrX | 152934527 | 152935063 | 1.78124 | 2.97843 | 1.672 |
| chr5 | 90521968 | 90522181 | 1.48908 | 2.48997 | 1.672 |
| chr11 | 61061554 | 61061845 | 0.425485 | 0.71156 | 1.672 |
| chr6 | 113201135 | 113201391 | 0.803796 | 1.34426 | 1.672 |
| chr8 | 142888912 | 142889361 | 0.645404 | 1.07941 | 1.672 |
| chr16 | 55977977 | 55978648 | 1.91332 | 3.19998 | 1.672 |
| chr1 | 54955124 | 54955408 | 1.53152 | 2.56146 | 1.673 |
| chr16 | 68554456 | 68554873 | 2.42049 | 4.04827 | 1.673 |
| chr1 | 43388691 | 43388984 | 1.27756 | 2.13675 | 1.673 |
| chr10 | 56763598 | 56763837 | 0.765279 | 1.27995 | 1.673 |
| chr5 | 77683507 | 77683719 | 0.522857 | 0.874499 | 1.673 |
| chr12 | 27030896 | 27031204 | 1.4134 | 2.36398 | 1.673 |
| chr1 | 181796651 | 181797235 | 0.458727 | 0.76728 | 1.673 |
| chr3 | 37000237 | 37000662 | 1.7396 | 2.90976 | 1.673 |
| chr10 | 2479055 | 2479299 | 0.558282 | 0.933853 | 1.673 |
| chr5 | 94525277 | 94525492 | 0.586431 | 0.980938 | 1.673 |
| chr17 | 6189469 | 6189896 | 0.933094 | 1.56084 | 1.673 |
| chr3 | 122661645 | 122662171 | 2.92188 | 4.88773 | 1.673 |
| chr7 | 141674736 | 141674950 | 1.20567 | 2.01688 | 1.673 |
| chr9 | 76893805 | 76894076 | 0.582569 | 0.974549 | 1.673 |
| chr8 | 49890531 | 49890815 | 0.510415 | 0.853853 | 1.673 |
| chr9 | 125215506 | 125215828 | 1.55735 | 2.60524 | 1.673 |
| chr22 | 37417962 | 37418390 | 1.81314 | 3.03326 | 1.673 |
| chr4 | 123638041 | 123638361 | 2.03621 | 3.40678 | 1.673 |
| chr15 | 83957483 | 83957725 | 0.686077 | 1.14789 | 1.673 |
| chr1 | 120428870 | 120429171 | 1.51171 | 2.52931 | 1.673 |
| chr14 | 100784515 | 100784948 | 1.75105 | 2.92977 | 1.673 |

| | | | | | |
|---|---|---|---|---|---|
| chr16 | 88850674 | 88850882 | 1.19318 | 1.99643 | 1.673 |
| chr19 | 58688021 | 58688259 | 0.674012 | 1.12776 | 1.673 |
| chr20 | 51150078 | 51150324 | 0.770187 | 1.2887 | 1.673 |
| chr15 | 73598006 | 73598771 | 1.28509 | 2.15026 | 1.673 |
| chr22 | 37595515 | 37595823 | 1.90605 | 3.1895 | 1.673 |
| chr17 | 62530369 | 62530609 | 1.47774 | 2.4728 | 1.673 |
| chr21 | 34185813 | 34186023 | 0.39821 | 0.666376 | 1.673 |
| chr15 | 40322529 | 40322822 | 1.27253 | 2.12953 | 1.673 |
| chr2 | 176944419 | 176945249 | 2.60551 | 4.36048 | 1.674 |
| chr5 | 128697022 | 128697360 | 0.577242 | 0.966048 | 1.674 |
| chr7 | 76876590 | 76876818 | 1.42998 | 2.39319 | 1.674 |
| chr7 | 15916377 | 15916609 | 0.62552 | 1.04687 | 1.674 |
| chr7 | 94829147 | 94829406 | 1.50689 | 2.52214 | 1.674 |
| chr1 | 85248130 | 85248480 | 2.12785 | 3.56164 | 1.674 |
| chr4 | 24689034 | 24689292 | 2.30486 | 3.85817 | 1.674 |
| chr4 | 1237229 | 1237473 | 0.895764 | 1.49954 | 1.674 |
| chr3 | 19214618 | 19214922 | 0.627626 | 1.05067 | 1.674 |
| chr15 | 33684667 | 33684869 | 0.726857 | 1.2168 | 1.674 |
| chr8 | 137132480 | 137132731 | 0.622086 | 1.04141 | 1.674 |
| chrX | 70199306 | 70199533 | 1.30241 | 2.18033 | 1.674 |
| chr1 | 12679916 | 12680158 | 1.64009 | 2.74566 | 1.674 |
| chr16 | 74851850 | 74852060 | 0.650501 | 1.089 | 1.674 |
| chr5 | 86175280 | 86175518 | 0.489505 | 0.819509 | 1.674 |
| chr2 | 38692943 | 38693289 | 2.05328 | 3.43757 | 1.674 |
| chr7 | 45620202 | 45620476 | 1.06361 | 1.78072 | 1.674 |
| chr1 | 229617618 | 229618138 | 0.900428 | 1.50759 | 1.674 |
| chr5 | 147237182 | 147237417 | 1.42557 | 2.38694 | 1.674 |
| chr9 | 19732567 | 19732804 | 0.589526 | 0.987112 | 1.674 |
| chr20 | 24924118 | 24924322 | 0.71055 | 1.18977 | 1.674 |
| chr12 | 48109109 | 48109430 | 2.13383 | 3.57313 | 1.675 |
| chr21 | 43678690 | 43679058 | 1.81975 | 3.04723 | 1.675 |
| chr19 | 45907393 | 45907735 | 1.90182 | 3.1847 | 1.675 |
| chr22 | 39264896 | 39265220 | 2.13041 | 3.56768 | 1.675 |
| chr12 | 68461664 | 68461881 | 1.38158 | 2.31366 | 1.675 |
| chr8 | 22021917 | 22022571 | 2.36125 | 3.95432 | 1.675 |
| chr2 | 169203209 | 169203483 | 1.24369 | 2.08281 | 1.675 |
| chr5 | 28426477 | 28426719 | 0.564104 | 0.944719 | 1.675 |
| chr7 | 106300374 | 106301123 | 2.98499 | 4.99908 | 1.675 |
| chr11 | 39885142 | 39885333 | 0.632941 | 1.06003 | 1.675 |
| chr11 | 83527680 | 83527926 | 1.20572 | 2.01963 | 1.675 |
| chr6 | 166492519 | 166492741 | 0.614761 | 1.02975 | 1.675 |
| chr4 | 178260999 | 178261229 | 1.30859 | 2.19199 | 1.675 |
| chr19 | 28568489 | 28568904 | 0.707198 | 1.18463 | 1.675 |
| chr3 | 132760183 | 132760429 | 1.38429 | 2.31896 | 1.675 |
| chr14 | 36287646 | 36287889 | 1.79437 | 3.00602 | 1.675 |
| chr1 | 180239364 | 180239662 | 1.42514 | 2.38748 | 1.675 |
| chr17 | 7340110 | 7340784 | 2.3982 | 4.01762 | 1.675 |
| chr3 | 163559990 | 163560267 | 0.327498 | 0.548657 | 1.675 |
| chr1 | 24716530 | 24716912 | 2.2838 | 3.82607 | 1.675 |
| chr10 | 111985745 | 111986046 | 2.0363 | 3.41178 | 1.675 |
| chr12 | 72940561 | 72940793 | 0.758907 | 1.27153 | 1.675 |
| chr20 | 38147274 | 38147493 | 0.525802 | 0.881024 | 1.676 |
| chr6 | 24856315 | 24856556 | 1.31265 | 2.19946 | 1.676 |
| chr19 | 45454740 | 45455180 | 1.98603 | 3.32796 | 1.676 |
| chr1 | 213750007 | 213750217 | 0.504815 | 0.845919 | 1.676 |
| chr2 | 10457729 | 10458041 | 0.6875 | 1.15208 | 1.676 |
| chr10 | 118208089 | 118208313 | 0.409267 | 0.685839 | 1.676 |
| chr14 | 55883605 | 55883979 | 1.05387 | 1.76608 | 1.676 |
| chr17 | 78900775 | 78901127 | 0.683934 | 1.14615 | 1.676 |
| chr8 | 94061615 | 94061976 | 0.534388 | 0.89557 | 1.676 |
| chr19 | 51661553 | 51662425 | 2.12031 | 3.55344 | 1.676 |

| | | | | | |
|---|---|---|---|---|---|
| chr12 | 113750325 | 113750614 | 2.0863 | 3.49654 | 1.676 |
| chr2 | 152893433 | 152893706 | 1.71363 | 2.87203 | 1.676 |
| chr12 | 105157176 | 105157456 | 0.573565 | 0.9613 | 1.676 |
| chr1 | 166853355 | 166853907 | 0.685416 | 1.14881 | 1.676 |
| chr4 | 107955266 | 107955655 | 0.891688 | 1.49458 | 1.676 |
| chr10 | 90694590 | 90695681 | 1.28794 | 2.15879 | 1.676 |
| chr14 | 97238702 | 97238984 | 0.56309 | 0.943846 | 1.676 |
| chr7 | 130581224 | 130581443 | 1.68987 | 2.83258 | 1.676 |
| chr15 | 51791645 | 51791923 | 1.67702 | 2.81115 | 1.676 |
| chr5 | 169861953 | 169862395 | 1.85841 | 3.11521 | 1.676 |
| chr2 | 190525765 | 190526078 | 1.66904 | 2.79781 | 1.676 |
| chr10 | 58694169 | 58694715 | 0.578587 | 0.969896 | 1.676 |
| chr9 | 112006838 | 112007325 | 1.98027 | 3.31957 | 1.676 |
| chr11 | 2215990 | 2216721 | 2.23023 | 3.73878 | 1.676 |
| chr20 | 5836819 | 5837085 | 0.746532 | 1.25153 | 1.676 |
| chr4 | 88014721 | 88015065 | 0.841173 | 1.41025 | 1.677 |
| chr15 | 77316260 | 77317014 | 2.28786 | 3.83597 | 1.677 |
| chr20 | 35184881 | 35185327 | 2.23953 | 3.75498 | 1.677 |
| chr5 | 55110538 | 55110835 | 0.652933 | 1.09476 | 1.677 |
| chr3 | 12871975 | 12872505 | 0.69305 | 1.16203 | 1.677 |
| chr18 | 74421787 | 74422122 | 1.79624 | 3.01177 | 1.677 |
| chr8 | 125561875 | 125562170 | 1.05671 | 1.77203 | 1.677 |
| chr16 | 10274531 | 10274891 | 0.924384 | 1.55017 | 1.677 |
| chr12 | 126170864 | 126171088 | 0.579687 | 0.972156 | 1.677 |
| chr18 | 45971909 | 45972164 | 1.87036 | 3.13676 | 1.677 |
| chr11 | 44958011 | 44958367 | 2.20904 | 3.70482 | 1.677 |
| chr10 | 20166505 | 20166892 | 0.556981 | 0.934143 | 1.677 |
| chr3 | 97801260 | 97801548 | 0.718437 | 1.20499 | 1.677 |
| chr12 | 1714434 | 1714840 | 1.66395 | 2.79104 | 1.677 |
| chr14 | 42201164 | 42201390 | 0.506289 | 0.849227 | 1.677 |
| chr3 | 50263521 | 50263854 | 1.91736 | 3.21613 | 1.677 |
| chr14 | 89270995 | 89271214 | 0.547834 | 0.918956 | 1.677 |
| chr8 | 142327226 | 142327537 | 0.472049 | 0.791839 | 1.677 |
| chr16 | 28560668 | 28560960 | 1.50396 | 2.52288 | 1.677 |
| chr12 | 73185888 | 73186287 | 0.544078 | 0.9127 | 1.678 |
| chr18 | 72369586 | 72369845 | 1.47427 | 2.47321 | 1.678 |
| chr16 | 21250608 | 21251017 | 0.441476 | 0.74062 | 1.678 |
| chr7 | 33787323 | 33787571 | 0.438902 | 0.736325 | 1.678 |
| chr16 | 89772318 | 89772705 | 2.10435 | 3.53044 | 1.678 |
| chr11 | 94495439 | 94495691 | 1.55335 | 2.60605 | 1.678 |
| chr19 | 15740060 | 15740451 | 2.59017 | 4.34559 | 1.678 |
| chr14 | 35141842 | 35142051 | 1.68801 | 2.83209 | 1.678 |
| chr5 | 8851158 | 8851423 | 0.621946 | 1.04369 | 1.678 |
| chr11 | 119599909 | 119600378 | 1.73992 | 2.91982 | 1.678 |
| chr19 | 16173565 | 16173895 | 2.42006 | 4.06127 | 1.678 |
| chr10 | 52643478 | 52643757 | 0.583233 | 0.978787 | 1.678 |
| chr9 | 84302038 | 84302267 | 1.09735 | 1.84178 | 1.678 |
| chr8 | 10131390 | 10131715 | 1.84204 | 3.09181 | 1.678 |
| chr17 | 61514162 | 61514390 | 1.15578 | 1.94003 | 1.679 |
| chr6 | 37451942 | 37452336 | 1.96377 | 3.29639 | 1.679 |
| chr6 | 166490553 | 166490802 | 0.557452 | 0.935776 | 1.679 |
| chr20 | 49262990 | 49263244 | 1.0793 | 1.81204 | 1.679 |
| chr19 | 49575894 | 49576153 | 0.845216 | 1.41914 | 1.679 |
| chr11 | 75243445 | 75243769 | 1.88405 | 3.16352 | 1.679 |
| chr2 | 71687888 | 71688306 | 1.90016 | 3.19062 | 1.679 |
| chr8 | 12726559 | 12726817 | 0.497437 | 0.835309 | 1.679 |
| chr13 | 63970877 | 63971082 | 0.629643 | 1.05734 | 1.679 |
| chr9 | 119423414 | 119423674 | 1.63533 | 2.74641 | 1.679 |
| chr10 | 92312080 | 92312286 | 0.631765 | 1.06105 | 1.680 |
| chr3 | 35220095 | 35220431 | 0.667286 | 1.12071 | 1.680 |
| chr20 | 17598073 | 17598623 | 1.79152 | 3.00891 | 1.680 |

| | | | | | |
|---|---|---|---|---|---|
| chr15 | 97516466 | 97516713 | 0.792932 | 1.33176 | 1.680 |
| chr15 | 94663173 | 94663423 | 1.46526 | 2.46113 | 1.680 |
| chr2 | 59049334 | 59049564 | 0.579529 | 0.973403 | 1.680 |
| chr11 | 32108170 | 32108395 | 0.868475 | 1.45876 | 1.680 |
| chr19 | 30237474 | 30237880 | 0.713022 | 1.19765 | 1.680 |
| chr7 | 137687095 | 137687424 | 1.85885 | 3.12229 | 1.680 |
| chr8 | 104169755 | 104169975 | 0.753953 | 1.26642 | 1.680 |
| chr1 | 103137180 | 103137440 | 0.641871 | 1.07817 | 1.680 |
| chr17 | 40591217 | 40591582 | 1.81217 | 3.044 | 1.680 |
| chr9 | 140196744 | 140197327 | 1.4248 | 2.39333 | 1.680 |
| chr1 | 236695090 | 236695360 | 1.87828 | 3.15535 | 1.680 |
| chr8 | 59372228 | 59372497 | 1.193 | 2.00413 | 1.680 |
| chr16 | 15871715 | 15872045 | 2.42953 | 4.08142 | 1.680 |
| chr12 | 24256329 | 24256556 | 0.67756 | 1.13825 | 1.680 |
| chr6 | 84224646 | 84224876 | 1.18335 | 1.98795 | 1.680 |
| chr16 | 27379472 | 27379823 | 2.0251 | 3.40206 | 1.680 |
| chr11 | 118798779 | 118799107 | 1.62211 | 2.72508 | 1.680 |
| chr17 | 50953916 | 50954131 | 0.929804 | 1.56204 | 1.680 |
| chr9 | 135897735 | 135898128 | 3.01003 | 5.05695 | 1.680 |
| chr19 | 47652834 | 47653057 | 0.919622 | 1.54503 | 1.680 |
| chr7 | 12663500 | 12663888 | 1.95893 | 3.29114 | 1.680 |
| chr5 | 151522331 | 151522843 | 1.64399 | 2.76208 | 1.680 |
| chr5 | 89902420 | 89902724 | 1.18041 | 1.98323 | 1.680 |
| chr12 | 2556713 | 2556966 | 0.42815 | 0.719349 | 1.680 |
| chr10 | 34944598 | 34944809 | 1.29455 | 2.17507 | 1.680 |
| chr1 | 201088376 | 201088740 | 2.04286 | 3.43242 | 1.680 |
| chr12 | 52316071 | 52316319 | 0.660517 | 1.10981 | 1.680 |
| chr11 | 61103679 | 61104044 | 2.38187 | 4.00208 | 1.680 |
| chr17 | 18248175 | 18248500 | 0.748082 | 1.25712 | 1.680 |
| chr3 | 73270331 | 73270578 | 0.623371 | 1.04755 | 1.680 |
| chr5 | 72750271 | 72750597 | 1.3499 | 2.26858 | 1.681 |
| chr17 | 35754691 | 35754994 | 1.69382 | 2.84668 | 1.681 |
| chr22 | 47193327 | 47193622 | 1.12001 | 1.88246 | 1.681 |
| chr3 | 76103959 | 76104210 | 0.5353 | 0.899717 | 1.681 |
| chr7 | 128763552 | 128763775 | 0.723102 | 1.21542 | 1.681 |
| chr6 | 41246835 | 41247055 | 1.28145 | 2.15398 | 1.681 |
| chr1 | 157939529 | 157939918 | 1.80731 | 3.03792 | 1.681 |
| chr3 | 192162254 | 192162636 | 2.71174 | 4.55882 | 1.681 |
| chr1 | 156095838 | 156096075 | 1.52404 | 2.56221 | 1.681 |
| chr2 | 36922808 | 36923186 | 0.670685 | 1.1276 | 1.681 |
| chr19 | 42794849 | 42795489 | 2.46723 | 4.1483 | 1.681 |
| chr14 | 23334734 | 23335022 | 1.18676 | 1.99541 | 1.681 |
| chr19 | 50238255 | 50238478 | 0.498113 | 0.837544 | 1.681 |
| chr5 | 172758985 | 172759325 | 2.17794 | 3.66205 | 1.681 |
| chr9 | 79548919 | 79549200 | 0.59468 | 0.999932 | 1.681 |
| chr16 | 238744 | 239005 | 1.06492 | 1.79063 | 1.681 |
| chr13 | 32820591 | 32820850 | 1.80058 | 3.02764 | 1.681 |
| chr21 | 41087821 | 41088035 | 1.15758 | 1.94649 | 1.682 |
| chr3 | 34677021 | 34677244 | 0.609271 | 1.02453 | 1.682 |
| chr1 | 247739862 | 247740097 | 0.925738 | 1.55682 | 1.682 |
| chr20 | 43496232 | 43496533 | 2.18291 | 3.67138 | 1.682 |
| chr12 | 133772031 | 133772473 | 0.539308 | 0.907053 | 1.682 |
| chr10 | 81689163 | 81689581 | 1.99274 | 3.3516 | 1.682 |
| chrX | 24389594 | 24389920 | 1.43516 | 2.41382 | 1.682 |
| chr5 | 134292780 | 134293031 | 0.560935 | 0.943463 | 1.682 |
| chr9 | 85147552 | 85147786 | 0.818783 | 1.37724 | 1.682 |
| chr9 | 135847863 | 135848400 | 1.95395 | 3.28683 | 1.682 |
| chr3 | 72107795 | 72108030 | 0.625271 | 1.05182 | 1.682 |
| chrX | 109656330 | 109656709 | 1.91046 | 3.21409 | 1.682 |
| chr4 | 156965656 | 156965869 | 0.571723 | 0.961855 | 1.682 |
| chr19 | 32679245 | 32679856 | 0.781435 | 1.31471 | 1.682 |

| | | | | | |
|---|---|---|---|---|---|
| chr4 | 8240141 | 8240524 | 1.16014 | 1.95185 | 1.682 |
| chr3 | 150583126 | 150583356 | 1.14007 | 1.91817 | 1.683 |
| chr11 | 119494820 | 119495322 | 1.41415 | 2.37941 | 1.683 |
| chr5 | 64372220 | 64372469 | 0.948841 | 1.59656 | 1.683 |
| chr10 | 102973291 | 102973745 | 2.00444 | 3.37288 | 1.683 |
| chr17 | 13616435 | 13616957 | 0.647057 | 1.08883 | 1.683 |
| chr16 | 31469277 | 31469496 | 0.981995 | 1.6525 | 1.683 |
| chr6 | 16331529 | 16331937 | 2.34844 | 3.95203 | 1.683 |
| chr21 | 41239132 | 41239377 | 1.61606 | 2.7196 | 1.683 |
| chr3 | 12537326 | 12537525 | 1.08712 | 1.82948 | 1.683 |
| chr10 | 52035500 | 52035817 | 1.77023 | 2.97945 | 1.683 |
| chr12 | 67742748 | 67743007 | 0.620919 | 1.04507 | 1.683 |
| chr17 | 12846691 | 12846897 | 0.715019 | 1.20347 | 1.683 |
| chr19 | 49522965 | 49523173 | 0.846337 | 1.42455 | 1.683 |
| chr19 | 47516232 | 47516626 | 3.42685 | 5.7681 | 1.683 |
| chr13 | 32005865 | 32006178 | 2.17098 | 3.65428 | 1.683 |
| chr3 | 10992497 | 10992840 | 1.53087 | 2.57689 | 1.683 |
| chr3 | 38537545 | 38537975 | 2.36086 | 3.97417 | 1.683 |
| chr8 | 57228574 | 57228784 | 0.583466 | 0.982235 | 1.683 |
| chr10 | 72221159 | 72221506 | 2.3084 | 3.88612 | 1.683 |
| chr11 | 90021474 | 90021702 | 1.27326 | 2.14356 | 1.684 |
| chr2 | 239276225 | 239276568 | 1.01613 | 1.71077 | 1.684 |
| chr2 | 38873358 | 38873569 | 0.476564 | 0.802362 | 1.684 |
| chr10 | 120480600 | 120480881 | 1.70402 | 2.86912 | 1.684 |
| chr15 | 91593144 | 91593338 | 1.69196 | 2.84884 | 1.684 |
| chr7 | 158575569 | 158575825 | 0.939982 | 1.58275 | 1.684 |
| chr18 | 60625227 | 60625484 | 1.02803 | 1.73105 | 1.684 |
| chr11 | 13356800 | 13357066 | 1.77947 | 2.99637 | 1.684 |
| chr7 | 30588463 | 30588793 | 1.68653 | 2.83996 | 1.684 |
| chr12 | 114728130 | 114728336 | 0.599358 | 1.00931 | 1.684 |
| chr3 | 163556994 | 163557258 | 0.686876 | 1.15674 | 1.684 |
| chr12 | 73405369 | 73405752 | 0.525009 | 0.884151 | 1.684 |
| chr5 | 156749899 | 156750422 | 0.767333 | 1.29232 | 1.684 |
| chr2 | 38542911 | 38543215 | 1.9123 | 3.22074 | 1.684 |
| chr10 | 105358311 | 105358860 | 1.20828 | 2.03503 | 1.684 |
| chr2 | 149431632 | 149431943 | 0.97795 | 1.6471 | 1.684 |
| chr6 | 147178511 | 147178922 | 2.32741 | 3.92015 | 1.684 |
| chr16 | 46707355 | 46707688 | 2.88575 | 4.86078 | 1.684 |
| chr3 | 52885366 | 52885660 | 1.47664 | 2.4873 | 1.684 |
| chr9 | 36248059 | 36248307 | 0.815272 | 1.37332 | 1.684 |
| chr20 | 20843651 | 20844024 | 0.849186 | 1.43045 | 1.685 |
| chr17 | 40833259 | 40833520 | 1.91327 | 3.22297 | 1.685 |
| chr3 | 73105569 | 73105804 | 0.974011 | 1.64076 | 1.685 |
| chr11 | 44680101 | 44680298 | 1.05379 | 1.77524 | 1.685 |
| chr15 | 60017134 | 60017456 | 1.46861 | 2.47423 | 1.685 |
| chr15 | 62533253 | 62533654 | 0.799629 | 1.3472 | 1.685 |
| chr11 | 124628266 | 124628840 | 1.05946 | 1.78504 | 1.685 |
| chr17 | 1968156 | 1968675 | 1.19583 | 2.01483 | 1.685 |
| chr7 | 150942391 | 150942986 | 2.0075 | 3.38239 | 1.685 |
| chr7 | 34799208 | 34799493 | 1.42503 | 2.40109 | 1.685 |
| chr7 | 44427124 | 44427350 | 0.719503 | 1.21237 | 1.685 |
| chr10 | 32236592 | 32236835 | 1.59472 | 2.68718 | 1.685 |
| chr7 | 128027680 | 128028080 | 2.38861 | 4.02494 | 1.685 |
| chr19 | 2956491 | 2956806 | 1.32356 | 2.23037 | 1.685 |
| chr2 | 7533167 | 7533418 | 0.430455 | 0.725386 | 1.685 |
| chr14 | 105669121 | 105669772 | 1.86524 | 3.14325 | 1.685 |
| chr19 | 39360399 | 39360900 | 2.14974 | 3.62283 | 1.685 |
| chr17 | 27442884 | 27443158 | 1.172 | 1.97514 | 1.685 |
| chr17 | 47317143 | 47317576 | 0.641737 | 1.0815 | 1.685 |
| chr2 | 7259110 | 7259658 | 1.0317 | 1.73869 | 1.685 |
| chr20 | 51337462 | 51337773 | 0.574824 | 0.968733 | 1.685 |

| | | | | | |
|---|---|---|---|---|---|
| chr10 | 92766018 | 92766387 | 1.887 | 3.18019 | 1.685 |
| chr18 | 60305754 | 60306018 | 1.09299 | 1.84204 | 1.685 |
| chr3 | 46746503 | 46746888 | 1.72781 | 2.91209 | 1.685 |
| chr11 | 62327994 | 62328314 | 1.58213 | 2.66666 | 1.685 |
| chr3 | 32185665 | 32185903 | 1.65828 | 2.79504 | 1.686 |
| chrX | 152820338 | 152820591 | 1.82562 | 3.07714 | 1.686 |
| chr15 | 52093597 | 52093944 | 2.67287 | 4.5053 | 1.686 |
| chr17 | 53730376 | 53730628 | 0.614132 | 1.0352 | 1.686 |
| chr7 | 141607316 | 141607538 | 0.63428 | 1.06926 | 1.686 |
| chr5 | 36578947 | 36579159 | 0.545483 | 0.919596 | 1.686 |
| chr12 | 109693172 | 109693492 | 0.801892 | 1.3519 | 1.686 |
| chr16 | 57126633 | 57127166 | 2.35828 | 3.9758 | 1.686 |
| chr7 | 115778256 | 115778606 | 1.65825 | 2.79583 | 1.686 |
| chr4 | 103967787 | 103968189 | 1.23592 | 2.08385 | 1.686 |
| chr2 | 43397170 | 43397410 | 2.07239 | 3.4944 | 1.686 |
| chr10 | 9868723 | 9868956 | 0.7193 | 1.21287 | 1.686 |
| chr15 | 60619026 | 60619386 | 2.67351 | 4.50805 | 1.686 |
| chr15 | 80061556 | 80061862 | 1.30284 | 2.19687 | 1.686 |
| chr5 | 172878123 | 172878347 | 1.49609 | 2.52273 | 1.686 |
| chr14 | 103571258 | 103571584 | 2.8103 | 4.73893 | 1.686 |
| chr12 | 2667690 | 2667984 | 0.468904 | 0.790726 | 1.686 |
| chr7 | 85831199 | 85831407 | 1.04381 | 1.76021 | 1.686 |
| chr20 | 17411829 | 17412833 | 1.41816 | 2.39164 | 1.686 |
| chr20 | 33150484 | 33150767 | 1.22335 | 2.06322 | 1.687 |
| chr9 | 38417054 | 38417491 | 1.423 | 2.39998 | 1.687 |
| chr20 | 5984053 | 5984287 | 1.07875 | 1.81939 | 1.687 |
| chr7 | 100702572 | 100703059 | 1.88984 | 3.18734 | 1.687 |
| chr6 | 89202576 | 89202784 | 1.0362 | 1.74763 | 1.687 |
| chr17 | 66773596 | 66773830 | 1.42885 | 2.40991 | 1.687 |
| chr3 | 107472378 | 107472669 | 1.49045 | 2.51382 | 1.687 |
| chr5 | 149213799 | 149214234 | 1.57086 | 2.64951 | 1.687 |
| chr3 | 129102035 | 129102454 | 1.30954 | 2.20878 | 1.687 |
| chr3 | 97377606 | 97377819 | 1.42346 | 2.40093 | 1.687 |
| chr18 | 24384490 | 24384711 | 1.22217 | 2.06145 | 1.687 |
| chr16 | 88810074 | 88810630 | 2.35224 | 3.96761 | 1.687 |
| chr7 | 145545918 | 145546122 | 0.747157 | 1.26026 | 1.687 |
| chr1 | 70953611 | 70953937 | 1.38145 | 2.3302 | 1.687 |
| chr4 | 106062109 | 106062371 | 1.92729 | 3.25091 | 1.687 |
| chr12 | 50232790 | 50233337 | 1.43122 | 2.4143 | 1.687 |
| chr14 | 100713255 | 100713479 | 1.41613 | 2.38899 | 1.687 |
| chr7 | 140489762 | 140490017 | 0.568829 | 0.959629 | 1.687 |
| chr3 | 13036425 | 13036817 | 2.32428 | 3.92119 | 1.687 |
| chr8 | 53024991 | 53025281 | 2.40227 | 4.05278 | 1.687 |
| chr6 | 1263347 | 1263598 | 0.625294 | 1.05492 | 1.687 |
| chr12 | 45235332 | 45235555 | 0.541806 | 0.914102 | 1.687 |
| chr5 | 32709889 | 32710101 | 1.45652 | 2.45741 | 1.687 |
| chr17 | 59857954 | 59858156 | 0.674 | 1.13718 | 1.687 |
| chr18 | 47385150 | 47385410 | 2.05919 | 3.47433 | 1.687 |
| chr5 | 131563761 | 131563975 | 0.743949 | 1.25525 | 1.687 |
| chr13 | 113098324 | 113098623 | 2.58257 | 4.35763 | 1.687 |
| chr15 | 100771638 | 100771841 | 0.591992 | 0.998887 | 1.687 |
| chr4 | 176031237 | 176031446 | 0.941543 | 1.58873 | 1.687 |
| chr6 | 28351431 | 28351657 | 1.20593 | 2.03501 | 1.688 |
| chr12 | 97456118 | 97456478 | 0.743534 | 1.25473 | 1.688 |
| chr14 | 46524779 | 46525016 | 0.583089 | 0.98398 | 1.688 |
| chr3 | 49332957 | 49333270 | 1.34878 | 2.27613 | 1.688 |
| chr2 | 42612387 | 42612605 | 1.21983 | 2.05866 | 1.688 |
| chr20 | 4766510 | 4766795 | 0.564336 | 0.952428 | 1.688 |
| chr1 | 183592414 | 183592638 | 1.69951 | 2.86829 | 1.688 |
| chr3 | 154493945 | 154494151 | 0.661147 | 1.1159 | 1.688 |
| chr20 | 36685929 | 36686294 | 1.92348 | 3.24661 | 1.688 |

| | | | | | |
|---|---|---|---|---|---|
| chr3 | 143877019 | 143877232 | 0.472265 | 0.797159 | 1.688 |
| chr1 | 156111019 | 156111422 | 1.53306 | 2.58776 | 1.688 |
| chr1 | 175696638 | 175696857 | 0.672226 | 1.1347 | 1.688 |
| chr1 | 18311087 | 18311358 | 1.29057 | 2.17847 | 1.688 |
| chr3 | 24040418 | 24040790 | 2.22422 | 3.75446 | 1.688 |
| chr5 | 136995995 | 136996418 | 1.97998 | 3.3423 | 1.688 |
| chr1 | 201528835 | 201529167 | 2.10859 | 3.55951 | 1.688 |
| chr8 | 133663686 | 133664048 | 1.5046 | 2.53996 | 1.688 |
| chr4 | 61066813 | 61067083 | 1.15127 | 1.94353 | 1.688 |
| chr17 | 77675880 | 77676103 | 1.0487 | 1.7704 | 1.688 |
| chr19 | 35837966 | 35838177 | 0.765795 | 1.29281 | 1.688 |
| chrX | 68353815 | 68354120 | 2.05412 | 3.46775 | 1.688 |
| chr4 | 7803908 | 7804177 | 1.94888 | 3.29025 | 1.688 |
| chr3 | 396137 | 396369 | 1.47837 | 2.49591 | 1.688 |
| chr17 | 80739641 | 80740087 | 4.10354 | 6.92821 | 1.688 |
| chr15 | 95095232 | 95095635 | 0.589103 | 0.994625 | 1.688 |
| chr8 | 54199288 | 54199681 | 1.61951 | 2.73454 | 1.689 |
| chr3 | 2447449 | 2447728 | 0.655189 | 1.10635 | 1.689 |
| chr8 | 13775046 | 13775416 | 0.779873 | 1.31689 | 1.689 |
| chr1 | 1279956 | 1280353 | 1.36376 | 2.30288 | 1.689 |
| chr1 | 94491520 | 94491773 | 1.10584 | 1.86737 | 1.689 |
| chr15 | 24102113 | 24102518 | 1.38419 | 2.33745 | 1.689 |
| chr10 | 86503461 | 86503790 | 0.551488 | 0.931317 | 1.689 |
| chr2 | 121766676 | 121766890 | 0.993571 | 1.67787 | 1.689 |
| chr20 | 21486730 | 21487262 | 2.27612 | 3.84384 | 1.689 |
| chr7 | 19309671 | 19309933 | 0.660958 | 1.11627 | 1.689 |
| chr10 | 96039881 | 96040269 | 2.88438 | 4.87136 | 1.689 |
| chr4 | 100729466 | 100729764 | 1.22782 | 2.07368 | 1.689 |
| chr17 | 75237937 | 75238159 | 0.54752 | 0.924738 | 1.689 |
| chr20 | 37678699 | 37679098 | 3.19662 | 5.39912 | 1.689 |
| chr1 | 10574352 | 10574769 | 1.01693 | 1.71764 | 1.689 |
| chr16 | 57509711 | 57510086 | 2.22004 | 3.74979 | 1.689 |
| chr7 | 111495382 | 111495620 | 1.74194 | 2.94235 | 1.689 |
| chr15 | 56111212 | 56111414 | 0.643967 | 1.08776 | 1.689 |
| chr10 | 111592770 | 111593019 | 0.64187 | 1.08423 | 1.689 |
| chr10 | 66354806 | 66355303 | 0.570056 | 0.962936 | 1.689 |
| chr8 | 58284400 | 58284620 | 0.528867 | 0.893371 | 1.689 |
| chr3 | 195621495 | 195622257 | 1.93462 | 3.26803 | 1.689 |
| chr20 | 54937961 | 54938214 | 0.579097 | 0.978309 | 1.689 |
| chr5 | 112183978 | 112184242 | 1.48803 | 2.51397 | 1.689 |
| chr5 | 145782671 | 145782900 | 0.562525 | 0.950366 | 1.689 |
| chr4 | 38186071 | 38186320 | 1.03095 | 1.74182 | 1.690 |
| chr14 | 38510026 | 38510295 | 0.558971 | 0.944402 | 1.690 |
| chr11 | 73071546 | 73071903 | 1.75806 | 2.97037 | 1.690 |
| chr19 | 31710112 | 31710376 | 0.547294 | 0.924731 | 1.690 |
| chr7 | 118815973 | 118816414 | 1.59476 | 2.69462 | 1.690 |
| chr17 | 67603283 | 67603643 | 2.52681 | 4.26962 | 1.690 |
| chr2 | 55395770 | 55395999 | 2.05731 | 3.47638 | 1.690 |
| chr11 | 78131347 | 78131853 | 1.58254 | 2.67415 | 1.690 |
| chr2 | 26785212 | 26785493 | 1.07735 | 1.8205 | 1.690 |
| chr15 | 80527071 | 80527879 | 1.00629 | 1.70044 | 1.690 |
| chr9 | 130567955 | 130568337 | 1.39295 | 2.35386 | 1.690 |
| chr6 | 121376759 | 121377023 | 1.21201 | 2.04819 | 1.690 |
| chr17 | 46956326 | 46956768 | 2.11294 | 3.57069 | 1.690 |
| chr17 | 76989119 | 76989693 | 1.93125 | 3.26366 | 1.690 |
| chr1 | 207368997 | 207369214 | 0.697768 | 1.17919 | 1.690 |
| chr9 | 121191175 | 121191432 | 0.710515 | 1.20079 | 1.690 |
| chr2 | 241505723 | 241506128 | 1.76565 | 2.98418 | 1.690 |
| chr12 | 103276944 | 103277366 | 2.07767 | 3.51173 | 1.690 |
| chr3 | 125965053 | 125965500 | 1.64833 | 2.78607 | 1.690 |
| chr17 | 79071199 | 79071665 | 1.77292 | 2.99675 | 1.690 |

| | | | | | |
|---|---|---|---|---|---|
| chr18 | 59850806 | 59851138 | 2.15886 | 3.64913 | 1.690 |
| chr1 | 95699567 | 95699846 | 2.12587 | 3.59348 | 1.690 |
| chr10 | 82423825 | 82424178 | 2.52084 | 4.26117 | 1.690 |
| chr19 | 4199726 | 4200198 | 0.794893 | 1.34367 | 1.690 |
| chr16 | 77633739 | 77634059 | 2.11809 | 3.58042 | 1.690 |
| chr17 | 52530209 | 52530498 | 0.50241 | 0.849281 | 1.690 |
| chr17 | 39844225 | 39844525 | 1.05818 | 1.78885 | 1.691 |
| chr12 | 13892560 | 13892838 | 1.69778 | 2.87012 | 1.691 |
| chr7 | 10767339 | 10767635 | 0.500421 | 0.846008 | 1.691 |
| chr10 | 17013953 | 17014171 | 0.763379 | 1.29057 | 1.691 |
| chr1 | 233088634 | 233088974 | 1.5492 | 2.61917 | 1.691 |
| chr10 | 36420452 | 36420675 | 0.560502 | 0.947622 | 1.691 |
| chr12 | 27886058 | 27886321 | 1.8946 | 3.20315 | 1.691 |
| chr10 | 64578311 | 64578716 | 0.632414 | 1.06921 | 1.691 |
| chr20 | 33680421 | 33680824 | 2.72985 | 4.61548 | 1.691 |
| chr7 | 15001037 | 15001271 | 0.930895 | 1.57391 | 1.691 |
| chr15 | 88326719 | 88326956 | 0.473538 | 0.800662 | 1.691 |
| chr20 | 44935706 | 44936211 | 2.02651 | 3.42644 | 1.691 |
| chr11 | 122451221 | 122451523 | 2.08583 | 3.52681 | 1.691 |
| chr15 | 42426079 | 42426503 | 1.68455 | 2.84842 | 1.691 |
| chr19 | 54769937 | 54770388 | 1.58831 | 2.68569 | 1.691 |
| chr6 | 46057915 | 46058173 | 1.57362 | 2.66096 | 1.691 |
| chr2 | 58798435 | 58798681 | 1.92765 | 3.25963 | 1.691 |
| chr17 | 45853962 | 45854522 | 1.59077 | 2.69006 | 1.691 |
| chr2 | 35468935 | 35469334 | 0.738495 | 1.24884 | 1.691 |
| chr5 | 11688794 | 11689027 | 0.590102 | 0.997906 | 1.691 |
| chr19 | 28971405 | 28971715 | 0.716344 | 1.21143 | 1.691 |
| chrX | 20009226 | 20009526 | 2.06167 | 3.48657 | 1.691 |
| chr8 | 140411961 | 140412223 | 0.66261 | 1.12057 | 1.691 |
| chr8 | 11424652 | 11424943 | 1.9841 | 3.35563 | 1.691 |
| chr9 | 71642233 | 71642855 | 1.06935 | 1.80856 | 1.691 |
| chr3 | 99488273 | 99488499 | 0.514228 | 0.869709 | 1.691 |
| chr20 | 32657722 | 32658009 | 1.57593 | 2.66538 | 1.691 |
| chr20 | 23321233 | 23321456 | 1.75114 | 2.9621 | 1.692 |
| chr14 | 105994642 | 105995104 | 2.45998 | 4.16141 | 1.692 |
| chr19 | 11939707 | 11939919 | 0.752131 | 1.27234 | 1.692 |
| chr16 | 30406811 | 30407246 | 1.3482 | 2.2807 | 1.692 |
| chr20 | 7907298 | 7907813 | 0.945035 | 1.59885 | 1.692 |
| chr8 | 55567184 | 55567386 | 0.698515 | 1.18182 | 1.692 |
| chr1 | 41882754 | 41883134 | 1.99147 | 3.36938 | 1.692 |
| chr15 | 97632377 | 97632599 | 0.666991 | 1.12849 | 1.692 |
| chr1 | 248028071 | 248028278 | 0.672804 | 1.13838 | 1.692 |
| chr11 | 92967824 | 92968110 | 2.56971 | 4.34798 | 1.692 |
| chr17 | 77041914 | 77042702 | 2.02597 | 3.42819 | 1.692 |
| chr7 | 41004549 | 41004800 | 0.700467 | 1.18533 | 1.692 |
| chr6 | 85725159 | 85725438 | 1.66237 | 2.8131 | 1.692 |
| chr10 | 32499362 | 32499958 | 1.22952 | 2.08068 | 1.692 |
| chr9 | 120545539 | 120545775 | 1.39912 | 2.36776 | 1.692 |
| chr3 | 48722925 | 48723653 | 1.87514 | 3.1734 | 1.692 |
| chrX | 139139138 | 139139477 | 1.85532 | 3.13986 | 1.692 |
| chr7 | 109096174 | 109096549 | 1.19359 | 2.02 | 1.692 |
| chr17 | 3551141 | 3551444 | 1.84078 | 3.1153 | 1.692 |
| chr22 | 45497708 | 45498031 | 1.12662 | 1.90674 | 1.692 |
| chr17 | 61575173 | 61575503 | 1.08214 | 1.83148 | 1.692 |
| chr19 | 45942708 | 45943282 | 3.23086 | 5.46814 | 1.692 |
| chr2 | 98357019 | 98357353 | 1.83678 | 3.10869 | 1.692 |
| chr8 | 74207113 | 74207437 | 1.04899 | 1.77541 | 1.692 |
| chr4 | 81287162 | 81287414 | 0.864337 | 1.46289 | 1.693 |
| chr10 | 60144843 | 60145106 | 1.81352 | 3.06942 | 1.693 |
| chr4 | 6136294 | 6136675 | 1.04768 | 1.77324 | 1.693 |
| chr1 | 201678584 | 201679008 | 2.4595 | 4.16284 | 1.693 |

| | | | | | |
|---|---|---|---|---|---|
| chr2 | 48333758 | 48333993 | 1.43733 | 2.43289 | 1.693 |
| chr17 | 47512411 | 47512763 | 2.12855 | 3.60294 | 1.693 |
| chr11 | 95787953 | 95788276 | 0.82332 | 1.39363 | 1.693 |
| chrX | 153627150 | 153627415 | 2.29632 | 3.88697 | 1.693 |
| chr2 | 48412734 | 48413108 | 0.520935 | 0.881791 | 1.693 |
| chr9 | 96725225 | 96725591 | 1.73317 | 2.93393 | 1.693 |
| chr9 | 106531353 | 106531657 | 0.56237 | 0.951995 | 1.693 |
| chr9 | 35051459 | 35051662 | 1.20427 | 2.03865 | 1.693 |
| chr9 | 93660557 | 93660811 | 0.893287 | 1.51241 | 1.693 |
| chr18 | 8654579 | 8654809 | 1.28122 | 2.16926 | 1.693 |
| chr5 | 127213921 | 127214165 | 1.20756 | 2.04454 | 1.693 |
| chr15 | 91361734 | 91362204 | 0.849644 | 1.43859 | 1.693 |
| chr17 | 32318379 | 32318603 | 0.767295 | 1.29917 | 1.693 |
| chr12 | 6667242 | 6667511 | 1.38769 | 2.34969 | 1.693 |
| chr17 | 78873190 | 78873454 | 2.00489 | 3.39498 | 1.693 |
| chr2 | 121433106 | 121433442 | 1.74302 | 2.95159 | 1.693 |
| chr20 | 47398088 | 47398388 | 0.612858 | 1.03781 | 1.693 |
| chr7 | 19752545 | 19752773 | 0.636821 | 1.0784 | 1.693 |
| chr1 | 207251169 | 207251482 | 3.19655 | 5.41333 | 1.693 |
| chr3 | 122880204 | 122880632 | 0.721361 | 1.22171 | 1.694 |
| chr12 | 111040086 | 111040299 | 1.47721 | 2.50198 | 1.694 |
| chr12 | 56478968 | 56479252 | 1.6913 | 2.86469 | 1.694 |
| chr8 | 82740977 | 82741219 | 0.481887 | 0.816211 | 1.694 |
| chr2 | 85906289 | 85906736 | 2.51235 | 4.25551 | 1.694 |
| chr7 | 102702444 | 102702667 | 1.17711 | 1.99384 | 1.694 |
| chr2 | 25517312 | 25517660 | 0.456105 | 0.772597 | 1.694 |
| chr17 | 45878118 | 45878324 | 0.672647 | 1.13943 | 1.694 |
| chr8 | 87631074 | 87631556 | 1.54659 | 2.61985 | 1.694 |
| chr12 | 131303338 | 131303945 | 2.09766 | 3.55345 | 1.694 |
| chr2 | 86991935 | 86992156 | 1.23301 | 2.08878 | 1.694 |
| chr16 | 66309773 | 66310070 | 1.89771 | 3.21485 | 1.694 |
| chr12 | 102545874 | 102546088 | 0.560981 | 0.95036 | 1.694 |
| chr9 | 125216789 | 125217020 | 1.19283 | 2.0209 | 1.694 |
| chr21 | 40760518 | 40760853 | 1.93791 | 3.28339 | 1.694 |
| chr5 | 132583508 | 132584016 | 2.02853 | 3.43724 | 1.694 |
| chr5 | 168729590 | 168729982 | 1.70647 | 2.89163 | 1.695 |
| chr3 | 128513939 | 128514149 | 1.58726 | 2.68966 | 1.695 |
| chr8 | 73722331 | 73722567 | 0.723083 | 1.22531 | 1.695 |
| chr3 | 120161901 | 120162202 | 1.15449 | 1.95641 | 1.695 |
| chr8 | 140493123 | 140493357 | 0.636955 | 1.0794 | 1.695 |
| chr8 | 126444170 | 126444568 | 2.48905 | 4.21813 | 1.695 |
| chr13 | 51860558 | 51860948 | 1.83521 | 3.11011 | 1.695 |
| chr20 | 32657181 | 32657411 | 1.27483 | 2.16046 | 1.695 |
| chr19 | 42486280 | 42486488 | 0.708615 | 1.2009 | 1.695 |
| chr16 | 67515077 | 67515408 | 2.06509 | 3.49978 | 1.695 |
| chr16 | 20326263 | 20326479 | 0.656373 | 1.11246 | 1.695 |
| chr10 | 67161383 | 67161926 | 0.494728 | 0.838507 | 1.695 |
| chr9 | 5425108 | 5425331 | 1.31949 | 2.2364 | 1.695 |
| chr9 | 111433576 | 111433781 | 1.41746 | 2.40248 | 1.695 |
| chr20 | 16477827 | 16478032 | 0.717257 | 1.2157 | 1.695 |
| chr1 | 11301830 | 11302108 | 1.21189 | 2.05417 | 1.695 |
| chr1 | 1235477 | 1235808 | 0.804514 | 1.36375 | 1.695 |
| chrX | 135659791 | 135660057 | 1.7899 | 3.03411 | 1.695 |
| chr13 | 46895822 | 46896117 | 1.40222 | 2.377 | 1.695 |
| chr10 | 95048841 | 95049096 | 1.38851 | 2.35381 | 1.695 |
| chr5 | 38193252 | 38193521 | 2.45317 | 4.15882 | 1.695 |
| chr22 | 19617420 | 19617763 | 1.80798 | 3.06509 | 1.695 |
| chr9 | 5886051 | 5886496 | 1.82308 | 3.09078 | 1.695 |
| chr15 | 93631029 | 93631597 | 2.3824 | 4.03909 | 1.695 |
| chr4 | 4432341 | 4432649 | 1.75852 | 2.98146 | 1.695 |
| chr8 | 23149273 | 23149475 | 1.52589 | 2.58706 | 1.695 |

| | | | | | |
|---|---|---|---|---|---|
| chr8 | 91764045 | 91764262 | 0.470637 | 0.797939 | 1.695 |
| chr1 | 182755941 | 182756127 | 1.88974 | 3.20406 | 1.696 |
| chr3 | 185960473 | 185960655 | 1.13882 | 1.93093 | 1.696 |
| chr8 | 130793755 | 130794041 | 0.589891 | 1.00022 | 1.696 |
| chr11 | 69059826 | 69060179 | 2.24562 | 3.8077 | 1.696 |
| chr9 | 21749436 | 21749722 | 0.565265 | 0.958521 | 1.696 |
| chr2 | 27069208 | 27069498 | 2.63826 | 4.47376 | 1.696 |
| chr10 | 113170385 | 113170750 | 0.48996 | 0.830896 | 1.696 |
| chr2 | 10975417 | 10975807 | 2.58097 | 4.3769 | 1.696 |
| chrX | 53100746 | 53101018 | 1.52938 | 2.59372 | 1.696 |
| chr8 | 143808252 | 143808688 | 2.44869 | 4.15332 | 1.696 |
| chrX | 30753938 | 30754260 | 1.88455 | 3.19648 | 1.696 |
| chr10 | 81198307 | 81198747 | 0.647474 | 1.09822 | 1.696 |
| chr17 | 48075565 | 48075925 | 2.12245 | 3.60003 | 1.696 |
| chr5 | 153900527 | 153900879 | 2.32461 | 3.94311 | 1.696 |
| chr19 | 1781902 | 1782354 | 0.950912 | 1.61301 | 1.696 |
| chr2 | 11213434 | 11213702 | 1.51437 | 2.56882 | 1.696 |
| chr16 | 10275325 | 10275867 | 1.95294 | 3.31326 | 1.697 |
| chr9 | 12215104 | 12215442 | 0.598984 | 1.01627 | 1.697 |
| chr2 | 232382301 | 232382648 | 1.2594 | 2.13684 | 1.697 |
| chr3 | 126471092 | 126471456 | 1.56364 | 2.65321 | 1.697 |
| chr20 | 48440453 | 48440708 | 0.67614 | 1.14734 | 1.697 |
| chr9 | 12626708 | 12626946 | 0.368366 | 0.625111 | 1.697 |
| chr3 | 6770204 | 6770428 | 0.871608 | 1.47913 | 1.697 |
| chr1 | 46813801 | 46814083 | 2.23937 | 3.80029 | 1.697 |
| chr10 | 2419766 | 2419948 | 0.668459 | 1.13441 | 1.697 |
| chr2 | 53650113 | 53650432 | 1.54884 | 2.62851 | 1.697 |
| chr10 | 129652216 | 129652589 | 2.20222 | 3.73746 | 1.697 |
| chr11 | 15444966 | 15445192 | 0.559982 | 0.950372 | 1.697 |
| chr12 | 82014992 | 82015397 | 0.691758 | 1.17403 | 1.697 |
| chr14 | 81102546 | 81102775 | 1.02404 | 1.73802 | 1.697 |
| chr19 | 1813038 | 1813255 | 1.23044 | 2.08845 | 1.697 |
| chr12 | 108003978 | 108004560 | 1.74989 | 2.97017 | 1.697 |
| chr1 | 45958830 | 45959093 | 1.13262 | 1.92247 | 1.697 |
| chr19 | 18390191 | 18390534 | 1.91649 | 3.25327 | 1.698 |
| chr19 | 42455656 | 42456187 | 2.97674 | 5.05327 | 1.698 |
| chrX | 11551282 | 11551637 | 1.30393 | 2.21362 | 1.698 |
| chr10 | 1411110 | 1411373 | 2.04319 | 3.46872 | 1.698 |
| chr13 | 75946692 | 75946897 | 0.926354 | 1.57269 | 1.698 |
| chr22 | 37663116 | 37663416 | 1.60697 | 2.72828 | 1.698 |
| chr12 | 77015263 | 77015682 | 1.66634 | 2.82919 | 1.698 |
| chr19 | 8643475 | 8644175 | 1.24039 | 2.1061 | 1.698 |
| chr1 | 180147933 | 180148295 | 2.20525 | 3.74444 | 1.698 |
| chr1 | 55410793 | 55411084 | 1.6435 | 2.7909 | 1.698 |
| chr17 | 55183963 | 55184261 | 2.01148 | 3.41582 | 1.698 |
| chr14 | 105914885 | 105915709 | 1.48202 | 2.51672 | 1.698 |
| chr19 | 11909540 | 11909751 | 0.64922 | 1.10252 | 1.698 |
| chr2 | 26744067 | 26744489 | 1.4849 | 2.52172 | 1.698 |
| chr7 | 135099681 | 135099910 | 1.33032 | 2.25922 | 1.698 |
| chr2 | 223803977 | 223804290 | 1.52359 | 2.58749 | 1.698 |
| chr1 | 235578712 | 235578948 | 1.59596 | 2.71049 | 1.698 |
| chr1 | 151416405 | 151416626 | 1.84636 | 3.1359 | 1.698 |
| chr8 | 138845984 | 138846334 | 0.738727 | 1.25476 | 1.699 |
| chr2 | 240063 | 240321 | 0.673918 | 1.14469 | 1.699 |
| chr20 | 1193615 | 1193904 | 2.16117 | 3.67088 | 1.699 |
| chrX | 130760972 | 130761222 | 1.22394 | 2.07916 | 1.699 |
| chr3 | 81959435 | 81959983 | 1.50292 | 2.55309 | 1.699 |
| chr9 | 14577233 | 14577491 | 0.622796 | 1.05809 | 1.699 |
| chr17 | 78056397 | 78056769 | 1.64474 | 2.79441 | 1.699 |
| chr17 | 65411362 | 65411582 | 0.768105 | 1.30503 | 1.699 |
| chr22 | 21984280 | 21984570 | 1.50331 | 2.55441 | 1.699 |

| | | | | | |
|---|---|---|---|---|---|
| chr9 | 90299444 | 90299847 | 1.59752 | 2.71462 | 1.699 |
| chr20 | 40809817 | 40810042 | 0.56123 | 0.953744 | 1.699 |
| chr7 | 35077167 | 35077858 | 1.43295 | 2.43513 | 1.699 |
| chr15 | 66803241 | 66803641 | 0.59376 | 1.00903 | 1.699 |
| chr3 | 13921810 | 13922475 | 1.85974 | 3.16044 | 1.699 |
| chr17 | 34303004 | 34303416 | 1.53245 | 2.6043 | 1.699 |
| chr1 | 199432580 | 199433002 | 1.34881 | 2.29224 | 1.699 |
| chr16 | 71455866 | 71456285 | 1.32278 | 2.24816 | 1.700 |
| chr16 | 66750955 | 66751253 | 1.12581 | 1.9134 | 1.700 |
| chr22 | 24917791 | 24918047 | 1.31394 | 2.23333 | 1.700 |
| chr1 | 175822094 | 175822494 | 0.634212 | 1.07801 | 1.700 |
| chr14 | 61923133 | 61923729 | 1.66088 | 2.82328 | 1.700 |
| chr15 | 63034318 | 63034956 | 2.01261 | 3.4212 | 1.700 |
| chr17 | 70556417 | 70556791 | 1.90538 | 3.23898 | 1.700 |
| chr2 | 25015702 | 25015929 | 1.56603 | 2.6622 | 1.700 |
| chr3 | 72314639 | 72314923 | 0.890767 | 1.51428 | 1.700 |
| chr9 | 123817870 | 123818409 | 1.34861 | 2.29266 | 1.700 |
| chr14 | 65377679 | 65378027 | 1.38744 | 2.3587 | 1.700 |
| chr1 | 67677115 | 67677465 | 0.462973 | 0.787078 | 1.700 |
| chr15 | 73419646 | 73420016 | 1.87265 | 3.18363 | 1.700 |
| chr11 | 131543829 | 131544182 | 2.05825 | 3.49918 | 1.700 |
| chr6 | 73995504 | 73995721 | 0.754674 | 1.28301 | 1.700 |
| chr12 | 8085766 | 8086195 | 1.68224 | 2.86001 | 1.700 |
| chr11 | 73104617 | 73104939 | 1.92621 | 3.27496 | 1.700 |
| chr19 | 4730762 | 4730974 | 0.869724 | 1.47871 | 1.700 |
| chr7 | 28277082 | 28277340 | 0.573819 | 0.975691 | 1.700 |
| chr20 | 16936435 | 16936732 | 0.655196 | 1.11407 | 1.700 |
| chr8 | 110396487 | 110396701 | 0.672557 | 1.14363 | 1.700 |
| chr12 | 51663357 | 51663655 | 1.47027 | 2.50012 | 1.700 |
| chr1 | 27372360 | 27372740 | 1.95269 | 3.32074 | 1.701 |
| chr2 | 135249435 | 135249663 | 1.04931 | 1.78446 | 1.701 |
| chr2 | 99265843 | 99266241 | 1.89414 | 3.22136 | 1.701 |
| chr1 | 90098364 | 90098737 | 1.38696 | 2.35885 | 1.701 |
| chr11 | 5903578 | 5903832 | 0.712431 | 1.21165 | 1.701 |
| chr5 | 88291791 | 88292061 | 0.664708 | 1.13049 | 1.701 |
| chr17 | 72086025 | 72086465 | 1.9274 | 3.27807 | 1.701 |
| chr16 | 56735957 | 56736318 | 2.56242 | 4.35832 | 1.701 |
| chr20 | 45167858 | 45168066 | 1.09458 | 1.86173 | 1.701 |
| chr19 | 49925566 | 49925994 | 1.54307 | 2.62463 | 1.701 |
| chr12 | 126797924 | 126798166 | 0.530665 | 0.902621 | 1.701 |
| chr6 | 52151698 | 52152149 | 1.26311 | 2.14845 | 1.701 |
| chr10 | 107929104 | 107929450 | 0.563289 | 0.958119 | 1.701 |
| chr17 | 26087094 | 26087600 | 2.25225 | 3.831 | 1.701 |
| chr5 | 114910302 | 114910552 | 1.25771 | 2.13935 | 1.701 |
| chr4 | 181749640 | 181749962 | 0.97186 | 1.65322 | 1.701 |
| chr8 | 75278383 | 75278718 | 1.19622 | 2.03489 | 1.701 |
| chr2 | 7154594 | 7154810 | 1.37435 | 2.33795 | 1.701 |
| chr7 | 50417675 | 50417966 | 0.631658 | 1.07453 | 1.701 |
| chr12 | 18293024 | 18293242 | 0.978499 | 1.66457 | 1.701 |
| chr9 | 764031 | 764477 | 2.82721 | 4.80957 | 1.701 |
| chr8 | 61145643 | 61146276 | 1.27247 | 2.16472 | 1.701 |
| chr6 | 5136077 | 5136306 | 1.56305 | 2.65908 | 1.701 |
| chr12 | 32465043 | 32465348 | 1.28843 | 2.19194 | 1.701 |
| chr5 | 156989112 | 156989465 | 0.633715 | 1.07821 | 1.701 |
| chr7 | 105819266 | 105819497 | 0.867621 | 1.47618 | 1.701 |
| chr19 | 38586387 | 38586742 | 2.18175 | 3.71228 | 1.702 |
| chr3 | 55519061 | 55519455 | 2.02506 | 3.44566 | 1.702 |
| chr20 | 4065870 | 4066286 | 3.47231 | 5.90826 | 1.702 |
| chr2 | 160778343 | 160778632 | 1.78731 | 3.04126 | 1.702 |
| chr8 | 5662770 | 5663072 | 0.484533 | 0.824492 | 1.702 |
| chr1 | 156695590 | 156695916 | 2.00564 | 3.41301 | 1.702 |

| | | | | | |
|---|---|---|---|---|---|
| chr18 | 13003328 | 13003989 | 1.08853 | 1.85242 | 1.702 |
| chr3 | 142399647 | 142399854 | 1.0054 | 1.71096 | 1.702 |
| chr3 | 136751272 | 136751768 | 2.11064 | 3.59193 | 1.702 |
| chr3 | 134513646 | 134514112 | 1.86232 | 3.16947 | 1.702 |
| chr15 | 73536692 | 73537035 | 1.35831 | 2.31176 | 1.702 |
| chr22 | 44477032 | 44477510 | 2.0639 | 3.51263 | 1.702 |
| chr5 | 157167308 | 157167629 | 1.35549 | 2.30707 | 1.702 |
| chr1 | 54547366 | 54547650 | 1.60356 | 2.72939 | 1.702 |
| chr15 | 89770373 | 89770579 | 0.680939 | 1.15902 | 1.702 |
| chr14 | 41254072 | 41254488 | 0.800905 | 1.3633 | 1.702 |
| chr20 | 12264421 | 12264640 | 0.617574 | 1.05124 | 1.702 |
| chr14 | 85005101 | 85005347 | 0.47263 | 0.80452 | 1.702 |
| chr1 | 207106703 | 207107100 | 1.4548 | 2.47644 | 1.702 |
| chr12 | 23285789 | 23285996 | 0.571346 | 0.972583 | 1.702 |
| chr3 | 9617265 | 9617476 | 0.462927 | 0.78811 | 1.702 |
| chr11 | 41121656 | 41121877 | 0.500793 | 0.852586 | 1.702 |
| chr19 | 48002436 | 48002756 | 0.437822 | 0.745378 | 1.702 |
| chr1 | 23543243 | 23543644 | 1.9618 | 3.33994 | 1.702 |
| chr11 | 90013539 | 90013776 | 0.980013 | 1.66847 | 1.703 |
| chr8 | 91657141 | 91657389 | 1.73036 | 2.94598 | 1.703 |
| chr8 | 26548196 | 26548931 | 1.45928 | 2.48464 | 1.703 |
| chr19 | 46852181 | 46852424 | 0.731702 | 1.24584 | 1.703 |
| chr1 | 180502657 | 180502977 | 2.32316 | 3.95561 | 1.703 |
| chr7 | 75948018 | 75948282 | 0.674645 | 1.14878 | 1.703 |
| chr6 | 43139007 | 43139346 | 1.43256 | 2.43937 | 1.703 |
| chr12 | 925584 | 925883 | 1.25296 | 2.13356 | 1.703 |
| chr8 | 9656968 | 9657230 | 0.615384 | 1.04792 | 1.703 |
| chr3 | 108943212 | 108943596 | 1.21207 | 2.06406 | 1.703 |
| chr6 | 37621746 | 37622009 | 1.38842 | 2.36449 | 1.703 |
| chr14 | 61763715 | 61763952 | 1.38502 | 2.35878 | 1.703 |
| chr16 | 72183987 | 72184240 | 0.721806 | 1.2293 | 1.703 |
| chr14 | 22128272 | 22128685 | 0.592531 | 1.00914 | 1.703 |
| chr14 | 57476185 | 57476413 | 0.58821 | 1.00178 | 1.703 |
| chr8 | 70366701 | 70366911 | 0.383353 | 0.652889 | 1.703 |
| chr14 | 25620174 | 25620688 | 0.982444 | 1.67331 | 1.703 |
| chr2 | 241377632 | 241378087 | 2.01096 | 3.42509 | 1.703 |
| chr12 | 22488490 | 22489043 | 1.57272 | 2.67877 | 1.703 |
| chr9 | 138460860 | 138461258 | 0.545661 | 0.929412 | 1.703 |
| chr19 | 50233895 | 50234136 | 0.733814 | 1.24993 | 1.703 |
| chr5 | 80096044 | 80096302 | 0.748545 | 1.27507 | 1.703 |
| chr2 | 109605498 | 109605876 | 1.49659 | 2.54932 | 1.703 |
| chr20 | 34790704 | 34791181 | 3.72864 | 6.35153 | 1.703 |
| chr8 | 142356972 | 142357500 | 3.10179 | 5.28376 | 1.703 |
| chr11 | 119978847 | 119979253 | 2.80456 | 4.77753 | 1.703 |
| chr11 | 18749143 | 18749529 | 2.07984 | 3.54305 | 1.704 |
| chr8 | 143379413 | 143379809 | 0.931286 | 1.58651 | 1.704 |
| chr15 | 58143671 | 58144023 | 0.883803 | 1.50569 | 1.704 |
| chr12 | 124816644 | 124816889 | 1.31718 | 2.24404 | 1.704 |
| chr10 | 126190761 | 126190976 | 0.40221 | 0.685244 | 1.704 |
| chr5 | 168758417 | 168758656 | 0.539084 | 0.918482 | 1.704 |
| chrX | 103810474 | 103810839 | 2.14715 | 3.65829 | 1.704 |
| chr5 | 160091636 | 160091900 | 0.542189 | 0.923784 | 1.704 |
| chr7 | 99331097 | 99331459 | 1.07562 | 1.83264 | 1.704 |
| chr12 | 67265727 | 67266049 | 0.638395 | 1.08771 | 1.704 |
| chr22 | 20960498 | 20960895 | 1.87195 | 3.18962 | 1.704 |
| chr6 | 141242812 | 141243091 | 1.26914 | 2.16249 | 1.704 |
| chr3 | 157250280 | 157250696 | 1.45331 | 2.47636 | 1.704 |
| chr5 | 67272486 | 67272819 | 0.526077 | 0.896407 | 1.704 |
| chr1 | 155293083 | 155293456 | 2.19002 | 3.73177 | 1.704 |
| chr8 | 135917794 | 135918065 | 2.05274 | 3.49793 | 1.704 |
| chr2 | 4186967 | 4187212 | 0.720282 | 1.2275 | 1.704 |

| | | | | | |
|---|---|---|---|---|---|
| chr2 | 62144996 | 62145306 | 1.5902 | 2.71 | 1.704 |
| chr8 | 129579612 | 129579837 | 0.79375 | 1.35279 | 1.704 |
| chr19 | 55978984 | 55979595 | 2.45548 | 4.18492 | 1.704 |
| chr15 | 100672102 | 100672472 | 1.47727 | 2.51783 | 1.704 |
| chr19 | 17457058 | 17457312 | 1.91466 | 3.2633 | 1.704 |
| chr7 | 120497589 | 120497915 | 1.35146 | 2.3034 | 1.704 |
| chr19 | 14982581 | 14982804 | 0.836838 | 1.42634 | 1.704 |
| chr1 | 8978204 | 8978491 | 1.25924 | 2.14646 | 1.705 |
| chr11 | 64216248 | 64216726 | 2.74676 | 4.68224 | 1.705 |
| chr13 | 22199608 | 22199838 | 1.47691 | 2.51761 | 1.705 |
| chr20 | 53096294 | 53096649 | 0.851336 | 1.45124 | 1.705 |
| chr2 | 176068443 | 176068670 | 1.50876 | 2.57203 | 1.705 |
| chr1 | 178971755 | 178972224 | 1.85885 | 3.16887 | 1.705 |
| chr15 | 94566759 | 94567061 | 0.691899 | 1.17955 | 1.705 |
| chr1 | 205888002 | 205888388 | 1.72905 | 2.94785 | 1.705 |
| chr3 | 21903993 | 21904255 | 0.684499 | 1.16701 | 1.705 |
| chr9 | 130705966 | 130706383 | 1.80894 | 3.08412 | 1.705 |
| chr15 | 24318564 | 24318808 | 0.703589 | 1.19959 | 1.705 |
| chr9 | 35955013 | 35955272 | 1.91046 | 3.25728 | 1.705 |
| chr11 | 72309974 | 72310543 | 1.33624 | 2.27826 | 1.705 |
| chr5 | 150902992 | 150903324 | 1.25667 | 2.1426 | 1.705 |
| chr1 | 231493648 | 231493868 | 1.69812 | 2.89528 | 1.705 |
| chr7 | 26416694 | 26417028 | 3.29065 | 5.61064 | 1.705 |
| chr8 | 15018469 | 15018673 | 0.523076 | 0.891894 | 1.705 |
| chr1 | 8621193 | 8621434 | 1.15547 | 1.97035 | 1.705 |
| chr1 | 9606804 | 9607022 | 0.433388 | 0.739029 | 1.705 |
| chr8 | 70141987 | 70142192 | 0.532614 | 0.908324 | 1.705 |
| chr9 | 72161232 | 72161514 | 0.624357 | 1.06481 | 1.705 |
| chr10 | 103296631 | 103296961 | 1.06382 | 1.81431 | 1.705 |
| chr19 | 6632076 | 6632593 | 1.54431 | 2.6339 | 1.706 |
| chr9 | 124506556 | 124506963 | 1.10554 | 1.88556 | 1.706 |
| chr12 | 121021146 | 121021606 | 2.30176 | 3.92586 | 1.706 |
| chr15 | 61806846 | 61807077 | 0.612134 | 1.04406 | 1.706 |
| chr10 | 124039243 | 124039512 | 1.54625 | 2.6374 | 1.706 |
| chr7 | 64774406 | 64774638 | 0.414403 | 0.706852 | 1.706 |
| chr5 | 166055450 | 166055898 | 0.570948 | 0.973887 | 1.706 |
| chr18 | 59916742 | 59917023 | 1.66737 | 2.84428 | 1.706 |
| chr8 | 80997323 | 80997579 | 2.06387 | 3.52076 | 1.706 |
| chr2 | 24632697 | 24633244 | 1.33887 | 2.28409 | 1.706 |
| chr2 | 236649646 | 236650090 | 0.605405 | 1.03292 | 1.706 |
| chr15 | 51699697 | 51700038 | 0.800255 | 1.36543 | 1.706 |
| chr11 | 75918583 | 75919118 | 2.57165 | 4.38791 | 1.706 |
| chr5 | 7746080 | 7746582 | 1.54842 | 2.64201 | 1.706 |
| chr8 | 37247988 | 37248296 | 2.08953 | 3.56562 | 1.706 |
| chr10 | 106109942 | 106110291 | 1.46378 | 2.49786 | 1.706 |
| chr8 | 91798024 | 91798641 | 0.488576 | 0.833761 | 1.707 |
| chr19 | 36489901 | 36490323 | 2.38069 | 4.06306 | 1.707 |
| chr21 | 45724850 | 45725140 | 1.79269 | 3.05954 | 1.707 |
| chr5 | 140348690 | 140348908 | 0.635995 | 1.08544 | 1.707 |
| chrX | 38034319 | 38034739 | 1.68046 | 2.86804 | 1.707 |
| chr3 | 71924370 | 71924901 | 1.77429 | 3.02834 | 1.707 |
| chr5 | 132652127 | 132652362 | 1.13205 | 1.93217 | 1.707 |
| chr2 | 40485049 | 40485309 | 0.637136 | 1.08748 | 1.707 |
| chr2 | 65058863 | 65059224 | 2.82293 | 4.81842 | 1.707 |
| chr8 | 145704335 | 145704793 | 1.70815 | 2.91567 | 1.707 |
| chr14 | 69737046 | 69737281 | 0.447773 | 0.764364 | 1.707 |
| chr3 | 26839894 | 26840190 | 0.628958 | 1.07367 | 1.707 |
| chr8 | 15242919 | 15243135 | 0.558122 | 0.952746 | 1.707 |
| chr11 | 1314904 | 1315139 | 1.512 | 2.58114 | 1.707 |
| chr16 | 3108730 | 3109603 | 2.24671 | 3.83542 | 1.707 |
| chr19 | 43100437 | 43101097 | 3.03566 | 5.18228 | 1.707 |

| | | | | | |
|---|---|---|---|---|---|
| chr3 | 29049375 | 29049579 | 0.634976 | 1.08401 | 1.707 |
| chr9 | 107322590 | 107322853 | 0.61761 | 1.05439 | 1.707 |
| chr2 | 7752011 | 7752312 | 0.468953 | 0.800613 | 1.707 |
| chr17 | 13846868 | 13847404 | 0.642015 | 1.0961 | 1.707 |
| chr11 | 91036538 | 91036771 | 0.576343 | 0.983991 | 1.707 |
| chr11 | 46404732 | 46405224 | 1.90042 | 3.24484 | 1.707 |
| chr14 | 42041977 | 42042540 | 0.629119 | 1.0742 | 1.707 |
| chr4 | 75719000 | 75719237 | 1.08961 | 1.86048 | 1.707 |
| chr15 | 75321529 | 75321915 | 1.99161 | 3.40091 | 1.708 |
| chr15 | 78533275 | 78533700 | 1.76515 | 3.01433 | 1.708 |
| chr11 | 2023788 | 2024488 | 0.912887 | 1.55903 | 1.708 |
| chr15 | 83339739 | 83340129 | 0.625778 | 1.06877 | 1.708 |
| chr7 | 2683025 | 2683318 | 0.960843 | 1.64104 | 1.708 |
| chr6 | 132236253 | 132236622 | 1.5387 | 2.62799 | 1.708 |
| chr9 | 137035788 | 137036369 | 2.55447 | 4.36291 | 1.708 |
| chr19 | 40811245 | 40811564 | 2.31317 | 3.95089 | 1.708 |
| chrX | 36926891 | 36927251 | 1.18502 | 2.02401 | 1.708 |
| chr7 | 141354315 | 141354604 | 1.86463 | 3.18492 | 1.708 |
| chr11 | 66549020 | 66549306 | 0.707725 | 1.20886 | 1.708 |
| chr2 | 238231272 | 238231502 | 1.33166 | 2.27471 | 1.708 |
| chr7 | 130756523 | 130756923 | 1.03967 | 1.7762 | 1.708 |
| chr11 | 64780996 | 64782034 | 2.26456 | 3.86914 | 1.709 |
| chr14 | 75608818 | 75609041 | 1.06183 | 1.81428 | 1.709 |
| chr11 | 6464650 | 6465366 | 1.45733 | 2.49024 | 1.709 |
| chr12 | 108384187 | 108384530 | 0.974481 | 1.66516 | 1.709 |
| chr7 | 3937019 | 3937411 | 1.89993 | 3.2467 | 1.709 |
| chr2 | 69064382 | 69064770 | 1.93526 | 3.30708 | 1.709 |
| chr7 | 1265108 | 1265682 | 2.27107 | 3.88094 | 1.709 |
| chr2 | 52741376 | 52741612 | 0.56259 | 0.961396 | 1.709 |
| chr11 | 61500901 | 61501280 | 3.09454 | 5.28838 | 1.709 |
| chr3 | 32217040 | 32217399 | 1.65193 | 2.82306 | 1.709 |
| chr2 | 106162148 | 106162562 | 1.87372 | 3.20224 | 1.709 |
| chr22 | 43456193 | 43456629 | 1.88424 | 3.2203 | 1.709 |
| chr9 | 87569012 | 87569352 | 2.71367 | 4.63787 | 1.709 |
| chr1 | 246166473 | 246166815 | 2.37644 | 4.06194 | 1.709 |
| chr2 | 18701819 | 18702166 | 2.3072 | 3.94363 | 1.709 |
| chr9 | 108788724 | 108789031 | 0.64975 | 1.11063 | 1.709 |
| chr16 | 11808471 | 11808800 | 1.89431 | 3.2383 | 1.709 |
| chr19 | 19754625 | 19754986 | 1.93207 | 3.3029 | 1.710 |
| chr5 | 81470455 | 81470808 | 1.97868 | 3.38272 | 1.710 |
| chr10 | 99831878 | 99832102 | 0.447751 | 0.765476 | 1.710 |
| chr12 | 54996997 | 54997362 | 1.46198 | 2.49953 | 1.710 |
| chr14 | 95827718 | 95828328 | 1.33729 | 2.28637 | 1.710 |
| chr1 | 71595320 | 71595530 | 0.671156 | 1.14748 | 1.710 |
| chr14 | 76819065 | 76819553 | 1.91211 | 3.26937 | 1.710 |
| chr14 | 50328700 | 50329176 | 4.95588 | 8.47383 | 1.710 |
| chr16 | 65898549 | 65898806 | 0.688538 | 1.17731 | 1.710 |
| chr3 | 158249985 | 158250189 | 1.30559 | 2.23239 | 1.710 |
| chr5 | 143526154 | 143526411 | 0.694935 | 1.1883 | 1.710 |
| chr2 | 19866926 | 19867141 | 0.933255 | 1.59583 | 1.710 |
| chr18 | 55297487 | 55297702 | 1.66185 | 2.84173 | 1.710 |
| chr3 | 107361921 | 107362172 | 1.04099 | 1.78007 | 1.710 |
| chr22 | 38658112 | 38658437 | 1.61622 | 2.76375 | 1.710 |
| chr14 | 91789842 | 91790318 | 2.50486 | 4.28337 | 1.710 |
| chr3 | 13301876 | 13302097 | 0.948531 | 1.62201 | 1.710 |
| chr3 | 76960887 | 76961108 | 0.819867 | 1.40204 | 1.710 |
| chr14 | 89259015 | 89259506 | 0.707745 | 1.21042 | 1.710 |
| chr12 | 25612662 | 25613183 | 0.682567 | 1.16737 | 1.710 |
| chr3 | 96532048 | 96532351 | 1.52847 | 2.61421 | 1.710 |
| chr7 | 144988460 | 144988768 | 0.621661 | 1.06328 | 1.710 |
| chr5 | 170440116 | 170440326 | 0.594639 | 1.01707 | 1.710 |

| | | | | | |
|---|---|---|---|---|---|
| chr2 | 11618075 | 11618454 | 1.79445 | 3.06927 | 1.710 |
| chr7 | 119648586 | 119649013 | 0.493663 | 0.844375 | 1.710 |
| chr1 | 174186903 | 174187157 | 1.37983 | 2.36019 | 1.710 |
| chr9 | 79060343 | 79060593 | 0.517224 | 0.884734 | 1.711 |
| chr19 | 49345870 | 49346408 | 2.26699 | 3.87789 | 1.711 |
| chr6 | 150250592 | 150250807 | 1.19457 | 2.04342 | 1.711 |
| chr7 | 33578785 | 33579023 | 0.857023 | 1.46604 | 1.711 |
| chr7 | 361986 | 362540 | 1.86725 | 3.19419 | 1.711 |
| chr2 | 232092048 | 232092343 | 1.675 | 2.86545 | 1.711 |
| chr17 | 73901037 | 73901427 | 2.59233 | 4.43481 | 1.711 |
| chr3 | 155761717 | 155761942 | 1.24673 | 2.13284 | 1.711 |
| chr10 | 116037588 | 116037803 | 0.852992 | 1.4593 | 1.711 |
| chr14 | 101370771 | 101370973 | 1.28588 | 2.1999 | 1.711 |
| chr10 | 62784543 | 62784957 | 1.48308 | 2.5373 | 1.711 |
| chr14 | 51750146 | 51750437 | 1.35376 | 2.31612 | 1.711 |
| chr15 | 72892273 | 72892531 | 2.41071 | 4.12446 | 1.711 |
| chr18 | 61007198 | 61007466 | 0.881541 | 1.50822 | 1.711 |
| chr4 | 8887616 | 8887836 | 1.30953 | 2.24051 | 1.711 |
| chr8 | 36396257 | 36396493 | 0.600786 | 1.0279 | 1.711 |
| chr3 | 79546723 | 79546940 | 1.26308 | 2.16105 | 1.711 |
| chr11 | 20694211 | 20694486 | 1.42722 | 2.44193 | 1.711 |
| chrX | 19024088 | 19024335 | 1.10231 | 1.88603 | 1.711 |
| chr1 | 111323472 | 111323780 | 1.37184 | 2.34741 | 1.711 |
| chr10 | 124326528 | 124326960 | 1.44407 | 2.47101 | 1.711 |
| chr17 | 55230805 | 55231083 | 0.652187 | 1.11601 | 1.711 |
| chr9 | 34702004 | 34702597 | 2.26229 | 3.87127 | 1.711 |
| chr22 | 32662177 | 32662440 | 1.27401 | 2.18023 | 1.711 |
| chr9 | 112083187 | 112083589 | 2.45965 | 4.20937 | 1.711 |
| chr5 | 169611072 | 169611478 | 1.777 | 3.04114 | 1.711 |
| chr1 | 232574778 | 232575011 | 0.669927 | 1.14653 | 1.711 |
| chr1 | 6464809 | 6465120 | 2.25121 | 3.8528 | 1.711 |
| chr5 | 14399195 | 14399446 | 1.1653 | 1.99444 | 1.712 |
| chr15 | 26844556 | 26844767 | 0.769346 | 1.31682 | 1.712 |
| chr1 | 45301516 | 45301863 | 1.43699 | 2.4596 | 1.712 |
| chr7 | 142273787 | 142273992 | 1.49032 | 2.55111 | 1.712 |
| chr22 | 22936599 | 22936801 | 1.37262 | 2.34965 | 1.712 |
| chr5 | 140474014 | 140474289 | 1.61736 | 2.76862 | 1.712 |
| chr7 | 128519988 | 128520457 | 2.42428 | 4.14998 | 1.712 |
| chr16 | 30124048 | 30124690 | 1.59951 | 2.73814 | 1.712 |
| chr10 | 11557646 | 11557901 | 1.50694 | 2.57995 | 1.712 |
| chr1 | 20113762 | 20114025 | 0.473716 | 0.811092 | 1.712 |
| chr2 | 191237113 | 191237432 | 2.12306 | 3.63518 | 1.712 |
| chrX | 66775527 | 66775770 | 1.68528 | 2.88568 | 1.712 |
| chr14 | 61491986 | 61492265 | 1.60278 | 2.74446 | 1.712 |
| chr12 | 51610932 | 51611274 | 1.4477 | 2.47914 | 1.712 |
| chr6 | 99287728 | 99287992 | 1.02437 | 1.75433 | 1.713 |
| chr12 | 89444853 | 89445198 | 1.43004 | 2.44912 | 1.713 |
| chr2 | 224835033 | 224835312 | 1.58838 | 2.72045 | 1.713 |
| chr12 | 123815309 | 123815550 | 0.78461 | 1.34385 | 1.713 |
| chr5 | 164806713 | 164806924 | 0.920323 | 1.57632 | 1.713 |
| chr18 | 47745250 | 47745517 | 1.36315 | 2.33491 | 1.713 |
| chr7 | 25022553 | 25022820 | 0.911472 | 1.56126 | 1.713 |
| chr11 | 130764326 | 130764597 | 1.83565 | 3.14444 | 1.713 |
| chr9 | 12221105 | 12221423 | 0.707568 | 1.21207 | 1.713 |
| chr10 | 45214598 | 45214916 | 0.533407 | 0.913743 | 1.713 |
| chr7 | 95500363 | 95500763 | 1.19012 | 2.03873 | 1.713 |
| chr3 | 131725536 | 131725751 | 1.56855 | 2.68713 | 1.713 |
| chr19 | 18383244 | 18383547 | 1.60434 | 2.74849 | 1.713 |
| chr12 | 73797298 | 73797682 | 0.838387 | 1.43633 | 1.713 |
| chr12 | 50482380 | 50482777 | 1.47043 | 2.51918 | 1.713 |
| chr2 | 79560150 | 79560414 | 0.538407 | 0.922423 | 1.713 |

| | | | | | |
|---|---|---|---|---|---|
| chr12 | 102563748 | 102564077 | 1.39801 | 2.39519 | 1.713 |
| chr7 | 116166148 | 116166465 | 2.43241 | 4.16756 | 1.713 |
| chr7 | 40031712 | 40031981 | 0.992626 | 1.70074 | 1.713 |
| chr11 | 6724757 | 6725019 | 1.3223 | 2.26561 | 1.713 |
| chr17 | 15785465 | 15785677 | 1.0338 | 1.77148 | 1.714 |
| chr11 | 126944055 | 126944321 | 1.54127 | 2.64107 | 1.714 |
| chr19 | 43951065 | 43951495 | 1.97082 | 3.37731 | 1.714 |
| chr7 | 73082016 | 73082358 | 2.02407 | 3.46868 | 1.714 |
| chr10 | 112306998 | 112307224 | 1.75113 | 3.00104 | 1.714 |
| chr14 | 42997230 | 42997505 | 0.573909 | 0.983555 | 1.714 |
| chr20 | 12652610 | 12652938 | 0.493799 | 0.846273 | 1.714 |
| chr12 | 54773389 | 54773905 | 4.10335 | 7.03238 | 1.714 |
| chr2 | 128211070 | 128211425 | 1.17478 | 2.01335 | 1.714 |
| chr5 | 177978259 | 177978702 | 0.669611 | 1.14764 | 1.714 |
| chr5 | 132050487 | 132050712 | 0.92038 | 1.57744 | 1.714 |
| chr7 | 54907312 | 54907675 | 2.414 | 4.13749 | 1.714 |
| chr18 | 25179203 | 25179467 | 1.90375 | 3.26297 | 1.714 |
| chr12 | 7791256 | 7791637 | 1.53132 | 2.62468 | 1.714 |
| chr2 | 11805225 | 11805454 | 0.847845 | 1.45322 | 1.714 |
| chr8 | 30027722 | 30028077 | 1.60093 | 2.74404 | 1.714 |
| chr1 | 208155292 | 208155640 | 1.01741 | 1.74389 | 1.714 |
| chr14 | 53899078 | 53899310 | 1.55948 | 2.67309 | 1.714 |
| chr20 | 42721313 | 42721964 | 0.882602 | 1.51291 | 1.714 |
| chr19 | 29679721 | 29680367 | 0.634347 | 1.08739 | 1.714 |
| chr5 | 110493857 | 110494192 | 2.56153 | 4.39114 | 1.714 |
| chr4 | 111553082 | 111553303 | 1.16672 | 2.0001 | 1.714 |
| chr7 | 107027593 | 107027842 | 0.536769 | 0.920176 | 1.714 |
| chr3 | 130257910 | 130258252 | 1.70579 | 2.9243 | 1.714 |
| chr11 | 1974701 | 1975080 | 1.91612 | 3.28495 | 1.714 |
| chr15 | 40882044 | 40882364 | 1.54954 | 2.65651 | 1.714 |
| chr11 | 57244497 | 57244714 | 1.15206 | 1.97517 | 1.714 |
| chr14 | 103590141 | 103590515 | 1.46158 | 2.50592 | 1.715 |
| chr5 | 41520034 | 41520355 | 0.743207 | 1.27428 | 1.715 |
| chr14 | 74292337 | 74292599 | 1.86261 | 3.19372 | 1.715 |
| chr1 | 3827398 | 3827665 | 1.81655 | 3.11487 | 1.715 |
| chr8 | 145634634 | 145635189 | 2.0905 | 3.58471 | 1.715 |
| chr8 | 145692983 | 145693464 | 1.99791 | 3.42598 | 1.715 |
| chr17 | 52968660 | 52968893 | 1.22409 | 2.09908 | 1.715 |
| chr19 | 32656300 | 32656571 | 0.64781 | 1.11088 | 1.715 |
| chr6 | 75284611 | 75284834 | 1.23934 | 2.12527 | 1.715 |
| chr11 | 5510468 | 5510818 | 0.782615 | 1.3422 | 1.715 |
| chr20 | 43313857 | 43314095 | 1.46056 | 2.50492 | 1.715 |
| chr8 | 125042699 | 125042936 | 1.34577 | 2.30812 | 1.715 |
| chr2 | 173330243 | 173330877 | 1.04082 | 1.78515 | 1.715 |
| chr7 | 34470080 | 34470434 | 1.06948 | 1.83437 | 1.715 |
| chr12 | 2792157 | 2792476 | 1.02918 | 1.76527 | 1.715 |
| chr19 | 13990867 | 13991270 | 2.72686 | 4.6773 | 1.715 |
| chr11 | 65133769 | 65134035 | 2.04541 | 3.50851 | 1.715 |
| chr17 | 35596620 | 35597016 | 1.23823 | 2.12398 | 1.715 |
| chr9 | 79407844 | 79408067 | 0.436238 | 0.748302 | 1.715 |
| chr19 | 45895160 | 45895430 | 0.535418 | 0.918456 | 1.715 |
| chr12 | 5940452 | 5940683 | 0.683716 | 1.17287 | 1.715 |
| chr8 | 36716475 | 36716702 | 1.53249 | 2.62891 | 1.715 |
| chr9 | 88225007 | 88225325 | 1.64019 | 2.81375 | 1.716 |
| chr12 | 68325650 | 68325984 | 0.851558 | 1.46089 | 1.716 |
| chr8 | 118095382 | 118095605 | 0.777601 | 1.33403 | 1.716 |
| chr6 | 166142878 | 166143176 | 1.4071 | 2.41409 | 1.716 |
| chr9 | 28043088 | 28043358 | 0.431187 | 0.739797 | 1.716 |
| chr5 | 71078778 | 71079103 | 1.87109 | 3.2104 | 1.716 |
| chrX | 118252708 | 118253096 | 1.64046 | 2.81468 | 1.716 |
| chr14 | 22446473 | 22446852 | 1.35837 | 2.33075 | 1.716 |

| | | | | | |
|---|---|---|---|---|---|
| chr7 | 142219078 | 142219318 | 1.25236 | 2.14889 | 1.716 |
| chr2 | 173478421 | 173478639 | 0.871966 | 1.49625 | 1.716 |
| chr2 | 21611014 | 21611318 | 0.618659 | 1.0616 | 1.716 |
| chr5 | 89145423 | 89145652 | 0.750033 | 1.28709 | 1.716 |
| chr10 | 82942361 | 82942798 | 0.580006 | 0.995351 | 1.716 |
| chr7 | 10366295 | 10366562 | 0.613997 | 1.05372 | 1.716 |
| chr2 | 36549338 | 36549560 | 1.42643 | 2.44801 | 1.716 |
| chr8 | 123507618 | 123507839 | 0.740252 | 1.27041 | 1.716 |
| chr10 | 49497742 | 49498071 | 1.73118 | 2.97106 | 1.716 |
| chr1 | 212004040 | 212004372 | 1.67462 | 2.87407 | 1.716 |
| chr14 | 71412417 | 71412680 | 0.556226 | 0.954677 | 1.716 |
| chr14 | 77176504 | 77176784 | 1.18203 | 2.02881 | 1.716 |
| chr6 | 42011987 | 42012277 | 1.94978 | 3.34678 | 1.716 |
| chr16 | 86553635 | 86554029 | 2.62966 | 4.51388 | 1.717 |
| chr15 | 89750036 | 89750525 | 2.43699 | 4.18322 | 1.717 |
| chr2 | 29746925 | 29747141 | 0.568828 | 0.976421 | 1.717 |
| chr3 | 131753817 | 131754153 | 2.07664 | 3.56465 | 1.717 |
| chr5 | 140762094 | 140762483 | 1.13249 | 1.94401 | 1.717 |
| chr13 | 103452400 | 103452804 | 1.72735 | 2.96522 | 1.717 |
| chr16 | 64123672 | 64123944 | 1.00975 | 1.73338 | 1.717 |
| chr5 | 159772237 | 159772740 | 1.17037 | 2.00928 | 1.717 |
| chr13 | 39837116 | 39837363 | 1.29152 | 2.2173 | 1.717 |
| chr19 | 10209727 | 10209973 | 1.34247 | 2.30477 | 1.717 |
| chr5 | 134241203 | 134241606 | 1.21259 | 2.08183 | 1.717 |
| chr10 | 120183393 | 120183838 | 1.52234 | 2.61394 | 1.717 |
| chr12 | 26767193 | 26767433 | 1.19244 | 2.04751 | 1.717 |
| chr14 | 24028652 | 24029232 | 2.16439 | 3.71644 | 1.717 |
| chr15 | 71395480 | 71395737 | 1.61497 | 2.77308 | 1.717 |
| chr7 | 133804759 | 133804983 | 0.480016 | 0.824244 | 1.717 |
| chr11 | 75514304 | 75514584 | 1.91322 | 3.28535 | 1.717 |
| chr4 | 89140162 | 89140467 | 1.04519 | 1.79479 | 1.717 |
| chr15 | 69945373 | 69945595 | 0.696236 | 1.19561 | 1.717 |
| chr5 | 71053779 | 71054096 | 1.94965 | 3.34804 | 1.717 |
| chrX | 81903595 | 81903804 | 1.10581 | 1.89895 | 1.717 |
| chr20 | 17434247 | 17434486 | 0.688628 | 1.18256 | 1.717 |
| chr7 | 126848391 | 126848705 | 0.51006 | 0.875963 | 1.717 |
| chr17 | 41560915 | 41561337 | 5.05174 | 8.67605 | 1.717 |
| chr1 | 15241873 | 15242262 | 1.72465 | 2.96208 | 1.718 |
| chr7 | 120014786 | 120015060 | 0.883626 | 1.51764 | 1.718 |
| chr1 | 248772953 | 248773223 | 0.670196 | 1.15111 | 1.718 |
| chr7 | 84930569 | 84930785 | 0.586435 | 1.00725 | 1.718 |
| chr8 | 104687404 | 104687648 | 0.525643 | 0.902832 | 1.718 |
| chr5 | 7697202 | 7697446 | 0.575868 | 0.989149 | 1.718 |
| chr11 | 7097043 | 7097317 | 0.581251 | 0.998407 | 1.718 |
| chr14 | 58968018 | 58968237 | 1.77495 | 3.04888 | 1.718 |
| chr17 | 11666353 | 11666639 | 0.612276 | 1.05176 | 1.718 |
| chr7 | 50700547 | 50700990 | 2.66335 | 4.57565 | 1.718 |
| chrX | 110177013 | 110177267 | 0.504663 | 0.86707 | 1.718 |
| chr10 | 43886211 | 43886540 | 1.5182 | 2.60853 | 1.718 |
| chr19 | 44249450 | 44249791 | 2.14905 | 3.69259 | 1.718 |
| chrX | 3880516 | 3880869 | 0.845106 | 1.4521 | 1.718 |
| chr22 | 45605017 | 45605368 | 1.4633 | 2.51438 | 1.718 |
| chr20 | 35824035 | 35824292 | 1.45365 | 2.49781 | 1.718 |
| chr8 | 138101794 | 138102109 | 0.65383 | 1.12351 | 1.718 |
| chr16 | 12065977 | 12066253 | 0.888176 | 1.52623 | 1.718 |
| chr11 | 45392780 | 45393146 | 2.54687 | 4.37657 | 1.718 |
| chr16 | 84507818 | 84508028 | 0.583079 | 1.00198 | 1.718 |
| chr16 | 89282923 | 89283171 | 1.24332 | 2.13656 | 1.718 |
| chr10 | 16851369 | 16851626 | 1.30802 | 2.2478 | 1.718 |
| chr16 | 9872782 | 9873081 | 0.688384 | 1.18302 | 1.719 |
| chr12 | 50432888 | 50433446 | 2.50013 | 4.29687 | 1.719 |

| | | | | | |
|---|---|---|---|---|---|
| chr5 | 89241499 | 89241709 | 0.682394 | 1.1728 | 1.719 |
| chr6 | 133144172 | 133144409 | 1.71758 | 2.95193 | 1.719 |
| chr8 | 123628776 | 123628981 | 0.980017 | 1.68432 | 1.719 |
| chr16 | 67280068 | 67280348 | 1.30387 | 2.24092 | 1.719 |
| chr11 | 120672952 | 120673295 | 1.55788 | 2.67754 | 1.719 |
| chr6 | 157893222 | 157893461 | 0.506787 | 0.871133 | 1.719 |
| chr9 | 34633807 | 34634362 | 1.83874 | 3.16067 | 1.719 |
| chr8 | 20233734 | 20234112 | 2.45964 | 4.22814 | 1.719 |
| chr7 | 130753992 | 130754279 | 1.64772 | 2.8325 | 1.719 |
| chr11 | 19332841 | 19333220 | 1.64227 | 2.82322 | 1.719 |
| chr15 | 78423619 | 78424114 | 1.65278 | 2.84137 | 1.719 |
| chr10 | 73329751 | 73330117 | 1.85947 | 3.19675 | 1.719 |
| chr10 | 116655205 | 116655452 | 0.525992 | 0.904286 | 1.719 |
| chr7 | 148573193 | 148573406 | 0.618054 | 1.06258 | 1.719 |
| chr8 | 71314761 | 71315063 | 1.15794 | 1.99081 | 1.719 |
| chr17 | 78074946 | 78075439 | 2.54875 | 4.38207 | 1.719 |
| chr17 | 74178269 | 74178563 | 2.38334 | 4.09788 | 1.719 |
| chr2 | 70870861 | 70871163 | 1.69279 | 2.91057 | 1.719 |
| chr22 | 37373468 | 37373797 | 1.35824 | 2.33536 | 1.719 |
| chr17 | 55969790 | 55970180 | 2.86433 | 4.92501 | 1.719 |
| chr4 | 55084022 | 55084277 | 1.70189 | 2.92629 | 1.719 |
| chr14 | 47566079 | 47566376 | 0.800702 | 1.37677 | 1.719 |
| chr4 | 78306578 | 78306864 | 1.56373 | 2.68875 | 1.719 |
| chr8 | 65487671 | 65487950 | 1.45142 | 2.49567 | 1.719 |
| chr2 | 131678031 | 131678313 | 1.97779 | 3.4009 | 1.720 |
| chr5 | 140729403 | 140729763 | 1.80638 | 3.10619 | 1.720 |
| chr1 | 167829720 | 167829960 | 0.430746 | 0.740735 | 1.720 |
| chr20 | 20574127 | 20574637 | 1.22231 | 2.102 | 1.720 |
| chr2 | 78426177 | 78426468 | 0.53754 | 0.924414 | 1.720 |
| chr7 | 131543972 | 131544326 | 2.52493 | 4.34224 | 1.720 |
| chr3 | 126701279 | 126701943 | 2.44608 | 4.20672 | 1.720 |
| chr11 | 64283839 | 64284045 | 1.01985 | 1.75404 | 1.720 |
| chr10 | 56119999 | 56120228 | 0.574284 | 0.98785 | 1.720 |
| chr9 | 93851889 | 93852244 | 1.52463 | 2.62265 | 1.720 |
| chr5 | 125874948 | 125875236 | 1.41392 | 2.43223 | 1.720 |
| chr8 | 124553950 | 124554233 | 0.620735 | 1.06783 | 1.720 |
| chr9 | 119622849 | 119623091 | 1.82869 | 3.14641 | 1.721 |
| chr4 | 114169494 | 114169765 | 0.837815 | 1.44156 | 1.721 |
| chr6 | 11044528 | 11044993 | 2.9486 | 5.07349 | 1.721 |
| chr16 | 75798045 | 75798392 | 1.64868 | 2.83681 | 1.721 |
| chr8 | 30776445 | 30776844 | 1.63314 | 2.81008 | 1.721 |
| chr18 | 46862168 | 46862373 | 0.923964 | 1.58986 | 1.721 |
| chr5 | 11749016 | 11749436 | 0.611953 | 1.05298 | 1.721 |
| chr1 | 9065234 | 9065543 | 1.99081 | 3.42558 | 1.721 |
| chr4 | 152020509 | 152020776 | 1.69401 | 2.9149 | 1.721 |
| chr10 | 121251390 | 121251604 | 0.570653 | 0.981955 | 1.721 |
| chr10 | 115761901 | 115762116 | 0.671676 | 1.1558 | 1.721 |
| chr6 | 108436969 | 108437250 | 1.47341 | 2.53549 | 1.721 |
| chr1 | 219986011 | 219986272 | 1.42888 | 2.45901 | 1.721 |
| chr10 | 48585930 | 48586313 | 0.705985 | 1.21496 | 1.721 |
| chr2 | 178354518 | 178354724 | 0.90041 | 1.54962 | 1.721 |
| chr9 | 138173092 | 138173305 | 0.959401 | 1.6512 | 1.721 |
| chr10 | 59609566 | 59609832 | 0.857341 | 1.47555 | 1.721 |
| chr11 | 57335328 | 57336112 | 2.23402 | 3.84493 | 1.721 |
| chr4 | 102651769 | 102651977 | 1.3703 | 2.35853 | 1.721 |
| chr12 | 109605706 | 109606083 | 0.578468 | 0.995663 | 1.721 |
| chr14 | 68249714 | 68250117 | 0.500895 | 0.862149 | 1.721 |
| chr11 | 46890168 | 46890396 | 0.483182 | 0.831705 | 1.721 |
| chr2 | 149846500 | 149846752 | 1.49093 | 2.56639 | 1.721 |
| chr1 | 240150512 | 240150719 | 1.51034 | 2.59986 | 1.721 |
| chr8 | 121274001 | 121274269 | 0.654357 | 1.1264 | 1.721 |

| | | | | | |
|---|---|---|---|---|---|
| chr12 | 55671114 | 55671381 | 0.64209 | 1.10537 | 1.722 |
| chr7 | 156742098 | 156742562 | 3.09797 | 5.33394 | 1.722 |
| chr19 | 50366166 | 50366443 | 1.79722 | 3.09438 | 1.722 |
| chr8 | 131761876 | 131762235 | 2.42125 | 4.169 | 1.722 |
| chr19 | 13373264 | 13373478 | 0.901364 | 1.55202 | 1.722 |
| chr17 | 61512291 | 61512579 | 0.791373 | 1.36266 | 1.722 |
| chr2 | 3923929 | 3924230 | 0.579388 | 0.997668 | 1.722 |
| chr7 | 98429213 | 98429801 | 1.59928 | 2.75391 | 1.722 |
| chr17 | 37395790 | 37396221 | 1.42054 | 2.44614 | 1.722 |
| chr8 | 61659545 | 61659780 | 0.742083 | 1.27786 | 1.722 |
| chr11 | 61284272 | 61284916 | 2.04917 | 3.52867 | 1.722 |
| chr15 | 102042053 | 102042509 | 1.94333 | 3.34642 | 1.722 |
| chr7 | 120721466 | 120721723 | 1.29458 | 2.22928 | 1.722 |
| chr11 | 119537144 | 119537475 | 1.92118 | 3.30836 | 1.722 |
| chr19 | 46386492 | 46386920 | 3.4582 | 5.95535 | 1.722 |
| chr9 | 78371307 | 78371522 | 0.47664 | 0.820849 | 1.722 |
| chr9 | 86875976 | 86876417 | 2.19691 | 3.78369 | 1.722 |
| chr10 | 61874443 | 61874894 | 1.06961 | 1.84224 | 1.722 |
| chr20 | 35441162 | 35441629 | 0.561475 | 0.967084 | 1.722 |
| chr7 | 150516370 | 150516574 | 0.592708 | 1.02088 | 1.722 |
| chr9 | 130605130 | 130605379 | 0.639703 | 1.10184 | 1.722 |
| chr17 | 37225692 | 37226003 | 1.3126 | 2.26092 | 1.722 |
| chr14 | 55556329 | 55556697 | 1.64588 | 2.83503 | 1.723 |
| chr17 | 56072840 | 56073082 | 1.43583 | 2.47328 | 1.723 |
| chr14 | 23494661 | 23494885 | 0.636935 | 1.09716 | 1.723 |
| chr8 | 69263849 | 69264077 | 0.427387 | 0.736198 | 1.723 |
| chr16 | 4303983 | 4304472 | 3.65422 | 6.29508 | 1.723 |
| chr17 | 43312065 | 43312287 | 1.58496 | 2.73045 | 1.723 |
| chr15 | 68598600 | 68599381 | 1.94022 | 3.34248 | 1.723 |
| chr17 | 13772823 | 13773050 | 0.640221 | 1.10295 | 1.723 |
| chr12 | 121078117 | 121078430 | 1.36787 | 2.35656 | 1.723 |
| chr12 | 121084609 | 121085037 | 1.68255 | 2.89872 | 1.723 |
| chrX | 79540958 | 79541216 | 1.34951 | 2.32501 | 1.723 |
| chr15 | 41758059 | 41758353 | 1.83844 | 3.16746 | 1.723 |
| chr13 | 68130051 | 68130273 | 0.53715 | 0.925488 | 1.723 |
| chr21 | 38226844 | 38227101 | 1.4007 | 2.41352 | 1.723 |
| chr10 | 89416297 | 89416576 | 1.49328 | 2.57308 | 1.723 |
| chr9 | 139962186 | 139962580 | 2.00816 | 3.46032 | 1.723 |
| chr16 | 10603341 | 10603791 | 1.67667 | 2.88916 | 1.723 |
| chr5 | 160217734 | 160218073 | 0.833435 | 1.43622 | 1.723 |
| chr10 | 12391147 | 12391350 | 1.01739 | 1.75323 | 1.723 |
| chr1 | 27995746 | 27996099 | 1.84442 | 3.17848 | 1.723 |
| chr12 | 120895040 | 120895241 | 1.24835 | 2.15132 | 1.723 |
| chr5 | 180586285 | 180586833 | 1.78578 | 3.07773 | 1.723 |
| chr12 | 109870554 | 109871010 | 2.33352 | 4.02179 | 1.723 |
| chr17 | 7997239 | 7997773 | 1.73441 | 2.98927 | 1.724 |
| chr4 | 89594224 | 89594436 | 1.06106 | 1.82878 | 1.724 |
| chr7 | 142438470 | 142438924 | 0.618979 | 1.06686 | 1.724 |
| chr1 | 208441256 | 208441490 | 0.493344 | 0.850357 | 1.724 |
| chr7 | 77669354 | 77669898 | 1.90195 | 3.27838 | 1.724 |
| chr14 | 84652264 | 84652490 | 0.7213 | 1.24331 | 1.724 |
| chr5 | 115166900 | 115167173 | 1.57004 | 2.70639 | 1.724 |
| chr8 | 48909799 | 48910199 | 2.2183 | 3.82392 | 1.724 |
| chr2 | 119884238 | 119884473 | 1.53624 | 2.64835 | 1.724 |
| chr19 | 49540074 | 49540363 | 1.12963 | 1.94742 | 1.724 |
| chr20 | 6167282 | 6167741 | 0.542793 | 0.935759 | 1.724 |
| chr2 | 48827454 | 48827708 | 1.5865 | 2.73526 | 1.724 |
| chr7 | 73043606 | 73043843 | 1.24803 | 2.15172 | 1.724 |
| chr8 | 11759752 | 11760083 | 2.44984 | 4.22375 | 1.724 |
| chr12 | 122296424 | 122296829 | 0.507798 | 0.875581 | 1.724 |
| chr7 | 156442843 | 156443112 | 1.03562 | 1.78578 | 1.724 |

| | | | | | |
|---|---|---|---|---|---|
| chr17 | 5610165 | 5610550 | 1.69631 | 2.92511 | 1.724 |
| chr5 | 180361401 | 180361637 | 0.721989 | 1.24508 | 1.725 |
| chr13 | 20756443 | 20756723 | 1.94884 | 3.36083 | 1.725 |
| chr14 | 69709348 | 69709762 | 1.07829 | 1.85954 | 1.725 |
| chr6 | 35520077 | 35520443 | 1.44472 | 2.49147 | 1.725 |
| chr16 | 146827 | 147271 | 3.17486 | 5.47518 | 1.725 |
| chr8 | 1926224 | 1926674 | 1.0413 | 1.79582 | 1.725 |
| chr12 | 115157000 | 115157207 | 0.486383 | 0.838819 | 1.725 |
| chr7 | 116853078 | 116853384 | 1.80483 | 3.11266 | 1.725 |
| chr14 | 99762905 | 99763122 | 1.21628 | 2.0977 | 1.725 |
| chr8 | 21095874 | 21096261 | 0.588733 | 1.01539 | 1.725 |
| chr9 | 71808111 | 71808364 | 1.24387 | 2.14531 | 1.725 |
| chr3 | 168867147 | 168867454 | 1.78713 | 3.08231 | 1.725 |
| chr17 | 38115219 | 38115540 | 2.19788 | 3.79092 | 1.725 |
| chr1 | 160804649 | 160805070 | 1.4025 | 2.4191 | 1.725 |
| chr4 | 27049980 | 27050226 | 1.00692 | 1.73689 | 1.725 |
| chr11 | 5744355 | 5744706 | 0.467651 | 0.80669 | 1.725 |
| chr17 | 50465150 | 50465417 | 0.605634 | 1.04471 | 1.725 |
| chr12 | 49656626 | 49656865 | 1.17206 | 2.0219 | 1.725 |
| chr7 | 4055159 | 4055366 | 0.561146 | 0.968057 | 1.725 |
| chr9 | 78633356 | 78633563 | 0.600401 | 1.03579 | 1.725 |
| chr15 | 65378715 | 65379071 | 1.60475 | 2.76852 | 1.725 |
| chr8 | 82214876 | 82215130 | 1.03095 | 1.77867 | 1.725 |
| chr19 | 51215291 | 51215590 | 0.538644 | 0.929324 | 1.725 |
| chr17 | 37264399 | 37264629 | 0.662821 | 1.14358 | 1.725 |
| chr11 | 74991218 | 74991605 | 2.94217 | 5.07633 | 1.725 |
| chr12 | 122941778 | 122942053 | 0.423984 | 0.731561 | 1.725 |
| chr5 | 98103861 | 98104196 | 2.04403 | 3.52689 | 1.725 |
| chr11 | 59304729 | 59305008 | 1.71385 | 2.95721 | 1.725 |
| chr1 | 155248185 | 155248470 | 1.01896 | 1.75828 | 1.726 |
| chr2 | 166150275 | 166150515 | 1.6222 | 2.79923 | 1.726 |
| chr17 | 1050781 | 1051150 | 1.95334 | 3.37073 | 1.726 |
| chr17 | 74710913 | 74711374 | 1.99654 | 3.44548 | 1.726 |
| chr5 | 10046909 | 10047372 | 0.530495 | 0.915497 | 1.726 |
| chr1 | 9241334 | 9241723 | 1.96205 | 3.38635 | 1.726 |
| chr17 | 37874973 | 37875307 | 1.52566 | 2.63318 | 1.726 |
| chr18 | 29738019 | 29738278 | 1.34113 | 2.31474 | 1.726 |
| chr19 | 40886496 | 40886722 | 0.577629 | 0.996977 | 1.726 |
| chr17 | 72919539 | 72920090 | 3.25486 | 5.61801 | 1.726 |
| chr5 | 135547849 | 135548310 | 2.19761 | 3.79322 | 1.726 |
| chr4 | 38070858 | 38071146 | 1.24934 | 2.15647 | 1.726 |
| chr22 | 24571132 | 24571506 | 2.70453 | 4.66843 | 1.726 |
| chr1 | 91187031 | 91187313 | 2.19499 | 3.78896 | 1.726 |
| chr1 | 156627249 | 156627597 | 2.60821 | 4.5023 | 1.726 |
| chr19 | 50175423 | 50176005 | 2.79334 | 4.82197 | 1.726 |
| chr8 | 128738009 | 128738331 | 1.45156 | 2.50576 | 1.726 |
| chr16 | 28857760 | 28857983 | 1.65185 | 2.85152 | 1.726 |
| chr16 | 3988988 | 3989536 | 3.12222 | 5.38995 | 1.726 |
| chr14 | 71605750 | 71606115 | 2.40658 | 4.15455 | 1.726 |
| chr5 | 172405813 | 172406105 | 1.78518 | 3.08183 | 1.726 |
| chr10 | 111449536 | 111449769 | 0.516904 | 0.892379 | 1.726 |
| chr19 | 59030876 | 59031652 | 1.43022 | 2.46913 | 1.726 |
| chr7 | 98050278 | 98050779 | 2.83568 | 4.89552 | 1.726 |
| chr6 | 100526450 | 100526728 | 1.12833 | 1.94798 | 1.726 |
| chr22 | 36697319 | 36697654 | 1.71093 | 2.95383 | 1.726 |
| chr15 | 32223439 | 32223658 | 0.6655 | 1.14897 | 1.726 |
| chr15 | 60472011 | 60472616 | 1.31352 | 2.26782 | 1.727 |
| chr7 | 134440237 | 134440491 | 0.994254 | 1.71664 | 1.727 |
| chr7 | 128482029 | 128482617 | 2.00646 | 3.46429 | 1.727 |
| chr3 | 81351175 | 81351440 | 1.16318 | 2.00834 | 1.727 |
| chr10 | 82203278 | 82203640 | 1.31756 | 2.27491 | 1.727 |

| | | | | | |
|---|---|---|---|---|---|
| chr9 | 125597495 | 125597722 | 1.00921 | 1.7426 | 1.727 |
| chr2 | 201577745 | 201577948 | 1.93217 | 3.33639 | 1.727 |
| chr3 | 133646296 | 133646777 | 1.66902 | 2.88201 | 1.727 |
| chr17 | 4690604 | 4690946 | 1.47802 | 2.55227 | 1.727 |
| chr11 | 26437907 | 26438137 | 0.626113 | 1.0812 | 1.727 |
| chr10 | 13441324 | 13441619 | 0.951268 | 1.6427 | 1.727 |
| chr19 | 6481990 | 6482261 | 1.50529 | 2.59948 | 1.727 |
| chr16 | 71679441 | 71679686 | 1.44599 | 2.49715 | 1.727 |
| chr16 | 25138092 | 25138400 | 1.30233 | 2.24909 | 1.727 |
| chr8 | 39020186 | 39020531 | 0.522599 | 0.902535 | 1.727 |
| chr8 | 702197 | 702497 | 0.753945 | 1.30208 | 1.727 |
| chr2 | 22695514 | 22695900 | 0.56734 | 0.979811 | 1.727 |
| chr11 | 66153652 | 66154052 | 1.72463 | 2.97859 | 1.727 |
| chr8 | 143705261 | 143705882 | 2.15794 | 3.72716 | 1.727 |
| chr1 | 2493430 | 2493692 | 1.1023 | 1.904 | 1.727 |
| chr7 | 20784575 | 20784834 | 1.14555 | 1.97875 | 1.727 |
| chr10 | 127030531 | 127030764 | 0.51656 | 0.89228 | 1.727 |
| chr1 | 237032080 | 237032501 | 1.7383 | 3.00276 | 1.727 |
| chr15 | 89959418 | 89960085 | 3.12241 | 5.3937 | 1.727 |
| chr17 | 11547594 | 11547997 | 1.53308 | 2.64836 | 1.727 |
| chr15 | 52424158 | 52424435 | 0.501068 | 0.86559 | 1.727 |
| chr7 | 127675793 | 127676074 | 0.840229 | 1.4515 | 1.728 |
| chr15 | 40566712 | 40567276 | 3.16289 | 5.46419 | 1.728 |
| chr2 | 34939330 | 34939706 | 0.62318 | 1.07662 | 1.728 |
| chr17 | 77879665 | 77879919 | 1.23032 | 2.1256 | 1.728 |
| chr3 | 13714771 | 13715090 | 1.27867 | 2.20914 | 1.728 |
| chr8 | 27323147 | 27323422 | 2.09192 | 3.61418 | 1.728 |
| chr1 | 165570874 | 165571142 | 1.54525 | 2.66978 | 1.728 |
| chr19 | 40220811 | 40221201 | 0.659203 | 1.13894 | 1.728 |
| chr14 | 65231423 | 65231807 | 2.58787 | 4.4715 | 1.728 |
| chr20 | 21517523 | 21517759 | 0.744214 | 1.28592 | 1.728 |
| chr11 | 19926852 | 19927148 | 1.39594 | 2.41216 | 1.728 |
| chr12 | 7830064 | 7830287 | 0.903267 | 1.5609 | 1.728 |
| chr10 | 101307805 | 101308002 | 1.1994 | 2.07267 | 1.728 |
| chr7 | 149389657 | 149389903 | 0.670471 | 1.15865 | 1.728 |
| chr6 | 108371163 | 108371379 | 1.36313 | 2.35571 | 1.728 |
| chr16 | 21095106 | 21095414 | 1.39293 | 2.40729 | 1.728 |
| chr9 | 23297124 | 23297412 | 0.655564 | 1.13296 | 1.728 |
| chr2 | 232406336 | 232406544 | 0.784767 | 1.35626 | 1.728 |
| chr9 | 17794253 | 17794522 | 1.12932 | 1.95172 | 1.728 |
| chr7 | 26318480 | 26318873 | 1.0553 | 1.82383 | 1.728 |
| chr14 | 34454307 | 34454568 | 0.697008 | 1.20462 | 1.728 |
| chr1 | 202284607 | 202285094 | 2.30831 | 3.98942 | 1.728 |
| chr10 | 55424314 | 55424746 | 0.506155 | 0.874801 | 1.728 |
| chr12 | 70272982 | 70273497 | 1.6543 | 2.85933 | 1.728 |
| chr4 | 122228998 | 122229235 | 1.58502 | 2.73979 | 1.729 |
| chr3 | 62187190 | 62187407 | 0.965529 | 1.66924 | 1.729 |
| chr8 | 31042780 | 31043106 | 1.52785 | 2.64149 | 1.729 |
| chr17 | 57082313 | 57082545 | 2.11227 | 3.65198 | 1.729 |
| chr4 | 155480675 | 155480913 | 1.23853 | 2.14135 | 1.729 |
| chr19 | 59030372 | 59030599 | 1.50154 | 2.59611 | 1.729 |
| chr16 | 65446879 | 65447112 | 0.495905 | 0.857435 | 1.729 |
| chr5 | 168692988 | 168693195 | 0.954327 | 1.65009 | 1.729 |
| chr22 | 42253637 | 42253861 | 1.74806 | 3.02258 | 1.729 |
| chr20 | 35067877 | 35068442 | 2.12314 | 3.67124 | 1.729 |
| chr12 | 100532208 | 100532420 | 1.34176 | 2.32015 | 1.729 |
| chr1 | 153793755 | 153794123 | 1.35876 | 2.34958 | 1.729 |
| chrX | 12585965 | 12586259 | 1.32076 | 2.28388 | 1.729 |
| chr5 | 164636184 | 164636386 | 0.83591 | 1.44551 | 1.729 |
| chr9 | 12224311 | 12224682 | 0.683647 | 1.18221 | 1.729 |
| chr19 | 31140925 | 31141317 | 2.89278 | 5.00244 | 1.729 |

| | | | | | |
|---|---|---|---|---|---|
| chrX | 71497026 | 71497275 | 1.7249 | 2.98285 | 1.729 |
| chr1 | 180157285 | 180157677 | 2.28768 | 3.9561 | 1.729 |
| chr5 | 52614230 | 52614434 | 1.02185 | 1.76711 | 1.729 |
| chrX | 43885840 | 43886044 | 1.37886 | 2.38455 | 1.729 |
| chr14 | 105881303 | 105881542 | 2.21854 | 3.83671 | 1.729 |
| chr16 | 19459179 | 19459461 | 0.836293 | 1.44634 | 1.729 |
| chr9 | 104431493 | 104431736 | 0.769027 | 1.33016 | 1.730 |
| chr5 | 180594856 | 180595169 | 2.43326 | 4.2088 | 1.730 |
| chr8 | 139429445 | 139429737 | 0.613081 | 1.06046 | 1.730 |
| chr11 | 6727076 | 6727474 | 2.06085 | 3.56488 | 1.730 |
| chr15 | 96897079 | 96897398 | 2.32379 | 4.01971 | 1.730 |
| chr6 | 43196985 | 43197401 | 1.80841 | 3.1282 | 1.730 |
| chr15 | 76102345 | 76102728 | 1.1137 | 1.92652 | 1.730 |
| chr19 | 49330077 | 49330341 | 1.07244 | 1.85515 | 1.730 |
| chr19 | 17814724 | 17815038 | 1.21125 | 2.0953 | 1.730 |
| chr11 | 134738512 | 134738848 | 2.15205 | 3.72286 | 1.730 |
| chr15 | 62941182 | 62941452 | 1.39318 | 2.41012 | 1.730 |
| chr19 | 41025253 | 41025792 | 1.07098 | 1.85273 | 1.730 |
| chr5 | 65221503 | 65221776 | 1.40583 | 2.432 | 1.730 |
| chr12 | 65524946 | 65525253 | 0.850094 | 1.47063 | 1.730 |
| chr2 | 239067350 | 239067651 | 1.68935 | 2.92259 | 1.730 |
| chr16 | 56352127 | 56352601 | 2.28427 | 3.95184 | 1.730 |
| chr14 | 65878971 | 65879282 | 1.26933 | 2.19598 | 1.730 |
| chr3 | 161126866 | 161127070 | 1.11276 | 1.92517 | 1.730 |
| chr19 | 58861794 | 58862011 | 0.5981 | 1.03482 | 1.730 |
| chr12 | 107842070 | 107842382 | 0.837171 | 1.44848 | 1.730 |
| chr2 | 69786875 | 69787122 | 1.36085 | 2.35463 | 1.730 |
| chr16 | 87357982 | 87358852 | 1.23989 | 2.1454 | 1.730 |
| chr19 | 51911025 | 51911333 | 1.53431 | 2.65493 | 1.730 |
| chr19 | 33210417 | 33210874 | 1.45921 | 2.52499 | 1.730 |
| chr7 | 93041487 | 93041902 | 0.687865 | 1.19029 | 1.730 |
| chr16 | 71780227 | 71780669 | 1.36346 | 2.35939 | 1.730 |
| chr16 | 75505707 | 75505961 | 1.766 | 3.05612 | 1.731 |
| chr1 | 195713275 | 195713522 | 0.778648 | 1.34763 | 1.731 |
| chr17 | 4806654 | 4807237 | 1.63881 | 2.83634 | 1.731 |
| chr5 | 38268428 | 38268659 | 0.635271 | 1.09951 | 1.731 |
| chr11 | 1409168 | 1409459 | 1.10113 | 1.90583 | 1.731 |
| chr6 | 119450455 | 119450836 | 1.50054 | 2.59712 | 1.731 |
| chr4 | 171180760 | 171181111 | 1.41285 | 2.44542 | 1.731 |
| chr5 | 135398327 | 135398687 | 1.28726 | 2.2281 | 1.731 |
| chr8 | 62722594 | 62722852 | 1.40755 | 2.43631 | 1.731 |
| chr7 | 1223154 | 1223456 | 1.46794 | 2.54088 | 1.731 |
| chr17 | 38218866 | 38219244 | 1.34767 | 2.33276 | 1.731 |
| chr1 | 1891504 | 1891859 | 2.58994 | 4.48322 | 1.731 |
| chr12 | 20799196 | 20799477 | 1.5725 | 2.72204 | 1.731 |
| chr6 | 117638391 | 117638652 | 0.59001 | 1.02133 | 1.731 |
| chr10 | 13124641 | 13125124 | 0.541801 | 0.937921 | 1.731 |
| chr20 | 20905750 | 20905992 | 0.781412 | 1.35274 | 1.731 |
| chr20 | 9893522 | 9893853 | 2.06086 | 3.56771 | 1.731 |
| chr1 | 151032052 | 151032288 | 1.47252 | 2.54924 | 1.731 |
| chr8 | 136670134 | 136670339 | 0.550504 | 0.953095 | 1.731 |
| chr2 | 206768497 | 206768764 | 1.19491 | 2.06884 | 1.731 |
| chr14 | 102701557 | 102701974 | 1.67858 | 2.90627 | 1.731 |
| chr3 | 46965020 | 46965423 | 1.90025 | 3.29013 | 1.731 |
| chr8 | 131687765 | 131688104 | 1.98819 | 3.44245 | 1.731 |
| chr12 | 54833478 | 54833880 | 1.26065 | 2.18279 | 1.731 |
| chr4 | 178030027 | 178030304 | 0.63392 | 1.09773 | 1.732 |
| chr1 | 36348470 | 36348877 | 2.3703 | 4.10457 | 1.732 |
| chr12 | 115915939 | 115916147 | 0.835536 | 1.44691 | 1.732 |
| chr1 | 162673556 | 162673816 | 0.527745 | 0.913954 | 1.732 |
| chr7 | 41736069 | 41736390 | 1.4029 | 2.42955 | 1.732 |

| | | | | | |
|---|---|---|---|---|---|
| chr11 | 114133430 | 114133753 | 1.6249 | 2.81409 | 1.732 |
| chr19 | 54735533 | 54735926 | 2.29563 | 3.97585 | 1.732 |
| chr10 | 62576391 | 62576632 | 1.44215 | 2.49773 | 1.732 |
| chr8 | 140238320 | 140238526 | 0.789875 | 1.36807 | 1.732 |
| chr6 | 15441551 | 15441801 | 1.77179 | 3.0689 | 1.732 |
| chr15 | 73992649 | 73992935 | 1.96509 | 3.40377 | 1.732 |
| chr1 | 204732906 | 204733276 | 2.07306 | 3.59081 | 1.732 |
| chr21 | 47563479 | 47563718 | 1.84499 | 3.19577 | 1.732 |
| chr12 | 60766748 | 60767025 | 0.736762 | 1.2762 | 1.732 |
| chr4 | 103953674 | 103954037 | 1.44657 | 2.5057 | 1.732 |
| chr7 | 75943550 | 75943966 | 3.35685 | 5.81465 | 1.732 |
| chr2 | 173729003 | 173729246 | 1.11968 | 1.93949 | 1.732 |
| chr4 | 119693319 | 119693526 | 0.869399 | 1.50596 | 1.732 |
| chr14 | 75411927 | 75412395 | 1.86076 | 3.22324 | 1.732 |
| chr1 | 204749565 | 204749811 | 0.444856 | 0.770606 | 1.732 |
| chr12 | 54399745 | 54400090 | 1.85009 | 3.20484 | 1.732 |
| chr22 | 42869574 | 42869822 | 0.854012 | 1.47946 | 1.732 |
| chr20 | 36621429 | 36621642 | 1.62966 | 2.82317 | 1.732 |
| chr5 | 11318172 | 11318436 | 0.737642 | 1.27788 | 1.732 |
| chr3 | 119098701 | 119099001 | 0.572214 | 0.991305 | 1.732 |
| chr5 | 143521896 | 143522155 | 1.6983 | 2.94215 | 1.732 |
| chr2 | 239693599 | 239694330 | 1.10237 | 1.90977 | 1.732 |
| chr19 | 51227574 | 51228160 | 1.98781 | 3.44375 | 1.732 |
| chr6 | 36985988 | 36986689 | 1.31278 | 2.27435 | 1.732 |
| chr1 | 156675682 | 156676298 | 1.84512 | 3.19673 | 1.733 |
| chr14 | 92325078 | 92325444 | 0.827215 | 1.43319 | 1.733 |
| chr1 | 228447567 | 228447968 | 1.03976 | 1.80145 | 1.733 |
| chr3 | 78406466 | 78406795 | 0.576592 | 0.998991 | 1.733 |
| chr9 | 97943600 | 97943830 | 1.35919 | 2.35498 | 1.733 |
| chr22 | 36682563 | 36682973 | 1.98147 | 3.43342 | 1.733 |
| chr5 | 138119757 | 138119964 | 0.679004 | 1.17655 | 1.733 |
| chr11 | 75039728 | 75040021 | 1.56016 | 2.70346 | 1.733 |
| chr9 | 119456179 | 119456424 | 0.587447 | 1.01805 | 1.733 |
| chr11 | 17410589 | 17411088 | 2.21003 | 3.83031 | 1.733 |
| chr10 | 76868467 | 76868771 | 1.44277 | 2.50065 | 1.733 |
| chr22 | 30682815 | 30683169 | 2.40849 | 4.17456 | 1.733 |
| chr1 | 91630740 | 91631121 | 1.48858 | 2.58013 | 1.733 |
| chr6 | 154135351 | 154135639 | 0.510222 | 0.884361 | 1.733 |
| chr3 | 42271316 | 42271655 | 1.12337 | 1.9472 | 1.733 |
| chr7 | 128065779 | 128066245 | 1.97205 | 3.41836 | 1.733 |
| chr10 | 129322603 | 129322864 | 0.804374 | 1.39432 | 1.733 |
| chr5 | 84478297 | 84478534 | 0.62948 | 1.09116 | 1.733 |
| chr8 | 144025020 | 144025433 | 1.56407 | 2.71139 | 1.734 |
| chr4 | 170731578 | 170731800 | 1.36887 | 2.37308 | 1.734 |
| chr9 | 85949200 | 85949502 | 2.50812 | 4.34818 | 1.734 |
| chr1 | 231163776 | 231163983 | 0.641044 | 1.11137 | 1.734 |
| chr4 | 114199509 | 114199862 | 1.7598 | 3.05108 | 1.734 |
| chr5 | 8864254 | 8864471 | 0.743317 | 1.28878 | 1.734 |
| chr8 | 125938422 | 125938649 | 0.679043 | 1.17734 | 1.734 |
| chr7 | 27275014 | 27275707 | 0.925817 | 1.60526 | 1.734 |
| chr9 | 132374397 | 132374723 | 0.652755 | 1.1318 | 1.734 |
| chr5 | 107038296 | 107038581 | 0.674444 | 1.16943 | 1.734 |
| chr5 | 170058909 | 170060377 | 1.41073 | 2.44615 | 1.734 |
| chr12 | 7265867 | 7266302 | 1.55808 | 2.70166 | 1.734 |
| chr3 | 66872732 | 66872992 | 1.29714 | 2.2494 | 1.734 |
| chr9 | 95264240 | 95264650 | 1.62996 | 2.82676 | 1.734 |
| chr1 | 55416356 | 55416574 | 1.05169 | 1.8239 | 1.734 |
| chr14 | 23369433 | 23369703 | 1.55087 | 2.68971 | 1.734 |
| chr1 | 238064675 | 238064877 | 1.51526 | 2.62801 | 1.734 |
| chr8 | 55260193 | 55260533 | 0.580171 | 1.00623 | 1.734 |
| chr6 | 5115468 | 5115701 | 0.792054 | 1.37377 | 1.734 |

| | | | | | |
|---|---|---|---|---|---|
| chr1 | 67813417 | 67813699 | 1.12635 | 1.95361 | 1.734 |
| chr17 | 7080587 | 7081085 | 2.09398 | 3.63212 | 1.735 |
| chr1 | 32254908 | 32255164 | 1.43449 | 2.48831 | 1.735 |
| chr5 | 142260028 | 142260441 | 1.53537 | 2.66354 | 1.735 |
| chr15 | 98964773 | 98965214 | 2.12616 | 3.6887 | 1.735 |
| chr6 | 142887808 | 142888299 | 1.7773 | 3.08352 | 1.735 |
| chr11 | 11683086 | 11683295 | 0.649724 | 1.12732 | 1.735 |
| chr15 | 77032281 | 77032487 | 0.668449 | 1.15982 | 1.735 |
| chr5 | 5497308 | 5497613 | 1.56886 | 2.72214 | 1.735 |
| chr19 | 51102596 | 51102985 | 0.717575 | 1.24512 | 1.735 |
| chr2 | 192431043 | 192431393 | 1.64463 | 2.8538 | 1.735 |
| chr5 | 114645502 | 114645725 | 0.789247 | 1.36953 | 1.735 |
| chr5 | 22415444 | 22415715 | 0.646017 | 1.12102 | 1.735 |
| chr5 | 153559763 | 153559991 | 1.53028 | 2.6555 | 1.735 |
| chr10 | 24906635 | 24907050 | 2.52292 | 4.37808 | 1.735 |
| chr2 | 137142333 | 137142557 | 1.17711 | 2.04271 | 1.735 |
| chr12 | 5378004 | 5378298 | 0.603215 | 1.04686 | 1.735 |
| chr15 | 101547163 | 101547469 | 2.38326 | 4.13611 | 1.735 |
| chr5 | 74345666 | 74346022 | 1.73583 | 3.01265 | 1.736 |
| chr14 | 53833099 | 53833375 | 0.664022 | 1.15252 | 1.736 |
| chr15 | 74189802 | 74190147 | 1.95157 | 3.38731 | 1.736 |
| chr3 | 133649238 | 133649473 | 0.999838 | 1.7354 | 1.736 |
| chr7 | 43878426 | 43878915 | 3.80963 | 6.6123 | 1.736 |
| chr8 | 26513030 | 26513313 | 1.70956 | 2.96725 | 1.736 |
| chr3 | 48193048 | 48193329 | 1.09959 | 1.90855 | 1.736 |
| chr12 | 81487063 | 81487277 | 0.668385 | 1.16023 | 1.736 |
| chr10 | 71186678 | 71187141 | 0.93068 | 1.61557 | 1.736 |
| chr9 | 132373355 | 132373779 | 1.71334 | 2.97418 | 1.736 |
| chr1 | 158094816 | 158095104 | 1.57274 | 2.73016 | 1.736 |
| chr12 | 133100408 | 133100900 | 1.65613 | 2.87504 | 1.736 |
| chr6 | 138608395 | 138608712 | 1.53309 | 2.66145 | 1.736 |
| chr2 | 121310491 | 121310783 | 2.06335 | 3.58204 | 1.736 |
| chr10 | 21567713 | 21567987 | 1.54851 | 2.68835 | 1.736 |
| chr19 | 47930407 | 47930830 | 2.08041 | 3.61178 | 1.736 |
| chr4 | 84173954 | 84174308 | 1.57404 | 2.73272 | 1.736 |
| chr10 | 76859657 | 76860097 | 2.48143 | 4.30822 | 1.736 |
| chr19 | 55652455 | 55652911 | 1.41585 | 2.45821 | 1.736 |
| chr5 | 58484352 | 58484569 | 1.60023 | 2.77833 | 1.736 |
| chrX | 150528601 | 150529166 | 1.35476 | 2.35215 | 1.736 |
| chr5 | 35926458 | 35927093 | 0.958896 | 1.66488 | 1.736 |
| chr8 | 92968446 | 92968682 | 0.593646 | 1.03074 | 1.736 |
| chr10 | 89494956 | 89495297 | 1.78227 | 3.09466 | 1.736 |
| chr11 | 74855382 | 74856018 | 1.25371 | 2.17689 | 1.736 |
| chr9 | 124754994 | 124755208 | 1.29715 | 2.25232 | 1.736 |
| chr4 | 48965134 | 48965451 | 1.96098 | 3.40506 | 1.736 |
| chr6 | 147712210 | 147712419 | 0.678401 | 1.17798 | 1.736 |
| chr6 | 44205662 | 44206024 | 2.31031 | 4.01182 | 1.736 |
| chr5 | 81134420 | 81134699 | 1.02642 | 1.78244 | 1.737 |
| chr8 | 138032441 | 138032753 | 0.5971 | 1.0369 | 1.737 |
| chr5 | 14270359 | 14270657 | 1.75418 | 3.04626 | 1.737 |
| chr16 | 2888809 | 2889228 | 2.16058 | 3.75215 | 1.737 |
| chr9 | 134053710 | 134054079 | 1.96626 | 3.41477 | 1.737 |
| chr1 | 228703022 | 228703385 | 1.20109 | 2.08602 | 1.737 |
| chr2 | 166984475 | 166984682 | 1.25436 | 2.17864 | 1.737 |
| chr5 | 37723602 | 37723922 | 1.15839 | 2.01201 | 1.737 |
| chr2 | 60484377 | 60484689 | 0.677096 | 1.17606 | 1.737 |
| chr1 | 220363173 | 220363418 | 1.19162 | 2.06982 | 1.737 |
| chr5 | 162634388 | 162635101 | 0.553412 | 0.961263 | 1.737 |
| chr4 | 25182275 | 25182507 | 0.888106 | 1.54263 | 1.737 |
| chr7 | 128313028 | 128313273 | 1.38352 | 2.40318 | 1.737 |
| chr14 | 105442135 | 105442789 | 1.96957 | 3.42122 | 1.737 |

| | | | | | |
|---|---|---|---|---|---|
| chr13 | 73414098 | 73414285 | 1.51486 | 2.63138 | 1.737 |
| chr5 | 32489884 | 32490171 | 1.66567 | 2.89339 | 1.737 |
| chr9 | 11855852 | 11856124 | 0.603317 | 1.04801 | 1.737 |
| chr16 | 57682876 | 57683358 | 4.22772 | 7.34402 | 1.737 |
| chr15 | 65677105 | 65677588 | 2.77856 | 4.82698 | 1.737 |
| chr7 | 49329516 | 49329842 | 0.662393 | 1.15075 | 1.737 |
| chr18 | 56735213 | 56735514 | 0.876219 | 1.52229 | 1.737 |
| chr3 | 158031196 | 158031429 | 1.45424 | 2.52662 | 1.737 |
| chr7 | 6566018 | 6566529 | 1.55279 | 2.69787 | 1.737 |
| chr8 | 2229559 | 2229784 | 0.693486 | 1.20494 | 1.738 |
| chr3 | 50158926 | 50159305 | 1.24634 | 2.1656 | 1.738 |
| chr3 | 56521403 | 56521636 | 0.700292 | 1.21687 | 1.738 |
| chr18 | 2906673 | 2906879 | 1.03855 | 1.80467 | 1.738 |
| chr4 | 141516199 | 141516608 | 1.85331 | 3.22046 | 1.738 |
| chr16 | 62457373 | 62457736 | 0.478153 | 0.830913 | 1.738 |
| chr10 | 86297212 | 86297538 | 1.31565 | 2.28634 | 1.738 |
| chr8 | 89033334 | 89033539 | 0.70326 | 1.22218 | 1.738 |
| chr11 | 58483486 | 58483833 | 0.649732 | 1.12919 | 1.738 |
| chr17 | 12883456 | 12883666 | 0.506418 | 0.880124 | 1.738 |
| chr5 | 39826714 | 39826951 | 0.525418 | 0.913163 | 1.738 |
| chr12 | 103548669 | 103549068 | 0.644634 | 1.12036 | 1.738 |
| chr5 | 93654709 | 93654916 | 0.63261 | 1.09948 | 1.738 |
| chr5 | 65379912 | 65380142 | 0.807685 | 1.40377 | 1.738 |
| chr17 | 6323532 | 6323998 | 1.09737 | 1.90733 | 1.738 |
| chr9 | 4315654 | 4316060 | 2.46228 | 4.27987 | 1.738 |
| chr7 | 15929190 | 15929434 | 1.09704 | 1.90691 | 1.738 |
| chr2 | 38342424 | 38342720 | 2.88854 | 5.02104 | 1.738 |
| rUn_gl0002 | 87248 | 87534 | 0.706653 | 1.22836 | 1.738 |
| chr1 | 166407245 | 166407611 | 0.563467 | 0.979472 | 1.738 |
| chr5 | 103971459 | 103971738 | 0.462443 | 0.803871 | 1.738 |
| chr9 | 20977126 | 20977425 | 0.600326 | 1.04358 | 1.738 |
| chr3 | 116583244 | 116583528 | 1.11418 | 1.93686 | 1.738 |
| chr7 | 39422119 | 39422401 | 0.458123 | 0.796405 | 1.738 |
| chr8 | 52926950 | 52927195 | 1.40785 | 2.44745 | 1.738 |
| chr3 | 169780517 | 169780799 | 1.15074 | 2.00063 | 1.739 |
| chr19 | 14616274 | 14616589 | 1.98487 | 3.45087 | 1.739 |
| chr1 | 245316371 | 245316623 | 1.19976 | 2.08593 | 1.739 |
| chr9 | 124402277 | 124402707 | 1.78535 | 3.1042 | 1.739 |
| chr7 | 5515867 | 5516109 | 0.478104 | 0.831385 | 1.739 |
| chr15 | 53591426 | 53591735 | 0.909962 | 1.58244 | 1.739 |
| chr9 | 82946192 | 82946529 | 0.47536 | 0.82666 | 1.739 |
| chr11 | 27180380 | 27180704 | 1.95823 | 3.40545 | 1.739 |
| chr12 | 108631128 | 108631330 | 1.01301 | 1.76172 | 1.739 |
| chr10 | 94608361 | 94608573 | 0.827566 | 1.43923 | 1.739 |
| chr2 | 74477181 | 74477655 | 1.5113 | 2.62843 | 1.739 |
| chr3 | 55214976 | 55215555 | 0.861027 | 1.49748 | 1.739 |
| chr9 | 25658577 | 25658832 | 0.635174 | 1.10473 | 1.739 |
| chr1 | 231942478 | 231942714 | 1.38609 | 2.41081 | 1.739 |
| chr5 | 111482634 | 111482936 | 1.157 | 2.0124 | 1.739 |
| chr11 | 68868687 | 68869263 | 2.39277 | 4.16184 | 1.739 |
| chr8 | 142919545 | 142919903 | 0.610002 | 1.06101 | 1.739 |
| chr4 | 34424186 | 34424416 | 0.744891 | 1.29566 | 1.739 |
| chr6 | 118228110 | 118228437 | 1.09251 | 1.90035 | 1.739 |
| chr12 | 118287784 | 118288091 | 1.97284 | 3.43186 | 1.740 |
| chr2 | 138224915 | 138225145 | 0.842584 | 1.46575 | 1.740 |
| chr13 | 49855269 | 49855542 | 1.45767 | 2.53579 | 1.740 |
| chr12 | 117346911 | 117347205 | 0.804271 | 1.39913 | 1.740 |
| chr12 | 9043087 | 9043649 | 1.87994 | 3.27044 | 1.740 |
| chr3 | 159334087 | 159334308 | 1.70825 | 2.972 | 1.740 |
| chr9 | 87873720 | 87874027 | 2.21667 | 3.85686 | 1.740 |
| chr10 | 126092113 | 126092340 | 1.43865 | 2.5032 | 1.740 |

| | | | | | |
|---|---|---|---|---|---|
| chr11 | 59590488 | 59590805 | 1.26043 | 2.19312 | 1.740 |
| chr1 | 27895046 | 27895434 | 1.72056 | 2.99375 | 1.740 |
| chr17 | 45928290 | 45928503 | 0.813286 | 1.41512 | 1.740 |
| chr16 | 58160667 | 58161105 | 0.655949 | 1.14138 | 1.740 |
| chr2 | 86595411 | 86595779 | 1.63822 | 2.85058 | 1.740 |
| chr16 | 71644913 | 71645310 | 3.01714 | 5.25003 | 1.740 |
| chr14 | 67886816 | 67887211 | 1.55867 | 2.7123 | 1.740 |
| chrX | 153044281 | 153044581 | 1.29903 | 2.26053 | 1.740 |
| chr9 | 79611703 | 79611956 | 1.59705 | 2.77938 | 1.740 |
| chr3 | 130478594 | 130478897 | 1.97151 | 3.43126 | 1.740 |
| chr20 | 24506080 | 24506315 | 0.624809 | 1.08744 | 1.740 |
| chr6 | 119215107 | 119215366 | 1.62441 | 2.8273 | 1.741 |
| chr14 | 87139729 | 87140068 | 0.70182 | 1.22156 | 1.741 |
| chr3 | 125156667 | 125157027 | 2.3156 | 4.0306 | 1.741 |
| chr4 | 7337545 | 7337808 | 2.38769 | 4.15635 | 1.741 |
| chr3 | 58554323 | 58554814 | 1.8901 | 3.29032 | 1.741 |
| chr7 | 99588494 | 99588893 | 2.65406 | 4.62024 | 1.741 |
| chr9 | 95553071 | 95553386 | 1.68676 | 2.93634 | 1.741 |
| chr1 | 18071628 | 18071938 | 0.733067 | 1.27619 | 1.741 |
| chr14 | 33180380 | 33180598 | 1.27385 | 2.21768 | 1.741 |
| chr8 | 50208196 | 50208411 | 0.752146 | 1.30951 | 1.741 |
| chr20 | 19783355 | 19783761 | 1.14405 | 1.99192 | 1.741 |
| chr1 | 6761779 | 6762170 | 2.26717 | 3.94746 | 1.741 |
| chr16 | 29859778 | 29860195 | 2.74246 | 4.77511 | 1.741 |
| chr4 | 54670911 | 54671229 | 1.56839 | 2.73085 | 1.741 |
| chr14 | 101176078 | 101176351 | 1.3041 | 2.27069 | 1.741 |
| chr12 | 115982203 | 115982609 | 0.637942 | 1.1108 | 1.741 |
| chr15 | 100889780 | 100890681 | 2.66058 | 4.63269 | 1.741 |
| chr11 | 3058443 | 3058926 | 2.39812 | 4.17584 | 1.741 |
| chr7 | 138330070 | 138330280 | 0.805973 | 1.40356 | 1.741 |
| chr17 | 6316434 | 6316701 | 1.46293 | 2.54767 | 1.741 |
| chr5 | 15003580 | 15003876 | 1.4955 | 2.6044 | 1.741 |
| chr11 | 70329088 | 70329368 | 0.720725 | 1.25515 | 1.742 |
| chr4 | 17474014 | 17474289 | 1.41821 | 2.46984 | 1.742 |
| chr9 | 130656625 | 130656891 | 0.671291 | 1.16909 | 1.742 |
| chr1 | 170323418 | 170323639 | 0.661466 | 1.15198 | 1.742 |
| chr8 | 37782093 | 37782671 | 1.31606 | 2.29202 | 1.742 |
| chr3 | 19904827 | 19905044 | 0.666057 | 1.16013 | 1.742 |
| chr6 | 21181193 | 21181396 | 0.587839 | 1.02391 | 1.742 |
| chr12 | 50333539 | 50333755 | 0.670091 | 1.16729 | 1.742 |
| chr11 | 36287570 | 36287878 | 1.55161 | 2.703 | 1.742 |
| chr20 | 36793618 | 36794023 | 2.20486 | 3.84109 | 1.742 |
| chr9 | 85814172 | 85814524 | 1.49138 | 2.59813 | 1.742 |
| chr3 | 130049059 | 130049291 | 0.56578 | 0.985741 | 1.742 |
| chr11 | 2326508 | 2326974 | 2.22574 | 3.87787 | 1.742 |
| chr17 | 10261153 | 10261363 | 1.28468 | 2.23829 | 1.742 |
| chr12 | 111374873 | 111375253 | 3.21893 | 5.60833 | 1.742 |
| chr10 | 54404491 | 54404745 | 1.52441 | 2.656 | 1.742 |
| chr17 | 40558004 | 40558451 | 2.9925 | 5.21401 | 1.742 |
| chr12 | 62248854 | 62249114 | 0.779722 | 1.35863 | 1.742 |
| chr16 | 56345227 | 56345444 | 0.932787 | 1.62534 | 1.742 |
| chr5 | 50452853 | 50453070 | 0.715097 | 1.24606 | 1.743 |
| chr19 | 38875158 | 38875722 | 1.62778 | 2.83643 | 1.743 |
| chr5 | 136410899 | 136411203 | 1.94136 | 3.38301 | 1.743 |
| chr19 | 29468832 | 29469188 | 0.870374 | 1.51681 | 1.743 |
| chr1 | 237963240 | 237963516 | 1.16316 | 2.02708 | 1.743 |
| chr10 | 2981172 | 2981445 | 0.896667 | 1.56266 | 1.743 |
| chr12 | 107011645 | 107011861 | 0.471207 | 0.821191 | 1.743 |
| chr12 | 112130731 | 112130999 | 1.31372 | 2.28947 | 1.743 |
| chr17 | 72433841 | 72434138 | 1.86906 | 3.25733 | 1.743 |
| chr5 | 36178140 | 36178361 | 0.908879 | 1.58398 | 1.743 |

| | | | | | |
|---|---|---|---|---|---|
| chr2 | 196943093 | 196943345 | 0.927023 | 1.61571 | 1.743 |
| chr11 | 119076547 | 119076822 | 1.75392 | 3.05699 | 1.743 |
| chr17 | 41653957 | 41654392 | 1.55431 | 2.7092 | 1.743 |
| chr16 | 67461893 | 67462464 | 2.03007 | 3.5386 | 1.743 |
| chr4 | 42373551 | 42373761 | 1.36184 | 2.37388 | 1.743 |
| chr20 | 62073515 | 62073824 | 1.75072 | 3.05183 | 1.743 |
| chr15 | 27262367 | 27262884 | 2.30585 | 4.0196 | 1.743 |
| chr5 | 164654411 | 164654762 | 0.6191 | 1.07924 | 1.743 |
| chr19 | 48316669 | 48317343 | 1.49597 | 2.60786 | 1.743 |
| chr12 | 85818672 | 85818949 | 1.29816 | 2.26308 | 1.743 |
| chr3 | 71779066 | 71779317 | 2.21698 | 3.86489 | 1.743 |
| chr9 | 95396950 | 95397816 | 1.87538 | 3.26938 | 1.743 |
| chr12 | 6647028 | 6647395 | 3.22859 | 5.62862 | 1.743 |
| chr16 | 49890970 | 49891229 | 0.981444 | 1.71108 | 1.743 |
| chr7 | 73466039 | 73466314 | 1.60594 | 2.79984 | 1.743 |
| chr20 | 48755120 | 48755330 | 0.842627 | 1.46912 | 1.744 |
| chr1 | 167370546 | 167370860 | 1.40725 | 2.45363 | 1.744 |
| chr1 | 183679119 | 183679349 | 0.505052 | 0.880586 | 1.744 |
| chr3 | 142840248 | 142840469 | 1.47398 | 2.56997 | 1.744 |
| chrX | 152915185 | 152915430 | 1.76979 | 3.08582 | 1.744 |
| chr4 | 134067805 | 134068192 | 2.2583 | 3.93765 | 1.744 |
| chr15 | 94034656 | 94035021 | 0.569247 | 0.992571 | 1.744 |
| chr1 | 11538760 | 11539009 | 1.65117 | 2.87918 | 1.744 |
| chr3 | 50283243 | 50284007 | 2.42821 | 4.23421 | 1.744 |
| chr11 | 36514303 | 36514619 | 1.76557 | 3.07874 | 1.744 |
| chr8 | 59447379 | 59447892 | 1.83954 | 3.20783 | 1.744 |
| chr20 | 56754588 | 56755004 | 2.62578 | 4.57899 | 1.744 |
| chr9 | 85210583 | 85210795 | 0.765415 | 1.33481 | 1.744 |
| chr16 | 51371689 | 51371901 | 0.848605 | 1.47997 | 1.744 |
| chr13 | 100089099 | 100089505 | 3.71136 | 6.47269 | 1.744 |
| chr8 | 107844179 | 107844382 | 0.69663 | 1.21496 | 1.744 |
| chr11 | 102217770 | 102218015 | 1.91004 | 3.33123 | 1.744 |
| chr7 | 12558740 | 12559110 | 1.13894 | 1.98639 | 1.744 |
| chr7 | 25019067 | 25019311 | 1.28791 | 2.24622 | 1.744 |
| chr9 | 82962732 | 82962994 | 0.530877 | 0.925902 | 1.744 |
| chr8 | 92703671 | 92703917 | 0.476643 | 0.831344 | 1.744 |
| chr11 | 40677007 | 40677211 | 0.432831 | 0.75495 | 1.744 |
| chr14 | 74100128 | 74100453 | 1.90879 | 3.32939 | 1.744 |
| chr15 | 101389789 | 101390031 | 1.76421 | 3.0772 | 1.744 |
| chr7 | 133701477 | 133701893 | 1.65798 | 2.89202 | 1.744 |
| chr9 | 74421758 | 74422074 | 2.28885 | 3.99258 | 1.744 |
| chr12 | 23319097 | 23319403 | 0.630544 | 1.0999 | 1.744 |
| chr16 | 11312403 | 11312664 | 1.2714 | 2.2178 | 1.744 |
| chr9 | 131865386 | 131865816 | 1.85624 | 3.23803 | 1.744 |
| chr2 | 65144829 | 65145098 | 2.35019 | 4.0997 | 1.744 |
| chr5 | 71852472 | 71852931 | 0.797897 | 1.39188 | 1.744 |
| chr5 | 159448890 | 159449155 | 1.6893 | 2.94688 | 1.744 |
| chr2 | 70439788 | 70440010 | 0.643531 | 1.12263 | 1.744 |
| chr17 | 38183210 | 38183720 | 2.55379 | 4.45533 | 1.745 |
| chr8 | 82620875 | 82621179 | 1.31799 | 2.29939 | 1.745 |
| chr11 | 30688678 | 30688929 | 0.635231 | 1.10824 | 1.745 |
| chr15 | 56210115 | 56210367 | 1.51154 | 2.63711 | 1.745 |
| chr10 | 834969 | 835679 | 2.0919 | 3.64967 | 1.745 |
| chr1 | 1376173 | 1376776 | 1.47179 | 2.56788 | 1.745 |
| chr16 | 66358751 | 66359057 | 1.17989 | 2.05863 | 1.745 |
| chr5 | 137078776 | 137078981 | 1.47962 | 2.58166 | 1.745 |
| chr15 | 67145715 | 67146070 | 2.28754 | 3.99139 | 1.745 |
| chr5 | 95116808 | 95117087 | 1.71227 | 2.98772 | 1.745 |
| chr17 | 42856411 | 42857039 | 2.95547 | 5.15704 | 1.745 |
| chr11 | 64455812 | 64456416 | 2.2733 | 3.96672 | 1.745 |
| chr1 | 242183166 | 242183462 | 1.05087 | 1.8337 | 1.745 |

| | | | | | |
|---|---|---|---|---|---|
| chr8 | 124232836 | 124233148 | 1.40588 | 2.45324 | 1.745 |
| chr10 | 85714468 | 85714680 | 0.533515 | 0.930996 | 1.745 |
| chr6 | 161154117 | 161154384 | 1.34551 | 2.34821 | 1.745 |
| chr17 | 46740407 | 46740729 | 0.718826 | 1.25452 | 1.745 |
| chr17 | 77015644 | 77015917 | 1.43714 | 2.50823 | 1.745 |
| chr8 | 142070070 | 142070533 | 1.27549 | 2.2261 | 1.745 |
| chr11 | 29590086 | 29590340 | 0.62257 | 1.08661 | 1.745 |
| chr1 | 228870788 | 228871288 | 1.6876 | 2.94552 | 1.745 |
| chr1 | 236006754 | 236006993 | 1.54004 | 2.68797 | 1.745 |
| chr17 | 41843737 | 41844154 | 2.57552 | 4.49545 | 1.745 |
| chrX | 69353104 | 69353396 | 1.91277 | 3.33872 | 1.745 |
| chr12 | 57881520 | 57881739 | 1.03415 | 1.80513 | 1.746 |
| chr1 | 160959943 | 160960445 | 2.22564 | 3.885 | 1.746 |
| chr2 | 144016983 | 144017380 | 1.17237 | 2.04646 | 1.746 |
| chr8 | 138851073 | 138851287 | 0.59105 | 1.03177 | 1.746 |
| chr16 | 67917598 | 67918205 | 2.38495 | 4.16334 | 1.746 |
| chr16 | 58528908 | 58529189 | 1.34775 | 2.35278 | 1.746 |
| chr15 | 68126188 | 68126528 | 1.15601 | 2.01815 | 1.746 |
| chr11 | 61021803 | 61022238 | 2.39351 | 4.17859 | 1.746 |
| chr8 | 134232653 | 134232902 | 0.675491 | 1.17928 | 1.746 |
| chr11 | 66445932 | 66446329 | 1.8725 | 3.26926 | 1.746 |
| chr12 | 66652509 | 66652726 | 0.53133 | 0.927688 | 1.746 |
| chr15 | 67238958 | 67239345 | 1.4903 | 2.60206 | 1.746 |
| chr7 | 95141054 | 95141387 | 2.01912 | 3.52547 | 1.746 |
| chr15 | 79561401 | 79561619 | 0.494889 | 0.86413 | 1.746 |
| chr11 | 24216421 | 24216621 | 0.753302 | 1.31539 | 1.746 |
| chr8 | 60855432 | 60855699 | 0.677454 | 1.18297 | 1.746 |
| chr7 | 37467230 | 37467483 | 0.548665 | 0.958132 | 1.746 |
| chr20 | 48729224 | 48729429 | 0.925395 | 1.61607 | 1.746 |
| chr4 | 122657950 | 122658181 | 1.06111 | 1.85321 | 1.746 |
| chr12 | 124339483 | 124339826 | 0.74563 | 1.30229 | 1.747 |
| chr5 | 133437599 | 133438056 | 3.00716 | 5.25246 | 1.747 |
| chrX | 37684489 | 37684802 | 1.60068 | 2.79583 | 1.747 |
| chr17 | 77848262 | 77848507 | 1.19024 | 2.07896 | 1.747 |
| chr14 | 25026272 | 25026493 | 0.578228 | 1.00998 | 1.747 |
| chr3 | 37855987 | 37856485 | 0.686592 | 1.19928 | 1.747 |
| chr9 | 132918936 | 132919309 | 1.41196 | 2.4664 | 1.747 |
| chr9 | 122621457 | 122621668 | 0.624547 | 1.09097 | 1.747 |
| chr8 | 120429027 | 120429369 | 1.73302 | 3.02729 | 1.747 |
| chr7 | 134922868 | 134923282 | 1.76906 | 3.0903 | 1.747 |
| chr17 | 16776506 | 16776941 | 2.18036 | 3.80886 | 1.747 |
| chr17 | 81048728 | 81048951 | 0.500002 | 0.873454 | 1.747 |
| chr8 | 30524392 | 30524782 | 1.26299 | 2.20632 | 1.747 |
| chr7 | 72756592 | 72756877 | 1.2759 | 2.22888 | 1.747 |
| chr2 | 44619565 | 44619784 | 1.03154 | 1.80202 | 1.747 |
| chr8 | 139069760 | 139069976 | 0.492087 | 0.859661 | 1.747 |
| chrX | 18969263 | 18969558 | 0.887026 | 1.54962 | 1.747 |
| chr8 | 72930783 | 72931256 | 1.93339 | 3.37765 | 1.747 |
| chr14 | 47980966 | 47981251 | 0.741576 | 1.29563 | 1.747 |
| chr20 | 2627352 | 2627768 | 1.2335 | 2.15508 | 1.747 |
| chr19 | 15261222 | 15261607 | 1.94319 | 3.39505 | 1.747 |
| chr19 | 42134906 | 42135109 | 0.623497 | 1.08937 | 1.747 |
| chr1 | 182038151 | 182038604 | 2.6615 | 4.65017 | 1.747 |
| chr11 | 32830791 | 32831166 | 1.77804 | 3.10661 | 1.747 |
| chr20 | 34035034 | 34035375 | 2.99497 | 5.23295 | 1.747 |
| chr9 | 928496 | 928727 | 1.20589 | 2.10717 | 1.747 |
| chr3 | 111434746 | 111435092 | 1.31864 | 2.30424 | 1.747 |
| chr16 | 55876799 | 55877040 | 1.43646 | 2.51019 | 1.747 |
| chr12 | 99392555 | 99392894 | 1.24959 | 2.18368 | 1.748 |
| chr10 | 135143269 | 135143642 | 2.40756 | 4.20745 | 1.748 |
| chr7 | 45961746 | 45962122 | 2.27476 | 3.97545 | 1.748 |

| | | | | | |
|---|---|---|---|---|---|
| chr17 | 47109563 | 47110305 | 1.9211 | 3.35755 | 1.748 |
| chr7 | 49581967 | 49582237 | 0.433926 | 0.758386 | 1.748 |
| chr9 | 137670633 | 137671138 | 2.38327 | 4.16539 | 1.748 |
| chr8 | 82448064 | 82448307 | 0.706503 | 1.23483 | 1.748 |
| chr1 | 233866400 | 233866948 | 0.661182 | 1.15563 | 1.748 |
| chr1 | 46347328 | 46347553 | 1.56835 | 2.74131 | 1.748 |
| chr1 | 207084374 | 207084757 | 1.46303 | 2.55722 | 1.748 |
| chr2 | 25923074 | 25923390 | 1.5982 | 2.79348 | 1.748 |
| chr8 | 11104507 | 11104711 | 1.16168 | 2.03049 | 1.748 |
| chr5 | 175173388 | 175173614 | 0.906905 | 1.58518 | 1.748 |
| chr6 | 131406224 | 131406460 | 0.809809 | 1.41548 | 1.748 |
| chr19 | 5987392 | 5987701 | 1.87828 | 3.28316 | 1.748 |
| chr21 | 34978103 | 34978374 | 1.60773 | 2.81024 | 1.748 |
| chr7 | 128523526 | 128524114 | 2.08631 | 3.64702 | 1.748 |
| chr2 | 54334130 | 54334436 | 1.51504 | 2.64842 | 1.748 |
| chr8 | 6167141 | 6167352 | 0.696268 | 1.21714 | 1.748 |
| chr11 | 70601544 | 70601966 | 2.91402 | 5.09403 | 1.748 |
| chr15 | 76478071 | 76478303 | 2.17726 | 3.80615 | 1.748 |
| chr9 | 140195772 | 140195991 | 0.928781 | 1.62383 | 1.748 |
| chr17 | 17291336 | 17291730 | 1.49409 | 2.61228 | 1.748 |
| chr19 | 13541810 | 13542107 | 1.73901 | 3.04053 | 1.748 |
| chr5 | 14364361 | 14365049 | 1.55585 | 2.72031 | 1.748 |
| chr14 | 99882077 | 99882371 | 1.87453 | 3.27752 | 1.748 |
| chr5 | 142042412 | 142042688 | 0.489953 | 0.856686 | 1.749 |
| chr15 | 70877557 | 70877910 | 2.21046 | 3.86511 | 1.749 |
| chr19 | 48292052 | 48292502 | 1.84436 | 3.22499 | 1.749 |
| chr5 | 54522216 | 54522609 | 1.94935 | 3.4088 | 1.749 |
| chr3 | 147795206 | 147795410 | 1.01559 | 1.77597 | 1.749 |
| chr3 | 154516861 | 154517130 | 0.629054 | 1.10004 | 1.749 |
| chr12 | 4926107 | 4926461 | 0.575186 | 1.00586 | 1.749 |
| chr18 | 56431510 | 56431888 | 1.49071 | 2.60693 | 1.749 |
| chr1 | 26471647 | 26471901 | 1.95276 | 3.41502 | 1.749 |
| chr16 | 10006665 | 10006948 | 0.520096 | 0.909554 | 1.749 |
| chr19 | 44064846 | 44065157 | 0.958561 | 1.67635 | 1.749 |
| chr7 | 134628634 | 134628842 | 1.1983 | 2.09561 | 1.749 |
| chr12 | 3150791 | 3151082 | 1.5949 | 2.78927 | 1.749 |
| chr5 | 150618846 | 150619113 | 1.98634 | 3.47387 | 1.749 |
| chr3 | 129118438 | 129118722 | 1.7007 | 2.97435 | 1.749 |
| rUn_gl0002 | 133934 | 134537 | 1.49352 | 2.61205 | 1.749 |
| chr2 | 41724103 | 41724314 | 0.988142 | 1.7282 | 1.749 |
| chr2 | 64276885 | 64277156 | 1.76974 | 3.09521 | 1.749 |
| chr12 | 120043332 | 120043793 | 0.496879 | 0.869092 | 1.749 |
| chr14 | 61026175 | 61026405 | 1.04782 | 1.83278 | 1.749 |
| chr8 | 69118875 | 69119095 | 0.794432 | 1.3896 | 1.749 |
| chr5 | 143337141 | 143337366 | 0.85486 | 1.49535 | 1.749 |
| chr8 | 121580445 | 121580682 | 0.632565 | 1.10653 | 1.749 |
| chr17 | 29716398 | 29716776 | 2.38274 | 4.16813 | 1.749 |
| chr4 | 129730711 | 129730922 | 1.05683 | 1.84882 | 1.749 |
| chr15 | 55940616 | 55940907 | 1.99345 | 3.48749 | 1.749 |
| chr4 | 77669952 | 77670181 | 1.03392 | 1.80885 | 1.750 |
| chr6 | 39197308 | 39197698 | 1.91828 | 3.35652 | 1.750 |
| chr11 | 35893881 | 35894093 | 0.509984 | 0.892359 | 1.750 |
| chr11 | 30097039 | 30097259 | 0.512408 | 0.896607 | 1.750 |
| chr11 | 46559392 | 46559780 | 1.99639 | 3.49338 | 1.750 |
| chr7 | 150398929 | 150399214 | 0.661658 | 1.15787 | 1.750 |
| chr17 | 73628936 | 73629511 | 2.29368 | 4.01408 | 1.750 |
| chr3 | 81715720 | 81716004 | 1.4777 | 2.58607 | 1.750 |
| chr15 | 101566191 | 101566603 | 0.591238 | 1.03473 | 1.750 |
| chr1 | 88646259 | 88646567 | 2.09956 | 3.67448 | 1.750 |
| chr4 | 178171277 | 178171643 | 3.28045 | 5.74117 | 1.750 |
| chr19 | 42532441 | 42532768 | 2.23374 | 3.90933 | 1.750 |

| | | | | | |
|---|---|---|---|---|---|
| chr11 | 73371924 | 73372658 | 1.64685 | 2.88226 | 1.750 |
| chr9 | 19926001 | 19926297 | 1.95109 | 3.41479 | 1.750 |
| chr1 | 172419049 | 172419356 | 2.00183 | 3.50365 | 1.750 |
| chr1 | 230410194 | 230410416 | 0.668711 | 1.17043 | 1.750 |
| chr19 | 54836387 | 54837000 | 2.00785 | 3.51432 | 1.750 |
| chr1 | 204415075 | 204415860 | 1.70524 | 2.98468 | 1.750 |
| chr16 | 19992273 | 19992497 | 1.38867 | 2.43066 | 1.750 |
| chr9 | 103938709 | 103938945 | 0.77792 | 1.36171 | 1.750 |
| chr2 | 113699304 | 113699657 | 2.05946 | 3.60502 | 1.750 |
| chr12 | 83325437 | 83325656 | 0.769736 | 1.34748 | 1.751 |
| chr12 | 111709848 | 111710189 | 1.22826 | 2.15016 | 1.751 |
| chr3 | 119107406 | 119107691 | 0.732683 | 1.28267 | 1.751 |
| chr19 | 5803962 | 5804439 | 3.22096 | 5.63879 | 1.751 |
| chr15 | 53132202 | 53132536 | 0.581349 | 1.01776 | 1.751 |
| chr7 | 144604799 | 144605019 | 0.418287 | 0.732303 | 1.751 |
| chrX | 79834065 | 79834306 | 1.73879 | 3.04418 | 1.751 |
| chrX | 125063789 | 125064004 | 0.442955 | 0.775527 | 1.751 |
| chr20 | 59251693 | 59251927 | 0.591012 | 1.03475 | 1.751 |
| chr14 | 49743814 | 49744072 | 0.52769 | 0.923908 | 1.751 |
| chr7 | 101007159 | 101007605 | 2.1499 | 3.76418 | 1.751 |
| chr1 | 220469562 | 220470003 | 1.80068 | 3.15283 | 1.751 |
| chr18 | 11163718 | 11163967 | 1.39496 | 2.44246 | 1.751 |
| chr10 | 95456979 | 95457330 | 0.964657 | 1.68909 | 1.751 |
| chr8 | 20049934 | 20050182 | 0.437328 | 0.765762 | 1.751 |
| chr13 | 99828772 | 99829161 | 1.83004 | 3.20446 | 1.751 |
| chr9 | 129464650 | 129465146 | 2.02135 | 3.53949 | 1.751 |
| chr14 | 50727351 | 50727595 | 0.92196 | 1.61442 | 1.751 |
| chr16 | 84780544 | 84780807 | 1.04168 | 1.82411 | 1.751 |
| chr19 | 1929315 | 1930040 | 1.45718 | 2.55169 | 1.751 |
| chr14 | 74684279 | 74684726 | 2.27865 | 3.99039 | 1.751 |
| chr8 | 3780610 | 3780824 | 0.502159 | 0.879449 | 1.751 |
| chr16 | 76687268 | 76687629 | 1.28073 | 2.24304 | 1.751 |
| chr8 | 94995251 | 94995688 | 1.6361 | 2.86547 | 1.751 |
| chr14 | 105952161 | 105952442 | 1.41647 | 2.48102 | 1.752 |
| chr4 | 22390926 | 22391236 | 1.28275 | 2.24684 | 1.752 |
| chr5 | 134801606 | 134802146 | 1.71183 | 2.9985 | 1.752 |
| chr5 | 109304731 | 109305064 | 0.542786 | 0.950778 | 1.752 |
| chr20 | 46341229 | 46341540 | 1.54352 | 2.70382 | 1.752 |
| chr7 | 38583946 | 38584174 | 0.557369 | 0.97636 | 1.752 |
| chr19 | 10230479 | 10230810 | 2.37445 | 4.15941 | 1.752 |
| chr20 | 2654796 | 2655017 | 1.6126 | 2.82491 | 1.752 |
| chr5 | 156769864 | 156770204 | 2.20427 | 3.8615 | 1.752 |
| chr1 | 155140356 | 155140667 | 2.2317 | 3.90964 | 1.752 |
| chr1 | 207672838 | 207673134 | 1.68276 | 2.948 | 1.752 |
| chr10 | 62127454 | 62127733 | 0.640799 | 1.12262 | 1.752 |
| chr7 | 154801109 | 154801636 | 0.974137 | 1.70663 | 1.752 |
| chr2 | 76674159 | 76674364 | 0.792345 | 1.38815 | 1.752 |
| chr13 | 29370801 | 29371153 | 1.44787 | 2.53661 | 1.752 |
| chr8 | 1082578 | 1082808 | 0.757516 | 1.32717 | 1.752 |
| chr8 | 69408757 | 69408981 | 0.503922 | 0.882926 | 1.752 |
| chr15 | 63758832 | 63759308 | 1.6235 | 2.84457 | 1.752 |
| chr5 | 64254718 | 64254951 | 0.974661 | 1.70777 | 1.752 |
| chr17 | 34296545 | 34296870 | 2.22394 | 3.89674 | 1.752 |
| chr1 | 156046642 | 156046967 | 0.417007 | 0.730769 | 1.752 |
| chr10 | 74096898 | 74097291 | 2.71692 | 4.7612 | 1.752 |
| chr3 | 10385697 | 10386394 | 2.17925 | 3.81903 | 1.752 |
| chr15 | 99125046 | 99125314 | 1.02488 | 1.7961 | 1.753 |
| chr1 | 44513389 | 44513621 | 1.71313 | 3.00228 | 1.753 |
| chr7 | 101915489 | 101915833 | 2.18446 | 3.82843 | 1.753 |
| chr2 | 78910011 | 78910438 | 0.601965 | 1.055 | 1.753 |
| chr7 | 8504965 | 8505292 | 0.541329 | 0.948739 | 1.753 |

| | | | | | |
|---|---|---|---|---|---|
| chr12 | 124538464 | 124538707 | 1.31827 | 2.31046 | 1.753 |
| chr12 | 81528511 | 81528804 | 0.78184 | 1.37032 | 1.753 |
| chr7 | 66843220 | 66843774 | 1.35557 | 2.37607 | 1.753 |
| chr12 | 54804243 | 54804488 | 0.691643 | 1.21234 | 1.753 |
| chr15 | 55731912 | 55732260 | 1.59231 | 2.79106 | 1.753 |
| chr17 | 77161831 | 77162054 | 0.680558 | 1.19292 | 1.753 |
| chr6 | 158447290 | 158447598 | 1.69227 | 2.96638 | 1.753 |
| chr1 | 113346839 | 113347155 | 1.98634 | 3.4819 | 1.753 |
| chr12 | 99349186 | 99349410 | 1.05709 | 1.85309 | 1.753 |
| chr4 | 155947274 | 155947485 | 1.1971 | 2.09863 | 1.753 |
| chr6 | 14046797 | 14046987 | 1.79152 | 3.1407 | 1.753 |
| chrX | 18372088 | 18372406 | 0.853925 | 1.49701 | 1.753 |
| chr15 | 95474121 | 95474393 | 0.604887 | 1.06047 | 1.753 |
| chr4 | 168949826 | 168950059 | 0.750972 | 1.31659 | 1.753 |
| chr9 | 128393626 | 128393967 | 1.14525 | 2.00787 | 1.753 |
| chr12 | 69548425 | 69548763 | 2.53988 | 4.45334 | 1.753 |
| chrX | 37604666 | 37604936 | 2.06904 | 3.6278 | 1.753 |
| chr2 | 197714726 | 197715022 | 0.945048 | 1.65713 | 1.753 |
| chr14 | 70046943 | 70047223 | 1.31467 | 2.30529 | 1.754 |
| chr15 | 89547262 | 89547656 | 1.11901 | 1.96224 | 1.754 |
| chr19 | 30417627 | 30417851 | 0.709318 | 1.24383 | 1.754 |
| chr3 | 186258294 | 186258663 | 1.11181 | 1.94963 | 1.754 |
| chr1 | 27003985 | 27004378 | 1.62615 | 2.85187 | 1.754 |
| chr17 | 4443616 | 4443827 | 1.50454 | 2.63862 | 1.754 |
| chr8 | 14438302 | 14438689 | 0.553455 | 0.970712 | 1.754 |
| chr7 | 33374069 | 33374589 | 0.8989 | 1.5766 | 1.754 |
| chr17 | 79486336 | 79486707 | 1.59248 | 2.7931 | 1.754 |
| chr20 | 21216166 | 21216973 | 2.52402 | 4.42698 | 1.754 |
| chr6 | 33712163 | 33712586 | 1.82832 | 3.20679 | 1.754 |
| chr3 | 127092881 | 127093198 | 1.30467 | 2.28839 | 1.754 |
| chr13 | 46064350 | 46064583 | 1.19209 | 2.09097 | 1.754 |
| chr14 | 52510466 | 52510768 | 2.03699 | 3.57297 | 1.754 |
| chr6 | 110712095 | 110712390 | 1.34136 | 2.35299 | 1.754 |
| chr7 | 147360296 | 147360509 | 0.595597 | 1.0448 | 1.754 |
| chr16 | 537522 | 538025 | 1.24613 | 2.18613 | 1.754 |
| chr1 | 68808468 | 68808842 | 1.79992 | 3.15776 | 1.754 |
| chr20 | 45386766 | 45387098 | 2.04736 | 3.59192 | 1.754 |
| chr4 | 183994790 | 183995014 | 1.20328 | 2.11111 | 1.754 |
| chr4 | 177044914 | 177045259 | 1.56045 | 2.73777 | 1.754 |
| chr20 | 37709595 | 37710140 | 1.67115 | 2.93209 | 1.755 |
| chr1 | 151566630 | 151567037 | 2.36487 | 4.14944 | 1.755 |
| chr12 | 7699263 | 7699516 | 0.517446 | 0.907937 | 1.755 |
| chr8 | 23239159 | 23239411 | 1.55436 | 2.72739 | 1.755 |
| chr7 | 100066823 | 100067202 | 2.15288 | 3.77765 | 1.755 |
| chr7 | 1163181 | 1163742 | 1.01573 | 1.78234 | 1.755 |
| chr8 | 42660487 | 42660801 | 1.45436 | 2.55203 | 1.755 |
| chr17 | 27966557 | 27966907 | 2.22731 | 3.90844 | 1.755 |
| chr8 | 42948445 | 42949259 | 1.76931 | 3.10475 | 1.755 |
| chr15 | 73997138 | 73997508 | 1.88391 | 3.30589 | 1.755 |
| chr1 | 20088712 | 20089293 | 1.39005 | 2.43928 | 1.755 |
| chr3 | 59561159 | 59561593 | 1.61004 | 2.82534 | 1.755 |
| chr5 | 60679070 | 60679285 | 0.799256 | 1.40268 | 1.755 |
| chr1 | 146966749 | 146967190 | 2.0065 | 3.52149 | 1.755 |
| chr1 | 15058470 | 15058680 | 1.857 | 3.2592 | 1.755 |
| chr1 | 109370744 | 109371197 | 1.34981 | 2.36904 | 1.755 |
| chr5 | 66160288 | 66160632 | 1.24672 | 2.18821 | 1.755 |
| chr5 | 24642182 | 24642438 | 0.73277 | 1.28616 | 1.755 |
| chr7 | 28136283 | 28136570 | 1.35542 | 2.37908 | 1.755 |
| chr7 | 73441052 | 73441485 | 2.62405 | 4.60612 | 1.755 |
| chr17 | 33866411 | 33866845 | 2.43932 | 4.28199 | 1.755 |
| chr6 | 41336527 | 41337016 | 3.06686 | 5.38358 | 1.755 |

| | | | | | |
|---|---|---|---|---|---|
| chr2 | 102439746 | 102440246 | 1.10176 | 1.93405 | 1.755 |
| chr20 | 1279961 | 1280444 | 0.722703 | 1.26868 | 1.755 |
| chr14 | 73938905 | 73939123 | 1.22324 | 2.1475 | 1.756 |
| chr9 | 129657566 | 129657797 | 0.462402 | 0.811819 | 1.756 |
| chr15 | 52873143 | 52873347 | 0.537236 | 0.943218 | 1.756 |
| chr12 | 51646294 | 51646586 | 1.20331 | 2.11265 | 1.756 |
| chr13 | 101808389 | 101808685 | 1.14353 | 2.00771 | 1.756 |
| chr8 | 37712197 | 37712416 | 0.521953 | 0.916402 | 1.756 |
| chr19 | 39443285 | 39443550 | 1.40942 | 2.47473 | 1.756 |
| chr5 | 32590322 | 32590533 | 1.06393 | 1.86824 | 1.756 |
| chr19 | 41937428 | 41937738 | 1.09293 | 1.91922 | 1.756 |
| chr10 | 109111992 | 109112218 | 0.492562 | 0.864972 | 1.756 |
| chr11 | 15848457 | 15848738 | 1.99147 | 3.49723 | 1.756 |
| chr14 | 65453360 | 65453626 | 1.97397 | 3.46653 | 1.756 |
| chr19 | 55185443 | 55185850 | 2.03789 | 3.57878 | 1.756 |
| chr17 | 48696276 | 48696574 | 0.685008 | 1.20298 | 1.756 |
| chr19 | 58918327 | 58918670 | 1.14915 | 2.01809 | 1.756 |
| chr6 | 1604554 | 1604966 | 2.24105 | 3.93587 | 1.756 |
| chr4 | 174222811 | 174223169 | 0.91824 | 1.61271 | 1.756 |
| chr10 | 101470955 | 101471259 | 1.23784 | 2.17406 | 1.756 |
| chr15 | 56081258 | 56081618 | 1.81206 | 3.18276 | 1.756 |
| chr1 | 153438202 | 153438635 | 1.66366 | 2.92214 | 1.756 |
| chr2 | 170646383 | 170646771 | 1.08021 | 1.89733 | 1.756 |
| chr12 | 7795511 | 7795754 | 1.10115 | 1.93416 | 1.756 |
| chr10 | 35415690 | 35416019 | 2.39808 | 4.21231 | 1.757 |
| chr7 | 106433589 | 106433887 | 1.19225 | 2.09423 | 1.757 |
| chr9 | 4987964 | 4988222 | 1.65549 | 2.90801 | 1.757 |
| chr12 | 101659948 | 101660287 | 0.527911 | 0.92735 | 1.757 |
| chr15 | 23712266 | 23712472 | 0.745824 | 1.31017 | 1.757 |
| chr12 | 31096585 | 31096844 | 0.422575 | 0.742349 | 1.757 |
| chr8 | 18938452 | 18938701 | 1.18342 | 2.07895 | 1.757 |
| chr7 | 150715124 | 150715599 | 2.78522 | 4.89297 | 1.757 |
| chr18 | 56940666 | 56940995 | 0.935627 | 1.64368 | 1.757 |
| chr5 | 7897187 | 7897404 | 1.23988 | 2.17821 | 1.757 |
| chr17 | 76996984 | 76997227 | 0.773825 | 1.35952 | 1.757 |
| chr14 | 94359300 | 94359778 | 2.10733 | 3.70237 | 1.757 |
| chr1 | 27743056 | 27743364 | 1.67968 | 2.95105 | 1.757 |
| chr19 | 33467112 | 33467542 | 2.05364 | 3.60833 | 1.757 |
| chr1 | 154321755 | 154322047 | 0.543836 | 0.95555 | 1.757 |
| chr14 | 61076843 | 61077313 | 0.604456 | 1.06209 | 1.757 |
| chr15 | 79515525 | 79515777 | 0.484062 | 0.850573 | 1.757 |
| chr18 | 11655366 | 11655656 | 1.57254 | 2.76323 | 1.757 |
| chr9 | 6584694 | 6585398 | 0.719961 | 1.26511 | 1.757 |
| chr8 | 36860996 | 36861257 | 0.482201 | 0.847346 | 1.757 |
| chr12 | 8810028 | 8810236 | 1.45386 | 2.55482 | 1.757 |
| chr1 | 212185330 | 212185622 | 3.00798 | 5.28588 | 1.757 |
| chr3 | 45252074 | 45252405 | 1.82724 | 3.21105 | 1.757 |
| chr19 | 56289023 | 56289228 | 0.736585 | 1.29449 | 1.757 |
| chrX | 70398815 | 70399284 | 0.888259 | 1.56115 | 1.758 |
| chr1 | 83241119 | 83241411 | 1.16825 | 2.05326 | 1.758 |
| chr9 | 16131356 | 16131667 | 0.652971 | 1.14765 | 1.758 |
| chr2 | 55290800 | 55291055 | 1.04027 | 1.82837 | 1.758 |
| chrX | 49042751 | 49043007 | 1.13178 | 1.98924 | 1.758 |
| chr17 | 53603717 | 53604106 | 2.13997 | 3.76144 | 1.758 |
| chr11 | 76802195 | 76802680 | 2.42458 | 4.26181 | 1.758 |
| chr12 | 30935638 | 30935944 | 0.639797 | 1.12464 | 1.758 |
| chr10 | 6150988 | 6151195 | 1.2598 | 2.21468 | 1.758 |
| chr20 | 36386837 | 36387122 | 1.0938 | 1.92287 | 1.758 |
| chr2 | 44314347 | 44314682 | 2.09022 | 3.67457 | 1.758 |
| chr20 | 15876312 | 15876532 | 0.519505 | 0.913291 | 1.758 |
| chr5 | 148509984 | 148510227 | 1.06315 | 1.86903 | 1.758 |

| | | | | | |
|---|---|---|---|---|---|
| chrX | 129194513 | 129194912 | 1.75335 | 3.08248 | 1.758 |
| chr2 | 6712646 | 6712939 | 0.547801 | 0.963107 | 1.758 |
| chr3 | 30735078 | 30735334 | 1.45703 | 2.56165 | 1.758 |
| chr8 | 78521481 | 78521770 | 0.571776 | 1.00531 | 1.758 |
| chr1 | 242630949 | 242631161 | 1.54463 | 2.7159 | 1.758 |
| chr1 | 241648972 | 241649313 | 1.33898 | 2.35433 | 1.758 |
| chr5 | 94450461 | 94450677 | 0.666487 | 1.17189 | 1.758 |
| chr16 | 49706794 | 49707027 | 0.787731 | 1.38508 | 1.758 |
| chr3 | 158970648 | 158970952 | 1.03851 | 1.82604 | 1.758 |
| chr5 | 149681918 | 149682252 | 1.50198 | 2.64105 | 1.758 |
| chr10 | 129976630 | 129976934 | 2.09502 | 3.68389 | 1.758 |
| chr11 | 63688554 | 63688922 | 1.56161 | 2.74595 | 1.758 |
| chr2 | 175613670 | 175613994 | 1.34418 | 2.36363 | 1.758 |
| chr22 | 46409333 | 46409559 | 1.37507 | 2.41797 | 1.758 |
| chr16 | 26271507 | 26271770 | 0.652411 | 1.14724 | 1.758 |
| chr4 | 13600888 | 13601115 | 0.710817 | 1.24995 | 1.758 |
| chr19 | 42339559 | 42340433 | 1.46318 | 2.57297 | 1.758 |
| chr11 | 70913747 | 70913962 | 0.556747 | 0.979032 | 1.758 |
| chr10 | 32000695 | 32001165 | 0.587544 | 1.0332 | 1.759 |
| chr7 | 104793325 | 104793591 | 0.994472 | 1.74882 | 1.759 |
| chr12 | 7293769 | 7294024 | 0.6321 | 1.11161 | 1.759 |
| chr14 | 89643884 | 89644114 | 1.57944 | 2.77761 | 1.759 |
| chrX | 109129442 | 109129645 | 1.78705 | 3.14274 | 1.759 |
| chr5 | 163098902 | 163099108 | 0.777944 | 1.36812 | 1.759 |
| chr17 | 74553582 | 74553801 | 1.75613 | 3.08857 | 1.759 |
| chr15 | 92119973 | 92120175 | 0.694267 | 1.22106 | 1.759 |
| chr9 | 92904934 | 92905285 | 1.2575 | 2.21169 | 1.759 |
| chr8 | 125452672 | 125452991 | 1.63744 | 2.88 | 1.759 |
| chr8 | 84784998 | 84785230 | 0.629965 | 1.10802 | 1.759 |
| chr1 | 194883175 | 194883403 | 0.663402 | 1.16688 | 1.759 |
| chr3 | 141435164 | 141435382 | 0.645613 | 1.13561 | 1.759 |
| chr1 | 248128639 | 248129028 | 0.668301 | 1.17553 | 1.759 |
| chr2 | 206750312 | 206750654 | 2.30905 | 4.06167 | 1.759 |
| chr10 | 118441847 | 118442227 | 2.23961 | 3.94016 | 1.759 |
| chr17 | 37363251 | 37363587 | 2.1738 | 3.82445 | 1.759 |
| chr10 | 76025863 | 76026097 | 1.52444 | 2.6822 | 1.759 |
| chr21 | 34733234 | 34733843 | 1.14123 | 2.00805 | 1.760 |
| chr13 | 106062957 | 106063289 | 1.57578 | 2.77276 | 1.760 |
| chr14 | 96741837 | 96742205 | 2.18387 | 3.84301 | 1.760 |
| chr7 | 85183604 | 85183898 | 0.993823 | 1.74885 | 1.760 |
| chr12 | 9000209 | 9000446 | 0.559386 | 0.984384 | 1.760 |
| chr10 | 108210171 | 108210375 | 0.554342 | 0.975521 | 1.760 |
| chr8 | 25910755 | 25911061 | 1.68671 | 2.9683 | 1.760 |
| chr11 | 120530925 | 120531226 | 0.924927 | 1.62777 | 1.760 |
| chr1 | 120351643 | 120352019 | 1.88683 | 3.32073 | 1.760 |
| chr1 | 61919094 | 61919335 | 1.53789 | 2.70662 | 1.760 |
| chr15 | 61969242 | 61969456 | 0.669707 | 1.17868 | 1.760 |
| chr17 | 49399699 | 49400050 | 1.12907 | 1.98727 | 1.760 |
| chr10 | 27150099 | 27150368 | 1.68797 | 2.97115 | 1.760 |
| chr1 | 33833383 | 33833766 | 2.04492 | 3.59946 | 1.760 |
| chr2 | 75103516 | 75103739 | 1.07305 | 1.88879 | 1.760 |
| chr1 | 239008716 | 239008939 | 0.380884 | 0.670444 | 1.760 |
| chr6 | 52769426 | 52769628 | 0.723959 | 1.27437 | 1.760 |
| chr2 | 10237462 | 10237888 | 2.40475 | 4.23314 | 1.760 |
| chr12 | 59558928 | 59559181 | 1.01439 | 1.78567 | 1.760 |
| chr15 | 69351400 | 69351809 | 1.59937 | 2.81558 | 1.760 |
| chr15 | 91875011 | 91875257 | 0.823914 | 1.45044 | 1.760 |
| chr15 | 29130450 | 29131039 | 0.546245 | 0.961645 | 1.760 |
| chr20 | 35376513 | 35376903 | 0.682678 | 1.20184 | 1.760 |
| chr10 | 101371966 | 101372455 | 2.03482 | 3.58252 | 1.761 |
| chr11 | 8268812 | 8269210 | 2.27063 | 3.99771 | 1.761 |

| | | | | | |
|---|---|---|---|---|---|
| chr20 | 61640403 | 61640765 | 1.96131 | 3.45314 | 1.761 |
| chr7 | 5717647 | 5718095 | 1.06499 | 1.87505 | 1.761 |
| chr14 | 37618855 | 37619179 | 0.722846 | 1.2727 | 1.761 |
| chr12 | 3312076 | 3312451 | 2.88377 | 5.07785 | 1.761 |
| chr9 | 37913615 | 37914011 | 2.45999 | 4.33173 | 1.761 |
| chr11 | 46941454 | 46941835 | 1.48199 | 2.60966 | 1.761 |
| chr7 | 26101312 | 26101704 | 2.03438 | 3.58241 | 1.761 |
| chr2 | 670263 | 670491 | 0.640175 | 1.12731 | 1.761 |
| chr14 | 94369941 | 94370149 | 0.896177 | 1.57816 | 1.761 |
| chr8 | 146223115 | 146223343 | 0.840543 | 1.4802 | 1.761 |
| chr10 | 66911187 | 66911481 | 0.880366 | 1.55039 | 1.761 |
| chr7 | 134043042 | 134043286 | 1.61397 | 2.84244 | 1.761 |
| chr1 | 184141448 | 184141680 | 0.722035 | 1.27165 | 1.761 |
| chr11 | 62264625 | 62264994 | 1.16949 | 2.05972 | 1.761 |
| chr9 | 24454253 | 24454573 | 0.623818 | 1.09869 | 1.761 |
| chr20 | 38565460 | 38565701 | 0.725432 | 1.27769 | 1.761 |
| chr19 | 45690657 | 45691285 | 1.36597 | 2.40588 | 1.761 |
| chr5 | 178194527 | 178194866 | 0.772708 | 1.36102 | 1.761 |
| chr11 | 68657628 | 68657926 | 1.87843 | 3.30861 | 1.761 |
| chr8 | 145575080 | 145575507 | 1.13022 | 1.99076 | 1.761 |
| chr12 | 56099560 | 56099931 | 1.16307 | 2.04873 | 1.761 |
| chr12 | 124805500 | 124806471 | 1.37919 | 2.42946 | 1.762 |
| chr16 | 66994857 | 66995356 | 1.97016 | 3.4705 | 1.762 |
| chr20 | 35905076 | 35905328 | 0.923034 | 1.62604 | 1.762 |
| chr15 | 86914991 | 86915294 | 0.512503 | 0.902891 | 1.762 |
| chr10 | 13098539 | 13099049 | 1.02928 | 1.81336 | 1.762 |
| chr19 | 33864905 | 33865216 | 1.48562 | 2.61739 | 1.762 |
| chr5 | 140578938 | 140579232 | 1.25993 | 2.21999 | 1.762 |
| chrX | 77375422 | 77375670 | 0.945799 | 1.66654 | 1.762 |
| chr14 | 103821188 | 103821416 | 1.24631 | 2.19609 | 1.762 |
| chr1 | 246175593 | 246176019 | 1.21123 | 2.13444 | 1.762 |
| chr8 | 8594463 | 8594824 | 1.50961 | 2.66041 | 1.762 |
| chr10 | 99372262 | 99372499 | 0.975812 | 1.71982 | 1.762 |
| chr3 | 79484472 | 79484785 | 0.502638 | 0.885924 | 1.763 |
| chr16 | 78039656 | 78040072 | 0.775001 | 1.36606 | 1.763 |
| chr15 | 57430503 | 57430802 | 1.81089 | 3.19201 | 1.763 |
| chr5 | 1667447 | 1667653 | 2.73819 | 4.82655 | 1.763 |
| chr17 | 79919082 | 79919339 | 1.32039 | 2.32746 | 1.763 |
| chr5 | 151031595 | 151031992 | 1.65429 | 2.91604 | 1.763 |
| chr3 | 131245613 | 131245916 | 1.21689 | 2.14515 | 1.763 |
| chr1 | 21861329 | 21861606 | 1.39346 | 2.45644 | 1.763 |
| chr3 | 72413120 | 72413416 | 0.859388 | 1.515 | 1.763 |
| chr15 | 101693123 | 101693413 | 1.39906 | 2.46641 | 1.763 |
| chr14 | 102131913 | 102132152 | 1.52352 | 2.68591 | 1.763 |
| chr21 | 17702755 | 17703016 | 0.775649 | 1.36746 | 1.763 |
| chr4 | 113242955 | 113243295 | 1.70486 | 3.00565 | 1.763 |
| chrX | 55187282 | 55187526 | 1.72367 | 3.03884 | 1.763 |
| chr7 | 27735047 | 27735360 | 0.514382 | 0.906897 | 1.763 |
| chr9 | 105289102 | 105289305 | 0.642381 | 1.1326 | 1.763 |
| chr17 | 8906254 | 8906477 | 0.926613 | 1.63375 | 1.763 |
| chr6 | 152873785 | 152874048 | 1.60835 | 2.83575 | 1.763 |
| chr11 | 33351290 | 33351609 | 1.69273 | 2.98459 | 1.763 |
| chr7 | 40801560 | 40801807 | 0.774411 | 1.36544 | 1.763 |
| chr2 | 48793731 | 48793974 | 1.35578 | 2.39056 | 1.763 |
| chrX | 15164680 | 15165008 | 1.51245 | 2.66684 | 1.763 |
| chr7 | 24615854 | 24616068 | 1.46117 | 2.57652 | 1.763 |
| chr13 | 50243918 | 50244195 | 2.04361 | 3.60366 | 1.763 |
| chr16 | 81657017 | 81657330 | 1.31151 | 2.31285 | 1.764 |
| chr9 | 4549143 | 4549466 | 2.21306 | 3.90304 | 1.764 |
| chrX | 114461573 | 114461848 | 1.20315 | 2.12195 | 1.764 |
| chr7 | 48194404 | 48194710 | 1.85401 | 3.26987 | 1.764 |

| | | | | | |
|---|---|---|---|---|---|
| chr10 | 23015627 | 23015880 | 0.551675 | 0.972979 | 1.764 |
| chr17 | 4103192 | 4103487 | 2.00962 | 3.54444 | 1.764 |
| chr3 | 1731227 | 1731501 | 0.71628 | 1.26336 | 1.764 |
| chr5 | 99483542 | 99483768 | 0.586905 | 1.03518 | 1.764 |
| chr1 | 205242590 | 205243026 | 2.05379 | 3.62251 | 1.764 |
| rUn_gl0002 | 118363 | 119057 | 0.796482 | 1.40489 | 1.764 |
| chr6 | 160362770 | 160363042 | 0.765729 | 1.35081 | 1.764 |
| chr6 | 148785893 | 148786208 | 1.45116 | 2.56005 | 1.764 |
| chr10 | 88447994 | 88448630 | 1.2497 | 2.20469 | 1.764 |
| chr17 | 27546443 | 27546845 | 2.38324 | 4.20451 | 1.764 |
| chr14 | 103857203 | 103857728 | 1.83248 | 3.23288 | 1.764 |
| chr15 | 79202129 | 79202523 | 1.81394 | 3.20023 | 1.764 |
| chr15 | 31782233 | 31782666 | 1.73553 | 3.06214 | 1.764 |
| chr4 | 128321953 | 128322235 | 1.54732 | 2.73014 | 1.764 |
| chr4 | 48111774 | 48112012 | 1.30106 | 2.29566 | 1.764 |
| chr5 | 4396244 | 4396473 | 0.446831 | 0.788409 | 1.764 |
| chr5 | 84330250 | 84330502 | 0.463031 | 0.817025 | 1.765 |
| chr8 | 31825089 | 31825330 | 0.571339 | 1.00814 | 1.765 |
| chr6 | 52099086 | 52099297 | 1.36111 | 2.40173 | 1.765 |
| chr14 | 23908656 | 23908956 | 0.47593 | 0.839855 | 1.765 |
| chr17 | 54804663 | 54805091 | 0.608492 | 1.07378 | 1.765 |
| chr17 | 13828744 | 13829138 | 0.691237 | 1.21981 | 1.765 |
| chr12 | 44469018 | 44469258 | 0.728764 | 1.28605 | 1.765 |
| chr9 | 4762556 | 4762878 | 1.29178 | 2.27972 | 1.765 |
| chr1 | 154738522 | 154738760 | 1.26553 | 2.23347 | 1.765 |
| chr10 | 104106997 | 104107457 | 1.49429 | 2.63726 | 1.765 |
| chr9 | 130887277 | 130887742 | 2.56945 | 4.53485 | 1.765 |
| chr3 | 6920360 | 6920692 | 0.609981 | 1.07657 | 1.765 |
| chr6 | 85836699 | 85836982 | 2.17376 | 3.83653 | 1.765 |
| chr5 | 165272644 | 165272850 | 0.544083 | 0.960287 | 1.765 |
| chr5 | 54062559 | 54062800 | 0.527824 | 0.931623 | 1.765 |
| chr8 | 11411082 | 11411698 | 1.61911 | 2.85782 | 1.765 |
| chr2 | 16246992 | 16247197 | 0.49564 | 0.874851 | 1.765 |
| chr3 | 128564781 | 128565246 | 1.23921 | 2.18744 | 1.765 |
| chr5 | 157145522 | 157145798 | 1.15056 | 2.03096 | 1.765 |
| chr9 | 138495729 | 138496057 | 2.48755 | 4.39113 | 1.765 |
| chr10 | 82459146 | 82459755 | 2.03346 | 3.58978 | 1.765 |
| chr1 | 207178391 | 207178611 | 1.61516 | 2.85144 | 1.765 |
| chr9 | 33517434 | 33517745 | 1.17893 | 2.08133 | 1.765 |
| chr12 | 126006042 | 126006384 | 0.642947 | 1.13512 | 1.766 |
| chr12 | 49482042 | 49482418 | 2.35631 | 4.1602 | 1.766 |
| chr19 | 16494138 | 16494424 | 1.632 | 2.8814 | 1.766 |
| chr12 | 110792992 | 110793202 | 1.04579 | 1.84644 | 1.766 |
| chr6 | 169114132 | 169114358 | 1.03786 | 1.83246 | 1.766 |
| chr5 | 137515964 | 137516179 | 1.48684 | 2.62521 | 1.766 |
| chr11 | 5632964 | 5633171 | 1.1222 | 1.98143 | 1.766 |
| chr17 | 49613184 | 49613645 | 1.58634 | 2.80097 | 1.766 |
| chr1 | 29572357 | 29572636 | 1.52844 | 2.69886 | 1.766 |
| chr11 | 36669478 | 36669685 | 0.558511 | 0.986204 | 1.766 |
| chr4 | 93308691 | 93308916 | 1.60179 | 2.82844 | 1.766 |
| chr1 | 160398343 | 160398689 | 1.18058 | 2.08469 | 1.766 |
| chr14 | 24950782 | 24951586 | 1.10888 | 1.95808 | 1.766 |
| chr16 | 30043414 | 30043874 | 1.56365 | 2.76115 | 1.766 |
| chr14 | 28069639 | 28069892 | 0.672378 | 1.18732 | 1.766 |
| chrX | 118616630 | 118616978 | 2.22263 | 3.92484 | 1.766 |
| chr19 | 33630064 | 33630331 | 0.826333 | 1.45919 | 1.766 |
| chr2 | 112655449 | 112655708 | 0.969643 | 1.71233 | 1.766 |
| chr16 | 62727516 | 62727780 | 0.477461 | 0.843181 | 1.766 |
| chr15 | 31492945 | 31493183 | 1.67179 | 2.95239 | 1.766 |
| chr17 | 10212430 | 10212768 | 0.592904 | 1.04708 | 1.766 |
| chr12 | 64195869 | 64196092 | 1.21016 | 2.13721 | 1.766 |

| | | | | | |
|---|---|---|---|---|---|
| chr13 | 78701422 | 78701651 | 0.560176 | 0.989322 | 1.766 |
| chr22 | 49442346 | 49442684 | 1.7767 | 3.1379 | 1.766 |
| chr3 | 776773 | 776988 | 0.648164 | 1.14475 | 1.766 |
| chr17 | 62935000 | 62935373 | 1.89052 | 3.33896 | 1.766 |
| chr3 | 178865036 | 178865251 | 0.993814 | 1.75527 | 1.766 |
| chr17 | 40115023 | 40115349 | 1.99376 | 3.52149 | 1.766 |
| chr12 | 111471085 | 111471313 | 1.47609 | 2.60725 | 1.766 |
| chr20 | 32888987 | 32889218 | 0.676133 | 1.19427 | 1.766 |
| chr9 | 21789484 | 21789924 | 1.25301 | 2.21325 | 1.766 |
| chr8 | 119132898 | 119133119 | 1.26642 | 2.23695 | 1.766 |
| chr16 | 10615500 | 10615929 | 1.61697 | 2.85639 | 1.767 |
| chr5 | 8410810 | 8411055 | 0.64538 | 1.1401 | 1.767 |
| chr1 | 160924047 | 160924339 | 2.21596 | 3.91465 | 1.767 |
| chrX | 75499462 | 75499800 | 1.34367 | 2.37374 | 1.767 |
| chr18 | 33357043 | 33357248 | 1.07864 | 1.9056 | 1.767 |
| chr16 | 28338207 | 28338582 | 2.48652 | 4.3929 | 1.767 |
| chr5 | 33578220 | 33578449 | 0.548273 | 0.968642 | 1.767 |
| chr15 | 90764528 | 90764910 | 2.7747 | 4.90226 | 1.767 |
| chr15 | 86699398 | 86699629 | 0.5311 | 0.938356 | 1.767 |
| chr14 | 89248937 | 89249285 | 0.542334 | 0.958213 | 1.767 |
| chr15 | 35288036 | 35288239 | 0.619379 | 1.09438 | 1.767 |
| chr12 | 52317143 | 52317611 | 2.5767 | 4.55283 | 1.767 |
| chr15 | 56242545 | 56242959 | 0.548865 | 0.969799 | 1.767 |
| chr12 | 76979315 | 76979551 | 0.669253 | 1.18253 | 1.767 |
| chr9 | 115626122 | 115626449 | 0.447372 | 0.790526 | 1.767 |
| chr9 | 113913993 | 113914231 | 0.720796 | 1.27368 | 1.767 |
| chr15 | 78251754 | 78252080 | 2.81241 | 4.96974 | 1.767 |
| chr6 | 159112441 | 159112724 | 1.09855 | 1.9413 | 1.767 |
| chr6 | 33557454 | 33557811 | 1.27309 | 2.24978 | 1.767 |
| chr7 | 134035052 | 134035277 | 0.794152 | 1.40342 | 1.767 |
| chr3 | 46599872 | 46600102 | 1.84583 | 3.26198 | 1.767 |
| chr16 | 5006037 | 5006503 | 2.10847 | 3.7262 | 1.767 |
| chr19 | 39260329 | 39260764 | 2.14179 | 3.78528 | 1.767 |
| chr5 | 119296445 | 119296723 | 0.561522 | 0.992402 | 1.767 |
| chr19 | 5892513 | 5892915 | 1.7529 | 3.09799 | 1.767 |
| chr1 | 20042117 | 20042466 | 1.69036 | 2.98748 | 1.767 |
| chr17 | 48034203 | 48034469 | 0.486894 | 0.860533 | 1.767 |
| chr5 | 71662271 | 71662531 | 1.44563 | 2.55502 | 1.767 |
| chr11 | 84509790 | 84510130 | 0.66487 | 1.17511 | 1.767 |
| chr1 | 205756580 | 205756851 | 0.716634 | 1.26665 | 1.768 |
| chr20 | 17346607 | 17346821 | 0.694517 | 1.22762 | 1.768 |
| chr12 | 109865849 | 109866217 | 3.94993 | 6.98253 | 1.768 |
| chr7 | 2700504 | 2700830 | 1.08159 | 1.91231 | 1.768 |
| chr8 | 25550170 | 25550377 | 0.516152 | 0.912603 | 1.768 |
| chr3 | 149654554 | 149654832 | 1.7198 | 3.04092 | 1.768 |
| chr17 | 29339750 | 29340052 | 0.641954 | 1.13513 | 1.768 |
| chrX | 48802983 | 48803437 | 1.70925 | 3.02238 | 1.768 |
| chr12 | 23336874 | 23337122 | 0.583098 | 1.03107 | 1.768 |
| chr7 | 73269846 | 73270274 | 3.08102 | 5.44822 | 1.768 |
| chr3 | 184186178 | 184186474 | 1.64345 | 2.90618 | 1.768 |
| chr7 | 1680031 | 1680400 | 1.76225 | 3.11625 | 1.768 |
| chr11 | 86121375 | 86121592 | 0.422057 | 0.746347 | 1.768 |
| chr5 | 151947723 | 151947928 | 0.707384 | 1.25093 | 1.768 |
| chr10 | 120610020 | 120610293 | 1.3337 | 2.3586 | 1.768 |
| chr3 | 50264784 | 50265061 | 1.54836 | 2.73821 | 1.768 |
| chr9 | 122448672 | 122449005 | 0.585409 | 1.03528 | 1.768 |
| chr5 | 19902438 | 19902670 | 0.814203 | 1.43994 | 1.769 |
| chr2 | 3183087 | 3183308 | 1.1037 | 1.95205 | 1.769 |
| chr3 | 151402818 | 151403058 | 0.629918 | 1.11411 | 1.769 |
| chrX | 114252744 | 114252976 | 0.757721 | 1.34018 | 1.769 |
| chr5 | 175145828 | 175146155 | 1.85158 | 3.275 | 1.769 |

| | | | | | |
|---|---|---|---|---|---|
| chr20 | 7863019 | 7863407 | 0.601934 | 1.06468 | 1.769 |
| chr17 | 79493857 | 79494087 | 0.620371 | 1.09734 | 1.769 |
| chr19 | 55459833 | 55460088 | 0.494236 | 0.874232 | 1.769 |
| chr7 | 129252612 | 129252910 | 1.11653 | 1.97504 | 1.769 |
| chr19 | 4624808 | 4625110 | 1.36672 | 2.41772 | 1.769 |
| chr20 | 49934730 | 49935080 | 1.62389 | 2.87264 | 1.769 |
| chr17 | 78549029 | 78549542 | 4.77521 | 8.44738 | 1.769 |
| chr1 | 22432171 | 22432438 | 1.48114 | 2.62019 | 1.769 |
| chr10 | 54644072 | 54644400 | 1.40672 | 2.48857 | 1.769 |
| chr7 | 45556253 | 45556528 | 0.62828 | 1.11149 | 1.769 |
| chr15 | 59914587 | 59914878 | 1.7322 | 3.06458 | 1.769 |
| chr8 | 144048659 | 144048895 | 0.376024 | 0.665264 | 1.769 |
| chr10 | 28917047 | 28917361 | 1.60141 | 2.83327 | 1.769 |
| chr4 | 68259419 | 68259699 | 0.944569 | 1.67116 | 1.769 |
| chr4 | 166052405 | 166052729 | 1.41471 | 2.50298 | 1.769 |
| chr17 | 16290020 | 16290242 | 1.38036 | 2.44221 | 1.769 |
| chr1 | 42039767 | 42040093 | 1.76571 | 3.12415 | 1.769 |
| chr10 | 64004793 | 64005120 | 1.57775 | 2.79178 | 1.769 |
| chr16 | 21719915 | 21720137 | 1.48763 | 2.63231 | 1.769 |
| chr1 | 160078593 | 160079167 | 1.01633 | 1.79838 | 1.769 |
| chr10 | 50615415 | 50615998 | 1.81729 | 3.21565 | 1.769 |
| chr2 | 4312116 | 4312373 | 0.586657 | 1.03808 | 1.769 |
| chr2 | 239172099 | 239172517 | 1.90627 | 3.37314 | 1.770 |
| chr5 | 78551773 | 78552125 | 1.85146 | 3.27622 | 1.770 |
| chr1 | 157743479 | 157743716 | 0.781219 | 1.3824 | 1.770 |
| chr9 | 22470002 | 22470288 | 1.67384 | 2.96192 | 1.770 |
| chr15 | 40602220 | 40602668 | 1.63733 | 2.89734 | 1.770 |
| chr3 | 48777395 | 48777694 | 1.45137 | 2.5686 | 1.770 |
| chr10 | 128437596 | 128437985 | 1.86054 | 3.29277 | 1.770 |
| chr7 | 80546129 | 80546445 | 1.01625 | 1.79856 | 1.770 |
| chr12 | 2061541 | 2061929 | 0.706882 | 1.25108 | 1.770 |
| chr5 | 160992689 | 160992955 | 0.482672 | 0.854283 | 1.770 |
| chr19 | 46092849 | 46093105 | 0.695191 | 1.23051 | 1.770 |
| chr3 | 56889955 | 56890219 | 1.48815 | 2.63407 | 1.770 |
| chr4 | 111440113 | 111440373 | 1.12244 | 1.98681 | 1.770 |
| chr15 | 76020658 | 76020920 | 1.25772 | 2.2263 | 1.770 |
| chr1 | 27932872 | 27933181 | 1.57222 | 2.78304 | 1.770 |
| chr13 | 99097474 | 99097960 | 1.92239 | 3.40289 | 1.770 |
| chr15 | 81511775 | 81512093 | 0.637096 | 1.12778 | 1.770 |
| chr3 | 185701297 | 185701634 | 1.39785 | 2.47448 | 1.770 |
| chr7 | 85602119 | 85602343 | 0.618037 | 1.09407 | 1.770 |
| chr8 | 42547534 | 42547962 | 1.90596 | 3.37398 | 1.770 |
| chr13 | 51486184 | 51486612 | 1.56625 | 2.77289 | 1.770 |
| chr6 | 110499839 | 110500157 | 0.899645 | 1.59277 | 1.770 |
| chr2 | 26407536 | 26407934 | 1.87029 | 3.31148 | 1.771 |
| chr3 | 89091431 | 89091687 | 0.606403 | 1.07375 | 1.771 |
| chr3 | 186471020 | 186471253 | 1.53174 | 2.71223 | 1.771 |
| chr9 | 36597502 | 36597778 | 1.19642 | 2.11849 | 1.771 |
| chr9 | 79721501 | 79721728 | 0.402874 | 0.713369 | 1.771 |
| chr10 | 14578939 | 14579284 | 1.7249 | 3.0544 | 1.771 |
| chr11 | 1858391 | 1859420 | 1.37579 | 2.43621 | 1.771 |
| chr18 | 37321784 | 37322032 | 0.463464 | 0.820726 | 1.771 |
| chr20 | 50929958 | 50930209 | 1.43639 | 2.54373 | 1.771 |
| chr10 | 19777892 | 19778153 | 0.646547 | 1.14499 | 1.771 |
| chr17 | 27064743 | 27065162 | 0.657205 | 1.16387 | 1.771 |
| chr19 | 2611221 | 2611548 | 2.29089 | 4.05704 | 1.771 |
| chr8 | 22480366 | 22480942 | 1.93916 | 3.43413 | 1.771 |
| chr3 | 137690033 | 137690415 | 1.79308 | 3.17545 | 1.771 |
| chr16 | 2523930 | 2524391 | 1.13151 | 2.00396 | 1.771 |
| chr10 | 101130769 | 101130973 | 1.26203 | 2.23516 | 1.771 |
| chr12 | 96312200 | 96312675 | 2.04366 | 3.61956 | 1.771 |

| | | | | | |
|---|---|---|---|---|---|
| chrX | 129059625 | 129059968 | 1.81997 | 3.22351 | 1.771 |
| chr4 | 184171070 | 184171302 | 0.887367 | 1.57171 | 1.771 |
| chr16 | 21652092 | 21652300 | 1.27618 | 2.26046 | 1.771 |
| chr1 | 203840180 | 203840663 | 1.60645 | 2.84552 | 1.771 |
| chr15 | 33992898 | 33993761 | 1.17924 | 2.08891 | 1.771 |
| chr6 | 99778022 | 99778256 | 1.51761 | 2.68837 | 1.771 |
| chr1 | 94339729 | 94339955 | 1.32465 | 2.34659 | 1.771 |
| chr7 | 142340084 | 142340385 | 1.25747 | 2.22772 | 1.772 |
| chr19 | 46850515 | 46850892 | 3.00336 | 5.32092 | 1.772 |
| chr3 | 53229799 | 53230311 | 1.79204 | 3.17497 | 1.772 |
| chr20 | 16019485 | 16019780 | 0.659221 | 1.16799 | 1.772 |
| chr5 | 93422405 | 93422630 | 1.15422 | 2.04516 | 1.772 |
| chr8 | 27172851 | 27173068 | 1.90046 | 3.36749 | 1.772 |
| chr9 | 139379458 | 139379833 | 2.60102 | 4.60896 | 1.772 |
| chr2 | 9916358 | 9916606 | 0.574545 | 1.0181 | 1.772 |
| chr17 | 75082612 | 75082968 | 2.50044 | 4.43082 | 1.772 |
| chr12 | 95730221 | 95730601 | 3.098 | 5.48977 | 1.772 |
| chr7 | 131231342 | 131231828 | 2.5938 | 4.5964 | 1.772 |
| chr7 | 128731510 | 128731897 | 2.69367 | 4.77346 | 1.772 |
| chr17 | 77979549 | 77979989 | 3.26185 | 5.78042 | 1.772 |
| chr9 | 74639903 | 74640265 | 1.76849 | 3.1344 | 1.772 |
| chr3 | 63335306 | 63335565 | 1.12438 | 1.99285 | 1.772 |
| chr1 | 226411376 | 226411655 | 0.820979 | 1.45512 | 1.772 |
| chr1 | 235247089 | 235247402 | 2.09898 | 3.72041 | 1.772 |
| chr12 | 82777740 | 82778018 | 1.17214 | 2.07762 | 1.773 |
| chr3 | 89181854 | 89182102 | 0.637357 | 1.12973 | 1.773 |
| chr20 | 52171839 | 52172044 | 0.574675 | 1.01872 | 1.773 |
| chr11 | 798130 | 798629 | 2.07985 | 3.687 | 1.773 |
| chr2 | 66141589 | 66141804 | 0.514301 | 0.911779 | 1.773 |
| chr3 | 151000788 | 151001017 | 1.30631 | 2.31589 | 1.773 |
| chr10 | 119237921 | 119238130 | 0.944511 | 1.67461 | 1.773 |
| chr1 | 45259413 | 45259690 | 1.22542 | 2.17272 | 1.773 |
| chr16 | 18015344 | 18015745 | 0.743877 | 1.31895 | 1.773 |
| chrX | 94145194 | 94145429 | 0.579725 | 1.02793 | 1.773 |
| chrX | 27701964 | 27702267 | 1.5474 | 2.74383 | 1.773 |
| chr20 | 22002409 | 22002661 | 0.592249 | 1.05022 | 1.773 |
| chr11 | 45315233 | 45315447 | 1.6301 | 2.89076 | 1.773 |
| chr8 | 141477343 | 141477598 | 2.12892 | 3.77534 | 1.773 |
| chr14 | 94993366 | 94993654 | 0.542969 | 0.962895 | 1.773 |
| chr11 | 38432847 | 38433053 | 0.755311 | 1.33947 | 1.773 |
| chr10 | 32895050 | 32895454 | 0.62232 | 1.10367 | 1.773 |
| chr14 | 70784578 | 70784854 | 1.10111 | 1.95282 | 1.774 |
| chr16 | 72133273 | 72133501 | 1.52697 | 2.70808 | 1.774 |
| chr16 | 47918190 | 47918489 | 0.589008 | 1.04462 | 1.774 |
| chr1 | 204131541 | 204131956 | 3.06393 | 5.43455 | 1.774 |
| chr2 | 220295501 | 220295812 | 0.809987 | 1.43674 | 1.774 |
| chr8 | 20898560 | 20898993 | 2.1332 | 3.78386 | 1.774 |
| chr2 | 17900908 | 17901126 | 1.46623 | 2.60082 | 1.774 |
| chr15 | 61789729 | 61789973 | 0.631658 | 1.12047 | 1.774 |
| chr15 | 31151479 | 31151816 | 1.5509 | 2.75119 | 1.774 |
| chr20 | 39974300 | 39974563 | 0.489551 | 0.868437 | 1.774 |
| chr3 | 12864559 | 12864822 | 1.17153 | 2.07828 | 1.774 |
| chr15 | 34902792 | 34903216 | 0.689846 | 1.22383 | 1.774 |
| chr5 | 13987935 | 13988263 | 1.30532 | 2.31571 | 1.774 |
| chr3 | 7209969 | 7210262 | 0.530147 | 0.940535 | 1.774 |
| chr15 | 64338610 | 64339018 | 1.91602 | 3.39925 | 1.774 |
| chr19 | 34827689 | 34828072 | 1.73859 | 3.08463 | 1.774 |
| chr17 | 49806738 | 49806964 | 0.563794 | 1.0003 | 1.774 |
| chr8 | 98834275 | 98834570 | 2.40718 | 4.27092 | 1.774 |
| chr1 | 225039134 | 225039550 | 2.11907 | 3.75983 | 1.774 |
| chr20 | 30101956 | 30102306 | 2.34302 | 4.15723 | 1.774 |

| | | | | | |
|---|---|---|---|---|---|
| chr19 | 44560144 | 44560408 | 1.06207 | 1.88449 | 1.774 |
| chr10 | 70373399 | 70373609 | 0.482727 | 0.856554 | 1.774 |
| chr14 | 92377985 | 92378221 | 1.77252 | 3.14521 | 1.774 |
| chr6 | 132231154 | 132231378 | 1.17813 | 2.09051 | 1.774 |
| chr1 | 166669905 | 166670133 | 0.426594 | 0.756969 | 1.774 |
| chr15 | 56285696 | 56286025 | 1.8987 | 3.36919 | 1.774 |
| chr17 | 39119923 | 39120411 | 1.42832 | 2.53472 | 1.775 |
| chr4 | 107423273 | 107423544 | 1.35952 | 2.41273 | 1.775 |
| chr10 | 102822325 | 102822716 | 1.29884 | 2.3051 | 1.775 |
| chr12 | 54069868 | 54070140 | 3.13188 | 5.55841 | 1.775 |
| chr15 | 52198120 | 52198325 | 1.55722 | 2.7638 | 1.775 |
| chr1 | 195345494 | 195345713 | 0.59699 | 1.05956 | 1.775 |
| chr15 | 33424981 | 33425244 | 1.64406 | 2.918 | 1.775 |
| chr5 | 139731330 | 139731665 | 1.91644 | 3.40143 | 1.775 |
| chr5 | 61229905 | 61230221 | 1.59819 | 2.8366 | 1.775 |
| chr8 | 28968007 | 28968279 | 0.725099 | 1.28697 | 1.775 |
| chr1 | 184951328 | 184951625 | 1.55638 | 2.76251 | 1.775 |
| chr12 | 122354744 | 122355141 | 0.886706 | 1.57392 | 1.775 |
| chr11 | 44473545 | 44473758 | 0.717094 | 1.27291 | 1.775 |
| chr20 | 3800725 | 3801112 | 3.10149 | 5.50556 | 1.775 |
| chr5 | 118092311 | 118092573 | 0.5495 | 0.975452 | 1.775 |
| chr4 | 178601390 | 178601628 | 0.471136 | 0.836353 | 1.775 |
| chr11 | 69177193 | 69177426 | 1.03459 | 1.83666 | 1.775 |
| chr2 | 45115654 | 45115871 | 0.497528 | 0.883297 | 1.775 |
| chr8 | 136072506 | 136072715 | 0.508026 | 0.901964 | 1.775 |
| chr9 | 9352801 | 9353174 | 0.881512 | 1.5651 | 1.775 |
| chr1 | 156100165 | 156100518 | 2.0119 | 3.57223 | 1.776 |
| chr14 | 29649319 | 29649620 | 0.427547 | 0.759139 | 1.776 |
| chr16 | 63564177 | 63564412 | 0.856884 | 1.52146 | 1.776 |
| gl000202_r | 10249 | 10497 | 1.41746 | 2.51684 | 1.776 |
| chr7 | 98470017 | 98470257 | 1.24449 | 2.20973 | 1.776 |
| chr16 | 2346577 | 2346920 | 1.06735 | 1.89522 | 1.776 |
| chr1 | 201975050 | 201975421 | 0.768255 | 1.36415 | 1.776 |
| chr12 | 56696102 | 56696482 | 1.50413 | 2.67089 | 1.776 |
| chr17 | 80202239 | 80202487 | 2.12283 | 3.7695 | 1.776 |
| chr12 | 110235680 | 110236090 | 1.12851 | 2.00392 | 1.776 |
| chr10 | 93129795 | 93130102 | 0.908202 | 1.61279 | 1.776 |
| chr1 | 214105793 | 214106024 | 0.74364 | 1.32058 | 1.776 |
| chr3 | 77864641 | 77865015 | 0.978055 | 1.7369 | 1.776 |
| chr4 | 6722007 | 6722320 | 1.59176 | 2.82681 | 1.776 |
| chr15 | 89985926 | 89986432 | 2.17066 | 3.85489 | 1.776 |
| chr1 | 101822945 | 101823274 | 1.70565 | 3.02913 | 1.776 |
| chr5 | 79940289 | 79940581 | 0.591013 | 1.04962 | 1.776 |
| chr1 | 86939781 | 86940172 | 1.40403 | 2.49363 | 1.776 |
| chr4 | 2980186 | 2980502 | 1.25091 | 2.22168 | 1.776 |
| chr20 | 21596171 | 21596487 | 2.87192 | 5.10084 | 1.776 |
| chr22 | 22763661 | 22763927 | 1.61242 | 2.86397 | 1.776 |
| chr12 | 42690392 | 42690639 | 1.74699 | 3.10301 | 1.776 |
| chr8 | 95625091 | 95625481 | 1.83868 | 3.26594 | 1.776 |
| chr9 | 72932839 | 72933184 | 1.02815 | 1.82625 | 1.776 |
| chr17 | 19669745 | 19670175 | 3.62836 | 6.44491 | 1.776 |
| chr7 | 90955895 | 90956121 | 0.642518 | 1.14142 | 1.776 |
| chr10 | 93449785 | 93450152 | 0.618398 | 1.09858 | 1.776 |
| chr16 | 79636111 | 79636347 | 1.74548 | 3.10084 | 1.777 |
| chr8 | 84405072 | 84405355 | 0.55038 | 0.97779 | 1.777 |
| chr18 | 20284343 | 20284591 | 1.27039 | 2.25711 | 1.777 |
| chr11 | 96126950 | 96127280 | 0.904309 | 1.60673 | 1.777 |
| chr8 | 79419814 | 79420148 | 2.00004 | 3.5536 | 1.777 |
| chr20 | 36192766 | 36193277 | 2.06965 | 3.67732 | 1.777 |
| chr14 | 83176692 | 83176966 | 1.34668 | 2.39279 | 1.777 |
| chr2 | 21756228 | 21756623 | 0.704017 | 1.25093 | 1.777 |

| | | | | | |
|---|---|---|---|---|---|
| chr10 | 126681104 | 126681421 | 1.52962 | 2.71794 | 1.777 |
| chr7 | 157225652 | 157225975 | 1.52261 | 2.70562 | 1.777 |
| chr3 | 191906099 | 191906416 | 1.32016 | 2.34592 | 1.777 |
| chr5 | 94980092 | 94980303 | 1.44026 | 2.55936 | 1.777 |
| chr5 | 114405119 | 114405441 | 1.45302 | 2.58203 | 1.777 |
| chr8 | 107773388 | 107773621 | 0.712085 | 1.26538 | 1.777 |
| chr16 | 67647798 | 67648062 | 1.47515 | 2.62146 | 1.777 |
| chr9 | 17811051 | 17811329 | 2.10167 | 3.73484 | 1.777 |
| chr2 | 17091213 | 17091628 | 0.554917 | 0.986148 | 1.777 |
| chr1 | 215482602 | 215482865 | 0.644475 | 1.14533 | 1.777 |
| chr12 | 48577440 | 48577679 | 0.991847 | 1.76269 | 1.777 |
| chr17 | 43312992 | 43313498 | 1.74393 | 3.09933 | 1.777 |
| chr15 | 93961551 | 93961818 | 0.836756 | 1.48711 | 1.777 |
| chr20 | 4207685 | 4207978 | 1.74287 | 3.0975 | 1.777 |
| chr15 | 101569440 | 101569711 | 0.790608 | 1.40521 | 1.777 |
| chr16 | 6427990 | 6428285 | 0.443984 | 0.789137 | 1.777 |
| chr2 | 222438231 | 222438590 | 2.03602 | 3.61882 | 1.777 |
| chr1 | 158596441 | 158596813 | 0.629227 | 1.11844 | 1.777 |
| chr22 | 35550903 | 35551181 | 1.59484 | 2.83479 | 1.777 |
| chr1 | 119818000 | 119818421 | 1.51097 | 2.68597 | 1.778 |
| chr3 | 99263663 | 99263885 | 0.529577 | 0.941411 | 1.778 |
| chr10 | 13413819 | 13414294 | 1.83163 | 3.25631 | 1.778 |
| chr10 | 107734586 | 107734973 | 0.55937 | 0.994488 | 1.778 |
| chr4 | 57410682 | 57411085 | 2.15258 | 3.82721 | 1.778 |
| chr12 | 2419041 | 2419254 | 0.61947 | 1.10141 | 1.778 |
| chr16 | 49525954 | 49526518 | 2.56738 | 4.56479 | 1.778 |
| chr17 | 80308197 | 80308440 | 1.14374 | 2.03358 | 1.778 |
| chr8 | 2449864 | 2450119 | 0.5806 | 1.03232 | 1.778 |
| chr4 | 124319322 | 124319657 | 1.29342 | 2.29975 | 1.778 |
| chr3 | 163381346 | 163381610 | 0.446846 | 0.794516 | 1.778 |
| chr9 | 120538049 | 120538604 | 0.586193 | 1.04237 | 1.778 |
| chr19 | 3719086 | 3719409 | 1.68267 | 2.9924 | 1.778 |
| chr8 | 8800215 | 8800431 | 0.950661 | 1.69064 | 1.778 |
| chr19 | 33668385 | 33668673 | 1.2699 | 2.25839 | 1.778 |
| chr3 | 103680084 | 103680286 | 0.487943 | 0.867786 | 1.778 |
| chr21 | 38003086 | 38003366 | 2.00939 | 3.57372 | 1.779 |
| chr9 | 127242571 | 127243185 | 1.31411 | 2.33737 | 1.779 |
| chr10 | 88699170 | 88699541 | 5.18251 | 9.21835 | 1.779 |
| chr10 | 1402236 | 1402528 | 1.07658 | 1.915 | 1.779 |
| chr16 | 28956758 | 28957237 | 2.42141 | 4.3073 | 1.779 |
| chr5 | 154347068 | 154347419 | 1.07366 | 1.9099 | 1.779 |
| chr17 | 58678539 | 58678852 | 2.22251 | 3.95358 | 1.779 |
| chr5 | 13479027 | 13479331 | 1.89681 | 3.37436 | 1.779 |
| chr12 | 110950290 | 110950504 | 1.1347 | 2.01861 | 1.779 |
| chr19 | 47628481 | 47628801 | 1.42322 | 2.5319 | 1.779 |
| chr19 | 44039647 | 44040128 | 1.59878 | 2.8443 | 1.779 |
| chr5 | 154996407 | 154996757 | 2.00675 | 3.57008 | 1.779 |
| chr1 | 160055407 | 160055788 | 2.17255 | 3.86508 | 1.779 |
| chr14 | 36352976 | 36353230 | 1.36616 | 2.43048 | 1.779 |
| chr17 | 17763497 | 17763887 | 3.33476 | 5.93284 | 1.779 |
| chr8 | 82692729 | 82693154 | 2.19094 | 3.89788 | 1.779 |
| chr11 | 8588086 | 8588496 | 1.09322 | 1.94495 | 1.779 |
| chr17 | 72206505 | 72207052 | 2.57978 | 4.5897 | 1.779 |
| chr16 | 54971209 | 54971697 | 2.21049 | 3.93272 | 1.779 |
| chr12 | 82986109 | 82986359 | 0.67564 | 1.20207 | 1.779 |
| chr8 | 91561673 | 91561876 | 0.681902 | 1.21326 | 1.779 |
| chr15 | 59249605 | 59249896 | 1.88231 | 3.34912 | 1.779 |
| chr5 | 1140365 | 1140898 | 2.51486 | 4.47467 | 1.779 |
| chr16 | 74069895 | 74070108 | 0.91338 | 1.62528 | 1.779 |
| chr12 | 49520959 | 49521191 | 0.6766 | 1.20396 | 1.779 |
| chr16 | 54667845 | 54668071 | 1.17897 | 2.09791 | 1.779 |

| | | | | | |
|---|---|---|---|---|---|
| chr12 | 121821380 | 121821589 | 1.55348 | 2.76435 | 1.779 |
| chr5 | 164420555 | 164420831 | 0.657233 | 1.16952 | 1.779 |
| chr1 | 112298191 | 112298460 | 1.67513 | 2.98096 | 1.780 |
| chr3 | 129543810 | 129544117 | 1.66692 | 2.96641 | 1.780 |
| chr18 | 9951327 | 9951606 | 1.42062 | 2.52813 | 1.780 |
| chr1 | 166171038 | 166171240 | 0.431374 | 0.767676 | 1.780 |
| chr22 | 50585241 | 50585512 | 2.75484 | 4.9027 | 1.780 |
| chr1 | 163098871 | 163099078 | 0.841941 | 1.49843 | 1.780 |
| chr19 | 33675064 | 33675521 | 2.41448 | 4.29715 | 1.780 |
| chr10 | 86111758 | 86112384 | 1.8503 | 3.29311 | 1.780 |
| chr1 | 113261217 | 113261750 | 2.02746 | 3.60843 | 1.780 |
| chr2 | 71228135 | 71228659 | 2.73129 | 4.86113 | 1.780 |
| chr17 | 25926918 | 25927192 | 1.65348 | 2.94297 | 1.780 |
| chr5 | 150785907 | 150786289 | 1.28897 | 2.2942 | 1.780 |
| chr15 | 94199369 | 94199645 | 0.615601 | 1.0957 | 1.780 |
| chr1 | 68577580 | 68577825 | 1.18163 | 2.10319 | 1.780 |
| chr3 | 12761565 | 12761862 | 1.79826 | 3.2008 | 1.780 |
| chr6 | 3748531 | 3748897 | 1.71465 | 3.052 | 1.780 |
| chr15 | 81398840 | 81399162 | 1.40135 | 2.49437 | 1.780 |
| chr20 | 17012235 | 17012520 | 0.727585 | 1.2951 | 1.780 |
| chr5 | 39570571 | 39570975 | 1.00427 | 1.7876 | 1.780 |
| chr4 | 78279462 | 78279746 | 1.54649 | 2.75285 | 1.780 |
| chr16 | 78513578 | 78513793 | 0.708392 | 1.261 | 1.780 |
| chr18 | 8478624 | 8479075 | 1.36821 | 2.43556 | 1.780 |
| chr3 | 192185026 | 192185371 | 1.58446 | 2.82067 | 1.780 |
| chr3 | 162397764 | 162398138 | 0.508475 | 0.90521 | 1.780 |
| chr17 | 20024820 | 20025431 | 1.76697 | 3.14576 | 1.780 |
| chr11 | 83313178 | 83313468 | 0.412066 | 0.733609 | 1.780 |
| chr9 | 114837095 | 114837430 | 2.5862 | 4.60436 | 1.780 |
| chr19 | 34131680 | 34131952 | 0.824717 | 1.46832 | 1.780 |
| chr20 | 39318580 | 39319168 | 1.77609 | 3.1622 | 1.780 |
| chr12 | 29428822 | 29429055 | 0.59718 | 1.0633 | 1.781 |
| chr4 | 10556321 | 10556568 | 1.26949 | 2.26054 | 1.781 |
| chr2 | 29809834 | 29810083 | 0.606901 | 1.08071 | 1.781 |
| chr1 | 157125768 | 157126215 | 2.48784 | 4.43016 | 1.781 |
| chr16 | 22012269 | 22012713 | 2.91801 | 5.19621 | 1.781 |
| chr8 | 95654489 | 95654844 | 1.59623 | 2.84248 | 1.781 |
| chr2 | 72998084 | 72998296 | 1.60375 | 2.85596 | 1.781 |
| chr12 | 6149206 | 6149530 | 2.71442 | 4.83405 | 1.781 |
| chr17 | 7259329 | 7259762 | 2.45069 | 4.36447 | 1.781 |
| chr17 | 36891310 | 36891794 | 2.66894 | 4.75331 | 1.781 |
| chr9 | 10165011 | 10165233 | 0.645434 | 1.1495 | 1.781 |
| chr1 | 2071186 | 2071671 | 1.71996 | 3.06322 | 1.781 |
| chr5 | 53530352 | 53530668 | 0.891591 | 1.58791 | 1.781 |
| chr12 | 78188634 | 78188851 | 0.591676 | 1.05381 | 1.781 |
| chr2 | 20050916 | 20051162 | 0.832505 | 1.48276 | 1.781 |
| chr19 | 44270647 | 44271417 | 1.60535 | 2.85941 | 1.781 |
| chr11 | 40315625 | 40315952 | 1.78934 | 3.18716 | 1.781 |
| chr8 | 19902277 | 19902487 | 0.423528 | 0.754382 | 1.781 |
| chr18 | 39608243 | 39608515 | 2.23709 | 3.98473 | 1.781 |
| chr20 | 30311672 | 30311997 | 1.94267 | 3.46033 | 1.781 |
| chr22 | 21457117 | 21457457 | 1.99643 | 3.55608 | 1.781 |
| chr1 | 22643784 | 22644420 | 1.20391 | 2.14445 | 1.781 |
| chr14 | 32724055 | 32724354 | 1.37162 | 2.44335 | 1.781 |
| chr15 | 61753417 | 61753714 | 0.763162 | 1.35949 | 1.781 |
| chr8 | 71851037 | 71851431 | 0.588812 | 1.04895 | 1.781 |
| chr8 | 143421206 | 143421500 | 0.508815 | 0.906456 | 1.782 |
| chr17 | 9648201 | 9648602 | 1.88482 | 3.35783 | 1.782 |
| chr19 | 6218496 | 6218939 | 1.87015 | 3.33175 | 1.782 |
| chr7 | 97795461 | 97795818 | 2.35153 | 4.18936 | 1.782 |
| chr6 | 47381810 | 47382027 | 1.31303 | 2.33946 | 1.782 |

| | | | | | |
|---|---|---|---|---|---|
| chr12 | 15304515 | 15304727 | 1.36917 | 2.4395 | 1.782 |
| chr5 | 102811425 | 102811682 | 1.65358 | 2.94634 | 1.782 |
| chr8 | 127082749 | 127083019 | 0.753496 | 1.34258 | 1.782 |
| chr12 | 56472721 | 56473407 | 2.15995 | 3.84861 | 1.782 |
| chr10 | 69236013 | 69236408 | 0.98159 | 1.74905 | 1.782 |
| chr5 | 134563019 | 134563329 | 0.553043 | 0.985446 | 1.782 |
| chr20 | 30048332 | 30048645 | 1.45127 | 2.5861 | 1.782 |
| chr13 | 110436906 | 110437324 | 1.35587 | 2.41618 | 1.782 |
| chr15 | 102012450 | 102012723 | 0.741853 | 1.32202 | 1.782 |
| chr11 | 1406417 | 1406654 | 0.545591 | 0.972294 | 1.782 |
| chr19 | 35738724 | 35738947 | 1.42873 | 2.54612 | 1.782 |
| chr2 | 22500262 | 22500509 | 0.514608 | 0.917091 | 1.782 |
| chr21 | 45864354 | 45864652 | 1.28595 | 2.29174 | 1.782 |
| chr2 | 15239173 | 15239409 | 0.671754 | 1.19719 | 1.782 |
| chr13 | 42002342 | 42002610 | 1.7168 | 3.05977 | 1.782 |
| chr14 | 72427743 | 72428331 | 0.529047 | 0.942894 | 1.782 |
| chr18 | 45938791 | 45939071 | 1.33857 | 2.3858 | 1.782 |
| chr15 | 58335441 | 58335973 | 1.47847 | 2.63517 | 1.782 |
| chr8 | 85810328 | 85810539 | 0.573673 | 1.02251 | 1.782 |
| chr6 | 159638608 | 159638839 | 1.32736 | 2.36588 | 1.782 |
| chr12 | 122791079 | 122791314 | 0.883953 | 1.57561 | 1.782 |
| chr14 | 33045345 | 33045606 | 1.71755 | 3.06159 | 1.783 |
| chr8 | 66977201 | 66977465 | 1.54152 | 2.74782 | 1.783 |
| chr11 | 41349863 | 41350201 | 0.584393 | 1.04171 | 1.783 |
| chr5 | 176513914 | 176514292 | 2.2182 | 3.95408 | 1.783 |
| chr8 | 137987518 | 137987735 | 0.663283 | 1.18236 | 1.783 |
| chr7 | 88620735 | 88621017 | 0.545122 | 0.97178 | 1.783 |
| chr7 | 155531399 | 155531639 | 1.33045 | 2.37191 | 1.783 |
| chr15 | 64044391 | 64044629 | 1.4145 | 2.52182 | 1.783 |
| chr9 | 98388594 | 98388900 | 0.424553 | 0.756904 | 1.783 |
| chr10 | 88726347 | 88726740 | 1.49557 | 2.66636 | 1.783 |
| chr7 | 155949465 | 155949824 | 1.59647 | 2.84632 | 1.783 |
| chr16 | 86021155 | 86021467 | 1.6193 | 2.88714 | 1.783 |
| chr17 | 42406431 | 42406637 | 1.23796 | 2.20725 | 1.783 |
| chr10 | 93453995 | 93454383 | 1.22696 | 2.18766 | 1.783 |
| chr11 | 60954979 | 60955357 | 1.59083 | 2.83652 | 1.783 |
| chr17 | 64723959 | 64724352 | 1.56462 | 2.7898 | 1.783 |
| chr1 | 205601141 | 205601484 | 1.14207 | 2.0365 | 1.783 |
| chr2 | 143791885 | 143792311 | 2.25474 | 4.02061 | 1.783 |
| chr7 | 100835606 | 100836129 | 1.89659 | 3.38208 | 1.783 |
| chr4 | 112864920 | 112865291 | 1.18102 | 2.10613 | 1.783 |
| chr17 | 8313646 | 8313948 | 2.02543 | 3.61202 | 1.783 |
| chr7 | 113493851 | 113494165 | 0.519147 | 0.925808 | 1.783 |
| chr15 | 31445568 | 31445776 | 0.773213 | 1.37895 | 1.783 |
| chr11 | 20375466 | 20375796 | 1.31327 | 2.34213 | 1.783 |
| chr8 | 130315626 | 130316167 | 1.53377 | 2.73548 | 1.784 |
| chr12 | 117146732 | 117147072 | 1.55601 | 2.77528 | 1.784 |
| chr10 | 112316730 | 112317153 | 1.97913 | 3.52997 | 1.784 |
| chr1 | 184341204 | 184341408 | 0.506681 | 0.903725 | 1.784 |
| chr20 | 23300990 | 23301299 | 1.28292 | 2.28827 | 1.784 |
| chr10 | 78925489 | 78925813 | 1.41556 | 2.52489 | 1.784 |
| chr14 | 69964189 | 69964391 | 0.578317 | 1.03153 | 1.784 |
| chr5 | 116079530 | 116079980 | 0.475259 | 0.847717 | 1.784 |
| chr3 | 139072814 | 139073132 | 1.58984 | 2.83593 | 1.784 |
| chr1 | 152707625 | 152707865 | 0.517521 | 0.923178 | 1.784 |
| chr10 | 105614854 | 105615377 | 1.09693 | 1.95685 | 1.784 |
| chr14 | 92848946 | 92849469 | 1.51818 | 2.70835 | 1.784 |
| chr11 | 124616534 | 124616759 | 1.09394 | 1.95155 | 1.784 |
| chr5 | 114370843 | 114371108 | 0.512726 | 0.914724 | 1.784 |
| chr19 | 28935334 | 28935702 | 0.602971 | 1.07574 | 1.784 |
| chr17 | 3163213 | 3163418 | 0.664828 | 1.18617 | 1.784 |

| | | | | | |
|---|---|---|---|---|---|
| chr9 | 121358105 | 121358355 | 1.76055 | 3.14118 | 1.784 |
| chr1 | 175109396 | 175109640 | 0.466452 | 0.8323 | 1.784 |
| chr11 | 64122608 | 64122937 | 2.60004 | 4.63932 | 1.784 |
| chr11 | 30989624 | 30989870 | 0.587799 | 1.04893 | 1.785 |
| chr2 | 85152805 | 85153364 | 2.69861 | 4.81573 | 1.785 |
| chr15 | 101212250 | 101212542 | 2.57542 | 4.59592 | 1.785 |
| chr19 | 3075968 | 3076401 | 0.923542 | 1.64809 | 1.785 |
| chr6 | 75940248 | 75940535 | 1.19396 | 2.13067 | 1.785 |
| chr12 | 51441568 | 51441917 | 2.26607 | 4.04392 | 1.785 |
| chr5 | 8473181 | 8473457 | 0.933148 | 1.66526 | 1.785 |
| chr17 | 70819725 | 70819947 | 0.867166 | 1.54757 | 1.785 |
| chr17 | 77384504 | 77385072 | 2.1495 | 3.8361 | 1.785 |
| chr5 | 84056647 | 84056917 | 0.63328 | 1.1302 | 1.785 |
| chr20 | 56229383 | 56229829 | 1.40333 | 2.50451 | 1.785 |
| chr2 | 219157113 | 219157515 | 1.91703 | 3.42156 | 1.785 |
| chr2 | 216795965 | 216796542 | 1.61038 | 2.87428 | 1.785 |
| chr7 | 130644694 | 130644944 | 1.37123 | 2.44748 | 1.785 |
| chr2 | 9013059 | 9013268 | 1.35032 | 2.41019 | 1.785 |
| chr12 | 78864299 | 78864751 | 1.24945 | 2.23015 | 1.785 |
| chr15 | 72072777 | 72073140 | 1.86387 | 3.32693 | 1.785 |
| chr8 | 109367926 | 109368265 | 0.485701 | 0.866958 | 1.785 |
| chr19 | 56117546 | 56117865 | 2.30559 | 4.11542 | 1.785 |
| chr11 | 2058122 | 2058460 | 1.37214 | 2.44951 | 1.785 |
| chr12 | 108372616 | 108372837 | 0.55606 | 0.992665 | 1.785 |
| chr2 | 178281004 | 178281267 | 1.22618 | 2.18913 | 1.785 |
| chr8 | 10873851 | 10874280 | 1.63225 | 2.91416 | 1.785 |
| chr9 | 91307904 | 91308108 | 0.705574 | 1.25972 | 1.785 |
| chr10 | 88109307 | 88109553 | 1.16016 | 2.07134 | 1.785 |
| chr11 | 61250024 | 61250334 | 1.97163 | 3.52016 | 1.785 |
| chr5 | 79153811 | 79154283 | 1.53035 | 2.73234 | 1.785 |
| chr9 | 94489581 | 94489862 | 1.55913 | 2.78374 | 1.785 |
| chr21 | 38382222 | 38382504 | 1.71495 | 3.06208 | 1.786 |
| chr15 | 44511899 | 44512169 | 1.65884 | 2.96194 | 1.786 |
| chr3 | 52265485 | 52265916 | 2.20514 | 3.93743 | 1.786 |
| chr16 | 83249075 | 83249400 | 1.61275 | 2.87979 | 1.786 |
| chr14 | 106714143 | 106714441 | 1.60354 | 2.86359 | 1.786 |
| chr8 | 121270286 | 121270617 | 0.571984 | 1.02145 | 1.786 |
| chr2 | 59142092 | 59142321 | 0.515556 | 0.920706 | 1.786 |
| chr5 | 43556929 | 43557386 | 1.85435 | 3.31169 | 1.786 |
| chr17 | 15151466 | 15151801 | 1.09121 | 1.94883 | 1.786 |
| chr17 | 46671234 | 46671608 | 2.59566 | 4.63584 | 1.786 |
| chr1 | 219759177 | 219759479 | 0.659561 | 1.17798 | 1.786 |
| chr19 | 10044821 | 10045099 | 1.86143 | 3.32456 | 1.786 |
| chr10 | 57606979 | 57607187 | 0.595339 | 1.06336 | 1.786 |
| chr14 | 21458483 | 21458703 | 0.918289 | 1.64019 | 1.786 |
| chr17 | 61484142 | 61484420 | 2.58072 | 4.60958 | 1.786 |
| chr8 | 5189635 | 5189845 | 0.742734 | 1.32666 | 1.786 |
| chr7 | 39493445 | 39493719 | 1.35785 | 2.42538 | 1.786 |
| chr19 | 19097513 | 19097898 | 1.55606 | 2.77943 | 1.786 |
| chr4 | 83206373 | 83206686 | 1.76375 | 3.15084 | 1.786 |
| chr7 | 44896576 | 44896861 | 1.48309 | 2.64945 | 1.786 |
| chr1 | 35147115 | 35147430 | 1.3639 | 2.43666 | 1.787 |
| chr15 | 101798240 | 101798764 | 1.2971 | 2.31732 | 1.787 |
| chr19 | 5523202 | 5523487 | 1.69305 | 3.0247 | 1.787 |
| chr10 | 122235643 | 122235865 | 0.469852 | 0.839427 | 1.787 |
| chr7 | 37026744 | 37027266 | 1.38447 | 2.47354 | 1.787 |
| chr14 | 48386653 | 48386863 | 0.934471 | 1.66973 | 1.787 |
| chr9 | 94537748 | 94538078 | 2.2154 | 3.95851 | 1.787 |
| chr6 | 3457373 | 3457610 | 2.33929 | 4.18017 | 1.787 |
| chr12 | 1227155 | 1227516 | 2.25211 | 4.02444 | 1.787 |
| chr3 | 127678790 | 127679046 | 1.05651 | 1.88795 | 1.787 |

| | | | | | |
|---|---|---|---|---|---|
| chr6 | 54582715 | 54582987 | 1.37529 | 2.45769 | 1.787 |
| chr3 | 103931892 | 103932096 | 0.54897 | 0.981099 | 1.787 |
| chr19 | 1438194 | 1438439 | 2.01572 | 3.60243 | 1.787 |
| chr17 | 38883260 | 38883511 | 1.26633 | 2.26322 | 1.787 |
| chr8 | 96400526 | 96400849 | 0.42006 | 0.750757 | 1.787 |
| chr1 | 31656842 | 31657231 | 1.81675 | 3.2471 | 1.787 |
| chr12 | 49330055 | 49330510 | 1.76953 | 3.1627 | 1.787 |
| chr11 | 64974423 | 64974634 | 0.638385 | 1.14103 | 1.787 |
| chr15 | 94990325 | 94990537 | 0.541788 | 0.968375 | 1.787 |
| chr12 | 3409060 | 3409432 | 2.63403 | 4.70835 | 1.788 |
| chr21 | 39047544 | 39047926 | 1.962 | 3.50717 | 1.788 |
| chr15 | 93237301 | 93237628 | 1.4681 | 2.62437 | 1.788 |
| chr17 | 46292990 | 46293236 | 1.76309 | 3.15173 | 1.788 |
| chr2 | 38059799 | 38060064 | 1.30626 | 2.3351 | 1.788 |
| chr8 | 105771902 | 105772307 | 2.93539 | 5.24791 | 1.788 |
| chr1 | 161135506 | 161135852 | 1.14976 | 2.05557 | 1.788 |
| chr5 | 134527045 | 134527475 | 2.31696 | 4.14239 | 1.788 |
| chr2 | 220394893 | 220395240 | 3.34259 | 5.97627 | 1.788 |
| chr13 | 44378653 | 44378858 | 0.431517 | 0.771574 | 1.788 |
| chr3 | 4332032 | 4332343 | 1.64721 | 2.94534 | 1.788 |
| chr6 | 166298382 | 166298675 | 1.68225 | 3.008 | 1.788 |
| chr17 | 76172612 | 76173037 | 3.29653 | 5.89477 | 1.788 |
| chr17 | 41957695 | 41958222 | 1.95111 | 3.48921 | 1.788 |
| chr9 | 112108086 | 112108424 | 1.40573 | 2.51389 | 1.788 |
| chr11 | 6778662 | 6778928 | 1.33001 | 2.37859 | 1.788 |
| chr5 | 177572937 | 177573376 | 2.16947 | 3.88006 | 1.788 |
| chr12 | 59198692 | 59198909 | 0.884278 | 1.58155 | 1.789 |
| chr11 | 65393343 | 65393821 | 1.44027 | 2.57597 | 1.789 |
| chr16 | 14013238 | 14013884 | 1.34988 | 2.4144 | 1.789 |
| chr9 | 114360110 | 114360473 | 1.47913 | 2.64557 | 1.789 |
| chr20 | 49196530 | 49196815 | 1.71575 | 3.0688 | 1.789 |
| chr1 | 212764586 | 212764808 | 0.424312 | 0.758964 | 1.789 |
| chr12 | 113568380 | 113568886 | 1.09413 | 1.95707 | 1.789 |
| chr17 | 6476099 | 6476557 | 0.690472 | 1.23508 | 1.789 |
| chr14 | 57999964 | 58000235 | 1.00753 | 1.80223 | 1.789 |
| chr5 | 148110516 | 148110879 | 1.84957 | 3.30844 | 1.789 |
| chr19 | 11314484 | 11314789 | 1.86986 | 3.34476 | 1.789 |
| chr8 | 22446433 | 22446909 | 2.13925 | 3.82665 | 1.789 |
| chr9 | 107444853 | 107445125 | 0.978134 | 1.74988 | 1.789 |
| chr12 | 30272058 | 30272313 | 0.548336 | 0.980985 | 1.789 |
| chr8 | 130738544 | 130738969 | 2.27938 | 4.07785 | 1.789 |
| chr3 | 150863656 | 150864415 | 1.43335 | 2.56431 | 1.789 |
| chr5 | 7851122 | 7851396 | 0.732988 | 1.31136 | 1.789 |
| chr19 | 15870849 | 15871102 | 1.28782 | 2.30401 | 1.789 |
| chr18 | 51654225 | 51654494 | 1.31015 | 2.34403 | 1.789 |
| chr3 | 3102316 | 3102898 | 0.496381 | 0.888091 | 1.789 |
| chr5 | 149903381 | 149903638 | 1.80137 | 3.22315 | 1.789 |
| chr10 | 52385004 | 52385217 | 0.983722 | 1.76023 | 1.789 |
| chr8 | 102149488 | 102149753 | 1.31586 | 2.35471 | 1.789 |
| chr9 | 117249834 | 117250230 | 1.95659 | 3.5013 | 1.789 |
| chr1 | 32403900 | 32404180 | 0.709023 | 1.26883 | 1.790 |
| chr8 | 142044757 | 142044964 | 1.14168 | 2.04315 | 1.790 |
| chr5 | 42750541 | 42750758 | 0.497432 | 0.890215 | 1.790 |
| chr3 | 118333240 | 118333467 | 0.429106 | 0.767949 | 1.790 |
| chr21 | 45066538 | 45066870 | 1.93095 | 3.45582 | 1.790 |
| chr2 | 88890465 | 88890911 | 1.53401 | 2.74556 | 1.790 |
| chr20 | 4721397 | 4722015 | 1.55228 | 2.77832 | 1.790 |
| chr5 | 116063191 | 116063457 | 0.652847 | 1.16852 | 1.790 |
| chr9 | 103085257 | 103085617 | 1.41641 | 2.53525 | 1.790 |
| chr5 | 62697337 | 62697684 | 1.51602 | 2.71358 | 1.790 |
| chr8 | 9122828 | 9123117 | 0.574421 | 1.02818 | 1.790 |

| | | | | | |
|---|---|---|---|---|---|
| chr9 | 129029821 | 129030055 | 0.732109 | 1.3105 | 1.790 |
| chr3 | 7120075 | 7120507 | 0.529501 | 0.947854 | 1.790 |
| chr15 | 88680617 | 88680893 | 0.584172 | 1.04574 | 1.790 |
| chr3 | 19613340 | 19613592 | 0.583217 | 1.04405 | 1.790 |
| chr10 | 15461316 | 15461674 | 1.35523 | 2.42609 | 1.790 |
| chr10 | 85160646 | 85160910 | 0.7026 | 1.25778 | 1.790 |
| chr7 | 83565481 | 83565785 | 2.25075 | 4.02931 | 1.790 |
| chr9 | 107256851 | 107257086 | 0.485465 | 0.869086 | 1.790 |
| chr8 | 36615619 | 36616053 | 0.610388 | 1.09282 | 1.790 |
| chr2 | 55276116 | 55276394 | 2.13491 | 3.82236 | 1.790 |
| chr7 | 133826043 | 133826433 | 2.43359 | 4.35719 | 1.790 |
| chr10 | 4945040 | 4945244 | 0.552022 | 0.988386 | 1.790 |
| chr6 | 165658836 | 165659102 | 0.677978 | 1.21392 | 1.791 |
| chr12 | 48153217 | 48153513 | 1.90785 | 3.41604 | 1.791 |
| chr8 | 95339640 | 95339881 | 1.01212 | 1.81223 | 1.791 |
| chr12 | 56551604 | 56551818 | 1.51481 | 2.71234 | 1.791 |
| chr5 | 15278119 | 15278375 | 0.533339 | 0.955018 | 1.791 |
| chr1 | 167028405 | 167028717 | 1.99634 | 3.57478 | 1.791 |
| chrX | 98652107 | 98652335 | 0.498626 | 0.892885 | 1.791 |
| chr17 | 1572615 | 1573166 | 1.45323 | 2.6023 | 1.791 |
| chr11 | 69458087 | 69458428 | 2.06342 | 3.69515 | 1.791 |
| chr5 | 113559562 | 113559894 | 1.09261 | 1.95667 | 1.791 |
| chr17 | 11060788 | 11061111 | 0.578959 | 1.03688 | 1.791 |
| chr15 | 68487459 | 68487778 | 0.728778 | 1.30521 | 1.791 |
| chr1 | 167525357 | 167525854 | 1.42482 | 2.55191 | 1.791 |
| chr9 | 101558712 | 101559119 | 1.58974 | 2.84733 | 1.791 |
| chrX | 102412975 | 102413273 | 1.20829 | 2.16416 | 1.791 |
| chr7 | 90496994 | 90497252 | 0.598624 | 1.0722 | 1.791 |
| chr8 | 22287881 | 22288093 | 1.29513 | 2.31976 | 1.791 |
| chr10 | 54326762 | 54327052 | 1.30264 | 2.33325 | 1.791 |
| chr17 | 36833285 | 36833724 | 2.16201 | 3.87253 | 1.791 |
| chr5 | 158743743 | 158744189 | 1.73824 | 3.1136 | 1.791 |
| chr12 | 106612780 | 106613017 | 0.65715 | 1.17714 | 1.791 |
| chr1 | 152842307 | 152842649 | 1.6222 | 2.9059 | 1.791 |
| chr19 | 4233415 | 4233888 | 1.3357 | 2.39269 | 1.791 |
| chr9 | 75207720 | 75207990 | 1.5599 | 2.79442 | 1.791 |
| chr15 | 61487388 | 61487712 | 1.31309 | 2.35232 | 1.791 |
| chr19 | 30718494 | 30718873 | 2.59471 | 4.64828 | 1.791 |
| chr2 | 20293393 | 20293794 | 1.17504 | 2.10502 | 1.791 |
| chr12 | 101271686 | 101272023 | 0.96348 | 1.72607 | 1.792 |
| chr2 | 73278851 | 73279131 | 0.428639 | 0.767947 | 1.792 |
| chr3 | 196729555 | 196730179 | 1.34998 | 2.41861 | 1.792 |
| chr5 | 126115968 | 126116283 | 1.29941 | 2.32807 | 1.792 |
| chr12 | 121445417 | 121445634 | 0.47191 | 0.845545 | 1.792 |
| chr12 | 122211135 | 122211531 | 2.55994 | 4.58681 | 1.792 |
| chr19 | 48199348 | 48199560 | 0.78676 | 1.40971 | 1.792 |
| chr9 | 1005133 | 1005428 | 1.20258 | 2.15481 | 1.792 |
| chr17 | 27090640 | 27091042 | 1.56339 | 2.80144 | 1.792 |
| chr3 | 33116003 | 33116267 | 2.06717 | 3.70426 | 1.792 |
| chr12 | 54354979 | 54355399 | 0.801892 | 1.43701 | 1.792 |
| chr8 | 73067494 | 73067710 | 1.10895 | 1.98733 | 1.792 |
| chr11 | 63916017 | 63916508 | 2.2841 | 4.09336 | 1.792 |
| chr2 | 179779060 | 179779279 | 1.47786 | 2.64852 | 1.792 |
| chr17 | 38242675 | 38243088 | 2.44201 | 4.3766 | 1.792 |
| chr11 | 120100451 | 120100843 | 0.818128 | 1.46634 | 1.792 |
| chr6 | 159525135 | 159525495 | 1.99208 | 3.5705 | 1.792 |
| chr19 | 47939895 | 47940313 | 1.88544 | 3.37941 | 1.792 |
| chr5 | 118670701 | 118671021 | 1.25815 | 2.25509 | 1.792 |
| chr5 | 144644064 | 144644326 | 0.567004 | 1.01635 | 1.792 |
| chr3 | 47841342 | 47841550 | 0.904415 | 1.62116 | 1.793 |
| chr11 | 66197388 | 66197868 | 1.96789 | 3.52755 | 1.793 |

| | | | | | |
|---|---|---|---|---|---|
| chr12 | 52755924 | 52756173 | 1.02207 | 1.83211 | 1.793 |
| chr9 | 135450424 | 135450753 | 2.43194 | 4.35946 | 1.793 |
| chr17 | 7255687 | 7255914 | 1.43733 | 2.57667 | 1.793 |
| chr11 | 118560526 | 118561363 | 1.97544 | 3.54144 | 1.793 |
| chr8 | 4566996 | 4567268 | 0.749813 | 1.3443 | 1.793 |
| chr5 | 6780976 | 6781356 | 1.34659 | 2.4143 | 1.793 |
| chr15 | 66791215 | 66791464 | 1.29007 | 2.31311 | 1.793 |
| chr1 | 62054388 | 62054635 | 1.85667 | 3.32906 | 1.793 |
| chr12 | 10653679 | 10653952 | 1.45102 | 2.60174 | 1.793 |
| chr15 | 35736884 | 35737148 | 1.17854 | 2.11318 | 1.793 |
| chr8 | 33751588 | 33751846 | 0.526775 | 0.94458 | 1.793 |
| chr3 | 15599057 | 15599396 | 1.12263 | 2.01305 | 1.793 |
| chr6 | 131840814 | 131841034 | 1.21527 | 2.1792 | 1.793 |
| chr12 | 71243264 | 71243511 | 0.84565 | 1.51641 | 1.793 |
| chr17 | 62770475 | 62770840 | 1.54106 | 2.76343 | 1.793 |
| chr15 | 78931039 | 78931382 | 1.49968 | 2.68925 | 1.793 |
| chr4 | 103178955 | 103179192 | 1.26967 | 2.27691 | 1.793 |
| chr19 | 35760351 | 35761000 | 3.02618 | 5.42705 | 1.793 |
| chr10 | 73533092 | 73533532 | 1.90473 | 3.41615 | 1.794 |
| chr10 | 2661788 | 2662004 | 1.0354 | 1.85704 | 1.794 |
| chr22 | 46518298 | 46518672 | 1.89416 | 3.3974 | 1.794 |
| chr18 | 43302860 | 43303148 | 1.51844 | 2.72364 | 1.794 |
| chr17 | 50408744 | 50409022 | 0.623449 | 1.1183 | 1.794 |
| chr7 | 155589571 | 155589962 | 2.63005 | 4.71768 | 1.794 |
| chr20 | 59421615 | 59421822 | 0.641875 | 1.15139 | 1.794 |
| chr3 | 75208400 | 75208627 | 0.71644 | 1.28516 | 1.794 |
| chr2 | 70530942 | 70531364 | 1.55083 | 2.78198 | 1.794 |
| chr19 | 30591938 | 30592162 | 0.885004 | 1.58762 | 1.794 |
| chr3 | 111686541 | 111686920 | 2.07838 | 3.72851 | 1.794 |
| chr16 | 83987685 | 83987958 | 1.29832 | 2.32917 | 1.794 |
| chr2 | 120399158 | 120399531 | 1.91744 | 3.4399 | 1.794 |
| chr12 | 64219873 | 64220108 | 1.31628 | 2.36145 | 1.794 |
| chr13 | 76591024 | 76591395 | 1.00045 | 1.79497 | 1.794 |
| chr3 | 141173837 | 141174056 | 1.33741 | 2.39954 | 1.794 |
| chr19 | 13172109 | 13172514 | 1.60869 | 2.8863 | 1.794 |
| chr20 | 1447409 | 1447875 | 3.7419 | 6.7138 | 1.794 |
| chrX | 45181660 | 45181893 | 1.03751 | 1.86157 | 1.794 |
| chr22 | 38039342 | 38039682 | 2.14603 | 3.8506 | 1.794 |
| chr12 | 127147838 | 127148122 | 0.655285 | 1.17578 | 1.794 |
| chr5 | 149232438 | 149233055 | 1.76817 | 3.17262 | 1.794 |
| chr14 | 63775486 | 63775754 | 1.82032 | 3.26624 | 1.794 |
| chr1 | 24181027 | 24181322 | 0.610365 | 1.0952 | 1.794 |
| chrX | 133055949 | 133056193 | 1.43762 | 2.57958 | 1.794 |
| chr15 | 51982295 | 51982637 | 0.632028 | 1.1341 | 1.794 |
| chr18 | 72020471 | 72020735 | 1.26499 | 2.26988 | 1.794 |
| chr11 | 108484023 | 108484372 | 0.525778 | 0.943454 | 1.794 |
| chr2 | 197876802 | 197877152 | 1.56826 | 2.8141 | 1.794 |
| chr15 | 58153271 | 58153657 | 1.50368 | 2.69825 | 1.794 |
| chr10 | 72300258 | 72300696 | 2.21078 | 3.96726 | 1.795 |
| chr11 | 90097275 | 90097521 | 0.502247 | 0.901285 | 1.795 |
| chr20 | 4072488 | 4073031 | 2.06317 | 3.7024 | 1.795 |
| chr15 | 28091171 | 28091381 | 0.413422 | 0.74191 | 1.795 |
| chr9 | 138756303 | 138756708 | 2.5945 | 4.656 | 1.795 |
| chr15 | 28074562 | 28074803 | 0.715453 | 1.284 | 1.795 |
| chr17 | 14715235 | 14715463 | 0.74142 | 1.33061 | 1.795 |
| chr1 | 153544991 | 153545278 | 2.25223 | 4.04207 | 1.795 |
| chr3 | 194054442 | 194054788 | 2.3392 | 4.19817 | 1.795 |
| chr17 | 49699195 | 49699624 | 0.637817 | 1.14472 | 1.795 |
| chr1 | 228353309 | 228353681 | 3.10431 | 5.57148 | 1.795 |
| chrX | 150320106 | 150320558 | 1.22333 | 2.1956 | 1.795 |
| chr15 | 41196216 | 41196539 | 1.89933 | 3.409 | 1.795 |

| | | | | | |
|---|---|---|---|---|---|
| chr15 | 91445504 | 91445757 | 1.75515 | 3.15024 | 1.795 |
| chr15 | 80145829 | 80146222 | 2.77345 | 4.97827 | 1.795 |
| chr6 | 56179556 | 56179795 | 1.33296 | 2.39264 | 1.795 |
| chr6 | 138322972 | 138323191 | 1.1899 | 2.13587 | 1.795 |
| chr11 | 6776261 | 6776974 | 1.32959 | 2.38666 | 1.795 |
| chr7 | 155597699 | 155598519 | 1.88295 | 3.37998 | 1.795 |
| chr19 | 39408720 | 39408979 | 1.04231 | 1.87106 | 1.795 |
| chr13 | 96276235 | 96276601 | 0.915511 | 1.64345 | 1.795 |
| chr3 | 50361967 | 50362333 | 1.53354 | 2.75299 | 1.795 |
| chr7 | 20817533 | 20817993 | 2.12579 | 3.81627 | 1.795 |
| chr7 | 26130613 | 26130880 | 1.46187 | 2.62449 | 1.795 |
| chr10 | 110938096 | 110938484 | 0.612653 | 1.09991 | 1.795 |
| chr3 | 131929590 | 131929973 | 1.7647 | 3.16838 | 1.795 |
| chr20 | 22293417 | 22293714 | 1.83746 | 3.2992 | 1.796 |
| chr14 | 81364192 | 81364443 | 0.595463 | 1.06923 | 1.796 |
| chr2 | 238384041 | 238384389 | 2.23943 | 4.02119 | 1.796 |
| chr12 | 95510983 | 95511228 | 1.18442 | 2.1268 | 1.796 |
| chr5 | 32916802 | 32917062 | 1.4595 | 2.62075 | 1.796 |
| chr8 | 2876917 | 2877262 | 0.598891 | 1.07546 | 1.796 |
| chr1 | 2126587 | 2126977 | 1.43997 | 2.58589 | 1.796 |
| chr10 | 61666202 | 61666842 | 1.33654 | 2.40037 | 1.796 |
| chr2 | 237936102 | 237936363 | 1.45572 | 2.6146 | 1.796 |
| chr2 | 231865071 | 231865267 | 1.25423 | 2.25272 | 1.796 |
| chr2 | 76129462 | 76129693 | 0.802766 | 1.44187 | 1.796 |
| chr3 | 18878777 | 18878993 | 0.534261 | 0.959616 | 1.796 |
| chr12 | 22682273 | 22682785 | 0.924497 | 1.66064 | 1.796 |
| chr2 | 170078359 | 170078576 | 1.27819 | 2.29596 | 1.796 |
| chr9 | 34861138 | 34861425 | 0.494498 | 0.888336 | 1.796 |
| chr1 | 228566262 | 228566634 | 1.49494 | 2.68559 | 1.796 |
| chr13 | 114048228 | 114048696 | 1.46705 | 2.63553 | 1.796 |
| chr19 | 35633797 | 35634247 | 2.95895 | 5.31575 | 1.797 |
| chr1 | 225965194 | 225965526 | 1.97927 | 3.55587 | 1.797 |
| chr11 | 18797103 | 18797612 | 2.04248 | 3.66945 | 1.797 |
| chr2 | 4669628 | 4669846 | 0.664744 | 1.19426 | 1.797 |
| chr10 | 13116814 | 13117268 | 2.03748 | 3.66068 | 1.797 |
| chr6 | 24581317 | 24581642 | 1.65642 | 2.97611 | 1.797 |
| chr12 | 85390033 | 85390235 | 0.492539 | 0.884969 | 1.797 |
| chr19 | 45668111 | 45668363 | 0.645909 | 1.16055 | 1.797 |
| chr7 | 157021621 | 157021899 | 1.29525 | 2.32727 | 1.797 |
| chr12 | 49681977 | 49682484 | 1.54066 | 2.76828 | 1.797 |
| chr9 | 104364467 | 104364820 | 1.36732 | 2.4569 | 1.797 |
| chr19 | 35047496 | 35048027 | 1.098 | 1.973 | 1.797 |
| chr4 | 85425027 | 85425354 | 1.52252 | 2.73583 | 1.797 |
| chr7 | 37241206 | 37241544 | 2.00298 | 3.59921 | 1.797 |
| chr7 | 133675773 | 133676026 | 1.44654 | 2.59941 | 1.797 |
| chr9 | 116499758 | 116500062 | 2.47034 | 4.43915 | 1.797 |
| chr7 | 153351925 | 153352242 | 0.431602 | 0.775616 | 1.797 |
| chr4 | 160367498 | 160367988 | 1.17306 | 2.10809 | 1.797 |
| chr2 | 58259816 | 58260146 | 0.715778 | 1.28637 | 1.797 |
| chr21 | 37792237 | 37792558 | 1.51991 | 2.73174 | 1.797 |
| chr19 | 46167464 | 46167763 | 0.614998 | 1.10536 | 1.797 |
| chr10 | 35484987 | 35485297 | 1.10935 | 1.9939 | 1.797 |
| chr6 | 112318401 | 112318643 | 1.64306 | 2.95326 | 1.797 |
| chr1 | 159906237 | 159906619 | 2.01223 | 3.61692 | 1.797 |
| chr8 | 133887797 | 133888058 | 1.58591 | 2.85066 | 1.797 |
| chr17 | 27470898 | 27471269 | 1.00213 | 1.80134 | 1.798 |
| chr7 | 124708477 | 124708687 | 0.500159 | 0.899052 | 1.798 |
| chr7 | 128938166 | 128938464 | 1.68715 | 3.03272 | 1.798 |
| chr15 | 66895564 | 66896374 | 1.67389 | 3.0089 | 1.798 |
| chr2 | 80937007 | 80937227 | 0.504367 | 0.906625 | 1.798 |
| chr12 | 34500315 | 34500607 | 1.94156 | 3.49021 | 1.798 |

| | | | | | |
|---|---|---|---|---|---|
| chr15 | 97225738 | 97226006 | 0.940835 | 1.6913 | 1.798 |
| chr17 | 10130099 | 10130788 | 2.00884 | 3.61126 | 1.798 |
| chr8 | 40851288 | 40851593 | 2.32422 | 4.17831 | 1.798 |
| chr8 | 86196403 | 86196662 | 0.543602 | 0.977252 | 1.798 |
| chr2 | 22023359 | 22023595 | 0.567008 | 1.01934 | 1.798 |
| chr8 | 143897694 | 143898114 | 1.67551 | 3.01216 | 1.798 |
| chr10 | 93518039 | 93518635 | 0.818546 | 1.4716 | 1.798 |
| chr14 | 92759745 | 92759963 | 0.481947 | 0.866499 | 1.798 |
| chr1 | 162702719 | 162702931 | 0.6673 | 1.19975 | 1.798 |
| chr3 | 4697686 | 4698058 | 1.78714 | 3.21325 | 1.798 |
| chr3 | 50690274 | 50690498 | 0.935369 | 1.68181 | 1.798 |
| chr4 | 120971051 | 120971324 | 1.52972 | 2.75047 | 1.798 |
| chr17 | 47296986 | 47297336 | 1.41719 | 2.54816 | 1.798 |
| chr10 | 104661243 | 104661493 | 1.37896 | 2.47949 | 1.798 |
| chr12 | 129785664 | 129786118 | 0.432111 | 0.777035 | 1.798 |
| chr8 | 127219102 | 127219504 | 0.597444 | 1.07436 | 1.798 |
| chr7 | 5517304 | 5517520 | 0.963245 | 1.73217 | 1.798 |
| chr15 | 80952525 | 80952774 | 1.28736 | 2.31513 | 1.798 |
| chr3 | 15093088 | 15093405 | 1.98565 | 3.57098 | 1.798 |
| chr5 | 142035235 | 142035601 | 3.22594 | 5.80202 | 1.799 |
| chr5 | 176795973 | 176796518 | 1.13884 | 2.04836 | 1.799 |
| chr19 | 32421477 | 32421741 | 0.783432 | 1.40912 | 1.799 |
| chr1 | 209791891 | 209792158 | 0.597535 | 1.07476 | 1.799 |
| chr14 | 24217806 | 24218055 | 0.538766 | 0.969097 | 1.799 |
| chr1 | 150253701 | 150253931 | 1.44534 | 2.59981 | 1.799 |
| chr11 | 65183823 | 65184238 | 1.96325 | 3.53141 | 1.799 |
| chr17 | 42423202 | 42423560 | 0.885221 | 1.59231 | 1.799 |
| chr10 | 35924450 | 35924754 | 1.76599 | 3.17666 | 1.799 |
| chr11 | 96197972 | 96198214 | 1.46521 | 2.63571 | 1.799 |
| chr1 | 9551122 | 9551440 | 1.56465 | 2.81463 | 1.799 |
| chr17 | 19707247 | 19707498 | 0.681415 | 1.22587 | 1.799 |
| chr9 | 36328891 | 36329105 | 1.3294 | 2.39161 | 1.799 |
| chr14 | 24900893 | 24901357 | 2.3518 | 4.23107 | 1.799 |
| chr22 | 36617827 | 36618178 | 1.71891 | 3.0925 | 1.799 |
| chr4 | 39471060 | 39471335 | 1.26326 | 2.27276 | 1.799 |
| chrX | 41593877 | 41594121 | 1.15554 | 2.07895 | 1.799 |
| chr17 | 68982187 | 68982538 | 0.590644 | 1.06266 | 1.799 |
| chr2 | 202725324 | 202725627 | 1.52447 | 2.74275 | 1.799 |
| chr12 | 125382952 | 125383387 | 2.56289 | 4.61113 | 1.799 |
| chr8 | 4263154 | 4263548 | 0.574279 | 1.03339 | 1.799 |
| chr5 | 155118721 | 155118955 | 0.574909 | 1.03461 | 1.800 |
| chr5 | 155370853 | 155371196 | 2.08707 | 3.75596 | 1.800 |
| chr12 | 96608987 | 96609273 | 2.11417 | 3.80486 | 1.800 |
| chr3 | 113006913 | 113007264 | 2.77139 | 4.98767 | 1.800 |
| chr17 | 77024707 | 77024920 | 1.16742 | 2.10106 | 1.800 |
| chr7 | 154744735 | 154745292 | 1.17469 | 2.11436 | 1.800 |
| chr12 | 9903391 | 9903644 | 1.65566 | 2.98009 | 1.800 |
| chr16 | 1463646 | 1463952 | 1.97943 | 3.56288 | 1.800 |
| chr16 | 6269249 | 6269514 | 0.474558 | 0.854189 | 1.800 |
| chr16 | 48029388 | 48029607 | 0.796002 | 1.43279 | 1.800 |
| chr14 | 49705805 | 49706062 | 0.605576 | 1.09003 | 1.800 |
| chr14 | 93673826 | 93674101 | 1.16977 | 2.10557 | 1.800 |
| chr8 | 121820812 | 121821327 | 1.98916 | 3.58047 | 1.800 |
| chr3 | 132254908 | 132255137 | 0.955975 | 1.72076 | 1.800 |
| chr3 | 80762963 | 80763212 | 0.503969 | 0.90717 | 1.800 |
| chr6 | 33600336 | 33600627 | 1.20848 | 2.17535 | 1.800 |
| chr20 | 59827873 | 59828667 | 0.6513 | 1.17239 | 1.800 |
| chr1 | 200108813 | 200109085 | 1.3556 | 2.44021 | 1.800 |
| chr9 | 130587659 | 130588229 | 1.77757 | 3.19982 | 1.800 |
| chr20 | 39370215 | 39370518 | 1.37493 | 2.47505 | 1.800 |
| chr3 | 160131486 | 160131760 | 1.76449 | 3.17634 | 1.800 |

| | | | | | |
|---|---|---|---|---|---|
| chr17 | 17258198 | 17258669 | 1.99456 | 3.59064 | 1.800 |
| chr7 | 100063442 | 100063769 | 1.56129 | 2.81073 | 1.800 |
| chr21 | 17375269 | 17375471 | 0.899671 | 1.61974 | 1.800 |
| chr17 | 15312984 | 15313203 | 0.728031 | 1.31075 | 1.800 |
| chr16 | 25719359 | 25719650 | 0.916792 | 1.6506 | 1.800 |
| chr2 | 63002208 | 63002567 | 1.48435 | 2.67245 | 1.800 |
| chr17 | 17565247 | 17565454 | 1.04959 | 1.88971 | 1.800 |
| chr18 | 72160912 | 72161186 | 1.38088 | 2.48622 | 1.800 |
| chr18 | 8988337 | 8988595 | 1.84029 | 3.31343 | 1.800 |
| chr16 | 49955792 | 49956158 | 1.45638 | 2.62222 | 1.801 |
| chr19 | 47343749 | 47344112 | 2.21015 | 3.97941 | 1.801 |
| chr17 | 16208148 | 16208487 | 1.64488 | 2.96174 | 1.801 |
| chr16 | 84818958 | 84819169 | 0.807715 | 1.45437 | 1.801 |
| chr20 | 19609250 | 19609506 | 2.14755 | 3.86715 | 1.801 |
| chr3 | 172195131 | 172195399 | 1.3478 | 2.42707 | 1.801 |
| chr13 | 28400497 | 28401256 | 1.42506 | 2.56621 | 1.801 |
| chr12 | 16234682 | 16235009 | 0.698908 | 1.25859 | 1.801 |
| chr17 | 48439164 | 48439736 | 1.65072 | 2.97279 | 1.801 |
| chr17 | 78840089 | 78840338 | 1.82813 | 3.29256 | 1.801 |
| chr15 | 38982987 | 38983278 | 1.16314 | 2.09491 | 1.801 |
| chr1 | 45825531 | 45825753 | 1.30018 | 2.34177 | 1.801 |
| chr6 | 41747618 | 41747971 | 2.78929 | 5.02388 | 1.801 |
| chr1 | 167026523 | 167026823 | 1.7061 | 3.07307 | 1.801 |
| chr7 | 92462306 | 92462711 | 2.67232 | 4.81347 | 1.801 |
| chr4 | 88760563 | 88760776 | 1.20717 | 2.1744 | 1.801 |
| chr17 | 9742775 | 9742992 | 1.04286 | 1.87859 | 1.801 |
| chr5 | 172143407 | 172144128 | 1.23103 | 2.21755 | 1.801 |
| chr10 | 74100180 | 74100641 | 2.25114 | 4.05524 | 1.801 |
| chr19 | 40476669 | 40476954 | 2.48449 | 4.47566 | 1.801 |
| chr10 | 53080203 | 53080432 | 0.544366 | 0.980646 | 1.801 |
| chr11 | 91766610 | 91766852 | 0.51056 | 0.919775 | 1.802 |
| chr10 | 36024344 | 36024812 | 0.557059 | 1.00357 | 1.802 |
| chr17 | 12199236 | 12199906 | 0.501024 | 0.902662 | 1.802 |
| chr11 | 45354012 | 45354269 | 1.43212 | 2.58017 | 1.802 |
| chr2 | 29461426 | 29461689 | 1.75248 | 3.15744 | 1.802 |
| chr17 | 74381843 | 74382249 | 2.11318 | 3.80754 | 1.802 |
| chr14 | 23442731 | 23443080 | 1.09134 | 1.96651 | 1.802 |
| chr1 | 182053359 | 182053804 | 1.07967 | 1.94568 | 1.802 |
| chr5 | 5886697 | 5887529 | 1.39729 | 2.51811 | 1.802 |
| chr16 | 29703313 | 29703688 | 2.97312 | 5.35804 | 1.802 |
| chr1 | 26946738 | 26947097 | 1.51702 | 2.73394 | 1.802 |
| chr8 | 4632239 | 4632460 | 0.663232 | 1.19531 | 1.802 |
| chr1 | 223302617 | 223303018 | 2.66911 | 4.81047 | 1.802 |
| chr17 | 36489569 | 36490083 | 1.67424 | 3.01753 | 1.802 |
| chr20 | 1417916 | 1418428 | 2.06286 | 3.71795 | 1.802 |
| chr3 | 20081235 | 20081614 | 1.46831 | 2.64642 | 1.802 |
| chr18 | 44530128 | 44530341 | 1.35303 | 2.43868 | 1.802 |
| chr17 | 21142997 | 21143504 | 1.23331 | 2.22313 | 1.803 |
| chr4 | 154128749 | 154128998 | 1.23074 | 2.21853 | 1.803 |
| chr22 | 49364069 | 49364348 | 1.39638 | 2.51725 | 1.803 |
| chr10 | 72309603 | 72309950 | 2.45923 | 4.43327 | 1.803 |
| chr2 | 192403871 | 192404115 | 1.40144 | 2.52645 | 1.803 |
| chr21 | 39282254 | 39282584 | 0.718024 | 1.2947 | 1.803 |
| chr7 | 21277030 | 21277276 | 0.591673 | 1.06687 | 1.803 |
| chr12 | 15680519 | 15680734 | 1.36984 | 2.47009 | 1.803 |
| chr15 | 102204617 | 102204863 | 1.42764 | 2.57433 | 1.803 |
| chr2 | 86009835 | 86010038 | 0.651489 | 1.17477 | 1.803 |
| chr8 | 121150042 | 121150247 | 0.679798 | 1.22582 | 1.803 |
| chr15 | 69815760 | 69816071 | 1.90771 | 3.44003 | 1.803 |
| chr7 | 139665224 | 139665448 | 0.64996 | 1.17203 | 1.803 |
| chr11 | 73046533 | 73046827 | 3.20191 | 5.77385 | 1.803 |

| | | | | | |
|---|---|---|---|---|---|
| chr6 | 17086719 | 17086931 | 1.12522 | 2.02905 | 1.803 |
| chr10 | 128286593 | 128286936 | 1.46396 | 2.63998 | 1.803 |
| chr5 | 117899111 | 117899343 | 0.455669 | 0.821732 | 1.803 |
| chr8 | 132567954 | 132568236 | 0.617033 | 1.11274 | 1.803 |
| chr17 | 36819378 | 36819771 | 2.13573 | 3.85162 | 1.803 |
| chr5 | 117435959 | 117436294 | 0.533333 | 0.961847 | 1.803 |
| chr5 | 4700315 | 4700582 | 0.516115 | 0.930822 | 1.804 |
| chr3 | 12851508 | 12851797 | 1.53859 | 2.77495 | 1.804 |
| chr22 | 23035287 | 23035572 | 1.64172 | 2.96103 | 1.804 |
| chr9 | 100796847 | 100797313 | 1.82352 | 3.28895 | 1.804 |
| chr4 | 110971170 | 110971420 | 1.29752 | 2.34029 | 1.804 |
| chr15 | 79303913 | 79304124 | 0.649779 | 1.17201 | 1.804 |
| chr4 | 28688192 | 28688551 | 0.954967 | 1.72248 | 1.804 |
| chr19 | 42924133 | 42924558 | 2.67847 | 4.83124 | 1.804 |
| chr11 | 67976636 | 67976967 | 2.14231 | 3.86416 | 1.804 |
| chr10 | 114449128 | 114449352 | 1.26876 | 2.28858 | 1.804 |
| chr17 | 65713765 | 65714080 | 2.69076 | 4.85356 | 1.804 |
| chr1 | 156658797 | 156659667 | 1.60354 | 2.89252 | 1.804 |
| chr10 | 106340614 | 106340912 | 1.62512 | 2.93148 | 1.804 |
| chr19 | 51762639 | 51762947 | 1.62454 | 2.93047 | 1.804 |
| chr17 | 46152550 | 46152889 | 0.605274 | 1.09189 | 1.804 |
| chr9 | 14431844 | 14432207 | 0.533927 | 0.963185 | 1.804 |
| chr1 | 177915049 | 177915534 | 1.96123 | 3.53803 | 1.804 |
| chr8 | 11816367 | 11816728 | 2.46361 | 4.44433 | 1.804 |
| chr17 | 42610878 | 42611238 | 2.81229 | 5.07343 | 1.804 |
| chr7 | 140623983 | 140624351 | 1.12278 | 2.02552 | 1.804 |
| chr5 | 42653767 | 42654110 | 0.542483 | 0.978672 | 1.804 |
| chr7 | 51435627 | 51435869 | 0.678388 | 1.2239 | 1.804 |
| chr19 | 55966043 | 55966621 | 2.12967 | 3.84223 | 1.804 |
| chr1 | 154850907 | 154851423 | 1.62971 | 2.9403 | 1.804 |
| chr10 | 45219879 | 45220280 | 0.554712 | 1.00082 | 1.804 |
| chr10 | 69066431 | 69066635 | 1.40276 | 2.53092 | 1.804 |
| chr20 | 37360084 | 37360657 | 2.00677 | 3.62069 | 1.804 |
| chr4 | 6112019 | 6112231 | 1.0952 | 1.97602 | 1.804 |
| chr19 | 42937732 | 42938263 | 2.79829 | 5.04884 | 1.804 |
| chr2 | 166810602 | 166810851 | 1.42224 | 2.56613 | 1.804 |
| chr5 | 63978529 | 63978786 | 0.899206 | 1.62244 | 1.804 |
| chr16 | 30856059 | 30856436 | 2.46554 | 4.44889 | 1.804 |
| chr2 | 49575684 | 49576217 | 0.539459 | 0.97343 | 1.804 |
| chr10 | 91811142 | 91811487 | 0.686427 | 1.23865 | 1.804 |
| chr5 | 131400241 | 131401349 | 1.44542 | 2.60835 | 1.805 |
| chr4 | 155427420 | 155427649 | 0.930904 | 1.6799 | 1.805 |
| chr4 | 2950708 | 2950946 | 1.07162 | 1.93385 | 1.805 |
| chr15 | 51169172 | 51169434 | 2.36015 | 4.25929 | 1.805 |
| chr17 | 46451910 | 46452325 | 1.46159 | 2.6378 | 1.805 |
| chr3 | 477827 | 478091 | 0.576835 | 1.04105 | 1.805 |
| chr7 | 148663807 | 148664305 | 2.55366 | 4.60883 | 1.805 |
| chr2 | 104695273 | 104695636 | 1.10222 | 1.98929 | 1.805 |
| chr17 | 53795513 | 53795892 | 1.67759 | 3.02777 | 1.805 |
| chr10 | 43693339 | 43693725 | 0.887429 | 1.60175 | 1.805 |
| chr17 | 64670009 | 64670232 | 1.55329 | 2.8036 | 1.805 |
| chr3 | 45634680 | 45634963 | 2.04327 | 3.68824 | 1.805 |
| chr1 | 207080843 | 207081278 | 1.15016 | 2.07616 | 1.805 |
| chr1 | 52467940 | 52468232 | 1.24613 | 2.24946 | 1.805 |
| chr15 | 27185479 | 27185718 | 0.461858 | 0.833746 | 1.805 |
| chr10 | 102102101 | 102102492 | 1.50014 | 2.70808 | 1.805 |
| chr7 | 90258712 | 90258960 | 1.06867 | 1.92919 | 1.805 |
| chr10 | 7147686 | 7148050 | 0.738021 | 1.33232 | 1.805 |
| chr14 | 81053461 | 81053683 | 1.46604 | 2.64659 | 1.805 |
| chr4 | 89377894 | 89378103 | 1.12177 | 2.02515 | 1.805 |
| chr20 | 51566219 | 51566555 | 0.630056 | 1.13751 | 1.805 |

| | | | | | |
|---|---|---|---|---|---|
| chr6 | 117290409 | 117290690 | 1.01977 | 1.84114 | 1.805 |
| chr11 | 73062169 | 73062642 | 2.225 | 4.01716 | 1.805 |
| chr19 | 38633041 | 38633514 | 2.45254 | 4.42831 | 1.806 |
| chr5 | 40904808 | 40905198 | 2.78558 | 5.02971 | 1.806 |
| chr5 | 43105511 | 43106159 | 2.80855 | 5.07165 | 1.806 |
| chr4 | 183839027 | 183839418 | 0.883119 | 1.59479 | 1.806 |
| chr16 | 57553699 | 57554058 | 2.69482 | 4.86654 | 1.806 |
| chr1 | 174027721 | 174027999 | 0.470642 | 0.849936 | 1.806 |
| chr12 | 21322845 | 21323138 | 1.70389 | 3.0771 | 1.806 |
| chr14 | 77111266 | 77111634 | 1.64017 | 2.96231 | 1.806 |
| chr15 | 69121373 | 69121591 | 0.670704 | 1.2114 | 1.806 |
| chr17 | 65374790 | 65375254 | 3.00634 | 5.43029 | 1.806 |
| chr2 | 52375362 | 52375665 | 0.537595 | 0.971107 | 1.806 |
| chr20 | 646617 | 646842 | 0.753024 | 1.36027 | 1.806 |
| chr8 | 112730675 | 112730988 | 0.706151 | 1.27562 | 1.806 |
| chr11 | 31663059 | 31663417 | 1.23735 | 2.23522 | 1.806 |
| chr18 | 7568500 | 7568821 | 0.908887 | 1.6419 | 1.807 |
| chr2 | 175206222 | 175206502 | 2.49684 | 4.51054 | 1.807 |
| chr14 | 24134874 | 24135404 | 1.31705 | 2.3794 | 1.807 |
| chr16 | 87669046 | 87669491 | 2.53132 | 4.57311 | 1.807 |
| chr11 | 65278279 | 65278542 | 1.71146 | 3.09195 | 1.807 |
| chr17 | 8735887 | 8736530 | 2.50949 | 4.53372 | 1.807 |
| chr15 | 84105312 | 84105632 | 1.49999 | 2.70998 | 1.807 |
| chr19 | 43446665 | 43447116 | 0.900532 | 1.62703 | 1.807 |
| chr16 | 66906978 | 66907421 | 2.4894 | 4.49773 | 1.807 |
| chr7 | 148436026 | 148436283 | 0.436829 | 0.789265 | 1.807 |
| chr8 | 39566289 | 39566495 | 0.624897 | 1.12911 | 1.807 |
| chr15 | 59821136 | 59821389 | 1.63939 | 2.96249 | 1.807 |
| chr11 | 113644641 | 113644961 | 1.29337 | 2.33725 | 1.807 |
| chr5 | 139348286 | 139348797 | 3.01707 | 5.45217 | 1.807 |
| chr5 | 94511241 | 94511498 | 0.636033 | 1.14941 | 1.807 |
| chr20 | 47863084 | 47863380 | 1.37718 | 2.4888 | 1.807 |
| chr4 | 174673674 | 174674074 | 1.24187 | 2.24434 | 1.807 |
| chr10 | 65479760 | 65480071 | 1.70224 | 3.07637 | 1.807 |
| chr2 | 84722724 | 84723022 | 1.19719 | 2.16365 | 1.807 |
| chr5 | 56653984 | 56654331 | 0.529172 | 0.956361 | 1.807 |
| chr8 | 70854448 | 70854904 | 1.36813 | 2.47268 | 1.807 |
| chr2 | 43358698 | 43359020 | 1.67894 | 3.03444 | 1.807 |
| chr3 | 128594497 | 128594739 | 1.34514 | 2.43116 | 1.807 |
| chr9 | 71081547 | 71081879 | 0.627259 | 1.13372 | 1.807 |
| chr12 | 22697129 | 22697331 | 1.27944 | 2.31255 | 1.807 |
| chr15 | 96918617 | 96918885 | 0.625327 | 1.13027 | 1.807 |
| chr12 | 45366698 | 45366906 | 1.76417 | 3.1891 | 1.808 |
| chr14 | 97888724 | 97888950 | 0.696769 | 1.25956 | 1.808 |
| chr3 | 111583681 | 111583943 | 0.957222 | 1.73043 | 1.808 |
| chr3 | 20553375 | 20553598 | 0.949752 | 1.71708 | 1.808 |
| chr11 | 34862360 | 34862708 | 1.29516 | 2.34162 | 1.808 |
| chr17 | 2479313 | 2479527 | 0.499867 | 0.903749 | 1.808 |
| chr14 | 55912960 | 55913216 | 1.43717 | 2.59845 | 1.808 |
| chr16 | 4363816 | 4364173 | 2.64514 | 4.78254 | 1.808 |
| chr7 | 43850237 | 43850468 | 1.18196 | 2.13708 | 1.808 |
| chr16 | 65151824 | 65152219 | 2.27426 | 4.11218 | 1.808 |
| chr3 | 13541561 | 13542213 | 1.51566 | 2.74065 | 1.808 |
| chrX | 48489756 | 48489992 | 0.749501 | 1.35528 | 1.808 |
| chr15 | 74595182 | 74595557 | 1.59758 | 2.88889 | 1.808 |
| chr16 | 8012209 | 8012622 | 0.641817 | 1.16059 | 1.808 |
| chr5 | 36303847 | 36304057 | 0.644051 | 1.1647 | 1.808 |
| chr1 | 41898411 | 41898771 | 1.70056 | 3.07533 | 1.808 |
| chr18 | 55710806 | 55711073 | 1.44424 | 2.61179 | 1.808 |
| chr16 | 68297727 | 68298298 | 2.09212 | 3.78348 | 1.808 |
| chr7 | 2143856 | 2144216 | 1.55782 | 2.8173 | 1.808 |

| | | | | | |
|---|---|---|---|---|---|
| chr17 | 2319318 | 2319747 | 1.53277 | 2.77201 | 1.809 |
| chr1 | 26611541 | 26611923 | 1.7003 | 3.07502 | 1.809 |
| chr16 | 75267616 | 75268034 | 1.90593 | 3.44712 | 1.809 |
| chr6 | 168472282 | 168472587 | 0.968774 | 1.75222 | 1.809 |
| chr2 | 20787298 | 20787626 | 0.669009 | 1.21005 | 1.809 |
| chr5 | 140000287 | 140000497 | 0.624486 | 1.12952 | 1.809 |
| chr2 | 231663595 | 231663852 | 1.46829 | 2.65582 | 1.809 |
| chr10 | 30648645 | 30649093 | 1.39636 | 2.52573 | 1.809 |
| chr20 | 10655313 | 10655762 | 2.17751 | 3.93871 | 1.809 |
| chr1 | 20820315 | 20820591 | 1.8584 | 3.36156 | 1.809 |
| chr7 | 124712633 | 124712897 | 0.512593 | 0.927245 | 1.809 |
| chr12 | 68788429 | 68788767 | 0.721344 | 1.30487 | 1.809 |
| chr1 | 1057493 | 1057839 | 2.25922 | 4.08681 | 1.809 |
| chr11 | 75258584 | 75258962 | 1.75345 | 3.17213 | 1.809 |
| chr11 | 87088594 | 87088834 | 0.635411 | 1.14952 | 1.809 |
| chr2 | 232057217 | 232057450 | 1.58636 | 2.87022 | 1.809 |
| chr12 | 89411012 | 89411256 | 1.69352 | 3.06418 | 1.809 |
| chr10 | 112506187 | 112506448 | 1.73371 | 3.13708 | 1.809 |
| chr8 | 119515789 | 119516084 | 1.1923 | 2.15745 | 1.809 |
| chr3 | 181427395 | 181427635 | 1.14877 | 2.0787 | 1.810 |
| chr9 | 110187550 | 110187999 | 1.71017 | 3.09466 | 1.810 |
| chr7 | 125019484 | 125019719 | 0.468569 | 0.847911 | 1.810 |
| chr19 | 46898768 | 46899404 | 2.578 | 4.66514 | 1.810 |
| chr2 | 37641605 | 37641873 | 1.97265 | 3.56973 | 1.810 |
| chr10 | 94833870 | 94834125 | 1.07839 | 1.9515 | 1.810 |
| chr5 | 164154257 | 164154465 | 0.448779 | 0.812127 | 1.810 |
| chr3 | 888689 | 888933 | 0.506501 | 0.916589 | 1.810 |
| chr7 | 26267881 | 26268113 | 0.467449 | 0.845941 | 1.810 |
| chr19 | 52281167 | 52281448 | 1.37683 | 2.49167 | 1.810 |
| chr17 | 26971074 | 26971283 | 0.978679 | 1.77122 | 1.810 |
| chr3 | 10266653 | 10266967 | 1.451 | 2.62603 | 1.810 |
| chr12 | 114404166 | 114404494 | 2.5741 | 4.6587 | 1.810 |
| chr13 | 78325669 | 78326345 | 1.14626 | 2.07462 | 1.810 |
| chr9 | 95449306 | 95449637 | 0.869087 | 1.57297 | 1.810 |
| chr17 | 56076872 | 56077249 | 1.37984 | 2.49741 | 1.810 |
| chr2 | 173842138 | 173842380 | 1.32451 | 2.39727 | 1.810 |
| chr20 | 49242659 | 49243030 | 2.39783 | 4.33999 | 1.810 |
| chr2 | 110472958 | 110473198 | 0.885076 | 1.60198 | 1.810 |
| chr18 | 8449970 | 8450220 | 1.64373 | 2.97517 | 1.810 |
| chr2 | 98326386 | 98326669 | 1.90481 | 3.44786 | 1.810 |
| chr15 | 42103681 | 42103995 | 1.73737 | 3.1448 | 1.810 |
| chr8 | 122261075 | 122261373 | 1.92094 | 3.47727 | 1.810 |
| chr2 | 171829838 | 171830268 | 2.24505 | 4.06398 | 1.810 |
| chr10 | 105644480 | 105644770 | 1.69226 | 3.06335 | 1.810 |
| chr15 | 45722412 | 45722899 | 2.79637 | 5.06209 | 1.810 |
| chr2 | 179565660 | 179565953 | 1.65961 | 3.00435 | 1.810 |
| chr3 | 157260477 | 157260779 | 1.74161 | 3.15287 | 1.810 |
| chr14 | 23764233 | 23764663 | 1.90971 | 3.45729 | 1.810 |
| chr3 | 69359035 | 69359315 | 1.18277 | 2.14133 | 1.810 |
| chr17 | 35774346 | 35774551 | 1.06856 | 1.93463 | 1.811 |
| chr1 | 228318720 | 228319062 | 1.80682 | 3.2713 | 1.811 |
| chr12 | 114593622 | 114593874 | 0.638814 | 1.15661 | 1.811 |
| chr4 | 187647011 | 187647351 | 1.41965 | 2.57047 | 1.811 |
| chr17 | 1162358 | 1162855 | 1.86474 | 3.37652 | 1.811 |
| chr15 | 73406455 | 73406762 | 0.628945 | 1.13888 | 1.811 |
| chr19 | 51228454 | 51228726 | 0.558419 | 1.01119 | 1.811 |
| chr8 | 49838091 | 49838480 | 1.2848 | 2.32666 | 1.811 |
| chr7 | 80676611 | 80676827 | 0.664435 | 1.20324 | 1.811 |
| chr5 | 17189571 | 17189955 | 1.69159 | 3.06347 | 1.811 |
| chr18 | 21078718 | 21079177 | 1.98573 | 3.59625 | 1.811 |
| chr12 | 91372002 | 91372259 | 1.39277 | 2.52246 | 1.811 |

| | | | | | |
|---|---|---|---|---|---|
| chr11 | 5312526 | 5312936 | 1.49448 | 2.70684 | 1.811 |
| chr8 | 81910561 | 81910832 | 2.19489 | 3.97555 | 1.811 |
| chr10 | 109625491 | 109625837 | 0.59964 | 1.08614 | 1.811 |
| chr6 | 35295292 | 35295536 | 1.54939 | 2.80645 | 1.811 |
| chr1 | 36615146 | 36615456 | 1.55137 | 2.81007 | 1.811 |
| chr8 | 98800143 | 98800442 | 1.42812 | 2.58683 | 1.811 |
| chr5 | 147558914 | 147559160 | 1.07123 | 1.9404 | 1.811 |
| chr7 | 148906586 | 148907016 | 1.2309 | 2.22963 | 1.811 |
| chr3 | 47040078 | 47040357 | 2.90617 | 5.26445 | 1.811 |
| chr16 | 60992274 | 60992485 | 0.571227 | 1.03481 | 1.812 |
| chr11 | 119194303 | 119194656 | 1.49915 | 2.71582 | 1.812 |
| chr3 | 71211378 | 71211661 | 0.606783 | 1.09923 | 1.812 |
| chr10 | 110088268 | 110088554 | 0.618262 | 1.12008 | 1.812 |
| chr19 | 33408135 | 33408390 | 0.717676 | 1.30022 | 1.812 |
| chr1 | 147283663 | 147283913 | 0.512028 | 0.927705 | 1.812 |
| chr1 | 113423498 | 113423866 | 2.06585 | 3.74297 | 1.812 |
| chr17 | 12440282 | 12440669 | 0.519334 | 0.940945 | 1.812 |
| chr16 | 1350739 | 1351301 | 2.08862 | 3.78432 | 1.812 |
| chr3 | 66866628 | 66866862 | 1.40861 | 2.55223 | 1.812 |
| chr3 | 3265625 | 3265962 | 0.537074 | 0.973146 | 1.812 |
| chr12 | 2450478 | 2450751 | 0.55773 | 1.01059 | 1.812 |
| chr9 | 136741354 | 136741603 | 1.22552 | 2.22066 | 1.812 |
| chr7 | 134199393 | 134199717 | 1.87618 | 3.39971 | 1.812 |
| chr15 | 93595807 | 93596082 | 1.10237 | 1.99755 | 1.812 |
| chr1 | 17988143 | 17988500 | 1.77118 | 3.2095 | 1.812 |
| chr12 | 3336443 | 3336960 | 0.569033 | 1.03117 | 1.812 |
| chr17 | 74366466 | 74367055 | 1.21966 | 2.2102 | 1.812 |
| chr12 | 6136308 | 6136570 | 0.560524 | 1.01576 | 1.812 |
| chr1 | 228800112 | 228800533 | 1.61656 | 2.92949 | 1.812 |
| chrX | 103029084 | 103029299 | 1.02591 | 1.85914 | 1.812 |
| chr16 | 85751447 | 85751862 | 2.3597 | 4.27631 | 1.812 |
| chr7 | 4923160 | 4923729 | 2.17002 | 3.93269 | 1.812 |
| chr1 | 2105735 | 2106171 | 2.20316 | 3.99284 | 1.812 |
| chr16 | 4556338 | 4556825 | 1.62528 | 2.9456 | 1.812 |
| chr11 | 75863191 | 75863617 | 2.53661 | 4.59744 | 1.812 |
| chr7 | 30840388 | 30840745 | 1.12857 | 2.04547 | 1.812 |
| chr19 | 1864813 | 1865049 | 1.08023 | 1.95786 | 1.812 |
| chr17 | 32747226 | 32747629 | 2.0762 | 3.76303 | 1.812 |
| chr12 | 129325379 | 129325585 | 0.619479 | 1.12281 | 1.813 |
| chr3 | 144001138 | 144001358 | 0.743854 | 1.34824 | 1.813 |
| chr5 | 98086182 | 98086726 | 0.778104 | 1.41036 | 1.813 |
| chr15 | 28163322 | 28163544 | 0.404383 | 0.732973 | 1.813 |
| chr3 | 10210502 | 10210861 | 1.0726 | 1.94431 | 1.813 |
| chr10 | 129291119 | 129291506 | 1.51901 | 2.75363 | 1.813 |
| chr11 | 65704620 | 65704906 | 0.507394 | 0.919811 | 1.813 |
| chr17 | 21323228 | 21323620 | 0.717767 | 1.30122 | 1.813 |
| chr9 | 33415428 | 33415809 | 1.96735 | 3.56664 | 1.813 |
| chr7 | 89927548 | 89927750 | 1.51958 | 2.75497 | 1.813 |
| chr5 | 142128208 | 142128512 | 1.43775 | 2.60699 | 1.813 |
| chr21 | 35274123 | 35274339 | 1.02583 | 1.8601 | 1.813 |
| chr9 | 29212143 | 29212569 | 1.80858 | 3.27942 | 1.813 |
| chr3 | 23001943 | 23002195 | 1.15714 | 2.09826 | 1.813 |
| chr22 | 23744877 | 23745163 | 2.32276 | 4.21192 | 1.813 |
| chr8 | 139711076 | 139711555 | 1.50615 | 2.7312 | 1.813 |
| chr10 | 54964230 | 54964440 | 0.378631 | 0.686607 | 1.813 |
| chr1 | 18626151 | 18626551 | 1.46296 | 2.65302 | 1.813 |
| chr1 | 52000738 | 52000974 | 1.44815 | 2.6262 | 1.813 |
| chr19 | 50511805 | 50512098 | 1.15978 | 2.10328 | 1.814 |
| chr8 | 146052079 | 146052355 | 1.15192 | 2.08907 | 1.814 |
| chrX | 117973259 | 117973653 | 2.54151 | 4.60946 | 1.814 |
| chr1 | 222628504 | 222628793 | 2.14487 | 3.89013 | 1.814 |

| | | | | | |
|---|---|---|---|---|---|
| chr8 | 101586246 | 101586560 | 1.4167 | 2.56949 | 1.814 |
| chr9 | 12096073 | 12096283 | 0.629085 | 1.14102 | 1.814 |
| chr3 | 67432164 | 67432353 | 1.18086 | 2.14189 | 1.814 |
| chr13 | 40351633 | 40351871 | 1.06477 | 1.93134 | 1.814 |
| chr12 | 110537964 | 110538658 | 1.3558 | 2.4593 | 1.814 |
| chr12 | 121647711 | 121647930 | 1.0856 | 1.96921 | 1.814 |
| chr12 | 127135579 | 127136020 | 0.574112 | 1.04142 | 1.814 |
| chr5 | 141436958 | 141437177 | 0.550443 | 0.998489 | 1.814 |
| chr6 | 36853445 | 36853673 | 1.5895 | 2.88333 | 1.814 |
| chr1 | 110088960 | 110089175 | 1.3722 | 2.48917 | 1.814 |
| chr19 | 30101271 | 30101851 | 2.04854 | 3.71611 | 1.814 |
| chr13 | 99495192 | 99495491 | 1.65869 | 3.00894 | 1.814 |
| chr18 | 19192035 | 19192348 | 1.98104 | 3.5938 | 1.814 |
| chr5 | 173411346 | 173411548 | 1.36255 | 2.47185 | 1.814 |
| chr17 | 55824930 | 55825287 | 1.71908 | 3.11868 | 1.814 |
| chr9 | 137335108 | 137335522 | 2.26329 | 4.10598 | 1.814 |
| chr1 | 176145652 | 176145896 | 0.555711 | 1.00817 | 1.814 |
| chr17 | 75426003 | 75426521 | 1.80226 | 3.26972 | 1.814 |
| chr17 | 80758054 | 80758305 | 0.624516 | 1.13302 | 1.814 |
| chrX | 24168448 | 24168751 | 0.796875 | 1.44572 | 1.814 |
| chr14 | 107091862 | 107092088 | 0.457717 | 0.830425 | 1.814 |
| chr5 | 150080058 | 150080361 | 1.24588 | 2.26046 | 1.814 |
| chr17 | 33923705 | 33923992 | 1.17629 | 2.13427 | 1.814 |
| chr13 | 82531275 | 82531489 | 0.749555 | 1.36003 | 1.814 |
| chr9 | 108175890 | 108176109 | 1.47744 | 2.6808 | 1.814 |
| chr17 | 43131948 | 43132173 | 0.430624 | 0.781408 | 1.815 |
| chr11 | 33829828 | 33830158 | 2.22126 | 4.03082 | 1.815 |
| chr5 | 122759109 | 122759340 | 2.27616 | 4.13051 | 1.815 |
| chr17 | 78835637 | 78835980 | 2.37437 | 4.30878 | 1.815 |
| chr10 | 101605630 | 101606072 | 1.41881 | 2.5748 | 1.815 |
| chr14 | 105478477 | 105479021 | 2.01557 | 3.65778 | 1.815 |
| chr19 | 45988443 | 45989421 | 2.05868 | 3.73606 | 1.815 |
| chr19 | 1445163 | 1445790 | 1.73981 | 3.15741 | 1.815 |
| chr11 | 57226070 | 57226518 | 2.17859 | 3.95374 | 1.815 |
| chr15 | 62546473 | 62546926 | 2.07852 | 3.77213 | 1.815 |
| chr17 | 26940258 | 26940631 | 1.824 | 3.31049 | 1.815 |
| chr14 | 49795862 | 49796114 | 0.467556 | 0.848612 | 1.815 |
| chr14 | 21560557 | 21561079 | 4.97364 | 9.02734 | 1.815 |
| chr17 | 25563766 | 25564107 | 1.32603 | 2.40688 | 1.815 |
| chr19 | 38625204 | 38625601 | 2.67216 | 4.85034 | 1.815 |
| chr11 | 45332739 | 45333064 | 0.529741 | 0.96156 | 1.815 |
| chr1 | 96143736 | 96143999 | 0.389008 | 0.70611 | 1.815 |
| chr10 | 59416942 | 59417220 | 1.9932 | 3.61797 | 1.815 |
| chr2 | 6637732 | 6637961 | 0.492979 | 0.894854 | 1.815 |
| chr17 | 20896168 | 20896572 | 2.39842 | 4.35364 | 1.815 |
| chr10 | 113376968 | 113377389 | 0.880295 | 1.59794 | 1.815 |
| chr10 | 24017810 | 24018064 | 0.60517 | 1.0986 | 1.815 |
| chr20 | 51820115 | 51820373 | 0.619145 | 1.12398 | 1.815 |
| chr3 | 129209097 | 129209379 | 2.1813 | 3.95998 | 1.815 |
| chr6 | 162258274 | 162258529 | 1.37065 | 2.4886 | 1.816 |
| chr16 | 7067114 | 7067335 | 0.660862 | 1.19993 | 1.816 |
| chr15 | 90613978 | 90614244 | 0.635965 | 1.15485 | 1.816 |
| chr11 | 47201060 | 47201435 | 1.75272 | 3.18278 | 1.816 |
| chr11 | 40126747 | 40126977 | 0.758852 | 1.37802 | 1.816 |
| chr7 | 143002586 | 143002902 | 2.16796 | 3.93687 | 1.816 |
| chr18 | 6543441 | 6543684 | 0.958668 | 1.74091 | 1.816 |
| chr1 | 45265404 | 45265685 | 2.2345 | 4.05789 | 1.816 |
| chr10 | 103057602 | 103058017 | 2.7769 | 5.043 | 1.816 |
| chr16 | 12189028 | 12189429 | 1.75942 | 3.19524 | 1.816 |
| chr1 | 97603884 | 97604068 | 1.00088 | 1.81775 | 1.816 |
| chr11 | 57045362 | 57045777 | 1.73881 | 3.15802 | 1.816 |

| | | | | | |
|---|---|---|---|---|---|
| chr3 | 58515205 | 58515563 | 1.18515 | 2.15247 | 1.816 |
| chr2 | 76707176 | 76707402 | 0.620553 | 1.12706 | 1.816 |
| chr2 | 56357603 | 56357929 | 1.53966 | 2.79649 | 1.816 |
| chr20 | 30081895 | 30082277 | 1.06355 | 1.93175 | 1.816 |
| chr5 | 135418127 | 135418612 | 1.98871 | 3.61218 | 1.816 |
| chr13 | 110522068 | 110522310 | 1.6283 | 2.95757 | 1.816 |
| chr19 | 49895039 | 49895489 | 1.75935 | 3.19571 | 1.816 |
| chr5 | 87076951 | 87077302 | 2.11395 | 3.83995 | 1.816 |
| chrX | 103381154 | 103381434 | 1.21018 | 2.19835 | 1.817 |
| chr2 | 74374901 | 74375272 | 2.86827 | 5.21039 | 1.817 |
| chr12 | 57632575 | 57633296 | 3.30537 | 6.00468 | 1.817 |
| chr7 | 149007363 | 149007632 | 1.35435 | 2.46037 | 1.817 |
| chr15 | 45691862 | 45692168 | 1.3679 | 2.48519 | 1.817 |
| chr5 | 109527408 | 109527634 | 0.593636 | 1.07852 | 1.817 |
| chr17 | 1546737 | 1547158 | 1.10497 | 2.00753 | 1.817 |
| chr3 | 10305659 | 10306008 | 1.19439 | 2.16999 | 1.817 |
| chr11 | 20376774 | 20377104 | 1.10621 | 2.00979 | 1.817 |
| chr11 | 67150049 | 67150374 | 2.65047 | 4.81553 | 1.817 |
| chrX | 127969487 | 127969742 | 0.501193 | 0.910609 | 1.817 |
| chr11 | 38433310 | 38433568 | 0.5731 | 1.04126 | 1.817 |
| chr8 | 75256895 | 75257214 | 0.875137 | 1.59003 | 1.817 |
| chr5 | 79582136 | 79582401 | 0.661068 | 1.20111 | 1.817 |
| chr11 | 61058034 | 61058356 | 0.548715 | 0.997001 | 1.817 |
| chr1 | 176256682 | 176256914 | 0.60975 | 1.10792 | 1.817 |
| chrX | 70188205 | 70188430 | 0.469236 | 0.852614 | 1.817 |
| chr8 | 31050007 | 31050282 | 0.390634 | 0.709804 | 1.817 |
| chr7 | 147959633 | 147960287 | 1.62203 | 2.94742 | 1.817 |
| chr3 | 118073690 | 118074246 | 0.481516 | 0.874978 | 1.817 |
| chr6 | 47748556 | 47748805 | 1.17088 | 2.12775 | 1.817 |
| chrX | 83984813 | 83985129 | 1.40109 | 2.54614 | 1.817 |
| chrX | 119648121 | 119648401 | 1.24924 | 2.27029 | 1.817 |
| chr3 | 169755397 | 169755932 | 1.98882 | 3.61468 | 1.818 |
| chr11 | 9769980 | 9770424 | 1.95109 | 3.54618 | 1.818 |
| chr16 | 66440350 | 66440638 | 1.33081 | 2.41895 | 1.818 |
| chr8 | 134152883 | 134153174 | 0.572656 | 1.04103 | 1.818 |
| chr3 | 150945463 | 150945918 | 1.54139 | 2.80217 | 1.818 |
| chr7 | 150145823 | 150146122 | 2.20554 | 4.00963 | 1.818 |
| chr14 | 67618127 | 67618352 | 0.968879 | 1.76156 | 1.818 |
| chr12 | 51128410 | 51128824 | 1.53465 | 2.79022 | 1.818 |
| chr17 | 76364531 | 76364746 | 0.717348 | 1.30425 | 1.818 |
| chr2 | 27026901 | 27027161 | 0.67811 | 1.23294 | 1.818 |
| chr14 | 105106155 | 105106741 | 2.52089 | 4.58358 | 1.818 |
| chr17 | 77808629 | 77809165 | 2.6437 | 4.80725 | 1.818 |
| chr18 | 4106378 | 4106719 | 1.20605 | 2.19307 | 1.818 |
| chr1 | 89948828 | 89949198 | 1.48128 | 2.69359 | 1.818 |
| chr17 | 65775125 | 65775472 | 2.75274 | 5.00567 | 1.818 |
| chr5 | 176918319 | 176918922 | 2.98405 | 5.42637 | 1.818 |
| chr14 | 96771346 | 96771578 | 1.35565 | 2.46534 | 1.819 |
| chr7 | 146943436 | 146943679 | 0.56666 | 1.03059 | 1.819 |
| chr14 | 67993349 | 67993795 | 1.93708 | 3.5231 | 1.819 |
| chr5 | 50282973 | 50283446 | 0.551714 | 1.00351 | 1.819 |
| chr3 | 149181777 | 149182046 | 0.534191 | 0.971668 | 1.819 |
| chr14 | 78270999 | 78271246 | 1.52537 | 2.77463 | 1.819 |
| chr1 | 200994901 | 200995140 | 1.11616 | 2.03029 | 1.819 |
| chr15 | 72565104 | 72565380 | 1.98208 | 3.6055 | 1.819 |
| chr20 | 34802704 | 34802915 | 1.14966 | 2.09133 | 1.819 |
| chr10 | 127004076 | 127004443 | 0.683644 | 1.24363 | 1.819 |
| chr11 | 68103573 | 68103862 | 0.49582 | 0.901968 | 1.819 |
| chr11 | 93904913 | 93905191 | 1.65294 | 3.00695 | 1.819 |
| chr20 | 30410611 | 30411004 | 2.20489 | 4.01103 | 1.819 |
| chr1 | 39724348 | 39724659 | 1.78082 | 3.23962 | 1.819 |

| | | | | | |
|---|---|---|---|---|---|
| chr3 | 72353136 | 72353371 | 0.427242 | 0.777228 | 1.819 |
| chr8 | 65608218 | 65608468 | 1.88267 | 3.42505 | 1.819 |
| chr11 | 67141600 | 67141915 | 1.00994 | 1.83734 | 1.819 |
| chr10 | 124420265 | 124420504 | 0.801296 | 1.45777 | 1.819 |
| chr1 | 110554303 | 110554512 | 0.955882 | 1.73903 | 1.819 |
| chr5 | 12346109 | 12346409 | 0.602006 | 1.09526 | 1.819 |
| chr5 | 79777028 | 79777453 | 1.6147 | 2.93786 | 1.819 |
| chr11 | 70004345 | 70004653 | 0.933636 | 1.69876 | 1.820 |
| chr5 | 172957534 | 172957841 | 0.780868 | 1.42085 | 1.820 |
| chr8 | 131983801 | 131984070 | 0.576748 | 1.04944 | 1.820 |
| chr16 | 14623053 | 14623280 | 1.43582 | 2.61261 | 1.820 |
| chr1 | 111942479 | 111942765 | 1.19792 | 2.17979 | 1.820 |
| chr14 | 57268052 | 57268299 | 0.473732 | 0.862109 | 1.820 |
| chr14 | 61752954 | 61753187 | 0.894694 | 1.62825 | 1.820 |
| chr8 | 130788722 | 130788996 | 1.57175 | 2.86042 | 1.820 |
| chr7 | 27176057 | 27176510 | 1.70336 | 3.10006 | 1.820 |
| chr5 | 1737856 | 1738266 | 0.923496 | 1.68076 | 1.820 |
| chr13 | 67192973 | 67193198 | 0.562592 | 1.02393 | 1.820 |
| chr17 | 75327471 | 75327842 | 3.32966 | 6.06007 | 1.820 |
| chr2 | 128028314 | 128028496 | 0.899672 | 1.63745 | 1.820 |
| chr10 | 56110455 | 56110857 | 0.634731 | 1.15527 | 1.820 |
| chr10 | 23676914 | 23677230 | 0.612508 | 1.11483 | 1.820 |
| chr22 | 22526049 | 22526265 | 0.766739 | 1.39555 | 1.820 |
| chr5 | 72251644 | 72251968 | 1.68737 | 3.07127 | 1.820 |
| chr15 | 74927490 | 74927954 | 1.18705 | 2.16064 | 1.820 |
| chr9 | 75472076 | 75472285 | 1.21536 | 2.21224 | 1.820 |
| chr6 | 108025658 | 108025957 | 1.19508 | 2.17541 | 1.820 |
| chr9 | 139218929 | 139219249 | 0.61351 | 1.11679 | 1.820 |
| chr3 | 185695237 | 185695561 | 1.65063 | 3.00509 | 1.821 |
| chr12 | 124918065 | 124918290 | 0.729967 | 1.32896 | 1.821 |
| chr17 | 9480430 | 9480783 | 1.45233 | 2.64412 | 1.821 |
| chr8 | 32081037 | 32081362 | 1.2651 | 2.30343 | 1.821 |
| chr10 | 78933558 | 78934037 | 2.10623 | 3.83509 | 1.821 |
| chr4 | 160408453 | 160408657 | 1.06726 | 1.9433 | 1.821 |
| chr17 | 34074501 | 34074876 | 1.95122 | 3.55286 | 1.821 |
| chr12 | 119344628 | 119344859 | 0.618675 | 1.12654 | 1.821 |
| chr5 | 157985816 | 157986023 | 0.801878 | 1.46013 | 1.821 |
| chr2 | 70874966 | 70875351 | 3.04132 | 5.53811 | 1.821 |
| chr4 | 37604681 | 37604960 | 2.222 | 4.04642 | 1.821 |
| chr20 | 33460526 | 33460917 | 2.14353 | 3.90359 | 1.821 |
| chr8 | 126742298 | 126742730 | 0.649805 | 1.18336 | 1.821 |
| chr13 | 32738842 | 32739162 | 1.54693 | 2.81726 | 1.821 |
| chr8 | 62008697 | 62008974 | 1.66722 | 3.03644 | 1.821 |
| chr7 | 135674457 | 135674667 | 0.469188 | 0.854529 | 1.821 |
| chrX | 95627817 | 95628042 | 1.71583 | 3.12508 | 1.821 |
| chr3 | 65306537 | 65306836 | 0.482346 | 0.878536 | 1.821 |
| chr22 | 45216497 | 45216845 | 2.00089 | 3.64441 | 1.821 |
| chr1 | 39876618 | 39876949 | 1.58538 | 2.88767 | 1.821 |
| chr1 | 156578871 | 156579161 | 1.32782 | 2.41856 | 1.821 |
| chr16 | 70285652 | 70286133 | 2.64889 | 4.82483 | 1.821 |
| chrX | 53122347 | 53122559 | 1.29752 | 2.36341 | 1.821 |
| chr11 | 67351000 | 67351306 | 1.87615 | 3.41743 | 1.822 |
| chr8 | 1683682 | 1683934 | 2.3089 | 4.20583 | 1.822 |
| chr5 | 36693238 | 36693662 | 1.55269 | 2.82836 | 1.822 |
| chr7 | 55317953 | 55318266 | 1.98888 | 3.62292 | 1.822 |
| chr16 | 64844181 | 64844387 | 0.613773 | 1.11809 | 1.822 |
| chr11 | 10955353 | 10955657 | 1.67291 | 3.04758 | 1.822 |
| chr8 | 10834558 | 10834976 | 1.11399 | 2.02944 | 1.822 |
| chr14 | 38060692 | 38061184 | 1.34381 | 2.44841 | 1.822 |
| chr7 | 51209191 | 51209403 | 1.4059 | 2.56166 | 1.822 |
| chr1 | 201515290 | 201515503 | 0.85415 | 1.55637 | 1.822 |

| | | | | | |
|---|---|---|---|---|---|
| chr17 | 5342387 | 5342793 | 4.01998 | 7.32506 | 1.822 |
| chr19 | 33121259 | 33121558 | 0.95582 | 1.74166 | 1.822 |
| chr17 | 80915157 | 80915500 | 0.47904 | 0.872895 | 1.822 |
| chr14 | 101701572 | 101701910 | 2.14065 | 3.90081 | 1.822 |
| chr10 | 77644969 | 77645277 | 0.552003 | 1.00596 | 1.822 |
| chr12 | 72740192 | 72740464 | 1.18028 | 2.15097 | 1.822 |
| chr21 | 30117430 | 30117695 | 1.60804 | 2.93056 | 1.822 |
| chr7 | 108570709 | 108570946 | 0.421681 | 0.76856 | 1.823 |
| chr17 | 42453423 | 42453767 | 1.76748 | 3.22158 | 1.823 |
| chr2 | 30387764 | 30388227 | 1.64748 | 3.00287 | 1.823 |
| chr11 | 72953448 | 72954039 | 2.09084 | 3.81108 | 1.823 |
| chr5 | 38809957 | 38810248 | 2.35632 | 4.29518 | 1.823 |
| chr10 | 124892521 | 124892962 | 1.74721 | 3.18489 | 1.823 |
| chr11 | 72889113 | 72889590 | 1.33749 | 2.43808 | 1.823 |
| chr9 | 20259966 | 20260279 | 0.607251 | 1.10697 | 1.823 |
| chr4 | 126238773 | 126239019 | 1.34408 | 2.45027 | 1.823 |
| chr11 | 109963368 | 109963938 | 0.960362 | 1.75076 | 1.823 |
| chr5 | 23978203 | 23978414 | 1.07511 | 1.95999 | 1.823 |
| chr8 | 56567013 | 56567265 | 0.542757 | 0.989525 | 1.823 |
| chr16 | 57118179 | 57118553 | 2.99157 | 5.45414 | 1.823 |
| chr3 | 31592230 | 31592544 | 1.33496 | 2.43404 | 1.823 |
| chr6 | 4957764 | 4958145 | 0.780484 | 1.42307 | 1.823 |
| chr17 | 42829405 | 42829713 | 0.671711 | 1.22476 | 1.823 |
| chr10 | 103539655 | 103540190 | 2.31 | 4.21195 | 1.823 |
| chr5 | 165755510 | 165755831 | 0.523156 | 0.953908 | 1.823 |
| chr17 | 6932337 | 6932597 | 0.548092 | 0.999398 | 1.823 |
| chr8 | 128176053 | 128176267 | 0.849026 | 1.54816 | 1.823 |
| chr7 | 134853887 | 134854114 | 1.51255 | 2.75807 | 1.823 |
| chr10 | 32910447 | 32910664 | 0.737159 | 1.34426 | 1.824 |
| chr12 | 111699889 | 111700095 | 0.490648 | 0.894765 | 1.824 |
| chr17 | 17045158 | 17045602 | 2.10493 | 3.83889 | 1.824 |
| chr14 | 23322200 | 23323032 | 2.01801 | 3.68043 | 1.824 |
| chr9 | 124457682 | 124458039 | 1.50342 | 2.74195 | 1.824 |
| chr12 | 110282984 | 110283485 | 3.05692 | 5.57526 | 1.824 |
| chr10 | 121049957 | 121050169 | 0.834327 | 1.52167 | 1.824 |
| chr3 | 184521096 | 184521402 | 3.15642 | 5.75683 | 1.824 |
| chr3 | 103682663 | 103682903 | 0.749683 | 1.36735 | 1.824 |
| chr8 | 140380919 | 140381296 | 0.566794 | 1.03378 | 1.824 |
| chr17 | 38555482 | 38555699 | 1.242 | 2.26548 | 1.824 |
| chr6 | 27156662 | 27156902 | 1.07311 | 1.95747 | 1.824 |
| chr8 | 123584565 | 123584882 | 1.89048 | 3.44848 | 1.824 |
| chr1 | 38440771 | 38441014 | 1.52507 | 2.78197 | 1.824 |
| chr3 | 9116573 | 9116882 | 0.509638 | 0.929661 | 1.824 |
| chr16 | 81462549 | 81462972 | 1.36501 | 2.49002 | 1.824 |
| chr20 | 31057957 | 31058391 | 1.37636 | 2.51082 | 1.824 |
| chr9 | 111152486 | 111152712 | 0.535434 | 0.97679 | 1.824 |
| chr16 | 30932930 | 30933320 | 1.65081 | 3.01159 | 1.824 |
| chr16 | 80403565 | 80403798 | 1.18572 | 2.16312 | 1.824 |
| chr10 | 43600825 | 43601330 | 2.55631 | 4.66366 | 1.824 |
| chr1 | 188097783 | 188098241 | 0.639991 | 1.16768 | 1.825 |
| chrX | 109100180 | 109100488 | 1.36891 | 2.49767 | 1.825 |
| chr3 | 152189786 | 152190073 | 1.19389 | 2.17857 | 1.825 |
| chr14 | 106410731 | 106411210 | 2.6211 | 4.7831 | 1.825 |
| chr7 | 90953294 | 90953572 | 0.621751 | 1.1346 | 1.825 |
| chr17 | 2752054 | 2752480 | 3.14403 | 5.73794 | 1.825 |
| chr9 | 134269281 | 134269524 | 2.35216 | 4.29286 | 1.825 |
| chr17 | 53045490 | 53045742 | 1.59956 | 2.91945 | 1.825 |
| chr7 | 38358463 | 38358717 | 1.29365 | 2.36114 | 1.825 |
| chr17 | 7982535 | 7982904 | 1.97255 | 3.60043 | 1.825 |
| chr5 | 125828637 | 125828848 | 0.905864 | 1.65345 | 1.825 |
| chr21 | 44212355 | 44212564 | 1.04094 | 1.90001 | 1.825 |

| | | | | | |
|---|---|---|---|---|---|
| chr16 | 68074393 | 68074927 | 2.10272 | 3.83822 | 1.825 |
| chr12 | 11706927 | 11707214 | 1.62326 | 2.96313 | 1.825 |
| chr19 | 28440972 | 28441232 | 0.699775 | 1.27739 | 1.825 |
| chr1 | 217229125 | 217229387 | 0.518061 | 0.945698 | 1.825 |
| chr10 | 56851584 | 56851841 | 1.12091 | 2.04626 | 1.826 |
| chr2 | 109618461 | 109618693 | 1.20726 | 2.20402 | 1.826 |
| chr17 | 46748990 | 46749349 | 2.17142 | 3.96437 | 1.826 |
| chr11 | 520726 | 521401 | 0.814856 | 1.48771 | 1.826 |
| chr19 | 11607207 | 11607486 | 2.05721 | 3.75597 | 1.826 |
| chr19 | 10582657 | 10582873 | 0.688673 | 1.2574 | 1.826 |
| chr5 | 149730140 | 149730369 | 1.27873 | 2.33481 | 1.826 |
| chr1 | 204760856 | 204761286 | 2.47305 | 4.51584 | 1.826 |
| chr12 | 98791266 | 98791619 | 1.68867 | 3.08356 | 1.826 |
| chr17 | 37683449 | 37683865 | 1.56719 | 2.86177 | 1.826 |
| chr3 | 123005109 | 123005467 | 2.73255 | 4.98979 | 1.826 |
| chr10 | 115464893 | 115465171 | 1.30828 | 2.38902 | 1.826 |
| chr17 | 55017286 | 55017548 | 1.48881 | 2.71871 | 1.826 |
| chr19 | 54533260 | 54533513 | 2.34203 | 4.27682 | 1.826 |
| chr12 | 45255201 | 45255529 | 0.587185 | 1.07228 | 1.826 |
| chr8 | 42529275 | 42529522 | 1.0379 | 1.89544 | 1.826 |
| chr16 | 49910475 | 49911023 | 2.46657 | 4.50458 | 1.826 |
| chr2 | 40405317 | 40405598 | 0.968369 | 1.76855 | 1.826 |
| chrX | 68313859 | 68314108 | 1.42468 | 2.60198 | 1.826 |
| chr20 | 43238547 | 43238954 | 2.03334 | 3.71369 | 1.826 |
| chr20 | 4176252 | 4176581 | 1.67233 | 3.05455 | 1.827 |
| chr2 | 119129143 | 119129398 | 1.4785 | 2.70052 | 1.827 |
| chr16 | 74343719 | 74344026 | 1.67379 | 3.05771 | 1.827 |
| chr20 | 15980971 | 15981342 | 0.523536 | 0.956478 | 1.827 |
| chr9 | 107148793 | 107149076 | 1.89695 | 3.46566 | 1.827 |
| chr2 | 118760944 | 118761282 | 1.71955 | 3.14156 | 1.827 |
| chr10 | 6709689 | 6709935 | 1.37653 | 2.51489 | 1.827 |
| chr8 | 57699905 | 57700418 | 1.55254 | 2.83652 | 1.827 |
| chr4 | 90148179 | 90148418 | 1.30054 | 2.37618 | 1.827 |
| chr3 | 33828900 | 33829197 | 1.61831 | 2.95683 | 1.827 |
| chr7 | 111515563 | 111515797 | 0.616373 | 1.1262 | 1.827 |
| chr12 | 41670874 | 41671188 | 1.50117 | 2.74296 | 1.827 |
| chr8 | 95369326 | 95369668 | 2.58445 | 4.72241 | 1.827 |
| chr15 | 76904496 | 76904845 | 0.750149 | 1.37071 | 1.827 |
| chr8 | 57160871 | 57161133 | 1.08465 | 1.98196 | 1.827 |
| chr7 | 24295921 | 24296131 | 0.54328 | 0.992748 | 1.827 |
| chr16 | 71916374 | 71916598 | 1.55835 | 2.84764 | 1.827 |
| chr8 | 82192463 | 82192739 | 0.869964 | 1.58972 | 1.827 |
| chr12 | 56984128 | 56984332 | 1.08016 | 1.97383 | 1.827 |
| chr8 | 100872025 | 100872445 | 1.78574 | 3.26321 | 1.827 |
| chr9 | 85818445 | 85818812 | 0.59091 | 1.07981 | 1.827 |
| chr2 | 203043155 | 203043364 | 1.11514 | 2.03779 | 1.827 |
| chr17 | 79116506 | 79116947 | 1.49608 | 2.73394 | 1.827 |
| chr2 | 46768905 | 46769261 | 2.01954 | 3.69051 | 1.827 |
| chr8 | 133338331 | 133338559 | 0.636423 | 1.16301 | 1.827 |
| chr4 | 7103768 | 7104038 | 1.90831 | 3.48771 | 1.828 |
| chr3 | 116275868 | 116276153 | 0.46131 | 0.843112 | 1.828 |
| chr6 | 138437080 | 138437409 | 1.22741 | 2.24328 | 1.828 |
| chr7 | 38550857 | 38551061 | 0.405561 | 0.741233 | 1.828 |
| chr3 | 67022274 | 67022615 | 1.56444 | 2.85945 | 1.828 |
| chr12 | 52900769 | 52901261 | 1.08643 | 1.9859 | 1.828 |
| chr2 | 207648542 | 207648803 | 1.30888 | 2.39264 | 1.828 |
| chr16 | 1651824 | 1652254 | 0.519641 | 0.949971 | 1.828 |
| chr3 | 183484855 | 183485144 | 1.62822 | 2.97659 | 1.828 |
| chr18 | 49866277 | 49866527 | 1.01514 | 1.85591 | 1.828 |
| chr16 | 86744000 | 86744295 | 0.496511 | 0.907758 | 1.828 |
| chr12 | 130916541 | 130916755 | 0.577944 | 1.05666 | 1.828 |

| | | | | | |
|---|---|---|---|---|---|
| chr11 | 35843331 | 35844008 | 2.029 | 3.70974 | 1.828 |
| chr1 | 41469472 | 41469778 | 1.12596 | 2.05872 | 1.828 |
| chr5 | 34474630 | 34475013 | 2.18982 | 4.00395 | 1.828 |
| chr18 | 62903507 | 62903744 | 0.558303 | 1.02083 | 1.828 |
| chr12 | 20573396 | 20573610 | 0.719717 | 1.31602 | 1.829 |
| chr10 | 121403985 | 121404258 | 1.26406 | 2.31141 | 1.829 |
| chr11 | 102811647 | 102812026 | 1.04749 | 1.9154 | 1.829 |
| chr16 | 64902956 | 64903214 | 0.502152 | 0.918223 | 1.829 |
| chr9 | 95571203 | 95571610 | 0.631926 | 1.15553 | 1.829 |
| chr19 | 12868384 | 12868778 | 2.12682 | 3.88909 | 1.829 |
| chr5 | 32057726 | 32057983 | 0.716114 | 1.30948 | 1.829 |
| chr21 | 35734144 | 35734473 | 1.76075 | 3.21972 | 1.829 |
| chr7 | 142564087 | 142564367 | 1.68453 | 3.08038 | 1.829 |
| chr19 | 4207244 | 4207471 | 1.14422 | 2.09238 | 1.829 |
| chr4 | 85450428 | 85450638 | 1.03146 | 1.88627 | 1.829 |
| chr2 | 27308442 | 27308690 | 1.1877 | 2.17203 | 1.829 |
| chr5 | 111770441 | 111770706 | 0.639891 | 1.17023 | 1.829 |
| chr20 | 37037248 | 37037700 | 1.79748 | 3.28725 | 1.829 |
| chr5 | 179720457 | 179720862 | 1.89209 | 3.46029 | 1.829 |
| chr20 | 30221996 | 30222388 | 2.2025 | 4.02801 | 1.829 |
| chr5 | 62790948 | 62791320 | 0.536133 | 0.980511 | 1.829 |
| chr11 | 23332942 | 23333178 | 0.666818 | 1.21954 | 1.829 |
| chr12 | 54185601 | 54185887 | 1.54152 | 2.81928 | 1.829 |
| chr7 | 25109629 | 25109895 | 1.93743 | 3.54343 | 1.829 |
| chr3 | 99859911 | 99860247 | 0.65518 | 1.19831 | 1.829 |
| chr8 | 106218018 | 106218280 | 0.395211 | 0.722832 | 1.829 |
| chr3 | 187457703 | 187458045 | 2.16784 | 3.96503 | 1.829 |
| chr12 | 92537862 | 92538100 | 1.6717 | 3.0576 | 1.829 |
| chr11 | 11811309 | 11811789 | 1.32331 | 2.42043 | 1.829 |
| chr2 | 230580636 | 230580926 | 0.936424 | 1.71286 | 1.829 |
| chr9 | 117266542 | 117266989 | 1.07976 | 1.97504 | 1.829 |
| chr16 | 27092317 | 27092725 | 1.28282 | 2.34655 | 1.829 |
| chr19 | 30215044 | 30215327 | 0.755203 | 1.38145 | 1.829 |
| chr1 | 225960038 | 225960336 | 1.75495 | 3.21029 | 1.829 |
| chr6 | 111378563 | 111378781 | 1.0738 | 1.96428 | 1.829 |
| chr9 | 86198095 | 86198453 | 1.567 | 2.8665 | 1.829 |
| chr9 | 128965645 | 128965938 | 1.00852 | 1.84492 | 1.829 |
| chr11 | 78774074 | 78774579 | 1.20334 | 2.20155 | 1.830 |
| chr8 | 57791373 | 57791640 | 0.526401 | 0.963077 | 1.830 |
| chr5 | 50469133 | 50469370 | 0.691779 | 1.26568 | 1.830 |
| chr5 | 145806785 | 145807063 | 1.095 | 2.00351 | 1.830 |
| chrX | 106515619 | 106516149 | 1.18964 | 2.17672 | 1.830 |
| chr9 | 74653863 | 74654192 | 1.68747 | 3.08786 | 1.830 |
| chr17 | 72957699 | 72957992 | 1.61323 | 2.95203 | 1.830 |
| chr1 | 241051346 | 241051699 | 2.13912 | 3.91438 | 1.830 |
| chr14 | 93250246 | 93250485 | 1.26063 | 2.30684 | 1.830 |
| chr10 | 53274407 | 53274689 | 0.463579 | 0.848328 | 1.830 |
| chr19 | 4911857 | 4912101 | 1.40338 | 2.56811 | 1.830 |
| chr5 | 75698829 | 75699269 | 1.81161 | 3.31521 | 1.830 |
| chr14 | 94083465 | 94083677 | 0.531228 | 0.972153 | 1.830 |
| chr10 | 83761827 | 83762122 | 0.614276 | 1.1242 | 1.830 |
| chr22 | 31544985 | 31545307 | 2.61963 | 4.79427 | 1.830 |
| chr1 | 239568876 | 239569279 | 1.08095 | 1.9783 | 1.830 |
| chr8 | 142171020 | 142171399 | 2.38699 | 4.36858 | 1.830 |
| chr5 | 31638939 | 31639630 | 0.806776 | 1.47655 | 1.830 |
| chr4 | 88954540 | 88954769 | 1.77012 | 3.2397 | 1.830 |
| chr11 | 16653811 | 16654086 | 1.2626 | 2.3109 | 1.830 |
| chr14 | 100462157 | 100462438 | 1.5648 | 2.864 | 1.830 |
| chr9 | 20524818 | 20525035 | 1.388 | 2.54041 | 1.830 |
| chr7 | 6475168 | 6475628 | 1.74931 | 3.20183 | 1.830 |
| chr7 | 4448408 | 4448750 | 0.438253 | 0.802156 | 1.830 |

| | | | | | |
|---|---|---|---|---|---|
| chr7 | 48141787 | 48141993 | 1.45512 | 2.66346 | 1.830 |
| chr1 | 968396 | 968610 | 1.04885 | 1.91993 | 1.831 |
| chr22 | 24204307 | 24204637 | 1.78734 | 3.27184 | 1.831 |
| chr20 | 11844264 | 11844495 | 0.62973 | 1.15282 | 1.831 |
| chr9 | 35096062 | 35096871 | 2.17132 | 3.97504 | 1.831 |
| chr7 | 150869532 | 150870065 | 4.75674 | 8.70882 | 1.831 |
| chr2 | 8875534 | 8876042 | 0.501535 | 0.918251 | 1.831 |
| chr1 | 146821773 | 146821982 | 0.519359 | 0.950914 | 1.831 |
| chr11 | 118481643 | 118482043 | 1.72548 | 3.1593 | 1.831 |
| chr5 | 104493479 | 104493737 | 0.586572 | 1.07403 | 1.831 |
| chr11 | 34514425 | 34514709 | 1.29869 | 2.378 | 1.831 |
| chr5 | 139619312 | 139619782 | 1.40532 | 2.57325 | 1.831 |
| chr2 | 136498866 | 136499147 | 1.31291 | 2.40406 | 1.831 |
| chr12 | 64943379 | 64943706 | 1.86589 | 3.41671 | 1.831 |
| chr19 | 4140429 | 4140702 | 1.51685 | 2.77758 | 1.831 |
| chr5 | 15558513 | 15559111 | 0.475949 | 0.871532 | 1.831 |
| chr15 | 49148103 | 49148337 | 1.13068 | 2.07066 | 1.831 |
| chr9 | 33738167 | 33738455 | 1.22271 | 2.2392 | 1.831 |
| chr15 | 24123098 | 24123456 | 1.68738 | 3.09034 | 1.831 |
| chr17 | 35872429 | 35872807 | 0.628699 | 1.15143 | 1.831 |
| chr1 | 160963446 | 160963681 | 1.0161 | 1.86096 | 1.831 |
| chr3 | 52254473 | 52254799 | 2.7395 | 5.01769 | 1.832 |
| chr18 | 77404075 | 77404461 | 2.1548 | 3.94678 | 1.832 |
| chr5 | 71690584 | 71690888 | 1.07029 | 1.96039 | 1.832 |
| chr2 | 70298007 | 70298233 | 1.79144 | 3.28129 | 1.832 |
| chr1 | 26743483 | 26743828 | 1.89488 | 3.47079 | 1.832 |
| chr3 | 73890882 | 73891100 | 0.602821 | 1.10419 | 1.832 |
| chr10 | 129948288 | 129948647 | 3.37239 | 6.17731 | 1.832 |
| chr3 | 164540675 | 164540878 | 0.497785 | 0.911844 | 1.832 |
| chr17 | 39705093 | 39705845 | 2.36846 | 4.33864 | 1.832 |
| chr19 | 38375979 | 38376229 | 0.850762 | 1.55847 | 1.832 |
| chr6 | 16129456 | 16129870 | 1.05304 | 1.92905 | 1.832 |
| chr12 | 68739301 | 68739544 | 0.587161 | 1.07564 | 1.832 |
| chr5 | 71051508 | 71051754 | 0.580525 | 1.06349 | 1.832 |
| chr16 | 69914535 | 69914753 | 0.580115 | 1.06282 | 1.832 |
| chr5 | 115625063 | 115625352 | 0.747255 | 1.36913 | 1.832 |
| chr9 | 15873651 | 15873879 | 1.39134 | 2.54935 | 1.832 |
| chr17 | 8041922 | 8042453 | 1.98948 | 3.64549 | 1.832 |
| chr3 | 14194186 | 14194538 | 0.838147 | 1.5358 | 1.832 |
| chr8 | 37637291 | 37637723 | 1.79957 | 3.29758 | 1.832 |
| chr11 | 129620755 | 129621017 | 2.76805 | 5.0724 | 1.832 |
| chr15 | 100663207 | 100663447 | 0.458472 | 0.840174 | 1.833 |
| chr2 | 233309146 | 233309566 | 1.8264 | 3.34703 | 1.833 |
| chr16 | 53544331 | 53544708 | 2.31926 | 4.25034 | 1.833 |
| chr7 | 66846244 | 66846458 | 0.602015 | 1.10327 | 1.833 |
| chr21 | 42655239 | 42655625 | 0.948275 | 1.73785 | 1.833 |
| chr17 | 65196718 | 65197168 | 0.947091 | 1.73572 | 1.833 |
| chr9 | 97833004 | 97833232 | 0.902718 | 1.65441 | 1.833 |
| chr7 | 53626596 | 53626862 | 0.559286 | 1.02502 | 1.833 |
| chr3 | 70113431 | 70113651 | 1.54965 | 2.84023 | 1.833 |
| chr16 | 25244832 | 25245356 | 1.43465 | 2.62956 | 1.833 |
| chr9 | 101925073 | 101925438 | 1.20257 | 2.20431 | 1.833 |
| chr7 | 51228681 | 51229091 | 2.35469 | 4.31617 | 1.833 |
| chr8 | 85094461 | 85094675 | 0.746016 | 1.3675 | 1.833 |
| chr12 | 120218568 | 120218981 | 1.72799 | 3.16762 | 1.833 |
| chr7 | 142705052 | 142705347 | 1.84749 | 3.38672 | 1.833 |
| chr7 | 98634491 | 98634801 | 1.14803 | 2.10452 | 1.833 |
| chr10 | 22745661 | 22746044 | 0.670873 | 1.22983 | 1.833 |
| chrX | 2737566 | 2737995 | 1.24434 | 2.28111 | 1.833 |
| chr17 | 73974933 | 73975353 | 0.769465 | 1.41058 | 1.833 |
| chr1 | 11702487 | 11702781 | 1.97555 | 3.62159 | 1.833 |

| | | | | | |
|---|---|---|---|---|---|
| chr12 | 56586820 | 56587035 | 1.51777 | 2.78245 | 1.833 |
| chr8 | 136686518 | 136686774 | 0.493639 | 0.905047 | 1.833 |
| chr2 | 97246579 | 97246855 | 0.897818 | 1.64616 | 1.834 |
| chrX | 106781147 | 106781404 | 1.1984 | 2.19729 | 1.834 |
| chr9 | 80456800 | 80457093 | 0.767961 | 1.40808 | 1.834 |
| chr2 | 158890855 | 158891089 | 1.10014 | 2.01715 | 1.834 |
| chr17 | 41923657 | 41923987 | 2.81451 | 5.1607 | 1.834 |
| chr4 | 152533764 | 152534051 | 0.893916 | 1.63911 | 1.834 |
| chr11 | 67499732 | 67499994 | 2.41087 | 4.42068 | 1.834 |
| chr4 | 4490185 | 4490487 | 1.32494 | 2.42951 | 1.834 |
| chr12 | 89413338 | 89413694 | 1.96024 | 3.59453 | 1.834 |
| chr9 | 10609920 | 10610189 | 0.528898 | 0.969854 | 1.834 |
| chr10 | 134174223 | 134174697 | 1.80827 | 3.31605 | 1.834 |
| chr16 | 88593842 | 88594240 | 2.2128 | 4.05806 | 1.834 |
| chr14 | 96874432 | 96874671 | 0.449511 | 0.824367 | 1.834 |
| chr16 | 89641024 | 89641356 | 1.81228 | 3.32359 | 1.834 |
| chr7 | 35470216 | 35470626 | 1.32602 | 2.43192 | 1.834 |
| chr15 | 43218844 | 43219256 | 1.45602 | 2.67035 | 1.834 |
| chr10 | 73143922 | 73144503 | 1.90236 | 3.48904 | 1.834 |
| chr8 | 61385907 | 61386149 | 0.895426 | 1.6423 | 1.834 |
| chr10 | 6193012 | 6193760 | 0.992873 | 1.82105 | 1.834 |
| chr11 | 1828929 | 1829305 | 0.800965 | 1.46909 | 1.834 |
| chr14 | 65103428 | 65103708 | 1.76595 | 3.23902 | 1.834 |
| chr8 | 130623805 | 130624022 | 0.646682 | 1.18614 | 1.834 |
| chr15 | 27112010 | 27112378 | 2.11034 | 3.87082 | 1.834 |
| chr3 | 111056980 | 111057284 | 1.75616 | 3.22119 | 1.834 |
| chr10 | 63738748 | 63738993 | 0.779089 | 1.42905 | 1.834 |
| chr9 | 122327579 | 122327876 | 0.464969 | 0.852879 | 1.834 |
| chr3 | 147352932 | 147353175 | 0.601976 | 1.10427 | 1.834 |
| chr20 | 55248216 | 55248544 | 0.609515 | 1.11812 | 1.834 |
| chr21 | 43546009 | 43546295 | 0.698688 | 1.28171 | 1.834 |
| chr20 | 37489048 | 37489276 | 1.87787 | 3.44489 | 1.834 |
| chr1 | 155126282 | 155126485 | 0.612952 | 1.12445 | 1.834 |
| chr20 | 17632560 | 17632770 | 0.625824 | 1.14809 | 1.835 |
| chr12 | 100785585 | 100785933 | 0.748658 | 1.37349 | 1.835 |
| chr17 | 16866188 | 16866886 | 0.731133 | 1.34136 | 1.835 |
| chr1 | 18838009 | 18838257 | 1.30431 | 2.39297 | 1.835 |
| chr8 | 63628351 | 63628697 | 0.550562 | 1.01011 | 1.835 |
| chr9 | 131124367 | 131124707 | 1.59607 | 2.92844 | 1.835 |
| chr8 | 91960574 | 91960803 | 1.03453 | 1.89818 | 1.835 |
| chr1 | 160921112 | 160921555 | 1.29708 | 2.37992 | 1.835 |
| chr12 | 79169282 | 79169494 | 0.931433 | 1.70905 | 1.835 |
| chr2 | 100877246 | 100877669 | 1.02806 | 1.88638 | 1.835 |
| chr5 | 38208851 | 38209211 | 1.33843 | 2.45594 | 1.835 |
| chr12 | 112608215 | 112608440 | 1.12171 | 2.05832 | 1.835 |
| chr2 | 73496129 | 73496636 | 0.755631 | 1.38669 | 1.835 |
| chr7 | 151456615 | 151456982 | 2.77721 | 5.09679 | 1.835 |
| chr12 | 129670552 | 129670926 | 0.707495 | 1.29842 | 1.835 |
| chr20 | 16492751 | 16492975 | 0.779939 | 1.43139 | 1.835 |
| chr20 | 49631693 | 49632051 | 1.66823 | 3.0617 | 1.835 |
| chr17 | 37074776 | 37075135 | 0.783541 | 1.43813 | 1.835 |
| chr11 | 24085814 | 24086066 | 0.658478 | 1.20866 | 1.836 |
| chr20 | 5509430 | 5509887 | 1.78949 | 3.28469 | 1.836 |
| chr9 | 7837134 | 7837347 | 0.6309 | 1.15806 | 1.836 |
| chr19 | 49946495 | 49947303 | 2.97481 | 5.46066 | 1.836 |
| chr19 | 29750182 | 29750394 | 1.13705 | 2.08731 | 1.836 |
| chr12 | 5294203 | 5294856 | 0.598189 | 1.09812 | 1.836 |
| chr5 | 153564814 | 153565027 | 0.60928 | 1.11851 | 1.836 |
| chr12 | 58225183 | 58225531 | 1.72273 | 3.16281 | 1.836 |
| chr9 | 140473272 | 140473687 | 2.82456 | 5.1857 | 1.836 |
| chr9 | 38432988 | 38433209 | 1.19572 | 2.19538 | 1.836 |

| | | | | | |
|---|---|---|---|---|---|
| chr9 | 111534680 | 111534883 | 0.622337 | 1.14265 | 1.836 |
| chr16 | 81038356 | 81038565 | 1.29795 | 2.38323 | 1.836 |
| chrX | 123442733 | 123442995 | 0.550993 | 1.01171 | 1.836 |
| chr5 | 154778170 | 154778392 | 0.414724 | 0.761547 | 1.836 |
| chrX | 113579323 | 113579559 | 1.07271 | 1.96989 | 1.836 |
| chr17 | 10657486 | 10657937 | 1.31059 | 2.40676 | 1.836 |
| chr18 | 45756905 | 45757165 | 2.06806 | 3.79784 | 1.836 |
| chr4 | 82388618 | 82388974 | 1.21967 | 2.23985 | 1.836 |
| chr11 | 20693414 | 20693859 | 0.98588 | 1.81057 | 1.837 |
| chr15 | 70557791 | 70558214 | 1.03562 | 1.90193 | 1.837 |
| chr7 | 140192938 | 140193168 | 1.23625 | 2.27041 | 1.837 |
| chr11 | 67138931 | 67139167 | 0.978693 | 1.79741 | 1.837 |
| chr14 | 25148119 | 25148711 | 1.93144 | 3.54717 | 1.837 |
| chr5 | 124563135 | 124563370 | 0.918545 | 1.68698 | 1.837 |
| chr2 | 60808915 | 60809302 | 2.51526 | 4.61949 | 1.837 |
| chr5 | 108804540 | 108804800 | 1.33479 | 2.45146 | 1.837 |
| chr10 | 16035249 | 16035459 | 0.592071 | 1.0874 | 1.837 |
| chr14 | 45614125 | 45614371 | 0.551024 | 1.01202 | 1.837 |
| chr5 | 18394366 | 18394661 | 1.76967 | 3.25024 | 1.837 |
| chrX | 23920055 | 23920284 | 1.1291 | 2.07379 | 1.837 |
| chrX | 152200985 | 152201368 | 2.06536 | 3.79373 | 1.837 |
| chr6 | 139013257 | 139013815 | 1.31806 | 2.42109 | 1.837 |
| chr22 | 41042289 | 41042543 | 1.1105 | 2.03991 | 1.837 |
| chr14 | 94519324 | 94519552 | 0.480927 | 0.883446 | 1.837 |
| chr11 | 36724129 | 36724520 | 1.28607 | 2.36248 | 1.837 |
| chr5 | 137839412 | 137839894 | 1.54874 | 2.84501 | 1.837 |
| chr7 | 91762966 | 91763516 | 2.14218 | 3.93521 | 1.837 |
| chr20 | 43765473 | 43765696 | 0.700371 | 1.28661 | 1.837 |
| chr7 | 36361282 | 36361590 | 0.574883 | 1.0561 | 1.837 |
| chr17 | 8157162 | 8157482 | 1.39213 | 2.55745 | 1.837 |
| chr10 | 57851177 | 57851535 | 0.630983 | 1.15917 | 1.837 |
| chr8 | 24655856 | 24656080 | 0.766833 | 1.40876 | 1.837 |
| chr4 | 72579455 | 72579773 | 0.991011 | 1.82073 | 1.837 |
| chr19 | 48610563 | 48610951 | 2.39492 | 4.40006 | 1.837 |
| chr6 | 138910026 | 138910258 | 1.6306 | 2.99586 | 1.837 |
| chr7 | 40321953 | 40322203 | 0.465579 | 0.855441 | 1.837 |
| chr2 | 70369794 | 70370126 | 2.56055 | 4.70489 | 1.837 |
| chr11 | 64490809 | 64491415 | 2.6887 | 4.9404 | 1.837 |
| chr19 | 11557552 | 11557936 | 1.55838 | 2.86354 | 1.838 |
| chr1 | 227636903 | 227637160 | 0.795674 | 1.46214 | 1.838 |
| chr2 | 69240033 | 69240256 | 2.13019 | 3.91462 | 1.838 |
| chr5 | 63759792 | 63760034 | 0.436832 | 0.802763 | 1.838 |
| chr1 | 211307316 | 211307883 | 3.1695 | 5.82469 | 1.838 |
| chr4 | 95311998 | 95312281 | 1.13548 | 2.08671 | 1.838 |
| chr7 | 79648462 | 79648863 | 1.70588 | 3.13496 | 1.838 |
| chr14 | 75555183 | 75555748 | 1.59928 | 2.93927 | 1.838 |
| chr11 | 8027932 | 8028246 | 1.96868 | 3.61827 | 1.838 |
| chr15 | 56224134 | 56224380 | 0.598576 | 1.10014 | 1.838 |
| chr12 | 54613897 | 54614321 | 1.89725 | 3.48705 | 1.838 |
| chr2 | 57565378 | 57565629 | 0.577467 | 1.06136 | 1.838 |
| chr15 | 90043846 | 90044244 | 1.29825 | 2.38618 | 1.838 |
| chr10 | 113232311 | 113232569 | 0.645994 | 1.18742 | 1.838 |
| chr17 | 43210596 | 43210884 | 2.52784 | 4.64659 | 1.838 |
| chr2 | 69664502 | 69664765 | 1.02478 | 1.88374 | 1.838 |
| chr19 | 31545564 | 31546167 | 0.667769 | 1.22767 | 1.838 |
| chr17 | 46048026 | 46048379 | 2.26604 | 4.16622 | 1.839 |
| chr19 | 36438199 | 36438586 | 2.23453 | 4.10829 | 1.839 |
| chr17 | 41755275 | 41755748 | 3.22169 | 5.92334 | 1.839 |
| chr2 | 171785326 | 171785839 | 2.62635 | 4.82876 | 1.839 |
| chr4 | 25314086 | 25314493 | 3.28506 | 6.03985 | 1.839 |
| chr7 | 149504316 | 149504611 | 1.74556 | 3.20943 | 1.839 |

| | | | | | |
|---|---|---|---|---|---|
| chr1 | 155057126 | 155057501 | 1.98026 | 3.64104 | 1.839 |
| chr11 | 78125156 | 78125659 | 0.941554 | 1.73127 | 1.839 |
| chr17 | 46882255 | 46882539 | 2.60377 | 4.7877 | 1.839 |
| chr15 | 88327636 | 88327931 | 0.557112 | 1.02448 | 1.839 |
| chr10 | 57617361 | 57617686 | 0.603877 | 1.11049 | 1.839 |
| chr9 | 121803596 | 121803850 | 0.551954 | 1.01502 | 1.839 |
| chr14 | 38455048 | 38455581 | 1.31551 | 2.41918 | 1.839 |
| chr14 | 91163951 | 91164317 | 2.56711 | 4.72096 | 1.839 |
| chr14 | 82524624 | 82524870 | 0.524085 | 0.963838 | 1.839 |
| chr22 | 46929406 | 46929655 | 1.38869 | 2.55399 | 1.839 |
| chr11 | 18743582 | 18744057 | 1.42075 | 2.61298 | 1.839 |
| chr2 | 27041810 | 27042019 | 0.664677 | 1.22251 | 1.839 |
| chr17 | 7963549 | 7963902 | 1.71435 | 3.15323 | 1.839 |
| chr2 | 66107890 | 66108109 | 0.572898 | 1.0538 | 1.839 |
| chr15 | 89019797 | 89020234 | 1.23483 | 2.2715 | 1.840 |
| chr4 | 75186299 | 75186543 | 0.930736 | 1.71212 | 1.840 |
| chr12 | 132991470 | 132991867 | 2.63318 | 4.8439 | 1.840 |
| chr6 | 72129828 | 72130038 | 0.688605 | 1.26674 | 1.840 |
| chr15 | 84107477 | 84107825 | 2.02706 | 3.72901 | 1.840 |
| chr17 | 39852212 | 39852582 | 2.005 | 3.68871 | 1.840 |
| chr11 | 18431831 | 18432318 | 1.10881 | 2.04002 | 1.840 |
| chr1 | 206785717 | 206786040 | 2.96996 | 5.46424 | 1.840 |
| chr16 | 8289935 | 8290155 | 0.609091 | 1.12063 | 1.840 |
| chr15 | 79724318 | 79724784 | 0.876688 | 1.61299 | 1.840 |
| chr7 | 64083 | 64347 | 1.27103 | 2.33854 | 1.840 |
| chr14 | 31978496 | 31978710 | 0.710165 | 1.30671 | 1.840 |
| chr12 | 3422783 | 3423134 | 1.24748 | 2.29539 | 1.840 |
| chr7 | 130987910 | 130988182 | 1.70527 | 3.13775 | 1.840 |
| chr20 | 32610007 | 32610231 | 1.13176 | 2.08252 | 1.840 |
| chr14 | 20401702 | 20402097 | 0.783534 | 1.44185 | 1.840 |
| chr17 | 17794302 | 17794663 | 2.52233 | 4.64156 | 1.840 |
| chr6 | 107318730 | 107319151 | 1.46768 | 2.70086 | 1.840 |
| chr3 | 42785732 | 42786119 | 1.45517 | 2.67786 | 1.840 |
| chrX | 9956863 | 9957166 | 1.18616 | 2.18283 | 1.840 |
| chr7 | 120685221 | 120685424 | 0.537048 | 0.988334 | 1.840 |
| chr6 | 111887160 | 111887508 | 0.503554 | 0.926763 | 1.840 |
| chr12 | 114079197 | 114079403 | 0.535458 | 0.985504 | 1.840 |
| chr9 | 81882550 | 81882786 | 0.556846 | 1.02489 | 1.841 |
| chrX | 19551366 | 19551570 | 1.12134 | 2.06386 | 1.841 |
| chr17 | 1785983 | 1786484 | 1.20921 | 2.22562 | 1.841 |
| chr17 | 21414923 | 21415251 | 2.08935 | 3.8456 | 1.841 |
| chr12 | 101320516 | 101320755 | 1.03655 | 1.9079 | 1.841 |
| chr5 | 112485644 | 112485918 | 0.530722 | 0.976861 | 1.841 |
| chr11 | 9635629 | 9635932 | 1.84813 | 3.40176 | 1.841 |
| chr19 | 8223374 | 8223713 | 0.882285 | 1.62398 | 1.841 |
| chr10 | 68711709 | 68711922 | 0.703194 | 1.29435 | 1.841 |
| chr10 | 84614728 | 84615113 | 0.44534 | 0.819725 | 1.841 |
| chr14 | 76373040 | 76373431 | 1.51929 | 2.79661 | 1.841 |
| chr3 | 147657513 | 147657804 | 0.562151 | 1.03477 | 1.841 |
| chr18 | 13813342 | 13813607 | 1.19231 | 2.19475 | 1.841 |
| chr12 | 133101585 | 133101833 | 0.610422 | 1.12365 | 1.841 |
| chr1 | 181899564 | 181899809 | 0.533489 | 0.982043 | 1.841 |
| chr3 | 73992914 | 73993183 | 0.661562 | 1.21781 | 1.841 |
| chr12 | 13814259 | 13814625 | 1.93964 | 3.57056 | 1.841 |
| chr14 | 99855503 | 99856060 | 2.84966 | 5.24586 | 1.841 |
| chr19 | 14185874 | 14186188 | 2.03628 | 3.74865 | 1.841 |
| chr19 | 45646877 | 45647286 | 1.2672 | 2.33283 | 1.841 |
| chr12 | 52721284 | 52721616 | 2.37919 | 4.37998 | 1.841 |
| chr3 | 13549315 | 13549623 | 0.642021 | 1.18197 | 1.841 |
| chr17 | 69019563 | 69019785 | 0.598055 | 1.10104 | 1.841 |
| chr7 | 39559581 | 39559831 | 1.08173 | 1.99152 | 1.841 |

| | | | | | |
|---|---|---|---|---|---|
| chr20 | 7094499 | 7094740 | 1.10093 | 2.0269 | 1.841 |
| chr11 | 30355026 | 30355257 | 1.11633 | 2.05531 | 1.841 |
| chr6 | 38002184 | 38002392 | 0.952034 | 1.75286 | 1.841 |
| chr1 | 163331002 | 163331298 | 1.17803 | 2.16897 | 1.841 |
| chr8 | 66618215 | 66618595 | 1.16165 | 2.13882 | 1.841 |
| chr1 | 222942558 | 222942878 | 1.081 | 1.99048 | 1.841 |
| chr8 | 75083556 | 75083976 | 1.0888 | 2.00487 | 1.841 |
| chr2 | 167241205 | 167241498 | 0.610273 | 1.12377 | 1.841 |
| chr17 | 58498594 | 58499276 | 3.1442 | 5.78999 | 1.841 |
| chr2 | 218653910 | 218654305 | 1.93372 | 3.56096 | 1.842 |
| chr14 | 103655336 | 103655625 | 1.13905 | 2.0976 | 1.842 |
| chr11 | 109809987 | 109810205 | 1.09426 | 2.01514 | 1.842 |
| chr19 | 15288260 | 15288677 | 1.93812 | 3.56919 | 1.842 |
| chr5 | 98104711 | 98105162 | 1.53392 | 2.82494 | 1.842 |
| chr1 | 179555441 | 179555933 | 2.27679 | 4.19341 | 1.842 |
| chr1 | 61803937 | 61804298 | 1.05795 | 1.94862 | 1.842 |
| chr2 | 76672140 | 76672389 | 0.70513 | 1.29878 | 1.842 |
| chr7 | 152847868 | 152848089 | 0.569718 | 1.04937 | 1.842 |
| chr3 | 170217378 | 170217718 | 1.44152 | 2.65516 | 1.842 |
| chr9 | 135111658 | 135112247 | 2.85952 | 5.26722 | 1.842 |
| chr12 | 39139244 | 39139518 | 0.581322 | 1.0708 | 1.842 |
| chr19 | 1064923 | 1065214 | 1.70247 | 3.13597 | 1.842 |
| chr19 | 6047917 | 6048301 | 1.82059 | 3.35384 | 1.842 |
| chr2 | 636888 | 637571 | 1.4916 | 2.7479 | 1.842 |
| chr1 | 207432799 | 207433012 | 1.0173 | 1.87413 | 1.842 |
| chr16 | 19178962 | 19179486 | 1.99453 | 3.67456 | 1.842 |
| chrX | 46716697 | 46717030 | 0.505701 | 0.931685 | 1.842 |
| chr15 | 90431453 | 90431797 | 1.27555 | 2.35013 | 1.842 |
| chr17 | 79885879 | 79886330 | 0.969947 | 1.78709 | 1.842 |
| chr19 | 3692566 | 3692813 | 1.88557 | 3.47417 | 1.843 |
| chr5 | 14326373 | 14326798 | 1.12602 | 2.07469 | 1.843 |
| chr10 | 87233256 | 87233513 | 0.590086 | 1.08724 | 1.843 |
| chr3 | 5847982 | 5848207 | 1.37651 | 2.53623 | 1.843 |
| chr3 | 4762405 | 4762625 | 1.22702 | 2.26088 | 1.843 |
| chr4 | 17449399 | 17449632 | 1.12722 | 2.07703 | 1.843 |
| chr16 | 81919586 | 81919974 | 1.81555 | 3.3455 | 1.843 |
| chr7 | 56147119 | 56147510 | 1.31688 | 2.42685 | 1.843 |
| chr12 | 48135970 | 48136444 | 2.6642 | 4.90995 | 1.843 |
| chr14 | 69316208 | 69316417 | 0.460998 | 0.849612 | 1.843 |
| chr19 | 30701019 | 30701342 | 0.719287 | 1.32566 | 1.843 |
| chrX | 118622773 | 118623021 | 1.20854 | 2.22739 | 1.843 |
| chr19 | 46991057 | 46991441 | 2.27757 | 4.19769 | 1.843 |
| chr1 | 159744014 | 159744261 | 0.664015 | 1.22385 | 1.843 |
| chr14 | 21482860 | 21483180 | 2.3637 | 4.35663 | 1.843 |
| chr1 | 182054091 | 182054460 | 1.67926 | 3.09519 | 1.843 |
| chr17 | 40219136 | 40219909 | 2.19805 | 4.05154 | 1.843 |
| chr7 | 97797990 | 97798201 | 1.70788 | 3.14855 | 1.844 |
| chr19 | 58439651 | 58439870 | 0.524311 | 0.966655 | 1.844 |
| chr2 | 242640714 | 242640923 | 1.3485 | 2.48622 | 1.844 |
| chr8 | 24772339 | 24772614 | 1.3151 | 2.42466 | 1.844 |
| chr14 | 47250460 | 47250750 | 0.474014 | 0.873997 | 1.844 |
| chr20 | 16784392 | 16785027 | 2.09545 | 3.86364 | 1.844 |
| chr13 | 73217587 | 73217845 | 1.03625 | 1.91067 | 1.844 |
| chr1 | 26796861 | 26797217 | 1.2911 | 2.38061 | 1.844 |
| chr3 | 184894624 | 184894916 | 0.871315 | 1.60658 | 1.844 |
| chr7 | 88481764 | 88482040 | 0.569657 | 1.05038 | 1.844 |
| chr5 | 152654429 | 152654801 | 0.50205 | 0.92574 | 1.844 |
| chr6 | 136871491 | 136871980 | 1.67879 | 3.09574 | 1.844 |
| chr3 | 70139498 | 70139787 | 1.38017 | 2.5451 | 1.844 |
| chr3 | 12883272 | 12883515 | 1.62228 | 2.99185 | 1.844 |
| chr1 | 159041883 | 159042086 | 0.957677 | 1.76619 | 1.844 |

| | | | | | |
|---|---|---|---|---|---|
| chr11 | 61666546 | 61667041 | 3.05853 | 5.6407 | 1.844 |
| chr2 | 121802123 | 121802353 | 0.971695 | 1.79211 | 1.844 |
| chr10 | 134210056 | 134210266 | 0.650485 | 1.1997 | 1.844 |
| chr8 | 26484296 | 26484580 | 0.511263 | 0.942942 | 1.844 |
| chr17 | 81034868 | 81035178 | 1.51932 | 2.80223 | 1.844 |
| chr8 | 127396011 | 127396229 | 1.40554 | 2.59244 | 1.844 |
| chr11 | 45894259 | 45894573 | 1.47384 | 2.71842 | 1.844 |
| chr12 | 127119153 | 127119377 | 0.738441 | 1.36203 | 1.844 |
| chr1 | 45180673 | 45181023 | 1.65616 | 3.05486 | 1.845 |
| chr14 | 37294121 | 37294387 | 0.583544 | 1.07637 | 1.845 |
| chr1 | 202206240 | 202206606 | 1.64598 | 3.03615 | 1.845 |
| chr12 | 21760681 | 21760933 | 0.977261 | 1.80264 | 1.845 |
| chr2 | 3584368 | 3584630 | 1.65464 | 3.05211 | 1.845 |
| chr10 | 87021969 | 87022240 | 0.423186 | 0.780614 | 1.845 |
| chrX | 19611839 | 19612066 | 0.923215 | 1.70297 | 1.845 |
| chr7 | 4793463 | 4793905 | 2.80449 | 5.17335 | 1.845 |
| chr3 | 159602621 | 159602928 | 1.11632 | 2.05932 | 1.845 |
| chr8 | 79632620 | 79632944 | 1.28498 | 2.37046 | 1.845 |
| chr14 | 31960089 | 31960489 | 0.605267 | 1.11663 | 1.845 |
| chr9 | 103881479 | 103881908 | 2.19655 | 4.05252 | 1.845 |
| chr5 | 149338972 | 149339339 | 1.07708 | 1.98725 | 1.845 |
| chr22 | 36866250 | 36866650 | 1.85904 | 3.4301 | 1.845 |
| chr20 | 43672748 | 43673161 | 1.74668 | 3.2228 | 1.845 |
| chr14 | 52480803 | 52481068 | 1.45833 | 2.69084 | 1.845 |
| chr15 | 98300045 | 98300256 | 0.80563 | 1.48652 | 1.845 |
| chr1 | 228026043 | 228026254 | 0.651237 | 1.20168 | 1.845 |
| chr12 | 14771990 | 14772383 | 0.936541 | 1.72815 | 1.845 |
| chr7 | 5526486 | 5527015 | 1.92189 | 3.54638 | 1.845 |
| chr1 | 35252089 | 35252479 | 1.75009 | 3.2295 | 1.845 |
| chr10 | 43759028 | 43759292 | 0.874163 | 1.61313 | 1.845 |
| chr16 | 27120938 | 27121700 | 2.44232 | 4.50713 | 1.845 |
| chr8 | 95604567 | 95604986 | 1.40237 | 2.58797 | 1.845 |
| chr4 | 128460031 | 128460277 | 1.48888 | 2.74768 | 1.845 |
| chr5 | 131746613 | 131746984 | 1.87662 | 3.46326 | 1.845 |
| chr3 | 67164296 | 67164553 | 2.08951 | 3.85628 | 1.846 |
| chr12 | 99522874 | 99523143 | 0.633523 | 1.16922 | 1.846 |
| chr19 | 42817491 | 42817812 | 0.875393 | 1.61562 | 1.846 |
| chr8 | 11446892 | 11447337 | 1.8288 | 3.37534 | 1.846 |
| chr1 | 77999433 | 77999674 | 2.03181 | 3.75006 | 1.846 |
| chr1 | 42612569 | 42612828 | 1.59151 | 2.93747 | 1.846 |
| chr2 | 163747218 | 163747545 | 1.26806 | 2.34051 | 1.846 |
| chr14 | 34770408 | 34770667 | 0.943627 | 1.74171 | 1.846 |
| chr12 | 108311794 | 108312374 | 1.38603 | 2.55833 | 1.846 |
| chr16 | 2301602 | 2301834 | 0.937009 | 1.72953 | 1.846 |
| chr5 | 139955019 | 139955291 | 1.22643 | 2.26388 | 1.846 |
| chr17 | 62946904 | 62947207 | 0.525391 | 0.969836 | 1.846 |
| chr1 | 207010722 | 207010967 | 0.546144 | 1.00816 | 1.846 |
| chr10 | 62900572 | 62900838 | 0.94797 | 1.74991 | 1.846 |
| chr20 | 25381393 | 25381641 | 0.507392 | 0.936623 | 1.846 |
| chr15 | 62126291 | 62126685 | 2.40566 | 4.44083 | 1.846 |
| chr3 | 9868945 | 9869313 | 1.36967 | 2.5284 | 1.846 |
| chr19 | 45492401 | 45492824 | 1.95769 | 3.61395 | 1.846 |
| chr11 | 33564468 | 33564900 | 2.02112 | 3.73119 | 1.846 |
| chr17 | 36055386 | 36055613 | 0.652557 | 1.20469 | 1.846 |
| chr17 | 71706209 | 71706650 | 1.53382 | 2.83162 | 1.846 |
| chr8 | 86375434 | 86375853 | 1.67096 | 3.08483 | 1.846 |
| chr22 | 50690624 | 50690917 | 1.34885 | 2.49028 | 1.846 |
| chr11 | 18265336 | 18265700 | 2.12074 | 3.91546 | 1.846 |
| chrX | 132804513 | 132804729 | 1.28878 | 2.37945 | 1.846 |
| chr16 | 24991589 | 24991799 | 0.577261 | 1.0658 | 1.846 |
| chr16 | 5666138 | 5666449 | 1.59557 | 2.94597 | 1.846 |

| | | | | | |
|---|---|---|---|---|---|
| chr9 | 13034224 | 13034514 | 1.75278 | 3.23627 | 1.846 |
| chr12 | 5944548 | 5945224 | 0.565242 | 1.04368 | 1.846 |
| chr22 | 26756324 | 26756730 | 1.19965 | 2.21518 | 1.847 |
| chr7 | 43548570 | 43548833 | 0.567259 | 1.04748 | 1.847 |
| chr17 | 57598863 | 57599320 | 2.31097 | 4.26737 | 1.847 |
| gl000205_r | 117962 | 118389 | 2.49553 | 4.60823 | 1.847 |
| chr5 | 169245142 | 169245408 | 0.535747 | 0.989357 | 1.847 |
| chr12 | 27258567 | 27258902 | 0.510022 | 0.941864 | 1.847 |
| chr1 | 104057637 | 104057843 | 1.11132 | 2.0523 | 1.847 |
| chr6 | 97634351 | 97634613 | 1.99739 | 3.6887 | 1.847 |
| chr1 | 113683225 | 113683452 | 1.2507 | 2.30979 | 1.847 |
| chr13 | 24534698 | 24535019 | 1.30032 | 2.40143 | 1.847 |
| chr19 | 13895760 | 13896062 | 1.65934 | 3.06455 | 1.847 |
| chr5 | 45590554 | 45590812 | 0.739634 | 1.36599 | 1.847 |
| chr15 | 95159891 | 95160181 | 0.726122 | 1.34105 | 1.847 |
| chr14 | 82456539 | 82456998 | 1.64007 | 3.02903 | 1.847 |
| chr2 | 20800636 | 20801133 | 0.94258 | 1.74085 | 1.847 |
| chr3 | 143318426 | 143318629 | 0.49845 | 0.920591 | 1.847 |
| chr1 | 64400919 | 64401142 | 0.985084 | 1.81937 | 1.847 |
| chr15 | 75242693 | 75243062 | 2.83976 | 5.24502 | 1.847 |
| chr17 | 48474766 | 48475140 | 4.19977 | 7.757 | 1.847 |
| chr12 | 57859631 | 57859862 | 0.665506 | 1.2292 | 1.847 |
| chr20 | 15995180 | 15995582 | 0.623075 | 1.15085 | 1.847 |
| chr8 | 56712032 | 56712414 | 0.507146 | 0.936749 | 1.847 |
| chr11 | 123172375 | 123172700 | 1.92034 | 3.54714 | 1.847 |
| chr6 | 49590789 | 49591224 | 0.985306 | 1.82002 | 1.847 |
| chr18 | 39610570 | 39611007 | 1.33039 | 2.4575 | 1.847 |
| chr10 | 12256698 | 12256968 | 1.43009 | 2.64167 | 1.847 |
| chr8 | 144818713 | 144819013 | 1.34299 | 2.48081 | 1.847 |
| chr16 | 86050015 | 86050321 | 1.81525 | 3.35325 | 1.847 |
| chr17 | 29276136 | 29276343 | 0.657559 | 1.21469 | 1.847 |
| chr9 | 137673856 | 137674172 | 0.965161 | 1.78291 | 1.847 |
| chr17 | 61689608 | 61689977 | 1.83307 | 3.3863 | 1.847 |
| chr9 | 122318359 | 122318754 | 1.95916 | 3.61927 | 1.847 |
| chr15 | 99202469 | 99202822 | 2.11753 | 3.91189 | 1.847 |
| chr15 | 66934104 | 66934366 | 0.697833 | 1.28921 | 1.847 |
| chr7 | 112500687 | 112501025 | 0.382927 | 0.707457 | 1.848 |
| chr9 | 123834391 | 123834679 | 1.25444 | 2.31759 | 1.848 |
| chr15 | 83679614 | 83680002 | 2.42417 | 4.47877 | 1.848 |
| chr12 | 118842714 | 118843060 | 1.72662 | 3.19005 | 1.848 |
| chr5 | 101114788 | 101115063 | 0.471915 | 0.871971 | 1.848 |
| chr7 | 53944549 | 53944850 | 0.571631 | 1.05622 | 1.848 |
| chr2 | 42890254 | 42890801 | 2.51388 | 4.64536 | 1.848 |
| chr5 | 145282656 | 145282989 | 0.869727 | 1.60716 | 1.848 |
| chr1 | 168572798 | 168573149 | 1.40505 | 2.59643 | 1.848 |
| chr7 | 135897548 | 135897790 | 0.513207 | 0.948375 | 1.848 |
| chr17 | 8656964 | 8657542 | 1.01877 | 1.88264 | 1.848 |
| chr14 | 100883823 | 100884089 | 1.66243 | 3.07215 | 1.848 |
| chr7 | 142246080 | 142246354 | 0.629982 | 1.16424 | 1.848 |
| chr4 | 16291587 | 16291824 | 1.10595 | 2.04393 | 1.848 |
| chr12 | 106751273 | 106751601 | 2.27009 | 4.19546 | 1.848 |
| chr9 | 140777555 | 140777875 | 0.527239 | 0.974483 | 1.848 |
| chr6 | 161188434 | 161188859 | 1.51678 | 2.80347 | 1.848 |
| chr2 | 86597720 | 86597957 | 0.884039 | 1.63398 | 1.848 |
| chr1 | 221047623 | 221047874 | 1.45825 | 2.69535 | 1.848 |
| chr9 | 100616148 | 100616484 | 1.82056 | 3.36511 | 1.848 |
| chr19 | 54311883 | 54312289 | 1.56858 | 2.89938 | 1.848 |
| chr22 | 25598326 | 25598740 | 1.61241 | 2.98049 | 1.848 |
| chr4 | 112775673 | 112775945 | 1.18036 | 2.18187 | 1.848 |
| chr1 | 89266145 | 89266429 | 1.30617 | 2.4146 | 1.849 |
| chr17 | 20841975 | 20842206 | 0.915439 | 1.69231 | 1.849 |

| | | | | | |
|---|---|---|---|---|---|
| chr19 | 42891181 | 42891618 | 1.43128 | 2.64594 | 1.849 |
| chr8 | 67089093 | 67089706 | 1.78477 | 3.29944 | 1.849 |
| chr2 | 182295263 | 182295592 | 0.574888 | 1.0628 | 1.849 |
| chr1 | 203307563 | 203308090 | 2.01763 | 3.73036 | 1.849 |
| chr5 | 54989948 | 54990289 | 1.18474 | 2.19046 | 1.849 |
| chr7 | 49212690 | 49212919 | 0.546625 | 1.01071 | 1.849 |
| chr1 | 237501225 | 237501433 | 0.629314 | 1.16362 | 1.849 |
| chr19 | 56587860 | 56588150 | 1.3796 | 2.55099 | 1.849 |
| chrX | 152804884 | 152805304 | 2.27211 | 4.20137 | 1.849 |
| chr15 | 76431827 | 76432203 | 1.81042 | 3.3477 | 1.849 |
| chr8 | 13800634 | 13800890 | 0.449418 | 0.831049 | 1.849 |
| chr17 | 12454835 | 12455039 | 0.559736 | 1.03505 | 1.849 |
| chr3 | 188674717 | 188674954 | 1.43976 | 2.66249 | 1.849 |
| chr16 | 56350465 | 56350744 | 0.550602 | 1.01822 | 1.849 |
| chr5 | 138728215 | 138728777 | 2.03386 | 3.76119 | 1.849 |
| chr5 | 153451914 | 153452186 | 1.90198 | 3.51767 | 1.849 |
| chr22 | 22676436 | 22676653 | 2.46985 | 4.56799 | 1.850 |
| chr10 | 11749250 | 11749468 | 0.823685 | 1.52346 | 1.850 |
| chr12 | 113573444 | 113573854 | 2.82079 | 5.21747 | 1.850 |
| chr1 | 246810563 | 246810854 | 0.846423 | 1.56561 | 1.850 |
| chr20 | 22433918 | 22434320 | 2.02454 | 3.74479 | 1.850 |
| chr12 | 93381134 | 93381367 | 1.05388 | 1.94937 | 1.850 |
| chr5 | 168214233 | 168214458 | 0.586266 | 1.08447 | 1.850 |
| chr1 | 215501083 | 215501374 | 0.640494 | 1.18482 | 1.850 |
| chr19 | 34077021 | 34077428 | 0.650603 | 1.20353 | 1.850 |
| chr22 | 36239086 | 36239308 | 1.55953 | 2.88497 | 1.850 |
| chr7 | 67642059 | 67642301 | 1.6397 | 3.03331 | 1.850 |
| chr17 | 49449003 | 49449289 | 1.89047 | 3.49727 | 1.850 |
| chr14 | 56174111 | 56174409 | 1.1586 | 2.14336 | 1.850 |
| chr2 | 156233737 | 156233982 | 0.720446 | 1.33281 | 1.850 |
| chr9 | 91836497 | 91836758 | 0.496567 | 0.918668 | 1.850 |
| chr8 | 22876709 | 22877128 | 2.38055 | 4.40422 | 1.850 |
| chr16 | 31393781 | 31394240 | 2.18312 | 4.03922 | 1.850 |
| chr8 | 81492139 | 81492472 | 1.46231 | 2.70559 | 1.850 |
| chr17 | 34965309 | 34966174 | 0.89334 | 1.65302 | 1.850 |
| chr17 | 31666227 | 31666479 | 0.588026 | 1.08811 | 1.850 |
| chr17 | 76418109 | 76418507 | 2.25153 | 4.16655 | 1.851 |
| chr8 | 119431305 | 119431609 | 1.7936 | 3.31918 | 1.851 |
| chr13 | 33041527 | 33041779 | 0.544591 | 1.00781 | 1.851 |
| chr8 | 124172688 | 124173066 | 3.98585 | 7.37613 | 1.851 |
| chr15 | 64237402 | 64237880 | 2.08134 | 3.85171 | 1.851 |
| chr22 | 38749114 | 38749534 | 2.30494 | 4.26555 | 1.851 |
| chr6 | 34122535 | 34123009 | 1.99527 | 3.69275 | 1.851 |
| chr8 | 103664123 | 103664333 | 1.08921 | 2.01589 | 1.851 |
| chr11 | 87836825 | 87837208 | 2.59793 | 4.80823 | 1.851 |
| chr1 | 201083284 | 201083582 | 1.78779 | 3.30889 | 1.851 |
| chr2 | 178197157 | 178197389 | 1.19864 | 2.21851 | 1.851 |
| chr17 | 79075458 | 79076052 | 4.22289 | 7.81608 | 1.851 |
| chr11 | 46330144 | 46330530 | 1.9255 | 3.5639 | 1.851 |
| chr17 | 60143323 | 60143659 | 2.18055 | 4.03604 | 1.851 |
| chr9 | 95871208 | 95871897 | 1.40657 | 2.60357 | 1.851 |
| chr19 | 2819622 | 2819992 | 2.19075 | 4.05521 | 1.851 |
| chr4 | 107250266 | 107250478 | 1.16701 | 2.16034 | 1.851 |
| chr11 | 123942114 | 123942439 | 1.11843 | 2.07045 | 1.851 |
| chr16 | 85286788 | 85287374 | 3.61984 | 6.7014 | 1.851 |
| chr22 | 46938237 | 46938661 | 1.9445 | 3.59994 | 1.851 |
| chr2 | 8665387 | 8665812 | 0.706461 | 1.30792 | 1.851 |
| chr8 | 18422440 | 18422776 | 1.08848 | 2.0152 | 1.851 |
| chr2 | 174904662 | 174904951 | 1.14755 | 2.12459 | 1.851 |
| chr19 | 14685042 | 14685314 | 1.17929 | 2.18353 | 1.852 |
| chr2 | 62220573 | 62220890 | 1.25738 | 2.32812 | 1.852 |

| | | | | | |
|---|---|---|---|---|---|
| chr8 | 108706748 | 108707028 | 0.656951 | 1.21644 | 1.852 |
| chr3 | 28957675 | 28957887 | 0.929397 | 1.72092 | 1.852 |
| chr22 | 27041310 | 27041628 | 1.1841 | 2.19255 | 1.852 |
| chr13 | 49343313 | 49343599 | 1.6743 | 3.10025 | 1.852 |
| chr5 | 61434477 | 61434712 | 0.817765 | 1.51424 | 1.852 |
| chr1 | 60225145 | 60225480 | 2.49104 | 4.61275 | 1.852 |
| chr1 | 70203267 | 70203567 | 1.29059 | 2.38992 | 1.852 |
| chr13 | 31618104 | 31618854 | 1.46786 | 2.71821 | 1.852 |
| chr16 | 67554758 | 67555764 | 2.19561 | 4.06607 | 1.852 |
| chr19 | 42298878 | 42299318 | 1.40537 | 2.60271 | 1.852 |
| chr22 | 42486920 | 42487257 | 1.88658 | 3.4941 | 1.852 |
| chr22 | 41913765 | 41914038 | 1.5992 | 2.96192 | 1.852 |
| chr19 | 29825347 | 29825550 | 0.479675 | 0.888454 | 1.852 |
| chr11 | 898624 | 898874 | 1.20347 | 2.22909 | 1.852 |
| chrX | 117593554 | 117593791 | 1.12725 | 2.08793 | 1.852 |
| chr16 | 56827580 | 56827825 | 1.24334 | 2.30297 | 1.852 |
| chr12 | 125670384 | 125670957 | 1.16457 | 2.1571 | 1.852 |
| chr5 | 121571834 | 121572196 | 0.470875 | 0.87224 | 1.852 |
| chr21 | 37400801 | 37401117 | 1.99965 | 3.7042 | 1.852 |
| chr15 | 57632867 | 57633274 | 2.14243 | 3.96885 | 1.853 |
| chr8 | 132250808 | 132251048 | 0.594395 | 1.10121 | 1.853 |
| chr1 | 246614164 | 246614410 | 1.24758 | 2.31145 | 1.853 |
| chr8 | 103639713 | 103639939 | 1.08686 | 2.01368 | 1.853 |
| chr11 | 65356184 | 65356527 | 1.36766 | 2.53398 | 1.853 |
| chr9 | 128137025 | 128137407 | 1.43214 | 2.6535 | 1.853 |
| chr1 | 169898154 | 169898406 | 1.14186 | 2.1157 | 1.853 |
| chr7 | 135329999 | 135330367 | 1.20714 | 2.23668 | 1.853 |
| chrX | 69884699 | 69884960 | 1.03433 | 1.91649 | 1.853 |
| chr5 | 125930986 | 125931253 | 1.59353 | 2.95264 | 1.853 |
| chr19 | 48201247 | 48201773 | 2.47091 | 4.5784 | 1.853 |
| chr12 | 4807130 | 4807401 | 0.455724 | 0.844451 | 1.853 |
| chr12 | 52386883 | 52387274 | 1.92625 | 3.56936 | 1.853 |
| chr14 | 67633120 | 67633519 | 1.43711 | 2.66315 | 1.853 |
| chr6 | 16184446 | 16184677 | 0.863448 | 1.6001 | 1.853 |
| chr12 | 31138698 | 31139116 | 1.8749 | 3.47463 | 1.853 |
| chr17 | 73597279 | 73597723 | 2.24475 | 4.16008 | 1.853 |
| chr12 | 6382722 | 6382950 | 0.565684 | 1.04837 | 1.853 |
| chr12 | 13687637 | 13688125 | 1.99073 | 3.68941 | 1.853 |
| chrX | 22003378 | 22003729 | 1.66628 | 3.08818 | 1.853 |
| chr17 | 47113263 | 47113771 | 1.77497 | 3.28964 | 1.853 |
| chr1 | 145427462 | 145427708 | 1.39422 | 2.58416 | 1.853 |
| chr12 | 16318438 | 16318702 | 1.49296 | 2.7673 | 1.854 |
| chr2 | 62952420 | 62952717 | 1.5838 | 2.9357 | 1.854 |
| chr20 | 17635608 | 17635990 | 1.77035 | 3.28151 | 1.854 |
| chr7 | 136107470 | 136107857 | 0.492801 | 0.913508 | 1.854 |
| chr13 | 95327163 | 95327469 | 1.34617 | 2.49542 | 1.854 |
| chr1 | 238723659 | 238723970 | 0.549028 | 1.01775 | 1.854 |
| chr10 | 72685347 | 72685874 | 2.27632 | 4.21969 | 1.854 |
| chr3 | 51536092 | 51536340 | 1.72704 | 3.20149 | 1.854 |
| chr12 | 121722478 | 121722831 | 0.492149 | 0.912323 | 1.854 |
| chr2 | 20215297 | 20215562 | 1.41936 | 2.63119 | 1.854 |
| chr7 | 42746337 | 42746559 | 0.492957 | 0.913847 | 1.854 |
| chr22 | 37957756 | 37958116 | 1.97909 | 3.66887 | 1.854 |
| chr6 | 123047781 | 123047987 | 1.36615 | 2.53264 | 1.854 |
| chr11 | 32906297 | 32906632 | 1.28327 | 2.37908 | 1.854 |
| chr17 | 36662646 | 36663022 | 0.666221 | 1.23512 | 1.854 |
| chr14 | 30739849 | 30740307 | 2.0819 | 3.8599 | 1.854 |
| chr6 | 107349171 | 107349481 | 2.28611 | 4.23892 | 1.854 |
| chr6 | 155093610 | 155093889 | 0.435797 | 0.808073 | 1.854 |
| chr11 | 57409014 | 57409267 | 1.51434 | 2.80814 | 1.854 |
| chr20 | 35233806 | 35234117 | 2.26463 | 4.19946 | 1.854 |

| | | | | | |
|---|---|---|---|---|---|
| chr7 | 152033884 | 152034151 | 1.51424 | 2.80796 | 1.854 |
| chr11 | 56032757 | 56033081 | 0.712002 | 1.32032 | 1.854 |
| chr8 | 140590887 | 140591119 | 0.685099 | 1.27046 | 1.854 |
| chr10 | 21399505 | 21399838 | 0.435697 | 0.807969 | 1.854 |
| chr11 | 125218613 | 125219006 | 2.21268 | 4.10328 | 1.854 |
| chr7 | 113350347 | 113350706 | 1.82288 | 3.38067 | 1.855 |
| chr6 | 36726931 | 36727329 | 2.45461 | 4.55234 | 1.855 |
| chr5 | 103769903 | 103770124 | 0.671951 | 1.24623 | 1.855 |
| chr19 | 1279575 | 1280097 | 1.9783 | 3.66908 | 1.855 |
| chr4 | 1811559 | 1812299 | 2.31054 | 4.2853 | 1.855 |
| chr11 | 113767004 | 113767348 | 1.48916 | 2.76193 | 1.855 |
| chr20 | 35636086 | 35636294 | 0.504713 | 0.936107 | 1.855 |
| chr10 | 135321390 | 135321595 | 0.51949 | 0.96356 | 1.855 |
| chr17 | 42428040 | 42428685 | 2.03269 | 3.7704 | 1.855 |
| chr11 | 8935974 | 8936410 | 1.34537 | 2.49559 | 1.855 |
| chr1 | 110572274 | 110572535 | 1.43676 | 2.66518 | 1.855 |
| chr10 | 53642309 | 53642532 | 0.544309 | 1.00969 | 1.855 |
| chr5 | 1468290 | 1468751 | 1.64254 | 3.04698 | 1.855 |
| chr2 | 20306541 | 20306827 | 0.977938 | 1.81418 | 1.855 |
| chr11 | 73336787 | 73337120 | 1.89824 | 3.52149 | 1.855 |
| chr5 | 141862690 | 141862944 | 0.517648 | 0.960362 | 1.855 |
| chrX | 24413918 | 24414192 | 1.35063 | 2.50584 | 1.855 |
| chr3 | 184988341 | 184988580 | 0.901623 | 1.67284 | 1.855 |
| chr1 | 182740747 | 182740961 | 0.603252 | 1.11927 | 1.855 |
| chr2 | 202023074 | 202023382 | 1.43202 | 2.65698 | 1.855 |
| chr5 | 176239813 | 176240224 | 1.58567 | 2.94221 | 1.856 |
| chr17 | 37400922 | 37401463 | 2.03686 | 3.77941 | 1.856 |
| chr20 | 9959092 | 9959381 | 1.66943 | 3.09768 | 1.856 |
| chr15 | 76564723 | 76564955 | 0.486767 | 0.903248 | 1.856 |
| chr3 | 53409053 | 53409429 | 1.96945 | 3.65465 | 1.856 |
| chr7 | 104065014 | 104065221 | 0.589675 | 1.09426 | 1.856 |
| chr15 | 83386864 | 83387361 | 0.577577 | 1.07185 | 1.856 |
| chr19 | 23196257 | 23196537 | 0.462651 | 0.858582 | 1.856 |
| chr5 | 17062246 | 17062695 | 1.59916 | 2.96776 | 1.856 |
| chr1 | 6694724 | 6695020 | 1.36371 | 2.53082 | 1.856 |
| chr7 | 73115752 | 73116078 | 1.73807 | 3.22565 | 1.856 |
| chr3 | 50609606 | 50609876 | 1.79697 | 3.33502 | 1.856 |
| chr3 | 76209224 | 76209469 | 0.605012 | 1.12286 | 1.856 |
| chr2 | 85081884 | 85082146 | 0.6323 | 1.17352 | 1.856 |
| chr3 | 34527211 | 34527524 | 0.528545 | 0.981024 | 1.856 |
| chr16 | 34492694 | 34493000 | 0.69236 | 1.28508 | 1.856 |
| chr14 | 68050495 | 68051154 | 1.50895 | 2.80086 | 1.856 |
| chr17 | 25928755 | 25929103 | 2.43682 | 4.52336 | 1.856 |
| chr10 | 49571656 | 49571883 | 1.66168 | 3.0847 | 1.856 |
| chr1 | 233500500 | 233500889 | 0.729954 | 1.3551 | 1.856 |
| chr2 | 52565862 | 52566072 | 0.482473 | 0.895687 | 1.856 |
| chr8 | 143820113 | 143820516 | 1.72153 | 3.19611 | 1.857 |
| chr1 | 873510 | 873714 | 1.08819 | 2.02032 | 1.857 |
| chr1 | 181901135 | 181901509 | 0.56065 | 1.0409 | 1.857 |
| chrX | 16854601 | 16854868 | 1.3213 | 2.45314 | 1.857 |
| chr8 | 3904458 | 3904899 | 0.556308 | 1.03285 | 1.857 |
| chr14 | 57761014 | 57761228 | 0.979816 | 1.81919 | 1.857 |
| chr17 | 28431564 | 28431865 | 2.18554 | 4.05806 | 1.857 |
| chr3 | 169379905 | 169380567 | 1.53045 | 2.84181 | 1.857 |
| chr8 | 74947485 | 74947710 | 0.482711 | 0.896331 | 1.857 |
| chr17 | 15116925 | 15117289 | 1.086 | 2.01658 | 1.857 |
| chr8 | 20081195 | 20081618 | 1.55222 | 2.88267 | 1.857 |
| chr13 | 96056565 | 96056887 | 1.69413 | 3.14623 | 1.857 |
| chr5 | 88618820 | 88619136 | 0.422259 | 0.784238 | 1.857 |
| chr17 | 27472347 | 27472639 | 1.20262 | 2.23356 | 1.857 |
| chr14 | 37702147 | 37702377 | 1.56062 | 2.89867 | 1.857 |

| | | | | | |
|---|---|---|---|---|---|
| chr17 | 6287845 | 6288084 | 0.581508 | 1.08012 | 1.857 |
| chr7 | 150683022 | 150683476 | 1.21315 | 2.25338 | 1.857 |
| chr3 | 23184601 | 23184865 | 0.594863 | 1.10494 | 1.857 |
| chr8 | 19171788 | 19172083 | 1.0434 | 1.93808 | 1.857 |
| chr19 | 29776863 | 29777191 | 0.859586 | 1.59671 | 1.858 |
| chr20 | 3730270 | 3730893 | 2.93979 | 5.46091 | 1.858 |
| chr20 | 10057765 | 10058078 | 1.56829 | 2.91324 | 1.858 |
| chr4 | 130283864 | 130284139 | 1.20942 | 2.24661 | 1.858 |
| chr10 | 95752325 | 95752633 | 0.707705 | 1.31463 | 1.858 |
| chr2 | 30020159 | 30020416 | 0.563238 | 1.04629 | 1.858 |
| chr17 | 40580492 | 40580796 | 1.84194 | 3.42168 | 1.858 |
| chr10 | 75403932 | 75404243 | 1.37124 | 2.54733 | 1.858 |
| chr1 | 33764247 | 33764602 | 1.66384 | 3.09091 | 1.858 |
| chr6 | 124123703 | 124124070 | 1.6627 | 3.08884 | 1.858 |
| chr9 | 116420283 | 116420604 | 2.36723 | 4.39769 | 1.858 |
| chr5 | 55610430 | 55611129 | 1.14044 | 2.11873 | 1.858 |
| chr3 | 55439630 | 55439887 | 0.651752 | 1.21091 | 1.858 |
| chr10 | 103600406 | 103600857 | 2.11972 | 3.93831 | 1.858 |
| chr10 | 130008368 | 130009553 | 1.13292 | 2.10499 | 1.858 |
| chr6 | 7239051 | 7239320 | 1.4543 | 2.70214 | 1.858 |
| chr14 | 103551971 | 103552248 | 1.05186 | 1.95441 | 1.858 |
| chr10 | 43957410 | 43957698 | 1.16515 | 2.16493 | 1.858 |
| chr14 | 77536733 | 77537289 | 0.880016 | 1.63515 | 1.858 |
| chr7 | 99707287 | 99707846 | 0.513858 | 0.954807 | 1.858 |
| chr7 | 116570386 | 116570824 | 1.33591 | 2.48235 | 1.858 |
| chr12 | 120241933 | 120242442 | 1.33739 | 2.48515 | 1.858 |
| chr16 | 25842157 | 25842371 | 0.52104 | 0.968241 | 1.858 |
| chr1 | 35667761 | 35668057 | 1.7032 | 3.16515 | 1.858 |
| chr3 | 123084694 | 123084954 | 1.6949 | 3.14984 | 1.858 |
| chr11 | 36063627 | 36063913 | 1.18519 | 2.20259 | 1.858 |
| chr6 | 99915495 | 99915777 | 0.813514 | 1.51193 | 1.859 |
| chr10 | 64314428 | 64314757 | 1.2128 | 2.25426 | 1.859 |
| chr2 | 28789624 | 28789998 | 3.8192 | 7.099 | 1.859 |
| chr16 | 53807116 | 53807424 | 1.89124 | 3.5156 | 1.859 |
| chr9 | 139929634 | 139930230 | 1.79097 | 3.32935 | 1.859 |
| chr1 | 212923241 | 212923538 | 1.61188 | 2.9965 | 1.859 |
| chr1 | 27435642 | 27435928 | 1.24115 | 2.30733 | 1.859 |
| chr16 | 67697180 | 67697752 | 0.671251 | 1.24791 | 1.859 |
| chr17 | 77337097 | 77337308 | 0.546066 | 1.0152 | 1.859 |
| chr22 | 23130661 | 23131100 | 1.68772 | 3.13786 | 1.859 |
| chr10 | 71841052 | 71841280 | 0.638087 | 1.18636 | 1.859 |
| chr12 | 23021524 | 23022165 | 0.596551 | 1.10916 | 1.859 |
| chr15 | 90303988 | 90304395 | 2.68241 | 4.98741 | 1.859 |
| chr21 | 43429430 | 43429731 | 0.672705 | 1.25076 | 1.859 |
| chr10 | 56232347 | 56232561 | 0.607257 | 1.12911 | 1.859 |
| chr9 | 136271892 | 136272476 | 2.06671 | 3.84282 | 1.859 |
| chr3 | 23705618 | 23705871 | 0.710058 | 1.32029 | 1.859 |
| chr8 | 111992951 | 111993168 | 0.718052 | 1.33516 | 1.859 |
| chr11 | 122451809 | 122452118 | 1.42321 | 2.64664 | 1.860 |
| chr19 | 4677185 | 4677447 | 1.90927 | 3.55056 | 1.860 |
| chr4 | 22899179 | 22899506 | 1.72222 | 3.20272 | 1.860 |
| chr8 | 9008119 | 9008468 | 2.65867 | 4.94449 | 1.860 |
| chr20 | 34582930 | 34583242 | 2.43226 | 4.52345 | 1.860 |
| chr20 | 45039588 | 45039813 | 1.73991 | 3.23584 | 1.860 |
| chr11 | 76750519 | 76751011 | 1.23549 | 2.29774 | 1.860 |
| chr9 | 123239641 | 123240031 | 2.16261 | 4.022 | 1.860 |
| chr6 | 126112760 | 126113045 | 1.14074 | 2.1216 | 1.860 |
| chr17 | 73775000 | 73775487 | 1.37855 | 2.56399 | 1.860 |
| chr20 | 56009495 | 56009708 | 1.16204 | 2.16151 | 1.860 |
| chr9 | 4048557 | 4048775 | 1.09794 | 2.04229 | 1.860 |
| chr21 | 36065546 | 36065942 | 1.37238 | 2.55283 | 1.860 |

| | | | | | |
|---|---|---|---|---|---|
| chr15 | 94089857 | 94090110 | 0.535856 | 0.996777 | 1.860 |
| chr9 | 96074934 | 96075141 | 1.49691 | 2.78454 | 1.860 |
| chr12 | 110050785 | 110051122 | 1.12174 | 2.08678 | 1.860 |
| chr20 | 48100655 | 48101045 | 2.06523 | 3.84201 | 1.860 |
| chr3 | 47830636 | 47830874 | 1.78415 | 3.31917 | 1.860 |
| chr3 | 36833911 | 36834128 | 1.33098 | 2.47617 | 1.860 |
| chr5 | 53823152 | 53823373 | 0.541768 | 1.00791 | 1.860 |
| chr17 | 45899121 | 45899411 | 1.66343 | 3.09468 | 1.860 |
| chr11 | 87802357 | 87802592 | 0.548134 | 1.01982 | 1.861 |
| chr17 | 19616842 | 19617054 | 0.97703 | 1.8178 | 1.861 |
| chr5 | 96237076 | 96237367 | 0.78054 | 1.45223 | 1.861 |
| chr14 | 91909680 | 91909932 | 0.533604 | 0.992852 | 1.861 |
| chr20 | 59695754 | 59696151 | 1.19888 | 2.23074 | 1.861 |
| chr20 | 34726621 | 34726872 | 1.23889 | 2.30522 | 1.861 |
| chr10 | 64353770 | 64354092 | 1.83318 | 3.41111 | 1.861 |
| chr17 | 7620213 | 7620849 | 1.14156 | 2.12417 | 1.861 |
| chr17 | 38195124 | 38195493 | 1.77109 | 3.29559 | 1.861 |
| chr8 | 141558860 | 141559093 | 0.527775 | 0.982141 | 1.861 |
| chr11 | 72635690 | 72635938 | 1.19735 | 2.22819 | 1.861 |
| chr22 | 39702704 | 39703010 | 1.30125 | 2.42159 | 1.861 |
| chr8 | 25493870 | 25494166 | 0.593529 | 1.10454 | 1.861 |
| chr8 | 124166886 | 124167330 | 1.71831 | 3.19775 | 1.861 |
| chr15 | 83916973 | 83917493 | 1.32087 | 2.45814 | 1.861 |
| chr1 | 181393588 | 181394469 | 1.93692 | 3.60462 | 1.861 |
| chr5 | 54640277 | 54640562 | 1.53411 | 2.85508 | 1.861 |
| chr1 | 203830387 | 203830720 | 2.53494 | 4.71782 | 1.861 |
| chr6 | 24431267 | 24431551 | 0.971079 | 1.80729 | 1.861 |
| chr8 | 25904090 | 25904318 | 1.42265 | 2.64773 | 1.861 |
| chr7 | 44121869 | 44122072 | 0.748734 | 1.39352 | 1.861 |
| chr11 | 71040073 | 71040360 | 1.05314 | 1.96011 | 1.861 |
| chr16 | 89665831 | 89666161 | 1.57316 | 2.92805 | 1.861 |
| chr1 | 205876975 | 205877267 | 1.64966 | 3.07066 | 1.861 |
| chr2 | 26800691 | 26801059 | 3.04018 | 5.65895 | 1.861 |
| chr15 | 95802733 | 95803008 | 1.48333 | 2.7611 | 1.861 |
| chr1 | 45480187 | 45480472 | 1.60377 | 2.98539 | 1.861 |
| chr1 | 212933468 | 212933776 | 1.42759 | 2.65744 | 1.861 |
| chr15 | 94867392 | 94867771 | 1.61419 | 3.00498 | 1.862 |
| chr11 | 47956040 | 47956339 | 1.51945 | 2.82872 | 1.862 |
| chr5 | 38515824 | 38516053 | 1.29426 | 2.40953 | 1.862 |
| chr6 | 10031295 | 10031513 | 0.75408 | 1.4039 | 1.862 |
| chr1 | 47213442 | 47213651 | 1.25292 | 2.33263 | 1.862 |
| chr19 | 18319167 | 18319603 | 1.24127 | 2.31099 | 1.862 |
| chr8 | 56870410 | 56870728 | 0.826715 | 1.53921 | 1.862 |
| chr2 | 231577343 | 231577638 | 1.68911 | 3.14499 | 1.862 |
| chr18 | 55104787 | 55105412 | 1.07578 | 2.00305 | 1.862 |
| chr11 | 1853610 | 1854129 | 2.13663 | 3.97836 | 1.862 |
| chr2 | 234766059 | 234766776 | 1.5509 | 2.88778 | 1.862 |
| chr14 | 102783250 | 102783467 | 1.58744 | 2.95585 | 1.862 |
| chr12 | 60962517 | 60962771 | 0.670359 | 1.24827 | 1.862 |
| chr20 | 17457370 | 17457575 | 0.62504 | 1.16391 | 1.862 |
| chr19 | 4851580 | 4852166 | 1.78617 | 3.32623 | 1.862 |
| chr4 | 111236654 | 111236969 | 1.28307 | 2.38946 | 1.862 |
| chr14 | 100574632 | 100575046 | 2.3835 | 4.43884 | 1.862 |
| chr7 | 73140105 | 73140418 | 2.39772 | 4.46541 | 1.862 |
| chr20 | 35242413 | 35242659 | 0.525303 | 0.978358 | 1.862 |
| chr11 | 61849550 | 61849826 | 2.08374 | 3.88091 | 1.862 |
| chr1 | 24717808 | 24718080 | 1.92907 | 3.59299 | 1.863 |
| chr6 | 107435464 | 107435788 | 1.49616 | 2.78668 | 1.863 |
| chr3 | 161318252 | 161318455 | 0.678664 | 1.26409 | 1.863 |
| chr5 | 96869802 | 96870032 | 0.916607 | 1.70734 | 1.863 |
| chr12 | 80119216 | 80119453 | 0.519947 | 0.968504 | 1.863 |

| | | | | | |
|---|---|---|---|---|---|
| chr15 | 85712010 | 85712362 | 1.30552 | 2.4318 | 1.863 |
| chr12 | 88873927 | 88874229 | 0.531025 | 0.989197 | 1.863 |
| chr3 | 27399229 | 27399494 | 1.68345 | 3.13601 | 1.863 |
| chr10 | 117253772 | 117253978 | 0.656606 | 1.22317 | 1.863 |
| chr22 | 43017434 | 43017913 | 1.82507 | 3.39997 | 1.863 |
| chr6 | 90502611 | 90502907 | 1.15192 | 2.14596 | 1.863 |
| chr1 | 32281585 | 32281909 | 2.44451 | 4.55402 | 1.863 |
| chr3 | 110919500 | 110919798 | 0.512945 | 0.955601 | 1.863 |
| chr1 | 8035374 | 8035652 | 1.99416 | 3.71538 | 1.863 |
| chr2 | 133304742 | 133304980 | 1.3783 | 2.56803 | 1.863 |
| chr3 | 176532103 | 176532363 | 1.3299 | 2.47786 | 1.863 |
| chr6 | 1382016 | 1382417 | 1.08363 | 2.01904 | 1.863 |
| chr9 | 14442273 | 14442586 | 1.01874 | 1.89819 | 1.863 |
| chr20 | 19516187 | 19516471 | 0.563741 | 1.05041 | 1.863 |
| chr1 | 234622671 | 234623038 | 1.50406 | 2.80251 | 1.863 |
| chr5 | 111828053 | 111828307 | 1.757 | 3.27383 | 1.863 |
| chr20 | 32717563 | 32717769 | 1.42777 | 2.6604 | 1.863 |
| chr1 | 6546566 | 6546870 | 2.26593 | 4.22223 | 1.863 |
| chr15 | 74604169 | 74604395 | 1.60061 | 2.98252 | 1.863 |
| chr18 | 3594358 | 3594563 | 2.18237 | 4.06661 | 1.863 |
| chr9 | 126803195 | 126803820 | 2.23628 | 4.16746 | 1.864 |
| chr10 | 103031998 | 103032343 | 1.4665 | 2.73299 | 1.864 |
| chr3 | 48586212 | 48586524 | 2.24691 | 4.18756 | 1.864 |
| chr12 | 53681499 | 53682043 | 1.66503 | 3.10332 | 1.864 |
| chr7 | 49396663 | 49396881 | 0.508366 | 0.947504 | 1.864 |
| chr7 | 8476372 | 8476876 | 1.67625 | 3.12429 | 1.864 |
| chr5 | 140891853 | 140892268 | 2.76898 | 5.1612 | 1.864 |
| chr17 | 65289609 | 65290226 | 1.73181 | 3.2281 | 1.864 |
| chr16 | 1823308 | 1823731 | 2.09545 | 3.90595 | 1.864 |
| chr19 | 1826000 | 1826465 | 1.74061 | 3.24458 | 1.864 |
| chr5 | 73632171 | 73632392 | 0.842241 | 1.56999 | 1.864 |
| chr4 | 1608345 | 1608712 | 1.12625 | 2.09948 | 1.864 |
| chr7 | 112190533 | 112190882 | 2.53535 | 4.72622 | 1.864 |
| chr22 | 44222244 | 44222599 | 1.38759 | 2.58672 | 1.864 |
| chr17 | 10521398 | 10521887 | 3.49587 | 6.51698 | 1.864 |
| chr19 | 33616773 | 33617285 | 1.9386 | 3.61397 | 1.864 |
| chr1 | 195628272 | 195628461 | 0.757759 | 1.41269 | 1.864 |
| chr7 | 129723825 | 129724269 | 1.05672 | 1.97006 | 1.864 |
| chr10 | 37177982 | 37178236 | 1.62641 | 3.03222 | 1.864 |
| chr3 | 119041576 | 119041961 | 0.678285 | 1.26457 | 1.864 |
| chr2 | 6253622 | 6253833 | 0.551125 | 1.0275 | 1.864 |
| chr12 | 4964632 | 4965006 | 1.32687 | 2.4738 | 1.864 |
| chr15 | 30163675 | 30163916 | 0.710566 | 1.32477 | 1.864 |
| chr17 | 5095130 | 5095480 | 1.71234 | 3.19273 | 1.865 |
| chr17 | 56788307 | 56788642 | 1.35588 | 2.52834 | 1.865 |
| chr7 | 107371594 | 107371912 | 0.548422 | 1.02271 | 1.865 |
| chr20 | 1292796 | 1293111 | 0.544534 | 1.01547 | 1.865 |
| chr3 | 160437981 | 160438450 | 0.653721 | 1.2191 | 1.865 |
| chr9 | 75815662 | 75815918 | 1.12016 | 2.08895 | 1.865 |
| chr17 | 2270374 | 2270713 | 1.61935 | 3.01991 | 1.865 |
| chr1 | 238444348 | 238444565 | 0.605158 | 1.12865 | 1.865 |
| chr11 | 95582305 | 95582542 | 0.789262 | 1.47201 | 1.865 |
| chr17 | 74502427 | 74502825 | 2.0337 | 3.79334 | 1.865 |
| chr10 | 3085816 | 3086061 | 1.58891 | 2.96372 | 1.865 |
| chr3 | 171284664 | 171284903 | 1.20866 | 2.25446 | 1.865 |
| chr19 | 33061068 | 33061435 | 1.68291 | 3.13915 | 1.865 |
| chr1 | 43692977 | 43693319 | 1.68683 | 3.14648 | 1.865 |
| chr11 | 10754001 | 10754240 | 0.521867 | 0.973458 | 1.865 |
| chr1 | 6264982 | 6265648 | 1.7172 | 3.20319 | 1.865 |
| chr8 | 33448491 | 33448733 | 1.36645 | 2.54896 | 1.865 |
| chr14 | 34697215 | 34697558 | 1.83869 | 3.43001 | 1.865 |

| | | | | | |
|---|---|---|---|---|---|
| chr17 | 9549248 | 9549506 | 0.953029 | 1.77786 | 1.865 |
| chr17 | 43296230 | 43296566 | 1.81034 | 3.37717 | 1.865 |
| chr21 | 32881980 | 32882231 | 1.23757 | 2.30872 | 1.866 |
| chr8 | 134594325 | 134594540 | 0.991915 | 1.85046 | 1.866 |
| chr19 | 39061429 | 39061634 | 0.599015 | 1.1175 | 1.866 |
| chr9 | 35489427 | 35490187 | 0.984386 | 1.83647 | 1.866 |
| chr10 | 117983724 | 117983947 | 0.529212 | 0.987311 | 1.866 |
| chr1 | 227318970 | 227319273 | 1.19538 | 2.23024 | 1.866 |
| chr1 | 229796934 | 229797353 | 2.22054 | 4.14305 | 1.866 |
| chr2 | 165873329 | 165873627 | 1.28673 | 2.4008 | 1.866 |
| chr9 | 94717045 | 94717432 | 2.22412 | 4.14983 | 1.866 |
| chr14 | 21536624 | 21537023 | 1.24825 | 2.32909 | 1.866 |
| chr11 | 10646021 | 10646396 | 1.33264 | 2.48664 | 1.866 |
| chr3 | 80823895 | 80824124 | 0.640722 | 1.19558 | 1.866 |
| chr5 | 4653119 | 4653337 | 0.50011 | 0.933213 | 1.866 |
| chr15 | 89207769 | 89207973 | 0.860502 | 1.60574 | 1.866 |
| chr2 | 85493623 | 85494083 | 1.65785 | 3.09363 | 1.866 |
| chr19 | 10589560 | 10589905 | 1.42195 | 2.65354 | 1.866 |
| chr7 | 71959033 | 71959463 | 1.72433 | 3.21785 | 1.866 |
| chr12 | 116384917 | 116385285 | 1.88506 | 3.51791 | 1.866 |
| chr14 | 29243654 | 29243871 | 1.48428 | 2.77002 | 1.866 |
| chr18 | 30294605 | 30294832 | 1.14782 | 2.14218 | 1.866 |
| chr14 | 82352558 | 82352937 | 0.450706 | 0.841159 | 1.866 |
| chr19 | 34268778 | 34269398 | 5.78931 | 10.8051 | 1.866 |
| chr9 | 36788469 | 36788868 | 0.881934 | 1.64604 | 1.866 |
| chr1 | 154298764 | 154298980 | 1.68012 | 3.1358 | 1.866 |
| chr6 | 159572309 | 159572631 | 1.14495 | 2.13695 | 1.866 |
| chr4 | 76579087 | 76579330 | 1.14603 | 2.13898 | 1.866 |
| chr7 | 102031107 | 102031570 | 2.74768 | 5.12843 | 1.866 |
| chr12 | 30960425 | 30960712 | 1.25786 | 2.34781 | 1.867 |
| chr6 | 16965106 | 16965495 | 1.79534 | 3.35106 | 1.867 |
| chr6 | 83777115 | 83777495 | 2.41664 | 4.51083 | 1.867 |
| chr2 | 28675319 | 28675681 | 2.51707 | 4.69841 | 1.867 |
| chr11 | 110334014 | 110334222 | 0.570693 | 1.06528 | 1.867 |
| chr17 | 46306183 | 46306437 | 0.943022 | 1.76031 | 1.867 |
| chr8 | 136464581 | 136464959 | 2.06903 | 3.86229 | 1.867 |
| chr7 | 95319747 | 95320182 | 1.0742 | 2.00529 | 1.867 |
| chr5 | 153801428 | 153801708 | 1.35129 | 2.52258 | 1.867 |
| chr14 | 34318361 | 34318793 | 2.17147 | 4.05374 | 1.867 |
| chr9 | 8507393 | 8507699 | 0.474806 | 0.886375 | 1.867 |
| chrX | 153176555 | 153176877 | 0.939003 | 1.75295 | 1.867 |
| chr12 | 69582785 | 69583046 | 0.522486 | 0.975441 | 1.867 |
| chr10 | 9424790 | 9425053 | 0.635653 | 1.18675 | 1.867 |
| chr10 | 126833700 | 126833939 | 1.44966 | 2.70651 | 1.867 |
| chr1 | 48176172 | 48176507 | 2.35318 | 4.39342 | 1.867 |
| chr16 | 62129252 | 62129456 | 0.419922 | 0.784007 | 1.867 |
| chr14 | 76044016 | 76044327 | 1.61925 | 3.02323 | 1.867 |
| chr10 | 124058256 | 124058647 | 1.4513 | 2.70973 | 1.867 |
| chr8 | 107777523 | 107777811 | 1.58498 | 2.95961 | 1.867 |
| chr11 | 6526923 | 6527216 | 1.32867 | 2.48107 | 1.867 |
| chr14 | 64292989 | 64293193 | 0.599552 | 1.11956 | 1.867 |
| chr12 | 14134991 | 14135380 | 2.16959 | 4.0514 | 1.867 |
| chr15 | 72634812 | 72635065 | 1.38778 | 2.5915 | 1.867 |
| chrX | 67819342 | 67819734 | 1.10088 | 2.05576 | 1.867 |
| chr15 | 70655158 | 70655597 | 1.27662 | 2.384 | 1.867 |
| chr15 | 79299376 | 79299761 | 1.21206 | 2.26345 | 1.867 |
| chr2 | 46870718 | 46870924 | 1.12377 | 2.09867 | 1.868 |
| chr3 | 101292438 | 101292728 | 2.06406 | 3.85475 | 1.868 |
| chr2 | 11552117 | 11552470 | 1.7082 | 3.19026 | 1.868 |
| chrX | 150250691 | 150251165 | 1.6947 | 3.16506 | 1.868 |
| chr10 | 3204859 | 3205149 | 1.0791 | 2.01544 | 1.868 |

| | | | | | |
|---|---|---|---|---|---|
| chr7 | 130125893 | 130126176 | 2.18479 | 4.08067 | 1.868 |
| chr2 | 10124405 | 10124892 | 1.34239 | 2.50728 | 1.868 |
| chr1 | 113615286 | 113615724 | 1.64801 | 3.07844 | 1.868 |
| chr6 | 166648780 | 166648998 | 0.692847 | 1.29423 | 1.868 |
| chr13 | 97865080 | 97865418 | 1.38268 | 2.58287 | 1.868 |
| chr1 | 206663382 | 206663823 | 2.88907 | 5.3971 | 1.868 |
| chr9 | 117001299 | 117001544 | 0.564833 | 1.0552 | 1.868 |
| chr11 | 26161803 | 26162222 | 0.558522 | 1.04342 | 1.868 |
| chr1 | 236706225 | 236706641 | 0.64404 | 1.20319 | 1.868 |
| chr8 | 124077885 | 124078203 | 1.35965 | 2.54018 | 1.868 |
| rUn_gl0002 | 92912 | 93189 | 1.27046 | 2.37364 | 1.868 |
| chr19 | 19663619 | 19663833 | 1.28701 | 2.40459 | 1.868 |
| chr6 | 27280540 | 27280781 | 1.8016 | 3.36608 | 1.868 |
| chr6 | 106808262 | 106808773 | 2.23408 | 4.17417 | 1.868 |
| chr6 | 12844983 | 12845271 | 1.11942 | 2.0916 | 1.868 |
| chr22 | 31741732 | 31742134 | 2.36109 | 4.41164 | 1.868 |
| chr14 | 104145807 | 104146027 | 1.51733 | 2.83513 | 1.869 |
| chr8 | 36439828 | 36440034 | 0.508377 | 0.949907 | 1.869 |
| chr19 | 2579438 | 2579803 | 1.91188 | 3.5725 | 1.869 |
| chr2 | 242254661 | 242254958 | 1.71189 | 3.19903 | 1.869 |
| chr1 | 240161067 | 240161428 | 1.16905 | 2.18475 | 1.869 |
| chr19 | 48269065 | 48269546 | 2.33142 | 4.35704 | 1.869 |
| chr20 | 55125736 | 55126030 | 2.09629 | 3.91775 | 1.869 |
| chr7 | 10094709 | 10094939 | 0.597565 | 1.11679 | 1.869 |
| chr14 | 96075747 | 96076132 | 0.920654 | 1.72065 | 1.869 |
| chr2 | 227850126 | 227850367 | 1.33539 | 2.49576 | 1.869 |
| chr9 | 139664392 | 139664642 | 2.16076 | 4.03857 | 1.869 |
| chr5 | 58686012 | 58686236 | 0.922004 | 1.72329 | 1.869 |
| chr6 | 88101107 | 88101369 | 0.732023 | 1.36824 | 1.869 |
| chr5 | 161476041 | 161476302 | 0.541848 | 1.01279 | 1.869 |
| chr16 | 22207228 | 22207582 | 2.21601 | 4.14207 | 1.869 |
| chr1 | 157659684 | 157660003 | 0.548067 | 1.02447 | 1.869 |
| chr12 | 29453839 | 29454128 | 0.476104 | 0.889956 | 1.869 |
| chr9 | 129064328 | 129064622 | 1.51857 | 2.83867 | 1.869 |
| chr5 | 140187132 | 140187362 | 1.56997 | 2.93478 | 1.869 |
| chr8 | 49231286 | 49231578 | 1.70485 | 3.18708 | 1.869 |
| chr19 | 42043934 | 42044637 | 1.31185 | 2.45251 | 1.870 |
| chr10 | 2088098 | 2088335 | 0.606271 | 1.13347 | 1.870 |
| chr8 | 8608173 | 8608656 | 2.2791 | 4.26097 | 1.870 |
| chr1 | 224521038 | 224521346 | 1.02317 | 1.91306 | 1.870 |
| chr7 | 100449680 | 100450037 | 2.2933 | 4.28787 | 1.870 |
| chr20 | 61454194 | 61454454 | 1.53446 | 2.86905 | 1.870 |
| chr7 | 155981481 | 155981822 | 0.505767 | 0.945686 | 1.870 |
| chr5 | 166076608 | 166076886 | 0.653019 | 1.22105 | 1.870 |
| chr2 | 20155642 | 20155959 | 1.7898 | 3.34681 | 1.870 |
| chr19 | 45414538 | 45414769 | 1.10784 | 2.07161 | 1.870 |
| chr12 | 122469511 | 122469839 | 1.72993 | 3.23493 | 1.870 |
| chr12 | 7527069 | 7527462 | 1.72388 | 3.22372 | 1.870 |
| chr16 | 80958440 | 80958852 | 1.52942 | 2.86011 | 1.870 |
| chr7 | 143077309 | 143077883 | 1.06061 | 1.98345 | 1.870 |
| chr4 | 83189765 | 83190049 | 1.97306 | 3.68995 | 1.870 |
| chr15 | 75660047 | 75660428 | 1.7571 | 3.28621 | 1.870 |
| chr15 | 27136525 | 27136901 | 1.66848 | 3.12051 | 1.870 |
| chr10 | 27014963 | 27015436 | 1.75857 | 3.28907 | 1.870 |
| chr16 | 85693295 | 85693885 | 1.72953 | 3.23482 | 1.870 |
| chr1 | 27714971 | 27715244 | 1.32821 | 2.48437 | 1.870 |
| chr16 | 30996206 | 30996629 | 2.17787 | 4.07371 | 1.871 |
| chr5 | 179897929 | 179898491 | 1.85588 | 3.47142 | 1.871 |
| chrX | 102431268 | 102431600 | 1.42493 | 2.66577 | 1.871 |
| chr14 | 56529836 | 56530076 | 0.664506 | 1.24319 | 1.871 |
| chr2 | 68238687 | 68238926 | 0.662109 | 1.23871 | 1.871 |

| | | | | | |
|---|---|---|---|---|---|
| chr6 | 145349433 | 145349695 | 1.08428 | 2.02862 | 1.871 |
| chr18 | 43753329 | 43754062 | 2.04217 | 3.82086 | 1.871 |
| chr2 | 65780365 | 65780706 | 1.76448 | 3.30146 | 1.871 |
| chr2 | 44787568 | 44787897 | 0.899327 | 1.68273 | 1.871 |
| chr16 | 56092860 | 56093083 | 0.409996 | 0.767148 | 1.871 |
| chr2 | 110493782 | 110494082 | 1.14596 | 2.14426 | 1.871 |
| chr10 | 110678806 | 110679106 | 0.690252 | 1.29159 | 1.871 |
| chr2 | 51262133 | 51262488 | 0.617441 | 1.15539 | 1.871 |
| chr4 | 5636406 | 5636744 | 1.03009 | 1.92757 | 1.871 |
| chr10 | 129188976 | 129189316 | 0.511191 | 0.956594 | 1.871 |
| chr1 | 156023626 | 156023884 | 1.36027 | 2.54557 | 1.871 |
| chr2 | 40195186 | 40195396 | 0.553204 | 1.03525 | 1.871 |
| chr2 | 60782109 | 60782371 | 0.826241 | 1.54623 | 1.871 |
| chr8 | 137333055 | 137333266 | 0.753381 | 1.40989 | 1.871 |
| chr1 | 19422762 | 19423096 | 1.27037 | 2.37743 | 1.871 |
| chr16 | 84958733 | 84959101 | 2.02136 | 3.783 | 1.872 |
| chr20 | 30207464 | 30207841 | 0.574324 | 1.07492 | 1.872 |
| chr16 | 30012184 | 30012388 | 0.928351 | 1.73761 | 1.872 |
| chr6 | 16409977 | 16410237 | 1.2296 | 2.30149 | 1.872 |
| chr12 | 66079420 | 66079626 | 1.47183 | 2.75505 | 1.872 |
| chr5 | 126828606 | 126828977 | 1.12633 | 2.10834 | 1.872 |
| chr8 | 123248514 | 123248759 | 0.905651 | 1.69529 | 1.872 |
| chr3 | 72973279 | 72973708 | 0.649134 | 1.21518 | 1.872 |
| chr14 | 105116639 | 105117141 | 2.73466 | 5.11936 | 1.872 |
| chr12 | 86266358 | 86266565 | 1.29955 | 2.43288 | 1.872 |
| chr12 | 54610900 | 54611226 | 1.49093 | 2.79118 | 1.872 |
| chr1 | 155033921 | 155034606 | 1.8424 | 3.4492 | 1.872 |
| chr10 | 125043649 | 125043929 | 0.739476 | 1.38439 | 1.872 |
| chr11 | 82313530 | 82313786 | 0.51363 | 0.961599 | 1.872 |
| chr14 | 68026373 | 68026611 | 0.748826 | 1.40196 | 1.872 |
| chr21 | 43527901 | 43528265 | 1.70972 | 3.20108 | 1.872 |
| chr2 | 31360869 | 31361143 | 1.32063 | 2.47272 | 1.872 |
| chrX | 134572367 | 134572634 | 1.41713 | 2.65349 | 1.872 |
| chr10 | 129876722 | 129877118 | 1.92739 | 3.60904 | 1.873 |
| chr16 | 84118387 | 84118792 | 1.95263 | 3.6564 | 1.873 |
| chr2 | 85977541 | 85977985 | 1.50628 | 2.82062 | 1.873 |
| chr2 | 46828608 | 46828824 | 1.00759 | 1.88692 | 1.873 |
| chr19 | 40257135 | 40257696 | 1.849 | 3.4627 | 1.873 |
| chr2 | 25451684 | 25451990 | 1.97291 | 3.69488 | 1.873 |
| chr1 | 52607582 | 52607870 | 1.84134 | 3.44849 | 1.873 |
| chr11 | 1224221 | 1224557 | 2.08206 | 3.89936 | 1.873 |
| chr17 | 61523341 | 61523635 | 1.14885 | 2.15164 | 1.873 |
| chr20 | 34535360 | 34535696 | 1.35263 | 2.53328 | 1.873 |
| chr5 | 111510774 | 111511052 | 2.12198 | 3.97433 | 1.873 |
| chr3 | 33773966 | 33774187 | 0.987622 | 1.84988 | 1.873 |
| chr16 | 23739241 | 23739491 | 1.50711 | 2.82297 | 1.873 |
| chr5 | 1380444 | 1380865 | 2.20243 | 4.12538 | 1.873 |
| chr12 | 103163291 | 103163695 | 1.70297 | 3.18994 | 1.873 |
| chr7 | 114062596 | 114062828 | 0.84351 | 1.58007 | 1.873 |
| chr1 | 46908000 | 46908678 | 1.70949 | 3.20231 | 1.873 |
| chr15 | 70752390 | 70752873 | 2.40291 | 4.50139 | 1.873 |
| chr11 | 62160848 | 62161136 | 1.0599 | 1.98553 | 1.873 |
| chr17 | 55749150 | 55749642 | 2.90207 | 5.43659 | 1.873 |
| chr5 | 6811126 | 6811354 | 0.465146 | 0.871395 | 1.873 |
| chr2 | 196531162 | 196531401 | 0.446175 | 0.835883 | 1.873 |
| chr1 | 32014550 | 32015029 | 1.23026 | 2.30483 | 1.873 |
| chr11 | 126860014 | 126860604 | 1.43514 | 2.68872 | 1.873 |
| chr15 | 75193880 | 75194369 | 2.20086 | 4.12328 | 1.873 |
| chr10 | 97776999 | 97777331 | 1.51144 | 2.8317 | 1.874 |
| chr19 | 46876171 | 46876501 | 1.46174 | 2.73869 | 1.874 |
| chr11 | 124615803 | 124616015 | 1.13184 | 2.12062 | 1.874 |

| | | | | | |
|---|---|---|---|---|---|
| chr16 | 12667635 | 12667995 | 1.77578 | 3.32723 | 1.874 |
| chrX | 107408718 | 107409022 | 0.966501 | 1.811 | 1.874 |
| chr1 | 213418010 | 213418254 | 1.42627 | 2.67256 | 1.874 |
| chr19 | 42942625 | 42942987 | 2.61406 | 4.89827 | 1.874 |
| chr19 | 42325165 | 42325491 | 1.78835 | 3.35111 | 1.874 |
| chr2 | 157446195 | 157446432 | 1.04342 | 1.95535 | 1.874 |
| chr14 | 58862442 | 58862869 | 1.78803 | 3.35082 | 1.874 |
| chr22 | 33445148 | 33445542 | 1.78328 | 3.34196 | 1.874 |
| chr20 | 54975098 | 54975450 | 2.09177 | 3.92012 | 1.874 |
| chr3 | 63834787 | 63835143 | 0.473792 | 0.887935 | 1.874 |
| chr10 | 68668493 | 68668736 | 0.610945 | 1.14503 | 1.874 |
| chr15 | 75940187 | 75940448 | 1.73388 | 3.24963 | 1.874 |
| chr19 | 50073390 | 50073812 | 2.9833 | 5.59151 | 1.874 |
| chr5 | 142396407 | 142396644 | 1.14588 | 2.14769 | 1.874 |
| chr4 | 162341101 | 162341447 | 0.694155 | 1.30104 | 1.874 |
| chr5 | 177662026 | 177662465 | 1.60488 | 3.00808 | 1.874 |
| chr5 | 56594617 | 56595031 | 1.10876 | 2.0782 | 1.874 |
| chr19 | 43837288 | 43837869 | 1.4162 | 2.6545 | 1.874 |
| chr3 | 135794090 | 135794365 | 1.08866 | 2.04056 | 1.874 |
| chr17 | 72653595 | 72653933 | 2.07446 | 3.88853 | 1.874 |
| chr19 | 48210205 | 48210883 | 1.51242 | 2.83512 | 1.875 |
| chr22 | 30651852 | 30652332 | 2.07579 | 3.89125 | 1.875 |
| chr2 | 14697708 | 14697947 | 0.514746 | 0.964979 | 1.875 |
| chr8 | 10783852 | 10784139 | 0.525261 | 0.984736 | 1.875 |
| chr15 | 64973649 | 64973973 | 2.07195 | 3.88465 | 1.875 |
| chr3 | 118351180 | 118351441 | 0.460928 | 0.86422 | 1.875 |
| chr12 | 26240077 | 26240286 | 0.511813 | 0.959634 | 1.875 |
| chr16 | 66958734 | 66959262 | 1.04764 | 1.96435 | 1.875 |
| chr6 | 26521938 | 26522210 | 0.576385 | 1.08075 | 1.875 |
| chr15 | 90514796 | 90515100 | 1.68049 | 3.15103 | 1.875 |
| chr18 | 68599534 | 68599746 | 0.955303 | 1.79134 | 1.875 |
| chr12 | 117140960 | 117141428 | 1.84523 | 3.46019 | 1.875 |
| chr16 | 67189130 | 67189729 | 2.44121 | 4.57796 | 1.875 |
| chr16 | 66638641 | 66639050 | 1.58775 | 2.97753 | 1.875 |
| chr2 | 48790280 | 48790520 | 0.454038 | 0.851513 | 1.875 |
| chr16 | 49441888 | 49442231 | 2.62231 | 4.91834 | 1.876 |
| chr10 | 70803015 | 70803224 | 0.406715 | 0.762829 | 1.876 |
| chr12 | 123749211 | 123749433 | 1.50299 | 2.81899 | 1.876 |
| chr1 | 31807943 | 31808196 | 0.874994 | 1.64116 | 1.876 |
| chr15 | 57853176 | 57853584 | 2.41264 | 4.52526 | 1.876 |
| chr17 | 3730494 | 3730798 | 1.96113 | 3.6787 | 1.876 |
| chr10 | 113897125 | 113897545 | 2.57581 | 4.83183 | 1.876 |
| chrX | 149852098 | 149852435 | 1.35784 | 2.5471 | 1.876 |
| chr10 | 46167954 | 46168248 | 1.37824 | 2.58542 | 1.876 |
| chr19 | 33351664 | 33351962 | 0.58198 | 1.09176 | 1.876 |
| chr8 | 30670183 | 30670659 | 2.5004 | 4.69065 | 1.876 |
| chr8 | 65377088 | 65377416 | 0.472454 | 0.886335 | 1.876 |
| chr15 | 66149213 | 66149660 | 1.73429 | 3.25361 | 1.876 |
| chr5 | 39283513 | 39283811 | 1.2714 | 2.38524 | 1.876 |
| chr1 | 207793707 | 207793953 | 1.29003 | 2.4202 | 1.876 |
| chr15 | 74257077 | 74257732 | 1.55488 | 2.91711 | 1.876 |
| chr14 | 101293498 | 101293809 | 1.84378 | 3.45913 | 1.876 |
| chr9 | 85677752 | 85678040 | 0.710621 | 1.33322 | 1.876 |
| chr19 | 45192443 | 45192826 | 3.54605 | 6.65296 | 1.876 |
| chr10 | 55283180 | 55283472 | 0.648293 | 1.21634 | 1.876 |
| chr15 | 75280350 | 75280711 | 1.29736 | 2.43422 | 1.876 |
| chr3 | 32775182 | 32775397 | 1.02892 | 1.93057 | 1.876 |
| chr12 | 6672655 | 6673132 | 2.32702 | 4.36656 | 1.876 |
| chr2 | 192386395 | 192386667 | 1.62579 | 3.0508 | 1.877 |
| chr17 | 6075041 | 6075272 | 0.699889 | 1.31342 | 1.877 |
| chr2 | 20850703 | 20851127 | 1.927 | 3.61627 | 1.877 |

| | | | | | |
|---|---|---|---|---|---|
| chr12 | 71932024 | 71932257 | 0.793398 | 1.48892 | 1.877 |
| chr7 | 51332890 | 51333186 | 0.49098 | 0.921397 | 1.877 |
| chr2 | 237252278 | 237252522 | 1.31989 | 2.47699 | 1.877 |
| chr22 | 19941113 | 19941568 | 2.05153 | 3.85007 | 1.877 |
| chr8 | 17433589 | 17433932 | 0.544053 | 1.02102 | 1.877 |
| chr15 | 65504475 | 65504725 | 1.41448 | 2.65456 | 1.877 |
| chrX | 8783529 | 8783783 | 1.23571 | 2.31906 | 1.877 |
| chr3 | 53144243 | 53144534 | 1.23494 | 2.31764 | 1.877 |
| chr17 | 75052170 | 75052766 | 2.64081 | 4.95617 | 1.877 |
| chr6 | 63481425 | 63481631 | 0.537232 | 1.00828 | 1.877 |
| chr11 | 32369513 | 32369747 | 0.573178 | 1.07581 | 1.877 |
| chr12 | 110662158 | 110662376 | 1.29609 | 2.43266 | 1.877 |
| chr4 | 5826458 | 5826684 | 0.987039 | 1.85263 | 1.877 |
| chr15 | 33584969 | 33585254 | 1.97108 | 3.6997 | 1.877 |
| chr3 | 103581671 | 103582029 | 0.599906 | 1.12602 | 1.877 |
| chrX | 9963812 | 9964139 | 2.22446 | 4.17537 | 1.877 |
| chr7 | 116960544 | 116960760 | 0.668232 | 1.2544 | 1.877 |
| chr3 | 95932308 | 95932534 | 0.964255 | 1.81017 | 1.877 |
| chr4 | 84561126 | 84561674 | 0.816818 | 1.5334 | 1.877 |
| chrX | 12968982 | 12969294 | 2.08638 | 3.91672 | 1.877 |
| chr1 | 226612241 | 226612676 | 1.27458 | 2.39276 | 1.877 |
| chr5 | 16690395 | 16690980 | 1.59561 | 2.99553 | 1.877 |
| chr15 | 43806664 | 43806974 | 1.01563 | 1.90672 | 1.877 |
| chr12 | 49374444 | 49374978 | 0.923665 | 1.7341 | 1.877 |
| chr20 | 43540060 | 43540319 | 1.86831 | 3.50765 | 1.877 |
| chr1 | 202540859 | 202541236 | 1.91611 | 3.59771 | 1.878 |
| chr8 | 10656323 | 10656545 | 1.13956 | 2.13967 | 1.878 |
| chr1 | 223899949 | 223900183 | 1.09743 | 2.06068 | 1.878 |
| chr19 | 40645741 | 40646003 | 1.70541 | 3.2023 | 1.878 |
| chr2 | 238330203 | 238330485 | 2.46019 | 4.61958 | 1.878 |
| chr2 | 171181067 | 171181538 | 1.75587 | 3.2972 | 1.878 |
| chr10 | 134271127 | 134271736 | 1.07281 | 2.01457 | 1.878 |
| chr2 | 240609803 | 240610138 | 1.20802 | 2.2685 | 1.878 |
| chr5 | 54095226 | 54095556 | 0.543001 | 1.0197 | 1.878 |
| chr12 | 9966735 | 9967316 | 2.20869 | 4.14809 | 1.878 |
| chr7 | 119912223 | 119912607 | 1.37369 | 2.57991 | 1.878 |
| chr2 | 56188494 | 56188746 | 1.64403 | 3.08765 | 1.878 |
| chr1 | 164756577 | 164756883 | 2.02758 | 3.80821 | 1.878 |
| chr3 | 172050252 | 172050630 | 1.77957 | 3.34239 | 1.878 |
| chr17 | 47957877 | 47958269 | 2.62042 | 4.92192 | 1.878 |
| chr2 | 19946443 | 19946742 | 1.1486 | 2.1574 | 1.878 |
| chr8 | 142204095 | 142204516 | 0.750346 | 1.40939 | 1.878 |
| chr5 | 133476032 | 133476540 | 1.47993 | 2.77986 | 1.878 |
| chr7 | 71648963 | 71649196 | 0.919085 | 1.72639 | 1.878 |
| chr2 | 62020483 | 62020767 | 1.44044 | 2.70612 | 1.879 |
| chr13 | 98462470 | 98462722 | 0.963615 | 1.81034 | 1.879 |
| chr8 | 144462653 | 144462887 | 0.424404 | 0.797327 | 1.879 |
| chr20 | 31377680 | 31378355 | 2.0919 | 3.93017 | 1.879 |
| chr9 | 132998450 | 132998887 | 2.93676 | 5.51785 | 1.879 |
| chr19 | 34262326 | 34263024 | 2.14385 | 4.0281 | 1.879 |
| chr8 | 133452460 | 133452742 | 0.509199 | 0.956773 | 1.879 |
| chr8 | 20547484 | 20547702 | 0.642506 | 1.20726 | 1.879 |
| chr6 | 4806318 | 4806671 | 1.79439 | 3.37167 | 1.879 |
| chr4 | 74595376 | 74595590 | 1.47879 | 2.77875 | 1.879 |
| chr9 | 129452593 | 129453454 | 1.21306 | 2.27943 | 1.879 |
| chr12 | 122492236 | 122492648 | 2.59631 | 4.87868 | 1.879 |
| chr19 | 45080166 | 45080624 | 1.61066 | 3.02658 | 1.879 |
| chr12 | 2962770 | 2963192 | 2.05724 | 3.86584 | 1.879 |
| chr17 | 47293892 | 47294133 | 0.56228 | 1.05664 | 1.879 |
| chr5 | 5819513 | 5819835 | 0.541383 | 1.01744 | 1.879 |
| chr3 | 172179404 | 172179645 | 1.32456 | 2.48934 | 1.879 |

| | | | | | |
|---|---|---|---|---|---|
| chr1 | 225110274 | 225110605 | 1.78951 | 3.3632 | 1.879 |
| chr12 | 12716994 | 12717284 | 2.04409 | 3.84181 | 1.879 |
| chr11 | 69476897 | 69477429 | 0.593603 | 1.11569 | 1.880 |
| chr12 | 44836116 | 44836510 | 0.654869 | 1.23096 | 1.880 |
| chr10 | 27541375 | 27541626 | 1.39772 | 2.62734 | 1.880 |
| chr2 | 79910523 | 79910727 | 0.489927 | 0.920933 | 1.880 |
| chr7 | 86020019 | 86020231 | 0.392857 | 0.738473 | 1.880 |
| chr9 | 7725206 | 7725448 | 0.521791 | 0.980905 | 1.880 |
| chr19 | 2613804 | 2614200 | 1.95244 | 3.67052 | 1.880 |
| chr17 | 74812672 | 74813016 | 1.98922 | 3.73977 | 1.880 |
| chr1 | 23030871 | 23031314 | 1.35465 | 2.54681 | 1.880 |
| chr2 | 74710358 | 74710614 | 1.95988 | 3.68467 | 1.880 |
| chr12 | 118025445 | 118025825 | 2.34908 | 4.41659 | 1.880 |
| chr5 | 180211536 | 180211835 | 1.75104 | 3.29237 | 1.880 |
| chr13 | 24627585 | 24627916 | 1.22264 | 2.29888 | 1.880 |
| chr11 | 11177630 | 11177874 | 0.94147 | 1.77023 | 1.880 |
| chr2 | 172455309 | 172455550 | 1.5627 | 2.93831 | 1.880 |
| chr18 | 45280922 | 45281239 | 1.91321 | 3.59741 | 1.880 |
| chr18 | 3062958 | 3063348 | 1.39014 | 2.61391 | 1.880 |
| chr22 | 43532253 | 43532610 | 1.63154 | 3.0679 | 1.880 |
| chr8 | 91863195 | 91863406 | 0.595596 | 1.11995 | 1.880 |
| chr8 | 62051394 | 62052060 | 2.11845 | 3.98358 | 1.880 |
| chr7 | 2594839 | 2595130 | 1.27071 | 2.38963 | 1.881 |
| chr12 | 56858143 | 56858437 | 1.97248 | 3.70937 | 1.881 |
| chr4 | 107971723 | 107972010 | 1.14721 | 2.1574 | 1.881 |
| chr7 | 108093201 | 108093438 | 1.06805 | 2.00853 | 1.881 |
| chr8 | 78374975 | 78375347 | 0.631775 | 1.18822 | 1.881 |
| chr5 | 135223034 | 135223380 | 1.67166 | 3.144 | 1.881 |
| chr4 | 185021457 | 185021745 | 1.42163 | 2.67383 | 1.881 |
| chr3 | 78166205 | 78166411 | 0.508805 | 0.95698 | 1.881 |
| chr7 | 8230031 | 8230299 | 1.18518 | 2.22913 | 1.881 |
| chr12 | 56324496 | 56324868 | 1.50771 | 2.83578 | 1.881 |
| chr6 | 153345611 | 153345863 | 1.33012 | 2.50176 | 1.881 |
| chr15 | 67066687 | 67067074 | 2.73183 | 5.13823 | 1.881 |
| chr7 | 155629429 | 155629706 | 0.587754 | 1.10552 | 1.881 |
| chr11 | 59740116 | 59740330 | 0.689043 | 1.29607 | 1.881 |
| chr9 | 119409549 | 119409898 | 1.46059 | 2.74741 | 1.881 |
| chr12 | 115473169 | 115473549 | 0.674828 | 1.26941 | 1.881 |
| chr5 | 149924220 | 149924689 | 2.81435 | 5.29405 | 1.881 |
| chr17 | 62487225 | 62487683 | 1.43603 | 2.70131 | 1.881 |
| chr11 | 27727453 | 27727762 | 1.39993 | 2.63358 | 1.881 |
| chr1 | 212782657 | 212783000 | 1.5206 | 2.86062 | 1.881 |
| chr18 | 74919490 | 74919739 | 1.78136 | 3.35122 | 1.881 |
| chr1 | 32211164 | 32211573 | 1.72117 | 3.23804 | 1.881 |
| chr3 | 22268788 | 22269048 | 0.469046 | 0.882425 | 1.881 |
| chr22 | 43784311 | 43784744 | 2.5899 | 4.87253 | 1.881 |
| chr8 | 2027601 | 2027982 | 1.58424 | 2.98061 | 1.881 |
| chr2 | 9337088 | 9337348 | 0.587059 | 1.10452 | 1.881 |
| chr3 | 12180741 | 12181043 | 1.12007 | 2.10736 | 1.881 |
| chr11 | 75236832 | 75237164 | 1.27239 | 2.39397 | 1.881 |
| chr20 | 37740685 | 37741023 | 1.54282 | 2.90288 | 1.882 |
| chr14 | 103557901 | 103558182 | 1.26989 | 2.38944 | 1.882 |
| chrX | 133654939 | 133655240 | 1.79233 | 3.37265 | 1.882 |
| chr7 | 144134680 | 144135017 | 0.716397 | 1.34809 | 1.882 |
| chr2 | 71191986 | 71192452 | 2.19906 | 4.13814 | 1.882 |
| chr12 | 14922492 | 14923166 | 1.48432 | 2.7932 | 1.882 |
| chr1 | 209971427 | 209971775 | 0.589823 | 1.10996 | 1.882 |
| chr19 | 45930863 | 45931233 | 2.15617 | 4.05783 | 1.882 |
| chr10 | 13701893 | 13702473 | 2.5668 | 4.83077 | 1.882 |
| chr14 | 72768173 | 72768384 | 0.505342 | 0.951111 | 1.882 |
| chr17 | 38589676 | 38590239 | 1.24186 | 2.33743 | 1.882 |

| | | | | | |
|---|---|---|---|---|---|
| chr5 | 148696757 | 148697014 | 1.06879 | 2.01177 | 1.882 |
| chr1 | 24136208 | 24136547 | 1.921 | 3.61606 | 1.882 |
| chr2 | 31954850 | 31955163 | 0.924141 | 1.7396 | 1.882 |
| chr15 | 67143276 | 67143858 | 1.85285 | 3.4879 | 1.882 |
| chr11 | 96197042 | 96197324 | 1.33241 | 2.50827 | 1.883 |
| chr11 | 1714365 | 1714884 | 2.51563 | 4.7358 | 1.883 |
| chr5 | 72466449 | 72466703 | 1.39957 | 2.63478 | 1.883 |
| chr1 | 95509007 | 95509329 | 1.68068 | 3.16407 | 1.883 |
| chr6 | 135410467 | 135410737 | 0.945073 | 1.77922 | 1.883 |
| chr11 | 32892170 | 32892569 | 1.18098 | 2.2234 | 1.883 |
| chr1 | 65289889 | 65290190 | 1.1904 | 2.24148 | 1.883 |
| chr16 | 85597197 | 85597611 | 0.852112 | 1.60449 | 1.883 |
| chr5 | 10452874 | 10453085 | 1.39761 | 2.63168 | 1.883 |
| chr2 | 30468915 | 30469134 | 0.639419 | 1.20414 | 1.883 |
| chr15 | 66968669 | 66968997 | 0.493347 | 0.929081 | 1.883 |
| chr14 | 83766865 | 83767117 | 1.33788 | 2.51953 | 1.883 |
| chr14 | 68038017 | 68038408 | 1.86339 | 3.50929 | 1.883 |
| chr2 | 8417826 | 8418073 | 0.636953 | 1.19958 | 1.883 |
| chr14 | 37117192 | 37117448 | 1.46663 | 2.76225 | 1.883 |
| chr19 | 42167731 | 42168123 | 0.528788 | 0.995947 | 1.883 |
| chr9 | 23196861 | 23197205 | 0.627987 | 1.18278 | 1.883 |
| chrX | 118646514 | 118646769 | 2.61427 | 4.92417 | 1.884 |
| chr17 | 11751193 | 11751429 | 0.569886 | 1.07344 | 1.884 |
| chr7 | 151267532 | 151267803 | 1.21229 | 2.2835 | 1.884 |
| chr3 | 5068604 | 5068859 | 1.56915 | 2.95573 | 1.884 |
| chr3 | 155523620 | 155524253 | 1.72729 | 3.25379 | 1.884 |
| chr1 | 937157 | 937594 | 2.40498 | 4.53055 | 1.884 |
| chr12 | 116435918 | 116436272 | 1.17439 | 2.21236 | 1.884 |
| chr16 | 77820118 | 77820340 | 0.53956 | 1.01646 | 1.884 |
| chr17 | 4207660 | 4208090 | 1.82418 | 3.43674 | 1.884 |
| chr17 | 42811107 | 42811529 | 1.21094 | 2.28151 | 1.884 |
| chr7 | 82440050 | 82440289 | 1.34773 | 2.53923 | 1.884 |
| chr16 | 87312261 | 87312687 | 2.10602 | 3.96798 | 1.884 |
| chr9 | 99507187 | 99507555 | 0.647435 | 1.21988 | 1.884 |
| chr7 | 1851971 | 1852709 | 1.19125 | 2.24469 | 1.884 |
| chr2 | 202968988 | 202969374 | 2.25675 | 4.25247 | 1.884 |
| chr12 | 122640175 | 122640404 | 0.87205 | 1.64326 | 1.884 |
| chr14 | 74225035 | 74225406 | 1.87872 | 3.54025 | 1.884 |
| chr19 | 17413984 | 17414313 | 0.800094 | 1.5077 | 1.884 |
| chr1 | 6168969 | 6169414 | 1.61089 | 3.03559 | 1.884 |
| chr1 | 30558831 | 30559095 | 1.59973 | 3.01471 | 1.885 |
| chr2 | 7237354 | 7237975 | 1.69274 | 3.19014 | 1.885 |
| chr3 | 115295865 | 115296089 | 0.509095 | 0.959471 | 1.885 |
| chrX | 12719818 | 12720127 | 1.47086 | 2.77208 | 1.885 |
| chr8 | 103612904 | 103613162 | 1.0726 | 2.0216 | 1.885 |
| chr16 | 70365501 | 70365730 | 1.17987 | 2.22379 | 1.885 |
| chr5 | 148292347 | 148292575 | 1.17057 | 2.20634 | 1.885 |
| chr6 | 12606109 | 12606338 | 1.4358 | 2.70632 | 1.885 |
| chr16 | 74044368 | 74044629 | 0.474892 | 0.895143 | 1.885 |
| gl000202_r | 16275 | 16887 | 1.62689 | 3.06663 | 1.885 |
| chr20 | 45395759 | 45396140 | 2.05209 | 3.86817 | 1.885 |
| chr1 | 219898435 | 219898968 | 0.608153 | 1.14638 | 1.885 |
| chr8 | 110338139 | 110338362 | 0.908392 | 1.71235 | 1.885 |
| chr18 | 19173941 | 19174146 | 0.641456 | 1.20917 | 1.885 |
| chr8 | 76824050 | 76824324 | 0.632175 | 1.19169 | 1.885 |
| chr17 | 56288344 | 56288673 | 0.985097 | 1.857 | 1.885 |
| chr5 | 88484797 | 88485060 | 0.44635 | 0.841415 | 1.885 |
| chr9 | 127177726 | 127178078 | 1.51504 | 2.856 | 1.885 |
| chr3 | 159724363 | 159724689 | 2.04865 | 3.862 | 1.885 |
| chr7 | 139330713 | 139331001 | 1.82216 | 3.43505 | 1.885 |
| chr18 | 29399630 | 29399932 | 1.59247 | 3.00208 | 1.885 |

| | | | | | |
|---|---|---|---|---|---|
| chr4 | 111132503 | 111132737 | 1.49648 | 2.82121 | 1.885 |
| chr1 | 47639509 | 47639822 | 1.14588 | 2.16031 | 1.885 |
| chr11 | 6742898 | 6743117 | 0.546054 | 1.02947 | 1.885 |
| chr1 | 172592866 | 172593161 | 0.629467 | 1.18674 | 1.885 |
| chr19 | 19164616 | 19164958 | 1.24974 | 2.35616 | 1.885 |
| chr9 | 74061759 | 74062065 | 0.87804 | 1.6554 | 1.885 |
| chr4 | 89580560 | 89580867 | 1.22817 | 2.31553 | 1.885 |
| chr9 | 140400557 | 140401224 | 0.57896 | 1.09155 | 1.885 |
| chr22 | 31481208 | 31481499 | 1.24133 | 2.34038 | 1.885 |
| chr2 | 192083095 | 192083301 | 1.0198 | 1.92272 | 1.885 |
| chr3 | 72704926 | 72705217 | 0.978329 | 1.84458 | 1.885 |
| chr9 | 104150772 | 104151054 | 1.07473 | 2.02634 | 1.885 |
| chr8 | 33192303 | 33192567 | 1.73138 | 3.26449 | 1.885 |
| chr17 | 8280001 | 8280263 | 1.14458 | 2.15816 | 1.886 |
| chr12 | 54095526 | 54096106 | 1.96014 | 3.69598 | 1.886 |
| chr12 | 43818962 | 43819399 | 0.554344 | 1.04529 | 1.886 |
| chr9 | 101538125 | 101538493 | 1.8576 | 3.50287 | 1.886 |
| chr7 | 157043315 | 157043702 | 2.05954 | 3.88377 | 1.886 |
| chr12 | 105673298 | 105673539 | 1.19003 | 2.24411 | 1.886 |
| chr8 | 32412259 | 32412524 | 1.86357 | 3.51428 | 1.886 |
| chr12 | 27856628 | 27856906 | 1.97395 | 3.72252 | 1.886 |
| chr2 | 221018973 | 221019248 | 0.949825 | 1.79122 | 1.886 |
| chr3 | 51426814 | 51427182 | 1.81539 | 3.42359 | 1.886 |
| chr3 | 132071334 | 132071769 | 1.22297 | 2.30644 | 1.886 |
| chr20 | 44882144 | 44882356 | 1.42505 | 2.68774 | 1.886 |
| chr12 | 69550793 | 69551186 | 3.12385 | 5.89184 | 1.886 |
| chr14 | 105174872 | 105175434 | 1.72429 | 3.25219 | 1.886 |
| chr22 | 50418072 | 50418424 | 1.25443 | 2.36603 | 1.886 |
| chr8 | 109631707 | 109631911 | 0.691558 | 1.30439 | 1.886 |
| chr19 | 51941545 | 51941889 | 1.71261 | 3.23042 | 1.886 |
| chr22 | 23152455 | 23152888 | 1.72409 | 3.25217 | 1.886 |
| chr5 | 98597709 | 98597931 | 0.480445 | 0.906297 | 1.886 |
| chr15 | 67532602 | 67532904 | 1.11329 | 2.10015 | 1.886 |
| chr1 | 9686987 | 9687434 | 4.57001 | 8.62134 | 1.887 |
| chr3 | 149945629 | 149945956 | 1.42001 | 2.67887 | 1.887 |
| chr3 | 34342100 | 34342392 | 0.431332 | 0.813743 | 1.887 |
| chr19 | 39612668 | 39613046 | 2.34664 | 4.42765 | 1.887 |
| chr17 | 7537712 | 7537949 | 1.74401 | 3.29069 | 1.887 |
| chr16 | 1813980 | 1814342 | 1.25759 | 2.37301 | 1.887 |
| chr19 | 36551337 | 36551707 | 1.74884 | 3.29997 | 1.887 |
| chr17 | 38175838 | 38176059 | 0.810932 | 1.53022 | 1.887 |
| chr10 | 61808021 | 61808223 | 1.15441 | 2.17846 | 1.887 |
| chr2 | 193614426 | 193614639 | 0.915515 | 1.72767 | 1.887 |
| chr10 | 6526034 | 6526253 | 0.602698 | 1.13736 | 1.887 |
| chr7 | 52541562 | 52542082 | 0.553313 | 1.04418 | 1.887 |
| chr1 | 222807834 | 222808096 | 1.73841 | 3.28069 | 1.887 |
| chr2 | 220267563 | 220267990 | 1.34547 | 2.53917 | 1.887 |
| chr17 | 3907864 | 3908310 | 2.391 | 4.51242 | 1.887 |
| chr10 | 94799546 | 94799770 | 1.1728 | 2.21338 | 1.887 |
| chr10 | 15013884 | 15014264 | 1.79387 | 3.38564 | 1.887 |
| chr3 | 122747396 | 122747653 | 1.21106 | 2.28568 | 1.887 |
| chr9 | 133741786 | 133742207 | 2.57125 | 4.85312 | 1.887 |
| chr6 | 144334530 | 144334903 | 1.91431 | 3.61321 | 1.887 |
| chr1 | 159283366 | 159283762 | 0.479222 | 0.904525 | 1.887 |
| chr10 | 134266628 | 134266992 | 2.15885 | 4.07482 | 1.888 |
| chr16 | 31484043 | 31484374 | 2.20988 | 4.17115 | 1.888 |
| chr19 | 44957107 | 44957436 | 0.600904 | 1.13421 | 1.888 |
| chr10 | 7166971 | 7167244 | 0.556236 | 1.04992 | 1.888 |
| chr2 | 38104680 | 38104932 | 1.12933 | 2.13167 | 1.888 |
| chr3 | 22252874 | 22253273 | 0.447251 | 0.844225 | 1.888 |
| chr20 | 1881013 | 1881323 | 2.29765 | 4.33706 | 1.888 |

| | | | | | |
|---|---|---|---|---|---|
| chr1 | 151128334 | 151128816 | 1.53012 | 2.88835 | 1.888 |
| chr10 | 104356249 | 104356623 | 1.79342 | 3.38537 | 1.888 |
| chr9 | 504388 | 504648 | 0.741817 | 1.40034 | 1.888 |
| chr5 | 6423416 | 6423637 | 0.524201 | 0.989647 | 1.888 |
| chr19 | 1244311 | 1244707 | 1.85126 | 3.49526 | 1.888 |
| chr14 | 100732640 | 100732856 | 0.552233 | 1.04265 | 1.888 |
| chr22 | 39164880 | 39165119 | 2.62546 | 4.95705 | 1.888 |
| chr1 | 226312850 | 226313167 | 1.88027 | 3.55012 | 1.888 |
| chr9 | 140429404 | 140429610 | 1.53997 | 2.90785 | 1.888 |
| chr17 | 56310308 | 56310520 | 0.619219 | 1.16925 | 1.888 |
| chr15 | 83680300 | 83680592 | 2.14924 | 4.05837 | 1.888 |
| chr15 | 90768915 | 90769120 | 0.621557 | 1.17368 | 1.888 |
| chr2 | 25745002 | 25745248 | 1.24567 | 2.35218 | 1.888 |
| chr3 | 169122440 | 169122644 | 0.749783 | 1.41581 | 1.888 |
| chr15 | 55582661 | 55583037 | 1.75429 | 3.31262 | 1.888 |
| chr10 | 88159545 | 88159912 | 2.33658 | 4.41218 | 1.888 |
| chr11 | 32395794 | 32396155 | 1.70538 | 3.22031 | 1.888 |
| chr19 | 40741573 | 40741850 | 1.735 | 3.2763 | 1.888 |
| chr1 | 36765553 | 36765806 | 1.97324 | 3.72623 | 1.888 |
| chr19 | 19296718 | 19297249 | 1.54793 | 2.92319 | 1.888 |
| chr16 | 2957206 | 2957445 | 0.813749 | 1.53673 | 1.888 |
| chr21 | 39288667 | 39289394 | 1.30982 | 2.47361 | 1.889 |
| chr4 | 39136922 | 39137238 | 1.63477 | 3.08728 | 1.889 |
| chr1 | 244462762 | 244463112 | 2.09225 | 3.9513 | 1.889 |
| chr9 | 29285913 | 29286178 | 1.03801 | 1.96047 | 1.889 |
| chr14 | 73544937 | 73545172 | 0.386847 | 0.730704 | 1.889 |
| chr20 | 55956768 | 55957179 | 1.8146 | 3.42758 | 1.889 |
| chr3 | 158981542 | 158981865 | 1.77892 | 3.36025 | 1.889 |
| chr3 | 47365836 | 47366048 | 1.58572 | 2.99545 | 1.889 |
| chr13 | 42684771 | 42685018 | 1.20201 | 2.27063 | 1.889 |
| chr1 | 15353907 | 15354312 | 1.44783 | 2.73507 | 1.889 |
| chr3 | 70393712 | 70393947 | 0.67431 | 1.27384 | 1.889 |
| chr10 | 877339 | 877613 | 0.721146 | 1.36241 | 1.889 |
| chr3 | 182622180 | 182622533 | 0.450916 | 0.851886 | 1.889 |
| chr7 | 150671926 | 150672386 | 0.862277 | 1.62908 | 1.889 |
| chr17 | 30169539 | 30169930 | 2.46893 | 4.66464 | 1.889 |
| chr20 | 17663228 | 17663590 | 1.46645 | 2.77066 | 1.889 |
| chr3 | 30054483 | 30054763 | 1.53132 | 2.89341 | 1.889 |
| chr1 | 206635147 | 206635548 | 1.79425 | 3.39045 | 1.890 |
| chr19 | 53693415 | 53693871 | 0.630772 | 1.19192 | 1.890 |
| chrX | 128860381 | 128860716 | 1.41315 | 2.67046 | 1.890 |
| chr7 | 26324109 | 26324317 | 0.561907 | 1.06196 | 1.890 |
| chr12 | 29462318 | 29462539 | 0.851755 | 1.60976 | 1.890 |
| chr1 | 76559684 | 76559892 | 0.438667 | 0.829057 | 1.890 |
| chr2 | 33620796 | 33621026 | 1.66531 | 3.14735 | 1.890 |
| chr2 | 22832417 | 22832694 | 0.655833 | 1.23952 | 1.890 |
| chr19 | 54534439 | 54534825 | 1.19982 | 2.26769 | 1.890 |
| chr3 | 39489356 | 39489706 | 1.68648 | 3.18756 | 1.890 |
| chr15 | 24313132 | 24313418 | 0.48786 | 0.922141 | 1.890 |
| chr7 | 149580528 | 149580873 | 1.61327 | 3.04948 | 1.890 |
| chr19 | 45347904 | 45348393 | 2.72563 | 5.15235 | 1.890 |
| chr1 | 159130799 | 159131133 | 2.78341 | 5.26179 | 1.890 |
| chr17 | 42201260 | 42201528 | 1.22569 | 2.31711 | 1.890 |
| chr7 | 50359014 | 50359282 | 0.540428 | 1.02167 | 1.890 |
| chr11 | 1254568 | 1254900 | 1.02432 | 1.9366 | 1.891 |
| chr4 | 173946278 | 173946521 | 1.07244 | 2.0276 | 1.891 |
| chr7 | 126422884 | 126423136 | 0.657449 | 1.24303 | 1.891 |
| chr7 | 86985267 | 86985513 | 0.538284 | 1.01775 | 1.891 |
| chr16 | 6587021 | 6587318 | 0.48078 | 0.909029 | 1.891 |
| chr20 | 741506 | 742044 | 2.52716 | 4.77826 | 1.891 |
| chr12 | 101717448 | 101717809 | 1.86831 | 3.5326 | 1.891 |

| | | | | | |
|---|---|---|---|---|---|
| chr7 | 23095229 | 23095585 | 2.32092 | 4.38839 | 1.891 |
| chr12 | 2297034 | 2297276 | 0.599833 | 1.13421 | 1.891 |
| chr8 | 111432044 | 111432376 | 0.586628 | 1.10926 | 1.891 |
| chr11 | 89170377 | 89170768 | 0.619233 | 1.17093 | 1.891 |
| chr10 | 18503799 | 18504206 | 1.57575 | 2.97973 | 1.891 |
| chr14 | 24058236 | 24058538 | 2.06687 | 3.90883 | 1.891 |
| chr22 | 39955771 | 39956051 | 1.78315 | 3.37228 | 1.891 |
| chr16 | 88503660 | 88504200 | 2.31577 | 4.37966 | 1.891 |
| chr5 | 167696359 | 167696740 | 1.82948 | 3.45999 | 1.891 |
| chr14 | 75327489 | 75328070 | 1.73832 | 3.28771 | 1.891 |
| chr3 | 118582827 | 118583128 | 1.08527 | 2.05258 | 1.891 |
| chr15 | 58362958 | 58363385 | 2.04476 | 3.86752 | 1.891 |
| chr8 | 124034654 | 124034936 | 1.72364 | 3.26042 | 1.892 |
| chr11 | 55798206 | 55798512 | 0.555367 | 1.05053 | 1.892 |
| chr5 | 173075783 | 173076098 | 0.470605 | 0.890197 | 1.892 |
| chr9 | 129976702 | 129977023 | 2.03684 | 3.85294 | 1.892 |
| chr2 | 96782698 | 96783045 | 1.48414 | 2.80752 | 1.892 |
| chr19 | 9958752 | 9959022 | 1.03863 | 1.96478 | 1.892 |
| chr9 | 105083502 | 105083735 | 0.648327 | 1.22651 | 1.892 |
| chr7 | 141233156 | 141233522 | 2.09748 | 3.96828 | 1.892 |
| chr7 | 36168078 | 36168398 | 1.18303 | 2.23827 | 1.892 |
| chr5 | 66445840 | 66446048 | 1.11768 | 2.11464 | 1.892 |
| chr1 | 26431405 | 26431764 | 1.32962 | 2.51576 | 1.892 |
| chr1 | 45060755 | 45061164 | 2.06869 | 3.91416 | 1.892 |
| chr16 | 3020159 | 3020467 | 1.12386 | 2.12654 | 1.892 |
| chr11 | 4208163 | 4209207 | 2.0072 | 3.79808 | 1.892 |
| chr12 | 120739827 | 120740139 | 2.00491 | 3.79375 | 1.892 |
| chr9 | 123975522 | 123975981 | 2.38344 | 4.51026 | 1.892 |
| chr1 | 205312517 | 205313137 | 1.20328 | 2.27705 | 1.892 |
| chr15 | 43663609 | 43663954 | 0.809536 | 1.53197 | 1.892 |
| chr7 | 953841 | 954142 | 0.483509 | 0.915038 | 1.892 |
| chr6 | 2750135 | 2750545 | 1.61463 | 3.05581 | 1.893 |
| chr20 | 31366231 | 31366628 | 2.42461 | 4.58885 | 1.893 |
| chr6 | 105535513 | 105535795 | 1.22862 | 2.32535 | 1.893 |
| chr4 | 8404939 | 8405228 | 1.991 | 3.76833 | 1.893 |
| chr8 | 145671770 | 145672002 | 0.574673 | 1.08774 | 1.893 |
| chr6 | 52589265 | 52589540 | 1.58987 | 3.00934 | 1.893 |
| chr17 | 56212317 | 56212558 | 0.666898 | 1.26234 | 1.893 |
| chr12 | 119402023 | 119402319 | 0.473341 | 0.895995 | 1.893 |
| chr10 | 21341113 | 21341374 | 0.616995 | 1.16795 | 1.893 |
| chr2 | 12753403 | 12753795 | 0.565859 | 1.07116 | 1.893 |
| chr3 | 194064668 | 194064989 | 2.16668 | 4.10154 | 1.893 |
| chr1 | 163041694 | 163041940 | 1.54866 | 2.93166 | 1.893 |
| chr12 | 54601530 | 54601978 | 2.17403 | 4.1155 | 1.893 |
| chr8 | 80067780 | 80067993 | 0.589364 | 1.11569 | 1.893 |
| chr10 | 105241729 | 105242450 | 1.0677 | 2.02126 | 1.893 |
| chr8 | 69046352 | 69046559 | 0.602548 | 1.14069 | 1.893 |
| chr16 | 74743286 | 74743649 | 2.20799 | 4.18027 | 1.893 |
| chr3 | 79884840 | 79885054 | 0.716946 | 1.35739 | 1.893 |
| chr19 | 48196808 | 48197082 | 0.68452 | 1.29602 | 1.893 |
| chr14 | 36156574 | 36156831 | 1.59746 | 3.0246 | 1.893 |
| chr19 | 8645518 | 8646001 | 3.54156 | 6.70589 | 1.893 |
| chr15 | 58474691 | 58475159 | 2.08647 | 3.9507 | 1.893 |
| chr1 | 31188074 | 31188350 | 1.3586 | 2.57261 | 1.894 |
| chr14 | 46813634 | 46813839 | 0.618923 | 1.17206 | 1.894 |
| chr12 | 104271733 | 104272027 | 1.65186 | 3.12816 | 1.894 |
| chr10 | 127625316 | 127625621 | 0.505509 | 0.957313 | 1.894 |
| chr11 | 67020212 | 67020571 | 1.08347 | 2.05185 | 1.894 |
| chr12 | 130770667 | 130770964 | 0.94524 | 1.7901 | 1.894 |
| chr9 | 85086753 | 85087081 | 0.534743 | 1.0127 | 1.894 |
| chr17 | 3920749 | 3921053 | 1.37442 | 2.60295 | 1.894 |

| | | | | | |
|---|---|---|---|---|---|
| chr18 | 74635501 | 74635730 | 1.42272 | 2.69444 | 1.894 |
| chr20 | 62406473 | 62407109 | 1.4139 | 2.67777 | 1.894 |
| chr12 | 103889822 | 103890268 | 1.13932 | 2.1578 | 1.894 |
| chr18 | 12394503 | 12394749 | 1.24091 | 2.35036 | 1.894 |
| chr12 | 7074224 | 7074546 | 2.33838 | 4.42917 | 1.894 |
| chr9 | 94506981 | 94507200 | 0.522824 | 0.990321 | 1.894 |
| chr12 | 71086459 | 71086782 | 1.28302 | 2.43032 | 1.894 |
| chr12 | 105993060 | 105993426 | 0.510415 | 0.966838 | 1.894 |
| chr19 | 39452769 | 39453094 | 2.40364 | 4.55308 | 1.894 |
| chr7 | 100787341 | 100787692 | 1.551 | 2.93809 | 1.894 |
| chr1 | 227669250 | 227669543 | 1.71523 | 3.24929 | 1.894 |
| chr5 | 113402557 | 113402785 | 0.436594 | 0.82709 | 1.894 |
| chr2 | 102622777 | 102623042 | 1.5319 | 2.90218 | 1.895 |
| chr8 | 29034490 | 29034839 | 2.43583 | 4.61471 | 1.895 |
| chr15 | 88103079 | 88103364 | 0.487588 | 0.923762 | 1.895 |
| chr19 | 35781400 | 35781746 | 0.568973 | 1.07795 | 1.895 |
| chr17 | 18144750 | 18145066 | 0.708078 | 1.34166 | 1.895 |
| chr19 | 40162705 | 40162940 | 1.58961 | 3.012 | 1.895 |
| chr4 | 141174342 | 141174644 | 1.8354 | 3.47776 | 1.895 |
| chr5 | 132623253 | 132623475 | 1.56538 | 2.96612 | 1.895 |
| chr12 | 6981427 | 6981957 | 2.25831 | 4.27916 | 1.895 |
| chr5 | 7927899 | 7928289 | 1.69687 | 3.21536 | 1.895 |
| chr19 | 15742634 | 15742924 | 2.44262 | 4.6286 | 1.895 |
| chr13 | 43543794 | 43544021 | 1.42381 | 2.69807 | 1.895 |
| chr19 | 40926509 | 40926929 | 2.61563 | 4.9566 | 1.895 |
| chr15 | 67096533 | 67096756 | 0.581716 | 1.10237 | 1.895 |
| chr1 | 159915416 | 159915815 | 1.15922 | 2.1968 | 1.895 |
| chr12 | 115310194 | 115310512 | 0.566774 | 1.07412 | 1.895 |
| chr14 | 88958570 | 88958896 | 1.46602 | 2.77884 | 1.896 |
| chr5 | 74729436 | 74729668 | 0.610061 | 1.15637 | 1.896 |
| chr7 | 44516987 | 44517280 | 1.4729 | 2.79188 | 1.896 |
| chr5 | 148414907 | 148415159 | 1.90016 | 3.60187 | 1.896 |
| chr9 | 88649772 | 88650139 | 1.84629 | 3.49989 | 1.896 |
| chr16 | 66517666 | 66518360 | 1.41932 | 2.69062 | 1.896 |
| chr5 | 43215599 | 43215856 | 0.747519 | 1.41717 | 1.896 |
| chr20 | 11185826 | 11186219 | 1.95979 | 3.71544 | 1.896 |
| chr1 | 245319763 | 245320195 | 1.80783 | 3.42768 | 1.896 |
| chr12 | 68288724 | 68289101 | 0.623324 | 1.18192 | 1.896 |
| chr15 | 99640757 | 99641135 | 2.39606 | 4.54332 | 1.896 |
| chr5 | 132082596 | 132082975 | 1.22015 | 2.3136 | 1.896 |
| chr17 | 37732476 | 37732872 | 1.77906 | 3.37345 | 1.896 |
| chr12 | 78540163 | 78540388 | 1.6979 | 3.21959 | 1.896 |
| chr12 | 92089737 | 92089941 | 0.806743 | 1.52982 | 1.896 |
| chr6 | 90081754 | 90082200 | 2.26181 | 4.28914 | 1.896 |
| chr19 | 33921537 | 33922007 | 2.44033 | 4.62769 | 1.896 |
| chr14 | 77342201 | 77342721 | 1.56327 | 2.96452 | 1.896 |
| chr5 | 19734798 | 19735222 | 0.482244 | 0.914513 | 1.896 |
| chr7 | 138701782 | 138702128 | 1.54986 | 2.93914 | 1.896 |
| chr9 | 6907815 | 6908128 | 2.25805 | 4.28285 | 1.897 |
| chr5 | 131343576 | 131343910 | 1.75899 | 3.33651 | 1.897 |
| chr8 | 3071976 | 3072195 | 0.550858 | 1.04489 | 1.897 |
| chr8 | 132070824 | 132071068 | 0.703525 | 1.33449 | 1.897 |
| chr1 | 40998598 | 40998809 | 1.07328 | 2.03591 | 1.897 |
| chrX | 22766606 | 22766938 | 1.54142 | 2.92395 | 1.897 |
| chr8 | 27618001 | 27618333 | 0.956464 | 1.81435 | 1.897 |
| chr20 | 58530250 | 58530569 | 0.992026 | 1.88193 | 1.897 |
| chr16 | 67527886 | 67528231 | 1.93003 | 3.66141 | 1.897 |
| chr11 | 43304451 | 43304873 | 2.30398 | 4.37083 | 1.897 |
| chr15 | 86159244 | 86159517 | 1.28692 | 2.44141 | 1.897 |
| chr18 | 21202282 | 21202544 | 2.36033 | 4.47778 | 1.897 |
| chr9 | 4077534 | 4077797 | 2.03864 | 3.86756 | 1.897 |

| | | | | | |
|---|---|---|---|---|---|
| chr14 | 61123896 | 61124228 | 1.2752 | 2.41923 | 1.897 |
| chr1 | 200479309 | 200479597 | 1.45429 | 2.7593 | 1.897 |
| chr5 | 16057110 | 16057335 | 0.584642 | 1.10928 | 1.897 |
| chr8 | 144918560 | 144918851 | 1.89754 | 3.60075 | 1.898 |
| chr19 | 59025173 | 59025627 | 1.82912 | 3.47098 | 1.898 |
| chr7 | 88643949 | 88644291 | 0.514177 | 0.97579 | 1.898 |
| chr3 | 99550922 | 99551145 | 0.822532 | 1.56102 | 1.898 |
| chr12 | 1900347 | 1900579 | 2.29306 | 4.35188 | 1.898 |
| chr17 | 30810518 | 30810772 | 0.929537 | 1.76416 | 1.898 |
| chr5 | 130752979 | 130753336 | 1.48558 | 2.8195 | 1.898 |
| chr1 | 155618381 | 155618676 | 1.26806 | 2.40673 | 1.898 |
| chr13 | 57102429 | 57102679 | 0.408404 | 0.775154 | 1.898 |
| chr3 | 161089594 | 161089810 | 0.951403 | 1.80582 | 1.898 |
| chr16 | 49626747 | 49627201 | 1.40696 | 2.67065 | 1.898 |
| chr2 | 208632663 | 208632878 | 1.5687 | 2.97777 | 1.898 |
| chr7 | 76961241 | 76961479 | 1.49209 | 2.83242 | 1.898 |
| chr4 | 158049680 | 158049887 | 1.20992 | 2.29679 | 1.898 |
| chr15 | 66274673 | 66274881 | 1.06424 | 2.02027 | 1.898 |
| chr14 | 24578063 | 24578349 | 2.50043 | 4.74697 | 1.898 |
| chr11 | 128775882 | 128776085 | 1.11969 | 2.1257 | 1.898 |
| chr12 | 71118278 | 71118596 | 1.01043 | 1.91833 | 1.899 |
| chr1 | 1976300 | 1976626 | 1.72932 | 3.2832 | 1.899 |
| chr10 | 49816195 | 49816664 | 1.66602 | 3.16308 | 1.899 |
| chr14 | 63121846 | 63122102 | 0.627421 | 1.19124 | 1.899 |
| chr22 | 50911724 | 50911927 | 1.01393 | 1.92508 | 1.899 |
| chr14 | 88109523 | 88109737 | 0.540585 | 1.02641 | 1.899 |
| chr14 | 94900134 | 94900570 | 1.53858 | 2.92133 | 1.899 |
| chr13 | 27948882 | 27949114 | 1.26716 | 2.406 | 1.899 |
| chr19 | 44314024 | 44314273 | 1.2221 | 2.32065 | 1.899 |
| chr11 | 26671379 | 26671707 | 1.63263 | 3.10034 | 1.899 |
| chr11 | 76521348 | 76521615 | 1.17637 | 2.23393 | 1.899 |
| chr1 | 205627920 | 205628541 | 1.16104 | 2.20483 | 1.899 |
| chr14 | 22508596 | 22508805 | 1.21599 | 2.30936 | 1.899 |
| chr12 | 116421054 | 116421377 | 1.47375 | 2.79897 | 1.899 |
| chr1 | 249135948 | 249136256 | 1.43337 | 2.72233 | 1.899 |
| chr3 | 98787309 | 98787513 | 0.429561 | 0.815909 | 1.899 |
| chr19 | 897517 | 897719 | 0.816224 | 1.55037 | 1.899 |
| chr5 | 148747035 | 148747449 | 2.37401 | 4.50949 | 1.900 |
| chr4 | 81911176 | 81911410 | 1.34545 | 2.55587 | 1.900 |
| chr10 | 121445968 | 121446357 | 2.06141 | 3.916 | 1.900 |
| chr15 | 34635298 | 34635625 | 2.83357 | 5.38293 | 1.900 |
| chr7 | 105517035 | 105517422 | 1.65341 | 3.14107 | 1.900 |
| chr7 | 129882401 | 129882780 | 2.11892 | 4.02555 | 1.900 |
| chr2 | 182942384 | 182942766 | 1.54632 | 2.93775 | 1.900 |
| chr2 | 194828320 | 194828555 | 1.13754 | 2.16124 | 1.900 |
| chr20 | 49545921 | 49546271 | 3.03605 | 5.76844 | 1.900 |
| chr3 | 129346813 | 129347080 | 1.6982 | 3.22666 | 1.900 |
| chr22 | 45022037 | 45022417 | 1.95487 | 3.71442 | 1.900 |
| chr4 | 7772153 | 7772488 | 0.912695 | 1.73421 | 1.900 |
| chr14 | 45756320 | 45756668 | 0.480713 | 0.913464 | 1.900 |
| chr9 | 134637804 | 134638260 | 1.58792 | 3.0175 | 1.900 |
| chr12 | 111332176 | 111332539 | 2.24985 | 4.27541 | 1.900 |
| chr22 | 24643601 | 24643900 | 1.74207 | 3.31049 | 1.900 |
| chr22 | 27231946 | 27232194 | 1.40346 | 2.66702 | 1.900 |
| chr6 | 142528583 | 142528928 | 1.54802 | 2.94174 | 1.900 |
| chr11 | 79629340 | 79629688 | 0.654193 | 1.24318 | 1.900 |
| chr2 | 27630985 | 27631208 | 1.16444 | 2.21286 | 1.900 |
| chr10 | 43700336 | 43700841 | 1.72267 | 3.27375 | 1.900 |
| chr9 | 100360951 | 100361402 | 2.09721 | 3.98551 | 1.900 |
| chr17 | 42749933 | 42750164 | 1.13645 | 2.15977 | 1.900 |
| chr9 | 79236572 | 79236851 | 0.547035 | 1.03961 | 1.900 |

| | | | | | |
|---|---|---|---|---|---|
| chr14 | 26134097 | 26134339 | 0.547497 | 1.04053 | 1.901 |
| rUn_gl0002 | 96758 | 97324 | 2.19121 | 4.16456 | 1.901 |
| chr19 | 45197589 | 45197817 | 1.19424 | 2.27001 | 1.901 |
| chr22 | 30841190 | 30841532 | 1.33094 | 2.52992 | 1.901 |
| chr15 | 70392820 | 70393031 | 0.860983 | 1.63661 | 1.901 |
| chr1 | 156289344 | 156289677 | 0.971859 | 1.84738 | 1.901 |
| chr7 | 54909443 | 54909772 | 1.21819 | 2.31562 | 1.901 |
| chr7 | 43739618 | 43739877 | 1.55226 | 2.95072 | 1.901 |
| chr17 | 53373312 | 53373642 | 1.92925 | 3.66741 | 1.901 |
| chr1 | 13992209 | 13992730 | 0.941854 | 1.79058 | 1.901 |
| chr20 | 33548361 | 33548947 | 1.26854 | 2.41173 | 1.901 |
| chr16 | 75525833 | 75526123 | 0.863739 | 1.64214 | 1.901 |
| chr10 | 93311546 | 93311955 | 1.63371 | 3.10603 | 1.901 |
| chr7 | 99510778 | 99511114 | 1.86842 | 3.55233 | 1.901 |
| chr2 | 227971749 | 227972176 | 1.55982 | 2.96567 | 1.901 |
| chr3 | 138665067 | 138665318 | 1.34934 | 2.56558 | 1.901 |
| chr11 | 130793250 | 130793543 | 1.54884 | 2.94493 | 1.901 |
| chr1 | 120154225 | 120154594 | 2.10306 | 3.99874 | 1.901 |
| chr13 | 91519611 | 91519854 | 1.13036 | 2.14925 | 1.901 |
| chr16 | 85461808 | 85462177 | 1.39228 | 2.64741 | 1.901 |
| chr12 | 62368392 | 62368645 | 0.699993 | 1.33104 | 1.902 |
| chr13 | 91697776 | 91697986 | 1.3301 | 2.52922 | 1.902 |
| chr11 | 23387712 | 23388059 | 0.643063 | 1.22282 | 1.902 |
| chr1 | 167656901 | 167657463 | 1.14409 | 2.17575 | 1.902 |
| chr7 | 157017420 | 157017770 | 0.800956 | 1.52326 | 1.902 |
| chr11 | 119067683 | 119067921 | 1.23367 | 2.34622 | 1.902 |
| chr3 | 81631190 | 81631525 | 1.60567 | 3.05381 | 1.902 |
| chr17 | 28663437 | 28663966 | 1.37694 | 2.61891 | 1.902 |
| chr3 | 51753183 | 51753520 | 1.92741 | 3.66599 | 1.902 |
| chr10 | 11514170 | 11514499 | 1.70108 | 3.23562 | 1.902 |
| chr2 | 233191896 | 233192196 | 2.12487 | 4.04174 | 1.902 |
| chr11 | 10675872 | 10676141 | 1.31532 | 2.50189 | 1.902 |
| chrX | 19035428 | 19035794 | 1.52437 | 2.89955 | 1.902 |
| chr9 | 25626220 | 25626490 | 0.582928 | 1.10888 | 1.902 |
| chr6 | 158088205 | 158088571 | 1.83201 | 3.48507 | 1.902 |
| chr5 | 40423483 | 40423765 | 0.548233 | 1.04299 | 1.902 |
| chr13 | 52389236 | 52389588 | 1.3549 | 2.57785 | 1.903 |
| chr8 | 141644053 | 141644432 | 0.738479 | 1.40509 | 1.903 |
| chr3 | 112453620 | 112453871 | 1.34044 | 2.55044 | 1.903 |
| chr12 | 45577092 | 45577294 | 0.821 | 1.56217 | 1.903 |
| chr16 | 28328311 | 28328830 | 1.75701 | 3.34323 | 1.903 |
| chr2 | 188318680 | 188318907 | 0.57601 | 1.09611 | 1.903 |
| chr11 | 82128416 | 82128622 | 0.588154 | 1.11925 | 1.903 |
| chr17 | 73823243 | 73824196 | 1.40421 | 2.67227 | 1.903 |
| chr11 | 82029259 | 82029474 | 0.409437 | 0.779181 | 1.903 |
| chr19 | 5578216 | 5578563 | 2.864 | 5.4504 | 1.903 |
| chr20 | 5325629 | 5325837 | 0.691667 | 1.3163 | 1.903 |
| chr5 | 172689727 | 172690046 | 0.501425 | 0.954257 | 1.903 |
| chr4 | 17638457 | 17638818 | 1.66406 | 3.16688 | 1.903 |
| chr1 | 23892718 | 23893070 | 1.01195 | 1.92589 | 1.903 |
| chr5 | 121549009 | 121549249 | 0.494764 | 0.94162 | 1.903 |
| chr13 | 50510318 | 50510598 | 2.25168 | 4.28539 | 1.903 |
| chr20 | 33110935 | 33111230 | 1.97308 | 3.75519 | 1.903 |
| chr12 | 94373000 | 94373236 | 1.04131 | 1.98184 | 1.903 |
| chr6 | 43742470 | 43742830 | 1.48509 | 2.82651 | 1.903 |
| chr12 | 53605569 | 53605916 | 1.62265 | 3.08844 | 1.903 |
| chr17 | 76113399 | 76113900 | 1.61669 | 3.07735 | 1.903 |
| chr1 | 94947041 | 94947440 | 1.44448 | 2.74959 | 1.904 |
| chr20 | 42684730 | 42685001 | 1.20388 | 2.29169 | 1.904 |
| chr2 | 103777946 | 103778187 | 2.11983 | 4.0353 | 1.904 |
| chr3 | 2555779 | 2556037 | 0.723133 | 1.37665 | 1.904 |

| | | | | | |
|---|---|---|---|---|---|
| chr1 | 25157242 | 25157505 | 1.55239 | 2.95551 | 1.904 |
| chr3 | 99508295 | 99508531 | 1.05863 | 2.01571 | 1.904 |
| chr10 | 6720179 | 6720479 | 1.49813 | 2.85266 | 1.904 |
| chr4 | 71370865 | 71371210 | 1.84722 | 3.5174 | 1.904 |
| chr5 | 34898375 | 34898742 | 1.3981 | 2.66223 | 1.904 |
| chr2 | 38143775 | 38144072 | 0.711623 | 1.35517 | 1.904 |
| chr5 | 156903999 | 156904271 | 1.84927 | 3.52165 | 1.904 |
| chr21 | 30118061 | 30118320 | 1.35326 | 2.57723 | 1.904 |
| chr19 | 33345027 | 33345229 | 0.57406 | 1.09334 | 1.905 |
| chr6 | 39329799 | 39330199 | 1.92277 | 3.66217 | 1.905 |
| chr15 | 23755703 | 23756013 | 0.776951 | 1.47982 | 1.905 |
| chr7 | 98957304 | 98957597 | 0.57409 | 1.09346 | 1.905 |
| chr10 | 61564867 | 61565132 | 1.13614 | 2.1641 | 1.905 |
| chr12 | 41772436 | 41772665 | 0.692671 | 1.31939 | 1.905 |
| chr11 | 93654915 | 93655275 | 1.05051 | 2.00106 | 1.905 |
| chr5 | 145432433 | 145432743 | 0.899951 | 1.71428 | 1.905 |
| chr20 | 12266930 | 12267231 | 0.628829 | 1.19801 | 1.905 |
| chr6 | 168518709 | 168518926 | 1.07716 | 2.05216 | 1.905 |
| chr15 | 71487656 | 71487887 | 0.978812 | 1.86484 | 1.905 |
| chr9 | 112642897 | 112643229 | 1.05058 | 2.00158 | 1.905 |
| chr15 | 90776485 | 90776848 | 1.17258 | 2.23407 | 1.905 |
| chr3 | 187456687 | 187456899 | 1.47118 | 2.803 | 1.905 |
| chr2 | 26476224 | 26476503 | 0.518507 | 0.987935 | 1.905 |
| chr8 | 5646007 | 5646255 | 0.681046 | 1.29764 | 1.905 |
| chr15 | 82303240 | 82303459 | 0.691359 | 1.3173 | 1.905 |
| chr1 | 35325324 | 35325656 | 1.69552 | 3.23065 | 1.905 |
| chr16 | 85022065 | 85022582 | 0.88425 | 1.68486 | 1.905 |
| chr5 | 172717730 | 172717941 | 0.388079 | 0.73952 | 1.906 |
| chr16 | 10462953 | 10463252 | 0.784234 | 1.49448 | 1.906 |
| chr9 | 103609479 | 103609834 | 0.546279 | 1.04102 | 1.906 |
| chr17 | 40180616 | 40180942 | 1.29781 | 2.47334 | 1.906 |
| chr17 | 76670758 | 76671106 | 0.770719 | 1.46884 | 1.906 |
| chr9 | 133894073 | 133894519 | 0.803886 | 1.53207 | 1.906 |
| chr17 | 5697291 | 5697584 | 0.864151 | 1.64694 | 1.906 |
| chr4 | 7930925 | 7931145 | 0.740074 | 1.41047 | 1.906 |
| chr16 | 82095527 | 82095796 | 1.03698 | 1.97639 | 1.906 |
| chr11 | 117924176 | 117924435 | 1.36577 | 2.6033 | 1.906 |
| chr13 | 21517762 | 21518009 | 1.86968 | 3.56382 | 1.906 |
| chr17 | 38231741 | 38232082 | 1.99721 | 3.80694 | 1.906 |
| chr17 | 79783806 | 79784020 | 0.926263 | 1.76559 | 1.906 |
| chr12 | 108523302 | 108523531 | 0.738546 | 1.40782 | 1.906 |
| chr2 | 236460561 | 236460820 | 1.22699 | 2.33891 | 1.906 |
| chr5 | 139012835 | 139013439 | 1.99091 | 3.79513 | 1.906 |
| chr1 | 205240596 | 205240809 | 0.876402 | 1.67072 | 1.906 |
| chr10 | 50583700 | 50583997 | 0.568717 | 1.08419 | 1.906 |
| chr1 | 204329134 | 204329418 | 1.27469 | 2.43005 | 1.906 |
| chr9 | 107267991 | 107268294 | 0.481267 | 0.917594 | 1.907 |
| chr6 | 43097631 | 43097874 | 1.14408 | 2.18138 | 1.907 |
| chr2 | 28509145 | 28509377 | 0.643637 | 1.22723 | 1.907 |
| chr20 | 6057709 | 6058376 | 1.17718 | 2.24457 | 1.907 |
| chr17 | 8385466 | 8385704 | 0.482953 | 0.920928 | 1.907 |
| chr5 | 60579357 | 60579577 | 0.580588 | 1.10711 | 1.907 |
| chrX | 10584058 | 10584304 | 1.43681 | 2.73994 | 1.907 |
| chr14 | 96523039 | 96523373 | 1.1626 | 2.2172 | 1.907 |
| chr10 | 84556354 | 84556572 | 0.728367 | 1.38908 | 1.907 |
| chr12 | 19099878 | 19100339 | 0.646282 | 1.23255 | 1.907 |
| chr15 | 90535506 | 90535739 | 0.559249 | 1.06658 | 1.907 |
| chr18 | 13159546 | 13159810 | 1.48521 | 2.83257 | 1.907 |
| chr8 | 29580552 | 29580886 | 1.38319 | 2.63813 | 1.907 |
| chr16 | 2244884 | 2245169 | 1.40216 | 2.67437 | 1.907 |
| chr10 | 102819728 | 102820200 | 1.55195 | 2.96014 | 1.907 |

| | | | | | |
|---|---|---|---|---|---|
| chr3 | 137833855 | 137834082 | 0.777864 | 1.48372 | 1.907 |
| chr15 | 50608230 | 50608511 | 1.07564 | 2.05182 | 1.908 |
| chr8 | 75648858 | 75649074 | 0.437339 | 0.834247 | 1.908 |
| chr5 | 80502731 | 80503113 | 0.785834 | 1.49913 | 1.908 |
| chr12 | 84035627 | 84035928 | 0.559742 | 1.06784 | 1.908 |
| chr16 | 3145991 | 3146595 | 0.574227 | 1.09548 | 1.908 |
| chr20 | 15217480 | 15217683 | 1.16095 | 2.2148 | 1.908 |
| chr7 | 825992 | 826329 | 2.09769 | 4.00192 | 1.908 |
| chr3 | 72004571 | 72004967 | 1.49503 | 2.8523 | 1.908 |
| chr11 | 1990044 | 1990680 | 1.28403 | 2.44986 | 1.908 |
| chr11 | 89938055 | 89938317 | 1.88236 | 3.59171 | 1.908 |
| chr6 | 97663057 | 97663326 | 1.29429 | 2.46977 | 1.908 |
| chr2 | 26074858 | 26075063 | 1.09182 | 2.08346 | 1.908 |
| chr10 | 66985139 | 66985566 | 0.5547 | 1.05853 | 1.908 |
| chr17 | 72199586 | 72199843 | 2.95478 | 5.63883 | 1.908 |
| chr5 | 132682794 | 132683065 | 1.08981 | 2.07978 | 1.908 |
| chr9 | 72535562 | 72535779 | 0.44024 | 0.840232 | 1.909 |
| chr6 | 1313886 | 1314283 | 1.06558 | 2.0338 | 1.909 |
| chr1 | 27659280 | 27659731 | 1.93421 | 3.69178 | 1.909 |
| chr16 | 599534 | 599830 | 0.898928 | 1.71577 | 1.909 |
| chr20 | 21459883 | 21460137 | 0.512068 | 0.977394 | 1.909 |
| chr12 | 47701640 | 47701919 | 1.93714 | 3.69755 | 1.909 |
| chr5 | 168106465 | 168106848 | 1.65614 | 3.16121 | 1.909 |
| chr22 | 37252285 | 37252781 | 3.55328 | 6.78257 | 1.909 |
| chr3 | 24216492 | 24216728 | 0.584483 | 1.1157 | 1.909 |
| chr3 | 72937484 | 72937687 | 0.706482 | 1.34873 | 1.909 |
| chr17 | 73319077 | 73319662 | 1.78328 | 3.40446 | 1.909 |
| chr10 | 127622889 | 127623223 | 1.71727 | 3.27848 | 1.909 |
| chr3 | 146821738 | 146822010 | 0.52328 | 0.999072 | 1.909 |
| chr12 | 42332968 | 42333212 | 0.864333 | 1.65036 | 1.909 |
| chr17 | 75999864 | 76000130 | 0.713637 | 1.36264 | 1.909 |
| chr9 | 128991184 | 128991547 | 1.98458 | 3.78983 | 1.910 |
| chr12 | 1743784 | 1744136 | 2.42269 | 4.62649 | 1.910 |
| chr10 | 13313584 | 13313844 | 1.17443 | 2.24278 | 1.910 |
| chr6 | 37140014 | 37140229 | 1.09952 | 2.09974 | 1.910 |
| chr12 | 112929545 | 112929789 | 0.563134 | 1.07547 | 1.910 |
| chr15 | 86374773 | 86375109 | 1.8317 | 3.49818 | 1.910 |
| chr16 | 30198608 | 30199063 | 1.76597 | 3.3727 | 1.910 |
| chr19 | 47120305 | 47120892 | 1.59059 | 3.0378 | 1.910 |
| chr11 | 30843851 | 30844107 | 0.460648 | 0.879795 | 1.910 |
| chrX | 3858088 | 3858308 | 1.02037 | 1.94883 | 1.910 |
| chr3 | 101426115 | 101426523 | 1.15936 | 2.21445 | 1.910 |
| chr12 | 60341061 | 60341575 | 0.595191 | 1.13689 | 1.910 |
| chr17 | 58217231 | 58217537 | 1.68089 | 3.21076 | 1.910 |
| chr12 | 53292396 | 53292683 | 1.03589 | 1.97886 | 1.910 |
| chr19 | 46905492 | 46906003 | 1.88273 | 3.59661 | 1.910 |
| chr7 | 129702866 | 129703208 | 2.06083 | 3.93689 | 1.910 |
| chr15 | 90441941 | 90442243 | 1.60948 | 3.07467 | 1.910 |
| chr3 | 2160450 | 2160991 | 0.522353 | 0.997884 | 1.910 |
| chr14 | 77206789 | 77207019 | 0.480884 | 0.91867 | 1.910 |
| chr3 | 97100287 | 97100550 | 0.578254 | 1.1047 | 1.910 |
| chr1 | 151507187 | 151507534 | 2.88778 | 5.51688 | 1.910 |
| chr10 | 7868958 | 7869175 | 0.762018 | 1.45579 | 1.910 |
| chr2 | 27047559 | 27047763 | 0.456053 | 0.871272 | 1.910 |
| chr1 | 36808309 | 36808661 | 1.6657 | 3.18231 | 1.910 |
| chr19 | 19051944 | 19052203 | 2.20075 | 4.20451 | 1.910 |
| chr17 | 73267170 | 73267578 | 1.96818 | 3.76034 | 1.911 |
| chr11 | 119192303 | 119192653 | 1.81665 | 3.4709 | 1.911 |
| chr8 | 91368615 | 91368908 | 0.435091 | 0.831288 | 1.911 |
| chr11 | 90953913 | 90954210 | 0.569934 | 1.08894 | 1.911 |
| chr12 | 96843092 | 96843344 | 0.567854 | 1.08497 | 1.911 |

| | | | | | |
|---|---|---|---|---|---|
| chr16 | 48198782 | 48199110 | 1.41162 | 2.6973 | 1.911 |
| chr11 | 60897164 | 60897678 | 2.82954 | 5.40669 | 1.911 |
| chr11 | 67396530 | 67397251 | 1.57022 | 3.00038 | 1.911 |
| chr11 | 67973599 | 67973984 | 1.25514 | 2.39835 | 1.911 |
| chr16 | 4250200 | 4250514 | 1.48562 | 2.83878 | 1.911 |
| chr6 | 24991888 | 24992391 | 1.04289 | 1.99285 | 1.911 |
| chr17 | 30427484 | 30427838 | 0.49569 | 0.94726 | 1.911 |
| chr10 | 29654057 | 29654271 | 0.374525 | 0.715757 | 1.911 |
| chr19 | 51976207 | 51976522 | 1.95904 | 3.74398 | 1.911 |
| chr7 | 130725249 | 130725537 | 1.80522 | 3.45008 | 1.911 |
| chr8 | 145982563 | 145982857 | 1.43222 | 2.73722 | 1.911 |
| chr18 | 67863161 | 67863384 | 0.992812 | 1.89752 | 1.911 |
| chr11 | 66025590 | 66026167 | 1.80794 | 3.45548 | 1.911 |
| chr8 | 145597439 | 145597796 | 2.87845 | 5.50164 | 1.911 |
| chr15 | 25939066 | 25939433 | 1.36548 | 2.61 | 1.911 |
| chr14 | 38034852 | 38035108 | 1.11249 | 2.12645 | 1.911 |
| chr20 | 2647280 | 2647570 | 0.500159 | 0.956048 | 1.911 |
| chr2 | 113956560 | 113956986 | 1.60274 | 3.0637 | 1.912 |
| chr3 | 155796989 | 155797288 | 0.555338 | 1.06155 | 1.912 |
| chr19 | 51599870 | 51600100 | 1.25079 | 2.39095 | 1.912 |
| chr9 | 71669047 | 71669502 | 1.54012 | 2.94401 | 1.912 |
| chr9 | 26103282 | 26103526 | 0.56637 | 1.08266 | 1.912 |
| chr11 | 82798148 | 82798369 | 0.709037 | 1.35546 | 1.912 |
| chr4 | 130562300 | 130562767 | 1.14984 | 2.19822 | 1.912 |
| chr2 | 21741438 | 21741741 | 0.592373 | 1.1325 | 1.912 |
| chr5 | 148998289 | 148998619 | 1.35316 | 2.58704 | 1.912 |
| chr12 | 71796879 | 71797125 | 0.818774 | 1.56538 | 1.912 |
| chr3 | 115641526 | 115641830 | 1.11333 | 2.12864 | 1.912 |
| chr4 | 5401534 | 5401981 | 1.82239 | 3.48439 | 1.912 |
| chr2 | 143868455 | 143868869 | 1.45897 | 2.78958 | 1.912 |
| chr4 | 20249782 | 20250022 | 1.28014 | 2.44771 | 1.912 |
| chr12 | 4863782 | 4864099 | 0.384395 | 0.734995 | 1.912 |
| chr8 | 140782358 | 140782575 | 0.587435 | 1.12322 | 1.912 |
| chr18 | 55989377 | 55989593 | 0.991313 | 1.89548 | 1.912 |
| chrX | 134185335 | 134185608 | 0.654245 | 1.25102 | 1.912 |
| chr1 | 154531971 | 154532319 | 1.77813 | 3.40025 | 1.912 |
| chr12 | 122607765 | 122608035 | 1.33999 | 2.5625 | 1.912 |
| chr2 | 131722119 | 131722442 | 1.1004 | 2.10433 | 1.912 |
| chr12 | 27788328 | 27788672 | 1.38668 | 2.65192 | 1.912 |
| chr12 | 34449809 | 34450019 | 0.463005 | 0.885465 | 1.912 |
| chr11 | 76899422 | 76899817 | 2.02099 | 3.86519 | 1.913 |
| chr3 | 155728641 | 155728972 | 1.18186 | 2.26036 | 1.913 |
| chr10 | 60174372 | 60174615 | 1.22746 | 2.34768 | 1.913 |
| chr4 | 164602492 | 164602696 | 0.667183 | 1.27608 | 1.913 |
| chr1 | 46912496 | 46912846 | 1.89118 | 3.61721 | 1.913 |
| chr14 | 33029519 | 33029786 | 1.48565 | 2.84169 | 1.913 |
| chr10 | 99930158 | 99930586 | 1.2553 | 2.40111 | 1.913 |
| chr3 | 146875213 | 146875576 | 0.488161 | 0.933796 | 1.913 |
| chr12 | 14238840 | 14239118 | 0.433491 | 0.829277 | 1.913 |
| chr17 | 38606440 | 38606805 | 1.76125 | 3.36932 | 1.913 |
| chr5 | 64106908 | 64107201 | 1.29704 | 2.48132 | 1.913 |
| chr3 | 45802719 | 45802923 | 1.5432 | 2.95255 | 1.913 |
| chr3 | 159130448 | 159130677 | 0.588091 | 1.12519 | 1.913 |
| chr16 | 84217556 | 84217890 | 2.26554 | 4.33493 | 1.913 |
| chr2 | 87035498 | 87035799 | 1.96251 | 3.75511 | 1.913 |
| chr2 | 238480581 | 238480990 | 2.07098 | 3.96277 | 1.913 |
| chr12 | 68244705 | 68244925 | 0.54275 | 1.03855 | 1.914 |
| chr10 | 23447006 | 23447612 | 1.23714 | 2.36728 | 1.914 |
| chr5 | 10628053 | 10628323 | 1.76799 | 3.38312 | 1.914 |
| chr17 | 21247251 | 21247477 | 0.632412 | 1.21015 | 1.914 |
| chr12 | 10078435 | 10078655 | 1.14705 | 2.195 | 1.914 |

| | | | | | |
|---|---|---|---|---|---|
| chr13 | 114909272 | 114909495 | 1.24505 | 2.38255 | 1.914 |
| chr1 | 165912627 | 165913032 | 0.607749 | 1.16304 | 1.914 |
| chr14 | 71570654 | 71571015 | 1.64114 | 3.14066 | 1.914 |
| chr15 | 101034867 | 101035407 | 0.707912 | 1.35474 | 1.914 |
| chr17 | 76704620 | 76704929 | 2.39007 | 4.57396 | 1.914 |
| chr12 | 9901327 | 9901714 | 1.40707 | 2.6928 | 1.914 |
| chr3 | 96365558 | 96365796 | 0.633253 | 1.21192 | 1.914 |
| chr16 | 1107408 | 1107963 | 2.3859 | 4.56624 | 1.914 |
| chr17 | 62772930 | 62773288 | 2.80296 | 5.36442 | 1.914 |
| chr7 | 4426362 | 4426617 | 0.534065 | 1.0222 | 1.914 |
| chr9 | 139839169 | 139839515 | 2.55925 | 4.89841 | 1.914 |
| chr9 | 36632956 | 36633319 | 0.981771 | 1.87913 | 1.914 |
| chr9 | 131901130 | 131901785 | 1.44148 | 2.75904 | 1.914 |
| chr22 | 45659716 | 45660057 | 1.29304 | 2.47495 | 1.914 |
| chr3 | 46901818 | 46902107 | 1.538 | 2.944 | 1.914 |
| chr17 | 59467163 | 59467771 | 2.61356 | 5.00283 | 1.914 |
| chr20 | 4587925 | 4588263 | 1.88014 | 3.59918 | 1.914 |
| chr1 | 59248093 | 59248361 | 0.632119 | 1.21008 | 1.914 |
| chr20 | 311233 | 311611 | 1.90563 | 3.64799 | 1.914 |
| chr5 | 176540854 | 176541330 | 2.48725 | 4.76139 | 1.914 |
| chr2 | 169630118 | 169630518 | 1.46997 | 2.81403 | 1.914 |
| chr20 | 43346714 | 43346944 | 0.983522 | 1.88288 | 1.914 |
| chr11 | 22884135 | 22884357 | 0.506837 | 0.970307 | 1.914 |
| chr7 | 53082603 | 53082829 | 0.529475 | 1.01366 | 1.914 |
| chr17 | 8399556 | 8399926 | 0.604474 | 1.15725 | 1.914 |
| chr7 | 20824521 | 20824760 | 1.38605 | 2.65359 | 1.915 |
| chr1 | 206807621 | 206807944 | 1.53885 | 2.94626 | 1.915 |
| chr19 | 31828994 | 31829316 | 1.04209 | 1.99521 | 1.915 |
| chr1 | 203311494 | 203311904 | 1.84207 | 3.52689 | 1.915 |
| chr1 | 223254695 | 223255086 | 2.34902 | 4.4978 | 1.915 |
| chr17 | 72740693 | 72741144 | 3.10452 | 5.94443 | 1.915 |
| chr6 | 129052377 | 129052625 | 1.29681 | 2.4833 | 1.915 |
| chr1 | 41315687 | 41316015 | 1.74396 | 3.3396 | 1.915 |
| chr4 | 120548326 | 120548760 | 1.38394 | 2.65024 | 1.915 |
| chr12 | 117682704 | 117682913 | 0.536581 | 1.02757 | 1.915 |
| chr19 | 44059314 | 44059748 | 2.12406 | 4.06767 | 1.915 |
| chr19 | 47494447 | 47494806 | 1.83772 | 3.5193 | 1.915 |
| chr10 | 33624065 | 33624300 | 1.01688 | 1.94741 | 1.915 |
| chr12 | 25589298 | 25589596 | 1.68035 | 3.21802 | 1.915 |
| chr12 | 86500188 | 86500410 | 0.650106 | 1.24503 | 1.915 |
| chr20 | 6557592 | 6557930 | 0.567029 | 1.08593 | 1.915 |
| chr3 | 45980320 | 45980683 | 1.65656 | 3.17266 | 1.915 |
| chr7 | 134386072 | 134386295 | 0.972452 | 1.86252 | 1.915 |
| chr15 | 71185406 | 71185610 | 1.32726 | 2.54219 | 1.915 |
| chr11 | 86319285 | 86319747 | 1.75294 | 3.35769 | 1.915 |
| chr11 | 61665143 | 61665632 | 2.25861 | 4.32638 | 1.916 |
| chr11 | 125811900 | 125812292 | 1.69257 | 3.24219 | 1.916 |
| chr20 | 58786180 | 58786527 | 2.50721 | 4.80269 | 1.916 |
| chr15 | 100860914 | 100861206 | 0.524993 | 1.00575 | 1.916 |
| chr12 | 69684729 | 69685158 | 1.88242 | 3.60627 | 1.916 |
| chr15 | 62358900 | 62359184 | 0.886783 | 1.69889 | 1.916 |
| chr19 | 3987817 | 3988139 | 1.66012 | 3.18051 | 1.916 |
| chr17 | 72472480 | 72472853 | 0.874191 | 1.67487 | 1.916 |
| chr6 | 170190254 | 170190924 | 1.20697 | 2.31245 | 1.916 |
| chr5 | 17178733 | 17178993 | 0.43611 | 0.835558 | 1.916 |
| chr7 | 77287143 | 77287380 | 1.07929 | 2.06787 | 1.916 |
| chr1 | 32059625 | 32059905 | 1.77249 | 3.3961 | 1.916 |
| chr14 | 79106364 | 79106709 | 0.544744 | 1.04375 | 1.916 |
| chr16 | 18990698 | 18990977 | 1.3422 | 2.57173 | 1.916 |
| chr11 | 20875974 | 20876318 | 0.646799 | 1.23932 | 1.916 |
| chr11 | 45238940 | 45239216 | 0.467591 | 0.896007 | 1.916 |

| | | | | | |
|---|---|---|---|---|---|
| chr1 | 147362334 | 147362699 | 1.66642 | 3.19327 | 1.916 |
| chr5 | 176830749 | 176831173 | 1.65733 | 3.1759 | 1.916 |
| chr16 | 84771913 | 84772521 | 1.59765 | 3.06161 | 1.916 |
| chr16 | 57450929 | 57451305 | 1.97995 | 3.79439 | 1.916 |
| chr9 | 86853646 | 86853965 | 0.978422 | 1.87506 | 1.916 |
| chr1 | 37169307 | 37169643 | 2.16252 | 4.14444 | 1.916 |
| chr1 | 238637606 | 238637923 | 0.670813 | 1.28561 | 1.917 |
| chr3 | 129548495 | 129548733 | 1.02034 | 1.95549 | 1.917 |
| chr16 | 62422870 | 62423116 | 0.538798 | 1.03268 | 1.917 |
| chr3 | 23583182 | 23583469 | 0.551232 | 1.05653 | 1.917 |
| chr12 | 84665763 | 84666021 | 0.474973 | 0.910425 | 1.917 |
| chr4 | 47507407 | 47507620 | 0.532146 | 1.02001 | 1.917 |
| chrX | 100023686 | 100024131 | 3.34614 | 6.41388 | 1.917 |
| chr2 | 121837808 | 121838163 | 1.10236 | 2.11303 | 1.917 |
| chr9 | 31893900 | 31894102 | 0.839236 | 1.60872 | 1.917 |
| chr6 | 130072962 | 130073276 | 1.00747 | 1.9313 | 1.917 |
| chr3 | 97512344 | 97512590 | 1.17814 | 2.25859 | 1.917 |
| chr10 | 37068265 | 37068513 | 1.36362 | 2.61427 | 1.917 |
| chr2 | 131130065 | 131130404 | 2.14894 | 4.11989 | 1.917 |
| chr12 | 21850908 | 21851152 | 0.851176 | 1.63186 | 1.917 |
| chr5 | 141531514 | 141531767 | 1.45215 | 2.78403 | 1.917 |
| chr5 | 67277653 | 67277870 | 0.375202 | 0.719352 | 1.917 |
| chr6 | 52288807 | 52289024 | 0.491568 | 0.942458 | 1.917 |
| chr19 | 20636533 | 20636755 | 0.55055 | 1.05555 | 1.917 |
| chr10 | 59115026 | 59115324 | 0.512793 | 0.983169 | 1.917 |
| chr10 | 47627347 | 47627670 | 1.67913 | 3.21951 | 1.917 |
| chr20 | 44591681 | 44592049 | 1.38132 | 2.6486 | 1.917 |
| chr22 | 20226120 | 20226361 | 1.42712 | 2.73642 | 1.917 |
| chrX | 118315510 | 118315783 | 1.43786 | 2.75701 | 1.917 |
| chr1 | 65210156 | 65210397 | 0.998256 | 1.91421 | 1.918 |
| chr2 | 12942249 | 12942538 | 0.484947 | 0.929939 | 1.918 |
| chr6 | 152677916 | 152678183 | 0.604774 | 1.15972 | 1.918 |
| chr7 | 138759848 | 138760173 | 1.62668 | 3.11933 | 1.918 |
| chr2 | 221965548 | 221965807 | 0.996516 | 1.91106 | 1.918 |
| chr7 | 814064 | 814519 | 1.66616 | 3.19529 | 1.918 |
| chr1 | 21661266 | 21661474 | 1.64562 | 3.15594 | 1.918 |
| chrX | 3569413 | 3569650 | 0.82646 | 1.58499 | 1.918 |
| chr20 | 18256489 | 18256897 | 0.618156 | 1.18552 | 1.918 |
| chr2 | 178469262 | 178469548 | 1.45076 | 2.78236 | 1.918 |
| chr2 | 239422064 | 239422371 | 1.38678 | 2.65969 | 1.918 |
| chr2 | 178997121 | 178997347 | 1.1308 | 2.16878 | 1.918 |
| chr12 | 52545844 | 52546223 | 2.0183 | 3.87099 | 1.918 |
| chr3 | 11197168 | 11197491 | 2.11863 | 4.06349 | 1.918 |
| chr18 | 74398711 | 74398970 | 1.55667 | 2.98567 | 1.918 |
| chr9 | 116916071 | 116916273 | 1.06901 | 2.05037 | 1.918 |
| chr8 | 90860499 | 90860758 | 1.38253 | 2.65172 | 1.918 |
| chr21 | 36060506 | 36060930 | 1.22834 | 2.35599 | 1.918 |
| chr17 | 39653011 | 39653373 | 2.27953 | 4.37246 | 1.918 |
| chr16 | 30497901 | 30498196 | 0.471632 | 0.904665 | 1.918 |
| chr12 | 42876424 | 42876948 | 2.02371 | 3.88185 | 1.918 |
| chr6 | 18789259 | 18789509 | 0.524469 | 1.00604 | 1.918 |
| chr15 | 91025669 | 91026019 | 1.44 | 2.76238 | 1.918 |
| chr8 | 80596907 | 80597132 | 1.67213 | 3.20769 | 1.918 |
| chr1 | 152346420 | 152346626 | 0.549822 | 1.05475 | 1.918 |
| chrX | 117670724 | 117670980 | 1.00733 | 1.93241 | 1.918 |
| chr20 | 44640523 | 44640995 | 1.39787 | 2.68165 | 1.918 |
| chr6 | 152589321 | 152589662 | 0.825751 | 1.58412 | 1.918 |
| chr13 | 36400860 | 36401124 | 1.06128 | 2.03597 | 1.918 |
| chr19 | 8571032 | 8571402 | 2.16189 | 4.14751 | 1.918 |
| chr8 | 58630647 | 58630859 | 0.557969 | 1.07047 | 1.919 |
| chr11 | 33278692 | 33279015 | 1.06916 | 2.0512 | 1.919 |

| | | | | | |
|---|---|---|---|---|---|
| chr19 | 30890795 | 30891019 | 1.2769 | 2.44978 | 1.919 |
| chr15 | 77695237 | 77695444 | 0.524856 | 1.007 | 1.919 |
| chr20 | 2435420 | 2435785 | 2.12518 | 4.07747 | 1.919 |
| chr3 | 86294427 | 86294791 | 0.853319 | 1.63723 | 1.919 |
| chr6 | 44398946 | 44399161 | 1.27827 | 2.45257 | 1.919 |
| chr12 | 77214728 | 77215003 | 1.76357 | 3.38375 | 1.919 |
| chr20 | 16782611 | 16782992 | 1.52808 | 2.93194 | 1.919 |
| chr7 | 123134310 | 123134583 | 1.05155 | 2.01762 | 1.919 |
| chr2 | 172991854 | 172992327 | 2.03897 | 3.91224 | 1.919 |
| chr8 | 27621427 | 27621666 | 1.13867 | 2.18483 | 1.919 |
| chr14 | 106803922 | 106804175 | 0.885168 | 1.69848 | 1.919 |
| chr7 | 149439214 | 149439996 | 1.2258 | 2.35236 | 1.919 |
| chr14 | 77591261 | 77591811 | 1.7584 | 3.37451 | 1.919 |
| chr1 | 3531695 | 3532000 | 1.7675 | 3.39216 | 1.919 |
| chr2 | 85765693 | 85766044 | 2.00705 | 3.8521 | 1.919 |
| chr16 | 77440638 | 77440861 | 1.07999 | 2.07283 | 1.919 |
| chr8 | 37185910 | 37186128 | 1.41848 | 2.72268 | 1.919 |
| chr4 | 151164017 | 151164398 | 1.6891 | 3.24228 | 1.920 |
| chr16 | 12576331 | 12576566 | 0.659915 | 1.26673 | 1.920 |
| chr6 | 159139490 | 159139843 | 1.53493 | 2.94636 | 1.920 |
| chr1 | 204535012 | 204535223 | 1.07918 | 2.07167 | 1.920 |
| chr17 | 43361401 | 43362021 | 1.19479 | 2.29394 | 1.920 |
| chr11 | 38364208 | 38364468 | 0.486945 | 0.93493 | 1.920 |
| chr11 | 58949906 | 58950163 | 0.635094 | 1.21939 | 1.920 |
| chr16 | 47721635 | 47722080 | 0.544473 | 1.04541 | 1.920 |
| chr13 | 22051169 | 22051501 | 2.92019 | 5.60701 | 1.920 |
| chr7 | 84950372 | 84950612 | 0.579888 | 1.11345 | 1.920 |
| chr14 | 64883828 | 64884085 | 0.502796 | 0.965431 | 1.920 |
| chr10 | 32058982 | 32059710 | 1.20835 | 2.32024 | 1.920 |
| chr10 | 53196432 | 53196687 | 0.528024 | 1.01396 | 1.920 |
| chr1 | 37074919 | 37075157 | 1.75647 | 3.37307 | 1.920 |
| chr17 | 48556880 | 48557120 | 1.57645 | 3.02753 | 1.920 |
| chr12 | 52998768 | 52998981 | 0.574513 | 1.10334 | 1.920 |
| chr16 | 16281260 | 16281661 | 1.65588 | 3.1802 | 1.921 |
| chr9 | 76509946 | 76510297 | 0.630785 | 1.21148 | 1.921 |
| chr12 | 1909851 | 1910084 | 0.87767 | 1.68565 | 1.921 |
| chr5 | 12510000 | 12510254 | 0.568852 | 1.09257 | 1.921 |
| chr7 | 2471075 | 2471484 | 1.38688 | 2.66416 | 1.921 |
| chr6 | 34214379 | 34214612 | 1.16205 | 2.23228 | 1.921 |
| chr15 | 70929212 | 70929455 | 1.09207 | 2.09793 | 1.921 |
| chr19 | 42461904 | 42462246 | 2.05688 | 3.9514 | 1.921 |
| chr11 | 31066929 | 31067268 | 0.595138 | 1.14332 | 1.921 |
| chr19 | 42194134 | 42194403 | 0.724221 | 1.39133 | 1.921 |
| chr11 | 86586365 | 86586643 | 0.502502 | 0.965388 | 1.921 |
| chr2 | 35635480 | 35635810 | 1.64457 | 3.15957 | 1.921 |
| chr15 | 78014900 | 78015285 | 0.556685 | 1.06952 | 1.921 |
| chr1 | 246764629 | 246764850 | 0.788716 | 1.51532 | 1.921 |
| chr16 | 82263623 | 82264052 | 2.31944 | 4.45623 | 1.921 |
| chr2 | 30249347 | 30249657 | 1.2827 | 2.46453 | 1.921 |
| chr9 | 105680232 | 105680671 | 0.535965 | 1.02978 | 1.921 |
| chr1 | 210484429 | 210484737 | 1.33275 | 2.56071 | 1.921 |
| chr19 | 39312010 | 39312220 | 0.70795 | 1.36032 | 1.921 |
| chr14 | 75380738 | 75381005 | 2.09299 | 4.02184 | 1.922 |
| chr2 | 86448293 | 86448879 | 1.2456 | 2.3936 | 1.922 |
| chr10 | 25401830 | 25402473 | 1.86135 | 3.57697 | 1.922 |
| chr19 | 54527140 | 54527488 | 2.19507 | 4.21832 | 1.922 |
| chr3 | 156227489 | 156227779 | 1.01826 | 1.95681 | 1.922 |
| chr3 | 111436424 | 111436724 | 2.15001 | 4.13182 | 1.922 |
| chr3 | 158720611 | 158720823 | 0.459794 | 0.88363 | 1.922 |
| chr5 | 36337376 | 36337632 | 0.77801 | 1.49518 | 1.922 |
| chr16 | 793887 | 794099 | 0.577267 | 1.10943 | 1.922 |

| | | | | | |
|---|---|---|---|---|---|
| chr9 | 139601604 | 139601913 | 0.433009 | 0.832202 | 1.922 |
| chr2 | 227661637 | 227661874 | 1.09236 | 2.09942 | 1.922 |
| chr1 | 113258926 | 113259217 | 1.99341 | 3.83124 | 1.922 |
| chr1 | 108293081 | 108293296 | 1.15773 | 2.2252 | 1.922 |
| chr11 | 96074916 | 96075124 | 0.958599 | 1.84248 | 1.922 |
| chr15 | 68447263 | 68447643 | 1.98398 | 3.81331 | 1.922 |
| chr16 | 29740520 | 29740900 | 1.13686 | 2.18514 | 1.922 |
| chr2 | 236688099 | 236688461 | 2.0868 | 4.01117 | 1.922 |
| chr5 | 81461772 | 81462067 | 1.2818 | 2.46387 | 1.922 |
| chr10 | 69102690 | 69102965 | 0.491154 | 0.94412 | 1.922 |
| chr8 | 79410913 | 79411230 | 0.639161 | 1.22867 | 1.922 |
| chr9 | 19466283 | 19466536 | 0.495064 | 0.951682 | 1.922 |
| chr10 | 59495364 | 59495744 | 0.431905 | 0.830292 | 1.922 |
| chr11 | 30092779 | 30093089 | 0.511102 | 0.982545 | 1.922 |
| chr19 | 49496464 | 49496839 | 2.11666 | 4.06914 | 1.922 |
| chr15 | 90501309 | 90501523 | 0.664424 | 1.27738 | 1.923 |
| chr8 | 139712583 | 139713025 | 1.63083 | 3.13533 | 1.923 |
| chr12 | 39970035 | 39970242 | 0.602728 | 1.15886 | 1.923 |
| chr7 | 130666947 | 130667255 | 1.47704 | 2.83995 | 1.923 |
| chr8 | 63616998 | 63617200 | 0.599106 | 1.15196 | 1.923 |
| chr17 | 9599989 | 9600257 | 1.15774 | 2.22613 | 1.923 |
| chr7 | 130499569 | 130499926 | 0.844964 | 1.62477 | 1.923 |
| chr1 | 27371020 | 27371352 | 1.4948 | 2.87443 | 1.923 |
| chr20 | 9340244 | 9340799 | 1.32447 | 2.54692 | 1.923 |
| chr1 | 178313663 | 178313876 | 0.757231 | 1.45619 | 1.923 |
| chr3 | 143360378 | 143360664 | 0.382867 | 0.736275 | 1.923 |
| chr17 | 13275848 | 13276052 | 0.583584 | 1.12228 | 1.923 |
| chr1 | 156326658 | 156327119 | 2.60302 | 5.00587 | 1.923 |
| chr16 | 12322509 | 12322986 | 1.4993 | 2.88338 | 1.923 |
| chr19 | 14169743 | 14170188 | 1.99083 | 3.82883 | 1.923 |
| chr7 | 34360554 | 34360932 | 2.04925 | 3.9412 | 1.923 |
| chr10 | 95575873 | 95576264 | 2.13352 | 4.10339 | 1.923 |
| chr17 | 10640409 | 10640788 | 1.59365 | 3.06508 | 1.923 |
| chr8 | 66472571 | 66472955 | 0.582335 | 1.12003 | 1.923 |
| chr11 | 44749376 | 44749818 | 2.44948 | 4.71121 | 1.923 |
| chr12 | 74809518 | 74809734 | 1.13491 | 2.18285 | 1.923 |
| chr12 | 40501332 | 40501679 | 1.76179 | 3.38867 | 1.923 |
| chr4 | 6623370 | 6623597 | 0.550526 | 1.0589 | 1.923 |
| chr11 | 9037181 | 9037471 | 1.84861 | 3.55595 | 1.924 |
| chr14 | 60210358 | 60210598 | 0.542556 | 1.04368 | 1.924 |
| chr20 | 33713595 | 33713849 | 0.645188 | 1.24113 | 1.924 |
| chr9 | 105750883 | 105751162 | 0.468289 | 0.900874 | 1.924 |
| chr17 | 43099275 | 43099716 | 1.88135 | 3.6193 | 1.924 |
| chr10 | 124739762 | 124740046 | 1.9194 | 3.69263 | 1.924 |
| chr17 | 10287369 | 10287617 | 0.508029 | 0.977372 | 1.924 |
| chr5 | 18584801 | 18585035 | 0.538333 | 1.03567 | 1.924 |
| chr12 | 48936099 | 48936443 | 2.26542 | 4.35843 | 1.924 |
| chr5 | 129107029 | 129107414 | 1.66041 | 3.19451 | 1.924 |
| chr4 | 71978935 | 71979249 | 0.796539 | 1.53253 | 1.924 |
| chr6 | 83299531 | 83299742 | 0.680035 | 1.3084 | 1.924 |
| chr17 | 52706831 | 52707367 | 0.479247 | 0.922104 | 1.924 |
| chr9 | 127299037 | 127299425 | 1.51996 | 2.92457 | 1.924 |
| chr9 | 16192199 | 16192471 | 2.46388 | 4.74082 | 1.924 |
| chr3 | 127263092 | 127263479 | 1.71022 | 3.29099 | 1.924 |
| chr9 | 100836607 | 100837025 | 3.14241 | 6.04698 | 1.924 |
| chr16 | 58034501 | 58034837 | 2.1618 | 4.16008 | 1.924 |
| chr15 | 89973866 | 89974310 | 2.26152 | 4.3521 | 1.924 |
| chr3 | 151139844 | 151140102 | 0.722896 | 1.39118 | 1.924 |
| chr1 | 52093446 | 52093662 | 1.04293 | 2.00712 | 1.925 |
| chr8 | 141403903 | 141404316 | 2.24349 | 4.31776 | 1.925 |
| chr7 | 3129119 | 3129474 | 1.52545 | 2.93598 | 1.925 |

| | | | | | |
|---|---|---|---|---|---|
| chr9 | 135889860 | 135890180 | 0.466725 | 0.898314 | 1.925 |
| chr18 | 42159014 | 42159259 | 1.36832 | 2.63364 | 1.925 |
| chr15 | 49153954 | 49154335 | 1.37557 | 2.64765 | 1.925 |
| chr7 | 99771387 | 99771764 | 0.659636 | 1.26968 | 1.925 |
| chr2 | 162129641 | 162130040 | 1.02913 | 1.98096 | 1.925 |
| chr11 | 129862338 | 129862564 | 0.679663 | 1.30832 | 1.925 |
| chr8 | 144065547 | 144066190 | 1.60707 | 3.09359 | 1.925 |
| chr17 | 42071962 | 42072616 | 1.51925 | 2.92457 | 1.925 |
| chr19 | 13094427 | 13094747 | 2.7479 | 5.2898 | 1.925 |
| chr11 | 11809292 | 11809793 | 1.40654 | 2.70765 | 1.925 |
| chr9 | 2859829 | 2860154 | 1.89221 | 3.64265 | 1.925 |
| chr18 | 5267732 | 5267959 | 0.434258 | 0.836002 | 1.925 |
| chr9 | 24177104 | 24177368 | 0.502477 | 0.967356 | 1.925 |
| chr2 | 95781116 | 95781447 | 2.74894 | 5.29229 | 1.925 |
| chr14 | 34800868 | 34801224 | 1.86903 | 3.59836 | 1.925 |
| chr14 | 106397742 | 106397949 | 0.801722 | 1.54354 | 1.925 |
| chr1 | 16321045 | 16321363 | 1.57772 | 3.03757 | 1.925 |
| chr13 | 98507829 | 98508074 | 1.77514 | 3.41768 | 1.925 |
| chr14 | 54396644 | 54396982 | 1.6976 | 3.26844 | 1.925 |
| chr20 | 55952466 | 55952674 | 0.520773 | 1.00267 | 1.925 |
| chr5 | 149312254 | 149312650 | 1.62853 | 3.13551 | 1.925 |
| chr2 | 121100972 | 121101355 | 1.53217 | 2.95 | 1.925 |
| chr8 | 76395645 | 76395891 | 0.996953 | 1.91952 | 1.925 |
| chr1 | 230256287 | 230256912 | 1.04584 | 2.01375 | 1.925 |
| chr8 | 128441667 | 128442020 | 0.488086 | 0.939854 | 1.926 |
| chr14 | 81462651 | 81463008 | 1.31241 | 2.52732 | 1.926 |
| chr20 | 56169356 | 56169899 | 1.44633 | 2.78525 | 1.926 |
| chr1 | 157581101 | 157581329 | 0.374887 | 0.721947 | 1.926 |
| chr14 | 23525361 | 23525752 | 1.6639 | 3.2043 | 1.926 |
| chr19 | 8210966 | 8211208 | 0.663174 | 1.27713 | 1.926 |
| chr10 | 14865213 | 14865505 | 1.68625 | 3.2475 | 1.926 |
| chr10 | 18959291 | 18959529 | 0.5936 | 1.1432 | 1.926 |
| chr10 | 55681227 | 55681466 | 0.448864 | 0.864468 | 1.926 |
| chr1 | 117117306 | 117117558 | 1.73182 | 3.33534 | 1.926 |
| chr9 | 13351854 | 13352067 | 1.15079 | 2.21633 | 1.926 |
| chr2 | 208490191 | 208490383 | 1.71769 | 3.30824 | 1.926 |
| chr9 | 135287343 | 135287862 | 0.830651 | 1.59987 | 1.926 |
| chr20 | 54025242 | 54025469 | 0.545362 | 1.0504 | 1.926 |
| chr8 | 22004314 | 22004759 | 1.78287 | 3.43395 | 1.926 |
| chr8 | 126741402 | 126741869 | 1.80628 | 3.4791 | 1.926 |
| chr7 | 155561027 | 155561377 | 1.67931 | 3.23475 | 1.926 |
| chr14 | 35891624 | 35892027 | 1.52954 | 2.94632 | 1.926 |
| chr11 | 46550706 | 46551228 | 1.81832 | 3.50265 | 1.926 |
| chr11 | 94887707 | 94887924 | 1.13257 | 2.18168 | 1.926 |
| chr10 | 52207624 | 52207919 | 0.515688 | 0.993403 | 1.926 |
| chr14 | 88362775 | 88363073 | 1.21183 | 2.33458 | 1.926 |
| chr16 | 86106380 | 86106766 | 1.71036 | 3.29513 | 1.927 |
| chr7 | 145155842 | 145156121 | 0.654548 | 1.26108 | 1.927 |
| chr6 | 6724718 | 6725049 | 1.13847 | 2.19346 | 1.927 |
| chrX | 153762913 | 153763441 | 2.82424 | 5.44148 | 1.927 |
| chr7 | 110729959 | 110730177 | 1.2519 | 2.41206 | 1.927 |
| chr6 | 34605906 | 34606138 | 0.953122 | 1.83645 | 1.927 |
| chr3 | 43741499 | 43741710 | 0.831597 | 1.60232 | 1.927 |
| chr20 | 1782944 | 1783278 | 1.66605 | 3.21018 | 1.927 |
| chr9 | 136647263 | 136647657 | 2.12036 | 4.08556 | 1.927 |
| chr8 | 19203805 | 19204333 | 1.43865 | 2.77204 | 1.927 |
| chr17 | 8901201 | 8901867 | 1.80328 | 3.47463 | 1.927 |
| chr15 | 76626265 | 76626502 | 1.67235 | 3.22248 | 1.927 |
| chr14 | 26258434 | 26258705 | 0.619323 | 1.19348 | 1.927 |
| chr6 | 119469671 | 119470037 | 1.08044 | 2.08211 | 1.927 |
| chr1 | 115655366 | 115655813 | 1.78599 | 3.442 | 1.927 |

| | | | | | |
|---|---|---|---|---|---|
| chr14 | 68005216 | 68005540 | 0.435725 | 0.839749 | 1.927 |
| chr15 | 90418215 | 90418588 | 2.49267 | 4.80402 | 1.927 |
| chr16 | 67701790 | 67702692 | 1.17578 | 2.2661 | 1.927 |
| chr11 | 62310105 | 62310627 | 1.8053 | 3.4795 | 1.927 |
| chr1 | 183250618 | 183250926 | 1.87519 | 3.6143 | 1.927 |
| chr1 | 155149039 | 155149591 | 1.85248 | 3.57055 | 1.927 |
| chr7 | 70929113 | 70929318 | 0.432745 | 0.83409 | 1.927 |
| chr3 | 6841066 | 6841330 | 1.23514 | 2.38067 | 1.927 |
| chr5 | 174362475 | 174363492 | 1.20902 | 2.33032 | 1.927 |
| chr5 | 66794142 | 66794437 | 0.546387 | 1.05315 | 1.927 |
| chr1 | 23961528 | 23961730 | 1.60638 | 3.09631 | 1.928 |
| chr17 | 68548627 | 68548934 | 1.45607 | 2.80667 | 1.928 |
| chr17 | 33860362 | 33860680 | 1.92032 | 3.70158 | 1.928 |
| chr20 | 57022914 | 57023132 | 1.51964 | 2.92926 | 1.928 |
| chr11 | 28719546 | 28719785 | 1.39022 | 2.67981 | 1.928 |
| chr1 | 33177747 | 33178146 | 3.01376 | 5.8094 | 1.928 |
| chr2 | 45663518 | 45663969 | 0.481982 | 0.929088 | 1.928 |
| chr19 | 47074814 | 47075097 | 0.96861 | 1.86715 | 1.928 |
| chr1 | 85809925 | 85810228 | 1.69077 | 3.25937 | 1.928 |
| chr12 | 55119012 | 55119452 | 1.04287 | 2.01039 | 1.928 |
| chr11 | 90026954 | 90027236 | 0.552282 | 1.06469 | 1.928 |
| chr7 | 138550679 | 138550921 | 1.02583 | 1.97768 | 1.928 |
| chr15 | 98628214 | 98628546 | 1.02773 | 1.98141 | 1.928 |
| chr8 | 19233560 | 19233801 | 0.977306 | 1.88422 | 1.928 |
| chr3 | 31652278 | 31652509 | 0.984391 | 1.89794 | 1.928 |
| chr17 | 72357184 | 72358014 | 1.66313 | 3.2066 | 1.928 |
| chr2 | 23728176 | 23728379 | 1.04172 | 2.00849 | 1.928 |
| chr4 | 65579027 | 65579233 | 0.695089 | 1.34029 | 1.928 |
| chr3 | 115538664 | 115539263 | 1.52741 | 2.94522 | 1.928 |
| chr18 | 5890437 | 5890892 | 1.2566 | 2.42312 | 1.928 |
| chr4 | 178476212 | 178476457 | 1.46591 | 2.82678 | 1.928 |
| chr8 | 133289571 | 133289799 | 0.549496 | 1.05965 | 1.928 |
| chr8 | 135397312 | 135397594 | 0.595829 | 1.14909 | 1.929 |
| chr2 | 196933051 | 196933295 | 1.69041 | 3.26007 | 1.929 |
| chr3 | 127409728 | 127410114 | 1.59983 | 3.08559 | 1.929 |
| chr15 | 52500526 | 52500898 | 2.41409 | 4.65609 | 1.929 |
| chr9 | 83814355 | 83814591 | 0.441764 | 0.852093 | 1.929 |
| chr15 | 68900369 | 68900659 | 0.527109 | 1.01672 | 1.929 |
| chr7 | 56072495 | 56072903 | 1.79316 | 3.45885 | 1.929 |
| chr1 | 20959699 | 20960062 | 2.50534 | 4.8326 | 1.929 |
| chr11 | 64972205 | 64972656 | 2.91019 | 5.6136 | 1.929 |
| chr19 | 29856861 | 29857305 | 0.808586 | 1.55976 | 1.929 |
| chr1 | 154518978 | 154519271 | 2.11346 | 4.07691 | 1.929 |
| chr1 | 231827391 | 231827777 | 2.01647 | 3.88984 | 1.929 |
| chr2 | 25829845 | 25830100 | 1.27102 | 2.45184 | 1.929 |
| chr7 | 37006967 | 37007265 | 2.42447 | 4.67693 | 1.929 |
| chrX | 15705246 | 15705557 | 1.86493 | 3.59755 | 1.929 |
| chr1 | 911513 | 911740 | 0.852295 | 1.64414 | 1.929 |
| chr10 | 103329919 | 103330357 | 1.90454 | 3.67402 | 1.929 |
| chr12 | 57636569 | 57637849 | 1.66787 | 3.21751 | 1.929 |
| chr14 | 62593649 | 62593890 | 1.17869 | 2.27384 | 1.929 |
| chr9 | 94195525 | 94195766 | 1.04599 | 2.01784 | 1.929 |
| chr11 | 69058304 | 69058567 | 0.56073 | 1.08174 | 1.929 |
| chr12 | 33419818 | 33420106 | 0.470058 | 0.90688 | 1.929 |
| chr5 | 131553526 | 131553755 | 0.472121 | 0.91086 | 1.929 |
| chr6 | 52704458 | 52704708 | 0.827016 | 1.59555 | 1.929 |
| chr5 | 169105747 | 169105956 | 0.700078 | 1.35069 | 1.929 |
| chr16 | 2835695 | 2835913 | 0.781876 | 1.50861 | 1.929 |
| gl000205_r | 89387 | 89774 | 2.07101 | 3.99598 | 1.929 |
| chr1 | 85725246 | 85725614 | 2.3103 | 4.45783 | 1.930 |
| chr17 | 79228733 | 79228956 | 1.41875 | 2.73756 | 1.930 |

| | | | | | |
|---|---|---|---|---|---|
| chr10 | 95573228 | 95573619 | 2.0465 | 3.94889 | 1.930 |
| chr6 | 122813109 | 122813389 | 1.33867 | 2.58322 | 1.930 |
| chr10 | 90447688 | 90447896 | 1.57985 | 3.04893 | 1.930 |
| chrX | 73235260 | 73235600 | 1.61208 | 3.11116 | 1.930 |
| chr2 | 55998861 | 55999064 | 1.1485 | 2.21651 | 1.930 |
| chr3 | 107379305 | 107379519 | 1.30101 | 2.51087 | 1.930 |
| chr3 | 48315150 | 48315387 | 1.80113 | 3.47611 | 1.930 |
| chr3 | 74545651 | 74546121 | 0.655382 | 1.2649 | 1.930 |
| chr5 | 75897268 | 75897601 | 1.07468 | 2.07418 | 1.930 |
| chr19 | 28464848 | 28465126 | 0.887556 | 1.71304 | 1.930 |
| chr16 | 85330568 | 85331053 | 2.563 | 4.94691 | 1.930 |
| chr5 | 1209013 | 1209447 | 1.98238 | 3.82624 | 1.930 |
| chr10 | 95681133 | 95681493 | 1.11638 | 2.15477 | 1.930 |
| chr5 | 141193627 | 141193993 | 1.7731 | 3.42238 | 1.930 |
| chr11 | 131557901 | 131558297 | 1.67658 | 3.23625 | 1.930 |
| chr5 | 160971231 | 160971674 | 1.4219 | 2.74467 | 1.930 |
| chr1 | 199514949 | 199515183 | 0.918624 | 1.77323 | 1.930 |
| chr2 | 48338296 | 48338815 | 1.80243 | 3.47929 | 1.930 |
| chr2 | 79446594 | 79446833 | 1.14642 | 2.213 | 1.930 |
| chr8 | 105738058 | 105738349 | 0.483657 | 0.933647 | 1.930 |
| chr9 | 97944955 | 97945237 | 0.54385 | 1.04985 | 1.930 |
| chr17 | 62598587 | 62598959 | 0.812738 | 1.5691 | 1.931 |
| chrX | 57004562 | 57004976 | 1.24624 | 2.40608 | 1.931 |
| chr12 | 94389080 | 94389355 | 0.500647 | 0.966595 | 1.931 |
| chr8 | 2448120 | 2448447 | 0.444717 | 0.858635 | 1.931 |
| chr8 | 135187762 | 135188083 | 0.541372 | 1.04538 | 1.931 |
| chr11 | 2552612 | 2553503 | 1.15372 | 2.22787 | 1.931 |
| chr22 | 47154744 | 47155021 | 1.94346 | 3.75299 | 1.931 |
| chr3 | 36878720 | 36878930 | 0.498477 | 0.962668 | 1.931 |
| chr19 | 5226016 | 5226360 | 1.18826 | 2.29483 | 1.931 |
| chr7 | 158364929 | 158365244 | 1.16121 | 2.2427 | 1.931 |
| chr3 | 38038382 | 38038643 | 2.09036 | 4.03726 | 1.931 |
| chr9 | 116639132 | 116639341 | 0.984503 | 1.90145 | 1.931 |
| chr20 | 6673554 | 6673758 | 0.565995 | 1.09316 | 1.931 |
| chr16 | 30988721 | 30989207 | 1.26529 | 2.44392 | 1.932 |
| chr19 | 42256956 | 42257538 | 2.15322 | 4.15896 | 1.932 |
| chr16 | 85449973 | 85450388 | 1.75391 | 3.38781 | 1.932 |
| chr13 | 36994492 | 36994887 | 1.44181 | 2.78499 | 1.932 |
| chr5 | 175128087 | 175128363 | 1.55053 | 2.99514 | 1.932 |
| chr14 | 52118613 | 52118944 | 2.14499 | 4.14351 | 1.932 |
| chr6 | 137635488 | 137635762 | 1.06937 | 2.06576 | 1.932 |
| chr7 | 147962126 | 147962484 | 1.1246 | 2.17245 | 1.932 |
| chr2 | 55065666 | 55066056 | 1.79542 | 3.46832 | 1.932 |
| chr2 | 7066298 | 7066554 | 0.498009 | 0.962039 | 1.932 |
| chr8 | 26771049 | 26771271 | 0.43747 | 0.845102 | 1.932 |
| chr8 | 56294006 | 56294250 | 0.659239 | 1.27353 | 1.932 |
| chr14 | 24881128 | 24881567 | 1.14646 | 2.21477 | 1.932 |
| chr17 | 46338715 | 46338967 | 2.1223 | 4.09993 | 1.932 |
| chr1 | 220983502 | 220983866 | 2.3592 | 4.55773 | 1.932 |
| chr16 | 24700476 | 24700772 | 1.80529 | 3.48765 | 1.932 |
| chr20 | 43378825 | 43379518 | 1.49688 | 2.89184 | 1.932 |
| chr13 | 28558449 | 28558688 | 1.2007 | 2.31969 | 1.932 |
| chr12 | 80663817 | 80664138 | 1.31865 | 2.5477 | 1.932 |
| chr12 | 101631225 | 101631449 | 0.484221 | 0.935546 | 1.932 |
| chr12 | 74415663 | 74416016 | 1.65216 | 3.19224 | 1.932 |
| chr2 | 7251868 | 7252156 | 1.00634 | 1.94445 | 1.932 |
| chr7 | 45641520 | 45641816 | 0.545985 | 1.05503 | 1.932 |
| chr16 | 80626175 | 80626379 | 0.574834 | 1.11088 | 1.933 |
| chr20 | 58781343 | 58781577 | 1.39816 | 2.702 | 1.933 |
| chr17 | 1617159 | 1617393 | 1.64309 | 3.17536 | 1.933 |
| chr5 | 176084828 | 176085460 | 1.64134 | 3.17211 | 1.933 |

| | | | | | |
|---|---|---|---|---|---|
| chr14 | 51181311 | 51181768 | 0.601185 | 1.1619 | 1.933 |
| chr7 | 28995278 | 28996671 | 1.02921 | 1.98918 | 1.933 |
| chrX | 150918205 | 150918625 | 1.38073 | 2.66861 | 1.933 |
| chr20 | 58358104 | 58358513 | 2.50654 | 4.84458 | 1.933 |
| chr1 | 110955160 | 110955490 | 1.96632 | 3.80055 | 1.933 |
| chr14 | 54006257 | 54006598 | 1.91193 | 3.69549 | 1.933 |
| chr17 | 73913718 | 73914167 | 0.483955 | 0.935442 | 1.933 |
| chr2 | 70923372 | 70923623 | 0.516407 | 0.998192 | 1.933 |
| chr2 | 191247694 | 191247942 | 1.47335 | 2.84793 | 1.933 |
| chr5 | 6933825 | 6934153 | 0.533359 | 1.03099 | 1.933 |
| chr10 | 97360938 | 97361213 | 2.21467 | 4.2811 | 1.933 |
| chr12 | 307285 | 307513 | 0.422166 | 0.816076 | 1.933 |
| chr15 | 85839792 | 85840103 | 1.37874 | 2.66521 | 1.933 |
| chr3 | 15309403 | 15309768 | 2.1269 | 4.11153 | 1.933 |
| chr15 | 42429985 | 42430440 | 0.854423 | 1.65172 | 1.933 |
| chr1 | 235895018 | 235895246 | 1.3298 | 2.57074 | 1.933 |
| chr14 | 46261954 | 46262168 | 0.571482 | 1.10478 | 1.933 |
| chr11 | 111435897 | 111436186 | 1.21892 | 2.35641 | 1.933 |
| chr7 | 108403091 | 108403392 | 0.815748 | 1.57709 | 1.933 |
| chr1 | 111308492 | 111308989 | 1.41452 | 2.73486 | 1.933 |
| chr5 | 59964614 | 59964887 | 1.00321 | 1.93963 | 1.933 |
| chr21 | 44238377 | 44238738 | 1.6072 | 3.1075 | 1.933 |
| chr7 | 39620031 | 39620326 | 1.40494 | 2.7166 | 1.934 |
| chr11 | 46830389 | 46830642 | 1.15813 | 2.23938 | 1.934 |
| chr5 | 169338748 | 169339108 | 0.70436 | 1.36197 | 1.934 |
| chr19 | 48812372 | 48812811 | 0.743653 | 1.43797 | 1.934 |
| chr17 | 7314772 | 7315041 | 0.485633 | 0.939109 | 1.934 |
| chr19 | 30537338 | 30537690 | 2.11321 | 4.08648 | 1.934 |
| chr22 | 39910444 | 39910692 | 1.80677 | 3.49389 | 1.934 |
| chr4 | 108104441 | 108104633 | 1.41867 | 2.74339 | 1.934 |
| chr12 | 81358029 | 81358256 | 0.817692 | 1.58126 | 1.934 |
| chr4 | 56814780 | 56815092 | 2.06742 | 3.99799 | 1.934 |
| chr8 | 78302237 | 78302519 | 0.634871 | 1.22778 | 1.934 |
| chr12 | 108221448 | 108221966 | 0.420971 | 0.814169 | 1.934 |
| chr5 | 44935469 | 44935887 | 0.669212 | 1.29429 | 1.934 |
| chr3 | 155174326 | 155174753 | 0.506584 | 0.979773 | 1.934 |
| chrX | 16463274 | 16463667 | 1.67261 | 3.23508 | 1.934 |
| chr3 | 160814601 | 160814906 | 1.16491 | 2.25318 | 1.934 |
| chr8 | 103646514 | 103646747 | 1.15832 | 2.24046 | 1.934 |
| chr3 | 188759291 | 188759618 | 1.19502 | 2.31177 | 1.935 |
| chr19 | 4267605 | 4268008 | 1.36409 | 2.63884 | 1.935 |
| chr18 | 12271716 | 12272147 | 2.26211 | 4.37635 | 1.935 |
| chr3 | 38339559 | 38340000 | 1.31172 | 2.53777 | 1.935 |
| chr7 | 70321882 | 70322282 | 1.451 | 2.80724 | 1.935 |
| chr15 | 72022775 | 72023541 | 1.27597 | 2.46867 | 1.935 |
| chr3 | 48477275 | 48477730 | 2.27163 | 4.39505 | 1.935 |
| chr7 | 55482934 | 55483188 | 1.76331 | 3.41164 | 1.935 |
| chr17 | 59488153 | 59488554 | 1.32982 | 2.57297 | 1.935 |
| chrX | 103515519 | 103515881 | 1.09755 | 2.1236 | 1.935 |
| chrX | 47243892 | 47244252 | 1.54557 | 2.99057 | 1.935 |
| chr14 | 46258106 | 46258416 | 0.580319 | 1.1229 | 1.935 |
| chr2 | 232535723 | 232535954 | 0.936603 | 1.81231 | 1.935 |
| chr19 | 51425551 | 51425923 | 2.03321 | 3.93425 | 1.935 |
| chr10 | 69064582 | 69064801 | 0.700849 | 1.35622 | 1.935 |
| chr8 | 127881658 | 127881925 | 1.09488 | 2.11872 | 1.935 |
| chr9 | 137977892 | 137978302 | 1.92971 | 3.7343 | 1.935 |
| chr10 | 123646546 | 123646980 | 1.40557 | 2.72038 | 1.935 |
| chr10 | 128576701 | 128576918 | 0.528178 | 1.02225 | 1.935 |
| chr3 | 163277403 | 163277840 | 0.537954 | 1.04123 | 1.936 |
| chr5 | 141928253 | 141928693 | 2.11493 | 4.09358 | 1.936 |
| chr2 | 19569218 | 19569462 | 1.31195 | 2.5394 | 1.936 |

| | | | | | |
|---|---|---|---|---|---|
| chr17 | 42643284 | 42643867 | 1.1146 | 2.15744 | 1.936 |
| chr5 | 54162964 | 54163761 | 1.09516 | 2.11985 | 1.936 |
| chr20 | 32379938 | 32380308 | 1.98536 | 3.84304 | 1.936 |
| chr2 | 161775476 | 161775795 | 1.22616 | 2.37355 | 1.936 |
| chr15 | 55727509 | 55727767 | 1.29378 | 2.50455 | 1.936 |
| chr3 | 35828347 | 35828560 | 0.543721 | 1.05258 | 1.936 |
| chr16 | 49156278 | 49156496 | 0.50755 | 0.982565 | 1.936 |
| chr2 | 190563086 | 190563298 | 0.926375 | 1.79337 | 1.936 |
| chr20 | 59838149 | 59838558 | 2.33443 | 4.5193 | 1.936 |
| chr2 | 176975962 | 176976279 | 2.08142 | 4.0296 | 1.936 |
| chr8 | 102768556 | 102768816 | 1.64454 | 3.18381 | 1.936 |
| chr15 | 85326911 | 85327186 | 0.787651 | 1.52493 | 1.936 |
| chr14 | 70193584 | 70194038 | 2.6891 | 5.2063 | 1.936 |
| chr10 | 62493190 | 62493434 | 0.86059 | 1.66618 | 1.936 |
| chr3 | 115859031 | 115859235 | 1.35316 | 2.61984 | 1.936 |
| chr18 | 9792641 | 9792872 | 0.939241 | 1.81846 | 1.936 |
| chr1 | 249120730 | 249121012 | 1.99061 | 3.85412 | 1.936 |
| chr12 | 49284459 | 49284844 | 2.33892 | 4.52854 | 1.936 |
| chr4 | 119902860 | 119903243 | 1.09431 | 2.11878 | 1.936 |
| chr3 | 39463267 | 39463584 | 0.503849 | 0.975547 | 1.936 |
| chr6 | 90383992 | 90384230 | 0.548245 | 1.06152 | 1.936 |
| chr17 | 9671896 | 9672178 | 0.939495 | 1.81909 | 1.936 |
| chr10 | 44448483 | 44448770 | 0.469059 | 0.908246 | 1.936 |
| chr1 | 29111581 | 29111878 | 1.13684 | 2.20134 | 1.936 |
| chr8 | 79446731 | 79447096 | 0.630768 | 1.22143 | 1.936 |
| chr1 | 204944742 | 204945302 | 1.48711 | 2.87968 | 1.936 |
| chr10 | 72518871 | 72519132 | 2.44453 | 4.73373 | 1.936 |
| chr13 | 96356791 | 96357101 | 1.08857 | 2.10798 | 1.936 |
| chr10 | 33297673 | 33298014 | 1.96817 | 3.8114 | 1.937 |
| chr4 | 736400 | 736842 | 2.28033 | 4.416 | 1.937 |
| chrX | 122556800 | 122557036 | 1.28009 | 2.47903 | 1.937 |
| chr6 | 39271036 | 39271294 | 1.74099 | 3.37164 | 1.937 |
| chr11 | 22457578 | 22457818 | 0.542708 | 1.05103 | 1.937 |
| chr12 | 112635895 | 112636331 | 1.28035 | 2.47964 | 1.937 |
| chr12 | 42270114 | 42270472 | 0.701315 | 1.35824 | 1.937 |
| chr5 | 446900 | 447179 | 1.32397 | 2.56418 | 1.937 |
| chr10 | 98094341 | 98095093 | 1.5686 | 3.03799 | 1.937 |
| chr14 | 55359841 | 55360191 | 1.03495 | 2.00444 | 1.937 |
| chr12 | 78575403 | 78575647 | 1.49403 | 2.89361 | 1.937 |
| chr3 | 30884812 | 30885133 | 0.655469 | 1.2695 | 1.937 |
| chr15 | 65809960 | 65810266 | 2.97447 | 5.76117 | 1.937 |
| chr8 | 142193547 | 142193908 | 1.87388 | 3.6295 | 1.937 |
| chr12 | 65190111 | 65190317 | 1.51811 | 2.94047 | 1.937 |
| chr19 | 47224859 | 47225285 | 0.881202 | 1.70684 | 1.937 |
| chr12 | 15685428 | 15685677 | 1.10609 | 2.14245 | 1.937 |
| chr12 | 7126128 | 7126430 | 1.96435 | 3.80511 | 1.937 |
| chr14 | 24483209 | 24483473 | 1.61476 | 3.12792 | 1.937 |
| chr6 | 6315950 | 6316190 | 1.30685 | 2.53147 | 1.937 |
| chr9 | 33925187 | 33925487 | 1.79081 | 3.46899 | 1.937 |
| chr3 | 93952949 | 93953415 | 1.28177 | 2.48294 | 1.937 |
| chr11 | 67983908 | 67984158 | 0.540566 | 1.04715 | 1.937 |
| chr19 | 57348531 | 57349178 | 0.758578 | 1.46963 | 1.937 |
| chr6 | 133282830 | 133283074 | 0.768031 | 1.48795 | 1.937 |
| chr19 | 34961711 | 34962218 | 1.30356 | 2.5256 | 1.937 |
| chr1 | 55137729 | 55138048 | 2.04817 | 3.96829 | 1.937 |
| chr3 | 103494603 | 103494853 | 0.726693 | 1.40795 | 1.937 |
| chr8 | 18671707 | 18671921 | 1.16491 | 2.25702 | 1.938 |
| chr15 | 52538347 | 52538681 | 0.60657 | 1.17527 | 1.938 |
| chr6 | 70492279 | 70492487 | 0.686695 | 1.33064 | 1.938 |
| chr10 | 99313173 | 99313628 | 1.79408 | 3.4765 | 1.938 |
| chr12 | 54426398 | 54426723 | 2.85435 | 5.53155 | 1.938 |

| | | | | | |
|---|---|---|---|---|---|
| chr17 | 17584422 | 17584800 | 2.44416 | 4.73664 | 1.938 |
| chr5 | 131608422 | 131608857 | 2.14778 | 4.16226 | 1.938 |
| chr2 | 85527435 | 85527916 | 2.27658 | 4.41201 | 1.938 |
| chr5 | 175123552 | 175123907 | 1.63253 | 3.16384 | 1.938 |
| chr1 | 20031048 | 20031378 | 2.42588 | 4.7014 | 1.938 |
| chr1 | 151273262 | 151273772 | 1.81399 | 3.51575 | 1.938 |
| chr7 | 31748462 | 31748751 | 0.457097 | 0.886053 | 1.938 |
| chr7 | 155205211 | 155205897 | 1.7322 | 3.35793 | 1.939 |
| chr1 | 179162974 | 179163292 | 1.84256 | 3.57194 | 1.939 |
| chr1 | 173224408 | 173224723 | 1.69664 | 3.28914 | 1.939 |
| chr14 | 99859386 | 99860039 | 1.77327 | 3.43784 | 1.939 |
| chr12 | 26677848 | 26678136 | 1.75226 | 3.39712 | 1.939 |
| chr5 | 138598293 | 138598707 | 2.84173 | 5.50937 | 1.939 |
| chr16 | 3181896 | 3182125 | 1.03349 | 2.00375 | 1.939 |
| chr1 | 95065042 | 95065308 | 1.21167 | 2.34922 | 1.939 |
| chr15 | 73565575 | 73566062 | 1.4482 | 2.80791 | 1.939 |
| chr7 | 31376145 | 31376369 | 1.20277 | 2.3321 | 1.939 |
| chr8 | 68751362 | 68751614 | 0.517526 | 1.00345 | 1.939 |
| chr3 | 34354152 | 34354488 | 0.45471 | 0.881674 | 1.939 |
| chr14 | 63743562 | 63743954 | 0.901649 | 1.74837 | 1.939 |
| chr10 | 43751268 | 43751777 | 1.84199 | 3.57178 | 1.939 |
| chr15 | 99424774 | 99425116 | 1.19087 | 2.30925 | 1.939 |
| chrX | 153976986 | 153977315 | 1.56094 | 3.02687 | 1.939 |
| chr17 | 65217266 | 65217517 | 1.36063 | 2.63852 | 1.939 |
| chr4 | 18595198 | 18595500 | 1.18712 | 2.30217 | 1.939 |
| chr12 | 2893046 | 2893565 | 2.10202 | 4.07644 | 1.939 |
| chr15 | 34502223 | 34502550 | 2.06202 | 3.99905 | 1.939 |
| chr11 | 61153052 | 61153369 | 2.0643 | 4.00358 | 1.939 |
| chr11 | 66601759 | 66602041 | 1.44604 | 2.80454 | 1.939 |
| chr1 | 167035833 | 167036281 | 2.47407 | 4.79842 | 1.939 |
| chr11 | 18874905 | 18875234 | 1.11862 | 2.16957 | 1.940 |
| chr19 | 34336898 | 34337134 | 0.862608 | 1.67304 | 1.940 |
| chr16 | 22089033 | 22089576 | 1.41564 | 2.7457 | 1.940 |
| chr10 | 47080932 | 47081309 | 1.75644 | 3.40689 | 1.940 |
| chr1 | 226033518 | 226033922 | 1.55244 | 3.01123 | 1.940 |
| chr18 | 56197039 | 56197286 | 1.26914 | 2.46174 | 1.940 |
| chr14 | 87124487 | 87124919 | 0.661666 | 1.28351 | 1.940 |
| chr10 | 91020100 | 91020426 | 1.86379 | 3.6155 | 1.940 |
| chr7 | 71464487 | 71464831 | 0.582969 | 1.13089 | 1.940 |
| chr2 | 68412747 | 68413071 | 2.23592 | 4.33743 | 1.940 |
| chr15 | 64441507 | 64441896 | 1.02679 | 1.99188 | 1.940 |
| chr10 | 32635082 | 32635404 | 3.27487 | 6.35319 | 1.940 |
| chr2 | 86560944 | 86561160 | 0.482036 | 0.935145 | 1.940 |
| chr19 | 1946028 | 1946254 | 2.18578 | 4.24056 | 1.940 |
| chr15 | 81567281 | 81567680 | 1.79124 | 3.4752 | 1.940 |
| chr3 | 113049748 | 113050072 | 0.468245 | 0.908462 | 1.940 |
| chr1 | 209381002 | 209381395 | 2.21318 | 4.2941 | 1.940 |
| chr1 | 47658574 | 47658946 | 1.4695 | 2.85124 | 1.940 |
| chr5 | 112579391 | 112579593 | 0.75768 | 1.47012 | 1.940 |
| chr16 | 21200018 | 21200347 | 1.83907 | 3.56849 | 1.940 |
| chr9 | 130470072 | 130470288 | 0.812335 | 1.57624 | 1.940 |
| chrX | 71465336 | 71465711 | 1.98176 | 3.8454 | 1.940 |
| chr9 | 138689526 | 138689890 | 1.33829 | 2.59689 | 1.940 |
| chr16 | 55735644 | 55736019 | 1.2999 | 2.5224 | 1.940 |
| chr5 | 85975412 | 85975656 | 0.854296 | 1.65776 | 1.941 |
| chr6 | 46459593 | 46459832 | 1.28179 | 2.48732 | 1.941 |
| chr14 | 80484078 | 80484478 | 0.612821 | 1.18922 | 1.941 |
| chr1 | 229650489 | 229650784 | 1.5552 | 3.01801 | 1.941 |
| chr18 | 11310532 | 11310833 | 1.9485 | 3.78154 | 1.941 |
| chrX | 53473001 | 53473261 | 1.61129 | 3.1271 | 1.941 |
| chr17 | 27944163 | 27944505 | 1.29829 | 2.51969 | 1.941 |

| | | | | | |
|---|---|---|---|---|---|
| chr18 | 3998751 | 3999008 | 1.00127 | 1.94333 | 1.941 |
| chr20 | 1166388 | 1166792 | 1.87658 | 3.6422 | 1.941 |
| chr11 | 36694201 | 36694657 | 0.905859 | 1.75816 | 1.941 |
| chr8 | 71580659 | 71580909 | 1.4951 | 2.90186 | 1.941 |
| chr16 | 68263347 | 68263717 | 1.79057 | 3.47539 | 1.941 |
| chr8 | 61350549 | 61350995 | 0.470006 | 0.91226 | 1.941 |
| chr19 | 6417753 | 6418016 | 1.55479 | 3.01781 | 1.941 |
| chr17 | 7249138 | 7249439 | 0.909804 | 1.76592 | 1.941 |
| chr1 | 97432145 | 97432566 | 1.31642 | 2.5552 | 1.941 |
| chr10 | 30324880 | 30325117 | 0.455665 | 0.884503 | 1.941 |
| chr12 | 66377455 | 66377708 | 1.6816 | 3.26429 | 1.941 |
| chr6 | 97345698 | 97345992 | 2.50411 | 4.86113 | 1.941 |
| chr1 | 150159627 | 150159979 | 0.939768 | 1.82435 | 1.941 |
| chr5 | 132219031 | 132219233 | 1.33402 | 2.58979 | 1.941 |
| chr13 | 111333194 | 111333492 | 0.967848 | 1.87899 | 1.941 |
| chr12 | 20938935 | 20939168 | 0.531504 | 1.03191 | 1.941 |
| chr4 | 68854204 | 68854538 | 1.0288 | 1.99742 | 1.942 |
| chr21 | 39877661 | 39877968 | 1.138 | 2.20947 | 1.942 |
| chr19 | 50183908 | 50184408 | 2.97858 | 5.78309 | 1.942 |
| chr20 | 9373863 | 9374308 | 1.76081 | 3.41878 | 1.942 |
| chr17 | 46337071 | 46337363 | 1.48373 | 2.88085 | 1.942 |
| chr17 | 11729308 | 11729628 | 0.385768 | 0.749048 | 1.942 |
| chr10 | 106051126 | 106051540 | 3.78884 | 7.35706 | 1.942 |
| chr14 | 54960067 | 54960302 | 0.817521 | 1.58752 | 1.942 |
| chr10 | 88136914 | 88137397 | 2.17866 | 4.23087 | 1.942 |
| chr16 | 30565207 | 30565423 | 0.457337 | 0.88817 | 1.942 |
| chr2 | 23683190 | 23683429 | 1.36066 | 2.64249 | 1.942 |
| chr11 | 58397430 | 58397705 | 1.11731 | 2.16993 | 1.942 |
| chr1 | 153769640 | 153770056 | 2.32205 | 4.50968 | 1.942 |
| chr11 | 74450840 | 74451132 | 1.941 | 3.76968 | 1.942 |
| chr7 | 676129 | 676471 | 1.32643 | 2.5761 | 1.942 |
| chr12 | 114423187 | 114423456 | 0.515123 | 1.00044 | 1.942 |
| chr1 | 63782573 | 63782989 | 1.69341 | 3.28888 | 1.942 |
| chr21 | 46369613 | 46370022 | 1.47635 | 2.86737 | 1.942 |
| chr12 | 69188270 | 69188752 | 0.521399 | 1.01268 | 1.942 |
| chr19 | 54989855 | 54990176 | 1.41972 | 2.75747 | 1.942 |
| chr4 | 55837607 | 55838062 | 1.07754 | 2.09286 | 1.942 |
| chr7 | 103645708 | 103645950 | 0.508547 | 0.987778 | 1.942 |
| chr17 | 76441473 | 76442150 | 1.81749 | 3.53025 | 1.942 |
| chr20 | 30249366 | 30249694 | 1.81948 | 3.53423 | 1.942 |
| chr17 | 25715445 | 25715650 | 1.19559 | 2.32241 | 1.942 |
| chr2 | 165352191 | 165352463 | 0.957644 | 1.86027 | 1.943 |
| chr22 | 44280835 | 44281161 | 2.40551 | 4.67292 | 1.943 |
| chrX | 55084139 | 55084416 | 1.26677 | 2.46083 | 1.943 |
| chr14 | 22382010 | 22382238 | 1.21995 | 2.36989 | 1.943 |
| chr15 | 66919105 | 66919406 | 0.646219 | 1.25536 | 1.943 |
| chr5 | 126365245 | 126365556 | 0.865455 | 1.68125 | 1.943 |
| chr10 | 66050444 | 66050793 | 0.457906 | 0.889559 | 1.943 |
| chr14 | 60273568 | 60273855 | 0.369691 | 0.718195 | 1.943 |
| chr7 | 147244504 | 147244740 | 0.934632 | 1.81571 | 1.943 |
| chr22 | 29225412 | 29225714 | 2.06412 | 4.01 | 1.943 |
| chr12 | 53432558 | 53432769 | 1.27366 | 2.47455 | 1.943 |
| chr7 | 130021555 | 130021865 | 1.348 | 2.61904 | 1.943 |
| chr15 | 76443261 | 76443728 | 2.06057 | 4.00353 | 1.943 |
| chr3 | 115247943 | 115248221 | 0.471687 | 0.916497 | 1.943 |
| chrX | 135282699 | 135283032 | 0.894663 | 1.73836 | 1.943 |
| chr15 | 71167392 | 71167708 | 1.51639 | 2.94642 | 1.943 |
| chr6 | 53593382 | 53593756 | 1.89944 | 3.69087 | 1.943 |
| chr4 | 116794267 | 116794610 | 0.733711 | 1.42573 | 1.943 |
| chr19 | 49804456 | 49804939 | 2.56927 | 4.99261 | 1.943 |
| chr22 | 46402205 | 46402703 | 1.50174 | 2.9184 | 1.943 |

| | | | | | |
|---|---|---|---|---|---|
| chr9 | 32322271 | 32322512 | 0.450685 | 0.875845 | 1.943 |
| chr7 | 42980776 | 42981348 | 1.76 | 3.42057 | 1.944 |
| chr12 | 71610765 | 71611067 | 0.975824 | 1.89654 | 1.944 |
| chr5 | 120191635 | 120191951 | 0.497362 | 0.966659 | 1.944 |
| chr10 | 101006533 | 101006815 | 1.4064 | 2.73345 | 1.944 |
| chr12 | 125721551 | 125721970 | 1.625 | 3.1584 | 1.944 |
| chr12 | 40158805 | 40159106 | 1.60969 | 3.12879 | 1.944 |
| chr8 | 8637209 | 8637525 | 2.07233 | 4.02808 | 1.944 |
| chr7 | 124970856 | 124971085 | 0.520362 | 1.0115 | 1.944 |
| chr2 | 23859403 | 23859866 | 0.674317 | 1.31078 | 1.944 |
| chr21 | 38311758 | 38312061 | 1.79604 | 3.49134 | 1.944 |
| chr12 | 113338156 | 113338417 | 2.00951 | 3.90632 | 1.944 |
| chr15 | 39873947 | 39874154 | 1.31617 | 2.55855 | 1.944 |
| chr8 | 56464838 | 56465077 | 0.509816 | 0.991071 | 1.944 |
| chr14 | 53456939 | 53457166 | 0.519384 | 1.00971 | 1.944 |
| chr2 | 69253134 | 69253389 | 1.11981 | 2.17697 | 1.944 |
| chr6 | 52528513 | 52528748 | 1.54287 | 2.99956 | 1.944 |
| chr16 | 55266577 | 55266892 | 0.466012 | 0.906002 | 1.944 |
| chr8 | 37659202 | 37659593 | 0.931496 | 1.81109 | 1.944 |
| chr9 | 98994746 | 98995250 | 1.6749 | 3.25651 | 1.944 |
| chr5 | 87047371 | 87047630 | 0.46543 | 0.904982 | 1.944 |
| chr9 | 131397522 | 131397815 | 2.26676 | 4.40758 | 1.944 |
| chr17 | 1125759 | 1126069 | 0.535911 | 1.04206 | 1.944 |
| chr9 | 126116316 | 126116838 | 1.91491 | 3.72354 | 1.945 |
| chr14 | 66731352 | 66731556 | 0.615656 | 1.19719 | 1.945 |
| chr3 | 8979782 | 8980002 | 1.09403 | 2.12744 | 1.945 |
| chr12 | 24503155 | 24503405 | 0.654538 | 1.27286 | 1.945 |
| chr4 | 2443259 | 2443815 | 1.58059 | 3.07385 | 1.945 |
| chr1 | 26865262 | 26865499 | 1.26312 | 2.45658 | 1.945 |
| chr18 | 29291001 | 29291210 | 0.786315 | 1.52927 | 1.945 |
| chr3 | 13689653 | 13690143 | 2.56523 | 4.98903 | 1.945 |
| chr12 | 26998049 | 26998371 | 2.14873 | 4.17934 | 1.945 |
| chr7 | 139705605 | 139705941 | 1.90091 | 3.69735 | 1.945 |
| chr15 | 90630039 | 90630373 | 2.28302 | 4.44062 | 1.945 |
| chr2 | 61886018 | 61886326 | 1.31316 | 2.55425 | 1.945 |
| chr1 | 234099608 | 234099985 | 0.486527 | 0.946372 | 1.945 |
| chr11 | 92255166 | 92255421 | 0.660029 | 1.28388 | 1.945 |
| chr7 | 142608369 | 142608610 | 0.495672 | 0.964197 | 1.945 |
| chr16 | 9232862 | 9233336 | 2.07986 | 4.04586 | 1.945 |
| chr18 | 30812013 | 30812299 | 1.26695 | 2.46457 | 1.945 |
| chr1 | 40439186 | 40439594 | 1.76318 | 3.43026 | 1.946 |
| chr1 | 93459713 | 93459957 | 1.28089 | 2.49217 | 1.946 |
| chr2 | 165286734 | 165286977 | 1.44184 | 2.80544 | 1.946 |
| chr6 | 75918709 | 75918985 | 1.34524 | 2.6175 | 1.946 |
| chr8 | 72043223 | 72043496 | 0.592384 | 1.15263 | 1.946 |
| chr5 | 102560345 | 102560614 | 1.13726 | 2.21289 | 1.946 |
| chr5 | 141524934 | 141525240 | 1.61002 | 3.1328 | 1.946 |
| chr11 | 45038949 | 45039165 | 0.806776 | 1.56987 | 1.946 |
| chr10 | 88470891 | 88471170 | 1.7262 | 3.35903 | 1.946 |
| chr7 | 96645907 | 96646366 | 0.648592 | 1.26221 | 1.946 |
| chr11 | 61312867 | 61313108 | 1.19686 | 2.32925 | 1.946 |
| chr12 | 122249706 | 122250292 | 2.18626 | 4.25485 | 1.946 |
| chr7 | 151708341 | 151708573 | 0.555182 | 1.08051 | 1.946 |
| chr8 | 130382896 | 130383137 | 0.535858 | 1.04299 | 1.946 |
| chr10 | 75841025 | 75841285 | 1.92912 | 3.75508 | 1.947 |
| chr7 | 112135988 | 112136233 | 1.2685 | 2.46916 | 1.947 |
| chr12 | 55830612 | 55830954 | 0.612518 | 1.19233 | 1.947 |
| chr5 | 137968022 | 137968264 | 0.519982 | 1.01222 | 1.947 |
| chr12 | 76004747 | 76005094 | 0.678132 | 1.3201 | 1.947 |
| chr12 | 105777193 | 105777478 | 1.12985 | 2.19948 | 1.947 |
| chr18 | 74712426 | 74712748 | 2.88596 | 5.6181 | 1.947 |

| | | | | | |
|---|---|---|---|---|---|
| chr10 | 88499663 | 88499893 | 0.453676 | 0.883206 | 1.947 |
| chr15 | 28371116 | 28371538 | 1.1764 | 2.29024 | 1.947 |
| chr1 | 95550193 | 95550428 | 1.31813 | 2.56618 | 1.947 |
| chr17 | 49909279 | 49909541 | 0.565953 | 1.10183 | 1.947 |
| chr5 | 174163016 | 174163333 | 1.50045 | 2.92128 | 1.947 |
| chr8 | 58333089 | 58333481 | 1.88567 | 3.6714 | 1.947 |
| chr3 | 11247634 | 11247946 | 1.21258 | 2.36091 | 1.947 |
| chr5 | 145420849 | 145421318 | 1.05982 | 2.06361 | 1.947 |
| chr13 | 101780152 | 101780500 | 1.07613 | 2.09544 | 1.947 |
| chr5 | 62765226 | 62765502 | 0.483671 | 0.941806 | 1.947 |
| chr7 | 152591384 | 152592030 | 1.77514 | 3.4566 | 1.947 |
| chr12 | 708816 | 709022 | 0.456629 | 0.889178 | 1.947 |
| chr9 | 97545403 | 97545628 | 1.30919 | 2.54943 | 1.947 |
| chr16 | 22199937 | 22200209 | 1.73212 | 3.37304 | 1.947 |
| chr11 | 45254883 | 45255500 | 1.70086 | 3.31219 | 1.947 |
| chr17 | 71185701 | 71186079 | 2.387 | 4.64836 | 1.947 |
| chr5 | 122232704 | 122233001 | 1.2241 | 2.38377 | 1.947 |
| chr1 | 186274139 | 186274403 | 1.32595 | 2.58216 | 1.947 |
| chr8 | 65397841 | 65398055 | 0.373955 | 0.728258 | 1.947 |
| chr6 | 132851526 | 132851862 | 1.11349 | 2.16852 | 1.948 |
| chr12 | 29984936 | 29985312 | 0.503031 | 0.979741 | 1.948 |
| chrX | 135110604 | 135110826 | 1.18412 | 2.30629 | 1.948 |
| chr16 | 52946520 | 52947051 | 1.64715 | 3.20813 | 1.948 |
| chr22 | 28200695 | 28201091 | 1.28611 | 2.50531 | 1.948 |
| chr4 | 41216592 | 41216976 | 1.66985 | 3.25301 | 1.948 |
| chr3 | 195923508 | 195923840 | 2.04111 | 3.97667 | 1.948 |
| chr14 | 103569740 | 103570001 | 0.913471 | 1.77979 | 1.948 |
| chr20 | 58094480 | 58094848 | 1.28203 | 2.49791 | 1.948 |
| chr17 | 75497163 | 75497628 | 1.73163 | 3.374 | 1.948 |
| chr12 | 117908651 | 117909112 | 1.37839 | 2.68582 | 1.949 |
| chr14 | 81735772 | 81735998 | 0.973434 | 1.8968 | 1.949 |
| chr17 | 41918171 | 41918399 | 0.910992 | 1.77515 | 1.949 |
| chr8 | 101153322 | 101153535 | 0.486025 | 0.947062 | 1.949 |
| chr2 | 85618283 | 85618706 | 2.04483 | 3.98468 | 1.949 |
| chr17 | 25659913 | 25660263 | 2.0398 | 3.97504 | 1.949 |
| chr5 | 149597669 | 149598222 | 1.23489 | 2.40663 | 1.949 |
| chr8 | 118341244 | 118341516 | 1.46752 | 2.86013 | 1.949 |
| chr20 | 25512843 | 25513117 | 0.620282 | 1.20892 | 1.949 |
| chr16 | 587791 | 588015 | 0.956213 | 1.86365 | 1.949 |
| chr1 | 155908004 | 155908317 | 1.32473 | 2.58196 | 1.949 |
| chr11 | 17526095 | 17526461 | 2.26615 | 4.41688 | 1.949 |
| chr21 | 39084433 | 39084750 | 2.69758 | 5.25778 | 1.949 |
| chr5 | 54532657 | 54532867 | 0.60488 | 1.17899 | 1.949 |
| chr12 | 115236234 | 115236489 | 0.525735 | 1.02474 | 1.949 |
| chr19 | 56844938 | 56845187 | 1.56305 | 3.04669 | 1.949 |
| chr6 | 37027095 | 37027300 | 1.34132 | 2.61453 | 1.949 |
| chr9 | 120067049 | 120067460 | 0.446949 | 0.871204 | 1.949 |
| chr10 | 117548705 | 117549022 | 0.48627 | 0.947873 | 1.949 |
| chr12 | 44351611 | 44351926 | 0.868602 | 1.69317 | 1.949 |
| chr3 | 143393154 | 143393586 | 2.13304 | 4.15795 | 1.949 |
| chr12 | 107528491 | 107528725 | 1.112 | 2.16769 | 1.949 |
| chr12 | 127061454 | 127061762 | 0.550951 | 1.07408 | 1.950 |
| chr3 | 16219794 | 16220253 | 2.50143 | 4.87658 | 1.950 |
| chr8 | 103819698 | 103820106 | 2.62066 | 5.1091 | 1.950 |
| chr7 | 30114103 | 30114352 | 1.04601 | 2.03926 | 1.950 |
| chr12 | 10902771 | 10903081 | 1.98404 | 3.86824 | 1.950 |
| chr17 | 26874961 | 26875569 | 2.59933 | 5.06789 | 1.950 |
| chr19 | 41598918 | 41599493 | 0.58281 | 1.13632 | 1.950 |
| chr15 | 50818216 | 50818451 | 0.77886 | 1.51859 | 1.950 |
| chr3 | 95471125 | 95471341 | 1.14987 | 2.24198 | 1.950 |
| chr1 | 225416595 | 225416940 | 0.430915 | 0.840195 | 1.950 |

| | | | | | |
|---|---|---|---|---|---|
| chr8 | 142147989 | 142148223 | 0.873882 | 1.70401 | 1.950 |
| chr14 | 30474455 | 30474659 | 1.03375 | 2.01583 | 1.950 |
| chr21 | 45930337 | 45930766 | 1.9034 | 3.71167 | 1.950 |
| chr7 | 20687871 | 20688125 | 1.42779 | 2.78422 | 1.950 |
| chr2 | 30937011 | 30937290 | 1.5208 | 2.96567 | 1.950 |
| chr17 | 17216917 | 17217297 | 1.58102 | 3.08311 | 1.950 |
| chr5 | 50418219 | 50418457 | 0.598988 | 1.16814 | 1.950 |
| chr7 | 154705429 | 154705818 | 1.92158 | 3.74758 | 1.950 |
| chr9 | 22244775 | 22244996 | 1.21419 | 2.36802 | 1.950 |
| chr7 | 65878684 | 65879008 | 1.08796 | 2.1219 | 1.950 |
| chr9 | 94900440 | 94900734 | 1.41816 | 2.76597 | 1.950 |
| chr4 | 148905190 | 148905521 | 1.45304 | 2.83401 | 1.950 |
| chr6 | 74448888 | 74449146 | 1.35221 | 2.63738 | 1.950 |
| chrX | 25162234 | 25162439 | 1.10434 | 2.15396 | 1.950 |
| chr15 | 86617583 | 86617978 | 0.65616 | 1.27983 | 1.950 |
| chr8 | 88962206 | 88962430 | 0.396196 | 0.772799 | 1.951 |
| chr15 | 97392773 | 97392984 | 0.625418 | 1.21994 | 1.951 |
| chr20 | 23917984 | 23918330 | 1.22366 | 2.38694 | 1.951 |
| chr20 | 34472196 | 34472451 | 2.02684 | 3.95367 | 1.951 |
| chr17 | 34334835 | 34335060 | 0.632537 | 1.23388 | 1.951 |
| chr20 | 42469230 | 42469494 | 1.78445 | 3.48102 | 1.951 |
| chr2 | 218007446 | 218007740 | 1.06128 | 2.0704 | 1.951 |
| chr3 | 96857974 | 96858273 | 1.04734 | 2.04322 | 1.951 |
| chr19 | 33802238 | 33802582 | 1.47661 | 2.88071 | 1.951 |
| chr3 | 102248412 | 102248715 | 0.778665 | 1.51915 | 1.951 |
| chr3 | 150127765 | 150128122 | 1.35749 | 2.64845 | 1.951 |
| chr17 | 75482796 | 75483087 | 0.742649 | 1.44897 | 1.951 |
| chr5 | 32852819 | 32853031 | 1.39787 | 2.72746 | 1.951 |
| chr5 | 8742802 | 8743009 | 0.548732 | 1.07067 | 1.951 |
| chr20 | 58319185 | 58319527 | 0.885011 | 1.72687 | 1.951 |
| chr1 | 112050885 | 112051180 | 1.2023 | 2.34601 | 1.951 |
| chr16 | 85679943 | 85680166 | 0.477325 | 0.931397 | 1.951 |
| chr2 | 30909011 | 30909241 | 0.967594 | 1.88806 | 1.951 |
| chr5 | 134896452 | 134896820 | 1.66172 | 3.2425 | 1.951 |
| chr1 | 110739542 | 110739977 | 1.49053 | 2.90847 | 1.951 |
| chr2 | 20861231 | 20861617 | 2.25676 | 4.40374 | 1.951 |
| chr9 | 131893484 | 131894325 | 1.41359 | 2.75846 | 1.951 |
| chr18 | 58352222 | 58352637 | 0.612754 | 1.19583 | 1.952 |
| chr19 | 46320373 | 46320863 | 3.07539 | 6.0021 | 1.952 |
| chr12 | 47069526 | 47069778 | 1.35321 | 2.641 | 1.952 |
| chr11 | 64948325 | 64948948 | 1.1536 | 2.25147 | 1.952 |
| chr4 | 119875407 | 119875740 | 0.638794 | 1.24673 | 1.952 |
| chr12 | 818781 | 819134 | 0.468849 | 0.915065 | 1.952 |
| chr17 | 47591700 | 47591998 | 0.887717 | 1.73261 | 1.952 |
| chr8 | 72012462 | 72012675 | 0.777274 | 1.51705 | 1.952 |
| chr1 | 18152633 | 18152897 | 1.49553 | 2.91895 | 1.952 |
| chr22 | 26763490 | 26763928 | 1.50924 | 2.94573 | 1.952 |
| chr5 | 159343288 | 159343562 | 2.27846 | 4.44727 | 1.952 |
| chr6 | 142847583 | 142847786 | 0.537708 | 1.04955 | 1.952 |
| chr11 | 10746428 | 10746782 | 0.51703 | 1.00922 | 1.952 |
| chr1 | 154642205 | 154642485 | 2.00217 | 3.90832 | 1.952 |
| chr12 | 66722987 | 66723360 | 0.756729 | 1.47717 | 1.952 |
| chr14 | 62374272 | 62374808 | 0.492696 | 0.961788 | 1.952 |
| chr2 | 18664 | 19253 | 1.15305 | 2.25087 | 1.952 |
| chr15 | 28275357 | 28275594 | 0.673896 | 1.31557 | 1.952 |
| chr11 | 109816973 | 109817208 | 1.35202 | 2.63947 | 1.952 |
| chrX | 136036259 | 136036671 | 1.32609 | 2.58889 | 1.952 |
| chr11 | 2207857 | 2208286 | 1.57968 | 3.08404 | 1.952 |
| chr2 | 15896286 | 15896536 | 0.483899 | 0.944734 | 1.952 |
| chr10 | 2881226 | 2881756 | 1.16992 | 2.28409 | 1.952 |
| chr19 | 48655717 | 48656136 | 1.43824 | 2.80801 | 1.952 |

| | | | | | |
|---|---|---|---|---|---|
| chr11 | 19575369 | 19575582 | 1.81086 | 3.53553 | 1.952 |
| chr1 | 205557861 | 205558225 | 1.04491 | 2.04016 | 1.952 |
| chr5 | 176883332 | 176883669 | 2.95741 | 5.77441 | 1.953 |
| chr5 | 70825208 | 70825416 | 0.529415 | 1.0337 | 1.953 |
| chr9 | 121606316 | 121606544 | 0.548795 | 1.07158 | 1.953 |
| chrX | 107467993 | 107468220 | 1.38029 | 2.69517 | 1.953 |
| chr19 | 11241660 | 11241918 | 1.27983 | 2.49906 | 1.953 |
| chr6 | 84221581 | 84221873 | 1.9655 | 3.83797 | 1.953 |
| chr14 | 100448820 | 100449113 | 1.45658 | 2.84423 | 1.953 |
| chr22 | 41929215 | 41929631 | 1.46985 | 2.87014 | 1.953 |
| chr19 | 584381 | 584787 | 1.99531 | 3.89624 | 1.953 |
| chr8 | 17009896 | 17010327 | 2.02758 | 3.95936 | 1.953 |
| chr7 | 135395558 | 135395947 | 1.84988 | 3.61239 | 1.953 |
| chr13 | 47548120 | 47548360 | 1.24859 | 2.43872 | 1.953 |
| chr4 | 88140818 | 88141022 | 0.834545 | 1.63002 | 1.953 |
| chr2 | 71012481 | 71012764 | 1.5806 | 3.08723 | 1.953 |
| chr18 | 43245127 | 43245430 | 1.57991 | 3.08589 | 1.953 |
| chr10 | 111268746 | 111269021 | 0.390671 | 0.763153 | 1.953 |
| chr16 | 78804495 | 78804891 | 1.01975 | 1.99206 | 1.953 |
| chr10 | 10377679 | 10377987 | 0.499148 | 0.975081 | 1.953 |
| chr17 | 74519139 | 74519696 | 3.97822 | 7.77178 | 1.954 |
| chr19 | 23623665 | 23623895 | 1.47956 | 2.89048 | 1.954 |
| chr12 | 119982842 | 119983383 | 1.20148 | 2.34732 | 1.954 |
| chr1 | 160016395 | 160016805 | 1.29325 | 2.52663 | 1.954 |
| chr3 | 32338021 | 32338575 | 1.66376 | 3.25054 | 1.954 |
| chr10 | 96961553 | 96961778 | 1.0102 | 1.97368 | 1.954 |
| chr5 | 151098888 | 151099157 | 1.52169 | 2.97303 | 1.954 |
| chr7 | 122827130 | 122827332 | 0.437676 | 0.855126 | 1.954 |
| chr11 | 49156706 | 49156971 | 0.531379 | 1.03821 | 1.954 |
| chr9 | 110905296 | 110905520 | 0.559596 | 1.09336 | 1.954 |
| chr4 | 169838457 | 169838818 | 0.618989 | 1.20942 | 1.954 |
| chr14 | 47141454 | 47141767 | 0.556401 | 1.08717 | 1.954 |
| chr10 | 34370953 | 34371202 | 0.63252 | 1.23594 | 1.954 |
| chr2 | 216898715 | 216899182 | 1.74354 | 3.40686 | 1.954 |
| chr15 | 23061104 | 23061367 | 1.52889 | 2.98749 | 1.954 |
| chr1 | 168408971 | 168409368 | 1.42574 | 2.78598 | 1.954 |
| chr10 | 31072817 | 31073139 | 1.99005 | 3.88869 | 1.954 |
| chr19 | 33705494 | 33705716 | 1.0764 | 2.1035 | 1.954 |
| chr7 | 22822285 | 22822627 | 1.7178 | 3.35706 | 1.954 |
| chr15 | 100879208 | 100879471 | 0.82436 | 1.61104 | 1.954 |
| chr1 | 156036130 | 156036477 | 2.65456 | 5.18786 | 1.954 |
| chr2 | 194939778 | 194939998 | 0.554873 | 1.0844 | 1.954 |
| chr12 | 1626288 | 1626633 | 1.41627 | 2.76794 | 1.954 |
| chr19 | 54315545 | 54315991 | 0.808382 | 1.58007 | 1.955 |
| chr2 | 3025830 | 3026181 | 0.508554 | 0.994038 | 1.955 |
| chr2 | 31456621 | 31456912 | 1.99302 | 3.89571 | 1.955 |
| chr11 | 37939271 | 37939803 | 0.493336 | 0.964371 | 1.955 |
| chr6 | 136547384 | 136547704 | 3.16435 | 6.18571 | 1.955 |
| chr9 | 134211647 | 134212074 | 1.75895 | 3.43848 | 1.955 |
| chr20 | 35283048 | 35283297 | 0.56903 | 1.11239 | 1.955 |
| chr4 | 81046627 | 81046879 | 1.23441 | 2.41317 | 1.955 |
| chr7 | 2249088 | 2249456 | 1.81183 | 3.54214 | 1.955 |
| chr9 | 135042071 | 135042449 | 1.15895 | 2.26576 | 1.955 |
| chr17 | 44919122 | 44919638 | 2.65793 | 5.19637 | 1.955 |
| chr9 | 101946918 | 101947229 | 1.77081 | 3.46204 | 1.955 |
| chr3 | 10825074 | 10825453 | 1.69744 | 3.3189 | 1.955 |
| chr5 | 10373526 | 10373813 | 0.665978 | 1.30215 | 1.955 |
| chr15 | 99190708 | 99191124 | 1.73907 | 3.40039 | 1.955 |
| chr19 | 32831162 | 32831569 | 1.98639 | 3.88399 | 1.955 |
| chr13 | 30909167 | 30909490 | 0.767881 | 1.50145 | 1.955 |
| chr22 | 42062660 | 42062988 | 2.31563 | 4.52788 | 1.955 |

| | | | | | |
|---|---|---|---|---|---|
| chr7 | 92509617 | 92510030 | 0.634364 | 1.24041 | 1.955 |
| chr13 | 52509998 | 52510216 | 0.801179 | 1.5666 | 1.955 |
| chr17 | 34446232 | 34446496 | 0.627205 | 1.22642 | 1.955 |
| chr6 | 143658797 | 143659071 | 1.64214 | 3.21112 | 1.955 |
| chr3 | 138830683 | 138830978 | 1.48512 | 2.90433 | 1.956 |
| chr8 | 143084040 | 143084604 | 1.90359 | 3.72287 | 1.956 |
| chr9 | 125572677 | 125572888 | 1.11283 | 2.17642 | 1.956 |
| chr1 | 19772222 | 19772480 | 0.853027 | 1.66836 | 1.956 |
| chr7 | 124405766 | 124406166 | 1.18481 | 2.31736 | 1.956 |
| chr13 | 99100586 | 99100952 | 1.92388 | 3.76302 | 1.956 |
| chr11 | 18046955 | 18047211 | 0.511019 | 0.999539 | 1.956 |
| chr20 | 8640971 | 8641203 | 0.588137 | 1.15038 | 1.956 |
| chr2 | 59137979 | 59138224 | 0.506801 | 0.9913 | 1.956 |
| chr3 | 111258613 | 111258924 | 1.69246 | 3.31043 | 1.956 |
| chr7 | 22526041 | 22526280 | 0.524587 | 1.02609 | 1.956 |
| chr2 | 1773552 | 1773917 | 1.62168 | 3.17202 | 1.956 |
| chr1 | 175115717 | 175116295 | 1.86109 | 3.64041 | 1.956 |
| chr8 | 30085229 | 30085711 | 1.04963 | 2.05319 | 1.956 |
| chr7 | 47859019 | 47859297 | 1.30635 | 2.5554 | 1.956 |
| chr4 | 71553729 | 71554111 | 1.90451 | 3.7255 | 1.956 |
| chr8 | 79365960 | 79366212 | 0.629527 | 1.23146 | 1.956 |
| chr5 | 88836064 | 88836309 | 0.67718 | 1.32472 | 1.956 |
| chr11 | 60312907 | 60313218 | 0.606389 | 1.18624 | 1.956 |
| chr13 | 47468114 | 47468612 | 2.14558 | 4.19736 | 1.956 |
| chr4 | 6333735 | 6334195 | 1.71358 | 3.35233 | 1.956 |
| chr15 | 95173934 | 95174157 | 0.604028 | 1.18174 | 1.956 |
| chr2 | 188853207 | 188853417 | 0.390771 | 0.764517 | 1.956 |
| chr13 | 100269614 | 100269972 | 1.72937 | 3.38353 | 1.957 |
| chr10 | 117559003 | 117559297 | 1.34907 | 2.63967 | 1.957 |
| chr5 | 174243934 | 174244193 | 0.881292 | 1.7244 | 1.957 |
| chr5 | 177502969 | 177503454 | 1.92865 | 3.77374 | 1.957 |
| chr9 | 132331922 | 132332129 | 1.26899 | 2.483 | 1.957 |
| chr3 | 58400389 | 58400650 | 1.24844 | 2.44298 | 1.957 |
| chr1 | 154836123 | 154836708 | 1.15988 | 2.2698 | 1.957 |
| chr5 | 7816294 | 7816754 | 2.61197 | 5.11144 | 1.957 |
| chr5 | 177547859 | 177548241 | 1.57066 | 3.07374 | 1.957 |
| chr13 | 32989241 | 32989484 | 1.06637 | 2.08692 | 1.957 |
| chr7 | 8335785 | 8335997 | 0.972785 | 1.90377 | 1.957 |
| chr9 | 139958414 | 139958946 | 1.91457 | 3.74689 | 1.957 |
| chr17 | 36175975 | 36176244 | 1.28002 | 2.50507 | 1.957 |
| chr5 | 7415998 | 7416635 | 0.518979 | 1.01576 | 1.957 |
| chrX | 67458488 | 67458800 | 1.57191 | 3.07667 | 1.957 |
| chr1 | 246660712 | 246660919 | 1.29444 | 2.5336 | 1.957 |
| chr15 | 60651865 | 60652173 | 1.235 | 2.41738 | 1.957 |
| chr15 | 68951047 | 68951280 | 0.714705 | 1.39902 | 1.957 |
| chr12 | 44170390 | 44170595 | 0.845679 | 1.65553 | 1.958 |
| chr12 | 109361590 | 109361879 | 1.50844 | 2.95297 | 1.958 |
| chr1 | 200179179 | 200179411 | 0.699093 | 1.36858 | 1.958 |
| chr11 | 35327390 | 35327613 | 1.38027 | 2.70211 | 1.958 |
| chr10 | 97134421 | 97134713 | 2.10586 | 4.12275 | 1.958 |
| chr3 | 136471068 | 136471396 | 2.08671 | 4.0854 | 1.958 |
| chr21 | 26979810 | 26980053 | 1.21561 | 2.37997 | 1.958 |
| chr12 | 25842951 | 25843177 | 0.609837 | 1.19397 | 1.958 |
| chr5 | 151283710 | 151283914 | 0.646763 | 1.26633 | 1.958 |
| chr17 | 27918618 | 27919065 | 3.28211 | 6.42632 | 1.958 |
| chr10 | 72968728 | 72969198 | 2.46591 | 4.8283 | 1.958 |
| chr11 | 115041016 | 115041299 | 1.42583 | 2.79181 | 1.958 |
| chr2 | 56150590 | 56150872 | 0.610375 | 1.19523 | 1.958 |
| chr8 | 84643632 | 84643849 | 0.449653 | 0.880508 | 1.958 |
| chr12 | 34446168 | 34446515 | 1.19456 | 2.33919 | 1.958 |
| chr4 | 122113637 | 122113940 | 1.53878 | 3.01325 | 1.958 |

| | | | | | |
|---|---|---|---|---|---|
| chr20 | 8098294 | 8098564 | 0.591211 | 1.15772 | 1.958 |
| chr7 | 50485751 | 50486127 | 2.70172 | 5.29068 | 1.958 |
| chr15 | 81896293 | 81896547 | 0.511911 | 1.00246 | 1.958 |
| chr11 | 115195354 | 115195596 | 1.17886 | 2.30863 | 1.958 |
| chr19 | 19453457 | 19453958 | 1.50311 | 2.94386 | 1.959 |
| chr4 | 30154625 | 30155251 | 0.515791 | 1.01019 | 1.959 |
| chr7 | 8257191 | 8257409 | 0.891348 | 1.74573 | 1.959 |
| chr3 | 11876503 | 11876823 | 0.863377 | 1.69096 | 1.959 |
| chr1 | 197149405 | 197149749 | 1.05792 | 2.07199 | 1.959 |
| chr8 | 108008375 | 108008739 | 0.520172 | 1.01879 | 1.959 |
| chr10 | 104159098 | 104159983 | 2.0257 | 3.9676 | 1.959 |
| chr10 | 97190061 | 97190576 | 1.79766 | 3.52099 | 1.959 |
| chr9 | 132552578 | 132552956 | 2.03714 | 3.99007 | 1.959 |
| chr16 | 20861229 | 20861581 | 1.90581 | 3.7329 | 1.959 |
| chr19 | 15530034 | 15530265 | 1.18315 | 2.31746 | 1.959 |
| chr3 | 140998150 | 140998479 | 1.11097 | 2.17608 | 1.959 |
| chr6 | 25882213 | 25882745 | 0.831986 | 1.62977 | 1.959 |
| chr12 | 4462961 | 4463215 | 1.49378 | 2.9262 | 1.959 |
| chr11 | 111299815 | 111300060 | 1.03761 | 2.03262 | 1.959 |
| chr9 | 35405590 | 35405793 | 0.404848 | 0.793092 | 1.959 |
| chr17 | 36714344 | 36714749 | 2.32915 | 4.56291 | 1.959 |
| chr2 | 89039503 | 89039908 | 2.009 | 3.93576 | 1.959 |
| chr16 | 4275066 | 4275372 | 0.546932 | 1.07149 | 1.959 |
| chr15 | 24923703 | 24924144 | 1.06865 | 2.09359 | 1.959 |
| chr3 | 172554299 | 172554501 | 1.7593 | 3.44691 | 1.959 |
| chr7 | 126851634 | 126851878 | 0.466128 | 0.913264 | 1.959 |
| chr7 | 131521570 | 131521793 | 0.827213 | 1.62082 | 1.959 |
| chr12 | 42963111 | 42963349 | 1.1567 | 2.26646 | 1.959 |
| chr17 | 17897688 | 17897929 | 1.12298 | 2.20041 | 1.959 |
| chr2 | 88486459 | 88486992 | 1.58857 | 3.11272 | 1.959 |
| chr22 | 50160855 | 50161264 | 1.03659 | 2.03124 | 1.960 |
| chr7 | 156129335 | 156129594 | 0.460214 | 0.901809 | 1.960 |
| chr16 | 72158765 | 72159151 | 1.57868 | 3.09354 | 1.960 |
| chr3 | 44240194 | 44240426 | 1.48869 | 2.9172 | 1.960 |
| chr16 | 90014352 | 90014708 | 0.439958 | 0.862178 | 1.960 |
| chr3 | 43811014 | 43811257 | 1.30234 | 2.55226 | 1.960 |
| chr8 | 11137042 | 11137321 | 0.814495 | 1.59621 | 1.960 |
| chr2 | 230138504 | 230138784 | 1.32298 | 2.59272 | 1.960 |
| chrX | 107382612 | 107382914 | 1.24638 | 2.44261 | 1.960 |
| chr5 | 156449504 | 156449708 | 0.549826 | 1.07753 | 1.960 |
| chr20 | 50745043 | 50745327 | 1.45473 | 2.85108 | 1.960 |
| chr17 | 72776731 | 72777347 | 1.25448 | 2.45871 | 1.960 |
| chr12 | 98581416 | 98581650 | 0.425539 | 0.834058 | 1.960 |
| chr5 | 54319770 | 54320056 | 0.629586 | 1.23409 | 1.960 |
| chr6 | 158018658 | 158019159 | 1.32158 | 2.59054 | 1.960 |
| chr12 | 18107772 | 18108055 | 1.02177 | 2.00288 | 1.960 |
| chr2 | 44271354 | 44271710 | 2.67469 | 5.24298 | 1.960 |
| chr11 | 17251686 | 17251955 | 1.43764 | 2.81811 | 1.960 |
| chr3 | 139146397 | 139146721 | 1.21596 | 2.38359 | 1.960 |
| chr3 | 36007457 | 36007666 | 1.03391 | 2.02674 | 1.960 |
| chr15 | 89899831 | 89900329 | 1.62386 | 3.18334 | 1.960 |
| chr1 | 228261609 | 228261842 | 1.36556 | 2.67703 | 1.960 |
| chr12 | 51815093 | 51815387 | 1.89108 | 3.70733 | 1.960 |
| chr10 | 102773972 | 102774388 | 1.60171 | 3.14009 | 1.960 |
| chr7 | 137846330 | 137846541 | 1.32665 | 2.6009 | 1.961 |
| chr11 | 65548681 | 65549048 | 1.12681 | 2.20917 | 1.961 |
| chr15 | 44358505 | 44358729 | 1.12425 | 2.20417 | 1.961 |
| chr10 | 115144911 | 115145250 | 1.64569 | 3.22675 | 1.961 |
| chr19 | 38970852 | 38971122 | 0.529547 | 1.03831 | 1.961 |
| chr10 | 24707573 | 24707796 | 1.09365 | 2.14441 | 1.961 |
| chr15 | 63695096 | 63695313 | 0.974896 | 1.9116 | 1.961 |

| | | | | | |
|---|---|---|---|---|---|
| chr19 | 46142325 | 46142574 | 0.824438 | 1.61664 | 1.961 |
| chr1 | 152969235 | 152969601 | 0.524491 | 1.0285 | 1.961 |
| chr10 | 101152232 | 101152643 | 2.37941 | 4.66613 | 1.961 |
| chr7 | 100048025 | 100048468 | 2.15384 | 4.22378 | 1.961 |
| chr16 | 27616885 | 27617215 | 1.7002 | 3.3342 | 1.961 |
| chr7 | 76142336 | 76142645 | 0.783746 | 1.53706 | 1.961 |
| chr10 | 131769729 | 131770341 | 1.20629 | 2.36579 | 1.961 |
| chr20 | 44399810 | 44400104 | 1.77583 | 3.48278 | 1.961 |
| chr11 | 45273932 | 45274314 | 2.4963 | 4.89591 | 1.961 |
| chr12 | 85268323 | 85268663 | 1.05412 | 2.06746 | 1.961 |
| chr17 | 8689119 | 8689552 | 1.70926 | 3.35241 | 1.961 |
| chr7 | 118837909 | 118838361 | 1.50048 | 2.94303 | 1.961 |
| chr7 | 116759552 | 116759830 | 1.63185 | 3.20087 | 1.962 |
| chr20 | 17114607 | 17115069 | 0.453704 | 0.889974 | 1.962 |
| chr3 | 10800257 | 10800628 | 1.56068 | 3.0614 | 1.962 |
| chr11 | 65835628 | 65836053 | 1.99625 | 3.91584 | 1.962 |
| chr15 | 61343700 | 61343987 | 1.39774 | 2.74181 | 1.962 |
| chr5 | 117833037 | 117833507 | 0.484987 | 0.951357 | 1.962 |
| chr9 | 34675045 | 34675542 | 2.45271 | 4.81132 | 1.962 |
| chr16 | 66642246 | 66642788 | 2.40346 | 4.71484 | 1.962 |
| chr16 | 30640061 | 30640429 | 1.5401 | 3.02125 | 1.962 |
| chr9 | 87091678 | 87091950 | 0.467248 | 0.916634 | 1.962 |
| chrX | 130936876 | 130937345 | 1.61475 | 3.16779 | 1.962 |
| chr3 | 97164869 | 97165158 | 0.571369 | 1.12092 | 1.962 |
| chr8 | 94834619 | 94834909 | 1.54756 | 3.03602 | 1.962 |
| chr19 | 45629268 | 45629743 | 1.8953 | 3.71828 | 1.962 |
| chr15 | 27733062 | 27733367 | 0.61926 | 1.2149 | 1.962 |
| chr5 | 98963397 | 98963673 | 0.498414 | 0.977839 | 1.962 |
| chr17 | 47645171 | 47645977 | 1.69009 | 3.31586 | 1.962 |
| chr3 | 108208646 | 108208859 | 1.08678 | 2.13222 | 1.962 |
| chr5 | 177982098 | 177982345 | 0.457245 | 0.897156 | 1.962 |
| chr1 | 202122613 | 202122831 | 0.539311 | 1.05819 | 1.962 |
| chr1 | 75199472 | 75199711 | 1.19469 | 2.34413 | 1.962 |
| chr17 | 79226930 | 79227407 | 1.27472 | 2.50117 | 1.962 |
| chr22 | 29550401 | 29550792 | 2.09644 | 4.11351 | 1.962 |
| chr7 | 51891368 | 51891633 | 0.627474 | 1.2312 | 1.962 |
| chr5 | 6435507 | 6435876 | 1.43183 | 2.80964 | 1.962 |
| chr17 | 73283921 | 73284283 | 0.513822 | 1.00828 | 1.962 |
| chr16 | 88571854 | 88572295 | 1.71879 | 3.3729 | 1.962 |
| chr3 | 114137577 | 114138030 | 1.2188 | 2.39175 | 1.962 |
| chr9 | 104096442 | 104096772 | 0.495453 | 0.972267 | 1.962 |
| chr11 | 63855649 | 63856190 | 1.03194 | 2.02508 | 1.962 |
| chr16 | 81974351 | 81974745 | 0.992207 | 1.94715 | 1.962 |
| chr7 | 43494603 | 43494907 | 0.941449 | 1.84771 | 1.963 |
| chr7 | 31483039 | 31483320 | 1.52347 | 2.99013 | 1.963 |
| chr6 | 116589764 | 116590056 | 1.4485 | 2.84314 | 1.963 |
| chr4 | 71517965 | 71518397 | 0.623269 | 1.2234 | 1.963 |
| chr2 | 55242149 | 55242377 | 1.50209 | 2.94843 | 1.963 |
| chr1 | 145524642 | 145525049 | 1.53305 | 3.00967 | 1.963 |
| chr15 | 86397617 | 86397895 | 1.75367 | 3.44279 | 1.963 |
| chr15 | 59453221 | 59453727 | 1.06613 | 2.09305 | 1.963 |
| chr4 | 103370960 | 103371217 | 1.01028 | 1.98365 | 1.963 |
| chr9 | 20885071 | 20885326 | 1.25314 | 2.46059 | 1.964 |
| chr17 | 69003056 | 69003262 | 0.677374 | 1.33007 | 1.964 |
| chr13 | 76593565 | 76593859 | 1.11333 | 2.18612 | 1.964 |
| chr8 | 141442059 | 141442368 | 1.56587 | 3.07492 | 1.964 |
| chr9 | 131580518 | 131580826 | 1.65139 | 3.24302 | 1.964 |
| chr2 | 68073622 | 68073904 | 1.87219 | 3.67671 | 1.964 |
| chr1 | 109288991 | 109289327 | 1.97233 | 3.87354 | 1.964 |
| chr3 | 24721968 | 24722389 | 2.03046 | 3.98779 | 1.964 |
| chr12 | 57701575 | 57701909 | 1.69536 | 3.3297 | 1.964 |

| | | | | | |
|---|---|---|---|---|---|
| chr3 | 26151922 | 26152166 | 0.652855 | 1.28228 | 1.964 |
| chr4 | 17810175 | 17810377 | 1.00907 | 1.98192 | 1.964 |
| chr12 | 58499635 | 58499883 | 1.01092 | 1.98566 | 1.964 |
| chr7 | 140932567 | 140932811 | 0.502457 | 0.98693 | 1.964 |
| chr17 | 56598201 | 56598475 | 0.918075 | 1.80332 | 1.964 |
| chr20 | 53598885 | 53599090 | 0.569667 | 1.11899 | 1.964 |
| chr1 | 210621670 | 210622121 | 0.919124 | 1.80548 | 1.964 |
| chr10 | 34973434 | 34973675 | 1.15792 | 2.27469 | 1.964 |
| chr8 | 91911178 | 91911446 | 0.729214 | 1.43256 | 1.965 |
| chr4 | 11721999 | 11722226 | 1.32685 | 2.60668 | 1.965 |
| chr17 | 74117427 | 74117884 | 2.96606 | 5.82704 | 1.965 |
| chr9 | 130110463 | 130110669 | 1.36704 | 2.68566 | 1.965 |
| chr10 | 99555508 | 99555884 | 0.513994 | 1.00983 | 1.965 |
| chr2 | 219894399 | 219894796 | 2.49201 | 4.89609 | 1.965 |
| chr5 | 174961090 | 174961587 | 1.97939 | 3.88898 | 1.965 |
| chr6 | 24583697 | 24584015 | 1.56341 | 3.07181 | 1.965 |
| chr15 | 79100883 | 79101128 | 1.38136 | 2.71415 | 1.965 |
| chr14 | 103384249 | 103384462 | 1.36093 | 2.67408 | 1.965 |
| chr16 | 49924782 | 49925029 | 0.857421 | 1.68476 | 1.965 |
| chr19 | 46538899 | 46539149 | 0.32103 | 0.630811 | 1.965 |
| chr16 | 12161768 | 12162065 | 1.70235 | 3.34507 | 1.965 |
| chr9 | 119499323 | 119499697 | 2.90221 | 5.70276 | 1.965 |
| chr7 | 2404444 | 2404790 | 1.46062 | 2.87009 | 1.965 |
| chr20 | 36967195 | 36967524 | 0.759241 | 1.49197 | 1.965 |
| chr6 | 106892852 | 106893081 | 1.34303 | 2.6392 | 1.965 |
| chr16 | 2551027 | 2551632 | 2.3608 | 4.63946 | 1.965 |
| chr17 | 4118625 | 4118984 | 1.9313 | 3.79541 | 1.965 |
| chr2 | 238074096 | 238074337 | 1.39203 | 2.73571 | 1.965 |
| chr3 | 138634182 | 138635122 | 1.73275 | 3.40533 | 1.965 |
| chr5 | 72044512 | 72044729 | 0.534992 | 1.05142 | 1.965 |
| chr11 | 72436296 | 72436587 | 1.88917 | 3.71283 | 1.965 |
| chr1 | 4645170 | 4645647 | 1.3812 | 2.71476 | 1.966 |
| chr17 | 9195410 | 9196038 | 1.68297 | 3.30819 | 1.966 |
| chr5 | 7138051 | 7138285 | 0.535057 | 1.0518 | 1.966 |
| chr16 | 65528401 | 65528628 | 0.418501 | 0.822704 | 1.966 |
| chr11 | 119244903 | 119245186 | 1.40538 | 2.76281 | 1.966 |
| chr18 | 11872570 | 11872842 | 0.951708 | 1.87095 | 1.966 |
| chr7 | 66022357 | 66022934 | 1.47764 | 2.90498 | 1.966 |
| chr22 | 25506942 | 25507289 | 1.07211 | 2.10777 | 1.966 |
| chr5 | 139525378 | 139525778 | 1.47773 | 2.90526 | 1.966 |
| chr17 | 72562294 | 72562829 | 1.89786 | 3.73127 | 1.966 |
| chr6 | 153269363 | 153269626 | 1.00586 | 1.97774 | 1.966 |
| chr6 | 26066149 | 26066448 | 1.42774 | 2.80731 | 1.966 |
| chr1 | 229694261 | 229694843 | 1.59638 | 3.13892 | 1.966 |
| chr19 | 17535833 | 17536129 | 1.44633 | 2.8439 | 1.966 |
| chr2 | 197457411 | 197457952 | 0.795899 | 1.56499 | 1.966 |
| chr4 | 114992268 | 114992488 | 1.36368 | 2.68147 | 1.966 |
| chr22 | 22292379 | 22293223 | 2.00052 | 3.93438 | 1.967 |
| chr3 | 79097581 | 79097830 | 0.443973 | 0.873167 | 1.967 |
| chr6 | 137982801 | 137983043 | 1.30921 | 2.57484 | 1.967 |
| chr1 | 199212581 | 199212899 | 1.77001 | 3.48112 | 1.967 |
| chr16 | 23937240 | 23937557 | 0.519554 | 1.02182 | 1.967 |
| chr17 | 59713372 | 59713619 | 1.13807 | 2.23828 | 1.967 |
| chr12 | 6978566 | 6978981 | 2.47382 | 4.86541 | 1.967 |
| chr9 | 140995446 | 140995667 | 0.626446 | 1.23208 | 1.967 |
| chr3 | 183543469 | 183543809 | 0.756962 | 1.4888 | 1.967 |
| chr3 | 119963056 | 119963363 | 1.1289 | 2.22057 | 1.967 |
| chr20 | 9947700 | 9947979 | 1.02187 | 2.01005 | 1.967 |
| chr11 | 46260607 | 46261031 | 2.58745 | 5.08981 | 1.967 |
| chr20 | 48598779 | 48599607 | 0.846243 | 1.66465 | 1.967 |
| chrX | 39590024 | 39590315 | 2.34559 | 4.61413 | 1.967 |

| | | | | | |
|---|---|---|---|---|---|
| chr2 | 71558621 | 71558930 | 2.44312 | 4.80638 | 1.967 |
| chr14 | 23410129 | 23410419 | 1.92971 | 3.79647 | 1.967 |
| chr8 | 90701487 | 90701703 | 1.44654 | 2.84597 | 1.967 |
| chr12 | 55462040 | 55462558 | 1.33478 | 2.62615 | 1.967 |
| chr13 | 42033093 | 42033326 | 1.41755 | 2.78905 | 1.968 |
| chr20 | 44002544 | 44002921 | 2.05607 | 4.0455 | 1.968 |
| chr5 | 1167775 | 1168683 | 1.83938 | 3.61935 | 1.968 |
| chr3 | 86405373 | 86405613 | 0.742993 | 1.46203 | 1.968 |
| chr3 | 150505571 | 150505935 | 1.85982 | 3.65974 | 1.968 |
| chr16 | 77232217 | 77232426 | 0.457677 | 0.900621 | 1.968 |
| chr2 | 120510976 | 120511297 | 1.61304 | 3.17418 | 1.968 |
| chr10 | 88122954 | 88123233 | 1.42449 | 2.80322 | 1.968 |
| chr10 | 73024398 | 73024637 | 0.624431 | 1.22881 | 1.968 |
| chr12 | 130057589 | 130057801 | 0.662406 | 1.30354 | 1.968 |
| chrX | 57555116 | 57555592 | 1.02025 | 2.00784 | 1.968 |
| chr18 | 12038905 | 12039286 | 1.71307 | 3.3714 | 1.968 |
| chr7 | 82234650 | 82234861 | 1.24806 | 2.45632 | 1.968 |
| chr11 | 79083023 | 79083369 | 0.455318 | 0.896153 | 1.968 |
| chr10 | 83027208 | 83027417 | 0.492687 | 0.969706 | 1.968 |
| chr1 | 202088128 | 202088426 | 1.46274 | 2.879 | 1.968 |
| chr20 | 24975428 | 24975807 | 2.31929 | 4.5649 | 1.968 |
| chr9 | 92897154 | 92897432 | 0.523205 | 1.0299 | 1.968 |
| chr12 | 4541025 | 4541242 | 0.487363 | 0.959353 | 1.968 |
| chr5 | 23209101 | 23209492 | 1.11047 | 2.18593 | 1.968 |
| chr5 | 102769813 | 102770153 | 0.602837 | 1.18671 | 1.969 |
| chr12 | 60816654 | 60816906 | 0.614684 | 1.21004 | 1.969 |
| chr6 | 35551639 | 35551904 | 1.17014 | 2.30352 | 1.969 |
| chr8 | 22530548 | 22530801 | 2.46348 | 4.84965 | 1.969 |
| chr3 | 170131943 | 170132335 | 2.56499 | 5.04972 | 1.969 |
| chr8 | 71026902 | 71027229 | 0.933784 | 1.83835 | 1.969 |
| chr5 | 87743374 | 87743596 | 0.593436 | 1.16836 | 1.969 |
| chr16 | 53580872 | 53581082 | 1.43929 | 2.83372 | 1.969 |
| chr3 | 28517767 | 28518019 | 1.20605 | 2.37453 | 1.969 |
| chr8 | 144302434 | 144302893 | 1.06263 | 2.09217 | 1.969 |
| chr9 | 14428777 | 14429099 | 1.37581 | 2.70887 | 1.969 |
| chr9 | 130547214 | 130547907 | 1.57104 | 3.09333 | 1.969 |
| chr5 | 116499406 | 116499671 | 0.523892 | 1.03154 | 1.969 |
| chr1 | 174713172 | 174713383 | 1.05339 | 2.07432 | 1.969 |
| chr1 | 8731340 | 8731634 | 0.989742 | 1.949 | 1.969 |
| chr15 | 65903473 | 65903757 | 1.82969 | 3.60313 | 1.969 |
| chr17 | 27053835 | 27054205 | 3.4888 | 6.87053 | 1.969 |
| chr1 | 201482096 | 201482458 | 2.16805 | 4.26964 | 1.969 |
| chr12 | 57395187 | 57395606 | 2.16916 | 4.27207 | 1.969 |
| chr1 | 201055027 | 201055373 | 1.89899 | 3.74009 | 1.970 |
| chr10 | 90979437 | 90979657 | 0.725133 | 1.42822 | 1.970 |
| chr3 | 21898939 | 21899224 | 0.585026 | 1.15235 | 1.970 |
| chr1 | 116350429 | 116350685 | 1.25815 | 2.47832 | 1.970 |
| chr20 | 33917393 | 33917694 | 1.58611 | 3.12434 | 1.970 |
| chr4 | 168543808 | 168544279 | 1.56288 | 3.0786 | 1.970 |
| chr17 | 70385219 | 70385428 | 1.11866 | 2.20362 | 1.970 |
| chr2 | 85647768 | 85648176 | 2.34174 | 4.61294 | 1.970 |
| chr5 | 109926105 | 109926380 | 0.539402 | 1.06257 | 1.970 |
| chr1 | 110602695 | 110603111 | 1.8181 | 3.58149 | 1.970 |
| chr1 | 9934882 | 9935258 | 1.83003 | 3.60504 | 1.970 |
| chr18 | 42336554 | 42336778 | 0.499251 | 0.983511 | 1.970 |
| chr20 | 49918449 | 49918656 | 0.647037 | 1.27465 | 1.970 |
| chrX | 24770524 | 24770803 | 0.979805 | 1.93026 | 1.970 |
| chr5 | 171478705 | 171478918 | 0.541274 | 1.06635 | 1.970 |
| chr7 | 55617751 | 55618104 | 2.385 | 4.69861 | 1.970 |
| chr2 | 169615826 | 169616060 | 0.840701 | 1.65627 | 1.970 |
| chr10 | 120219073 | 120219444 | 2.07669 | 4.09135 | 1.970 |

| | | | | | |
|---|---|---|---|---|---|
| chr16 | 863466 | 863992 | 1.05051 | 2.06964 | 1.970 |
| chr5 | 36875987 | 36876213 | 0.957712 | 1.88683 | 1.970 |
| chr10 | 1119557 | 1120246 | 0.52955 | 1.0433 | 1.970 |
| chr17 | 14205833 | 14206226 | 2.50633 | 4.93795 | 1.970 |
| chr5 | 101489133 | 101489365 | 0.651235 | 1.28306 | 1.970 |
| chr1 | 248847031 | 248847307 | 0.506206 | 0.99733 | 1.970 |
| chr2 | 42964204 | 42964566 | 0.498341 | 0.981849 | 1.970 |
| chr12 | 130327164 | 130327422 | 0.722733 | 1.42402 | 1.970 |
| chr3 | 65422341 | 65422601 | 1.61876 | 3.1895 | 1.970 |
| chr20 | 49604241 | 49604577 | 1.543 | 3.04048 | 1.971 |
| chr12 | 112226009 | 112226299 | 1.11182 | 2.19099 | 1.971 |
| chr11 | 121462182 | 121462417 | 1.01857 | 2.00728 | 1.971 |
| chr9 | 92904172 | 92904378 | 0.969629 | 1.91083 | 1.971 |
| chr5 | 61408123 | 61408400 | 0.501557 | 0.988413 | 1.971 |
| chr19 | 36297404 | 36297681 | 0.681133 | 1.34233 | 1.971 |
| chr7 | 48213397 | 48213739 | 0.399336 | 0.786991 | 1.971 |
| chr6 | 141398873 | 141399121 | 0.88184 | 1.73799 | 1.971 |
| chr3 | 138047959 | 138048322 | 1.32046 | 2.60249 | 1.971 |
| chr6 | 86033198 | 86033455 | 1.3127 | 2.58725 | 1.971 |
| chr15 | 71418352 | 71418813 | 0.851881 | 1.67913 | 1.971 |
| chr2 | 64715143 | 64715473 | 2.42233 | 4.7747 | 1.971 |
| chr1 | 223809847 | 223810080 | 0.542458 | 1.06928 | 1.971 |
| chr17 | 39894242 | 39894567 | 1.804 | 3.55613 | 1.971 |
| chr2 | 45812460 | 45812696 | 1.73558 | 3.42134 | 1.971 |
| chr10 | 30147371 | 30147662 | 1.79842 | 3.54533 | 1.971 |
| chr7 | 149119391 | 149119817 | 1.968 | 3.87964 | 1.971 |
| chr13 | 27949894 | 27950207 | 1.61589 | 3.18558 | 1.971 |
| chr1 | 27932195 | 27932538 | 0.475546 | 0.937502 | 1.971 |
| chr5 | 171880239 | 171880491 | 0.813336 | 1.60345 | 1.971 |
| chr1 | 29310203 | 29310424 | 0.653125 | 1.28764 | 1.972 |
| chr13 | 114112651 | 114113105 | 0.794661 | 1.5667 | 1.972 |
| chr11 | 88285632 | 88285931 | 1.72784 | 3.40651 | 1.972 |
| chr12 | 68434269 | 68434619 | 1.9019 | 3.74977 | 1.972 |
| chr11 | 38502903 | 38503127 | 0.529229 | 1.04343 | 1.972 |
| chr15 | 98110787 | 98111019 | 0.60195 | 1.18681 | 1.972 |
| chr11 | 32180367 | 32180642 | 0.627291 | 1.23682 | 1.972 |
| chr15 | 86303694 | 86303980 | 1.50222 | 2.96192 | 1.972 |
| chr2 | 134965236 | 134965474 | 1.38025 | 2.72152 | 1.972 |
| chr8 | 27449744 | 27450191 | 2.12692 | 4.19398 | 1.972 |
| chr11 | 124745942 | 124746419 | 1.10533 | 2.17959 | 1.972 |
| chr6 | 46749266 | 46749495 | 1.39269 | 2.74627 | 1.972 |
| chr11 | 127215799 | 127216119 | 1.12166 | 2.21184 | 1.972 |
| chr12 | 23981969 | 23982293 | 0.576993 | 1.13779 | 1.972 |
| chr7 | 1105383 | 1105820 | 2.05707 | 4.05645 | 1.972 |
| chr5 | 92335289 | 92335533 | 0.494383 | 0.974964 | 1.972 |
| chr17 | 46891838 | 46892162 | 2.01799 | 3.9797 | 1.972 |
| chr7 | 37354467 | 37354677 | 0.562963 | 1.11023 | 1.972 |
| chr17 | 9432331 | 9432651 | 1.38482 | 2.73104 | 1.972 |
| chr14 | 49928564 | 49928835 | 1.95084 | 3.84734 | 1.972 |
| chr3 | 37254480 | 37254807 | 1.72244 | 3.397 | 1.972 |
| chr10 | 1245867 | 1246072 | 0.500313 | 0.98672 | 1.972 |
| chr5 | 22709000 | 22709312 | 0.563797 | 1.11193 | 1.972 |
| chr2 | 47615192 | 47615808 | 1.28708 | 2.53847 | 1.972 |
| chr14 | 88273818 | 88274162 | 0.493515 | 0.973399 | 1.972 |
| chr1 | 11710459 | 11710825 | 1.47477 | 2.90883 | 1.972 |
| chr6 | 79519960 | 79520190 | 1.15133 | 2.27104 | 1.973 |
| chr12 | 1692689 | 1692969 | 1.26538 | 2.49606 | 1.973 |
| chr6 | 127125419 | 127125767 | 1.69764 | 3.34879 | 1.973 |
| chr12 | 14291513 | 14291717 | 0.980278 | 1.93376 | 1.973 |
| chr19 | 31692737 | 31692940 | 0.622724 | 1.22843 | 1.973 |
| chr11 | 74733924 | 74734348 | 1.77003 | 3.49186 | 1.973 |

| | | | | | |
|---|---|---|---|---|---|
| chrX | 41640835 | 41641059 | 1.03193 | 2.03576 | 1.973 |
| chr1 | 214961573 | 214961895 | 0.512308 | 1.0107 | 1.973 |
| chr7 | 31853874 | 31854154 | 1.12054 | 2.21065 | 1.973 |
| chr3 | 72919308 | 72919570 | 0.53392 | 1.05335 | 1.973 |
| chr6 | 132600661 | 132601211 | 1.5526 | 3.06326 | 1.973 |
| chr3 | 51703287 | 51703649 | 1.72904 | 3.41153 | 1.973 |
| chr8 | 143039973 | 143040226 | 0.573106 | 1.13082 | 1.973 |
| chr17 | 52485041 | 52485299 | 0.667214 | 1.31651 | 1.973 |
| chr5 | 149940003 | 149940821 | 1.03969 | 2.05146 | 1.973 |
| chr8 | 16384984 | 16385509 | 0.59653 | 1.17716 | 1.973 |
| chr13 | 102568788 | 102568992 | 0.834655 | 1.64708 | 1.973 |
| chr7 | 37238431 | 37238936 | 1.60418 | 3.16577 | 1.973 |
| chr15 | 73088985 | 73089467 | 1.71696 | 3.38835 | 1.973 |
| chr12 | 129404673 | 129404905 | 0.654477 | 1.29171 | 1.974 |
| chr1 | 63793256 | 63793651 | 1.47209 | 2.90551 | 1.974 |
| chr20 | 60877619 | 60877969 | 1.70208 | 3.35956 | 1.974 |
| chr7 | 108678868 | 108679081 | 0.667754 | 1.31802 | 1.974 |
| chr12 | 48160987 | 48161279 | 1.80388 | 3.56056 | 1.974 |
| chr5 | 144433299 | 144433560 | 0.579936 | 1.14472 | 1.974 |
| chr16 | 87622332 | 87622807 | 1.19007 | 2.34915 | 1.974 |
| chr12 | 124277720 | 124277944 | 0.930921 | 1.83764 | 1.974 |
| chr20 | 36944723 | 36944929 | 0.46755 | 0.923039 | 1.974 |
| chr5 | 42842363 | 42842571 | 0.556105 | 1.09788 | 1.974 |
| chr17 | 65821206 | 65821701 | 1.72965 | 3.41477 | 1.974 |
| chr2 | 161826811 | 161827025 | 1.12635 | 2.22371 | 1.974 |
| chr13 | 98283515 | 98283746 | 0.664848 | 1.31263 | 1.974 |
| chr18 | 29265149 | 29265456 | 0.733013 | 1.44725 | 1.974 |
| chr10 | 97205630 | 97205926 | 1.33443 | 2.63473 | 1.974 |
| chr14 | 55249501 | 55249744 | 0.766965 | 1.5144 | 1.975 |
| chr13 | 65173516 | 65173733 | 0.505949 | 0.999029 | 1.975 |
| chrX | 96940872 | 96941101 | 1.27958 | 2.52665 | 1.975 |
| chr15 | 37175902 | 37176171 | 1.5931 | 3.14576 | 1.975 |
| chr21 | 43565639 | 43566044 | 1.5392 | 3.0394 | 1.975 |
| chr12 | 2391865 | 2392121 | 0.485654 | 0.959027 | 1.975 |
| chr8 | 38544654 | 38544873 | 0.615101 | 1.21465 | 1.975 |
| chr16 | 87604729 | 87605223 | 1.04934 | 2.07216 | 1.975 |
| chr17 | 1171652 | 1171905 | 0.978928 | 1.93325 | 1.975 |
| chr9 | 136658103 | 136658481 | 2.45932 | 4.8569 | 1.975 |
| chr7 | 86866172 | 86866622 | 1.28848 | 2.54464 | 1.975 |
| chr12 | 52071884 | 52072242 | 1.95391 | 3.85895 | 1.975 |
| chr7 | 158187644 | 158187979 | 1.67466 | 3.30754 | 1.975 |
| chr6 | 131855382 | 131855615 | 1.08337 | 2.13974 | 1.975 |
| chr17 | 38948064 | 38948422 | 1.87323 | 3.69996 | 1.975 |
| chr1 | 216608900 | 216609194 | 0.447017 | 0.882945 | 1.975 |
| chr19 | 31268895 | 31269153 | 0.571895 | 1.12963 | 1.975 |
| chr17 | 56277461 | 56277787 | 1.43639 | 2.83725 | 1.975 |
| chr13 | 77900683 | 77900960 | 1.05946 | 2.09274 | 1.975 |
| chr15 | 67725907 | 67726203 | 1.45764 | 2.87938 | 1.975 |
| chr8 | 6698110 | 6698508 | 2.3784 | 4.69823 | 1.975 |
| chr17 | 61926511 | 61926935 | 2.28532 | 4.5144 | 1.975 |
| chr10 | 11861866 | 11862105 | 1.12455 | 2.22144 | 1.975 |
| chr14 | 72172197 | 72172540 | 0.690222 | 1.36347 | 1.975 |
| chr10 | 32125719 | 32126078 | 1.90736 | 3.76789 | 1.975 |
| chr5 | 31313440 | 31313698 | 0.612201 | 1.20939 | 1.975 |
| chr15 | 69557696 | 69558034 | 1.45956 | 2.88334 | 1.975 |
| chr1 | 237970113 | 237970504 | 1.20584 | 2.38214 | 1.976 |
| chr19 | 46196547 | 46197027 | 2.27472 | 4.49376 | 1.976 |
| chr1 | 23963970 | 23964274 | 1.70059 | 3.35956 | 1.976 |
| chr4 | 37385241 | 37385543 | 0.79564 | 1.57183 | 1.976 |
| chr7 | 13316658 | 13316922 | 1.04833 | 2.07112 | 1.976 |
| chr12 | 62399804 | 62400303 | 0.984188 | 1.94442 | 1.976 |

| | | | | | |
|---|---|---|---|---|---|
| chr7 | 150709911 | 150710392 | 1.39844 | 2.7629 | 1.976 |
| chr1 | 242161703 | 242162184 | 1.51619 | 2.99567 | 1.976 |
| chr17 | 7976920 | 7977480 | 2.16963 | 4.28678 | 1.976 |
| chr6 | 3056351 | 3056570 | 0.498013 | 0.984 | 1.976 |
| chr3 | 162900555 | 162900818 | 0.423855 | 0.837486 | 1.976 |
| chr7 | 51584559 | 51584772 | 0.443733 | 0.876761 | 1.976 |
| chr17 | 42600551 | 42600825 | 1.67992 | 3.31935 | 1.976 |
| chr22 | 37153641 | 37154082 | 1.76499 | 3.48747 | 1.976 |
| chrX | 15511669 | 15512043 | 0.889566 | 1.75776 | 1.976 |
| chr20 | 52720014 | 52720306 | 2.02768 | 4.00676 | 1.976 |
| chr20 | 30606940 | 30607229 | 0.590182 | 1.16627 | 1.976 |
| chr1 | 31635548 | 31635849 | 1.77455 | 3.50682 | 1.976 |
| chr9 | 34403775 | 34404236 | 1.88417 | 3.72353 | 1.976 |
| chr12 | 91461947 | 91462229 | 0.495137 | 0.978516 | 1.976 |
| chr13 | 95841683 | 95841889 | 1.12688 | 2.22711 | 1.976 |
| chr20 | 33124778 | 33125207 | 1.47907 | 2.92317 | 1.976 |
| chr1 | 215893546 | 215893869 | 0.457858 | 0.904923 | 1.976 |
| chr21 | 45717544 | 45717895 | 1.61336 | 3.18869 | 1.976 |
| chr4 | 56212444 | 56212983 | 1.47706 | 2.91937 | 1.976 |
| chr2 | 47589367 | 47589593 | 0.593699 | 1.17347 | 1.977 |
| chr2 | 17827398 | 17827675 | 1.89116 | 3.73798 | 1.977 |
| chr7 | 23610224 | 23610602 | 1.42473 | 2.81606 | 1.977 |
| chr16 | 23566667 | 23566917 | 0.685717 | 1.35537 | 1.977 |
| chr16 | 88851788 | 88852275 | 2.06148 | 4.07473 | 1.977 |
| chr20 | 30605622 | 30605965 | 0.733565 | 1.45003 | 1.977 |
| chr3 | 5772691 | 5773085 | 0.75818 | 1.49872 | 1.977 |
| chr13 | 30046476 | 30046890 | 2.60909 | 5.15749 | 1.977 |
| chr5 | 144064764 | 144064996 | 0.644906 | 1.27481 | 1.977 |
| chr3 | 164924522 | 164924914 | 2.05324 | 4.059 | 1.977 |
| chr11 | 29365222 | 29365678 | 2.33582 | 4.61776 | 1.977 |
| chr14 | 90048403 | 90048722 | 0.440239 | 0.870339 | 1.977 |
| chr3 | 42485320 | 42485704 | 2.01884 | 3.99123 | 1.977 |
| chr12 | 120375849 | 120376385 | 1.26749 | 2.50584 | 1.977 |
| chr1 | 233530666 | 233530921 | 1.68213 | 3.32561 | 1.977 |
| chr1 | 6240463 | 6240675 | 0.718866 | 1.42128 | 1.977 |
| chr14 | 76522559 | 76522880 | 1.01669 | 2.01011 | 1.977 |
| chr5 | 3856214 | 3856475 | 0.650675 | 1.2866 | 1.977 |
| chr11 | 66246404 | 66246753 | 1.42531 | 2.81833 | 1.977 |
| chr11 | 22869853 | 22870093 | 1.22811 | 2.42854 | 1.977 |
| chr20 | 4153097 | 4153519 | 4.02986 | 7.96952 | 1.978 |
| chr7 | 48376689 | 48376909 | 1.054 | 2.08443 | 1.978 |
| chr10 | 94840038 | 94840559 | 1.7483 | 3.45752 | 1.978 |
| chr5 | 77763606 | 77763974 | 1.57094 | 3.10677 | 1.978 |
| chr12 | 52374117 | 52374490 | 2.52048 | 4.9848 | 1.978 |
| chr7 | 97182523 | 97182856 | 0.500067 | 0.98903 | 1.978 |
| chr8 | 47457016 | 47457367 | 0.555356 | 1.0984 | 1.978 |
| chr17 | 49990450 | 49990658 | 0.595316 | 1.17744 | 1.978 |
| chr15 | 70539532 | 70539847 | 0.678893 | 1.34275 | 1.978 |
| chr9 | 115912994 | 115913243 | 0.573737 | 1.13477 | 1.978 |
| chr16 | 24053313 | 24053763 | 2.11932 | 4.19192 | 1.978 |
| chr19 | 38716804 | 38717349 | 2.8867 | 5.71004 | 1.978 |
| chr8 | 28692742 | 28692948 | 1.08967 | 2.15557 | 1.978 |
| chr19 | 42103843 | 42104412 | 2.78061 | 5.50061 | 1.978 |
| chr10 | 57345183 | 57345414 | 0.492234 | 0.973772 | 1.978 |
| chr11 | 68411104 | 68411329 | 0.961018 | 1.90118 | 1.978 |
| chr5 | 137087584 | 137087863 | 1.5771 | 3.11998 | 1.978 |
| chr20 | 44761262 | 44761581 | 2.18105 | 4.3151 | 1.978 |
| chr16 | 30825620 | 30825946 | 2.13231 | 4.21873 | 1.978 |
| chr14 | 40031308 | 40031686 | 0.521812 | 1.03248 | 1.979 |
| chr1 | 156023077 | 156023295 | 0.837638 | 1.65742 | 1.979 |
| chr16 | 78269120 | 78269578 | 2.06204 | 4.08011 | 1.979 |

| | | | | | |
|---|---|---|---|---|---|
| chr17 | 40995368 | 40995746 | 1.90126 | 3.76198 | 1.979 |
| chr17 | 71307036 | 71307472 | 1.81579 | 3.59321 | 1.979 |
| chr1 | 176042767 | 176042998 | 1.1318 | 2.23973 | 1.979 |
| chr14 | 94372985 | 94373376 | 0.839127 | 1.66056 | 1.979 |
| chr10 | 98315821 | 98316137 | 0.892098 | 1.76545 | 1.979 |
| chr17 | 13110262 | 13110479 | 0.509164 | 1.00763 | 1.979 |
| chr10 | 103255233 | 103255461 | 0.870983 | 1.7237 | 1.979 |
| chr2 | 206631530 | 206631834 | 1.48689 | 2.94261 | 1.979 |
| chr17 | 43324321 | 43324596 | 1.03355 | 2.04545 | 1.979 |
| chr8 | 56923149 | 56923514 | 2.76811 | 5.47832 | 1.979 |
| chr3 | 36582421 | 36582841 | 2.09622 | 4.14893 | 1.979 |
| chr12 | 113487923 | 113488193 | 1.67348 | 3.31228 | 1.979 |
| chr15 | 78400592 | 78401103 | 2.39837 | 4.74715 | 1.979 |
| chr5 | 68303580 | 68303956 | 1.79779 | 3.55841 | 1.979 |
| chr19 | 41055094 | 41055351 | 2.05476 | 4.06723 | 1.979 |
| chr8 | 139555394 | 139555826 | 0.507117 | 1.00382 | 1.979 |
| chr12 | 113293261 | 113293495 | 1.03683 | 2.05253 | 1.980 |
| chr11 | 117714689 | 117715022 | 1.42805 | 2.82705 | 1.980 |
| chr11 | 56769198 | 56769470 | 1.01509 | 2.00954 | 1.980 |
| chrX | 107315332 | 107315602 | 1.43583 | 2.84248 | 1.980 |
| chr13 | 39719127 | 39719356 | 1.21984 | 2.41515 | 1.980 |
| chr1 | 206837495 | 206838220 | 1.15911 | 2.29494 | 1.980 |
| chr2 | 46290088 | 46290417 | 2.1631 | 4.28308 | 1.980 |
| chr11 | 74588241 | 74588536 | 1.26021 | 2.49536 | 1.980 |
| chr18 | 22067685 | 22067932 | 1.17385 | 2.32439 | 1.980 |
| chr9 | 347730 | 348067 | 2.05145 | 4.06222 | 1.980 |
| chr10 | 59825070 | 59825321 | 2.19625 | 4.34897 | 1.980 |
| chr2 | 57516840 | 57517042 | 0.541275 | 1.07183 | 1.980 |
| chr16 | 1413399 | 1413874 | 2.51787 | 4.98599 | 1.980 |
| chr14 | 61719544 | 61719894 | 1.48959 | 2.9498 | 1.980 |
| chr8 | 41559212 | 41559593 | 2.06819 | 4.09593 | 1.980 |
| chr9 | 34074226 | 34074576 | 2.16852 | 4.29471 | 1.980 |
| chr14 | 61000262 | 61000524 | 0.545411 | 1.0802 | 1.981 |
| chr16 | 87495065 | 87495273 | 1.05496 | 2.08938 | 1.981 |
| chr3 | 140911058 | 140911503 | 2.0821 | 4.12376 | 1.981 |
| chr14 | 59848214 | 59848463 | 0.508039 | 1.00622 | 1.981 |
| chr4 | 157889703 | 157889970 | 1.49135 | 2.95386 | 1.981 |
| chr5 | 7105527 | 7105775 | 0.795288 | 1.57522 | 1.981 |
| chr15 | 93105846 | 93106050 | 0.637894 | 1.26354 | 1.981 |
| chr1 | 2479411 | 2479829 | 1.75948 | 3.48525 | 1.981 |
| chr5 | 109289195 | 109289524 | 0.539399 | 1.06851 | 1.981 |
| chr1 | 202104338 | 202104615 | 1.66326 | 3.29491 | 1.981 |
| chr2 | 225866747 | 225867057 | 1.21243 | 2.4019 | 1.981 |
| chr9 | 27247542 | 27248039 | 1.12914 | 2.23693 | 1.981 |
| chr18 | 35117800 | 35118179 | 1.52642 | 3.02404 | 1.981 |
| chr19 | 57139199 | 57139550 | 0.565358 | 1.12012 | 1.981 |
| chr7 | 156609752 | 156609974 | 1.10405 | 2.18741 | 1.981 |
| chr22 | 29168968 | 29169172 | 0.814341 | 1.61343 | 1.981 |
| chr9 | 23580034 | 23580322 | 0.524247 | 1.03869 | 1.981 |
| chr11 | 116146894 | 116147651 | 2.39463 | 4.74493 | 1.981 |
| chr6 | 39590463 | 39590697 | 0.468391 | 0.928157 | 1.982 |
| chr12 | 56660527 | 56660753 | 1.3213 | 2.61829 | 1.982 |
| chr5 | 2017770 | 2018085 | 0.969873 | 1.9219 | 1.982 |
| chr12 | 13069189 | 13069542 | 2.10385 | 4.16909 | 1.982 |
| chr6 | 44067880 | 44068099 | 0.816716 | 1.61851 | 1.982 |
| chr6 | 108840197 | 108840530 | 0.951778 | 1.88618 | 1.982 |
| chr5 | 156898634 | 156899116 | 1.32168 | 2.61924 | 1.982 |
| chr3 | 72715350 | 72715745 | 1.30588 | 2.58798 | 1.982 |
| chr7 | 142491109 | 142491438 | 2.02941 | 4.02187 | 1.982 |
| chr14 | 55030798 | 55031005 | 0.869048 | 1.72228 | 1.982 |
| chr17 | 47573538 | 47574240 | 1.88104 | 3.72785 | 1.982 |

| | | | | | |
|---|---|---|---|---|---|
| chr8 | 143453558 | 143453967 | 0.588867 | 1.16705 | 1.982 |
| chr16 | 1234554 | 1234842 | 1.00496 | 1.9917 | 1.982 |
| chr11 | 18210606 | 18210842 | 0.404825 | 0.802319 | 1.982 |
| chr2 | 59283444 | 59283656 | 0.383385 | 0.759857 | 1.982 |
| chr15 | 100604601 | 100604962 | 0.536234 | 1.06281 | 1.982 |
| chr9 | 101761052 | 101761570 | 1.72932 | 3.42754 | 1.982 |
| chr9 | 35826008 | 35826572 | 1.27847 | 2.53396 | 1.982 |
| chr10 | 133782400 | 133782642 | 0.477248 | 0.94596 | 1.982 |
| chrX | 68378341 | 68378590 | 1.27012 | 2.51754 | 1.982 |
| chr8 | 97507429 | 97507667 | 0.628931 | 1.24663 | 1.982 |
| chr5 | 176763174 | 176763475 | 1.85532 | 3.67758 | 1.982 |
| chr17 | 48238837 | 48239228 | 3.00562 | 5.95775 | 1.982 |
| chr11 | 71949078 | 71949683 | 1.54168 | 3.05604 | 1.982 |
| chr5 | 142097151 | 142097418 | 1.80504 | 3.57817 | 1.982 |
| chr3 | 48989980 | 48990320 | 0.781698 | 1.5496 | 1.982 |
| chr2 | 118593558 | 118593853 | 1.99406 | 3.95302 | 1.982 |
| chr13 | 99325157 | 99325577 | 1.15753 | 2.29476 | 1.982 |
| chr9 | 3181675 | 3181877 | 1.02527 | 2.03259 | 1.982 |
| chr3 | 11130481 | 11130766 | 0.433847 | 0.860101 | 1.983 |
| chr7 | 85093523 | 85093788 | 0.672475 | 1.33319 | 1.983 |
| chr16 | 67548295 | 67548589 | 1.63177 | 3.23527 | 1.983 |
| chr6 | 52859935 | 52860241 | 1.40848 | 2.79266 | 1.983 |
| chr17 | 30451902 | 30452311 | 0.477945 | 0.94768 | 1.983 |
| chr15 | 100540093 | 100540380 | 0.405999 | 0.80503 | 1.983 |
| chr11 | 64216989 | 64217539 | 2.47042 | 4.89861 | 1.983 |
| chr11 | 448218 | 448581 | 2.52058 | 4.99814 | 1.983 |
| chr19 | 45055535 | 45055986 | 1.65481 | 3.28146 | 1.983 |
| chr1 | 100151353 | 100151588 | 1.27411 | 2.52655 | 1.983 |
| chr4 | 65040116 | 65040468 | 0.638728 | 1.26661 | 1.983 |
| chr9 | 120359802 | 120360005 | 0.428696 | 0.850113 | 1.983 |
| chr2 | 161067170 | 161067529 | 2.15066 | 4.2649 | 1.983 |
| chr15 | 93155238 | 93155808 | 0.776938 | 1.54075 | 1.983 |
| chr5 | 12731387 | 12731639 | 0.522631 | 1.03648 | 1.983 |
| chr20 | 48225185 | 48225899 | 2.26078 | 4.48371 | 1.983 |
| chr3 | 37585419 | 37585727 | 0.596737 | 1.18349 | 1.983 |
| chr9 | 115739995 | 115740286 | 0.564305 | 1.11917 | 1.983 |
| chr5 | 13019954 | 13020231 | 0.531691 | 1.05449 | 1.983 |
| chr16 | 47178172 | 47178577 | 1.06504 | 2.11229 | 1.983 |
| chr9 | 37262489 | 37262736 | 1.60144 | 3.17624 | 1.983 |
| chr7 | 29917734 | 29918045 | 1.62619 | 3.22534 | 1.983 |
| chr20 | 52560296 | 52560610 | 1.26203 | 2.50312 | 1.983 |
| chr8 | 4284547 | 4284792 | 0.556401 | 1.10357 | 1.983 |
| chr7 | 33925827 | 33926098 | 1.08561 | 2.15322 | 1.983 |
| chr1 | 156890200 | 156890548 | 2.42094 | 4.80187 | 1.983 |
| chr11 | 118777780 | 118778327 | 0.923882 | 1.83249 | 1.983 |
| chr10 | 65527329 | 65527585 | 1.02378 | 2.03065 | 1.983 |
| chr12 | 110906904 | 110907294 | 2.63128 | 5.21912 | 1.983 |
| chr15 | 71735852 | 71736119 | 1.91214 | 3.79273 | 1.984 |
| chr4 | 36401481 | 36401700 | 1.20247 | 2.38512 | 1.984 |
| chr15 | 75074121 | 75074544 | 2.4535 | 4.86682 | 1.984 |
| chr10 | 54074098 | 54074551 | 2.37646 | 4.7141 | 1.984 |
| chr3 | 148775968 | 148776185 | 0.56258 | 1.11597 | 1.984 |
| chr8 | 10682826 | 10683223 | 1.33253 | 2.64331 | 1.984 |
| chr3 | 47563649 | 47563914 | 1.87971 | 3.72886 | 1.984 |
| chr14 | 95826473 | 95826964 | 0.62508 | 1.24003 | 1.984 |
| chr11 | 118478570 | 118478823 | 0.963311 | 1.91105 | 1.984 |
| chr1 | 231153297 | 231153517 | 0.382701 | 0.759287 | 1.984 |
| chr11 | 12936073 | 12936519 | 1.96921 | 3.90711 | 1.984 |
| chr12 | 24048720 | 24048936 | 0.538075 | 1.0676 | 1.984 |
| chr9 | 140339724 | 140340067 | 2.02075 | 4.00948 | 1.984 |
| chr7 | 129794886 | 129795340 | 1.10242 | 2.18739 | 1.984 |

| | | | | | |
|---|---|---|---|---|---|
| chr3 | 74940522 | 74940733 | 0.753122 | 1.49433 | 1.984 |
| chr3 | 169906953 | 169907241 | 1.83657 | 3.6441 | 1.984 |
| chr5 | 114433782 | 114434096 | 1.11083 | 2.20415 | 1.984 |
| chr14 | 84987344 | 84987547 | 0.617006 | 1.22432 | 1.984 |
| chr18 | 47294459 | 47294676 | 1.16114 | 2.30411 | 1.984 |
| chr12 | 48137783 | 48138167 | 2.02067 | 4.00983 | 1.984 |
| chr15 | 52475819 | 52476148 | 1.6385 | 3.2515 | 1.984 |
| chr16 | 11625988 | 11626369 | 2.16668 | 4.2998 | 1.985 |
| chr11 | 74345752 | 74345974 | 1.21021 | 2.40176 | 1.985 |
| chr17 | 48275438 | 48275999 | 1.95655 | 3.88311 | 1.985 |
| chr17 | 36648314 | 36648528 | 2.35252 | 4.66906 | 1.985 |
| chr20 | 55667107 | 55667672 | 1.92632 | 3.82325 | 1.985 |
| chr17 | 72375483 | 72375968 | 2.37927 | 4.72225 | 1.985 |
| chr16 | 6184531 | 6184739 | 0.41143 | 0.81659 | 1.985 |
| chr18 | 34267992 | 34268359 | 1.52398 | 3.02484 | 1.985 |
| chr2 | 85934322 | 85934662 | 1.43936 | 2.85703 | 1.985 |
| chr7 | 150725748 | 150725981 | 1.17863 | 2.33951 | 1.985 |
| chr3 | 8225341 | 8225686 | 0.505377 | 1.00315 | 1.985 |
| chr9 | 140589632 | 140589992 | 1.60825 | 3.19233 | 1.985 |
| chr12 | 112802309 | 112802513 | 0.508052 | 1.0085 | 1.985 |
| chr3 | 152242377 | 152242655 | 1.27925 | 2.53935 | 1.985 |
| chr9 | 36205285 | 36205567 | 0.999142 | 1.98333 | 1.985 |
| chr11 | 94884691 | 94884977 | 1.50969 | 2.99681 | 1.985 |
| chr2 | 27946225 | 27946581 | 1.58364 | 3.14361 | 1.985 |
| chr11 | 27507469 | 27507798 | 1.96284 | 3.89664 | 1.985 |
| chr6 | 36818763 | 36819222 | 1.26396 | 2.50924 | 1.985 |
| chr11 | 58825968 | 58826272 | 1.76585 | 3.5058 | 1.985 |
| chr19 | 11353802 | 11354283 | 1.42078 | 2.82072 | 1.985 |
| chr10 | 51565005 | 51565224 | 1.26571 | 2.51295 | 1.985 |
| chr5 | 158338100 | 158338325 | 0.448197 | 0.889857 | 1.985 |
| chr14 | 77534543 | 77534909 | 1.82571 | 3.62498 | 1.986 |
| chr20 | 52816213 | 52816496 | 2.90033 | 5.75873 | 1.986 |
| chr7 | 36676928 | 36677379 | 0.492173 | 0.977234 | 1.986 |
| chr14 | 21325812 | 21326222 | 1.73189 | 3.4388 | 1.986 |
| chr4 | 91236304 | 91236542 | 1.99453 | 3.96043 | 1.986 |
| chr7 | 41920105 | 41920361 | 0.412582 | 0.819254 | 1.986 |
| chr3 | 34640836 | 34641151 | 0.512652 | 1.01798 | 1.986 |
| chr13 | 77576763 | 77577044 | 1.07084 | 2.12646 | 1.986 |
| chr8 | 142215937 | 142216246 | 0.59237 | 1.17633 | 1.986 |
| chr12 | 58015380 | 58015908 | 2.63224 | 5.22738 | 1.986 |
| chr12 | 109956574 | 109956994 | 1.39641 | 2.77335 | 1.986 |
| chr2 | 165871517 | 165871909 | 3.05518 | 6.06779 | 1.986 |
| chr1 | 231451589 | 231451951 | 1.39544 | 2.77151 | 1.986 |
| chr19 | 41480448 | 41480960 | 1.89127 | 3.75633 | 1.986 |
| chr16 | 69893569 | 69893812 | 1.16819 | 2.32024 | 1.986 |
| chr20 | 18361838 | 18362055 | 0.534767 | 1.06215 | 1.986 |
| chr2 | 41217995 | 41218236 | 0.417609 | 0.829482 | 1.986 |
| chr2 | 25104689 | 25104910 | 0.86087 | 1.70992 | 1.986 |
| chr7 | 44237488 | 44237703 | 0.3905 | 0.775639 | 1.986 |
| chr5 | 4214514 | 4214764 | 0.374578 | 0.744053 | 1.986 |
| chr8 | 30810678 | 30810927 | 0.508825 | 1.01072 | 1.986 |
| chr15 | 38991881 | 38992160 | 1.38033 | 2.7419 | 1.986 |
| chr14 | 104372330 | 104372548 | 0.888818 | 1.76559 | 1.986 |
| chr19 | 733996 | 734406 | 1.50201 | 2.98369 | 1.986 |
| chr13 | 96130825 | 96131293 | 1.12198 | 2.22887 | 1.987 |
| chr19 | 23348351 | 23348573 | 0.499744 | 0.992768 | 1.987 |
| chr12 | 96213970 | 96214251 | 1.14337 | 2.27142 | 1.987 |
| chr1 | 236272617 | 236273017 | 1.12061 | 2.22621 | 1.987 |
| chr6 | 149753116 | 149753437 | 1.46706 | 2.9145 | 1.987 |
| chr1 | 51810661 | 51810895 | 0.919241 | 1.82621 | 1.987 |
| chr8 | 28213109 | 28213372 | 1.56859 | 3.11645 | 1.987 |

| | | | | | |
|---|---|---|---|---|---|
| chr5 | 137404479 | 137404891 | 1.548 | 3.07558 | 1.987 |
| chr17 | 51511434 | 51511722 | 0.697047 | 1.38494 | 1.987 |
| chr7 | 4650456 | 4650822 | 1.73148 | 3.44026 | 1.987 |
| chr7 | 35946026 | 35946412 | 1.40234 | 2.78641 | 1.987 |
| chr8 | 72203365 | 72203609 | 0.682222 | 1.3556 | 1.987 |
| chr12 | 128554031 | 128554236 | 1.19458 | 2.37375 | 1.987 |
| chr16 | 596829 | 597462 | 1.51054 | 3.00159 | 1.987 |
| chr19 | 46273340 | 46273794 | 1.96291 | 3.90085 | 1.987 |
| chr11 | 29350537 | 29350807 | 1.09089 | 2.16791 | 1.987 |
| chr19 | 3545075 | 3545652 | 2.26383 | 4.49903 | 1.987 |
| chr15 | 75101362 | 75101595 | 0.676673 | 1.3448 | 1.987 |
| chr8 | 23145333 | 23145819 | 3.37228 | 6.70231 | 1.987 |
| chr20 | 33357414 | 33357860 | 0.997911 | 1.98343 | 1.988 |
| chr17 | 80452581 | 80452889 | 2.88304 | 5.7303 | 1.988 |
| chr9 | 136055329 | 136055770 | 1.85155 | 3.68013 | 1.988 |
| chr19 | 6862815 | 6863079 | 1.46582 | 2.91347 | 1.988 |
| chr4 | 78161707 | 78161928 | 0.921399 | 1.83155 | 1.988 |
| chr17 | 40474775 | 40475175 | 2.6034 | 5.17511 | 1.988 |
| chr22 | 19949680 | 19950088 | 1.50662 | 2.99491 | 1.988 |
| chr7 | 45194236 | 45194643 | 1.36412 | 2.71164 | 1.988 |
| chr8 | 2222871 | 2223260 | 0.893541 | 1.77626 | 1.988 |
| chr5 | 102894417 | 102894838 | 0.525822 | 1.04528 | 1.988 |
| chr17 | 77719925 | 77720270 | 2.04623 | 4.06785 | 1.988 |
| chr7 | 67090169 | 67090549 | 1.90608 | 3.78924 | 1.988 |
| chr15 | 29395585 | 29396302 | 2.17822 | 4.33031 | 1.988 |
| chr17 | 51004172 | 51004428 | 0.58812 | 1.16918 | 1.988 |
| chr17 | 5490825 | 5491123 | 1.81333 | 3.60491 | 1.988 |
| chr16 | 89234577 | 89234852 | 0.57313 | 1.1394 | 1.988 |
| chr8 | 61759993 | 61760288 | 0.930257 | 1.84938 | 1.988 |
| chr6 | 149881916 | 149882189 | 0.564243 | 1.12174 | 1.988 |
| chr3 | 124153137 | 124153420 | 0.659159 | 1.31045 | 1.988 |
| chr2 | 67878013 | 67878277 | 1.3368 | 2.65772 | 1.988 |
| chr3 | 17454487 | 17454756 | 1.66666 | 3.31357 | 1.988 |
| chr9 | 131625825 | 131626245 | 2.54863 | 5.06714 | 1.988 |
| chr7 | 2517216 | 2517803 | 1.96722 | 3.91122 | 1.988 |
| chr5 | 982611 | 982936 | 1.9228 | 3.82304 | 1.988 |
| chr8 | 99182227 | 99182664 | 1.84977 | 3.67795 | 1.988 |
| chr14 | 100849908 | 100850425 | 1.72832 | 3.43664 | 1.988 |
| chr19 | 1671914 | 1672622 | 1.97667 | 3.93055 | 1.988 |
| chr1 | 163207020 | 163207514 | 0.509589 | 1.01333 | 1.989 |
| chr14 | 46959574 | 46959911 | 0.568765 | 1.13102 | 1.989 |
| chr5 | 139267836 | 139268162 | 0.697895 | 1.3878 | 1.989 |
| chr9 | 38160591 | 38160977 | 2.29858 | 4.57097 | 1.989 |
| chr10 | 2301243 | 2301495 | 0.387935 | 0.771454 | 1.989 |
| chr6 | 52861510 | 52861828 | 1.56707 | 3.11633 | 1.989 |
| chr12 | 23883976 | 23884241 | 0.633736 | 1.26029 | 1.989 |
| chr3 | 127402587 | 127402844 | 0.836405 | 1.66333 | 1.989 |
| chr7 | 11383426 | 11383735 | 1.26233 | 2.5104 | 1.989 |
| chr20 | 5891951 | 5892233 | 1.09523 | 2.17815 | 1.989 |
| chr16 | 22195335 | 22195592 | 0.911754 | 1.81332 | 1.989 |
| chr3 | 25090350 | 25090553 | 0.519677 | 1.03358 | 1.989 |
| chr1 | 225999368 | 225999677 | 1.27073 | 2.52742 | 1.989 |
| chr16 | 89374826 | 89375130 | 1.16384 | 2.31486 | 1.989 |
| chr7 | 101151026 | 101151514 | 2.11149 | 4.19986 | 1.989 |
| chr12 | 33586762 | 33586998 | 0.411968 | 0.819452 | 1.989 |
| chr11 | 107879430 | 107879747 | 1.20621 | 2.39931 | 1.989 |
| chr12 | 95128516 | 95129019 | 0.558636 | 1.11122 | 1.989 |
| chr19 | 47165287 | 47165640 | 2.4149 | 4.8037 | 1.989 |
| chr7 | 54732012 | 54732326 | 1.17464 | 2.33664 | 1.989 |
| chr5 | 154191267 | 154191930 | 1.74972 | 3.48067 | 1.989 |
| chr4 | 84692287 | 84692543 | 1.18201 | 2.35136 | 1.989 |

| | | | | | |
|---|---|---|---|---|---|
| chr20 | 46365769 | 46365987 | 0.470164 | 0.935308 | 1.989 |
| chr7 | 154244022 | 154244264 | 0.523875 | 1.04216 | 1.989 |
| chr1 | 15827987 | 15828254 | 1.67201 | 3.32642 | 1.989 |
| chr14 | 95897570 | 95897946 | 1.07545 | 2.13962 | 1.990 |
| chr12 | 3113916 | 3114150 | 1.06276 | 2.11439 | 1.990 |
| chr3 | 2661805 | 2662178 | 0.551745 | 1.09776 | 1.990 |
| chr5 | 12234206 | 12234424 | 0.600322 | 1.19442 | 1.990 |
| chr5 | 153128282 | 153128494 | 0.618994 | 1.23159 | 1.990 |
| chr12 | 44409232 | 44409486 | 1.61378 | 3.21097 | 1.990 |
| chr7 | 93918341 | 93918652 | 3.20826 | 6.38362 | 1.990 |
| chr8 | 98321985 | 98322188 | 0.676928 | 1.34695 | 1.990 |
| chr20 | 22228752 | 22229004 | 1.61379 | 3.21134 | 1.990 |
| chr7 | 155859964 | 155860235 | 0.484725 | 0.964594 | 1.990 |
| chr19 | 35959572 | 35959991 | 1.90903 | 3.79901 | 1.990 |
| chr3 | 103505174 | 103505451 | 0.70346 | 1.3999 | 1.990 |
| chr10 | 25369890 | 25370266 | 1.58527 | 3.15477 | 1.990 |
| chr12 | 52615845 | 52616180 | 2.14273 | 4.26434 | 1.990 |
| chr7 | 88123787 | 88124080 | 0.447277 | 0.890159 | 1.990 |
| chr9 | 33722097 | 33722454 | 2.73754 | 5.44821 | 1.990 |
| chr17 | 64251249 | 64251465 | 1.28523 | 2.55792 | 1.990 |
| chr15 | 82545366 | 82545684 | 0.59117 | 1.17658 | 1.990 |
| chr14 | 92035009 | 92035318 | 0.904758 | 1.80089 | 1.990 |
| chr4 | 1727970 | 1728336 | 1.69607 | 3.37617 | 1.991 |
| chrX | 114781754 | 114782081 | 1.03427 | 2.05881 | 1.991 |
| chrX | 58052499 | 58052704 | 1.05534 | 2.10082 | 1.991 |
| chr20 | 197016 | 197262 | 0.623853 | 1.24192 | 1.991 |
| chr14 | 91885119 | 91885518 | 2.65777 | 5.29095 | 1.991 |
| chr11 | 72896270 | 72896715 | 1.81961 | 3.62249 | 1.991 |
| chr6 | 150740205 | 150740498 | 2.0946 | 4.17008 | 1.991 |
| chr17 | 77785141 | 77785618 | 1.85883 | 3.70079 | 1.991 |
| chr14 | 37586824 | 37587103 | 1.50713 | 3.00079 | 1.991 |
| chr18 | 23985390 | 23985672 | 1.23022 | 2.44946 | 1.991 |
| chr17 | 11518027 | 11518287 | 0.54047 | 1.07616 | 1.991 |
| chr11 | 72542496 | 72543026 | 2.28228 | 4.54459 | 1.991 |
| chr12 | 120877495 | 120877818 | 1.33556 | 2.65944 | 1.991 |
| chr8 | 33663261 | 33663783 | 1.06918 | 2.129 | 1.991 |
| chr12 | 122906626 | 122907030 | 1.11946 | 2.22915 | 1.991 |
| chr9 | 127087590 | 127087806 | 0.963615 | 1.91882 | 1.991 |
| chr14 | 74744968 | 74745442 | 0.484273 | 0.964392 | 1.991 |
| chr19 | 10501355 | 10501708 | 2.06396 | 4.1104 | 1.992 |
| chr12 | 24382982 | 24383287 | 0.529934 | 1.05538 | 1.992 |
| chr19 | 41304061 | 41304526 | 0.988947 | 1.96955 | 1.992 |
| chr3 | 149956868 | 149957080 | 1.39852 | 2.78536 | 1.992 |
| chr10 | 115713355 | 115713567 | 0.841222 | 1.67543 | 1.992 |
| chr6 | 123159341 | 123159545 | 0.848808 | 1.69059 | 1.992 |
| chr7 | 8405208 | 8405452 | 1.15619 | 2.30288 | 1.992 |
| chr22 | 27801835 | 27802227 | 1.53198 | 3.05142 | 1.992 |
| chr5 | 177591154 | 177591559 | 2.59623 | 5.17143 | 1.992 |
| chr16 | 85336874 | 85337346 | 2.58061 | 5.14038 | 1.992 |
| chr20 | 21985815 | 21986227 | 0.622946 | 1.24088 | 1.992 |
| chr13 | 52423809 | 52424127 | 1.28899 | 2.56773 | 1.992 |
| chr17 | 11783372 | 11783634 | 0.723536 | 1.44133 | 1.992 |
| chr2 | 39455210 | 39455596 | 2.21879 | 4.42005 | 1.992 |
| chr22 | 23881796 | 23882054 | 1.62368 | 3.23462 | 1.992 |
| chr1 | 3236552 | 3236828 | 1.29699 | 2.58381 | 1.992 |
| chr12 | 1695743 | 1695993 | 0.400913 | 0.79873 | 1.992 |
| chr5 | 36690304 | 36690763 | 1.28074 | 2.55187 | 1.993 |
| chr19 | 18699401 | 18699746 | 2.32757 | 4.6377 | 1.993 |
| chr17 | 31203444 | 31204551 | 1.59549 | 3.17904 | 1.993 |
| chr5 | 170845676 | 170846084 | 1.70135 | 3.39001 | 1.993 |
| chr14 | 94402622 | 94402921 | 1.24439 | 2.47951 | 1.993 |

| | | | | | |
|---|---|---|---|---|---|
| chr10 | 110318897 | 110319112 | 0.473697 | 0.943889 | 1.993 |
| chr11 | 36768512 | 36769126 | 2.37556 | 4.73354 | 1.993 |
| chr5 | 13977080 | 13977298 | 1.34633 | 2.68282 | 1.993 |
| chr17 | 60770473 | 60770894 | 1.19655 | 2.38451 | 1.993 |
| chr9 | 73993160 | 73993554 | 0.636774 | 1.26903 | 1.993 |
| chr10 | 73656125 | 73656605 | 2.11702 | 4.21908 | 1.993 |
| chr5 | 139927205 | 139927429 | 0.762981 | 1.52057 | 1.993 |
| chr15 | 67736816 | 67737096 | 0.545424 | 1.087 | 1.993 |
| chr5 | 36744753 | 36745062 | 1.55842 | 3.10599 | 1.993 |
| chr17 | 7199715 | 7200203 | 2.22378 | 4.43232 | 1.993 |
| chr11 | 65308280 | 65308742 | 2.01746 | 4.02114 | 1.993 |
| chr9 | 893731 | 894110 | 1.59543 | 3.17997 | 1.993 |
| chr1 | 24137317 | 24137598 | 1.57162 | 3.13263 | 1.993 |
| chr8 | 39910484 | 39910696 | 0.842863 | 1.68022 | 1.993 |
| chr7 | 98476500 | 98476959 | 2.73706 | 5.45628 | 1.993 |
| chr1 | 50798639 | 50799668 | 1.54968 | 3.08936 | 1.994 |
| chrX | 17393064 | 17393345 | 2.1661 | 4.31828 | 1.994 |
| chr3 | 108742566 | 108742823 | 1.29528 | 2.58232 | 1.994 |
| chr10 | 102921488 | 102921826 | 1.64604 | 3.28168 | 1.994 |
| chr19 | 41536625 | 41536831 | 0.725669 | 1.44675 | 1.994 |
| chr19 | 57996483 | 57996883 | 0.537858 | 1.07233 | 1.994 |
| chr9 | 132382373 | 132382592 | 0.766937 | 1.52906 | 1.994 |
| chr11 | 128729617 | 128730127 | 1.60391 | 3.19779 | 1.994 |
| chr8 | 9226317 | 9226853 | 1.75094 | 3.49112 | 1.994 |
| chr17 | 37828243 | 37828936 | 1.34684 | 2.68548 | 1.994 |
| chr17 | 38769624 | 38769909 | 0.637977 | 1.27207 | 1.994 |
| chr7 | 129937405 | 129937750 | 0.409503 | 0.816516 | 1.994 |
| chr16 | 81479100 | 81479308 | 1.2173 | 2.42725 | 1.994 |
| chr3 | 120501354 | 120501577 | 0.492426 | 0.981913 | 1.994 |
| chr17 | 33416353 | 33416705 | 2.41656 | 4.81877 | 1.994 |
| chr5 | 11523872 | 11524304 | 0.390726 | 0.779222 | 1.994 |
| chr2 | 84910060 | 84910266 | 0.566578 | 1.12998 | 1.994 |
| chr6 | 2938921 | 2939248 | 1.24971 | 2.4925 | 1.994 |
| chr4 | 139792266 | 139792504 | 1.09471 | 2.18337 | 1.994 |
| chr19 | 51281061 | 51281329 | 0.558926 | 1.11489 | 1.995 |
| chr16 | 3078189 | 3078643 | 1.85637 | 3.70293 | 1.995 |
| chr1 | 210534214 | 210534510 | 1.36574 | 2.72431 | 1.995 |
| chr17 | 3026167 | 3026413 | 0.499109 | 0.995614 | 1.995 |
| chr12 | 102694929 | 102695172 | 0.955488 | 1.90606 | 1.995 |
| chr3 | 183100275 | 183100543 | 1.18317 | 2.36026 | 1.995 |
| chr20 | 34099721 | 34100016 | 1.16422 | 2.32263 | 1.995 |
| chr3 | 196564770 | 196565014 | 1.22707 | 2.44809 | 1.995 |
| chr3 | 24965055 | 24965266 | 0.411294 | 0.820564 | 1.995 |
| chr4 | 115728976 | 115729262 | 1.25597 | 2.5058 | 1.995 |
| chr12 | 69809513 | 69809888 | 1.57464 | 3.14176 | 1.995 |
| chr18 | 39694389 | 39694620 | 1.33969 | 2.6733 | 1.995 |
| chr14 | 22518457 | 22518774 | 0.49366 | 0.985085 | 1.995 |
| chr1 | 21544390 | 21544691 | 1.34608 | 2.6861 | 1.996 |
| chr2 | 10212697 | 10213023 | 2.45118 | 4.89146 | 1.996 |
| chr7 | 65619532 | 65619747 | 0.919116 | 1.83415 | 1.996 |
| chr1 | 180723680 | 180723939 | 0.499454 | 0.996693 | 1.996 |
| chr7 | 44253995 | 44254268 | 0.730877 | 1.45852 | 1.996 |
| chr6 | 38997251 | 38997546 | 1.56694 | 3.12702 | 1.996 |
| chr14 | 38652715 | 38653026 | 0.619642 | 1.23661 | 1.996 |
| chr10 | 65626167 | 65626939 | 1.3292 | 2.6527 | 1.996 |
| chrX | 49684294 | 49684699 | 1.5052 | 3.00395 | 1.996 |
| chr10 | 96080073 | 96080369 | 2.10556 | 4.20212 | 1.996 |
| chr4 | 134404819 | 134405067 | 0.965619 | 1.92712 | 1.996 |
| chr14 | 71373856 | 71374316 | 1.77758 | 3.54761 | 1.996 |
| chr19 | 39702917 | 39703258 | 1.44368 | 2.88127 | 1.996 |
| chr3 | 176656830 | 176657110 | 1.1484 | 2.29195 | 1.996 |

| | | | | | |
|---|---|---|---|---|---|
| chr18 | 7272405 | 7272937 | 0.616896 | 1.23123 | 1.996 |
| chr7 | 33652093 | 33652513 | 0.521435 | 1.04071 | 1.996 |
| chr12 | 44050422 | 44050661 | 0.595333 | 1.18827 | 1.996 |
| chr1 | 43539304 | 43539561 | 0.821745 | 1.6402 | 1.996 |
| chr1 | 117027798 | 117028293 | 1.32662 | 2.64801 | 1.996 |
| chr17 | 33469181 | 33469611 | 3.01978 | 6.02784 | 1.996 |
| chr14 | 55679852 | 55680093 | 1.30308 | 2.60116 | 1.996 |
| chr17 | 12053428 | 12053691 | 0.603999 | 1.20571 | 1.996 |
| chr1 | 18561309 | 18561566 | 1.44415 | 2.88294 | 1.996 |
| chr12 | 126832062 | 126832277 | 0.714727 | 1.42686 | 1.996 |
| chr1 | 205225313 | 205225651 | 2.67101 | 5.33243 | 1.996 |
| chr5 | 81521329 | 81521751 | 1.87991 | 3.75308 | 1.996 |
| chr6 | 167555697 | 167556034 | 1.38799 | 2.77103 | 1.996 |
| chr17 | 71653660 | 71654003 | 1.9338 | 3.86082 | 1.996 |
| chr7 | 40985679 | 40985993 | 0.434344 | 0.867189 | 1.997 |
| chr3 | 78203254 | 78203470 | 0.556941 | 1.11198 | 1.997 |
| chr9 | 95913413 | 95913635 | 0.74578 | 1.48905 | 1.997 |
| chr20 | 37179014 | 37179220 | 0.920506 | 1.83799 | 1.997 |
| chrX | 9925891 | 9926101 | 0.873506 | 1.74421 | 1.997 |
| chr15 | 95567755 | 95567998 | 0.996566 | 1.98994 | 1.997 |
| chr8 | 100307751 | 100307980 | 0.627218 | 1.25245 | 1.997 |
| chr2 | 48816095 | 48816658 | 1.17627 | 2.34885 | 1.997 |
| chr8 | 78975767 | 78976023 | 0.588877 | 1.17592 | 1.997 |
| chr1 | 19000535 | 19000756 | 0.859195 | 1.71573 | 1.997 |
| chr6 | 19812789 | 19813091 | 1.21509 | 2.42651 | 1.997 |
| chr4 | 20424180 | 20424395 | 0.839632 | 1.67675 | 1.997 |
| chr17 | 52956814 | 52957020 | 0.780598 | 1.55889 | 1.997 |
| chr3 | 39419567 | 39419918 | 1.42848 | 2.85274 | 1.997 |
| chr5 | 33944546 | 33945103 | 0.453413 | 0.905491 | 1.997 |
| chr4 | 129231520 | 129231729 | 0.670992 | 1.34002 | 1.997 |
| chr7 | 122679158 | 122679762 | 0.519714 | 1.03791 | 1.997 |
| chr19 | 18057159 | 18057463 | 1.645 | 3.28523 | 1.997 |
| chr12 | 65901126 | 65901341 | 0.498143 | 0.994853 | 1.997 |
| chr14 | 77251122 | 77251526 | 2.54883 | 5.09035 | 1.997 |
| chr2 | 25913182 | 25913605 | 0.647212 | 1.29258 | 1.997 |
| chr20 | 56073869 | 56074137 | 1.01972 | 2.03654 | 1.997 |
| chr1 | 44683355 | 44683764 | 1.38089 | 2.75789 | 1.997 |
| chr15 | 40613147 | 40613383 | 2.06108 | 4.1166 | 1.997 |
| chr14 | 70480047 | 70480305 | 1.963 | 3.92077 | 1.997 |
| chr1 | 221766027 | 221766307 | 0.643321 | 1.28494 | 1.997 |
| chr7 | 16197280 | 16197615 | 0.665758 | 1.3298 | 1.997 |
| chr12 | 31402715 | 31402941 | 0.549892 | 1.0984 | 1.997 |
| chr6 | 169284776 | 169285068 | 1.57217 | 3.1404 | 1.997 |
| chr3 | 4753113 | 4753396 | 1.76272 | 3.52111 | 1.998 |
| chr7 | 91481795 | 91482054 | 0.905185 | 1.80818 | 1.998 |
| chr1 | 209422059 | 209422419 | 1.76317 | 3.52209 | 1.998 |
| chr19 | 14159369 | 14159750 | 1.6773 | 3.35075 | 1.998 |
| chr5 | 170911420 | 170911859 | 1.56622 | 3.12887 | 1.998 |
| chr8 | 55081940 | 55082175 | 1.36438 | 2.72567 | 1.998 |
| chr1 | 235257152 | 235257512 | 1.96196 | 3.91951 | 1.998 |
| chr16 | 49773332 | 49773629 | 0.679181 | 1.35685 | 1.998 |
| chr1 | 182753767 | 182754018 | 2.07501 | 4.14551 | 1.998 |
| chr12 | 3053637 | 3054121 | 1.52669 | 3.05006 | 1.998 |
| chr11 | 16635232 | 16635545 | 1.65166 | 3.29992 | 1.998 |
| chr6 | 101562544 | 101562804 | 0.927134 | 1.85237 | 1.998 |
| chr16 | 31117013 | 31117361 | 2.64394 | 5.28285 | 1.998 |
| chr14 | 103984804 | 103985367 | 2.26632 | 4.52835 | 1.998 |
| chr5 | 158942909 | 158943194 | 1.73231 | 3.46138 | 1.998 |
| chr6 | 27046648 | 27046932 | 1.05634 | 2.11074 | 1.998 |
| chr17 | 959021 | 959657 | 1.7465 | 3.48987 | 1.998 |
| chr12 | 118292108 | 118292587 | 1.90868 | 3.81418 | 1.998 |

| | | | | | |
|---|---|---|---|---|---|
| chr5 | 66452829 | 66453173 | 1.37924 | 2.75617 | 1.998 |
| chr5 | 7857071 | 7857322 | 1.58761 | 3.17261 | 1.998 |
| chr5 | 106179540 | 106179855 | 0.723864 | 1.44665 | 1.999 |
| chr5 | 170769412 | 170769680 | 1.28455 | 2.56721 | 1.999 |
| chr3 | 13926811 | 13927298 | 1.97164 | 3.9404 | 1.999 |
| chr15 | 38184034 | 38184266 | 1.30547 | 2.60913 | 1.999 |
| chr17 | 75276979 | 75277431 | 1.54682 | 3.0915 | 1.999 |
| chr14 | 86990611 | 86990941 | 0.530811 | 1.0609 | 1.999 |
| chr7 | 101514171 | 101514419 | 0.843963 | 1.68679 | 1.999 |
| chr12 | 106542947 | 106543275 | 1.65851 | 3.31484 | 1.999 |
| chr2 | 216814712 | 216814973 | 1.06477 | 2.12814 | 1.999 |
| chr14 | 22940535 | 22940792 | 1.4281 | 2.8544 | 1.999 |
| chr7 | 16460614 | 16461097 | 0.595944 | 1.19115 | 1.999 |
| chr10 | 90595398 | 90595681 | 1.74065 | 3.47933 | 1.999 |
| chr17 | 9832215 | 9832532 | 1.58094 | 3.1601 | 1.999 |
| chr8 | 136641041 | 136641496 | 0.461275 | 0.922034 | 1.999 |
| chr7 | 107519888 | 107520090 | 1.52212 | 3.04257 | 1.999 |
| chr2 | 37738752 | 37739028 | 0.518107 | 1.03566 | 1.999 |
| chr19 | 40887282 | 40887598 | 0.672019 | 1.34333 | 1.999 |
| chr3 | 106954790 | 106955008 | 1.39405 | 2.78664 | 1.999 |
| chr17 | 80123252 | 80123563 | 0.896377 | 1.79184 | 1.999 |
| chr3 | 133382577 | 133382952 | 1.16654 | 2.33189 | 1.999 |
| chr15 | 88384738 | 88384951 | 0.633728 | 1.26683 | 1.999 |
| chr2 | 201374690 | 201374983 | 2.02705 | 4.05214 | 1.999 |
| chr1 | 38325787 | 38326067 | 2.13526 | 4.26848 | 1.999 |
| chr6 | 136929374 | 136929655 | 1.75261 | 3.50367 | 1.999 |
| chr19 | 41814326 | 41814809 | 2.04002 | 4.07842 | 1.999 |
| chr5 | 148822046 | 148822434 | 1.99478 | 3.98809 | 1.999 |
| chr1 | 55537218 | 55537447 | 1.20262 | 2.40446 | 1.999 |
| chr8 | 117727998 | 117728200 | 1.41498 | 2.82939 | 2.000 |
| chr20 | 61478751 | 61479120 | 0.74024 | 1.48019 | 2.000 |
| chr17 | 11805267 | 11805547 | 0.554959 | 1.10972 | 2.000 |
| chr1 | 159976140 | 159976490 | 2.58032 | 5.15981 | 2.000 |
| chr12 | 89923593 | 89923886 | 1.22027 | 2.44022 | 2.000 |
| chr19 | 32053885 | 32054232 | 0.680915 | 1.36171 | 2.000 |
| chr3 | 16475591 | 16475810 | 0.635802 | 1.27152 | 2.000 |
| chr18 | 32586045 | 32586258 | 1.20184 | 2.40368 | 2.000 |
| chr8 | 39619308 | 39619746 | 0.567936 | 1.13587 | 2.000 |
| chr9 | 129424804 | 129425184 | 0.554146 | 1.1083 | 2.000 |
| chr21 | 40510094 | 40510410 | 2.05851 | 4.11718 | 2.000 |
| chr12 | 114795947 | 114796153 | 0.524163 | 1.04842 | 2.000 |
| chr7 | 131198152 | 131198838 | 1.57864 | 3.15756 | 2.000 |
| chr6 | 62872911 | 62873127 | 0.400911 | 0.801913 | 2.000 |
| chr2 | 230872890 | 230873217 | 2.18162 | 4.36394 | 2.000 |
| chr10 | 90973005 | 90973345 | 1.67087 | 3.34235 | 2.000 |
| chr16 | 21312813 | 21313238 | 0.786431 | 1.57321 | 2.000 |
| chr5 | 95672167 | 95672423 | 1.72717 | 3.45521 | 2.001 |
| chr20 | 35465134 | 35465378 | 1.53288 | 3.06665 | 2.001 |
| chr11 | 130934359 | 130934665 | 1.63743 | 3.27594 | 2.001 |
| chr1 | 178279820 | 178280082 | 1.01349 | 2.02766 | 2.001 |
| chr4 | 36424471 | 36424870 | 0.419054 | 0.838405 | 2.001 |
| chr8 | 86380740 | 86380991 | 0.430291 | 0.860903 | 2.001 |
| chr20 | 21700301 | 21700611 | 1.36484 | 2.73074 | 2.001 |
| chr9 | 140181427 | 140181753 | 1.44184 | 2.88482 | 2.001 |
| chr16 | 69146515 | 69147063 | 2.01195 | 4.02554 | 2.001 |
| chr1 | 227080361 | 227080643 | 0.440082 | 0.880585 | 2.001 |
| chr2 | 9697861 | 9698069 | 0.890356 | 1.78168 | 2.001 |
| chr12 | 90496884 | 90497096 | 1.08551 | 2.17227 | 2.001 |
| chr3 | 96099651 | 96099930 | 0.416648 | 0.83381 | 2.001 |
| chr12 | 56664942 | 56665253 | 1.20019 | 2.40192 | 2.001 |
| chr11 | 112150569 | 112151058 | 1.57803 | 3.15838 | 2.001 |

| | | | | | |
|---|---|---|---|---|---|
| chr5 | 133897581 | 133897798 | 1.11148 | 2.22463 | 2.002 |
| chr12 | 123278154 | 123278568 | 1.26185 | 2.52561 | 2.002 |
| chr18 | 44238173 | 44238551 | 1.30119 | 2.60436 | 2.002 |
| chr16 | 4635426 | 4635809 | 2.12469 | 4.25274 | 2.002 |
| chr18 | 60786901 | 60787149 | 1.15934 | 2.32051 | 2.002 |
| chr9 | 137311602 | 137311904 | 0.98434 | 1.97029 | 2.002 |
| chr20 | 48737374 | 48737699 | 2.0442 | 4.0918 | 2.002 |
| chr7 | 69142109 | 69142312 | 0.937288 | 1.8762 | 2.002 |
| chr19 | 6412354 | 6412726 | 1.40456 | 2.81163 | 2.002 |
| chr11 | 74959562 | 74960122 | 3.38223 | 6.77055 | 2.002 |
| chr14 | 23527205 | 23527660 | 2.43915 | 4.88268 | 2.002 |
| chr7 | 134966275 | 134966604 | 1.81439 | 3.63204 | 2.002 |
| chr11 | 119039301 | 119039637 | 1.56849 | 3.13988 | 2.002 |
| chr4 | 26030452 | 26030705 | 1.01004 | 2.02195 | 2.002 |
| chr1 | 41134482 | 41134949 | 1.46462 | 2.93197 | 2.002 |
| chr12 | 48288479 | 48288697 | 0.504188 | 1.00932 | 2.002 |
| chr17 | 40270690 | 40270960 | 1.51637 | 3.03576 | 2.002 |
| chr12 | 56711625 | 56711967 | 0.679616 | 1.36063 | 2.002 |
| chr2 | 113017056 | 113017363 | 2.25099 | 4.50664 | 2.002 |
| chr1 | 40648677 | 40648964 | 1.01232 | 2.0268 | 2.002 |
| chr9 | 92962833 | 92963046 | 0.380252 | 0.761343 | 2.002 |
| chr8 | 101571900 | 101572212 | 2.61425 | 5.23443 | 2.002 |
| chr12 | 86260973 | 86261337 | 1.34265 | 2.68838 | 2.002 |
| chr11 | 63278863 | 63279075 | 0.742376 | 1.48646 | 2.002 |
| chr1 | 208001968 | 208002242 | 1.27823 | 2.55941 | 2.002 |
| chr5 | 50046920 | 50047265 | 1.41798 | 2.83925 | 2.002 |
| chr4 | 58029778 | 58030549 | 1.46883 | 2.9412 | 2.002 |
| chr10 | 130169156 | 130169393 | 0.522295 | 1.04588 | 2.002 |
| chr10 | 103061217 | 103061524 | 0.969432 | 1.94134 | 2.003 |
| chr3 | 89163664 | 89164204 | 1.29017 | 2.58364 | 2.003 |
| chr9 | 81436105 | 81436330 | 1.09173 | 2.18629 | 2.003 |
| chr1 | 27684012 | 27684404 | 1.94714 | 3.89945 | 2.003 |
| chr11 | 69413044 | 69413267 | 0.753151 | 1.50831 | 2.003 |
| chr8 | 142248696 | 142248916 | 0.617595 | 1.23688 | 2.003 |
| chr7 | 116624940 | 116625222 | 1.45645 | 2.91704 | 2.003 |
| chr15 | 88086765 | 88087069 | 0.545849 | 1.09327 | 2.003 |
| chr11 | 16807041 | 16807401 | 1.47077 | 2.94586 | 2.003 |
| chr5 | 67363540 | 67363853 | 0.623521 | 1.24889 | 2.003 |
| chr6 | 35383042 | 35383291 | 0.892614 | 1.78793 | 2.003 |
| chr10 | 102902038 | 102902371 | 1.99605 | 3.99827 | 2.003 |
| chr10 | 70853108 | 70853372 | 1.68097 | 3.36717 | 2.003 |
| chr14 | 93181088 | 93181407 | 1.67152 | 3.34835 | 2.003 |
| chr12 | 69257754 | 69257990 | 1.37716 | 2.75872 | 2.003 |
| chr5 | 12590178 | 12590397 | 0.42364 | 0.848636 | 2.003 |
| chr15 | 99826663 | 99826947 | 1.45781 | 2.9203 | 2.003 |
| chr6 | 162615863 | 162616215 | 0.43826 | 0.87794 | 2.003 |
| chr16 | 74808386 | 74808759 | 1.15147 | 2.3067 | 2.003 |
| chr9 | 100685810 | 100686269 | 1.28033 | 2.56496 | 2.003 |
| chr19 | 43139047 | 43139734 | 1.61637 | 3.23828 | 2.003 |
| chr19 | 33370863 | 33371161 | 0.669302 | 1.34092 | 2.003 |
| chr12 | 52774040 | 52774362 | 1.2582 | 2.52087 | 2.004 |
| chr3 | 74834288 | 74834505 | 0.603619 | 1.20939 | 2.004 |
| chr17 | 72838996 | 72839577 | 1.22394 | 2.4524 | 2.004 |
| chr6 | 20126831 | 20127346 | 1.90249 | 3.812 | 2.004 |
| chr7 | 12726321 | 12727191 | 1.67679 | 3.35978 | 2.004 |
| chr10 | 82778902 | 82779196 | 1.47345 | 2.95236 | 2.004 |
| chr2 | 219844633 | 219845121 | 2.56166 | 5.13284 | 2.004 |
| chr2 | 102500371 | 102500576 | 0.949218 | 1.9021 | 2.004 |
| chr19 | 39270827 | 39271365 | 0.632906 | 1.26826 | 2.004 |
| chr15 | 92991484 | 92991898 | 0.600191 | 1.20271 | 2.004 |
| chr5 | 169694861 | 169695213 | 1.96004 | 3.92787 | 2.004 |

| | | | | | |
|---|---|---|---|---|---|
| chr10 | 118547518 | 118547919 | 2.49996 | 5.0102 | 2.004 |
| chr10 | 130574377 | 130574619 | 0.390475 | 0.782558 | 2.004 |
| chr7 | 28966751 | 28966987 | 1.49109 | 2.98837 | 2.004 |
| chr16 | 68738213 | 68738735 | 1.86008 | 3.72798 | 2.004 |
| chr16 | 47292853 | 47293095 | 1.50964 | 3.02568 | 2.004 |
| chr2 | 109544886 | 109545162 | 1.86452 | 3.73712 | 2.004 |
| chr6 | 135144118 | 135144401 | 2.33761 | 4.68538 | 2.004 |
| chr20 | 60989059 | 60989454 | 1.03595 | 2.07644 | 2.004 |
| chr2 | 11272721 | 11273064 | 1.82227 | 3.65265 | 2.004 |
| chr9 | 132404318 | 132404876 | 2.63735 | 5.28672 | 2.005 |
| chrX | 152989184 | 152989447 | 1.21885 | 2.44339 | 2.005 |
| chr12 | 98568166 | 98568439 | 0.770574 | 1.54476 | 2.005 |
| chr5 | 173400231 | 173400469 | 1.37897 | 2.76454 | 2.005 |
| chr19 | 11646917 | 11647199 | 1.62654 | 3.26091 | 2.005 |
| chr10 | 117031412 | 117031809 | 0.5658 | 1.13434 | 2.005 |
| chr7 | 78124172 | 78124460 | 0.4351 | 0.872317 | 2.005 |
| chr16 | 80825273 | 80825504 | 0.592883 | 1.18868 | 2.005 |
| chr11 | 58291582 | 58292384 | 1.8875 | 3.78434 | 2.005 |
| chr17 | 698101 | 698405 | 1.45004 | 2.90727 | 2.005 |
| chr14 | 91643751 | 91644014 | 1.90488 | 3.81926 | 2.005 |
| chr17 | 16266831 | 16267512 | 1.5801 | 3.16823 | 2.005 |
| chr2 | 21059303 | 21059534 | 0.641688 | 1.28664 | 2.005 |
| chr5 | 37775039 | 37775484 | 1.11345 | 2.23257 | 2.005 |
| chr11 | 86572972 | 86573216 | 0.505341 | 1.01326 | 2.005 |
| chr15 | 52371109 | 52371401 | 2.73159 | 5.47725 | 2.005 |
| chr9 | 135661694 | 135661969 | 0.450492 | 0.903303 | 2.005 |
| chr5 | 7853867 | 7854116 | 0.703477 | 1.41059 | 2.005 |
| chr4 | 90149340 | 90149547 | 0.544482 | 1.09182 | 2.005 |
| chr15 | 81923974 | 81924547 | 0.516335 | 1.03542 | 2.005 |
| chr5 | 104048327 | 104048685 | 0.573002 | 1.14906 | 2.005 |
| chr20 | 1450712 | 1451041 | 2.00441 | 4.01952 | 2.005 |
| chr22 | 29436720 | 29437052 | 1.32128 | 2.64985 | 2.006 |
| chr1 | 156647867 | 156648154 | 2.16351 | 4.33901 | 2.006 |
| chr8 | 130628200 | 130628624 | 0.486668 | 0.976118 | 2.006 |
| chr15 | 29534895 | 29535304 | 0.461421 | 0.925503 | 2.006 |
| chr1 | 159181253 | 159181666 | 1.82501 | 3.6606 | 2.006 |
| chr15 | 65102037 | 65102524 | 2.23145 | 4.47586 | 2.006 |
| chr3 | 153231094 | 153231409 | 0.868754 | 1.74257 | 2.006 |
| chr5 | 112299739 | 112299956 | 1.07868 | 2.16365 | 2.006 |
| chr16 | 70713848 | 70714262 | 2.15903 | 4.33071 | 2.006 |
| chr7 | 33580527 | 33580946 | 0.464637 | 0.932066 | 2.006 |
| chr3 | 147579153 | 147579770 | 0.460334 | 0.923448 | 2.006 |
| chr10 | 107442817 | 107443120 | 0.47629 | 0.955468 | 2.006 |
| chr20 | 12139264 | 12139628 | 0.533126 | 1.0695 | 2.006 |
| chr12 | 39539095 | 39539545 | 2.36084 | 4.73614 | 2.006 |
| chr19 | 13075712 | 13076522 | 2.08844 | 4.18972 | 2.006 |
| chr8 | 86202409 | 86202679 | 1.0791 | 2.16488 | 2.006 |
| chr5 | 94073509 | 94074003 | 1.25423 | 2.51627 | 2.006 |
| chr3 | 46671411 | 46671776 | 1.99634 | 4.00516 | 2.006 |
| chr14 | 58371856 | 58372516 | 0.54035 | 1.08411 | 2.006 |
| chr11 | 65236339 | 65236657 | 1.58272 | 3.1757 | 2.006 |
| chr2 | 101226791 | 101227083 | 1.74287 | 3.49705 | 2.006 |
| chr1 | 204335955 | 204336209 | 1.74917 | 3.5098 | 2.007 |
| chr5 | 55496996 | 55497293 | 1.52139 | 3.05275 | 2.007 |
| chr20 | 15959876 | 15960278 | 0.626413 | 1.25697 | 2.007 |
| chr7 | 116860994 | 116861212 | 0.63612 | 1.27645 | 2.007 |
| chr18 | 7926613 | 7926820 | 1.0616 | 2.13028 | 2.007 |
| chr5 | 135253971 | 135254213 | 0.451053 | 0.905142 | 2.007 |
| chr21 | 37703211 | 37703486 | 0.721385 | 1.44764 | 2.007 |
| chr13 | 75793874 | 75794246 | 1.86436 | 3.74143 | 2.007 |
| chr9 | 131057580 | 131058024 | 1.50808 | 3.02645 | 2.007 |

| | | | | | |
|---|---|---|---|---|---|
| chr19 | 14516974 | 14517358 | 2.9693 | 5.95921 | 2.007 |
| chr6 | 44119525 | 44119999 | 2.71774 | 5.45463 | 2.007 |
| chr11 | 71274594 | 71275258 | 0.816532 | 1.63886 | 2.007 |
| chr17 | 11096497 | 11096759 | 0.620701 | 1.24588 | 2.007 |
| chr16 | 19386928 | 19387295 | 1.54308 | 3.09739 | 2.007 |
| chr15 | 98854855 | 98855180 | 1.51131 | 3.03372 | 2.007 |
| chr5 | 139536712 | 139537008 | 1.85645 | 3.72652 | 2.007 |
| chr7 | 107710024 | 107710283 | 0.566646 | 1.13745 | 2.007 |
| chr12 | 103567255 | 103567491 | 0.520558 | 1.04499 | 2.007 |
| chr20 | 9598586 | 9598913 | 2.00069 | 4.01632 | 2.007 |
| chr17 | 38518855 | 38519198 | 1.8462 | 3.70642 | 2.008 |
| chr2 | 5928184 | 5928410 | 0.582881 | 1.1702 | 2.008 |
| chr5 | 179228809 | 179229184 | 2.15072 | 4.31783 | 2.008 |
| chr1 | 66708332 | 66708706 | 2.74017 | 5.50125 | 2.008 |
| chr10 | 29280729 | 29281023 | 1.68599 | 3.38489 | 2.008 |
| chr20 | 36950352 | 36950748 | 1.66505 | 3.3429 | 2.008 |
| chr12 | 118294493 | 118294726 | 1.43509 | 2.88125 | 2.008 |
| chr12 | 45083260 | 45083589 | 0.536148 | 1.07646 | 2.008 |
| chr12 | 133131385 | 133131837 | 1.50818 | 3.02827 | 2.008 |
| chr17 | 9809183 | 9809568 | 1.5254 | 3.06309 | 2.008 |
| chr10 | 133955885 | 133956346 | 2.45419 | 4.92847 | 2.008 |
| chr17 | 75062837 | 75063256 | 2.37426 | 4.76799 | 2.008 |
| chr8 | 131366917 | 131367247 | 0.619343 | 1.24385 | 2.008 |
| chr12 | 91501754 | 91502024 | 0.594946 | 1.19487 | 2.008 |
| chr13 | 34651319 | 34651560 | 1.20423 | 2.41864 | 2.008 |
| chr7 | 32101778 | 32102084 | 1.36363 | 2.73889 | 2.009 |
| chr10 | 103832576 | 103833036 | 2.03127 | 4.07991 | 2.009 |
| chr17 | 29885645 | 29886061 | 2.08743 | 4.19272 | 2.009 |
| chr17 | 49134200 | 49134471 | 1.16434 | 2.33865 | 2.009 |
| chr15 | 68966249 | 68966819 | 2.24699 | 4.51326 | 2.009 |
| chr4 | 122871343 | 122871669 | 1.6099 | 3.23378 | 2.009 |
| chr15 | 81702700 | 81702933 | 0.999461 | 2.00768 | 2.009 |
| chr6 | 1555267 | 1555584 | 1.73337 | 3.48198 | 2.009 |
| chr4 | 58057158 | 58057378 | 1.37991 | 2.77258 | 2.009 |
| chr11 | 62673421 | 62673835 | 2.58702 | 5.19817 | 2.009 |
| chr7 | 37221580 | 37221947 | 2.46196 | 4.9469 | 2.009 |
| chr11 | 108519062 | 108519336 | 1.5823 | 3.17941 | 2.009 |
| chr17 | 19912428 | 19912865 | 2.37298 | 4.76817 | 2.009 |
| chr1 | 156931548 | 156931764 | 0.831149 | 1.67009 | 2.009 |
| chr17 | 76861835 | 76862232 | 1.79715 | 3.61121 | 2.009 |
| chr7 | 33225386 | 33225605 | 1.38536 | 2.78383 | 2.009 |
| chr8 | 19229595 | 19229963 | 2.11242 | 4.24503 | 2.010 |
| chr3 | 84524720 | 84524955 | 0.55262 | 1.11055 | 2.010 |
| chr20 | 35170476 | 35170733 | 0.707251 | 1.42133 | 2.010 |
| chr1 | 110074895 | 110075242 | 1.94978 | 3.91839 | 2.010 |
| chr17 | 46095748 | 46096094 | 1.93982 | 3.89837 | 2.010 |
| chr1 | 160651322 | 160651583 | 1.4223 | 2.85837 | 2.010 |
| chr10 | 104488963 | 104489383 | 2.10496 | 4.23034 | 2.010 |
| chr6 | 116882643 | 116882885 | 0.984249 | 1.97807 | 2.010 |
| chr19 | 495627 | 495989 | 1.55563 | 3.12642 | 2.010 |
| chr5 | 60487863 | 60488100 | 0.469089 | 0.942828 | 2.010 |
| chr12 | 18520812 | 18521117 | 0.536058 | 1.07745 | 2.010 |
| chr7 | 158118866 | 158119138 | 0.975649 | 1.96104 | 2.010 |
| chr4 | 89300020 | 89300292 | 1.24734 | 2.50715 | 2.010 |
| chr14 | 70014672 | 70015192 | 1.2447 | 2.50186 | 2.010 |
| chr11 | 58281294 | 58281687 | 0.932935 | 1.87522 | 2.010 |
| chr1 | 180748002 | 180748260 | 1.20814 | 2.42849 | 2.010 |
| chr3 | 129047513 | 129047957 | 1.5222 | 3.05981 | 2.010 |
| chr12 | 69201793 | 69202129 | 1.24909 | 2.51087 | 2.010 |
| chr10 | 35283555 | 35283999 | 2.00965 | 4.04012 | 2.010 |
| chr7 | 149484564 | 149484892 | 1.03411 | 2.07907 | 2.010 |

| | | | | | |
|---|---|---|---|---|---|
| chr14 | 76120438 | 76120850 | 2.53509 | 5.09703 | 2.011 |
| chr12 | 10305891 | 10306492 | 1.31635 | 2.6467 | 2.011 |
| chr12 | 109730756 | 109731106 | 0.499324 | 1.00398 | 2.011 |
| chr22 | 33606145 | 33606508 | 1.64715 | 3.3119 | 2.011 |
| chr7 | 34863328 | 34863624 | 2.00498 | 4.03139 | 2.011 |
| chr2 | 174313951 | 174314245 | 1.66296 | 3.34374 | 2.011 |
| chr6 | 143748353 | 143748671 | 2.28586 | 4.59634 | 2.011 |
| chr4 | 25441600 | 25441883 | 1.88621 | 3.79275 | 2.011 |
| chr1 | 152162356 | 152162643 | 1.64387 | 3.30555 | 2.011 |
| chr15 | 89265937 | 89266204 | 0.524688 | 1.05509 | 2.011 |
| chr10 | 114216395 | 114216633 | 0.673193 | 1.35374 | 2.011 |
| chr5 | 139607961 | 139608390 | 0.500066 | 1.00563 | 2.011 |
| chr5 | 130925281 | 130925612 | 1.81215 | 3.64424 | 2.011 |
| chr22 | 38448213 | 38448650 | 1.67663 | 3.37173 | 2.011 |
| chr1 | 6403511 | 6403794 | 1.97006 | 3.96192 | 2.011 |
| chr1 | 224788150 | 224788638 | 0.579589 | 1.16563 | 2.011 |
| chr2 | 40065009 | 40065393 | 2.00434 | 4.031 | 2.011 |
| chr19 | 45311256 | 45311564 | 1.59243 | 3.20263 | 2.011 |
| chr5 | 112253233 | 112253526 | 1.06829 | 2.14855 | 2.011 |
| chr17 | 262050 | 262380 | 1.45473 | 2.92577 | 2.011 |
| chr2 | 18474078 | 18474638 | 0.624638 | 1.25635 | 2.011 |
| chr7 | 64772601 | 64773056 | 1.47401 | 2.96473 | 2.011 |
| chr5 | 111546214 | 111546431 | 1.60465 | 3.22754 | 2.011 |
| chr9 | 2857636 | 2857915 | 1.62329 | 3.26509 | 2.011 |
| chr21 | 34499481 | 34500010 | 1.37099 | 2.75762 | 2.011 |
| chr8 | 34287218 | 34287503 | 0.437688 | 0.880378 | 2.011 |
| chr10 | 11716478 | 11716708 | 0.478147 | 0.961778 | 2.011 |
| chr11 | 69703347 | 69703715 | 1.85791 | 3.73717 | 2.011 |
| chr2 | 59450052 | 59450332 | 0.593353 | 1.19355 | 2.012 |
| chr1 | 232067635 | 232067975 | 1.31496 | 2.64513 | 2.012 |
| chr12 | 131600913 | 131601295 | 1.27235 | 2.55944 | 2.012 |
| chr8 | 10282232 | 10283155 | 1.13288 | 2.27888 | 2.012 |
| chr1 | 237152550 | 237152846 | 0.580303 | 1.16733 | 2.012 |
| chr19 | 50917906 | 50918366 | 2.78949 | 5.61139 | 2.012 |
| chr11 | 62107725 | 62108148 | 1.24068 | 2.4958 | 2.012 |
| chr22 | 37934817 | 37935177 | 2.10234 | 4.22962 | 2.012 |
| chr12 | 41755554 | 41755880 | 0.655414 | 1.31863 | 2.012 |
| chr1 | 181135742 | 181136032 | 1.79774 | 3.6169 | 2.012 |
| chr15 | 50295964 | 50296248 | 2.25419 | 4.53534 | 2.012 |
| chr22 | 33454248 | 33454550 | 1.39542 | 2.80753 | 2.012 |
| chr2 | 71504686 | 71504889 | 0.96732 | 1.94623 | 2.012 |
| chr6 | 128058155 | 128058413 | 1.49507 | 3.00814 | 2.012 |
| chr14 | 106760914 | 106761251 | 0.45625 | 0.918011 | 2.012 |
| chr19 | 6733445 | 6733710 | 1.1238 | 2.26129 | 2.012 |
| chr20 | 2504983 | 2505461 | 2.58246 | 5.19638 | 2.012 |
| chr8 | 29940403 | 29940659 | 1.04393 | 2.10069 | 2.012 |
| chr19 | 12871281 | 12871795 | 1.24627 | 2.50787 | 2.012 |
| chr10 | 94032128 | 94032660 | 0.438526 | 0.882463 | 2.012 |
| chr10 | 104677140 | 104677573 | 2.43549 | 4.90103 | 2.012 |
| chr12 | 110318204 | 110318575 | 1.67139 | 3.36348 | 2.012 |
| chr2 | 48904625 | 48904953 | 0.596514 | 1.20052 | 2.013 |
| chr13 | 102409889 | 102410139 | 1.06853 | 2.15051 | 2.013 |
| chr17 | 77389304 | 77389700 | 2.92423 | 5.88548 | 2.013 |
| chr10 | 100227230 | 100227499 | 1.75387 | 3.53 | 2.013 |
| chr1 | 183208555 | 183208772 | 0.44052 | 0.8867 | 2.013 |
| chr12 | 102835956 | 102836163 | 0.47832 | 0.962792 | 2.013 |
| chr3 | 62318595 | 62318950 | 0.669911 | 1.34844 | 2.013 |
| chr4 | 4996805 | 4997213 | 1.80242 | 3.62807 | 2.013 |
| chr17 | 17050078 | 17050401 | 1.60152 | 3.2237 | 2.013 |
| chr16 | 57698282 | 57698526 | 0.981946 | 1.97662 | 2.013 |
| chr16 | 22692045 | 22692297 | 1.25611 | 2.52853 | 2.013 |

| | | | | | |
|---|---|---|---|---|---|
| chr16 | 26087468 | 26087774 | 0.452377 | 0.910664 | 2.013 |
| chr19 | 41945832 | 41946134 | 1.89453 | 3.81389 | 2.013 |
| chr4 | 147163158 | 147163595 | 0.677862 | 1.36464 | 2.013 |
| chr16 | 50038254 | 50038647 | 0.458167 | 0.922367 | 2.013 |
| chr16 | 87454178 | 87454589 | 1.06102 | 2.13602 | 2.013 |
| chr8 | 137042935 | 137043227 | 0.796253 | 1.60314 | 2.013 |
| chr5 | 165897543 | 165897797 | 0.533127 | 1.07341 | 2.013 |
| chr5 | 31929345 | 31929724 | 1.37847 | 2.77558 | 2.014 |
| chr5 | 134905705 | 134906253 | 1.29588 | 2.60929 | 2.014 |
| chr1 | 150759052 | 150759293 | 0.521127 | 1.04934 | 2.014 |
| chr11 | 18000291 | 18000629 | 2.21659 | 4.46332 | 2.014 |
| chr12 | 127302404 | 127302614 | 0.555014 | 1.11765 | 2.014 |
| chr11 | 95569236 | 95569479 | 1.4243 | 2.86826 | 2.014 |
| chr20 | 36985091 | 36985318 | 1.35924 | 2.73734 | 2.014 |
| chr1 | 152464113 | 152464322 | 1.33974 | 2.69812 | 2.014 |
| chr8 | 82410065 | 82410331 | 0.495645 | 0.99821 | 2.014 |
| chr20 | 45925688 | 45925939 | 0.517686 | 1.04264 | 2.014 |
| chr9 | 138593781 | 138594218 | 2.29786 | 4.62809 | 2.014 |
| chr12 | 78157414 | 78157880 | 0.69054 | 1.39083 | 2.014 |
| chr2 | 148958960 | 148959363 | 1.68938 | 3.4027 | 2.014 |
| chr12 | 132108285 | 132108519 | 1.20708 | 2.43129 | 2.014 |
| chr5 | 56516853 | 56517081 | 1.16449 | 2.34552 | 2.014 |
| chr11 | 56194081 | 56194309 | 0.543126 | 1.09398 | 2.014 |
| chr10 | 29683237 | 29683614 | 0.469333 | 0.945408 | 2.014 |
| chr8 | 10226780 | 10227003 | 0.974532 | 1.96314 | 2.014 |
| chr12 | 123344587 | 123345245 | 1.53163 | 3.08541 | 2.014 |
| chr5 | 126616288 | 126616510 | 0.70397 | 1.41822 | 2.015 |
| chr3 | 12179578 | 12179861 | 1.01794 | 2.05099 | 2.015 |
| chr8 | 103420088 | 103420295 | 1.29108 | 2.60132 | 2.015 |
| chr5 | 95228555 | 95228828 | 1.29899 | 2.61729 | 2.015 |
| chr5 | 102307505 | 102307725 | 0.910418 | 1.83442 | 2.015 |
| chr1 | 44399988 | 44400778 | 1.42881 | 2.87909 | 2.015 |
| chr18 | 19180821 | 19181142 | 2.15181 | 4.33601 | 2.015 |
| chr7 | 150865255 | 150865686 | 2.63423 | 5.30836 | 2.015 |
| chr9 | 101077644 | 101078236 | 2.14779 | 4.32863 | 2.015 |
| chr1 | 111441955 | 111442185 | 0.445754 | 0.898376 | 2.015 |
| chr21 | 27173819 | 27174208 | 2.27112 | 4.5773 | 2.015 |
| chr9 | 15211419 | 15211853 | 2.01375 | 4.05877 | 2.016 |
| chr12 | 98354827 | 98355111 | 1.38686 | 2.79528 | 2.016 |
| chr1 | 17980273 | 17980659 | 1.97701 | 3.98479 | 2.016 |
| chr5 | 66461938 | 66462385 | 2.20006 | 4.43437 | 2.016 |
| chr16 | 66308042 | 66308668 | 1.04074 | 2.09771 | 2.016 |
| chrX | 110924199 | 110924466 | 1.73879 | 3.50478 | 2.016 |
| chr6 | 41904098 | 41904405 | 1.42121 | 2.86467 | 2.016 |
| chr17 | 34274174 | 34274412 | 2.12303 | 4.27937 | 2.016 |
| chr9 | 38130054 | 38130257 | 0.334027 | 0.673299 | 2.016 |
| chr1 | 27692969 | 27693233 | 1.0571 | 2.13089 | 2.016 |
| chr10 | 64577632 | 64578107 | 1.55331 | 3.13115 | 2.016 |
| chr3 | 127550714 | 127551151 | 1.01476 | 2.04557 | 2.016 |
| chr12 | 55153559 | 55153856 | 1.19679 | 2.41252 | 2.016 |
| chr17 | 8798263 | 8798596 | 1.36323 | 2.74806 | 2.016 |
| chr8 | 106041605 | 106041896 | 0.657618 | 1.32568 | 2.016 |
| chr11 | 17549626 | 17550065 | 1.81651 | 3.66195 | 2.016 |
| chr10 | 6056141 | 6056723 | 0.99755 | 2.01107 | 2.016 |
| chr19 | 34711380 | 34711769 | 2.20107 | 4.43752 | 2.016 |
| chr2 | 32551628 | 32551834 | 1.10822 | 2.23428 | 2.016 |
| chr8 | 92462532 | 92462873 | 0.506875 | 1.02193 | 2.016 |
| chr12 | 49328194 | 49328408 | 1.44006 | 2.90337 | 2.016 |
| chr19 | 10514889 | 10515111 | 2.67315 | 5.38946 | 2.016 |
| chr3 | 194193066 | 194193399 | 2.6722 | 5.38773 | 2.016 |
| chr3 | 118603494 | 118604081 | 1.94776 | 3.92714 | 2.016 |

| | | | | | |
|---|---|---|---|---|---|
| chr10 | 81749262 | 81749517 | 0.419437 | 0.8457 | 2.016 |
| chr2 | 74488702 | 74488992 | 1.6455 | 3.31781 | 2.016 |
| chr1 | 228912274 | 228912570 | 0.493786 | 0.995619 | 2.016 |
| chr17 | 45019738 | 45020293 | 2.02732 | 4.08775 | 2.016 |
| chr1 | 46872027 | 46872349 | 1.89116 | 3.81343 | 2.016 |
| chr1 | 226384673 | 226385032 | 1.74383 | 3.51637 | 2.016 |
| chr10 | 71577896 | 71578098 | 0.375023 | 0.75626 | 2.017 |
| chr16 | 20751597 | 20751927 | 1.16692 | 2.35318 | 2.017 |
| chr16 | 69006648 | 69006888 | 1.08828 | 2.19461 | 2.017 |
| chr17 | 4064512 | 4064724 | 0.785053 | 1.58318 | 2.017 |
| chr2 | 232161796 | 232162097 | 1.71236 | 3.45335 | 2.017 |
| chr8 | 53396559 | 53396781 | 0.954489 | 1.92506 | 2.017 |
| chr12 | 125440247 | 125440570 | 0.570914 | 1.15152 | 2.017 |
| chr14 | 20415172 | 20415583 | 0.461188 | 0.930222 | 2.017 |
| chr12 | 101134823 | 101135272 | 1.15867 | 2.33712 | 2.017 |
| chr1 | 11845191 | 11845629 | 1.78681 | 3.60416 | 2.017 |
| chr1 | 160514539 | 160514758 | 0.915464 | 1.8466 | 2.017 |
| chr18 | 3995581 | 3995803 | 1.46082 | 2.94665 | 2.017 |
| chr14 | 72187683 | 72188092 | 1.12178 | 2.26279 | 2.017 |
| chr5 | 136411430 | 136411803 | 0.560603 | 1.13086 | 2.017 |
| chr10 | 5336136 | 5336416 | 1.23479 | 2.49089 | 2.017 |
| chr10 | 105302536 | 105302793 | 0.820466 | 1.6551 | 2.017 |
| chr8 | 65459165 | 65459508 | 0.465867 | 0.939778 | 2.017 |
| chr15 | 58043404 | 58043708 | 1.0593 | 2.137 | 2.017 |
| chr1 | 113008735 | 113009049 | 1.48771 | 3.00132 | 2.017 |
| chr20 | 22151550 | 22151769 | 0.473153 | 0.954548 | 2.017 |
| chr12 | 67835571 | 67835890 | 2.06503 | 4.16606 | 2.017 |
| chr9 | 71822463 | 71822710 | 0.825545 | 1.66552 | 2.017 |
| chr8 | 121617962 | 121618238 | 1.39286 | 2.81017 | 2.018 |
| chr3 | 190529658 | 190529897 | 1.12851 | 2.27692 | 2.018 |
| chr2 | 85470870 | 85471102 | 1.31514 | 2.65349 | 2.018 |
| chr11 | 30367748 | 30367953 | 0.751833 | 1.51701 | 2.018 |
| chrX | 103410967 | 103411225 | 1.16977 | 2.36048 | 2.018 |
| chr20 | 48595519 | 48595985 | 2.54909 | 5.14406 | 2.018 |
| chr3 | 157324383 | 157324715 | 2.03776 | 4.11223 | 2.018 |
| chr5 | 133913465 | 133913854 | 1.85947 | 3.75254 | 2.018 |
| chr12 | 39036972 | 39037277 | 0.503244 | 1.01559 | 2.018 |
| chr5 | 139422796 | 139423334 | 1.26451 | 2.55192 | 2.018 |
| chr14 | 97752676 | 97752944 | 0.474827 | 0.958257 | 2.018 |
| chr1 | 222966181 | 222966401 | 0.807307 | 1.62927 | 2.018 |
| chr20 | 53107135 | 53107651 | 0.515658 | 1.04068 | 2.018 |
| chr12 | 2143962 | 2144427 | 2.07761 | 4.19327 | 2.018 |
| chr18 | 839350 | 839758 | 2.08448 | 4.20718 | 2.018 |
| chr4 | 1580795 | 1581205 | 1.67882 | 3.38849 | 2.018 |
| chrX | 113845552 | 113845844 | 1.62244 | 3.27502 | 2.019 |
| chr8 | 138738319 | 138738699 | 0.621596 | 1.25478 | 2.019 |
| chr11 | 82486811 | 82487158 | 1.62035 | 3.27106 | 2.019 |
| chr3 | 21617348 | 21617618 | 0.575087 | 1.16096 | 2.019 |
| chr1 | 22602305 | 22602685 | 1.80115 | 3.63617 | 2.019 |
| chr3 | 10024656 | 10024940 | 1.39817 | 2.82269 | 2.019 |
| chr3 | 81539644 | 81540027 | 1.39647 | 2.81929 | 2.019 |
| chr2 | 246574 | 246798 | 1.74901 | 3.53133 | 2.019 |
| chr17 | 79265565 | 79265873 | 2.11158 | 4.26362 | 2.019 |
| chr5 | 42912372 | 42912614 | 1.34058 | 2.7069 | 2.019 |
| chr4 | 20775075 | 20775315 | 0.977785 | 1.97435 | 2.019 |
| chr14 | 105197624 | 105198082 | 1.23252 | 2.48875 | 2.019 |
| chr19 | 28824037 | 28824463 | 0.631978 | 1.27616 | 2.019 |
| chr16 | 70804587 | 70805369 | 1.87698 | 3.79028 | 2.019 |
| chr8 | 48078998 | 48079311 | 1.95007 | 3.93797 | 2.019 |
| chr2 | 29314549 | 29314870 | 0.489282 | 0.988061 | 2.019 |
| chr6 | 151996905 | 151997379 | 1.53354 | 3.09692 | 2.019 |

| | | | | | |
|---|---|---|---|---|---|
| chr10 | 24448487 | 24448735 | 0.59757 | 1.20682 | 2.020 |
| chr12 | 117125666 | 117126047 | 1.34135 | 2.70892 | 2.020 |
| chr16 | 63237513 | 63237729 | 0.477951 | 0.965276 | 2.020 |
| chr15 | 92258770 | 92259048 | 0.459727 | 0.928482 | 2.020 |
| chr19 | 30415416 | 30415726 | 0.660778 | 1.33457 | 2.020 |
| chr16 | 57180736 | 57181102 | 0.525981 | 1.06233 | 2.020 |
| chr16 | 66864744 | 66865099 | 2.611 | 5.2735 | 2.020 |
| chr17 | 73408723 | 73409013 | 2.03373 | 4.10779 | 2.020 |
| chr6 | 78172114 | 78172478 | 0.889522 | 1.79668 | 2.020 |
| chr17 | 41089805 | 41090292 | 1.35582 | 2.73862 | 2.020 |
| chr21 | 32470871 | 32471247 | 1.53341 | 3.09742 | 2.020 |
| chr12 | 121382301 | 121382960 | 1.39746 | 2.82284 | 2.020 |
| chr8 | 133607237 | 133607441 | 0.59662 | 1.20516 | 2.020 |
| chr1 | 153650123 | 153650574 | 1.92318 | 3.88517 | 2.020 |
| chr11 | 62651798 | 62652036 | 1.12975 | 2.28232 | 2.020 |
| chr6 | 90280561 | 90280772 | 0.334537 | 0.675834 | 2.020 |
| chr14 | 74828985 | 74829378 | 1.50912 | 3.04876 | 2.020 |
| chr5 | 88478261 | 88478504 | 0.415895 | 0.840225 | 2.020 |
| chr10 | 115214097 | 115214666 | 1.54165 | 3.11475 | 2.020 |
| chr1 | 182992742 | 182993352 | 1.01975 | 2.06034 | 2.020 |
| chr9 | 124857550 | 124857908 | 1.71868 | 3.47255 | 2.020 |
| chr3 | 154476100 | 154476379 | 0.713645 | 1.44191 | 2.020 |
| chr15 | 29623222 | 29623498 | 0.397485 | 0.803161 | 2.021 |
| chr21 | 43870087 | 43870358 | 1.33765 | 2.70289 | 2.021 |
| chr1 | 999138 | 999572 | 1.73982 | 3.51556 | 2.021 |
| chr7 | 1597351 | 1598401 | 0.698385 | 1.4112 | 2.021 |
| chr7 | 38145554 | 38145834 | 1.30557 | 2.63822 | 2.021 |
| chr12 | 56118425 | 56118707 | 1.37584 | 2.78023 | 2.021 |
| chr12 | 127636077 | 127636299 | 0.511276 | 1.03321 | 2.021 |
| chr2 | 20835500 | 20835903 | 1.9444 | 3.92934 | 2.021 |
| chr15 | 92940167 | 92940462 | 0.456838 | 0.923302 | 2.021 |
| chr9 | 613752 | 614157 | 1.40118 | 2.83196 | 2.021 |
| chr19 | 32985450 | 32985695 | 0.747616 | 1.51104 | 2.021 |
| chr20 | 48506364 | 48506826 | 0.944191 | 1.90839 | 2.021 |
| chr15 | 35384784 | 35384989 | 0.5416 | 1.09471 | 2.021 |
| chr3 | 193323718 | 193324039 | 1.14546 | 2.31533 | 2.021 |
| chr16 | 68470572 | 68471037 | 0.840884 | 1.69985 | 2.022 |
| chr4 | 87816949 | 87817229 | 1.16898 | 2.36316 | 2.022 |
| chr12 | 84829342 | 84829635 | 0.57538 | 1.16321 | 2.022 |
| chr20 | 30157072 | 30157723 | 2.06372 | 4.17211 | 2.022 |
| chr13 | 26760730 | 26760995 | 1.21035 | 2.447 | 2.022 |
| chr9 | 80070179 | 80070586 | 1.40641 | 2.84344 | 2.022 |
| chr1 | 20521698 | 20522047 | 1.67883 | 3.39427 | 2.022 |
| chr1 | 145421713 | 145421935 | 1.45728 | 2.94639 | 2.022 |
| chr4 | 45553569 | 45553829 | 0.519154 | 1.0497 | 2.022 |
| chr8 | 140571374 | 140571592 | 0.445847 | 0.901475 | 2.022 |
| chr15 | 29717281 | 29717537 | 0.466196 | 0.942632 | 2.022 |
| chr19 | 8009158 | 8009420 | 1.86513 | 3.77123 | 2.022 |
| chr5 | 108017098 | 108017324 | 1.59435 | 3.22397 | 2.022 |
| chr20 | 38402056 | 38402595 | 0.453268 | 0.916565 | 2.022 |
| chr10 | 125332649 | 125333114 | 1.89826 | 3.83856 | 2.022 |
| chr15 | 41165827 | 41166179 | 1.12743 | 2.27987 | 2.022 |
| chr11 | 55710366 | 55710568 | 0.94927 | 1.91968 | 2.022 |
| chr2 | 142073 | 142577 | 0.6911 | 1.3976 | 2.022 |
| chr6 | 153833351 | 153833618 | 1.19023 | 2.40699 | 2.022 |
| chr2 | 43037021 | 43037223 | 0.727869 | 1.47211 | 2.022 |
| chr17 | 46103753 | 46104129 | 2.20841 | 4.46661 | 2.023 |
| chr19 | 48572883 | 48573109 | 0.815027 | 1.64843 | 2.023 |
| chr9 | 4073446 | 4073699 | 1.4166 | 2.86527 | 2.023 |
| chr1 | 228191936 | 228192499 | 1.46713 | 2.96753 | 2.023 |
| chr21 | 16855554 | 16855855 | 2.1682 | 4.38604 | 2.023 |

| | | | | | |
|---|---|---|---|---|---|
| chr10 | 128192569 | 128193023 | 1.19232 | 2.41195 | 2.023 |
| chr12 | 125235418 | 125235635 | 1.86355 | 3.76987 | 2.023 |
| chr10 | 17292805 | 17293056 | 0.870506 | 1.76104 | 2.023 |
| chr10 | 43187693 | 43187957 | 0.384593 | 0.778034 | 2.023 |
| chr9 | 105871877 | 105872227 | 0.491608 | 0.994574 | 2.023 |
| chr2 | 220217156 | 220217456 | 1.12866 | 2.2834 | 2.023 |
| chr12 | 656074 | 656434 | 2.33519 | 4.72437 | 2.023 |
| chr15 | 82121334 | 82121581 | 0.56219 | 1.13742 | 2.023 |
| chr7 | 105172391 | 105172763 | 1.80948 | 3.66112 | 2.023 |
| chr9 | 127656899 | 127657146 | 1.21008 | 2.44836 | 2.023 |
| chr2 | 165714746 | 165714975 | 0.466073 | 0.943016 | 2.023 |
| chrX | 53858470 | 53858692 | 1.08464 | 2.19468 | 2.023 |
| chr14 | 74037618 | 74037859 | 0.675288 | 1.36642 | 2.023 |
| chr17 | 56199338 | 56199535 | 1.23477 | 2.49851 | 2.023 |
| chr12 | 85030107 | 85030329 | 0.598868 | 1.21179 | 2.023 |
| chr11 | 126873219 | 126873810 | 1.28709 | 2.60442 | 2.023 |
| chr9 | 135383063 | 135383480 | 1.89642 | 3.83755 | 2.024 |
| chr7 | 149186556 | 149186841 | 1.27423 | 2.57859 | 2.024 |
| chr17 | 31816634 | 31816845 | 0.438939 | 0.888271 | 2.024 |
| chr1 | 62317555 | 62318150 | 0.866075 | 1.75273 | 2.024 |
| chr12 | 49752488 | 49752773 | 1.2867 | 2.60401 | 2.024 |
| chr7 | 50916955 | 50917215 | 0.651865 | 1.31928 | 2.024 |
| chr3 | 31725958 | 31726206 | 0.405215 | 0.820108 | 2.024 |
| chr7 | 137583922 | 137584264 | 2.33343 | 4.72284 | 2.024 |
| chr20 | 61687694 | 61688127 | 1.88699 | 3.81939 | 2.024 |
| chr9 | 84716516 | 84716924 | 0.803547 | 1.62647 | 2.024 |
| chr1 | 222478200 | 222478445 | 0.574487 | 1.16286 | 2.024 |
| chr1 | 89920782 | 89921007 | 1.34755 | 2.72774 | 2.024 |
| chr5 | 41708274 | 41708685 | 0.718935 | 1.45529 | 2.024 |
| chr8 | 6637104 | 6637667 | 1.97526 | 3.99855 | 2.024 |
| chr8 | 57700652 | 57700913 | 0.823821 | 1.66772 | 2.024 |
| chr7 | 96130074 | 96130344 | 0.57978 | 1.17372 | 2.024 |
| chr11 | 8054758 | 8055096 | 1.7718 | 3.58704 | 2.025 |
| chr5 | 110399375 | 110399672 | 0.597398 | 1.20949 | 2.025 |
| chr2 | 113466662 | 113466900 | 0.571557 | 1.15718 | 2.025 |
| chr10 | 74087898 | 74088402 | 1.12682 | 2.28139 | 2.025 |
| chr1 | 92295887 | 92296134 | 0.767146 | 1.55325 | 2.025 |
| chr12 | 9950687 | 9950931 | 1.26582 | 2.56292 | 2.025 |
| chr15 | 60903950 | 60904199 | 0.441961 | 0.894866 | 2.025 |
| chr9 | 127181840 | 127182305 | 1.78413 | 3.61254 | 2.025 |
| chr3 | 50648923 | 50649201 | 1.05049 | 2.12711 | 2.025 |
| chr12 | 50569281 | 50569577 | 1.04414 | 2.11433 | 2.025 |
| chr1 | 154118498 | 154118761 | 0.822341 | 1.66535 | 2.025 |
| chr5 | 5419501 | 5419820 | 1.55894 | 3.15708 | 2.025 |
| chrX | 153212736 | 153213153 | 3.14234 | 6.36396 | 2.025 |
| chr7 | 86722848 | 86723126 | 0.439775 | 0.890651 | 2.025 |
| chr12 | 109802206 | 109802418 | 0.595695 | 1.20659 | 2.026 |
| chr17 | 55779551 | 55779758 | 0.621758 | 1.25944 | 2.026 |
| chr3 | 13535927 | 13536346 | 1.79246 | 3.63092 | 2.026 |
| chr1 | 239273934 | 239274206 | 0.576771 | 1.16835 | 2.026 |
| chr12 | 118620054 | 118620374 | 1.77471 | 3.595 | 2.026 |
| chr2 | 27321421 | 27321824 | 2.02604 | 4.10414 | 2.026 |
| chr3 | 163439359 | 163439669 | 0.768247 | 1.55624 | 2.026 |
| chr6 | 26003388 | 26003700 | 1.24911 | 2.53033 | 2.026 |
| chr9 | 100173909 | 100174410 | 3.79291 | 7.68344 | 2.026 |
| chr5 | 116926080 | 116926313 | 0.359563 | 0.728432 | 2.026 |
| chr16 | 2918315 | 2918743 | 1.72531 | 3.4953 | 2.026 |
| chr17 | 30916002 | 30916294 | 0.65147 | 1.31982 | 2.026 |
| chr20 | 12316191 | 12316412 | 0.459746 | 0.93142 | 2.026 |
| chr20 | 9597753 | 9598022 | 1.44124 | 2.92009 | 2.026 |
| chr12 | 110888101 | 110888431 | 2.63047 | 5.32971 | 2.026 |

| | | | | | |
|---|---|---|---|---|---|
| chr12 | 6172274 | 6172727 | 2.18557 | 4.42831 | 2.026 |
| chr14 | 73415229 | 73415612 | 1.42112 | 2.87954 | 2.026 |
| chr12 | 98378780 | 98378995 | 0.475669 | 0.963874 | 2.026 |
| chr10 | 29824450 | 29824785 | 2.13241 | 4.32106 | 2.026 |
| chr7 | 73815531 | 73815932 | 2.34318 | 4.74839 | 2.026 |
| chr15 | 99602226 | 99602540 | 3.03075 | 6.142 | 2.027 |
| chr10 | 121204789 | 121205334 | 2.23614 | 4.53187 | 2.027 |
| chr10 | 71275366 | 71275720 | 2.30987 | 4.68132 | 2.027 |
| chr10 | 92803082 | 92803348 | 0.980223 | 1.98659 | 2.027 |
| chr7 | 44713344 | 44713569 | 0.567138 | 1.14943 | 2.027 |
| chr19 | 49521206 | 49521495 | 1.36109 | 2.75857 | 2.027 |
| chr7 | 36406728 | 36407076 | 1.65798 | 3.36039 | 2.027 |
| chr14 | 99738478 | 99738764 | 1.34975 | 2.7358 | 2.027 |
| chr12 | 3244637 | 3244995 | 1.7035 | 3.45288 | 2.027 |
| chr12 | 132845295 | 132845555 | 0.664712 | 1.34735 | 2.027 |
| chr7 | 15344711 | 15344990 | 1.58798 | 3.21878 | 2.027 |
| chr19 | 38703969 | 38704402 | 0.558157 | 1.13142 | 2.027 |
| chr12 | 68498176 | 68498396 | 0.636929 | 1.2911 | 2.027 |
| chr9 | 28774539 | 28774761 | 1.32829 | 2.6926 | 2.027 |
| chr12 | 55143026 | 55143488 | 1.3072 | 2.64989 | 2.027 |
| chr19 | 40723142 | 40723656 | 2.20087 | 4.46156 | 2.027 |
| chr5 | 74230812 | 74231060 | 1.08719 | 2.20396 | 2.027 |
| chr15 | 101648662 | 101648891 | 0.603179 | 1.22278 | 2.027 |
| chr9 | 7610449 | 7610669 | 0.309808 | 0.628054 | 2.027 |
| chr6 | 145535685 | 145535937 | 0.72636 | 1.47254 | 2.027 |
| chr7 | 158553767 | 158553974 | 1.1963 | 2.42536 | 2.027 |
| chr17 | 73649028 | 73649452 | 2.33992 | 4.74397 | 2.027 |
| chr5 | 53837068 | 53837357 | 0.537623 | 1.09002 | 2.027 |
| chr1 | 109736302 | 109736585 | 1.82644 | 3.70311 | 2.028 |
| chr17 | 12114110 | 12114323 | 0.452696 | 0.917896 | 2.028 |
| chr1 | 99259912 | 99260210 | 1.43506 | 2.90977 | 2.028 |
| chr2 | 100170785 | 100171000 | 0.602137 | 1.22092 | 2.028 |
| chr17 | 56664971 | 56665415 | 2.38064 | 4.82727 | 2.028 |
| chr5 | 110261090 | 110261347 | 0.686469 | 1.39197 | 2.028 |
| chrX | 84339866 | 84340128 | 1.40248 | 2.84386 | 2.028 |
| chr15 | 27834409 | 27834625 | 0.644163 | 1.3062 | 2.028 |
| chr17 | 49298414 | 49298623 | 0.648902 | 1.31582 | 2.028 |
| chr8 | 17557000 | 17557439 | 1.73046 | 3.50917 | 2.028 |
| chr15 | 99394179 | 99394493 | 1.77974 | 3.60933 | 2.028 |
| chr3 | 157336020 | 157336294 | 0.818816 | 1.66062 | 2.028 |
| chr19 | 49729268 | 49729958 | 1.28458 | 2.60524 | 2.028 |
| chr13 | 100418892 | 100419105 | 1.09266 | 2.21604 | 2.028 |
| chr7 | 110003414 | 110003712 | 0.776039 | 1.5739 | 2.028 |
| chr3 | 24227259 | 24227798 | 1.32407 | 2.68538 | 2.028 |
| chr1 | 206679275 | 206679671 | 1.02392 | 2.07667 | 2.028 |
| chr1 | 201941784 | 201942069 | 0.940425 | 1.90736 | 2.028 |
| chr6 | 100449773 | 100450182 | 1.50822 | 3.05895 | 2.028 |
| chr9 | 130496539 | 130497870 | 1.27244 | 2.58077 | 2.028 |
| chr16 | 50699199 | 50699609 | 2.02291 | 4.10312 | 2.028 |
| chr17 | 288431 | 288761 | 0.630529 | 1.27893 | 2.028 |
| chr12 | 106739658 | 106739920 | 1.35566 | 2.74975 | 2.028 |
| chr10 | 58340313 | 58340600 | 0.584848 | 1.18629 | 2.028 |
| chrX | 46763152 | 46763696 | 1.67754 | 3.4027 | 2.028 |
| chr20 | 50317345 | 50317565 | 1.49685 | 3.03623 | 2.028 |
| chr8 | 88856927 | 88857147 | 0.547554 | 1.11074 | 2.029 |
| chr2 | 158354161 | 158354419 | 0.93729 | 1.90137 | 2.029 |
| chr18 | 57364442 | 57364940 | 2.06797 | 4.19519 | 2.029 |
| chr4 | 151219042 | 151219337 | 0.980818 | 1.98977 | 2.029 |
| chr9 | 96776403 | 96776709 | 0.967384 | 1.96268 | 2.029 |
| chr10 | 35887318 | 35887676 | 0.988868 | 2.00628 | 2.029 |
| chr5 | 923422 | 923889 | 2.49774 | 5.06765 | 2.029 |

| | | | | | |
|---|---|---|---|---|---|
| chr16 | 84170722 | 84171005 | 0.519851 | 1.05479 | 2.029 |
| chr10 | 43570856 | 43571373 | 1.05943 | 2.14964 | 2.029 |
| chr6 | 44064440 | 44064755 | 1.12913 | 2.29106 | 2.029 |
| chr6 | 116418227 | 116418493 | 0.839671 | 1.70377 | 2.029 |
| chr3 | 126241090 | 126241455 | 1.92164 | 3.89932 | 2.029 |
| chr17 | 12424675 | 12424957 | 0.797777 | 1.61885 | 2.029 |
| chr3 | 10987370 | 10987885 | 1.77301 | 3.59782 | 2.029 |
| chr6 | 166689130 | 166689516 | 2.47832 | 5.02909 | 2.029 |
| chr13 | 76374751 | 76374963 | 0.625629 | 1.26956 | 2.029 |
| chr2 | 25388950 | 25389215 | 1.79162 | 3.63571 | 2.029 |
| chr12 | 39058492 | 39058829 | 1.42471 | 2.89125 | 2.029 |
| chr16 | 83920793 | 83920997 | 1.20624 | 2.44796 | 2.029 |
| chr19 | 14590759 | 14591261 | 2.2254 | 4.51625 | 2.029 |
| chr2 | 78301190 | 78301469 | 0.432753 | 0.878245 | 2.029 |
| chr11 | 69319533 | 69319750 | 0.461116 | 0.93587 | 2.030 |
| chr13 | 103523003 | 103523285 | 1.29815 | 2.63474 | 2.030 |
| chr14 | 64194750 | 64195052 | 0.643576 | 1.30631 | 2.030 |
| chr8 | 144465654 | 144466268 | 2.50719 | 5.08902 | 2.030 |
| chr12 | 22095022 | 22095291 | 1.14242 | 2.31897 | 2.030 |
| chr8 | 143690478 | 143690783 | 1.00447 | 2.03898 | 2.030 |
| chr20 | 62605248 | 62605579 | 2.94117 | 5.97038 | 2.030 |
| chr4 | 165982046 | 165982301 | 0.615771 | 1.2501 | 2.030 |
| chr12 | 7959505 | 7959811 | 2.06752 | 4.1974 | 2.030 |
| chr1 | 160645287 | 160645713 | 2.03492 | 4.13163 | 2.030 |
| chr5 | 134668336 | 134668541 | 0.975032 | 1.97969 | 2.030 |
| chr2 | 238380163 | 238380437 | 1.87606 | 3.80932 | 2.030 |
| chr7 | 81585941 | 81586157 | 1.50278 | 3.05138 | 2.030 |
| chr8 | 68867703 | 68867980 | 0.411052 | 0.834657 | 2.031 |
| chr10 | 43843926 | 43844262 | 1.76587 | 3.58569 | 2.031 |
| chr19 | 2648526 | 2648765 | 1.39465 | 2.83217 | 2.031 |
| chr1 | 113585337 | 113585578 | 1.09834 | 2.23052 | 2.031 |
| chr14 | 70494552 | 70494888 | 2.18003 | 4.42737 | 2.031 |
| chr7 | 100813004 | 100813536 | 1.53172 | 3.11082 | 2.031 |
| chr13 | 110899905 | 110900114 | 1.13206 | 2.29919 | 2.031 |
| chr13 | 67568372 | 67568636 | 1.34516 | 2.73204 | 2.031 |
| chr1 | 29208566 | 29208891 | 1.62235 | 3.29502 | 2.031 |
| chr19 | 57977140 | 57977436 | 1.10464 | 2.24358 | 2.031 |
| chr12 | 111751412 | 111752107 | 1.47256 | 2.99087 | 2.031 |
| chr8 | 104111936 | 104112146 | 0.494588 | 1.00456 | 2.031 |
| chr12 | 106780002 | 106780355 | 1.28303 | 2.6061 | 2.031 |
| chr12 | 121511837 | 121512146 | 0.491034 | 0.997398 | 2.031 |
| chr7 | 21139218 | 21139440 | 0.541448 | 1.09985 | 2.031 |
| chr1 | 211687335 | 211687638 | 1.62685 | 3.30467 | 2.031 |
| chr17 | 65370749 | 65371149 | 2.39401 | 4.86308 | 2.031 |
| chr1 | 214434821 | 214435235 | 2.07658 | 4.2184 | 2.031 |
| chr8 | 23789644 | 23789854 | 1.25007 | 2.53945 | 2.031 |
| chr2 | 53037638 | 53037842 | 0.992896 | 2.01703 | 2.031 |
| chr12 | 31789595 | 31789916 | 1.49466 | 3.03635 | 2.031 |
| chr20 | 55664648 | 55665281 | 1.77646 | 3.60894 | 2.032 |
| chr6 | 155589302 | 155589541 | 0.993258 | 2.0179 | 2.032 |
| chr10 | 123747305 | 123747517 | 0.518687 | 1.05377 | 2.032 |
| chr5 | 140772090 | 140772541 | 0.671715 | 1.36467 | 2.032 |
| chr22 | 40744140 | 40744342 | 1.21751 | 2.47367 | 2.032 |
| chr12 | 129986279 | 129986645 | 0.50273 | 1.02145 | 2.032 |
| chr3 | 123965920 | 123966167 | 1.01279 | 2.05781 | 2.032 |
| chr9 | 26856467 | 26856710 | 1.12962 | 2.29518 | 2.032 |
| chr17 | 73825983 | 73826707 | 1.4255 | 2.89641 | 2.032 |
| chr13 | 26083942 | 26084256 | 1.23683 | 2.5131 | 2.032 |
| chr17 | 37320895 | 37321208 | 2.31214 | 4.69828 | 2.032 |
| chr2 | 115572370 | 115572729 | 0.634069 | 1.28846 | 2.032 |
| chr8 | 30765741 | 30766163 | 0.731665 | 1.48678 | 2.032 |

| | | | | | |
|---|---|---|---|---|---|
| chr2 | 47214154 | 47214593 | 3.33701 | 6.78111 | 2.032 |
| chr11 | 59447898 | 59448474 | 1.16998 | 2.37759 | 2.032 |
| chr9 | 109224668 | 109224981 | 0.545646 | 1.10885 | 2.032 |
| chr12 | 44183924 | 44184189 | 0.502325 | 1.02085 | 2.032 |
| chr9 | 140107020 | 140107251 | 0.602366 | 1.2242 | 2.032 |
| chr1 | 211604536 | 211604778 | 0.557976 | 1.13399 | 2.032 |
| chr10 | 94334144 | 94334473 | 2.46677 | 5.01349 | 2.032 |
| chr1 | 174200716 | 174201098 | 0.570664 | 1.15985 | 2.032 |
| chr10 | 24262368 | 24262592 | 0.466327 | 0.947829 | 2.033 |
| chr16 | 66344493 | 66344810 | 0.439291 | 0.892883 | 2.033 |
| chr8 | 99793207 | 99793480 | 0.448772 | 0.912183 | 2.033 |
| chr5 | 10863932 | 10864238 | 1.62727 | 3.30778 | 2.033 |
| chr9 | 139240330 | 139240645 | 1.47145 | 2.99113 | 2.033 |
| chr15 | 90800972 | 90801277 | 0.67217 | 1.3664 | 2.033 |
| chr1 | 11124052 | 11124326 | 0.86919 | 1.76703 | 2.033 |
| chr15 | 42349202 | 42349557 | 1.81957 | 3.69927 | 2.033 |
| chr14 | 55712454 | 55712712 | 1.42457 | 2.89623 | 2.033 |
| chr14 | 53427896 | 53428123 | 0.962466 | 1.95683 | 2.033 |
| chr17 | 76130261 | 76130807 | 1.71332 | 3.48344 | 2.033 |
| chr12 | 51159071 | 51159434 | 1.33856 | 2.72158 | 2.033 |
| chr1 | 26560305 | 26560592 | 1.59526 | 3.24354 | 2.033 |
| chr7 | 98439368 | 98439799 | 2.47643 | 5.03539 | 2.033 |
| chr5 | 337945 | 338241 | 1.43554 | 2.91904 | 2.033 |
| chrX | 109388984 | 109389335 | 0.984434 | 2.0019 | 2.034 |
| chr1 | 151438911 | 151439310 | 0.479875 | 0.975867 | 2.034 |
| chr5 | 154383881 | 154384249 | 1.79469 | 3.64966 | 2.034 |
| chr16 | 71474534 | 71474853 | 1.3913 | 2.82941 | 2.034 |
| chr12 | 85967741 | 85967943 | 0.793569 | 1.61384 | 2.034 |
| chr12 | 38797201 | 38797422 | 0.433925 | 0.882463 | 2.034 |
| chr11 | 18785819 | 18786207 | 1.51488 | 3.08081 | 2.034 |
| chr3 | 94243968 | 94244180 | 1.09539 | 2.22769 | 2.034 |
| chr4 | 150817631 | 150817891 | 0.998698 | 2.03111 | 2.034 |
| chrX | 118744078 | 118744404 | 1.16341 | 2.36612 | 2.034 |
| chr3 | 48235963 | 48236298 | 1.76626 | 3.59227 | 2.034 |
| chr7 | 81863840 | 81864060 | 1.14592 | 2.33061 | 2.034 |
| chr17 | 20946807 | 20947201 | 2.02997 | 4.12863 | 2.034 |
| chr5 | 55274080 | 55274318 | 0.785786 | 1.59818 | 2.034 |
| chr17 | 1533930 | 1534178 | 0.553725 | 1.12621 | 2.034 |
| chr7 | 158488641 | 158489063 | 1.17961 | 2.39921 | 2.034 |
| chr9 | 107274816 | 107275046 | 0.421464 | 0.857217 | 2.034 |
| chr19 | 42150452 | 42150886 | 1.89234 | 3.84915 | 2.034 |
| chr11 | 36949850 | 36950242 | 0.464425 | 0.944677 | 2.034 |
| chr14 | 73698597 | 73699027 | 1.10576 | 2.24923 | 2.034 |
| chr9 | 102569614 | 102569897 | 1.61302 | 3.28126 | 2.034 |
| chr4 | 153688395 | 153688683 | 0.725653 | 1.47617 | 2.034 |
| chr4 | 57826418 | 57826740 | 1.42677 | 2.90247 | 2.034 |
| chr10 | 26176067 | 26176362 | 0.602038 | 1.22477 | 2.034 |
| chr15 | 58358433 | 58358915 | 2.89124 | 5.88194 | 2.034 |
| chr11 | 66462178 | 66462639 | 2.95455 | 6.01082 | 2.034 |
| chr17 | 57297755 | 57298006 | 1.57584 | 3.206 | 2.034 |
| chr2 | 10549232 | 10549682 | 2.43724 | 4.95852 | 2.034 |
| chr16 | 4359738 | 4360098 | 2.30971 | 4.69912 | 2.035 |
| chr8 | 90554657 | 90554887 | 0.649098 | 1.32065 | 2.035 |
| chr12 | 50495478 | 50495698 | 0.93728 | 1.90701 | 2.035 |
| chr19 | 56577660 | 56578321 | 2.11239 | 4.29809 | 2.035 |
| chr14 | 24505977 | 24506216 | 2.04042 | 4.15186 | 2.035 |
| chr7 | 132106901 | 132107134 | 0.493946 | 1.00511 | 2.035 |
| chr8 | 89015454 | 89015713 | 0.634936 | 1.29207 | 2.035 |
| chr9 | 4339365 | 4339683 | 2.49779 | 5.083 | 2.035 |
| chr1 | 165045052 | 165045315 | 0.982902 | 2.00023 | 2.035 |
| chr16 | 629074 | 629415 | 2.25399 | 4.58701 | 2.035 |

| | | | | | |
|---|---|---|---|---|---|
| chr8 | 31896226 | 31896463 | 1.50846 | 3.0698 | 2.035 |
| chr15 | 35261908 | 35262251 | 2.26059 | 4.60052 | 2.035 |
| chr12 | 24384556 | 24384758 | 0.489031 | 0.995263 | 2.035 |
| chr8 | 11269033 | 11269460 | 1.46111 | 2.97363 | 2.035 |
| chr1 | 240139681 | 240140111 | 1.88305 | 3.83238 | 2.035 |
| chr10 | 4891826 | 4892206 | 2.2317 | 4.54214 | 2.035 |
| chr3 | 35060387 | 35060589 | 0.470426 | 0.957456 | 2.035 |
| chr4 | 152283903 | 152284155 | 1.73228 | 3.52576 | 2.035 |
| chr7 | 10518755 | 10519151 | 0.583526 | 1.18769 | 2.035 |
| chr12 | 114418280 | 114418670 | 2.2929 | 4.66693 | 2.035 |
| chr22 | 24503037 | 24503242 | 0.597947 | 1.21706 | 2.035 |
| chr7 | 87799118 | 87799330 | 1.14984 | 2.34041 | 2.035 |
| chr16 | 71682870 | 71683203 | 1.00313 | 2.0418 | 2.035 |
| chr7 | 37956692 | 37957028 | 1.53044 | 3.11515 | 2.035 |
| chr1 | 156595380 | 156596075 | 1.22827 | 2.50034 | 2.036 |
| chr7 | 104166326 | 104166578 | 0.500077 | 1.01803 | 2.036 |
| chr8 | 143626826 | 143627444 | 1.71296 | 3.4873 | 2.036 |
| chr19 | 48510109 | 48510363 | 1.34259 | 2.73331 | 2.036 |
| chr3 | 38521339 | 38521672 | 1.69625 | 3.45339 | 2.036 |
| chr5 | 72539271 | 72539554 | 0.407375 | 0.82939 | 2.036 |
| chr19 | 11492437 | 11492923 | 0.88894 | 1.80998 | 2.036 |
| chr12 | 14817493 | 14817872 | 0.793033 | 1.61472 | 2.036 |
| chr7 | 122168995 | 122169226 | 0.969136 | 1.97329 | 2.036 |
| chr7 | 138423485 | 138423811 | 1.56113 | 3.17869 | 2.036 |
| chr5 | 109529751 | 109530079 | 0.605589 | 1.23308 | 2.036 |
| chr5 | 111755879 | 111756342 | 1.82917 | 3.72466 | 2.036 |
| chr20 | 17517634 | 17517982 | 2.19042 | 4.46032 | 2.036 |
| chr11 | 48272131 | 48272383 | 0.543273 | 1.10631 | 2.036 |
| chr1 | 233440275 | 233440610 | 1.16139 | 2.36504 | 2.036 |
| chr11 | 102651301 | 102651558 | 0.99141 | 2.01898 | 2.036 |
| chr19 | 33963866 | 33964089 | 0.499333 | 1.01688 | 2.036 |
| chr8 | 101859391 | 101859984 | 1.19202 | 2.4276 | 2.037 |
| chr6 | 127963156 | 127963462 | 1.24957 | 2.54481 | 2.037 |
| chr2 | 136231215 | 136231468 | 1.13244 | 2.30629 | 2.037 |
| chr6 | 34954506 | 34954974 | 1.18922 | 2.42216 | 2.037 |
| chr17 | 6814402 | 6814627 | 1.06911 | 2.17754 | 2.037 |
| chr8 | 110294248 | 110294713 | 1.04535 | 2.12916 | 2.037 |
| chr11 | 77757199 | 77757651 | 2.60259 | 5.30097 | 2.037 |
| chr17 | 20943362 | 20943774 | 1.17209 | 2.38732 | 2.037 |
| chr3 | 186334867 | 186335156 | 1.70093 | 3.46448 | 2.037 |
| chr1 | 22992315 | 22992544 | 0.982293 | 2.00081 | 2.037 |
| chr5 | 52211091 | 52211562 | 1.28757 | 2.62262 | 2.037 |
| chr11 | 13365669 | 13366015 | 1.50928 | 3.07427 | 2.037 |
| chr7 | 33515990 | 33516200 | 0.531439 | 1.0825 | 2.037 |
| chr10 | 123368098 | 123368563 | 2.65461 | 5.40734 | 2.037 |
| chr11 | 18241813 | 18242053 | 0.477004 | 0.971636 | 2.037 |
| chr17 | 78258475 | 78258802 | 0.819648 | 1.6696 | 2.037 |
| chr20 | 37554587 | 37554961 | 2.46347 | 5.01816 | 2.037 |
| chr14 | 105189757 | 105190402 | 2.44752 | 4.98572 | 2.037 |
| chr19 | 31996426 | 31996777 | 0.50877 | 1.03639 | 2.037 |
| chr9 | 10776114 | 10776388 | 0.581942 | 1.18545 | 2.037 |
| chr6 | 35172868 | 35173148 | 1.92862 | 3.92876 | 2.037 |
| chr2 | 147844679 | 147844885 | 0.647139 | 1.3184 | 2.037 |
| chr22 | 29251644 | 29251948 | 1.31803 | 2.68518 | 2.037 |
| chr17 | 66978844 | 66979087 | 0.462082 | 0.941402 | 2.037 |
| chr2 | 219717552 | 219717852 | 1.31391 | 2.67691 | 2.037 |
| chr15 | 73757037 | 73757240 | 0.856166 | 1.74434 | 2.037 |
| chr16 | 88814341 | 88814705 | 2.03837 | 4.15305 | 2.037 |
| chr11 | 37346092 | 37346360 | 0.571326 | 1.16406 | 2.037 |
| chr1 | 155163259 | 155163611 | 2.52776 | 5.15081 | 2.038 |
| chr3 | 152743861 | 152744084 | 1.15442 | 2.35239 | 2.038 |

| | | | | | |
|---|---|---|---|---|---|
| chr4 | 169549526 | 169549786 | 0.596177 | 1.21485 | 2.038 |
| chr15 | 96870494 | 96870737 | 1.28342 | 2.61534 | 2.038 |
| chr19 | 42400086 | 42400528 | 1.2885 | 2.6257 | 2.038 |
| chr14 | 52246791 | 52247115 | 1.8454 | 3.76058 | 2.038 |
| chr1 | 157459150 | 157459498 | 1.65756 | 3.37782 | 2.038 |
| chr12 | 93813092 | 93813499 | 0.94231 | 1.92036 | 2.038 |
| chr1 | 213081133 | 213081388 | 0.513427 | 1.04636 | 2.038 |
| chr4 | 102711817 | 102712224 | 0.671991 | 1.36952 | 2.038 |
| chr22 | 19897526 | 19897839 | 1.89377 | 3.85974 | 2.038 |
| chr17 | 68618523 | 68618853 | 1.52693 | 3.11215 | 2.038 |
| chr8 | 58255729 | 58256359 | 1.84142 | 3.75316 | 2.038 |
| chr20 | 47603542 | 47603747 | 1.12833 | 2.29976 | 2.038 |
| chr12 | 126590530 | 126590741 | 0.516794 | 1.05337 | 2.038 |
| chr17 | 38020478 | 38020709 | 0.94257 | 1.92124 | 2.038 |
| chr5 | 125936248 | 125937249 | 1.46005 | 2.97616 | 2.038 |
| chr10 | 101245055 | 101245291 | 1.12518 | 2.29369 | 2.039 |
| chr4 | 156708308 | 156708602 | 1.30345 | 2.65723 | 2.039 |
| chr12 | 82768980 | 82769259 | 0.607887 | 1.2393 | 2.039 |
| chr11 | 56438971 | 56439272 | 0.50477 | 1.02908 | 2.039 |
| chr3 | 142547534 | 142547863 | 1.80979 | 3.68973 | 2.039 |
| chr10 | 94713049 | 94713252 | 0.569612 | 1.16134 | 2.039 |
| chr15 | 27588185 | 27588421 | 0.576309 | 1.175 | 2.039 |
| chr14 | 98349653 | 98349861 | 0.559943 | 1.14169 | 2.039 |
| chr1 | 153800087 | 153800350 | 0.831908 | 1.69626 | 2.039 |
| chr17 | 41188317 | 41188637 | 0.826317 | 1.68488 | 2.039 |
| chr17 | 11357648 | 11357979 | 0.570051 | 1.16236 | 2.039 |
| chr2 | 57133057 | 57133304 | 0.616772 | 1.25764 | 2.039 |
| chr10 | 79010996 | 79011340 | 1.59705 | 3.25656 | 2.039 |
| chr16 | 90148351 | 90148735 | 3.24105 | 6.60912 | 2.039 |
| chr2 | 239015887 | 239016117 | 0.743504 | 1.51616 | 2.039 |
| chr5 | 88761534 | 88761882 | 0.588604 | 1.20031 | 2.039 |
| chr11 | 122666881 | 122667154 | 1.01742 | 2.07488 | 2.039 |
| chr12 | 6574724 | 6575014 | 2.12019 | 4.32401 | 2.039 |
| chr11 | 46344287 | 46344569 | 0.748187 | 1.52589 | 2.039 |
| chr17 | 21219823 | 21220255 | 2.5392 | 5.17862 | 2.039 |
| chr12 | 31424892 | 31425223 | 1.27118 | 2.59261 | 2.040 |
| chr2 | 161850961 | 161851215 | 1.00824 | 2.05635 | 2.040 |
| chr1 | 21022895 | 21023662 | 1.16823 | 2.38268 | 2.040 |
| chr9 | 5186431 | 5186780 | 1.96293 | 4.00358 | 2.040 |
| chr20 | 45848768 | 45849112 | 1.51517 | 3.0904 | 2.040 |
| chr19 | 36046559 | 36046910 | 1.5804 | 3.22372 | 2.040 |
| chr6 | 80579299 | 80579692 | 1.99481 | 4.06906 | 2.040 |
| chr7 | 151065152 | 151065558 | 1.77144 | 3.61379 | 2.040 |
| chr12 | 49147289 | 49147540 | 1.38146 | 2.81825 | 2.040 |
| chr2 | 85154034 | 85154504 | 1.84649 | 3.76729 | 2.040 |
| chr20 | 18257354 | 18257734 | 1.76392 | 3.59895 | 2.040 |
| chrX | 129326251 | 129326586 | 1.74402 | 3.55842 | 2.040 |
| chr7 | 44236034 | 44236349 | 1.0348 | 2.11137 | 2.040 |
| chrX | 46838012 | 46838298 | 1.26712 | 2.58541 | 2.040 |
| chr10 | 114480938 | 114481154 | 1.39637 | 2.84914 | 2.040 |
| chr8 | 135215125 | 135215344 | 0.631988 | 1.28951 | 2.040 |
| chr8 | 127064192 | 127064423 | 0.660618 | 1.34794 | 2.040 |
| chr14 | 66962205 | 66962586 | 1.55973 | 3.18255 | 2.040 |
| chr17 | 47647329 | 47647801 | 2.90029 | 5.91792 | 2.040 |
| chrX | 139815955 | 139816259 | 1.46926 | 2.99805 | 2.041 |
| chr9 | 136019758 | 136020252 | 2.51473 | 5.13137 | 2.041 |
| chr10 | 23991760 | 23992265 | 1.64785 | 3.36251 | 2.041 |
| chr10 | 20296049 | 20296570 | 0.432906 | 0.88337 | 2.041 |
| chr3 | 85946605 | 85946829 | 0.686114 | 1.40008 | 2.041 |
| chr21 | 35739685 | 35739988 | 0.982227 | 2.00436 | 2.041 |
| chr8 | 40471024 | 40471397 | 0.70581 | 1.44038 | 2.041 |

| | | | | | |
|---|---|---|---|---|---|
| chr7 | 77670497 | 77670750 | 0.436348 | 0.890542 | 2.041 |
| chr15 | 99949228 | 99949608 | 1.46771 | 2.99549 | 2.041 |
| chr5 | 134540072 | 134540289 | 1.0377 | 2.1179 | 2.041 |
| chr19 | 10527174 | 10527919 | 2.29741 | 4.68897 | 2.041 |
| chr5 | 113957600 | 113957813 | 0.600558 | 1.22573 | 2.041 |
| chr8 | 48051513 | 48051899 | 0.511576 | 1.04419 | 2.041 |
| chr7 | 134143896 | 134144191 | 1.83406 | 3.74363 | 2.041 |
| chr20 | 51934359 | 51934624 | 0.540741 | 1.10382 | 2.041 |
| chr21 | 16236688 | 16237243 | 1.04939 | 2.14213 | 2.041 |
| chr9 | 129263100 | 129263687 | 2.13414 | 4.35667 | 2.041 |
| chr6 | 151656056 | 151656259 | 1.05058 | 2.14471 | 2.041 |
| chr7 | 13218744 | 13218982 | 1.13145 | 2.30982 | 2.041 |
| chr2 | 31577784 | 31578040 | 1.04172 | 2.12665 | 2.041 |
| chr11 | 250793 | 251135 | 1.63438 | 3.33667 | 2.042 |
| chr3 | 118320126 | 118320354 | 0.399182 | 0.814967 | 2.042 |
| chr3 | 54887808 | 54888191 | 1.91682 | 3.91359 | 2.042 |
| chr17 | 58716064 | 58716441 | 1.91588 | 3.91176 | 2.042 |
| chr10 | 86300264 | 86300687 | 1.69638 | 3.46363 | 2.042 |
| chrX | 128674064 | 128674349 | 1.48857 | 3.03933 | 2.042 |
| chr11 | 120039687 | 120040233 | 1.12241 | 2.29172 | 2.042 |
| chr15 | 43577758 | 43578049 | 1.33692 | 2.72971 | 2.042 |
| chr11 | 10388156 | 10388600 | 1.93335 | 3.94756 | 2.042 |
| chr1 | 106909437 | 106909642 | 0.556679 | 1.13666 | 2.042 |
| chr5 | 171720421 | 171720884 | 1.87532 | 3.82923 | 2.042 |
| chr11 | 128718761 | 128719056 | 1.09936 | 2.24484 | 2.042 |
| chr10 | 93058030 | 93058380 | 2.55879 | 5.22495 | 2.042 |
| chr5 | 50189047 | 50189305 | 0.594904 | 1.21478 | 2.042 |
| chrX | 42300943 | 42301183 | 0.481863 | 0.983955 | 2.042 |
| chr6 | 116324067 | 116324296 | 0.352917 | 0.720673 | 2.042 |
| chr7 | 18705769 | 18706108 | 2.07763 | 4.24288 | 2.042 |
| chr6 | 15491611 | 15491961 | 1.89181 | 3.86344 | 2.042 |
| chr19 | 54356126 | 54356563 | 2.46851 | 5.04136 | 2.042 |
| chr9 | 117197050 | 117197492 | 1.13642 | 2.32096 | 2.042 |
| chr8 | 32759507 | 32759721 | 0.37002 | 0.755714 | 2.042 |
| chr2 | 73119832 | 73120169 | 2.2277 | 4.54991 | 2.042 |
| chr20 | 1089055 | 1089497 | 2.10106 | 4.29135 | 2.042 |
| chr17 | 25896322 | 25896694 | 1.88651 | 3.85332 | 2.043 |
| chr14 | 53942502 | 53942885 | 0.652482 | 1.33282 | 2.043 |
| chr5 | 76788212 | 76788542 | 1.98569 | 4.05619 | 2.043 |
| chr3 | 72867298 | 72867555 | 1.32876 | 2.7143 | 2.043 |
| chr12 | 16538795 | 16539377 | 0.898969 | 1.83637 | 2.043 |
| chr19 | 3484964 | 3485313 | 1.63262 | 3.33513 | 2.043 |
| chr7 | 75887343 | 75887599 | 0.745453 | 1.52286 | 2.043 |
| chr9 | 25210513 | 25210770 | 0.605321 | 1.23659 | 2.043 |
| chr12 | 109344908 | 109345319 | 2.25842 | 4.61406 | 2.043 |
| chr20 | 57966017 | 57966290 | 0.579039 | 1.18301 | 2.043 |
| chr20 | 335055 | 335515 | 2.42761 | 4.96069 | 2.043 |
| chr1 | 53642471 | 53642776 | 1.33017 | 2.71817 | 2.043 |
| chr10 | 10499917 | 10500276 | 0.796853 | 1.62836 | 2.043 |
| chr1 | 21696985 | 21697454 | 2.45828 | 5.02392 | 2.044 |
| chr12 | 117592992 | 117593441 | 2.6354 | 5.38605 | 2.044 |
| chr19 | 24043085 | 24043291 | 0.360231 | 0.736221 | 2.044 |
| chr11 | 123937041 | 123937345 | 0.507937 | 1.03816 | 2.044 |
| chr10 | 32496353 | 32496731 | 1.37652 | 2.81357 | 2.044 |
| chr8 | 133085951 | 133086154 | 0.711127 | 1.45353 | 2.044 |
| chr19 | 46935151 | 46935443 | 1.31695 | 2.69203 | 2.044 |
| chr15 | 92525747 | 92525975 | 0.977601 | 1.99847 | 2.044 |
| chr19 | 55031147 | 55031431 | 0.399629 | 0.817019 | 2.044 |
| chr6 | 85138047 | 85138406 | 1.55066 | 3.17023 | 2.044 |
| chr12 | 48152724 | 48152989 | 0.883034 | 1.80537 | 2.045 |
| chr11 | 86282660 | 86282983 | 1.30392 | 2.66598 | 2.045 |

| | | | | | |
|---|---|---|---|---|---|
| chr9 | 11016012 | 11016251 | 0.569265 | 1.16395 | 2.045 |
| chr5 | 22494003 | 22494285 | 0.489725 | 1.00132 | 2.045 |
| chr2 | 241853705 | 241854207 | 1.66507 | 3.4046 | 2.045 |
| chr17 | 66168591 | 66169082 | 1.48569 | 3.03796 | 2.045 |
| chr19 | 17403698 | 17403916 | 1.4266 | 2.91713 | 2.045 |
| chr9 | 139061906 | 139062704 | 1.43482 | 2.93393 | 2.045 |
| chr12 | 85862473 | 85862771 | 1.29943 | 2.65715 | 2.045 |
| chr1 | 118438185 | 118438480 | 1.40578 | 2.87476 | 2.045 |
| chr5 | 142150814 | 142151028 | 1.13801 | 2.32719 | 2.045 |
| chrX | 154444970 | 154445306 | 1.12251 | 2.29558 | 2.045 |
| chr19 | 36342148 | 36342520 | 1.06604 | 2.18014 | 2.045 |
| chr7 | 125525436 | 125525676 | 0.48645 | 0.994847 | 2.045 |
| chr1 | 74970863 | 74971146 | 0.618902 | 1.26578 | 2.045 |
| chr11 | 45204498 | 45204982 | 1.54206 | 3.15407 | 2.045 |
| chr8 | 89105126 | 89105353 | 0.674681 | 1.38008 | 2.046 |
| chr11 | 7534265 | 7534820 | 1.58766 | 3.24764 | 2.046 |
| chr22 | 37594546 | 37594817 | 1.91769 | 3.92289 | 2.046 |
| chr1 | 17662583 | 17663061 | 1.58874 | 3.25001 | 2.046 |
| chr7 | 40174312 | 40174728 | 1.88114 | 3.84818 | 2.046 |
| chr12 | 49437344 | 49437934 | 1.56395 | 3.19942 | 2.046 |
| chr11 | 8816057 | 8816662 | 0.683171 | 1.39759 | 2.046 |
| chr8 | 110574015 | 110574201 | 1.05455 | 2.15735 | 2.046 |
| chr18 | 48804524 | 48804771 | 1.1095 | 2.26981 | 2.046 |
| chr11 | 1536697 | 1537167 | 2.11509 | 4.32747 | 2.046 |
| chr14 | 105556774 | 105557171 | 2.46327 | 5.04004 | 2.046 |
| chrX | 7909782 | 7910059 | 1.16288 | 2.37936 | 2.046 |
| chr1 | 198125265 | 198125543 | 2.13927 | 4.37726 | 2.046 |
| chr11 | 44998982 | 44999191 | 1.16839 | 2.39074 | 2.046 |
| chr5 | 96785095 | 96785359 | 0.540611 | 1.10622 | 2.046 |
| chr4 | 90175986 | 90176226 | 1.5356 | 3.14236 | 2.046 |
| chr12 | 121498316 | 121498737 | 1.33191 | 2.72555 | 2.046 |
| chr2 | 30295156 | 30295478 | 0.934092 | 1.91148 | 2.046 |
| chr19 | 40013699 | 40013902 | 0.639485 | 1.30867 | 2.046 |
| chr4 | 980923 | 981264 | 1.12597 | 2.30427 | 2.046 |
| chr8 | 133502919 | 133503366 | 0.504015 | 1.03146 | 2.046 |
| chr1 | 201154142 | 201154453 | 0.664882 | 1.36073 | 2.047 |
| chr5 | 134094238 | 134094539 | 1.58824 | 3.25049 | 2.047 |
| chr16 | 50357602 | 50357819 | 1.00826 | 2.06351 | 2.047 |
| chr9 | 11674311 | 11674526 | 0.480213 | 0.982811 | 2.047 |
| chr11 | 67382821 | 67383213 | 1.82392 | 3.73287 | 2.047 |
| chr2 | 46521388 | 46521598 | 1.26386 | 2.58672 | 2.047 |
| chr17 | 79353226 | 79353597 | 0.480636 | 0.983771 | 2.047 |
| chr6 | 100194174 | 100194482 | 1.11789 | 2.28812 | 2.047 |
| chr3 | 9542082 | 9542762 | 1.38757 | 2.84015 | 2.047 |
| chr10 | 6448276 | 6448491 | 0.913897 | 1.87066 | 2.047 |
| chr11 | 84275694 | 84275968 | 1.96287 | 4.01793 | 2.047 |
| chr1 | 192775015 | 192775224 | 1.07689 | 2.20437 | 2.047 |
| chr10 | 120327827 | 120328104 | 1.1795 | 2.41457 | 2.047 |
| chr11 | 30607847 | 30608107 | 0.733695 | 1.50196 | 2.047 |
| chr1 | 15105654 | 15105989 | 1.78234 | 3.64868 | 2.047 |
| chr1 | 244436320 | 244436637 | 1.43972 | 2.94748 | 2.047 |
| chr17 | 40700140 | 40700497 | 1.87726 | 3.84348 | 2.047 |
| chr22 | 42564986 | 42565375 | 1.9558 | 4.00435 | 2.047 |
| chr12 | 94333956 | 94334245 | 0.549848 | 1.12586 | 2.048 |
| chrX | 135005605 | 135005815 | 0.857216 | 1.75535 | 2.048 |
| chr14 | 105956866 | 105957273 | 1.6791 | 3.43852 | 2.048 |
| chr22 | 24817471 | 24817775 | 0.632962 | 1.29621 | 2.048 |
| chr1 | 156195246 | 156195522 | 2.04218 | 4.1821 | 2.048 |
| chr9 | 26139707 | 26139916 | 0.516403 | 1.05754 | 2.048 |
| chr1 | 20703387 | 20703733 | 1.26204 | 2.58469 | 2.048 |
| chr8 | 99853157 | 99853381 | 1.53312 | 3.13989 | 2.048 |

| | | | | | |
|---|---|---|---|---|---|
| chr11 | 36585994 | 36586201 | 1.07605 | 2.20381 | 2.048 |
| chr7 | 148659877 | 148660092 | 1.61611 | 3.30997 | 2.048 |
| chr1 | 2345796 | 2346224 | 1.84299 | 3.77472 | 2.048 |
| chr12 | 133191464 | 133191822 | 2.31795 | 4.74767 | 2.048 |
| chr2 | 59332512 | 59332965 | 0.613583 | 1.25675 | 2.048 |
| chr1 | 155022852 | 155023280 | 2.15978 | 4.42381 | 2.048 |
| chr6 | 41559788 | 41560120 | 1.537 | 3.1482 | 2.048 |
| chr2 | 13003953 | 13004196 | 0.422319 | 0.86504 | 2.048 |
| chr7 | 146704262 | 146704534 | 0.626043 | 1.28235 | 2.048 |
| chr18 | 24878376 | 24878630 | 0.482995 | 0.989344 | 2.048 |
| chr2 | 237980887 | 237981359 | 1.92671 | 3.94685 | 2.048 |
| chr14 | 27712163 | 27712405 | 0.680451 | 1.39392 | 2.049 |
| chr4 | 147016312 | 147016590 | 1.19692 | 2.45209 | 2.049 |
| chr5 | 104840327 | 104840686 | 0.572696 | 1.17327 | 2.049 |
| chr7 | 28839792 | 28840138 | 0.470689 | 0.964305 | 2.049 |
| chr17 | 65272265 | 65272585 | 1.20635 | 2.4715 | 2.049 |
| chr19 | 39035089 | 39035320 | 1.30854 | 2.68086 | 2.049 |
| chr5 | 132946639 | 132947002 | 1.37923 | 2.82578 | 2.049 |
| chr4 | 167515563 | 167515798 | 1.40793 | 2.88465 | 2.049 |
| chr9 | 25636062 | 25636347 | 0.548365 | 1.12353 | 2.049 |
| chr17 | 25981617 | 25982372 | 2.30518 | 4.72311 | 2.049 |
| chr5 | 175267343 | 175267586 | 1.33456 | 2.73449 | 2.049 |
| chr6 | 105872086 | 105872417 | 1.10893 | 2.2722 | 2.049 |
| chr3 | 38065556 | 38065962 | 0.493987 | 1.01219 | 2.049 |
| chr7 | 134231935 | 134232442 | 1.64958 | 3.38005 | 2.049 |
| chr1 | 85401922 | 85402224 | 1.58421 | 3.24612 | 2.049 |
| chr10 | 63510700 | 63511052 | 1.64613 | 3.37307 | 2.049 |
| chr12 | 4855845 | 4856173 | 0.426907 | 0.874805 | 2.049 |
| chr17 | 7738158 | 7738556 | 2.21582 | 4.54063 | 2.049 |
| chr2 | 169998283 | 169998656 | 1.50106 | 3.07597 | 2.049 |
| chr1 | 163454083 | 163454296 | 0.521277 | 1.06826 | 2.049 |
| chr12 | 57977045 | 57977603 | 2.04544 | 4.19184 | 2.049 |
| chr9 | 81744831 | 81745115 | 2.0564 | 4.21435 | 2.049 |
| chr11 | 75259832 | 75260074 | 0.458898 | 0.940466 | 2.049 |
| chr19 | 3879778 | 3880011 | 1.13749 | 2.33122 | 2.049 |
| chr12 | 10186447 | 10186694 | 0.979772 | 2.0081 | 2.050 |
| chr9 | 116844827 | 116845085 | 1.76233 | 3.61206 | 2.050 |
| chr3 | 111707986 | 111708283 | 1.39301 | 2.85513 | 2.050 |
| chr13 | 31377027 | 31377375 | 1.65903 | 3.40041 | 2.050 |
| chr2 | 183531748 | 183532049 | 0.871895 | 1.78707 | 2.050 |
| chr4 | 159588863 | 159589147 | 1.23797 | 2.53745 | 2.050 |
| chrX | 96025419 | 96025647 | 0.730082 | 1.49645 | 2.050 |
| chr19 | 17564230 | 17564610 | 1.64422 | 3.37018 | 2.050 |
| chr18 | 6886233 | 6886566 | 0.966142 | 1.98039 | 2.050 |
| chr5 | 140207458 | 140208250 | 0.513971 | 1.05354 | 2.050 |
| chr4 | 183846445 | 183846799 | 1.53115 | 3.13868 | 2.050 |
| chr10 | 80063176 | 80063447 | 1.18637 | 2.43195 | 2.050 |
| chr17 | 36130889 | 36131241 | 2.02922 | 4.15978 | 2.050 |
| chr14 | 34967980 | 34968343 | 1.69291 | 3.47046 | 2.050 |
| chr1 | 226735373 | 226735767 | 3.18685 | 6.53311 | 2.050 |
| chr20 | 61531582 | 61532024 | 2.17956 | 4.46823 | 2.050 |
| chr12 | 74329540 | 74329875 | 1.58163 | 3.2425 | 2.050 |
| chr4 | 90870957 | 90871278 | 0.974491 | 1.99784 | 2.050 |
| chr11 | 66156384 | 66156859 | 1.84067 | 3.77367 | 2.050 |
| chr3 | 194827140 | 194827547 | 1.7893 | 3.66836 | 2.050 |
| chr8 | 55182099 | 55182353 | 0.5425 | 1.11222 | 2.050 |
| chr16 | 368700 | 368938 | 0.801784 | 1.64381 | 2.050 |
| gl000202_r | 12726 | 13179 | 2.62872 | 5.38939 | 2.050 |
| chr19 | 47747441 | 47747845 | 2.40442 | 4.92988 | 2.050 |
| chr4 | 79873124 | 79873376 | 1.15433 | 2.36688 | 2.050 |
| chr5 | 40688295 | 40688599 | 1.32804 | 2.72312 | 2.050 |

| | | | | | |
|---|---|---|---|---|---|
| chr2 | 110914788 | 110915050 | 1.3352 | 2.73792 | 2.051 |
| chr1 | 205680250 | 205680613 | 1.44465 | 2.96239 | 2.051 |
| chr7 | 150941363 | 150941785 | 1.68636 | 3.45804 | 2.051 |
| chr5 | 14405372 | 14405729 | 1.33931 | 2.74641 | 2.051 |
| chr17 | 71948902 | 71949199 | 1.52789 | 3.13316 | 2.051 |
| chr9 | 129373906 | 129374176 | 2.41684 | 4.95614 | 2.051 |
| chr2 | 64558036 | 64558280 | 1.35446 | 2.77763 | 2.051 |
| chr3 | 98520285 | 98520556 | 1.02531 | 2.10264 | 2.051 |
| chr5 | 27523749 | 27524098 | 0.52701 | 1.08078 | 2.051 |
| chr4 | 153516942 | 153517147 | 0.468363 | 0.960514 | 2.051 |
| chr16 | 66901813 | 66902141 | 1.77939 | 3.64919 | 2.051 |
| chr11 | 69883153 | 69883358 | 0.639763 | 1.31206 | 2.051 |
| chr16 | 9426211 | 9426438 | 0.48275 | 0.990083 | 2.051 |
| chr5 | 148970685 | 148970987 | 0.588679 | 1.20739 | 2.051 |
| chrX | 103588361 | 103588700 | 1.34276 | 2.75403 | 2.051 |
| chr8 | 1009162 | 1009408 | 0.516088 | 1.05858 | 2.051 |
| chr3 | 146757569 | 146757905 | 0.502475 | 1.03066 | 2.051 |
| chr10 | 95836138 | 95836365 | 0.929611 | 1.90682 | 2.051 |
| chr10 | 80529385 | 80529602 | 0.404587 | 0.8299 | 2.051 |
| chr5 | 165306741 | 165306948 | 0.347169 | 0.712152 | 2.051 |
| chr8 | 63057232 | 63057469 | 0.457431 | 0.938362 | 2.051 |
| chr12 | 129266934 | 129267120 | 1.46818 | 3.01219 | 2.052 |
| chr3 | 32219384 | 32219758 | 1.83152 | 3.758 | 2.052 |
| chr8 | 71465288 | 71465515 | 0.522081 | 1.07129 | 2.052 |
| chr1 | 245242364 | 245242671 | 1.48142 | 3.03988 | 2.052 |
| chr7 | 98374825 | 98375433 | 1.45211 | 2.97973 | 2.052 |
| chr22 | 37656892 | 37657300 | 1.28015 | 2.62689 | 2.052 |
| chr15 | 94252081 | 94252336 | 0.541433 | 1.11108 | 2.052 |
| chr15 | 87487280 | 87487558 | 0.464895 | 0.954023 | 2.052 |
| chr9 | 33287063 | 33287311 | 1.4928 | 3.06349 | 2.052 |
| chr7 | 18630471 | 18630730 | 0.892155 | 1.83087 | 2.052 |
| chr2 | 18416852 | 18417059 | 0.469592 | 0.963703 | 2.052 |
| chr4 | 186017299 | 186017741 | 0.713055 | 1.46337 | 2.052 |
| chr17 | 33611111 | 33611409 | 0.502489 | 1.03125 | 2.052 |
| chr13 | 48701127 | 48701340 | 0.882316 | 1.81083 | 2.052 |
| chr12 | 122393646 | 122394148 | 1.13508 | 2.32962 | 2.052 |
| chr17 | 64563332 | 64563676 | 2.08339 | 4.2759 | 2.052 |
| chr10 | 94194790 | 94195045 | 1.45584 | 2.98795 | 2.052 |
| chr15 | 65692974 | 65693986 | 1.62978 | 3.345 | 2.052 |
| chr12 | 97379199 | 97379661 | 2.81137 | 5.77025 | 2.052 |
| chr15 | 100417289 | 100417519 | 0.569192 | 1.16826 | 2.052 |
| chr22 | 37963366 | 37963628 | 1.57333 | 3.22927 | 2.053 |
| chr5 | 139448063 | 139448265 | 0.532248 | 1.09245 | 2.053 |
| chr20 | 1293981 | 1294512 | 2.05401 | 4.21603 | 2.053 |
| chr3 | 88229638 | 88229850 | 0.885334 | 1.81722 | 2.053 |
| chr11 | 130849380 | 130849582 | 0.808889 | 1.66034 | 2.053 |
| chr8 | 140717977 | 140718418 | 1.85632 | 3.81033 | 2.053 |
| chr15 | 96900082 | 96900520 | 2.38231 | 4.89002 | 2.053 |
| chr16 | 30381212 | 30381467 | 0.553298 | 1.13573 | 2.053 |
| chr9 | 34646872 | 34647109 | 1.11867 | 2.29625 | 2.053 |
| chr10 | 73808289 | 73808733 | 2.50872 | 5.14972 | 2.053 |
| chr18 | 29869562 | 29869804 | 1.19962 | 2.4625 | 2.053 |
| chr16 | 49316057 | 49316353 | 1.19815 | 2.4595 | 2.053 |
| chr6 | 7243490 | 7244014 | 1.06982 | 2.19611 | 2.053 |
| chr10 | 20240392 | 20240623 | 0.522881 | 1.07339 | 2.053 |
| chr2 | 232537593 | 232537963 | 1.0757 | 2.20832 | 2.053 |
| chr19 | 8418883 | 8419086 | 1.26085 | 2.58846 | 2.053 |
| chr6 | 34131667 | 34131901 | 1.76657 | 3.62674 | 2.053 |
| chr2 | 208491914 | 208492207 | 1.70925 | 3.50933 | 2.053 |
| chr9 | 15837098 | 15837416 | 1.25701 | 2.58105 | 2.053 |
| chr5 | 81608214 | 81608612 | 1.25206 | 2.57105 | 2.053 |

| | | | | | |
|---|---|---|---|---|---|
| chr7 | 103871924 | 103872215 | 1.45334 | 2.98437 | 2.053 |
| chr4 | 141444961 | 141445242 | 2.12139 | 4.35646 | 2.054 |
| chr8 | 14914200 | 14914636 | 0.490007 | 1.00629 | 2.054 |
| chr17 | 79174611 | 79175183 | 1.06688 | 2.19098 | 2.054 |
| chr15 | 31822678 | 31823044 | 0.407257 | 0.836361 | 2.054 |
| chr15 | 90630720 | 90630922 | 0.858942 | 1.76398 | 2.054 |
| chr17 | 48940758 | 48941049 | 1.72745 | 3.54762 | 2.054 |
| chr8 | 79788587 | 79788824 | 0.575432 | 1.18177 | 2.054 |
| chr3 | 154200042 | 154200269 | 0.523235 | 1.07464 | 2.054 |
| chr17 | 30334162 | 30334498 | 2.73751 | 5.62254 | 2.054 |
| chr4 | 71570193 | 71570450 | 0.853127 | 1.75225 | 2.054 |
| chrX | 151800941 | 151801273 | 1.35479 | 2.78266 | 2.054 |
| chr7 | 50490871 | 50491081 | 0.453052 | 0.930547 | 2.054 |
| chr12 | 44952571 | 44952847 | 0.405887 | 0.833683 | 2.054 |
| chr15 | 82400784 | 82401243 | 3.14273 | 6.45527 | 2.054 |
| chr7 | 151479673 | 151480412 | 2.1761 | 4.46986 | 2.054 |
| chr4 | 40926595 | 40926873 | 1.72162 | 3.53638 | 2.054 |
| chr19 | 17178284 | 17178584 | 1.74761 | 3.58982 | 2.054 |
| chr19 | 42944473 | 42945038 | 2.27662 | 4.67664 | 2.054 |
| chr3 | 8178237 | 8178488 | 0.472253 | 0.970168 | 2.054 |
| chr7 | 25949812 | 25950088 | 0.522578 | 1.07358 | 2.054 |
| chr18 | 47794821 | 47795160 | 2.07586 | 4.26468 | 2.054 |
| chr14 | 78506857 | 78507361 | 0.594342 | 1.22104 | 2.054 |
| chr20 | 49404584 | 49404915 | 2.05501 | 4.22191 | 2.054 |
| chr10 | 44210260 | 44210489 | 0.499514 | 1.02623 | 2.054 |
| chr10 | 123902250 | 123902657 | 1.66985 | 3.43073 | 2.055 |
| chr10 | 30291392 | 30291692 | 2.35527 | 4.83913 | 2.055 |
| chr9 | 806593 | 806888 | 1.54619 | 3.1768 | 2.055 |
| chr4 | 83383591 | 83383812 | 1.47115 | 3.02268 | 2.055 |
| chr1 | 178000963 | 178001309 | 1.90247 | 3.90894 | 2.055 |
| chr7 | 51135305 | 51135886 | 1.07758 | 2.21419 | 2.055 |
| chr3 | 195577753 | 195578138 | 2.0365 | 4.18467 | 2.055 |
| chr18 | 45557408 | 45557691 | 1.54971 | 3.18444 | 2.055 |
| chr5 | 95296871 | 95297351 | 2.09939 | 4.31402 | 2.055 |
| chr8 | 30413308 | 30413936 | 1.76843 | 3.63404 | 2.055 |
| chr16 | 70504787 | 70505159 | 1.58993 | 3.26728 | 2.055 |
| chr7 | 19748635 | 19748915 | 1.64551 | 3.38153 | 2.055 |
| chr9 | 139922738 | 139924412 | 1.45188 | 2.98369 | 2.055 |
| chr20 | 1726097 | 1726361 | 0.607099 | 1.24774 | 2.055 |
| chrX | 102118624 | 102118967 | 1.21111 | 2.48913 | 2.055 |
| chr3 | 186494 | 186730 | 0.761246 | 1.56456 | 2.055 |
| chr17 | 11024276 | 11024648 | 0.585212 | 1.20279 | 2.055 |
| chr1 | 189189980 | 189190229 | 1.19148 | 2.44894 | 2.055 |
| chr1 | 216484249 | 216484668 | 0.643945 | 1.32357 | 2.055 |
| chr19 | 11404457 | 11404764 | 2.23021 | 4.58402 | 2.055 |
| chr20 | 45328349 | 45328752 | 1.29158 | 2.65477 | 2.055 |
| chr15 | 91429173 | 91429753 | 2.84516 | 5.84815 | 2.055 |
| chr2 | 47550283 | 47550730 | 2.06259 | 4.23985 | 2.056 |
| chr8 | 144366812 | 144367112 | 1.63375 | 3.35836 | 2.056 |
| chr14 | 70070322 | 70070638 | 3.45385 | 7.09981 | 2.056 |
| chr19 | 33211720 | 33212155 | 0.80683 | 1.65857 | 2.056 |
| chr3 | 36440447 | 36440692 | 0.45381 | 0.932902 | 2.056 |
| chr14 | 107020159 | 107020412 | 0.902456 | 1.85522 | 2.056 |
| chr5 | 43498711 | 43498965 | 0.45806 | 0.941659 | 2.056 |
| chr10 | 35852198 | 35852659 | 2.56106 | 5.26517 | 2.056 |
| chr10 | 71442467 | 71442721 | 0.459599 | 0.944878 | 2.056 |
| chr9 | 82899040 | 82899424 | 0.665794 | 1.36879 | 2.056 |
| chr17 | 2289021 | 2289320 | 0.503354 | 1.03488 | 2.056 |
| chr9 | 129320399 | 129320966 | 2.27364 | 4.67465 | 2.056 |
| chr3 | 134396088 | 134396361 | 0.887623 | 1.82498 | 2.056 |
| chr3 | 184231486 | 184231861 | 1.82678 | 3.75603 | 2.056 |

| | | | | | |
|---|---|---|---|---|---|
| chr19 | 18272136 | 18272432 | 1.78126 | 3.66244 | 2.056 |
| chr20 | 13283292 | 13283584 | 2.14802 | 4.41662 | 2.056 |
| chr9 | 117443872 | 117444110 | 1.45802 | 2.99802 | 2.056 |
| chr6 | 36308626 | 36309101 | 1.38564 | 2.84923 | 2.056 |
| chr9 | 136022788 | 136023261 | 2.19938 | 4.5225 | 2.056 |
| chr19 | 7982615 | 7983018 | 1.1419 | 2.34809 | 2.056 |
| chr16 | 11499440 | 11499844 | 1.13511 | 2.33415 | 2.056 |
| chr5 | 176758861 | 176759240 | 0.475149 | 0.977131 | 2.056 |
| chr12 | 45662969 | 45663213 | 1.28345 | 2.63949 | 2.057 |
| chr16 | 71528014 | 71528243 | 0.915302 | 1.8824 | 2.057 |
| chr7 | 113648644 | 113648989 | 0.555566 | 1.14259 | 2.057 |
| chr10 | 119202365 | 119202762 | 1.76286 | 3.62555 | 2.057 |
| chr20 | 34787589 | 34787964 | 1.95408 | 4.0189 | 2.057 |
| chr16 | 67596530 | 67596900 | 1.57421 | 3.23769 | 2.057 |
| chr3 | 125800442 | 125800722 | 1.14103 | 2.34677 | 2.057 |
| chr15 | 29407185 | 29407741 | 1.15255 | 2.37049 | 2.057 |
| chr4 | 152038559 | 152038852 | 1.38702 | 2.85275 | 2.057 |
| chr6 | 130342253 | 130342529 | 1.18803 | 2.44349 | 2.057 |
| chr16 | 15616900 | 15617150 | 0.964944 | 1.98467 | 2.057 |
| chr1 | 196233963 | 196234185 | 0.467711 | 0.961998 | 2.057 |
| chr12 | 99646896 | 99647248 | 0.935183 | 1.92357 | 2.057 |
| chr11 | 124663433 | 124663870 | 0.717116 | 1.47504 | 2.057 |
| chr8 | 11324999 | 11325429 | 1.58882 | 3.26822 | 2.057 |
| chr14 | 105837154 | 105837568 | 1.80523 | 3.71373 | 2.057 |
| chr11 | 55265575 | 55265803 | 0.465152 | 0.956953 | 2.057 |
| chr15 | 100271990 | 100272223 | 0.559876 | 1.15183 | 2.057 |
| chr3 | 141998940 | 141999175 | 0.881943 | 1.81443 | 2.057 |
| chr7 | 125531570 | 125531778 | 0.502667 | 1.03416 | 2.057 |
| chr8 | 15278772 | 15278996 | 0.417985 | 0.859954 | 2.057 |
| chr9 | 77815873 | 77816090 | 0.9223 | 1.8976 | 2.057 |
| chr22 | 48888260 | 48888476 | 1.02511 | 2.10915 | 2.057 |
| chr8 | 138600961 | 138601367 | 0.562244 | 1.15684 | 2.058 |
| chrX | 154285433 | 154285967 | 0.933478 | 1.92077 | 2.058 |
| chr19 | 30364281 | 30364844 | 1.87449 | 3.85707 | 2.058 |
| chr10 | 67663157 | 67663398 | 0.629834 | 1.296 | 2.058 |
| chr2 | 100175530 | 100175906 | 2.50433 | 5.15329 | 2.058 |
| chr11 | 37993254 | 37993457 | 0.624802 | 1.2857 | 2.058 |
| chr12 | 123827518 | 123827769 | 0.509415 | 1.04826 | 2.058 |
| chr7 | 107515104 | 107515388 | 1.50156 | 3.08996 | 2.058 |
| chr14 | 69754760 | 69754970 | 0.814132 | 1.67539 | 2.058 |
| chr3 | 58291698 | 58292032 | 1.79077 | 3.68527 | 2.058 |
| chr14 | 21077530 | 21077838 | 1.71837 | 3.53629 | 2.058 |
| chr17 | 32189957 | 32190183 | 0.5406 | 1.11253 | 2.058 |
| chr7 | 124425329 | 124425568 | 1.29999 | 2.67533 | 2.058 |
| chr9 | 2359924 | 2360134 | 0.593021 | 1.22042 | 2.058 |
| chr10 | 9486845 | 9487323 | 0.517753 | 1.06563 | 2.058 |
| chr20 | 62873587 | 62873844 | 0.486133 | 1.00059 | 2.058 |
| chr7 | 78422315 | 78422522 | 0.524195 | 1.07893 | 2.058 |
| chr16 | 71392325 | 71392774 | 1.84675 | 3.80122 | 2.058 |
| chr9 | 130953594 | 130954124 | 1.78606 | 3.67638 | 2.058 |
| chr7 | 99679379 | 99679797 | 1.62545 | 3.34581 | 2.058 |
| chr1 | 47805167 | 47805526 | 0.967623 | 1.99177 | 2.058 |
| chr6 | 111963838 | 111964053 | 1.49787 | 3.08358 | 2.059 |
| chr7 | 148823306 | 148823704 | 2.04322 | 4.20637 | 2.059 |
| chr11 | 44796980 | 44797312 | 1.56184 | 3.21541 | 2.059 |
| chr17 | 39804026 | 39805237 | 1.88078 | 3.87201 | 2.059 |
| chr5 | 171354400 | 171354661 | 0.816732 | 1.68147 | 2.059 |
| chr18 | 47654821 | 47655048 | 0.35117 | 0.722994 | 2.059 |
| chr7 | 55708924 | 55709273 | 1.09166 | 2.24754 | 2.059 |
| chr7 | 88097294 | 88097643 | 0.497837 | 1.02497 | 2.059 |
| chr17 | 77696152 | 77696500 | 1.23487 | 2.54251 | 2.059 |

| | | | | | |
|---|---|---|---|---|---|
| chr12 | 5986520 | 5986808 | 0.515864 | 1.06215 | 2.059 |
| chr12 | 2994485 | 2994793 | 2.22287 | 4.57692 | 2.059 |
| chr3 | 14962642 | 14963087 | 2.11405 | 4.35285 | 2.059 |
| chr12 | 25490390 | 25490759 | 0.637125 | 1.31187 | 2.059 |
| chr17 | 40086522 | 40086761 | 0.708429 | 1.4587 | 2.059 |
| chr20 | 33166962 | 33167229 | 1.18987 | 2.45009 | 2.059 |
| chr11 | 4018492 | 4018835 | 0.562021 | 1.15729 | 2.059 |
| chr20 | 61922587 | 61922932 | 1.79214 | 3.69042 | 2.059 |
| chr8 | 134215146 | 134215560 | 1.60008 | 3.29497 | 2.059 |
| chr14 | 57126704 | 57127070 | 1.26863 | 2.6125 | 2.059 |
| chr17 | 72947227 | 72947475 | 0.492982 | 1.01524 | 2.059 |
| chr20 | 2384200 | 2384545 | 0.532282 | 1.09619 | 2.059 |
| chr11 | 20153892 | 20154298 | 1.54609 | 3.18428 | 2.060 |
| chr7 | 1543989 | 1544322 | 1.91689 | 3.94811 | 2.060 |
| chr11 | 64071808 | 64072079 | 2.22102 | 4.57467 | 2.060 |
| chr12 | 60174594 | 60174844 | 0.941981 | 1.94028 | 2.060 |
| chr20 | 34861441 | 34861652 | 0.96046 | 1.97838 | 2.060 |
| chr12 | 21916559 | 21916786 | 0.590537 | 1.21642 | 2.060 |
| chr7 | 137084171 | 137084703 | 0.432987 | 0.89191 | 2.060 |
| chr5 | 139144816 | 139145596 | 2.7582 | 5.68211 | 2.060 |
| chr7 | 2286704 | 2287207 | 1.65959 | 3.41893 | 2.060 |
| chr11 | 46028004 | 46028303 | 1.31035 | 2.69948 | 2.060 |
| chr11 | 47395453 | 47395850 | 3.01046 | 6.20196 | 2.060 |
| chr6 | 86137577 | 86137785 | 0.497523 | 1.02502 | 2.060 |
| chr12 | 124876932 | 124877197 | 1.43415 | 2.95482 | 2.060 |
| chr1 | 183512806 | 183513036 | 1.71324 | 3.52989 | 2.060 |
| chr5 | 159392575 | 159392875 | 1.34703 | 2.77542 | 2.060 |
| chr3 | 27588536 | 27588738 | 0.504624 | 1.03975 | 2.060 |
| chr6 | 18716106 | 18716317 | 0.464754 | 0.957618 | 2.060 |
| chr11 | 125744434 | 125745009 | 1.05548 | 2.17481 | 2.060 |
| chr3 | 163108809 | 163109052 | 0.647579 | 1.33437 | 2.061 |
| chr2 | 238595291 | 238595564 | 1.04092 | 2.14492 | 2.061 |
| chr7 | 136910604 | 136910862 | 0.539966 | 1.11268 | 2.061 |
| chr7 | 25132981 | 25133273 | 2.08484 | 4.29635 | 2.061 |
| chr11 | 7715909 | 7716218 | 1.39387 | 2.87248 | 2.061 |
| chr11 | 34388149 | 34388470 | 1.0072 | 2.07568 | 2.061 |
| chr4 | 105781760 | 105781997 | 1.15077 | 2.37157 | 2.061 |
| chr6 | 148498754 | 148499176 | 1.00748 | 2.0763 | 2.061 |
| chr15 | 35581298 | 35581575 | 1.59711 | 3.29149 | 2.061 |
| chr17 | 41751113 | 41751521 | 2.64294 | 5.44724 | 2.061 |
| chr15 | 77952049 | 77952387 | 2.10533 | 4.33937 | 2.061 |
| chr15 | 53072771 | 53073051 | 1.8953 | 3.90684 | 2.061 |
| chr13 | 91669610 | 91669880 | 1.42535 | 2.93813 | 2.061 |
| chr3 | 126378908 | 126379238 | 0.724019 | 1.4925 | 2.061 |
| chr14 | 95899446 | 95899703 | 0.652557 | 1.34523 | 2.061 |
| chr15 | 74200609 | 74200968 | 1.73258 | 3.57171 | 2.062 |
| chr4 | 164465553 | 164465900 | 1.68727 | 3.4783 | 2.062 |
| chr2 | 60763689 | 60763909 | 0.648518 | 1.33695 | 2.062 |
| chr5 | 176881425 | 176881753 | 2.05979 | 4.24637 | 2.062 |
| chr18 | 56161069 | 56161376 | 1.46784 | 3.02607 | 2.062 |
| chr2 | 68452419 | 68452645 | 1.24396 | 2.56456 | 2.062 |
| chr12 | 57608090 | 57608634 | 1.63454 | 3.36981 | 2.062 |
| chr5 | 83222917 | 83223420 | 1.66266 | 3.42785 | 2.062 |
| chr20 | 49609493 | 49609853 | 1.73856 | 3.58438 | 2.062 |
| chr2 | 40164862 | 40165127 | 0.504273 | 1.03969 | 2.062 |
| chr2 | 26394371 | 26394716 | 1.56984 | 3.23674 | 2.062 |
| chr10 | 1887705 | 1888027 | 0.529249 | 1.09124 | 2.062 |
| chr8 | 67854996 | 67855208 | 0.461311 | 0.951188 | 2.062 |
| chr1 | 210048937 | 210049177 | 0.838459 | 1.72894 | 2.062 |
| chr12 | 72410827 | 72411188 | 1.64811 | 3.3985 | 2.062 |
| chr22 | 24899127 | 24899371 | 1.41071 | 2.90906 | 2.062 |

| | | | | | |
|---|---|---|---|---|---|
| chr8 | 21772801 | 21773140 | 1.07761 | 2.22224 | 2.062 |
| chr7 | 33937661 | 33937915 | 0.401952 | 0.828904 | 2.062 |
| chr15 | 99753369 | 99753852 | 1.63916 | 3.3804 | 2.062 |
| chr19 | 1401476 | 1401801 | 1.3284 | 2.73959 | 2.062 |
| chr2 | 160663982 | 160664305 | 1.80469 | 3.72184 | 2.062 |
| chr20 | 5914473 | 5914783 | 1.47265 | 3.03719 | 2.062 |
| chr8 | 10447455 | 10448118 | 1.99174 | 4.10786 | 2.062 |
| chr8 | 24920768 | 24920970 | 0.659059 | 1.35929 | 2.062 |
| chr3 | 141160275 | 141160662 | 1.31855 | 2.7196 | 2.063 |
| chr7 | 40092118 | 40092372 | 1.14645 | 2.36471 | 2.063 |
| chr3 | 134319640 | 134320059 | 2.37805 | 4.90508 | 2.063 |
| chr2 | 46592037 | 46592329 | 1.3341 | 2.75186 | 2.063 |
| chr10 | 20285122 | 20285401 | 0.530861 | 1.09504 | 2.063 |
| chr20 | 21888824 | 21889151 | 0.522149 | 1.07707 | 2.063 |
| chr19 | 7004386 | 7004667 | 1.49964 | 3.09362 | 2.063 |
| chr14 | 94889878 | 94890152 | 1.57943 | 3.25831 | 2.063 |
| chr1 | 19908783 | 19909061 | 1.35758 | 2.80072 | 2.063 |
| chr5 | 87976191 | 87976625 | 3.17855 | 6.55751 | 2.063 |
| chr19 | 45287680 | 45287981 | 2.27679 | 4.69733 | 2.063 |
| chr2 | 194448805 | 194449025 | 1.1088 | 2.28761 | 2.063 |
| chr1 | 157333403 | 157333682 | 1.36008 | 2.8061 | 2.063 |
| chr20 | 35220838 | 35221079 | 1.23898 | 2.55625 | 2.063 |
| chr11 | 85904638 | 85904910 | 1.09154 | 2.25235 | 2.063 |
| chr4 | 100081002 | 100081333 | 0.550801 | 1.13658 | 2.064 |
| chr9 | 105241733 | 105241952 | 0.582058 | 1.2011 | 2.064 |
| chr11 | 2951736 | 2952397 | 2.15713 | 4.45154 | 2.064 |
| chr15 | 44794080 | 44794290 | 0.741682 | 1.53061 | 2.064 |
| chr17 | 10532016 | 10532421 | 1.54218 | 3.18273 | 2.064 |
| chr15 | 101548583 | 101549364 | 1.86474 | 3.84866 | 2.064 |
| chr5 | 25197698 | 25197983 | 0.419725 | 0.866293 | 2.064 |
| chr8 | 117560609 | 117560824 | 0.820056 | 1.69263 | 2.064 |
| chr1 | 67998089 | 67998372 | 0.989587 | 2.04259 | 2.064 |
| chr6 | 126277743 | 126278152 | 2.11442 | 4.36446 | 2.064 |
| chr1 | 31895648 | 31895985 | 1.6697 | 3.44652 | 2.064 |
| chr19 | 2273647 | 2274088 | 1.87198 | 3.86415 | 2.064 |
| chr17 | 27507262 | 27507673 | 1.87161 | 3.86355 | 2.064 |
| chr19 | 1218131 | 1218360 | 1.17575 | 2.42712 | 2.064 |
| chr10 | 71875847 | 71876372 | 1.71385 | 3.53802 | 2.064 |
| chr18 | 19400320 | 19400677 | 0.666075 | 1.37503 | 2.064 |
| chr16 | 1979562 | 1980081 | 2.35598 | 4.8637 | 2.064 |
| chr12 | 132406901 | 132407104 | 0.443609 | 0.915891 | 2.065 |
| chr1 | 162025764 | 162026154 | 1.74635 | 3.60561 | 2.065 |
| chr11 | 45233367 | 45233750 | 0.911384 | 1.88172 | 2.065 |
| chr11 | 4203296 | 4203734 | 1.76197 | 3.63797 | 2.065 |
| chr19 | 49298342 | 49298755 | 2.6176 | 5.40467 | 2.065 |
| chr12 | 95928180 | 95928570 | 1.37048 | 2.82995 | 2.065 |
| chr17 | 1026325 | 1026630 | 2.52723 | 5.21856 | 2.065 |
| chr9 | 74434075 | 74434356 | 1.76337 | 3.64133 | 2.065 |
| chr4 | 6976177 | 6976423 | 1.58386 | 3.27072 | 2.065 |
| chr4 | 138030400 | 138030604 | 0.435146 | 0.898597 | 2.065 |
| chr7 | 101457435 | 101457775 | 1.1625 | 2.40074 | 2.065 |
| chr9 | 91003363 | 91003783 | 1.82539 | 3.76987 | 2.065 |
| chr13 | 100568835 | 100569097 | 1.95126 | 4.02994 | 2.065 |
| chr17 | 30228770 | 30228991 | 1.28431 | 2.65283 | 2.066 |
| chr14 | 79163805 | 79164034 | 0.718469 | 1.48416 | 2.066 |
| chr20 | 20941944 | 20942177 | 0.416774 | 0.860947 | 2.066 |
| chr6 | 42629368 | 42629632 | 1.39086 | 2.87316 | 2.066 |
| chr4 | 88191398 | 88191629 | 1.30405 | 2.69388 | 2.066 |
| chr19 | 40161811 | 40162048 | 0.596605 | 1.2325 | 2.066 |
| chr7 | 117904178 | 117904402 | 0.65194 | 1.34685 | 2.066 |
| chr12 | 77443620 | 77443884 | 1.52808 | 3.15697 | 2.066 |

| | | | | | |
|---|---|---|---|---|---|
| chr10 | 77796070 | 77796497 | 1.64863 | 3.40617 | 2.066 |
| chr15 | 27395711 | 27395927 | 0.452618 | 0.935201 | 2.066 |
| chr8 | 85538348 | 85538731 | 2.52495 | 5.21713 | 2.066 |
| chr21 | 35800918 | 35801206 | 0.9705 | 2.0053 | 2.066 |
| chr22 | 37712945 | 37713272 | 1.37194 | 2.83496 | 2.066 |
| chr17 | 47031623 | 47031980 | 1.93145 | 3.99114 | 2.066 |
| chr9 | 33374908 | 33375246 | 1.41509 | 2.92415 | 2.066 |
| chr10 | 99111129 | 99111358 | 0.567732 | 1.17319 | 2.066 |
| chr1 | 36030224 | 36030541 | 1.25756 | 2.5987 | 2.066 |
| chr12 | 55052477 | 55052721 | 0.952363 | 1.96802 | 2.066 |
| chr11 | 58903376 | 58903783 | 2.18478 | 4.5152 | 2.067 |
| chr11 | 364500 | 364716 | 1.44171 | 2.97966 | 2.067 |
| chr3 | 183197850 | 183198434 | 0.83457 | 1.72505 | 2.067 |
| chr20 | 53984055 | 53984286 | 0.402735 | 0.83249 | 2.067 |
| chr12 | 78118208 | 78118440 | 1.31585 | 2.72004 | 2.067 |
| chr17 | 27907263 | 27907689 | 1.83577 | 3.79485 | 2.067 |
| chr2 | 201440754 | 201441116 | 0.846357 | 1.74962 | 2.067 |
| chr17 | 70982240 | 70982849 | 1.68263 | 3.47848 | 2.067 |
| chr17 | 68140480 | 68140790 | 0.929831 | 1.92233 | 2.067 |
| chr12 | 111712621 | 111712921 | 1.43269 | 2.96196 | 2.067 |
| chr5 | 167913225 | 167913548 | 2.08043 | 4.30115 | 2.067 |
| chr2 | 118571694 | 118572312 | 2.00565 | 4.14659 | 2.067 |
| chr1 | 14137771 | 14137999 | 1.18806 | 2.45631 | 2.068 |
| chr21 | 46975946 | 46976326 | 1.60501 | 3.31838 | 2.068 |
| chr15 | 31183727 | 31183933 | 0.434035 | 0.89744 | 2.068 |
| chr12 | 109011877 | 109012324 | 2.20213 | 4.55329 | 2.068 |
| chr17 | 54669745 | 54670247 | 2.58603 | 5.34707 | 2.068 |
| chr12 | 65700274 | 65700726 | 1.63101 | 3.37265 | 2.068 |
| chr3 | 39231324 | 39231583 | 1.42124 | 2.93888 | 2.068 |
| chr2 | 73439641 | 73440006 | 2.21629 | 4.58304 | 2.068 |
| chr10 | 134257143 | 134257420 | 1.90246 | 3.93424 | 2.068 |
| chr8 | 73856221 | 73856689 | 1.87763 | 3.88291 | 2.068 |
| chr11 | 110306065 | 110306451 | 0.810898 | 1.67693 | 2.068 |
| chr2 | 13147032 | 13147366 | 1.6911 | 3.49741 | 2.068 |
| chr14 | 100844221 | 100844498 | 1.58589 | 3.2799 | 2.068 |
| chr11 | 133797519 | 133797847 | 1.2683 | 2.62308 | 2.068 |
| chr11 | 86002828 | 86003147 | 1.3894 | 2.87358 | 2.068 |
| chr11 | 99610291 | 99610513 | 0.960681 | 1.98709 | 2.068 |
| chr17 | 1477639 | 1478054 | 2.47912 | 5.12796 | 2.068 |
| chr12 | 11400729 | 11401001 | 0.700556 | 1.4491 | 2.069 |
| chr9 | 77800539 | 77800896 | 1.62223 | 3.3556 | 2.069 |
| chr10 | 126749613 | 126750120 | 1.85812 | 3.84362 | 2.069 |
| chr9 | 133313941 | 133314226 | 0.462614 | 0.956946 | 2.069 |
| chr6 | 47599127 | 47599381 | 1.22714 | 2.53842 | 2.069 |
| chr6 | 12151104 | 12151391 | 2.21874 | 4.58999 | 2.069 |
| chr1 | 160070815 | 160071169 | 1.68559 | 3.48706 | 2.069 |
| chr11 | 8008467 | 8008724 | 1.92599 | 3.98446 | 2.069 |
| chr3 | 71056018 | 71056294 | 1.2873 | 2.66323 | 2.069 |
| chr7 | 49076662 | 49076901 | 0.581013 | 1.20219 | 2.069 |
| chr13 | 30082220 | 30082640 | 2.14698 | 4.44244 | 2.069 |
| chr20 | 50240583 | 50241178 | 1.39314 | 2.88274 | 2.069 |
| chr10 | 54686031 | 54686245 | 0.460943 | 0.953827 | 2.069 |
| chr7 | 30603840 | 30604256 | 1.33295 | 2.75826 | 2.069 |
| chr2 | 207558414 | 207558723 | 0.842111 | 1.74265 | 2.069 |
| chr3 | 112012252 | 112012508 | 1.18044 | 2.44278 | 2.069 |
| chr5 | 64509732 | 64510125 | 1.23008 | 2.54553 | 2.069 |
| chr7 | 90733541 | 90733752 | 0.485763 | 1.0053 | 2.070 |
| chr20 | 39597258 | 39597891 | 1.64622 | 3.40692 | 2.070 |
| chr7 | 99728611 | 99728992 | 1.76079 | 3.64402 | 2.070 |
| chrX | 129464718 | 129465002 | 1.17222 | 2.42602 | 2.070 |
| chr5 | 171889207 | 171889605 | 2.19477 | 4.54231 | 2.070 |

| | | | | | |
|---|---|---|---|---|---|
| chr10 | 34955296 | 34955511 | 1.35106 | 2.79618 | 2.070 |
| chr2 | 50094537 | 50094768 | 0.579507 | 1.19937 | 2.070 |
| chr7 | 126296199 | 126296458 | 0.477058 | 0.987343 | 2.070 |
| chr4 | 73161756 | 73162079 | 0.902121 | 1.86708 | 2.070 |
| chr5 | 149217212 | 149217638 | 2.69613 | 5.58017 | 2.070 |
| chrX | 38079302 | 38079505 | 0.939729 | 1.94507 | 2.070 |
| chr12 | 126195124 | 126195349 | 0.54818 | 1.13465 | 2.070 |
| chrX | 109651383 | 109651670 | 1.15391 | 2.38846 | 2.070 |
| chr11 | 61468857 | 61469081 | 1.06529 | 2.20522 | 2.070 |
| chr3 | 30810197 | 30810585 | 1.83176 | 3.79209 | 2.070 |
| chr11 | 8287282 | 8287500 | 0.841118 | 1.74128 | 2.070 |
| chr1 | 17746356 | 17746723 | 2.31009 | 4.78246 | 2.070 |
| chr2 | 42067831 | 42068146 | 2.74264 | 5.67794 | 2.070 |
| chr1 | 207985292 | 207985515 | 1.53191 | 3.1715 | 2.070 |
| chr21 | 35975383 | 35975683 | 1.95394 | 4.04525 | 2.070 |
| chr12 | 85969416 | 85969808 | 1.38539 | 2.86818 | 2.070 |
| chr2 | 7573189 | 7573412 | 0.644493 | 1.33431 | 2.070 |
| chr5 | 116990466 | 116990701 | 0.562663 | 1.16491 | 2.070 |
| chr14 | 23452558 | 23452793 | 1.23713 | 2.56144 | 2.070 |
| chr4 | 89627682 | 89627916 | 0.947359 | 1.96152 | 2.071 |
| chr19 | 45157692 | 45158030 | 1.80323 | 3.73384 | 2.071 |
| chr3 | 83110208 | 83110518 | 0.575363 | 1.19137 | 2.071 |
| chr7 | 134959211 | 134959572 | 1.47669 | 3.05775 | 2.071 |
| chr11 | 63604754 | 63605025 | 1.7436 | 3.61052 | 2.071 |
| chr5 | 142092655 | 142093357 | 2.84809 | 5.89762 | 2.071 |
| chr14 | 69362883 | 69363165 | 1.29011 | 2.6715 | 2.071 |
| chr3 | 39583218 | 39583729 | 0.852285 | 1.76489 | 2.071 |
| chr3 | 13653067 | 13653535 | 0.463787 | 0.960427 | 2.071 |
| chr1 | 111325954 | 111326251 | 1.93029 | 3.99734 | 2.071 |
| chr9 | 135923897 | 135924633 | 0.821263 | 1.70082 | 2.071 |
| chr16 | 5792890 | 5793131 | 0.49465 | 1.02458 | 2.071 |
| chr19 | 47251723 | 47252182 | 1.98391 | 4.10934 | 2.071 |
| chr1 | 110033297 | 110033591 | 1.5413 | 3.19256 | 2.071 |
| chr17 | 36823755 | 36824213 | 2.97006 | 6.152 | 2.071 |
| chr19 | 30185361 | 30185971 | 2.79471 | 5.78884 | 2.071 |
| chr1 | 160324227 | 160324473 | 1.06085 | 2.19744 | 2.071 |
| chr7 | 135694293 | 135694563 | 1.05075 | 2.17657 | 2.071 |
| chr8 | 135898689 | 135899203 | 1.22589 | 2.53946 | 2.072 |
| chr12 | 96392439 | 96392794 | 1.32433 | 2.74353 | 2.072 |
| chr5 | 37917472 | 37917689 | 0.415268 | 0.86029 | 2.072 |
| chr16 | 69788760 | 69789053 | 1.82388 | 3.77856 | 2.072 |
| chr7 | 44240569 | 44240786 | 0.667198 | 1.38226 | 2.072 |
| chr15 | 84381720 | 84381992 | 0.613636 | 1.27131 | 2.072 |
| chr2 | 65800328 | 65800569 | 0.97507 | 2.02012 | 2.072 |
| chr20 | 48911016 | 48911535 | 0.715501 | 1.48236 | 2.072 |
| chr9 | 90293684 | 90293919 | 1.4934 | 3.09405 | 2.072 |
| chr9 | 123822384 | 123822603 | 1.40465 | 2.9102 | 2.072 |
| chr7 | 145787013 | 145787449 | 0.952777 | 1.974 | 2.072 |
| chr11 | 6689170 | 6689407 | 1.05014 | 2.17576 | 2.072 |
| chr11 | 71083717 | 71084298 | 0.984006 | 2.03876 | 2.072 |
| chr2 | 98423234 | 98423517 | 1.98831 | 4.11983 | 2.072 |
| chr9 | 133826093 | 133826496 | 0.703406 | 1.45749 | 2.072 |
| chr5 | 50415270 | 50415680 | 0.483444 | 1.00173 | 2.072 |
| chr20 | 53285077 | 53285569 | 1.57041 | 3.25415 | 2.072 |
| chr4 | 38730600 | 38730816 | 1.08191 | 2.24193 | 2.072 |
| chr7 | 81894035 | 81894493 | 0.428404 | 0.887758 | 2.072 |
| chr11 | 118530121 | 118530552 | 1.77707 | 3.68254 | 2.072 |
| chr8 | 23817920 | 23818201 | 1.23319 | 2.55548 | 2.072 |
| chr20 | 24435651 | 24435883 | 0.579784 | 1.20148 | 2.072 |
| chr4 | 55094249 | 55094535 | 1.57927 | 3.27288 | 2.072 |
| chr4 | 24737758 | 24737960 | 0.735462 | 1.52429 | 2.073 |

| | | | | | |
|---|---|---|---|---|---|
| chr17 | 14789572 | 14789800 | 0.951751 | 1.97267 | 2.073 |
| chr2 | 85512563 | 85513023 | 2.15537 | 4.46752 | 2.073 |
| chr14 | 64977766 | 64978176 | 1.47339 | 3.05412 | 2.073 |
| chr8 | 136794697 | 136794918 | 0.504069 | 1.0449 | 2.073 |
| chr5 | 147191323 | 147191543 | 0.561661 | 1.16429 | 2.073 |
| chr17 | 38347835 | 38348231 | 1.89713 | 3.93277 | 2.073 |
| chr19 | 41520047 | 41520352 | 0.518801 | 1.07549 | 2.073 |
| chr7 | 127849791 | 127850169 | 2.38138 | 4.93683 | 2.073 |
| chr8 | 63077964 | 63078198 | 0.959151 | 1.98843 | 2.073 |
| chr6 | 133120117 | 133120424 | 1.30321 | 2.70184 | 2.073 |
| chr14 | 72331090 | 72331292 | 0.513736 | 1.06514 | 2.073 |
| chr15 | 88850358 | 88850590 | 0.53556 | 1.11041 | 2.073 |
| chr17 | 58308576 | 58308800 | 0.958514 | 1.98736 | 2.073 |
| chr6 | 36128811 | 36129018 | 0.510091 | 1.05762 | 2.073 |
| chr8 | 102644815 | 102645028 | 1.60421 | 3.32622 | 2.073 |
| chr9 | 72164379 | 72164661 | 1.89845 | 3.93644 | 2.074 |
| chr13 | 31773822 | 31774625 | 1.40476 | 2.91283 | 2.074 |
| chr1 | 236150857 | 236151087 | 0.64233 | 1.33197 | 2.074 |
| chr3 | 133478188 | 133478420 | 0.508387 | 1.05422 | 2.074 |
| chr8 | 130744074 | 130744318 | 1.63769 | 3.39606 | 2.074 |
| chr19 | 49338986 | 49339448 | 1.59218 | 3.30175 | 2.074 |
| chr3 | 81986193 | 81986509 | 0.503087 | 1.04331 | 2.074 |
| chr10 | 106065302 | 106065628 | 1.53821 | 3.19019 | 2.074 |
| chr2 | 79679161 | 79679441 | 0.56084 | 1.16319 | 2.074 |
| chr9 | 116353927 | 116354305 | 2.29958 | 4.76949 | 2.074 |
| chr10 | 15413262 | 15413574 | 1.8813 | 3.90209 | 2.074 |
| chr17 | 44223822 | 44224202 | 0.576753 | 1.19635 | 2.074 |
| chr5 | 138897542 | 138897893 | 2.55324 | 5.29617 | 2.074 |
| chr12 | 75920699 | 75920995 | 1.11878 | 2.32071 | 2.074 |
| chr3 | 74864488 | 74864696 | 0.532345 | 1.10427 | 2.074 |
| chr16 | 30461105 | 30461502 | 1.95385 | 4.05319 | 2.074 |
| chr17 | 42296460 | 42297148 | 0.814403 | 1.68945 | 2.074 |
| chr8 | 55371960 | 55372300 | 0.89492 | 1.85648 | 2.074 |
| chr6 | 44424061 | 44424364 | 1.62052 | 3.36188 | 2.075 |
| chr10 | 65594780 | 65594990 | 0.374325 | 0.776569 | 2.075 |
| chr2 | 206678278 | 206678560 | 1.43277 | 2.97262 | 2.075 |
| chrX | 135014078 | 135014373 | 1.67646 | 3.47836 | 2.075 |
| chr1 | 33592828 | 33593095 | 2.08461 | 4.32534 | 2.075 |
| chr11 | 61532064 | 61532366 | 0.522043 | 1.08318 | 2.075 |
| chr9 | 80379800 | 80380162 | 1.18915 | 2.46736 | 2.075 |
| chr1 | 209376107 | 209376673 | 1.23118 | 2.55464 | 2.075 |
| chr11 | 80195017 | 80195315 | 0.419821 | 0.87115 | 2.075 |
| chr8 | 8345108 | 8345361 | 0.602399 | 1.25004 | 2.075 |
| chr22 | 37894352 | 37894706 | 1.37823 | 2.86001 | 2.075 |
| chr10 | 85907857 | 85908177 | 1.32453 | 2.7487 | 2.075 |
| chr8 | 124805174 | 124805394 | 0.664485 | 1.37896 | 2.075 |
| chr22 | 39566842 | 39567165 | 2.2031 | 4.57197 | 2.075 |
| chr7 | 44837023 | 44837275 | 1.29521 | 2.68788 | 2.075 |
| chr5 | 137558446 | 137558820 | 1.16099 | 2.40936 | 2.075 |
| chr6 | 26322381 | 26322597 | 0.949279 | 1.97006 | 2.075 |
| chr9 | 86930241 | 86930507 | 1.09435 | 2.27113 | 2.075 |
| chr4 | 10539634 | 10539882 | 1.16488 | 2.41762 | 2.075 |
| chr10 | 19939670 | 19940048 | 0.493936 | 1.02516 | 2.075 |
| chr7 | 48339230 | 48339497 | 1.49804 | 3.10916 | 2.075 |
| chr6 | 81047967 | 81048313 | 1.57301 | 3.26487 | 2.076 |
| chr7 | 4837788 | 4838255 | 2.56648 | 5.32696 | 2.076 |
| chr16 | 85004188 | 85004485 | 1.99267 | 4.13601 | 2.076 |
| chr1 | 247488222 | 247488438 | 0.445848 | 0.925523 | 2.076 |
| chr21 | 38289845 | 38290184 | 1.40088 | 2.90805 | 2.076 |
| chr5 | 140248972 | 140249327 | 1.17572 | 2.44065 | 2.076 |
| chr19 | 51172581 | 51173031 | 2.11637 | 4.39343 | 2.076 |

| | | | | | |
|---|---|---|---|---|---|
| chr8 | 65769522 | 65769738 | 0.704159 | 1.46179 | 2.076 |
| chr10 | 52321539 | 52321813 | 1.04777 | 2.17513 | 2.076 |
| chr8 | 72398537 | 72398905 | 0.522172 | 1.08405 | 2.076 |
| chr10 | 107987803 | 107988076 | 0.448511 | 0.931134 | 2.076 |
| chr3 | 156258904 | 156259175 | 1.41499 | 2.93774 | 2.076 |
| chr13 | 105030612 | 105030816 | 0.634871 | 1.31811 | 2.076 |
| chr17 | 6757039 | 6757262 | 1.15093 | 2.38966 | 2.076 |
| chr11 | 85753012 | 85753252 | 2.06361 | 4.2847 | 2.076 |
| chr4 | 76837198 | 76837482 | 1.89877 | 3.94245 | 2.076 |
| chr9 | 108604400 | 108604761 | 1.31306 | 2.72638 | 2.076 |
| chr2 | 113896040 | 113896249 | 1.3389 | 2.78005 | 2.076 |
| chr14 | 103763357 | 103763562 | 0.787615 | 1.63539 | 2.076 |
| chr5 | 1672675 | 1672879 | 0.593171 | 1.23165 | 2.076 |
| chr6 | 73121807 | 73122355 | 2.90466 | 6.03147 | 2.076 |
| chr8 | 72024202 | 72024514 | 0.549222 | 1.14047 | 2.077 |
| chr5 | 94307905 | 94308137 | 0.515899 | 1.07134 | 2.077 |
| chr16 | 56459386 | 56459674 | 1.18634 | 2.46366 | 2.077 |
| chr16 | 3033643 | 3034004 | 2.34028 | 4.86027 | 2.077 |
| chr7 | 55661260 | 55662395 | 1.12272 | 2.33179 | 2.077 |
| chr3 | 151901314 | 151901533 | 0.829268 | 1.72236 | 2.077 |
| chr1 | 201978696 | 201978959 | 1.28841 | 2.67599 | 2.077 |
| chr3 | 140950403 | 140950819 | 1.95639 | 4.06352 | 2.077 |
| chr5 | 176079739 | 176080050 | 0.616393 | 1.28028 | 2.077 |
| chr16 | 21289364 | 21289900 | 1.99812 | 4.15026 | 2.077 |
| chr5 | 131755346 | 131755691 | 2.21552 | 4.60185 | 2.077 |
| chr1 | 45978259 | 45978576 | 1.32868 | 2.75981 | 2.077 |
| chr12 | 78078955 | 78079224 | 1.77522 | 3.68732 | 2.077 |
| chr12 | 551449 | 551767 | 2.02825 | 4.21299 | 2.077 |
| chr9 | 93720843 | 93721112 | 0.405295 | 0.841885 | 2.077 |
| chr9 | 113659593 | 113659812 | 0.524653 | 1.08984 | 2.077 |
| chr17 | 27077061 | 27077518 | 2.64318 | 5.49067 | 2.077 |
| chr17 | 76261222 | 76261435 | 0.414046 | 0.860121 | 2.077 |
| chr9 | 136200387 | 136200639 | 0.479177 | 0.995457 | 2.077 |
| chr3 | 14332187 | 14332406 | 0.507915 | 1.05519 | 2.077 |
| chr8 | 48114866 | 48115184 | 1.73067 | 3.59549 | 2.078 |
| chr15 | 87538471 | 87538879 | 0.495134 | 1.02865 | 2.078 |
| chr2 | 209118959 | 209119318 | 1.99277 | 4.14008 | 2.078 |
| chr19 | 39378046 | 39378500 | 2.38385 | 4.95265 | 2.078 |
| chr10 | 13141661 | 13141953 | 1.45599 | 3.02503 | 2.078 |
| chr4 | 42570683 | 42570933 | 0.899745 | 1.86936 | 2.078 |
| chr19 | 921644 | 922049 | 2.17009 | 4.50874 | 2.078 |
| chr12 | 118619385 | 118619710 | 1.26823 | 2.63514 | 2.078 |
| chr16 | 84430872 | 84431079 | 1.1205 | 2.32825 | 2.078 |
| chr2 | 224171185 | 224171413 | 1.43683 | 2.9856 | 2.078 |
| chr1 | 226806910 | 226807183 | 0.355854 | 0.739441 | 2.078 |
| chr3 | 114811102 | 114811632 | 1.09381 | 2.27288 | 2.078 |
| chr3 | 19243943 | 19244152 | 0.467375 | 0.971216 | 2.078 |
| chr17 | 25929912 | 25930584 | 2.87076 | 5.96565 | 2.078 |
| chr9 | 79075027 | 79075568 | 0.695658 | 1.44563 | 2.078 |
| chr2 | 228749132 | 228749517 | 2.00962 | 4.17624 | 2.078 |
| chr1 | 34100497 | 34100708 | 1.00346 | 2.08535 | 2.078 |
| chr3 | 49709083 | 49709426 | 1.28269 | 2.66567 | 2.078 |
| chr1 | 33338231 | 33338583 | 1.88133 | 3.90982 | 2.078 |
| chr10 | 101972777 | 101973133 | 0.519026 | 1.07865 | 2.078 |
| chr7 | 144733777 | 144734359 | 0.872421 | 1.81316 | 2.078 |
| chr17 | 37866451 | 37866692 | 0.587307 | 1.22061 | 2.078 |
| chr7 | 26118578 | 26118822 | 1.62056 | 3.36818 | 2.078 |
| chr19 | 42653996 | 42654247 | 0.464413 | 0.965293 | 2.079 |
| chr10 | 82762041 | 82762312 | 0.533641 | 1.1092 | 2.079 |
| chr1 | 186775691 | 186775912 | 1.13363 | 2.35632 | 2.079 |
| chr3 | 141024328 | 141024572 | 0.867583 | 1.80344 | 2.079 |

| | | | | | |
|---|---|---|---|---|---|
| chr15 | 58562137 | 58562508 | 1.7988 | 3.73916 | 2.079 |
| chr17 | 78623662 | 78624206 | 1.72133 | 3.57816 | 2.079 |
| chr17 | 76638379 | 76638658 | 0.604579 | 1.25676 | 2.079 |
| chr3 | 97527159 | 97527417 | 0.622935 | 1.29497 | 2.079 |
| chr17 | 27200111 | 27200337 | 0.513015 | 1.06647 | 2.079 |
| chr17 | 19266129 | 19266594 | 2.14699 | 4.4633 | 2.079 |
| chr16 | 26193749 | 26193979 | 0.381804 | 0.793766 | 2.079 |
| chr8 | 139791666 | 139791884 | 0.570238 | 1.18555 | 2.079 |
| chr17 | 33152752 | 33152960 | 0.460486 | 0.957373 | 2.079 |
| chr1 | 154083184 | 154083650 | 0.54934 | 1.14211 | 2.079 |
| chr13 | 24152814 | 24153159 | 1.07773 | 2.24071 | 2.079 |
| chr1 | 150022621 | 150022909 | 1.8849 | 3.91904 | 2.079 |
| chr20 | 36701884 | 36702204 | 1.7883 | 3.71831 | 2.079 |
| chr4 | 4337062 | 4337281 | 1.2666 | 2.63356 | 2.079 |
| chr3 | 27320722 | 27321006 | 1.19188 | 2.47822 | 2.079 |
| chr17 | 17489600 | 17489833 | 1.39503 | 2.90064 | 2.079 |
| chr5 | 77596312 | 77596801 | 1.08536 | 2.25688 | 2.079 |
| chr8 | 83490841 | 83491153 | 0.967633 | 2.01208 | 2.079 |
| chr1 | 241110217 | 241110587 | 1.12332 | 2.33584 | 2.079 |
| chr3 | 158390148 | 158390700 | 0.950603 | 1.97671 | 2.079 |
| chr1 | 201450998 | 201451558 | 1.7476 | 3.63411 | 2.079 |
| chr6 | 150390407 | 150390662 | 1.06762 | 2.22015 | 2.080 |
| chr9 | 36452640 | 36452857 | 0.401244 | 0.834398 | 2.080 |
| chr16 | 82744562 | 82744830 | 0.628491 | 1.307 | 2.080 |
| chr8 | 15235740 | 15236384 | 1.7925 | 3.72776 | 2.080 |
| chr11 | 46373576 | 46374242 | 1.68644 | 3.50727 | 2.080 |
| chr5 | 131619171 | 131619428 | 1.23952 | 2.57784 | 2.080 |
| chr1 | 18882907 | 18883184 | 1.05549 | 2.19527 | 2.080 |
| chr5 | 141330867 | 141331199 | 1.11661 | 2.32242 | 2.080 |
| chr6 | 41889028 | 41889301 | 1.79472 | 3.73282 | 2.080 |
| chr10 | 29666350 | 29666712 | 1.51996 | 3.16141 | 2.080 |
| chr14 | 95908746 | 95909213 | 1.82523 | 3.79636 | 2.080 |
| chr11 | 23057068 | 23057394 | 0.545835 | 1.13534 | 2.080 |
| chr7 | 25256436 | 25256757 | 1.46245 | 3.04207 | 2.080 |
| chr16 | 31345064 | 31345363 | 1.3588 | 2.82655 | 2.080 |
| chr12 | 2848811 | 2849334 | 1.05053 | 2.18543 | 2.080 |
| chr7 | 128509669 | 128510022 | 0.585446 | 1.21793 | 2.080 |
| chr5 | 148761378 | 148761835 | 1.81616 | 3.77842 | 2.080 |
| chr12 | 62604168 | 62604495 | 1.96834 | 4.09508 | 2.080 |
| chr8 | 130013587 | 130013897 | 0.577979 | 1.20252 | 2.081 |
| chr1 | 27623304 | 27623731 | 1.4458 | 3.00811 | 2.081 |
| chr10 | 4698824 | 4699156 | 2.15401 | 4.48162 | 2.081 |
| chr19 | 31570609 | 31571099 | 0.506412 | 1.05367 | 2.081 |
| chr1 | 40004883 | 40005198 | 1.76759 | 3.67779 | 2.081 |
| chr2 | 59066694 | 59066921 | 0.497491 | 1.0352 | 2.081 |
| chr16 | 58028741 | 58028966 | 1.10604 | 2.30152 | 2.081 |
| chr19 | 47582513 | 47582915 | 2.08815 | 4.34519 | 2.081 |
| chr12 | 52288783 | 52289110 | 2.004 | 4.17017 | 2.081 |
| chr7 | 26002480 | 26002699 | 0.450821 | 0.938147 | 2.081 |
| chr9 | 13088772 | 13089379 | 1.26001 | 2.62214 | 2.081 |
| chr4 | 7846022 | 7846291 | 1.01121 | 2.10447 | 2.081 |
| chr3 | 137837798 | 137838042 | 1.66502 | 3.46515 | 2.081 |
| chr3 | 120136037 | 120136353 | 1.92583 | 4.00803 | 2.081 |
| chr7 | 107738184 | 107738433 | 1.281 | 2.66602 | 2.081 |
| chr17 | 80103427 | 80103729 | 0.531662 | 1.10654 | 2.081 |
| chr15 | 75063355 | 75063828 | 3.03982 | 6.32674 | 2.081 |
| chr6 | 91143828 | 91144042 | 0.657837 | 1.36919 | 2.081 |
| chr15 | 91475582 | 91476055 | 2.08177 | 4.33301 | 2.081 |
| chr1 | 224033784 | 224034034 | 1.42149 | 2.95884 | 2.082 |
| chr7 | 111403321 | 111403837 | 0.94891 | 1.97522 | 2.082 |
| chr1 | 50814378 | 50814700 | 1.31991 | 2.74758 | 2.082 |

| | | | | | |
|---|---|---|---|---|---|
| chr21 | 21132039 | 21132292 | 0.507517 | 1.05653 | 2.082 |
| chr7 | 118978251 | 118978476 | 0.402266 | 0.837461 | 2.082 |
| chr7 | 51650181 | 51650616 | 0.457735 | 0.952965 | 2.082 |
| chr12 | 72174058 | 72174421 | 1.32441 | 2.75738 | 2.082 |
| chr5 | 158583787 | 158584139 | 1.6893 | 3.5171 | 2.082 |
| chr1 | 33936033 | 33936362 | 1.12107 | 2.33408 | 2.082 |
| chr7 | 55212094 | 55212357 | 1.00351 | 2.08942 | 2.082 |
| chr3 | 128303279 | 128303631 | 1.59379 | 3.31849 | 2.082 |
| chr4 | 57167703 | 57167954 | 1.80374 | 3.75564 | 2.082 |
| chr7 | 2648656 | 2648943 | 0.975458 | 2.03105 | 2.082 |
| chr7 | 37960097 | 37960827 | 0.832351 | 1.73311 | 2.082 |
| chr16 | 49417007 | 49417223 | 0.713916 | 1.48653 | 2.082 |
| chr2 | 105986291 | 105986596 | 2.22311 | 4.62903 | 2.082 |
| chr15 | 86383820 | 86384138 | 2.60964 | 5.434 | 2.082 |
| chr12 | 114946360 | 114946720 | 1.14255 | 2.37915 | 2.082 |
| chr16 | 75515465 | 75515823 | 0.890982 | 1.85533 | 2.082 |
| chr1 | 188922649 | 188922891 | 0.559781 | 1.16566 | 2.082 |
| chr6 | 13615340 | 13615681 | 1.60585 | 3.34394 | 2.082 |
| chr12 | 26170539 | 26170882 | 1.36954 | 2.85202 | 2.082 |
| chr14 | 104187742 | 104187981 | 0.58297 | 1.21402 | 2.082 |
| chr19 | 43116675 | 43116889 | 0.546481 | 1.13807 | 2.083 |
| chr1 | 231347004 | 231347416 | 2.12661 | 4.42887 | 2.083 |
| chr8 | 103597463 | 103597714 | 1.41284 | 2.94253 | 2.083 |
| chr5 | 121698858 | 121699079 | 0.436686 | 0.909495 | 2.083 |
| chr16 | 4033013 | 4033435 | 2.38125 | 4.95949 | 2.083 |
| chr20 | 56784791 | 56785315 | 3.68189 | 7.66838 | 2.083 |
| chr7 | 45299567 | 45300261 | 1.62012 | 3.37445 | 2.083 |
| chr4 | 77299230 | 77299435 | 1.1259 | 2.34519 | 2.083 |
| chr8 | 10948471 | 10948717 | 1.31533 | 2.73984 | 2.083 |
| chr15 | 45030664 | 45030891 | 1.0917 | 2.27403 | 2.083 |
| chr7 | 30266513 | 30266791 | 1.36671 | 2.84705 | 2.083 |
| chr11 | 5705898 | 5706119 | 1.51638 | 3.15904 | 2.083 |
| chr5 | 112027316 | 112027968 | 0.848523 | 1.76777 | 2.083 |
| chr9 | 73864674 | 73864973 | 1.20523 | 2.5111 | 2.084 |
| chr13 | 111567473 | 111567757 | 1.82415 | 3.80095 | 2.084 |
| chr8 | 121135754 | 121136158 | 1.77332 | 3.69506 | 2.084 |
| chr12 | 45187989 | 45188238 | 0.491104 | 1.02332 | 2.084 |
| chr3 | 123047614 | 123048295 | 1.58673 | 3.30629 | 2.084 |
| chr19 | 56764904 | 56765396 | 0.692381 | 1.4428 | 2.084 |
| chr9 | 5265758 | 5266200 | 0.650344 | 1.35523 | 2.084 |
| chr1 | 8931619 | 8931949 | 1.02612 | 2.13833 | 2.084 |
| chr8 | 71520455 | 71520880 | 1.8847 | 3.92756 | 2.084 |
| chr11 | 74413368 | 74413873 | 2.01149 | 4.1922 | 2.084 |
| chr19 | 34258827 | 34259054 | 0.843859 | 1.75873 | 2.084 |
| chr8 | 22452684 | 22453105 | 1.66284 | 3.46571 | 2.084 |
| chr14 | 65741517 | 65741830 | 0.446457 | 0.930556 | 2.084 |
| chr16 | 67193930 | 67194216 | 0.955386 | 1.99134 | 2.084 |
| chr9 | 124885242 | 124885663 | 3.02701 | 6.30936 | 2.084 |
| chr5 | 126015285 | 126015509 | 1.18476 | 2.46969 | 2.085 |
| chr6 | 107639385 | 107639668 | 1.38373 | 2.88446 | 2.085 |
| chr9 | 12191600 | 12191918 | 0.675613 | 1.4084 | 2.085 |
| chr1 | 32180000 | 32180243 | 1.00526 | 2.09572 | 2.085 |
| chr16 | 64158420 | 64158681 | 0.429045 | 0.894463 | 2.085 |
| chr4 | 88157026 | 88157263 | 1.1275 | 2.35059 | 2.085 |
| chr15 | 101829191 | 101829456 | 1.54674 | 3.22464 | 2.085 |
| chr3 | 67160189 | 67160428 | 1.01568 | 2.11763 | 2.085 |
| chr5 | 58187616 | 58187848 | 1.3791 | 2.87534 | 2.085 |
| chr19 | 49938929 | 49939189 | 0.573203 | 1.1951 | 2.085 |
| chr4 | 131484681 | 131484990 | 1.41157 | 2.94315 | 2.085 |
| chr5 | 99014912 | 99015188 | 0.489486 | 1.02059 | 2.085 |
| chr8 | 143820773 | 143821292 | 2.06722 | 4.31033 | 2.085 |

| | | | | | |
|---|---|---|---|---|---|
| chr4 | 109486819 | 109487022 | 0.418173 | 0.871932 | 2.085 |
| chr16 | 83859674 | 83860051 | 1.52003 | 3.16943 | 2.085 |
| chr12 | 130820186 | 130820542 | 0.755312 | 1.57496 | 2.085 |
| chr16 | 8999675 | 9000085 | 0.859172 | 1.79162 | 2.085 |
| chr12 | 104595018 | 104595407 | 1.47161 | 3.06879 | 2.085 |
| chr8 | 97340139 | 97340586 | 1.8297 | 3.81554 | 2.085 |
| chr2 | 55301242 | 55301515 | 0.487926 | 1.0175 | 2.085 |
| chr1 | 177331696 | 177332069 | 0.426246 | 0.888901 | 2.085 |
| chr1 | 159750314 | 159750600 | 1.61706 | 3.37235 | 2.085 |
| chr1 | 109806161 | 109806671 | 1.81984 | 3.79538 | 2.086 |
| chr1 | 192755919 | 192756125 | 0.563766 | 1.17577 | 2.086 |
| chr15 | 45459651 | 45459869 | 1.59551 | 3.32755 | 2.086 |
| chr11 | 19228386 | 19228788 | 0.56271 | 1.1736 | 2.086 |
| chr21 | 47016592 | 47016960 | 1.8547 | 3.8682 | 2.086 |
| chr5 | 180296032 | 180296322 | 2.2508 | 4.69434 | 2.086 |
| chr13 | 30687120 | 30687446 | 2.10879 | 4.39826 | 2.086 |
| chr7 | 35664127 | 35664539 | 1.99006 | 4.15066 | 2.086 |
| chr19 | 33090411 | 33090926 | 1.72511 | 3.59824 | 2.086 |
| chr16 | 49677904 | 49678245 | 0.503426 | 1.05006 | 2.086 |
| chr7 | 128067232 | 128067661 | 0.979133 | 2.04234 | 2.086 |
| chr8 | 23331797 | 23332336 | 1.94889 | 4.06518 | 2.086 |
| chr19 | 18117772 | 18118212 | 1.99588 | 4.1633 | 2.086 |
| chr6 | 150161714 | 150162027 | 1.41274 | 2.94693 | 2.086 |
| chr10 | 72982451 | 72982911 | 0.50979 | 1.06341 | 2.086 |
| chr19 | 18977100 | 18977752 | 2.00805 | 4.18905 | 2.086 |
| chr2 | 119640255 | 119640652 | 1.84633 | 3.85169 | 2.086 |
| chr16 | 67580881 | 67581262 | 2.29591 | 4.78975 | 2.086 |
| chr1 | 156897576 | 156897979 | 1.64877 | 3.43981 | 2.086 |
| chr7 | 99577395 | 99577634 | 1.17821 | 2.45813 | 2.086 |
| chrX | 154377257 | 154377531 | 1.03131 | 2.15166 | 2.086 |
| chr16 | 84096296 | 84096851 | 2.05552 | 4.28871 | 2.086 |
| chr11 | 81170941 | 81171201 | 0.472346 | 0.985589 | 2.087 |
| chr20 | 34026842 | 34027170 | 1.89096 | 3.9457 | 2.087 |
| chr5 | 45696369 | 45696656 | 2.17173 | 4.53168 | 2.087 |
| chr4 | 12634762 | 12635016 | 1.43215 | 2.98848 | 2.087 |
| chr12 | 56389951 | 56390281 | 1.82213 | 3.80231 | 2.087 |
| chr8 | 8105403 | 8105628 | 1.29865 | 2.70995 | 2.087 |
| chr10 | 133748668 | 133748997 | 0.558239 | 1.16492 | 2.087 |
| chr19 | 33907022 | 33907538 | 2.37439 | 4.95485 | 2.087 |
| chr14 | 22314554 | 22314760 | 0.810581 | 1.69153 | 2.087 |
| chr5 | 7656868 | 7657140 | 0.384223 | 0.801836 | 2.087 |
| chr12 | 30907891 | 30908094 | 0.803213 | 1.67625 | 2.087 |
| chrX | 68047642 | 68048008 | 1.58484 | 3.30754 | 2.087 |
| chr1 | 37642949 | 37643156 | 1.0534 | 2.19845 | 2.087 |
| chr15 | 61141909 | 61142155 | 0.501133 | 1.04587 | 2.087 |
| chr7 | 107922054 | 107922313 | 0.749662 | 1.56456 | 2.087 |
| chr1 | 47903396 | 47903847 | 1.88757 | 3.93953 | 2.087 |
| chr7 | 126326651 | 126326875 | 0.504365 | 1.05267 | 2.087 |
| chr19 | 41827683 | 41828213 | 1.38924 | 2.89954 | 2.087 |
| chr9 | 135045187 | 135045674 | 1.96827 | 4.1081 | 2.087 |
| chr1 | 218694884 | 218695225 | 1.03222 | 2.15443 | 2.087 |
| chr5 | 104466676 | 104466932 | 0.516431 | 1.07789 | 2.087 |
| chr11 | 65439175 | 65439391 | 0.438855 | 0.916038 | 2.087 |
| chr19 | 31111037 | 31111413 | 0.604725 | 1.26227 | 2.087 |
| chr7 | 127348853 | 127349204 | 1.29639 | 2.70618 | 2.087 |
| chr1 | 93527793 | 93528010 | 0.608278 | 1.26982 | 2.088 |
| chr5 | 144793697 | 144793911 | 0.475863 | 0.9934 | 2.088 |
| chr3 | 74990763 | 74991193 | 0.5645 | 1.17855 | 2.088 |
| chr8 | 116237300 | 116237509 | 0.487843 | 1.01851 | 2.088 |
| chr11 | 103553105 | 103553335 | 1.26988 | 2.65126 | 2.088 |
| chr12 | 58026342 | 58027155 | 1.48809 | 3.10685 | 2.088 |

| | | | | | |
|---|---|---|---|---|---|
| chr14 | 64810772 | 64811221 | 1.21679 | 2.54045 | 2.088 |
| chr11 | 66234705 | 66234996 | 0.597693 | 1.24795 | 2.088 |
| chr5 | 158713535 | 158713747 | 0.944213 | 1.97151 | 2.088 |
| chr17 | 1624099 | 1624510 | 2.72317 | 5.6863 | 2.088 |
| chr2 | 26214483 | 26214685 | 1.14086 | 2.38232 | 2.088 |
| chr2 | 159321221 | 159321552 | 0.940954 | 1.96495 | 2.088 |
| chr8 | 39749591 | 39749831 | 1.36484 | 2.85022 | 2.088 |
| chr1 | 238674635 | 238675109 | 0.519243 | 1.08442 | 2.088 |
| chr17 | 73511674 | 73512283 | 2.69362 | 5.6258 | 2.089 |
| chr6 | 111470192 | 111470430 | 0.373286 | 0.779646 | 2.089 |
| chr17 | 80163659 | 80164067 | 2.23708 | 4.6724 | 2.089 |
| chr1 | 246862490 | 246862766 | 1.65878 | 3.46489 | 2.089 |
| chr17 | 36570947 | 36571380 | 1.88541 | 3.93834 | 2.089 |
| chr6 | 107739438 | 107739681 | 1.1348 | 2.37048 | 2.089 |
| chr6 | 70506788 | 70507093 | 2.60428 | 5.44039 | 2.089 |
| chr9 | 116555435 | 116555809 | 0.431552 | 0.901554 | 2.089 |
| chr7 | 139702127 | 139702481 | 2.12128 | 4.43158 | 2.089 |
| chr6 | 6677108 | 6677390 | 1.98767 | 4.15297 | 2.089 |
| chr14 | 44008506 | 44008735 | 0.908939 | 1.89919 | 2.089 |
| chr1 | 112257555 | 112257834 | 1.15776 | 2.41932 | 2.090 |
| chr12 | 106617855 | 106618345 | 3.16053 | 6.60451 | 2.090 |
| chr3 | 126679146 | 126679494 | 0.721841 | 1.50842 | 2.090 |
| chr11 | 27318000 | 27318258 | 1.49823 | 3.13094 | 2.090 |
| chr2 | 850197 | 850477 | 1.39645 | 2.91843 | 2.090 |
| chr12 | 52789570 | 52790015 | 1.00603 | 2.10263 | 2.090 |
| chr3 | 170408220 | 170408493 | 1.70214 | 3.55756 | 2.090 |
| chr19 | 52968136 | 52968616 | 1.06614 | 2.22832 | 2.090 |
| chr2 | 4563968 | 4564253 | 0.710709 | 1.48544 | 2.090 |
| chr2 | 172430588 | 172430949 | 2.41478 | 5.04722 | 2.090 |
| chr9 | 93633228 | 93633663 | 1.74169 | 3.6405 | 2.090 |
| chr14 | 76448830 | 76449097 | 2.05045 | 4.28589 | 2.090 |
| chr9 | 15135231 | 15135493 | 1.26377 | 2.64157 | 2.090 |
| chr12 | 98754158 | 98754416 | 0.406825 | 0.850415 | 2.090 |
| chr19 | 28457306 | 28457585 | 1.1391 | 2.38118 | 2.090 |
| chr11 | 113403443 | 113403799 | 1.52996 | 3.19833 | 2.090 |
| chr17 | 68763305 | 68763541 | 1.24893 | 2.61097 | 2.091 |
| chr15 | 51675071 | 51675282 | 1.33388 | 2.78881 | 2.091 |
| chr11 | 45986441 | 45987129 | 1.34471 | 2.81153 | 2.091 |
| chr3 | 88052358 | 88052579 | 1.15579 | 2.41655 | 2.091 |
| chr20 | 43015104 | 43015483 | 1.44551 | 3.02234 | 2.091 |
| chr10 | 71866819 | 71867356 | 1.97586 | 4.13126 | 2.091 |
| chr7 | 1032604 | 1033293 | 3.31064 | 6.92215 | 2.091 |
| chr17 | 45434044 | 45434246 | 0.825456 | 1.72594 | 2.091 |
| chr8 | 129701353 | 129701587 | 1.13608 | 2.37548 | 2.091 |
| chr14 | 100506730 | 100506989 | 0.569221 | 1.19021 | 2.091 |
| chr17 | 75243024 | 75243576 | 2.04894 | 4.2843 | 2.091 |
| chr15 | 69868279 | 69868668 | 2.32717 | 4.86617 | 2.091 |
| chr18 | 20711692 | 20712040 | 1.24824 | 2.61009 | 2.091 |
| chrX | 41926313 | 41926518 | 0.38297 | 0.800815 | 2.091 |
| chr14 | 50100903 | 50101229 | 2.45239 | 5.12819 | 2.091 |
| chr10 | 99166888 | 99167538 | 1.58735 | 3.31949 | 2.091 |
| chr1 | 234445695 | 234446020 | 2.0237 | 4.23225 | 2.091 |
| chr1 | 182049824 | 182050450 | 0.960054 | 2.0079 | 2.091 |
| chr22 | 30005223 | 30005449 | 0.641192 | 1.34104 | 2.091 |
| chr12 | 18081973 | 18082181 | 1.06329 | 2.22386 | 2.091 |
| chr3 | 42588831 | 42589275 | 2.13961 | 4.475 | 2.092 |
| chrX | 150891011 | 150891251 | 1.16954 | 2.44614 | 2.092 |
| chr20 | 3746678 | 3746882 | 0.438953 | 0.918105 | 2.092 |
| chr14 | 71180798 | 71181153 | 1.14251 | 2.38988 | 2.092 |
| chr11 | 26306455 | 26306743 | 1.39996 | 2.92844 | 2.092 |
| chrX | 22210813 | 22211122 | 1.63902 | 3.42854 | 2.092 |

| | | | | | |
|---|---|---|---|---|---|
| chr12 | 44276407 | 44276687 | 1.70827 | 3.57365 | 2.092 |
| chrX | 71131052 | 71131449 | 1.15442 | 2.41509 | 2.092 |
| chr15 | 91489693 | 91489959 | 0.699178 | 1.46275 | 2.092 |
| chr11 | 10757249 | 10757765 | 2.34222 | 4.90021 | 2.092 |
| chr12 | 114986331 | 114986760 | 0.462154 | 0.966947 | 2.092 |
| chr6 | 74695792 | 74696036 | 0.896816 | 1.8764 | 2.092 |
| chr16 | 64336156 | 64336463 | 0.525316 | 1.09917 | 2.092 |
| chr7 | 12725220 | 12725451 | 1.36686 | 2.86002 | 2.092 |
| chr20 | 597054 | 597315 | 0.78159 | 1.63552 | 2.093 |
| chr3 | 5178415 | 5178656 | 1.12441 | 2.35293 | 2.093 |
| chr19 | 39818749 | 39819153 | 2.87857 | 6.02428 | 2.093 |
| chr5 | 1235466 | 1235790 | 0.627515 | 1.31334 | 2.093 |
| chr12 | 55387100 | 55387627 | 1.14784 | 2.4024 | 2.093 |
| chr12 | 56617777 | 56618440 | 2.33033 | 4.87739 | 2.093 |
| chr2 | 134986068 | 134986374 | 1.22082 | 2.55526 | 2.093 |
| chr4 | 184642429 | 184642760 | 1.30427 | 2.72995 | 2.093 |
| chr12 | 62829445 | 62829779 | 1.41019 | 2.9517 | 2.093 |
| chr16 | 67427188 | 67427469 | 1.11272 | 2.32922 | 2.093 |
| chr15 | 41624250 | 41624545 | 1.12001 | 2.34454 | 2.093 |
| chr11 | 10503131 | 10503345 | 0.421822 | 0.883012 | 2.093 |
| chr3 | 74573697 | 74574073 | 0.587755 | 1.23038 | 2.093 |
| chr9 | 106939297 | 106939581 | 0.994879 | 2.08264 | 2.093 |
| chr9 | 112945569 | 112945841 | 1.45983 | 3.05596 | 2.093 |
| chr6 | 151645394 | 151645736 | 1.72614 | 3.61388 | 2.094 |
| chr12 | 55596985 | 55597193 | 0.417802 | 0.874732 | 2.094 |
| chr15 | 53676673 | 53676963 | 1.47734 | 3.09304 | 2.094 |
| chr18 | 13103523 | 13103807 | 0.760508 | 1.59225 | 2.094 |
| chr10 | 6622018 | 6622562 | 1.13445 | 2.37521 | 2.094 |
| chr8 | 90273710 | 90273944 | 0.397229 | 0.831702 | 2.094 |
| chr1 | 110855209 | 110855587 | 1.50127 | 3.14331 | 2.094 |
| chr1 | 150504762 | 150505616 | 1.18721 | 2.48576 | 2.094 |
| chr9 | 96190366 | 96190595 | 1.13456 | 2.37565 | 2.094 |
| chr11 | 64845653 | 64845956 | 1.48903 | 3.11794 | 2.094 |
| chr11 | 87749645 | 87749881 | 1.21046 | 2.53476 | 2.094 |
| chr17 | 52012115 | 52012433 | 0.602876 | 1.26245 | 2.094 |
| chr20 | 1215445 | 1215706 | 0.687398 | 1.43945 | 2.094 |
| chr6 | 27759971 | 27760348 | 1.00299 | 2.10052 | 2.094 |
| chr2 | 71221900 | 71222343 | 1.97339 | 4.13301 | 2.094 |
| chr12 | 130531910 | 130532264 | 1.47679 | 3.09301 | 2.094 |
| chr10 | 85259262 | 85259568 | 0.5662 | 1.18586 | 2.094 |
| chr4 | 57666331 | 57666620 | 1.75491 | 3.67554 | 2.094 |
| chr12 | 53298802 | 53299142 | 1.84743 | 3.86945 | 2.095 |
| chr11 | 4514424 | 4514846 | 1.13858 | 2.38478 | 2.095 |
| chr18 | 43728765 | 43729081 | 1.14453 | 2.39733 | 2.095 |
| chr8 | 125250306 | 125250519 | 0.446128 | 0.934471 | 2.095 |
| chr5 | 142246644 | 142246864 | 1.0517 | 2.20293 | 2.095 |
| chr12 | 24587969 | 24588216 | 0.530152 | 1.11059 | 2.095 |
| chr12 | 49514712 | 49515195 | 1.14164 | 2.39161 | 2.095 |
| chr9 | 139383032 | 139383474 | 2.27435 | 4.76467 | 2.095 |
| chr8 | 139800408 | 139800631 | 0.332121 | 0.695791 | 2.095 |
| chr2 | 39457072 | 39457284 | 1.62278 | 3.39973 | 2.095 |
| chr9 | 97431144 | 97431502 | 2.17902 | 4.56507 | 2.095 |
| chr1 | 47898094 | 47898404 | 1.35499 | 2.83874 | 2.095 |
| chr4 | 7877811 | 7878093 | 0.918006 | 1.92329 | 2.095 |
| chr13 | 45945034 | 45945452 | 2.24207 | 4.69732 | 2.095 |
| chr21 | 37497852 | 37498161 | 1.83375 | 3.84198 | 2.095 |
| chr4 | 15233150 | 15233438 | 1.13419 | 2.37632 | 2.095 |
| chr6 | 56457764 | 56458031 | 1.48251 | 3.10637 | 2.095 |
| chr10 | 84925111 | 84925386 | 0.649087 | 1.3601 | 2.095 |
| chr1 | 181053435 | 181053726 | 1.05107 | 2.2026 | 2.096 |
| chr14 | 54795906 | 54796201 | 1.71035 | 3.58418 | 2.096 |

| | | | | | |
|---|---|---|---|---|---|
| chr5 | 176817461 | 176817918 | 2.01888 | 4.23079 | 2.096 |
| chr16 | 928423 | 928730 | 1.42647 | 2.98939 | 2.096 |
| chr3 | 134027259 | 134027551 | 2.05702 | 4.31084 | 2.096 |
| chr15 | 83669151 | 83669484 | 1.32454 | 2.77589 | 2.096 |
| chr7 | 37541204 | 37541509 | 0.59071 | 1.23798 | 2.096 |
| chr7 | 95156771 | 95157072 | 1.98725 | 4.16489 | 2.096 |
| chr22 | 29755555 | 29755834 | 1.39408 | 2.9218 | 2.096 |
| chr14 | 94405824 | 94406588 | 1.55315 | 3.2552 | 2.096 |
| chr17 | 72915514 | 72915871 | 1.23917 | 2.59714 | 2.096 |
| chr13 | 52703030 | 52703398 | 1.2684 | 2.65844 | 2.096 |
| chrX | 134099412 | 134099733 | 1.1923 | 2.49896 | 2.096 |
| chr16 | 284906 | 285195 | 1.03172 | 2.16247 | 2.096 |
| chr17 | 63247513 | 63247767 | 0.599365 | 1.25627 | 2.096 |
| chr20 | 55255089 | 55255321 | 0.561089 | 1.17608 | 2.096 |
| chr1 | 205110666 | 205111088 | 1.25849 | 2.63798 | 2.096 |
| chr8 | 1949415 | 1949627 | 0.734728 | 1.54015 | 2.096 |
| chr20 | 62417613 | 62418010 | 1.96101 | 4.11073 | 2.096 |
| chr5 | 11972234 | 11972441 | 0.586762 | 1.23011 | 2.096 |
| chr14 | 47050214 | 47050683 | 1.29609 | 2.71719 | 2.096 |
| chr22 | 36520050 | 36520407 | 1.68527 | 3.53321 | 2.097 |
| chr8 | 110877776 | 110878067 | 0.444916 | 0.932778 | 2.097 |
| chr1 | 233119249 | 233119515 | 0.466115 | 0.977239 | 2.097 |
| chr7 | 28312032 | 28312352 | 1.15644 | 2.42456 | 2.097 |
| chr16 | 50023841 | 50024090 | 0.493809 | 1.03532 | 2.097 |
| chr12 | 53553017 | 53553545 | 1.90507 | 3.99433 | 2.097 |
| chr22 | 22562171 | 22562561 | 1.18078 | 2.47579 | 2.097 |
| chr8 | 139525728 | 139526002 | 0.525913 | 1.10273 | 2.097 |
| chr8 | 142396841 | 142397308 | 3.02808 | 6.34947 | 2.097 |
| chr10 | 3035070 | 3035284 | 0.723718 | 1.51755 | 2.097 |
| chr17 | 6817196 | 6817541 | 1.36529 | 2.86292 | 2.097 |
| chr7 | 53450903 | 53451358 | 0.445584 | 0.934368 | 2.097 |
| chr16 | 23359907 | 23360334 | 0.792978 | 1.66289 | 2.097 |
| chr1 | 28867270 | 28867490 | 0.94415 | 1.97995 | 2.097 |
| chr14 | 91893664 | 91893934 | 0.833529 | 1.74797 | 2.097 |
| chr1 | 153483341 | 153483765 | 2.16086 | 4.53175 | 2.097 |
| chr11 | 1595822 | 1596300 | 1.84716 | 3.87405 | 2.097 |
| chr10 | 24807947 | 24808482 | 1.38466 | 2.90406 | 2.097 |
| chr7 | 36024769 | 36025205 | 1.80888 | 3.79397 | 2.097 |
| chr9 | 114384186 | 114384581 | 1.69424 | 3.55353 | 2.097 |
| chr6 | 170231713 | 170232106 | 1.86109 | 3.90354 | 2.097 |
| chr17 | 47652871 | 47653534 | 1.88695 | 3.9578 | 2.097 |
| chr12 | 12857457 | 12857953 | 1.95433 | 4.09914 | 2.097 |
| chr6 | 43726269 | 43726614 | 1.6118 | 3.38075 | 2.098 |
| chr19 | 45160399 | 45160827 | 2.18263 | 4.57808 | 2.098 |
| chr6 | 18596503 | 18596771 | 1.53192 | 3.21322 | 2.098 |
| chr11 | 44087566 | 44087936 | 1.7334 | 3.63609 | 2.098 |
| chr20 | 37277981 | 37278235 | 1.42839 | 2.99627 | 2.098 |
| chr8 | 13665223 | 13665572 | 0.605997 | 1.27119 | 2.098 |
| chr5 | 165264329 | 165264550 | 0.38921 | 0.816443 | 2.098 |
| chr4 | 5461867 | 5462192 | 1.91489 | 4.01698 | 2.098 |
| chr8 | 110328474 | 110328704 | 1.02868 | 2.158 | 2.098 |
| chr5 | 85796903 | 85797217 | 0.524733 | 1.10081 | 2.098 |
| chr12 | 115106424 | 115106662 | 1.18865 | 2.49362 | 2.098 |
| chr8 | 15932376 | 15932589 | 0.639169 | 1.3409 | 2.098 |
| chr3 | 4356421 | 4356648 | 0.983926 | 2.06418 | 2.098 |
| chr9 | 35882175 | 35882622 | 2.06356 | 4.32919 | 2.098 |
| chr17 | 17400014 | 17400568 | 2.43034 | 5.09868 | 2.098 |
| chr1 | 155959070 | 155959550 | 2.14393 | 4.4981 | 2.098 |
| chr7 | 92961204 | 92961469 | 0.604928 | 1.26923 | 2.098 |
| chr17 | 47856453 | 47857349 | 0.738879 | 1.55032 | 2.098 |
| chr16 | 67318781 | 67319152 | 1.28883 | 2.70426 | 2.098 |

| | | | | | |
|---|---|---|---|---|---|
| chr22 | 51001199 | 51001530 | 1.63949 | 3.44004 | 2.098 |
| chrX | 19553619 | 19553932 | 1.18183 | 2.47977 | 2.098 |
| chr11 | 43203993 | 43204203 | 0.553521 | 1.16147 | 2.098 |
| chr11 | 124632953 | 124633437 | 1.19458 | 2.50665 | 2.098 |
| chr2 | 208610769 | 208611091 | 1.32525 | 2.78086 | 2.098 |
| chr5 | 129516268 | 129516485 | 0.448253 | 0.94062 | 2.098 |
| chr2 | 172290356 | 172290576 | 0.819105 | 1.71883 | 2.098 |
| chr15 | 86530061 | 86530295 | 0.658662 | 1.38216 | 2.098 |
| chr12 | 801588 | 801798 | 0.564917 | 1.18546 | 2.098 |
| chr16 | 30016725 | 30017294 | 2.00403 | 4.20539 | 2.098 |
| chr22 | 50323168 | 50323577 | 1.68384 | 3.53348 | 2.098 |
| chr10 | 90746753 | 90747130 | 1.51503 | 3.17929 | 2.099 |
| chrX | 25038968 | 25039353 | 1.92046 | 4.03008 | 2.099 |
| chr7 | 124022870 | 124023243 | 0.649566 | 1.36312 | 2.099 |
| chr6 | 112484519 | 112484996 | 1.97179 | 4.1379 | 2.099 |
| chr11 | 94630478 | 94630872 | 1.34561 | 2.82407 | 2.099 |
| chr16 | 49024066 | 49024277 | 0.622813 | 1.30713 | 2.099 |
| chr2 | 201567953 | 201568158 | 0.948799 | 1.99135 | 2.099 |
| chr1 | 27901749 | 27902083 | 0.82516 | 1.73189 | 2.099 |
| chr14 | 81919604 | 81919900 | 1.52588 | 3.2027 | 2.099 |
| chr4 | 100384122 | 100384362 | 1.09626 | 2.30103 | 2.099 |
| chr3 | 122426571 | 122426950 | 0.532671 | 1.11808 | 2.099 |
| chr15 | 57933112 | 57933868 | 0.872546 | 1.83149 | 2.099 |
| chr11 | 47174561 | 47174955 | 1.86036 | 3.90501 | 2.099 |
| chr6 | 37534066 | 37534493 | 1.93323 | 4.05825 | 2.099 |
| chr5 | 112331678 | 112332072 | 1.27155 | 2.66926 | 2.099 |
| chr5 | 179544092 | 179544481 | 1.39012 | 2.91825 | 2.099 |
| chr4 | 24990321 | 24990614 | 0.905544 | 1.90111 | 2.099 |
| chr6 | 20291978 | 20292232 | 1.62045 | 3.40209 | 2.099 |
| chr1 | 196446509 | 196446747 | 0.745313 | 1.56483 | 2.100 |
| chr5 | 137114799 | 137115031 | 0.875478 | 1.83816 | 2.100 |
| chr11 | 76630154 | 76630434 | 1.46698 | 3.08013 | 2.100 |
| chr5 | 153833414 | 153833672 | 0.552416 | 1.15989 | 2.100 |
| chr9 | 37789841 | 37790147 | 0.502937 | 1.056 | 2.100 |
| chr12 | 9903944 | 9904159 | 1.47679 | 3.10096 | 2.100 |
| chrX | 56274912 | 56275245 | 0.79396 | 1.66719 | 2.100 |
| chr10 | 85289839 | 85290047 | 0.456667 | 0.958949 | 2.100 |
| chrX | 104166807 | 104167229 | 2.11578 | 4.44346 | 2.100 |
| chr2 | 62758778 | 62759019 | 0.682772 | 1.43398 | 2.100 |
| chr9 | 72027355 | 72028095 | 1.43509 | 3.01408 | 2.100 |
| chr12 | 103433405 | 103433623 | 0.506747 | 1.06434 | 2.100 |
| chr15 | 89600722 | 89601109 | 2.93833 | 6.17162 | 2.100 |
| chr8 | 26428747 | 26429166 | 1.99052 | 4.18085 | 2.100 |
| chr1 | 233016148 | 233016521 | 1.80809 | 3.79794 | 2.101 |
| chr17 | 1768780 | 1769127 | 1.81238 | 3.80703 | 2.101 |
| chr16 | 75233253 | 75233554 | 1.91837 | 4.03013 | 2.101 |
| chr2 | 205124844 | 205125157 | 0.943222 | 1.98154 | 2.101 |
| chr12 | 121548088 | 121548653 | 1.66657 | 3.50142 | 2.101 |
| chr5 | 39376427 | 39376728 | 1.56107 | 3.27977 | 2.101 |
| chr13 | 113597046 | 113597365 | 1.52304 | 3.19991 | 2.101 |
| chr12 | 53591365 | 53591864 | 1.77905 | 3.73794 | 2.101 |
| chr12 | 124389885 | 124390285 | 1.79127 | 3.76369 | 2.101 |
| chr2 | 74051104 | 74051433 | 1.5174 | 3.18832 | 2.101 |
| chr10 | 102757393 | 102757792 | 1.72187 | 3.61796 | 2.101 |
| chr5 | 6526136 | 6526372 | 0.977133 | 2.05315 | 2.101 |
| chr6 | 144426079 | 144426411 | 0.793889 | 1.6682 | 2.101 |
| chr2 | 71327661 | 71327911 | 1.51133 | 3.17581 | 2.101 |
| chr1 | 249152826 | 249153030 | 1.87993 | 3.95039 | 2.101 |
| chr4 | 7196230 | 7196415 | 1.2731 | 2.67543 | 2.102 |
| chr5 | 24233158 | 24233406 | 0.569889 | 1.19764 | 2.102 |
| chr13 | 60970933 | 60971209 | 1.23296 | 2.59116 | 2.102 |

| | | | | | |
|---|---|---|---|---|---|
| chr8 | 75306603 | 75306922 | 0.431168 | 0.906138 | 2.102 |
| chr2 | 66430237 | 66430481 | 0.694581 | 1.45975 | 2.102 |
| chr1 | 36626702 | 36627133 | 0.922574 | 1.93905 | 2.102 |
| chr1 | 146714080 | 146714972 | 1.75079 | 3.67984 | 2.102 |
| chr4 | 142228379 | 142228664 | 1.33902 | 2.81449 | 2.102 |
| chr9 | 139560381 | 139560623 | 0.770224 | 1.61895 | 2.102 |
| chr15 | 91599085 | 91599450 | 1.69744 | 3.56807 | 2.102 |
| chr20 | 6359108 | 6359319 | 0.445156 | 0.93577 | 2.102 |
| chr22 | 20000987 | 20001212 | 1.12379 | 2.36242 | 2.102 |
| chr20 | 5456012 | 5456259 | 0.541553 | 1.13853 | 2.102 |
| chr6 | 41034653 | 41034877 | 1.17474 | 2.4697 | 2.102 |
| chr6 | 40862580 | 40862913 | 1.29583 | 2.7243 | 2.102 |
| chr1 | 201377326 | 201377730 | 2.25747 | 4.7461 | 2.102 |
| chr5 | 168624388 | 168624691 | 1.15862 | 2.4359 | 2.102 |
| chr12 | 23166252 | 23166494 | 0.373433 | 0.785141 | 2.102 |
| chr12 | 125827170 | 125827374 | 0.456469 | 0.959788 | 2.103 |
| chr6 | 146025391 | 146025599 | 0.99794 | 2.09836 | 2.103 |
| chr10 | 114235453 | 114235660 | 0.845954 | 1.77885 | 2.103 |
| chr17 | 75210560 | 75211135 | 1.29807 | 2.72955 | 2.103 |
| chr9 | 139525917 | 139526183 | 1.90726 | 4.01069 | 2.103 |
| chr1 | 17803860 | 17804221 | 1.28803 | 2.70861 | 2.103 |
| chr17 | 37309116 | 37309489 | 1.67598 | 3.5245 | 2.103 |
| chr1 | 150594060 | 150594294 | 1.29884 | 2.73151 | 2.103 |
| chr19 | 54822644 | 54822857 | 0.729065 | 1.5333 | 2.103 |
| chr4 | 103266600 | 103266914 | 1.52433 | 3.20588 | 2.103 |
| chr19 | 32816968 | 32817175 | 0.556182 | 1.16974 | 2.103 |
| chr2 | 219762901 | 219763329 | 2.47058 | 5.19609 | 2.103 |
| chr5 | 1011017 | 1011586 | 1.22351 | 2.57343 | 2.103 |
| chr14 | 21558446 | 21558804 | 1.58694 | 3.33801 | 2.103 |
| chr17 | 74677261 | 74677691 | 1.75633 | 3.69433 | 2.103 |
| chr5 | 6837695 | 6837943 | 1.56876 | 3.29992 | 2.104 |
| chr4 | 140524795 | 140525069 | 0.738201 | 1.55283 | 2.104 |
| chr17 | 73422264 | 73422724 | 1.3934 | 2.93107 | 2.104 |
| chr2 | 55659161 | 55659448 | 1.4391 | 3.0272 | 2.104 |
| chr14 | 93813416 | 93813750 | 0.900028 | 1.89339 | 2.104 |
| chr7 | 101879972 | 101880278 | 2.19486 | 4.61739 | 2.104 |
| chr20 | 52163838 | 52164802 | 0.826017 | 1.73773 | 2.104 |
| chr15 | 85587591 | 85587823 | 1.34518 | 2.82997 | 2.104 |
| chr1 | 220286537 | 220286890 | 0.978177 | 2.058 | 2.104 |
| chr19 | 49111966 | 49112409 | 1.98594 | 4.17828 | 2.104 |
| chr19 | 47505339 | 47505735 | 1.44079 | 3.03138 | 2.104 |
| chr3 | 14898858 | 14899280 | 0.878013 | 1.84739 | 2.104 |
| chr15 | 42677749 | 42677985 | 1.88243 | 3.96084 | 2.104 |
| chr8 | 105524846 | 105525049 | 0.790424 | 1.66319 | 2.104 |
| chr10 | 66916955 | 66917169 | 0.534179 | 1.12401 | 2.104 |
| chr9 | 35599955 | 35600160 | 0.66088 | 1.39062 | 2.104 |
| chr8 | 141646119 | 141646518 | 2.61442 | 5.50153 | 2.104 |
| chr14 | 94522006 | 94522210 | 0.38699 | 0.814354 | 2.104 |
| chr1 | 94801493 | 94801767 | 1.42582 | 3.00045 | 2.104 |
| chr14 | 103651769 | 103652014 | 0.99986 | 2.1041 | 2.104 |
| chr12 | 120868181 | 120868615 | 2.43491 | 5.12415 | 2.104 |
| chr5 | 117893096 | 117893345 | 0.477715 | 1.00536 | 2.105 |
| chr3 | 187396743 | 187396972 | 1.2368 | 2.60291 | 2.105 |
| chrX | 15779720 | 15780123 | 1.64579 | 3.46379 | 2.105 |
| chr1 | 244766792 | 244767055 | 0.976079 | 2.05438 | 2.105 |
| chr12 | 85451193 | 85451499 | 1.57459 | 3.31412 | 2.105 |
| chr1 | 59755889 | 59756164 | 0.993623 | 2.09138 | 2.105 |
| chr12 | 119766078 | 119766367 | 0.701731 | 1.4771 | 2.105 |
| chr6 | 162598913 | 162599143 | 0.535457 | 1.12711 | 2.105 |
| chr1 | 12178712 | 12179009 | 2.29985 | 4.84116 | 2.105 |
| chr3 | 123166727 | 123167120 | 1.19889 | 2.52375 | 2.105 |

| | | | | | |
|---|---|---|---|---|---|
| chrX | 49680156 | 49680702 | 1.25598 | 2.64401 | 2.105 |
| chr1 | 208136857 | 208137306 | 1.85348 | 3.90218 | 2.105 |
| chr19 | 36618561 | 36618970 | 2.59788 | 5.46952 | 2.105 |
| chr3 | 62174588 | 62174802 | 1.25883 | 2.65044 | 2.105 |
| chr5 | 179223020 | 179223514 | 1.33274 | 2.80606 | 2.105 |
| chr16 | 87093594 | 87093945 | 0.540876 | 1.13881 | 2.105 |
| chr4 | 15884281 | 15884585 | 1.37404 | 2.8931 | 2.106 |
| chr3 | 63966606 | 63966876 | 0.623769 | 1.3134 | 2.106 |
| chr2 | 88971250 | 88971750 | 2.22326 | 4.68153 | 2.106 |
| chr6 | 52463026 | 52463383 | 1.30102 | 2.73977 | 2.106 |
| chr11 | 20669440 | 20669860 | 1.34153 | 2.82511 | 2.106 |
| chr20 | 33064037 | 33064436 | 2.30697 | 4.85831 | 2.106 |
| chr10 | 2537694 | 2537921 | 0.472665 | 0.995417 | 2.106 |
| chr5 | 41925226 | 41925545 | 1.14127 | 2.40349 | 2.106 |
| chr1 | 120413525 | 120413971 | 1.91581 | 4.03467 | 2.106 |
| chr15 | 93028576 | 93028834 | 0.450036 | 0.947771 | 2.106 |
| chr17 | 49513003 | 49513329 | 2.26974 | 4.78009 | 2.106 |
| chr15 | 92634333 | 92634725 | 2.41252 | 5.08111 | 2.106 |
| chr7 | 69466774 | 69466988 | 0.554401 | 1.16767 | 2.106 |
| chr19 | 6260891 | 6261156 | 0.799287 | 1.68349 | 2.106 |
| chr15 | 67195949 | 67196395 | 1.91887 | 4.04166 | 2.106 |
| chr6 | 43806107 | 43806322 | 0.806087 | 1.69784 | 2.106 |
| chr20 | 46340017 | 46340263 | 0.465722 | 0.980946 | 2.106 |
| chr5 | 175917338 | 175917586 | 0.613236 | 1.29168 | 2.106 |
| chr15 | 45997908 | 45998151 | 1.2597 | 2.65339 | 2.106 |
| chr16 | 56998358 | 56998892 | 2.77219 | 5.83935 | 2.106 |
| chrX | 114924975 | 114925276 | 1.52041 | 3.20261 | 2.106 |
| chr14 | 100851552 | 100851775 | 0.933521 | 1.96651 | 2.107 |
| chr17 | 40443792 | 40444086 | 0.753811 | 1.58805 | 2.107 |
| chr18 | 12237117 | 12237560 | 1.99379 | 4.20045 | 2.107 |
| chr19 | 3104014 | 3104581 | 1.57944 | 3.32757 | 2.107 |
| chr15 | 76411022 | 76411237 | 0.417895 | 0.880464 | 2.107 |
| chr5 | 6821019 | 6821351 | 1.93518 | 4.07766 | 2.107 |
| chr5 | 76011744 | 76012219 | 1.573 | 3.31468 | 2.107 |
| chr20 | 36888795 | 36889066 | 0.403688 | 0.850681 | 2.107 |
| chr7 | 117633531 | 117633766 | 1.19367 | 2.51539 | 2.107 |
| chr5 | 178701323 | 178701812 | 0.609054 | 1.28345 | 2.107 |
| chr13 | 34116870 | 34117157 | 1.40107 | 2.95252 | 2.107 |
| chr11 | 33167248 | 33167478 | 0.591415 | 1.24634 | 2.107 |
| chr14 | 51327302 | 51327525 | 1.26567 | 2.6673 | 2.107 |
| chr3 | 14302638 | 14303030 | 2.08671 | 4.39778 | 2.108 |
| chr12 | 49389271 | 49389525 | 1.53997 | 3.24557 | 2.108 |
| chr21 | 43032305 | 43032601 | 1.09831 | 2.31474 | 2.108 |
| chr3 | 184904001 | 184904221 | 0.529359 | 1.11566 | 2.108 |
| chr3 | 123798287 | 123798552 | 1.28865 | 2.71596 | 2.108 |
| chr3 | 77443323 | 77443629 | 0.71953 | 1.51651 | 2.108 |
| chr5 | 58969326 | 58969597 | 1.89976 | 4.00406 | 2.108 |
| chr2 | 6690732 | 6690952 | 0.635141 | 1.33871 | 2.108 |
| chr7 | 22765938 | 22766303 | 1.06248 | 2.23967 | 2.108 |
| chr7 | 92250367 | 92250616 | 1.4015 | 2.95434 | 2.108 |
| chr10 | 47600438 | 47600856 | 1.5309 | 3.22718 | 2.108 |
| chr1 | 12335428 | 12335821 | 1.58184 | 3.33465 | 2.108 |
| chr7 | 145402301 | 145402605 | 0.718834 | 1.51537 | 2.108 |
| chr12 | 53845529 | 53845776 | 1.86275 | 3.92687 | 2.108 |
| chr9 | 130348782 | 130349166 | 1.81875 | 3.83422 | 2.108 |
| chr16 | 14404526 | 14404736 | 0.960226 | 2.02432 | 2.108 |
| chr12 | 57399934 | 57400690 | 1.99527 | 4.20638 | 2.108 |
| chr8 | 128746199 | 128746499 | 1.51791 | 3.20008 | 2.108 |
| chr8 | 21867769 | 21868100 | 0.746954 | 1.57474 | 2.108 |
| chr17 | 8649237 | 8649627 | 2.1633 | 4.56109 | 2.108 |
| chr12 | 95728259 | 95728494 | 1.37997 | 2.90956 | 2.108 |

| | | | | | |
|---|---|---|---|---|---|
| chr8 | 134465509 | 134465896 | 2.34885 | 4.95243 | 2.108 |
| chr9 | 117167849 | 117168197 | 1.43322 | 3.0219 | 2.108 |
| chr15 | 63003926 | 63004137 | 0.519982 | 1.09637 | 2.108 |
| chr3 | 88574618 | 88574868 | 0.473562 | 0.998494 | 2.108 |
| chr17 | 34890154 | 34890524 | 1.75855 | 3.70791 | 2.109 |
| chr22 | 38510446 | 38510913 | 1.80441 | 3.80475 | 2.109 |
| chr5 | 39425118 | 39425365 | 1.28216 | 2.70354 | 2.109 |
| chr12 | 50898451 | 50899106 | 1.74162 | 3.67245 | 2.109 |
| chr14 | 100659170 | 100659754 | 1.71629 | 3.6193 | 2.109 |
| chr10 | 45956593 | 45956817 | 0.428283 | 0.903173 | 2.109 |
| chr19 | 14225548 | 14225912 | 1.31722 | 2.77786 | 2.109 |
| chr3 | 197632827 | 197633177 | 1.65789 | 3.4963 | 2.109 |
| chr3 | 117245371 | 117245812 | 1.05885 | 2.23314 | 2.109 |
| chr19 | 5794237 | 5794549 | 0.801733 | 1.69096 | 2.109 |
| chr8 | 62381388 | 62381700 | 1.84363 | 3.8885 | 2.109 |
| chr17 | 58798072 | 58798452 | 1.62434 | 3.42602 | 2.109 |
| chr15 | 44870749 | 44871051 | 1.08882 | 2.29659 | 2.109 |
| chr3 | 99562163 | 99562383 | 1.21021 | 2.55266 | 2.109 |
| chr8 | 146209477 | 146209755 | 0.646041 | 1.36271 | 2.109 |
| chr17 | 9305971 | 9306292 | 1.49581 | 3.1553 | 2.109 |
| chr2 | 27926680 | 27926958 | 2.16635 | 4.56986 | 2.109 |
| chr12 | 98419782 | 98420025 | 0.685321 | 1.44569 | 2.110 |
| chr20 | 46292440 | 46292977 | 2.06925 | 4.36512 | 2.110 |
| chr6 | 41754876 | 41755450 | 1.34845 | 2.8447 | 2.110 |
| chr22 | 20918569 | 20919034 | 3.48371 | 7.34959 | 2.110 |
| chr9 | 138366817 | 138367066 | 1.12231 | 2.36781 | 2.110 |
| chr7 | 147862729 | 147863072 | 0.39128 | 0.825514 | 2.110 |
| chr10 | 107951745 | 107951967 | 0.475167 | 1.00251 | 2.110 |
| chr16 | 71779426 | 71779684 | 0.480891 | 1.01461 | 2.110 |
| chr10 | 91149085 | 91149300 | 1.08136 | 2.28174 | 2.110 |
| chr17 | 42659295 | 42659537 | 1.33214 | 2.81108 | 2.110 |
| chr2 | 55914448 | 55914665 | 1.17045 | 2.46992 | 2.110 |
| chr3 | 28841392 | 28841667 | 0.511368 | 1.07915 | 2.110 |
| chr3 | 38043030 | 38043326 | 1.20219 | 2.53708 | 2.110 |
| chr17 | 66201371 | 66202035 | 2.35984 | 4.98028 | 2.110 |
| chr5 | 140201622 | 140202103 | 1.72214 | 3.6346 | 2.111 |
| chr16 | 56922057 | 56922390 | 0.478775 | 1.01047 | 2.111 |
| chr16 | 68562366 | 68562903 | 1.76238 | 3.71956 | 2.111 |
| chr4 | 106442091 | 106442398 | 1.14163 | 2.40947 | 2.111 |
| chr2 | 1554334 | 1554951 | 1.93301 | 4.0799 | 2.111 |
| chr11 | 9528322 | 9528674 | 1.73386 | 3.65975 | 2.111 |
| chr17 | 49841734 | 49842173 | 1.75011 | 3.6941 | 2.111 |
| chr14 | 67893874 | 67894249 | 1.80245 | 3.8048 | 2.111 |
| chr3 | 76333890 | 76334148 | 0.62653 | 1.3226 | 2.111 |
| chr6 | 53583756 | 53583978 | 1.20249 | 2.53852 | 2.111 |
| chr7 | 92076478 | 92076809 | 1.19594 | 2.52474 | 2.111 |
| chr8 | 118486619 | 118486851 | 1.30791 | 2.76111 | 2.111 |
| chr6 | 132482511 | 132482733 | 1.22555 | 2.58736 | 2.111 |
| chr17 | 4469974 | 4470445 | 2.46921 | 5.21318 | 2.111 |
| chr9 | 12775991 | 12776590 | 0.99121 | 2.09271 | 2.111 |
| chr14 | 76517163 | 76517507 | 1.19789 | 2.52917 | 2.111 |
| chr17 | 72766812 | 72767246 | 2.43485 | 5.14081 | 2.111 |
| chr19 | 45016222 | 45017345 | 1.18855 | 2.50965 | 2.112 |
| chr12 | 89814995 | 89815232 | 1.03406 | 2.1835 | 2.112 |
| chr3 | 93968320 | 93968784 | 1.28983 | 2.72358 | 2.112 |
| chr1 | 37008604 | 37008828 | 0.712335 | 1.50421 | 2.112 |
| chr4 | 4254527 | 4254745 | 1.55499 | 3.28371 | 2.112 |
| chr10 | 105023006 | 105023261 | 0.511123 | 1.07937 | 2.112 |
| chr5 | 179518241 | 179518545 | 1.11095 | 2.34607 | 2.112 |
| chr11 | 86071883 | 86072130 | 0.793007 | 1.67493 | 2.112 |
| chr14 | 104836627 | 104836961 | 1.21743 | 2.57138 | 2.112 |

| | | | | | |
|---|---|---|---|---|---|
| chr8 | 145174273 | 145174484 | 0.532367 | 1.12445 | 2.112 |
| chr6 | 135466192 | 135466401 | 0.906645 | 1.91503 | 2.112 |
| chr17 | 37778094 | 37778706 | 3.64962 | 7.70907 | 2.112 |
| chr20 | 18581775 | 18582129 | 0.658543 | 1.39108 | 2.112 |
| chr16 | 85364470 | 85364688 | 1.07085 | 2.26206 | 2.112 |
| chr19 | 17607087 | 17607521 | 2.75662 | 5.82311 | 2.112 |
| chr3 | 146307771 | 146307996 | 1.36279 | 2.87898 | 2.113 |
| chr1 | 199923899 | 199924157 | 1.05114 | 2.22061 | 2.113 |
| chr8 | 39191984 | 39192191 | 0.439585 | 0.928655 | 2.113 |
| chr12 | 9102594 | 9102803 | 1.36261 | 2.87866 | 2.113 |
| chr2 | 75867686 | 75868111 | 2.39564 | 5.06106 | 2.113 |
| chr5 | 140145545 | 140145770 | 0.600325 | 1.26825 | 2.113 |
| chr18 | 29321114 | 29321449 | 1.16483 | 2.46089 | 2.113 |
| chr2 | 95622395 | 95622910 | 1.88568 | 3.98382 | 2.113 |
| chr7 | 105297415 | 105297749 | 1.4088 | 2.97634 | 2.113 |
| chr5 | 63443690 | 63444032 | 1.43225 | 3.02595 | 2.113 |
| chr1 | 21558536 | 21558753 | 1.27274 | 2.68906 | 2.113 |
| chr7 | 151773422 | 151773715 | 1.00146 | 2.11589 | 2.113 |
| chr10 | 134468165 | 134468689 | 1.92042 | 4.05757 | 2.113 |
| chr20 | 23255489 | 23255824 | 0.641863 | 1.3562 | 2.113 |
| chr1 | 226150018 | 226150477 | 2.00576 | 4.23817 | 2.113 |
| chrX | 53313764 | 53314210 | 1.62534 | 3.43437 | 2.113 |
| chr10 | 2369632 | 2369858 | 0.645897 | 1.3648 | 2.113 |
| chr12 | 14809826 | 14810075 | 0.776602 | 1.641 | 2.113 |
| chr12 | 56216099 | 56216366 | 0.93656 | 1.979 | 2.113 |
| chr5 | 78496468 | 78496726 | 1.31514 | 2.77902 | 2.113 |
| chr9 | 24788157 | 24788448 | 0.457107 | 0.965936 | 2.113 |
| chr15 | 89440661 | 89441015 | 0.842503 | 1.78034 | 2.113 |
| chr19 | 18108999 | 18109580 | 1.05088 | 2.22072 | 2.113 |
| chr1 | 165083598 | 165083807 | 1.25914 | 2.66083 | 2.113 |
| chr19 | 17886235 | 17886448 | 0.999549 | 2.11228 | 2.113 |
| chr4 | 88565448 | 88565653 | 0.541326 | 1.14395 | 2.113 |
| chr1 | 151707533 | 151707978 | 1.45524 | 3.07533 | 2.113 |
| chr15 | 84415839 | 84416079 | 0.791132 | 1.67188 | 2.113 |
| chr2 | 164039319 | 164039521 | 0.773641 | 1.63492 | 2.113 |
| chr12 | 78278127 | 78278360 | 0.565026 | 1.19407 | 2.113 |
| chr11 | 7686918 | 7687303 | 1.87569 | 3.96411 | 2.113 |
| chr10 | 29759258 | 29759665 | 1.16721 | 2.46698 | 2.114 |
| chr8 | 24272476 | 24272892 | 1.436 | 3.03513 | 2.114 |
| chr18 | 45523904 | 45524205 | 1.03082 | 2.17881 | 2.114 |
| chrY | 14645157 | 14645362 | 1.00599 | 2.12637 | 2.114 |
| chr13 | 111471746 | 111472136 | 1.36581 | 2.8871 | 2.114 |
| chr16 | 11907630 | 11907870 | 1.11128 | 2.34914 | 2.114 |
| chr15 | 33299546 | 33299758 | 1.07375 | 2.26982 | 2.114 |
| chr11 | 81726823 | 81727025 | 0.582513 | 1.23142 | 2.114 |
| chr8 | 138809649 | 138809856 | 0.517002 | 1.09293 | 2.114 |
| chr2 | 1821190 | 1821636 | 2.30678 | 4.87651 | 2.114 |
| chr12 | 76017624 | 76017857 | 0.981594 | 2.07509 | 2.114 |
| chr11 | 70264324 | 70264573 | 0.56764 | 1.20005 | 2.114 |
| chr12 | 25763492 | 25763739 | 0.346925 | 0.733447 | 2.114 |
| chr7 | 3688663 | 3689113 | 0.71882 | 1.51969 | 2.114 |
| chr2 | 110267704 | 110268055 | 2.08225 | 4.40221 | 2.114 |
| chr16 | 29874802 | 29875069 | 2.02666 | 4.28472 | 2.114 |
| chr12 | 46156382 | 46156590 | 1.29354 | 2.73481 | 2.114 |
| chr19 | 4186485 | 4186757 | 1.6738 | 3.53878 | 2.114 |
| chr19 | 38389346 | 38389664 | 1.6395 | 3.46644 | 2.114 |
| chr12 | 57463262 | 57463549 | 1.35995 | 2.87542 | 2.114 |
| chr15 | 86709548 | 86709769 | 0.5831 | 1.23292 | 2.114 |
| chr22 | 31847744 | 31848005 | 1.31734 | 2.78541 | 2.114 |
| chr6 | 41651326 | 41651706 | 1.6148 | 3.41443 | 2.114 |
| chr10 | 70997317 | 70997694 | 2.13884 | 4.52261 | 2.115 |

| | | | | | |
|---|---|---|---|---|---|
| chr3 | 52016502 | 52016765 | 1.14002 | 2.41078 | 2.115 |
| chr8 | 3263142 | 3263598 | 0.66229 | 1.40057 | 2.115 |
| chr16 | 56823412 | 56823660 | 1.07167 | 2.26638 | 2.115 |
| chr7 | 100781559 | 100781885 | 1.38075 | 2.92004 | 2.115 |
| chr15 | 74265266 | 74265612 | 1.17673 | 2.4887 | 2.115 |
| chr5 | 161862612 | 161862882 | 0.447434 | 0.946321 | 2.115 |
| chr8 | 281029 | 281250 | 0.478183 | 1.01136 | 2.115 |
| chr3 | 143581407 | 143581616 | 0.398047 | 0.841914 | 2.115 |
| chr3 | 3847677 | 3847904 | 0.493783 | 1.04442 | 2.115 |
| chr1 | 240312856 | 240313146 | 1.04561 | 2.21165 | 2.115 |
| chr8 | 51330422 | 51330633 | 0.596709 | 1.26216 | 2.115 |
| chr8 | 141678372 | 141678758 | 1.7819 | 3.76921 | 2.115 |
| chr9 | 127534353 | 127534639 | 1.91878 | 4.05876 | 2.115 |
| chr11 | 44058989 | 44059239 | 1.19233 | 2.52224 | 2.115 |
| chr7 | 22528445 | 22529041 | 1.30806 | 2.76715 | 2.115 |
| chr12 | 132638383 | 132638936 | 1.97701 | 4.18244 | 2.116 |
| chr2 | 44942813 | 44943039 | 1.19044 | 2.51843 | 2.116 |
| chr3 | 9757539 | 9757783 | 1.86891 | 3.95387 | 2.116 |
| chrX | 130767576 | 130767877 | 1.98582 | 4.20121 | 2.116 |
| chr22 | 44279017 | 44279389 | 1.67825 | 3.5506 | 2.116 |
| chr2 | 35185662 | 35185916 | 0.381195 | 0.806535 | 2.116 |
| chr7 | 44605063 | 44606046 | 1.54538 | 3.26986 | 2.116 |
| chr8 | 91013047 | 91013621 | 1.07885 | 2.28283 | 2.116 |
| chr11 | 67236797 | 67237096 | 0.7567 | 1.60127 | 2.116 |
| chr8 | 18537751 | 18538129 | 2.03844 | 4.31358 | 2.116 |
| chr20 | 34744134 | 34744907 | 1.06496 | 2.25362 | 2.116 |
| chr9 | 101610833 | 101611209 | 2.47128 | 5.22977 | 2.116 |
| chr9 | 100472985 | 100473255 | 1.42778 | 3.02155 | 2.116 |
| chr18 | 11519000 | 11519509 | 1.56564 | 3.31376 | 2.117 |
| chr5 | 43627187 | 43627465 | 0.97292 | 2.05933 | 2.117 |
| chr14 | 69941960 | 69942302 | 0.953805 | 2.01892 | 2.117 |
| chr4 | 126234608 | 126234926 | 1.03683 | 2.19475 | 2.117 |
| chr2 | 109561302 | 109561614 | 0.687723 | 1.4558 | 2.117 |
| chr1 | 51762411 | 51762880 | 1.75905 | 3.7237 | 2.117 |
| chr3 | 32218783 | 32219169 | 1.56118 | 3.30504 | 2.117 |
| chr7 | 33148892 | 33149376 | 1.4592 | 3.08917 | 2.117 |
| chr2 | 236044563 | 236044940 | 1.10923 | 2.3483 | 2.117 |
| chr20 | 34542339 | 34542695 | 2.26559 | 4.79636 | 2.117 |
| chr13 | 20139011 | 20139243 | 0.750932 | 1.58977 | 2.117 |
| chr1 | 223306891 | 223307354 | 1.89949 | 4.02147 | 2.117 |
| chr7 | 54841896 | 54842157 | 1.83324 | 3.88156 | 2.117 |
| chr18 | 47085752 | 47085976 | 0.941738 | 1.994 | 2.117 |
| chr3 | 118212921 | 118213213 | 0.661492 | 1.40063 | 2.117 |
| chr11 | 108525754 | 108526043 | 1.40663 | 2.9784 | 2.117 |
| chr2 | 99134610 | 99134845 | 1.09135 | 2.31086 | 2.117 |
| chr7 | 4281495 | 4281798 | 0.920647 | 1.94948 | 2.118 |
| chr22 | 50780703 | 50781127 | 1.64554 | 3.48451 | 2.118 |
| chr1 | 112901439 | 112901828 | 1.61664 | 3.42336 | 2.118 |
| chr11 | 132952675 | 132952995 | 2.82989 | 5.99252 | 2.118 |
| chr16 | 83178297 | 83178581 | 0.606901 | 1.28518 | 2.118 |
| chr20 | 44074679 | 44074988 | 1.48758 | 3.15034 | 2.118 |
| chr1 | 176613079 | 176613327 | 0.481843 | 1.02045 | 2.118 |
| chr8 | 85312226 | 85312664 | 0.507409 | 1.07463 | 2.118 |
| chr20 | 61940718 | 61941435 | 1.77066 | 3.75011 | 2.118 |
| chr17 | 10861395 | 10861650 | 0.48864 | 1.03491 | 2.118 |
| chr5 | 2038543 | 2039009 | 3.17228 | 6.71892 | 2.118 |
| chr5 | 180432908 | 180433120 | 0.540413 | 1.1446 | 2.118 |
| chr12 | 75899504 | 75899836 | 1.11625 | 2.3643 | 2.118 |
| chr3 | 64384880 | 64385163 | 1.35916 | 2.87881 | 2.118 |
| chr12 | 52269045 | 52269377 | 2.0297 | 4.29911 | 2.118 |
| chr14 | 96930493 | 96930789 | 0.543239 | 1.15069 | 2.118 |

| | | | | | |
|---|---|---|---|---|---|
| chr1 | 33684304 | 33684525 | 1.07758 | 2.28254 | 2.118 |
| chr19 | 31039890 | 31040130 | 0.525136 | 1.1124 | 2.118 |
| chr7 | 89840973 | 89841293 | 1.48957 | 3.15538 | 2.118 |
| chr19 | 35727485 | 35727728 | 0.832644 | 1.76387 | 2.118 |
| chr9 | 2743875 | 2744090 | 0.6053 | 1.28233 | 2.119 |
| chr14 | 90505656 | 90505889 | 0.419448 | 0.888659 | 2.119 |
| chr8 | 99176006 | 99176257 | 1.27347 | 2.69825 | 2.119 |
| chr1 | 223946965 | 223947189 | 0.410762 | 0.870339 | 2.119 |
| chr7 | 125575213 | 125575468 | 0.332493 | 0.704522 | 2.119 |
| chr9 | 133308658 | 133309168 | 1.32417 | 2.80585 | 2.119 |
| chrX | 16901027 | 16901666 | 1.18288 | 2.50648 | 2.119 |
| chr5 | 144542170 | 144542408 | 0.478615 | 1.01429 | 2.119 |
| chr5 | 126011789 | 126012137 | 1.40576 | 2.97923 | 2.119 |
| chr19 | 47019034 | 47019649 | 1.04882 | 2.22287 | 2.119 |
| chr16 | 21566218 | 21566428 | 2.04071 | 4.32511 | 2.119 |
| chr3 | 42543236 | 42543453 | 0.983869 | 2.08531 | 2.120 |
| chr11 | 73000482 | 73000865 | 2.11082 | 4.4741 | 2.120 |
| chr1 | 215879056 | 215879274 | 0.410321 | 0.86975 | 2.120 |
| chr19 | 42148173 | 42148678 | 2.30302 | 4.88168 | 2.120 |
| chr6 | 154996726 | 154997033 | 2.49355 | 5.28561 | 2.120 |
| chr1 | 180221723 | 180222145 | 1.51282 | 3.20698 | 2.120 |
| chr8 | 56602039 | 56602328 | 0.657686 | 1.39423 | 2.120 |
| chr13 | 86190371 | 86190594 | 0.566557 | 1.20107 | 2.120 |
| chr3 | 4392561 | 4392788 | 1.32955 | 2.8186 | 2.120 |
| chr16 | 115637 | 116050 | 2.20732 | 4.67948 | 2.120 |
| chr8 | 86304289 | 86304511 | 0.447818 | 0.949388 | 2.120 |
| chr12 | 59507538 | 59507976 | 2.71725 | 5.76079 | 2.120 |
| chr5 | 118771124 | 118771510 | 1.52803 | 3.23959 | 2.120 |
| chr4 | 139497976 | 139498232 | 1.02235 | 2.1675 | 2.120 |
| chr11 | 30982022 | 30982262 | 1.53531 | 3.25508 | 2.120 |
| chr5 | 153955366 | 153955721 | 1.0535 | 2.23375 | 2.120 |
| chr17 | 12849128 | 12849347 | 0.583914 | 1.23812 | 2.120 |
| chr19 | 41039460 | 41039726 | 1.39988 | 2.96835 | 2.120 |
| chr16 | 2393396 | 2393626 | 0.535257 | 1.13501 | 2.121 |
| chr12 | 124516699 | 124517210 | 1.70953 | 3.62515 | 2.121 |
| chr4 | 20546403 | 20546613 | 0.860672 | 1.8251 | 2.121 |
| chr8 | 32403930 | 32404191 | 0.956011 | 2.02727 | 2.121 |
| chr10 | 103227102 | 103227413 | 1.53157 | 3.24781 | 2.121 |
| chr2 | 25887646 | 25887903 | 1.81407 | 3.84716 | 2.121 |
| chr14 | 93079403 | 93079763 | 1.85161 | 3.92694 | 2.121 |
| chr16 | 53088683 | 53089166 | 1.55594 | 3.29994 | 2.121 |
| chr22 | 35979055 | 35979512 | 1.9159 | 4.0634 | 2.121 |
| chr5 | 65519536 | 65519793 | 1.16233 | 2.46523 | 2.121 |
| chr7 | 84815993 | 84816404 | 2.0515 | 4.35129 | 2.121 |
| chr6 | 131949324 | 131949701 | 2.6773 | 5.67867 | 2.121 |
| chr11 | 74581573 | 74581813 | 1.00223 | 2.12578 | 2.121 |
| chr5 | 11617054 | 11617275 | 0.464379 | 0.984979 | 2.121 |
| chr12 | 31391671 | 31392128 | 1.64532 | 3.49009 | 2.121 |
| chr17 | 62063097 | 62063331 | 0.603932 | 1.28112 | 2.121 |
| chr11 | 124514265 | 124514662 | 1.29306 | 2.7431 | 2.121 |
| chr2 | 47881531 | 47881898 | 0.699512 | 1.484 | 2.121 |
| chr5 | 90795812 | 90796055 | 0.660081 | 1.40037 | 2.122 |
| chr17 | 19241433 | 19241745 | 1.32761 | 2.81655 | 2.122 |
| chr8 | 37551971 | 37552182 | 1.36745 | 2.90114 | 2.122 |
| chr20 | 47842305 | 47842773 | 0.678723 | 1.43997 | 2.122 |
| chr19 | 17010446 | 17010815 | 1.7638 | 3.74227 | 2.122 |
| chr7 | 147144326 | 147144746 | 0.444046 | 0.942209 | 2.122 |
| chr19 | 52160133 | 52160577 | 2.1675 | 4.59924 | 2.122 |
| chr13 | 46085657 | 46086067 | 0.854875 | 1.81398 | 2.122 |
| chr3 | 195589973 | 195590315 | 2.04469 | 4.339 | 2.122 |
| chr1 | 51657957 | 51658197 | 1.99238 | 4.22808 | 2.122 |

| | | | | | |
|---|---|---|---|---|---|
| chr8 | 801160 | 801411 | 0.454523 | 0.964565 | 2.122 |
| chr22 | 35826809 | 35827151 | 2.2613 | 4.7989 | 2.122 |
| chr2 | 192366326 | 192366597 | 0.802428 | 1.70303 | 2.122 |
| chr3 | 179276595 | 179276944 | 1.30162 | 2.76253 | 2.122 |
| chr20 | 33200841 | 33201052 | 0.509161 | 1.08064 | 2.122 |
| chr7 | 32434635 | 32435167 | 0.509171 | 1.08066 | 2.122 |
| chr2 | 70233229 | 70233457 | 0.611132 | 1.29708 | 2.122 |
| chr9 | 90626423 | 90626634 | 0.720945 | 1.53019 | 2.122 |
| chr5 | 145076891 | 145077113 | 1.39432 | 2.95954 | 2.123 |
| chrX | 30326342 | 30326995 | 1.88591 | 4.00298 | 2.123 |
| chr3 | 11845507 | 11845744 | 1.4413 | 3.05931 | 2.123 |
| chr15 | 24112030 | 24112308 | 1.12298 | 2.38396 | 2.123 |
| chr1 | 86042685 | 86043009 | 1.18519 | 2.51604 | 2.123 |
| chr5 | 137157921 | 137158154 | 1.77518 | 3.76862 | 2.123 |
| chrX | 70218380 | 70218618 | 0.904726 | 1.92072 | 2.123 |
| chr1 | 201607926 | 201608140 | 0.54768 | 1.16273 | 2.123 |
| chr4 | 104836425 | 104836700 | 0.790579 | 1.67841 | 2.123 |
| chr4 | 156363498 | 156363818 | 1.77421 | 3.7668 | 2.123 |
| chr21 | 38034212 | 38034525 | 1.07082 | 2.27354 | 2.123 |
| chr12 | 26587233 | 26587656 | 1.34183 | 2.84907 | 2.123 |
| chr10 | 81052827 | 81053100 | 1.92893 | 4.09571 | 2.123 |
| chr19 | 47260518 | 47260952 | 2.6489 | 5.62458 | 2.123 |
| chr12 | 108154903 | 108155111 | 0.941761 | 1.99974 | 2.123 |
| chr22 | 39518088 | 39518294 | 1.10374 | 2.34375 | 2.123 |
| chr9 | 138396448 | 138396856 | 1.95306 | 4.14726 | 2.123 |
| chr19 | 22675829 | 22676342 | 0.809862 | 1.71973 | 2.123 |
| chr7 | 147350486 | 147350728 | 0.448385 | 0.95216 | 2.124 |
| chr5 | 134901878 | 134902211 | 2.46367 | 5.23184 | 2.124 |
| chr2 | 54433809 | 54434031 | 0.856319 | 1.81853 | 2.124 |
| chr14 | 35908057 | 35908370 | 1.451 | 3.08157 | 2.124 |
| chr15 | 28257478 | 28257702 | 0.482323 | 1.02434 | 2.124 |
| chr2 | 171221449 | 171221662 | 0.50601 | 1.07467 | 2.124 |
| chr3 | 196255653 | 196255956 | 1.88245 | 3.99837 | 2.124 |
| chr22 | 31607922 | 31608289 | 2.01829 | 4.28717 | 2.124 |
| chr7 | 95025379 | 95025869 | 1.37999 | 2.9315 | 2.124 |
| chr10 | 55361094 | 55361435 | 1.18945 | 2.52676 | 2.124 |
| chr10 | 133252306 | 133253051 | 1.18783 | 2.5234 | 2.124 |
| chr5 | 68338719 | 68338983 | 1.18312 | 2.51342 | 2.124 |
| chr14 | 88480066 | 88480447 | 3.41682 | 7.25877 | 2.124 |
| chr11 | 48131532 | 48131793 | 1.18153 | 2.51008 | 2.124 |
| chr7 | 78083904 | 78084186 | 0.363633 | 0.772615 | 2.125 |
| chrX | 38627398 | 38627602 | 0.779705 | 1.65668 | 2.125 |
| chr20 | 48423983 | 48424189 | 0.683708 | 1.45276 | 2.125 |
| chr17 | 9885446 | 9885872 | 1.21674 | 2.58538 | 2.125 |
| chr10 | 3257594 | 3257998 | 0.560501 | 1.19098 | 2.125 |
| chr11 | 68856222 | 68856570 | 1.67882 | 3.56725 | 2.125 |
| chr9 | 115873752 | 115874085 | 1.6417 | 3.48841 | 2.125 |
| chr14 | 104798208 | 104798495 | 0.799189 | 1.69825 | 2.125 |
| chr11 | 28378371 | 28378605 | 0.55275 | 1.17461 | 2.125 |
| chr12 | 773725 | 773963 | 0.559594 | 1.18919 | 2.125 |
| chr19 | 38724388 | 38724880 | 2.04188 | 4.33917 | 2.125 |
| chr1 | 23823489 | 23823868 | 1.21958 | 2.59174 | 2.125 |
| chr3 | 131757159 | 131757376 | 1.1638 | 2.47323 | 2.125 |
| chr8 | 6786106 | 6786403 | 1.87392 | 3.98252 | 2.125 |
| chr7 | 11380997 | 11381299 | 0.771582 | 1.63981 | 2.125 |
| chr16 | 50244480 | 50244743 | 1.61027 | 3.42249 | 2.125 |
| chr1 | 3370810 | 3371030 | 0.868577 | 1.84625 | 2.126 |
| chr17 | 38121900 | 38122179 | 0.461438 | 0.980855 | 2.126 |
| chr16 | 78261772 | 78262040 | 1.06751 | 2.26918 | 2.126 |
| chr2 | 107199006 | 107199374 | 1.74018 | 3.69914 | 2.126 |
| chr8 | 11161948 | 11162151 | 1.09318 | 2.32392 | 2.126 |

| | | | | | |
|---|---|---|---|---|---|
| chr8 | 120886307 | 120887012 | 1.03177 | 2.1934 | 2.126 |
| chr1 | 17386215 | 17386434 | 1.29807 | 2.75954 | 2.126 |
| chr7 | 157066198 | 157066658 | 2.13265 | 4.53381 | 2.126 |
| chr9 | 109447174 | 109447384 | 0.567284 | 1.20599 | 2.126 |
| chr4 | 180949952 | 180950179 | 0.715784 | 1.52173 | 2.126 |
| chr1 | 204098157 | 204098568 | 2.56523 | 5.45382 | 2.126 |
| chr18 | 21756660 | 21757115 | 1.65718 | 3.52327 | 2.126 |
| chr5 | 107978004 | 107978361 | 1.82982 | 3.89037 | 2.126 |
| chr5 | 17132519 | 17133047 | 0.901488 | 1.91668 | 2.126 |
| chr15 | 37426884 | 37427111 | 1.04417 | 2.22005 | 2.126 |
| chr1 | 154970678 | 154970909 | 0.696769 | 1.48145 | 2.126 |
| chr19 | 6022293 | 6022643 | 1.20644 | 2.56513 | 2.126 |
| chr8 | 37217812 | 37218035 | 0.891549 | 1.89573 | 2.126 |
| chr8 | 41998195 | 41998513 | 1.4563 | 3.09658 | 2.126 |
| chr15 | 89577492 | 89577763 | 1.15776 | 2.46184 | 2.126 |
| chr10 | 115438815 | 115439136 | 1.59801 | 3.39799 | 2.126 |
| chrX | 122589860 | 122590112 | 0.459213 | 0.976475 | 2.126 |
| chr17 | 9828098 | 9828383 | 1.34855 | 2.8676 | 2.126 |
| chr11 | 1673903 | 1674316 | 1.80384 | 3.836 | 2.127 |
| chr22 | 20772393 | 20772751 | 1.62978 | 3.46607 | 2.127 |
| chr9 | 118286797 | 118287100 | 1.2737 | 2.70887 | 2.127 |
| chr1 | 201762147 | 201762513 | 1.99525 | 4.24381 | 2.127 |
| chr2 | 30146008 | 30146368 | 1.73292 | 3.68589 | 2.127 |
| chr7 | 111370643 | 111370861 | 1.24222 | 2.64222 | 2.127 |
| chr7 | 149558498 | 149558815 | 1.16801 | 2.48439 | 2.127 |
| chr14 | 90439614 | 90440066 | 1.30942 | 2.78523 | 2.127 |
| chr1 | 157918354 | 157918632 | 1.42649 | 3.03446 | 2.127 |
| chr12 | 81988574 | 81988863 | 0.556486 | 1.18383 | 2.127 |
| chr16 | 19022539 | 19022748 | 0.815603 | 1.73511 | 2.127 |
| chr9 | 74033170 | 74033375 | 0.48633 | 1.03462 | 2.127 |
| chr11 | 90882124 | 90882359 | 0.501779 | 1.06753 | 2.127 |
| chr10 | 11291124 | 11291358 | 0.443266 | 0.94305 | 2.128 |
| chr20 | 62733284 | 62733840 | 1.95814 | 4.16621 | 2.128 |
| chr10 | 49390909 | 49391207 | 0.537133 | 1.14297 | 2.128 |
| chr14 | 36843293 | 36843603 | 1.63808 | 3.48577 | 2.128 |
| chr1 | 145713517 | 145713980 | 2.28324 | 4.85869 | 2.128 |
| chr16 | 81907416 | 81907943 | 1.70009 | 3.61805 | 2.128 |
| chr9 | 7823453 | 7823727 | 0.587185 | 1.24962 | 2.128 |
| chr17 | 42245784 | 42246171 | 2.30005 | 4.89495 | 2.128 |
| chr1 | 113249885 | 113250265 | 1.80794 | 3.84786 | 2.128 |
| chr10 | 22590178 | 22590446 | 0.762386 | 1.62261 | 2.128 |
| chr3 | 27270291 | 27270571 | 0.650122 | 1.3837 | 2.128 |
| chr10 | 30840106 | 30840372 | 0.48481 | 1.03186 | 2.128 |
| chr3 | 167458865 | 167459121 | 0.800367 | 1.7035 | 2.128 |
| chr2 | 192419857 | 192420146 | 1.01282 | 2.15575 | 2.128 |
| chr2 | 80300161 | 80300575 | 0.821301 | 1.74813 | 2.128 |
| chr6 | 10404256 | 10404470 | 0.896537 | 1.90827 | 2.128 |
| chr7 | 30315667 | 30316077 | 1.59211 | 3.38886 | 2.129 |
| chr11 | 44559976 | 44560408 | 1.55338 | 3.30665 | 2.129 |
| chr5 | 4512190 | 4512413 | 0.955623 | 2.03424 | 2.129 |
| chr9 | 97012693 | 97012964 | 0.425396 | 0.905577 | 2.129 |
| chr3 | 53333645 | 53333881 | 0.99406 | 2.11621 | 2.129 |
| chr5 | 140743622 | 140743838 | 0.958747 | 2.0411 | 2.129 |
| chr14 | 98972941 | 98973178 | 0.461796 | 0.983154 | 2.129 |
| chr11 | 26966492 | 26966783 | 0.473351 | 1.00777 | 2.129 |
| chr20 | 39707053 | 39707315 | 1.18851 | 2.53049 | 2.129 |
| chr4 | 26460884 | 26461092 | 0.893479 | 1.90237 | 2.129 |
| chr7 | 100425466 | 100425679 | 1.2262 | 2.61083 | 2.129 |
| chr5 | 4049362 | 4049705 | 0.459973 | 0.979377 | 2.129 |
| chr12 | 128464986 | 128465411 | 0.526183 | 1.12042 | 2.129 |
| chr15 | 70861512 | 70861807 | 1.88911 | 4.02264 | 2.129 |

| | | | | | |
|---|---|---|---|---|---|
| chr1 | 12027278 | 12027555 | 1.00948 | 2.14959 | 2.129 |
| chr2 | 86025138 | 86025368 | 0.9326 | 1.98588 | 2.129 |
| chr10 | 1339886 | 1340108 | 0.619882 | 1.31999 | 2.129 |
| chr12 | 46122591 | 46123007 | 2.39574 | 5.10158 | 2.129 |
| chr1 | 244344424 | 244344678 | 0.427454 | 0.910266 | 2.130 |
| chr8 | 14055677 | 14055919 | 0.38526 | 0.820419 | 2.130 |
| chr3 | 119354238 | 119354458 | 0.444327 | 0.94624 | 2.130 |
| chr5 | 67250334 | 67250631 | 0.549863 | 1.171 | 2.130 |
| chr11 | 57294542 | 57294930 | 1.67694 | 3.57129 | 2.130 |
| chr14 | 80345716 | 80346028 | 0.524561 | 1.11714 | 2.130 |
| chr19 | 14429687 | 14429987 | 0.819683 | 1.74566 | 2.130 |
| chr16 | 67313149 | 67313549 | 2.16015 | 4.60048 | 2.130 |
| chr2 | 79802706 | 79802985 | 0.782721 | 1.66696 | 2.130 |
| chr7 | 65479035 | 65479267 | 0.91407 | 1.94685 | 2.130 |
| chr12 | 7000300 | 7000630 | 2.19815 | 4.68183 | 2.130 |
| chr11 | 76630930 | 76631423 | 2.14249 | 4.56342 | 2.130 |
| chr3 | 37821072 | 37821383 | 1.39734 | 2.97638 | 2.130 |
| chr17 | 16894795 | 16895099 | 2.5202 | 5.36833 | 2.130 |
| chr11 | 47429570 | 47430015 | 1.41399 | 3.01207 | 2.130 |
| chr4 | 169104899 | 169105153 | 0.815151 | 1.73663 | 2.130 |
| chr12 | 10880524 | 10880735 | 0.577022 | 1.22932 | 2.130 |
| chr14 | 24740620 | 24740981 | 2.09306 | 4.45951 | 2.131 |
| chr7 | 158286830 | 158287053 | 1.11547 | 2.37666 | 2.131 |
| chr5 | 21686232 | 21686450 | 0.474828 | 1.01174 | 2.131 |
| chr5 | 111228395 | 111228654 | 0.437179 | 0.931544 | 2.131 |
| chr14 | 55369612 | 55370116 | 1.14295 | 2.43561 | 2.131 |
| chr14 | 95003244 | 95003453 | 0.81281 | 1.73209 | 2.131 |
| chr4 | 186518741 | 186519189 | 2.00914 | 4.28152 | 2.131 |
| chr13 | 107857678 | 107858116 | 1.73718 | 3.70207 | 2.131 |
| chr5 | 179911511 | 179911862 | 1.22572 | 2.61216 | 2.131 |
| chr2 | 10328411 | 10328706 | 1.18076 | 2.51635 | 2.131 |
| chr15 | 55460547 | 55461013 | 1.47178 | 3.13657 | 2.131 |
| chr11 | 115375683 | 115375918 | 0.716328 | 1.52662 | 2.131 |
| chr8 | 92796762 | 92796994 | 0.412644 | 0.879422 | 2.131 |
| chr16 | 29711814 | 29712132 | 2.21735 | 4.72573 | 2.131 |
| chr4 | 7763436 | 7763729 | 1.32722 | 2.8287 | 2.131 |
| chr19 | 44156354 | 44156858 | 1.43221 | 3.05253 | 2.131 |
| chr20 | 43362037 | 43362385 | 2.28688 | 4.8742 | 2.131 |
| chr9 | 72129514 | 72129903 | 0.506851 | 1.08029 | 2.131 |
| chr17 | 55044837 | 55045106 | 0.531895 | 1.13373 | 2.131 |
| chr12 | 8147917 | 8148137 | 0.655758 | 1.39775 | 2.132 |
| chr20 | 18984449 | 18984846 | 0.524788 | 1.11859 | 2.132 |
| chr3 | 150784794 | 150785152 | 0.772771 | 1.64725 | 2.132 |
| chr7 | 88433048 | 88433725 | 0.450207 | 0.959675 | 2.132 |
| chr3 | 172428876 | 172429172 | 2.10398 | 4.48498 | 2.132 |
| chr18 | 25757442 | 25757703 | 0.46792 | 0.997466 | 2.132 |
| chr13 | 25569325 | 25569681 | 1.13457 | 2.41863 | 2.132 |
| chr16 | 87712086 | 87712500 | 1.15042 | 2.45242 | 2.132 |
| chr10 | 24283451 | 24283799 | 1.37402 | 2.92939 | 2.132 |
| chr7 | 34967610 | 34967857 | 0.667706 | 1.42354 | 2.132 |
| chr12 | 62024785 | 62025042 | 0.549856 | 1.17231 | 2.132 |
| chr3 | 157211468 | 157211718 | 1.17119 | 2.49705 | 2.132 |
| chr15 | 81148838 | 81149042 | 1.22641 | 2.61481 | 2.132 |
| chr17 | 15901847 | 15902096 | 1.49182 | 3.18073 | 2.132 |
| chr7 | 29801054 | 29801339 | 0.549421 | 1.17143 | 2.132 |
| chr11 | 34534427 | 34534761 | 1.65784 | 3.53474 | 2.132 |
| chr15 | 89560771 | 89561176 | 1.81124 | 3.8619 | 2.132 |
| chr18 | 45275470 | 45275959 | 2.8115 | 5.99488 | 2.132 |
| chr10 | 102321682 | 102322257 | 3.20957 | 6.84384 | 2.132 |
| chr7 | 147182855 | 147183330 | 0.483292 | 1.03056 | 2.132 |
| chr11 | 67118631 | 67118981 | 2.17731 | 4.64291 | 2.132 |

| | | | | | |
|---|---|---|---|---|---|
| chr12 | 30403012 | 30403216 | 0.366055 | 0.780622 | 2.133 |
| chr20 | 15524485 | 15524719 | 0.69276 | 1.47747 | 2.133 |
| chr4 | 60165826 | 60166057 | 0.513094 | 1.0943 | 2.133 |
| chr12 | 13365067 | 13365316 | 1.83916 | 3.92254 | 2.133 |
| chr4 | 99977278 | 99977481 | 1.18836 | 2.53454 | 2.133 |
| chr12 | 61781866 | 61782149 | 0.520937 | 1.11109 | 2.133 |
| chr7 | 26893366 | 26893690 | 1.0715 | 2.28538 | 2.133 |
| chr8 | 107717334 | 107717566 | 1.23211 | 2.62797 | 2.133 |
| chr7 | 128348162 | 128348531 | 0.372289 | 0.794075 | 2.133 |
| chr17 | 40171816 | 40172165 | 1.65609 | 3.53253 | 2.133 |
| chr7 | 92864636 | 92864844 | 0.525688 | 1.12133 | 2.133 |
| chr12 | 122214016 | 122214491 | 2.25033 | 4.80013 | 2.133 |
| chr15 | 98796836 | 98797210 | 0.556359 | 1.18676 | 2.133 |
| chr19 | 30262639 | 30263082 | 1.9866 | 4.23757 | 2.133 |
| chr9 | 81054386 | 81054749 | 1.48372 | 3.16512 | 2.133 |
| chr5 | 178688285 | 178688654 | 0.960938 | 2.04999 | 2.133 |
| chr8 | 13761825 | 13762038 | 0.680577 | 1.45189 | 2.133 |
| chr2 | 232277063 | 232277427 | 2.40737 | 5.13574 | 2.133 |
| chr12 | 58129662 | 58129894 | 0.829934 | 1.77055 | 2.133 |
| chr1 | 175121139 | 175121576 | 1.80662 | 3.85424 | 2.133 |
| chr15 | 88043579 | 88043783 | 0.73362 | 1.56512 | 2.133 |
| chr22 | 18267946 | 18268397 | 1.10374 | 2.35474 | 2.133 |
| chr7 | 155137512 | 155137947 | 2.52567 | 5.38835 | 2.133 |
| chr1 | 23256825 | 23257156 | 1.73885 | 3.70982 | 2.133 |
| chr7 | 367317 | 367872 | 0.449043 | 0.958057 | 2.134 |
| chr9 | 134639067 | 134639285 | 0.501338 | 1.06966 | 2.134 |
| chr17 | 56385639 | 56386027 | 1.38576 | 2.95683 | 2.134 |
| chr1 | 13843735 | 13844000 | 1.1478 | 2.4491 | 2.134 |
| chr8 | 109829212 | 109829453 | 0.44462 | 0.948713 | 2.134 |
| chrX | 117737928 | 117738192 | 2.08519 | 4.44952 | 2.134 |
| chrX | 57182914 | 57183287 | 0.864489 | 1.84483 | 2.134 |
| chr1 | 225937521 | 225937724 | 0.306316 | 0.65369 | 2.134 |
| chr2 | 27906316 | 27906728 | 2.59961 | 5.54795 | 2.134 |
| chr7 | 47984885 | 47985158 | 1.15113 | 2.45672 | 2.134 |
| chr20 | 25159766 | 25160662 | 1.5579 | 3.32487 | 2.134 |
| chr3 | 127794424 | 127794752 | 1.60237 | 3.4198 | 2.134 |
| chr17 | 8274491 | 8274707 | 0.485388 | 1.03594 | 2.134 |
| chr8 | 144976584 | 144977043 | 2.52181 | 5.38221 | 2.134 |
| chr8 | 134668232 | 134668489 | 0.840312 | 1.79345 | 2.134 |
| chr15 | 57819759 | 57820004 | 1.09988 | 2.3475 | 2.134 |
| chr11 | 71726746 | 71726976 | 1.11727 | 2.38465 | 2.134 |
| chr19 | 8429374 | 8429608 | 0.743405 | 1.58671 | 2.134 |
| chr5 | 103730211 | 103730506 | 0.390648 | 0.833796 | 2.134 |
| chr18 | 55899274 | 55899497 | 0.940781 | 2.00801 | 2.134 |
| chr7 | 105697419 | 105697750 | 1.42915 | 3.05049 | 2.134 |
| chr10 | 17031858 | 17032136 | 0.576404 | 1.23035 | 2.135 |
| chr1 | 12061712 | 12062078 | 1.31168 | 2.79983 | 2.135 |
| chr14 | 62304380 | 62304672 | 2.34847 | 5.01295 | 2.135 |
| chr12 | 98884794 | 98885082 | 1.77982 | 3.7993 | 2.135 |
| chr7 | 37498719 | 37498998 | 0.538302 | 1.14919 | 2.135 |
| chr2 | 230931714 | 230932059 | 2.15458 | 4.59973 | 2.135 |
| chr11 | 67085004 | 67085341 | 1.61267 | 3.44286 | 2.135 |
| chrX | 65948624 | 65949145 | 1.50977 | 3.22325 | 2.135 |
| chr7 | 50159451 | 50159721 | 0.921269 | 1.96698 | 2.135 |
| chr12 | 41315417 | 41315679 | 1.03824 | 2.21681 | 2.135 |
| chr12 | 39314956 | 39315216 | 2.03112 | 4.33681 | 2.135 |
| chr10 | 98021718 | 98021972 | 0.494612 | 1.05611 | 2.135 |
| chr5 | 4899860 | 4900065 | 0.42868 | 0.915353 | 2.135 |
| chr6 | 25265944 | 25266270 | 1.20741 | 2.57823 | 2.135 |
| chr7 | 107383936 | 107384164 | 1.55424 | 3.31897 | 2.135 |
| chr3 | 150886045 | 150886456 | 1.83926 | 3.92763 | 2.135 |

| | | | | | |
|---|---|---|---|---|---|
| chr9 | 84911279 | 84911538 | 0.581981 | 1.2428 | 2.135 |
| chr11 | 39018826 | 39019054 | 0.4333 | 0.925338 | 2.136 |
| chr5 | 117336640 | 117337023 | 0.466912 | 0.997121 | 2.136 |
| chr11 | 44116971 | 44117356 | 1.90439 | 4.06703 | 2.136 |
| chr8 | 128750014 | 128750266 | 1.12927 | 2.41169 | 2.136 |
| chr19 | 49831158 | 49831536 | 1.93719 | 4.13718 | 2.136 |
| chr5 | 83087434 | 83087705 | 0.464926 | 0.99295 | 2.136 |
| chr14 | 105030215 | 105030773 | 1.66211 | 3.54987 | 2.136 |
| chr12 | 10954434 | 10954644 | 0.67381 | 1.43932 | 2.136 |
| chr1 | 94511163 | 94511651 | 1.37869 | 2.94525 | 2.136 |
| chr5 | 135170154 | 135170511 | 1.62442 | 3.47023 | 2.136 |
| chr15 | 65346608 | 65347020 | 2.61842 | 5.59379 | 2.136 |
| chr12 | 101421442 | 101421711 | 0.575596 | 1.22968 | 2.136 |
| chr5 | 90099304 | 90099517 | 0.815003 | 1.74114 | 2.136 |
| chr11 | 71688850 | 71689365 | 1.05027 | 2.24385 | 2.136 |
| chr7 | 77550160 | 77550381 | 0.478664 | 1.02268 | 2.137 |
| chr20 | 33296307 | 33296703 | 0.999867 | 2.13642 | 2.137 |
| chr12 | 72764555 | 72764895 | 0.538719 | 1.1511 | 2.137 |
| chr9 | 26723436 | 26723826 | 1.52759 | 3.26416 | 2.137 |
| chr16 | 59103543 | 59103795 | 0.457826 | 0.978291 | 2.137 |
| chr11 | 12310128 | 12310695 | 1.73317 | 3.70358 | 2.137 |
| chr11 | 100447209 | 100447458 | 1.31274 | 2.80517 | 2.137 |
| chr2 | 26173867 | 26174221 | 1.9423 | 4.15055 | 2.137 |
| chr4 | 22238853 | 22239166 | 1.6034 | 3.42635 | 2.137 |
| chr8 | 13074076 | 13074378 | 0.484764 | 1.03591 | 2.137 |
| chr1 | 178349010 | 178349333 | 1.50608 | 3.21852 | 2.137 |
| chr20 | 47273327 | 47273822 | 1.32506 | 2.83181 | 2.137 |
| chr14 | 75774562 | 75774835 | 2.17485 | 4.64811 | 2.137 |
| chr5 | 113665509 | 113666023 | 0.546378 | 1.16777 | 2.137 |
| chr11 | 6460275 | 6460512 | 1.44896 | 3.0969 | 2.137 |
| chr7 | 148958913 | 148959396 | 1.72612 | 3.68941 | 2.137 |
| chr1 | 152371641 | 152372000 | 1.42999 | 3.05652 | 2.137 |
| chr12 | 54793058 | 54793511 | 2.53587 | 5.42031 | 2.137 |
| chr16 | 11057810 | 11058238 | 1.90238 | 4.06634 | 2.138 |
| chr1 | 189694927 | 189695181 | 0.591352 | 1.2641 | 2.138 |
| chr15 | 72604883 | 72605407 | 2.11364 | 4.51823 | 2.138 |
| chr14 | 59106627 | 59106975 | 1.31858 | 2.81868 | 2.138 |
| chr19 | 14542715 | 14543067 | 1.71972 | 3.67625 | 2.138 |
| chr12 | 74586812 | 74587097 | 0.454015 | 0.970559 | 2.138 |
| chr5 | 141044520 | 141044908 | 2.2135 | 4.73187 | 2.138 |
| chr10 | 99115522 | 99115977 | 0.754535 | 1.613 | 2.138 |
| chr19 | 14014050 | 14014326 | 1.30182 | 2.78297 | 2.138 |
| chr8 | 80104223 | 80104456 | 0.674818 | 1.4427 | 2.138 |
| chr20 | 41019945 | 41020272 | 0.53445 | 1.14265 | 2.138 |
| chr12 | 94948204 | 94948457 | 0.568463 | 1.21539 | 2.138 |
| chr12 | 45465403 | 45465622 | 0.535131 | 1.14416 | 2.138 |
| chr2 | 10444247 | 10444478 | 0.882229 | 1.8863 | 2.138 |
| chr6 | 80382432 | 80382744 | 1.19943 | 2.56454 | 2.138 |
| chr11 | 33709493 | 33709729 | 1.47329 | 3.15015 | 2.138 |
| chr17 | 32633508 | 32633747 | 0.898251 | 1.92062 | 2.138 |
| chr3 | 138642287 | 138642649 | 1.76589 | 3.7759 | 2.138 |
| chr12 | 8979029 | 8979493 | 0.585524 | 1.25201 | 2.138 |
| chr20 | 21825819 | 21826092 | 0.529995 | 1.1333 | 2.138 |
| chr5 | 79806532 | 79806817 | 0.454406 | 0.971669 | 2.138 |
| chr9 | 34396526 | 34396887 | 0.62906 | 1.34522 | 2.138 |
| chr2 | 239048174 | 239048550 | 1.5934 | 3.40755 | 2.139 |
| chr10 | 129492916 | 129493183 | 2.16543 | 4.63096 | 2.139 |
| chr12 | 1046893 | 1047310 | 0.566817 | 1.21223 | 2.139 |
| chr9 | 36905617 | 36905866 | 0.566023 | 1.21055 | 2.139 |
| chr2 | 208556986 | 208557291 | 1.45755 | 3.1173 | 2.139 |
| chr4 | 53411669 | 53412009 | 1.69931 | 3.63436 | 2.139 |

| | | | | | |
|---|---|---|---|---|---|
| chr5 | 107211761 | 107212003 | 0.742936 | 1.58901 | 2.139 |
| chr16 | 57625322 | 57625636 | 2.16876 | 4.6386 | 2.139 |
| chr2 | 24835937 | 24836645 | 0.85043 | 1.819 | 2.139 |
| chr5 | 4096292 | 4096612 | 0.469585 | 1.00444 | 2.139 |
| chr21 | 37794598 | 37794881 | 1.93129 | 4.13109 | 2.139 |
| chr10 | 80955175 | 80955589 | 1.83977 | 3.93535 | 2.139 |
| chr16 | 10608551 | 10608780 | 1.20246 | 2.57215 | 2.139 |
| chr7 | 98638024 | 98638283 | 1.43619 | 3.07214 | 2.139 |
| chr8 | 15670283 | 15670654 | 0.658933 | 1.40958 | 2.139 |
| chr4 | 81283834 | 81284052 | 1.12682 | 2.41051 | 2.139 |
| chr1 | 211854079 | 211854378 | 1.11803 | 2.39175 | 2.139 |
| chr12 | 46736364 | 46736663 | 1.12203 | 2.40032 | 2.139 |
| chr7 | 128858307 | 128858744 | 1.33498 | 2.85592 | 2.139 |
| chr20 | 38915884 | 38916087 | 0.55179 | 1.18046 | 2.139 |
| chr16 | 81294630 | 81294982 | 1.35183 | 2.89204 | 2.139 |
| chr19 | 41208431 | 41208834 | 2.41339 | 5.16312 | 2.139 |
| chr1 | 245392349 | 245392655 | 0.607558 | 1.29985 | 2.139 |
| chr16 | 87291764 | 87291976 | 0.469545 | 1.00461 | 2.140 |
| chr20 | 35806652 | 35806866 | 1.27873 | 2.73601 | 2.140 |
| chr11 | 128027578 | 128028100 | 1.50313 | 3.21618 | 2.140 |
| chr8 | 95825778 | 95826117 | 1.19956 | 2.56669 | 2.140 |
| chr7 | 39080593 | 39080858 | 0.494747 | 1.05861 | 2.140 |
| chr10 | 73079304 | 73079645 | 1.87727 | 4.01682 | 2.140 |
| chr17 | 2833311 | 2833892 | 1.54128 | 3.29805 | 2.140 |
| chr18 | 45972986 | 45973326 | 1.35612 | 2.90187 | 2.140 |
| chr15 | 95213739 | 95214012 | 1.04618 | 2.23869 | 2.140 |
| chr6 | 2791435 | 2791699 | 1.03598 | 2.21705 | 2.140 |
| chr18 | 25265863 | 25266116 | 1.39373 | 2.98275 | 2.140 |
| chr20 | 46415278 | 46415509 | 1.22391 | 2.61947 | 2.140 |
| chr15 | 76483077 | 76483450 | 2.1886 | 4.68443 | 2.140 |
| chr10 | 32097608 | 32097880 | 0.996613 | 2.13314 | 2.140 |
| chr15 | 51673406 | 51673813 | 1.83909 | 3.93662 | 2.141 |
| chr16 | 63011357 | 63011611 | 0.419417 | 0.897782 | 2.141 |
| chr2 | 49849586 | 49849860 | 0.540538 | 1.15712 | 2.141 |
| chr14 | 105399689 | 105399964 | 1.75944 | 3.76652 | 2.141 |
| chr7 | 23586466 | 23587107 | 1.39114 | 2.9781 | 2.141 |
| chr1 | 27423927 | 27424295 | 1.66637 | 3.56738 | 2.141 |
| chr8 | 87774616 | 87774831 | 0.437874 | 0.937406 | 2.141 |
| chr14 | 105947111 | 105947541 | 2.1051 | 4.50697 | 2.141 |
| chr12 | 111294663 | 111295001 | 0.42483 | 0.909562 | 2.141 |
| chr21 | 43198273 | 43198742 | 1.23135 | 2.63642 | 2.141 |
| chr15 | 41234101 | 41234438 | 1.72707 | 3.69784 | 2.141 |
| chr10 | 16540805 | 16541029 | 0.49107 | 1.05146 | 2.141 |
| chr12 | 65517277 | 65517573 | 1.38636 | 2.96845 | 2.141 |
| chr2 | 158184043 | 158184415 | 2.21435 | 4.7415 | 2.141 |
| chr7 | 33211488 | 33211699 | 0.452247 | 0.968421 | 2.141 |
| chr1 | 201278711 | 201278950 | 0.989557 | 2.11908 | 2.141 |
| chr16 | 19896064 | 19896280 | 1.05906 | 2.26792 | 2.141 |
| chr16 | 4594425 | 4594646 | 0.544772 | 1.16663 | 2.142 |
| chr11 | 9556639 | 9557027 | 2.07453 | 4.44271 | 2.142 |
| chr9 | 134630989 | 134631339 | 2.94157 | 6.29979 | 2.142 |
| chr9 | 10136029 | 10136304 | 0.436708 | 0.935296 | 2.142 |
| chr1 | 153195437 | 153195827 | 0.88697 | 1.89963 | 2.142 |
| chr1 | 7535016 | 7535513 | 0.887507 | 1.90081 | 2.142 |
| chr3 | 32785049 | 32785327 | 0.393101 | 0.841929 | 2.142 |
| chr4 | 113525843 | 113526052 | 0.973413 | 2.08494 | 2.142 |
| chr14 | 105999070 | 105999434 | 1.02731 | 2.20049 | 2.142 |
| chr8 | 60117650 | 60117865 | 0.519847 | 1.11353 | 2.142 |
| chr7 | 140217749 | 140218098 | 1.67272 | 3.58314 | 2.142 |
| chr22 | 38131199 | 38131443 | 1.04559 | 2.23987 | 2.142 |
| chr4 | 6000065 | 6000514 | 2.77638 | 5.94762 | 2.142 |

| | | | | | |
|---|---|---|---|---|---|
| chr7 | 105794256 | 105794660 | 2.18866 | 4.68864 | 2.142 |
| chr12 | 99137180 | 99137459 | 2.31705 | 4.96371 | 2.142 |
| chr1 | 173389521 | 173389794 | 1.30143 | 2.78822 | 2.142 |
| chr1 | 226821788 | 226822231 | 2.26591 | 4.85491 | 2.143 |
| chr15 | 75801361 | 75801593 | 1.11142 | 2.38135 | 2.143 |
| chr7 | 18650028 | 18650322 | 1.15808 | 2.48134 | 2.143 |
| chr17 | 29403793 | 29404007 | 0.50123 | 1.07401 | 2.143 |
| chr5 | 20366624 | 20366876 | 0.481544 | 1.03185 | 2.143 |
| chr3 | 136110518 | 136110741 | 1.2371 | 2.65089 | 2.143 |
| chr16 | 32518192 | 32518570 | 0.42992 | 0.921252 | 2.143 |
| chr4 | 77822830 | 77823073 | 0.490489 | 1.05106 | 2.143 |
| chr4 | 76816891 | 76817229 | 1.27262 | 2.72708 | 2.143 |
| chr16 | 402583 | 402817 | 1.60951 | 3.44907 | 2.143 |
| chr22 | 43058059 | 43058273 | 1.5993 | 3.42723 | 2.143 |
| chr4 | 118998417 | 118998765 | 1.40688 | 3.01503 | 2.143 |
| chr17 | 4965686 | 4966058 | 2.11873 | 4.54063 | 2.143 |
| chr5 | 34882919 | 34883156 | 0.531191 | 1.13839 | 2.143 |
| chr2 | 160568776 | 160569020 | 1.47895 | 3.16966 | 2.143 |
| chr14 | 64770203 | 64770495 | 1.1061 | 2.37063 | 2.143 |
| chr19 | 5170391 | 5170738 | 1.20961 | 2.5925 | 2.143 |
| chr6 | 133089125 | 133089615 | 1.42006 | 3.04355 | 2.143 |
| chr8 | 109657679 | 109658020 | 0.56807 | 1.21757 | 2.143 |
| chr11 | 9779674 | 9779987 | 1.99085 | 4.26721 | 2.143 |
| chr10 | 49864149 | 49864684 | 2.6928 | 5.77189 | 2.143 |
| chr4 | 4670860 | 4671276 | 1.70722 | 3.65937 | 2.143 |
| chr3 | 159522975 | 159523282 | 1.83418 | 3.93167 | 2.144 |
| chr1 | 68001114 | 68001437 | 1.66242 | 3.56352 | 2.144 |
| chr18 | 45282784 | 45283184 | 2.0295 | 4.35041 | 2.144 |
| chr15 | 92625472 | 92625842 | 1.66851 | 3.57669 | 2.144 |
| chr1 | 41239854 | 41240241 | 1.61528 | 3.46274 | 2.144 |
| chr3 | 157624538 | 157624834 | 0.6586 | 1.41192 | 2.144 |
| chr8 | 144594698 | 144595364 | 2.15751 | 4.62546 | 2.144 |
| chr2 | 61982263 | 61982483 | 0.506283 | 1.08543 | 2.144 |
| chr12 | 48583345 | 48583677 | 1.66014 | 3.55936 | 2.144 |
| chr14 | 77059594 | 77059816 | 0.49571 | 1.06282 | 2.144 |
| chrX | 19533338 | 19533904 | 0.988469 | 2.11932 | 2.144 |
| chr11 | 86343861 | 86344170 | 1.02083 | 2.18873 | 2.144 |
| chr4 | 55185088 | 55185414 | 1.45705 | 3.12404 | 2.144 |
| chr19 | 510546 | 510953 | 2.65501 | 5.69274 | 2.144 |
| chr1 | 154317691 | 154318032 | 1.89813 | 4.06993 | 2.144 |
| chr18 | 59992264 | 59992878 | 1.0751 | 2.30524 | 2.144 |
| chr12 | 96209477 | 96209730 | 1.28959 | 2.76532 | 2.144 |
| chr17 | 884339 | 884658 | 1.2321 | 2.64223 | 2.144 |
| chr2 | 26993878 | 26994391 | 1.00668 | 2.15888 | 2.145 |
| chrX | 77379828 | 77380101 | 0.831987 | 1.78432 | 2.145 |
| chr22 | 41939789 | 41940131 | 1.39887 | 3.00017 | 2.145 |
| chr11 | 18866341 | 18866649 | 1.27681 | 2.73843 | 2.145 |
| chr1 | 238737000 | 238737307 | 0.549327 | 1.17817 | 2.145 |
| chr10 | 102809964 | 102810355 | 1.97226 | 4.23025 | 2.145 |
| chr2 | 86054469 | 86054863 | 1.82698 | 3.9189 | 2.145 |
| chr12 | 54762435 | 54762805 | 1.91222 | 4.10183 | 2.145 |
| chr9 | 71682062 | 71682310 | 0.662983 | 1.42217 | 2.145 |
| chr14 | 25869807 | 25870123 | 1.05322 | 2.25944 | 2.145 |
| chr7 | 19481879 | 19482229 | 0.98227 | 2.1074 | 2.145 |
| chr4 | 27012056 | 27012342 | 1.23316 | 2.64571 | 2.145 |
| chr19 | 41953848 | 41954105 | 0.535602 | 1.14915 | 2.146 |
| chr15 | 76440193 | 76440676 | 2.91372 | 6.25153 | 2.146 |
| chr7 | 79351180 | 79351496 | 0.452079 | 0.970124 | 2.146 |
| chr10 | 29291146 | 29291380 | 0.515734 | 1.10675 | 2.146 |
| chr11 | 41985935 | 41986186 | 0.376477 | 0.80792 | 2.146 |
| chr12 | 119843514 | 119843809 | 0.676741 | 1.45239 | 2.146 |

| | | | | | |
|---|---|---|---|---|---|
| chr17 | 77397154 | 77397619 | 2.47348 | 5.3085 | 2.146 |
| chr5 | 153508898 | 153509328 | 0.46563 | 0.999343 | 2.146 |
| chr14 | 104376713 | 104377004 | 1.76521 | 3.78857 | 2.146 |
| chr8 | 111844670 | 111844952 | 1.16376 | 2.49771 | 2.146 |
| chr11 | 60534756 | 60535131 | 1.64728 | 3.53556 | 2.146 |
| chr2 | 19549835 | 19550279 | 1.7102 | 3.67062 | 2.146 |
| chr1 | 158368852 | 158369133 | 0.709132 | 1.52207 | 2.146 |
| chr17 | 73857323 | 73857746 | 2.44618 | 5.25054 | 2.146 |
| chr2 | 71091696 | 71092070 | 1.45748 | 3.12836 | 2.146 |
| chr4 | 88906732 | 88907099 | 1.88524 | 4.04671 | 2.147 |
| chr4 | 140418978 | 140419235 | 1.03163 | 2.21442 | 2.147 |
| chr12 | 81240799 | 81241079 | 0.672712 | 1.44405 | 2.147 |
| chr11 | 75308719 | 75309070 | 1.49729 | 3.2143 | 2.147 |
| chr11 | 119404492 | 119405106 | 2.11304 | 4.53616 | 2.147 |
| chrX | 119923962 | 119924175 | 1.24325 | 2.66899 | 2.147 |
| chr3 | 3221202 | 3221648 | 2.61837 | 5.62108 | 2.147 |
| chr17 | 73236689 | 73237091 | 2.63964 | 5.66697 | 2.147 |
| chr20 | 10209098 | 10209374 | 0.476445 | 1.02287 | 2.147 |
| chr15 | 100243980 | 100244188 | 1.11742 | 2.39932 | 2.147 |
| chr5 | 176538834 | 176539256 | 2.51011 | 5.38988 | 2.147 |
| chr10 | 135127385 | 135127589 | 0.967894 | 2.07835 | 2.147 |
| chr3 | 46484107 | 46484410 | 1.07513 | 2.30865 | 2.147 |
| chr9 | 76789491 | 76789733 | 0.69653 | 1.49571 | 2.147 |
| chr5 | 141137025 | 141137267 | 1.0381 | 2.2292 | 2.147 |
| chr3 | 54864766 | 54864988 | 1.2127 | 2.60425 | 2.147 |
| chr11 | 62980179 | 62980435 | 1.10394 | 2.37077 | 2.148 |
| chr11 | 32334476 | 32334779 | 1.26491 | 2.71651 | 2.148 |
| chr12 | 3994776 | 3995012 | 1.59983 | 3.43592 | 2.148 |
| chr16 | 8885772 | 8886106 | 1.70964 | 3.6718 | 2.148 |
| chr18 | 77006539 | 77006911 | 1.97258 | 4.23661 | 2.148 |
| chr6 | 34216842 | 34217133 | 1.39254 | 2.99085 | 2.148 |
| chr15 | 78113797 | 78114184 | 1.94719 | 4.18245 | 2.148 |
| chr1 | 11771192 | 11771430 | 0.987208 | 2.12053 | 2.148 |
| chr7 | 82201562 | 82201882 | 2.03734 | 4.3763 | 2.148 |
| chr10 | 74918849 | 74919168 | 1.34867 | 2.89703 | 2.148 |
| chr12 | 129743489 | 129743858 | 0.464869 | 0.998571 | 2.148 |
| chr3 | 2912285 | 2912502 | 0.382718 | 0.82211 | 2.148 |
| chr8 | 22225307 | 22225596 | 1.78907 | 3.84306 | 2.148 |
| chr11 | 63513142 | 63513378 | 0.860676 | 1.84884 | 2.148 |
| chr5 | 100125864 | 100126141 | 0.909923 | 1.95466 | 2.148 |
| chr12 | 54140361 | 54140813 | 3.27767 | 7.04118 | 2.148 |
| chr12 | 99401213 | 99401619 | 0.55371 | 1.18954 | 2.148 |
| chrX | 49846320 | 49846589 | 0.783589 | 1.68342 | 2.148 |
| chr17 | 2952014 | 2952316 | 1.75213 | 3.76449 | 2.149 |
| chr8 | 74086519 | 74086903 | 2.77722 | 5.96692 | 2.149 |
| chr8 | 17770168 | 17770414 | 1.23584 | 2.65527 | 2.149 |
| chr2 | 3852722 | 3853004 | 1.65196 | 3.54953 | 2.149 |
| chrX | 135350648 | 135350958 | 1.33975 | 2.8789 | 2.149 |
| chr16 | 7309082 | 7309454 | 0.530872 | 1.14076 | 2.149 |
| chr12 | 62586534 | 62586775 | 1.30171 | 2.79725 | 2.149 |
| chr6 | 167764924 | 167765209 | 1.58812 | 3.41279 | 2.149 |
| chr18 | 34409082 | 34409350 | 1.0079 | 2.166 | 2.149 |
| chr1 | 213930937 | 213931226 | 0.545812 | 1.17297 | 2.149 |
| chr10 | 132424284 | 132424578 | 0.37852 | 0.813495 | 2.149 |
| chr8 | 146245331 | 146245762 | 1.17622 | 2.52788 | 2.149 |
| chr12 | 18403328 | 18403547 | 1.18108 | 2.53843 | 2.149 |
| chr3 | 112301114 | 112301419 | 1.37933 | 2.96458 | 2.149 |
| chr6 | 158368178 | 158368548 | 0.641834 | 1.37949 | 2.149 |
| chr10 | 7020053 | 7020311 | 0.600991 | 1.29172 | 2.149 |
| chr4 | 134075903 | 134076145 | 0.903443 | 1.9418 | 2.149 |
| chr3 | 67840571 | 67840773 | 1.25924 | 2.70654 | 2.149 |

| | | | | | |
|---|---|---|---|---|---|
| chr3 | 72036268 | 72036512 | 0.874156 | 1.87886 | 2.149 |
| chr1 | 110036436 | 110036847 | 1.60879 | 3.45788 | 2.149 |
| chr12 | 57233656 | 57234026 | 1.55337 | 3.33891 | 2.149 |
| chr7 | 135485893 | 135486194 | 1.28433 | 2.76064 | 2.149 |
| chr18 | 7038282 | 7038600 | 0.76119 | 1.63629 | 2.150 |
| chr16 | 56618229 | 56618456 | 0.49907 | 1.07284 | 2.150 |
| chr10 | 23613379 | 23613610 | 0.567416 | 1.21983 | 2.150 |
| chr4 | 52762034 | 52762325 | 1.34604 | 2.89401 | 2.150 |
| chr12 | 95611109 | 95611504 | 1.27749 | 2.74665 | 2.150 |
| chr10 | 61435556 | 61435846 | 0.833281 | 1.79175 | 2.150 |
| chr17 | 5696554 | 5696803 | 1.03788 | 2.23171 | 2.150 |
| chr12 | 80807352 | 80807575 | 0.973951 | 2.09428 | 2.150 |
| gl000204_r | 67540 | 67800 | 0.755163 | 1.62386 | 2.150 |
| chr8 | 95143010 | 95143260 | 0.467573 | 1.00544 | 2.150 |
| chr12 | 107498545 | 107498786 | 1.28964 | 2.77331 | 2.150 |
| chr20 | 19023870 | 19024320 | 0.706059 | 1.51847 | 2.151 |
| chr19 | 13305398 | 13305673 | 0.991871 | 2.13317 | 2.151 |
| chr2 | 30934425 | 30934927 | 3.8326 | 8.2427 | 2.151 |
| chr3 | 13234400 | 13234786 | 1.18183 | 2.54177 | 2.151 |
| chr15 | 93994245 | 93994606 | 0.571734 | 1.22966 | 2.151 |
| chr12 | 101493600 | 101493817 | 0.465876 | 1.002 | 2.151 |
| chr20 | 60980355 | 60980635 | 1.26339 | 2.71736 | 2.151 |
| chr8 | 33278656 | 33279079 | 1.17164 | 2.52002 | 2.151 |
| chr10 | 77826662 | 77826985 | 0.455291 | 0.979299 | 2.151 |
| chr1 | 203242180 | 203242666 | 2.51442 | 5.40837 | 2.151 |
| chr12 | 61531658 | 61532170 | 0.485251 | 1.04378 | 2.151 |
| chr9 | 73099486 | 73099709 | 1.08403 | 2.33178 | 2.151 |
| chr7 | 4143001 | 4143467 | 0.652597 | 1.40385 | 2.151 |
| chr14 | 45722354 | 45722663 | 1.67877 | 3.6114 | 2.151 |
| chr16 | 71571608 | 71571938 | 1.43705 | 3.09155 | 2.151 |
| chr10 | 73062896 | 73063497 | 1.1365 | 2.44504 | 2.151 |
| chr20 | 22955072 | 22955346 | 0.997635 | 2.14631 | 2.151 |
| chr1 | 174952371 | 174952660 | 0.922591 | 1.98493 | 2.151 |
| chr9 | 130266234 | 130266616 | 2.39845 | 5.16034 | 2.152 |
| chr16 | 84992745 | 84993054 | 1.47338 | 3.1701 | 2.152 |
| chr1 | 108668224 | 108668447 | 1.49815 | 3.22359 | 2.152 |
| chr16 | 441838 | 442169 | 1.98158 | 4.26391 | 2.152 |
| chr1 | 44843829 | 44844219 | 1.28445 | 2.76387 | 2.152 |
| chr1 | 147364860 | 147365216 | 1.81331 | 3.90187 | 2.152 |
| chr22 | 18038173 | 18038512 | 1.52997 | 3.29221 | 2.152 |
| chr13 | 103244252 | 103244601 | 1.05822 | 2.27721 | 2.152 |
| chr10 | 71404169 | 71404949 | 1.63205 | 3.51206 | 2.152 |
| chr14 | 64108165 | 64108388 | 1.38994 | 2.99107 | 2.152 |
| chr7 | 155149390 | 155149788 | 1.21042 | 2.60477 | 2.152 |
| chr5 | 140420869 | 140421117 | 1.42732 | 3.07157 | 2.152 |
| chr12 | 105184258 | 105184690 | 1.79112 | 3.85462 | 2.152 |
| chr12 | 89936774 | 89937020 | 2.01002 | 4.32587 | 2.152 |
| chr4 | 12726175 | 12726446 | 0.846109 | 1.82099 | 2.152 |
| chr10 | 15184256 | 15184569 | 0.554603 | 1.19366 | 2.152 |
| chr8 | 41160431 | 41160807 | 0.875149 | 1.88363 | 2.152 |
| chr8 | 115858914 | 115859141 | 1.02335 | 2.20261 | 2.152 |
| chr19 | 45408053 | 45408280 | 1.02531 | 2.20687 | 2.152 |
| chr10 | 3292422 | 3292648 | 0.487466 | 1.04923 | 2.152 |
| chr4 | 168998167 | 168998625 | 1.37697 | 2.96386 | 2.152 |
| chr17 | 40831672 | 40832191 | 1.91771 | 4.12785 | 2.152 |
| chr8 | 139424020 | 139424242 | 0.466931 | 1.00508 | 2.153 |
| chr8 | 56972633 | 56972929 | 0.865276 | 1.86255 | 2.153 |
| chr4 | 21770075 | 21770287 | 1.42464 | 3.06666 | 2.153 |
| chr10 | 29676320 | 29676613 | 0.493068 | 1.06143 | 2.153 |
| chr7 | 14497453 | 14497788 | 1.46491 | 3.15355 | 2.153 |
| chr15 | 66859402 | 66859756 | 1.74442 | 3.75543 | 2.153 |

| | | | | | |
|---|---|---|---|---|---|
| chr15 | 38940011 | 38940376 | 0.564541 | 1.21546 | 2.153 |
| chr6 | 54528752 | 54529006 | 0.403209 | 0.868114 | 2.153 |
| chr17 | 5142850 | 5143110 | 0.425038 | 0.915123 | 2.153 |
| chr5 | 10284072 | 10284473 | 2.28834 | 4.92699 | 2.153 |
| chr8 | 71060238 | 71060693 | 0.773363 | 1.66514 | 2.153 |
| chr13 | 108994796 | 108995021 | 1.07242 | 2.30906 | 2.153 |
| chr1 | 156329847 | 156330083 | 0.805076 | 1.73354 | 2.153 |
| chr2 | 71150559 | 71150822 | 1.05368 | 2.26886 | 2.153 |
| chr4 | 72050006 | 72050259 | 1.04939 | 2.25964 | 2.153 |
| chr11 | 35259571 | 35260059 | 1.17113 | 2.5218 | 2.153 |
| chr12 | 7167778 | 7168153 | 1.75507 | 3.77923 | 2.153 |
| chr8 | 18725931 | 18726207 | 1.63326 | 3.51698 | 2.153 |
| chr5 | 94290706 | 94290918 | 0.58558 | 1.26098 | 2.153 |
| chr17 | 48255733 | 48256178 | 1.69908 | 3.65881 | 2.153 |
| chr12 | 113096283 | 113096559 | 0.43531 | 0.937406 | 2.153 |
| chr20 | 47833666 | 47834014 | 2.19917 | 4.73596 | 2.154 |
| chr1 | 68582370 | 68582645 | 1.34205 | 2.89018 | 2.154 |
| chr14 | 26632990 | 26633205 | 0.769918 | 1.6581 | 2.154 |
| chr12 | 9860609 | 9861121 | 1.2255 | 2.63938 | 2.154 |
| chr3 | 11888309 | 11888633 | 2.38838 | 5.14404 | 2.154 |
| chr8 | 29197707 | 29198043 | 1.61657 | 3.48195 | 2.154 |
| chr3 | 149205448 | 149205702 | 0.92092 | 1.98362 | 2.154 |
| chr9 | 79755450 | 79755665 | 0.462865 | 0.997006 | 2.154 |
| chr12 | 122699969 | 122700291 | 1.19181 | 2.56726 | 2.154 |
| chr5 | 56795243 | 56795614 | 1.35705 | 2.9234 | 2.154 |
| chr8 | 145048871 | 145049397 | 2.17995 | 4.69639 | 2.154 |
| chr5 | 151038495 | 151038881 | 2.41937 | 5.2125 | 2.154 |
| chr18 | 70429834 | 70430038 | 0.900257 | 1.93968 | 2.155 |
| chr9 | 97807251 | 97807636 | 2.01077 | 4.33243 | 2.155 |
| chr4 | 7297644 | 7298099 | 1.22885 | 2.64775 | 2.155 |
| chr6 | 71079091 | 71079383 | 1.63447 | 3.5218 | 2.155 |
| chr7 | 47879005 | 47879359 | 0.579149 | 1.2479 | 2.155 |
| chr7 | 108253676 | 108254021 | 1.26364 | 2.72282 | 2.155 |
| chr12 | 60887968 | 60888354 | 0.89645 | 1.93166 | 2.155 |
| chr16 | 19566463 | 19566794 | 1.78304 | 3.84229 | 2.155 |
| chr10 | 92511013 | 92511594 | 0.969624 | 2.08954 | 2.155 |
| chr12 | 132464975 | 132465306 | 0.628237 | 1.35387 | 2.155 |
| chr5 | 89825924 | 89826184 | 1.21223 | 2.61239 | 2.155 |
| chr3 | 14411176 | 14411451 | 0.969177 | 2.08878 | 2.155 |
| chr17 | 65685682 | 65685947 | 1.06133 | 2.28743 | 2.155 |
| chr14 | 24379734 | 24380155 | 1.85821 | 4.00497 | 2.155 |
| chr2 | 211367687 | 211368134 | 1.90534 | 4.10656 | 2.155 |
| chr19 | 11146638 | 11146933 | 1.75468 | 3.78189 | 2.155 |
| chr10 | 99551698 | 99552181 | 2.26152 | 4.87481 | 2.156 |
| chr2 | 161180775 | 161180983 | 0.964984 | 2.08008 | 2.156 |
| chr4 | 53777900 | 53778243 | 1.79841 | 3.87658 | 2.156 |
| chr1 | 234462385 | 234462753 | 2.01844 | 4.35107 | 2.156 |
| chr10 | 4554886 | 4555325 | 1.09026 | 2.35023 | 2.156 |
| chr2 | 119622774 | 119623027 | 0.954502 | 2.05759 | 2.156 |
| chr6 | 71594113 | 71594429 | 1.44612 | 3.11753 | 2.156 |
| rUn_gl0002 | 36323 | 36666 | 1.56618 | 3.37636 | 2.156 |
| chr2 | 31913276 | 31913834 | 0.983452 | 2.12014 | 2.156 |
| chr11 | 69008253 | 69008459 | 1.41895 | 3.0591 | 2.156 |
| chr15 | 84049345 | 84049703 | 1.21801 | 2.62589 | 2.156 |
| chr16 | 88747539 | 88747910 | 1.64065 | 3.53707 | 2.156 |
| chr8 | 57716570 | 57716897 | 0.517311 | 1.11529 | 2.156 |
| chr8 | 141015448 | 141015796 | 0.602484 | 1.299 | 2.156 |
| chr1 | 27324097 | 27324498 | 1.35573 | 2.92315 | 2.156 |
| chr15 | 34058613 | 34058962 | 0.632509 | 1.36382 | 2.156 |
| chr2 | 49224930 | 49225182 | 0.635046 | 1.36932 | 2.156 |
| chr20 | 48785961 | 48786240 | 1.33515 | 2.8792 | 2.156 |

| | | | | | |
|---|---|---|---|---|---|
| chr9 | 122052115 | 122052403 | 1.0859 | 2.34172 | 2.156 |
| chr14 | 26300427 | 26300738 | 0.520476 | 1.12242 | 2.157 |
| chr11 | 46563549 | 46563827 | 0.76835 | 1.65698 | 2.157 |
| chr10 | 57390857 | 57391312 | 1.24368 | 2.68213 | 2.157 |
| chr1 | 156862081 | 156862381 | 1.74639 | 3.76637 | 2.157 |
| chr7 | 50863502 | 50863754 | 0.902582 | 1.94658 | 2.157 |
| chr5 | 79378648 | 79379024 | 1.78565 | 3.8513 | 2.157 |
| chr19 | 3761672 | 3761909 | 0.936074 | 2.01894 | 2.157 |
| chr12 | 108070021 | 108070398 | 1.72201 | 3.71408 | 2.157 |
| chr5 | 89735629 | 89735832 | 0.771048 | 1.66304 | 2.157 |
| chr8 | 65415738 | 65416016 | 0.577362 | 1.24529 | 2.157 |
| chr3 | 103467112 | 103467321 | 0.414416 | 0.893845 | 2.157 |
| chr9 | 101737162 | 101737603 | 2.45661 | 5.29868 | 2.157 |
| chr6 | 4210912 | 4211140 | 1.13935 | 2.45756 | 2.157 |
| chr11 | 74427195 | 74427545 | 0.450101 | 0.970925 | 2.157 |
| chr7 | 150784479 | 150784916 | 0.790267 | 1.70471 | 2.157 |
| chr2 | 52819367 | 52819610 | 0.538387 | 1.16138 | 2.157 |
| chr11 | 10098294 | 10098737 | 1.31137 | 2.82888 | 2.157 |
| chrX | 41301466 | 41301784 | 1.18334 | 2.55283 | 2.157 |
| chr14 | 57046109 | 57046326 | 1.72308 | 3.71728 | 2.157 |
| chr15 | 50914630 | 50914889 | 1.50279 | 3.24217 | 2.157 |
| chr3 | 182628198 | 182628480 | 0.331512 | 0.715217 | 2.157 |
| chr1 | 59247492 | 59247699 | 0.814351 | 1.75708 | 2.158 |
| chr3 | 184592866 | 184593137 | 1.06404 | 2.29581 | 2.158 |
| chr14 | 95754475 | 95754714 | 0.38319 | 0.826814 | 2.158 |
| chr7 | 36358663 | 36358993 | 1.57472 | 3.39812 | 2.158 |
| chr1 | 25224624 | 25224957 | 1.46567 | 3.16289 | 2.158 |
| chr5 | 36688891 | 36689093 | 1.05929 | 2.28594 | 2.158 |
| chr15 | 60564797 | 60565062 | 0.65687 | 1.41756 | 2.158 |
| chr5 | 92221701 | 92221961 | 0.8482 | 1.83046 | 2.158 |
| chr7 | 147781253 | 147781544 | 0.890796 | 1.92252 | 2.158 |
| chr19 | 15272226 | 15272790 | 1.13453 | 2.44861 | 2.158 |
| chr1 | 64041034 | 64041374 | 0.979447 | 2.11391 | 2.158 |
| chr1 | 231892504 | 231892867 | 2.36489 | 5.10418 | 2.158 |
| chr10 | 43977445 | 43977658 | 0.442382 | 0.954808 | 2.158 |
| chr11 | 43846807 | 43847177 | 2.13002 | 4.59735 | 2.158 |
| chr6 | 138724601 | 138724844 | 0.853928 | 1.84312 | 2.158 |
| chr8 | 74516862 | 74517184 | 0.719676 | 1.55338 | 2.158 |
| chr12 | 58299109 | 58299524 | 2.77916 | 5.99869 | 2.158 |
| chr3 | 27295383 | 27295683 | 1.50056 | 3.23889 | 2.158 |
| chr14 | 31343507 | 31343719 | 0.712951 | 1.53888 | 2.158 |
| chr19 | 48293620 | 48294124 | 2.75953 | 5.95649 | 2.159 |
| chr16 | 71995360 | 71995661 | 2.16683 | 4.67723 | 2.159 |
| chr15 | 64230465 | 64230975 | 0.606139 | 1.30844 | 2.159 |
| chr3 | 172800392 | 172800729 | 0.53538 | 1.15571 | 2.159 |
| chr2 | 220424989 | 220425545 | 1.34551 | 2.90458 | 2.159 |
| chr7 | 21024794 | 21025010 | 0.502036 | 1.08377 | 2.159 |
| chrX | 11788370 | 11788754 | 1.85346 | 4.00141 | 2.159 |
| chr22 | 43781752 | 43782064 | 1.83827 | 3.96865 | 2.159 |
| chr21 | 42440969 | 42441319 | 1.8918 | 4.08436 | 2.159 |
| chr9 | 84647048 | 84647255 | 0.490222 | 1.0584 | 2.159 |
| chr20 | 33857055 | 33857445 | 1.5091 | 3.25825 | 2.159 |
| chr12 | 122794249 | 122794488 | 0.60362 | 1.30329 | 2.159 |
| chr4 | 10020345 | 10020712 | 2.28944 | 4.94332 | 2.159 |
| chr1 | 197325979 | 197326422 | 1.17731 | 2.54206 | 2.159 |
| chr6 | 104973594 | 104973998 | 1.44629 | 3.12287 | 2.159 |
| chrX | 130809590 | 130810019 | 1.22692 | 2.64927 | 2.159 |
| chr12 | 119760253 | 119760468 | 0.558596 | 1.20624 | 2.159 |
| chr10 | 92547644 | 92548019 | 1.45441 | 3.14077 | 2.159 |
| chr7 | 158794516 | 158794726 | 0.438559 | 0.94707 | 2.160 |
| chr5 | 4172550 | 4172805 | 0.441657 | 0.953864 | 2.160 |

| | | | | | |
|---|---|---|---|---|---|
| chr11 | 128754662 | 128754933 | 1.29961 | 2.80683 | 2.160 |
| chr13 | 27858335 | 27858614 | 0.753861 | 1.6282 | 2.160 |
| chr11 | 22266572 | 22266897 | 1.72836 | 3.73297 | 2.160 |
| chr4 | 31676009 | 31676222 | 0.935339 | 2.02026 | 2.160 |
| chr7 | 140510930 | 140511310 | 1.45895 | 3.15125 | 2.160 |
| chr6 | 7615756 | 7615944 | 1.34152 | 2.89761 | 2.160 |
| chr10 | 19725119 | 19725352 | 0.49559 | 1.07048 | 2.160 |
| chr1 | 204715925 | 204716288 | 1.06234 | 2.29468 | 2.160 |
| chr19 | 44173324 | 44173636 | 1.47374 | 3.18338 | 2.160 |
| chr2 | 190847637 | 190847907 | 1.11274 | 2.40371 | 2.160 |
| chr3 | 4040604 | 4040891 | 0.455824 | 0.984708 | 2.160 |
| chr16 | 89707629 | 89707962 | 1.43689 | 3.10413 | 2.160 |
| chr5 | 164214122 | 164214376 | 0.392312 | 0.84752 | 2.160 |
| chr3 | 104941636 | 104941840 | 1.03123 | 2.22788 | 2.160 |
| chr17 | 79674396 | 79674625 | 0.974651 | 2.10582 | 2.161 |
| chr19 | 48823261 | 48823731 | 2.01252 | 4.34854 | 2.161 |
| chr8 | 105800142 | 105800401 | 0.971948 | 2.10013 | 2.161 |
| chr1 | 226756264 | 226756501 | 0.832084 | 1.79798 | 2.161 |
| chr7 | 97735831 | 97736232 | 1.96303 | 4.24176 | 2.161 |
| chr10 | 43186045 | 43186299 | 0.91004 | 1.96644 | 2.161 |
| chr22 | 40420847 | 40421187 | 1.19536 | 2.58322 | 2.161 |
| chr1 | 155058359 | 155058973 | 1.80039 | 3.8908 | 2.161 |
| chr15 | 71258562 | 71258877 | 0.524976 | 1.13458 | 2.161 |
| chr14 | 70724281 | 70724597 | 0.922565 | 1.99389 | 2.161 |
| chr14 | 23656301 | 23656598 | 1.37937 | 2.9812 | 2.161 |
| chr12 | 80779356 | 80779568 | 0.451708 | 0.976334 | 2.161 |
| chr4 | 40899685 | 40900016 | 0.851195 | 1.83981 | 2.161 |
| chr16 | 1755905 | 1756342 | 2.18751 | 4.72825 | 2.161 |
| chr2 | 37623042 | 37623367 | 1.81298 | 3.91875 | 2.162 |
| chr19 | 45996784 | 45997248 | 1.62618 | 3.51503 | 2.162 |
| chr15 | 55010111 | 55010317 | 0.563029 | 1.21701 | 2.162 |
| chr4 | 88600635 | 88600885 | 1.30154 | 2.8138 | 2.162 |
| chr16 | 16221517 | 16222080 | 1.36812 | 2.9578 | 2.162 |
| chr2 | 79754622 | 79754826 | 0.446675 | 0.96569 | 2.162 |
| chr15 | 40396974 | 40397432 | 2.435 | 5.26449 | 2.162 |
| chr5 | 78429475 | 78429760 | 1.96622 | 4.25111 | 2.162 |
| chr14 | 100029480 | 100029791 | 1.65657 | 3.58209 | 2.162 |
| chr6 | 122851672 | 122851921 | 1.23769 | 2.67634 | 2.162 |
| chr15 | 42376598 | 42376952 | 1.62236 | 3.50831 | 2.162 |
| chr12 | 54990200 | 54990685 | 1.68355 | 3.64082 | 2.163 |
| chr5 | 172720944 | 172721298 | 1.51043 | 3.26649 | 2.163 |
| chr22 | 24552895 | 24553265 | 1.50149 | 3.24718 | 2.163 |
| chr12 | 94875639 | 94876567 | 0.503414 | 1.08875 | 2.163 |
| chr11 | 8214078 | 8214737 | 1.38904 | 3.00414 | 2.163 |
| chr20 | 55921488 | 55921777 | 1.36842 | 2.95961 | 2.163 |
| chr16 | 57334506 | 57334822 | 1.58593 | 3.43009 | 2.163 |
| chr7 | 139119243 | 139119744 | 2.3193 | 5.01641 | 2.163 |
| chr8 | 49386498 | 49386700 | 0.475846 | 1.02922 | 2.163 |
| chr6 | 37271774 | 37272056 | 0.961456 | 2.07973 | 2.163 |
| chr12 | 118058056 | 118058304 | 0.442263 | 0.956667 | 2.163 |
| chr5 | 128830094 | 128830318 | 0.523586 | 1.1326 | 2.163 |
| chr9 | 131398377 | 131398635 | 1.31517 | 2.84525 | 2.163 |
| chr4 | 100870114 | 100870423 | 1.47293 | 3.18661 | 2.163 |
| chr9 | 4662146 | 4662375 | 1.44561 | 3.12755 | 2.163 |
| chr1 | 53951154 | 53951490 | 2.0591 | 4.45513 | 2.164 |
| chr5 | 4398696 | 4398944 | 0.603537 | 1.30587 | 2.164 |
| chr1 | 32681208 | 32681603 | 1.42741 | 3.08871 | 2.164 |
| chr9 | 74383842 | 74384514 | 0.857157 | 1.85477 | 2.164 |
| chr12 | 56320553 | 56320887 | 2.11276 | 4.57173 | 2.164 |
| chr11 | 38004644 | 38004928 | 0.556036 | 1.2032 | 2.164 |
| chr11 | 130082971 | 130083317 | 1.86815 | 4.04288 | 2.164 |

| | | | | | |
|---|---|---|---|---|---|
| chr7 | 8075581 | 8075924 | 1.30402 | 2.82205 | 2.164 |
| chr3 | 5347135 | 5347432 | 1.2947 | 2.80197 | 2.164 |
| chr3 | 105237736 | 105237976 | 1.19221 | 2.58016 | 2.164 |
| chr2 | 172015837 | 172016050 | 0.91718 | 1.98507 | 2.164 |
| chr8 | 145657719 | 145657965 | 0.444813 | 0.962717 | 2.164 |
| chr17 | 47304652 | 47305262 | 2.0872 | 4.51759 | 2.164 |
| chr5 | 33087434 | 33087736 | 0.566311 | 1.22578 | 2.165 |
| chr12 | 116354935 | 116355412 | 1.82132 | 3.94233 | 2.165 |
| chr14 | 100639762 | 100639977 | 0.721342 | 1.56142 | 2.165 |
| chr10 | 104343647 | 104343858 | 1.47896 | 3.20138 | 2.165 |
| chr1 | 164902965 | 164903172 | 0.495644 | 1.07289 | 2.165 |
| chr14 | 84144462 | 84144672 | 0.493931 | 1.0692 | 2.165 |
| chr18 | 8426490 | 8426784 | 0.952575 | 2.06207 | 2.165 |
| chr22 | 42840441 | 42840743 | 1.7595 | 3.80895 | 2.165 |
| chr4 | 190123482 | 190123730 | 1.43281 | 3.10174 | 2.165 |
| chr5 | 160663866 | 160664196 | 0.349963 | 0.757615 | 2.165 |
| chr12 | 106392262 | 106392473 | 0.509385 | 1.10276 | 2.165 |
| chr20 | 7118740 | 7118980 | 0.495558 | 1.07288 | 2.165 |
| chr3 | 179164318 | 179164597 | 0.879535 | 1.90429 | 2.165 |
| chr10 | 71981320 | 71981549 | 1.04657 | 2.26598 | 2.165 |
| chr12 | 7341002 | 7341431 | 1.59913 | 3.46235 | 2.165 |
| chr14 | 107012741 | 107012989 | 1.06021 | 2.29554 | 2.165 |
| chr10 | 133040356 | 133040620 | 1.30925 | 2.83483 | 2.165 |
| chr10 | 76951557 | 76951762 | 1.13899 | 2.46635 | 2.165 |
| chr8 | 19946065 | 19946375 | 0.583558 | 1.26363 | 2.165 |
| chr13 | 21872024 | 21872304 | 1.40922 | 3.05163 | 2.165 |
| chr9 | 32579760 | 32580080 | 2.19158 | 4.74593 | 2.166 |
| chr12 | 1772230 | 1772640 | 1.59973 | 3.46457 | 2.166 |
| chr8 | 143553045 | 143553687 | 1.87418 | 4.0591 | 2.166 |
| chr7 | 77532974 | 77533235 | 0.83487 | 1.80821 | 2.166 |
| chr3 | 27722756 | 27722964 | 1.06228 | 2.30076 | 2.166 |
| chr8 | 145911123 | 145911463 | 2.11375 | 4.5781 | 2.166 |
| chr20 | 36013132 | 36013489 | 2.14544 | 4.64684 | 2.166 |
| chr8 | 136323053 | 136323300 | 0.483605 | 1.04745 | 2.166 |
| chr4 | 165653006 | 165653325 | 1.13126 | 2.45026 | 2.166 |
| chr4 | 185926163 | 185926720 | 1.12378 | 2.43409 | 2.166 |
| chr9 | 137979137 | 137979559 | 1.84839 | 4.00372 | 2.166 |
| chr8 | 140805544 | 140805847 | 0.421697 | 0.913443 | 2.166 |
| chr21 | 16893193 | 16893721 | 0.611626 | 1.32489 | 2.166 |
| chr17 | 14342 | 14686 | 1.33726 | 2.89679 | 2.166 |
| chr11 | 27286632 | 27286915 | 0.519525 | 1.12547 | 2.166 |
| chr9 | 32432010 | 32432301 | 1.45017 | 3.14161 | 2.166 |
| chr16 | 85148461 | 85149127 | 1.55619 | 3.37136 | 2.166 |
| chr20 | 49407113 | 49407489 | 1.41826 | 3.07258 | 2.166 |
| chr9 | 8978513 | 8978887 | 0.691217 | 1.49754 | 2.167 |
| chr17 | 77818559 | 77818998 | 1.95371 | 4.23289 | 2.167 |
| chr8 | 33519939 | 33520231 | 0.429415 | 0.930405 | 2.167 |
| chr6 | 84302567 | 84302800 | 1.02277 | 2.21634 | 2.167 |
| chr3 | 52007958 | 52008275 | 2.10603 | 4.56386 | 2.167 |
| chr1 | 2246518 | 2246909 | 2.32281 | 5.03368 | 2.167 |
| chr14 | 22293398 | 22293670 | 1.32363 | 2.86853 | 2.167 |
| chrX | 19352283 | 19352712 | 1.1455 | 2.48255 | 2.167 |
| chr12 | 6888126 | 6888466 | 2.23508 | 4.84416 | 2.167 |
| chr4 | 3320322 | 3320566 | 0.649944 | 1.40867 | 2.167 |
| chr9 | 71650386 | 71650853 | 3.1985 | 6.9324 | 2.167 |
| chr5 | 143474636 | 143474902 | 0.612051 | 1.32656 | 2.167 |
| chr19 | 33710321 | 33710687 | 1.54314 | 3.34472 | 2.167 |
| chr12 | 105629532 | 105629777 | 0.7373 | 1.5981 | 2.168 |
| chr16 | 58877715 | 58877951 | 0.944674 | 2.04764 | 2.168 |
| chr17 | 11884194 | 11884437 | 0.715038 | 1.5499 | 2.168 |
| chr17 | 28258398 | 28258690 | 1.99721 | 4.32911 | 2.168 |

| | | | | | |
|---|---|---|---|---|---|
| chr13 | 29253700 | 29254028 | 0.56937 | 1.2342 | 2.168 |
| chr8 | 57154480 | 57154865 | 2.09901 | 4.55014 | 2.168 |
| chr17 | 80372363 | 80372832 | 1.73935 | 3.77056 | 2.168 |
| chr6 | 159362346 | 159362548 | 0.358982 | 0.778244 | 2.168 |
| chr5 | 169847844 | 169848126 | 1.40707 | 3.05049 | 2.168 |
| chr8 | 67372083 | 67372344 | 0.933359 | 2.02349 | 2.168 |
| chr10 | 6880251 | 6880530 | 1.04096 | 2.25714 | 2.168 |
| chr3 | 155181769 | 155182008 | 0.462563 | 1.003 | 2.168 |
| chr17 | 74028383 | 74028863 | 0.726339 | 1.575 | 2.168 |
| chr9 | 15532989 | 15533609 | 1.00845 | 2.18675 | 2.168 |
| chr16 | 17894130 | 17894346 | 0.61032 | 1.32345 | 2.168 |
| chr18 | 24127666 | 24128078 | 1.22758 | 2.662 | 2.168 |
| chr15 | 58505178 | 58505441 | 1.59106 | 3.45024 | 2.169 |
| chr8 | 28645279 | 28645612 | 1.43332 | 3.10822 | 2.169 |
| chr1 | 16408814 | 16409479 | 1.2141 | 2.63305 | 2.169 |
| chr8 | 22766503 | 22766920 | 1.59874 | 3.46727 | 2.169 |
| chr6 | 149068467 | 149068815 | 0.49884 | 1.08194 | 2.169 |
| chr19 | 770270 | 770580 | 1.80592 | 3.91719 | 2.169 |
| chr1 | 22534836 | 22535044 | 1.12981 | 2.45066 | 2.169 |
| chr8 | 125384873 | 125385199 | 0.840217 | 1.82256 | 2.169 |
| chr13 | 104535497 | 104535699 | 0.752386 | 1.63216 | 2.169 |
| chr14 | 94952569 | 94953036 | 2.37022 | 5.14177 | 2.169 |
| chr14 | 90865871 | 90866234 | 2.76903 | 6.00696 | 2.169 |
| chr9 | 16154624 | 16154888 | 1.02193 | 2.21691 | 2.169 |
| chr8 | 72528977 | 72529264 | 0.646624 | 1.40279 | 2.169 |
| chr20 | 48158261 | 48158608 | 1.92365 | 4.17321 | 2.169 |
| chr1 | 37041593 | 37042048 | 1.75277 | 3.80252 | 2.169 |
| chr9 | 107382028 | 107382345 | 0.450789 | 0.977971 | 2.169 |
| chr1 | 17933367 | 17933755 | 1.59922 | 3.46947 | 2.169 |
| chr11 | 133920411 | 133920750 | 1.38481 | 3.00434 | 2.170 |
| chr12 | 95831401 | 95832006 | 0.540083 | 1.17175 | 2.170 |
| chr17 | 1551677 | 1552442 | 1.75287 | 3.80299 | 2.170 |
| chr10 | 134957482 | 134957684 | 1.39994 | 3.03753 | 2.170 |
| chr11 | 41785581 | 41785796 | 0.632958 | 1.37339 | 2.170 |
| chr20 | 3994608 | 3994943 | 1.4261 | 3.09436 | 2.170 |
| chr2 | 119931828 | 119932401 | 1.51451 | 3.28621 | 2.170 |
| chr18 | 48636508 | 48637107 | 1.434 | 3.11157 | 2.170 |
| chr20 | 37075030 | 37075419 | 2.18505 | 4.74123 | 2.170 |
| chr5 | 167474316 | 167474679 | 1.13105 | 2.45423 | 2.170 |
| chr2 | 20303300 | 20303621 | 1.05681 | 2.29317 | 2.170 |
| chr16 | 15521495 | 15521742 | 1.26752 | 2.7504 | 2.170 |
| chr8 | 130619313 | 130619553 | 0.607681 | 1.31868 | 2.170 |
| chr10 | 45503970 | 45504373 | 1.37772 | 2.9897 | 2.170 |
| chr17 | 235807 | 236301 | 2.47387 | 5.36845 | 2.170 |
| chr9 | 116917513 | 116917873 | 2.62187 | 5.68961 | 2.170 |
| chr2 | 29436686 | 29436928 | 0.686374 | 1.48953 | 2.170 |
| chr12 | 103210614 | 103210956 | 1.43177 | 3.10717 | 2.170 |
| chr17 | 3438819 | 3439147 | 1.39512 | 3.02765 | 2.170 |
| chr4 | 78980796 | 78981006 | 1.78648 | 3.87698 | 2.170 |
| chr2 | 27137174 | 27137387 | 0.476712 | 1.03459 | 2.170 |
| chr10 | 128918232 | 128918507 | 0.56105 | 1.2177 | 2.170 |
| chr15 | 63902403 | 63902754 | 1.38707 | 3.01053 | 2.170 |
| chr14 | 81636974 | 81637298 | 2.26654 | 4.9194 | 2.170 |
| chr1 | 185689838 | 185690111 | 1.2116 | 2.62972 | 2.170 |
| chr1 | 154714360 | 154714704 | 0.910552 | 1.97632 | 2.170 |
| chr2 | 223966510 | 223966828 | 1.25422 | 2.72223 | 2.170 |
| chr17 | 26132478 | 26132772 | 1.34225 | 2.91346 | 2.171 |
| chr1 | 78610739 | 78611102 | 1.54146 | 3.34599 | 2.171 |
| chr4 | 154006300 | 154006687 | 1.68515 | 3.65789 | 2.171 |
| chr3 | 52968634 | 52968960 | 1.25292 | 2.7197 | 2.171 |
| chr5 | 72733362 | 72733710 | 1.48858 | 3.23132 | 2.171 |

| | | | | | |
|---|---|---|---|---|---|
| chr22 | 32774700 | 32774990 | 1.38635 | 3.00943 | 2.171 |
| chr4 | 74207182 | 74207414 | 1.04243 | 2.26293 | 2.171 |
| chr14 | 31151009 | 31151257 | 1.1971 | 2.59871 | 2.171 |
| chr1 | 223871974 | 223872228 | 1.88204 | 4.08578 | 2.171 |
| chr10 | 81321883 | 81322320 | 2.33388 | 5.06669 | 2.171 |
| chr3 | 138698645 | 138698874 | 1.29692 | 2.81559 | 2.171 |
| chr17 | 46679974 | 46680288 | 2.03101 | 4.40971 | 2.171 |
| chr12 | 4492609 | 4492812 | 0.533726 | 1.15889 | 2.171 |
| chr3 | 152639940 | 152640241 | 0.498626 | 1.08268 | 2.171 |
| chr1 | 24913797 | 24914008 | 1.10453 | 2.39834 | 2.171 |
| chr20 | 4052705 | 4053078 | 1.81451 | 3.94008 | 2.171 |
| chr19 | 49702094 | 49702500 | 1.60941 | 3.49484 | 2.172 |
| chr2 | 71204566 | 71204770 | 1.28042 | 2.78044 | 2.172 |
| chr1 | 7729602 | 7729970 | 1.71083 | 3.71519 | 2.172 |
| chr9 | 101955513 | 101956044 | 0.550583 | 1.19567 | 2.172 |
| chr13 | 27913481 | 27913771 | 1.59753 | 3.46927 | 2.172 |
| chr7 | 33972507 | 33972888 | 2.03803 | 4.42592 | 2.172 |
| chr17 | 17822577 | 17822808 | 1.53642 | 3.33662 | 2.172 |
| chr8 | 5216512 | 5216780 | 0.458408 | 0.99552 | 2.172 |
| chr6 | 44235748 | 44236093 | 2.05787 | 4.46938 | 2.172 |
| chr11 | 8625524 | 8625982 | 1.17912 | 2.5609 | 2.172 |
| chr12 | 48111324 | 48111629 | 1.9699 | 4.27842 | 2.172 |
| chr18 | 60822618 | 60823061 | 1.94908 | 4.23326 | 2.172 |
| chr20 | 61075087 | 61075504 | 1.64878 | 3.58123 | 2.172 |
| chr4 | 7044910 | 7045123 | 1.11311 | 2.41776 | 2.172 |
| chr19 | 3146605 | 3146941 | 1.34625 | 2.92447 | 2.172 |
| chrX | 152127502 | 152127737 | 1.24697 | 2.70908 | 2.173 |
| chr6 | 89609260 | 89609524 | 0.42236 | 0.917599 | 2.173 |
| chrX | 80376377 | 80376610 | 1.21484 | 2.63931 | 2.173 |
| chr3 | 46908198 | 46908551 | 1.70093 | 3.69542 | 2.173 |
| chr5 | 180145796 | 180146150 | 1.17968 | 2.563 | 2.173 |
| chr12 | 102186369 | 102186623 | 1.04063 | 2.26092 | 2.173 |
| chr17 | 80279276 | 80279820 | 2.59764 | 5.64392 | 2.173 |
| chr5 | 56086317 | 56086561 | 0.460876 | 1.00135 | 2.173 |
| chr10 | 26839773 | 26840067 | 0.631885 | 1.37291 | 2.173 |
| chr16 | 22308454 | 22308818 | 2.0676 | 4.49239 | 2.173 |
| chr18 | 74513920 | 74514356 | 1.29937 | 2.82338 | 2.173 |
| chr6 | 126492093 | 126492336 | 0.951861 | 2.06829 | 2.173 |
| chr12 | 104020335 | 104020585 | 1.1084 | 2.4085 | 2.173 |
| chr1 | 56167822 | 56168034 | 0.904631 | 1.96573 | 2.173 |
| chr7 | 156686095 | 156686380 | 0.739423 | 1.60678 | 2.173 |
| chr1 | 209742753 | 209743178 | 1.52883 | 3.32223 | 2.173 |
| chr13 | 40516920 | 40517253 | 1.85319 | 4.02731 | 2.173 |
| chr7 | 141437777 | 141438148 | 1.77125 | 3.84925 | 2.173 |
| chr10 | 1463063 | 1463573 | 1.87925 | 4.08406 | 2.173 |
| chr7 | 28882063 | 28882268 | 0.670171 | 1.45645 | 2.173 |
| chr2 | 35339850 | 35340177 | 1.08276 | 2.35323 | 2.173 |
| chr19 | 12886412 | 12886702 | 2.82305 | 6.13621 | 2.174 |
| chr16 | 761675 | 762146 | 2.18804 | 4.75606 | 2.174 |
| chr5 | 139737233 | 139737547 | 1.82082 | 3.95785 | 2.174 |
| chr3 | 69170051 | 69170367 | 1.74437 | 3.79168 | 2.174 |
| chr9 | 6143673 | 6144059 | 0.733599 | 1.59467 | 2.174 |
| chr3 | 21226625 | 21226869 | 0.391117 | 0.850218 | 2.174 |
| chr1 | 64386531 | 64386790 | 0.792612 | 1.72317 | 2.174 |
| chr10 | 129705120 | 129705456 | 1.85683 | 4.03722 | 2.174 |
| chr12 | 108881405 | 108881675 | 0.410123 | 0.891735 | 2.174 |
| chr17 | 37886614 | 37887016 | 2.07158 | 4.50433 | 2.174 |
| chr8 | 22094782 | 22095091 | 1.77146 | 3.85197 | 2.174 |
| chr19 | 47207588 | 47208104 | 2.99341 | 6.50928 | 2.175 |
| chr19 | 24345484 | 24345772 | 0.515878 | 1.12182 | 2.175 |
| chr1 | 47644867 | 47645132 | 1.65321 | 3.59506 | 2.175 |

| | | | | | |
|---|---|---|---|---|---|
| gl000204_r | 68134 | 68434 | 0.450535 | 0.979741 | 2.175 |
| chr17 | 30438485 | 30438737 | 0.891911 | 1.93966 | 2.175 |
| chr8 | 105142401 | 105142817 | 1.32363 | 2.87852 | 2.175 |
| chr7 | 142042959 | 142043277 | 0.433465 | 0.942668 | 2.175 |
| chr20 | 52824185 | 52824490 | 1.88398 | 4.09776 | 2.175 |
| chr5 | 172134253 | 172134548 | 2.19492 | 4.77405 | 2.175 |
| chr7 | 106787988 | 106788235 | 1.1058 | 2.40518 | 2.175 |
| chr12 | 71974207 | 71974521 | 0.466927 | 1.01562 | 2.175 |
| chr8 | 60834135 | 60834412 | 1.17897 | 2.5644 | 2.175 |
| chr11 | 119950476 | 119950797 | 0.889956 | 1.93577 | 2.175 |
| chr21 | 38299857 | 38300272 | 1.43898 | 3.13007 | 2.175 |
| chr20 | 42378363 | 42378843 | 2.0796 | 4.52358 | 2.175 |
| chr11 | 5844087 | 5844344 | 0.463679 | 1.00863 | 2.175 |
| chr1 | 20208592 | 20209057 | 0.907592 | 1.97433 | 2.175 |
| chr1 | 177049930 | 177050280 | 0.503617 | 1.09559 | 2.175 |
| chr6 | 5422638 | 5422913 | 1.04531 | 2.27411 | 2.176 |
| chr8 | 55585906 | 55586484 | 0.574052 | 1.24888 | 2.176 |
| chr19 | 31067384 | 31067645 | 0.537458 | 1.16928 | 2.176 |
| chr5 | 149509534 | 149509892 | 1.99667 | 4.34393 | 2.176 |
| chr7 | 137311249 | 137311652 | 1.27734 | 2.779 | 2.176 |
| chr1 | 40075303 | 40075612 | 1.48298 | 3.2264 | 2.176 |
| chr19 | 39677804 | 39678322 | 2.58045 | 5.61409 | 2.176 |
| chr3 | 58202975 | 58203517 | 1.07412 | 2.33704 | 2.176 |
| chr9 | 90449311 | 90449660 | 0.644933 | 1.40325 | 2.176 |
| chr12 | 108905649 | 108906090 | 1.72334 | 3.74972 | 2.176 |
| chr9 | 36407051 | 36407738 | 1.37312 | 2.98776 | 2.176 |
| chr5 | 1204891 | 1205111 | 0.864546 | 1.8812 | 2.176 |
| chr11 | 64736987 | 64737385 | 0.537314 | 1.16919 | 2.176 |
| chr22 | 31001545 | 31001849 | 1.18979 | 2.58898 | 2.176 |
| chr2 | 234357719 | 234358044 | 1.71101 | 3.72317 | 2.176 |
| chr16 | 19612454 | 19613051 | 0.896101 | 1.94997 | 2.176 |
| chr10 | 97625025 | 97625360 | 1.51655 | 3.30014 | 2.176 |
| chr17 | 76080997 | 76081443 | 2.45168 | 5.33504 | 2.176 |
| chr20 | 12035581 | 12035788 | 0.522811 | 1.13769 | 2.176 |
| chr15 | 92395902 | 92396730 | 1.86146 | 4.05077 | 2.176 |
| chr3 | 179367440 | 179367675 | 2.0447 | 4.4496 | 2.176 |
| chr16 | 7016273 | 7016481 | 0.480703 | 1.0461 | 2.176 |
| chr20 | 3767384 | 3767729 | 2.52654 | 5.49849 | 2.176 |
| chr11 | 9883724 | 9884041 | 1.40219 | 3.05177 | 2.176 |
| chr11 | 2906265 | 2906565 | 0.685299 | 1.49158 | 2.177 |
| chr15 | 79001754 | 79002093 | 1.85409 | 4.03566 | 2.177 |
| chr17 | 54440507 | 54440748 | 0.450478 | 0.980524 | 2.177 |
| chr10 | 86024334 | 86024733 | 2.08063 | 4.52887 | 2.177 |
| chr14 | 41824436 | 41824729 | 0.437212 | 0.951677 | 2.177 |
| chr12 | 50171609 | 50172350 | 1.56005 | 3.39599 | 2.177 |
| chr22 | 41922689 | 41923168 | 2.08929 | 4.54809 | 2.177 |
| chr22 | 17555940 | 17556188 | 0.549033 | 1.19521 | 2.177 |
| chr17 | 57517866 | 57518624 | 1.15785 | 2.52077 | 2.177 |
| chr15 | 27871897 | 27872315 | 0.498355 | 1.08498 | 2.177 |
| chr3 | 133537726 | 133538033 | 1.79916 | 3.91704 | 2.177 |
| chr11 | 8787910 | 8788115 | 0.56549 | 1.23118 | 2.177 |
| chr2 | 75801302 | 75801679 | 0.896116 | 1.95103 | 2.177 |
| chr10 | 82204471 | 82204822 | 2.47915 | 5.39816 | 2.177 |
| chr2 | 1135992 | 1136456 | 2.31146 | 5.03317 | 2.177 |
| chr9 | 111539356 | 111539780 | 0.541233 | 1.17854 | 2.178 |
| chr14 | 104224723 | 104224936 | 0.463883 | 1.01013 | 2.178 |
| chr12 | 57487044 | 57487438 | 1.56021 | 3.39747 | 2.178 |
| chr1 | 215448846 | 215449094 | 0.544686 | 1.18611 | 2.178 |
| chr7 | 129650690 | 129650975 | 1.96513 | 4.2793 | 2.178 |
| chr9 | 91208773 | 91209054 | 1.32231 | 2.87952 | 2.178 |
| chr12 | 80589020 | 80589277 | 1.07416 | 2.33914 | 2.178 |

| | | | | | |
|---|---|---|---|---|---|
| chr10 | 88861406 | 88861668 | 0.466001 | 1.01483 | 2.178 |
| chr5 | 167849544 | 167849829 | 0.928306 | 2.02163 | 2.178 |
| chr5 | 118747570 | 118748055 | 1.36416 | 2.97088 | 2.178 |
| chr17 | 48703471 | 48703883 | 1.41141 | 3.07384 | 2.178 |
| chr11 | 76391864 | 76392256 | 1.52042 | 3.31127 | 2.178 |
| chr15 | 77789125 | 77789522 | 2.54393 | 5.54036 | 2.178 |
| chr11 | 32345060 | 32345403 | 1.24517 | 2.71191 | 2.178 |
| chr11 | 64257316 | 64257855 | 1.79681 | 3.91334 | 2.178 |
| chr18 | 2083711 | 2083965 | 0.808807 | 1.76153 | 2.178 |
| chr7 | 145892992 | 145893217 | 0.544607 | 1.18612 | 2.178 |
| chr7 | 137588373 | 137588712 | 1.36446 | 2.97212 | 2.178 |
| chrX | 24072348 | 24072685 | 1.84208 | 4.01249 | 2.178 |
| chr9 | 100161449 | 100161785 | 2.81547 | 6.13287 | 2.178 |
| chr16 | 2190600 | 2191006 | 1.69986 | 3.70321 | 2.179 |
| chr9 | 123505234 | 123505590 | 1.45973 | 3.18008 | 2.179 |
| chr7 | 159068960 | 159069165 | 0.545648 | 1.18876 | 2.179 |
| chr11 | 130697747 | 130697994 | 0.914213 | 1.99179 | 2.179 |
| chr6 | 43741062 | 43741459 | 1.7405 | 3.79214 | 2.179 |
| chr2 | 69066367 | 69066870 | 1.4243 | 3.10323 | 2.179 |
| chr1 | 53859021 | 53859356 | 1.51656 | 3.30441 | 2.179 |
| chr13 | 45621437 | 45621800 | 1.96782 | 4.28781 | 2.179 |
| chr17 | 7944721 | 7944996 | 1.45826 | 3.17761 | 2.179 |
| chr2 | 27526365 | 27526665 | 1.61368 | 3.51634 | 2.179 |
| chr9 | 134622857 | 134623114 | 1.72991 | 3.76982 | 2.179 |
| chr3 | 25873163 | 25873403 | 0.543005 | 1.18336 | 2.179 |
| chr7 | 75272211 | 75272522 | 1.68478 | 3.67168 | 2.179 |
| chr10 | 104153663 | 104154060 | 2.0717 | 4.51505 | 2.179 |
| chrX | 8706706 | 8706957 | 0.978143 | 2.13178 | 2.179 |
| chr6 | 17584482 | 17584905 | 0.976119 | 2.12739 | 2.179 |
| chr4 | 168912513 | 168912751 | 0.485858 | 1.05892 | 2.179 |
| chr7 | 5867444 | 5867766 | 1.06185 | 2.31432 | 2.180 |
| chr7 | 103177819 | 103178253 | 0.449766 | 0.980282 | 2.180 |
| chr4 | 55189049 | 55189343 | 1.22516 | 2.67042 | 2.180 |
| chr8 | 12698038 | 12698510 | 1.28351 | 2.79763 | 2.180 |
| chr17 | 26743802 | 26744194 | 2.11949 | 4.62001 | 2.180 |
| chr20 | 1149989 | 1150204 | 0.453017 | 0.987508 | 2.180 |
| chr8 | 41852830 | 41853061 | 1.29918 | 2.83204 | 2.180 |
| chr17 | 38069897 | 38070122 | 0.561885 | 1.22486 | 2.180 |
| chr1 | 116012311 | 116012624 | 2.15962 | 4.70784 | 2.180 |
| chr2 | 219166019 | 219166336 | 1.98221 | 4.32115 | 2.180 |
| chr8 | 42178132 | 42178368 | 0.540843 | 1.17904 | 2.180 |
| chr1 | 243852932 | 243853152 | 0.460418 | 1.00372 | 2.180 |
| chr14 | 78603785 | 78604077 | 0.580604 | 1.26575 | 2.180 |
| chr12 | 123352231 | 123352555 | 1.08615 | 2.36809 | 2.180 |
| chr10 | 90313001 | 90313244 | 1.24306 | 2.71027 | 2.180 |
| chr4 | 22240476 | 22240717 | 1.07567 | 2.34536 | 2.180 |
| chr9 | 77259845 | 77260269 | 1.05282 | 2.29568 | 2.181 |
| chr19 | 48027004 | 48027233 | 0.51889 | 1.13146 | 2.181 |
| chr7 | 110738731 | 110739072 | 1.02318 | 2.2311 | 2.181 |
| chr15 | 87167833 | 87168166 | 1.37337 | 2.99483 | 2.181 |
| chr20 | 5866628 | 5866930 | 0.502941 | 1.09674 | 2.181 |
| chr1 | 211432251 | 211432634 | 1.77468 | 3.87011 | 2.181 |
| chr20 | 21485780 | 21486168 | 2.33746 | 5.09741 | 2.181 |
| chr14 | 61848493 | 61848717 | 0.444556 | 0.969485 | 2.181 |
| chr9 | 105125649 | 105125886 | 0.44826 | 0.977644 | 2.181 |
| chr2 | 226533403 | 226533670 | 1.56233 | 3.40749 | 2.181 |
| chr7 | 139286896 | 139287424 | 1.21749 | 2.65546 | 2.181 |
| chr18 | 56435665 | 56436017 | 1.60168 | 3.49345 | 2.181 |
| chr1 | 202821476 | 202821947 | 1.24906 | 2.72452 | 2.181 |
| chr12 | 128771181 | 128771478 | 0.619191 | 1.35068 | 2.181 |
| chr5 | 88025490 | 88025711 | 1.05546 | 2.30234 | 2.181 |

| | | | | | |
|---|---|---|---|---|---|
| chr11 | 56392115 | 56392324 | 0.741083 | 1.61658 | 2.181 |
| chr16 | 2106103 | 2106540 | 0.918635 | 2.0039 | 2.181 |
| chr17 | 72888113 | 72888427 | 0.743634 | 1.62216 | 2.181 |
| chr5 | 172366068 | 172366656 | 0.986529 | 2.15209 | 2.181 |
| chr7 | 307072 | 307337 | 0.974891 | 2.12675 | 2.182 |
| chr7 | 128702958 | 128703278 | 0.55997 | 1.22159 | 2.182 |
| chr1 | 183578448 | 183578738 | 1.27244 | 2.77593 | 2.182 |
| chr1 | 160053845 | 160054547 | 1.4571 | 3.17882 | 2.182 |
| chr11 | 70357611 | 70357844 | 0.845037 | 1.84358 | 2.182 |
| chr1 | 178470363 | 178471234 | 1.55804 | 3.39915 | 2.182 |
| chr5 | 32942615 | 32942856 | 1.39042 | 3.03353 | 2.182 |
| chr5 | 176127305 | 176127587 | 0.995985 | 2.17309 | 2.182 |
| chr10 | 15469884 | 15470151 | 1.01588 | 2.21657 | 2.182 |
| chr8 | 19616028 | 19616357 | 1.3658 | 2.98008 | 2.182 |
| chr10 | 53639309 | 53639658 | 0.535912 | 1.16936 | 2.182 |
| chr22 | 21316103 | 21316454 | 1.32601 | 2.89338 | 2.182 |
| chr3 | 32281543 | 32281855 | 0.829411 | 1.80984 | 2.182 |
| chr1 | 154916295 | 154916594 | 1.49545 | 3.2632 | 2.182 |
| chr11 | 27545836 | 27546108 | 0.886986 | 1.93556 | 2.182 |
| chr10 | 22021090 | 22021443 | 2.26568 | 4.94452 | 2.182 |
| chr17 | 47785603 | 47785936 | 3.05806 | 6.6738 | 2.182 |
| chr16 | 4598822 | 4599140 | 1.69278 | 3.69432 | 2.182 |
| chr7 | 99080867 | 99081326 | 0.562825 | 1.22831 | 2.182 |
| chr16 | 83373171 | 83373417 | 0.464932 | 1.01468 | 2.182 |
| chrX | 45661074 | 45661302 | 0.915976 | 1.99908 | 2.182 |
| chr7 | 26540426 | 26540673 | 1.21493 | 2.65159 | 2.183 |
| chr7 | 121036258 | 121036624 | 1.69015 | 3.68882 | 2.183 |
| chr9 | 97365570 | 97365892 | 1.32043 | 2.88197 | 2.183 |
| chr8 | 142179118 | 142179647 | 1.78448 | 3.89519 | 2.183 |
| chr16 | 53926043 | 53926289 | 1.57624 | 3.44067 | 2.183 |
| chr11 | 19171306 | 19171607 | 0.448756 | 0.979642 | 2.183 |
| chr17 | 48073844 | 48074455 | 1.86092 | 4.06255 | 2.183 |
| chr12 | 6074894 | 6075435 | 1.56859 | 3.4245 | 2.183 |
| chr20 | 48236808 | 48237060 | 0.465868 | 1.01708 | 2.183 |
| chr2 | 60461562 | 60461903 | 0.490415 | 1.07069 | 2.183 |
| chr3 | 188672420 | 188672877 | 1.38501 | 3.02402 | 2.183 |
| chrX | 131156172 | 131156400 | 0.86224 | 1.88262 | 2.183 |
| chr16 | 3068159 | 3068382 | 1.13819 | 2.48519 | 2.183 |
| chr16 | 1862674 | 1862987 | 1.6563 | 3.61673 | 2.184 |
| chr7 | 31017498 | 31017905 | 1.82357 | 3.98204 | 2.184 |
| chr16 | 74846941 | 74847426 | 2.834 | 6.18852 | 2.184 |
| chr17 | 15160580 | 15161058 | 2.13725 | 4.66718 | 2.184 |
| chr9 | 134406051 | 134406335 | 0.689619 | 1.50603 | 2.184 |
| chr12 | 101884168 | 101884752 | 0.8173 | 1.78488 | 2.184 |
| chr1 | 27189208 | 27189511 | 1.00318 | 2.19088 | 2.184 |
| chr1 | 189828572 | 189828870 | 1.43779 | 3.14024 | 2.184 |
| chr22 | 23728075 | 23728451 | 2.20467 | 4.81529 | 2.184 |
| chr16 | 1152916 | 1153272 | 1.24365 | 2.71636 | 2.184 |
| chr12 | 52463273 | 52463541 | 2.29176 | 5.00571 | 2.184 |
| chr8 | 20089239 | 20089639 | 1.90918 | 4.1704 | 2.184 |
| chrX | 135245023 | 135245478 | 1.0372 | 2.26569 | 2.184 |
| chr5 | 58330315 | 58330519 | 1.0771 | 2.35289 | 2.184 |
| chr8 | 70023767 | 70024036 | 0.476372 | 1.04063 | 2.184 |
| chr10 | 1757911 | 1758318 | 1.78188 | 3.89257 | 2.185 |
| chr18 | 73968853 | 73969206 | 1.07538 | 2.34932 | 2.185 |
| chr13 | 97670367 | 97670709 | 1.26234 | 2.75777 | 2.185 |
| chr6 | 74513534 | 74513804 | 1.19055 | 2.60093 | 2.185 |
| chr8 | 3848408 | 3848651 | 0.451572 | 0.986567 | 2.185 |
| chr17 | 33325139 | 33325599 | 2.41849 | 5.28381 | 2.185 |
| chr17 | 65599191 | 65599417 | 0.992502 | 2.16839 | 2.185 |
| chr4 | 166414528 | 166414816 | 1.01441 | 2.21629 | 2.185 |

| | | | | | |
|---|---|---|---|---|---|
| chr17 | 39747722 | 39747970 | 0.551245 | 1.20439 | 2.185 |
| chr15 | 80142177 | 80142458 | 1.61456 | 3.52771 | 2.185 |
| chr10 | 97453813 | 97454150 | 2.28479 | 4.99219 | 2.185 |
| chr5 | 83014053 | 83014396 | 1.57637 | 3.44432 | 2.185 |
| chr11 | 124632169 | 124632399 | 0.56053 | 1.22476 | 2.185 |
| chr10 | 121925410 | 121925721 | 0.514329 | 1.12382 | 2.185 |
| chr11 | 45288537 | 45288868 | 1.32573 | 2.89676 | 2.185 |
| chr19 | 51881271 | 51881486 | 0.391266 | 0.854963 | 2.185 |
| chr14 | 24423169 | 24423507 | 1.51673 | 3.3143 | 2.185 |
| chr2 | 33171238 | 33171687 | 1.7779 | 3.88506 | 2.185 |
| chr9 | 114454132 | 114454340 | 0.820285 | 1.79249 | 2.185 |
| chr15 | 62363948 | 62364289 | 1.11158 | 2.42931 | 2.185 |
| chr19 | 48763442 | 48763651 | 1.79955 | 3.93285 | 2.185 |
| chr8 | 93022441 | 93022755 | 0.595163 | 1.30072 | 2.185 |
| chr1 | 168051434 | 168051854 | 1.92262 | 4.20191 | 2.186 |
| chr14 | 35911445 | 35911963 | 0.60375 | 1.31951 | 2.186 |
| chr8 | 10589999 | 10590302 | 1.00835 | 2.20391 | 2.186 |
| chr19 | 37430172 | 37430416 | 1.06143 | 2.32011 | 2.186 |
| chr1 | 229556547 | 229556819 | 0.589837 | 1.28931 | 2.186 |
| chr15 | 34659544 | 34660139 | 2.0941 | 4.57753 | 2.186 |
| chr7 | 44347043 | 44347290 | 0.722852 | 1.58012 | 2.186 |
| chr15 | 100705295 | 100705782 | 0.492634 | 1.07696 | 2.186 |
| chr1 | 31192103 | 31192402 | 1.21004 | 2.64568 | 2.186 |
| chr19 | 10683070 | 10683407 | 2.21495 | 4.84335 | 2.187 |
| chrX | 27204767 | 27205112 | 1.15372 | 2.52279 | 2.187 |
| chr3 | 45698315 | 45698563 | 2.434 | 5.32236 | 2.187 |
| chr19 | 1315655 | 1316016 | 1.27523 | 2.78856 | 2.187 |
| chr4 | 40843045 | 40843394 | 1.49884 | 3.27754 | 2.187 |
| chr17 | 13126395 | 13126667 | 0.509299 | 1.11372 | 2.187 |
| chrX | 134730518 | 134730827 | 1.05275 | 2.30219 | 2.187 |
| chr16 | 85309164 | 85309682 | 2.4445 | 5.34575 | 2.187 |
| chr5 | 160534001 | 160534245 | 0.438728 | 0.959431 | 2.187 |
| chr17 | 68368642 | 68368969 | 0.59401 | 1.29902 | 2.187 |
| chr17 | 57394303 | 57394673 | 1.03661 | 2.26705 | 2.187 |
| chr2 | 80162918 | 80163203 | 0.950095 | 2.07785 | 2.187 |
| chr3 | 42872787 | 42873195 | 0.962559 | 2.10519 | 2.187 |
| chr5 | 137687993 | 137688953 | 3.04454 | 6.65888 | 2.187 |
| chr2 | 17120479 | 17120777 | 0.414952 | 0.907608 | 2.187 |
| chr11 | 14730991 | 14731236 | 0.50072 | 1.09522 | 2.187 |
| chr12 | 120904387 | 120904644 | 1.31727 | 2.88128 | 2.187 |
| chr19 | 45227163 | 45227571 | 2.09949 | 4.59242 | 2.187 |
| chr2 | 3619502 | 3619728 | 0.621988 | 1.36054 | 2.187 |
| chr2 | 46843102 | 46843346 | 1.09729 | 2.40024 | 2.187 |
| chr12 | 3678720 | 3678953 | 0.402779 | 0.881099 | 2.188 |
| chr19 | 45634242 | 45634709 | 1.11062 | 2.42962 | 2.188 |
| chr13 | 80604736 | 80604967 | 0.823683 | 1.80193 | 2.188 |
| chr2 | 27745300 | 27745732 | 0.515576 | 1.12797 | 2.188 |
| chr5 | 32232511 | 32232894 | 1.61976 | 3.54372 | 2.188 |
| chr15 | 89714648 | 89715057 | 1.58844 | 3.47558 | 2.188 |
| chr19 | 50440876 | 50441339 | 2.31745 | 5.07069 | 2.188 |
| chrY | 7212792 | 7213006 | 0.805073 | 1.76156 | 2.188 |
| chr8 | 143528729 | 143529279 | 2.10013 | 4.59536 | 2.188 |
| chr10 | 75973371 | 75973633 | 1.24416 | 2.72242 | 2.188 |
| chr2 | 13334073 | 13334313 | 1.00099 | 2.19034 | 2.188 |
| chr20 | 23587189 | 23587539 | 1.85636 | 4.06209 | 2.188 |
| chr16 | 46887866 | 46888200 | 1.86301 | 4.07689 | 2.188 |
| chr5 | 67957176 | 67957393 | 0.6266 | 1.37122 | 2.188 |
| chr17 | 8897565 | 8898028 | 1.81597 | 3.97401 | 2.188 |
| chr1 | 224645259 | 224645498 | 0.953783 | 2.08727 | 2.188 |
| chr11 | 36398794 | 36399177 | 1.78909 | 3.9154 | 2.188 |
| chr7 | 23736610 | 23736989 | 1.56162 | 3.41779 | 2.189 |

| | | | | | |
|---|---|---|---|---|---|
| chr20 | 25057405 | 25057658 | 1.11036 | 2.43022 | 2.189 |
| chr20 | 54775502 | 54775841 | 0.963377 | 2.10864 | 2.189 |
| chr8 | 14740124 | 14740374 | 0.547273 | 1.19789 | 2.189 |
| chr18 | 70803434 | 70803776 | 0.957255 | 2.09534 | 2.189 |
| chr5 | 122291317 | 122291605 | 0.490333 | 1.07331 | 2.189 |
| chr20 | 62451099 | 62451335 | 1.08425 | 2.37345 | 2.189 |
| chr12 | 128292603 | 128292914 | 0.570762 | 1.24945 | 2.189 |
| chr10 | 121066209 | 121066432 | 1.02622 | 2.24651 | 2.189 |
| chr12 | 97515027 | 97515285 | 0.473874 | 1.0374 | 2.189 |
| chr14 | 107210698 | 107211014 | 1.26368 | 2.76644 | 2.189 |
| chr18 | 33336495 | 33336785 | 1.026 | 2.24626 | 2.189 |
| chr17 | 50603764 | 50603989 | 0.342405 | 0.749647 | 2.189 |
| chr17 | 79864796 | 79865022 | 0.794957 | 1.74046 | 2.189 |
| chr5 | 86266754 | 86267053 | 0.577768 | 1.26496 | 2.189 |
| chr9 | 132488893 | 132489273 | 3.57143 | 7.81949 | 2.189 |
| chr14 | 65266795 | 65267106 | 1.5406 | 3.37317 | 2.190 |
| chr2 | 159580762 | 159581070 | 1.35043 | 2.9568 | 2.190 |
| chr8 | 72887441 | 72887866 | 1.63693 | 3.58423 | 2.190 |
| chr3 | 187465819 | 187466152 | 2.22925 | 4.88121 | 2.190 |
| chr10 | 28063061 | 28063333 | 0.447373 | 0.979586 | 2.190 |
| chrX | 150921828 | 150922237 | 1.78259 | 3.90345 | 2.190 |
| chr11 | 75434223 | 75434693 | 1.49621 | 3.27636 | 2.190 |
| chr17 | 9458187 | 9458411 | 1.6084 | 3.5223 | 2.190 |
| chr12 | 6387856 | 6388298 | 2.00615 | 4.39338 | 2.190 |
| chr1 | 161733363 | 161733636 | 1.50939 | 3.30564 | 2.190 |
| chr1 | 78149089 | 78149418 | 1.64956 | 3.61267 | 2.190 |
| chr16 | 13734912 | 13735135 | 0.388617 | 0.851113 | 2.190 |
| chr18 | 71510079 | 71510394 | 2.0785 | 4.55216 | 2.190 |
| chr8 | 135855914 | 135856322 | 0.793658 | 1.73831 | 2.190 |
| chr13 | 110761310 | 110761550 | 1.89156 | 4.14317 | 2.190 |
| chr11 | 32802461 | 32802727 | 0.503001 | 1.10176 | 2.190 |
| chr1 | 181111636 | 181111938 | 1.38085 | 3.02461 | 2.190 |
| chr4 | 146155330 | 146155568 | 1.30063 | 2.84897 | 2.190 |
| chr2 | 180568 | 180797 | 0.590285 | 1.29302 | 2.191 |
| chr12 | 128588903 | 128589233 | 0.547224 | 1.19874 | 2.191 |
| chr1 | 221651383 | 221651700 | 1.49702 | 3.27939 | 2.191 |
| chr6 | 164377693 | 164377943 | 1.62356 | 3.55661 | 2.191 |
| chr3 | 81839725 | 81840071 | 0.493847 | 1.08187 | 2.191 |
| chr5 | 154011422 | 154011647 | 0.69608 | 1.52492 | 2.191 |
| chr8 | 117863450 | 117863779 | 1.00265 | 2.19654 | 2.191 |
| chr3 | 133103255 | 133103643 | 1.03629 | 2.27024 | 2.191 |
| chr2 | 27309337 | 27309681 | 2.13122 | 4.669 | 2.191 |
| chr19 | 18460378 | 18460712 | 1.51933 | 3.32871 | 2.191 |
| chr1 | 16405419 | 16405709 | 1.2089 | 2.64863 | 2.191 |
| chr11 | 125439080 | 125439312 | 0.860869 | 1.88611 | 2.191 |
| chr2 | 192623664 | 192623935 | 1.10833 | 2.42837 | 2.191 |
| chr19 | 29884714 | 29884927 | 0.388408 | 0.851025 | 2.191 |
| chr15 | 63720304 | 63720539 | 0.585666 | 1.28325 | 2.191 |
| chr7 | 147574003 | 147574258 | 0.459529 | 1.00693 | 2.191 |
| chr17 | 69671390 | 69671825 | 0.658114 | 1.44209 | 2.191 |
| chr15 | 53879031 | 53879341 | 1.52148 | 3.33408 | 2.191 |
| chr10 | 124543430 | 124543724 | 0.515569 | 1.12982 | 2.191 |
| chr1 | 22458325 | 22458782 | 1.9028 | 4.16989 | 2.191 |
| chr10 | 73456394 | 73456820 | 2.27327 | 4.98176 | 2.191 |
| chr7 | 46128035 | 46128357 | 0.332221 | 0.728065 | 2.192 |
| chr22 | 23219674 | 23219937 | 2.1382 | 4.68599 | 2.192 |
| chr13 | 27546266 | 27546647 | 1.2604 | 2.76236 | 2.192 |
| chr12 | 125815416 | 125816113 | 2.28027 | 4.99761 | 2.192 |
| chr7 | 1620166 | 1620398 | 0.535704 | 1.17413 | 2.192 |
| chr3 | 49059317 | 49059630 | 1.34002 | 2.9372 | 2.192 |
| chr10 | 1146937 | 1147212 | 0.68133 | 1.49343 | 2.192 |

| | | | | | |
|---|---|---|---|---|---|
| chr12 | 116268567 | 116268814 | 0.449703 | 0.985728 | 2.192 |
| chr7 | 114505683 | 114505953 | 1.82152 | 3.99271 | 2.192 |
| chr5 | 157421909 | 157422263 | 1.34862 | 2.95622 | 2.192 |
| chr9 | 131768875 | 131769575 | 1.87785 | 4.11655 | 2.192 |
| chrX | 51203977 | 51204224 | 1.08468 | 2.37792 | 2.192 |
| chr3 | 42232646 | 42233046 | 1.58445 | 3.4737 | 2.192 |
| chr20 | 24776215 | 24776419 | 0.393574 | 0.862885 | 2.192 |
| chr6 | 36107334 | 36107556 | 0.616015 | 1.35057 | 2.192 |
| chr10 | 56176813 | 56177025 | 0.478621 | 1.04936 | 2.192 |
| chr11 | 45792774 | 45793836 | 1.37444 | 3.01343 | 2.192 |
| chr1 | 10848706 | 10849182 | 0.886452 | 1.94366 | 2.193 |
| chr16 | 84705850 | 84706290 | 2.29859 | 5.03998 | 2.193 |
| chr19 | 42750341 | 42750666 | 1.50584 | 3.30178 | 2.193 |
| chr7 | 149194723 | 149195157 | 1.16733 | 2.55961 | 2.193 |
| chr2 | 217552442 | 217552699 | 1.26008 | 2.76314 | 2.193 |
| chr8 | 141554162 | 141554415 | 0.441016 | 0.967089 | 2.193 |
| chr14 | 73371595 | 73371970 | 1.47185 | 3.22763 | 2.193 |
| chr1 | 67191705 | 67192095 | 1.80415 | 3.95663 | 2.193 |
| chrX | 47171881 | 47172084 | 1.21294 | 2.66009 | 2.193 |
| chr11 | 64320871 | 64321369 | 1.79852 | 3.94448 | 2.193 |
| chr1 | 76619801 | 76620153 | 1.70637 | 3.74244 | 2.193 |
| chr5 | 60275521 | 60275725 | 0.482915 | 1.05915 | 2.193 |
| chr14 | 58041722 | 58041947 | 1.07655 | 2.36117 | 2.193 |
| chr11 | 108605723 | 108606048 | 0.842531 | 1.84792 | 2.193 |
| chr16 | 21579920 | 21580185 | 0.526938 | 1.15578 | 2.193 |
| chr14 | 46057897 | 46058220 | 0.474336 | 1.04043 | 2.193 |
| chr7 | 5534229 | 5534653 | 2.05437 | 4.50628 | 2.194 |
| chr1 | 87430943 | 87431239 | 1.33995 | 2.93925 | 2.194 |
| chr1 | 156426209 | 156426533 | 1.7685 | 3.87931 | 2.194 |
| chr5 | 158122558 | 158122877 | 1.67185 | 3.66735 | 2.194 |
| chr3 | 190418870 | 190419142 | 1.35658 | 2.97599 | 2.194 |
| chr9 | 34701245 | 34701756 | 2.15609 | 4.72997 | 2.194 |
| chr13 | 44624695 | 44624922 | 1.34419 | 2.94888 | 2.194 |
| chr19 | 48854514 | 48855003 | 1.85656 | 4.07298 | 2.194 |
| chr7 | 28936729 | 28937001 | 0.375911 | 0.824685 | 2.194 |
| chr21 | 34025712 | 34025966 | 1.29189 | 2.83423 | 2.194 |
| chr5 | 164667651 | 164667962 | 0.468114 | 1.0271 | 2.194 |
| chr12 | 66526748 | 66526975 | 0.6471 | 1.41985 | 2.194 |
| chr10 | 6684681 | 6684949 | 0.45569 | 0.999887 | 2.194 |
| chr3 | 40761436 | 40761644 | 0.580932 | 1.27472 | 2.194 |
| chr11 | 72125183 | 72125535 | 1.83822 | 4.03357 | 2.194 |
| chr3 | 152679069 | 152679292 | 0.950577 | 2.0859 | 2.194 |
| chr10 | 109824444 | 109824750 | 0.452953 | 0.993945 | 2.194 |
| chr7 | 134117277 | 134117773 | 1.11976 | 2.45723 | 2.194 |
| chr2 | 62684085 | 62684467 | 1.75976 | 3.86204 | 2.195 |
| chr1 | 37920462 | 37920823 | 1.88011 | 4.12626 | 2.195 |
| chr12 | 48099277 | 48099614 | 1.61506 | 3.54455 | 2.195 |
| chr10 | 134938194 | 134938626 | 2.05298 | 4.50586 | 2.195 |
| chr12 | 57518036 | 57518429 | 2.14998 | 4.71876 | 2.195 |
| chr5 | 17951502 | 17951723 | 0.686667 | 1.50715 | 2.195 |
| chr6 | 79149207 | 79149414 | 0.461493 | 1.01293 | 2.195 |
| chr3 | 122473762 | 122474089 | 1.39372 | 3.05924 | 2.195 |
| chr6 | 12894081 | 12894391 | 0.891577 | 1.95704 | 2.195 |
| chr7 | 32467390 | 32467760 | 0.551315 | 1.21017 | 2.195 |
| chr9 | 20941097 | 20941412 | 1.54444 | 3.39039 | 2.195 |
| chr10 | 73576886 | 73577174 | 0.513661 | 1.12765 | 2.195 |
| chr22 | 47194435 | 47194841 | 1.65056 | 3.6236 | 2.195 |
| chr7 | 151048157 | 151048454 | 1.46368 | 3.21347 | 2.195 |
| chr1 | 89073897 | 89074106 | 0.805026 | 1.76759 | 2.196 |
| chr5 | 142529849 | 142530062 | 0.591656 | 1.29921 | 2.196 |
| chr11 | 31213883 | 31214159 | 0.564832 | 1.24035 | 2.196 |

| | | | | | |
|---|---|---|---|---|---|
| chr11 | 125215628 | 125215944 | 1.60987 | 3.53527 | 2.196 |
| chr2 | 86041353 | 86041705 | 2.18684 | 4.80249 | 2.196 |
| chr1 | 194435909 | 194436144 | 0.934843 | 2.05304 | 2.196 |
| chr7 | 68868246 | 68868482 | 0.928356 | 2.0388 | 2.196 |
| chr10 | 59069203 | 59069419 | 0.501035 | 1.10038 | 2.196 |
| chr14 | 22409181 | 22409416 | 1.00657 | 2.21066 | 2.196 |
| chr3 | 187036708 | 187036995 | 0.921657 | 2.02419 | 2.196 |
| chr17 | 79865659 | 79865918 | 0.672288 | 1.47655 | 2.196 |
| chr12 | 129775316 | 129775721 | 0.425914 | 0.935449 | 2.196 |
| chr12 | 53493245 | 53493523 | 2.12558 | 4.66868 | 2.196 |
| chr19 | 46902511 | 46902721 | 1.09448 | 2.40401 | 2.196 |
| chr14 | 69926881 | 69927139 | 1.62215 | 3.56305 | 2.197 |
| chr3 | 88662441 | 88662686 | 0.584848 | 1.28463 | 2.197 |
| chr12 | 103942450 | 103942725 | 1.61707 | 3.55194 | 2.197 |
| chr17 | 40202361 | 40202760 | 1.6018 | 3.51854 | 2.197 |
| chr5 | 139069553 | 139069919 | 0.899244 | 1.97534 | 2.197 |
| chr3 | 15321485 | 15321778 | 0.478504 | 1.05114 | 2.197 |
| chr6 | 169726538 | 169726784 | 0.864338 | 1.89879 | 2.197 |
| chr8 | 51816497 | 51816713 | 0.506783 | 1.11336 | 2.197 |
| chr22 | 21983550 | 21983830 | 1.24967 | 2.74548 | 2.197 |
| chr10 | 26624039 | 26624370 | 1.44893 | 3.18326 | 2.197 |
| chr6 | 113101762 | 113102009 | 1.25374 | 2.75443 | 2.197 |
| chr1 | 15650062 | 15650356 | 1.937 | 4.25567 | 2.197 |
| chr19 | 19477657 | 19478170 | 1.81501 | 3.98776 | 2.197 |
| chr17 | 25676085 | 25676586 | 2.05574 | 4.51676 | 2.197 |
| chr3 | 183187208 | 183187444 | 1.34586 | 2.95707 | 2.197 |
| chr5 | 10819018 | 10819275 | 0.424405 | 0.932489 | 2.197 |
| chr17 | 54857513 | 54857993 | 2.86104 | 6.2866 | 2.197 |
| chr4 | 79533600 | 79533845 | 0.740758 | 1.62768 | 2.197 |
| chr16 | 81817520 | 81817843 | 0.433863 | 0.9534 | 2.197 |
| chr9 | 89704279 | 89704623 | 0.526541 | 1.15706 | 2.197 |
| chr14 | 91030837 | 91031056 | 0.46247 | 1.01628 | 2.198 |
| chr11 | 9050879 | 9051182 | 0.450185 | 0.989289 | 2.198 |
| chr9 | 128190301 | 128190951 | 0.961552 | 2.11327 | 2.198 |
| chr10 | 4193638 | 4193950 | 0.943992 | 2.07484 | 2.198 |
| chr3 | 112754293 | 112754583 | 1.53178 | 3.36679 | 2.198 |
| chr11 | 68024691 | 68025148 | 1.29703 | 2.85096 | 2.198 |
| chr19 | 45949557 | 45949935 | 0.720339 | 1.58342 | 2.198 |
| chr10 | 100802521 | 100802807 | 0.874485 | 1.92232 | 2.198 |
| chr17 | 28422768 | 28423017 | 0.87237 | 1.91774 | 2.198 |
| chr4 | 87419769 | 87420053 | 1.46546 | 3.22163 | 2.198 |
| chr4 | 169631567 | 169631773 | 1.13806 | 2.50199 | 2.198 |
| chr9 | 22244038 | 22244445 | 1.07895 | 2.37207 | 2.199 |
| chr8 | 108854890 | 108855227 | 0.295421 | 0.649489 | 2.199 |
| chr9 | 94422060 | 94422447 | 0.760419 | 1.67187 | 2.199 |
| chr16 | 2283817 | 2284047 | 0.586439 | 1.28937 | 2.199 |
| chr12 | 110035457 | 110035874 | 2.87984 | 6.33196 | 2.199 |
| chr15 | 98418792 | 98419065 | 1.02288 | 2.24906 | 2.199 |
| chr9 | 17964283 | 17964486 | 0.49546 | 1.08943 | 2.199 |
| chr1 | 9170870 | 9171227 | 1.53672 | 3.37905 | 2.199 |
| chr17 | 56326892 | 56327530 | 2.07052 | 4.55284 | 2.199 |
| chr14 | 69291509 | 69292066 | 1.59832 | 3.51455 | 2.199 |
| chr17 | 48772531 | 48772754 | 1.36516 | 3.00191 | 2.199 |
| chr8 | 127711292 | 127711509 | 1.51743 | 3.33674 | 2.199 |
| chr1 | 79962897 | 79963183 | 1.42077 | 3.12426 | 2.199 |
| chr14 | 20999143 | 20999473 | 1.7505 | 3.8496 | 2.199 |
| chrX | 109134189 | 109134460 | 1.50309 | 3.30555 | 2.199 |
| chr22 | 30683821 | 30684228 | 0.748567 | 1.64626 | 2.199 |
| chr12 | 94523153 | 94523589 | 0.567483 | 1.24806 | 2.199 |
| chr19 | 56164938 | 56165327 | 3.2001 | 7.03822 | 2.199 |
| chr7 | 106655286 | 106655569 | 0.349958 | 0.76969 | 2.199 |

| | | | | | |
|---|---|---|---|---|---|
| chr12 | 26211314 | 26211535 | 0.7791 | 1.7136 | 2.199 |
| chr9 | 82761423 | 82761726 | 0.914877 | 2.01234 | 2.200 |
| chr1 | 212489252 | 212489457 | 0.775637 | 1.7061 | 2.200 |
| chr14 | 37125068 | 37125658 | 1.95578 | 4.30242 | 2.200 |
| chr8 | 105504558 | 105504760 | 0.542947 | 1.19442 | 2.200 |
| chr14 | 59950482 | 59951026 | 0.965178 | 2.12336 | 2.200 |
| chr15 | 99062329 | 99062642 | 1.32311 | 2.91081 | 2.200 |
| chr10 | 15148390 | 15148735 | 1.82715 | 4.01988 | 2.200 |
| chr12 | 53345950 | 53346279 | 1.17789 | 2.59157 | 2.200 |
| chr3 | 190102093 | 190102377 | 1.79684 | 3.95338 | 2.200 |
| chr15 | 38706561 | 38706938 | 0.955289 | 2.10182 | 2.200 |
| chr4 | 68193041 | 68193293 | 1.35028 | 2.97087 | 2.200 |
| chr17 | 46088727 | 46088985 | 1.23315 | 2.71325 | 2.200 |
| chr15 | 29267918 | 29268197 | 1.47803 | 3.25225 | 2.200 |
| chr4 | 160096756 | 160096972 | 1.34161 | 2.95215 | 2.200 |
| chr20 | 60951592 | 60951839 | 0.9556 | 2.10282 | 2.201 |
| chr3 | 33414340 | 33414686 | 1.8027 | 3.96693 | 2.201 |
| chr4 | 129495864 | 129496116 | 0.937307 | 2.06259 | 2.201 |
| chr11 | 61715097 | 61715475 | 2.17955 | 4.79633 | 2.201 |
| chr3 | 88839056 | 88839299 | 0.52721 | 1.16019 | 2.201 |
| chr19 | 29471115 | 29471455 | 0.772501 | 1.70003 | 2.201 |
| chr1 | 161162221 | 161162659 | 2.03315 | 4.47457 | 2.201 |
| chr11 | 12915670 | 12915951 | 1.47663 | 3.24983 | 2.201 |
| chr14 | 96731250 | 96731561 | 1.24874 | 2.74828 | 2.201 |
| chr20 | 38406265 | 38406508 | 0.454794 | 1.00093 | 2.201 |
| chr14 | 21572653 | 21572958 | 1.40111 | 3.08363 | 2.201 |
| chr11 | 46269172 | 46269458 | 1.05819 | 2.32894 | 2.201 |
| chr5 | 104889484 | 104889799 | 0.51748 | 1.13892 | 2.201 |
| chr5 | 13335787 | 13336071 | 0.526308 | 1.15839 | 2.201 |
| chr1 | 179783841 | 179784194 | 1.43412 | 3.15652 | 2.201 |
| chr13 | 26758870 | 26759244 | 1.37023 | 3.01593 | 2.201 |
| chr2 | 210288343 | 210288830 | 2.14886 | 4.72984 | 2.201 |
| chr9 | 96299204 | 96299426 | 0.581719 | 1.28043 | 2.201 |
| chr9 | 115649569 | 115649911 | 0.590913 | 1.30069 | 2.201 |
| chr5 | 157389645 | 157389864 | 0.318364 | 0.700771 | 2.201 |
| chr10 | 61590439 | 61590838 | 1.1236 | 2.47336 | 2.201 |
| chr12 | 61353682 | 61353969 | 0.497469 | 1.09509 | 2.201 |
| chr8 | 103045061 | 103045375 | 0.854097 | 1.88014 | 2.201 |
| chr11 | 66481064 | 66481306 | 0.598931 | 1.31848 | 2.201 |
| chr9 | 97858803 | 97859013 | 0.948014 | 2.08699 | 2.201 |
| chr12 | 57849944 | 57850353 | 1.32212 | 2.91063 | 2.201 |
| chr3 | 112770091 | 112770417 | 1.61695 | 3.55976 | 2.202 |
| chr9 | 118355840 | 118356122 | 1.15188 | 2.536 | 2.202 |
| chr6 | 42844851 | 42845117 | 0.837571 | 1.84402 | 2.202 |
| chr2 | 25419273 | 25419725 | 1.75203 | 3.85742 | 2.202 |
| chr18 | 20657423 | 20657625 | 0.892274 | 1.96457 | 2.202 |
| chr8 | 89428592 | 89428812 | 0.537595 | 1.18366 | 2.202 |
| chr19 | 48005934 | 48006759 | 0.630051 | 1.38725 | 2.202 |
| chr2 | 39957914 | 39958160 | 1.30724 | 2.87832 | 2.202 |
| chr13 | 27447037 | 27447322 | 1.18736 | 2.61448 | 2.202 |
| chr10 | 120966161 | 120966587 | 1.55635 | 3.42699 | 2.202 |
| chrX | 107978384 | 107978654 | 0.797251 | 1.7555 | 2.202 |
| chr5 | 131722315 | 131722545 | 1.58205 | 3.484 | 2.202 |
| chr7 | 51434763 | 51435073 | 1.07095 | 2.35849 | 2.202 |
| chr15 | 78469661 | 78470191 | 2.20011 | 4.84528 | 2.202 |
| chr5 | 133204890 | 133205316 | 1.18596 | 2.61189 | 2.202 |
| chr17 | 42193033 | 42193424 | 1.89042 | 4.16338 | 2.202 |
| chr5 | 96845052 | 96845306 | 0.618314 | 1.36178 | 2.202 |
| chr5 | 103778073 | 103778317 | 0.56362 | 1.24132 | 2.202 |
| chrX | 132973181 | 132973428 | 1.84575 | 4.06517 | 2.202 |
| chr12 | 23880967 | 23881269 | 0.466168 | 1.02673 | 2.202 |

| | | | | | |
|---|---|---|---|---|---|
| chr3 | 108800670 | 108801245 | 1.41526 | 3.11709 | 2.202 |
| chr6 | 35208845 | 35209254 | 1.65704 | 3.64963 | 2.203 |
| chr7 | 158009849 | 158010252 | 0.583449 | 1.28509 | 2.203 |
| chr17 | 3749464 | 3749793 | 2.56234 | 5.64425 | 2.203 |
| chr5 | 17216274 | 17216525 | 1.42886 | 3.14757 | 2.203 |
| chr13 | 95948801 | 95949138 | 1.40249 | 3.08952 | 2.203 |
| chr6 | 160141734 | 160141940 | 0.674959 | 1.48686 | 2.203 |
| chr5 | 6745751 | 6746171 | 1.42428 | 3.13754 | 2.203 |
| chr8 | 70827160 | 70827443 | 0.44852 | 0.988078 | 2.203 |
| chr9 | 27529651 | 27530055 | 2.05181 | 4.52007 | 2.203 |
| chr6 | 163602211 | 163602578 | 2.87423 | 6.33187 | 2.203 |
| chr8 | 75977410 | 75977619 | 0.445714 | 0.981901 | 2.203 |
| chr11 | 18824237 | 18824568 | 1.54447 | 3.40245 | 2.203 |
| chr20 | 41816256 | 41816761 | 1.76539 | 3.88923 | 2.203 |
| chr7 | 13302656 | 13302863 | 1.14644 | 2.52568 | 2.203 |
| chr7 | 20638270 | 20638473 | 1.40851 | 3.10309 | 2.203 |
| chr7 | 157668167 | 157668420 | 1.96744 | 4.33447 | 2.203 |
| chr19 | 56457234 | 56457441 | 0.942922 | 2.07744 | 2.203 |
| chr5 | 72114751 | 72115102 | 1.50758 | 3.32151 | 2.203 |
| chr9 | 3331902 | 3332123 | 1.10935 | 2.44429 | 2.203 |
| chr9 | 139652742 | 139653172 | 2.52224 | 5.55765 | 2.203 |
| chr10 | 77780387 | 77780659 | 0.478918 | 1.05532 | 2.204 |
| chr1 | 227917984 | 227918361 | 1.17086 | 2.58006 | 2.204 |
| chr16 | 66349135 | 66349523 | 2.51598 | 5.54435 | 2.204 |
| chr20 | 58560911 | 58561254 | 0.662576 | 1.46012 | 2.204 |
| chr16 | 31708282 | 31708552 | 1.39667 | 3.07795 | 2.204 |
| chr1 | 195909010 | 195909338 | 0.642102 | 1.41514 | 2.204 |
| chr15 | 60812999 | 60813319 | 2.2437 | 4.94535 | 2.204 |
| chr17 | 10887001 | 10887408 | 0.510678 | 1.12563 | 2.204 |
| chr2 | 238766972 | 238767332 | 1.5597 | 3.43796 | 2.204 |
| chr10 | 71152256 | 71152626 | 2.22284 | 4.89988 | 2.204 |
| chr10 | 65832981 | 65833204 | 0.711309 | 1.568 | 2.204 |
| chr13 | 30706702 | 30707002 | 1.45177 | 3.20027 | 2.204 |
| chr12 | 58153137 | 58153509 | 1.26749 | 2.79408 | 2.204 |
| chrX | 106916147 | 106916429 | 1.68143 | 3.70657 | 2.204 |
| chr7 | 81646741 | 81646952 | 0.993893 | 2.19098 | 2.204 |
| chr5 | 137799706 | 137800332 | 1.32787 | 2.92739 | 2.205 |
| chr7 | 104118844 | 104119091 | 0.483613 | 1.06619 | 2.205 |
| chr2 | 181845356 | 181845577 | 1.39498 | 3.07544 | 2.205 |
| chr7 | 760771 | 761009 | 0.476828 | 1.05126 | 2.205 |
| chr7 | 12283756 | 12284105 | 1.1956 | 2.63596 | 2.205 |
| chr12 | 111325756 | 111326317 | 2.43415 | 5.36667 | 2.205 |
| chr12 | 21154851 | 21155110 | 0.624637 | 1.37717 | 2.205 |
| chr22 | 26760455 | 26760749 | 1.08053 | 2.3824 | 2.205 |
| chr20 | 3762147 | 3762447 | 1.3907 | 3.06635 | 2.205 |
| chr16 | 87849248 | 87849550 | 1.03496 | 2.28202 | 2.205 |
| chr15 | 69976415 | 69976773 | 0.46593 | 1.0275 | 2.205 |
| chr6 | 52747207 | 52747410 | 1.46973 | 3.24128 | 2.205 |
| chr17 | 15312230 | 15312439 | 1.1436 | 2.52206 | 2.205 |
| chr8 | 22613572 | 22614010 | 1.61367 | 3.55877 | 2.205 |
| chr7 | 95405305 | 95405564 | 0.421835 | 0.930334 | 2.205 |
| chr20 | 4201825 | 4202347 | 1.81212 | 3.99656 | 2.205 |
| chr19 | 18631147 | 18631355 | 1.33765 | 2.95023 | 2.206 |
| chr5 | 176882239 | 176882637 | 1.28209 | 2.82772 | 2.206 |
| chr14 | 93156707 | 93157124 | 1.60142 | 3.53211 | 2.206 |
| chr16 | 31463171 | 31463469 | 1.00473 | 2.21604 | 2.206 |
| chr11 | 10750573 | 10750817 | 0.432208 | 0.953289 | 2.206 |
| chr2 | 49728278 | 49728515 | 0.591316 | 1.30423 | 2.206 |
| chr2 | 208898909 | 208899134 | 0.933303 | 2.05861 | 2.206 |
| chr22 | 39399044 | 39399399 | 1.33552 | 2.94583 | 2.206 |
| chr8 | 57261814 | 57262027 | 0.402329 | 0.887483 | 2.206 |

| | | | | | |
|---|---|---|---|---|---|
| chr7 | 129465582 | 129465930 | 2.22695 | 4.91236 | 2.206 |
| chr17 | 54245219 | 54245524 | 0.601149 | 1.32615 | 2.206 |
| chr10 | 10240700 | 10240917 | 0.545314 | 1.20303 | 2.206 |
| chr11 | 40629351 | 40629572 | 0.64056 | 1.41315 | 2.206 |
| chr17 | 45918573 | 45918989 | 1.9359 | 4.27118 | 2.206 |
| chr8 | 34342593 | 34342839 | 0.500323 | 1.10387 | 2.206 |
| chr20 | 17210957 | 17211159 | 0.415995 | 0.917836 | 2.206 |
| chr16 | 82271565 | 82271786 | 0.601115 | 1.3263 | 2.206 |
| chr17 | 73629820 | 73630248 | 1.67357 | 3.6927 | 2.206 |
| chr2 | 86252467 | 86252701 | 0.838593 | 1.85038 | 2.207 |
| chr1 | 117267318 | 117267811 | 1.01759 | 2.24536 | 2.207 |
| chr3 | 195586129 | 195586632 | 1.90674 | 4.20731 | 2.207 |
| chr10 | 120958850 | 120959270 | 1.64123 | 3.62161 | 2.207 |
| chr7 | 87849220 | 87849610 | 1.09759 | 2.42205 | 2.207 |
| chr8 | 142309530 | 142309782 | 0.863331 | 1.90515 | 2.207 |
| chrX | 40974341 | 40974662 | 1.59652 | 3.52312 | 2.207 |
| chr9 | 90589689 | 90589916 | 0.934364 | 2.06197 | 2.207 |
| chr1 | 165236785 | 165237085 | 0.437572 | 0.965647 | 2.207 |
| chr11 | 64661693 | 64662106 | 2.21671 | 4.89204 | 2.207 |
| chr7 | 128038583 | 128039067 | 1.7349 | 3.82883 | 2.207 |
| chr5 | 36307535 | 36307768 | 0.466438 | 1.02941 | 2.207 |
| chr3 | 8343622 | 8344218 | 1.11045 | 2.45073 | 2.207 |
| chr1 | 153721905 | 153722304 | 1.9155 | 4.22749 | 2.207 |
| chr2 | 190663544 | 190663764 | 0.65143 | 1.43774 | 2.207 |
| chr4 | 48946701 | 48947012 | 1.151 | 2.54055 | 2.207 |
| chr15 | 81127649 | 81127914 | 0.656969 | 1.4501 | 2.207 |
| chr3 | 131975879 | 131976113 | 0.983782 | 2.17147 | 2.207 |
| chr19 | 42377216 | 42377616 | 1.77413 | 3.91602 | 2.207 |
| chr2 | 73404480 | 73404946 | 2.78534 | 6.14809 | 2.207 |
| chr14 | 24712113 | 24712390 | 1.31031 | 2.89238 | 2.207 |
| chr11 | 68151428 | 68151783 | 1.93294 | 4.26682 | 2.207 |
| chr13 | 62932310 | 62932574 | 0.871778 | 1.92441 | 2.207 |
| chr11 | 1661299 | 1661730 | 2.1757 | 4.80298 | 2.208 |
| chr11 | 63972056 | 63972386 | 1.37274 | 3.03042 | 2.208 |
| chr16 | 68544017 | 68544449 | 2.61746 | 5.77822 | 2.208 |
| chr16 | 62215767 | 62216003 | 0.601525 | 1.32794 | 2.208 |
| chr9 | 123582788 | 123583273 | 0.482365 | 1.06508 | 2.208 |
| chr12 | 105865683 | 105866012 | 0.586913 | 1.29594 | 2.208 |
| chr2 | 20578645 | 20579147 | 1.92394 | 4.24818 | 2.208 |
| chr20 | 42876772 | 42877033 | 1.04524 | 2.30795 | 2.208 |
| chr19 | 4167587 | 4167850 | 0.934851 | 2.06423 | 2.208 |
| chr20 | 41642891 | 41643183 | 0.534346 | 1.17989 | 2.208 |
| chr8 | 98861903 | 98862185 | 0.985963 | 2.17716 | 2.208 |
| chr5 | 71576559 | 71576903 | 0.765387 | 1.69013 | 2.208 |
| chr2 | 84530069 | 84530394 | 2.04563 | 4.51724 | 2.208 |
| chr8 | 10275291 | 10275654 | 2.07409 | 4.58008 | 2.208 |
| chr10 | 53354821 | 53355103 | 0.581185 | 1.28342 | 2.208 |
| chr2 | 40426567 | 40426812 | 0.557822 | 1.23185 | 2.208 |
| chrX | 83301457 | 83301736 | 1.66358 | 3.67384 | 2.208 |
| chr10 | 10428093 | 10428332 | 0.385422 | 0.851172 | 2.208 |
| chr19 | 3618480 | 3618955 | 2.24831 | 4.96526 | 2.208 |
| chr15 | 100244642 | 100244880 | 1.3102 | 2.89354 | 2.208 |
| chr5 | 31809228 | 31809525 | 1.3437 | 2.96759 | 2.209 |
| chr9 | 104227613 | 104228044 | 1.0251 | 2.26399 | 2.209 |
| chr1 | 220853436 | 220853738 | 1.48073 | 3.27029 | 2.209 |
| chr7 | 22883693 | 22884047 | 0.439231 | 0.970112 | 2.209 |
| chr17 | 73892528 | 73892880 | 0.826729 | 1.82603 | 2.209 |
| chr17 | 49034082 | 49034531 | 1.56671 | 3.46048 | 2.209 |
| chr18 | 62883329 | 62883550 | 0.439006 | 0.9697 | 2.209 |
| chr14 | 24685122 | 24685414 | 1.25801 | 2.77906 | 2.209 |
| chr1 | 233376844 | 233377145 | 1.8179 | 4.01593 | 2.209 |

| | | | | | |
|---|---|---|---|---|---|
| chr10 | 130829808 | 130830176 | 1.14541 | 2.53038 | 2.209 |
| chr1 | 36554233 | 36554568 | 2.98011 | 6.58367 | 2.209 |
| chr4 | 131257869 | 131258084 | 0.713023 | 1.57524 | 2.209 |
| chr15 | 71104466 | 71104713 | 1.16205 | 2.56731 | 2.209 |
| chrX | 53360247 | 53360501 | 0.865463 | 1.91212 | 2.209 |
| chr1 | 153505627 | 153505950 | 2.23392 | 4.9358 | 2.209 |
| chr20 | 57212148 | 57212375 | 0.49762 | 1.09951 | 2.210 |
| chrX | 152909489 | 152909828 | 1.46156 | 3.22939 | 2.210 |
| chr5 | 71314007 | 71314470 | 1.79581 | 3.968 | 2.210 |
| chr1 | 160189316 | 160189686 | 1.1339 | 2.50546 | 2.210 |
| chr11 | 36077950 | 36078246 | 1.06442 | 2.35205 | 2.210 |
| chr9 | 124861443 | 124861966 | 1.1801 | 2.60769 | 2.210 |
| chr10 | 9762164 | 9762367 | 0.493929 | 1.09145 | 2.210 |
| chr6 | 26075630 | 26075902 | 0.565766 | 1.25026 | 2.210 |
| chrX | 52014440 | 52014674 | 0.726775 | 1.60607 | 2.210 |
| chr5 | 160468318 | 160468624 | 0.318901 | 0.70475 | 2.210 |
| chr20 | 17976531 | 17977313 | 1.42959 | 3.1596 | 2.210 |
| chr5 | 105718357 | 105718599 | 0.597144 | 1.31977 | 2.210 |
| chr6 | 49574894 | 49575114 | 1.05663 | 2.33531 | 2.210 |
| chr12 | 115653000 | 115653458 | 0.487313 | 1.07705 | 2.210 |
| chr17 | 8061905 | 8062423 | 1.59401 | 3.52316 | 2.210 |
| chr3 | 69904646 | 69904873 | 1.29624 | 2.86503 | 2.210 |
| chr17 | 74462054 | 74462470 | 2.16748 | 4.79079 | 2.210 |
| chr1 | 55606228 | 55606506 | 1.4284 | 3.15729 | 2.210 |
| chrX | 133887690 | 133887944 | 0.903615 | 1.99732 | 2.210 |
| chr17 | 46713796 | 46714307 | 2.42208 | 5.35391 | 2.210 |
| chr7 | 2710525 | 2710975 | 2.52501 | 5.58146 | 2.210 |
| chr5 | 32329482 | 32329775 | 2.11371 | 4.67233 | 2.210 |
| chr4 | 170644437 | 170644670 | 0.868105 | 1.91895 | 2.211 |
| chr8 | 27454661 | 27455000 | 1.79496 | 3.96813 | 2.211 |
| chr10 | 127031746 | 127032067 | 1.11145 | 2.4571 | 2.211 |
| chr11 | 87383096 | 87383433 | 0.576476 | 1.27443 | 2.211 |
| chr1 | 155066943 | 155067176 | 0.901347 | 1.99269 | 2.211 |
| chr3 | 191956043 | 191956256 | 0.657579 | 1.45377 | 2.211 |
| chr12 | 62124351 | 62124563 | 0.487878 | 1.0786 | 2.211 |
| chr8 | 22431833 | 22432356 | 3.4652 | 7.66098 | 2.211 |
| chr7 | 912875 | 913287 | 1.70801 | 3.77618 | 2.211 |
| chr6 | 37175827 | 37176102 | 1.29028 | 2.85271 | 2.211 |
| chr2 | 44706459 | 44706918 | 1.83754 | 4.06273 | 2.211 |
| chr9 | 14406117 | 14406544 | 0.508277 | 1.1238 | 2.211 |
| chr12 | 114085004 | 114085281 | 0.782496 | 1.73017 | 2.211 |
| chr10 | 98832518 | 98833090 | 1.94539 | 4.30159 | 2.211 |
| chr11 | 119512735 | 119513116 | 0.888517 | 1.96466 | 2.211 |
| chr16 | 31750703 | 31751039 | 0.641433 | 1.41832 | 2.211 |
| chr10 | 99461142 | 99461560 | 1.74026 | 3.84816 | 2.211 |
| chr20 | 47607192 | 47607403 | 0.476758 | 1.05427 | 2.211 |
| chr12 | 707198 | 707434 | 0.350195 | 0.774404 | 2.211 |
| chr17 | 67519475 | 67519755 | 1.51856 | 3.35826 | 2.211 |
| chr4 | 85404086 | 85404365 | 0.990708 | 2.19101 | 2.212 |
| chr8 | 17434339 | 17435206 | 0.821768 | 1.81743 | 2.212 |
| chr9 | 130865943 | 130866213 | 1.02461 | 2.26605 | 2.212 |
| chr3 | 155100224 | 155100527 | 1.89644 | 4.19453 | 2.212 |
| chr10 | 102690152 | 102690419 | 0.627222 | 1.38735 | 2.212 |
| chr16 | 4674133 | 4674588 | 2.21007 | 4.88901 | 2.212 |
| chr15 | 98838391 | 98838704 | 1.20303 | 2.66139 | 2.212 |
| chr9 | 18255101 | 18255436 | 1.51948 | 3.36161 | 2.212 |
| chr7 | 35837243 | 35837447 | 0.507589 | 1.12307 | 2.213 |
| chr19 | 51433119 | 51433447 | 2.01402 | 4.45621 | 2.213 |
| chr22 | 47199180 | 47199653 | 1.92423 | 4.25765 | 2.213 |
| chr20 | 50416619 | 50417079 | 2.81154 | 6.22112 | 2.213 |
| chr6 | 116332038 | 116332253 | 1.20082 | 2.65721 | 2.213 |

| | | | | | |
|---|---|---|---|---|---|
| chr12 | 41016097 | 41016376 | 1.41259 | 3.12584 | 2.213 |
| chr6 | 160524375 | 160524696 | 1.2604 | 2.78906 | 2.213 |
| chr10 | 21367596 | 21368162 | 1.31214 | 2.90358 | 2.213 |
| chr5 | 65621612 | 65622227 | 0.441148 | 0.976207 | 2.213 |
| chr11 | 25704864 | 25705398 | 0.536481 | 1.18718 | 2.213 |
| chr17 | 49924319 | 49924541 | 0.425215 | 0.940959 | 2.213 |
| chr13 | 28633416 | 28633680 | 1.2511 | 2.76859 | 2.213 |
| chr11 | 17036624 | 17036902 | 1.78553 | 3.95161 | 2.213 |
| chr19 | 30019205 | 30019662 | 2.05259 | 4.54288 | 2.213 |
| chr7 | 152191471 | 152191722 | 0.470238 | 1.0408 | 2.213 |
| chr11 | 105958594 | 105958930 | 1.74164 | 3.85515 | 2.214 |
| chrX | 10087416 | 10088054 | 2.8035 | 6.20561 | 2.214 |
| chr17 | 78451884 | 78452708 | 0.75462 | 1.67041 | 2.214 |
| chr16 | 28989063 | 28989380 | 1.47243 | 3.25981 | 2.214 |
| chrX | 43817806 | 43818011 | 1.02852 | 2.27715 | 2.214 |
| chr11 | 116976761 | 116976976 | 1.41695 | 3.13737 | 2.214 |
| chrX | 68441973 | 68442313 | 1.83273 | 4.05801 | 2.214 |
| chr10 | 98479599 | 98480094 | 0.921173 | 2.03968 | 2.214 |
| chr8 | 64843004 | 64843534 | 0.537458 | 1.19009 | 2.214 |
| chr5 | 65603479 | 65603876 | 1.32547 | 2.935 | 2.214 |
| chr3 | 105405182 | 105405461 | 1.72086 | 3.81062 | 2.214 |
| chr15 | 71117539 | 71117800 | 1.62067 | 3.58882 | 2.214 |
| chr8 | 70279290 | 70279533 | 0.551278 | 1.22076 | 2.214 |
| chr16 | 86607808 | 86608473 | 0.428929 | 0.949878 | 2.215 |
| chr8 | 142184924 | 142185614 | 1.88663 | 4.17805 | 2.215 |
| chr20 | 1158456 | 1158661 | 0.547701 | 1.21297 | 2.215 |
| chr4 | 142296977 | 142297294 | 1.25081 | 2.7702 | 2.215 |
| chr8 | 56286889 | 56287278 | 0.407983 | 0.903572 | 2.215 |
| chr10 | 86904738 | 86905079 | 0.694894 | 1.53903 | 2.215 |
| chr20 | 5203546 | 5204109 | 1.79619 | 3.97817 | 2.215 |
| chr12 | 19910290 | 19910594 | 1.41754 | 3.13961 | 2.215 |
| chr10 | 4216423 | 4216706 | 1.17448 | 2.60131 | 2.215 |
| chr9 | 19464399 | 19465366 | 1.7849 | 3.95336 | 2.215 |
| chr17 | 27475854 | 27476233 | 1.74121 | 3.85683 | 2.215 |
| chr10 | 9384859 | 9385154 | 1.20336 | 2.66554 | 2.215 |
| chr7 | 73482191 | 73482483 | 1.76872 | 3.91804 | 2.215 |
| chr15 | 51200527 | 51200984 | 2.09359 | 4.63776 | 2.215 |
| chr10 | 88345227 | 88345614 | 2.18698 | 4.84482 | 2.215 |
| chr3 | 101527869 | 101528356 | 1.01317 | 2.24459 | 2.215 |
| chr16 | 53308213 | 53308453 | 0.492282 | 1.09063 | 2.215 |
| chr6 | 53850999 | 53851276 | 1.23791 | 2.74255 | 2.215 |
| chr2 | 235166796 | 235167068 | 1.31709 | 2.91798 | 2.215 |
| chr11 | 57412040 | 57412769 | 1.56206 | 3.46117 | 2.216 |
| chr10 | 81921487 | 81921703 | 0.721324 | 1.59835 | 2.216 |
| chr11 | 66968067 | 66968277 | 0.517935 | 1.1477 | 2.216 |
| chr4 | 47707672 | 47707905 | 0.621086 | 1.37629 | 2.216 |
| chr17 | 75490970 | 75491408 | 1.94284 | 4.30533 | 2.216 |
| chr14 | 21439010 | 21439928 | 2.70483 | 5.99416 | 2.216 |
| chr9 | 117053241 | 117053612 | 1.74198 | 3.86045 | 2.216 |
| chr15 | 74073668 | 74073887 | 0.926803 | 2.05402 | 2.216 |
| chr16 | 12863478 | 12863694 | 0.518733 | 1.14964 | 2.216 |
| chr17 | 77083486 | 77084141 | 2.00767 | 4.44954 | 2.216 |
| chr4 | 155849818 | 155850122 | 1.1986 | 2.65663 | 2.216 |
| chr10 | 9932761 | 9933002 | 0.436817 | 0.968204 | 2.217 |
| chr7 | 48279395 | 48279615 | 0.480415 | 1.06486 | 2.217 |
| chr7 | 123556524 | 123556746 | 0.753859 | 1.67099 | 2.217 |
| chr11 | 107434420 | 107434647 | 0.936552 | 2.07595 | 2.217 |
| chr6 | 52281428 | 52281693 | 0.974031 | 2.15908 | 2.217 |
| chr6 | 150401816 | 150402322 | 1.09918 | 2.43651 | 2.217 |
| chr6 | 44140490 | 44140947 | 1.25813 | 2.78894 | 2.217 |
| chr14 | 100751511 | 100752396 | 2.17814 | 4.82864 | 2.217 |

| | | | | | |
|---|---|---|---|---|---|
| chr1 | 6787571 | 6787989 | 1.90948 | 4.23327 | 2.217 |
| chr11 | 66821287 | 66821711 | 1.37604 | 3.0508 | 2.217 |
| chr1 | 233160881 | 233161235 | 1.12999 | 2.50534 | 2.217 |
| chr9 | 79273566 | 79273942 | 0.910286 | 2.01825 | 2.217 |
| chr9 | 129087949 | 129088181 | 0.499363 | 1.10717 | 2.217 |
| chr1 | 186181795 | 186182082 | 1.56719 | 3.47472 | 2.217 |
| chr14 | 63329289 | 63329656 | 0.536231 | 1.18898 | 2.217 |
| chr1 | 46645265 | 46645542 | 1.85133 | 4.10498 | 2.217 |
| chr7 | 115898810 | 115899012 | 0.818978 | 1.81593 | 2.217 |
| chr2 | 68067251 | 68068051 | 1.21666 | 2.69793 | 2.217 |
| chr16 | 51796007 | 51796346 | 2.13347 | 4.73108 | 2.218 |
| chrY | 2668426 | 2668732 | 0.679678 | 1.50727 | 2.218 |
| chr10 | 16158425 | 16158671 | 1.10085 | 2.44146 | 2.218 |
| chr11 | 15671556 | 15671898 | 1.56294 | 3.46668 | 2.218 |
| chr15 | 76882045 | 76882515 | 0.502402 | 1.11439 | 2.218 |
| chr9 | 28337523 | 28337789 | 0.505936 | 1.12232 | 2.218 |
| chr15 | 69323799 | 69324177 | 1.85114 | 4.10662 | 2.218 |
| chr1 | 40042486 | 40042792 | 1.33662 | 2.96523 | 2.218 |
| chr11 | 66650585 | 66650981 | 1.52579 | 3.38492 | 2.218 |
| chr10 | 32151253 | 32151466 | 0.765243 | 1.69775 | 2.219 |
| chr2 | 43354230 | 43354667 | 1.45517 | 3.22844 | 2.219 |
| chr7 | 5405565 | 5406034 | 1.67235 | 3.7103 | 2.219 |
| chr12 | 70023981 | 70024266 | 0.827834 | 1.83667 | 2.219 |
| chr2 | 201600470 | 201600732 | 0.833297 | 1.8488 | 2.219 |
| chr8 | 37559224 | 37559466 | 0.636975 | 1.41326 | 2.219 |
| chr11 | 4846423 | 4846658 | 0.511037 | 1.13385 | 2.219 |
| chrX | 119347496 | 119347929 | 1.37798 | 3.05737 | 2.219 |
| chr17 | 2731684 | 2732132 | 2.09502 | 4.64834 | 2.219 |
| chr11 | 10203462 | 10203684 | 0.726646 | 1.6123 | 2.219 |
| chr6 | 8743467 | 8743684 | 0.68796 | 1.52657 | 2.219 |
| chr14 | 49931658 | 49931933 | 0.663217 | 1.47169 | 2.219 |
| chr5 | 151807470 | 151807720 | 0.465021 | 1.03195 | 2.219 |
| chr12 | 26618409 | 26618656 | 0.905542 | 2.00965 | 2.219 |
| chr16 | 30075414 | 30075616 | 0.731334 | 1.62313 | 2.219 |
| chr5 | 167002518 | 167002900 | 0.572736 | 1.27116 | 2.219 |
| chr20 | 24353476 | 24354023 | 0.469268 | 1.04152 | 2.219 |
| chr5 | 42583337 | 42583762 | 1.1849 | 2.62989 | 2.220 |
| chr17 | 70721654 | 70721980 | 1.98328 | 4.40217 | 2.220 |
| chr20 | 58220883 | 58221179 | 0.74267 | 1.64849 | 2.220 |
| chr12 | 32909145 | 32909444 | 1.12234 | 2.49126 | 2.220 |
| chr3 | 8664490 | 8664941 | 2.10221 | 4.66628 | 2.220 |
| chr7 | 55891161 | 55891386 | 0.424218 | 0.941647 | 2.220 |
| chr21 | 32976991 | 32977284 | 1.03794 | 2.30408 | 2.220 |
| chr9 | 9424244 | 9424491 | 0.536018 | 1.18991 | 2.220 |
| chr6 | 152659685 | 152659979 | 1.61105 | 3.57647 | 2.220 |
| chr3 | 148027256 | 148027517 | 0.979436 | 2.17444 | 2.220 |
| chr4 | 183859205 | 183859656 | 1.61526 | 3.58603 | 2.220 |
| chr10 | 22875210 | 22875519 | 1.35508 | 3.00841 | 2.220 |
| chr19 | 56143711 | 56144100 | 1.51087 | 3.35432 | 2.220 |
| chr18 | 29851757 | 29851979 | 1.23617 | 2.74456 | 2.220 |
| chr11 | 36531497 | 36532086 | 1.56289 | 3.46998 | 2.220 |
| chr20 | 13752595 | 13752818 | 0.83847 | 1.86172 | 2.220 |
| chr1 | 1307289 | 1307816 | 1.50087 | 3.33256 | 2.220 |
| chr8 | 81152808 | 81153224 | 2.98023 | 6.61788 | 2.221 |
| chr3 | 118960366 | 118960648 | 1.79211 | 3.97972 | 2.221 |
| chr10 | 92535134 | 92535353 | 0.647528 | 1.43805 | 2.221 |
| chr1 | 158831274 | 158831591 | 0.48417 | 1.07528 | 2.221 |
| chr17 | 74456641 | 74456966 | 1.61664 | 3.59059 | 2.221 |
| chr14 | 39049353 | 39049949 | 0.480683 | 1.06763 | 2.221 |
| chr2 | 239628763 | 239629192 | 1.34933 | 2.99703 | 2.221 |
| chr9 | 80981130 | 80981346 | 1.16034 | 2.57732 | 2.221 |

| | | | | | |
|---|---|---|---|---|---|
| chr6 | 134499621 | 134499903 | 1.4681 | 3.26093 | 2.221 |
| chr17 | 27003045 | 27003275 | 1.41717 | 3.14783 | 2.221 |
| chr2 | 201766254 | 201766520 | 1.00748 | 2.23782 | 2.221 |
| chr8 | 35838904 | 35839126 | 0.401064 | 0.890863 | 2.221 |
| chr2 | 106505194 | 106505588 | 0.673807 | 1.4967 | 2.221 |
| chr8 | 33621520 | 33621751 | 0.403521 | 0.89635 | 2.221 |
| chrX | 117735531 | 117735770 | 1.00878 | 2.2409 | 2.221 |
| chr11 | 36280258 | 36280639 | 1.4354 | 3.18867 | 2.221 |
| chr7 | 28873986 | 28874241 | 0.344784 | 0.765921 | 2.221 |
| chr12 | 31142806 | 31143145 | 0.97395 | 2.16365 | 2.222 |
| chr1 | 21922060 | 21922313 | 1.57838 | 3.50647 | 2.222 |
| chr19 | 33793108 | 33793781 | 1.70168 | 3.78038 | 2.222 |
| chr12 | 31003589 | 31003931 | 0.583079 | 1.29538 | 2.222 |
| chr2 | 30431273 | 30431587 | 1.92465 | 4.27595 | 2.222 |
| chr3 | 141170924 | 141171159 | 1.44973 | 3.22088 | 2.222 |
| chr1 | 90308838 | 90309112 | 1.64128 | 3.6466 | 2.222 |
| chr5 | 148277842 | 148278065 | 0.439412 | 0.976286 | 2.222 |
| chr1 | 16335477 | 16335839 | 1.3583 | 3.0179 | 2.222 |
| chr11 | 123278095 | 123278405 | 1.15536 | 2.56705 | 2.222 |
| chr9 | 74895826 | 74896149 | 1.71146 | 3.80274 | 2.222 |
| chr15 | 86732852 | 86733175 | 0.527667 | 1.17245 | 2.222 |
| chr9 | 116163624 | 116163955 | 1.92403 | 4.27517 | 2.222 |
| chr19 | 34992339 | 34992907 | 3.35974 | 7.46552 | 2.222 |
| chr12 | 106099549 | 106099833 | 1.39607 | 3.10224 | 2.222 |
| chr7 | 75807386 | 75807872 | 2.55162 | 5.67014 | 2.222 |
| chr8 | 144869020 | 144869271 | 1.20091 | 2.66868 | 2.222 |
| chr3 | 30877090 | 30877434 | 0.603793 | 1.34176 | 2.222 |
| chr10 | 58738238 | 58738469 | 0.364586 | 0.810193 | 2.222 |
| chr8 | 84708314 | 84708789 | 0.500348 | 1.11191 | 2.222 |
| chr16 | 27468421 | 27468867 | 1.71926 | 3.82068 | 2.222 |
| chr1 | 227949903 | 227950483 | 1.45946 | 3.24335 | 2.222 |
| chr19 | 47359404 | 47359700 | 1.71236 | 3.80538 | 2.222 |
| chr17 | 62936839 | 62937082 | 0.448029 | 0.995663 | 2.222 |
| chr7 | 126084902 | 126085216 | 0.498236 | 1.10724 | 2.222 |
| chr5 | 35627599 | 35627804 | 0.537296 | 1.19406 | 2.222 |
| chr12 | 129810395 | 129810604 | 0.370959 | 0.824415 | 2.222 |
| chr12 | 124439251 | 124439535 | 0.621399 | 1.38101 | 2.222 |
| chr9 | 34994413 | 34994746 | 1.03394 | 2.29786 | 2.222 |
| chr2 | 74814797 | 74815089 | 1.26687 | 2.81576 | 2.223 |
| chr4 | 143274291 | 143274601 | 1.28904 | 2.86513 | 2.223 |
| chr16 | 74760659 | 74761002 | 1.33852 | 2.97525 | 2.223 |
| chr4 | 48129784 | 48130120 | 1.18081 | 2.62471 | 2.223 |
| chr16 | 89976581 | 89977207 | 1.98374 | 4.40954 | 2.223 |
| chr20 | 8062715 | 8062994 | 0.931651 | 2.07093 | 2.223 |
| chr7 | 89577975 | 89578183 | 0.471898 | 1.04898 | 2.223 |
| chr12 | 119551086 | 119551349 | 0.456579 | 1.01506 | 2.223 |
| chr13 | 41959310 | 41959525 | 0.739392 | 1.64384 | 2.223 |
| chr7 | 90084760 | 90085085 | 1.06014 | 2.35702 | 2.223 |
| chr16 | 18980620 | 18980977 | 2.36737 | 5.26342 | 2.223 |
| chr22 | 50026206 | 50026495 | 1.39192 | 3.09493 | 2.224 |
| chr7 | 36123934 | 36124472 | 0.763857 | 1.69846 | 2.224 |
| chr7 | 141433993 | 141434230 | 0.504743 | 1.12236 | 2.224 |
| chr13 | 88323433 | 88323808 | 1.78026 | 3.95889 | 2.224 |
| chr11 | 15792426 | 15792698 | 1.61679 | 3.5954 | 2.224 |
| chr1 | 158057040 | 158057548 | 2.43019 | 5.40425 | 2.224 |
| chr10 | 111477240 | 111477700 | 0.463985 | 1.03183 | 2.224 |
| chr7 | 24019757 | 24019980 | 0.632043 | 1.40559 | 2.224 |
| chr17 | 28667179 | 28667451 | 1.20736 | 2.68503 | 2.224 |
| chr1 | 64023182 | 64023475 | 1.1216 | 2.49434 | 2.224 |
| chr7 | 97308182 | 97308405 | 0.375525 | 0.835173 | 2.224 |
| chr15 | 49438245 | 49438497 | 1.03388 | 2.29943 | 2.224 |

| | | | | | |
|---|---|---|---|---|---|
| chr2 | 13721593 | 13721969 | 0.877011 | 1.95061 | 2.224 |
| chr11 | 16627730 | 16628113 | 1.64532 | 3.65947 | 2.224 |
| chr17 | 7816349 | 7816618 | 1.42339 | 3.16586 | 2.224 |
| chr15 | 100296000 | 100296377 | 1.82144 | 4.05122 | 2.224 |
| chr1 | 151986140 | 151986452 | 1.23664 | 2.75055 | 2.224 |
| chrX | 39511687 | 39511919 | 1.08658 | 2.41684 | 2.224 |
| chr12 | 103272836 | 103273315 | 1.53756 | 3.42003 | 2.224 |
| chr12 | 114305043 | 114305479 | 1.2298 | 2.73551 | 2.224 |
| chr13 | 36920850 | 36921166 | 1.51037 | 3.35962 | 2.224 |
| chr12 | 110043653 | 110043872 | 0.562099 | 1.25033 | 2.224 |
| chr16 | 71917669 | 71918050 | 0.772318 | 1.71798 | 2.224 |
| chr5 | 42829445 | 42829703 | 0.420551 | 0.935513 | 2.224 |
| chr20 | 49942784 | 49943124 | 1.59659 | 3.55164 | 2.225 |
| chr3 | 30240627 | 30240942 | 1.51194 | 3.36336 | 2.225 |
| chr16 | 14343418 | 14343772 | 2.06059 | 4.58387 | 2.225 |
| chr17 | 47392855 | 47393179 | 1.63187 | 3.63018 | 2.225 |
| chr12 | 130399258 | 130399493 | 1.10895 | 2.46693 | 2.225 |
| chr11 | 3876057 | 3876484 | 2.30511 | 5.12787 | 2.225 |
| chr8 | 126708962 | 126709243 | 1.34767 | 2.998 | 2.225 |
| chr11 | 76445040 | 76445255 | 1.77257 | 3.94327 | 2.225 |
| chr22 | 45093246 | 45093679 | 1.51556 | 3.37172 | 2.225 |
| chr11 | 57259200 | 57259538 | 1.99863 | 4.44648 | 2.225 |
| chr3 | 138313444 | 138313814 | 0.760072 | 1.69099 | 2.225 |
| chr5 | 67582201 | 67582475 | 1.10517 | 2.45881 | 2.225 |
| chr11 | 8114183 | 8114654 | 1.35038 | 3.00438 | 2.225 |
| chr9 | 130632172 | 130632528 | 2.21427 | 4.92656 | 2.225 |
| chr2 | 26726105 | 26726669 | 2.65935 | 5.91683 | 2.225 |
| chr15 | 59131680 | 59131959 | 1.13071 | 2.51575 | 2.225 |
| chr19 | 54296175 | 54296644 | 2.81856 | 6.27118 | 2.225 |
| chr10 | 54638860 | 54639070 | 0.50244 | 1.11794 | 2.225 |
| chr15 | 76622507 | 76622945 | 1.05722 | 2.35237 | 2.225 |
| chr10 | 127318343 | 127318628 | 0.792816 | 1.76417 | 2.225 |
| chr4 | 155624329 | 155624582 | 0.4711 | 1.04834 | 2.225 |
| chr5 | 128367931 | 128368341 | 0.478194 | 1.06413 | 2.225 |
| chr5 | 139523805 | 139524185 | 1.62872 | 3.62457 | 2.225 |
| chr17 | 13369896 | 13370125 | 0.677745 | 1.50827 | 2.225 |
| chr4 | 13550512 | 13550885 | 1.30095 | 2.89531 | 2.226 |
| chr6 | 28464016 | 28464275 | 0.582942 | 1.29738 | 2.226 |
| chr12 | 120105306 | 120105741 | 1.89084 | 4.20836 | 2.226 |
| chr2 | 164328513 | 164328779 | 1.62388 | 3.61427 | 2.226 |
| chr10 | 6244459 | 6244940 | 2.56126 | 5.70072 | 2.226 |
| chr21 | 47567369 | 47567664 | 1.83025 | 4.07369 | 2.226 |
| chr11 | 62448354 | 62448556 | 0.981849 | 2.18543 | 2.226 |
| chr10 | 101769237 | 101770071 | 1.63346 | 3.63591 | 2.226 |
| chr5 | 93137030 | 93137355 | 1.07014 | 2.38242 | 2.226 |
| chr14 | 104013107 | 104013560 | 1.48995 | 3.31705 | 2.226 |
| chr6 | 85484376 | 85484806 | 1.59787 | 3.55737 | 2.226 |
| chr12 | 62354783 | 62355088 | 0.498766 | 1.11052 | 2.227 |
| chr12 | 29772239 | 29772445 | 0.495272 | 1.1028 | 2.227 |
| chr5 | 104822050 | 104822258 | 0.479523 | 1.06777 | 2.227 |
| chr14 | 103312848 | 103313196 | 0.428221 | 0.953569 | 2.227 |
| chr14 | 22191707 | 22191979 | 0.711282 | 1.58397 | 2.227 |
| chr1 | 183204049 | 183204452 | 1.45627 | 3.24302 | 2.227 |
| chr17 | 27944896 | 27945479 | 1.99187 | 4.4359 | 2.227 |
| chr12 | 71809491 | 71809750 | 1.17389 | 2.61427 | 2.227 |
| chr4 | 12131813 | 12132062 | 0.513639 | 1.14388 | 2.227 |
| chr2 | 14277830 | 14278088 | 0.764385 | 1.70231 | 2.227 |
| chr13 | 40186247 | 40186494 | 1.14264 | 2.54495 | 2.227 |
| chr16 | 77077094 | 77077323 | 0.89586 | 1.99533 | 2.227 |
| chr1 | 179220856 | 179221139 | 1.00712 | 2.24331 | 2.227 |
| chr1 | 188324333 | 188324581 | 0.499358 | 1.11235 | 2.228 |

| | | | | | |
|---|---|---|---|---|---|
| chr3 | 123370982 | 123371564 | 1.79312 | 3.99429 | 2.228 |
| chr3 | 128399534 | 128399892 | 1.99292 | 4.43952 | 2.228 |
| chr20 | 2723632 | 2724163 | 2.69987 | 6.01451 | 2.228 |
| chr6 | 34023020 | 34023596 | 1.45207 | 3.23483 | 2.228 |
| chr5 | 103289770 | 103289984 | 0.499769 | 1.1134 | 2.228 |
| chr2 | 220024708 | 220025191 | 1.87776 | 4.18362 | 2.228 |
| chr14 | 105234777 | 105235547 | 1.53586 | 3.42201 | 2.228 |
| chr5 | 78852042 | 78852311 | 1.45513 | 3.24215 | 2.228 |
| chr18 | 46974250 | 46974531 | 1.78427 | 3.97578 | 2.228 |
| chr5 | 111827036 | 111827310 | 0.971847 | 2.16557 | 2.228 |
| chr19 | 8461559 | 8461868 | 1.78321 | 3.97356 | 2.228 |
| chr20 | 61810607 | 61810980 | 1.46487 | 3.26444 | 2.228 |
| chr8 | 142610201 | 142610569 | 0.443527 | 0.988396 | 2.228 |
| chr11 | 122925041 | 122925285 | 1.13256 | 2.52425 | 2.229 |
| chr12 | 65996289 | 65996624 | 1.54707 | 3.44811 | 2.229 |
| chr19 | 2069693 | 2070129 | 2.42014 | 5.39404 | 2.229 |
| chr6 | 14144836 | 14145165 | 1.54212 | 3.43724 | 2.229 |
| chr15 | 27666074 | 27666370 | 0.658753 | 1.46834 | 2.229 |
| chr1 | 242149158 | 242149504 | 0.886968 | 1.97712 | 2.229 |
| chr14 | 104808269 | 104808623 | 1.36494 | 3.04257 | 2.229 |
| chr16 | 75555382 | 75555591 | 0.721661 | 1.60865 | 2.229 |
| chr1 | 145542804 | 145543212 | 1.6175 | 3.60569 | 2.229 |
| chr3 | 102255414 | 102255735 | 0.652707 | 1.4551 | 2.229 |
| chr7 | 111392227 | 111392582 | 2.29685 | 5.12047 | 2.229 |
| chr12 | 130723223 | 130723507 | 0.539824 | 1.20347 | 2.229 |
| chr19 | 10110884 | 10111331 | 1.56586 | 3.49089 | 2.229 |
| chr10 | 102778714 | 102779158 | 1.96934 | 4.39051 | 2.229 |
| chr9 | 132629916 | 132630304 | 0.735492 | 1.63977 | 2.229 |
| chr17 | 31236940 | 31237212 | 1.60962 | 3.58864 | 2.230 |
| chr10 | 35226918 | 35227202 | 1.29819 | 2.89433 | 2.230 |
| chr4 | 114015975 | 114016215 | 1.19341 | 2.66078 | 2.230 |
| chr7 | 122710794 | 122711052 | 0.763253 | 1.70178 | 2.230 |
| chr6 | 112364895 | 112365219 | 1.41539 | 3.15585 | 2.230 |
| chr14 | 85881659 | 85881883 | 1.1906 | 2.65471 | 2.230 |
| chr6 | 137397651 | 137398000 | 1.34915 | 3.00822 | 2.230 |
| chr6 | 88506285 | 88506538 | 1.09467 | 2.44083 | 2.230 |
| chr7 | 1209557 | 1209888 | 2.48468 | 5.54018 | 2.230 |
| chr9 | 77642680 | 77643344 | 0.940442 | 2.09697 | 2.230 |
| chr11 | 7108570 | 7108802 | 0.416126 | 0.927914 | 2.230 |
| chr22 | 23112614 | 23113014 | 1.6934 | 3.77609 | 2.230 |
| chr3 | 1171991 | 1172340 | 0.560504 | 1.24986 | 2.230 |
| chr9 | 124492422 | 124492803 | 0.361796 | 0.806803 | 2.230 |
| chr9 | 21110881 | 21111271 | 1.38522 | 3.0894 | 2.230 |
| chr2 | 180990233 | 180990557 | 1.3818 | 3.0818 | 2.230 |
| chr2 | 223183863 | 223184155 | 1.11082 | 2.47748 | 2.230 |
| chr1 | 235339964 | 235340187 | 0.624463 | 1.39286 | 2.230 |
| chr1 | 217060131 | 217060419 | 0.417732 | 0.93178 | 2.231 |
| chr4 | 75600974 | 75601212 | 1.0185 | 2.27198 | 2.231 |
| chr18 | 12893801 | 12894118 | 1.44019 | 3.21282 | 2.231 |
| chr4 | 170929648 | 170929900 | 0.584125 | 1.30311 | 2.231 |
| chr8 | 102345147 | 102345486 | 1.20515 | 2.68854 | 2.231 |
| chr5 | 53513782 | 53514044 | 0.869863 | 1.94069 | 2.231 |
| chr6 | 53349477 | 53349742 | 1.41793 | 3.16345 | 2.231 |
| chr5 | 96612805 | 96613052 | 0.387731 | 0.865068 | 2.231 |
| chr1 | 217458483 | 217458783 | 0.465298 | 1.03817 | 2.231 |
| chr20 | 62752941 | 62753144 | 0.425279 | 0.94888 | 2.231 |
| chr10 | 98970230 | 98970616 | 2.02862 | 4.52641 | 2.231 |
| chr17 | 45810368 | 45810676 | 1.04139 | 2.32369 | 2.231 |
| chr1 | 4716421 | 4716669 | 1.50324 | 3.35436 | 2.231 |
| chr5 | 115764759 | 115764974 | 0.582924 | 1.30083 | 2.232 |
| chr19 | 12996326 | 12996597 | 1.44232 | 3.21883 | 2.232 |

| | | | | | |
|---|---|---|---|---|---|
| chr19 | 50964451 | 50964831 | 1.83481 | 4.09481 | 2.232 |
| chr5 | 179049772 | 179050159 | 1.30665 | 2.91624 | 2.232 |
| chr9 | 139820619 | 139821000 | 0.804146 | 1.79474 | 2.232 |
| chr6 | 40973930 | 40974223 | 1.16347 | 2.59672 | 2.232 |
| chr13 | 28572037 | 28572453 | 1.35644 | 3.02743 | 2.232 |
| chr7 | 102090686 | 102091047 | 0.869154 | 1.93987 | 2.232 |
| chr1 | 68585960 | 68586187 | 0.988256 | 2.20584 | 2.232 |
| chr13 | 111596701 | 111597025 | 1.17125 | 2.61435 | 2.232 |
| chr16 | 67850373 | 67850838 | 3.71528 | 8.29356 | 2.232 |
| chr5 | 61488945 | 61489181 | 0.369399 | 0.824635 | 2.232 |
| chr1 | 201865038 | 201865450 | 2.74151 | 6.12008 | 2.232 |
| chr3 | 141178878 | 141179302 | 1.7801 | 3.97404 | 2.232 |
| chr3 | 48545128 | 48545376 | 0.880288 | 1.96524 | 2.233 |
| chr7 | 5013428 | 5013826 | 1.95731 | 4.36975 | 2.233 |
| chr15 | 75127493 | 75128389 | 1.39691 | 3.11867 | 2.233 |
| chr6 | 109848238 | 109848511 | 1.02049 | 2.27835 | 2.233 |
| chr8 | 19637052 | 19637357 | 1.57126 | 3.50801 | 2.233 |
| chr10 | 86134733 | 86135059 | 1.30038 | 2.9033 | 2.233 |
| chr8 | 81143118 | 81143333 | 1.51566 | 3.38403 | 2.233 |
| chr9 | 135490749 | 135491066 | 0.607371 | 1.35612 | 2.233 |
| chr14 | 68264904 | 68265122 | 0.422589 | 0.943553 | 2.233 |
| chr19 | 21465953 | 21466254 | 0.53676 | 1.19849 | 2.233 |
| chr4 | 1066709 | 1066962 | 1.05244 | 2.34992 | 2.233 |
| chr4 | 3785548 | 3785816 | 1.48271 | 3.31067 | 2.233 |
| chr5 | 173846592 | 173846909 | 1.89359 | 4.22827 | 2.233 |
| chr11 | 10658523 | 10658795 | 0.411705 | 0.919367 | 2.233 |
| chr6 | 116422122 | 116422351 | 0.5687 | 1.26998 | 2.233 |
| chr3 | 50587489 | 50588039 | 1.00134 | 2.23623 | 2.233 |
| chr16 | 70404030 | 70404263 | 1.18413 | 2.64451 | 2.233 |
| chr1 | 235746979 | 235747319 | 1.09486 | 2.44516 | 2.233 |
| chr12 | 128954641 | 128954848 | 0.604153 | 1.34929 | 2.233 |
| chr12 | 48680529 | 48680770 | 0.82507 | 1.84269 | 2.233 |
| chr11 | 14926349 | 14927280 | 1.14025 | 2.54679 | 2.234 |
| chr2 | 165456545 | 165456808 | 1.66333 | 3.71518 | 2.234 |
| chr2 | 36891342 | 36891577 | 1.06223 | 2.37262 | 2.234 |
| chr10 | 129752599 | 129752976 | 0.402871 | 0.899889 | 2.234 |
| chr14 | 90405771 | 90406360 | 1.72739 | 3.8585 | 2.234 |
| chr3 | 48442331 | 48442644 | 1.53486 | 3.42853 | 2.234 |
| chr10 | 33405174 | 33405502 | 1.16132 | 2.59414 | 2.234 |
| chrX | 133160804 | 133161018 | 1.13827 | 2.54266 | 2.234 |
| chr9 | 136275419 | 136275809 | 0.534386 | 1.19371 | 2.234 |
| chr19 | 46073148 | 46073549 | 2.42462 | 5.41621 | 2.234 |
| chr12 | 94940000 | 94940280 | 1.51022 | 3.3738 | 2.234 |
| chr15 | 86408190 | 86408683 | 1.32895 | 2.96888 | 2.234 |
| chr15 | 28027806 | 28028089 | 0.54577 | 1.2193 | 2.234 |
| chr8 | 136615222 | 136615424 | 0.647161 | 1.44582 | 2.234 |
| chr14 | 24801305 | 24801710 | 1.31665 | 2.94158 | 2.234 |
| chr7 | 1798331 | 1798571 | 1.23836 | 2.76701 | 2.234 |
| chr17 | 6569304 | 6569681 | 2.13118 | 4.76202 | 2.234 |
| chr12 | 55537088 | 55537407 | 1.1938 | 2.66753 | 2.234 |
| chr11 | 82429128 | 82429741 | 1.40325 | 3.13571 | 2.235 |
| chr14 | 100654761 | 100655043 | 0.483493 | 1.08042 | 2.235 |
| chr7 | 100397063 | 100397419 | 1.48527 | 3.31909 | 2.235 |
| chr6 | 3849403 | 3849744 | 1.90604 | 4.25952 | 2.235 |
| chr14 | 23133851 | 23134146 | 0.509438 | 1.13847 | 2.235 |
| chr10 | 103540727 | 103541235 | 2.07445 | 4.63645 | 2.235 |
| chr19 | 30236244 | 30236810 | 0.951241 | 2.12612 | 2.235 |
| chr10 | 3162111 | 3162635 | 1.22143 | 2.73006 | 2.235 |
| chr12 | 95467357 | 95467637 | 2.03898 | 4.55772 | 2.235 |
| chr12 | 75831376 | 75831671 | 0.999802 | 2.23487 | 2.235 |
| chr12 | 122120004 | 122120340 | 2.13021 | 4.76169 | 2.235 |

| | | | | | |
|---|---|---|---|---|---|
| chr7 | 23694103 | 23694530 | 1.05029 | 2.34773 | 2.235 |
| chr4 | 103930326 | 103930604 | 1.19313 | 2.66715 | 2.235 |
| chr15 | 67749268 | 67749739 | 2.11238 | 4.7224 | 2.236 |
| chr2 | 64822624 | 64822887 | 0.963264 | 2.15346 | 2.236 |
| chr1 | 202938148 | 202938441 | 1.22686 | 2.74277 | 2.236 |
| chr1 | 239927150 | 239927521 | 0.724667 | 1.62007 | 2.236 |
| chr7 | 27645353 | 27645584 | 1.34705 | 3.01155 | 2.236 |
| chr5 | 176498897 | 176499163 | 1.5519 | 3.46963 | 2.236 |
| chr10 | 128276489 | 128276842 | 0.371335 | 0.830207 | 2.236 |
| chr11 | 133817285 | 133817544 | 0.744457 | 1.66443 | 2.236 |
| chr7 | 92722186 | 92722603 | 0.860472 | 1.92385 | 2.236 |
| chr19 | 12787155 | 12787546 | 1.65382 | 3.69765 | 2.236 |
| chr11 | 126586360 | 126586620 | 1.03144 | 2.30612 | 2.236 |
| chr19 | 36001212 | 36001885 | 2.18248 | 4.87969 | 2.236 |
| chr4 | 96046120 | 96046342 | 0.887924 | 1.98531 | 2.236 |
| chr10 | 68981394 | 68981612 | 0.482524 | 1.07889 | 2.236 |
| chr3 | 28359005 | 28359272 | 1.69791 | 3.79645 | 2.236 |
| chr9 | 105718445 | 105718760 | 0.467999 | 1.04644 | 2.236 |
| chr9 | 79906706 | 79906975 | 0.980645 | 2.19273 | 2.236 |
| chr12 | 97092393 | 97092684 | 1.1067 | 2.47461 | 2.236 |
| chr4 | 41695767 | 41695971 | 0.553064 | 1.23677 | 2.236 |
| chr10 | 32013205 | 32013471 | 0.374259 | 0.836936 | 2.236 |
| chr2 | 40471778 | 40472115 | 1.19794 | 2.67893 | 2.236 |
| chr22 | 40432679 | 40432960 | 1.59389 | 3.56444 | 2.236 |
| chr17 | 76493203 | 76493524 | 1.45377 | 3.2511 | 2.236 |
| chr17 | 42768267 | 42768609 | 0.956151 | 2.13827 | 2.236 |
| chr9 | 139682113 | 139682414 | 1.40707 | 3.14706 | 2.237 |
| chr3 | 148097629 | 148097869 | 0.806127 | 1.80303 | 2.237 |
| chr1 | 154971519 | 154972111 | 1.79461 | 4.01395 | 2.237 |
| chr2 | 228523036 | 228523348 | 1.1165 | 2.49729 | 2.237 |
| chr6 | 7723325 | 7723707 | 1.13924 | 2.54818 | 2.237 |
| chr7 | 55500958 | 55501194 | 0.817781 | 1.82921 | 2.237 |
| chr14 | 102426559 | 102426881 | 1.56878 | 3.5092 | 2.237 |
| chr1 | 228528498 | 228528772 | 0.881937 | 1.97286 | 2.237 |
| chr2 | 36687018 | 36687319 | 1.84374 | 4.12457 | 2.237 |
| chr1 | 17955986 | 17956254 | 0.835758 | 1.86975 | 2.237 |
| chr2 | 198175501 | 198175736 | 1.11201 | 2.48783 | 2.237 |
| chr17 | 72453652 | 72454070 | 2.3386 | 5.23208 | 2.237 |
| chr3 | 27763008 | 27764030 | 1.59763 | 3.57436 | 2.237 |
| chr5 | 140479859 | 140480111 | 1.31968 | 2.9526 | 2.237 |
| chr1 | 145589169 | 145589631 | 2.84105 | 6.35657 | 2.237 |
| chr16 | 30066234 | 30066751 | 1.61261 | 3.60813 | 2.237 |
| chr9 | 2728045 | 2728475 | 1.18741 | 2.65678 | 2.237 |
| chr4 | 185172001 | 185172291 | 1.26655 | 2.83387 | 2.237 |
| chr11 | 15151387 | 15151607 | 0.546978 | 1.22385 | 2.237 |
| chr1 | 183901658 | 183902149 | 0.452772 | 1.0131 | 2.238 |
| chr16 | 67706702 | 67706925 | 1.2914 | 2.88967 | 2.238 |
| chr2 | 241580458 | 241580700 | 1.62815 | 3.64355 | 2.238 |
| chr12 | 45937649 | 45937969 | 1.58773 | 3.55314 | 2.238 |
| chr1 | 193224021 | 193224227 | 0.780352 | 1.74638 | 2.238 |
| chr9 | 27276468 | 27276738 | 1.40575 | 3.146 | 2.238 |
| chr8 | 86132843 | 86133193 | 2.43687 | 5.4539 | 2.238 |
| chr3 | 74249013 | 74249330 | 0.512044 | 1.14602 | 2.238 |
| chr3 | 46951065 | 46951404 | 1.53007 | 3.42471 | 2.238 |
| chr3 | 129995060 | 129995332 | 1.28153 | 2.86851 | 2.238 |
| chr3 | 43585755 | 43586009 | 0.733294 | 1.64141 | 2.238 |
| chr13 | 58199071 | 58199302 | 0.946608 | 2.11894 | 2.238 |
| chr2 | 153073901 | 153074170 | 0.360207 | 0.806316 | 2.238 |
| chr1 | 109782088 | 109782355 | 1.39738 | 3.12804 | 2.239 |
| chr17 | 73400223 | 73400542 | 1.51945 | 3.40148 | 2.239 |
| chr2 | 65291911 | 65292404 | 0.957442 | 2.14336 | 2.239 |

| | | | | | |
|---|---|---|---|---|---|
| chr6 | 74679793 | 74680015 | 1.25566 | 2.81098 | 2.239 |
| chr19 | 16152331 | 16152707 | 1.7669 | 3.95565 | 2.239 |
| chr3 | 99527010 | 99527354 | 1.42643 | 3.19347 | 2.239 |
| chr14 | 104601486 | 104601731 | 0.999467 | 2.2376 | 2.239 |
| chr9 | 22452502 | 22452753 | 1.54623 | 3.46173 | 2.239 |
| chr10 | 53419005 | 53419575 | 0.440701 | 0.98668 | 2.239 |
| chr17 | 49943487 | 49943870 | 0.494585 | 1.10749 | 2.239 |
| chr12 | 41510165 | 41510476 | 0.647858 | 1.45075 | 2.239 |
| chr13 | 105426119 | 105426419 | 1.00405 | 2.24837 | 2.239 |
| chr4 | 4727809 | 4728113 | 1.06986 | 2.39574 | 2.239 |
| chr6 | 82750328 | 82750550 | 1.19084 | 2.66671 | 2.239 |
| chr3 | 8649248 | 8649540 | 1.17913 | 2.64053 | 2.239 |
| chr14 | 77462825 | 77463143 | 0.505068 | 1.13105 | 2.239 |
| chr1 | 211496539 | 211497133 | 1.46642 | 3.28399 | 2.239 |
| chr4 | 41663510 | 41663780 | 1.71099 | 3.83174 | 2.239 |
| chr11 | 105908500 | 105908742 | 0.46882 | 1.04998 | 2.240 |
| chr19 | 35535164 | 35535673 | 2.20457 | 4.93741 | 2.240 |
| chr4 | 27862909 | 27863220 | 0.806799 | 1.80696 | 2.240 |
| chr16 | 83469091 | 83469415 | 0.530851 | 1.18895 | 2.240 |
| chr4 | 156128767 | 156128976 | 0.91705 | 2.05399 | 2.240 |
| chr4 | 90032213 | 90032593 | 1.67405 | 3.74978 | 2.240 |
| chr10 | 66791553 | 66791759 | 0.444246 | 0.995107 | 2.240 |
| chr14 | 34261916 | 34262207 | 2.07975 | 4.65862 | 2.240 |
| chr8 | 127839040 | 127839367 | 0.639817 | 1.43328 | 2.240 |
| chr9 | 38017421 | 38017630 | 1.20741 | 2.70493 | 2.240 |
| chr8 | 61936036 | 61936448 | 2.54726 | 5.70658 | 2.240 |
| chr7 | 31776025 | 31776329 | 1.34931 | 3.02286 | 2.240 |
| chr19 | 52740896 | 52741402 | 1.42593 | 3.19457 | 2.240 |
| chr14 | 86829821 | 86830040 | 0.465726 | 1.0434 | 2.240 |
| chr9 | 130797408 | 130797817 | 1.35588 | 3.03774 | 2.240 |
| chr17 | 39822157 | 39822499 | 2.83673 | 6.3555 | 2.240 |
| chr12 | 8198583 | 8198972 | 1.75627 | 3.93481 | 2.240 |
| chr8 | 22941587 | 22941997 | 1.71447 | 3.84119 | 2.240 |
| chr12 | 26249963 | 26250423 | 1.70512 | 3.82039 | 2.241 |
| chr2 | 202096975 | 202097279 | 1.8956 | 4.24726 | 2.241 |
| chr13 | 98227202 | 98227546 | 1.1959 | 2.67959 | 2.241 |
| chr6 | 1424624 | 1425025 | 1.05138 | 2.35585 | 2.241 |
| chr12 | 96910587 | 96910792 | 0.8099 | 1.81479 | 2.241 |
| chr10 | 35900815 | 35901040 | 0.718482 | 1.6101 | 2.241 |
| chr12 | 107407867 | 107408146 | 0.978876 | 2.19364 | 2.241 |
| chr17 | 71994710 | 71995149 | 1.63928 | 3.67381 | 2.241 |
| chr15 | 58826257 | 58826482 | 0.575568 | 1.28996 | 2.241 |
| chr15 | 74532316 | 74532572 | 1.29501 | 2.90253 | 2.241 |
| chr10 | 32479933 | 32480193 | 0.750017 | 1.68104 | 2.241 |
| chr3 | 141456884 | 141457329 | 1.58878 | 3.56107 | 2.241 |
| chr9 | 92208758 | 92209166 | 1.97894 | 4.43564 | 2.241 |
| chr7 | 51230455 | 51230824 | 1.67557 | 3.75568 | 2.241 |
| chr7 | 53090652 | 53090855 | 0.409476 | 0.91781 | 2.241 |
| chr11 | 21972358 | 21972575 | 0.466365 | 1.04535 | 2.241 |
| chr6 | 44598235 | 44598469 | 0.941668 | 2.11075 | 2.242 |
| chr19 | 36554525 | 36554810 | 0.711059 | 1.59388 | 2.242 |
| chr8 | 38824027 | 38824294 | 1.34232 | 3.00889 | 2.242 |
| chr17 | 39485596 | 39485919 | 1.50631 | 3.37657 | 2.242 |
| chr5 | 169253708 | 169253913 | 0.948779 | 2.12685 | 2.242 |
| chr4 | 163566619 | 163566854 | 1.59736 | 3.58088 | 2.242 |
| chr17 | 41771332 | 41772129 | 1.20667 | 2.70506 | 2.242 |
| chr2 | 97136289 | 97136780 | 1.30922 | 2.93504 | 2.242 |
| chr3 | 154647122 | 154647382 | 0.529472 | 1.18706 | 2.242 |
| chr5 | 138758052 | 138758278 | 1.24588 | 2.79329 | 2.242 |
| chr15 | 78916401 | 78916606 | 0.558125 | 1.25135 | 2.242 |
| chr3 | 184258913 | 184259281 | 1.36119 | 3.05209 | 2.242 |

| | | | | | |
|---|---|---|---|---|---|
| chr18 | 10560510 | 10561000 | 1.07631 | 2.4134 | 2.242 |
| chr17 | 73050788 | 73051010 | 0.917641 | 2.05767 | 2.242 |
| chr19 | 45406807 | 45407419 | 2.8281 | 6.3418 | 2.242 |
| chr15 | 60209689 | 60210092 | 0.997272 | 2.23633 | 2.242 |
| chr6 | 160955220 | 160955629 | 1.44502 | 3.2406 | 2.243 |
| chr3 | 159756689 | 159756978 | 0.603654 | 1.35376 | 2.243 |
| chr12 | 52607175 | 52607683 | 2.11964 | 4.75367 | 2.243 |
| chr2 | 95652216 | 95652429 | 1.16901 | 2.62173 | 2.243 |
| chr16 | 56637528 | 56637813 | 0.915629 | 2.05362 | 2.243 |
| chrX | 41417822 | 41418074 | 1.06453 | 2.38764 | 2.243 |
| chr17 | 79407745 | 79408223 | 2.22164 | 4.98354 | 2.243 |
| chr15 | 94628853 | 94629132 | 0.467399 | 1.04847 | 2.243 |
| chr11 | 120042971 | 120043267 | 1.57443 | 3.53199 | 2.243 |
| chr1 | 27293435 | 27293741 | 2.24192 | 5.02947 | 2.243 |
| chrX | 75473960 | 75474198 | 0.848101 | 1.90262 | 2.243 |
| chr7 | 150053304 | 150053919 | 1.19741 | 2.68632 | 2.243 |
| chr3 | 70114919 | 70115181 | 1.05067 | 2.35723 | 2.244 |
| chr17 | 49376214 | 49376467 | 1.34225 | 3.01149 | 2.244 |
| chr14 | 100351691 | 100352007 | 1.12746 | 2.52978 | 2.244 |
| chr7 | 23469943 | 23470270 | 1.24076 | 2.78416 | 2.244 |
| chr3 | 22534302 | 22534560 | 0.355806 | 0.798426 | 2.244 |
| chr7 | 69750989 | 69751279 | 0.606104 | 1.36018 | 2.244 |
| chr3 | 52480973 | 52481425 | 1.99896 | 4.48601 | 2.244 |
| chrX | 24099773 | 24100369 | 1.34121 | 3.00991 | 2.244 |
| chrX | 43419026 | 43419262 | 1.06196 | 2.38327 | 2.244 |
| chr12 | 107472986 | 107473230 | 1.22215 | 2.743 | 2.244 |
| chr20 | 61439551 | 61440028 | 1.69708 | 3.80903 | 2.244 |
| chr7 | 139184645 | 139185116 | 1.77844 | 3.99169 | 2.244 |
| chr8 | 137296864 | 137297120 | 0.543964 | 1.22093 | 2.245 |
| chr10 | 56342371 | 56342631 | 0.553931 | 1.24341 | 2.245 |
| chr1 | 169461360 | 169461570 | 0.563506 | 1.26493 | 2.245 |
| chr16 | 84984659 | 84984936 | 1.11235 | 2.49697 | 2.245 |
| chr13 | 110351977 | 110352223 | 0.564655 | 1.26754 | 2.245 |
| chr11 | 55623632 | 55623903 | 0.452341 | 1.01542 | 2.245 |
| chrX | 153601730 | 153602287 | 1.70882 | 3.83612 | 2.245 |
| chr8 | 51739350 | 51739625 | 0.696553 | 1.5637 | 2.245 |
| chr15 | 98682121 | 98682482 | 0.471247 | 1.05792 | 2.245 |
| chr17 | 58513309 | 58513764 | 1.36168 | 3.05696 | 2.245 |
| chr12 | 53374637 | 53374982 | 2.39449 | 5.37594 | 2.245 |
| chr4 | 6329027 | 6329420 | 1.58613 | 3.56125 | 2.245 |
| chr3 | 155701661 | 155701871 | 0.532126 | 1.19478 | 2.245 |
| chr9 | 133922087 | 133922548 | 2.26498 | 5.08563 | 2.245 |
| chr17 | 40915426 | 40915903 | 2.20428 | 4.94949 | 2.245 |
| chr5 | 133375175 | 133375729 | 2.43606 | 5.47035 | 2.246 |
| chr18 | 61173348 | 61173630 | 1.01691 | 2.28356 | 2.246 |
| chr17 | 41471396 | 41471673 | 1.6864 | 3.78717 | 2.246 |
| chr1 | 244816519 | 244816782 | 0.861789 | 1.93537 | 2.246 |
| chr8 | 127313173 | 127313554 | 1.33212 | 2.99162 | 2.246 |
| chr18 | 19477738 | 19478028 | 1.02987 | 2.3129 | 2.246 |
| chr14 | 25659990 | 25660329 | 0.460435 | 1.03406 | 2.246 |
| chr20 | 4716173 | 4716432 | 0.433808 | 0.974258 | 2.246 |
| chr17 | 72350168 | 72350838 | 2.14502 | 4.81765 | 2.246 |
| chr1 | 20848824 | 20849141 | 1.76003 | 3.95301 | 2.246 |
| chr3 | 127889812 | 127890041 | 0.857437 | 1.9259 | 2.246 |
| chr9 | 3240582 | 3241065 | 1.38967 | 3.12141 | 2.246 |
| chr11 | 76518437 | 76518895 | 1.55056 | 3.48294 | 2.246 |
| chr14 | 96411849 | 96412052 | 0.546208 | 1.22696 | 2.246 |
| chr5 | 158139021 | 158139300 | 0.562492 | 1.26356 | 2.246 |
| chr12 | 99400315 | 99400517 | 0.568129 | 1.27633 | 2.247 |
| chr5 | 57756109 | 57756904 | 1.15901 | 2.60386 | 2.247 |
| chr5 | 173891611 | 173891990 | 0.655415 | 1.47249 | 2.247 |

| | | | | | |
|---|---|---|---|---|---|
| chr19 | 41250423 | 41250807 | 0.813305 | 1.82727 | 2.247 |
| chr14 | 36875357 | 36875697 | 0.387937 | 0.871616 | 2.247 |
| chr11 | 42339229 | 42339546 | 0.442728 | 0.994732 | 2.247 |
| chr7 | 150864309 | 150864734 | 0.620039 | 1.39312 | 2.247 |
| chr17 | 29852292 | 29853706 | 1.51171 | 3.39678 | 2.247 |
| chr4 | 29972194 | 29972469 | 0.619555 | 1.39213 | 2.247 |
| chrX | 117896262 | 117896495 | 0.405621 | 0.911451 | 2.247 |
| chr4 | 122184088 | 122184332 | 0.4238 | 0.952327 | 2.247 |
| chr11 | 1567702 | 1568174 | 1.54795 | 3.47845 | 2.247 |
| chr8 | 99643977 | 99644243 | 0.390583 | 0.877703 | 2.247 |
| chr8 | 107093316 | 107093532 | 1.26127 | 2.83438 | 2.247 |
| chr1 | 156829809 | 156830209 | 1.89709 | 4.26328 | 2.247 |
| chr12 | 126182276 | 126182552 | 0.581456 | 1.30676 | 2.247 |
| chr17 | 80669618 | 80669976 | 1.22013 | 2.7422 | 2.247 |
| chr12 | 92406841 | 92407117 | 0.562831 | 1.26503 | 2.248 |
| chr6 | 44443444 | 44443793 | 1.74556 | 3.92354 | 2.248 |
| chr9 | 131200798 | 131201291 | 2.02475 | 4.55111 | 2.248 |
| chr18 | 24229706 | 24230022 | 1.14904 | 2.58275 | 2.248 |
| chr11 | 15286398 | 15286614 | 0.372925 | 0.838245 | 2.248 |
| chr19 | 46520121 | 46520450 | 2.94368 | 6.617 | 2.248 |
| chr17 | 18832068 | 18832403 | 0.707028 | 1.58941 | 2.248 |
| chr11 | 18834458 | 18834679 | 0.471704 | 1.06041 | 2.248 |
| chr10 | 91434510 | 91434837 | 0.491078 | 1.10399 | 2.248 |
| chr2 | 159602220 | 159602449 | 1.43987 | 3.23697 | 2.248 |
| chr4 | 110343597 | 110343877 | 1.35227 | 3.04006 | 2.248 |
| chr1 | 167021696 | 167022040 | 1.52546 | 3.42944 | 2.248 |
| chr11 | 15027698 | 15028125 | 0.874067 | 1.96506 | 2.248 |
| chr7 | 134105331 | 134105607 | 1.54802 | 3.48023 | 2.248 |
| chr4 | 177968469 | 177968801 | 1.22977 | 2.76476 | 2.248 |
| chr10 | 123811164 | 123811453 | 1.73878 | 3.90953 | 2.248 |
| chr9 | 124312760 | 124313210 | 2.18474 | 4.91238 | 2.249 |
| chr12 | 56114741 | 56115230 | 1.21941 | 2.74187 | 2.249 |
| chr15 | 56783343 | 56783704 | 1.97611 | 4.44332 | 2.249 |
| chr11 | 62356827 | 62357035 | 0.810871 | 1.82343 | 2.249 |
| chr7 | 138411027 | 138411426 | 1.06599 | 2.39719 | 2.249 |
| chr1 | 6498101 | 6498525 | 1.23592 | 2.77944 | 2.249 |
| chr17 | 50547978 | 50548284 | 0.641382 | 1.4424 | 2.249 |
| chr15 | 34256359 | 34256594 | 1.94641 | 4.37728 | 2.249 |
| chr4 | 128508182 | 128508464 | 0.844883 | 1.90013 | 2.249 |
| chr5 | 4182360 | 4182653 | 0.638397 | 1.43583 | 2.249 |
| chr12 | 130516135 | 130516531 | 2.10761 | 4.74093 | 2.249 |
| chr16 | 47175601 | 47175952 | 2.34988 | 5.28597 | 2.249 |
| chr9 | 108188326 | 108188638 | 0.881363 | 1.98259 | 2.249 |
| chr2 | 97616858 | 97617252 | 2.34086 | 5.26572 | 2.249 |
| chr1 | 38455716 | 38456036 | 1.85387 | 4.17026 | 2.249 |
| chr12 | 22596361 | 22596691 | 1.45826 | 3.28043 | 2.250 |
| chr11 | 61555507 | 61555994 | 1.67835 | 3.77563 | 2.250 |
| chr2 | 187993025 | 187993271 | 0.390466 | 0.878467 | 2.250 |
| chr8 | 41400171 | 41400485 | 1.32787 | 2.98744 | 2.250 |
| chr14 | 50999200 | 50999625 | 2.17507 | 4.8935 | 2.250 |
| chr2 | 43832288 | 43832571 | 1.00441 | 2.25974 | 2.250 |
| chr8 | 24923054 | 24923467 | 1.5045 | 3.3851 | 2.250 |
| chr13 | 107185434 | 107185720 | 1.16126 | 2.61293 | 2.250 |
| chr18 | 63014881 | 63015144 | 0.404055 | 0.909193 | 2.250 |
| chr19 | 46580817 | 46581230 | 2.83065 | 6.36972 | 2.250 |
| chr1 | 100925110 | 100925425 | 1.18197 | 2.65995 | 2.250 |
| chr1 | 156717164 | 156717396 | 0.788331 | 1.77411 | 2.250 |
| chr16 | 8223196 | 8223504 | 1.56804 | 3.52887 | 2.251 |
| chr22 | 37941226 | 37941462 | 2.08588 | 4.69429 | 2.251 |
| chr9 | 93080647 | 93080866 | 1.18015 | 2.65607 | 2.251 |
| chr3 | 17073977 | 17074236 | 1.14139 | 2.56895 | 2.251 |

| | | | | | |
|---|---|---|---|---|---|
| chr3 | 77019138 | 77019413 | 0.417128 | 0.938845 | 2.251 |
| chr6 | 99293717 | 99293964 | 1.80201 | 4.05594 | 2.251 |
| chr8 | 127836688 | 127837002 | 1.62677 | 3.66152 | 2.251 |
| chr20 | 17999807 | 18000105 | 0.85453 | 1.9234 | 2.251 |
| chr5 | 77304232 | 77304526 | 1.87653 | 4.22385 | 2.251 |
| chr16 | 88343519 | 88343894 | 0.876917 | 1.97384 | 2.251 |
| chr3 | 79027269 | 79027535 | 0.997394 | 2.24507 | 2.251 |
| chr17 | 74467248 | 74467490 | 0.574935 | 1.29415 | 2.251 |
| chr15 | 40574658 | 40575024 | 1.08531 | 2.443 | 2.251 |
| chr8 | 146159749 | 146160173 | 0.79143 | 1.78151 | 2.251 |
| chr10 | 77154882 | 77155455 | 2.00727 | 4.51843 | 2.251 |
| chr7 | 95174337 | 95174616 | 1.23619 | 2.78287 | 2.251 |
| chr12 | 103035824 | 103036048 | 0.469821 | 1.05767 | 2.251 |
| chr10 | 101860348 | 101860699 | 1.11668 | 2.51399 | 2.251 |
| chr17 | 58766607 | 58766834 | 0.490579 | 1.10445 | 2.251 |
| chr4 | 89436865 | 89437106 | 1.16898 | 2.63203 | 2.252 |
| chr2 | 177518610 | 177518880 | 1.20897 | 2.72244 | 2.252 |
| chr16 | 75855087 | 75855324 | 0.539567 | 1.21514 | 2.252 |
| chr8 | 143381201 | 143381542 | 1.28757 | 2.89977 | 2.252 |
| chr9 | 34899550 | 34899969 | 1.30837 | 2.94677 | 2.252 |
| chr13 | 101854959 | 101855255 | 0.604447 | 1.36143 | 2.252 |
| chr15 | 30072575 | 30072784 | 0.708134 | 1.59503 | 2.252 |
| chr1 | 220164440 | 220164657 | 0.918665 | 2.06935 | 2.253 |
| chr8 | 143863637 | 143864388 | 1.31636 | 2.96521 | 2.253 |
| chr6 | 131019116 | 131019468 | 1.51322 | 3.40892 | 2.253 |
| chr9 | 137552315 | 137552607 | 1.64149 | 3.698 | 2.253 |
| chr12 | 76474540 | 76474751 | 0.859401 | 1.93614 | 2.253 |
| chr19 | 46145382 | 46145695 | 2.33888 | 5.26928 | 2.253 |
| chr14 | 61188465 | 61188694 | 0.822674 | 1.85353 | 2.253 |
| chr17 | 56214047 | 56214458 | 1.57036 | 3.53811 | 2.253 |
| chr3 | 156391729 | 156392239 | 1.39518 | 3.14353 | 2.253 |
| chr4 | 117101288 | 117101496 | 0.833883 | 1.87888 | 2.253 |
| chr3 | 17686622 | 17686943 | 1.33005 | 2.99684 | 2.253 |
| chr2 | 45870774 | 45871171 | 1.72892 | 3.89565 | 2.253 |
| chr11 | 123425594 | 123425970 | 1.31056 | 2.95306 | 2.253 |
| chr12 | 18473006 | 18473262 | 1.09986 | 2.47829 | 2.253 |
| chr12 | 68752635 | 68752918 | 1.31106 | 2.95428 | 2.253 |
| chr6 | 110026378 | 110026781 | 1.89794 | 4.27674 | 2.253 |
| chr18 | 21095904 | 21096273 | 1.57667 | 3.55293 | 2.253 |
| chr6 | 45890582 | 45890810 | 1.62712 | 3.66667 | 2.253 |
| chr10 | 72639784 | 72640165 | 2.21389 | 4.989 | 2.254 |
| chr15 | 60981069 | 60981271 | 0.546641 | 1.23196 | 2.254 |
| chr1 | 241581588 | 241581870 | 0.534267 | 1.20408 | 2.254 |
| chr1 | 196692546 | 196692898 | 1.04906 | 2.3643 | 2.254 |
| chr9 | 119922598 | 119922803 | 0.625006 | 1.40864 | 2.254 |
| chr14 | 41319483 | 41319759 | 0.441368 | 0.99476 | 2.254 |
| chr14 | 90985153 | 90985563 | 1.09407 | 2.46594 | 2.254 |
| chr17 | 1533223 | 1533713 | 1.51944 | 3.42486 | 2.254 |
| chr10 | 65800965 | 65801309 | 2.58655 | 5.83029 | 2.254 |
| chr2 | 68440966 | 68441175 | 0.415698 | 0.937041 | 2.254 |
| chr14 | 37411461 | 37411789 | 1.85569 | 4.18323 | 2.254 |
| chr5 | 140624748 | 140625181 | 1.02118 | 2.30202 | 2.254 |
| chr10 | 35882929 | 35883242 | 0.467358 | 1.05356 | 2.254 |
| chr10 | 107503765 | 107504030 | 0.549826 | 1.23947 | 2.254 |
| chr9 | 135852927 | 135853437 | 2.47498 | 5.57932 | 2.254 |
| chrX | 149849519 | 149849726 | 0.884174 | 1.99329 | 2.254 |
| chr12 | 122675404 | 122675934 | 1.62632 | 3.66642 | 2.254 |
| chr12 | 78393872 | 78394168 | 1.27037 | 2.86404 | 2.254 |
| chr20 | 247484 | 248030 | 2.12988 | 4.80217 | 2.255 |
| chr17 | 79917469 | 79918020 | 2.74414 | 6.18715 | 2.255 |
| chr2 | 232316717 | 232316942 | 1.2389 | 2.79342 | 2.255 |

| | | | | | |
|---|---|---|---|---|---|
| chr11 | 61519726 | 61520267 | 2.02925 | 4.57549 | 2.255 |
| chr10 | 60712396 | 60712783 | 1.19152 | 2.68665 | 2.255 |
| chr5 | 70771633 | 70772016 | 1.19616 | 2.69727 | 2.255 |
| chr6 | 127442496 | 127442892 | 1.52413 | 3.43686 | 2.255 |
| chr15 | 77411149 | 77411551 | 1.50331 | 3.38996 | 2.255 |
| chr17 | 30241739 | 30241977 | 1.02767 | 2.31742 | 2.255 |
| chr1 | 168421389 | 168421915 | 0.924537 | 2.08487 | 2.255 |
| chr8 | 120339809 | 120340055 | 0.500654 | 1.129 | 2.255 |
| chr14 | 39179972 | 39180217 | 0.585457 | 1.32024 | 2.255 |
| chr10 | 72575386 | 72575814 | 2.95718 | 6.66891 | 2.255 |
| chr1 | 37952988 | 37953375 | 1.51371 | 3.41378 | 2.255 |
| chr6 | 159125202 | 159125594 | 2.43019 | 5.4807 | 2.255 |
| chr8 | 97192363 | 97192704 | 1.56108 | 3.52064 | 2.255 |
| chr1 | 110691627 | 110691866 | 0.787299 | 1.77557 | 2.255 |
| chr7 | 49774293 | 49774523 | 0.624379 | 1.40824 | 2.255 |
| chr11 | 20631653 | 20632050 | 1.99682 | 4.50371 | 2.255 |
| chr12 | 53583866 | 53584271 | 1.9674 | 4.43745 | 2.255 |
| chr3 | 9394844 | 9395212 | 1.68202 | 3.79386 | 2.256 |
| chr7 | 127218311 | 127218702 | 0.847497 | 1.91157 | 2.256 |
| chr11 | 44832210 | 44832440 | 0.395398 | 0.891866 | 2.256 |
| chr2 | 31046759 | 31047017 | 0.361923 | 0.816383 | 2.256 |
| chr7 | 148224743 | 148225088 | 1.53023 | 3.45185 | 2.256 |
| chr3 | 3071770 | 3072070 | 0.54521 | 1.23 | 2.256 |
| chr14 | 69404547 | 69404825 | 1.05925 | 2.38969 | 2.256 |
| chr1 | 5569949 | 5570297 | 1.92373 | 4.34003 | 2.256 |
| chr7 | 134236748 | 134237213 | 0.73799 | 1.66498 | 2.256 |
| chr8 | 68251741 | 68252011 | 1.41906 | 3.20163 | 2.256 |
| chr14 | 82720867 | 82721082 | 0.478956 | 1.08063 | 2.256 |
| chr14 | 105511953 | 105512420 | 3.36763 | 7.59826 | 2.256 |
| chr11 | 61348153 | 61348966 | 0.89698 | 2.02391 | 2.256 |
| chr8 | 39184922 | 39185268 | 2.15962 | 4.8729 | 2.256 |
| chr3 | 133142458 | 133142722 | 1.38691 | 3.1294 | 2.256 |
| chr8 | 79712481 | 79712697 | 0.376672 | 0.849958 | 2.256 |
| chr2 | 232513329 | 232513643 | 1.59539 | 3.60004 | 2.257 |
| chr8 | 55078197 | 55079046 | 1.05634 | 2.38367 | 2.257 |
| chr18 | 13530945 | 13531167 | 0.869883 | 1.96307 | 2.257 |
| chr14 | 29226362 | 29226644 | 1.44166 | 3.25343 | 2.257 |
| chr10 | 112480949 | 112481235 | 0.510727 | 1.15259 | 2.257 |
| chr17 | 53957133 | 53957366 | 0.525422 | 1.18577 | 2.257 |
| chr13 | 69438075 | 69438348 | 0.635845 | 1.43506 | 2.257 |
| chr7 | 140353211 | 140353595 | 1.23851 | 2.79535 | 2.257 |
| chr12 | 115656895 | 115657122 | 0.416143 | 0.939251 | 2.257 |
| chr17 | 17633050 | 17633317 | 1.34517 | 3.03619 | 2.257 |
| chr12 | 133137558 | 133138160 | 1.94716 | 4.39508 | 2.257 |
| chr8 | 136378716 | 136378938 | 0.467055 | 1.05423 | 2.257 |
| chr3 | 43003191 | 43003424 | 0.829812 | 1.87309 | 2.257 |
| chr11 | 42923771 | 42924123 | 0.451858 | 1.01999 | 2.257 |
| chr12 | 49974260 | 49974896 | 1.31516 | 2.96883 | 2.257 |
| chr7 | 50535749 | 50536079 | 1.80201 | 4.06791 | 2.257 |
| chr3 | 186938023 | 186938268 | 1.17828 | 2.65994 | 2.257 |
| chr12 | 24696272 | 24696527 | 0.593702 | 1.34031 | 2.258 |
| chr16 | 72501106 | 72501429 | 1.25405 | 2.83119 | 2.258 |
| chr18 | 48662196 | 48662456 | 0.838812 | 1.89374 | 2.258 |
| chr3 | 46120181 | 46120636 | 1.49265 | 3.36994 | 2.258 |
| chr16 | 67980567 | 67980769 | 0.555124 | 1.25332 | 2.258 |
| chr4 | 98074291 | 98074500 | 0.714146 | 1.61239 | 2.258 |
| chr7 | 150820971 | 150821452 | 2.59937 | 5.86892 | 2.258 |
| chr11 | 3238374 | 3238863 | 2.10158 | 4.74504 | 2.258 |
| chr21 | 34198994 | 34199214 | 0.98345 | 2.22047 | 2.258 |
| chr8 | 62295224 | 62295461 | 0.525832 | 1.18728 | 2.258 |
| chr16 | 89927199 | 89927576 | 2.65963 | 6.00526 | 2.258 |

| | | | | | |
|---|---|---|---|---|---|
| chr18 | 72912171 | 72912439 | 0.518668 | 1.17112 | 2.258 |
| chr10 | 116308521 | 116308815 | 0.973609 | 2.19843 | 2.258 |
| chr12 | 76849623 | 76849830 | 0.987584 | 2.23006 | 2.258 |
| chr9 | 31449085 | 31449336 | 0.46454 | 1.04898 | 2.258 |
| chr12 | 83079502 | 83079808 | 1.66292 | 3.75515 | 2.258 |
| chr12 | 52417460 | 52417893 | 2.03808 | 4.60238 | 2.258 |
| chr10 | 111085959 | 111086177 | 0.381403 | 0.861287 | 2.258 |
| chr12 | 42227941 | 42228195 | 0.487727 | 1.10145 | 2.258 |
| chr11 | 64099336 | 64099683 | 1.24703 | 2.81642 | 2.259 |
| chr11 | 2913223 | 2913686 | 1.95681 | 4.41964 | 2.259 |
| chr17 | 16492596 | 16493075 | 2.41882 | 5.46313 | 2.259 |
| chr1 | 33761038 | 33761483 | 1.96776 | 4.44442 | 2.259 |
| chr7 | 137263055 | 137263278 | 0.504129 | 1.13864 | 2.259 |
| chr19 | 41768166 | 41768472 | 2.48042 | 5.60248 | 2.259 |
| chr8 | 41560326 | 41560543 | 0.428623 | 0.96814 | 2.259 |
| chr8 | 992018 | 992285 | 0.464327 | 1.04882 | 2.259 |
| chr3 | 100345257 | 100345465 | 0.474839 | 1.07257 | 2.259 |
| chr4 | 25443218 | 25443445 | 1.27109 | 2.87116 | 2.259 |
| chr19 | 39083692 | 39084109 | 0.691638 | 1.56229 | 2.259 |
| chr17 | 79185021 | 79185390 | 0.390541 | 0.882171 | 2.259 |
| chr1 | 47902106 | 47902637 | 1.79805 | 4.06152 | 2.259 |
| chr6 | 93523367 | 93523598 | 0.392667 | 0.886986 | 2.259 |
| chr1 | 101300644 | 101300890 | 1.00384 | 2.26765 | 2.259 |
| chr19 | 44023672 | 44023995 | 1.92708 | 4.35334 | 2.259 |
| chr2 | 121856906 | 121857217 | 1.16916 | 2.64117 | 2.259 |
| chr2 | 162283203 | 162283563 | 0.873537 | 1.97351 | 2.259 |
| chr3 | 126108084 | 126108472 | 2.01555 | 4.55385 | 2.259 |
| chr7 | 153616255 | 153616482 | 0.327027 | 0.738872 | 2.259 |
| chr14 | 102789409 | 102789634 | 1.20151 | 2.71466 | 2.259 |
| chr16 | 69830449 | 69830723 | 1.51391 | 3.42048 | 2.259 |
| chr5 | 15933789 | 15934044 | 0.486807 | 1.09993 | 2.259 |
| chr8 | 134898292 | 134898752 | 1.68173 | 3.79989 | 2.260 |
| chr21 | 48059719 | 48060010 | 0.778415 | 1.75886 | 2.260 |
| chr3 | 76388510 | 76388760 | 0.554354 | 1.25265 | 2.260 |
| chr1 | 154244714 | 154244962 | 1.34944 | 3.0495 | 2.260 |
| chr8 | 142183278 | 142183787 | 1.47895 | 3.34224 | 2.260 |
| chr1 | 205479462 | 205479737 | 0.47419 | 1.07164 | 2.260 |
| chr22 | 32242748 | 32242966 | 0.685395 | 1.54897 | 2.260 |
| chr18 | 51970751 | 51970992 | 1.65275 | 3.73539 | 2.260 |
| chr17 | 47781287 | 47781536 | 1.46436 | 3.30965 | 2.260 |
| chr10 | 118386974 | 118387275 | 0.887577 | 2.00606 | 2.260 |
| chr1 | 60437118 | 60437391 | 2.02439 | 4.57549 | 2.260 |
| chr10 | 134391173 | 134391409 | 1.16616 | 2.63576 | 2.260 |
| chr13 | 61228496 | 61228898 | 0.734084 | 1.6592 | 2.260 |
| chr11 | 58717683 | 58718194 | 0.487411 | 1.10171 | 2.260 |
| chr8 | 60964729 | 60964986 | 1.00257 | 2.26626 | 2.260 |
| chr1 | 199653843 | 199654137 | 1.48264 | 3.35155 | 2.261 |
| chr10 | 34079728 | 34079965 | 0.552146 | 1.24819 | 2.261 |
| chr3 | 132061909 | 132062177 | 1.59305 | 3.60132 | 2.261 |
| chr2 | 208936994 | 208937438 | 1.4104 | 3.18843 | 2.261 |
| chr12 | 6793778 | 6794123 | 1.2089 | 2.73296 | 2.261 |
| chr11 | 6827669 | 6827884 | 0.407361 | 0.920949 | 2.261 |
| chr7 | 101386527 | 101386851 | 1.36657 | 3.08953 | 2.261 |
| chr12 | 61793268 | 61793479 | 0.461274 | 1.04288 | 2.261 |
| chr17 | 10534342 | 10534761 | 2.46828 | 5.58058 | 2.261 |
| chr20 | 8771110 | 8771498 | 2.41973 | 5.47094 | 2.261 |
| chr5 | 147329328 | 147329533 | 0.729007 | 1.64829 | 2.261 |
| chr3 | 65736720 | 65737141 | 1.18348 | 2.676 | 2.261 |
| chr16 | 70749437 | 70749650 | 0.466808 | 1.05555 | 2.261 |
| chr17 | 34091057 | 34091985 | 1.9483 | 4.4057 | 2.261 |
| chr11 | 65585549 | 65586153 | 2.6925 | 6.08971 | 2.262 |

| | | | | | |
|---|---|---|---|---|---|
| chr11 | 14844329 | 14844554 | 0.445382 | 1.00737 | 2.262 |
| chr7 | 2749244 | 2749775 | 2.46643 | 5.57863 | 2.262 |
| chr19 | 49979231 | 49979874 | 1.0564 | 2.38949 | 2.262 |
| chr8 | 108783042 | 108783355 | 0.606243 | 1.37128 | 2.262 |
| chr17 | 17888890 | 17889300 | 0.944933 | 2.1374 | 2.262 |
| chr11 | 59686569 | 59686785 | 1.06643 | 2.41223 | 2.262 |
| chr4 | 153856435 | 153856673 | 0.552852 | 1.25058 | 2.262 |
| chr1 | 242352954 | 242353266 | 1.12238 | 2.53894 | 2.262 |
| chr2 | 165773874 | 165774095 | 1.09846 | 2.48485 | 2.262 |
| chr19 | 48685723 | 48685956 | 0.870682 | 1.96974 | 2.262 |
| chr3 | 123847526 | 123847729 | 0.390818 | 0.884184 | 2.262 |
| chr20 | 9388785 | 9389126 | 1.34398 | 3.0408 | 2.263 |
| chr16 | 58155824 | 58156215 | 1.29305 | 2.92567 | 2.263 |
| chr1 | 2313185 | 2313600 | 2.03609 | 4.60715 | 2.263 |
| chr12 | 18312175 | 18312464 | 1.28227 | 2.90144 | 2.263 |
| chr12 | 48343194 | 48343615 | 2.08817 | 4.72502 | 2.263 |
| chr5 | 144466651 | 144466859 | 0.358975 | 0.812298 | 2.263 |
| chr8 | 1562826 | 1563185 | 0.500705 | 1.13302 | 2.263 |
| chr6 | 141747545 | 141747761 | 0.587174 | 1.32874 | 2.263 |
| chr5 | 18305509 | 18305837 | 1.2534 | 2.83646 | 2.263 |
| chr15 | 93461228 | 93461465 | 1.56668 | 3.54561 | 2.263 |
| chr8 | 118000524 | 118000738 | 0.428748 | 0.970336 | 2.263 |
| chr16 | 85524571 | 85525020 | 2.62401 | 5.93864 | 2.263 |
| chr20 | 1484272 | 1484629 | 2.33585 | 5.28662 | 2.263 |
| chr1 | 9762335 | 9762586 | 0.913026 | 2.06647 | 2.263 |
| chr8 | 87883348 | 87883558 | 0.787627 | 1.78276 | 2.263 |
| chrX | 10222796 | 10223028 | 0.825864 | 1.86932 | 2.263 |
| chr10 | 80417334 | 80417664 | 1.59307 | 3.60598 | 2.264 |
| chr12 | 124247717 | 124248076 | 0.601983 | 1.36266 | 2.264 |
| chr17 | 16424614 | 16425029 | 1.34814 | 3.05167 | 2.264 |
| chr22 | 39266265 | 39266625 | 1.61235 | 3.64979 | 2.264 |
| chr14 | 103059584 | 103059852 | 0.607756 | 1.37575 | 2.264 |
| chr2 | 173037131 | 173037498 | 1.25437 | 2.83948 | 2.264 |
| chr6 | 86192770 | 86193219 | 1.71679 | 3.88628 | 2.264 |
| chr16 | 717818 | 718159 | 1.56694 | 3.54731 | 2.264 |
| chr1 | 53166660 | 53166894 | 0.994551 | 2.25158 | 2.264 |
| chr2 | 6183259 | 6183462 | 0.634009 | 1.43536 | 2.264 |
| chr17 | 31003473 | 31003854 | 1.93582 | 4.38266 | 2.264 |
| chr7 | 99214086 | 99214422 | 1.09047 | 2.46887 | 2.264 |
| chr2 | 150954590 | 150954841 | 0.565073 | 1.27939 | 2.264 |
| chr6 | 85259730 | 85259945 | 0.663928 | 1.50325 | 2.264 |
| chr8 | 66948814 | 66949047 | 1.12348 | 2.54377 | 2.264 |
| chr21 | 44013351 | 44013744 | 1.08233 | 2.45061 | 2.264 |
| chr14 | 20715861 | 20716132 | 1.25898 | 2.8507 | 2.264 |
| chr3 | 149230065 | 149230463 | 1.73931 | 3.93878 | 2.265 |
| chr10 | 103892240 | 103892616 | 1.60792 | 3.64134 | 2.265 |
| chr3 | 29806198 | 29806482 | 1.61096 | 3.64831 | 2.265 |
| chr8 | 98852189 | 98852398 | 0.89438 | 2.02552 | 2.265 |
| chr14 | 21323859 | 21324180 | 1.33031 | 3.01284 | 2.265 |
| chr15 | 61775984 | 61776261 | 0.616941 | 1.39723 | 2.265 |
| chr1 | 177917533 | 177917806 | 1.48121 | 3.35472 | 2.265 |
| chr2 | 45251898 | 45252200 | 0.793972 | 1.79836 | 2.265 |
| chr8 | 89863563 | 89863783 | 0.528681 | 1.19751 | 2.265 |
| chr18 | 61603366 | 61603764 | 1.16504 | 2.63906 | 2.265 |
| chr17 | 64447683 | 64447944 | 1.3932 | 3.15596 | 2.265 |
| chr3 | 170529794 | 170530128 | 1.50625 | 3.41211 | 2.265 |
| chr8 | 49084665 | 49085062 | 2.08113 | 4.71449 | 2.265 |
| chr3 | 115411789 | 115412060 | 1.70739 | 3.86802 | 2.265 |
| chr1 | 7727612 | 7727965 | 2.15169 | 4.87459 | 2.265 |
| chr20 | 2832688 | 2833130 | 1.44709 | 3.27849 | 2.266 |
| chr8 | 53001667 | 53001916 | 1.08417 | 2.45627 | 2.266 |

| | | | | | |
|---|---|---|---|---|---|
| chr12 | 15925408 | 15925616 | 0.874127 | 1.98046 | 2.266 |
| chr5 | 179553978 | 179554456 | 1.72576 | 3.90999 | 2.266 |
| chr9 | 75289378 | 75289640 | 0.518032 | 1.1737 | 2.266 |
| chr18 | 11553334 | 11553673 | 1.52554 | 3.45646 | 2.266 |
| chr11 | 102212524 | 102212823 | 1.1047 | 2.50302 | 2.266 |
| chr5 | 98964278 | 98964518 | 0.535242 | 1.21275 | 2.266 |
| chr17 | 14763729 | 14763964 | 0.441768 | 1.00106 | 2.266 |
| chr20 | 3657240 | 3657855 | 1.61144 | 3.65168 | 2.266 |
| chr17 | 65624655 | 65625116 | 1.10523 | 2.50461 | 2.266 |
| chr20 | 40506772 | 40507039 | 0.495182 | 1.12217 | 2.266 |
| chr20 | 61854685 | 61855165 | 1.29433 | 2.93319 | 2.266 |
| chr7 | 38255208 | 38255683 | 1.8824 | 4.26605 | 2.266 |
| chr7 | 126893925 | 126894309 | 1.8166 | 4.11696 | 2.266 |
| chr2 | 45721865 | 45722116 | 1.24851 | 2.82961 | 2.266 |
| chr15 | 85478110 | 85478607 | 1.27672 | 2.89356 | 2.266 |
| chr6 | 143348235 | 143348455 | 0.952863 | 2.15959 | 2.266 |
| chr11 | 77723072 | 77723340 | 0.477164 | 1.08147 | 2.266 |
| chr6 | 165645280 | 165645603 | 1.03177 | 2.33845 | 2.266 |
| chr9 | 21694500 | 21694778 | 0.47221 | 1.07024 | 2.266 |
| chr2 | 85359977 | 85360341 | 1.98075 | 4.48945 | 2.267 |
| chr3 | 87399826 | 87400101 | 1.57071 | 3.5601 | 2.267 |
| chr9 | 128755041 | 128755256 | 0.420693 | 0.953526 | 2.267 |
| chr14 | 106598138 | 106598476 | 0.542069 | 1.22865 | 2.267 |
| chr14 | 74958678 | 74958913 | 0.620275 | 1.40592 | 2.267 |
| chr16 | 31053379 | 31053816 | 0.872077 | 1.97671 | 2.267 |
| chr5 | 142078660 | 142079015 | 1.93879 | 4.39477 | 2.267 |
| chr3 | 122397969 | 122398359 | 1.34163 | 3.0412 | 2.267 |
| chr17 | 21032422 | 21032646 | 1.03818 | 2.35335 | 2.267 |
| chr10 | 103565971 | 103566218 | 0.450928 | 1.02217 | 2.267 |
| chr12 | 119882313 | 119882617 | 1.28893 | 2.92176 | 2.267 |
| chr15 | 81818550 | 81818756 | 0.549721 | 1.24612 | 2.267 |
| chr18 | 51645909 | 51646139 | 1.04782 | 2.37524 | 2.267 |
| chr8 | 51725678 | 51726122 | 0.661638 | 1.49988 | 2.267 |
| chr2 | 171669237 | 171669557 | 1.02025 | 2.31284 | 2.267 |
| chr5 | 94999897 | 95000355 | 1.00145 | 2.2703 | 2.267 |
| chr4 | 15964714 | 15965120 | 1.02186 | 2.31665 | 2.267 |
| chr8 | 2370383 | 2370626 | 0.402348 | 0.912213 | 2.267 |
| chr7 | 84820824 | 84821049 | 0.796978 | 1.80702 | 2.267 |
| chr15 | 78933427 | 78933828 | 0.457087 | 1.03646 | 2.268 |
| chr20 | 33461884 | 33462138 | 1.70187 | 3.85915 | 2.268 |
| chr6 | 169356896 | 169357338 | 0.934353 | 2.11882 | 2.268 |
| chr12 | 22254025 | 22254230 | 0.623784 | 1.41457 | 2.268 |
| chr17 | 42218967 | 42219358 | 2.02523 | 4.59283 | 2.268 |
| chr7 | 73504994 | 73505235 | 0.475904 | 1.07927 | 2.268 |
| chr12 | 62975534 | 62975885 | 1.19245 | 2.70431 | 2.268 |
| chr5 | 87380265 | 87380538 | 0.430126 | 0.975497 | 2.268 |
| chr14 | 61295385 | 61295631 | 0.550073 | 1.24756 | 2.268 |
| chr8 | 95586299 | 95586818 | 1.26345 | 2.8655 | 2.268 |
| chr8 | 21911597 | 21911904 | 1.78406 | 4.04628 | 2.268 |
| chr19 | 48774464 | 48774728 | 1.45003 | 3.28875 | 2.268 |
| chr9 | 139117798 | 139118394 | 1.64369 | 3.72837 | 2.268 |
| chr15 | 65481493 | 65481795 | 1.57472 | 3.57199 | 2.268 |
| chr10 | 83679980 | 83680270 | 1.17874 | 2.67408 | 2.269 |
| chr7 | 100230594 | 100230905 | 1.41985 | 3.22126 | 2.269 |
| chr13 | 101167132 | 101167396 | 1.22243 | 2.77338 | 2.269 |
| chr16 | 62747788 | 62748047 | 0.383504 | 0.870112 | 2.269 |
| chr8 | 144987012 | 144987290 | 0.708215 | 1.60687 | 2.269 |
| chr10 | 90570056 | 90570448 | 1.81755 | 4.1241 | 2.269 |
| chr18 | 32723975 | 32724427 | 1.26295 | 2.86574 | 2.269 |
| chr17 | 48919268 | 48919554 | 1.79897 | 4.08206 | 2.269 |
| chr1 | 166134148 | 166134398 | 1.20744 | 2.73983 | 2.269 |

| | | | | | |
|---|---|---|---|---|---|
| chr8 | 97810436 | 97810836 | 0.519162 | 1.17804 | 2.269 |
| chr1 | 183624639 | 183624939 | 1.51676 | 3.44175 | 2.269 |
| chr2 | 25264214 | 25264447 | 0.955466 | 2.1681 | 2.269 |
| chr15 | 76025158 | 76025435 | 1.65563 | 3.75694 | 2.269 |
| chr7 | 136694975 | 136695324 | 0.446362 | 1.01291 | 2.269 |
| chrX | 106203189 | 106203459 | 1.17332 | 2.66263 | 2.269 |
| chr5 | 21912768 | 21912981 | 0.440397 | 0.999438 | 2.269 |
| chr3 | 159040144 | 159040553 | 1.597 | 3.62437 | 2.269 |
| chr2 | 176231576 | 176231868 | 1.04874 | 2.38029 | 2.270 |
| chr14 | 75489381 | 75489701 | 1.17262 | 2.66157 | 2.270 |
| chr19 | 47917745 | 47918048 | 1.53367 | 3.48133 | 2.270 |
| chr14 | 100149050 | 100149588 | 2.26887 | 5.15042 | 2.270 |
| chr13 | 21402736 | 21402967 | 0.732233 | 1.66223 | 2.270 |
| chr12 | 103576919 | 103577170 | 0.437141 | 0.992361 | 2.270 |
| chr5 | 7415534 | 7415738 | 0.418776 | 0.950711 | 2.270 |
| chr11 | 115373769 | 115374063 | 0.58915 | 1.33756 | 2.270 |
| chr8 | 23606522 | 23607040 | 1.13949 | 2.58703 | 2.270 |
| chr15 | 90853255 | 90853532 | 0.718243 | 1.63066 | 2.270 |
| chr19 | 18169117 | 18169547 | 1.67842 | 3.81064 | 2.270 |
| chr2 | 228336596 | 228337002 | 2.46571 | 5.59826 | 2.270 |
| chr7 | 36061648 | 36062070 | 0.647389 | 1.46989 | 2.270 |
| chr5 | 173352527 | 173352867 | 0.999273 | 2.26889 | 2.271 |
| chr12 | 80280786 | 80281058 | 1.27253 | 2.88961 | 2.271 |
| chr12 | 124066673 | 124067048 | 1.50955 | 3.42792 | 2.271 |
| chr5 | 139752499 | 139752796 | 1.74442 | 3.96129 | 2.271 |
| chr1 | 32181240 | 32181548 | 1.28354 | 2.91484 | 2.271 |
| chr20 | 49411269 | 49411684 | 2.0266 | 4.60271 | 2.271 |
| chr5 | 102568168 | 102568410 | 1.39045 | 3.15792 | 2.271 |
| chr9 | 124952927 | 124953176 | 0.386297 | 0.877345 | 2.271 |
| chr1 | 87299298 | 87299603 | 1.1158 | 2.53423 | 2.271 |
| chr12 | 67102491 | 67102697 | 0.843734 | 1.91632 | 2.271 |
| chr12 | 94975502 | 94975727 | 0.564827 | 1.28287 | 2.271 |
| chr15 | 23786147 | 23786586 | 0.56358 | 1.28012 | 2.271 |
| chr5 | 57794210 | 57794507 | 1.19156 | 2.70659 | 2.271 |
| chr20 | 33894790 | 33895176 | 1.78132 | 4.04629 | 2.272 |
| chr12 | 53264238 | 53264676 | 1.68808 | 3.83451 | 2.272 |
| chr18 | 8389596 | 8389958 | 1.50233 | 3.41259 | 2.272 |
| chr19 | 4732638 | 4732978 | 1.24096 | 2.81891 | 2.272 |
| chr19 | 54816421 | 54816660 | 0.502072 | 1.14053 | 2.272 |
| chr15 | 93726219 | 93726544 | 0.493499 | 1.12107 | 2.272 |
| chr2 | 9939752 | 9940174 | 2.34309 | 5.32282 | 2.272 |
| chr9 | 73072786 | 73072995 | 0.51663 | 1.17367 | 2.272 |
| chr1 | 7707345 | 7707630 | 1.85143 | 4.20628 | 2.272 |
| chr5 | 148494325 | 148494569 | 1.47361 | 3.3482 | 2.272 |
| chr18 | 12402083 | 12402315 | 0.365534 | 0.830556 | 2.272 |
| chr1 | 179764477 | 179764859 | 0.423759 | 0.962904 | 2.272 |
| chr11 | 47475798 | 47476112 | 1.71015 | 3.88604 | 2.272 |
| chr12 | 58131268 | 58131934 | 0.632462 | 1.43722 | 2.272 |
| chr7 | 30424349 | 30424559 | 0.434486 | 0.987351 | 2.272 |
| chr11 | 91611227 | 91611496 | 0.5545 | 1.26019 | 2.273 |
| chr15 | 89421934 | 89422362 | 1.89205 | 4.3 | 2.273 |
| chr10 | 73074081 | 73074425 | 1.69844 | 3.86037 | 2.273 |
| chr9 | 120932380 | 120932694 | 1.29076 | 2.93392 | 2.273 |
| chr8 | 69367519 | 69367733 | 0.476132 | 1.08226 | 2.273 |
| chr22 | 27428120 | 27428378 | 0.539412 | 1.22614 | 2.273 |
| chr5 | 171533882 | 171534221 | 2.02848 | 4.61106 | 2.273 |
| chr15 | 62841692 | 62842000 | 1.26669 | 2.87941 | 2.273 |
| chr3 | 43897144 | 43897466 | 1.13912 | 2.58942 | 2.273 |
| chr19 | 7445550 | 7445940 | 0.840137 | 1.9099 | 2.273 |
| chr20 | 3691418 | 3692083 | 1.75002 | 3.97859 | 2.273 |
| chr15 | 90039479 | 90039790 | 1.65899 | 3.77168 | 2.273 |

| | | | | | |
|---|---|---|---|---|---|
| chr12 | 94922787 | 94923026 | 0.617419 | 1.40377 | 2.274 |
| chr8 | 10787772 | 10788034 | 0.360563 | 0.8198 | 2.274 |
| chr12 | 7434448 | 7434710 | 0.345882 | 0.786425 | 2.274 |
| chr3 | 111762542 | 111762823 | 1.77814 | 4.04294 | 2.274 |
| chr15 | 59430368 | 59430734 | 2.41636 | 5.49407 | 2.274 |
| chr1 | 52234736 | 52234939 | 0.908781 | 2.06637 | 2.274 |
| chr4 | 157966695 | 157967057 | 1.09641 | 2.49303 | 2.274 |
| chr19 | 42048408 | 42048792 | 2.1447 | 4.87669 | 2.274 |
| chr11 | 90938596 | 90938836 | 0.419202 | 0.953222 | 2.274 |
| chr19 | 267242 | 267731 | 1.72901 | 3.93172 | 2.274 |
| chr16 | 25390747 | 25390980 | 0.369413 | 0.840036 | 2.274 |
| chr2 | 6579922 | 6580169 | 0.476968 | 1.08463 | 2.274 |
| chrX | 152704012 | 152704474 | 0.903018 | 2.05352 | 2.274 |
| chr7 | 79371048 | 79371322 | 0.919907 | 2.09194 | 2.274 |
| chr21 | 46712296 | 46712777 | 2.26541 | 5.15189 | 2.274 |
| chr10 | 73030145 | 73030442 | 1.90696 | 4.33693 | 2.274 |
| chr13 | 103233007 | 103233242 | 1.02272 | 2.32595 | 2.274 |
| chr1 | 171282880 | 171283254 | 1.30026 | 2.95725 | 2.274 |
| chr5 | 150180671 | 150181351 | 0.964817 | 2.19439 | 2.274 |
| chrX | 34937032 | 34937278 | 0.884353 | 2.01142 | 2.274 |
| chr15 | 38856860 | 38857077 | 0.59803 | 1.3602 | 2.274 |
| chr1 | 169778827 | 169779064 | 1.03165 | 2.34655 | 2.275 |
| chr4 | 138741090 | 138741450 | 0.806071 | 1.83352 | 2.275 |
| chr8 | 124475046 | 124475364 | 0.948872 | 2.15841 | 2.275 |
| chr5 | 58182459 | 58182788 | 1.90985 | 4.34439 | 2.275 |
| chr3 | 56836194 | 56836405 | 1.02382 | 2.32901 | 2.275 |
| chr8 | 25002319 | 25002544 | 0.461218 | 1.0492 | 2.275 |
| chr7 | 93616812 | 93617031 | 0.439838 | 1.00061 | 2.275 |
| chr20 | 2652676 | 2652928 | 0.753793 | 1.71486 | 2.275 |
| chr8 | 73888450 | 73888657 | 0.414076 | 0.942029 | 2.275 |
| chr12 | 22274241 | 22274554 | 0.871493 | 1.98267 | 2.275 |
| chr3 | 111410247 | 111410478 | 0.71461 | 1.62579 | 2.275 |
| chr13 | 43052761 | 43053021 | 0.873783 | 1.98815 | 2.275 |
| chr14 | 23455688 | 23456206 | 1.53352 | 3.48928 | 2.275 |
| chr8 | 57350185 | 57350733 | 1.39606 | 3.17655 | 2.275 |
| chrX | 37764977 | 37765249 | 1.22112 | 2.7785 | 2.275 |
| chr19 | 54950494 | 54950972 | 1.4116 | 3.21193 | 2.275 |
| chr9 | 101553404 | 101553847 | 1.93791 | 4.40954 | 2.275 |
| chr5 | 150677710 | 150678077 | 0.901042 | 2.0503 | 2.275 |
| chr14 | 105818596 | 105818848 | 2.80994 | 6.39414 | 2.276 |
| chr20 | 19100613 | 19101087 | 0.429753 | 0.977953 | 2.276 |
| chr10 | 81106921 | 81107377 | 1.90058 | 4.32523 | 2.276 |
| chr4 | 33062595 | 33062889 | 1.1404 | 2.59533 | 2.276 |
| chr15 | 53958167 | 53958531 | 0.915066 | 2.08261 | 2.276 |
| chr10 | 135060583 | 135060785 | 1.21827 | 2.77278 | 2.276 |
| chr11 | 65618789 | 65619011 | 1.01508 | 2.31044 | 2.276 |
| chr1 | 177726384 | 177726672 | 0.407296 | 0.927077 | 2.276 |
| chr2 | 73114937 | 73115379 | 2.42891 | 5.52877 | 2.276 |
| chr7 | 135306049 | 135306357 | 1.75509 | 3.99502 | 2.276 |
| chr9 | 3027421 | 3027650 | 0.509619 | 1.16002 | 2.276 |
| chr9 | 90427735 | 90428183 | 2.00362 | 4.56089 | 2.276 |
| chrX | 46772451 | 46772791 | 1.32225 | 3.00997 | 2.276 |
| chr8 | 127074239 | 127074457 | 0.471209 | 1.07268 | 2.276 |
| chr16 | 72410194 | 72410426 | 0.391292 | 0.890769 | 2.276 |
| chr1 | 203186493 | 203186711 | 0.704143 | 1.60299 | 2.277 |
| chr15 | 80465528 | 80465997 | 1.28174 | 2.91793 | 2.277 |
| chr3 | 196336148 | 196336514 | 1.88447 | 4.29015 | 2.277 |
| chr12 | 109663081 | 109663381 | 0.473867 | 1.07883 | 2.277 |
| chr11 | 119209071 | 119209361 | 1.81622 | 4.13494 | 2.277 |
| chr18 | 19069045 | 19069417 | 1.18053 | 2.68781 | 2.277 |
| chr17 | 43229399 | 43229781 | 1.79286 | 4.08201 | 2.277 |

| | | | | | |
|---|---|---|---|---|---|
| chr15 | 29885683 | 29886008 | 0.517143 | 1.17745 | 2.277 |
| chr6 | 127837890 | 127838174 | 2.24209 | 5.10488 | 2.277 |
| chr1 | 67966128 | 67966495 | 2.37877 | 5.41634 | 2.277 |
| chr15 | 58416671 | 58417038 | 0.489089 | 1.11368 | 2.277 |
| chr15 | 69767914 | 69768474 | 1.32462 | 3.01638 | 2.277 |
| chr7 | 137277061 | 137277339 | 1.6304 | 3.71277 | 2.277 |
| chr9 | 134225427 | 134225776 | 2.13964 | 4.87278 | 2.277 |
| chr12 | 56050034 | 56050583 | 1.25987 | 2.8694 | 2.278 |
| chr8 | 97176587 | 97176938 | 1.33765 | 3.04663 | 2.278 |
| chr12 | 25023904 | 25024139 | 0.400074 | 0.911224 | 2.278 |
| chr21 | 46859330 | 46859670 | 1.51016 | 3.43972 | 2.278 |
| chr9 | 89654839 | 89655104 | 0.494316 | 1.12593 | 2.278 |
| chr7 | 78937628 | 78937818 | 1.28731 | 2.93221 | 2.278 |
| chr8 | 107604032 | 107604340 | 1.21357 | 2.76433 | 2.278 |
| chr1 | 55487720 | 55488105 | 1.9053 | 4.34007 | 2.278 |
| chr5 | 114309454 | 114309714 | 0.431628 | 0.983199 | 2.278 |
| chr17 | 62503500 | 62503837 | 0.995565 | 2.26808 | 2.278 |
| chr9 | 120713608 | 120713823 | 0.472636 | 1.07681 | 2.278 |
| chr15 | 44174236 | 44174499 | 1.10535 | 2.51839 | 2.278 |
| chr21 | 27162439 | 27162735 | 0.759422 | 1.73035 | 2.279 |
| chr22 | 42510855 | 42511126 | 0.779445 | 1.77606 | 2.279 |
| chr16 | 63323803 | 63324171 | 1.63231 | 3.71955 | 2.279 |
| chr6 | 36733878 | 36734283 | 1.13885 | 2.59511 | 2.279 |
| chrX | 153402079 | 153402593 | 1.80394 | 4.11071 | 2.279 |
| chr16 | 6221979 | 6222318 | 0.346559 | 0.789776 | 2.279 |
| chr17 | 47269235 | 47269482 | 0.6734 | 1.53462 | 2.279 |
| chr11 | 102261452 | 102261763 | 1.72547 | 3.93229 | 2.279 |
| chr1 | 183654355 | 183654596 | 1.27414 | 2.90384 | 2.279 |
| chr14 | 26138424 | 26138751 | 0.495752 | 1.12985 | 2.279 |
| chr5 | 6583095 | 6583459 | 0.97517 | 2.22258 | 2.279 |
| chr8 | 71833282 | 71833610 | 0.492301 | 1.12204 | 2.279 |
| chr20 | 4230156 | 4230554 | 1.78468 | 4.06764 | 2.279 |
| chr17 | 68626142 | 68626402 | 1.27785 | 2.91271 | 2.279 |
| chr12 | 102333624 | 102333863 | 1.33178 | 3.03565 | 2.279 |
| chr14 | 103391504 | 103391716 | 0.432497 | 0.985835 | 2.279 |
| chr20 | 31039480 | 31039688 | 0.634008 | 1.44516 | 2.279 |
| chr10 | 111738235 | 111738539 | 0.926577 | 2.11207 | 2.279 |
| chr14 | 99732892 | 99733257 | 2.08331 | 4.74909 | 2.280 |
| chr19 | 29930911 | 29931212 | 0.486343 | 1.10867 | 2.280 |
| chr3 | 3237834 | 3238193 | 1.20955 | 2.75731 | 2.280 |
| chr9 | 123631447 | 123631755 | 1.08967 | 2.48403 | 2.280 |
| chr3 | 31608552 | 31608811 | 1.08891 | 2.48233 | 2.280 |
| chr1 | 92757007 | 92757226 | 1.10741 | 2.52472 | 2.280 |
| chr15 | 69240454 | 69240662 | 0.411343 | 0.93782 | 2.280 |
| chr10 | 94312103 | 94312382 | 1.20858 | 2.7556 | 2.280 |
| chr17 | 54793242 | 54793580 | 1.21752 | 2.77607 | 2.280 |
| chr20 | 32307207 | 32307459 | 0.401345 | 0.915117 | 2.280 |
| chr14 | 28844601 | 28844842 | 1.24601 | 2.84114 | 2.280 |
| chr10 | 66133543 | 66133754 | 0.368304 | 0.839845 | 2.280 |
| chr7 | 134442552 | 134442846 | 0.388777 | 0.886548 | 2.280 |
| chr12 | 7984637 | 7985002 | 1.15498 | 2.63377 | 2.280 |
| chr1 | 247391556 | 247391774 | 0.952098 | 2.17122 | 2.280 |
| chr12 | 110365335 | 110365715 | 1.87282 | 4.27107 | 2.281 |
| chr10 | 127136945 | 127137478 | 1.05495 | 2.40597 | 2.281 |
| chr10 | 77101871 | 77102084 | 0.547555 | 1.24879 | 2.281 |
| chr2 | 237085088 | 237085436 | 1.98442 | 4.52584 | 2.281 |
| chr8 | 76560438 | 76560666 | 0.823709 | 1.87863 | 2.281 |
| chr8 | 42010863 | 42011218 | 1.84729 | 4.21317 | 2.281 |
| chr13 | 70439258 | 70439462 | 0.444105 | 1.01292 | 2.281 |
| chr17 | 54707270 | 54707519 | 0.861686 | 1.96536 | 2.281 |
| chr11 | 86808595 | 86808941 | 1.3217 | 3.01469 | 2.281 |

| | | | | | |
|---|---|---|---|---|---|
| chr7 | 127987656 | 127988016 | 1.59546 | 3.63937 | 2.281 |
| chr15 | 89437448 | 89437962 | 1.5433 | 3.52051 | 2.281 |
| chr19 | 54276188 | 54276443 | 1.00652 | 2.29608 | 2.281 |
| chr9 | 135674436 | 135674676 | 0.671747 | 1.53242 | 2.281 |
| chr1 | 40105556 | 40105807 | 1.28528 | 2.93206 | 2.281 |
| chr14 | 55215426 | 55215796 | 0.529575 | 1.20819 | 2.281 |
| chr3 | 179485863 | 179486164 | 1.33027 | 3.035 | 2.281 |
| chr14 | 67091079 | 67091392 | 1.26082 | 2.87657 | 2.282 |
| chr10 | 36210952 | 36211233 | 0.528279 | 1.20532 | 2.282 |
| chr5 | 164418891 | 164419210 | 0.477376 | 1.08937 | 2.282 |
| chr1 | 8786533 | 8786856 | 0.599061 | 1.3671 | 2.282 |
| chr7 | 35577011 | 35577417 | 1.94762 | 4.44461 | 2.282 |
| chr4 | 155539920 | 155540180 | 1.00313 | 2.28923 | 2.282 |
| chr3 | 38016179 | 38016497 | 1.61795 | 3.69239 | 2.282 |
| chr22 | 50765406 | 50765700 | 1.0484 | 2.39262 | 2.282 |
| chr20 | 36280296 | 36280570 | 1.07362 | 2.45019 | 2.282 |
| chr10 | 34927440 | 34927710 | 1.44463 | 3.29699 | 2.282 |
| chr16 | 719227 | 719465 | 0.454893 | 1.03824 | 2.282 |
| gl000204_r | 58623 | 59112 | 2.20441 | 5.03174 | 2.283 |
| chr7 | 49925448 | 49925798 | 0.617121 | 1.40866 | 2.283 |
| chr8 | 62704690 | 62704921 | 1.35316 | 3.08877 | 2.283 |
| chr18 | 7452355 | 7452687 | 0.92198 | 2.10455 | 2.283 |
| chr1 | 31274494 | 31275001 | 1.35895 | 3.10223 | 2.283 |
| chr1 | 159347561 | 159347834 | 0.471831 | 1.07711 | 2.283 |
| chr8 | 125438245 | 125438631 | 1.13222 | 2.58469 | 2.283 |
| chr5 | 82726230 | 82726556 | 1.71962 | 3.92566 | 2.283 |
| chr15 | 51482703 | 51482963 | 0.551223 | 1.25842 | 2.283 |
| chr17 | 7068509 | 7068831 | 0.626039 | 1.42922 | 2.283 |
| chr20 | 31592872 | 31593392 | 0.724242 | 1.65342 | 2.283 |
| chr10 | 93668495 | 93668866 | 1.44776 | 3.3053 | 2.283 |
| chrX | 17178860 | 17179403 | 0.601938 | 1.37426 | 2.283 |
| chr17 | 2578327 | 2579038 | 0.908159 | 2.07347 | 2.283 |
| chr9 | 20356337 | 20356628 | 1.72792 | 3.94538 | 2.283 |
| chr11 | 91570947 | 91571166 | 0.449121 | 1.02552 | 2.283 |
| chr8 | 80677426 | 80677836 | 2.16527 | 4.94449 | 2.284 |
| chr22 | 23088140 | 23088531 | 1.3105 | 2.99292 | 2.284 |
| chr14 | 74971594 | 74971800 | 0.388966 | 0.888333 | 2.284 |
| chr20 | 17661679 | 17662174 | 1.94123 | 4.43344 | 2.284 |
| chrX | 12912167 | 12912583 | 1.15387 | 2.63528 | 2.284 |
| chr12 | 132434257 | 132434563 | 1.38405 | 3.161 | 2.284 |
| chr15 | 67155195 | 67155519 | 2.8044 | 6.40526 | 2.284 |
| chr12 | 129900009 | 129900281 | 0.435999 | 0.995878 | 2.284 |
| chr10 | 99538306 | 99538555 | 0.501776 | 1.14614 | 2.284 |
| chr14 | 101121064 | 101121553 | 1.79686 | 4.10454 | 2.284 |
| chr13 | 95363468 | 95363783 | 0.874537 | 1.99774 | 2.284 |
| chr15 | 77399912 | 77400349 | 1.4321 | 3.27148 | 2.284 |
| chr19 | 29466673 | 29466950 | 0.853885 | 1.95072 | 2.285 |
| chr8 | 146034284 | 146034789 | 0.358527 | 0.819075 | 2.285 |
| chr17 | 77297453 | 77297819 | 0.488863 | 1.11685 | 2.285 |
| chr15 | 51454872 | 51455463 | 2.29115 | 5.2344 | 2.285 |
| chr17 | 36740585 | 36741020 | 2.78821 | 6.37004 | 2.285 |
| chr18 | 13801015 | 13801357 | 1.26715 | 2.89519 | 2.285 |
| chr20 | 62151363 | 62151616 | 0.63352 | 1.44749 | 2.285 |
| chr16 | 68098621 | 68099051 | 1.4037 | 3.20729 | 2.285 |
| chr20 | 2853277 | 2853536 | 1.55007 | 3.54194 | 2.285 |
| chr2 | 151405153 | 151405372 | 1.26054 | 2.88037 | 2.285 |
| chr5 | 87160546 | 87160822 | 0.377825 | 0.863347 | 2.285 |
| chr3 | 129661096 | 129661427 | 1.39639 | 3.19086 | 2.285 |
| chr14 | 74968854 | 74969429 | 1.06548 | 2.43477 | 2.285 |
| chr1 | 186451810 | 186452180 | 1.74212 | 3.98104 | 2.285 |
| chr8 | 135393719 | 135393960 | 0.486938 | 1.11275 | 2.285 |

| | | | | | |
|---|---|---|---|---|---|
| chr9 | 35477271 | 35477601 | 1.71383 | 3.91647 | 2.285 |
| chr3 | 53744155 | 53744501 | 1.43739 | 3.2848 | 2.285 |
| chr12 | 129736197 | 129736528 | 0.396536 | 0.90619 | 2.285 |
| chr4 | 140540876 | 140541099 | 1.14647 | 2.62021 | 2.285 |
| chr12 | 131232638 | 131233041 | 1.81216 | 4.14174 | 2.286 |
| chr10 | 53555311 | 53555536 | 0.566039 | 1.29377 | 2.286 |
| chr10 | 115999136 | 115999385 | 0.69441 | 1.58719 | 2.286 |
| chr1 | 221173348 | 221173595 | 0.372723 | 0.851944 | 2.286 |
| chr1 | 20539308 | 20539668 | 1.51195 | 3.45609 | 2.286 |
| chr3 | 1190524 | 1190760 | 0.580728 | 1.32758 | 2.286 |
| chr5 | 31270627 | 31270856 | 0.777071 | 1.77658 | 2.286 |
| chr4 | 158899972 | 158900240 | 1.32877 | 3.03797 | 2.286 |
| chr19 | 36184421 | 36184983 | 3.05759 | 6.99061 | 2.286 |
| chr5 | 133481504 | 133481889 | 1.59068 | 3.63681 | 2.286 |
| chr12 | 121151324 | 121152031 | 1.17075 | 2.67674 | 2.286 |
| chr5 | 72591481 | 72591775 | 0.5257 | 1.20198 | 2.286 |
| chr16 | 67051750 | 67052035 | 1.60296 | 3.66521 | 2.287 |
| chr5 | 105672236 | 105672512 | 0.452231 | 1.03405 | 2.287 |
| chr12 | 63187692 | 63188071 | 1.75349 | 4.01013 | 2.287 |
| chr2 | 72380231 | 72380453 | 0.36257 | 0.829198 | 2.287 |
| chr11 | 13204628 | 13204886 | 0.716857 | 1.63949 | 2.287 |
| chr11 | 45201677 | 45202049 | 0.783072 | 1.79094 | 2.287 |
| chr7 | 111381864 | 111382078 | 1.14102 | 2.60965 | 2.287 |
| chr9 | 32379330 | 32379595 | 1.48534 | 3.39731 | 2.287 |
| chr12 | 58074267 | 58074480 | 0.448628 | 1.02614 | 2.287 |
| chr11 | 124736150 | 124736540 | 1.91549 | 4.38156 | 2.287 |
| chr12 | 3318573 | 3318865 | 0.374948 | 0.857678 | 2.287 |
| chr10 | 73885965 | 73886279 | 0.864936 | 1.97855 | 2.288 |
| chr12 | 53815128 | 53815667 | 1.48461 | 3.39615 | 2.288 |
| chr5 | 56633945 | 56634297 | 1.3653 | 3.12337 | 2.288 |
| chr8 | 99483314 | 99483708 | 1.18671 | 2.71489 | 2.288 |
| chr3 | 153486612 | 153486996 | 1.43434 | 3.28144 | 2.288 |
| chr9 | 98031137 | 98031357 | 0.901629 | 2.06275 | 2.288 |
| chr14 | 102562796 | 102563303 | 1.417 | 3.24203 | 2.288 |
| chr3 | 146795900 | 146796102 | 0.595194 | 1.36183 | 2.288 |
| chr7 | 48324880 | 48325091 | 0.464975 | 1.06395 | 2.288 |
| chr6 | 34360507 | 34360775 | 0.919181 | 2.1033 | 2.288 |
| chr11 | 117492049 | 117492301 | 0.908941 | 2.0799 | 2.288 |
| chr20 | 20308599 | 20308858 | 0.651808 | 1.4917 | 2.289 |
| chr12 | 98926545 | 98926837 | 1.1259 | 2.5767 | 2.289 |
| chr5 | 92932790 | 92933044 | 0.610313 | 1.39681 | 2.289 |
| chr3 | 147180881 | 147181105 | 0.496671 | 1.13673 | 2.289 |
| chr1 | 247309762 | 247310248 | 0.859901 | 1.96806 | 2.289 |
| chr14 | 49936929 | 49937152 | 0.615756 | 1.40933 | 2.289 |
| chr20 | 43077936 | 43078152 | 0.775396 | 1.77477 | 2.289 |
| chr10 | 97002843 | 97003095 | 1.64692 | 3.76959 | 2.289 |
| chr16 | 75823092 | 75823410 | 1.1729 | 2.68467 | 2.289 |
| chr20 | 61325394 | 61325679 | 0.492555 | 1.12753 | 2.289 |
| chr11 | 60655029 | 60655604 | 0.762012 | 1.74446 | 2.289 |
| chr4 | 15651667 | 15651883 | 1.03627 | 2.37231 | 2.289 |
| chr5 | 81152820 | 81153071 | 0.722072 | 1.65318 | 2.289 |
| chr15 | 72492039 | 72492378 | 2.08266 | 4.76827 | 2.290 |
| chr3 | 40358605 | 40358868 | 1.13855 | 2.60673 | 2.290 |
| chr17 | 37391937 | 37392315 | 2.41646 | 5.53266 | 2.290 |
| chr17 | 48104030 | 48104774 | 1.31336 | 3.00713 | 2.290 |
| chr20 | 43312186 | 43312522 | 1.60636 | 3.67807 | 2.290 |
| chr11 | 46368717 | 46369132 | 1.70595 | 3.90618 | 2.290 |
| chr2 | 21149599 | 21149887 | 0.395222 | 0.90496 | 2.290 |
| chr20 | 45191102 | 45191607 | 1.79305 | 4.10568 | 2.290 |
| chr1 | 203208912 | 203209157 | 0.480574 | 1.10045 | 2.290 |
| chr7 | 25219888 | 25220386 | 2.90595 | 6.65481 | 2.290 |

| | | | | | |
|---|---|---|---|---|---|
| chr1 | 227445702 | 227445918 | 0.918156 | 2.10268 | 2.290 |
| chr14 | 35235232 | 35235627 | 1.16122 | 2.65932 | 2.290 |
| chr4 | 1900979 | 1901291 | 1.37314 | 3.14464 | 2.290 |
| chr12 | 125968754 | 125969261 | 0.408072 | 0.934585 | 2.290 |
| chr11 | 103408653 | 103408924 | 1.21014 | 2.77156 | 2.290 |
| chr12 | 94404191 | 94404435 | 0.560246 | 1.28315 | 2.290 |
| chr1 | 249200459 | 249200776 | 1.67877 | 3.84507 | 2.290 |
| chr5 | 154022069 | 154022395 | 1.89048 | 4.32999 | 2.290 |
| chr4 | 153700617 | 153700881 | 0.619765 | 1.41972 | 2.291 |
| chr15 | 65197654 | 65198072 | 1.20196 | 2.75344 | 2.291 |
| chr17 | 65383078 | 65383560 | 1.81681 | 4.16209 | 2.291 |
| chr15 | 33248228 | 33248523 | 1.45504 | 3.3334 | 2.291 |
| chr7 | 13399339 | 13399547 | 0.914231 | 2.09453 | 2.291 |
| chr10 | 50720150 | 50720582 | 1.1012 | 2.52292 | 2.291 |
| chr17 | 7983285 | 7983672 | 2.35591 | 5.39786 | 2.291 |
| chr19 | 46692791 | 46693007 | 1.65737 | 3.79752 | 2.291 |
| chr11 | 43966242 | 43966776 | 2.01919 | 4.62684 | 2.291 |
| chr11 | 121322846 | 121323214 | 1.59964 | 3.66557 | 2.292 |
| chr8 | 49834208 | 49834706 | 0.841339 | 1.92793 | 2.292 |
| chr1 | 229026204 | 229026448 | 0.43329 | 0.992916 | 2.292 |
| chr1 | 19977659 | 19977952 | 1.69773 | 3.8905 | 2.292 |
| chr3 | 113187044 | 113187369 | 0.454592 | 1.04174 | 2.292 |
| chr12 | 77002715 | 77002972 | 0.451568 | 1.03484 | 2.292 |
| chr14 | 97050552 | 97050931 | 1.33167 | 3.05173 | 2.292 |
| chr18 | 28821945 | 28822238 | 1.35098 | 3.096 | 2.292 |
| chr7 | 92053955 | 92054198 | 1.49654 | 3.42958 | 2.292 |
| chr20 | 60070234 | 60070494 | 0.430831 | 0.987345 | 2.292 |
| chr10 | 28682811 | 28683262 | 1.98125 | 4.54056 | 2.292 |
| chr17 | 39621835 | 39622165 | 0.405363 | 0.929003 | 2.292 |
| chr2 | 38830579 | 38830902 | 1.92331 | 4.40781 | 2.292 |
| chr1 | 85086350 | 85086946 | 2.25813 | 5.17544 | 2.292 |
| chr7 | 125298141 | 125298384 | 0.299409 | 0.686221 | 2.292 |
| chr12 | 58087390 | 58087692 | 1.3886 | 3.18264 | 2.292 |
| chr4 | 109792727 | 109792973 | 0.370614 | 0.849467 | 2.292 |
| chr4 | 119274007 | 119274422 | 1.69685 | 3.88929 | 2.292 |
| chrX | 84366426 | 84366684 | 1.57919 | 3.61964 | 2.292 |
| chr10 | 6215187 | 6215444 | 0.702053 | 1.60919 | 2.292 |
| chr7 | 90663895 | 90664131 | 0.982137 | 2.25122 | 2.292 |
| chr14 | 23464034 | 23464269 | 1.00208 | 2.29698 | 2.292 |
| chr9 | 139588620 | 139588978 | 1.74429 | 3.99828 | 2.292 |
| chr1 | 218442349 | 218442602 | 0.435456 | 0.998181 | 2.292 |
| chr8 | 108920594 | 108920865 | 0.873146 | 2.00154 | 2.292 |
| chr19 | 33623181 | 33623852 | 1.52936 | 3.50581 | 2.292 |
| chr5 | 156739474 | 156739760 | 1.59983 | 3.6674 | 2.292 |
| chr6 | 90319729 | 90320110 | 1.82231 | 4.17745 | 2.292 |
| chr19 | 54409948 | 54410389 | 2.18408 | 5.00687 | 2.292 |
| chr1 | 30853278 | 30853558 | 2.03149 | 4.65723 | 2.293 |
| chr2 | 138596099 | 138596355 | 0.970274 | 2.2244 | 2.293 |
| chr12 | 8761579 | 8762109 | 1.1684 | 2.67871 | 2.293 |
| chr11 | 33087061 | 33087559 | 2.10023 | 4.81525 | 2.293 |
| chr3 | 184261711 | 184262076 | 1.28618 | 2.949 | 2.293 |
| chr12 | 53902305 | 53902750 | 2.1046 | 4.82562 | 2.293 |
| chrX | 67952722 | 67953053 | 1.55785 | 3.57198 | 2.293 |
| chr5 | 16552852 | 16553304 | 1.71521 | 3.9328 | 2.293 |
| chr1 | 210023828 | 210024074 | 1.67763 | 3.84682 | 2.293 |
| chr14 | 69408536 | 69408858 | 1.48149 | 3.3972 | 2.293 |
| chr2 | 239266002 | 239266333 | 1.1091 | 2.54333 | 2.293 |
| chr16 | 57452708 | 57453098 | 1.54842 | 3.55083 | 2.293 |
| chr7 | 30467545 | 30467911 | 0.417995 | 0.958558 | 2.293 |
| chr17 | 7576297 | 7576912 | 0.807425 | 1.85173 | 2.293 |
| chr10 | 106870665 | 106871120 | 0.482451 | 1.10645 | 2.293 |

| | | | | | |
|---|---|---|---|---|---|
| chr6 | 133561735 | 133562071 | 0.479722 | 1.10032 | 2.294 |
| chr1 | 31181031 | 31181290 | 1.11596 | 2.55966 | 2.294 |
| chr17 | 32688396 | 32688907 | 3.67389 | 8.42678 | 2.294 |
| chr5 | 139840204 | 139840416 | 0.605952 | 1.38992 | 2.294 |
| chr13 | 20737543 | 20737859 | 1.57627 | 3.6159 | 2.294 |
| chr5 | 80147979 | 80148351 | 0.607462 | 1.39353 | 2.294 |
| chr14 | 106003979 | 106004608 | 1.83521 | 4.21052 | 2.294 |
| chr12 | 122615598 | 122615955 | 1.50528 | 3.45363 | 2.294 |
| chr3 | 182971924 | 182972156 | 0.677319 | 1.55424 | 2.295 |
| chr3 | 190040094 | 190040516 | 1.88902 | 4.33483 | 2.295 |
| chr1 | 208218916 | 208219349 | 1.63044 | 3.74157 | 2.295 |
| chr11 | 45126936 | 45127446 | 1.98343 | 4.55186 | 2.295 |
| chr20 | 45149706 | 45149973 | 1.35199 | 3.10291 | 2.295 |
| chr3 | 137816239 | 137816705 | 0.902512 | 2.07136 | 2.295 |
| chr8 | 125784350 | 125784694 | 0.51914 | 1.19149 | 2.295 |
| chr11 | 123305029 | 123305276 | 0.911591 | 2.09237 | 2.295 |
| chr1 | 24243313 | 24243606 | 1.19358 | 2.73974 | 2.295 |
| chr1 | 120286665 | 120287104 | 1.59407 | 3.65912 | 2.295 |
| chr5 | 7534830 | 7535117 | 0.467311 | 1.07271 | 2.295 |
| chr11 | 119005064 | 119005418 | 1.26669 | 2.9077 | 2.296 |
| chr11 | 11870807 | 11871106 | 1.57464 | 3.61482 | 2.296 |
| chr9 | 14768541 | 14768848 | 1.21081 | 2.77974 | 2.296 |
| chr21 | 33973392 | 33973666 | 1.9895 | 4.56758 | 2.296 |
| chr19 | 5377421 | 5377811 | 1.21133 | 2.78123 | 2.296 |
| chr1 | 173400340 | 173400722 | 2.15553 | 4.94925 | 2.296 |
| chr3 | 196411764 | 196412219 | 1.01781 | 2.33697 | 2.296 |
| chr17 | 54756587 | 54756915 | 1.16319 | 2.67083 | 2.296 |
| chr20 | 60970475 | 60971075 | 0.930143 | 2.13573 | 2.296 |
| chr5 | 135509749 | 135510100 | 1.32597 | 3.04465 | 2.296 |
| chr16 | 58848154 | 58848473 | 1.58259 | 3.63393 | 2.296 |
| chr8 | 4483567 | 4483777 | 0.407394 | 0.935458 | 2.296 |
| chr11 | 32356288 | 32356810 | 1.32331 | 3.03885 | 2.296 |
| chr10 | 134753817 | 134754228 | 2.58828 | 5.94377 | 2.296 |
| chr14 | 102990815 | 102991186 | 2.53352 | 5.81809 | 2.296 |
| chr5 | 157282492 | 157282751 | 1.40525 | 3.22709 | 2.296 |
| chr12 | 25015011 | 25015345 | 0.543892 | 1.24903 | 2.296 |
| chr5 | 16268972 | 16269574 | 0.351328 | 0.80682 | 2.296 |
| chr1 | 20478808 | 20479190 | 1.62985 | 3.74332 | 2.297 |
| chr14 | 23849119 | 23849511 | 2.25469 | 5.17875 | 2.297 |
| chr17 | 15163978 | 15164726 | 1.56265 | 3.5893 | 2.297 |
| chr1 | 161197010 | 161197564 | 1.24717 | 2.86487 | 2.297 |
| chr5 | 16282751 | 16283007 | 0.542638 | 1.24652 | 2.297 |
| chr2 | 39174891 | 39175193 | 1.26638 | 2.90917 | 2.297 |
| chr15 | 82339478 | 82339841 | 1.75782 | 4.03824 | 2.297 |
| chr1 | 147112550 | 147112955 | 1.94353 | 4.4649 | 2.297 |
| chr14 | 94154563 | 94154765 | 0.404832 | 0.930024 | 2.297 |
| chr12 | 124883984 | 124884548 | 1.36822 | 3.14339 | 2.297 |
| chr8 | 17191560 | 17191783 | 1.22912 | 2.82414 | 2.298 |
| chr1 | 212118058 | 212118410 | 2.09724 | 4.8189 | 2.298 |
| chr10 | 98797034 | 98797513 | 1.60675 | 3.69201 | 2.298 |
| chr5 | 89561111 | 89561427 | 2.08963 | 4.80162 | 2.298 |
| chr16 | 23607539 | 23607922 | 2.13962 | 4.91683 | 2.298 |
| chr6 | 56819965 | 56820274 | 2.42921 | 5.58236 | 2.298 |
| chr9 | 123698796 | 123699161 | 1.6767 | 3.85318 | 2.298 |
| chr21 | 46974846 | 46975061 | 1.1699 | 2.68852 | 2.298 |
| chr10 | 107710046 | 107710507 | 0.78797 | 1.81092 | 2.298 |
| chr10 | 12209952 | 12210166 | 0.412801 | 0.948722 | 2.298 |
| chr11 | 68565533 | 68566122 | 2.08121 | 4.7832 | 2.298 |
| chrX | 9677042 | 9677434 | 1.21856 | 2.80063 | 2.298 |
| chr3 | 53080585 | 53080940 | 1.0308 | 2.36911 | 2.298 |
| chr16 | 52540193 | 52540484 | 0.391282 | 0.899297 | 2.298 |

| | | | | | |
|---|---|---|---|---|---|
| chr9 | 25348677 | 25348887 | 0.432697 | 0.994497 | 2.298 |
| chr11 | 107419951 | 107420281 | 0.675618 | 1.55287 | 2.298 |
| chr3 | 26607942 | 26608269 | 0.63798 | 1.46636 | 2.298 |
| chr9 | 80876427 | 80876643 | 0.959759 | 2.20597 | 2.298 |
| chr4 | 39437626 | 39437836 | 0.579134 | 1.33116 | 2.299 |
| chr17 | 46868423 | 46868714 | 1.84823 | 4.24825 | 2.299 |
| chr8 | 70375705 | 70375944 | 0.375373 | 0.862818 | 2.299 |
| chr12 | 50478757 | 50479026 | 1.68214 | 3.86661 | 2.299 |
| chr6 | 129573301 | 129573592 | 1.37265 | 3.15539 | 2.299 |
| chr6 | 62993556 | 62994017 | 0.571161 | 1.31298 | 2.299 |
| chr9 | 75814841 | 75815186 | 1.41081 | 3.24321 | 2.299 |
| chr17 | 12823839 | 12824079 | 0.558949 | 1.28502 | 2.299 |
| chr8 | 143210196 | 143210577 | 0.894896 | 2.0575 | 2.299 |
| chr20 | 48526393 | 48526609 | 0.657949 | 1.51274 | 2.299 |
| chr6 | 109509055 | 109509395 | 1.49294 | 3.43264 | 2.299 |
| chr9 | 21812987 | 21813301 | 1.41409 | 3.25144 | 2.299 |
| chr10 | 31040727 | 31041171 | 2.01437 | 4.63195 | 2.299 |
| chr12 | 47973122 | 47973618 | 0.486802 | 1.11938 | 2.299 |
| chr12 | 112190949 | 112191377 | 1.89712 | 4.36243 | 2.300 |
| chr12 | 55942596 | 55942852 | 0.420284 | 0.966447 | 2.300 |
| chr5 | 10857191 | 10857500 | 2.16951 | 4.98882 | 2.300 |
| chr6 | 74442884 | 74443114 | 1.41668 | 3.25776 | 2.300 |
| chr12 | 63670511 | 63670953 | 1.4215 | 3.26911 | 2.300 |
| chr15 | 68851929 | 68852373 | 2.14075 | 4.92335 | 2.300 |
| chr1 | 180137895 | 180138226 | 1.35782 | 3.12289 | 2.300 |
| chr16 | 10296701 | 10296917 | 0.363108 | 0.835126 | 2.300 |
| chr19 | 41312716 | 41313179 | 2.40372 | 5.52846 | 2.300 |
| chrX | 152875248 | 152875657 | 1.6759 | 3.85452 | 2.300 |
| chr9 | 19486516 | 19486836 | 0.374513 | 0.861386 | 2.300 |
| chr11 | 116706799 | 116707049 | 1.42677 | 3.28173 | 2.300 |
| chr5 | 39074993 | 39075203 | 0.880378 | 2.02505 | 2.300 |
| chr4 | 87843543 | 87843888 | 1.91058 | 4.39493 | 2.300 |
| chr5 | 149666470 | 149666805 | 1.06916 | 2.45986 | 2.301 |
| chr11 | 34009545 | 34009856 | 1.47898 | 3.40283 | 2.301 |
| chr3 | 158509170 | 158509430 | 1.40386 | 3.23004 | 2.301 |
| chr17 | 37168504 | 37168785 | 1.00202 | 2.30549 | 2.301 |
| chrX | 67930695 | 67931031 | 0.966265 | 2.22322 | 2.301 |
| chr6 | 56886005 | 56886244 | 0.549297 | 1.26387 | 2.301 |
| chr11 | 83641292 | 83641541 | 0.576975 | 1.32757 | 2.301 |
| chr19 | 45074192 | 45074500 | 1.44434 | 3.32335 | 2.301 |
| chr15 | 89901784 | 89902629 | 3.21431 | 7.39605 | 2.301 |
| chr1 | 42638832 | 42639367 | 2.11205 | 4.86058 | 2.301 |
| chr15 | 74782088 | 74782434 | 1.17106 | 2.69503 | 2.301 |
| chr6 | 25279161 | 25279429 | 1.17287 | 2.69923 | 2.301 |
| chr8 | 145501515 | 145501956 | 2.42043 | 5.57039 | 2.301 |
| chr2 | 206890727 | 206890997 | 0.289832 | 0.667027 | 2.301 |
| chr12 | 7741370 | 7741701 | 1.18899 | 2.7364 | 2.301 |
| chr12 | 51611808 | 51612072 | 0.785079 | 1.80688 | 2.302 |
| chr3 | 120341924 | 120342136 | 0.899086 | 2.06934 | 2.302 |
| chr5 | 77943903 | 77944115 | 0.564367 | 1.29897 | 2.302 |
| chr7 | 16931023 | 16931261 | 1.09034 | 2.50962 | 2.302 |
| chr7 | 99527182 | 99527430 | 1.02451 | 2.35811 | 2.302 |
| chr14 | 98852064 | 98852534 | 0.397364 | 0.91463 | 2.302 |
| chr12 | 82634957 | 82635165 | 0.580876 | 1.33715 | 2.302 |
| chr7 | 86941641 | 86941935 | 2.14414 | 4.93586 | 2.302 |
| chr8 | 96423185 | 96423428 | 0.439008 | 1.01062 | 2.302 |
| chr9 | 94188218 | 94188676 | 0.868607 | 1.99959 | 2.302 |
| chr17 | 5433861 | 5434301 | 0.627614 | 1.44482 | 2.302 |
| chr17 | 17304571 | 17305049 | 0.590178 | 1.35867 | 2.302 |
| chr17 | 28347601 | 28347908 | 0.596654 | 1.37361 | 2.302 |
| chr5 | 140018799 | 140019073 | 0.848359 | 1.95313 | 2.302 |

| | | | | | |
|---|---|---|---|---|---|
| chr1 | 146861728 | 146862066 | 1.15912 | 2.66859 | 2.302 |
| chr17 | 34959391 | 34959683 | 2.0824 | 4.79435 | 2.302 |
| chr2 | 234352190 | 234352544 | 1.59546 | 3.67335 | 2.302 |
| chr1 | 237205113 | 237205424 | 1.08505 | 2.49821 | 2.302 |
| chr2 | 11460888 | 11461130 | 0.757868 | 1.74491 | 2.302 |
| chr16 | 84220507 | 84220890 | 1.77798 | 4.09369 | 2.302 |
| chr6 | 148880899 | 148881316 | 1.85022 | 4.26032 | 2.303 |
| chr22 | 19667021 | 19667518 | 1.63828 | 3.77236 | 2.303 |
| chr8 | 134467956 | 134468363 | 2.02853 | 4.671 | 2.303 |
| chr2 | 8825792 | 8826296 | 2.8485 | 6.55929 | 2.303 |
| chr6 | 123110411 | 123111266 | 0.861154 | 1.98312 | 2.303 |
| chr6 | 46816738 | 46816946 | 1.29921 | 2.99191 | 2.303 |
| chr17 | 9065956 | 9066265 | 1.25957 | 2.90073 | 2.303 |
| chr12 | 2049228 | 2049514 | 1.81307 | 4.17549 | 2.303 |
| chr16 | 80956766 | 80957114 | 0.424679 | 0.978058 | 2.303 |
| chr9 | 112282770 | 112283167 | 1.58972 | 3.66129 | 2.303 |
| chr9 | 13121666 | 13121995 | 0.458474 | 1.05594 | 2.303 |
| chr7 | 103047307 | 103047660 | 1.17355 | 2.7029 | 2.303 |
| chr4 | 81973667 | 81973933 | 0.918783 | 2.11621 | 2.303 |
| chr16 | 68626478 | 68626686 | 0.945827 | 2.17853 | 2.303 |
| chr1 | 209739462 | 209739716 | 1.53034 | 3.52487 | 2.303 |
| chr12 | 97542719 | 97543257 | 1.36905 | 3.15337 | 2.303 |
| chr22 | 50095400 | 50095708 | 1.29153 | 2.97486 | 2.303 |
| chr7 | 11251767 | 11252034 | 1.60049 | 3.68656 | 2.303 |
| chr7 | 94065362 | 94065638 | 1.41053 | 3.24901 | 2.303 |
| chr3 | 110033400 | 110033678 | 1.23328 | 2.84076 | 2.303 |
| chr5 | 159935817 | 159936127 | 1.52001 | 3.50125 | 2.303 |
| chr7 | 45930751 | 45931007 | 1.22331 | 2.81811 | 2.304 |
| chr2 | 180205495 | 180205790 | 1.17967 | 2.71763 | 2.304 |
| chr7 | 4587351 | 4587626 | 0.473182 | 1.09009 | 2.304 |
| chr17 | 5140780 | 5141203 | 1.75767 | 4.04933 | 2.304 |
| chr7 | 127228912 | 127229276 | 1.29439 | 2.98206 | 2.304 |
| chr5 | 94574608 | 94574925 | 0.578312 | 1.33239 | 2.304 |
| chr19 | 664885 | 665303 | 1.29057 | 2.97339 | 2.304 |
| chr11 | 102604043 | 102604429 | 1.42137 | 3.27479 | 2.304 |
| chr2 | 20881960 | 20882309 | 1.17672 | 2.71114 | 2.304 |
| chr9 | 136454387 | 136454887 | 1.1236 | 2.58875 | 2.304 |
| chr11 | 112151325 | 112151579 | 1.45526 | 3.35293 | 2.304 |
| chr10 | 17299769 | 17300019 | 1.2616 | 2.9068 | 2.304 |
| chr6 | 114180123 | 114180478 | 1.91638 | 4.41553 | 2.304 |
| chr1 | 176650244 | 176650551 | 0.504605 | 1.1627 | 2.304 |
| chr18 | 74118026 | 74118283 | 1.14192 | 2.63119 | 2.304 |
| chr9 | 129728082 | 129728434 | 1.5946 | 3.67449 | 2.304 |
| chr1 | 210580555 | 210580936 | 2.59701 | 5.98468 | 2.304 |
| chr3 | 24536582 | 24536890 | 0.909301 | 2.09554 | 2.305 |
| chr11 | 57408448 | 57408716 | 0.941515 | 2.16985 | 2.305 |
| chr7 | 103213720 | 103214185 | 0.607328 | 1.39969 | 2.305 |
| chr22 | 50838055 | 50838492 | 0.921057 | 2.12274 | 2.305 |
| chr12 | 18705818 | 18706095 | 0.995556 | 2.29445 | 2.305 |
| chr19 | 51924958 | 51925578 | 2.20421 | 5.08014 | 2.305 |
| chr6 | 43636027 | 43636364 | 1.23513 | 2.84665 | 2.305 |
| chr8 | 90996332 | 90997122 | 2.23295 | 5.14637 | 2.305 |
| chr7 | 304091 | 304326 | 0.465056 | 1.07186 | 2.305 |
| chr8 | 51869342 | 51869547 | 0.497421 | 1.14652 | 2.305 |
| chr12 | 3182543 | 3182986 | 2.81783 | 6.49521 | 2.305 |
| chr6 | 3126633 | 3126931 | 1.97498 | 4.55245 | 2.305 |
| chr4 | 38677170 | 38677448 | 1.39827 | 3.22323 | 2.305 |
| chr3 | 171547697 | 171548077 | 1.60733 | 3.70542 | 2.305 |
| chr8 | 138175882 | 138176086 | 0.391998 | 0.903697 | 2.305 |
| chr13 | 100308456 | 100308671 | 0.900566 | 2.07618 | 2.305 |
| chr5 | 72063477 | 72063689 | 0.77444 | 1.78548 | 2.306 |

| | | | | | |
|---|---|---|---|---|---|
| chr2 | 26538956 | 26539318 | 0.853401 | 1.96757 | 2.306 |
| chr11 | 19764032 | 19764584 | 1.21431 | 2.79968 | 2.306 |
| chr1 | 18023040 | 18023283 | 1.17611 | 2.71165 | 2.306 |
| chr15 | 76349757 | 76350105 | 1.07181 | 2.47125 | 2.306 |
| chr17 | 46955552 | 46955861 | 1.20588 | 2.78046 | 2.306 |
| chr7 | 104351787 | 104352015 | 0.477032 | 1.09998 | 2.306 |
| chr12 | 52537298 | 52537628 | 2.03417 | 4.69063 | 2.306 |
| chr2 | 151230091 | 151230306 | 0.371143 | 0.855902 | 2.306 |
| chr5 | 169652777 | 169653196 | 1.76004 | 4.05887 | 2.306 |
| chr11 | 9634754 | 9635085 | 1.7051 | 3.93227 | 2.306 |
| chr8 | 126656849 | 126657101 | 1.63007 | 3.75928 | 2.306 |
| chr11 | 105947633 | 105947976 | 0.697928 | 1.60959 | 2.306 |
| chr6 | 167188763 | 167189000 | 1.51233 | 3.48784 | 2.306 |
| chr8 | 19234556 | 19234891 | 0.691815 | 1.59551 | 2.306 |
| chr16 | 80832041 | 80832283 | 0.509782 | 1.17572 | 2.306 |
| chr1 | 113044497 | 113044785 | 1.32643 | 3.05929 | 2.306 |
| chr7 | 142571032 | 142571321 | 0.878019 | 2.02508 | 2.306 |
| chr2 | 19065409 | 19065729 | 1.27538 | 2.94168 | 2.307 |
| chr5 | 119120561 | 119120876 | 0.489718 | 1.12954 | 2.307 |
| chr9 | 138386815 | 138387019 | 0.445352 | 1.02721 | 2.307 |
| chr19 | 36542193 | 36542467 | 1.27533 | 2.94161 | 2.307 |
| chr3 | 141871219 | 141871542 | 1.08087 | 2.49313 | 2.307 |
| chr2 | 20611134 | 20611505 | 1.80834 | 4.17117 | 2.307 |
| chr10 | 55479255 | 55479472 | 0.479606 | 1.1063 | 2.307 |
| chr19 | 40661603 | 40661933 | 1.40046 | 3.23048 | 2.307 |
| chr6 | 42515310 | 42515689 | 2.42871 | 5.60241 | 2.307 |
| chr10 | 102827616 | 102827952 | 1.82854 | 4.21802 | 2.307 |
| chr9 | 131764840 | 131765239 | 0.451455 | 1.04144 | 2.307 |
| chr9 | 7721484 | 7721697 | 0.427102 | 0.985312 | 2.307 |
| chr15 | 78609597 | 78609854 | 1.12663 | 2.59911 | 2.307 |
| chr17 | 899775 | 900119 | 2.15447 | 4.97039 | 2.307 |
| chr5 | 18056773 | 18057042 | 0.429235 | 0.990251 | 2.307 |
| chr5 | 29698014 | 29698296 | 0.521588 | 1.20334 | 2.307 |
| chr2 | 150454126 | 150454330 | 0.997822 | 2.30221 | 2.307 |
| chr16 | 80560362 | 80560775 | 1.38253 | 3.18984 | 2.307 |
| chr14 | 70051758 | 70052070 | 1.49029 | 3.43855 | 2.307 |
| chr11 | 44856462 | 44856722 | 0.587738 | 1.35612 | 2.307 |
| chr16 | 68150243 | 68150457 | 0.928812 | 2.14309 | 2.307 |
| chr4 | 19793084 | 19793287 | 0.629064 | 1.45149 | 2.307 |
| chr7 | 158526607 | 158526824 | 0.64399 | 1.48593 | 2.307 |
| chr7 | 53471588 | 53471796 | 0.524244 | 1.20966 | 2.307 |
| chr11 | 892131 | 892481 | 0.71911 | 1.65933 | 2.307 |
| chr19 | 28419052 | 28419256 | 0.484437 | 1.11785 | 2.308 |
| chr1 | 18907922 | 18908227 | 1.50284 | 3.46788 | 2.308 |
| chr5 | 97671778 | 97672122 | 0.486429 | 1.12252 | 2.308 |
| chr1 | 29063128 | 29063396 | 2.17907 | 5.02861 | 2.308 |
| chr14 | 95918560 | 95919073 | 0.903782 | 2.08569 | 2.308 |
| chr2 | 56899213 | 56899702 | 0.646519 | 1.492 | 2.308 |
| chr17 | 72353782 | 72354033 | 0.479313 | 1.10621 | 2.308 |
| chr16 | 15736525 | 15736794 | 0.951505 | 2.19601 | 2.308 |
| chr1 | 219378053 | 219378287 | 0.847223 | 1.95535 | 2.308 |
| chr17 | 36715817 | 36716042 | 0.631164 | 1.45678 | 2.308 |
| chr13 | 53024606 | 53025069 | 2.45154 | 5.65859 | 2.308 |
| chr4 | 129784630 | 129784973 | 1.23692 | 2.85511 | 2.308 |
| chr4 | 141280920 | 141281573 | 1.08989 | 2.51578 | 2.308 |
| chr5 | 166771817 | 166772060 | 0.438332 | 1.01186 | 2.308 |
| chr2 | 62137767 | 62137984 | 0.987517 | 2.27968 | 2.309 |
| chr12 | 98396772 | 98396976 | 1.42507 | 3.28982 | 2.309 |
| chr6 | 27863315 | 27863564 | 1.0567 | 2.43948 | 2.309 |
| chr20 | 49548142 | 49548364 | 0.834113 | 1.92568 | 2.309 |
| chr16 | 82049971 | 82050178 | 0.96738 | 2.23342 | 2.309 |

| | | | | | |
|---|---|---|---|---|---|
| chr8 | 120098759 | 120098970 | 0.556129 | 1.28396 | 2.309 |
| chr3 | 122024763 | 122025009 | 0.886634 | 2.04706 | 2.309 |
| chr19 | 42108230 | 42108453 | 0.665627 | 1.53681 | 2.309 |
| chr3 | 63336828 | 63337231 | 0.950779 | 2.19519 | 2.309 |
| chr15 | 67081447 | 67081728 | 1.15108 | 2.65771 | 2.309 |
| chr14 | 82122041 | 82122319 | 0.488642 | 1.12823 | 2.309 |
| chr15 | 93064792 | 93065058 | 1.25492 | 2.89751 | 2.309 |
| chr11 | 76309382 | 76309921 | 1.2497 | 2.88584 | 2.309 |
| chr12 | 45390355 | 45390578 | 0.504918 | 1.16598 | 2.309 |
| chr9 | 98103809 | 98104282 | 0.679971 | 1.57033 | 2.309 |
| chr9 | 103989278 | 103989563 | 0.516848 | 1.19365 | 2.309 |
| chr12 | 31593242 | 31593475 | 0.426659 | 0.985364 | 2.309 |
| chr20 | 50928744 | 50929056 | 1.21984 | 2.81721 | 2.309 |
| chr10 | 128989369 | 128989584 | 0.653666 | 1.50964 | 2.310 |
| chr15 | 65611139 | 65611526 | 2.12783 | 4.91424 | 2.310 |
| chr3 | 65658586 | 65658890 | 1.2956 | 2.99223 | 2.310 |
| chr10 | 91005729 | 91005951 | 0.419738 | 0.969405 | 2.310 |
| chr20 | 5173044 | 5173281 | 1.37087 | 3.16613 | 2.310 |
| chr20 | 21697769 | 21698152 | 0.536485 | 1.23908 | 2.310 |
| chr4 | 185666932 | 185667262 | 1.20297 | 2.77855 | 2.310 |
| chr19 | 39329372 | 39329605 | 0.592197 | 1.36786 | 2.310 |
| chr3 | 134192823 | 134193056 | 0.893273 | 2.0633 | 2.310 |
| chr8 | 27283679 | 27284194 | 1.58659 | 3.66476 | 2.310 |
| chr12 | 6934740 | 6935186 | 2.15454 | 4.97671 | 2.310 |
| chr16 | 52225471 | 52225767 | 1.68681 | 3.89634 | 2.310 |
| chr19 | 46278071 | 46278315 | 0.80826 | 1.86702 | 2.310 |
| chr8 | 8130765 | 8131001 | 1.39016 | 3.21124 | 2.310 |
| chr1 | 27152931 | 27153207 | 1.1674 | 2.69672 | 2.310 |
| chr13 | 36295119 | 36295453 | 1.76407 | 4.07506 | 2.310 |
| chr7 | 158200714 | 158200985 | 0.853672 | 1.97205 | 2.310 |
| chr1 | 156355364 | 156355669 | 0.996439 | 2.30186 | 2.310 |
| chr2 | 11051790 | 11052202 | 0.547304 | 1.26452 | 2.310 |
| chr4 | 154239907 | 154240140 | 1.0677 | 2.46689 | 2.310 |
| chr1 | 9256458 | 9256794 | 1.18909 | 2.74774 | 2.311 |
| chr3 | 30821222 | 30821492 | 1.04971 | 2.42567 | 2.311 |
| chr10 | 60059331 | 60059648 | 1.42497 | 3.29292 | 2.311 |
| chr11 | 2187300 | 2187886 | 1.75093 | 4.04622 | 2.311 |
| chr8 | 133064172 | 133064513 | 1.53419 | 3.54554 | 2.311 |
| chr8 | 9252405 | 9252629 | 0.354927 | 0.820264 | 2.311 |
| chr9 | 127602857 | 127603224 | 1.05643 | 2.44167 | 2.311 |
| chr1 | 39843945 | 39844244 | 1.09042 | 2.52028 | 2.311 |
| chr17 | 37824683 | 37825047 | 1.48879 | 3.44109 | 2.311 |
| chr9 | 22935382 | 22935611 | 0.428743 | 0.990977 | 2.311 |
| chr1 | 16067804 | 16068226 | 2.26072 | 5.22539 | 2.311 |
| chr10 | 7343443 | 7343666 | 0.987053 | 2.28154 | 2.311 |
| chr12 | 69464093 | 69464543 | 1.85235 | 4.28167 | 2.311 |
| chr19 | 59011014 | 59011250 | 0.649191 | 1.50061 | 2.312 |
| chr5 | 2437891 | 2438257 | 0.467039 | 1.07959 | 2.312 |
| chr16 | 85513842 | 85514116 | 1.98178 | 4.58103 | 2.312 |
| chr22 | 30822548 | 30822788 | 1.39492 | 3.22445 | 2.312 |
| chr11 | 15357563 | 15357830 | 0.485569 | 1.12243 | 2.312 |
| chr10 | 112257958 | 112258160 | 0.561254 | 1.2974 | 2.312 |
| chr18 | 43554052 | 43554369 | 0.932848 | 2.15659 | 2.312 |
| chr1 | 116138495 | 116138780 | 1.6453 | 3.8038 | 2.312 |
| chr5 | 36593588 | 36594040 | 0.446679 | 1.03272 | 2.312 |
| chr8 | 8978177 | 8978411 | 0.546385 | 1.26324 | 2.312 |
| chr9 | 139655315 | 139655660 | 1.70412 | 3.93997 | 2.312 |
| chr12 | 40404951 | 40405265 | 2.21063 | 5.11115 | 2.312 |
| chr5 | 140792423 | 140792679 | 0.69104 | 1.59779 | 2.312 |
| chr12 | 25540449 | 25540715 | 1.38999 | 3.2139 | 2.312 |
| chr10 | 6128386 | 6128720 | 1.83513 | 4.24335 | 2.312 |

| | | | | | |
|---|---|---|---|---|---|
| chr11 | 109567954 | 109568295 | 0.856 | 1.97971 | 2.313 |
| chr3 | 79895724 | 79896040 | 0.636914 | 1.47313 | 2.313 |
| chr12 | 25036744 | 25037014 | 0.630703 | 1.45877 | 2.313 |
| chr8 | 103540958 | 103541314 | 0.961378 | 2.22361 | 2.313 |
| chr8 | 121963461 | 121963744 | 1.19555 | 2.76539 | 2.313 |
| chr16 | 48560482 | 48560930 | 0.966822 | 2.23635 | 2.313 |
| chr5 | 140165598 | 140165816 | 0.771848 | 1.78536 | 2.313 |
| chr6 | 53158910 | 53159291 | 0.980603 | 2.26823 | 2.313 |
| chr2 | 220253232 | 220253448 | 1.37776 | 3.18697 | 2.313 |
| chr8 | 26527780 | 26528408 | 1.47192 | 3.40483 | 2.313 |
| chr1 | 151976617 | 151976835 | 0.772227 | 1.7864 | 2.313 |
| chr9 | 26385070 | 26385278 | 0.355036 | 0.82135 | 2.313 |
| chr1 | 26124528 | 26124791 | 1.41845 | 3.28172 | 2.314 |
| chr1 | 23000689 | 23001003 | 1.33015 | 3.07751 | 2.314 |
| chr10 | 81389241 | 81389500 | 0.48444 | 1.12083 | 2.314 |
| chr1 | 230123247 | 230123460 | 0.455073 | 1.05291 | 2.314 |
| chr3 | 1847288 | 1847490 | 0.396433 | 0.917244 | 2.314 |
| chr15 | 69770527 | 69770932 | 2.32936 | 5.38964 | 2.314 |
| chr15 | 59825043 | 59825258 | 1.25703 | 2.90853 | 2.314 |
| chr1 | 183155339 | 183155595 | 0.968737 | 2.24149 | 2.314 |
| chr13 | 101247072 | 101247530 | 0.923895 | 2.13787 | 2.314 |
| chr1 | 215319670 | 215319922 | 0.880384 | 2.03732 | 2.314 |
| chr9 | 130879808 | 130880345 | 3.819 | 8.83774 | 2.314 |
| chr9 | 75545741 | 75545953 | 0.866859 | 2.00611 | 2.314 |
| chr11 | 70042168 | 70042650 | 1.95038 | 4.51378 | 2.314 |
| chr17 | 63793055 | 63793500 | 0.506928 | 1.17322 | 2.314 |
| chr11 | 120555623 | 120556015 | 1.06057 | 2.45459 | 2.314 |
| chr1 | 51215740 | 51216012 | 1.15694 | 2.6779 | 2.315 |
| chr3 | 12024286 | 12024726 | 0.397739 | 0.92063 | 2.315 |
| chr9 | 98027287 | 98027492 | 0.986255 | 2.28286 | 2.315 |
| chr12 | 124261702 | 124261963 | 0.446882 | 1.03442 | 2.315 |
| chr8 | 37593803 | 37594154 | 1.54645 | 3.57974 | 2.315 |
| chr19 | 50420020 | 50420355 | 0.955656 | 2.21219 | 2.315 |
| chr19 | 51111342 | 51111890 | 1.27428 | 2.94987 | 2.315 |
| chr14 | 85478100 | 85478399 | 0.406421 | 0.940865 | 2.315 |
| chr8 | 140398476 | 140398719 | 0.557405 | 1.29043 | 2.315 |
| chr19 | 6165992 | 6166257 | 1.28821 | 2.98237 | 2.315 |
| chr7 | 148844275 | 148844685 | 2.13265 | 4.93744 | 2.315 |
| chr3 | 13324712 | 13324928 | 0.82118 | 1.9012 | 2.315 |
| chr8 | 136638611 | 136638848 | 0.478558 | 1.10798 | 2.315 |
| chr16 | 88887801 | 88888153 | 0.413505 | 0.95737 | 2.315 |
| chr14 | 54833508 | 54833798 | 1.15987 | 2.68542 | 2.315 |
| chr17 | 36204258 | 36204693 | 1.56066 | 3.61342 | 2.315 |
| chr10 | 30681408 | 30681797 | 2.15739 | 4.99546 | 2.316 |
| chr17 | 37790549 | 37791142 | 1.37907 | 3.1933 | 2.316 |
| chr1 | 228659291 | 228659834 | 1.03324 | 2.39253 | 2.316 |
| chr10 | 31320460 | 31320788 | 1.48987 | 3.44988 | 2.316 |
| chr12 | 114720298 | 114720505 | 0.491679 | 1.13851 | 2.316 |
| chr15 | 95835194 | 95835465 | 1.09454 | 2.53449 | 2.316 |
| chr7 | 142912110 | 142912568 | 2.78378 | 6.44606 | 2.316 |
| chr9 | 104221268 | 104221633 | 1.43682 | 3.32715 | 2.316 |
| chr5 | 167646855 | 167647178 | 1.62068 | 3.75299 | 2.316 |
| chr3 | 16090294 | 16090521 | 1.38334 | 3.20355 | 2.316 |
| chr16 | 23193590 | 23194250 | 2.5825 | 5.9807 | 2.316 |
| chr1 | 117112956 | 117113259 | 1.60865 | 3.72553 | 2.316 |
| chr16 | 2969621 | 2969984 | 1.3032 | 3.01818 | 2.316 |
| chr17 | 29024947 | 29025207 | 2.0991 | 4.86148 | 2.316 |
| chr3 | 14591011 | 14591389 | 0.787331 | 1.82345 | 2.316 |
| chr18 | 55113588 | 55113905 | 1.18746 | 2.75023 | 2.316 |
| chr14 | 103438974 | 103439257 | 1.22644 | 2.84056 | 2.316 |
| chr5 | 177888157 | 177888508 | 1.56021 | 3.61365 | 2.316 |

| | | | | | |
|---|---|---|---|---|---|
| chr11 | 63807922 | 63808496 | 1.38091 | 3.1986 | 2.316 |
| chr5 | 168419734 | 168419972 | 1.00373 | 2.32499 | 2.316 |
| chr22 | 47826510 | 47826766 | 1.34109 | 3.10659 | 2.316 |
| chr14 | 42693522 | 42693844 | 1.88299 | 4.36191 | 2.316 |
| chr14 | 51009695 | 51009920 | 1.15146 | 2.66745 | 2.317 |
| chr15 | 23887518 | 23888121 | 0.973959 | 2.25625 | 2.317 |
| chr15 | 32943401 | 32943785 | 1.68578 | 3.90525 | 2.317 |
| chr17 | 40976224 | 40976606 | 2.18789 | 5.0685 | 2.317 |
| chr17 | 65471550 | 65471900 | 2.02635 | 4.69429 | 2.317 |
| chr1 | 154558796 | 154559016 | 0.534133 | 1.2374 | 2.317 |
| chr16 | 75655066 | 75655355 | 1.06869 | 2.47582 | 2.317 |
| chr15 | 50124822 | 50125212 | 1.37493 | 3.1853 | 2.317 |
| chr11 | 121526206 | 121526495 | 1.67205 | 3.87373 | 2.317 |
| chr19 | 6516875 | 6517274 | 1.88595 | 4.36943 | 2.317 |
| chr21 | 26934458 | 26934813 | 0.395024 | 0.915232 | 2.317 |
| chr14 | 66291470 | 66291755 | 1.83967 | 4.26238 | 2.317 |
| chr22 | 37727744 | 37728168 | 0.821279 | 1.90285 | 2.317 |
| chr16 | 2820469 | 2820759 | 0.684242 | 1.5854 | 2.317 |
| chr12 | 4073476 | 4073859 | 0.817922 | 1.89515 | 2.317 |
| chr13 | 80916941 | 80917342 | 1.96438 | 4.55156 | 2.317 |
| chr11 | 39387461 | 39387694 | 0.623926 | 1.44568 | 2.317 |
| chr5 | 170764016 | 170764254 | 1.25818 | 2.91535 | 2.317 |
| chr1 | 183287730 | 183287972 | 1.16002 | 2.68799 | 2.317 |
| chr12 | 72140805 | 72141075 | 1.36674 | 3.16701 | 2.317 |
| chr7 | 73981000 | 73981486 | 2.32759 | 5.39361 | 2.317 |
| chr19 | 43281077 | 43281425 | 0.533169 | 1.23549 | 2.317 |
| chr9 | 7809820 | 7810055 | 0.583122 | 1.35132 | 2.317 |
| chr1 | 52593080 | 52593493 | 0.918282 | 2.12811 | 2.317 |
| chr12 | 31556098 | 31556411 | 1.97702 | 4.58182 | 2.318 |
| chr8 | 47395061 | 47395299 | 0.502645 | 1.16494 | 2.318 |
| chr14 | 23287348 | 23287542 | 1.16283 | 2.69502 | 2.318 |
| chr6 | 132805292 | 132805576 | 0.788223 | 1.82683 | 2.318 |
| chr9 | 139334192 | 139334594 | 2.43414 | 5.64167 | 2.318 |
| chr5 | 3230414 | 3230687 | 0.446782 | 1.03555 | 2.318 |
| chr14 | 21575026 | 21575350 | 1.89203 | 4.38536 | 2.318 |
| chr8 | 132471785 | 132471994 | 0.510458 | 1.18316 | 2.318 |
| chr6 | 114302204 | 114302438 | 0.963024 | 2.23221 | 2.318 |
| chr11 | 126350600 | 126350931 | 1.30759 | 3.03092 | 2.318 |
| chr16 | 9030349 | 9030698 | 1.2931 | 2.99741 | 2.318 |
| chr12 | 65515202 | 65515558 | 1.83764 | 4.2597 | 2.318 |
| chr12 | 130498517 | 130499362 | 1.01436 | 2.35145 | 2.318 |
| chr18 | 21656301 | 21656570 | 1.25769 | 2.91604 | 2.319 |
| chr10 | 65632133 | 65632476 | 2.06453 | 4.78683 | 2.319 |
| chr17 | 10748210 | 10748832 | 1.21532 | 2.81786 | 2.319 |
| chr8 | 134157375 | 134157628 | 0.494754 | 1.14715 | 2.319 |
| chr9 | 97491565 | 97491855 | 0.575907 | 1.33536 | 2.319 |
| chr3 | 119408983 | 119409346 | 1.09441 | 2.53765 | 2.319 |
| chr6 | 44272529 | 44272871 | 1.44397 | 3.34829 | 2.319 |
| chr5 | 31086888 | 31087122 | 0.423718 | 0.982543 | 2.319 |
| chr17 | 17259343 | 17259822 | 3.00115 | 6.95954 | 2.319 |
| chr7 | 64096824 | 64097137 | 0.923554 | 2.14173 | 2.319 |
| chr22 | 39925210 | 39925631 | 2.29431 | 5.32063 | 2.319 |
| chr8 | 14114615 | 14114911 | 0.591768 | 1.37237 | 2.319 |
| chr11 | 62078547 | 62078997 | 2.21667 | 5.14075 | 2.319 |
| chr16 | 18995007 | 18995466 | 2.46485 | 5.71641 | 2.319 |
| chr22 | 50421840 | 50422183 | 1.33041 | 3.08545 | 2.319 |
| chr4 | 8268941 | 8269232 | 2.2625 | 5.24719 | 2.319 |
| chr17 | 73065923 | 73066263 | 1.00131 | 2.32229 | 2.319 |
| chr7 | 75244360 | 75244639 | 0.899912 | 2.08724 | 2.319 |
| chr9 | 97930258 | 97930573 | 1.35583 | 3.14478 | 2.319 |
| chr3 | 51409472 | 51409814 | 1.01697 | 2.35886 | 2.320 |

| | | | | | |
|---|---|---|---|---|---|
| chr6 | 11575654 | 11576102 | 1.32724 | 3.07859 | 2.320 |
| chr15 | 51178073 | 51178421 | 3.06139 | 7.10127 | 2.320 |
| chr4 | 668034 | 668336 | 1.62408 | 3.7676 | 2.320 |
| chr10 | 75863381 | 75863595 | 0.842551 | 1.95466 | 2.320 |
| chr5 | 130500294 | 130500670 | 0.725509 | 1.68314 | 2.320 |
| chr15 | 98298426 | 98298649 | 0.794955 | 1.84442 | 2.320 |
| chr9 | 36166289 | 36166888 | 1.64604 | 3.81913 | 2.320 |
| chr10 | 71718042 | 71718410 | 1.94882 | 4.52191 | 2.320 |
| chr17 | 42227967 | 42228364 | 1.26932 | 2.94535 | 2.320 |
| chr22 | 32809818 | 32810022 | 0.9459 | 2.19509 | 2.321 |
| chr12 | 120524682 | 120525038 | 2.46141 | 5.71208 | 2.321 |
| chr19 | 19256787 | 19257206 | 2.16587 | 5.02652 | 2.321 |
| chr10 | 86436582 | 86436799 | 0.379103 | 0.879837 | 2.321 |
| chr9 | 4197618 | 4197941 | 1.60544 | 3.72604 | 2.321 |
| chr1 | 175587821 | 175588251 | 0.441191 | 1.02396 | 2.321 |
| chr1 | 237976302 | 237976571 | 0.505575 | 1.17341 | 2.321 |
| chr1 | 182112297 | 182112623 | 2.14339 | 4.97475 | 2.321 |
| chr4 | 106392734 | 106393020 | 1.08882 | 2.52715 | 2.321 |
| chr15 | 41149783 | 41150202 | 1.74134 | 4.04172 | 2.321 |
| chr3 | 45701557 | 45701891 | 1.52699 | 3.54431 | 2.321 |
| chr1 | 150228282 | 150228678 | 1.11679 | 2.59221 | 2.321 |
| chr20 | 45901614 | 45901844 | 1.25312 | 2.90868 | 2.321 |
| chr6 | 41754101 | 41754491 | 1.46053 | 3.39012 | 2.321 |
| chr19 | 11698531 | 11698817 | 1.28032 | 2.97198 | 2.321 |
| chr17 | 31886204 | 31886470 | 0.473287 | 1.09864 | 2.321 |
| chr22 | 29699173 | 29699391 | 0.335828 | 0.779557 | 2.321 |
| chr7 | 147394778 | 147394990 | 0.449808 | 1.04419 | 2.321 |
| chr5 | 75976344 | 75976575 | 0.606582 | 1.40821 | 2.322 |
| chr5 | 156755294 | 156755841 | 1.4975 | 3.47662 | 2.322 |
| chr13 | 63200993 | 63201214 | 1.14766 | 2.66444 | 2.322 |
| chr16 | 87110835 | 87111289 | 1.99007 | 4.62026 | 2.322 |
| chr4 | 101014628 | 101014884 | 1.06538 | 2.47348 | 2.322 |
| chr12 | 112993749 | 112994039 | 0.505694 | 1.17409 | 2.322 |
| chr12 | 58934277 | 58934641 | 1.80509 | 4.19153 | 2.322 |
| chr16 | 50361019 | 50361292 | 1.2752 | 2.96123 | 2.322 |
| chr5 | 148750879 | 148751319 | 1.76611 | 4.10139 | 2.322 |
| chr8 | 27445686 | 27446106 | 1.65729 | 3.84867 | 2.322 |
| chr3 | 194354076 | 194354369 | 0.979106 | 2.27381 | 2.322 |
| chr16 | 78537491 | 78537790 | 1.4503 | 3.36809 | 2.322 |
| chr2 | 77313679 | 77313924 | 0.477062 | 1.1079 | 2.322 |
| chr16 | 61009539 | 61009918 | 0.484093 | 1.12424 | 2.322 |
| chr2 | 15686656 | 15687044 | 0.963297 | 2.23713 | 2.322 |
| chr16 | 58545747 | 58546440 | 1.63505 | 3.7972 | 2.322 |
| chr9 | 71564154 | 71564377 | 0.502903 | 1.16794 | 2.322 |
| chr8 | 37853757 | 37854162 | 2.20639 | 5.12421 | 2.322 |
| chr19 | 43067571 | 43067851 | 1.33523 | 3.10103 | 2.322 |
| chr10 | 44482867 | 44483256 | 0.447679 | 1.03973 | 2.322 |
| chr16 | 86674859 | 86675204 | 0.461837 | 1.07273 | 2.323 |
| chr19 | 6280163 | 6280481 | 1.71573 | 3.98561 | 2.323 |
| chr16 | 10666086 | 10666292 | 0.458549 | 1.06525 | 2.323 |
| chr7 | 95468393 | 95468654 | 0.293673 | 0.68224 | 2.323 |
| chr9 | 126462618 | 126462830 | 0.480722 | 1.11681 | 2.323 |
| chr13 | 65104083 | 65104345 | 0.942288 | 2.18934 | 2.323 |
| chr3 | 11183687 | 11183901 | 0.642917 | 1.49378 | 2.323 |
| chr2 | 61777789 | 61778069 | 0.442895 | 1.02905 | 2.323 |
| chr11 | 103072875 | 103073086 | 1.00566 | 2.33676 | 2.324 |
| chr8 | 146214238 | 146214517 | 0.65227 | 1.51564 | 2.324 |
| chr14 | 22515414 | 22515632 | 1.07448 | 2.49682 | 2.324 |
| chr19 | 52215959 | 52216262 | 1.28679 | 2.99027 | 2.324 |
| chr20 | 41067388 | 41067682 | 0.443423 | 1.03052 | 2.324 |
| chr16 | 47052158 | 47052368 | 1.05154 | 2.44385 | 2.324 |

| | | | | | |
|---|---|---|---|---|---|
| chr3 | 128964569 | 128964966 | 1.7963 | 4.17479 | 2.324 |
| chr15 | 42358793 | 42359055 | 0.705404 | 1.63947 | 2.324 |
| chr10 | 89293454 | 89293671 | 1.14511 | 2.66147 | 2.324 |
| chr9 | 134351328 | 134351692 | 1.46978 | 3.41609 | 2.324 |
| chr6 | 161574030 | 161574297 | 1.54586 | 3.59302 | 2.324 |
| chr16 | 85644089 | 85644734 | 1.71603 | 3.9886 | 2.324 |
| chr16 | 85090839 | 85091095 | 1.17132 | 2.72279 | 2.325 |
| chr19 | 42688259 | 42688857 | 2.1312 | 4.95414 | 2.325 |
| chr13 | 109549220 | 109549689 | 0.932636 | 2.16802 | 2.325 |
| chr1 | 84326105 | 84326488 | 1.7492 | 4.06626 | 2.325 |
| chr12 | 103488998 | 103489224 | 0.63268 | 1.47081 | 2.325 |
| chr10 | 120957998 | 120958259 | 1.4384 | 3.34397 | 2.325 |
| chr17 | 13241117 | 13241451 | 1.82422 | 4.24097 | 2.325 |
| chr20 | 51605286 | 51605497 | 0.51188 | 1.19005 | 2.325 |
| chr3 | 185071929 | 185072188 | 1.25794 | 2.92453 | 2.325 |
| chr5 | 94686550 | 94686847 | 0.577465 | 1.34254 | 2.325 |
| chr3 | 48031507 | 48031726 | 1.27799 | 2.97124 | 2.325 |
| chr19 | 51040734 | 51041406 | 1.8859 | 4.38464 | 2.325 |
| chr6 | 3732810 | 3733108 | 1.82073 | 4.23336 | 2.325 |
| chr16 | 21595364 | 21595839 | 0.652012 | 1.51599 | 2.325 |
| chr8 | 10696384 | 10696778 | 1.62164 | 3.77055 | 2.325 |
| chr15 | 65677870 | 65678316 | 2.04518 | 4.75543 | 2.325 |
| chr9 | 136302906 | 136303335 | 1.01161 | 2.35221 | 2.325 |
| chr14 | 31957370 | 31957604 | 0.542813 | 1.26216 | 2.325 |
| chr7 | 129275383 | 129275721 | 1.60165 | 3.72422 | 2.325 |
| chr20 | 16053748 | 16053953 | 0.554792 | 1.29003 | 2.325 |
| chr5 | 141348361 | 141348737 | 1.91729 | 4.4582 | 2.325 |
| chr20 | 30150625 | 30150980 | 2.13045 | 4.95398 | 2.325 |
| chr1 | 60412934 | 60413467 | 0.980039 | 2.27909 | 2.326 |
| chr16 | 9504447 | 9504760 | 0.358379 | 0.833488 | 2.326 |
| chr1 | 223884056 | 223884469 | 1.57482 | 3.66261 | 2.326 |
| chr11 | 15378853 | 15379188 | 0.411634 | 0.957349 | 2.326 |
| chr7 | 28893739 | 28894180 | 1.89961 | 4.41802 | 2.326 |
| chr6 | 43823918 | 43824199 | 1.52527 | 3.54743 | 2.326 |
| chr12 | 129781293 | 129781679 | 0.416605 | 0.968971 | 2.326 |
| chr15 | 30124568 | 30124781 | 0.483326 | 1.12418 | 2.326 |
| chr2 | 233034172 | 233034403 | 0.411992 | 0.958266 | 2.326 |
| chr10 | 95136197 | 95136437 | 1.63468 | 3.80223 | 2.326 |
| chr20 | 8112596 | 8112844 | 2.00241 | 4.65781 | 2.326 |
| chr12 | 46384413 | 46384670 | 2.05094 | 4.77081 | 2.326 |
| chr7 | 44887244 | 44887632 | 1.61471 | 3.75628 | 2.326 |
| chr2 | 183649507 | 183649711 | 0.575117 | 1.33792 | 2.326 |
| chr5 | 157078633 | 157078949 | 0.588837 | 1.36985 | 2.326 |
| chr5 | 8174021 | 8174259 | 0.468254 | 1.08935 | 2.326 |
| chr5 | 153216442 | 153216783 | 0.522146 | 1.21476 | 2.326 |
| chr5 | 101313517 | 101313752 | 1.15613 | 2.68973 | 2.326 |
| chr10 | 9373517 | 9373855 | 1.20503 | 2.80358 | 2.327 |
| chr17 | 45018170 | 45018560 | 2.19149 | 5.09871 | 2.327 |
| chr2 | 105796252 | 105796575 | 1.37682 | 3.20348 | 2.327 |
| chr10 | 116546258 | 116546533 | 0.942746 | 2.19355 | 2.327 |
| chr17 | 42440884 | 42441709 | 1.84118 | 4.28403 | 2.327 |
| chr5 | 154519170 | 154519556 | 1.57131 | 3.65629 | 2.327 |
| chr1 | 223316017 | 223316574 | 1.28926 | 3.00003 | 2.327 |
| chr20 | 58146220 | 58147065 | 1.82114 | 4.23784 | 2.327 |
| chr7 | 79763827 | 79764095 | 1.40548 | 3.27061 | 2.327 |
| chr7 | 75988670 | 75988892 | 1.42559 | 3.3175 | 2.327 |
| chr3 | 69474503 | 69474706 | 0.766082 | 1.7828 | 2.327 |
| chr8 | 17220318 | 17220525 | 0.405784 | 0.944328 | 2.327 |
| chr5 | 31232430 | 31232751 | 1.11104 | 2.58562 | 2.327 |
| chr12 | 57030437 | 57031090 | 1.3841 | 3.22112 | 2.327 |
| chr11 | 78604031 | 78604279 | 0.439692 | 1.0233 | 2.327 |

| | | | | | |
|---|---|---|---|---|---|
| chr7 | 71588400 | 71588612 | 0.440341 | 1.02481 | 2.327 |
| chr12 | 103897788 | 103898010 | 0.966397 | 2.24927 | 2.327 |
| chr11 | 46911032 | 46911629 | 1.20139 | 2.79637 | 2.328 |
| chr10 | 82939626 | 82939889 | 1.06452 | 2.47792 | 2.328 |
| chr1 | 38148083 | 38148420 | 1.11707 | 2.60043 | 2.328 |
| chr12 | 57028784 | 57029057 | 1.42738 | 3.32297 | 2.328 |
| chr12 | 76552565 | 76552910 | 1.59355 | 3.70982 | 2.328 |
| chr16 | 53726099 | 53726334 | 1.11561 | 2.59735 | 2.328 |
| chr15 | 100284428 | 100284820 | 0.854714 | 1.98996 | 2.328 |
| chr12 | 120086987 | 120087352 | 1.37092 | 3.19184 | 2.328 |
| chr7 | 151007625 | 151008058 | 0.842091 | 1.96062 | 2.328 |
| chr7 | 82369500 | 82369918 | 1.51118 | 3.51874 | 2.328 |
| chr10 | 13248469 | 13249149 | 0.67502 | 1.57181 | 2.329 |
| chr9 | 120009693 | 120009912 | 0.884504 | 2.05968 | 2.329 |
| chr12 | 26986269 | 26986494 | 1.42448 | 3.31713 | 2.329 |
| chrX | 149131029 | 149131409 | 1.20699 | 2.81077 | 2.329 |
| chr14 | 62361230 | 62361774 | 1.17082 | 2.72664 | 2.329 |
| chr19 | 50982308 | 50982828 | 2.30851 | 5.37618 | 2.329 |
| chr9 | 13580811 | 13581097 | 0.617862 | 1.43891 | 2.329 |
| chr15 | 85471383 | 85471731 | 2.47926 | 5.77392 | 2.329 |
| chr9 | 100652289 | 100652706 | 1.22156 | 2.84499 | 2.329 |
| chr9 | 89695830 | 89696032 | 0.638072 | 1.48614 | 2.329 |
| chr13 | 20531458 | 20531791 | 0.7901 | 1.8403 | 2.329 |
| chr10 | 26661914 | 26662324 | 0.405046 | 0.943453 | 2.329 |
| chr14 | 100095177 | 100095652 | 1.97982 | 4.61152 | 2.329 |
| chr13 | 20534327 | 20534587 | 1.37347 | 3.1993 | 2.329 |
| chr14 | 58338253 | 58338536 | 1.2621 | 2.93993 | 2.329 |
| chr9 | 22447322 | 22447636 | 0.890499 | 2.07438 | 2.329 |
| chr19 | 6863669 | 6863906 | 0.93446 | 2.17686 | 2.330 |
| chr17 | 79346310 | 79346666 | 1.31678 | 3.06751 | 2.330 |
| chr9 | 14353385 | 14353848 | 1.57214 | 3.66238 | 2.330 |
| chr14 | 25850812 | 25851026 | 0.75191 | 1.75164 | 2.330 |
| chr17 | 15357655 | 15357998 | 0.478517 | 1.11484 | 2.330 |
| chr20 | 59454964 | 59455203 | 0.412533 | 0.961178 | 2.330 |
| chr9 | 2662535 | 2662783 | 0.913624 | 2.12879 | 2.330 |
| chr8 | 30458264 | 30458575 | 1.12926 | 2.63125 | 2.330 |
| chr14 | 35698696 | 35699000 | 0.856561 | 1.99588 | 2.330 |
| chr19 | 56666534 | 56667042 | 0.963604 | 2.24538 | 2.330 |
| chr6 | 159081311 | 159081587 | 1.23675 | 2.88193 | 2.330 |
| chr17 | 19372632 | 19373045 | 1.64211 | 3.82653 | 2.330 |
| chr17 | 1268739 | 1269106 | 2.02217 | 4.71223 | 2.330 |
| chr3 | 128336332 | 128336919 | 1.73232 | 4.03695 | 2.330 |
| chr6 | 111531525 | 111531779 | 1.01484 | 2.36497 | 2.330 |
| chr10 | 29785346 | 29785699 | 1.65035 | 3.84629 | 2.331 |
| chrX | 148593843 | 148594194 | 1.20718 | 2.81367 | 2.331 |
| chr19 | 10223745 | 10224026 | 2.12931 | 4.963 | 2.331 |
| chr13 | 21278092 | 21278426 | 0.649697 | 1.51439 | 2.331 |
| chr9 | 28105751 | 28106109 | 0.41138 | 0.958904 | 2.331 |
| chr20 | 49192433 | 49192794 | 1.65494 | 3.85761 | 2.331 |
| chr6 | 5303524 | 5303838 | 1.03305 | 2.40811 | 2.331 |
| chr15 | 86433307 | 86433525 | 0.335597 | 0.782315 | 2.331 |
| chr9 | 22792556 | 22792800 | 0.509324 | 1.18735 | 2.331 |
| chr12 | 52476997 | 52477592 | 2.15334 | 5.02005 | 2.331 |
| chr11 | 83839852 | 83840285 | 0.476843 | 1.1118 | 2.332 |
| chr15 | 100411825 | 100412152 | 0.405865 | 0.946326 | 2.332 |
| chr11 | 63044880 | 63045191 | 0.824959 | 1.92351 | 2.332 |
| chr19 | 41452048 | 41452275 | 0.570128 | 1.32937 | 2.332 |
| chr14 | 96566917 | 96567278 | 3.06954 | 7.15794 | 2.332 |
| chr7 | 20691679 | 20691983 | 0.692174 | 1.61416 | 2.332 |
| chr6 | 130096252 | 130096656 | 2.05632 | 4.79539 | 2.332 |
| chr11 | 14558595 | 14558881 | 1.21415 | 2.83148 | 2.332 |

| | | | | | |
|---|---|---|---|---|---|
| chr17 | 29153016 | 29153274 | 1.29286 | 3.01508 | 2.332 |
| chr2 | 88558867 | 88559167 | 0.459483 | 1.07164 | 2.332 |
| chr11 | 58974181 | 58974436 | 1.37669 | 3.21091 | 2.332 |
| chr5 | 154708065 | 154708333 | 0.370257 | 0.863607 | 2.332 |
| chr17 | 57612090 | 57612293 | 0.42427 | 0.989623 | 2.333 |
| chr15 | 72640660 | 72640864 | 0.541769 | 1.26373 | 2.333 |
| chr2 | 19474375 | 19474722 | 0.553396 | 1.29089 | 2.333 |
| chr20 | 61912469 | 61912879 | 1.12005 | 2.61276 | 2.333 |
| chr9 | 77027724 | 77028011 | 0.439329 | 1.02483 | 2.333 |
| chr2 | 227664492 | 227664835 | 1.26223 | 2.94454 | 2.333 |
| chr5 | 133922280 | 133923190 | 1.49372 | 3.48461 | 2.333 |
| chr2 | 3060785 | 3060992 | 0.500622 | 1.16791 | 2.333 |
| chr18 | 19536068 | 19536341 | 1.64554 | 3.839 | 2.333 |
| chrX | 82890342 | 82890597 | 1.2401 | 2.89311 | 2.333 |
| chr16 | 2079903 | 2080196 | 0.536483 | 1.25161 | 2.333 |
| chr7 | 154207493 | 154207806 | 0.510348 | 1.19069 | 2.333 |
| chr12 | 50017262 | 50017521 | 1.75912 | 4.10426 | 2.333 |
| chr3 | 38570912 | 38571508 | 0.529131 | 1.23483 | 2.334 |
| chrX | 102884943 | 102885192 | 1.18863 | 2.77412 | 2.334 |
| chr5 | 3401060 | 3401308 | 0.540364 | 1.26118 | 2.334 |
| chr1 | 59012316 | 59012734 | 1.67424 | 3.90771 | 2.334 |
| chr11 | 4647483 | 4647808 | 1.10672 | 2.58311 | 2.334 |
| chr6 | 38639679 | 38639945 | 1.19887 | 2.7982 | 2.334 |
| chr15 | 90524492 | 90524731 | 0.459364 | 1.07221 | 2.334 |
| chr2 | 46005754 | 46006098 | 1.83025 | 4.27203 | 2.334 |
| chr5 | 65994068 | 65994512 | 0.445614 | 1.04014 | 2.334 |
| chr18 | 25769336 | 25769637 | 1.22021 | 2.84849 | 2.334 |
| chr14 | 102904395 | 102904662 | 1.34886 | 3.149 | 2.335 |
| chr15 | 41198565 | 41199216 | 1.06417 | 2.48467 | 2.335 |
| chr12 | 82752193 | 82752842 | 2.78795 | 6.50945 | 2.335 |
| chr1 | 8919491 | 8919784 | 1.19987 | 2.80166 | 2.335 |
| chr11 | 69532912 | 69533247 | 1.26735 | 2.9594 | 2.335 |
| chr11 | 92969763 | 92970218 | 0.833574 | 1.9465 | 2.335 |
| chr9 | 112820697 | 112820944 | 1.14424 | 2.67196 | 2.335 |
| chr5 | 135313101 | 135313451 | 1.85814 | 4.3391 | 2.335 |
| chr17 | 79995024 | 79995488 | 2.29488 | 5.35903 | 2.335 |
| chr3 | 147404548 | 147404829 | 0.436565 | 1.01947 | 2.335 |
| chr3 | 167758377 | 167758692 | 1.15286 | 2.69222 | 2.335 |
| chr15 | 27960182 | 27960468 | 0.605136 | 1.41315 | 2.335 |
| chr20 | 49988789 | 49989149 | 2.01061 | 4.69529 | 2.335 |
| chr14 | 55272319 | 55272919 | 1.92968 | 4.50642 | 2.335 |
| chr7 | 98239850 | 98240126 | 0.403895 | 0.943228 | 2.335 |
| chr19 | 49449388 | 49449834 | 0.691193 | 1.61422 | 2.335 |
| chr3 | 9974314 | 9975028 | 1.28972 | 3.01216 | 2.336 |
| chr6 | 123737063 | 123737273 | 0.869955 | 2.03188 | 2.336 |
| chr8 | 144313597 | 144314020 | 1.44301 | 3.37034 | 2.336 |
| chr5 | 14727449 | 14727851 | 1.79144 | 4.18418 | 2.336 |
| chr1 | 181732917 | 181733168 | 0.387222 | 0.904518 | 2.336 |
| chr15 | 59837797 | 59838016 | 0.516407 | 1.20631 | 2.336 |
| chr20 | 15689637 | 15690028 | 1.54384 | 3.60649 | 2.336 |
| chr19 | 28265894 | 28266118 | 1.01049 | 2.36064 | 2.336 |
| chr5 | 158791850 | 158792151 | 0.557273 | 1.30192 | 2.336 |
| chr22 | 17652125 | 17653074 | 1.79121 | 4.18478 | 2.336 |
| chr5 | 153569506 | 153569881 | 0.9414 | 2.19947 | 2.336 |
| chr12 | 56048478 | 56049133 | 0.531682 | 1.24224 | 2.336 |
| chr15 | 78774707 | 78775048 | 1.62259 | 3.7913 | 2.337 |
| chr9 | 71456965 | 71457195 | 0.616678 | 1.44093 | 2.337 |
| chr9 | 95798396 | 95798644 | 0.645886 | 1.50923 | 2.337 |
| chr1 | 237503122 | 237503381 | 0.562577 | 1.31458 | 2.337 |
| chr17 | 3838321 | 3838738 | 2.07612 | 4.85129 | 2.337 |
| chr12 | 70033182 | 70033607 | 1.78061 | 4.1608 | 2.337 |

| | | | | | |
|---|---|---|---|---|---|
| chr14 | 44367933 | 44368156 | 0.510366 | 1.19262 | 2.337 |
| chr1 | 1875520 | 1875838 | 0.491012 | 1.14741 | 2.337 |
| chr18 | 46777305 | 46777624 | 0.834547 | 1.9503 | 2.337 |
| chr8 | 127154694 | 127154898 | 0.456445 | 1.06671 | 2.337 |
| chr16 | 3929926 | 3930226 | 1.18293 | 2.76471 | 2.337 |
| chr5 | 45797132 | 45797346 | 1.03967 | 2.42995 | 2.337 |
| chr3 | 13052265 | 13052489 | 0.440359 | 1.0293 | 2.337 |
| chr12 | 12808907 | 12809221 | 1.54055 | 3.601 | 2.337 |
| chr1 | 166688987 | 166689216 | 0.404231 | 0.944915 | 2.338 |
| chr11 | 72145027 | 72145603 | 3.44927 | 8.0632 | 2.338 |
| chr1 | 214315046 | 214315278 | 0.610675 | 1.42765 | 2.338 |
| chr18 | 12327846 | 12328278 | 1.83627 | 4.29318 | 2.338 |
| chr1 | 9070056 | 9070440 | 1.21345 | 2.83707 | 2.338 |
| chr4 | 73885219 | 73885481 | 1.01141 | 2.3647 | 2.338 |
| chr6 | 107976473 | 107976715 | 0.659223 | 1.54147 | 2.338 |
| chr14 | 80002129 | 80002334 | 0.510275 | 1.19331 | 2.339 |
| chr12 | 56440911 | 56441288 | 1.30752 | 3.0578 | 2.339 |
| chr10 | 61963887 | 61964105 | 1.25445 | 2.93371 | 2.339 |
| chr6 | 167120570 | 167120811 | 1.06656 | 2.49435 | 2.339 |
| chr2 | 17652190 | 17652468 | 0.798568 | 1.86763 | 2.339 |
| chr12 | 66663640 | 66663872 | 1.27989 | 2.99335 | 2.339 |
| chr1 | 16514282 | 16514566 | 2.40892 | 5.63396 | 2.339 |
| chr1 | 244615175 | 244615637 | 0.816554 | 1.90979 | 2.339 |
| chr7 | 82358795 | 82359030 | 0.922558 | 2.1578 | 2.339 |
| chr1 | 90270240 | 90270580 | 1.72525 | 4.03527 | 2.339 |
| chr7 | 134197952 | 134198282 | 1.6351 | 3.82441 | 2.339 |
| chr17 | 75725710 | 75725943 | 0.516228 | 1.20745 | 2.339 |
| chr3 | 10780292 | 10780964 | 2.17281 | 5.08223 | 2.339 |
| chr12 | 55465841 | 55466247 | 1.49793 | 3.50394 | 2.339 |
| chr4 | 1350058 | 1350368 | 1.79738 | 4.20458 | 2.339 |
| chr1 | 157056313 | 157056555 | 0.496845 | 1.16228 | 2.339 |
| chr5 | 3373092 | 3373322 | 0.46217 | 1.08117 | 2.339 |
| chr13 | 111875816 | 111876053 | 1.3172 | 3.08146 | 2.339 |
| chr3 | 93755796 | 93756045 | 1.26119 | 2.95045 | 2.339 |
| chr20 | 56478105 | 56478338 | 0.460601 | 1.07755 | 2.339 |
| chr8 | 146252106 | 146252434 | 1.71359 | 4.00902 | 2.340 |
| chr16 | 412146 | 412520 | 2.44699 | 5.72501 | 2.340 |
| chr1 | 27611168 | 27611520 | 1.16807 | 2.73291 | 2.340 |
| chr20 | 31879578 | 31880043 | 2.52933 | 5.91787 | 2.340 |
| chr11 | 65307723 | 65308023 | 0.674197 | 1.57744 | 2.340 |
| chr11 | 45864737 | 45865005 | 1.72184 | 4.02868 | 2.340 |
| chr8 | 99097861 | 99098109 | 1.29016 | 3.01866 | 2.340 |
| chr4 | 140216786 | 140217165 | 1.71998 | 4.02441 | 2.340 |
| chrX | 10737190 | 10737539 | 0.987637 | 2.31091 | 2.340 |
| chr4 | 41694913 | 41695234 | 2.08621 | 4.88142 | 2.340 |
| chr19 | 13539201 | 13539521 | 1.5768 | 3.68955 | 2.340 |
| chr20 | 31490181 | 31490665 | 2.06066 | 4.82188 | 2.340 |
| chr10 | 13482423 | 13482853 | 2.22707 | 5.21137 | 2.340 |
| chr6 | 130686493 | 130686917 | 1.84995 | 4.32922 | 2.340 |
| chr1 | 94979773 | 94979958 | 1.21513 | 2.84365 | 2.340 |
| chr5 | 82904046 | 82904249 | 1.16497 | 2.72635 | 2.340 |
| chr11 | 110300631 | 110301114 | 1.18062 | 2.76298 | 2.340 |
| chr11 | 128042266 | 128042524 | 0.956576 | 2.23868 | 2.340 |
| chr1 | 177150819 | 177151137 | 2.26668 | 5.30476 | 2.340 |
| chr19 | 11491644 | 11491972 | 1.06234 | 2.48626 | 2.340 |
| chr4 | 126663089 | 126663311 | 1.01032 | 2.36451 | 2.340 |
| chr5 | 77833690 | 77833953 | 0.982871 | 2.30027 | 2.340 |
| chr12 | 58386877 | 58387102 | 1.20598 | 2.82248 | 2.340 |
| chr1 | 176088819 | 176089078 | 0.840883 | 1.96801 | 2.340 |
| chr22 | 32914841 | 32915071 | 0.899197 | 2.10454 | 2.340 |
| chr7 | 111375817 | 111376078 | 1.41046 | 3.30148 | 2.341 |

| | | | | | |
|---|---|---|---|---|---|
| chr20 | 22366690 | 22366961 | 1.45659 | 3.40955 | 2.341 |
| chr3 | 151045324 | 151045797 | 0.846712 | 1.982 | 2.341 |
| chr2 | 46133199 | 46133434 | 0.581975 | 1.36233 | 2.341 |
| chr18 | 22306650 | 22307082 | 1.11286 | 2.60509 | 2.341 |
| chr22 | 22929469 | 22929724 | 1.55707 | 3.64494 | 2.341 |
| chr3 | 113871298 | 113871654 | 1.72647 | 4.0418 | 2.341 |
| chr8 | 95006956 | 95007180 | 1.03357 | 2.41973 | 2.341 |
| chr4 | 113687650 | 113687935 | 0.86402 | 2.02282 | 2.341 |
| chr5 | 50657639 | 50657897 | 0.47344 | 1.10846 | 2.341 |
| chr7 | 95590863 | 95591087 | 0.469322 | 1.09883 | 2.341 |
| chr12 | 6210402 | 6210674 | 0.410757 | 0.96175 | 2.341 |
| chr17 | 17173313 | 17173539 | 1.07541 | 2.51798 | 2.341 |
| chr1 | 211526270 | 211526578 | 1.57785 | 3.69441 | 2.341 |
| chr22 | 25843770 | 25844092 | 1.61703 | 3.78614 | 2.341 |
| chr3 | 86156769 | 86157032 | 0.434442 | 1.01722 | 2.341 |
| chr1 | 166879625 | 166879921 | 1.00934 | 2.36338 | 2.342 |
| chr15 | 49714155 | 49714415 | 0.937143 | 2.19435 | 2.342 |
| chr5 | 154407674 | 154407891 | 0.735087 | 1.72123 | 2.342 |
| chr20 | 17705442 | 17705774 | 0.496865 | 1.16345 | 2.342 |
| chr11 | 2291175 | 2291597 | 0.845047 | 1.97876 | 2.342 |
| chr7 | 152925692 | 152925954 | 0.495811 | 1.16099 | 2.342 |
| chr12 | 123755347 | 123755953 | 1.45687 | 3.41143 | 2.342 |
| chr8 | 1729408 | 1729736 | 1.2152 | 2.84566 | 2.342 |
| chr2 | 31084958 | 31085184 | 1.6146 | 3.78107 | 2.342 |
| chr17 | 79283092 | 79283586 | 0.844514 | 1.9779 | 2.342 |
| chr3 | 185943336 | 185943640 | 0.334158 | 0.782673 | 2.342 |
| chr16 | 30996871 | 30997078 | 0.797012 | 1.8668 | 2.342 |
| chr10 | 118925285 | 118925597 | 1.441 | 3.37521 | 2.342 |
| chr5 | 166671669 | 166671930 | 0.712654 | 1.66929 | 2.342 |
| chr15 | 80462836 | 80463068 | 0.399177 | 0.935094 | 2.343 |
| chrX | 106106180 | 106106705 | 1.1353 | 2.65953 | 2.343 |
| chr19 | 3304980 | 3305344 | 1.65277 | 3.87182 | 2.343 |
| chr3 | 183220640 | 183220869 | 0.617771 | 1.4472 | 2.343 |
| chr16 | 2009254 | 2009562 | 1.95193 | 4.57276 | 2.343 |
| chr17 | 44859600 | 44859931 | 1.95143 | 4.57194 | 2.343 |
| chr9 | 36862064 | 36862543 | 1.89384 | 4.43709 | 2.343 |
| chr5 | 23800249 | 23800604 | 0.385002 | 0.902032 | 2.343 |
| chr1 | 233654164 | 233654366 | 0.976272 | 2.2874 | 2.343 |
| chr15 | 33900973 | 33901251 | 0.729473 | 1.70921 | 2.343 |
| chr5 | 4256294 | 4256511 | 0.555545 | 1.30172 | 2.343 |
| chr8 | 132081981 | 132082289 | 0.475689 | 1.11462 | 2.343 |
| chr15 | 57506102 | 57506335 | 1.05656 | 2.47583 | 2.343 |
| chr7 | 80310650 | 80310887 | 1.24943 | 2.92783 | 2.343 |
| chr18 | 18759660 | 18759912 | 1.31693 | 3.08601 | 2.343 |
| chr16 | 76673944 | 76674162 | 0.812299 | 1.90351 | 2.343 |
| chr13 | 51695056 | 51695364 | 1.25263 | 2.93537 | 2.343 |
| chr16 | 58712770 | 58712995 | 0.610771 | 1.43127 | 2.343 |
| chr14 | 21296168 | 21296431 | 0.780851 | 1.82986 | 2.343 |
| chr17 | 2916296 | 2916681 | 1.85696 | 4.35164 | 2.343 |
| chr1 | 16142061 | 16142395 | 1.10312 | 2.58526 | 2.344 |
| chr6 | 74098390 | 74098697 | 1.16339 | 2.72655 | 2.344 |
| chr10 | 88453558 | 88453886 | 1.34338 | 3.14849 | 2.344 |
| chr14 | 64527390 | 64527749 | 1.22887 | 2.88029 | 2.344 |
| chr1 | 43703773 | 43704128 | 1.20738 | 2.82999 | 2.344 |
| chr7 | 158813740 | 158814021 | 1.02319 | 2.39828 | 2.344 |
| chr1 | 26490630 | 26491046 | 1.12414 | 2.63491 | 2.344 |
| chr10 | 119297031 | 119297498 | 1.81484 | 4.25386 | 2.344 |
| chr8 | 36206370 | 36207023 | 0.426675 | 1.00014 | 2.344 |
| chr6 | 75053594 | 75053797 | 0.936753 | 2.19586 | 2.344 |
| chr12 | 723567 | 723775 | 1.28506 | 3.01239 | 2.344 |
| chr9 | 127099469 | 127099730 | 0.494624 | 1.15957 | 2.344 |

| | | | | | |
|---|---|---|---|---|---|
| chr11 | 64068097 | 64068839 | 1.59116 | 3.73028 | 2.344 |
| chr9 | 124462460 | 124462794 | 1.68032 | 3.93937 | 2.344 |
| chr12 | 64563640 | 64563976 | 0.620493 | 1.4547 | 2.344 |
| chr17 | 73760599 | 73761093 | 1.19644 | 2.80505 | 2.345 |
| chr11 | 19623825 | 19624176 | 0.357243 | 0.837593 | 2.345 |
| chr5 | 174220762 | 174221016 | 1.5066 | 3.53243 | 2.345 |
| chr3 | 113898489 | 113898781 | 0.957642 | 2.24536 | 2.345 |
| chr3 | 77423894 | 77424210 | 0.46047 | 1.07972 | 2.345 |
| chr17 | 63984208 | 63984438 | 1.09145 | 2.5595 | 2.345 |
| chr3 | 170767665 | 170767940 | 0.968937 | 2.27229 | 2.345 |
| chr1 | 163498596 | 163498858 | 1.2155 | 2.85055 | 2.345 |
| chr16 | 84544934 | 84545365 | 1.75852 | 4.12414 | 2.345 |
| chr11 | 36328551 | 36328754 | 0.816227 | 1.91426 | 2.345 |
| chr17 | 78095757 | 78096000 | 0.396436 | 0.929764 | 2.345 |
| chr3 | 172942874 | 172943273 | 0.543239 | 1.27407 | 2.345 |
| chr19 | 54345374 | 54345673 | 2.03242 | 4.76676 | 2.345 |
| chr20 | 57445347 | 57445642 | 2.06771 | 4.85 | 2.346 |
| chr19 | 29334665 | 29334891 | 0.766865 | 1.79883 | 2.346 |
| chr11 | 65537365 | 65537609 | 1.00041 | 2.34666 | 2.346 |
| chr12 | 7047167 | 7047491 | 2.17271 | 5.09668 | 2.346 |
| chr15 | 82490388 | 82490658 | 1.24998 | 2.93216 | 2.346 |
| chr3 | 142575975 | 142576238 | 1.06143 | 2.48997 | 2.346 |
| chr11 | 21238204 | 21238509 | 0.333181 | 0.781615 | 2.346 |
| chr3 | 120226737 | 120227258 | 0.417551 | 0.979553 | 2.346 |
| chr8 | 144089477 | 144089905 | 0.945254 | 2.21752 | 2.346 |
| chr10 | 7565727 | 7565947 | 0.754729 | 1.77058 | 2.346 |
| chr5 | 59692405 | 59692630 | 0.516459 | 1.2116 | 2.346 |
| chr10 | 110560188 | 110560442 | 0.628077 | 1.47347 | 2.346 |
| chr14 | 79080310 | 79080551 | 0.391525 | 0.918516 | 2.346 |
| chr11 | 61638498 | 61639003 | 1.81534 | 4.25906 | 2.346 |
| chr11 | 74742959 | 74743439 | 1.61402 | 3.7873 | 2.347 |
| chr3 | 170881043 | 170881282 | 0.894924 | 2.09994 | 2.347 |
| chr11 | 61439066 | 61439461 | 1.12993 | 2.65149 | 2.347 |
| chrX | 55161680 | 55161891 | 1.2202 | 2.86334 | 2.347 |
| chr3 | 157261099 | 157261353 | 0.831979 | 1.95236 | 2.347 |
| chr17 | 72787619 | 72788048 | 3.21898 | 7.55443 | 2.347 |
| chr3 | 185959645 | 185959956 | 1.21781 | 2.85806 | 2.347 |
| chr7 | 52989055 | 52989332 | 0.561778 | 1.31843 | 2.347 |
| chr1 | 45390782 | 45391011 | 0.712831 | 1.67296 | 2.347 |
| chr10 | 3235090 | 3235429 | 1.81993 | 4.27178 | 2.347 |
| chr12 | 118312644 | 118312986 | 2.22473 | 5.22193 | 2.347 |
| chr2 | 216784924 | 216785171 | 1.19295 | 2.80021 | 2.347 |
| chr12 | 39928844 | 39929089 | 0.909385 | 2.13465 | 2.347 |
| chr11 | 43968694 | 43969060 | 0.699135 | 1.64125 | 2.348 |
| chr14 | 57784035 | 57784268 | 1.39111 | 3.2657 | 2.348 |
| chr9 | 132869419 | 132869755 | 1.35858 | 3.18936 | 2.348 |
| chr17 | 69356724 | 69356932 | 0.489495 | 1.14918 | 2.348 |
| chr1 | 149238821 | 149239029 | 0.530879 | 1.24635 | 2.348 |
| chr1 | 46290356 | 46290572 | 0.74131 | 1.74045 | 2.348 |
| chr13 | 76056582 | 76056928 | 1.79807 | 4.22171 | 2.348 |
| chr14 | 59593951 | 59594176 | 0.882693 | 2.07249 | 2.348 |
| chr13 | 46863069 | 46863424 | 1.12259 | 2.63585 | 2.348 |
| chr5 | 73321950 | 73322288 | 0.414072 | 0.972298 | 2.348 |
| chr4 | 157893036 | 157893312 | 0.720894 | 1.69302 | 2.349 |
| chr17 | 33639753 | 33640101 | 1.84294 | 4.32841 | 2.349 |
| chr12 | 49730774 | 49731250 | 2.52094 | 5.92125 | 2.349 |
| chr14 | 83573609 | 83573818 | 0.377151 | 0.885878 | 2.349 |
| chr7 | 3713169 | 3713434 | 1.2435 | 2.92086 | 2.349 |
| chr7 | 1233449 | 1233827 | 1.41142 | 3.3154 | 2.349 |
| chr7 | 85455452 | 85455769 | 0.514027 | 1.20744 | 2.349 |
| chr6 | 138430600 | 138430882 | 1.46657 | 3.44498 | 2.349 |

| | | | | | |
|---|---|---|---|---|---|
| chr12 | 120666172 | 120666750 | 1.55889 | 3.66213 | 2.349 |
| chr8 | 19615318 | 19615573 | 1.19839 | 2.81524 | 2.349 |
| chr9 | 77193012 | 77193245 | 0.992601 | 2.33183 | 2.349 |
| chr4 | 26083595 | 26083983 | 1.08068 | 2.53884 | 2.349 |
| chr11 | 107560393 | 107560688 | 1.16882 | 2.74595 | 2.349 |
| chr15 | 41222663 | 41223144 | 1.38451 | 3.25273 | 2.349 |
| chr19 | 10534944 | 10535372 | 2.0412 | 4.79554 | 2.349 |
| chr16 | 80031210 | 80031590 | 1.08042 | 2.53854 | 2.350 |
| chr7 | 146942429 | 146942734 | 1.34368 | 3.15724 | 2.350 |
| chr1 | 165840731 | 165840962 | 0.965018 | 2.26762 | 2.350 |
| chr17 | 79568569 | 79568907 | 1.72992 | 4.065 | 2.350 |
| chr18 | 61646592 | 61646992 | 1.18532 | 2.78533 | 2.350 |
| chr1 | 36565267 | 36565870 | 1.63007 | 3.83054 | 2.350 |
| chr3 | 141184116 | 141184404 | 0.690428 | 1.62247 | 2.350 |
| chr3 | 188466787 | 188467093 | 0.976845 | 2.29563 | 2.350 |
| chr2 | 160855475 | 160855737 | 1.21995 | 2.867 | 2.350 |
| chr14 | 102173688 | 102173968 | 1.48103 | 3.48072 | 2.350 |
| chr15 | 100521496 | 100521767 | 0.607961 | 1.42884 | 2.350 |
| chr4 | 113859617 | 113859917 | 1.03529 | 2.43317 | 2.350 |
| chr8 | 25991147 | 25991587 | 2.10959 | 4.95827 | 2.350 |
| chr2 | 79595803 | 79596227 | 0.385847 | 0.906879 | 2.350 |
| chr12 | 20219620 | 20219902 | 0.418733 | 0.984178 | 2.350 |
| chr6 | 114315983 | 114316394 | 1.4926 | 3.50823 | 2.350 |
| chr20 | 52029684 | 52029995 | 1.3688 | 3.21733 | 2.350 |
| chr11 | 10792221 | 10792510 | 1.85423 | 4.35839 | 2.351 |
| chr10 | 46983352 | 46984039 | 2.39774 | 5.63602 | 2.351 |
| chr3 | 157787952 | 157788209 | 0.331701 | 0.779685 | 2.351 |
| chr4 | 8243917 | 8244337 | 2.30682 | 5.42248 | 2.351 |
| chr8 | 96069123 | 96069468 | 1.36306 | 3.20409 | 2.351 |
| chr19 | 11236772 | 11237218 | 1.23566 | 2.90471 | 2.351 |
| chr10 | 96122683 | 96122935 | 1.22278 | 2.87462 | 2.351 |
| chr12 | 120132949 | 120133233 | 1.58712 | 3.7313 | 2.351 |
| chr22 | 36461951 | 36462527 | 3.14804 | 7.40103 | 2.351 |
| chr7 | 4204306 | 4204509 | 0.554888 | 1.30455 | 2.351 |
| chr8 | 74416724 | 74416988 | 1.28733 | 3.02652 | 2.351 |
| chr1 | 228581165 | 228581510 | 1.37081 | 3.2228 | 2.351 |
| chr19 | 8650592 | 8651128 | 1.37918 | 3.24254 | 2.351 |
| chr12 | 75954016 | 75954255 | 0.763526 | 1.79514 | 2.351 |
| chr1 | 110319524 | 110319773 | 1.45581 | 3.4228 | 2.351 |
| chr5 | 29115317 | 29115599 | 0.563872 | 1.32575 | 2.351 |
| chr20 | 35436929 | 35437385 | 2.47765 | 5.82565 | 2.351 |
| chr20 | 38533137 | 38533385 | 0.662027 | 1.55669 | 2.351 |
| chr5 | 32343033 | 32343293 | 0.400353 | 0.941389 | 2.351 |
| chr8 | 31497606 | 31498006 | 1.16431 | 2.73782 | 2.351 |
| chr10 | 62723925 | 62724208 | 1.34019 | 3.15143 | 2.351 |
| chr1 | 169703102 | 169703328 | 0.460013 | 1.08173 | 2.352 |
| chr20 | 15911026 | 15911290 | 1.05733 | 2.48634 | 2.352 |
| chr6 | 53599829 | 53600105 | 1.00389 | 2.3607 | 2.352 |
| chr7 | 85374584 | 85374813 | 1.05425 | 2.47914 | 2.352 |
| chr19 | 50646118 | 50646705 | 1.15107 | 2.70686 | 2.352 |
| chr11 | 65652064 | 65652580 | 2.30129 | 5.41183 | 2.352 |
| chr1 | 116710899 | 116711304 | 1.49661 | 3.51956 | 2.352 |
| chr12 | 52565875 | 52566243 | 1.2766 | 3.00219 | 2.352 |
| chr14 | 102983028 | 102983583 | 1.44239 | 3.39211 | 2.352 |
| chr3 | 149938985 | 149939406 | 2.47912 | 5.83027 | 2.352 |
| chr9 | 86153475 | 86153937 | 1.2868 | 3.0263 | 2.352 |
| chr10 | 19739707 | 19739958 | 0.360882 | 0.848761 | 2.352 |
| chr5 | 317257 | 317587 | 1.9677 | 4.6279 | 2.352 |
| chr1 | 182570431 | 182570830 | 1.34713 | 3.16861 | 2.352 |
| chr15 | 54100300 | 54100596 | 0.662491 | 1.55827 | 2.352 |
| chr5 | 118806599 | 118806846 | 1.07604 | 2.53109 | 2.352 |

| | | | | | |
|---|---|---|---|---|---|
| chr1 | 211000191 | 211000472 | 0.451347 | 1.06168 | 2.352 |
| chr8 | 130740868 | 130741151 | 0.985294 | 2.31771 | 2.352 |
| chr9 | 12884837 | 12885098 | 0.982201 | 2.31045 | 2.352 |
| chr20 | 8188329 | 8188579 | 0.425084 | 0.999987 | 2.352 |
| chr1 | 235677074 | 235677446 | 2.08841 | 4.91293 | 2.352 |
| chr12 | 10826664 | 10827159 | 2.31166 | 5.43812 | 2.352 |
| chr10 | 132774203 | 132774516 | 0.426571 | 1.0035 | 2.352 |
| chr13 | 26918590 | 26918805 | 0.374344 | 0.880641 | 2.352 |
| chr2 | 86745709 | 86745946 | 1.19444 | 2.80993 | 2.353 |
| chr19 | 54672168 | 54672519 | 2.72604 | 6.4134 | 2.353 |
| chr7 | 121376328 | 121376652 | 0.465602 | 1.09541 | 2.353 |
| chr17 | 32277367 | 32277786 | 0.46521 | 1.09454 | 2.353 |
| chr16 | 67962189 | 67962761 | 1.55201 | 3.65157 | 2.353 |
| chr20 | 43922327 | 43922739 | 1.76706 | 4.15777 | 2.353 |
| chr10 | 13820026 | 13820320 | 1.30613 | 3.0733 | 2.353 |
| chr5 | 112257880 | 112258327 | 1.86254 | 4.38282 | 2.353 |
| chr14 | 50234587 | 50234885 | 1.40905 | 3.31575 | 2.353 |
| chr4 | 57904177 | 57904516 | 2.35197 | 5.53463 | 2.353 |
| chr9 | 117776331 | 117776638 | 1.54601 | 3.63809 | 2.353 |
| chr8 | 27713025 | 27713326 | 1.89954 | 4.47006 | 2.353 |
| chr16 | 30649327 | 30649603 | 1.16293 | 2.7367 | 2.353 |
| chr17 | 77387777 | 77388255 | 2.11963 | 4.98814 | 2.353 |
| chr9 | 18090345 | 18090591 | 0.434694 | 1.023 | 2.353 |
| chr1 | 11705701 | 11706046 | 1.2865 | 3.02772 | 2.353 |
| chr14 | 104563396 | 104563598 | 0.746637 | 1.75718 | 2.353 |
| chr3 | 164027135 | 164027344 | 0.491267 | 1.1562 | 2.354 |
| chr6 | 143554100 | 143554310 | 0.832268 | 1.9588 | 2.354 |
| chr18 | 60990343 | 60990569 | 0.769385 | 1.81107 | 2.354 |
| chr11 | 31286348 | 31286555 | 0.475147 | 1.11861 | 2.354 |
| chr2 | 235314839 | 235315042 | 0.738678 | 1.73903 | 2.354 |
| chr12 | 125667658 | 125668095 | 2.63003 | 6.19177 | 2.354 |
| chr9 | 140189707 | 140190221 | 1.91558 | 4.50982 | 2.354 |
| chr8 | 127189525 | 127189768 | 0.49081 | 1.15556 | 2.354 |
| chr15 | 77835659 | 77836158 | 2.67982 | 6.30939 | 2.354 |
| chr12 | 95043696 | 95043953 | 1.07305 | 2.52652 | 2.355 |
| chr1 | 20543093 | 20543416 | 0.881934 | 2.07661 | 2.355 |
| chr6 | 52128533 | 52128886 | 1.58239 | 3.72603 | 2.355 |
| chr15 | 87286579 | 87286863 | 1.54624 | 3.64132 | 2.355 |
| chr7 | 16138231 | 16138599 | 2.01414 | 4.74334 | 2.355 |
| chr2 | 28378488 | 28378833 | 1.84872 | 4.3538 | 2.355 |
| chr3 | 27286059 | 27286270 | 0.499324 | 1.17596 | 2.355 |
| chr19 | 14153914 | 14154137 | 0.918243 | 2.16275 | 2.355 |
| chr21 | 35882916 | 35883192 | 1.19462 | 2.81378 | 2.355 |
| chr8 | 71657833 | 71658180 | 0.991489 | 2.33541 | 2.355 |
| chr3 | 9988810 | 9989313 | 1.12133 | 2.64138 | 2.356 |
| chr19 | 16497445 | 16497810 | 0.941025 | 2.21685 | 2.356 |
| chr20 | 55549418 | 55549807 | 1.30328 | 3.07056 | 2.356 |
| chr2 | 5506058 | 5506394 | 1.24437 | 2.93179 | 2.356 |
| chr8 | 98608041 | 98608408 | 2.1113 | 4.9743 | 2.356 |
| chr11 | 5145324 | 5145617 | 1.34466 | 3.16833 | 2.356 |
| chr14 | 48867935 | 48868243 | 1.46606 | 3.45447 | 2.356 |
| chr12 | 32479530 | 32479759 | 0.480805 | 1.13294 | 2.356 |
| chr5 | 150842981 | 150843323 | 1.15729 | 2.72697 | 2.356 |
| chr11 | 61938675 | 61939109 | 1.42278 | 3.35269 | 2.356 |
| chr1 | 23871260 | 23871517 | 1.22587 | 2.88873 | 2.356 |
| chr7 | 89712755 | 89712974 | 0.333031 | 0.784819 | 2.357 |
| chr3 | 13271063 | 13271410 | 0.909933 | 2.14451 | 2.357 |
| chr11 | 1248105 | 1248410 | 0.944324 | 2.22569 | 2.357 |
| chr19 | 1848653 | 1849011 | 1.44924 | 3.41575 | 2.357 |
| chr7 | 37940926 | 37941135 | 1.12566 | 2.65315 | 2.357 |
| chr12 | 23221448 | 23221770 | 0.49059 | 1.15631 | 2.357 |

| | | | | | |
|---|---|---|---|---|---|
| chr6 | 23204765 | 23204972 | 0.32502 | 0.766122 | 2.357 |
| chr19 | 18263410 | 18263617 | 0.867653 | 2.04527 | 2.357 |
| chr14 | 69378310 | 69378733 | 2.21184 | 5.21389 | 2.357 |
| chr19 | 1220456 | 1220880 | 0.623396 | 1.46953 | 2.357 |
| chr6 | 10465953 | 10466178 | 1.43892 | 3.39201 | 2.357 |
| chr5 | 12243273 | 12243618 | 0.38848 | 0.915781 | 2.357 |
| chr10 | 125727138 | 125727569 | 1.70247 | 4.01361 | 2.358 |
| chr5 | 1763795 | 1764083 | 1.35278 | 3.18925 | 2.358 |
| chr10 | 74820885 | 74821113 | 1.05501 | 2.48725 | 2.358 |
| chr12 | 10714982 | 10715238 | 0.984466 | 2.32098 | 2.358 |
| chr16 | 35043531 | 35043766 | 0.660374 | 1.55698 | 2.358 |
| chr19 | 47538985 | 47539398 | 1.42707 | 3.36479 | 2.358 |
| chr10 | 51533023 | 51533312 | 0.436101 | 1.02836 | 2.358 |
| chr3 | 187037457 | 187037856 | 1.57761 | 3.7202 | 2.358 |
| chr16 | 156974 | 157430 | 1.49209 | 3.51854 | 2.358 |
| chr15 | 56538234 | 56538638 | 4.35308 | 10.2655 | 2.358 |
| chr16 | 23072089 | 23072441 | 1.51666 | 3.577 | 2.358 |
| chr8 | 109900012 | 109900314 | 0.839539 | 1.98005 | 2.359 |
| chr4 | 1750354 | 1750693 | 2.03325 | 4.79544 | 2.359 |
| chr12 | 92801884 | 92802180 | 1.44318 | 3.40407 | 2.359 |
| chr18 | 3380531 | 3380836 | 0.439319 | 1.03625 | 2.359 |
| chr7 | 152728694 | 152728946 | 0.458943 | 1.08256 | 2.359 |
| chr6 | 18824022 | 18824287 | 0.71727 | 1.69191 | 2.359 |
| chr20 | 61846470 | 61846676 | 1.28166 | 3.02326 | 2.359 |
| chr1 | 248129318 | 248129638 | 0.576425 | 1.35972 | 2.359 |
| chr14 | 102529562 | 102529779 | 0.790402 | 1.86446 | 2.359 |
| chr20 | 31850429 | 31850787 | 1.96764 | 4.64162 | 2.359 |
| chr19 | 17393486 | 17393797 | 1.76168 | 4.15583 | 2.359 |
| chr12 | 54378685 | 54378926 | 3.13357 | 7.39233 | 2.359 |
| chr17 | 13498109 | 13498419 | 1.74654 | 4.12032 | 2.359 |
| chr11 | 76902977 | 76903535 | 2.292 | 5.40737 | 2.359 |
| chr9 | 99204195 | 99204463 | 1.09463 | 2.58251 | 2.359 |
| chr7 | 92219601 | 92219917 | 2.37784 | 5.60994 | 2.359 |
| chr22 | 31503675 | 31503996 | 1.81512 | 4.28236 | 2.359 |
| chr8 | 28602634 | 28602899 | 1.71203 | 4.03943 | 2.359 |
| chr17 | 3817003 | 3817501 | 1.87009 | 4.41239 | 2.359 |
| chr19 | 38826124 | 38826440 | 2.95813 | 6.97959 | 2.359 |
| chr14 | 95147683 | 95147960 | 1.86941 | 4.41081 | 2.359 |
| chr14 | 20528390 | 20528602 | 0.392568 | 0.926318 | 2.360 |
| chr17 | 48171670 | 48172000 | 1.80083 | 4.24943 | 2.360 |
| chr12 | 109005900 | 109006254 | 0.980472 | 2.31364 | 2.360 |
| chr15 | 78600722 | 78601118 | 2.16393 | 5.10639 | 2.360 |
| chr17 | 79827203 | 79827723 | 2.11682 | 4.99524 | 2.360 |
| chr21 | 47390440 | 47390980 | 1.73281 | 4.08928 | 2.360 |
| chr16 | 72586606 | 72586878 | 0.426707 | 1.00701 | 2.360 |
| chr9 | 95922164 | 95922668 | 2.44254 | 5.76441 | 2.360 |
| chr15 | 74665967 | 74666333 | 1.65601 | 3.90829 | 2.360 |
| chr1 | 46502683 | 46503142 | 1.52598 | 3.60149 | 2.360 |
| chr11 | 62676685 | 62677005 | 1.13353 | 2.67531 | 2.360 |
| chr3 | 50297431 | 50297719 | 2.1653 | 5.11055 | 2.360 |
| chr11 | 30939484 | 30939757 | 0.9879 | 2.33166 | 2.360 |
| chr15 | 31507541 | 31508020 | 1.80004 | 4.24853 | 2.360 |
| chr5 | 40958308 | 40958550 | 0.443594 | 1.04705 | 2.360 |
| chrX | 139844342 | 139844547 | 1.71093 | 4.03845 | 2.360 |
| chr15 | 81693059 | 81693317 | 1.31056 | 3.0937 | 2.361 |
| chr8 | 26231515 | 26231942 | 1.38268 | 3.26449 | 2.361 |
| chr9 | 96648372 | 96648808 | 0.467128 | 1.10289 | 2.361 |
| chr3 | 138553183 | 138553412 | 0.630479 | 1.48862 | 2.361 |
| chr13 | 28107164 | 28107646 | 2.00731 | 4.73964 | 2.361 |
| chr3 | 143806035 | 143806307 | 0.521473 | 1.23131 | 2.361 |
| chr14 | 65268565 | 65268861 | 1.4574 | 3.44125 | 2.361 |

| | | | | | |
|---|---|---|---|---|---|
| chr2 | 187506520 | 187506765 | 1.34096 | 3.16638 | 2.361 |
| chr14 | 103987433 | 103987783 | 1.2653 | 2.98787 | 2.361 |
| chr15 | 56633539 | 56633789 | 0.460133 | 1.08659 | 2.361 |
| chr8 | 51989771 | 51990083 | 0.412338 | 0.973729 | 2.361 |
| chr14 | 87174374 | 87174650 | 0.418214 | 0.987614 | 2.362 |
| chr2 | 31522568 | 31522792 | 0.405509 | 0.957654 | 2.362 |
| chr9 | 117669276 | 117669552 | 1.35996 | 3.2117 | 2.362 |
| chr10 | 91122167 | 91122546 | 0.459208 | 1.08455 | 2.362 |
| chr12 | 31432479 | 31432713 | 0.627659 | 1.4824 | 2.362 |
| chr7 | 125054097 | 125054317 | 0.467364 | 1.10384 | 2.362 |
| chrX | 28677435 | 28677782 | 0.800223 | 1.89002 | 2.362 |
| chr9 | 5442503 | 5442835 | 0.929027 | 2.19433 | 2.362 |
| chr12 | 26182127 | 26182524 | 1.2487 | 2.94942 | 2.362 |
| chr18 | 61630916 | 61631207 | 1.39372 | 3.29199 | 2.362 |
| chr20 | 49922953 | 49923698 | 1.07574 | 2.54096 | 2.362 |
| chr9 | 121591479 | 121591918 | 0.534094 | 1.26166 | 2.362 |
| chr21 | 45905046 | 45905315 | 1.16953 | 2.76311 | 2.363 |
| chr20 | 5537731 | 5538084 | 0.950329 | 2.24531 | 2.363 |
| chr4 | 140430998 | 140431287 | 0.494542 | 1.16853 | 2.363 |
| chr10 | 83598803 | 83599008 | 0.74704 | 1.76515 | 2.363 |
| chr2 | 232660259 | 232660549 | 1.12349 | 2.65483 | 2.363 |
| chr7 | 5569171 | 5569574 | 1.85617 | 4.3864 | 2.363 |
| chr17 | 59489683 | 59490153 | 2.62067 | 6.1934 | 2.363 |
| chr10 | 118556395 | 118556965 | 1.44795 | 3.42218 | 2.363 |
| chr3 | 112046272 | 112046477 | 0.804853 | 1.9024 | 2.364 |
| chr11 | 22152266 | 22152531 | 1.07832 | 2.54882 | 2.364 |
| chr16 | 57079753 | 57079994 | 1.02549 | 2.42399 | 2.364 |
| chr3 | 10234547 | 10234840 | 0.73933 | 1.74762 | 2.364 |
| chr3 | 58518463 | 58518808 | 1.66891 | 3.94509 | 2.364 |
| chr16 | 78133805 | 78134196 | 2.73183 | 6.45771 | 2.364 |
| chr8 | 123197950 | 123198194 | 0.774243 | 1.83028 | 2.364 |
| chr14 | 75762769 | 75763174 | 2.46032 | 5.81627 | 2.364 |
| chr8 | 74585258 | 74585538 | 1.12309 | 2.65503 | 2.364 |
| chrX | 153545832 | 153546091 | 1.07332 | 2.53741 | 2.364 |
| chr8 | 143154845 | 143155057 | 1.00031 | 2.36492 | 2.364 |
| chr22 | 45080540 | 45080960 | 1.57092 | 3.71397 | 2.364 |
| chr11 | 102180957 | 102181190 | 1.19049 | 2.81471 | 2.364 |
| chr1 | 55841297 | 55841651 | 1.29046 | 3.05117 | 2.364 |
| chr10 | 44882022 | 44882512 | 2.06125 | 4.87363 | 2.364 |
| chr1 | 61515576 | 61515958 | 1.91077 | 4.51799 | 2.364 |
| chr8 | 74217802 | 74218097 | 1.75117 | 4.14065 | 2.365 |
| chr20 | 55889988 | 55890228 | 0.626849 | 1.48219 | 2.365 |
| chr12 | 50763259 | 50763489 | 0.293695 | 0.694448 | 2.365 |
| chr8 | 27472100 | 27472509 | 1.20232 | 2.84304 | 2.365 |
| chr11 | 2177681 | 2178136 | 1.64611 | 3.89245 | 2.365 |
| chr3 | 153970806 | 153971222 | 1.5838 | 3.74511 | 2.365 |
| chr16 | 47177238 | 47177693 | 1.58255 | 3.74231 | 2.365 |
| chr14 | 100338116 | 100338367 | 0.570515 | 1.34922 | 2.365 |
| chr7 | 38262155 | 38262403 | 1.35473 | 3.20395 | 2.365 |
| chr12 | 68337141 | 68337482 | 1.11579 | 2.63893 | 2.365 |
| chr6 | 35704081 | 35704406 | 1.62378 | 3.84038 | 2.365 |
| chr22 | 46763504 | 46763871 | 2.08078 | 4.92157 | 2.365 |
| chr1 | 226175183 | 226175619 | 1.29636 | 3.06628 | 2.365 |
| chr11 | 33879415 | 33879822 | 1.74001 | 4.11587 | 2.365 |
| chr5 | 126899693 | 126900123 | 0.566763 | 1.34065 | 2.365 |
| chr20 | 13527191 | 13527504 | 1.67668 | 3.96628 | 2.366 |
| chr20 | 40040065 | 40040310 | 0.843183 | 1.99476 | 2.366 |
| chr8 | 31354835 | 31355145 | 1.42175 | 3.36362 | 2.366 |
| chr5 | 71651621 | 71651856 | 1.16331 | 2.75226 | 2.366 |
| chr6 | 126274268 | 126274493 | 1.013 | 2.3967 | 2.366 |
| chr1 | 167029750 | 167030028 | 0.631576 | 1.49428 | 2.366 |

| | | | | | |
|---|---|---|---|---|---|
| chr19 | 15264867 | 15265106 | 1.03225 | 2.44226 | 2.366 |
| chr12 | 5704217 | 5704611 | 0.486586 | 1.15128 | 2.366 |
| chr17 | 11369864 | 11370221 | 0.52123 | 1.23328 | 2.366 |
| chr2 | 206540736 | 206540939 | 0.952601 | 2.25397 | 2.366 |
| chr14 | 64100536 | 64100990 | 0.657997 | 1.55705 | 2.366 |
| chr18 | 74515715 | 74516040 | 1.69601 | 4.01343 | 2.366 |
| chr7 | 119805431 | 119805686 | 0.427788 | 1.01237 | 2.367 |
| chr10 | 124102192 | 124102469 | 1.31886 | 3.12118 | 2.367 |
| chr19 | 57610290 | 57610958 | 1.53501 | 3.63301 | 2.367 |
| chr12 | 121671101 | 121671354 | 0.519844 | 1.23038 | 2.367 |
| chr12 | 119763375 | 119763831 | 0.44912 | 1.063 | 2.367 |
| chr6 | 41286567 | 41286937 | 1.755 | 4.15422 | 2.367 |
| chr12 | 67254937 | 67255344 | 1.54078 | 3.64742 | 2.367 |
| chr7 | 29846978 | 29847338 | 0.885821 | 2.0971 | 2.367 |
| chrX | 22099601 | 22099993 | 1.67515 | 3.96584 | 2.367 |
| chr4 | 128400274 | 128400485 | 0.965528 | 2.28587 | 2.367 |
| chr4 | 138101591 | 138101857 | 0.650434 | 1.53993 | 2.368 |
| chr7 | 120327974 | 120328222 | 0.539028 | 1.27619 | 2.368 |
| chr5 | 25853456 | 25853758 | 0.529005 | 1.25248 | 2.368 |
| chr10 | 10095162 | 10095386 | 0.371458 | 0.879474 | 2.368 |
| chr6 | 37992162 | 37992445 | 1.03281 | 2.44537 | 2.368 |
| chr15 | 71266410 | 71266624 | 1.0066 | 2.38334 | 2.368 |
| chr6 | 27125367 | 27126005 | 0.471316 | 1.11599 | 2.368 |
| chr15 | 32350503 | 32350735 | 0.550463 | 1.30344 | 2.368 |
| chr8 | 3434443 | 3434730 | 0.467449 | 1.10689 | 2.368 |
| chr9 | 129467542 | 129467848 | 1.44549 | 3.42313 | 2.368 |
| chr14 | 100549137 | 100549349 | 0.374407 | 0.886689 | 2.368 |
| chr5 | 149016421 | 149016687 | 1.42296 | 3.36997 | 2.368 |
| chr12 | 38532499 | 38532857 | 2.09629 | 4.96508 | 2.369 |
| chr10 | 7798314 | 7798700 | 1.30755 | 3.09708 | 2.369 |
| chr1 | 230703796 | 230704017 | 0.731421 | 1.73252 | 2.369 |
| chr11 | 61299609 | 61300010 | 2.89932 | 6.86761 | 2.369 |
| chr20 | 6248440 | 6248659 | 0.433592 | 1.02706 | 2.369 |
| chr15 | 90863234 | 90863633 | 2.47066 | 5.85233 | 2.369 |
| chr12 | 127268922 | 127269185 | 1.64119 | 3.88755 | 2.369 |
| chr7 | 43994149 | 43994399 | 0.851913 | 2.01805 | 2.369 |
| chr12 | 112825178 | 112825445 | 1.78742 | 4.23443 | 2.369 |
| chr20 | 13768903 | 13769148 | 0.916348 | 2.17092 | 2.369 |
| chr19 | 2261737 | 2262101 | 1.27311 | 3.01616 | 2.369 |
| chr12 | 56040230 | 56040538 | 0.571296 | 1.35349 | 2.369 |
| chr19 | 42211482 | 42211749 | 0.918221 | 2.17552 | 2.369 |
| chr1 | 220976972 | 220977297 | 1.22474 | 2.90191 | 2.369 |
| chr4 | 6682910 | 6683282 | 1.76762 | 4.18831 | 2.369 |
| chr9 | 8217337 | 8217558 | 0.415974 | 0.985674 | 2.370 |
| chr3 | 13938020 | 13938410 | 1.41669 | 3.35702 | 2.370 |
| chr12 | 60138156 | 60138421 | 0.420326 | 0.996024 | 2.370 |
| chr5 | 7526054 | 7526286 | 0.496814 | 1.17733 | 2.370 |
| chr1 | 246655947 | 246656297 | 2.05463 | 4.86921 | 2.370 |
| chr14 | 96027212 | 96027658 | 2.144 | 5.08109 | 2.370 |
| chr7 | 49892317 | 49892666 | 0.932089 | 2.20912 | 2.370 |
| chr7 | 42019427 | 42019817 | 1.0532 | 2.49629 | 2.370 |
| chr5 | 159414381 | 159414715 | 1.96044 | 4.64679 | 2.370 |
| chr3 | 119539743 | 119540173 | 1.50032 | 3.5565 | 2.370 |
| chr19 | 41897142 | 41897433 | 0.580754 | 1.37672 | 2.371 |
| chr1 | 178063541 | 178063783 | 0.865525 | 2.05192 | 2.371 |
| chr5 | 2208129 | 2208465 | 1.19543 | 2.83406 | 2.371 |
| chr19 | 6126432 | 6126690 | 1.35028 | 3.2012 | 2.371 |
| chr5 | 68266633 | 68266871 | 0.438639 | 1.04004 | 2.371 |
| chr1 | 226251956 | 226252329 | 1.83708 | 4.35586 | 2.371 |
| chr8 | 144663768 | 144664003 | 0.466609 | 1.10645 | 2.371 |
| chr15 | 93352292 | 93352749 | 2.2083 | 5.23675 | 2.371 |

| | | | | | |
|---|---|---|---|---|---|
| chr10 | 75639355 | 75639786 | 1.82335 | 4.32391 | 2.371 |
| chr8 | 74961160 | 74961480 | 1.11524 | 2.64473 | 2.371 |
| chr2 | 198224638 | 198224891 | 0.791686 | 1.87745 | 2.371 |
| chr3 | 120067743 | 120068144 | 0.66494 | 1.57688 | 2.371 |
| chr15 | 50792620 | 50792879 | 0.466524 | 1.10635 | 2.371 |
| chr2 | 30358635 | 30359005 | 0.611936 | 1.45124 | 2.372 |
| chr11 | 3845117 | 3845335 | 0.45818 | 1.08665 | 2.372 |
| chr13 | 99305715 | 99305986 | 1.09233 | 2.59068 | 2.372 |
| chr10 | 72108270 | 72108508 | 1.18004 | 2.79872 | 2.372 |
| chr17 | 79977825 | 79978441 | 2.2966 | 5.44694 | 2.372 |
| chr4 | 110327585 | 110327867 | 0.958156 | 2.2725 | 2.372 |
| chr6 | 157756023 | 157756247 | 0.535634 | 1.27039 | 2.372 |
| chr10 | 15675462 | 15675675 | 0.414406 | 0.982884 | 2.372 |
| chr15 | 41896070 | 41896412 | 1.42338 | 3.376 | 2.372 |
| chr11 | 117052006 | 117052382 | 1.18172 | 2.80287 | 2.372 |
| chr14 | 61747221 | 61747853 | 0.965613 | 2.29036 | 2.372 |
| chr3 | 119610706 | 119611106 | 1.71493 | 4.06781 | 2.372 |
| chr15 | 59539647 | 59539912 | 1.30728 | 3.10094 | 2.372 |
| chr15 | 91499750 | 91500200 | 3.23402 | 7.67125 | 2.372 |
| chr22 | 24236170 | 24236410 | 1.35965 | 3.22523 | 2.372 |
| chr3 | 70867272 | 70867624 | 0.40452 | 0.959597 | 2.372 |
| chr9 | 139148056 | 139148276 | 0.770084 | 1.82681 | 2.372 |
| chr8 | 97696622 | 97696885 | 0.558373 | 1.32468 | 2.372 |
| chr4 | 23137508 | 23137720 | 1.03123 | 2.44659 | 2.373 |
| chr11 | 3845894 | 3846406 | 1.62944 | 3.8659 | 2.373 |
| chr19 | 31952488 | 31952826 | 0.539391 | 1.27978 | 2.373 |
| chr12 | 54224355 | 54224660 | 0.549739 | 1.3044 | 2.373 |
| chr2 | 71775817 | 71776116 | 1.56874 | 3.72225 | 2.373 |
| chr19 | 982719 | 983064 | 1.32392 | 3.14148 | 2.373 |
| chr17 | 38959792 | 38960079 | 1.45793 | 3.45988 | 2.373 |
| chr20 | 30144990 | 30145608 | 1.22611 | 2.90974 | 2.373 |
| chr19 | 42017785 | 42018078 | 0.458566 | 1.08825 | 2.373 |
| chr8 | 112248619 | 112248846 | 0.925775 | 2.19701 | 2.373 |
| chr5 | 133279960 | 133280330 | 1.14618 | 2.72015 | 2.373 |
| chr10 | 57771992 | 57772376 | 0.567927 | 1.34783 | 2.373 |
| chr10 | 102021824 | 102022055 | 0.403249 | 0.95701 | 2.373 |
| chr5 | 150427344 | 150427787 | 1.73952 | 4.12835 | 2.373 |
| chr19 | 54249585 | 54249907 | 2.3937 | 5.68094 | 2.373 |
| chr8 | 127407303 | 127407517 | 0.747531 | 1.77412 | 2.373 |
| chr3 | 25635526 | 25635828 | 0.783262 | 1.859 | 2.373 |
| chr10 | 90149265 | 90149527 | 0.886875 | 2.10504 | 2.374 |
| chr11 | 12055177 | 12055424 | 0.529739 | 1.25736 | 2.374 |
| chr6 | 156885530 | 156885867 | 1.98968 | 4.72314 | 2.374 |
| chr11 | 839968 | 840179 | 0.862914 | 2.0486 | 2.374 |
| chr5 | 73423820 | 73424043 | 0.749096 | 1.77841 | 2.374 |
| chr10 | 126228716 | 126229120 | 1.29408 | 3.0723 | 2.374 |
| chrX | 40507188 | 40507407 | 0.517416 | 1.22841 | 2.374 |
| chr2 | 236504150 | 236504392 | 1.31522 | 3.12255 | 2.374 |
| chr7 | 54564908 | 54565193 | 0.480132 | 1.13994 | 2.374 |
| chr12 | 57657584 | 57657921 | 1.27662 | 3.0311 | 2.374 |
| chr11 | 74032501 | 74033027 | 1.23508 | 2.93251 | 2.374 |
| chr21 | 40890256 | 40890520 | 1.18235 | 2.80738 | 2.374 |
| chr15 | 30218011 | 30218398 | 2.15871 | 5.12572 | 2.374 |
| chr11 | 87132373 | 87132729 | 1.07367 | 2.54969 | 2.375 |
| chr11 | 35413835 | 35414062 | 1.30137 | 3.09054 | 2.375 |
| chrX | 149533634 | 149534039 | 1.20924 | 2.87181 | 2.375 |
| chr17 | 6325296 | 6325687 | 2.25274 | 5.3503 | 2.375 |
| chr12 | 132559981 | 132560238 | 0.971975 | 2.30859 | 2.375 |
| chr13 | 92001205 | 92001485 | 1.76096 | 4.18278 | 2.375 |
| chr11 | 85375147 | 85375433 | 1.2901 | 3.06437 | 2.375 |
| chr6 | 151335476 | 151335826 | 1.67539 | 3.97957 | 2.375 |

| | | | | | |
|---|---|---|---|---|---|
| chr16 | 451928 | 452181 | 0.702246 | 1.66816 | 2.375 |
| chr15 | 73590681 | 73590943 | 0.517646 | 1.22968 | 2.376 |
| chr16 | 34666383 | 34666813 | 1.17733 | 2.79692 | 2.376 |
| chr20 | 21846637 | 21846882 | 0.463705 | 1.10165 | 2.376 |
| chr7 | 118694658 | 118694877 | 0.464681 | 1.10397 | 2.376 |
| chr19 | 33856713 | 33856959 | 0.673 | 1.59889 | 2.376 |
| chr14 | 104213236 | 104213450 | 0.367411 | 0.872935 | 2.376 |
| chr19 | 8764031 | 8764378 | 2.3217 | 5.51664 | 2.376 |
| chr1 | 174002948 | 174003168 | 0.450681 | 1.07089 | 2.376 |
| chr8 | 131823123 | 131823361 | 1.34265 | 3.19038 | 2.376 |
| chr4 | 11430463 | 11430819 | 1.34097 | 3.18661 | 2.376 |
| chr5 | 83171196 | 83171459 | 0.523186 | 1.24335 | 2.377 |
| chr2 | 27868675 | 27868917 | 0.472626 | 1.1232 | 2.377 |
| chr21 | 47573169 | 47573562 | 1.48787 | 3.53599 | 2.377 |
| chr5 | 22429357 | 22429666 | 0.407444 | 0.96839 | 2.377 |
| chr10 | 71267592 | 71267928 | 2.51913 | 5.9874 | 2.377 |
| chr11 | 61447578 | 61447938 | 2.24943 | 5.34655 | 2.377 |
| chr17 | 49929212 | 49929611 | 0.369737 | 0.878854 | 2.377 |
| chr10 | 88146981 | 88147410 | 0.765941 | 1.8208 | 2.377 |
| chr10 | 98135070 | 98135495 | 1.49497 | 3.55387 | 2.377 |
| chr17 | 15406131 | 15406433 | 1.96645 | 4.67469 | 2.377 |
| chr13 | 103166626 | 103166916 | 0.414581 | 0.985585 | 2.377 |
| chr10 | 56336788 | 56337201 | 0.422243 | 1.00383 | 2.377 |
| chr4 | 186953506 | 186953724 | 0.382282 | 0.908869 | 2.377 |
| chr11 | 58347098 | 58347336 | 1.52607 | 3.62829 | 2.378 |
| chr5 | 57773677 | 57773937 | 0.837842 | 1.992 | 2.378 |
| chr5 | 173744275 | 173744691 | 1.72967 | 4.11236 | 2.378 |
| chr1 | 111765526 | 111765840 | 1.61286 | 3.83468 | 2.378 |
| chr1 | 116694615 | 116695020 | 1.32505 | 3.1504 | 2.378 |
| chr12 | 48255320 | 48255560 | 0.58901 | 1.40042 | 2.378 |
| chr6 | 11094022 | 11094548 | 0.969569 | 2.30529 | 2.378 |
| chr15 | 38364785 | 38365136 | 0.494454 | 1.17564 | 2.378 |
| chr20 | 37063857 | 37064297 | 2.51109 | 5.97061 | 2.378 |
| chr5 | 173754292 | 173754684 | 1.79553 | 4.26928 | 2.378 |
| chr17 | 77813522 | 77813929 | 1.88242 | 4.47595 | 2.378 |
| chr6 | 89490564 | 89490835 | 1.56342 | 3.71745 | 2.378 |
| chr10 | 79401648 | 79401870 | 1.22227 | 2.90653 | 2.378 |
| chr1 | 246893497 | 246893795 | 0.389543 | 0.926353 | 2.378 |
| chr15 | 60632786 | 60633103 | 0.663358 | 1.5775 | 2.378 |
| chr13 | 33213738 | 33214022 | 1.0658 | 2.53455 | 2.378 |
| chr9 | 14982451 | 14982757 | 0.354983 | 0.844185 | 2.378 |
| chr19 | 44227831 | 44228043 | 0.944289 | 2.24564 | 2.378 |
| chr7 | 109770710 | 109770984 | 0.699488 | 1.66349 | 2.378 |
| chr11 | 21535923 | 21536188 | 1.20684 | 2.87008 | 2.378 |
| chr14 | 105734123 | 105734483 | 0.745591 | 1.77315 | 2.378 |
| chr3 | 9882269 | 9882744 | 0.467052 | 1.11082 | 2.378 |
| chr9 | 136580878 | 136581238 | 1.01721 | 2.41931 | 2.378 |
| chr19 | 43934209 | 43934714 | 0.56175 | 1.33615 | 2.379 |
| chr12 | 14115716 | 14115967 | 1.05465 | 2.50863 | 2.379 |
| chr10 | 62997444 | 62997753 | 0.557768 | 1.32674 | 2.379 |
| chr3 | 184495500 | 184495866 | 0.939568 | 2.23496 | 2.379 |
| chr2 | 134875914 | 134876129 | 1.22098 | 2.90469 | 2.379 |
| chr4 | 15807304 | 15807560 | 1.28747 | 3.06314 | 2.379 |
| chr5 | 167513410 | 167513630 | 0.961794 | 2.28837 | 2.379 |
| chr17 | 48156905 | 48157464 | 3.20222 | 7.61897 | 2.379 |
| chr17 | 34093975 | 34094297 | 1.22078 | 2.90466 | 2.379 |
| chr13 | 83054590 | 83054814 | 0.500336 | 1.19049 | 2.379 |
| chr2 | 80315158 | 80315455 | 0.392962 | 0.935088 | 2.380 |
| chr11 | 57247827 | 57248119 | 1.88172 | 4.47795 | 2.380 |
| chr5 | 58238668 | 58238887 | 0.51962 | 1.23655 | 2.380 |
| chr5 | 53525062 | 53525343 | 2.01321 | 4.791 | 2.380 |

| | | | | | |
|---|---|---|---|---|---|
| chr5 | 162279124 | 162279359 | 0.531015 | 1.26373 | 2.380 |
| chr9 | 126306333 | 126306636 | 1.49799 | 3.56504 | 2.380 |
| chr12 | 29883264 | 29883501 | 0.384104 | 0.91413 | 2.380 |
| chr20 | 55636338 | 55636640 | 0.758523 | 1.80526 | 2.380 |
| chr10 | 79789273 | 79789811 | 1.43327 | 3.41143 | 2.380 |
| chr3 | 16288798 | 16289206 | 2.02753 | 4.82596 | 2.380 |
| chr12 | 50360689 | 50361670 | 1.87208 | 4.45618 | 2.380 |
| chr12 | 27006766 | 27007045 | 1.6063 | 3.8239 | 2.381 |
| chr6 | 113410291 | 113410553 | 0.569778 | 1.3564 | 2.381 |
| chr2 | 24387211 | 24387591 | 1.14928 | 2.73602 | 2.381 |
| chr1 | 205497885 | 205498479 | 1.99283 | 4.74436 | 2.381 |
| chr2 | 28976292 | 28976540 | 1.19767 | 2.85139 | 2.381 |
| chr19 | 46181354 | 46181558 | 0.548665 | 1.30635 | 2.381 |
| chr11 | 111472704 | 111473014 | 1.98459 | 4.72557 | 2.381 |
| chr4 | 129301073 | 129301354 | 1.09816 | 2.61493 | 2.381 |
| chr19 | 47257042 | 47257372 | 1.16916 | 2.78432 | 2.381 |
| chr17 | 16404271 | 16404500 | 0.355002 | 0.845429 | 2.381 |
| chr19 | 38747001 | 38747521 | 2.62868 | 6.26034 | 2.382 |
| chr9 | 80512302 | 80512577 | 1.00535 | 2.39448 | 2.382 |
| chr14 | 104079930 | 104080236 | 0.860757 | 2.05011 | 2.382 |
| chr16 | 28081952 | 28082522 | 1.17279 | 2.7933 | 2.382 |
| chr7 | 133029091 | 133029328 | 0.695867 | 1.65746 | 2.382 |
| chr2 | 72972986 | 72973210 | 0.980307 | 2.33501 | 2.382 |
| chr3 | 42141336 | 42141643 | 1.98977 | 4.73977 | 2.382 |
| chr5 | 24110194 | 24110433 | 0.799135 | 1.90364 | 2.382 |
| chr2 | 102035576 | 102035801 | 1.44509 | 3.44243 | 2.382 |
| chr9 | 112231938 | 112232289 | 1.86849 | 4.45122 | 2.382 |
| chr2 | 100997597 | 100997954 | 1.22745 | 2.92415 | 2.382 |
| chrX | 153029361 | 153029712 | 2.26943 | 5.40742 | 2.383 |
| chr1 | 69543506 | 69543723 | 0.352246 | 0.839343 | 2.383 |
| chr16 | 4338328 | 4338936 | 2.18302 | 5.20194 | 2.383 |
| chr9 | 140567305 | 140567558 | 1.27545 | 3.03942 | 2.383 |
| chr15 | 83994971 | 83995206 | 0.679467 | 1.61922 | 2.383 |
| chr6 | 135770313 | 135770527 | 0.420139 | 1.00123 | 2.383 |
| chr10 | 48520422 | 48520631 | 0.367561 | 0.875934 | 2.383 |
| chr20 | 20923905 | 20924211 | 0.525585 | 1.25253 | 2.383 |
| chr5 | 160793637 | 160793929 | 0.571014 | 1.36082 | 2.383 |
| chr12 | 19796604 | 19796896 | 1.46891 | 3.501 | 2.383 |
| chr12 | 57846646 | 57847073 | 1.9497 | 4.64707 | 2.383 |
| chr16 | 28985327 | 28985534 | 1.45283 | 3.46281 | 2.383 |
| chr2 | 61863012 | 61863295 | 0.852223 | 2.03129 | 2.384 |
| chr8 | 20238356 | 20238630 | 1.48358 | 3.53615 | 2.384 |
| chr17 | 72195727 | 72196165 | 2.97144 | 7.08321 | 2.384 |
| chr20 | 749401 | 749682 | 1.37007 | 3.266 | 2.384 |
| chr1 | 116681729 | 116682091 | 1.30735 | 3.11661 | 2.384 |
| chr19 | 3163249 | 3163655 | 1.62948 | 3.88479 | 2.384 |
| chr20 | 47823711 | 47823978 | 0.942495 | 2.24698 | 2.384 |
| chr5 | 7392472 | 7392825 | 1.21332 | 2.89279 | 2.384 |
| chr9 | 136036347 | 136036714 | 1.36263 | 3.24893 | 2.384 |
| chr7 | 55322500 | 55322973 | 1.83115 | 4.36624 | 2.384 |
| chr2 | 61062873 | 61063130 | 0.517155 | 1.23316 | 2.385 |
| chr3 | 137665258 | 137665499 | 1.0881 | 2.59462 | 2.385 |
| chr3 | 17586103 | 17586392 | 0.648225 | 1.54573 | 2.385 |
| chr12 | 15175217 | 15175433 | 0.680639 | 1.62303 | 2.385 |
| chr17 | 73487394 | 73487943 | 0.373117 | 0.889728 | 2.385 |
| chr3 | 150439151 | 150439493 | 1.45974 | 3.48088 | 2.385 |
| chr8 | 8528111 | 8528389 | 1.62885 | 3.88416 | 2.385 |
| chr19 | 46302258 | 46302656 | 1.34325 | 3.20331 | 2.385 |
| chr16 | 48265828 | 48266073 | 1.16749 | 2.78422 | 2.385 |
| chr9 | 21466010 | 21466227 | 0.979346 | 2.33553 | 2.385 |
| chr14 | 65462900 | 65463143 | 0.471556 | 1.12458 | 2.385 |

| | | | | | |
|---|---|---|---|---|---|
| chr10 | 127849773 | 127850103 | 0.721961 | 1.72179 | 2.385 |
| chr8 | 16511021 | 16511243 | 0.507772 | 1.21102 | 2.385 |
| chr8 | 23020861 | 23021234 | 1.40984 | 3.36245 | 2.385 |
| chr19 | 56683214 | 56683643 | 1.68329 | 4.0148 | 2.385 |
| chr3 | 79369505 | 79369729 | 0.679438 | 1.62054 | 2.385 |
| chr7 | 91017451 | 91017727 | 0.351243 | 0.837761 | 2.385 |
| chrX | 152665492 | 152665975 | 2.5032 | 5.97062 | 2.385 |
| chr8 | 140988893 | 140989149 | 0.370494 | 0.883715 | 2.385 |
| chr1 | 57050643 | 57050904 | 0.927999 | 2.21358 | 2.385 |
| chr10 | 104375138 | 104375403 | 0.818729 | 1.95297 | 2.385 |
| chr6 | 150186126 | 150186375 | 1.13382 | 2.70459 | 2.385 |
| chr8 | 97072486 | 97072694 | 0.543106 | 1.29552 | 2.385 |
| chr8 | 62637362 | 62637624 | 0.966455 | 2.30553 | 2.386 |
| chr19 | 10652125 | 10652527 | 1.20376 | 2.87171 | 2.386 |
| chr1 | 33551981 | 33552257 | 1.72336 | 4.11134 | 2.386 |
| chr5 | 122657724 | 122657926 | 0.708113 | 1.68938 | 2.386 |
| chrX | 120442444 | 120442674 | 1.28761 | 3.07217 | 2.386 |
| chr1 | 225470000 | 225470276 | 0.403232 | 0.96211 | 2.386 |
| chr17 | 34838773 | 34839202 | 2.02428 | 4.83007 | 2.386 |
| chr16 | 46883138 | 46883462 | 0.369977 | 0.882829 | 2.386 |
| chr4 | 38895263 | 38895566 | 1.00216 | 2.39136 | 2.386 |
| chr14 | 98183147 | 98183455 | 0.483038 | 1.15269 | 2.386 |
| chr12 | 13355945 | 13356217 | 1.15989 | 2.76792 | 2.386 |
| chr16 | 57634300 | 57634666 | 1.81046 | 4.32064 | 2.386 |
| chr22 | 23190484 | 23190952 | 2.05024 | 4.89298 | 2.387 |
| chr8 | 127234136 | 127234339 | 1.07692 | 2.57011 | 2.387 |
| chr11 | 45676789 | 45677237 | 1.65604 | 3.95228 | 2.387 |
| chr11 | 66072395 | 66072611 | 0.524127 | 1.2509 | 2.387 |
| chr21 | 15594016 | 15594311 | 0.407078 | 0.971566 | 2.387 |
| chr20 | 3799057 | 3799373 | 1.26966 | 3.03029 | 2.387 |
| chr11 | 83487539 | 83487792 | 0.458788 | 1.09503 | 2.387 |
| chr8 | 137187544 | 137187848 | 0.624777 | 1.49124 | 2.387 |
| chr5 | 35753821 | 35754055 | 0.504447 | 1.20404 | 2.387 |
| chr6 | 118939964 | 118940266 | 1.58258 | 3.77746 | 2.387 |
| chrX | 154365850 | 154366069 | 1.15212 | 2.75002 | 2.387 |
| chr6 | 108813390 | 108813754 | 1.03913 | 2.48039 | 2.387 |
| chrX | 71238434 | 71239013 | 1.70338 | 4.06617 | 2.387 |
| chr16 | 1982313 | 1982775 | 1.05369 | 2.51532 | 2.387 |
| chrX | 12964894 | 12965246 | 2.65569 | 6.33963 | 2.387 |
| chr1 | 93324482 | 93324815 | 1.24035 | 2.96106 | 2.387 |
| chr4 | 40910489 | 40911051 | 1.14283 | 2.72842 | 2.387 |
| chr5 | 19460697 | 19460916 | 0.5133 | 1.22551 | 2.388 |
| chr11 | 64611747 | 64612110 | 1.31612 | 3.14266 | 2.388 |
| chr15 | 34089432 | 34089659 | 0.502706 | 1.20039 | 2.388 |
| chr19 | 5369936 | 5370208 | 1.245 | 2.97289 | 2.388 |
| chr9 | 83098992 | 83099213 | 0.494074 | 1.17981 | 2.388 |
| chr8 | 98881533 | 98882050 | 1.998 | 4.77122 | 2.388 |
| chr2 | 63416060 | 63416459 | 1.53065 | 3.65529 | 2.388 |
| chr16 | 84655807 | 84656065 | 1.5794 | 3.77177 | 2.388 |
| chr12 | 58570125 | 58570347 | 1.38165 | 3.2998 | 2.388 |
| chr5 | 153471462 | 153471706 | 0.492138 | 1.17539 | 2.388 |
| chr2 | 62138675 | 62139003 | 0.737933 | 1.76254 | 2.388 |
| chr17 | 41837710 | 41838149 | 2.73836 | 6.54066 | 2.389 |
| chr8 | 121951339 | 121951657 | 1.14628 | 2.73793 | 2.389 |
| chr12 | 12529644 | 12529848 | 0.933482 | 2.22967 | 2.389 |
| chr5 | 171285298 | 171285651 | 1.50486 | 3.59447 | 2.389 |
| chr17 | 57525507 | 57525841 | 0.36616 | 0.87466 | 2.389 |
| chr12 | 12186290 | 12186573 | 1.43144 | 3.41947 | 2.389 |
| chr1 | 222340419 | 222340710 | 1.18941 | 2.84135 | 2.389 |
| chr7 | 36961353 | 36961571 | 0.527778 | 1.26083 | 2.389 |
| chr12 | 126168385 | 126168633 | 0.598724 | 1.43032 | 2.389 |

| | | | | | |
|---|---|---|---|---|---|
| chr19 | 42101550 | 42102174 | 1.31765 | 3.14783 | 2.389 |
| chr22 | 32366644 | 32366970 | 1.86469 | 4.45518 | 2.389 |
| chr15 | 83696155 | 83696487 | 0.560995 | 1.3404 | 2.389 |
| chr4 | 142575254 | 142575483 | 1.01138 | 2.41659 | 2.389 |
| chr7 | 92238063 | 92238310 | 0.610977 | 1.45995 | 2.390 |
| chr3 | 14567786 | 14567990 | 0.399093 | 0.953663 | 2.390 |
| chr1 | 15455033 | 15455392 | 1.58756 | 3.79364 | 2.390 |
| chr1 | 212650485 | 212650838 | 1.75605 | 4.19627 | 2.390 |
| chr4 | 141805648 | 141805907 | 1.45288 | 3.47183 | 2.390 |
| chr5 | 87155378 | 87155639 | 0.478183 | 1.14271 | 2.390 |
| chrX | 122967250 | 122967511 | 1.12974 | 2.69992 | 2.390 |
| chr3 | 38841831 | 38842099 | 0.494065 | 1.18075 | 2.390 |
| chr12 | 78058694 | 78058917 | 0.482395 | 1.15289 | 2.390 |
| chr5 | 151204965 | 151205367 | 1.21405 | 2.90153 | 2.390 |
| chr11 | 2449283 | 2449798 | 1.55759 | 3.72269 | 2.390 |
| chr7 | 139475666 | 139475921 | 1.06267 | 2.53983 | 2.390 |
| chr20 | 57414509 | 57414834 | 1.22756 | 2.93408 | 2.390 |
| chr5 | 14142993 | 14143263 | 1.03929 | 2.48413 | 2.390 |
| chr17 | 10777214 | 10777421 | 0.496722 | 1.18738 | 2.390 |
| chr1 | 6483441 | 6483854 | 1.54247 | 3.68728 | 2.391 |
| chr10 | 103769601 | 103769939 | 2.13115 | 5.09452 | 2.391 |
| chr22 | 48972546 | 48972972 | 1.63003 | 3.89675 | 2.391 |
| chr12 | 116082094 | 116082408 | 0.606574 | 1.4502 | 2.391 |
| chr3 | 124487725 | 124488050 | 1.01893 | 2.4362 | 2.391 |
| chr8 | 62188597 | 62188831 | 0.501607 | 1.19932 | 2.391 |
| chr9 | 104061394 | 104061644 | 0.544715 | 1.30248 | 2.391 |
| chr3 | 42791528 | 42791745 | 1.10256 | 2.63645 | 2.391 |
| chr12 | 122820603 | 122820906 | 2.09144 | 5.00116 | 2.391 |
| chr16 | 28991774 | 28992015 | 0.849296 | 2.03093 | 2.391 |
| chr9 | 34437416 | 34437684 | 0.977285 | 2.33699 | 2.391 |
| chr10 | 78532250 | 78532464 | 0.482605 | 1.15415 | 2.392 |
| chr4 | 188337913 | 188338160 | 0.661553 | 1.58223 | 2.392 |
| chr4 | 152065927 | 152066155 | 0.412415 | 0.986372 | 2.392 |
| chr1 | 151665567 | 151665872 | 1.22997 | 2.94187 | 2.392 |
| chr9 | 131410612 | 131410833 | 1.00338 | 2.4001 | 2.392 |
| chr9 | 133336585 | 133337148 | 1.35669 | 3.24549 | 2.392 |
| chr11 | 60598455 | 60598904 | 2.01904 | 4.82998 | 2.392 |
| chr14 | 68551107 | 68551363 | 1.18344 | 2.83106 | 2.392 |
| chr2 | 66547020 | 66547227 | 0.369966 | 0.885055 | 2.392 |
| chr19 | 47852260 | 47852569 | 2.79714 | 6.69158 | 2.392 |
| chr1 | 90228437 | 90228906 | 1.62024 | 3.87617 | 2.392 |
| chr19 | 46160558 | 46161009 | 0.375909 | 0.899366 | 2.393 |
| chr17 | 4692975 | 4693740 | 1.19312 | 2.85462 | 2.393 |
| chr7 | 1595925 | 1596242 | 1.91308 | 4.5774 | 2.393 |
| chr10 | 6458825 | 6459218 | 1.02898 | 2.46204 | 2.393 |
| chr11 | 2019497 | 2020435 | 1.00672 | 2.40887 | 2.393 |
| chr14 | 56198681 | 56198953 | 1.22846 | 2.93949 | 2.393 |
| chr7 | 42054245 | 42054575 | 1.67834 | 4.016 | 2.393 |
| chr16 | 19918773 | 19919420 | 1.55096 | 3.71173 | 2.393 |
| chr17 | 42988438 | 42988768 | 1.29019 | 3.08773 | 2.393 |
| chrX | 106903014 | 106903279 | 0.969695 | 2.32073 | 2.393 |
| chr17 | 47625833 | 47626359 | 1.46219 | 3.49941 | 2.393 |
| chr10 | 9347050 | 9347328 | 0.877571 | 2.10029 | 2.393 |
| chr7 | 44283903 | 44284212 | 1.00966 | 2.41646 | 2.393 |
| chr20 | 3749280 | 3749563 | 1.30559 | 3.1249 | 2.393 |
| chr12 | 119687234 | 119687490 | 0.409436 | 0.979997 | 2.394 |
| chr8 | 6224848 | 6225064 | 0.375572 | 0.898943 | 2.394 |
| chr5 | 105318223 | 105318476 | 0.441783 | 1.05745 | 2.394 |
| chr7 | 77649955 | 77650218 | 0.757551 | 1.81329 | 2.394 |
| chr4 | 57702817 | 57703090 | 1.20248 | 2.87834 | 2.394 |
| chr15 | 38287943 | 38288228 | 0.918898 | 2.19968 | 2.394 |

| | | | | | |
|---|---|---|---|---|---|
| chr17 | 80111729 | 80112159 | 1.86589 | 4.46681 | 2.394 |
| chr12 | 53606908 | 53607471 | 2.08739 | 4.99711 | 2.394 |
| chr10 | 98590338 | 98590625 | 1.13634 | 2.72043 | 2.394 |
| chr1 | 227625451 | 227625736 | 0.789631 | 1.89042 | 2.394 |
| chr14 | 38343904 | 38344196 | 1.40161 | 3.35553 | 2.394 |
| chr5 | 10724878 | 10725137 | 1.26418 | 3.02652 | 2.394 |
| chr3 | 155266052 | 155266278 | 0.455643 | 1.09086 | 2.394 |
| chr5 | 86402251 | 86402509 | 0.414278 | 0.991839 | 2.394 |
| chr22 | 45969486 | 45969739 | 1.05346 | 2.52237 | 2.394 |
| chr15 | 88617745 | 88617982 | 0.486769 | 1.16553 | 2.394 |
| chr22 | 45492175 | 45492460 | 1.63873 | 3.92394 | 2.395 |
| chr9 | 15642405 | 15642667 | 0.892616 | 2.13749 | 2.395 |
| chrY | 15548485 | 15548710 | 1.25383 | 3.00261 | 2.395 |
| chr1 | 223377890 | 223378149 | 0.407219 | 0.975238 | 2.395 |
| chr8 | 83219811 | 83220115 | 1.62444 | 3.89037 | 2.395 |
| chr6 | 150739348 | 150739606 | 1.17888 | 2.82339 | 2.395 |
| chr10 | 99829974 | 99830393 | 0.353412 | 0.846452 | 2.395 |
| chr5 | 36696167 | 36696462 | 1.18761 | 2.84444 | 2.395 |
| chr5 | 50254758 | 50254966 | 0.689691 | 1.65194 | 2.395 |
| chr15 | 64445701 | 64446176 | 2.74841 | 6.58309 | 2.395 |
| chr19 | 42954613 | 42955021 | 1.81392 | 4.34478 | 2.395 |
| chr20 | 51017600 | 51017865 | 0.561419 | 1.3448 | 2.395 |
| chr1 | 8469242 | 8469573 | 0.586113 | 1.40398 | 2.395 |
| chr5 | 63345777 | 63345988 | 0.416517 | 0.997728 | 2.395 |
| chr17 | 80724957 | 80725564 | 1.21164 | 2.90239 | 2.395 |
| chr5 | 133747430 | 133747866 | 1.41505 | 3.38982 | 2.396 |
| chr12 | 55074260 | 55074479 | 0.862257 | 2.06566 | 2.396 |
| chr14 | 71422697 | 71422909 | 1.01925 | 2.44176 | 2.396 |
| chr13 | 31248283 | 31248633 | 1.45364 | 3.48256 | 2.396 |
| chr20 | 4936882 | 4937216 | 1.33201 | 3.19131 | 2.396 |
| chr6 | 161814919 | 161815136 | 0.967036 | 2.31694 | 2.396 |
| chr20 | 44220996 | 44221576 | 1.59076 | 3.81141 | 2.396 |
| chr19 | 35503800 | 35504241 | 2.45022 | 5.87116 | 2.396 |
| chr1 | 33358437 | 33359169 | 1.58479 | 3.79766 | 2.396 |
| chr1 | 46689130 | 46689587 | 1.13712 | 2.72495 | 2.396 |
| chr12 | 103342583 | 103343077 | 1.36904 | 3.28073 | 2.396 |
| chr7 | 156280781 | 156280983 | 0.560011 | 1.34202 | 2.396 |
| chr19 | 11545655 | 11546044 | 1.09719 | 2.62943 | 2.397 |
| chr18 | 68449148 | 68449426 | 1.32937 | 3.18589 | 2.397 |
| chr10 | 99185766 | 99186012 | 1.86882 | 4.47885 | 2.397 |
| chr4 | 150902097 | 150902308 | 0.829141 | 1.98714 | 2.397 |
| chr5 | 149208005 | 149208319 | 1.61669 | 3.87508 | 2.397 |
| chr17 | 47937625 | 47937995 | 0.833165 | 1.99705 | 2.397 |
| chr1 | 227730134 | 227730625 | 1.79704 | 4.30742 | 2.397 |
| chr1 | 240855374 | 240855752 | 1.59971 | 3.83442 | 2.397 |
| chr16 | 58370586 | 58370892 | 2.17396 | 5.21102 | 2.397 |
| chr5 | 33298020 | 33298277 | 1.00642 | 2.41242 | 2.397 |
| chr2 | 9922674 | 9922877 | 0.971902 | 2.32973 | 2.397 |
| chr12 | 76741736 | 76742163 | 1.17682 | 2.82098 | 2.397 |
| chrX | 46241107 | 46241434 | 0.922294 | 2.21124 | 2.398 |
| chr22 | 49115796 | 49116078 | 0.946218 | 2.2689 | 2.398 |
| chr2 | 100106612 | 100107208 | 1.09342 | 2.62194 | 2.398 |
| chr12 | 123962094 | 123962315 | 0.487107 | 1.16815 | 2.398 |
| chr14 | 29624959 | 29625286 | 0.433018 | 1.03844 | 2.398 |
| chr22 | 37440135 | 37440498 | 1.46811 | 3.52098 | 2.398 |
| chr13 | 51796387 | 51796768 | 0.586893 | 1.40756 | 2.398 |
| chr1 | 78366426 | 78366692 | 1.01351 | 2.43073 | 2.398 |
| chr3 | 113213029 | 113213310 | 1.01887 | 2.44372 | 2.398 |
| chr12 | 14797491 | 14797859 | 0.37735 | 0.905061 | 2.398 |
| chr14 | 51240286 | 51240698 | 1.35736 | 3.2556 | 2.398 |
| chr6 | 107506896 | 107507138 | 1.13597 | 2.72461 | 2.398 |

| | | | | | |
|---|---|---|---|---|---|
| chr12 | 45524614 | 45524972 | 1.4887 | 3.57064 | 2.399 |
| chr7 | 26659907 | 26660403 | 0.432788 | 1.03804 | 2.399 |
| chr17 | 54145536 | 54145912 | 0.629269 | 1.50957 | 2.399 |
| chr5 | 30040680 | 30040934 | 1.52246 | 3.65227 | 2.399 |
| chr6 | 8983428 | 8983730 | 1.52464 | 3.65756 | 2.399 |
| chr9 | 7922284 | 7922497 | 0.468942 | 1.12498 | 2.399 |
| chrX | 19244098 | 19244417 | 0.746249 | 1.79026 | 2.399 |
| chr5 | 80256323 | 80256636 | 0.847757 | 2.03381 | 2.399 |
| chr2 | 71088903 | 71089406 | 2.51081 | 6.02362 | 2.399 |
| chr14 | 60431778 | 60432026 | 1.26748 | 3.04085 | 2.399 |
| chrX | 43808909 | 43809222 | 1.27144 | 3.05036 | 2.399 |
| chr1 | 6474397 | 6474694 | 1.55506 | 3.73087 | 2.399 |
| chrX | 99353772 | 99354171 | 0.346901 | 0.832302 | 2.399 |
| chr12 | 30848353 | 30848630 | 0.762787 | 1.83015 | 2.399 |
| chr12 | 122231994 | 122233246 | 1.79956 | 4.31781 | 2.399 |
| chr5 | 171615717 | 171616030 | 1.60909 | 3.86081 | 2.399 |
| chr10 | 73821264 | 73821524 | 1.28958 | 3.09436 | 2.400 |
| chr1 | 149192680 | 149193119 | 1.71547 | 4.11634 | 2.400 |
| chr1 | 26496137 | 26496473 | 1.84731 | 4.43275 | 2.400 |
| chr2 | 237261728 | 237262010 | 0.866467 | 2.07926 | 2.400 |
| chr5 | 11611545 | 11611792 | 0.481136 | 1.15462 | 2.400 |
| chr8 | 11801140 | 11801539 | 1.79576 | 4.30989 | 2.400 |
| chr14 | 101550251 | 101550605 | 0.579583 | 1.39105 | 2.400 |
| chr10 | 75157819 | 75158095 | 0.924176 | 2.21812 | 2.400 |
| chr7 | 146903339 | 146903591 | 0.576915 | 1.38467 | 2.400 |
| chr4 | 46771690 | 46771961 | 0.398774 | 0.957159 | 2.400 |
| chr12 | 111967190 | 111967544 | 1.42511 | 3.42069 | 2.400 |
| chr20 | 1905711 | 1905951 | 0.503854 | 1.2094 | 2.400 |
| chr17 | 38716531 | 38716881 | 2.07431 | 4.97933 | 2.400 |
| chr12 | 123436870 | 123437233 | 1.64944 | 3.9596 | 2.401 |
| chr5 | 140114044 | 140114247 | 0.800813 | 1.92253 | 2.401 |
| chr5 | 23413110 | 23413339 | 0.349361 | 0.838753 | 2.401 |
| chr8 | 59249083 | 59249503 | 0.431845 | 1.0368 | 2.401 |
| chr21 | 37212575 | 37212870 | 1.06705 | 2.56186 | 2.401 |
| chr19 | 48018685 | 48019035 | 2.12558 | 5.1033 | 2.401 |
| chr1 | 178511648 | 178512383 | 2.13169 | 5.11819 | 2.401 |
| chr9 | 1632533 | 1632779 | 0.440451 | 1.05753 | 2.401 |
| chr9 | 101903046 | 101903337 | 1.62409 | 3.89951 | 2.401 |
| chr4 | 41884694 | 41885145 | 2.26514 | 5.43871 | 2.401 |
| chr21 | 37582018 | 37582235 | 1.12092 | 2.69142 | 2.401 |
| chr3 | 40233340 | 40233673 | 1.50473 | 3.61311 | 2.401 |
| chr17 | 39823245 | 39823572 | 1.71267 | 4.11257 | 2.401 |
| chr5 | 146833426 | 146833888 | 1.89879 | 4.5598 | 2.401 |
| chrX | 106137613 | 106137929 | 1.58107 | 3.79681 | 2.401 |
| chr10 | 12122464 | 12122684 | 0.405934 | 0.974828 | 2.401 |
| chr1 | 156415625 | 156415942 | 1.64062 | 3.93993 | 2.401 |
| chr19 | 51021671 | 51022108 | 1.27094 | 3.05217 | 2.402 |
| chr19 | 857103 | 857322 | 1.52302 | 3.65758 | 2.402 |
| chr10 | 134430811 | 134431027 | 0.541233 | 1.29983 | 2.402 |
| chr1 | 43231586 | 43231874 | 0.77121 | 1.85217 | 2.402 |
| chr11 | 61515102 | 61515551 | 2.10761 | 5.06174 | 2.402 |
| chr19 | 29532509 | 29532725 | 0.59709 | 1.43403 | 2.402 |
| chr16 | 2033423 | 2034078 | 0.865926 | 2.07981 | 2.402 |
| chr21 | 33950431 | 33950684 | 1.23945 | 2.97704 | 2.402 |
| chr8 | 896763 | 897204 | 1.59137 | 3.82232 | 2.402 |
| chr6 | 85677201 | 85677419 | 1.06513 | 2.5584 | 2.402 |
| chr16 | 74141586 | 74142064 | 1.56145 | 3.75055 | 2.402 |
| chr3 | 28771937 | 28772244 | 0.507772 | 1.21966 | 2.402 |
| chr20 | 6290144 | 6290400 | 0.480582 | 1.15438 | 2.402 |
| chr7 | 131934520 | 131934788 | 1.0752 | 2.58269 | 2.402 |
| chr8 | 121968912 | 121969309 | 1.7808 | 4.27764 | 2.402 |

| | | | | | |
|---|---|---|---|---|---|
| chr14 | 77607412 | 77607842 | 0.898318 | 2.15786 | 2.402 |
| chr17 | 66756035 | 66756454 | 1.53345 | 3.68352 | 2.402 |
| chr18 | 43459954 | 43460293 | 2.11841 | 5.08865 | 2.402 |
| chr13 | 81553232 | 81553529 | 1.19454 | 2.86949 | 2.402 |
| chr16 | 86597394 | 86597879 | 3.4413 | 8.26671 | 2.402 |
| chr5 | 175875076 | 175875411 | 2.00891 | 4.82586 | 2.402 |
| chr2 | 234741538 | 234741744 | 0.980157 | 2.35464 | 2.402 |
| chr1 | 156629637 | 156630190 | 2.76114 | 6.63315 | 2.402 |
| chr10 | 135333597 | 135333997 | 2.87574 | 6.90851 | 2.402 |
| chr6 | 41274057 | 41274329 | 0.990426 | 2.37943 | 2.402 |
| chr5 | 66103627 | 66103862 | 1.42774 | 3.43012 | 2.402 |
| chrX | 128957205 | 128957412 | 0.913893 | 2.19564 | 2.403 |
| chr16 | 72870904 | 72871167 | 0.700616 | 1.68328 | 2.403 |
| chr14 | 105633896 | 105634198 | 0.813367 | 1.95445 | 2.403 |
| chr12 | 9202218 | 9202431 | 0.830688 | 1.99615 | 2.403 |
| chr10 | 56239877 | 56240099 | 0.395309 | 0.949991 | 2.403 |
| chr20 | 35898126 | 35898387 | 2.02693 | 4.8712 | 2.403 |
| chr7 | 156352837 | 156353120 | 0.613908 | 1.47547 | 2.403 |
| chr12 | 49907896 | 49908154 | 1.3632 | 3.27635 | 2.403 |
| chr8 | 37037448 | 37037660 | 1.21279 | 2.91496 | 2.404 |
| chr8 | 80328377 | 80328648 | 0.317199 | 0.762408 | 2.404 |
| chr14 | 105553113 | 105553395 | 1.39653 | 3.35666 | 2.404 |
| chr2 | 18604293 | 18604591 | 0.44378 | 1.06669 | 2.404 |
| chr16 | 81248311 | 81248720 | 1.06858 | 2.56854 | 2.404 |
| chr19 | 1057268 | 1057509 | 1.24394 | 2.99006 | 2.404 |
| chr14 | 91227048 | 91227383 | 1.27417 | 3.06275 | 2.404 |
| chr11 | 60737737 | 60738378 | 1.72981 | 4.15802 | 2.404 |
| chr10 | 66113191 | 66113415 | 0.427949 | 1.02872 | 2.404 |
| chr1 | 201354401 | 201354851 | 2.06251 | 4.95815 | 2.404 |
| chr15 | 74600172 | 74600515 | 1.00897 | 2.42553 | 2.404 |
| chr5 | 173019047 | 173019401 | 1.35821 | 3.26512 | 2.404 |
| chr19 | 35605445 | 35606020 | 2.67672 | 6.43484 | 2.404 |
| chr17 | 42118822 | 42119084 | 1.07903 | 2.59403 | 2.404 |
| chr1 | 42065664 | 42065871 | 1.22774 | 2.95156 | 2.404 |
| chr3 | 149901979 | 149902279 | 1.18064 | 2.83847 | 2.404 |
| chr12 | 54730282 | 54730672 | 1.55289 | 3.73348 | 2.404 |
| chr20 | 48685031 | 48685243 | 0.351474 | 0.845023 | 2.404 |
| chr8 | 82802721 | 82803125 | 1.18631 | 2.8522 | 2.404 |
| chr6 | 136638236 | 136638502 | 0.984374 | 2.3667 | 2.404 |
| chr12 | 131400742 | 131401136 | 3.16424 | 7.60791 | 2.404 |
| chr10 | 125755216 | 125755661 | 0.810073 | 1.94771 | 2.404 |
| chr5 | 153099963 | 153100428 | 0.504105 | 1.21214 | 2.405 |
| chr9 | 76282688 | 76283020 | 0.719117 | 1.72918 | 2.405 |
| chr7 | 68271146 | 68271587 | 0.395217 | 0.95034 | 2.405 |
| chr6 | 97876014 | 97876228 | 0.309236 | 0.743616 | 2.405 |
| chr8 | 52041257 | 52041627 | 0.359475 | 0.864463 | 2.405 |
| chr2 | 207048696 | 207048909 | 0.395226 | 0.950477 | 2.405 |
| chr17 | 30646406 | 30646937 | 2.41363 | 5.8046 | 2.405 |
| chr20 | 17384640 | 17384876 | 0.574637 | 1.38198 | 2.405 |
| chr15 | 22921381 | 22921648 | 0.384862 | 0.925581 | 2.405 |
| chr22 | 27075038 | 27075307 | 1.21193 | 2.91509 | 2.405 |
| chr11 | 125741394 | 125741680 | 0.951168 | 2.28814 | 2.406 |
| chr14 | 58574671 | 58575022 | 1.36623 | 3.2867 | 2.406 |
| chr3 | 29194008 | 29194256 | 0.63065 | 1.51716 | 2.406 |
| chr13 | 95014722 | 95014940 | 0.758033 | 1.82365 | 2.406 |
| chr12 | 56987926 | 56988231 | 1.82623 | 4.39353 | 2.406 |
| chr7 | 142700554 | 142700901 | 1.28046 | 3.08069 | 2.406 |
| chr20 | 10160248 | 10160596 | 1.19877 | 2.88421 | 2.406 |
| chr9 | 104457972 | 104458277 | 0.47799 | 1.15005 | 2.406 |
| chr11 | 1846124 | 1846574 | 1.74347 | 4.19487 | 2.406 |
| chr11 | 2320527 | 2320876 | 0.992504 | 2.38809 | 2.406 |

| | | | | | |
|---|---|---|---|---|---|
| chr12 | 11352520 | 11352933 | 1.06803 | 2.56986 | 2.406 |
| chr11 | 40252034 | 40252331 | 0.411625 | 0.990519 | 2.406 |
| chr5 | 170030956 | 170031357 | 1.53535 | 3.69468 | 2.406 |
| chr9 | 36995767 | 36996257 | 1.51344 | 3.64195 | 2.406 |
| chr9 | 137273565 | 137273809 | 1.10231 | 2.65272 | 2.407 |
| chr5 | 8700596 | 8700884 | 0.439151 | 1.057 | 2.407 |
| chr8 | 51021447 | 51021847 | 0.450154 | 1.08353 | 2.407 |
| chr2 | 178974651 | 178974950 | 1.00804 | 2.42642 | 2.407 |
| chr5 | 179628299 | 179628655 | 0.933368 | 2.24687 | 2.407 |
| chr17 | 74789832 | 74790262 | 1.27949 | 3.08017 | 2.407 |
| chr2 | 158050592 | 158050899 | 0.873072 | 2.10181 | 2.407 |
| chr9 | 123374568 | 123374841 | 1.04072 | 2.50556 | 2.408 |
| chr3 | 69021600 | 69021845 | 1.28031 | 3.08248 | 2.408 |
| chr3 | 168783289 | 168783530 | 0.596311 | 1.43575 | 2.408 |
| chr2 | 233963421 | 233963638 | 1.13524 | 2.73344 | 2.408 |
| chr8 | 132049173 | 132049466 | 1.55365 | 3.74092 | 2.408 |
| chrX | 32280629 | 32280841 | 0.305411 | 0.735396 | 2.408 |
| chr9 | 72645921 | 72646154 | 0.498071 | 1.19931 | 2.408 |
| chr1 | 93434062 | 93434273 | 0.962687 | 2.31808 | 2.408 |
| chr5 | 90397634 | 90397890 | 1.10904 | 2.6707 | 2.408 |
| chr8 | 18871748 | 18872190 | 1.46711 | 3.53316 | 2.408 |
| chr8 | 142105576 | 142105807 | 0.527785 | 1.27113 | 2.408 |
| chr2 | 235790399 | 235790680 | 1.52808 | 3.6803 | 2.408 |
| chr22 | 39706873 | 39707226 | 1.76033 | 4.23995 | 2.409 |
| chr15 | 62568604 | 62568870 | 0.406182 | 0.978369 | 2.409 |
| chr9 | 99215944 | 99216208 | 0.883771 | 2.12892 | 2.409 |
| chr12 | 65091832 | 65092140 | 1.49809 | 3.60876 | 2.409 |
| chr20 | 626032 | 626545 | 2.18785 | 5.27038 | 2.409 |
| chr8 | 80732359 | 80732635 | 0.489083 | 1.17824 | 2.409 |
| chr9 | 71838434 | 71838809 | 1.74384 | 4.20141 | 2.409 |
| chr18 | 44626650 | 44626930 | 1.17678 | 2.83524 | 2.409 |
| chr10 | 94822376 | 94822748 | 1.01025 | 2.4341 | 2.409 |
| chr10 | 124487102 | 124487331 | 0.493478 | 1.18901 | 2.409 |
| chr1 | 209920860 | 209921682 | 1.88868 | 4.55077 | 2.410 |
| chr7 | 142171967 | 142172444 | 1.17218 | 2.82445 | 2.410 |
| chr1 | 110881024 | 110881284 | 0.722517 | 1.74096 | 2.410 |
| chr2 | 18308694 | 18308956 | 0.409234 | 0.986098 | 2.410 |
| chr20 | 7266552 | 7266903 | 1.18033 | 2.84419 | 2.410 |
| chr2 | 158866887 | 158867114 | 0.797036 | 1.92071 | 2.410 |
| chr4 | 6335646 | 6335947 | 1.6015 | 3.85948 | 2.410 |
| chr9 | 93220853 | 93221152 | 1.03086 | 2.48432 | 2.410 |
| chr16 | 84539973 | 84540360 | 2.13629 | 5.14856 | 2.410 |
| chr16 | 11065362 | 11065670 | 1.12757 | 2.71763 | 2.410 |
| chr18 | 33876631 | 33876843 | 0.986738 | 2.37823 | 2.410 |
| chr1 | 115681565 | 115681860 | 1.53623 | 3.70263 | 2.410 |
| chr11 | 19098508 | 19098875 | 1.8999 | 4.5793 | 2.410 |
| chr1 | 173701176 | 173701401 | 0.47784 | 1.15174 | 2.410 |
| chr5 | 178922989 | 178923208 | 0.974375 | 2.34864 | 2.410 |
| chr17 | 6652983 | 6653210 | 0.604436 | 1.45696 | 2.410 |
| chr2 | 102972009 | 102972396 | 1.77083 | 4.26853 | 2.410 |
| chr15 | 66978025 | 66978431 | 1.04087 | 2.50908 | 2.411 |
| chrX | 130897781 | 130898026 | 1.35371 | 3.2632 | 2.411 |
| chr1 | 200457924 | 200458179 | 1.2544 | 3.02397 | 2.411 |
| chr21 | 33153228 | 33153524 | 0.637188 | 1.53607 | 2.411 |
| chr13 | 40643373 | 40643632 | 1.38082 | 3.32882 | 2.411 |
| chr1 | 25174371 | 25174806 | 1.43628 | 3.46256 | 2.411 |
| chr19 | 32757393 | 32757620 | 0.819501 | 1.97566 | 2.411 |
| chr7 | 85379640 | 85380117 | 1.28799 | 3.1052 | 2.411 |
| chr6 | 88102805 | 88103042 | 0.781559 | 1.88432 | 2.411 |
| chr1 | 164423953 | 164424155 | 0.43446 | 1.04749 | 2.411 |
| chr1 | 170043877 | 170044088 | 1.14451 | 2.7595 | 2.411 |

| | | | | | |
|---|---|---|---|---|---|
| chr9 | 73698238 | 73698468 | 0.445992 | 1.07533 | 2.411 |
| chr10 | 131926085 | 131926424 | 1.31621 | 3.17358 | 2.411 |
| chr20 | 412008 | 412553 | 2.00831 | 4.84247 | 2.411 |
| chr17 | 7591287 | 7591651 | 2.40342 | 5.79538 | 2.411 |
| chr8 | 6643051 | 6643339 | 0.679572 | 1.63874 | 2.411 |
| chr8 | 22552499 | 22552790 | 0.77132 | 1.86016 | 2.412 |
| chr17 | 70557181 | 70557422 | 1.09503 | 2.64089 | 2.412 |
| chr19 | 1837591 | 1838067 | 1.76631 | 4.25986 | 2.412 |
| chr20 | 33683987 | 33684375 | 1.31306 | 3.1668 | 2.412 |
| chr17 | 46208105 | 46208927 | 1.28329 | 3.09508 | 2.412 |
| chr15 | 94447380 | 94447693 | 0.289654 | 0.698599 | 2.412 |
| chr5 | 176797849 | 176798152 | 1.06731 | 2.57424 | 2.412 |
| chr1 | 22743063 | 22743462 | 1.26009 | 3.03937 | 2.412 |
| chr12 | 2841865 | 2842084 | 0.660552 | 1.59331 | 2.412 |
| chr3 | 42613881 | 42614084 | 1.41021 | 3.40156 | 2.412 |
| chr20 | 3034470 | 3034843 | 1.95228 | 4.70917 | 2.412 |
| chr7 | 93429136 | 93429377 | 0.712039 | 1.7176 | 2.412 |
| chr19 | 12958376 | 12958847 | 1.96104 | 4.73072 | 2.412 |
| chr3 | 97151435 | 97151684 | 0.422327 | 1.01882 | 2.412 |
| chr8 | 143775501 | 143775748 | 0.511405 | 1.23374 | 2.412 |
| chr16 | 30441493 | 30442046 | 1.1614 | 2.80185 | 2.412 |
| chr4 | 99860819 | 99861252 | 0.912664 | 2.20189 | 2.413 |
| chr21 | 35574326 | 35574551 | 1.10414 | 2.6641 | 2.413 |
| chr10 | 50801057 | 50801480 | 1.66597 | 4.01976 | 2.413 |
| chr18 | 60798358 | 60798623 | 0.95116 | 2.29508 | 2.413 |
| chr4 | 186942689 | 186943072 | 1.77035 | 4.27173 | 2.413 |
| chr7 | 99998350 | 99998708 | 0.43764 | 1.05604 | 2.413 |
| chr6 | 2932361 | 2932763 | 1.7949 | 4.33148 | 2.413 |
| chr12 | 127459329 | 127459597 | 0.511859 | 1.2353 | 2.413 |
| chr7 | 115763080 | 115763390 | 1.15603 | 2.79007 | 2.413 |
| chr15 | 26189558 | 26189800 | 0.520521 | 1.2563 | 2.414 |
| chr11 | 26353318 | 26353744 | 1.05771 | 2.55289 | 2.414 |
| chr5 | 166688754 | 166689065 | 0.880238 | 2.12464 | 2.414 |
| chr1 | 45696307 | 45696539 | 0.53733 | 1.29698 | 2.414 |
| chr11 | 44642662 | 44643058 | 2.49721 | 6.02809 | 2.414 |
| chr5 | 39217833 | 39218226 | 1.36832 | 3.30339 | 2.414 |
| chr20 | 60731873 | 60732388 | 1.9758 | 4.77006 | 2.414 |
| chr13 | 95068769 | 95068994 | 0.741359 | 1.78995 | 2.414 |
| chr6 | 149364654 | 149364881 | 0.656417 | 1.5849 | 2.414 |
| chr7 | 7143094 | 7143310 | 1.18882 | 2.87043 | 2.415 |
| chr9 | 123382102 | 123382339 | 1.29037 | 3.11608 | 2.415 |
| chr15 | 35212460 | 35212747 | 1.21236 | 2.9279 | 2.415 |
| chr3 | 64009103 | 64009464 | 2.27372 | 5.49138 | 2.415 |
| chr8 | 61048631 | 61049011 | 1.47406 | 3.56012 | 2.415 |
| chr7 | 158769790 | 158770202 | 1.22999 | 2.9707 | 2.415 |
| chr9 | 28951984 | 28952274 | 0.62 | 1.49756 | 2.415 |
| chr8 | 85273441 | 85273684 | 0.476792 | 1.15166 | 2.415 |
| chr16 | 57940377 | 57940710 | 0.908727 | 2.19499 | 2.415 |
| chr12 | 15374137 | 15374629 | 2.38757 | 5.76743 | 2.416 |
| chr8 | 68402594 | 68402837 | 1.58524 | 3.82954 | 2.416 |
| chr8 | 21994697 | 21995295 | 1.96866 | 4.75583 | 2.416 |
| chr15 | 89786933 | 89787206 | 0.943568 | 2.27945 | 2.416 |
| chr5 | 146414724 | 146415010 | 0.444602 | 1.07409 | 2.416 |
| chr7 | 71391030 | 71391400 | 0.418719 | 1.01159 | 2.416 |
| chr4 | 186427652 | 186427906 | 0.3542 | 0.855777 | 2.416 |
| chr17 | 36061414 | 36061827 | 1.78784 | 4.31962 | 2.416 |
| chr12 | 59923652 | 59923900 | 0.509554 | 1.23115 | 2.416 |
| chr5 | 28014964 | 28015493 | 0.834736 | 2.01694 | 2.416 |
| chr12 | 30131310 | 30131516 | 0.396474 | 0.958016 | 2.416 |
| chr2 | 84304669 | 84304876 | 0.61949 | 1.49698 | 2.416 |
| chr6 | 25992727 | 25993138 | 2.1582 | 5.21522 | 2.416 |

| | | | | | |
|---|---|---|---|---|---|
| chr6 | 35490620 | 35490964 | 2.00225 | 4.83838 | 2.416 |
| chr7 | 73831598 | 73832097 | 1.98037 | 4.78554 | 2.416 |
| chr7 | 139778887 | 139779158 | 1.09108 | 2.63669 | 2.417 |
| chr1 | 245134033 | 245134277 | 1.1099 | 2.68267 | 2.417 |
| chr1 | 93425966 | 93426172 | 0.980907 | 2.37094 | 2.417 |
| chr1 | 24044982 | 24045286 | 1.36642 | 3.30286 | 2.417 |
| chr8 | 23605370 | 23605670 | 1.16298 | 2.81119 | 2.417 |
| chr10 | 126408101 | 126408343 | 1.24413 | 3.00741 | 2.417 |
| chr4 | 171011320 | 171011593 | 1.18364 | 2.86122 | 2.417 |
| chr4 | 141348474 | 141349003 | 0.689432 | 1.66658 | 2.417 |
| chr4 | 12676557 | 12676816 | 0.440256 | 1.06428 | 2.417 |
| chr7 | 94078514 | 94078734 | 1.02227 | 2.47134 | 2.418 |
| chr20 | 8834406 | 8834820 | 1.45414 | 3.51541 | 2.418 |
| chr15 | 66121593 | 66121826 | 0.939913 | 2.27234 | 2.418 |
| chr9 | 86263992 | 86264332 | 2.16692 | 5.2389 | 2.418 |
| chr22 | 42337295 | 42337589 | 1.50476 | 3.63809 | 2.418 |
| chr11 | 58701032 | 58701318 | 0.418379 | 1.01153 | 2.418 |
| chr6 | 138188103 | 138188408 | 1.3858 | 3.35056 | 2.418 |
| chr9 | 132862007 | 132862209 | 0.603836 | 1.45996 | 2.418 |
| chr5 | 22697265 | 22697467 | 0.426669 | 1.03162 | 2.418 |
| chr10 | 7450224 | 7450802 | 1.52571 | 3.68895 | 2.418 |
| chr7 | 116426822 | 116427111 | 1.03296 | 2.49761 | 2.418 |
| chr1 | 35916689 | 35916983 | 1.37123 | 3.31555 | 2.418 |
| chr20 | 18924595 | 18925068 | 0.38856 | 0.939524 | 2.418 |
| chr6 | 7992293 | 7992539 | 0.868054 | 2.09893 | 2.418 |
| chr2 | 70528354 | 70529522 | 1.34051 | 3.24134 | 2.418 |
| chr1 | 12393415 | 12393641 | 0.476852 | 1.15303 | 2.418 |
| chr8 | 127907632 | 127907869 | 0.417765 | 1.01016 | 2.418 |
| chr12 | 121772573 | 121772991 | 1.98934 | 4.81027 | 2.418 |
| chr1 | 111158322 | 111158720 | 1.40607 | 3.39993 | 2.418 |
| chr14 | 23938338 | 23938609 | 1.3332 | 3.22375 | 2.418 |
| chrX | 17609301 | 17609612 | 1.34935 | 3.2628 | 2.418 |
| chr4 | 22131154 | 22131426 | 0.935567 | 2.26228 | 2.418 |
| chr18 | 33530284 | 33530683 | 1.12588 | 2.72248 | 2.418 |
| chr18 | 73881840 | 73882050 | 0.749798 | 1.81318 | 2.418 |
| chr14 | 105337038 | 105337418 | 1.15717 | 2.79831 | 2.418 |
| chr4 | 4329451 | 4329702 | 0.846606 | 2.04745 | 2.418 |
| chr8 | 62191696 | 62191912 | 0.498363 | 1.20529 | 2.419 |
| chrX | 55744525 | 55744938 | 1.07968 | 2.61122 | 2.419 |
| chr10 | 89619952 | 89620217 | 0.679638 | 1.64373 | 2.419 |
| chr2 | 715942 | 716150 | 0.850688 | 2.05744 | 2.419 |
| chr14 | 68085392 | 68085721 | 1.00001 | 2.41863 | 2.419 |
| chr1 | 216394533 | 216394860 | 1.18807 | 2.87356 | 2.419 |
| chr19 | 34532648 | 34533399 | 1.9584 | 4.73703 | 2.419 |
| chr8 | 53436079 | 53436282 | 0.927635 | 2.24381 | 2.419 |
| chr6 | 2952750 | 2953240 | 2.20906 | 5.34373 | 2.419 |
| chr4 | 109493741 | 109493947 | 1.27391 | 3.08162 | 2.419 |
| chr2 | 106720888 | 106721223 | 1.37453 | 3.32505 | 2.419 |
| chr2 | 101805171 | 101805450 | 1.23396 | 2.98516 | 2.419 |
| chr5 | 76941325 | 76941741 | 1.41891 | 3.43273 | 2.419 |
| chr15 | 59785136 | 59785527 | 1.26113 | 3.0511 | 2.419 |
| chr18 | 8882617 | 8882829 | 1.02621 | 2.48291 | 2.420 |
| chr3 | 76148038 | 76148378 | 0.557489 | 1.34888 | 2.420 |
| chr15 | 79718639 | 79718966 | 0.513008 | 1.24128 | 2.420 |
| chr12 | 122301204 | 122301423 | 0.426877 | 1.03295 | 2.420 |
| chr15 | 99066566 | 99066791 | 0.825411 | 1.99739 | 2.420 |
| chr16 | 52909823 | 52910182 | 0.487824 | 1.18053 | 2.420 |
| chr3 | 98376274 | 98376641 | 1.15356 | 2.79165 | 2.420 |
| chr8 | 140064224 | 140064502 | 0.478092 | 1.15706 | 2.420 |
| chr11 | 35462356 | 35462762 | 0.441038 | 1.06741 | 2.420 |
| chr3 | 125116676 | 125116946 | 0.318497 | 0.770853 | 2.420 |

| | | | | | |
|---|---|---|---|---|---|
| chr6 | 139034943 | 139035150 | 0.370862 | 0.897589 | 2.420 |
| chr17 | 20130068 | 20130474 | 2.00293 | 4.84775 | 2.420 |
| chr11 | 88277573 | 88277883 | 1.89573 | 4.58837 | 2.420 |
| chr12 | 121880789 | 121881397 | 1.75841 | 4.25625 | 2.421 |
| chr20 | 35820674 | 35821074 | 1.63204 | 3.9504 | 2.421 |
| chr22 | 30401234 | 30401592 | 1.61111 | 3.89978 | 2.421 |
| chr11 | 15958409 | 15958658 | 0.358853 | 0.86863 | 2.421 |
| chr5 | 159507863 | 159508221 | 1.38886 | 3.36184 | 2.421 |
| chr6 | 6547344 | 6547586 | 1.25136 | 3.02903 | 2.421 |
| chr12 | 50468884 | 50469374 | 1.9179 | 4.64251 | 2.421 |
| chr1 | 179074548 | 179074867 | 1.1999 | 2.90451 | 2.421 |
| chr19 | 33915066 | 33915317 | 0.462832 | 1.12043 | 2.421 |
| chr6 | 141255593 | 141255797 | 0.418641 | 1.01351 | 2.421 |
| chr1 | 226098876 | 226099425 | 2.36947 | 5.73641 | 2.421 |
| chr20 | 35383038 | 35383446 | 1.01039 | 2.44625 | 2.421 |
| chr17 | 13592048 | 13592393 | 0.378178 | 0.91561 | 2.421 |
| chr8 | 94053397 | 94053621 | 0.440767 | 1.06715 | 2.421 |
| chr1 | 68027343 | 68027719 | 1.82073 | 4.4085 | 2.421 |
| chr5 | 158533817 | 158534217 | 1.7466 | 4.22901 | 2.421 |
| chr13 | 95272618 | 95272917 | 0.887981 | 2.15034 | 2.422 |
| chr8 | 10444800 | 10445264 | 1.10953 | 2.68714 | 2.422 |
| chr8 | 86211517 | 86211720 | 0.596167 | 1.44386 | 2.422 |
| chr3 | 133213434 | 133213642 | 0.755612 | 1.83004 | 2.422 |
| chr17 | 20816643 | 20816898 | 0.537696 | 1.30231 | 2.422 |
| chr8 | 118316417 | 118316760 | 1.47587 | 3.57458 | 2.422 |
| chr1 | 7177557 | 7177773 | 0.636367 | 1.54141 | 2.422 |
| chr2 | 75652377 | 75652640 | 0.963479 | 2.3338 | 2.422 |
| chr5 | 51937013 | 51937238 | 0.361825 | 0.876485 | 2.422 |
| chr5 | 40755129 | 40755432 | 1.74747 | 4.23325 | 2.423 |
| chr12 | 99506552 | 99507051 | 0.441414 | 1.06933 | 2.423 |
| chr6 | 163975797 | 163976062 | 1.10427 | 2.6753 | 2.423 |
| chr16 | 27180857 | 27181188 | 0.698903 | 1.69342 | 2.423 |
| chr11 | 35426064 | 35426505 | 1.11605 | 2.70439 | 2.423 |
| chr16 | 10830899 | 10831256 | 1.48243 | 3.59219 | 2.423 |
| chr18 | 72335434 | 72335662 | 0.850011 | 2.05974 | 2.423 |
| chr6 | 44623266 | 44623667 | 1.84342 | 4.46696 | 2.423 |
| chr7 | 154499670 | 154499873 | 0.379426 | 0.91946 | 2.423 |
| chr14 | 107250836 | 107251249 | 1.72578 | 4.18233 | 2.423 |
| chr15 | 71015051 | 71015268 | 0.984516 | 2.38604 | 2.424 |
| chr11 | 128500348 | 128500565 | 1.06632 | 2.58443 | 2.424 |
| chr8 | 87249720 | 87249991 | 0.51753 | 1.25436 | 2.424 |
| chr15 | 96394148 | 96394384 | 1.12441 | 2.72536 | 2.424 |
| chr3 | 106990616 | 106990945 | 0.982773 | 2.3821 | 2.424 |
| chr12 | 22142789 | 22143040 | 1.02029 | 2.47313 | 2.424 |
| chr13 | 25257959 | 25258299 | 1.46947 | 3.56201 | 2.424 |
| chr16 | 48721453 | 48721670 | 1.4556 | 3.52842 | 2.424 |
| chr17 | 25707954 | 25708431 | 3.72215 | 9.02379 | 2.424 |
| chr20 | 52229523 | 52229904 | 0.584123 | 1.41615 | 2.424 |
| chr1 | 209996487 | 209997000 | 1.01718 | 2.46624 | 2.425 |
| chr17 | 27417887 | 27418104 | 0.358382 | 0.868944 | 2.425 |
| chr2 | 88969235 | 88969489 | 1.02992 | 2.49721 | 2.425 |
| chr21 | 43032861 | 43033147 | 1.48751 | 3.60679 | 2.425 |
| chr19 | 10981907 | 10982267 | 1.92279 | 4.66249 | 2.425 |
| chr11 | 12433808 | 12434182 | 2.08416 | 5.05423 | 2.425 |
| chr12 | 61909290 | 61909497 | 0.53575 | 1.29924 | 2.425 |
| chr9 | 122170956 | 122171176 | 0.504569 | 1.22367 | 2.425 |
| chr2 | 7812145 | 7812612 | 1.37039 | 3.32356 | 2.425 |
| chr5 | 53813209 | 53813628 | 2.0831 | 5.05214 | 2.425 |
| chr16 | 11351146 | 11351482 | 1.20049 | 2.91157 | 2.425 |
| chr21 | 46557025 | 46557391 | 1.58527 | 3.84516 | 2.426 |
| chr17 | 75084678 | 75084978 | 1.43925 | 3.49102 | 2.426 |

| | | | | | |
|---|---|---|---|---|---|
| chr8 | 121136539 | 121136750 | 1.19551 | 2.89982 | 2.426 |
| chr1 | 62996258 | 62996569 | 1.54145 | 3.73903 | 2.426 |
| chr2 | 54311803 | 54312043 | 1.12743 | 2.7348 | 2.426 |
| chr9 | 129957518 | 129957779 | 0.74169 | 1.79913 | 2.426 |
| chr10 | 82137929 | 82138280 | 1.05833 | 2.56741 | 2.426 |
| chr3 | 164807554 | 164807766 | 0.451495 | 1.09538 | 2.426 |
| chr6 | 27486672 | 27486972 | 0.944518 | 2.29153 | 2.426 |
| chr10 | 69968264 | 69968591 | 1.29164 | 3.13372 | 2.426 |
| chr5 | 58154618 | 58154832 | 0.556925 | 1.3512 | 2.426 |
| chr1 | 202182578 | 202182803 | 0.940028 | 2.28081 | 2.426 |
| chr20 | 2327843 | 2328209 | 1.57951 | 3.83242 | 2.426 |
| chr3 | 9544860 | 9545274 | 1.99106 | 4.83107 | 2.426 |
| chr10 | 71399016 | 71399298 | 1.45152 | 3.52195 | 2.426 |
| chr2 | 213727898 | 213728182 | 1.28818 | 3.12585 | 2.427 |
| chr11 | 118436630 | 118436910 | 1.13426 | 2.75237 | 2.427 |
| chr8 | 142094324 | 142094921 | 2.009 | 4.87511 | 2.427 |
| chr1 | 27989682 | 27990002 | 0.851301 | 2.06583 | 2.427 |
| chr11 | 117097161 | 117097532 | 1.7224 | 4.17984 | 2.427 |
| chr12 | 45895248 | 45895586 | 0.499541 | 1.21232 | 2.427 |
| chr10 | 34570062 | 34570294 | 1.07676 | 2.61317 | 2.427 |
| chr1 | 228651470 | 228651811 | 0.592387 | 1.43774 | 2.427 |
| chr12 | 51818791 | 51819183 | 2.07921 | 5.04633 | 2.427 |
| chr8 | 144461844 | 144462244 | 2.22421 | 5.39857 | 2.427 |
| chr8 | 110986566 | 110986916 | 1.58039 | 3.83607 | 2.427 |
| chr15 | 57438657 | 57438901 | 1.11466 | 2.70571 | 2.427 |
| chr7 | 136011570 | 136011827 | 0.465372 | 1.12964 | 2.427 |
| chr21 | 43030533 | 43030915 | 1.64974 | 4.00458 | 2.427 |
| chr12 | 43226455 | 43226786 | 0.460116 | 1.11697 | 2.428 |
| chr17 | 17343370 | 17343630 | 1.59369 | 3.86891 | 2.428 |
| chr7 | 69479991 | 69480205 | 0.387317 | 0.940275 | 2.428 |
| chr11 | 46410725 | 46411361 | 1.73139 | 4.20334 | 2.428 |
| chr8 | 407821 | 408106 | 0.796666 | 1.93409 | 2.428 |
| chr8 | 102534029 | 102534243 | 0.984825 | 2.39102 | 2.428 |
| chr17 | 49717506 | 49717692 | 1.37856 | 3.34698 | 2.428 |
| chr6 | 160940891 | 160941226 | 1.90783 | 4.63198 | 2.428 |
| chr10 | 33482263 | 33482569 | 1.44829 | 3.51635 | 2.428 |
| chr16 | 46981798 | 46982065 | 1.41242 | 3.42932 | 2.428 |
| chr19 | 58414712 | 58415088 | 0.560341 | 1.36049 | 2.428 |
| chr14 | 104602368 | 104602684 | 0.662396 | 1.60829 | 2.428 |
| chr1 | 242618138 | 242618405 | 1.91989 | 4.66176 | 2.428 |
| chr12 | 99333232 | 99333435 | 1.09115 | 2.64948 | 2.428 |
| chr9 | 13950555 | 13950794 | 0.470215 | 1.14176 | 2.428 |
| chrX | 10604459 | 10604727 | 1.68759 | 4.09807 | 2.428 |
| chr5 | 81660374 | 81660577 | 0.826447 | 2.00698 | 2.428 |
| chr14 | 65486711 | 65487033 | 1.42114 | 3.45139 | 2.429 |
| chr5 | 134761845 | 134762121 | 1.21024 | 2.9392 | 2.429 |
| chr11 | 19281315 | 19281673 | 1.54632 | 3.75545 | 2.429 |
| chrX | 120388207 | 120388465 | 0.687786 | 1.67047 | 2.429 |
| chr9 | 5799139 | 5799597 | 1.52559 | 3.70561 | 2.429 |
| chr20 | 60174078 | 60174311 | 0.724023 | 1.75869 | 2.429 |
| chr14 | 58654693 | 58654953 | 0.458851 | 1.11464 | 2.429 |
| chr22 | 31485097 | 31485504 | 1.53016 | 3.71728 | 2.429 |
| chr11 | 74952617 | 74953044 | 2.46739 | 5.99423 | 2.429 |
| chr11 | 64555058 | 64555468 | 2.19286 | 5.32741 | 2.429 |
| chr8 | 127822153 | 127822504 | 1.19031 | 2.89184 | 2.429 |
| chr16 | 30669328 | 30669591 | 1.62923 | 3.95833 | 2.430 |
| chr7 | 3522104 | 3522365 | 1.55065 | 3.76754 | 2.430 |
| chr4 | 188527708 | 188528015 | 1.704 | 4.14026 | 2.430 |
| chr9 | 80366436 | 80366659 | 1.22744 | 2.98252 | 2.430 |
| chr17 | 5972190 | 5972630 | 2.2499 | 5.46721 | 2.430 |
| chr12 | 5973301 | 5973709 | 0.431003 | 1.04734 | 2.430 |

| | | | | | |
|---|---|---|---|---|---|
| chr17 | 2285861 | 2286230 | 1.7447 | 4.23969 | 2.430 |
| chr1 | 9781801 | 9782268 | 1.00434 | 2.44068 | 2.430 |
| chr4 | 175143493 | 175143774 | 1.0128 | 2.4613 | 2.430 |
| chr1 | 179038277 | 179038535 | 0.734369 | 1.78466 | 2.430 |
| chr5 | 172710785 | 172711212 | 1.46161 | 3.55213 | 2.430 |
| chr5 | 112979219 | 112979594 | 0.362886 | 0.881954 | 2.430 |
| chr6 | 43474270 | 43474760 | 1.25724 | 3.05561 | 2.430 |
| chr10 | 70834934 | 70835488 | 0.448744 | 1.09067 | 2.430 |
| chr11 | 13295398 | 13295827 | 0.446783 | 1.08591 | 2.431 |
| chr17 | 20528251 | 20528655 | 0.479913 | 1.16652 | 2.431 |
| chr6 | 33553451 | 33553751 | 1.6322 | 3.96746 | 2.431 |
| chr6 | 8670888 | 8671144 | 0.340895 | 0.828642 | 2.431 |
| chr17 | 47472394 | 47472712 | 1.85373 | 4.50621 | 2.431 |
| chr3 | 141027539 | 141027778 | 1.21652 | 2.95765 | 2.431 |
| chr14 | 52739145 | 52739411 | 0.942695 | 2.29198 | 2.431 |
| chr11 | 18753336 | 18753927 | 1.23688 | 3.00733 | 2.431 |
| chr22 | 42069172 | 42069423 | 0.846641 | 2.05855 | 2.431 |
| chr5 | 34523264 | 34523538 | 1.1322 | 2.75292 | 2.431 |
| chr14 | 86521571 | 86521828 | 0.555893 | 1.35174 | 2.432 |
| chr17 | 42598989 | 42599356 | 0.414191 | 1.00722 | 2.432 |
| chr3 | 124407713 | 124408056 | 1.41194 | 3.43383 | 2.432 |
| chr7 | 44584528 | 44584748 | 0.606527 | 1.4751 | 2.432 |
| chr2 | 15437600 | 15437854 | 0.544712 | 1.32477 | 2.432 |
| chr10 | 110922434 | 110922788 | 0.420053 | 1.02161 | 2.432 |
| chr20 | 22567258 | 22567753 | 2.02724 | 4.93051 | 2.432 |
| chr12 | 128482238 | 128482488 | 0.594869 | 1.44692 | 2.432 |
| chr19 | 41354498 | 41354808 | 0.909868 | 2.2131 | 2.432 |
| chr8 | 22046377 | 22046654 | 1.13143 | 2.75202 | 2.432 |
| chr19 | 28268740 | 28268996 | 0.911896 | 2.21815 | 2.432 |
| chr1 | 101393693 | 101393968 | 0.509817 | 1.24016 | 2.433 |
| chrX | 129213769 | 129214094 | 1.25152 | 3.04461 | 2.433 |
| chr22 | 21158248 | 21158642 | 0.978437 | 2.3805 | 2.433 |
| chr17 | 56296609 | 56296859 | 1.17893 | 2.86835 | 2.433 |
| chr4 | 167173991 | 167174236 | 0.782284 | 1.90332 | 2.433 |
| chr2 | 169231795 | 169232018 | 0.917915 | 2.23333 | 2.433 |
| chr19 | 17501952 | 17502185 | 1.16332 | 2.83048 | 2.433 |
| chr13 | 113689347 | 113689589 | 1.20557 | 2.93358 | 2.433 |
| chrX | 132083070 | 132083359 | 0.517165 | 1.25845 | 2.433 |
| chr12 | 14998277 | 14998509 | 1.39511 | 3.39484 | 2.433 |
| chr5 | 172603707 | 172603937 | 0.440988 | 1.07312 | 2.433 |
| chr10 | 77159620 | 77159956 | 1.19335 | 2.90402 | 2.434 |
| chr12 | 58641532 | 58641923 | 0.761807 | 1.85387 | 2.434 |
| chr1 | 120218227 | 120218469 | 1.07982 | 2.62793 | 2.434 |
| chr18 | 68159015 | 68159334 | 1.32512 | 3.22505 | 2.434 |
| chr4 | 49034507 | 49034790 | 1.18237 | 2.87779 | 2.434 |
| chr8 | 136264964 | 136265306 | 0.497254 | 1.21036 | 2.434 |
| chr20 | 14016661 | 14016880 | 1.03225 | 2.5126 | 2.434 |
| chr21 | 45358912 | 45359336 | 1.97368 | 4.80428 | 2.434 |
| chr3 | 169581230 | 169581577 | 1.46638 | 3.56942 | 2.434 |
| chr12 | 70294442 | 70294738 | 1.51308 | 3.68312 | 2.434 |
| chr21 | 35296295 | 35296657 | 1.58362 | 3.85523 | 2.434 |
| chr1 | 9310110 | 9310369 | 1.04584 | 2.54622 | 2.435 |
| chr3 | 176590586 | 176590930 | 1.16057 | 2.82563 | 2.435 |
| chr20 | 61856117 | 61856335 | 0.795573 | 1.93699 | 2.435 |
| chr17 | 53082950 | 53083212 | 1.08175 | 2.6339 | 2.435 |
| chr18 | 65183116 | 65183476 | 1.23942 | 3.01787 | 2.435 |
| chr5 | 74199253 | 74199577 | 1.59538 | 3.8846 | 2.435 |
| chr6 | 101789725 | 101789947 | 0.358255 | 0.872322 | 2.435 |
| chr12 | 113536156 | 113536415 | 1.48542 | 3.61691 | 2.435 |
| chr2 | 47003889 | 47004217 | 1.45088 | 3.5331 | 2.435 |
| chr3 | 38159542 | 38159900 | 1.46967 | 3.57886 | 2.435 |

| | | | | | |
|---|---|---|---|---|---|
| chr4 | 56178770 | 56179043 | 0.972709 | 2.36869 | 2.435 |
| chr10 | 114325291 | 114325527 | 1.52774 | 3.72033 | 2.435 |
| chr14 | 86670307 | 86670573 | 0.418439 | 1.01899 | 2.435 |
| gl000205_r | 151487 | 151714 | 0.36486 | 0.888519 | 2.435 |
| chr14 | 61655694 | 61656050 | 1.44966 | 3.53034 | 2.435 |
| chr5 | 150505684 | 150506034 | 1.82885 | 4.45386 | 2.435 |
| chr4 | 32464589 | 32464892 | 0.867702 | 2.11326 | 2.435 |
| chr7 | 136980601 | 136980808 | 0.39574 | 0.963851 | 2.436 |
| chr4 | 63575463 | 63575674 | 0.514253 | 1.25253 | 2.436 |
| chr17 | 31785822 | 31786336 | 0.465655 | 1.13417 | 2.436 |
| chr16 | 24516083 | 24516379 | 0.96448 | 2.34916 | 2.436 |
| chr1 | 215779996 | 215780258 | 0.971061 | 2.36538 | 2.436 |
| chr10 | 12967825 | 12968114 | 0.412364 | 1.00447 | 2.436 |
| chrX | 20085383 | 20085587 | 0.88369 | 2.15259 | 2.436 |
| chr6 | 52516919 | 52517206 | 1.56371 | 3.80928 | 2.436 |
| chr19 | 44580271 | 44580521 | 0.44712 | 1.08924 | 2.436 |
| chr7 | 93906219 | 93906443 | 1.29057 | 3.14399 | 2.436 |
| chr6 | 108169412 | 108169782 | 1.55 | 3.77604 | 2.436 |
| chr20 | 50255813 | 50256169 | 1.42548 | 3.47295 | 2.436 |
| chr7 | 35749120 | 35749450 | 1.35018 | 3.28994 | 2.437 |
| chr12 | 112266429 | 112266671 | 0.503734 | 1.22748 | 2.437 |
| chr1 | 231752677 | 231752906 | 0.35648 | 0.868737 | 2.437 |
| chr10 | 24672361 | 24672855 | 0.799655 | 1.94885 | 2.437 |
| chr5 | 67296902 | 67297287 | 0.421541 | 1.02736 | 2.437 |
| chr16 | 22311809 | 22312102 | 1.5483 | 3.7737 | 2.437 |
| chr14 | 55239964 | 55240206 | 0.959941 | 2.33975 | 2.437 |
| chr10 | 92631039 | 92631265 | 1.0178 | 2.48088 | 2.437 |
| chr20 | 23059695 | 23059982 | 1.22518 | 2.98674 | 2.438 |
| chr12 | 95158083 | 95158315 | 1.27489 | 3.10794 | 2.438 |
| chr8 | 27319028 | 27319437 | 0.708065 | 1.72639 | 2.438 |
| chr14 | 73101501 | 73102057 | 0.528015 | 1.28745 | 2.438 |
| chr5 | 52244362 | 52244644 | 1.47154 | 3.5881 | 2.438 |
| chr9 | 120891563 | 120891786 | 0.349401 | 0.851967 | 2.438 |
| chr8 | 17352749 | 17352957 | 0.926097 | 2.25817 | 2.438 |
| chr18 | 52346884 | 52347117 | 1.01815 | 2.48279 | 2.439 |
| chr7 | 27183367 | 27183709 | 1.65017 | 4.02399 | 2.439 |
| chr8 | 35446083 | 35446327 | 0.485113 | 1.18298 | 2.439 |
| chr11 | 30793672 | 30793998 | 0.460137 | 1.12209 | 2.439 |
| chr6 | 10622091 | 10622361 | 0.599651 | 1.46233 | 2.439 |
| chr8 | 54500659 | 54500909 | 0.82113 | 2.00251 | 2.439 |
| chr8 | 142098615 | 142098848 | 1.07848 | 2.63018 | 2.439 |
| chr11 | 73966901 | 73967134 | 1.21271 | 2.95758 | 2.439 |
| chr12 | 55582858 | 55583164 | 0.981809 | 2.39459 | 2.439 |
| chr15 | 26179085 | 26179418 | 1.46802 | 3.58077 | 2.439 |
| chr19 | 18499245 | 18499625 | 1.851 | 4.5151 | 2.439 |
| chr11 | 107988622 | 107988847 | 0.769693 | 1.87755 | 2.439 |
| chr8 | 145024115 | 145024318 | 0.710642 | 1.73353 | 2.439 |
| chr3 | 100309420 | 100309705 | 0.921104 | 2.24711 | 2.440 |
| chr1 | 197838102 | 197838409 | 1.07838 | 2.63088 | 2.440 |
| chr11 | 26308057 | 26308370 | 0.989551 | 2.41417 | 2.440 |
| chr5 | 171486555 | 171486977 | 1.89744 | 4.62925 | 2.440 |
| chr19 | 39832542 | 39832868 | 1.98884 | 4.85265 | 2.440 |
| chr6 | 136231003 | 136231220 | 0.939071 | 2.29131 | 2.440 |
| chr3 | 145110965 | 145111204 | 0.488974 | 1.19316 | 2.440 |
| chr10 | 27898877 | 27899347 | 0.369114 | 0.900739 | 2.440 |
| chr14 | 41513170 | 41513561 | 0.393625 | 0.960571 | 2.440 |
| chr7 | 100933786 | 100934063 | 1.15768 | 2.82518 | 2.440 |
| chr3 | 99787584 | 99787973 | 0.976976 | 2.38421 | 2.440 |
| chr11 | 44373570 | 44373876 | 0.661469 | 1.61429 | 2.440 |
| chr20 | 5446107 | 5446395 | 0.6149 | 1.50064 | 2.440 |
| chr19 | 7294827 | 7295119 | 1.17665 | 2.8716 | 2.440 |

| | | | | | |
|---|---|---|---|---|---|
| chr3 | 32509083 | 32509788 | 1.82048 | 4.44308 | 2.441 |
| chr2 | 63528115 | 63528368 | 0.900661 | 2.19821 | 2.441 |
| chr4 | 184328226 | 184328584 | 1.8562 | 4.53056 | 2.441 |
| chr2 | 210421170 | 210421382 | 0.758553 | 1.85146 | 2.441 |
| chr17 | 52503676 | 52503951 | 0.942074 | 2.2995 | 2.441 |
| chr17 | 15340041 | 15340286 | 0.814419 | 1.98797 | 2.441 |
| chr10 | 112440835 | 112441120 | 1.36997 | 3.34431 | 2.441 |
| chr2 | 216979431 | 216979708 | 1.64705 | 4.02091 | 2.441 |
| chr5 | 141071632 | 141072059 | 1.82043 | 4.44425 | 2.441 |
| chr4 | 154265491 | 154265881 | 1.70204 | 4.15526 | 2.441 |
| chr5 | 105028494 | 105028967 | 0.517455 | 1.26331 | 2.441 |
| chr5 | 176005381 | 176005680 | 1.01752 | 2.48417 | 2.441 |
| chr3 | 194992247 | 194992625 | 1.77625 | 4.33683 | 2.442 |
| chr17 | 46695976 | 46696273 | 1.82855 | 4.46455 | 2.442 |
| chr10 | 67535809 | 67536079 | 0.514788 | 1.2569 | 2.442 |
| chr11 | 26318826 | 26319034 | 1.22496 | 2.99086 | 2.442 |
| chr12 | 111700718 | 111700934 | 0.542817 | 1.32539 | 2.442 |
| chr2 | 227663129 | 227663405 | 1.14183 | 2.788 | 2.442 |
| chr18 | 19958678 | 19958993 | 1.03619 | 2.53018 | 2.442 |
| chr3 | 128689103 | 128689444 | 1.4407 | 3.51802 | 2.442 |
| chr8 | 58986792 | 58987146 | 1.33093 | 3.25019 | 2.442 |
| chr17 | 80626337 | 80626822 | 1.82281 | 4.45161 | 2.442 |
| chr3 | 197236907 | 197237417 | 1.75067 | 4.27589 | 2.442 |
| chr10 | 95506457 | 95506876 | 0.940558 | 2.29728 | 2.442 |
| chr12 | 11659044 | 11659285 | 0.495167 | 1.2095 | 2.443 |
| chr12 | 63545862 | 63546192 | 1.77217 | 4.32887 | 2.443 |
| chr5 | 151777099 | 151777383 | 0.398714 | 0.973933 | 2.443 |
| chr14 | 25780885 | 25781306 | 0.406697 | 0.993497 | 2.443 |
| chr7 | 112538787 | 112539083 | 1.95079 | 4.76576 | 2.443 |
| chr20 | 35402615 | 35403020 | 1.63234 | 3.98781 | 2.443 |
| chr8 | 50558565 | 50558904 | 0.525265 | 1.28323 | 2.443 |
| chr2 | 36899413 | 36899654 | 1.28681 | 3.14377 | 2.443 |
| chr1 | 75155538 | 75155760 | 1.0681 | 2.6096 | 2.443 |
| chr5 | 178893010 | 178893279 | 0.423089 | 1.0337 | 2.443 |
| chr17 | 33815273 | 33815562 | 0.796373 | 1.94581 | 2.443 |
| chr7 | 6362642 | 6362978 | 2.12471 | 5.19147 | 2.443 |
| chr1 | 23536825 | 23537099 | 0.570658 | 1.39436 | 2.443 |
| chr1 | 177623661 | 177623867 | 0.407156 | 0.994926 | 2.444 |
| chr3 | 195980489 | 195980722 | 1.10277 | 2.69474 | 2.444 |
| chr8 | 27373376 | 27373622 | 0.637744 | 1.55854 | 2.444 |
| chr1 | 46879372 | 46879584 | 0.612192 | 1.4961 | 2.444 |
| chr11 | 24176859 | 24177187 | 0.394619 | 0.964394 | 2.444 |
| chr1 | 36751128 | 36751395 | 1.07351 | 2.62373 | 2.444 |
| chr7 | 6144334 | 6144641 | 1.43436 | 3.50569 | 2.444 |
| chr1 | 32782116 | 32782374 | 1.15218 | 2.81627 | 2.444 |
| chr5 | 116915970 | 116916227 | 0.347602 | 0.849676 | 2.444 |
| chr12 | 56013306 | 56013814 | 1.35896 | 3.32189 | 2.444 |
| chr9 | 137701029 | 137701234 | 0.382089 | 0.934044 | 2.445 |
| chr2 | 103422767 | 103423051 | 1.44141 | 3.52381 | 2.445 |
| chr12 | 23924222 | 23924451 | 0.517861 | 1.26609 | 2.445 |
| chr4 | 8689879 | 8690256 | 1.5265 | 3.73208 | 2.445 |
| chr2 | 33666329 | 33666636 | 1.61415 | 3.94642 | 2.445 |
| chr20 | 35241134 | 35241420 | 1.09166 | 2.66907 | 2.445 |
| chr12 | 52217793 | 52218066 | 1.34284 | 3.28326 | 2.445 |
| chr9 | 110249807 | 110250149 | 1.46854 | 3.59077 | 2.445 |
| chr14 | 103449297 | 103449625 | 0.585842 | 1.4327 | 2.446 |
| chr10 | 33444834 | 33445073 | 1.33739 | 3.27079 | 2.446 |
| chr3 | 106517896 | 106518170 | 0.548587 | 1.34176 | 2.446 |
| chr12 | 111736571 | 111737202 | 1.79492 | 4.39033 | 2.446 |
| chr15 | 33136416 | 33136630 | 1.31895 | 3.22646 | 2.446 |
| chr18 | 74766986 | 74767268 | 1.09442 | 2.67744 | 2.446 |

| | | | | | |
|---|---|---|---|---|---|
| chr18 | 74771582 | 74771806 | 0.889484 | 2.17611 | 2.446 |
| chr17 | 47633592 | 47634096 | 2.46599 | 6.03317 | 2.447 |
| chr20 | 44615827 | 44616532 | 1.34778 | 3.29743 | 2.447 |
| chr11 | 102483736 | 102483975 | 1.01202 | 2.476 | 2.447 |
| chr7 | 1633164 | 1633553 | 1.10253 | 2.69747 | 2.447 |
| chrX | 46501594 | 46501878 | 0.563191 | 1.37797 | 2.447 |
| chr9 | 101299182 | 101299416 | 0.844005 | 2.06517 | 2.447 |
| chr15 | 65326110 | 65326362 | 1.1904 | 2.91278 | 2.447 |
| chr12 | 128266791 | 128266997 | 0.449862 | 1.10078 | 2.447 |
| chr19 | 13983515 | 13983761 | 1.07449 | 2.62939 | 2.447 |
| chr8 | 97152527 | 97152877 | 0.347487 | 0.85035 | 2.447 |
| chr13 | 96672410 | 96672688 | 1.03388 | 2.5302 | 2.447 |
| chr10 | 70310381 | 70310840 | 0.833034 | 2.03869 | 2.447 |
| chr11 | 102678780 | 102679107 | 1.21129 | 2.96453 | 2.447 |
| chr10 | 81818540 | 81818998 | 1.26769 | 3.10264 | 2.447 |
| chr3 | 159706286 | 159706806 | 2.04661 | 5.00914 | 2.448 |
| chr1 | 178006612 | 178007214 | 0.80584 | 1.97237 | 2.448 |
| chr16 | 67433247 | 67433846 | 1.6583 | 4.05906 | 2.448 |
| chrX | 18904593 | 18904878 | 1.43257 | 3.50658 | 2.448 |
| chr20 | 22419588 | 22419849 | 0.994141 | 2.43359 | 2.448 |
| chr1 | 38492360 | 38492592 | 1.45963 | 3.57344 | 2.448 |
| chr17 | 5582052 | 5582256 | 0.429887 | 1.05244 | 2.448 |
| chr15 | 81880761 | 81881010 | 0.474087 | 1.16072 | 2.448 |
| chr12 | 57156798 | 57157166 | 0.92224 | 2.25801 | 2.448 |
| chr22 | 20167784 | 20168098 | 0.565934 | 1.38571 | 2.449 |
| chr5 | 73944659 | 73944985 | 1.29154 | 3.16267 | 2.449 |
| chr13 | 98452586 | 98452935 | 1.05031 | 2.57197 | 2.449 |
| chr22 | 23009075 | 23009283 | 1.03336 | 2.53046 | 2.449 |
| chr1 | 193402962 | 193403215 | 0.467992 | 1.14603 | 2.449 |
| chr13 | 59384289 | 59384496 | 0.343346 | 0.8408 | 2.449 |
| chr14 | 55240696 | 55241097 | 1.89307 | 4.63602 | 2.449 |
| chr7 | 78636203 | 78636669 | 0.546404 | 1.33813 | 2.449 |
| chr7 | 77388660 | 77388924 | 1.85303 | 4.53822 | 2.449 |
| chr19 | 51841969 | 51842357 | 1.5754 | 3.85844 | 2.449 |
| chr11 | 47574669 | 47574991 | 1.6057 | 3.9327 | 2.449 |
| chr7 | 141891820 | 141892119 | 0.572442 | 1.40214 | 2.449 |
| chr20 | 43940199 | 43940619 | 1.81734 | 4.45151 | 2.449 |
| chr11 | 76510253 | 76510670 | 1.76481 | 4.32285 | 2.449 |
| chrX | 46656049 | 46656391 | 1.72233 | 4.21934 | 2.450 |
| chr11 | 1970191 | 1970516 | 0.780033 | 1.91116 | 2.450 |
| chr6 | 64512578 | 64512788 | 0.822528 | 2.01538 | 2.450 |
| chr2 | 47261272 | 47261528 | 1.13143 | 2.77274 | 2.451 |
| chr1 | 240772083 | 240772450 | 1.11171 | 2.72461 | 2.451 |
| chr3 | 32195736 | 32195944 | 0.853364 | 2.09156 | 2.451 |
| chr2 | 127729598 | 127730015 | 1.44964 | 3.55311 | 2.451 |
| chr5 | 140235382 | 140235631 | 0.576493 | 1.41309 | 2.451 |
| chr16 | 57201970 | 57202207 | 1.30593 | 3.20121 | 2.451 |
| chr1 | 33560475 | 33560977 | 1.60543 | 3.93543 | 2.451 |
| chr12 | 49736054 | 49736764 | 2.59631 | 6.36443 | 2.451 |
| chrX | 8716929 | 8717218 | 1.24278 | 3.04659 | 2.451 |
| chr12 | 85306555 | 85306826 | 1.10494 | 2.70879 | 2.452 |
| chr1 | 114889123 | 114889544 | 3.40089 | 8.33753 | 2.452 |
| chr19 | 18043386 | 18043626 | 1.80992 | 4.43724 | 2.452 |
| chr6 | 36390997 | 36391208 | 1.0169 | 2.49314 | 2.452 |
| chr1 | 224701395 | 224701754 | 1.4605 | 3.58076 | 2.452 |
| chr2 | 65498970 | 65499182 | 0.455832 | 1.11758 | 2.452 |
| chr12 | 57381277 | 57381715 | 2.32432 | 5.69895 | 2.452 |
| chr19 | 57259860 | 57260220 | 0.841106 | 2.06234 | 2.452 |
| chr7 | 148936520 | 148936809 | 2.01348 | 4.93702 | 2.452 |
| chr20 | 33732929 | 33733274 | 1.14533 | 2.80843 | 2.452 |
| chr14 | 22386044 | 22386344 | 1.35803 | 3.33002 | 2.452 |

| | | | | | |
|---|---|---|---|---|---|
| chr11 | 81021421 | 81021753 | 0.444446 | 1.08984 | 2.452 |
| chr19 | 17733166 | 17733414 | 1.03459 | 2.53698 | 2.452 |
| chr11 | 91038674 | 91039060 | 0.385926 | 0.946369 | 2.452 |
| chr2 | 36873448 | 36873719 | 1.07053 | 2.62515 | 2.452 |
| chr10 | 13141120 | 13141350 | 1.01563 | 2.49054 | 2.452 |
| chr2 | 178199768 | 178200053 | 1.32902 | 3.25903 | 2.452 |
| chr14 | 29409784 | 29410034 | 0.314585 | 0.77146 | 2.452 |
| chr8 | 104669374 | 104669660 | 0.446856 | 1.09585 | 2.452 |
| chr11 | 117103077 | 117103410 | 1.40617 | 3.44848 | 2.452 |
| chr2 | 149873972 | 149874318 | 1.58037 | 3.8758 | 2.452 |
| chr19 | 50871911 | 50872211 | 0.942217 | 2.31076 | 2.452 |
| chr7 | 16958002 | 16958227 | 0.916014 | 2.24659 | 2.453 |
| chr6 | 41743614 | 41744016 | 1.44393 | 3.54138 | 2.453 |
| chr12 | 113194601 | 113195161 | 0.434044 | 1.06456 | 2.453 |
| chr1 | 95212675 | 95212941 | 0.335464 | 0.822795 | 2.453 |
| chr3 | 7111252 | 7111548 | 0.384639 | 0.943425 | 2.453 |
| chr5 | 53715558 | 53715830 | 1.00465 | 2.46438 | 2.453 |
| chr5 | 93230488 | 93230803 | 0.912255 | 2.23776 | 2.453 |
| chr1 | 228856385 | 228856804 | 0.557164 | 1.36677 | 2.453 |
| chr13 | 27849798 | 27850043 | 0.330648 | 0.811116 | 2.453 |
| chr11 | 8388334 | 8388664 | 0.431797 | 1.05926 | 2.453 |
| chr8 | 69087490 | 69087708 | 0.627749 | 1.54 | 2.453 |
| chr1 | 172711752 | 172712075 | 1.68883 | 4.14308 | 2.453 |
| chr12 | 72354482 | 72354712 | 0.435095 | 1.06748 | 2.453 |
| chr12 | 64233249 | 64233557 | 1.26403 | 3.10136 | 2.454 |
| chr11 | 65990122 | 65990558 | 2.78006 | 6.82113 | 2.454 |
| chr16 | 54946301 | 54946521 | 0.470014 | 1.15323 | 2.454 |
| chr1 | 230033032 | 230033270 | 0.512724 | 1.25807 | 2.454 |
| chr17 | 46114258 | 46114725 | 1.56737 | 3.84597 | 2.454 |
| chr10 | 13113461 | 13113752 | 0.488026 | 1.19751 | 2.454 |
| chr19 | 10966869 | 10967155 | 1.06628 | 2.61644 | 2.454 |
| chr11 | 61334925 | 61335385 | 1.73236 | 4.25091 | 2.454 |
| chr6 | 158641892 | 158642193 | 2.3842 | 5.85058 | 2.454 |
| chr11 | 41854085 | 41854504 | 0.478988 | 1.17541 | 2.454 |
| chr8 | 4091932 | 4092342 | 0.544922 | 1.33723 | 2.454 |
| chr1 | 47000783 | 47001125 | 1.12693 | 2.76554 | 2.454 |
| chr15 | 85874375 | 85874687 | 2.13208 | 5.23229 | 2.454 |
| chr13 | 45968145 | 45968402 | 1.04049 | 2.55346 | 2.454 |
| chr9 | 90788321 | 90788683 | 2.05285 | 5.03791 | 2.454 |
| chr3 | 57896806 | 57897018 | 0.929545 | 2.28129 | 2.454 |
| chrX | 9853148 | 9853508 | 1.28252 | 3.14777 | 2.454 |
| chr1 | 165753382 | 165753639 | 0.482238 | 1.18363 | 2.454 |
| chr5 | 109745863 | 109746129 | 0.390028 | 0.957355 | 2.455 |
| chr14 | 75079071 | 75079672 | 2.47577 | 6.07701 | 2.455 |
| chr11 | 66935764 | 66935999 | 0.880762 | 2.16192 | 2.455 |
| chr15 | 53076038 | 53076526 | 2.00845 | 4.92998 | 2.455 |
| chr10 | 115439339 | 115439691 | 0.842521 | 2.06809 | 2.455 |
| chr14 | 69970457 | 69970911 | 1.14711 | 2.8159 | 2.455 |
| chr9 | 131905227 | 131905593 | 1.90588 | 4.67864 | 2.455 |
| chr11 | 2023326 | 2023544 | 0.902906 | 2.21654 | 2.455 |
| chr16 | 51494956 | 51495450 | 0.495689 | 1.21688 | 2.455 |
| chr12 | 111264469 | 111264855 | 1.48959 | 3.65693 | 2.455 |
| chr22 | 31478680 | 31479070 | 0.953452 | 2.34074 | 2.455 |
| chr1 | 62973906 | 62974164 | 1.06598 | 2.61709 | 2.455 |
| chr12 | 50917698 | 50917916 | 0.689489 | 1.69278 | 2.455 |
| chr15 | 89089863 | 89090127 | 2.12125 | 5.20811 | 2.455 |
| chr3 | 134097273 | 134097497 | 1.01273 | 2.48676 | 2.456 |
| chr13 | 32884143 | 32884597 | 0.738893 | 1.81442 | 2.456 |
| chr8 | 15098031 | 15098250 | 0.487032 | 1.19604 | 2.456 |
| chr4 | 150835704 | 150836021 | 1.44347 | 3.54486 | 2.456 |
| chr21 | 33942092 | 33942424 | 1.44911 | 3.55876 | 2.456 |

| | | | | | |
|---|---|---|---|---|---|
| chr10 | 17472756 | 17473109 | 1.90348 | 4.67465 | 2.456 |
| chr7 | 127775688 | 127776096 | 1.77474 | 4.35862 | 2.456 |
| chr7 | 94712570 | 94712834 | 0.479894 | 1.1786 | 2.456 |
| chr1 | 176932845 | 176933164 | 0.366073 | 0.899127 | 2.456 |
| chr8 | 66746811 | 66747139 | 1.49658 | 3.67609 | 2.456 |
| chr8 | 102513706 | 102513935 | 1.2927 | 3.17533 | 2.456 |
| chr11 | 66636160 | 66636678 | 1.27649 | 3.13558 | 2.456 |
| chr11 | 105110387 | 105110688 | 0.404303 | 0.993153 | 2.456 |
| chr3 | 9012645 | 9013068 | 2.09965 | 5.15779 | 2.457 |
| chr5 | 98396064 | 98396403 | 1.37472 | 3.37713 | 2.457 |
| chr13 | 21754913 | 21755139 | 0.884766 | 2.17352 | 2.457 |
| chr16 | 47867654 | 47867865 | 0.424283 | 1.04231 | 2.457 |
| chr8 | 76284723 | 76285008 | 1.01176 | 2.48556 | 2.457 |
| chr8 | 140035333 | 140035747 | 1.98944 | 4.8874 | 2.457 |
| chr7 | 1408722 | 1408930 | 1.01964 | 2.50494 | 2.457 |
| chr7 | 115820280 | 115820597 | 0.933102 | 2.29239 | 2.457 |
| chr10 | 44178947 | 44179215 | 1.11901 | 2.7492 | 2.457 |
| chr3 | 43226724 | 43226955 | 0.349653 | 0.859051 | 2.457 |
| chr12 | 6663269 | 6663691 | 1.23067 | 3.02361 | 2.457 |
| chr17 | 18886841 | 18887044 | 0.36394 | 0.894157 | 2.457 |
| chr1 | 240190301 | 240190659 | 1.61428 | 3.96611 | 2.457 |
| chr5 | 147836793 | 147837115 | 0.871271 | 2.14094 | 2.457 |
| chr7 | 2140006 | 2140246 | 0.423856 | 1.04155 | 2.457 |
| chr9 | 137341050 | 137341738 | 1.73665 | 4.26751 | 2.457 |
| chr14 | 60631935 | 60632379 | 1.10315 | 2.71103 | 2.458 |
| chr12 | 110348681 | 110349104 | 1.02585 | 2.5211 | 2.458 |
| chr4 | 84031610 | 84031870 | 1.26788 | 3.11591 | 2.458 |
| chr17 | 16189204 | 16189413 | 1.31966 | 3.24342 | 2.458 |
| chr7 | 106102931 | 106103157 | 0.619724 | 1.52326 | 2.458 |
| chr3 | 141030649 | 141030884 | 1.23977 | 3.04734 | 2.458 |
| chr4 | 103531195 | 103531500 | 0.916773 | 2.25369 | 2.458 |
| chr19 | 5823608 | 5823850 | 1.75888 | 4.32394 | 2.458 |
| chr12 | 55799438 | 55799719 | 0.409711 | 1.00729 | 2.459 |
| chr20 | 45381240 | 45381625 | 1.39314 | 3.42539 | 2.459 |
| chr20 | 16681741 | 16682366 | 0.70913 | 1.74364 | 2.459 |
| chr17 | 10675668 | 10676151 | 1.45429 | 3.57594 | 2.459 |
| chr1 | 238751189 | 238751427 | 0.364901 | 0.8973 | 2.459 |
| chr20 | 15957061 | 15957291 | 0.34211 | 0.841261 | 2.459 |
| chr20 | 61610773 | 61610984 | 0.412471 | 1.0143 | 2.459 |
| chr2 | 122353823 | 122354117 | 1.13127 | 2.7819 | 2.459 |
| chr7 | 140714719 | 140714985 | 1.40577 | 3.45699 | 2.459 |
| chr9 | 19471851 | 19472200 | 1.69346 | 4.1645 | 2.459 |
| chr6 | 82463617 | 82463918 | 1.45385 | 3.57528 | 2.459 |
| chr19 | 18049064 | 18049432 | 1.72909 | 4.25223 | 2.459 |
| chr19 | 5029691 | 5030037 | 1.8221 | 4.48101 | 2.459 |
| chr15 | 43030080 | 43030382 | 1.52085 | 3.74021 | 2.459 |
| chr3 | 36770635 | 36771296 | 0.696364 | 1.71266 | 2.459 |
| chr14 | 59911007 | 59911278 | 1.02404 | 2.51876 | 2.460 |
| chr11 | 17129309 | 17129745 | 1.12012 | 2.75515 | 2.460 |
| chr14 | 50309856 | 50310180 | 0.519245 | 1.27719 | 2.460 |
| chr7 | 110953075 | 110953281 | 0.524283 | 1.28959 | 2.460 |
| chr3 | 156192395 | 156192683 | 0.583125 | 1.43442 | 2.460 |
| chr14 | 74769672 | 74770030 | 1.35202 | 3.32585 | 2.460 |
| chr14 | 23113244 | 23113481 | 0.761467 | 1.87323 | 2.460 |
| chr12 | 123722677 | 123723003 | 1.78863 | 4.40016 | 2.460 |
| chr17 | 47772811 | 47773124 | 1.87579 | 4.61465 | 2.460 |
| chr2 | 42683371 | 42683574 | 0.460598 | 1.13313 | 2.460 |
| chr7 | 24421544 | 24421886 | 0.725771 | 1.78554 | 2.460 |
| chr12 | 50170125 | 50170490 | 1.50103 | 3.69301 | 2.460 |
| chr12 | 99563496 | 99563828 | 0.566302 | 1.3933 | 2.460 |
| chr13 | 49323897 | 49324173 | 1.48343 | 3.64975 | 2.460 |

| | | | | | |
|---|---|---|---|---|---|
| chr16 | 9694257 | 9694495 | 0.313158 | 0.770477 | 2.460 |
| chr4 | 148846340 | 148846571 | 1.08276 | 2.66406 | 2.460 |
| chr9 | 73209513 | 73209788 | 0.450107 | 1.10746 | 2.460 |
| chr16 | 75584202 | 75584416 | 0.973374 | 2.39496 | 2.460 |
| chr8 | 17722107 | 17722359 | 1.4279 | 3.51346 | 2.461 |
| chr5 | 79387636 | 79387872 | 0.938295 | 2.3089 | 2.461 |
| chr5 | 76733083 | 76733384 | 0.83871 | 2.06386 | 2.461 |
| chr17 | 4323878 | 4324435 | 1.41223 | 3.47526 | 2.461 |
| chr17 | 66684992 | 66685444 | 0.462502 | 1.13819 | 2.461 |
| chr16 | 87811158 | 87811895 | 1.64587 | 4.05064 | 2.461 |
| chr19 | 56015336 | 56015800 | 2.39102 | 5.88456 | 2.461 |
| chr12 | 13128644 | 13128956 | 1.55938 | 3.83788 | 2.461 |
| chr20 | 50912542 | 50912823 | 0.443207 | 1.09091 | 2.461 |
| chr10 | 14074471 | 14074907 | 1.64765 | 4.05561 | 2.461 |
| chr8 | 61325139 | 61325406 | 1.56875 | 3.86163 | 2.462 |
| chr17 | 27189080 | 27189296 | 0.464164 | 1.14259 | 2.462 |
| chr19 | 32004581 | 32004902 | 0.465068 | 1.1449 | 2.462 |
| chr7 | 73689606 | 73690053 | 1.45204 | 3.57473 | 2.462 |
| chr19 | 40306739 | 40307074 | 0.623582 | 1.53522 | 2.462 |
| chr15 | 100846598 | 100846868 | 0.607212 | 1.49493 | 2.462 |
| chr20 | 5549780 | 5550039 | 1.03641 | 2.55171 | 2.462 |
| chr6 | 16763316 | 16763565 | 1.19272 | 2.9367 | 2.462 |
| chr17 | 62106270 | 62106553 | 1.6106 | 3.96591 | 2.462 |
| chr10 | 112065996 | 112066221 | 0.883191 | 2.17485 | 2.462 |
| chr2 | 36364531 | 36364763 | 0.387657 | 0.954619 | 2.463 |
| chr20 | 200148 | 200449 | 0.810316 | 1.99576 | 2.463 |
| chr11 | 7168001 | 7168242 | 0.491712 | 1.21108 | 2.463 |
| chr9 | 12075174 | 12075412 | 0.521308 | 1.28402 | 2.463 |
| chr1 | 204332799 | 204333148 | 1.59704 | 3.93372 | 2.463 |
| chr10 | 71396716 | 71397129 | 0.887646 | 2.18639 | 2.463 |
| chr11 | 119252212 | 119252541 | 1.86252 | 4.58764 | 2.463 |
| chr6 | 797608 | 797836 | 0.932615 | 2.29718 | 2.463 |
| chr17 | 2947018 | 2947688 | 1.32006 | 3.25167 | 2.463 |
| chr2 | 7567468 | 7567753 | 0.858844 | 2.11558 | 2.463 |
| chr5 | 63259019 | 63259262 | 1.10332 | 2.71781 | 2.463 |
| chrX | 128812146 | 128812510 | 0.900012 | 2.21715 | 2.463 |
| chr2 | 33748458 | 33748693 | 0.456868 | 1.1255 | 2.464 |
| chr19 | 31408549 | 31408767 | 0.675995 | 1.66542 | 2.464 |
| chr7 | 70277451 | 70277833 | 0.739777 | 1.82261 | 2.464 |
| chr2 | 46691667 | 46691978 | 1.35578 | 3.34032 | 2.464 |
| chr4 | 152966946 | 152967241 | 1.38272 | 3.40725 | 2.464 |
| chr15 | 90283180 | 90283636 | 1.59719 | 3.93603 | 2.464 |
| chr7 | 16604540 | 16604989 | 1.10148 | 2.71452 | 2.464 |
| chr11 | 50382473 | 50382708 | 0.815786 | 2.01047 | 2.464 |
| chr15 | 74913376 | 74913794 | 1.93375 | 4.76587 | 2.465 |
| chr10 | 38041317 | 38041521 | 0.532531 | 1.31261 | 2.465 |
| chr1 | 156066380 | 156066744 | 1.94077 | 4.78377 | 2.465 |
| chr17 | 8286495 | 8287107 | 2.64588 | 6.5218 | 2.465 |
| chr2 | 26552920 | 26553239 | 1.88129 | 4.63721 | 2.465 |
| chr14 | 95898489 | 95898890 | 2.0105 | 4.95608 | 2.465 |
| chr9 | 80157854 | 80158068 | 0.71593 | 1.76493 | 2.465 |
| chr10 | 107592526 | 107592760 | 0.292325 | 0.720659 | 2.465 |
| chr17 | 782838 | 783235 | 1.10295 | 2.71909 | 2.465 |
| chr3 | 113693481 | 113693781 | 0.446424 | 1.10058 | 2.465 |
| chr6 | 104425265 | 104425475 | 0.355126 | 0.875507 | 2.465 |
| chr9 | 37415198 | 37415547 | 1.07993 | 2.66253 | 2.465 |
| chr11 | 134230163 | 134230519 | 1.76651 | 4.35531 | 2.465 |
| chr1 | 110042626 | 110042871 | 1.14506 | 2.82321 | 2.466 |
| chr11 | 19442622 | 19442862 | 0.457993 | 1.12922 | 2.466 |
| chr9 | 21624033 | 21624270 | 0.425194 | 1.04836 | 2.466 |
| chr7 | 141485658 | 141486200 | 1.3182 | 3.25017 | 2.466 |

| | | | | | |
|---|---|---|---|---|---|
| chr8 | 41386284 | 41386687 | 1.53143 | 3.77592 | 2.466 |
| chr17 | 15564992 | 15565207 | 0.377805 | 0.931585 | 2.466 |
| chr1 | 115817920 | 115818148 | 0.941346 | 2.32127 | 2.466 |
| chr17 | 39957877 | 39958186 | 1.76545 | 4.3536 | 2.466 |
| chr10 | 93616444 | 93616692 | 1.0073 | 2.48403 | 2.466 |
| chr15 | 72518975 | 72519266 | 0.905755 | 2.23368 | 2.466 |
| chr8 | 28479715 | 28480057 | 1.71762 | 4.23598 | 2.466 |
| chr11 | 34018554 | 34018889 | 1.1289 | 2.78418 | 2.466 |
| chrX | 118603010 | 118603391 | 1.26216 | 3.11293 | 2.466 |
| chr11 | 40328546 | 40328841 | 1.02483 | 2.52781 | 2.467 |
| chr10 | 25989735 | 25990090 | 1.07876 | 2.66108 | 2.467 |
| chr16 | 46588333 | 46588542 | 0.917365 | 2.26296 | 2.467 |
| chr7 | 22599521 | 22599864 | 1.63747 | 4.03938 | 2.467 |
| chr16 | 30053151 | 30053538 | 0.747612 | 1.84433 | 2.467 |
| chr8 | 140290168 | 140290410 | 0.680584 | 1.67909 | 2.467 |
| chr6 | 166511404 | 166511714 | 1.77115 | 4.36976 | 2.467 |
| chr16 | 56366559 | 56366943 | 1.53093 | 3.77718 | 2.467 |
| chr1 | 181825381 | 181825611 | 0.40933 | 1.00999 | 2.467 |
| chr13 | 20702807 | 20703119 | 0.850007 | 2.09735 | 2.467 |
| chr10 | 131090462 | 131090762 | 0.492657 | 1.21568 | 2.468 |
| chr20 | 37380070 | 37380362 | 0.910254 | 2.2462 | 2.468 |
| chr14 | 20344435 | 20344835 | 0.722543 | 1.78308 | 2.468 |
| chr2 | 46698469 | 46698676 | 0.478636 | 1.18122 | 2.468 |
| chr22 | 23297958 | 23298431 | 1.08755 | 2.68396 | 2.468 |
| chr12 | 107798823 | 107799378 | 1.43415 | 3.53976 | 2.468 |
| chr13 | 24289946 | 24290333 | 1.05716 | 2.6093 | 2.468 |
| chr16 | 24048073 | 24048370 | 1.13638 | 2.80508 | 2.468 |
| chr1 | 85237994 | 85238392 | 1.4551 | 3.59226 | 2.469 |
| chr10 | 115280847 | 115281420 | 2.337 | 5.76955 | 2.469 |
| chr1 | 147196063 | 147196393 | 0.639036 | 1.57766 | 2.469 |
| chr15 | 35380586 | 35380847 | 0.410948 | 1.01464 | 2.469 |
| chr1 | 21815276 | 21815550 | 0.662328 | 1.63536 | 2.469 |
| chr10 | 101390287 | 101390578 | 0.752972 | 1.85921 | 2.469 |
| chr22 | 32266892 | 32267102 | 0.67893 | 1.67643 | 2.469 |
| chr12 | 49367878 | 49368317 | 1.07462 | 2.6535 | 2.469 |
| chr20 | 40321863 | 40322250 | 0.849216 | 2.09692 | 2.469 |
| chr12 | 42078930 | 42079142 | 0.765822 | 1.89106 | 2.469 |
| chr8 | 48651463 | 48651767 | 1.41787 | 3.50123 | 2.469 |
| chr12 | 50449520 | 50450344 | 1.43498 | 3.54381 | 2.470 |
| chr5 | 8689930 | 8690206 | 0.443654 | 1.09565 | 2.470 |
| chr5 | 169643166 | 169643647 | 1.73388 | 4.28208 | 2.470 |
| chr1 | 218524946 | 218525175 | 0.708648 | 1.75024 | 2.470 |
| chr6 | 168551501 | 168551915 | 1.28158 | 3.16532 | 2.470 |
| chr5 | 51787090 | 51787387 | 0.499746 | 1.23436 | 2.470 |
| chr12 | 130578786 | 130579029 | 0.504255 | 1.24556 | 2.470 |
| chr18 | 54230815 | 54231145 | 0.662196 | 1.63569 | 2.470 |
| chr3 | 53168775 | 53169051 | 1.68146 | 4.15353 | 2.470 |
| chr1 | 203117322 | 203117569 | 0.609036 | 1.50452 | 2.470 |
| chr5 | 43306127 | 43306424 | 0.978199 | 2.41655 | 2.470 |
| chr1 | 209775374 | 209775720 | 1.0632 | 2.62655 | 2.470 |
| chr16 | 21113370 | 21113661 | 1.15071 | 2.8428 | 2.470 |
| chr17 | 38771760 | 38772232 | 1.47697 | 3.64907 | 2.471 |
| chr1 | 161179611 | 161179933 | 2.13016 | 5.26317 | 2.471 |
| chr1 | 231923438 | 231923666 | 0.359838 | 0.889135 | 2.471 |
| chr10 | 116636898 | 116637133 | 1.5628 | 3.86158 | 2.471 |
| chr14 | 47578794 | 47579030 | 0.315822 | 0.780423 | 2.471 |
| chr7 | 98477382 | 98477592 | 0.587112 | 1.45085 | 2.471 |
| chr9 | 72547865 | 72548148 | 1.24187 | 3.06897 | 2.471 |
| chr20 | 43709554 | 43709768 | 0.923825 | 2.28313 | 2.471 |
| chr19 | 43389092 | 43389494 | 1.60222 | 3.95976 | 2.471 |
| chr20 | 43767526 | 43767817 | 1.14862 | 2.83874 | 2.471 |

| | | | | | |
|---|---|---|---|---|---|
| chr1 | 212208337 | 212208825 | 1.85687 | 4.58935 | 2.472 |
| chr19 | 34232594 | 34232810 | 0.515869 | 1.27508 | 2.472 |
| chr7 | 88102511 | 88102725 | 0.394316 | 0.974636 | 2.472 |
| chr8 | 77080180 | 77080390 | 0.543477 | 1.34334 | 2.472 |
| chr3 | 147944172 | 147944411 | 0.578706 | 1.43042 | 2.472 |
| chr13 | 66559843 | 66560125 | 0.862833 | 2.13273 | 2.472 |
| chr15 | 75871545 | 75871927 | 1.24157 | 3.06896 | 2.472 |
| chr14 | 102094787 | 102095293 | 1.42856 | 3.53122 | 2.472 |
| chr9 | 37594554 | 37594759 | 1.26127 | 3.11787 | 2.472 |
| chr5 | 68389482 | 68389813 | 1.58421 | 3.91637 | 2.472 |
| chr5 | 114970416 | 114970682 | 1.91807 | 4.74187 | 2.472 |
| chr12 | 21449687 | 21450108 | 1.24682 | 3.08252 | 2.472 |
| chr11 | 3862753 | 3863123 | 1.52129 | 3.76112 | 2.472 |
| chr2 | 74942158 | 74942547 | 1.99609 | 4.93504 | 2.472 |
| chr11 | 128701111 | 128701387 | 1.07339 | 2.65386 | 2.472 |
| chr1 | 238451763 | 238452007 | 0.727194 | 1.79806 | 2.473 |
| chr1 | 248159360 | 248159585 | 0.390067 | 0.96448 | 2.473 |
| chr7 | 38375171 | 38375639 | 1.4413 | 3.56421 | 2.473 |
| chrX | 20271017 | 20271346 | 1.42804 | 3.53147 | 2.473 |
| chr8 | 59634291 | 59634497 | 0.7466 | 1.84647 | 2.473 |
| chr12 | 88446518 | 88446853 | 1.46501 | 3.62336 | 2.473 |
| chr17 | 77944500 | 77944854 | 1.28417 | 3.17611 | 2.473 |
| chr11 | 74111798 | 74112192 | 0.450719 | 1.1148 | 2.473 |
| chr3 | 182902452 | 182902736 | 0.741797 | 1.8348 | 2.473 |
| chr3 | 65034194 | 65034400 | 0.448917 | 1.1104 | 2.474 |
| chr4 | 85663811 | 85664060 | 0.311309 | 0.770035 | 2.474 |
| chr12 | 107480330 | 107480666 | 0.47412 | 1.17276 | 2.474 |
| chr19 | 49291323 | 49291731 | 2.02849 | 5.01759 | 2.474 |
| chr19 | 41650209 | 41650733 | 1.86577 | 4.61527 | 2.474 |
| chr3 | 160889503 | 160889734 | 0.587344 | 1.45292 | 2.474 |
| chr5 | 59351990 | 59352505 | 0.472418 | 1.16865 | 2.474 |
| chr12 | 125391230 | 125391644 | 1.90999 | 4.72498 | 2.474 |
| chr16 | 87905257 | 87905485 | 1.03505 | 2.5606 | 2.474 |
| chr9 | 114522405 | 114522663 | 1.00011 | 2.47416 | 2.474 |
| chr14 | 52462908 | 52463169 | 1.14696 | 2.83783 | 2.474 |
| chr10 | 79479363 | 79479694 | 2.45221 | 6.06767 | 2.474 |
| chr4 | 88769869 | 88770444 | 0.805406 | 1.99291 | 2.474 |
| chr16 | 58340241 | 58340560 | 1.07899 | 2.66997 | 2.475 |
| chr9 | 116169220 | 116169574 | 1.76588 | 4.36981 | 2.475 |
| chr5 | 134889408 | 134889915 | 0.900796 | 2.22912 | 2.475 |
| chr15 | 52180884 | 52181093 | 1.16061 | 2.87216 | 2.475 |
| chr11 | 65756453 | 65757042 | 1.62104 | 4.01217 | 2.475 |
| chr17 | 36579887 | 36580427 | 2.28786 | 5.66289 | 2.475 |
| chr2 | 84541945 | 84542337 | 2.11421 | 5.23314 | 2.475 |
| chr7 | 16840795 | 16841025 | 1.06258 | 2.63033 | 2.475 |
| chr8 | 130036224 | 130036465 | 1.27764 | 3.16278 | 2.475 |
| chr6 | 112364178 | 112364534 | 1.84976 | 4.57916 | 2.476 |
| chr15 | 101628908 | 101629421 | 2.61815 | 6.48146 | 2.476 |
| chr8 | 42177215 | 42177505 | 1.57119 | 3.88976 | 2.476 |
| chr10 | 29136326 | 29136582 | 0.554817 | 1.37373 | 2.476 |
| chr10 | 11476445 | 11476700 | 1.18178 | 2.92637 | 2.476 |
| chr1 | 41077948 | 41078306 | 1.6019 | 3.96685 | 2.476 |
| chr3 | 117716308 | 117716715 | 1.58495 | 3.92491 | 2.476 |
| chr1 | 225741984 | 225742199 | 0.738943 | 1.82998 | 2.476 |
| chr17 | 42021306 | 42021935 | 1.79344 | 4.44174 | 2.477 |
| chr13 | 25861490 | 25861895 | 1.69217 | 4.19125 | 2.477 |
| chr2 | 202228627 | 202228991 | 1.35199 | 3.34867 | 2.477 |
| chr5 | 21810880 | 21811091 | 0.754444 | 1.86865 | 2.477 |
| chr3 | 182335958 | 182336220 | 1.22543 | 3.03524 | 2.477 |
| chr10 | 47665597 | 47665853 | 0.934123 | 2.31376 | 2.477 |
| chr9 | 114784543 | 114784752 | 0.987205 | 2.44533 | 2.477 |

| | | | | | |
|---|---|---|---|---|---|
| chr17 | 42287618 | 42288186 | 2.69519 | 6.6761 | 2.477 |
| chr1 | 41733837 | 41734186 | 1.74259 | 4.31673 | 2.477 |
| chr12 | 104609352 | 104609729 | 1.59669 | 3.95535 | 2.477 |
| chr5 | 67907317 | 67907578 | 0.467609 | 1.15841 | 2.477 |
| chr2 | 72006103 | 72006306 | 0.742561 | 1.83968 | 2.477 |
| chr18 | 12886557 | 12886834 | 1.07997 | 2.67573 | 2.478 |
| chr8 | 79885015 | 79885224 | 0.647671 | 1.60471 | 2.478 |
| chr6 | 153185021 | 153185380 | 0.810708 | 2.00869 | 2.478 |
| chr7 | 125451893 | 125452290 | 0.4226 | 1.04712 | 2.478 |
| chr3 | 190602070 | 190602320 | 1.15349 | 2.85813 | 2.478 |
| chr20 | 23042789 | 23043099 | 1.46024 | 3.61832 | 2.478 |
| chr7 | 116238406 | 116238653 | 0.653799 | 1.62018 | 2.478 |
| chr14 | 23027445 | 23027811 | 0.63944 | 1.58461 | 2.478 |
| chr6 | 42182565 | 42182844 | 0.871362 | 2.15937 | 2.478 |
| chr1 | 80683132 | 80683334 | 0.880025 | 2.18088 | 2.478 |
| chr6 | 2989226 | 2989581 | 1.51762 | 3.76109 | 2.478 |
| chr12 | 66136044 | 66136293 | 0.636981 | 1.57865 | 2.478 |
| chr15 | 98971549 | 98972246 | 1.52764 | 3.78617 | 2.478 |
| chr11 | 35394978 | 35395242 | 0.547992 | 1.35828 | 2.479 |
| chr3 | 167764988 | 167765386 | 1.07687 | 2.6692 | 2.479 |
| chr16 | 31027401 | 31027662 | 0.461256 | 1.14331 | 2.479 |
| chr17 | 74404682 | 74405036 | 2.2391 | 5.55015 | 2.479 |
| chr19 | 51831007 | 51831281 | 0.67492 | 1.67308 | 2.479 |
| chr15 | 49267924 | 49268329 | 1.48896 | 3.69124 | 2.479 |
| chr6 | 138599468 | 138599898 | 0.888692 | 2.20316 | 2.479 |
| chr2 | 7177881 | 7178232 | 2.14498 | 5.31796 | 2.479 |
| chr12 | 56583233 | 56583707 | 2.42034 | 6.00112 | 2.479 |
| chr10 | 36025744 | 36026027 | 0.517632 | 1.28353 | 2.480 |
| chr19 | 42951007 | 42951309 | 0.383363 | 0.950621 | 2.480 |
| chrX | 107244711 | 107245070 | 0.803241 | 1.9918 | 2.480 |
| chr5 | 99854187 | 99854401 | 0.787044 | 1.95169 | 2.480 |
| chr1 | 17600693 | 17600995 | 1.66852 | 4.1376 | 2.480 |
| chr2 | 173796742 | 173796966 | 1.23094 | 3.05254 | 2.480 |
| chr11 | 29181202 | 29181462 | 1.30916 | 3.24664 | 2.480 |
| chr12 | 101941840 | 101942116 | 0.385121 | 0.95509 | 2.480 |
| chr2 | 239165127 | 239165643 | 2.25966 | 5.60389 | 2.480 |
| chr1 | 238177925 | 238178141 | 0.430607 | 1.06792 | 2.480 |
| chr1 | 79083269 | 79083681 | 1.32619 | 3.28907 | 2.480 |
| chr6 | 4844422 | 4844737 | 1.01317 | 2.51276 | 2.480 |
| chr10 | 107562979 | 107563384 | 0.541625 | 1.3433 | 2.480 |
| chrX | 135793408 | 135793612 | 0.864961 | 2.14529 | 2.480 |
| chr14 | 58633037 | 58633453 | 1.39754 | 3.46672 | 2.481 |
| chr18 | 19577430 | 19577947 | 0.908854 | 2.25453 | 2.481 |
| chr1 | 160749752 | 160750278 | 1.08277 | 2.68599 | 2.481 |
| chr14 | 94471789 | 94472235 | 0.755876 | 1.87513 | 2.481 |
| chr7 | 103199940 | 103200171 | 0.403984 | 1.00222 | 2.481 |
| chr10 | 60051363 | 60051679 | 1.33175 | 3.30392 | 2.481 |
| chr15 | 71740962 | 71741292 | 1.7343 | 4.30278 | 2.481 |
| chr20 | 47645168 | 47645583 | 0.646231 | 1.60329 | 2.481 |
| chr1 | 65896811 | 65897121 | 0.930663 | 2.30898 | 2.481 |
| chr19 | 41104451 | 41104867 | 2.72067 | 6.75009 | 2.481 |
| chr22 | 43176079 | 43176409 | 1.53256 | 3.80246 | 2.481 |
| chr1 | 155017433 | 155017705 | 1.41477 | 3.51042 | 2.481 |
| chr7 | 22800227 | 22800447 | 0.513931 | 1.27525 | 2.481 |
| chr12 | 53455559 | 53455859 | 0.972781 | 2.41386 | 2.481 |
| chr18 | 32241285 | 32241889 | 1.2157 | 3.01688 | 2.482 |
| chr19 | 3108674 | 3109057 | 1.68513 | 4.18235 | 2.482 |
| chr17 | 59378716 | 59378974 | 0.519873 | 1.29032 | 2.482 |
| chr17 | 53434083 | 53434478 | 1.66238 | 4.12654 | 2.482 |
| chr10 | 84950415 | 84950708 | 0.42692 | 1.05976 | 2.482 |
| chr2 | 236015840 | 236016051 | 1.57367 | 3.90641 | 2.482 |

| | | | | | |
|---|---|---|---|---|---|
| chr20 | 46924413 | 46924620 | 1.09141 | 2.70932 | 2.482 |
| chr3 | 34021854 | 34022114 | 0.942096 | 2.3387 | 2.482 |
| chr16 | 27701945 | 27702265 | 0.959317 | 2.38183 | 2.483 |
| chr14 | 77605942 | 77606388 | 1.86298 | 4.62569 | 2.483 |
| chr8 | 131713931 | 131714293 | 0.545681 | 1.35492 | 2.483 |
| chr5 | 134240213 | 134240688 | 1.69471 | 4.20814 | 2.483 |
| chr2 | 11558124 | 11558446 | 1.36311 | 3.38475 | 2.483 |
| chr4 | 81990932 | 81991294 | 1.57339 | 3.90724 | 2.483 |
| chr7 | 43147940 | 43148339 | 1.7736 | 4.40451 | 2.483 |
| chr3 | 112361782 | 112362067 | 1.94041 | 4.81888 | 2.483 |
| chr8 | 53320262 | 53320485 | 1.46108 | 3.62853 | 2.483 |
| chr19 | 35464043 | 35464441 | 2.30658 | 5.72834 | 2.483 |
| chr9 | 139155811 | 139156117 | 0.677533 | 1.68265 | 2.484 |
| chr1 | 10010369 | 10010750 | 1.60199 | 3.97868 | 2.484 |
| chr17 | 69773050 | 69773369 | 0.438938 | 1.09014 | 2.484 |
| chr14 | 91752865 | 91753225 | 1.37846 | 3.42368 | 2.484 |
| chr15 | 68545296 | 68545596 | 1.57324 | 3.90749 | 2.484 |
| chr7 | 104524102 | 104524462 | 1.87218 | 4.6501 | 2.484 |
| chr17 | 77697091 | 77697324 | 0.558156 | 1.38637 | 2.484 |
| chr5 | 131841872 | 131842139 | 1.03644 | 2.5745 | 2.484 |
| chr9 | 35071946 | 35072223 | 0.656119 | 1.6298 | 2.484 |
| chr3 | 50311736 | 50312186 | 1.73935 | 4.32113 | 2.484 |
| chr5 | 71604225 | 71604569 | 1.78435 | 4.43299 | 2.484 |
| chr1 | 919542 | 919901 | 2.4012 | 5.96571 | 2.484 |
| chr11 | 25966889 | 25967105 | 0.483997 | 1.20248 | 2.484 |
| chr17 | 6182922 | 6183369 | 0.980592 | 2.43629 | 2.485 |
| chr11 | 19138943 | 19139209 | 1.24592 | 3.09555 | 2.485 |
| chr17 | 72846174 | 72846544 | 0.724781 | 1.80083 | 2.485 |
| chr7 | 32530492 | 32530720 | 0.673247 | 1.67282 | 2.485 |
| chr1 | 247553398 | 247553748 | 2.12415 | 5.27797 | 2.485 |
| chr12 | 7296920 | 7297407 | 0.879374 | 2.18502 | 2.485 |
| chr17 | 6339242 | 6339696 | 1.59191 | 3.95635 | 2.485 |
| chr9 | 125554750 | 125555210 | 0.553687 | 1.37612 | 2.485 |
| chr15 | 71407496 | 71407957 | 2.39385 | 5.94988 | 2.485 |
| chr16 | 51789048 | 51789255 | 0.448477 | 1.11471 | 2.486 |
| chr19 | 4139760 | 4140055 | 1.83432 | 4.55944 | 2.486 |
| chr4 | 77973414 | 77973801 | 1.1478 | 2.85312 | 2.486 |
| chrX | 11962688 | 11962912 | 1.0568 | 2.62706 | 2.486 |
| chrX | 48828727 | 48829167 | 1.19927 | 2.98123 | 2.486 |
| chr15 | 70198398 | 70198815 | 0.385117 | 0.9574 | 2.486 |
| chr3 | 158152270 | 158152568 | 1.01076 | 2.51281 | 2.486 |
| chr16 | 13154846 | 13155051 | 0.367302 | 0.91315 | 2.486 |
| chr6 | 144263289 | 144263683 | 1.32592 | 3.29668 | 2.486 |
| chr6 | 126978719 | 126979006 | 1.13293 | 2.81689 | 2.486 |
| chr18 | 30510745 | 30510979 | 0.861435 | 2.14186 | 2.486 |
| chr1 | 158741811 | 158742080 | 0.390503 | 0.97099 | 2.487 |
| chr12 | 7315197 | 7315611 | 1.9119 | 4.75397 | 2.487 |
| chr11 | 60666813 | 60667190 | 1.85702 | 4.6179 | 2.487 |
| chr11 | 27325979 | 27326379 | 0.902402 | 2.24408 | 2.487 |
| chr14 | 83094404 | 83094641 | 0.434846 | 1.08138 | 2.487 |
| chr2 | 187168378 | 187168608 | 0.42008 | 1.04469 | 2.487 |
| chr20 | 36919732 | 36920057 | 1.73362 | 4.31143 | 2.487 |
| chr9 | 22422450 | 22422652 | 0.303832 | 0.755745 | 2.487 |
| chr5 | 171906014 | 171906418 | 1.61385 | 4.01436 | 2.487 |
| chr2 | 197615746 | 197616119 | 1.52 | 3.78101 | 2.488 |
| chr21 | 43729815 | 43730158 | 1.25946 | 3.13295 | 2.488 |
| chr1 | 166327416 | 166327824 | 0.446635 | 1.11109 | 2.488 |
| chr5 | 157502234 | 157502599 | 1.72231 | 4.28458 | 2.488 |
| chr12 | 51380758 | 51381123 | 1.37391 | 3.41789 | 2.488 |
| chr15 | 59467020 | 59467320 | 1.22891 | 3.05725 | 2.488 |
| chr7 | 82073516 | 82073762 | 0.533699 | 1.32774 | 2.488 |

| | | | | | |
|---|---|---|---|---|---|
| chr12 | 57670776 | 57671244 | 1.2736 | 3.16849 | 2.488 |
| chr7 | 93717041 | 93717405 | 0.968726 | 2.41008 | 2.488 |
| chr14 | 75412742 | 75413479 | 1.16567 | 2.90017 | 2.488 |
| chr12 | 103424307 | 103424527 | 0.39676 | 0.987265 | 2.488 |
| chr17 | 77078607 | 77079147 | 2.23635 | 5.56479 | 2.488 |
| chr7 | 104794168 | 104794443 | 0.393499 | 0.979189 | 2.488 |
| chr22 | 25778833 | 25779047 | 0.828043 | 2.06056 | 2.488 |
| chr11 | 17994632 | 17994899 | 1.25181 | 3.11524 | 2.489 |
| chr10 | 110967160 | 110967391 | 0.438232 | 1.0906 | 2.489 |
| chr17 | 72404138 | 72404714 | 0.765641 | 1.90553 | 2.489 |
| chr2 | 38118013 | 38118435 | 1.81028 | 4.50544 | 2.489 |
| chr9 | 97886893 | 97887212 | 1.02694 | 2.55589 | 2.489 |
| chrX | 100150432 | 100150650 | 0.934646 | 2.3262 | 2.489 |
| chr16 | 3740460 | 3740798 | 1.79622 | 4.47078 | 2.489 |
| chr11 | 67407427 | 67407812 | 1.65765 | 4.12596 | 2.489 |
| chr11 | 111712202 | 111712509 | 1.62509 | 4.04499 | 2.489 |
| chr8 | 142394626 | 142395067 | 1.07954 | 2.68723 | 2.489 |
| chr21 | 45230686 | 45231022 | 2.13843 | 5.32335 | 2.489 |
| chr4 | 122405747 | 122406122 | 0.782048 | 1.94685 | 2.489 |
| chr6 | 53275513 | 53275784 | 1.61012 | 4.00851 | 2.490 |
| chr4 | 117253301 | 117253506 | 0.381656 | 0.950232 | 2.490 |
| chr12 | 107796154 | 107796542 | 1.54074 | 3.83616 | 2.490 |
| chr21 | 34312798 | 34313214 | 1.38852 | 3.45721 | 2.490 |
| chr3 | 38704156 | 38704457 | 1.06355 | 2.64815 | 2.490 |
| chr3 | 197669858 | 197670210 | 0.985266 | 2.45325 | 2.490 |
| chr9 | 71779766 | 71780170 | 0.471502 | 1.17401 | 2.490 |
| chr3 | 65718053 | 65718423 | 1.18992 | 2.96294 | 2.490 |
| chr17 | 27570810 | 27571040 | 1.18383 | 2.94791 | 2.490 |
| chr15 | 49103736 | 49103988 | 0.41647 | 1.03713 | 2.490 |
| chr8 | 86291623 | 86291835 | 0.487799 | 1.21479 | 2.490 |
| chr10 | 112360327 | 112360590 | 1.44718 | 3.60423 | 2.491 |
| chr2 | 30784086 | 30784311 | 0.428679 | 1.06768 | 2.491 |
| chr16 | 57829325 | 57829612 | 0.97161 | 2.41995 | 2.491 |
| chr18 | 45550220 | 45550508 | 1.23703 | 3.08116 | 2.491 |
| chr14 | 24196826 | 24197047 | 1.32712 | 3.30569 | 2.491 |
| chr1 | 159074778 | 159075332 | 0.379443 | 0.945166 | 2.491 |
| chr15 | 68155658 | 68156168 | 1.70948 | 4.2582 | 2.491 |
| chr1 | 26602055 | 26602553 | 0.950923 | 2.36883 | 2.491 |
| chr4 | 1731533 | 1731799 | 1.43726 | 3.58047 | 2.491 |
| chrX | 45684051 | 45684273 | 1.33589 | 3.32801 | 2.491 |
| chr7 | 107786655 | 107787297 | 1.42825 | 3.55824 | 2.491 |
| chr6 | 158635241 | 158635489 | 1.37632 | 3.42893 | 2.491 |
| chr22 | 28405306 | 28405735 | 1.76009 | 4.38512 | 2.491 |
| chr12 | 53775718 | 53775953 | 1.27827 | 3.18473 | 2.491 |
| chr8 | 55344318 | 55344590 | 1.53191 | 3.81677 | 2.492 |
| chr1 | 48559398 | 48559811 | 1.61549 | 4.02505 | 2.492 |
| chr17 | 59486754 | 59487322 | 1.52043 | 3.78828 | 2.492 |
| chr9 | 34985313 | 34985664 | 1.76315 | 4.39323 | 2.492 |
| chr19 | 46367283 | 46367719 | 1.80015 | 4.48548 | 2.492 |
| chr5 | 166604507 | 166605123 | 0.379254 | 0.945006 | 2.492 |
| chr1 | 87791716 | 87792100 | 0.933059 | 2.32504 | 2.492 |
| chr1 | 32800769 | 32801222 | 0.970711 | 2.41889 | 2.492 |
| chr1 | 215380555 | 215380787 | 0.380322 | 0.947862 | 2.492 |
| chr1 | 22029820 | 22030064 | 0.860145 | 2.14377 | 2.492 |
| chr9 | 6054561 | 6054854 | 1.56031 | 3.88882 | 2.492 |
| chr10 | 22392660 | 22392958 | 0.689703 | 1.71903 | 2.492 |
| chr11 | 58938824 | 58939651 | 1.7209 | 4.28926 | 2.492 |
| chr1 | 18573055 | 18573331 | 0.753233 | 1.8775 | 2.493 |
| chr20 | 20064424 | 20064791 | 0.832732 | 2.07569 | 2.493 |
| chr2 | 1746817 | 1747076 | 0.740256 | 1.84519 | 2.493 |
| chr15 | 72094795 | 72095394 | 0.88445 | 2.20485 | 2.493 |

| | | | | | |
|---|---|---|---|---|---|
| chr10 | 118934373 | 118934799 | 1.50632 | 3.75552 | 2.493 |
| chr14 | 29943844 | 29944182 | 0.44953 | 1.12076 | 2.493 |
| chr16 | 67597137 | 67597442 | 0.515952 | 1.28638 | 2.493 |
| chr22 | 50675651 | 50676024 | 0.998904 | 2.49069 | 2.493 |
| chr1 | 5697331 | 5697573 | 1.38298 | 3.44837 | 2.493 |
| chr16 | 30913456 | 30913902 | 1.58596 | 3.9545 | 2.493 |
| chr6 | 53302431 | 53302652 | 0.864853 | 2.15648 | 2.493 |
| chr1 | 26928912 | 26929168 | 1.10989 | 2.76748 | 2.493 |
| chrX | 14050571 | 14050836 | 1.07229 | 2.67372 | 2.493 |
| chr12 | 44386128 | 44386452 | 0.495914 | 1.23664 | 2.494 |
| chr4 | 39697272 | 39697617 | 1.32519 | 3.30463 | 2.494 |
| chr17 | 68243241 | 68243534 | 0.623918 | 1.55591 | 2.494 |
| chrX | 94213041 | 94213282 | 0.336184 | 0.83841 | 2.494 |
| chr8 | 144467138 | 144467717 | 1.56791 | 3.91024 | 2.494 |
| chr17 | 50206026 | 50206323 | 0.453003 | 1.12978 | 2.494 |
| chr14 | 82084405 | 82084669 | 0.388125 | 0.968009 | 2.494 |
| chr10 | 120909282 | 120909484 | 0.638668 | 1.59291 | 2.494 |
| chr4 | 174415501 | 174415893 | 1.37652 | 3.43372 | 2.494 |
| chr1 | 19717853 | 19718087 | 0.646149 | 1.61184 | 2.495 |
| chr5 | 131573973 | 131574436 | 1.33025 | 3.31835 | 2.495 |
| chr3 | 196696722 | 196697159 | 1.45712 | 3.63498 | 2.495 |
| chr18 | 60253475 | 60253776 | 1.16151 | 2.89755 | 2.495 |
| chr22 | 22001051 | 22001299 | 1.30879 | 3.26503 | 2.495 |
| chr5 | 16075904 | 16076132 | 0.411521 | 1.02663 | 2.495 |
| chr1 | 70598675 | 70598961 | 1.07467 | 2.6811 | 2.495 |
| chr5 | 151941055 | 151941482 | 0.479695 | 1.19679 | 2.495 |
| chr5 | 11642489 | 11642759 | 0.402128 | 1.00333 | 2.495 |
| chr4 | 97091345 | 97091668 | 0.637928 | 1.59167 | 2.495 |
| chr10 | 99294507 | 99295010 | 1.40756 | 3.51199 | 2.495 |
| chr22 | 32598408 | 32598724 | 0.797172 | 1.98904 | 2.495 |
| chr20 | 35515591 | 35515936 | 1.81384 | 4.5259 | 2.495 |
| chr9 | 136063598 | 136063919 | 0.777117 | 1.93919 | 2.495 |
| chr5 | 2151248 | 2151782 | 1.14339 | 2.85323 | 2.495 |
| chr6 | 53820939 | 53821230 | 1.54664 | 3.85956 | 2.495 |
| chr7 | 51422712 | 51422927 | 0.963146 | 2.40378 | 2.496 |
| chr14 | 94603019 | 94603364 | 1.74612 | 4.35792 | 2.496 |
| chr11 | 64571793 | 64572048 | 0.479317 | 1.19628 | 2.496 |
| chr15 | 27083726 | 27084342 | 1.30067 | 3.24632 | 2.496 |
| chr11 | 112325020 | 112325232 | 0.648497 | 1.61861 | 2.496 |
| chr11 | 65789872 | 65790283 | 2.38349 | 5.94927 | 2.496 |
| chr14 | 55737792 | 55738204 | 2.15676 | 5.38341 | 2.496 |
| chr11 | 8981163 | 8981440 | 1.10666 | 2.76232 | 2.496 |
| chr16 | 27527080 | 27527780 | 0.930675 | 2.32315 | 2.496 |
| chr8 | 47034645 | 47034855 | 0.516869 | 1.29022 | 2.496 |
| chr2 | 153032463 | 153032803 | 1.36125 | 3.39805 | 2.496 |
| chr6 | 593203 | 593456 | 0.82141 | 2.05057 | 2.496 |
| chr17 | 40464246 | 40464858 | 1.39713 | 3.48788 | 2.496 |
| chr7 | 91347397 | 91347681 | 0.432971 | 1.08092 | 2.497 |
| chr17 | 57469189 | 57469482 | 0.358526 | 0.895075 | 2.497 |
| chr16 | 21668530 | 21668896 | 1.18555 | 2.95985 | 2.497 |
| chr14 | 54430381 | 54430672 | 1.09568 | 2.73554 | 2.497 |
| chr5 | 111347637 | 111347894 | 0.465017 | 1.16111 | 2.497 |
| chr10 | 19082909 | 19083158 | 0.693638 | 1.73218 | 2.497 |
| chr8 | 82147542 | 82147816 | 1.21595 | 3.03681 | 2.497 |
| chrX | 23926059 | 23926402 | 1.60226 | 4.00185 | 2.498 |
| chr3 | 122913391 | 122913666 | 1.14671 | 2.86417 | 2.498 |
| chr5 | 93817411 | 93817617 | 0.545155 | 1.36165 | 2.498 |
| chr3 | 10206321 | 10206733 | 1.7033 | 4.25449 | 2.498 |
| chr14 | 106403495 | 106403784 | 0.488829 | 1.22102 | 2.498 |
| chr1 | 8908894 | 8909241 | 1.35974 | 3.39649 | 2.498 |
| chr1 | 186649314 | 186649618 | 0.891278 | 2.22656 | 2.498 |

| | | | | | |
|---|---|---|---|---|---|
| chr10 | 108848115 | 108848413 | 0.384381 | 0.960315 | 2.498 |
| chr10 | 30702970 | 30703235 | 1.35553 | 3.38692 | 2.499 |
| chr7 | 64174967 | 64175245 | 0.524136 | 1.30962 | 2.499 |
| chr16 | 19191199 | 19191574 | 1.48671 | 3.71488 | 2.499 |
| chr20 | 23028425 | 23028669 | 0.719686 | 1.79839 | 2.499 |
| chr14 | 88598456 | 88598673 | 0.462198 | 1.15497 | 2.499 |
| chr16 | 67481535 | 67481831 | 1.16453 | 2.91003 | 2.499 |
| chr12 | 2955115 | 2955388 | 1.08938 | 2.72235 | 2.499 |
| chr19 | 49617438 | 49617670 | 0.807956 | 2.01908 | 2.499 |
| chr8 | 141090523 | 141090815 | 0.453592 | 1.13353 | 2.499 |
| chr12 | 54973409 | 54973876 | 2.67305 | 6.68052 | 2.499 |
| chr4 | 90562288 | 90562665 | 1.15118 | 2.87734 | 2.499 |
| chr5 | 107009399 | 107009711 | 1.59267 | 3.98086 | 2.499 |
| chr14 | 57541486 | 57541724 | 0.360816 | 0.901903 | 2.500 |
| chr8 | 50765360 | 50765648 | 0.410685 | 1.02659 | 2.500 |
| chr18 | 7022278 | 7022564 | 1.09217 | 2.73035 | 2.500 |
| chr4 | 148440013 | 148440359 | 1.52011 | 3.80027 | 2.500 |
| chr4 | 78539813 | 78540113 | 1.11872 | 2.79684 | 2.500 |
| chr16 | 67233421 | 67233650 | 0.769411 | 1.9236 | 2.500 |
| chr10 | 76947708 | 76948031 | 1.16216 | 2.90567 | 2.500 |
| chr9 | 110103420 | 110103888 | 2.28843 | 5.72175 | 2.500 |
| chr14 | 57544305 | 57544527 | 0.330138 | 0.825469 | 2.500 |
| chr2 | 149362093 | 149362324 | 0.653966 | 1.63517 | 2.500 |
| chr16 | 70429091 | 70429637 | 1.49004 | 3.72575 | 2.500 |
| chr17 | 66736997 | 66737600 | 0.862786 | 2.15752 | 2.501 |
| chr12 | 123338235 | 123338479 | 0.397393 | 0.99376 | 2.501 |
| chr3 | 50310402 | 50310682 | 1.62875 | 4.07315 | 2.501 |
| chr8 | 24857425 | 24857852 | 2.41052 | 6.02825 | 2.501 |
| chr9 | 79765744 | 79766052 | 1.67692 | 4.19371 | 2.501 |
| chr1 | 175288726 | 175289115 | 1.02351 | 2.55981 | 2.501 |
| chr8 | 86152823 | 86153056 | 0.493961 | 1.23547 | 2.501 |
| chr13 | 76450550 | 76450863 | 1.50003 | 3.75205 | 2.501 |
| chr10 | 82359247 | 82359574 | 1.2231 | 3.05954 | 2.501 |
| chr5 | 76373028 | 76373797 | 1.55763 | 3.89648 | 2.502 |
| chr12 | 131355422 | 131355783 | 0.813929 | 2.0361 | 2.502 |
| chr15 | 68664741 | 68665214 | 0.771478 | 1.92999 | 2.502 |
| chr17 | 80258570 | 80259071 | 2.0641 | 5.16379 | 2.502 |
| chr11 | 28695961 | 28696194 | 0.795056 | 1.98912 | 2.502 |
| chr15 | 88998072 | 88998594 | 1.53984 | 3.85256 | 2.502 |
| chr2 | 175561856 | 175562259 | 1.3309 | 3.32981 | 2.502 |
| chr8 | 427550 | 427926 | 2.32454 | 5.81582 | 2.502 |
| chr8 | 133535933 | 133536203 | 0.499308 | 1.24925 | 2.502 |
| chr20 | 32054692 | 32054905 | 0.347792 | 0.87018 | 2.502 |
| chr5 | 4193231 | 4193523 | 0.434676 | 1.08758 | 2.502 |
| chr5 | 140181080 | 140181547 | 1.37786 | 3.44753 | 2.502 |
| chr12 | 13431023 | 13431499 | 2.29528 | 5.74302 | 2.502 |
| chr5 | 148507531 | 148507954 | 1.5854 | 3.96686 | 2.502 |
| chr9 | 34155658 | 34155899 | 1.00978 | 2.5266 | 2.502 |
| chr14 | 74005744 | 74006002 | 0.646334 | 1.61727 | 2.502 |
| chr12 | 97455035 | 97455334 | 0.589332 | 1.47469 | 2.502 |
| chr8 | 141647427 | 141647871 | 2.50355 | 6.2647 | 2.502 |
| chr14 | 23578343 | 23578747 | 0.861159 | 2.15499 | 2.502 |
| chr12 | 16009383 | 16009595 | 0.941757 | 2.35705 | 2.503 |
| chr7 | 33147365 | 33147608 | 1.18446 | 2.96456 | 2.503 |
| chr9 | 77098864 | 77099066 | 0.433131 | 1.08414 | 2.503 |
| chr8 | 64227500 | 64227754 | 0.560367 | 1.40271 | 2.503 |
| chr7 | 157091142 | 157091435 | 1.0325 | 2.58461 | 2.503 |
| chr13 | 110262976 | 110263234 | 0.564301 | 1.4126 | 2.503 |
| chr1 | 157460427 | 157460782 | 1.05729 | 2.64672 | 2.503 |
| chr15 | 55933888 | 55934138 | 1.10255 | 2.76003 | 2.503 |
| chr11 | 4404393 | 4404676 | 0.444021 | 1.11154 | 2.503 |

| | | | | | |
|---|---|---|---|---|---|
| chr5 | 90228002 | 90228271 | 1.42803 | 3.57486 | 2.503 |
| chr8 | 102313621 | 102313978 | 1.78713 | 4.474 | 2.503 |
| chr8 | 22832054 | 22832388 | 1.25682 | 3.14676 | 2.504 |
| chr5 | 173116800 | 173117109 | 0.546244 | 1.36768 | 2.504 |
| chr5 | 159865413 | 159866558 | 1.58684 | 3.97322 | 2.504 |
| chr10 | 134201462 | 134201688 | 1.28629 | 3.22096 | 2.504 |
| chr4 | 75857990 | 75858404 | 1.51035 | 3.78223 | 2.504 |
| chr14 | 59495337 | 59495616 | 1.61132 | 4.03519 | 2.504 |
| chr10 | 14541011 | 14541271 | 1.11203 | 2.78486 | 2.504 |
| chr3 | 51834001 | 51834229 | 1.47423 | 3.69199 | 2.504 |
| chr9 | 119351604 | 119352115 | 1.319 | 3.30339 | 2.504 |
| chr4 | 129349051 | 129349341 | 1.35349 | 3.39002 | 2.505 |
| chr18 | 57069644 | 57070095 | 1.62855 | 4.0793 | 2.505 |
| chr5 | 63545074 | 63545363 | 0.386059 | 0.96704 | 2.505 |
| chr4 | 83723115 | 83723469 | 0.902176 | 2.25988 | 2.505 |
| chrX | 11792721 | 11793008 | 0.917427 | 2.29821 | 2.505 |
| chrX | 8837062 | 8837286 | 0.6969 | 1.74595 | 2.505 |
| chr12 | 4949734 | 4949969 | 0.518018 | 1.29782 | 2.505 |
| chr19 | 44251589 | 44252038 | 0.57098 | 1.43051 | 2.505 |
| chrX | 40523477 | 40523776 | 0.793377 | 1.98773 | 2.505 |
| chr3 | 108110434 | 108110819 | 0.438696 | 1.09914 | 2.505 |
| chr8 | 34012719 | 34013052 | 0.386338 | 0.967979 | 2.506 |
| chr6 | 134432142 | 134432601 | 1.59323 | 3.99189 | 2.506 |
| chr5 | 80461340 | 80461622 | 1.4065 | 3.52408 | 2.506 |
| chr17 | 48194567 | 48194984 | 1.1799 | 2.95635 | 2.506 |
| chr12 | 56967217 | 56967450 | 1.28063 | 3.20875 | 2.506 |
| chr12 | 57566724 | 57567191 | 1.66563 | 4.17341 | 2.506 |
| chr5 | 167718427 | 167718881 | 0.64849 | 1.62488 | 2.506 |
| chr9 | 94896184 | 94896553 | 1.33606 | 3.34769 | 2.506 |
| chr20 | 41288475 | 41288690 | 0.51166 | 1.28205 | 2.506 |
| chr11 | 65547195 | 65547838 | 2.45263 | 6.14569 | 2.506 |
| chr15 | 77925712 | 77926247 | 3.96846 | 9.94443 | 2.506 |
| chr22 | 45944933 | 45945246 | 0.896512 | 2.24654 | 2.506 |
| chr7 | 36015533 | 36015882 | 1.06998 | 2.68136 | 2.506 |
| chr20 | 15472210 | 15472718 | 0.969459 | 2.42967 | 2.506 |
| chr19 | 4352611 | 4352992 | 1.49086 | 3.73645 | 2.506 |
| chr8 | 92553354 | 92553587 | 0.48152 | 1.20699 | 2.507 |
| chr17 | 72280863 | 72281156 | 1.39085 | 3.48636 | 2.507 |
| chr5 | 55565420 | 55565694 | 1.20014 | 3.00834 | 2.507 |
| chr17 | 6334224 | 6334464 | 0.854678 | 2.14244 | 2.507 |
| chr12 | 98556643 | 98556852 | 0.427027 | 1.07046 | 2.507 |
| chr10 | 105512512 | 105512825 | 1.28445 | 3.21988 | 2.507 |
| chr19 | 48044014 | 48044560 | 0.690427 | 1.73089 | 2.507 |
| chr10 | 56047781 | 56048056 | 0.444828 | 1.11518 | 2.507 |
| chr7 | 24561876 | 24562105 | 0.462945 | 1.16066 | 2.507 |
| chr12 | 126633376 | 126633701 | 0.571383 | 1.43257 | 2.507 |
| chr15 | 41522735 | 41523193 | 2.038 | 5.11005 | 2.507 |
| chr15 | 63779491 | 63779885 | 2.29705 | 5.75961 | 2.507 |
| chr6 | 62860013 | 62860261 | 1.05824 | 2.65358 | 2.508 |
| chr22 | 46307472 | 46307744 | 0.841396 | 2.10988 | 2.508 |
| chr5 | 96333044 | 96333246 | 1.09427 | 2.74407 | 2.508 |
| chr1 | 153327967 | 153328268 | 1.21898 | 3.05744 | 2.508 |
| chr17 | 45693267 | 45693485 | 1.1335 | 2.84307 | 2.508 |
| chr9 | 101127409 | 101127785 | 0.792947 | 1.98902 | 2.508 |
| chr11 | 121075384 | 121075603 | 0.618346 | 1.55113 | 2.509 |
| chr15 | 28343963 | 28344229 | 0.941797 | 2.36252 | 2.509 |
| chr15 | 96303780 | 96304034 | 1.1946 | 2.99681 | 2.509 |
| chr20 | 30679646 | 30680073 | 2.04433 | 5.12851 | 2.509 |
| chr2 | 116342138 | 116342375 | 0.447535 | 1.12282 | 2.509 |
| chr20 | 45834368 | 45834608 | 1.01923 | 2.55733 | 2.509 |
| chrX | 13664212 | 13664527 | 1.14041 | 2.86144 | 2.509 |

| | | | | | |
|---|---|---|---|---|---|
| chr2 | 188365653 | 188365868 | 0.708413 | 1.77762 | 2.509 |
| chr1 | 203436011 | 203436244 | 0.345019 | 0.865764 | 2.509 |
| chr5 | 55479998 | 55480204 | 0.314948 | 0.790304 | 2.509 |
| chr10 | 47594351 | 47594674 | 0.84004 | 2.10811 | 2.510 |
| chr6 | 157695680 | 157696136 | 1.09843 | 2.75674 | 2.510 |
| chr10 | 124194843 | 124195049 | 0.918394 | 2.30492 | 2.510 |
| chr19 | 18223840 | 18224223 | 1.35736 | 3.40685 | 2.510 |
| chr2 | 33899118 | 33899336 | 0.552349 | 1.38636 | 2.510 |
| chr5 | 32613196 | 32613531 | 1.75719 | 4.41042 | 2.510 |
| chr7 | 139874998 | 139875289 | 1.55884 | 3.91264 | 2.510 |
| chr3 | 54904468 | 54904691 | 0.777401 | 1.95127 | 2.510 |
| chr7 | 39155652 | 39155911 | 0.49425 | 1.24062 | 2.510 |
| chr4 | 126672318 | 126672687 | 0.805436 | 2.02175 | 2.510 |
| chr14 | 22370780 | 22371077 | 1.0691 | 2.68359 | 2.510 |
| chr11 | 56884105 | 56884456 | 0.823652 | 2.06749 | 2.510 |
| chr9 | 88817130 | 88817561 | 1.27644 | 3.20406 | 2.510 |
| chr5 | 145061176 | 145061463 | 1.57489 | 3.95327 | 2.510 |
| chr20 | 6121711 | 6121935 | 0.835543 | 2.0975 | 2.510 |
| chr12 | 117028433 | 117028637 | 0.576816 | 1.44806 | 2.510 |
| chr9 | 119605007 | 119605337 | 0.582926 | 1.4635 | 2.511 |
| chr1 | 220011095 | 220011456 | 0.82164 | 2.06285 | 2.511 |
| chrX | 10565948 | 10566226 | 1.10173 | 2.7661 | 2.511 |
| chr12 | 52680755 | 52680985 | 0.760111 | 1.90849 | 2.511 |
| chr19 | 49149679 | 49150079 | 2.12327 | 5.33114 | 2.511 |
| chr7 | 149296168 | 149296510 | 0.414582 | 1.04096 | 2.511 |
| chr13 | 21684940 | 21685275 | 1.40716 | 3.53327 | 2.511 |
| chr6 | 18504243 | 18504607 | 1.30211 | 3.26954 | 2.511 |
| chr17 | 55208350 | 55208608 | 1.55209 | 3.89745 | 2.511 |
| chr7 | 42437411 | 42437630 | 0.639781 | 1.60666 | 2.511 |
| chr19 | 43071034 | 43071514 | 3.10787 | 7.80476 | 2.511 |
| chr11 | 76706843 | 76707074 | 0.421403 | 1.05835 | 2.511 |
| chr2 | 66180435 | 66180715 | 0.443836 | 1.11469 | 2.511 |
| chr12 | 50464280 | 50464776 | 1.68364 | 4.22851 | 2.512 |
| chr13 | 41927180 | 41927781 | 0.678176 | 1.70327 | 2.512 |
| chr19 | 1513759 | 1514221 | 1.32473 | 3.32729 | 2.512 |
| chr1 | 175123956 | 175124441 | 1.41272 | 3.54833 | 2.512 |
| chr7 | 94333801 | 94334010 | 0.442082 | 1.11039 | 2.512 |
| chr2 | 158485028 | 158485300 | 0.680045 | 1.70816 | 2.512 |
| chrX | 154892047 | 154892336 | 0.845856 | 2.12481 | 2.512 |
| chr12 | 32810440 | 32810796 | 0.873558 | 2.19443 | 2.512 |
| chr17 | 3822066 | 3822336 | 0.439918 | 1.1052 | 2.512 |
| chr15 | 50440619 | 50440909 | 1.5821 | 3.97478 | 2.512 |
| chrX | 151985649 | 151985885 | 0.749449 | 1.88292 | 2.512 |
| chr10 | 102800972 | 102801307 | 1.81641 | 4.56366 | 2.512 |
| chr20 | 50211571 | 50211829 | 1.38477 | 3.4793 | 2.513 |
| chr12 | 125887705 | 125888085 | 1.04194 | 2.61796 | 2.513 |
| chr3 | 171269954 | 171270172 | 0.388688 | 0.976643 | 2.513 |
| chr8 | 71025798 | 71026276 | 1.21216 | 3.04579 | 2.513 |
| chr20 | 42077661 | 42077933 | 1.58741 | 3.9887 | 2.513 |
| chr17 | 33446695 | 33447130 | 1.92824 | 4.84514 | 2.513 |
| chr5 | 129060114 | 129060376 | 0.358427 | 0.900645 | 2.513 |
| chr1 | 211822394 | 211822676 | 1.28136 | 3.21987 | 2.513 |
| chr22 | 29641411 | 29641905 | 1.52484 | 3.83172 | 2.513 |
| chr7 | 98362347 | 98362802 | 0.857271 | 2.15421 | 2.513 |
| chr3 | 124186985 | 124187450 | 1.81297 | 4.55654 | 2.513 |
| chr17 | 3766907 | 3767377 | 2.09672 | 5.26975 | 2.513 |
| chr12 | 67273663 | 67273925 | 0.492038 | 1.23667 | 2.513 |
| chr3 | 792808 | 793188 | 0.40217 | 1.0108 | 2.513 |
| chr21 | 44270937 | 44271281 | 1.55789 | 3.91567 | 2.513 |
| chr19 | 12888358 | 12888709 | 1.70468 | 4.28466 | 2.513 |
| chr14 | 59989426 | 59989680 | 0.462092 | 1.16146 | 2.513 |

| | | | | | |
|---|---|---|---|---|---|
| chr2 | 25584761 | 25585162 | 1.93186 | 4.85574 | 2.514 |
| chr12 | 121723326 | 121723635 | 0.521572 | 1.31099 | 2.514 |
| chr7 | 99229321 | 99229584 | 1.09385 | 2.74946 | 2.514 |
| chr12 | 5390711 | 5390971 | 0.34867 | 0.876437 | 2.514 |
| chr19 | 38893415 | 38894073 | 2.04171 | 5.13227 | 2.514 |
| chr19 | 55671867 | 55672260 | 3.0247 | 7.60324 | 2.514 |
| chr20 | 60876863 | 60877183 | 2.49151 | 6.26346 | 2.514 |
| chr4 | 48292434 | 48292690 | 1.16698 | 2.93375 | 2.514 |
| chr3 | 65425307 | 65425606 | 0.82622 | 2.07716 | 2.514 |
| chr21 | 37632881 | 37633109 | 0.91332 | 2.29627 | 2.514 |
| chr3 | 5021991 | 5022328 | 2.03058 | 5.10537 | 2.514 |
| chr8 | 135883230 | 135883484 | 0.412119 | 1.03626 | 2.514 |
| chr20 | 43089476 | 43089777 | 0.974638 | 2.45094 | 2.515 |
| chr19 | 16528542 | 16528857 | 1.55512 | 3.91086 | 2.515 |
| chr22 | 31030312 | 31030706 | 1.37205 | 3.4505 | 2.515 |
| chr1 | 75237680 | 75237919 | 1.25747 | 3.16242 | 2.515 |
| chr9 | 106027890 | 106028150 | 0.36978 | 0.929991 | 2.515 |
| chr15 | 41184076 | 41184353 | 1.47318 | 3.70507 | 2.515 |
| chr9 | 37482350 | 37482652 | 0.994655 | 2.50175 | 2.515 |
| chr2 | 42997693 | 42997996 | 1.35454 | 3.40694 | 2.515 |
| chr2 | 143919887 | 143920107 | 1.15742 | 2.91118 | 2.515 |
| chr21 | 40852316 | 40852536 | 0.333095 | 0.837832 | 2.515 |
| chr1 | 16382571 | 16382932 | 1.9587 | 4.92671 | 2.515 |
| chr8 | 58026284 | 58026570 | 1.15714 | 2.91059 | 2.515 |
| chr16 | 1396936 | 1397322 | 0.643285 | 1.61837 | 2.516 |
| chr14 | 91639498 | 91639793 | 1.88448 | 4.74098 | 2.516 |
| chr10 | 100424104 | 100424584 | 1.02197 | 2.57112 | 2.516 |
| chrX | 48749049 | 48749253 | 0.936929 | 2.35728 | 2.516 |
| chr15 | 75497106 | 75497508 | 2.50725 | 6.30829 | 2.516 |
| chr19 | 43033351 | 43033989 | 1.56663 | 3.94186 | 2.516 |
| chr17 | 56682900 | 56683148 | 0.466777 | 1.17452 | 2.516 |
| chr5 | 14352117 | 14352573 | 1.69045 | 4.25356 | 2.516 |
| chr12 | 122326876 | 122327190 | 1.12481 | 2.83038 | 2.516 |
| chr1 | 117610470 | 117610709 | 0.996464 | 2.5075 | 2.516 |
| chr4 | 7153581 | 7153939 | 0.983256 | 2.47431 | 2.516 |
| chr16 | 1478562 | 1478958 | 1.36989 | 3.44754 | 2.517 |
| chr22 | 39162151 | 39162574 | 2.0967 | 5.27686 | 2.517 |
| chr12 | 33686835 | 33687095 | 0.464821 | 1.16988 | 2.517 |
| chr2 | 231968915 | 231969183 | 1.67005 | 4.20361 | 2.517 |
| chr11 | 123132209 | 123132536 | 1.8825 | 4.73864 | 2.517 |
| chr9 | 90656383 | 90656717 | 0.834534 | 2.10075 | 2.517 |
| chr8 | 88916209 | 88916664 | 0.454823 | 1.14493 | 2.517 |
| chr10 | 22052636 | 22052933 | 0.398269 | 1.00264 | 2.517 |
| chr3 | 133930748 | 133930986 | 0.741077 | 1.86582 | 2.518 |
| chr10 | 83704714 | 83704942 | 0.428556 | 1.07899 | 2.518 |
| chr17 | 47929051 | 47929508 | 1.92529 | 4.848 | 2.518 |
| chr5 | 89717068 | 89717312 | 0.816416 | 2.05585 | 2.518 |
| chr7 | 157190509 | 157191026 | 1.66061 | 4.18168 | 2.518 |
| chr14 | 101368691 | 101368908 | 0.775402 | 1.95266 | 2.518 |
| chr22 | 23053150 | 23053587 | 1.7389 | 4.37907 | 2.518 |
| gl000195_ra | 103275 | 103676 | 1.7494 | 4.4056 | 2.518 |
| chr12 | 121668000 | 121668536 | 1.92803 | 4.85548 | 2.518 |
| chr13 | 108972663 | 108972895 | 0.260131 | 0.655116 | 2.518 |
| chr12 | 52645903 | 52646201 | 1.1752 | 2.95972 | 2.518 |
| chr4 | 21767496 | 21767712 | 1.24667 | 3.13993 | 2.519 |
| chr12 | 117588927 | 117589239 | 1.65059 | 4.15739 | 2.519 |
| chr5 | 150764284 | 150764514 | 0.798759 | 2.01196 | 2.519 |
| chr22 | 30085786 | 30086218 | 1.06548 | 2.68396 | 2.519 |
| chr1 | 192475859 | 192476174 | 0.92215 | 2.32295 | 2.519 |
| chr9 | 35646741 | 35647168 | 1.74646 | 4.39972 | 2.519 |
| chr8 | 85930210 | 85930491 | 0.452416 | 1.13975 | 2.519 |

| | | | | | |
|---|---|---|---|---|---|
| chr15 | 66208970 | 66209326 | 1.46792 | 3.69813 | 2.519 |
| chrX | 129527471 | 129527832 | 1.40373 | 3.53643 | 2.519 |
| chr1 | 216774481 | 216774825 | 1.28369 | 3.2341 | 2.519 |
| chr7 | 19794177 | 19794463 | 0.972561 | 2.45026 | 2.519 |
| chr12 | 99437102 | 99437462 | 1.63203 | 4.11187 | 2.519 |
| chr21 | 46518794 | 46519057 | 0.546295 | 1.3764 | 2.520 |
| chr17 | 55935293 | 55935629 | 1.43227 | 3.60888 | 2.520 |
| chr8 | 2034501 | 2034794 | 1.98368 | 4.9985 | 2.520 |
| chr10 | 95505302 | 95505534 | 0.932718 | 2.35035 | 2.520 |
| chr12 | 53441117 | 53441474 | 1.46125 | 3.68224 | 2.520 |
| chr15 | 33228603 | 33228827 | 0.893454 | 2.25161 | 2.520 |
| chr14 | 24249725 | 24249986 | 0.446983 | 1.12646 | 2.520 |
| chr12 | 91775860 | 91776144 | 1.50826 | 3.80122 | 2.520 |
| chr5 | 68703305 | 68703552 | 0.932838 | 2.35106 | 2.520 |
| chr15 | 81395880 | 81396118 | 0.561004 | 1.41393 | 2.520 |
| chr8 | 67424393 | 67424747 | 1.43548 | 3.61821 | 2.521 |
| chr7 | 81099391 | 81099715 | 1.37832 | 3.47446 | 2.521 |
| chr17 | 38597253 | 38597605 | 1.27873 | 3.22344 | 2.521 |
| chr2 | 192734056 | 192734261 | 0.795667 | 2.00581 | 2.521 |
| chr4 | 20360790 | 20361034 | 0.476887 | 1.20223 | 2.521 |
| chr10 | 10853022 | 10853303 | 0.355754 | 0.896913 | 2.521 |
| chr1 | 186344515 | 186344905 | 1.63307 | 4.11727 | 2.521 |
| chr10 | 27361423 | 27361738 | 0.683598 | 1.7235 | 2.521 |
| chr12 | 62277070 | 62277332 | 0.391234 | 0.986427 | 2.521 |
| chr12 | 49491634 | 49492233 | 1.47529 | 3.71995 | 2.522 |
| chr18 | 46568292 | 46568622 | 1.09644 | 2.76473 | 2.522 |
| chr9 | 124082355 | 124082745 | 2.53163 | 6.38367 | 2.522 |
| chrX | 17139650 | 17139893 | 0.835182 | 2.10611 | 2.522 |
| chr20 | 4740975 | 4741576 | 1.55867 | 3.93065 | 2.522 |
| chr15 | 33287036 | 33287353 | 0.826779 | 2.08519 | 2.522 |
| chr2 | 31248172 | 31248456 | 0.516751 | 1.30329 | 2.522 |
| chr17 | 46569846 | 46570227 | 2.63706 | 6.65103 | 2.522 |
| chr7 | 85459232 | 85459475 | 0.703491 | 1.77433 | 2.522 |
| chr15 | 88048582 | 88048999 | 0.371917 | 0.938116 | 2.522 |
| chr12 | 120794379 | 120794654 | 1.74078 | 4.39125 | 2.523 |
| chr16 | 18119089 | 18119393 | 0.422877 | 1.06675 | 2.523 |
| chr1 | 240061475 | 240061731 | 1.55326 | 3.91846 | 2.523 |
| chr11 | 86529596 | 86529830 | 0.897592 | 2.26443 | 2.523 |
| chr3 | 5288460 | 5288731 | 0.912845 | 2.30296 | 2.523 |
| chr4 | 174911321 | 174911618 | 0.892632 | 2.25199 | 2.523 |
| chr16 | 52895567 | 52895813 | 0.981552 | 2.47638 | 2.523 |
| chr6 | 114417756 | 114418106 | 1.15541 | 2.91502 | 2.523 |
| chr8 | 35016782 | 35017035 | 0.550863 | 1.38983 | 2.523 |
| chr16 | 1380427 | 1380905 | 1.52646 | 3.85166 | 2.523 |
| chr5 | 4188952 | 4189161 | 0.445602 | 1.12439 | 2.523 |
| chr2 | 7057162 | 7057684 | 1.74195 | 4.39552 | 2.523 |
| chr1 | 172010942 | 172011231 | 0.617677 | 1.55867 | 2.523 |
| chr20 | 36568548 | 36568793 | 0.314138 | 0.792708 | 2.523 |
| chr3 | 187459859 | 187460152 | 1.53837 | 3.88221 | 2.524 |
| chr14 | 76177508 | 76177791 | 1.69367 | 4.27423 | 2.524 |
| chrX | 53346656 | 53347111 | 1.90368 | 4.80433 | 2.524 |
| chr15 | 86343756 | 86343962 | 0.833649 | 2.10395 | 2.524 |
| chr2 | 136555063 | 136555389 | 1.75012 | 4.41697 | 2.524 |
| chr5 | 94322375 | 94322578 | 0.484741 | 1.22341 | 2.524 |
| chr1 | 47993350 | 47993644 | 0.811324 | 2.0477 | 2.524 |
| chr18 | 48346302 | 48346569 | 1.63271 | 4.1208 | 2.524 |
| chr17 | 42160764 | 42161242 | 1.80341 | 4.55217 | 2.524 |
| chr14 | 66281887 | 66282136 | 0.57027 | 1.43958 | 2.524 |
| chr12 | 62517598 | 62517839 | 1.55059 | 3.91462 | 2.525 |
| chrX | 10294715 | 10294945 | 0.826301 | 2.08623 | 2.525 |
| chr10 | 63798479 | 63798742 | 1.07443 | 2.71274 | 2.525 |

| | | | | | |
|---|---|---|---|---|---|
| chr12 | 34276685 | 34276909 | 0.413791 | 1.0448 | 2.525 |
| chr7 | 140338312 | 140338682 | 0.548344 | 1.38461 | 2.525 |
| chr2 | 120963856 | 120964212 | 1.27279 | 3.21398 | 2.525 |
| chr4 | 74439414 | 74439834 | 1.78481 | 4.50749 | 2.525 |
| chr8 | 41685832 | 41686706 | 1.17451 | 2.96627 | 2.526 |
| chrX | 152736324 | 152736595 | 1.62674 | 4.10857 | 2.526 |
| chr4 | 120550038 | 120550421 | 1.17642 | 2.97154 | 2.526 |
| chr9 | 121978412 | 121978690 | 0.383852 | 0.969585 | 2.526 |
| chr16 | 27190797 | 27191206 | 1.78029 | 4.49704 | 2.526 |
| chr6 | 80577957 | 80578181 | 0.844578 | 2.13347 | 2.526 |
| chr8 | 143420467 | 143420776 | 0.479735 | 1.21186 | 2.526 |
| chr12 | 116259385 | 116259609 | 0.409479 | 1.03442 | 2.526 |
| chr10 | 119256108 | 119256443 | 1.91623 | 4.84151 | 2.527 |
| chr20 | 48162184 | 48162469 | 1.06122 | 2.68129 | 2.527 |
| chr3 | 108212796 | 108213042 | 1.22022 | 3.08336 | 2.527 |
| chr12 | 64237543 | 64238053 | 2.14103 | 5.41021 | 2.527 |
| chr2 | 27392694 | 27393090 | 2.33297 | 5.89522 | 2.527 |
| chrX | 99668266 | 99668567 | 0.675195 | 1.70621 | 2.527 |
| chr11 | 77268676 | 77268884 | 0.816958 | 2.06452 | 2.527 |
| chr12 | 14338864 | 14339286 | 1.40539 | 3.55164 | 2.527 |
| chr11 | 10813939 | 10814376 | 2.31878 | 5.85997 | 2.527 |
| chr9 | 26189222 | 26189465 | 0.44107 | 1.11474 | 2.527 |
| chr12 | 110213002 | 110213468 | 2.33 | 5.88876 | 2.527 |
| chr16 | 16288674 | 16289036 | 1.73746 | 4.39128 | 2.527 |
| chr7 | 101331554 | 101332023 | 2.26695 | 5.72959 | 2.527 |
| chr12 | 111136120 | 111136367 | 1.0337 | 2.61262 | 2.527 |
| chr10 | 98877216 | 98877471 | 0.963416 | 2.43509 | 2.528 |
| chr3 | 185378222 | 185378625 | 1.37569 | 3.47723 | 2.528 |
| chr19 | 30250925 | 30251572 | 2.27988 | 5.76279 | 2.528 |
| chr2 | 172386499 | 172386774 | 1.10986 | 2.80546 | 2.528 |
| chr12 | 120339852 | 120340247 | 1.43815 | 3.6354 | 2.528 |
| chr19 | 1221464 | 1221947 | 1.2906 | 3.26258 | 2.528 |
| chr14 | 76027278 | 76027657 | 1.84333 | 4.65989 | 2.528 |
| chr11 | 25115169 | 25115407 | 0.498542 | 1.26036 | 2.528 |
| chr21 | 40497919 | 40498188 | 0.958497 | 2.42321 | 2.528 |
| chr10 | 55639453 | 55639745 | 0.505068 | 1.2769 | 2.528 |
| chr10 | 60994622 | 60994871 | 1.1694 | 2.95654 | 2.528 |
| chr8 | 3758918 | 3759176 | 0.419856 | 1.06152 | 2.528 |
| chr2 | 47043847 | 47044090 | 0.323123 | 0.816967 | 2.528 |
| chrX | 75776282 | 75776484 | 0.664383 | 1.67982 | 2.528 |
| chr16 | 50388771 | 50389110 | 0.9025 | 2.28193 | 2.528 |
| chr5 | 100235800 | 100236198 | 0.711735 | 1.79961 | 2.528 |
| chr11 | 119352046 | 119352395 | 1.62838 | 4.11747 | 2.529 |
| chr22 | 32373258 | 32373497 | 1.09874 | 2.77835 | 2.529 |
| chr5 | 171879480 | 171879758 | 0.907689 | 2.29531 | 2.529 |
| chrX | 117388676 | 117389006 | 0.910121 | 2.30173 | 2.529 |
| chr7 | 50670762 | 50671033 | 0.533206 | 1.34851 | 2.529 |
| chr14 | 32851971 | 32852237 | 1.59838 | 4.04255 | 2.529 |
| chr12 | 52419169 | 52419786 | 1.89196 | 4.78521 | 2.529 |
| chr13 | 50571418 | 50571776 | 1.54164 | 3.89948 | 2.529 |
| chr12 | 52709596 | 52709915 | 0.697788 | 1.76502 | 2.529 |
| chr1 | 61508636 | 61508977 | 1.4362 | 3.63295 | 2.530 |
| chr18 | 22068884 | 22069156 | 0.777057 | 1.96565 | 2.530 |
| chr19 | 15212386 | 15212642 | 0.89724 | 2.26977 | 2.530 |
| chr10 | 130010153 | 130010592 | 2.20605 | 5.58072 | 2.530 |
| chr18 | 7221444 | 7221853 | 1.37403 | 3.476 | 2.530 |
| chrX | 102719737 | 102719925 | 1.1716 | 2.96416 | 2.530 |
| chr9 | 124308281 | 124308616 | 2.40864 | 6.0941 | 2.530 |
| chr12 | 43838795 | 43839108 | 1.41189 | 3.57224 | 2.530 |
| chr17 | 48504822 | 48505087 | 1.34944 | 3.41431 | 2.530 |
| chr9 | 109388364 | 109388659 | 0.444716 | 1.12524 | 2.530 |

| | | | | | |
|---|---|---|---|---|---|
| chr1 | 112312248 | 112312570 | 1.31121 | 3.31776 | 2.530 |
| chr15 | 100945736 | 100945974 | 0.419789 | 1.0622 | 2.530 |
| chr3 | 16290895 | 16291209 | 1.27441 | 3.22484 | 2.530 |
| chr19 | 5720589 | 5720854 | 1.73641 | 4.39393 | 2.530 |
| chr19 | 1223084 | 1223543 | 0.982824 | 2.48719 | 2.531 |
| chr14 | 98109836 | 98110137 | 0.412116 | 1.04293 | 2.531 |
| chr12 | 117538064 | 117538442 | 0.849754 | 2.15061 | 2.531 |
| chr19 | 45751903 | 45752287 | 1.47581 | 3.73508 | 2.531 |
| chr9 | 95978133 | 95978511 | 0.805815 | 2.03945 | 2.531 |
| chr1 | 110316289 | 110316646 | 1.49081 | 3.77315 | 2.531 |
| chr11 | 32771168 | 32771439 | 0.763339 | 1.93199 | 2.531 |
| chr16 | 58056071 | 58056312 | 0.416711 | 1.05471 | 2.531 |
| chr14 | 51422568 | 51422772 | 1.21435 | 3.07361 | 2.531 |
| chr3 | 13968779 | 13969189 | 0.8006 | 2.02641 | 2.531 |
| chr1 | 221608768 | 221609106 | 1.19726 | 3.03062 | 2.531 |
| chr12 | 43935772 | 43936067 | 0.425829 | 1.07793 | 2.531 |
| chr3 | 148032899 | 148033130 | 0.594499 | 1.50493 | 2.531 |
| chr12 | 68963691 | 68963895 | 0.464129 | 1.17492 | 2.531 |
| chr1 | 220392056 | 220392261 | 1.10466 | 2.79642 | 2.531 |
| chr8 | 11593904 | 11594216 | 1.89404 | 4.79472 | 2.531 |
| chr12 | 63025604 | 63025843 | 0.763528 | 1.93286 | 2.531 |
| chr8 | 36692943 | 36693219 | 0.457542 | 1.1583 | 2.532 |
| chr5 | 139224003 | 139224395 | 1.72492 | 4.36704 | 2.532 |
| chr1 | 150436905 | 150437150 | 1.35251 | 3.42425 | 2.532 |
| chr17 | 655419 | 655828 | 2.33831 | 5.92079 | 2.532 |
| chr10 | 70092483 | 70092754 | 0.682736 | 1.7288 | 2.532 |
| chr6 | 133361159 | 133361466 | 1.13755 | 2.88051 | 2.532 |
| chr12 | 99784396 | 99784815 | 0.96705 | 2.44891 | 2.532 |
| chr14 | 37715623 | 37715857 | 0.843918 | 2.13719 | 2.532 |
| chr12 | 56435338 | 56435557 | 1.2326 | 3.12154 | 2.532 |
| chr7 | 107641821 | 107642083 | 1.0822 | 2.7407 | 2.533 |
| chr12 | 103096347 | 103096753 | 1.32695 | 3.36057 | 2.533 |
| chr5 | 129103541 | 129104027 | 2.40923 | 6.10216 | 2.533 |
| chr8 | 76692575 | 76692852 | 0.47008 | 1.19064 | 2.533 |
| chr3 | 36742518 | 36742819 | 1.39142 | 3.5244 | 2.533 |
| chr12 | 54585353 | 54585704 | 1.86147 | 4.71515 | 2.533 |
| chr1 | 214331376 | 214331680 | 0.509698 | 1.29114 | 2.533 |
| chr5 | 142783137 | 142783770 | 0.986773 | 2.49966 | 2.533 |
| chr20 | 16959658 | 16959906 | 0.26068 | 0.660349 | 2.533 |
| chr9 | 124887379 | 124887850 | 1.75691 | 4.45089 | 2.533 |
| chr8 | 133204154 | 133204712 | 2.31932 | 5.87593 | 2.533 |
| chr1 | 10969989 | 10970364 | 1.43313 | 3.63105 | 2.534 |
| chr19 | 22306336 | 22306540 | 0.543921 | 1.37823 | 2.534 |
| chr15 | 45497172 | 45497659 | 1.33206 | 3.37537 | 2.534 |
| chr22 | 49700784 | 49701632 | 1.14115 | 2.89164 | 2.534 |
| chr19 | 50220721 | 50221121 | 1.59944 | 4.05295 | 2.534 |
| chr5 | 149540660 | 149541657 | 1.82394 | 4.62183 | 2.534 |
| chr9 | 98207886 | 98208126 | 0.36174 | 0.916664 | 2.534 |
| chr3 | 28740525 | 28740932 | 0.401306 | 1.01694 | 2.534 |
| chrX | 70872402 | 70872628 | 0.969397 | 2.4567 | 2.534 |
| chr17 | 64711876 | 64712185 | 1.27777 | 3.23821 | 2.534 |
| chr12 | 66501204 | 66501522 | 1.09001 | 2.76241 | 2.534 |
| chr20 | 10831882 | 10832179 | 0.540783 | 1.37054 | 2.534 |
| chr14 | 69329991 | 69330272 | 1.25583 | 3.18278 | 2.534 |
| chr6 | 13575513 | 13575730 | 0.905887 | 2.2959 | 2.534 |
| chr1 | 37899903 | 37900228 | 1.62321 | 4.11425 | 2.535 |
| chr7 | 68483187 | 68483585 | 0.941405 | 2.38621 | 2.535 |
| chr12 | 130101983 | 130102213 | 0.445209 | 1.1285 | 2.535 |
| chr21 | 39591048 | 39591303 | 1.52647 | 3.86936 | 2.535 |
| chr12 | 23209364 | 23209644 | 0.558971 | 1.41703 | 2.535 |
| chr20 | 3098609 | 3098887 | 1.51389 | 3.83806 | 2.535 |

| | | | | | |
|---|---|---|---|---|---|
| chr2 | 178013695 | 178013992 | 1.65501 | 4.19586 | 2.535 |
| chr17 | 1956725 | 1957205 | 1.36489 | 3.46037 | 2.535 |
| chr20 | 8646833 | 8647162 | 0.350282 | 0.888075 | 2.535 |
| chr21 | 17980826 | 17981035 | 0.950336 | 2.4094 | 2.535 |
| chr1 | 10092561 | 10092855 | 1.77569 | 4.5021 | 2.535 |
| chr17 | 35531254 | 35531529 | 0.937976 | 2.37818 | 2.535 |
| chr20 | 18609005 | 18609574 | 1.36705 | 3.46631 | 2.536 |
| chr17 | 57824271 | 57824534 | 1.14141 | 2.89432 | 2.536 |
| chr7 | 29649404 | 29649700 | 1.24791 | 3.16445 | 2.536 |
| chr12 | 57506155 | 57506398 | 1.42826 | 3.62224 | 2.536 |
| chr12 | 118935420 | 118935685 | 0.368405 | 0.934361 | 2.536 |
| chr12 | 122771565 | 122771852 | 0.392887 | 0.996479 | 2.536 |
| chr7 | 95862162 | 95862386 | 0.421068 | 1.06799 | 2.536 |
| chr1 | 155294983 | 155295234 | 1.00793 | 2.55658 | 2.536 |
| chr17 | 35730480 | 35730851 | 2.57819 | 6.53963 | 2.537 |
| chr7 | 151323386 | 151323992 | 1.04783 | 2.65791 | 2.537 |
| chr11 | 13689688 | 13690041 | 0.80935 | 2.05318 | 2.537 |
| chr12 | 13361460 | 13361680 | 1.54742 | 3.92565 | 2.537 |
| chr9 | 4604588 | 4604881 | 1.1978 | 3.03878 | 2.537 |
| chr3 | 79566984 | 79567249 | 0.784374 | 1.98997 | 2.537 |
| chr12 | 52766314 | 52766804 | 1.49118 | 3.78321 | 2.537 |
| chr13 | 28055979 | 28056330 | 0.902642 | 2.2901 | 2.537 |
| chr16 | 2066504 | 2067009 | 0.854601 | 2.16824 | 2.537 |
| chr1 | 210033666 | 210033991 | 1.39718 | 3.54489 | 2.537 |
| chr10 | 20825964 | 20826323 | 0.336093 | 0.852776 | 2.537 |
| chr17 | 63843760 | 63844113 | 0.465366 | 1.1808 | 2.537 |
| chr6 | 12289640 | 12289883 | 2.01311 | 5.1084 | 2.538 |
| chr8 | 127888777 | 127889264 | 2.1631 | 5.48935 | 2.538 |
| chr1 | 41314236 | 41314523 | 1.4252 | 3.61683 | 2.538 |
| chr6 | 110374923 | 110375138 | 0.765807 | 1.94349 | 2.538 |
| chr6 | 90996613 | 90996942 | 1.32018 | 3.3504 | 2.538 |
| chr1 | 15757541 | 15757933 | 1.56913 | 3.98245 | 2.538 |
| chr17 | 41376354 | 41376561 | 0.462629 | 1.17417 | 2.538 |
| chr2 | 121453155 | 121453565 | 1.16173 | 2.94861 | 2.538 |
| chr22 | 22901526 | 22901891 | 1.98737 | 5.04419 | 2.538 |
| chr12 | 123528028 | 123528371 | 1.87931 | 4.76994 | 2.538 |
| chr1 | 154378154 | 154378480 | 0.995418 | 2.52663 | 2.538 |
| chrX | 69674218 | 69674595 | 2.36001 | 5.99036 | 2.538 |
| chr1 | 180484994 | 180485281 | 0.96328 | 2.44509 | 2.538 |
| chrX | 152726296 | 152726729 | 1.67234 | 4.24494 | 2.538 |
| chr19 | 50715064 | 50715273 | 0.590627 | 1.49928 | 2.538 |
| chr9 | 17593285 | 17593541 | 0.459871 | 1.16738 | 2.538 |
| chr12 | 14409309 | 14409763 | 0.80843 | 2.05221 | 2.539 |
| chr14 | 94773760 | 94774104 | 1.01769 | 2.58348 | 2.539 |
| chr9 | 130352166 | 130352525 | 1.40457 | 3.56575 | 2.539 |
| chr10 | 108534718 | 108534930 | 0.519389 | 1.31861 | 2.539 |
| chr9 | 34562079 | 34562582 | 1.41487 | 3.59204 | 2.539 |
| chr3 | 13503030 | 13503498 | 1.81418 | 4.60603 | 2.539 |
| chr7 | 64079429 | 64079778 | 0.949122 | 2.40986 | 2.539 |
| chr11 | 61058721 | 61059138 | 0.928525 | 2.35762 | 2.539 |
| chr3 | 2054232 | 2054525 | 0.430398 | 1.09283 | 2.539 |
| chr19 | 54865524 | 54865855 | 1.57344 | 3.99516 | 2.539 |
| chr5 | 74532583 | 74532886 | 1.75585 | 4.45863 | 2.539 |
| chr7 | 105042076 | 105042339 | 1.11629 | 2.83484 | 2.540 |
| chr7 | 100674443 | 100674663 | 0.691378 | 1.75589 | 2.540 |
| chr18 | 37421441 | 37421808 | 1.02882 | 2.61292 | 2.540 |
| chr12 | 27805137 | 27805430 | 1.3396 | 3.40226 | 2.540 |
| chr9 | 111576322 | 111576566 | 0.932419 | 2.36827 | 2.540 |
| chr12 | 71435154 | 71435654 | 0.457099 | 1.16101 | 2.540 |
| chrX | 10690866 | 10691131 | 1.21076 | 3.07562 | 2.540 |
| chr14 | 95982476 | 95982682 | 0.489528 | 1.24353 | 2.540 |

| | | | | | |
|---|---|---|---|---|---|
| chr1 | 172104290 | 172104526 | 0.948216 | 2.40876 | 2.540 |
| chr18 | 50906529 | 50906766 | 0.623754 | 1.58453 | 2.540 |
| chr10 | 109376675 | 109377078 | 0.367525 | 0.933763 | 2.541 |
| chr7 | 70163254 | 70163539 | 1.44833 | 3.67977 | 2.541 |
| chrX | 114885331 | 114885567 | 1.06059 | 2.69479 | 2.541 |
| chr14 | 40896538 | 40896771 | 0.377289 | 0.958666 | 2.541 |
| chr3 | 57134212 | 57134434 | 1.05805 | 2.68847 | 2.541 |
| chr11 | 112090451 | 112090782 | 0.985663 | 2.50462 | 2.541 |
| chr4 | 96201061 | 96201269 | 0.486887 | 1.23721 | 2.541 |
| chr8 | 82553747 | 82554017 | 1.59878 | 4.06263 | 2.541 |
| chr10 | 7153990 | 7154192 | 0.421031 | 1.0699 | 2.541 |
| chr2 | 45878044 | 45878303 | 0.8027 | 2.0398 | 2.541 |
| chr3 | 53106724 | 53107103 | 1.27405 | 3.23766 | 2.541 |
| chr1 | 41981717 | 41982155 | 1.39578 | 3.54712 | 2.541 |
| chr5 | 12176261 | 12176479 | 0.45041 | 1.14465 | 2.541 |
| chr10 | 1422002 | 1422311 | 0.625647 | 1.59002 | 2.541 |
| chr2 | 72193960 | 72194215 | 0.457715 | 1.16324 | 2.541 |
| chr6 | 36997077 | 36997394 | 1.60332 | 4.07493 | 2.542 |
| chr5 | 72497737 | 72498156 | 1.8373 | 4.66971 | 2.542 |
| chr8 | 140502248 | 140502486 | 0.553493 | 1.40678 | 2.542 |
| chr22 | 30484920 | 30485224 | 1.15901 | 2.9458 | 2.542 |
| chr3 | 101396131 | 101396398 | 1.06639 | 2.71043 | 2.542 |
| chr10 | 133294678 | 133294886 | 0.515111 | 1.30927 | 2.542 |
| chr12 | 108766986 | 108767231 | 0.61003 | 1.55057 | 2.542 |
| chr20 | 33891616 | 33891963 | 1.86665 | 4.74465 | 2.542 |
| chr1 | 36886453 | 36886747 | 1.44323 | 3.66841 | 2.542 |
| chr6 | 137592758 | 137593060 | 1.15676 | 2.94039 | 2.542 |
| chr17 | 57243308 | 57243598 | 1.20426 | 3.0612 | 2.542 |
| chr5 | 131435650 | 131435937 | 0.344486 | 0.875686 | 2.542 |
| chr2 | 406988 | 407236 | 0.424177 | 1.07832 | 2.542 |
| chr12 | 112328804 | 112329045 | 0.424219 | 1.07846 | 2.542 |
| chr20 | 42744700 | 42745715 | 1.212 | 3.0812 | 2.542 |
| chr19 | 47329432 | 47329644 | 0.423731 | 1.07726 | 2.542 |
| chr3 | 112057340 | 112057577 | 0.550056 | 1.39848 | 2.542 |
| chr10 | 65389175 | 65389485 | 1.89809 | 4.82579 | 2.542 |
| chr7 | 39545222 | 39545431 | 0.392122 | 0.996951 | 2.542 |
| chr4 | 78520605 | 78520861 | 1.02866 | 2.61536 | 2.542 |
| chr4 | 155547829 | 155548263 | 1.90907 | 4.85381 | 2.543 |
| chr11 | 14322308 | 14322613 | 1.24196 | 3.1577 | 2.543 |
| chr14 | 104544260 | 104544717 | 3.11053 | 7.90855 | 2.543 |
| chr1 | 172413162 | 172413441 | 2.13896 | 5.43854 | 2.543 |
| chr7 | 56160667 | 56160970 | 1.08048 | 2.74724 | 2.543 |
| chr9 | 37904104 | 37904620 | 1.30244 | 3.31177 | 2.543 |
| chr3 | 163634697 | 163634910 | 0.692385 | 1.76066 | 2.543 |
| chr20 | 30090869 | 30091288 | 2.01042 | 5.11236 | 2.543 |
| chr11 | 65812517 | 65813037 | 2.26053 | 5.74839 | 2.543 |
| chr11 | 62037185 | 62037539 | 0.544491 | 1.38465 | 2.543 |
| chr14 | 95465189 | 95465409 | 0.480193 | 1.22115 | 2.543 |
| chr2 | 20297958 | 20298296 | 2.49185 | 6.33691 | 2.543 |
| chr18 | 29536952 | 29537192 | 1.089 | 2.7694 | 2.543 |
| chr11 | 77939470 | 77939704 | 1.16466 | 2.96185 | 2.543 |
| chr2 | 21813198 | 21813417 | 0.437658 | 1.11301 | 2.543 |
| chr6 | 49578628 | 49578893 | 0.323261 | 0.822179 | 2.543 |
| chr11 | 45105121 | 45105395 | 1.22825 | 3.1241 | 2.544 |
| chr2 | 4627235 | 4627522 | 0.398095 | 1.01257 | 2.544 |
| chr11 | 6656917 | 6657120 | 0.334145 | 0.849928 | 2.544 |
| chr22 | 20850766 | 20851100 | 1.44726 | 3.68145 | 2.544 |
| chr12 | 111263378 | 111263636 | 0.426062 | 1.08394 | 2.544 |
| chr10 | 131675914 | 131676471 | 0.903834 | 2.29965 | 2.544 |
| chr5 | 7821741 | 7822159 | 0.455075 | 1.15789 | 2.544 |
| chr15 | 89942567 | 89942867 | 1.43821 | 3.65943 | 2.544 |

| | | | | | |
|---|---|---|---|---|---|
| chr19 | 31568372 | 31568636 | 0.409321 | 1.04149 | 2.544 |
| chr5 | 10424304 | 10424524 | 1.24625 | 3.17099 | 2.544 |
| chr11 | 37781386 | 37781639 | 0.370618 | 0.94304 | 2.545 |
| chr5 | 176927920 | 176928554 | 1.63094 | 4.15001 | 2.545 |
| chr2 | 67957369 | 67957650 | 0.38381 | 0.976685 | 2.545 |
| chr7 | 125533914 | 125534139 | 0.430957 | 1.09671 | 2.545 |
| chr10 | 6487661 | 6487967 | 1.42164 | 3.61783 | 2.545 |
| chr1 | 208305503 | 208305935 | 1.41464 | 3.60004 | 2.545 |
| chr7 | 95704748 | 95705083 | 1.67082 | 4.25216 | 2.545 |
| chr2 | 236845161 | 236845519 | 1.2665 | 3.2232 | 2.545 |
| chr4 | 78619418 | 78619724 | 0.935116 | 2.37989 | 2.545 |
| chr10 | 50076191 | 50076557 | 0.449569 | 1.14429 | 2.545 |
| chr9 | 128320777 | 128321020 | 1.5447 | 3.93174 | 2.545 |
| chr9 | 130672978 | 130673433 | 0.750889 | 1.91135 | 2.545 |
| chr7 | 98030342 | 98030892 | 1.56626 | 3.98691 | 2.546 |
| chr3 | 16925515 | 16925894 | 1.40739 | 3.58253 | 2.546 |
| chr12 | 52513603 | 52514151 | 1.95618 | 4.97967 | 2.546 |
| chr11 | 727886 | 728424 | 1.81907 | 4.63074 | 2.546 |
| chr1 | 150785605 | 150785999 | 1.30151 | 3.31321 | 2.546 |
| chr5 | 140213715 | 140214150 | 0.508231 | 1.29392 | 2.546 |
| chr5 | 80490836 | 80491235 | 1.74985 | 4.45516 | 2.546 |
| chr14 | 103430517 | 103430736 | 0.582019 | 1.48184 | 2.546 |
| chr13 | 48652324 | 48652543 | 0.763373 | 1.9437 | 2.546 |
| chr1 | 21581765 | 21582105 | 1.40688 | 3.58227 | 2.546 |
| chr1 | 221706068 | 221706276 | 0.432288 | 1.10091 | 2.547 |
| chr7 | 140375134 | 140375354 | 0.775573 | 1.97524 | 2.547 |
| chr9 | 34603556 | 34604213 | 1.33308 | 3.39512 | 2.547 |
| chr4 | 122924616 | 122924849 | 1.1069 | 2.8191 | 2.547 |
| chr14 | 35876176 | 35876456 | 1.15084 | 2.93113 | 2.547 |
| chr11 | 68065190 | 68065592 | 1.88884 | 4.81081 | 2.547 |
| chr6 | 41392270 | 41392608 | 1.06214 | 2.70531 | 2.547 |
| chr10 | 126218641 | 126219050 | 2.76299 | 7.03778 | 2.547 |
| chr7 | 24760506 | 24760772 | 0.435339 | 1.10891 | 2.547 |
| chr9 | 36547392 | 36547641 | 0.432364 | 1.10142 | 2.547 |
| chr1 | 172398481 | 172398818 | 1.64746 | 4.19684 | 2.547 |
| chrX | 149654319 | 149654642 | 1.45955 | 3.71825 | 2.548 |
| chr12 | 77404602 | 77405154 | 1.89373 | 4.82441 | 2.548 |
| chr5 | 39716662 | 39716892 | 1.08285 | 2.75865 | 2.548 |
| chr2 | 3307398 | 3307845 | 1.84034 | 4.68858 | 2.548 |
| chr8 | 64483113 | 64483470 | 0.43011 | 1.09593 | 2.548 |
| chr8 | 110237589 | 110237824 | 0.441666 | 1.12538 | 2.548 |
| chr9 | 116417701 | 116418100 | 1.55888 | 3.97209 | 2.548 |
| chr12 | 118274984 | 118275474 | 2.18862 | 5.57683 | 2.548 |
| chr13 | 101240858 | 101241312 | 1.54506 | 3.93699 | 2.548 |
| chr14 | 86378258 | 86378514 | 0.385659 | 0.98272 | 2.548 |
| chr1 | 94706446 | 94706678 | 0.723758 | 1.84426 | 2.548 |
| chr5 | 162226348 | 162226550 | 0.495652 | 1.26301 | 2.548 |
| chr1 | 116088819 | 116089110 | 1.24038 | 3.16085 | 2.548 |
| chr19 | 31092541 | 31092759 | 0.505106 | 1.28719 | 2.548 |
| chr17 | 43309098 | 43309402 | 1.62082 | 4.13053 | 2.548 |
| chr19 | 56616534 | 56616805 | 1.56194 | 3.98061 | 2.549 |
| chr7 | 90089321 | 90089575 | 0.643018 | 1.63881 | 2.549 |
| chr16 | 50279785 | 50280090 | 0.831193 | 2.11842 | 2.549 |
| chr7 | 35311023 | 35311243 | 0.40268 | 1.02629 | 2.549 |
| chrX | 77285236 | 77285472 | 1.16244 | 2.96267 | 2.549 |
| chr20 | 2383358 | 2383646 | 0.996731 | 2.54035 | 2.549 |
| chrX | 120712950 | 120713255 | 0.758042 | 1.93201 | 2.549 |
| chr12 | 24510285 | 24510692 | 0.44675 | 1.13864 | 2.549 |
| chr8 | 92863833 | 92864343 | 0.408259 | 1.04054 | 2.549 |
| chr12 | 107036894 | 107037115 | 0.466969 | 1.19024 | 2.549 |
| chr13 | 71585420 | 71585671 | 0.504375 | 1.28559 | 2.549 |

| | | | | | |
|---|---|---|---|---|---|
| chrX | 93313710 | 93314045 | 0.684701 | 1.74541 | 2.549 |
| chr10 | 81016934 | 81017196 | 0.372141 | 0.94871 | 2.549 |
| chr7 | 130130737 | 130131019 | 0.883613 | 2.25271 | 2.549 |
| chr20 | 19765585 | 19765875 | 1.45294 | 3.70432 | 2.550 |
| chr1 | 218344752 | 218344960 | 1.0285 | 2.62223 | 2.550 |
| chr20 | 17485468 | 17485930 | 0.860431 | 2.19378 | 2.550 |
| chr1 | 9675675 | 9676015 | 1.51536 | 3.86374 | 2.550 |
| chr19 | 4791058 | 4791431 | 1.5385 | 3.92315 | 2.550 |
| chr6 | 11172766 | 11173068 | 1.75231 | 4.46835 | 2.550 |
| chr20 | 48687768 | 48688223 | 1.82288 | 4.64878 | 2.550 |
| chr11 | 7377630 | 7377846 | 0.442072 | 1.12747 | 2.550 |
| chr7 | 30805034 | 30805336 | 0.416081 | 1.06119 | 2.550 |
| chr1 | 218433195 | 218433653 | 0.596699 | 1.52192 | 2.551 |
| chrX | 11162221 | 11162686 | 1.07562 | 2.74347 | 2.551 |
| chr12 | 114659993 | 114660541 | 0.379459 | 0.967852 | 2.551 |
| chr2 | 99029372 | 99029665 | 1.58967 | 4.05478 | 2.551 |
| chr2 | 49573570 | 49573890 | 0.475266 | 1.21228 | 2.551 |
| chr17 | 35111894 | 35112296 | 1.44703 | 3.69102 | 2.551 |
| chr1 | 177368821 | 177369028 | 0.473848 | 1.20868 | 2.551 |
| chr17 | 71425343 | 71426033 | 1.86511 | 4.75759 | 2.551 |
| chr1 | 210184732 | 210184951 | 0.392584 | 1.00144 | 2.551 |
| chr10 | 68685603 | 68685846 | 1.06375 | 2.71356 | 2.551 |
| chr19 | 45593288 | 45593560 | 0.471125 | 1.20181 | 2.551 |
| chr14 | 84915572 | 84915822 | 0.438009 | 1.11736 | 2.551 |
| chr8 | 107542536 | 107542788 | 0.498809 | 1.2725 | 2.551 |
| chr11 | 71802317 | 71802623 | 1.2824 | 3.27152 | 2.551 |
| chr10 | 134699955 | 134700273 | 0.807034 | 2.05893 | 2.551 |
| chr17 | 4812421 | 4812819 | 1.5713 | 4.00884 | 2.551 |
| chr18 | 76635478 | 76635874 | 1.15777 | 2.95388 | 2.551 |
| chr19 | 2749190 | 2749624 | 1.95481 | 4.98747 | 2.551 |
| chr17 | 72745127 | 72745363 | 0.560453 | 1.42995 | 2.551 |
| chr3 | 169790649 | 169790950 | 0.437112 | 1.11526 | 2.551 |
| chr2 | 6986129 | 6986345 | 0.934662 | 2.38482 | 2.552 |
| chr7 | 118600081 | 118600292 | 0.355702 | 0.90764 | 2.552 |
| chr20 | 55201621 | 55202065 | 1.05209 | 2.68463 | 2.552 |
| chr10 | 106670590 | 106671111 | 0.408307 | 1.04189 | 2.552 |
| chr11 | 61062177 | 61062390 | 0.832293 | 2.12396 | 2.552 |
| chr16 | 19897526 | 19897946 | 2.40346 | 6.13436 | 2.552 |
| chr12 | 108278894 | 108279208 | 1.46184 | 3.73118 | 2.552 |
| chr10 | 33619482 | 33619722 | 1.61494 | 4.12211 | 2.552 |
| chr5 | 179742538 | 179743110 | 1.42897 | 3.64753 | 2.553 |
| chr14 | 21304900 | 21305142 | 1.08944 | 2.78094 | 2.553 |
| chr10 | 98525457 | 98525834 | 1.41809 | 3.61987 | 2.553 |
| chr4 | 57687711 | 57687979 | 0.743306 | 1.89743 | 2.553 |
| chr1 | 120491290 | 120491598 | 1.54336 | 3.93981 | 2.553 |
| chr2 | 193150894 | 193151119 | 0.845597 | 2.15861 | 2.553 |
| chr10 | 121249215 | 121249626 | 1.78451 | 4.55561 | 2.553 |
| chr4 | 113156267 | 113156519 | 0.97897 | 2.4992 | 2.553 |
| chr19 | 22529356 | 22529825 | 0.663771 | 1.6946 | 2.553 |
| chr4 | 139416177 | 139416402 | 0.750203 | 1.91538 | 2.553 |
| chr3 | 32551541 | 32551918 | 0.935309 | 2.38821 | 2.553 |
| chr10 | 98273438 | 98273733 | 0.956243 | 2.44225 | 2.554 |
| chr17 | 58114447 | 58114900 | 2.03369 | 5.19442 | 2.554 |
| chr10 | 79403673 | 79404223 | 1.35999 | 3.47378 | 2.554 |
| chr12 | 103605182 | 103605411 | 0.539544 | 1.37817 | 2.554 |
| chr12 | 3384419 | 3385036 | 1.71171 | 4.37235 | 2.554 |
| chr14 | 74718213 | 74718650 | 1.4929 | 3.81348 | 2.554 |
| chr20 | 55623333 | 55623636 | 1.3179 | 3.36648 | 2.554 |
| chr8 | 56110070 | 56110447 | 0.301356 | 0.7698 | 2.554 |
| chr14 | 97351089 | 97351319 | 0.473911 | 1.21062 | 2.555 |
| chr7 | 139183384 | 139183609 | 0.411144 | 1.05028 | 2.555 |

| | | | | | |
|---|---|---|---|---|---|
| chr1 | 66719095 | 66719328 | 0.955967 | 2.4421 | 2.555 |
| chr9 | 5784058 | 5784339 | 0.503373 | 1.28595 | 2.555 |
| chr6 | 53036655 | 53037104 | 3.25286 | 8.3104 | 2.555 |
| chr4 | 72136705 | 72136971 | 0.59172 | 1.51175 | 2.555 |
| chr1 | 189677790 | 189678084 | 1.02509 | 2.61909 | 2.555 |
| chr14 | 101367276 | 101367515 | 1.04741 | 2.67629 | 2.555 |
| chr7 | 77960542 | 77960756 | 0.449391 | 1.1483 | 2.555 |
| chr6 | 47701937 | 47702216 | 0.623877 | 1.59422 | 2.555 |
| chr19 | 50006408 | 50006676 | 1.11075 | 2.83838 | 2.555 |
| chr17 | 56190363 | 56190899 | 1.55446 | 3.97234 | 2.555 |
| chr6 | 165341033 | 165341398 | 2.18549 | 5.58496 | 2.555 |
| chr5 | 1349437 | 1349700 | 0.711933 | 1.81934 | 2.555 |
| chr10 | 96943237 | 96943893 | 2.4262 | 6.20051 | 2.556 |
| chr6 | 116217789 | 116218034 | 0.992944 | 2.53762 | 2.556 |
| chr11 | 102139510 | 102139785 | 0.763495 | 1.95123 | 2.556 |
| chr2 | 42423908 | 42424155 | 1.57561 | 4.02686 | 2.556 |
| chr20 | 3835713 | 3836104 | 1.81457 | 4.6379 | 2.556 |
| chr5 | 118323740 | 118324413 | 2.62655 | 6.71327 | 2.556 |
| chr1 | 950852 | 951127 | 1.0596 | 2.70845 | 2.556 |
| chr5 | 172329941 | 172330247 | 1.5397 | 3.93608 | 2.556 |
| chr19 | 19642993 | 19643418 | 1.4869 | 3.80137 | 2.557 |
| chr5 | 175274496 | 175274759 | 1.04661 | 2.67584 | 2.557 |
| chr12 | 99590700 | 99590984 | 0.487383 | 1.24612 | 2.557 |
| chr5 | 154352083 | 154352422 | 1.34146 | 3.42991 | 2.557 |
| chr19 | 50550016 | 50550658 | 0.96938 | 2.47863 | 2.557 |
| chr12 | 116715474 | 116715829 | 1.25228 | 3.20199 | 2.557 |
| chr15 | 70021513 | 70021841 | 1.35806 | 3.47252 | 2.557 |
| chr3 | 125931561 | 125932183 | 0.89094 | 2.27811 | 2.557 |
| chr3 | 13698196 | 13698411 | 0.316668 | 0.809727 | 2.557 |
| chr20 | 39968549 | 39969006 | 1.82745 | 4.67336 | 2.557 |
| chr15 | 75092320 | 75092999 | 2.16375 | 5.53343 | 2.557 |
| chr9 | 115096014 | 115096394 | 1.36807 | 3.49865 | 2.557 |
| chr15 | 69686904 | 69687368 | 1.64879 | 4.21658 | 2.557 |
| chr3 | 118601271 | 118601539 | 1.32518 | 3.38904 | 2.557 |
| chr16 | 71533690 | 71533981 | 1.0152 | 2.59633 | 2.557 |
| chr3 | 16558583 | 16558991 | 1.07686 | 2.75405 | 2.557 |
| chr14 | 69310623 | 69311053 | 1.58427 | 4.05175 | 2.557 |
| chr8 | 114501474 | 114501716 | 0.798307 | 2.04177 | 2.558 |
| chr5 | 114064302 | 114064531 | 0.409995 | 1.04862 | 2.558 |
| chr5 | 151769644 | 151769852 | 0.363581 | 0.929938 | 2.558 |
| chr11 | 1285660 | 1285922 | 0.96023 | 2.45628 | 2.558 |
| chr15 | 102118738 | 102119197 | 1.40449 | 3.59308 | 2.558 |
| chr15 | 86659579 | 86659825 | 0.44663 | 1.14264 | 2.558 |
| chr1 | 182931628 | 182931942 | 1.78175 | 4.55861 | 2.559 |
| chr5 | 145945380 | 145945602 | 1.03856 | 2.6573 | 2.559 |
| chr9 | 71131325 | 71131537 | 0.510094 | 1.30515 | 2.559 |
| chr18 | 34862454 | 34862739 | 0.967396 | 2.47528 | 2.559 |
| chr21 | 18533876 | 18534117 | 0.31291 | 0.80067 | 2.559 |
| chr15 | 92620555 | 92620994 | 1.98439 | 5.07766 | 2.559 |
| chr13 | 70111642 | 70111875 | 0.503376 | 1.28805 | 2.559 |
| chr14 | 105906302 | 105906934 | 1.04365 | 2.67053 | 2.559 |
| chr17 | 30469246 | 30469576 | 1.72566 | 4.41585 | 2.559 |
| chr1 | 3643876 | 3644177 | 1.42382 | 3.64347 | 2.559 |
| chr20 | 61169276 | 61169538 | 0.752228 | 1.92503 | 2.559 |
| chr21 | 26980634 | 26980897 | 1.25182 | 3.20354 | 2.559 |
| chr12 | 116127998 | 116128439 | 0.426466 | 1.09144 | 2.559 |
| chr6 | 10429917 | 10430144 | 0.840044 | 2.14994 | 2.559 |
| chr6 | 56708994 | 56709333 | 1.00368 | 2.56881 | 2.559 |
| chr5 | 160326830 | 160327111 | 0.610731 | 1.56312 | 2.559 |
| chr17 | 49339447 | 49339759 | 0.419198 | 1.07291 | 2.559 |
| chr15 | 31734795 | 31735055 | 0.575407 | 1.47288 | 2.560 |

| | | | | | |
|---|---|---|---|---|---|
| chr17 | 74275524 | 74276019 | 2.27844 | 5.83225 | 2.560 |
| chrX | 12788454 | 12788852 | 0.799576 | 2.04688 | 2.560 |
| chr1 | 230108809 | 230109049 | 1.02023 | 2.61185 | 2.560 |
| chr16 | 31471999 | 31472292 | 1.10631 | 2.83254 | 2.560 |
| chr19 | 2156066 | 2156295 | 1.28896 | 3.30028 | 2.560 |
| chr14 | 64266412 | 64266663 | 0.513871 | 1.31579 | 2.561 |
| chr14 | 67076858 | 67077206 | 0.887729 | 2.27309 | 2.561 |
| chr5 | 176946971 | 176947279 | 0.481657 | 1.2334 | 2.561 |
| chr18 | 25330126 | 25330331 | 0.72417 | 1.85447 | 2.561 |
| chr2 | 163795035 | 163795272 | 0.78343 | 2.00625 | 2.561 |
| chr1 | 15373039 | 15373536 | 1.12898 | 2.89124 | 2.561 |
| chr12 | 131246180 | 131246592 | 0.937433 | 2.40071 | 2.561 |
| chr8 | 38578930 | 38579177 | 1.07057 | 2.74173 | 2.561 |
| chr12 | 109825160 | 109825595 | 1.14627 | 2.9358 | 2.561 |
| chr19 | 5048303 | 5048615 | 1.68279 | 4.30994 | 2.561 |
| chr8 | 144354678 | 144355011 | 1.01817 | 2.60775 | 2.561 |
| chr6 | 157505452 | 157505709 | 1.48208 | 3.796 | 2.561 |
| chr2 | 41504811 | 41505064 | 0.53072 | 1.35943 | 2.561 |
| chr16 | 23814013 | 23814304 | 0.488597 | 1.25155 | 2.562 |
| chr3 | 22674545 | 22674771 | 0.262648 | 0.6728 | 2.562 |
| chr4 | 131280769 | 131280985 | 0.834155 | 2.13684 | 2.562 |
| chr10 | 86187604 | 86187832 | 0.477953 | 1.22439 | 2.562 |
| chr14 | 24630187 | 24630548 | 1.65202 | 4.23245 | 2.562 |
| chr3 | 27753563 | 27753929 | 1.52089 | 3.89649 | 2.562 |
| chr1 | 230404674 | 230404958 | 1.35407 | 3.46922 | 2.562 |
| chr8 | 102339215 | 102339516 | 1.7652 | 4.52278 | 2.562 |
| chr12 | 64758176 | 64758391 | 0.441044 | 1.13009 | 2.562 |
| chr5 | 177706948 | 177707318 | 1.71545 | 4.3961 | 2.563 |
| chr8 | 99953134 | 99953574 | 1.70193 | 4.3615 | 2.563 |
| chrX | 47691243 | 47691527 | 1.29152 | 3.30988 | 2.563 |
| chr17 | 38571646 | 38571899 | 1.02201 | 2.61925 | 2.563 |
| chr1 | 9161641 | 9161983 | 0.912633 | 2.33899 | 2.563 |
| chr7 | 152266855 | 152267192 | 0.383977 | 0.984155 | 2.563 |
| chr15 | 31421447 | 31421885 | 2.18579 | 5.60233 | 2.563 |
| chr16 | 230144 | 230973 | 1.11904 | 2.86819 | 2.563 |
| chr8 | 25602694 | 25602982 | 0.468592 | 1.20107 | 2.563 |
| chr11 | 4560871 | 4561157 | 1.26769 | 3.2493 | 2.563 |
| chr15 | 40660099 | 40660611 | 1.53868 | 3.94399 | 2.563 |
| chr22 | 27727911 | 27728124 | 1.1065 | 2.83621 | 2.563 |
| chr12 | 7071062 | 7071491 | 1.04183 | 2.67051 | 2.563 |
| chr22 | 29920478 | 29920720 | 0.81426 | 2.0872 | 2.563 |
| chr7 | 91506134 | 91506386 | 0.744688 | 1.90888 | 2.563 |
| chr11 | 76906853 | 76907160 | 0.920588 | 2.35991 | 2.563 |
| chr3 | 196756627 | 196757006 | 1.85577 | 4.75734 | 2.564 |
| chr1 | 170995433 | 170995717 | 0.354282 | 0.908227 | 2.564 |
| chr17 | 7183827 | 7184258 | 2.51869 | 6.45685 | 2.564 |
| chr6 | 56767746 | 56768040 | 1.58088 | 4.05271 | 2.564 |
| chr15 | 40337310 | 40337647 | 1.64713 | 4.22288 | 2.564 |
| chr1 | 158091501 | 158091811 | 1.17025 | 3.00027 | 2.564 |
| chr8 | 27459597 | 27459890 | 1.51705 | 3.8894 | 2.564 |
| chr12 | 119417501 | 119417908 | 1.54546 | 3.96273 | 2.564 |
| chr7 | 158947033 | 158947320 | 0.713865 | 1.83049 | 2.564 |
| chr5 | 72511881 | 72512474 | 1.61225 | 4.13436 | 2.564 |
| chr17 | 56249846 | 56250127 | 1.03933 | 2.66522 | 2.564 |
| chr17 | 55841082 | 55841288 | 0.447516 | 1.14763 | 2.564 |
| chr1 | 220050065 | 220050479 | 0.92502 | 2.37219 | 2.564 |
| chr14 | 29332727 | 29332942 | 0.373641 | 0.958208 | 2.565 |
| chr7 | 85566851 | 85567061 | 0.528506 | 1.35542 | 2.565 |
| chr7 | 91808145 | 91808573 | 2.03792 | 5.22697 | 2.565 |
| chr8 | 108792092 | 108792296 | 0.964207 | 2.47336 | 2.565 |
| chr6 | 27719996 | 27720357 | 1.3098 | 3.35991 | 2.565 |

| | | | | | |
|---|---|---|---|---|---|
| chr10 | 129041797 | 129042183 | 1.54904 | 3.9737 | 2.565 |
| chr8 | 82426316 | 82426545 | 0.41836 | 1.07321 | 2.565 |
| chr10 | 94602878 | 94603186 | 1.50294 | 3.85562 | 2.565 |
| chr7 | 94024820 | 94025059 | 0.889806 | 2.28284 | 2.566 |
| chr9 | 136313605 | 136313841 | 0.397654 | 1.02023 | 2.566 |
| chr8 | 144363747 | 144364094 | 2.27216 | 5.82953 | 2.566 |
| chr11 | 13306145 | 13306370 | 0.391096 | 1.00341 | 2.566 |
| chr19 | 10380128 | 10380521 | 1.82389 | 4.67961 | 2.566 |
| chr22 | 21239763 | 21240266 | 1.63116 | 4.18517 | 2.566 |
| chr5 | 37837402 | 37837801 | 1.06382 | 2.72958 | 2.566 |
| chr1 | 247536031 | 247536335 | 1.63202 | 4.18762 | 2.566 |
| chr16 | 75551274 | 75551696 | 1.73056 | 4.4409 | 2.566 |
| chr1 | 11380244 | 11380584 | 1.31937 | 3.38586 | 2.566 |
| chr5 | 165508961 | 165509189 | 0.405539 | 1.04076 | 2.566 |
| chr5 | 7373264 | 7373574 | 0.423342 | 1.08655 | 2.567 |
| chr12 | 110989899 | 110990404 | 1.15552 | 2.9658 | 2.567 |
| chr19 | 47210452 | 47210729 | 0.483445 | 1.24084 | 2.567 |
| chr1 | 238463250 | 238463453 | 0.425552 | 1.09226 | 2.567 |
| chr13 | 110764796 | 110765066 | 1.20804 | 3.10073 | 2.567 |
| chr1 | 160747311 | 160747573 | 0.97288 | 2.49732 | 2.567 |
| chr15 | 90099934 | 90100190 | 1.222 | 3.1369 | 2.567 |
| chr15 | 69217171 | 69217552 | 2.07237 | 5.32025 | 2.567 |
| chr10 | 7302443 | 7302645 | 0.973038 | 2.49807 | 2.567 |
| chrX | 48901268 | 48901620 | 1.3338 | 3.42428 | 2.567 |
| chr1 | 158083185 | 158083659 | 1.05588 | 2.71089 | 2.567 |
| chr15 | 50474223 | 50474619 | 0.987786 | 2.53616 | 2.568 |
| chr14 | 105667819 | 105668075 | 0.586639 | 1.50627 | 2.568 |
| chr16 | 55055815 | 55056156 | 0.803773 | 2.06383 | 2.568 |
| chr12 | 115100398 | 115100736 | 1.40427 | 3.60584 | 2.568 |
| chr1 | 159009208 | 159009495 | 0.329547 | 0.846232 | 2.568 |
| chr20 | 42709367 | 42709819 | 1.57812 | 4.05239 | 2.568 |
| chr8 | 59521079 | 59521555 | 0.950269 | 2.44018 | 2.568 |
| chr11 | 60733917 | 60734551 | 0.324399 | 0.833034 | 2.568 |
| chr4 | 24474597 | 24475018 | 1.50662 | 3.86891 | 2.568 |
| chr1 | 204777234 | 204777542 | 1.05835 | 2.71795 | 2.568 |
| chr15 | 98861018 | 98861310 | 0.4283 | 1.09995 | 2.568 |
| chr14 | 51702848 | 51703137 | 0.532873 | 1.36852 | 2.568 |
| chr17 | 1958081 | 1958607 | 2.35311 | 6.04337 | 2.568 |
| chr5 | 105151643 | 105151882 | 0.353962 | 0.909245 | 2.569 |
| chr20 | 18612831 | 18613196 | 3.25063 | 8.35015 | 2.569 |
| chr3 | 5468093 | 5468397 | 1.87373 | 4.81331 | 2.569 |
| chr11 | 46389002 | 46389735 | 1.94637 | 5.00004 | 2.569 |
| chr4 | 44450932 | 44451182 | 1.06091 | 2.72548 | 2.569 |
| chr14 | 64236976 | 64237204 | 1.33256 | 3.4234 | 2.569 |
| chr3 | 2046537 | 2046817 | 0.522741 | 1.34296 | 2.569 |
| chr5 | 149370311 | 149370693 | 1.60624 | 4.12662 | 2.569 |
| chr17 | 2689455 | 2689959 | 2.21816 | 5.69874 | 2.569 |
| chr14 | 100259505 | 100259721 | 0.820352 | 2.10761 | 2.569 |
| chr16 | 19097770 | 19098141 | 2.09491 | 5.38221 | 2.569 |
| chr5 | 137469176 | 137469448 | 0.795446 | 2.04366 | 2.569 |
| chr8 | 24749298 | 24749655 | 1.33043 | 3.41839 | 2.569 |
| chr1 | 112841161 | 112841539 | 0.902835 | 2.31978 | 2.569 |
| chr2 | 24149837 | 24150211 | 1.79403 | 4.60979 | 2.570 |
| chr13 | 87691265 | 87691544 | 0.371013 | 0.953363 | 2.570 |
| chr16 | 1873911 | 1874124 | 0.85531 | 2.1979 | 2.570 |
| chr3 | 194315532 | 194315738 | 0.795836 | 2.04508 | 2.570 |
| chr2 | 224858439 | 224858665 | 0.769769 | 1.97811 | 2.570 |
| chr8 | 48796320 | 48796555 | 0.606906 | 1.55959 | 2.570 |
| chrX | 15516036 | 15516332 | 1.17256 | 3.0135 | 2.570 |
| chr4 | 118955414 | 118955667 | 1.22554 | 3.14988 | 2.570 |
| chr4 | 75973661 | 75973889 | 0.398298 | 1.02374 | 2.570 |

| | | | | | |
|---|---|---|---|---|---|
| chr5 | 18386562 | 18386851 | 0.591008 | 1.51929 | 2.571 |
| chr12 | 41761892 | 41762105 | 0.961796 | 2.47254 | 2.571 |
| chrX | 129673675 | 129673984 | 1.05632 | 2.71555 | 2.571 |
| chr1 | 27883280 | 27883608 | 1.40486 | 3.61186 | 2.571 |
| chr15 | 88183731 | 88184039 | 0.562006 | 1.44494 | 2.571 |
| chr14 | 70041347 | 70041863 | 2.3658 | 6.08282 | 2.571 |
| chr19 | 19026573 | 19026794 | 0.699416 | 1.79836 | 2.571 |
| chr11 | 44578205 | 44578576 | 1.53135 | 3.93762 | 2.571 |
| chr14 | 22479063 | 22479279 | 0.44112 | 1.13427 | 2.571 |
| chr4 | 167942783 | 167943045 | 0.924393 | 2.37739 | 2.572 |
| chr5 | 109782408 | 109782666 | 0.326412 | 0.839494 | 2.572 |
| chr1 | 11397574 | 11397981 | 1.72447 | 4.4355 | 2.572 |
| chr3 | 153891976 | 153892196 | 0.492787 | 1.26753 | 2.572 |
| chr17 | 55038286 | 55038648 | 2.12557 | 5.46773 | 2.572 |
| chr1 | 38356138 | 38356345 | 0.635559 | 1.63497 | 2.572 |
| chr16 | 60494626 | 60494897 | 0.484419 | 1.24623 | 2.573 |
| chr11 | 61584314 | 61584642 | 1.66663 | 4.28771 | 2.573 |
| chr15 | 99045655 | 99045899 | 0.456703 | 1.17498 | 2.573 |
| chr6 | 14711327 | 14711620 | 0.736437 | 1.89488 | 2.573 |
| chr20 | 49204553 | 49205092 | 2.26604 | 5.83077 | 2.573 |
| chr10 | 37552329 | 37552563 | 0.817938 | 2.10477 | 2.573 |
| chr16 | 51823328 | 51823596 | 0.420804 | 1.0829 | 2.573 |
| chr2 | 237051927 | 237052167 | 0.967224 | 2.48908 | 2.573 |
| chr3 | 53253960 | 53254329 | 1.45168 | 3.73587 | 2.573 |
| chr22 | 31957326 | 31957587 | 1.40824 | 3.62409 | 2.573 |
| chr4 | 142054601 | 142054889 | 1.37395 | 3.5359 | 2.574 |
| chr11 | 76757042 | 76757525 | 2.23066 | 5.74087 | 2.574 |
| chr11 | 46511575 | 46511833 | 1.08006 | 2.7797 | 2.574 |
| chr4 | 78394787 | 78395001 | 0.570357 | 1.46797 | 2.574 |
| chr9 | 11135926 | 11136156 | 0.350663 | 0.902571 | 2.574 |
| chr16 | 84759598 | 84759980 | 1.86437 | 4.79875 | 2.574 |
| chr15 | 98985609 | 98986013 | 1.70085 | 4.37795 | 2.574 |
| chrX | 118201054 | 118201271 | 1.12331 | 2.89142 | 2.574 |
| chr19 | 58883744 | 58884022 | 0.516347 | 1.32914 | 2.574 |
| chr5 | 101807618 | 101807839 | 0.412729 | 1.06253 | 2.574 |
| chr12 | 96409571 | 96409779 | 0.899954 | 2.31688 | 2.574 |
| chr8 | 135941720 | 135942153 | 0.437163 | 1.12545 | 2.574 |
| chr8 | 104776581 | 104776827 | 0.952945 | 2.45347 | 2.575 |
| chr6 | 166869895 | 166870106 | 0.813233 | 2.09379 | 2.575 |
| chr16 | 55090455 | 55090692 | 0.810862 | 2.08772 | 2.575 |
| chr1 | 805200 | 805580 | 0.493422 | 1.2706 | 2.575 |
| chr16 | 27983813 | 27984168 | 1.07775 | 2.77557 | 2.575 |
| chr2 | 99539694 | 99539998 | 0.920387 | 2.37033 | 2.575 |
| chr12 | 8671816 | 8672116 | 0.395152 | 1.01769 | 2.575 |
| chr3 | 188865882 | 188866108 | 0.812195 | 2.09188 | 2.576 |
| chr2 | 240301630 | 240301885 | 0.611532 | 1.57506 | 2.576 |
| chr11 | 71054412 | 71054955 | 1.37862 | 3.55081 | 2.576 |
| chr1 | 9555291 | 9555707 | 1.80654 | 4.65301 | 2.576 |
| chr5 | 92943477 | 92943818 | 1.03069 | 2.65473 | 2.576 |
| chr6 | 146864436 | 146864767 | 0.894652 | 2.3044 | 2.576 |
| chr11 | 84648966 | 84649238 | 0.416011 | 1.07158 | 2.576 |
| chr11 | 10329350 | 10329724 | 1.77723 | 4.57797 | 2.576 |
| chr20 | 48505022 | 48505404 | 1.99545 | 5.14013 | 2.576 |
| chr3 | 183757816 | 183758103 | 1.48702 | 3.83061 | 2.576 |
| chr9 | 129535381 | 129535633 | 1.08937 | 2.80638 | 2.576 |
| chr17 | 66462226 | 66462699 | 1.31803 | 3.39556 | 2.576 |
| chr17 | 73779465 | 73779718 | 1.05277 | 2.71251 | 2.577 |
| chr5 | 80667209 | 80667580 | 0.444256 | 1.14465 | 2.577 |
| chr8 | 91341311 | 91341529 | 0.463108 | 1.19324 | 2.577 |
| chr12 | 128028978 | 128029251 | 0.464911 | 1.19791 | 2.577 |
| chr20 | 62751833 | 62752331 | 1.11076 | 2.86204 | 2.577 |

| | | | | | |
|---|---|---|---|---|---|
| chr12 | 48345678 | 48346007 | 0.443329 | 1.1425 | 2.577 |
| chr14 | 90701538 | 90702040 | 1.87522 | 4.83265 | 2.577 |
| chr4 | 131729703 | 131729961 | 0.877416 | 2.26134 | 2.577 |
| chr9 | 136242188 | 136242543 | 0.960966 | 2.47669 | 2.577 |
| chr3 | 181976393 | 181976699 | 1.60976 | 4.14925 | 2.578 |
| chr19 | 40727729 | 40728120 | 2.08955 | 5.38601 | 2.578 |
| chr17 | 7490054 | 7490489 | 1.61987 | 4.17538 | 2.578 |
| chr19 | 47525033 | 47525258 | 0.329495 | 0.849306 | 2.578 |
| chr20 | 2332497 | 2332935 | 1.70926 | 4.40587 | 2.578 |
| chr1 | 235416513 | 235416842 | 1.44906 | 3.73542 | 2.578 |
| chr19 | 13193233 | 13193531 | 1.10433 | 2.84685 | 2.578 |
| chr17 | 13501128 | 13501412 | 1.19372 | 3.07734 | 2.578 |
| chr9 | 107688766 | 107689081 | 1.66042 | 4.2805 | 2.578 |
| chr16 | 56346556 | 56346831 | 0.396291 | 1.02163 | 2.578 |
| chr6 | 126354897 | 126355099 | 0.679096 | 1.75095 | 2.578 |
| chr11 | 118068986 | 118069316 | 1.43168 | 3.6915 | 2.578 |
| chr17 | 4017755 | 4018450 | 0.430997 | 1.11132 | 2.578 |
| chr5 | 179517447 | 179517831 | 2.55714 | 6.59364 | 2.579 |
| chr1 | 23866298 | 23866621 | 1.59434 | 4.11131 | 2.579 |
| chr10 | 102998554 | 102998827 | 1.0476 | 2.70161 | 2.579 |
| chr16 | 88695148 | 88695523 | 0.956468 | 2.46665 | 2.579 |
| chr10 | 21506913 | 21507226 | 0.6046 | 1.55925 | 2.579 |
| chr19 | 44136559 | 44137001 | 0.504182 | 1.30032 | 2.579 |
| chr19 | 8477715 | 8478075 | 0.924739 | 2.38502 | 2.579 |
| chr4 | 25793920 | 25794199 | 1.11813 | 2.88389 | 2.579 |
| chr2 | 179279445 | 179279787 | 1.21342 | 3.12974 | 2.579 |
| chr12 | 32213574 | 32213816 | 1.25731 | 3.24298 | 2.579 |
| chr12 | 60202156 | 60202415 | 1.34803 | 3.47705 | 2.579 |
| chr2 | 79541209 | 79541502 | 0.452575 | 1.16738 | 2.579 |
| chr17 | 63290187 | 63290762 | 1.5551 | 4.01148 | 2.580 |
| chr20 | 38153039 | 38153419 | 0.396131 | 1.02191 | 2.580 |
| chr6 | 107996313 | 107996892 | 1.98403 | 5.11837 | 2.580 |
| chr17 | 77982178 | 77982577 | 2.39319 | 6.17404 | 2.580 |
| chr2 | 43251642 | 43251987 | 1.69493 | 4.37298 | 2.580 |
| chr3 | 145907401 | 145907668 | 1.15125 | 2.97033 | 2.580 |
| chr2 | 20490743 | 20491133 | 1.49082 | 3.84654 | 2.580 |
| chr3 | 100322408 | 100322717 | 0.794674 | 2.05039 | 2.580 |
| chr10 | 23550903 | 23551222 | 0.407329 | 1.05103 | 2.580 |
| chr6 | 54132696 | 54132928 | 0.774608 | 1.99872 | 2.580 |
| chr8 | 124936918 | 124937155 | 1.09953 | 2.83713 | 2.580 |
| chr9 | 82873192 | 82873416 | 0.358716 | 0.925612 | 2.580 |
| chr1 | 1977885 | 1978234 | 1.52234 | 3.92821 | 2.580 |
| chr2 | 242882487 | 242882823 | 0.837111 | 2.1602 | 2.581 |
| chr14 | 93418222 | 93418632 | 1.73758 | 4.48395 | 2.581 |
| chr16 | 11890939 | 11891301 | 1.49186 | 3.84985 | 2.581 |
| chr16 | 4986259 | 4986717 | 0.782421 | 2.01911 | 2.581 |
| chr13 | 29210915 | 29211203 | 1.57689 | 4.06934 | 2.581 |
| chr15 | 63891757 | 63892091 | 1.20552 | 3.11098 | 2.581 |
| chr4 | 86965167 | 86965460 | 1.14296 | 2.94972 | 2.581 |
| chr9 | 140731342 | 140731635 | 0.565572 | 1.45969 | 2.581 |
| chr11 | 125322609 | 125322852 | 1.43397 | 3.701 | 2.581 |
| chr15 | 93027079 | 93027321 | 0.418851 | 1.08105 | 2.581 |
| chr16 | 2140786 | 2141051 | 0.847397 | 2.18729 | 2.581 |
| chr20 | 2188136 | 2188558 | 1.97775 | 5.10503 | 2.581 |
| chr2 | 29044293 | 29044497 | 1.13792 | 2.93724 | 2.581 |
| chr19 | 2637072 | 2637311 | 1.84725 | 4.76876 | 2.582 |
| chr1 | 206652770 | 206653255 | 1.13916 | 2.94097 | 2.582 |
| chr7 | 5704943 | 5705372 | 1.27379 | 3.28854 | 2.582 |
| chr6 | 37178295 | 37178519 | 1.19178 | 3.07698 | 2.582 |
| chr16 | 48190651 | 48191054 | 1.93623 | 4.9994 | 2.582 |
| chrX | 42537444 | 42537652 | 0.805736 | 2.08045 | 2.582 |

| | | | | | |
|---|---|---|---|---|---|
| chr12 | 54718394 | 54718688 | 1.46117 | 3.77287 | 2.582 |
| chr15 | 102192534 | 102192736 | 0.858676 | 2.21722 | 2.582 |
| chr12 | 26244850 | 26245065 | 0.57204 | 1.47714 | 2.582 |
| chr19 | 35685044 | 35685372 | 0.610186 | 1.57571 | 2.582 |
| chr5 | 38484116 | 38484403 | 0.671434 | 1.73395 | 2.582 |
| chr14 | 78495875 | 78496088 | 0.6194 | 1.59975 | 2.583 |
| chr19 | 43188081 | 43188491 | 1.45059 | 3.74651 | 2.583 |
| chr8 | 36634135 | 36634475 | 0.503661 | 1.30083 | 2.583 |
| chr6 | 145252543 | 145252821 | 0.564961 | 1.45923 | 2.583 |
| chr14 | 37809659 | 37809889 | 0.902141 | 2.33015 | 2.583 |
| chr18 | 27454623 | 27454942 | 0.651915 | 1.68384 | 2.583 |
| chr8 | 26323870 | 26324079 | 0.756032 | 1.95284 | 2.583 |
| chrX | 108876223 | 108876426 | 0.825814 | 2.13318 | 2.583 |
| chr14 | 68171215 | 68171574 | 1.20803 | 3.12075 | 2.583 |
| chrX | 38141403 | 38141670 | 0.76524 | 1.97699 | 2.583 |
| chr8 | 96114927 | 96115151 | 1.2123 | 3.13201 | 2.584 |
| chr15 | 79888419 | 79888756 | 2.19318 | 5.66618 | 2.584 |
| chr20 | 12849872 | 12850264 | 0.503664 | 1.30132 | 2.584 |
| chr16 | 88625500 | 88625827 | 1.07503 | 2.77782 | 2.584 |
| chr10 | 93505040 | 93505256 | 0.985712 | 2.54713 | 2.584 |
| chr18 | 9516495 | 9516746 | 1.35583 | 3.50363 | 2.584 |
| chr20 | 32255666 | 32256072 | 1.21463 | 3.13875 | 2.584 |
| chr19 | 8317809 | 8318229 | 1.4752 | 3.81212 | 2.584 |
| chr4 | 1160932 | 1161339 | 0.966762 | 2.49826 | 2.584 |
| chr9 | 104292423 | 104292736 | 1.32042 | 3.41216 | 2.584 |
| chr11 | 27153543 | 27153757 | 0.394195 | 1.01868 | 2.584 |
| chr2 | 99953556 | 99953873 | 1.33485 | 3.44965 | 2.584 |
| chr19 | 16889679 | 16889977 | 1.04141 | 2.6914 | 2.584 |
| chr20 | 56197501 | 56197921 | 1.88232 | 4.86485 | 2.585 |
| chr8 | 24763213 | 24763763 | 0.675914 | 1.74695 | 2.585 |
| chr10 | 127557036 | 127557250 | 1.2167 | 3.14469 | 2.585 |
| chr20 | 23029110 | 23029718 | 1.19594 | 3.09118 | 2.585 |
| chr15 | 52432032 | 52432615 | 1.79702 | 4.64496 | 2.585 |
| chr5 | 33683161 | 33683381 | 0.421343 | 1.08913 | 2.585 |
| chr12 | 91755596 | 91755899 | 1.2252 | 3.16703 | 2.585 |
| chr19 | 46094017 | 46094746 | 1.34527 | 3.4774 | 2.585 |
| chr16 | 27246293 | 27246702 | 1.69706 | 4.38678 | 2.585 |
| chr14 | 30522359 | 30522666 | 0.745017 | 1.92586 | 2.585 |
| chr17 | 75515324 | 75515557 | 0.476651 | 1.23222 | 2.585 |
| chr18 | 45549090 | 45549313 | 1.18146 | 3.05429 | 2.585 |
| chr10 | 129809555 | 129809911 | 1.75622 | 4.54019 | 2.585 |
| chr3 | 46950416 | 46950711 | 0.453746 | 1.17308 | 2.585 |
| chr8 | 123567173 | 123567384 | 0.318342 | 0.823034 | 2.585 |
| chr1 | 33238614 | 33238996 | 1.88545 | 4.87495 | 2.586 |
| chr2 | 122113650 | 122113979 | 1.33562 | 3.45347 | 2.586 |
| chr3 | 88252700 | 88253208 | 1.00201 | 2.59092 | 2.586 |
| chrX | 65206170 | 65206401 | 1.04172 | 2.69362 | 2.586 |
| chr17 | 76809491 | 76809693 | 0.443973 | 1.14803 | 2.586 |
| chr19 | 11493428 | 11493822 | 1.1644 | 3.01093 | 2.586 |
| chr15 | 83518375 | 83518760 | 1.34871 | 3.48759 | 2.586 |
| chr10 | 21254365 | 21254691 | 0.524811 | 1.35714 | 2.586 |
| chr1 | 1227274 | 1227594 | 1.40212 | 3.62596 | 2.586 |
| chr7 | 128099449 | 128099670 | 1.04137 | 2.69308 | 2.586 |
| chr7 | 143042483 | 143042813 | 1.38242 | 3.57509 | 2.586 |
| chr1 | 32083706 | 32084159 | 1.29652 | 3.35298 | 2.586 |
| chr3 | 8483750 | 8484000 | 1.338 | 3.46038 | 2.586 |
| chr8 | 145800820 | 145801183 | 1.48199 | 3.83287 | 2.586 |
| chr17 | 79644374 | 79644725 | 0.442123 | 1.14352 | 2.586 |
| chr1 | 85410448 | 85410911 | 1.40982 | 3.64649 | 2.586 |
| chr11 | 3112891 | 3113402 | 2.48544 | 6.42901 | 2.587 |
| chr7 | 90057042 | 90057401 | 0.856246 | 2.21493 | 2.587 |

| | | | | | |
|---|---|---|---|---|---|
| chr17 | 36886130 | 36886568 | 1.65295 | 4.2759 | 2.587 |
| chr8 | 25224626 | 25224961 | 1.28315 | 3.31932 | 2.587 |
| chr14 | 94502904 | 94503237 | 1.58805 | 4.10812 | 2.587 |
| chr12 | 127865528 | 127866062 | 1.77693 | 4.59709 | 2.587 |
| chr5 | 68382046 | 68382284 | 0.385261 | 0.996722 | 2.587 |
| chr17 | 6376426 | 6376858 | 1.69757 | 4.39214 | 2.587 |
| chr5 | 162197209 | 162197424 | 0.548005 | 1.4179 | 2.587 |
| chr1 | 40904168 | 40904436 | 1.14009 | 2.94999 | 2.588 |
| chr3 | 194033384 | 194033920 | 1.34774 | 3.48737 | 2.588 |
| chr4 | 174153090 | 174153322 | 0.609125 | 1.57621 | 2.588 |
| chr3 | 101355147 | 101355377 | 1.38589 | 3.58659 | 2.588 |
| chr5 | 7857566 | 7857809 | 0.554685 | 1.43571 | 2.588 |
| chr11 | 64877743 | 64878125 | 1.73942 | 4.50245 | 2.588 |
| chr12 | 96575966 | 96576179 | 0.486052 | 1.2582 | 2.589 |
| chr16 | 61162806 | 61163026 | 0.515422 | 1.3343 | 2.589 |
| chr19 | 3783687 | 3783922 | 0.684246 | 1.77147 | 2.589 |
| chr15 | 23819997 | 23820211 | 0.649771 | 1.68223 | 2.589 |
| chr9 | 127314069 | 127314385 | 1.40752 | 3.64412 | 2.589 |
| chr2 | 73383541 | 73384097 | 1.81104 | 4.6889 | 2.589 |
| chr7 | 143018897 | 143019191 | 1.0126 | 2.62173 | 2.589 |
| chr1 | 230933656 | 230933975 | 1.36482 | 3.53385 | 2.589 |
| chr15 | 43496569 | 43496793 | 0.880984 | 2.28112 | 2.589 |
| chr4 | 65899731 | 65899953 | 0.685958 | 1.77618 | 2.589 |
| chr7 | 54559279 | 54559544 | 0.410443 | 1.06278 | 2.589 |
| chr16 | 73703995 | 73704248 | 0.515213 | 1.33408 | 2.589 |
| chr12 | 360905 | 361201 | 0.41582 | 1.07676 | 2.589 |
| chr14 | 56331569 | 56331836 | 0.757017 | 1.96048 | 2.590 |
| chr5 | 140501203 | 140501516 | 0.798827 | 2.06883 | 2.590 |
| chr20 | 62427302 | 62427889 | 1.59241 | 4.1243 | 2.590 |
| chr5 | 148810607 | 148810906 | 0.90946 | 2.35547 | 2.590 |
| chr20 | 61334857 | 61335063 | 0.822839 | 2.13128 | 2.590 |
| chr12 | 22189755 | 22189957 | 0.433647 | 1.12323 | 2.590 |
| rUn_gl0002 | 77545 | 77774 | 0.826507 | 2.14121 | 2.591 |
| chr7 | 75495783 | 75496191 | 2.325 | 6.02333 | 2.591 |
| chr14 | 57420675 | 57420890 | 0.368134 | 0.953801 | 2.591 |
| chr1 | 168311954 | 168312225 | 1.06347 | 2.75549 | 2.591 |
| chr18 | 70211561 | 70211883 | 2.14137 | 5.54865 | 2.591 |
| chr7 | 38407791 | 38408565 | 1.06155 | 2.75084 | 2.591 |
| chr9 | 23473038 | 23473401 | 0.479115 | 1.24155 | 2.591 |
| chr13 | 108715497 | 108715715 | 0.562844 | 1.45861 | 2.592 |
| chr12 | 28325445 | 28325702 | 1.23851 | 3.20976 | 2.592 |
| chr15 | 78405565 | 78405966 | 1.52829 | 3.96077 | 2.592 |
| chr1 | 201499935 | 201500258 | 1.72009 | 4.45808 | 2.592 |
| chr14 | 88676762 | 88677214 | 0.404453 | 1.04833 | 2.592 |
| chr13 | 28404061 | 28404448 | 1.7814 | 4.61754 | 2.592 |
| chr14 | 75016133 | 75016500 | 0.96171 | 2.49295 | 2.592 |
| chr9 | 136060427 | 136060642 | 0.909415 | 2.35741 | 2.592 |
| chr5 | 141990602 | 141991009 | 1.55588 | 4.03321 | 2.592 |
| chr14 | 21508280 | 21508639 | 0.982702 | 2.54746 | 2.592 |
| chr15 | 40571624 | 40571992 | 1.64468 | 4.2635 | 2.592 |
| chr17 | 48555881 | 48556449 | 2.56795 | 6.657 | 2.592 |
| chr8 | 79420683 | 79420894 | 0.465964 | 1.20795 | 2.592 |
| chr8 | 119427229 | 119427501 | 1.45949 | 3.78424 | 2.593 |
| chr11 | 42004414 | 42004675 | 0.465269 | 1.20641 | 2.593 |
| chr12 | 27295104 | 27295350 | 1.86827 | 4.84432 | 2.593 |
| chr20 | 49094816 | 49095092 | 1.26744 | 3.28639 | 2.593 |
| chr1 | 111573208 | 111573443 | 1.01113 | 2.62181 | 2.593 |
| chr14 | 91224810 | 91225087 | 1.73293 | 4.49341 | 2.593 |
| chr19 | 47141717 | 47142255 | 2.16378 | 5.61074 | 2.593 |
| chr5 | 150826880 | 150827388 | 1.96287 | 5.08992 | 2.593 |
| chr12 | 97798351 | 97798625 | 1.50686 | 3.9075 | 2.593 |

| | | | | | |
|---|---|---|---|---|---|
| chr15 | 27286276 | 27286571 | 0.422346 | 1.09521 | 2.593 |
| chr12 | 15675598 | 15675852 | 1.10407 | 2.86331 | 2.593 |
| chr1 | 240114868 | 240115173 | 1.30427 | 3.38261 | 2.593 |
| chr19 | 29472406 | 29472624 | 0.563209 | 1.46074 | 2.594 |
| chr3 | 27884770 | 27884976 | 0.478661 | 1.24151 | 2.594 |
| chr3 | 45742304 | 45742523 | 0.444135 | 1.15198 | 2.594 |
| chr11 | 123451317 | 123451668 | 2.35228 | 6.10155 | 2.594 |
| chr9 | 110227853 | 110228223 | 1.97973 | 5.13536 | 2.594 |
| chr20 | 48403257 | 48403491 | 0.686229 | 1.78027 | 2.594 |
| chr4 | 1630773 | 1631408 | 1.48729 | 3.85849 | 2.594 |
| chr20 | 45779331 | 45779586 | 1.46827 | 3.80935 | 2.594 |
| chr8 | 38269147 | 38269412 | 1.17091 | 3.03788 | 2.594 |
| chr13 | 52355425 | 52355669 | 0.883609 | 2.29271 | 2.595 |
| chr6 | 1524118 | 1524555 | 1.28489 | 3.33392 | 2.595 |
| chr17 | 26568142 | 26568375 | 0.397071 | 1.03029 | 2.595 |
| chr10 | 129122183 | 129122402 | 0.381174 | 0.989042 | 2.595 |
| chr12 | 7053782 | 7054080 | 1.36116 | 3.53187 | 2.595 |
| chr12 | 81787526 | 81787809 | 0.394649 | 1.02404 | 2.595 |
| chr12 | 95968679 | 95968960 | 1.36326 | 3.53769 | 2.595 |
| chr9 | 35611204 | 35611575 | 1.30549 | 3.38795 | 2.595 |
| chr7 | 38467593 | 38467903 | 0.353553 | 0.91755 | 2.595 |
| chr12 | 77650890 | 77651208 | 0.50875 | 1.32035 | 2.595 |
| chr12 | 116504440 | 116504797 | 1.78644 | 4.63655 | 2.595 |
| chr9 | 38672419 | 38672643 | 0.8898 | 2.30949 | 2.596 |
| chrX | 110688834 | 110689353 | 0.67139 | 1.74264 | 2.596 |
| chr10 | 13578732 | 13579201 | 1.57885 | 4.09808 | 2.596 |
| chr17 | 73110166 | 73110584 | 1.69496 | 4.39953 | 2.596 |
| chr2 | 230665950 | 230666242 | 1.27201 | 3.30175 | 2.596 |
| chr2 | 65492214 | 65492483 | 0.856741 | 2.22402 | 2.596 |
| chr12 | 4867290 | 4867567 | 1.01857 | 2.64413 | 2.596 |
| chr5 | 72892576 | 72892810 | 1.41035 | 3.66115 | 2.596 |
| chr9 | 123739815 | 123740024 | 0.363675 | 0.944146 | 2.596 |
| chr8 | 139205996 | 139206251 | 0.425217 | 1.10395 | 2.596 |
| chr11 | 108288388 | 108288659 | 0.612725 | 1.59078 | 2.596 |
| chr17 | 40599373 | 40599607 | 0.877574 | 2.27841 | 2.596 |
| chr19 | 49087555 | 49087869 | 1.5821 | 4.10778 | 2.596 |
| chr16 | 24448700 | 24448904 | 0.393631 | 1.02207 | 2.597 |
| chr6 | 15464555 | 15464823 | 1.60453 | 4.1662 | 2.597 |
| chr15 | 58158294 | 58158717 | 1.50065 | 3.89659 | 2.597 |
| chr3 | 44277505 | 44277747 | 1.19748 | 3.10963 | 2.597 |
| chr12 | 52784808 | 52785132 | 1.47478 | 3.82975 | 2.597 |
| chr12 | 117804863 | 117805241 | 1.65767 | 4.30478 | 2.597 |
| chr7 | 90042262 | 90042499 | 1.14557 | 2.97501 | 2.597 |
| chr16 | 30032886 | 30033222 | 1.88721 | 4.90125 | 2.597 |
| chr19 | 39850410 | 39850658 | 0.532753 | 1.38366 | 2.597 |
| chr14 | 45495861 | 45496117 | 0.687251 | 1.78499 | 2.597 |
| chr14 | 70659738 | 70659965 | 0.431573 | 1.12101 | 2.598 |
| chr14 | 61736301 | 61736579 | 1.08631 | 2.82172 | 2.598 |
| chr10 | 23603656 | 23603952 | 0.683872 | 1.77639 | 2.598 |
| chr12 | 119742962 | 119743213 | 0.570109 | 1.48092 | 2.598 |
| chr8 | 58487789 | 58488187 | 1.79734 | 4.66888 | 2.598 |
| chr12 | 69781904 | 69782456 | 0.578756 | 1.50348 | 2.598 |
| chr1 | 225625838 | 225626052 | 0.914232 | 2.37499 | 2.598 |
| chr6 | 100564590 | 100564807 | 0.899399 | 2.33652 | 2.598 |
| chr15 | 97447532 | 97447765 | 0.367656 | 0.955151 | 2.598 |
| chr3 | 44955318 | 44955737 | 1.3315 | 3.45924 | 2.598 |
| chr17 | 10549683 | 10550112 | 1.86872 | 4.85506 | 2.598 |
| chr17 | 43173573 | 43173904 | 1.50791 | 3.91833 | 2.599 |
| chr12 | 121879179 | 121879807 | 0.934803 | 2.42921 | 2.599 |
| chr1 | 25176238 | 25176500 | 0.951427 | 2.47249 | 2.599 |
| chr7 | 158235000 | 158235500 | 0.573112 | 1.4894 | 2.599 |

| | | | | | |
|---|---|---|---|---|---|
| chr1 | 54647393 | 54647699 | 1.28565 | 3.34151 | 2.599 |
| chr7 | 100878422 | 100878842 | 0.781962 | 2.03255 | 2.599 |
| chr12 | 131587854 | 131588172 | 0.524074 | 1.36225 | 2.599 |
| chr1 | 237925690 | 237925911 | 0.479348 | 1.24616 | 2.600 |
| chr4 | 14873802 | 14874176 | 0.656249 | 1.70608 | 2.600 |
| chr13 | 109160646 | 109160880 | 0.765351 | 1.98972 | 2.600 |
| chr5 | 178157687 | 178157999 | 1.68224 | 4.37349 | 2.600 |
| chrX | 154913612 | 154913819 | 0.379795 | 0.987429 | 2.600 |
| chr5 | 130056055 | 130056342 | 1.17364 | 3.05145 | 2.600 |
| chr9 | 87282491 | 87283592 | 0.923292 | 2.40061 | 2.600 |
| chr1 | 157670064 | 157670364 | 0.435333 | 1.13195 | 2.600 |
| chr5 | 87804641 | 87804973 | 0.430928 | 1.12053 | 2.600 |
| chr2 | 47116231 | 47116534 | 1.24147 | 3.22847 | 2.601 |
| chr13 | 106689300 | 106689585 | 0.897869 | 2.33495 | 2.601 |
| chr15 | 95569388 | 95569737 | 1.18136 | 3.07234 | 2.601 |
| chr12 | 120804950 | 120805233 | 0.953974 | 2.48124 | 2.601 |
| chr3 | 163208000 | 163208338 | 1.21986 | 3.17295 | 2.601 |
| chr4 | 119890518 | 119890736 | 0.84531 | 2.19875 | 2.601 |
| chr8 | 23329885 | 23330267 | 2.19584 | 5.71169 | 2.601 |
| chr1 | 48963759 | 48964461 | 1.13119 | 2.94249 | 2.601 |
| chr1 | 153541035 | 153541410 | 1.61601 | 4.20368 | 2.601 |
| chr14 | 23770636 | 23770975 | 1.82421 | 4.74526 | 2.601 |
| chr1 | 19986381 | 19986730 | 1.06046 | 2.75876 | 2.601 |
| chr9 | 33044793 | 33045005 | 1.02156 | 2.65772 | 2.602 |
| chr6 | 41570827 | 41571160 | 1.30954 | 3.40697 | 2.602 |
| chr1 | 118296614 | 118297051 | 1.20515 | 3.13555 | 2.602 |
| chr21 | 23298002 | 23298235 | 0.458579 | 1.19316 | 2.602 |
| chr5 | 172631235 | 172631564 | 1.13656 | 2.9573 | 2.602 |
| chr16 | 58231977 | 58232229 | 1.87073 | 4.86764 | 2.602 |
| chr20 | 14245555 | 14245836 | 0.964962 | 2.51102 | 2.602 |
| chr8 | 57360439 | 57360907 | 1.59405 | 4.14812 | 2.602 |
| chr21 | 41160268 | 41160536 | 1.29486 | 3.36956 | 2.602 |
| chr2 | 121437880 | 121438088 | 1.373 | 3.57301 | 2.602 |
| chr1 | 34098635 | 34099064 | 1.83276 | 4.76949 | 2.602 |
| chr2 | 2848015 | 2848243 | 0.822567 | 2.14066 | 2.602 |
| chr8 | 63290031 | 63290370 | 0.653796 | 1.70152 | 2.603 |
| chr12 | 29822148 | 29822425 | 0.485769 | 1.26428 | 2.603 |
| chr17 | 16900862 | 16901296 | 1.79357 | 4.66806 | 2.603 |
| chr3 | 94581625 | 94581877 | 1.08433 | 2.82235 | 2.603 |
| chr11 | 66360170 | 66360504 | 1.44495 | 3.76109 | 2.603 |
| chr12 | 52207848 | 52208397 | 1.21396 | 3.15987 | 2.603 |
| chr8 | 10190252 | 10190605 | 1.65973 | 4.32021 | 2.603 |
| chr1 | 86046807 | 86047297 | 1.38613 | 3.60806 | 2.603 |
| chr13 | 52701933 | 52702190 | 1.2293 | 3.2 | 2.603 |
| chr2 | 196996112 | 196996349 | 0.793334 | 2.06519 | 2.603 |
| chr20 | 6809499 | 6809765 | 1.20997 | 3.14986 | 2.603 |
| chr2 | 12003745 | 12004154 | 1.66148 | 4.32554 | 2.603 |
| chr10 | 86016839 | 86017125 | 1.56717 | 4.08005 | 2.603 |
| chr16 | 6487257 | 6487503 | 0.293714 | 0.76468 | 2.603 |
| chr8 | 34266076 | 34266366 | 0.570915 | 1.48638 | 2.604 |
| chr4 | 56915104 | 56915382 | 0.628387 | 1.63608 | 2.604 |
| chr17 | 2920392 | 2920669 | 0.505907 | 1.31723 | 2.604 |
| chr9 | 18213281 | 18213597 | 1.0842 | 2.82308 | 2.604 |
| chr6 | 83767467 | 83767814 | 1.36001 | 3.54134 | 2.604 |
| chr3 | 146979466 | 146979676 | 0.460693 | 1.19961 | 2.604 |
| chr19 | 5563165 | 5563368 | 0.561268 | 1.46151 | 2.604 |
| chr5 | 28883388 | 28883604 | 0.379237 | 0.98751 | 2.604 |
| chr20 | 31073063 | 31073272 | 1.16084 | 3.02278 | 2.604 |
| chr7 | 30549773 | 30550063 | 1.08016 | 2.81312 | 2.604 |
| chr7 | 6036872 | 6037151 | 1.03938 | 2.70692 | 2.604 |
| chr12 | 127851344 | 127851629 | 0.522088 | 1.35972 | 2.604 |

| | | | | | |
|---|---|---|---|---|---|
| chr7 | 4885565 | 4885768 | 0.769738 | 2.00473 | 2.604 |
| chr6 | 75913992 | 75914221 | 0.976013 | 2.54205 | 2.605 |
| chr8 | 130903945 | 130904198 | 1.14495 | 2.98205 | 2.605 |
| chr17 | 71152321 | 71152672 | 1.6403 | 4.27224 | 2.605 |
| chr6 | 161919570 | 161919857 | 1.02994 | 2.68266 | 2.605 |
| chr15 | 40636829 | 40637101 | 1.41982 | 3.69837 | 2.605 |
| chr11 | 61543984 | 61544267 | 2.03864 | 5.31062 | 2.605 |
| chr10 | 96106909 | 96107129 | 0.440703 | 1.14808 | 2.605 |
| chr2 | 82250925 | 82251129 | 0.900658 | 2.34646 | 2.605 |
| chr17 | 14204488 | 14204708 | 0.686422 | 1.78843 | 2.605 |
| chr10 | 115870352 | 115870631 | 1.0327 | 2.69069 | 2.605 |
| chr10 | 132417598 | 132417807 | 0.366642 | 0.955306 | 2.606 |
| chr22 | 24819646 | 24820033 | 1.89133 | 4.92823 | 2.606 |
| chr9 | 131176768 | 131177027 | 0.74088 | 1.93059 | 2.606 |
| chr10 | 99469278 | 99469751 | 1.00434 | 2.61714 | 2.606 |
| chr17 | 40679276 | 40679607 | 2.0451 | 5.32924 | 2.606 |
| chr10 | 105365458 | 105365824 | 1.50564 | 3.92354 | 2.606 |
| chr3 | 102211245 | 102211478 | 0.573805 | 1.49531 | 2.606 |
| chr3 | 48693549 | 48694026 | 1.04436 | 2.72164 | 2.606 |
| chr22 | 37178096 | 37178484 | 1.51068 | 3.93744 | 2.606 |
| chr18 | 53051821 | 53052084 | 0.803396 | 2.09398 | 2.606 |
| chr5 | 175843502 | 175843892 | 1.77723 | 4.63231 | 2.606 |
| chr15 | 70780366 | 70780748 | 1.51533 | 3.94981 | 2.607 |
| chr5 | 162415034 | 162415269 | 0.331356 | 0.863711 | 2.607 |
| chr12 | 133757827 | 133758099 | 1.96426 | 5.12045 | 2.607 |
| chr7 | 73149061 | 73149329 | 1.46584 | 3.82132 | 2.607 |
| chr22 | 31668819 | 31669097 | 0.954065 | 2.48722 | 2.607 |
| chr12 | 77579102 | 77579332 | 0.500197 | 1.30416 | 2.607 |
| chr14 | 56806818 | 56807195 | 1.43378 | 3.73837 | 2.607 |
| chr11 | 102164800 | 102165038 | 1.25834 | 3.28105 | 2.607 |
| chr11 | 76909718 | 76909926 | 1.15829 | 3.02026 | 2.608 |
| chr3 | 48970432 | 48970642 | 0.600532 | 1.56592 | 2.608 |
| chr5 | 143639821 | 143640025 | 0.495036 | 1.29083 | 2.608 |
| chr9 | 37747117 | 37747579 | 1.55533 | 4.0556 | 2.608 |
| chr9 | 88040031 | 88040410 | 1.35124 | 3.52342 | 2.608 |
| chr8 | 104383524 | 104383782 | 0.405746 | 1.05804 | 2.608 |
| chr5 | 11273658 | 11273903 | 0.382912 | 0.998511 | 2.608 |
| chr19 | 41168630 | 41169039 | 2.56224 | 6.6817 | 2.608 |
| chr2 | 74423179 | 74423445 | 0.381654 | 0.99529 | 2.608 |
| chrX | 47382802 | 47383086 | 1.6873 | 4.4003 | 2.608 |
| chr7 | 28059964 | 28060316 | 1.39652 | 3.64198 | 2.608 |
| chr3 | 51976078 | 51976593 | 1.53862 | 4.01272 | 2.608 |
| chr8 | 82503146 | 82503493 | 1.09021 | 2.84327 | 2.608 |
| chr19 | 47016687 | 47017114 | 2.61034 | 6.80782 | 2.608 |
| chr5 | 177488921 | 177489361 | 1.42302 | 3.71134 | 2.608 |
| chr10 | 126077851 | 126078175 | 1.79437 | 4.68001 | 2.608 |
| chrX | 47503430 | 47503698 | 0.688957 | 1.79693 | 2.608 |
| chr13 | 59907303 | 59907523 | 0.385835 | 1.00634 | 2.608 |
| chr1 | 247657802 | 247658075 | 0.314671 | 0.820782 | 2.608 |
| chr2 | 233305279 | 233305653 | 1.91969 | 5.00734 | 2.608 |
| chr12 | 24992246 | 24992497 | 0.537633 | 1.40241 | 2.608 |
| chr8 | 144516907 | 144517258 | 1.01499 | 2.64783 | 2.609 |
| chr12 | 19048704 | 19048917 | 0.306558 | 0.799729 | 2.609 |
| chr12 | 132335603 | 132336179 | 1.85773 | 4.84681 | 2.609 |
| chr4 | 75548364 | 75548588 | 0.751276 | 1.96018 | 2.609 |
| chr8 | 87899074 | 87899288 | 0.448929 | 1.17135 | 2.609 |
| chr11 | 93628592 | 93628886 | 1.20953 | 3.156 | 2.609 |
| chr13 | 33243213 | 33243479 | 1.05993 | 2.76572 | 2.609 |
| chr1 | 6780100 | 6780382 | 1.58163 | 4.12737 | 2.610 |
| chr8 | 123691122 | 123691484 | 1.65064 | 4.30746 | 2.610 |
| chr6 | 134444610 | 134444967 | 0.435586 | 1.13674 | 2.610 |

| | | | | | |
|---|---|---|---|---|---|
| chr19 | 34732362 | 34732680 | 1.67235 | 4.36447 | 2.610 |
| chrX | 64815224 | 64815639 | 2.12252 | 5.53968 | 2.610 |
| chr8 | 61694337 | 61694542 | 0.734592 | 1.9174 | 2.610 |
| chr8 | 25473283 | 25473604 | 1.74097 | 4.54441 | 2.610 |
| chr10 | 118084199 | 118084455 | 0.46046 | 1.20196 | 2.610 |
| chr10 | 103929032 | 103929522 | 1.28515 | 3.35499 | 2.611 |
| chr16 | 88739250 | 88739501 | 1.00268 | 2.61791 | 2.611 |
| chrX | 40433046 | 40433354 | 1.34902 | 3.52218 | 2.611 |
| chr5 | 38781141 | 38781472 | 0.63407 | 1.65551 | 2.611 |
| chr12 | 48558859 | 48559295 | 1.55475 | 4.05947 | 2.611 |
| chr17 | 47257619 | 47258000 | 1.87778 | 4.90312 | 2.611 |
| chr10 | 108082794 | 108083059 | 0.480174 | 1.25381 | 2.611 |
| chr9 | 130287907 | 130288437 | 1.09144 | 2.84993 | 2.611 |
| chr17 | 41505935 | 41506187 | 0.761273 | 1.98784 | 2.611 |
| chr22 | 37442053 | 37442313 | 0.974164 | 2.54382 | 2.611 |
| chr17 | 28804179 | 28804545 | 1.32549 | 3.46131 | 2.611 |
| chr1 | 84588537 | 84588829 | 1.19084 | 3.10986 | 2.611 |
| chr16 | 6610957 | 6611188 | 0.34246 | 0.894348 | 2.612 |
| chr3 | 40626027 | 40626381 | 1.88865 | 4.93229 | 2.612 |
| chr10 | 134752189 | 134752483 | 1.36454 | 3.56377 | 2.612 |
| chr12 | 97056029 | 97056318 | 0.310849 | 0.811856 | 2.612 |
| chr16 | 66649282 | 66649519 | 0.506751 | 1.3236 | 2.612 |
| chr6 | 131437973 | 131438193 | 0.816501 | 2.13268 | 2.612 |
| chrX | 134246228 | 134246455 | 1.11991 | 2.92519 | 2.612 |
| chr21 | 45209183 | 45209589 | 1.70972 | 4.46582 | 2.612 |
| chr12 | 114369629 | 114369869 | 0.410811 | 1.07314 | 2.612 |
| chr7 | 156779789 | 156780118 | 0.594889 | 1.55401 | 2.612 |
| chr3 | 138136398 | 138136733 | 1.95211 | 5.09997 | 2.613 |
| chr14 | 33057750 | 33057965 | 1.14012 | 2.97877 | 2.613 |
| chr12 | 119488972 | 119489287 | 0.366827 | 0.958516 | 2.613 |
| chr8 | 95725627 | 95725956 | 1.02971 | 2.69071 | 2.613 |
| chr12 | 6097347 | 6098168 | 1.45366 | 3.7988 | 2.613 |
| chr5 | 41007545 | 41007794 | 0.614705 | 1.60649 | 2.613 |
| chr11 | 72404683 | 72404928 | 0.619787 | 1.61978 | 2.613 |
| chr19 | 11593387 | 11593770 | 2.03888 | 5.32858 | 2.613 |
| chr15 | 92194953 | 92195193 | 0.422585 | 1.10442 | 2.613 |
| chr5 | 3287575 | 3287842 | 0.498462 | 1.30278 | 2.614 |
| chr6 | 13769979 | 13770245 | 1.01129 | 2.64314 | 2.614 |
| chr15 | 29743431 | 29743806 | 1.91948 | 5.0172 | 2.614 |
| chr13 | 61047758 | 61048054 | 2.77593 | 7.25589 | 2.614 |
| chr20 | 56812140 | 56812520 | 1.65617 | 4.3291 | 2.614 |
| chr5 | 53483012 | 53483337 | 0.46943 | 1.22707 | 2.614 |
| chr12 | 24745631 | 24745842 | 0.483556 | 1.26415 | 2.614 |
| chr12 | 10519566 | 10519781 | 1.17591 | 3.07418 | 2.614 |
| chr3 | 12909463 | 12909754 | 1.94002 | 5.07206 | 2.614 |
| chr11 | 102158769 | 102159130 | 0.642496 | 1.67978 | 2.614 |
| chr9 | 33229302 | 33229625 | 1.80234 | 4.7127 | 2.615 |
| chr5 | 106211529 | 106211911 | 0.961112 | 2.5131 | 2.615 |
| chr11 | 72395525 | 72395976 | 1.30923 | 3.4234 | 2.615 |
| chr15 | 34485599 | 34485854 | 0.936882 | 2.4498 | 2.615 |
| chr14 | 75770189 | 75770535 | 1.4808 | 3.87214 | 2.615 |
| chr6 | 126179952 | 126180168 | 1.223 | 3.19802 | 2.615 |
| chr3 | 76433070 | 76433313 | 0.545117 | 1.42544 | 2.615 |
| chr15 | 45906495 | 45906700 | 0.96786 | 2.53089 | 2.615 |
| chr8 | 128748373 | 128748603 | 1.12563 | 2.94347 | 2.615 |
| chr1 | 110523188 | 110523581 | 1.45434 | 3.80309 | 2.615 |
| chr1 | 241133686 | 241134068 | 0.523047 | 1.3678 | 2.615 |
| chr12 | 13779059 | 13779284 | 0.415903 | 1.08763 | 2.615 |
| chr16 | 80167120 | 80167334 | 0.481553 | 1.25939 | 2.615 |
| chr15 | 62683949 | 62684225 | 0.897127 | 2.34651 | 2.616 |
| chr5 | 75552133 | 75552463 | 1.10988 | 2.90308 | 2.616 |

| | | | | | |
|---|---|---|---|---|---|
| chr5 | 137371693 | 137372016 | 1.5613 | 4.08421 | 2.616 |
| chr6 | 88032032 | 88032415 | 2.52374 | 6.60206 | 2.616 |
| chr19 | 40283599 | 40284051 | 2.58815 | 6.77072 | 2.616 |
| chrX | 35908840 | 35909151 | 1.49763 | 3.91788 | 2.616 |
| chr11 | 65042520 | 65042805 | 1.49095 | 3.90052 | 2.616 |
| chr20 | 4573197 | 4573640 | 1.50153 | 3.92824 | 2.616 |
| chr11 | 85405527 | 85405756 | 0.798647 | 2.08945 | 2.616 |
| chr5 | 175300125 | 175300379 | 1.4096 | 3.68814 | 2.616 |
| chr19 | 48246880 | 48247183 | 1.76188 | 4.60996 | 2.617 |
| chr12 | 56513212 | 56513439 | 1.54218 | 4.03514 | 2.617 |
| chr1 | 65541035 | 65541419 | 1.16846 | 3.05746 | 2.617 |
| chr11 | 9994442 | 9994745 | 1.24894 | 3.26813 | 2.617 |
| chr21 | 37312675 | 37312912 | 1.23862 | 3.24121 | 2.617 |
| chr7 | 151630406 | 151630618 | 0.871017 | 2.2794 | 2.617 |
| chr2 | 163190548 | 163190805 | 1.31871 | 3.4511 | 2.617 |
| chr12 | 7261767 | 7262088 | 1.97563 | 5.17047 | 2.617 |
| chrX | 103100089 | 103100301 | 0.84209 | 2.20408 | 2.617 |
| chr19 | 51850047 | 51850274 | 0.650605 | 1.70311 | 2.618 |
| chr8 | 115183061 | 115183340 | 0.670608 | 1.75556 | 2.618 |
| chr15 | 44067726 | 44068025 | 0.771104 | 2.01873 | 2.618 |
| chr22 | 42142631 | 42142927 | 1.30685 | 3.42133 | 2.618 |
| chr3 | 14649865 | 14650151 | 0.662037 | 1.73326 | 2.618 |
| chr17 | 34385060 | 34385331 | 0.469155 | 1.22829 | 2.618 |
| chr3 | 8027589 | 8028068 | 0.437016 | 1.14415 | 2.618 |
| chr20 | 12905784 | 12906127 | 1.65746 | 4.33961 | 2.618 |
| chr5 | 116107238 | 116107456 | 0.328783 | 0.860831 | 2.618 |
| chr22 | 41972937 | 41973266 | 1.20443 | 3.15351 | 2.618 |
| chr22 | 47848015 | 47848257 | 0.796886 | 2.08648 | 2.618 |
| chr2 | 112879190 | 112879523 | 0.63414 | 1.6605 | 2.619 |
| chr15 | 95427243 | 95427550 | 0.494609 | 1.29516 | 2.619 |
| chr20 | 17879165 | 17879523 | 2.17581 | 5.69762 | 2.619 |
| chr1 | 7267943 | 7268255 | 0.965502 | 2.52829 | 2.619 |
| chr14 | 78017757 | 78017980 | 1.34509 | 3.52237 | 2.619 |
| chr7 | 151710001 | 151710329 | 0.334774 | 0.876692 | 2.619 |
| chr8 | 131428855 | 131429137 | 0.874917 | 2.2912 | 2.619 |
| chr1 | 26605998 | 26606371 | 0.729508 | 1.91042 | 2.619 |
| chr9 | 117673536 | 117673882 | 1.5 | 3.92875 | 2.619 |
| chr5 | 117548602 | 117548813 | 0.288275 | 0.755087 | 2.619 |
| chr5 | 161020905 | 161021180 | 0.422527 | 1.1068 | 2.619 |
| chr5 | 173950561 | 173950786 | 0.891158 | 2.3344 | 2.620 |
| chr11 | 93916939 | 93917298 | 1.2921 | 3.38479 | 2.620 |
| chr8 | 139465822 | 139466085 | 0.499362 | 1.30818 | 2.620 |
| chr3 | 50226708 | 50227096 | 1.37743 | 3.60855 | 2.620 |
| chr3 | 104702865 | 104703162 | 0.385717 | 1.01057 | 2.620 |
| chr1 | 234201620 | 234201895 | 1.51991 | 3.98223 | 2.620 |
| chr19 | 56150390 | 56150828 | 1.20479 | 3.15685 | 2.620 |
| chr14 | 36539680 | 36540045 | 1.47778 | 3.87237 | 2.620 |
| chr7 | 97063555 | 97063878 | 0.400653 | 1.04996 | 2.621 |
| chr7 | 96694256 | 96694652 | 1.32254 | 3.46615 | 2.621 |
| chr12 | 113749691 | 113749909 | 0.503081 | 1.31854 | 2.621 |
| chr15 | 86511784 | 86512126 | 0.436322 | 1.14369 | 2.621 |
| chr17 | 79296296 | 79296929 | 1.02953 | 2.69873 | 2.621 |
| chr16 | 3729598 | 3729913 | 0.418072 | 1.09592 | 2.621 |
| chr12 | 83991257 | 83991503 | 0.419569 | 1.0999 | 2.622 |
| chr1 | 232766295 | 232766752 | 1.38934 | 3.64234 | 2.622 |
| chr4 | 159093717 | 159093934 | 0.948328 | 2.48618 | 2.622 |
| chr8 | 71000782 | 71001225 | 1.77635 | 4.65697 | 2.622 |
| chr9 | 104211390 | 104211938 | 1.8022 | 4.72479 | 2.622 |
| chr3 | 38409870 | 38410141 | 0.774726 | 2.03141 | 2.622 |
| chr6 | 57086427 | 57086638 | 0.802485 | 2.10425 | 2.622 |
| chr14 | 23291960 | 23292336 | 1.29312 | 3.3908 | 2.622 |

| | | | | | |
|---|---|---|---|---|---|
| chr1 | 29508456 | 29508670 | 0.669751 | 1.75626 | 2.622 |
| chr12 | 80580377 | 80580764 | 1.26426 | 3.31533 | 2.622 |
| chr10 | 75818616 | 75818859 | 0.363092 | 0.952157 | 2.622 |
| chr1 | 183971241 | 183971568 | 0.422011 | 1.10672 | 2.622 |
| chr9 | 19048980 | 19049327 | 1.21996 | 3.19937 | 2.623 |
| chr4 | 55098993 | 55099267 | 1.36927 | 3.59127 | 2.623 |
| chr3 | 41244504 | 41244769 | 0.914077 | 2.39743 | 2.623 |
| chr15 | 75930626 | 75930954 | 1.28482 | 3.37021 | 2.623 |
| chr10 | 116840874 | 116841115 | 0.45238 | 1.18672 | 2.623 |
| chr7 | 100926186 | 100926705 | 1.6502 | 4.32895 | 2.623 |
| chr7 | 131229774 | 131229989 | 0.714232 | 1.87367 | 2.623 |
| chr8 | 92695292 | 92695512 | 0.583749 | 1.53137 | 2.623 |
| chr1 | 30578932 | 30579305 | 1.02594 | 2.69144 | 2.623 |
| chr3 | 130611801 | 130612056 | 1.18624 | 3.11198 | 2.623 |
| chr12 | 32824894 | 32825195 | 0.945586 | 2.48066 | 2.623 |
| chr8 | 49033770 | 49034184 | 1.10724 | 2.90475 | 2.623 |
| chr7 | 123792974 | 123793193 | 0.646924 | 1.69716 | 2.623 |
| chr6 | 108863265 | 108863490 | 1.128 | 2.95933 | 2.624 |
| chr17 | 49027712 | 49028040 | 1.71761 | 4.5062 | 2.624 |
| chr12 | 14876667 | 14877259 | 0.932774 | 2.44721 | 2.624 |
| chr14 | 37022910 | 37023177 | 1.01779 | 2.6705 | 2.624 |
| chrX | 118708677 | 118708914 | 0.564815 | 1.48201 | 2.624 |
| chr8 | 97325459 | 97325778 | 0.601699 | 1.57891 | 2.624 |
| chr16 | 9391407 | 9391658 | 1.10621 | 2.90314 | 2.624 |
| chr12 | 54595020 | 54595538 | 1.94438 | 5.10344 | 2.625 |
| chr15 | 77133255 | 77133506 | 1.18526 | 3.11097 | 2.625 |
| chr17 | 36504990 | 36505375 | 2.07747 | 5.45281 | 2.625 |
| chr3 | 159204414 | 159204675 | 0.7073 | 1.85652 | 2.625 |
| chr12 | 131310770 | 131311300 | 1.76433 | 4.63121 | 2.625 |
| chr12 | 94926558 | 94926918 | 0.935935 | 2.45683 | 2.625 |
| chr9 | 35665212 | 35665447 | 0.93245 | 2.44793 | 2.625 |
| chr15 | 63140138 | 63140419 | 1.39895 | 3.67276 | 2.625 |
| chr14 | 22941977 | 22942296 | 0.886614 | 2.32777 | 2.625 |
| chr15 | 101810145 | 101810358 | 1.17453 | 3.08393 | 2.626 |
| chr10 | 43761304 | 43761634 | 1.29624 | 3.40357 | 2.626 |
| chr12 | 102233565 | 102233991 | 1.19649 | 3.14181 | 2.626 |
| chr9 | 79319582 | 79319837 | 0.872328 | 2.29071 | 2.626 |
| chr12 | 29025604 | 29025997 | 1.29007 | 3.38782 | 2.626 |
| chr8 | 2387932 | 2388140 | 0.487821 | 1.28119 | 2.626 |
| chr7 | 127238494 | 127238719 | 0.907751 | 2.38417 | 2.626 |
| chr1 | 28535386 | 28535667 | 0.919973 | 2.41629 | 2.626 |
| chr11 | 110044843 | 110045226 | 1.30363 | 3.42396 | 2.626 |
| chr4 | 120134197 | 120134437 | 0.790929 | 2.07739 | 2.627 |
| chr15 | 93226744 | 93227104 | 1.65105 | 4.33661 | 2.627 |
| chr15 | 39109882 | 39110253 | 2.08677 | 5.48113 | 2.627 |
| chr19 | 19005813 | 19006130 | 1.12606 | 2.95776 | 2.627 |
| chr10 | 108223386 | 108223591 | 0.370573 | 0.973389 | 2.627 |
| chr9 | 79034770 | 79035017 | 1.11302 | 2.92364 | 2.627 |
| chr10 | 21625401 | 21625659 | 0.683988 | 1.79678 | 2.627 |
| chr1 | 231739825 | 231740068 | 1.24711 | 3.2761 | 2.627 |
| chr14 | 102427419 | 102427797 | 2.03889 | 5.35606 | 2.627 |
| chr17 | 38358071 | 38358420 | 1.48341 | 3.89688 | 2.627 |
| chr18 | 71707923 | 71708246 | 1.46728 | 3.85467 | 2.627 |
| chr2 | 1914252 | 1914723 | 0.691756 | 1.8174 | 2.627 |
| chr11 | 122974006 | 122974249 | 1.1948 | 3.13904 | 2.627 |
| chr16 | 58768087 | 58768433 | 1.71002 | 4.49267 | 2.627 |
| chr7 | 5744119 | 5744505 | 1.38477 | 3.63837 | 2.627 |
| chr12 | 112103508 | 112103810 | 0.441877 | 1.1611 | 2.628 |
| chr7 | 35464459 | 35464836 | 1.35193 | 3.55248 | 2.628 |
| chr11 | 5829683 | 5830010 | 1.09829 | 2.88603 | 2.628 |
| chr8 | 16246596 | 16246858 | 0.398971 | 1.04844 | 2.628 |

| | | | | | |
|---|---|---|---|---|---|
| chr9 | 92981764 | 92982256 | 1.2189 | 3.20325 | 2.628 |
| chr5 | 40737180 | 40737435 | 0.53976 | 1.41857 | 2.628 |
| chr4 | 185185362 | 185185597 | 0.996094 | 2.61815 | 2.628 |
| chr4 | 37972465 | 37972721 | 0.998183 | 2.62381 | 2.629 |
| chr3 | 99809269 | 99809568 | 0.814146 | 2.14007 | 2.629 |
| chr5 | 28473702 | 28473936 | 0.714052 | 1.87702 | 2.629 |
| chr21 | 39498858 | 39499071 | 1.26198 | 3.31756 | 2.629 |
| chr7 | 110082364 | 110082713 | 1.10707 | 2.91054 | 2.629 |
| chr7 | 147810662 | 147810875 | 0.42813 | 1.12559 | 2.629 |
| chr16 | 10805302 | 10805620 | 0.348022 | 0.915029 | 2.629 |
| chr17 | 16331402 | 16331766 | 2.03778 | 5.35783 | 2.629 |
| chr7 | 42871260 | 42871602 | 1.15002 | 3.02369 | 2.629 |
| chr9 | 16874338 | 16874592 | 0.427308 | 1.12356 | 2.629 |
| rUn_gl0002 | 40160 | 40490 | 1.74102 | 4.57786 | 2.629 |
| chr6 | 126274761 | 126275161 | 0.977686 | 2.57082 | 2.629 |
| chr7 | 77068947 | 77069207 | 1.3011 | 3.42144 | 2.630 |
| chr3 | 148445138 | 148445355 | 0.893089 | 2.34856 | 2.630 |
| chr7 | 25133866 | 25134166 | 1.47459 | 3.87786 | 2.630 |
| chr7 | 150808122 | 150808557 | 2.16692 | 5.69857 | 2.630 |
| chr8 | 113813408 | 113813623 | 0.995704 | 2.61857 | 2.630 |
| chr8 | 22423505 | 22423744 | 0.419423 | 1.10304 | 2.630 |
| chrX | 45046245 | 45046459 | 1.02827 | 2.70426 | 2.630 |
| chr7 | 94988816 | 94989314 | 0.38609 | 1.01541 | 2.630 |
| chr3 | 89215965 | 89216174 | 0.617689 | 1.62452 | 2.630 |
| chr22 | 43583492 | 43583740 | 1.46782 | 3.86044 | 2.630 |
| chr12 | 104775556 | 104776156 | 1.15969 | 3.05016 | 2.630 |
| chr5 | 169277504 | 169277777 | 0.897022 | 2.35958 | 2.630 |
| chr12 | 82112097 | 82112415 | 0.414629 | 1.09073 | 2.631 |
| chr12 | 109831506 | 109831950 | 1.75146 | 4.60757 | 2.631 |
| chr15 | 34880241 | 34880567 | 0.857571 | 2.25636 | 2.631 |
| chr12 | 114992557 | 114992909 | 0.37148 | 0.977469 | 2.631 |
| chr6 | 126317130 | 126317517 | 1.40804 | 3.7053 | 2.632 |
| chr12 | 29029493 | 29029772 | 0.46843 | 1.23277 | 2.632 |
| chr3 | 14503247 | 14503707 | 1.22417 | 3.22166 | 2.632 |
| chr17 | 78950199 | 78950417 | 0.373964 | 0.98425 | 2.632 |
| chr17 | 46755699 | 46756149 | 1.94887 | 5.13003 | 2.632 |
| chrX | 102465698 | 102466278 | 0.95236 | 2.50692 | 2.632 |
| chr12 | 46547123 | 46547443 | 0.958075 | 2.52233 | 2.633 |
| chr20 | 39147946 | 39148159 | 0.40963 | 1.07849 | 2.633 |
| chr10 | 67092366 | 67092617 | 0.322904 | 0.850188 | 2.633 |
| chr18 | 74587186 | 74587654 | 1.32318 | 3.48405 | 2.633 |
| chr9 | 11439941 | 11440327 | 0.353204 | 0.930128 | 2.633 |
| chr20 | 56662302 | 56662682 | 1.84086 | 4.84808 | 2.634 |
| chr15 | 67215761 | 67216082 | 1.27198 | 3.34992 | 2.634 |
| chr11 | 72504642 | 72505069 | 1.83218 | 4.82593 | 2.634 |
| chr11 | 118016054 | 118016327 | 1.01709 | 2.67903 | 2.634 |
| chr10 | 96259891 | 96260142 | 0.912908 | 2.40467 | 2.634 |
| chr17 | 8085313 | 8085560 | 1.06561 | 2.80692 | 2.634 |
| chr16 | 74463986 | 74464417 | 1.23028 | 3.24074 | 2.634 |
| chr11 | 59898064 | 59898304 | 0.767068 | 2.0206 | 2.634 |
| chr15 | 101557497 | 101557898 | 1.1511 | 3.03237 | 2.634 |
| chr10 | 91186091 | 91186386 | 0.94685 | 2.49437 | 2.634 |
| chr1 | 153324915 | 153325308 | 1.32761 | 3.49757 | 2.634 |
| chr9 | 102396029 | 102396260 | 0.269453 | 0.709881 | 2.635 |
| chr17 | 9868896 | 9869319 | 2.26809 | 5.97539 | 2.635 |
| chr1 | 158589984 | 158590312 | 0.483544 | 1.27398 | 2.635 |
| chr17 | 26973917 | 26974185 | 1.08069 | 2.84729 | 2.635 |
| chr10 | 121510148 | 121510500 | 2.05553 | 5.41587 | 2.635 |
| chr6 | 52680373 | 52680631 | 0.777352 | 2.0482 | 2.635 |
| chr7 | 53389642 | 53390007 | 0.431425 | 1.13674 | 2.635 |
| chr13 | 111064613 | 111064906 | 1.30904 | 3.44959 | 2.635 |

| | | | | | |
|---|---|---|---|---|---|
| chr1 | 207866561 | 207866824 | 0.427034 | 1.12536 | 2.635 |
| chr10 | 129635487 | 129635788 | 0.434178 | 1.14425 | 2.635 |
| chr1 | 171220752 | 171221177 | 1.23692 | 3.25998 | 2.636 |
| chr1 | 50834177 | 50834536 | 1.77638 | 4.6822 | 2.636 |
| chr1 | 118301472 | 118301842 | 1.27715 | 3.36639 | 2.636 |
| chr5 | 78316863 | 78317183 | 1.56325 | 4.1206 | 2.636 |
| chr17 | 54731219 | 54731636 | 0.803004 | 2.11703 | 2.636 |
| chr1 | 4838061 | 4838327 | 1.25093 | 3.29836 | 2.637 |
| chr7 | 82702293 | 82702564 | 1.03382 | 2.72644 | 2.637 |
| chr17 | 80486692 | 80486982 | 1.25115 | 3.30019 | 2.638 |
| chr19 | 19665203 | 19665530 | 0.99336 | 2.62026 | 2.638 |
| chrX | 152772963 | 152773387 | 1.79267 | 4.7289 | 2.638 |
| chr7 | 37889882 | 37890098 | 0.345284 | 0.910851 | 2.638 |
| chr12 | 56519971 | 56520309 | 1.90996 | 5.03859 | 2.638 |
| chr19 | 4204581 | 4204918 | 1.7579 | 4.63758 | 2.638 |
| chr6 | 126090840 | 126091220 | 0.549957 | 1.45093 | 2.638 |
| chr7 | 77627431 | 77627807 | 0.490362 | 1.29371 | 2.638 |
| chr6 | 37349383 | 37349687 | 1.12387 | 2.96519 | 2.638 |
| chr16 | 89766334 | 89766863 | 0.982048 | 2.59102 | 2.638 |
| chr10 | 101844627 | 101844890 | 0.462684 | 1.22074 | 2.638 |
| chr12 | 7244956 | 7245358 | 1.51564 | 3.99888 | 2.638 |
| chr7 | 47892069 | 47892612 | 0.680337 | 1.79513 | 2.639 |
| chr5 | 148608646 | 148608952 | 1.22309 | 3.22726 | 2.639 |
| chr4 | 8178575 | 8178947 | 1.92488 | 5.0801 | 2.639 |
| chr9 | 119835512 | 119836171 | 1.08637 | 2.86727 | 2.639 |
| chr12 | 63328873 | 63329245 | 1.90221 | 5.02063 | 2.639 |
| chr17 | 71192700 | 71193080 | 0.894435 | 2.36076 | 2.639 |
| chr5 | 90098541 | 90098847 | 1.2679 | 3.34665 | 2.640 |
| chr10 | 122164618 | 122165067 | 0.395389 | 1.04364 | 2.640 |
| chr15 | 93257883 | 93258399 | 1.64876 | 4.35201 | 2.640 |
| chr8 | 38327648 | 38327945 | 0.839798 | 2.21674 | 2.640 |
| chr3 | 121315026 | 121315379 | 0.378544 | 0.999241 | 2.640 |
| chr6 | 109214699 | 109214935 | 0.588862 | 1.55442 | 2.640 |
| chr2 | 223555292 | 223555667 | 2.963 | 7.82171 | 2.640 |
| chrX | 18950191 | 18950408 | 0.701535 | 1.85193 | 2.640 |
| chr1 | 242687366 | 242687754 | 0.859226 | 2.26824 | 2.640 |
| chr5 | 66541983 | 66542190 | 0.882885 | 2.33073 | 2.640 |
| chr8 | 138827172 | 138827375 | 0.32197 | 0.849973 | 2.640 |
| chr17 | 41597911 | 41598309 | 1.45971 | 3.8546 | 2.641 |
| chr2 | 18076875 | 18077179 | 0.381228 | 1.00672 | 2.641 |
| chr2 | 85519919 | 85520175 | 0.388223 | 1.02521 | 2.641 |
| chr14 | 60944379 | 60944707 | 0.569312 | 1.50347 | 2.641 |
| chr12 | 123365311 | 123365669 | 1.17992 | 3.11616 | 2.641 |
| chr10 | 105004453 | 105004826 | 1.51256 | 3.99492 | 2.641 |
| chr3 | 58153090 | 58153548 | 1.3193 | 3.48456 | 2.641 |
| chr1 | 31384212 | 31384551 | 1.52894 | 4.03829 | 2.641 |
| chr3 | 118781 | 118996 | 0.458464 | 1.21095 | 2.641 |
| chr12 | 118602875 | 118603171 | 1.48131 | 3.91278 | 2.641 |
| chr14 | 100906580 | 100906908 | 1.70716 | 4.50997 | 2.642 |
| chr20 | 39724010 | 39724383 | 1.47473 | 3.89601 | 2.642 |
| chr17 | 49538818 | 49539321 | 2.35398 | 6.21893 | 2.642 |
| chr20 | 62714773 | 62715412 | 1.24115 | 3.279 | 2.642 |
| chr6 | 24521219 | 24521586 | 1.06376 | 2.81037 | 2.642 |
| chr1 | 43396773 | 43397023 | 0.854726 | 2.25824 | 2.642 |
| chr22 | 32869398 | 32869635 | 0.885947 | 2.34074 | 2.642 |
| chr5 | 176037157 | 176037361 | 0.594051 | 1.56954 | 2.642 |
| chr14 | 77114030 | 77114422 | 1.35035 | 3.56787 | 2.642 |
| chr3 | 159304401 | 159304760 | 1.43569 | 3.79336 | 2.642 |
| chr22 | 31627215 | 31627423 | 0.848567 | 2.24226 | 2.642 |
| chr10 | 61624907 | 61625243 | 0.804422 | 2.12562 | 2.642 |
| chr9 | 95766058 | 95766398 | 1.46074 | 3.85993 | 2.642 |

| | | | | | |
|---|---|---|---|---|---|
| chr8 | 123685735 | 123686029 | 1.1826 | 3.12498 | 2.642 |
| chr8 | 142085750 | 142086216 | 0.801414 | 2.11783 | 2.643 |
| chr4 | 7083759 | 7084109 | 0.93668 | 2.47543 | 2.643 |
| chr7 | 148762862 | 148763302 | 2.29708 | 6.07133 | 2.643 |
| chr1 | 208209636 | 208209851 | 0.573606 | 1.51608 | 2.643 |
| chr11 | 61405830 | 61406257 | 1.74295 | 4.60705 | 2.643 |
| chr12 | 75728102 | 75728496 | 2.10383 | 5.56099 | 2.643 |
| chr18 | 43200958 | 43201467 | 1.26374 | 3.3406 | 2.643 |
| chr3 | 52178303 | 52178682 | 1.09567 | 2.89643 | 2.644 |
| chr8 | 73413082 | 73413337 | 0.356166 | 0.94157 | 2.644 |
| chr19 | 57275706 | 57276060 | 0.895147 | 2.36652 | 2.644 |
| chr17 | 72599498 | 72599724 | 0.390374 | 1.03207 | 2.644 |
| chr9 | 14781941 | 14782261 | 0.491883 | 1.30056 | 2.644 |
| chr10 | 116454477 | 116454838 | 1.22636 | 3.24278 | 2.644 |
| chr1 | 246859777 | 246860033 | 1.76726 | 4.6733 | 2.644 |
| chr12 | 132647057 | 132647338 | 0.480959 | 1.27187 | 2.644 |
| chr19 | 19285783 | 19286244 | 1.78221 | 4.71308 | 2.645 |
| chr20 | 9196273 | 9196529 | 0.949382 | 2.51065 | 2.645 |
| chr13 | 36204769 | 36205130 | 1.49662 | 3.95804 | 2.645 |
| chr11 | 113677185 | 113677423 | 1.01373 | 2.68102 | 2.645 |
| chr14 | 93347107 | 93347476 | 0.655453 | 1.73356 | 2.645 |
| chr5 | 93398361 | 93398643 | 0.400354 | 1.05894 | 2.645 |
| chr11 | 47376651 | 47377249 | 2.89107 | 7.64693 | 2.645 |
| chr22 | 26168932 | 26169246 | 1.15458 | 3.05402 | 2.645 |
| chr4 | 75236654 | 75236893 | 1.14801 | 3.03677 | 2.645 |
| chr16 | 74841169 | 74841463 | 0.443259 | 1.17254 | 2.645 |
| chr12 | 56333982 | 56334424 | 1.91221 | 5.05848 | 2.645 |
| chr9 | 3441268 | 3441520 | 1.04995 | 2.77755 | 2.645 |
| chr21 | 32552919 | 32553239 | 0.688521 | 1.82145 | 2.645 |
| chr6 | 40555310 | 40555965 | 1.02502 | 2.7117 | 2.646 |
| chr9 | 78129361 | 78129641 | 0.394812 | 1.04456 | 2.646 |
| chr13 | 96032981 | 96033267 | 1.22187 | 3.23303 | 2.646 |
| chr4 | 1695086 | 1695454 | 1.46397 | 3.87372 | 2.646 |
| chr12 | 52515421 | 52516080 | 1.5672 | 4.1471 | 2.646 |
| chr12 | 84084185 | 84084403 | 0.504688 | 1.33555 | 2.646 |
| chr6 | 135203599 | 135203956 | 2.08975 | 5.53018 | 2.646 |
| chr17 | 767812 | 768014 | 0.367788 | 0.973353 | 2.647 |
| chr12 | 4958300 | 4958568 | 1.46379 | 3.87407 | 2.647 |
| chr17 | 75538735 | 75539115 | 1.81457 | 4.80257 | 2.647 |
| chr15 | 41230237 | 41230606 | 1.69762 | 4.49326 | 2.647 |
| chr9 | 104782437 | 104782741 | 0.398005 | 1.05356 | 2.647 |
| chr4 | 18239751 | 18240121 | 0.409119 | 1.08303 | 2.647 |
| chr13 | 66500947 | 66501275 | 0.966751 | 2.55949 | 2.648 |
| chr12 | 110474470 | 110474720 | 0.921494 | 2.43986 | 2.648 |
| chr3 | 141280307 | 141280655 | 1.85261 | 4.9052 | 2.648 |
| chr19 | 1248849 | 1249254 | 1.10634 | 2.9293 | 2.648 |
| chr1 | 60164968 | 60165211 | 0.810672 | 2.14654 | 2.648 |
| chr16 | 6623538 | 6623920 | 0.449524 | 1.19043 | 2.648 |
| chrX | 40653298 | 40653576 | 1.0275 | 2.72112 | 2.648 |
| chr17 | 41056149 | 41056448 | 0.529025 | 1.40103 | 2.648 |
| chr3 | 165416574 | 165416804 | 0.497252 | 1.31691 | 2.648 |
| chr8 | 76202087 | 76202294 | 0.482996 | 1.27933 | 2.649 |
| chr9 | 107372551 | 107372795 | 0.369993 | 0.980203 | 2.649 |
| chr6 | 17200604 | 17200981 | 0.885257 | 2.34539 | 2.649 |
| chr18 | 76746212 | 76746523 | 0.978801 | 2.59324 | 2.649 |
| chr12 | 30099388 | 30099664 | 0.392139 | 1.03899 | 2.650 |
| chr8 | 86059720 | 86059967 | 1.05387 | 2.79231 | 2.650 |
| chr3 | 194722659 | 194723001 | 1.2995 | 3.44347 | 2.650 |
| chr4 | 7805734 | 7806095 | 1.90107 | 5.0383 | 2.650 |
| chr15 | 56310278 | 56310653 | 0.535825 | 1.42007 | 2.650 |
| chr19 | 41036141 | 41036509 | 1.23398 | 3.27091 | 2.651 |

| | | | | | |
|---|---|---|---|---|---|
| chr1 | 17452257 | 17452550 | 1.39025 | 3.68528 | 2.651 |
| chr7 | 42610241 | 42610557 | 0.402566 | 1.06719 | 2.651 |
| chr10 | 45428112 | 45428541 | 2.20953 | 5.8578 | 2.651 |
| chr3 | 79135638 | 79136028 | 1.29012 | 3.4203 | 2.651 |
| chr11 | 45361606 | 45362002 | 1.69168 | 4.48495 | 2.651 |
| chr19 | 17416236 | 17416561 | 1.42179 | 3.76949 | 2.651 |
| chr12 | 47055198 | 47055493 | 1.07344 | 2.84595 | 2.651 |
| chr12 | 70017880 | 70018112 | 0.458457 | 1.21554 | 2.651 |
| chr5 | 131721107 | 131721399 | 1.42102 | 3.76769 | 2.651 |
| chrX | 130926463 | 130926678 | 0.982795 | 2.60597 | 2.652 |
| chr14 | 20525821 | 20526035 | 0.450235 | 1.19385 | 2.652 |
| chr7 | 131838906 | 131839311 | 0.437706 | 1.16063 | 2.652 |
| chr8 | 131776898 | 131777100 | 0.737424 | 1.9554 | 2.652 |
| chr10 | 98009368 | 98009620 | 0.431831 | 1.14511 | 2.652 |
| chr3 | 89178275 | 89178485 | 0.50634 | 1.34276 | 2.652 |
| chr3 | 77020862 | 77021097 | 0.439017 | 1.16426 | 2.652 |
| chr2 | 42323150 | 42323371 | 0.855267 | 2.26815 | 2.652 |
| chrX | 140372530 | 140372732 | 0.961397 | 2.54961 | 2.652 |
| chr4 | 3420211 | 3420465 | 0.826928 | 2.19307 | 2.652 |
| chr7 | 158146584 | 158146981 | 1.17008 | 3.10366 | 2.653 |
| chr11 | 12782692 | 12783009 | 0.765022 | 2.02929 | 2.653 |
| chr9 | 131017530 | 131017864 | 1.7196 | 4.56154 | 2.653 |
| chr12 | 61579607 | 61579901 | 0.378084 | 1.00295 | 2.653 |
| chr12 | 124018360 | 124018733 | 0.964214 | 2.55804 | 2.653 |
| chr12 | 69349569 | 69349958 | 0.411717 | 1.09228 | 2.653 |
| chr20 | 8826784 | 8827010 | 0.38481 | 1.02093 | 2.653 |
| chr12 | 113226046 | 113226452 | 0.778493 | 2.06541 | 2.653 |
| chr8 | 134476944 | 134477260 | 0.410644 | 1.08952 | 2.653 |
| chr15 | 32954879 | 32955148 | 1.73637 | 4.60723 | 2.653 |
| chr12 | 15609997 | 15610267 | 0.470534 | 1.24853 | 2.653 |
| chr16 | 49142645 | 49142896 | 0.429403 | 1.13939 | 2.653 |
| chr7 | 71566575 | 71566907 | 0.393368 | 1.0438 | 2.653 |
| chr8 | 101839231 | 101839545 | 1.08151 | 2.86982 | 2.654 |
| chr15 | 71303882 | 71304217 | 1.4921 | 3.95936 | 2.654 |
| chr9 | 111175055 | 111175313 | 0.78908 | 2.09392 | 2.654 |
| chr1 | 229984853 | 229985222 | 0.741839 | 1.96871 | 2.654 |
| chr11 | 17848655 | 17848882 | 0.748878 | 1.98751 | 2.654 |
| chr4 | 138654339 | 138654592 | 1.13412 | 3.00999 | 2.654 |
| chr11 | 30316606 | 30316846 | 0.448974 | 1.19167 | 2.654 |
| chr3 | 161278143 | 161278380 | 0.992922 | 2.63542 | 2.654 |
| chr17 | 76450330 | 76450654 | 0.400173 | 1.06219 | 2.654 |
| chr12 | 4032248 | 4032487 | 0.406573 | 1.07919 | 2.654 |
| chr16 | 2073130 | 2073450 | 2.02668 | 5.37958 | 2.654 |
| chr12 | 71113571 | 71113958 | 1.56034 | 4.14176 | 2.654 |
| chr14 | 103056921 | 103057407 | 1.38325 | 3.67178 | 2.654 |
| chr5 | 67109592 | 67110087 | 0.427353 | 1.13448 | 2.655 |
| chr1 | 221961165 | 221961379 | 0.359503 | 0.95437 | 2.655 |
| chr11 | 23561685 | 23562004 | 1.06652 | 2.83134 | 2.655 |
| chr20 | 25046373 | 25046713 | 0.825696 | 2.19217 | 2.655 |
| chr14 | 99991273 | 99991492 | 0.906858 | 2.40792 | 2.655 |
| chr14 | 49836790 | 49837024 | 0.455618 | 1.20981 | 2.655 |
| chr12 | 132320763 | 132321336 | 0.802154 | 2.13016 | 2.656 |
| chr4 | 677754 | 678202 | 1.59265 | 4.22936 | 2.656 |
| chr12 | 63207039 | 63207396 | 1.25929 | 3.34434 | 2.656 |
| chrX | 41039514 | 41039788 | 1.49761 | 3.97734 | 2.656 |
| chr11 | 1715183 | 1715573 | 1.67653 | 4.45256 | 2.656 |
| chr7 | 130560062 | 130560355 | 1.8663 | 4.95748 | 2.656 |
| chr19 | 45273980 | 45274221 | 1.37582 | 3.65477 | 2.656 |
| chr2 | 36008880 | 36009110 | 0.483418 | 1.2842 | 2.657 |
| chr8 | 16637379 | 16637677 | 0.622541 | 1.65378 | 2.657 |
| chr22 | 49827877 | 49828158 | 0.924008 | 2.45468 | 2.657 |

| | | | | | |
|---|---|---|---|---|---|
| chr11 | 113160328 | 113160651 | 1.88251 | 5.00161 | 2.657 |
| chr11 | 34195841 | 34196394 | 2.8554 | 7.58705 | 2.657 |
| chr17 | 79526105 | 79526454 | 1.47214 | 3.91186 | 2.657 |
| chr7 | 24627499 | 24627778 | 1.0968 | 2.91454 | 2.657 |
| chr4 | 110920033 | 110920250 | 0.863924 | 2.29596 | 2.658 |
| chr6 | 142795987 | 142796201 | 1.03655 | 2.75497 | 2.658 |
| chr7 | 15697539 | 15697883 | 0.601064 | 1.59753 | 2.658 |
| chr4 | 145277671 | 145277874 | 0.576949 | 1.53349 | 2.658 |
| chr1 | 209363951 | 209364358 | 1.10809 | 2.94527 | 2.658 |
| chr3 | 119487127 | 119487534 | 0.397801 | 1.05736 | 2.658 |
| chr10 | 56310663 | 56310867 | 0.420848 | 1.11864 | 2.658 |
| chr13 | 111335556 | 111335879 | 0.846433 | 2.25 | 2.658 |
| chr8 | 143649988 | 143650208 | 0.410132 | 1.09022 | 2.658 |
| chr3 | 102417122 | 102417481 | 0.486156 | 1.29232 | 2.658 |
| chr15 | 72578253 | 72578651 | 1.7603 | 4.67963 | 2.658 |
| chr3 | 149206911 | 149207259 | 1.77611 | 4.72194 | 2.659 |
| chr17 | 21416226 | 21416588 | 2.4115 | 6.41122 | 2.659 |
| chr1 | 54007440 | 54007737 | 0.687861 | 1.82883 | 2.659 |
| chr15 | 32254239 | 32254515 | 0.477135 | 1.26876 | 2.659 |
| chr11 | 68899050 | 68899346 | 1.84403 | 4.90405 | 2.659 |
| chr19 | 50650733 | 50650944 | 1.30091 | 3.46036 | 2.660 |
| chr7 | 147235460 | 147235693 | 0.483573 | 1.28635 | 2.660 |
| chr15 | 72036277 | 72036745 | 1.15222 | 3.06503 | 2.660 |
| chr7 | 126002772 | 126003049 | 0.366487 | 0.974898 | 2.660 |
| chr19 | 35378863 | 35379266 | 0.570546 | 1.51775 | 2.660 |
| chr5 | 130790816 | 130791098 | 0.564926 | 1.5028 | 2.660 |
| chr15 | 66544658 | 66545122 | 1.41599 | 3.76692 | 2.660 |
| chr19 | 48202501 | 48203063 | 2.01681 | 5.36547 | 2.660 |
| chr18 | 60093009 | 60093275 | 0.763421 | 2.03115 | 2.661 |
| chr12 | 125603117 | 125603323 | 0.370351 | 0.985381 | 2.661 |
| chr11 | 122652138 | 122652540 | 1.2398 | 3.29895 | 2.661 |
| chr3 | 1040580 | 1040950 | 0.430229 | 1.14491 | 2.661 |
| chr16 | 85454160 | 85454911 | 2.70313 | 7.19398 | 2.661 |
| chr8 | 146125500 | 146125871 | 2.06963 | 5.50804 | 2.661 |
| chr17 | 40176913 | 40177443 | 1.99332 | 5.30496 | 2.661 |
| chr11 | 93276466 | 93276749 | 1.18527 | 3.1545 | 2.661 |
| chr3 | 14930699 | 14931070 | 0.3104 | 0.826105 | 2.661 |
| chr2 | 118960300 | 118960611 | 1.78595 | 4.75322 | 2.661 |
| chr19 | 13915695 | 13915945 | 0.847685 | 2.25608 | 2.661 |
| chr4 | 96077338 | 96077672 | 1.04117 | 2.77106 | 2.661 |
| chr20 | 62689057 | 62689495 | 2.309 | 6.14572 | 2.662 |
| chr17 | 7893451 | 7893898 | 2.24013 | 5.9633 | 2.662 |
| chr3 | 46419904 | 46420173 | 1.59247 | 4.23937 | 2.662 |
| chr17 | 77782529 | 77782785 | 1.03292 | 2.74981 | 2.662 |
| chr3 | 179040479 | 179040831 | 1.47826 | 3.93545 | 2.662 |
| chr9 | 131182414 | 131182813 | 1.49404 | 3.97754 | 2.662 |
| chr19 | 55723833 | 55724179 | 1.38549 | 3.68858 | 2.662 |
| chr5 | 139769681 | 139769912 | 0.345031 | 0.918574 | 2.662 |
| chr17 | 70026628 | 70026867 | 0.990761 | 2.63781 | 2.662 |
| chr17 | 70536413 | 70536874 | 1.8781 | 5.00034 | 2.662 |
| chr11 | 79847791 | 79848167 | 0.434632 | 1.15719 | 2.662 |
| chr2 | 45179217 | 45179893 | 1.70723 | 4.54554 | 2.663 |
| chr5 | 138954671 | 138954936 | 1.24307 | 3.30983 | 2.663 |
| chr18 | 59798626 | 59798861 | 1.08661 | 2.89335 | 2.663 |
| chr14 | 29602431 | 29602668 | 0.335551 | 0.893484 | 2.663 |
| chr11 | 111648078 | 111648280 | 0.920157 | 2.45015 | 2.663 |
| chr12 | 8012580 | 8012803 | 0.51501 | 1.37144 | 2.663 |
| chr2 | 139572183 | 139572405 | 0.616387 | 1.6414 | 2.663 |
| chr1 | 85070327 | 85070552 | 0.756723 | 2.01513 | 2.663 |
| chr9 | 104329574 | 104329845 | 1.07141 | 2.85326 | 2.663 |
| chr5 | 155079872 | 155080186 | 0.897811 | 2.39117 | 2.663 |

| | | | | | |
|---|---|---|---|---|---|
| chr6 | 151797080 | 151797351 | 1.14337 | 3.04521 | 2.663 |
| chr17 | 73585150 | 73585744 | 1.11191 | 2.96159 | 2.664 |
| chr10 | 14244671 | 14244958 | 0.466588 | 1.24281 | 2.664 |
| chr14 | 70512683 | 70513099 | 1.6364 | 4.35941 | 2.664 |
| chr11 | 44844640 | 44844869 | 1.17239 | 3.12328 | 2.664 |
| chr10 | 29419018 | 29419441 | 1.47827 | 3.93826 | 2.664 |
| chr1 | 226595677 | 226596065 | 1.91532 | 5.10313 | 2.664 |
| chr4 | 75800037 | 75800341 | 1.03236 | 2.75068 | 2.664 |
| chr11 | 469753 | 470061 | 0.586059 | 1.56156 | 2.665 |
| chr5 | 52239192 | 52239550 | 0.738687 | 1.96826 | 2.665 |
| chr16 | 78005535 | 78005767 | 1.20856 | 3.22049 | 2.665 |
| chr17 | 33939980 | 33940321 | 1.5161 | 4.0402 | 2.665 |
| chr3 | 149388519 | 149388730 | 0.403052 | 1.07409 | 2.665 |
| chr20 | 45336786 | 45337072 | 0.829433 | 2.2106 | 2.665 |
| chr4 | 81105143 | 81105625 | 2.01336 | 5.36613 | 2.665 |
| chr5 | 10201701 | 10201934 | 0.676446 | 1.80315 | 2.666 |
| chr1 | 84348617 | 84348862 | 0.740169 | 1.97326 | 2.666 |
| chr12 | 25308050 | 25308319 | 1.15895 | 3.08974 | 2.666 |
| chr13 | 102167062 | 102167383 | 0.644382 | 1.71797 | 2.666 |
| chr1 | 226377194 | 226377418 | 0.682325 | 1.8193 | 2.666 |
| chr6 | 107453645 | 107453928 | 0.95382 | 2.54322 | 2.666 |
| chr7 | 4603441 | 4603673 | 1.03069 | 2.7482 | 2.666 |
| chr3 | 24130364 | 24130634 | 0.798147 | 2.12838 | 2.667 |
| chr3 | 128745983 | 128746327 | 1.76024 | 4.69396 | 2.667 |
| chr17 | 46174877 | 46175209 | 1.02085 | 2.72228 | 2.667 |
| chr17 | 41804476 | 41804752 | 0.497998 | 1.32809 | 2.667 |
| chr2 | 11885302 | 11885555 | 1.05627 | 2.81695 | 2.667 |
| chr3 | 158144706 | 158144943 | 0.418848 | 1.11709 | 2.667 |
| chr3 | 48701490 | 48701823 | 1.32526 | 3.53489 | 2.667 |
| chr20 | 24549574 | 24549781 | 0.38512 | 1.0274 | 2.668 |
| chr6 | 27577105 | 27577392 | 1.41888 | 3.78523 | 2.668 |
| chr8 | 37888390 | 37888626 | 0.824089 | 2.19848 | 2.668 |
| chr15 | 45002822 | 45003052 | 0.485711 | 1.2958 | 2.668 |
| chr15 | 62599123 | 62599473 | 1.01271 | 2.70182 | 2.668 |
| chr14 | 78121370 | 78121881 | 1.07525 | 2.86892 | 2.668 |
| chr8 | 37695080 | 37695324 | 0.348808 | 0.930734 | 2.668 |
| chr8 | 6635713 | 6635968 | 0.400396 | 1.06843 | 2.668 |
| chr7 | 35841453 | 35841789 | 1.46831 | 3.91816 | 2.668 |
| chr12 | 60272527 | 60272774 | 0.490091 | 1.30781 | 2.669 |
| chr14 | 101869752 | 101869954 | 0.522563 | 1.39448 | 2.669 |
| chr15 | 89223747 | 89223995 | 0.577202 | 1.54034 | 2.669 |
| chr7 | 41982544 | 41983258 | 1.60922 | 4.29474 | 2.669 |
| chr12 | 133165596 | 133165815 | 0.388631 | 1.0372 | 2.669 |
| chr11 | 36069638 | 36069900 | 0.391079 | 1.04379 | 2.669 |
| chr8 | 126606295 | 126606595 | 0.60959 | 1.62713 | 2.669 |
| chr5 | 8791488 | 8791815 | 0.474148 | 1.26568 | 2.669 |
| chr5 | 132165590 | 132166016 | 1.73956 | 4.64376 | 2.670 |
| chr17 | 7232234 | 7232441 | 0.496593 | 1.32567 | 2.670 |
| chr5 | 150158749 | 150159060 | 1.015 | 2.70963 | 2.670 |
| chr12 | 45571265 | 45571497 | 0.493986 | 1.31877 | 2.670 |
| chr18 | 31689197 | 31689420 | 0.243637 | 0.650453 | 2.670 |
| chr16 | 9352384 | 9352693 | 1.14592 | 3.05941 | 2.670 |
| chr14 | 71088889 | 71089144 | 1.10003 | 2.93693 | 2.670 |
| chr7 | 20480258 | 20480708 | 1.12225 | 2.99645 | 2.670 |
| chr1 | 55755427 | 55755651 | 0.889393 | 2.37485 | 2.670 |
| chr6 | 41649965 | 41650268 | 1.20964 | 3.22999 | 2.670 |
| chr5 | 143571661 | 143571903 | 0.979711 | 2.61609 | 2.670 |
| chr6 | 52027790 | 52028034 | 1.0285 | 2.74637 | 2.670 |
| chr1 | 230777858 | 230778338 | 2.0451 | 5.46101 | 2.670 |
| chr12 | 49212812 | 49213044 | 0.485219 | 1.29568 | 2.670 |
| chr12 | 53259111 | 53259510 | 1.54165 | 4.11667 | 2.670 |

| | | | | | |
|---|---|---|---|---|---|
| chr1 | 151043516 | 151044006 | 1.99258 | 5.32113 | 2.670 |
| chr6 | 168461209 | 168461521 | 1.0377 | 2.77122 | 2.671 |
| chr1 | 54965212 | 54965428 | 0.828627 | 2.21293 | 2.671 |
| chr20 | 31683698 | 31684183 | 1.76307 | 4.70909 | 2.671 |
| chr10 | 112431821 | 112432355 | 1.32796 | 3.54701 | 2.671 |
| chr12 | 5977956 | 5978223 | 0.50047 | 1.33678 | 2.671 |
| chr8 | 80649436 | 80649732 | 0.487688 | 1.30264 | 2.671 |
| chr17 | 1520902 | 1521143 | 0.80351 | 2.14636 | 2.671 |
| chr12 | 61497963 | 61498217 | 0.628093 | 1.67797 | 2.672 |
| chr19 | 40030814 | 40031072 | 1.79531 | 4.79628 | 2.672 |
| chr5 | 136556197 | 136556428 | 1.10463 | 2.95127 | 2.672 |
| chr5 | 67010440 | 67010755 | 0.441625 | 1.17991 | 2.672 |
| chr1 | 27115682 | 27116039 | 1.48036 | 3.95524 | 2.672 |
| chr7 | 99730274 | 99730620 | 1.68273 | 4.49603 | 2.672 |
| chr3 | 107148813 | 107149059 | 1.82511 | 4.87661 | 2.672 |
| chr17 | 47976539 | 47976825 | 0.823899 | 2.20154 | 2.672 |
| chr7 | 129222695 | 129222923 | 0.931562 | 2.48929 | 2.672 |
| chr3 | 150021605 | 150021928 | 0.955792 | 2.5542 | 2.672 |
| chr13 | 28527770 | 28528350 | 1.45855 | 3.89805 | 2.673 |
| chr7 | 131376789 | 131377210 | 2.02767 | 5.4191 | 2.673 |
| chr8 | 29504893 | 29505153 | 0.685351 | 1.83171 | 2.673 |
| chr5 | 67548545 | 67548840 | 1.38479 | 3.70133 | 2.673 |
| chr20 | 39801725 | 39802304 | 1.5728 | 4.20412 | 2.673 |
| chr17 | 31263725 | 31264340 | 1.66554 | 4.45217 | 2.673 |
| chr12 | 111843057 | 111843424 | 0.97739 | 2.61327 | 2.674 |
| chr3 | 58645095 | 58645303 | 0.584922 | 1.56393 | 2.674 |
| chr11 | 130786187 | 130786571 | 1.25341 | 3.3514 | 2.674 |
| chr8 | 85027737 | 85028053 | 0.404978 | 1.08285 | 2.674 |
| chr20 | 30313506 | 30313710 | 0.722872 | 1.93312 | 2.674 |
| chr15 | 94347188 | 94347423 | 0.455887 | 1.21931 | 2.675 |
| chr19 | 56214955 | 56215579 | 1.56254 | 4.17931 | 2.675 |
| chr9 | 90341350 | 90341596 | 0.867363 | 2.32004 | 2.675 |
| chr12 | 34371402 | 34371667 | 1.19612 | 3.19946 | 2.675 |
| chr12 | 114602022 | 114602226 | 0.525056 | 1.40456 | 2.675 |
| chr15 | 65372030 | 65372276 | 0.594521 | 1.59047 | 2.675 |
| chrX | 85106510 | 85106772 | 1.20868 | 3.23356 | 2.675 |
| chr4 | 129375473 | 129375747 | 1.24552 | 3.33264 | 2.676 |
| chrX | 49012560 | 49012872 | 1.45424 | 3.89121 | 2.676 |
| chr12 | 127847978 | 127848295 | 0.532354 | 1.42449 | 2.676 |
| chr11 | 15122651 | 15122954 | 1.65451 | 4.42737 | 2.676 |
| chr20 | 3676925 | 3677164 | 1.41583 | 3.7887 | 2.676 |
| chr12 | 14319952 | 14320293 | 1.46419 | 3.91815 | 2.676 |
| chr4 | 15781044 | 15781253 | 0.628258 | 1.68122 | 2.676 |
| chr12 | 33072118 | 33072427 | 0.93038 | 2.48974 | 2.676 |
| chr12 | 45276652 | 45276870 | 0.354627 | 0.949016 | 2.676 |
| chr5 | 163322688 | 163322935 | 0.549534 | 1.47066 | 2.676 |
| chr8 | 41393412 | 41393710 | 1.16098 | 3.10704 | 2.676 |
| chr3 | 104746323 | 104746625 | 0.394605 | 1.05607 | 2.676 |
| chr10 | 63663014 | 63663318 | 0.501705 | 1.34276 | 2.676 |
| chr12 | 117910855 | 117911289 | 1.56592 | 4.19108 | 2.676 |
| chr3 | 13343793 | 13344009 | 0.850604 | 2.2766 | 2.676 |
| chr17 | 29151436 | 29151954 | 1.25375 | 3.35564 | 2.676 |
| chr22 | 39323133 | 39323673 | 1.66512 | 4.4569 | 2.677 |
| chr10 | 556715 | 557362 | 1.67659 | 4.48772 | 2.677 |
| chr6 | 24283270 | 24283706 | 0.983528 | 2.6326 | 2.677 |
| chr17 | 73874467 | 73874894 | 0.446677 | 1.19568 | 2.677 |
| chr6 | 1397227 | 1397570 | 2.07031 | 5.54237 | 2.677 |
| chr6 | 110663246 | 110663543 | 0.793793 | 2.12505 | 2.677 |
| chr9 | 215039 | 215302 | 0.811244 | 2.17192 | 2.677 |
| chr9 | 132302048 | 132302278 | 0.505809 | 1.35438 | 2.678 |
| chr16 | 29827462 | 29827924 | 1.92007 | 5.14162 | 2.678 |

| | | | | | |
|---|---|---|---|---|---|
| chr8 | 131433794 | 131434050 | 0.766025 | 2.05132 | 2.678 |
| chr3 | 65001121 | 65001343 | 0.3542 | 0.948572 | 2.678 |
| chr1 | 37980277 | 37980540 | 0.973912 | 2.60835 | 2.678 |
| chr10 | 2326222 | 2326442 | 0.348044 | 0.932162 | 2.678 |
| chr5 | 26395828 | 26396096 | 0.384858 | 1.03082 | 2.678 |
| chr11 | 30555113 | 30555358 | 0.402767 | 1.07884 | 2.679 |
| chr14 | 66343967 | 66344186 | 0.469333 | 1.25716 | 2.679 |
| chr17 | 7418071 | 7418411 | 1.24794 | 3.34285 | 2.679 |
| chr1 | 159861321 | 159861788 | 2.55204 | 6.83619 | 2.679 |
| chr6 | 16128932 | 16129237 | 1.13415 | 3.0384 | 2.679 |
| chr14 | 50296913 | 50297333 | 0.642799 | 1.72207 | 2.679 |
| chr9 | 114854157 | 114854401 | 0.821992 | 2.20224 | 2.679 |
| chr12 | 55505100 | 55505341 | 0.403242 | 1.0804 | 2.679 |
| chr14 | 105457875 | 105458155 | 0.482087 | 1.29171 | 2.679 |
| chr1 | 218454477 | 218454803 | 0.459274 | 1.23063 | 2.680 |
| chr11 | 10562667 | 10562987 | 1.70417 | 4.56653 | 2.680 |
| chr6 | 133505984 | 133506302 | 1.02455 | 2.74545 | 2.680 |
| chr9 | 37582646 | 37582888 | 0.826697 | 2.21537 | 2.680 |
| chr15 | 40714124 | 40714558 | 1.48294 | 3.97406 | 2.680 |
| chrX | 46246003 | 46246227 | 0.904337 | 2.4235 | 2.680 |
| chr4 | 73168878 | 73169145 | 0.895186 | 2.39902 | 2.680 |
| chr15 | 74284566 | 74284893 | 1.32582 | 3.55323 | 2.680 |
| chr7 | 7164126 | 7164379 | 0.529217 | 1.41838 | 2.680 |
| chr12 | 31388295 | 31389092 | 0.650419 | 1.74323 | 2.680 |
| chr19 | 45086125 | 45086486 | 1.61845 | 4.3378 | 2.680 |
| chr1 | 99214745 | 99214984 | 0.813903 | 2.18155 | 2.680 |
| chr6 | 26026536 | 26026805 | 1.14121 | 3.059 | 2.680 |
| chr12 | 13408660 | 13409012 | 2.16022 | 5.79057 | 2.681 |
| chr1 | 160800360 | 160800698 | 1.33905 | 3.58982 | 2.681 |
| chr3 | 101100781 | 101101000 | 0.619645 | 1.66122 | 2.681 |
| chr17 | 28353754 | 28354001 | 1.06838 | 2.86429 | 2.681 |
| chr10 | 86101217 | 86101432 | 0.467068 | 1.25226 | 2.681 |
| chr10 | 105352360 | 105352881 | 1.78427 | 4.78395 | 2.681 |
| chrX | 73353316 | 73353675 | 1.04109 | 2.79139 | 2.681 |
| chr5 | 54523086 | 54523323 | 0.816506 | 2.18947 | 2.682 |
| chr3 | 188913263 | 188913533 | 0.68666 | 1.84148 | 2.682 |
| chr12 | 5789392 | 5789646 | 0.364098 | 0.976451 | 2.682 |
| chr1 | 152020375 | 152020649 | 1.22175 | 3.27692 | 2.682 |
| chr3 | 45079161 | 45079570 | 1.4569 | 3.90762 | 2.682 |
| chr9 | 120576816 | 120577099 | 0.446597 | 1.19785 | 2.682 |
| chr16 | 31880007 | 31880220 | 0.900664 | 2.41574 | 2.682 |
| chr9 | 131095858 | 131096180 | 0.363815 | 0.975905 | 2.682 |
| chr14 | 104603802 | 104604282 | 1.81364 | 4.86509 | 2.683 |
| chr3 | 147589999 | 147590211 | 0.368441 | 0.988377 | 2.683 |
| chr12 | 74505554 | 74505768 | 0.770813 | 2.06794 | 2.683 |
| chr8 | 63160593 | 63160904 | 0.955295 | 2.56293 | 2.683 |
| chr13 | 74861127 | 74861341 | 0.965382 | 2.59013 | 2.683 |
| chr9 | 11470251 | 11470499 | 0.372642 | 0.999969 | 2.683 |
| chr7 | 145199179 | 145199448 | 0.482985 | 1.2961 | 2.684 |
| chr7 | 23301464 | 23301846 | 1.52942 | 4.10424 | 2.684 |
| chr20 | 56809014 | 56809293 | 1.65327 | 4.43678 | 2.684 |
| chr6 | 52926553 | 52926848 | 1.0428 | 2.79859 | 2.684 |
| chr12 | 120638985 | 120639369 | 1.48213 | 3.97775 | 2.684 |
| chr17 | 7919981 | 7920377 | 1.36906 | 3.67444 | 2.684 |
| chr16 | 69887217 | 69887501 | 0.429773 | 1.15348 | 2.684 |
| chr9 | 124040975 | 124041441 | 1.36803 | 3.6717 | 2.684 |
| chr22 | 48022778 | 48022982 | 0.554535 | 1.48834 | 2.684 |
| chr12 | 15743317 | 15743647 | 1.35638 | 3.6405 | 2.684 |
| chr10 | 11799957 | 11800266 | 1.76497 | 4.73717 | 2.684 |
| chr11 | 57224808 | 57225179 | 2.12237 | 5.69642 | 2.684 |
| chr11 | 122893880 | 122894155 | 1.50948 | 4.05143 | 2.684 |

| | | | | | |
|---|---|---|---|---|---|
| chr1 | 179338979 | 179339194 | 0.832429 | 2.23425 | 2.684 |
| chr14 | 95565335 | 95565683 | 1.39875 | 3.75495 | 2.685 |
| chr9 | 124260425 | 124260884 | 1.00507 | 2.69843 | 2.685 |
| chr12 | 4386057 | 4386497 | 1.69695 | 4.55602 | 2.685 |
| chr14 | 91440813 | 91441054 | 0.304953 | 0.81881 | 2.685 |
| chr1 | 40367752 | 40368229 | 0.814636 | 2.18741 | 2.685 |
| chr8 | 92600115 | 92600340 | 0.507926 | 1.36386 | 2.685 |
| chr13 | 88608919 | 88609127 | 0.22573 | 0.606151 | 2.685 |
| chr6 | 5402700 | 5402904 | 0.509878 | 1.36927 | 2.685 |
| chr3 | 101522052 | 101522366 | 1.3102 | 3.51855 | 2.686 |
| chr17 | 44953778 | 44954192 | 1.44089 | 3.8697 | 2.686 |
| chr1 | 194264880 | 194265123 | 1.54037 | 4.13727 | 2.686 |
| chr8 | 21600351 | 21600611 | 0.331472 | 0.890297 | 2.686 |
| chr5 | 5492385 | 5492843 | 1.22179 | 3.28174 | 2.686 |
| chr11 | 29355868 | 29356078 | 0.841003 | 2.25923 | 2.686 |
| chr3 | 23371094 | 23371382 | 0.434618 | 1.16757 | 2.686 |
| chr11 | 27720512 | 27720823 | 1.53496 | 4.12395 | 2.687 |
| chr16 | 54316004 | 54316579 | 1.07525 | 2.88893 | 2.687 |
| chr6 | 12234265 | 12234532 | 1.03635 | 2.78478 | 2.687 |
| chr3 | 100612536 | 100612762 | 0.905182 | 2.43235 | 2.687 |
| chr19 | 8134250 | 8134591 | 1.14534 | 3.0777 | 2.687 |
| chr15 | 56064647 | 56064907 | 0.547384 | 1.47091 | 2.687 |
| chr4 | 86717860 | 86718401 | 1.42947 | 3.8415 | 2.687 |
| chr17 | 30530462 | 30530755 | 0.86003 | 2.31149 | 2.688 |
| chr5 | 177034300 | 177034904 | 1.68793 | 4.53674 | 2.688 |
| chr1 | 29839861 | 29840134 | 1.01303 | 2.7228 | 2.688 |
| chr10 | 121115461 | 121115749 | 0.965949 | 2.59636 | 2.688 |
| chr17 | 50552460 | 50552673 | 0.349932 | 0.940582 | 2.688 |
| chr1 | 160248865 | 160249221 | 0.42856 | 1.15195 | 2.688 |
| chr14 | 103759187 | 103759578 | 1.61063 | 4.32957 | 2.688 |
| chr11 | 64420482 | 64420947 | 1.97757 | 5.31613 | 2.688 |
| chr16 | 22188268 | 22188728 | 0.304123 | 0.817567 | 2.688 |
| chr11 | 62742583 | 62742937 | 1.51308 | 4.06767 | 2.688 |
| chr19 | 57580462 | 57581074 | 0.584396 | 1.57106 | 2.688 |
| chr6 | 137663058 | 137663331 | 0.824355 | 2.21653 | 2.689 |
| chr12 | 73809408 | 73809674 | 0.497295 | 1.33723 | 2.689 |
| chr8 | 143376736 | 143377196 | 0.878148 | 2.3616 | 2.689 |
| chr12 | 111461917 | 111462401 | 2.16305 | 5.81727 | 2.689 |
| chr2 | 65267815 | 65268225 | 1.49479 | 4.02007 | 2.689 |
| chr8 | 131704525 | 131704822 | 0.821714 | 2.20991 | 2.689 |
| chr7 | 26581209 | 26581457 | 0.320716 | 0.862534 | 2.689 |
| chr10 | 103860969 | 103861202 | 0.828153 | 2.22729 | 2.689 |
| chr9 | 23378066 | 23378278 | 0.379173 | 1.01985 | 2.690 |
| chr4 | 39978025 | 39978385 | 1.60089 | 4.30595 | 2.690 |
| chr19 | 18701449 | 18701814 | 0.74757 | 2.01076 | 2.690 |
| chr9 | 111328120 | 111328400 | 1.29852 | 3.49286 | 2.690 |
| chr2 | 120165366 | 120165595 | 1.56222 | 4.20252 | 2.690 |
| chr7 | 9700501 | 9700905 | 0.528231 | 1.42106 | 2.690 |
| chr12 | 58196997 | 58197524 | 1.08172 | 2.91016 | 2.690 |
| chr9 | 86159377 | 86159677 | 1.10554 | 2.97432 | 2.690 |
| chr10 | 98100045 | 98100423 | 1.33343 | 3.58764 | 2.691 |
| chr6 | 12638276 | 12638608 | 1.40959 | 3.79256 | 2.691 |
| chr2 | 9910518 | 9910914 | 1.56698 | 4.21629 | 2.691 |
| chr22 | 37562604 | 37562913 | 1.5404 | 4.14486 | 2.691 |
| chr1 | 154879091 | 154879515 | 1.06015 | 2.8527 | 2.691 |
| chr14 | 89761422 | 89761631 | 0.440506 | 1.18542 | 2.691 |
| chr19 | 14247673 | 14248443 | 0.773799 | 2.08234 | 2.691 |
| chr3 | 70128587 | 70128819 | 0.374349 | 1.0074 | 2.691 |
| chr22 | 22697120 | 22697625 | 1.38729 | 3.7336 | 2.691 |
| chr16 | 790929 | 791450 | 1.17161 | 3.15323 | 2.691 |
| chr12 | 128010570 | 128010774 | 0.474733 | 1.27777 | 2.692 |

| | | | | | |
|---|---|---|---|---|---|
| chr8 | 143953251 | 143953511 | 0.597536 | 1.60838 | 2.692 |
| chr15 | 86029232 | 86029544 | 1.65218 | 4.44743 | 2.692 |
| chr1 | 36981903 | 36982320 | 2.09084 | 5.62871 | 2.692 |
| chr6 | 13795359 | 13795639 | 1.24594 | 3.35428 | 2.692 |
| chr6 | 155661623 | 155661849 | 0.744157 | 2.00344 | 2.692 |
| chr1 | 164509844 | 164510105 | 0.48298 | 1.30035 | 2.692 |
| chr3 | 180397842 | 180398112 | 0.747773 | 2.01336 | 2.692 |
| chr8 | 38571793 | 38572158 | 2.13024 | 5.73573 | 2.693 |
| chr5 | 30791273 | 30791507 | 0.414473 | 1.11599 | 2.693 |
| chr7 | 133427544 | 133427843 | 0.873073 | 2.35095 | 2.693 |
| chr11 | 8449557 | 8449806 | 0.437573 | 1.17829 | 2.693 |
| chr2 | 82509260 | 82509620 | 0.981767 | 2.64393 | 2.693 |
| chr10 | 33846374 | 33846678 | 0.369448 | 0.994968 | 2.693 |
| chr7 | 143113717 | 143113978 | 1.32351 | 3.56451 | 2.693 |
| chr6 | 139694384 | 139694656 | 0.600519 | 1.61747 | 2.693 |
| chr7 | 134204980 | 134205190 | 0.688966 | 1.85577 | 2.694 |
| chr1 | 29667500 | 29667715 | 0.90162 | 2.42865 | 2.694 |
| chr1 | 211688734 | 211689090 | 1.96317 | 5.28916 | 2.694 |
| chr1 | 170928330 | 170928557 | 0.312264 | 0.841306 | 2.694 |
| chr12 | 52550575 | 52550915 | 1.89282 | 5.09969 | 2.694 |
| chr6 | 27094874 | 27095132 | 1.14723 | 3.09102 | 2.694 |
| chr2 | 225987555 | 225987830 | 0.689312 | 1.85747 | 2.695 |
| chrY | 2803981 | 2804259 | 1.16497 | 3.13929 | 2.695 |
| chr5 | 74062188 | 74062525 | 0.612022 | 1.64925 | 2.695 |
| chr17 | 17854192 | 17854456 | 0.958243 | 2.58225 | 2.695 |
| chr5 | 36079241 | 36079839 | 0.820374 | 2.21075 | 2.695 |
| chr10 | 80208431 | 80208681 | 0.257361 | 0.69364 | 2.695 |
| chr19 | 11373181 | 11373396 | 0.897075 | 2.41833 | 2.696 |
| chr20 | 46483508 | 46483883 | 1.33921 | 3.61023 | 2.696 |
| chr19 | 51524252 | 51524583 | 1.03705 | 2.79573 | 2.696 |
| chr3 | 32989238 | 32989606 | 1.04751 | 2.82394 | 2.696 |
| chr10 | 50910963 | 50911205 | 0.336289 | 0.906642 | 2.696 |
| chrX | 40327626 | 40327933 | 0.779353 | 2.10124 | 2.696 |
| chr12 | 49371661 | 49372254 | 1.62634 | 4.38504 | 2.696 |
| chr6 | 147827735 | 147828093 | 1.14137 | 3.07747 | 2.696 |
| chr10 | 84571461 | 84571697 | 0.69456 | 1.87286 | 2.696 |
| chr6 | 38461762 | 38462041 | 0.872173 | 2.35182 | 2.697 |
| chr12 | 13515175 | 13515546 | 1.59759 | 4.30799 | 2.697 |
| chr7 | 156825514 | 156825754 | 0.354965 | 0.957233 | 2.697 |
| chr17 | 65154271 | 65154529 | 0.929697 | 2.50717 | 2.697 |
| chr16 | 52016709 | 52016971 | 1.16753 | 3.14857 | 2.697 |
| chr1 | 227635362 | 227635747 | 1.14522 | 3.08843 | 2.697 |
| chr11 | 3870493 | 3870811 | 1.11777 | 3.01446 | 2.697 |
| chr18 | 3771329 | 3771635 | 1.56478 | 4.21998 | 2.697 |
| chr7 | 39950634 | 39950919 | 0.357722 | 0.964733 | 2.697 |
| chr11 | 120983345 | 120983582 | 0.978751 | 2.6396 | 2.697 |
| chr1 | 17901167 | 17901393 | 0.778731 | 2.10039 | 2.697 |
| chr1 | 150141128 | 150141517 | 1.8688 | 5.04068 | 2.697 |
| chr12 | 102878222 | 102878731 | 1.54755 | 4.17421 | 2.697 |
| chr6 | 16511389 | 16511700 | 0.885612 | 2.38876 | 2.697 |
| chr18 | 644480 | 644719 | 0.529056 | 1.4271 | 2.697 |
| chr15 | 97789068 | 97789516 | 0.934446 | 2.52066 | 2.697 |
| chr19 | 10025374 | 10025620 | 0.90343 | 2.43717 | 2.698 |
| chr7 | 151991017 | 151991280 | 0.557182 | 1.50313 | 2.698 |
| chr6 | 159127491 | 159127730 | 0.948048 | 2.55796 | 2.698 |
| chr20 | 33297664 | 33298018 | 0.996078 | 2.68776 | 2.698 |
| chr21 | 42505935 | 42506138 | 0.995653 | 2.68666 | 2.698 |
| chr16 | 53450665 | 53450954 | 0.515467 | 1.39099 | 2.699 |
| chr2 | 194560167 | 194560370 | 0.706289 | 1.90592 | 2.699 |
| chr3 | 509959 | 510193 | 0.603638 | 1.62897 | 2.699 |
| chr12 | 83146186 | 83146448 | 1.41706 | 3.82442 | 2.699 |

| | | | | | |
|---|---|---|---|---|---|
| chr20 | 54565263 | 54565729 | 0.40012 | 1.07991 | 2.699 |
| chr1 | 163549778 | 163550114 | 0.412969 | 1.11461 | 2.699 |
| chr5 | 66985244 | 66985454 | 0.379079 | 1.02316 | 2.699 |
| chr10 | 49681813 | 49682075 | 0.51151 | 1.38074 | 2.699 |
| chr3 | 100035477 | 100035737 | 1.11201 | 3.00176 | 2.699 |
| chr22 | 36023300 | 36023506 | 0.963511 | 2.601 | 2.700 |
| chr11 | 19347992 | 19348413 | 0.445239 | 1.20193 | 2.700 |
| chr5 | 170050511 | 170050941 | 1.83628 | 4.95721 | 2.700 |
| chr17 | 42700545 | 42700765 | 0.824838 | 2.22674 | 2.700 |
| chr12 | 20702149 | 20702382 | 1.2659 | 3.41752 | 2.700 |
| chr2 | 230296138 | 230296355 | 0.81208 | 2.19239 | 2.700 |
| chr13 | 99083399 | 99083722 | 1.3523 | 3.65091 | 2.700 |
| chr3 | 100789093 | 100789460 | 1.56859 | 4.23487 | 2.700 |
| chr18 | 11040888 | 11041136 | 0.789296 | 2.13095 | 2.700 |
| chr9 | 35949652 | 35949937 | 0.458101 | 1.23679 | 2.700 |
| chr17 | 60781022 | 60781490 | 1.9028 | 5.13742 | 2.700 |
| chr5 | 111276358 | 111276567 | 0.356777 | 0.963325 | 2.700 |
| chr9 | 100377847 | 100378198 | 1.04239 | 2.81462 | 2.700 |
| chr6 | 106782987 | 106783205 | 0.889542 | 2.40221 | 2.701 |
| chr6 | 76644680 | 76644987 | 1.01211 | 2.73325 | 2.701 |
| chr19 | 49092093 | 49092522 | 2.35853 | 6.36973 | 2.701 |
| chr1 | 31687678 | 31687943 | 0.90307 | 2.43901 | 2.701 |
| chr8 | 106672738 | 106672989 | 0.440349 | 1.18935 | 2.701 |
| chr16 | 53583894 | 53584445 | 0.796554 | 2.15147 | 2.701 |
| chr8 | 71036093 | 71036320 | 0.476012 | 1.2857 | 2.701 |
| chr19 | 295013 | 295693 | 1.2762 | 3.44748 | 2.701 |
| chr14 | 77782297 | 77782581 | 1.05102 | 2.83923 | 2.701 |
| chr15 | 91648927 | 91649139 | 0.483913 | 1.30732 | 2.702 |
| chr17 | 6616450 | 6617165 | 0.845058 | 2.283 | 2.702 |
| chr16 | 16240291 | 16240511 | 0.372906 | 1.00749 | 2.702 |
| chr21 | 33238384 | 33238816 | 1.7082 | 4.61511 | 2.702 |
| chr4 | 83350502 | 83350865 | 1.33144 | 3.59745 | 2.702 |
| chr12 | 119553103 | 119553443 | 1.63686 | 4.42274 | 2.702 |
| chr10 | 82422485 | 82422797 | 1.79494 | 4.84994 | 2.702 |
| chr8 | 55745607 | 55745864 | 0.434453 | 1.17398 | 2.702 |
| chr3 | 131899073 | 131899505 | 1.03205 | 2.7892 | 2.703 |
| chr17 | 32098993 | 32099297 | 0.457299 | 1.23601 | 2.703 |
| chr1 | 156060547 | 156060848 | 1.62124 | 4.38204 | 2.703 |
| chr3 | 159721312 | 159721666 | 1.2467 | 3.36971 | 2.703 |
| chr13 | 59965772 | 59966098 | 0.841984 | 2.2761 | 2.703 |
| chr2 | 200850313 | 200850716 | 1.001 | 2.70604 | 2.703 |
| chr15 | 101203348 | 101203576 | 0.888362 | 2.40172 | 2.704 |
| chr1 | 55803843 | 55804075 | 0.831159 | 2.24745 | 2.704 |
| chr19 | 34286123 | 34286620 | 2.41034 | 6.51766 | 2.704 |
| chr12 | 81536835 | 81537039 | 0.514099 | 1.39021 | 2.704 |
| chr22 | 17539135 | 17539479 | 1.82266 | 4.92916 | 2.704 |
| chr10 | 124467471 | 124467724 | 0.36758 | 0.994155 | 2.705 |
| chr5 | 163081118 | 163081681 | 0.335825 | 0.908329 | 2.705 |
| chr16 | 28992630 | 28993393 | 1.17951 | 3.19033 | 2.705 |
| chr20 | 10517147 | 10517426 | 1.19702 | 3.23771 | 2.705 |
| chr7 | 96734002 | 96734403 | 1.36698 | 3.69742 | 2.705 |
| chr12 | 118813910 | 118814231 | 2.26489 | 6.12647 | 2.705 |
| chr12 | 104227767 | 104227986 | 0.66674 | 1.80359 | 2.705 |
| chr17 | 61556269 | 61556639 | 2.29839 | 6.21735 | 2.705 |
| chr20 | 50808180 | 50808711 | 1.31842 | 3.56658 | 2.705 |
| chr10 | 133181553 | 133181833 | 0.355743 | 0.962407 | 2.705 |
| chr15 | 91474910 | 91475257 | 1.36418 | 3.69058 | 2.705 |
| chr4 | 162173678 | 162174040 | 0.96201 | 2.60297 | 2.706 |
| chr1 | 234247246 | 234247578 | 1.03337 | 2.79624 | 2.706 |
| chr15 | 74731042 | 74731467 | 1.45435 | 3.93548 | 2.706 |
| chr17 | 31435620 | 31435832 | 0.619029 | 1.67513 | 2.706 |

| | | | | | |
|---|---|---|---|---|---|
| chr12 | 125946310 | 125946670 | 0.385949 | 1.04444 | 2.706 |
| chr11 | 32197806 | 32198088 | 1.21889 | 3.29852 | 2.706 |
| chr16 | 11012606 | 11012949 | 2.30994 | 6.25113 | 2.706 |
| chr5 | 63897754 | 63898044 | 0.488708 | 1.32264 | 2.706 |
| chr16 | 54991274 | 54991539 | 1.40463 | 3.80191 | 2.707 |
| chr9 | 79073852 | 79074068 | 0.72411 | 1.95999 | 2.707 |
| chr4 | 140682720 | 140682925 | 0.722469 | 1.95561 | 2.707 |
| chr2 | 53203521 | 53203831 | 1.10045 | 2.97875 | 2.707 |
| chr9 | 87287053 | 87287300 | 0.492366 | 1.33276 | 2.707 |
| chr17 | 73197824 | 73198179 | 1.45732 | 3.94519 | 2.707 |
| chr1 | 26123350 | 26123609 | 2.18825 | 5.92426 | 2.707 |
| chr12 | 108890207 | 108890483 | 0.380766 | 1.03086 | 2.707 |
| chr6 | 37759288 | 37759549 | 1.22141 | 3.30682 | 2.707 |
| chr1 | 228141667 | 228141938 | 1.57036 | 4.25223 | 2.708 |
| chr17 | 19350438 | 19350699 | 1.13284 | 3.06756 | 2.708 |
| chr5 | 67542566 | 67542887 | 0.430304 | 1.16521 | 2.708 |
| chr16 | 65539120 | 65539354 | 0.545226 | 1.47642 | 2.708 |
| chr2 | 44518868 | 44519129 | 1.35551 | 3.67079 | 2.708 |
| chrX | 128771531 | 128771845 | 1.36411 | 3.69417 | 2.708 |
| chr3 | 12747961 | 12748345 | 1.30966 | 3.54684 | 2.708 |
| chr10 | 31454873 | 31455160 | 0.472287 | 1.27908 | 2.708 |
| chrX | 54479451 | 54479658 | 0.4065 | 1.10092 | 2.708 |
| chr4 | 39979198 | 39979787 | 1.3026 | 3.52792 | 2.708 |
| chr16 | 75974979 | 75975367 | 0.404535 | 1.09565 | 2.708 |
| chr10 | 43574269 | 43574716 | 1.51256 | 4.09671 | 2.708 |
| chr2 | 8767781 | 8768102 | 0.789649 | 2.13875 | 2.708 |
| chr4 | 10015340 | 10015690 | 0.891496 | 2.41479 | 2.709 |
| chr11 | 65512015 | 65512268 | 1.14995 | 3.11508 | 2.709 |
| chr17 | 47276196 | 47276440 | 0.38848 | 1.05241 | 2.709 |
| chr10 | 102772590 | 102773240 | 1.88885 | 5.11706 | 2.709 |
| chr12 | 55780932 | 55781156 | 0.456341 | 1.23627 | 2.709 |
| chr1 | 206898993 | 206899245 | 1.30976 | 3.54827 | 2.709 |
| chr1 | 44412021 | 44412367 | 0.950274 | 2.57453 | 2.709 |
| chr1 | 16482244 | 16482447 | 0.308685 | 0.836312 | 2.709 |
| chr8 | 86985627 | 86985923 | 0.95012 | 2.57414 | 2.709 |
| chr12 | 103134472 | 103134719 | 0.601054 | 1.62846 | 2.709 |
| chr3 | 148748434 | 148748682 | 0.516093 | 1.39827 | 2.709 |
| chr17 | 41312089 | 41312357 | 0.319431 | 0.865471 | 2.709 |
| chr3 | 79441697 | 79441908 | 0.446742 | 1.21048 | 2.710 |
| chr13 | 110874574 | 110874892 | 1.40723 | 3.81306 | 2.710 |
| chr9 | 35658503 | 35658830 | 1.30099 | 3.52525 | 2.710 |
| chr7 | 105405553 | 105405947 | 1.30819 | 3.54496 | 2.710 |
| chr14 | 103541836 | 103542248 | 1.3726 | 3.71963 | 2.710 |
| chr7 | 93180455 | 93180694 | 0.969784 | 2.6281 | 2.710 |
| chr14 | 106986402 | 106987068 | 0.732343 | 1.9847 | 2.710 |
| chr12 | 576565 | 576864 | 0.58594 | 1.58798 | 2.710 |
| chr17 | 17997308 | 17997812 | 1.04418 | 2.82993 | 2.710 |
| chr10 | 26164551 | 26164833 | 1.74958 | 4.74222 | 2.710 |
| chr12 | 87308576 | 87308848 | 1.21682 | 3.29852 | 2.711 |
| chr7 | 19787416 | 19787660 | 1.07521 | 2.91471 | 2.711 |
| chr20 | 46012752 | 46013118 | 2.21283 | 5.9987 | 2.711 |
| chr5 | 60139930 | 60140163 | 0.772387 | 2.09387 | 2.711 |
| chr18 | 864765 | 865143 | 2.03917 | 5.52808 | 2.711 |
| chr5 | 134252787 | 134253136 | 1.05739 | 2.86656 | 2.711 |
| chr10 | 17272595 | 17273000 | 1.49229 | 4.04558 | 2.711 |
| chr3 | 138654918 | 138655459 | 1.05384 | 2.85695 | 2.711 |
| chr12 | 52397601 | 52398154 | 1.01484 | 2.75142 | 2.711 |
| chr16 | 2053687 | 2054221 | 1.90467 | 5.16398 | 2.711 |
| chr2 | 135415318 | 135415568 | 1.62246 | 4.39914 | 2.711 |
| chr10 | 7108273 | 7108686 | 0.346071 | 0.938419 | 2.712 |
| chr16 | 75730172 | 75730396 | 0.368829 | 1.00015 | 2.712 |

| | | | | | |
|---|---|---|---|---|---|
| chr12 | 108587510 | 108587988 | 1.06656 | 2.89234 | 2.712 |
| chr2 | 68423299 | 68423574 | 1.05633 | 2.86464 | 2.712 |
| chr5 | 56378324 | 56378596 | 0.75535 | 2.04846 | 2.712 |
| chr22 | 38077574 | 38077897 | 1.37666 | 3.73345 | 2.712 |
| chr8 | 21826700 | 21826930 | 0.616207 | 1.67123 | 2.712 |
| chr1 | 156597185 | 156597517 | 1.37862 | 3.73922 | 2.712 |
| chr20 | 37274692 | 37275157 | 1.7625 | 4.78126 | 2.713 |
| chr7 | 120632862 | 120633148 | 0.452633 | 1.22789 | 2.713 |
| chr14 | 101649651 | 101650015 | 0.763088 | 2.07012 | 2.713 |
| chr17 | 80958026 | 80958317 | 0.481098 | 1.30516 | 2.713 |
| chr7 | 65495931 | 65496192 | 0.799187 | 2.16819 | 2.713 |
| chr18 | 3297446 | 3297716 | 1.21806 | 3.30471 | 2.713 |
| chr12 | 111834649 | 111835102 | 2.13518 | 5.79343 | 2.713 |
| chr3 | 191848244 | 191848669 | 1.00968 | 2.73966 | 2.713 |
| chr19 | 23065687 | 23065937 | 0.630897 | 1.71189 | 2.713 |
| chr17 | 71512498 | 71512720 | 0.370711 | 1.00599 | 2.714 |
| chr1 | 214502126 | 214502539 | 2.17533 | 5.90339 | 2.714 |
| chr11 | 69500612 | 69501029 | 2.06573 | 5.60609 | 2.714 |
| chr3 | 11815780 | 11816074 | 1.27104 | 3.44941 | 2.714 |
| chrX | 67403704 | 67404016 | 0.792325 | 2.15033 | 2.714 |
| chr9 | 21709518 | 21709788 | 1.33837 | 3.63247 | 2.714 |
| chr2 | 206428421 | 206428679 | 0.607447 | 1.64878 | 2.714 |
| chr6 | 65215298 | 65215600 | 0.559544 | 1.51885 | 2.714 |
| chr15 | 40630286 | 40630664 | 1.59478 | 4.32903 | 2.715 |
| chr10 | 70166806 | 70167280 | 2.23123 | 6.05708 | 2.715 |
| chr17 | 77965390 | 77965889 | 1.31025 | 3.55692 | 2.715 |
| chr22 | 44320049 | 44320355 | 0.532197 | 1.44476 | 2.715 |
| chr10 | 6711302 | 6711517 | 0.858009 | 2.32941 | 2.715 |
| chr11 | 65319525 | 65319950 | 1.41443 | 3.84026 | 2.715 |
| chr12 | 132155522 | 132155750 | 0.332646 | 0.903166 | 2.715 |
| chr20 | 55570685 | 55571045 | 1.15236 | 3.12889 | 2.715 |
| chr3 | 128327115 | 128327579 | 1.53897 | 4.17874 | 2.715 |
| chr6 | 16206860 | 16207107 | 1.16857 | 3.17304 | 2.715 |
| chr8 | 107858178 | 107858485 | 0.954899 | 2.59296 | 2.715 |
| chr1 | 157140589 | 157140875 | 1.5129 | 4.1082 | 2.715 |
| chr7 | 100317588 | 100317950 | 0.890506 | 2.4185 | 2.716 |
| chr10 | 105881425 | 105881639 | 1.33426 | 3.6239 | 2.716 |
| chr9 | 139622586 | 139622908 | 1.64888 | 4.47868 | 2.716 |
| chr12 | 21466426 | 21466731 | 1.36703 | 3.71324 | 2.716 |
| chr8 | 56987047 | 56987365 | 1.67046 | 4.53812 | 2.717 |
| chr10 | 103843868 | 103844074 | 0.363835 | 0.988435 | 2.717 |
| chr1 | 206804493 | 206804748 | 1.02549 | 2.78597 | 2.717 |
| chr8 | 126712952 | 126713237 | 1.0837 | 2.94419 | 2.717 |
| chr6 | 125475990 | 125476285 | 1.10385 | 2.99911 | 2.717 |
| chr12 | 58033178 | 58033426 | 0.390249 | 1.06047 | 2.717 |
| chr1 | 42780174 | 42780419 | 0.989535 | 2.68903 | 2.717 |
| chr6 | 117771109 | 117771316 | 0.665894 | 1.8096 | 2.718 |
| chr3 | 156799506 | 156799984 | 1.55537 | 4.22691 | 2.718 |
| chr5 | 134332450 | 134332658 | 0.344788 | 0.937004 | 2.718 |
| chr7 | 47621618 | 47621851 | 0.630024 | 1.71232 | 2.718 |
| chr22 | 49178948 | 49179216 | 1.05202 | 2.85951 | 2.718 |
| chr2 | 41161501 | 41161713 | 0.550584 | 1.49661 | 2.718 |
| chr19 | 39420802 | 39421341 | 2.9703 | 8.07412 | 2.718 |
| chr9 | 115473418 | 115473701 | 0.917832 | 2.49515 | 2.719 |
| chr10 | 45960401 | 45960821 | 1.99647 | 5.42764 | 2.719 |
| chr2 | 149315675 | 149315897 | 0.815424 | 2.21697 | 2.719 |
| chr12 | 111284800 | 111285086 | 1.33064 | 3.61789 | 2.719 |
| chr16 | 59446198 | 59446467 | 1.23184 | 3.34939 | 2.719 |
| chr3 | 167587551 | 167587792 | 0.235384 | 0.640013 | 2.719 |
| chr10 | 82820898 | 82821118 | 0.429565 | 1.168 | 2.719 |
| chr1 | 27721915 | 27722209 | 1.4175 | 3.85427 | 2.719 |

| | | | | | |
|---|---|---|---|---|---|
| chr15 | 29147590 | 29147817 | 0.734284 | 1.99661 | 2.719 |
| chr9 | 138304337 | 138304866 | 2.41239 | 6.56005 | 2.719 |
| chr6 | 146847387 | 146847638 | 1.0658 | 2.89827 | 2.719 |
| chr21 | 37876527 | 37876887 | 0.865474 | 2.35355 | 2.719 |
| chr3 | 99973021 | 99973431 | 1.18358 | 3.21863 | 2.719 |
| chr12 | 113821786 | 113821990 | 0.524832 | 1.42727 | 2.719 |
| chr6 | 21935870 | 21936166 | 0.770066 | 2.09418 | 2.719 |
| chr15 | 59704805 | 59705170 | 1.3029 | 3.54323 | 2.719 |
| chr22 | 37374104 | 37374440 | 1.31378 | 3.57287 | 2.720 |
| chr10 | 131988312 | 131988853 | 2.54481 | 6.92072 | 2.720 |
| chr13 | 46302954 | 46303216 | 1.00649 | 2.73725 | 2.720 |
| chr3 | 5625723 | 5625928 | 0.62698 | 1.70528 | 2.720 |
| chr9 | 36008734 | 36009259 | 2.705 | 7.35743 | 2.720 |
| chr4 | 89739829 | 89740216 | 0.964825 | 2.62428 | 2.720 |
| chr4 | 124694007 | 124694341 | 1.13009 | 3.07384 | 2.720 |
| chr12 | 132022211 | 132022629 | 0.448784 | 1.22073 | 2.720 |
| chr6 | 53357806 | 53358037 | 0.770261 | 2.09521 | 2.720 |
| chr17 | 39480464 | 39480801 | 1.27214 | 3.4605 | 2.720 |
| chr10 | 75645843 | 75646227 | 1.41624 | 3.85327 | 2.721 |
| chr7 | 111201978 | 111202203 | 0.697589 | 1.89812 | 2.721 |
| chr3 | 111830665 | 111830973 | 1.31268 | 3.57225 | 2.721 |
| chr20 | 3613313 | 3613520 | 0.785775 | 2.13844 | 2.721 |
| chr20 | 52556475 | 52556756 | 1.05872 | 2.8813 | 2.721 |
| chr5 | 109024278 | 109024644 | 0.986284 | 2.68453 | 2.722 |
| chr5 | 95717299 | 95717514 | 0.700485 | 1.90665 | 2.722 |
| chr12 | 76755356 | 76755741 | 0.298449 | 0.812415 | 2.722 |
| chr1 | 228248310 | 228248908 | 0.709478 | 1.9313 | 2.722 |
| chr10 | 12698596 | 12698884 | 1.13322 | 3.08499 | 2.722 |
| chr7 | 54745136 | 54745419 | 0.404383 | 1.10095 | 2.723 |
| chr17 | 53591382 | 53591742 | 1.4199 | 3.86577 | 2.723 |
| chr1 | 69065746 | 69066089 | 1.21771 | 3.31536 | 2.723 |
| chr7 | 71200861 | 71201366 | 1.42537 | 3.88077 | 2.723 |
| chr4 | 148787906 | 148788352 | 1.19439 | 3.25197 | 2.723 |
| chr9 | 84815173 | 84815382 | 0.477062 | 1.29892 | 2.723 |
| chr11 | 29905859 | 29906150 | 0.28508 | 0.776312 | 2.723 |
| chr2 | 208337945 | 208338163 | 1.08269 | 2.94832 | 2.723 |
| chrX | 7893879 | 7894236 | 1.54372 | 4.20379 | 2.723 |
| chr12 | 113634705 | 113635068 | 1.3764 | 3.74824 | 2.723 |
| chr3 | 11420735 | 11421013 | 0.692476 | 1.88577 | 2.723 |
| chr20 | 25317216 | 25317448 | 0.940797 | 2.56209 | 2.723 |
| chr16 | 22110387 | 22110655 | 1.06667 | 2.90499 | 2.723 |
| chr1 | 47889999 | 47890392 | 1.26189 | 3.43675 | 2.723 |
| chr13 | 27056672 | 27057015 | 0.858487 | 2.33811 | 2.724 |
| chr14 | 48004469 | 48004783 | 0.346754 | 0.944428 | 2.724 |
| chr7 | 80886390 | 80886592 | 0.857247 | 2.33488 | 2.724 |
| chr22 | 23933242 | 23933708 | 1.62025 | 4.41343 | 2.724 |
| chr5 | 177801967 | 177802202 | 0.4281 | 1.16611 | 2.724 |
| chr8 | 27721368 | 27721688 | 1.63133 | 4.44367 | 2.724 |
| chr7 | 125358272 | 125358491 | 0.422569 | 1.15111 | 2.724 |
| chr12 | 49741318 | 49741797 | 2.08019 | 5.66683 | 2.724 |
| chr12 | 124944815 | 124945109 | 0.405323 | 1.10422 | 2.724 |
| chrX | 140000623 | 140000881 | 1.28555 | 3.50246 | 2.724 |
| chr4 | 37454662 | 37454933 | 0.747229 | 2.03586 | 2.725 |
| chr16 | 89623864 | 89624455 | 1.1795 | 3.21403 | 2.725 |
| chr22 | 39727207 | 39727482 | 0.589667 | 1.60691 | 2.725 |
| chr12 | 121106195 | 121106441 | 0.362022 | 0.986562 | 2.725 |
| chr20 | 55955262 | 55955562 | 0.356258 | 0.970894 | 2.725 |
| chr6 | 161413844 | 161414089 | 0.887172 | 2.41784 | 2.725 |
| chr4 | 184393433 | 184393678 | 1.18406 | 3.22716 | 2.726 |
| chr8 | 96036967 | 96037268 | 1.26705 | 3.45345 | 2.726 |
| chr17 | 14717086 | 14717306 | 1.10318 | 3.00696 | 2.726 |

| | | | | | |
|---|---|---|---|---|---|
| chr19 | 56320864 | 56321222 | 1.07048 | 2.91786 | 2.726 |
| chr2 | 64555102 | 64555331 | 0.859207 | 2.34198 | 2.726 |
| chr5 | 16975341 | 16975559 | 0.377503 | 1.02903 | 2.726 |
| chr1 | 24528339 | 24528853 | 0.78639 | 2.14379 | 2.726 |
| chr5 | 156575884 | 156576278 | 1.54574 | 4.21388 | 2.726 |
| chr3 | 21525759 | 21526167 | 1.11948 | 3.05211 | 2.726 |
| chr6 | 17825803 | 17826052 | 0.347503 | 0.947432 | 2.726 |
| chr10 | 92457829 | 92458052 | 0.400862 | 1.09299 | 2.727 |
| chr5 | 62619849 | 62620133 | 0.402606 | 1.09777 | 2.727 |
| chr3 | 185977311 | 185977676 | 1.64492 | 4.48537 | 2.727 |
| chr12 | 112849447 | 112849671 | 0.694617 | 1.89425 | 2.727 |
| chr19 | 43929865 | 43930112 | 1.02589 | 2.79775 | 2.727 |
| chr17 | 76794590 | 76795482 | 0.747452 | 2.03853 | 2.727 |
| chr10 | 35893926 | 35894227 | 0.325517 | 0.887802 | 2.727 |
| chr1 | 87170067 | 87170812 | 0.953565 | 2.60073 | 2.727 |
| chr3 | 6892189 | 6892427 | 0.52386 | 1.42881 | 2.727 |
| chr15 | 79632346 | 79632712 | 2.26039 | 6.16526 | 2.728 |
| chr10 | 90977485 | 90977744 | 1.25821 | 3.43244 | 2.728 |
| chr13 | 61192248 | 61192485 | 0.690897 | 1.88482 | 2.728 |
| chr14 | 52482415 | 52482728 | 0.84301 | 2.29983 | 2.728 |
| chr12 | 112383097 | 112383337 | 1.06834 | 2.91459 | 2.728 |
| chr4 | 188301276 | 188301512 | 0.715866 | 1.95307 | 2.728 |
| chr12 | 102685741 | 102686123 | 0.418434 | 1.14164 | 2.728 |
| chr12 | 29735530 | 29735778 | 0.418571 | 1.14203 | 2.728 |
| chr9 | 102773401 | 102773668 | 1.04871 | 2.86172 | 2.729 |
| chr7 | 103402812 | 103403034 | 0.3579 | 0.97669 | 2.729 |
| chr10 | 73155925 | 73156308 | 0.943856 | 2.57589 | 2.729 |
| chr5 | 16934194 | 16934563 | 2.26047 | 6.16937 | 2.729 |
| chr8 | 128508686 | 128509029 | 1.20533 | 3.2898 | 2.729 |
| chr3 | 124505567 | 124505828 | 0.495621 | 1.35276 | 2.729 |
| chr16 | 68936287 | 68936604 | 1.40732 | 3.84126 | 2.729 |
| chr12 | 130765692 | 130766095 | 2.03776 | 5.56307 | 2.730 |
| chr17 | 17328367 | 17328701 | 0.742968 | 2.02838 | 2.730 |
| chrX | 20324891 | 20325190 | 0.333115 | 0.909455 | 2.730 |
| chr15 | 68675756 | 68676146 | 1.20849 | 3.29937 | 2.730 |
| chr1 | 9048124 | 9048519 | 2.06 | 5.62437 | 2.730 |
| chr17 | 46662742 | 46663051 | 1.51493 | 4.13618 | 2.730 |
| chr11 | 87541829 | 87542099 | 0.40758 | 1.11281 | 2.730 |
| chr15 | 41120899 | 41121158 | 0.865019 | 2.36182 | 2.730 |
| chr4 | 187867846 | 187868160 | 1.35309 | 3.69489 | 2.731 |
| chr17 | 34985661 | 34986129 | 2.68278 | 7.32666 | 2.731 |
| chr5 | 29132714 | 29132932 | 0.504227 | 1.37721 | 2.731 |
| chr2 | 27184436 | 27184902 | 1.23188 | 3.3647 | 2.731 |
| chr10 | 115323180 | 115323675 | 2.17527 | 5.94155 | 2.731 |
| chr10 | 128075604 | 128075920 | 1.46963 | 4.01472 | 2.732 |
| chr12 | 93861319 | 93861671 | 1.51344 | 4.13454 | 2.732 |
| chr19 | 55939139 | 55939344 | 0.327069 | 0.893634 | 2.732 |
| chr7 | 146130172 | 146130455 | 0.440033 | 1.20233 | 2.732 |
| chr18 | 12391690 | 12391960 | 1.02036 | 2.78814 | 2.733 |
| chr1 | 3434715 | 3435149 | 1.28676 | 3.51612 | 2.733 |
| chr11 | 46336754 | 46337027 | 1.54054 | 4.20997 | 2.733 |
| chr1 | 203236412 | 203236779 | 1.53966 | 4.20808 | 2.733 |
| chr10 | 36999203 | 36999405 | 0.48731 | 1.3319 | 2.733 |
| chr12 | 131939289 | 131939666 | 1.82714 | 4.99569 | 2.734 |
| chr16 | 1903320 | 1903577 | 0.320056 | 0.875101 | 2.734 |
| chr7 | 15428726 | 15429010 | 0.543488 | 1.48613 | 2.734 |
| chr2 | 163183724 | 163183954 | 0.70027 | 1.91485 | 2.734 |
| chr3 | 14187389 | 14187645 | 0.475925 | 1.30149 | 2.735 |
| chr3 | 127174412 | 127174746 | 1.77971 | 4.8669 | 2.735 |
| chr14 | 89026286 | 89026588 | 0.82847 | 2.26561 | 2.735 |
| chrX | 9167439 | 9167663 | 0.487577 | 1.33351 | 2.735 |

| | | | | | |
|---|---|---|---|---|---|
| chr12 | 76430949 | 76431250 | 1.29521 | 3.54237 | 2.735 |
| chrX | 34318946 | 34319282 | 0.535972 | 1.46589 | 2.735 |
| chr17 | 4934714 | 4935026 | 1.66891 | 4.56452 | 2.735 |
| chr5 | 168983307 | 168983660 | 0.337658 | 0.923614 | 2.735 |
| chr2 | 46877705 | 46878043 | 1.55554 | 4.25509 | 2.735 |
| chr3 | 5295421 | 5295627 | 0.492983 | 1.34856 | 2.736 |
| chr15 | 70870690 | 70871028 | 0.429179 | 1.17403 | 2.736 |
| chr14 | 52070681 | 52070889 | 0.477504 | 1.30623 | 2.736 |
| chr17 | 42295604 | 42296021 | 1.06308 | 2.90811 | 2.736 |
| chr4 | 20315278 | 20315588 | 1.12481 | 3.07736 | 2.736 |
| chr10 | 56828666 | 56828989 | 0.496484 | 1.35838 | 2.736 |
| chr19 | 4809613 | 4809924 | 1.47548 | 4.03743 | 2.736 |
| chr1 | 211643369 | 211643743 | 1.74027 | 4.762 | 2.736 |
| chr2 | 96011221 | 96011664 | 2.37567 | 6.50069 | 2.736 |
| chr17 | 8791806 | 8792236 | 1.23256 | 3.37286 | 2.736 |
| chr18 | 8696777 | 8697056 | 1.71836 | 4.7024 | 2.737 |
| chr19 | 56582473 | 56582760 | 1.19255 | 3.26356 | 2.737 |
| chr12 | 65950517 | 65950847 | 1.69769 | 4.64646 | 2.737 |
| chr10 | 79925544 | 79925825 | 0.690353 | 1.88953 | 2.737 |
| chr14 | 104367298 | 104367932 | 2.15747 | 5.90516 | 2.737 |
| chr19 | 49992973 | 49993348 | 0.989331 | 2.70789 | 2.737 |
| chr8 | 118627665 | 118627914 | 0.739185 | 2.02353 | 2.738 |
| chr3 | 186332645 | 186333006 | 1.72111 | 4.71166 | 2.738 |
| chr17 | 20704974 | 20705298 | 0.945885 | 2.58969 | 2.738 |
| chr10 | 69408754 | 69409064 | 0.291578 | 0.798326 | 2.738 |
| chr3 | 15160126 | 15160509 | 2.19454 | 6.00865 | 2.738 |
| chrX | 70775083 | 70775305 | 0.860224 | 2.35533 | 2.738 |
| chr21 | 37485107 | 37485431 | 1.62891 | 4.46012 | 2.738 |
| chr20 | 42818810 | 42819200 | 1.2951 | 3.54626 | 2.738 |
| chr6 | 43969064 | 43969325 | 1.1088 | 3.03622 | 2.738 |
| chr7 | 118070753 | 118070992 | 0.347563 | 0.951767 | 2.738 |
| chr3 | 37783203 | 37783449 | 0.89775 | 2.45882 | 2.739 |
| chr2 | 43451346 | 43451620 | 1.0945 | 2.99799 | 2.739 |
| chrX | 40737228 | 40737448 | 0.578678 | 1.58517 | 2.739 |
| chr10 | 73804151 | 73804370 | 1.33409 | 3.65452 | 2.739 |
| chr12 | 38778508 | 38778780 | 0.783894 | 2.14782 | 2.740 |
| chr4 | 1124333 | 1124736 | 1.8369 | 5.03322 | 2.740 |
| chr12 | 110152097 | 110152516 | 0.73607 | 2.01704 | 2.740 |
| chr5 | 90734770 | 90735004 | 0.668466 | 1.83184 | 2.740 |
| chr12 | 131741869 | 131742100 | 0.403612 | 1.10608 | 2.740 |
| chr10 | 118998726 | 118998961 | 0.504912 | 1.38369 | 2.740 |
| chr9 | 96444328 | 96444914 | 1.05474 | 2.89055 | 2.741 |
| chr4 | 31964644 | 31964846 | 0.422497 | 1.15788 | 2.741 |
| chr12 | 109701782 | 109702316 | 1.4819 | 4.06138 | 2.741 |
| chr22 | 25758111 | 25758461 | 1.83629 | 5.0327 | 2.741 |
| chr21 | 45557820 | 45558340 | 1.14742 | 3.14486 | 2.741 |
| chr4 | 139951806 | 139952109 | 1.22328 | 3.35296 | 2.741 |
| chr5 | 140998649 | 140999038 | 1.77572 | 4.86742 | 2.741 |
| chr8 | 146127353 | 146127734 | 1.60406 | 4.3972 | 2.741 |
| chr10 | 112370115 | 112370466 | 1.58001 | 4.33137 | 2.741 |
| chr14 | 52766502 | 52766708 | 0.900728 | 2.46922 | 2.741 |
| chr15 | 72143226 | 72143508 | 1.07057 | 2.93506 | 2.742 |
| chr11 | 6704594 | 6704996 | 1.61255 | 4.42128 | 2.742 |
| chr3 | 128772274 | 128772539 | 0.993529 | 2.72409 | 2.742 |
| chr10 | 17659369 | 17659652 | 0.701222 | 1.92306 | 2.742 |
| chr15 | 29431780 | 29432459 | 1.60496 | 4.40169 | 2.743 |
| chr7 | 105022333 | 105022585 | 0.236139 | 0.647643 | 2.743 |
| chr2 | 122055083 | 122055386 | 1.59373 | 4.37112 | 2.743 |
| chr1 | 246852638 | 246853219 | 1.37068 | 3.75955 | 2.743 |
| chr20 | 52195374 | 52195590 | 1.43969 | 3.94909 | 2.743 |
| chr20 | 33796220 | 33796548 | 1.36618 | 3.74797 | 2.743 |

| | | | | | |
|---|---|---|---|---|---|
| chr4 | 84203252 | 84203462 | 0.858395 | 2.35507 | 2.744 |
| chr1 | 201798030 | 201798414 | 2.55263 | 7.00354 | 2.744 |
| chr15 | 92462328 | 92462530 | 0.417439 | 1.14532 | 2.744 |
| chr5 | 84860974 | 84861214 | 0.342187 | 0.938852 | 2.744 |
| chr19 | 39274102 | 39274349 | 0.519592 | 1.42569 | 2.744 |
| chr19 | 41047082 | 41047525 | 1.39435 | 3.82611 | 2.744 |
| chr20 | 62612026 | 62612512 | 1.39535 | 3.82895 | 2.744 |
| chr9 | 19248883 | 19249101 | 0.295429 | 0.810686 | 2.744 |
| chr6 | 42042690 | 42043579 | 0.661488 | 1.81526 | 2.744 |
| chr10 | 91898736 | 91898940 | 0.615673 | 1.68954 | 2.744 |
| chr17 | 25680820 | 25681160 | 1.85469 | 5.09038 | 2.745 |
| chr1 | 229363190 | 229363565 | 1.42791 | 3.91922 | 2.745 |
| chr14 | 44961285 | 44961602 | 0.465865 | 1.27868 | 2.745 |
| chr13 | 79965443 | 79965940 | 1.09792 | 3.01375 | 2.745 |
| chr13 | 25823360 | 25823588 | 0.710513 | 1.95034 | 2.745 |
| chr14 | 45382879 | 45383268 | 1.27145 | 3.49041 | 2.745 |
| chr1 | 236922584 | 236922848 | 0.95637 | 2.6257 | 2.745 |
| chr11 | 131274092 | 131274312 | 0.365544 | 1.00365 | 2.746 |
| chr10 | 93677686 | 93677952 | 0.434208 | 1.19219 | 2.746 |
| chr7 | 32982564 | 32982969 | 1.10128 | 3.02428 | 2.746 |
| chr1 | 232595803 | 232596040 | 0.358346 | 0.984227 | 2.747 |
| chr12 | 122326310 | 122326546 | 0.68042 | 1.86904 | 2.747 |
| chr1 | 175706889 | 175707092 | 0.35245 | 0.968204 | 2.747 |
| chr16 | 85793427 | 85793764 | 1.73413 | 4.76441 | 2.747 |
| chr11 | 44970593 | 44970956 | 1.25819 | 3.45685 | 2.747 |
| chr3 | 128742897 | 128743201 | 0.557458 | 1.53168 | 2.748 |
| chr19 | 11488944 | 11489294 | 1.32283 | 3.63485 | 2.748 |
| chr20 | 16414724 | 16415127 | 1.76628 | 4.85356 | 2.748 |
| chr2 | 219260629 | 219260990 | 1.18811 | 3.26482 | 2.748 |
| chr4 | 75000061 | 75000276 | 1.07142 | 2.94419 | 2.748 |
| chr10 | 123734637 | 123735047 | 1.46111 | 4.01565 | 2.748 |
| chrX | 43417143 | 43417362 | 1.18789 | 3.26481 | 2.748 |
| chr1 | 41910034 | 41910283 | 0.736008 | 2.02314 | 2.749 |
| chr7 | 87872616 | 87873053 | 1.02846 | 2.82712 | 2.749 |
| chr14 | 51597595 | 51597907 | 0.357213 | 0.981962 | 2.749 |
| chr20 | 43229805 | 43230026 | 0.50404 | 1.38558 | 2.749 |
| chr10 | 13432827 | 13433111 | 0.349337 | 0.960316 | 2.749 |
| chr10 | 102586559 | 102586946 | 0.616029 | 1.6935 | 2.749 |
| chr5 | 96478431 | 96478645 | 1.02643 | 2.82181 | 2.749 |
| chr12 | 57569453 | 57570021 | 1.52243 | 4.1854 | 2.749 |
| chr20 | 32899406 | 32899663 | 0.570585 | 1.56866 | 2.749 |
| chr17 | 54671790 | 54672228 | 0.500444 | 1.37592 | 2.749 |
| chr16 | 21311888 | 21312201 | 1.24402 | 3.42034 | 2.749 |
| chr14 | 49640573 | 49640795 | 1.03283 | 2.83972 | 2.749 |
| chr8 | 95003613 | 95003860 | 1.24764 | 3.43047 | 2.750 |
| chr15 | 39156460 | 39156740 | 0.667002 | 1.83413 | 2.750 |
| chr15 | 90293604 | 90294250 | 0.874176 | 2.40393 | 2.750 |
| chr1 | 169555576 | 169555896 | 1.07332 | 2.95166 | 2.750 |
| chr14 | 82025931 | 82026137 | 0.457973 | 1.25945 | 2.750 |
| chr1 | 173475698 | 173475900 | 0.7497 | 2.06187 | 2.750 |
| chr20 | 44190751 | 44190998 | 0.408495 | 1.12349 | 2.750 |
| chr11 | 41653933 | 41654218 | 0.31631 | 0.869982 | 2.750 |
| chr12 | 107525627 | 107525900 | 1.20295 | 3.30873 | 2.751 |
| chr1 | 162583493 | 162583835 | 1.14711 | 3.15527 | 2.751 |
| chr6 | 147543734 | 147543953 | 0.553033 | 1.52122 | 2.751 |
| chr5 | 188699 | 189067 | 1.34406 | 3.69741 | 2.751 |
| chr12 | 3409668 | 3410049 | 1.75178 | 4.81963 | 2.751 |
| chr16 | 88509408 | 88509848 | 2.1026 | 5.7849 | 2.751 |
| chr1 | 160967122 | 160967511 | 1.49893 | 4.12433 | 2.752 |
| chr11 | 82393281 | 82393486 | 0.822833 | 2.26417 | 2.752 |
| chr15 | 40811940 | 40812288 | 1.59723 | 4.39518 | 2.752 |

| | | | | | |
|---|---|---|---|---|---|
| chr20 | 59804172 | 59804589 | 2.31328 | 6.36592 | 2.752 |
| chr5 | 72832657 | 72833076 | 1.30073 | 3.57956 | 2.752 |
| chr12 | 45610092 | 45610804 | 0.763222 | 2.10043 | 2.752 |
| chr18 | 8480584 | 8480821 | 0.985287 | 2.71159 | 2.752 |
| chr5 | 140699720 | 140700115 | 2.02713 | 5.579 | 2.752 |
| chr8 | 38625815 | 38626151 | 1.64022 | 4.51427 | 2.752 |
| chr10 | 49423490 | 49423941 | 1.3543 | 3.7274 | 2.752 |
| chr4 | 15807851 | 15808087 | 0.576747 | 1.58744 | 2.752 |
| chr12 | 19725821 | 19726193 | 1.33381 | 3.67123 | 2.752 |
| chr9 | 89682489 | 89682763 | 0.416386 | 1.14611 | 2.753 |
| chr17 | 30714946 | 30715362 | 0.477479 | 1.31428 | 2.753 |
| chr4 | 150931724 | 150931977 | 0.623008 | 1.7149 | 2.753 |
| chr3 | 12236119 | 12236414 | 1.43781 | 3.95774 | 2.753 |
| chr10 | 59098743 | 59099226 | 0.706737 | 1.94539 | 2.753 |
| chr2 | 74759439 | 74759649 | 0.852208 | 2.34585 | 2.753 |
| chr10 | 30880182 | 30880438 | 0.936966 | 2.57919 | 2.753 |
| chr4 | 112563328 | 112563597 | 1.41314 | 3.88998 | 2.753 |
| chr1 | 175591354 | 175591573 | 0.399678 | 1.10031 | 2.753 |
| chr5 | 149942504 | 149942749 | 0.705129 | 1.94129 | 2.753 |
| chr18 | 20143151 | 20143366 | 0.780279 | 2.14822 | 2.753 |
| chr2 | 53336796 | 53337105 | 0.45725 | 1.25894 | 2.753 |
| chr11 | 75417257 | 75417818 | 1.858 | 5.116 | 2.754 |
| chr12 | 74482014 | 74482389 | 0.511907 | 1.40955 | 2.754 |
| chr18 | 18822110 | 18822351 | 0.770955 | 2.12295 | 2.754 |
| chr12 | 126554107 | 126554429 | 1.15893 | 3.1915 | 2.754 |
| chr1 | 16766954 | 16767260 | 1.75137 | 4.823 | 2.754 |
| chr6 | 52441308 | 52441692 | 1.24553 | 3.4301 | 2.754 |
| chr4 | 3882299 | 3882622 | 1.0832 | 2.98332 | 2.754 |
| chr19 | 14682835 | 14683111 | 1.62742 | 4.48225 | 2.754 |
| chr11 | 69536653 | 69536876 | 0.797888 | 2.19756 | 2.754 |
| chr6 | 25071076 | 25071413 | 1.04508 | 2.87844 | 2.754 |
| chr11 | 23060514 | 23060740 | 0.334162 | 0.920556 | 2.755 |
| chr7 | 138499267 | 138499470 | 0.409206 | 1.12739 | 2.755 |
| chr8 | 48137174 | 48137632 | 0.638286 | 1.7588 | 2.756 |
| chr20 | 13242686 | 13243032 | 1.76658 | 4.86784 | 2.756 |
| chr5 | 21669888 | 21670093 | 0.356544 | 0.982469 | 2.756 |
| chr9 | 5263642 | 5263931 | 0.527991 | 1.45497 | 2.756 |
| chr1 | 178737469 | 178737925 | 1.3202 | 3.63812 | 2.756 |
| chr1 | 32029707 | 32030237 | 1.0808 | 2.97857 | 2.756 |
| chr9 | 137630917 | 137631328 | 2.09968 | 5.78693 | 2.756 |
| chr3 | 42160338 | 42160553 | 0.895016 | 2.46695 | 2.756 |
| chr9 | 17402435 | 17402660 | 0.560652 | 1.54535 | 2.756 |
| chr8 | 3190572 | 3190797 | 0.434327 | 1.19721 | 2.756 |
| chr2 | 197141823 | 197142029 | 0.827952 | 2.28223 | 2.756 |
| chrX | 38186771 | 38187108 | 1.60644 | 4.42852 | 2.757 |
| chr7 | 114045569 | 114045788 | 0.767698 | 2.11646 | 2.757 |
| chrX | 4465327 | 4465747 | 1.07867 | 2.97403 | 2.757 |
| chr1 | 19980196 | 19980518 | 1.4859 | 4.09694 | 2.757 |
| chr13 | 77554189 | 77554705 | 1.99219 | 5.49296 | 2.757 |
| chr16 | 22990403 | 22990701 | 0.445571 | 1.22855 | 2.757 |
| chr13 | 53824485 | 53824728 | 1.04754 | 2.8884 | 2.757 |
| chr17 | 79014542 | 79015054 | 1.39973 | 3.85984 | 2.758 |
| chr8 | 8527266 | 8527530 | 1.83438 | 5.05855 | 2.758 |
| chr15 | 85498620 | 85498914 | 1.48647 | 4.09944 | 2.758 |
| chr4 | 164253947 | 164254192 | 1.26007 | 3.4753 | 2.758 |
| chr1 | 18795193 | 18795412 | 0.518509 | 1.43008 | 2.758 |
| chr12 | 48591306 | 48591597 | 0.829518 | 2.28786 | 2.758 |
| chr14 | 23797201 | 23797566 | 1.4161 | 3.90575 | 2.758 |
| chr6 | 116857382 | 116857631 | 0.756413 | 2.08639 | 2.758 |
| chr11 | 73018282 | 73018621 | 1.66646 | 4.59674 | 2.758 |
| chr5 | 5512135 | 5512364 | 0.485726 | 1.33992 | 2.759 |

| | | | | | |
|---|---|---|---|---|---|
| chr9 | 121821904 | 121822113 | 0.506887 | 1.39841 | 2.759 |
| chr7 | 3194607 | 3194844 | 1.09308 | 3.01598 | 2.759 |
| chr19 | 39888222 | 39888674 | 2.50487 | 6.91139 | 2.759 |
| chr1 | 157961724 | 157962157 | 2.05624 | 5.67387 | 2.759 |
| chr8 | 143682066 | 143682308 | 0.448445 | 1.23751 | 2.760 |
| chr11 | 105731982 | 105732244 | 0.73299 | 2.02283 | 2.760 |
| chr2 | 66656182 | 66656502 | 1.92032 | 5.30012 | 2.760 |
| chr19 | 29969480 | 29970334 | 0.928575 | 2.56295 | 2.760 |
| chr2 | 103320275 | 103320554 | 0.746572 | 2.06083 | 2.760 |
| chr3 | 118434081 | 118434299 | 0.564397 | 1.55842 | 2.761 |
| chr9 | 123111220 | 123111541 | 1.24778 | 3.44569 | 2.761 |
| chr14 | 35873725 | 35874047 | 0.82533 | 2.27932 | 2.762 |
| chr2 | 136576848 | 136577143 | 0.952726 | 2.63125 | 2.762 |
| chr11 | 62682193 | 62682425 | 0.40607 | 1.12154 | 2.762 |
| chr10 | 74856502 | 74856735 | 1.13416 | 3.1325 | 2.762 |
| chr11 | 106190223 | 106190598 | 0.748172 | 2.06649 | 2.762 |
| chr14 | 102780549 | 102780937 | 1.92962 | 5.32973 | 2.762 |
| chr22 | 38361007 | 38361498 | 0.748407 | 2.0672 | 2.762 |
| chr16 | 19722367 | 19722602 | 0.588592 | 1.62589 | 2.762 |
| chr7 | 101537543 | 101537891 | 0.400151 | 1.10538 | 2.762 |
| chr12 | 39300353 | 39300772 | 2.00109 | 5.52796 | 2.762 |
| chr3 | 178154553 | 178154812 | 0.638695 | 1.76448 | 2.763 |
| chr6 | 42228652 | 42228990 | 1.07064 | 2.95783 | 2.763 |
| chr12 | 57852688 | 57853176 | 1.55768 | 4.30362 | 2.763 |
| chr10 | 123813532 | 123813923 | 1.39915 | 3.86574 | 2.763 |
| chr11 | 76824385 | 76824886 | 0.976395 | 2.69796 | 2.763 |
| chr21 | 40084103 | 40084324 | 0.724343 | 2.00158 | 2.763 |
| chr4 | 129486623 | 129487090 | 0.698952 | 1.93145 | 2.763 |
| chrX | 73603524 | 73603744 | 0.882913 | 2.43986 | 2.763 |
| chr10 | 25971136 | 25971869 | 0.991968 | 2.74219 | 2.764 |
| chr7 | 21113999 | 21114212 | 0.326779 | 0.903365 | 2.764 |
| chr5 | 94553174 | 94553379 | 0.490063 | 1.35476 | 2.764 |
| chr17 | 22193930 | 22194350 | 1.70632 | 4.71716 | 2.765 |
| chr1 | 176407264 | 176407553 | 0.40417 | 1.11747 | 2.765 |
| chr18 | 72608439 | 72608664 | 0.912007 | 2.5216 | 2.765 |
| chr19 | 2951097 | 2951356 | 0.826489 | 2.28519 | 2.765 |
| chr19 | 3852805 | 3853025 | 0.644998 | 1.78338 | 2.765 |
| chr15 | 98467381 | 98467611 | 0.579456 | 1.60224 | 2.765 |
| chr18 | 72219209 | 72219508 | 1.49198 | 4.12547 | 2.765 |
| chrX | 130881271 | 130881493 | 0.972 | 2.68769 | 2.765 |
| chr1 | 93914760 | 93914962 | 0.705157 | 1.94985 | 2.765 |
| chr10 | 27017775 | 27018210 | 0.638535 | 1.76582 | 2.765 |
| chr6 | 24802851 | 24803135 | 0.3314 | 0.916461 | 2.765 |
| chr14 | 100145567 | 100145814 | 0.429441 | 1.18774 | 2.766 |
| chr8 | 122723160 | 122723479 | 1.08302 | 2.99553 | 2.766 |
| chr10 | 62950405 | 62950653 | 1.00043 | 2.76712 | 2.766 |
| chr7 | 118224326 | 118224559 | 0.40773 | 1.12777 | 2.766 |
| chr11 | 67465713 | 67466071 | 0.828646 | 2.29251 | 2.767 |
| chr18 | 30050898 | 30051164 | 0.984836 | 2.72463 | 2.767 |
| chr17 | 27229810 | 27230466 | 2.77164 | 7.66853 | 2.767 |
| chr19 | 56861809 | 56862154 | 1.15766 | 3.20335 | 2.767 |
| chr8 | 131840639 | 131840938 | 0.948756 | 2.62542 | 2.767 |
| chr12 | 27037783 | 27038282 | 1.92602 | 5.32984 | 2.767 |
| chrX | 40595047 | 40595265 | 0.642345 | 1.77758 | 2.767 |
| chr9 | 23899005 | 23899382 | 0.428636 | 1.18624 | 2.767 |
| chr12 | 39497379 | 39497692 | 1.3901 | 3.84736 | 2.768 |
| chr17 | 28281549 | 28281832 | 0.839398 | 2.32342 | 2.768 |
| chrX | 71346464 | 71346829 | 1.34852 | 3.73294 | 2.768 |
| chr17 | 17516773 | 17517457 | 0.789725 | 2.18613 | 2.768 |
| chr22 | 32785746 | 32786001 | 0.816127 | 2.25932 | 2.768 |
| chr3 | 124647449 | 124647806 | 1.38247 | 3.82726 | 2.768 |

| | | | | | |
|---|---|---|---|---|---|
| chr1 | 53387187 | 53387556 | 0.812733 | 2.25002 | 2.768 |
| chr14 | 23538062 | 23538555 | 1.96016 | 5.42671 | 2.769 |
| chr21 | 30872045 | 30872501 | 1.63595 | 4.52925 | 2.769 |
| chr1 | 170746429 | 170746700 | 0.307774 | 0.852156 | 2.769 |
| chr19 | 35986247 | 35986855 | 2.23059 | 6.17598 | 2.769 |
| chr5 | 158758349 | 158758701 | 2.30679 | 6.38748 | 2.769 |
| chr12 | 49390510 | 49390862 | 1.37447 | 3.80625 | 2.769 |
| chr12 | 117463370 | 117463732 | 1.12718 | 3.12148 | 2.769 |
| chr19 | 10662388 | 10662635 | 0.576035 | 1.59525 | 2.769 |
| chrX | 21498011 | 21498387 | 1.27982 | 3.54454 | 2.770 |
| chr16 | 62259638 | 62259900 | 1.09558 | 3.0343 | 2.770 |
| chr6 | 53590603 | 53590961 | 1.39317 | 3.85876 | 2.770 |
| chr12 | 116453332 | 116453670 | 0.847301 | 2.3471 | 2.770 |
| chr15 | 35272722 | 35273122 | 1.18203 | 3.27449 | 2.770 |
| chr16 | 16236943 | 16237209 | 1.02662 | 2.84413 | 2.770 |
| chr5 | 153784723 | 153785102 | 1.21453 | 3.36473 | 2.770 |
| chr6 | 39277706 | 39278083 | 1.53237 | 4.24528 | 2.770 |
| chr13 | 37310312 | 37310566 | 0.875114 | 2.42443 | 2.770 |
| chr17 | 42429860 | 42430130 | 0.423496 | 1.17347 | 2.771 |
| chr4 | 103220808 | 103221138 | 1.16706 | 3.23392 | 2.771 |
| chr20 | 58100472 | 58100900 | 2.08797 | 5.78662 | 2.771 |
| chr5 | 141394626 | 141394953 | 0.769374 | 2.13256 | 2.772 |
| chr18 | 64163164 | 64163448 | 0.701561 | 1.94469 | 2.772 |
| chr11 | 77711205 | 77711479 | 1.14859 | 3.1839 | 2.772 |
| chr9 | 90565729 | 90566021 | 0.962154 | 2.66724 | 2.772 |
| chr10 | 10136663 | 10136915 | 0.41349 | 1.14632 | 2.772 |
| chr12 | 33609241 | 33609629 | 0.473909 | 1.31382 | 2.772 |
| chr22 | 39739547 | 39739882 | 1.19269 | 3.30656 | 2.772 |
| chr19 | 58666430 | 58666833 | 1.55705 | 4.31685 | 2.772 |
| chr6 | 40989703 | 40989947 | 0.886034 | 2.45654 | 2.773 |
| chr8 | 13595359 | 13595785 | 0.423345 | 1.17377 | 2.773 |
| chr1 | 41414617 | 41414938 | 1.16773 | 3.23784 | 2.773 |
| chr3 | 121234229 | 121234432 | 0.359093 | 0.9957 | 2.773 |
| chr1 | 151168818 | 151169043 | 0.979007 | 2.71475 | 2.773 |
| chr12 | 94135979 | 94136375 | 1.17083 | 3.24675 | 2.773 |
| chr14 | 102204431 | 102204670 | 0.791997 | 2.1963 | 2.773 |
| chr19 | 6299810 | 6300081 | 1.2629 | 3.50227 | 2.773 |
| chr4 | 163880292 | 163880539 | 0.897493 | 2.48908 | 2.773 |
| chr20 | 39861088 | 39861316 | 1.05233 | 2.91858 | 2.773 |
| chr7 | 148383302 | 148383579 | 0.447481 | 1.24113 | 2.774 |
| chr19 | 12876776 | 12877174 | 1.77678 | 4.92879 | 2.774 |
| chr8 | 103875326 | 103875705 | 0.801773 | 2.22412 | 2.774 |
| chr19 | 28816288 | 28816588 | 0.757874 | 2.10235 | 2.774 |
| chr11 | 19217548 | 19217802 | 0.319674 | 0.886818 | 2.774 |
| chr12 | 100948381 | 100948711 | 1.84616 | 5.12169 | 2.774 |
| chr8 | 133118863 | 133119080 | 0.524841 | 1.45614 | 2.774 |
| chr6 | 133258560 | 133258829 | 1.09004 | 3.02439 | 2.775 |
| chr11 | 13260451 | 13260791 | 1.40049 | 3.88629 | 2.775 |
| chr11 | 57117277 | 57117755 | 1.83453 | 5.09084 | 2.775 |
| chr3 | 36910314 | 36910695 | 1.18082 | 3.27683 | 2.775 |
| chr5 | 122039996 | 122040283 | 0.849583 | 2.35768 | 2.775 |
| chr9 | 38620522 | 38621078 | 1.17386 | 3.25775 | 2.775 |
| chr14 | 102821708 | 102821996 | 0.300904 | 0.83514 | 2.775 |
| chr3 | 146358680 | 146358947 | 1.01194 | 2.80858 | 2.775 |
| chr17 | 26578160 | 26579020 | 1.61869 | 4.49323 | 2.776 |
| chr1 | 27136581 | 27136820 | 1.42523 | 3.95624 | 2.776 |
| chr11 | 72929376 | 72929720 | 1.32766 | 3.68574 | 2.776 |
| chr5 | 179190201 | 179190465 | 1.64002 | 4.55294 | 2.776 |
| chr9 | 72287790 | 72288159 | 1.35405 | 3.75906 | 2.776 |
| chr6 | 133151415 | 133151646 | 0.602577 | 1.67296 | 2.776 |
| chr14 | 103241314 | 103241558 | 0.987756 | 2.74242 | 2.776 |

| | | | | | |
|---|---|---|---|---|---|
| chr14 | 68400826 | 68401067 | 0.885856 | 2.45961 | 2.777 |
| chr10 | 20144802 | 20145018 | 0.243969 | 0.677395 | 2.777 |
| chr12 | 13386990 | 13387304 | 1.61143 | 4.47444 | 2.777 |
| chr18 | 60382083 | 60382352 | 1.12225 | 3.11626 | 2.777 |
| chr12 | 6862781 | 6863146 | 2.17003 | 6.02598 | 2.777 |
| chr9 | 35945803 | 35946042 | 1.4222 | 3.94941 | 2.777 |
| chr17 | 41034686 | 41034893 | 0.665289 | 1.84757 | 2.777 |
| chr10 | 103990710 | 103990958 | 0.676035 | 1.87744 | 2.777 |
| chr15 | 75080131 | 75080465 | 1.58388 | 4.39868 | 2.777 |
| chr16 | 31107178 | 31107488 | 1.38729 | 3.85368 | 2.778 |
| chr7 | 57217283 | 57217670 | 0.353841 | 0.982919 | 2.778 |
| chr12 | 54608470 | 54609007 | 2.39137 | 6.64301 | 2.778 |
| chr8 | 28558884 | 28559157 | 0.807049 | 2.24221 | 2.778 |
| chr14 | 93147378 | 93147846 | 2.14713 | 5.96545 | 2.778 |
| chr12 | 81493739 | 81493981 | 0.526725 | 1.46347 | 2.778 |
| chr5 | 31358975 | 31359343 | 0.404746 | 1.12458 | 2.778 |
| chr14 | 99786362 | 99786784 | 1.85066 | 5.14283 | 2.779 |
| chr8 | 4346294 | 4346542 | 0.333704 | 0.927355 | 2.779 |
| chr3 | 164804875 | 164805124 | 0.349949 | 0.972523 | 2.779 |
| chr1 | 237883842 | 237884081 | 0.445769 | 1.23917 | 2.780 |
| chr7 | 87193644 | 87194006 | 1.3465 | 3.74307 | 2.780 |
| chr12 | 2764411 | 2764780 | 0.361963 | 1.00629 | 2.780 |
| chr8 | 105288315 | 105288531 | 0.367717 | 1.0223 | 2.780 |
| chr5 | 132954818 | 132955024 | 0.410417 | 1.14106 | 2.780 |
| chr19 | 10138351 | 10138749 | 1.93169 | 5.37092 | 2.780 |
| chr11 | 7678664 | 7679107 | 0.799443 | 2.2229 | 2.781 |
| chr11 | 55850778 | 55850986 | 0.567964 | 1.57928 | 2.781 |
| chr19 | 22545541 | 22545752 | 0.390556 | 1.08599 | 2.781 |
| chr15 | 65589968 | 65590398 | 1.3291 | 3.69624 | 2.781 |
| chr5 | 40798342 | 40798624 | 0.98136 | 2.72919 | 2.781 |
| chr14 | 29078486 | 29078755 | 1.3124 | 3.64994 | 2.781 |
| chr6 | 16286274 | 16286690 | 1.44084 | 4.00721 | 2.781 |
| chr3 | 118864270 | 118864555 | 1.24285 | 3.45672 | 2.781 |
| chr10 | 108165928 | 108166133 | 0.363643 | 1.01144 | 2.781 |
| chr10 | 6013964 | 6014204 | 0.345434 | 0.960818 | 2.781 |
| chr4 | 101232347 | 101232568 | 0.544765 | 1.51526 | 2.781 |
| chr16 | 4732846 | 4733340 | 1.16115 | 3.22975 | 2.782 |
| chr12 | 6394141 | 6394541 | 1.48327 | 4.12581 | 2.782 |
| chr19 | 5690067 | 5690392 | 1.71039 | 4.75789 | 2.782 |
| chr1 | 150824793 | 150825019 | 0.382238 | 1.06333 | 2.782 |
| chr20 | 5769054 | 5769308 | 1.47242 | 4.09614 | 2.782 |
| chr12 | 27459244 | 27459460 | 0.74389 | 2.06962 | 2.782 |
| chr18 | 61662035 | 61662444 | 0.884208 | 2.46002 | 2.782 |
| chr20 | 10297344 | 10297601 | 0.964566 | 2.68369 | 2.782 |
| chr12 | 60691019 | 60691257 | 0.464322 | 1.29206 | 2.783 |
| chr2 | 88991104 | 88991419 | 1.14915 | 3.19813 | 2.783 |
| chr10 | 29528861 | 29529137 | 1.14648 | 3.19083 | 2.783 |
| chr7 | 158148211 | 158148678 | 0.969634 | 2.69894 | 2.783 |
| chr22 | 39129245 | 39129662 | 1.28397 | 3.57389 | 2.783 |
| chr11 | 67113341 | 67113649 | 1.72598 | 4.80492 | 2.784 |
| chr1 | 117664728 | 117665067 | 1.34421 | 3.74228 | 2.784 |
| chr13 | 68310713 | 68311065 | 0.64433 | 1.79383 | 2.784 |
| chr13 | 28534425 | 28534706 | 1.20891 | 3.36593 | 2.784 |
| chr10 | 4671167 | 4671508 | 0.418699 | 1.16586 | 2.784 |
| chr12 | 114383475 | 114383701 | 0.390746 | 1.08828 | 2.785 |
| chr3 | 111732358 | 111732719 | 1.36211 | 3.79373 | 2.785 |
| chr19 | 41310670 | 41310995 | 0.8694 | 2.42149 | 2.785 |
| chr8 | 101637859 | 101638206 | 1.12908 | 3.14495 | 2.785 |
| chr14 | 53117542 | 53117906 | 1.11176 | 3.09685 | 2.786 |
| chr8 | 27584778 | 27585194 | 1.29813 | 3.61622 | 2.786 |
| chr10 | 102977230 | 102977513 | 1.1858 | 3.30364 | 2.786 |

| | | | | | |
|---|---|---|---|---|---|
| chr5 | 95182702 | 95183036 | 1.28598 | 3.58301 | 2.786 |
| chr14 | 78283676 | 78284000 | 0.751594 | 2.09416 | 2.786 |
| chr7 | 138482744 | 138482948 | 0.411821 | 1.14758 | 2.787 |
| chr6 | 34497425 | 34497709 | 0.525364 | 1.46407 | 2.787 |
| chr12 | 52557689 | 52558101 | 1.88953 | 5.2661 | 2.787 |
| chr5 | 36265579 | 36265857 | 1.19187 | 3.3218 | 2.787 |
| chrX | 40270681 | 40270887 | 0.467811 | 1.30392 | 2.787 |
| chr16 | 56547842 | 56548100 | 0.961996 | 2.6818 | 2.788 |
| chr6 | 145955173 | 145955461 | 1.15745 | 3.22668 | 2.788 |
| chr6 | 43422240 | 43422588 | 1.08419 | 3.02248 | 2.788 |
| chr1 | 85219214 | 85219610 | 1.77054 | 4.93596 | 2.788 |
| chr9 | 97385566 | 97385867 | 1.03554 | 2.88698 | 2.788 |
| chr1 | 31658524 | 31658846 | 0.998978 | 2.78507 | 2.788 |
| chr3 | 52833109 | 52833410 | 0.849809 | 2.3692 | 2.788 |
| chr10 | 116639335 | 116639667 | 1.40716 | 3.92344 | 2.788 |
| chr3 | 108125430 | 108125754 | 1.07328 | 2.99261 | 2.788 |
| chr11 | 89872119 | 89872368 | 0.365522 | 1.01919 | 2.788 |
| chr22 | 45224111 | 45224361 | 1.10175 | 3.07208 | 2.788 |
| chr1 | 228186972 | 228187421 | 1.83381 | 5.11338 | 2.788 |
| chr6 | 110360701 | 110360971 | 1.6682 | 4.65205 | 2.789 |
| chr4 | 4261999 | 4262236 | 0.647245 | 1.80502 | 2.789 |
| chr1 | 223936812 | 223937202 | 2.09921 | 5.85469 | 2.789 |
| chr14 | 77875610 | 77875847 | 0.73033 | 2.03709 | 2.789 |
| chr5 | 44428170 | 44428435 | 0.582359 | 1.62454 | 2.790 |
| chr6 | 158065463 | 158065724 | 0.970082 | 2.70627 | 2.790 |
| chr8 | 135703407 | 135703910 | 1.70862 | 4.76669 | 2.790 |
| chrX | 53323214 | 53323572 | 0.717865 | 2.00287 | 2.790 |
| chr21 | 44072600 | 44072882 | 1.16806 | 3.25905 | 2.790 |
| chr1 | 12289820 | 12290265 | 1.63505 | 4.56222 | 2.790 |
| chr3 | 194991451 | 194991833 | 0.814694 | 2.27349 | 2.791 |
| chr12 | 105107545 | 105107795 | 0.442106 | 1.23376 | 2.791 |
| chr7 | 66836790 | 66837112 | 1.1176 | 3.11905 | 2.791 |
| chr10 | 3239597 | 3239902 | 1.41143 | 3.9393 | 2.791 |
| chr11 | 65479206 | 65479618 | 1.80468 | 5.03731 | 2.791 |
| chr6 | 8499835 | 8500122 | 0.712533 | 1.98888 | 2.791 |
| chrX | 45374476 | 45374715 | 1.03558 | 2.89075 | 2.791 |
| chr14 | 33035022 | 33035231 | 0.420912 | 1.17495 | 2.791 |
| chr11 | 2441964 | 2442385 | 1.52881 | 4.26759 | 2.791 |
| chr12 | 52955849 | 52956099 | 0.768436 | 2.14512 | 2.792 |
| chr1 | 30385362 | 30385623 | 0.503626 | 1.40598 | 2.792 |
| chr11 | 124539090 | 124539408 | 0.965564 | 2.69558 | 2.792 |
| chr10 | 71389708 | 71390246 | 1.52662 | 4.26196 | 2.792 |
| chr8 | 35288084 | 35288510 | 0.444752 | 1.24167 | 2.792 |
| chr8 | 56941424 | 56941741 | 1.20484 | 3.36393 | 2.792 |
| chr12 | 53718523 | 53718794 | 1.48936 | 4.1587 | 2.792 |
| chr3 | 185451768 | 185452030 | 0.938247 | 2.62027 | 2.793 |
| chr12 | 84219924 | 84220310 | 1.19127 | 3.32712 | 2.793 |
| chr11 | 57017248 | 57017542 | 1.1652 | 3.25455 | 2.793 |
| chr17 | 1722776 | 1723004 | 0.416341 | 1.16291 | 2.793 |
| chr15 | 58513820 | 58514186 | 1.08758 | 3.03793 | 2.793 |
| chr4 | 160400343 | 160400586 | 0.37119 | 1.03693 | 2.794 |
| chr12 | 129184044 | 129184329 | 0.408366 | 1.14083 | 2.794 |
| chr19 | 16487497 | 16487809 | 0.906785 | 2.53338 | 2.794 |
| chr20 | 36851050 | 36851482 | 1.77938 | 4.97143 | 2.794 |
| chr12 | 61641648 | 61641922 | 0.559472 | 1.56312 | 2.794 |
| chr15 | 81068740 | 81069118 | 1.85571 | 5.18509 | 2.794 |
| chr7 | 101500254 | 101500770 | 0.65985 | 1.8438 | 2.794 |
| chr21 | 35802099 | 35802346 | 1.08626 | 3.03543 | 2.794 |
| chr10 | 96723629 | 96723975 | 1.02967 | 2.87737 | 2.794 |
| chr1 | 86601741 | 86602001 | 0.339146 | 0.947745 | 2.795 |
| chr8 | 116140134 | 116140369 | 0.689035 | 1.92555 | 2.795 |

| | | | | | |
|---|---|---|---|---|---|
| chr6 | 134495167 | 134495407 | 0.91676 | 2.5622 | 2.795 |
| chr15 | 29696029 | 29696268 | 0.751944 | 2.10171 | 2.795 |
| chr6 | 127875643 | 127875929 | 1.00529 | 2.80983 | 2.795 |
| chr19 | 35329159 | 35329965 | 1.37819 | 3.85216 | 2.795 |
| chr5 | 41754968 | 41755249 | 0.77527 | 2.16703 | 2.795 |
| chr8 | 70858465 | 70858686 | 0.843714 | 2.35842 | 2.795 |
| chr7 | 150705737 | 150706222 | 1.71349 | 4.79023 | 2.796 |
| chr11 | 86706408 | 86706658 | 1.03295 | 2.8881 | 2.796 |
| chr1 | 147172001 | 147172423 | 1.73628 | 4.85491 | 2.796 |
| chr18 | 61616359 | 61616601 | 0.63917 | 1.78724 | 2.796 |
| chr16 | 77269815 | 77270360 | 1.72907 | 4.83524 | 2.796 |
| chr18 | 32142997 | 32143285 | 0.398663 | 1.11507 | 2.797 |
| chr19 | 741662 | 741875 | 1.012 | 2.83091 | 2.797 |
| chr17 | 62645254 | 62645584 | 1.29716 | 3.62884 | 2.798 |
| chr11 | 72064997 | 72065200 | 0.764687 | 2.13943 | 2.798 |
| chr6 | 130614996 | 130615243 | 1.0949 | 3.0633 | 2.798 |
| chr17 | 37080239 | 37080544 | 1.24875 | 3.49389 | 2.798 |
| chr16 | 30609490 | 30609838 | 1.39358 | 3.89919 | 2.798 |
| chr19 | 9957690 | 9958066 | 1.20827 | 3.38101 | 2.798 |
| chr10 | 23357146 | 23357440 | 1.26647 | 3.54407 | 2.798 |
| chr10 | 10903148 | 10903361 | 0.499075 | 1.39664 | 2.798 |
| chr4 | 125183418 | 125183725 | 1.08007 | 3.02264 | 2.799 |
| chr2 | 133426043 | 133426447 | 0.762728 | 2.13455 | 2.799 |
| chr12 | 118492931 | 118493293 | 1.25767 | 3.51994 | 2.799 |
| chr12 | 115307365 | 115307567 | 0.444473 | 1.24402 | 2.799 |
| chr11 | 48138425 | 48138697 | 0.847415 | 2.37182 | 2.799 |
| chr22 | 25246295 | 25246647 | 1.73531 | 4.85739 | 2.799 |
| chr11 | 61826302 | 61826545 | 0.295528 | 0.827309 | 2.799 |
| chr10 | 104263398 | 104263724 | 1.8014 | 5.04292 | 2.799 |
| chr12 | 57606961 | 57607560 | 1.70604 | 4.77598 | 2.799 |
| chr19 | 29958744 | 29959111 | 1.40939 | 3.94556 | 2.799 |
| chr3 | 106815274 | 106815519 | 0.391412 | 1.09575 | 2.799 |
| chrX | 19922979 | 19923247 | 1.0455 | 2.92694 | 2.800 |
| chr8 | 12523023 | 12523255 | 1.14366 | 3.20202 | 2.800 |
| chr1 | 79818979 | 79819238 | 0.783579 | 2.19393 | 2.800 |
| chr17 | 36692638 | 36692884 | 0.340401 | 0.953107 | 2.800 |
| chr9 | 35553966 | 35554300 | 1.72202 | 4.8217 | 2.800 |
| chr3 | 160555305 | 160555549 | 0.3371 | 0.943926 | 2.800 |
| chr11 | 18672716 | 18673014 | 1.23316 | 3.45304 | 2.800 |
| chr1 | 100239432 | 100239666 | 0.760505 | 2.12973 | 2.800 |
| chr19 | 45089072 | 45089303 | 0.816794 | 2.28743 | 2.801 |
| chr20 | 56172006 | 56172411 | 1.3418 | 3.75846 | 2.801 |
| chr11 | 9159293 | 9159665 | 2.02815 | 5.68098 | 2.801 |
| chr6 | 8905096 | 8905395 | 0.646499 | 1.81099 | 2.801 |
| chr10 | 5991368 | 5991653 | 1.02553 | 2.87281 | 2.801 |
| chr3 | 132361135 | 132361374 | 0.898433 | 2.51678 | 2.801 |
| chr1 | 23984193 | 23984507 | 1.49035 | 4.17494 | 2.801 |
| chr9 | 103357902 | 103358344 | 1.52844 | 4.28178 | 2.801 |
| chr3 | 183602599 | 183602921 | 1.9511 | 5.46746 | 2.802 |
| chr5 | 38338705 | 38338933 | 0.337907 | 0.94691 | 2.802 |
| chr11 | 72158052 | 72158389 | 1.26043 | 3.5321 | 2.802 |
| chr19 | 53522932 | 53523277 | 1.30754 | 3.66412 | 2.802 |
| chr5 | 175156819 | 175157199 | 2.15627 | 6.04287 | 2.802 |
| chr14 | 95993773 | 95994077 | 1.04789 | 2.93686 | 2.803 |
| chr14 | 62237310 | 62237567 | 0.348264 | 0.976081 | 2.803 |
| chr17 | 7961004 | 7961286 | 1.37546 | 3.8552 | 2.803 |
| chr8 | 107282363 | 107282759 | 1.74714 | 4.89704 | 2.803 |
| chr3 | 121714459 | 121714711 | 1.35076 | 3.78658 | 2.803 |
| chrX | 48963838 | 48964142 | 0.893119 | 2.50376 | 2.803 |
| chr12 | 132093609 | 132093890 | 1.39316 | 3.90565 | 2.803 |
| chr8 | 33101788 | 33102248 | 0.756041 | 2.11966 | 2.804 |

| | | | | | |
|---|---|---|---|---|---|
| chr1 | 101774874 | 101775121 | 1.65272 | 4.63373 | 2.804 |
| chr12 | 131260616 | 131261048 | 0.369632 | 1.03642 | 2.804 |
| chr12 | 48356992 | 48357435 | 2.3053 | 6.46415 | 2.804 |
| chr12 | 29557374 | 29557670 | 0.462854 | 1.29791 | 2.804 |
| chr5 | 172386159 | 172386524 | 2.08307 | 5.84159 | 2.804 |
| chr1 | 53151935 | 53152237 | 1.42311 | 3.99087 | 2.804 |
| chr7 | 90225144 | 90225789 | 1.08885 | 3.05398 | 2.805 |
| chr20 | 23216063 | 23216408 | 1.40843 | 3.95116 | 2.805 |
| chr5 | 37188382 | 37188664 | 0.413347 | 1.15959 | 2.805 |
| chr18 | 46941844 | 46942070 | 0.633786 | 1.77817 | 2.806 |
| chr12 | 27222317 | 27222523 | 0.381365 | 1.06999 | 2.806 |
| chr17 | 44920218 | 44920585 | 1.57633 | 4.42298 | 2.806 |
| chr4 | 6703155 | 6703481 | 1.17479 | 3.29663 | 2.806 |
| chr4 | 175240382 | 175240653 | 1.18984 | 3.33889 | 2.806 |
| chr16 | 85474749 | 85475016 | 0.68405 | 1.9196 | 2.806 |
| chr7 | 5820981 | 5821515 | 1.06043 | 2.97589 | 2.806 |
| chr20 | 39591383 | 39592063 | 1.68884 | 4.73991 | 2.807 |
| chr7 | 87974645 | 87974868 | 0.863483 | 2.42363 | 2.807 |
| chr8 | 31047866 | 31048111 | 0.37613 | 1.05582 | 2.807 |
| chr3 | 153008909 | 153009174 | 1.3727 | 3.85369 | 2.807 |
| chr17 | 42408631 | 42409009 | 1.35118 | 3.79392 | 2.808 |
| chr18 | 57504825 | 57505103 | 1.17549 | 3.30074 | 2.808 |
| chr19 | 45291253 | 45291621 | 0.399838 | 1.12276 | 2.808 |
| chr1 | 91631712 | 91632098 | 1.35269 | 3.79861 | 2.808 |
| chr15 | 53027737 | 53028007 | 1.02129 | 2.86803 | 2.808 |
| chr16 | 86551325 | 86551667 | 1.45364 | 4.08233 | 2.808 |
| chr4 | 133254154 | 133254424 | 0.888247 | 2.49463 | 2.808 |
| chr12 | 16342841 | 16343210 | 1.19324 | 3.35127 | 2.809 |
| chr11 | 111169866 | 111170150 | 1.00406 | 2.82047 | 2.809 |
| chr11 | 67397989 | 67398414 | 1.36478 | 3.83396 | 2.809 |
| chr4 | 5981773 | 5982044 | 0.888048 | 2.49504 | 2.810 |
| chr2 | 64115935 | 64116317 | 1.40756 | 3.95484 | 2.810 |
| chr17 | 76183123 | 76183381 | 1.22684 | 3.44712 | 2.810 |
| chr8 | 143069909 | 143070348 | 1.21263 | 3.40722 | 2.810 |
| chr22 | 38329914 | 38330276 | 1.76699 | 4.96523 | 2.810 |
| chr6 | 51952526 | 51952800 | 0.697102 | 1.95903 | 2.810 |
| chr6 | 43302264 | 43302565 | 0.655149 | 1.84117 | 2.810 |
| chr14 | 44010286 | 44010515 | 0.589226 | 1.65601 | 2.810 |
| chr5 | 139167052 | 139167962 | 1.32403 | 3.72141 | 2.811 |
| chr14 | 90912861 | 90913115 | 0.375949 | 1.05673 | 2.811 |
| chr12 | 40874111 | 40874565 | 0.418352 | 1.17596 | 2.811 |
| chr9 | 91233512 | 91233817 | 0.269562 | 0.757766 | 2.811 |
| chr5 | 174337218 | 174337555 | 2.07546 | 5.83447 | 2.811 |
| chr15 | 57734863 | 57735089 | 0.441774 | 1.24197 | 2.811 |
| chr3 | 73149545 | 73149817 | 0.355996 | 1.00084 | 2.811 |
| chr11 | 662664 | 663240 | 1.22772 | 3.4517 | 2.811 |
| chr7 | 128911670 | 128912021 | 0.920135 | 2.58695 | 2.811 |
| chr9 | 32572339 | 32572601 | 1.15194 | 3.23905 | 2.812 |
| chr19 | 24342601 | 24342870 | 0.975027 | 2.7417 | 2.812 |
| chr1 | 165027863 | 165028086 | 0.283968 | 0.798564 | 2.812 |
| chr10 | 114231694 | 114232077 | 0.999731 | 2.81161 | 2.812 |
| chr1 | 159768581 | 159769028 | 1.24784 | 3.51014 | 2.813 |
| chr2 | 228190150 | 228190455 | 0.531808 | 1.49597 | 2.813 |
| chr5 | 42889469 | 42889766 | 1.93 | 5.42907 | 2.813 |
| chr7 | 105817124 | 105817499 | 1.65626 | 4.65921 | 2.813 |
| chr9 | 96328646 | 96328957 | 1.77264 | 4.98729 | 2.813 |
| chr14 | 21924460 | 21924698 | 1.00159 | 2.81799 | 2.814 |
| chr21 | 36117827 | 36118154 | 1.50603 | 4.23733 | 2.814 |
| chr15 | 33452330 | 33452536 | 0.485046 | 1.36472 | 2.814 |
| chr5 | 43064983 | 43065300 | 1.74644 | 4.91414 | 2.814 |
| chr3 | 120513953 | 120514262 | 1.27464 | 3.58664 | 2.814 |

| | | | | | |
|---|---|---|---|---|---|
| chr22 | 42353590 | 42354070 | 2.0783 | 5.84824 | 2.814 |
| chr7 | 148892433 | 148892824 | 1.21141 | 3.40896 | 2.814 |
| chr1 | 17313530 | 17313878 | 1.09876 | 3.09212 | 2.814 |
| chr8 | 67211336 | 67211641 | 0.434028 | 1.2215 | 2.814 |
| chr20 | 57456188 | 57456546 | 0.494307 | 1.39118 | 2.814 |
| chr16 | 5075186 | 5075654 | 0.951999 | 2.67966 | 2.815 |
| chr5 | 99490024 | 99490244 | 0.449612 | 1.26556 | 2.815 |
| chr11 | 130002173 | 130002484 | 1.09548 | 3.08356 | 2.815 |
| chr1 | 213145949 | 213146261 | 0.344045 | 0.968458 | 2.815 |
| chr3 | 58285908 | 58286113 | 0.800027 | 2.2521 | 2.815 |
| chr14 | 74723918 | 74724530 | 2.07175 | 5.83254 | 2.815 |
| chr8 | 47156886 | 47157101 | 0.715296 | 2.01384 | 2.815 |
| chr8 | 69195190 | 69195397 | 0.319803 | 0.900389 | 2.815 |
| chr16 | 85279228 | 85279560 | 1.11769 | 3.14704 | 2.816 |
| chr13 | 108974649 | 108974864 | 0.730463 | 2.05733 | 2.816 |
| chr8 | 40809097 | 40809524 | 1.36674 | 3.84963 | 2.817 |
| chr3 | 122025251 | 122025540 | 0.804094 | 2.2649 | 2.817 |
| chr10 | 93598882 | 93599148 | 0.351206 | 0.989261 | 2.817 |
| chr17 | 25782879 | 25783152 | 0.875169 | 2.46515 | 2.817 |
| chr19 | 9900896 | 9901120 | 1.05569 | 2.97375 | 2.817 |
| chr3 | 19815896 | 19816240 | 0.363329 | 1.0235 | 2.817 |
| chr5 | 74632507 | 74632724 | 0.512018 | 1.44249 | 2.817 |
| chr17 | 7801155 | 7801457 | 0.86229 | 2.42944 | 2.817 |
| chr2 | 199161452 | 199161856 | 1.28704 | 3.62617 | 2.817 |
| chr17 | 40268709 | 40269108 | 1.32807 | 3.74185 | 2.818 |
| chr1 | 16570848 | 16571254 | 0.967191 | 2.72522 | 2.818 |
| chr16 | 26453039 | 26453283 | 0.335857 | 0.946336 | 2.818 |
| chr20 | 43376211 | 43376437 | 1.16322 | 3.2776 | 2.818 |
| chr19 | 50308302 | 50308757 | 0.888892 | 2.50476 | 2.818 |
| chr14 | 44337925 | 44338235 | 0.477608 | 1.34598 | 2.818 |
| chr8 | 118050216 | 118050419 | 0.774514 | 2.18279 | 2.818 |
| chr14 | 75955145 | 75955484 | 2.26022 | 6.37004 | 2.818 |
| chr16 | 66391439 | 66391712 | 0.99705 | 2.81015 | 2.818 |
| chr4 | 89181491 | 89181706 | 0.353475 | 0.996261 | 2.818 |
| chr12 | 51421957 | 51422201 | 0.828279 | 2.33463 | 2.819 |
| chr11 | 2194180 | 2194698 | 0.967716 | 2.72778 | 2.819 |
| chr2 | 97523796 | 97524194 | 1.07724 | 3.03657 | 2.819 |
| chr19 | 33736850 | 33737305 | 1.58657 | 4.47251 | 2.819 |
| chr16 | 51847513 | 51847811 | 1.13325 | 3.19465 | 2.819 |
| chr8 | 94706361 | 94706679 | 1.15422 | 3.25405 | 2.819 |
| chr6 | 83775429 | 83775785 | 1.63848 | 4.61955 | 2.819 |
| chr11 | 17084865 | 17085353 | 1.64793 | 4.64624 | 2.819 |
| chr5 | 168133617 | 168134073 | 1.05135 | 2.96444 | 2.820 |
| chr7 | 2760666 | 2760874 | 0.540699 | 1.52462 | 2.820 |
| chr3 | 127275504 | 127275737 | 0.654185 | 1.84484 | 2.820 |
| chr1 | 32866449 | 32866827 | 1.13308 | 3.19583 | 2.820 |
| chr11 | 1526766 | 1527154 | 0.592433 | 1.67098 | 2.821 |
| chr12 | 20951574 | 20951984 | 0.710485 | 2.004 | 2.821 |
| chr1 | 200276587 | 200276953 | 2.36803 | 6.67984 | 2.821 |
| chr4 | 104986217 | 104986489 | 1.07523 | 3.0336 | 2.821 |
| chr3 | 142305657 | 142305960 | 0.938734 | 2.6488 | 2.822 |
| chr5 | 112980516 | 112980807 | 0.268589 | 0.757887 | 2.822 |
| chr4 | 129112519 | 129112785 | 1.2381 | 3.49382 | 2.822 |
| chr12 | 112910962 | 112911171 | 0.920255 | 2.5972 | 2.822 |
| chr13 | 85470328 | 85470590 | 0.748191 | 2.11171 | 2.822 |
| chr17 | 70030882 | 70031107 | 0.409852 | 1.1568 | 2.822 |
| chr1 | 48191911 | 48192239 | 1.28633 | 3.63072 | 2.823 |
| chr2 | 205190177 | 205190390 | 0.92037 | 2.59781 | 2.823 |
| chr11 | 41453340 | 41453638 | 0.806726 | 2.27707 | 2.823 |
| chr12 | 84896895 | 84897332 | 1.24146 | 3.5043 | 2.823 |
| chr20 | 44065490 | 44065859 | 1.69538 | 4.78581 | 2.823 |

| | | | | | |
|---|---|---|---|---|---|
| chr1 | 28967334 | 28967578 | 0.45398 | 1.28158 | 2.823 |
| chr19 | 46184698 | 46185201 | 1.84435 | 5.20665 | 2.823 |
| chr2 | 78603407 | 78603623 | 0.397458 | 1.12204 | 2.823 |
| chr18 | 71749734 | 71750056 | 1.3006 | 3.67169 | 2.823 |
| chr12 | 44022189 | 44022393 | 0.441969 | 1.24786 | 2.823 |
| chr9 | 13596567 | 13596895 | 0.418746 | 1.18231 | 2.823 |
| chr19 | 51520002 | 51520604 | 1.58256 | 4.46833 | 2.823 |
| chr14 | 53716094 | 53716322 | 0.472292 | 1.33352 | 2.824 |
| chr13 | 107172940 | 107173257 | 0.831824 | 2.34867 | 2.824 |
| chr3 | 39521610 | 39521917 | 0.371074 | 1.0478 | 2.824 |
| chr5 | 176915048 | 176915266 | 0.850718 | 2.40217 | 2.824 |
| chr3 | 17935996 | 17936349 | 0.95417 | 2.69434 | 2.824 |
| chr4 | 38063656 | 38063951 | 0.938061 | 2.64892 | 2.824 |
| chr5 | 143204692 | 143205094 | 1.73536 | 4.90046 | 2.824 |
| chr18 | 47037035 | 47037363 | 1.24492 | 3.51559 | 2.824 |
| chr12 | 70893982 | 70894348 | 0.426596 | 1.20471 | 2.824 |
| chr14 | 58829395 | 58829776 | 0.426437 | 1.20428 | 2.824 |
| chr7 | 71217045 | 71217708 | 1.05719 | 2.98568 | 2.824 |
| chr18 | 8899066 | 8899309 | 0.487513 | 1.37684 | 2.824 |
| chr8 | 125965993 | 125966331 | 0.835762 | 2.36046 | 2.824 |
| chr19 | 11876607 | 11876824 | 0.34708 | 0.980279 | 2.824 |
| chr12 | 7613118 | 7613566 | 1.28836 | 3.63893 | 2.824 |
| chr1 | 29447741 | 29448052 | 1.17917 | 3.33065 | 2.825 |
| chrX | 86838748 | 86838987 | 0.794665 | 2.2448 | 2.825 |
| chr3 | 47462577 | 47462978 | 0.984726 | 2.78192 | 2.825 |
| chr6 | 133890053 | 133890311 | 1.05266 | 2.97387 | 2.825 |
| chr1 | 45769700 | 45769932 | 1.43401 | 4.05124 | 2.825 |
| chr9 | 21586422 | 21586642 | 0.858683 | 2.42599 | 2.825 |
| chr9 | 33617497 | 33617818 | 0.7474 | 2.11159 | 2.825 |
| chr12 | 50101288 | 50101736 | 1.60743 | 4.54168 | 2.825 |
| chr1 | 2361527 | 2361868 | 1.25947 | 3.55871 | 2.826 |
| chr12 | 31093034 | 31093317 | 0.368469 | 1.04145 | 2.826 |
| chr3 | 113251640 | 113251856 | 0.497008 | 1.40478 | 2.826 |
| chr1 | 1720550 | 1720796 | 0.895172 | 2.53029 | 2.827 |
| chr15 | 66911476 | 66911868 | 1.8118 | 5.12139 | 2.827 |
| chr20 | 4001829 | 4002146 | 1.11117 | 3.14124 | 2.827 |
| chrX | 30731088 | 30731475 | 1.59895 | 4.52025 | 2.827 |
| chr20 | 49077259 | 49077606 | 2.0515 | 5.79988 | 2.827 |
| chr17 | 75092598 | 75092895 | 1.48222 | 4.19092 | 2.827 |
| chr9 | 113798839 | 113799071 | 0.883848 | 2.49921 | 2.828 |
| chr12 | 114540482 | 114540757 | 0.365694 | 1.03416 | 2.828 |
| chr3 | 13494186 | 13494528 | 0.879573 | 2.48743 | 2.828 |
| chr2 | 230451808 | 230452165 | 1.5255 | 4.31441 | 2.828 |
| chr9 | 93808420 | 93808646 | 0.421211 | 1.19147 | 2.829 |
| chr3 | 160498225 | 160498538 | 0.40211 | 1.13754 | 2.829 |
| chr1 | 12184840 | 12185204 | 1.6026 | 4.53408 | 2.829 |
| chr22 | 19705431 | 19705900 | 1.48841 | 4.21111 | 2.829 |
| chr1 | 6949917 | 6950308 | 1.28074 | 3.62374 | 2.829 |
| chr10 | 134036556 | 134037023 | 2.52483 | 7.1441 | 2.830 |
| chr20 | 17629809 | 17630240 | 1.19304 | 3.37575 | 2.830 |
| chr7 | 3539363 | 3539604 | 0.348102 | 0.984969 | 2.830 |
| chr3 | 87560272 | 87560631 | 0.329268 | 0.931696 | 2.830 |
| chr15 | 88878780 | 88879122 | 0.509139 | 1.44083 | 2.830 |
| chr4 | 77172809 | 77173100 | 0.670236 | 1.89677 | 2.830 |
| chr19 | 51135472 | 51135977 | 1.02814 | 2.90986 | 2.830 |
| chr20 | 15634838 | 15635076 | 1.1333 | 3.20749 | 2.830 |
| chr13 | 27504326 | 27504884 | 1.5687 | 4.44027 | 2.831 |
| chr12 | 13138996 | 13139354 | 0.817364 | 2.31391 | 2.831 |
| chr17 | 22176460 | 22176869 | 0.379735 | 1.07501 | 2.831 |
| chr18 | 75612012 | 75612467 | 2.11917 | 5.99947 | 2.831 |
| chr14 | 99896889 | 99897209 | 0.72792 | 2.06085 | 2.831 |

| | | | | | |
|---|---|---|---|---|---|
| chr19 | 43695840 | 43696162 | 0.656843 | 1.85977 | 2.831 |
| chr5 | 85511731 | 85511981 | 0.319911 | 0.905826 | 2.831 |
| chr5 | 111496281 | 111496615 | 2.3865 | 6.76034 | 2.833 |
| chr7 | 23978631 | 23978870 | 0.912525 | 2.58504 | 2.833 |
| chr3 | 69831047 | 69831258 | 0.386427 | 1.09476 | 2.833 |
| chr10 | 115740235 | 115740472 | 0.781749 | 2.21501 | 2.833 |
| chr21 | 46213276 | 46213533 | 1.10936 | 3.14326 | 2.833 |
| chr1 | 199082472 | 199082738 | 1.33063 | 3.7704 | 2.834 |
| chr17 | 67565198 | 67565506 | 1.14945 | 3.25747 | 2.834 |
| chr22 | 33698016 | 33698398 | 1.52196 | 4.31345 | 2.834 |
| chr19 | 44166718 | 44166987 | 1.27745 | 3.6205 | 2.834 |
| chr16 | 89520293 | 89520615 | 0.680218 | 1.92809 | 2.835 |
| chr5 | 149341715 | 149341943 | 0.871703 | 2.47096 | 2.835 |
| chr13 | 90347972 | 90348232 | 1.32694 | 3.76141 | 2.835 |
| chr8 | 18138228 | 18138435 | 0.368624 | 1.04492 | 2.835 |
| chr1 | 151020931 | 151021197 | 1.18411 | 3.35693 | 2.835 |
| chr7 | 134395454 | 134395692 | 0.309042 | 0.876158 | 2.835 |
| chr4 | 8888782 | 8889159 | 1.08751 | 3.08331 | 2.835 |
| chr15 | 100468659 | 100468923 | 1.10732 | 3.13949 | 2.835 |
| chr17 | 33001219 | 33001597 | 0.384783 | 1.09101 | 2.835 |
| chr4 | 72924451 | 72924691 | 0.385887 | 1.09419 | 2.836 |
| chr1 | 68319289 | 68319638 | 1.32036 | 3.74417 | 2.836 |
| chr14 | 107183137 | 107183502 | 1.1386 | 3.22883 | 2.836 |
| chr17 | 41149997 | 41150289 | 1.0483 | 2.97276 | 2.836 |
| chr1 | 26826687 | 26827030 | 1.43589 | 4.07217 | 2.836 |
| chr7 | 72385727 | 72386071 | 0.487303 | 1.38212 | 2.836 |
| chr14 | 62763726 | 62764110 | 0.428737 | 1.21608 | 2.836 |
| chr15 | 56189941 | 56190243 | 1.42359 | 4.03818 | 2.837 |
| chr1 | 197446819 | 197447095 | 0.600261 | 1.70272 | 2.837 |
| chr9 | 118456317 | 118456552 | 0.802054 | 2.27513 | 2.837 |
| chr6 | 160509525 | 160509807 | 1.01045 | 2.86631 | 2.837 |
| chr1 | 36240334 | 36240568 | 1.23756 | 3.51069 | 2.837 |
| chr9 | 74886849 | 74887118 | 1.31214 | 3.72236 | 2.837 |
| chr4 | 140626290 | 140626546 | 0.827102 | 2.34672 | 2.837 |
| chr12 | 69590444 | 69590785 | 1.02193 | 2.89954 | 2.837 |
| chrX | 38051270 | 38051568 | 1.15355 | 3.27318 | 2.837 |
| chr7 | 26707155 | 26707516 | 1.36963 | 3.88685 | 2.838 |
| chr3 | 49577105 | 49577510 | 1.27153 | 3.60897 | 2.838 |
| chr8 | 52671487 | 52671786 | 0.367151 | 1.04213 | 2.838 |
| chr14 | 40567080 | 40567308 | 0.361871 | 1.02717 | 2.839 |
| chr9 | 23813332 | 23813627 | 0.324654 | 0.921533 | 2.839 |
| chr12 | 14607093 | 14607379 | 1.29358 | 3.67187 | 2.839 |
| chr12 | 25394160 | 25394449 | 1.161 | 3.29592 | 2.839 |
| chr14 | 54492362 | 54492850 | 0.7603 | 2.15842 | 2.839 |
| chr8 | 73508252 | 73508682 | 0.374069 | 1.06196 | 2.839 |
| chr1 | 68907774 | 68908005 | 0.482283 | 1.36934 | 2.839 |
| chr8 | 2453392 | 2453805 | 0.353645 | 1.00412 | 2.839 |
| chr14 | 107280730 | 107281577 | 1.07093 | 3.04082 | 2.839 |
| chr6 | 160054320 | 160054612 | 1.00672 | 2.85853 | 2.839 |
| chr11 | 106908308 | 106908600 | 0.730447 | 2.07417 | 2.840 |
| chr1 | 156450538 | 156450867 | 1.47055 | 4.17602 | 2.840 |
| chr12 | 120393962 | 120394264 | 0.794653 | 2.25678 | 2.840 |
| chr2 | 68690503 | 68690713 | 0.950202 | 2.69881 | 2.840 |
| chr11 | 66049905 | 66050394 | 1.77494 | 5.04164 | 2.840 |
| chr13 | 45855240 | 45855512 | 1.09255 | 3.10334 | 2.840 |
| chr3 | 123751895 | 123752371 | 1.33988 | 3.80603 | 2.841 |
| chr20 | 17817269 | 17817577 | 0.458607 | 1.30272 | 2.841 |
| chr1 | 152598202 | 152598524 | 0.326885 | 0.928581 | 2.841 |
| chr3 | 13025673 | 13026124 | 1.27949 | 3.63477 | 2.841 |
| chr1 | 211828574 | 211828870 | 1.80942 | 5.14023 | 2.841 |
| chr8 | 61577585 | 61577801 | 0.370829 | 1.05365 | 2.841 |

| | | | | | |
|---|---|---|---|---|---|
| chr9 | 21696732 | 21696945 | 0.658004 | 1.86962 | 2.841 |
| chr10 | 90453234 | 90453441 | 0.499096 | 1.41817 | 2.841 |
| chrX | 77347766 | 77348233 | 0.999982 | 2.84202 | 2.842 |
| chr8 | 112265335 | 112265665 | 0.767801 | 2.18217 | 2.842 |
| chr7 | 42225759 | 42226210 | 1.09807 | 3.12092 | 2.842 |
| chr15 | 34402311 | 34402591 | 0.965156 | 2.74321 | 2.842 |
| chr12 | 128586389 | 128586663 | 0.358525 | 1.01904 | 2.842 |
| chr15 | 101413351 | 101413603 | 1.59451 | 4.53294 | 2.843 |
| chrX | 52950309 | 52950630 | 1.76211 | 5.00951 | 2.843 |
| chr19 | 6125561 | 6125958 | 1.28254 | 3.64626 | 2.843 |
| chr14 | 31886760 | 31887077 | 0.636904 | 1.81077 | 2.843 |
| chr1 | 160464441 | 160464692 | 1.15875 | 3.29449 | 2.843 |
| chr5 | 145267584 | 145267877 | 1.15829 | 3.29357 | 2.843 |
| chr19 | 40980453 | 40980659 | 0.755729 | 2.14892 | 2.844 |
| chr19 | 9098885 | 9099152 | 0.791042 | 2.24969 | 2.844 |
| chr11 | 73750103 | 73750383 | 0.418002 | 1.18889 | 2.844 |
| chr10 | 73675802 | 73676018 | 1.03693 | 2.94944 | 2.844 |
| chr11 | 1968379 | 1968763 | 1.30211 | 3.70395 | 2.845 |
| chr17 | 3760400 | 3760616 | 0.422502 | 1.2019 | 2.845 |
| chr10 | 69072149 | 69072353 | 0.370372 | 1.05361 | 2.845 |
| chr16 | 57456277 | 57456595 | 1.61476 | 4.59363 | 2.845 |
| chr17 | 2273631 | 2273944 | 0.350142 | 0.996076 | 2.845 |
| chr15 | 80859993 | 80860391 | 1.46309 | 4.16222 | 2.845 |
| chr16 | 67232145 | 67232578 | 1.67316 | 4.76006 | 2.845 |
| chr2 | 232542370 | 232542669 | 1.19407 | 3.39748 | 2.845 |
| chr12 | 60039155 | 60039370 | 0.547585 | 1.55809 | 2.845 |
| chr12 | 125906865 | 125907081 | 0.585771 | 1.66704 | 2.846 |
| chr12 | 16229555 | 16229844 | 1.1762 | 3.34744 | 2.846 |
| chr11 | 117296921 | 117297260 | 1.97761 | 5.6283 | 2.846 |
| chr19 | 13944570 | 13944790 | 0.646091 | 1.83888 | 2.846 |
| chr19 | 51799686 | 51799926 | 0.54122 | 1.54064 | 2.847 |
| chr7 | 96661700 | 96661914 | 0.483658 | 1.37684 | 2.847 |
| chr6 | 3157745 | 3158084 | 1.11896 | 3.18547 | 2.847 |
| chr9 | 71773094 | 71773403 | 0.733176 | 2.08724 | 2.847 |
| chr13 | 30287953 | 30288337 | 0.825543 | 2.35022 | 2.847 |
| chr19 | 30652144 | 30652380 | 0.43519 | 1.23895 | 2.847 |
| chr6 | 135827462 | 135827677 | 0.746523 | 2.12532 | 2.847 |
| chr19 | 3821233 | 3821585 | 1.56087 | 4.44404 | 2.847 |
| chr12 | 16929623 | 16929922 | 0.4638 | 1.32057 | 2.847 |
| chr19 | 1001630 | 1002055 | 1.70763 | 4.86244 | 2.847 |
| chr6 | 54485910 | 54486368 | 0.443891 | 1.26402 | 2.848 |
| chr19 | 47750603 | 47751082 | 1.6619 | 4.73242 | 2.848 |
| chr8 | 118342390 | 118342709 | 1.7602 | 5.01238 | 2.848 |
| chr17 | 48860440 | 48860925 | 2.39705 | 6.82684 | 2.848 |
| chr6 | 117996367 | 117996642 | 1.27101 | 3.62029 | 2.848 |
| chr12 | 544830 | 545081 | 0.480259 | 1.36802 | 2.849 |
| chr12 | 1761987 | 1762274 | 1.53574 | 4.37486 | 2.849 |
| chr15 | 79317622 | 79317853 | 0.784777 | 2.2358 | 2.849 |
| chr7 | 146794788 | 146795034 | 0.289609 | 0.825175 | 2.849 |
| chr14 | 64319501 | 64319776 | 0.724806 | 2.0654 | 2.850 |
| chr2 | 84645139 | 84645369 | 0.466907 | 1.33051 | 2.850 |
| chr5 | 161131808 | 161132030 | 0.322136 | 0.917977 | 2.850 |
| chr1 | 241998109 | 241998388 | 1.00761 | 2.87147 | 2.850 |
| chr10 | 102278549 | 102278864 | 1.04743 | 2.98502 | 2.850 |
| chr7 | 102045139 | 102045422 | 0.688092 | 1.96101 | 2.850 |
| chr14 | 104387771 | 104388277 | 1.87965 | 5.35717 | 2.850 |
| chr12 | 41584255 | 41584550 | 1.22824 | 3.50089 | 2.850 |
| chr12 | 58176337 | 58176614 | 1.47753 | 4.21155 | 2.850 |
| chr1 | 200941020 | 200941523 | 1.58585 | 4.52038 | 2.850 |
| chr7 | 37636410 | 37636627 | 0.503504 | 1.43532 | 2.851 |
| chr22 | 29791032 | 29791274 | 1.16829 | 3.33056 | 2.851 |

| | | | | | |
|---|---|---|---|---|---|
| chr7 | 94876232 | 94876475 | 1.43906 | 4.1025 | 2.851 |
| chr1 | 203141226 | 203141861 | 2.04679 | 5.83547 | 2.851 |
| chr5 | 60306705 | 60306941 | 0.717982 | 2.04703 | 2.851 |
| chr18 | 28069627 | 28069859 | 0.287052 | 0.818501 | 2.851 |
| chr22 | 25631839 | 25632135 | 1.36602 | 3.89515 | 2.851 |
| chr3 | 181666134 | 181666419 | 1.22032 | 3.4797 | 2.851 |
| chr9 | 94626894 | 94627186 | 0.370491 | 1.05649 | 2.852 |
| chr11 | 133800631 | 133800951 | 1.26945 | 3.62014 | 2.852 |
| chr7 | 104503232 | 104503439 | 0.86933 | 2.47915 | 2.852 |
| chr8 | 100180421 | 100180686 | 1.04315 | 2.97489 | 2.852 |
| chr9 | 100565896 | 100566183 | 0.507709 | 1.44799 | 2.852 |
| chr6 | 36591140 | 36591545 | 0.830717 | 2.36929 | 2.852 |
| chr12 | 53273162 | 53274012 | 1.734 | 4.94582 | 2.852 |
| chr16 | 85854968 | 85855370 | 2.0088 | 5.7308 | 2.853 |
| chr12 | 3913071 | 3913401 | 1.33224 | 3.80084 | 2.853 |
| chr15 | 44876395 | 44876693 | 1.20164 | 3.42828 | 2.853 |
| chr20 | 54620888 | 54621150 | 0.413789 | 1.18056 | 2.853 |
| chr11 | 79597531 | 79597829 | 0.509489 | 1.45361 | 2.853 |
| chr12 | 82633867 | 82634097 | 0.466302 | 1.33054 | 2.853 |
| chr12 | 56132662 | 56132888 | 0.747403 | 2.1327 | 2.853 |
| chr18 | 2766292 | 2766588 | 0.636172 | 1.81552 | 2.854 |
| chr4 | 103541846 | 103542061 | 0.882016 | 2.51725 | 2.854 |
| chr17 | 55675180 | 55675604 | 1.95121 | 5.56889 | 2.854 |
| chr8 | 28220954 | 28221232 | 0.926194 | 2.64342 | 2.854 |
| chr12 | 48045453 | 48045728 | 1.07472 | 3.06763 | 2.854 |
| chr4 | 119786283 | 119786590 | 1.37076 | 3.9127 | 2.854 |
| chr14 | 100075766 | 100076163 | 1.23131 | 3.51478 | 2.855 |
| chr6 | 136347132 | 136347396 | 0.367501 | 1.04908 | 2.855 |
| chr9 | 38727410 | 38727718 | 1.43932 | 4.10909 | 2.855 |
| chr5 | 1010167 | 1010549 | 0.639463 | 1.8256 | 2.855 |
| chr18 | 21939763 | 21939974 | 0.610919 | 1.74432 | 2.855 |
| chr12 | 130395874 | 130396365 | 0.938428 | 2.67978 | 2.856 |
| chr9 | 35603862 | 35604336 | 1.74114 | 4.97205 | 2.856 |
| chr17 | 19657192 | 19657494 | 1.31264 | 3.7485 | 2.856 |
| chr8 | 106730285 | 106730562 | 1.06514 | 3.04221 | 2.856 |
| chr7 | 5279875 | 5280154 | 1.5583 | 4.45082 | 2.856 |
| chr5 | 138446505 | 138446707 | 0.312383 | 0.892329 | 2.857 |
| chr1 | 211123390 | 211123641 | 1.48369 | 4.23857 | 2.857 |
| chr8 | 127568542 | 127569058 | 1.37226 | 3.92041 | 2.857 |
| chrY | 15850589 | 15850852 | 1.01763 | 2.90745 | 2.857 |
| chr18 | 45564059 | 45564477 | 0.592093 | 1.69168 | 2.857 |
| chr14 | 24765997 | 24766404 | 1.23315 | 3.52334 | 2.857 |
| chr12 | 65196022 | 65196404 | 1.45707 | 4.16342 | 2.857 |
| chr5 | 138738254 | 138738596 | 1.29876 | 3.71136 | 2.858 |
| chr1 | 47915650 | 47915900 | 0.981637 | 2.80519 | 2.858 |
| chr12 | 49208254 | 49208878 | 2.15141 | 6.14854 | 2.858 |
| chr4 | 25989745 | 25990126 | 1.82745 | 5.22282 | 2.858 |
| chr6 | 721995 | 722390 | 1.0597 | 3.02868 | 2.858 |
| chr3 | 186831122 | 186831384 | 0.651032 | 1.86077 | 2.858 |
| chr2 | 227080661 | 227081031 | 0.848964 | 2.42684 | 2.859 |
| chr16 | 86609376 | 86609578 | 0.873686 | 2.49753 | 2.859 |
| chr21 | 32831790 | 32832178 | 1.3179 | 3.76784 | 2.859 |
| chr8 | 59720458 | 59720839 | 1.61025 | 4.60385 | 2.859 |
| chr1 | 236901444 | 236901687 | 0.347122 | 0.992512 | 2.859 |
| chr3 | 15419799 | 15420082 | 1.675 | 4.79023 | 2.860 |
| chr7 | 73438814 | 73439040 | 0.443742 | 1.26904 | 2.860 |
| chr7 | 48914942 | 48915277 | 0.97514 | 2.78904 | 2.860 |
| chr5 | 120267671 | 120268002 | 0.409315 | 1.17076 | 2.860 |
| chr20 | 56084710 | 56085181 | 1.10568 | 3.16271 | 2.860 |
| chr20 | 5494807 | 5495166 | 0.413649 | 1.18326 | 2.861 |
| chr10 | 5325517 | 5325781 | 1.54146 | 4.40965 | 2.861 |

| | | | | | |
|---|---|---|---|---|---|
| chr16 | 71760171 | 71760373 | 0.358386 | 1.02529 | 2.861 |
| chr11 | 19585552 | 19585886 | 1.54726 | 4.42654 | 2.861 |
| chr4 | 56504406 | 56504666 | 0.561404 | 1.60627 | 2.861 |
| chr9 | 136061442 | 136061746 | 1.48177 | 4.23961 | 2.861 |
| chr4 | 48548406 | 48548652 | 1.02082 | 2.92085 | 2.861 |
| chr2 | 73626387 | 73626589 | 0.294493 | 0.842644 | 2.861 |
| chr3 | 4146097 | 4146379 | 0.38794 | 1.11009 | 2.862 |
| chr12 | 30131776 | 30132038 | 0.41227 | 1.17974 | 2.862 |
| chr14 | 95155766 | 95156211 | 2.38059 | 6.81227 | 2.862 |
| chr12 | 104851104 | 104851468 | 0.924075 | 2.64436 | 2.862 |
| chr20 | 2367825 | 2368366 | 1.01623 | 2.90844 | 2.862 |
| chr11 | 86167167 | 86167433 | 1.23287 | 3.52858 | 2.862 |
| chr14 | 27997588 | 27997836 | 0.89069 | 2.54928 | 2.862 |
| chr3 | 58173468 | 58173778 | 2.04261 | 5.84638 | 2.862 |
| chr3 | 71119074 | 71119291 | 0.953934 | 2.7306 | 2.862 |
| chr9 | 110594525 | 110594773 | 0.406991 | 1.16504 | 2.863 |
| chr10 | 44806398 | 44806835 | 1.76429 | 5.05071 | 2.863 |
| chr16 | 50711329 | 50711647 | 0.654338 | 1.8732 | 2.863 |
| chr11 | 77531677 | 77531996 | 1.29219 | 3.69957 | 2.863 |
| chr1 | 100202663 | 100202867 | 1.28628 | 3.68267 | 2.863 |
| chr8 | 120464189 | 120464426 | 0.997874 | 2.85716 | 2.863 |
| chr5 | 95194441 | 95194782 | 2.43432 | 6.9702 | 2.863 |
| chr13 | 31978965 | 31979203 | 0.446069 | 1.27729 | 2.863 |
| chr20 | 42939789 | 42940108 | 1.90739 | 5.46191 | 2.864 |
| chr12 | 43732808 | 43733078 | 0.986681 | 2.82559 | 2.864 |
| chr4 | 88274738 | 88275008 | 0.854267 | 2.44643 | 2.864 |
| chr20 | 35548180 | 35548455 | 1.14909 | 3.29081 | 2.864 |
| chr1 | 176821902 | 176822207 | 0.279891 | 0.801734 | 2.864 |
| chr12 | 15334707 | 15334968 | 0.761614 | 2.18172 | 2.865 |
| chr1 | 202569287 | 202569666 | 1.67521 | 4.79962 | 2.865 |
| chr12 | 18144888 | 18145630 | 0.855883 | 2.45219 | 2.865 |
| chr12 | 69292954 | 69293269 | 1.23986 | 3.55241 | 2.865 |
| chr19 | 43936998 | 43937392 | 1.98456 | 5.68651 | 2.865 |
| chr1 | 216526248 | 216526531 | 0.393653 | 1.12797 | 2.865 |
| chr20 | 62169193 | 62169909 | 1.14451 | 3.27967 | 2.866 |
| chr2 | 88491021 | 88491322 | 1.07014 | 3.06663 | 2.866 |
| chr12 | 82864941 | 82865143 | 0.469008 | 1.34401 | 2.866 |
| chr20 | 1207476 | 1207841 | 0.678684 | 1.94507 | 2.866 |
| chr12 | 66538952 | 66539280 | 1.18852 | 3.40626 | 2.866 |
| chr20 | 45221326 | 45221534 | 0.630975 | 1.80843 | 2.866 |
| chr3 | 27700710 | 27701049 | 1.43789 | 4.12139 | 2.866 |
| chr12 | 3196217 | 3196433 | 0.347097 | 0.9949 | 2.866 |
| chr17 | 79779680 | 79779980 | 1.31857 | 3.77948 | 2.866 |
| chr10 | 28952747 | 28953129 | 1.77517 | 5.08838 | 2.866 |
| chr19 | 1624473 | 1624766 | 1.18299 | 3.39096 | 2.866 |
| chr20 | 60135565 | 60135804 | 0.450184 | 1.2905 | 2.867 |
| chr1 | 119750032 | 119750280 | 0.34677 | 0.994127 | 2.867 |
| chr7 | 6529305 | 6529560 | 0.579413 | 1.66111 | 2.867 |
| chr20 | 12793199 | 12793441 | 0.425948 | 1.22124 | 2.867 |
| chr19 | 42928860 | 42929192 | 1.18683 | 3.40296 | 2.867 |
| chr3 | 170027925 | 170028308 | 1.43178 | 4.10602 | 2.868 |
| chr17 | 58663675 | 58663946 | 1.01244 | 2.90387 | 2.868 |
| chr12 | 126853283 | 126853640 | 0.469683 | 1.34726 | 2.868 |
| chr5 | 16550459 | 16550863 | 1.80927 | 5.19054 | 2.869 |
| chr12 | 4416905 | 4417378 | 1.57935 | 4.53122 | 2.869 |
| chr11 | 42435204 | 42435420 | 0.367858 | 1.05541 | 2.869 |
| chr5 | 50258801 | 50259230 | 2.06113 | 5.91372 | 2.869 |
| chr16 | 2924885 | 2925240 | 1.56388 | 4.48751 | 2.869 |
| chr3 | 33173130 | 33173502 | 1.30351 | 3.74039 | 2.869 |
| chr9 | 35813284 | 35813782 | 0.894468 | 2.56673 | 2.870 |
| chr10 | 50405358 | 50405641 | 0.429986 | 1.23394 | 2.870 |

| | | | | | |
|---|---|---|---|---|---|
| chr3 | 134032064 | 134032336 | 0.960184 | 2.75547 | 2.870 |
| chrX | 120715870 | 120716092 | 0.883297 | 2.5349 | 2.870 |
| chr1 | 110527033 | 110527371 | 1.31604 | 3.77696 | 2.870 |
| chr6 | 41703062 | 41703563 | 1.74426 | 5.00674 | 2.870 |
| chr9 | 7186235 | 7186522 | 1.33109 | 3.8208 | 2.870 |
| chr9 | 36168636 | 36168880 | 0.395238 | 1.13452 | 2.870 |
| chr18 | 47254949 | 47255337 | 0.85987 | 2.46856 | 2.871 |
| chr5 | 1772809 | 1773161 | 0.852205 | 2.44656 | 2.871 |
| chrX | 69417165 | 69417422 | 0.831198 | 2.3867 | 2.871 |
| chr6 | 8059379 | 8059641 | 0.513792 | 1.47531 | 2.871 |
| chr17 | 61775175 | 61775617 | 1.64364 | 4.71975 | 2.872 |
| chr4 | 188550142 | 188550366 | 0.468225 | 1.34452 | 2.872 |
| chr5 | 431095 | 431430 | 1.22816 | 3.5267 | 2.872 |
| chr6 | 71665580 | 71666020 | 1.63534 | 4.69592 | 2.872 |
| chr3 | 183968005 | 183968382 | 1.3651 | 3.92001 | 2.872 |
| chr12 | 108957346 | 108957675 | 1.24063 | 3.56272 | 2.872 |
| chr18 | 34337155 | 34337402 | 0.804919 | 2.31158 | 2.872 |
| chr9 | 35661266 | 35661544 | 0.783641 | 2.25048 | 2.872 |
| chrX | 19353066 | 19353901 | 0.8403 | 2.41321 | 2.872 |
| chr7 | 48031420 | 48031729 | 1.27797 | 3.67016 | 2.872 |
| chr14 | 50787780 | 50788082 | 1.68025 | 4.82549 | 2.872 |
| chr1 | 79725537 | 79725761 | 0.552821 | 1.58765 | 2.872 |
| chr12 | 13227475 | 13227799 | 0.546516 | 1.56957 | 2.872 |
| chr19 | 14667411 | 14667795 | 1.62477 | 4.66645 | 2.872 |
| chr3 | 49131943 | 49132237 | 0.712681 | 2.04698 | 2.872 |
| chr5 | 17203525 | 17203793 | 0.872653 | 2.50663 | 2.872 |
| chr19 | 49242198 | 49242484 | 1.15188 | 3.30932 | 2.873 |
| chr10 | 134717691 | 134718190 | 1.75522 | 5.04295 | 2.873 |
| chr16 | 11075653 | 11075892 | 1.24188 | 3.56831 | 2.873 |
| chr20 | 32003897 | 32004130 | 0.715683 | 2.05638 | 2.873 |
| chr22 | 41079157 | 41079418 | 1.76124 | 5.06083 | 2.873 |
| chr12 | 126867903 | 126868323 | 0.307545 | 0.883872 | 2.874 |
| chr18 | 55789133 | 55789494 | 1.20462 | 3.46239 | 2.874 |
| chr6 | 21597249 | 21597484 | 0.625062 | 1.7966 | 2.874 |
| chr5 | 59457198 | 59457460 | 0.738904 | 2.12382 | 2.874 |
| chr17 | 73101472 | 73102203 | 0.814022 | 2.33976 | 2.874 |
| chr9 | 132499897 | 132500264 | 1.82504 | 5.24599 | 2.874 |
| chr2 | 162270849 | 162271134 | 0.418207 | 1.20212 | 2.874 |
| chr5 | 138677342 | 138677574 | 1.08071 | 3.10692 | 2.875 |
| chr12 | 120789009 | 120789285 | 0.320144 | 0.920493 | 2.875 |
| chr20 | 50677677 | 50677955 | 1.1038 | 3.17381 | 2.875 |
| chr5 | 11094375 | 11094590 | 0.306787 | 0.882135 | 2.875 |
| chr21 | 33775515 | 33775722 | 0.906944 | 2.60785 | 2.875 |
| chr1 | 206310370 | 206310757 | 1.61571 | 4.64608 | 2.876 |
| chr5 | 436506 | 436749 | 0.842167 | 2.42174 | 2.876 |
| chr10 | 72147429 | 72147661 | 0.61568 | 1.7705 | 2.876 |
| chr8 | 98594770 | 98595177 | 1.00284 | 2.88388 | 2.876 |
| chr12 | 75784782 | 75785167 | 1.53306 | 4.40974 | 2.876 |
| chrX | 118138857 | 118139274 | 1.7817 | 5.12507 | 2.877 |
| chr6 | 10954782 | 10955080 | 0.91867 | 2.64263 | 2.877 |
| chr7 | 135711466 | 135711769 | 0.33148 | 0.953555 | 2.877 |
| chr7 | 100720769 | 100721174 | 1.61401 | 4.64341 | 2.877 |
| chr3 | 109128717 | 109129123 | 0.69329 | 1.9947 | 2.877 |
| chr3 | 138051416 | 138051618 | 0.673624 | 1.93823 | 2.877 |
| chr9 | 14465588 | 14465886 | 0.96747 | 2.7838 | 2.877 |
| chr11 | 10900238 | 10900546 | 1.16486 | 3.35178 | 2.877 |
| chr4 | 142782213 | 142782441 | 0.674119 | 1.93972 | 2.877 |
| chr12 | 48950729 | 48951126 | 0.424355 | 1.22112 | 2.878 |
| chr5 | 23821023 | 23821308 | 0.699336 | 2.0125 | 2.878 |
| chr12 | 59797154 | 59797392 | 1.08971 | 3.13599 | 2.878 |
| chr19 | 14472608 | 14472859 | 0.697488 | 2.00729 | 2.878 |

| | | | | | |
|---|---|---|---|---|---|
| chr14 | 21571750 | 21571982 | 1.50114 | 4.32013 | 2.878 |
| chr19 | 52164326 | 52164573 | 1.31269 | 3.77785 | 2.878 |
| chr7 | 86688961 | 86689290 | 0.706115 | 2.03231 | 2.878 |
| chr12 | 20616916 | 20617195 | 0.975058 | 2.80685 | 2.879 |
| chr1 | 41418130 | 41418530 | 1.25378 | 3.60927 | 2.879 |
| chr15 | 33990599 | 33990815 | 0.401995 | 1.15728 | 2.879 |
| chr6 | 33736373 | 33736712 | 1.73344 | 4.99065 | 2.879 |
| chr15 | 75437899 | 75438253 | 1.1512 | 3.3145 | 2.879 |
| chr7 | 42566608 | 42566865 | 0.340705 | 0.981055 | 2.879 |
| chr1 | 9648503 | 9648810 | 1.06517 | 3.06716 | 2.880 |
| chr9 | 78180110 | 78180365 | 0.344518 | 0.992086 | 2.880 |
| chr8 | 1978739 | 1979165 | 1.32943 | 3.82881 | 2.880 |
| chr19 | 42363625 | 42363913 | 0.767929 | 2.21168 | 2.880 |
| chr12 | 121937763 | 121938078 | 2.82656 | 8.14217 | 2.881 |
| chr7 | 150660060 | 150660548 | 1.33568 | 3.8476 | 2.881 |
| chr17 | 34131709 | 34132126 | 1.99226 | 5.73906 | 2.881 |
| chr9 | 4924219 | 4924529 | 0.813206 | 2.34272 | 2.881 |
| chr7 | 43300880 | 43301304 | 1.43123 | 4.12346 | 2.881 |
| chr17 | 63133272 | 63133529 | 2.81634 | 8.11481 | 2.881 |
| chr6 | 78175727 | 78176005 | 1.1591 | 3.33977 | 2.881 |
| chr6 | 9540794 | 9541032 | 0.95278 | 2.74552 | 2.882 |
| chr4 | 146137772 | 146138053 | 0.649114 | 1.8706 | 2.882 |
| chr19 | 5687140 | 5687480 | 1.57141 | 4.5287 | 2.882 |
| chr3 | 139048199 | 139048592 | 2.20778 | 6.36281 | 2.882 |
| chr10 | 99491551 | 99491880 | 1.2094 | 3.48556 | 2.882 |
| chr14 | 20774039 | 20774315 | 1.53891 | 4.43573 | 2.882 |
| chr4 | 87417254 | 87417617 | 0.904 | 2.60569 | 2.882 |
| chr10 | 23324619 | 23324969 | 0.594293 | 1.71302 | 2.882 |
| chr5 | 10656614 | 10656821 | 0.325126 | 0.937167 | 2.882 |
| chr6 | 131815628 | 131815851 | 0.496131 | 1.43019 | 2.883 |
| chr2 | 85358290 | 85358509 | 0.518019 | 1.49336 | 2.883 |
| chr10 | 104163582 | 104163963 | 1.82087 | 5.24985 | 2.883 |
| chr3 | 50651505 | 50651850 | 1.797 | 5.18145 | 2.883 |
| chr5 | 140971632 | 140971883 | 1.38249 | 3.98683 | 2.884 |
| chr10 | 69548197 | 69548596 | 1.14723 | 3.30904 | 2.884 |
| chr3 | 58449426 | 58449760 | 1.31948 | 3.80623 | 2.885 |
| chr3 | 130950576 | 130950946 | 0.501479 | 1.44674 | 2.885 |
| chr7 | 35559872 | 35560181 | 1.10652 | 3.19253 | 2.885 |
| chr5 | 23710113 | 23710320 | 0.318452 | 0.918811 | 2.885 |
| chr16 | 28006370 | 28006572 | 0.514328 | 1.48422 | 2.886 |
| chr10 | 35012660 | 35013113 | 1.10483 | 3.18867 | 2.886 |
| chr5 | 4862722 | 4862954 | 0.72947 | 2.10539 | 2.886 |
| chr2 | 234389178 | 234389426 | 0.637505 | 1.84 | 2.886 |
| chr10 | 71168309 | 71168896 | 1.78452 | 5.1506 | 2.886 |
| chr1 | 199560685 | 199560939 | 0.445489 | 1.28583 | 2.886 |
| chr9 | 4382781 | 4383019 | 0.906965 | 2.6182 | 2.887 |
| chr2 | 232571709 | 232572095 | 1.20654 | 3.48333 | 2.887 |
| chr10 | 88021681 | 88021987 | 1.01491 | 2.93019 | 2.887 |
| chr12 | 52533254 | 52533667 | 1.75902 | 5.0786 | 2.887 |
| chr10 | 115953191 | 115953396 | 0.343063 | 0.990493 | 2.887 |
| chr19 | 39493770 | 39494104 | 0.608043 | 1.7556 | 2.887 |
| chr6 | 41149896 | 41150172 | 0.874412 | 2.52478 | 2.887 |
| chr12 | 39836922 | 39837288 | 0.694081 | 2.00414 | 2.887 |
| chr9 | 21813905 | 21814183 | 1.20486 | 3.47925 | 2.888 |
| chr3 | 110731575 | 110731777 | 0.886839 | 2.56097 | 2.888 |
| chr11 | 17494988 | 17495326 | 1.05855 | 3.0569 | 2.888 |
| chr8 | 32405341 | 32406929 | 1.16895 | 3.37576 | 2.888 |
| chr14 | 41479789 | 41479993 | 0.319649 | 0.923166 | 2.888 |
| chr16 | 11368644 | 11368891 | 0.256127 | 0.739736 | 2.888 |
| chr7 | 155326525 | 155326902 | 1.50305 | 4.34109 | 2.888 |
| chr5 | 113353908 | 113354260 | 0.564928 | 1.63163 | 2.888 |

| | | | | | |
|---|---|---|---|---|---|
| chr12 | 102167376 | 102167709 | 1.32672 | 3.83254 | 2.889 |
| chr14 | 100621371 | 100621750 | 1.91789 | 5.54093 | 2.889 |
| chr9 | 77388859 | 77389218 | 1.25059 | 3.61357 | 2.889 |
| chr1 | 84271573 | 84271832 | 0.351064 | 1.01442 | 2.890 |
| chr9 | 96064896 | 96065357 | 1.69496 | 4.89782 | 2.890 |
| chr7 | 37738587 | 37739180 | 0.323425 | 0.934618 | 2.890 |
| chr7 | 154982343 | 154982720 | 2.02574 | 5.85465 | 2.890 |
| chr4 | 17513649 | 17514177 | 1.03886 | 3.00249 | 2.890 |
| chr10 | 121390567 | 121390802 | 0.247089 | 0.714164 | 2.890 |
| chr6 | 2483108 | 2483334 | 0.587015 | 1.69682 | 2.891 |
| chr16 | 89355775 | 89356215 | 2.30524 | 6.66362 | 2.891 |
| chr7 | 64082079 | 64082348 | 0.69884 | 2.02051 | 2.891 |
| chr2 | 234823901 | 234824171 | 0.636264 | 1.83959 | 2.891 |
| chr12 | 92972591 | 92972813 | 0.262211 | 0.758154 | 2.891 |
| chr11 | 30571896 | 30572258 | 0.419465 | 1.21286 | 2.891 |
| chr18 | 35065228 | 35065586 | 1.24765 | 3.60764 | 2.892 |
| chr12 | 78319300 | 78319784 | 1.56947 | 4.5384 | 2.892 |
| chr7 | 101054419 | 101054874 | 1.84074 | 5.32315 | 2.892 |
| chr12 | 97222050 | 97222339 | 0.595563 | 1.72239 | 2.892 |
| chr17 | 56674099 | 56674351 | 0.413918 | 1.19733 | 2.893 |
| chr19 | 10691592 | 10692038 | 1.28989 | 3.73158 | 2.893 |
| chr10 | 23787478 | 23787916 | 0.381019 | 1.10229 | 2.893 |
| chr3 | 83572753 | 83572959 | 0.911192 | 2.63613 | 2.893 |
| chr7 | 1222635 | 1222939 | 1.19576 | 3.45954 | 2.893 |
| chr3 | 172313960 | 172314221 | 0.822003 | 2.37867 | 2.894 |
| chr7 | 134855496 | 134855771 | 1.0575 | 3.06031 | 2.894 |
| chr16 | 7666524 | 7667097 | 0.385347 | 1.11519 | 2.894 |
| chr12 | 101127675 | 101128007 | 1.21007 | 3.50198 | 2.894 |
| chr1 | 96823715 | 96823953 | 0.783193 | 2.26682 | 2.894 |
| chr19 | 30363115 | 30363355 | 0.792101 | 2.29274 | 2.895 |
| chr22 | 45837087 | 45837433 | 1.28564 | 3.72132 | 2.895 |
| chr14 | 56889252 | 56889645 | 1.1851 | 3.4306 | 2.895 |
| chrX | 118918881 | 118919175 | 1.19349 | 3.45492 | 2.895 |
| chr5 | 73088560 | 73088910 | 2.26517 | 6.55747 | 2.895 |
| chr16 | 66512637 | 66512982 | 1.45812 | 4.22136 | 2.895 |
| chr10 | 71310691 | 71310894 | 0.355714 | 1.02987 | 2.895 |
| chr8 | 24086867 | 24087218 | 1.32977 | 3.84998 | 2.895 |
| chr17 | 29445900 | 29446283 | 0.391961 | 1.13494 | 2.896 |
| chr9 | 130217535 | 130218049 | 1.04601 | 3.02931 | 2.896 |
| chr20 | 58325995 | 58326532 | 1.42907 | 4.13875 | 2.896 |
| chr12 | 77718478 | 77718744 | 1.1166 | 3.23401 | 2.896 |
| chr5 | 26009861 | 26010181 | 0.786017 | 2.27671 | 2.897 |
| chr9 | 20058686 | 20058990 | 0.418714 | 1.21282 | 2.897 |
| chr1 | 994634 | 995033 | 0.879274 | 2.54688 | 2.897 |
| chr14 | 77334999 | 77335390 | 0.504616 | 1.46169 | 2.897 |
| chr14 | 50332112 | 50332464 | 1.62583 | 4.70955 | 2.897 |
| chr16 | 25679588 | 25679839 | 0.338174 | 0.979594 | 2.897 |
| chr7 | 152964670 | 152964915 | 0.518504 | 1.50214 | 2.897 |
| chr9 | 137033850 | 137034151 | 0.944217 | 2.73554 | 2.897 |
| chr7 | 121436754 | 121437035 | 0.391884 | 1.13535 | 2.897 |
| chr10 | 118935480 | 118935814 | 1.10309 | 3.19639 | 2.898 |
| chr15 | 90905242 | 90905604 | 1.06834 | 3.09579 | 2.898 |
| chr5 | 54367180 | 54367406 | 0.456387 | 1.32253 | 2.898 |
| chr9 | 76599850 | 76600181 | 0.428757 | 1.24249 | 2.898 |
| chr10 | 69972488 | 69972928 | 0.948657 | 2.74918 | 2.898 |
| chr12 | 23338368 | 23338787 | 0.358151 | 1.03792 | 2.898 |
| chr14 | 32214943 | 32215276 | 0.804933 | 2.33294 | 2.898 |
| chr11 | 64764329 | 64764598 | 0.93017 | 2.69614 | 2.899 |
| chr5 | 176916393 | 176916631 | 0.578101 | 1.6757 | 2.899 |
| chr3 | 147663898 | 147664149 | 1.09551 | 3.17553 | 2.899 |
| chr3 | 38387957 | 38388417 | 1.20304 | 3.48745 | 2.899 |

| | | | | | |
|---|---|---|---|---|---|
| chr2 | 44270620 | 44270851 | 0.402008 | 1.16537 | 2.899 |
| chr19 | 39965944 | 39966298 | 0.73011 | 2.11653 | 2.899 |
| chr11 | 103456481 | 103456789 | 0.324295 | 0.940127 | 2.899 |
| chrX | 53462980 | 53463279 | 1.14731 | 3.32625 | 2.899 |
| chr2 | 82461500 | 82461821 | 0.951741 | 2.75954 | 2.899 |
| chr12 | 20521041 | 20521284 | 0.385681 | 1.1183 | 2.900 |
| chr3 | 69210807 | 69211077 | 1.28228 | 3.71817 | 2.900 |
| chr4 | 23910981 | 23911207 | 0.298453 | 0.865467 | 2.900 |
| chr7 | 5603221 | 5603603 | 1.98338 | 5.75176 | 2.900 |
| chr16 | 11774237 | 11774566 | 0.32164 | 0.9328 | 2.900 |
| chr1 | 72116061 | 72116286 | 0.34882 | 1.01176 | 2.901 |
| chr14 | 31020051 | 31020257 | 0.306392 | 0.888736 | 2.901 |
| chr4 | 38321482 | 38321736 | 1.45192 | 4.21303 | 2.902 |
| chr17 | 52914521 | 52914730 | 0.443203 | 1.28607 | 2.902 |
| chr1 | 232002371 | 232002774 | 1.30758 | 3.79442 | 2.902 |
| chr6 | 35995191 | 35995424 | 1.51654 | 4.40087 | 2.902 |
| chr17 | 63452144 | 63452429 | 0.85074 | 2.46903 | 2.902 |
| chr14 | 100120533 | 100120812 | 0.441169 | 1.28038 | 2.902 |
| chr7 | 37873118 | 37873516 | 1.114 | 3.23323 | 2.902 |
| chr16 | 67312571 | 67312808 | 0.903495 | 2.62247 | 2.903 |
| chr9 | 139233540 | 139234091 | 2.09652 | 6.08592 | 2.903 |
| chr1 | 238499739 | 238500035 | 0.440609 | 1.27909 | 2.903 |
| chr5 | 178017558 | 178017942 | 1.43578 | 4.16808 | 2.903 |
| chr5 | 45653199 | 45653401 | 0.413017 | 1.19901 | 2.903 |
| chr12 | 27577609 | 27577960 | 0.870181 | 2.52681 | 2.904 |
| chr6 | 36091618 | 36091884 | 0.800477 | 2.32447 | 2.904 |
| chr5 | 16020653 | 16020886 | 0.339048 | 0.984559 | 2.904 |
| chr20 | 30667504 | 30667749 | 0.570924 | 1.65794 | 2.904 |
| chr6 | 85489189 | 85489405 | 0.552903 | 1.6057 | 2.904 |
| chr7 | 106779702 | 106779913 | 0.366642 | 1.06487 | 2.904 |
| chr7 | 118417072 | 118417286 | 0.329109 | 0.955923 | 2.905 |
| chrX | 19562833 | 19563129 | 1.3005 | 3.7775 | 2.905 |
| chr12 | 126951182 | 126951392 | 0.432459 | 1.25617 | 2.905 |
| chr2 | 169193353 | 169193707 | 0.778815 | 2.26223 | 2.905 |
| chr7 | 156701608 | 156701960 | 1.63762 | 4.75683 | 2.905 |
| chr20 | 57662887 | 57663354 | 1.47582 | 4.28686 | 2.905 |
| chr15 | 98447357 | 98447774 | 1.39517 | 4.05263 | 2.905 |
| chr1 | 208047444 | 208047680 | 0.335419 | 0.974338 | 2.905 |
| chr18 | 32810997 | 32811268 | 1.40529 | 4.08244 | 2.905 |
| chr10 | 26014872 | 26015283 | 1.38216 | 4.01527 | 2.905 |
| chr16 | 63736989 | 63737210 | 0.737649 | 2.14298 | 2.905 |
| chr10 | 93847428 | 93847855 | 1.42037 | 4.1264 | 2.905 |
| chr15 | 73432492 | 73432739 | 0.569079 | 1.6533 | 2.905 |
| chr1 | 55556739 | 55557008 | 0.414829 | 1.2052 | 2.905 |
| chr7 | 97200464 | 97200674 | 0.376303 | 1.09333 | 2.905 |
| chr18 | 26771302 | 26771525 | 0.634413 | 1.84327 | 2.905 |
| chr3 | 30975727 | 30975990 | 0.723758 | 2.10297 | 2.906 |
| chr3 | 73142847 | 73143466 | 0.421356 | 1.22434 | 2.906 |
| chr19 | 970688 | 971009 | 1.21674 | 3.53555 | 2.906 |
| chr20 | 19098172 | 19098437 | 0.355366 | 1.03261 | 2.906 |
| chr6 | 109557669 | 109557901 | 0.527133 | 1.53174 | 2.906 |
| chr1 | 171952776 | 171953037 | 0.816403 | 2.37231 | 2.906 |
| chr9 | 16032900 | 16033206 | 0.681288 | 1.97971 | 2.906 |
| chr6 | 150656342 | 150656632 | 1.07208 | 3.11546 | 2.906 |
| chr12 | 109646947 | 109647188 | 0.465155 | 1.35181 | 2.906 |
| chr8 | 143777616 | 143777957 | 0.474574 | 1.37932 | 2.906 |
| chr13 | 76123415 | 76123674 | 1.11497 | 3.24095 | 2.907 |
| chr4 | 103652348 | 103652607 | 0.503378 | 1.46321 | 2.907 |
| chr1 | 150894940 | 150895284 | 1.1319 | 3.29022 | 2.907 |
| chr7 | 151145417 | 151146106 | 1.03328 | 3.00394 | 2.907 |
| chr15 | 70271693 | 70271911 | 0.561793 | 1.63325 | 2.907 |

| | | | | | |
|---|---|---|---|---|---|
| chr2 | 129226824 | 129227113 | 0.94778 | 2.7554 | 2.907 |
| chr16 | 23534699 | 23534913 | 0.42525 | 1.23647 | 2.908 |
| chr8 | 103823076 | 103823481 | 1.74552 | 5.07545 | 2.908 |
| chr4 | 94120786 | 94121027 | 0.836559 | 2.43249 | 2.908 |
| chr12 | 15145991 | 15146217 | 0.433383 | 1.26026 | 2.908 |
| chr9 | 96442832 | 96443106 | 0.552598 | 1.60695 | 2.908 |
| chr7 | 136362170 | 136362411 | 0.456255 | 1.32697 | 2.908 |
| chr16 | 90014966 | 90015230 | 1.96273 | 5.70917 | 2.909 |
| chr1 | 227015732 | 227016102 | 1.60215 | 4.66076 | 2.909 |
| chr12 | 28093759 | 28094076 | 1.29935 | 3.77995 | 2.909 |
| chr15 | 67793632 | 67793885 | 0.546813 | 1.59081 | 2.909 |
| chr15 | 62352649 | 62353011 | 1.13991 | 3.31628 | 2.909 |
| chr10 | 13344373 | 13344659 | 0.920651 | 2.67891 | 2.910 |
| chr17 | 7036814 | 7037113 | 1.01192 | 2.94453 | 2.910 |
| chr1 | 77366868 | 77367237 | 1.07382 | 3.12473 | 2.910 |
| chr12 | 127287747 | 127288050 | 0.345875 | 1.00647 | 2.910 |
| chr9 | 75722482 | 75722733 | 1.41243 | 4.11006 | 2.910 |
| chr1 | 117279976 | 117280190 | 1.10514 | 3.21598 | 2.910 |
| chr7 | 154719244 | 154719569 | 1.11555 | 3.24633 | 2.910 |
| chr3 | 94055241 | 94055450 | 0.883558 | 2.57126 | 2.910 |
| chr5 | 117755548 | 117755822 | 0.472452 | 1.37491 | 2.910 |
| chrX | 106117391 | 106117643 | 0.608212 | 1.77034 | 2.911 |
| chr15 | 90701168 | 90701422 | 1.64678 | 4.79336 | 2.911 |
| chr6 | 37670025 | 37670315 | 0.431325 | 1.2555 | 2.911 |
| chr13 | 20771541 | 20771792 | 0.946999 | 2.75722 | 2.912 |
| chr11 | 107769106 | 107769321 | 0.811893 | 2.36399 | 2.912 |
| chr2 | 11878631 | 11879018 | 1.35676 | 3.95077 | 2.912 |
| chr3 | 31489994 | 31490294 | 1.22303 | 3.56147 | 2.912 |
| chr12 | 113778630 | 113779109 | 0.335322 | 0.976468 | 2.912 |
| chr5 | 137883634 | 137883982 | 0.859437 | 2.50271 | 2.912 |
| chr4 | 65404019 | 65404252 | 0.646763 | 1.88362 | 2.912 |
| chr8 | 140353096 | 140353460 | 1.63708 | 4.76824 | 2.913 |
| chr6 | 100036482 | 100036801 | 1.09879 | 3.20049 | 2.913 |
| chr5 | 121961454 | 121961688 | 0.380034 | 1.10704 | 2.913 |
| chr19 | 34836554 | 34837168 | 1.45095 | 4.22744 | 2.914 |
| chr7 | 95353001 | 95353207 | 0.756465 | 2.20407 | 2.914 |
| chr12 | 117650363 | 117650721 | 0.961575 | 2.80169 | 2.914 |
| chr7 | 47128446 | 47128740 | 1.58311 | 4.61265 | 2.914 |
| chr2 | 231729373 | 231729678 | 1.0009 | 2.91631 | 2.914 |
| chr19 | 22728642 | 22728899 | 1.083 | 3.15554 | 2.914 |
| chrX | 152901772 | 152902096 | 1.42732 | 4.15934 | 2.914 |
| chr12 | 128739810 | 128740075 | 0.340476 | 0.99226 | 2.914 |
| chr7 | 140026066 | 140026460 | 1.57168 | 4.58049 | 2.914 |
| chr14 | 77413513 | 77413764 | 0.735987 | 2.14507 | 2.915 |
| chr8 | 60386062 | 60386271 | 0.349982 | 1.02006 | 2.915 |
| chr1 | 226521112 | 226521527 | 1.25082 | 3.64577 | 2.915 |
| chr1 | 23303003 | 23303456 | 0.983122 | 2.86552 | 2.915 |
| chr3 | 184714201 | 184714503 | 0.38596 | 1.12504 | 2.915 |
| chr5 | 136190080 | 136190313 | 0.528075 | 1.53939 | 2.915 |
| chr10 | 79606365 | 79606792 | 1.64401 | 4.79265 | 2.915 |
| chr18 | 72675870 | 72676099 | 0.68001 | 1.98245 | 2.915 |
| chr5 | 26482876 | 26483299 | 0.808906 | 2.35827 | 2.915 |
| chr4 | 3482211 | 3482640 | 1.74007 | 5.07304 | 2.915 |
| chr4 | 184761431 | 184761749 | 0.987003 | 2.87791 | 2.916 |
| chr8 | 142736996 | 142737524 | 1.92944 | 5.62621 | 2.916 |
| chr20 | 46249958 | 46250310 | 1.1092 | 3.23465 | 2.916 |
| chr12 | 52214151 | 52214817 | 1.54907 | 4.51753 | 2.916 |
| chr10 | 61469548 | 61469801 | 0.845624 | 2.46647 | 2.917 |
| chr11 | 46639162 | 46639597 | 1.30385 | 3.8031 | 2.917 |
| chr16 | 19371419 | 19371699 | 0.390814 | 1.14009 | 2.917 |
| chr8 | 28181811 | 28182234 | 1.82615 | 5.32749 | 2.917 |

| | | | | | |
|---|---|---|---|---|---|
| chr5 | 91854109 | 91854342 | 1.16087 | 3.38684 | 2.918 |
| chr5 | 158868445 | 158868900 | 1.02253 | 2.98335 | 2.918 |
| chrX | 109403743 | 109404034 | 0.917364 | 2.67652 | 2.918 |
| chr6 | 155495917 | 155496147 | 0.858267 | 2.50416 | 2.918 |
| chrX | 119021596 | 119022021 | 1.19165 | 3.4771 | 2.918 |
| chr8 | 135550473 | 135550749 | 0.922628 | 2.69254 | 2.918 |
| chr12 | 62396897 | 62397306 | 0.443592 | 1.29466 | 2.919 |
| chr10 | 29802022 | 29802277 | 1.24422 | 3.63159 | 2.919 |
| chr12 | 17748299 | 17748536 | 0.45591 | 1.33085 | 2.919 |
| chr20 | 49280057 | 49280270 | 0.747385 | 2.18184 | 2.919 |
| chr9 | 44227351 | 44227577 | 1.38402 | 4.04048 | 2.919 |
| chr9 | 136268945 | 136269401 | 1.37518 | 4.01482 | 2.919 |
| chr5 | 92014363 | 92014675 | 0.8042 | 2.34789 | 2.920 |
| chr7 | 5179617 | 5179945 | 0.590272 | 1.72343 | 2.920 |
| chr10 | 6828452 | 6828835 | 0.967462 | 2.82486 | 2.920 |
| chr17 | 45815629 | 45815876 | 0.372783 | 1.08852 | 2.920 |
| chr14 | 31159863 | 31160103 | 0.301403 | 0.880103 | 2.920 |
| chr6 | 43634966 | 43635273 | 1.50182 | 4.38567 | 2.920 |
| chr5 | 32425691 | 32425925 | 0.484259 | 1.41463 | 2.921 |
| chr21 | 44559151 | 44559401 | 1.14307 | 3.3393 | 2.921 |
| chr8 | 52391163 | 52391641 | 0.505228 | 1.47616 | 2.922 |
| chr9 | 127536691 | 127537096 | 1.10823 | 3.23808 | 2.922 |
| chr12 | 7351492 | 7351763 | 0.444454 | 1.29866 | 2.922 |
| chr15 | 74419850 | 74420217 | 2.09191 | 6.11255 | 2.922 |
| chr22 | 31316267 | 31316600 | 0.993812 | 2.90398 | 2.922 |
| chr1 | 202968302 | 202968820 | 1.07459 | 3.14018 | 2.922 |
| chr18 | 65666860 | 65667122 | 0.394004 | 1.15138 | 2.922 |
| chr8 | 85472858 | 85473090 | 0.353909 | 1.03422 | 2.922 |
| chr3 | 49878623 | 49878935 | 0.983287 | 2.87357 | 2.922 |
| chr11 | 20132313 | 20132798 | 1.67652 | 4.89973 | 2.923 |
| chr17 | 66768046 | 66768340 | 1.3332 | 3.89638 | 2.923 |
| chr3 | 23930120 | 23930402 | 0.824054 | 2.40843 | 2.923 |
| chr7 | 69359024 | 69359230 | 1.17497 | 3.43426 | 2.923 |
| chr12 | 130681474 | 130681697 | 0.498291 | 1.45663 | 2.923 |
| chr12 | 56423525 | 56423849 | 1.72448 | 5.04176 | 2.924 |
| chr12 | 96017376 | 96017736 | 1.48054 | 4.32874 | 2.924 |
| chr13 | 30982514 | 30982716 | 1.04148 | 3.0451 | 2.924 |
| chr16 | 66981109 | 66981391 | 1.28207 | 3.7487 | 2.924 |
| chr15 | 31117039 | 31117290 | 0.319056 | 0.932905 | 2.924 |
| chr16 | 81619519 | 81619730 | 0.905647 | 2.64826 | 2.924 |
| chr3 | 144510170 | 144510435 | 0.28839 | 0.84331 | 2.924 |
| chr18 | 11398788 | 11398998 | 0.705027 | 2.06198 | 2.925 |
| chr4 | 122383895 | 122384162 | 1.11738 | 3.26824 | 2.925 |
| chr2 | 49190469 | 49190727 | 0.613509 | 1.79451 | 2.925 |
| chr22 | 46127605 | 46127903 | 0.632887 | 1.85121 | 2.925 |
| chr12 | 101490065 | 101490279 | 0.298916 | 0.874349 | 2.925 |
| chr10 | 102728839 | 102729167 | 1.74197 | 5.09596 | 2.925 |
| chr5 | 171033515 | 171033720 | 0.889016 | 2.6008 | 2.925 |
| chr7 | 29628716 | 29629208 | 1.15238 | 3.37158 | 2.926 |
| chr1 | 53935755 | 53936082 | 0.988305 | 2.89156 | 2.926 |
| chr10 | 59476734 | 59476991 | 0.801575 | 2.34528 | 2.926 |
| chr12 | 114172184 | 114172390 | 0.266415 | 0.779493 | 2.926 |
| chr12 | 3654236 | 3654521 | 0.284374 | 0.832048 | 2.926 |
| chr11 | 56130408 | 56130669 | 0.507997 | 1.48643 | 2.926 |
| chr15 | 45560365 | 45560588 | 0.660289 | 1.93215 | 2.926 |
| chr9 | 133326877 | 133327196 | 0.65409 | 1.91405 | 2.926 |
| chr16 | 5106425 | 5106688 | 0.869346 | 2.54396 | 2.926 |
| chr3 | 24681868 | 24682124 | 1.08324 | 3.1701 | 2.927 |
| chr2 | 84485095 | 84485376 | 0.680629 | 1.99201 | 2.927 |
| chr19 | 8399379 | 8399681 | 0.411029 | 1.20298 | 2.927 |
| chr12 | 27167148 | 27167633 | 1.76431 | 5.16406 | 2.927 |

| | | | | | |
|---|---|---|---|---|---|
| chr7 | 22768138 | 22768354 | 1.10371 | 3.23073 | 2.927 |
| chr16 | 9102526 | 9103029 | 1.83849 | 5.38182 | 2.927 |
| chr14 | 90234003 | 90234429 | 0.775637 | 2.27085 | 2.928 |
| chrX | 100872292 | 100872519 | 0.813088 | 2.38062 | 2.928 |
| chr12 | 11639541 | 11639963 | 1.42262 | 4.16544 | 2.928 |
| chr16 | 25236501 | 25236929 | 1.32175 | 3.8702 | 2.928 |
| chr8 | 63015391 | 63015671 | 1.09422 | 3.20414 | 2.928 |
| chr19 | 48661739 | 48662042 | 0.648278 | 1.8984 | 2.928 |
| chr15 | 36793358 | 36793602 | 1.61494 | 4.7292 | 2.928 |
| chr18 | 596999 | 597203 | 1.08483 | 3.17684 | 2.928 |
| chr22 | 29985934 | 29986263 | 1.29868 | 3.80314 | 2.928 |
| chr6 | 786963 | 787296 | 0.899979 | 2.63574 | 2.929 |
| chr9 | 118912180 | 118912446 | 1.06826 | 3.12873 | 2.929 |
| chr10 | 135252432 | 135252650 | 0.551318 | 1.61471 | 2.929 |
| chr15 | 91814864 | 91815140 | 1.08242 | 3.17049 | 2.929 |
| chr3 | 47866232 | 47866602 | 1.5743 | 4.61151 | 2.929 |
| chr4 | 127525483 | 127525729 | 0.809497 | 2.37143 | 2.930 |
| chr6 | 35435900 | 35436196 | 1.51534 | 4.4392 | 2.930 |
| chr3 | 51671872 | 51672348 | 1.13252 | 3.31774 | 2.930 |
| chr3 | 63979301 | 63979687 | 1.01752 | 2.98087 | 2.930 |
| chr4 | 177711110 | 177711353 | 0.895009 | 2.6221 | 2.930 |
| chr11 | 69659755 | 69660297 | 0.268633 | 0.787042 | 2.930 |
| chr6 | 130643834 | 130644146 | 1.13014 | 3.31125 | 2.930 |
| chr3 | 50556926 | 50557331 | 1.5934 | 4.66868 | 2.930 |
| chr3 | 34632478 | 34632983 | 0.317065 | 0.929026 | 2.930 |
| chr2 | 24766091 | 24766402 | 0.805613 | 2.36058 | 2.930 |
| chr7 | 11467385 | 11467592 | 0.853115 | 2.49988 | 2.930 |
| chr8 | 37924559 | 37925019 | 1.16416 | 3.41187 | 2.931 |
| chr2 | 60901635 | 60902096 | 2.15362 | 6.31206 | 2.931 |
| chr11 | 2986045 | 2986308 | 0.381366 | 1.11791 | 2.931 |
| chr19 | 46849234 | 46849658 | 0.827505 | 2.42602 | 2.932 |
| chr16 | 466687 | 466915 | 0.356472 | 1.04511 | 2.932 |
| chr8 | 54016230 | 54016434 | 0.368374 | 1.08006 | 2.932 |
| chr5 | 129534653 | 129534894 | 0.911613 | 2.67295 | 2.932 |
| chr3 | 100915007 | 100915308 | 1.18324 | 3.46949 | 2.932 |
| chr10 | 104325606 | 104325859 | 0.489009 | 1.43405 | 2.933 |
| chr1 | 86078800 | 86079056 | 1.09652 | 3.21563 | 2.933 |
| chr1 | 146644077 | 146644507 | 1.55929 | 4.573 | 2.933 |
| chr6 | 25591331 | 25591550 | 0.308629 | 0.905182 | 2.933 |
| chr20 | 10652122 | 10652343 | 0.909098 | 2.66644 | 2.933 |
| chr9 | 87506183 | 87506387 | 0.302269 | 0.886599 | 2.933 |
| chr1 | 67461768 | 67462097 | 0.974889 | 2.85974 | 2.933 |
| chr15 | 75227639 | 75227847 | 0.680035 | 1.99499 | 2.934 |
| chr16 | 62169693 | 62170006 | 0.358986 | 1.05327 | 2.934 |
| chr9 | 6107099 | 6107417 | 0.798864 | 2.34389 | 2.934 |
| chr4 | 7187821 | 7188087 | 0.791759 | 2.32308 | 2.934 |
| chr7 | 27219343 | 27219677 | 1.36615 | 4.00877 | 2.934 |
| chr5 | 43040446 | 43040757 | 1.40386 | 4.11979 | 2.935 |
| chr12 | 56074594 | 56075594 | 0.754477 | 2.21429 | 2.935 |
| chrX | 48506612 | 48506901 | 0.486236 | 1.42707 | 2.935 |
| chr12 | 101538207 | 101538531 | 0.344145 | 1.01011 | 2.935 |
| chr15 | 32038793 | 32039004 | 0.49605 | 1.45608 | 2.935 |
| chr22 | 51170555 | 51170951 | 1.50147 | 4.40738 | 2.935 |
| chrX | 107535406 | 107535659 | 0.630107 | 1.84963 | 2.935 |
| chr14 | 97319200 | 97319464 | 0.950709 | 2.79087 | 2.936 |
| chr3 | 70169535 | 70169847 | 1.4434 | 4.23723 | 2.936 |
| chr14 | 104038341 | 104038651 | 0.836108 | 2.45457 | 2.936 |
| chr22 | 30786229 | 30786529 | 0.4788 | 1.40567 | 2.936 |
| chr14 | 50778562 | 50778804 | 1.28975 | 3.78664 | 2.936 |
| chr11 | 90625294 | 90625571 | 0.424825 | 1.24727 | 2.936 |
| chr13 | 110768400 | 110768717 | 1.83106 | 5.37603 | 2.936 |

| | | | | | |
|---|---|---|---|---|---|
| chr20 | 32900498 | 32900886 | 2.44089 | 7.1668 | 2.936 |
| chr7 | 12251003 | 12251347 | 0.465906 | 1.36798 | 2.936 |
| chr11 | 122499663 | 122499988 | 1.41608 | 4.15805 | 2.936 |
| chr13 | 36857058 | 36857316 | 1.04167 | 3.05882 | 2.936 |
| chrX | 11805632 | 11806106 | 1.06979 | 3.14143 | 2.936 |
| chr12 | 29054094 | 29054324 | 0.400748 | 1.17681 | 2.937 |
| chr4 | 113683894 | 113684280 | 0.972681 | 2.85637 | 2.937 |
| chr15 | 90798193 | 90798429 | 0.375541 | 1.10303 | 2.937 |
| chr12 | 118295560 | 118295817 | 1.14859 | 3.37378 | 2.937 |
| chr15 | 91846045 | 91846398 | 1.47382 | 4.32918 | 2.937 |
| chr17 | 80781005 | 80781260 | 0.988136 | 2.90255 | 2.937 |
| chr13 | 25611418 | 25611639 | 1.06602 | 3.13142 | 2.937 |
| chr5 | 80802865 | 80803138 | 1.11784 | 3.28374 | 2.938 |
| chrX | 69653675 | 69653973 | 1.06386 | 3.12555 | 2.938 |
| chr2 | 10972465 | 10972959 | 1.72537 | 5.06909 | 2.938 |
| chr11 | 111307388 | 111307705 | 0.583236 | 1.71366 | 2.938 |
| chr12 | 65002022 | 65002300 | 1.20892 | 3.55208 | 2.938 |
| chr9 | 124300471 | 124300796 | 0.540875 | 1.58922 | 2.938 |
| chr21 | 47240499 | 47240794 | 1.2736 | 3.74224 | 2.938 |
| chr6 | 45898080 | 45898491 | 1.54235 | 4.53213 | 2.938 |
| chr14 | 72188370 | 72188575 | 0.407867 | 1.19851 | 2.938 |
| chr1 | 219559098 | 219559502 | 1.15965 | 3.40841 | 2.939 |
| chr11 | 80110703 | 80110969 | 0.425215 | 1.2499 | 2.939 |
| chr14 | 21945132 | 21945471 | 2.47654 | 7.28033 | 2.940 |
| chr1 | 32429717 | 32430165 | 2.21405 | 6.5091 | 2.940 |
| chr1 | 24194623 | 24195123 | 0.853398 | 2.50914 | 2.940 |
| chr6 | 43919038 | 43919283 | 0.826406 | 2.42978 | 2.940 |
| chr20 | 8156966 | 8157188 | 0.428561 | 1.2601 | 2.940 |
| chr9 | 38371571 | 38371840 | 0.844711 | 2.48373 | 2.940 |
| chr19 | 29506027 | 29506258 | 0.659978 | 1.94063 | 2.940 |
| chr17 | 64286279 | 64286495 | 0.486118 | 1.42943 | 2.941 |
| chr8 | 14744299 | 14744544 | 0.466267 | 1.37112 | 2.941 |
| chr8 | 140641357 | 140641606 | 0.552858 | 1.626 | 2.941 |
| chr10 | 19159934 | 19160144 | 0.500791 | 1.47293 | 2.941 |
| chr9 | 140121758 | 140122317 | 1.30993 | 3.85302 | 2.941 |
| chr9 | 3732663 | 3732909 | 1.04455 | 3.07244 | 2.941 |
| chr3 | 196249583 | 196249802 | 0.701962 | 2.06481 | 2.941 |
| chr19 | 588096 | 588438 | 0.894634 | 2.63164 | 2.942 |
| chr1 | 64239474 | 64239720 | 1.24892 | 3.67405 | 2.942 |
| chr1 | 151313353 | 151313687 | 0.912997 | 2.68587 | 2.942 |
| chr13 | 109797909 | 109798222 | 1.08121 | 3.18094 | 2.942 |
| chr4 | 113444959 | 113445391 | 1.44191 | 4.24222 | 2.942 |
| chr3 | 112841689 | 112841902 | 0.279273 | 0.821682 | 2.942 |
| chr6 | 11144672 | 11144875 | 1.13578 | 3.34173 | 2.942 |
| chr10 | 75700328 | 75700677 | 1.86681 | 5.49271 | 2.942 |
| chr14 | 45272458 | 45272801 | 0.876577 | 2.57933 | 2.943 |
| chr7 | 67102008 | 67102280 | 0.914033 | 2.6896 | 2.943 |
| chr7 | 67960938 | 67961155 | 0.346707 | 1.02023 | 2.943 |
| chr16 | 72100661 | 72100918 | 1.4872 | 4.37677 | 2.943 |
| chr5 | 134887524 | 134887953 | 1.3265 | 3.90383 | 2.943 |
| chr1 | 51086249 | 51086560 | 0.763624 | 2.24739 | 2.943 |
| chr1 | 62641327 | 62641533 | 0.737432 | 2.17061 | 2.943 |
| chr1 | 23425216 | 23425546 | 1.10522 | 3.25332 | 2.944 |
| chr13 | 102438272 | 102438539 | 0.822696 | 2.4217 | 2.944 |
| chr12 | 122056776 | 122056986 | 0.397256 | 1.16942 | 2.944 |
| chr10 | 18641138 | 18641412 | 0.327598 | 0.964369 | 2.944 |
| chr3 | 57022949 | 57023243 | 1.12535 | 3.31293 | 2.944 |
| chrX | 110090596 | 110090852 | 0.692958 | 2.04019 | 2.944 |
| chr19 | 31157080 | 31157612 | 2.44135 | 7.18863 | 2.945 |
| chr8 | 118874324 | 118874670 | 1.62178 | 4.77591 | 2.945 |
| chr11 | 1897677 | 1897954 | 1.1558 | 3.4037 | 2.945 |

| | | | | | |
|---|---|---|---|---|---|
| chr21 | 39531416 | 39531679 | 0.924288 | 2.72208 | 2.945 |
| chr14 | 96460601 | 96461047 | 1.45661 | 4.28996 | 2.945 |
| chr9 | 139030520 | 139030858 | 1.07346 | 3.16159 | 2.945 |
| chr8 | 69764522 | 69764816 | 0.495345 | 1.45893 | 2.945 |
| chr22 | 36019395 | 36019687 | 1.12575 | 3.31569 | 2.945 |
| chr1 | 152157986 | 152158355 | 1.1131 | 3.27908 | 2.946 |
| chr11 | 83889395 | 83889626 | 0.430193 | 1.26736 | 2.946 |
| chr20 | 52486412 | 52486654 | 0.982025 | 2.89324 | 2.946 |
| chr11 | 85652333 | 85652832 | 0.646043 | 1.90378 | 2.947 |
| chr18 | 7567055 | 7567487 | 0.933632 | 2.75129 | 2.947 |
| chr3 | 8811003 | 8811391 | 2.07795 | 6.12381 | 2.947 |
| chr12 | 125256362 | 125256819 | 1.48915 | 4.38893 | 2.947 |
| chr4 | 160376836 | 160377090 | 0.874424 | 2.57716 | 2.947 |
| chr17 | 59914638 | 59914894 | 1.33369 | 3.93106 | 2.948 |
| chr6 | 169890473 | 169890684 | 0.315748 | 0.930691 | 2.948 |
| chr4 | 170534343 | 170534633 | 1.13742 | 3.35287 | 2.948 |
| chr3 | 148990932 | 148991221 | 1.25958 | 3.71319 | 2.948 |
| chr9 | 90498920 | 90499227 | 0.716972 | 2.1137 | 2.948 |
| chr1 | 24077779 | 24078129 | 1.49215 | 4.39979 | 2.949 |
| chr3 | 193498591 | 193498885 | 1.21453 | 3.58145 | 2.949 |
| chr22 | 28387062 | 28387345 | 0.694233 | 2.04728 | 2.949 |
| chr21 | 44775756 | 44775996 | 0.699234 | 2.06228 | 2.949 |
| chr7 | 155191718 | 155192034 | 1.05404 | 3.10873 | 2.949 |
| chr13 | 49794463 | 49794775 | 1.01245 | 2.9861 | 2.949 |
| chr12 | 12990010 | 12990239 | 1.00661 | 2.96925 | 2.950 |
| chr15 | 99644925 | 99645354 | 2.02849 | 5.98532 | 2.951 |
| chr1 | 169681215 | 169681512 | 1.40817 | 4.15502 | 2.951 |
| chr7 | 100769871 | 100770117 | 0.578993 | 1.70845 | 2.951 |
| chr11 | 62242342 | 62242545 | 0.541181 | 1.59691 | 2.951 |
| chr5 | 178570519 | 178570774 | 0.729695 | 2.15319 | 2.951 |
| chr6 | 41159369 | 41159597 | 1.1184 | 3.30036 | 2.951 |
| chr9 | 8768250 | 8768521 | 0.3075 | 0.907429 | 2.951 |
| chr5 | 33712832 | 33713097 | 0.454938 | 1.34261 | 2.951 |
| chr6 | 3887772 | 3888003 | 0.519515 | 1.53321 | 2.951 |
| chr9 | 126196985 | 126197191 | 0.689756 | 2.03579 | 2.951 |
| chr8 | 23612799 | 23613035 | 1.13408 | 3.34723 | 2.951 |
| chr9 | 139258072 | 139258482 | 2.14081 | 6.31888 | 2.952 |
| chr1 | 241432465 | 241432760 | 0.876297 | 2.58736 | 2.953 |
| chr4 | 2415069 | 2415531 | 0.774352 | 2.28645 | 2.953 |
| chr6 | 136965291 | 136965711 | 1.49427 | 4.4122 | 2.953 |
| chr3 | 142647976 | 142648280 | 1.32102 | 3.90071 | 2.953 |
| chr3 | 73551918 | 73552168 | 1.07212 | 3.16577 | 2.953 |
| chr14 | 91369441 | 91369707 | 1.19621 | 3.5323 | 2.953 |
| chr3 | 47517404 | 47517676 | 1.32198 | 3.90406 | 2.953 |
| chr11 | 120815813 | 120816089 | 0.898776 | 2.65432 | 2.953 |
| chr17 | 41867676 | 41867992 | 1.17228 | 3.46219 | 2.953 |
| chr7 | 27190454 | 27190795 | 1.70084 | 5.02337 | 2.953 |
| chr7 | 32481603 | 32481844 | 0.339361 | 1.00229 | 2.953 |
| chr12 | 119594243 | 119594487 | 0.486577 | 1.43724 | 2.954 |
| chr10 | 32735015 | 32735252 | 0.899914 | 2.65839 | 2.954 |
| chr1 | 112912299 | 112912737 | 1.6168 | 4.77658 | 2.954 |
| chr6 | 131781137 | 131781339 | 0.807838 | 2.38663 | 2.954 |
| chr2 | 238030893 | 238031190 | 1.17066 | 3.45874 | 2.955 |
| chr5 | 79412218 | 79412545 | 0.371006 | 1.09621 | 2.955 |
| chr10 | 72139374 | 72139735 | 1.3457 | 3.97653 | 2.955 |
| chr3 | 115502865 | 115503705 | 1.01912 | 3.01158 | 2.955 |
| chr12 | 56821702 | 56821912 | 0.311537 | 0.920642 | 2.955 |
| chr16 | 69345083 | 69345445 | 1.80352 | 5.32989 | 2.955 |
| chr5 | 68630655 | 68630978 | 2.08581 | 6.16416 | 2.955 |
| chr4 | 131975875 | 131976118 | 0.689616 | 2.03802 | 2.955 |
| chr6 | 73890811 | 73891067 | 0.94108 | 2.78121 | 2.955 |

| | | | | | |
|---|---|---|---|---|---|
| chr17 | 62359393 | 62359647 | 1.68365 | 4.97578 | 2.955 |
| chr5 | 42949456 | 42949664 | 0.764821 | 2.26041 | 2.955 |
| chr8 | 10648032 | 10648275 | 0.529839 | 1.56607 | 2.956 |
| chr10 | 30236682 | 30236979 | 0.357804 | 1.05761 | 2.956 |
| chr2 | 227086477 | 227086688 | 0.353238 | 1.0443 | 2.956 |
| chr15 | 34286186 | 34286551 | 1.24635 | 3.68529 | 2.957 |
| chr8 | 123479324 | 123479803 | 1.37042 | 4.05243 | 2.957 |
| chr1 | 35318232 | 35318625 | 1.85007 | 5.47084 | 2.957 |
| chr12 | 125248858 | 125249236 | 1.2374 | 3.65948 | 2.957 |
| chr2 | 239867401 | 239867822 | 1.3027 | 3.85307 | 2.958 |
| chr7 | 45911658 | 45912076 | 1.14243 | 3.37957 | 2.958 |
| chr3 | 49025295 | 49025577 | 0.992511 | 2.9361 | 2.958 |
| chr20 | 52679344 | 52679662 | 0.901801 | 2.66796 | 2.958 |
| chr4 | 74864574 | 74864785 | 0.467788 | 1.38407 | 2.959 |
| chr5 | 21588935 | 21589311 | 0.40352 | 1.19431 | 2.960 |
| chr7 | 3202019 | 3202223 | 0.592709 | 1.75426 | 2.960 |
| chr11 | 58442956 | 58443442 | 2.16182 | 6.3991 | 2.960 |
| chr1 | 1980881 | 1981192 | 0.568076 | 1.68191 | 2.961 |
| chr22 | 49059137 | 49059433 | 0.872895 | 2.58453 | 2.961 |
| chr12 | 8059806 | 8060013 | 0.826156 | 2.44626 | 2.961 |
| chr5 | 171797620 | 171798034 | 0.99238 | 2.93847 | 2.961 |
| chr22 | 29598638 | 29598953 | 0.709294 | 2.10027 | 2.961 |
| chr12 | 40719152 | 40719509 | 0.954907 | 2.82788 | 2.961 |
| chr20 | 43325052 | 43325330 | 0.930837 | 2.75663 | 2.961 |
| chr8 | 102135470 | 102135884 | 2.13916 | 6.33521 | 2.962 |
| chr8 | 93614009 | 93614287 | 0.336725 | 0.997297 | 2.962 |
| chr10 | 79734221 | 79734496 | 0.800822 | 2.3719 | 2.962 |
| chr17 | 64621454 | 64621690 | 1.00972 | 2.99094 | 2.962 |
| chr20 | 633809 | 634187 | 1.2641 | 3.74495 | 2.963 |
| chr21 | 47343125 | 47343426 | 1.39536 | 4.13404 | 2.963 |
| chr10 | 14271701 | 14271929 | 0.334785 | 0.992054 | 2.963 |
| chr1 | 101704308 | 101704682 | 1.7853 | 5.29056 | 2.963 |
| chr17 | 25936669 | 25937108 | 0.957868 | 2.83855 | 2.963 |
| chr10 | 23529293 | 23529498 | 0.2681 | 0.794527 | 2.964 |
| chr5 | 147716608 | 147716961 | 0.676238 | 2.00409 | 2.964 |
| chr5 | 109884934 | 109885160 | 0.363005 | 1.07615 | 2.965 |
| chr10 | 50802610 | 50802898 | 1.25671 | 3.72607 | 2.965 |
| chr19 | 17970266 | 17970579 | 1.04937 | 3.11156 | 2.965 |
| chr2 | 232551964 | 232552281 | 1.03851 | 3.07943 | 2.965 |
| chr5 | 66570050 | 66570396 | 0.281821 | 0.835759 | 2.966 |
| chr16 | 65752398 | 65752724 | 0.345936 | 1.02594 | 2.966 |
| chr19 | 2474538 | 2474904 | 1.40109 | 4.15531 | 2.966 |
| chr3 | 145062798 | 145063001 | 0.463311 | 1.37409 | 2.966 |
| chr19 | 57237350 | 57237785 | 0.908708 | 2.69525 | 2.966 |
| chr14 | 21165946 | 21166198 | 1.19326 | 3.5393 | 2.966 |
| chr12 | 112617836 | 112618122 | 0.48202 | 1.42978 | 2.966 |
| chr10 | 7729188 | 7729597 | 1.88353 | 5.58726 | 2.966 |
| chr7 | 157257361 | 157257585 | 0.350756 | 1.04049 | 2.966 |
| chr10 | 38265290 | 38265883 | 0.97077 | 2.88008 | 2.967 |
| chr8 | 37788958 | 37789188 | 0.73974 | 2.19483 | 2.967 |
| chr10 | 93147140 | 93147360 | 0.492983 | 1.46285 | 2.967 |
| chr9 | 4366560 | 4367070 | 1.06286 | 3.15424 | 2.968 |
| chr1 | 98519651 | 98520023 | 1.05223 | 3.12278 | 2.968 |
| chr10 | 28721087 | 28721375 | 1.02299 | 3.03601 | 2.968 |
| chr12 | 110683296 | 110683639 | 1.45299 | 4.31215 | 2.968 |
| chr20 | 37355916 | 37356171 | 0.829163 | 2.46082 | 2.968 |
| chr6 | 44189298 | 44189586 | 1.36317 | 4.04594 | 2.968 |
| chr12 | 111742393 | 111742781 | 1.94724 | 5.78076 | 2.969 |
| chr14 | 23590008 | 23590383 | 2.75629 | 8.18338 | 2.969 |
| chr2 | 223824067 | 223824416 | 0.443663 | 1.31739 | 2.969 |
| chr16 | 14383632 | 14383965 | 1.26662 | 3.76144 | 2.970 |

| | | | | | |
|---|---|---|---|---|---|
| chr19 | 12086411 | 12086700 | 0.477859 | 1.4191 | 2.970 |
| chr1 | 152458862 | 152459245 | 0.805031 | 2.39077 | 2.970 |
| chr4 | 7852174 | 7852480 | 0.492041 | 1.46142 | 2.970 |
| chr5 | 149059647 | 149059986 | 0.388472 | 1.15389 | 2.970 |
| chr5 | 7076006 | 7076330 | 0.348847 | 1.03627 | 2.971 |
| chr9 | 114132175 | 114132410 | 0.500415 | 1.48664 | 2.971 |
| chr5 | 163048263 | 163048556 | 0.334125 | 0.992624 | 2.971 |
| chr12 | 71127182 | 71127452 | 0.794094 | 2.35974 | 2.972 |
| chr1 | 95507195 | 95507446 | 1.17876 | 3.50329 | 2.972 |
| chr15 | 87787450 | 87787679 | 0.471145 | 1.40038 | 2.972 |
| chr1 | 59234952 | 59235210 | 1.24876 | 3.71183 | 2.972 |
| chr15 | 66680174 | 66680389 | 0.965058 | 2.86927 | 2.973 |
| chr12 | 102774490 | 102774709 | 0.560946 | 1.66779 | 2.973 |
| chr16 | 11311460 | 11311703 | 0.315684 | 0.938621 | 2.973 |
| chr12 | 101670388 | 101670618 | 0.386376 | 1.14895 | 2.974 |
| chr3 | 172468253 | 172468483 | 1.31993 | 3.92508 | 2.974 |
| chr3 | 126896161 | 126896416 | 0.706299 | 2.10034 | 2.974 |
| chr5 | 10695189 | 10695546 | 1.46875 | 4.36821 | 2.974 |
| chr7 | 64348435 | 64348802 | 1.21063 | 3.60101 | 2.974 |
| chr21 | 28104812 | 28105162 | 0.683783 | 2.03391 | 2.975 |
| chr12 | 133176979 | 133177396 | 1.14349 | 3.4014 | 2.975 |
| chr10 | 36424357 | 36424641 | 0.352447 | 1.04843 | 2.975 |
| chr5 | 104041690 | 104041981 | 0.484405 | 1.44102 | 2.975 |
| chr16 | 2873505 | 2873781 | 0.532238 | 1.58346 | 2.975 |
| chr18 | 60649902 | 60650227 | 0.830022 | 2.4695 | 2.975 |
| chr12 | 49951586 | 49951878 | 0.57739 | 1.71798 | 2.975 |
| chr19 | 22610992 | 22611457 | 0.503932 | 1.49958 | 2.976 |
| chr22 | 37382211 | 37382629 | 0.798308 | 2.37564 | 2.976 |
| chr9 | 5437639 | 5437842 | 0.497867 | 1.4817 | 2.976 |
| chr1 | 45196933 | 45197313 | 1.24832 | 3.71517 | 2.976 |
| chr16 | 6734057 | 6734426 | 0.607173 | 1.80719 | 2.976 |
| chr13 | 44765611 | 44765853 | 0.652193 | 1.94128 | 2.977 |
| chr11 | 114151343 | 114151621 | 1.12398 | 3.34605 | 2.977 |
| chr19 | 59028469 | 59028694 | 0.688989 | 2.05111 | 2.977 |
| chr22 | 31343836 | 31344108 | 0.703702 | 2.09502 | 2.977 |
| chr8 | 108006458 | 108006730 | 0.411244 | 1.22444 | 2.977 |
| chr16 | 20758837 | 20759189 | 0.34398 | 1.02437 | 2.978 |
| chr1 | 202531285 | 202531522 | 0.52846 | 1.57395 | 2.978 |
| chr14 | 74926359 | 74926598 | 1.09425 | 3.25921 | 2.978 |
| chr6 | 140762494 | 140762752 | 0.670239 | 1.9963 | 2.978 |
| chr2 | 83423422 | 83423638 | 0.676555 | 2.01526 | 2.979 |
| chr6 | 3991061 | 3991432 | 1.1323 | 3.37282 | 2.979 |
| chr14 | 26724525 | 26724749 | 0.757491 | 2.25646 | 2.979 |
| chr12 | 49457716 | 49458060 | 1.60218 | 4.77296 | 2.979 |
| chr10 | 43593023 | 43593376 | 0.426195 | 1.26979 | 2.979 |
| chr1 | 33520787 | 33521508 | 0.729413 | 2.17324 | 2.979 |
| chr19 | 4400450 | 4400840 | 2.21044 | 6.586 | 2.980 |
| chr5 | 78320725 | 78321267 | 1.72287 | 5.13478 | 2.980 |
| chr6 | 56234897 | 56235155 | 0.91054 | 2.7138 | 2.980 |
| chr1 | 6417243 | 6417656 | 0.646956 | 1.92823 | 2.980 |
| chr16 | 2203516 | 2203861 | 1.61835 | 4.82413 | 2.981 |
| chrX | 73563894 | 73564170 | 1.0272 | 3.06228 | 2.981 |
| chr12 | 123122418 | 123122641 | 0.680082 | 2.02752 | 2.981 |
| chr4 | 128544004 | 128544234 | 0.68248 | 2.03469 | 2.981 |
| chr12 | 1058984 | 1059485 | 1.55063 | 4.62324 | 2.982 |
| chr8 | 94987351 | 94987785 | 1.71333 | 5.10861 | 2.982 |
| chr1 | 51339101 | 51339580 | 1.20644 | 3.59769 | 2.982 |
| chr12 | 46119505 | 46119854 | 1.41842 | 4.2299 | 2.982 |
| chr19 | 54312780 | 54313136 | 1.1117 | 3.31525 | 2.982 |
| chr3 | 42158665 | 42158993 | 1.1984 | 3.57401 | 2.982 |
| chr4 | 159174556 | 159174853 | 0.951125 | 2.83657 | 2.982 |

| | | | | | |
|---|---|---|---|---|---|
| chr14 | 69618190 | 69618604 | 1.16691 | 3.48037 | 2.983 |
| chr9 | 116269569 | 116269928 | 0.403658 | 1.20404 | 2.983 |
| chr9 | 86329345 | 86329678 | 0.600254 | 1.79051 | 2.983 |
| chr11 | 66619677 | 66619939 | 0.772377 | 2.30402 | 2.983 |
| chr8 | 15170273 | 15170507 | 0.347452 | 1.03646 | 2.983 |
| chr6 | 169920315 | 169920574 | 0.470598 | 1.40408 | 2.984 |
| chr3 | 36964971 | 36965369 | 0.875808 | 2.61317 | 2.984 |
| chr12 | 5311009 | 5311356 | 0.398336 | 1.18863 | 2.984 |
| chr11 | 15841231 | 15841621 | 1.96892 | 5.87534 | 2.984 |
| chr2 | 88607419 | 88607732 | 1.05726 | 3.15537 | 2.984 |
| chr11 | 67168922 | 67169577 | 0.723589 | 2.15956 | 2.985 |
| chr4 | 114468732 | 114469017 | 1.30718 | 3.90135 | 2.985 |
| chr10 | 70883343 | 70883603 | 0.727894 | 2.17262 | 2.985 |
| chr1 | 24397381 | 24397748 | 1.54349 | 4.60745 | 2.985 |
| chr9 | 38708055 | 38708268 | 1.11678 | 3.33409 | 2.985 |
| chr3 | 12176516 | 12176920 | 1.75956 | 5.25316 | 2.986 |
| chr4 | 169844923 | 169845197 | 1.21689 | 3.63306 | 2.986 |
| chr17 | 66011974 | 66012207 | 0.921246 | 2.75058 | 2.986 |
| chr17 | 43354564 | 43354980 | 1.67619 | 5.00496 | 2.986 |
| chr12 | 12188957 | 12189329 | 1.23389 | 3.68434 | 2.986 |
| chr16 | 31870739 | 31871077 | 1.09325 | 3.26467 | 2.986 |
| chr9 | 72546220 | 72546579 | 0.984203 | 2.93908 | 2.986 |
| chr3 | 36739185 | 36739437 | 0.868849 | 2.59472 | 2.986 |
| chr8 | 53649692 | 53649974 | 0.975718 | 2.91491 | 2.987 |
| chr17 | 52114283 | 52114641 | 0.470392 | 1.40529 | 2.987 |
| chr6 | 34383069 | 34383396 | 0.633482 | 1.89252 | 2.987 |
| chr14 | 49005554 | 49005806 | 0.656121 | 1.96029 | 2.988 |
| chr3 | 101947916 | 101948300 | 1.38386 | 4.13496 | 2.988 |
| chr19 | 35644591 | 35644952 | 1.15151 | 3.44091 | 2.988 |
| chr3 | 24306205 | 24306499 | 1.18167 | 3.53137 | 2.988 |
| chr6 | 125513535 | 125513799 | 0.778808 | 2.32762 | 2.989 |
| chr19 | 47713982 | 47714341 | 1.4043 | 4.19707 | 2.989 |
| chr12 | 88061270 | 88061474 | 0.796863 | 2.38191 | 2.989 |
| chr12 | 58119784 | 58120171 | 1.90587 | 5.6977 | 2.990 |
| chr10 | 106113362 | 106113627 | 0.954598 | 2.85392 | 2.990 |
| chr8 | 60032130 | 60032515 | 1.45051 | 4.33671 | 2.990 |
| chr3 | 113296788 | 113297046 | 1.26434 | 3.78034 | 2.990 |
| chr15 | 22946787 | 22947011 | 0.385202 | 1.15175 | 2.990 |
| chr12 | 79814017 | 79814269 | 0.903695 | 2.70214 | 2.990 |
| chr5 | 150336157 | 150336368 | 0.463737 | 1.38677 | 2.990 |
| chr3 | 133206807 | 133207097 | 0.888249 | 2.65627 | 2.990 |
| chr3 | 145516393 | 145516634 | 0.401082 | 1.19955 | 2.991 |
| chr17 | 60755701 | 60756007 | 0.350522 | 1.04837 | 2.991 |
| chr8 | 32411633 | 32411930 | 0.466944 | 1.39673 | 2.991 |
| chr19 | 45303434 | 45304026 | 1.4998 | 4.48641 | 2.991 |
| chr8 | 72756093 | 72756323 | 0.430352 | 1.28736 | 2.991 |
| chr8 | 136452710 | 136453038 | 0.389417 | 1.16495 | 2.992 |
| chr8 | 21481513 | 21481717 | 0.786006 | 2.35139 | 2.992 |
| chr12 | 59740100 | 59740320 | 0.903865 | 2.70437 | 2.992 |
| chr22 | 36673743 | 36674248 | 0.88698 | 2.654 | 2.992 |
| chr19 | 53238320 | 53238570 | 0.76367 | 2.28544 | 2.993 |
| chr1 | 157506073 | 157506283 | 0.319715 | 0.956936 | 2.993 |
| chr17 | 1811995 | 1812364 | 1.36914 | 4.0984 | 2.993 |
| chr20 | 3696937 | 3697304 | 1.53076 | 4.58228 | 2.993 |
| chr10 | 105541023 | 105541296 | 1.13874 | 3.40907 | 2.994 |
| chr11 | 70023382 | 70023774 | 1.89042 | 5.65958 | 2.994 |
| chr7 | 25932456 | 25932758 | 0.997151 | 2.98532 | 2.994 |
| chr5 | 174008547 | 174008787 | 1.19878 | 3.58919 | 2.994 |
| chr14 | 88929735 | 88930030 | 0.72702 | 2.17687 | 2.994 |
| chr14 | 79587376 | 79587613 | 0.409568 | 1.22641 | 2.994 |
| chr7 | 15055756 | 15056032 | 0.99476 | 2.97891 | 2.995 |

| | | | | | |
|---|---|---|---|---|---|
| chr1 | 161708616 | 161708845 | 0.778616 | 2.33209 | 2.995 |
| chr18 | 8367474 | 8367725 | 0.817629 | 2.44933 | 2.996 |
| chr20 | 56788270 | 56788480 | 0.803809 | 2.40812 | 2.996 |
| chr8 | 35406788 | 35407032 | 0.379931 | 1.13829 | 2.996 |
| chr21 | 38954897 | 38955190 | 1.16499 | 3.49082 | 2.996 |
| chr4 | 48277252 | 48277602 | 0.686586 | 2.05733 | 2.996 |
| chr1 | 222269394 | 222269929 | 1.06754 | 3.19896 | 2.997 |
| chr12 | 42523471 | 42523746 | 1.10979 | 3.32562 | 2.997 |
| chr1 | 11809285 | 11809605 | 1.12827 | 3.38178 | 2.997 |
| chr12 | 56652059 | 56652447 | 1.42454 | 4.27015 | 2.998 |
| chr15 | 74426596 | 74427265 | 0.720752 | 2.16054 | 2.998 |
| chr17 | 42115578 | 42115802 | 0.400519 | 1.20065 | 2.998 |
| chr3 | 102464828 | 102465053 | 0.642327 | 1.9256 | 2.998 |
| chr11 | 14062678 | 14062891 | 0.754169 | 2.26093 | 2.998 |
| chr21 | 36262246 | 36262727 | 0.982592 | 2.94578 | 2.998 |
| chr19 | 28237887 | 28238251 | 0.891137 | 2.67164 | 2.998 |
| chr7 | 55347611 | 55347934 | 0.874302 | 2.62122 | 2.998 |
| chr10 | 100022304 | 100022791 | 0.994239 | 2.98081 | 2.998 |
| chr5 | 149155374 | 149155695 | 0.830763 | 2.49077 | 2.998 |
| chr17 | 53426145 | 53426470 | 2.18777 | 6.55939 | 2.998 |
| chr17 | 79163533 | 79163748 | 0.78483 | 2.35321 | 2.998 |
| chr1 | 177995292 | 177995499 | 0.462072 | 1.38548 | 2.998 |
| chr17 | 64902815 | 64903119 | 1.7204 | 5.1592 | 2.999 |
| chr13 | 46700764 | 46701036 | 1.29721 | 3.89057 | 2.999 |
| chr19 | 8804664 | 8805013 | 1.30941 | 3.92723 | 2.999 |
| chr12 | 46465910 | 46466260 | 2.33582 | 7.0057 | 2.999 |
| chr18 | 72753561 | 72753840 | 0.738209 | 2.21422 | 2.999 |
| chr6 | 78167509 | 78167831 | 0.693338 | 2.07968 | 3.000 |
| chr1 | 176581310 | 176581713 | 0.304536 | 0.913478 | 3.000 |
| chr8 | 99032197 | 99032427 | 0.628091 | 1.884 | 3.000 |
| chr7 | 151689409 | 151689685 | 0.643228 | 1.92951 | 3.000 |
| chr9 | 5450450 | 5450730 | 1.29049 | 3.87116 | 3.000 |
| chr15 | 55818718 | 55819073 | 1.01999 | 3.06006 | 3.000 |
| chr10 | 116231189 | 116231398 | 1.27149 | 3.81466 | 3.000 |
| chr17 | 75453964 | 75454271 | 0.401412 | 1.20468 | 3.001 |
| chr14 | 54805449 | 54805767 | 1.22815 | 3.68601 | 3.001 |
| chr1 | 201673320 | 201673726 | 2.37187 | 7.11865 | 3.001 |
| chr3 | 123270941 | 123271378 | 0.978635 | 2.93722 | 3.001 |
| chr9 | 120249753 | 120250185 | 2.12199 | 6.36911 | 3.001 |
| chr3 | 148661408 | 148661696 | 0.896964 | 2.69231 | 3.002 |
| chr12 | 126678789 | 126679005 | 0.458429 | 1.37602 | 3.002 |
| chr12 | 64660184 | 64660416 | 0.635117 | 1.9066 | 3.002 |
| chr1 | 207494438 | 207494649 | 0.799856 | 2.40138 | 3.002 |
| chr17 | 34336199 | 34336911 | 0.95084 | 2.85483 | 3.002 |
| chr20 | 6138075 | 6138350 | 0.77752 | 2.33469 | 3.003 |
| chr16 | 12549054 | 12549563 | 2.22648 | 6.68565 | 3.003 |
| chr20 | 31109558 | 31109887 | 1.88968 | 5.67508 | 3.003 |
| chr16 | 49383610 | 49383820 | 0.382371 | 1.14846 | 3.004 |
| chr15 | 101458155 | 101458544 | 1.60775 | 4.82909 | 3.004 |
| chr20 | 57525620 | 57526220 | 0.319099 | 0.958509 | 3.004 |
| chr16 | 3011548 | 3012150 | 1.39656 | 4.19517 | 3.004 |
| chr4 | 2187902 | 2188383 | 0.695004 | 2.08796 | 3.004 |
| chr7 | 144151616 | 144151839 | 0.611886 | 1.83852 | 3.005 |
| chr8 | 87905167 | 87905432 | 0.980283 | 2.94557 | 3.005 |
| chr15 | 65596583 | 65596852 | 1.44001 | 4.32728 | 3.005 |
| chr1 | 19147212 | 19147635 | 1.54243 | 4.63517 | 3.005 |
| chr10 | 28591545 | 28591873 | 0.6388 | 1.91968 | 3.005 |
| chr8 | 132907184 | 132907429 | 0.29027 | 0.872377 | 3.005 |
| chr6 | 100315998 | 100316272 | 1.0265 | 3.08541 | 3.006 |
| chr1 | 111038656 | 111039028 | 1.60903 | 4.83637 | 3.006 |
| chr12 | 69864681 | 69864892 | 0.98032 | 2.94673 | 3.006 |

| | | | | | |
|---|---|---|---|---|---|
| chr5 | 39236860 | 39237132 | 0.326255 | 0.98069 | 3.006 |
| chr19 | 8333727 | 8334134 | 1.04261 | 3.13403 | 3.006 |
| chr22 | 40018921 | 40019172 | 1.10713 | 3.32803 | 3.006 |
| chr6 | 35310022 | 35310385 | 1.75557 | 5.27788 | 3.006 |
| chr5 | 16438203 | 16438441 | 0.504907 | 1.51798 | 3.006 |
| chr10 | 21202830 | 21203100 | 0.404884 | 1.21735 | 3.007 |
| chr10 | 23277822 | 23278120 | 1.05834 | 3.18217 | 3.007 |
| chr10 | 60290880 | 60291227 | 0.824793 | 2.48022 | 3.007 |
| chr12 | 2750105 | 2750628 | 0.447715 | 1.34637 | 3.007 |
| chr11 | 75162470 | 75162889 | 1.53165 | 4.60623 | 3.007 |
| chr3 | 184278999 | 184279347 | 1.73826 | 5.22832 | 3.008 |
| chr8 | 15420174 | 15420439 | 0.782401 | 2.35338 | 3.008 |
| chr17 | 29972309 | 29972623 | 0.707518 | 2.12818 | 3.008 |
| chr5 | 111575787 | 111576001 | 0.822367 | 2.47387 | 3.008 |
| chr7 | 147939372 | 147939779 | 1.16874 | 3.51614 | 3.008 |
| chr9 | 129361399 | 129361626 | 0.681287 | 2.05016 | 3.009 |
| chr21 | 45909895 | 45910187 | 0.511326 | 1.53876 | 3.009 |
| chr8 | 145669344 | 145669823 | 1.05261 | 3.16769 | 3.009 |
| chr19 | 7188036 | 7188247 | 0.350751 | 1.05566 | 3.010 |
| chr15 | 81071224 | 81071787 | 1.56174 | 4.70042 | 3.010 |
| chr10 | 34236060 | 34236266 | 0.319402 | 0.961413 | 3.010 |
| chr5 | 78760589 | 78760979 | 0.852572 | 2.56649 | 3.010 |
| chr14 | 38063435 | 38063769 | 1.03032 | 3.10162 | 3.010 |
| chr10 | 5476038 | 5476291 | 1.34075 | 4.03636 | 3.011 |
| chr12 | 45941567 | 45941773 | 0.440226 | 1.32532 | 3.011 |
| chr10 | 97742632 | 97742991 | 0.298907 | 0.899934 | 3.011 |
| chr9 | 35116831 | 35117221 | 1.6844 | 5.07133 | 3.011 |
| chr7 | 71132895 | 71133140 | 0.381464 | 1.14858 | 3.011 |
| chr14 | 31676908 | 31677277 | 1.34952 | 4.06362 | 3.011 |
| chr9 | 116810033 | 116810326 | 0.982153 | 2.95749 | 3.011 |
| chr11 | 66032438 | 66032723 | 0.987854 | 2.97469 | 3.011 |
| chr15 | 81611116 | 81611562 | 0.396611 | 1.19444 | 3.012 |
| chr22 | 45713609 | 45713813 | 0.543083 | 1.63589 | 3.012 |
| chr11 | 46884574 | 46884940 | 0.362032 | 1.09053 | 3.012 |
| chr8 | 134221462 | 134221920 | 1.61786 | 4.87378 | 3.012 |
| chr5 | 990216 | 990505 | 0.475828 | 1.43353 | 3.013 |
| chr12 | 82720755 | 82721167 | 0.79698 | 2.40118 | 3.013 |
| chr22 | 47392572 | 47392786 | 0.593658 | 1.78872 | 3.013 |
| chr12 | 84096503 | 84096752 | 0.409808 | 1.23484 | 3.013 |
| chr12 | 89431469 | 89431733 | 0.443017 | 1.33494 | 3.013 |
| chr1 | 199391413 | 199391658 | 0.708595 | 2.13532 | 3.013 |
| chr12 | 88974076 | 88974351 | 0.870271 | 2.62255 | 3.013 |
| chr12 | 101045742 | 101046075 | 0.543238 | 1.63706 | 3.014 |
| chr14 | 88211931 | 88212221 | 0.932408 | 2.81008 | 3.014 |
| chr22 | 23470624 | 23470887 | 0.825927 | 2.48937 | 3.014 |
| chr4 | 10016889 | 10017158 | 0.821386 | 2.47593 | 3.014 |
| chr3 | 109115322 | 109115543 | 0.764469 | 2.30438 | 3.014 |
| chr11 | 95701865 | 95702078 | 0.658873 | 1.98619 | 3.015 |
| chr2 | 149684235 | 149684558 | 1.33696 | 4.03043 | 3.015 |
| chr8 | 146030485 | 146030797 | 1.09581 | 3.30478 | 3.016 |
| chr8 | 94973148 | 94973515 | 1.09862 | 3.3133 | 3.016 |
| chr7 | 23125666 | 23126017 | 1.29663 | 3.91081 | 3.016 |
| chr11 | 111317361 | 111317652 | 1.15211 | 3.47494 | 3.016 |
| chr3 | 195800126 | 195800462 | 1.01988 | 3.07611 | 3.016 |
| chr5 | 14277931 | 14278154 | 0.391693 | 1.18144 | 3.016 |
| chr17 | 41955823 | 41956361 | 2.07199 | 6.24979 | 3.016 |
| chrX | 5154295 | 5154573 | 0.727333 | 2.19417 | 3.017 |
| chr3 | 194980719 | 194981088 | 0.933475 | 2.81627 | 3.017 |
| chrX | 119586042 | 119586304 | 0.799905 | 2.41349 | 3.017 |
| chr22 | 22785884 | 22786090 | 1.21441 | 3.66421 | 3.017 |
| chr9 | 122541439 | 122541723 | 0.319622 | 0.964563 | 3.018 |

| | | | | | |
|---|---|---|---|---|---|
| chr5 | 21269948 | 21270239 | 0.349764 | 1.05559 | 3.018 |
| chr1 | 184774728 | 184775016 | 1.50174 | 4.53235 | 3.018 |
| chr11 | 125298994 | 125299321 | 0.992911 | 2.99667 | 3.018 |
| chr1 | 15480318 | 15480884 | 0.730611 | 2.20509 | 3.018 |
| chr1 | 11424321 | 11424539 | 0.669028 | 2.01944 | 3.018 |
| chr12 | 50505261 | 50505657 | 1.3948 | 4.21044 | 3.019 |
| chr4 | 52697381 | 52697606 | 0.996402 | 3.00857 | 3.019 |
| chr13 | 82061373 | 82061643 | 0.694436 | 2.09683 | 3.019 |
| chr9 | 35258870 | 35259158 | 1.66519 | 5.02932 | 3.020 |
| chr7 | 114179709 | 114179936 | 0.617943 | 1.8665 | 3.021 |
| chr1 | 204400432 | 204400780 | 1.57878 | 4.76944 | 3.021 |
| chr1 | 55257588 | 55257942 | 0.865866 | 2.61585 | 3.021 |
| chr1 | 171685052 | 171685440 | 0.693245 | 2.09441 | 3.021 |
| chr16 | 32639586 | 32639807 | 0.383046 | 1.15728 | 3.021 |
| chr17 | 66687900 | 66688140 | 0.504537 | 1.52435 | 3.021 |
| chr13 | 23314467 | 23314689 | 0.799344 | 2.41533 | 3.022 |
| chr7 | 156266129 | 156266480 | 1.46087 | 4.41447 | 3.022 |
| chr10 | 24931854 | 24932093 | 0.619843 | 1.87317 | 3.022 |
| chr10 | 11895514 | 11895778 | 0.942267 | 2.84776 | 3.022 |
| chr9 | 100672308 | 100672521 | 0.488784 | 1.47731 | 3.022 |
| chr14 | 95015707 | 95016252 | 1.37492 | 4.15625 | 3.023 |
| chr16 | 56707808 | 56708194 | 0.899983 | 2.72059 | 3.023 |
| chr13 | 32575468 | 32575797 | 1.13744 | 3.43841 | 3.023 |
| chr11 | 131564694 | 131565059 | 1.80027 | 5.44244 | 3.023 |
| chr13 | 61201163 | 61201528 | 0.657781 | 1.98856 | 3.023 |
| chr16 | 10762192 | 10762621 | 1.21613 | 3.67692 | 3.023 |
| chr10 | 107316192 | 107316481 | 0.403527 | 1.22005 | 3.023 |
| chr12 | 12044995 | 12045240 | 0.767606 | 2.32095 | 3.024 |
| chr11 | 69612073 | 69612351 | 0.629396 | 1.90306 | 3.024 |
| chr21 | 46560241 | 46560576 | 0.952172 | 2.87957 | 3.024 |
| chr1 | 45195009 | 45195300 | 1.34418 | 4.0651 | 3.024 |
| chr7 | 37699904 | 37700140 | 0.385313 | 1.16533 | 3.024 |
| chr5 | 154708745 | 154708971 | 0.370431 | 1.12046 | 3.025 |
| chr5 | 89357009 | 89357475 | 0.36437 | 1.10214 | 3.025 |
| chr1 | 229390767 | 229390972 | 0.805118 | 2.4354 | 3.025 |
| chr20 | 11534294 | 11534543 | 0.871295 | 2.63559 | 3.025 |
| chr17 | 39967803 | 39968246 | 1.40284 | 4.24442 | 3.026 |
| chr14 | 54130432 | 54130700 | 0.936057 | 2.83213 | 3.026 |
| chr20 | 51390769 | 51390992 | 0.454072 | 1.37385 | 3.026 |
| chr1 | 110648767 | 110649095 | 1.66511 | 5.0381 | 3.026 |
| chr1 | 113351820 | 113352254 | 1.35675 | 4.10537 | 3.026 |
| chr12 | 126218835 | 126219272 | 0.742298 | 2.24629 | 3.026 |
| chr13 | 40176659 | 40176972 | 0.569669 | 1.72392 | 3.026 |
| chr17 | 49553152 | 49553394 | 0.385348 | 1.16615 | 3.026 |
| chr1 | 193090897 | 193091604 | 1.12604 | 3.40867 | 3.027 |
| chr8 | 23238350 | 23238676 | 0.538916 | 1.63139 | 3.027 |
| chr9 | 75423145 | 75423354 | 0.693971 | 2.10115 | 3.028 |
| chr7 | 75590599 | 75590871 | 0.6409 | 1.94063 | 3.028 |
| chr4 | 96126077 | 96126321 | 0.871551 | 2.63909 | 3.028 |
| chr18 | 10877851 | 10878171 | 0.417472 | 1.26424 | 3.028 |
| chr14 | 93579161 | 93579451 | 1.05446 | 3.19331 | 3.028 |
| chr15 | 57113958 | 57114281 | 1.16337 | 3.52324 | 3.028 |
| chr4 | 165700947 | 165701250 | 1.3066 | 3.95725 | 3.029 |
| chr9 | 73900073 | 73900434 | 1.78502 | 5.40632 | 3.029 |
| chr17 | 27041473 | 27041681 | 0.681653 | 2.06475 | 3.029 |
| chr20 | 43017214 | 43017541 | 1.549 | 4.69203 | 3.029 |
| chr14 | 88634839 | 88635261 | 1.55009 | 4.69576 | 3.029 |
| chr19 | 5535014 | 5535315 | 1.00996 | 3.05969 | 3.030 |
| chr5 | 43484277 | 43484701 | 1.2599 | 3.81703 | 3.030 |
| chr18 | 18772020 | 18772320 | 1.30095 | 3.94171 | 3.030 |
| chr1 | 229732887 | 229733158 | 0.527405 | 1.59805 | 3.030 |

| | | | | | |
|---|---|---|---|---|---|
| chr4 | 146478786 | 146479163 | 0.960564 | 2.91053 | 3.030 |
| chr19 | 6551979 | 6552331 | 0.566281 | 1.71597 | 3.030 |
| chr22 | 30787161 | 30787396 | 1.38771 | 4.20514 | 3.030 |
| chr19 | 5967941 | 5968396 | 1.04688 | 3.17286 | 3.031 |
| chr15 | 32195969 | 32196176 | 0.420448 | 1.27444 | 3.031 |
| chr3 | 131748106 | 131748452 | 1.59948 | 4.84859 | 3.031 |
| chr1 | 204472368 | 204472587 | 0.776228 | 2.35313 | 3.031 |
| chr1 | 204721004 | 204721386 | 0.611672 | 1.85442 | 3.032 |
| chr10 | 67526290 | 67526494 | 0.432875 | 1.3124 | 3.032 |
| chr17 | 53930189 | 53930438 | 0.668511 | 2.02681 | 3.032 |
| chr19 | 14134271 | 14134599 | 1.17616 | 3.56605 | 3.032 |
| chr3 | 138418097 | 138418333 | 0.768314 | 2.32968 | 3.032 |
| chr12 | 73451331 | 73451567 | 0.590599 | 1.79098 | 3.032 |
| chr15 | 81288616 | 81288977 | 1.05451 | 3.19796 | 3.033 |
| chr1 | 177154345 | 177154661 | 0.645392 | 1.95741 | 3.033 |
| chr2 | 44471265 | 44471491 | 1.02829 | 3.11871 | 3.033 |
| chr12 | 4253112 | 4253481 | 0.6451 | 1.95658 | 3.033 |
| chr16 | 74903609 | 74903868 | 0.622688 | 1.88867 | 3.033 |
| chr6 | 35417221 | 35417458 | 1.00941 | 3.06209 | 3.034 |
| chr12 | 29959107 | 29959395 | 0.282895 | 0.858414 | 3.034 |
| chr9 | 92864641 | 92864977 | 0.918754 | 2.788 | 3.035 |
| chr4 | 54663746 | 54663949 | 0.367946 | 1.11682 | 3.035 |
| chr9 | 140502945 | 140503182 | 0.339853 | 1.03157 | 3.035 |
| chr22 | 18507281 | 18507489 | 1.37553 | 4.17576 | 3.036 |
| chr10 | 124789264 | 124789506 | 0.417161 | 1.26645 | 3.036 |
| chr19 | 2316002 | 2316315 | 1.02935 | 3.12531 | 3.036 |
| chr12 | 12965855 | 12966396 | 1.50461 | 4.56835 | 3.036 |
| chr3 | 146876903 | 146877193 | 0.365307 | 1.10921 | 3.036 |
| chr4 | 65024439 | 65024743 | 0.875972 | 2.65987 | 3.036 |
| chr4 | 84283730 | 84284105 | 0.449065 | 1.36368 | 3.037 |
| chrX | 152917277 | 152917491 | 0.703144 | 2.13533 | 3.037 |
| chr5 | 44536849 | 44537099 | 0.839212 | 2.54856 | 3.037 |
| chr16 | 88751598 | 88751927 | 0.93231 | 2.83132 | 3.037 |
| chr22 | 26898922 | 26899326 | 0.702962 | 2.13483 | 3.037 |
| chr14 | 66943878 | 66944260 | 0.333934 | 1.01413 | 3.037 |
| chr14 | 69595990 | 69596378 | 1.91455 | 5.81437 | 3.037 |
| chr7 | 22143345 | 22143685 | 1.1558 | 3.51021 | 3.037 |
| chr11 | 86581054 | 86581376 | 1.44078 | 4.37583 | 3.037 |
| chr9 | 21251699 | 21251959 | 0.429417 | 1.30426 | 3.037 |
| chr14 | 77745231 | 77745501 | 0.400817 | 1.21752 | 3.038 |
| chr16 | 56390242 | 56390668 | 1.78557 | 5.42409 | 3.038 |
| chr12 | 120645053 | 120645332 | 0.416873 | 1.26649 | 3.038 |
| chr19 | 8412611 | 8412933 | 0.93941 | 2.85409 | 3.038 |
| chr21 | 46690154 | 46690451 | 0.844578 | 2.56647 | 3.039 |
| chr12 | 12480793 | 12481057 | 0.867308 | 2.63579 | 3.039 |
| chr1 | 241354158 | 241354377 | 0.35209 | 1.07005 | 3.039 |
| chr5 | 149838408 | 149838655 | 2.35096 | 7.14491 | 3.039 |
| chr14 | 39886270 | 39886516 | 0.752504 | 2.28701 | 3.039 |
| chr9 | 25716319 | 25716570 | 0.640315 | 1.9461 | 3.039 |
| chr3 | 108994567 | 108994932 | 1.55883 | 4.73783 | 3.039 |
| chr3 | 5017854 | 5018153 | 1.56817 | 4.76634 | 3.039 |
| chr11 | 22646561 | 22646871 | 1.00186 | 3.04552 | 3.040 |
| chr18 | 10520142 | 10520392 | 1.08861 | 3.30933 | 3.040 |
| chr11 | 2229992 | 2230267 | 0.814052 | 2.47474 | 3.040 |
| chr7 | 32886324 | 32886589 | 0.889017 | 2.70269 | 3.040 |
| chr14 | 105559965 | 105560336 | 0.942655 | 2.86577 | 3.040 |
| chr12 | 57856442 | 57856845 | 1.77341 | 5.39159 | 3.040 |
| chr1 | 92775602 | 92775949 | 0.546088 | 1.66051 | 3.041 |
| chr1 | 25015037 | 25015367 | 1.40341 | 4.26745 | 3.041 |
| chr3 | 196213181 | 196213502 | 0.837892 | 2.54809 | 3.041 |
| chr16 | 79055767 | 79056057 | 0.350848 | 1.06697 | 3.041 |

| | | | | | |
|---|---|---|---|---|---|
| chr7 | 115388299 | 115388531 | 0.642909 | 1.95535 | 3.041 |
| chr15 | 74614934 | 74615335 | 1.40928 | 4.28639 | 3.042 |
| chr9 | 78309853 | 78310100 | 0.340319 | 1.03512 | 3.042 |
| chr16 | 29721622 | 29722036 | 0.987138 | 3.0026 | 3.042 |
| chr14 | 91470412 | 91470781 | 0.29764 | 0.905343 | 3.042 |
| chr5 | 61113569 | 61113898 | 1.05509 | 3.20948 | 3.042 |
| chr10 | 65225135 | 65225358 | 0.443342 | 1.3488 | 3.042 |
| chr17 | 78226658 | 78227049 | 1.31771 | 4.00893 | 3.042 |
| chr6 | 122901995 | 122902288 | 0.409671 | 1.24642 | 3.042 |
| chr15 | 66990605 | 66991126 | 1.52269 | 4.63281 | 3.043 |
| chr12 | 104752172 | 104752466 | 1.13391 | 3.45002 | 3.043 |
| chr5 | 19544780 | 19545053 | 0.529247 | 1.61033 | 3.043 |
| chr7 | 15047196 | 15047422 | 0.830844 | 2.52805 | 3.043 |
| chr5 | 56359114 | 56359327 | 0.344455 | 1.04818 | 3.043 |
| chr11 | 125034314 | 125034612 | 0.827392 | 2.51787 | 3.043 |
| chr4 | 7072456 | 7072881 | 1.50605 | 4.58406 | 3.044 |
| chr2 | 71211942 | 71212401 | 1.83135 | 5.57438 | 3.044 |
| chr7 | 103987771 | 103988010 | 0.348828 | 1.06182 | 3.044 |
| chr19 | 48837134 | 48837558 | 1.4013 | 4.26558 | 3.044 |
| chr2 | 40160404 | 40160644 | 0.409517 | 1.24666 | 3.044 |
| chr6 | 6057758 | 6058003 | 0.569604 | 1.73403 | 3.044 |
| chr16 | 89250578 | 89250908 | 0.391177 | 1.19088 | 3.044 |
| chr12 | 45111577 | 45111836 | 0.579469 | 1.76412 | 3.044 |
| chr1 | 87597371 | 87597812 | 1.34378 | 4.09138 | 3.045 |
| chr8 | 9618063 | 9618316 | 1.06698 | 3.24861 | 3.045 |
| chr5 | 180097583 | 180097924 | 1.40241 | 4.26995 | 3.045 |
| chr9 | 136591543 | 136591754 | 1.40398 | 4.27521 | 3.045 |
| chr12 | 2170605 | 2170909 | 0.451211 | 1.37399 | 3.045 |
| chr1 | 87490946 | 87491178 | 0.585721 | 1.78399 | 3.046 |
| chr3 | 21631569 | 21631831 | 0.341979 | 1.04166 | 3.046 |
| chr16 | 72536038 | 72536417 | 1.20442 | 3.66915 | 3.046 |
| chr15 | 43803822 | 43804030 | 0.699874 | 2.13212 | 3.046 |
| chr12 | 56130853 | 56131297 | 1.88837 | 5.75323 | 3.047 |
| chr5 | 148520891 | 148521245 | 2.37243 | 7.2282 | 3.047 |
| chr3 | 106592091 | 106592335 | 0.297372 | 0.906057 | 3.047 |
| chr3 | 169339009 | 169339333 | 0.328823 | 1.00189 | 3.047 |
| chr8 | 36694170 | 36694448 | 0.363281 | 1.10692 | 3.047 |
| chr13 | 23309647 | 23309855 | 0.459913 | 1.40143 | 3.047 |
| chr7 | 127121095 | 127121340 | 0.281332 | 0.857265 | 3.047 |
| chr9 | 111713389 | 111713766 | 1.08736 | 3.31345 | 3.047 |
| chr9 | 131790756 | 131791121 | 2.04074 | 6.22015 | 3.048 |
| chr17 | 77899070 | 77899445 | 0.555446 | 1.69306 | 3.048 |
| chr1 | 178501263 | 178501792 | 0.899468 | 2.74184 | 3.048 |
| chr12 | 129629812 | 129630116 | 0.318788 | 0.971804 | 3.048 |
| chr5 | 129477431 | 129477803 | 0.975091 | 2.97259 | 3.049 |
| chr14 | 60981650 | 60982123 | 1.16667 | 3.55666 | 3.049 |
| chr5 | 158841826 | 158842065 | 0.380792 | 1.16088 | 3.049 |
| chr6 | 135857113 | 135857390 | 0.682634 | 2.08142 | 3.049 |
| chr14 | 90807551 | 90807836 | 0.289313 | 0.882237 | 3.049 |
| chr3 | 46530152 | 46530603 | 1.83446 | 5.5943 | 3.050 |
| chr15 | 95663737 | 95663943 | 0.399498 | 1.2183 | 3.050 |
| chr15 | 72335215 | 72335461 | 0.662489 | 2.02049 | 3.050 |
| chr7 | 3240164 | 3240500 | 1.2847 | 3.91816 | 3.050 |
| chr7 | 75920897 | 75921264 | 1.37769 | 4.20176 | 3.050 |
| chr21 | 30848808 | 30849108 | 1.5234 | 4.64675 | 3.050 |
| chr11 | 22809151 | 22809401 | 1.09987 | 3.35535 | 3.051 |
| chr3 | 125917083 | 125917405 | 1.19665 | 3.65069 | 3.051 |
| chr11 | 124737706 | 124737937 | 0.613613 | 1.87206 | 3.051 |
| chr1 | 202153847 | 202154155 | 1.64837 | 5.02916 | 3.051 |
| chr17 | 25821247 | 25821600 | 0.93968 | 2.86697 | 3.051 |
| chr1 | 76098091 | 76098390 | 0.886621 | 2.70543 | 3.051 |

| | | | | | |
|---|---|---|---|---|---|
| chr3 | 158362063 | 158362395 | 2.11406 | 6.45112 | 3.052 |
| chr19 | 58535439 | 58535656 | 0.64949 | 1.98227 | 3.052 |
| chr6 | 80487742 | 80488297 | 0.938037 | 2.86295 | 3.052 |
| chr6 | 137335043 | 137335307 | 0.72612 | 2.21631 | 3.052 |
| chr15 | 102148376 | 102148788 | 0.403229 | 1.23078 | 3.052 |
| chr8 | 11603880 | 11604290 | 0.955338 | 2.91602 | 3.052 |
| chr12 | 65333704 | 65334029 | 0.459718 | 1.40322 | 3.052 |
| chr1 | 210465740 | 210466182 | 0.93491 | 2.85372 | 3.052 |
| chr19 | 34849995 | 34850235 | 1.20658 | 3.68301 | 3.052 |
| chr7 | 102937658 | 102937987 | 1.58779 | 4.84727 | 3.053 |
| chr4 | 177198814 | 177199025 | 1.1805 | 3.60401 | 3.053 |
| chr12 | 56600278 | 56600659 | 1.3514 | 4.12584 | 3.053 |
| chr12 | 113900112 | 113900541 | 2.02822 | 6.19232 | 3.053 |
| chr3 | 173040089 | 173040374 | 0.351658 | 1.07368 | 3.053 |
| chr8 | 24245191 | 24245472 | 1.3066 | 3.98943 | 3.053 |
| chr8 | 145873470 | 145873736 | 0.619274 | 1.89101 | 3.054 |
| chr15 | 55509792 | 55510129 | 1.0322 | 3.15202 | 3.054 |
| chr5 | 180038229 | 180038461 | 0.897266 | 2.74026 | 3.054 |
| chr2 | 102868127 | 102868454 | 1.15695 | 3.53364 | 3.054 |
| chr11 | 71894387 | 71894840 | 1.5085 | 4.60768 | 3.054 |
| chr16 | 83960242 | 83960640 | 2.07311 | 6.33251 | 3.055 |
| chr19 | 6504872 | 6505104 | 0.891112 | 2.7223 | 3.055 |
| chr4 | 142632483 | 142632697 | 0.567958 | 1.73519 | 3.055 |
| chr7 | 143598797 | 143599092 | 0.648282 | 1.9807 | 3.055 |
| chr9 | 139294560 | 139294957 | 1.98678 | 6.07073 | 3.056 |
| chr5 | 55777620 | 55777855 | 0.634476 | 1.93881 | 3.056 |
| chr7 | 88601125 | 88601334 | 0.36871 | 1.12684 | 3.056 |
| chr1 | 236670442 | 236670696 | 0.375554 | 1.14776 | 3.056 |
| chr12 | 129463742 | 129463960 | 0.279775 | 0.855086 | 3.056 |
| chr20 | 5081416 | 5081720 | 1.77414 | 5.42249 | 3.056 |
| chr19 | 10360751 | 10361107 | 1.60561 | 4.90756 | 3.057 |
| chr4 | 90823038 | 90823315 | 1.08926 | 3.32937 | 3.057 |
| chr3 | 156949167 | 156949380 | 0.309318 | 0.945755 | 3.058 |
| chr1 | 182004320 | 182004892 | 0.612111 | 1.8716 | 3.058 |
| chr12 | 32414628 | 32414867 | 0.565139 | 1.72818 | 3.058 |
| chr15 | 76013370 | 76013766 | 0.993972 | 3.03967 | 3.058 |
| chr10 | 131651906 | 131652117 | 0.725307 | 2.21808 | 3.058 |
| chr16 | 9253922 | 9254358 | 1.37369 | 4.20124 | 3.058 |
| chr5 | 96497694 | 96497975 | 1.62779 | 4.97861 | 3.059 |
| chr17 | 71370022 | 71370604 | 0.351296 | 1.07448 | 3.059 |
| chr11 | 13487476 | 13487758 | 0.816248 | 2.49662 | 3.059 |
| chr19 | 45004509 | 45004894 | 0.605264 | 1.85148 | 3.059 |
| chr5 | 142295445 | 142295706 | 1.31896 | 4.03465 | 3.059 |
| chr7 | 69626691 | 69626917 | 0.882446 | 2.70055 | 3.060 |
| chr14 | 51686612 | 51686891 | 0.802868 | 2.45707 | 3.060 |
| chr3 | 50294452 | 50294793 | 0.890145 | 2.72427 | 3.060 |
| chr10 | 112563642 | 112563936 | 1.45573 | 4.45543 | 3.061 |
| chr2 | 85999581 | 85999858 | 0.801834 | 2.45412 | 3.061 |
| chr6 | 83924927 | 83925180 | 1.05937 | 3.2426 | 3.061 |
| chr14 | 105292730 | 105293092 | 1.64127 | 5.02437 | 3.061 |
| chr12 | 45564294 | 45564594 | 1.51732 | 4.64578 | 3.062 |
| chr5 | 133847953 | 133848206 | 0.693312 | 2.12283 | 3.062 |
| chr14 | 74693998 | 74694310 | 0.388866 | 1.19067 | 3.062 |
| chrX | 4076597 | 4076818 | 0.367568 | 1.12547 | 3.062 |
| chr8 | 100955463 | 100955735 | 0.363107 | 1.11185 | 3.062 |
| chr7 | 19161659 | 19161951 | 1.11579 | 3.41673 | 3.062 |
| chr20 | 6800112 | 6800444 | 0.393713 | 1.20566 | 3.062 |
| chr17 | 13640401 | 13640604 | 0.331488 | 1.01515 | 3.062 |
| chr19 | 56764131 | 56764604 | 1.33196 | 4.07995 | 3.063 |
| chr6 | 156983081 | 156983490 | 1.4635 | 4.48316 | 3.063 |
| chr3 | 5099454 | 5099923 | 0.73412 | 2.24907 | 3.064 |

| | | | | | |
|---|---|---|---|---|---|
| chr7 | 96415751 | 96416008 | 0.303679 | 0.930387 | 3.064 |
| chr12 | 3317112 | 3317362 | 0.675361 | 2.06915 | 3.064 |
| chr15 | 38295053 | 38295440 | 0.845896 | 2.59195 | 3.064 |
| chr7 | 15365757 | 15366077 | 0.79578 | 2.43843 | 3.064 |
| chr21 | 44839419 | 44839681 | 0.835705 | 2.56089 | 3.064 |
| chr12 | 34453080 | 34453367 | 0.651156 | 1.99539 | 3.064 |
| chr11 | 73675384 | 73675744 | 1.56134 | 4.78496 | 3.065 |
| chr5 | 150880248 | 150880652 | 1.59216 | 4.88021 | 3.065 |
| chr19 | 12355290 | 12355519 | 0.641018 | 1.96483 | 3.065 |
| chr4 | 170712949 | 170713208 | 1.04988 | 3.21889 | 3.066 |
| chr8 | 41287589 | 41288106 | 1.01654 | 3.11702 | 3.066 |
| chr9 | 12702206 | 12702459 | 0.901233 | 2.76364 | 3.067 |
| chr10 | 16733642 | 16733917 | 0.558298 | 1.71219 | 3.067 |
| chr12 | 71094381 | 71094697 | 0.498307 | 1.52825 | 3.067 |
| chr5 | 73079671 | 73079957 | 0.723314 | 2.21837 | 3.067 |
| chr17 | 59379455 | 59379683 | 0.458582 | 1.40648 | 3.067 |
| chr9 | 126164244 | 126164507 | 1.37295 | 4.21094 | 3.067 |
| chr7 | 54371612 | 54371972 | 0.407853 | 1.25147 | 3.068 |
| chr10 | 38043649 | 38043977 | 0.941479 | 2.8891 | 3.069 |
| chr6 | 149269036 | 149269354 | 1.08284 | 3.32411 | 3.070 |
| chr3 | 149052167 | 149052399 | 1.08752 | 3.33879 | 3.070 |
| chr4 | 173123689 | 173123912 | 1.07087 | 3.28768 | 3.070 |
| chr8 | 46941079 | 46941468 | 0.866185 | 2.65969 | 3.071 |
| chrX | 41152447 | 41152855 | 0.576827 | 1.77122 | 3.071 |
| chr17 | 43094317 | 43094537 | 0.701052 | 2.15279 | 3.071 |
| chr17 | 2148021 | 2148473 | 2.1659 | 6.65134 | 3.071 |
| chr5 | 122964798 | 122965013 | 0.365205 | 1.12172 | 3.071 |
| chr9 | 130705008 | 130705444 | 1.99312 | 6.12347 | 3.072 |
| chr11 | 72353968 | 72354523 | 1.60017 | 4.91654 | 3.073 |
| chr2 | 47995001 | 47995526 | 2.26509 | 6.95998 | 3.073 |
| chr1 | 163622022 | 163622311 | 0.466139 | 1.43232 | 3.073 |
| chr9 | 33778786 | 33779011 | 1.1268 | 3.46266 | 3.073 |
| chr11 | 6319090 | 6319359 | 0.818823 | 2.51625 | 3.073 |
| chr4 | 86669082 | 86669322 | 0.903618 | 2.77752 | 3.074 |
| chr9 | 19457431 | 19457755 | 1.82192 | 5.60027 | 3.074 |
| chr7 | 98910554 | 98910850 | 1.5538 | 4.77682 | 3.074 |
| chr9 | 140916659 | 140917076 | 0.896266 | 2.75551 | 3.074 |
| chr5 | 172359415 | 172359934 | 1.36419 | 4.19431 | 3.075 |
| chr12 | 121990499 | 121990958 | 1.73757 | 5.3429 | 3.075 |
| chr10 | 8095171 | 8095554 | 1.84156 | 5.66326 | 3.075 |
| chr14 | 94125705 | 94125957 | 0.307701 | 0.946269 | 3.075 |
| chr3 | 173163335 | 173163588 | 0.323548 | 0.995034 | 3.075 |
| chr1 | 150592247 | 150592594 | 1.00784 | 3.09951 | 3.075 |
| chr9 | 135519774 | 135519989 | 0.776776 | 2.389 | 3.076 |
| chr11 | 22851450 | 22851685 | 1.2575 | 3.86752 | 3.076 |
| chr3 | 183735099 | 183735558 | 1.35102 | 4.15537 | 3.076 |
| chr14 | 61663585 | 61663797 | 0.785992 | 2.41759 | 3.076 |
| chr17 | 80382349 | 80382752 | 1.47615 | 4.54073 | 3.076 |
| chr19 | 17877476 | 17877802 | 1.48752 | 4.57581 | 3.076 |
| chr20 | 31799692 | 31799911 | 1.22788 | 3.77722 | 3.076 |
| chr11 | 102485222 | 102485455 | 0.64914 | 1.99717 | 3.077 |
| chr16 | 23560633 | 23560904 | 0.326162 | 1.00349 | 3.077 |
| chr15 | 69755018 | 69755333 | 1.4919 | 4.59023 | 3.077 |
| chr3 | 150804444 | 150804746 | 0.717705 | 2.20833 | 3.077 |
| chr3 | 73042599 | 73042885 | 0.589168 | 1.81292 | 3.077 |
| chr8 | 33482218 | 33482430 | 0.71261 | 2.1929 | 3.077 |
| chr7 | 45453996 | 45454245 | 0.439983 | 1.35412 | 3.078 |
| chr19 | 18551276 | 18551522 | 0.853389 | 2.62653 | 3.078 |
| chr3 | 10814025 | 10814735 | 0.933663 | 2.8737 | 3.078 |
| chr21 | 33732063 | 33732302 | 0.953327 | 2.93444 | 3.078 |
| chr11 | 103472020 | 103472257 | 1.33684 | 4.11509 | 3.078 |

| | | | | | |
|---|---|---|---|---|---|
| chr5 | 171334734 | 171335176 | 0.949026 | 2.92154 | 3.078 |
| chr8 | 98565717 | 98565945 | 0.919318 | 2.83034 | 3.079 |
| chr4 | 151881991 | 151882366 | 1.21221 | 3.73247 | 3.079 |
| chr6 | 124609183 | 124609427 | 0.342985 | 1.05608 | 3.079 |
| chr3 | 64016382 | 64016645 | 1.05131 | 3.23744 | 3.079 |
| chr11 | 6303204 | 6303609 | 0.757893 | 2.33447 | 3.080 |
| chr4 | 131751203 | 131751473 | 0.297796 | 0.917398 | 3.081 |
| chr6 | 116608907 | 116609140 | 0.803398 | 2.47509 | 3.081 |
| chr5 | 138000923 | 138001255 | 0.287872 | 0.886883 | 3.081 |
| chr12 | 74335722 | 74335930 | 0.42599 | 1.31241 | 3.081 |
| chr7 | 91509974 | 91510342 | 1.73192 | 5.336 | 3.081 |
| chr10 | 125343248 | 125343558 | 1.51265 | 4.66116 | 3.081 |
| chr15 | 55700226 | 55700755 | 0.880176 | 2.71235 | 3.082 |
| chr16 | 23166820 | 23167042 | 0.784632 | 2.41809 | 3.082 |
| chr6 | 159173698 | 159173918 | 0.677064 | 2.08681 | 3.082 |
| chr3 | 30789588 | 30789805 | 0.896181 | 2.76217 | 3.082 |
| chr9 | 98981846 | 98982346 | 1.73742 | 5.35514 | 3.082 |
| chr8 | 96260963 | 96261280 | 1.42816 | 4.40216 | 3.082 |
| chr8 | 69081267 | 69081549 | 0.318927 | 0.983227 | 3.083 |
| chr16 | 337381 | 337692 | 0.558234 | 1.721 | 3.083 |
| chr19 | 49206110 | 49206347 | 0.346845 | 1.06933 | 3.083 |
| chr12 | 101659112 | 101659352 | 0.46447 | 1.43204 | 3.083 |
| chr6 | 168308753 | 168309080 | 0.853928 | 2.63302 | 3.083 |
| chr9 | 89939619 | 89939821 | 1.06328 | 3.27863 | 3.084 |
| chr16 | 54328876 | 54329111 | 0.401314 | 1.23746 | 3.084 |
| chr3 | 163503575 | 163503824 | 0.488329 | 1.50593 | 3.084 |
| chr1 | 212838537 | 212838964 | 1.63124 | 5.03094 | 3.084 |
| chr19 | 5031596 | 5031981 | 1.18855 | 3.66575 | 3.084 |
| chr9 | 24967786 | 24968025 | 0.320728 | 0.989252 | 3.084 |
| chr17 | 38473670 | 38473943 | 0.999269 | 3.08267 | 3.085 |
| chr10 | 30024440 | 30025099 | 1.58711 | 4.89654 | 3.085 |
| chr14 | 65239200 | 65239419 | 0.840203 | 2.59258 | 3.086 |
| chr19 | 5773829 | 5774055 | 0.914321 | 2.82155 | 3.086 |
| chr10 | 134606717 | 134607116 | 0.674741 | 2.08256 | 3.086 |
| chr15 | 88789947 | 88790469 | 1.98689 | 6.13321 | 3.087 |
| chr3 | 25574316 | 25574588 | 1.15207 | 3.55688 | 3.087 |
| chr11 | 134595941 | 134596191 | 0.746493 | 2.30501 | 3.088 |
| chr2 | 4652185 | 4652647 | 0.50146 | 1.54851 | 3.088 |
| chr17 | 41186347 | 41186843 | 0.430456 | 1.32926 | 3.088 |
| chr3 | 156848461 | 156848678 | 0.994257 | 3.07032 | 3.088 |
| chr2 | 132170829 | 132171151 | 0.884086 | 2.73015 | 3.088 |
| chr4 | 139011708 | 139011923 | 1.22491 | 3.78303 | 3.088 |
| chr12 | 48336091 | 48336389 | 0.91663 | 2.83105 | 3.089 |
| chr2 | 207256746 | 207257217 | 0.818851 | 2.52909 | 3.089 |
| chr7 | 115316194 | 115316459 | 0.792725 | 2.44841 | 3.089 |
| chr10 | 95676382 | 95676615 | 0.747675 | 2.30929 | 3.089 |
| chr10 | 65393548 | 65393765 | 0.752172 | 2.32328 | 3.089 |
| chr9 | 107922039 | 107922324 | 0.987221 | 3.0496 | 3.089 |
| chr12 | 98910065 | 98910469 | 1.06651 | 3.29455 | 3.089 |
| chr14 | 22860259 | 22860624 | 1.44084 | 4.45152 | 3.090 |
| chr20 | 19642996 | 19643432 | 2.07605 | 6.41422 | 3.090 |
| chr6 | 1436902 | 1437190 | 0.661524 | 2.04406 | 3.090 |
| chr10 | 87833116 | 87833382 | 0.294508 | 0.910024 | 3.090 |
| chr17 | 43212594 | 43213059 | 3.00953 | 9.29943 | 3.090 |
| chr19 | 56904462 | 56905034 | 1.09333 | 3.37843 | 3.090 |
| chr6 | 42134221 | 42134724 | 1.22519 | 3.78589 | 3.090 |
| chr17 | 28563845 | 28564143 | 1.69018 | 5.22283 | 3.090 |
| chr22 | 30021599 | 30021974 | 1.10901 | 3.42738 | 3.090 |
| chr7 | 136008997 | 136009301 | 0.348373 | 1.07665 | 3.091 |
| chr9 | 6354180 | 6354419 | 1.07522 | 3.32341 | 3.091 |
| chr1 | 29706042 | 29706334 | 1.03894 | 3.21136 | 3.091 |

| | | | | | |
|---|---|---|---|---|---|
| chr20 | 38173077 | 38173281 | 0.34689 | 1.07228 | 3.091 |
| chr1 | 44471265 | 44471753 | 1.52872 | 4.72547 | 3.091 |
| chr16 | 24845761 | 24846131 | 1.27441 | 3.94038 | 3.092 |
| chr17 | 19998667 | 19998978 | 0.930762 | 2.87819 | 3.092 |
| chr5 | 158946397 | 158946618 | 0.360844 | 1.11607 | 3.093 |
| chr6 | 3251733 | 3252048 | 1.30332 | 4.03117 | 3.093 |
| chr16 | 5624495 | 5624704 | 0.306816 | 0.949262 | 3.094 |
| chr16 | 29022502 | 29022817 | 0.658808 | 2.03875 | 3.095 |
| chr15 | 85878536 | 85878755 | 0.458531 | 1.41901 | 3.095 |
| chr16 | 65988054 | 65988308 | 0.367835 | 1.13835 | 3.095 |
| chr3 | 127730110 | 127730382 | 1.04877 | 3.24588 | 3.095 |
| chr5 | 92023814 | 92024080 | 0.917409 | 2.83951 | 3.095 |
| chr9 | 7234174 | 7234492 | 1.19565 | 3.70106 | 3.095 |
| chr9 | 127563724 | 127563954 | 0.330115 | 1.02195 | 3.096 |
| chr4 | 6682438 | 6682656 | 0.755259 | 2.33822 | 3.096 |
| chr12 | 6844796 | 6845210 | 1.09805 | 3.39983 | 3.096 |
| chr20 | 44803139 | 44803611 | 1.10696 | 3.42751 | 3.096 |
| chr15 | 97926033 | 97926242 | 0.504246 | 1.56137 | 3.096 |
| chr2 | 28830134 | 28830345 | 0.613548 | 1.90057 | 3.098 |
| chr5 | 26301693 | 26301964 | 0.354185 | 1.09737 | 3.098 |
| chr10 | 82295597 | 82296201 | 1.45935 | 4.52152 | 3.098 |
| chr1 | 240352720 | 240353036 | 1.13734 | 3.52392 | 3.098 |
| chr6 | 85278768 | 85279102 | 0.652477 | 2.02165 | 3.098 |
| chr16 | 31454340 | 31454778 | 1.94195 | 6.01772 | 3.099 |
| chr2 | 37449040 | 37449360 | 0.3278 | 1.01586 | 3.099 |
| chr4 | 104917207 | 104917437 | 0.675112 | 2.09219 | 3.099 |
| chr9 | 127474216 | 127474557 | 1.60779 | 4.98279 | 3.099 |
| chr10 | 104182388 | 104182813 | 1.53951 | 4.77119 | 3.099 |
| chr10 | 43471093 | 43471342 | 1.0094 | 3.12838 | 3.099 |
| chr4 | 20471400 | 20471702 | 0.312574 | 0.968754 | 3.099 |
| chr8 | 139481060 | 139481267 | 0.469569 | 1.45536 | 3.099 |
| chr7 | 6629354 | 6629874 | 1.25149 | 3.87902 | 3.100 |
| chr16 | 13819095 | 13819450 | 0.381479 | 1.18246 | 3.100 |
| chr5 | 98563472 | 98563678 | 0.376172 | 1.16607 | 3.100 |
| chr4 | 106395099 | 106395476 | 1.64048 | 5.08562 | 3.100 |
| chr2 | 205009910 | 205010275 | 1.67207 | 5.18395 | 3.100 |
| chr9 | 133646398 | 133646665 | 1.25857 | 3.9021 | 3.100 |
| chr18 | 3441387 | 3441611 | 0.582168 | 1.80498 | 3.100 |
| chr12 | 50339279 | 50339714 | 2.01277 | 6.2408 | 3.101 |
| chr6 | 54934035 | 54934292 | 0.725518 | 2.24956 | 3.101 |
| chr12 | 15148841 | 15149054 | 0.550555 | 1.7073 | 3.101 |
| chr22 | 35420058 | 35420428 | 1.38136 | 4.28452 | 3.102 |
| chr12 | 129453970 | 129454206 | 0.696893 | 2.16171 | 3.102 |
| chr14 | 76445537 | 76445910 | 0.785671 | 2.43779 | 3.103 |
| chr9 | 102446511 | 102446964 | 0.429514 | 1.33272 | 3.103 |
| chr6 | 134962142 | 134962445 | 0.829211 | 2.57329 | 3.103 |
| chr8 | 116286362 | 116286576 | 0.767733 | 2.38254 | 3.103 |
| chr11 | 68400274 | 68400593 | 1.11719 | 3.46711 | 3.103 |
| chr12 | 56898463 | 56899137 | 0.402416 | 1.24893 | 3.104 |
| chr10 | 15287593 | 15287861 | 0.679276 | 2.1082 | 3.104 |
| chr15 | 56345020 | 56345263 | 1.36767 | 4.2447 | 3.104 |
| chr20 | 25295729 | 25296056 | 1.66637 | 5.17257 | 3.104 |
| chr8 | 79608758 | 79608988 | 0.302842 | 0.940183 | 3.105 |
| chr1 | 233576619 | 233576951 | 0.983781 | 3.05468 | 3.105 |
| chr18 | 14086626 | 14086886 | 0.900654 | 2.79665 | 3.105 |
| chr22 | 37853240 | 37853789 | 1.41265 | 4.38646 | 3.105 |
| chr18 | 47536192 | 47536437 | 0.52713 | 1.63729 | 3.106 |
| chr19 | 36681362 | 36681671 | 1.03087 | 3.20199 | 3.106 |
| chr5 | 66505051 | 66505355 | 0.995418 | 3.092 | 3.106 |
| chr8 | 127755449 | 127755698 | 0.423098 | 1.31435 | 3.106 |
| chr22 | 30783322 | 30783791 | 1.49614 | 4.64796 | 3.107 |

| | | | | | |
|---|---|---|---|---|---|
| chr22 | 27152611 | 27153125 | 0.73057 | 2.26971 | 3.107 |
| chr20 | 5591440 | 5591684 | 0.691398 | 2.14812 | 3.107 |
| chr8 | 37010696 | 37011130 | 1.13505 | 3.52661 | 3.107 |
| chr3 | 39702777 | 39703121 | 0.351483 | 1.09208 | 3.107 |
| chr22 | 22080976 | 22081190 | 0.566087 | 1.75896 | 3.107 |
| chrX | 47400559 | 47400809 | 0.947114 | 2.94295 | 3.107 |
| chr4 | 87144702 | 87145088 | 0.861785 | 2.67789 | 3.107 |
| chr16 | 48657075 | 48657540 | 2.11119 | 6.56067 | 3.108 |
| chr3 | 42881221 | 42881448 | 0.680679 | 2.11527 | 3.108 |
| chr20 | 57978612 | 57979200 | 1.51324 | 4.70258 | 3.108 |
| chr21 | 40177007 | 40177249 | 1.02127 | 3.17414 | 3.108 |
| chr12 | 10484329 | 10484533 | 0.966644 | 3.00467 | 3.108 |
| chr19 | 43820739 | 43821399 | 1.27516 | 3.96367 | 3.108 |
| chr3 | 122746554 | 122746977 | 0.974058 | 3.02779 | 3.108 |
| chr11 | 2218422 | 2218914 | 1.44746 | 4.49955 | 3.109 |
| chr16 | 57933614 | 57933976 | 1.5073 | 4.68564 | 3.109 |
| chr8 | 145758151 | 145758411 | 0.679422 | 2.11207 | 3.109 |
| chr5 | 55368225 | 55368558 | 0.75124 | 2.33546 | 3.109 |
| chr12 | 104124890 | 104125111 | 0.51821 | 1.61104 | 3.109 |
| chr7 | 85625130 | 85625348 | 0.377074 | 1.17249 | 3.109 |
| chr17 | 46429229 | 46429534 | 0.839696 | 2.6112 | 3.110 |
| chr6 | 41060328 | 41060636 | 0.637765 | 1.98353 | 3.110 |
| chr3 | 151008512 | 151008804 | 0.829748 | 2.58071 | 3.110 |
| chr8 | 141954304 | 141954550 | 0.911167 | 2.83422 | 3.111 |
| chr4 | 187071175 | 187071387 | 0.887117 | 2.7596 | 3.111 |
| chr1 | 85575134 | 85575494 | 1.12978 | 3.51461 | 3.111 |
| chr12 | 75197094 | 75197349 | 0.913668 | 2.84232 | 3.111 |
| chr15 | 45749791 | 45750394 | 0.645565 | 2.00833 | 3.111 |
| chr1 | 39657351 | 39657555 | 0.887037 | 2.75972 | 3.111 |
| chr9 | 36910881 | 36911668 | 0.469164 | 1.45968 | 3.111 |
| chr11 | 7592887 | 7593304 | 1.60278 | 4.98727 | 3.112 |
| chr2 | 198056250 | 198056455 | 1.00304 | 3.12115 | 3.112 |
| chr1 | 35208266 | 35208663 | 1.41657 | 4.40819 | 3.112 |
| chr16 | 81756664 | 81757070 | 0.912736 | 2.8404 | 3.112 |
| chr1 | 176542372 | 176542578 | 0.360099 | 1.12076 | 3.112 |
| chr9 | 140083489 | 140083802 | 0.592733 | 1.84482 | 3.112 |
| chr11 | 70790847 | 70791077 | 0.747637 | 2.32696 | 3.112 |
| chr7 | 158823305 | 158823521 | 0.625457 | 1.94672 | 3.112 |
| chr2 | 29419647 | 29419990 | 1.55791 | 4.84909 | 3.113 |
| chr2 | 131113601 | 131113896 | 0.820334 | 2.55384 | 3.113 |
| chr5 | 14108730 | 14108948 | 1.025 | 3.19167 | 3.114 |
| chr8 | 36419016 | 36419289 | 0.337729 | 1.05165 | 3.114 |
| chr4 | 167090384 | 167090613 | 0.570663 | 1.777 | 3.114 |
| chr9 | 139035099 | 139035460 | 0.742333 | 2.31157 | 3.114 |
| chr12 | 114569688 | 114570267 | 0.315286 | 0.982011 | 3.115 |
| chr2 | 209121196 | 209121482 | 0.476967 | 1.48579 | 3.115 |
| chr5 | 86696210 | 86696451 | 0.628051 | 1.95658 | 3.115 |
| chr3 | 50384749 | 50384962 | 0.90116 | 2.80742 | 3.115 |
| chr5 | 145912763 | 145913161 | 1.43506 | 4.47157 | 3.116 |
| chr14 | 91912812 | 91913020 | 0.383713 | 1.1957 | 3.116 |
| chr21 | 43188861 | 43189068 | 0.508782 | 1.58554 | 3.116 |
| chr2 | 121194798 | 121195155 | 1.20887 | 3.76789 | 3.117 |
| chr14 | 65307990 | 65308212 | 0.671519 | 2.09331 | 3.117 |
| chr4 | 155903504 | 155903976 | 0.930514 | 2.90085 | 3.117 |
| chr9 | 139940983 | 139941325 | 2.16343 | 6.74526 | 3.118 |
| chr20 | 55756105 | 55756327 | 0.390051 | 1.21628 | 3.118 |
| chr12 | 23722394 | 23722606 | 0.417054 | 1.30055 | 3.118 |
| chr1 | 92351272 | 92351624 | 0.372584 | 1.16198 | 3.119 |
| chr5 | 149668582 | 149668860 | 0.315908 | 0.985297 | 3.119 |
| chr1 | 240552977 | 240553180 | 0.302241 | 0.942799 | 3.119 |
| chr14 | 34792187 | 34792556 | 1.25057 | 3.90124 | 3.120 |

| | | | | | |
|---|---|---|---|---|---|
| chr8 | 24102518 | 24102818 | 0.905725 | 2.82563 | 3.120 |
| chr18 | 3585833 | 3586133 | 1.12253 | 3.50219 | 3.120 |
| chr1 | 172059378 | 172059712 | 1.37183 | 4.28027 | 3.120 |
| chr12 | 21099767 | 21100190 | 0.726335 | 2.26642 | 3.120 |
| chr11 | 25626961 | 25627208 | 0.350311 | 1.09352 | 3.122 |
| chr7 | 126581243 | 126581672 | 0.352755 | 1.1013 | 3.122 |
| chr20 | 61506895 | 61507151 | 1.0806 | 3.37372 | 3.122 |
| chr6 | 151727264 | 151727508 | 0.994468 | 3.10492 | 3.122 |
| chr12 | 68616928 | 68617273 | 1.67151 | 5.21885 | 3.122 |
| chr16 | 6967932 | 6968172 | 0.289866 | 0.905053 | 3.122 |
| chr19 | 22060423 | 22060635 | 0.484691 | 1.5135 | 3.123 |
| chr4 | 37803783 | 37804034 | 0.674353 | 2.10613 | 3.123 |
| chr20 | 39619206 | 39619550 | 1.15489 | 3.60705 | 3.123 |
| chr4 | 174772796 | 174773009 | 0.353214 | 1.1032 | 3.123 |
| chr11 | 118971054 | 118971291 | 0.586795 | 1.83285 | 3.123 |
| chrX | 150882372 | 150882700 | 1.14231 | 3.56809 | 3.124 |
| chr16 | 30420695 | 30421149 | 1.67649 | 5.23734 | 3.124 |
| chr4 | 111437547 | 111437879 | 1.17437 | 3.66943 | 3.125 |
| chr5 | 76070394 | 76070757 | 1.45318 | 4.54146 | 3.125 |
| chr5 | 178220077 | 178220557 | 0.671608 | 2.09903 | 3.125 |
| chr19 | 15695449 | 15695857 | 2.39132 | 7.47455 | 3.126 |
| chr10 | 100275833 | 100276068 | 0.24378 | 0.762041 | 3.126 |
| chr11 | 114210978 | 114211310 | 1.14882 | 3.59158 | 3.126 |
| chr7 | 52884158 | 52884435 | 0.475456 | 1.48644 | 3.126 |
| chr16 | 68624380 | 68624732 | 1.46613 | 4.58377 | 3.126 |
| chr5 | 176102663 | 176103015 | 0.885848 | 2.76973 | 3.127 |
| chr14 | 32403954 | 32404191 | 0.793917 | 2.48237 | 3.127 |
| chr15 | 29987754 | 29987997 | 0.849405 | 2.65601 | 3.127 |
| chr1 | 206729363 | 206729629 | 0.388815 | 1.21586 | 3.127 |
| chr12 | 114909309 | 114909630 | 0.41859 | 1.30902 | 3.127 |
| chr12 | 124328973 | 124329176 | 0.36491 | 1.14117 | 3.127 |
| chr4 | 180228707 | 180229010 | 1.03897 | 3.2494 | 3.128 |
| chr1 | 1727985 | 1728208 | 0.66612 | 2.08382 | 3.128 |
| chr8 | 86157766 | 86158046 | 0.646371 | 2.02224 | 3.129 |
| chr8 | 8923194 | 8923623 | 1.04336 | 3.26456 | 3.129 |
| chr7 | 23902893 | 23903152 | 1.46451 | 4.58263 | 3.129 |
| chr12 | 51925705 | 51926052 | 1.70369 | 5.33131 | 3.129 |
| chr1 | 33966243 | 33966458 | 0.601875 | 1.88374 | 3.130 |
| chr8 | 75686757 | 75686987 | 0.32357 | 1.01275 | 3.130 |
| chr6 | 8540575 | 8540777 | 0.458176 | 1.43423 | 3.130 |
| chr1 | 212244119 | 212244370 | 0.880895 | 2.75749 | 3.130 |
| chr8 | 110965998 | 110966347 | 0.689905 | 2.15967 | 3.130 |
| chr13 | 52419278 | 52419575 | 1.33663 | 4.18449 | 3.131 |
| chrX | 119619923 | 119620281 | 1.07654 | 3.37043 | 3.131 |
| chr13 | 24463565 | 24463789 | 1.12702 | 3.5287 | 3.131 |
| chr3 | 32822816 | 32823356 | 1.10765 | 3.46808 | 3.131 |
| chr17 | 64851380 | 64851696 | 1.15057 | 3.60258 | 3.131 |
| chr18 | 55549073 | 55549472 | 1.24903 | 3.91159 | 3.132 |
| chr18 | 9683419 | 9683706 | 1.30975 | 4.10195 | 3.132 |
| chr11 | 117692705 | 117693127 | 0.973048 | 3.04746 | 3.132 |
| chr16 | 27876613 | 27876833 | 0.864365 | 2.70717 | 3.132 |
| chr10 | 120899010 | 120899325 | 1.19946 | 3.75676 | 3.132 |
| chr11 | 8120914 | 8121192 | 0.999107 | 3.12955 | 3.132 |
| chr6 | 109917215 | 109917587 | 0.700938 | 2.19581 | 3.133 |
| chr5 | 57183744 | 57184034 | 1.65428 | 5.18234 | 3.133 |
| chrX | 53509125 | 53509400 | 1.27456 | 3.99285 | 3.133 |
| chr12 | 2097234 | 2097447 | 0.369222 | 1.15682 | 3.133 |
| chr8 | 137769080 | 137769397 | 0.332717 | 1.04254 | 3.133 |
| chr11 | 7432205 | 7432408 | 0.436681 | 1.36833 | 3.133 |
| chr13 | 32842062 | 32842280 | 0.693639 | 2.17381 | 3.134 |
| chr10 | 13203176 | 13203546 | 1.37434 | 4.30733 | 3.134 |

| | | | | | |
|---|---|---|---|---|---|
| chr12 | 79738666 | 79738969 | 1.12166 | 3.51581 | 3.134 |
| chr8 | 101950880 | 101951136 | 1.12843 | 3.53708 | 3.135 |
| chr22 | 32061672 | 32061921 | 0.890804 | 2.79243 | 3.135 |
| chr9 | 117777347 | 117777655 | 1.06071 | 3.32504 | 3.135 |
| chr15 | 42066711 | 42066961 | 1.00359 | 3.14605 | 3.135 |
| chr19 | 35632288 | 35632608 | 1.5611 | 4.8939 | 3.135 |
| chr19 | 56701260 | 56701584 | 0.400027 | 1.25411 | 3.135 |
| chr8 | 78059736 | 78060069 | 0.589885 | 1.84939 | 3.135 |
| chr8 | 145164581 | 145164832 | 0.742953 | 2.32955 | 3.136 |
| chr4 | 140020528 | 140021053 | 1.06572 | 3.34179 | 3.136 |
| chr16 | 49743743 | 49743996 | 0.659051 | 2.06664 | 3.136 |
| chrX | 12789569 | 12790189 | 1.04372 | 3.27291 | 3.136 |
| chr5 | 125519209 | 125519504 | 0.339531 | 1.06471 | 3.136 |
| chr2 | 36584980 | 36585287 | 0.641441 | 2.01192 | 3.137 |
| chr1 | 234733716 | 234733981 | 0.798832 | 2.50567 | 3.137 |
| chr9 | 103363431 | 103363855 | 1.23042 | 3.85986 | 3.137 |
| chr5 | 63575623 | 63575992 | 0.53698 | 1.68477 | 3.137 |
| chr1 | 39564342 | 39564644 | 1.04693 | 3.28491 | 3.138 |
| chr2 | 86435635 | 86435878 | 0.465594 | 1.46107 | 3.138 |
| chr1 | 16359357 | 16359685 | 1.40119 | 4.39723 | 3.138 |
| chr5 | 118357745 | 118357974 | 0.760317 | 2.38619 | 3.138 |
| chr1 | 177435571 | 177435921 | 0.399456 | 1.25378 | 3.139 |
| chr8 | 35503174 | 35503395 | 0.335893 | 1.05438 | 3.139 |
| chr8 | 12963987 | 12964498 | 0.306964 | 0.96369 | 3.139 |
| chr9 | 36974787 | 36975070 | 1.02484 | 3.21795 | 3.140 |
| chr3 | 53955404 | 53955607 | 0.742402 | 2.33118 | 3.140 |
| chr17 | 75007226 | 75007703 | 1.23555 | 3.88094 | 3.141 |
| chr3 | 47555239 | 47555464 | 0.798934 | 2.50952 | 3.141 |
| chr9 | 127939514 | 127939762 | 0.4161 | 1.3071 | 3.141 |
| chr3 | 149526077 | 149526431 | 1.16884 | 3.67203 | 3.142 |
| chr5 | 178266390 | 178266669 | 0.576577 | 1.81139 | 3.142 |
| chr11 | 7906223 | 7906434 | 0.329233 | 1.03435 | 3.142 |
| chr5 | 130744351 | 130744730 | 0.34193 | 1.0743 | 3.142 |
| chr17 | 57936876 | 57937106 | 0.867545 | 2.72602 | 3.142 |
| chr11 | 79438227 | 79438431 | 0.337852 | 1.06172 | 3.143 |
| chr9 | 124767941 | 124768197 | 1.0298 | 3.23658 | 3.143 |
| chr8 | 22144770 | 22145108 | 1.04483 | 3.28388 | 3.143 |
| chr12 | 40546585 | 40546947 | 1.03199 | 3.24365 | 3.143 |
| chr5 | 149611890 | 149612106 | 0.418072 | 1.31405 | 3.143 |
| chr3 | 122075383 | 122075629 | 0.976887 | 3.07089 | 3.144 |
| chr10 | 22549174 | 22549456 | 0.312195 | 0.98144 | 3.144 |
| chr13 | 90577405 | 90577623 | 0.682662 | 2.14609 | 3.144 |
| chr6 | 14280994 | 14281283 | 0.742784 | 2.3351 | 3.144 |
| chr16 | 56402082 | 56402288 | 0.415166 | 1.30525 | 3.144 |
| chr13 | 30345440 | 30345645 | 0.545148 | 1.71409 | 3.144 |
| chr1 | 206982885 | 206983257 | 0.993524 | 3.12412 | 3.144 |
| chr3 | 151186118 | 151186423 | 1.46302 | 4.60123 | 3.145 |
| chr19 | 53093963 | 53094282 | 0.776179 | 2.44113 | 3.145 |
| chr1 | 151940441 | 151940795 | 1.1381 | 3.57982 | 3.145 |
| chr5 | 173447349 | 173447685 | 0.560853 | 1.76414 | 3.145 |
| chr18 | 77138971 | 77139354 | 1.40424 | 4.41735 | 3.146 |
| chr9 | 135569680 | 135570041 | 0.635973 | 2.00081 | 3.146 |
| chr5 | 20556340 | 20556558 | 0.38069 | 1.19771 | 3.146 |
| chr4 | 141348014 | 141348257 | 0.417441 | 1.31335 | 3.146 |
| chr12 | 52309845 | 52310491 | 0.308779 | 0.9715 | 3.146 |
| chr14 | 100185624 | 100185853 | 0.327746 | 1.03125 | 3.146 |
| chr16 | 46891525 | 46891746 | 0.548544 | 1.72608 | 3.147 |
| chrX | 100645773 | 100646027 | 1.51347 | 4.76337 | 3.147 |
| chr12 | 113590819 | 113591483 | 0.665793 | 2.09553 | 3.147 |
| chr7 | 63781475 | 63781699 | 0.333417 | 1.04945 | 3.148 |
| chr19 | 17346430 | 17346675 | 0.541372 | 1.7041 | 3.148 |

| | | | | | |
|---|---|---|---|---|---|
| chr10 | 120902646 | 120903034 | 0.426357 | 1.34208 | 3.148 |
| chr15 | 75600396 | 75600716 | 1.44376 | 4.54467 | 3.148 |
| chr7 | 140004380 | 140004670 | 0.738612 | 2.32545 | 3.148 |
| chr17 | 18069912 | 18070565 | 0.62292 | 1.9613 | 3.149 |
| chr3 | 119187850 | 119188083 | 0.491234 | 1.54671 | 3.149 |
| chr1 | 39300982 | 39301218 | 0.881529 | 2.77575 | 3.149 |
| chr14 | 65332058 | 65332268 | 0.367095 | 1.15606 | 3.149 |
| chr3 | 68925365 | 68925654 | 1.01779 | 3.20524 | 3.149 |
| chr12 | 69647817 | 69648050 | 1.16797 | 3.67879 | 3.150 |
| chr12 | 91353371 | 91353747 | 1.00055 | 3.15186 | 3.150 |
| chr6 | 144261180 | 144261465 | 0.903524 | 2.8463 | 3.150 |
| chr20 | 25398202 | 25398482 | 0.396098 | 1.24781 | 3.150 |
| chr1 | 39558069 | 39558400 | 1.30656 | 4.11662 | 3.151 |
| chr1 | 225662581 | 225663065 | 2.69261 | 8.48395 | 3.151 |
| chr9 | 98980301 | 98980809 | 1.67013 | 5.26299 | 3.151 |
| chr3 | 111856027 | 111856235 | 0.864482 | 2.72427 | 3.151 |
| chr6 | 10510804 | 10511089 | 1.1723 | 3.69434 | 3.151 |
| chr5 | 178639972 | 178640225 | 0.657047 | 2.07068 | 3.151 |
| chr10 | 101058948 | 101059233 | 0.313183 | 0.987025 | 3.152 |
| chr10 | 73815818 | 73816142 | 0.582681 | 1.83648 | 3.152 |
| chr1 | 76500808 | 76501087 | 0.752512 | 2.37183 | 3.152 |
| chr15 | 76096936 | 76097155 | 0.367815 | 1.1594 | 3.152 |
| chr7 | 40780645 | 40780872 | 0.378451 | 1.19295 | 3.152 |
| chr15 | 76405786 | 76406112 | 0.644273 | 2.03121 | 3.153 |
| chr12 | 131616842 | 131617420 | 1.50446 | 4.74315 | 3.153 |
| chr1 | 212464906 | 212465166 | 1.01285 | 3.19331 | 3.153 |
| chr3 | 39009151 | 39009435 | 0.24831 | 0.782886 | 3.153 |
| chr4 | 139840342 | 139840601 | 0.861372 | 2.71581 | 3.153 |
| chr7 | 143589782 | 143590035 | 0.709463 | 2.23704 | 3.153 |
| chr12 | 113430533 | 113431033 | 1.72224 | 5.43065 | 3.153 |
| chr1 | 1610334 | 1610572 | 0.75204 | 2.37145 | 3.153 |
| chr1 | 156389771 | 156389974 | 0.678179 | 2.1388 | 3.154 |
| chr13 | 100094581 | 100094892 | 1.05402 | 3.32431 | 3.154 |
| chr8 | 89048577 | 89048831 | 0.234998 | 0.741265 | 3.154 |
| chr1 | 157742079 | 157742283 | 0.360921 | 1.13865 | 3.155 |
| chrX | 118863903 | 118864189 | 0.953215 | 3.00728 | 3.155 |
| chr8 | 4177074 | 4177355 | 0.575322 | 1.81509 | 3.155 |
| chr22 | 25333158 | 25333385 | 0.694833 | 2.1922 | 3.155 |
| chr1 | 234382670 | 234382907 | 0.262744 | 0.829041 | 3.155 |
| chr3 | 36165660 | 36165948 | 1.18107 | 3.72682 | 3.155 |
| chr4 | 10065385 | 10065653 | 0.796356 | 2.51298 | 3.156 |
| chrX | 23890171 | 23890375 | 0.520093 | 1.6413 | 3.156 |
| chr4 | 24490321 | 24490706 | 1.11693 | 3.52483 | 3.156 |
| chr10 | 24446808 | 24447257 | 0.39832 | 1.25719 | 3.156 |
| chr12 | 122332664 | 122333085 | 0.503229 | 1.58837 | 3.156 |
| chr8 | 18066468 | 18066883 | 0.748564 | 2.36306 | 3.157 |
| chr8 | 11026353 | 11026759 | 0.459645 | 1.4511 | 3.157 |
| chr6 | 9413251 | 9413510 | 1.136 | 3.58719 | 3.158 |
| chr20 | 22832401 | 22832766 | 1.79218 | 5.65971 | 3.158 |
| chr2 | 33587666 | 33587968 | 0.417104 | 1.31733 | 3.158 |
| chr19 | 10880844 | 10881066 | 0.547481 | 1.72966 | 3.159 |
| chr10 | 30823540 | 30823771 | 1.14595 | 3.62062 | 3.159 |
| chr8 | 74361373 | 74361707 | 1.01753 | 3.21515 | 3.160 |
| chr13 | 78052615 | 78052839 | 0.926367 | 2.92744 | 3.160 |
| chr3 | 97540939 | 97541167 | 0.559711 | 1.76885 | 3.160 |
| chr20 | 39756280 | 39756515 | 0.384074 | 1.21387 | 3.161 |
| chr7 | 157267220 | 157267748 | 1.01012 | 3.19298 | 3.161 |
| chr19 | 52157996 | 52158444 | 0.700695 | 2.21494 | 3.161 |
| chr7 | 25267523 | 25267842 | 0.343367 | 1.08554 | 3.161 |
| chr4 | 166300050 | 166300329 | 0.634933 | 2.0075 | 3.162 |
| chr12 | 103509711 | 103509913 | 0.338101 | 1.069 | 3.162 |

| | | | | | |
|---|---|---|---|---|---|
| chr16 | 60832790 | 60833018 | 0.376152 | 1.18932 | 3.162 |
| chr10 | 90551763 | 90551970 | 0.934395 | 2.95455 | 3.162 |
| chr14 | 68066732 | 68067032 | 0.655716 | 2.07337 | 3.162 |
| chr10 | 22862996 | 22863274 | 0.872646 | 2.75948 | 3.162 |
| chr3 | 33957316 | 33957638 | 0.817536 | 2.5857 | 3.163 |
| chr19 | 49934077 | 49934432 | 1.0199 | 3.22579 | 3.163 |
| chr11 | 1051034 | 1051446 | 1.38921 | 4.39413 | 3.163 |
| chr10 | 111643875 | 111644343 | 1.55586 | 4.92138 | 3.163 |
| chr15 | 97015804 | 97016236 | 0.358991 | 1.13554 | 3.163 |
| chr4 | 137867443 | 137867724 | 0.353777 | 1.11907 | 3.163 |
| chr11 | 118800972 | 118801368 | 1.05978 | 3.35255 | 3.163 |
| chr1 | 81880146 | 81880491 | 0.811207 | 2.56631 | 3.164 |
| chr17 | 65955219 | 65955449 | 0.363398 | 1.14968 | 3.164 |
| chr10 | 133850287 | 133850682 | 0.628703 | 1.98908 | 3.164 |
| chr19 | 57122775 | 57123029 | 0.516241 | 1.6333 | 3.164 |
| chr7 | 126987692 | 126988138 | 1.08073 | 3.41928 | 3.164 |
| chr6 | 57132950 | 57133190 | 0.731873 | 2.31559 | 3.164 |
| chr10 | 90342856 | 90343294 | 1.14784 | 3.63183 | 3.164 |
| chr7 | 89568945 | 89569214 | 0.4777 | 1.51162 | 3.164 |
| chr17 | 55888405 | 55888631 | 0.421766 | 1.33473 | 3.165 |
| chr17 | 8456091 | 8456299 | 0.491066 | 1.55444 | 3.165 |
| chr12 | 133345042 | 133345348 | 1.44574 | 4.5766 | 3.166 |
| chr1 | 17766123 | 17766333 | 0.885372 | 2.8029 | 3.166 |
| chr1 | 3692518 | 3692839 | 0.964662 | 3.05404 | 3.166 |
| chr8 | 42010081 | 42010400 | 1.76614 | 5.59152 | 3.166 |
| chr9 | 76180824 | 76181139 | 1.16984 | 3.70382 | 3.166 |
| chr20 | 14318264 | 14318558 | 0.970632 | 3.07319 | 3.166 |
| chr4 | 180788528 | 180788809 | 1.22514 | 3.87962 | 3.167 |
| chrX | 64671538 | 64671849 | 0.928138 | 2.93917 | 3.167 |
| chr8 | 144822498 | 144822955 | 1.10014 | 3.48388 | 3.167 |
| chr17 | 14418390 | 14418667 | 0.682987 | 2.16299 | 3.167 |
| chr19 | 56028213 | 56028691 | 1.46758 | 4.64818 | 3.167 |
| chr4 | 111510648 | 111510852 | 0.339607 | 1.07563 | 3.167 |
| chr5 | 61649950 | 61650182 | 0.653801 | 2.07114 | 3.168 |
| chr14 | 98477952 | 98478164 | 0.308882 | 0.978677 | 3.168 |
| chr1 | 209250120 | 209250394 | 1.22725 | 3.88888 | 3.169 |
| chr8 | 80525408 | 80525795 | 0.752496 | 2.38456 | 3.169 |
| chr11 | 76812257 | 76812628 | 0.805828 | 2.55358 | 3.169 |
| chr2 | 5927413 | 5927669 | 0.396402 | 1.25618 | 3.169 |
| chr7 | 92965840 | 92966071 | 0.3565 | 1.12979 | 3.169 |
| chr6 | 108188145 | 108188664 | 0.873899 | 2.76963 | 3.169 |
| chr3 | 108196421 | 108196715 | 1.08201 | 3.42964 | 3.170 |
| chr3 | 44574315 | 44574561 | 0.536425 | 1.70031 | 3.170 |
| chr17 | 62028162 | 62028645 | 1.20339 | 3.81446 | 3.170 |
| chr14 | 39503988 | 39504227 | 1.05699 | 3.35043 | 3.170 |
| chr3 | 122111758 | 122112061 | 0.791627 | 2.50935 | 3.170 |
| chr5 | 2711797 | 2712056 | 0.295206 | 0.935776 | 3.170 |
| chr12 | 74710854 | 74711097 | 0.573137 | 1.81691 | 3.170 |
| chr6 | 134477991 | 134478201 | 0.673044 | 2.13376 | 3.170 |
| chr12 | 62653888 | 62654155 | 1.37275 | 4.35222 | 3.170 |
| chr6 | 159589662 | 159589894 | 0.678101 | 2.14988 | 3.170 |
| chr7 | 151560554 | 151560784 | 0.450221 | 1.42749 | 3.171 |
| chr12 | 68444958 | 68445162 | 0.459213 | 1.45606 | 3.171 |
| chr19 | 50990123 | 50990557 | 1.45992 | 4.6294 | 3.171 |
| chr2 | 60175280 | 60175523 | 0.409037 | 1.29712 | 3.171 |
| chr15 | 31727488 | 31727857 | 1.23064 | 3.90272 | 3.171 |
| chr12 | 12965106 | 12965429 | 1.32709 | 4.20865 | 3.171 |
| chr9 | 131214172 | 131214411 | 0.377289 | 1.19666 | 3.172 |
| chrX | 129709831 | 129710148 | 1.72965 | 5.48723 | 3.172 |
| chr7 | 83576607 | 83576910 | 0.999633 | 3.17176 | 3.173 |
| chr12 | 103969071 | 103969429 | 1.59981 | 5.07635 | 3.173 |

| | | | | | |
|---|---|---|---|---|---|
| chr17 | 21071913 | 21072136 | 0.456992 | 1.45008 | 3.173 |
| chr5 | 124133494 | 124133732 | 0.451012 | 1.43116 | 3.173 |
| chr10 | 87307853 | 87308082 | 0.942039 | 2.98991 | 3.174 |
| chr1 | 186124802 | 186125090 | 0.793315 | 2.51806 | 3.174 |
| chr6 | 44280925 | 44281298 | 1.45897 | 4.63094 | 3.174 |
| chr12 | 13154180 | 13154599 | 0.667 | 2.11744 | 3.175 |
| chr1 | 92012532 | 92012813 | 0.837404 | 2.65856 | 3.175 |
| chr15 | 52509298 | 52509722 | 1.08582 | 3.44724 | 3.175 |
| chr7 | 135392776 | 135392978 | 0.371543 | 1.17968 | 3.175 |
| chrX | 141023985 | 141024196 | 0.400998 | 1.27337 | 3.176 |
| chr11 | 103196761 | 103197002 | 0.967402 | 3.07221 | 3.176 |
| chr4 | 146540426 | 146540861 | 1.21257 | 3.85093 | 3.176 |
| chr7 | 152404051 | 152404413 | 1.28885 | 4.09321 | 3.176 |
| chr12 | 83783132 | 83783436 | 0.823844 | 2.61654 | 3.176 |
| chr5 | 22078592 | 22078939 | 0.377113 | 1.19773 | 3.176 |
| chr4 | 154004654 | 154004954 | 0.613615 | 1.94889 | 3.176 |
| chr17 | 27243359 | 27243760 | 1.24212 | 3.94549 | 3.176 |
| chrX | 114913873 | 114914080 | 0.538506 | 1.71068 | 3.177 |
| chr15 | 85897764 | 85898072 | 0.400518 | 1.27258 | 3.177 |
| chr17 | 46534613 | 46535014 | 2.42127 | 7.69323 | 3.177 |
| chr17 | 48680782 | 48680989 | 0.599069 | 1.90379 | 3.178 |
| chr14 | 87280075 | 87280464 | 0.411594 | 1.30811 | 3.178 |
| chr16 | 4380269 | 4380723 | 1.89907 | 6.03577 | 3.178 |
| chr8 | 75467116 | 75467330 | 0.40921 | 1.30075 | 3.179 |
| chr11 | 68873226 | 68873472 | 0.243359 | 0.773813 | 3.180 |
| chr7 | 77428176 | 77428384 | 0.653344 | 2.07752 | 3.180 |
| chr5 | 127082504 | 127082840 | 0.961974 | 3.05906 | 3.180 |
| chr17 | 57930629 | 57930885 | 1.34992 | 4.29362 | 3.181 |
| chr3 | 127813952 | 127814356 | 1.07998 | 3.4353 | 3.181 |
| chr17 | 59652282 | 59652539 | 1.19882 | 3.81369 | 3.181 |
| chr4 | 14789609 | 14789887 | 0.742889 | 2.36364 | 3.182 |
| chr12 | 110480926 | 110481330 | 1.03396 | 3.28987 | 3.182 |
| chr7 | 131630123 | 131630546 | 1.22482 | 3.89773 | 3.182 |
| chr19 | 1774879 | 1775270 | 1.26736 | 4.03408 | 3.183 |
| chr6 | 149067718 | 149068050 | 0.64387 | 2.0497 | 3.183 |
| chr12 | 19660907 | 19661270 | 0.450529 | 1.43437 | 3.184 |
| chrX | 57480408 | 57480714 | 0.74244 | 2.36391 | 3.184 |
| chr8 | 134461735 | 134462031 | 1.50714 | 4.7987 | 3.184 |
| chr5 | 33058421 | 33058785 | 0.334062 | 1.06368 | 3.184 |
| chr15 | 80659493 | 80659767 | 1.18316 | 3.76748 | 3.184 |
| chr7 | 151008467 | 151008876 | 0.95965 | 3.0558 | 3.184 |
| chr1 | 202091671 | 202092120 | 1.20964 | 3.85202 | 3.184 |
| chr10 | 4309683 | 4309920 | 0.654228 | 2.08394 | 3.185 |
| chr12 | 121533749 | 121534142 | 1.79018 | 5.70264 | 3.186 |
| chr14 | 76964192 | 76964396 | 0.519099 | 1.65366 | 3.186 |
| chr3 | 2773821 | 2774030 | 0.253564 | 0.807776 | 3.186 |
| chr9 | 96975843 | 96976123 | 0.748736 | 2.38563 | 3.186 |
| chr8 | 54775888 | 54776201 | 0.631243 | 2.01155 | 3.187 |
| chr22 | 39369645 | 39369979 | 1.16279 | 3.70551 | 3.187 |
| chr20 | 57797334 | 57797966 | 1.55335 | 4.95073 | 3.187 |
| chr3 | 14528984 | 14529215 | 0.272764 | 0.869357 | 3.187 |
| chr12 | 16722095 | 16722350 | 0.411564 | 1.31209 | 3.188 |
| chr5 | 1641698 | 1642045 | 1.82854 | 5.82973 | 3.188 |
| chr3 | 38697249 | 38697491 | 1.10234 | 3.51477 | 3.188 |
| chr10 | 119208969 | 119209391 | 0.72821 | 2.32214 | 3.189 |
| chr17 | 18000181 | 18000449 | 0.389565 | 1.24228 | 3.189 |
| chr12 | 53715316 | 53715627 | 2.09243 | 6.67254 | 3.189 |
| chr15 | 101475136 | 101475347 | 0.585443 | 1.86751 | 3.190 |
| chr17 | 18015144 | 18015348 | 0.350166 | 1.11708 | 3.190 |
| chr4 | 182064471 | 182064729 | 0.92658 | 2.95613 | 3.190 |
| chr11 | 6319726 | 6320143 | 1.29924 | 4.14509 | 3.190 |

| | | | | | |
|---|---|---|---|---|---|
| chr1 | 181158950 | 181159472 | 2.60414 | 8.30843 | 3.190 |
| chr6 | 111332888 | 111333119 | 0.724327 | 2.31127 | 3.191 |
| chr5 | 109843672 | 109843947 | 0.347801 | 1.10988 | 3.191 |
| chr17 | 60378241 | 60378524 | 0.374221 | 1.19431 | 3.191 |
| chr4 | 97273465 | 97273689 | 0.595592 | 1.90088 | 3.192 |
| chr14 | 86002856 | 86003078 | 0.670307 | 2.13942 | 3.192 |
| chr11 | 83004022 | 83004384 | 1.11549 | 3.56033 | 3.192 |
| chr9 | 125261092 | 125261480 | 0.887476 | 2.83292 | 3.192 |
| chr17 | 49244266 | 49244809 | 1.27549 | 4.07185 | 3.192 |
| chr1 | 200948669 | 200949097 | 1.75809 | 5.61259 | 3.192 |
| chr9 | 34612798 | 34613223 | 1.28164 | 4.09257 | 3.193 |
| chr16 | 49991867 | 49992101 | 1.77411 | 5.66589 | 3.194 |
| chr12 | 51984447 | 51984824 | 2.13476 | 6.81787 | 3.194 |
| chr12 | 66672931 | 66673286 | 1.11401 | 3.55803 | 3.194 |
| chr6 | 139540314 | 139540753 | 1.06324 | 3.39601 | 3.194 |
| chrX | 41619176 | 41619450 | 0.675549 | 2.15779 | 3.194 |
| chr8 | 47484588 | 47484808 | 0.541138 | 1.72856 | 3.194 |
| chr12 | 6532742 | 6533070 | 1.11614 | 3.56543 | 3.194 |
| chr4 | 112996364 | 112996618 | 0.843265 | 2.69381 | 3.195 |
| chr6 | 5057415 | 5057677 | 0.939437 | 3.00104 | 3.195 |
| chr11 | 91290174 | 91290446 | 0.269967 | 0.862512 | 3.195 |
| chr4 | 53848825 | 53849051 | 0.721038 | 2.30368 | 3.195 |
| chr9 | 105160569 | 105160807 | 0.273378 | 0.873486 | 3.195 |
| chr7 | 141853243 | 141853480 | 0.764002 | 2.44117 | 3.195 |
| chr1 | 230717464 | 230717801 | 0.792049 | 2.53105 | 3.196 |
| chr4 | 86677904 | 86678439 | 0.602392 | 1.92503 | 3.196 |
| chr17 | 53626567 | 53626857 | 0.834404 | 2.66701 | 3.196 |
| chr6 | 100456889 | 100457107 | 0.374109 | 1.19584 | 3.197 |
| chr14 | 71633184 | 71633503 | 1.25928 | 4.02558 | 3.197 |
| chr17 | 30849579 | 30849865 | 0.395236 | 1.26351 | 3.197 |
| chr8 | 124778458 | 124778814 | 1.10215 | 3.52358 | 3.197 |
| chr12 | 83398481 | 83398750 | 0.906803 | 2.89985 | 3.198 |
| chr1 | 44502231 | 44502597 | 0.969346 | 3.10011 | 3.198 |
| chr12 | 130071651 | 130071960 | 0.387854 | 1.2405 | 3.198 |
| chr10 | 47696308 | 47696572 | 0.705464 | 2.25735 | 3.200 |
| chr13 | 20710892 | 20711257 | 1.31316 | 4.20191 | 3.200 |
| chr3 | 135797691 | 135797918 | 0.698333 | 2.23497 | 3.200 |
| chr4 | 159651141 | 159651465 | 1.13349 | 3.62785 | 3.201 |
| chr9 | 91356304 | 91356564 | 0.865976 | 2.77166 | 3.201 |
| chr3 | 189838266 | 189838898 | 0.457469 | 1.4643 | 3.201 |
| chr17 | 19771048 | 19771417 | 0.494998 | 1.58455 | 3.201 |
| chr6 | 40995682 | 40996124 | 1.30373 | 4.17385 | 3.201 |
| chr15 | 23894114 | 23894356 | 0.695986 | 2.22822 | 3.202 |
| chr9 | 98230434 | 98230649 | 0.451659 | 1.44601 | 3.202 |
| chr19 | 50758385 | 50758689 | 1.06611 | 3.41362 | 3.202 |
| chr10 | 127650368 | 127650600 | 0.610891 | 1.95626 | 3.202 |
| chr20 | 47291037 | 47291343 | 0.338868 | 1.08565 | 3.204 |
| chr3 | 119957355 | 119957614 | 0.322894 | 1.03452 | 3.204 |
| chr7 | 112923262 | 112923467 | 0.477025 | 1.52869 | 3.205 |
| chr12 | 58158743 | 58159309 | 0.823314 | 2.63881 | 3.205 |
| chr16 | 30547148 | 30547457 | 0.347558 | 1.11396 | 3.205 |
| chr12 | 56349806 | 56350136 | 0.993817 | 3.18532 | 3.205 |
| chr1 | 59118296 | 59118590 | 1.42392 | 4.56427 | 3.205 |
| chr9 | 136294756 | 136295030 | 0.800584 | 2.56644 | 3.206 |
| chr3 | 181418264 | 181418600 | 1.17599 | 3.77002 | 3.206 |
| chr3 | 10909116 | 10909446 | 0.358707 | 1.14997 | 3.206 |
| chr17 | 7892108 | 7892345 | 0.788341 | 2.52772 | 3.206 |
| chr10 | 7571677 | 7571910 | 0.808885 | 2.59386 | 3.207 |
| chr15 | 54009221 | 54009451 | 0.613296 | 1.96676 | 3.207 |
| chr4 | 170414093 | 170414361 | 0.843308 | 2.70454 | 3.207 |
| chr3 | 109450804 | 109451079 | 0.963702 | 3.09087 | 3.207 |

| | | | | | |
|---|---|---|---|---|---|
| chr10 | 66138147 | 66138369 | 0.434023 | 1.3922 | 3.208 |
| chr12 | 132860900 | 132861288 | 0.835982 | 2.68219 | 3.208 |
| chr1 | 12268953 | 12269338 | 1.18737 | 3.80999 | 3.209 |
| chr7 | 82243619 | 82243851 | 0.37793 | 1.21269 | 3.209 |
| chr10 | 26895423 | 26895640 | 0.445244 | 1.4288 | 3.209 |
| chr17 | 33096677 | 33096919 | 0.514118 | 1.64985 | 3.209 |
| chr14 | 54642113 | 54642355 | 1.26247 | 4.05187 | 3.209 |
| chr14 | 91604757 | 91605124 | 1.06102 | 3.40566 | 3.210 |
| chr10 | 107523779 | 107524035 | 0.296665 | 0.952269 | 3.210 |
| chr5 | 36682694 | 36682998 | 0.261497 | 0.839391 | 3.210 |
| chr15 | 75986993 | 75987290 | 0.859477 | 2.75889 | 3.210 |
| chr10 | 56616311 | 56616528 | 0.341676 | 1.09691 | 3.210 |
| chr3 | 124705437 | 124705689 | 1.35513 | 4.35119 | 3.211 |
| chr10 | 76060206 | 76060423 | 0.400411 | 1.28574 | 3.211 |
| chr5 | 12578735 | 12578971 | 0.391882 | 1.25858 | 3.212 |
| chr5 | 39997751 | 39997967 | 0.38922 | 1.25007 | 3.212 |
| chr5 | 64678532 | 64678792 | 0.434882 | 1.39734 | 3.213 |
| chr4 | 30716056 | 30716374 | 0.79381 | 2.55073 | 3.213 |
| chr15 | 93174952 | 93175276 | 0.871601 | 2.80075 | 3.213 |
| chr4 | 148082150 | 148082385 | 0.860639 | 2.76553 | 3.213 |
| chr7 | 72217000 | 72217271 | 0.64593 | 2.07567 | 3.213 |
| chr9 | 112203058 | 112203323 | 1.16467 | 3.74265 | 3.213 |
| chr7 | 87308203 | 87308501 | 0.350164 | 1.12548 | 3.214 |
| chr12 | 5994184 | 5994415 | 0.3686 | 1.18479 | 3.214 |
| chr19 | 19250465 | 19250831 | 1.22393 | 3.93421 | 3.214 |
| chrX | 131226536 | 131226810 | 0.969739 | 3.11735 | 3.215 |
| chr10 | 93640173 | 93640426 | 0.400043 | 1.286 | 3.215 |
| chr19 | 42829265 | 42829617 | 1.9315 | 6.20957 | 3.215 |
| chr20 | 43858231 | 43858477 | 1.0562 | 3.396 | 3.215 |
| chr14 | 27453806 | 27454111 | 0.974687 | 3.13418 | 3.216 |
| chr12 | 12941820 | 12942039 | 1.02114 | 3.28392 | 3.216 |
| chr8 | 32809115 | 32809317 | 0.799007 | 2.56977 | 3.216 |
| chr8 | 50679438 | 50679642 | 0.338751 | 1.08949 | 3.216 |
| chr4 | 155558905 | 155559142 | 0.656008 | 2.1099 | 3.216 |
| chr5 | 149815577 | 149816040 | 1.24377 | 4.0003 | 3.216 |
| chr14 | 20693192 | 20693469 | 0.316917 | 1.01932 | 3.216 |
| chr18 | 56936417 | 56936711 | 1.14718 | 3.68996 | 3.217 |
| chr10 | 42924053 | 42924257 | 0.732385 | 2.35585 | 3.217 |
| chr12 | 57495837 | 57496053 | 0.459733 | 1.47892 | 3.217 |
| chr16 | 67975993 | 67976436 | 1.2442 | 4.0026 | 3.217 |
| chr14 | 67826666 | 67827011 | 1.2518 | 4.02728 | 3.217 |
| chr17 | 47331337 | 47331713 | 0.708837 | 2.28052 | 3.217 |
| chr18 | 61245681 | 61245912 | 0.84691 | 2.72581 | 3.219 |
| chr11 | 13174320 | 13174574 | 0.782904 | 2.52027 | 3.219 |
| chr14 | 23723368 | 23723598 | 0.536073 | 1.72582 | 3.219 |
| chr12 | 108015740 | 108016104 | 1.25921 | 4.05407 | 3.220 |
| chr11 | 113176447 | 113176827 | 1.59459 | 5.13407 | 3.220 |
| chr10 | 93170022 | 93170243 | 0.657232 | 2.11643 | 3.220 |
| chr6 | 10887140 | 10887411 | 1.14439 | 3.68589 | 3.221 |
| chr1 | 238336400 | 238336672 | 0.381239 | 1.228 | 3.221 |
| chr13 | 80938115 | 80938353 | 0.766419 | 2.46877 | 3.221 |
| chr14 | 95288894 | 95289171 | 0.342476 | 1.10323 | 3.221 |
| chr8 | 11141866 | 11142205 | 0.889333 | 2.86496 | 3.221 |
| chr9 | 137300330 | 137300581 | 0.826678 | 2.66316 | 3.222 |
| chr7 | 22388101 | 22388366 | 0.316072 | 1.01825 | 3.222 |
| chr8 | 23103684 | 23104606 | 0.972866 | 3.13446 | 3.222 |
| chr2 | 83743714 | 83743920 | 0.458426 | 1.47711 | 3.222 |
| chr5 | 171761219 | 171761531 | 0.570394 | 1.83796 | 3.222 |
| chr12 | 107485775 | 107486199 | 0.941318 | 3.03344 | 3.223 |
| chr8 | 47579865 | 47580155 | 1.33156 | 4.29105 | 3.223 |
| chr5 | 57185559 | 57185790 | 0.945235 | 3.04625 | 3.223 |

| | | | | | |
|---|---|---|---|---|---|
| chr16 | 57701880 | 57702167 | 0.875544 | 2.82183 | 3.223 |
| chr5 | 144201368 | 144201681 | 0.288185 | 0.928802 | 3.223 |
| chr13 | 34298069 | 34298372 | 0.75897 | 2.44647 | 3.223 |
| chr17 | 62141149 | 62141495 | 1.13855 | 3.67001 | 3.223 |
| chr14 | 67920138 | 67920344 | 0.27047 | 0.872034 | 3.224 |
| chr5 | 1511867 | 1512134 | 0.862199 | 2.78036 | 3.225 |
| chr6 | 33679489 | 33679750 | 0.692868 | 2.23435 | 3.225 |
| chr15 | 74667572 | 74667915 | 1.1018 | 3.55373 | 3.225 |
| chr11 | 13894763 | 13895068 | 1.35778 | 4.37993 | 3.226 |
| chr16 | 21168119 | 21168321 | 0.83186 | 2.68347 | 3.226 |
| chr1 | 221437202 | 221437427 | 0.804124 | 2.59411 | 3.226 |
| chr7 | 18610588 | 18610845 | 0.614025 | 1.98087 | 3.226 |
| chr12 | 5499323 | 5499532 | 0.508658 | 1.64131 | 3.227 |
| chr17 | 12429772 | 12430006 | 0.426591 | 1.37653 | 3.227 |
| chr10 | 12294913 | 12295145 | 0.728654 | 2.35126 | 3.227 |
| chr11 | 66370032 | 66370236 | 0.363406 | 1.17272 | 3.227 |
| chr16 | 69432275 | 69432492 | 0.930002 | 3.00121 | 3.227 |
| chr11 | 3828988 | 3829417 | 1.24928 | 4.03167 | 3.227 |
| chr9 | 100684613 | 100684825 | 0.999875 | 3.22748 | 3.228 |
| chr20 | 43556255 | 43556506 | 0.398635 | 1.28686 | 3.228 |
| chr12 | 115858186 | 115858999 | 0.450929 | 1.45576 | 3.228 |
| chr13 | 71894665 | 71894900 | 0.635392 | 2.05136 | 3.229 |
| chr7 | 149157606 | 149157891 | 1.23355 | 3.98323 | 3.229 |
| chr7 | 99591724 | 99592153 | 1.16227 | 3.75328 | 3.229 |
| chr8 | 88707460 | 88707773 | 0.623684 | 2.01477 | 3.230 |
| chr16 | 48576038 | 48576388 | 1.24342 | 4.01717 | 3.231 |
| chr1 | 33352188 | 33352530 | 1.29713 | 4.19082 | 3.231 |
| chr5 | 50270746 | 50271027 | 0.427038 | 1.37972 | 3.231 |
| chr7 | 100231545 | 100231751 | 1.03481 | 3.34346 | 3.231 |
| chr5 | 92917095 | 92917391 | 1.81163 | 5.85449 | 3.232 |
| chr4 | 164471498 | 164471759 | 1.20502 | 3.89428 | 3.232 |
| chr14 | 24905944 | 24906258 | 1.38121 | 4.4645 | 3.232 |
| chr7 | 139024800 | 139025154 | 1.28132 | 4.14178 | 3.232 |
| chr9 | 2971692 | 2972034 | 0.408989 | 1.32222 | 3.233 |
| chr16 | 52102382 | 52102594 | 0.824574 | 2.66587 | 3.233 |
| chr14 | 31739374 | 31739803 | 1.29479 | 4.18646 | 3.233 |
| chr8 | 64305483 | 64305719 | 0.289515 | 0.936095 | 3.233 |
| chr1 | 24439112 | 24439340 | 0.868904 | 2.80949 | 3.233 |
| chrX | 130887239 | 130887444 | 0.661956 | 2.14036 | 3.233 |
| chr8 | 53853280 | 53854515 | 1.13618 | 3.67388 | 3.234 |
| chr12 | 96794159 | 96794532 | 1.53127 | 4.95205 | 3.234 |
| chr10 | 43951124 | 43951444 | 1.15639 | 3.74049 | 3.235 |
| chr6 | 55806546 | 55806822 | 0.474023 | 1.53334 | 3.235 |
| chr19 | 52179196 | 52179476 | 0.961932 | 3.11165 | 3.235 |
| chr5 | 1053070 | 1053513 | 0.513596 | 1.66159 | 3.235 |
| chr12 | 49523988 | 49524365 | 2.25842 | 7.30745 | 3.236 |
| chr17 | 38599880 | 38600375 | 0.724229 | 2.34374 | 3.236 |
| chr2 | 1595722 | 1596036 | 0.828478 | 2.68134 | 3.236 |
| chr9 | 125141652 | 125142017 | 1.06613 | 3.45054 | 3.237 |
| chr9 | 78468392 | 78468651 | 0.282826 | 0.915424 | 3.237 |
| chr12 | 55068001 | 55068219 | 0.373522 | 1.20927 | 3.237 |
| chr16 | 73906066 | 73906334 | 0.388646 | 1.25849 | 3.238 |
| chr1 | 227930139 | 227930413 | 0.297935 | 0.964801 | 3.238 |
| chr2 | 11970095 | 11970321 | 1.05812 | 3.42652 | 3.238 |
| chr16 | 61267345 | 61267668 | 0.524114 | 1.69772 | 3.239 |
| chr10 | 95547189 | 95547476 | 0.288232 | 0.933649 | 3.239 |
| chr5 | 173713367 | 173713643 | 0.378613 | 1.22653 | 3.240 |
| chr9 | 94520627 | 94521053 | 1.02162 | 3.31009 | 3.240 |
| chr20 | 51180477 | 51180703 | 0.365794 | 1.1853 | 3.240 |
| chr10 | 132484761 | 132485225 | 0.45122 | 1.46245 | 3.241 |
| chr5 | 108774594 | 108774874 | 0.610699 | 1.97953 | 3.241 |

| | | | | | |
|---|---|---|---|---|---|
| chr11 | 91727677 | 91727890 | 0.3626 | 1.17578 | 3.243 |
| chr11 | 126310764 | 126311130 | 1.26381 | 4.09821 | 3.243 |
| chr3 | 154124105 | 154124314 | 0.309494 | 1.00362 | 3.243 |
| chr10 | 12753092 | 12753495 | 1.38683 | 4.49729 | 3.243 |
| chr10 | 96935889 | 96936180 | 0.960823 | 3.11629 | 3.243 |
| chr10 | 11767202 | 11767516 | 0.971954 | 3.15241 | 3.243 |
| chr7 | 27200515 | 27200997 | 1.44503 | 4.68722 | 3.244 |
| chr7 | 139415404 | 139415707 | 0.321591 | 1.04317 | 3.244 |
| chrX | 46618700 | 46619059 | 1.4577 | 4.7287 | 3.244 |
| chr3 | 85616279 | 85616498 | 0.84566 | 2.74384 | 3.245 |
| chr17 | 26220204 | 26220677 | 2.30025 | 7.46353 | 3.245 |
| chr14 | 23246749 | 23246977 | 0.62547 | 2.0295 | 3.245 |
| chr1 | 3612235 | 3612589 | 1.34535 | 4.36564 | 3.245 |
| chr11 | 4088197 | 4088712 | 0.367109 | 1.19131 | 3.245 |
| chr8 | 22280082 | 22280435 | 1.09614 | 3.55783 | 3.246 |
| chr16 | 67450301 | 67450600 | 1.41057 | 4.57856 | 3.246 |
| chrX | 68165756 | 68166079 | 0.56184 | 1.82394 | 3.246 |
| chr12 | 48299113 | 48299349 | 0.847176 | 2.75067 | 3.247 |
| chr16 | 55677135 | 55677366 | 0.415653 | 1.34965 | 3.247 |
| chr1 | 105323198 | 105323402 | 0.779859 | 2.53328 | 3.248 |
| chr20 | 21468855 | 21469058 | 0.410477 | 1.33345 | 3.249 |
| chr15 | 25746996 | 25747344 | 0.515562 | 1.67497 | 3.249 |
| chr6 | 119097766 | 119098039 | 0.811404 | 2.6363 | 3.249 |
| chr3 | 188988080 | 188988301 | 1.12576 | 3.65803 | 3.249 |
| chr10 | 4524409 | 4524617 | 0.437302 | 1.42109 | 3.250 |
| chr5 | 125674915 | 125675246 | 1.01149 | 3.28703 | 3.250 |
| chr7 | 41355462 | 41355727 | 0.674523 | 2.1925 | 3.250 |
| chr3 | 97438098 | 97438410 | 1.69888 | 5.52254 | 3.251 |
| chr11 | 113534003 | 113534210 | 0.302781 | 0.984257 | 3.251 |
| chr7 | 105292761 | 105293155 | 1.64685 | 5.35351 | 3.251 |
| chr1 | 245374482 | 245374704 | 0.455622 | 1.48155 | 3.252 |
| chr18 | 57663687 | 57663956 | 0.830491 | 2.70138 | 3.253 |
| chr1 | 91316825 | 91317359 | 0.661745 | 2.15341 | 3.254 |
| chr15 | 51484874 | 51485289 | 2.01407 | 6.55488 | 3.255 |
| chr3 | 32943559 | 32943836 | 0.803269 | 2.6149 | 3.255 |
| chr11 | 67415686 | 67415904 | 0.587055 | 1.91125 | 3.256 |
| chr14 | 69283418 | 69283769 | 1.9662 | 6.4019 | 3.256 |
| chr6 | 53409175 | 53409783 | 0.749306 | 2.43983 | 3.256 |
| chr12 | 130737259 | 130737688 | 0.331624 | 1.08005 | 3.257 |
| chr8 | 143374957 | 143375337 | 0.694893 | 2.26319 | 3.257 |
| chr12 | 77359176 | 77359520 | 0.287577 | 0.936707 | 3.257 |
| chr19 | 30341605 | 30341822 | 0.349996 | 1.14006 | 3.257 |
| chr8 | 47829206 | 47829699 | 1.83781 | 5.98647 | 3.257 |
| chr11 | 33857031 | 33857241 | 0.442332 | 1.44093 | 3.258 |
| chrX | 23800910 | 23801219 | 1.16461 | 3.79406 | 3.258 |
| chr20 | 7177596 | 7177820 | 0.412138 | 1.34289 | 3.258 |
| chr11 | 122073644 | 122074022 | 1.55628 | 5.07093 | 3.258 |
| chr11 | 10664355 | 10664595 | 0.268512 | 0.874948 | 3.259 |
| chr16 | 66549660 | 66549997 | 0.853969 | 2.78338 | 3.259 |
| chr3 | 146577682 | 146577905 | 0.54713 | 1.78378 | 3.260 |
| chr9 | 136858060 | 136858367 | 0.926032 | 3.02075 | 3.262 |
| chr11 | 518616 | 519094 | 2.35678 | 7.68814 | 3.262 |
| chr19 | 35342394 | 35342647 | 0.772505 | 2.52009 | 3.262 |
| chr1 | 239549887 | 239550131 | 0.456331 | 1.48901 | 3.263 |
| chr15 | 53052152 | 53052396 | 0.397061 | 1.29563 | 3.263 |
| chr19 | 11641385 | 11641604 | 0.699199 | 2.28163 | 3.263 |
| chr1 | 244964500 | 244964837 | 1.06643 | 3.48054 | 3.264 |
| chr11 | 17743604 | 17743881 | 0.603638 | 1.97029 | 3.264 |
| chr2 | 28619717 | 28619928 | 0.843531 | 2.75353 | 3.264 |
| chr2 | 67549831 | 67550127 | 1.16072 | 3.78905 | 3.264 |
| chr12 | 66859770 | 66860216 | 0.281036 | 0.917542 | 3.265 |

| | | | | | |
|---|---|---|---|---|---|
| chr12 | 9531241 | 9531457 | 0.715688 | 2.33668 | 3.265 |
| chr10 | 98994201 | 98994523 | 0.800068 | 2.61231 | 3.265 |
| chr1 | 25427950 | 25428247 | 1.47546 | 4.81962 | 3.267 |
| chrY | 15841482 | 15841736 | 0.564906 | 1.84549 | 3.267 |
| chr14 | 107287354 | 107287817 | 1.32516 | 4.32927 | 3.267 |
| chr11 | 98886502 | 98886925 | 0.96431 | 3.15073 | 3.267 |
| chr12 | 27192026 | 27192653 | 0.495034 | 1.61753 | 3.268 |
| chr8 | 144876718 | 144877055 | 0.473879 | 1.5487 | 3.268 |
| chr6 | 138838733 | 138839100 | 0.863875 | 2.82345 | 3.268 |
| chr16 | 49789257 | 49789468 | 0.697151 | 2.27863 | 3.268 |
| chr14 | 39484599 | 39484912 | 0.873612 | 2.8554 | 3.269 |
| chr12 | 110059577 | 110059851 | 0.361991 | 1.18319 | 3.269 |
| chr5 | 128010582 | 128010869 | 1.36184 | 4.45127 | 3.269 |
| chr8 | 72819243 | 72819551 | 0.309006 | 1.01002 | 3.269 |
| chr17 | 38136717 | 38137002 | 0.920921 | 3.0104 | 3.269 |
| chr12 | 106529966 | 106530313 | 1.0781 | 3.52422 | 3.269 |
| chr12 | 123144685 | 123145042 | 0.809646 | 2.64683 | 3.269 |
| chr3 | 188951054 | 188951374 | 0.982093 | 3.21148 | 3.270 |
| chr16 | 55302489 | 55302708 | 0.468271 | 1.53187 | 3.271 |
| chr5 | 164999566 | 164999772 | 0.399462 | 1.30682 | 3.271 |
| chr12 | 59543808 | 59544141 | 1.70023 | 5.56462 | 3.273 |
| chr15 | 66678883 | 66679197 | 1.27344 | 4.16795 | 3.273 |
| chr16 | 88878781 | 88879246 | 0.458784 | 1.50165 | 3.273 |
| chr11 | 2421417 | 2421839 | 1.77098 | 5.79745 | 3.274 |
| chr5 | 131690792 | 131691106 | 0.370624 | 1.2133 | 3.274 |
| chr14 | 86946838 | 86947058 | 0.414377 | 1.35655 | 3.274 |
| chr14 | 36414076 | 36414381 | 0.943144 | 3.08764 | 3.274 |
| chr9 | 27333188 | 27333440 | 0.907059 | 2.9701 | 3.274 |
| chr15 | 34742288 | 34742684 | 0.407064 | 1.33312 | 3.275 |
| chr21 | 38362308 | 38362693 | 0.900285 | 2.94925 | 3.276 |
| chr10 | 94875576 | 94875794 | 1.23842 | 4.05722 | 3.276 |
| chr18 | 58493506 | 58493778 | 0.410817 | 1.34601 | 3.276 |
| chr18 | 24393071 | 24393295 | 0.805892 | 2.64086 | 3.277 |
| chr7 | 81877155 | 81877443 | 0.893081 | 2.92684 | 3.277 |
| chr1 | 9144811 | 9145196 | 0.306192 | 1.00362 | 3.278 |
| chr12 | 38554003 | 38554272 | 0.748303 | 2.45283 | 3.278 |
| chr6 | 2354169 | 2354451 | 1.06625 | 3.49515 | 3.278 |
| chr2 | 122159908 | 122160214 | 1.46221 | 4.79388 | 3.279 |
| chr12 | 13136404 | 13136694 | 1.01021 | 3.31212 | 3.279 |
| chr18 | 32586553 | 32586755 | 0.268689 | 0.881135 | 3.279 |
| chr13 | 50443740 | 50443986 | 0.963579 | 3.16009 | 3.280 |
| chr1 | 164409544 | 164409773 | 0.540084 | 1.77128 | 3.280 |
| chr5 | 123379000 | 123379468 | 0.753537 | 2.47135 | 3.280 |
| chr9 | 108418347 | 108418678 | 0.640998 | 2.10238 | 3.280 |
| chr12 | 62283955 | 62284296 | 0.398273 | 1.30633 | 3.280 |
| rUn_gl0002 | 83086 | 83296 | 0.339798 | 1.11454 | 3.280 |
| chr12 | 20777729 | 20777956 | 1.16347 | 3.8162 | 3.280 |
| chr6 | 134158932 | 134159353 | 1.26466 | 4.14873 | 3.281 |
| chr5 | 159626296 | 159626715 | 2.08375 | 6.83722 | 3.281 |
| chr22 | 31286599 | 31286963 | 0.997707 | 3.2738 | 3.281 |
| chr8 | 106907801 | 106908067 | 0.359394 | 1.1793 | 3.281 |
| chr4 | 171296013 | 171296348 | 0.347852 | 1.14149 | 3.282 |
| chr10 | 17068495 | 17068747 | 0.304258 | 0.998453 | 3.282 |
| chr3 | 33376260 | 33376471 | 0.369768 | 1.2136 | 3.282 |
| chr14 | 101174660 | 101174955 | 0.47065 | 1.54476 | 3.282 |
| chr6 | 137962697 | 137962903 | 0.617066 | 2.02558 | 3.283 |
| chr8 | 123617242 | 123617473 | 0.388786 | 1.27631 | 3.283 |
| chr1 | 98516693 | 98516934 | 0.841066 | 2.76153 | 3.283 |
| chr13 | 95087887 | 95088289 | 0.599893 | 1.96972 | 3.283 |
| chr4 | 2464507 | 2464755 | 0.798338 | 2.62186 | 3.284 |
| chrY | 23899158 | 23899394 | 0.750984 | 2.46641 | 3.284 |

| | | | | | |
|---|---|---|---|---|---|
| chr12 | 49359327 | 49359553 | 0.79217 | 2.60237 | 3.285 |
| chr4 | 121992370 | 121992730 | 0.452368 | 1.48612 | 3.285 |
| chr17 | 2135013 | 2135294 | 1.05064 | 3.4516 | 3.285 |
| chr18 | 20510086 | 20510376 | 0.954252 | 3.13497 | 3.285 |
| chr13 | 103046512 | 103046793 | 0.636255 | 2.09037 | 3.285 |
| chr13 | 49169837 | 49170060 | 0.972708 | 3.19596 | 3.286 |
| chr11 | 111802565 | 111802877 | 0.607253 | 1.99523 | 3.286 |
| chr4 | 127700961 | 127701322 | 0.419519 | 1.37841 | 3.286 |
| chr19 | 13832113 | 13832367 | 0.993747 | 3.26553 | 3.286 |
| chr3 | 11278146 | 11278650 | 1.48359 | 4.87529 | 3.286 |
| chr16 | 85061661 | 85061950 | 0.612697 | 2.01346 | 3.286 |
| chr5 | 77923077 | 77923384 | 0.872351 | 2.8669 | 3.286 |
| chr8 | 42315597 | 42315848 | 1.01956 | 3.35089 | 3.287 |
| chr15 | 85388805 | 85389114 | 1.6231 | 5.33493 | 3.287 |
| chr4 | 107819864 | 107820094 | 0.592812 | 1.94859 | 3.287 |
| chr3 | 169176011 | 169176303 | 0.393589 | 1.29386 | 3.287 |
| chrX | 131765730 | 131766032 | 0.776007 | 2.55116 | 3.288 |
| chr1 | 150186133 | 150186396 | 0.611195 | 2.00934 | 3.288 |
| chr9 | 93571024 | 93571448 | 0.343125 | 1.12816 | 3.288 |
| chr2 | 30931747 | 30932048 | 0.85361 | 2.80709 | 3.288 |
| chr20 | 25291043 | 25291356 | 0.9052 | 2.97677 | 3.289 |
| chr15 | 93307941 | 93308178 | 0.374796 | 1.23259 | 3.289 |
| chr17 | 64911256 | 64911630 | 1.14775 | 3.7747 | 3.289 |
| chr5 | 140201056 | 140201320 | 0.553303 | 1.81994 | 3.289 |
| chr12 | 298231 | 298620 | 1.30639 | 4.29742 | 3.290 |
| chr8 | 215685 | 216025 | 1.67907 | 5.5234 | 3.290 |
| chr17 | 1687307 | 1687701 | 0.723439 | 2.37997 | 3.290 |
| chr16 | 17770088 | 17770294 | 0.32702 | 1.07595 | 3.290 |
| chr3 | 158839751 | 158840050 | 0.274151 | 0.902052 | 3.290 |
| chr7 | 45978576 | 45978784 | 0.879748 | 2.89515 | 3.291 |
| chrX | 81473856 | 81474084 | 0.695333 | 2.28842 | 3.291 |
| chr1 | 6305161 | 6305431 | 1.14144 | 3.75688 | 3.291 |
| chr20 | 42073551 | 42074044 | 0.670819 | 2.208 | 3.292 |
| chr7 | 16110931 | 16111248 | 0.740041 | 2.43627 | 3.292 |
| chr1 | 202895877 | 202896119 | 0.854167 | 2.81212 | 3.292 |
| chr1 | 29668464 | 29668708 | 0.572658 | 1.88557 | 3.293 |
| chr15 | 44231327 | 44231676 | 1.07964 | 3.55499 | 3.293 |
| chr3 | 5163688 | 5164142 | 1.52099 | 5.00833 | 3.293 |
| chr2 | 95897426 | 95897739 | 1.26581 | 4.16841 | 3.293 |
| chr5 | 172197558 | 172197989 | 0.713093 | 2.34829 | 3.293 |
| chr12 | 70970863 | 70971117 | 1.12336 | 3.69988 | 3.294 |
| chr18 | 831011 | 831234 | 0.687975 | 2.26611 | 3.294 |
| chr1 | 7599599 | 7600120 | 0.661957 | 2.18103 | 3.295 |
| chr3 | 195501843 | 195502087 | 1.41029 | 4.64671 | 3.295 |
| chr5 | 40160620 | 40160829 | 0.375685 | 1.23793 | 3.295 |
| chr7 | 304608 | 304856 | 0.289304 | 0.953349 | 3.295 |
| chr13 | 37573395 | 37573715 | 1.68882 | 5.56542 | 3.295 |
| chr19 | 11337834 | 11338123 | 0.772423 | 2.54552 | 3.296 |
| chr10 | 80962286 | 80962621 | 1.22953 | 4.05204 | 3.296 |
| chr12 | 100040034 | 100040279 | 0.398126 | 1.31208 | 3.296 |
| chr19 | 17981143 | 17981388 | 1.08048 | 3.561 | 3.296 |
| chr22 | 25477376 | 25477684 | 0.753325 | 2.48336 | 3.297 |
| chr8 | 99847479 | 99847717 | 0.318062 | 1.04862 | 3.297 |
| chr19 | 23603635 | 23603905 | 0.701749 | 2.31411 | 3.298 |
| chr15 | 42942705 | 42942947 | 0.695821 | 2.29476 | 3.298 |
| chr12 | 262329 | 262779 | 1.78635 | 5.89126 | 3.298 |
| chr12 | 131937149 | 131937552 | 1.4242 | 4.69691 | 3.298 |
| chr4 | 189379539 | 189379867 | 0.796959 | 2.62847 | 3.298 |
| chr9 | 99429005 | 99429405 | 1.11829 | 3.68837 | 3.298 |
| chr6 | 74019971 | 74020270 | 0.869059 | 2.86665 | 3.299 |
| chr11 | 74437572 | 74437966 | 1.66708 | 5.49956 | 3.299 |

| | | | | | |
|---|---|---|---|---|---|
| chr19 | 12167112 | 12167428 | 0.562668 | 1.85653 | 3.300 |
| chr5 | 149145469 | 149145779 | 0.637591 | 2.10413 | 3.300 |
| chr19 | 10930956 | 10931176 | 0.584345 | 1.92844 | 3.300 |
| chr3 | 42822368 | 42822623 | 0.657318 | 2.16946 | 3.300 |
| chr4 | 74730145 | 74730428 | 1.02034 | 3.36782 | 3.301 |
| chr16 | 57790864 | 57791518 | 1.21284 | 4.00356 | 3.301 |
| chr17 | 79766730 | 79767000 | 0.872363 | 2.87979 | 3.301 |
| chr15 | 78395563 | 78395806 | 0.416414 | 1.37465 | 3.301 |
| chr1 | 235667840 | 235668174 | 1.07663 | 3.55416 | 3.301 |
| chr19 | 40334635 | 40334921 | 0.97907 | 3.23325 | 3.302 |
| chr15 | 91878913 | 91879132 | 0.435143 | 1.43743 | 3.303 |
| chr12 | 103603368 | 103603638 | 0.364773 | 1.205 | 3.303 |
| chr5 | 121412395 | 121412716 | 1.67362 | 5.52879 | 3.303 |
| chr11 | 72193285 | 72193756 | 0.960948 | 3.1754 | 3.304 |
| chr7 | 102558169 | 102558401 | 0.508925 | 1.68179 | 3.305 |
| chr16 | 31725915 | 31726143 | 0.842118 | 2.78294 | 3.305 |
| chr17 | 71751227 | 71751563 | 0.359543 | 1.18835 | 3.305 |
| chr1 | 161195386 | 161195622 | 0.681435 | 2.25246 | 3.305 |
| chr7 | 55462090 | 55462317 | 0.363447 | 1.20153 | 3.306 |
| chr14 | 93626644 | 93627059 | 1.06743 | 3.52939 | 3.306 |
| chr8 | 52921025 | 52921422 | 1.93514 | 6.40039 | 3.307 |
| chr11 | 111101099 | 111101431 | 1.14688 | 3.79406 | 3.308 |
| chr19 | 15363276 | 15363579 | 0.799113 | 2.64375 | 3.308 |
| chr17 | 10568079 | 10568281 | 0.423916 | 1.40249 | 3.308 |
| chr19 | 46009678 | 46009912 | 0.775207 | 2.56547 | 3.309 |
| chr1 | 52762031 | 52762234 | 1.29148 | 4.27418 | 3.310 |
| chr14 | 87691312 | 87691566 | 0.239831 | 0.793778 | 3.310 |
| chr5 | 172337831 | 172338163 | 0.952851 | 3.15376 | 3.310 |
| chr1 | 175349470 | 175349750 | 0.250421 | 0.828888 | 3.310 |
| chr12 | 48848517 | 48848811 | 0.795179 | 2.63203 | 3.310 |
| chr1 | 101071944 | 101072405 | 0.961517 | 3.18317 | 3.311 |
| chr12 | 3427496 | 3427710 | 0.308766 | 1.02261 | 3.312 |
| rUn_gl0002 | 99687 | 99895 | 1.16489 | 3.85816 | 3.312 |
| chr2 | 192096715 | 192097022 | 0.637655 | 2.11227 | 3.313 |
| chr1 | 30179955 | 30180165 | 1.4677 | 4.86228 | 3.313 |
| chr11 | 65017865 | 65018233 | 0.948251 | 3.14162 | 3.313 |
| chr1 | 168204556 | 168204829 | 1.10888 | 3.67463 | 3.314 |
| chr5 | 1004710 | 1005082 | 1.12994 | 3.74543 | 3.315 |
| chr11 | 64655278 | 64655722 | 2.13552 | 7.07951 | 3.315 |
| chr5 | 59632107 | 59632357 | 0.44172 | 1.46451 | 3.315 |
| chr8 | 6281495 | 6281701 | 0.755352 | 2.50438 | 3.316 |
| chr3 | 58980504 | 58980751 | 0.90591 | 3.00429 | 3.316 |
| chr20 | 19252702 | 19252926 | 0.335157 | 1.1115 | 3.316 |
| chr10 | 85933414 | 85933686 | 0.343394 | 1.13883 | 3.316 |
| chr5 | 157375795 | 157376176 | 1.03964 | 3.44826 | 3.317 |
| chr8 | 19932236 | 19932466 | 0.372987 | 1.23758 | 3.318 |
| chr18 | 56975570 | 56976054 | 1.01228 | 3.35895 | 3.318 |
| chr12 | 860420 | 860734 | 1.26401 | 4.19486 | 3.319 |
| chr14 | 89479392 | 89479613 | 0.828313 | 2.74897 | 3.319 |
| chr11 | 63185293 | 63185525 | 0.318123 | 1.05595 | 3.319 |
| chr17 | 80891704 | 80892133 | 0.421302 | 1.39854 | 3.320 |
| chr4 | 6676363 | 6676685 | 0.537616 | 1.78467 | 3.320 |
| chr4 | 7193988 | 7194335 | 1.3253 | 4.39999 | 3.320 |
| chr1 | 67678941 | 67679159 | 0.945289 | 3.1386 | 3.320 |
| chr1 | 152452646 | 152453047 | 1.13141 | 3.75675 | 3.320 |
| chr16 | 74132682 | 74132913 | 0.310534 | 1.03151 | 3.322 |
| chr4 | 147837677 | 147837945 | 0.817288 | 2.71535 | 3.322 |
| chr3 | 19597389 | 19597637 | 0.287171 | 0.954196 | 3.323 |
| chr16 | 87236794 | 87237217 | 1.76569 | 5.86723 | 3.323 |
| chr3 | 44956397 | 44956714 | 0.523614 | 1.73995 | 3.323 |
| chr6 | 14211357 | 14211728 | 0.940285 | 3.12473 | 3.323 |

| | | | | | |
|---|---|---|---|---|---|
| chr9 | 95491730 | 95492024 | 0.603927 | 2.00708 | 3.323 |
| chr5 | 98646578 | 98646835 | 0.28084 | 0.933403 | 3.324 |
| chr1 | 230303131 | 230303626 | 1.16958 | 3.88779 | 3.324 |
| chr17 | 51110170 | 51110439 | 0.68806 | 2.28733 | 3.324 |
| chr1 | 35764205 | 35764411 | 0.782351 | 2.60083 | 3.324 |
| chr5 | 29330990 | 29331306 | 0.430692 | 1.43199 | 3.325 |
| chr4 | 657590 | 657849 | 0.453828 | 1.50947 | 3.326 |
| chr1 | 1837665 | 1837983 | 1.29243 | 4.29885 | 3.326 |
| chr17 | 4316514 | 4316804 | 2.15427 | 7.16601 | 3.326 |
| chr6 | 5004217 | 5004549 | 1.11442 | 3.70746 | 3.327 |
| chr9 | 116615653 | 116615928 | 1.01059 | 3.36247 | 3.327 |
| chr11 | 696479 | 696716 | 1.35628 | 4.51291 | 3.327 |
| chr9 | 4776281 | 4776635 | 0.361474 | 1.20293 | 3.328 |
| chr7 | 122516330 | 122516596 | 0.374824 | 1.24738 | 3.328 |
| chr7 | 97112711 | 97112956 | 0.314755 | 1.04761 | 3.328 |
| chr8 | 56946065 | 56946317 | 0.3255 | 1.08338 | 3.328 |
| chr7 | 16965959 | 16966227 | 1.07216 | 3.56859 | 3.328 |
| chr15 | 57959512 | 57959785 | 0.987258 | 3.28622 | 3.329 |
| chr7 | 91677811 | 91678021 | 0.734816 | 2.44651 | 3.329 |
| chr14 | 91853785 | 91854153 | 0.236109 | 0.786322 | 3.330 |
| chr1 | 243488671 | 243488934 | 0.937882 | 3.12381 | 3.331 |
| chr17 | 67605711 | 67606087 | 1.2468 | 4.1528 | 3.331 |
| chr16 | 27878789 | 27879145 | 1.26054 | 4.19922 | 3.331 |
| chr12 | 85443801 | 85444026 | 0.366656 | 1.2215 | 3.331 |
| chr1 | 156783528 | 156783891 | 1.22664 | 4.08685 | 3.332 |
| chr3 | 169462490 | 169462741 | 0.520192 | 1.73324 | 3.332 |
| chr16 | 31153736 | 31154127 | 1.72965 | 5.76333 | 3.332 |
| chr16 | 53164224 | 53164667 | 1.04803 | 3.49242 | 3.332 |
| chr13 | 92979240 | 92979449 | 0.383524 | 1.27819 | 3.333 |
| chr12 | 53568803 | 53569102 | 0.923999 | 3.08088 | 3.334 |
| chr10 | 94886269 | 94886530 | 0.952739 | 3.17675 | 3.334 |
| chr8 | 145566724 | 145566996 | 0.695634 | 2.31954 | 3.334 |
| chr11 | 40743788 | 40744119 | 0.947016 | 3.15786 | 3.335 |
| chr22 | 45831588 | 45831868 | 1.46656 | 4.89216 | 3.336 |
| chr12 | 82920998 | 82921307 | 1.41888 | 4.73348 | 3.336 |
| chr7 | 13564801 | 13565033 | 0.621049 | 2.07233 | 3.337 |
| chr7 | 108233172 | 108233467 | 0.646887 | 2.15855 | 3.337 |
| chr16 | 53317312 | 53317534 | 0.81304 | 2.71311 | 3.337 |
| chrX | 69482831 | 69483173 | 1.69243 | 5.64803 | 3.337 |
| chr21 | 43801464 | 43801913 | 1.43556 | 4.79083 | 3.337 |
| chrX | 28765680 | 28765886 | 0.407897 | 1.36133 | 3.337 |
| chr3 | 181419913 | 181420278 | 1.53775 | 5.13216 | 3.337 |
| chr22 | 42322073 | 42322401 | 1.56221 | 5.21589 | 3.339 |
| chr3 | 112060330 | 112060546 | 0.379791 | 1.26809 | 3.339 |
| chr12 | 4359960 | 4360188 | 0.381452 | 1.27366 | 3.339 |
| chr11 | 69498646 | 69499019 | 1.28872 | 4.30342 | 3.339 |
| chr19 | 36984121 | 36984573 | 0.697808 | 2.33033 | 3.340 |
| chr13 | 78139892 | 78140094 | 0.665743 | 2.22372 | 3.340 |
| chr4 | 25368667 | 25368910 | 0.55878 | 1.86693 | 3.341 |
| chr5 | 127093270 | 127093530 | 1.19409 | 3.99052 | 3.342 |
| chr19 | 5137877 | 5138127 | 0.621997 | 2.07881 | 3.342 |
| chr5 | 140255395 | 140255717 | 1.10633 | 3.69766 | 3.342 |
| chr19 | 13140823 | 13141121 | 0.864335 | 2.88913 | 3.343 |
| chr18 | 1914447 | 1914805 | 0.694533 | 2.32162 | 3.343 |
| chr4 | 183127398 | 183127630 | 0.343697 | 1.14889 | 3.343 |
| chr5 | 91806538 | 91806787 | 0.609554 | 2.03764 | 3.343 |
| chr19 | 15661780 | 15662106 | 0.519682 | 1.73725 | 3.343 |
| chr19 | 49895742 | 49896034 | 0.347894 | 1.163 | 3.343 |
| chr11 | 124961144 | 124961424 | 0.589846 | 1.97271 | 3.344 |
| chr7 | 99045607 | 99045898 | 1.0804 | 3.61344 | 3.345 |
| chr7 | 136745128 | 136745445 | 0.445801 | 1.49103 | 3.345 |

| | | | | | |
|---|---|---|---|---|---|
| chr12 | 27970595 | 27970871 | 0.412033 | 1.37814 | 3.345 |
| chr5 | 103849027 | 103849301 | 0.498263 | 1.66662 | 3.345 |
| chr3 | 93723237 | 93723504 | 1.165 | 3.89684 | 3.345 |
| chr20 | 10071231 | 10071571 | 1.12591 | 3.76616 | 3.345 |
| chr21 | 35267125 | 35267359 | 1.17404 | 3.9278 | 3.346 |
| chr9 | 126723332 | 126723571 | 0.471406 | 1.57718 | 3.346 |
| chr3 | 100515514 | 100515718 | 0.63316 | 2.11886 | 3.346 |
| chr5 | 178614178 | 178614503 | 0.413018 | 1.38225 | 3.347 |
| chr10 | 49652360 | 49652721 | 0.519448 | 1.73891 | 3.348 |
| chr7 | 141524197 | 141524457 | 0.336399 | 1.12619 | 3.348 |
| chr20 | 34192356 | 34192640 | 0.761468 | 2.54936 | 3.348 |
| chr3 | 106259606 | 106259905 | 0.499309 | 1.67233 | 3.349 |
| chr4 | 54796249 | 54796545 | 0.772702 | 2.58846 | 3.350 |
| chr10 | 83173742 | 83174102 | 0.591347 | 1.98107 | 3.350 |
| chr5 | 113181097 | 113181327 | 0.36103 | 1.20963 | 3.351 |
| chr8 | 124664501 | 124664727 | 0.272469 | 0.912944 | 3.351 |
| chr6 | 168770944 | 168771369 | 1.24628 | 4.17599 | 3.351 |
| chr12 | 96485666 | 96485992 | 0.711524 | 2.38417 | 3.351 |
| chr5 | 154429390 | 154429678 | 0.747034 | 2.50326 | 3.351 |
| chr4 | 185089590 | 185089934 | 1.49648 | 5.01468 | 3.351 |
| chr1 | 208042349 | 208042732 | 1.41173 | 4.73129 | 3.351 |
| chr13 | 45768704 | 45768991 | 0.664439 | 2.22689 | 3.352 |
| chrX | 36975412 | 36975656 | 0.690736 | 2.31537 | 3.352 |
| chr1 | 183997633 | 183997855 | 0.378557 | 1.26898 | 3.352 |
| chr4 | 186392481 | 186392765 | 0.984903 | 3.30213 | 3.353 |
| chr7 | 73226053 | 73226449 | 1.1606 | 3.89188 | 3.353 |
| chr8 | 109105348 | 109105598 | 0.323133 | 1.08367 | 3.354 |
| chr1 | 110466009 | 110466437 | 0.764476 | 2.56388 | 3.354 |
| chr2 | 48243786 | 48244068 | 0.649356 | 2.17797 | 3.354 |
| chr6 | 132244591 | 132244869 | 0.797161 | 2.67465 | 3.355 |
| chr21 | 47977909 | 47978127 | 0.301438 | 1.01161 | 3.356 |
| chr5 | 140753311 | 140753686 | 0.718212 | 2.41096 | 3.357 |
| chr6 | 141884013 | 141884284 | 1.01763 | 3.41654 | 3.357 |
| chr16 | 49642684 | 49643063 | 0.645826 | 2.16837 | 3.358 |
| chr11 | 58925921 | 58926316 | 1.05594 | 3.54546 | 3.358 |
| chr11 | 130894994 | 130895300 | 0.773159 | 2.59643 | 3.358 |
| chr12 | 115105952 | 115106206 | 1.08566 | 3.64592 | 3.358 |
| chr9 | 90640304 | 90640657 | 0.43012 | 1.4445 | 3.358 |
| chr1 | 37789366 | 37789737 | 1.33367 | 4.47901 | 3.358 |
| chr4 | 144480054 | 144480373 | 1.12968 | 3.79439 | 3.359 |
| chr12 | 99038233 | 99038542 | 0.417182 | 1.40127 | 3.359 |
| chr1 | 181392722 | 181393032 | 0.873006 | 2.93277 | 3.359 |
| chr1 | 197584318 | 197584606 | 0.761469 | 2.55818 | 3.360 |
| chr12 | 2465555 | 2465870 | 0.324806 | 1.09128 | 3.360 |
| chr8 | 127616130 | 127616407 | 0.781307 | 2.62507 | 3.360 |
| chr14 | 99147328 | 99147552 | 0.390229 | 1.31113 | 3.360 |
| chr12 | 65694656 | 65694869 | 0.848707 | 2.85158 | 3.360 |
| chr11 | 65689729 | 65689947 | 0.285089 | 0.958032 | 3.360 |
| chr16 | 15968662 | 15969035 | 0.89891 | 3.02085 | 3.361 |
| chr3 | 50231976 | 50232330 | 1.32543 | 4.45494 | 3.361 |
| chr16 | 89307211 | 89307464 | 1.56829 | 5.27142 | 3.361 |
| chr5 | 122071853 | 122072248 | 0.372797 | 1.25339 | 3.362 |
| chr13 | 47125623 | 47125924 | 1.21737 | 4.09346 | 3.363 |
| chr17 | 4634961 | 4635282 | 1.14496 | 3.85004 | 3.363 |
| chr1 | 231229334 | 231229656 | 0.828137 | 2.7849 | 3.363 |
| chr11 | 64109899 | 64110160 | 0.682934 | 2.29667 | 3.363 |
| chr6 | 118830206 | 118830468 | 1.19099 | 4.00558 | 3.363 |
| chr12 | 13251001 | 13251251 | 1.19594 | 4.0223 | 3.363 |
| chr13 | 89719233 | 89719527 | 0.735748 | 2.47486 | 3.364 |
| chr1 | 95582223 | 95582501 | 1.16208 | 3.9091 | 3.364 |
| chr8 | 117715416 | 117715667 | 0.841574 | 2.83102 | 3.364 |

| | | | | | |
|---|---|---|---|---|---|
| chr19 | 29083913 | 29084156 | 0.339779 | 1.14302 | 3.364 |
| chr7 | 102663586 | 102663818 | 0.247563 | 0.832823 | 3.364 |
| chr10 | 81233505 | 81233765 | 0.42467 | 1.4288 | 3.364 |
| chr21 | 43430709 | 43430918 | 0.376674 | 1.26739 | 3.365 |
| chr8 | 143734953 | 143735217 | 0.809553 | 2.72427 | 3.365 |
| chr21 | 27464062 | 27464312 | 0.526747 | 1.77315 | 3.366 |
| chr13 | 93880157 | 93880517 | 0.47937 | 1.61405 | 3.367 |
| chr3 | 141079710 | 141080046 | 1.26009 | 4.24333 | 3.367 |
| chr17 | 60186762 | 60187171 | 2.07097 | 6.97572 | 3.368 |
| chr20 | 49252372 | 49252851 | 2.03864 | 6.86748 | 3.369 |
| chr14 | 64307745 | 64308006 | 0.61127 | 2.05933 | 3.369 |
| chr12 | 84217443 | 84217816 | 1.16912 | 3.93885 | 3.369 |
| chr12 | 53242723 | 53242989 | 0.257392 | 0.867189 | 3.369 |
| chr10 | 44162744 | 44162982 | 1.02265 | 3.4458 | 3.369 |
| chr5 | 154650039 | 154650267 | 0.341646 | 1.15119 | 3.370 |
| chr8 | 36390282 | 36390526 | 0.286027 | 0.963869 | 3.370 |
| chr11 | 85929565 | 85929843 | 1.24272 | 4.18842 | 3.370 |
| chr1 | 32098746 | 32099029 | 0.584523 | 1.97043 | 3.371 |
| chr4 | 5379790 | 5380004 | 0.587723 | 1.98124 | 3.371 |
| chr7 | 45702900 | 45703273 | 1.43012 | 4.82217 | 3.372 |
| chr17 | 67390888 | 67391215 | 1.169 | 3.94209 | 3.372 |
| chr21 | 33797911 | 33798253 | 0.407946 | 1.37586 | 3.373 |
| chr15 | 77711894 | 77712148 | 0.665493 | 2.24483 | 3.373 |
| chr16 | 87051078 | 87051376 | 0.625148 | 2.10913 | 3.374 |
| chr10 | 91226334 | 91226563 | 0.758257 | 2.55863 | 3.374 |
| chr19 | 54385386 | 54385782 | 0.847414 | 2.85954 | 3.374 |
| chr16 | 86711542 | 86711768 | 0.42786 | 1.44381 | 3.374 |
| chr10 | 27179861 | 27180148 | 0.587918 | 1.98435 | 3.375 |
| chr11 | 96076345 | 96076678 | 1.33375 | 4.50196 | 3.375 |
| chr9 | 131936296 | 131936937 | 1.23439 | 4.16715 | 3.376 |
| chr3 | 136477381 | 136477647 | 0.887805 | 2.99726 | 3.376 |
| chr11 | 13797762 | 13798009 | 0.650216 | 2.19516 | 3.376 |
| chr11 | 124994141 | 124994384 | 0.714974 | 2.41381 | 3.376 |
| chr8 | 135984543 | 135984754 | 0.345757 | 1.16733 | 3.376 |
| chr8 | 101158024 | 101158290 | 0.806221 | 2.72204 | 3.376 |
| chr9 | 91193388 | 91193799 | 1.43524 | 4.84623 | 3.377 |
| chr20 | 60471297 | 60471727 | 1.42638 | 4.81634 | 3.377 |
| chr18 | 52314500 | 52314717 | 0.637465 | 2.1525 | 3.377 |
| chr8 | 133415974 | 133416193 | 0.3198 | 1.07988 | 3.377 |
| chr14 | 53772950 | 53773337 | 0.930371 | 3.14177 | 3.377 |
| chr3 | 52419251 | 52419645 | 1.02334 | 3.45574 | 3.377 |
| chr14 | 66299619 | 66299845 | 0.937234 | 3.165 | 3.377 |
| chr6 | 56224119 | 56224337 | 0.699326 | 2.36179 | 3.377 |
| chr1 | 48175253 | 48175524 | 0.619683 | 2.09296 | 3.377 |
| chr14 | 25501696 | 25502017 | 0.872724 | 2.9478 | 3.378 |
| chr19 | 51208755 | 51209127 | 1.17316 | 3.96278 | 3.378 |
| chr6 | 7425250 | 7425515 | 0.745047 | 2.51692 | 3.378 |
| chr8 | 919242 | 919456 | 0.270797 | 0.91517 | 3.380 |
| chr2 | 71837128 | 71837381 | 0.615385 | 2.08005 | 3.380 |
| chr6 | 2634708 | 2635035 | 1.41717 | 4.79042 | 3.380 |
| chr1 | 171954849 | 171955078 | 0.811354 | 2.74267 | 3.380 |
| chr1 | 201346013 | 201346362 | 0.753651 | 2.5478 | 3.381 |
| chr17 | 73610146 | 73610451 | 1.00566 | 3.40003 | 3.381 |
| chr14 | 30396278 | 30396528 | 0.663683 | 2.24474 | 3.382 |
| chr3 | 20329630 | 20329866 | 0.333358 | 1.12781 | 3.383 |
| chr13 | 66230942 | 66231172 | 0.590255 | 1.99698 | 3.383 |
| chr11 | 101731577 | 101731807 | 0.894177 | 3.02656 | 3.385 |
| chr19 | 19226807 | 19227131 | 1.05756 | 3.57977 | 3.385 |
| chr9 | 9719965 | 9720167 | 0.40765 | 1.37987 | 3.385 |
| chr20 | 58090686 | 58091058 | 1.38311 | 4.68328 | 3.386 |
| chr11 | 60941083 | 60941286 | 0.597899 | 2.02483 | 3.387 |

| | | | | | |
|---|---|---|---|---|---|
| chr12 | 1654188 | 1654475 | 0.691994 | 2.34354 | 3.387 |
| chrX | 10693179 | 10693411 | 0.637962 | 2.16073 | 3.387 |
| chr12 | 42417936 | 42418381 | 1.34619 | 4.56025 | 3.388 |
| chr14 | 64011454 | 64011695 | 0.784617 | 2.65791 | 3.388 |
| chr12 | 53278583 | 53278933 | 1.26286 | 4.27869 | 3.388 |
| chr10 | 86189734 | 86190025 | 0.913571 | 3.09531 | 3.388 |
| chr12 | 73908387 | 73908769 | 0.736326 | 2.49481 | 3.388 |
| chr6 | 139309332 | 139309714 | 1.60626 | 5.44272 | 3.388 |
| chr7 | 25268736 | 25269065 | 0.424741 | 1.43926 | 3.389 |
| chr5 | 58746101 | 58746436 | 0.96394 | 3.26693 | 3.389 |
| chrX | 99940733 | 99941110 | 0.833081 | 2.82422 | 3.390 |
| chr10 | 80826152 | 80826458 | 0.741934 | 2.51548 | 3.390 |
| chr4 | 156680043 | 156680341 | 0.416522 | 1.41265 | 3.392 |
| chr11 | 6106363 | 6106668 | 0.362154 | 1.22852 | 3.392 |
| chr7 | 79212124 | 79212357 | 0.234435 | 0.795362 | 3.393 |
| chr17 | 22202539 | 22203665 | 1.31265 | 4.45362 | 3.393 |
| chr2 | 174432830 | 174433117 | 0.74599 | 2.53126 | 3.393 |
| chr1 | 178158969 | 178159321 | 0.708594 | 2.4047 | 3.394 |
| chr9 | 26975676 | 26975992 | 1.16622 | 3.95786 | 3.394 |
| chr11 | 133705095 | 133705472 | 0.901838 | 3.06072 | 3.394 |
| chr5 | 16076887 | 16077130 | 0.403003 | 1.36777 | 3.394 |
| chr1 | 167183963 | 167184233 | 0.41485 | 1.40799 | 3.394 |
| chr7 | 94536802 | 94537085 | 1.1775 | 3.99655 | 3.394 |
| chr12 | 96455643 | 96455932 | 0.792271 | 2.68916 | 3.394 |
| chr3 | 127945631 | 127946013 | 1.35 | 4.58249 | 3.394 |
| chr6 | 41431083 | 41431376 | 0.828215 | 2.81143 | 3.395 |
| chr15 | 92309109 | 92309352 | 0.273012 | 0.92697 | 3.395 |
| chr15 | 36043889 | 36044143 | 0.965162 | 3.27719 | 3.395 |
| chr11 | 10012256 | 10012544 | 0.5775 | 1.96111 | 3.396 |
| chr1 | 21866406 | 21866632 | 0.696627 | 2.36567 | 3.396 |
| chr2 | 21980587 | 21980797 | 0.343835 | 1.1677 | 3.396 |
| chr12 | 125083658 | 125083882 | 0.778897 | 2.64556 | 3.397 |
| chr9 | 109462575 | 109462837 | 0.235412 | 0.799702 | 3.397 |
| chr13 | 63086351 | 63086640 | 1.09762 | 3.72882 | 3.397 |
| chr10 | 88444022 | 88444399 | 0.935146 | 3.1774 | 3.398 |
| chr9 | 131464597 | 131465269 | 1.62867 | 5.534 | 3.398 |
| chr7 | 10086075 | 10086393 | 0.337401 | 1.14648 | 3.398 |
| chr5 | 64589929 | 64590177 | 0.323286 | 1.09853 | 3.398 |
| chr6 | 139446068 | 139446302 | 0.476059 | 1.61771 | 3.398 |
| chr3 | 169480762 | 169481002 | 1.31285 | 4.46129 | 3.398 |
| chr22 | 25164747 | 25165106 | 0.833145 | 2.83177 | 3.399 |
| chr5 | 147079337 | 147079559 | 0.27108 | 0.921377 | 3.399 |
| chr15 | 96370123 | 96370442 | 0.905368 | 3.07738 | 3.399 |
| chr6 | 83075113 | 83075366 | 0.405144 | 1.37722 | 3.399 |
| chr7 | 151047412 | 151047695 | 1.05537 | 3.58948 | 3.401 |
| chr14 | 92923049 | 92923412 | 1.43537 | 4.88233 | 3.401 |
| chr6 | 114200743 | 114200974 | 0.874119 | 2.97398 | 3.402 |
| chr9 | 134224034 | 134224293 | 0.666428 | 2.26748 | 3.402 |
| chrX | 123522474 | 123522720 | 1.32869 | 4.52079 | 3.402 |
| chr12 | 123237016 | 123237444 | 1.31892 | 4.48821 | 3.403 |
| chr22 | 22067270 | 22067862 | 0.786792 | 2.67763 | 3.403 |
| chr11 | 60665829 | 60666070 | 0.42657 | 1.45191 | 3.404 |
| chrX | 82671408 | 82671636 | 0.712164 | 2.42405 | 3.404 |
| chr8 | 128906940 | 128907189 | 0.810778 | 2.7598 | 3.404 |
| chr11 | 82972330 | 82972671 | 0.412704 | 1.40509 | 3.405 |
| chr11 | 8256121 | 8256436 | 0.81604 | 2.77832 | 3.405 |
| chr1 | 113498406 | 113498624 | 0.634917 | 2.162 | 3.405 |
| chr19 | 42748965 | 42749238 | 0.887802 | 3.02497 | 3.407 |
| chr5 | 127211246 | 127211482 | 0.652394 | 2.22362 | 3.408 |
| chr17 | 72508416 | 72508629 | 0.496413 | 1.69198 | 3.408 |
| chr8 | 109902687 | 109902889 | 0.621924 | 2.11984 | 3.409 |

| | | | | | |
|---|---|---|---|---|---|
| chr20 | 30310627 | 30311251 | 0.576801 | 1.96626 | 3.409 |
| chr18 | 60761277 | 60761641 | 0.28368 | 0.967076 | 3.409 |
| chr1 | 109631796 | 109632193 | 1.3216 | 4.50554 | 3.409 |
| chr11 | 58335411 | 58335755 | 1.12752 | 3.84415 | 3.409 |
| chr19 | 54683721 | 54684112 | 0.721257 | 2.45919 | 3.410 |
| chr1 | 178055492 | 178055710 | 0.439172 | 1.49784 | 3.411 |
| chr12 | 85835554 | 85835844 | 0.821908 | 2.80334 | 3.411 |
| chr15 | 71867849 | 71868168 | 0.741501 | 2.52914 | 3.411 |
| chr1 | 66153401 | 66153727 | 1.00562 | 3.43019 | 3.411 |
| chr10 | 47599757 | 47600150 | 1.25112 | 4.26784 | 3.411 |
| chr5 | 113641357 | 113641600 | 0.688954 | 2.35166 | 3.413 |
| chr2 | 121493287 | 121493655 | 1.62953 | 5.56328 | 3.414 |
| chr2 | 101346781 | 101347075 | 1.17586 | 4.01541 | 3.415 |
| chr12 | 78623339 | 78623649 | 0.909801 | 3.10702 | 3.415 |
| chr16 | 87812709 | 87813075 | 1.23517 | 4.21831 | 3.415 |
| chr3 | 14903313 | 14903518 | 0.340265 | 1.1625 | 3.416 |
| chr8 | 39891516 | 39891826 | 0.921163 | 3.14821 | 3.418 |
| chr2 | 27593082 | 27593402 | 1.68731 | 5.76702 | 3.418 |
| chr5 | 122327160 | 122327391 | 0.299859 | 1.0249 | 3.418 |
| chr8 | 33538973 | 33539345 | 1.56596 | 5.35242 | 3.418 |
| chr5 | 140734430 | 140734824 | 0.664475 | 2.2714 | 3.418 |
| chr12 | 4918770 | 4918974 | 0.28896 | 0.98798 | 3.419 |
| chr11 | 44115841 | 44116162 | 1.17244 | 4.00883 | 3.419 |
| chr9 | 137791641 | 137791916 | 0.690928 | 2.36254 | 3.419 |
| chr6 | 27740168 | 27740582 | 1.03932 | 3.5544 | 3.420 |
| chr3 | 128145118 | 128145598 | 1.1066 | 3.78462 | 3.420 |
| chr18 | 53590926 | 53591134 | 0.292472 | 1.00029 | 3.420 |
| chr12 | 66745979 | 66746318 | 0.810421 | 2.77224 | 3.421 |
| chr4 | 115101715 | 115101961 | 0.950136 | 3.2503 | 3.421 |
| chr3 | 35949881 | 35950206 | 1.51802 | 5.19354 | 3.421 |
| chr18 | 58081331 | 58081719 | 0.756664 | 2.5888 | 3.421 |
| chr3 | 45874228 | 45874504 | 0.7493 | 2.56363 | 3.421 |
| chr12 | 28390452 | 28390693 | 0.663668 | 2.27085 | 3.422 |
| chr17 | 9407058 | 9407311 | 0.303569 | 1.03882 | 3.422 |
| chr12 | 53321270 | 53321621 | 1.0701 | 3.66217 | 3.422 |
| chr15 | 33378743 | 33379144 | 0.62724 | 2.14676 | 3.423 |
| chr16 | 85665729 | 85665950 | 0.54302 | 1.85858 | 3.423 |
| chr19 | 52168260 | 52168615 | 0.532961 | 1.8252 | 3.425 |
| chr17 | 28927069 | 28927480 | 0.855432 | 2.92969 | 3.425 |
| chr5 | 135585413 | 135585719 | 1.22379 | 4.19125 | 3.425 |
| chr22 | 50468681 | 50469063 | 1.65273 | 5.6604 | 3.425 |
| chr14 | 65007031 | 65007286 | 0.771065 | 2.64095 | 3.425 |
| chr4 | 179007542 | 179007779 | 0.624208 | 2.1383 | 3.426 |
| chr1 | 205392387 | 205392612 | 0.427898 | 1.46603 | 3.426 |
| chr15 | 61459585 | 61460046 | 0.773377 | 2.64991 | 3.426 |
| chr10 | 116581264 | 116581825 | 1.25402 | 4.29688 | 3.426 |
| chr12 | 109197298 | 109197631 | 1.42586 | 4.88612 | 3.427 |
| chr9 | 134501564 | 134501795 | 0.409878 | 1.40469 | 3.427 |
| chr5 | 79450842 | 79451107 | 0.356847 | 1.22303 | 3.427 |
| chr5 | 59010262 | 59010556 | 0.539337 | 1.84857 | 3.427 |
| chr19 | 23586561 | 23586863 | 0.823754 | 2.82343 | 3.428 |
| chr4 | 45455354 | 45455577 | 0.624315 | 2.13998 | 3.428 |
| chr10 | 93056338 | 93056570 | 0.809314 | 2.77494 | 3.429 |
| chr8 | 73654736 | 73655006 | 0.266803 | 0.915084 | 3.430 |
| chr20 | 10534056 | 10534335 | 1.39673 | 4.79133 | 3.430 |
| chr1 | 155176606 | 155176983 | 1.36139 | 4.67018 | 3.430 |
| chr5 | 105009573 | 105009929 | 0.322277 | 1.10565 | 3.431 |
| chr5 | 150226728 | 150227032 | 0.840576 | 2.88418 | 3.431 |
| chr1 | 219997600 | 219997808 | 0.336657 | 1.15532 | 3.432 |
| chr6 | 47755825 | 47756153 | 1.21783 | 4.17993 | 3.432 |
| chr7 | 27249983 | 27250519 | 1.02787 | 3.52836 | 3.433 |

| | | | | | |
|---|---|---|---|---|---|
| chr12 | 57542579 | 57542784 | 0.351685 | 1.20729 | 3.433 |
| chr16 | 3550740 | 3551026 | 1.33244 | 4.5746 | 3.433 |
| chr17 | 4613244 | 4613466 | 1.16625 | 4.00404 | 3.433 |
| chr9 | 112415287 | 112415590 | 0.725473 | 2.49081 | 3.433 |
| chr18 | 11732950 | 11733325 | 1.63624 | 5.61817 | 3.434 |
| chr12 | 52976828 | 52977128 | 1.12409 | 3.85976 | 3.434 |
| chr9 | 135421886 | 135422173 | 0.521382 | 1.79077 | 3.435 |
| chr5 | 95170682 | 95170959 | 1.06084 | 3.64383 | 3.435 |
| chr19 | 3700443 | 3700766 | 1.03573 | 3.55763 | 3.435 |
| chrX | 82216181 | 82216550 | 0.379285 | 1.30288 | 3.435 |
| chr5 | 14685872 | 14686163 | 0.809634 | 2.78146 | 3.435 |
| chr5 | 168938869 | 168939118 | 0.349863 | 1.20221 | 3.436 |
| chr7 | 38491567 | 38491771 | 0.438343 | 1.50636 | 3.436 |
| chr2 | 105028928 | 105029214 | 0.684427 | 2.35209 | 3.437 |
| chrX | 106898349 | 106898766 | 1.4374 | 4.94048 | 3.437 |
| chr4 | 65211923 | 65212132 | 0.28568 | 0.982071 | 3.438 |
| chr20 | 44223936 | 44224335 | 0.358461 | 1.23238 | 3.438 |
| chr3 | 37845594 | 37845814 | 0.365325 | 1.25623 | 3.439 |
| chr3 | 13611978 | 13612352 | 0.617361 | 2.12368 | 3.440 |
| chr14 | 56875990 | 56876210 | 0.393587 | 1.35418 | 3.441 |
| chr4 | 141247644 | 141247870 | 0.698644 | 2.4038 | 3.441 |
| chr5 | 36870631 | 36870871 | 0.790066 | 2.71897 | 3.441 |
| chr8 | 47868502 | 47869040 | 1.06676 | 3.67149 | 3.442 |
| chr22 | 21096091 | 21096405 | 0.789277 | 2.71678 | 3.442 |
| chr11 | 65361164 | 65361762 | 1.08465 | 3.73385 | 3.442 |
| chr14 | 37256070 | 37256276 | 0.424307 | 1.46081 | 3.443 |
| chr5 | 176831381 | 176831750 | 1.0808 | 3.72129 | 3.443 |
| chr12 | 34273565 | 34273786 | 0.241295 | 0.831323 | 3.445 |
| chr15 | 99447735 | 99448072 | 1.377 | 4.74485 | 3.446 |
| chr21 | 39727983 | 39728270 | 1.51484 | 5.21986 | 3.446 |
| chr11 | 62276001 | 62276299 | 1.03001 | 3.55005 | 3.447 |
| chr11 | 22213829 | 22214112 | 1.33114 | 4.5881 | 3.447 |
| chr11 | 5116635 | 5116920 | 0.273387 | 0.942356 | 3.447 |
| chr12 | 23604392 | 23604645 | 0.414963 | 1.43045 | 3.447 |
| chr9 | 79062444 | 79062781 | 0.932918 | 3.21666 | 3.448 |
| chr9 | 18208220 | 18208548 | 1.13988 | 3.93155 | 3.449 |
| chr20 | 61175432 | 61175757 | 0.628318 | 2.16744 | 3.450 |
| chr6 | 146209233 | 146209473 | 0.395447 | 1.36449 | 3.451 |
| chr16 | 12267981 | 12268217 | 0.403058 | 1.39078 | 3.451 |
| chr22 | 46840588 | 46840791 | 0.53247 | 1.83757 | 3.451 |
| chr5 | 15686136 | 15686373 | 0.406735 | 1.40372 | 3.451 |
| chr11 | 12309180 | 12309421 | 1.25737 | 4.33943 | 3.451 |
| chrX | 120181233 | 120181478 | 0.716165 | 2.47174 | 3.451 |
| chr22 | 43988464 | 43988679 | 0.498619 | 1.72093 | 3.451 |
| chr15 | 45927607 | 45927912 | 0.828883 | 2.86157 | 3.452 |
| chr8 | 95370060 | 95370273 | 0.945592 | 3.26488 | 3.453 |
| chr7 | 86958229 | 86958451 | 0.468234 | 1.61672 | 3.453 |
| chr3 | 96419828 | 96420148 | 0.455098 | 1.57137 | 3.453 |
| chr7 | 43518651 | 43519017 | 0.465708 | 1.60843 | 3.454 |
| chr4 | 140452034 | 140452347 | 0.852077 | 2.94337 | 3.454 |
| chr14 | 59313703 | 59313924 | 0.36343 | 1.25549 | 3.455 |
| chr13 | 97762160 | 97762512 | 1.47991 | 5.11284 | 3.455 |
| chr6 | 137893038 | 137893406 | 0.726378 | 2.50951 | 3.455 |
| chr3 | 127293859 | 127294168 | 1.26385 | 4.36673 | 3.455 |
| chr16 | 6698591 | 6698801 | 0.301431 | 1.04148 | 3.455 |
| chr2 | 45396649 | 45396872 | 0.432913 | 1.4958 | 3.455 |
| chr1 | 41057875 | 41058148 | 0.655399 | 2.26457 | 3.455 |
| chr12 | 98850733 | 98851063 | 0.367823 | 1.27109 | 3.456 |
| chr5 | 140954480 | 140954743 | 0.373306 | 1.29005 | 3.456 |
| chr19 | 8680602 | 8680929 | 1.00192 | 3.46256 | 3.456 |
| chrX | 46889822 | 46890030 | 0.539484 | 1.86448 | 3.456 |

| | | | | | |
|---|---|---|---|---|---|
| chr3 | 178733390 | 178733787 | 0.759456 | 2.62526 | 3.457 |
| chr6 | 70100106 | 70100332 | 0.306416 | 1.05928 | 3.457 |
| chr16 | 58333601 | 58334093 | 1.13086 | 3.91009 | 3.458 |
| chr10 | 121878829 | 121879155 | 0.300137 | 1.03783 | 3.458 |
| chr10 | 97789653 | 97789875 | 0.342264 | 1.18357 | 3.458 |
| chr16 | 89985063 | 89985290 | 0.768991 | 2.65988 | 3.459 |
| chrX | 68042535 | 68042869 | 0.937414 | 3.24306 | 3.460 |
| chr19 | 7745486 | 7745775 | 1.33413 | 4.61556 | 3.460 |
| chr3 | 18336753 | 18337111 | 1.22089 | 4.22406 | 3.460 |
| chr1 | 229850764 | 229851049 | 1.02044 | 3.53134 | 3.461 |
| chr9 | 38630627 | 38630837 | 0.246423 | 0.852845 | 3.461 |
| chr4 | 40810639 | 40810883 | 0.341348 | 1.18138 | 3.461 |
| chr12 | 45981996 | 45982201 | 0.551485 | 1.9088 | 3.461 |
| chr1 | 233696879 | 233697122 | 0.446846 | 1.54664 | 3.461 |
| chr2 | 38763162 | 38763620 | 1.76703 | 6.11749 | 3.462 |
| chr3 | 77486072 | 77486351 | 0.336456 | 1.16491 | 3.462 |
| chr3 | 139199508 | 139199787 | 0.39378 | 1.3634 | 3.462 |
| chr9 | 114162296 | 114162641 | 1.05671 | 3.65934 | 3.463 |
| chr3 | 84597089 | 84597404 | 0.584334 | 2.02442 | 3.464 |
| chr18 | 3666290 | 3666588 | 1.16461 | 4.03494 | 3.465 |
| chr17 | 59628607 | 59628828 | 0.747675 | 2.59052 | 3.465 |
| chr11 | 19975540 | 19975791 | 0.869673 | 3.01323 | 3.465 |
| chr1 | 40780950 | 40781370 | 1.26137 | 4.37054 | 3.465 |
| chr11 | 55474550 | 55474798 | 0.635584 | 2.20235 | 3.465 |
| chr19 | 9316246 | 9316504 | 0.902913 | 3.1292 | 3.466 |
| chr10 | 133495604 | 133496094 | 0.328386 | 1.13827 | 3.466 |
| chr12 | 56790953 | 56791158 | 0.392998 | 1.36224 | 3.466 |
| chr11 | 113667185 | 113667674 | 0.616042 | 2.13541 | 3.466 |
| chr19 | 10369715 | 10370019 | 0.940946 | 3.26166 | 3.466 |
| chr20 | 36966059 | 36966539 | 1.67702 | 5.81373 | 3.467 |
| chr7 | 3439597 | 3439895 | 0.717395 | 2.48742 | 3.467 |
| chr14 | 78690282 | 78690485 | 0.333437 | 1.15616 | 3.467 |
| chr11 | 62192214 | 62192609 | 1.55054 | 5.37667 | 3.468 |
| chr5 | 43402018 | 43402279 | 0.696031 | 2.41422 | 3.469 |
| chr12 | 52713009 | 52713358 | 0.480585 | 1.66704 | 3.469 |
| chr7 | 56031930 | 56032365 | 1.12517 | 3.90431 | 3.470 |
| chr7 | 62476052 | 62476394 | 1.17821 | 4.08849 | 3.470 |
| chr7 | 139526884 | 139527122 | 0.578808 | 2.0089 | 3.471 |
| chr17 | 33882790 | 33883020 | 0.686818 | 2.38384 | 3.471 |
| chr4 | 100504046 | 100504428 | 0.281463 | 0.977177 | 3.472 |
| chr8 | 106436976 | 106437432 | 0.603557 | 2.09613 | 3.473 |
| chr12 | 55514554 | 55514843 | 0.301148 | 1.04597 | 3.473 |
| chr3 | 79510474 | 79510679 | 0.293707 | 1.02016 | 3.473 |
| chr11 | 70660832 | 70661286 | 1.89138 | 6.5697 | 3.474 |
| chr1 | 190342870 | 190343209 | 0.800145 | 2.77937 | 3.474 |
| chr6 | 138725140 | 138725705 | 0.863149 | 2.99842 | 3.474 |
| chr5 | 143583317 | 143583638 | 1.00161 | 3.48012 | 3.475 |
| chr17 | 69114290 | 69114497 | 0.616781 | 2.14376 | 3.476 |
| chr19 | 3224370 | 3224691 | 1.24521 | 4.32885 | 3.476 |
| chr7 | 36710472 | 36710770 | 1.8218 | 6.33359 | 3.477 |
| chr8 | 63882679 | 63882914 | 0.296145 | 1.02961 | 3.477 |
| chr12 | 128506023 | 128506382 | 0.374624 | 1.30254 | 3.477 |
| chr20 | 19389073 | 19389347 | 0.248473 | 0.864578 | 3.480 |
| chr19 | 9542204 | 9542418 | 0.551356 | 1.91859 | 3.480 |
| chr10 | 33355762 | 33356076 | 0.532612 | 1.85343 | 3.480 |
| chr5 | 140705586 | 140705848 | 1.02973 | 3.58356 | 3.480 |
| chr10 | 111319161 | 111319399 | 0.316668 | 1.10213 | 3.480 |
| chr19 | 11908258 | 11908587 | 1.26009 | 4.38566 | 3.480 |
| chr11 | 123117971 | 123118280 | 1.18812 | 4.13523 | 3.480 |
| chr3 | 109827938 | 109828537 | 0.359681 | 1.25192 | 3.481 |
| chr3 | 46759322 | 46759625 | 0.729303 | 2.53853 | 3.481 |

| | | | | | |
|---|---|---|---|---|---|
| chr12 | 17092745 | 17093157 | 1.86861 | 6.5053 | 3.481 |
| chr3 | 187432182 | 187432613 | 0.975154 | 3.39497 | 3.481 |
| chr10 | 100183383 | 100183883 | 0.803335 | 2.79691 | 3.482 |
| chr9 | 96067118 | 96067508 | 1.36042 | 4.73654 | 3.482 |
| chr13 | 52470414 | 52470811 | 1.01735 | 3.54308 | 3.483 |
| chr15 | 71944094 | 71944418 | 1.1657 | 4.0598 | 3.483 |
| chr14 | 44078030 | 44078246 | 0.309901 | 1.07944 | 3.483 |
| chr4 | 31372505 | 31372812 | 0.73346 | 2.55519 | 3.484 |
| chr18 | 43419448 | 43419704 | 0.885454 | 3.08593 | 3.485 |
| chr13 | 74409375 | 74409602 | 0.585395 | 2.04091 | 3.486 |
| chr6 | 161794949 | 161795254 | 1.21386 | 4.23269 | 3.487 |
| chr10 | 74124050 | 74124494 | 1.36056 | 4.74483 | 3.487 |
| chr12 | 69964605 | 69964947 | 0.935661 | 3.26303 | 3.487 |
| chr14 | 106873307 | 106873903 | 1.63024 | 5.68585 | 3.488 |
| chr3 | 158512740 | 158513094 | 0.959915 | 3.34805 | 3.488 |
| chr1 | 192443315 | 192443612 | 0.536899 | 1.87298 | 3.489 |
| chr10 | 90078534 | 90078858 | 0.906883 | 3.16466 | 3.490 |
| chr3 | 150874497 | 150874741 | 0.943839 | 3.29415 | 3.490 |
| chrX | 47004352 | 47004649 | 0.909542 | 3.17506 | 3.491 |
| chr6 | 40792075 | 40792440 | 0.819203 | 2.85976 | 3.491 |
| chr16 | 31170960 | 31171348 | 0.932945 | 3.25684 | 3.491 |
| chr19 | 23620690 | 23620985 | 0.728228 | 2.54223 | 3.491 |
| chr19 | 57615392 | 57615679 | 1.15171 | 4.02102 | 3.491 |
| chr11 | 37967812 | 37968065 | 0.770739 | 2.69119 | 3.492 |
| chr12 | 121294869 | 121295137 | 0.780976 | 2.72744 | 3.492 |
| chr14 | 100889780 | 100890021 | 0.302734 | 1.05734 | 3.493 |
| chr5 | 28074552 | 28074794 | 0.358682 | 1.25279 | 3.493 |
| chr14 | 74129169 | 74129386 | 0.71582 | 2.50058 | 3.493 |
| chr12 | 127593180 | 127593570 | 1.14769 | 4.00939 | 3.493 |
| chr7 | 26721505 | 26721748 | 1.1765 | 4.1114 | 3.495 |
| chr12 | 52935361 | 52935572 | 0.427291 | 1.49357 | 3.495 |
| chr8 | 57114829 | 57115156 | 0.818992 | 2.86315 | 3.496 |
| chr7 | 151843044 | 151843470 | 1.19913 | 4.19263 | 3.496 |
| chr4 | 178801100 | 178801302 | 0.513415 | 1.79536 | 3.497 |
| chr11 | 123582028 | 123582313 | 1.61664 | 5.65346 | 3.497 |
| chr5 | 94229543 | 94229757 | 0.646963 | 2.26273 | 3.497 |
| chr5 | 175218015 | 175218262 | 0.396575 | 1.3871 | 3.498 |
| chr8 | 89182066 | 89182302 | 0.375203 | 1.31252 | 3.498 |
| chr12 | 17181280 | 17181600 | 1.42579 | 4.98797 | 3.498 |
| chr3 | 16870387 | 16870636 | 0.533561 | 1.8669 | 3.499 |
| chr12 | 57206824 | 57207079 | 0.304575 | 1.06581 | 3.499 |
| chr4 | 101770105 | 101770311 | 0.416504 | 1.4575 | 3.499 |
| chr6 | 149347702 | 149347917 | 0.761067 | 2.66407 | 3.500 |
| chr19 | 19335786 | 19336217 | 0.705142 | 2.4685 | 3.501 |
| chr13 | 82214190 | 82214430 | 0.620544 | 2.17244 | 3.501 |
| chr12 | 67863275 | 67863557 | 0.310888 | 1.08876 | 3.502 |
| chr16 | 8834898 | 8835249 | 0.293779 | 1.02887 | 3.502 |
| chr16 | 74800650 | 74800987 | 0.649394 | 2.27434 | 3.502 |
| chr17 | 45330967 | 45331462 | 0.983171 | 3.44344 | 3.502 |
| chr2 | 236511081 | 236511324 | 0.315658 | 1.10579 | 3.503 |
| chr5 | 125879816 | 125880163 | 1.13545 | 3.97795 | 3.503 |
| chr10 | 29593525 | 29593839 | 0.966827 | 3.38847 | 3.505 |
| chr1 | 40282522 | 40282744 | 1.05346 | 3.69214 | 3.505 |
| chr4 | 150185359 | 150185806 | 0.465822 | 1.63301 | 3.506 |
| chr12 | 42631233 | 42631555 | 0.62369 | 2.18682 | 3.506 |
| chr14 | 23127179 | 23127422 | 0.528928 | 1.8547 | 3.507 |
| chr9 | 118353746 | 118354105 | 1.14447 | 4.01312 | 3.507 |
| chr12 | 6726884 | 6727139 | 0.943316 | 3.30847 | 3.507 |
| chr7 | 31023167 | 31023590 | 1.47792 | 5.18352 | 3.507 |
| chr11 | 134009732 | 134009947 | 1.17964 | 4.13746 | 3.507 |
| chr14 | 55039090 | 55039378 | 1.23732 | 4.33984 | 3.507 |

| | | | | | |
|---|---|---|---|---|---|
| chrX | 10877585 | 10877871 | 0.324001 | 1.13643 | 3.507 |
| chr18 | 39632163 | 39632394 | 0.695509 | 2.43963 | 3.508 |
| chr6 | 82715922 | 82716169 | 0.804995 | 2.82368 | 3.508 |
| chr7 | 20578871 | 20579111 | 0.626742 | 2.19843 | 3.508 |
| chr16 | 30456739 | 30456944 | 0.6242 | 2.19041 | 3.509 |
| chr8 | 129334975 | 129335264 | 1.25513 | 4.40458 | 3.509 |
| chr5 | 161853878 | 161854155 | 0.442535 | 1.55321 | 3.510 |
| chr12 | 11659517 | 11659748 | 0.79404 | 2.78732 | 3.510 |
| chr4 | 53454328 | 53454660 | 0.642231 | 2.25448 | 3.510 |
| chr16 | 58634675 | 58634941 | 0.310113 | 1.08867 | 3.511 |
| chr8 | 28277042 | 28277415 | 0.885543 | 3.10876 | 3.511 |
| chr7 | 70181353 | 70181722 | 0.627802 | 2.20408 | 3.511 |
| chr5 | 173008658 | 173009012 | 1.24884 | 4.38543 | 3.512 |
| chr10 | 37996862 | 37997154 | 0.576474 | 2.02523 | 3.513 |
| chr12 | 109531205 | 109531558 | 2.25894 | 7.93662 | 3.513 |
| chr13 | 48669175 | 48669577 | 1.27557 | 4.48275 | 3.514 |
| chr14 | 22689468 | 22689688 | 0.776829 | 2.73022 | 3.515 |
| chr16 | 6916519 | 6916901 | 0.292066 | 1.02678 | 3.516 |
| chr15 | 94205832 | 94206077 | 0.483951 | 1.70204 | 3.517 |
| chr12 | 119407575 | 119407780 | 0.373542 | 1.31381 | 3.517 |
| chr8 | 32964195 | 32964483 | 0.676188 | 2.37907 | 3.518 |
| chr3 | 170736291 | 170736523 | 0.70514 | 2.48169 | 3.519 |
| chr18 | 3703716 | 3703927 | 0.864217 | 3.04168 | 3.520 |
| chrX | 49969210 | 49969468 | 0.626699 | 2.20577 | 3.520 |
| chr5 | 140652292 | 140652780 | 0.611753 | 2.15335 | 3.520 |
| chr12 | 28123042 | 28123394 | 0.94542 | 3.32795 | 3.520 |
| chr10 | 105883248 | 105883518 | 0.685947 | 2.41503 | 3.521 |
| chr12 | 57863156 | 57863475 | 0.304032 | 1.07073 | 3.522 |
| chr8 | 18244441 | 18244994 | 0.622876 | 2.19405 | 3.522 |
| chr11 | 1903601 | 1904109 | 1.83859 | 6.47643 | 3.523 |
| chr8 | 143799551 | 143799856 | 1.25215 | 4.41073 | 3.523 |
| chr6 | 27219568 | 27219811 | 0.819362 | 2.88631 | 3.523 |
| chr12 | 132918624 | 132919225 | 1.01767 | 3.58518 | 3.523 |
| chr16 | 18045193 | 18045411 | 0.327532 | 1.15432 | 3.524 |
| chrX | 136007272 | 136007650 | 1.29318 | 4.55874 | 3.525 |
| chr3 | 153313005 | 153313321 | 1.22397 | 4.31512 | 3.526 |
| chr12 | 56885608 | 56885874 | 0.268351 | 0.946492 | 3.527 |
| chr15 | 41074833 | 41075205 | 1.47728 | 5.2109 | 3.527 |
| chr12 | 44247066 | 44247395 | 0.414914 | 1.4641 | 3.529 |
| chr9 | 115707870 | 115708281 | 1.29308 | 4.56292 | 3.529 |
| chr16 | 48631836 | 48632042 | 0.369609 | 1.30427 | 3.529 |
| chr19 | 46907204 | 46907525 | 0.509479 | 1.79813 | 3.529 |
| chr10 | 21784616 | 21784946 | 1.67561 | 5.91532 | 3.530 |
| chr12 | 1175455 | 1175790 | 0.902919 | 3.18808 | 3.531 |
| chr15 | 101017044 | 101017386 | 0.403305 | 1.42402 | 3.531 |
| chr22 | 32924974 | 32925286 | 0.909754 | 3.21265 | 3.531 |
| chr12 | 113468629 | 113469067 | 1.43457 | 5.0664 | 3.532 |
| chr8 | 36948390 | 36948709 | 1.04564 | 3.69297 | 3.532 |
| chr9 | 22307092 | 22307327 | 1.08865 | 3.84561 | 3.532 |
| chr15 | 22833071 | 22833359 | 0.351424 | 1.24149 | 3.533 |
| chr12 | 131298003 | 131298216 | 0.411943 | 1.45538 | 3.533 |
| chr14 | 69122748 | 69123089 | 0.664821 | 2.34901 | 3.533 |
| chr19 | 13829630 | 13829859 | 0.536976 | 1.89803 | 3.535 |
| chr14 | 55518103 | 55518517 | 2.07592 | 7.33924 | 3.535 |
| chr20 | 31332656 | 31332901 | 0.542589 | 1.91837 | 3.536 |
| chr22 | 44481644 | 44481904 | 0.877361 | 3.10221 | 3.536 |
| chr1 | 41090826 | 41091039 | 0.636421 | 2.25031 | 3.536 |
| chr12 | 116673968 | 116674258 | 1.04655 | 3.70129 | 3.537 |
| chr12 | 126789792 | 126790027 | 0.309104 | 1.09325 | 3.537 |
| chr2 | 62650380 | 62650630 | 0.602972 | 2.13273 | 3.537 |
| chr8 | 27930331 | 27930565 | 1.27999 | 4.52819 | 3.538 |

| | | | | | |
|---|---|---|---|---|---|
| chr12 | 32908082 | 32908385 | 1.48712 | 5.26206 | 3.538 |
| chr11 | 94474602 | 94474804 | 0.61706 | 2.18466 | 3.540 |
| chr12 | 5427583 | 5427899 | 0.333416 | 1.1805 | 3.541 |
| chrX | 115028664 | 115029016 | 0.439112 | 1.55484 | 3.541 |
| chr6 | 160274000 | 160274224 | 0.410993 | 1.45595 | 3.543 |
| chr6 | 112350414 | 112350751 | 1.09458 | 3.87933 | 3.544 |
| chr1 | 8375922 | 8376168 | 0.661989 | 2.34636 | 3.544 |
| chr1 | 9785085 | 9785472 | 1.61912 | 5.74089 | 3.546 |
| chr8 | 51689400 | 51689603 | 0.301573 | 1.06944 | 3.546 |
| chr20 | 19669001 | 19669378 | 1.16802 | 4.14288 | 3.547 |
| chr14 | 89506619 | 89507079 | 1.43198 | 5.07936 | 3.547 |
| chr2 | 175564690 | 175565010 | 0.874259 | 3.10234 | 3.549 |
| chr15 | 74436811 | 74437023 | 0.580197 | 2.05896 | 3.549 |
| chr4 | 181370522 | 181370864 | 0.896736 | 3.18271 | 3.549 |
| chr12 | 85250801 | 85251041 | 0.263452 | 0.935092 | 3.549 |
| chr6 | 158177668 | 158178038 | 1.3056 | 4.63431 | 3.550 |
| chr4 | 104012868 | 104013147 | 0.870173 | 3.08889 | 3.550 |
| chr15 | 86402035 | 86402422 | 1.82838 | 6.49066 | 3.550 |
| chr10 | 77863184 | 77863638 | 0.344653 | 1.22383 | 3.551 |
| chr7 | 120445969 | 120446311 | 1.1855 | 4.21 | 3.551 |
| chr22 | 37269661 | 37270001 | 0.748911 | 2.65989 | 3.552 |
| chr12 | 105219230 | 105219542 | 0.637903 | 2.26571 | 3.552 |
| chr6 | 137289251 | 137289705 | 0.812687 | 2.88667 | 3.552 |
| chr19 | 23528720 | 23529053 | 0.865822 | 3.07553 | 3.552 |
| chr17 | 76170693 | 76171322 | 0.642608 | 2.28283 | 3.552 |
| chr12 | 118109434 | 118109860 | 1.41608 | 5.03057 | 3.552 |
| chr4 | 119811346 | 119811600 | 0.374262 | 1.32957 | 3.553 |
| chr12 | 127232028 | 127232538 | 1.85235 | 6.58095 | 3.553 |
| chr19 | 52207240 | 52207491 | 0.636551 | 2.26151 | 3.553 |
| chr18 | 74138672 | 74139047 | 1.17746 | 4.18372 | 3.553 |
| chr2 | 75468876 | 75469188 | 0.712102 | 2.53057 | 3.554 |
| chr9 | 104625812 | 104626070 | 0.28884 | 1.02645 | 3.554 |
| chr13 | 108194096 | 108194346 | 0.229747 | 0.816457 | 3.554 |
| chr16 | 3317011 | 3317368 | 1.00094 | 3.55733 | 3.554 |
| chr6 | 37828035 | 37828300 | 0.524685 | 1.86475 | 3.554 |
| chr1 | 175767628 | 175767858 | 0.267155 | 0.949564 | 3.554 |
| chr5 | 136435497 | 136435720 | 0.351108 | 1.24809 | 3.555 |
| chr17 | 30465933 | 30466307 | 1.27332 | 4.52682 | 3.555 |
| chr12 | 133037222 | 133037521 | 0.822253 | 2.92324 | 3.555 |
| chr11 | 106004347 | 106004603 | 0.551462 | 1.96056 | 3.555 |
| chr1 | 158087539 | 158087978 | 0.953609 | 3.39185 | 3.557 |
| chr10 | 121367793 | 121368205 | 1.36523 | 4.85639 | 3.557 |
| chr8 | 116599184 | 116599483 | 0.989867 | 3.52119 | 3.557 |
| chr20 | 61804163 | 61804417 | 0.564575 | 2.00852 | 3.558 |
| chr12 | 104502975 | 104503384 | 1.36173 | 4.84481 | 3.558 |
| chr3 | 83977587 | 83977848 | 0.611504 | 2.176 | 3.558 |
| chr1 | 42642645 | 42642967 | 1.28646 | 4.57814 | 3.559 |
| chr7 | 150574247 | 150574533 | 0.867812 | 3.08859 | 3.559 |
| chr3 | 25851722 | 25851932 | 0.287725 | 1.02403 | 3.559 |
| chr2 | 169102915 | 169103159 | 0.559189 | 1.99037 | 3.559 |
| chr14 | 73509847 | 73510111 | 0.465107 | 1.65558 | 3.560 |
| chr20 | 31097456 | 31097673 | 0.636943 | 2.2676 | 3.560 |
| chr18 | 29261258 | 29261474 | 0.440544 | 1.56844 | 3.560 |
| chr6 | 34494996 | 34495231 | 0.802865 | 2.85872 | 3.561 |
| chr14 | 88259887 | 88260144 | 0.407809 | 1.45247 | 3.562 |
| chr8 | 11266900 | 11267513 | 0.643718 | 2.29271 | 3.562 |
| chr21 | 46352562 | 46352912 | 0.650388 | 2.31665 | 3.562 |
| chr12 | 41139307 | 41139512 | 0.684299 | 2.4377 | 3.562 |
| chr12 | 130505491 | 130505784 | 0.3397 | 1.21054 | 3.564 |
| chr9 | 97013880 | 97014148 | 0.326616 | 1.16414 | 3.564 |
| chr11 | 107447278 | 107447557 | 1.15655 | 4.12332 | 3.565 |

| | | | | | |
|---|---|---|---|---|---|
| chr11 | 62148287 | 62148501 | 0.960634 | 3.425 | 3.565 |
| chr15 | 99901831 | 99902141 | 0.660874 | 2.35626 | 3.565 |
| chr17 | 72786252 | 72786503 | 0.310512 | 1.10711 | 3.565 |
| chr13 | 31998264 | 31998588 | 1.13857 | 4.06113 | 3.567 |
| chr11 | 100726280 | 100726529 | 0.510828 | 1.82214 | 3.567 |
| chr12 | 45567123 | 45567511 | 1.05038 | 3.74733 | 3.568 |
| chr15 | 73448747 | 73448996 | 0.431263 | 1.5386 | 3.568 |
| chr5 | 88148584 | 88148898 | 0.477144 | 1.70239 | 3.568 |
| chr7 | 42606771 | 42607004 | 0.191728 | 0.684083 | 3.568 |
| chr15 | 52074584 | 52074925 | 0.976064 | 3.48338 | 3.569 |
| chr7 | 75386415 | 75386677 | 0.797463 | 2.84608 | 3.569 |
| chr3 | 31531062 | 31531272 | 0.230588 | 0.8231 | 3.570 |
| chr12 | 49116001 | 49116207 | 0.392196 | 1.39998 | 3.570 |
| chr20 | 6123017 | 6123440 | 1.0295 | 3.67592 | 3.571 |
| chr14 | 36497493 | 36497743 | 0.621304 | 2.21878 | 3.571 |
| chr15 | 97436614 | 97436818 | 0.59728 | 2.13354 | 3.572 |
| chr8 | 17612570 | 17612863 | 0.748331 | 2.67345 | 3.573 |
| chr3 | 57568602 | 57568906 | 0.632418 | 2.2594 | 3.573 |
| chr20 | 5510282 | 5510643 | 1.82088 | 6.50563 | 3.573 |
| chr1 | 99645897 | 99646134 | 0.77608 | 2.77278 | 3.573 |
| chr16 | 87670043 | 87670328 | 0.853436 | 3.04926 | 3.573 |
| chr1 | 154326161 | 154326365 | 0.547254 | 1.95615 | 3.574 |
| chr3 | 38008730 | 38009196 | 1.12527 | 4.02228 | 3.575 |
| chr10 | 125947770 | 125948077 | 0.662074 | 2.36687 | 3.575 |
| chr8 | 110463208 | 110463493 | 1.02097 | 3.65053 | 3.576 |
| chr19 | 1000075 | 1000442 | 0.99451 | 3.55624 | 3.576 |
| chr5 | 178594038 | 178594490 | 1.34072 | 4.79424 | 3.576 |
| chr7 | 2607350 | 2607624 | 0.6854 | 2.45134 | 3.577 |
| chr11 | 3216385 | 3216755 | 0.783243 | 2.8017 | 3.577 |
| chr15 | 92392966 | 92393236 | 1.1155 | 3.99227 | 3.579 |
| chr3 | 118523883 | 118524215 | 0.288434 | 1.03247 | 3.580 |
| chr5 | 128166450 | 128166694 | 0.284959 | 1.02004 | 3.580 |
| chr13 | 95844013 | 95844342 | 1.26838 | 4.54179 | 3.581 |
| chr17 | 14200461 | 14201001 | 0.80522 | 2.88449 | 3.582 |
| chr18 | 32367931 | 32368211 | 0.792066 | 2.83756 | 3.582 |
| chrX | 51603875 | 51604137 | 0.625969 | 2.24278 | 3.583 |
| chr4 | 32776773 | 32776977 | 0.547405 | 1.96186 | 3.584 |
| chr16 | 12434317 | 12434679 | 0.654587 | 2.3477 | 3.587 |
| chr17 | 66395183 | 66395552 | 1.40881 | 5.05401 | 3.587 |
| chr8 | 67341046 | 67341386 | 1.104 | 3.9622 | 3.589 |
| chr17 | 80020422 | 80020695 | 0.476589 | 1.71057 | 3.589 |
| chr1 | 62107413 | 62107793 | 1.22852 | 4.41077 | 3.590 |
| chr12 | 94997229 | 94997603 | 1.70515 | 6.12205 | 3.590 |
| chr12 | 57151273 | 57151598 | 1.12754 | 4.04883 | 3.591 |
| chr19 | 43427174 | 43427520 | 0.451938 | 1.62294 | 3.591 |
| chr9 | 77247902 | 77248345 | 0.347809 | 1.24912 | 3.591 |
| chr15 | 69702625 | 69702981 | 1.08617 | 3.90096 | 3.591 |
| chr19 | 19471919 | 19472155 | 1.08367 | 3.89218 | 3.592 |
| chr13 | 50422105 | 50422408 | 0.870024 | 3.12852 | 3.596 |
| chr12 | 49628178 | 49628608 | 1.66277 | 5.9807 | 3.597 |
| chr20 | 51690689 | 51690913 | 0.297164 | 1.06902 | 3.597 |
| chr10 | 25516005 | 25516374 | 1.10796 | 3.98649 | 3.598 |
| chr19 | 17213998 | 17214311 | 1.10331 | 3.96976 | 3.598 |
| chr14 | 70001951 | 70002330 | 1.13002 | 4.06591 | 3.598 |
| chr13 | 96865730 | 96865971 | 0.236342 | 0.850505 | 3.599 |
| chr7 | 134012710 | 134012915 | 1.06787 | 3.84354 | 3.599 |
| chr21 | 46728944 | 46729298 | 1.38756 | 4.99429 | 3.599 |
| chr18 | 32987929 | 32988155 | 0.64991 | 2.33975 | 3.600 |
| chr7 | 10757573 | 10757840 | 0.284637 | 1.02493 | 3.601 |
| chr11 | 63275452 | 63275812 | 1.10147 | 3.96624 | 3.601 |
| chr12 | 59766822 | 59767036 | 0.770115 | 2.77364 | 3.602 |

| | | | | | |
|---|---|---|---|---|---|
| chr5 | 115911773 | 115912116 | 1.63892 | 5.90331 | 3.602 |
| chr9 | 90691144 | 90691392 | 0.364023 | 1.31183 | 3.604 |
| chr17 | 38514764 | 38515204 | 1.72998 | 6.23515 | 3.604 |
| chr17 | 54382547 | 54383052 | 0.375818 | 1.35453 | 3.604 |
| chr8 | 114384663 | 114384907 | 0.555811 | 2.00349 | 3.605 |
| chr14 | 64016771 | 64016989 | 0.224629 | 0.809828 | 3.605 |
| chr20 | 43991433 | 43991803 | 1.43462 | 5.17328 | 3.606 |
| chr1 | 28184571 | 28184963 | 1.36104 | 4.90887 | 3.607 |
| chr3 | 79081899 | 79082157 | 0.635572 | 2.2925 | 3.607 |
| chr1 | 153377209 | 153377564 | 0.804145 | 2.90141 | 3.608 |
| chr3 | 156919455 | 156919990 | 0.807726 | 2.91452 | 3.608 |
| chr6 | 35114204 | 35114583 | 0.861641 | 3.10916 | 3.608 |
| chr16 | 71598876 | 71599151 | 0.755455 | 2.72615 | 3.609 |
| chr14 | 29454656 | 29454920 | 0.275719 | 0.995021 | 3.609 |
| chr20 | 60449793 | 60450019 | 0.513385 | 1.85322 | 3.610 |
| chr16 | 6693067 | 6693510 | 0.572334 | 2.06646 | 3.611 |
| chr10 | 45420700 | 45420932 | 0.815002 | 2.94264 | 3.611 |
| chr9 | 90282200 | 90282505 | 0.938106 | 3.38744 | 3.611 |
| chr12 | 685248 | 685682 | 1.80734 | 6.52625 | 3.611 |
| chr11 | 14665300 | 14665507 | 0.353527 | 1.2766 | 3.611 |
| chr4 | 26475550 | 26475831 | 0.37248 | 1.34524 | 3.612 |
| chr12 | 129060523 | 129060737 | 0.255096 | 0.92135 | 3.612 |
| chr17 | 1090372 | 1090958 | 1.68476 | 6.09041 | 3.615 |
| chr14 | 107198670 | 107199064 | 1.22127 | 4.41591 | 3.616 |
| chrX | 106242228 | 106242553 | 1.13755 | 4.11433 | 3.617 |
| chr13 | 92409666 | 92409941 | 0.378799 | 1.37008 | 3.617 |
| chr19 | 9906889 | 9907162 | 0.593066 | 2.14524 | 3.617 |
| chr14 | 93055560 | 93055937 | 0.673594 | 2.43729 | 3.618 |
| chr8 | 12709648 | 12709939 | 0.323527 | 1.17077 | 3.619 |
| chr4 | 149116402 | 149116668 | 0.513071 | 1.85685 | 3.619 |
| chr18 | 2223972 | 2224350 | 0.36904 | 1.33562 | 3.619 |
| chr1 | 198057543 | 198057813 | 0.956091 | 3.46037 | 3.619 |
| chr5 | 134539260 | 134539570 | 0.827746 | 2.99596 | 3.619 |
| chr10 | 120560469 | 120560733 | 0.86249 | 3.12184 | 3.620 |
| chr1 | 178722606 | 178722845 | 0.849962 | 3.07652 | 3.620 |
| chr9 | 83718853 | 83719055 | 0.671114 | 2.42963 | 3.620 |
| chr1 | 160912689 | 160913140 | 1.05118 | 3.80562 | 3.620 |
| chr14 | 104760287 | 104760815 | 0.425407 | 1.54037 | 3.621 |
| chr11 | 72378481 | 72378868 | 0.705358 | 2.5543 | 3.621 |
| chr19 | 51166777 | 51166990 | 0.700899 | 2.53896 | 3.622 |
| chr16 | 69971336 | 69971748 | 0.734338 | 2.66017 | 3.623 |
| chr11 | 128646916 | 128647263 | 0.782387 | 2.83428 | 3.623 |
| chr18 | 61185778 | 61186005 | 0.769984 | 2.78936 | 3.623 |
| chr3 | 178684456 | 178684674 | 0.449846 | 1.62966 | 3.623 |
| chr14 | 24527500 | 24527788 | 0.95714 | 3.46851 | 3.624 |
| chr11 | 44545139 | 44545388 | 0.983114 | 3.5637 | 3.625 |
| chr11 | 75479039 | 75479356 | 0.637852 | 2.31218 | 3.625 |
| chrX | 107263807 | 107264148 | 1.04619 | 3.79368 | 3.626 |
| chr6 | 44225130 | 44225565 | 1.30071 | 4.71769 | 3.627 |
| chr9 | 16257619 | 16257874 | 0.948134 | 3.43892 | 3.627 |
| chr4 | 92886393 | 92886649 | 0.477142 | 1.73087 | 3.628 |
| chr9 | 93747548 | 93747809 | 0.914408 | 3.31958 | 3.630 |
| chr8 | 32885505 | 32885891 | 0.639542 | 2.32263 | 3.632 |
| chr19 | 10112632 | 10112873 | 0.526926 | 1.91369 | 3.632 |
| chr8 | 64212687 | 64212918 | 0.316402 | 1.14918 | 3.632 |
| chr19 | 48894677 | 48895141 | 1.58808 | 5.76899 | 3.633 |
| chr1 | 62601493 | 62601758 | 1.00615 | 3.65524 | 3.633 |
| chr20 | 36871306 | 36871620 | 1.28187 | 4.65797 | 3.634 |
| chr12 | 101520221 | 101520512 | 0.348839 | 1.26798 | 3.635 |
| chr15 | 76221003 | 76221316 | 1.07473 | 3.90668 | 3.635 |
| chr11 | 3362553 | 3362838 | 0.685881 | 2.49343 | 3.635 |

| | | | | | |
|---|---|---|---|---|---|
| chr8 | 40059324 | 40059597 | 1.04688 | 3.80613 | 3.636 |
| chr9 | 4375397 | 4375673 | 0.997062 | 3.6252 | 3.636 |
| chr16 | 88677773 | 88677997 | 0.28067 | 1.0207 | 3.637 |
| chr19 | 48216266 | 48216852 | 1.49514 | 5.4374 | 3.637 |
| chr8 | 37684063 | 37684283 | 0.524026 | 1.90586 | 3.637 |
| chr2 | 10640051 | 10640280 | 0.320155 | 1.1644 | 3.637 |
| chr5 | 164927741 | 164927973 | 0.269196 | 0.979085 | 3.637 |
| chr11 | 129205860 | 129206068 | 0.270313 | 0.983214 | 3.637 |
| chr10 | 124899867 | 124900163 | 0.542843 | 1.97485 | 3.638 |
| chr5 | 140262145 | 140262581 | 0.841565 | 3.06165 | 3.638 |
| chr14 | 55158252 | 55158648 | 1.62538 | 5.91384 | 3.638 |
| chr3 | 174839102 | 174839353 | 0.431559 | 1.57043 | 3.639 |
| chr5 | 180515273 | 180515586 | 0.732019 | 2.66393 | 3.639 |
| chr15 | 90686335 | 90686764 | 1.44847 | 5.27177 | 3.640 |
| chr15 | 84228114 | 84228527 | 1.35331 | 4.92687 | 3.641 |
| chr16 | 9481928 | 9482133 | 0.328715 | 1.19673 | 3.641 |
| chr12 | 46686705 | 46687039 | 0.372959 | 1.35791 | 3.641 |
| chr12 | 93427285 | 93427579 | 0.311234 | 1.1332 | 3.641 |
| chr3 | 79410990 | 79411211 | 0.917033 | 3.33943 | 3.642 |
| chr15 | 51187079 | 51187293 | 0.432694 | 1.57587 | 3.642 |
| chr5 | 29053327 | 29053567 | 0.442836 | 1.6131 | 3.643 |
| chr5 | 38189871 | 38190083 | 0.284289 | 1.03579 | 3.643 |
| chr4 | 6345623 | 6345923 | 0.411935 | 1.50092 | 3.644 |
| chr19 | 38074274 | 38074574 | 0.917833 | 3.34422 | 3.644 |
| chr7 | 17809542 | 17809756 | 0.946771 | 3.44968 | 3.644 |
| chr14 | 59437074 | 59437276 | 0.302195 | 1.10129 | 3.644 |
| chr7 | 1479280 | 1479572 | 0.581668 | 2.11985 | 3.644 |
| chr13 | 105792447 | 105792687 | 0.858603 | 3.12951 | 3.645 |
| chr16 | 31722468 | 31722815 | 0.355027 | 1.29405 | 3.645 |
| chr19 | 480161 | 480447 | 1.03788 | 3.78445 | 3.646 |
| chr13 | 51824309 | 51824566 | 0.713705 | 2.60311 | 3.647 |
| chr9 | 92736415 | 92736724 | 0.252572 | 0.921339 | 3.648 |
| chr3 | 157485510 | 157485722 | 0.335399 | 1.2236 | 3.648 |
| chr5 | 56946148 | 56946482 | 0.828188 | 3.02155 | 3.648 |
| chr4 | 35936124 | 35936326 | 0.656967 | 2.39761 | 3.650 |
| chr1 | 220390714 | 220391026 | 0.85348 | 3.11598 | 3.651 |
| chr5 | 140248342 | 140248590 | 0.43701 | 1.59558 | 3.651 |
| chr19 | 52996316 | 52996678 | 0.631333 | 2.30509 | 3.651 |
| chr3 | 182906728 | 182907036 | 0.606371 | 2.21453 | 3.652 |
| chr18 | 53448914 | 53449184 | 0.661008 | 2.41431 | 3.652 |
| chr6 | 47065158 | 47065394 | 0.75687 | 2.76467 | 3.653 |
| chr15 | 89943165 | 89943408 | 0.609697 | 2.22716 | 3.653 |
| chr7 | 21565705 | 21565970 | 0.734099 | 2.6831 | 3.655 |
| chr7 | 104563350 | 104563653 | 1.13064 | 4.1348 | 3.657 |
| chr8 | 98656369 | 98656622 | 0.903849 | 3.3059 | 3.658 |
| chr6 | 154477185 | 154477506 | 0.807932 | 2.95522 | 3.658 |
| chr8 | 119824099 | 119824469 | 0.617267 | 2.25889 | 3.660 |
| chr19 | 13345333 | 13345704 | 1.13198 | 4.143 | 3.660 |
| chr11 | 59980917 | 59981122 | 0.252209 | 0.923241 | 3.661 |
| chr11 | 65105978 | 65106184 | 0.380437 | 1.39269 | 3.661 |
| chr8 | 95069217 | 95069419 | 0.45757 | 1.67506 | 3.661 |
| chr9 | 115875015 | 115875363 | 1.2453 | 4.55969 | 3.662 |
| chr9 | 74219381 | 74219741 | 0.978964 | 3.58454 | 3.662 |
| chr4 | 89212101 | 89212341 | 0.214774 | 0.786508 | 3.662 |
| chr1 | 154377512 | 154377907 | 0.6981 | 2.55673 | 3.662 |
| chr12 | 44470691 | 44470933 | 0.344251 | 1.26096 | 3.663 |
| chr11 | 32456016 | 32456379 | 0.657235 | 2.40748 | 3.663 |
| chr4 | 140572808 | 140573078 | 1.06847 | 3.91411 | 3.663 |
| chr1 | 6706562 | 6706861 | 0.566396 | 2.07507 | 3.664 |
| chr12 | 129885602 | 129885811 | 0.2754 | 1.00902 | 3.664 |
| chr16 | 71641598 | 71641967 | 1.04798 | 3.84039 | 3.665 |

| | | | | | |
|---|---|---|---|---|---|
| chr11 | 44807887 | 44808210 | 0.577237 | 2.11589 | 3.666 |
| chr16 | 87428121 | 87428351 | 0.972522 | 3.56531 | 3.666 |
| chr6 | 37282701 | 37282989 | 0.982765 | 3.60313 | 3.666 |
| chr20 | 653726 | 654000 | 0.677901 | 2.48549 | 3.666 |
| chr6 | 132359589 | 132359852 | 0.290962 | 1.06707 | 3.667 |
| chr10 | 87551960 | 87552198 | 0.536681 | 1.96945 | 3.670 |
| chr2 | 23847290 | 23847624 | 1.09746 | 4.02791 | 3.670 |
| chr7 | 126602200 | 126602417 | 0.272458 | 1.00016 | 3.671 |
| chr15 | 84811537 | 84811885 | 1.1081 | 4.06897 | 3.672 |
| chr2 | 47104042 | 47104281 | 0.735825 | 2.70269 | 3.673 |
| chr15 | 89309035 | 89309322 | 0.341261 | 1.25347 | 3.673 |
| chr3 | 56528736 | 56529103 | 1.49419 | 5.48886 | 3.673 |
| chr5 | 912922 | 913180 | 0.889368 | 3.26734 | 3.674 |
| chr7 | 129112258 | 129112488 | 0.69258 | 2.54478 | 3.674 |
| chr12 | 101381665 | 101381876 | 0.725205 | 2.66472 | 3.674 |
| chrX | 41112904 | 41113299 | 0.982565 | 3.61089 | 3.675 |
| chr1 | 177639033 | 177639286 | 0.33065 | 1.21519 | 3.675 |
| chr12 | 119224541 | 119224797 | 0.343474 | 1.26235 | 3.675 |
| chr19 | 50755462 | 50756093 | 0.901144 | 3.31214 | 3.675 |
| chr11 | 118081174 | 118081435 | 0.28229 | 1.03765 | 3.676 |
| chr4 | 170540647 | 170540857 | 0.840546 | 3.08974 | 3.676 |
| chr10 | 64251856 | 64252059 | 0.594721 | 2.18645 | 3.676 |
| chr1 | 197369210 | 197369533 | 1.31554 | 4.83676 | 3.677 |
| chr3 | 24239364 | 24239609 | 0.471506 | 1.734 | 3.678 |
| chrX | 69869280 | 69869507 | 0.312269 | 1.14907 | 3.680 |
| chr12 | 50134882 | 50135224 | 1.51623 | 5.58154 | 3.681 |
| chr9 | 33170879 | 33171210 | 0.722695 | 2.66043 | 3.681 |
| chr2 | 204680074 | 204680318 | 0.868018 | 3.19628 | 3.682 |
| chr10 | 114886332 | 114886797 | 1.30774 | 4.81696 | 3.683 |
| chr9 | 124497299 | 124497593 | 0.92552 | 3.40951 | 3.684 |
| chr10 | 126849990 | 126850291 | 0.699733 | 2.5787 | 3.685 |
| chr19 | 15284630 | 15284861 | 0.580205 | 2.1387 | 3.686 |
| chr1 | 114031423 | 114031639 | 0.484926 | 1.78765 | 3.686 |
| chrX | 71214618 | 71214827 | 0.298858 | 1.10186 | 3.687 |
| chr3 | 141632100 | 141632419 | 0.990083 | 3.65149 | 3.688 |
| chr16 | 69139382 | 69139598 | 0.733964 | 2.707 | 3.688 |
| chr2 | 95980816 | 95981163 | 1.33031 | 4.90939 | 3.690 |
| chr21 | 36423552 | 36423865 | 1.08596 | 4.00785 | 3.691 |
| chr7 | 73393454 | 73393799 | 1.51951 | 5.6083 | 3.691 |
| chr15 | 73052032 | 73052449 | 0.321238 | 1.18568 | 3.691 |
| chrX | 133895875 | 133896127 | 0.657739 | 2.42924 | 3.693 |
| chr19 | 9247196 | 9247468 | 0.925233 | 3.41896 | 3.695 |
| chr16 | 19533275 | 19533482 | 0.711519 | 2.6294 | 3.695 |
| chr1 | 205091349 | 205091655 | 1.44546 | 5.34364 | 3.697 |
| chr3 | 115705434 | 115705713 | 0.935996 | 3.46069 | 3.697 |
| chr9 | 140126682 | 140126893 | 0.328438 | 1.21461 | 3.698 |
| chr14 | 74998189 | 74998492 | 0.53963 | 1.99633 | 3.699 |
| chr12 | 112250900 | 112251373 | 1.04923 | 3.88194 | 3.700 |
| chr3 | 577717 | 577983 | 0.33456 | 1.238 | 3.700 |
| chr16 | 89008208 | 89008671 | 1.5545 | 5.75403 | 3.702 |
| chr19 | 9697445 | 9697727 | 0.622017 | 2.30243 | 3.702 |
| chr22 | 21310681 | 21310895 | 0.856065 | 3.16883 | 3.702 |
| chr3 | 143199388 | 143199624 | 0.249391 | 0.923167 | 3.702 |
| chrX | 114382342 | 114382570 | 0.303081 | 1.12224 | 3.703 |
| chr10 | 121176361 | 121176757 | 0.920285 | 3.4079 | 3.703 |
| chr8 | 190417 | 190631 | 0.677563 | 2.50988 | 3.704 |
| chrX | 9319918 | 9320189 | 0.558899 | 2.07037 | 3.704 |
| chr20 | 61053252 | 61053478 | 0.353285 | 1.30966 | 3.707 |
| chr1 | 178252039 | 178252281 | 0.685044 | 2.53957 | 3.707 |
| chr8 | 13691059 | 13691295 | 0.35297 | 1.30863 | 3.707 |
| chr5 | 140943695 | 140944122 | 1.60268 | 5.94483 | 3.709 |

| | | | | | |
|---|---|---|---|---|---|
| chr8 | 19612167 | 19612452 | 1.44137 | 5.34924 | 3.711 |
| chr1 | 27728853 | 27729096 | 0.587694 | 2.18148 | 3.712 |
| chr3 | 98710337 | 98710569 | 0.713928 | 2.65046 | 3.713 |
| chr10 | 6443831 | 6444085 | 1.01951 | 3.78543 | 3.713 |
| chr5 | 49615919 | 49616327 | 0.934814 | 3.47281 | 3.715 |
| chr13 | 25245556 | 25245825 | 0.798446 | 2.96628 | 3.715 |
| chr10 | 26496185 | 26496387 | 0.32413 | 1.2044 | 3.716 |
| chr10 | 37723123 | 37723338 | 0.412704 | 1.53358 | 3.716 |
| chr14 | 84174895 | 84175131 | 0.375867 | 1.39704 | 3.717 |
| chr3 | 177873308 | 177873611 | 0.35216 | 1.30895 | 3.717 |
| chr16 | 66730711 | 66731039 | 0.478063 | 1.77704 | 3.717 |
| chr3 | 188382681 | 188382887 | 0.639288 | 2.37727 | 3.719 |
| chr15 | 58519945 | 58520258 | 1.3239 | 4.92376 | 3.719 |
| chr13 | 29943333 | 29943589 | 0.737097 | 2.74171 | 3.720 |
| chr1 | 208794684 | 208794956 | 0.231248 | 0.860488 | 3.721 |
| chr5 | 140220606 | 140221559 | 0.719086 | 2.67602 | 3.721 |
| chr8 | 10785923 | 10786386 | 2.01338 | 7.49377 | 3.722 |
| chr8 | 107016664 | 107016905 | 1.04055 | 3.87325 | 3.722 |
| chr5 | 140709862 | 140710080 | 0.743943 | 2.7696 | 3.723 |
| chr14 | 35250620 | 35250845 | 0.354017 | 1.31812 | 3.723 |
| chr12 | 56190927 | 56191158 | 0.849316 | 3.16239 | 3.723 |
| chr9 | 6701466 | 6701678 | 0.309515 | 1.15285 | 3.725 |
| chr4 | 185084984 | 185085355 | 1.22546 | 4.56492 | 3.725 |
| chr13 | 75455184 | 75455461 | 0.350968 | 1.30776 | 3.726 |
| chr1 | 27718785 | 27719037 | 0.910397 | 3.3941 | 3.728 |
| chr1 | 163626090 | 163626360 | 0.270976 | 1.01056 | 3.729 |
| chr12 | 14782938 | 14783458 | 1.85419 | 6.91676 | 3.730 |
| chr5 | 176934315 | 176934540 | 0.505237 | 1.88494 | 3.731 |
| chr9 | 82182779 | 82183086 | 0.862924 | 3.21978 | 3.731 |
| chr17 | 16487541 | 16487954 | 1.30918 | 4.88522 | 3.732 |
| chr3 | 104282943 | 104283235 | 0.767071 | 2.86295 | 3.732 |
| chr22 | 25568164 | 25568539 | 0.765741 | 2.85849 | 3.733 |
| chr12 | 46696864 | 46697145 | 0.600707 | 2.24262 | 3.733 |
| chrX | 148368348 | 148368707 | 0.79824 | 2.98148 | 3.735 |
| chr1 | 7278387 | 7278589 | 0.908892 | 3.39486 | 3.735 |
| chr12 | 101590557 | 101590812 | 0.339236 | 1.26737 | 3.736 |
| chr18 | 29557256 | 29557583 | 0.869012 | 3.24667 | 3.736 |
| chr4 | 83660419 | 83660896 | 1.58573 | 5.92441 | 3.736 |
| chr3 | 197228875 | 197229428 | 0.68545 | 2.56103 | 3.736 |
| chrX | 12919634 | 12919900 | 0.944338 | 3.52865 | 3.737 |
| chr1 | 93250245 | 93250485 | 0.35468 | 1.32616 | 3.739 |
| chr22 | 19166933 | 19167280 | 0.913611 | 3.41657 | 3.740 |
| chr8 | 29873625 | 29873907 | 0.78381 | 2.93193 | 3.741 |
| chr17 | 46151700 | 46152017 | 1.01806 | 3.80822 | 3.741 |
| chr17 | 63575458 | 63575681 | 0.379385 | 1.41983 | 3.742 |
| chr14 | 51709391 | 51709615 | 0.502723 | 1.8817 | 3.743 |
| chr12 | 93043779 | 93044099 | 0.991266 | 3.71041 | 3.743 |
| chr9 | 31959861 | 31960090 | 0.268382 | 1.00463 | 3.743 |
| chr19 | 22095266 | 22095809 | 0.576363 | 2.15756 | 3.743 |
| chr11 | 3131324 | 3131836 | 1.00586 | 3.76588 | 3.744 |
| chr9 | 136328269 | 136328686 | 0.777804 | 2.91277 | 3.745 |
| chr13 | 113260695 | 113261071 | 1.24849 | 4.67731 | 3.746 |
| chr3 | 48274150 | 48274458 | 0.834792 | 3.12781 | 3.747 |
| chr12 | 54754635 | 54755022 | 2.28439 | 8.56026 | 3.747 |
| chr1 | 201795980 | 201796224 | 0.274873 | 1.03065 | 3.750 |
| chr1 | 95391886 | 95392199 | 0.80647 | 3.0244 | 3.750 |
| chr18 | 74713999 | 74714262 | 0.38639 | 1.44965 | 3.752 |
| chr4 | 162175618 | 162175883 | 0.531743 | 1.99512 | 3.752 |
| chr7 | 26331357 | 26331559 | 0.718751 | 2.6969 | 3.752 |
| chr1 | 20693701 | 20693927 | 1.20233 | 4.51209 | 3.753 |
| chr6 | 126682229 | 126682451 | 0.503512 | 1.89041 | 3.754 |

| | | | | | |
|---|---|---|---|---|---|
| chr5 | 140797027 | 140797301 | 0.876212 | 3.28999 | 3.755 |
| chr12 | 53738840 | 53739377 | 0.78719 | 2.95705 | 3.756 |
| chr5 | 133362337 | 133362624 | 0.967483 | 3.63456 | 3.757 |
| chr12 | 76425233 | 76425542 | 1.00259 | 3.76684 | 3.757 |
| chr1 | 18081603 | 18081917 | 1.03541 | 3.89182 | 3.759 |
| chr15 | 32093304 | 32093599 | 0.355803 | 1.33759 | 3.759 |
| chr12 | 61002345 | 61002598 | 0.291815 | 1.09743 | 3.761 |
| chr12 | 75040773 | 75040985 | 0.516395 | 1.94205 | 3.761 |
| chr9 | 86140584 | 86140795 | 0.258731 | 0.97306 | 3.761 |
| chr2 | 224854021 | 224854233 | 0.660597 | 2.48514 | 3.762 |
| chr19 | 52759687 | 52759959 | 0.834794 | 3.14084 | 3.762 |
| chr5 | 35611596 | 35611954 | 0.366222 | 1.37794 | 3.763 |
| chr5 | 35617716 | 35618061 | 1.47274 | 5.54316 | 3.764 |
| chrX | 134220523 | 134220809 | 0.790129 | 2.97437 | 3.764 |
| chr12 | 107141223 | 107141450 | 0.319954 | 1.20461 | 3.765 |
| chr12 | 49297754 | 49297973 | 1.7719 | 6.67149 | 3.765 |
| chr1 | 109102472 | 109102823 | 1.16304 | 4.37908 | 3.765 |
| chr4 | 117624192 | 117624454 | 0.718536 | 2.70552 | 3.765 |
| chr7 | 97037015 | 97037256 | 0.219682 | 0.827201 | 3.765 |
| chr3 | 9988127 | 9988469 | 1.39841 | 5.2657 | 3.765 |
| chr19 | 56425101 | 56425529 | 0.874689 | 3.29409 | 3.766 |
| chr1 | 219860285 | 219860623 | 0.346236 | 1.30397 | 3.766 |
| chr8 | 16619977 | 16620240 | 0.326862 | 1.23106 | 3.766 |
| chr2 | 67808858 | 67809239 | 0.513581 | 1.93477 | 3.767 |
| chr18 | 32309237 | 32309559 | 0.665885 | 2.50867 | 3.767 |
| chr7 | 2234671 | 2234879 | 0.613393 | 2.3114 | 3.768 |
| chr4 | 110495623 | 110495876 | 0.711433 | 2.6823 | 3.770 |
| chr18 | 9612298 | 9612513 | 1.12331 | 4.23521 | 3.770 |
| chrX | 7050541 | 7050756 | 0.803723 | 3.03039 | 3.770 |
| chr2 | 6067241 | 6067460 | 0.36727 | 1.38498 | 3.771 |
| chr1 | 43243583 | 43243836 | 0.758729 | 2.8612 | 3.771 |
| chr11 | 4819228 | 4819448 | 0.341833 | 1.28933 | 3.772 |
| chr4 | 76862137 | 76862369 | 0.56949 | 2.14821 | 3.772 |
| chr1 | 26362427 | 26362840 | 1.3165 | 4.96679 | 3.773 |
| chr14 | 61892937 | 61893145 | 0.251214 | 0.947825 | 3.773 |
| chr4 | 82272790 | 82273203 | 1.14824 | 4.33314 | 3.774 |
| chr12 | 89918464 | 89918787 | 1.06064 | 4.00268 | 3.774 |
| chr13 | 25268251 | 25268550 | 1.13673 | 4.2905 | 3.774 |
| chr14 | 100491606 | 100491886 | 0.71522 | 2.69971 | 3.775 |
| chr14 | 69218759 | 69218988 | 0.361324 | 1.36427 | 3.776 |
| chr9 | 118646065 | 118646357 | 1.34221 | 5.06813 | 3.776 |
| chr15 | 40799257 | 40799692 | 0.853771 | 3.22398 | 3.776 |
| chr7 | 30452741 | 30452959 | 0.752442 | 2.84154 | 3.776 |
| chr1 | 66912154 | 66912395 | 0.819182 | 3.09414 | 3.777 |
| chr19 | 42681230 | 42681554 | 0.875181 | 3.30636 | 3.778 |
| chr11 | 59317840 | 59318095 | 0.83375 | 3.15153 | 3.780 |
| chr2 | 122476288 | 122476637 | 0.936795 | 3.54253 | 3.782 |
| chr4 | 33781664 | 33781878 | 0.291539 | 1.10258 | 3.782 |
| chr3 | 75252545 | 75252779 | 0.305903 | 1.15727 | 3.783 |
| chr8 | 67976373 | 67976662 | 2.2891 | 8.66106 | 3.784 |
| chr12 | 129484261 | 129484479 | 0.275311 | 1.04216 | 3.785 |
| chr17 | 74454685 | 74455016 | 1.18002 | 4.46793 | 3.786 |
| chr18 | 45534508 | 45535008 | 0.718144 | 2.71953 | 3.787 |
| chr20 | 34189499 | 34189809 | 0.530397 | 2.00856 | 3.787 |
| chr14 | 82096216 | 82096479 | 0.381664 | 1.44541 | 3.787 |
| chr5 | 173494798 | 173495004 | 0.313696 | 1.18834 | 3.788 |
| chr13 | 24803018 | 24803302 | 1.08241 | 4.10095 | 3.789 |
| chr12 | 131589190 | 131589614 | 1.74346 | 6.6061 | 3.789 |
| chr5 | 91270790 | 91271064 | 0.414604 | 1.57155 | 3.790 |
| chr20 | 35492111 | 35492393 | 0.835856 | 3.16929 | 3.792 |
| chr11 | 65627647 | 65628053 | 1.33768 | 5.07376 | 3.793 |

| | | | | | |
|---|---|---|---|---|---|
| chr8 | 41435336 | 41435766 | 0.717757 | 2.72327 | 3.794 |
| chr1 | 94311758 | 94312012 | 0.431587 | 1.63809 | 3.796 |
| chr8 | 69588306 | 69588617 | 0.342987 | 1.30213 | 3.796 |
| chr15 | 100639042 | 100639245 | 0.399763 | 1.51785 | 3.797 |
| chr17 | 54116031 | 54116421 | 0.361066 | 1.37094 | 3.797 |
| chr11 | 35571956 | 35572167 | 0.251291 | 0.954583 | 3.799 |
| chr19 | 12201700 | 12202056 | 0.591276 | 2.24663 | 3.800 |
| chr16 | 34760557 | 34760951 | 0.567295 | 2.15686 | 3.802 |
| chr19 | 52642489 | 52642835 | 1.00584 | 3.8248 | 3.803 |
| chrX | 117724609 | 117724872 | 0.302948 | 1.15215 | 3.803 |
| chr1 | 228401211 | 228401470 | 0.889208 | 3.3826 | 3.804 |
| chr9 | 115733774 | 115733996 | 0.753617 | 2.86722 | 3.805 |
| chr1 | 147817762 | 147818003 | 0.364957 | 1.38854 | 3.805 |
| chr3 | 105085466 | 105085725 | 0.9506 | 3.61686 | 3.805 |
| chr9 | 139559790 | 139560053 | 0.65769 | 2.50247 | 3.805 |
| chr12 | 95793439 | 95793789 | 0.27571 | 1.04919 | 3.805 |
| chr5 | 77591543 | 77591789 | 0.739928 | 2.81589 | 3.806 |
| chr4 | 34955074 | 34955345 | 0.605747 | 2.30566 | 3.806 |
| chr9 | 120869436 | 120869667 | 0.229289 | 0.872897 | 3.807 |
| chr16 | 58060287 | 58060505 | 0.265513 | 1.01114 | 3.808 |
| chr9 | 5715143 | 5715421 | 0.915793 | 3.48829 | 3.809 |
| chr2 | 151344174 | 151344507 | 1.16519 | 4.4396 | 3.810 |
| chr22 | 39239247 | 39239617 | 0.438421 | 1.6705 | 3.810 |
| chr7 | 91015705 | 91016016 | 0.260658 | 0.99325 | 3.811 |
| chr5 | 146270483 | 146270704 | 0.225409 | 0.858959 | 3.811 |
| chr19 | 52405165 | 52405432 | 0.637128 | 2.42832 | 3.811 |
| chr11 | 57016505 | 57016860 | 0.927641 | 3.53572 | 3.812 |
| chr16 | 35143414 | 35143641 | 0.539298 | 2.05582 | 3.812 |
| chr6 | 21589466 | 21589804 | 1.58839 | 6.0554 | 3.812 |
| chr1 | 26648077 | 26648345 | 1.1932 | 4.54913 | 3.813 |
| chr12 | 49247037 | 49247384 | 0.812469 | 3.09782 | 3.813 |
| chr5 | 156751299 | 156751595 | 0.881683 | 3.36263 | 3.814 |
| chr12 | 48030931 | 48031350 | 0.710094 | 2.7092 | 3.815 |
| chr10 | 133521425 | 133521716 | 0.977521 | 3.7299 | 3.816 |
| chr5 | 145287871 | 145288125 | 0.506843 | 1.934 | 3.816 |
| chr3 | 3242359 | 3242696 | 0.855427 | 3.26597 | 3.818 |
| chr1 | 59307250 | 59307513 | 0.531003 | 2.02768 | 3.819 |
| chr4 | 163418783 | 163419078 | 0.658151 | 2.51415 | 3.820 |
| chr12 | 50163736 | 50163971 | 0.327105 | 1.25008 | 3.822 |
| chr5 | 145826520 | 145826789 | 1.32076 | 5.04809 | 3.822 |
| chr19 | 52854810 | 52855084 | 0.866027 | 3.31079 | 3.823 |
| chr14 | 61092146 | 61092423 | 0.846199 | 3.23513 | 3.823 |
| chr16 | 34794743 | 34794988 | 0.454092 | 1.73608 | 3.823 |
| chr3 | 149435325 | 149435552 | 0.560996 | 2.14488 | 3.823 |
| chr19 | 57917933 | 57918263 | 1.10724 | 4.23597 | 3.826 |
| chr10 | 118525169 | 118525420 | 0.666469 | 2.55076 | 3.827 |
| chr9 | 87047733 | 87047964 | 0.280079 | 1.07241 | 3.829 |
| chr14 | 91721426 | 91721662 | 0.400153 | 1.53272 | 3.830 |
| chr19 | 53580241 | 53580575 | 0.927633 | 3.55329 | 3.830 |
| chr12 | 22964229 | 22964609 | 0.281493 | 1.07829 | 3.831 |
| chr7 | 41744723 | 41745063 | 1.37531 | 5.26936 | 3.831 |
| chr16 | 25430488 | 25430765 | 0.309105 | 1.18445 | 3.832 |
| chr10 | 34071753 | 34072005 | 0.313196 | 1.20018 | 3.832 |
| chr5 | 147376008 | 147376267 | 0.322993 | 1.23794 | 3.833 |
| chr19 | 57058951 | 57059310 | 0.792427 | 3.03853 | 3.834 |
| chr22 | 42362591 | 42362932 | 0.495908 | 1.90217 | 3.836 |
| chr13 | 80253483 | 80253723 | 0.577786 | 2.2166 | 3.836 |
| chr18 | 68944800 | 68945046 | 0.2521 | 0.967319 | 3.837 |
| chr1 | 99745021 | 99745243 | 0.284773 | 1.09363 | 3.840 |
| chr3 | 117617637 | 117617891 | 0.715464 | 2.74772 | 3.840 |
| chr6 | 165824003 | 165824244 | 0.2681 | 1.02978 | 3.841 |

| | | | | | |
|---|---|---|---|---|---|
| chr19 | 49982107 | 49982445 | 1.0291 | 3.95362 | 3.842 |
| chr3 | 15373992 | 15374239 | 0.684176 | 2.62917 | 3.843 |
| chr20 | 20111095 | 20111467 | 0.887233 | 3.41006 | 3.843 |
| chr15 | 57904245 | 57904619 | 0.793837 | 3.05155 | 3.844 |
| chr2 | 147170292 | 147170515 | 0.809106 | 3.11135 | 3.845 |
| chr6 | 108434561 | 108434927 | 0.454123 | 1.74696 | 3.847 |
| chr22 | 32519733 | 32519967 | 0.493722 | 1.89942 | 3.847 |
| chr3 | 180407921 | 180408139 | 0.394793 | 1.51901 | 3.848 |
| chr20 | 44144530 | 44144756 | 0.248379 | 0.955774 | 3.848 |
| chr22 | 19744112 | 19744326 | 1.12988 | 4.34809 | 3.848 |
| chr21 | 45907712 | 45908023 | 0.894372 | 3.44189 | 3.848 |
| chr16 | 78739605 | 78739895 | 0.677126 | 2.60589 | 3.848 |
| chr1 | 30265895 | 30266165 | 1.36223 | 5.24435 | 3.850 |
| chr16 | 85132970 | 85133291 | 0.504334 | 1.94167 | 3.850 |
| chr12 | 106599255 | 106599557 | 0.945138 | 3.63897 | 3.850 |
| chr9 | 11414314 | 11414529 | 0.265899 | 1.02392 | 3.851 |
| chr14 | 105237140 | 105237680 | 0.826815 | 3.18554 | 3.853 |
| chr10 | 118481717 | 118481962 | 0.330651 | 1.27407 | 3.853 |
| chr14 | 93439573 | 93439946 | 1.11227 | 4.28626 | 3.854 |
| chr14 | 42958211 | 42958485 | 0.316786 | 1.2212 | 3.855 |
| chr1 | 68786809 | 68787117 | 1.28801 | 4.96574 | 3.855 |
| chr9 | 33447413 | 33447725 | 0.620202 | 2.39149 | 3.856 |
| chr5 | 177920596 | 177920852 | 0.371616 | 1.43304 | 3.856 |
| chr11 | 57386824 | 57387143 | 0.630784 | 2.43337 | 3.858 |
| chrX | 42574603 | 42575015 | 0.910459 | 3.51423 | 3.860 |
| chr4 | 190017991 | 190018199 | 0.6556 | 2.53069 | 3.860 |
| chr12 | 23515035 | 23515246 | 0.358519 | 1.38395 | 3.860 |
| chr22 | 29702197 | 29702418 | 0.791234 | 3.05452 | 3.860 |
| chr3 | 38109826 | 38110129 | 0.220958 | 0.8532 | 3.861 |
| chr19 | 58554308 | 58554695 | 0.929745 | 3.59028 | 3.862 |
| chr11 | 34912461 | 34912682 | 0.59201 | 2.28618 | 3.862 |
| chr5 | 140514624 | 140514904 | 1.03775 | 4.00812 | 3.862 |
| chr13 | 45451075 | 45451403 | 0.852245 | 3.29183 | 3.863 |
| chr11 | 106526089 | 106526316 | 0.322217 | 1.2448 | 3.863 |
| chr11 | 69993813 | 69994061 | 0.271502 | 1.04894 | 3.863 |
| chr7 | 48149719 | 48150128 | 1.53994 | 5.95282 | 3.866 |
| chr9 | 95748517 | 95748828 | 0.670565 | 2.59232 | 3.866 |
| chr5 | 78796620 | 78796881 | 0.294955 | 1.14082 | 3.868 |
| chr8 | 26519356 | 26519635 | 1.07887 | 4.17409 | 3.869 |
| chr16 | 58874183 | 58874438 | 0.281237 | 1.08862 | 3.871 |
| chr12 | 122089186 | 122089718 | 0.316895 | 1.22702 | 3.872 |
| chr19 | 37407270 | 37407524 | 0.758286 | 2.93665 | 3.873 |
| chr19 | 49818425 | 49818717 | 0.475347 | 1.84115 | 3.873 |
| chr11 | 60683030 | 60683280 | 0.51782 | 2.00587 | 3.874 |
| chr12 | 116360793 | 116361116 | 0.934553 | 3.62059 | 3.874 |
| chr4 | 139443661 | 139443908 | 0.580821 | 2.2506 | 3.875 |
| chr8 | 129871436 | 129871773 | 1.11032 | 4.30242 | 3.875 |
| chr8 | 144614197 | 144614712 | 1.41729 | 5.49325 | 3.876 |
| chr16 | 19400569 | 19400843 | 1.05472 | 4.08803 | 3.876 |
| chr1 | 48463252 | 48463687 | 0.998212 | 3.86902 | 3.876 |
| chr9 | 91933797 | 91934055 | 0.918145 | 3.55924 | 3.877 |
| chr16 | 71475317 | 71475819 | 0.742308 | 2.8783 | 3.878 |
| chr6 | 120398836 | 120399111 | 0.298058 | 1.15594 | 3.878 |
| chr2 | 228270866 | 228271127 | 0.780003 | 3.02529 | 3.879 |
| chr10 | 43716837 | 43717195 | 0.68947 | 2.67573 | 3.881 |
| chr3 | 194756092 | 194756356 | 0.494953 | 1.92238 | 3.884 |
| chr11 | 82952465 | 82952686 | 0.89301 | 3.46889 | 3.884 |
| chr21 | 30536758 | 30537011 | 0.564168 | 2.19178 | 3.885 |
| chr20 | 50328997 | 50329395 | 1.00822 | 3.9182 | 3.886 |
| chr18 | 74187486 | 74187743 | 0.677115 | 2.633 | 3.889 |
| chr4 | 105372767 | 105373056 | 0.543527 | 2.11356 | 3.889 |

| | | | | | |
|---|---|---|---|---|---|
| chr5 | 37199879 | 37200108 | 0.421828 | 1.64053 | 3.889 |
| chr1 | 97061876 | 97062121 | 1.00582 | 3.91257 | 3.890 |
| chr11 | 90813527 | 90813743 | 0.337394 | 1.31277 | 3.891 |
| chr12 | 48166729 | 48167112 | 1.17511 | 4.57235 | 3.891 |
| chr9 | 87992906 | 87993154 | 0.232345 | 0.904082 | 3.891 |
| chr1 | 203941898 | 203942103 | 0.217677 | 0.847264 | 3.892 |
| chr11 | 69460065 | 69460422 | 0.519876 | 2.024 | 3.893 |
| chr1 | 3333631 | 3334010 | 0.558973 | 2.17661 | 3.894 |
| chr1 | 2040078 | 2040389 | 0.645304 | 2.51306 | 3.894 |
| chr7 | 825104 | 825316 | 0.363093 | 1.41408 | 3.895 |
| chr8 | 42994181 | 42994546 | 0.620821 | 2.41805 | 3.895 |
| chr5 | 122335660 | 122335993 | 0.963437 | 3.75265 | 3.895 |
| chr4 | 48945367 | 48945695 | 0.496988 | 1.93625 | 3.896 |
| chr7 | 51068629 | 51069051 | 0.909084 | 3.54265 | 3.897 |
| chr12 | 114607705 | 114607928 | 0.282681 | 1.10167 | 3.897 |
| chr3 | 125803095 | 125803322 | 0.530885 | 2.06898 | 3.897 |
| chr3 | 109088395 | 109088690 | 0.648358 | 2.52763 | 3.899 |
| chr12 | 52672746 | 52673129 | 1.92169 | 7.49208 | 3.899 |
| chr7 | 91210891 | 91211177 | 0.275648 | 1.07473 | 3.899 |
| chr5 | 140588094 | 140588441 | 0.623184 | 2.43031 | 3.900 |
| chr12 | 13964615 | 13964828 | 0.399757 | 1.55922 | 3.900 |
| chr6 | 81785681 | 81786044 | 0.51524 | 2.00989 | 3.901 |
| chr2 | 121214592 | 121214835 | 0.570052 | 2.22374 | 3.901 |
| chr4 | 114605954 | 114606243 | 0.512109 | 1.99788 | 3.901 |
| chr7 | 97325650 | 97325884 | 0.290932 | 1.13517 | 3.902 |
| chr7 | 71198096 | 71198685 | 1.25841 | 4.91088 | 3.902 |
| chr7 | 158766220 | 158766474 | 0.644763 | 2.51663 | 3.903 |
| chr21 | 30671030 | 30671251 | 0.475593 | 1.85649 | 3.904 |
| chr11 | 65297557 | 65298025 | 0.609119 | 2.37792 | 3.904 |
| chr11 | 101723079 | 101723326 | 0.677576 | 2.64539 | 3.904 |
| chr11 | 66206509 | 66206818 | 0.930192 | 3.63329 | 3.906 |
| chr8 | 48105686 | 48105945 | 0.69464 | 2.71324 | 3.906 |
| chr5 | 565994 | 566201 | 0.402625 | 1.57289 | 3.907 |
| chr8 | 71191047 | 71191263 | 0.30406 | 1.188 | 3.907 |
| chr5 | 132446941 | 132447222 | 1.20489 | 4.70771 | 3.907 |
| chr3 | 8396457 | 8396660 | 0.287929 | 1.12504 | 3.907 |
| chr13 | 27936090 | 27936458 | 1.29099 | 5.04474 | 3.908 |
| chrX | 100670542 | 100670797 | 0.972074 | 3.79867 | 3.908 |
| chr1 | 197884925 | 197885216 | 1.21543 | 4.74982 | 3.908 |
| chr10 | 97050532 | 97050791 | 0.669411 | 2.61602 | 3.908 |
| chr15 | 76380237 | 76380509 | 0.260719 | 1.01888 | 3.908 |
| chr12 | 110869094 | 110869524 | 1.9598 | 7.66053 | 3.909 |
| chr4 | 189892395 | 189892600 | 0.622103 | 2.43197 | 3.909 |
| chr20 | 3057490 | 3057819 | 0.825793 | 3.22915 | 3.910 |
| chr6 | 134138600 | 134138993 | 0.99268 | 3.88259 | 3.911 |
| chr15 | 79165849 | 79166148 | 1.06148 | 4.15171 | 3.911 |
| chr7 | 23053879 | 23054108 | 0.620389 | 2.42705 | 3.912 |
| chr5 | 60179085 | 60179376 | 0.890999 | 3.48612 | 3.913 |
| chr2 | 76011846 | 76012106 | 0.292176 | 1.14331 | 3.913 |
| chr20 | 1757979 | 1758223 | 1.5094 | 5.90992 | 3.915 |
| chr13 | 68015094 | 68015344 | 0.238453 | 0.933927 | 3.917 |
| chr11 | 5621479 | 5621866 | 1.07738 | 4.21987 | 3.917 |
| chr5 | 174979463 | 174979688 | 0.261982 | 1.02616 | 3.917 |
| chr13 | 27180081 | 27180299 | 0.403012 | 1.57874 | 3.917 |
| chr5 | 21803040 | 21803296 | 0.673675 | 2.64031 | 3.919 |
| chr1 | 188768667 | 188768957 | 0.266223 | 1.04343 | 3.919 |
| chr19 | 3129155 | 3129423 | 0.608906 | 2.38661 | 3.920 |
| chr5 | 119970060 | 119970316 | 0.349536 | 1.37006 | 3.920 |
| chr22 | 27333255 | 27333485 | 0.536423 | 2.10286 | 3.920 |
| chr9 | 25686574 | 25686779 | 0.347304 | 1.36152 | 3.920 |
| chr12 | 17368701 | 17369047 | 0.570703 | 2.23752 | 3.921 |

| | | | | | |
|---|---|---|---|---|---|
| chr10 | 55779208 | 55779436 | 0.316795 | 1.24219 | 3.921 |
| chr3 | 113685044 | 113685251 | 0.479841 | 1.88183 | 3.922 |
| chrX | 62470237 | 62470470 | 1.02994 | 4.03998 | 3.923 |
| chr19 | 56820858 | 56821442 | 1.65761 | 6.50556 | 3.925 |
| chr1 | 200248978 | 200249245 | 1.37779 | 5.40849 | 3.925 |
| chr9 | 116444494 | 116444863 | 1.40358 | 5.51061 | 3.926 |
| chr21 | 39131353 | 39131588 | 1.1579 | 4.5466 | 3.927 |
| chr17 | 72081595 | 72081917 | 0.729722 | 2.86584 | 3.927 |
| chr19 | 11896076 | 11896282 | 0.828916 | 3.2558 | 3.928 |
| chr16 | 29801648 | 29801989 | 0.915318 | 3.59716 | 3.930 |
| chr10 | 50472064 | 50472277 | 0.216786 | 0.852072 | 3.930 |
| chr9 | 131641590 | 131641881 | 0.675506 | 2.6556 | 3.931 |
| chr11 | 11688666 | 11688953 | 0.458799 | 1.80379 | 3.932 |
| chr1 | 227422664 | 227423007 | 1.11455 | 4.38377 | 3.933 |
| chr9 | 106236023 | 106236454 | 0.23056 | 0.906999 | 3.934 |
| chrX | 7089041 | 7089287 | 1.06075 | 4.17323 | 3.934 |
| chr9 | 100955382 | 100955648 | 0.680269 | 2.67667 | 3.935 |
| chr12 | 132010321 | 132010559 | 0.253145 | 0.996371 | 3.936 |
| chr8 | 145955381 | 145955712 | 1.22495 | 4.82186 | 3.936 |
| chr16 | 67465158 | 67465435 | 0.311529 | 1.22636 | 3.937 |
| chr9 | 120842208 | 120842432 | 0.204076 | 0.803433 | 3.937 |
| chr12 | 63322304 | 63322509 | 0.341413 | 1.34441 | 3.938 |
| chr4 | 173813539 | 173813845 | 0.819171 | 3.22586 | 3.938 |
| chr14 | 25532418 | 25532744 | 0.293237 | 1.15512 | 3.939 |
| chr19 | 5515766 | 5516083 | 0.805509 | 3.17311 | 3.939 |
| chrX | 53118510 | 53118756 | 1.03707 | 4.08545 | 3.939 |
| chr1 | 234225216 | 234225431 | 0.789111 | 3.109 | 3.940 |
| chr7 | 121150405 | 121150652 | 0.66731 | 2.62988 | 3.941 |
| chr7 | 26438933 | 26439154 | 0.446071 | 1.75885 | 3.943 |
| chr5 | 44862974 | 44863187 | 0.554545 | 2.18679 | 3.943 |
| chr10 | 119444294 | 119444556 | 0.621193 | 2.4499 | 3.944 |
| chr7 | 25138003 | 25138392 | 0.977018 | 3.85356 | 3.944 |
| chr5 | 176829555 | 176829792 | 0.999754 | 3.94353 | 3.945 |
| chr5 | 176046356 | 176046570 | 0.778579 | 3.07284 | 3.947 |
| chr14 | 50809524 | 50809738 | 0.72131 | 2.84682 | 3.947 |
| chr12 | 52444574 | 52444862 | 0.360608 | 1.42336 | 3.947 |
| chr19 | 37718935 | 37719247 | 1.13832 | 4.49323 | 3.947 |
| chr12 | 33684191 | 33684469 | 0.231202 | 0.912713 | 3.948 |
| chr12 | 13232308 | 13232547 | 0.650488 | 2.56812 | 3.948 |
| chr1 | 208425802 | 208426129 | 1.04802 | 4.13762 | 3.948 |
| chr3 | 52462589 | 52462872 | 0.58639 | 2.31618 | 3.950 |
| chr10 | 5057210 | 5057489 | 0.56063 | 2.21521 | 3.951 |
| chr15 | 74243482 | 74243769 | 1.04468 | 4.12813 | 3.952 |
| chr12 | 6875896 | 6876303 | 1.08343 | 4.28176 | 3.952 |
| chr15 | 39872352 | 39872716 | 1.33801 | 5.28881 | 3.953 |
| chr19 | 58428066 | 58428295 | 0.764525 | 3.0249 | 3.957 |
| chr7 | 27784749 | 27785024 | 1.12539 | 4.45328 | 3.957 |
| chr10 | 116463383 | 116463745 | 1.22591 | 4.85473 | 3.960 |
| chr17 | 73717430 | 73717795 | 0.402963 | 1.59579 | 3.960 |
| chr3 | 150735992 | 150736214 | 0.23465 | 0.92934 | 3.961 |
| chr14 | 107150503 | 107150877 | 0.766172 | 3.03535 | 3.962 |
| chr9 | 20175965 | 20176188 | 0.257663 | 1.02124 | 3.963 |
| chr10 | 19457564 | 19457809 | 0.713166 | 2.82714 | 3.964 |
| chr8 | 127287586 | 127287833 | 0.524469 | 2.07969 | 3.965 |
| chr22 | 33599257 | 33599551 | 0.825676 | 3.27599 | 3.968 |
| chr4 | 6666352 | 6666654 | 0.719328 | 2.85479 | 3.969 |
| chr8 | 61972235 | 61972458 | 0.317472 | 1.26054 | 3.971 |
| chr1 | 54587282 | 54587564 | 1.14485 | 4.54574 | 3.971 |
| chr6 | 148466130 | 148466390 | 0.990472 | 3.93323 | 3.971 |
| chr19 | 11923018 | 11923271 | 0.713199 | 2.83252 | 3.972 |
| chr4 | 88976099 | 88976407 | 1.07181 | 4.25695 | 3.972 |

| | | | | | |
|---|---|---|---|---|---|
| chr3 | 102460071 | 102460566 | 0.391198 | 1.55463 | 3.974 |
| chr17 | 62368581 | 62369024 | 0.895869 | 3.56021 | 3.974 |
| chr8 | 50811693 | 50811911 | 0.341045 | 1.35537 | 3.974 |
| chr5 | 100683486 | 100683748 | 0.596261 | 2.37007 | 3.975 |
| chr9 | 104784497 | 104784747 | 0.358582 | 1.4254 | 3.975 |
| chr12 | 48211128 | 48211430 | 0.490681 | 1.95056 | 3.975 |
| chr19 | 50050254 | 50050524 | 0.791401 | 3.14673 | 3.976 |
| chr7 | 771786 | 772169 | 0.540037 | 2.14735 | 3.976 |
| chr5 | 109026736 | 109027013 | 0.915465 | 3.64079 | 3.977 |
| chr18 | 18714891 | 18715160 | 0.564624 | 2.24634 | 3.978 |
| chr19 | 18519071 | 18519334 | 0.974177 | 3.87701 | 3.980 |
| chr22 | 48233480 | 48233763 | 0.641638 | 2.55383 | 3.980 |
| chr4 | 88465077 | 88465283 | 0.49565 | 1.97342 | 3.981 |
| chr4 | 127548500 | 127548765 | 0.461345 | 1.83701 | 3.982 |
| chr5 | 132460533 | 132460781 | 0.280463 | 1.11689 | 3.982 |
| chr16 | 11750973 | 11751192 | 0.292638 | 1.16616 | 3.985 |
| chr7 | 1749586 | 1749891 | 0.654171 | 2.60714 | 3.985 |
| chr16 | 27439865 | 27440140 | 0.563056 | 2.24468 | 3.987 |
| chr20 | 38531146 | 38531357 | 0.249196 | 0.993527 | 3.987 |
| chr5 | 3852801 | 3853045 | 0.545313 | 2.17429 | 3.987 |
| chr10 | 88108576 | 88108910 | 0.76608 | 3.05462 | 3.987 |
| chr6 | 34163978 | 34164421 | 1.35871 | 5.41902 | 3.988 |
| chr4 | 184284454 | 184284669 | 0.52078 | 2.07747 | 3.989 |
| chr16 | 67360511 | 67360936 | 1.17556 | 4.69067 | 3.990 |
| chr5 | 140777291 | 140777549 | 0.502229 | 2.00407 | 3.990 |
| chr15 | 42916093 | 42916331 | 0.151782 | 0.605706 | 3.991 |
| chr12 | 91347641 | 91348088 | 1.13139 | 4.51506 | 3.991 |
| chr9 | 5306716 | 5306963 | 0.300494 | 1.20039 | 3.995 |
| chr5 | 134073913 | 134074237 | 0.938405 | 3.7498 | 3.996 |
| chr10 | 93395213 | 93395475 | 0.953126 | 3.80924 | 3.997 |
| chr4 | 10159427 | 10159711 | 0.83462 | 3.3357 | 3.997 |
| chr20 | 25555140 | 25555516 | 0.595666 | 2.38099 | 3.997 |
| chr6 | 25490338 | 25490647 | 0.960936 | 3.84153 | 3.998 |
| chr6 | 50524685 | 50524993 | 0.192467 | 0.769464 | 3.998 |
| chr12 | 100013629 | 100014046 | 0.909596 | 3.63657 | 3.998 |
| chr9 | 5832886 | 5833300 | 1.57581 | 6.30064 | 3.998 |
| chr17 | 10798998 | 10799220 | 0.336208 | 1.34436 | 3.999 |
| chr5 | 35116050 | 35116262 | 0.267341 | 1.06904 | 3.999 |
| chr12 | 109058606 | 109058910 | 0.961344 | 3.84688 | 4.002 |
| chr7 | 54078963 | 54079214 | 0.324976 | 1.30083 | 4.003 |
| chr6 | 84410557 | 84410795 | 0.231293 | 0.926021 | 4.004 |
| chrX | 47885956 | 47886466 | 0.957786 | 3.83519 | 4.004 |
| chr3 | 157473198 | 157473459 | 0.359902 | 1.44114 | 4.004 |
| chr7 | 151451187 | 151451527 | 1.85521 | 7.42881 | 4.004 |
| chr10 | 92613796 | 92614027 | 0.316895 | 1.2694 | 4.006 |
| chr7 | 40206192 | 40206488 | 0.240732 | 0.964744 | 4.008 |
| chr9 | 77385831 | 77386250 | 1.84442 | 7.39322 | 4.008 |
| chr8 | 61573118 | 61573417 | 0.827998 | 3.31905 | 4.009 |
| chr8 | 43291143 | 43291502 | 0.533045 | 2.13731 | 4.010 |
| chr12 | 1905105 | 1905601 | 1.50654 | 6.04127 | 4.010 |
| chr4 | 4482398 | 4482600 | 0.617568 | 2.47668 | 4.010 |
| chr3 | 5829523 | 5829917 | 0.82317 | 3.30132 | 4.011 |
| chr1 | 229588968 | 229589370 | 0.852163 | 3.41902 | 4.012 |
| chr9 | 127527170 | 127527374 | 0.683926 | 2.74423 | 4.012 |
| chr12 | 133498341 | 133498684 | 0.804387 | 3.22767 | 4.013 |
| chr12 | 6176831 | 6177062 | 0.93717 | 3.76134 | 4.014 |
| chr10 | 13457047 | 13457319 | 0.531595 | 2.13376 | 4.014 |
| chr4 | 115580593 | 115580909 | 1.02707 | 4.12298 | 4.014 |
| chr6 | 137910530 | 137910847 | 0.843004 | 3.3855 | 4.016 |
| chr19 | 57171749 | 57172405 | 0.810094 | 3.25348 | 4.016 |
| chr19 | 628163 | 628532 | 0.813845 | 3.2689 | 4.017 |

| | | | | | |
|---|---|---|---|---|---|
| chr16 | 12692810 | 12693057 | 0.276684 | 1.11138 | 4.017 |
| chr1 | 157157608 | 157157887 | 0.277878 | 1.11637 | 4.017 |
| chr13 | 74423471 | 74423687 | 0.491504 | 1.97472 | 4.018 |
| chr12 | 121086967 | 121087379 | 1.27299 | 5.11554 | 4.019 |
| chrY | 15040039 | 15040269 | 0.515655 | 2.07259 | 4.019 |
| chr10 | 95462084 | 95462315 | 0.513054 | 2.06328 | 4.022 |
| chr17 | 26674061 | 26674287 | 0.671582 | 2.7013 | 4.022 |
| chr20 | 5799776 | 5799990 | 0.857333 | 3.44924 | 4.023 |
| chr1 | 90246260 | 90246555 | 0.481483 | 1.93773 | 4.025 |
| chr12 | 14017010 | 14017480 | 1.56206 | 6.28849 | 4.026 |
| chr3 | 94435551 | 94435762 | 0.68809 | 2.7704 | 4.026 |
| chrX | 119384489 | 119384759 | 0.935002 | 3.76506 | 4.027 |
| chr15 | 69453970 | 69454230 | 0.823858 | 3.31855 | 4.028 |
| chr5 | 170731073 | 170731317 | 0.245548 | 0.989651 | 4.030 |
| chr2 | 76826881 | 76827190 | 0.678452 | 2.73462 | 4.031 |
| chr17 | 32395721 | 32395928 | 0.326589 | 1.31657 | 4.031 |
| chr1 | 94375025 | 94375475 | 1.35261 | 5.45295 | 4.031 |
| chr3 | 36885221 | 36885427 | 0.672671 | 2.71184 | 4.031 |
| chr20 | 42037909 | 42038360 | 0.781663 | 3.15146 | 4.032 |
| chr6 | 151213073 | 151213325 | 1.00813 | 4.0655 | 4.033 |
| chr12 | 45858371 | 45858705 | 0.404465 | 1.63189 | 4.035 |
| chr8 | 99713010 | 99713381 | 1.02105 | 4.1221 | 4.037 |
| chr15 | 68831398 | 68831760 | 1.09041 | 4.40293 | 4.038 |
| chr19 | 22785473 | 22785685 | 0.402731 | 1.62634 | 4.038 |
| chr15 | 45491028 | 45491364 | 0.748023 | 3.02135 | 4.039 |
| chr9 | 10612977 | 10613184 | 0.57176 | 2.3104 | 4.041 |
| chr20 | 37361745 | 37362053 | 0.289172 | 1.16896 | 4.042 |
| chrX | 29496808 | 29497125 | 1.22778 | 4.9633 | 4.043 |
| chr10 | 80184579 | 80184795 | 0.499324 | 2.01903 | 4.044 |
| chr15 | 32082216 | 32082471 | 0.323941 | 1.31012 | 4.044 |
| chr16 | 88320906 | 88321268 | 0.948056 | 3.83472 | 4.045 |
| chrX | 134133570 | 134133789 | 0.744153 | 3.0102 | 4.045 |
| chr16 | 65703949 | 65704225 | 0.247221 | 1.00007 | 4.045 |
| chr19 | 35797598 | 35797999 | 0.925134 | 3.74429 | 4.047 |
| chr12 | 13623213 | 13623419 | 0.342333 | 1.38558 | 4.047 |
| chr18 | 11937927 | 11938248 | 1.12777 | 4.56523 | 4.048 |
| chr10 | 14946334 | 14946566 | 0.693548 | 2.80782 | 4.048 |
| chrY | 14733702 | 14733998 | 0.603253 | 2.44333 | 4.050 |
| chr12 | 53447427 | 53447712 | 0.721881 | 2.92409 | 4.051 |
| chr15 | 52069160 | 52069673 | 1.12993 | 4.57721 | 4.051 |
| chr11 | 117009485 | 117009718 | 0.601351 | 2.43684 | 4.052 |
| chr19 | 11811045 | 11811278 | 0.728215 | 2.95108 | 4.052 |
| chr4 | 3454214 | 3454547 | 0.814519 | 3.30171 | 4.054 |
| chr4 | 3347824 | 3348219 | 0.740648 | 3.00462 | 4.057 |
| chrX | 133898290 | 133898521 | 0.595408 | 2.41595 | 4.058 |
| chr14 | 23821449 | 23821672 | 0.248376 | 1.00788 | 4.058 |
| chr12 | 96883122 | 96883440 | 1.96192 | 7.96304 | 4.059 |
| chr22 | 50982661 | 50982973 | 0.818982 | 3.32646 | 4.062 |
| chr16 | 67294599 | 67294931 | 1.18089 | 4.798 | 4.063 |
| chr14 | 40202324 | 40202593 | 0.194393 | 0.78999 | 4.064 |
| chr11 | 78745963 | 78746168 | 0.261503 | 1.06303 | 4.065 |
| chr5 | 42995047 | 42995322 | 1.30047 | 5.29049 | 4.068 |
| chr9 | 83789018 | 83789285 | 0.640189 | 2.60451 | 4.068 |
| chr9 | 3044655 | 3044970 | 0.295461 | 1.20218 | 4.069 |
| chr3 | 51008890 | 51009240 | 1.0302 | 4.19451 | 4.072 |
| chr10 | 31372630 | 31372951 | 0.650222 | 2.64764 | 4.072 |
| chr12 | 130526665 | 130527172 | 1.19297 | 4.85854 | 4.073 |
| chr2 | 66159281 | 66159485 | 0.342976 | 1.39685 | 4.073 |
| chr20 | 51995526 | 51995748 | 0.263231 | 1.07215 | 4.073 |
| chr22 | 29949764 | 29950027 | 0.733873 | 2.98992 | 4.074 |
| chr14 | 71885309 | 71885546 | 0.320887 | 1.30766 | 4.075 |

| | | | | | |
|---|---|---|---|---|---|
| chr3 | 189679393 | 189679648 | 1.02309 | 4.17035 | 4.076 |
| chr19 | 47634015 | 47634230 | 0.464516 | 1.89403 | 4.077 |
| chr3 | 111335443 | 111335715 | 0.459092 | 1.87231 | 4.078 |
| chr17 | 60256996 | 60257244 | 0.966807 | 3.944 | 4.079 |
| chr10 | 35274839 | 35275094 | 0.954759 | 3.89788 | 4.083 |
| chr14 | 65512340 | 65512576 | 0.303185 | 1.23792 | 4.083 |
| chr5 | 171550230 | 171550454 | 0.435703 | 1.77963 | 4.085 |
| chr16 | 55681807 | 55682115 | 0.374375 | 1.52915 | 4.085 |
| chr4 | 30382077 | 30382345 | 0.857759 | 3.50376 | 4.085 |
| chr6 | 18538413 | 18538666 | 0.22844 | 0.933364 | 4.086 |
| chr1 | 221916080 | 221916372 | 0.751971 | 3.07287 | 4.086 |
| chr19 | 12313122 | 12313401 | 0.595444 | 2.43348 | 4.087 |
| chr3 | 37850385 | 37850661 | 0.198194 | 0.810138 | 4.088 |
| chr10 | 31757219 | 31757571 | 0.773336 | 3.16153 | 4.088 |
| chr4 | 99916256 | 99916970 | 0.616674 | 2.52108 | 4.088 |
| chr4 | 41283110 | 41283337 | 0.520875 | 2.13185 | 4.093 |
| chr2 | 106586079 | 106586369 | 0.809046 | 3.31241 | 4.094 |
| chr12 | 8674110 | 8674334 | 0.231634 | 0.948462 | 4.095 |
| chr19 | 11248444 | 11248681 | 0.644969 | 2.64217 | 4.097 |
| chr7 | 145108533 | 145108738 | 0.320794 | 1.31476 | 4.098 |
| chr7 | 50437191 | 50437472 | 0.757837 | 3.1067 | 4.099 |
| chr5 | 44952469 | 44952704 | 0.880171 | 3.60825 | 4.099 |
| chr11 | 99858898 | 99859107 | 0.218653 | 0.896482 | 4.100 |
| chr4 | 56210081 | 56210348 | 0.265066 | 1.08683 | 4.100 |
| chr15 | 85405522 | 85405745 | 0.602177 | 2.46958 | 4.101 |
| chr19 | 16436086 | 16436312 | 0.545295 | 2.23652 | 4.101 |
| chr3 | 10983735 | 10984128 | 0.993532 | 4.07496 | 4.101 |
| chr15 | 70805087 | 70805464 | 1.5238 | 6.25226 | 4.103 |
| chr3 | 40350804 | 40351151 | 0.942292 | 3.86727 | 4.104 |
| chr4 | 148103317 | 148103548 | 0.196944 | 0.808495 | 4.105 |
| chr12 | 125196083 | 125196302 | 0.333117 | 1.36775 | 4.106 |
| chr17 | 40333329 | 40333702 | 1.21013 | 4.96875 | 4.106 |
| chr8 | 47130609 | 47130911 | 0.454806 | 1.86744 | 4.106 |
| chr8 | 62669092 | 62669363 | 1.21228 | 4.97829 | 4.107 |
| chr17 | 32257521 | 32257802 | 0.369078 | 1.51597 | 4.107 |
| chr19 | 22397248 | 22397549 | 0.377836 | 1.55205 | 4.108 |
| chr5 | 100726118 | 100726334 | 0.739934 | 3.03955 | 4.108 |
| chr21 | 35916364 | 35916739 | 0.453099 | 1.86179 | 4.109 |
| chr12 | 32552445 | 32553022 | 0.692389 | 2.84631 | 4.111 |
| chr4 | 8943042 | 8943262 | 0.802559 | 3.29986 | 4.112 |
| chr12 | 89189620 | 89189837 | 0.283448 | 1.16578 | 4.113 |
| chr13 | 82446746 | 82446971 | 0.727773 | 2.99384 | 4.114 |
| chr10 | 122283009 | 122283251 | 0.243754 | 1.00331 | 4.116 |
| chr4 | 23457203 | 23457414 | 0.402811 | 1.65897 | 4.118 |
| chr8 | 35403494 | 35403734 | 0.334842 | 1.37923 | 4.119 |
| chr10 | 95117472 | 95117757 | 0.637868 | 2.62771 | 4.120 |
| chr7 | 4831641 | 4832090 | 0.997256 | 4.1085 | 4.120 |
| chr7 | 155638275 | 155638583 | 0.220107 | 0.906896 | 4.120 |
| chr20 | 57056577 | 57056937 | 0.750791 | 3.0936 | 4.120 |
| chr20 | 21117638 | 21117845 | 0.282478 | 1.16524 | 4.125 |
| chr12 | 122117478 | 122117693 | 0.23348 | 0.963724 | 4.128 |
| chr14 | 33520139 | 33520343 | 0.156498 | 0.646094 | 4.128 |
| chr19 | 5561573 | 5561854 | 0.770701 | 3.18236 | 4.129 |
| chr16 | 661061 | 661475 | 0.905286 | 3.73906 | 4.130 |
| chr7 | 96360902 | 96361111 | 0.251691 | 1.0397 | 4.131 |
| chr4 | 90881279 | 90881497 | 0.240889 | 0.995683 | 4.133 |
| chr1 | 32713757 | 32714162 | 1.14322 | 4.72618 | 4.134 |
| chr14 | 26410689 | 26410930 | 0.259512 | 1.07287 | 4.134 |
| chrX | 36916389 | 36916810 | 0.702324 | 2.90635 | 4.138 |
| chrX | 131353360 | 131353743 | 1.00444 | 4.15773 | 4.139 |
| chr2 | 104802977 | 104803210 | 0.408507 | 1.69122 | 4.140 |

| | | | | | |
|---|---|---|---|---|---|
| chrX | 81931908 | 81932174 | 0.507197 | 2.10014 | 4.141 |
| chr4 | 190011422 | 190011688 | 0.917565 | 3.80011 | 4.142 |
| chr12 | 124086294 | 124086641 | 0.357901 | 1.48304 | 4.144 |
| chr14 | 36989861 | 36990140 | 0.741694 | 3.07408 | 4.145 |
| chr9 | 14322531 | 14322737 | 0.406238 | 1.68373 | 4.145 |
| chr9 | 120127780 | 120128052 | 0.297021 | 1.23113 | 4.145 |
| chr14 | 90381698 | 90381956 | 0.65711 | 2.72532 | 4.147 |
| chrX | 83100417 | 83100720 | 0.711407 | 2.95278 | 4.151 |
| chr18 | 30129221 | 30129555 | 1.08935 | 4.5256 | 4.154 |
| chr5 | 140174900 | 140175136 | 0.649473 | 2.69879 | 4.155 |
| chr17 | 16127442 | 16127724 | 0.338335 | 1.4061 | 4.156 |
| chr12 | 67087690 | 67088040 | 0.873239 | 3.63078 | 4.158 |
| chr6 | 166519584 | 166519869 | 0.684792 | 2.84765 | 4.158 |
| chr12 | 131832656 | 131832943 | 0.230693 | 0.959353 | 4.159 |
| chr6 | 12890491 | 12890741 | 0.613474 | 2.55216 | 4.160 |
| chr3 | 81267263 | 81267501 | 0.546696 | 2.27479 | 4.161 |
| chr3 | 177621042 | 177621250 | 0.264081 | 1.09965 | 4.164 |
| chr5 | 179234867 | 179235194 | 0.471174 | 1.96336 | 4.167 |
| chr4 | 16207286 | 16207495 | 0.644099 | 2.68409 | 4.167 |
| chrX | 27573690 | 27573944 | 0.521816 | 2.17496 | 4.168 |
| chr17 | 16256713 | 16257041 | 1.04467 | 4.35488 | 4.169 |
| chr13 | 30689185 | 30689474 | 1.35955 | 5.67038 | 4.171 |
| chr12 | 17741974 | 17742235 | 0.75378 | 3.14447 | 4.172 |
| chr16 | 85982037 | 85982430 | 0.205494 | 0.857239 | 4.172 |
| chr2 | 232274933 | 232275349 | 0.922917 | 3.8505 | 4.172 |
| chr14 | 38155029 | 38155260 | 0.289227 | 1.20747 | 4.175 |
| chr8 | 115466712 | 115467049 | 1.08864 | 4.54633 | 4.176 |
| chr14 | 50346442 | 50346816 | 1.05149 | 4.39511 | 4.180 |
| chr6 | 3933326 | 3933675 | 1.07816 | 4.50725 | 4.181 |
| chr5 | 23097840 | 23098085 | 0.277495 | 1.16011 | 4.181 |
| chr1 | 41788599 | 41788889 | 0.682454 | 2.85379 | 4.182 |
| chr4 | 189702904 | 189703131 | 0.394993 | 1.65205 | 4.182 |
| chr19 | 20740899 | 20741137 | 0.688455 | 2.88022 | 4.184 |
| chr11 | 111749651 | 111750007 | 1.12634 | 4.71273 | 4.184 |
| chr10 | 73423843 | 73424154 | 0.401624 | 1.68053 | 4.184 |
| chr5 | 3603070 | 3603415 | 0.536527 | 2.24519 | 4.185 |
| chr8 | 37851135 | 37851572 | 0.98185 | 4.10895 | 4.185 |
| chr5 | 96868041 | 96868297 | 0.256021 | 1.07188 | 4.187 |
| chr11 | 35507246 | 35507659 | 1.31379 | 5.50236 | 4.188 |
| chr18 | 46439598 | 46439867 | 0.596223 | 2.49891 | 4.191 |
| chr1 | 25846997 | 25847235 | 0.645286 | 2.70586 | 4.193 |
| chr10 | 57694154 | 57694366 | 0.428077 | 1.79522 | 4.194 |
| chr4 | 117279826 | 117280069 | 0.75979 | 3.1865 | 4.194 |
| chr4 | 61329935 | 61330146 | 0.519339 | 2.17857 | 4.195 |
| chr4 | 159593391 | 159593642 | 0.845593 | 3.55088 | 4.199 |
| chr9 | 99448939 | 99449182 | 0.876674 | 3.68156 | 4.199 |
| chr10 | 87189747 | 87190032 | 0.241404 | 1.01387 | 4.200 |
| chr13 | 103499931 | 103500147 | 0.652047 | 2.73915 | 4.201 |
| chr12 | 94496046 | 94496351 | 0.983853 | 4.13341 | 4.201 |
| chr11 | 126477210 | 126477502 | 1.08123 | 4.54855 | 4.207 |
| chr5 | 77908593 | 77908817 | 0.323123 | 1.35951 | 4.207 |
| chr6 | 48733067 | 48733321 | 0.683249 | 2.87489 | 4.208 |
| chr3 | 111614035 | 111614244 | 0.455227 | 1.91615 | 4.209 |
| chr16 | 84498593 | 84498933 | 0.343592 | 1.44681 | 4.211 |
| chr10 | 133995640 | 133995878 | 0.299532 | 1.26167 | 4.212 |
| chr13 | 93900843 | 93901093 | 0.506705 | 2.13479 | 4.213 |
| chr11 | 117689845 | 117690144 | 1.01749 | 4.28759 | 4.214 |
| chr8 | 42129208 | 42129553 | 0.410793 | 1.73131 | 4.215 |
| chr12 | 42877375 | 42877591 | 0.347756 | 1.46627 | 4.216 |
| chr12 | 94161054 | 94161306 | 0.854364 | 3.60241 | 4.216 |
| chr8 | 13272171 | 13272483 | 0.248486 | 1.04804 | 4.218 |

| | | | | | |
|---|---|---|---|---|---|
| chr4 | 88427121 | 88427383 | 0.447873 | 1.88912 | 4.218 |
| chr5 | 29812039 | 29812300 | 0.721875 | 3.04639 | 4.220 |
| chr5 | 60653653 | 60653882 | 0.778444 | 3.28559 | 4.221 |
| chr13 | 46721139 | 46721373 | 0.343242 | 1.44884 | 4.221 |
| chr17 | 74581803 | 74582452 | 0.412301 | 1.74056 | 4.222 |
| chr4 | 42403384 | 42403632 | 0.561771 | 2.37223 | 4.223 |
| chr10 | 111775839 | 111776139 | 0.781199 | 3.29961 | 4.224 |
| chr18 | 34522297 | 34522541 | 0.546089 | 2.30801 | 4.226 |
| chrX | 38577204 | 38577426 | 0.514401 | 2.17432 | 4.227 |
| chr5 | 45038516 | 45038843 | 0.829216 | 3.50577 | 4.228 |
| chr4 | 36352794 | 36353021 | 0.550015 | 2.3262 | 4.229 |
| chr16 | 51235588 | 51235844 | 0.485648 | 2.05492 | 4.231 |
| chr17 | 27206025 | 27206428 | 0.762842 | 3.22872 | 4.232 |
| chr6 | 91189085 | 91189367 | 0.679534 | 2.87796 | 4.235 |
| chr12 | 20373344 | 20373586 | 0.626316 | 2.65271 | 4.235 |
| chr5 | 149036169 | 149036566 | 1.10833 | 4.69432 | 4.235 |
| chr17 | 26661746 | 26661963 | 0.282162 | 1.1952 | 4.236 |
| chr3 | 119759140 | 119759430 | 0.725103 | 3.07192 | 4.237 |
| chrY | 15455515 | 15455847 | 0.596742 | 2.52829 | 4.237 |
| chr6 | 74385233 | 74385532 | 1.17715 | 4.99386 | 4.242 |
| chr4 | 185662163 | 185662405 | 0.281719 | 1.19537 | 4.243 |
| chr4 | 115646869 | 115647167 | 0.148328 | 0.629458 | 4.244 |
| chr5 | 97121183 | 97121456 | 0.289476 | 1.22861 | 4.244 |
| chr15 | 79603448 | 79603668 | 0.82664 | 3.50925 | 4.245 |
| chr16 | 2904731 | 2905234 | 0.862933 | 3.6634 | 4.245 |
| chrX | 47504199 | 47504433 | 0.922683 | 3.91896 | 4.247 |
| chr12 | 66563309 | 66563850 | 0.663314 | 2.8176 | 4.248 |
| chr14 | 35879755 | 35880016 | 0.264721 | 1.12464 | 4.248 |
| chr10 | 99524908 | 99525239 | 0.274436 | 1.16592 | 4.248 |
| chr6 | 131081231 | 131081452 | 0.685714 | 2.9133 | 4.249 |
| chr12 | 70393425 | 70393637 | 0.38144 | 1.62154 | 4.251 |
| chr17 | 63093588 | 63093802 | 0.387734 | 1.64867 | 4.252 |
| chr11 | 124706949 | 124707178 | 0.48672 | 2.07055 | 4.254 |
| chr7 | 68676841 | 68677117 | 0.487649 | 2.07639 | 4.258 |
| chr1 | 205560935 | 205561428 | 1.31378 | 5.5948 | 4.259 |
| chr7 | 28714052 | 28714274 | 0.378748 | 1.61331 | 4.260 |
| chr22 | 50431853 | 50432088 | 1.23435 | 5.25906 | 4.261 |
| chr4 | 33452206 | 33452415 | 0.217685 | 0.927611 | 4.261 |
| chr5 | 67196349 | 67196589 | 0.331156 | 1.41122 | 4.262 |
| chr6 | 64284310 | 64284528 | 0.508044 | 2.16639 | 4.264 |
| chr3 | 185792477 | 185792705 | 0.64535 | 2.75191 | 4.264 |
| chr12 | 123431267 | 123431552 | 0.602327 | 2.56866 | 4.265 |
| chr4 | 114682906 | 114683136 | 1.0059 | 4.2904 | 4.265 |
| chr12 | 17795047 | 17795297 | 1.42302 | 6.06969 | 4.265 |
| chr4 | 187123011 | 187123336 | 1.10919 | 4.73198 | 4.266 |
| chr16 | 64085697 | 64086172 | 0.406445 | 1.73458 | 4.268 |
| chr10 | 126220660 | 126221005 | 0.722007 | 3.08169 | 4.268 |
| chr16 | 4767368 | 4767667 | 0.514094 | 2.19443 | 4.269 |
| chr12 | 78978535 | 78978804 | 0.695798 | 2.97096 | 4.270 |
| chr14 | 105654243 | 105654601 | 0.905972 | 3.8686 | 4.270 |
| chr9 | 36793340 | 36793612 | 0.896013 | 3.8264 | 4.270 |
| chr1 | 242654527 | 242654747 | 0.27968 | 1.19469 | 4.272 |
| chr5 | 123001585 | 123001965 | 0.442623 | 1.891 | 4.272 |
| chr20 | 31756965 | 31757437 | 0.851289 | 3.63881 | 4.274 |
| chr18 | 61409098 | 61409342 | 0.529134 | 2.26297 | 4.277 |
| chr17 | 13923078 | 13923310 | 0.325069 | 1.39027 | 4.277 |
| chr22 | 44399695 | 44399925 | 0.684042 | 2.92573 | 4.277 |
| chr2 | 72595039 | 72595353 | 0.838377 | 3.58842 | 4.280 |
| chr12 | 132239854 | 132240137 | 0.703187 | 3.01125 | 4.282 |
| chr9 | 136634122 | 136634366 | 0.463007 | 1.98279 | 4.282 |
| chr1 | 2023084 | 2023429 | 0.748293 | 3.20798 | 4.287 |

| | | | | | |
|---|---|---|---|---|---|
| chr1 | 93297351 | 93297631 | 1.10458 | 4.73634 | 4.288 |
| chr3 | 126194228 | 126194621 | 0.54282 | 2.32793 | 4.289 |
| chr20 | 46149798 | 46150163 | 0.735073 | 3.15257 | 4.289 |
| chr7 | 134432362 | 134432758 | 0.23596 | 1.01202 | 4.289 |
| chr7 | 121366656 | 121366888 | 0.159053 | 0.682673 | 4.292 |
| chr3 | 13570004 | 13570237 | 0.453034 | 1.94459 | 4.292 |
| chr19 | 53463009 | 53463268 | 0.305533 | 1.31182 | 4.294 |
| chr19 | 58740740 | 58741013 | 0.685327 | 2.94318 | 4.295 |
| chr20 | 61866183 | 61866400 | 0.751783 | 3.22924 | 4.295 |
| chr8 | 122050134 | 122050340 | 0.294121 | 1.26417 | 4.298 |
| chr6 | 75609176 | 75609400 | 0.62275 | 2.67831 | 4.301 |
| chr19 | 56592055 | 56592360 | 0.751533 | 3.23419 | 4.303 |
| chr14 | 102356378 | 102356603 | 0.305766 | 1.31668 | 4.306 |
| chr4 | 125667478 | 125667758 | 0.291484 | 1.25548 | 4.307 |
| chr6 | 85395784 | 85396011 | 0.764548 | 3.29436 | 4.309 |
| chr1 | 155277258 | 155277477 | 0.437405 | 1.88475 | 4.309 |
| chr3 | 191134845 | 191135181 | 0.706468 | 3.04432 | 4.309 |
| chr1 | 46799297 | 46799586 | 0.341761 | 1.47275 | 4.309 |
| chr12 | 107958825 | 107959107 | 0.689503 | 2.97506 | 4.315 |
| chr12 | 107235580 | 107235814 | 0.738935 | 3.18852 | 4.315 |
| chr3 | 150481668 | 150482030 | 0.71653 | 3.09329 | 4.317 |
| chr19 | 8751618 | 8751824 | 0.336241 | 1.45205 | 4.318 |
| chr4 | 187641173 | 187641381 | 0.459274 | 1.98439 | 4.321 |
| chr14 | 81439149 | 81439436 | 0.276447 | 1.19455 | 4.321 |
| chr12 | 122484095 | 122484585 | 0.263873 | 1.14025 | 4.321 |
| chr5 | 135292942 | 135293168 | 0.244662 | 1.05729 | 4.321 |
| chr19 | 56859671 | 56859918 | 0.747766 | 3.2321 | 4.322 |
| chr16 | 80508027 | 80508247 | 0.265996 | 1.15006 | 4.324 |
| chr16 | 56677427 | 56677711 | 0.582196 | 2.51737 | 4.324 |
| chr4 | 167423011 | 167423315 | 0.831505 | 3.59623 | 4.325 |
| chr16 | 28565451 | 28565811 | 0.588442 | 2.54601 | 4.327 |
| chr12 | 26739105 | 26739319 | 0.294311 | 1.27354 | 4.327 |
| chr1 | 30560358 | 30560643 | 0.422514 | 1.82833 | 4.327 |
| chr5 | 71433686 | 71433940 | 0.617485 | 2.67217 | 4.328 |
| chr1 | 212631102 | 212631450 | 0.924075 | 4.00275 | 4.332 |
| chr16 | 1139724 | 1139972 | 0.488728 | 2.11709 | 4.332 |
| chr8 | 47329580 | 47329812 | 0.408779 | 1.77165 | 4.334 |
| chrX | 43821005 | 43821217 | 0.463218 | 2.00833 | 4.336 |
| chr12 | 102224591 | 102225078 | 1.18412 | 5.13447 | 4.336 |
| chr6 | 47277585 | 47277921 | 1.18359 | 5.13351 | 4.337 |
| chr5 | 4437861 | 4438163 | 0.218074 | 0.946269 | 4.339 |
| chr12 | 9208547 | 9208771 | 0.307983 | 1.33723 | 4.342 |
| chr20 | 5107100 | 5107637 | 0.73113 | 3.17537 | 4.343 |
| chr5 | 6577039 | 6577355 | 0.884508 | 3.84287 | 4.345 |
| chr1 | 156306161 | 156306423 | 0.572369 | 2.48736 | 4.346 |
| chr5 | 23851950 | 23852221 | 0.290841 | 1.26396 | 4.346 |
| chr17 | 72655750 | 72656032 | 0.269032 | 1.1692 | 4.346 |
| chr12 | 130677492 | 130677825 | 0.741038 | 3.22166 | 4.348 |
| chrY | 2725697 | 2725960 | 0.338133 | 1.47066 | 4.349 |
| chr12 | 47791546 | 47791820 | 0.792716 | 3.44922 | 4.351 |
| chr20 | 49080873 | 49081080 | 0.323423 | 1.40792 | 4.353 |
| chr4 | 140374238 | 140374471 | 0.449785 | 1.95862 | 4.355 |
| chr15 | 74612983 | 74613337 | 0.702667 | 3.05991 | 4.355 |
| chr19 | 56061435 | 56061640 | 0.278116 | 1.21117 | 4.355 |
| chr4 | 14751625 | 14751865 | 0.522581 | 2.27596 | 4.355 |
| chr14 | 56405050 | 56405259 | 0.267385 | 1.16469 | 4.356 |
| chr21 | 46331546 | 46331865 | 0.503878 | 2.19518 | 4.357 |
| chr4 | 7556725 | 7556946 | 0.492908 | 2.14762 | 4.357 |
| chr1 | 41588857 | 41589060 | 0.502934 | 2.19194 | 4.358 |
| chr12 | 114625612 | 114625823 | 0.312998 | 1.36789 | 4.370 |
| chr11 | 63506263 | 63506561 | 1.01182 | 4.42274 | 4.371 |

| | | | | | |
|---|---|---|---|---|---|
| chr20 | 41844178 | 41844381 | 0.172975 | 0.75612 | 4.371 |
| chr13 | 25688352 | 25688713 | 0.755295 | 3.30243 | 4.372 |
| chr12 | 72494540 | 72494852 | 0.26772 | 1.17073 | 4.373 |
| chr3 | 25607264 | 25607539 | 0.238449 | 1.04292 | 4.374 |
| chr2 | 81551202 | 81551469 | 0.667413 | 2.91922 | 4.374 |
| chr15 | 85166603 | 85166851 | 0.507514 | 2.21991 | 4.374 |
| chr6 | 100061279 | 100061573 | 1.06711 | 4.66772 | 4.374 |
| chr1 | 156323549 | 156323767 | 0.271117 | 1.18601 | 4.375 |
| chr14 | 102237341 | 102237563 | 0.974018 | 4.26233 | 4.376 |
| chr12 | 93963358 | 93963672 | 0.986346 | 4.31786 | 4.378 |
| chr19 | 37853584 | 37854045 | 0.853121 | 3.7347 | 4.378 |
| chr9 | 130717658 | 130717914 | 0.415662 | 1.81974 | 4.378 |
| chr2 | 159892634 | 159892842 | 0.675606 | 2.95807 | 4.378 |
| chr3 | 68761282 | 68761520 | 0.22935 | 1.00425 | 4.379 |
| chr10 | 83580216 | 83580515 | 0.69893 | 3.06113 | 4.380 |
| chr1 | 14024355 | 14024557 | 0.351525 | 1.53976 | 4.380 |
| chr20 | 8740288 | 8740681 | 1.02507 | 4.49008 | 4.380 |
| chr10 | 71993527 | 71993744 | 0.418884 | 1.83635 | 4.384 |
| chr10 | 18666993 | 18667275 | 0.187624 | 0.82272 | 4.385 |
| chr9 | 29830859 | 29831155 | 0.676759 | 2.96774 | 4.385 |
| chr3 | 4763742 | 4764093 | 0.688506 | 3.02109 | 4.388 |
| chr15 | 80878062 | 80878328 | 0.584203 | 2.56387 | 4.389 |
| chr18 | 27860910 | 27861126 | 0.586898 | 2.57604 | 4.389 |
| chr12 | 44719654 | 44719869 | 0.276387 | 1.21375 | 4.391 |
| chr9 | 115128860 | 115129155 | 0.632917 | 2.78462 | 4.400 |
| chr10 | 114632509 | 114632873 | 0.72805 | 3.20479 | 4.402 |
| chr1 | 89985461 | 89985675 | 0.464741 | 2.0467 | 4.404 |
| chr20 | 540461 | 540811 | 0.231491 | 1.01951 | 4.404 |
| chr12 | 115852886 | 115853201 | 0.351488 | 1.54872 | 4.406 |
| chr6 | 166570501 | 166570800 | 0.557339 | 2.45791 | 4.410 |
| chr12 | 64575535 | 64575787 | 0.911586 | 4.02087 | 4.411 |
| chr4 | 151881303 | 151881555 | 0.629278 | 2.77847 | 4.415 |
| chr1 | 86965745 | 86966094 | 0.577891 | 2.55353 | 4.419 |
| chr21 | 39739609 | 39739838 | 0.216733 | 0.958039 | 4.420 |
| chr11 | 69459176 | 69459588 | 0.524091 | 2.31813 | 4.423 |
| chr10 | 80828805 | 80829068 | 0.508854 | 2.25079 | 4.423 |
| chr3 | 134349374 | 134349629 | 0.651597 | 2.88275 | 4.424 |
| chr4 | 113153297 | 113153518 | 0.490617 | 2.1712 | 4.425 |
| chr1 | 183439262 | 183439513 | 0.738935 | 3.27115 | 4.427 |
| chr18 | 45299014 | 45299349 | 1.21947 | 5.39881 | 4.427 |
| chr13 | 114782542 | 114782780 | 0.641165 | 2.839 | 4.428 |
| chr16 | 34625741 | 34626026 | 0.626243 | 2.77415 | 4.430 |
| chr22 | 49998561 | 49998779 | 0.356644 | 1.57989 | 4.430 |
| chr7 | 155025877 | 155026080 | 0.590468 | 2.61617 | 4.431 |
| chr12 | 114224064 | 114224282 | 0.668736 | 2.96395 | 4.432 |
| chr15 | 101409724 | 101410093 | 1.16627 | 5.17234 | 4.435 |
| chr3 | 138660100 | 138660504 | 1.00822 | 4.48479 | 4.448 |
| chr7 | 81130661 | 81130875 | 0.38294 | 1.70434 | 4.451 |
| chr1 | 227980453 | 227980680 | 0.531221 | 2.36855 | 4.459 |
| chr6 | 5379876 | 5380195 | 0.631859 | 2.81741 | 4.459 |
| chr16 | 60874828 | 60875090 | 0.248934 | 1.10999 | 4.459 |
| chr1 | 237592730 | 237592970 | 0.223698 | 0.997753 | 4.460 |
| chr12 | 42982738 | 42983125 | 1.00338 | 4.47704 | 4.462 |
| chr10 | 83909822 | 83910099 | 0.392842 | 1.75382 | 4.464 |
| chr9 | 81848499 | 81848723 | 0.423281 | 1.89235 | 4.471 |
| chr16 | 29796225 | 29796699 | 0.681981 | 3.04918 | 4.471 |
| chr17 | 57388393 | 57388685 | 0.861625 | 3.85262 | 4.471 |
| chr10 | 38277484 | 38277698 | 0.468666 | 2.09898 | 4.479 |
| chr1 | 200591507 | 200591743 | 0.505553 | 2.26446 | 4.479 |
| chr17 | 52250091 | 52250401 | 0.829482 | 3.71747 | 4.482 |
| chr3 | 5610175 | 5610441 | 0.621724 | 2.78648 | 4.482 |

| | | | | | |
|---|---|---|---|---|---|
| chr3 | 9300647 | 9300855 | 0.377819 | 1.69337 | 4.482 |
| chr7 | 148787778 | 148788092 | 0.951056 | 4.26361 | 4.483 |
| chr3 | 96050262 | 96050527 | 0.578747 | 2.59585 | 4.485 |
| chr1 | 24718865 | 24719206 | 0.953345 | 4.27688 | 4.486 |
| chr5 | 140535195 | 140535538 | 0.740735 | 3.32327 | 4.486 |
| chrX | 150970517 | 150970778 | 0.91331 | 4.1007 | 4.490 |
| chr3 | 103168837 | 103169102 | 0.260404 | 1.16921 | 4.490 |
| chr11 | 29041945 | 29042156 | 0.579559 | 2.60404 | 4.493 |
| chr5 | 121451896 | 121452113 | 0.323519 | 1.45438 | 4.496 |
| chr7 | 50200146 | 50200361 | 0.223003 | 1.00313 | 4.498 |
| chr12 | 119285617 | 119285833 | 0.228456 | 1.02808 | 4.500 |
| chr3 | 121188844 | 121189083 | 0.211175 | 0.952077 | 4.508 |
| chr3 | 27183234 | 27183485 | 0.702391 | 3.16715 | 4.509 |
| chr5 | 178831398 | 178831673 | 0.49499 | 2.23256 | 4.510 |
| chr22 | 36882747 | 36883094 | 0.677706 | 3.05716 | 4.511 |
| chr1 | 227849285 | 227849495 | 0.527032 | 2.37773 | 4.512 |
| chr18 | 54662957 | 54663226 | 0.264806 | 1.19587 | 4.516 |
| chr5 | 140561750 | 140562002 | 0.506174 | 2.28654 | 4.517 |
| chr8 | 16299086 | 16299300 | 0.209658 | 0.948808 | 4.526 |
| chr7 | 137282616 | 137282864 | 0.260276 | 1.17795 | 4.526 |
| chr11 | 72619893 | 72620132 | 0.206075 | 0.932944 | 4.527 |
| chr5 | 140174281 | 140174624 | 0.483228 | 2.18945 | 4.531 |
| chr6 | 134418358 | 134418562 | 0.47455 | 2.1504 | 4.531 |
| chr8 | 51896435 | 51896650 | 0.251807 | 1.14113 | 4.532 |
| chr4 | 167245568 | 167245783 | 0.472918 | 2.14349 | 4.532 |
| chr10 | 53162528 | 53162800 | 0.240368 | 1.08951 | 4.533 |
| chr1 | 219491165 | 219491442 | 0.703262 | 3.18864 | 4.534 |
| chr10 | 66268880 | 66269210 | 0.226718 | 1.02814 | 4.535 |
| chr7 | 11260233 | 11260438 | 0.198737 | 0.902129 | 4.539 |
| chr13 | 99681180 | 99681415 | 0.752894 | 3.42021 | 4.543 |
| chr7 | 31528278 | 31528567 | 1.45902 | 6.63103 | 4.545 |
| chr14 | 101294265 | 101294664 | 0.558022 | 2.53639 | 4.545 |
| chr12 | 7294675 | 7295114 | 1.22691 | 5.57725 | 4.546 |
| chr7 | 68532401 | 68532751 | 1.15753 | 5.26612 | 4.549 |
| chr11 | 83816253 | 83816481 | 0.28064 | 1.2775 | 4.552 |
| chr5 | 140552038 | 140552382 | 0.720956 | 3.28275 | 4.553 |
| chr19 | 12758985 | 12759217 | 0.477541 | 2.17492 | 4.554 |
| chr5 | 40756003 | 40756299 | 1.06199 | 4.83759 | 4.555 |
| chr19 | 13637393 | 13637690 | 0.446997 | 2.03643 | 4.556 |
| chr15 | 25513382 | 25513622 | 0.401452 | 1.8298 | 4.558 |
| chr12 | 66481202 | 66481460 | 0.901937 | 4.11203 | 4.559 |
| chr10 | 31780122 | 31780437 | 0.738106 | 3.36583 | 4.560 |
| chr16 | 90095724 | 90095976 | 0.211756 | 0.965994 | 4.562 |
| chr11 | 125293230 | 125293520 | 0.761166 | 3.47303 | 4.563 |
| chr15 | 69896422 | 69896627 | 0.534698 | 2.44031 | 4.564 |
| chr12 | 29474178 | 29474404 | 0.263152 | 1.20117 | 4.565 |
| chr8 | 25946234 | 25946483 | 1.34063 | 6.12078 | 4.566 |
| chr3 | 30176699 | 30177022 | 0.864821 | 3.94849 | 4.566 |
| chr1 | 43540029 | 43540239 | 0.445868 | 2.03572 | 4.566 |
| chr7 | 45614260 | 45614498 | 0.285653 | 1.30466 | 4.567 |
| chrX | 8847766 | 8848163 | 0.593649 | 2.71288 | 4.570 |
| chr12 | 49947334 | 49947704 | 0.768905 | 3.51514 | 4.572 |
| chr8 | 126889657 | 126889869 | 0.334314 | 1.53043 | 4.578 |
| chr11 | 118796629 | 118796848 | 0.594171 | 2.72268 | 4.582 |
| chr19 | 37820640 | 37820860 | 0.678495 | 3.11023 | 4.584 |
| chr12 | 125053076 | 125053338 | 0.922455 | 4.23213 | 4.588 |
| chr18 | 48188597 | 48188863 | 0.429827 | 1.97267 | 4.589 |
| chr12 | 2211479 | 2211744 | 0.255731 | 1.17414 | 4.591 |
| chr16 | 84001643 | 84001927 | 0.602168 | 2.76489 | 4.592 |
| chr1 | 109542465 | 109542712 | 0.243977 | 1.12057 | 4.593 |
| chr6 | 2524401 | 2524720 | 0.599202 | 2.75241 | 4.593 |

| | | | | | |
|---|---|---|---|---|---|
| chr16 | 15561120 | 15561353 | 0.562457 | 2.58365 | 4.594 |
| chr19 | 8160866 | 8161292 | 0.605622 | 2.78287 | 4.595 |
| chr7 | 41085159 | 41085445 | 0.671427 | 3.08593 | 4.596 |
| chr4 | 22436584 | 22436795 | 0.652863 | 3.00288 | 4.600 |
| chrX | 151358081 | 151358389 | 0.27669 | 1.27275 | 4.600 |
| chr2 | 191184651 | 191184903 | 0.828786 | 3.81318 | 4.601 |
| chr17 | 5100277 | 5100623 | 0.956594 | 4.40195 | 4.602 |
| chr8 | 80009645 | 80009865 | 0.548077 | 2.52235 | 4.602 |
| chr1 | 154618322 | 154618542 | 0.22365 | 1.02947 | 4.603 |
| chr14 | 89651247 | 89651477 | 0.184261 | 0.848217 | 4.603 |
| chr8 | 24052063 | 24052328 | 0.783075 | 3.60606 | 4.605 |
| chr10 | 43362187 | 43362391 | 0.642202 | 2.95746 | 4.605 |
| chr12 | 110290879 | 110291262 | 0.581753 | 2.68254 | 4.611 |
| chr12 | 84891654 | 84891918 | 0.640721 | 2.95465 | 4.611 |
| chrX | 76708886 | 76709114 | 0.236305 | 1.08992 | 4.612 |
| chr6 | 154936875 | 154937113 | 0.428629 | 1.97751 | 4.614 |
| chr7 | 128434225 | 128434541 | 0.355299 | 1.63974 | 4.615 |
| chr8 | 43553932 | 43554249 | 0.508644 | 2.34763 | 4.615 |
| chr15 | 61029440 | 61029679 | 0.413331 | 1.90817 | 4.617 |
| chr8 | 38415053 | 38415498 | 0.44146 | 2.03852 | 4.618 |
| chr6 | 128841713 | 128841950 | 0.415721 | 1.92199 | 4.623 |
| chr5 | 26306672 | 26306909 | 0.32278 | 1.49311 | 4.626 |
| chr14 | 107074545 | 107074842 | 0.340517 | 1.5754 | 4.626 |
| chr12 | 127358880 | 127359515 | 1.04391 | 4.83259 | 4.629 |
| chr3 | 31173699 | 31173966 | 0.533443 | 2.47073 | 4.632 |
| chr9 | 76964403 | 76964641 | 0.172129 | 0.797308 | 4.632 |
| chr6 | 9852301 | 9852616 | 0.617558 | 2.86076 | 4.632 |
| chr12 | 26427387 | 26427707 | 0.542531 | 2.51652 | 4.638 |
| chr14 | 52613237 | 52613501 | 0.411583 | 1.90945 | 4.639 |
| chr22 | 30953271 | 30953545 | 0.270935 | 1.25734 | 4.641 |
| chr16 | 50705616 | 50706033 | 0.576822 | 2.67739 | 4.642 |
| chr12 | 105073908 | 105074170 | 0.672388 | 3.12249 | 4.644 |
| chr8 | 55589761 | 55589991 | 0.215034 | 0.99865 | 4.644 |
| chr10 | 102470127 | 102470411 | 0.214611 | 0.9976 | 4.648 |
| chr1 | 247134568 | 247134923 | 0.324855 | 1.51012 | 4.649 |
| chr20 | 18110338 | 18110718 | 0.234552 | 1.09048 | 4.649 |
| chrX | 81213432 | 81213635 | 0.614201 | 2.85572 | 4.649 |
| chr20 | 35200456 | 35200670 | 0.285531 | 1.32761 | 4.650 |
| chr1 | 183939477 | 183939690 | 0.276133 | 1.28399 | 4.650 |
| chr20 | 49434960 | 49435281 | 1.12261 | 5.22002 | 4.650 |
| chr2 | 127954570 | 127954813 | 0.751795 | 3.49976 | 4.655 |
| chr1 | 111002946 | 111003192 | 0.962268 | 4.4806 | 4.656 |
| chr6 | 76077824 | 76078145 | 0.938195 | 4.37334 | 4.661 |
| chr4 | 25927474 | 25927727 | 0.767474 | 3.57876 | 4.663 |
| chr3 | 172007415 | 172007699 | 0.387968 | 1.81005 | 4.665 |
| chr15 | 99067025 | 99067229 | 0.331247 | 1.5468 | 4.670 |
| chrX | 65091814 | 65092052 | 0.309249 | 1.4452 | 4.673 |
| chr12 | 95825053 | 95825406 | 0.38775 | 1.8124 | 4.674 |
| chr3 | 152760882 | 152761129 | 0.862483 | 4.03175 | 4.675 |
| chr8 | 125391363 | 125391586 | 0.251043 | 1.17416 | 4.677 |
| chr11 | 122891730 | 122892029 | 1.02124 | 4.77686 | 4.678 |
| chr11 | 67056402 | 67056619 | 0.352518 | 1.64903 | 4.678 |
| chr14 | 54970249 | 54970504 | 0.548157 | 2.56467 | 4.679 |
| chr9 | 18320074 | 18320290 | 0.367459 | 1.71956 | 4.680 |
| chr21 | 37987614 | 37987840 | 0.581502 | 2.72321 | 4.683 |
| chr3 | 18486550 | 18486927 | 0.504849 | 2.36637 | 4.687 |
| chr7 | 57485562 | 57486001 | 0.766668 | 3.59551 | 4.690 |
| chr19 | 23253022 | 23253428 | 0.647739 | 3.03834 | 4.691 |
| chr17 | 32674131 | 32674359 | 0.385654 | 1.80915 | 4.691 |
| chr5 | 140809941 | 140810222 | 0.572109 | 2.69178 | 4.705 |
| chr11 | 34489829 | 34490123 | 0.856869 | 4.03212 | 4.706 |

| | | | | | |
|---|---|---|---|---|---|
| chr10 | 13829779 | 13830060 | 0.689812 | 3.2474 | 4.708 |
| chr9 | 118925828 | 118926085 | 0.56428 | 2.65684 | 4.708 |
| chrX | 141528862 | 141529112 | 0.182565 | 0.860076 | 4.711 |
| chr6 | 2209541 | 2209757 | 0.7339 | 3.45753 | 4.711 |
| chr4 | 19488214 | 19488432 | 0.354335 | 1.67072 | 4.715 |
| chr5 | 52391796 | 52392110 | 1.05518 | 4.97615 | 4.716 |
| chr12 | 82326722 | 82326978 | 0.24993 | 1.17895 | 4.717 |
| chr19 | 58533236 | 58533607 | 0.734871 | 3.46762 | 4.719 |
| chr19 | 21439536 | 21439912 | 0.528078 | 2.49278 | 4.720 |
| chr9 | 98228281 | 98228494 | 0.261125 | 1.23434 | 4.727 |
| chr5 | 30401623 | 30401905 | 0.544186 | 2.57324 | 4.729 |
| chr7 | 65876284 | 65876550 | 0.157005 | 0.742465 | 4.729 |
| chr8 | 129281803 | 129282087 | 0.708519 | 3.35084 | 4.729 |
| chr18 | 20958832 | 20959110 | 0.719364 | 3.40493 | 4.733 |
| chr18 | 12539915 | 12540164 | 0.812296 | 3.84913 | 4.739 |
| chr3 | 112156565 | 112156794 | 0.430691 | 2.0423 | 4.742 |
| chr5 | 10741847 | 10742085 | 0.610241 | 2.89463 | 4.743 |
| chr12 | 121022390 | 121022596 | 0.614203 | 2.91391 | 4.744 |
| chr1 | 15389086 | 15389475 | 0.534787 | 2.53935 | 4.748 |
| chr12 | 110249172 | 110249742 | 0.433887 | 2.06229 | 4.753 |
| chr5 | 157170372 | 157170895 | 1.79355 | 8.5338 | 4.758 |
| chr3 | 43197298 | 43197513 | 0.688198 | 3.27455 | 4.758 |
| chr22 | 19132231 | 19132551 | 0.713022 | 3.39514 | 4.762 |
| chr3 | 64470129 | 64470443 | 0.564805 | 2.68953 | 4.762 |
| chr19 | 53497235 | 53497475 | 0.384008 | 1.83047 | 4.767 |
| chr8 | 101156244 | 101156460 | 0.937898 | 4.47567 | 4.772 |
| chr20 | 44021502 | 44021720 | 0.298795 | 1.42604 | 4.773 |
| chr7 | 42289498 | 42289737 | 0.242394 | 1.15689 | 4.773 |
| chr5 | 38571249 | 38571472 | 0.317743 | 1.51729 | 4.775 |
| chr7 | 13782393 | 13782673 | 0.588251 | 2.80939 | 4.776 |
| chr16 | 12913276 | 12913565 | 1.25488 | 5.99611 | 4.778 |
| chr12 | 13423832 | 13424038 | 0.313371 | 1.49737 | 4.778 |
| chr3 | 138286306 | 138286521 | 0.270332 | 1.29305 | 4.783 |
| chr1 | 7058373 | 7058635 | 0.566926 | 2.71189 | 4.784 |
| chr5 | 5340461 | 5340737 | 0.792426 | 3.7942 | 4.788 |
| chr5 | 19553807 | 19554048 | 0.301185 | 1.44241 | 4.789 |
| chr11 | 78545390 | 78545609 | 0.201687 | 0.966169 | 4.790 |
| chr19 | 17494880 | 17495161 | 0.480889 | 2.30438 | 4.792 |
| chr15 | 41038750 | 41039002 | 0.699836 | 3.35461 | 4.793 |
| chr15 | 43648648 | 43648855 | 0.580443 | 2.78468 | 4.798 |
| chr7 | 158021736 | 158022084 | 0.70071 | 3.36391 | 4.801 |
| chr2 | 166731017 | 166731235 | 0.612446 | 2.94204 | 4.804 |
| chr11 | 70788740 | 70789047 | 0.817357 | 3.92684 | 4.804 |
| chr20 | 46444622 | 46444856 | 0.15429 | 0.741318 | 4.805 |
| chr12 | 112681285 | 112681508 | 0.403991 | 1.94248 | 4.808 |
| chr5 | 31664692 | 31665159 | 0.507839 | 2.44329 | 4.811 |
| chr5 | 134346731 | 134347033 | 0.54587 | 2.62671 | 4.812 |
| chr7 | 158288599 | 158288841 | 0.393461 | 1.89364 | 4.813 |
| chr10 | 93403458 | 93403828 | 0.737044 | 3.54746 | 4.813 |
| chr22 | 49944660 | 49944939 | 0.451901 | 2.17563 | 4.814 |
| chr17 | 75795116 | 75795360 | 0.274714 | 1.3227 | 4.815 |
| chr1 | 25236036 | 25236342 | 0.890791 | 4.29153 | 4.818 |
| chr19 | 24050782 | 24050997 | 0.681285 | 3.28227 | 4.818 |
| chrX | 27354660 | 27354975 | 0.745124 | 3.59143 | 4.820 |
| chr22 | 36635584 | 36635792 | 0.417488 | 2.01338 | 4.823 |
| chr5 | 82181904 | 82182158 | 0.517066 | 2.49376 | 4.823 |
| chr6 | 150247375 | 150247654 | 0.763682 | 3.68364 | 4.824 |
| chr13 | 84354422 | 84354701 | 0.610243 | 2.94798 | 4.831 |
| chr6 | 55482124 | 55482379 | 0.660753 | 3.19211 | 4.831 |
| chr4 | 178588934 | 178589197 | 0.869305 | 4.19976 | 4.831 |
| chrX | 13065793 | 13066077 | 0.207356 | 1.0023 | 4.834 |

| | | | | | |
|---|---|---|---|---|---|
| chr12 | 98766771 | 98767120 | 0.690246 | 3.33689 | 4.834 |
| chr1 | 209604748 | 209605031 | 0.160577 | 0.776341 | 4.835 |
| chr20 | 23031430 | 23031714 | 1.25215 | 6.05723 | 4.837 |
| chrX | 8751170 | 8751464 | 0.863556 | 4.17804 | 4.838 |
| chr19 | 52751432 | 52751952 | 0.718563 | 3.47657 | 4.838 |
| chr1 | 10488171 | 10488425 | 0.901425 | 4.36476 | 4.842 |
| chr17 | 18528994 | 18529309 | 0.464159 | 2.24755 | 4.842 |
| chr6 | 37980364 | 37980714 | 0.563342 | 2.72945 | 4.845 |
| chr22 | 47609363 | 47609754 | 0.975235 | 4.72894 | 4.849 |
| chr5 | 112784828 | 112785111 | 0.233588 | 1.13373 | 4.854 |
| chr1 | 222344470 | 222344815 | 0.611308 | 2.97233 | 4.862 |
| chrX | 149705202 | 149705444 | 0.652436 | 3.17301 | 4.863 |
| chr19 | 53030350 | 53030569 | 0.550734 | 2.68024 | 4.867 |
| chr20 | 36569745 | 36570159 | 1.03621 | 5.047 | 4.871 |
| chr9 | 30065940 | 30066242 | 0.322076 | 1.56883 | 4.871 |
| chr14 | 91830281 | 91830491 | 0.233218 | 1.13742 | 4.877 |
| chr4 | 94685818 | 94686133 | 0.703173 | 3.43106 | 4.879 |
| chr5 | 140571713 | 140572102 | 0.679903 | 3.32067 | 4.884 |
| chr5 | 86042585 | 86042882 | 0.293672 | 1.4367 | 4.892 |
| chr11 | 48231760 | 48231996 | 0.370669 | 1.81673 | 4.901 |
| chr20 | 30442186 | 30442519 | 0.228616 | 1.1206 | 4.902 |
| chr4 | 184319837 | 184320192 | 0.836606 | 4.10655 | 4.909 |
| chr9 | 100249492 | 100249718 | 0.617342 | 3.03508 | 4.916 |
| chr19 | 56732933 | 56733147 | 0.46866 | 2.30623 | 4.921 |
| chr5 | 115367880 | 115368125 | 0.644094 | 3.17133 | 4.924 |
| chr12 | 132678634 | 132678996 | 0.996093 | 4.90603 | 4.925 |
| chr3 | 7432809 | 7433030 | 0.492138 | 2.42576 | 4.929 |
| chr14 | 87985071 | 87985306 | 0.333891 | 1.64708 | 4.933 |
| chr6 | 142783459 | 142783669 | 0.257694 | 1.27142 | 4.934 |
| chr4 | 65137280 | 65137637 | 0.662356 | 3.26856 | 4.935 |
| chr5 | 37555052 | 37555307 | 0.77189 | 3.81038 | 4.936 |
| chr8 | 121133131 | 121133375 | 0.223353 | 1.10306 | 4.939 |
| chr6 | 26341871 | 26342091 | 0.680558 | 3.36142 | 4.939 |
| chrX | 40883619 | 40883941 | 0.787448 | 3.89201 | 4.943 |
| chr14 | 59345214 | 59345450 | 0.324724 | 1.60646 | 4.947 |
| chr3 | 152192436 | 152192711 | 0.842849 | 4.17231 | 4.950 |
| chr1 | 236029450 | 236029726 | 0.576368 | 2.85467 | 4.953 |
| chr16 | 21295141 | 21295519 | 0.780772 | 3.87016 | 4.957 |
| chr4 | 13655751 | 13656431 | 0.750346 | 3.71968 | 4.957 |
| chr12 | 18583376 | 18583719 | 0.814846 | 4.03996 | 4.958 |
| chr17 | 74337471 | 74337769 | 0.241406 | 1.19804 | 4.963 |
| chr8 | 60127709 | 60127946 | 0.530011 | 2.63066 | 4.963 |
| chr10 | 82841692 | 82842082 | 0.492644 | 2.44675 | 4.967 |
| chr3 | 181444403 | 181444703 | 0.509187 | 2.52901 | 4.967 |
| chr10 | 29408383 | 29408678 | 1.07084 | 5.31963 | 4.968 |
| chr5 | 20972516 | 20972767 | 0.640324 | 3.18123 | 4.968 |
| chr6 | 82175653 | 82175891 | 0.412581 | 2.05044 | 4.970 |
| chr19 | 5547810 | 5548162 | 0.781846 | 3.88655 | 4.971 |
| chrX | 125132512 | 125132759 | 0.388502 | 1.93129 | 4.971 |
| chr19 | 53698507 | 53698876 | 0.698052 | 3.47356 | 4.976 |
| chr19 | 771280 | 771516 | 1.02248 | 5.0913 | 4.979 |
| chr2 | 159816998 | 159817208 | 0.149962 | 0.749081 | 4.995 |
| chr1 | 153187022 | 153187231 | 0.215046 | 1.07432 | 4.996 |
| chr10 | 96919624 | 96919844 | 0.216136 | 1.08062 | 5.000 |
| chr11 | 113601711 | 113601982 | 0.616904 | 3.08539 | 5.001 |
| chr8 | 89855832 | 89856055 | 0.373017 | 1.86802 | 5.008 |
| chr1 | 231558249 | 231558473 | 0.404024 | 2.0236 | 5.009 |
| chr18 | 56338869 | 56339259 | 0.250861 | 1.25725 | 5.012 |
| chr7 | 131225588 | 131225802 | 0.39775 | 1.9939 | 5.013 |
| chr21 | 44866540 | 44866813 | 0.600504 | 3.01043 | 5.013 |
| chr12 | 14869860 | 14870150 | 0.806053 | 4.04184 | 5.014 |

| | | | | | |
|---|---|---|---|---|---|
| chr2 | 104575730 | 104575957 | 0.618019 | 3.09996 | 5.016 |
| chr5 | 23352596 | 23352843 | 0.510607 | 2.56183 | 5.017 |
| chr12 | 8549260 | 8549483 | 0.555518 | 2.78723 | 5.017 |
| chr4 | 596452 | 596784 | 0.771332 | 3.87122 | 5.019 |
| chr8 | 93126067 | 93126312 | 0.250089 | 1.2558 | 5.021 |
| chr1 | 247282277 | 247282552 | 0.509371 | 2.55834 | 5.023 |
| chr19 | 23466784 | 23467097 | 0.731823 | 3.67943 | 5.028 |
| chr5 | 105647006 | 105647259 | 0.280091 | 1.40943 | 5.032 |
| chr1 | 185355547 | 185355837 | 0.191665 | 0.966213 | 5.041 |
| chr12 | 93678016 | 93678355 | 0.848514 | 4.27776 | 5.041 |
| chr12 | 75141197 | 75141409 | 0.42716 | 2.15455 | 5.044 |
| chr5 | 159200123 | 159200435 | 0.699513 | 3.52978 | 5.046 |
| chr1 | 183252338 | 183252668 | 0.21845 | 1.10316 | 5.050 |
| chr8 | 28491294 | 28491556 | 0.697139 | 3.52298 | 5.053 |
| chr4 | 41727385 | 41727588 | 0.537425 | 2.71816 | 5.058 |
| chr12 | 104026272 | 104026967 | 1.25998 | 6.37402 | 5.059 |
| chr10 | 26217228 | 26217438 | 0.237439 | 1.20384 | 5.070 |
| chr5 | 26460757 | 26461165 | 0.314145 | 1.59287 | 5.070 |
| chr3 | 13187153 | 13187419 | 0.291863 | 1.48005 | 5.071 |
| chr8 | 115242256 | 115242620 | 0.451456 | 2.28953 | 5.071 |
| chr12 | 59135071 | 59135305 | 0.562575 | 2.85516 | 5.075 |
| chr11 | 102979946 | 102980167 | 0.508623 | 2.58798 | 5.088 |
| chr6 | 75304177 | 75304455 | 0.364448 | 1.85791 | 5.098 |
| chr7 | 63641415 | 63641691 | 0.559142 | 2.8507 | 5.098 |
| chr4 | 142526882 | 142527088 | 0.271057 | 1.38471 | 5.109 |
| chr8 | 119522743 | 119523063 | 0.644791 | 3.29399 | 5.109 |
| chr10 | 70748485 | 70748844 | 0.366853 | 1.87451 | 5.110 |
| chr3 | 13380661 | 13381025 | 0.265526 | 1.35718 | 5.111 |
| chr7 | 4869687 | 4870214 | 1.13694 | 5.81269 | 5.113 |
| chr11 | 109736584 | 109736861 | 0.769664 | 3.93882 | 5.118 |
| chr5 | 318715 | 319027 | 0.26306 | 1.35137 | 5.137 |
| chr19 | 58730366 | 58730689 | 0.557883 | 2.86735 | 5.140 |
| chr8 | 47144345 | 47144591 | 0.61586 | 3.16538 | 5.140 |
| chr19 | 24038514 | 24038799 | 0.614134 | 3.15755 | 5.141 |
| chr12 | 41658166 | 41658418 | 0.421757 | 2.16947 | 5.144 |
| chr1 | 20760642 | 20760864 | 0.319044 | 1.64151 | 5.145 |
| chr1 | 224294528 | 224294750 | 0.741206 | 3.81587 | 5.148 |
| chr6 | 15642575 | 15642872 | 0.436416 | 2.24925 | 5.154 |
| chr4 | 23338995 | 23339220 | 0.502448 | 2.59017 | 5.155 |
| chr22 | 20098653 | 20099128 | 0.328482 | 1.69422 | 5.158 |
| chr12 | 116864152 | 116864372 | 0.589668 | 3.0419 | 5.159 |
| chrX | 54666337 | 54666604 | 0.75995 | 3.92496 | 5.165 |
| chr5 | 144073286 | 144073669 | 0.19733 | 1.01968 | 5.167 |
| chr6 | 155851405 | 155851675 | 0.40508 | 2.09365 | 5.168 |
| chr8 | 49314968 | 49315214 | 0.285295 | 1.47637 | 5.175 |
| chr19 | 35911301 | 35911623 | 1.01622 | 5.26182 | 5.178 |
| chr12 | 120891769 | 120892058 | 0.984267 | 5.10778 | 5.189 |
| chr1 | 33921486 | 33921722 | 0.679616 | 3.52803 | 5.191 |
| chr9 | 117609434 | 117609652 | 0.199169 | 1.03533 | 5.198 |
| chr12 | 117769418 | 117769785 | 0.508522 | 2.64719 | 5.206 |
| chr8 | 114183947 | 114184294 | 0.464613 | 2.41948 | 5.208 |
| chr15 | 41186264 | 41186637 | 0.999114 | 5.20403 | 5.209 |
| chr14 | 58249941 | 58250182 | 0.451732 | 2.35305 | 5.209 |
| chr18 | 37739438 | 37739717 | 0.568364 | 2.96289 | 5.213 |
| chr3 | 47335741 | 47336004 | 0.1903 | 0.993071 | 5.218 |
| chr14 | 77907227 | 77907452 | 0.236366 | 1.23458 | 5.223 |
| chr11 | 94891916 | 94892245 | 0.480666 | 2.51741 | 5.237 |
| chr14 | 105631534 | 105631764 | 0.21351 | 1.11854 | 5.239 |
| chr1 | 21817369 | 21817658 | 0.77804 | 4.07782 | 5.241 |
| chr4 | 96756896 | 96757126 | 0.547252 | 2.87162 | 5.247 |
| chr7 | 89636514 | 89636767 | 0.266103 | 1.39689 | 5.249 |

| | | | | | |
|---|---|---|---|---|---|
| chr7 | 105349395 | 105349614 | 0.175134 | 0.92249 | 5.267 |
| chr3 | 96582012 | 96582269 | 0.66859 | 3.52224 | 5.268 |
| chr16 | 86053063 | 86053392 | 0.867499 | 4.57378 | 5.272 |
| chr19 | 50254404 | 50254689 | 0.691096 | 3.64491 | 5.274 |
| chr19 | 52140667 | 52140961 | 0.665422 | 3.51052 | 5.276 |
| chr18 | 77958106 | 77958425 | 0.493052 | 2.60596 | 5.285 |
| chr14 | 107147517 | 107148590 | 0.696715 | 3.68358 | 5.287 |
| chr5 | 179684031 | 179684274 | 0.481644 | 2.5475 | 5.289 |
| chr12 | 48191228 | 48191581 | 0.665139 | 3.51888 | 5.290 |
| chr6 | 15893888 | 15894118 | 0.458818 | 2.42974 | 5.296 |
| chr7 | 4681694 | 4681947 | 0.668961 | 3.54716 | 5.302 |
| chr11 | 84433686 | 84434004 | 0.320802 | 1.70336 | 5.310 |
| chr6 | 2765593 | 2765813 | 0.50371 | 2.68128 | 5.323 |
| chr7 | 5128163 | 5128407 | 0.300941 | 1.60238 | 5.325 |
| chr5 | 174442053 | 174442306 | 0.21384 | 1.14004 | 5.331 |
| chr3 | 81810060 | 81810282 | 0.493443 | 2.63239 | 5.335 |
| chrX | 77154637 | 77154852 | 0.467878 | 2.49675 | 5.336 |
| chr3 | 133845109 | 133845324 | 0.387677 | 2.07118 | 5.343 |
| chr12 | 62069419 | 62069708 | 0.202359 | 1.08178 | 5.346 |
| chr3 | 185025983 | 185026195 | 0.554914 | 2.96674 | 5.346 |
| chr12 | 4697561 | 4697775 | 0.240394 | 1.28968 | 5.365 |
| chr12 | 39207565 | 39207822 | 0.23177 | 1.24538 | 5.373 |
| chr7 | 97656698 | 97656971 | 0.330569 | 1.77672 | 5.375 |
| chr9 | 128771403 | 128771619 | 0.502865 | 2.70601 | 5.381 |
| chr10 | 17563819 | 17564034 | 0.20108 | 1.08243 | 5.383 |
| chr1 | 172696976 | 172697506 | 1.16078 | 6.25785 | 5.391 |
| chr11 | 69796143 | 69796621 | 0.932066 | 5.02801 | 5.394 |
| chr6 | 76272757 | 76272995 | 0.391022 | 2.12062 | 5.423 |
| chr18 | 56565809 | 56566272 | 0.179409 | 0.973014 | 5.423 |
| chr12 | 9216774 | 9217009 | 0.54464 | 2.95565 | 5.427 |
| chr10 | 52550663 | 52550950 | 0.256893 | 1.39587 | 5.434 |
| chr1 | 92083112 | 92083354 | 0.655074 | 3.56164 | 5.437 |
| chr22 | 27218376 | 27218602 | 0.282356 | 1.53767 | 5.446 |
| chr7 | 31247110 | 31247315 | 0.227271 | 1.23861 | 5.450 |
| chr12 | 21103229 | 21103525 | 0.688513 | 3.75742 | 5.457 |
| chrX | 27416807 | 27417374 | 0.626338 | 3.41979 | 5.460 |
| chr12 | 133095899 | 133096310 | 0.73453 | 4.01569 | 5.467 |
| chr2 | 1588939 | 1589203 | 0.317578 | 1.73746 | 5.471 |
| chr5 | 115089575 | 115089799 | 0.285577 | 1.5637 | 5.476 |
| chr11 | 64570715 | 64570971 | 0.379448 | 2.08014 | 5.482 |
| chr13 | 55004318 | 55004522 | 0.153172 | 0.840968 | 5.490 |
| chr20 | 62495586 | 62495860 | 0.673634 | 3.70286 | 5.497 |
| chr9 | 34869438 | 34869718 | 0.333654 | 1.83612 | 5.503 |
| chr14 | 73356287 | 73356492 | 0.204807 | 1.12808 | 5.508 |
| chr19 | 54015080 | 54015535 | 0.808543 | 4.45417 | 5.509 |
| chr8 | 2854436 | 2854680 | 0.225311 | 1.24127 | 5.509 |
| chr2 | 235519918 | 235520172 | 0.408188 | 2.25075 | 5.514 |
| chr16 | 447819 | 448060 | 0.504118 | 2.78077 | 5.516 |
| chr19 | 12632870 | 12633140 | 0.390466 | 2.15637 | 5.523 |
| chr3 | 68981748 | 68982052 | 0.512032 | 2.83087 | 5.529 |
| chr7 | 36977350 | 36977583 | 0.306606 | 1.69591 | 5.531 |
| chr19 | 37991263 | 37991481 | 0.384582 | 2.12772 | 5.533 |
| chr18 | 10851804 | 10852055 | 0.617088 | 3.41469 | 5.534 |
| chr7 | 70272675 | 70272933 | 0.267609 | 1.48448 | 5.547 |
| chr3 | 138739804 | 138740105 | 0.330506 | 1.83418 | 5.550 |
| chr12 | 124953666 | 124954138 | 1.06967 | 5.93832 | 5.552 |
| chr12 | 133481317 | 133481676 | 0.519436 | 2.88403 | 5.552 |
| chr16 | 22217347 | 22217644 | 0.92647 | 5.15582 | 5.565 |
| chr19 | 46785726 | 46785966 | 0.544855 | 3.03531 | 5.571 |
| chr13 | 80547686 | 80547891 | 0.33622 | 1.87336 | 5.572 |
| chr12 | 20639241 | 20639489 | 0.492678 | 2.74547 | 5.573 |

| | | | | | |
|---|---|---|---|---|---|
| chr18 | 19749394 | 19749617 | 0.335357 | 1.86881 | 5.573 |
| chr10 | 43857836 | 43858120 | 0.530265 | 2.95533 | 5.573 |
| chr3 | 85392170 | 85392440 | 0.509154 | 2.83833 | 5.575 |
| chr5 | 154134502 | 154134735 | 0.431059 | 2.40603 | 5.582 |
| chr8 | 47399695 | 47400114 | 0.647255 | 3.61287 | 5.582 |
| chr16 | 67532978 | 67533211 | 0.162855 | 0.90974 | 5.586 |
| chr5 | 140613713 | 140614015 | 0.635295 | 3.55062 | 5.589 |
| chr14 | 106465837 | 106466355 | 0.299557 | 1.67425 | 5.589 |
| chr19 | 54162664 | 54162898 | 0.818844 | 4.57695 | 5.590 |
| chr19 | 12248338 | 12248650 | 0.510011 | 2.8526 | 5.593 |
| chr8 | 141349233 | 141349482 | 0.384763 | 2.15309 | 5.596 |
| chr10 | 71327725 | 71327929 | 0.205932 | 1.15428 | 5.605 |
| chr4 | 101663266 | 101663468 | 0.439929 | 2.46953 | 5.613 |
| chr5 | 140787332 | 140787645 | 0.931144 | 5.23001 | 5.617 |
| chr10 | 123872185 | 123872449 | 0.655008 | 3.68246 | 5.622 |
| chr6 | 145829381 | 145829604 | 0.449747 | 2.52997 | 5.625 |
| chr17 | 4161807 | 4162134 | 0.489639 | 2.75491 | 5.626 |
| chr17 | 72717241 | 72717449 | 0.260574 | 1.4681 | 5.634 |
| chr13 | 72890899 | 72891113 | 0.23845 | 1.34414 | 5.637 |
| chr15 | 58200609 | 58200903 | 0.201991 | 1.13874 | 5.638 |
| chr15 | 71825324 | 71825605 | 0.87577 | 4.95555 | 5.659 |
| chr14 | 79110010 | 79110224 | 0.187468 | 1.0619 | 5.664 |
| chr12 | 108696194 | 108696416 | 0.462847 | 2.62477 | 5.671 |
| chr14 | 103246209 | 103246433 | 0.191884 | 1.08829 | 5.672 |
| chr11 | 70295084 | 70295507 | 0.790925 | 4.48786 | 5.674 |
| chr11 | 36344818 | 36345078 | 0.169746 | 0.965144 | 5.686 |
| chr5 | 140800426 | 140800705 | 0.599295 | 3.40793 | 5.687 |
| chr3 | 14138313 | 14138573 | 0.455127 | 2.59476 | 5.701 |
| chr1 | 231629656 | 231629887 | 0.201005 | 1.14681 | 5.705 |
| chrX | 84544466 | 84544670 | 0.337374 | 1.92574 | 5.708 |
| chr11 | 27205544 | 27205805 | 0.569187 | 3.24927 | 5.709 |
| chr10 | 105969427 | 105969645 | 0.444902 | 2.54444 | 5.719 |
| chr1 | 246424993 | 246425256 | 0.410867 | 2.35159 | 5.723 |
| chr2 | 173065340 | 173065591 | 0.514443 | 2.95032 | 5.735 |
| chr7 | 64090897 | 64091120 | 0.477819 | 2.74455 | 5.744 |
| chr19 | 23276679 | 23276906 | 0.260734 | 1.49945 | 5.751 |
| chr1 | 89202308 | 89202640 | 0.493907 | 2.84157 | 5.753 |
| chrY | 14533410 | 14533632 | 0.457444 | 2.63735 | 5.765 |
| chr19 | 58518704 | 58518973 | 0.67727 | 3.90493 | 5.766 |
| chr20 | 43594838 | 43595065 | 0.347277 | 2.00534 | 5.774 |
| chr16 | 57989636 | 57989880 | 0.23746 | 1.37182 | 5.777 |
| chr12 | 121282585 | 121282790 | 0.179865 | 1.04067 | 5.786 |
| chr19 | 44455101 | 44455348 | 0.378509 | 2.19251 | 5.792 |
| chr6 | 43921524 | 43921793 | 0.425024 | 2.46305 | 5.795 |
| chr11 | 64645935 | 64646223 | 0.585406 | 3.39892 | 5.806 |
| chr20 | 32535475 | 32535900 | 0.77552 | 4.50566 | 5.810 |
| chr7 | 88169207 | 88169490 | 0.207453 | 1.20544 | 5.811 |
| chr6 | 104524511 | 104524728 | 0.134634 | 0.782411 | 5.811 |
| chr19 | 24006779 | 24007016 | 0.610077 | 3.54609 | 5.813 |
| chr4 | 93084618 | 93084939 | 0.603409 | 3.50892 | 5.815 |
| chr9 | 26326518 | 26326830 | 0.639818 | 3.72449 | 5.821 |
| chr11 | 13508681 | 13509055 | 1.31148 | 7.63684 | 5.823 |
| chr3 | 185995394 | 185995598 | 0.467213 | 2.7242 | 5.831 |
| chr4 | 98908264 | 98908511 | 0.43507 | 2.54863 | 5.858 |
| chr4 | 8170029 | 8170244 | 0.361137 | 2.11879 | 5.867 |
| chrX | 96775932 | 96776154 | 0.103294 | 0.606585 | 5.872 |
| chr5 | 177711101 | 177711505 | 0.527615 | 3.10166 | 5.879 |
| chr11 | 98674844 | 98675076 | 0.173729 | 1.02182 | 5.882 |
| chr4 | 6909535 | 6909832 | 0.618006 | 3.64168 | 5.893 |
| chr1 | 31867469 | 31867834 | 0.865548 | 5.10196 | 5.894 |
| chr2 | 43249434 | 43249664 | 0.753766 | 4.44469 | 5.897 |

| | | | | | |
|---|---|---|---|---|---|
| chr3 | 174864084 | 174864325 | 0.244592 | 1.44453 | 5.906 |
| chr6 | 22737695 | 22737925 | 0.533036 | 3.15208 | 5.913 |
| chrX | 116877951 | 116878269 | 0.334631 | 1.98129 | 5.921 |
| chr3 | 96532788 | 96533111 | 0.361875 | 2.14362 | 5.924 |
| chr19 | 52111156 | 52111584 | 1.14566 | 6.78893 | 5.926 |
| chr6 | 15650558 | 15650826 | 0.181238 | 1.07749 | 5.945 |
| chr3 | 22275604 | 22275863 | 0.169567 | 1.01265 | 5.972 |
| chr3 | 85409958 | 85410282 | 0.463627 | 2.77948 | 5.995 |
| chr4 | 96614800 | 96615097 | 0.482254 | 2.89285 | 5.999 |
| chr9 | 121284364 | 121284706 | 0.537373 | 3.22612 | 6.004 |
| chr19 | 53811620 | 53812145 | 0.721554 | 4.33774 | 6.012 |
| chr14 | 93700434 | 93700642 | 0.340814 | 2.05066 | 6.017 |
| chr5 | 150369311 | 150369556 | 0.603095 | 3.6342 | 6.026 |
| chr4 | 164113232 | 164113454 | 0.407383 | 2.46348 | 6.047 |
| chr19 | 4256571 | 4256922 | 0.640974 | 3.88314 | 6.058 |
| chr8 | 25574304 | 25574528 | 0.113119 | 0.688655 | 6.088 |
| chr19 | 10184961 | 10185363 | 0.419315 | 2.55822 | 6.101 |
| chr10 | 17466125 | 17466355 | 0.172723 | 1.05839 | 6.128 |
| chr12 | 6754203 | 6754408 | 0.341618 | 2.10181 | 6.153 |
| chr19 | 21651672 | 21651942 | 0.32986 | 2.03316 | 6.164 |
| chr12 | 116844371 | 116844681 | 0.363346 | 2.24129 | 6.168 |
| chr7 | 63643385 | 63643906 | 0.599605 | 3.70003 | 6.171 |
| chr9 | 29691918 | 29692366 | 0.487864 | 3.01539 | 6.181 |
| chr19 | 12674888 | 12675184 | 0.591373 | 3.65931 | 6.188 |
| chr14 | 107238801 | 107240021 | 1.00188 | 6.20048 | 6.189 |
| chr11 | 103480258 | 103480499 | 0.444383 | 2.75912 | 6.209 |
| chr3 | 154960228 | 154960510 | 0.177205 | 1.104 | 6.230 |
| chr19 | 21666758 | 21666968 | 0.650444 | 4.05321 | 6.231 |
| chr12 | 13701266 | 13701468 | 0.220483 | 1.37432 | 6.233 |
| chr9 | 80371296 | 80371662 | 0.588077 | 3.67343 | 6.247 |
| chr8 | 90083919 | 90084242 | 0.590363 | 3.70805 | 6.281 |
| chr7 | 64085544 | 64085775 | 0.498239 | 3.14038 | 6.303 |
| chr6 | 28072482 | 28072720 | 0.234082 | 1.48483 | 6.343 |
| chr12 | 57769726 | 57770125 | 0.177554 | 1.1271 | 6.348 |
| chr19 | 53928133 | 53928647 | 0.458964 | 2.91617 | 6.354 |
| chr10 | 83544173 | 83544449 | 0.437905 | 2.78548 | 6.361 |
| chr6 | 153633792 | 153634112 | 0.517635 | 3.3069 | 6.388 |
| chr4 | 5806367 | 5806748 | 0.468699 | 3.00771 | 6.417 |
| chr12 | 84902363 | 84902612 | 0.362491 | 2.33376 | 6.438 |
| chr19 | 58693299 | 58693762 | 0.704468 | 4.54446 | 6.451 |
| chr14 | 106765059 | 106765686 | 0.710365 | 4.60571 | 6.484 |
| chr19 | 37464267 | 37464543 | 0.735515 | 4.78075 | 6.500 |
| chr19 | 53718290 | 53718528 | 0.633004 | 4.12781 | 6.521 |
| chr13 | 105223976 | 105224349 | 0.598641 | 3.90459 | 6.522 |
| chr4 | 121730692 | 121730994 | 0.504512 | 3.29232 | 6.526 |
| chr1 | 21629258 | 21629465 | 0.275828 | 1.80099 | 6.529 |
| chrY | 19542443 | 19542852 | 0.632165 | 4.12828 | 6.530 |
| chr19 | 56048320 | 56048710 | 0.578471 | 3.78819 | 6.549 |
| chr7 | 21547286 | 21547531 | 0.539499 | 3.53671 | 6.556 |
| chr19 | 52846800 | 52847222 | 0.484173 | 3.18873 | 6.586 |
| chr19 | 37800777 | 37801038 | 0.49752 | 3.28795 | 6.609 |
| chr11 | 134945496 | 134946025 | 0.95283 | 6.30122 | 6.613 |
| chr16 | 86907764 | 86908141 | 0.615005 | 4.06918 | 6.617 |
| chrY | 21665024 | 21665346 | 0.451329 | 2.99176 | 6.629 |
| chr5 | 127791415 | 127791648 | 0.207105 | 1.37885 | 6.658 |
| chr4 | 96016595 | 96016821 | 0.38235 | 2.55934 | 6.694 |
| chr19 | 12750879 | 12751354 | 0.403627 | 2.70279 | 6.696 |
| chr6 | 107699713 | 107699923 | 0.169015 | 1.15021 | 6.805 |
| chr4 | 1280013 | 1280278 | 0.561164 | 3.8208 | 6.809 |
| chr19 | 24000362 | 24000645 | 0.708458 | 4.8339 | 6.823 |
| chr14 | 41691475 | 41691705 | 0.175838 | 1.20059 | 6.828 |

| | | | | | |
|---|---|---|---|---|---|
| chrX | 152094854 | 152095087 | 0.543533 | 3.71164 | 6.829 |
| chr12 | 77706158 | 77706465 | 0.189367 | 1.2935 | 6.831 |
| chr4 | 16066866 | 16067079 | 0.165474 | 1.13032 | 6.831 |
| chr7 | 57483570 | 57483810 | 0.279436 | 1.91954 | 6.869 |
| chr13 | 66541102 | 66541398 | 0.584962 | 4.0205 | 6.873 |
| chr3 | 16183147 | 16183404 | 0.349253 | 2.41358 | 6.911 |
| chr6 | 148569002 | 148569238 | 0.132599 | 0.916492 | 6.912 |
| chr20 | 35420845 | 35421098 | 0.355999 | 2.46618 | 6.927 |
| chr7 | 57376318 | 57376662 | 0.465493 | 3.23 | 6.939 |
| chr12 | 5327346 | 5327558 | 0.17972 | 1.2488 | 6.949 |
| chr3 | 84126662 | 84127018 | 0.406668 | 2.83233 | 6.965 |
| chrX | 9830598 | 9830897 | 0.548562 | 3.84008 | 7.000 |
| chr12 | 32259521 | 32259793 | 0.354938 | 2.48798 | 7.010 |
| chr5 | 30324269 | 30324538 | 0.31023 | 2.18024 | 7.028 |
| chr19 | 12074729 | 12075063 | 0.493559 | 3.48392 | 7.059 |
| chr4 | 31694120 | 31694435 | 0.264496 | 1.86886 | 7.066 |
| chr20 | 6103441 | 6103843 | 0.202068 | 1.43209 | 7.087 |
| chr8 | 83628979 | 83629238 | 0.468433 | 3.32545 | 7.099 |
| chr3 | 18015775 | 18015980 | 0.365549 | 2.5979 | 7.107 |
| chr4 | 116011745 | 116011954 | 0.466276 | 3.31474 | 7.109 |
| chr1 | 18085587 | 18085938 | 0.691886 | 4.92028 | 7.111 |
| chr12 | 52195799 | 52196072 | 0.129948 | 0.928822 | 7.148 |
| chr19 | 12457800 | 12458023 | 0.339097 | 2.43671 | 7.186 |
| chr10 | 74003873 | 74004093 | 0.437538 | 3.15372 | 7.208 |
| chr19 | 2661690 | 2661986 | 0.488356 | 3.52144 | 7.211 |
| chr20 | 62776837 | 62777173 | 0.466779 | 3.36889 | 7.217 |
| chr19 | 3302831 | 3303035 | 0.361095 | 2.6313 | 7.287 |
| chr12 | 54686878 | 54687238 | 0.172142 | 1.25455 | 7.288 |
| chr17 | 75229816 | 75230031 | 0.502102 | 3.66192 | 7.293 |
| chr1 | 228424566 | 228424886 | 0.443831 | 3.24425 | 7.310 |
| chr5 | 89466979 | 89467233 | 0.163164 | 1.19773 | 7.341 |
| chr19 | 20471197 | 20471424 | 0.368101 | 2.70401 | 7.346 |
| chr2 | 122505808 | 122506096 | 0.435391 | 3.20051 | 7.351 |
| chrY | 21594978 | 21595213 | 0.212819 | 1.57098 | 7.382 |
| chr16 | 33618857 | 33619488 | 0.941277 | 6.98017 | 7.416 |
| chr3 | 180630007 | 180630273 | 0.36562 | 2.7157 | 7.428 |
| chr6 | 137454346 | 137454574 | 0.2253 | 1.68706 | 7.488 |
| chr5 | 140566401 | 140566856 | 0.395989 | 2.97579 | 7.515 |
| chr8 | 111072521 | 111072853 | 0.259586 | 1.95286 | 7.523 |
| chr14 | 69392885 | 69393428 | 0.596072 | 4.53005 | 7.600 |
| chr7 | 127087398 | 127087788 | 0.717221 | 5.45658 | 7.608 |
| chr8 | 83355771 | 83356059 | 0.479312 | 3.64872 | 7.612 |
| chr2 | 11352 | 12111 | 1.21474 | 9.28766 | 7.646 |
| chr19 | 12052581 | 12052844 | 0.274025 | 2.09832 | 7.657 |
| chr19 | 12237561 | 12237829 | 0.31708 | 2.45152 | 7.732 |
| chr14 | 42579393 | 42579735 | 0.624599 | 4.8333 | 7.738 |
| chr19 | 54096153 | 54096566 | 0.479218 | 3.70867 | 7.739 |
| chr11 | 112484125 | 112484329 | 0.107021 | 0.830026 | 7.756 |
| chr17 | 16440205 | 16440543 | 0.646 | 5.01511 | 7.763 |
| chr15 | 67198228 | 67198709 | 0.583746 | 4.53261 | 7.765 |
| chr1 | 51524475 | 51524753 | 0.439418 | 3.49725 | 7.959 |
| chr10 | 42970661 | 42970864 | 0.298138 | 2.38364 | 7.995 |
| chr4 | 236081 | 236363 | 0.63326 | 5.09838 | 8.051 |
| chr19 | 20108509 | 20108818 | 0.374361 | 3.02187 | 8.072 |
| chr3 | 175658765 | 175659002 | 0.271876 | 2.22361 | 8.179 |
| chr8 | 43556411 | 43556615 | 0.340176 | 2.80338 | 8.241 |
| chr9 | 30774521 | 30774791 | 0.297903 | 2.47556 | 8.310 |
| chr10 | 102464982 | 102465192 | 0.191239 | 1.59447 | 8.338 |
| chr6 | 56502446 | 56502651 | 0.342934 | 2.87492 | 8.383 |
| chr1 | 206994805 | 206995035 | 0.159383 | 1.38763 | 8.706 |
| chr19 | 13358691 | 13358904 | 0.342376 | 2.98972 | 8.732 |

| | | | | | |
|---|---|---|---|---|---|
| chrX | 65330016 | 65330291 | 0.373102 | 3.28937 | 8.816 |
| chr15 | 25092206 | 25092485 | 0.271638 | 2.42297 | 8.920 |
| chr4 | 3550751 | 3551037 | 0.0527912 | 0.472333 | 8.947 |
| chr1 | 87246679 | 87246900 | 0.304278 | 2.7318 | 8.978 |
| chr4 | 129141503 | 129141730 | 0.148833 | 1.3371 | 8.984 |
| chr5 | 140755059 | 140755300 | 0.283072 | 2.57494 | 9.096 |
| chr4 | 7240073 | 7240363 | 0.356966 | 3.31181 | 9.278 |
| chr19 | 12251960 | 12252312 | 0.385195 | 3.59142 | 9.324 |
| chr19 | 21385720 | 21386026 | 0.481395 | 4.54204 | 9.435 |
| chr19 | 12095765 | 12096086 | 0.290449 | 2.7417 | 9.440 |
| chr12 | 50216694 | 50216904 | 0.139036 | 1.31712 | 9.473 |
| chr6 | 26550645 | 26550904 | 0.0864302 | 0.830789 | 9.612 |
| chr19 | 12170290 | 12170735 | 0.376227 | 3.65204 | 9.707 |
| chr19 | 20357869 | 20358154 | 0.316913 | 3.11827 | 9.840 |
| chr6 | 75781243 | 75781486 | 0.12016 | 1.24902 | 10.395 |
| chr18 | 58193003 | 58193276 | 0.108892 | 1.14363 | 10.502 |
| chr5 | 140723229 | 140723530 | 0.362935 | 3.88225 | 10.697 |
| chr4 | 30723456 | 30723838 | 0.376648 | 4.12096 | 10.941 |
| chr3 | 56741521 | 56741749 | 0.185914 | 2.07094 | 11.139 |
| chr5 | 66784050 | 66784306 | 0.107614 | 1.22386 | 11.373 |
| chr19 | 11784361 | 11784715 | 0.298114 | 3.72447 | 12.493 |
| chr19 | 21665544 | 21665930 | 0.33774 | 4.26364 | 12.624 |
| chr4 | 87574442 | 87574790 | 0.324652 | 4.60585 | 14.187 |
| chr19 | 12034451 | 12034783 | 0.21281 | 3.6132 | 16.979 |
| chr4 | 189983249 | 189983514 | 0.11908 | 2.34077 | 19.657 |

**Supplementary Table 3, Gene ontology analysis of genes associated with H3K27ac increased peaks, Related to Figure 3D**

| GO term name | GO term ID | FDR | -log10(FDR) | GO term size | Intersection size |
|---|---|---|---|---|---|
| cell adhesion | GO:0007155 | 1.56E-08 | 7.808 | 1135 | 416 |
| biological adhesion | GO:0022610 | 1.56E-08 | 7.808 | 1142 | 418 |
| multicellular organismal homeostasis | GO:0048871 | 1.95E-07 | 6.710 | 377 | 162 |
| animal organ morphogenesis | GO:0009887 | 2.03E-07 | 6.693 | 624 | 245 |
| cell morphogenesis | GO:0000902 | 1.35814E-06 | 5.867 | 756 | 284 |
| renal system development | GO:0072001 | 2.63552E-06 | 5.579 | 150 | 76 |
| urogenital system development | GO:0001655 | 3.17457E-06 | 5.498 | 159 | 79 |
| kidney development | GO:0001822 | 8.03588E-06 | 5.095 | 146 | 73 |
| positive regulation of multicellular organismal process | GO:0051240 | 8.4887E-06 | 5.071 | 1375 | 471 |
| sodium ion transport | GO:0006814 | 8.4887E-06 | 5.071 | 136 | 69 |
| cell-cell signaling | GO:0007267 | 9.61525E-06 | 5.017 | 1499 | 508 |
| cell-cell adhesion | GO:0098609 | 9.65439E-06 | 5.015 | 666 | 249 |
| positive regulation of RNA metabolic process | GO:0051254 | 1.26828E-05 | 4.897 | 1395 | 475 |
| positive regulation of transcription, DNA-templated | GO:0045893 | 1.38217E-05 | 4.859 | 1308 | 448 |
| positive regulation of RNA biosynthetic process | GO:1902680 | 1.38217E-05 | 4.859 | 1312 | 449 |
| positive regulation of nucleic acid-templated transcription | GO:1903508 | 1.67951E-05 | 4.775 | 1311 | 448 |
| mesenchymal cell differentiation | GO:0048762 | 1.8226E-05 | 4.739 | 175 | 82 |
| cellular component morphogenesis | GO:0032989 | 2.29732E-05 | 4.639 | 566 | 214 |
| heart morphogenesis | GO:0003007 | 3.23736E-05 | 4.490 | 169 | 79 |
| mesenchyme development | GO:0060485 | 3.32468E-05 | 4.478 | 208 | 93 |
| heart development | GO:0007507 | 3.97316E-05 | 4.401 | 342 | 139 |
| positive regulation of transcription by RNA polymerase II | GO:0045944 | 4.33346E-05 | 4.363 | 987 | 345 |
| sodium ion transmembrane transport | GO:0035725 | 4.62203E-05 | 4.335 | 104 | 54 |
| cell development | GO:0048468 | 5.27507E-05 | 4.278 | 1369 | 461 |
| nephron epithelium development | GO:0072009 | 5.67028E-05 | 4.246 | 65 | 38 |
| temperature homeostasis | GO:0001659 | 5.7751E-05 | 4.238 | 158 | 74 |
| kidney epithelium development | GO:0072073 | 9.58413E-05 | 4.018 | 71 | 40 |
| homeostatic process | GO:0042592 | 9.80708E-05 | 4.008 | 1500 | 498 |
| cold-induced thermogenesis | GO:0106106 | 0.000104229 | 3.982 | 144 | 68 |
| regulation of cold-induced thermogenesis | GO:0120161 | 0.000104229 | 3.982 | 144 | 68 |
| adaptive thermogenesis | GO:1990845 | 0.00014176 | 3.848 | 151 | 70 |
| renal tubule development | GO:0061326 | 0.000155991 | 3.807 | 51 | 31 |
| cell part morphogenesis | GO:0032990 | 0.000185124 | 3.733 | 494 | 185 |
| positive regulation of molecular function | GO:0044093 | 0.000198901 | 3.701 | 1342 | 447 |
| mesonephros development | GO:0001823 | 0.000207812 | 3.682 | 47 | 29 |
| cell projection morphogenesis | GO:0048858 | 0.000214781 | 3.668 | 483 | 181 |
| negative regulation of transcription by RNA polymerase II | GO:0000122 | 0.000254697 | 3.594 | 664 | 238 |
| skeletal system development | GO:0001501 | 0.000275733 | 3.560 | 296 | 119 |
| nephron tubule development | GO:0072080 | 0.000275733 | 3.560 | 50 | 30 |
| plasma membrane bounded cell projection morphogenesis | GO:0120039 | 0.000300393 | 3.522 | 480 | 179 |
| outflow tract morphogenesis | GO:0003151 | 0.000301233 | 3.521 | 55 | 32 |
| cell morphogenesis involved in neuron differentiation | GO:0048667 | 0.0003425 | 3.465 | 410 | 156 |
| actin filament-based process | GO:0030029 | 0.000344506 | 3.463 | 690 | 245 |

| | | | | | |
|---|---|---|---|---|---|
| neuron differentiation | GO:0030182 | 0.000376079 | 3.425 | 904 | 311 |
| morphogenesis of a branching structure | GO:0001763 | 0.000401884 | 3.396 | 78 | 41 |
| positive regulation of catalytic activity | GO:0043085 | 0.000402564 | 3.395 | 1049 | 355 |
| nephron development | GO:0072006 | 0.000403594 | 3.394 | 91 | 46 |
| positive regulation of cell-cell adhesion | GO:0022409 | 0.000403594 | 3.394 | 222 | 93 |
| neuron projection morphogenesis | GO:0048812 | 0.000438614 | 3.358 | 468 | 174 |
| ureteric bud development | GO:0001657 | 0.000470072 | 3.328 | 42 | 26 |
| positive regulation of cell differentiation | GO:0045597 | 0.000497084 | 3.304 | 723 | 254 |
| cellular response to oxygen-containing compound | GO:1901701 | 0.000497084 | 3.304 | 843 | 291 |
| circulatory system development | GO:0072359 | 0.000497084 | 3.304 | 814 | 282 |
| generation of neurons | GO:0048699 | 0.000497084 | 3.304 | 1013 | 343 |
| negative regulation of nucleic acid-templated transcription | GO:1903507 | 0.000497084 | 3.304 | 1046 | 353 |
| neurogenesis | GO:0022008 | 0.000497084 | 3.304 | 1069 | 360 |
| cardiac chamber morphogenesis | GO:0003206 | 0.000503579 | 3.298 | 79 | 41 |
| negative regulation of transcription, DNA-templated | GO:0045892 | 0.000503579 | 3.298 | 1043 | 352 |
| negative regulation of RNA biosynthetic process | GO:1902679 | 0.000523112 | 3.281 | 1047 | 353 |
| tube development | GO:0035295 | 0.000550574 | 3.259 | 696 | 245 |
| axonogenesis | GO:0007409 | 0.000559991 | 3.252 | 335 | 130 |
| gland development | GO:0048732 | 0.000638926 | 3.195 | 139 | 63 |
| positive regulation of developmental process | GO:0051094 | 0.00068792 | 3.162 | 1034 | 348 |
| mesonephric tubule development | GO:0072164 | 0.00071185 | 3.148 | 43 | 26 |
| mesonephric epithelium development | GO:0072163 | 0.00071185 | 3.148 | 43 | 26 |
| regulation of sodium ion transport | GO:0002028 | 0.000751431 | 3.124 | 70 | 37 |
| mesonephric tubule morphogenesis | GO:0072171 | 0.000751431 | 3.124 | 34 | 22 |
| anatomical structure formation involved in morphogenesis | GO:0048646 | 0.000820707 | 3.086 | 807 | 278 |
| renal tubule morphogenesis | GO:0061333 | 0.000833935 | 3.079 | 41 | 25 |
| cellular response to lipid | GO:0071396 | 0.000880092 | 3.055 | 406 | 152 |
| positive regulation of leukocyte cell-cell adhesion | GO:1903039 | 0.000880092 | 3.055 | 186 | 79 |
| morphogenesis of a branching epithelium | GO:0061138 | 0.00090193 | 3.045 | 68 | 36 |
| O-glycan processing | GO:0016266 | 0.00090193 | 3.045 | 58 | 32 |
| regulation of hormone levels | GO:0010817 | 0.000912135 | 3.040 | 357 | 136 |
| sensory organ morphogenesis | GO:0090596 | 0.000947077 | 3.024 | 105 | 50 |
| actin cytoskeleton organization | GO:0030036 | 0.001057025 | 2.976 | 597 | 212 |
| circulatory system process | GO:0003013 | 0.001060012 | 2.975 | 417 | 155 |
| positive regulation of T cell activation | GO:0050870 | 0.001084967 | 2.965 | 167 | 72 |
| cardiac chamber development | GO:0003205 | 0.001166438 | 2.933 | 95 | 46 |
| response to oxygen-containing compound | GO:1901700 | 0.001166438 | 2.933 | 1038 | 347 |
| axon development | GO:0061564 | 0.001169705 | 2.932 | 359 | 136 |
| regulation of transporter activity | GO:0032409 | 0.001252255 | 2.902 | 250 | 100 |
| negative regulation of RNA metabolic process | GO:0051253 | 0.001252255 | 2.902 | 1132 | 375 |
| protein O-linked glycosylation | GO:0006493 | 0.001318297 | 2.880 | 109 | 51 |
| ion transport | GO:0006811 | 0.001350294 | 2.870 | 1330 | 434 |
| regulation of transmembrane transport | GO:0034762 | 0.001410231 | 2.851 | 379 | 142 |
| glycoprotein biosynthetic process | GO:0009101 | 0.001410231 | 2.851 | 290 | 113 |
| cell morphogenesis involved in differentiation | GO:0000904 | 0.001411897 | 2.850 | 530 | 190 |
| branching morphogenesis of an epithelial tube | GO:0048754 | 0.001894795 | 2.722 | 65 | 34 |
| blood circulation | GO:0008015 | 0.001908295 | 2.719 | 403 | 149 |
| embryo development | GO:0009790 | 0.001947339 | 2.711 | 463 | 168 |
| regulation of cation transmembrane transport | GO:1904062 | 0.001947339 | 2.711 | 280 | 109 |
| cell surface receptor signaling pathway involved in cell-cell signaling | GO:1905114 | 0.001947339 | 2.711 | 482 | 174 |
| sensory organ development | GO:0007423 | 0.001980524 | 2.703 | 226 | 91 |
| positive regulation of cold-induced thermogenesis | GO:0120162 | 0.001992517 | 2.701 | 97 | 46 |
| cellular response to endogenous stimulus | GO:0071495 | 0.002013361 | 2.696 | 1069 | 354 |
| positive regulation of lymphocyte proliferation | GO:0050671 | 0.002149007 | 2.668 | 100 | 47 |
| action potential | GO:0001508 | 0.002149007 | 2.668 | 111 | 51 |
| cellular chemical homeostasis | GO:0055082 | 0.002200186 | 2.658 | 650 | 226 |
| regulation of transmembrane transporter activity | GO:0022898 | 0.002294692 | 2.639 | 236 | 94 |
| cell junction organization | GO:0034330 | 0.002294692 | 2.639 | 487 | 175 |

| | | | | | |
|---|---|---|---|---|---|
| multicellular organismal signaling | GO:0035637 | 0.002294692 | 2.639 | 171 | 72 |
| regulation of cardiac conduction | GO:1903779 | 0.002352634 | 2.628 | 71 | 36 |
| regulation of cell morphogenesis | GO:0022604 | 0.002429601 | 2.614 | 365 | 136 |
| regulation of transport | GO:0051049 | 0.002430087 | 2.614 | 1478 | 475 |
| positive regulation of transferase activity | GO:0051347 | 0.002430087 | 2.614 | 574 | 202 |
| positive regulation of cell adhesion | GO:0045785 | 0.002486256 | 2.604 | 325 | 123 |
| chemical homeostasis | GO:0048878 | 0.002505028 | 2.601 | 940 | 314 |
| regulation of ion transmembrane transport | GO:0034765 | 0.002505028 | 2.601 | 319 | 121 |
| epithelial to mesenchymal transition | GO:0001837 | 0.002505028 | 2.601 | 123 | 55 |
| positive regulation of mononuclear cell proliferation | GO:0032946 | 0.002631957 | 2.580 | 101 | 47 |
| ureteric bud morphogenesis | GO:0060675 | 0.00273385 | 2.563 | 32 | 20 |
| regulation of ion transport | GO:0043269 | 0.002772164 | 2.557 | 464 | 167 |
| neuron development | GO:0048666 | 0.002938403 | 2.532 | 732 | 250 |
| cellular response to organic cyclic compound | GO:0071407 | 0.003068368 | 2.513 | 361 | 134 |
| regulation of cell adhesion | GO:0030155 | 0.003206747 | 2.494 | 574 | 201 |
| cell-cell signaling by wnt | GO:0198738 | 0.003289441 | 2.483 | 437 | 158 |
| alpha-beta T cell differentiation | GO:0046632 | 0.003289441 | 2.483 | 67 | 34 |
| carbohydrate derivative biosynthetic process | GO:1901137 | 0.00333995 | 2.476 | 581 | 203 |
| regulation of ion transmembrane transporter activity | GO:0032412 | 0.004183173 | 2.378 | 231 | 91 |
| calcium ion homeostasis | GO:0055074 | 0.004190333 | 2.378 | 376 | 138 |
| response to lipid | GO:0033993 | 0.004190333 | 2.378 | 522 | 184 |
| cell migration | GO:0016477 | 0.004190333 | 2.378 | 1342 | 432 |
| Wnt signaling pathway | GO:0016055 | 0.004217116 | 2.375 | 433 | 156 |
| cation transmembrane transport | GO:0098655 | 0.004217116 | 2.375 | 619 | 214 |
| regulation of lymphocyte proliferation | GO:0050670 | 0.004217116 | 2.375 | 160 | 67 |
| regulation of protein phosphorylation | GO:0001932 | 0.004217116 | 2.375 | 1238 | 401 |
| tissue development | GO:0009888 | 0.0043539 | 2.361 | 1414 | 453 |
| cellular homeostasis | GO:0019725 | 0.004447652 | 2.352 | 760 | 257 |
| stem cell development | GO:0048864 | 0.004468436 | 2.350 | 50 | 27 |
| tissue homeostasis | GO:0001894 | 0.004606009 | 2.337 | 146 | 62 |
| heart contraction | GO:0060047 | 0.004646766 | 2.333 | 229 | 90 |
| axon guidance | GO:0007411 | 0.00467267 | 2.330 | 211 | 84 |
| cell projection organization | GO:0030030 | 0.004811977 | 2.318 | 1180 | 383 |
| localization of cell | GO:0051674 | 0.004902145 | 2.310 | 1474 | 470 |
| cell motility | GO:0048870 | 0.004902145 | 2.310 | 1474 | 470 |
| nephron tubule morphogenesis | GO:0072078 | 0.004964265 | 2.304 | 38 | 22 |
| positive regulation of animal organ morphogenesis | GO:0110110 | 0.005220773 | 2.282 | 53 | 28 |
| limb morphogenesis | GO:0035108 | 0.005220773 | 2.282 | 48 | 26 |
| kidney morphogenesis | GO:0060993 | 0.005220773 | 2.282 | 48 | 26 |
| cardiac muscle cell contraction | GO:0086003 | 0.005220773 | 2.282 | 66 | 33 |
| regulation of blood circulation | GO:1903522 | 0.005220773 | 2.282 | 233 | 91 |
| appendage morphogenesis | GO:0035107 | 0.005220773 | 2.282 | 48 | 26 |
| regulation of heart contraction | GO:0008016 | 0.005274455 | 2.278 | 206 | 82 |
| regulation of peptidyl-tyrosine phosphorylation | GO:0050730 | 0.005274455 | 2.278 | 203 | 81 |
| neuron projection guidance | GO:0097485 | 0.005274455 | 2.278 | 212 | 84 |
| positive regulation of leukocyte proliferation | GO:0070665 | 0.005301848 | 2.276 | 107 | 48 |
| cellular ion homeostasis | GO:0006873 | 0.005402388 | 2.267 | 561 | 195 |
| positive regulation of kinase activity | GO:0033674 | 0.005581598 | 2.253 | 497 | 175 |
| regulation of mononuclear cell proliferation | GO:0032944 | 0.005632624 | 2.249 | 162 | 67 |
| cellular calcium ion homeostasis | GO:0006874 | 0.005632624 | 2.249 | 370 | 135 |
| cellular cation homeostasis | GO:0030003 | 0.005647093 | 2.248 | 552 | 192 |
| embryo development ending in birth or egg hatching | GO:0009792 | 0.006029028 | 2.220 | 195 | 78 |
| eye morphogenesis | GO:0048592 | 0.006212913 | 2.207 | 72 | 35 |
| positive regulation of cell population proliferation | GO:0008284 | 0.006255663 | 2.204 | 680 | 231 |
| positive regulation of peptidyl-tyrosine phosphorylation | GO:0050731 | 0.006255663 | 2.204 | 148 | 62 |
| endocrine system development | GO:0035270 | 0.006531487 | 2.185 | 59 | 30 |
| cardiac ventricle development | GO:0003231 | 0.006677724 | 2.175 | 75 | 36 |
| heart process | GO:0003015 | 0.006677724 | 2.175 | 235 | 91 |
| chordate embryonic development | GO:0043009 | 0.006730877 | 2.172 | 181 | 73 |
| regulation of sodium ion transmembrane transport | GO:1902305 | 0.006730877 | 2.172 | 54 | 28 |
| response to endogenous stimulus | GO:0009719 | 0.006730877 | 2.172 | 1177 | 380 |

| | | | | | |
|---|---|---|---|---|---|
| positive regulation of protein kinase activity | GO:0045860 | 0.006730877 | 2.172 | 432 | 154 |
| neural crest cell differentiation | GO:0014033 | 0.006730877 | 2.172 | 54 | 28 |
| cell-cell adhesion via plasma-membrane adhesion molecules | GO:0098742 | 0.006730877 | 2.172 | 184 | 74 |
| striated muscle tissue development | GO:0014706 | 0.006730877 | 2.172 | 196 | 78 |
| insulin secretion | GO:0030073 | 0.006730877 | 2.172 | 134 | 57 |
| cardiac conduction | GO:0061337 | 0.00680183 | 2.167 | 140 | 59 |
| cell differentiation involved in kidney development | GO:0061005 | 0.00680183 | 2.167 | 39 | 22 |
| neuron projection development | GO:0031175 | 0.00680183 | 2.167 | 672 | 228 |
| actin filament bundle assembly | GO:0051017 | 0.00680183 | 2.167 | 140 | 59 |
| cation homeostasis | GO:0055080 | 0.006817875 | 2.166 | 597 | 205 |
| ion transmembrane transport | GO:0034220 | 0.006928524 | 2.159 | 837 | 278 |
| appendage development | GO:0048736 | 0.006963701 | 2.157 | 70 | 34 |
| glycosaminoglycan biosynthetic process | GO:0006024 | 0.006963701 | 2.157 | 103 | 46 |
| limb development | GO:0060173 | 0.006963701 | 2.157 | 70 | 34 |
| potassium ion transport | GO:0006813 | 0.007593609 | 2.120 | 215 | 84 |
| positive regulation of ossification | GO:0045778 | 0.007725561 | 2.112 | 65 | 32 |
| inorganic cation transmembrane transport | GO:0098662 | 0.007725561 | 2.112 | 553 | 191 |
| inorganic ion homeostasis | GO:0098771 | 0.007725561 | 2.112 | 605 | 207 |
| import across plasma membrane | GO:0098739 | 0.007848924 | 2.105 | 112 | 49 |
| inorganic ion transmembrane transport | GO:0098660 | 0.00788738 | 2.103 | 589 | 202 |
| regulation of small GTPase mediated signal transduction | GO:0051056 | 0.008004421 | 2.097 | 249 | 95 |
| regulation of phosphorylation | GO:0042325 | 0.008051441 | 2.094 | 1380 | 439 |
| regulation of interleukin-2 production | GO:0032663 | 0.008221316 | 2.085 | 42 | 23 |
| calcium-dependent cell-cell adhesion via plasma membrane cell adhesion molecules | GO:0016339 | 0.008221316 | 2.085 | 42 | 23 |
| positive regulation of phosphate metabolic process | GO:0045937 | 0.008264702 | 2.083 | 983 | 321 |
| positive regulation of phosphorus metabolic process | GO:0010562 | 0.008264702 | 2.083 | 983 | 321 |
| regulation of cellular component movement | GO:0051270 | 0.008264702 | 2.083 | 953 | 312 |
| response to purine-containing compound | GO:0014074 | 0.008264702 | 2.083 | 68 | 33 |
| positive regulation of secretion | GO:0051047 | 0.008466457 | 2.072 | 296 | 110 |
| T cell differentiation | GO:0030217 | 0.008476479 | 2.072 | 159 | 65 |
| branching involved in ureteric bud morphogenesis | GO:0001658 | 0.008526316 | 2.069 | 30 | 18 |
| cation transport | GO:0006812 | 0.008940354 | 2.049 | 881 | 290 |
| monovalent inorganic cation transport | GO:0015672 | 0.008940354 | 2.049 | 372 | 134 |
| chondrocyte differentiation | GO:0002062 | 0.008940354 | 2.049 | 71 | 34 |
| leukocyte cell-cell adhesion | GO:0007159 | 0.009299574 | 2.032 | 297 | 110 |
| negative chemotaxis | GO:0050919 | 0.009299574 | 2.032 | 35 | 20 |
| positive regulation of bone mineralization | GO:0030501 | 0.009299574 | 2.032 | 35 | 20 |
| regulation of kidney development | GO:0090183 | 0.009494021 | 2.023 | 40 | 22 |
| nephron epithelium morphogenesis | GO:0072088 | 0.009494021 | 2.023 | 40 | 22 |
| nephron morphogenesis | GO:0072028 | 0.009494021 | 2.023 | 40 | 22 |
| regulation of cell development | GO:0060284 | 0.009494021 | 2.023 | 618 | 210 |
| embryonic organ development | GO:0048568 | 0.00975308 | 2.011 | 190 | 75 |
| regulation of leukocyte proliferation | GO:0070663 | 0.00975308 | 2.011 | 172 | 69 |
| biomineralization | GO:0110148 | 0.00975308 | 2.011 | 102 | 45 |
| ion homeostasis | GO:0050801 | 0.00975308 | 2.011 | 661 | 223 |
| hormone metabolic process | GO:0042445 | 0.00975308 | 2.011 | 163 | 66 |
| regulation of cell-cell adhesion | GO:0022407 | 0.00975308 | 2.011 | 351 | 127 |
| biomineral tissue development | GO:0031214 | 0.00975308 | 2.011 | 102 | 45 |
| plasma membrane bounded cell projection organization | GO:0120036 | 0.010007771 | 2.000 | 1165 | 374 |
| regulation of Wnt signaling pathway | GO:0030111 | 0.010142101 | 1.994 | 323 | 118 |
| metal ion transport | GO:0030001 | 0.010159634 | 1.993 | 688 | 231 |
| regulation of DNA binding | GO:0051101 | 0.010304529 | 1.987 | 91 | 41 |
| aminoglycan biosynthetic process | GO:0006023 | 0.010382757 | 1.984 | 108 | 47 |
| regulation of kinase activity | GO:0043549 | 0.010556282 | 1.976 | 738 | 246 |
| supramolecular fiber organization | GO:0097435 | 0.01061209 | 1.974 | 600 | 204 |
| alpha-beta T cell differentiation involved in immune response | GO:0002293 | 0.010653281 | 1.973 | 48 | 25 |
| alpha-beta T cell activation involved in immune response | GO:0002287 | 0.010653281 | 1.973 | 48 | 25 |
| neural crest cell development | GO:0014032 | 0.010653281 | 1.973 | 48 | 25 |

| | | | | | |
|---|---|---|---|---|---|
| metanephros development | GO:0001656 | 0.010653281 | 1.973 | 48 | 25 |
| actin filament bundle organization | GO:0061572 | 0.010654825 | 1.972 | 143 | 59 |
| muscle tissue development | GO:0060537 | 0.010803304 | 1.966 | 209 | 81 |
| positive regulation of cellular component organization | GO:0051130 | 0.010887417 | 1.963 | 965 | 314 |
| regulation of metal ion transport | GO:0010959 | 0.010934668 | 1.961 | 305 | 112 |
| potassium ion transmembrane transport | GO:0071805 | 0.011015812 | 1.958 | 194 | 76 |
| regulation of insulin secretion | GO:0050796 | 0.011015812 | 1.958 | 117 | 50 |
| CD4-positive, alpha-beta T cell differentiation | GO:0043367 | 0.01118567 | 1.951 | 56 | 28 |
| regulation of transferase activity | GO:0051338 | 0.011403502 | 1.943 | 839 | 276 |
| regulation of leukocyte cell-cell adhesion | GO:1903037 | 0.011787114 | 1.929 | 262 | 98 |
| regulation of cytosolic calcium ion concentration | GO:0051480 | 0.011899256 | 1.924 | 287 | 106 |
| regulation of neurogenesis | GO:0050767 | 0.01227701 | 1.911 | 514 | 177 |
| divalent inorganic cation homeostasis | GO:0072507 | 0.01227701 | 1.911 | 414 | 146 |
| regulation of cell migration | GO:0030334 | 0.012542968 | 1.902 | 827 | 272 |
| cellular response to cAMP | GO:0071320 | 0.012581569 | 1.900 | 31 | 18 |
| positive regulation of phosphorylation | GO:0042327 | 0.012632172 | 1.899 | 934 | 304 |
| interleukin-2 production | GO:0032623 | 0.012703335 | 1.896 | 46 | 24 |
| eye development | GO:0001654 | 0.012892829 | 1.890 | 165 | 66 |
| sensory system development | GO:0048880 | 0.012892829 | 1.890 | 174 | 69 |
| regulation of protein kinase activity | GO:0045859 | 0.01313003 | 1.882 | 649 | 218 |
| positive regulation of protein modification process | GO:0031401 | 0.01313003 | 1.882 | 1059 | 341 |
| peptidyl-tyrosine modification | GO:0018212 | 0.013963647 | 1.855 | 251 | 94 |
| peptidyl-tyrosine phosphorylation | GO:0018108 | 0.014056426 | 1.852 | 248 | 93 |
| glycoprotein metabolic process | GO:0009100 | 0.014056426 | 1.852 | 358 | 128 |
| positive regulation of binding | GO:0051099 | 0.014056426 | 1.852 | 130 | 54 |
| connective tissue development | GO:0061448 | 0.014224419 | 1.847 | 133 | 55 |
| cardiac septum morphogenesis | GO:0060411 | 0.014427059 | 1.841 | 49 | 25 |
| tube morphogenesis | GO:0035239 | 0.014551174 | 1.837 | 634 | 213 |
| regulation of cell projection organization | GO:0031344 | 0.014927062 | 1.826 | 484 | 167 |
| positive regulation of protein phosphorylation | GO:0001934 | 0.015319549 | 1.815 | 867 | 283 |
| organic anion transport | GO:0015711 | 0.015319549 | 1.815 | 378 | 134 |
| regulation of cell death | GO:0010941 | 0.015451078 | 1.811 | 1316 | 416 |
| regulation of neuron differentiation | GO:0045664 | 0.015488726 | 1.810 | 407 | 143 |
| regulation of locomotion | GO:0040012 | 0.015799575 | 1.801 | 911 | 296 |
| positive regulation of transport | GO:0051050 | 0.016019022 | 1.795 | 771 | 254 |
| cardiac ventricle morphogenesis | GO:0003208 | 0.016151671 | 1.792 | 52 | 26 |
| mesenchymal cell development | GO:0014031 | 0.016151671 | 1.792 | 52 | 26 |
| cellular metal ion homeostasis | GO:0006875 | 0.016159368 | 1.792 | 485 | 167 |
| central nervous system neuron differentiation | GO:0021953 | 0.016282241 | 1.788 | 82 | 37 |
| alpha-beta T cell activation | GO:0046631 | 0.016317002 | 1.787 | 99 | 43 |
| CD4-positive, alpha-beta T cell activation | GO:0035710 | 0.016619836 | 1.779 | 71 | 33 |
| positive regulation of neurogenesis | GO:0050769 | 0.016657779 | 1.778 | 287 | 105 |
| forebrain development | GO:0030900 | 0.016826194 | 1.774 | 140 | 57 |
| positive regulation of secretion by cell | GO:1903532 | 0.016826194 | 1.774 | 284 | 104 |
| positive regulation of DNA binding | GO:0043388 | 0.017147539 | 1.766 | 47 | 24 |
| CD4-positive, alpha-beta T cell differentiation involved in immune response | GO:0002294 | 0.017147539 | 1.766 | 47 | 24 |
| cardiac septum development | GO:0003279 | 0.017163488 | 1.765 | 63 | 30 |
| energy reserve metabolic process | GO:0006112 | 0.017163488 | 1.765 | 63 | 30 |
| ear development | GO:0043583 | 0.017163488 | 1.765 | 63 | 30 |
| cell communication by electrical coupling | GO:0010644 | 0.017163488 | 1.765 | 27 | 16 |
| cellular divalent inorganic cation homeostasis | GO:0072503 | 0.017275207 | 1.763 | 402 | 141 |
| second-messenger-mediated signaling | GO:0019932 | 0.017345148 | 1.761 | 370 | 131 |
| telencephalon development | GO:0021537 | 0.017928839 | 1.746 | 91 | 40 |
| positive regulation of biomineralization | GO:0110151 | 0.017928839 | 1.746 | 42 | 22 |
| positive regulation of biomineral tissue development | GO:0070169 | 0.017928839 | 1.746 | 42 | 22 |
| regulation of nervous system development | GO:0051960 | 0.017928839 | 1.746 | 578 | 195 |
| roof of mouth development | GO:0060021 | 0.018530686 | 1.732 | 37 | 20 |
| camera-type eye development | GO:0043010 | 0.019256157 | 1.715 | 129 | 53 |
| glycosaminoglycan metabolic process | GO:0030203 | 0.019406046 | 1.712 | 132 | 54 |

| | | | | | |
|---|---|---|---|---|---|
| negative regulation of nucleobase-containing compound metabolic process | GO:0045934 | 0.019600759 | 1.708 | 1219 | 386 |
| transmembrane transport | GO:0055085 | 0.019600759 | 1.708 | 1124 | 358 |
| adenylate cyclase-inhibiting G protein-coupled receptor signaling pathway | GO:0007193 | 0.019661925 | 1.706 | 69 | 32 |
| visual system development | GO:0150063 | 0.019661925 | 1.706 | 168 | 66 |
| aminoglycan metabolic process | GO:0006022 | 0.019664776 | 1.706 | 144 | 58 |
| regulation of plasma membrane bounded cell projection organization | GO:0120035 | 0.019804771 | 1.703 | 478 | 164 |
| regulation of cell motility | GO:2000145 | 0.019804771 | 1.703 | 885 | 287 |
| regulation of epithelial to mesenchymal transition | GO:0010717 | 0.019804771 | 1.703 | 83 | 37 |
| chemotaxis | GO:0006935 | 0.020016519 | 1.699 | 514 | 175 |
| taxis | GO:0042330 | 0.020016519 | 1.699 | 514 | 175 |
| animal organ formation | GO:0048645 | 0.020075317 | 1.697 | 25 | 15 |
| macromolecule glycosylation | GO:0043413 | 0.020155979 | 1.696 | 211 | 80 |
| positive regulation of epithelial to mesenchymal transition | GO:0010718 | 0.020220278 | 1.694 | 45 | 23 |
| T-helper cell differentiation | GO:0042093 | 0.020220278 | 1.694 | 45 | 23 |
| regulation of sodium ion transmembrane transporter activity | GO:2000649 | 0.020220278 | 1.694 | 45 | 23 |
| lipid metabolic process | GO:0006629 | 0.020429821 | 1.690 | 1193 | 378 |
| regulation of cell differentiation | GO:0045595 | 0.021173277 | 1.674 | 1388 | 435 |
| regulation of biomineral tissue development | GO:0070167 | 0.021380709 | 1.670 | 75 | 34 |
| regulation of biomineralization | GO:0110149 | 0.021380709 | 1.670 | 75 | 34 |
| epithelial cell differentiation involved in kidney development | GO:0035850 | 0.021881165 | 1.660 | 35 | 19 |
| neuron apoptotic process | GO:0051402 | 0.022252598 | 1.653 | 130 | 53 |
| regulation of action potential | GO:0098900 | 0.022601274 | 1.646 | 48 | 24 |
| cell fate commitment | GO:0045165 | 0.022601274 | 1.646 | 160 | 63 |
| muscle organ development | GO:0007517 | 0.022601274 | 1.646 | 212 | 80 |
| regulation of T cell activation | GO:0050863 | 0.023814946 | 1.623 | 256 | 94 |
| cartilage development | GO:0051216 | 0.023814946 | 1.623 | 104 | 44 |
| anion transport | GO:0006820 | 0.023911432 | 1.621 | 474 | 162 |
| regulation of neuron projection development | GO:0010975 | 0.024935068 | 1.603 | 326 | 116 |
| regulation of membrane potential | GO:0042391 | 0.024935068 | 1.603 | 326 | 116 |
| T cell differentiation involved in immune response | GO:0002292 | 0.024997337 | 1.602 | 51 | 25 |
| actin filament organization | GO:0007015 | 0.025285038 | 1.597 | 371 | 130 |
| metal ion homeostasis | GO:0055065 | 0.025285038 | 1.597 | 527 | 178 |
| activation of protein kinase activity | GO:0032147 | 0.025300502 | 1.597 | 266 | 97 |
| endocrine pancreas development | GO:0031018 | 0.025468922 | 1.594 | 28 | 16 |
| peptide hormone secretion | GO:0030072 | 0.025468922 | 1.594 | 167 | 65 |
| ossification | GO:0001503 | 0.025468922 | 1.594 | 241 | 89 |
| regulation of neural precursor cell proliferation | GO:2000177 | 0.025468922 | 1.594 | 38 | 20 |
| cardiac muscle cell action potential | GO:0086001 | 0.025468922 | 1.594 | 73 | 33 |
| regulation of cell shape | GO:0008360 | 0.025471713 | 1.594 | 125 | 51 |
| protein glycosylation | GO:0006486 | 0.025597296 | 1.592 | 210 | 79 |
| regulation of secretion | GO:0051046 | 0.025667801 | 1.591 | 613 | 204 |
| polysaccharide biosynthetic process | GO:0000271 | 0.025680332 | 1.590 | 62 | 29 |
| regulation of peptide hormone secretion | GO:0090276 | 0.025869774 | 1.587 | 143 | 57 |
| retinoid metabolic process | GO:0001523 | 0.02589129 | 1.587 | 93 | 40 |
| response to metal ion | GO:0010038 | 0.02605217 | 1.584 | 204 | 77 |
| bone mineralization | GO:0030282 | 0.026198489 | 1.582 | 76 | 34 |
| regulation of potassium ion transmembrane transport | GO:1901379 | 0.026198489 | 1.582 | 76 | 34 |
| T cell activation | GO:0042110 | 0.026328659 | 1.580 | 346 | 122 |
| regulation of anatomical structure morphogenesis | GO:0022603 | 0.027030304 | 1.568 | 938 | 301 |
| regulation of MAP kinase activity | GO:0043405 | 0.027328063 | 1.563 | 270 | 98 |
| cell junction assembly | GO:0034329 | 0.027328063 | 1.563 | 324 | 115 |
| terpenoid metabolic process | GO:0006721 | 0.027392242 | 1.562 | 102 | 43 |
| positive regulation of cell development | GO:0010720 | 0.027784665 | 1.556 | 353 | 124 |
| positive regulation of nervous system development | GO:0051962 | 0.027784665 | 1.556 | 321 | 114 |
| odontogenesis | GO:0042476 | 0.028680909 | 1.542 | 57 | 27 |
| proteoglycan biosynthetic process | GO:0030166 | 0.028680909 | 1.542 | 57 | 27 |

| | | | | | |
|---|---|---|---|---|---|
| camera-type eye morphogenesis | GO:0048593 | 0.028680909 | 1.542 | 57 | 27 |
| cellular polysaccharide biosynthetic process | GO:0033692 | 0.028680909 | 1.542 | 57 | 27 |
| regulation of neuron apoptotic process | GO:0043523 | 0.029121783 | 1.536 | 117 | 48 |
| head development | GO:0060322 | 0.029225081 | 1.534 | 360 | 126 |
| regulation of binding | GO:0051098 | 0.030948474 | 1.509 | 268 | 97 |
| regulation of bone remodeling | GO:0046850 | 0.031118018 | 1.507 | 31 | 17 |
| glycosylation | GO:0070085 | 0.03120179 | 1.506 | 221 | 82 |
| canonical Wnt signaling pathway | GO:0060070 | 0.032666492 | 1.486 | 297 | 106 |
| positive regulation of T cell proliferation | GO:0042102 | 0.032843667 | 1.484 | 77 | 34 |
| phenol-containing compound metabolic process | GO:0018958 | 0.032903742 | 1.483 | 63 | 29 |
| extracellular matrix organization | GO:0030198 | 0.033224532 | 1.479 | 294 | 105 |
| skeletal system morphogenesis | GO:0048705 | 0.033717567 | 1.472 | 106 | 44 |
| epithelial tube morphogenesis | GO:0060562 | 0.034114043 | 1.467 | 163 | 63 |
| regulation of protein tyrosine kinase activity | GO:0061097 | 0.034869776 | 1.458 | 66 | 30 |
| inorganic ion import across plasma membrane | GO:0099587 | 0.035055726 | 1.455 | 83 | 36 |
| inorganic cation import across plasma membrane | GO:0098659 | 0.035055726 | 1.455 | 83 | 36 |
| cardiac muscle tissue development | GO:0048738 | 0.035373998 | 1.451 | 136 | 54 |
| developmental growth involved in morphogenesis | GO:0060560 | 0.035373998 | 1.451 | 130 | 52 |
| membrane repolarization | GO:0086009 | 0.036114783 | 1.442 | 47 | 23 |
| positive regulation of signal transduction | GO:0009967 | 0.036267803 | 1.440 | 1393 | 433 |
| neural precursor cell proliferation | GO:0061351 | 0.036267803 | 1.440 | 69 | 31 |
| extracellular structure organization | GO:0043062 | 0.036568602 | 1.437 | 295 | 105 |
| regulation of bone mineralization | GO:0030500 | 0.037190089 | 1.430 | 58 | 27 |
| ventricular septum morphogenesis | GO:0060412 | 0.037190089 | 1.430 | 29 | 16 |
| positive regulation of kidney development | GO:0090184 | 0.037190089 | 1.430 | 29 | 16 |
| regulation of MAPK cascade | GO:0043408 | 0.037258022 | 1.429 | 612 | 202 |
| diterpenoid metabolic process | GO:0016101 | 0.037404125 | 1.427 | 95 | 40 |
| autophagy of mitochondrion | GO:0000422 | 0.039090521 | 1.408 | 61 | 28 |
| mitochondrion disassembly | GO:0061726 | 0.039090521 | 1.408 | 61 | 28 |
| positive regulation of small molecule metabolic process | GO:0062013 | 0.039139308 | 1.407 | 104 | 43 |
| developmental growth | GO:0048589 | 0.03916806 | 1.407 | 299 | 106 |
| cellular response to organonitrogen compound | GO:0071417 | 0.039756651 | 1.401 | 448 | 152 |
| cell-cell junction assembly | GO:0007043 | 0.040588611 | 1.392 | 119 | 48 |
| positive regulation of cell motility | GO:2000147 | 0.041074618 | 1.386 | 491 | 165 |
| tissue morphogenesis | GO:0048729 | 0.041122177 | 1.386 | 419 | 143 |
| cell communication involved in cardiac conduction | GO:0086065 | 0.041695307 | 1.380 | 53 | 25 |
| response to organic cyclic compound | GO:0014070 | 0.04191115 | 1.378 | 452 | 153 |
| mesenchyme morphogenesis | GO:0072132 | 0.04191115 | 1.378 | 37 | 19 |
| regulation of cardiac muscle cell contraction | GO:0086004 | 0.04191115 | 1.378 | 37 | 19 |
| activation of adenylate cyclase activity | GO:0007190 | 0.04191115 | 1.378 | 37 | 19 |
| positive regulation of cell migration | GO:0030335 | 0.042941511 | 1.367 | 472 | 159 |
| positive regulation of neuron differentiation | GO:0045666 | 0.042941511 | 1.367 | 227 | 83 |
| regulation of potassium ion transport | GO:0043266 | 0.043636602 | 1.360 | 90 | 38 |
| lymphocyte proliferation | GO:0046651 | 0.043733761 | 1.359 | 199 | 74 |
| actomyosin structure organization | GO:0031032 | 0.043796193 | 1.359 | 165 | 63 |
| positive regulation of cytosolic calcium ion concentration | GO:0007204 | 0.044080762 | 1.356 | 262 | 94 |
| synaptic signaling | GO:0099536 | 0.044643108 | 1.350 | 512 | 171 |
| cytosolic calcium ion transport | GO:0060401 | 0.045006722 | 1.347 | 156 | 60 |
| apoptotic process | GO:0006915 | 0.045127411 | 1.346 | 1443 | 446 |
| synapse organization | GO:0050808 | 0.046572486 | 1.332 | 253 | 91 |
| bone development | GO:0060348 | 0.046719514 | 1.331 | 111 | 45 |
| respiratory system development | GO:0060541 | 0.046719514 | 1.331 | 59 | 27 |
| T cell proliferation | GO:0042098 | 0.046719514 | 1.331 | 135 | 53 |
| ventricular septum development | GO:0003281 | 0.046773135 | 1.330 | 40 | 20 |
| regulation of T cell proliferation | GO:0042129 | 0.046999315 | 1.328 | 123 | 49 |
| homophilic cell adhesion via plasma membrane adhesion molecules | GO:0007156 | 0.047046484 | 1.327 | 76 | 33 |
| enzyme linked receptor protein signaling pathway | GO:0007167 | 0.048217609 | 1.317 | 915 | 291 |
| positive regulation of locomotion | GO:0040017 | 0.048437036 | 1.315 | 500 | 167 |
| cellular polysaccharide metabolic process | GO:0044264 | 0.048437036 | 1.315 | 79 | 34 |
| monocarboxylic acid metabolic process | GO:0032787 | 0.048793346 | 1.312 | 553 | 183 |

| | | | | | |
|---|---|---|---|---|---|
| oxoacid metabolic process | GO:0043436 | 0.049774948 | 1.303 | 997 | 315 |
| glycogen biosynthetic process | GO:0005978 | 0.049918671 | 1.302 | 35 | 18 |
| glucan biosynthetic process | GO:0009250 | 0.049918671 | 1.302 | 35 | 18 |
| ventricular cardiac muscle tissue development | GO:0003229 | 0.049918671 | 1.302 | 35 | 18 |

**Supplementary Table 3, Gene ontology analysis of genes associated with H3K27ac decreased peaks, Related to Figure 3D**

| GO term name | GO term ID | FDR | -log10(FDR) | GO term size | Intersection size |
|---|---|---|---|---|---|
| neuron differentiation | GO:0030182 | 1.12841E-05 | 4.947534985 | 904 | 171 |
| mesenchyme development | GO:0060485 | 1.52669E-05 | 4.816249128 | 208 | 56 |
| embryo development | GO:0009790 | 2.61539E-05 | 4.582463164 | 463 | 99 |
| generation of neurons | GO:0048699 | 2.76546E-05 | 4.558231929 | 1013 | 183 |
| mesenchymal cell differentiation | GO:0048762 | 3.31435E-05 | 4.479601216 | 175 | 48 |
| smooth muscle contraction | GO:0006939 | 5.9913E-05 | 4.222478654 | 62 | 24 |
| neuron development | GO:0048666 | 6.81713E-05 | 4.166398258 | 732 | 138 |
| neurogenesis | GO:0022008 | 6.84206E-05 | 4.164812985 | 1069 | 188 |
| cell junction organization | GO:0034330 | 0.000104782 | 3.979711533 | 487 | 99 |
| neuron projection development | GO:0031175 | 0.000147206 | 3.832073108 | 672 | 127 |
| circulatory system development | GO:0072359 | 0.000162801 | 3.788344179 | 814 | 148 |
| tube morphogenesis | GO:0035239 | 0.000217042 | 3.663455504 | 634 | 120 |
| positive regulation of multicellular organismal process | GO:0051240 | 0.000238949 | 3.621694945 | 1375 | 228 |
| regulation of nervous system development | GO:0051960 | 0.000251373 | 3.599682202 | 578 | 111 |
| tube development | GO:0035295 | 0.000251373 | 3.599682202 | 696 | 129 |
| nephron epithelium development | GO:0072009 | 0.000317761 | 3.497898776 | 65 | 23 |
| kidney epithelium development | GO:0072073 | 0.000428085 | 3.368469841 | 71 | 24 |
| positive regulation of molecular function | GO:0044093 | 0.000454608 | 3.342363031 | 1342 | 221 |
| regulation of smooth muscle contraction | GO:0006940 | 0.000468945 | 3.328878138 | 33 | 15 |
| embryonic morphogenesis | GO:0048598 | 0.000521363 | 3.282859929 | 305 | 66 |
| cell development | GO:0048468 | 0.000527613 | 3.277684774 | 1369 | 224 |
| animal organ morphogenesis | GO:0009887 | 0.000527613 | 3.277684774 | 624 | 116 |
| cellular response to endogenous stimulus | GO:0071495 | 0.00058667 | 3.231606397 | 1069 | 181 |
| tissue morphogenesis | GO:0048729 | 0.000598968 | 3.222596393 | 419 | 84 |
| cell junction assembly | GO:0034329 | 0.000892506 | 3.049388985 | 324 | 68 |
| response to endogenous stimulus | GO:0009719 | 0.000892506 | 3.049388985 | 1177 | 195 |
| positive regulation of transferase activity | GO:0051347 | 0.000893367 | 3.048969963 | 574 | 107 |
| response to oxygen-containing compound | GO:1901700 | 0.00092269 | 3.034944219 | 1038 | 175 |
| blood vessel development | GO:0001568 | 0.00092269 | 3.034944219 | 555 | 104 |
| regulation of transferase activity | GO:0051338 | 0.001011191 | 2.995166815 | 839 | 146 |
| enzyme linked receptor protein signaling pathway | GO:0007167 | 0.001014834 | 2.99360495 | 915 | 157 |
| polysaccharide metabolic process | GO:0005976 | 0.001058352 | 2.975369814 | 86 | 26 |
| cardiovascular system development | GO:0072358 | 0.001253811 | 2.901767886 | 586 | 108 |
| positive regulation of signal transduction | GO:0009967 | 0.001324012 | 2.87810796 | 1393 | 224 |
| positive regulation of developmental process | GO:0051094 | 0.001513662 | 2.819971089 | 1034 | 173 |
| positive regulation of protein kinase activity | GO:0045860 | 0.001529632 | 2.815413009 | 432 | 84 |
| blood vessel morphogenesis | GO:0048514 | 0.001529632 | 2.815413009 | 523 | 98 |
| response to organonitrogen compound | GO:0010243 | 0.001598729 | 2.796225273 | 637 | 115 |
| Wnt signaling pathway, calcium modulating pathway | GO:0007223 | 0.00161147 | 2.792777655 | 37 | 15 |
| cellular polysaccharide metabolic process | GO:0044264 | 0.00186383 | 2.729593815 | 79 | 24 |
| cell morphogenesis | GO:0000902 | 0.001978318 | 2.703703893 | 756 | 132 |
| embryonic limb morphogenesis | GO:0030326 | 0.002196991 | 2.65817169 | 38 | 15 |
| embryonic appendage morphogenesis | GO:0035113 | 0.002196991 | 2.65817169 | 38 | 15 |
| vasculature development | GO:0001944 | 0.002280366 | 2.641995518 | 583 | 106 |

| | | | | | |
|---|---|---|---|---|---|
| regulation of neuron differentiation | GO:0045664 | 0.002501612 | 2.601780034 | 407 | 79 |
| nephron development | GO:0072006 | 0.002549485 | 2.593547485 | 91 | 26 |
| epithelial to mesenchymal transition | GO:0001837 | 0.00258311 | 2.58785715 | 123 | 32 |
| cation transport | GO:0006812 | 0.002794446 | 2.553704314 | 881 | 149 |
| myoblast differentiation | GO:0045445 | 0.003365213 | 2.472987506 | 53 | 18 |
| cellular response to oxygen-containing compound | GO:1901701 | 0.003377078 | 2.4714589 | 843 | 143 |
| positive regulation of kinase activity | GO:0033674 | 0.003486391 | 2.457623929 | 497 | 92 |
| cellular response to hormone stimulus | GO:0032870 | 0.003559809 | 2.448573294 | 471 | 88 |
| energy reserve metabolic process | GO:0006112 | 0.003671746 | 2.435127414 | 63 | 20 |
| positive regulation of catalytic activity | GO:0043085 | 0.003704526 | 2.431267352 | 1049 | 172 |
| regulation of kinase activity | GO:0043549 | 0.004296038 | 2.366931857 | 738 | 127 |
| regulation of neurogenesis | GO:0050767 | 0.004296038 | 2.366931857 | 514 | 94 |
| cellular response to organonitrogen compound | GO:0071417 | 0.004296038 | 2.366931857 | 448 | 84 |
| regulation of cell junction assembly | GO:1901888 | 0.004296038 | 2.366931857 | 138 | 34 |
| neuron projection morphogenesis | GO:0048812 | 0.004315862 | 2.364932428 | 468 | 87 |
| renal system development | GO:0072001 | 0.004549783 | 2.342009354 | 150 | 36 |
| nephron tubule development | GO:0072080 | 0.004654879 | 2.332091639 | 50 | 17 |
| regulation of ion transport | GO:0043269 | 0.005073461 | 2.29469565 | 464 | 86 |
| synapse organization | GO:0050808 | 0.005345022 | 2.272050472 | 253 | 53 |
| kidney development | GO:0001822 | 0.005574742 | 2.253775245 | 146 | 35 |
| dorsal/ventral pattern formation | GO:0009953 | 0.005821574 | 2.234959577 | 33 | 13 |
| renal tubule development | GO:0061326 | 0.005876379 | 2.230890234 | 51 | 17 |
| response to nitrogen compound | GO:1901698 | 0.005876503 | 2.230881005 | 717 | 123 |
| carbohydrate derivative biosynthetic process | GO:1901137 | 0.006063513 | 2.217275656 | 581 | 103 |
| regulation of protein kinase activity | GO:0045859 | 0.006063513 | 2.217275656 | 649 | 113 |
| ureteric bud development | GO:0001657 | 0.006071526 | 2.216702164 | 42 | 15 |
| cell part morphogenesis | GO:0032990 | 0.006134128 | 2.212247191 | 494 | 90 |
| developmental growth | GO:0048589 | 0.006271781 | 2.202609139 | 299 | 60 |
| urogenital system development | GO:0001655 | 0.006396595 | 2.194051144 | 159 | 37 |
| regulation of protein phosphorylation | GO:0001932 | 0.007088898 | 2.149421269 | 1238 | 196 |
| mesonephric tubule morphogenesis | GO:0072171 | 0.007492981 | 2.125345379 | 34 | 13 |
| embryonic organ development | GO:0048568 | 0.007492981 | 2.125345379 | 190 | 42 |
| mesonephric epithelium development | GO:0072163 | 0.007593919 | 2.119534023 | 43 | 15 |
| mesonephric tubule development | GO:0072164 | 0.007593919 | 2.119534023 | 43 | 15 |
| morphogenesis | GO:0048646 | 0.007756367 | 2.110341669 | 807 | 135 |
| regulation of phosphorylation | GO:0042325 | 0.008098349 | 2.091603485 | 1380 | 215 |
| limb morphogenesis | GO:0035108 | 0.00817728 | 2.087391122 | 48 | 16 |
| appendage morphogenesis | GO:0035107 | 0.00817728 | 2.087391122 | 48 | 16 |
| cellular response to alcohol | GO:0097306 | 0.008534828 | 2.068805221 | 53 | 17 |
| morphogenesis | GO:0120039 | 0.008534828 | 2.068805221 | 480 | 87 |
| cell differentiation involved in kidney development | GO:0061005 | 0.008534828 | 2.068805221 | 39 | 14 |
| regulation of anatomical structure morphogenesis | GO:0022603 | 0.00928504 | 2.032216231 | 938 | 153 |
| cellular response to nitrogen compound | GO:1901699 | 0.009557456 | 2.019657697 | 502 | 90 |
| axon guidance | GO:0007411 | 0.009562316 | 2.019436923 | 211 | 45 |
| muscle contraction | GO:0006936 | 0.009623258 | 2.016677892 | 286 | 57 |
| localization of cell | GO:0051674 | 0.009623258 | 2.016677892 | 1474 | 227 |
| cardiac chamber development | GO:0003205 | 0.009623258 | 2.016677892 | 95 | 25 |
| cell motility | GO:0048870 | 0.009623258 | 2.016677892 | 1474 | 227 |
| cardiac chamber morphogenesis | GO:0003206 | 0.009623258 | 2.016677892 | 79 | 22 |
| cell projection morphogenesis | GO:0048858 | 0.009646735 | 2.015619645 | 483 | 87 |
| actin filament bundle assembly | GO:0051017 | 0.009646735 | 2.015619645 | 140 | 33 |
| cell morphogenesis involved in differentiation | GO:0000904 | 0.009646735 | 2.015619645 | 530 | 94 |
| metal ion transport | GO:0030001 | 0.009694611 | 2.013469595 | 688 | 117 |
| cell morphogenesis involved in neuron differentiation | GO:0048667 | 0.009694611 | 2.013469595 | 410 | 76 |
| neuron projection guidance | GO:0097485 | 0.009786486 | 2.009373213 | 212 | 45 |
| regulation of cell differentiation | GO:0045595 | 0.009786486 | 2.009373213 | 1388 | 215 |
| glycogen metabolic process | GO:0005977 | 0.009786486 | 2.009373213 | 54 | 17 |
| ion transport | GO:0006811 | 0.009828043 | 2.007532945 | 1330 | 207 |
| regulation of neuron projection development | GO:0010975 | 0.010087034 | 1.996236517 | 326 | 63 |

| | | | | | |
|---|---|---|---|---|---|
| nephron epithelium morphogenesis | GO:0072088 | 0.010087034 | 1.996236517 | 40 | 14 |
| nephron morphogenesis | GO:0072028 | 0.010087034 | 1.996236517 | 40 | 14 |
| cellular component morphogenesis | GO:0032989 | 0.010276339 | 1.988161584 | 566 | 99 |
| response to hormone | GO:0009725 | 0.011378905 | 1.943899513 | 561 | 98 |
| cell migration | GO:0016477 | 0.011448243 | 1.94126117 | 1342 | 208 |
| regulation of gastrulation | GO:0010470 | 0.01149104 | 1.939640674 | 36 | 13 |
| cellular glucan metabolic process | GO:0006073 | 0.011676587 | 1.932684072 | 55 | 17 |
| glucan metabolic process | GO:0044042 | 0.011676587 | 1.932684072 | 55 | 17 |
| glycoprotein biosynthetic process | GO:0009101 | 0.011959499 | 1.922287025 | 290 | 57 |
| renal tubule morphogenesis | GO:0061333 | 0.012651576 | 1.897855378 | 41 | 14 |
| morphogenesis of an epithelium | GO:0002009 | 0.012651576 | 1.897855378 | 323 | 62 |
| vasoconstriction | GO:0042310 | 0.012651576 | 1.897855378 | 41 | 14 |
| actin filament bundle organization | GO:0061572 | 0.012651576 | 1.897855378 | 143 | 33 |
| ureteric bud morphogenesis | GO:0060675 | 0.012688576 | 1.896587119 | 32 | 12 |
| regulation of transport | GO:0051049 | 0.012851935 | 1.891031486 | 1478 | 226 |
| protein localization to plasma membrane | GO:0072659 | 0.013910613 | 1.856653717 | 235 | 48 |
| regulation of cell-substrate junction organization | GO:0150116 | 0.014120238 | 1.850157991 | 56 | 17 |
| glycoprotein metabolic process | GO:0009100 | 0.014889835 | 1.827110113 | 358 | 67 |
| stress fiber assembly | GO:0043149 | 0.015594334 | 1.807033176 | 88 | 23 |
| contractile actin filament bundle assembly | GO:0030038 | 0.015594334 | 1.807033176 | 88 | 23 |
| epithelial tube morphogenesis | GO:0060562 | 0.016137843 | 1.792154517 | 163 | 36 |
| regulation of BMP signaling pathway | GO:0030510 | 0.01636568 | 1.786065946 | 67 | 19 |
| mesonephros development | GO:0001823 | 0.016734247 | 1.77639383 | 47 | 15 |
| regulation of focal adhesion assembly | GO:0051893 | 0.016983421 | 1.769974821 | 52 | 16 |
| regulation of cell-substrate junction assembly | GO:0090109 | 0.016983421 | 1.769974821 | 52 | 16 |
| positive regulation of catabolic process | GO:0009896 | 0.017010133 | 1.769292296 | 380 | 70 |
| regulation of cell development | GO:0060284 | 0.017392292 | 1.759643172 | 618 | 105 |
| positive regulation of cell differentiation | GO:0045597 | 0.017657939 | 1.753059981 | 723 | 120 |
| proteoglycan metabolic process | GO:0006029 | 0.018271492 | 1.738225998 | 73 | 20 |
| nephron tubule morphogenesis | GO:0072078 | 0.018412871 | 1.734878499 | 38 | 13 |
| heart development | GO:0007507 | 0.018691694 | 1.728351343 | 342 | 64 |
| cell-substrate junction organization | GO:0150115 | 0.018870785 | 1.724210029 | 84 | 22 |
| peptidyl-threonine modification | GO:0018210 | 0.019092056 | 1.719147301 | 112 | 27 |
| actomyosin structure organization | GO:0031032 | 0.01919226 | 1.716873884 | 165 | 36 |
| positive regulation of muscle contraction | GO:0045933 | 0.019991876 | 1.699146456 | 25 | 10 |
| positive regulation of protein metabolic process | GO:0051247 | 0.019991876 | 1.699146456 | 1485 | 225 |
| kidney morphogenesis | GO:0060993 | 0.019997046 | 1.699034145 | 48 | 15 |
| second-messenger-mediated signaling | GO:0019932 | 0.020495112 | 1.688349703 | 370 | 68 |
| activation of protein kinase activity | GO:0032147 | 0.020981049 | 1.678172794 | 266 | 52 |
| regulation of embryonic development | GO:0045995 | 0.021453279 | 1.668506314 | 85 | 22 |
| response to peptide hormone | GO:0043434 | 0.021573421 | 1.666080978 | 260 | 51 |
| signaling pathway | GO:0007178 | 0.022899379 | 1.640176302 | 280 | 54 |
| cellular response to peptide hormone stimulus | GO:0071375 | 0.022935913 | 1.639483965 | 229 | 46 |
| taxis | GO:0042330 | 0.022935913 | 1.639483965 | 514 | 89 |
| chemotaxis | GO:0006935 | 0.022935913 | 1.639483965 | 514 | 89 |
| positive regulation of RNA metabolic process | GO:0051254 | 0.02344449 | 1.629959204 | 1395 | 212 |
| branching involved in ureteric bud morphogenesis | GO:0001658 | 0.023495195 | 1.629020943 | 30 | 11 |
| glomerulus development | GO:0032835 | 0.023495195 | 1.629020943 | 44 | 14 |
| cell-cell signaling | GO:0007267 | 0.0237114 | 1.625042795 | 1499 | 226 |
| ameboidal-type cell migration | GO:0001667 | 0.024482031 | 1.611152565 | 400 | 72 |
| muscle system process | GO:0003012 | 0.025246512 | 1.59779862 | 354 | 65 |
| positive regulation of nervous system development | GO:0051962 | 0.025246512 | 1.59779862 | 321 | 60 |
| appendage development | GO:0048736 | 0.025246512 | 1.59779862 | 70 | 19 |
| actin filament-based process | GO:0030029 | 0.025246512 | 1.59779862 | 690 | 114 |
| limb development | GO:0060173 | 0.025246512 | 1.59779862 | 70 | 19 |
| positive regulation of intracellular signal transduction | GO:1902533 | 0.025447962 | 1.594346984 | 889 | 142 |
| development | GO:0035850 | 0.025911869 | 1.586501267 | 35 | 12 |
| cell projection organization | GO:0030030 | 0.026988044 | 1.568828586 | 1180 | 182 |
| positive regulation of calcium ion transport | GO:0051928 | 0.027109308 | 1.566881564 | 87 | 22 |
| response to BMP | GO:0071772 | 0.027109308 | 1.566881564 | 133 | 30 |
| cellular response to BMP stimulus | GO:0071773 | 0.027109308 | 1.566881564 | 133 | 30 |

| | | | | | |
|---|---|---|---|---|---|
| cellular response to external stimulus | GO:0071496 | 0.027109308 | 1.566881564 | 200 | 41 |
| negative regulation of ossification | GO:0030279 | 0.027490129 | 1.560823219 | 55 | 16 |
| locomotory behavior | GO:0007626 | 0.027753438 | 1.556683203 | 45 | 14 |
| cell-cell signaling by wnt | GO:0198738 | 0.028459915 | 1.5457664 | 437 | 77 |
| protein localization to cell periphery | GO:1990778 | 0.028602218 | 1.543600281 | 271 | 52 |
| integrin-mediated signaling pathway | GO:0007229 | 0.028704594 | 1.542048589 | 82 | 21 |
| actin cytoskeleton organization | GO:0030036 | 0.031003006 | 1.508596196 | 597 | 100 |
| serine/threonine kinase signaling pathway | GO:0090092 | 0.032459605 | 1.488656769 | 196 | 40 |
| positive regulation of ion transport | GO:0043270 | 0.033049535 | 1.480834648 | 190 | 39 |
| Wnt signaling pathway | GO:0016055 | 0.033170098 | 1.479253242 | 433 | 76 |
| negative regulation of blood vessel diameter | GO:0097756 | 0.033170098 | 1.479253242 | 46 | 14 |
| plasma membrane bounded cell projection organization | GO:0120036 | 0.033170098 | 1.479253242 | 1165 | 179 |
| cellular response to ketone | GO:1901655 | 0.033170098 | 1.479253242 | 41 | 13 |
| cell signaling | GO:1905114 | 0.034311007 | 1.464566539 | 482 | 83 |
| focal adhesion assembly | GO:0048041 | 0.034311007 | 1.464566539 | 67 | 18 |
| positive regulation of neurogenesis | GO:0050769 | 0.034311007 | 1.464566539 | 287 | 54 |
| receptor internalization | GO:0031623 | 0.034311007 | 1.464566539 | 89 | 22 |
| regulation of cell migration | GO:0030334 | 0.035203365 | 1.453415823 | 827 | 132 |
| polysaccharide biosynthetic process | GO:0000271 | 0.035871358 | 1.445252176 | 62 | 17 |
| regulation of small GTPase mediated signal transduction | GO:0051056 | 0.036586246 | 1.436682154 | 249 | 48 |
| proteoglycan biosynthetic process | GO:0030166 | 0.037615157 | 1.424637116 | 57 | 16 |
| purine-containing compound biosynthetic process | GO:0072522 | 0.03787238 | 1.421677402 | 161 | 34 |
| renal system process | GO:0003014 | 0.038468092 | 1.414899355 | 90 | 22 |
| cardiac ventricle morphogenesis | GO:0003208 | 0.038488055 | 1.414674035 | 52 | 15 |
| embryonic organ morphogenesis | GO:0048562 | 0.038639355 | 1.412970133 | 131 | 29 |
| response to ketone | GO:1901654 | 0.038861643 | 1.410478844 | 68 | 18 |
| axis specification | GO:0009798 | 0.038861643 | 1.410478844 | 47 | 14 |
| cell-substrate junction assembly | GO:0007044 | 0.038877736 | 1.410299029 | 79 | 20 |
| signaling pathway | GO:0007169 | 0.039850228 | 1.399569194 | 646 | 106 |
| positive regulation of protein phosphorylation | GO:0001934 | 0.039850228 | 1.399569194 | 867 | 137 |
| import into cell | GO:0098657 | 0.040520983 | 1.392320031 | 162 | 34 |
| regulation of actomyosin structure organization | GO:0110020 | 0.040806995 | 1.389265386 | 85 | 21 |
| regulation of actin filament bundle assembly | GO:0032231 | 0.040806995 | 1.389265386 | 85 | 21 |
| heparan sulfate proteoglycan metabolic process | GO:0030201 | 0.040999192 | 1.387224697 | 28 | 10 |
| purine nucleotide biosynthetic process | GO:0006164 | 0.041231526 | 1.384770588 | 150 | 32 |
| regulation of muscle contraction | GO:0006937 | 0.042021133 | 1.376532243 | 126 | 28 |
| cell-substrate adhesion | GO:0031589 | 0.042125314 | 1.375456851 | 271 | 51 |
| actin filament organization | GO:0007015 | 0.042666491 | 1.369913077 | 371 | 66 |
| positive regulation of embryonic development | GO:0040019 | 0.044768467 | 1.349027781 | 33 | 11 |
| growth | GO:0040007 | 0.045652676 | 1.340533757 | 530 | 89 |
| regulation of synapse structure or activity | GO:0050803 | 0.046722859 | 1.330470588 | 133 | 29 |
| cardiac ventricle development | GO:0003231 | 0.048099426 | 1.317860109 | 75 | 19 |
| signaling pathway | GO:0007200 | 0.048291327 | 1.316130864 | 92 | 22 |
| cell-matrix adhesion | GO:0007160 | 0.049183062 | 1.308184439 | 183 | 37 |
| regionalization | GO:0003002 | 0.049183062 | 1.308184439 | 158 | 33 |
